# A draft genome of the ascomycotal fungal species *Pseudopithomyces maydicus* (family *Didymosphaeriaceae*)

**DOI:** 10.1101/2022.11.26.518055

**Authors:** Krithika Arumugam, Sherilyn Ho, Irina Bessarab, Falicia Q. Y. Goh, Mindia A. S. Haryono, Ezequiel Santillan, Stefan Wuertz, Yvonne Chow, Rohan B. H. Williams

## Abstract

We report a draft genome of the ascomycotal fungal species Pseudopithomyces maydicus (isolate name SBW1) obtained using a culture isolate from brewery wastewater. From a 22 contig assembly, we predict 13502 protein coding gene models, of which 4389 (32.5%) were annotated to KEGG Orthology and identify 39 biosynthetic gene clusters.

## Announcement

*Pseudopithomyces maydicus* is a fungal species within phylum *Ascomycota* (order *Pleosporales*), previously named *Pithomyces maydicus* and recently renamed with the introduction of genus *Pseudopithomyces* into family *Didymosphaeriaceae* (Ariyawansa *et al.*, 2015). Members of this species have been identified as potential human pathogens, based on identification from several clinical specimens (da Cunha *et al*., 2014), and natural products from this species have recently been characterized, some of which hold antimicrobial activity (Ningsih *et al*., 2021). Neither *Pseudopithomyces maydicus* nor any member of genus *Pseudopithomyces* have a reference or draft genome available, with the most relevant related genome sequence data being 11 draft genomes collected from other genera in family *Didymosphaeriaceae* (NCBI Assembly, accessed 2022/03/31).

We report a draft genome of *Pseudopithomyces maydicus* (isolate name SBW1) obtained from a culture isolate from brewery wastewater in Singapore. Genomic characterization of microbes isolated from food-processing wastewater can provide foundational data for biotechnological applications in the circular economy, such as the production of microbial protein (Vethathirri *et al*. 2021).

We obtained an isolate from brewery wastewater by culturing on solid Yeast Extract– Peptone–Dextrose (YPD) agar media at 30°C for 2 days. The colony was isolated and streaked out on a new YPD plate. Taxonomic classification was made via Sanger sequencing of the D1/D2 domain of the large-subunit (28S) ribosomal DNA (NCBI BLASTN webserver against the NCBI nr/nt database in megablast mode with top hit annotated to a *P. maydicus* partial 28S sequence, MF919633.1, at 99% identity; see **Supplementary Figure 1** for alignment; https://blast.ncbi.nlm.nih.gov/Blast.cgi, executed on 06/31/2022). Genomic DNA was extracted using liquid nitrogen and mechanical grinding of the fungal culture followed by application of the Qiagen DNeasy PowerSoil Pro Kit. 1.5μg of input DNA was subjected to shearing (Megaruptor3, Diagenode Inc, Denville, NJ, USA; operated for 20kb target at speed 35) and then 800ng sheared DNA was used to construct a sequencing library using the SQK-LSK109 ligation sequencing kit (Oxord Nanopore Technologies Ltd, Oxford, UK), barcoded using the EXP-NDB104 native barcoding kit (Oxford Nanopore Technologies; barcode 10). Following construction, 200 ng of the library was sequenced on a GridION instrument (Oxford Nanopore Technologies; release 21.05.20) for 72 hours. Basecalling was performed using Guppy 5.0.13 (Oxford Nanopore Technologies) in SUP mode.

The run generated 233,209 raw reads from the cognate barcode (232,644 reads following the application of Porechop version 0.2.4 using default parameters except --discard_middle, -t 20) (Porechop, 2018), comprising a total of 1.53 Gbp of sequence. Genome assembly was performed using Flye version 2.9 (using parameters --nano-hq, -t 44) (Kolmogorov *et al*., 2019). A total of 36 contigs were obtained with a total sequence length of 39,781,613 bp.

Based on visualization of per-contig GC content, mean coverage and length, we determined a working draft of the genome to be comprised of 22 contigs (mean length 1,792,464 bp, range: 81,297-3,886,452 bp), with an N50 of 2,331,148 bp and a total sequence length of 39,434,212 bp. The mean GC content was 0.5 (range: 0.48-0.51) and mean coverage was 35 (range: 33-36). (**Supplementary Figure 2** and **Supplementary Data File 1**). One high coverage circular contig (37,662 bp; contig 40) was aligned to the mitochondrial genome of the closely related species *Pseudopithomyces chartarum* (97% nucleotide identity with 80% query coverage to Genbank KY792993.1; annotated to *Pithomyces chartarum*, but refer Ariyawansa *et al*., 2015, for discussion of reassignment to genus *Pseudopithomyces*). The remaining 13 contigs held substantially lower mean GC content values than those observed from the draft genome. These 13 contigs accounted for 309,739 bp of sequence (mean: 23,826 bp, median: 8,979 bp, range: 2897-126,848 bp), and were considered to arise from potentially mis-assembled telomeric or repeat sequences and/or sequences from intra-plate contaminants (**Supplementary Figure 2)**.

The quality of the draft genome was examined using gene-level analysis with BUSCO package (v5.3.2; exeuted in genome mode using the lineage dataset for *Pleosporales;*pleosporales_odb10) (Manni *et al*., 2021). BUSCO identified 5637 complete marker genes (of 6641 searched), of which 5610 were complete and single copy, 27 complete and duplicated, 231 were fragmented and 733 missing (BUSCO notation: C:84.9% [S:84.5%, D:0.4%], F:3.5%, M:11.6%, *n*:6641).

From the draft genome, five 18S SSU-rRNA genes were predicted with RNAmmer (Lagesen *et al*., 2007) all of which annotated to order *Pleosporales* using the SILVA Alignment, Classification and Tree Service (ACT) (Pruesse *et al*., 2012) (**Supplementary Data File 2**). Three 28S LSU-rRNA genes were recovered from the assembly, to which the partial 28S sequence obtained above aligned with 99% identity (BLASTN, run with default settings; Camacho *et al*., 2009; alignments provided in **Supplementary Data File 3**). We note these ribosomal gene numbers are less than expected based on recent estimates (Lofgren *et al*., 2019) made within ensembles of complete fungal genomes and may be related to limited reconstructability of closely related DNA fragments harbouring ribosomal operons. Further taxonomic analysis was undertaken using sourmash (Pierce *et al*., 2019), comparing the draft genome against all 9563 fungal genomes downloadable from the NCBI (2022/03/29), with the 7 most similar genomes observed to be members of family *Didymosphaeriaceae* (**Supplementary Data File 4**). Collectively these results are consistent with the original taxonomic assignment, within the limits of fungal genome availability for closely related fungal groups.

To gain some insight into possible chromosomal structures, all assembled contig sequences were searched for more than three or more repeat units of exact matches to CCCTAA and TTAGGG motifs (Rahnama et al., 2021) using the find function in a text editor to identify possible telomeric regions. Of the 22 contigs in the draft genome, 4 contigs have multiple repeat units of CCCTAA and 8 contigs have multiple repeat units of TTAGGG at the 5’ and 3’ end respectively. 3 contigs have both CCCTAA and TTAGGG at the 5’ and 3’ end (**Supplementary Data File 1)**suggesting these sequences may represent distinct chromosomes.

An initial catalogue of gene models, predicted using GeneMark-ES (run with --fungus flag set) (Ter-Hovhannisyan *et al*., 2008), was comprised of 13502 protein coding genes, of which 4389 (32.5%) were annotated to one or more KEGG Orthology identifiers using BlastKOALA (Kanehisa *et al*., 2016; **Supplementary Data File 5**). A total of 162 tRNA encoding genes were predicted using EuFindtRNA search algorithm in tRNAscan-SE (version 2.0, running default parameters) (Lowe and Eddy, 1997).

Recently Ningsih *et al*. (2021) isolated and characterized seven natural product compounds from an isolate of *P. maydicus* isolated from marine bryozoan (genus *Schizoporella)*. To identify potentially-related biosynthetic gene clusters, we analysed the recovered genome sequence using the biosynthetic gene cluster (BGC) finder antiSMASH6 (Blin *et al*., 2021). In total we identified 39 BGCs, comprised of 19 Type 1 polyketide synthase (T1PKS) clusters and 14 non-ribosomal peptide synthetase (NRPS) or NRPS-like clusters, 4 terpene encoding clusters and two indole encoding clusters (**Supplementary Results**). Further examination of the relationships between these detected BGCs and the compounds defined by Ningsih *et al*. (2021) may provide insight into the relevant biosynthesis pathways for these specalised metabolites, as recently highlighted by Louwen and van der Hooft (2021).

## Supporting information

Supplementary Results

## Acknowledgements

We thank colleagues at the Integrated Genomics Platform at the Genome Institute of Singapore for performing DNA sequencing and basecalling. This research was supported by the Singapore National Research Foundation (NRF) and Ministry of Education under the Research Centre of Excellence Program and the NRF Competitive Research Programme (NRF-CRP21-2018-0006) *“Recovery and microbial synthesis of high-value aquaculture feed additives from food-processing wastewater”*. We thank Dr Chang Ying (National University of Singapore) for her critical reading of the manuscript.

## Author contributions

Y.C, S.W and R.B.H.W each conceptualized separate components of this work as part of a broader collaborative research programme. E.S. coordinated and project managed the broader project, and obtained and characterized source wastewater samples. S.H, F.Q.Y.G and Y.C isolated and cultured the microbe, performed taxonomic classification and extracted genomic DNA. I.B. performed further characterization of genomic DNA and coordinated sequencing. K.A., M.A.S.H and R.B.H.W performed data analysis. All authors were involved in data interpretation. The manuscript was written by R.B.H.W with inputs from other authors.

## Data Availability Statement

The draft genome sequence has been deposited at DDBJ/ENA/GenBank under the accession JANTUC000000000. The version described in this paper is version JANTUC010000000. Raw sequence data are pending release at the time of writing.

## List of Supplementary Material

**Supplementary Figure 1**

Sequence alignment between Sanger-sequenced partial 28S LSU-rRNA sequence and the top ranked BLASTN hit from NCBI nr/nt database.

**Supplementary Figure 2**

Pairs plot for contig GC-content, contig coverage and contig length from the *P. maydicus assembly*.

**Supplementary Data File 1**

Table listing properties of contigs from the *P. maydicus* assembly.

**Supplementary Data File 2**

Summary of taxonomic classification analysis of recovered 18S SSU-rRNA sequences to the SILVA 138 database.

**Supplementary Data File 3**

Alignment of Sanger-sequenced partial 28S LSU-rRNA sequence against three 28S LSU-rRNA gene sequences recovered from the *P. maydicus* long read genome assembly and a set of 62 28S LSU-rRNA sequences from members of genus *Psuedopithomyces* (NCBI Nucleotide searched for “Pseudopithomyces AND 28S” on 30th May 2022).

**Supplementary Data File 4**

MASH similarity statistics obtained by comparing the *P. maydicus* long read genome assembly sequence to 9563 fungal genomes obtained from NCBI. The reference genomes from NCBI were downloaded using the NCBI ‘dataset’ (version 13.6.0) command line tool (datasets_13.6.0 download genome taxon 4751 --filename fungi.zip --assembly-level complete_genome,chromosome,scaffold,contig --exclude-gff3 --exclude-protein --exclude-rna).

**Supplementary Data File 5**

BlastKOALA annotation data for all proteins predicted from *P. maydicus* long read assembly.

**Supplementary Results**

Complete output from the antiSMASH6 analysis of the *P. maydicus* long read assembly.

