## Supplementary Data File 3 for "A draft genome of the ascomycotal fungal species *Pseudopithomyces maydicus* (family *Didymosphaeriaceae*)"

```
BLASTN 2.4.0+


Reference: Zheng Zhang, Scott Schwartz, Lukas Wagner, and Webb
Miller (2000), "A greedy algorithm for aligning DNA sequences", J
Comput Biol 2000; 7(1-2):203-14.



Database: combined28S.fna
           66 sequences; 66,483 total letters



Query= YPD_A_1_partial_28S_Sanger

Length=551
                                                                      Score     E
Sequences producing significant alignments:                          (Bits)  Value

  YPD_A_1_partial_28S_Sanger                                          1018    0.0
  HG933820.1 Pithomyces maydicus genomic DNA containing, 28S rRNA...  1013    0.0
  HG933821.1 Pithomyces maydicus genomic DNA containing, 28S rRNA...  1009    0.0
  HG933822.1 Pithomyces maydicus genomic DNA containing, 28S rRNA...  1009    0.0
  LC158628.1 Pseudopithomyces maydicus gene for 28S ribosomal RNA...  1005    0.0
  NG_066305.1 Pseudopithomyces entadae MFLUCC 17-0917 28S rRNA ge...  1002    0.0
  rRNA_contig_5_11377-17618_DIR- /molecule=28s_rRNA /score=3291.0     1000    0.0
  rRNA_contig_5_3179-9420_DIR- /molecule=28s_rRNA /score=3291.0       1000    0.0
  KU554627.1 Pseudopithomyces palmicola strain UC32 28S ribosomal...  979     0.0
  KU554628.1 Pseudopithomyces palmicola strain UC33 28S ribosomal...  979     0.0
  KU554629.1 Pseudopithomyces palmicola strain UC15 28S ribosomal...  979     0.0
  KU554630.1 Pseudopithomyces palmicola strain UC22 28S ribosomal...  979     0.0
  MF804533.1 Pseudopithomyces palmicola strain PP4 28S ribosomal ...  979     0.0
  MF173605.1 Pseudopithomyces kunmingensis strain MFLUCC 17-0314 ...  979     0.0
  MH376738.1 Pseudopithomyces pandanicola voucher MFLUCC 18-0116 ...  979     0.0
  MK047478.1 Pseudopithomyces angolensis culture CBS:145056 28S r...  979     0.0
  NG_070067.1 Pseudopithomyces angolensis 28S rRNA gene, partial ...  979     0.0
  MG829064.1 Pseudopithomyces rosae strain MFLUCC 15-0035 28S rib...  979     0.0
  NG_059876.1 Pseudopithomyces rosae MFLU 18-0109 28S rRNA gene, ...  979     0.0
  MF804532.1 Pseudopithomyces palmicola strain L12 28S ribosomal ...  974     0.0
  MF804534.1 Pseudopithomyces palmicola strain PP28B 28S ribosoma...  974     0.0
  MF804535.1 Pseudopithomyces palmicola strain AC4 28S ribosomal ...  974     0.0
  HG933828.1 Pseudopithomyces karoo genomic DNA containing, 28S r...  968     0.0
  HG933829.1 Pseudopithomyces karoo genomic DNA containing, 28S r...  968     0.0
  NG_057865.1 Pseudopithomyces karoo CBS 804.72 28S rRNA gene, pa...  968     0.0
  LT671616.1 Pseudopithomyces atro-olivaceus genomic DNA sequence...  963     0.0
  LT671617.1 Pseudopithomyces atro-olivaceus genomic DNA sequence...  963     0.0
  LT671618.1 Pseudopithomyces atro-olivaceus genomic DNA sequence...  963     0.0
  HG933818.1 Pithomyces maydicus genomic DNA containing, 28S rRNA...  955     0.0
  HG933819.1 Pithomyces maydicus genomic DNA containing, 28S rRNA...  953     0.0
  LK936378.1 Pithomyces sacchari partial 28S rRNA gene, isolate C...  946     0.0
  LK936379.1 Pithomyces sacchari partial 28S rRNA gene, isolate C...  946     0.0
  HG933824.1 Pithomyces sp. I 2014-KDC genomic DNA containing, 28...  904     0.0
  HG933813.1 Pithomyces sacchari genomic DNA containing, 28S rRNA...  894     0.0
  LK936380.1 Pithomyces sp. 1 ALT-2014 partial 28S rRNA gene, str...  893     0.0
  HG933816.1 Pithomyces sacchari genomic DNA containing, 28S rRNA...  883     0.0
  HG933823.1 Pithomyces sp. I 2014-KDC genomic DNA containing, 28...  881     0.0
  HG933825.1 Pithomyces sp. I 2014-KDC genomic DNA containing, 28...  869     0.0
  HG933826.1 Pithomyces sp. I 2014-KDC genomic DNA containing, 28...  856     0.0
  HG933817.1 Pithomyces sacchari genomic DNA containing, 28S rRNA...  852     0.0
  LK936381.1 Pithomyces sp. 2 ALT-2014 partial 28S rRNA gene, str...  852     0.0
  HG933815.1 Pithomyces sacchari genomic DNA containing, 28S rRNA...  850     0.0
  HG933814.1 Pithomyces sacchari genomic DNA containing, 28S rRNA...  833     0.0
  HG933827.1 Pseudopithomyces diversisporus genomic DNA containin...  824     0.0
```

```
  KX034666.1 Pseudopithomyces maydicus strain MFLUCC 14 0391 28S ...  606     7e-176
  KX034665.1 Pseudopithomyces palmicola strain MFLUCC 14-0392 28S...  577     6e-167


> YPD_A_1_partial_28S_Sanger
Length=551

 Score = 1018 bits (551),  Expect = 0.0
 Identities = 551/551 (100%), Gaps = 0/551 (0%)
 Strand=Plus/Plus

Query  1    CGGCGAGTGAGCGGCTACAGCTCAAATTTGAAATCTGGCCTCCTTTGGTGGTCCGAGTTG  60
            ||||||||||||||||||||||||||||||||||||||||||||||||||||||||||||
Sbjct  1    CGGCGAGTGAGCGGCTACAGCTCAAATTTGAAATCTGGCCTCCTTTGGTGGTCCGAGTTG  60

Query  61   TAATTTGCAGAGGATGCTTTGGCATTGGCGGCGGTCTAAGTTCCTTGGAACAGGACATCG  120
            ||||||||||||||||||||||||||||||||||||||||||||||||||||||||||||
Sbjct  61   TAATTTGCAGAGGATGCTTTGGCATTGGCGGCGGTCTAAGTTCCTTGGAACAGGACATCG  120

Query  121  CAGAGGGTGAGAATCCCGTACGTGGGCGCCTGCCTTTGCCGTGTAAAGCTCCTTCGACGA  180
            ||||||||||||||||||||||||||||||||||||||||||||||||||||||||||||
Sbjct  121  CAGAGGGTGAGAATCCCGTACGTGGGCGCCTGCCTTTGCCGTGTAAAGCTCCTTCGACGA  180

Query  181  GTCGAGTTGTTTGGGAATGCAGCTCTAAATGGGAGGTAAATTTCTCCTAAAGCTAAATAC  240
            ||||||||||||||||||||||||||||||||||||||||||||||||||||||||||||
Sbjct  181  GTCGAGTTGTTTGGGAATGCAGCTCTAAATGGGAGGTAAATTTCTCCTAAAGCTAAATAC  240

Query  241  CGGCCAGAGACCGATAGCGCACAAGTAGAGTGATCGAAAGATGAAAAGTACTTTGGAAAG  300
            ||||||||||||||||||||||||||||||||||||||||||||||||||||||||||||
Sbjct  241  CGGCCAGAGACCGATAGCGCACAAGTAGAGTGATCGAAAGATGAAAAGTACTTTGGAAAG  300

Query  301  AGAGTCAAATAGCACGTGAAATTGTTGAAAGGGAAGCGCTTGCAGCCAGACTTGCCCGCA  360
            ||||||||||||||||||||||||||||||||||||||||||||||||||||||||||||
Sbjct  301  AGAGTCAAATAGCACGTGAAATTGTTGAAAGGGAAGCGCTTGCAGCCAGACTTGCCCGCA  360

Query  361  GTTGCTCACCCAGGCTCTCGCCTGGGGCACTCTTCTGCGGGCAGGCCAGCATCAGTTTGG  420
            ||||||||||||||||||||||||||||||||||||||||||||||||||||||||||||
Sbjct  361  GTTGCTCACCCAGGCTCTCGCCTGGGGCACTCTTCTGCGGGCAGGCCAGCATCAGTTTGG  420

Query  421  GCGGTCGGATAAAGGCTCCTGTCATGTACCACCCCTCGGGGTGGCCTTATAggggggggCG  480
            ||||||||||||||||||||||||||||||||||||||||||||||||||||||||||||
Sbjct  421  GCGGTCGGATAAAGGCTCCTGTCATGTACCACCCCTCGGGGTGGCCTTATAGGGGGGGCG  480

Query  481  TAATGCGACCAGCCCGGACTGAGGTCCGCGCATCTGCTAGGATGCTGGCGTAATGGCTGT  540
            ||||||||||||||||||||||||||||||||||||||||||||||||||||||||||||
Sbjct  481  TAATGCGACCAGCCCGGACTGAGGTCCGCGCATCTGCTAGGATGCTGGCGTAATGGCTGT  540

Query  541  AAGCGGCCCGT  551
            |||||||||||
Sbjct  541  AAGCGGCCCGT  551


> HG933820.1 Pithomyces maydicus genomic DNA containing, 28S rRNA
gene
Length=1055

 Score = 1013 bits (548),  Expect = 0.0
 Identities = 551/552 (99%), Gaps = 1/552 (0%)
 Strand=Plus/Plus

Query  1    CGGCGAGTG-AGCGGCTACAGCTCAAATTTGAAATCTGGCCTCCTTTGGTGGTCCGAGTT  59
            ||||||||| ||||||||||||||||||||||||||||||||||||||||||||||||||
Sbjct  38   CGGCGAGTGAAGCGGCTACAGCTCAAATTTGAAATCTGGCCTCCTTTGGTGGTCCGAGTT  97
```

```
Query  60    GTAATTTGCAGAGGATGCTTTGGCATTGGCGGCGGTCTAAGTTCCTTGGAACAGGACATC  119
             ||||||||||||||||||||||||||||||||||||||||||||||||||||||||||||
Sbjct  98    GTAATTTGCAGAGGATGCTTTGGCATTGGCGGCGGTCTAAGTTCCTTGGAACAGGACATC  157

Query  120   GCAGAGGGTGAGAATCCCGTACGTGGGCGCCTGCCTTTGCCGTGTAAAGCTCCTTCGACG  179
             ||||||||||||||||||||||||||||||||||||||||||||||||||||||||||||
Sbjct  158   GCAGAGGGTGAGAATCCCGTACGTGGGCGCCTGCCTTTGCCGTGTAAAGCTCCTTCGACG  217

Query  180   AGTCGAGTTGTTTGGGAATGCAGCTCTAAATGGGAGGTAAATTTCTCCTAAAGCTAAATA  239
             ||||||||||||||||||||||||||||||||||||||||||||||||||||||||||||
Sbjct  218   AGTCGAGTTGTTTGGGAATGCAGCTCTAAATGGGAGGTAAATTTCTCCTAAAGCTAAATA  277

Query  240   CCGGCCAGAGACCGATAGCGCACAAGTAGAGTGATCGAAAGATGAAAAGTACTTTGGAAA  299
             ||||||||||||||||||||||||||||||||||||||||||||||||||||||||||||
Sbjct  278   CCGGCCAGAGACCGATAGCGCACAAGTAGAGTGATCGAAAGATGAAAAGTACTTTGGAAA  337

Query  300   GAGAGTCAAATAGCACGTGAAATTGTTGAAAGGGAAGCGCTTGCAGCCAGACTTGCCCGC  359
             ||||||||||||||||||||||||||||||||||||||||||||||||||||||||||||
Sbjct  338   GAGAGTCAAATAGCACGTGAAATTGTTGAAAGGGAAGCGCTTGCAGCCAGACTTGCCCGC  397

Query  360   AGTTGCTCACCCAGGCTCTCGCCTGGGGCACTCTTCTGCGGGCAGGCCAGCATCAGTTTG  419
             ||||||||||||||||||||||||||||||||||||||||||||||||||||||||||||
Sbjct  398   AGTTGCTCACCCAGGCTCTCGCCTGGGGCACTCTTCTGCGGGCAGGCCAGCATCAGTTTG  457

Query  420   GGCGGTCGGATAAAGGCTCCTGTCATGTACCACCCCTCGGGGTGGCCTTATAgggggggC  479
             ||||||||||||||||||||||||||||||||||||||||||||||||||||||||||||
Sbjct  458   GGCGGTCGGATAAAGGCTCCTGTCATGTACCACCCCTCGGGGTGGCCTTATAGGGGGGGC  517

Query  480   GTAATGCGACCAGCCCGGACTGAGGTCCGCGCATCTGCTAGGATGCTGGCGTAATGGCTG  539
             ||||||||||||||||||||||||||||||||||||||||||||||||||||||||||||
Sbjct  518   GTAATGCGACCAGCCCGGACTGAGGTCCGCGCATCTGCTAGGATGCTGGCGTAATGGCTG  577

Query  540   TAAGCGGCCCGT  551
             ||||||||||||
Sbjct  578   TAAGCGGCCCGT  589


> HG933821.1 Pithomyces maydicus genomic DNA containing, 28S rRNA
gene
Length=1022

 Score = 1009 bits (546),  Expect = 0.0
 Identities = 551/553 (99%), Gaps = 2/553 (0%)
 Strand=Plus/Plus

Query  1     CGGCGAGTG-AGCGGCTACAGCTCAAATTTGAAATCTGGCCTCCTTTGGTGGTCCGAGTT  59
             ||||||||| ||||||||||||||||||||||||||||||||||||||||||||||||||
Sbjct  38    CGGCGAGTGAAGCGGCTACAGCTCAAATTTGAAATCTGGCCTCCTTTGGTGGTCCGAGTT  97

Query  60    GTAATTTGCAGAGGATGCTTTGGCATTGGCGGCGGTCTAAGTTCCTTGGAACAGGACATC  119
             ||||||||||||||||||||||||||||||||||||||||||||||||||||||||||||
Sbjct  98    GTAATTTGCAGAGGATGCTTTGGCATTGGCGGCGGTCTAAGTTCCTTGGAACAGGACATC  157

Query  120   GCAGAGGGTGAGAATCCCGTACGTGGGCGCCTGCCTTTGCCGTGTAAAGCTCCTTCGACG  179
             ||||||||||||||||||||||||||||||||||||||||||||||||||||||||||||
Sbjct  158   GCAGAGGGTGAGAATCCCGTACGTGGGCGCCTGCCTTTGCCGTGTAAAGCTCCTTCGACG  217

Query  180   AGTCGAGTTGTTTGGGAATGCAGCTCTAAATGGGAGGTAAATTTCTCCTAAAGCTAAATA  239
             ||||||||||||||||||||||||||||||||||||||||||||||||||||||||||||
Sbjct  218   AGTCGAGTTGTTTGGGAATGCAGCTCTAAATGGGAGGTAAATTTCTCCTAAAGCTAAATA  277

Query  240   CCGGCCAGAGACCGATAGCGCACAAGTAGAGTGATCGAAAGATGAAAAGTACTTTGGAAA  299
             ||||||||||||||||||||||||||||||||||||||||||||||||||||||||||||
Sbjct  278   CCGGCCAGAGACCGATAGCGCACAAGTAGAGTGATCGAAAGATGAAAAGTACTTTGGAAA  337
```

```
Query  300   GAGAGTCAAATAGCACGTGAAATTGTTGAAAGGGAAGCGCTTGCAGCCAGACTTGCCCGC  359
             ||||||||||||||||||||||||||||||||||||||||||||||||||||||||||||
Sbjct  338   GAGAGTCAAATAGCACGTGAAATTGTTGAAAGGGAAGCGCTTGCAGCCAGACTTGCCCGC  397

Query  360   AGTTGCTCACCCAGGCTCTCGCCTGGGGCACTCTTCTGCGGGCAGGCCAGCATCAGTTTG  419
             ||||||||||||||||||||||||||||||||||||||||||||||||||||||||||||
Sbjct  398   AGTTGCTCACCCAGGCTCTCGCCTGGGGCACTCTTCTGCGGGCAGGCCAGCATCAGTTTG  457

Query  420   GGCGGTCGGATAAAGGCTCCTGTCATGTACCACCCCTCGGGGTGGCCTTATA-ggggggg  478
             |||||||||||||||||||||||||||||||||||||||||||||||||||| |||||||
Sbjct  458   GGCGGTCGGATAAAGGCTCCTGTCATGTACCACCCCTCGGGGTGGCCTTATAGGGGGGGG  517

Query  479   CGTAATGCGACCAGCCCGGACTGAGGTCCGCGCATCTGCTAGGATGCTGGCGTAATGGCT  538
             ||||||||||||||||||||||||||||||||||||||||||||||||||||||||||||
Sbjct  518   CGTAATGCGACCAGCCCGGACTGAGGTCCGCGCATCTGCTAGGATGCTGGCGTAATGGCT  577

Query  539   GTAAGCGGCCCGT  551
             |||||||||||||
Sbjct  578   GTAAGCGGCCCGT  590


> HG933822.1 Pithomyces maydicus genomic DNA containing, 28S rRNA
gene
Length=1006

 Score = 1009 bits (546),  Expect = 0.0
 Identities = 551/553 (99%), Gaps = 2/553 (0%)
 Strand=Plus/Plus

Query  1     CGGCGAGTG-AGCGGCTACAGCTCAAATTTGAAATCTGGCCTCCTTTGGTGGTCCGAGTT  59
             ||||||||| ||||||||||||||||||||||||||||||||||||||||||||||||||
Sbjct  38    CGGCGAGTGAAGCGGCTACAGCTCAAATTTGAAATCTGGCCTCCTTTGGTGGTCCGAGTT  97

Query  60    GTAATTTGCAGAGGATGCTTTGGCATTGGCGGCGGTCTAAGTTCCTTGGAACAGGACATC  119
             ||||||||||||||||||||||||||||||||||||||||||||||||||||||||||||
Sbjct  98    GTAATTTGCAGAGGATGCTTTGGCATTGGCGGCGGTCTAAGTTCCTTGGAACAGGACATC  157

Query  120   GCAGAGGGTGAGAATCCCGTACGTGGGCGCCTGCCTTTGCCGTGTAAAGCTCCTTCGACG  179
             ||||||||||||||||||||||||||||||||||||||||||||||||||||||||||||
Sbjct  158   GCAGAGGGTGAGAATCCCGTACGTGGGCGCCTGCCTTTGCCGTGTAAAGCTCCTTCGACG  217

Query  180   AGTCGAGTTGTTTGGGAATGCAGCTCTAAATGGGAGGTAAATTTCTCCTAAAGCTAAATA  239
             ||||||||||||||||||||||||||||||||||||||||||||||||||||||||||||
Sbjct  218   AGTCGAGTTGTTTGGGAATGCAGCTCTAAATGGGAGGTAAATTTCTCCTAAAGCTAAATA  277

Query  240   CCGGCCAGAGACCGATAGCGCACAAGTAGAGTGATCGAAAGATGAAAAGTACTTTGGAAA  299
             ||||||||||||||||||||||||||||||||||||||||||||||||||||||||||||
Sbjct  278   CCGGCCAGAGACCGATAGCGCACAAGTAGAGTGATCGAAAGATGAAAAGTACTTTGGAAA  337

Query  300   GAGAGTCAAATAGCACGTGAAATTGTTGAAAGGGAAGCGCTTGCAGCCAGACTTGCCCGC  359
             ||||||||||||||||||||||||||||||||||||||||||||||||||||||||||||
Sbjct  338   GAGAGTCAAATAGCACGTGAAATTGTTGAAAGGGAAGCGCTTGCAGCCAGACTTGCCCGC  397

Query  360   AGTTGCTCACCCAGGCTCTCGCCTGGGGCACTCTTCTGCGGGCAGGCCAGCATCAGTTTG  419
             ||||||||||||||||||||||||||||||||||||||||||||||||||||||||||||
Sbjct  398   AGTTGCTCACCCAGGCTCTCGCCTGGGGCACTCTTCTGCGGGCAGGCCAGCATCAGTTTG  457

Query  420   GGCGGTCGGATAAAGGCTCCTGTCATGTACCACCCCTCGGGGTGGCCTTATA-ggggggg  478
             |||||||||||||||||||||||||||||||||||||||||||||||||||| |||||||
Sbjct  458   GGCGGTCGGATAAAGGCTCCTGTCATGTACCACCCCTCGGGGTGGCCTTATAGGGGGGGG  517

Query  479   CGTAATGCGACCAGCCCGGACTGAGGTCCGCGCATCTGCTAGGATGCTGGCGTAATGGCT  538
             ||||||||||||||||||||||||||||||||||||||||||||||||||||||||||||
```

```
Sbjct  518  CGTAATGCGACCAGCCCGGACTGAGGTCCGCGCATCTGCTAGGATGCTGGCGTAATGGCT  577

Query  539  GTAAGCGGCCCGT  551
            |||||||||||||
Sbjct  578  GTAAGCGGCCCGT  590


> LC158628.1 Pseudopithomyces maydicus gene for 28S ribosomal RNA,
partial sequence, strain: PW2861
Length=573

 Score = 1005 bits (544),  Expect = 0.0
 Identities = 547/548 (99%), Gaps = 1/548 (0%)
 Strand=Plus/Plus

Query  1    CGGCGAGTG-AGCGGCTACAGCTCAAATTTGAAATCTGGCCTCCTTTGGTGGTCCGAGTT  59
            ||||||||| ||||||||||||||||||||||||||||||||||||||||||||||||||
Sbjct  26   CGGCGAGTGAAGCGGCTACAGCTCAAATTTGAAATCTGGCCTCCTTTGGTGGTCCGAGTT  85

Query  60   GTAATTTGCAGAGGATGCTTTGGCATTGGCGGCGGTCTAAGTTCCTTGGAACAGGACATC  119
            ||||||||||||||||||||||||||||||||||||||||||||||||||||||||||||
Sbjct  86   GTAATTTGCAGAGGATGCTTTGGCATTGGCGGCGGTCTAAGTTCCTTGGAACAGGACATC  145

Query  120  GCAGAGGGTGAGAATCCCGTACGTGGGCGCCTGCCTTTGCCGTGTAAAGCTCCTTCGACG  179
            ||||||||||||||||||||||||||||||||||||||||||||||||||||||||||||
Sbjct  146  GCAGAGGGTGAGAATCCCGTACGTGGGCGCCTGCCTTTGCCGTGTAAAGCTCCTTCGACG  205

Query  180  AGTCGAGTTGTTTGGGAATGCAGCTCTAAATGGGAGGTAAATTTCTCCTAAAGCTAAATA  239
            ||||||||||||||||||||||||||||||||||||||||||||||||||||||||||||
Sbjct  206  AGTCGAGTTGTTTGGGAATGCAGCTCTAAATGGGAGGTAAATTTCTCCTAAAGCTAAATA  265

Query  240  CCGGCCAGAGACCGATAGCGCACAAGTAGAGTGATCGAAAGATGAAAAGTACTTTGGAAA  299
            ||||||||||||||||||||||||||||||||||||||||||||||||||||||||||||
Sbjct  266  CCGGCCAGAGACCGATAGCGCACAAGTAGAGTGATCGAAAGATGAAAAGTACTTTGGAAA  325

Query  300  GAGAGTCAAATAGCACGTGAAATTGTTGAAAGGGAAGCGCTTGCAGCCAGACTTGCCCGC  359
            ||||||||||||||||||||||||||||||||||||||||||||||||||||||||||||
Sbjct  326  GAGAGTCAAATAGCACGTGAAATTGTTGAAAGGGAAGCGCTTGCAGCCAGACTTGCCCGC  385

Query  360  AGTTGCTCACCCAGGCTCTCGCCTGGGGCACTCTTCTGCGGGCAGGCCAGCATCAGTTTG  419
            ||||||||||||||||||||||||||||||||||||||||||||||||||||||||||||
Sbjct  386  AGTTGCTCACCCAGGCTCTCGCCTGGGGCACTCTTCTGCGGGCAGGCCAGCATCAGTTTG  445

Query  420  GGCGGTCGGATAAAGGCTCCTGTCATGTACCACCCCTCGGGGTGGCCTTATAgggggggC  479
            ||||||||||||||||||||||||||||||||||||||||||||||||||||||||||||
Sbjct  446  GGCGGTCGGATAAAGGCTCCTGTCATGTACCACCCCTCGGGGTGGCCTTATAGGGGGGGC  505

Query  480  GTAATGCGACCAGCCCGGACTGAGGTCCGCGCATCTGCTAGGATGCTGGCGTAATGGCTG  539
            ||||||||||||||||||||||||||||||||||||||||||||||||||||||||||||
Sbjct  506  GTAATGCGACCAGCCCGGACTGAGGTCCGCGCATCTGCTAGGATGCTGGCGTAATGGCTG  565

Query  540  TAAGCGGC  547
            ||||||||
Sbjct  566  TAAGCGGC  573


> NG_066305.1 Pseudopithomyces entadae MFLUCC 17-0917 28S rRNA
gene, partial sequence; from TYPE material
Length=846

 Score = 1002 bits (542),  Expect = 0.0
 Identities = 549/552 (99%), Gaps = 1/552 (0%)
 Strand=Plus/Plus
```

```
Query  1    CGGCGAGTG-AGCGGCTACAGCTCAAATTTGAAATCTGGCCTCCTTTGGTGGTCCGAGTT  59
            ||||||||| ||||||||||||||||||||||||||||||||||||||||||||||||||
Sbjct  14   CGGCGAGTGAAGCGGCTACAGCTCAAATTTGAAATCTGGCCTCCTTTGGTGGTCCGAGTT  73

Query  60   GTAATTTGCAGAGGATGCTTTGGCATTGGCGGCGGTCTAAGTTCCTTGGAACAGGACATC  119
            ||||||||||||||||||||||||||||||||||||||||||||||||||||||||||||
Sbjct  74   GTAATTTGCAGAGGATGCTTTGGCATTGGCGGCGGTCTAAGTTCCTTGGAACAGGACATC  133

Query  120  GCAGAGGGTGAGAATCCCGTACGTGGGCGCCTGCCTTTGCCGTGTAAAGCTCCTTCGACG  179
            ||||||||||||||||||||||||||||||||||||||||||||||||||||||||||||
Sbjct  134  GCAGAGGGTGAGAATCCCGTACGTGGGCGCCTGCCTTTGCCGTGTAAAGCTCCTTCGACG  193

Query  180  AGTCGAGTTGTTTGGGAATGCAGCTCTAAATGGGAGGTAAATTTCTCCTAAAGCTAAATA  239
            ||||||||||||||||||||||||||||||||||||||||||||||||||||||||||||
Sbjct  194  AGTCGAGTTGTTTGGGAATGCAGCTCTAAATGGGAGGTAAATTTCTCCTAAAGCTAAATA  253

Query  240  CCGGCCAGAGACCGATAGCGCACAAGTAGAGTGATCGAAAGATGAAAAGTACTTTGGAAA  299
            ||||||||||||||||||||||||||||||||||||||||||||||||||||||||||||
Sbjct  254  CCGGCCAGAGACCGATAGCGCACAAGTAGAGTGATCGAAAGATGAAAAGTACTTTGGAAA  313

Query  300  GAGAGTCAAATAGCACGTGAAATTGTTGAAAGGGAAGCGCTTGCAGCCAGACTTGCCCGC  359
            ||||||||||||||||||||||||||||||||||||||||||||||||||||||||||||
Sbjct  314  GAGAGTCAAATAGCACGTGAAATTGTTGAAAGGGAAGCGCTTGCAGCCAGACTTGCCCGC  373

Query  360  AGTTGCTCACCCAGGCTCTCGCCTGGGGCACTCTTCTGCGGGCAGGCCAGCATCAGTTTG  419
            ||||||||||||||||||||||||||||||||||||||||||||||||||||||||||||
Sbjct  374  AGTTGCTCACCCAGGCTCTCGCCTGGGGCACTCTTCTGCGGGCAGGCCAGCATCAGTTTG  433

Query  420  GGCGGTCGGATAAAGGCTCCTGTCATGTACCACCCCTCGGGGTGGCCTTATAgggggggC  479
            ||||||||||||||||||||||||||||||||||||||||||||||||||||||||| ||
Sbjct  434  GGCGGTCGGATAAAGGCTCCTGTCATGTACCACCCCTCGGGGTGGCCTTATAGGGGGAGC  493

Query  480  GTAATGCGACCAGCCCGGACTGAGGTCCGCGCATCTGCTAGGATGCTGGCGTAATGGCTG  539
            | ||||||||||||||||||||||||||||||||||||||||||||||||||||||||||
Sbjct  494  GCAATGCGACCAGCCCGGACTGAGGTCCGCGCATCTGCTAGGATGCTGGCGTAATGGCTG  553

Query  540  TAAGCGGCCCGT  551
            ||||||||||||
Sbjct  554  TAAGCGGCCCGT  565


> rRNA_contig_5_11377-17618_DIR- /molecule=28s_rRNA /score=3291.0
Length=6242

 Score = 1000 bits (541),  Expect = 0.0
 Identities = 549/552 (99%), Gaps = 3/552 (1%)
 Strand=Plus/Plus

Query  1    CGGCGAGTG-AGCGGCTACAGCTCAAATTTGAAATCTGGCCTCCTTTGGTGGTCCGAGTT  59
            ||||||||| ||||||||||||||||||||||||||||||||||||||||||||||||||
Sbjct  399  CGGCGAGTGAAGCGGCTACAGCTCAAATTTGAAATCTGGCCTCCTTTGGTGGTCCGAGTT  458

Query  60   GTAATTTGCAGAGGATGCTTTGGCATTGGCGGCGGTCTAAGTTCCTTGGAACAGGACATC  119
            ||||||||||||||||||||||||||||||||||||||||||||||||||||||||||||
Sbjct  459  GTAATTTGCAGAGGATGCTTTGGCATTGGCGGCGGTCTAAGTTCCTTGGAACAGGACATC  518

Query  120  GCAGAGGGTGAGAATCCCGTACGTGGGCGCCTGCCTTTGCCGTGTAAAGCTCCTTCGACG  179
            ||||||||||||||||||||||||||||||||||||||||||||||||||||||||||||
Sbjct  519  GCAGAGGGTGAGAATCCCGTACGTGGGCGCCTGCCTTTGCCGTGTAAAGCTCCTTCGACG  578

Query  180  AGTCGAGTTGTTTGGGAATGCAGCTCTAAATGGGAGGTAAATTTCTCCTAAAGCTAAATA  239
            ||||||||||||||||||||||||||||||||||||||||||||||||||||||||||||
Sbjct  579  AGTCGAGTTGTTTGGGAATGCAGCTCTAAATGGGAGGTAAATTTCTCCTAAAGCTAAATA  638
```

```
Query  240   CCGGCCAGAGACCGATAGCGCACAAGTAGAGTGATCGAAAGATGAAAAGTACTTTGGAAA  299
             ||||||||||||||||||||||||||||||||||||||||||||||||||||||||||||
Sbjct  639   CCGGCCAGAGACCGATAGCGCACAAGTAGAGTGATCGAAAGATGAAAAGTACTTTGGAAA  698

Query  300   GAGAGTCAAATAGCACGTGAAATTGTTGAAAGGGAAGCGCTTGCAGCCAGACTTGCCCGC  359
             ||||||||||||||||||||||||||||||||||||||||||||||||||||||||||||
Sbjct  699   GAGAGTCAAATAGCACGTGAAATTGTTGAAAGGGAAGCGCTTGCAGCCAGACTTGCCCGC  758

Query  360   AGTTGCTCACCCAGGCTCTCGCCTGGGGCACTCTTCTGCGGGCAGGCCAGCATCAGTTTG  419
             ||||||||||||||||||||||||||||||||||||||||||||||||||||||||||||
Sbjct  759   AGTTGCTCACCCAGGCTCTCGCCTGGGGCACTCTTCTGCGGGCAGGCCAGCATCAGTTTG  818

Query  420   GGCGGTCGGATAAAGGCTCCTGTCATGTACCACCCCTCGGGGTGGCCTTATAgggggggC  479
             ||||||||||||||||||||||||||||||||||||||||||||||||||||  ||||||
Sbjct  819   GGCGGTCGGATAAAGGCTCCTGTCATGTACCACCCCTCGGGGTGGCCTTATA--GGGGGC  876

Query  480   GTAATGCGACCAGCCCGGACTGAGGTCCGCGCATCTGCTAGGATGCTGGCGTAATGGCTG  539
             ||||||||||||||||||||||||||||||||||||||||||||||||||||||||||||
Sbjct  877   GTAATGCGACCAGCCCGGACTGAGGTCCGCGCATCTGCTAGGATGCTGGCGTAATGGCTG  936

Query  540   TAAGCGGCCCGT  551
             ||||||||||||
Sbjct  937   TAAGCGGCCCGT  948


> rRNA_contig_5_3179-9420_DIR- /molecule=28s_rRNA /score=3291.0
Length=6242

 Score = 1000 bits (541),  Expect = 0.0
 Identities = 549/552 (99%), Gaps = 3/552 (1%)
 Strand=Plus/Plus

Query  1     CGGCGAGTG-AGCGGCTACAGCTCAAATTTGAAATCTGGCCTCCTTTGGTGGTCCGAGTT  59
             ||||||||| ||||||||||||||||||||||||||||||||||||||||||||||||||
Sbjct  399   CGGCGAGTGAAGCGGCTACAGCTCAAATTTGAAATCTGGCCTCCTTTGGTGGTCCGAGTT  458

Query  60    GTAATTTGCAGAGGATGCTTTGGCATTGGCGGCGGTCTAAGTTCCTTGGAACAGGACATC  119
             ||||||||||||||||||||||||||||||||||||||||||||||||||||||||||||
Sbjct  459   GTAATTTGCAGAGGATGCTTTGGCATTGGCGGCGGTCTAAGTTCCTTGGAACAGGACATC  518

Query  120   GCAGAGGGTGAGAATCCCGTACGTGGGCGCCTGCCTTTGCCGTGTAAAGCTCCTTCGACG  179
             ||||||||||||||||||||||||||||||||||||||||||||||||||||||||||||
Sbjct  519   GCAGAGGGTGAGAATCCCGTACGTGGGCGCCTGCCTTTGCCGTGTAAAGCTCCTTCGACG  578

Query  180   AGTCGAGTTGTTTGGGAATGCAGCTCTAAATGGGAGGTAAATTTCTCCTAAAGCTAAATA  239
             ||||||||||||||||||||||||||||||||||||||||||||||||||||||||||||
Sbjct  579   AGTCGAGTTGTTTGGGAATGCAGCTCTAAATGGGAGGTAAATTTCTCCTAAAGCTAAATA  638

Query  240   CCGGCCAGAGACCGATAGCGCACAAGTAGAGTGATCGAAAGATGAAAAGTACTTTGGAAA  299
             ||||||||||||||||||||||||||||||||||||||||||||||||||||||||||||
Sbjct  639   CCGGCCAGAGACCGATAGCGCACAAGTAGAGTGATCGAAAGATGAAAAGTACTTTGGAAA  698

Query  300   GAGAGTCAAATAGCACGTGAAATTGTTGAAAGGGAAGCGCTTGCAGCCAGACTTGCCCGC  359
             ||||||||||||||||||||||||||||||||||||||||||||||||||||||||||||
Sbjct  699   GAGAGTCAAATAGCACGTGAAATTGTTGAAAGGGAAGCGCTTGCAGCCAGACTTGCCCGC  758

Query  360   AGTTGCTCACCCAGGCTCTCGCCTGGGGCACTCTTCTGCGGGCAGGCCAGCATCAGTTTG  419
             ||||||||||||||||||||||||||||||||||||||||||||||||||||||||||||
Sbjct  759   AGTTGCTCACCCAGGCTCTCGCCTGGGGCACTCTTCTGCGGGCAGGCCAGCATCAGTTTG  818

Query  420   GGCGGTCGGATAAAGGCTCCTGTCATGTACCACCCCTCGGGGTGGCCTTATAgggggggC  479
             ||||||||||||||||||||||||||||||||||||||||||||||||||||  ||||||
Sbjct  819   GGCGGTCGGATAAAGGCTCCTGTCATGTACCACCCCTCGGGGTGGCCTTATA--GGGGGC  876
```

```
Query  480  GTAATGCGACCAGCCCGGACTGAGGTCCGCGCATCTGCTAGGATGCTGGCGTAATGGCTG  539
            ||||||||||||||||||||||||||||||||||||||||||||||||||||||||||||
Sbjct  877  GTAATGCGACCAGCCCGGACTGAGGTCCGCGCATCTGCTAGGATGCTGGCGTAATGGCTG  936

Query  540  TAAGCGGCCCGT  551
            ||||||||||||
Sbjct  937  TAAGCGGCCCGT  948


> KU554627.1 Pseudopithomyces palmicola strain UC32 28S ribosomal
RNA gene, partial sequence
Length=1190

 Score = 979 bits (530),  Expect = 0.0
 Identities = 545/552 (99%), Gaps = 1/552 (0%)
 Strand=Plus/Plus

Query  1    CGGCGAGTG-AGCGGCTACAGCTCAAATTTGAAATCTGGCCTCCTTTGGTGGTCCGAGTT  59
            ||||||||| ||||||||||||||||||||||||||||||||||||||| ||||||||||
Sbjct  20   CGGCGAGTGAAGCGGCTACAGCTCAAATTTGAAATCTGGCCTCCTTTGGGGGTCCGAGTT  79

Query  60   GTAATTTGCAGAGGATGCTTTGGCATTGGCGGCGGTCTAAGTTCCTTGGAACAGGACATC  119
            ||||||||||||||||||||||||||||||||||||||||||||||||||||||||||||
Sbjct  80   GTAATTTGCAGAGGATGCTTTGGCATTGGCGGCGGTCTAAGTTCCTTGGAACAGGACATC  139

Query  120  GCAGAGGGTGAGAATCCCGTACGTGGGCGCCTGCCTTTGCCGTGTAAAGCTCCTTCGACG  179
            ||||||||||||||||||||||||||||||||||||||||||||||||||||||||||||
Sbjct  140  GCAGAGGGTGAGAATCCCGTACGTGGGCGCCTGCCTTTGCCGTGTAAAGCTCCTTCGACG  199

Query  180  AGTCGAGTTGTTTGGGAATGCAGCTCTAAATGGGAGGTAAATTTCTCCTAAAGCTAAATA  239
            ||||||||||||||||||||||||||||||||||||||||||||||||||||||||||||
Sbjct  200  AGTCGAGTTGTTTGGGAATGCAGCTCTAAATGGGAGGTAAATTTCTCCTAAAGCTAAATA  259

Query  240  CCGGCCAGAGACCGATAGCGCACAAGTAGAGTGATCGAAAGATGAAAAGTACTTTGGAAA  299
            ||||||||||||||||||||||||||||||||||||||||||||||||||||||||||||
Sbjct  260  CCGGCCAGAGACCGATAGCGCACAAGTAGAGTGATCGAAAGATGAAAAGTACTTTGGAAA  319

Query  300  GAGAGTCAAATAGCACGTGAAATTGTTGAAAGGGAAGCGCTTGCAGCCAGACTTGCCCGC  359
            ||||||||||||||||||||||||||||||||||||||||||||||||||||||||||||
Sbjct  320  GAGAGTCAAATAGCACGTGAAATTGTTGAAAGGGAAGCGCTTGCAGCCAGACTTGCCCGC  379

Query  360  AGTTGCTCACCCAGGCTCTCGCCTGGGGCACTCTTCTGCGGGCAGGCCAGCATCAGTTTG  419
            ||||||||||| ||||| ||||||||||||||||||||||||||||||||||||||||||
Sbjct  380  AGTTGCTCACCTAGGCTTTCGCCTGGGGCACTCTTCTGCGGGCAGGCCAGCATCAGTTTG  439

Query  420  GGCGGTCGGATAAAGGCTCCTGTCATGTACCACCCCTCGGGGTGGCCTTATAgggggggC  479
            |||||||||||||||||  ||||||||||||||||||||||||||||||||||||| |||
Sbjct  440  GGCGGTCGGATAAAGGCCTCTGTCATGTACCACCCCTCGGGGTGGCCTTATAGGGGAGGC  499

Query  480  GTAATGCGACCAGCCCGGACTGAGGTCCGCGCATCTGCTAGGATGCTGGCGTAATGGCTG  539
            ||||||||||||||||||||||||||||||||||||||||||||||||||||||||||||
Sbjct  500  GTAATGCGACCAGCCCGGACTGAGGTCCGCGCATCTGCTAGGATGCTGGCGTAATGGCTG  559

Query  540  TAAGCGGCCCGT  551
            ||||||||||||
Sbjct  560  TAAGCGGCCCGT  571


> KU554628.1 Pseudopithomyces palmicola strain UC33 28S ribosomal
RNA gene, partial sequence
Length=1191

 Score = 979 bits (530),  Expect = 0.0
 Identities = 545/552 (99%), Gaps = 1/552 (0%)
```

```
 Strand=Plus/Plus

Query  1    CGGCGAGTG-AGCGGCTACAGCTCAAATTTGAAATCTGGCCTCCTTTGGTGGTCCGAGTT  59
            ||||||||| ||||||||||||||||||||||||||||||||||||||| ||||||||||
Sbjct  20   CGGCGAGTGAAGCGGCTACAGCTCAAATTTGAAATCTGGCCTCCTTTGGGGGTCCGAGTT  79

Query  60   GTAATTTGCAGAGGATGCTTTGGCATTGGCGGCGGTCTAAGTTCCTTGGAACAGGACATC  119
            ||||||||||||||||||||||||||||||||||||||||||||||||||||||||||||
Sbjct  80   GTAATTTGCAGAGGATGCTTTGGCATTGGCGGCGGTCTAAGTTCCTTGGAACAGGACATC  139

Query  120  GCAGAGGGTGAGAATCCCGTACGTGGGCGCCTGCCTTTGCCGTGTAAAGCTCCTTCGACG  179
            ||||||||||||||||||||||||||||||||||||||||||||||||||||||||||||
Sbjct  140  GCAGAGGGTGAGAATCCCGTACGTGGGCGCCTGCCTTTGCCGTGTAAAGCTCCTTCGACG  199

Query  180  AGTCGAGTTGTTTGGGAATGCAGCTCTAAATGGGAGGTAAATTTCTCCTAAAGCTAAATA  239
            ||||||||||||||||||||||||||||||||||||||||||||||||||||||||||||
Sbjct  200  AGTCGAGTTGTTTGGGAATGCAGCTCTAAATGGGAGGTAAATTTCTCCTAAAGCTAAATA  259

Query  240  CCGGCCAGAGACCGATAGCGCACAAGTAGAGTGATCGAAAGATGAAAAGTACTTTGGAAA  299
            ||||||||||||||||||||||||||||||||||||||||||||||||||||||||||||
Sbjct  260  CCGGCCAGAGACCGATAGCGCACAAGTAGAGTGATCGAAAGATGAAAAGTACTTTGGAAA  319

Query  300  GAGAGTCAAATAGCACGTGAAATTGTTGAAAGGGAAGCGCTTGCAGCCAGACTTGCCCGC  359
            ||||||||||||||||||||||||||||||||||||||||||||||||||||||||||||
Sbjct  320  GAGAGTCAAATAGCACGTGAAATTGTTGAAAGGGAAGCGCTTGCAGCCAGACTTGCCCGC  379

Query  360  AGTTGCTCACCCAGGCTCTCGCCTGGGGCACTCTTCTGCGGGCAGGCCAGCATCAGTTTG  419
            ||||||||||| ||||| ||||||||||||||||||||||||||||||||||||||||||
Sbjct  380  AGTTGCTCACCTAGGCTTTCGCCTGGGGCACTCTTCTGCGGGCAGGCCAGCATCAGTTTG  439

Query  420  GGCGGTCGGATAAAGGCTCCTGTCATGTACCACCCCTCGGGGTGGCCTTATAgggggggC  479
            |||||||||||||||||  ||||||||||||||||||||||||||||||||||||| |||
Sbjct  440  GGCGGTCGGATAAAGGCCTCTGTCATGTACCACCCCTCGGGGTGGCCTTATAGGGGAGGC  499

Query  480  GTAATGCGACCAGCCCGGACTGAGGTCCGCGCATCTGCTAGGATGCTGGCGTAATGGCTG  539
            ||||||||||||||||||||||||||||||||||||||||||||||||||||||||||||
Sbjct  500  GTAATGCGACCAGCCCGGACTGAGGTCCGCGCATCTGCTAGGATGCTGGCGTAATGGCTG  559

Query  540  TAAGCGGCCCGT  551
            ||||||||||||
Sbjct  560  TAAGCGGCCCGT  571


> KU554629.1 Pseudopithomyces palmicola strain UC15 28S ribosomal
RNA gene, partial sequence
Length=1235

 Score = 979 bits (530),  Expect = 0.0
 Identities = 545/552 (99%), Gaps = 1/552 (0%)
 Strand=Plus/Plus

Query  1    CGGCGAGTG-AGCGGCTACAGCTCAAATTTGAAATCTGGCCTCCTTTGGTGGTCCGAGTT  59
            ||||||||| ||||||||||||||||||||||||||||||||||||||| ||||||||||
Sbjct  18   CGGCGAGTGAAGCGGCTACAGCTCAAATTTGAAATCTGGCCTCCTTTGGGGGTCCGAGTT  77

Query  60   GTAATTTGCAGAGGATGCTTTGGCATTGGCGGCGGTCTAAGTTCCTTGGAACAGGACATC  119
            ||||||||||||||||||||||||||||||||||||||||||||||||||||||||||||
Sbjct  78   GTAATTTGCAGAGGATGCTTTGGCATTGGCGGCGGTCTAAGTTCCTTGGAACAGGACATC  137

Query  120  GCAGAGGGTGAGAATCCCGTACGTGGGCGCCTGCCTTTGCCGTGTAAAGCTCCTTCGACG  179
            ||||||||||||||||||||||||||||||||||||||||||||||||||||||||||||
Sbjct  138  GCAGAGGGTGAGAATCCCGTACGTGGGCGCCTGCCTTTGCCGTGTAAAGCTCCTTCGACG  197

Query  180  AGTCGAGTTGTTTGGGAATGCAGCTCTAAATGGGAGGTAAATTTCTCCTAAAGCTAAATA  239
```

```
                ||||||||||||||||||||||||||||||||||||||||||||||||||||||||||||
Sbjct  198   AGTCGAGTTGTTTGGGAATGCAGCTCTAAATGGGAGGTAAATTTCTCCTAAAGCTAAATA  257

Query  240   CCGGCCAGAGACCGATAGCGCACAAGTAGAGTGATCGAAAGATGAAAAGTACTTTGGAAA  299
             ||||||||||||||||||||||||||||||||||||||||||||||||||||||||||||
Sbjct  258   CCGGCCAGAGACCGATAGCGCACAAGTAGAGTGATCGAAAGATGAAAAGTACTTTGGAAA  317

Query  300   GAGAGTCAAATAGCACGTGAAATTGTTGAAAGGGAAGCGCTTGCAGCCAGACTTGCCCGC  359
             ||||||||||||||||||||||||||||||||||||||||||||||||||||||||||||
Sbjct  318   GAGAGTCAAATAGCACGTGAAATTGTTGAAAGGGAAGCGCTTGCAGCCAGACTTGCCCGC  377

Query  360   AGTTGCTCACCCAGGCTCTCGCCTGGGGCACTCTTCTGCGGGCAGGCCAGCATCAGTTTG  419
             ||||||||||| ||||| ||||||||||||||||||||||||||||||||||||||||||
Sbjct  378   AGTTGCTCACCTAGGCTTTCGCCTGGGGCACTCTTCTGCGGGCAGGCCAGCATCAGTTTG  437

Query  420   GGCGGTCGGATAAAGGCTCCTGTCATGTACCACCCCTCGGGGTGGCCTTATAgggggggC  479
             |||||||||||||||||  ||||||||||||||||||||||||||||||||||||| |||
Sbjct  438   GGCGGTCGGATAAAGGCCTCTGTCATGTACCACCCCTCGGGGTGGCCTTATAGGGGAGGC  497

Query  480   GTAATGCGACCAGCCCGGACTGAGGTCCGCGCATCTGCTAGGATGCTGGCGTAATGGCTG  539
             ||||||||||||||||||||||||||||||||||||||||||||||||||||||||||||
Sbjct  498   GTAATGCGACCAGCCCGGACTGAGGTCCGCGCATCTGCTAGGATGCTGGCGTAATGGCTG  557

Query  540   TAAGCGGCCCGT  551
             ||||||||||||
Sbjct  558   TAAGCGGCCCGT  569


> KU554630.1 Pseudopithomyces palmicola strain UC22 28S ribosomal
RNA gene, partial sequence
Length=1235

 Score = 979 bits (530),  Expect = 0.0
 Identities = 545/552 (99%), Gaps = 1/552 (0%)
 Strand=Plus/Plus

Query  1     CGGCGAGTG-AGCGGCTACAGCTCAAATTTGAAATCTGGCCTCCTTTGGTGGTCCGAGTT  59
             ||||||||| ||||||||||||||||||||||||||||||||||||||| ||||||||||
Sbjct  18    CGGCGAGTGAAGCGGCTACAGCTCAAATTTGAAATCTGGCCTCCTTTGGGGGTCCGAGTT  77

Query  60    GTAATTTGCAGAGGATGCTTTGGCATTGGCGGCGGTCTAAGTTCCTTGGAACAGGACATC  119
             ||||||||||||||||||||||||||||||||||||||||||||||||||||||||||||
Sbjct  78    GTAATTTGCAGAGGATGCTTTGGCATTGGCGGCGGTCTAAGTTCCTTGGAACAGGACATC  137

Query  120   GCAGAGGGTGAGAATCCCGTACGTGGGCGCCTGCCTTTGCCGTGTAAAGCTCCTTCGACG  179
             ||||||||||||||||||||||||||||||||||||||||||||||||||||||||||||
Sbjct  138   GCAGAGGGTGAGAATCCCGTACGTGGGCGCCTGCCTTTGCCGTGTAAAGCTCCTTCGACG  197

Query  180   AGTCGAGTTGTTTGGGAATGCAGCTCTAAATGGGAGGTAAATTTCTCCTAAAGCTAAATA  239
             ||||||||||||||||||||||||||||||||||||||||||||||||||||||||||||
Sbjct  198   AGTCGAGTTGTTTGGGAATGCAGCTCTAAATGGGAGGTAAATTTCTCCTAAAGCTAAATA  257

Query  240   CCGGCCAGAGACCGATAGCGCACAAGTAGAGTGATCGAAAGATGAAAAGTACTTTGGAAA  299
             ||||||||||||||||||||||||||||||||||||||||||||||||||||||||||||
Sbjct  258   CCGGCCAGAGACCGATAGCGCACAAGTAGAGTGATCGAAAGATGAAAAGTACTTTGGAAA  317

Query  300   GAGAGTCAAATAGCACGTGAAATTGTTGAAAGGGAAGCGCTTGCAGCCAGACTTGCCCGC  359
             ||||||||||||||||||||||||||||||||||||||||||||||||||||||||||||
Sbjct  318   GAGAGTCAAATAGCACGTGAAATTGTTGAAAGGGAAGCGCTTGCAGCCAGACTTGCCCGC  377

Query  360   AGTTGCTCACCCAGGCTCTCGCCTGGGGCACTCTTCTGCGGGCAGGCCAGCATCAGTTTG  419
             ||||||||||| ||||| ||||||||||||||||||||||||||||||||||||||||||
Sbjct  378   AGTTGCTCACCTAGGCTTTCGCCTGGGGCACTCTTCTGCGGGCAGGCCAGCATCAGTTTG  437
```

```
Query  420   GGCGGTCGGATAAAGGCTCCTGTCATGTACCACCCCTCGGGGTGGCCTTATAgggggggC  479
             |||||||||||||||||  ||||||||||||||||||||||||||||||||||||| |||
Sbjct  438   GGCGGTCGGATAAAGGCCTCTGTCATGTACCACCCCTCGGGGTGGCCTTATAGGGGAGGC  497

Query  480   GTAATGCGACCAGCCCGGACTGAGGTCCGCGCATCTGCTAGGATGCTGGCGTAATGGCTG  539
             ||||||||||||||||||||||||||||||||||||||||||||||||||||||||||||
Sbjct  498   GTAATGCGACCAGCCCGGACTGAGGTCCGCGCATCTGCTAGGATGCTGGCGTAATGGCTG  557

Query  540   TAAGCGGCCCGT  551
             ||||||||||||
Sbjct  558   TAAGCGGCCCGT  569


> MF804533.1 Pseudopithomyces palmicola strain PP4 28S ribosomal
RNA gene, partial sequence
Length=867

 Score = 979 bits (530),  Expect = 0.0
 Identities = 545/552 (99%), Gaps = 1/552 (0%)
 Strand=Plus/Plus

Query  1     CGGCGAGTG-AGCGGCTACAGCTCAAATTTGAAATCTGGCCTCCTTTGGTGGTCCGAGTT  59
             ||||||||| ||||||||||||||||||||||||||||||||||||||| ||||||||||
Sbjct  17    CGGCGAGTGAAGCGGCTACAGCTCAAATTTGAAATCTGGCCTCCTTTGGGGGTCCGAGTT  76

Query  60    GTAATTTGCAGAGGATGCTTTGGCATTGGCGGCGGTCTAAGTTCCTTGGAACAGGACATC  119
             ||||||||||||||||||||||||||||||||||||||||||||||||||||||||||||
Sbjct  77    GTAATTTGCAGAGGATGCTTTGGCATTGGCGGCGGTCTAAGTTCCTTGGAACAGGACATC  136

Query  120   GCAGAGGGTGAGAATCCCGTACGTGGGCGCCTGCCTTTGCCGTGTAAAGCTCCTTCGACG  179
             ||||||||||||||||||||||||||||||||||||||||||||||||||||||||||||
Sbjct  137   GCAGAGGGTGAGAATCCCGTACGTGGGCGCCTGCCTTTGCCGTGTAAAGCTCCTTCGACG  196

Query  180   AGTCGAGTTGTTTGGGAATGCAGCTCTAAATGGGAGGTAAATTTCTCCTAAAGCTAAATA  239
             ||||||||||||||||||||||||||||||||||||||||||||||||||||||||||||
Sbjct  197   AGTCGAGTTGTTTGGGAATGCAGCTCTAAATGGGAGGTAAATTTCTCCTAAAGCTAAATA  256

Query  240   CCGGCCAGAGACCGATAGCGCACAAGTAGAGTGATCGAAAGATGAAAAGTACTTTGGAAA  299
             ||||||||||||||||||||||||||||||||||||||||||||||||||||||||||||
Sbjct  257   CCGGCCAGAGACCGATAGCGCACAAGTAGAGTGATCGAAAGATGAAAAGTACTTTGGAAA  316

Query  300   GAGAGTCAAATAGCACGTGAAATTGTTGAAAGGGAAGCGCTTGCAGCCAGACTTGCCCGC  359
             ||||||||||||||||||||||||||||||||||||||||||||||||||||||||||||
Sbjct  317   GAGAGTCAAATAGCACGTGAAATTGTTGAAAGGGAAGCGCTTGCAGCCAGACTTGCCCGC  376

Query  360   AGTTGCTCACCCAGGCTCTCGCCTGGGGCACTCTTCTGCGGGCAGGCCAGCATCAGTTTG  419
             ||||||||||| ||||| ||||||||||||||||||||||||||||||||||||||||||
Sbjct  377   AGTTGCTCACCTAGGCTTTCGCCTGGGGCACTCTTCTGCGGGCAGGCCAGCATCAGTTTG  436

Query  420   GGCGGTCGGATAAAGGCTCCTGTCATGTACCACCCCTCGGGGTGGCCTTATAgggggggC  479
             |||||||||||||||||  ||||||||||||||||||||||||||||||||||||| |||
Sbjct  437   GGCGGTCGGATAAAGGCCTCTGTCATGTACCACCCCTCGGGGTGGCCTTATAGGGGAGGC  496

Query  480   GTAATGCGACCAGCCCGGACTGAGGTCCGCGCATCTGCTAGGATGCTGGCGTAATGGCTG  539
             ||||||||||||||||||||||||||||||||||||||||||||||||||||||||||||
Sbjct  497   GTAATGCGACCAGCCCGGACTGAGGTCCGCGCATCTGCTAGGATGCTGGCGTAATGGCTG  556

Query  540   TAAGCGGCCCGT  551
             ||||||||||||
Sbjct  557   TAAGCGGCCCGT  568


> MF173605.1 Pseudopithomyces kunmingensis strain MFLUCC 17-0314
28S ribosomal RNA gene, partial sequence
```

```
Length=824

 Score = 979 bits (530),  Expect = 0.0
 Identities = 545/552 (99%), Gaps = 1/552 (0%)
 Strand=Plus/Plus

Query  1    CGGCGAGTG-AGCGGCTACAGCTCAAATTTGAAATCTGGCCTCCTTTGGTGGTCCGAGTT  59
            ||||||||| ||||||||||||||||||||||||||||||||||||||| ||||||||||
Sbjct  14   CGGCGAGTGAAGCGGCTACAGCTCAAATTTGAAATCTGGCCTCCTTTGGGGGTCCGAGTT  73

Query  60   GTAATTTGCAGAGGATGCTTTGGCATTGGCGGCGGTCTAAGTTCCTTGGAACAGGACATC  119
            ||||||||||||||||||||||||||||||||||||||||||||||||||||||||||||
Sbjct  74   GTAATTTGCAGAGGATGCTTTGGCATTGGCGGCGGTCTAAGTTCCTTGGAACAGGACATC  133

Query  120  GCAGAGGGTGAGAATCCCGTACGTGGGCGCCTGCCTTTGCCGTGTAAAGCTCCTTCGACG  179
            ||||||||||||||||||||||||||||||||||||||||||||||||||||||||||||
Sbjct  134  GCAGAGGGTGAGAATCCCGTACGTGGGCGCCTGCCTTTGCCGTGTAAAGCTCCTTCGACG  193

Query  180  AGTCGAGTTGTTTGGGAATGCAGCTCTAAATGGGAGGTAAATTTCTCCTAAAGCTAAATA  239
            ||||||||||||||||||||||||||||||||||||||||||||||||||||||||||||
Sbjct  194  AGTCGAGTTGTTTGGGAATGCAGCTCTAAATGGGAGGTAAATTTCTCCTAAAGCTAAATA  253

Query  240  CCGGCCAGAGACCGATAGCGCACAAGTAGAGTGATCGAAAGATGAAAAGTACTTTGGAAA  299
            ||||||||||||||||||||||||||||||||||||||||||||||||||||||||||||
Sbjct  254  CCGGCCAGAGACCGATAGCGCACAAGTAGAGTGATCGAAAGATGAAAAGTACTTTGGAAA  313

Query  300  GAGAGTCAAATAGCACGTGAAATTGTTGAAAGGGAAGCGCTTGCAGCCAGACTTGCCCGC  359
            ||||||||||||||||||||||||||||||||||||||||||||||||||||||||||||
Sbjct  314  GAGAGTCAAATAGCACGTGAAATTGTTGAAAGGGAAGCGCTTGCAGCCAGACTTGCCCGC  373

Query  360  AGTTGCTCACCCAGGCTCTCGCCTGGGGCACTCTTCTGCGGGCAGGCCAGCATCAGTTTG  419
            ||||||||||| ||||| ||||||||||||||||||||||||||||||||||||||||||
Sbjct  374  AGTTGCTCACCTAGGCTTTCGCCTGGGGCACTCTTCTGCGGGCAGGCCAGCATCAGTTTG  433

Query  420  GGCGGTCGGATAAAGGCTCCTGTCATGTACCACCCCTCGGGGTGGCCTTATAgggggggC  479
            |||||||||||||||||  ||||||||||||||||||||||||||||||||||||| |||
Sbjct  434  GGCGGTCGGATAAAGGCCTCTGTCATGTACCACCCCTCGGGGTGGCCTTATAGGGGAGGC  493

Query  480  GTAATGCGACCAGCCCGGACTGAGGTCCGCGCATCTGCTAGGATGCTGGCGTAATGGCTG  539
            ||||||||||||||||||||||||||||||||||||||||||||||||||||||||||||
Sbjct  494  GTAATGCGACCAGCCCGGACTGAGGTCCGCGCATCTGCTAGGATGCTGGCGTAATGGCTG  553

Query  540  TAAGCGGCCCGT  551
            ||||||||||||
Sbjct  554  TAAGCGGCCCGT  565


> MH376738.1 Pseudopithomyces pandanicola voucher MFLUCC 18-0116
28S large subunit ribosomal RNA gene, partial sequence
Length=861

 Score = 979 bits (530),  Expect = 0.0
 Identities = 545/552 (99%), Gaps = 1/552 (0%)
 Strand=Plus/Plus

Query  1    CGGCGAGTG-AGCGGCTACAGCTCAAATTTGAAATCTGGCCTCCTTTGGTGGTCCGAGTT  59
            ||||||||| ||||||||||||||||||||||||||||||||||||||| ||||||||||
Sbjct  17   CGGCGAGTGAAGCGGCTACAGCTCAAATTTGAAATCTGGCCTCCTTTGGGGGTCCGAGTT  76

Query  60   GTAATTTGCAGAGGATGCTTTGGCATTGGCGGCGGTCTAAGTTCCTTGGAACAGGACATC  119
            ||||||||||||||||||||||||||||||||||||||||||||||||||||||||||||
Sbjct  77   GTAATTTGCAGAGGATGCTTTGGCATTGGCGGCGGTCTAAGTTCCTTGGAACAGGACATC  136

Query  120  GCAGAGGGTGAGAATCCCGTACGTGGGCGCCTGCCTTTGCCGTGTAAAGCTCCTTCGACG  179
```

```
             ||||||||||||||||||||||||||||||||||||||||||||||||||||||||||||
Sbjct  137   GCAGAGGGTGAGAATCCCGTACGTGGGCGCCTGCCTTTGCCGTGTAAAGCTCCTTCGACG  196

Query  180   AGTCGAGTTGTTTGGGAATGCAGCTCTAAATGGGAGGTAAATTTCTCCTAAAGCTAAATA  239
             ||||||||||||||||||||||||||||||||||||||||||||||||||||||||||||
Sbjct  197   AGTCGAGTTGTTTGGGAATGCAGCTCTAAATGGGAGGTAAATTTCTCCTAAAGCTAAATA  256

Query  240   CCGGCCAGAGACCGATAGCGCACAAGTAGAGTGATCGAAAGATGAAAAGTACTTTGGAAA  299
             ||||||||||||||||||||||||||||||||||||||||||||||||||||||||||||
Sbjct  257   CCGGCCAGAGACCGATAGCGCACAAGTAGAGTGATCGAAAGATGAAAAGTACTTTGGAAA  316

Query  300   GAGAGTCAAATAGCACGTGAAATTGTTGAAAGGGAAGCGCTTGCAGCCAGACTTGCCCGC  359
             ||||||||||||||||||||||||||||||||||||||||||||||||||||||||||||
Sbjct  317   GAGAGTCAAATAGCACGTGAAATTGTTGAAAGGGAAGCGCTTGCAGCCAGACTTGCCCGC  376

Query  360   AGTTGCTCACCCAGGCTCTCGCCTGGGGCACTCTTCTGCGGGCAGGCCAGCATCAGTTTG  419
             ||||||||||| ||||| ||||||||||||||||||||||||||||||||||||||||||
Sbjct  377   AGTTGCTCACCTAGGCTTTCGCCTGGGGCACTCTTCTGCGGGCAGGCCAGCATCAGTTTG  436

Query  420   GGCGGTCGGATAAAGGCTCCTGTCATGTACCACCCCTCGGGGTGGCCTTATAgggggggC  479
             |||||||||||||||||  ||||||||||||||||||||||||||||||||||||| |||
Sbjct  437   GGCGGTCGGATAAAGGCCTCTGTCATGTACCACCCCTCGGGGTGGCCTTATAGGGGAGGC  496

Query  480   GTAATGCGACCAGCCCGGACTGAGGTCCGCGCATCTGCTAGGATGCTGGCGTAATGGCTG  539
             ||||||||||||||||||||||||||||||||||||||||||||||||||||||||||||
Sbjct  497   GTAATGCGACCAGCCCGGACTGAGGTCCGCGCATCTGCTAGGATGCTGGCGTAATGGCTG  556

Query  540   TAAGCGGCCCGT  551
             ||||||||||||
Sbjct  557   TAAGCGGCCCGT  568


> MK047478.1 Pseudopithomyces angolensis culture CBS:145056 28S
ribosomal RNA gene, partial sequence
Length=848

 Score = 979 bits (530),  Expect = 0.0
 Identities = 545/552 (99%), Gaps = 1/552 (0%)
 Strand=Plus/Plus

Query  1     CGGCGAGTG-AGCGGCTACAGCTCAAATTTGAAATCTGGCCTCCTTTGGTGGTCCGAGTT  59
             ||||||||| ||||||||||||||||||||||||||||||||||||||| ||||||||||
Sbjct  20    CGGCGAGTGAAGCGGCTACAGCTCAAATTTGAAATCTGGCCTCCTTTGGGGGTCCGAGTT  79

Query  60    GTAATTTGCAGAGGATGCTTTGGCATTGGCGGCGGTCTAAGTTCCTTGGAACAGGACATC  119
             ||||||||||||||||||||||||||||||||||||||||||||||||||||||||||||
Sbjct  80    GTAATTTGCAGAGGATGCTTTGGCATTGGCGGCGGTCTAAGTTCCTTGGAACAGGACATC  139

Query  120   GCAGAGGGTGAGAATCCCGTACGTGGGCGCCTGCCTTTGCCGTGTAAAGCTCCTTCGACG  179
             ||||||||||||||||||||||||||||||||||||||||||||||||||||||||||||
Sbjct  140   GCAGAGGGTGAGAATCCCGTACGTGGGCGCCTGCCTTTGCCGTGTAAAGCTCCTTCGACG  199

Query  180   AGTCGAGTTGTTTGGGAATGCAGCTCTAAATGGGAGGTAAATTTCTCCTAAAGCTAAATA  239
             ||||||||||||||||||||||||||||||||||||||||||||||||||||||||||||
Sbjct  200   AGTCGAGTTGTTTGGGAATGCAGCTCTAAATGGGAGGTAAATTTCTCCTAAAGCTAAATA  259

Query  240   CCGGCCAGAGACCGATAGCGCACAAGTAGAGTGATCGAAAGATGAAAAGTACTTTGGAAA  299
             ||||||||||||||||||||||||||||||||||||||||||||||||||||||||||||
Sbjct  260   CCGGCCAGAGACCGATAGCGCACAAGTAGAGTGATCGAAAGATGAAAAGTACTTTGGAAA  319

Query  300   GAGAGTCAAATAGCACGTGAAATTGTTGAAAGGGAAGCGCTTGCAGCCAGACTTGCCCGC  359
             ||||||||||||||||||||||||||||||||||||||||||||||||||||||||||||
Sbjct  320   GAGAGTCAAATAGCACGTGAAATTGTTGAAAGGGAAGCGCTTGCAGCCAGACTTGCCCGC  379
```

```
Query  360   AGTTGCTCACCCAGGCTCTCGCCTGGGGCACTCTTCTGCGGGCAGGCCAGCATCAGTTTG  419
             ||||||||||| ||||| ||||||||||||||||||||||||||||||||||||||||||
Sbjct  380   AGTTGCTCACCTAGGCTTTCGCCTGGGGCACTCTTCTGCGGGCAGGCCAGCATCAGTTTG  439

Query  420   GGCGGTCGGATAAAGGCTCCTGTCATGTACCACCCCTCGGGGTGGCCTTATAgggggggC  479
             |||||||||||||||||  ||||||||||||||||||||||||||||||||||||| |||
Sbjct  440   GGCGGTCGGATAAAGGCCTCTGTCATGTACCACCCCTCGGGGTGGCCTTATAGGGGAGGC  499

Query  480   GTAATGCGACCAGCCCGGACTGAGGTCCGCGCATCTGCTAGGATGCTGGCGTAATGGCTG  539
             ||||||||||||||||||||||||||||||||||||||||||||||||||||||||||||
Sbjct  500   GTAATGCGACCAGCCCGGACTGAGGTCCGCGCATCTGCTAGGATGCTGGCGTAATGGCTG  559

Query  540   TAAGCGGCCCGT  551
             ||||||||||||
Sbjct  560   TAAGCGGCCCGT  571


> NG_070067.1 Pseudopithomyces angolensis 28S rRNA gene, partial
sequence; from TYPE material
Length=848

 Score = 979 bits (530),  Expect = 0.0
 Identities = 545/552 (99%), Gaps = 1/552 (0%)
 Strand=Plus/Plus

Query  1     CGGCGAGTG-AGCGGCTACAGCTCAAATTTGAAATCTGGCCTCCTTTGGTGGTCCGAGTT  59
             ||||||||| ||||||||||||||||||||||||||||||||||||||| ||||||||||
Sbjct  20    CGGCGAGTGAAGCGGCTACAGCTCAAATTTGAAATCTGGCCTCCTTTGGGGGTCCGAGTT  79

Query  60    GTAATTTGCAGAGGATGCTTTGGCATTGGCGGCGGTCTAAGTTCCTTGGAACAGGACATC  119
             ||||||||||||||||||||||||||||||||||||||||||||||||||||||||||||
Sbjct  80    GTAATTTGCAGAGGATGCTTTGGCATTGGCGGCGGTCTAAGTTCCTTGGAACAGGACATC  139

Query  120   GCAGAGGGTGAGAATCCCGTACGTGGGCGCCTGCCTTTGCCGTGTAAAGCTCCTTCGACG  179
             ||||||||||||||||||||||||||||||||||||||||||||||||||||||||||||
Sbjct  140   GCAGAGGGTGAGAATCCCGTACGTGGGCGCCTGCCTTTGCCGTGTAAAGCTCCTTCGACG  199

Query  180   AGTCGAGTTGTTTGGGAATGCAGCTCTAAATGGGAGGTAAATTTCTCCTAAAGCTAAATA  239
             ||||||||||||||||||||||||||||||||||||||||||||||||||||||||||||
Sbjct  200   AGTCGAGTTGTTTGGGAATGCAGCTCTAAATGGGAGGTAAATTTCTCCTAAAGCTAAATA  259

Query  240   CCGGCCAGAGACCGATAGCGCACAAGTAGAGTGATCGAAAGATGAAAAGTACTTTGGAAA  299
             ||||||||||||||||||||||||||||||||||||||||||||||||||||||||||||
Sbjct  260   CCGGCCAGAGACCGATAGCGCACAAGTAGAGTGATCGAAAGATGAAAAGTACTTTGGAAA  319

Query  300   GAGAGTCAAATAGCACGTGAAATTGTTGAAAGGGAAGCGCTTGCAGCCAGACTTGCCCGC  359
             ||||||||||||||||||||||||||||||||||||||||||||||||||||||||||||
Sbjct  320   GAGAGTCAAATAGCACGTGAAATTGTTGAAAGGGAAGCGCTTGCAGCCAGACTTGCCCGC  379

Query  360   AGTTGCTCACCCAGGCTCTCGCCTGGGGCACTCTTCTGCGGGCAGGCCAGCATCAGTTTG  419
             ||||||||||| ||||| ||||||||||||||||||||||||||||||||||||||||||
Sbjct  380   AGTTGCTCACCTAGGCTTTCGCCTGGGGCACTCTTCTGCGGGCAGGCCAGCATCAGTTTG  439

Query  420   GGCGGTCGGATAAAGGCTCCTGTCATGTACCACCCCTCGGGGTGGCCTTATAgggggggC  479
             |||||||||||||||||  ||||||||||||||||||||||||||||||||||||| |||
Sbjct  440   GGCGGTCGGATAAAGGCCTCTGTCATGTACCACCCCTCGGGGTGGCCTTATAGGGGAGGC  499

Query  480   GTAATGCGACCAGCCCGGACTGAGGTCCGCGCATCTGCTAGGATGCTGGCGTAATGGCTG  539
             ||||||||||||||||||||||||||||||||||||||||||||||||||||||||||||
Sbjct  500   GTAATGCGACCAGCCCGGACTGAGGTCCGCGCATCTGCTAGGATGCTGGCGTAATGGCTG  559

Query  540   TAAGCGGCCCGT  551
             ||||||||||||
Sbjct  560   TAAGCGGCCCGT  571
```

```
> MG829064.1 Pseudopithomyces rosae strain MFLUCC 15-0035 28S ribosomal
RNA gene, partial sequence
Length=853

 Score = 979 bits (530),  Expect = 0.0
 Identities = 545/552 (99%), Gaps = 1/552 (0%)
 Strand=Plus/Plus

Query  1    CGGCGAGTG-AGCGGCTACAGCTCAAATTTGAAATCTGGCCTCCTTTGGTGGTCCGAGTT  59
            ||||||||| ||||||||||||||||||||||||||||||||||||||| ||||||||||
Sbjct  14   CGGCGAGTGAAGCGGCTACAGCTCAAATTTGAAATCTGGCCTCCTTTGGGGGTCCGAGTT  73

Query  60   GTAATTTGCAGAGGATGCTTTGGCATTGGCGGCGGTCTAAGTTCCTTGGAACAGGACATC  119
            ||||||||||||||||||||||||||||||||||||||||||||||||||||||||||||
Sbjct  74   GTAATTTGCAGAGGATGCTTTGGCATTGGCGGCGGTCTAAGTTCCTTGGAACAGGACATC  133

Query  120  GCAGAGGGTGAGAATCCCGTACGTGGGCGCCTGCCTTTGCCGTGTAAAGCTCCTTCGACG  179
            ||||||||||||||||||||||||||||||||||||||||||||||||||||||||||||
Sbjct  134  GCAGAGGGTGAGAATCCCGTACGTGGGCGCCTGCCTTTGCCGTGTAAAGCTCCTTCGACG  193

Query  180  AGTCGAGTTGTTTGGGAATGCAGCTCTAAATGGGAGGTAAATTTCTCCTAAAGCTAAATA  239
            ||||||||||||||||||||||||||||||||||||||||||||||||||||||||||||
Sbjct  194  AGTCGAGTTGTTTGGGAATGCAGCTCTAAATGGGAGGTAAATTTCTCCTAAAGCTAAATA  253

Query  240  CCGGCCAGAGACCGATAGCGCACAAGTAGAGTGATCGAAAGATGAAAAGTACTTTGGAAA  299
            ||||||||||||||||||||||||||||||||||||||||||||||||||||||||||||
Sbjct  254  CCGGCCAGAGACCGATAGCGCACAAGTAGAGTGATCGAAAGATGAAAAGTACTTTGGAAA  313

Query  300  GAGAGTCAAATAGCACGTGAAATTGTTGAAAGGGAAGCGCTTGCAGCCAGACTTGCCCGC  359
            ||||||||||||||||||||||||||||||||||||||||||||||||||||||||||||
Sbjct  314  GAGAGTCAAATAGCACGTGAAATTGTTGAAAGGGAAGCGCTTGCAGCCAGACTTGCCCGC  373

Query  360  AGTTGCTCACCCAGGCTCTCGCCTGGGGCACTCTTCTGCGGGCAGGCCAGCATCAGTTTG  419
            ||||||||||| ||||| ||||||||||||||||||||||||||||||||||||||||||
Sbjct  374  AGTTGCTCACCTAGGCTTTCGCCTGGGGCACTCTTCTGCGGGCAGGCCAGCATCAGTTTG  433

Query  420  GGCGGTCGGATAAAGGCTCCTGTCATGTACCACCCCTCGGGGTGGCCTTATAgggggggC  479
            |||||||||||||||||  ||||||||||||||||||||||||||||||||||||| |||
Sbjct  434  GGCGGTCGGATAAAGGCCTCTGTCATGTACCACCCCTCGGGGTGGCCTTATAGGGGAGGC  493

Query  480  GTAATGCGACCAGCCCGGACTGAGGTCCGCGCATCTGCTAGGATGCTGGCGTAATGGCTG  539
            ||||||||||||||||||||||||||||||||||||||||||||||||||||||||||||
Sbjct  494  GTAATGCGACCAGCCCGGACTGAGGTCCGCGCATCTGCTAGGATGCTGGCGTAATGGCTG  553

Query  540  TAAGCGGCCCGT  551
            ||||||||||||
Sbjct  554  TAAGCGGCCCGT  565


> NG_059876.1 Pseudopithomyces rosae MFLU 18-0109 28S rRNA gene,
partial sequence; from TYPE material
Length=853

 Score = 979 bits (530),  Expect = 0.0
 Identities = 545/552 (99%), Gaps = 1/552 (0%)
 Strand=Plus/Plus

Query  1    CGGCGAGTG-AGCGGCTACAGCTCAAATTTGAAATCTGGCCTCCTTTGGTGGTCCGAGTT  59
            ||||||||| ||||||||||||||||||||||||||||||||||||||| ||||||||||
Sbjct  14   CGGCGAGTGAAGCGGCTACAGCTCAAATTTGAAATCTGGCCTCCTTTGGGGGTCCGAGTT  73

Query  60   GTAATTTGCAGAGGATGCTTTGGCATTGGCGGCGGTCTAAGTTCCTTGGAACAGGACATC  119
```

```
            ||||||||||||||||||||||||||||||||||||||||||||||||||||||||||||
Sbjct  74   GTAATTTGCAGAGGATGCTTTGGCATTGGCGGCGGTCTAAGTTCCTTGGAACAGGACATC  133

Query  120  GCAGAGGGTGAGAATCCCGTACGTGGGCGCCTGCCTTTGCCGTGTAAAGCTCCTTCGACG  179
            ||||||||||||||||||||||||||||||||||||||||||||||||||||||||||||
Sbjct  134  GCAGAGGGTGAGAATCCCGTACGTGGGCGCCTGCCTTTGCCGTGTAAAGCTCCTTCGACG  193

Query  180  AGTCGAGTTGTTTGGGAATGCAGCTCTAAATGGGAGGTAAATTTCTCCTAAAGCTAAATA  239
            ||||||||||||||||||||||||||||||||||||||||||||||||||||||||||||
Sbjct  194  AGTCGAGTTGTTTGGGAATGCAGCTCTAAATGGGAGGTAAATTTCTCCTAAAGCTAAATA  253

Query  240  CCGGCCAGAGACCGATAGCGCACAAGTAGAGTGATCGAAAGATGAAAAGTACTTTGGAAA  299
            ||||||||||||||||||||||||||||||||||||||||||||||||||||||||||||
Sbjct  254  CCGGCCAGAGACCGATAGCGCACAAGTAGAGTGATCGAAAGATGAAAAGTACTTTGGAAA  313

Query  300  GAGAGTCAAATAGCACGTGAAATTGTTGAAAGGGAAGCGCTTGCAGCCAGACTTGCCCGC  359
            ||||||||||||||||||||||||||||||||||||||||||||||||||||||||||||
Sbjct  314  GAGAGTCAAATAGCACGTGAAATTGTTGAAAGGGAAGCGCTTGCAGCCAGACTTGCCCGC  373

Query  360  AGTTGCTCACCCAGGCTCTCGCCTGGGGCACTCTTCTGCGGGCAGGCCAGCATCAGTTTG  419
            ||||||||||| ||||| ||||||||||||||||||||||||||||||||||||||||||
Sbjct  374  AGTTGCTCACCTAGGCTTTCGCCTGGGGCACTCTTCTGCGGGCAGGCCAGCATCAGTTTG  433

Query  420  GGCGGTCGGATAAAGGCTCCTGTCATGTACCACCCCTCGGGGTGGCCTTATAgggggggC  479
            |||||||||||||||||  ||||||||||||||||||||||||||||||||||||| |||
Sbjct  434  GGCGGTCGGATAAAGGCCTCTGTCATGTACCACCCCTCGGGGTGGCCTTATAGGGGAGGC  493

Query  480  GTAATGCGACCAGCCCGGACTGAGGTCCGCGCATCTGCTAGGATGCTGGCGTAATGGCTG  539
            ||||||||||||||||||||||||||||||||||||||||||||||||||||||||||||
Sbjct  494  GTAATGCGACCAGCCCGGACTGAGGTCCGCGCATCTGCTAGGATGCTGGCGTAATGGCTG  553

Query  540  TAAGCGGCCCGT  551
            ||||||||||||
Sbjct  554  TAAGCGGCCCGT  565


> MF804532.1 Pseudopithomyces palmicola strain L12 28S ribosomal
RNA gene, partial sequence
Length=867

 Score = 974 bits (527),  Expect = 0.0
 Identities = 544/552 (99%), Gaps = 1/552 (0%)
 Strand=Plus/Plus

Query  1    CGGCGAGTG-AGCGGCTACAGCTCAAATTTGAAATCTGGCCTCCTTTGGTGGTCCGAGTT  59
            ||||||||| ||||||||||||||||||||||||||||||||||||||| ||||||||||
Sbjct  17   CGGCGAGTGAAGCGGCTACAGCTCAAATTTGAAATCTGGCCTCCTTTGGGGGTCCGAGTT  76

Query  60   GTAATTTGCAGAGGATGCTTTGGCATTGGCGGCGGTCTAAGTTCCTTGGAACAGGACATC  119
            ||||||||||||||||||||||||||||||||||||||||||||||||||||||||||||
Sbjct  77   GTAATTTGCAGAGGATGCTTTGGCATTGGCGGCGGTCTAAGTTCCTTGGAACAGGACATC  136

Query  120  GCAGAGGGTGAGAATCCCGTACGTGGGCGCCTGCCTTTGCCGTGTAAAGCTCCTTCGACG  179
            ||||||||||||||||||||||||||||||||||||||||||||||||||||||||||||
Sbjct  137  GCAGAGGGTGAGAATCCCGTACGTGGGCGCCTGCCTTTGCCGTGTAAAGCTCCTTCGACG  196

Query  180  AGTCGAGTTGTTTGGGAATGCAGCTCTAAATGGGAGGTAAATTTCTCCTAAAGCTAAATA  239
            ||||||||||||||||||||||||||||||||||||||||||||||||||||||||||||
Sbjct  197  AGTCGAGTTGTTTGGGAATGCAGCTCTAAATGGGAGGTAAATTTCTCCTAAAGCTAAATA  256

Query  240  CCGGCCAGAGACCGATAGCGCACAAGTAGAGTGATCGAAAGATGAAAAGTACTTTGGAAA  299
            ||||||||||||||||||||||||||||||||||||||||||||||||||||||||||||
Sbjct  257  CCGGCCAGAGACCGATAGCGCACAAGTAGAGTGATCGAAAGATGAAAAGTACTTTGGAAA  316
```

```
Query  300   GAGAGTCAAATAGCACGTGAAATTGTTGAAAGGGAAGCGCTTGCAGCCAGACTTGCCCGC  359
             ||||||||||||||||||||||||||||||||||||||||||||||||||||||||||||
Sbjct  317   GAGAGTCAAATAGCACGTGAAATTGTTGAAAGGGAAGCGCTTGCAGCCAGACTTGCCCGC  376

Query  360   AGTTGCTCACCCAGGCTCTCGCCTGGGGCACTCTTCTGCGGGCAGGCCAGCATCAGTTTG  419
             ||||||||||| ||||| ||||||||||||||||||||||||||||||||||||||||||
Sbjct  377   AGTTGCTCACCTAGGCTTTCGCCTGGGGCACTCTTCTGCGGGCAGGCCAGCATCAGTTTG  436

Query  420   GGCGGTCGGATAAAGGCTCCTGTCATGTACCACCCCTCGGGGTGGCCTTATAgggggggC  479
             |||||||||||||||||  ||||||||||||||||||||||||||||||||||||| || 
Sbjct  437   GGCGGTCGGATAAAGGCCTCTGTCATGTACCACCCCTCGGGGTGGCCTTATAGGGGAGGT  496

Query  480   GTAATGCGACCAGCCCGGACTGAGGTCCGCGCATCTGCTAGGATGCTGGCGTAATGGCTG  539
             ||||||||||||||||||||||||||||||||||||||||||||||||||||||||||||
Sbjct  497   GTAATGCGACCAGCCCGGACTGAGGTCCGCGCATCTGCTAGGATGCTGGCGTAATGGCTG  556

Query  540   TAAGCGGCCCGT  551
             ||||||||||||
Sbjct  557   TAAGCGGCCCGT  568


> MF804534.1 Pseudopithomyces palmicola strain PP28B 28S ribosomal
RNA gene, partial sequence
Length=867

 Score = 974 bits (527),  Expect = 0.0
 Identities = 544/552 (99%), Gaps = 1/552 (0%)
 Strand=Plus/Plus

Query  1     CGGCGAGTG-AGCGGCTACAGCTCAAATTTGAAATCTGGCCTCCTTTGGTGGTCCGAGTT  59
             ||||||||| ||||||||||||||||||||||||||||||||||||||| ||||||||||
Sbjct  17    CGGCGAGTGAAGCGGCTACAGCTCAAATTTGAAATCTGGCCTCCTTTGGGGGTCCGAGTT  76

Query  60    GTAATTTGCAGAGGATGCTTTGGCATTGGCGGCGGTCTAAGTTCCTTGGAACAGGACATC  119
             ||||||||||||||||||||||||||||||||||||||||||||||||||||||||||||
Sbjct  77    GTAATTTGCAGAGGATGCTTTGGCATTGGCGGCGGTCTAAGTTCCTTGGAACAGGACATC  136

Query  120   GCAGAGGGTGAGAATCCCGTACGTGGGCGCCTGCCTTTGCCGTGTAAAGCTCCTTCGACG  179
             ||||||||||||||||||||||||||||||||||||||||||||||||||||||||||||
Sbjct  137   GCAGAGGGTGAGAATCCCGTACGTGGGCGCCTGCCTTTGCCGTGTAAAGCTCCTTCGACG  196

Query  180   AGTCGAGTTGTTTGGGAATGCAGCTCTAAATGGGAGGTAAATTTCTCCTAAAGCTAAATA  239
             ||||||||||||||||||||||||||||||||||||||||||||||||||||||||||||
Sbjct  197   AGTCGAGTTGTTTGGGAATGCAGCTCTAAATGGGAGGTAAATTTCTCCTAAAGCTAAATA  256

Query  240   CCGGCCAGAGACCGATAGCGCACAAGTAGAGTGATCGAAAGATGAAAAGTACTTTGGAAA  299
             ||||||||||||||||||||||||||||||||||||||||||||||||||||||||||||
Sbjct  257   CCGGCCAGAGACCGATAGCGCACAAGTAGAGTGATCGAAAGATGAAAAGTACTTTGGAAA  316

Query  300   GAGAGTCAAATAGCACGTGAAATTGTTGAAAGGGAAGCGCTTGCAGCCAGACTTGCCCGC  359
             ||||||||||||||||||||||||||||||||||||||||||||||||||||||||||||
Sbjct  317   GAGAGTCAAATAGCACGTGAAATTGTTGAAAGGGAAGCGCTTGCAGCCAGACTTGCCCGC  376

Query  360   AGTTGCTCACCCAGGCTCTCGCCTGGGGCACTCTTCTGCGGGCAGGCCAGCATCAGTTTG  419
             ||||||||||| ||||| ||||||||||||||||||||||||||||||||||||||||||
Sbjct  377   AGTTGCTCACCTAGGCTTTCGCCTGGGGCACTCTTCTGCGGGCAGGCCAGCATCAGTTTG  436

Query  420   GGCGGTCGGATAAAGGCTCCTGTCATGTACCACCCCTCGGGGTGGCCTTATAgggggggC  479
             |||||||||||||||||  ||||||||||||||||||||||||||||||||||||| || 
Sbjct  437   GGCGGTCGGATAAAGGCCTCTGTCATGTACCACCCCTCGGGGTGGCCTTATAGGGGAGGT  496

Query  480   GTAATGCGACCAGCCCGGACTGAGGTCCGCGCATCTGCTAGGATGCTGGCGTAATGGCTG  539
             ||||||||||||||||||||||||||||||||||||||||||||||||||||||||||||
Sbjct  497   GTAATGCGACCAGCCCGGACTGAGGTCCGCGCATCTGCTAGGATGCTGGCGTAATGGCTG  556
```

```
Query  540   TAAGCGGCCCGT  551
             ||||||||||||
Sbjct  557   TAAGCGGCCCGT  568


> MF804535.1 Pseudopithomyces palmicola strain AC4 28S ribosomal
RNA gene, partial sequence
Length=867

 Score = 974 bits (527),  Expect = 0.0
 Identities = 544/552 (99%), Gaps = 1/552 (0%)
 Strand=Plus/Plus

Query  1     CGGCGAGTG-AGCGGCTACAGCTCAAATTTGAAATCTGGCCTCCTTTGGTGGTCCGAGTT  59
             ||||||||| ||||||||||||||||||||||||||||||||||| ||| ||||||||||
Sbjct  17    CGGCGAGTGAAGCGGCTACAGCTCAAATTTGAAATCTGGCCTCCTCTGGGGGTCCGAGTT  76

Query  60    GTAATTTGCAGAGGATGCTTTGGCATTGGCGGCGGTCTAAGTTCCTTGGAACAGGACATC  119
             ||||||||||||||||||||||||||||||||||||||||||||||||||||||||||||
Sbjct  77    GTAATTTGCAGAGGATGCTTTGGCATTGGCGGCGGTCTAAGTTCCTTGGAACAGGACATC  136

Query  120   GCAGAGGGTGAGAATCCCGTACGTGGGCGCCTGCCTTTGCCGTGTAAAGCTCCTTCGACG  179
             ||||||||||||||||||||||||||||||||||||||||||||||||||||||||||||
Sbjct  137   GCAGAGGGTGAGAATCCCGTACGTGGGCGCCTGCCTTTGCCGTGTAAAGCTCCTTCGACG  196

Query  180   AGTCGAGTTGTTTGGGAATGCAGCTCTAAATGGGAGGTAAATTTCTCCTAAAGCTAAATA  239
             ||||||||||||||||||||||||||||||||||||||||||||||||||||||||||||
Sbjct  197   AGTCGAGTTGTTTGGGAATGCAGCTCTAAATGGGAGGTAAATTTCTCCTAAAGCTAAATA  256

Query  240   CCGGCCAGAGACCGATAGCGCACAAGTAGAGTGATCGAAAGATGAAAAGTACTTTGGAAA  299
             ||||||||||||||||||||||||||||||||||||||||||||||||||||||||||||
Sbjct  257   CCGGCCAGAGACCGATAGCGCACAAGTAGAGTGATCGAAAGATGAAAAGTACTTTGGAAA  316

Query  300   GAGAGTCAAATAGCACGTGAAATTGTTGAAAGGGAAGCGCTTGCAGCCAGACTTGCCCGC  359
             ||||||||||||||||||||||||||||||||||||||||||||||||||||||||||||
Sbjct  317   GAGAGTCAAATAGCACGTGAAATTGTTGAAAGGGAAGCGCTTGCAGCCAGACTTGCCCGC  376

Query  360   AGTTGCTCACCCAGGCTCTCGCCTGGGGCACTCTTCTGCGGGCAGGCCAGCATCAGTTTG  419
             ||||||||||| ||||| ||||||||||||||||||||||||||||||||||||||||||
Sbjct  377   AGTTGCTCACCTAGGCTTTCGCCTGGGGCACTCTTCTGCGGGCAGGCCAGCATCAGTTTG  436

Query  420   GGCGGTCGGATAAAGGCTCCTGTCATGTACCACCCCTCGGGGTGGCCTTATAgggggggC  479
             |||||||||||||||||  ||||||||||||||||||||||||||||||||||||| |||
Sbjct  437   GGCGGTCGGATAAAGGCCTCTGTCATGTACCACCCCTCGGGGTGGCCTTATAGGGGAGGC  496

Query  480   GTAATGCGACCAGCCCGGACTGAGGTCCGCGCATCTGCTAGGATGCTGGCGTAATGGCTG  539
             ||||||||||||||||||||||||||||||||||||||||||||||||||||||||||||
Sbjct  497   GTAATGCGACCAGCCCGGACTGAGGTCCGCGCATCTGCTAGGATGCTGGCGTAATGGCTG  556

Query  540   TAAGCGGCCCGT  551
             ||||||||||||
Sbjct  557   TAAGCGGCCCGT  568


> HG933828.1 Pseudopithomyces karoo genomic DNA containing, 28S
rRNA gene
Length=955

 Score = 968 bits (524),  Expect = 0.0
 Identities = 544/553 (98%), Gaps = 3/553 (1%)
 Strand=Plus/Plus

Query  1     CGGCGAGTG-AGCGGCTACAGCTCAAATTTGAAATCTGGCCTCCTTTGGTGGTCCGAGTT  59
```

```
            ||||||||| ||||||||||||||||||||||||||||||| ||||||| ||||||||||
Sbjct  38   CGGCGAGTGAAGCGGCTACAGCTCAAATTTGAAATCTGGCCCCCTTTGGAGGTCCGAGTT  97

Query  60   GTAATTTGCAGAGGATGCTTTGGCATTGGCGGCGGTCTAAGTTCCTTGGAACAGGACATC  119
            ||||||||||||||||||||||||||||||||||||||||||||||||||||||||||||
Sbjct  98   GTAATTTGCAGAGGATGCTTTGGCATTGGCGGCGGTCTAAGTTCCTTGGAACAGGACATC  157

Query  120  GCAGAGGGTGAGAATCCCGTACGTGGGCGCCTGCCTTTGCCGTGTAAAGCTCCTTCGACG  179
            ||||||||||||||||||||||||||||||||||||||||||||||||||||||||||||
Sbjct  158  GCAGAGGGTGAGAATCCCGTACGTGGGCGCCTGCCTTTGCCGTGTAAAGCTCCTTCGACG  217

Query  180  AGTCGAGTTGTTTGGGAATGCAGCTCTAAATGGGAGGTAAATTTCTCCTAAAGCTAAATA  239
            ||||||||||||||||||||||||||||||||||||||||||||||||||||||||||||
Sbjct  218  AGTCGAGTTGTTTGGGAATGCAGCTCTAAATGGGAGGTAAATTTCTCCTAAAGCTAAATA  277

Query  240  CCGGCCAGAGACCGATAGCGCACAAGTAGAGTGATCGAAAGATGAAAAGTACTTTGGAAA  299
            ||||||||||||||||||||||||||||||||||||||||||||||||||||||||||||
Sbjct  278  CCGGCCAGAGACCGATAGCGCACAAGTAGAGTGATCGAAAGATGAAAAGTACTTTGGAAA  337

Query  300  GAGAGTCAAATAGCACGTGAAATTGTTGAAAGGGAAGCGCTTGCAGCCAGACTTGCCCGC  359
            ||||||||||||||||||||||||||||||||||||||||||||||||||||||||||||
Sbjct  338  GAGAGTCAAATAGCACGTGAAATTGTTGAAAGGGAAGCGCTTGCAGCCAGACTTGCCCGC  397

Query  360  AGTTGCTCACCCAGGCTCTCGCCTGGGGCACTCTTCTGCGGGCAGGCCAGCATCAGTTTG  419
            ||||||||||||||||||| ||||||||||||||||||||||||||||||||||||||||
Sbjct  398  AGTTGCTCACCCAGGCTCTTGCCTGGGGCACTCTTCTGCGGGCAGGCCAGCATCAGTTTG  457

Query  420  GGCGGTCGGATAAAGG-CTCCTGTCATGTACCACCCCTCGGGGTGGCCTTATAggggggg  478
            |||||||||||||||| || ||||||||||||||||||||||||||||||||||||| |
Sbjct  458  GGCGGTCGGATAAAGGTCT-CTGTCATGTACCACCCCTCGGGGTGGCCTTATAGGGGAGA  516

Query  479  CGTAATGCGACCAGCCCGGACTGAGGTCCGCGCATCTGCTAGGATGCTGGCGTAATGGCT  538
            || |||||||||||||||||||||||||||||||||||||||||||||||||||||||||
Sbjct  517  CGCAATGCGACCAGCCCGGACTGAGGTCCGCGCATCTGCTAGGATGCTGGCGTAATGGCT  576

Query  539  GTAAGCGGCCCGT  551
            |||||||||||||
Sbjct  577  GTAAGCGGCCCGT  589


> HG933829.1 Pseudopithomyces karoo genomic DNA containing, 28S
rRNA gene
Length=987

 Score = 968 bits (524),  Expect = 0.0
 Identities = 544/553 (98%), Gaps = 3/553 (1%)
 Strand=Plus/Plus

Query  1    CGGCGAGTG-AGCGGCTACAGCTCAAATTTGAAATCTGGCCTCCTTTGGTGGTCCGAGTT  59
            ||||||||| ||||||||||||||||||||||||||||||| ||||||| ||||||||||
Sbjct  38   CGGCGAGTGAAGCGGCTACAGCTCAAATTTGAAATCTGGCCCCCTTTGGAGGTCCGAGTT  97

Query  60   GTAATTTGCAGAGGATGCTTTGGCATTGGCGGCGGTCTAAGTTCCTTGGAACAGGACATC  119
            ||||||||||||||||||||||||||||||||||||||||||||||||||||||||||||
Sbjct  98   GTAATTTGCAGAGGATGCTTTGGCATTGGCGGCGGTCTAAGTTCCTTGGAACAGGACATC  157

Query  120  GCAGAGGGTGAGAATCCCGTACGTGGGCGCCTGCCTTTGCCGTGTAAAGCTCCTTCGACG  179
            ||||||||||||||||||||||||||||||||||||||||||||||||||||||||||||
Sbjct  158  GCAGAGGGTGAGAATCCCGTACGTGGGCGCCTGCCTTTGCCGTGTAAAGCTCCTTCGACG  217

Query  180  AGTCGAGTTGTTTGGGAATGCAGCTCTAAATGGGAGGTAAATTTCTCCTAAAGCTAAATA  239
            ||||||||||||||||||||||||||||||||||||||||||||||||||||||||||||
Sbjct  218  AGTCGAGTTGTTTGGGAATGCAGCTCTAAATGGGAGGTAAATTTCTCCTAAAGCTAAATA  277
```

```
Query  240  CCGGCCAGAGACCGATAGCGCACAAGTAGAGTGATCGAAAGATGAAAAGTACTTTGGAAA  299
            ||||||||||||||||||||||||||||||||||||||||||||||||||||||||||||
Sbjct  278  CCGGCCAGAGACCGATAGCGCACAAGTAGAGTGATCGAAAGATGAAAAGTACTTTGGAAA  337

Query  300  GAGAGTCAAATAGCACGTGAAATTGTTGAAAGGGAAGCGCTTGCAGCCAGACTTGCCCGC  359
            ||||||||||||||||||||||||||||||||||||||||||||||||||||||||||||
Sbjct  338  GAGAGTCAAATAGCACGTGAAATTGTTGAAAGGGAAGCGCTTGCAGCCAGACTTGCCCGC  397

Query  360  AGTTGCTCACCCAGGCTCTCGCCTGGGGCACTCTTCTGCGGGCAGGCCAGCATCAGTTTG  419
            ||||||||||||||||||| ||||||||||||||||||||||||||||||||||||||||
Sbjct  398  AGTTGCTCACCCAGGCTCTTGCCTGGGGCACTCTTCTGCGGGCAGGCCAGCATCAGTTTG  457

Query  420  GGCGGTCGGATAAAGG-CTCCTGTCATGTACCACCCCTCGGGGTGGCCTTATAggggggg  478
            |||||||||||||||| || |||||||||||||||||||||||||||||||||||| |
Sbjct  458  GGCGGTCGGATAAAGGTCT-CTGTCATGTACCACCCCTCGGGGTGGCCTTATAGGGGAGA  516

Query  479  CGTAATGCGACCAGCCCGGACTGAGGTCCGCGCATCTGCTAGGATGCTGGCGTAATGGCT  538
            || |||||||||||||||||||||||||||||||||||||||||||||||||||||||||
Sbjct  517  CGCAATGCGACCAGCCCGGACTGAGGTCCGCGCATCTGCTAGGATGCTGGCGTAATGGCT  576

Query  539  GTAAGCGGCCCGT  551
            |||||||||||||
Sbjct  577  GTAAGCGGCCCGT  589


> NG_057865.1 Pseudopithomyces karoo CBS 804.72 28S rRNA gene,
partial sequence; from TYPE material
Length=955

 Score = 968 bits (524),  Expect = 0.0
 Identities = 544/553 (98%), Gaps = 3/553 (1%)
 Strand=Plus/Plus

Query  1    CGGCGAGTG-AGCGGCTACAGCTCAAATTTGAAATCTGGCCTCCTTTGGTGGTCCGAGTT  59
            ||||||||| ||||||||||||||||||||||||||||||| ||||||| ||||||||||
Sbjct  38   CGGCGAGTGAAGCGGCTACAGCTCAAATTTGAAATCTGGCCCCCTTTGGAGGTCCGAGTT  97

Query  60   GTAATTTGCAGAGGATGCTTTGGCATTGGCGGCGGTCTAAGTTCCTTGGAACAGGACATC  119
            ||||||||||||||||||||||||||||||||||||||||||||||||||||||||||||
Sbjct  98   GTAATTTGCAGAGGATGCTTTGGCATTGGCGGCGGTCTAAGTTCCTTGGAACAGGACATC  157

Query  120  GCAGAGGGTGAGAATCCCGTACGTGGGCGCCTGCCTTTGCCGTGTAAAGCTCCTTCGACG  179
            ||||||||||||||||||||||||||||||||||||||||||||||||||||||||||||
Sbjct  158  GCAGAGGGTGAGAATCCCGTACGTGGGCGCCTGCCTTTGCCGTGTAAAGCTCCTTCGACG  217

Query  180  AGTCGAGTTGTTTGGGAATGCAGCTCTAAATGGGAGGTAAATTTCTCCTAAAGCTAAATA  239
            ||||||||||||||||||||||||||||||||||||||||||||||||||||||||||||
Sbjct  218  AGTCGAGTTGTTTGGGAATGCAGCTCTAAATGGGAGGTAAATTTCTCCTAAAGCTAAATA  277

Query  240  CCGGCCAGAGACCGATAGCGCACAAGTAGAGTGATCGAAAGATGAAAAGTACTTTGGAAA  299
            ||||||||||||||||||||||||||||||||||||||||||||||||||||||||||||
Sbjct  278  CCGGCCAGAGACCGATAGCGCACAAGTAGAGTGATCGAAAGATGAAAAGTACTTTGGAAA  337

Query  300  GAGAGTCAAATAGCACGTGAAATTGTTGAAAGGGAAGCGCTTGCAGCCAGACTTGCCCGC  359
            ||||||||||||||||||||||||||||||||||||||||||||||||||||||||||||
Sbjct  338  GAGAGTCAAATAGCACGTGAAATTGTTGAAAGGGAAGCGCTTGCAGCCAGACTTGCCCGC  397

Query  360  AGTTGCTCACCCAGGCTCTCGCCTGGGGCACTCTTCTGCGGGCAGGCCAGCATCAGTTTG  419
            ||||||||||||||||||| ||||||||||||||||||||||||||||||||||||||||
Sbjct  398  AGTTGCTCACCCAGGCTCTTGCCTGGGGCACTCTTCTGCGGGCAGGCCAGCATCAGTTTG  457

Query  420  GGCGGTCGGATAAAGG-CTCCTGTCATGTACCACCCCTCGGGGTGGCCTTATAggggggg  478
            |||||||||||||||| || |||||||||||||||||||||||||||||||||||| |
Sbjct  458  GGCGGTCGGATAAAGGTCT-CTGTCATGTACCACCCCTCGGGGTGGCCTTATAGGGGAGA  516
```

```
Query  479   CGTAATGCGACCAGCCCGGACTGAGGTCCGCGCATCTGCTAGGATGCTGGCGTAATGGCT  538
             || |||||||||||||||||||||||||||||||||||||||||||||||||||||||||
Sbjct  517   CGCAATGCGACCAGCCCGGACTGAGGTCCGCGCATCTGCTAGGATGCTGGCGTAATGGCT  576

Query  539   GTAAGCGGCCCGT  551
             |||||||||||||
Sbjct  577   GTAAGCGGCCCGT  589
```

> LT671616.1 Pseudopithomyces atro-olivaceus genomic DNA sequence
contains 28S rRNA gene, strain CBS 244.96
Length=1070

Score = 963 bits (521),  Expect = 0.0
Identities = 543/553 (98%), Gaps = 3/553 (1%)
Strand=Plus/Plus

```
Query  1     CGGCGAGTG-AGCGGCTACAGCTCAAATTTGAAATCTGGCCTCCTTTGGTGGTCCGAGTT  59
             ||||||||| ||||||||||||||||||||||||||||||||||||||||||||||||||
Sbjct  51    CGGCGAGTGAAGCGGCTACAGCTCAAATTTGAAATCTGGCCTCCTTTGGTGGTCCGAGTT  110

Query  60    GTAATTTGCAGAGGATGCTTTGGCATTGGCGGCGGTCTAAGTTCCTTGGAACAGGACATC  119
             ||||||||||||||||||||||||||||||||||||||||||||||||||||||||||||
Sbjct  111   GTAATTTGCAGAGGATGCTTTGGCATTGGCGGCGGTCTAAGTTCCTTGGAACAGGACATC  170

Query  120   GCAGAGGGTGAGAATCCCGTACGTGGGCGCCTGCCTTTGCCGTGTAAAGCTCCTTCGACG  179
             ||||||||||||||||||||||||||||||||||||||||||||||||||||||||||||
Sbjct  171   GCAGAGGGTGAGAATCCCGTACGTGGGCGCCTGCCTTTGCCGTGTAAAGCTCCTTCGACG  230

Query  180   AGTCGAGTTGTTTGGGAATGCAGCTCTAAATGGGAGGTAAATTTCTCCTAAAGCTAAATA  239
             ||||||||||||||||||||||||||||||||||||||||||||||||||||||||||||
Sbjct  231   AGTCGAGTTGTTTGGGAATGCAGCTCTAAATGGGAGGTAAATTTCTCCTAAAGCTAAATA  290

Query  240   CCGGCCAGAGACCGATAGCGCACAAGTAGAGTGATCGAAAGATGAAAAGTACTTTGGAAA  299
             ||||||||||||||||||||||||||||||||||||||||||||||||||||||||||||
Sbjct  291   CCGGCCAGAGACCGATAGCGCACAAGTAGAGTGATCGAAAGATGAAAAGTACTTTGGAAA  350

Query  300   GAGAGTCAAATAGCACGTGAAATTGTTGAAAGGGAAGCGCTTGCAGCCAGACTTGCCCGC  359
             ||||||||||||||||||||||||||||||||||||||||||||||||||||||||||||
Sbjct  351   GAGAGTCAAATAGCACGTGAAATTGTTGAAAGGGAAGCGCTTGCAGCCAGACTTGCCCGC  410

Query  360   AGTTGCTCACCCAGGCTCTCGCCTGGGGCACTCTTCTGCGGGCAGGCCAGCATCAGTTTG  419
             ||||||||||| ||||| | ||||||||||||||||||||||||||||||||||||||||
Sbjct  411   AGTTGCTCACCTAGGCTTTTGCCTGGGGCACTCTTCTGCGGGCAGGCCAGCATCAGTTTG  470

Query  420   GGCGGTCGGATAAAGG-CTCCTGTCATGTACCACCCCTCGGGGTGGCCTTATAggggggg  478
             |||||||||||||||| || |||||||||||||||| |||||||||||||||||||| |
Sbjct  471   GGCGGTCGGATAAAGGTCT-CTGTCATGTACCACCCTTCGGGGTGGCCTTATAGGGGAGA  529

Query  479   CGTAATGCGACCAGCCCGGACTGAGGTCCGCGCATCTGCTAGGATGCTGGCGTAATGGCT  538
             || |||||||||||||||||||||||||||||||||||||||||||||||||||||||||
Sbjct  530   CGCAATGCGACCAGCCCGGACTGAGGTCCGCGCATCTGCTAGGATGCTGGCGTAATGGCT  589

Query  539   GTAAGCGGCCCGT  551
             |||||||||||||
Sbjct  590   GTAAGCGGCCCGT  602
```

> LT671617.1 Pseudopithomyces atro-olivaceus genomic DNA sequence
contains 28S rRNA gene, strain MUCL 33112
Length=1025

Score = 963 bits (521),  Expect = 0.0

```
 Identities = 543/553 (98%), Gaps = 3/553 (1%)
 Strand=Plus/Plus

Query  1    CGGCGAGTG-AGCGGCTACAGCTCAAATTTGAAATCTGGCCTCCTTTGGTGGTCCGAGTT  59
            ||||||||| ||||||||||||||||||||||||||||||||||||||||||||||||||
Sbjct  23   CGGCGAGTGAAGCGGCTACAGCTCAAATTTGAAATCTGGCCTCCTTTGGTGGTCCGAGTT  82

Query  60   GTAATTTGCAGAGGATGCTTTGGCATTGGCGGCGGTCTAAGTTCCTTGGAACAGGACATC  119
            ||||||||||||||||||||||||||||||||||||||||||||||||||||||||||||
Sbjct  83   GTAATTTGCAGAGGATGCTTTGGCATTGGCGGCGGTCTAAGTTCCTTGGAACAGGACATC  142

Query  120  GCAGAGGGTGAGAATCCCGTACGTGGGCGCCTGCCTTTGCCGTGTAAAGCTCCTTCGACG  179
            ||||||||||||||||||||||||||||||||||||||||||||||||||||||||||||
Sbjct  143  GCAGAGGGTGAGAATCCCGTACGTGGGCGCCTGCCTTTGCCGTGTAAAGCTCCTTCGACG  202

Query  180  AGTCGAGTTGTTTGGGAATGCAGCTCTAAATGGGAGGTAAATTTCTCCTAAAGCTAAATA  239
            ||||||||||||||||||||||||||||||||||||||||||||||||||||||||||||
Sbjct  203  AGTCGAGTTGTTTGGGAATGCAGCTCTAAATGGGAGGTAAATTTCTCCTAAAGCTAAATA  262

Query  240  CCGGCCAGAGACCGATAGCGCACAAGTAGAGTGATCGAAAGATGAAAAGTACTTTGGAAA  299
            ||||||||||||||||||||||||||||||||||||||||||||||||||||||||||||
Sbjct  263  CCGGCCAGAGACCGATAGCGCACAAGTAGAGTGATCGAAAGATGAAAAGTACTTTGGAAA  322

Query  300  GAGAGTCAAATAGCACGTGAAATTGTTGAAAGGGAAGCGCTTGCAGCCAGACTTGCCCGC  359
            ||||||||||||||||||||||||||||||||||||||||||||||||||||||||||||
Sbjct  323  GAGAGTCAAATAGCACGTGAAATTGTTGAAAGGGAAGCGCTTGCAGCCAGACTTGCCCGC  382

Query  360  AGTTGCTCACCCAGGCTCTCGCCTGGGGCACTCTTCTGCGGGCAGGCCAGCATCAGTTTG  419
            ||||||||||| ||||| | ||||||||||||||||||||||||||||||||||||||||
Sbjct  383  AGTTGCTCACCTAGGCTTTTGCCTGGGGCACTCTTCTGCGGGCAGGCCAGCATCAGTTTG  442

Query  420  GGCGGTCGGATAAAGG-CTCCTGTCATGTACCACCCCTCGGGGTGGCCTTATAggggggg  478
            |||||||||||||||| || |||||||||||||||| |||||||||||||||||||| |
Sbjct  443  GGCGGTCGGATAAAGGTCT-CTGTCATGTACCACCCTTCGGGGTGGCCTTATAGGGGAGA  501

Query  479  CGTAATGCGACCAGCCCGGACTGAGGTCCGCGCATCTGCTAGGATGCTGGCGTAATGGCT  538
            || |||||||||||||||||||||||||||||||||||||||||||||||||||||||||
Sbjct  502  CGCAATGCGACCAGCCCGGACTGAGGTCCGCGCATCTGCTAGGATGCTGGCGTAATGGCT  561

Query  539  GTAAGCGGCCCGT  551
            |||||||||||||
Sbjct  562  GTAAGCGGCCCGT  574


> LT671618.1 Pseudopithomyces atro-olivaceus genomic DNA sequence
contains 28S rRNA gene, strain MUCL 50391
Length=1033

 Score = 963 bits (521),  Expect = 0.0
 Identities = 543/553 (98%), Gaps = 3/553 (1%)
 Strand=Plus/Plus

Query  1    CGGCGAGTG-AGCGGCTACAGCTCAAATTTGAAATCTGGCCTCCTTTGGTGGTCCGAGTT  59
            ||||||||| ||||||||||||||||||||||||||||||||||||||||||||||||||
Sbjct  25   CGGCGAGTGAAGCGGCTACAGCTCAAATTTGAAATCTGGCCTCCTTTGGTGGTCCGAGTT  84

Query  60   GTAATTTGCAGAGGATGCTTTGGCATTGGCGGCGGTCTAAGTTCCTTGGAACAGGACATC  119
            ||||||||||||||||||||||||||||||||||||||||||||||||||||||||||||
Sbjct  85   GTAATTTGCAGAGGATGCTTTGGCATTGGCGGCGGTCTAAGTTCCTTGGAACAGGACATC  144

Query  120  GCAGAGGGTGAGAATCCCGTACGTGGGCGCCTGCCTTTGCCGTGTAAAGCTCCTTCGACG  179
            ||||||||||||||||||||||||||||||||||||||||||||||||||||||||||||
Sbjct  145  GCAGAGGGTGAGAATCCCGTACGTGGGCGCCTGCCTTTGCCGTGTAAAGCTCCTTCGACG  204
```

```
Query  180  AGTCGAGTTGTTTGGGAATGCAGCTCTAAATGGGAGGTAAATTTCTCCTAAAGCTAAATA  239
            ||||||||||||||||||||||||||||||||||||||||||||||||||||||||||||
Sbjct  205  AGTCGAGTTGTTTGGGAATGCAGCTCTAAATGGGAGGTAAATTTCTCCTAAAGCTAAATA  264

Query  240  CCGGCCAGAGACCGATAGCGCACAAGTAGAGTGATCGAAAGATGAAAAGTACTTTGGAAA  299
            ||||||||||||||||||||||||||||||||||||||||||||||||||||||||||||
Sbjct  265  CCGGCCAGAGACCGATAGCGCACAAGTAGAGTGATCGAAAGATGAAAAGTACTTTGGAAA  324

Query  300  GAGAGTCAAATAGCACGTGAAATTGTTGAAAGGGAAGCGCTTGCAGCCAGACTTGCCCGC  359
            ||||||||||||||||||||||||||||||||||||||||||||||||||||||||||||
Sbjct  325  GAGAGTCAAATAGCACGTGAAATTGTTGAAAGGGAAGCGCTTGCAGCCAGACTTGCCCGC  384

Query  360  AGTTGCTCACCCAGGCTCTCGCCTGGGGCACTCTTCTGCGGGCAGGCCAGCATCAGTTTG  419
            ||||||||||| ||||| | ||||||||||||||||||||||||||||||||||||||||
Sbjct  385  AGTTGCTCACCTAGGCTTTTGCCTGGGGCACTCTTCTGCGGGCAGGCCAGCATCAGTTTG  444

Query  420  GGCGGTCGGATAAAGG-CTCCTGTCATGTACCACCCCTCGGGGTGGCCTTATAggggggg  478
            |||||||||||||||| || |||||||||||||||| |||||||||||||||||||| |
Sbjct  445  GGCGGTCGGATAAAGGTCT-CTGTCATGTACCACCCTTCGGGGTGGCCTTATAGGGGAGA  503

Query  479  CGTAATGCGACCAGCCCGGACTGAGGTCCGCGCATCTGCTAGGATGCTGGCGTAATGGCT  538
            || |||||||||||||||||||||||||||||||||||||||||||||||||||||||||
Sbjct  504  CGCAATGCGACCAGCCCGGACTGAGGTCCGCGCATCTGCTAGGATGCTGGCGTAATGGCT  563

Query  539  GTAAGCGGCCCGT  551
            |||||||||||||
Sbjct  564  GTAAGCGGCCCGT  576


> HG933818.1 Pithomyces maydicus genomic DNA containing, 28S rRNA
gene, strain UTHSC 06-1549
Length=558

 Score = 955 bits (517),  Expect = 0.0
 Identities = 520/521 (99%), Gaps = 1/521 (0%)
 Strand=Plus/Plus

Query  1    CGGCGAGTG-AGCGGCTACAGCTCAAATTTGAAATCTGGCCTCCTTTGGTGGTCCGAGTT  59
            ||||||||| ||||||||||||||||||||||||||||||||||||||||||||||||||
Sbjct  38   CGGCGAGTGAAGCGGCTACAGCTCAAATTTGAAATCTGGCCTCCTTTGGTGGTCCGAGTT  97

Query  60   GTAATTTGCAGAGGATGCTTTGGCATTGGCGGCGGTCTAAGTTCCTTGGAACAGGACATC  119
            ||||||||||||||||||||||||||||||||||||||||||||||||||||||||||||
Sbjct  98   GTAATTTGCAGAGGATGCTTTGGCATTGGCGGCGGTCTAAGTTCCTTGGAACAGGACATC  157

Query  120  GCAGAGGGTGAGAATCCCGTACGTGGGCGCCTGCCTTTGCCGTGTAAAGCTCCTTCGACG  179
            ||||||||||||||||||||||||||||||||||||||||||||||||||||||||||||
Sbjct  158  GCAGAGGGTGAGAATCCCGTACGTGGGCGCCTGCCTTTGCCGTGTAAAGCTCCTTCGACG  217

Query  180  AGTCGAGTTGTTTGGGAATGCAGCTCTAAATGGGAGGTAAATTTCTCCTAAAGCTAAATA  239
            ||||||||||||||||||||||||||||||||||||||||||||||||||||||||||||
Sbjct  218  AGTCGAGTTGTTTGGGAATGCAGCTCTAAATGGGAGGTAAATTTCTCCTAAAGCTAAATA  277

Query  240  CCGGCCAGAGACCGATAGCGCACAAGTAGAGTGATCGAAAGATGAAAAGTACTTTGGAAA  299
            ||||||||||||||||||||||||||||||||||||||||||||||||||||||||||||
Sbjct  278  CCGGCCAGAGACCGATAGCGCACAAGTAGAGTGATCGAAAGATGAAAAGTACTTTGGAAA  337

Query  300  GAGAGTCAAATAGCACGTGAAATTGTTGAAAGGGAAGCGCTTGCAGCCAGACTTGCCCGC  359
            ||||||||||||||||||||||||||||||||||||||||||||||||||||||||||||
Sbjct  338  GAGAGTCAAATAGCACGTGAAATTGTTGAAAGGGAAGCGCTTGCAGCCAGACTTGCCCGC  397

Query  360  AGTTGCTCACCCAGGCTCTCGCCTGGGGCACTCTTCTGCGGGCAGGCCAGCATCAGTTTG  419
            ||||||||||||||||||||||||||||||||||||||||||||||||||||||||||||
Sbjct  398  AGTTGCTCACCCAGGCTCTCGCCTGGGGCACTCTTCTGCGGGCAGGCCAGCATCAGTTTG  457
```

```
Query  420   GGCGGTCGGATAAAGGCTCCTGTCATGTACCACCCCTCGGGGTGGCCTTATAgggggggC  479
             ||||||||||||||||||||||||||||||||||||||||||||||||||||||||||||
Sbjct  458   GGCGGTCGGATAAAGGCTCCTGTCATGTACCACCCCTCGGGGTGGCCTTATAGGGGGGGC  517

Query  480   GTAATGCGACCAGCCCGGACTGAGGTCCGCGCATCTGCTAG  520
             |||||||||||||||||||||||||||||||||||||||||
Sbjct  518   GTAATGCGACCAGCCCGGACTGAGGTCCGCGCATCTGCTAG  558


> HG933819.1 Pithomyces maydicus genomic DNA containing, 28S rRNA
gene, strain UTHSC 06-3954
Length=557

 Score = 953 bits (516),  Expect = 0.0
 Identities = 519/520 (99%), Gaps = 1/520 (0%)
 Strand=Plus/Plus

Query  1     CGGCGAGTG-AGCGGCTACAGCTCAAATTTGAAATCTGGCCTCCTTTGGTGGTCCGAGTT  59
             ||||||||| ||||||||||||||||||||||||||||||||||||||||||||||||||
Sbjct  38    CGGCGAGTGAAGCGGCTACAGCTCAAATTTGAAATCTGGCCTCCTTTGGTGGTCCGAGTT  97

Query  60    GTAATTTGCAGAGGATGCTTTGGCATTGGCGGCGGTCTAAGTTCCTTGGAACAGGACATC  119
             ||||||||||||||||||||||||||||||||||||||||||||||||||||||||||||
Sbjct  98    GTAATTTGCAGAGGATGCTTTGGCATTGGCGGCGGTCTAAGTTCCTTGGAACAGGACATC  157

Query  120   GCAGAGGGTGAGAATCCCGTACGTGGGCGCCTGCCTTTGCCGTGTAAAGCTCCTTCGACG  179
             ||||||||||||||||||||||||||||||||||||||||||||||||||||||||||||
Sbjct  158   GCAGAGGGTGAGAATCCCGTACGTGGGCGCCTGCCTTTGCCGTGTAAAGCTCCTTCGACG  217

Query  180   AGTCGAGTTGTTTGGGAATGCAGCTCTAAATGGGAGGTAAATTTCTCCTAAAGCTAAATA  239
             ||||||||||||||||||||||||||||||||||||||||||||||||||||||||||||
Sbjct  218   AGTCGAGTTGTTTGGGAATGCAGCTCTAAATGGGAGGTAAATTTCTCCTAAAGCTAAATA  277

Query  240   CCGGCCAGAGACCGATAGCGCACAAGTAGAGTGATCGAAAGATGAAAAGTACTTTGGAAA  299
             ||||||||||||||||||||||||||||||||||||||||||||||||||||||||||||
Sbjct  278   CCGGCCAGAGACCGATAGCGCACAAGTAGAGTGATCGAAAGATGAAAAGTACTTTGGAAA  337

Query  300   GAGAGTCAAATAGCACGTGAAATTGTTGAAAGGGAAGCGCTTGCAGCCAGACTTGCCCGC  359
             ||||||||||||||||||||||||||||||||||||||||||||||||||||||||||||
Sbjct  338   GAGAGTCAAATAGCACGTGAAATTGTTGAAAGGGAAGCGCTTGCAGCCAGACTTGCCCGC  397

Query  360   AGTTGCTCACCCAGGCTCTCGCCTGGGGCACTCTTCTGCGGGCAGGCCAGCATCAGTTTG  419
             ||||||||||||||||||||||||||||||||||||||||||||||||||||||||||||
Sbjct  398   AGTTGCTCACCCAGGCTCTCGCCTGGGGCACTCTTCTGCGGGCAGGCCAGCATCAGTTTG  457

Query  420   GGCGGTCGGATAAAGGCTCCTGTCATGTACCACCCCTCGGGGTGGCCTTATAgggggggC  479
             ||||||||||||||||||||||||||||||||||||||||||||||||||||||||||||
Sbjct  458   GGCGGTCGGATAAAGGCTCCTGTCATGTACCACCCCTCGGGGTGGCCTTATAGGGGGGGC  517

Query  480   GTAATGCGACCAGCCCGGACTGAGGTCCGCGCATCTGCTA  519
             ||||||||||||||||||||||||||||||||||||||||
Sbjct  518   GTAATGCGACCAGCCCGGACTGAGGTCCGCGCATCTGCTA  557


> LK936378.1 Pithomyces sacchari partial 28S rRNA gene, isolate
CBS120504
Length=1011

 Score = 946 bits (512),  Expect = 0.0
 Identities = 540/553 (98%), Gaps = 3/553 (1%)
 Strand=Plus/Plus

Query  1     CGGCGAGTG-AGCGGCTACAGCTCAAATTTGAAATCTGGCCTCCTTTGGTGGTCCGAGTT  59
```

```
            ||||||||| ||||||||||||||||||||||||||||||  |||||||  |||||||||
Sbjct  52   CGGCGAGTGAAGCGGCTACAGCTCAAATTTGAAATCTGGCTCCCTTTGGGCGTCCGAGTT  111

Query  60   GTAATTTGCAGAGGATGCTTTGGCATTGGCGGCGGTCTAAGTTCCTTGGAACAGGACATC  119
            ||||||||||||||||||||||||||||||||||||||||||||||||||||||||||||
Sbjct  112  GTAATTTGCAGAGGATGCTTTGGCATTGGCGGCGGTCTAAGTTCCTTGGAACAGGACATC  171

Query  120  GCAGAGGGTGAGAATCCCGTACGTGGGCGCCTGCCTTTGCCGTGTAAAGCTCCTTCGACG  179
            ||||||||||||||||||||||||||||||||||||||||||||||||||||||||||||
Sbjct  172  GCAGAGGGTGAGAATCCCGTACGTGGGCGCCTGCCTTTGCCGTGTAAAGCTCCTTCGACG  231

Query  180  AGTCGAGTTGTTTGGGAATGCAGCTCTAAATGGGAGGTAAATTTCTCCTAAAGCTAAATA  239
            ||||||||||||||||||||||||||||||||||||||||||||||||||||||||||||
Sbjct  232  AGTCGAGTTGTTTGGGAATGCAGCTCTAAATGGGAGGTAAATTTCTCCTAAAGCTAAATA  291

Query  240  CCGGCCAGAGACCGATAGCGCACAAGTAGAGTGATCGAAAGATGAAAAGTACTTTGGAAA  299
            ||||||||||||||||||||||||||||||||||||||||||||||||||||||||||||
Sbjct  292  CCGGCCAGAGACCGATAGCGCACAAGTAGAGTGATCGAAAGATGAAAAGTACTTTGGAAA  351

Query  300  GAGAGTCAAATAGCACGTGAAATTGTTGAAAGGGAAGCGCTTGCAGCCAGACTTGCCCGC  359
            ||||||||||||||||||||||||||||||||||||||||||||||||||||||||||||
Sbjct  352  GAGAGTCAAATAGCACGTGAAATTGTTGAAAGGGAAGCGCTTGCAGCCAGACTTGCCCGC  411

Query  360  AGTTGCTCACCCAGGCTCTCGCCTGGGGCACTCTTCTGCGGGCAGGCCAGCATCAGTTTG  419
            ||||||||||| ||||| | ||||||||||||||||||||||||||||||||||||||||
Sbjct  412  AGTTGCTCACCTAGGCTTTTGCCTGGGGCACTCTTCTGCGGGCAGGCCAGCATCAGTTTG  471

Query  420  GGCGGTCGGATAAAGG-CTCCTGTCATGTACCACCCCTCGGGGTGGCCTTATAggggggg  478
            |||||||||||||||| || ||||||||||||||||||||||||||||||||||||| |
Sbjct  472  GGCGGTCGGATAAAGGTCT-CTGTCATGTACCACCCCTCGGGGTGGCCTTATAGGGGAGA  530

Query  479  CGTAATGCGACCAGCCCGGACTGAGGTCCGCGCATCTGCTAGGATGCTGGCGTAATGGCT  538
            || |||||||||||||||||||||||||||||||||||||||||||||||||||||||||
Sbjct  531  CGCAATGCGACCAGCCCGGACTGAGGTCCGCGCATCTGCTAGGATGCTGGCGTAATGGCT  590

Query  539  GTAAGCGGCCCGT  551
            |||||||||||||
Sbjct  591  GTAAGCGGCCCGT  603


> LK936379.1 Pithomyces sacchari partial 28S rRNA gene, isolate
CBS803.72
Length=1017

 Score = 946 bits (512),  Expect = 0.0
 Identities = 540/553 (98%), Gaps = 3/553 (1%)
 Strand=Plus/Plus

Query  1    CGGCGAGTG-AGCGGCTACAGCTCAAATTTGAAATCTGGCCTCCTTTGGTGGTCCGAGTT  59
            ||||||||| ||||||||||||||||||||||||||||||  |||||||  |||||||||
Sbjct  51   CGGCGAGTGAAGCGGCTACAGCTCAAATTTGAAATCTGGCTCCCTTTGGGCGTCCGAGTT  110

Query  60   GTAATTTGCAGAGGATGCTTTGGCATTGGCGGCGGTCTAAGTTCCTTGGAACAGGACATC  119
            ||||||||||||||||||||||||||||||||||||||||||||||||||||||||||||
Sbjct  111  GTAATTTGCAGAGGATGCTTTGGCATTGGCGGCGGTCTAAGTTCCTTGGAACAGGACATC  170

Query  120  GCAGAGGGTGAGAATCCCGTACGTGGGCGCCTGCCTTTGCCGTGTAAAGCTCCTTCGACG  179
            ||||||||||||||||||||||||||||||||||||||||||||||||||||||||||||
Sbjct  171  GCAGAGGGTGAGAATCCCGTACGTGGGCGCCTGCCTTTGCCGTGTAAAGCTCCTTCGACG  230

Query  180  AGTCGAGTTGTTTGGGAATGCAGCTCTAAATGGGAGGTAAATTTCTCCTAAAGCTAAATA  239
            ||||||||||||||||||||||||||||||||||||||||||||||||||||||||||||
Sbjct  231  AGTCGAGTTGTTTGGGAATGCAGCTCTAAATGGGAGGTAAATTTCTCCTAAAGCTAAATA  290
```

```
Query  240   CCGGCCAGAGACCGATAGCGCACAAGTAGAGTGATCGAAAGATGAAAAGTACTTTGGAAA  299
             ||||||||||||||||||||||||||||||||||||||||||||||||||||||||||||
Sbjct  291   CCGGCCAGAGACCGATAGCGCACAAGTAGAGTGATCGAAAGATGAAAAGTACTTTGGAAA  350

Query  300   GAGAGTCAAATAGCACGTGAAATTGTTGAAAGGGAAGCGCTTGCAGCCAGACTTGCCCGC  359
             ||||||||||||||||||||||||||||||||||||||||||||||||||||||||||||
Sbjct  351   GAGAGTCAAATAGCACGTGAAATTGTTGAAAGGGAAGCGCTTGCAGCCAGACTTGCCCGC  410

Query  360   AGTTGCTCACCCAGGCTCTCGCCTGGGGCACTCTTCTGCGGGCAGGCCAGCATCAGTTTG  419
             ||||||||||| ||||| | ||||||||||||||||||||||||||||||||||||||||
Sbjct  411   AGTTGCTCACCTAGGCTTTTGCCTGGGGCACTCTTCTGCGGGCAGGCCAGCATCAGTTTG  470

Query  420   GGCGGTCGGATAAAGG-CTCCTGTCATGTACCACCCCTCGGGGTGGCCTTATAggggggg  478
             |||||||||||||||| || ||||||||||||||||||||||||||||||||||||| | 
Sbjct  471   GGCGGTCGGATAAAGGTCT-CTGTCATGTACCACCCCTCGGGGTGGCCTTATAGGGGAGA  529

Query  479   CGTAATGCGACCAGCCCGGACTGAGGTCCGCGCATCTGCTAGGATGCTGGCGTAATGGCT  538
             || |||||||||||||||||||||||||||||||||||||||||||||||||||||||||
Sbjct  530   CGCAATGCGACCAGCCCGGACTGAGGTCCGCGCATCTGCTAGGATGCTGGCGTAATGGCT  589

Query  539   GTAAGCGGCCCGT  551
             |||||||||||||
Sbjct  590   GTAAGCGGCCCGT  602


> HG933824.1 Pithomyces sp. I 2014-KDC genomic DNA containing,
28S rRNA gene, strain UTHSC 05-3373
Length=557

 Score = 904 bits (489),  Expect = 0.0
 Identities = 511/521 (98%), Gaps = 3/521 (1%)
 Strand=Plus/Plus

Query  1     CGGCGAGTG-AGCGGCTACAGCTCAAATTTGAAATCTGGCCTCCTTTGGTGGTCCGAGTT  59
             ||||||||| ||||||||||||||||||||||||||||||||||||||||||||||||||
Sbjct  38    CGGCGAGTGAAGCGGCTACAGCTCAAATTTGAAATCTGGCCTCCTTTGGTGGTCCGAGTT  97

Query  60    GTAATTTGCAGAGGATGCTTTGGCATTGGCGGCGGTCTAAGTTCCTTGGAACAGGACATC  119
             ||||||||||||||||||||||||||||||||||||||||||||||||||||||||||||
Sbjct  98    GTAATTTGCAGAGGATGCTTTGGCATTGGCGGCGGTCTAAGTTCCTTGGAACAGGACATC  157

Query  120   GCAGAGGGTGAGAATCCCGTACGTGGGCGCCTGCCTTTGCCGTGTAAAGCTCCTTCGACG  179
             ||||||||||||||||||||||||||||||||||||||||||||||||||||||||||||
Sbjct  158   GCAGAGGGTGAGAATCCCGTACGTGGGCGCCTGCCTTTGCCGTGTAAAGCTCCTTCGACG  217

Query  180   AGTCGAGTTGTTTGGGAATGCAGCTCTAAATGGGAGGTAAATTTCTCCTAAAGCTAAATA  239
             ||||||||||||||||||||||||||||||||||||||||||||||||||||||||||||
Sbjct  218   AGTCGAGTTGTTTGGGAATGCAGCTCTAAATGGGAGGTAAATTTCTCCTAAAGCTAAATA  277

Query  240   CCGGCCAGAGACCGATAGCGCACAAGTAGAGTGATCGAAAGATGAAAAGTACTTTGGAAA  299
             ||||||||||||||||||||||||||||||||||||||||||||||||||||||||||||
Sbjct  278   CCGGCCAGAGACCGATAGCGCACAAGTAGAGTGATCGAAAGATGAAAAGTACTTTGGAAA  337

Query  300   GAGAGTCAAATAGCACGTGAAATTGTTGAAAGGGAAGCGCTTGCAGCCAGACTTGCCCGC  359
             ||||||||||||||||||||||||||||||||||||||||||||||||||||||||||||
Sbjct  338   GAGAGTCAAATAGCACGTGAAATTGTTGAAAGGGAAGCGCTTGCAGCCAGACTTGCCCGC  397

Query  360   AGTTGCTCACCCAGGCTCTCGCCTGGGGCACTCTTCTGCGGGCAGGCCAGCATCAGTTTG  419
             ||||||||||| ||||| | ||||||||||||||||||||||||||||||||||||||||
Sbjct  398   AGTTGCTCACCTAGGCTTTTGCCTGGGGCACTCTTCTGCGGGCAGGCCAGCATCAGTTTG  457

Query  420   GGCGGTCGGATAAAGG-CTCCTGTCATGTACCACCCCTCGGGGTGGCCTTATAggggggg  478
             |||||||||||||||| || |||||||||||||||| |||||||||||||||||||| | 
Sbjct  458   GGCGGTCGGATAAAGGTCT-CTGTCATGTACCACCCTTCGGGGTGGCCTTATAGGGGAGA  516
```

```
Query  479  CGTAATGCGACCAGCCCGGACTGAGGTCCGCGCATCTGCTA  519
            || ||||||||||||||||||||||||||||||||||||||
Sbjct  517  CGCAATGCGACCAGCCCGGACTGAGGTCCGCGCATCTGCTA  557
```

> HG933813.1 Pithomyces sacchari genomic DNA containing, 28S rRNA
gene, strain UTHSC 03-1337
Length=558

```
 Score = 894 bits (484),  Expect = 0.0
 Identities = 510/522 (98%), Gaps = 3/522 (1%)
 Strand=Plus/Plus

Query  1    CGGCGAGTG-AGCGGCTACAGCTCAAATTTGAAATCTGGCCTCCTTTGGTGGTCCGAGTT  59
            ||||||||| ||||||||||||||||||||||||||||||  |||||||  |||||||||
Sbjct  38   CGGCGAGTGAAGCGGCTACAGCTCAAATTTGAAATCTGGCTCCCTTTGGGCGTCCGAGTT  97

Query  60   GTAATTTGCAGAGGATGCTTTGGCATTGGCGGCGGTCTAAGTTCCTTGGAACAGGACATC  119
            ||||||||||||||||||||||||||||||||||||||||||||||||||||||||||||
Sbjct  98   GTAATTTGCAGAGGATGCTTTGGCATTGGCGGCGGTCTAAGTTCCTTGGAACAGGACATC  157

Query  120  GCAGAGGGTGAGAATCCCGTACGTGGGCGCCTGCCTTTGCCGTGTAAAGCTCCTTCGACG  179
            ||||||||||||||||||||||||||||||||||||||||||||||||||||||||||||
Sbjct  158  GCAGAGGGTGAGAATCCCGTACGTGGGCGCCTGCCTTTGCCGTGTAAAGCTCCTTCGACG  217

Query  180  AGTCGAGTTGTTTGGGAATGCAGCTCTAAATGGGAGGTAAATTTCTCCTAAAGCTAAATA  239
            ||||||||||||||||||||||||||||||||||||||||||||||||||||||||||||
Sbjct  218  AGTCGAGTTGTTTGGGAATGCAGCTCTAAATGGGAGGTAAATTTCTCCTAAAGCTAAATA  277

Query  240  CCGGCCAGAGACCGATAGCGCACAAGTAGAGTGATCGAAAGATGAAAAGTACTTTGGAAA  299
            ||||||||||||||||||||||||||||||||||||||||||||||||||||||||||||
Sbjct  278  CCGGCCAGAGACCGATAGCGCACAAGTAGAGTGATCGAAAGATGAAAAGTACTTTGGAAA  337

Query  300  GAGAGTCAAATAGCACGTGAAATTGTTGAAAGGGAAGCGCTTGCAGCCAGACTTGCCCGC  359
            ||||||||||||||||||||||||||||||||||||||||||||||||||||||||||||
Sbjct  338  GAGAGTCAAATAGCACGTGAAATTGTTGAAAGGGAAGCGCTTGCAGCCAGACTTGCCCGC  397

Query  360  AGTTGCTCACCCAGGCTCTCGCCTGGGGCACTCTTCTGCGGGCAGGCCAGCATCAGTTTG  419
            ||||||||||||||||| | ||||||||||||||||||||||||||||||||||||||||
Sbjct  398  AGTTGCTCACCCAGGCTTTTGCCTGGGGCACTCTTCTGCGGGCAGGCCAGCATCAGTTTG  457

Query  420  GGCGGTCGGATAAAGG-CTCCTGTCATGTACCACCCCTCGGGGTGGCCTTATAggggggg  478
            |||||||||||||||| || ||||||||||||||||||||||||||||||||||||| |
Sbjct  458  GGCGGTCGGATAAAGGTCT-CTGTCATGTACCACCCCTCGGGGTGGCCTTATAGGGGAGA  516

Query  479  CGTAATGCGACCAGCCCGGACTGAGGTCCGCGCATCTGCTAG  520
            || |||||||||||||||||||||||||||||||||||||||
Sbjct  517  CGCAATGCGACCAGCCCGGACTGAGGTCCGCGCATCTGCTAG  558
```

> LK936380.1 Pithomyces sp. 1 ALT-2014 partial 28S rRNA gene, strain
UTHSC07-995
Length=551

```
 Score = 893 bits (483),  Expect = 0.0
 Identities = 505/515 (98%), Gaps = 3/515 (1%)
 Strand=Plus/Plus

Query  1    CGGCGAGTG-AGCGGCTACAGCTCAAATTTGAAATCTGGCCTCCTTTGGTGGTCCGAGTT  59
            ||||||||| ||||||||||||||||||||||||||||||||||||||||||||||||||
Sbjct  38   CGGCGAGTGAAGCGGCTACAGCTCAAATTTGAAATCTGGCCTCCTTTGGTGGTCCGAGTT  97

Query  60   GTAATTTGCAGAGGATGCTTTGGCATTGGCGGCGGTCTAAGTTCCTTGGAACAGGACATC  119
```

```
               ||||||||||||||||||||||||||||||||||||||||||||||||||||||||||||
Sbjct  98      GTAATTTGCAGAGGATGCTTTGGCATTGGCGGCGGTCTAAGTTCCTTGGAACAGGACATC  157

Query  120     GCAGAGGGTGAGAATCCCGTACGTGGGCGCCTGCCTTTGCCGTGTAAAGCTCCTTCGACG  179
               ||||||||||||||||||||||||||||||||||||||||||||||||||||||||||||
Sbjct  158     GCAGAGGGTGAGAATCCCGTACGTGGGCGCCTGCCTTTGCCGTGTAAAGCTCCTTCGACG  217

Query  180     AGTCGAGTTGTTTGGGAATGCAGCTCTAAATGGGAGGTAAATTTCTCCTAAAGCTAAATA  239
               ||||||||||||||||||||||||||||||||||||||||||||||||||||||||||||
Sbjct  218     AGTCGAGTTGTTTGGGAATGCAGCTCTAAATGGGAGGTAAATTTCTCCTAAAGCTAAATA  277

Query  240     CCGGCCAGAGACCGATAGCGCACAAGTAGAGTGATCGAAAGATGAAAAGTACTTTGGAAA  299
               ||||||||||||||||||||||||||||||||||||||||||||||||||||||||||||
Sbjct  278     CCGGCCAGAGACCGATAGCGCACAAGTAGAGTGATCGAAAGATGAAAAGTACTTTGGAAA  337

Query  300     GAGAGTCAAATAGCACGTGAAATTGTTGAAAGGGAAGCGCTTGCAGCCAGACTTGCCCGC  359
               ||||||||||||||||||||||||||||||||||||||||||||||||||||||||||||
Sbjct  338     GAGAGTCAAATAGCACGTGAAATTGTTGAAAGGGAAGCGCTTGCAGCCAGACTTGCCCGC  397

Query  360     AGTTGCTCACCCAGGCTCTCGCCTGGGGCACTCTTCTGCGGGCAGGCCAGCATCAGTTTG  419
               ||||||||||| ||||| | ||||||||||||||||||||||||||||||||||||||||
Sbjct  398     AGTTGCTCACCTAGGCTTTTGCCTGGGGCACTCTTCTGCGGGCAGGCCAGCATCAGTTTG  457

Query  420     GGCGGTCGGATAAAGG-CTCCTGTCATGTACCACCCCTCGGGGTGGCCTTATAggggggg  478
               |||||||||||||||| || |||||||||||||||| |||||||||||||||||||| |
Sbjct  458     GGCGGTCGGATAAAGGTCT-CTGTCATGTACCACCCTTCGGGGTGGCCTTATAGGGGAGA  516

Query  479     CGTAATGCGACCAGCCCGGACTGAGGTCCGCGCAT  513
               || ||||||||||||||||||||||||||||||||
Sbjct  517     CGCAATGCGACCAGCCCGGACTGAGGTCCGCGCAT  551


> HG933816.1 Pithomyces sacchari genomic DNA containing, 28S rRNA
gene, strain UTHSC 04-2746
Length=555

 Score = 883 bits (478),  Expect = 0.0
 Identities = 506/519 (97%), Gaps = 3/519 (1%)
 Strand=Plus/Plus

Query  1       CGGCGAGTG-AGCGGCTACAGCTCAAATTTGAAATCTGGCCTCCTTTGGTGGTCCGAGTT  59
               ||||||||| ||||||||||||||||||||||||||||||  |||||||  |||||||||
Sbjct  38      CGGCGAGTGAAGCGGCTACAGCTCAAATTTGAAATCTGGCTCCCTTTGGGCGTCCGAGTT  97

Query  60      GTAATTTGCAGAGGATGCTTTGGCATTGGCGGCGGTCTAAGTTCCTTGGAACAGGACATC  119
               ||||||||||||||||||||||||||||||||||||||||||||||||||||||||||||
Sbjct  98      GTAATTTGCAGAGGATGCTTTGGCATTGGCGGCGGTCTAAGTTCCTTGGAACAGGACATC  157

Query  120     GCAGAGGGTGAGAATCCCGTACGTGGGCGCCTGCCTTTGCCGTGTAAAGCTCCTTCGACG  179
               ||||||||||||||||||||||||||||||||||||||||||||||||||||||||||||
Sbjct  158     GCAGAGGGTGAGAATCCCGTACGTGGGCGCCTGCCTTTGCCGTGTAAAGCTCCTTCGACG  217

Query  180     AGTCGAGTTGTTTGGGAATGCAGCTCTAAATGGGAGGTAAATTTCTCCTAAAGCTAAATA  239
               ||||||||||||||||||||||||||||||||||||||||||||||||||||||||||||
Sbjct  218     AGTCGAGTTGTTTGGGAATGCAGCTCTAAATGGGAGGTAAATTTCTCCTAAAGCTAAATA  277

Query  240     CCGGCCAGAGACCGATAGCGCACAAGTAGAGTGATCGAAAGATGAAAAGTACTTTGGAAA  299
               ||||||||||||||||||||||||||||||||||||||||||||||||||||||||||||
Sbjct  278     CCGGCCAGAGACCGATAGCGCACAAGTAGAGTGATCGAAAGATGAAAAGTACTTTGGAAA  337

Query  300     GAGAGTCAAATAGCACGTGAAATTGTTGAAAGGGAAGCGCTTGCAGCCAGACTTGCCCGC  359
               ||||||||||||||||||||||||||||||||||||||||||||||||||||||||||||
Sbjct  338     GAGAGTCAAATAGCACGTGAAATTGTTGAAAGGGAAGCGCTTGCAGCCAGACTTGCCCGC  397
```

```
Query  360   AGTTGCTCACCCAGGCTCTCGCCTGGGGCACTCTTCTGCGGGCAGGCCAGCATCAGTTTG  419
             ||||||||||| ||||| | ||||||||||||||||||||||||||||||||||||||||
Sbjct  398   AGTTGCTCACCTAGGCTTTTGCCTGGGGCACTCTTCTGCGGGCAGGCCAGCATCAGTTTG  457

Query  420   GGCGGTCGGATAAAGG-CTCCTGTCATGTACCACCCCTCGGGGTGGCCTTATAggggggg  478
             |||||||||||||||| || ||||||||||||||||||||||||||||||||||||| |
Sbjct  458   GGCGGTCGGATAAAGGTCT-CTGTCATGTACCACCCCTCGGGGTGGCCTTATAGGGGAGA  516

Query  479   CGTAATGCGACCAGCCCGGACTGAGGTCCGCGCATCTGC  517
             || ||||||||||||||||||||||||||||||||||||
Sbjct  517   CGCAATGCGACCAGCCCGGACTGAGGTCCGCGCATCTGC  555
```

> HG933823.1 Pithomyces sp. I 2014-KDC genomic DNA containing,
28S rRNA gene, strain UTHSC 05-3161
Length=545

Score = 881 bits (477),  Expect = 0.0
Identities = 499/509 (98%), Gaps = 3/509 (1%)
Strand=Plus/Plus

```
Query  1     CGGCGAGTG-AGCGGCTACAGCTCAAATTTGAAATCTGGCCTCCTTTGGTGGTCCGAGTT  59
             ||||||||| ||||||||||||||||||||||||||||||||||||||||||||||||||
Sbjct  38    CGGCGAGTGAAGCGGCTACAGCTCAAATTTGAAATCTGGCCTCCTTTGGTGGTCCGAGTT  97

Query  60    GTAATTTGCAGAGGATGCTTTGGCATTGGCGGCGGTCTAAGTTCCTTGGAACAGGACATC  119
             ||||||||||||||||||||||||||||||||||||||||||||||||||||||||||||
Sbjct  98    GTAATTTGCAGAGGATGCTTTGGCATTGGCGGCGGTCTAAGTTCCTTGGAACAGGACATC  157

Query  120   GCAGAGGGTGAGAATCCCGTACGTGGGCGCCTGCCTTTGCCGTGTAAAGCTCCTTCGACG  179
             ||||||||||||||||||||||||||||||||||||||||||||||||||||||||||||
Sbjct  158   GCAGAGGGTGAGAATCCCGTACGTGGGCGCCTGCCTTTGCCGTGTAAAGCTCCTTCGACG  217

Query  180   AGTCGAGTTGTTTGGGAATGCAGCTCTAAATGGGAGGTAAATTTCTCCTAAAGCTAAATA  239
             ||||||||||||||||||||||||||||||||||||||||||||||||||||||||||||
Sbjct  218   AGTCGAGTTGTTTGGGAATGCAGCTCTAAATGGGAGGTAAATTTCTCCTAAAGCTAAATA  277

Query  240   CCGGCCAGAGACCGATAGCGCACAAGTAGAGTGATCGAAAGATGAAAAGTACTTTGGAAA  299
             ||||||||||||||||||||||||||||||||||||||||||||||||||||||||||||
Sbjct  278   CCGGCCAGAGACCGATAGCGCACAAGTAGAGTGATCGAAAGATGAAAAGTACTTTGGAAA  337

Query  300   GAGAGTCAAATAGCACGTGAAATTGTTGAAAGGGAAGCGCTTGCAGCCAGACTTGCCCGC  359
             ||||||||||||||||||||||||||||||||||||||||||||||||||||||||||||
Sbjct  338   GAGAGTCAAATAGCACGTGAAATTGTTGAAAGGGAAGCGCTTGCAGCCAGACTTGCCCGC  397

Query  360   AGTTGCTCACCCAGGCTCTCGCCTGGGGCACTCTTCTGCGGGCAGGCCAGCATCAGTTTG  419
             ||||||||||| ||||| | ||||||||||||||||||||||||||||||||||||||||
Sbjct  398   AGTTGCTCACCTAGGCTTTTGCCTGGGGCACTCTTCTGCGGGCAGGCCAGCATCAGTTTG  457

Query  420   GGCGGTCGGATAAAGG-CTCCTGTCATGTACCACCCCTCGGGGTGGCCTTATAggggggg  478
             |||||||||||||||| || |||||||||||||||| |||||||||||||||||||| |
Sbjct  458   GGCGGTCGGATAAAGGTCT-CTGTCATGTACCACCCTTCGGGGTGGCCTTATAGGGGAGA  516

Query  479   CGTAATGCGACCAGCCCGGACTGAGGTCC  507
             || ||||||||||||||||||||||||||
Sbjct  517   CGCAATGCGACCAGCCCGGACTGAGGTCC  545
```

> HG933825.1 Pithomyces sp. I 2014-KDC genomic DNA containing,
28S rRNA gene, strain UTHSC 06-3492
Length=538

Score = 869 bits (470),  Expect = 0.0
Identities = 492/502 (98%), Gaps = 3/502 (1%)

```
 Strand=Plus/Plus

Query  1    CGGCGAGTG-AGCGGCTACAGCTCAAATTTGAAATCTGGCCTCCTTTGGTGGTCCGAGTT  59
            ||||||||| ||||||||||||||||||||||||||||||||||||||||||||||||||
Sbjct  38   CGGCGAGTGAAGCGGCTACAGCTCAAATTTGAAATCTGGCCTCCTTTGGTGGTCCGAGTT  97

Query  60   GTAATTTGCAGAGGATGCTTTGGCATTGGCGGCGGTCTAAGTTCCTTGGAACAGGACATC  119
            ||||||||||||||||||||||||||||||||||||||||||||||||||||||||||||
Sbjct  98   GTAATTTGCAGAGGATGCTTTGGCATTGGCGGCGGTCTAAGTTCCTTGGAACAGGACATC  157

Query  120  GCAGAGGGTGAGAATCCCGTACGTGGGCGCCTGCCTTTGCCGTGTAAAGCTCCTTCGACG  179
            ||||||||||||||||||||||||||||||||||||||||||||||||||||||||||||
Sbjct  158  GCAGAGGGTGAGAATCCCGTACGTGGGCGCCTGCCTTTGCCGTGTAAAGCTCCTTCGACG  217

Query  180  AGTCGAGTTGTTTGGGAATGCAGCTCTAAATGGGAGGTAAATTTCTCCTAAAGCTAAATA  239
            ||||||||||||||||||||||||||||||||||||||||||||||||||||||||||||
Sbjct  218  AGTCGAGTTGTTTGGGAATGCAGCTCTAAATGGGAGGTAAATTTCTCCTAAAGCTAAATA  277

Query  240  CCGGCCAGAGACCGATAGCGCACAAGTAGAGTGATCGAAAGATGAAAAGTACTTTGGAAA  299
            ||||||||||||||||||||||||||||||||||||||||||||||||||||||||||||
Sbjct  278  CCGGCCAGAGACCGATAGCGCACAAGTAGAGTGATCGAAAGATGAAAAGTACTTTGGAAA  337

Query  300  GAGAGTCAAATAGCACGTGAAATTGTTGAAAGGGAAGCGCTTGCAGCCAGACTTGCCCGC  359
            ||||||||||||||||||||||||||||||||||||||||||||||||||||||||||||
Sbjct  338  GAGAGTCAAATAGCACGTGAAATTGTTGAAAGGGAAGCGCTTGCAGCCAGACTTGCCCGC  397

Query  360  AGTTGCTCACCCAGGCTCTCGCCTGGGGCACTCTTCTGCGGGCAGGCCAGCATCAGTTTG  419
            ||||||||||| ||||| | ||||||||||||||||||||||||||||||||||||||||
Sbjct  398  AGTTGCTCACCTAGGCTTTTGCCTGGGGCACTCTTCTGCGGGCAGGCCAGCATCAGTTTG  457

Query  420  GGCGGTCGGATAAAGG-CTCCTGTCATGTACCACCCCTCGGGGTGGCCTTATAggggggg  478
            |||||||||||||||| || |||||||||||||||| |||||||||||||||||||| |
Sbjct  458  GGCGGTCGGATAAAGGTCT-CTGTCATGTACCACCCTTCGGGGTGGCCTTATAGGGGAGA  516

Query  479  CGTAATGCGACCAGCCCGGACT  500
            || |||||||||||||||||||
Sbjct  517  CGCAATGCGACCAGCCCGGACT  538


> HG933826.1 Pithomyces sp. I 2014-KDC genomic DNA containing,
28S rRNA gene, strain UTHSC 06-3706
Length=534

 Score = 856 bits (463),  Expect = 0.0
 Identities = 487/498 (98%), Gaps = 4/498 (1%)
 Strand=Plus/Plus

Query  1    CGGCGAGTG-AGCGGCTACAGCTCAAATTTGAAATCTGGCCTCCTTTGGTGGTCCGAGTT  59
            ||||||||| ||||||||||||||||||||||||||||||||||||||||||||||||||
Sbjct  38   CGGCGAGTGAAGCGGCTACAGCTCAAATTTGAAATCTGGCCTCCTTTGGTGGTCCGAGTT  97

Query  60   GTAATTTGCAGAGGATGCTTTGGCATTGGCGGCGGTCTAAGTTCCTTGGAACAGGACATC  119
            ||||||||||||||||||||||||||||||||||||||||||||||||||||||||||||
Sbjct  98   GTAATTTGCAGAGGATGCTTTGGCATTGGCGGCGGTCTAAGTTCCTTGGAACAGGACATC  157

Query  120  GCAGAGGGTGAGAATCCCGTACGTGGGCGCCTGCCTTTGCCGTGTAAAGCTCCTTCGACG  179
            ||||||||||||||||||||||||||||||||||||||||||||||||||||||||||||
Sbjct  158  GCAGAGGGTGAGAATCCCGTACGTGGGCGCCTGCCTTTGCCGTGTAAAGCTCCTTCGACG  217

Query  180  AGTCGAGTTGTTTGGGAATGCAGCTCTAAATGGGAGGTAAATTTCTCCTAAAGCTAAATA  239
            ||||||||||||||||||||||||||||||||||||||||||||||||||||||||||||
Sbjct  218  AGTCGAGTTGTTTGGGAATGCAGCTCTAAATGGGAGGTAAATTTCTCCTAAAGCTAAATA  277

Query  240  CCGGCCAGAGACCGATAGCGCACAAGTAGAGTGATCGAAAGATGAAAAGTACTTTGGAAA  299
```

```
            ||||||||||||||||||||||||||||||||||||||||||||||||||||||||||||
Sbjct  278  CCGGCCAGAGACCGATAGCGCACAAGTAGAGTGATCGAAAGATGAAAAGTACTTTGGAAA  337

Query  300  GAGAGTCAAATAGCACGTGAAATTGTTGAAAGGGAAGCGCTTGCAGCCAGACTTGCCCGC  359
            ||||||||||||||||||||||||||||||||||||||||||||||||||||||||||||
Sbjct  338  GAGAGTCAAATAGCACGTGAAATTGTTGAAAGGGAAGCGCTTGCAGCCAGACTTGCCCGC  397

Query  360  AGTTGCTCACCCAGGCTCTCGCCTGGGGCACTCTTCTGCGGGCAGGCCAGCATCAGTTTG  419
            ||||||||||| ||||| | ||||||||||||||||||||||||||||||||||||||||
Sbjct  398  AGTTGCTCACCTAGGCTTTTGCCTGGGGCACTCTTCTGCGGGCAGGCCAGCATCAGTTTG  457

Query  420  GGCGGTCGGATAAAGG-CTCCTGTCATGTACCACCCCTCGGGGTGGCCTTATAggggggg  478
            |||||||||||||||| || |||||||||||||||| |||||||||||||||||||| |
Sbjct  458  GGCGGTCGGATAAAGGTCT-CTGTCATGTACCACCCTTCGGGGTGGCCTTATAGGGGAGA  516

Query  479  CGTA-ATGCGACCAGCCC  495
            || | |||||||||||||
Sbjct  517  CGCATATGCGACCAGCCC  534


> HG933817.1 Pithomyces sacchari genomic DNA containing, 28S rRNA
gene, strain UTHSC 05-3251
Length=543

 Score = 852 bits (461),  Expect = 0.0
 Identities = 491/505 (97%), Gaps = 3/505 (1%)
 Strand=Plus/Plus

Query  1    CGGCGAGTG-AGCGGCTACAGCTCAAATTTGAAATCTGGCCTCCTTTGGTGGTCCGAGTT  59
            ||||||||| ||||||||||||||||||||||||||||||  |||||||  |||||||||
Sbjct  38   CGGCGAGTGAAGCGGCTACAGCTCAAATTTGAAATCTGGCTCCCTTTGGGCGTCCGAGTT  97

Query  60   GTAATTTGCAGAGGATGCTTTGGCATTGGCGGCGGTCTAAGTTCCTTGGAACAGGACATC  119
            ||||||||||||||||||||||||||||||||||||||||||||||||||||||||||||
Sbjct  98   GTAATTTGCAGAGGATGCTTTGGCATTGGCGGCGGTCTAAGTTCCTTGGAACAGGACATC  157

Query  120  GCAGAGGGTGAGAATCCCGTACGTGGGCGCCTGCCTTTGCCGTGTAAAGCTCCTTCGACG  179
            ||||||||||||||||||||||||||||||||||||||||||||||||||||||||||||
Sbjct  158  GCAGAGGGTGAGAATCCCGTACGTGGGCGCCTGCCTTTGCCGTGTAAAGCTCCTTCGACG  217

Query  180  AGTCGAGTTGTTTGGGAATGCAGCTCTAAATGGGAGGTAAATTTCTCCTAAAGCTAAATA  239
            ||||||||||||||||||||||||||||||||||||||||||||||||||||||||||||
Sbjct  218  AGTCGAGTTGTTTGGGAATGCAGCTCTAAATGGGAGGTAAATTTCTCCTAAAGCTAAATA  277

Query  240  CCGGCCAGAGACCGATAGCGCACAAGTAGAGTGATCGAAAGATGAAAAGTACTTTGGAAA  299
            ||||||||||||||||||||||||||||||||||||||||||||||||||||||||||||
Sbjct  278  CCGGCCAGAGACCGATAGCGCACAAGTAGAGTGATCGAAAGATGAAAAGTACTTTGGAAA  337

Query  300  GAGAGTCAAATAGCACGTGAAATTGTTGAAAGGGAAGCGCTTGCAGCCAGACTTGCCCGC  359
            ||||||||||||||||||||||||||||||||||||||||||||||||||||||||||||
Sbjct  338  GAGAGTCAAATAGCACGTGAAATTGTTGAAAGGGAAGCGCTTGCAGCCAGACTTGCCCGC  397

Query  360  AGTTGCTCACCCAGGCTCTCGCCTGGGGCACTCTTCTGCGGGCAGGCCAGCATCAGTTTG  419
            ||||||||||| ||||| | ||||||||||||||||||||||||||||||||||||||||
Sbjct  398  AGTTGCTCACCTAGGCTTTTGCCTGGGGCACTCTTCTGCGGGCAGGCCAGCATCAGTTTG  457

Query  420  GGCGGTCGGATAAAGG-CTCCTGTCATGTACCACCCCTCGGGGTGGCCTTATAggggggg  478
            |||||||||||||||| || ||||||||||||||||||| ||||||||||||||||| |
Sbjct  458  GGCGGTCGGATAAAGGTCT-CTGTCATGTACCACCCCTCAGGGTGGCCTTATAGGGGAGA  516

Query  479  CGTAATGCGACCAGCCCGGACTGAG  503
            || ||||||||||||||||||||||
Sbjct  517  CGCAATGCGACCAGCCCGGACTGAG  541
```

```
> LK936381.1 Pithomyces sp. 2 ALT-2014 partial 28S rRNA gene, strain
UTHSC07-578
Length=529

 Score = 852 bits (461),  Expect = 0.0
 Identities = 483/493 (98%), Gaps = 3/493 (1%)
 Strand=Plus/Plus

Query  1    CGGCGAGTG-AGCGGCTACAGCTCAAATTTGAAATCTGGCCTCCTTTGGTGGTCCGAGTT  59
            ||||||||| ||||||||||||||||||||||||||||||||||||||||||||||||||
Sbjct  38   CGGCGAGTGAAGCGGCTACAGCTCAAATTTGAAATCTGGCCTCCTTTGGTGGTCCGAGTT  97

Query  60   GTAATTTGCAGAGGATGCTTTGGCATTGGCGGCGGTCTAAGTTCCTTGGAACAGGACATC  119
            ||||||||||||||||||||||||||||||||||||||||||||||||||||||||||||
Sbjct  98   GTAATTTGCAGAGGATGCTTTGGCATTGGCGGCGGTCTAAGTTCCTTGGAACAGGACATC  157

Query  120  GCAGAGGGTGAGAATCCCGTACGTGGGCGCCTGCCTTTGCCGTGTAAAGCTCCTTCGACG  179
            ||||||||||||||||||||||||||||||||||||||||||||||||||||||||||||
Sbjct  158  GCAGAGGGTGAGAATCCCGTACGTGGGCGCCTGCCTTTGCCGTGTAAAGCTCCTTCGACG  217

Query  180  AGTCGAGTTGTTTGGGAATGCAGCTCTAAATGGGAGGTAAATTTCTCCTAAAGCTAAATA  239
            ||||||||||||||||||||||||||||||||||||||||||||||||||||||||||||
Sbjct  218  AGTCGAGTTGTTTGGGAATGCAGCTCTAAATGGGAGGTAAATTTCTCCTAAAGCTAAATA  277

Query  240  CCGGCCAGAGACCGATAGCGCACAAGTAGAGTGATCGAAAGATGAAAAGTACTTTGGAAA  299
            ||||||||||||||||||||||||||||||||||||||||||||||||||||||||||||
Sbjct  278  CCGGCCAGAGACCGATAGCGCACAAGTAGAGTGATCGAAAGATGAAAAGTACTTTGGAAA  337

Query  300  GAGAGTCAAATAGCACGTGAAATTGTTGAAAGGGAAGCGCTTGCAGCCAGACTTGCCCGC  359
            ||||||||||||||||||||||||||||||||||||||||||||||||||||||||||||
Sbjct  338  GAGAGTCAAATAGCACGTGAAATTGTTGAAAGGGAAGCGCTTGCAGCCAGACTTGCCCGC  397

Query  360  AGTTGCTCACCCAGGCTCTCGCCTGGGGCACTCTTCTGCGGGCAGGCCAGCATCAGTTTG  419
            ||||||||||| ||||| | ||||||||||||||||||||||||||||||||||||||||
Sbjct  398  AGTTGCTCACCTAGGCTTTTGCCTGGGGCACTCTTCTGCGGGCAGGCCAGCATCAGTTTG  457

Query  420  GGCGGTCGGATAAAGG-CTCCTGTCATGTACCACCCCTCGGGGTGGCCTTATAggggggg  478
            |||||||||||||||| || |||||||||||||||| |||||||||||||||||||| |
Sbjct  458  GGCGGTCGGATAAAGGTCT-CTGTCATGTACCACCCTTCGGGGTGGCCTTATAGGGGAGA  516

Query  479  CGTAATGCGACCA  491
            || ||||||||||
Sbjct  517  CGCAATGCGACCA  529


> HG933815.1 Pithomyces sacchari genomic DNA containing, 28S rRNA
gene, strain UTHSC 04-2483
Length=538

 Score = 850 bits (460),  Expect = 0.0
 Identities = 488/501 (97%), Gaps = 3/501 (1%)
 Strand=Plus/Plus

Query  1    CGGCGAGTG-AGCGGCTACAGCTCAAATTTGAAATCTGGCCTCCTTTGGTGGTCCGAGTT  59
            ||||||||| ||||||||||||||||||||||||||||||  |||||||  |||||||||
Sbjct  38   CGGCGAGTGAAGCGGCTACAGCTCAAATTTGAAATCTGGCTCCCTTTGGGCGTCCGAGTT  97

Query  60   GTAATTTGCAGAGGATGCTTTGGCATTGGCGGCGGTCTAAGTTCCTTGGAACAGGACATC  119
            ||||||||||||||||||||||||||||||||||||||||||||||||||||||||||||
Sbjct  98   GTAATTTGCAGAGGATGCTTTGGCATTGGCGGCGGTCTAAGTTCCTTGGAACAGGACATC  157

Query  120  GCAGAGGGTGAGAATCCCGTACGTGGGCGCCTGCCTTTGCCGTGTAAAGCTCCTTCGACG  179
            ||||||||||||||||||||||||||||||||||||||||||||||||||||||||||||
```

```
Sbjct  158   GCAGAGGGTGAGAATCCCGTACGTGGGCGCCTGCCTTTGCCGTGTAAAGCTCCTTCGACG  217

Query  180   AGTCGAGTTGTTTGGGAATGCAGCTCTAAATGGGAGGTAAATTTCTCCTAAAGCTAAATA  239
             ||||||||||||||||||||||||||||||||||||||||||||||||||||||||||||
Sbjct  218   AGTCGAGTTGTTTGGGAATGCAGCTCTAAATGGGAGGTAAATTTCTCCTAAAGCTAAATA  277

Query  240   CCGGCCAGAGACCGATAGCGCACAAGTAGAGTGATCGAAAGATGAAAAGTACTTTGGAAA  299
             ||||||||||||||||||||||||||||||||||||||||||||||||||||||||||||
Sbjct  278   CCGGCCAGAGACCGATAGCGCACAAGTAGAGTGATCGAAAGATGAAAAGTACTTTGGAAA  337

Query  300   GAGAGTCAAATAGCACGTGAAATTGTTGAAAGGGAAGCGCTTGCAGCCAGACTTGCCCGC  359
             ||||||||||||||||||||||||||||||||||||||||||||||||||||||||||||
Sbjct  338   GAGAGTCAAATAGCACGTGAAATTGTTGAAAGGGAAGCGCTTGCAGCCAGACTTGCCCGC  397

Query  360   AGTTGCTCACCCAGGCTCTCGCCTGGGGCACTCTTCTGCGGGCAGGCCAGCATCAGTTTG  419
             ||||||||||| ||||| | ||||||||||||||||||||||||||||||||||||||||
Sbjct  398   AGTTGCTCACCTAGGCTTTTGCCTGGGGCACTCTTCTGCGGGCAGGCCAGCATCAGTTTG  457

Query  420   GGCGGTCGGATAAAGG-CTCCTGTCATGTACCACCCCTCGGGGTGGCCTTATAggggggg  478
             |||||||||||||||| || |||||||||||||||||||||||||||||||||||| |
Sbjct  458   GGCGGTCGGATAAAGGTCT-CTGTCATGTACCACCCCTCGGGGTGGCCTTATAGGGGAGA  516

Query  479   CGTAATGCGACCAGCCCGGAC  499
             || ||||||||||||||||||
Sbjct  517   CGCAATGCGACCAGCCCGGAC  537


> HG933814.1 Pithomyces sacchari genomic DNA containing, 28S rRNA
gene, strain UTHSC 03-3221
Length=528

 Score = 833 bits (451),  Expect = 0.0
 Identities = 479/492 (97%), Gaps = 3/492 (1%)
 Strand=Plus/Plus

Query  1     CGGCGAGTG-AGCGGCTACAGCTCAAATTTGAAATCTGGCCTCCTTTGGTGGTCCGAGTT  59
             ||||||||| ||||||||||||||||||||||||||||||  |||||||  |||||||||
Sbjct  38    CGGCGAGTGAAGCGGCTACAGCTCAAATTTGAAATCTGGCTCCCTTTGGGCGTCCGAGTT  97

Query  60    GTAATTTGCAGAGGATGCTTTGGCATTGGCGGCGGTCTAAGTTCCTTGGAACAGGACATC  119
             ||||||||||||||||||||||||||||||||||||||||||||||||||||||||||||
Sbjct  98    GTAATTTGCAGAGGATGCTTTGGCATTGGCGGCGGTCTAAGTTCCTTGGAACAGGACATC  157

Query  120   GCAGAGGGTGAGAATCCCGTACGTGGGCGCCTGCCTTTGCCGTGTAAAGCTCCTTCGACG  179
             ||||||||||||||||||||||||||||||||||||||||||||||||||||||||||||
Sbjct  158   GCAGAGGGTGAGAATCCCGTACGTGGGCGCCTGCCTTTGCCGTGTAAAGCTCCTTCGACG  217

Query  180   AGTCGAGTTGTTTGGGAATGCAGCTCTAAATGGGAGGTAAATTTCTCCTAAAGCTAAATA  239
             ||||||||||||||||||||||||||||||||||||||||||||||||||||||||||||
Sbjct  218   AGTCGAGTTGTTTGGGAATGCAGCTCTAAATGGGAGGTAAATTTCTCCTAAAGCTAAATA  277

Query  240   CCGGCCAGAGACCGATAGCGCACAAGTAGAGTGATCGAAAGATGAAAAGTACTTTGGAAA  299
             ||||||||||||||||||||||||||||||||||||||||||||||||||||||||||||
Sbjct  278   CCGGCCAGAGACCGATAGCGCACAAGTAGAGTGATCGAAAGATGAAAAGTACTTTGGAAA  337

Query  300   GAGAGTCAAATAGCACGTGAAATTGTTGAAAGGGAAGCGCTTGCAGCCAGACTTGCCCGC  359
             ||||||||||||||||||||||||||||||||||||||||||||||||||||||||||||
Sbjct  338   GAGAGTCAAATAGCACGTGAAATTGTTGAAAGGGAAGCGCTTGCAGCCAGACTTGCCCGC  397

Query  360   AGTTGCTCACCCAGGCTCTCGCCTGGGGCACTCTTCTGCGGGCAGGCCAGCATCAGTTTG  419
             ||||||||||| ||||| | ||||||||||||||||||||||||||||||||||||||||
Sbjct  398   AGTTGCTCACCTAGGCTTTTGCCTGGGGCACTCTTCTGCGGGCAGGCCAGCATCAGTTTG  457

Query  420   GGCGGTCGGATAAAGG-CTCCTGTCATGTACCACCCCTCGGGGTGGCCTTATAggggggg  478
```

```
               |||||||||||||||| || ||||||||||||||||||||||||||||||||||||| | 
Sbjct  458   GGCGGTCGGATAAAGGTCT-CTGTCATGTACCACCCCTCGGGGTGGCCTTATAGGGGAGA  516

Query  479   CGTAATGCGACC  490
             || |||||||||
Sbjct  517   CGCAATGCGACC  528


> HG933827.1 Pseudopithomyces diversisporus genomic DNA containing,
28S rRNA gene, strain UTHSC 06-4528
Length=505

 Score = 824 bits (446),  Expect = 0.0
 Identities = 462/469 (99%), Gaps = 3/469 (1%)
 Strand=Plus/Plus

Query  1     CGGCGAGTG-AGCGGCTACAGCTCAAATTTGAAATCTGGCCTCCTTTGGTGGTCCGAGTT  59
             ||||||||| ||||||||||||||||||||||||||||||||||||||||||||||||||
Sbjct  38    CGGCGAGTGAAGCGGCTACAGCTCAAATTTGAAATCTGGCCTCCTTTGGTGGTCCGAGTT  97

Query  60    GTAATTTGCAGAGGATGCTTTGGCATTGGCGGCGGTCTAAGTTCCTTGGAACAGGACATC  119
             ||||||||||||||||||||||||||||||||||||||||||||||||||||||||||||
Sbjct  98    GTAATTTGCAGAGGATGCTTTGGCATTGGCGGCGGTCTAAGTTCCTTGGAACAGGACATC  157

Query  120   GCAGAGGGTGAGAATCCCGTACGTGGGCGCCTGCCTTTGCCGTGTAAAGCTCCTTCGACG  179
             ||||||||||||||||||||||||||||||||||||||||||||||||||||||||||||
Sbjct  158   GCAGAGGGTGAGAATCCCGTACGTGGGCGCCTGCCTTTGCCGTGTAAAGCTCCTTCGACG  217

Query  180   AGTCGAGTTGTTTGGGAATGCAGCTCTAAATGGGAGGTAAATTTCTCCTAAAGCTAAATA  239
             ||||||||||||||||||||||||||||||||||||||||||||||||||||||||||||
Sbjct  218   AGTCGAGTTGTTTGGGAATGCAGCTCTAAATGGGAGGTAAATTTCTCCTAAAGCTAAATA  277

Query  240   CCGGCCAGAGACCGATAGCGCACAAGTAGAGTGATCGAAAGATGAAAAGTACTTTGGAAA  299
             ||||||||||||||||||||||||||||||||||||||||||||||||||||||||||||
Sbjct  278   CCGGCCAGAGACCGATAGCGCACAAGTAGAGTGATCGAAAGATGAAAAGTACTTTGGAAA  337

Query  300   GAGAGTCAAATAGCACGTGAAATTGTTGAAAGGGAAGCGCTTGCAGCCAGACTTGCCCGC  359
             ||||||||||||||||||||||||||||||||||||||||||||||||||||||||||||
Sbjct  338   GAGAGTCAAATAGCACGTGAAATTGTTGAAAGGGAAGCGCTTGCAGCCAGACTTGCCCGC  397

Query  360   AGTTGCTCACCCAGGCTCTCGCCTGGGGCACTCTTCTGCGGGCAGGCCAGCATCAGTTTG  419
             ||||||||||| ||||| | ||||||||||||||||||||||||||||||||||||||||
Sbjct  398   AGTTGCTCACCTAGGCTTTTGCCTGGGGCACTCTTCTGCGGGCAGGCCAGCATCAGTTTG  457

Query  420   GGCGGTCGGATAAAGG-CTCCTGTCATGTACCACCCCTCGGGGTGGCCT  467
             |||||||||||||||| || ||||||||||||||| ||||||||||||
Sbjct  458   GGCGGTCGGATAAAGGTCT-CTGTCATGTACCACCCTTCGGGGTGGCCT  505


> KX034666.1 Pseudopithomyces maydicus strain MFLUCC 14 0391 28S
ribosomal RNA gene, partial sequence
Length=592

 Score = 606 bits (328),  Expect = 7e-176
 Identities = 328/328 (100%), Gaps = 0/328 (0%)
 Strand=Plus/Plus

Query  224   CTCCTAAAGCTAAATACCGGCCAGAGACCGATAGCGCACAAGTAGAGTGATCGAAAGATG  283
             ||||||||||||||||||||||||||||||||||||||||||||||||||||||||||||
Sbjct  1     CTCCTAAAGCTAAATACCGGCCAGAGACCGATAGCGCACAAGTAGAGTGATCGAAAGATG  60

Query  284   AAAAGTACTTTGGAAAGAGAGTCAAATAGCACGTGAAATTGTTGAAAGGGAAGCGCTTGC  343
             ||||||||||||||||||||||||||||||||||||||||||||||||||||||||||||
Sbjct  61    AAAAGTACTTTGGAAAGAGAGTCAAATAGCACGTGAAATTGTTGAAAGGGAAGCGCTTGC  120
```

```
Query  344  AGCCAGACTTGCCCGCAGTTGCTCACCCAGGCTCTCGCCTGGGGCACTCTTCTGCGGGCA  403
            ||||||||||||||||||||||||||||||||||||||||||||||||||||||||||||
Sbjct  121  AGCCAGACTTGCCCGCAGTTGCTCACCCAGGCTCTCGCCTGGGGCACTCTTCTGCGGGCA  180

Query  404  GGCCAGCATCAGTTTGGGCGGTCGGATAAAGGCTCCTGTCATGTACCACCCCTCGGGGTG  463
            ||||||||||||||||||||||||||||||||||||||||||||||||||||||||||||
Sbjct  181  GGCCAGCATCAGTTTGGGCGGTCGGATAAAGGCTCCTGTCATGTACCACCCCTCGGGGTG  240

Query  464  GCCTTATAgggggggCGTAATGCGACCAGCCCGGACTGAGGTCCGCGCATCTGCTAGGAT  523
            ||||||||||||||||||||||||||||||||||||||||||||||||||||||||||||
Sbjct  241  GCCTTATAGGGGGGGCGTAATGCGACCAGCCCGGACTGAGGTCCGCGCATCTGCTAGGAT  300

Query  524  GCTGGCGTAATGGCTGTAAGCGGCCCGT  551
            ||||||||||||||||||||||||||||
Sbjct  301  GCTGGCGTAATGGCTGTAAGCGGCCCGT  328


> KX034665.1 Pseudopithomyces palmicola strain MFLUCC 14-0392 28S
ribosomal RNA gene, partial sequence
Length=565

 Score = 577 bits (312),  Expect = 6e-167
 Identities = 312/312 (100%), Gaps = 0/312 (0%)
 Strand=Plus/Plus

Query  240  CCGGCCAGAGACCGATAGCGCACAAGTAGAGTGATCGAAAGATGAAAAGTACTTTGGAAA  299
            ||||||||||||||||||||||||||||||||||||||||||||||||||||||||||||
Sbjct  1    CCGGCCAGAGACCGATAGCGCACAAGTAGAGTGATCGAAAGATGAAAAGTACTTTGGAAA  60

Query  300  GAGAGTCAAATAGCACGTGAAATTGTTGAAAGGGAAGCGCTTGCAGCCAGACTTGCCCGC  359
            ||||||||||||||||||||||||||||||||||||||||||||||||||||||||||||
Sbjct  61   GAGAGTCAAATAGCACGTGAAATTGTTGAAAGGGAAGCGCTTGCAGCCAGACTTGCCCGC  120

Query  360  AGTTGCTCACCCAGGCTCTCGCCTGGGGCACTCTTCTGCGGGCAGGCCAGCATCAGTTTG  419
            ||||||||||||||||||||||||||||||||||||||||||||||||||||||||||||
Sbjct  121  AGTTGCTCACCCAGGCTCTCGCCTGGGGCACTCTTCTGCGGGCAGGCCAGCATCAGTTTG  180

Query  420  GGCGGTCGGATAAAGGCTCCTGTCATGTACCACCCCTCGGGGTGGCCTTATAgggggggC  479
            ||||||||||||||||||||||||||||||||||||||||||||||||||||||||||||
Sbjct  181  GGCGGTCGGATAAAGGCTCCTGTCATGTACCACCCCTCGGGGTGGCCTTATAGGGGGGGC  240

Query  480  GTAATGCGACCAGCCCGGACTGAGGTCCGCGCATCTGCTAGGATGCTGGCGTAATGGCTG  539
            ||||||||||||||||||||||||||||||||||||||||||||||||||||||||||||
Sbjct  241  GTAATGCGACCAGCCCGGACTGAGGTCCGCGCATCTGCTAGGATGCTGGCGTAATGGCTG  300

Query  540  TAAGCGGCCCGT  551
            ||||||||||||
Sbjct  301  TAAGCGGCCCGT  312



Lambda      K        H
    1.33    0.621     1.12

Gapped
Lambda      K        H
    1.28    0.460    0.850

Effective search space used: 34902774


  Database: combined28S.fna
    Posted date:  May 31, 2022  9:56 PM
```

```
  Number of letters in database: 66,483
  Number of sequences in database:  66



Matrix: blastn matrix 1 -2
Gap Penalties: Existence: 0, Extension: 2.5
```
