## Supplementary Figure 1 for "A draft genome of the ascomycotal fungal species *Pseudopithomyces maydicus* (family *Didymosphaeriaceae*)"

```
Query: YPD_A_1_partial_28S_Sanger Query ID: lcl|Query_55593 Length: 551


>Pseudopithomyces maydicus culture BCC&lt;THA&gt;:84332 large subunit ribosomal RNA gene,
 partial sequence
Sequence ID: MF919633.1 Length: 840
Range 1: 32 to 583

Score:1013 bits(548), Expect:0.0,
Identities:551/552(99%),  Gaps:1/552(0%), Strand: Plus/Plus

Query  1    CGGCGAGTG-AGCGGCTACAGCTCAAATTTGAAATCTGGCCTCCTTTGGTGGTCCGAGTT  59
            ||||||||| ||||||||||||||||||||||||||||||||||||||||||||||||||
Sbjct  32   CGGCGAGTGAAGCGGCTACAGCTCAAATTTGAAATCTGGCCTCCTTTGGTGGTCCGAGTT  91

Query  60   GTAATTTGCAGAGGATGCTTTGGCATTGGCGGCGGTCTAAGTTCCTTGGAACAGGACATC  119
            ||||||||||||||||||||||||||||||||||||||||||||||||||||||||||||
Sbjct  92   GTAATTTGCAGAGGATGCTTTGGCATTGGCGGCGGTCTAAGTTCCTTGGAACAGGACATC  151

Query  120  GCAGAGGGTGAGAATCCCGTACGTGGGCGCCTGCCTTTGCCGTGTAAAGCTCCTTCGACG  179
            ||||||||||||||||||||||||||||||||||||||||||||||||||||||||||||
Sbjct  152  GCAGAGGGTGAGAATCCCGTACGTGGGCGCCTGCCTTTGCCGTGTAAAGCTCCTTCGACG  211

Query  180  AGTCGAGTTGTTTGGGAATGCAGCTCTAAATGGGAGGTAAATTTCTCCTAAAGCTAAATA  239
            ||||||||||||||||||||||||||||||||||||||||||||||||||||||||||||
Sbjct  212  AGTCGAGTTGTTTGGGAATGCAGCTCTAAATGGGAGGTAAATTTCTCCTAAAGCTAAATA  271

Query  240  CCGGCCAGAGACCGATAGCGCACAAGTAGAGTGATCGAAAGATGAAAAGTACTTTGGAAA  299
            ||||||||||||||||||||||||||||||||||||||||||||||||||||||||||||
Sbjct  272  CCGGCCAGAGACCGATAGCGCACAAGTAGAGTGATCGAAAGATGAAAAGTACTTTGGAAA  331

Query  300  GAGAGTCAAATAGCACGTGAAATTGTTGAAAGGGAAGCGCTTGCAGCCAGACTTGCCCGC  359
            ||||||||||||||||||||||||||||||||||||||||||||||||||||||||||||
Sbjct  332  GAGAGTCAAATAGCACGTGAAATTGTTGAAAGGGAAGCGCTTGCAGCCAGACTTGCCCGC  391

Query  360  AGTTGCTCACCCAGGCTCTCGCCTGGGGCACTCTTCTGCGGGCAGGCCAGCATCAGTTTG  419
            ||||||||||||||||||||||||||||||||||||||||||||||||||||||||||||
Sbjct  392  AGTTGCTCACCCAGGCTCTCGCCTGGGGCACTCTTCTGCGGGCAGGCCAGCATCAGTTTG  451

Query  420  GGCGGTCGGATAAAGGCTCCTGTCATGTACCACCCCTCGGGGTGGCCTTATAgggggggC  479
            ||||||||||||||||||||||||||||||||||||||||||||||||||||||||||||
Sbjct  452  GGCGGTCGGATAAAGGCTCCTGTCATGTACCACCCCTCGGGGTGGCCTTATAGGGGGGGC  511

Query  480  GTAATGCGACCAGCCCGGACTGAGGTCCGCGCATCTGCTAGGATGCTGGCGTAATGGCTG  539
            ||||||||||||||||||||||||||||||||||||||||||||||||||||||||||||
Sbjct  512  GTAATGCGACCAGCCCGGACTGAGGTCCGCGCATCTGCTAGGATGCTGGCGTAATGGCTG  571

Query  540  TAAGCGGCCCGT  551
            ||||||||||||
Sbjct  572  TAAGCGGCCCGT  583
```
