## Supplementary figures and images for "A draft genome of the ascomycotal fungal species *Pseudopithomyces maydicus* (family *Didymosphaeriaceae*)"

### bacteria_about.png

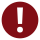

### bacteria_download.png

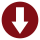

### bacteria_help.png

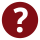

### bacteria_home.png

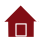

### bacteria_logo.png

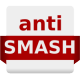

### fungi_about.png

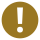

### fungi_download.png

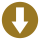

### fungi_help.png

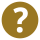

### fungi_home.png

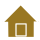

### fungi_logo.png

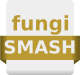

### mail.png

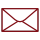

### nostructure_icon.png

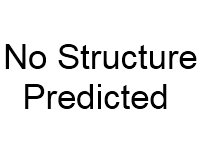

### Supplementary Figure 2

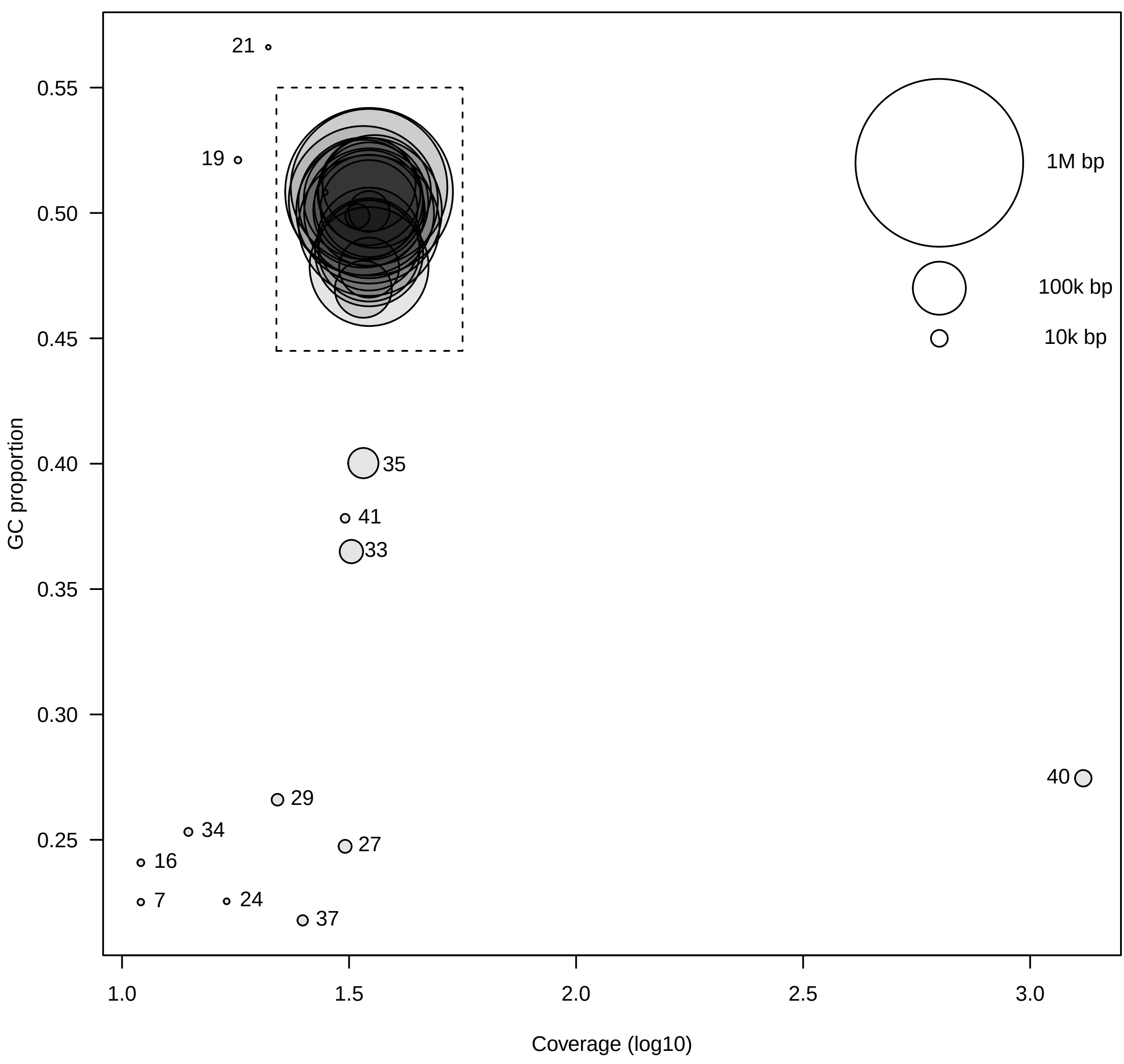
