## Supplementary Results for "A draft genome of the ascomycotal fungal species *Pseudopithomyces maydicus* (family *Didymosphaeriaceae*)": contig_3_c1.txt

ClusterBlast scores for contig_3

Table of genes, locations, strands and annotations of query cluster:
input.path1.gene130	486670	487111	+		
input.path1.gene131	487949	490047	+		
input.path1.gene132	494297	496062	+		
input.path1.gene133	496371	496704	-		
input.path1.gene134	497566	498757	-		
input.path1.gene135	499006	500476	-		
input.path1.gene136	501063	502619	+		
input.path1.gene137	506326	511102	+		
input.path1.gene138	512517	514516	-		
input.path1.gene139	520161	521156	+		
input.path1.gene140	521904	522071	+		
input.path1.gene141	523636	525575	-		
input.path1.gene142	527325	527967	+		
input.path1.gene143	528581	529545	+		


Significant hits: 
1. NW_006271975	Cordyceps militaris CM01 unplaced genomic scaffold CCM_S00007, whole genome shotgun sequence
2. NW_006917091	Pestalotiopsis fici W106-1 unplaced genomic scaffold PFICI_Scf_1, whole genome shotgun sequence
3. NW_003052497	Uncinocarpus reesii 1704 scaffold_4 genomic scaffold, whole genome shotgun sequence


Details:

>>
1. NW_006271975
Source: Cordyceps militaris CM01 unplaced genomic scaffold CCM_S00007, whole genome shotgun sequence
Type: T1PKS
Number of proteins with BLAST hits to this cluster: 2
Cumulative BLAST score: 759

Table of genes, locations, strands and annotations of subject cluster:

Table of Blast hits (query gene, subject gene, %identity, blast score, %coverage, e-value):
input.path1.gene137	CCM_09042	33	596	105.52147239263803	7.1e-168
input.path1.gene143	CCM_09038	42	163	78.3882783882784	3.2e-38


>>
2. NW_006917091
Source: Pestalotiopsis fici W106-1 unplaced genomic scaffold PFICI_Scf_1, whole genome shotgun sequence
Type: T1PKS
Number of proteins with BLAST hits to this cluster: 2
Cumulative BLAST score: 741

Table of genes, locations, strands and annotations of subject cluster:

Table of Blast hits (query gene, subject gene, %identity, blast score, %coverage, e-value):
input.path1.gene137	PFICI_00294	33	641	96.47239263803681	2.5e-181
input.path1.gene141	PFICI_00288	42	100	48.80952380952381	3.1e-19


>>
3. NW_003052497
Source: Uncinocarpus reesii 1704 scaffold_4 genomic scaffold, whole genome shotgun sequence
Type: NRPS,T1PKS
Number of proteins with BLAST hits to this cluster: 2
Cumulative BLAST score: 647

Table of genes, locations, strands and annotations of subject cluster:

Table of Blast hits (query gene, subject gene, %identity, blast score, %coverage, e-value):
input.path1.gene130	UREG_06505	51	75	87.65432098765432	2.6e-12
input.path1.gene137	UREG_06499	32	572	97.62269938650306	1.4e-160
