## Supplementary Results for "A draft genome of the ascomycotal fungal species *Pseudopithomyces maydicus* (family *Didymosphaeriaceae*)": contig_4_c1.txt

ClusterBlast scores for contig_4

Table of genes, locations, strands and annotations of query cluster:
input.path1.gene103	354413	355557	-		
input.path1.gene104	360322	361219	-		
input.path1.gene105	361306	362766	+		
input.path1.gene106	363258	364317	+		
input.path1.gene107	364516	365570	-		
input.path1.gene108	365875	366276	+		
input.path1.gene109	371338	373741	+		
input.path1.gene110	376034	377180	+		
input.path1.gene111	377444	379239	-		
input.path1.gene112	381789	382134	+		
input.path1.gene113	383003	384665	-		
input.path1.gene114	388890	389490	+		
input.path1.gene115	390223	392004	+		


Significant hits: 
1. NC_007199	Aspergillus fumigatus Af293 chromosome 6, whole genome shotgun sequence
2. NW_022984634	Aspergillus tanneri strain NIH1004 chromosome Unknown AapTan_7_pilon_pilon_pilon, whole genome shotgun sequence
3. NZ_FWXV01000013	Kibdelosporangium aridum strain DSM 43828, whole genome shotgun sequence
4. NZ_BBXF01000001	Herbidospora daliensis strain NBRC 106372, whole genome shotgun sequence
5. NZ_JXCA02000019	Scytonema tolypothrichoides VB-61278 scaffold_003, whole genome shotgun sequence
6. NZ_FNQS01000007	Lonsdalea quercina strain ATCC 29281, whole genome shotgun sequence
7. NZ_CP023009	Lonsdalea britannica strain 477 chromosome, complete genome
8. NZ_FO818637	Xenorhabdus bovienii strain CS03
9. NZ_CP036282	Rhodoferax sediminis strain Gr-4 chromosome, complete genome
10. NZ_CP015698	Curvibacter sp. AEP1-3 genome
11. NZ_KB290824	Corynebacterium durum F0235 Scfld189, whole genome shotgun sequence
12. NZ_CP009220	Corynebacterium deserti GIMN1.010, complete genome
13. NZ_JVVM01000022	Corynebacterium vitaeruminis strain 1285_CVIT 125_346261_8890490, whole genome shotgun sequence
14. NZ_CP004350	Corynebacterium casei LMG S-19264 chromosome, complete genome
15. NC_004369	Corynebacterium efficiens YS-314, complete genome
16. NZ_CP009244	Corynebacterium ammoniagenes DSM 20306 strain DSM 20306 = 9.6 chromosome, complete genome
17. NZ_CP059404	Corynebacterium sp. Marseille-Q3630 chromosome
18. NZ_PTJO01000001	Corynebacterium sp. CCUG 69366 69366_contig_0000001, whole genome shotgun sequence
19. NC_016783	Corynebacterium diphtheriae INCA 402, complete sequence
20. NC_016799	Corynebacterium diphtheriae 31A, complete sequence
21. NZ_CP039247	Corynebacterium endometrii strain LMM-1653 chromosome, complete genome
22. NZ_LR134538	Corynebacterium diphtheriae strain NCTC3529 chromosome 1
23. NC_016789	Corynebacterium diphtheriae PW8, complete sequence
24. NC_016785	Corynebacterium diphtheriae CDCE 8392, complete sequence
25. NZ_CP029644	Corynebacterium diphtheriae strain BQ11 chromosome, complete genome
26. NZ_CP031418	Nocardia farcinica strain W6977 chromosome, complete genome
27. NC_002935	Corynebacterium diphtheriae NCTC 13129, complete genome
28. NC_016801	Corynebacterium diphtheriae C7 (beta), complete sequence
29. NZ_CP018331	Corynebacterium diphtheriae strain B-D-16-78 chromosome, complete genome
30. NZ_CP038504	Corynebacterium diphtheriae strain TH1526 chromosome, complete genome
31. NZ_LR134537	Corynebacterium diphtheriae strain NCTC7838 chromosome 1
32. NC_016782	Corynebacterium diphtheriae 241, complete sequence
33. NC_016787	Corynebacterium diphtheriae HC03, complete sequence
34. NC_016790	Corynebacterium diphtheriae VA01, complete sequence
35. NC_016800	Corynebacterium diphtheriae BH8, complete sequence
36. NC_016802	Corynebacterium diphtheriae HC02, complete sequence
37. NZ_CP006764	Corynebacterium doosanense CAU 212 = DSM 45436 strain CAU 212(T) chromosome, complete genome
38. NZ_CP011541	Corynebacterium epidermidicanis strain DSM 45586 chromosome, complete genome
39. NZ_CP059399	Nocardia sp. WCH-YHL-001 chromosome, complete genome
40. NZ_FOET01000004	Streptomyces radiopugnans strain CGMCC 4.3519, whole genome shotgun sequence
41. NZ_CP020700	Streptomyces tsukubensis strain NRRL 18488 chromosome, complete genome
42. NZ_CP046172	Nocardia arthritidis strain AUSMDU00012717 chromosome, complete genome
43. NC_014158	Tsukamurella paurometabola DSM 20162, complete sequence
44. NZ_CP010519	Streptomyces albus strain DSM 41398 chromosome, complete genome
45. NZ_HE997626	Tsukamurella sp. 1534, whole genome shotgun sequence
46. NZ_CP034550	Saccharothrix syringae strain NRRL B-16468 chromosome, complete genome
47. NW_003345200	Nannizzia gypsea CBS 118893 supercont1.2 genomic scaffold, whole genome shotgun sequence
48. NW_003456424	Trichophyton rubrum CBS 118892 genomic scaffold supercont2.5, whole genome shotgun sequence
49. NC_036436	Aspergillus oryzae RIB40 DNA, chromosome 2
50. NW_013550604	Rhinocladiella mackenziei CBS 650.93 unplaced genomic scaffold supercont1.1, whole genome shotgun sequence


Details:

>>
1. NC_007199
Source: Aspergillus fumigatus Af293 chromosome 6, whole genome shotgun sequence
Type: NRPS
Number of proteins with BLAST hits to this cluster: 3
Cumulative BLAST score: 632

Table of genes, locations, strands and annotations of subject cluster:

Table of Blast hits (query gene, subject gene, %identity, blast score, %coverage, e-value):
input.path1.gene107	AFUA_6G09740	32	133	102.01612903225808	3.2e-29
input.path1.gene109	AFUA_6G09660	41	183	45.729537366548044	4.7e-44
input.path1.gene113	AFUA_6G09680	46	316	70.0	5.2e-84


>>
2. NW_022984634
Source: Aspergillus tanneri strain NIH1004 chromosome Unknown AapTan_7_pilon_pilon_pilon, whole genome shotgun sequence
Type: NRPS
Number of proteins with BLAST hits to this cluster: 3
Cumulative BLAST score: 607

Table of genes, locations, strands and annotations of subject cluster:

Table of Blast hits (query gene, subject gene, %identity, blast score, %coverage, e-value):
input.path1.gene107	ATNIH1004_006715	33	135	100.0	6.5e-30
input.path1.gene109	ATNIH1004_006709	32	128	50.88967971530249	1.8e-27
input.path1.gene113	ATNIH1004_006716	50	344	70.20833333333333	1.8e-92


>>
3. NZ_FWXV01000013
Source: Kibdelosporangium aridum strain DSM 43828, whole genome shotgun sequence
Type: NRPS
Number of proteins with BLAST hits to this cluster: 2
Cumulative BLAST score: 416

Table of genes, locations, strands and annotations of subject cluster:

Table of Blast hits (query gene, subject gene, %identity, blast score, %coverage, e-value):
input.path1.gene109	B9C43_RS47680	37	99	33.096085409252666	9e-19
input.path1.gene109	B9C43_RS47675	34	92	34.16370106761566	1.4e-16
input.path1.gene111	B9C43_RS47730	31	225	88.53615520282186	1.9e-56


>>
4. NZ_BBXF01000001
Source: Herbidospora daliensis strain NBRC 106372, whole genome shotgun sequence
Type: NRPS
Number of proteins with BLAST hits to this cluster: 2
Cumulative BLAST score: 305

Table of genes, locations, strands and annotations of subject cluster:

Table of Blast hits (query gene, subject gene, %identity, blast score, %coverage, e-value):
input.path1.gene109	AW274_RS01855	34	88	32.028469750889684	2.7e-15
input.path1.gene111	AW274_RS01915	31	217	89.0652557319224	3e-54


>>
5. NZ_JXCA02000019
Source: Scytonema tolypothrichoides VB-61278 scaffold_003, whole genome shotgun sequence
Type: NRPS,T1PKS
Number of proteins with BLAST hits to this cluster: 2
Cumulative BLAST score: 293

Table of genes, locations, strands and annotations of subject cluster:
SD80_RS18880	SD80_RS18880	503615	503837	+	hypothetical_protein

Table of Blast hits (query gene, subject gene, %identity, blast score, %coverage, e-value):
input.path1.gene109	SD80_RS18800	36	95	40.0355871886121	2.2e-17
input.path1.gene115	SD80_RS18720	32	198	97.15370018975332	1.3e-48


>>
6. NZ_FNQS01000007
Source: Lonsdalea quercina strain ATCC 29281, whole genome shotgun sequence
Type: NRPS
Number of proteins with BLAST hits to this cluster: 2
Cumulative BLAST score: 281

Table of genes, locations, strands and annotations of subject cluster:

Table of Blast hits (query gene, subject gene, %identity, blast score, %coverage, e-value):
input.path1.gene109	BLU64_RS11210	31	85	42.34875444839858	2.3e-14
input.path1.gene115	BLU64_RS11255	36	196	66.98292220113852	8.6e-48


>>
7. NZ_CP023009
Source: Lonsdalea britannica strain 477 chromosome, complete genome
Type: NRPS
Number of proteins with BLAST hits to this cluster: 2
Cumulative BLAST score: 280

Table of genes, locations, strands and annotations of subject cluster:
CKQ53_RS08300	CKQ53_RS08300	1757154	1758052	+	LysR_family_transcriptional_regulator
CKQ53_RS18535	CKQ53_RS18535	1776848	1778455	+	amino_acid_adenylation_domain-containing_protein
CKQ53_RS18540	CKQ53_RS18540	1778615	1780075	+	non-ribosomal_peptide_synthetase
CKQ53_RS18545	CKQ53_RS18545	1781525	1781851	+	hypothetical_protein
CKQ53_RS08365	CKQ53_RS08365	1792435	1793810	+	phosphomannomutase_CpsG

Table of Blast hits (query gene, subject gene, %identity, blast score, %coverage, e-value):
input.path1.gene109	CKQ53_RS18535	33	86	32.20640569395018	1.3e-14
input.path1.gene115	CKQ53_RS08280	36	194	66.98292220113852	2.5e-47


>>
8. NZ_FO818637
Source: Xenorhabdus bovienii strain CS03
Type: NRPS,T1PKS
Number of proteins with BLAST hits to this cluster: 2
Cumulative BLAST score: 277

Table of genes, locations, strands and annotations of subject cluster:
XBW1_RS24140	XBW1_RS24140	3629630	3630081	+	aspartyl_protease_family_protein
XBW1_RS24145	XBW1_RS24145	3630814	3630975	+	type_VI_secretion_system_tube_protein_Hcp
XBW1_RS24150	XBW1_RS24150	3634625	3635309	-	IS1_family_transposase
XBW1_RS17000	XBW1_RS17000	3636749	3636913	+	branched-chain_amino_acid_transport_system_II_carrier_protein
XBW1_RS17040	XBW1_RS17040	3643492	3645078	-	multidrug_efflux_MFS_transporter
XBW1_RS25205	XBW1_RS25205	3660006	3660155	+	transposase
XBW1_RS25210	XBW1_RS25210	3660110	3660345	+	transposase
XBW1_RS24850	XBW1_RS24850	3723348	3723458	+	IS6_family_transposase
XBW1_RS24160	XBW1_RS24160	3731918	3732608	-	IS1_family_transposase

Table of Blast hits (query gene, subject gene, %identity, blast score, %coverage, e-value):
input.path1.gene109	XBW1_RS17045	33	82	34.69750889679715	1.5e-13
input.path1.gene115	XBW1_RS17125	35	195	66.41366223908919	1.9e-47


>>
9. NZ_CP036282
Source: Rhodoferax sediminis strain Gr-4 chromosome, complete genome
Type: NRPS-like
Number of proteins with BLAST hits to this cluster: 2
Cumulative BLAST score: 276

Table of genes, locations, strands and annotations of subject cluster:

Table of Blast hits (query gene, subject gene, %identity, blast score, %coverage, e-value):
input.path1.gene109	EXZ61_RS15205	37	91	31.494661921708182	2.4e-16
input.path1.gene115	EXZ61_RS15175	34	185	67.93168880455408	2e-44


>>
10. NZ_CP015698
Source: Curvibacter sp. AEP1-3 genome
Type: NRPS-like
Number of proteins with BLAST hits to this cluster: 2
Cumulative BLAST score: 276

Table of genes, locations, strands and annotations of subject cluster:

Table of Blast hits (query gene, subject gene, %identity, blast score, %coverage, e-value):
input.path1.gene109	AEP_RS19195	36	87	29.715302491103202	6e-15
input.path1.gene115	AEP_RS19240	35	189	67.93168880455408	8.1e-46


>>
11. NZ_KB290824
Source: Corynebacterium durum F0235 Scfld189, whole genome shotgun sequence
Type: NRPS-like
Number of proteins with BLAST hits to this cluster: 2
Cumulative BLAST score: 254

Table of genes, locations, strands and annotations of subject cluster:

Table of Blast hits (query gene, subject gene, %identity, blast score, %coverage, e-value):
input.path1.gene109	HMPREF9997_RS04345	34	91	32.74021352313167	4.2e-16
input.path1.gene115	HMPREF9997_RS04450	34	163	59.96204933586338	8.1e-38


>>
12. NZ_CP009220
Source: Corynebacterium deserti GIMN1.010, complete genome
Type: NRPS-like
Number of proteins with BLAST hits to this cluster: 2
Cumulative BLAST score: 253

Table of genes, locations, strands and annotations of subject cluster:
CDES_RS11695	CDES_RS11695	2524225	2525943	+	hypothetical_protein
CDES_RS14700	CDES_RS14700	2542340	2542402	-	hypothetical_protein

Table of Blast hits (query gene, subject gene, %identity, blast score, %coverage, e-value):
input.path1.gene109	CDES_RS11795	35	86	29.715302491103202	1e-14
input.path1.gene115	CDES_RS11710	33	167	59.58254269449716	3.3e-39


>>
13. NZ_JVVM01000022
Source: Corynebacterium vitaeruminis strain 1285_CVIT 125_346261_8890490, whole genome shotgun sequence
Type: NRPS-like
Number of proteins with BLAST hits to this cluster: 2
Cumulative BLAST score: 249

Table of genes, locations, strands and annotations of subject cluster:

Table of Blast hits (query gene, subject gene, %identity, blast score, %coverage, e-value):
input.path1.gene109	ADA16_RS06945	32	86	36.654804270462634	1e-14
input.path1.gene115	ADA16_RS07035	32	163	59.39278937381404	8.1e-38


>>
14. NZ_CP004350
Source: Corynebacterium casei LMG S-19264 chromosome, complete genome
Type: NRPS-like,terpene
Number of proteins with BLAST hits to this cluster: 2
Cumulative BLAST score: 247

Table of genes, locations, strands and annotations of subject cluster:

Table of Blast hits (query gene, subject gene, %identity, blast score, %coverage, e-value):
input.path1.gene109	CCASEI_RS12625	36	91	29.715302491103202	4.2e-16
input.path1.gene115	CCASEI_RS12515	31	156	59.58254269449716	5.8e-36


>>
15. NC_004369
Source: Corynebacterium efficiens YS-314, complete genome
Type: NRPS-like
Number of proteins with BLAST hits to this cluster: 2
Cumulative BLAST score: 247

Table of genes, locations, strands and annotations of subject cluster:

Table of Blast hits (query gene, subject gene, %identity, blast score, %coverage, e-value):
input.path1.gene109	CE_RS12675	35	88	29.715302491103202	2.7e-15
input.path1.gene115	CE_RS12565	31	159	60.72106261859582	8.9e-37


>>
16. NZ_CP009244
Source: Corynebacterium ammoniagenes DSM 20306 strain DSM 20306 = 9.6 chromosome, complete genome
Type: NRPS-like,terpene
Number of proteins with BLAST hits to this cluster: 2
Cumulative BLAST score: 246

Table of genes, locations, strands and annotations of subject cluster:
CAMM_RS11295	CAMM_RS11295	2462616	2463802	-	glutamate--cysteine_ligase

Table of Blast hits (query gene, subject gene, %identity, blast score, %coverage, e-value):
input.path1.gene109	CAMM_RS11220	34	87	29.715302491103202	6e-15
input.path1.gene115	CAMM_RS11120	31	159	59.58254269449716	8.9e-37


>>
17. NZ_CP059404
Source: Corynebacterium sp. Marseille-Q3630 chromosome
Type: NRPS-like
Number of proteins with BLAST hits to this cluster: 2
Cumulative BLAST score: 246

Table of genes, locations, strands and annotations of subject cluster:

Table of Blast hits (query gene, subject gene, %identity, blast score, %coverage, e-value):
input.path1.gene109	H0194_RS04130	32	84	36.83274021352313	3e-14
input.path1.gene115	H0194_RS04030	32	162	59.58254269449716	1.4e-37


>>
18. NZ_PTJO01000001
Source: Corynebacterium sp. CCUG 69366 69366_contig_0000001, whole genome shotgun sequence
Type: NRPS-like
Number of proteins with BLAST hits to this cluster: 2
Cumulative BLAST score: 244

Table of genes, locations, strands and annotations of subject cluster:

Table of Blast hits (query gene, subject gene, %identity, blast score, %coverage, e-value):
input.path1.gene109	C5L39_RS00520	35	86	29.715302491103202	1.3e-14
input.path1.gene115	C5L39_RS00425	33	158	59.96204933586338	2e-36


>>
19. NC_016783
Source: Corynebacterium diphtheriae INCA 402, complete sequence
Type: LAP,NRPS-like,thiopeptide
Number of proteins with BLAST hits to this cluster: 2
Cumulative BLAST score: 243

Table of genes, locations, strands and annotations of subject cluster:
CDB402_RS12365	CDB402_RS12365	2003402	2003692	+	hypothetical_protein
CDB402_RS11935	CDB402_RS11935	2019032	2019268	+	IS110_family_transposase
CDB402_RS11940	CDB402_RS11940	2039935	2040825	+	lipase
CDB402_RS11945	CDB402_RS11945	2045974	2046159	-	IS256_family_transposase
CDB402_RS11950	CDB402_RS11950	2046700	2046990	+	Fic_family_protein
CDB402_RS12370	CDB402_RS12370	2047153	2047655	+	Fic_family_protein
CDB402_RS11960	CDB402_RS11960	2049618	2050684	-	IS256_family_transposase
CDB402_RS11965	CDB402_RS11965	2052887	2053158	-	transposase
CDB402_RS09670	CDB402_RS09670	2054770	2055975	+	IS1249_family_transposase

Table of Blast hits (query gene, subject gene, %identity, blast score, %coverage, e-value):
input.path1.gene109	CDB402_RS09585	31	88	39.679715302491104	2.7e-15
input.path1.gene115	CDB402_RS09440	32	155	59.58254269449716	1.3e-35


>>
20. NC_016799
Source: Corynebacterium diphtheriae 31A, complete sequence
Type: LAP,NRPS-like,thiopeptide
Number of proteins with BLAST hits to this cluster: 2
Cumulative BLAST score: 243

Table of genes, locations, strands and annotations of subject cluster:
CD31A_RS09880	CD31A_RS09880	2059963	2060978	+	IS110_family_transposase
CD31A_RS09985	CD31A_RS09985	2079765	2080313	-	hypothetical_protein
CD31A_RS09990	CD31A_RS09990	2080322	2081154	-	SpaH/EbpB_family_LPXTG-anchored_major_pilin
CD31A_RS13075	CD31A_RS13075	2088402	2088973	-	IS256_family_transposase
CD31A_RS10025	CD31A_RS10025	2091184	2092212	-	IS256_family_transposase
CD31A_RS10030	CD31A_RS10030	2092759	2093885	-	IS110_family_transposase
CD31A_RS11990	CD31A_RS11990	2096698	2097856	+	IS1249_family_transposase

Table of Blast hits (query gene, subject gene, %identity, blast score, %coverage, e-value):
input.path1.gene109	CD31A_RS09980	31	88	39.679715302491104	2.7e-15
input.path1.gene115	CD31A_RS09825	32	155	59.58254269449716	1.7e-35


>>
21. NZ_CP039247
Source: Corynebacterium endometrii strain LMM-1653 chromosome, complete genome
Type: NRPS-like,terpene
Number of proteins with BLAST hits to this cluster: 2
Cumulative BLAST score: 243

Table of genes, locations, strands and annotations of subject cluster:

Table of Blast hits (query gene, subject gene, %identity, blast score, %coverage, e-value):
input.path1.gene109	CENDO_RS09925	31	85	43.77224199288256	1.7e-14
input.path1.gene115	CENDO_RS09830	32	158	58.44402277039848	2e-36


>>
22. NZ_LR134538
Source: Corynebacterium diphtheriae strain NCTC3529 chromosome 1
Type: LAP,NRPS-like,thiopeptide
Number of proteins with BLAST hits to this cluster: 2
Cumulative BLAST score: 242

Table of genes, locations, strands and annotations of subject cluster:
EL331_RS06000	EL331_RS06000	1267401	1267598	+	transposase
EL331_RS06065	EL331_RS06065	1282919	1283155	+	IS110_family_transposase
EL331_RS06090	EL331_RS06090	1286551	1287173	-	hypothetical_protein
EL331_RS06180	EL331_RS06180	1303794	1304684	+	lipase
EL331_RS06200	EL331_RS06200	1309833	1310018	-	IS256_family_transposase
EL331_RS06210	EL331_RS06210	1310559	1310849	+	Fic_family_protein
EL331_RS06215	EL331_RS06215	1311012	1311514	+	Fic_family_protein
EL331_RS06230	EL331_RS06230	1313477	1314543	-	IS256_family_transposase
EL331_RS06245	EL331_RS06245	1316746	1317017	-	transposase
EL331_RS06255	EL331_RS06255	1318629	1319834	+	IS1249_family_transposase

Table of Blast hits (query gene, subject gene, %identity, blast score, %coverage, e-value):
input.path1.gene109	EL331_RS06165	31	88	39.679715302491104	2.7e-15
input.path1.gene115	EL331_RS06010	32	154	59.58254269449716	2.9e-35


>>
23. NC_016789
Source: Corynebacterium diphtheriae PW8, complete sequence
Type: LAP,NRPS-like,thiopeptide
Number of proteins with BLAST hits to this cluster: 2
Cumulative BLAST score: 242

Table of genes, locations, strands and annotations of subject cluster:
CDPW8_RS12480	CDPW8_RS12480	2071642	2072649	+	IS110_family_transposase
CDPW8_RS09970	CDPW8_RS09970	2087586	2091389	+	amino_acid_adenylation_domain-containing_protein
CDPW8_RS09975	CDPW8_RS09975	2091386	2096522	-	VWA_domain-containing_protein
CDPW8_RS12485	CDPW8_RS12485	2100879	2101570	+	lipase
CDPW8_RS10020	CDPW8_RS10020	2106765	2107335	-	IS256_family_transposase
CDPW8_RS10040	CDPW8_RS10040	2109342	2110572	-	IS110_family_transposase

Table of Blast hits (query gene, subject gene, %identity, blast score, %coverage, e-value):
input.path1.gene109	CDPW8_RS09970	31	87	39.679715302491104	3.5e-15
input.path1.gene115	CDPW8_RS09820	32	155	59.58254269449716	1.7e-35


>>
24. NC_016785
Source: Corynebacterium diphtheriae CDCE 8392, complete sequence
Type: LAP,NRPS-like,thiopeptide
Number of proteins with BLAST hits to this cluster: 2
Cumulative BLAST score: 242

Table of genes, locations, strands and annotations of subject cluster:
CDCE8392_RS09600	CDCE8392_RS09600	2003209	2004035	+	hypothetical_protein
CDCE8392_RS12405	CDCE8392_RS12405	2004255	2005046	+	hypothetical_protein
CDCE8392_RS12010	CDCE8392_RS12010	2010312	2010687	-	IS256_family_transposase
CDCE8392_RS09620	CDCE8392_RS09620	2011251	2011843	+	hypothetical_protein
CDCE8392_RS12015	CDCE8392_RS12015	2013179	2013415	-	transposase
CDCE8392_RS12020	CDCE8392_RS12020	2013525	2013662	+	IS3_family_transposase

Table of Blast hits (query gene, subject gene, %identity, blast score, %coverage, e-value):
input.path1.gene109	CDCE8392_RS09575	31	87	39.679715302491104	4.6e-15
input.path1.gene115	CDCE8392_RS09430	32	155	59.58254269449716	1.7e-35


>>
25. NZ_CP029644
Source: Corynebacterium diphtheriae strain BQ11 chromosome, complete genome
Type: LAP,NRPS-like,thiopeptide
Number of proteins with BLAST hits to this cluster: 2
Cumulative BLAST score: 241

Table of genes, locations, strands and annotations of subject cluster:
B11Q_RS10255	B11Q_RS10255	2066416	2067210	+	alpha/beta_hydrolase
B11Q_RS10340	B11Q_RS10340	2086430	2086666	+	IS110_family_transposase
B11Q_RS10470	B11Q_RS10470	2115552	2116765	+	IS110_family_transposase
B11Q_RS10475	B11Q_RS10475	2116965	2117738	+	lipase
B11Q_RS10495	B11Q_RS10495	2122972	2123393	-	IS256_family_transposase
B11Q_RS10500	B11Q_RS10500	2123623	2124848	+	IS30_family_transposase

Table of Blast hits (query gene, subject gene, %identity, blast score, %coverage, e-value):
input.path1.gene109	B11Q_RS10440	32	86	38.967971530249116	7.8e-15
input.path1.gene115	B11Q_RS10285	32	155	59.58254269449716	1.3e-35


>>
26. NZ_CP031418
Source: Nocardia farcinica strain W6977 chromosome, complete genome
Type: NRPS
Number of proteins with BLAST hits to this cluster: 2
Cumulative BLAST score: 240

Table of genes, locations, strands and annotations of subject cluster:
DXT66_RS11465	DXT66_RS11465	2422560	2422778	+	hypothetical_protein
DXT66_RS11505	DXT66_RS11505	2442384	2444419	-	MFS_transporter

Table of Blast hits (query gene, subject gene, %identity, blast score, %coverage, e-value):
input.path1.gene109	DXT66_RS11485	36	103	34.16370106761566	6.2e-20
input.path1.gene111	DXT66_RS11505	32	137	52.38095238095239	3.9e-30


>>
27. NC_002935
Source: Corynebacterium diphtheriae NCTC 13129, complete genome
Type: NRPS-like
Number of proteins with BLAST hits to this cluster: 2
Cumulative BLAST score: 240

Table of genes, locations, strands and annotations of subject cluster:
DIP_RS23945	DIP_RS23945	2040651	2040884	+	IS110_family_transposase
DIP_RS23625	DIP_RS23625	2075826	2076099	-	hypothetical_protein
DIP_RS21290	DIP_RS21290	2076133	2076546	-	hypothetical_protein
DIP_RS21295	DIP_RS21295	2076725	2077456	+	Fic_family_protein

Table of Blast hits (query gene, subject gene, %identity, blast score, %coverage, e-value):
input.path1.gene109	DIP_RS21235	31	85	39.679715302491104	1.7e-14
input.path1.gene115	DIP_RS21120	32	155	59.58254269449716	2.2e-35


>>
28. NC_016801
Source: Corynebacterium diphtheriae C7 (beta), complete sequence
Type: LAP,NRPS-like,thiopeptide
Number of proteins with BLAST hits to this cluster: 2
Cumulative BLAST score: 240

Table of genes, locations, strands and annotations of subject cluster:
CDC7B_RS12935	CDC7B_RS12935	2048478	2048714	+	IS110_family_transposase
CDC7B_RS12940	CDC7B_RS12940	2069362	2070210	+	lipase
CDC7B_RS10045	CDC7B_RS10045	2075379	2075879	-	IS256_family_transposase
CDC7B_RS12510	CDC7B_RS12510	2078590	2078825	-	transposase
CDC7B_RS12515	CDC7B_RS12515	2078932	2079072	+	IS3_family_transposase
CDC7B_RS12520	CDC7B_RS12520	2079248	2080452	+	IS1249_family_transposase

Table of Blast hits (query gene, subject gene, %identity, blast score, %coverage, e-value):
input.path1.gene109	CDC7B_RS10010	31	85	39.679715302491104	1.7e-14
input.path1.gene115	CDC7B_RS09865	32	155	59.58254269449716	1.7e-35


>>
29. NZ_CP018331
Source: Corynebacterium diphtheriae strain B-D-16-78 chromosome, complete genome
Type: LAP,NRPS-like,thiopeptide
Number of proteins with BLAST hits to this cluster: 2
Cumulative BLAST score: 240

Table of genes, locations, strands and annotations of subject cluster:
BS112_RS05945	BS112_RS05945	1235852	1241437	-	VWA_domain-containing_protein
BS112_RS05970	BS112_RS05970	1245784	1247165	-	IS3_family_transposase
BS112_RS12075	BS112_RS12075	1248606	1249246	+	hypothetical_protein
BS112_RS06005	BS112_RS06005	1254330	1254640	-	IS256_family_transposase

Table of Blast hits (query gene, subject gene, %identity, blast score, %coverage, e-value):
input.path1.gene109	BS112_RS05940	31	85	39.679715302491104	1.7e-14
input.path1.gene115	BS112_RS05780	32	155	59.58254269449716	1.7e-35


>>
30. NZ_CP038504
Source: Corynebacterium diphtheriae strain TH1526 chromosome, complete genome
Type: NRPS-like
Number of proteins with BLAST hits to this cluster: 2
Cumulative BLAST score: 240

Table of genes, locations, strands and annotations of subject cluster:
E4651_RS08965	E4651_RS08965	1848319	1848552	+	IS110_family_transposase
E4651_RS09120	E4651_RS09120	1883462	1883735	-	hypothetical_protein
E4651_RS09125	E4651_RS09125	1883769	1884182	-	hypothetical_protein
E4651_RS09130	E4651_RS09130	1884402	1885269	+	Fic_family_protein

Table of Blast hits (query gene, subject gene, %identity, blast score, %coverage, e-value):
input.path1.gene109	E4651_RS09065	31	85	39.679715302491104	1.7e-14
input.path1.gene115	E4651_RS08945	32	155	59.58254269449716	1.3e-35


>>
31. NZ_LR134537
Source: Corynebacterium diphtheriae strain NCTC7838 chromosome 1
Type: LAP,NRPS-like,thiopeptide
Number of proteins with BLAST hits to this cluster: 2
Cumulative BLAST score: 240

Table of genes, locations, strands and annotations of subject cluster:
EL333_RS09100	EL333_RS09100	1877636	1878674	+	IS110_family_transposase
EL333_RS09250	EL333_RS09250	1911587	1912929	-	IS3_family_transposase
EL333_RS09260	EL333_RS09260	1915018	1915439	-	IS256_family_transposase

Table of Blast hits (query gene, subject gene, %identity, blast score, %coverage, e-value):
input.path1.gene109	EL333_RS09200	31	85	39.679715302491104	1.7e-14
input.path1.gene115	EL333_RS09045	32	155	59.58254269449716	1.7e-35


>>
32. NC_016782
Source: Corynebacterium diphtheriae 241, complete sequence
Type: NRPS-like
Number of proteins with BLAST hits to this cluster: 2
Cumulative BLAST score: 239

Table of genes, locations, strands and annotations of subject cluster:
CD241_RS11875	CD241_RS11875	1997660	1997893	+	IS110_family_transposase
CD241_RS11880	CD241_RS11880	2030896	2032239	+	IS3_family_transposase
CD241_RS11885	CD241_RS11885	2032293	2033635	-	IS3_family_transposase
CD241_RS11890	CD241_RS11890	2035724	2036145	-	IS256_family_transposase

Table of Blast hits (query gene, subject gene, %identity, blast score, %coverage, e-value):
input.path1.gene109	CD241_RS09550	31	84	39.679715302491104	3e-14
input.path1.gene115	CD241_RS09435	32	155	59.58254269449716	2.2e-35


>>
33. NC_016787
Source: Corynebacterium diphtheriae HC03, complete sequence
Type: LAP,NRPS-like,thiopeptide
Number of proteins with BLAST hits to this cluster: 2
Cumulative BLAST score: 239

Table of genes, locations, strands and annotations of subject cluster:
CDHC03_RS09365	CDHC03_RS09365	1987111	1989710	+	lantibiotic_dehydratase
CDHC03_RS12320	CDHC03_RS12320	1995280	1995543	+	hypothetical_protein
CDHC03_RS11915	CDHC03_RS11915	2030429	2030702	-	hypothetical_protein
CDHC03_RS12325	CDHC03_RS12325	2031514	2031714	+	hypothetical_protein
CDHC03_RS11920	CDHC03_RS11920	2031877	2032379	+	Fic_family_protein

Table of Blast hits (query gene, subject gene, %identity, blast score, %coverage, e-value):
input.path1.gene109	CDHC03_RS09490	31	84	39.679715302491104	3e-14
input.path1.gene115	CDHC03_RS09345	32	155	59.58254269449716	1.3e-35


>>
34. NC_016790
Source: Corynebacterium diphtheriae VA01, complete sequence
Type: NRPS-like
Number of proteins with BLAST hits to this cluster: 2
Cumulative BLAST score: 239

Table of genes, locations, strands and annotations of subject cluster:
CDVA01_RS11935	CDVA01_RS11935	1961424	1961657	+	IS110_family_transposase
CDVA01_RS11525	CDVA01_RS11525	1996640	1996825	-	IS256_family_transposase
CDVA01_RS09380	CDVA01_RS09380	1996859	1997272	-	hypothetical_protein

Table of Blast hits (query gene, subject gene, %identity, blast score, %coverage, e-value):
input.path1.gene109	CDVA01_RS09325	31	84	38.967971530249116	3e-14
input.path1.gene115	CDVA01_RS09210	32	155	59.58254269449716	2.2e-35


>>
35. NC_016800
Source: Corynebacterium diphtheriae BH8, complete sequence
Type: LAP,NRPS-like,thiopeptide
Number of proteins with BLAST hits to this cluster: 2
Cumulative BLAST score: 239

Table of genes, locations, strands and annotations of subject cluster:
CDBH8_RS13005	CDBH8_RS13005	2042939	2043121	+	IS110_family_transposase
CDBH8_RS13010	CDBH8_RS13010	2071816	2072665	+	lipase
CDBH8_RS13015	CDBH8_RS13015	2077784	2078296	-	IS256_family_transposase
CDBH8_RS13025	CDBH8_RS13025	2080005	2081201	-	IS3_family_transposase

Table of Blast hits (query gene, subject gene, %identity, blast score, %coverage, e-value):
input.path1.gene109	CDBH8_RS09925	31	84	39.679715302491104	3e-14
input.path1.gene115	CDBH8_RS09775	32	155	59.58254269449716	1.7e-35


>>
36. NC_016802
Source: Corynebacterium diphtheriae HC02, complete sequence
Type: NRPS-like
Number of proteins with BLAST hits to this cluster: 2
Cumulative BLAST score: 239

Table of genes, locations, strands and annotations of subject cluster:
CDHC02_RS12710	CDHC02_RS12710	2013929	2014369	+	IS110_family_transposase
CDHC02_RS12485	CDHC02_RS12485	2016744	2016980	+	IS110_family_transposase
CDHC02_RS12115	CDHC02_RS12115	2049943	2051285	+	IS3_family_transposase
CDHC02_RS12490	CDHC02_RS12490	2053314	2053587	-	hypothetical_protein

Table of Blast hits (query gene, subject gene, %identity, blast score, %coverage, e-value):
input.path1.gene109	CDHC02_RS09670	31	84	38.967971530249116	5.1e-14
input.path1.gene115	CDHC02_RS09550	32	155	59.58254269449716	2.2e-35


>>
37. NZ_CP006764
Source: Corynebacterium doosanense CAU 212 = DSM 45436 strain CAU 212(T) chromosome, complete genome
Type: NRPS-like
Number of proteins with BLAST hits to this cluster: 2
Cumulative BLAST score: 236

Table of genes, locations, strands and annotations of subject cluster:
CDOO_RS11545	CDOO_RS11545	2360066	2361294	-	IS3_family_transposase

Table of Blast hits (query gene, subject gene, %identity, blast score, %coverage, e-value):
input.path1.gene109	CDOO_RS11530	33	84	33.985765124555165	3.9e-14
input.path1.gene115	CDOO_RS11430	31	152	59.58254269449716	1.4e-34


>>
38. NZ_CP011541
Source: Corynebacterium epidermidicanis strain DSM 45586 chromosome, complete genome
Type: NRPS-like
Number of proteins with BLAST hits to this cluster: 2
Cumulative BLAST score: 234

Table of genes, locations, strands and annotations of subject cluster:

Table of Blast hits (query gene, subject gene, %identity, blast score, %coverage, e-value):
input.path1.gene109	CEPID_RS10510	34	82	30.2491103202847	1.1e-13
input.path1.gene115	CEPID_RS10425	31	152	59.96204933586338	1.4e-34


>>
39. NZ_CP059399
Source: Nocardia sp. WCH-YHL-001 chromosome, complete genome
Type: NRPS
Number of proteins with BLAST hits to this cluster: 2
Cumulative BLAST score: 180

Table of genes, locations, strands and annotations of subject cluster:
H0264_RS09270	H0264_RS09270	2071658	2071939	+	DUF559_domain-containing_protein

Table of Blast hits (query gene, subject gene, %identity, blast score, %coverage, e-value):
input.path1.gene107	H0264_RS09385	32	64	39.11290322580645	2.4e-08
input.path1.gene109	H0264_RS09315	36	116	38.967971530249116	7.1e-24


>>
40. NZ_FOET01000004
Source: Streptomyces radiopugnans strain CGMCC 4.3519, whole genome shotgun sequence
Type: NRPS,NRPS-like,T2PKS,lanthipeptide,lassopeptide
Number of proteins with BLAST hits to this cluster: 2
Cumulative BLAST score: 180

Table of genes, locations, strands and annotations of subject cluster:
BMY40_RS10715	BMY40_RS10715	132607	132807	+	polyketide_cyclase
BMY40_RS10875	BMY40_RS10875	159608	159754	+	hemerythrin_domain-containing_protein

Table of Blast hits (query gene, subject gene, %identity, blast score, %coverage, e-value):
input.path1.gene109	BMY40_RS10595	38	88	31.672597864768683	2.1e-15
input.path1.gene113	BMY40_RS10865	31	92	61.25000000000001	1.6e-16


>>
41. NZ_CP020700
Source: Streptomyces tsukubensis strain NRRL 18488 chromosome, complete genome
Type: NRPS,T1PKS,butyrolactone,ectoine,other
Number of proteins with BLAST hits to this cluster: 2
Cumulative BLAST score: 163

Table of genes, locations, strands and annotations of subject cluster:
B7R87_RS00030	B7R87_RS00030	5929	6168	-	acyltransferase
B7R87_RS00045	B7R87_RS00045	8518	8900	-	hypothetical_protein
B7R87_RS00100	B7R87_RS00100	31199	31333	-	phytanoyl-CoA_dioxygenase_family_protein
B7R87_RS00170	B7R87_RS00170	63756	64256	+	antibiotic_biosynthesis_protein
B7R87_RS00490	B7R87_RS00490	153105	169037	-	SDR_family_NAD(P)-dependent_oxidoreductase
B7R87_RS00495	B7R87_RS00495	169974	172445	-	KR_domain-containing_protein
B7R87_RS00500	B7R87_RS00500	173316	176066	-	acyltransferase_domain-containing_protein
B7R87_RS00505	B7R87_RS00505	176397	190526	-	SDR_family_NAD(P)-dependent_oxidoreductase
B7R87_RS00510	B7R87_RS00510	190557	191306	-	KR_domain-containing_protein
B7R87_RS00515	B7R87_RS00515	191640	195098	-	acyltransferase_domain-containing_protein
B7R87_RS00520	B7R87_RS00520	196248	196397	-	hypothetical_protein
B7R87_RS00525	B7R87_RS00525	196440	197945	-	acyltransferase_domain-containing_protein
B7R87_RS00535	B7R87_RS00535	209273	210141	-	proline_iminopeptidase-family_hydrolase

Table of Blast hits (query gene, subject gene, %identity, blast score, %coverage, e-value):
input.path1.gene109	B7R87_RS00135	34	94	39.32384341637011	2.9e-17
input.path1.gene113	B7R87_RS00105	31	69	39.166666666666664	8.5e-10


>>
42. NZ_CP046172
Source: Nocardia arthritidis strain AUSMDU00012717 chromosome, complete genome
Type: NRPS,lanthipeptide
Number of proteins with BLAST hits to this cluster: 2
Cumulative BLAST score: 159

Table of genes, locations, strands and annotations of subject cluster:

Table of Blast hits (query gene, subject gene, %identity, blast score, %coverage, e-value):
input.path1.gene107	F5544_RS08915	31	68	48.38709677419355	1.3e-09
input.path1.gene109	F5544_RS08935	31	91	42.52669039145908	4.2e-16


>>
43. NC_014158
Source: Tsukamurella paurometabola DSM 20162, complete sequence
Type: NRPS
Number of proteins with BLAST hits to this cluster: 2
Cumulative BLAST score: 156

Table of genes, locations, strands and annotations of subject cluster:
TPAU_RS18505	TPAU_RS18505	3817333	3818139	+	helix-turn-helix_domain-containing_protein

Table of Blast hits (query gene, subject gene, %identity, blast score, %coverage, e-value):
input.path1.gene107	TPAU_RS18335	38	62	33.87096774193548	5.4e-08
input.path1.gene109	TPAU_RS18470	32	94	43.238434163701065	4.9e-17


>>
44. NZ_CP010519
Source: Streptomyces albus strain DSM 41398 chromosome, complete genome
Type: NRPS
Number of proteins with BLAST hits to this cluster: 2
Cumulative BLAST score: 155

Table of genes, locations, strands and annotations of subject cluster:
SLNWT_RS37405	SLNWT_RS37405	4062432	4062512	-	energy_transducer_TonB
SLNWT_RS15595	SLNWT_RS15595	4069819	4071693	+	ABC_transporter_ATP-binding_protein
SLNWT_RS37415	SLNWT_RS37415	4106256	4106531	+	FAD-binding_protein

Table of Blast hits (query gene, subject gene, %identity, blast score, %coverage, e-value):
input.path1.gene107	SLNWT_RS35350	36	69	36.29032258064516	4.4e-10
input.path1.gene109	SLNWT_RS15620	32	86	40.569395017793596	7.8e-15


>>
45. NZ_HE997626
Source: Tsukamurella sp. 1534, whole genome shotgun sequence
Type: NRPS
Number of proteins with BLAST hits to this cluster: 2
Cumulative BLAST score: 151

Table of genes, locations, strands and annotations of subject cluster:
BN39_RS09165	BN39_RS09165	1898718	1899786	-	non-ribosomal_peptide_synthetase

Table of Blast hits (query gene, subject gene, %identity, blast score, %coverage, e-value):
input.path1.gene107	BN39_RS09325	31	62	61.29032258064516	7e-08
input.path1.gene109	BN39_RS09195	32	89	37.36654804270463	1.6e-15


>>
46. NZ_CP034550
Source: Saccharothrix syringae strain NRRL B-16468 chromosome, complete genome
Type: NRPS,T2PKS
Number of proteins with BLAST hits to this cluster: 2
Cumulative BLAST score: 145

Table of genes, locations, strands and annotations of subject cluster:
EKG83_RS32120	EKG83_RS32120	7609229	7610120	+	TetR_family_transcriptional_regulator
EKG83_RS32320	EKG83_RS32320	7693327	7693569	-	monooxygenase

Table of Blast hits (query gene, subject gene, %identity, blast score, %coverage, e-value):
input.path1.gene107	EKG83_RS32265	33	63	41.935483870967744	3.1e-08
input.path1.gene109	EKG83_RS32175	33	82	36.47686832740214	1.9e-13


>>
47. NW_003345200
Source: Nannizzia gypsea CBS 118893 supercont1.2 genomic scaffold, whole genome shotgun sequence
Type: NRPS
Number of proteins with BLAST hits to this cluster: 4
Cumulative BLAST score: 1218

Table of genes, locations, strands and annotations of subject cluster:

Table of Blast hits (query gene, subject gene, %identity, blast score, %coverage, e-value):
input.path1.gene106	MGYG_03297	54	314	98.93992932862191	8.9e-84
input.path1.gene107	MGYG_03298	35	164	100.0	1.3e-38
input.path1.gene110	MGYG_03304	31	140	89.76377952755905	4.1e-31
input.path1.gene113	MGYG_09014	47	311	70.83333333333334	2.2e-82
input.path1.gene113	MGYG_03307	43	289	71.04166666666667	6.8e-76


>>
48. NW_003456424
Source: Trichophyton rubrum CBS 118892 genomic scaffold supercont2.5, whole genome shotgun sequence
Type: NRPS
Number of proteins with BLAST hits to this cluster: 4
Cumulative BLAST score: 1190

Table of genes, locations, strands and annotations of subject cluster:

Table of Blast hits (query gene, subject gene, %identity, blast score, %coverage, e-value):
input.path1.gene106	TERG_05131	53	310	98.93992932862191	2.2e-82
input.path1.gene107	TERG_05132	34	157	100.0	1.6e-36
input.path1.gene110	TERG_05139	31	135	83.2020997375328	1.3e-29
input.path1.gene113	TERG_05134	47	309	70.83333333333334	8.3e-82
input.path1.gene113	TERG_05143	41	279	71.04166666666667	5.4e-73


>>
49. NC_036436
Source: Aspergillus oryzae RIB40 DNA, chromosome 2
Type: NRPS
Number of proteins with BLAST hits to this cluster: 4
Cumulative BLAST score: 1097

Table of genes, locations, strands and annotations of subject cluster:

Table of Blast hits (query gene, subject gene, %identity, blast score, %coverage, e-value):
input.path1.gene106	AO090001000036	60	187	49.82332155477032	2.1e-45
input.path1.gene107	AO090001000037	35	156	100.40322580645163	3.6e-36
input.path1.gene110	AO090001000043	31	136	83.98950131233596	5.9e-30
input.path1.gene113	AO090001000039	48	342	71.04166666666667	5.2e-92
input.path1.gene113	AO090001000046	42	276	70.625	4.6e-72


>>
50. NW_013550604
Source: Rhinocladiella mackenziei CBS 650.93 unplaced genomic scaffold supercont1.1, whole genome shotgun sequence
Type: NRPS
Number of proteins with BLAST hits to this cluster: 4
Cumulative BLAST score: 1076

Table of genes, locations, strands and annotations of subject cluster:

Table of Blast hits (query gene, subject gene, %identity, blast score, %coverage, e-value):
input.path1.gene106	Z518_00668	43	229	98.58657243816255	3.8e-58
input.path1.gene110	Z518_00658	32	128	83.2020997375328	1.6e-27
input.path1.gene111	Z518_00652	41	402	91.53439153439153	6.5e-110
input.path1.gene113	Z518_00668	46	317	71.66666666666667	3e-84
