## Supplementary Results for "A draft genome of the ascomycotal fungal species *Pseudopithomyces maydicus* (family *Didymosphaeriaceae*)": contig_4_c2.txt

ClusterBlast scores for contig_4

Table of genes, locations, strands and annotations of query cluster:
input.path1.gene340	1284316	1284607	+		
input.path1.gene341	1286354	1288678	+		
input.path1.gene342	1288948	1289596	-		
input.path1.gene343	1294518	1295517	+		
input.path1.gene344	1295611	1297093	-		
input.path1.gene345	1299349	1304830	-		
input.path1.gene346	1307660	1309419	-		
input.path1.gene347	1309571	1310220	+		
input.path1.gene348	1311642	1312949	+		
input.path1.gene349	1313508	1315245	-		
input.path1.gene350	1315986	1317843	+		
input.path1.gene351	1320309	1321467	-		


Significant hits: 
1. NZ_CP034352	Streptomyces sp. W1SF4 plasmid p2, complete sequence
2. NZ_CP016824	Streptomyces sampsonii strain KJ40 chromosome, complete genome
3. NZ_CP056773	Streptomyces violascens strain YIM 100212 chromosome, complete genome
4. NZ_CM002271	Streptomyces sp. GBA 94-10 chromosome, whole genome shotgun sequence
5. NC_020990	Streptomyces albidoflavus, complete sequence
6. NZ_CP047147	Streptomyces sp. GF20 chromosome, complete genome
7. NZ_CP031742	Streptomyces koyangensis strain VK-A60T chromosome, complete genome
8. NZ_AMPN02000002	Streptomyces sp. SM8 Scaffold_2, whole genome shotgun sequence
9. NZ_CP024052	Micromonospora sp. WMMA2032 chromosome
10. NZ_SMBL01000011	Streptomyces sp. BK215 Ga0307669_111, whole genome shotgun sequence
11. NZ_JYNX01000025	Mycolicibacterium chubuense strain DSM 44219 MCHUDSM44219_contig000025, whole genome shotgun sequence
12. NZ_KQ948552	Streptomyces longwoodensis strain DSM 41677 PRJNA299227_s004, whole genome shotgun sequence
13. NZ_ML123127	Streptomyces sp. ADI98-12 scaffold002, whole genome shotgun sequence
14. NC_013440	Haliangium ochraceum DSM 14365, complete genome
15. NZ_LWLA01000004	Streptomyces sp. CB03911 scaffold12, whole genome shotgun sequence
16. NZ_CP023778	Nocardia terpenica strain NC_YFY_NT001 chromosome, complete genome
17. NZ_AUBB01000021	Pseudonocardia spinosispora DSM 44797 G406DRAFT_scaffold00011.11_C, whole genome shotgun sequence
18. NZ_KB907309	Nocardia sp. BMG111209 G361DRAFT_scaffold00003.3, whole genome shotgun sequence
19. NW_023336287	Aspergillus tubingensis WU-2223L DNA, scaffold_13, whole genome shotgun sequence
20. NZ_JACHJF010000011	Streptomyces eurocidicus strain CECT 3259 Ga0436944_11, whole genome shotgun sequence
21. NZ_CP014306	Burkholderia sp. PAMC 26561 chromosome 1, complete sequence
22. NZ_CP018082	Nocardia mangyaensis strain Y48 chromosome, complete genome
23. NC_036441	Aspergillus oryzae RIB40 DNA, chromosome 7
24. NZ_CP016438	Streptomyces lincolnensis strain NRRL 2936 chromosome
25. NZ_CP047146	Streptomyces sp. GS7 chromosome, complete genome
26. NZ_CP054569	Achromobacter denitrificans strain FDAARGOS_787 chromosome, complete genome
27. NZ_CP020917	Achromobacter denitrificans strain PR1 chromosome, complete genome
28. NZ_CP053986	Achromobacter denitrificans strain FDAARGOS_788 chromosome, complete genome
29. NZ_CP033730	Achromobacter denitrificans strain PheN1 chromosome, complete genome
30. NZ_MEBE01000001	Burkholderia ubonensis strain MSMB0245 Contig_1, whole genome shotgun sequence
31. NZ_KN150849	Burkholderia gladioli strain ATCC 25417 scaffold1, whole genome shotgun sequence
32. NZ_CADEXT010000003	Burkholderia gladioli strain BCC1700, whole genome shotgun sequence
33. NZ_CADEQD010000001	Burkholderia gladioli strain BCC1730, whole genome shotgun sequence
34. NZ_MECO01000028	Burkholderia ubonensis strain MSMB0803 Contig_7, whole genome shotgun sequence
35. NZ_CP009487	Burkholderia ubonensis MSMB22 chromosome III, complete sequence
36. NZ_CADEQR010000002	Burkholderia gladioli strain BCC1732, whole genome shotgun sequence
37. NZ_CADEYA010000006	Burkholderia gladioli strain BCC1733, whole genome shotgun sequence
38. NZ_CADEYO010000001	Burkholderia gladioli strain BCC1728, whole genome shotgun sequence
39. NZ_LNJU01000004	Burkholderia ubonensis strain MSMB1157 MSMB1157WGS_5, whole genome shotgun sequence
40. NZ_MEBC01000001	Burkholderia ubonensis strain MSMB0170 Contig_1, whole genome shotgun sequence
41. NZ_MEBF01000045	Burkholderia ubonensis strain MSMB0258b Contig_5, whole genome shotgun sequence
42. NZ_NPFX01000014	Rhodococcus sp. 14-2483-1-1 RS1B5_1, whole genome shotgun sequence
43. NZ_NSDZ01000001	Rhodococcus sp. ACS1 Contig_1, whole genome shotgun sequence
44. NZ_CP023701	Streptomyces subrutilus strain ATCC 27467 chromosome, complete genome
45. NZ_CP048641	Streptomyces aureoverticillatus strain HN6 chromosome, complete genome
46. NZ_CP008947	Rhodococcus opacus strain R7 chromosome
47. NC_012522	Rhodococcus opacus B4, complete genome
48. NZ_CP009111	Rhodococcus opacus strain 1CP, complete genome
49. NZ_CADEPP010000002	Burkholderia gladioli strain BCC1701, whole genome shotgun sequence
50. NZ_CP051855	Rhodococcus imtechensis strain XM24D chromosome 1, complete sequence


Details:

>>
1. NZ_CP034352
Source: Streptomyces sp. W1SF4 plasmid p2, complete sequence
Type: NRPS,NRPS-like,T1PKS,T2PKS,butyrolactone,melanin
Number of proteins with BLAST hits to this cluster: 2
Cumulative BLAST score: 2821

Table of genes, locations, strands and annotations of subject cluster:
D1J60_RS36170	D1J60_RS36170	1038	1672	+	TetR_family_transcriptional_regulator
D1J60_RS36175	D1J60_RS36175	1727	1906	-	hypothetical_protein
D1J60_RS36435	D1J60_RS36435	136120	141220	-	acyltransferase_domain-containing_protein
D1J60_RS37620	D1J60_RS37620	183563	183817	-	ATP-binding_protein
D1J60_RS36485	D1J60_RS36485	184321	185095	-	AfsR/SARP_family_transcriptional_regulator
D1J60_RS36505	D1J60_RS36505	189346	192303	+	acyltransferase_domain-containing_protein
D1J60_RS36675	D1J60_RS36675	227432	229621	+	MMPL_family_transporter
D1J60_RS36725	D1J60_RS36725	243784	245379	-	acyl--CoA_ligase
D1J60_RS36990	D1J60_RS36990	298697	299017	-	TetR_family_transcriptional_regulator

Table of Blast hits (query gene, subject gene, %identity, blast score, %coverage, e-value):
input.path1.gene345	D1J60_RS36425	33	495	58.14732142857143	1.8e-137
input.path1.gene345	D1J60_RS36430	32	464	57.70089285714286	4.4e-128
input.path1.gene345	D1J60_RS36450	32	445	57.75669642857143	2.8e-122
input.path1.gene345	D1J60_RS36445	32	441	58.92857142857143	5.2e-121
input.path1.gene345	D1J60_RS36440	33	404	50.55803571428571	5.4e-110
input.path1.gene345	D1J60_RS36505	34	394	44.754464285714285	4.3e-107
input.path1.gene350	D1J60_RS36750	33	178	50.48543689320388	2.8e-42


>>
2. NZ_CP016824
Source: Streptomyces sampsonii strain KJ40 chromosome, complete genome
Type: NRPS,NRPS-like,T1PKS,T2PKS,butyrolactone,ectoine,lanthipeptide,transAT-PKS
Number of proteins with BLAST hits to this cluster: 2
Cumulative BLAST score: 1603

Table of genes, locations, strands and annotations of subject cluster:
BEN82_RS30405	BEN82_RS30405	64728	66076	+	transposase
BEN82_RS30410	BEN82_RS30410	88873	89136	-	transposase
BEN82_RS00470	BEN82_RS00470	102837	103672	-	hypothetical_protein
BEN82_RS00485	BEN82_RS00485	106160	106854	+	TetR_family_transcriptional_regulator
BEN82_RS00490	BEN82_RS00490	106885	108331	-	hypothetical_protein
BEN82_RS00495	BEN82_RS00495	108480	109117	-	PadR_family_transcriptional_regulator
BEN82_RS00505	BEN82_RS00505	110261	111066	+	ABC_transporter_permease
BEN82_RS00525	BEN82_RS00525	115803	116493	+	IS701_family_transposase
BEN82_RS00915	BEN82_RS00915	290441	307060	-	SDR_family_NAD(P)-dependent_oxidoreductase

Table of Blast hits (query gene, subject gene, %identity, blast score, %coverage, e-value):
input.path1.gene345	BEN82_RS00925	32	478	57.86830357142857	3.9e-132
input.path1.gene345	BEN82_RS00930	34	456	51.171875	1.2e-125
input.path1.gene345	BEN82_RS00895	33	412	51.72991071428571	2.6e-112
input.path1.gene350	BEN82_RS00540	45	257	49.676375404530745	2.8e-66


>>
3. NZ_CP056773
Source: Streptomyces violascens strain YIM 100212 chromosome, complete genome
Type: NRPS,NRPS-like,T1PKS,lanthipeptide
Number of proteins with BLAST hits to this cluster: 2
Cumulative BLAST score: 1600

Table of genes, locations, strands and annotations of subject cluster:
HWN34_RS30010	HWN34_RS30010	6954131	6955383	-	IS3_family_transposase
HWN34_RS30015	HWN34_RS30015	6955434	6955916	-	TniB_family_NTP-binding_protein

Table of Blast hits (query gene, subject gene, %identity, blast score, %coverage, e-value):
input.path1.gene345	HWN34_RS29605	32	477	57.86830357142857	5e-132
input.path1.gene345	HWN34_RS29600	35	455	51.33928571428571	2.1e-125
input.path1.gene345	HWN34_RS29635	33	409	51.89732142857143	1.7e-111
input.path1.gene350	HWN34_RS29995	45	259	50.0	1.3e-66


>>
4. NZ_CM002271
Source: Streptomyces sp. GBA 94-10 chromosome, whole genome shotgun sequence
Type: NRPS,NRPS-like,T1PKS,T3PKS,lanthipeptide
Number of proteins with BLAST hits to this cluster: 2
Cumulative BLAST score: 1600

Table of genes, locations, strands and annotations of subject cluster:
B591_RS32375	B591_RS32375	6561762	6562382	-	MerR_family_DNA-binding_transcriptional_regulator
B591_RS28710	B591_RS28710	6563143	6564837	+	DHA2_family_efflux_MFS_transporter_permease_subunit
B591_RS33260	B591_RS33260	6565107	6565220	-	pirin_family_protein
B591_RS33265	B591_RS33265	6583759	6583929	-	kinase
B591_RS32385	B591_RS32385	6583905	6584282	-	DUF664_domain-containing_protein
B591_RS33270	B591_RS33270	6584868	6585010	-	alcohol_dehydrogenase
B591_RS29180	B591_RS29180	6813727	6822663	+	SDR_family_NAD(P)-dependent_oxidoreductase
B591_RS29185	B591_RS29185	6822702	6826045	+	amino_acid_adenylation_protein
B591_RS29275	B591_RS29275	6846847	6848680	-	DDE-type_integrase/transposase/recombinase

Table of Blast hits (query gene, subject gene, %identity, blast score, %coverage, e-value):
input.path1.gene345	B591_RS28870	32	473	57.86830357142857	9.5e-131
input.path1.gene345	B591_RS28865	34	455	50.66964285714286	3.5e-125
input.path1.gene345	B591_RS28900	33	409	51.89732142857143	1.7e-111
input.path1.gene350	B591_RS29255	46	263	49.51456310679612	5.2e-68


>>
5. NC_020990
Source: Streptomyces albidoflavus, complete sequence
Type: NRPS,NRPS-like,T1PKS,lanthipeptide
Number of proteins with BLAST hits to this cluster: 2
Cumulative BLAST score: 1599

Table of genes, locations, strands and annotations of subject cluster:
XNR_RS29035	XNR_RS29035	6661707	6682502	+	SDR_family_NAD(P)-dependent_oxidoreductase
XNR_RS30645	XNR_RS30645	6682626	6684770	+	type_I_polyketide_synthase
XNR_RS30545	XNR_RS30545	6731362	6732209	-	tryptophan_2,3-dioxygenase
XNR_RS29330	XNR_RS29330	6803548	6809793	+	SDR_family_NAD(P)-dependent_oxidoreductase
XNR_RS30555	XNR_RS30555	6810133	6810444	+	hypothetical_protein
XNR_RS30560	XNR_RS30560	6811201	6812472	+	SDR_family_NAD(P)-dependent_oxidoreductase

Table of Blast hits (query gene, subject gene, %identity, blast score, %coverage, e-value):
input.path1.gene345	XNR_RS29020	32	475	57.86830357142857	1.9e-131
input.path1.gene345	XNR_RS29015	35	456	51.33928571428571	9.2e-126
input.path1.gene345	XNR_RS29050	33	409	51.89732142857143	2.2e-111
input.path1.gene350	XNR_RS29400	46	259	48.86731391585761	9.7e-67


>>
6. NZ_CP047147
Source: Streptomyces sp. GF20 chromosome, complete genome
Type: NRPS,NRPS-like,T1PKS,lanthipeptide
Number of proteins with BLAST hits to this cluster: 2
Cumulative BLAST score: 1593

Table of genes, locations, strands and annotations of subject cluster:
GR131_RS00230	GR131_RS00230	63290	64222	+	ATP-dependent_DNA_ligase
GR131_RS00360	GR131_RS00360	112627	113098	+	hypothetical_protein
GR131_RS00660	GR131_RS00660	319311	320616	+	flavohemoprotein

Table of Blast hits (query gene, subject gene, %identity, blast score, %coverage, e-value):
input.path1.gene345	GR131_RS00635	32	475	57.86830357142857	1.9e-131
input.path1.gene345	GR131_RS00640	35	453	51.28348214285714	1e-124
input.path1.gene345	GR131_RS00605	33	410	51.72991071428571	1.3e-111
input.path1.gene350	GR131_RS00255	45	255	49.35275080906149	1.4e-65


>>
7. NZ_CP031742
Source: Streptomyces koyangensis strain VK-A60T chromosome, complete genome
Type: NRPS,NRPS-like,T1PKS
Number of proteins with BLAST hits to this cluster: 2
Cumulative BLAST score: 1582

Table of genes, locations, strands and annotations of subject cluster:
D0C37_RS31655	D0C37_RS31655	7131468	7132262	-	alpha/beta_fold_hydrolase
D0C37_RS31730	D0C37_RS31730	7156172	7156258	-	TetR/AcrR_family_transcriptional_regulator
D0C37_RS31765	D0C37_RS31765	7162683	7163491	+	IS5_family_transposase
D0C37_RS31795	D0C37_RS31795	7168589	7169197	+	IS110_family_transposase
D0C37_RS31825	D0C37_RS31825	7174805	7175050	+	transposase

Table of Blast hits (query gene, subject gene, %identity, blast score, %coverage, e-value):
input.path1.gene345	D0C37_RS31530	32	465	57.36607142857143	2e-128
input.path1.gene345	D0C37_RS31525	34	456	51.28348214285714	1.6e-125
input.path1.gene345	D0C37_RS31560	33	405	51.67410714285714	2.4e-110
input.path1.gene350	D0C37_RS31745	44	256	50.323624595469255	8.2e-66


>>
8. NZ_AMPN02000002
Source: Streptomyces sp. SM8 Scaffold_2, whole genome shotgun sequence
Type: NRPS,NRPS-like,T1PKS,lanthipeptide,transAT-PKS
Number of proteins with BLAST hits to this cluster: 2
Cumulative BLAST score: 1547

Table of genes, locations, strands and annotations of subject cluster:
SM8_RS31120	SM8_RS31120	6664429	6665119	-	cholesterol_esterase
SM8_RS31195	SM8_RS31195	6678278	6678512	+	hypothetical_protein
SM8_RS31200	SM8_RS31200	6678613	6679047	+	hypothetical_protein
SM8_RS31210	SM8_RS31210	6680780	6681884	+	beta-ketoacyl_synthase
SM8_RS31215	SM8_RS31215	6681985	6698753	+	SDR_family_NAD(P)-dependent_oxidoreductase
SM8_RS31245	SM8_RS31245	6719896	6725146	+	SDR_family_NAD(P)-dependent_oxidoreductase
SM8_RS31250	SM8_RS31250	6725249	6732084	+	acyltransferase_domain-containing_protein
SM8_RS31255	SM8_RS31255	6732122	6733405	+	hypothetical_protein
SM8_RS31265	SM8_RS31265	6738034	6754680	+	SDR_family_NAD(P)-dependent_oxidoreductase
SM8_RS31275	SM8_RS31275	6762230	6763171	+	beta-ketoacyl_synthase
SM8_RS31280	SM8_RS31280	6763274	6767068	+	SDR_family_NAD(P)-dependent_oxidoreductase
SM8_RS31285	SM8_RS31285	6767170	6774262	+	SDR_family_NAD(P)-dependent_oxidoreductase
SM8_RS31335	SM8_RS31335	6797755	6798823	-	cytochrome_P450
SM8_RS31350	SM8_RS31350	6801846	6803492	+	AAA_family_ATPase
SM8_RS31355	SM8_RS31355	6803593	6804549	+	LuxR_family_transcriptional_regulator
SM8_RS31360	SM8_RS31360	6804608	6806826	+	AAA_family_ATPase
SM8_RS31370	SM8_RS31370	6808930	6809589	+	LuxR_family_transcriptional_regulator
SM8_RS31390	SM8_RS31390	6813476	6814140	-	ATP-binding_cassette_domain-containing_protein
SM8_RS31400	SM8_RS31400	6814945	6816965	-	branched-chain_amino_acid_ABC_transporter_permease
SM8_RS31500	SM8_RS31500	6847086	6853025	-	amino_acid_adenylation_domain-containing_protein
SM8_RS31505	SM8_RS31505	6853293	6853880	-	MFS_transporter
SM8_RS31510	SM8_RS31510	6853983	6858239	-	acyltransferase_domain-containing_protein
SM8_RS31515	SM8_RS31515	6858238	6859023	-	MFS_transporter
SM8_RS31530	SM8_RS31530	6863053	6865822	+	AAA_family_ATPase
SM8_RS31535	SM8_RS31535	6865977	6867676	+	AfsR_family_transcriptional_regulator
SM8_RS31565	SM8_RS31565	6873794	6876729	+	AAA_family_ATPase
SM8_RS31580	SM8_RS31580	6878382	6878911	-	hypothetical_protein
SM8_RS31615	SM8_RS31615	6893773	6899043	+	SDR_family_NAD(P)-dependent_oxidoreductase
SM8_RS31620	SM8_RS31620	6899044	6902267	+	hypothetical_protein
SM8_RS31625	SM8_RS31625	6902306	6905404	+	amino_acid_adenylation_protein
SM8_RS31640	SM8_RS31640	6909292	6913033	+	aminotransferase_class_I/II-fold_pyridoxal_phosphate-dependent_enzyme
SM8_RS31710	SM8_RS31710	6925065	6925755	-	IS701_family_transposase

Table of Blast hits (query gene, subject gene, %identity, blast score, %coverage, e-value):
input.path1.gene345	SM8_RS31240	32	477	57.86830357142857	5e-132
input.path1.gene345	SM8_RS31315	33	406	51.89732142857143	1.4e-110
input.path1.gene345	SM8_RS31235	33	406	50.94866071428571	1.9e-110
input.path1.gene350	SM8_RS31695	46	258	49.676375404530745	1.7e-66


>>
9. NZ_CP024052
Source: Micromonospora sp. WMMA2032 chromosome
Type: NRPS-like,T1PKS
Number of proteins with BLAST hits to this cluster: 2
Cumulative BLAST score: 1517

Table of genes, locations, strands and annotations of subject cluster:
CO540_RS05160	CO540_RS05160	1134854	1139826	-	SDR_family_NAD(P)-dependent_oxidoreductase
CO540_RS05235	CO540_RS05235	1280835	1281329	-	zinc-binding_dehydrogenase
CO540_RS05240	CO540_RS05240	1281522	1281941	-	alcohol_dehydrogenase_catalytic_domain-containing_protein
CO540_RS05245	CO540_RS05245	1282242	1294178	-	SDR_family_NAD(P)-dependent_oxidoreductase

Table of Blast hits (query gene, subject gene, %identity, blast score, %coverage, e-value):
input.path1.gene345	CO540_RS05175	33	486	59.65401785714286	1.4e-134
input.path1.gene345	CO540_RS05170	35	477	50.11160714285714	5e-132
input.path1.gene345	CO540_RS05195	32	454	58.87276785714286	6e-125
input.path1.gene349	CO540_RS05080	35	100	39.285714285714285	5.2e-19


>>
10. NZ_SMBL01000011
Source: Streptomyces sp. BK215 Ga0307669_111, whole genome shotgun sequence
Type: T1PKS,lanthipeptide
Number of proteins with BLAST hits to this cluster: 2
Cumulative BLAST score: 1507

Table of genes, locations, strands and annotations of subject cluster:
EV288_RS20985	EV288_RS20985	177415	178259	-	IS5_family_transposase
EV288_RS20995	EV288_RS20995	178638	179484	-	IS5_family_transposase
EV288_RS21050	EV288_RS21050	185356	190200	+	acyltransferase_domain-containing_protein
EV288_RS21055	EV288_RS21055	191200	192069	+	SDR_family_NAD(P)-dependent_oxidoreductase
EV288_RS21060	EV288_RS21060	192079	194283	+	polyketide_synthase
EV288_RS21065	EV288_RS21065	194521	195372	+	hypothetical_protein
EV288_RS21070	EV288_RS21070	195511	196749	+	SDR_family_NAD(P)-dependent_oxidoreductase
EV288_RS21075	EV288_RS21075	197326	200391	+	acyltransferase_domain-containing_protein
EV288_RS21080	EV288_RS21080	201976	202791	+	SDR_family_NAD(P)-dependent_oxidoreductase
EV288_RS21090	EV288_RS21090	204139	207132	+	acyltransferase_domain-containing_protein
EV288_RS21100	EV288_RS21100	207490	210756	+	acyltransferase_domain-containing_protein
EV288_RS21105	EV288_RS21105	212374	213609	+	SDR_family_NAD(P)-dependent_oxidoreductase
EV288_RS21115	EV288_RS21115	214497	216797	+	polyketide_synthase
EV288_RS21120	EV288_RS21120	217683	218762	+	beta-ketoacyl_synthase
EV288_RS35540	EV288_RS35540	218799	220025	+	SDR_family_NAD(P)-dependent_oxidoreductase
EV288_RS35545	EV288_RS35545	221850	222236	+	hypothetical_protein
EV288_RS21130	EV288_RS21130	222279	223463	+	acyltransferase_domain-containing_protein
EV288_RS21135	EV288_RS21135	224763	225707	+	SDR_family_NAD(P)-dependent_oxidoreductase
EV288_RS21155	EV288_RS21155	228673	229439	-	ABC-F_family_ATP-binding_cassette_domain-containing_protein
EV288_RS21215	EV288_RS21215	240818	241598	+	IS5_family_transposase

Table of Blast hits (query gene, subject gene, %identity, blast score, %coverage, e-value):
input.path1.gene345	EV288_RS21050	34	436	51.78571428571429	9.9e-120
input.path1.gene345	EV288_RS21090	32	418	53.18080357142857	3.6e-114
input.path1.gene345	EV288_RS21075	32	398	51.45089285714286	3e-108
input.path1.gene350	EV288_RS20930	44	255	50.161812297734635	1.8e-65


>>
11. NZ_JYNX01000025
Source: Mycolicibacterium chubuense strain DSM 44219 MCHUDSM44219_contig000025, whole genome shotgun sequence
Type: NRPS,T1PKS
Number of proteins with BLAST hits to this cluster: 2
Cumulative BLAST score: 1482

Table of genes, locations, strands and annotations of subject cluster:
MCHUDSM44219_RS06690	MCHUDSM44219_RS06690	83851	85136	+	phosphotransferase_family_protein
MCHUDSM44219_RS06800	MCHUDSM44219_RS06800	120914	125560	-	acyltransferase_domain-containing_protein

Table of Blast hits (query gene, subject gene, %identity, blast score, %coverage, e-value):
input.path1.gene345	MCHUDSM44219_RS06790	34	498	52.34375	2.8e-138
input.path1.gene345	MCHUDSM44219_RS06785	34	465	51.28348214285714	3.4e-128
input.path1.gene345	MCHUDSM44219_RS06800	41	426	32.142857142857146	1.3e-116
input.path1.gene349	MCHUDSM44219_RS06670	31	93	37.5	8.4e-17


>>
12. NZ_KQ948552
Source: Streptomyces longwoodensis strain DSM 41677 PRJNA299227_s004, whole genome shotgun sequence
Type: PKS-like,T1PKS,T3PKS,ectoine
Number of proteins with BLAST hits to this cluster: 2
Cumulative BLAST score: 1460

Table of genes, locations, strands and annotations of subject cluster:
AQJ30_RS37870	AQJ30_RS37870	296227	296445	+	helix-turn-helix_transcriptional_regulator
AQJ30_RS13850	AQJ30_RS13850	307907	308350	-	transcriptional_regulator
AQJ30_RS14105	AQJ30_RS14105	415254	415574	-	MarR_family_transcriptional_regulator
AQJ30_RS14110	AQJ30_RS14110	415722	415874	-	MarR_family_transcriptional_regulator

Table of Blast hits (query gene, subject gene, %identity, blast score, %coverage, e-value):
input.path1.gene345	AQJ30_RS13895	37	488	50.0	2.2e-135
input.path1.gene345	AQJ30_RS13870	33	484	58.70535714285714	5.4e-134
input.path1.gene345	AQJ30_RS13865	31	393	51.50669642857143	1.2e-106
input.path1.gene350	AQJ30_RS13910	33	95	39.15857605177994	2.4e-17


>>
13. NZ_ML123127
Source: Streptomyces sp. ADI98-12 scaffold002, whole genome shotgun sequence
Type: NRPS,T1PKS,lanthipeptide
Number of proteins with BLAST hits to this cluster: 2
Cumulative BLAST score: 1139

Table of genes, locations, strands and annotations of subject cluster:
EES47_RS14315	EES47_RS14315	140375	140512	-	site-specific_integrase
EES47_RS30480	EES47_RS30480	140592	140782	+	hypothetical_protein
EES47_RS14355	EES47_RS14355	146381	146569	-	transposase_family_protein
EES47_RS14365	EES47_RS14365	146980	147826	-	IS5_family_transposase
EES47_RS14405	EES47_RS14405	168168	168860	-	transposase
EES47_RS14425	EES47_RS14425	171642	172349	+	TetR_family_transcriptional_regulator

Table of Blast hits (query gene, subject gene, %identity, blast score, %coverage, e-value):
input.path1.gene345	EES47_RS14380	37	454	47.20982142857143	4.6e-125
input.path1.gene345	EES47_RS14385	32	427	54.40848214285714	6e-117
input.path1.gene350	EES47_RS14520	44	258	50.323624595469255	1.7e-66


>>
14. NC_013440
Source: Haliangium ochraceum DSM 14365, complete genome
Type: NRPS,T1PKS,phenazine
Number of proteins with BLAST hits to this cluster: 2
Cumulative BLAST score: 1060

Table of genes, locations, strands and annotations of subject cluster:
HOCH_RS38735	HOCH_RS38735	4002239	4004815	+	DUF4157_domain-containing_protein
HOCH_RS38745	HOCH_RS38745	4008109	4008345	+	AAA_family_ATPase
HOCH_RS14540	HOCH_RS14540	4119152	4128711	+	cytochrome_P450
HOCH_RS40125	HOCH_RS40125	4135752	4135844	-	site-specific_integrase
HOCH_RS14585	HOCH_RS14585	4139211	4139527	+	hypothetical_protein

Table of Blast hits (query gene, subject gene, %identity, blast score, %coverage, e-value):
input.path1.gene345	HOCH_RS14530	34	505	53.23660714285714	2.9e-140
input.path1.gene345	HOCH_RS14440	35	466	51.11607142857143	1.5e-128
input.path1.gene350	HOCH_RS14425	31	89	35.760517799352755	1.7e-15


>>
15. NZ_LWLA01000004
Source: Streptomyces sp. CB03911 scaffold12, whole genome shotgun sequence
Type: T1PKS
Number of proteins with BLAST hits to this cluster: 2
Cumulative BLAST score: 1022

Table of genes, locations, strands and annotations of subject cluster:
A6A07_RS41780	A6A07_RS41780	311553	313975	-	hypothetical_protein
A6A07_RS41785	A6A07_RS41785	314223	314306	-	hypothetical_protein

Table of Blast hits (query gene, subject gene, %identity, blast score, %coverage, e-value):
input.path1.gene345	A6A07_RS07645	36	488	52.39955357142857	2.2e-135
input.path1.gene345	A6A07_RS07650	36	466	50.22321428571429	1.5e-128
input.path1.gene351	A6A07_RS07690	34	68	44.675324675324674	1.5e-09


>>
16. NZ_CP023778
Source: Nocardia terpenica strain NC_YFY_NT001 chromosome, complete genome
Type: T1PKS
Number of proteins with BLAST hits to this cluster: 2
Cumulative BLAST score: 906

Table of genes, locations, strands and annotations of subject cluster:
CRH09_RS40180	CRH09_RS40180	1743578	1743625	-	hypothetical_protein
CRH09_RS40185	CRH09_RS40185	1777333	1778892	+	hypothetical_protein
CRH09_RS40190	CRH09_RS40190	1779121	1779936	+	acyltransferase_domain-containing_protein
CRH09_RS40195	CRH09_RS40195	1780723	1781469	+	SDR_family_NAD(P)-dependent_oxidoreductase
CRH09_RS40200	CRH09_RS40200	1781860	1782756	+	polyketide_synthase
CRH09_RS40205	CRH09_RS40205	1783063	1783467	+	hypothetical_protein
CRH09_RS40210	CRH09_RS40210	1784284	1785936	+	SDR_family_NAD(P)-dependent_oxidoreductase
CRH09_RS40215	CRH09_RS40215	1786099	1786413	+	hypothetical_protein
CRH09_RS40220	CRH09_RS40220	1786504	1787940	+	hypothetical_protein
CRH09_RS40225	CRH09_RS40225	1788082	1789011	+	acyltransferase_domain-containing_protein
CRH09_RS40230	CRH09_RS40230	1789378	1790181	+	hypothetical_protein
CRH09_RS40235	CRH09_RS40235	1790542	1791408	+	polyketide_synthase
CRH09_RS40240	CRH09_RS40240	1791664	1792122	+	polyketide_synthase
CRH09_RS40245	CRH09_RS40245	1792294	1793199	+	acyltransferase_domain-containing_protein
CRH09_RS40250	CRH09_RS40250	1794145	1794732	+	SDR_family_NAD(P)-dependent_oxidoreductase
CRH09_RS40255	CRH09_RS40255	1795195	1796091	+	hypothetical_protein
CRH09_RS40260	CRH09_RS40260	1796407	1796820	+	hypothetical_protein
CRH09_RS40265	CRH09_RS40265	1796968	1797774	+	acyltransferase_domain-containing_protein
CRH09_RS40270	CRH09_RS40270	1798825	1800309	+	zinc-binding_dehydrogenase
CRH09_RS40290	CRH09_RS40290	1827070	1827948	+	polyketide_synthase
CRH09_RS40295	CRH09_RS40295	1828270	1828689	+	hypothetical_protein
CRH09_RS40300	CRH09_RS40300	1828810	1829598	+	acyltransferase_domain-containing_protein
CRH09_RS40305	CRH09_RS40305	1829995	1830807	+	SDR_family_NAD(P)-dependent_oxidoreductase
CRH09_RS40310	CRH09_RS40310	1831177	1831941	+	6-deoxyerythronolide-B_synthase
CRH09_RS40315	CRH09_RS40315	1832053	1832448	+	hypothetical_protein
CRH09_RS40320	CRH09_RS40320	1832650	1833843	+	acyltransferase_domain-containing_protein
CRH09_RS40325	CRH09_RS40325	1835026	1835730	+	alcohol_dehydrogenase_catalytic_domain-containing_protein
CRH09_RS40330	CRH09_RS40330	1836310	1836984	+	SDR_family_NAD(P)-dependent_oxidoreductase

Table of Blast hits (query gene, subject gene, %identity, blast score, %coverage, e-value):
input.path1.gene345	CRH09_RS08230	34	415	49.94419642857143	2.4e-113
input.path1.gene345	CRH09_RS08085	32	394	54.12946428571429	5.6e-107
input.path1.gene349	CRH09_RS08075	31	97	37.142857142857146	4.4e-18


>>
17. NZ_AUBB01000021
Source: Pseudonocardia spinosispora DSM 44797 G406DRAFT_scaffold00011.11_C, whole genome shotgun sequence
Type: NRPS,T1PKS
Number of proteins with BLAST hits to this cluster: 2
Cumulative BLAST score: 729

Table of genes, locations, strands and annotations of subject cluster:

Table of Blast hits (query gene, subject gene, %identity, blast score, %coverage, e-value):
input.path1.gene345	G406_RS53815	36	482	49.94419642857143	2.7e-133
input.path1.gene350	G406_RS0109750	44	247	48.86731391585761	2.9e-63


>>
18. NZ_KB907309
Source: Nocardia sp. BMG111209 G361DRAFT_scaffold00003.3, whole genome shotgun sequence
Type: T1PKS
Number of proteins with BLAST hits to this cluster: 2
Cumulative BLAST score: 643

Table of genes, locations, strands and annotations of subject cluster:

Table of Blast hits (query gene, subject gene, %identity, blast score, %coverage, e-value):
input.path1.gene345	G361_RS0135515	34	495	55.18973214285714	2.3e-137
input.path1.gene350	G361_RS0135580	44	148	29.61165048543689	1.8e-33


>>
19. NW_023336287
Source: Aspergillus tubingensis WU-2223L DNA, scaffold_13, whole genome shotgun sequence
Type: T1PKS
Number of proteins with BLAST hits to this cluster: 2
Cumulative BLAST score: 640

Table of genes, locations, strands and annotations of subject cluster:

Table of Blast hits (query gene, subject gene, %identity, blast score, %coverage, e-value):
input.path1.gene345	AtWU_10486	31	371	51.78571428571429	6.6e-100
input.path1.gene346	AtWU_10480	36	269	79.64912280701755	6.6e-70


>>
20. NZ_JACHJF010000011
Source: Streptomyces eurocidicus strain CECT 3259 Ga0436944_11, whole genome shotgun sequence
Type: NRPS,NRPS-like,T1PKS,T3PKS,fused,siderophore
Number of proteins with BLAST hits to this cluster: 2
Cumulative BLAST score: 632

Table of genes, locations, strands and annotations of subject cluster:
FHS36_RS18320	FHS36_RS18320	126019	126207	+	MerR_family_transcriptional_regulator
FHS36_RS18470	FHS36_RS18470	166410	167332	-	AfsR/SARP_family_transcriptional_regulator
FHS36_RS18660	FHS36_RS18660	213263	213884	-	DUF4166_domain-containing_protein

Table of Blast hits (query gene, subject gene, %identity, blast score, %coverage, e-value):
input.path1.gene345	FHS36_RS18550	32	402	50.0	2.7e-109
input.path1.gene350	FHS36_RS18210	41	230	49.35275080906149	6.3e-58


>>
21. NZ_CP014306
Source: Burkholderia sp. PAMC 26561 chromosome 1, complete sequence
Type: T1PKS
Number of proteins with BLAST hits to this cluster: 2
Cumulative BLAST score: 623

Table of genes, locations, strands and annotations of subject cluster:
AXG89_RS02585	AXG89_RS02585	563263	564069	+	MFS_transporter
AXG89_RS02595	AXG89_RS02595	564518	564788	-	type_II_toxin-antitoxin_system_RelE/ParE_family_toxin
AXG89_RS02745	AXG89_RS02745	604748	605592	+	glycosyltransferase_family_4_protein

Table of Blast hits (query gene, subject gene, %identity, blast score, %coverage, e-value):
input.path1.gene345	AXG89_RS02685	31	468	61.55133928571429	3.1e-129
input.path1.gene349	AXG89_RS02660	33	155	58.03571428571429	1.8e-35


>>
22. NZ_CP018082
Source: Nocardia mangyaensis strain Y48 chromosome, complete genome
Type: T1PKS
Number of proteins with BLAST hits to this cluster: 2
Cumulative BLAST score: 606

Table of genes, locations, strands and annotations of subject cluster:

Table of Blast hits (query gene, subject gene, %identity, blast score, %coverage, e-value):
input.path1.gene345	BOX37_RS32800	32	455	58.81696428571429	3.5e-125
input.path1.gene350	BOX37_RS32755	43	151	33.33333333333333	2.2e-34


>>
23. NC_036441
Source: Aspergillus oryzae RIB40 DNA, chromosome 7
Type: T1PKS
Number of proteins with BLAST hits to this cluster: 2
Cumulative BLAST score: 605

Table of genes, locations, strands and annotations of subject cluster:

Table of Blast hits (query gene, subject gene, %identity, blast score, %coverage, e-value):
input.path1.gene345	AO090011000015	31	394	55.13392857142857	4.3e-107
input.path1.gene351	AO090011000020	36	211	90.64935064935065	1.9e-52


>>
24. NZ_CP016438
Source: Streptomyces lincolnensis strain NRRL 2936 chromosome
Type: T1PKS
Number of proteins with BLAST hits to this cluster: 2
Cumulative BLAST score: 596

Table of genes, locations, strands and annotations of subject cluster:
SLINC_RS46955	SLINC_RS46955	10167861	10168082	+	NADP-dependent_oxidoreductase
SLINC_RS46970	SLINC_RS46970	10186173	10186352	-	IS256_family_transposase
SLINC_RS48705	SLINC_RS48705	10196082	10196888	+	shikimate_dehydrogenase

Table of Blast hits (query gene, subject gene, %identity, blast score, %coverage, e-value):
input.path1.gene345	SLINC_RS44855	32	392	50.44642857142857	2.1e-106
input.path1.gene350	SLINC_RS44810	36	204	49.676375404530745	2.8e-50


>>
25. NZ_CP047146
Source: Streptomyces sp. GS7 chromosome, complete genome
Type: T1PKS
Number of proteins with BLAST hits to this cluster: 2
Cumulative BLAST score: 587

Table of genes, locations, strands and annotations of subject cluster:
GR130_RS20715	GR130_RS20715	4644792	4645232	+	serine_hydrolase
GR130_RS20740	GR130_RS20740	4650725	4651521	-	IS5_family_transposase
GR130_RS20750	GR130_RS20750	4653270	4653542	-	IS1380_family_transposase
GR130_RS20755	GR130_RS20755	4653720	4654489	-	transposase
GR130_RS20820	GR130_RS20820	4678315	4679084	+	transposase

Table of Blast hits (query gene, subject gene, %identity, blast score, %coverage, e-value):
input.path1.gene345	GR130_RS20795	36	470	49.720982142857146	1.1e-129
input.path1.gene349	GR130_RS20760	33	117	46.964285714285715	5.4e-24


>>
26. NZ_CP054569
Source: Achromobacter denitrificans strain FDAARGOS_787 chromosome, complete genome
Type: T1PKS
Number of proteins with BLAST hits to this cluster: 2
Cumulative BLAST score: 540

Table of genes, locations, strands and annotations of subject cluster:

Table of Blast hits (query gene, subject gene, %identity, blast score, %coverage, e-value):
input.path1.gene345	FOC81_RS12390	33	458	57.75669642857143	2.4e-126
input.path1.gene349	FOC81_RS12460	32	82	36.607142857142854	1.9e-13


>>
27. NZ_CP020917
Source: Achromobacter denitrificans strain PR1 chromosome, complete genome
Type: T1PKS
Number of proteins with BLAST hits to this cluster: 2
Cumulative BLAST score: 539

Table of genes, locations, strands and annotations of subject cluster:

Table of Blast hits (query gene, subject gene, %identity, blast score, %coverage, e-value):
input.path1.gene345	B9P52_RS26690	33	457	57.75669642857143	5.4e-126
input.path1.gene349	B9P52_RS26755	32	82	36.607142857142854	1.9e-13


>>
28. NZ_CP053986
Source: Achromobacter denitrificans strain FDAARGOS_788 chromosome, complete genome
Type: T1PKS
Number of proteins with BLAST hits to this cluster: 2
Cumulative BLAST score: 537

Table of genes, locations, strands and annotations of subject cluster:

Table of Blast hits (query gene, subject gene, %identity, blast score, %coverage, e-value):
input.path1.gene345	FOC82_RS07295	33	455	57.75669642857143	2.7e-125
input.path1.gene349	FOC82_RS07225	32	82	36.607142857142854	1.9e-13


>>
29. NZ_CP033730
Source: Achromobacter denitrificans strain PheN1 chromosome, complete genome
Type: T1PKS
Number of proteins with BLAST hits to this cluster: 2
Cumulative BLAST score: 535

Table of genes, locations, strands and annotations of subject cluster:
EC609_RS27940	EC609_RS27940	5915296	5916583	+	Vi_polysaccharide_biosynthesis_UDP-N-acetylglucosamine_C-6_dehydrogenase_TviB
EC609_RS28055	EC609_RS28055	5948509	5949746	+	high-affinity_branched-chain_amino_acid_ABC_transporter_permease_LivM
EC609_RS28060	EC609_RS28060	5949743	5950509	+	ABC_transporter_ATP-binding_protein

Table of Blast hits (query gene, subject gene, %identity, blast score, %coverage, e-value):
input.path1.gene345	EC609_RS27970	32	453	57.75669642857143	7.8e-125
input.path1.gene349	EC609_RS28035	32	82	36.607142857142854	1.9e-13


>>
30. NZ_MEBE01000001
Source: Burkholderia ubonensis strain MSMB0245 Contig_1, whole genome shotgun sequence
Type: NRPS,T1PKS,other,transAT-PKS-like
Number of proteins with BLAST hits to this cluster: 2
Cumulative BLAST score: 534

Table of genes, locations, strands and annotations of subject cluster:
BGV51_RS34825	BGV51_RS34825	54291	54571	-	hypothetical_protein
BGV51_RS00230	BGV51_RS00230	54575	55018	-	(2Fe-2S)-binding_protein

Table of Blast hits (query gene, subject gene, %identity, blast score, %coverage, e-value):
input.path1.gene345	BGV51_RS00285	33	443	50.72544642857143	1.1e-121
input.path1.gene349	BGV51_RS00430	31	91	37.32142857142857	4.1e-16


>>
31. NZ_KN150849
Source: Burkholderia gladioli strain ATCC 25417 scaffold1, whole genome shotgun sequence
Type: NRPS,T1PKS
Number of proteins with BLAST hits to this cluster: 2
Cumulative BLAST score: 533

Table of genes, locations, strands and annotations of subject cluster:

Table of Blast hits (query gene, subject gene, %identity, blast score, %coverage, e-value):
input.path1.gene345	DM48_RS15660	32	387	51.22767857142857	6.9e-105
input.path1.gene350	DM48_RS15710	42	146	31.55339805825243	9.2e-33


>>
32. NZ_CADEXT010000003
Source: Burkholderia gladioli strain BCC1700, whole genome shotgun sequence
Type: NRPS,T1PKS
Number of proteins with BLAST hits to this cluster: 2
Cumulative BLAST score: 532

Table of genes, locations, strands and annotations of subject cluster:

Table of Blast hits (query gene, subject gene, %identity, blast score, %coverage, e-value):
input.path1.gene345	H9D60_RS17120	32	386	51.22767857142857	1.2e-104
input.path1.gene350	H9D60_RS17170	42	146	31.55339805825243	9.2e-33


>>
33. NZ_CADEQD010000001
Source: Burkholderia gladioli strain BCC1730, whole genome shotgun sequence
Type: NRPS,T1PKS
Number of proteins with BLAST hits to this cluster: 2
Cumulative BLAST score: 532

Table of genes, locations, strands and annotations of subject cluster:

Table of Blast hits (query gene, subject gene, %identity, blast score, %coverage, e-value):
input.path1.gene345	H9D02_RS00675	32	386	51.22767857142857	1.5e-104
input.path1.gene350	H9D02_RS00615	42	146	31.55339805825243	9.2e-33


>>
34. NZ_MECO01000028
Source: Burkholderia ubonensis strain MSMB0803 Contig_7, whole genome shotgun sequence
Type: NRPS,T1PKS,other,transAT-PKS-like
Number of proteins with BLAST hits to this cluster: 2
Cumulative BLAST score: 531

Table of genes, locations, strands and annotations of subject cluster:
BGV64_RS27190	BGV64_RS27190	68447	68890	-	(2Fe-2S)-binding_protein

Table of Blast hits (query gene, subject gene, %identity, blast score, %coverage, e-value):
input.path1.gene345	BGV64_RS27255	33	441	50.72544642857143	3.1e-121
input.path1.gene349	BGV64_RS27400	31	90	37.32142857142857	5.4e-16


>>
35. NZ_CP009487
Source: Burkholderia ubonensis MSMB22 chromosome III, complete sequence
Type: NRPS,T1PKS,other,transAT-PKS-like
Number of proteins with BLAST hits to this cluster: 2
Cumulative BLAST score: 531

Table of genes, locations, strands and annotations of subject cluster:

Table of Blast hits (query gene, subject gene, %identity, blast score, %coverage, e-value):
input.path1.gene345	BW23_RS12150	32	440	50.11160714285714	8.9e-121
input.path1.gene349	BW23_RS12300	31	91	37.32142857142857	4.1e-16


>>
36. NZ_CADEQR010000002
Source: Burkholderia gladioli strain BCC1732, whole genome shotgun sequence
Type: NRPS,T1PKS
Number of proteins with BLAST hits to this cluster: 2
Cumulative BLAST score: 531

Table of genes, locations, strands and annotations of subject cluster:

Table of Blast hits (query gene, subject gene, %identity, blast score, %coverage, e-value):
input.path1.gene345	H9D15_RS06130	32	385	51.22767857142857	3.4e-104
input.path1.gene350	H9D15_RS06075	42	146	31.55339805825243	9.2e-33


>>
37. NZ_CADEYA010000006
Source: Burkholderia gladioli strain BCC1733, whole genome shotgun sequence
Type: NRPS,T1PKS
Number of proteins with BLAST hits to this cluster: 2
Cumulative BLAST score: 530

Table of genes, locations, strands and annotations of subject cluster:

Table of Blast hits (query gene, subject gene, %identity, blast score, %coverage, e-value):
input.path1.gene345	H9D74_RS18145	32	384	51.22767857142857	5.8e-104
input.path1.gene350	H9D74_RS18200	42	146	31.55339805825243	9.2e-33


>>
38. NZ_CADEYO010000001
Source: Burkholderia gladioli strain BCC1728, whole genome shotgun sequence
Type: NRPS,T1PKS
Number of proteins with BLAST hits to this cluster: 2
Cumulative BLAST score: 530

Table of genes, locations, strands and annotations of subject cluster:

Table of Blast hits (query gene, subject gene, %identity, blast score, %coverage, e-value):
input.path1.gene345	H9D70_RS03215	32	384	51.22767857142857	5.8e-104
input.path1.gene350	H9D70_RS03270	42	146	31.55339805825243	9.2e-33


>>
39. NZ_LNJU01000004
Source: Burkholderia ubonensis strain MSMB1157 MSMB1157WGS_5, whole genome shotgun sequence
Type: NRPS,T1PKS,other,transAT-PKS-like
Number of proteins with BLAST hits to this cluster: 2
Cumulative BLAST score: 529

Table of genes, locations, strands and annotations of subject cluster:

Table of Blast hits (query gene, subject gene, %identity, blast score, %coverage, e-value):
input.path1.gene345	WK57_RS33175	32	439	50.72544642857143	1.5e-120
input.path1.gene349	WK57_RS33030	31	90	37.32142857142857	7.1e-16


>>
40. NZ_MEBC01000001
Source: Burkholderia ubonensis strain MSMB0170 Contig_1, whole genome shotgun sequence
Type: NRPS,T1PKS,other,transAT-PKS-like
Number of proteins with BLAST hits to this cluster: 2
Cumulative BLAST score: 528

Table of genes, locations, strands and annotations of subject cluster:

Table of Blast hits (query gene, subject gene, %identity, blast score, %coverage, e-value):
input.path1.gene345	BGV49_RS00265	33	438	50.11160714285714	3.4e-120
input.path1.gene349	BGV49_RS00415	31	90	37.32142857142857	5.4e-16


>>
41. NZ_MEBF01000045
Source: Burkholderia ubonensis strain MSMB0258b Contig_5, whole genome shotgun sequence
Type: NRPS,T1PKS,bacteriocin,transAT-PKS-like
Number of proteins with BLAST hits to this cluster: 2
Cumulative BLAST score: 525

Table of genes, locations, strands and annotations of subject cluster:
BGV52_RS25705	BGV52_RS25705	123214	123483	+	2Fe-2S_ferredoxin
BGV52_RS25855	BGV52_RS25855	199212	199298	+	activator_protein
BGV52_RS25890	BGV52_RS25890	203586	204014	-	acyl-CoA_dehydrogenase_family_protein

Table of Blast hits (query gene, subject gene, %identity, blast score, %coverage, e-value):
input.path1.gene345	BGV52_RS25645	32	435	50.11160714285714	2.2e-119
input.path1.gene349	BGV52_RS25775	31	90	37.32142857142857	7.1e-16


>>
42. NZ_NPFX01000014
Source: Rhodococcus sp. 14-2483-1-1 RS1B5_1, whole genome shotgun sequence
Type: PKS-like,T1PKS
Number of proteins with BLAST hits to this cluster: 2
Cumulative BLAST score: 518

Table of genes, locations, strands and annotations of subject cluster:

Table of Blast hits (query gene, subject gene, %identity, blast score, %coverage, e-value):
input.path1.gene345	CH294_RS11710	33	426	51.39508928571429	1.7e-116
input.path1.gene349	CH294_RS11780	33	92	35.535714285714285	1.1e-16


>>
43. NZ_NSDZ01000001
Source: Rhodococcus sp. ACS1 Contig_1, whole genome shotgun sequence
Type: T1PKS,terpene
Number of proteins with BLAST hits to this cluster: 2
Cumulative BLAST score: 517

Table of genes, locations, strands and annotations of subject cluster:
CJ179_RS02215	CJ179_RS02215	472511	472959	+	DUF1298_domain-containing_protein

Table of Blast hits (query gene, subject gene, %identity, blast score, %coverage, e-value):
input.path1.gene345	CJ179_RS02110	36	414	44.86607142857143	5.2e-113
input.path1.gene349	CJ179_RS02045	31	103	59.464285714285715	8.1e-20


>>
44. NZ_CP023701
Source: Streptomyces subrutilus strain ATCC 27467 chromosome, complete genome
Type: T1PKS
Number of proteins with BLAST hits to this cluster: 2
Cumulative BLAST score: 515

Table of genes, locations, strands and annotations of subject cluster:
CP968_RS11070	CP968_RS11070	2615228	2615698	+	hypothetical_protein
CP968_RS11115	CP968_RS11115	2625819	2626425	-	helix-turn-helix_transcriptional_regulator
CP968_RS11235	CP968_RS11235	2651881	2653329	-	penicillin-binding_protein_2

Table of Blast hits (query gene, subject gene, %identity, blast score, %coverage, e-value):
input.path1.gene345	CP968_RS11150	34	439	50.33482142857143	2e-120
input.path1.gene349	CP968_RS11075	31	76	33.57142857142857	8.1e-12


>>
45. NZ_CP048641
Source: Streptomyces aureoverticillatus strain HN6 chromosome, complete genome
Type: NRPS,T1PKS
Number of proteins with BLAST hits to this cluster: 2
Cumulative BLAST score: 508

Table of genes, locations, strands and annotations of subject cluster:
G3H79_RS08245	G3H79_RS08245	2333592	2334477	+	FAD-binding_molybdopterin_dehydrogenase
G3H79_RS08280	G3H79_RS08280	2343705	2345852	-	MMPL_family_transporter

Table of Blast hits (query gene, subject gene, %identity, blast score, %coverage, e-value):
input.path1.gene345	G3H79_RS08210	31	433	54.6875	8.3e-119
input.path1.gene349	G3H79_RS08155	31	75	40.89285714285714	1.8e-11


>>
46. NZ_CP008947
Source: Rhodococcus opacus strain R7 chromosome
Type: T1PKS,terpene
Number of proteins with BLAST hits to this cluster: 2
Cumulative BLAST score: 499

Table of genes, locations, strands and annotations of subject cluster:
EP51_RS04310	EP51_RS04310	940632	944941	+	aminotransferase_class_I/II-fold_pyridoxal_phosphate-dependent_enzyme

Table of Blast hits (query gene, subject gene, %identity, blast score, %coverage, e-value):
input.path1.gene345	EP51_RS04310	34	394	46.595982142857146	7.3e-107
input.path1.gene349	EP51_RS04245	34	105	44.82142857142858	1.6e-20


>>
47. NC_012522
Source: Rhodococcus opacus B4, complete genome
Type: T1PKS,terpene
Number of proteins with BLAST hits to this cluster: 2
Cumulative BLAST score: 496

Table of genes, locations, strands and annotations of subject cluster:
ROP_RS34850	ROP_RS34850	7660536	7662284	-	fatty_acyl-AMP_ligase

Table of Blast hits (query gene, subject gene, %identity, blast score, %coverage, e-value):
input.path1.gene345	ROP_RS34840	35	394	44.86607142857143	4.3e-107
input.path1.gene349	ROP_RS34905	33	102	44.642857142857146	1.8e-19


>>
48. NZ_CP009111
Source: Rhodococcus opacus strain 1CP, complete genome
Type: T1PKS,terpene
Number of proteins with BLAST hits to this cluster: 2
Cumulative BLAST score: 488

Table of genes, locations, strands and annotations of subject cluster:
R1CP_RS17005	R1CP_RS17005	3736315	3736742	+	hypothetical_protein
R1CP_RS17015	R1CP_RS17015	3737489	3738978	+	LuxR_family_transcriptional_regulator
R1CP_RS17025	R1CP_RS17025	3740669	3741445	+	LuxR_family_transcriptional_regulator

Table of Blast hits (query gene, subject gene, %identity, blast score, %coverage, e-value):
input.path1.gene345	R1CP_RS17125	34	382	46.372767857142854	2.2e-103
input.path1.gene349	R1CP_RS17190	35	106	45.357142857142854	9.5e-21


>>
49. NZ_CADEPP010000002
Source: Burkholderia gladioli strain BCC1701, whole genome shotgun sequence
Type: T1PKS,transAT-PKS-like
Number of proteins with BLAST hits to this cluster: 2
Cumulative BLAST score: 483

Table of genes, locations, strands and annotations of subject cluster:

Table of Blast hits (query gene, subject gene, %identity, blast score, %coverage, e-value):
input.path1.gene345	H9C75_RS05790	33	387	50.0	9e-105
input.path1.gene349	H9C75_RS05765	32	96	37.5	9.9e-18


>>
50. NZ_CP051855
Source: Rhodococcus imtechensis strain XM24D chromosome 1, complete sequence
Type: T1PKS,terpene
Number of proteins with BLAST hits to this cluster: 2
Cumulative BLAST score: 483

Table of genes, locations, strands and annotations of subject cluster:
HHX23_RS17305	HHX23_RS17305	3752479	3754702	+	LuxR_family_transcriptional_regulator

Table of Blast hits (query gene, subject gene, %identity, blast score, %coverage, e-value):
input.path1.gene345	HHX23_RS17405	34	379	46.372767857142854	1.4e-102
input.path1.gene349	HHX23_RS17470	34	104	45.357142857142854	4.7e-20
