## Supplementary Results for "A draft genome of the ascomycotal fungal species *Pseudopithomyces maydicus* (family *Didymosphaeriaceae*)": contig_4_c3.txt

ClusterBlast scores for contig_4

Table of genes, locations, strands and annotations of query cluster:
input.path1.gene494	1833627	1834165	+		
input.path1.gene495	1834940	1835906	-		
input.path1.gene496	1837251	1839309	+		
input.path1.gene497	1841891	1842339	+		
input.path1.gene498	1842677	1843676	-		
input.path1.gene499	1847056	1848403	-		
input.path1.gene500	1848604	1850314	+		
input.path1.gene501	1850600	1876090	-		
input.path1.gene502	1879655	1884675	+		
input.path1.gene503	1886992	1888560	-		
input.path1.gene504	1888972	1889613	+		
input.path1.gene505	1890258	1891226	-		
input.path1.gene506	1891455	1893435	-		


Significant hits: 
1. NC_036442	Aspergillus oryzae RIB40 DNA, chromosome 8
2. NW_001939244	Pyrenophora tritici-repentis Pt-1C-BFP supercont1.1, whole genome shotgun sequence
3. NW_003456428	Trichophyton rubrum CBS 118892 genomic scaffold supercont2.1, whole genome shotgun sequence
4. NW_003299167	Microsporum canis CBS 113480 supercont1.3 genomic scaffold, whole genome shotgun sequence
5. NW_022983863	Arthroderma uncinatum strain CBS 119779 chromosome Unknown scaffold1, whole genome shotgun sequence
6. NW_003316003	Coccidioides posadasii C735 delta SOWgp chromosome Unknown gcontig_1105601051733, whole genome shotgun sequence
7. NW_017971444	Talaromyces atroroseus strain IBT 11181 chromosome Unknown scaffold_12, whole genome shotgun sequence
8. NC_016458	Thermothielavioides terrestris NRRL 8126 chromosome 2, complete sequence
9. NC_016477	Thermothelomyces thermophilus ATCC 42464 chromosome 6, complete sequence
10. CM000574	Fusarium graminearum PH-1 chromosome 1, whole genome shotgun sequence
11. NC_026474	Fusarium graminearum PH-1 chromosome 1, whole genome shotgun sequence
12. NW_022194794	Fusarium proliferatum ET1 genome assembly, contig: FPRO_scaffold005, whole genome shotgun sequence
13. NC_036628	Fusarium fujikuroi IMI 58289 draft genome, chromosome FFUJ_chr07
14. NC_038012	Fusarium venenatum strain A3/5 genome assembly, chromosome: I
15. NT_165978	Chaetomium globosum CBS 148.51 scaffold_3 genomic scaffold, whole genome shotgun sequence
16. NC_036440	Aspergillus oryzae RIB40 DNA, chromosome 6
17. NW_023336283	Aspergillus tubingensis WU-2223L DNA, scaffold_9, whole genome shotgun sequence
18. CM004175	Drechmeria coniospora strain ARSEF 6962 chromosome 02, whole genome shotgun sequence
19. NW_022474210	Venustampulla echinocandica strain BP 5553 chromosome Unknown scaffold_5, whole genome shotgun sequence
20. NW_019716268	Ramularia collo-cygni strain URUG2 genome assembly, contig: RCC_scaffold14, whole genome shotgun sequence
21. NW_007361001	Glarea lozoyensis ATCC 20868 chromosome Unknown GLAREA6, whole genome shotgun sequence
22. NW_013550610	Fonsecaea pedrosoi CBS 271.37 unplaced genomic scaffold supercont1.6, whole genome shotgun sequence
23. NW_022474205	Venustampulla echinocandica strain BP 5553 chromosome Unknown scaffold_0, whole genome shotgun sequence
24. CM000597	Fusarium oxysporum f. sp. lycopersici 4287 chromosome 9, whole genome shotgun sequence
25. NC_030994	Fusarium oxysporum f. sp. lycopersici 4287 chromosome 9, whole genome shotgun sequence
26. NW_003052499	Uncinocarpus reesii 1704 scaffold_2 genomic scaffold, whole genome shotgun sequence
27. NW_004504309	Coccidioides immitis RS genomic scaffold supercont3.2, whole genome shotgun sequence
28. NW_003345201	Nannizzia gypsea CBS 118893 supercont1.1 genomic scaffold, whole genome shotgun sequence
29. NC_049561	Talaromyces rugulosus chromosome I, complete sequence
30. NW_001914850	Podospora anserina S mat+ genomic DNA chromosome 4, supercontig 4
31. NW_011942171	Metarhizium robertsii ARSEF 23 MAA_Scf_3, whole genome shotgun sequence
32. NW_014574611	Penicillium digitatum Pd1 unplaced genomic scaffold Scaffold_14, whole genome shotgun sequence
33. NW_014574578	Penicillium digitatum Pd1 unplaced genomic scaffold Scaffold_47, whole genome shotgun sequence
34. NW_023336279	Aspergillus tubingensis WU-2223L DNA, scaffold_5, whole genome shotgun sequence
35. NW_022194798	Fusarium proliferatum ET1 genome assembly, contig: FPRO_scaffold009, whole genome shotgun sequence
36. CM000574	Fusarium graminearum PH-1 chromosome 1, whole genome shotgun sequence
37. NC_026474	Fusarium graminearum PH-1 chromosome 1, whole genome shotgun sequence
38. NW_015971594	Cladophialophora bantiana CBS 173.52 unplaced genomic scaffold supercont1.20, whole genome shotgun sequence
39. NC_036435	Aspergillus oryzae RIB40 DNA, chromosome 1
40. NC_030961	Colletotrichum higginsianum IMI 349063 chromosome 9, whole genome shotgun sequence
41. NW_001914833	Podospora anserina S mat+ genomic DNA chromosome 3, supercontig 3
42. NW_015622507	Exophiala mesophila strain CBS 40295 unplaced genomic scaffold supercont1.4, whole genome shotgun sequence
43. NW_011942160	Metarhizium robertsii ARSEF 23 MAA_Scf_2, whole genome shotgun sequence
44. NW_014574716	Metarhizium brunneum ARSEF 3297 chromosome Unknown scaffold_1, whole genome shotgun sequence
45. NW_006271973	Cordyceps militaris CM01 unplaced genomic scaffold CCM_S00005, whole genome shotgun sequence
46. NC_035791	Pochonia chlamydosporia 170 chromosome 2, whole genome shotgun sequence
47. NW_006911257	Baudoinia panamericana UAMH 10762 unplaced genomic scaffold BAUCOscaffold_1, whole genome shotgun sequence
48. CM002799	Penicillium chrysogenum strain P2niaD18 chromosome II, whole genome shotgun sequence
49. NC_035790	Pochonia chlamydosporia 170 chromosome 1, whole genome shotgun sequence
50. NW_020290247	Acaromyces ingoldii strain MCA 4198 unplaced genomic scaffold FA10scaffold_1, whole genome shotgun sequence


Details:

>>
1. NC_036442
Source: Aspergillus oryzae RIB40 DNA, chromosome 8
Type: NRPS
Number of proteins with BLAST hits to this cluster: 3
Cumulative BLAST score: 2178

Table of genes, locations, strands and annotations of subject cluster:

Table of Blast hits (query gene, subject gene, %identity, blast score, %coverage, e-value):
input.path1.gene499	AO090103000359	35	131	82.33333333333334	1.5e-28
input.path1.gene501	AO090103000355	34	1404	37.00608978898173	0.0
input.path1.gene502	AO090103000356	33	643	92.48175182481752	6.9e-182


>>
2. NW_001939244
Source: Pyrenophora tritici-repentis Pt-1C-BFP supercont1.1, whole genome shotgun sequence
Type: NRPS
Number of proteins with BLAST hits to this cluster: 2
Cumulative BLAST score: 8804

Table of genes, locations, strands and annotations of subject cluster:

Table of Blast hits (query gene, subject gene, %identity, blast score, %coverage, e-value):
input.path1.gene501	PTRG_01800	53	7099	99.07945050276165	0.0
input.path1.gene502	PTRG_01799	68	1705	91.97080291970804	0.0


>>
3. NW_003456428
Source: Trichophyton rubrum CBS 118892 genomic scaffold supercont2.1, whole genome shotgun sequence
Type: NRPS
Number of proteins with BLAST hits to this cluster: 2
Cumulative BLAST score: 6699

Table of genes, locations, strands and annotations of subject cluster:

Table of Blast hits (query gene, subject gene, %identity, blast score, %coverage, e-value):
input.path1.gene501	TERG_01444	40	5417	111.37232686588301	0.0
input.path1.gene502	TERG_01443	53	1282	93.64963503649635	0.0


>>
4. NW_003299167
Source: Microsporum canis CBS 113480 supercont1.3 genomic scaffold, whole genome shotgun sequence
Type: NRPS
Number of proteins with BLAST hits to this cluster: 2
Cumulative BLAST score: 6681

Table of genes, locations, strands and annotations of subject cluster:

Table of Blast hits (query gene, subject gene, %identity, blast score, %coverage, e-value):
input.path1.gene501	MCYG_02927	40	5464	111.71222206486333	0.0
input.path1.gene502	MCYG_02928	51	1217	92.84671532846716	0.0


>>
5. NW_022983863
Source: Arthroderma uncinatum strain CBS 119779 chromosome Unknown scaffold1, whole genome shotgun sequence
Type: NRPS
Number of proteins with BLAST hits to this cluster: 2
Cumulative BLAST score: 6661

Table of genes, locations, strands and annotations of subject cluster:

Table of Blast hits (query gene, subject gene, %identity, blast score, %coverage, e-value):
input.path1.gene501	GIQ15_00338	40	5384	111.28735306613795	0.0
input.path1.gene502	GIQ15_00337	53	1277	94.08759124087591	0.0


>>
6. NW_003316003
Source: Coccidioides posadasii C735 delta SOWgp chromosome Unknown gcontig_1105601051733, whole genome shotgun sequence
Type: NRPS
Number of proteins with BLAST hits to this cluster: 2
Cumulative BLAST score: 6621

Table of genes, locations, strands and annotations of subject cluster:

Table of Blast hits (query gene, subject gene, %identity, blast score, %coverage, e-value):
input.path1.gene501	CPC735_023110	39	5310	113.25591276023226	0.0
input.path1.gene502	CPC735_023100	55	1311	92.4087591240876	0.0


>>
7. NW_017971444
Source: Talaromyces atroroseus strain IBT 11181 chromosome Unknown scaffold_12, whole genome shotgun sequence
Type: NRPS
Number of proteins with BLAST hits to this cluster: 2
Cumulative BLAST score: 6251

Table of genes, locations, strands and annotations of subject cluster:

Table of Blast hits (query gene, subject gene, %identity, blast score, %coverage, e-value):
input.path1.gene501	UA08_07809	37	4963	111.59892366520323	0.0
input.path1.gene502	UA08_07808	53	1288	93.13868613138686	0.0


>>
8. NC_016458
Source: Thermothielavioides terrestris NRRL 8126 chromosome 2, complete sequence
Type: NRPS
Number of proteins with BLAST hits to this cluster: 2
Cumulative BLAST score: 6081

Table of genes, locations, strands and annotations of subject cluster:

Table of Blast hits (query gene, subject gene, %identity, blast score, %coverage, e-value):
input.path1.gene501	THITE_112015	38	4813	113.35504885993485	0.0
input.path1.gene502	THITE_35966	54	1268	90.14598540145985	0.0


>>
9. NC_016477
Source: Thermothelomyces thermophilus ATCC 42464 chromosome 6, complete sequence
Type: NRPS
Number of proteins with BLAST hits to this cluster: 2
Cumulative BLAST score: 5859

Table of genes, locations, strands and annotations of subject cluster:

Table of Blast hits (query gene, subject gene, %identity, blast score, %coverage, e-value):
input.path1.gene501	MYCTH_75351	37	4628	112.972666761082	0.0
input.path1.gene502	MYCTH_61507	51	1231	91.75182481751824	0.0


>>
10. CM000574
Source: Fusarium graminearum PH-1 chromosome 1, whole genome shotgun sequence
Type: NRPS,T1PKS
Number of proteins with BLAST hits to this cluster: 2
Cumulative BLAST score: 5799

Table of genes, locations, strands and annotations of subject cluster:

Table of Blast hits (query gene, subject gene, %identity, blast score, %coverage, e-value):
input.path1.gene501	FGSG_02315	37	4693	108.66732757399802	0.0
input.path1.gene502	FGSG_02316	48	1106	92.62773722627738	0.0


>>
11. NC_026474
Source: Fusarium graminearum PH-1 chromosome 1, whole genome shotgun sequence
Type: NRPS,T1PKS
Number of proteins with BLAST hits to this cluster: 2
Cumulative BLAST score: 5799

Table of genes, locations, strands and annotations of subject cluster:

Table of Blast hits (query gene, subject gene, %identity, blast score, %coverage, e-value):
input.path1.gene501	FGSG_02315	37	4693	108.66732757399802	0.0
input.path1.gene502	FGSG_02316	48	1106	92.62773722627738	0.0


>>
12. NW_022194794
Source: Fusarium proliferatum ET1 genome assembly, contig: FPRO_scaffold005, whole genome shotgun sequence
Type: NRPS
Number of proteins with BLAST hits to this cluster: 2
Cumulative BLAST score: 5744

Table of genes, locations, strands and annotations of subject cluster:

Table of Blast hits (query gene, subject gene, %identity, blast score, %coverage, e-value):
input.path1.gene501	FPRO_11310	37	4626	108.22829627531512	0.0
input.path1.gene502	FPRO_11309	48	1118	92.11678832116789	0.0


>>
13. NC_036628
Source: Fusarium fujikuroi IMI 58289 draft genome, chromosome FFUJ_chr07
Type: NRPS
Number of proteins with BLAST hits to this cluster: 2
Cumulative BLAST score: 5732

Table of genes, locations, strands and annotations of subject cluster:

Table of Blast hits (query gene, subject gene, %identity, blast score, %coverage, e-value):
input.path1.gene501	FFUJ_08113	37	4616	108.27078317518766	0.0
input.path1.gene502	FFUJ_08114	47	1116	92.04379562043795	0.0


>>
14. NC_038012
Source: Fusarium venenatum strain A3/5 genome assembly, chromosome: I
Type: NRPS,T1PKS
Number of proteins with BLAST hits to this cluster: 2
Cumulative BLAST score: 5688

Table of genes, locations, strands and annotations of subject cluster:

Table of Blast hits (query gene, subject gene, %identity, blast score, %coverage, e-value):
input.path1.gene501	FVRRES_02708	36	4594	108.5398668743804	0.0
input.path1.gene502	FVRRES_02709	47	1094	92.62773722627738	0.0


>>
15. NT_165978
Source: Chaetomium globosum CBS 148.51 scaffold_3 genomic scaffold, whole genome shotgun sequence
Type: NRPS
Number of proteins with BLAST hits to this cluster: 2
Cumulative BLAST score: 5522

Table of genes, locations, strands and annotations of subject cluster:

Table of Blast hits (query gene, subject gene, %identity, blast score, %coverage, e-value):
input.path1.gene501	CHGG_06789	36	4614	112.67525846197421	0.0
input.path1.gene502	CHGG_06790	49	908	74.23357664233576	8.8e-262


>>
16. NC_036440
Source: Aspergillus oryzae RIB40 DNA, chromosome 6
Type: NRPS
Number of proteins with BLAST hits to this cluster: 2
Cumulative BLAST score: 3739

Table of genes, locations, strands and annotations of subject cluster:

Table of Blast hits (query gene, subject gene, %identity, blast score, %coverage, e-value):
input.path1.gene501	AO090038000390	38	2861	62.045036113864896	0.0
input.path1.gene502	AO090038000389	43	618	55.98540145985401	1.4e-174
input.path1.gene502	AO090038000388	50	260	19.34306569343066	7.4e-67


>>
17. NW_023336283
Source: Aspergillus tubingensis WU-2223L DNA, scaffold_9, whole genome shotgun sequence
Type: NRPS
Number of proteins with BLAST hits to this cluster: 2
Cumulative BLAST score: 3034

Table of genes, locations, strands and annotations of subject cluster:

Table of Blast hits (query gene, subject gene, %identity, blast score, %coverage, e-value):
input.path1.gene501	AtWU_08918	33	1800	51.35249964594251	0.0
input.path1.gene502	AtWU_08917	51	1234	92.04379562043795	0.0


>>
18. CM004175
Source: Drechmeria coniospora strain ARSEF 6962 chromosome 02, whole genome shotgun sequence
Type: NRPS
Number of proteins with BLAST hits to this cluster: 2
Cumulative BLAST score: 1791

Table of genes, locations, strands and annotations of subject cluster:

Table of Blast hits (query gene, subject gene, %identity, blast score, %coverage, e-value):
input.path1.gene501	DCS_03520	34	1036	29.245149412264553	1.1e-299
input.path1.gene502	DCS_03521	35	755	93.64963503649635	9.6e-216


>>
19. NW_022474210
Source: Venustampulla echinocandica strain BP 5553 chromosome Unknown scaffold_5, whole genome shotgun sequence
Type: NRPS
Number of proteins with BLAST hits to this cluster: 2
Cumulative BLAST score: 1566

Table of genes, locations, strands and annotations of subject cluster:

Table of Blast hits (query gene, subject gene, %identity, blast score, %coverage, e-value):
input.path1.gene501	BP5553_06962	36	994	24.67072652598782	6.3e-287
input.path1.gene502	BP5553_06963	33	572	84.81751824817519	1.1e-160


>>
20. NW_019716268
Source: Ramularia collo-cygni strain URUG2 genome assembly, contig: RCC_scaffold14, whole genome shotgun sequence
Type: NRPS
Number of proteins with BLAST hits to this cluster: 2
Cumulative BLAST score: 1491

Table of genes, locations, strands and annotations of subject cluster:

Table of Blast hits (query gene, subject gene, %identity, blast score, %coverage, e-value):
input.path1.gene501	RCC_08343	32	749	23.29698343010905	2.7e-213
input.path1.gene502	RCC_08344	35	742	91.67883211678833	8.4e-212


>>
21. NW_007361001
Source: Glarea lozoyensis ATCC 20868 chromosome Unknown GLAREA6, whole genome shotgun sequence
Type: NRPS
Number of proteins with BLAST hits to this cluster: 2
Cumulative BLAST score: 1256

Table of genes, locations, strands and annotations of subject cluster:

Table of Blast hits (query gene, subject gene, %identity, blast score, %coverage, e-value):
input.path1.gene501	GLAREA_02479	33	589	15.918425152244724	4.7e-165
input.path1.gene502	GLAREA_02480	34	667	94.52554744525547	2.6e-189


>>
22. NW_013550610
Source: Fonsecaea pedrosoi CBS 271.37 unplaced genomic scaffold supercont1.6, whole genome shotgun sequence
Type: NRPS
Number of proteins with BLAST hits to this cluster: 2
Cumulative BLAST score: 689

Table of genes, locations, strands and annotations of subject cluster:

Table of Blast hits (query gene, subject gene, %identity, blast score, %coverage, e-value):
input.path1.gene501	Z517_10058	34	581	15.63517915309446	1.3e-162
input.path1.gene502	Z517_10057	31	108	18.905109489051096	8e-21


>>
23. NW_022474205
Source: Venustampulla echinocandica strain BP 5553 chromosome Unknown scaffold_0, whole genome shotgun sequence
Type: NRPS
Number of proteins with BLAST hits to this cluster: 2
Cumulative BLAST score: 6798

Table of genes, locations, strands and annotations of subject cluster:

Table of Blast hits (query gene, subject gene, %identity, blast score, %coverage, e-value):
input.path1.gene501	BP5553_00199	41	4903	97.87565500637302	0.0
input.path1.gene501	BP5553_00198	39	564	13.085965160742106	2.1e-157
input.path1.gene502	BP5553_00200	56	1331	91.0948905109489	0.0


>>
24. CM000597
Source: Fusarium oxysporum f. sp. lycopersici 4287 chromosome 9, whole genome shotgun sequence
Type: NRPS
Number of proteins with BLAST hits to this cluster: 2
Cumulative BLAST score: 6702

Table of genes, locations, strands and annotations of subject cluster:

Table of Blast hits (query gene, subject gene, %identity, blast score, %coverage, e-value):
input.path1.gene501	FOXG_13024	37	4665	107.10947457867157	0.0
input.path1.gene502	FOXG_13025	47	1126	92.04379562043795	0.0
input.path1.gene502	FOXG_13025_KNB13553.1	47	911	74.96350364963503	8e-263


>>
25. NC_030994
Source: Fusarium oxysporum f. sp. lycopersici 4287 chromosome 9, whole genome shotgun sequence
Type: NRPS
Number of proteins with BLAST hits to this cluster: 2
Cumulative BLAST score: 6702

Table of genes, locations, strands and annotations of subject cluster:

Table of Blast hits (query gene, subject gene, %identity, blast score, %coverage, e-value):
input.path1.gene501	FOXG_13024	37	4665	107.10947457867157	0.0
input.path1.gene502	FOXG_13025	47	1126	92.04379562043795	0.0
input.path1.gene502	FOXG_13025_XP_018251598.1	47	911	74.96350364963503	8e-263


>>
26. NW_003052499
Source: Uncinocarpus reesii 1704 scaffold_2 genomic scaffold, whole genome shotgun sequence
Type: NRPS
Number of proteins with BLAST hits to this cluster: 2
Cumulative BLAST score: 6662

Table of genes, locations, strands and annotations of subject cluster:

Table of Blast hits (query gene, subject gene, %identity, blast score, %coverage, e-value):
input.path1.gene501	UREG_04677	43	4407	80.51267525846197	0.0
input.path1.gene501	UREG_04676	39	939	20.56365953830902	1.8e-270
input.path1.gene502	UREG_04679	55	1316	92.91970802919708	0.0


>>
27. NW_004504309
Source: Coccidioides immitis RS genomic scaffold supercont3.2, whole genome shotgun sequence
Type: NRPS
Number of proteins with BLAST hits to this cluster: 2
Cumulative BLAST score: 6642

Table of genes, locations, strands and annotations of subject cluster:

Table of Blast hits (query gene, subject gene, %identity, blast score, %coverage, e-value):
input.path1.gene501	CIMG_09750	39	5332	113.1851012604447	0.0
input.path1.gene502	CIMG_09753	55	1310	92.33576642335767	0.0


>>
28. NW_003345201
Source: Nannizzia gypsea CBS 118893 supercont1.1 genomic scaffold, whole genome shotgun sequence
Type: NRPS
Number of proteins with BLAST hits to this cluster: 2
Cumulative BLAST score: 6584

Table of genes, locations, strands and annotations of subject cluster:

Table of Blast hits (query gene, subject gene, %identity, blast score, %coverage, e-value):
input.path1.gene501	MGYG_00094	40	5301	111.44313836567059	0.0
input.path1.gene502	MGYG_00096	53	1283	93.64963503649635	0.0


>>
29. NC_049561
Source: Talaromyces rugulosus chromosome I, complete sequence
Type: NRPS
Number of proteins with BLAST hits to this cluster: 2
Cumulative BLAST score: 6331

Table of genes, locations, strands and annotations of subject cluster:

Table of Blast hits (query gene, subject gene, %identity, blast score, %coverage, e-value):
input.path1.gene501	TRUGW13939_01108	38	5068	111.1457300665628	0.0
input.path1.gene502	TRUGW13939_01110	53	1263	92.4087591240876	0.0


>>
30. NW_001914850
Source: Podospora anserina S mat+ genomic DNA chromosome 4, supercontig 4
Type: NRPS
Number of proteins with BLAST hits to this cluster: 2
Cumulative BLAST score: 5559

Table of genes, locations, strands and annotations of subject cluster:

Table of Blast hits (query gene, subject gene, %identity, blast score, %coverage, e-value):
input.path1.gene501	PODANS_4_4640	37	4704	113.12845206061463	0.0
input.path1.gene502	PODANS_4_4610	45	855	78.46715328467153	6.8e-246


>>
31. NW_011942171
Source: Metarhizium robertsii ARSEF 23 MAA_Scf_3, whole genome shotgun sequence
Type: NRPS,T1PKS
Number of proteins with BLAST hits to this cluster: 2
Cumulative BLAST score: 3815

Table of genes, locations, strands and annotations of subject cluster:

Table of Blast hits (query gene, subject gene, %identity, blast score, %coverage, e-value):
input.path1.gene501	MAA_09953	33	3009	85.4128310437615	0.0
input.path1.gene502	MAA_09951	38	806	91.0948905109489	4.7e-231


>>
32. NW_014574611
Source: Penicillium digitatum Pd1 unplaced genomic scaffold Scaffold_14, whole genome shotgun sequence
Type: NRPS
Number of proteins with BLAST hits to this cluster: 2
Cumulative BLAST score: 3168

Table of genes, locations, strands and annotations of subject cluster:

Table of Blast hits (query gene, subject gene, %identity, blast score, %coverage, e-value):
input.path1.gene501	PDIP_25130	31	2361	75.18765047443705	0.0
input.path1.gene502	PDIP_25150	38	807	88.17518248175182	2.8e-231


>>
33. NW_014574578
Source: Penicillium digitatum Pd1 unplaced genomic scaffold Scaffold_47, whole genome shotgun sequence
Type: NRPS
Number of proteins with BLAST hits to this cluster: 2
Cumulative BLAST score: 2623

Table of genes, locations, strands and annotations of subject cluster:

Table of Blast hits (query gene, subject gene, %identity, blast score, %coverage, e-value):
input.path1.gene501	PDIP_87660	35	1893	50.1628664495114	0.0
input.path1.gene502	PDIP_87640	39	730	81.75182481751825	4.3e-208


>>
34. NW_023336279
Source: Aspergillus tubingensis WU-2223L DNA, scaffold_5, whole genome shotgun sequence
Type: NRPS
Number of proteins with BLAST hits to this cluster: 2
Cumulative BLAST score: 2531

Table of genes, locations, strands and annotations of subject cluster:

Table of Blast hits (query gene, subject gene, %identity, blast score, %coverage, e-value):
input.path1.gene501	AtWU_03451	31	1764	59.623282821130154	0.0
input.path1.gene502	AtWU_03456	36	767	98.97810218978103	2.4e-219


>>
35. NW_022194798
Source: Fusarium proliferatum ET1 genome assembly, contig: FPRO_scaffold009, whole genome shotgun sequence
Type: NRPS,T1PKS,terpene
Number of proteins with BLAST hits to this cluster: 2
Cumulative BLAST score: 2259

Table of genes, locations, strands and annotations of subject cluster:

Table of Blast hits (query gene, subject gene, %identity, blast score, %coverage, e-value):
input.path1.gene501	FPRO_15719	32	1621	51.73488174479536	0.0
input.path1.gene502	FPRO_15733	32	638	90.87591240875912	1.7e-180


>>
36. CM000574
Source: Fusarium graminearum PH-1 chromosome 1, whole genome shotgun sequence
Type: NRPS
Number of proteins with BLAST hits to this cluster: 2
Cumulative BLAST score: 1703

Table of genes, locations, strands and annotations of subject cluster:

Table of Blast hits (query gene, subject gene, %identity, blast score, %coverage, e-value):
input.path1.gene501	FGSG_11659	31	1016	33.93287069820139	1.2e-293
input.path1.gene502	FGSG_00046	33	687	94.67153284671534	2.5e-195


>>
37. NC_026474
Source: Fusarium graminearum PH-1 chromosome 1, whole genome shotgun sequence
Type: NRPS
Number of proteins with BLAST hits to this cluster: 2
Cumulative BLAST score: 1703

Table of genes, locations, strands and annotations of subject cluster:

Table of Blast hits (query gene, subject gene, %identity, blast score, %coverage, e-value):
input.path1.gene501	FGSG_11659	31	1016	33.93287069820139	1.2e-293
input.path1.gene502	FGSG_00046	33	687	94.67153284671534	2.5e-195


>>
38. NW_015971594
Source: Cladophialophora bantiana CBS 173.52 unplaced genomic scaffold supercont1.20, whole genome shotgun sequence
Type: NRPS
Number of proteins with BLAST hits to this cluster: 2
Cumulative BLAST score: 801

Table of genes, locations, strands and annotations of subject cluster:

Table of Blast hits (query gene, subject gene, %identity, blast score, %coverage, e-value):
input.path1.gene500	Z519_11088	33	230	96.06126914660832	2.7e-58
input.path1.gene501	Z519_11086	33	571	15.776802152669594	1.7e-159


>>
39. NC_036435
Source: Aspergillus oryzae RIB40 DNA, chromosome 1
Type: NRPS
Number of proteins with BLAST hits to this cluster: 2
Cumulative BLAST score: 774

Table of genes, locations, strands and annotations of subject cluster:

Table of Blast hits (query gene, subject gene, %identity, blast score, %coverage, e-value):
input.path1.gene501	AO090005000993	32	588	16.640702450077892	1e-164
input.path1.gene502	AO090005000987	36	186	21.605839416058394	2.3e-44


>>
40. NC_030961
Source: Colletotrichum higginsianum IMI 349063 chromosome 9, whole genome shotgun sequence
Type: NRPS
Number of proteins with BLAST hits to this cluster: 2
Cumulative BLAST score: 742

Table of genes, locations, strands and annotations of subject cluster:

Table of Blast hits (query gene, subject gene, %identity, blast score, %coverage, e-value):
input.path1.gene501	CH63R_12697	36	626	15.84761365245716	2.6e-176
input.path1.gene502	CH63R_12704	31	116	18.613138686131386	2.9e-23


>>
41. NW_001914833
Source: Podospora anserina S mat+ genomic DNA chromosome 3, supercontig 3
Type: NRPS
Number of proteins with BLAST hits to this cluster: 2
Cumulative BLAST score: 712

Table of genes, locations, strands and annotations of subject cluster:

Table of Blast hits (query gene, subject gene, %identity, blast score, %coverage, e-value):
input.path1.gene501	PODANS_3_11200	35	606	16.683189349950432	3.7e-170
input.path1.gene502	PODANS_3_11220	31	106	20.21897810218978	1.8e-20


>>
42. NW_015622507
Source: Exophiala mesophila strain CBS 40295 unplaced genomic scaffold supercont1.4, whole genome shotgun sequence
Type: NRPS
Number of proteins with BLAST hits to this cluster: 2
Cumulative BLAST score: 701

Table of genes, locations, strands and annotations of subject cluster:

Table of Blast hits (query gene, subject gene, %identity, blast score, %coverage, e-value):
input.path1.gene501	PV10_05618	33	585	15.776802152669594	6.8e-164
input.path1.gene502	PV10_05615	34	116	16.4963503649635	1.7e-23


>>
43. NW_011942160
Source: Metarhizium robertsii ARSEF 23 MAA_Scf_2, whole genome shotgun sequence
Type: NRPS
Number of proteins with BLAST hits to this cluster: 2
Cumulative BLAST score: 697

Table of genes, locations, strands and annotations of subject cluster:

Table of Blast hits (query gene, subject gene, %identity, blast score, %coverage, e-value):
input.path1.gene501	MAA_05334	34	574	15.70599065288203	1.2e-160
input.path1.gene502	MAA_05338	31	123	21.021897810218977	1.4e-25


>>
44. NW_014574716
Source: Metarhizium brunneum ARSEF 3297 chromosome Unknown scaffold_1, whole genome shotgun sequence
Type: NRPS
Number of proteins with BLAST hits to this cluster: 2
Cumulative BLAST score: 694

Table of genes, locations, strands and annotations of subject cluster:

Table of Blast hits (query gene, subject gene, %identity, blast score, %coverage, e-value):
input.path1.gene501	MBR_01852	33	575	15.649341453051976	9.1e-161
input.path1.gene502	MBR_01848	31	119	19.48905109489051	2.7e-24


>>
45. NW_006271973
Source: Cordyceps militaris CM01 unplaced genomic scaffold CCM_S00005, whole genome shotgun sequence
Type: NRPS
Number of proteins with BLAST hits to this cluster: 2
Cumulative BLAST score: 690

Table of genes, locations, strands and annotations of subject cluster:

Table of Blast hits (query gene, subject gene, %identity, blast score, %coverage, e-value):
input.path1.gene501	CCM_06702	34	571	15.25279705424161	1.3e-159
input.path1.gene502	CCM_06697	32	119	18.83211678832117	3.5e-24


>>
46. NC_035791
Source: Pochonia chlamydosporia 170 chromosome 2, whole genome shotgun sequence
Type: NRPS
Number of proteins with BLAST hits to this cluster: 2
Cumulative BLAST score: 683

Table of genes, locations, strands and annotations of subject cluster:

Table of Blast hits (query gene, subject gene, %identity, blast score, %coverage, e-value):
input.path1.gene501	VFPPC_03467	34	568	15.63517915309446	8.6e-159
input.path1.gene502	VFPPC_03464	32	115	18.248175182481752	5e-23


>>
47. NW_006911257
Source: Baudoinia panamericana UAMH 10762 unplaced genomic scaffold BAUCOscaffold_1, whole genome shotgun sequence
Type: NRPS-like
Number of proteins with BLAST hits to this cluster: 2
Cumulative BLAST score: 688

Table of genes, locations, strands and annotations of subject cluster:

Table of Blast hits (query gene, subject gene, %identity, blast score, %coverage, e-value):
input.path1.gene498	BAUCODRAFT_135756	51	285	98.95833333333334	4.5e-75
input.path1.gene499	BAUCODRAFT_29396	74	403	88.0	2e-110


>>
48. CM002799
Source: Penicillium chrysogenum strain P2niaD18 chromosome II, whole genome shotgun sequence
Type: NRPS-like
Number of proteins with BLAST hits to this cluster: 2
Cumulative BLAST score: 597

Table of genes, locations, strands and annotations of subject cluster:

Table of Blast hits (query gene, subject gene, %identity, blast score, %coverage, e-value):
input.path1.gene498	EN45_053030	51	280	101.73611111111111	1.9e-73
input.path1.gene499	EN45_053020	55	317	88.0	1.1e-84


>>
49. NC_035790
Source: Pochonia chlamydosporia 170 chromosome 1, whole genome shotgun sequence
Type: NRPS
Number of proteins with BLAST hits to this cluster: 2
Cumulative BLAST score: 848

Table of genes, locations, strands and annotations of subject cluster:

Table of Blast hits (query gene, subject gene, %identity, blast score, %coverage, e-value):
input.path1.gene500	VFPPC_12684	31	200	100.87527352297595	3e-49
input.path1.gene502	VFPPC_00080	34	648	90.36496350364963	1.6e-183


>>
50. NW_020290247
Source: Acaromyces ingoldii strain MCA 4198 unplaced genomic scaffold FA10scaffold_1, whole genome shotgun sequence
Type: NRPS
Number of proteins with BLAST hits to this cluster: 2
Cumulative BLAST score: 585

Table of genes, locations, strands and annotations of subject cluster:

Table of Blast hits (query gene, subject gene, %identity, blast score, %coverage, e-value):
input.path1.gene500	FA10DRAFT_290725	31	192	97.59299781181619	8.3e-47
input.path1.gene502	FA10DRAFT_298914	31	393	66.35036496350365	1.2e-106
