## Supplementary Results for "A draft genome of the ascomycotal fungal species *Pseudopithomyces maydicus* (family *Didymosphaeriaceae*)": contig_6_c2.txt

ClusterBlast scores for contig_6

Table of genes, locations, strands and annotations of query cluster:
input.path1.gene162	595470	599488	+		
input.path1.gene163	601142	604218	+		
input.path1.gene164	610523	611109	-		
input.path1.gene165	611770	614747	-		
input.path1.gene166	615872	617148	-		


Significant hits: 
1. NC_049562	Talaromyces rugulosus chromosome II, complete sequence
2. NZ_KQ976354	Scytonema hofmannii PCC 7110 Scaffold1, whole genome shotgun sequence
3. NZ_FYDI01000001	Pseudomonas sp. Irchel 3A18 isolate Pseudomonas sp. Irchel 3A18, whole genome shotgun sequence
4. NZ_LN847264	Pseudomonas sp. CCOS 191 chromosome I
5. NZ_QPDJ01000011	Pseudomonas coronafaciens pv. garcae strain 462 Contig_11, whole genome shotgun sequence
6. NZ_CP026386	Pseudomonas sp. PONIH3 chromosome, complete genome
7. NZ_SLXS01000001	Tumebacillus sp. BK434 Ga0307714_101, whole genome shotgun sequence
8. NZ_CP007039	Pseudomonas cichorii JBC1 chromosome, complete genome
9. NZ_LT962481	Pseudomonas syringae pv. syringae isolate CFBP2118 chromosome 1
10. NZ_CP005969	Pseudomonas syringae pv. syringae B301D chromosome, complete genome
11. NZ_FOPR01000003	Pseudomonas syringae strain BS3829, whole genome shotgun sequence
12. NZ_KB316531	Pseudomonas syringae BRIP34881 scaffold1, whole genome shotgun sequence
13. NZ_CP019871	Pseudomonas syringae pv. tomato strain B13-200 chromosome, complete genome
14. NZ_CP027706	Pseudomonas sp. CMR12a chromosome, complete genome
15. NZ_JACHOW010000002	Longimicrobium terrae strain CECT 8660 Ga0372432_02, whole genome shotgun sequence
16. NZ_CP015367	Methylobacterium phyllosphaerae strain CBMB27 chromosome, complete genome
17. NZ_PCQC01000010	Pseudomonas sp. MYb2 MYb2_10, whole genome shotgun sequence
18. NZ_LTAQ01000023	Bacillus gaemokensis strain KCTC 13318 scaffold3.1, whole genome shotgun sequence
19. NZ_AP020337	Pseudomonas sp. KUIN-1
20. NZ_CP047267	Pseudomonas syringae UB303 chromosome, complete genome
21. NZ_FNJH01000001	Pseudomonas congelans strain DSM 14939, whole genome shotgun sequence
22. NZ_KI519271	Pseudomonas aeruginosa CF5 adgfB-supercont1.1, whole genome shotgun sequence
23. NZ_CP026681	Nostoc sp. 'Peltigera membranacea cyanobiont' N6 chromosome, complete genome
24. NZ_LDCN01000002	Brevibacillus formosus strain DSM 9885 Scaffold2, whole genome shotgun sequence
25. NC_015379	Pseudomonas brassicacearum subsp. brassicacearum NFM421, complete sequence
26. NZ_AP015029	Pseudomonas putida strain KF715 chromosome 1
27. NZ_LT222319	Pseudomonas cerasi isolate Sour cherry (Prunus cerasus) symptomatic leaf chromosome 1
28. NZ_CP045302	Azotobacter salinestris strain KACC 13899 chromosome, complete genome
29. NZ_CP008714	Xanthomonas translucens pv. undulosa strain Xtu 4699 chromosome, complete genome
30. NZ_KB900108	Pseudomonas sp. 35MFCvi1.1 F571DRAFT_scaffold00001.1, whole genome shotgun sequence
31. NZ_CP011567	Pseudomonas sp. GR 6-02 chromosome, complete genome
32. NC_022600	Gloeobacter kilaueensis JS1, complete sequence
33. NZ_LT963391	Pseudomonas syringae pv. cerasicola isolate CFBP6109 chromosome 1
34. NZ_FOPV01000006	Nitrosospira sp. Nsp14, whole genome shotgun sequence
35. NZ_CP023466	Pseudomonas frederiksbergensis strain KNU-15 chromosome, complete genome
36. NZ_MSDI01000042	Pseudomonas fluorescens strain E24 PF_E24_contig_26, whole genome shotgun sequence
37. NZ_CP013612	Pseudoalteromonas rubra strain SCSIO 6842 chromosome 2, complete sequence
38. NZ_CP046622	Variovorax paradoxus strain CSUSB chromosome, complete genome
39. NZ_CP007203	Nodularia spumigena CCY9414
40. NZ_CP020114	Nodularia spumigena UHCC 0039 chromosome, complete genome
41. NC_015563	Delftia sp. Cs1-4, complete sequence
42. NZ_NIBV01000002	Xenorhabdus szentirmaii DSM 16338 Xsze_2, whole genome shotgun sequence
43. NZ_AP019314	Microcystis viridis NIES-102
44. CM000574	Fusarium graminearum PH-1 chromosome 1, whole genome shotgun sequence
45. NC_026474	Fusarium graminearum PH-1 chromosome 1, whole genome shotgun sequence
46. NZ_CP043056	Dolichospermum sp. UHCC 0315A chromosome, complete genome
47. NZ_CP022561	Pseudomonas putida strain B4 chromosome, complete genome
48. NZ_QJRL01000019	Pseudomonas sp. LB-090624 NODE_19_length_108915_cov_19.230908, whole genome shotgun sequence
49. NZ_CP017938	Acinetobacter pittii strain YMC2010/8/T346 chromosome, complete genome
50. NZ_CP045551	Pseudomonas putida strain 15420352 chromosome, complete genome


Details:

>>
1. NC_049562
Source: Talaromyces rugulosus chromosome II, complete sequence
Type: NRPS
Number of proteins with BLAST hits to this cluster: 3
Cumulative BLAST score: 662

Table of genes, locations, strands and annotations of subject cluster:

Table of Blast hits (query gene, subject gene, %identity, blast score, %coverage, e-value):
input.path1.gene162	TRUGW13939_04796	33	213	56.72853828306265	6.5e-53
input.path1.gene163	TRUGW13939_04796	40	358	68.58789625360231	2.3e-96
input.path1.gene166	TRUGW13939_04797	33	91	47.83783783783784	2.7e-16


>>
2. NZ_KQ976354
Source: Scytonema hofmannii PCC 7110 Scaffold1, whole genome shotgun sequence
Type: NRPS,microviridin
Number of proteins with BLAST hits to this cluster: 2
Cumulative BLAST score: 1475

Table of genes, locations, strands and annotations of subject cluster:
WA1_RS15490	WA1_RS15490	3712567	3714927	-	hypothetical_protein
WA1_RS15505	WA1_RS15505	3719729	3741925	+	non-ribosomal_peptide_synthase/polyketide_synthase

Table of Blast hits (query gene, subject gene, %identity, blast score, %coverage, e-value):
input.path1.gene162	WA1_RS15365	32	153	32.59860788863109	8e-35
input.path1.gene162	WA1_RS15425	33	148	31.786542923433874	2.6e-33
input.path1.gene162	WA1_RS15395	32	135	31.322505800464036	2.3e-29
input.path1.gene162	WA1_RS15390	31	132	33.29466357308585	2.5e-28
input.path1.gene162	WA1_RS15420	31	131	31.90255220417633	3.3e-28
input.path1.gene162	WA1_RS15400	31	125	31.786542923433874	3.1e-26
input.path1.gene163	WA1_RS15395	33	241	67.0028818443804	2.4e-61
input.path1.gene163	WA1_RS15365	31	207	66.71469740634006	4.9e-51
input.path1.gene163	WA1_RS15360	32	203	67.0028818443804	9.3e-50


>>
3. NZ_FYDI01000001
Source: Pseudomonas sp. Irchel 3A18 isolate Pseudomonas sp. Irchel 3A18, whole genome shotgun sequence
Type: NRPS
Number of proteins with BLAST hits to this cluster: 2
Cumulative BLAST score: 905

Table of genes, locations, strands and annotations of subject cluster:
CKW23_RS01540	CKW23_RS01540	345248	345733	-	hypothetical_protein
CKW23_RS01545	CKW23_RS01545	345830	346119	-	hypothetical_protein

Table of Blast hits (query gene, subject gene, %identity, blast score, %coverage, e-value):
input.path1.gene162	CKW23_RS01575	33	136	31.786542923433874	1.3e-29
input.path1.gene162	CKW23_RS01580	31	121	31.206496519721576	3.4e-25
input.path1.gene162	CKW23_RS01570	32	119	31.90255220417633	1.7e-24
input.path1.gene162	CKW23_RS01565	33	114	27.1461716937355	4.1e-23
input.path1.gene163	CKW23_RS01560	33	210	62.824207492795395	4.4e-52
input.path1.gene163	CKW23_RS01475	32	205	68.44380403458213	2.4e-50


>>
4. NZ_LN847264
Source: Pseudomonas sp. CCOS 191 chromosome I
Type: NRPS
Number of proteins with BLAST hits to this cluster: 2
Cumulative BLAST score: 823

Table of genes, locations, strands and annotations of subject cluster:

Table of Blast hits (query gene, subject gene, %identity, blast score, %coverage, e-value):
input.path1.gene162	CCOS191_RS11050	31	128	33.75870069605568	4.7e-27
input.path1.gene162	CCOS191_RS27205	33	120	31.786542923433874	7.5e-25
input.path1.gene163	CCOS191_RS11050	32	196	63.976945244956774	8.7e-48
input.path1.gene163	CCOS191_RS11045	33	192	61.09510086455331	1.3e-46
input.path1.gene163	CCOS191_RS10855	31	187	61.527377521613836	4e-45


>>
5. NZ_QPDJ01000011
Source: Pseudomonas coronafaciens pv. garcae strain 462 Contig_11, whole genome shotgun sequence
Type: NRPS,acyl_amino_acids,hserlactone
Number of proteins with BLAST hits to this cluster: 2
Cumulative BLAST score: 821

Table of genes, locations, strands and annotations of subject cluster:
D2G28_RS15460	D2G28_RS15460	70105	71355	+	M20/M25/M40_family_metallo-hydrolase
D2G28_RS15595	D2G28_RS15595	96742	97277	-	RNA_polymerase_factor_sigma-70
D2G28_RS15680	D2G28_RS15680	138891	147518	+	amino_acid_adenylation_domain-containing_protein
D2G28_RS15685	D2G28_RS15685	147452	147645	-	DUF1534_domain-containing_protein

Table of Blast hits (query gene, subject gene, %identity, blast score, %coverage, e-value):
input.path1.gene162	D2G28_RS15675	32	120	31.670533642691417	7.5e-25
input.path1.gene162	D2G28_RS15670	34	113	24.94199535962877	1.2e-22
input.path1.gene163	D2G28_RS15665	32	200	66.4265129682997	6e-49
input.path1.gene163	D2G28_RS15605	31	198	68.01152737752162	2.3e-48
input.path1.gene163	D2G28_RS15675	31	190	71.75792507204612	4.8e-46


>>
6. NZ_CP026386
Source: Pseudomonas sp. PONIH3 chromosome, complete genome
Type: NRPS
Number of proteins with BLAST hits to this cluster: 2
Cumulative BLAST score: 803

Table of genes, locations, strands and annotations of subject cluster:
C3F42_RS29525	C3F42_RS29525	6153021	6153276	+	hypothetical_protein

Table of Blast hits (query gene, subject gene, %identity, blast score, %coverage, e-value):
input.path1.gene162	C3F42_RS28605	34	114	25.290023201856147	5.4e-23
input.path1.gene162	C3F42_RS28600	33	114	30.8584686774942	7e-23
input.path1.gene163	C3F42_RS28595	32	195	63.976945244956774	2.5e-47
input.path1.gene163	C3F42_RS28405	32	190	61.67146974063401	4.8e-46
input.path1.gene163	C3F42_RS28590	33	190	61.09510086455331	4.8e-46


>>
7. NZ_SLXS01000001
Source: Tumebacillus sp. BK434 Ga0307714_101, whole genome shotgun sequence
Type: NRPS,T1PKS
Number of proteins with BLAST hits to this cluster: 2
Cumulative BLAST score: 794

Table of genes, locations, strands and annotations of subject cluster:

Table of Blast hits (query gene, subject gene, %identity, blast score, %coverage, e-value):
input.path1.gene162	EV586_RS02480	32	133	31.438515081206496	8.6e-29
input.path1.gene162	EV586_RS02575	33	128	29.118329466357306	4.7e-27
input.path1.gene162	EV586_RS02600	31	124	31.90255220417633	4e-26
input.path1.gene163	EV586_RS02605	32	206	65.4178674351585	8.4e-51
input.path1.gene163	EV586_RS02610	31	203	65.70605187319885	9.3e-50


>>
8. NZ_CP007039
Source: Pseudomonas cichorii JBC1 chromosome, complete genome
Type: NRPS
Number of proteins with BLAST hits to this cluster: 2
Cumulative BLAST score: 788

Table of genes, locations, strands and annotations of subject cluster:

Table of Blast hits (query gene, subject gene, %identity, blast score, %coverage, e-value):
input.path1.gene162	BH81_RS15510	33	136	31.786542923433874	1.3e-29
input.path1.gene162	BH81_RS15505	31	121	31.206496519721576	3.4e-25
input.path1.gene162	BH81_RS15515	33	115	27.1461716937355	3.2e-23
input.path1.gene163	BH81_RS15520	32	211	63.40057636887608	3.4e-52
input.path1.gene163	BH81_RS15605	32	205	68.44380403458213	1.4e-50


>>
9. NZ_LT962481
Source: Pseudomonas syringae pv. syringae isolate CFBP2118 chromosome 1
Type: NRPS,hserlactone
Number of proteins with BLAST hits to this cluster: 2
Cumulative BLAST score: 762

Table of genes, locations, strands and annotations of subject cluster:
C6H33_RS17355	C6H33_RS17355	3887284	3887912	-	hypothetical_protein
C6H33_RS17360	C6H33_RS17360	3888006	3888801	-	DNA_adenine_methylase

Table of Blast hits (query gene, subject gene, %identity, blast score, %coverage, e-value):
input.path1.gene162	C6H33_RS17210	33	122	31.438515081206496	2.6e-25
input.path1.gene162	C6H33_RS17205	31	119	31.09048723897912	2.2e-24
input.path1.gene162	C6H33_RS17215	31	118	30.278422273781903	4.9e-24
input.path1.gene163	C6H33_RS17220	32	203	66.13832853025937	5.4e-50
input.path1.gene163	C6H33_RS17290	31	200	68.01152737752162	4.6e-49


>>
10. NZ_CP005969
Source: Pseudomonas syringae pv. syringae B301D chromosome, complete genome
Type: NRPS,hserlactone
Number of proteins with BLAST hits to this cluster: 2
Cumulative BLAST score: 756

Table of genes, locations, strands and annotations of subject cluster:
PSYRB_RS27040	PSYRB_RS27040	2111230	2111657	+	transposase
PSYRB_RS09395	PSYRB_RS09395	2116205	2116690	-	DNA_methyltransferase
PSYRB_RS09410	PSYRB_RS09410	2118424	2118812	-	chorismate_mutase
PSYRB_RS09440	PSYRB_RS09440	2123925	2125046	-	sensor_histidine_kinase
PSYRB_RS27060	PSYRB_RS27060	2150406	2150651	-	transposase
PSYRB_RS29300	PSYRB_RS29300	2177516	2177993	-	DUF1534_domain-containing_protein

Table of Blast hits (query gene, subject gene, %identity, blast score, %coverage, e-value):
input.path1.gene162	PSYRB_RS09530	32	121	31.206496519721576	5.8e-25
input.path1.gene162	PSYRB_RS09535	31	119	31.09048723897912	2.2e-24
input.path1.gene162	PSYRB_RS09525	31	114	30.278422273781903	4.1e-23
input.path1.gene163	PSYRB_RS09520	32	201	68.58789625360231	2.1e-49
input.path1.gene163	PSYRB_RS09460	31	201	67.86743515850145	3.5e-49


>>
11. NZ_FOPR01000003
Source: Pseudomonas syringae strain BS3829, whole genome shotgun sequence
Type: NRPS
Number of proteins with BLAST hits to this cluster: 2
Cumulative BLAST score: 755

Table of genes, locations, strands and annotations of subject cluster:
BM038_RS27580	BM038_RS27580	301290	301472	+	hypothetical_protein

Table of Blast hits (query gene, subject gene, %identity, blast score, %coverage, e-value):
input.path1.gene162	BM038_RS10190	31	122	31.206496519721576	2.6e-25
input.path1.gene162	BM038_RS10195	31	118	31.09048723897912	3.7e-24
input.path1.gene162	BM038_RS10185	31	115	30.510440835266824	2.4e-23
input.path1.gene163	BM038_RS10180	31	200	65.99423631123919	4.6e-49
input.path1.gene163	BM038_RS10115	31	200	68.29971181556196	6e-49


>>
12. NZ_KB316531
Source: Pseudomonas syringae BRIP34881 scaffold1, whole genome shotgun sequence
Type: NRPS
Number of proteins with BLAST hits to this cluster: 2
Cumulative BLAST score: 754

Table of genes, locations, strands and annotations of subject cluster:

Table of Blast hits (query gene, subject gene, %identity, blast score, %coverage, e-value):
input.path1.gene162	A987_RS23850	32	121	30.97447795823666	3.4e-25
input.path1.gene162	A987_RS23845	31	116	31.09048723897912	1.1e-23
input.path1.gene162	A987_RS23855	31	116	30.510440835266824	1.9e-23
input.path1.gene163	A987_RS23860	31	201	65.99423631123919	2.1e-49
input.path1.gene163	A987_RS23920	31	200	68.01152737752162	7.9e-49


>>
13. NZ_CP019871
Source: Pseudomonas syringae pv. tomato strain B13-200 chromosome, complete genome
Type: NRPS
Number of proteins with BLAST hits to this cluster: 2
Cumulative BLAST score: 738

Table of genes, locations, strands and annotations of subject cluster:
XJ28_RS18770	XJ28_RS18770	3916607	3916693	-	DNA_adenine_methylase
XJ28_RS18775	XJ28_RS18775	3916679	3917034	+	DNA_cytosine_methyltransferase
XJ28_RS32775	XJ28_RS32775	3917414	3917617	+	IS66_family_transposase

Table of Blast hits (query gene, subject gene, %identity, blast score, %coverage, e-value):
input.path1.gene162	XJ28_RS18625	31	117	31.554524361948953	6.4e-24
input.path1.gene162	XJ28_RS18620	31	116	30.8584686774942	1.4e-23
input.path1.gene162	XJ28_RS18630	32	111	30.62645011600928	3.5e-22
input.path1.gene163	XJ28_RS18635	31	198	69.164265129683	1.7e-48
input.path1.gene163	XJ28_RS18695	31	196	68.01152737752162	6.6e-48


>>
14. NZ_CP027706
Source: Pseudomonas sp. CMR12a chromosome, complete genome
Type: NRPS
Number of proteins with BLAST hits to this cluster: 2
Cumulative BLAST score: 732

Table of genes, locations, strands and annotations of subject cluster:

Table of Blast hits (query gene, subject gene, %identity, blast score, %coverage, e-value):
input.path1.gene162	C4K39_RS29750	31	121	31.322505800464036	3.4e-25
input.path1.gene162	C4K39_RS29755	35	116	24.593967517401392	1.9e-23
input.path1.gene162	C4K39_RS29745	31	113	32.36658932714617	9.2e-23
input.path1.gene163	C4K39_RS29750	31	192	63.25648414985591	1.6e-46
input.path1.gene163	C4K39_RS29755	31	190	64.26512968299711	4.8e-46


>>
15. NZ_JACHOW010000002
Source: Longimicrobium terrae strain CECT 8660 Ga0372432_02, whole genome shotgun sequence
Type: NRPS,T1PKS
Number of proteins with BLAST hits to this cluster: 2
Cumulative BLAST score: 699

Table of genes, locations, strands and annotations of subject cluster:

Table of Blast hits (query gene, subject gene, %identity, blast score, %coverage, e-value):
input.path1.gene162	FHS01_RS05110	34	114	26.218097447795824	5.4e-23
input.path1.gene163	FHS01_RS05115	32	207	67.86743515850145	3.8e-51
input.path1.gene163	FHS01_RS05245	32	190	60.95100864553314	6.2e-46
input.path1.gene163	FHS01_RS05240	32	188	65.27377521613833	1.8e-45


>>
16. NZ_CP015367
Source: Methylobacterium phyllosphaerae strain CBMB27 chromosome, complete genome
Type: NRPS
Number of proteins with BLAST hits to this cluster: 2
Cumulative BLAST score: 695

Table of genes, locations, strands and annotations of subject cluster:
MCBMB27_RS04035	MCBMB27_RS04035	873491	873982	+	IS66_family_transposase_zinc-finger_binding_domain-containing_protein
MCBMB27_RS04050	MCBMB27_RS04050	876202	876997	+	IS66_family_transposase
MCBMB27_RS29050	MCBMB27_RS29050	935736	936246	+	transposase
MCBMB27_RS04080	MCBMB27_RS04080	936253	937467	+	IS110_family_transposase
MCBMB27_RS28745	MCBMB27_RS28745	949326	950599	+	flagellin

Table of Blast hits (query gene, subject gene, %identity, blast score, %coverage, e-value):
input.path1.gene162	MCBMB27_RS04055	31	121	28.65429234338747	4.4e-25
input.path1.gene162	MCBMB27_RS04060	32	118	24.013921113689097	3.7e-24
input.path1.gene163	MCBMB27_RS04065	33	237	66.85878962536023	5.8e-60
input.path1.gene163	MCBMB27_RS04055	31	219	67.14697406340058	1.3e-54


>>
17. NZ_PCQC01000010
Source: Pseudomonas sp. MYb2 MYb2_10, whole genome shotgun sequence
Type: NRPS,terpene
Number of proteins with BLAST hits to this cluster: 2
Cumulative BLAST score: 653

Table of genes, locations, strands and annotations of subject cluster:
CQ009_RS18140	CQ009_RS18140	111777	112759	+	GlxA_family_transcriptional_regulator

Table of Blast hits (query gene, subject gene, %identity, blast score, %coverage, e-value):
input.path1.gene162	CQ009_RS18195	33	128	32.01856148491879	4.7e-27
input.path1.gene162	CQ009_RS18190	35	115	24.593967517401392	2.4e-23
input.path1.gene163	CQ009_RS18195	31	208	66.57060518731988	2.9e-51
input.path1.gene163	CQ009_RS18210	32	202	67.72334293948127	1.2e-49


>>
18. NZ_LTAQ01000023
Source: Bacillus gaemokensis strain KCTC 13318 scaffold3.1, whole genome shotgun sequence
Type: NRPS
Number of proteins with BLAST hits to this cluster: 2
Cumulative BLAST score: 648

Table of genes, locations, strands and annotations of subject cluster:
AZF08_RS18025	AZF08_RS18025	196933	197334	+	DUF3980_domain-containing_protein
AZF08_RS18050	AZF08_RS18050	202231	202527	-	transposase
AZF08_RS28955	AZF08_RS28955	231235	231420	-	DDE-type_integrase/transposase/recombinase
AZF08_RS18085	AZF08_RS18085	231493	231689	-	transposase
AZF08_RS18105	AZF08_RS18105	235122	235700	+	alpha/beta_hydrolase

Table of Blast hits (query gene, subject gene, %identity, blast score, %coverage, e-value):
input.path1.gene162	AZF08_RS18075	31	128	30.8584686774942	4.7e-27
input.path1.gene162	AZF08_RS18065	32	124	28.538283062645007	6.8e-26
input.path1.gene163	AZF08_RS18070	31	200	65.85014409221903	4.6e-49
input.path1.gene163	AZF08_RS18075	31	196	69.88472622478386	8.7e-48


>>
19. NZ_AP020337
Source: Pseudomonas sp. KUIN-1
Type: NRPS,hserlactone
Number of proteins with BLAST hits to this cluster: 2
Cumulative BLAST score: 648

Table of genes, locations, strands and annotations of subject cluster:
KUIN1_RS10050	KUIN1_RS10050	2128409	2129328	+	alpha/beta_fold_hydrolase

Table of Blast hits (query gene, subject gene, %identity, blast score, %coverage, e-value):
input.path1.gene162	KUIN1_RS10235	33	123	31.438515081206496	1.5e-25
input.path1.gene162	KUIN1_RS10230	31	119	30.278422273781903	1.7e-24
input.path1.gene163	KUIN1_RS10225	32	205	66.13832853025937	1.4e-50
input.path1.gene163	KUIN1_RS10150	31	201	67.86743515850145	3.5e-49


>>
20. NZ_CP047267
Source: Pseudomonas syringae UB303 chromosome, complete genome
Type: NRPS,hserlactone
Number of proteins with BLAST hits to this cluster: 2
Cumulative BLAST score: 644

Table of genes, locations, strands and annotations of subject cluster:
N026_RS09470	N026_RS09470	2056328	2057538	-	hypothetical_protein
N026_RS09585	N026_RS09585	2089715	2090039	-	DUF1534_domain-containing_protein
N026_RS09610	N026_RS09610	2116478	2116747	-	hypothetical_protein
N026_RS09650	N026_RS09650	2129543	2129959	+	DUF1534_domain-containing_protein

Table of Blast hits (query gene, subject gene, %identity, blast score, %coverage, e-value):
input.path1.gene162	N026_RS09600	33	121	31.438515081206496	3.4e-25
input.path1.gene162	N026_RS09595	31	118	30.278422273781903	3.7e-24
input.path1.gene163	N026_RS09590	31	204	65.99423631123919	4.2e-50
input.path1.gene163	N026_RS09515	31	201	67.86743515850145	2.7e-49


>>
21. NZ_FNJH01000001
Source: Pseudomonas congelans strain DSM 14939, whole genome shotgun sequence
Type: NRPS
Number of proteins with BLAST hits to this cluster: 2
Cumulative BLAST score: 641

Table of genes, locations, strands and annotations of subject cluster:
BLR20_RS26330	BLR20_RS26330	185506	185747	+	DUF1534_domain-containing_protein

Table of Blast hits (query gene, subject gene, %identity, blast score, %coverage, e-value):
input.path1.gene162	BLR20_RS00710	31	124	31.206496519721576	6.8e-26
input.path1.gene162	BLR20_RS00705	31	117	30.278422273781903	6.4e-24
input.path1.gene163	BLR20_RS00700	32	201	65.4178674351585	2.1e-49
input.path1.gene163	BLR20_RS00635	31	199	68.29971181556196	1e-48


>>
22. NZ_KI519271
Source: Pseudomonas aeruginosa CF5 adgfB-supercont1.1, whole genome shotgun sequence
Type: NRPS
Number of proteins with BLAST hits to this cluster: 2
Cumulative BLAST score: 634

Table of genes, locations, strands and annotations of subject cluster:
Q004_RS16110	Q004_RS16110	2062799	2063564	+	thioesterase
Q004_RS0106160	Q004_RS0106160	2095814	2096675	-	pirin_family_protein
Q004_RS0106165	Q004_RS0106165	2122939	2135897	+	amino_acid_adenylation_domain-containing_protein
Q004_RS0106170	Q004_RS0106170	2143994	2145353	-	membrane_dipeptidase

Table of Blast hits (query gene, subject gene, %identity, blast score, %coverage, e-value):
input.path1.gene162	Q004_RS0106165	31	117	30.62645011600928	6.4e-24
input.path1.gene162	Q004_RS16280	31	113	30.278422273781903	9.2e-23
input.path1.gene163	Q004_RS16275	33	207	67.43515850144092	4.9e-51
input.path1.gene163	Q004_RS16115	32	197	69.30835734870317	3.9e-48


>>
23. NZ_CP026681
Source: Nostoc sp. 'Peltigera membranacea cyanobiont' N6 chromosome, complete genome
Type: LAP,NRPS,PKS-like,T1PKS,T3PKS,bacteriocin,transAT-PKS
Number of proteins with BLAST hits to this cluster: 2
Cumulative BLAST score: 596

Table of genes, locations, strands and annotations of subject cluster:
NPM_RS09640	NPM_RS09640	2163044	2163652	+	DNA_polymerase_IV
NPM_RS09655	NPM_RS09655	2165651	2165803	-	low_molecular_weight_phosphatase_family_protein
NPM_RS09660	NPM_RS09660	2166032	2166565	+	DUF928_domain-containing_protein
NPM_RS09720	NPM_RS09720	2179150	2179521	+	hypothetical_protein
NPM_RS09730	NPM_RS09730	2181686	2182046	-	transposase
NPM_RS09745	NPM_RS09745	2208790	2209012	-	transposase
NPM_RS09770	NPM_RS09770	2212363	2213672	-	hypothetical_protein
NPM_RS09830	NPM_RS09830	2223971	2224245	+	hypothetical_protein
NPM_RS09960	NPM_RS09960	2237449	2237805	-	NmrA_family_NAD(P)-binding_protein
NPM_RS09965	NPM_RS09965	2237804	2238340	+	LysR_family_transcriptional_regulator
NPM_RS09975	NPM_RS09975	2240638	2242125	-	IS1380_family_transposase
NPM_RS10050	NPM_RS10050	2302249	2302470	-	IS4_family_transposase
NPM_RS10060	NPM_RS10060	2303242	2303487	+	transposase_family_protein
NPM_RS10085	NPM_RS10085	2306795	2307071	+	HU_family_DNA-binding_protein
NPM_RS10205	NPM_RS10205	2330539	2330865	+	hypothetical_protein
NPM_RS10235	NPM_RS10235	2345934	2346944	-	DNA_cytosine_methyltransferase
NPM_RS10300	NPM_RS10300	2355943	2356377	-	hypothetical_protein
NPM_RS10310	NPM_RS10310	2358423	2360234	-	hypothetical_protein
NPM_RS10320	NPM_RS10320	2360701	2360910	-	hypothetical_protein

Table of Blast hits (query gene, subject gene, %identity, blast score, %coverage, e-value):
input.path1.gene162	NPM_RS09740	31	139	36.542923433874705	2e-30
input.path1.gene163	NPM_RS09735	31	234	69.02017291066282	2.9e-59
input.path1.gene163	NPM_RS10230	31	223	62.39193083573487	5.1e-56


>>
24. NZ_LDCN01000002
Source: Brevibacillus formosus strain DSM 9885 Scaffold2, whole genome shotgun sequence
Type: NRPS
Number of proteins with BLAST hits to this cluster: 2
Cumulative BLAST score: 568

Table of genes, locations, strands and annotations of subject cluster:
AA984_RS09005	AA984_RS09005	883114	898581	+	non-ribosomal_peptide_synthase/polyketide_synthase

Table of Blast hits (query gene, subject gene, %identity, blast score, %coverage, e-value):
input.path1.gene162	AA984_RS09010	32	138	31.786542923433874	2.7e-30
input.path1.gene163	AA984_RS09015	31	219	69.164265129683	1.3e-54
input.path1.gene163	AA984_RS09005	31	211	63.25648414985591	2.6e-52


>>
25. NC_015379
Source: Pseudomonas brassicacearum subsp. brassicacearum NFM421, complete sequence
Type: NRPS
Number of proteins with BLAST hits to this cluster: 2
Cumulative BLAST score: 553

Table of genes, locations, strands and annotations of subject cluster:
PSEBR_RS20980	PSEBR_RS20980	4814463	4824674	-	amino_acid_adenylation_domain-containing_protein
PSEBR_RS20985	PSEBR_RS20985	4824910	4827258	-	AMP-binding_protein

Table of Blast hits (query gene, subject gene, %identity, blast score, %coverage, e-value):
input.path1.gene162	PSEBR_RS20975	32	122	33.17865429234338	2.6e-25
input.path1.gene162	PSEBR_RS20985	31	119	32.01856148491879	1.3e-24
input.path1.gene162	PSEBR_RS20980	35	114	24.825986078886313	4.1e-23
input.path1.gene163	PSEBR_RS20975	31	198	66.4265129682997	3e-48


>>
26. NZ_AP015029
Source: Pseudomonas putida strain KF715 chromosome 1
Type: NRPS
Number of proteins with BLAST hits to this cluster: 2
Cumulative BLAST score: 553

Table of genes, locations, strands and annotations of subject cluster:
KF715C_RS21360	KF715C_RS21360	4593459	4601231	-	amino_acid_adenylation_domain-containing_protein
KF715C_RS21380	KF715C_RS21380	4620417	4621793	-	aspartate_aminotransferase_family_protein

Table of Blast hits (query gene, subject gene, %identity, blast score, %coverage, e-value):
input.path1.gene162	KF715C_RS21360	31	120	30.8584686774942	7.5e-25
input.path1.gene162	KF715C_RS21355	31	112	27.7262180974478	2e-22
input.path1.gene162	KF715C_RS21365	33	112	26.450116009280745	2e-22
input.path1.gene163	KF715C_RS21360	31	209	68.01152737752162	1.3e-51


>>
27. NZ_LT222319
Source: Pseudomonas cerasi isolate Sour cherry (Prunus cerasus) symptomatic leaf chromosome 1
Type: NRPS
Number of proteins with BLAST hits to this cluster: 2
Cumulative BLAST score: 545

Table of genes, locations, strands and annotations of subject cluster:
PCPL58_RS18750	PCPL58_RS18750	3657003	3657753	-	SDR_family_oxidoreductase

Table of Blast hits (query gene, subject gene, %identity, blast score, %coverage, e-value):
input.path1.gene162	PCPL58_RS18805	32	119	31.09048723897912	2.2e-24
input.path1.gene162	PCPL58_RS18800	31	117	31.09048723897912	6.4e-24
input.path1.gene162	PCPL58_RS18810	31	112	30.510440835266824	2.7e-22
input.path1.gene163	PCPL58_RS18815	32	197	66.13832853025937	3.9e-48


>>
28. NZ_CP045302
Source: Azotobacter salinestris strain KACC 13899 chromosome, complete genome
Type: NRPS,T1PKS
Number of proteins with BLAST hits to this cluster: 2
Cumulative BLAST score: 532

Table of genes, locations, strands and annotations of subject cluster:
GCU53_RS13710	GCU53_RS13710	2964838	2965461	-	TolC_family_protein

Table of Blast hits (query gene, subject gene, %identity, blast score, %coverage, e-value):
input.path1.gene162	GCU53_RS13660	32	132	31.322505800464036	1.9e-28
input.path1.gene163	GCU53_RS13665	31	214	69.88472622478386	3.1e-53
input.path1.gene163	GCU53_RS13670	31	186	65.70605187319885	6.9e-45


>>
29. NZ_CP008714
Source: Xanthomonas translucens pv. undulosa strain Xtu 4699 chromosome, complete genome
Type: NRPS,T3PKS
Number of proteins with BLAST hits to this cluster: 2
Cumulative BLAST score: 531

Table of genes, locations, strands and annotations of subject cluster:
FD63_RS18990	FD63_RS18990	2218868	2219995	-	amino_acid_adenylation_domain-containing_protein
FD63_RS20930	FD63_RS20930	2220519	2221646	-	non-ribosomal_peptide_synthetase
FD63_RS19485	FD63_RS19485	2291476	2292249	-	IS5_family_transposase

Table of Blast hits (query gene, subject gene, %identity, blast score, %coverage, e-value):
input.path1.gene162	FD63_RS09435	37	117	24.709976798143853	8.3e-24
input.path1.gene163	FD63_RS09440	33	218	64.69740634005764	2.8e-54
input.path1.gene163	FD63_RS09430	33	196	62.824207492795395	1.1e-47


>>
30. NZ_KB900108
Source: Pseudomonas sp. 35MFCvi1.1 F571DRAFT_scaffold00001.1, whole genome shotgun sequence
Type: NRPS
Number of proteins with BLAST hits to this cluster: 2
Cumulative BLAST score: 530

Table of genes, locations, strands and annotations of subject cluster:
F571_RS0101795	F571_RS0101795	416704	416910	-	sigma-54-dependent_Fis_family_transcriptional_regulator

Table of Blast hits (query gene, subject gene, %identity, blast score, %coverage, e-value):
input.path1.gene162	F571_RS0101710	31	114	31.438515081206496	5.4e-23
input.path1.gene163	F571_RS0101705	31	209	65.12968299711815	9.9e-52
input.path1.gene163	F571_RS0101690	31	207	66.71469740634006	3.8e-51


>>
31. NZ_CP011567
Source: Pseudomonas sp. GR 6-02 chromosome, complete genome
Type: NRPS
Number of proteins with BLAST hits to this cluster: 2
Cumulative BLAST score: 525

Table of genes, locations, strands and annotations of subject cluster:
PGR6_RS19225	PGR6_RS19225	4397346	4397552	-	sigma-54-dependent_Fis_family_transcriptional_regulator

Table of Blast hits (query gene, subject gene, %identity, blast score, %coverage, e-value):
input.path1.gene162	PGR6_RS19140	31	119	32.59860788863109	2.2e-24
input.path1.gene163	PGR6_RS19135	32	205	65.12968299711815	2.4e-50
input.path1.gene163	PGR6_RS19120	32	201	65.85014409221903	2.7e-49


>>
32. NC_022600
Source: Gloeobacter kilaueensis JS1, complete sequence
Type: NRPS,NRPS-like,T1PKS
Number of proteins with BLAST hits to this cluster: 2
Cumulative BLAST score: 524

Table of genes, locations, strands and annotations of subject cluster:
GKIL_RS08965	GKIL_RS08965	1936886	1939507	-	glucosidase

Table of Blast hits (query gene, subject gene, %identity, blast score, %coverage, e-value):
input.path1.gene162	GKIL_RS08820	33	116	24.245939675174014	1.9e-23
input.path1.gene163	GKIL_RS08825	32	209	67.43515850144092	9.9e-52
input.path1.gene163	GKIL_RS08830	31	199	63.40057636887608	1e-48


>>
33. NZ_LT963391
Source: Pseudomonas syringae pv. cerasicola isolate CFBP6109 chromosome 1
Type: NRPS
Number of proteins with BLAST hits to this cluster: 2
Cumulative BLAST score: 523

Table of genes, locations, strands and annotations of subject cluster:
C6H44_RS07210	C6H44_RS07210	1477746	1478205	+	RulA_protein
C6H44_RS07260	C6H44_RS07260	1487783	1490119	-	PAS_domain_S-box_protein
C6H44_RS07290	C6H44_RS07290	1494461	1507474	+	non-ribosomal_peptide_synthetase
C6H44_RS30710	C6H44_RS30710	1509426	1509506	-	DUF1534_domain-containing_protein
C6H44_RS30715	C6H44_RS30715	1545172	1545273	+	DUF1534_domain-containing_protein
C6H44_RS07420	C6H44_RS07420	1552349	1552666	-	hypothetical_protein

Table of Blast hits (query gene, subject gene, %identity, blast score, %coverage, e-value):
input.path1.gene162	C6H44_RS07360	31	121	32.01856148491879	3.4e-25
input.path1.gene163	C6H44_RS07355	31	207	70.02881844380403	6.4e-51
input.path1.gene163	C6H44_RS07290	33	195	66.28242074927954	1.5e-47


>>
34. NZ_FOPV01000006
Source: Nitrosospira sp. Nsp14, whole genome shotgun sequence
Type: NRPS,T1PKS
Number of proteins with BLAST hits to this cluster: 2
Cumulative BLAST score: 523

Table of genes, locations, strands and annotations of subject cluster:
BM022_RS05420	BM022_RS05420	71778	71888	+	PEP-CTERM_sorting_domain-containing_protein

Table of Blast hits (query gene, subject gene, %identity, blast score, %coverage, e-value):
input.path1.gene162	BM022_RS05390	33	113	27.37819025522042	9.2e-23
input.path1.gene163	BM022_RS05385	33	211	66.57060518731988	2.6e-52
input.path1.gene163	BM022_RS05400	32	199	66.13832853025937	1.3e-48


>>
35. NZ_CP023466
Source: Pseudomonas frederiksbergensis strain KNU-15 chromosome, complete genome
Type: NRPS
Number of proteins with BLAST hits to this cluster: 2
Cumulative BLAST score: 521

Table of genes, locations, strands and annotations of subject cluster:
CNN82_RS10230	CNN82_RS10230	2152214	2152420	+	sigma-54-dependent_Fis_family_transcriptional_regulator

Table of Blast hits (query gene, subject gene, %identity, blast score, %coverage, e-value):
input.path1.gene162	CNN82_RS10320	31	117	31.438515081206496	8.3e-24
input.path1.gene163	CNN82_RS10325	32	205	65.12968299711815	1.9e-50
input.path1.gene163	CNN82_RS10340	31	199	66.71469740634006	1e-48


>>
36. NZ_MSDI01000042
Source: Pseudomonas fluorescens strain E24 PF_E24_contig_26, whole genome shotgun sequence
Type: NRPS
Number of proteins with BLAST hits to this cluster: 2
Cumulative BLAST score: 520

Table of genes, locations, strands and annotations of subject cluster:

Table of Blast hits (query gene, subject gene, %identity, blast score, %coverage, e-value):
input.path1.gene162	BFW91_RS28945	31	135	31.438515081206496	2.3e-29
input.path1.gene163	BFW91_RS28940	32	194	67.43515850144092	4.3e-47
input.path1.gene163	BFW91_RS28935	31	191	69.59654178674351	2.1e-46


>>
37. NZ_CP013612
Source: Pseudoalteromonas rubra strain SCSIO 6842 chromosome 2, complete sequence
Type: NRPS
Number of proteins with BLAST hits to this cluster: 2
Cumulative BLAST score: 519

Table of genes, locations, strands and annotations of subject cluster:
AT705_RS23445	AT705_RS23445	1012940	1019631	+	amino_acid_adenylation_domain-containing_protein

Table of Blast hits (query gene, subject gene, %identity, blast score, %coverage, e-value):
input.path1.gene162	AT705_RS23480	32	113	27.1461716937355	1.2e-22
input.path1.gene163	AT705_RS23475	32	212	64.98559077809799	2e-52
input.path1.gene163	AT705_RS23470	31	194	61.95965417867435	3.3e-47


>>
38. NZ_CP046622
Source: Variovorax paradoxus strain CSUSB chromosome, complete genome
Type: NRPS,T1PKS
Number of proteins with BLAST hits to this cluster: 2
Cumulative BLAST score: 518

Table of genes, locations, strands and annotations of subject cluster:
GOQ09_RS19030	GOQ09_RS19030	4075902	4080039	+	filamentous_hemagglutinin_N-terminal_domain-containing_protein
GOQ09_RS19160	GOQ09_RS19160	4122995	4123297	+	LysR_family_transcriptional_regulator

Table of Blast hits (query gene, subject gene, %identity, blast score, %coverage, e-value):
input.path1.gene162	GOQ09_RS19080	34	116	24.129930394431554	1.9e-23
input.path1.gene163	GOQ09_RS19075	33	207	70.46109510086455	6.4e-51
input.path1.gene163	GOQ09_RS19095	31	195	73.05475504322767	1.9e-47


>>
39. NZ_CP007203
Source: Nodularia spumigena CCY9414
Type: NRPS
Number of proteins with BLAST hits to this cluster: 2
Cumulative BLAST score: 500

Table of genes, locations, strands and annotations of subject cluster:

Table of Blast hits (query gene, subject gene, %identity, blast score, %coverage, e-value):
input.path1.gene162	NSP_RS22225	32	132	31.670533642691417	1.9e-28
input.path1.gene162	NSP_RS22220	32	124	32.36658932714617	6.8e-26
input.path1.gene163	NSP_RS22225	35	244	63.25648414985591	2.8e-62


>>
40. NZ_CP020114
Source: Nodularia spumigena UHCC 0039 chromosome, complete genome
Type: NRPS
Number of proteins with BLAST hits to this cluster: 2
Cumulative BLAST score: 497

Table of genes, locations, strands and annotations of subject cluster:
BMF81_RS19160	BMF81_RS19160	4277171	4277245	+	cytochrome
BMF81_RS19255	BMF81_RS19255	4307014	4307207	+	hypothetical_protein

Table of Blast hits (query gene, subject gene, %identity, blast score, %coverage, e-value):
input.path1.gene162	BMF81_RS19225	32	132	31.670533642691417	1.9e-28
input.path1.gene162	BMF81_RS19230	32	121	32.36658932714617	3.4e-25
input.path1.gene163	BMF81_RS19225	35	244	63.25648414985591	2.8e-62


>>
41. NC_015563
Source: Delftia sp. Cs1-4, complete sequence
Type: NRPS,T1PKS
Number of proteins with BLAST hits to this cluster: 2
Cumulative BLAST score: 494

Table of genes, locations, strands and annotations of subject cluster:

Table of Blast hits (query gene, subject gene, %identity, blast score, %coverage, e-value):
input.path1.gene162	DELCS14_RS10585	35	117	24.709976798143853	6.4e-24
input.path1.gene163	DELCS14_RS10590	33	189	65.27377521613833	1.1e-45
input.path1.gene163	DELCS14_RS10595	32	188	63.68876080691642	1.8e-45


>>
42. NZ_NIBV01000002
Source: Xenorhabdus szentirmaii DSM 16338 Xsze_2, whole genome shotgun sequence
Type: NRPS
Number of proteins with BLAST hits to this cluster: 2
Cumulative BLAST score: 486

Table of genes, locations, strands and annotations of subject cluster:
Xsze_RS14625	Xsze_RS14625	1046645	1046785	-	IS1_family_transposase

Table of Blast hits (query gene, subject gene, %identity, blast score, %coverage, e-value):
input.path1.gene162	Xsze_RS14615	32	140	30.97447795823666	9.2e-31
input.path1.gene162	Xsze_RS14620	33	131	31.438515081206496	5.6e-28
input.path1.gene163	Xsze_RS14615	31	215	64.84149855907782	2.4e-53


>>
43. NZ_AP019314
Source: Microcystis viridis NIES-102
Type: NRPS,microviridin
Number of proteins with BLAST hits to this cluster: 2
Cumulative BLAST score: 482

Table of genes, locations, strands and annotations of subject cluster:
myaer102_RS17575	myaer102_RS17575	3404725	3405567	+	HindVP_family_restriction_endonuclease
myaer102_RS17580	myaer102_RS17580	3405680	3406063	+	DNA_cytosine_methyltransferase
myaer102_RS17605	myaer102_RS17605	3409299	3412041	-	DNA_topoisomerase_4_subunit_A
myaer102_RS17725	myaer102_RS17725	3462931	3463486	-	transposase
myaer102_RS17740	myaer102_RS17740	3467520	3468214	+	hypothetical_protein

Table of Blast hits (query gene, subject gene, %identity, blast score, %coverage, e-value):
input.path1.gene162	myaer102_RS17655	32	134	31.670533642691417	3.9e-29
input.path1.gene162	myaer102_RS17650	31	133	31.438515081206496	1.5e-28
input.path1.gene163	myaer102_RS17655	31	215	63.54466858789626	2.4e-53


>>
44. CM000574
Source: Fusarium graminearum PH-1 chromosome 1, whole genome shotgun sequence
Type: NRPS
Number of proteins with BLAST hits to this cluster: 2
Cumulative BLAST score: 481

Table of genes, locations, strands and annotations of subject cluster:

Table of Blast hits (query gene, subject gene, %identity, blast score, %coverage, e-value):
input.path1.gene162	FGSG_11660	39	151	22.8538283062645	4e-34
input.path1.gene163	FGSG_11659	39	330	66.28242074927954	3.9e-88


>>
45. NC_026474
Source: Fusarium graminearum PH-1 chromosome 1, whole genome shotgun sequence
Type: NRPS
Number of proteins with BLAST hits to this cluster: 2
Cumulative BLAST score: 481

Table of genes, locations, strands and annotations of subject cluster:

Table of Blast hits (query gene, subject gene, %identity, blast score, %coverage, e-value):
input.path1.gene162	FGSG_11660	39	151	22.8538283062645	4e-34
input.path1.gene163	FGSG_11659	39	330	66.28242074927954	3.9e-88


>>
46. NZ_CP043056
Source: Dolichospermum sp. UHCC 0315A chromosome, complete genome
Type: NRPS
Number of proteins with BLAST hits to this cluster: 2
Cumulative BLAST score: 475

Table of genes, locations, strands and annotations of subject cluster:

Table of Blast hits (query gene, subject gene, %identity, blast score, %coverage, e-value):
input.path1.gene162	BMF77_RS03355	31	139	29.930394431554525	1.6e-30
input.path1.gene162	BMF77_RS03350	32	139	31.90255220417633	2e-30
input.path1.gene163	BMF77_RS03355	31	197	65.70605187319885	5.1e-48


>>
47. NZ_CP022561
Source: Pseudomonas putida strain B4 chromosome, complete genome
Type: NRPS
Number of proteins with BLAST hits to this cluster: 2
Cumulative BLAST score: 473

Table of genes, locations, strands and annotations of subject cluster:

Table of Blast hits (query gene, subject gene, %identity, blast score, %coverage, e-value):
input.path1.gene162	CHN49_RS08835	33	126	29.00232018561485	1.8e-26
input.path1.gene162	CHN49_RS08805	31	122	31.786542923433874	2.6e-25
input.path1.gene163	CHN49_RS08810	32	225	66.85878962536023	1.7e-56


>>
48. NZ_QJRL01000019
Source: Pseudomonas sp. LB-090624 NODE_19_length_108915_cov_19.230908, whole genome shotgun sequence
Type: NRPS
Number of proteins with BLAST hits to this cluster: 2
Cumulative BLAST score: 473

Table of genes, locations, strands and annotations of subject cluster:

Table of Blast hits (query gene, subject gene, %identity, blast score, %coverage, e-value):
input.path1.gene162	DMX11_RS17635	33	123	29.00232018561485	8.9e-26
input.path1.gene162	DMX11_RS17605	31	122	31.90255220417633	2e-25
input.path1.gene163	DMX11_RS17610	31	228	66.85878962536023	2.7e-57


>>
49. NZ_CP017938
Source: Acinetobacter pittii strain YMC2010/8/T346 chromosome, complete genome
Type: NRPS,T1PKS
Number of proteins with BLAST hits to this cluster: 2
Cumulative BLAST score: 472

Table of genes, locations, strands and annotations of subject cluster:
BMU11_RS19705	BMU11_RS19705	11807	12064	-	hypothetical_protein
BMU11_RS00075	BMU11_RS00075	12272	21229	-	Ig-like_domain_repeat_protein
BMU11_RS19545	BMU11_RS19545	21545	21742	-	BapA_prefix-like_domain-containing_protein
BMU11_RS00085	BMU11_RS00085	23940	24227	-	neutral_zinc_metallopeptidase
BMU11_RS00200	BMU11_RS00200	69241	69367	+	hypothetical_protein

Table of Blast hits (query gene, subject gene, %identity, blast score, %coverage, e-value):
input.path1.gene162	BMU11_RS00120	33	135	32.01856148491879	3e-29
input.path1.gene162	BMU11_RS00125	31	131	35.150812064965194	3.3e-28
input.path1.gene163	BMU11_RS00120	32	206	67.57925072046109	8.4e-51


>>
50. NZ_CP045551
Source: Pseudomonas putida strain 15420352 chromosome, complete genome
Type: NRPS
Number of proteins with BLAST hits to this cluster: 2
Cumulative BLAST score: 470

Table of genes, locations, strands and annotations of subject cluster:

Table of Blast hits (query gene, subject gene, %identity, blast score, %coverage, e-value):
input.path1.gene162	GEV41_RS09070	33	123	29.466357308584683	1.5e-25
input.path1.gene162	GEV41_RS09040	31	122	32.25058004640371	2e-25
input.path1.gene163	GEV41_RS09045	32	225	66.85878962536023	1.7e-56
