## Supplementary Results for "A draft genome of the ascomycotal fungal species *Pseudopithomyces maydicus* (family *Didymosphaeriaceae*)": contig_6_c3.txt

ClusterBlast scores for contig_6

Table of genes, locations, strands and annotations of query cluster:
input.path1.gene476	1815539	1815863	+		
input.path1.gene477	1822932	1824289	+		
input.path1.gene478	1824501	1825526	-		
input.path1.gene479	1826819	1827933	+		
input.path1.gene480	1828057	1831447	-		
input.path1.gene481	1833111	1834765	-		
input.path1.gene482	1834902	1841987	+		
input.path1.gene483	1843352	1846304	-		
input.path1.gene484	1846627	1848332	+		
input.path1.gene485	1851154	1852027	+		
input.path1.gene486	1853403	1854775	+		
input.path1.gene487	1856541	1857965	-		
input.path1.gene488	1858243	1858591	+		
input.path1.gene489	1858971	1860177	-		
input.path1.gene490	1860400	1861468	+		


Significant hits: 
1. NW_011942164	Metarhizium robertsii ARSEF 23 MAA_Scf_23, whole genome shotgun sequence
2. NW_014574700	Metarhizium brunneum ARSEF 3297 chromosome Unknown scaffold_17, whole genome shotgun sequence
3. NC_016459	Thermothielavioides terrestris NRRL 8126 chromosome 3, complete sequence
4. NW_008481799	Cyphellophora europaea CBS 101466 unplaced genomic scaffold supercont1.11, whole genome shotgun sequence
5. NW_013550604	Rhinocladiella mackenziei CBS 650.93 unplaced genomic scaffold supercont1.1, whole genome shotgun sequence
6. CM000593	Fusarium oxysporum f. sp. lycopersici 4287 chromosome 5, whole genome shotgun sequence
7. NC_030990	Fusarium oxysporum f. sp. lycopersici 4287 chromosome 5, whole genome shotgun sequence
8. NC_036624	Fusarium fujikuroi IMI 58289 draft genome, chromosome FFUJ_chr03
9. NW_022983866	Arthroderma uncinatum strain CBS 119779 chromosome Unknown scaffold4, whole genome shotgun sequence
10. NW_003345200	Nannizzia gypsea CBS 118893 supercont1.2 genomic scaffold, whole genome shotgun sequence
11. NZ_CP012159	Chondromyces crocatus strain Cm c5 chromosome, complete genome
12. NZ_FOKG01000022	Amycolatopsis marina strain CGMCC 4.3568, whole genome shotgun sequence
13. NZ_CP022434	Nocardiopsis dassonvillei strain HZNU_N_1 chromosome, complete genome
14. NZ_KB906854	Nocardiopsis sp. CNS-639 strain CNS639 G011DRAFT_scaffold00001.1, whole genome shotgun sequence
15. NW_003345193	Nannizzia gypsea CBS 118893 supercont1.9 genomic scaffold, whole genome shotgun sequence
16. NZ_FOIB01000013	Myxococcus fulvus strain DSM 16525, whole genome shotgun sequence
17. NZ_LWLA01000014	Streptomyces sp. CB03911 scaffold21, whole genome shotgun sequence
18. NZ_CP028490	Pseudomonas syringae pv. atrofaciens strain LMG5095 isolate Triticum aestivum (Aestivum Group) chromosome, complete genome
19. NZ_BAZE01000009	Nocardiopsis alba strain TP-A0876, whole genome shotgun sequence
20. NC_018524	Nocardiopsis alba ATCC BAA-2165, complete sequence
21. NZ_JACHJO010000007	Nocardiopsis algeriensis strain CECT 8712 Ga0436954_07, whole genome shotgun sequence
22. NZ_CP015515	Rathayibacter tritici strain NCPPB 1953 chromosome, complete genome
23. NZ_AUBI01000011	Acetobacter nitrogenifigens DSM 23921 = LMG 23498 strain DSM 23921 G442DRAFT_scaffold00010.10_C, whole genome shotgun sequence
24. NZ_AP019807	Microbulbifer sp. GL-2
25. NZ_VLHY01000008	Pseudomonas syringae strain P12854 CFBP1285408, whole genome shotgun sequence
26. NZ_CP006256	Pseudomonas syringae pv. syringae HS191 chromosome, complete genome
27. NW_001820823	Sclerotinia sclerotiorum 1980 UF-70 scaffold_13 genomic scaffold, whole genome shotgun sequence
28. NW_023336283	Aspergillus tubingensis WU-2223L DNA, scaffold_9, whole genome shotgun sequence
29. NW_001820834	Sclerotinia sclerotiorum 1980 UF-70 scaffold_2 genomic scaffold, whole genome shotgun sequence
30. NW_003299165	Microsporum canis CBS 113480 supercont1.5 genomic scaffold, whole genome shotgun sequence


Details:

>>
1. NW_011942164
Source: Metarhizium robertsii ARSEF 23 MAA_Scf_23, whole genome shotgun sequence
Type: NRPS-like,T1PKS
Number of proteins with BLAST hits to this cluster: 3
Cumulative BLAST score: 1381

Table of genes, locations, strands and annotations of subject cluster:

Table of Blast hits (query gene, subject gene, %identity, blast score, %coverage, e-value):
input.path1.gene480	MAA_10222	37	616	114.5077720207254	3.7e-174
input.path1.gene481	MAA_10221	36	74	72.62569832402235	9.8e-12
input.path1.gene482	MAA_10220	31	691	107.70688566013898	2.6e-196


>>
2. NW_014574700
Source: Metarhizium brunneum ARSEF 3297 chromosome Unknown scaffold_17, whole genome shotgun sequence
Type: NRPS-like,T1PKS
Number of proteins with BLAST hits to this cluster: 3
Cumulative BLAST score: 1377

Table of genes, locations, strands and annotations of subject cluster:

Table of Blast hits (query gene, subject gene, %identity, blast score, %coverage, e-value):
input.path1.gene480	MBR_08561	36	611	114.5077720207254	1.6e-172
input.path1.gene481	MBR_08562	35	74	72.62569832402235	1.3e-11
input.path1.gene482	MBR_08563	31	692	107.70688566013898	1.1e-196


>>
3. NC_016459
Source: Thermothielavioides terrestris NRRL 8126 chromosome 3, complete sequence
Type: NRPS-like,T1PKS
Number of proteins with BLAST hits to this cluster: 3
Cumulative BLAST score: 1483

Table of genes, locations, strands and annotations of subject cluster:

Table of Blast hits (query gene, subject gene, %identity, blast score, %coverage, e-value):
input.path1.gene480	THITE_2155249	37	646	112.84974093264249	4.4e-183
input.path1.gene481	THITE_76326	41	94	73.74301675977654	1.2e-17
input.path1.gene482	THITE_170170	35	743	89.07138344914719	3.3e-212


>>
4. NW_008481799
Source: Cyphellophora europaea CBS 101466 unplaced genomic scaffold supercont1.11, whole genome shotgun sequence
Type: NRPS-like,T1PKS
Number of proteins with BLAST hits to this cluster: 2
Cumulative BLAST score: 841

Table of genes, locations, strands and annotations of subject cluster:

Table of Blast hits (query gene, subject gene, %identity, blast score, %coverage, e-value):
input.path1.gene480	HMPREF1541_09825	31	459	111.29533678756476	7.6e-127
input.path1.gene482	HMPREF1541_09820	35	382	46.11497157296272	1.9e-103


>>
5. NW_013550604
Source: Rhinocladiella mackenziei CBS 650.93 unplaced genomic scaffold supercont1.1, whole genome shotgun sequence
Type: NRPS-like,T1PKS
Number of proteins with BLAST hits to this cluster: 2
Cumulative BLAST score: 813

Table of genes, locations, strands and annotations of subject cluster:

Table of Blast hits (query gene, subject gene, %identity, blast score, %coverage, e-value):
input.path1.gene480	Z518_00501	31	437	111.0880829015544	4.1e-120
input.path1.gene482	Z518_00509	34	376	46.304485154769424	1.4e-101


>>
6. CM000593
Source: Fusarium oxysporum f. sp. lycopersici 4287 chromosome 5, whole genome shotgun sequence
Type: NRPS-like,T1PKS
Number of proteins with BLAST hits to this cluster: 2
Cumulative BLAST score: 760

Table of genes, locations, strands and annotations of subject cluster:

Table of Blast hits (query gene, subject gene, %identity, blast score, %coverage, e-value):
input.path1.gene480	FOXG_15239	32	396	88.60103626943005	8e-108
input.path1.gene482	FOXG_15248	34	364	44.47252053063803	5.5e-98


>>
7. NC_030990
Source: Fusarium oxysporum f. sp. lycopersici 4287 chromosome 5, whole genome shotgun sequence
Type: NRPS-like,T1PKS
Number of proteins with BLAST hits to this cluster: 2
Cumulative BLAST score: 760

Table of genes, locations, strands and annotations of subject cluster:

Table of Blast hits (query gene, subject gene, %identity, blast score, %coverage, e-value):
input.path1.gene480	FOXG_15239	32	396	88.60103626943005	8e-108
input.path1.gene482	FOXG_15248	34	364	44.47252053063803	5.5e-98


>>
8. NC_036624
Source: Fusarium fujikuroi IMI 58289 draft genome, chromosome FFUJ_chr03
Type: NRPS-like,T1PKS
Number of proteins with BLAST hits to this cluster: 2
Cumulative BLAST score: 749

Table of genes, locations, strands and annotations of subject cluster:

Table of Blast hits (query gene, subject gene, %identity, blast score, %coverage, e-value):
input.path1.gene480	FFUJ_02115	32	388	88.60103626943005	1.7e-105
input.path1.gene482	FFUJ_02105	34	361	44.598862918509155	3.6e-97


>>
9. NW_022983866
Source: Arthroderma uncinatum strain CBS 119779 chromosome Unknown scaffold4, whole genome shotgun sequence
Type: T1PKS
Number of proteins with BLAST hits to this cluster: 2
Cumulative BLAST score: 732

Table of genes, locations, strands and annotations of subject cluster:

Table of Blast hits (query gene, subject gene, %identity, blast score, %coverage, e-value):
input.path1.gene482	GIQ15_05697	37	344	42.57738471257107	7.7e-92
input.path1.gene483	GIQ15_05703	33	388	72.63479145473042	1.7e-105


>>
10. NW_003345200
Source: Nannizzia gypsea CBS 118893 supercont1.2 genomic scaffold, whole genome shotgun sequence
Type: NRPS-like,T1PKS
Number of proteins with BLAST hits to this cluster: 2
Cumulative BLAST score: 690

Table of genes, locations, strands and annotations of subject cluster:

Table of Blast hits (query gene, subject gene, %identity, blast score, %coverage, e-value):
input.path1.gene480	MGYG_02804	31	393	92.53886010362694	8.8e-107
input.path1.gene482	MGYG_02807	33	297	41.88250157927985	1.1e-77


>>
11. NZ_CP012159
Source: Chondromyces crocatus strain Cm c5 chromosome, complete genome
Type: NRPS,T1PKS
Number of proteins with BLAST hits to this cluster: 2
Cumulative BLAST score: 687

Table of genes, locations, strands and annotations of subject cluster:

Table of Blast hits (query gene, subject gene, %identity, blast score, %coverage, e-value):
input.path1.gene482	CMC5_RS27400	34	305	38.97662665824384	2.3e-80
input.path1.gene483	CMC5_RS27420	37	382	63.377416073245165	1.2e-103


>>
12. NZ_FOKG01000022
Source: Amycolatopsis marina strain CGMCC 4.3568, whole genome shotgun sequence
Type: T1PKS
Number of proteins with BLAST hits to this cluster: 2
Cumulative BLAST score: 614

Table of genes, locations, strands and annotations of subject cluster:

Table of Blast hits (query gene, subject gene, %identity, blast score, %coverage, e-value):
input.path1.gene482	BM199_RS27525	38	230	22.109917877447884	1.6e-57
input.path1.gene483	BM199_RS27470	38	384	64.59816887080366	4.2e-104


>>
13. NZ_CP022434
Source: Nocardiopsis dassonvillei strain HZNU_N_1 chromosome, complete genome
Type: T1PKS
Number of proteins with BLAST hits to this cluster: 2
Cumulative BLAST score: 603

Table of genes, locations, strands and annotations of subject cluster:

Table of Blast hits (query gene, subject gene, %identity, blast score, %coverage, e-value):
input.path1.gene482	CGQ36_RS06160	31	272	43.52495262160455	2.8e-70
input.path1.gene483	CGQ36_RS06175	34	331	66.9379450661241	3.2e-88


>>
14. NZ_KB906854
Source: Nocardiopsis sp. CNS-639 strain CNS639 G011DRAFT_scaffold00001.1, whole genome shotgun sequence
Type: T1PKS
Number of proteins with BLAST hits to this cluster: 2
Cumulative BLAST score: 603

Table of genes, locations, strands and annotations of subject cluster:

Table of Blast hits (query gene, subject gene, %identity, blast score, %coverage, e-value):
input.path1.gene482	G011_RS0100355	31	272	43.52495262160455	2.8e-70
input.path1.gene483	G011_RS0100370	34	331	66.9379450661241	3.2e-88


>>
15. NW_003345193
Source: Nannizzia gypsea CBS 118893 supercont1.9 genomic scaffold, whole genome shotgun sequence
Type: NRPS-like,T1PKS
Number of proteins with BLAST hits to this cluster: 2
Cumulative BLAST score: 587

Table of genes, locations, strands and annotations of subject cluster:

Table of Blast hits (query gene, subject gene, %identity, blast score, %coverage, e-value):
input.path1.gene480	MGYG_08311	31	374	94.19689119170984	4.2e-101
input.path1.gene482	MGYG_08317	34	213	23.499684144030322	1.2e-52


>>
16. NZ_FOIB01000013
Source: Myxococcus fulvus strain DSM 16525, whole genome shotgun sequence
Type: NRPS,T1PKS
Number of proteins with BLAST hits to this cluster: 2
Cumulative BLAST score: 583

Table of genes, locations, strands and annotations of subject cluster:

Table of Blast hits (query gene, subject gene, %identity, blast score, %coverage, e-value):
input.path1.gene482	BMY20_RS37625	35	217	23.373341756159192	1.4e-53
input.path1.gene483	BMY20_RS37655	36	366	66.02238046795524	1.2e-98


>>
17. NZ_LWLA01000014
Source: Streptomyces sp. CB03911 scaffold21, whole genome shotgun sequence
Type: PKS-like,T1PKS
Number of proteins with BLAST hits to this cluster: 2
Cumulative BLAST score: 579

Table of genes, locations, strands and annotations of subject cluster:

Table of Blast hits (query gene, subject gene, %identity, blast score, %coverage, e-value):
input.path1.gene482	A6A07_RS19125	35	217	23.05748578648136	1.4e-53
input.path1.gene483	A6A07_RS19185	37	362	66.02238046795524	1.3e-97


>>
18. NZ_CP028490
Source: Pseudomonas syringae pv. atrofaciens strain LMG5095 isolate Triticum aestivum (Aestivum Group) chromosome, complete genome
Type: transAT-PKS
Number of proteins with BLAST hits to this cluster: 2
Cumulative BLAST score: 577

Table of genes, locations, strands and annotations of subject cluster:
DA456_RS24140	DA456_RS24140	5367893	5370046	+	ATP-dependent_helicase_HrpB
DA456_RS24145	DA456_RS24145	5370087	5370450	+	NAD-dependent_deacylase
DA456_RS24220	DA456_RS24220	5407095	5407316	+	TauD/TfdA_family_dioxygenase

Table of Blast hits (query gene, subject gene, %identity, blast score, %coverage, e-value):
input.path1.gene482	DA456_RS24185	35	208	24.19456727732154	3.9e-51
input.path1.gene483	DA456_RS24115	34	243	46.1851475076297	8.8e-62
input.path1.gene483	DA456_RS24140	31	126	31.943031536113935	1.6e-26


>>
19. NZ_BAZE01000009
Source: Nocardiopsis alba strain TP-A0876, whole genome shotgun sequence
Type: T1PKS
Number of proteins with BLAST hits to this cluster: 2
Cumulative BLAST score: 560

Table of genes, locations, strands and annotations of subject cluster:
TR33_RS17250	TR33_RS17250	105192	105766	+	ATP-binding_protein

Table of Blast hits (query gene, subject gene, %identity, blast score, %coverage, e-value):
input.path1.gene482	TR33_RS17260	34	211	22.236260265319014	7.8e-52
input.path1.gene483	TR33_RS17275	35	349	67.34486266531027	1.1e-93


>>
20. NC_018524
Source: Nocardiopsis alba ATCC BAA-2165, complete sequence
Type: T1PKS
Number of proteins with BLAST hits to this cluster: 2
Cumulative BLAST score: 559

Table of genes, locations, strands and annotations of subject cluster:
B005_RS15195	B005_RS15195	3485572	3487176	-	alpha/beta_fold_hydrolase
B005_RS15290	B005_RS15290	3523510	3524262	+	SDR_family_oxidoreductase

Table of Blast hits (query gene, subject gene, %identity, blast score, %coverage, e-value):
input.path1.gene482	B005_RS15260	34	210	22.236260265319014	1e-51
input.path1.gene483	B005_RS15275	35	349	67.34486266531027	1.1e-93


>>
21. NZ_JACHJO010000007
Source: Nocardiopsis algeriensis strain CECT 8712 Ga0436954_07, whole genome shotgun sequence
Type: T1PKS
Number of proteins with BLAST hits to this cluster: 2
Cumulative BLAST score: 556

Table of genes, locations, strands and annotations of subject cluster:
FHS13_RS13090	FHS13_RS13090	234749	236771	+	tetratricopeptide_repeat_protein

Table of Blast hits (query gene, subject gene, %identity, blast score, %coverage, e-value):
input.path1.gene482	FHS13_RS13105	38	215	21.857233101705624	4.1e-53
input.path1.gene483	FHS13_RS13080	36	341	66.9379450661241	4e-91


>>
22. NZ_CP015515
Source: Rathayibacter tritici strain NCPPB 1953 chromosome, complete genome
Type: NRPS,PKS-like,T3PKS,transAT-PKS,transAT-PKS-like
Number of proteins with BLAST hits to this cluster: 2
Cumulative BLAST score: 545

Table of genes, locations, strands and annotations of subject cluster:
A6122_RS16930	A6122_RS16930	915702	916007	-	hypothetical_protein
A6122_RS16935	A6122_RS16935	916125	916271	-	peptidoglycan-binding_protein
A6122_RS16940	A6122_RS16940	916377	916697	-	peptidoglycan-binding_protein

Table of Blast hits (query gene, subject gene, %identity, blast score, %coverage, e-value):
input.path1.gene482	A6122_RS16395	36	212	23.94188250157928	2.7e-52
input.path1.gene483	A6122_RS04035	34	333	70.09155645981689	6.4e-89


>>
23. NZ_AUBI01000011
Source: Acetobacter nitrogenifigens DSM 23921 = LMG 23498 strain DSM 23921 G442DRAFT_scaffold00010.10_C, whole genome shotgun sequence
Type: T1PKS
Number of proteins with BLAST hits to this cluster: 2
Cumulative BLAST score: 489

Table of genes, locations, strands and annotations of subject cluster:

Table of Blast hits (query gene, subject gene, %identity, blast score, %coverage, e-value):
input.path1.gene482	G442_RS0112720	31	293	45.35691724573594	1.6e-76
input.path1.gene483	G442_RS0112680	32	196	46.1851475076297	1.2e-47


>>
24. NZ_AP019807
Source: Microbulbifer sp. GL-2
Type: NRPS,T1PKS
Number of proteins with BLAST hits to this cluster: 2
Cumulative BLAST score: 452

Table of genes, locations, strands and annotations of subject cluster:

Table of Blast hits (query gene, subject gene, %identity, blast score, %coverage, e-value):
input.path1.gene482	GL2_RS16115	37	222	22.046746683512318	4.4e-55
input.path1.gene483	GL2_RS16105	33	230	46.69379450661241	5.9e-58


>>
25. NZ_VLHY01000008
Source: Pseudomonas syringae strain P12854 CFBP1285408, whole genome shotgun sequence
Type: transAT-PKS
Number of proteins with BLAST hits to this cluster: 2
Cumulative BLAST score: 406

Table of genes, locations, strands and annotations of subject cluster:
PspP12854CL_RS15295	PspP12854CL_RS15295	114282	114604	+	iron_dicitrate_transport_regulator_FecR
PspP12854CL_RS15365	PspP12854CL_RS15365	151291	151512	+	TauD/TfdA_family_dioxygenase

Table of Blast hits (query gene, subject gene, %identity, blast score, %coverage, e-value):
input.path1.gene482	PspP12854CL_RS15330	35	208	24.19456727732154	3.9e-51
input.path1.gene483	PspP12854CL_RS15290	31	198	44.557477110885046	3.2e-48


>>
26. NZ_CP006256
Source: Pseudomonas syringae pv. syringae HS191 chromosome, complete genome
Type: transAT-PKS
Number of proteins with BLAST hits to this cluster: 2
Cumulative BLAST score: 403

Table of genes, locations, strands and annotations of subject cluster:
PSYRH_RS04540	PSYRH_RS04540	993104	993426	+	NAD-dependent_deacylase
PSYRH_RS26410	PSYRH_RS26410	1030122	1030343	+	TauD/TfdA_family_dioxygenase

Table of Blast hits (query gene, subject gene, %identity, blast score, %coverage, e-value):
input.path1.gene482	PSYRH_RS04570	35	208	24.19456727732154	3.9e-51
input.path1.gene483	PSYRH_RS04535	31	195	44.557477110885046	2.7e-47


>>
27. NW_001820823
Source: Sclerotinia sclerotiorum 1980 UF-70 scaffold_13 genomic scaffold, whole genome shotgun sequence
Type: NRPS-like,T1PKS,other
Number of proteins with BLAST hits to this cluster: 2
Cumulative BLAST score: 852

Table of genes, locations, strands and annotations of subject cluster:

Table of Blast hits (query gene, subject gene, %identity, blast score, %coverage, e-value):
input.path1.gene480	SS1G_09449	40	723	106.6321243523316	3.7e-206
input.path1.gene481	SS1G_09448	45	129	91.06145251396647	3.4e-28


>>
28. NW_023336283
Source: Aspergillus tubingensis WU-2223L DNA, scaffold_9, whole genome shotgun sequence
Type: NRPS-like
Number of proteins with BLAST hits to this cluster: 2
Cumulative BLAST score: 706

Table of genes, locations, strands and annotations of subject cluster:

Table of Blast hits (query gene, subject gene, %identity, blast score, %coverage, e-value):
input.path1.gene480	AtWU_08710	37	639	110.77720207253886	4.1e-181
input.path1.gene481	AtWU_08709	32	67	81.00558659217877	2e-09


>>
29. NW_001820834
Source: Sclerotinia sclerotiorum 1980 UF-70 scaffold_2 genomic scaffold, whole genome shotgun sequence
Type: NRPS-like
Number of proteins with BLAST hits to this cluster: 2
Cumulative BLAST score: 648

Table of genes, locations, strands and annotations of subject cluster:

Table of Blast hits (query gene, subject gene, %identity, blast score, %coverage, e-value):
input.path1.gene480	SS1G_01217	33	584	111.50259067357513	2.7e-164
input.path1.gene481	SS1G_01219	33	64	63.687150837988824	1.7e-08


>>
30. NW_003299165
Source: Microsporum canis CBS 113480 supercont1.5 genomic scaffold, whole genome shotgun sequence
Type: NRPS,indole
Number of proteins with BLAST hits to this cluster: 2
Cumulative BLAST score: 494

Table of genes, locations, strands and annotations of subject cluster:

Table of Blast hits (query gene, subject gene, %identity, blast score, %coverage, e-value):
input.path1.gene484	MCYG_05764	54	435	95.70093457943926	6.6e-120
input.path1.gene485	MCYG_05766	54	59	26.359832635983267	5.7e-07
