## Supplementary Results for "A draft genome of the ascomycotal fungal species *Pseudopithomyces maydicus* (family *Didymosphaeriaceae*)": contig_9_c1.txt

ClusterBlast scores for contig_9

Table of genes, locations, strands and annotations of query cluster:
input.path1.gene386	1341733	1344667	-		
input.path1.gene387	1345508	1347838	-		
input.path1.gene388	1348257	1349376	-		
input.path1.gene389	1350289	1352207	-		
input.path1.gene390	1357441	1358404	+		
input.path1.gene391	1359897	1364011	-		
input.path1.gene392	1364647	1365403	-		
input.path1.gene393	1365823	1367967	-		
input.path1.gene394	1369122	1371124	+		
input.path1.gene395	1374591	1376148	+		
input.path1.gene396	1377489	1377751	-		
input.path1.gene397	1378137	1381941	+		
input.path1.gene398	1382563	1388922	+		
input.path1.gene399	1389309	1399185	+		
input.path1.gene400	1401788	1402819	-		
input.path1.gene401	1403089	1406156	-		
input.path1.gene402	1407443	1407775	-		
input.path1.gene403	1408109	1411452	-		
input.path1.gene404	1411775	1412992	+		
input.path1.gene405	1413825	1415634	-		


Significant hits: 
1. NW_017971440	Talaromyces atroroseus strain IBT 11181 chromosome Unknown scaffold_8, whole genome shotgun sequence
2. CM004174	Drechmeria coniospora strain ARSEF 6962 chromosome 01, whole genome shotgun sequence
3. NW_003315027	Verticillium alfalfae VaMs.102 supercont1.12 genomic scaffold, whole genome shotgun sequence
4. NT_165937	Aspergillus terreus NIH2624 scaffold_14 genomic scaffold, whole genome shotgun sequence
5. NT_165929	Aspergillus terreus NIH2624 scaffold_6 genomic scaffold, whole genome shotgun sequence
6. NC_035790	Pochonia chlamydosporia 170 chromosome 1, whole genome shotgun sequence
7. NC_038012	Fusarium venenatum strain A3/5 genome assembly, chromosome: I
8. NW_017387250	Fonsecaea erecta strain CBS 125763 chromosome Unknown Scaff7, whole genome shotgun sequence
9. NW_013550602	Rhinocladiella mackenziei CBS 650.93 unplaced genomic scaffold supercont1.3, whole genome shotgun sequence
10. NW_022984629	Aspergillus tanneri strain NIH1004 chromosome Unknown AapTan_2_pilon_pilon_pilon, whole genome shotgun sequence
11. NT_107012	Aspergillus nidulans FGSC A4 chromosome II map unlocalized scaffold_4, whole genome shotgun sequence
12. NC_007199	Aspergillus fumigatus Af293 chromosome 6, whole genome shotgun sequence
13. NW_022983863	Arthroderma uncinatum strain CBS 119779 chromosome Unknown scaffold1, whole genome shotgun sequence
14. NW_019716257	Ramularia collo-cygni strain URUG2 genome assembly, contig: RCC_scaffold03, whole genome shotgun sequence
15. NW_019716264	Ramularia collo-cygni strain URUG2 genome assembly, contig: RCC_scaffold10, whole genome shotgun sequence
16. CM000576	Fusarium graminearum PH-1 chromosome 3, whole genome shotgun sequence
17. NC_026476	Fusarium graminearum PH-1 chromosome 3, whole genome shotgun sequence
18. NW_003345200	Nannizzia gypsea CBS 118893 supercont1.2 genomic scaffold, whole genome shotgun sequence
19. NW_015971668	Fonsecaea multimorphosa CBS 102226 unplaced genomic scaffold supercont1.34, whole genome shotgun sequence
20. NW_022984633	Aspergillus tanneri strain NIH1004 chromosome Unknown AapTan_6_pilon_pilon_pilon, whole genome shotgun sequence
21. NW_011371358	Paracoccidioides brasiliensis Pb18 unplaced genomic scaffold supercont2.1, whole genome shotgun sequence
22. NW_014013634	Trichoderma atroviride IMI 206040 chromosome Unknown TRIATcontig_22, whole genome shotgun sequence
23. CM000601	Fusarium oxysporum f. sp. lycopersici 4287 chromosome 13, whole genome shotgun sequence
24. NC_030998	Fusarium oxysporum f. sp. lycopersici 4287 chromosome 13, whole genome shotgun sequence
25. NW_022983863	Arthroderma uncinatum strain CBS 119779 chromosome Unknown scaffold1, whole genome shotgun sequence
26. NW_001939250	Pyrenophora tritici-repentis Pt-1C-BFP supercont1.7, whole genome shotgun sequence
27. NW_011942151	Metarhizium robertsii ARSEF 23 MAA_Scf_11, whole genome shotgun sequence
28. NW_008481809	Cyphellophora europaea CBS 101466 unplaced genomic scaffold supercont1.3, whole genome shotgun sequence
29. CP042185	Venturia effusa strain albino chromosome 1, complete sequence
30. NT_165979	Chaetomium globosum CBS 148.51 scaffold_4 genomic scaffold, whole genome shotgun sequence
31. NC_036632	Fusarium fujikuroi IMI 58289 draft genome, chromosome FFUJ_chr11
32. NW_022194798	Fusarium proliferatum ET1 genome assembly, contig: FPRO_scaffold009, whole genome shotgun sequence
33. NC_018206	Zymoseptoria tritici IPO323 chromosome 13, whole genome shotgun sequence
34. CM004174	Drechmeria coniospora strain ARSEF 6962 chromosome 01, whole genome shotgun sequence
35. NW_006911260	Baudoinia panamericana UAMH 10762 unplaced genomic scaffold BAUCOscaffold_4, whole genome shotgun sequence
36. NW_007361004	Glarea lozoyensis ATCC 20868 chromosome Unknown GLAREA9, whole genome shotgun sequence
37. NW_019716255	Ramularia collo-cygni strain URUG2 genome assembly, contig: RCC_scaffold01, whole genome shotgun sequence
38. NW_006763039	Marssonina brunnea f. sp. 'multigermtubi' MB_m1 unplaced genomic scaffold M6_S00001, whole genome shotgun sequence
39. NC_038013	Fusarium venenatum strain A3/5 genome assembly, chromosome: II
40. CM000575	Fusarium graminearum PH-1 chromosome 2, whole genome shotgun sequence
41. NC_026475	Fusarium graminearum PH-1 chromosome 2, whole genome shotgun sequence
42. CP051141	Peltaster fructicola strain LNHT1506 chromosome 3
43. NW_017264202	Xylona heveae TC161 unplaced genomic scaffold L228scaffold_10, whole genome shotgun sequence
44. NC_016947	Mycobacterium intracellulare MOTT-02, complete sequence
45. NC_016946	Mycobacterium intracellulare ATCC 13950, complete sequence
46. NZ_CP023149	Mycobacterium intracellulare strain FLAC0181 chromosome, complete genome
47. NC_018612	Mycobacterium intracellulare subsp. intracellulare MTCC 9506, complete sequence
48. NC_016948	Mycobacterium paraintracellulare, complete sequence
49. NZ_KL446995	Mycobacterium sp. TKK-01-0059 supercont2.3, whole genome shotgun sequence
50. NW_022984635	Aspergillus tanneri strain NIH1004 chromosome Unknown AapTan_8_pilon_pilon_pilon, whole genome shotgun sequence


Details:

>>
1. NW_017971440
Source: Talaromyces atroroseus strain IBT 11181 chromosome Unknown scaffold_8, whole genome shotgun sequence
Type: NRPS
Number of proteins with BLAST hits to this cluster: 4
Cumulative BLAST score: 2548

Table of genes, locations, strands and annotations of subject cluster:

Table of Blast hits (query gene, subject gene, %identity, blast score, %coverage, e-value):
input.path1.gene394	UA08_06361	32	62	47.368421052631575	9.1e-08
input.path1.gene397	UA08_06365	32	371	99.74391805377721	2.2e-100
input.path1.gene398	UA08_06365	37	479	58.18553888130968	1.4e-132
input.path1.gene399	UA08_06365	36	1636	112.07635372029607	0.0


>>
2. CM004174
Source: Drechmeria coniospora strain ARSEF 6962 chromosome 01, whole genome shotgun sequence
Type: NRPS
Number of proteins with BLAST hits to this cluster: 4
Cumulative BLAST score: 1709

Table of genes, locations, strands and annotations of subject cluster:

Table of Blast hits (query gene, subject gene, %identity, blast score, %coverage, e-value):
input.path1.gene394	DCS_02866	37	58	28.34008097165992	1.3e-06
input.path1.gene397	DCS_02863	33	220	58.25864276568502	8.3e-55
input.path1.gene398	DCS_02860	31	381	65.89358799454298	5.3e-103
input.path1.gene398	DCS_02863	38	312	33.42428376534789	2.3e-82
input.path1.gene399	DCS_02860	33	738	55.940786910790806	1.7e-210


>>
3. NW_003315027
Source: Verticillium alfalfae VaMs.102 supercont1.12 genomic scaffold, whole genome shotgun sequence
Type: NRPS
Number of proteins with BLAST hits to this cluster: 3
Cumulative BLAST score: 3181

Table of genes, locations, strands and annotations of subject cluster:

Table of Blast hits (query gene, subject gene, %identity, blast score, %coverage, e-value):
input.path1.gene397	VDBG_08383	34	201	44.81434058898848	2.3e-49
input.path1.gene398	VDBG_08381	31	440	65.41609822646657	9.5e-121
input.path1.gene398	VDBG_08382	32	423	58.04911323328786	7.1e-116
input.path1.gene399	VDBG_08382	31	723	61.3167121153097	7.6e-206
input.path1.gene399	VDBG_08381	31	709	65.64082586677054	1.1e-201
input.path1.gene399	VDBG_08380	32	685	56.369302687962595	3e-194


>>
4. NT_165937
Source: Aspergillus terreus NIH2624 scaffold_14 genomic scaffold, whole genome shotgun sequence
Type: NRPS
Number of proteins with BLAST hits to this cluster: 3
Cumulative BLAST score: 2893

Table of genes, locations, strands and annotations of subject cluster:

Table of Blast hits (query gene, subject gene, %identity, blast score, %coverage, e-value):
input.path1.gene397	ATEG_09068	32	337	97.0550576184379	2.7e-90
input.path1.gene398	ATEG_09064	33	469	60.436562073669855	1.5e-129
input.path1.gene398	ATEG_09068	35	432	57.094133697135064	1.5e-118
input.path1.gene399	ATEG_09064	31	915	79.89871445266849	1.4e-263
input.path1.gene399	ATEG_09068	36	740	48.5391507596416	4.6e-211


>>
5. NT_165929
Source: Aspergillus terreus NIH2624 scaffold_6 genomic scaffold, whole genome shotgun sequence
Type: NRPS
Number of proteins with BLAST hits to this cluster: 3
Cumulative BLAST score: 2272

Table of genes, locations, strands and annotations of subject cluster:

Table of Blast hits (query gene, subject gene, %identity, blast score, %coverage, e-value):
input.path1.gene397	ATEG_04322	34	275	62.099871959026885	2.2e-71
input.path1.gene398	ATEG_04323	31	584	87.92633015006821	2.4e-164
input.path1.gene398	ATEG_04322	31	381	60.436562073669855	4e-103
input.path1.gene399	ATEG_04322	33	579	40.20257109466303	1.8e-162
input.path1.gene399	ATEG_04323	32	453	37.12504869497468	1.9e-124


>>
6. NC_035790
Source: Pochonia chlamydosporia 170 chromosome 1, whole genome shotgun sequence
Type: NRPS
Number of proteins with BLAST hits to this cluster: 3
Cumulative BLAST score: 3179

Table of genes, locations, strands and annotations of subject cluster:

Table of Blast hits (query gene, subject gene, %identity, blast score, %coverage, e-value):
input.path1.gene398	VFPPC_00082	32	412	59.276944065484315	1.6e-112
input.path1.gene398	VFPPC_17432	33	404	59.61800818553888	5.8e-110
input.path1.gene399	VFPPC_17432	31	1067	91.58550837553564	1.6e-309
input.path1.gene399	VFPPC_00082	32	1060	90.84534476042072	3.4e-307
input.path1.gene405	VFPPC_12683	35	236	94.10569105691057	7e-60


>>
7. NC_038012
Source: Fusarium venenatum strain A3/5 genome assembly, chromosome: I
Type: NRPS,T1PKS
Number of proteins with BLAST hits to this cluster: 3
Cumulative BLAST score: 1783

Table of genes, locations, strands and annotations of subject cluster:

Table of Blast hits (query gene, subject gene, %identity, blast score, %coverage, e-value):
input.path1.gene398	FVRRES_00020	34	428	58.663028649386085	2.2e-117
input.path1.gene399	FVRRES_00020	31	1231	113.75146084924035	0.0
input.path1.gene401	FVRRES_00024	32	124	74.23469387755102	1.8e-26


>>
8. NW_017387250
Source: Fonsecaea erecta strain CBS 125763 chromosome Unknown Scaff7, whole genome shotgun sequence
Type: terpene
Number of proteins with BLAST hits to this cluster: 2
Cumulative BLAST score: 544

Table of genes, locations, strands and annotations of subject cluster:

Table of Blast hits (query gene, subject gene, %identity, blast score, %coverage, e-value):
input.path1.gene388	AYL99_08261	40	199	102.8673835125448	4.1e-49
input.path1.gene389	AYL99_08260	50	345	96.55172413793103	8.8e-93


>>
9. NW_013550602
Source: Rhinocladiella mackenziei CBS 650.93 unplaced genomic scaffold supercont1.3, whole genome shotgun sequence
Type: terpene
Number of proteins with BLAST hits to this cluster: 2
Cumulative BLAST score: 534

Table of genes, locations, strands and annotations of subject cluster:

Table of Blast hits (query gene, subject gene, %identity, blast score, %coverage, e-value):
input.path1.gene388	Z518_04126	45	223	94.26523297491039	2.7e-56
input.path1.gene389	Z518_04127	46	311	103.94088669950739	1.8e-82


>>
10. NW_022984629
Source: Aspergillus tanneri strain NIH1004 chromosome Unknown AapTan_2_pilon_pilon_pilon, whole genome shotgun sequence
Type: NRPS,T1PKS,indole
Number of proteins with BLAST hits to this cluster: 2
Cumulative BLAST score: 4288

Table of genes, locations, strands and annotations of subject cluster:

Table of Blast hits (query gene, subject gene, %identity, blast score, %coverage, e-value):
input.path1.gene398	ATNIH1004_001011	31	668	102.387448840382	2.2e-189
input.path1.gene398	ATNIH1004_001010	31	619	101.09140518417463	8.8e-175
input.path1.gene398	ATNIH1004_001002	33	367	49.24965893587995	7.8e-99
input.path1.gene398	ATNIH1004_001005	31	345	48.158253751705324	2.4e-92
input.path1.gene398	ATNIH1004_001003	31	327	50.272851296043655	9e-87
input.path1.gene399	ATNIH1004_001011	33	755	58.16127775613557	1.4e-215
input.path1.gene399	ATNIH1004_001010	33	417	32.60615504479938	8.9e-114
input.path1.gene399	ATNIH1004_001002	34	398	28.74951305025321	7.3e-108
input.path1.gene399	ATNIH1004_001003	35	392	25.165562913907287	4e-106


>>
11. NT_107012
Source: Aspergillus nidulans FGSC A4 chromosome II map unlocalized scaffold_4, whole genome shotgun sequence
Type: NRPS
Number of proteins with BLAST hits to this cluster: 2
Cumulative BLAST score: 2005

Table of genes, locations, strands and annotations of subject cluster:

Table of Blast hits (query gene, subject gene, %identity, blast score, %coverage, e-value):
input.path1.gene398	AN3495.2	32	387	56.68485675306958	7.3e-105
input.path1.gene398	AN3496.2	32	377	57.094133697135064	7.6e-102
input.path1.gene399	AN3496.2	33	761	57.615894039735096	2.5e-217
input.path1.gene399	AN3495.2	33	480	34.35917413322945	1.5e-132


>>
12. NC_007199
Source: Aspergillus fumigatus Af293 chromosome 6, whole genome shotgun sequence
Type: NRPS
Number of proteins with BLAST hits to this cluster: 2
Cumulative BLAST score: 1403

Table of genes, locations, strands and annotations of subject cluster:

Table of Blast hits (query gene, subject gene, %identity, blast score, %coverage, e-value):
input.path1.gene398	AFUA_6G12050	33	329	50.0	1.8e-87
input.path1.gene399	AFUA_6G12080	32	727	56.641994546162834	5.2e-207
input.path1.gene399	AFUA_6G12050	35	347	25.204518893650175	1.5e-92


>>
13. NW_022983863
Source: Arthroderma uncinatum strain CBS 119779 chromosome Unknown scaffold1, whole genome shotgun sequence
Type: NRPS
Number of proteins with BLAST hits to this cluster: 2
Cumulative BLAST score: 1401

Table of genes, locations, strands and annotations of subject cluster:

Table of Blast hits (query gene, subject gene, %identity, blast score, %coverage, e-value):
input.path1.gene398	GIQ15_01912	31	366	58.79945429740792	1.7e-98
input.path1.gene398	GIQ15_01907	32	358	57.43519781718963	3.6e-96
input.path1.gene399	GIQ15_01907	32	677	56.758862485391504	6.2e-192


>>
14. NW_019716257
Source: Ramularia collo-cygni strain URUG2 genome assembly, contig: RCC_scaffold03, whole genome shotgun sequence
Type: NRPS
Number of proteins with BLAST hits to this cluster: 2
Cumulative BLAST score: 1066

Table of genes, locations, strands and annotations of subject cluster:

Table of Blast hits (query gene, subject gene, %identity, blast score, %coverage, e-value):
input.path1.gene398	RCC_02695	35	371	47.40791268758527	4.2e-100
input.path1.gene398	RCC_02693	31	305	48.0218281036835	2.1e-80
input.path1.gene399	RCC_02695	35	390	27.035449941566032	1.5e-105


>>
15. NW_019716264
Source: Ramularia collo-cygni strain URUG2 genome assembly, contig: RCC_scaffold10, whole genome shotgun sequence
Type: NRPS
Number of proteins with BLAST hits to this cluster: 2
Cumulative BLAST score: 1030

Table of genes, locations, strands and annotations of subject cluster:

Table of Blast hits (query gene, subject gene, %identity, blast score, %coverage, e-value):
input.path1.gene398	RCC_12189	31	337	51.637107776261935	5.1e-90
input.path1.gene398	RCC_06743	31	322	50.4774897680764	2.2e-85
input.path1.gene399	RCC_06743	35	371	24.464355278535255	9.5e-100


>>
16. CM000576
Source: Fusarium graminearum PH-1 chromosome 3, whole genome shotgun sequence
Type: NRPS
Number of proteins with BLAST hits to this cluster: 2
Cumulative BLAST score: 1009

Table of genes, locations, strands and annotations of subject cluster:

Table of Blast hits (query gene, subject gene, %identity, blast score, %coverage, e-value):
input.path1.gene398	FGSG_13878	31	658	101.09140518417463	2.2e-186
input.path1.gene399	FGSG_10990	34	351	25.67199065056486	7.8e-94


>>
17. NC_026476
Source: Fusarium graminearum PH-1 chromosome 3, whole genome shotgun sequence
Type: NRPS
Number of proteins with BLAST hits to this cluster: 2
Cumulative BLAST score: 1009

Table of genes, locations, strands and annotations of subject cluster:

Table of Blast hits (query gene, subject gene, %identity, blast score, %coverage, e-value):
input.path1.gene398	FGSG_13878	31	658	101.09140518417463	2.2e-186
input.path1.gene399	FGSG_10990	34	351	25.67199065056486	7.8e-94


>>
18. NW_003345200
Source: Nannizzia gypsea CBS 118893 supercont1.2 genomic scaffold, whole genome shotgun sequence
Type: NRPS,terpene
Number of proteins with BLAST hits to this cluster: 2
Cumulative BLAST score: 755

Table of genes, locations, strands and annotations of subject cluster:

Table of Blast hits (query gene, subject gene, %identity, blast score, %coverage, e-value):
input.path1.gene389	MGYG_02096	47	344	103.44827586206897	1.2e-92
input.path1.gene398	MGYG_02098	32	411	57.36698499317872	2.8e-112


>>
19. NW_015971668
Source: Fonsecaea multimorphosa CBS 102226 unplaced genomic scaffold supercont1.34, whole genome shotgun sequence
Type: NRPS
Number of proteins with BLAST hits to this cluster: 3
Cumulative BLAST score: 737

Table of genes, locations, strands and annotations of subject cluster:

Table of Blast hits (query gene, subject gene, %identity, blast score, %coverage, e-value):
input.path1.gene394	Z520_10940	42	87	42.51012145748988	2.6e-15
input.path1.gene398	Z520_10935	35	343	46.93042291950887	9.3e-92
input.path1.gene401	Z520_10940	51	307	73.46938775510205	2e-81


>>
20. NW_022984633
Source: Aspergillus tanneri strain NIH1004 chromosome Unknown AapTan_6_pilon_pilon_pilon, whole genome shotgun sequence
Type: NRPS
Number of proteins with BLAST hits to this cluster: 2
Cumulative BLAST score: 625

Table of genes, locations, strands and annotations of subject cluster:

Table of Blast hits (query gene, subject gene, %identity, blast score, %coverage, e-value):
input.path1.gene397	ATNIH1004_008076	31	347	102.8169014084507	3.4e-93
input.path1.gene398	ATNIH1004_008080	32	278	43.58799454297408	3.7e-72


>>
21. NW_011371358
Source: Paracoccidioides brasiliensis Pb18 unplaced genomic scaffold supercont2.1, whole genome shotgun sequence
Type: NRPS
Number of proteins with BLAST hits to this cluster: 3
Cumulative BLAST score: 588

Table of genes, locations, strands and annotations of subject cluster:

Table of Blast hits (query gene, subject gene, %identity, blast score, %coverage, e-value):
input.path1.gene394	PADG_00099	32	61	45.34412955465587	1.2e-07
input.path1.gene398	PADG_00102	34	350	46.24829467939973	9.9e-94
input.path1.gene400	PADG_00096	43	177	86.7704280155642	1.5e-42


>>
22. NW_014013634
Source: Trichoderma atroviride IMI 206040 chromosome Unknown TRIATcontig_22, whole genome shotgun sequence
Type: NRPS
Number of proteins with BLAST hits to this cluster: 3
Cumulative BLAST score: 584

Table of genes, locations, strands and annotations of subject cluster:

Table of Blast hits (query gene, subject gene, %identity, blast score, %coverage, e-value):
input.path1.gene394	TRIATDRAFT_90126	35	99	78.54251012145748	3.9e-19
input.path1.gene398	TRIATDRAFT_154060	33	342	45.975443383356065	2.1e-91
input.path1.gene401	TRIATDRAFT_90126	32	143	80.35714285714286	4.9e-32


>>
23. CM000601
Source: Fusarium oxysporum f. sp. lycopersici 4287 chromosome 13, whole genome shotgun sequence
Type: terpene
Number of proteins with BLAST hits to this cluster: 2
Cumulative BLAST score: 1218

Table of genes, locations, strands and annotations of subject cluster:

Table of Blast hits (query gene, subject gene, %identity, blast score, %coverage, e-value):
input.path1.gene388	FOXG_12142	57	322	106.4516129032258	5.5e-86
input.path1.gene389	FOXG_12144_KNB12574.1	56	448	104.67980295566504	7.4e-124
input.path1.gene389	FOXG_12144	56	448	104.67980295566504	7.4e-124


>>
24. NC_030998
Source: Fusarium oxysporum f. sp. lycopersici 4287 chromosome 13, whole genome shotgun sequence
Type: terpene
Number of proteins with BLAST hits to this cluster: 2
Cumulative BLAST score: 1218

Table of genes, locations, strands and annotations of subject cluster:

Table of Blast hits (query gene, subject gene, %identity, blast score, %coverage, e-value):
input.path1.gene388	FOXG_12142	57	322	106.4516129032258	5.5e-86
input.path1.gene389	FOXG_12144_XP_018250619.1	56	448	104.67980295566504	7.4e-124
input.path1.gene389	FOXG_12144	56	448	104.67980295566504	7.4e-124


>>
25. NW_022983863
Source: Arthroderma uncinatum strain CBS 119779 chromosome Unknown scaffold1, whole genome shotgun sequence
Type: NRPS
Number of proteins with BLAST hits to this cluster: 2
Cumulative BLAST score: 841

Table of genes, locations, strands and annotations of subject cluster:

Table of Blast hits (query gene, subject gene, %identity, blast score, %coverage, e-value):
input.path1.gene391	GIQ15_00312	36	361	105.6384742951907	2.3e-97
input.path1.gene399	GIQ15_00309	31	480	36.891312816517335	8.5e-133


>>
26. NW_001939250
Source: Pyrenophora tritici-repentis Pt-1C-BFP supercont1.7, whole genome shotgun sequence
Type: terpene
Number of proteins with BLAST hits to this cluster: 2
Cumulative BLAST score: 832

Table of genes, locations, strands and annotations of subject cluster:

Table of Blast hits (query gene, subject gene, %identity, blast score, %coverage, e-value):
input.path1.gene388	PTRG_07364	56	312	101.4336917562724	3.3e-83
input.path1.gene389	PTRG_07366	65	520	102.70935960591132	2e-145


>>
27. NW_011942151
Source: Metarhizium robertsii ARSEF 23 MAA_Scf_11, whole genome shotgun sequence
Type: NRPS,T1PKS,terpene
Number of proteins with BLAST hits to this cluster: 2
Cumulative BLAST score: 816

Table of genes, locations, strands and annotations of subject cluster:

Table of Blast hits (query gene, subject gene, %identity, blast score, %coverage, e-value):
input.path1.gene387	MAA_06580	65	604	87.05440900562851	1.1e-170
input.path1.gene397	MAA_06584	32	212	57.74647887323944	1.3e-52


>>
28. NW_008481809
Source: Cyphellophora europaea CBS 101466 unplaced genomic scaffold supercont1.3, whole genome shotgun sequence
Type: NRPS
Number of proteins with BLAST hits to this cluster: 2
Cumulative BLAST score: 799

Table of genes, locations, strands and annotations of subject cluster:

Table of Blast hits (query gene, subject gene, %identity, blast score, %coverage, e-value):
input.path1.gene398	HMPREF1541_03025	33	732	101.56889495225103	9.3e-209
input.path1.gene400	HMPREF1541_03028	41	67	40.07782101167315	1.7e-09


>>
29. CP042185
Source: Venturia effusa strain albino chromosome 1, complete sequence
Type: terpene
Number of proteins with BLAST hits to this cluster: 2
Cumulative BLAST score: 783

Table of genes, locations, strands and annotations of subject cluster:

Table of Blast hits (query gene, subject gene, %identity, blast score, %coverage, e-value):
input.path1.gene388	FKW77_005808	55	319	113.26164874551972	3.6e-85
input.path1.gene389	FKW77_005773	56	464	107.14285714285714	1.3e-128


>>
30. NT_165979
Source: Chaetomium globosum CBS 148.51 scaffold_4 genomic scaffold, whole genome shotgun sequence
Type: NRPS,T1PKS
Number of proteins with BLAST hits to this cluster: 2
Cumulative BLAST score: 772

Table of genes, locations, strands and annotations of subject cluster:

Table of Blast hits (query gene, subject gene, %identity, blast score, %coverage, e-value):
input.path1.gene394	CHGG_04474	31	92	97.97570850202429	6.3e-17
input.path1.gene399	CHGG_04477	32	680	61.12193221659524	9.6e-193


>>
31. NC_036632
Source: Fusarium fujikuroi IMI 58289 draft genome, chromosome FFUJ_chr11
Type: terpene
Number of proteins with BLAST hits to this cluster: 2
Cumulative BLAST score: 767

Table of genes, locations, strands and annotations of subject cluster:

Table of Blast hits (query gene, subject gene, %identity, blast score, %coverage, e-value):
input.path1.gene388	FFUJ_11804	56	319	106.4516129032258	2.7e-85
input.path1.gene389	FFUJ_11802	56	448	104.67980295566504	7.4e-124


>>
32. NW_022194798
Source: Fusarium proliferatum ET1 genome assembly, contig: FPRO_scaffold009, whole genome shotgun sequence
Type: terpene
Number of proteins with BLAST hits to this cluster: 2
Cumulative BLAST score: 764

Table of genes, locations, strands and annotations of subject cluster:

Table of Blast hits (query gene, subject gene, %identity, blast score, %coverage, e-value):
input.path1.gene388	FPRO_15514	57	321	106.4516129032258	1.2e-85
input.path1.gene389	FPRO_15512	55	443	104.67980295566504	1.8e-122


>>
33. NC_018206
Source: Zymoseptoria tritici IPO323 chromosome 13, whole genome shotgun sequence
Type: terpene
Number of proteins with BLAST hits to this cluster: 2
Cumulative BLAST score: 756

Table of genes, locations, strands and annotations of subject cluster:

Table of Blast hits (query gene, subject gene, %identity, blast score, %coverage, e-value):
input.path1.gene388	MYCGRDRAFT_106573	55	311	109.31899641577061	9.7e-83
input.path1.gene389	MYCGRDRAFT_51545	55	445	104.92610837438423	4.8e-123


>>
34. CM004174
Source: Drechmeria coniospora strain ARSEF 6962 chromosome 01, whole genome shotgun sequence
Type: NRPS
Number of proteins with BLAST hits to this cluster: 2
Cumulative BLAST score: 747

Table of genes, locations, strands and annotations of subject cluster:

Table of Blast hits (query gene, subject gene, %identity, blast score, %coverage, e-value):
input.path1.gene398	DCS_00047	32	688	102.31923601637108	1.5e-195
input.path1.gene400	DCS_00044	36	59	34.63035019455253	6.1e-07


>>
35. NW_006911260
Source: Baudoinia panamericana UAMH 10762 unplaced genomic scaffold BAUCOscaffold_4, whole genome shotgun sequence
Type: terpene
Number of proteins with BLAST hits to this cluster: 2
Cumulative BLAST score: 740

Table of genes, locations, strands and annotations of subject cluster:

Table of Blast hits (query gene, subject gene, %identity, blast score, %coverage, e-value):
input.path1.gene388	BAUCODRAFT_31870	58	312	102.8673835125448	4.4e-83
input.path1.gene389	BAUCODRAFT_66702	53	428	109.85221674876848	7.9e-118


>>
36. NW_007361004
Source: Glarea lozoyensis ATCC 20868 chromosome Unknown GLAREA9, whole genome shotgun sequence
Type: terpene
Number of proteins with BLAST hits to this cluster: 2
Cumulative BLAST score: 720

Table of genes, locations, strands and annotations of subject cluster:

Table of Blast hits (query gene, subject gene, %identity, blast score, %coverage, e-value):
input.path1.gene388	GLAREA_08042	49	285	111.8279569892473	4.4e-75
input.path1.gene389	GLAREA_08044	54	435	106.1576354679803	8.5e-120


>>
37. NW_019716255
Source: Ramularia collo-cygni strain URUG2 genome assembly, contig: RCC_scaffold01, whole genome shotgun sequence
Type: terpene
Number of proteins with BLAST hits to this cluster: 2
Cumulative BLAST score: 719

Table of genes, locations, strands and annotations of subject cluster:

Table of Blast hits (query gene, subject gene, %identity, blast score, %coverage, e-value):
input.path1.gene388	RCC_00418	56	310	111.4695340501792	2.2e-82
input.path1.gene389	RCC_00415	51	409	109.11330049261083	5e-112


>>
38. NW_006763039
Source: Marssonina brunnea f. sp. 'multigermtubi' MB_m1 unplaced genomic scaffold M6_S00001, whole genome shotgun sequence
Type: terpene
Number of proteins with BLAST hits to this cluster: 2
Cumulative BLAST score: 718

Table of genes, locations, strands and annotations of subject cluster:

Table of Blast hits (query gene, subject gene, %identity, blast score, %coverage, e-value):
input.path1.gene388	MBM_00657	52	258	99.6415770609319	7.5e-67
input.path1.gene389	MBM_00659	56	460	105.91133004926108	2.5e-127


>>
39. NC_038013
Source: Fusarium venenatum strain A3/5 genome assembly, chromosome: II
Type: terpene
Number of proteins with BLAST hits to this cluster: 2
Cumulative BLAST score: 691

Table of genes, locations, strands and annotations of subject cluster:

Table of Blast hits (query gene, subject gene, %identity, blast score, %coverage, e-value):
input.path1.gene388	FVRRES_05978	48	256	107.5268817204301	2.8e-66
input.path1.gene389	FVRRES_05980	55	435	104.43349753694582	5e-120


>>
40. CM000575
Source: Fusarium graminearum PH-1 chromosome 2, whole genome shotgun sequence
Type: terpene
Number of proteins with BLAST hits to this cluster: 2
Cumulative BLAST score: 679

Table of genes, locations, strands and annotations of subject cluster:

Table of Blast hits (query gene, subject gene, %identity, blast score, %coverage, e-value):
input.path1.gene388	FGSG_03064	46	245	107.5268817204301	8.6e-63
input.path1.gene389	FGSG_03066	54	434	106.1576354679803	1.4e-119


>>
41. NC_026475
Source: Fusarium graminearum PH-1 chromosome 2, whole genome shotgun sequence
Type: terpene
Number of proteins with BLAST hits to this cluster: 2
Cumulative BLAST score: 679

Table of genes, locations, strands and annotations of subject cluster:

Table of Blast hits (query gene, subject gene, %identity, blast score, %coverage, e-value):
input.path1.gene388	FGSG_03064	46	245	107.5268817204301	8.6e-63
input.path1.gene389	FGSG_03066	54	434	106.1576354679803	1.4e-119


>>
42. CP051141
Source: Peltaster fructicola strain LNHT1506 chromosome 3
Type: terpene
Number of proteins with BLAST hits to this cluster: 2
Cumulative BLAST score: 676

Table of genes, locations, strands and annotations of subject cluster:

Table of Blast hits (query gene, subject gene, %identity, blast score, %coverage, e-value):
input.path1.gene388	AMS68_004790	67	273	71.68458781362007	2.2e-71
input.path1.gene389	AMS68_004793	52	403	106.1576354679803	3.6e-110


>>
43. NW_017264202
Source: Xylona heveae TC161 unplaced genomic scaffold L228scaffold_10, whole genome shotgun sequence
Type: terpene
Number of proteins with BLAST hits to this cluster: 2
Cumulative BLAST score: 668

Table of genes, locations, strands and annotations of subject cluster:

Table of Blast hits (query gene, subject gene, %identity, blast score, %coverage, e-value):
input.path1.gene388	L228DRAFT_250153	52	280	97.4910394265233	2.4e-73
input.path1.gene389	L228DRAFT_285165	49	388	103.44827586206897	9.1e-106


>>
44. NC_016947
Source: Mycobacterium intracellulare MOTT-02, complete sequence
Type: NRPS
Number of proteins with BLAST hits to this cluster: 2
Cumulative BLAST score: 644

Table of genes, locations, strands and annotations of subject cluster:
OCO_RS15030	OCO_RS15030	3321728	3324620	+	RND_family_transporter
OCO_RS15035	OCO_RS15035	3324710	3327626	+	RND_family_transporter
OCO_RS15040	OCO_RS15040	3327800	3328744	+	daunorubicin_resistance_protein_DrrA_family_ABC_transporter_ATP-binding_protein

Table of Blast hits (query gene, subject gene, %identity, blast score, %coverage, e-value):
input.path1.gene394	OCO_RS14915	43	74	39.67611336032389	1.8e-11
input.path1.gene398	OCO_RS15010	31	288	48.84038199181446	3.5e-75
input.path1.gene398	OCO_RS15015	31	282	49.04502046384721	1.9e-73


>>
45. NC_016946
Source: Mycobacterium intracellulare ATCC 13950, complete sequence
Type: NRPS
Number of proteins with BLAST hits to this cluster: 2
Cumulative BLAST score: 643

Table of genes, locations, strands and annotations of subject cluster:
OCU_RS39950	OCU_RS39950	3323815	3324759	+	daunorubicin_resistance_protein_DrrA_family_ABC_transporter_ATP-binding_protein

Table of Blast hits (query gene, subject gene, %identity, blast score, %coverage, e-value):
input.path1.gene394	OCU_RS39825	43	74	39.67611336032389	1.8e-11
input.path1.gene398	OCU_RS39920	31	287	48.84038199181446	7.9e-75
input.path1.gene398	OCU_RS39925	31	282	49.04502046384721	1.9e-73


>>
46. NZ_CP023149
Source: Mycobacterium intracellulare strain FLAC0181 chromosome, complete genome
Type: NRPS
Number of proteins with BLAST hits to this cluster: 2
Cumulative BLAST score: 643

Table of genes, locations, strands and annotations of subject cluster:

Table of Blast hits (query gene, subject gene, %identity, blast score, %coverage, e-value):
input.path1.gene394	CKJ67_RS15625	42	73	39.67611336032389	3.9e-11
input.path1.gene398	CKJ67_RS15720	31	288	48.84038199181446	3.5e-75
input.path1.gene398	CKJ67_RS15725	31	282	49.04502046384721	1.9e-73


>>
47. NC_018612
Source: Mycobacterium intracellulare subsp. intracellulare MTCC 9506, complete sequence
Type: NRPS
Number of proteins with BLAST hits to this cluster: 2
Cumulative BLAST score: 643

Table of genes, locations, strands and annotations of subject cluster:
MIP_RS15260	MIP_RS15260	3432168	3432560	-	GDP-mannose_4,6-dehydratase
MIP_RS15275	MIP_RS15275	3433575	3434380	-	ATP-binding_protein

Table of Blast hits (query gene, subject gene, %identity, blast score, %coverage, e-value):
input.path1.gene394	MIP_RS15095	41	71	39.67611336032389	2e-10
input.path1.gene398	MIP_RS15190	31	290	48.84038199181446	7.1e-76
input.path1.gene398	MIP_RS15195	31	282	49.04502046384721	1.9e-73


>>
48. NC_016948
Source: Mycobacterium paraintracellulare, complete sequence
Type: NRPS
Number of proteins with BLAST hits to this cluster: 2
Cumulative BLAST score: 642

Table of genes, locations, strands and annotations of subject cluster:

Table of Blast hits (query gene, subject gene, %identity, blast score, %coverage, e-value):
input.path1.gene394	OCQ_RS15240	42	73	39.67611336032389	3.9e-11
input.path1.gene398	OCQ_RS15335	31	287	48.84038199181446	7.9e-75
input.path1.gene398	OCQ_RS15340	31	282	49.04502046384721	1.9e-73


>>
49. NZ_KL446995
Source: Mycobacterium sp. TKK-01-0059 supercont2.3, whole genome shotgun sequence
Type: NRPS
Number of proteins with BLAST hits to this cluster: 2
Cumulative BLAST score: 640

Table of genes, locations, strands and annotations of subject cluster:
K883_RS23335	K883_RS23335	276314	276720	+	group_1_truncated_hemoglobin
K883_RS23450	K883_RS23450	343713	344111	-	GDP-mannose_4,6-dehydratase
K883_RS23475	K883_RS23475	349652	350287	-	GDP-mannose_4,6-dehydratase

Table of Blast hits (query gene, subject gene, %identity, blast score, %coverage, e-value):
input.path1.gene394	K883_RS23295	41	71	39.67611336032389	2e-10
input.path1.gene398	K883_RS23390	31	286	48.84038199181446	1.3e-74
input.path1.gene398	K883_RS23395	31	283	49.04502046384721	1.5e-73


>>
50. NW_022984635
Source: Aspergillus tanneri strain NIH1004 chromosome Unknown AapTan_8_pilon_pilon_pilon, whole genome shotgun sequence
Type: NRPS
Number of proteins with BLAST hits to this cluster: 2
Cumulative BLAST score: 635

Table of genes, locations, strands and annotations of subject cluster:

Table of Blast hits (query gene, subject gene, %identity, blast score, %coverage, e-value):
input.path1.gene399	ATNIH1004_011596	34	420	29.80132450331126	1.4e-114
input.path1.gene405	ATNIH1004_011601	31	215	100.20325203252031	1.7e-53
