## Supplementary Results for "A draft genome of the ascomycotal fungal species *Pseudopithomyces maydicus* (family *Didymosphaeriaceae*)": contig_9_c2.txt

ClusterBlast scores for contig_9

Table of genes, locations, strands and annotations of query cluster:
input.path1.gene511	1787953	1788622	+		
input.path1.gene512	1790829	1792452	-		
input.path1.gene513	1793072	1793939	-		
input.path1.gene514	1795178	1796398	-		
input.path1.gene515	1803508	1805696	+		


Significant hits: 
1. NW_001939250	Pyrenophora tritici-repentis Pt-1C-BFP supercont1.7, whole genome shotgun sequence
2. CM000590	Fusarium oxysporum f. sp. lycopersici 4287 chromosome 2, whole genome shotgun sequence
3. NC_030987	Fusarium oxysporum f. sp. lycopersici 4287 chromosome 2, whole genome shotgun sequence
4. NC_026507	Neurospora crassa OR74A linkage group VII, whole genome shotgun sequence
5. NW_020194478	Amorphotheca resinae ATCC 22711 unplaced genomic scaffold M430scaffold_8, whole genome shotgun sequence
6. NW_013562494	Exophiala xenobiotica strain CBS 118157 unplaced genomic scaffold supercont1.1, whole genome shotgun sequence
7. NW_013550612	Fonsecaea pedrosoi CBS 271.37 unplaced genomic scaffold supercont1.4, whole genome shotgun sequence
8. NW_015971664	Fonsecaea multimorphosa CBS 102226 unplaced genomic scaffold supercont1.30, whole genome shotgun sequence
9. NW_015971575	Cladophialophora bantiana CBS 173.52 unplaced genomic scaffold supercont1.1, whole genome shotgun sequence
10. NW_006763066	Marssonina brunnea f. sp. 'multigermtubi' MB_m1 unplaced genomic scaffold M6_S00028, whole genome shotgun sequence
11. NW_017387246	Fonsecaea erecta strain CBS 125763 chromosome Unknown Scaff3, whole genome shotgun sequence
12. NW_015622515	Exophiala spinifera strain CBS 89968 unplaced genomic scaffold supercont1.3, whole genome shotgun sequence
13. NC_007200	Aspergillus fumigatus Af293 chromosome 7, whole genome shotgun sequence
14. NT_107013	Aspergillus nidulans FGSC A4 chromosome VI map unlocalized scaffold_3, whole genome shotgun sequence
15. NC_036442	Aspergillus oryzae RIB40 DNA, chromosome 8
16. NW_004504308	Coccidioides immitis RS genomic scaffold supercont3.3, whole genome shotgun sequence
17. NW_017264195	Xylona heveae TC161 unplaced genomic scaffold L228scaffold_3, whole genome shotgun sequence
18. NW_006912911	Capronia epimyces CBS 606.96 aczIQ-supercont1.8, whole genome shotgun sequence
19. NW_003345195	Nannizzia gypsea CBS 118893 supercont1.7 genomic scaffold, whole genome shotgun sequence
20. NW_006912893	Capronia coronata CBS 617.96 acBCG-supercont1.1, whole genome shotgun sequence
21. CP042186	Venturia effusa strain albino chromosome 2, complete sequence
22. NT_165936	Aspergillus terreus NIH2624 scaffold_13 genomic scaffold, whole genome shotgun sequence
23. NW_007360991	Glarea lozoyensis ATCC 20868 chromosome Unknown GLAREA17, whole genome shotgun sequence
24. NW_008481816	Cladophialophora carrionii CBS 160.54 unplaced genomic scaffold supercont1.1, whole genome shotgun sequence
25. NW_003456423	Trichophyton rubrum CBS 118892 genomic scaffold supercont2.6, whole genome shotgun sequence
26. NW_006913045	Cladophialophora yegresii CBS 114405 aczIN-supercont1.1, whole genome shotgun sequence
27. NW_008751646	Exophiala dermatitidis NIH/UT8656 unplaced genomic scaffold supercont1.1, whole genome shotgun sequence
28. NW_014574584	Penicillium digitatum Pd1 unplaced genomic scaffold Scaffold_41, whole genome shotgun sequence
29. NW_008481812	Cyphellophora europaea CBS 101466 unplaced genomic scaffold supercont1.6, whole genome shotgun sequence
30. NC_016476	Thermothelomyces thermophilus ATCC 42464 chromosome 5, complete sequence
31. NW_023336260	Colletotrichum scovillei strain TJNH1 chromosome Unknown Contig00002, whole genome shotgun sequence
32. NW_015622505	Exophiala mesophila strain CBS 40295 unplaced genomic scaffold supercont1.2, whole genome shotgun sequence
33. NW_006917094	Pestalotiopsis fici W106-1 unplaced genomic scaffold PFICI_Scf_4, whole genome shotgun sequence
34. NC_016459	Thermothielavioides terrestris NRRL 8126 chromosome 3, complete sequence
35. NC_030957	Colletotrichum higginsianum IMI 349063 chromosome 4, whole genome shotgun sequence
36. NW_003299168	Microsporum canis CBS 113480 supercont1.2 genomic scaffold, whole genome shotgun sequence
37. NW_011371380	Paracoccidioides brasiliensis Pb18 unplaced genomic scaffold supercont2.23, whole genome shotgun sequence
38. NW_001914847	Podospora anserina S mat+ genomic DNA chromosome 5, supercontig 5
39. NW_017971433	Talaromyces atroroseus strain IBT 11181 chromosome Unknown scaffold_1, whole genome shotgun sequence
40. NW_013550596	Rhinocladiella mackenziei CBS 650.93 unplaced genomic scaffold supercont1.9, whole genome shotgun sequence
41. NW_023336279	Aspergillus tubingensis WU-2223L DNA, scaffold_5, whole genome shotgun sequence
42. NW_022983866	Arthroderma uncinatum strain CBS 119779 chromosome Unknown scaffold4, whole genome shotgun sequence
43. NW_015971145	Sporothrix schenckii 1099-18 chromosome Unknown Cont44, whole genome shotgun sequence
44. NC_038015	Fusarium venenatum strain A3/5 genome assembly, chromosome: IIII
45. CM002799	Penicillium chrysogenum strain P2niaD18 chromosome II, whole genome shotgun sequence
46. NW_003315110	Trichophyton benhamiae CBS 112371 chromosome Unknown Abenha-G-c-322p01.r1, whole genome shotgun sequence
47. CM000577	Fusarium graminearum PH-1 chromosome 4, whole genome shotgun sequence
48. NC_026477	Fusarium graminearum PH-1 chromosome 4, whole genome shotgun sequence
49. NC_049566	Talaromyces rugulosus chromosome VI, complete sequence
50. NW_022984634	Aspergillus tanneri strain NIH1004 chromosome Unknown AapTan_7_pilon_pilon_pilon, whole genome shotgun sequence


Details:

>>
1. NW_001939250
Source: Pyrenophora tritici-repentis Pt-1C-BFP supercont1.7, whole genome shotgun sequence
Type: terpene
Number of proteins with BLAST hits to this cluster: 3
Cumulative BLAST score: 1666

Table of genes, locations, strands and annotations of subject cluster:

Table of Blast hits (query gene, subject gene, %identity, blast score, %coverage, e-value):
input.path1.gene512	PTRG_07460	83	580	95.76923076923077	2.1e-163
input.path1.gene513	PTRG_07461	86	457	100.0	1e-126
input.path1.gene514	PTRG_07462	79	629	97.9328165374677	1.7e-178


>>
2. CM000590
Source: Fusarium oxysporum f. sp. lycopersici 4287 chromosome 2, whole genome shotgun sequence
Type: terpene
Number of proteins with BLAST hits to this cluster: 2
Cumulative BLAST score: 2257

Table of genes, locations, strands and annotations of subject cluster:

Table of Blast hits (query gene, subject gene, %identity, blast score, %coverage, e-value):
input.path1.gene512	FOXG_06059	67	489	99.23076923076923	3.7e-136
input.path1.gene512	FOXG_06059_KNB03632.1	66	452	94.03846153846153	6.6e-125
input.path1.gene512	FOXG_06059_KNB03633.1	64	352	54.61538461538461	7.1e-95
input.path1.gene514	FOXG_06061_KNB03637.1	63	482	95.8656330749354	5.8e-134
input.path1.gene514	FOXG_06061	63	482	95.8656330749354	5.8e-134


>>
3. NC_030987
Source: Fusarium oxysporum f. sp. lycopersici 4287 chromosome 2, whole genome shotgun sequence
Type: terpene
Number of proteins with BLAST hits to this cluster: 2
Cumulative BLAST score: 2257

Table of genes, locations, strands and annotations of subject cluster:

Table of Blast hits (query gene, subject gene, %identity, blast score, %coverage, e-value):
input.path1.gene512	FOXG_06059	67	489	99.23076923076923	3.7e-136
input.path1.gene512	FOXG_06059_XP_018241677.1	66	452	94.03846153846153	6.6e-125
input.path1.gene512	FOXG_06059_XP_018241678.1	64	352	54.61538461538461	7.1e-95
input.path1.gene514	FOXG_06061_XP_018241682.1	63	482	95.8656330749354	5.8e-134
input.path1.gene514	FOXG_06061	63	482	95.8656330749354	5.8e-134


>>
4. NC_026507
Source: Neurospora crassa OR74A linkage group VII, whole genome shotgun sequence
Type: terpene
Number of proteins with BLAST hits to this cluster: 2
Cumulative BLAST score: 1584

Table of genes, locations, strands and annotations of subject cluster:

Table of Blast hits (query gene, subject gene, %identity, blast score, %coverage, e-value):
input.path1.gene512	NCU06052	63	508	96.73076923076923	5.9e-142
input.path1.gene514	NCU06054_XP_011395224.1	68	538	96.12403100775194	6.8e-151
input.path1.gene514	NCU06054	68	538	96.12403100775194	6.8e-151


>>
5. NW_020194478
Source: Amorphotheca resinae ATCC 22711 unplaced genomic scaffold M430scaffold_8, whole genome shotgun sequence
Type: terpene
Number of proteins with BLAST hits to this cluster: 2
Cumulative BLAST score: 1119

Table of genes, locations, strands and annotations of subject cluster:

Table of Blast hits (query gene, subject gene, %identity, blast score, %coverage, e-value):
input.path1.gene512	M430DRAFT_142656	62	549	95.57692307692308	4e-154
input.path1.gene514	M430DRAFT_123948	73	570	97.1576227390181	1.6e-160


>>
6. NW_013562494
Source: Exophiala xenobiotica strain CBS 118157 unplaced genomic scaffold supercont1.1, whole genome shotgun sequence
Type: terpene
Number of proteins with BLAST hits to this cluster: 2
Cumulative BLAST score: 1113

Table of genes, locations, strands and annotations of subject cluster:

Table of Blast hits (query gene, subject gene, %identity, blast score, %coverage, e-value):
input.path1.gene512	PV05_01916	72	557	95.57692307692308	1.1e-156
input.path1.gene514	PV05_01915	69	556	97.9328165374677	1.8e-156


>>
7. NW_013550612
Source: Fonsecaea pedrosoi CBS 271.37 unplaced genomic scaffold supercont1.4, whole genome shotgun sequence
Type: terpene
Number of proteins with BLAST hits to this cluster: 2
Cumulative BLAST score: 1104

Table of genes, locations, strands and annotations of subject cluster:

Table of Blast hits (query gene, subject gene, %identity, blast score, %coverage, e-value):
input.path1.gene512	Z517_07112	60	542	95.38461538461539	4.8e-152
input.path1.gene514	Z517_07111	70	562	96.64082687338501	3.4e-158


>>
8. NW_015971664
Source: Fonsecaea multimorphosa CBS 102226 unplaced genomic scaffold supercont1.30, whole genome shotgun sequence
Type: terpene
Number of proteins with BLAST hits to this cluster: 2
Cumulative BLAST score: 1103

Table of genes, locations, strands and annotations of subject cluster:

Table of Blast hits (query gene, subject gene, %identity, blast score, %coverage, e-value):
input.path1.gene512	Z520_10463	65	559	95.38461538461539	2.9e-157
input.path1.gene514	Z520_10462	67	544	97.9328165374677	9.5e-153


>>
9. NW_015971575
Source: Cladophialophora bantiana CBS 173.52 unplaced genomic scaffold supercont1.1, whole genome shotgun sequence
Type: terpene
Number of proteins with BLAST hits to this cluster: 2
Cumulative BLAST score: 1103

Table of genes, locations, strands and annotations of subject cluster:

Table of Blast hits (query gene, subject gene, %identity, blast score, %coverage, e-value):
input.path1.gene512	Z519_00478	60	554	95.38461538461539	9.4e-156
input.path1.gene514	Z519_00477	67	549	97.9328165374677	2.3e-154


>>
10. NW_006763066
Source: Marssonina brunnea f. sp. 'multigermtubi' MB_m1 unplaced genomic scaffold M6_S00028, whole genome shotgun sequence
Type: terpene
Number of proteins with BLAST hits to this cluster: 2
Cumulative BLAST score: 1099

Table of genes, locations, strands and annotations of subject cluster:

Table of Blast hits (query gene, subject gene, %identity, blast score, %coverage, e-value):
input.path1.gene512	MBM_09221	68	542	95.1923076923077	4.8e-152
input.path1.gene514	MBM_09225	69	557	97.1576227390181	1.4e-156


>>
11. NW_017387246
Source: Fonsecaea erecta strain CBS 125763 chromosome Unknown Scaff3, whole genome shotgun sequence
Type: terpene
Number of proteins with BLAST hits to this cluster: 2
Cumulative BLAST score: 1094

Table of genes, locations, strands and annotations of subject cluster:

Table of Blast hits (query gene, subject gene, %identity, blast score, %coverage, e-value):
input.path1.gene512	AYL99_04356	64	540	95.57692307692308	2.4e-151
input.path1.gene514	AYL99_04355	68	554	97.9328165374677	7e-156


>>
12. NW_015622515
Source: Exophiala spinifera strain CBS 89968 unplaced genomic scaffold supercont1.3, whole genome shotgun sequence
Type: terpene
Number of proteins with BLAST hits to this cluster: 2
Cumulative BLAST score: 1094

Table of genes, locations, strands and annotations of subject cluster:

Table of Blast hits (query gene, subject gene, %identity, blast score, %coverage, e-value):
input.path1.gene512	PV08_04840	68	537	95.57692307692308	1.6e-150
input.path1.gene514	PV08_04839	69	557	97.9328165374677	1.1e-156


>>
13. NC_007200
Source: Aspergillus fumigatus Af293 chromosome 7, whole genome shotgun sequence
Type: terpene
Number of proteins with BLAST hits to this cluster: 2
Cumulative BLAST score: 1093

Table of genes, locations, strands and annotations of subject cluster:

Table of Blast hits (query gene, subject gene, %identity, blast score, %coverage, e-value):
input.path1.gene512	AFUA_7G01230	70	557	96.15384615384616	1.1e-156
input.path1.gene514	AFUA_7G01220	68	536	95.8656330749354	2e-150


>>
14. NT_107013
Source: Aspergillus nidulans FGSC A4 chromosome VI map unlocalized scaffold_3, whole genome shotgun sequence
Type: terpene
Number of proteins with BLAST hits to this cluster: 2
Cumulative BLAST score: 1090

Table of genes, locations, strands and annotations of subject cluster:

Table of Blast hits (query gene, subject gene, %identity, blast score, %coverage, e-value):
input.path1.gene512	AN3375.2	66	541	95.96153846153847	8.2e-152
input.path1.gene514	AN3376.2	68	549	97.41602067183463	3.8e-154


>>
15. NC_036442
Source: Aspergillus oryzae RIB40 DNA, chromosome 8
Type: terpene
Number of proteins with BLAST hits to this cluster: 2
Cumulative BLAST score: 1089

Table of genes, locations, strands and annotations of subject cluster:

Table of Blast hits (query gene, subject gene, %identity, blast score, %coverage, e-value):
input.path1.gene512	AO090010000203	75	562	96.34615384615385	3.5e-158
input.path1.gene514	AO090010000204	66	527	96.64082687338501	1.2e-147


>>
16. NW_004504308
Source: Coccidioides immitis RS genomic scaffold supercont3.3, whole genome shotgun sequence
Type: terpene
Number of proteins with BLAST hits to this cluster: 2
Cumulative BLAST score: 1088

Table of genes, locations, strands and annotations of subject cluster:

Table of Blast hits (query gene, subject gene, %identity, blast score, %coverage, e-value):
input.path1.gene512	CIMG_08117	71	566	95.38461538461539	2.4e-159
input.path1.gene514	CIMG_08115	66	522	95.8656330749354	3.9e-146


>>
17. NW_017264195
Source: Xylona heveae TC161 unplaced genomic scaffold L228scaffold_3, whole genome shotgun sequence
Type: terpene
Number of proteins with BLAST hits to this cluster: 2
Cumulative BLAST score: 1086

Table of genes, locations, strands and annotations of subject cluster:

Table of Blast hits (query gene, subject gene, %identity, blast score, %coverage, e-value):
input.path1.gene512	L228DRAFT_237153	62	551	96.34615384615385	1e-154
input.path1.gene514	L228DRAFT_245095	65	535	97.9328165374677	5.7e-150


>>
18. NW_006912911
Source: Capronia epimyces CBS 606.96 aczIQ-supercont1.8, whole genome shotgun sequence
Type: terpene
Number of proteins with BLAST hits to this cluster: 2
Cumulative BLAST score: 1083

Table of genes, locations, strands and annotations of subject cluster:

Table of Blast hits (query gene, subject gene, %identity, blast score, %coverage, e-value):
input.path1.gene512	A1O3_08740	67	537	96.15384615384616	1.6e-150
input.path1.gene514	A1O3_08741	68	546	97.9328165374677	1.9e-153


>>
19. NW_003345195
Source: Nannizzia gypsea CBS 118893 supercont1.7 genomic scaffold, whole genome shotgun sequence
Type: terpene
Number of proteins with BLAST hits to this cluster: 2
Cumulative BLAST score: 1081

Table of genes, locations, strands and annotations of subject cluster:

Table of Blast hits (query gene, subject gene, %identity, blast score, %coverage, e-value):
input.path1.gene512	MGYG_07530	62	564	96.34615384615385	1.2e-158
input.path1.gene514	MGYG_07528	64	517	96.64082687338501	1.2e-144


>>
20. NW_006912893
Source: Capronia coronata CBS 617.96 acBCG-supercont1.1, whole genome shotgun sequence
Type: terpene
Number of proteins with BLAST hits to this cluster: 2
Cumulative BLAST score: 1079

Table of genes, locations, strands and annotations of subject cluster:

Table of Blast hits (query gene, subject gene, %identity, blast score, %coverage, e-value):
input.path1.gene512	A1O1_01159	73	534	96.15384615384616	1.3e-149
input.path1.gene514	A1O1_01160	68	545	97.9328165374677	5.6e-153


>>
21. CP042186
Source: Venturia effusa strain albino chromosome 2, complete sequence
Type: terpene
Number of proteins with BLAST hits to this cluster: 2
Cumulative BLAST score: 1077

Table of genes, locations, strands and annotations of subject cluster:

Table of Blast hits (query gene, subject gene, %identity, blast score, %coverage, e-value):
input.path1.gene512	FKW77_007280	75	538	97.11538461538461	7e-151
input.path1.gene514	FKW77_007286	68	539	97.41602067183463	3e-151


>>
22. NT_165936
Source: Aspergillus terreus NIH2624 scaffold_13 genomic scaffold, whole genome shotgun sequence
Type: NRPS-like,terpene
Number of proteins with BLAST hits to this cluster: 2
Cumulative BLAST score: 1075

Table of genes, locations, strands and annotations of subject cluster:

Table of Blast hits (query gene, subject gene, %identity, blast score, %coverage, e-value):
input.path1.gene512	ATEG_08675	74	571	95.96153846153847	1.3e-160
input.path1.gene514	ATEG_08674	67	504	91.98966408268734	8.3e-141


>>
23. NW_007360991
Source: Glarea lozoyensis ATCC 20868 chromosome Unknown GLAREA17, whole genome shotgun sequence
Type: terpene
Number of proteins with BLAST hits to this cluster: 2
Cumulative BLAST score: 1074

Table of genes, locations, strands and annotations of subject cluster:

Table of Blast hits (query gene, subject gene, %identity, blast score, %coverage, e-value):
input.path1.gene512	GLAREA_11899	69	529	95.57692307692308	3.2e-148
input.path1.gene514	GLAREA_11903	70	545	97.1576227390181	3.3e-153


>>
24. NW_008481816
Source: Cladophialophora carrionii CBS 160.54 unplaced genomic scaffold supercont1.1, whole genome shotgun sequence
Type: terpene
Number of proteins with BLAST hits to this cluster: 2
Cumulative BLAST score: 1073

Table of genes, locations, strands and annotations of subject cluster:

Table of Blast hits (query gene, subject gene, %identity, blast score, %coverage, e-value):
input.path1.gene512	G647_01370	72	526	95.38461538461539	3.6e-147
input.path1.gene514	G647_01369	67	547	97.9328165374677	8.6e-154


>>
25. NW_003456423
Source: Trichophyton rubrum CBS 118892 genomic scaffold supercont2.6, whole genome shotgun sequence
Type: terpene
Number of proteins with BLAST hits to this cluster: 2
Cumulative BLAST score: 1071

Table of genes, locations, strands and annotations of subject cluster:

Table of Blast hits (query gene, subject gene, %identity, blast score, %coverage, e-value):
input.path1.gene512	TERG_05885	67	557	96.34615384615385	1.1e-156
input.path1.gene514	TERG_05883	63	514	96.64082687338501	8e-144


>>
26. NW_006913045
Source: Cladophialophora yegresii CBS 114405 aczIN-supercont1.1, whole genome shotgun sequence
Type: terpene
Number of proteins with BLAST hits to this cluster: 2
Cumulative BLAST score: 1068

Table of genes, locations, strands and annotations of subject cluster:

Table of Blast hits (query gene, subject gene, %identity, blast score, %coverage, e-value):
input.path1.gene512	A1O7_01306	64	523	95.57692307692308	2.3e-146
input.path1.gene514	A1O7_01305	67	545	97.9328165374677	5.6e-153


>>
27. NW_008751646
Source: Exophiala dermatitidis NIH/UT8656 unplaced genomic scaffold supercont1.1, whole genome shotgun sequence
Type: terpene
Number of proteins with BLAST hits to this cluster: 2
Cumulative BLAST score: 1066

Table of genes, locations, strands and annotations of subject cluster:

Table of Blast hits (query gene, subject gene, %identity, blast score, %coverage, e-value):
input.path1.gene512	HMPREF1120_00254	64	524	95.57692307692308	1e-146
input.path1.gene514	HMPREF1120_00253	67	542	97.9328165374677	4.7e-152


>>
28. NW_014574584
Source: Penicillium digitatum Pd1 unplaced genomic scaffold Scaffold_41, whole genome shotgun sequence
Type: terpene
Number of proteins with BLAST hits to this cluster: 2
Cumulative BLAST score: 1053

Table of genes, locations, strands and annotations of subject cluster:

Table of Blast hits (query gene, subject gene, %identity, blast score, %coverage, e-value):
input.path1.gene512	PDIP_55110	57	517	96.34615384615385	1.3e-144
input.path1.gene514	PDIP_55100	68	536	95.8656330749354	2.6e-150


>>
29. NW_008481812
Source: Cyphellophora europaea CBS 101466 unplaced genomic scaffold supercont1.6, whole genome shotgun sequence
Type: terpene
Number of proteins with BLAST hits to this cluster: 2
Cumulative BLAST score: 1050

Table of genes, locations, strands and annotations of subject cluster:

Table of Blast hits (query gene, subject gene, %identity, blast score, %coverage, e-value):
input.path1.gene512	HMPREF1541_06269	61	495	96.34615384615385	6.8e-138
input.path1.gene514	HMPREF1541_06270	69	555	97.9328165374677	5.4e-156


>>
30. NC_016476
Source: Thermothelomyces thermophilus ATCC 42464 chromosome 5, complete sequence
Type: terpene
Number of proteins with BLAST hits to this cluster: 2
Cumulative BLAST score: 1045

Table of genes, locations, strands and annotations of subject cluster:

Table of Blast hits (query gene, subject gene, %identity, blast score, %coverage, e-value):
input.path1.gene512	MYCTH_2315997	60	510	96.53846153846153	2e-142
input.path1.gene514	MYCTH_112159	67	535	97.9328165374677	4.4e-150


>>
31. NW_023336260
Source: Colletotrichum scovillei strain TJNH1 chromosome Unknown Contig00002, whole genome shotgun sequence
Type: terpene
Number of proteins with BLAST hits to this cluster: 2
Cumulative BLAST score: 1044

Table of genes, locations, strands and annotations of subject cluster:

Table of Blast hits (query gene, subject gene, %identity, blast score, %coverage, e-value):
input.path1.gene512	HER10_EVM0006642	68	517	96.73076923076923	1.7e-144
input.path1.gene514	HER10_EVM0007108	68	527	95.8656330749354	1.2e-147


>>
32. NW_015622505
Source: Exophiala mesophila strain CBS 40295 unplaced genomic scaffold supercont1.2, whole genome shotgun sequence
Type: terpene
Number of proteins with BLAST hits to this cluster: 2
Cumulative BLAST score: 1044

Table of genes, locations, strands and annotations of subject cluster:

Table of Blast hits (query gene, subject gene, %identity, blast score, %coverage, e-value):
input.path1.gene512	PV10_02395	65	512	95.57692307692308	5.4e-143
input.path1.gene514	PV10_02396	67	532	96.64082687338501	2.9e-149


>>
33. NW_006917094
Source: Pestalotiopsis fici W106-1 unplaced genomic scaffold PFICI_Scf_4, whole genome shotgun sequence
Type: terpene
Number of proteins with BLAST hits to this cluster: 2
Cumulative BLAST score: 1042

Table of genes, locations, strands and annotations of subject cluster:

Table of Blast hits (query gene, subject gene, %identity, blast score, %coverage, e-value):
input.path1.gene512	PFICI_06796	72	498	96.34615384615385	6.1e-139
input.path1.gene514	PFICI_06798	68	544	97.9328165374677	7.2e-153


>>
34. NC_016459
Source: Thermothielavioides terrestris NRRL 8126 chromosome 3, complete sequence
Type: terpene
Number of proteins with BLAST hits to this cluster: 2
Cumulative BLAST score: 1038

Table of genes, locations, strands and annotations of subject cluster:

Table of Blast hits (query gene, subject gene, %identity, blast score, %coverage, e-value):
input.path1.gene512	THITE_2116916	65	495	96.34615384615385	6.8e-138
input.path1.gene514	THITE_76823	69	543	96.89922480620154	1.6e-152


>>
35. NC_030957
Source: Colletotrichum higginsianum IMI 349063 chromosome 4, whole genome shotgun sequence
Type: terpene
Number of proteins with BLAST hits to this cluster: 2
Cumulative BLAST score: 1034

Table of genes, locations, strands and annotations of subject cluster:

Table of Blast hits (query gene, subject gene, %identity, blast score, %coverage, e-value):
input.path1.gene512	CH63R_06221	74	510	96.73076923076923	1.6e-142
input.path1.gene514	CH63R_06219	68	524	95.8656330749354	1e-146


>>
36. NW_003299168
Source: Microsporum canis CBS 113480 supercont1.2 genomic scaffold, whole genome shotgun sequence
Type: terpene
Number of proteins with BLAST hits to this cluster: 2
Cumulative BLAST score: 1029

Table of genes, locations, strands and annotations of subject cluster:

Table of Blast hits (query gene, subject gene, %identity, blast score, %coverage, e-value):
input.path1.gene512	MCYG_02535	58	528	96.15384615384616	7.2e-148
input.path1.gene514	MCYG_02533	62	501	96.64082687338501	7e-140


>>
37. NW_011371380
Source: Paracoccidioides brasiliensis Pb18 unplaced genomic scaffold supercont2.23, whole genome shotgun sequence
Type: terpene
Number of proteins with BLAST hits to this cluster: 2
Cumulative BLAST score: 1026

Table of genes, locations, strands and annotations of subject cluster:

Table of Blast hits (query gene, subject gene, %identity, blast score, %coverage, e-value):
input.path1.gene512	PADG_08599	56	521	96.34615384615385	1.2e-145
input.path1.gene514	PADG_08600	67	505	91.98966408268734	6.4e-141


>>
38. NW_001914847
Source: Podospora anserina S mat+ genomic DNA chromosome 5, supercontig 5
Type: terpene
Number of proteins with BLAST hits to this cluster: 2
Cumulative BLAST score: 1024

Table of genes, locations, strands and annotations of subject cluster:

Table of Blast hits (query gene, subject gene, %identity, blast score, %coverage, e-value):
input.path1.gene512	PODANS_5_4940	65	477	95.76923076923077	1.5e-132
input.path1.gene514	PODANS_5_4960	68	547	97.9328165374677	8.6e-154


>>
39. NW_017971433
Source: Talaromyces atroroseus strain IBT 11181 chromosome Unknown scaffold_1, whole genome shotgun sequence
Type: terpene
Number of proteins with BLAST hits to this cluster: 2
Cumulative BLAST score: 1024

Table of genes, locations, strands and annotations of subject cluster:

Table of Blast hits (query gene, subject gene, %identity, blast score, %coverage, e-value):
input.path1.gene512	UA08_01225	63	472	95.76923076923077	4.7e-131
input.path1.gene514	UA08_01226	68	552	96.64082687338501	4.5e-155


>>
40. NW_013550596
Source: Rhinocladiella mackenziei CBS 650.93 unplaced genomic scaffold supercont1.9, whole genome shotgun sequence
Type: terpene
Number of proteins with BLAST hits to this cluster: 2
Cumulative BLAST score: 1020

Table of genes, locations, strands and annotations of subject cluster:

Table of Blast hits (query gene, subject gene, %identity, blast score, %coverage, e-value):
input.path1.gene512	Z518_10373	64	467	95.57692307692308	1.5e-129
input.path1.gene514	Z518_10372	68	553	97.9328165374677	2e-155


>>
41. NW_023336279
Source: Aspergillus tubingensis WU-2223L DNA, scaffold_5, whole genome shotgun sequence
Type: terpene
Number of proteins with BLAST hits to this cluster: 2
Cumulative BLAST score: 1019

Table of genes, locations, strands and annotations of subject cluster:

Table of Blast hits (query gene, subject gene, %identity, blast score, %coverage, e-value):
input.path1.gene512	AtWU_03592	64	497	96.15384615384616	1.8e-138
input.path1.gene514	AtWU_03593	66	522	95.8656330749354	3e-146


>>
42. NW_022983866
Source: Arthroderma uncinatum strain CBS 119779 chromosome Unknown scaffold4, whole genome shotgun sequence
Type: terpene
Number of proteins with BLAST hits to this cluster: 2
Cumulative BLAST score: 1018

Table of genes, locations, strands and annotations of subject cluster:

Table of Blast hits (query gene, subject gene, %identity, blast score, %coverage, e-value):
input.path1.gene512	GIQ15_05935	65	515	95.96153846153847	6.3e-144
input.path1.gene514	GIQ15_05936	62	503	96.64082687338501	1.4e-140


>>
43. NW_015971145
Source: Sporothrix schenckii 1099-18 chromosome Unknown Cont44, whole genome shotgun sequence
Type: terpene
Number of proteins with BLAST hits to this cluster: 2
Cumulative BLAST score: 1012

Table of genes, locations, strands and annotations of subject cluster:

Table of Blast hits (query gene, subject gene, %identity, blast score, %coverage, e-value):
input.path1.gene512	SPSK_09035	62	492	97.11538461538461	4.4e-137
input.path1.gene514	SPSK_09158	65	520	98.70801033591732	1.5e-145


>>
44. NC_038015
Source: Fusarium venenatum strain A3/5 genome assembly, chromosome: IIII
Type: terpene
Number of proteins with BLAST hits to this cluster: 2
Cumulative BLAST score: 1009

Table of genes, locations, strands and annotations of subject cluster:

Table of Blast hits (query gene, subject gene, %identity, blast score, %coverage, e-value):
input.path1.gene512	FVRRES_13379	72	524	96.34615384615385	1.8e-146
input.path1.gene514	FVRRES_13378	62	485	95.8656330749354	4e-135


>>
45. CM002799
Source: Penicillium chrysogenum strain P2niaD18 chromosome II, whole genome shotgun sequence
Type: terpene
Number of proteins with BLAST hits to this cluster: 2
Cumulative BLAST score: 1008

Table of genes, locations, strands and annotations of subject cluster:

Table of Blast hits (query gene, subject gene, %identity, blast score, %coverage, e-value):
input.path1.gene512	EN45_076160	65	466	91.53846153846153	3.4e-129
input.path1.gene514	EN45_076170	69	542	95.8656330749354	3.6e-152


>>
46. NW_003315110
Source: Trichophyton benhamiae CBS 112371 chromosome Unknown Abenha-G-c-322p01.r1, whole genome shotgun sequence
Type: terpene
Number of proteins with BLAST hits to this cluster: 2
Cumulative BLAST score: 1007

Table of genes, locations, strands and annotations of subject cluster:

Table of Blast hits (query gene, subject gene, %identity, blast score, %coverage, e-value):
input.path1.gene512	ARB_06260	64	545	100.38461538461539	5.7e-153
input.path1.gene514	ARB_06257	62	462	88.37209302325581	4.7e-128


>>
47. CM000577
Source: Fusarium graminearum PH-1 chromosome 4, whole genome shotgun sequence
Type: terpene
Number of proteins with BLAST hits to this cluster: 2
Cumulative BLAST score: 1005

Table of genes, locations, strands and annotations of subject cluster:

Table of Blast hits (query gene, subject gene, %identity, blast score, %coverage, e-value):
input.path1.gene512	FGSG_09380	72	520	96.73076923076923	2e-145
input.path1.gene514	FGSG_09381	63	485	95.8656330749354	5.2e-135


>>
48. NC_026477
Source: Fusarium graminearum PH-1 chromosome 4, whole genome shotgun sequence
Type: terpene
Number of proteins with BLAST hits to this cluster: 2
Cumulative BLAST score: 1005

Table of genes, locations, strands and annotations of subject cluster:

Table of Blast hits (query gene, subject gene, %identity, blast score, %coverage, e-value):
input.path1.gene512	FGSG_09380	72	520	96.73076923076923	2e-145
input.path1.gene514	FGSG_09381	63	485	95.8656330749354	5.2e-135


>>
49. NC_049566
Source: Talaromyces rugulosus chromosome VI, complete sequence
Type: terpene
Number of proteins with BLAST hits to this cluster: 2
Cumulative BLAST score: 1000

Table of genes, locations, strands and annotations of subject cluster:

Table of Blast hits (query gene, subject gene, %identity, blast score, %coverage, e-value):
input.path1.gene512	TRUGW13939_11747	62	470	95.57692307692308	1.8e-130
input.path1.gene514	TRUGW13939_11746	66	530	96.64082687338501	1.1e-148


>>
50. NW_022984634
Source: Aspergillus tanneri strain NIH1004 chromosome Unknown AapTan_7_pilon_pilon_pilon, whole genome shotgun sequence
Type: terpene
Number of proteins with BLAST hits to this cluster: 2
Cumulative BLAST score: 988

Table of genes, locations, strands and annotations of subject cluster:

Table of Blast hits (query gene, subject gene, %identity, blast score, %coverage, e-value):
input.path1.gene512	ATNIH1004_006810	63	465	96.15384615384616	9.8e-129
input.path1.gene514	ATNIH1004_006811	65	523	97.1576227390181	1.7e-146
