## Supplementary Results for "A draft genome of the ascomycotal fungal species *Pseudopithomyces maydicus* (family *Didymosphaeriaceae*)": contig_10_c1.txt

ClusterBlast scores for contig_10

Table of genes, locations, strands and annotations of query cluster:
input.path1.gene25	133009	134410	+		
input.path1.gene26	134565	136623	-		
input.path1.gene27	138570	144953	+		
input.path1.gene28	148133	148778	+		
input.path1.gene29	150420	154559	-		
input.path1.gene30	156618	157514	+		
input.path1.gene31	157797	158496	-		
input.path1.gene32	163864	164850	+		
input.path1.gene33	165304	166494	+		
input.path1.gene34	167003	170014	-		
input.path1.gene35	170499	170964	+		
input.path1.gene36	173054	174203	-		


Significant hits: 
1. NW_001820829	Sclerotinia sclerotiorum 1980 UF-70 scaffold_7 genomic scaffold, whole genome shotgun sequence
2. NW_009276941	Verticillium dahliae VdLs.17 supercont1.26 genomic scaffold, whole genome shotgun sequence
3. NW_022984633	Aspergillus tanneri strain NIH1004 chromosome Unknown AapTan_6_pilon_pilon_pilon, whole genome shotgun sequence
4. NC_030962	Colletotrichum higginsianum IMI 349063 chromosome 10, whole genome shotgun sequence
5. CM000595	Fusarium oxysporum f. sp. lycopersici 4287 chromosome 7, whole genome shotgun sequence
6. NC_030992	Fusarium oxysporum f. sp. lycopersici 4287 chromosome 7, whole genome shotgun sequence
7. NZ_JAAALO010000001	Micromonospora sp. NEAU-HG-1 Scaffold1, whole genome shotgun sequence
8. NW_003315103	Trichophyton benhamiae CBS 112371 chromosome Unknown contig00075.49, whole genome shotgun sequence
9. NW_003456426	Trichophyton rubrum CBS 118892 genomic scaffold supercont2.3, whole genome shotgun sequence
10. NW_003345192	Nannizzia gypsea CBS 118893 supercont1.10 genomic scaffold, whole genome shotgun sequence
11. NZ_CP023699	Streptomyces kanamyceticus strain ATCC 12853 chromosome, complete genome
12. NW_017264193	Xylona heveae TC161 unplaced genomic scaffold L228scaffold_1, whole genome shotgun sequence
13. NW_003299161	Microsporum canis CBS 113480 supercont1.9 genomic scaffold, whole genome shotgun sequence
14. NW_022984632	Aspergillus tanneri strain NIH1004 chromosome Unknown AapTan_5_pilon_pilon_pilon, whole genome shotgun sequence
15. NC_038015	Fusarium venenatum strain A3/5 genome assembly, chromosome: IIII
16. NC_030957	Colletotrichum higginsianum IMI 349063 chromosome 4, whole genome shotgun sequence
17. NW_022194791	Fusarium proliferatum ET1 genome assembly, contig: FPRO_scaffold002, whole genome shotgun sequence
18. NC_036623	Fusarium fujikuroi IMI 58289 draft genome, chromosome FFUJ_chr02
19. CM000574	Fusarium graminearum PH-1 chromosome 1, whole genome shotgun sequence
20. NC_026474	Fusarium graminearum PH-1 chromosome 1, whole genome shotgun sequence
21. NC_038012	Fusarium venenatum strain A3/5 genome assembly, chromosome: I
22. CM000577	Fusarium graminearum PH-1 chromosome 4, whole genome shotgun sequence
23. NC_026477	Fusarium graminearum PH-1 chromosome 4, whole genome shotgun sequence
24. NW_023336269	Colletotrichum scovillei strain TJNH1 chromosome Unknown Contig00011, whole genome shotgun sequence
25. NW_006271969	Cordyceps militaris CM01 unplaced genomic scaffold CCM_S00001, whole genome shotgun sequence
26. NC_049562	Talaromyces rugulosus chromosome II, complete sequence
27. NZ_RFFH01000003	Nocardia stercoris strain NEAU-LL90 Scaffold3, whole genome shotgun sequence
28. NW_022984633	Aspergillus tanneri strain NIH1004 chromosome Unknown AapTan_6_pilon_pilon_pilon, whole genome shotgun sequence
29. NW_023336282	Aspergillus tubingensis WU-2223L DNA, scaffold_8, whole genome shotgun sequence
30. NZ_CP048405	Streptacidiphilus sp. PB12-B1b chromosome, complete genome
31. CM000590	Fusarium oxysporum f. sp. lycopersici 4287 chromosome 2, whole genome shotgun sequence
32. NC_030987	Fusarium oxysporum f. sp. lycopersici 4287 chromosome 2, whole genome shotgun sequence
33. NC_036442	Aspergillus oryzae RIB40 DNA, chromosome 8
34. NT_165931	Aspergillus terreus NIH2624 scaffold_8 genomic scaffold, whole genome shotgun sequence
35. NZ_WOFH01000003	Actinomadura sp. NEAU-AAG5 Scaffold3, whole genome shotgun sequence
36. NZ_GL877879	Saccharopolyspora spinosa NRRL 18395 Scaffold000002, whole genome shotgun sequence
37. NW_022194796	Fusarium proliferatum ET1 genome assembly, contig: FPRO_scaffold007, whole genome shotgun sequence
38. NC_036626	Fusarium fujikuroi IMI 58289 draft genome, chromosome FFUJ_chr05
39. NW_017971433	Talaromyces atroroseus strain IBT 11181 chromosome Unknown scaffold_1, whole genome shotgun sequence
40. NW_013562491	Exophiala xenobiotica strain CBS 118157 unplaced genomic scaffold supercont1.4, whole genome shotgun sequence
41. NW_001914858	Podospora anserina S mat+ genomic DNA chromosome 2, supercontig 2
42. NC_049561	Talaromyces rugulosus chromosome I, complete sequence
43. NZ_CP023694	Streptomyces coeruleorubidus strain ATCC 13740 chromosome, complete genome
44. NZ_LT629775	Streptomyces sp. TLI_053 chromosome I
45. NZ_CP036534	Streptomyces sp. VN1 chromosome, complete genome
46. NZ_LJIW01000001	Streptomyces malaysiensis strain F913 SMF9131, whole genome shotgun sequence
47. NZ_CP054932	Actinomadura sp. NAK00032 chromosome, complete genome
48. NZ_CP040605	Saccharopolyspora sp. ASAGF58 chromosome, complete genome
49. NC_010612	Mycobacterium marinum M, complete genome
50. NC_049561	Talaromyces rugulosus chromosome I, complete sequence


Details:

>>
1. NW_001820829
Source: Sclerotinia sclerotiorum 1980 UF-70 scaffold_7 genomic scaffold, whole genome shotgun sequence
Type: T1PKS
Number of proteins with BLAST hits to this cluster: 6
Cumulative BLAST score: 1994

Table of genes, locations, strands and annotations of subject cluster:

Table of Blast hits (query gene, subject gene, %identity, blast score, %coverage, e-value):
input.path1.gene29	SS1G_05681	43	750	113.58695652173914	2.7e-214
input.path1.gene30	SS1G_05676	54	297	99.27272727272727	1.1e-78
input.path1.gene31	SS1G_05675	55	108	54.65116279069767	7.7e-22
input.path1.gene32	SS1G_05677	54	242	77.44360902255639	6.9e-62
input.path1.gene33	SS1G_05678	42	288	101.82291666666667	9.3e-76
input.path1.gene34	SS1G_05679	47	309	63.09963099630996	9.4e-82


>>
2. NW_009276941
Source: Verticillium dahliae VdLs.17 supercont1.26 genomic scaffold, whole genome shotgun sequence
Type: NRPS-like,T1PKS
Number of proteins with BLAST hits to this cluster: 3
Cumulative BLAST score: 555

Table of genes, locations, strands and annotations of subject cluster:

Table of Blast hits (query gene, subject gene, %identity, blast score, %coverage, e-value):
input.path1.gene29	VDAG_09534	36	374	71.52173913043478	3.1e-101
input.path1.gene33	VDAG_09529	37	84	36.19791666666667	2.7e-14
input.path1.gene34	VDAG_09526	32	97	37.45387453874539	4.3e-18


>>
3. NW_022984633
Source: Aspergillus tanneri strain NIH1004 chromosome Unknown AapTan_6_pilon_pilon_pilon, whole genome shotgun sequence
Type: T1PKS
Number of proteins with BLAST hits to this cluster: 3
Cumulative BLAST score: 553

Table of genes, locations, strands and annotations of subject cluster:

Table of Blast hits (query gene, subject gene, %identity, blast score, %coverage, e-value):
input.path1.gene29	ATNIH1004_008249	39	265	47.71739130434783	2e-68
input.path1.gene30	ATNIH1004_008244	31	98	88.72727272727273	9.8e-19
input.path1.gene32	ATNIH1004_008247	37	190	108.6466165413534	3.1e-46


>>
4. NC_030962
Source: Colletotrichum higginsianum IMI 349063 chromosome 10, whole genome shotgun sequence
Type: T1PKS,indole
Number of proteins with BLAST hits to this cluster: 3
Cumulative BLAST score: 541

Table of genes, locations, strands and annotations of subject cluster:

Table of Blast hits (query gene, subject gene, %identity, blast score, %coverage, e-value):
input.path1.gene29	CH63R_14371	43	282	42.71739130434782	1.6e-73
input.path1.gene30	CH63R_14368	31	94	84.0	2.4e-17
input.path1.gene32	CH63R_14376	34	165	109.39849624060149	1.1e-38


>>
5. CM000595
Source: Fusarium oxysporum f. sp. lycopersici 4287 chromosome 7, whole genome shotgun sequence
Type: T1PKS
Number of proteins with BLAST hits to this cluster: 2
Cumulative BLAST score: 1383

Table of genes, locations, strands and annotations of subject cluster:

Table of Blast hits (query gene, subject gene, %identity, blast score, %coverage, e-value):
input.path1.gene29	FOXG_04757	39	589	100.32608695652175	6.1e-166
input.path1.gene29	FOXG_04757_KNB01536.1	39	589	100.32608695652175	6.1e-166
input.path1.gene33	FOXG_04755	37	205	92.70833333333334	1.4e-50


>>
6. NC_030992
Source: Fusarium oxysporum f. sp. lycopersici 4287 chromosome 7, whole genome shotgun sequence
Type: T1PKS
Number of proteins with BLAST hits to this cluster: 2
Cumulative BLAST score: 1383

Table of genes, locations, strands and annotations of subject cluster:

Table of Blast hits (query gene, subject gene, %identity, blast score, %coverage, e-value):
input.path1.gene29	FOXG_04757	39	589	100.32608695652175	6.1e-166
input.path1.gene29	FOXG_04757_XP_018239581.1	39	589	100.32608695652175	6.1e-166
input.path1.gene33	FOXG_04755	37	205	92.70833333333334	1.4e-50


>>
7. NZ_JAAALO010000001
Source: Micromonospora sp. NEAU-HG-1 Scaffold1, whole genome shotgun sequence
Type: NRPS,T1PKS,bacteriocin,bottromycin,transAT-PKS,transAT-PKS-like
Number of proteins with BLAST hits to this cluster: 2
Cumulative BLAST score: 1046

Table of genes, locations, strands and annotations of subject cluster:
GVV04_RS03545	GVV04_RS03545	909458	909989	+	carbohydrate-binding_protein

Table of Blast hits (query gene, subject gene, %identity, blast score, %coverage, e-value):
input.path1.gene29	GVV04_RS03530	38	251	44.891304347826086	3.9e-64
input.path1.gene29	GVV04_RS03525	38	249	44.565217391304344	1.1e-63
input.path1.gene29	GVV04_RS03520	37	243	41.630434782608695	8.2e-62
input.path1.gene29	GVV04_RS03515	37	240	43.913043478260875	9.1e-61
input.path1.gene33	GVV04_RS03490	34	63	32.8125	4.9e-08


>>
8. NW_003315103
Source: Trichophyton benhamiae CBS 112371 chromosome Unknown contig00075.49, whole genome shotgun sequence
Type: T1PKS
Number of proteins with BLAST hits to this cluster: 2
Cumulative BLAST score: 1036

Table of genes, locations, strands and annotations of subject cluster:

Table of Blast hits (query gene, subject gene, %identity, blast score, %coverage, e-value):
input.path1.gene29	ARB_07994	40	770	122.93478260869566	1.5e-220
input.path1.gene33	ARB_07989	42	266	104.6875	3.8e-69


>>
9. NW_003456426
Source: Trichophyton rubrum CBS 118892 genomic scaffold supercont2.3, whole genome shotgun sequence
Type: T1PKS
Number of proteins with BLAST hits to this cluster: 2
Cumulative BLAST score: 1032

Table of genes, locations, strands and annotations of subject cluster:

Table of Blast hits (query gene, subject gene, %identity, blast score, %coverage, e-value):
input.path1.gene29	TERG_02850	40	767	122.93478260869566	2.1e-219
input.path1.gene33	TERG_02845	44	265	91.14583333333334	1.1e-68


>>
10. NW_003345192
Source: Nannizzia gypsea CBS 118893 supercont1.10 genomic scaffold, whole genome shotgun sequence
Type: T1PKS
Number of proteins with BLAST hits to this cluster: 2
Cumulative BLAST score: 1030

Table of genes, locations, strands and annotations of subject cluster:

Table of Blast hits (query gene, subject gene, %identity, blast score, %coverage, e-value):
input.path1.gene29	MGYG_08684	40	768	121.95652173913044	7.3e-220
input.path1.gene33	MGYG_08689	44	262	91.14583333333334	5.5e-68


>>
11. NZ_CP023699
Source: Streptomyces kanamyceticus strain ATCC 12853 chromosome, complete genome
Type: T1PKS,transAT-PKS
Number of proteins with BLAST hits to this cluster: 2
Cumulative BLAST score: 1029

Table of genes, locations, strands and annotations of subject cluster:
CP970_RS36570	CP970_RS36570	8385184	8386041	-	SDR_family_NAD(P)-dependent_oxidoreductase
CP970_RS36625	CP970_RS36625	8410567	8411358	+	acyltransferase_domain-containing_protein
CP970_RS36630	CP970_RS36630	8412391	8413578	+	SDR_family_NAD(P)-dependent_oxidoreductase
CP970_RS36635	CP970_RS36635	8414071	8414936	+	type_I_polyketide_synthase
CP970_RS36640	CP970_RS36640	8417718	8418905	+	SDR_family_NAD(P)-dependent_oxidoreductase
CP970_RS36645	CP970_RS36645	8418924	8420870	+	hypothetical_protein
CP970_RS36650	CP970_RS36650	8421222	8422013	+	acyltransferase_domain-containing_protein
CP970_RS36655	CP970_RS36655	8422041	8423066	+	hypothetical_protein
CP970_RS36670	CP970_RS36670	8429343	8429846	+	hypothetical_protein

Table of Blast hits (query gene, subject gene, %identity, blast score, %coverage, e-value):
input.path1.gene29	CP970_RS36685	34	242	46.19565217391305	2.4e-61
input.path1.gene29	CP970_RS36665	33	238	49.130434782608695	2e-60
input.path1.gene29	CP970_RS36645	35	237	45.65217391304348	4.5e-60
input.path1.gene29	CP970_RS36675	36	237	45.97826086956522	5.9e-60
input.path1.gene34	CP970_RS36695	32	75	32.84132841328413	2.3e-11


>>
12. NW_017264193
Source: Xylona heveae TC161 unplaced genomic scaffold L228scaffold_1, whole genome shotgun sequence
Type: T1PKS
Number of proteins with BLAST hits to this cluster: 2
Cumulative BLAST score: 1014

Table of genes, locations, strands and annotations of subject cluster:

Table of Blast hits (query gene, subject gene, %identity, blast score, %coverage, e-value):
input.path1.gene29	L228DRAFT_235437	40	771	125.0	1.1e-220
input.path1.gene33	L228DRAFT_235439	42	243	92.96875	4.5e-62


>>
13. NW_003299161
Source: Microsporum canis CBS 113480 supercont1.9 genomic scaffold, whole genome shotgun sequence
Type: T1PKS
Number of proteins with BLAST hits to this cluster: 2
Cumulative BLAST score: 970

Table of genes, locations, strands and annotations of subject cluster:

Table of Blast hits (query gene, subject gene, %identity, blast score, %coverage, e-value):
input.path1.gene29	MCYG_08535	40	743	118.4782608695652	3.3e-212
input.path1.gene33	MCYG_08540	43	227	76.04166666666666	3.3e-57


>>
14. NW_022984632
Source: Aspergillus tanneri strain NIH1004 chromosome Unknown AapTan_5_pilon_pilon_pilon, whole genome shotgun sequence
Type: T1PKS
Number of proteins with BLAST hits to this cluster: 2
Cumulative BLAST score: 968

Table of genes, locations, strands and annotations of subject cluster:

Table of Blast hits (query gene, subject gene, %identity, blast score, %coverage, e-value):
input.path1.gene29	ATNIH1004_009518	40	735	123.58695652173913	9e-210
input.path1.gene33	ATNIH1004_009523	34	233	100.0	2.7e-59


>>
15. NC_038015
Source: Fusarium venenatum strain A3/5 genome assembly, chromosome: IIII
Type: T1PKS
Number of proteins with BLAST hits to this cluster: 2
Cumulative BLAST score: 932

Table of genes, locations, strands and annotations of subject cluster:

Table of Blast hits (query gene, subject gene, %identity, blast score, %coverage, e-value):
input.path1.gene29	FVRRES_13609	39	711	121.52173913043478	8.2e-203
input.path1.gene33	FVRRES_13608	35	221	90.88541666666666	1.8e-55


>>
16. NC_030957
Source: Colletotrichum higginsianum IMI 349063 chromosome 4, whole genome shotgun sequence
Type: T1PKS
Number of proteins with BLAST hits to this cluster: 2
Cumulative BLAST score: 923

Table of genes, locations, strands and annotations of subject cluster:

Table of Blast hits (query gene, subject gene, %identity, blast score, %coverage, e-value):
input.path1.gene29	CH63R_06245	38	691	123.80434782608695	1.1e-196
input.path1.gene33	CH63R_06246	38	232	99.47916666666666	7.9e-59


>>
17. NW_022194791
Source: Fusarium proliferatum ET1 genome assembly, contig: FPRO_scaffold002, whole genome shotgun sequence
Type: T1PKS
Number of proteins with BLAST hits to this cluster: 2
Cumulative BLAST score: 915

Table of genes, locations, strands and annotations of subject cluster:

Table of Blast hits (query gene, subject gene, %identity, blast score, %coverage, e-value):
input.path1.gene29	FPRO_03539	38	696	121.52173913043478	4.6e-198
input.path1.gene33	FPRO_03538	36	219	90.625	5.3e-55


>>
18. NC_036623
Source: Fusarium fujikuroi IMI 58289 draft genome, chromosome FFUJ_chr02
Type: T1PKS
Number of proteins with BLAST hits to this cluster: 2
Cumulative BLAST score: 914

Table of genes, locations, strands and annotations of subject cluster:

Table of Blast hits (query gene, subject gene, %identity, blast score, %coverage, e-value):
input.path1.gene29	FFUJ_03984	38	691	121.52173913043478	1.1e-196
input.path1.gene33	FFUJ_03985	36	223	91.14583333333334	4.8e-56


>>
19. CM000574
Source: Fusarium graminearum PH-1 chromosome 1, whole genome shotgun sequence
Type: NRPS,T1PKS
Number of proteins with BLAST hits to this cluster: 2
Cumulative BLAST score: 898

Table of genes, locations, strands and annotations of subject cluster:

Table of Blast hits (query gene, subject gene, %identity, blast score, %coverage, e-value):
input.path1.gene29	FGSG_02324	41	661	100.8695652173913	1.3e-187
input.path1.gene33	FGSG_02326	40	237	85.67708333333334	3.2e-60


>>
20. NC_026474
Source: Fusarium graminearum PH-1 chromosome 1, whole genome shotgun sequence
Type: NRPS,T1PKS
Number of proteins with BLAST hits to this cluster: 2
Cumulative BLAST score: 898

Table of genes, locations, strands and annotations of subject cluster:

Table of Blast hits (query gene, subject gene, %identity, blast score, %coverage, e-value):
input.path1.gene29	FGSG_02324	41	661	100.8695652173913	1.3e-187
input.path1.gene33	FGSG_02326	40	237	85.67708333333334	3.2e-60


>>
21. NC_038012
Source: Fusarium venenatum strain A3/5 genome assembly, chromosome: I
Type: NRPS,T1PKS
Number of proteins with BLAST hits to this cluster: 2
Cumulative BLAST score: 896

Table of genes, locations, strands and annotations of subject cluster:

Table of Blast hits (query gene, subject gene, %identity, blast score, %coverage, e-value):
input.path1.gene29	FVRRES_02715	41	655	98.69565217391305	9.1e-186
input.path1.gene33	FVRRES_02717	40	241	86.97916666666666	1.3e-61


>>
22. CM000577
Source: Fusarium graminearum PH-1 chromosome 4, whole genome shotgun sequence
Type: T1PKS
Number of proteins with BLAST hits to this cluster: 2
Cumulative BLAST score: 895

Table of genes, locations, strands and annotations of subject cluster:

Table of Blast hits (query gene, subject gene, %identity, blast score, %coverage, e-value):
input.path1.gene29	FGSG_09182	38	685	122.28260869565217	6.3e-195
input.path1.gene33	FGSG_09183	38	210	83.85416666666666	4.2e-52


>>
23. NC_026477
Source: Fusarium graminearum PH-1 chromosome 4, whole genome shotgun sequence
Type: T1PKS
Number of proteins with BLAST hits to this cluster: 2
Cumulative BLAST score: 895

Table of genes, locations, strands and annotations of subject cluster:

Table of Blast hits (query gene, subject gene, %identity, blast score, %coverage, e-value):
input.path1.gene29	FGSG_09182	38	685	122.28260869565217	6.3e-195
input.path1.gene33	FGSG_09183	38	210	83.85416666666666	4.2e-52


>>
24. NW_023336269
Source: Colletotrichum scovillei strain TJNH1 chromosome Unknown Contig00011, whole genome shotgun sequence
Type: T1PKS
Number of proteins with BLAST hits to this cluster: 2
Cumulative BLAST score: 882

Table of genes, locations, strands and annotations of subject cluster:

Table of Blast hits (query gene, subject gene, %identity, blast score, %coverage, e-value):
input.path1.gene29	HER10_EVM0004425	38	680	123.80434782608695	2.6e-193
input.path1.gene33	HER10_EVM0004116	43	202	69.27083333333334	8.8e-50


>>
25. NW_006271969
Source: Cordyceps militaris CM01 unplaced genomic scaffold CCM_S00001, whole genome shotgun sequence
Type: T1PKS
Number of proteins with BLAST hits to this cluster: 2
Cumulative BLAST score: 873

Table of genes, locations, strands and annotations of subject cluster:

Table of Blast hits (query gene, subject gene, %identity, blast score, %coverage, e-value):
input.path1.gene29	CCM_01921	40	628	99.67391304347825	9.1e-178
input.path1.gene33	CCM_01923	41	245	91.40625	1.2e-62


>>
26. NC_049562
Source: Talaromyces rugulosus chromosome II, complete sequence
Type: T1PKS
Number of proteins with BLAST hits to this cluster: 2
Cumulative BLAST score: 866

Table of genes, locations, strands and annotations of subject cluster:

Table of Blast hits (query gene, subject gene, %identity, blast score, %coverage, e-value):
input.path1.gene29	TRUGW13939_04214	36	631	122.93478260869566	1.1e-178
input.path1.gene33	TRUGW13939_04213	40	235	93.48958333333334	9.3e-60


>>
27. NZ_RFFH01000003
Source: Nocardia stercoris strain NEAU-LL90 Scaffold3, whole genome shotgun sequence
Type: NRPS,NRPS-like,T1PKS
Number of proteins with BLAST hits to this cluster: 2
Cumulative BLAST score: 864

Table of genes, locations, strands and annotations of subject cluster:
EBN03_RS09775	EBN03_RS09775	297319	297710	-	nitroreductase_family_deazaflavin-dependent_oxidoreductase
EBN03_RS09790	EBN03_RS09790	299480	300307	+	helix-turn-helix_domain-containing_protein
EBN03_RS09805	EBN03_RS09805	302297	302948	-	methyltransferase_domain-containing_protein
EBN03_RS09810	EBN03_RS09810	303006	303536	+	TetR/AcrR_family_transcriptional_regulator

Table of Blast hits (query gene, subject gene, %identity, blast score, %coverage, e-value):
input.path1.gene29	EBN03_RS09630	40	281	44.891304347826086	2.7e-73
input.path1.gene29	EBN03_RS09770	36	251	47.71739130434783	3.9e-64
input.path1.gene29	EBN03_RS09635	37	241	44.565217391304344	3.1e-61
input.path1.gene30	EBN03_RS09785	31	91	84.0	1.2e-16


>>
28. NW_022984633
Source: Aspergillus tanneri strain NIH1004 chromosome Unknown AapTan_6_pilon_pilon_pilon, whole genome shotgun sequence
Type: T1PKS
Number of proteins with BLAST hits to this cluster: 2
Cumulative BLAST score: 852

Table of genes, locations, strands and annotations of subject cluster:

Table of Blast hits (query gene, subject gene, %identity, blast score, %coverage, e-value):
input.path1.gene29	ATNIH1004_008649	40	592	98.80434782608695	9.4e-167
input.path1.gene33	ATNIH1004_008651	44	260	93.75	2.7e-67


>>
29. NW_023336282
Source: Aspergillus tubingensis WU-2223L DNA, scaffold_8, whole genome shotgun sequence
Type: NRPS,T1PKS
Number of proteins with BLAST hits to this cluster: 2
Cumulative BLAST score: 830

Table of genes, locations, strands and annotations of subject cluster:

Table of Blast hits (query gene, subject gene, %identity, blast score, %coverage, e-value):
input.path1.gene29	AtWU_06971	35	523	98.15217391304348	5.4e-146
input.path1.gene29	AtWU_06980	35	240	45.76086956521739	9.1e-61
input.path1.gene34	AtWU_06979	31	67	36.715867158671585	3.6e-09


>>
30. NZ_CP048405
Source: Streptacidiphilus sp. PB12-B1b chromosome, complete genome
Type: NRPS,PKS-like,T1PKS,T2PKS,other,transAT-PKS
Number of proteins with BLAST hits to this cluster: 2
Cumulative BLAST score: 822

Table of genes, locations, strands and annotations of subject cluster:
GXW83_RS23155	GXW83_RS23155	5302976	5304168	+	DUF1205_domain-containing_protein
GXW83_RS23245	GXW83_RS23245	5323573	5324002	-	AraC_family_ligand_binding_domain-containing_protein
GXW83_RS23415	GXW83_RS23415	5359878	5361014	+	oleate_hydratase
GXW83_RS23440	GXW83_RS23440	5363877	5364622	-	MmgE/PrpD_family_protein
GXW83_RS23505	GXW83_RS23505	5377925	5378838	-	phytanoyl-CoA_dioxygenase_family_protein
GXW83_RS23615	GXW83_RS23615	5402867	5403112	+	aldo/keto_reductase
GXW83_RS23625	GXW83_RS23625	5403626	5403796	+	aldo/keto_reductase
GXW83_RS23780	GXW83_RS23780	5495658	5496320	+	4'-phosphopantetheinyl_transferase_superfamily_protein
GXW83_RS23935	GXW83_RS23935	5528603	5528897	+	carotenoid_oxygenase_family_protein

Table of Blast hits (query gene, subject gene, %identity, blast score, %coverage, e-value):
input.path1.gene29	GXW83_RS23740	36	253	45.0	6.1e-65
input.path1.gene29	GXW83_RS23735	40	242	41.52173913043478	1.4e-61
input.path1.gene29	GXW83_RS23760	36	236	46.19565217391305	1.3e-59
input.path1.gene34	GXW83_RS23230	32	91	38.00738007380074	4e-16


>>
31. CM000590
Source: Fusarium oxysporum f. sp. lycopersici 4287 chromosome 2, whole genome shotgun sequence
Type: T1PKS
Number of proteins with BLAST hits to this cluster: 2
Cumulative BLAST score: 820

Table of genes, locations, strands and annotations of subject cluster:

Table of Blast hits (query gene, subject gene, %identity, blast score, %coverage, e-value):
input.path1.gene29	FOXG_05816	41	638	97.28260869565217	1.5e-180
input.path1.gene33	FOXG_05817	38	182	68.75	9.4e-44


>>
32. NC_030987
Source: Fusarium oxysporum f. sp. lycopersici 4287 chromosome 2, whole genome shotgun sequence
Type: T1PKS
Number of proteins with BLAST hits to this cluster: 2
Cumulative BLAST score: 820

Table of genes, locations, strands and annotations of subject cluster:

Table of Blast hits (query gene, subject gene, %identity, blast score, %coverage, e-value):
input.path1.gene29	FOXG_05816	41	638	97.28260869565217	1.5e-180
input.path1.gene33	FOXG_05817	38	182	68.75	9.4e-44


>>
33. NC_036442
Source: Aspergillus oryzae RIB40 DNA, chromosome 8
Type: T1PKS
Number of proteins with BLAST hits to this cluster: 2
Cumulative BLAST score: 808

Table of genes, locations, strands and annotations of subject cluster:

Table of Blast hits (query gene, subject gene, %identity, blast score, %coverage, e-value):
input.path1.gene29	AO090010000048	39	583	98.04347826086956	3.3e-164
input.path1.gene33	AO090010000047	40	225	94.01041666666666	7.4e-57


>>
34. NT_165931
Source: Aspergillus terreus NIH2624 scaffold_8 genomic scaffold, whole genome shotgun sequence
Type: T1PKS
Number of proteins with BLAST hits to this cluster: 2
Cumulative BLAST score: 806

Table of genes, locations, strands and annotations of subject cluster:

Table of Blast hits (query gene, subject gene, %identity, blast score, %coverage, e-value):
input.path1.gene29	ATEG_06206	41	549	88.80434782608695	7e-154
input.path1.gene33	ATEG_06203	43	257	82.8125	3e-66


>>
35. NZ_WOFH01000003
Source: Actinomadura sp. NEAU-AAG5 Scaffold3, whole genome shotgun sequence
Type: NRPS,NRPS-like,T1PKS,blactam
Number of proteins with BLAST hits to this cluster: 2
Cumulative BLAST score: 804

Table of genes, locations, strands and annotations of subject cluster:
GNZ18_RS40765	GNZ18_RS40765	243139	244056	-	SDR_family_NAD(P)-dependent_oxidoreductase
GNZ18_RS40770	GNZ18_RS40770	244258	245247	-	polyketide_synthase_dehydratase_domain-containing_protein
GNZ18_RS40775	GNZ18_RS40775	246382	247647	-	polyketide_synthase
GNZ18_RS40780	GNZ18_RS40780	247690	248199	-	hypothetical_protein
GNZ18_RS40785	GNZ18_RS40785	248215	249246	-	SDR_family_NAD(P)-dependent_oxidoreductase
GNZ18_RS40790	GNZ18_RS40790	250207	251586	-	acyltransferase_domain-containing_protein
GNZ18_RS40795	GNZ18_RS40795	251701	252762	-	polyketide_synthase

Table of Blast hits (query gene, subject gene, %identity, blast score, %coverage, e-value):
input.path1.gene29	GNZ18_RS09130	36	243	45.869565217391305	6.3e-62
input.path1.gene29	GNZ18_RS09165	37	240	46.95652173913044	9.1e-61
input.path1.gene29	GNZ18_RS09170	36	238	48.80434782608696	2.6e-60
input.path1.gene34	GNZ18_RS09135	31	83	34.87084870848708	8.4e-14


>>
36. NZ_GL877879
Source: Saccharopolyspora spinosa NRRL 18395 Scaffold000002, whole genome shotgun sequence
Type: NRPS,T1PKS
Number of proteins with BLAST hits to this cluster: 2
Cumulative BLAST score: 800

Table of genes, locations, strands and annotations of subject cluster:
SSPN_RS51610	SSPN_RS51610	745795	746152	+	phage_terminase_small_subunit_P27_family
SSPN_RS45120	SSPN_RS45120	753971	755255	+	IS256_family_transposase
SSPN_RS55220	SSPN_RS55220	785451	791151	-	SDR_family_NAD(P)-dependent_oxidoreductase
SSPN_RS55225	SSPN_RS55225	793424	797587	-	SDR_family_NAD(P)-dependent_oxidoreductase
SSPN_RS45145	SSPN_RS45145	797915	805840	-	SDR_family_NAD(P)-dependent_oxidoreductase

Table of Blast hits (query gene, subject gene, %identity, blast score, %coverage, e-value):
input.path1.gene29	SSPN_RS45150	35	243	45.21739130434783	8.2e-62
input.path1.gene29	SSPN_RS51660	37	243	45.43478260869565	8.2e-62
input.path1.gene29	SSPN_RS0122785	35	242	45.869565217391305	2.4e-61
input.path1.gene34	SSPN_RS45175	32	72	32.84132841328413	1.1e-10


>>
37. NW_022194796
Source: Fusarium proliferatum ET1 genome assembly, contig: FPRO_scaffold007, whole genome shotgun sequence
Type: T1PKS
Number of proteins with BLAST hits to this cluster: 2
Cumulative BLAST score: 799

Table of genes, locations, strands and annotations of subject cluster:

Table of Blast hits (query gene, subject gene, %identity, blast score, %coverage, e-value):
input.path1.gene29	FPRO_07275	40	593	98.47826086956522	3.2e-167
input.path1.gene33	FPRO_07277	37	206	92.70833333333334	6.1e-51


>>
38. NC_036626
Source: Fusarium fujikuroi IMI 58289 draft genome, chromosome FFUJ_chr05
Type: T1PKS
Number of proteins with BLAST hits to this cluster: 2
Cumulative BLAST score: 795

Table of genes, locations, strands and annotations of subject cluster:

Table of Blast hits (query gene, subject gene, %identity, blast score, %coverage, e-value):
input.path1.gene29	FFUJ_06742	39	589	98.47826086956522	6.1e-166
input.path1.gene33	FFUJ_06744	37	206	92.70833333333334	4.6e-51


>>
39. NW_017971433
Source: Talaromyces atroroseus strain IBT 11181 chromosome Unknown scaffold_1, whole genome shotgun sequence
Type: T1PKS
Number of proteins with BLAST hits to this cluster: 2
Cumulative BLAST score: 728

Table of genes, locations, strands and annotations of subject cluster:

Table of Blast hits (query gene, subject gene, %identity, blast score, %coverage, e-value):
input.path1.gene29	UA08_01172	36	521	97.3913043478261	2e-145
input.path1.gene33	UA08_01176	38	207	85.41666666666666	2.1e-51


>>
40. NW_013562491
Source: Exophiala xenobiotica strain CBS 118157 unplaced genomic scaffold supercont1.4, whole genome shotgun sequence
Type: T1PKS
Number of proteins with BLAST hits to this cluster: 2
Cumulative BLAST score: 687

Table of genes, locations, strands and annotations of subject cluster:

Table of Blast hits (query gene, subject gene, %identity, blast score, %coverage, e-value):
input.path1.gene29	PV05_06233	39	561	97.28260869565217	1.4e-157
input.path1.gene33	PV05_06224	31	126	89.32291666666666	8e-27


>>
41. NW_001914858
Source: Podospora anserina S mat+ genomic DNA chromosome 2, supercontig 2
Type: T1PKS
Number of proteins with BLAST hits to this cluster: 2
Cumulative BLAST score: 677

Table of genes, locations, strands and annotations of subject cluster:

Table of Blast hits (query gene, subject gene, %identity, blast score, %coverage, e-value):
input.path1.gene29	PODANS_2_7410	39	591	97.93478260869564	2.1e-166
input.path1.gene33	PODANS_2_7430	31	86	56.25	7e-15


>>
42. NC_049561
Source: Talaromyces rugulosus chromosome I, complete sequence
Type: T1PKS
Number of proteins with BLAST hits to this cluster: 2
Cumulative BLAST score: 656

Table of genes, locations, strands and annotations of subject cluster:

Table of Blast hits (query gene, subject gene, %identity, blast score, %coverage, e-value):
input.path1.gene29	TRUGW13939_02416	42	300	44.45652173913043	7.4e-79
input.path1.gene29	TRUGW13939_02410	37	250	41.52173913043478	6.7e-64
input.path1.gene34	TRUGW13939_02411	35	106	38.56088560885609	1.2e-20


>>
43. NZ_CP023694
Source: Streptomyces coeruleorubidus strain ATCC 13740 chromosome, complete genome
Type: NRPS,T1PKS,siderophore,transAT-PKS,transAT-PKS-like
Number of proteins with BLAST hits to this cluster: 2
Cumulative BLAST score: 621

Table of genes, locations, strands and annotations of subject cluster:
CP976_RS38345	CP976_RS38345	8350275	8350792	+	GNAT_family_N-acetyltransferase
CP976_RS38520	CP976_RS38520	8451328	8452352	-	helix-turn-helix_transcriptional_regulator
CP976_RS38535	CP976_RS38535	8454105	8454338	-	DUF1275_domain-containing_protein
CP976_RS38600	CP976_RS38600	8473894	8474031	+	FAD-dependent_oxidoreductase
CP976_RS38710	CP976_RS38710	8497138	8497328	+	alpha/beta_hydrolase

Table of Blast hits (query gene, subject gene, %identity, blast score, %coverage, e-value):
input.path1.gene29	CP976_RS38415	40	271	44.23913043478261	3.7e-70
input.path1.gene29	CP976_RS38395	39	253	41.41304347826087	1e-64
input.path1.gene34	CP976_RS38705	31	97	35.79335793357934	3.3e-18


>>
44. NZ_LT629775
Source: Streptomyces sp. TLI_053 chromosome I
Type: NRPS,NRPS-like,T1PKS
Number of proteins with BLAST hits to this cluster: 2
Cumulative BLAST score: 599

Table of genes, locations, strands and annotations of subject cluster:
BLU95_RS37245	BLU95_RS37245	8949849	8950946	-	hypothetical_protein
BLU95_RS43135	BLU95_RS43135	8990001	8990090	+	DUF2662_domain-containing_protein
BLU95_RS43140	BLU95_RS43140	8990791	8991612	-	zinc-binding_dehydrogenase
BLU95_RS43145	BLU95_RS43145	8992627	8994342	-	polyketide_synthase

Table of Blast hits (query gene, subject gene, %identity, blast score, %coverage, e-value):
input.path1.gene29	BLU95_RS37320	40	277	44.565217391304344	3.9e-72
input.path1.gene29	BLU95_RS37330	39	259	44.78260869565218	1.4e-66
input.path1.gene33	BLU95_RS37345	33	63	36.19791666666667	4.9e-08


>>
45. NZ_CP036534
Source: Streptomyces sp. VN1 chromosome, complete genome
Type: NRPS,T1PKS,T3PKS,oligosaccharide,other
Number of proteins with BLAST hits to this cluster: 2
Cumulative BLAST score: 579

Table of genes, locations, strands and annotations of subject cluster:
EZV63_RS00065	EZV63_RS00065	14386	16827	-	helix-turn-helix_transcriptional_regulator
EZV63_RS00090	EZV63_RS00090	23641	23749	-	thioesterase
EZV63_RS00150	EZV63_RS00150	53965	56637	+	type_I_polyketide_synthase
EZV63_RS00155	EZV63_RS00155	57787	58674	+	KR_domain-containing_protein
EZV63_RS00160	EZV63_RS00160	58753	59187	+	hypothetical_protein
EZV63_RS00165	EZV63_RS00165	59272	60000	+	polyketide_synthase
EZV63_RS00170	EZV63_RS00170	60322	60717	+	hypothetical_protein
EZV63_RS00175	EZV63_RS00175	61651	62700	+	hypothetical_protein
EZV63_RS00180	EZV63_RS00180	63217	65358	+	SDR_family_NAD(P)-dependent_oxidoreductase
EZV63_RS00185	EZV63_RS00185	65563	65940	+	hypothetical_protein
EZV63_RS00190	EZV63_RS00190	66166	66975	+	acyltransferase_domain-containing_protein
EZV63_RS00195	EZV63_RS00195	67927	69399	+	SDR_family_NAD(P)-dependent_oxidoreductase
EZV63_RS00200	EZV63_RS00200	69610	72615	+	acyltransferase_domain-containing_protein
EZV63_RS00205	EZV63_RS00205	73447	74706	+	SDR_family_NAD(P)-dependent_oxidoreductase
EZV63_RS00210	EZV63_RS00210	74719	75225	+	hypothetical_protein
EZV63_RS00450	EZV63_RS00450	175527	176252	-	KR_domain-containing_protein
EZV63_RS00455	EZV63_RS00455	177255	179702	-	type_I_polyketide_synthase
EZV63_RS00460	EZV63_RS00460	179823	180350	-	hypothetical_protein
EZV63_RS00465	EZV63_RS00465	180360	180968	-	SDR_family_NAD(P)-dependent_oxidoreductase
EZV63_RS00470	EZV63_RS00470	181623	182228	-	alcohol_dehydrogenase_catalytic_domain-containing_protein
EZV63_RS00475	EZV63_RS00475	182517	183347	-	polyketide_synthase
EZV63_RS00480	EZV63_RS00480	183477	184385	-	acyltransferase_domain-containing_protein
EZV63_RS00485	EZV63_RS00485	184644	186137	-	polyketide_synthase
EZV63_RS00495	EZV63_RS00495	186973	187644	-	SDR_family_NAD(P)-dependent_oxidoreductase
EZV63_RS00500	EZV63_RS00500	188296	188715	-	alcohol_dehydrogenase_catalytic_domain-containing_protein
EZV63_RS00505	EZV63_RS00505	190135	192780	-	type_I_polyketide_synthase
EZV63_RS00510	EZV63_RS00510	192886	193260	-	acyl_carrier_protein
EZV63_RS00515	EZV63_RS00515	193417	194178	-	SDR_family_NAD(P)-dependent_oxidoreductase
EZV63_RS00520	EZV63_RS00520	194800	198006	-	acyltransferase_domain-containing_protein
EZV63_RS00525	EZV63_RS00525	198028	198996	-	SDR_family_NAD(P)-dependent_oxidoreductase
EZV63_RS00530	EZV63_RS00530	199183	200313	-	polyketide_synthase
EZV63_RS00535	EZV63_RS00535	201355	202782	-	polyketide_synthase
EZV63_RS00540	EZV63_RS00540	202933	203100	-	acyl_carrier_protein
EZV63_RS00545	EZV63_RS00545	203695	204063	-	acyltransferase_domain-containing_protein
EZV63_RS00550	EZV63_RS00550	204310	204786	-	hypothetical_protein
EZV63_RS00555	EZV63_RS00555	205057	205803	-	polyketide_synthase
EZV63_RS00575	EZV63_RS00575	209243	210280	+	LLM_class_flavin-dependent_oxidoreductase

Table of Blast hits (query gene, subject gene, %identity, blast score, %coverage, e-value):
input.path1.gene29	EZV63_RS00145	36	247	44.45652173913043	5.7e-63
input.path1.gene29	EZV63_RS00485	35	241	46.95652173913044	3.1e-61
input.path1.gene34	EZV63_RS00265	32	91	38.191881918819185	3.1e-16


>>
46. NZ_LJIW01000001
Source: Streptomyces malaysiensis strain F913 SMF9131, whole genome shotgun sequence
Type: T1PKS,T3PKS,nucleoside,transAT-PKS
Number of proteins with BLAST hits to this cluster: 2
Cumulative BLAST score: 578

Table of genes, locations, strands and annotations of subject cluster:
SMF913_RS00780	SMF913_RS00780	136718	136825	+	IS5/IS1182_family_transposase
SMF913_RS00800	SMF913_RS00800	139724	139840	+	IS5/IS1182_family_transposase
SMF913_RS00805	SMF913_RS00805	139882	140073	-	helix-turn-helix_transcriptional_regulator
SMF913_RS00810	SMF913_RS00810	140115	140512	+	SDR_family_NAD(P)-dependent_oxidoreductase
SMF913_RS00835	SMF913_RS00835	142522	142893	+	transposase
SMF913_RS00890	SMF913_RS00890	156788	157000	-	transposase
SMF913_RS01010	SMF913_RS01010	214453	216225	-	polyketide_synthase_dehydratase_domain-containing_protein
SMF913_RS01015	SMF913_RS01015	216352	218898	-	KR_domain-containing_protein
SMF913_RS48480	SMF913_RS48480	218902	219378	-	zinc-binding_dehydrogenase
SMF913_RS48485	SMF913_RS48485	219577	220368	-	alcohol_dehydrogenase_catalytic_domain-containing_protein
SMF913_RS48490	SMF913_RS48490	220396	222192	-	acyltransferase_domain-containing_protein
SMF913_RS01030	SMF913_RS01030	222229	223938	-	hypothetical_protein
SMF913_RS01055	SMF913_RS01055	241763	255700	-	SDR_family_NAD(P)-dependent_oxidoreductase
SMF913_RS48495	SMF913_RS48495	255842	256696	-	polyketide_synthase
SMF913_RS47930	SMF913_RS47930	256694	257680	-	type_I_polyketide_synthase
SMF913_RS47935	SMF913_RS47935	257822	262231	-	SDR_family_NAD(P)-dependent_oxidoreductase
SMF913_RS47940	SMF913_RS47940	262274	262378	-	polyketide_synthase_docking_domain-containing_protein
SMF913_RS01115	SMF913_RS01115	317275	317529	-	ribonuclease_PH

Table of Blast hits (query gene, subject gene, %identity, blast score, %coverage, e-value):
input.path1.gene29	SMF913_RS01055	38	243	40.869565217391305	1.1e-61
input.path1.gene29	SMF913_RS01065	35	238	44.45652173913043	3.5e-60
input.path1.gene30	SMF913_RS00970	31	97	83.63636363636363	2.8e-18


>>
47. NZ_CP054932
Source: Actinomadura sp. NAK00032 chromosome, complete genome
Type: NRPS,T1PKS,other
Number of proteins with BLAST hits to this cluster: 2
Cumulative BLAST score: 570

Table of genes, locations, strands and annotations of subject cluster:

Table of Blast hits (query gene, subject gene, %identity, blast score, %coverage, e-value):
input.path1.gene29	HUT06_RS29200	35	237	45.65217391304348	5.9e-60
input.path1.gene29	HUT06_RS29215	35	236	46.73913043478261	1e-59
input.path1.gene34	HUT06_RS29150	32	97	33.76383763837638	5.6e-18


>>
48. NZ_CP040605
Source: Saccharopolyspora sp. ASAGF58 chromosome, complete genome
Type: NRPS,T1PKS,lanthipeptide
Number of proteins with BLAST hits to this cluster: 2
Cumulative BLAST score: 557

Table of genes, locations, strands and annotations of subject cluster:
FDZ84_RS24220	FDZ84_RS24220	5119327	5119622	-	hypothetical_protein
FDZ84_RS24335	FDZ84_RS24335	5145731	5145966	+	IS110_family_transposase
FDZ84_RS24340	FDZ84_RS24340	5146682	5147634	-	tyrosine-type_recombinase/integrase
FDZ84_RS24425	FDZ84_RS24425	5173290	5175128	+	Gfo/Idh/MocA_family_oxidoreductase
FDZ84_RS24540	FDZ84_RS24540	5193069	5193302	+	NUDIX_hydrolase
FDZ84_RS24550	FDZ84_RS24550	5194153	5194705	-	kinase
FDZ84_RS24605	FDZ84_RS24605	5205379	5206534	-	hypothetical_protein
FDZ84_RS24620	FDZ84_RS24620	5208008	5209032	-	hypothetical_protein
FDZ84_RS24695	FDZ84_RS24695	5222453	5223239	-	IS982_family_transposase
FDZ84_RS24700	FDZ84_RS24700	5223317	5223517	+	alpha/beta_hydrolase

Table of Blast hits (query gene, subject gene, %identity, blast score, %coverage, e-value):
input.path1.gene29	FDZ84_RS24210	36	244	46.52173913043478	4.8e-62
input.path1.gene29	FDZ84_RS24200	36	243	44.45652173913043	8.2e-62
input.path1.gene34	FDZ84_RS24170	32	70	32.84132841328413	7.3e-10


>>
49. NC_010612
Source: Mycobacterium marinum M, complete genome
Type: NRPS,T1PKS
Number of proteins with BLAST hits to this cluster: 2
Cumulative BLAST score: 549

Table of genes, locations, strands and annotations of subject cluster:
MMAR_RS27975	MMAR_RS27975	77197	77486	+	serine_hydrolase
MMAR_RS27980	MMAR_RS27980	91634	91978	+	methyltransferase_domain-containing_protein
MMAR_RS00470	MMAR_RS00470	93220	93537	+	SAM-dependent_methyltransferase

Table of Blast hits (query gene, subject gene, %identity, blast score, %coverage, e-value):
input.path1.gene29	MMAR_RS00515	34	237	48.04347826086956	4.5e-60
input.path1.gene29	MMAR_RS00510	35	237	45.326086956521735	5.9e-60
input.path1.gene34	MMAR_RS00555	31	75	31.54981549815498	2.3e-11


>>
50. NC_049561
Source: Talaromyces rugulosus chromosome I, complete sequence
Type: T1PKS
Number of proteins with BLAST hits to this cluster: 2
Cumulative BLAST score: 476

Table of genes, locations, strands and annotations of subject cluster:

Table of Blast hits (query gene, subject gene, %identity, blast score, %coverage, e-value):
input.path1.gene29	TRUGW13939_01413	41	277	44.67391304347826	6.7e-72
input.path1.gene32	TRUGW13939_01410	41	199	109.39849624060149	3.9e-49
