## Supplementary Results for "A draft genome of the ascomycotal fungal species *Pseudopithomyces maydicus* (family *Didymosphaeriaceae*)": contig_10_c2.txt

ClusterBlast scores for contig_10

Table of genes, locations, strands and annotations of query cluster:
input.path1.gene173	638688	639604	-		
input.path1.gene174	640219	642139	+		
input.path1.gene175	644124	648112	-		
input.path1.gene176	651038	652466	+		
input.path1.gene177	653296	656734	-		
input.path1.gene178	657822	662907	+		
input.path1.gene179	663094	664224	-		
input.path1.gene180	666162	668145	+		
input.path1.gene181	668454	668627	+		
input.path1.gene182	669026	671092	-		
input.path1.gene183	672052	673122	-		
input.path1.gene184	674274	676506	+		
input.path1.gene185	678483	681267	+		
input.path1.gene186	681672	682178	+		


Significant hits: 
1. NW_022984631	Aspergillus tanneri strain NIH1004 chromosome Unknown AapTan_4_pilon_pilon_pilon, whole genome shotgun sequence
2. NC_007200	Aspergillus fumigatus Af293 chromosome 7, whole genome shotgun sequence
3. NW_003299166	Microsporum canis CBS 113480 supercont1.4 genomic scaffold, whole genome shotgun sequence
4. CM000595	Fusarium oxysporum f. sp. lycopersici 4287 chromosome 7, whole genome shotgun sequence
5. NC_030992	Fusarium oxysporum f. sp. lycopersici 4287 chromosome 7, whole genome shotgun sequence
6. NZ_CP030864	Streptomyces globosus strain LZH-48 plasmid unnamed2, complete sequence
7. NW_017971433	Talaromyces atroroseus strain IBT 11181 chromosome Unknown scaffold_1, whole genome shotgun sequence
8. NW_022984634	Aspergillus tanneri strain NIH1004 chromosome Unknown AapTan_7_pilon_pilon_pilon, whole genome shotgun sequence
9. NC_007197	Aspergillus fumigatus Af293 chromosome 4, whole genome shotgun sequence
10. NW_019154043	Pochonia chlamydosporia 170 chromosome Unknown PCv3seq00008, whole genome shotgun sequence
11. NT_165935	Aspergillus terreus NIH2624 scaffold_12 genomic scaffold, whole genome shotgun sequence
12. NW_023336287	Aspergillus tubingensis WU-2223L DNA, scaffold_13, whole genome shotgun sequence
13. CM000574	Fusarium graminearum PH-1 chromosome 1, whole genome shotgun sequence
14. NC_026474	Fusarium graminearum PH-1 chromosome 1, whole genome shotgun sequence
15. NC_049562	Talaromyces rugulosus chromosome II, complete sequence
16. NC_016109	Kitasatospora setae KM-6054, complete genome
17. NW_003345196	Nannizzia gypsea CBS 118893 supercont1.6 genomic scaffold, whole genome shotgun sequence
18. NZ_JACHJH010000002	Streptomyces olivoverticillatus strain CECT 3266 Ga0436946_02, whole genome shotgun sequence
19. NC_036626	Fusarium fujikuroi IMI 58289 draft genome, chromosome FFUJ_chr05
20. NW_003315100	Trichophyton benhamiae CBS 112371 chromosome Unknown contig01356.0, whole genome shotgun sequence
21. NC_038012	Fusarium venenatum strain A3/5 genome assembly, chromosome: I
22. NW_022984633	Aspergillus tanneri strain NIH1004 chromosome Unknown AapTan_6_pilon_pilon_pilon, whole genome shotgun sequence
23. NW_023336282	Aspergillus tubingensis WU-2223L DNA, scaffold_8, whole genome shotgun sequence
24. NT_107009	Aspergillus nidulans FGSC A4 chromosome I map unlocalized scaffold_7, whole genome shotgun sequence
25. NW_003299167	Microsporum canis CBS 113480 supercont1.3 genomic scaffold, whole genome shotgun sequence
26. NW_022194796	Fusarium proliferatum ET1 genome assembly, contig: FPRO_scaffold007, whole genome shotgun sequence
27. NC_036442	Aspergillus oryzae RIB40 DNA, chromosome 8
28. NW_017264193	Xylona heveae TC161 unplaced genomic scaffold L228scaffold_1, whole genome shotgun sequence
29. NW_003345192	Nannizzia gypsea CBS 118893 supercont1.10 genomic scaffold, whole genome shotgun sequence
30. NW_003315103	Trichophyton benhamiae CBS 112371 chromosome Unknown contig00075.49, whole genome shotgun sequence
31. NW_003456426	Trichophyton rubrum CBS 118892 genomic scaffold supercont2.3, whole genome shotgun sequence
32. NW_003299161	Microsporum canis CBS 113480 supercont1.9 genomic scaffold, whole genome shotgun sequence
33. CM000577	Fusarium graminearum PH-1 chromosome 4, whole genome shotgun sequence
34. NC_026477	Fusarium graminearum PH-1 chromosome 4, whole genome shotgun sequence
35. NW_022984632	Aspergillus tanneri strain NIH1004 chromosome Unknown AapTan_5_pilon_pilon_pilon, whole genome shotgun sequence
36. NW_001820829	Sclerotinia sclerotiorum 1980 UF-70 scaffold_7 genomic scaffold, whole genome shotgun sequence
37. NW_006912895	Capronia coronata CBS 617.96 acBCG-supercont1.3, whole genome shotgun sequence
38. NT_165931	Aspergillus terreus NIH2624 scaffold_8 genomic scaffold, whole genome shotgun sequence
39. NC_036623	Fusarium fujikuroi IMI 58289 draft genome, chromosome FFUJ_chr02
40. NW_022194791	Fusarium proliferatum ET1 genome assembly, contig: FPRO_scaffold002, whole genome shotgun sequence
41. NZ_FRCX01000002	Duganella sacchari strain Sac-22, whole genome shotgun sequence
42. CM000590	Fusarium oxysporum f. sp. lycopersici 4287 chromosome 2, whole genome shotgun sequence
43. NC_030987	Fusarium oxysporum f. sp. lycopersici 4287 chromosome 2, whole genome shotgun sequence
44. NW_001939246	Pyrenophora tritici-repentis Pt-1C-BFP supercont1.3, whole genome shotgun sequence
45. NW_006912904	Capronia epimyces CBS 606.96 aczIQ-supercont1.1, whole genome shotgun sequence
46. NW_019716258	Ramularia collo-cygni strain URUG2 genome assembly, contig: RCC_scaffold04, whole genome shotgun sequence
47. NC_036437	Aspergillus oryzae RIB40 DNA, chromosome 3
48. NW_017971438	Talaromyces atroroseus strain IBT 11181 chromosome Unknown scaffold_6, whole genome shotgun sequence
49. NW_020939749	Sparassis crispa DNA, contig_000002F, whole genome shotgun sequence
50. NW_009276941	Verticillium dahliae VdLs.17 supercont1.26 genomic scaffold, whole genome shotgun sequence


Details:

>>
1. NW_022984631
Source: Aspergillus tanneri strain NIH1004 chromosome Unknown AapTan_4_pilon_pilon_pilon, whole genome shotgun sequence
Type: T1PKS,indole
Number of proteins with BLAST hits to this cluster: 2
Cumulative BLAST score: 912

Table of genes, locations, strands and annotations of subject cluster:

Table of Blast hits (query gene, subject gene, %identity, blast score, %coverage, e-value):
input.path1.gene177	ATNIH1004_005530	33	74	31.610044313146236	6.3e-11
input.path1.gene178	ATNIH1004_005531	37	838	114.97105045492142	7.6e-241


>>
2. NC_007200
Source: Aspergillus fumigatus Af293 chromosome 7, whole genome shotgun sequence
Type: T1PKS,indole
Number of proteins with BLAST hits to this cluster: 2
Cumulative BLAST score: 911

Table of genes, locations, strands and annotations of subject cluster:

Table of Blast hits (query gene, subject gene, %identity, blast score, %coverage, e-value):
input.path1.gene177	AFUA_7G00150	33	71	25.849335302806498	5.4e-10
input.path1.gene178	AFUA_7G00160	37	840	116.21174524400331	2e-241


>>
3. NW_003299166
Source: Microsporum canis CBS 113480 supercont1.4 genomic scaffold, whole genome shotgun sequence
Type: T1PKS
Number of proteins with BLAST hits to this cluster: 3
Cumulative BLAST score: 1099

Table of genes, locations, strands and annotations of subject cluster:

Table of Blast hits (query gene, subject gene, %identity, blast score, %coverage, e-value):
input.path1.gene176	MCYG_04450	33	91	55.04087193460491	1.6e-16
input.path1.gene177	MCYG_04450	37	157	40.02954209748892	4.4e-36
input.path1.gene178	MCYG_04442	38	851	116.70802315963607	1.1e-244


>>
4. CM000595
Source: Fusarium oxysporum f. sp. lycopersici 4287 chromosome 7, whole genome shotgun sequence
Type: T1PKS
Number of proteins with BLAST hits to this cluster: 2
Cumulative BLAST score: 1485

Table of genes, locations, strands and annotations of subject cluster:

Table of Blast hits (query gene, subject gene, %identity, blast score, %coverage, e-value):
input.path1.gene176	FOXG_04755	34	239	111.44414168937328	4.7e-61
input.path1.gene178	FOXG_04757	38	623	82.96112489660877	3.8e-176
input.path1.gene178	FOXG_04757_KNB01536.1	38	623	82.96112489660877	3.8e-176


>>
5. NC_030992
Source: Fusarium oxysporum f. sp. lycopersici 4287 chromosome 7, whole genome shotgun sequence
Type: T1PKS
Number of proteins with BLAST hits to this cluster: 2
Cumulative BLAST score: 1485

Table of genes, locations, strands and annotations of subject cluster:

Table of Blast hits (query gene, subject gene, %identity, blast score, %coverage, e-value):
input.path1.gene176	FOXG_04755	34	239	111.44414168937328	4.7e-61
input.path1.gene178	FOXG_04757	38	623	82.96112489660877	3.8e-176
input.path1.gene178	FOXG_04757_XP_018239581.1	38	623	82.96112489660877	3.8e-176


>>
6. NZ_CP030864
Source: Streptomyces globosus strain LZH-48 plasmid unnamed2, complete sequence
Type: NRPS,NRPS-like,T2PKS,butyrolactone,ectoine,hglE-KS,linaridin,transAT-PKS
Number of proteins with BLAST hits to this cluster: 2
Cumulative BLAST score: 1169

Table of genes, locations, strands and annotations of subject cluster:
C0216_RS31350	C0216_RS31350	25068	25503	+	transposase
C0216_RS33995	C0216_RS33995	31513	32988	-	hypothetical_protein
C0216_RS34000	C0216_RS34000	33058	33558	-	hypothetical_protein
C0216_RS34005	C0216_RS34005	33571	34563	-	SDR_family_NAD(P)-dependent_oxidoreductase
C0216_RS34010	C0216_RS34010	34585	35604	-	polyketide_synthase_dehydratase_domain-containing_protein
C0216_RS34015	C0216_RS34015	36697	38229	-	polyketide_synthase_docking_domain-containing_protein
C0216_RS31715	C0216_RS31715	175226	175600	-	hypothetical_protein
C0216_RS31790	C0216_RS31790	191392	191821	-	DUF1772_domain-containing_protein
C0216_RS34025	C0216_RS34025	296568	297596	+	NAD-dependent_epimerase/dehydratase_family_protein
C0216_RS32170	C0216_RS32170	303897	303992	+	short-chain_dehydrogenase

Table of Blast hits (query gene, subject gene, %identity, blast score, %coverage, e-value):
input.path1.gene176	C0216_RS31470	32	74	43.596730245231605	3.4e-11
input.path1.gene178	C0216_RS31395	42	233	26.63358147229115	8.6e-59
input.path1.gene178	C0216_RS31380	36	220	31.679073614557485	1.3e-54
input.path1.gene178	C0216_RS31390	36	218	32.42349048800662	4.9e-54
input.path1.gene178	C0216_RS31385	39	214	28.03970223325062	5.4e-53
input.path1.gene178	C0216_RS34015	37	210	27.626137303556657	1.3e-51


>>
7. NW_017971433
Source: Talaromyces atroroseus strain IBT 11181 chromosome Unknown scaffold_1, whole genome shotgun sequence
Type: T1PKS
Number of proteins with BLAST hits to this cluster: 2
Cumulative BLAST score: 1115

Table of genes, locations, strands and annotations of subject cluster:

Table of Blast hits (query gene, subject gene, %identity, blast score, %coverage, e-value):
input.path1.gene176	UA08_01176	32	225	110.62670299727519	1.2e-56
input.path1.gene178	UA08_01172	38	890	114.72291149710504	2.2e-256


>>
8. NW_022984634
Source: Aspergillus tanneri strain NIH1004 chromosome Unknown AapTan_7_pilon_pilon_pilon, whole genome shotgun sequence
Type: T1PKS
Number of proteins with BLAST hits to this cluster: 2
Cumulative BLAST score: 1037

Table of genes, locations, strands and annotations of subject cluster:

Table of Blast hits (query gene, subject gene, %identity, blast score, %coverage, e-value):
input.path1.gene176	ATNIH1004_008025	39	115	47.68392370572207	1e-23
input.path1.gene178	ATNIH1004_008023	40	922	114.64019851116625	4e-266


>>
9. NC_007197
Source: Aspergillus fumigatus Af293 chromosome 4, whole genome shotgun sequence
Type: T1PKS
Number of proteins with BLAST hits to this cluster: 2
Cumulative BLAST score: 1030

Table of genes, locations, strands and annotations of subject cluster:

Table of Blast hits (query gene, subject gene, %identity, blast score, %coverage, e-value):
input.path1.gene176	AFUA_4G14580	34	107	49.31880108991826	3.7e-21
input.path1.gene178	AFUA_4G14560	40	923	115.55004135649295	2.3e-266


>>
10. NW_019154043
Source: Pochonia chlamydosporia 170 chromosome Unknown PCv3seq00008, whole genome shotgun sequence
Type: T1PKS
Number of proteins with BLAST hits to this cluster: 2
Cumulative BLAST score: 1019

Table of genes, locations, strands and annotations of subject cluster:

Table of Blast hits (query gene, subject gene, %identity, blast score, %coverage, e-value):
input.path1.gene175	VFPPC_06839	40	151	32.61802575107296	3.2e-34
input.path1.gene178	VFPPC_06844	37	868	116.62531017369726	8.9e-250


>>
11. NT_165935
Source: Aspergillus terreus NIH2624 scaffold_12 genomic scaffold, whole genome shotgun sequence
Type: NRPS,T1PKS
Number of proteins with BLAST hits to this cluster: 2
Cumulative BLAST score: 1007

Table of genes, locations, strands and annotations of subject cluster:

Table of Blast hits (query gene, subject gene, %identity, blast score, %coverage, e-value):
input.path1.gene176	ATEG_08449	36	111	47.13896457765668	2.5e-22
input.path1.gene178	ATEG_08451	39	896	115.79818031430935	3.1e-258


>>
12. NW_023336287
Source: Aspergillus tubingensis WU-2223L DNA, scaffold_13, whole genome shotgun sequence
Type: T1PKS
Number of proteins with BLAST hits to this cluster: 2
Cumulative BLAST score: 998

Table of genes, locations, strands and annotations of subject cluster:

Table of Blast hits (query gene, subject gene, %identity, blast score, %coverage, e-value):
input.path1.gene176	AtWU_10462	40	294	111.44414168937328	1.6e-77
input.path1.gene178	AtWU_10461	34	704	114.2266335814723	1.7e-200


>>
13. CM000574
Source: Fusarium graminearum PH-1 chromosome 1, whole genome shotgun sequence
Type: NRPS,T1PKS
Number of proteins with BLAST hits to this cluster: 2
Cumulative BLAST score: 954

Table of genes, locations, strands and annotations of subject cluster:

Table of Blast hits (query gene, subject gene, %identity, blast score, %coverage, e-value):
input.path1.gene176	FGSG_02326	31	203	112.80653950953679	4.9e-50
input.path1.gene178	FGSG_02324	35	751	115.05376344086022	1.6e-214


>>
14. NC_026474
Source: Fusarium graminearum PH-1 chromosome 1, whole genome shotgun sequence
Type: NRPS,T1PKS
Number of proteins with BLAST hits to this cluster: 2
Cumulative BLAST score: 954

Table of genes, locations, strands and annotations of subject cluster:

Table of Blast hits (query gene, subject gene, %identity, blast score, %coverage, e-value):
input.path1.gene176	FGSG_02326	31	203	112.80653950953679	4.9e-50
input.path1.gene178	FGSG_02324	35	751	115.05376344086022	1.6e-214


>>
15. NC_049562
Source: Talaromyces rugulosus chromosome II, complete sequence
Type: T1PKS
Number of proteins with BLAST hits to this cluster: 2
Cumulative BLAST score: 953

Table of genes, locations, strands and annotations of subject cluster:

Table of Blast hits (query gene, subject gene, %identity, blast score, %coverage, e-value):
input.path1.gene176	TRUGW13939_04213	33	248	110.89918256130791	1e-63
input.path1.gene178	TRUGW13939_04214	34	705	116.87344913151365	1e-200


>>
16. NC_016109
Source: Kitasatospora setae KM-6054, complete genome
Type: NRPS,T1PKS,transAT-PKS
Number of proteins with BLAST hits to this cluster: 2
Cumulative BLAST score: 949

Table of genes, locations, strands and annotations of subject cluster:
KSE_RS42480	KSE_RS42480	7957789	7957881	+	HEAT_repeat_domain-containing_protein
KSE_RS40580	KSE_RS40580	7958636	7958893	-	IS5/IS1182_family_transposase

Table of Blast hits (query gene, subject gene, %identity, blast score, %coverage, e-value):
input.path1.gene176	KSE_RS34840	31	76	52.043596730245234	9e-12
input.path1.gene178	KSE_RS34905	39	225	26.63358147229115	3e-56
input.path1.gene178	KSE_RS34910	38	219	27.04714640198511	1.7e-54
input.path1.gene178	KSE_RS34920	37	217	27.377998345740277	6.3e-54
input.path1.gene178	KSE_RS34915	39	212	26.881720430107524	3.5e-52


>>
17. NW_003345196
Source: Nannizzia gypsea CBS 118893 supercont1.6 genomic scaffold, whole genome shotgun sequence
Type: T1PKS,indole
Number of proteins with BLAST hits to this cluster: 2
Cumulative BLAST score: 940

Table of genes, locations, strands and annotations of subject cluster:

Table of Blast hits (query gene, subject gene, %identity, blast score, %coverage, e-value):
input.path1.gene177	MGYG_06590	32	92	31.610044313146236	1.3e-16
input.path1.gene178	MGYG_06588	38	848	116.62531017369726	9.6e-244


>>
18. NZ_JACHJH010000002
Source: Streptomyces olivoverticillatus strain CECT 3266 Ga0436946_02, whole genome shotgun sequence
Type: NRPS,NRPS-like,T1PKS,butyrolactone
Number of proteins with BLAST hits to this cluster: 2
Cumulative BLAST score: 939

Table of genes, locations, strands and annotations of subject cluster:
FHS39_RS06710	FHS39_RS06710	316604	317638	-	SDR_family_NAD(P)-dependent_oxidoreductase
FHS39_RS06715	FHS39_RS06715	318503	319534	-	acyltransferase_domain-containing_protein
FHS39_RS06720	FHS39_RS06720	319691	321565	-	beta-ketoacyl_synthase
FHS39_RS06725	FHS39_RS06725	321581	322669	-	SDR_family_NAD(P)-dependent_oxidoreductase
FHS39_RS06730	FHS39_RS06730	323510	324409	-	acyltransferase_domain-containing_protein
FHS39_RS06735	FHS39_RS06735	324719	326374	-	type_I_polyketide_synthase

Table of Blast hits (query gene, subject gene, %identity, blast score, %coverage, e-value):
input.path1.gene176	FHS39_RS06650	33	61	26.430517711171664	1.8e-07
input.path1.gene178	FHS39_RS06660	37	225	28.205128205128204	2.3e-56
input.path1.gene178	FHS39_RS06735	36	221	28.53598014888337	4.4e-55
input.path1.gene178	FHS39_RS06720	37	218	27.70885028949545	2.8e-54
input.path1.gene178	FHS39_RS06665	37	214	27.70885028949545	5.4e-53


>>
19. NC_036626
Source: Fusarium fujikuroi IMI 58289 draft genome, chromosome FFUJ_chr05
Type: T1PKS
Number of proteins with BLAST hits to this cluster: 2
Cumulative BLAST score: 938

Table of genes, locations, strands and annotations of subject cluster:

Table of Blast hits (query gene, subject gene, %identity, blast score, %coverage, e-value):
input.path1.gene176	FFUJ_06744	33	238	111.44414168937328	1.4e-60
input.path1.gene178	FFUJ_06742	34	700	114.97105045492142	3.2e-199


>>
20. NW_003315100
Source: Trichophyton benhamiae CBS 112371 chromosome Unknown contig01356.0, whole genome shotgun sequence
Type: T1PKS,indole
Number of proteins with BLAST hits to this cluster: 2
Cumulative BLAST score: 937

Table of genes, locations, strands and annotations of subject cluster:

Table of Blast hits (query gene, subject gene, %identity, blast score, %coverage, e-value):
input.path1.gene177	ARB_00536	33	83	29.68980797636632	8e-14
input.path1.gene178	ARB_00538	38	854	116.29445822994211	1e-245


>>
21. NC_038012
Source: Fusarium venenatum strain A3/5 genome assembly, chromosome: I
Type: NRPS,T1PKS
Number of proteins with BLAST hits to this cluster: 2
Cumulative BLAST score: 934

Table of genes, locations, strands and annotations of subject cluster:

Table of Blast hits (query gene, subject gene, %identity, blast score, %coverage, e-value):
input.path1.gene176	FVRRES_02717	31	196	112.80653950953679	4.6e-48
input.path1.gene178	FVRRES_02715	35	738	114.39205955334988	8.2e-211


>>
22. NW_022984633
Source: Aspergillus tanneri strain NIH1004 chromosome Unknown AapTan_6_pilon_pilon_pilon, whole genome shotgun sequence
Type: T1PKS
Number of proteins with BLAST hits to this cluster: 2
Cumulative BLAST score: 922

Table of genes, locations, strands and annotations of subject cluster:

Table of Blast hits (query gene, subject gene, %identity, blast score, %coverage, e-value):
input.path1.gene176	ATNIH1004_008651	32	238	110.89918256130791	1.4e-60
input.path1.gene178	ATNIH1004_008649	34	684	114.64019851116625	1.8e-194


>>
23. NW_023336282
Source: Aspergillus tubingensis WU-2223L DNA, scaffold_8, whole genome shotgun sequence
Type: NRPS,T1PKS
Number of proteins with BLAST hits to this cluster: 2
Cumulative BLAST score: 921

Table of genes, locations, strands and annotations of subject cluster:

Table of Blast hits (query gene, subject gene, %identity, blast score, %coverage, e-value):
input.path1.gene177	AtWU_06973	32	86	31.314623338257014	9.5e-15
input.path1.gene178	AtWU_06971	36	835	116.70802315963607	6.4e-240


>>
24. NT_107009
Source: Aspergillus nidulans FGSC A4 chromosome I map unlocalized scaffold_7, whole genome shotgun sequence
Type: T1PKS
Number of proteins with BLAST hits to this cluster: 2
Cumulative BLAST score: 921

Table of genes, locations, strands and annotations of subject cluster:

Table of Blast hits (query gene, subject gene, %identity, blast score, %coverage, e-value):
input.path1.gene177	AN6002.2	32	80	31.0192023633678	6.8e-13
input.path1.gene178	AN6000.2	37	841	116.45988420181969	1.5e-241


>>
25. NW_003299167
Source: Microsporum canis CBS 113480 supercont1.3 genomic scaffold, whole genome shotgun sequence
Type: T1PKS,indole
Number of proteins with BLAST hits to this cluster: 2
Cumulative BLAST score: 917

Table of genes, locations, strands and annotations of subject cluster:

Table of Blast hits (query gene, subject gene, %identity, blast score, %coverage, e-value):
input.path1.gene177	MCYG_03600	33	84	31.610044313146236	3.6e-14
input.path1.gene178	MCYG_03598	37	833	116.45988420181969	3.2e-239


>>
26. NW_022194796
Source: Fusarium proliferatum ET1 genome assembly, contig: FPRO_scaffold007, whole genome shotgun sequence
Type: T1PKS
Number of proteins with BLAST hits to this cluster: 2
Cumulative BLAST score: 908

Table of genes, locations, strands and annotations of subject cluster:

Table of Blast hits (query gene, subject gene, %identity, blast score, %coverage, e-value):
input.path1.gene176	FPRO_07277	33	210	102.17983651226159	2.4e-52
input.path1.gene178	FPRO_07275	34	698	114.97105045492142	1.2e-198


>>
27. NC_036442
Source: Aspergillus oryzae RIB40 DNA, chromosome 8
Type: T1PKS
Number of proteins with BLAST hits to this cluster: 2
Cumulative BLAST score: 905

Table of genes, locations, strands and annotations of subject cluster:

Table of Blast hits (query gene, subject gene, %identity, blast score, %coverage, e-value):
input.path1.gene176	AO090010000047	32	222	111.1716621253406	6e-56
input.path1.gene178	AO090010000048	34	683	115.88089330024813	4.1e-194


>>
28. NW_017264193
Source: Xylona heveae TC161 unplaced genomic scaffold L228scaffold_1, whole genome shotgun sequence
Type: T1PKS
Number of proteins with BLAST hits to this cluster: 2
Cumulative BLAST score: 902

Table of genes, locations, strands and annotations of subject cluster:

Table of Blast hits (query gene, subject gene, %identity, blast score, %coverage, e-value):
input.path1.gene176	L228DRAFT_235439	31	215	111.44414168937328	9.6e-54
input.path1.gene178	L228DRAFT_235437	33	687	115.88089330024813	2.8e-195


>>
29. NW_003345192
Source: Nannizzia gypsea CBS 118893 supercont1.10 genomic scaffold, whole genome shotgun sequence
Type: T1PKS
Number of proteins with BLAST hits to this cluster: 2
Cumulative BLAST score: 880

Table of genes, locations, strands and annotations of subject cluster:

Table of Blast hits (query gene, subject gene, %identity, blast score, %coverage, e-value):
input.path1.gene176	MGYG_08689	33	235	112.80653950953679	8.9e-60
input.path1.gene178	MGYG_08684	38	645	86.10421836228288	9.4e-183


>>
30. NW_003315103
Source: Trichophyton benhamiae CBS 112371 chromosome Unknown contig00075.49, whole genome shotgun sequence
Type: T1PKS
Number of proteins with BLAST hits to this cluster: 2
Cumulative BLAST score: 874

Table of genes, locations, strands and annotations of subject cluster:

Table of Blast hits (query gene, subject gene, %identity, blast score, %coverage, e-value):
input.path1.gene176	ARB_07989	32	232	112.80653950953679	9.9e-59
input.path1.gene178	ARB_07994	38	642	85.8560794044665	8e-182


>>
31. NW_003456426
Source: Trichophyton rubrum CBS 118892 genomic scaffold supercont2.3, whole genome shotgun sequence
Type: T1PKS
Number of proteins with BLAST hits to this cluster: 2
Cumulative BLAST score: 866

Table of genes, locations, strands and annotations of subject cluster:

Table of Blast hits (query gene, subject gene, %identity, blast score, %coverage, e-value):
input.path1.gene176	TERG_02845	32	228	112.26158038147138	1.4e-57
input.path1.gene178	TERG_02850	38	638	87.75847808105873	1.5e-180


>>
32. NW_003299161
Source: Microsporum canis CBS 113480 supercont1.9 genomic scaffold, whole genome shotgun sequence
Type: T1PKS
Number of proteins with BLAST hits to this cluster: 2
Cumulative BLAST score: 826

Table of genes, locations, strands and annotations of subject cluster:

Table of Blast hits (query gene, subject gene, %identity, blast score, %coverage, e-value):
input.path1.gene176	MCYG_08540	31	203	112.80653950953679	4.9e-50
input.path1.gene178	MCYG_08535	38	623	85.11166253101737	6.5e-176


>>
33. CM000577
Source: Fusarium graminearum PH-1 chromosome 4, whole genome shotgun sequence
Type: T1PKS
Number of proteins with BLAST hits to this cluster: 2
Cumulative BLAST score: 785

Table of genes, locations, strands and annotations of subject cluster:

Table of Blast hits (query gene, subject gene, %identity, blast score, %coverage, e-value):
input.path1.gene176	FGSG_09183	37	154	58.58310626702997	2.6e-35
input.path1.gene178	FGSG_09182	38	631	81.05872622001654	1.4e-178


>>
34. NC_026477
Source: Fusarium graminearum PH-1 chromosome 4, whole genome shotgun sequence
Type: T1PKS
Number of proteins with BLAST hits to this cluster: 2
Cumulative BLAST score: 785

Table of genes, locations, strands and annotations of subject cluster:

Table of Blast hits (query gene, subject gene, %identity, blast score, %coverage, e-value):
input.path1.gene176	FGSG_09183	37	154	58.58310626702997	2.6e-35
input.path1.gene178	FGSG_09182	38	631	81.05872622001654	1.4e-178


>>
35. NW_022984632
Source: Aspergillus tanneri strain NIH1004 chromosome Unknown AapTan_5_pilon_pilon_pilon, whole genome shotgun sequence
Type: T1PKS
Number of proteins with BLAST hits to this cluster: 2
Cumulative BLAST score: 774

Table of genes, locations, strands and annotations of subject cluster:

Table of Blast hits (query gene, subject gene, %identity, blast score, %coverage, e-value):
input.path1.gene176	ATNIH1004_009523	31	181	96.45776566757493	1.2e-43
input.path1.gene178	ATNIH1004_009518	37	593	82.46484698097602	5.6e-167


>>
36. NW_001820829
Source: Sclerotinia sclerotiorum 1980 UF-70 scaffold_7 genomic scaffold, whole genome shotgun sequence
Type: T1PKS
Number of proteins with BLAST hits to this cluster: 2
Cumulative BLAST score: 701

Table of genes, locations, strands and annotations of subject cluster:

Table of Blast hits (query gene, subject gene, %identity, blast score, %coverage, e-value):
input.path1.gene177	SS1G_05679	37	90	25.406203840472674	6.5e-16
input.path1.gene178	SS1G_05681	35	611	103.63937138130686	2.6e-172


>>
37. NW_006912895
Source: Capronia coronata CBS 617.96 acBCG-supercont1.3, whole genome shotgun sequence
Type: T1PKS
Number of proteins with BLAST hits to this cluster: 2
Cumulative BLAST score: 678

Table of genes, locations, strands and annotations of subject cluster:

Table of Blast hits (query gene, subject gene, %identity, blast score, %coverage, e-value):
input.path1.gene176	A1O1_04521	32	203	109.8092643051771	4.9e-50
input.path1.gene178	A1O1_04517	42	262	28.949545078577337	2.2e-67
input.path1.gene178	A1O1_04527	34	213	32.754342431761785	9.2e-53


>>
38. NT_165931
Source: Aspergillus terreus NIH2624 scaffold_8 genomic scaffold, whole genome shotgun sequence
Type: T1PKS
Number of proteins with BLAST hits to this cluster: 2
Cumulative BLAST score: 674

Table of genes, locations, strands and annotations of subject cluster:

Table of Blast hits (query gene, subject gene, %identity, blast score, %coverage, e-value):
input.path1.gene176	ATEG_06203	34	174	86.37602179836512	2.4e-41
input.path1.gene178	ATEG_06206	33	500	77.25392886683208	4.9e-139


>>
39. NC_036623
Source: Fusarium fujikuroi IMI 58289 draft genome, chromosome FFUJ_chr02
Type: T1PKS
Number of proteins with BLAST hits to this cluster: 2
Cumulative BLAST score: 572

Table of genes, locations, strands and annotations of subject cluster:

Table of Blast hits (query gene, subject gene, %identity, blast score, %coverage, e-value):
input.path1.gene176	FFUJ_03985	31	166	97.54768392370572	3.9e-39
input.path1.gene178	FFUJ_03984	59	406	27.791563275434246	1.3e-110


>>
40. NW_022194791
Source: Fusarium proliferatum ET1 genome assembly, contig: FPRO_scaffold002, whole genome shotgun sequence
Type: T1PKS
Number of proteins with BLAST hits to this cluster: 2
Cumulative BLAST score: 567

Table of genes, locations, strands and annotations of subject cluster:

Table of Blast hits (query gene, subject gene, %identity, blast score, %coverage, e-value):
input.path1.gene176	FPRO_03538	31	163	97.82016348773843	5.6e-38
input.path1.gene178	FPRO_03539	59	404	27.791563275434246	3.7e-110


>>
41. NZ_FRCX01000002
Source: Duganella sacchari strain Sac-22, whole genome shotgun sequence
Type: NRPS,hserlactone,transAT-PKS
Number of proteins with BLAST hits to this cluster: 2
Cumulative BLAST score: 563

Table of genes, locations, strands and annotations of subject cluster:

Table of Blast hits (query gene, subject gene, %identity, blast score, %coverage, e-value):
input.path1.gene177	BUA36_RS05195	31	105	51.994091580502214	2.6e-20
input.path1.gene178	BUA36_RS05370	38	231	27.46071133167907	5.5e-58
input.path1.gene178	BUA36_RS05365	38	227	29.03225806451613	8e-57


>>
42. CM000590
Source: Fusarium oxysporum f. sp. lycopersici 4287 chromosome 2, whole genome shotgun sequence
Type: T1PKS
Number of proteins with BLAST hits to this cluster: 2
Cumulative BLAST score: 560

Table of genes, locations, strands and annotations of subject cluster:

Table of Blast hits (query gene, subject gene, %identity, blast score, %coverage, e-value):
input.path1.gene176	FOXG_05817	38	155	58.58310626702997	1.5e-35
input.path1.gene178	FOXG_05816	59	405	27.791563275434246	2.1e-110


>>
43. NC_030987
Source: Fusarium oxysporum f. sp. lycopersici 4287 chromosome 2, whole genome shotgun sequence
Type: T1PKS
Number of proteins with BLAST hits to this cluster: 2
Cumulative BLAST score: 560

Table of genes, locations, strands and annotations of subject cluster:

Table of Blast hits (query gene, subject gene, %identity, blast score, %coverage, e-value):
input.path1.gene176	FOXG_05817	38	155	58.58310626702997	1.5e-35
input.path1.gene178	FOXG_05816	59	405	27.791563275434246	2.1e-110


>>
44. NW_001939246
Source: Pyrenophora tritici-repentis Pt-1C-BFP supercont1.3, whole genome shotgun sequence
Type: NRPS,T1PKS
Number of proteins with BLAST hits to this cluster: 2
Cumulative BLAST score: 443

Table of genes, locations, strands and annotations of subject cluster:

Table of Blast hits (query gene, subject gene, %identity, blast score, %coverage, e-value):
input.path1.gene177	PTRG_04242	40	225	48.005908419497786	1.7e-56
input.path1.gene178	PTRG_04244	38	218	27.377998345740277	3.7e-54


>>
45. NW_006912904
Source: Capronia epimyces CBS 606.96 aczIQ-supercont1.1, whole genome shotgun sequence
Type: T1PKS
Number of proteins with BLAST hits to this cluster: 2
Cumulative BLAST score: 441

Table of genes, locations, strands and annotations of subject cluster:

Table of Blast hits (query gene, subject gene, %identity, blast score, %coverage, e-value):
input.path1.gene176	A1O3_01769	31	176	95.91280653950953	6.4e-42
input.path1.gene178	A1O3_01765	44	265	28.453267162944584	2.7e-68


>>
46. NW_019716258
Source: Ramularia collo-cygni strain URUG2 genome assembly, contig: RCC_scaffold04, whole genome shotgun sequence
Type: NRPS,T1PKS
Number of proteins with BLAST hits to this cluster: 2
Cumulative BLAST score: 431

Table of genes, locations, strands and annotations of subject cluster:

Table of Blast hits (query gene, subject gene, %identity, blast score, %coverage, e-value):
input.path1.gene177	RCC_03310	40	220	48.301329394387	5.5e-55
input.path1.gene178	RCC_03313	37	211	28.453267162944584	4.5e-52


>>
47. NC_036437
Source: Aspergillus oryzae RIB40 DNA, chromosome 3
Type: T1PKS
Number of proteins with BLAST hits to this cluster: 2
Cumulative BLAST score: 421

Table of genes, locations, strands and annotations of subject cluster:

Table of Blast hits (query gene, subject gene, %identity, blast score, %coverage, e-value):
input.path1.gene176	AO090023000878	33	103	58.31062670299727	4.1e-20
input.path1.gene178	AO090023000877	31	318	67.99007444168734	2.6e-84


>>
48. NW_017971438
Source: Talaromyces atroroseus strain IBT 11181 chromosome Unknown scaffold_6, whole genome shotgun sequence
Type: T1PKS
Number of proteins with BLAST hits to this cluster: 2
Cumulative BLAST score: 414

Table of genes, locations, strands and annotations of subject cluster:

Table of Blast hits (query gene, subject gene, %identity, blast score, %coverage, e-value):
input.path1.gene176	UA08_04988	32	161	93.18801089918256	1.3e-37
input.path1.gene178	UA08_04992	41	253	29.114971050454923	8e-65


>>
49. NW_020939749
Source: Sparassis crispa DNA, contig_000002F, whole genome shotgun sequence
Type: NRPS-like,T1PKS
Number of proteins with BLAST hits to this cluster: 2
Cumulative BLAST score: 409

Table of genes, locations, strands and annotations of subject cluster:

Table of Blast hits (query gene, subject gene, %identity, blast score, %coverage, e-value):
input.path1.gene176	SCP_0306500	37	117	50.13623978201635	3.5e-24
input.path1.gene178	SCP_0306480	41	292	32.919768403639374	1.6e-76


>>
50. NW_009276941
Source: Verticillium dahliae VdLs.17 supercont1.26 genomic scaffold, whole genome shotgun sequence
Type: NRPS-like,T1PKS
Number of proteins with BLAST hits to this cluster: 2
Cumulative BLAST score: 401

Table of genes, locations, strands and annotations of subject cluster:

Table of Blast hits (query gene, subject gene, %identity, blast score, %coverage, e-value):
input.path1.gene176	VDAG_09529	36	110	56.94822888283378	4.3e-22
input.path1.gene178	VDAG_09534	32	291	56.07940446650124	4.5e-76
