## Supplementary Results for "A draft genome of the ascomycotal fungal species *Pseudopithomyces maydicus* (family *Didymosphaeriaceae*)": contig_12_c1.txt

ClusterBlast scores for contig_12

Table of genes, locations, strands and annotations of query cluster:
input.path1.gene385	1263495	1264447	+		
input.path1.gene386	1265341	1266587	+		
input.path1.gene387	1267172	1271073	-		
input.path1.gene388	1271535	1273114	+		
input.path1.gene389	1273834	1273969	+		
input.path1.gene390	1277277	1278624	+		
input.path1.gene391	1279972	1281248	-		
input.path1.gene392	1282186	1294181	+		
input.path1.gene393	1294284	1294541	-		
input.path1.gene394	1294822	1301524	-		
input.path1.gene395	1301935	1304132	+		
input.path1.gene396	1305158	1308096	+		
input.path1.gene397	1310405	1311834	+		


Significant hits: 
1. NW_022984629	Aspergillus tanneri strain NIH1004 chromosome Unknown AapTan_2_pilon_pilon_pilon, whole genome shotgun sequence
2. NW_022474207	Venustampulla echinocandica strain BP 5553 chromosome Unknown scaffold_2, whole genome shotgun sequence
3. NW_007360993	Glarea lozoyensis ATCC 20868 chromosome Unknown GLAREA19, whole genome shotgun sequence
4. NW_022983866	Arthroderma uncinatum strain CBS 119779 chromosome Unknown scaffold4, whole genome shotgun sequence
5. NW_006917091	Pestalotiopsis fici W106-1 unplaced genomic scaffold PFICI_Scf_1, whole genome shotgun sequence
6. NW_014574692	Metarhizium brunneum ARSEF 3297 chromosome Unknown scaffold_25, whole genome shotgun sequence
7. NW_011942153	Metarhizium robertsii ARSEF 23 MAA_Scf_13, whole genome shotgun sequence
8. NC_036436	Aspergillus oryzae RIB40 DNA, chromosome 2
9. NW_006917095	Pestalotiopsis fici W106-1 unplaced genomic scaffold PFICI_Scf_5, whole genome shotgun sequence
10. NW_022474213	Venustampulla echinocandica strain BP 5553 chromosome Unknown scaffold_8, whole genome shotgun sequence
11. NC_026507	Neurospora crassa OR74A linkage group VII, whole genome shotgun sequence
12. RRCJ01000024	Pyricularia sp. CBS 133598 strain NI919 Pyricularia_sp_LS_NI919_Scf_24, whole genome shotgun sequence
13. CM000593	Fusarium oxysporum f. sp. lycopersici 4287 chromosome 5, whole genome shotgun sequence
14. NC_030990	Fusarium oxysporum f. sp. lycopersici 4287 chromosome 5, whole genome shotgun sequence
15. NW_022194803	Fusarium proliferatum ET1 genome assembly, contig: FPRO_scaffold014, whole genome shotgun sequence
16. NW_017971438	Talaromyces atroroseus strain IBT 11181 chromosome Unknown scaffold_6, whole genome shotgun sequence
17. NC_036624	Fusarium fujikuroi IMI 58289 draft genome, chromosome FFUJ_chr03
18. NW_003345199	Nannizzia gypsea CBS 118893 supercont1.3 genomic scaffold, whole genome shotgun sequence
19. NW_017264196	Xylona heveae TC161 unplaced genomic scaffold L228scaffold_4, whole genome shotgun sequence
20. NW_009276923	Verticillium dahliae VdLs.17 supercont1.18 genomic scaffold, whole genome shotgun sequence
21. NW_017971435	Talaromyces atroroseus strain IBT 11181 chromosome Unknown scaffold_3, whole genome shotgun sequence
22. NW_007360988	Glarea lozoyensis ATCC 20868 chromosome Unknown GLAREA14, whole genome shotgun sequence
23. CM002798	Penicillium chrysogenum strain P2niaD18 chromosome I, whole genome shotgun sequence
24. NC_036629	Fusarium fujikuroi IMI 58289 draft genome, chromosome FFUJ_chr08
25. NW_022194804	Fusarium proliferatum ET1 genome assembly, contig: FPRO_scaffold015, whole genome shotgun sequence
26. NW_017387257	Fonsecaea erecta strain CBS 125763 chromosome Unknown Scaff14, whole genome shotgun sequence
27. NT_165976	Chaetomium globosum CBS 148.51 scaffold_1 genomic scaffold, whole genome shotgun sequence
28. NW_019716258	Ramularia collo-cygni strain URUG2 genome assembly, contig: RCC_scaffold04, whole genome shotgun sequence
29. NW_022474219	Venustampulla echinocandica strain BP 5553 chromosome Unknown scaffold_14, whole genome shotgun sequence
30. NC_030955	Colletotrichum higginsianum IMI 349063 chromosome 2, whole genome shotgun sequence
31. NW_013550613	Fonsecaea pedrosoi CBS 271.37 unplaced genomic scaffold supercont1.3, whole genome shotgun sequence
32. NW_015971579	Cladophialophora bantiana CBS 173.52 unplaced genomic scaffold supercont1.5, whole genome shotgun sequence
33. NW_023336267	Colletotrichum scovillei strain TJNH1 chromosome Unknown Contig00009, whole genome shotgun sequence
34. RRCJ01000011	Pyricularia sp. CBS 133598 strain NI919 Pyricularia_sp_LS_NI919_Scf_11, whole genome shotgun sequence
35. NT_165939	Aspergillus terreus NIH2624 scaffold_16 genomic scaffold, whole genome shotgun sequence
36. NC_030959	Colletotrichum higginsianum IMI 349063 chromosome 6, whole genome shotgun sequence
37. NW_007360994	Glarea lozoyensis ATCC 20868 chromosome Unknown GLAREA2, whole genome shotgun sequence
38. NC_035794	Pochonia chlamydosporia 170 chromosome 5, whole genome shotgun sequence
39. NC_030954	Colletotrichum higginsianum IMI 349063 chromosome 1, whole genome shotgun sequence
40. NT_107004	Aspergillus nidulans FGSC A4 chromosome Unknown scaffold_12, whole genome shotgun sequence
41. NW_011942237	Metarhizium robertsii ARSEF 23 MAA_Scf_9, whole genome shotgun sequence
42. CM000600	Fusarium oxysporum f. sp. lycopersici 4287 chromosome 12, whole genome shotgun sequence
43. NC_030997	Fusarium oxysporum f. sp. lycopersici 4287 chromosome 12, whole genome shotgun sequence
44. NW_022194799	Fusarium proliferatum ET1 genome assembly, contig: FPRO_scaffold010, whole genome shotgun sequence
45. NC_016457	Thermothielavioides terrestris NRRL 8126 chromosome 1, complete sequence
46. NC_030962	Colletotrichum higginsianum IMI 349063 chromosome 10, whole genome shotgun sequence
47. NC_036629	Fusarium fujikuroi IMI 58289 draft genome, chromosome FFUJ_chr08
48. NW_017263917	Colletotrichum higginsianum IMI 349063 chromosome 7 map unlocalized, whole genome shotgun sequence
49. CM000574	Fusarium graminearum PH-1 chromosome 1, whole genome shotgun sequence
50. NC_026474	Fusarium graminearum PH-1 chromosome 1, whole genome shotgun sequence


Details:

>>
1. NW_022984629
Source: Aspergillus tanneri strain NIH1004 chromosome Unknown AapTan_2_pilon_pilon_pilon, whole genome shotgun sequence
Type: NRPS,T1PKS
Number of proteins with BLAST hits to this cluster: 4
Cumulative BLAST score: 2097

Table of genes, locations, strands and annotations of subject cluster:

Table of Blast hits (query gene, subject gene, %identity, blast score, %coverage, e-value):
input.path1.gene387	ATNIH1004_001273	40	84	19.482758620689655	3.1e-14
input.path1.gene388	ATNIH1004_001273	50	259	82.7485380116959	4.1e-67
input.path1.gene390	ATNIH1004_001269	32	143	75.82697201017811	3.8e-32
input.path1.gene392	ATNIH1004_001274	31	1611	144.22417079679758	0.0


>>
2. NW_022474207
Source: Venustampulla echinocandica strain BP 5553 chromosome Unknown scaffold_2, whole genome shotgun sequence
Type: NRPS,T1PKS
Number of proteins with BLAST hits to this cluster: 3
Cumulative BLAST score: 2389

Table of genes, locations, strands and annotations of subject cluster:

Table of Blast hits (query gene, subject gene, %identity, blast score, %coverage, e-value):
input.path1.gene387	BP5553_05151	31	209	77.93103448275862	8.3e-52
input.path1.gene388	BP5553_05146	46	309	99.70760233918129	5.9e-82
input.path1.gene392	BP5553_05145	33	1871	144.2622950819672	0.0


>>
3. NW_007360993
Source: Glarea lozoyensis ATCC 20868 chromosome Unknown GLAREA19, whole genome shotgun sequence
Type: NRPS,T1PKS
Number of proteins with BLAST hits to this cluster: 3
Cumulative BLAST score: 1545

Table of genes, locations, strands and annotations of subject cluster:

Table of Blast hits (query gene, subject gene, %identity, blast score, %coverage, e-value):
input.path1.gene386	GLAREA_12718	37	224	104.33526011560694	1.9e-56
input.path1.gene388	GLAREA_12720	38	240	100.29239766081872	2.6e-61
input.path1.gene392	GLAREA_12721	39	1081	59.81700343118567	0.0


>>
4. NW_022983866
Source: Arthroderma uncinatum strain CBS 119779 chromosome Unknown scaffold4, whole genome shotgun sequence
Type: T1PKS
Number of proteins with BLAST hits to this cluster: 3
Cumulative BLAST score: 1523

Table of genes, locations, strands and annotations of subject cluster:

Table of Blast hits (query gene, subject gene, %identity, blast score, %coverage, e-value):
input.path1.gene386	GIQ15_05694	38	209	102.60115606936415	6.5e-52
input.path1.gene388	GIQ15_05696	38	241	101.16959064327486	1.2e-61
input.path1.gene392	GIQ15_05697	38	1073	64.5825390773923	3.4e-311


>>
5. NW_006917091
Source: Pestalotiopsis fici W106-1 unplaced genomic scaffold PFICI_Scf_1, whole genome shotgun sequence
Type: T1PKS
Number of proteins with BLAST hits to this cluster: 3
Cumulative BLAST score: 1490

Table of genes, locations, strands and annotations of subject cluster:

Table of Blast hits (query gene, subject gene, %identity, blast score, %coverage, e-value):
input.path1.gene386	PFICI_00299	38	182	89.30635838150289	8.4e-44
input.path1.gene388	PFICI_00296	38	231	100.87719298245614	1.6e-58
input.path1.gene392	PFICI_00294	38	1077	62.7906976744186	0.0


>>
6. NW_014574692
Source: Metarhizium brunneum ARSEF 3297 chromosome Unknown scaffold_25, whole genome shotgun sequence
Type: T1PKS
Number of proteins with BLAST hits to this cluster: 3
Cumulative BLAST score: 1438

Table of genes, locations, strands and annotations of subject cluster:

Table of Blast hits (query gene, subject gene, %identity, blast score, %coverage, e-value):
input.path1.gene386	MBR_09790	35	191	104.62427745664739	1.4e-46
input.path1.gene388	MBR_09792	37	233	102.92397660818713	4.1e-59
input.path1.gene392	MBR_09791	36	1014	62.256957682043456	1.7e-293


>>
7. NW_011942153
Source: Metarhizium robertsii ARSEF 23 MAA_Scf_13, whole genome shotgun sequence
Type: T1PKS
Number of proteins with BLAST hits to this cluster: 3
Cumulative BLAST score: 1422

Table of genes, locations, strands and annotations of subject cluster:

Table of Blast hits (query gene, subject gene, %identity, blast score, %coverage, e-value):
input.path1.gene386	MAA_01345	35	191	104.62427745664739	1.4e-46
input.path1.gene388	MAA_01347	37	233	102.92397660818713	3.2e-59
input.path1.gene392	MAA_01346	36	998	62.409454822722076	1.2e-288


>>
8. NC_036436
Source: Aspergillus oryzae RIB40 DNA, chromosome 2
Type: NRPS,T1PKS
Number of proteins with BLAST hits to this cluster: 2
Cumulative BLAST score: 1976

Table of genes, locations, strands and annotations of subject cluster:

Table of Blast hits (query gene, subject gene, %identity, blast score, %coverage, e-value):
input.path1.gene388	AO090001000287	41	245	92.98245614035088	6.2e-63
input.path1.gene392	AO090001000277	35	918	66.75562333206251	2.1e-264
input.path1.gene392	AO090001000293	34	813	58.52077773541746	5.7e-233


>>
9. NW_006917095
Source: Pestalotiopsis fici W106-1 unplaced genomic scaffold PFICI_Scf_5, whole genome shotgun sequence
Type: T1PKS
Number of proteins with BLAST hits to this cluster: 2
Cumulative BLAST score: 1689

Table of genes, locations, strands and annotations of subject cluster:

Table of Blast hits (query gene, subject gene, %identity, blast score, %coverage, e-value):
input.path1.gene388	PFICI_07939	39	246	98.53801169590643	4.7e-63
input.path1.gene392	PFICI_07941	39	1012	59.43576057948914	6.4e-293
input.path1.gene392	PFICI_07942	37	431	30.537552420892105	7.9e-118


>>
10. NW_022474213
Source: Venustampulla echinocandica strain BP 5553 chromosome Unknown scaffold_8, whole genome shotgun sequence
Type: NRPS,T1PKS
Number of proteins with BLAST hits to this cluster: 2
Cumulative BLAST score: 1658

Table of genes, locations, strands and annotations of subject cluster:

Table of Blast hits (query gene, subject gene, %identity, blast score, %coverage, e-value):
input.path1.gene388	BP5553_08694	34	193	104.09356725146199	3.6e-47
input.path1.gene392	BP5553_08690	38	1025	64.0106747998475	7.3e-297
input.path1.gene392	BP5553_08674	31	440	43.88105223027068	1.3e-120


>>
11. NC_026507
Source: Neurospora crassa OR74A linkage group VII, whole genome shotgun sequence
Type: T1PKS
Number of proteins with BLAST hits to this cluster: 2
Cumulative BLAST score: 1619

Table of genes, locations, strands and annotations of subject cluster:

Table of Blast hits (query gene, subject gene, %identity, blast score, %coverage, e-value):
input.path1.gene386	NCU08398	37	247	113.58381502890174	2.8e-63
input.path1.gene392	NCU08399_XP_011395280.1	39	686	40.83110941669844	1.4e-194
input.path1.gene392	NCU08399	39	686	40.83110941669844	1.4e-194


>>
12. RRCJ01000024
Source: Pyricularia sp. CBS 133598 strain NI919 Pyricularia_sp_LS_NI919_Scf_24, whole genome shotgun sequence
Type: NRPS,T1PKS
Number of proteins with BLAST hits to this cluster: 2
Cumulative BLAST score: 1496

Table of genes, locations, strands and annotations of subject cluster:

Table of Blast hits (query gene, subject gene, %identity, blast score, %coverage, e-value):
input.path1.gene388	PspLS_11937	36	215	99.41520467836257	1.2e-53
input.path1.gene392	PspLS_11933	34	858	62.48570339306138	1.5e-246
input.path1.gene392	PspLS_11937	36	423	31.414410979794127	2.2e-115


>>
13. CM000593
Source: Fusarium oxysporum f. sp. lycopersici 4287 chromosome 5, whole genome shotgun sequence
Type: NRPS,T1PKS
Number of proteins with BLAST hits to this cluster: 2
Cumulative BLAST score: 1331

Table of genes, locations, strands and annotations of subject cluster:

Table of Blast hits (query gene, subject gene, %identity, blast score, %coverage, e-value):
input.path1.gene388	FOXG_15298	42	276	100.0	4.2e-72
input.path1.gene392	FOXG_15296	37	1055	64.27754479603507	8.6e-306


>>
14. NC_030990
Source: Fusarium oxysporum f. sp. lycopersici 4287 chromosome 5, whole genome shotgun sequence
Type: NRPS,T1PKS
Number of proteins with BLAST hits to this cluster: 2
Cumulative BLAST score: 1331

Table of genes, locations, strands and annotations of subject cluster:

Table of Blast hits (query gene, subject gene, %identity, blast score, %coverage, e-value):
input.path1.gene388	FOXG_15298	42	276	100.0	4.2e-72
input.path1.gene392	FOXG_15296	37	1055	64.27754479603507	8.6e-306


>>
15. NW_022194803
Source: Fusarium proliferatum ET1 genome assembly, contig: FPRO_scaffold014, whole genome shotgun sequence
Type: NRPS,T1PKS
Number of proteins with BLAST hits to this cluster: 2
Cumulative BLAST score: 1324

Table of genes, locations, strands and annotations of subject cluster:

Table of Blast hits (query gene, subject gene, %identity, blast score, %coverage, e-value):
input.path1.gene388	FPRO_03978	43	280	100.0	2.9e-73
input.path1.gene392	FPRO_03976	37	1044	64.27754479603507	1.5e-302


>>
16. NW_017971438
Source: Talaromyces atroroseus strain IBT 11181 chromosome Unknown scaffold_6, whole genome shotgun sequence
Type: NRPS,T1PKS
Number of proteins with BLAST hits to this cluster: 2
Cumulative BLAST score: 1317

Table of genes, locations, strands and annotations of subject cluster:

Table of Blast hits (query gene, subject gene, %identity, blast score, %coverage, e-value):
input.path1.gene388	UA08_05016	38	243	102.92397660818713	4e-62
input.path1.gene392	UA08_05018	38	1074	64.7350362180709	1.7e-311


>>
17. NC_036624
Source: Fusarium fujikuroi IMI 58289 draft genome, chromosome FFUJ_chr03
Type: NRPS,T1PKS
Number of proteins with BLAST hits to this cluster: 2
Cumulative BLAST score: 1315

Table of genes, locations, strands and annotations of subject cluster:

Table of Blast hits (query gene, subject gene, %identity, blast score, %coverage, e-value):
input.path1.gene388	FFUJ_02221	42	271	100.0	1e-70
input.path1.gene392	FFUJ_02219	37	1044	63.743804803659934	1.5e-302


>>
18. NW_003345199
Source: Nannizzia gypsea CBS 118893 supercont1.3 genomic scaffold, whole genome shotgun sequence
Type: NRPS,T1PKS
Number of proteins with BLAST hits to this cluster: 2
Cumulative BLAST score: 1312

Table of genes, locations, strands and annotations of subject cluster:

Table of Blast hits (query gene, subject gene, %identity, blast score, %coverage, e-value):
input.path1.gene388	MGYG_04006	39	252	100.58479532163742	5e-65
input.path1.gene392	MGYG_04007	37	1060	64.69691193290126	3.5e-307


>>
19. NW_017264196
Source: Xylona heveae TC161 unplaced genomic scaffold L228scaffold_4, whole genome shotgun sequence
Type: NRPS,T1PKS
Number of proteins with BLAST hits to this cluster: 2
Cumulative BLAST score: 1295

Table of genes, locations, strands and annotations of subject cluster:

Table of Blast hits (query gene, subject gene, %identity, blast score, %coverage, e-value):
input.path1.gene388	L228DRAFT_246731	38	254	100.58479532163742	1.7e-65
input.path1.gene392	L228DRAFT_282560	38	1041	61.608844834159356	1.7e-301


>>
20. NW_009276923
Source: Verticillium dahliae VdLs.17 supercont1.18 genomic scaffold, whole genome shotgun sequence
Type: NRPS,T1PKS
Number of proteins with BLAST hits to this cluster: 2
Cumulative BLAST score: 1289

Table of genes, locations, strands and annotations of subject cluster:

Table of Blast hits (query gene, subject gene, %identity, blast score, %coverage, e-value):
input.path1.gene388	VDAG_07923	37	207	100.29239766081872	3.2e-51
input.path1.gene392	VDAG_07928	39	1082	65.1544033549371	0.0


>>
21. NW_017971435
Source: Talaromyces atroroseus strain IBT 11181 chromosome Unknown scaffold_3, whole genome shotgun sequence
Type: NRPS,T1PKS
Number of proteins with BLAST hits to this cluster: 2
Cumulative BLAST score: 1283

Table of genes, locations, strands and annotations of subject cluster:

Table of Blast hits (query gene, subject gene, %identity, blast score, %coverage, e-value):
input.path1.gene388	UA08_03200	36	206	102.92397660818713	4.1e-51
input.path1.gene392	UA08_03198	39	1077	60.160121997712544	0.0


>>
22. NW_007360988
Source: Glarea lozoyensis ATCC 20868 chromosome Unknown GLAREA14, whole genome shotgun sequence
Type: NRPS,T1PKS
Number of proteins with BLAST hits to this cluster: 2
Cumulative BLAST score: 1274

Table of genes, locations, strands and annotations of subject cluster:

Table of Blast hits (query gene, subject gene, %identity, blast score, %coverage, e-value):
input.path1.gene388	GLAREA_06758	34	218	103.21637426900585	1.1e-54
input.path1.gene392	GLAREA_06760	38	1056	63.97255051467785	5e-306


>>
23. CM002798
Source: Penicillium chrysogenum strain P2niaD18 chromosome I, whole genome shotgun sequence
Type: NRPS,T1PKS
Number of proteins with BLAST hits to this cluster: 2
Cumulative BLAST score: 1270

Table of genes, locations, strands and annotations of subject cluster:

Table of Blast hits (query gene, subject gene, %identity, blast score, %coverage, e-value):
input.path1.gene388	EN45_000080	37	221	100.0	1.2e-55
input.path1.gene392	EN45_000070	39	1049	60.808234845596644	8e-304


>>
24. NC_036629
Source: Fusarium fujikuroi IMI 58289 draft genome, chromosome FFUJ_chr08
Type: NRPS,T1PKS
Number of proteins with BLAST hits to this cluster: 2
Cumulative BLAST score: 1250

Table of genes, locations, strands and annotations of subject cluster:

Table of Blast hits (query gene, subject gene, %identity, blast score, %coverage, e-value):
input.path1.gene388	FFUJ_12708	36	216	95.02923976608187	4e-54
input.path1.gene392	FFUJ_12707	38	1034	61.30385055280213	1.6e-299


>>
25. NW_022194804
Source: Fusarium proliferatum ET1 genome assembly, contig: FPRO_scaffold015, whole genome shotgun sequence
Type: NRPS,T1PKS
Number of proteins with BLAST hits to this cluster: 2
Cumulative BLAST score: 1248

Table of genes, locations, strands and annotations of subject cluster:

Table of Blast hits (query gene, subject gene, %identity, blast score, %coverage, e-value):
input.path1.gene388	FPRO_12373	36	214	95.02923976608187	1.5e-53
input.path1.gene392	FPRO_12372	38	1034	61.30385055280213	2e-299


>>
26. NW_017387257
Source: Fonsecaea erecta strain CBS 125763 chromosome Unknown Scaff14, whole genome shotgun sequence
Type: T1PKS
Number of proteins with BLAST hits to this cluster: 2
Cumulative BLAST score: 1237

Table of genes, locations, strands and annotations of subject cluster:

Table of Blast hits (query gene, subject gene, %identity, blast score, %coverage, e-value):
input.path1.gene388	AYL99_11335	36	222	98.83040935672514	9.5e-56
input.path1.gene392	AYL99_11333	38	1015	59.35951200914983	1.3e-293


>>
27. NT_165976
Source: Chaetomium globosum CBS 148.51 scaffold_1 genomic scaffold, whole genome shotgun sequence
Type: NRPS,T1PKS
Number of proteins with BLAST hits to this cluster: 2
Cumulative BLAST score: 1236

Table of genes, locations, strands and annotations of subject cluster:

Table of Blast hits (query gene, subject gene, %identity, blast score, %coverage, e-value):
input.path1.gene388	CHGG_01240	38	227	101.75438596491229	1.7e-57
input.path1.gene392	CHGG_01239	39	1009	61.18947769729317	7e-292


>>
28. NW_019716258
Source: Ramularia collo-cygni strain URUG2 genome assembly, contig: RCC_scaffold04, whole genome shotgun sequence
Type: NRPS,T1PKS
Number of proteins with BLAST hits to this cluster: 2
Cumulative BLAST score: 1235

Table of genes, locations, strands and annotations of subject cluster:

Table of Blast hits (query gene, subject gene, %identity, blast score, %coverage, e-value):
input.path1.gene388	RCC_03309	37	225	102.92397660818713	1.1e-56
input.path1.gene392	RCC_03313	37	1010	64.65878764773161	4.1e-292


>>
29. NW_022474219
Source: Venustampulla echinocandica strain BP 5553 chromosome Unknown scaffold_14, whole genome shotgun sequence
Type: NRPS,T1PKS
Number of proteins with BLAST hits to this cluster: 2
Cumulative BLAST score: 1229

Table of genes, locations, strands and annotations of subject cluster:

Table of Blast hits (query gene, subject gene, %identity, blast score, %coverage, e-value):
input.path1.gene388	BP5553_10292	37	240	106.43274853801171	2.6e-61
input.path1.gene392	BP5553_10291	39	989	61.72321768966832	5.8e-286


>>
30. NC_030955
Source: Colletotrichum higginsianum IMI 349063 chromosome 2, whole genome shotgun sequence
Type: NRPS,T1PKS
Number of proteins with BLAST hits to this cluster: 2
Cumulative BLAST score: 1217

Table of genes, locations, strands and annotations of subject cluster:

Table of Blast hits (query gene, subject gene, %identity, blast score, %coverage, e-value):
input.path1.gene388	CH63R_03383	37	217	101.16959064327486	3.1e-54
input.path1.gene392	CH63R_03382	38	1000	61.15135341212352	4.3e-289


>>
31. NW_013550613
Source: Fonsecaea pedrosoi CBS 271.37 unplaced genomic scaffold supercont1.3, whole genome shotgun sequence
Type: T1PKS
Number of proteins with BLAST hits to this cluster: 2
Cumulative BLAST score: 1213

Table of genes, locations, strands and annotations of subject cluster:

Table of Blast hits (query gene, subject gene, %identity, blast score, %coverage, e-value):
input.path1.gene388	Z517_05664	35	215	98.83040935672514	1.2e-53
input.path1.gene392	Z517_05666	37	998	59.85512771635532	9.5e-289


>>
32. NW_015971579
Source: Cladophialophora bantiana CBS 173.52 unplaced genomic scaffold supercont1.5, whole genome shotgun sequence
Type: T1PKS
Number of proteins with BLAST hits to this cluster: 2
Cumulative BLAST score: 1206

Table of genes, locations, strands and annotations of subject cluster:

Table of Blast hits (query gene, subject gene, %identity, blast score, %coverage, e-value):
input.path1.gene388	Z519_04307	37	232	98.53801169590643	7e-59
input.path1.gene392	Z519_04309	37	974	59.24513915364087	1.9e-281


>>
33. NW_023336267
Source: Colletotrichum scovillei strain TJNH1 chromosome Unknown Contig00009, whole genome shotgun sequence
Type: NRPS,T1PKS
Number of proteins with BLAST hits to this cluster: 2
Cumulative BLAST score: 1206

Table of genes, locations, strands and annotations of subject cluster:

Table of Blast hits (query gene, subject gene, %identity, blast score, %coverage, e-value):
input.path1.gene388	HER10_EVM0003518	35	208	98.83040935672514	1.1e-51
input.path1.gene392	HER10_EVM0009395	37	998	61.608844834159356	9.5e-289


>>
34. RRCJ01000011
Source: Pyricularia sp. CBS 133598 strain NI919 Pyricularia_sp_LS_NI919_Scf_11, whole genome shotgun sequence
Type: T1PKS
Number of proteins with BLAST hits to this cluster: 2
Cumulative BLAST score: 1203

Table of genes, locations, strands and annotations of subject cluster:

Table of Blast hits (query gene, subject gene, %identity, blast score, %coverage, e-value):
input.path1.gene388	PspLS_09588	40	251	101.46198830409357	1.5e-64
input.path1.gene392	PspLS_09593	38	952	61.4944719786504	1e-274


>>
35. NT_165939
Source: Aspergillus terreus NIH2624 scaffold_16 genomic scaffold, whole genome shotgun sequence
Type: T1PKS
Number of proteins with BLAST hits to this cluster: 2
Cumulative BLAST score: 1199

Table of genes, locations, strands and annotations of subject cluster:

Table of Blast hits (query gene, subject gene, %identity, blast score, %coverage, e-value):
input.path1.gene388	ATEG_09963	39	236	100.58479532163742	4.9e-60
input.path1.gene392	ATEG_09961	36	963	64.46816622188334	5.8e-278


>>
36. NC_030959
Source: Colletotrichum higginsianum IMI 349063 chromosome 6, whole genome shotgun sequence
Type: T1PKS
Number of proteins with BLAST hits to this cluster: 2
Cumulative BLAST score: 1198

Table of genes, locations, strands and annotations of subject cluster:

Table of Blast hits (query gene, subject gene, %identity, blast score, %coverage, e-value):
input.path1.gene388	CH63R_09681	40	247	101.46198830409357	1.6e-63
input.path1.gene392	CH63R_09687	36	951	62.63820053374	1.7e-274


>>
37. NW_007360994
Source: Glarea lozoyensis ATCC 20868 chromosome Unknown GLAREA2, whole genome shotgun sequence
Type: NRPS,T1PKS
Number of proteins with BLAST hits to this cluster: 2
Cumulative BLAST score: 1194

Table of genes, locations, strands and annotations of subject cluster:

Table of Blast hits (query gene, subject gene, %identity, blast score, %coverage, e-value):
input.path1.gene388	GLAREA_03921	39	243	99.70760233918129	3.1e-62
input.path1.gene392	GLAREA_03913	36	951	60.160121997712544	1.7e-274


>>
38. NC_035794
Source: Pochonia chlamydosporia 170 chromosome 5, whole genome shotgun sequence
Type: NRPS,T1PKS
Number of proteins with BLAST hits to this cluster: 2
Cumulative BLAST score: 1193

Table of genes, locations, strands and annotations of subject cluster:

Table of Blast hits (query gene, subject gene, %identity, blast score, %coverage, e-value):
input.path1.gene388	VFPPC_10859	34	223	108.47953216374269	2.5e-56
input.path1.gene392	VFPPC_10857	38	970	60.083873427373234	2.8e-280


>>
39. NC_030954
Source: Colletotrichum higginsianum IMI 349063 chromosome 1, whole genome shotgun sequence
Type: NRPS,T1PKS
Number of proteins with BLAST hits to this cluster: 2
Cumulative BLAST score: 1183

Table of genes, locations, strands and annotations of subject cluster:

Table of Blast hits (query gene, subject gene, %identity, blast score, %coverage, e-value):
input.path1.gene388	CH63R_01265	33	188	102.63157894736842	1.2e-45
input.path1.gene392	CH63R_01271	38	995	61.45634769348075	1.4e-287


>>
40. NT_107004
Source: Aspergillus nidulans FGSC A4 chromosome Unknown scaffold_12, whole genome shotgun sequence
Type: NRPS,T1PKS
Number of proteins with BLAST hits to this cluster: 2
Cumulative BLAST score: 1181

Table of genes, locations, strands and annotations of subject cluster:

Table of Blast hits (query gene, subject gene, %identity, blast score, %coverage, e-value):
input.path1.gene388	AN8409.2	39	233	92.39766081871345	2.4e-59
input.path1.gene392	AN8412.2	37	948	60.57948913457872	1.9e-273


>>
41. NW_011942237
Source: Metarhizium robertsii ARSEF 23 MAA_Scf_9, whole genome shotgun sequence
Type: NRPS,T1PKS
Number of proteins with BLAST hits to this cluster: 2
Cumulative BLAST score: 1180

Table of genes, locations, strands and annotations of subject cluster:

Table of Blast hits (query gene, subject gene, %identity, blast score, %coverage, e-value):
input.path1.gene388	MAA_10116	34	179	101.75438596491229	7.1e-43
input.path1.gene392	MAA_10114	39	1001	59.81700343118567	1.9e-289


>>
42. CM000600
Source: Fusarium oxysporum f. sp. lycopersici 4287 chromosome 12, whole genome shotgun sequence
Type: NRPS,T1PKS
Number of proteins with BLAST hits to this cluster: 2
Cumulative BLAST score: 1175

Table of genes, locations, strands and annotations of subject cluster:

Table of Blast hits (query gene, subject gene, %identity, blast score, %coverage, e-value):
input.path1.gene388	FOXG_14588	42	154	49.707602339181285	2.4e-35
input.path1.gene392	FOXG_14587	38	1021	60.8463591307663	1.8e-295


>>
43. NC_030997
Source: Fusarium oxysporum f. sp. lycopersici 4287 chromosome 12, whole genome shotgun sequence
Type: NRPS,T1PKS
Number of proteins with BLAST hits to this cluster: 2
Cumulative BLAST score: 1175

Table of genes, locations, strands and annotations of subject cluster:

Table of Blast hits (query gene, subject gene, %identity, blast score, %coverage, e-value):
input.path1.gene388	FOXG_14588	42	154	49.707602339181285	2.4e-35
input.path1.gene392	FOXG_14587	38	1021	60.8463591307663	1.8e-295


>>
44. NW_022194799
Source: Fusarium proliferatum ET1 genome assembly, contig: FPRO_scaffold010, whole genome shotgun sequence
Type: T1PKS
Number of proteins with BLAST hits to this cluster: 2
Cumulative BLAST score: 1171

Table of genes, locations, strands and annotations of subject cluster:

Table of Blast hits (query gene, subject gene, %identity, blast score, %coverage, e-value):
input.path1.gene388	FPRO_11892	33	186	99.70760233918129	5.8e-45
input.path1.gene392	FPRO_11891	37	985	60.57948913457872	1.4e-284


>>
45. NC_016457
Source: Thermothielavioides terrestris NRRL 8126 chromosome 1, complete sequence
Type: NRPS,T1PKS
Number of proteins with BLAST hits to this cluster: 2
Cumulative BLAST score: 1168

Table of genes, locations, strands and annotations of subject cluster:

Table of Blast hits (query gene, subject gene, %identity, blast score, %coverage, e-value):
input.path1.gene388	THITE_2141905	33	202	99.70760233918129	6e-50
input.path1.gene392	THITE_115563	37	966	61.72321768966832	6.8e-279


>>
46. NC_030962
Source: Colletotrichum higginsianum IMI 349063 chromosome 10, whole genome shotgun sequence
Type: NRPS,T1PKS
Number of proteins with BLAST hits to this cluster: 2
Cumulative BLAST score: 1167

Table of genes, locations, strands and annotations of subject cluster:

Table of Blast hits (query gene, subject gene, %identity, blast score, %coverage, e-value):
input.path1.gene388	CH63R_14277	37	220	94.15204678362574	3.6e-55
input.path1.gene392	CH63R_14283	36	947	61.68509340449867	3.3e-273


>>
47. NC_036629
Source: Fusarium fujikuroi IMI 58289 draft genome, chromosome FFUJ_chr08
Type: T1PKS
Number of proteins with BLAST hits to this cluster: 2
Cumulative BLAST score: 1167

Table of genes, locations, strands and annotations of subject cluster:

Table of Blast hits (query gene, subject gene, %identity, blast score, %coverage, e-value):
input.path1.gene388	FFUJ_12240	33	185	99.70760233918129	1.3e-44
input.path1.gene392	FFUJ_12239	37	982	60.57948913457872	9.2e-284


>>
48. NW_017263917
Source: Colletotrichum higginsianum IMI 349063 chromosome 7 map unlocalized, whole genome shotgun sequence
Type: T1PKS
Number of proteins with BLAST hits to this cluster: 2
Cumulative BLAST score: 1164

Table of genes, locations, strands and annotations of subject cluster:

Table of Blast hits (query gene, subject gene, %identity, blast score, %coverage, e-value):
input.path1.gene388	CH63R_10652	39	224	95.32163742690058	1.9e-56
input.path1.gene392	CH63R_10646	36	940	60.61761341974838	5.2e-271


>>
49. CM000574
Source: Fusarium graminearum PH-1 chromosome 1, whole genome shotgun sequence
Type: NRPS-like,T1PKS
Number of proteins with BLAST hits to this cluster: 2
Cumulative BLAST score: 1164

Table of genes, locations, strands and annotations of subject cluster:

Table of Blast hits (query gene, subject gene, %identity, blast score, %coverage, e-value):
input.path1.gene388	FGSG_10460	35	192	84.21052631578947	6.2e-47
input.path1.gene392	FGSG_10464	36	972	65.26877621044605	9.5e-281


>>
50. NC_026474
Source: Fusarium graminearum PH-1 chromosome 1, whole genome shotgun sequence
Type: NRPS-like,T1PKS
Number of proteins with BLAST hits to this cluster: 2
Cumulative BLAST score: 1164

Table of genes, locations, strands and annotations of subject cluster:

Table of Blast hits (query gene, subject gene, %identity, blast score, %coverage, e-value):
input.path1.gene388	FGSG_10460	35	192	84.21052631578947	6.2e-47
input.path1.gene392	FGSG_10464	36	972	65.26877621044605	9.5e-281
