## Supplementary Results for "A draft genome of the ascomycotal fungal species *Pseudopithomyces maydicus* (family *Didymosphaeriaceae*)": contig_12_c2.txt

ClusterBlast scores for contig_12

Table of genes, locations, strands and annotations of query cluster:
input.path1.gene638	2325386	2326715	-		
input.path1.gene639	2329982	2331603	+		
input.path1.gene640	2331896	2333866	-		
input.path1.gene641	2336738	2343315	+		
input.path1.gene642	2345182	2349957	+		
input.path1.gene643	2350125	2351567	-		
input.path1.gene644	2352390	2353742	-		
input.path1.gene645	2363625	2365011	+		
input.path1.gene646	2365115	2367616	-		
input.path1.gene647	2367807	2369320	+		


Significant hits: 
1. NW_011371358	Paracoccidioides brasiliensis Pb18 unplaced genomic scaffold supercont2.1, whole genome shotgun sequence
2. NW_004504310	Coccidioides immitis RS genomic scaffold supercont3.1, whole genome shotgun sequence
3. NW_003052500	Uncinocarpus reesii 1704 scaffold_1 genomic scaffold, whole genome shotgun sequence
4. NW_003315979	Coccidioides posadasii C735 delta SOWgp chromosome Unknown gcontig_1105601051781, whole genome shotgun sequence
5. NW_015971594	Cladophialophora bantiana CBS 173.52 unplaced genomic scaffold supercont1.20, whole genome shotgun sequence
6. NW_013550610	Fonsecaea pedrosoi CBS 271.37 unplaced genomic scaffold supercont1.6, whole genome shotgun sequence
7. NW_022474213	Venustampulla echinocandica strain BP 5553 chromosome Unknown scaffold_8, whole genome shotgun sequence
8. NW_008481827	Cladophialophora carrionii CBS 160.54 unplaced genomic scaffold supercont1.2, whole genome shotgun sequence
9. NW_015971668	Fonsecaea multimorphosa CBS 102226 unplaced genomic scaffold supercont1.34, whole genome shotgun sequence
10. NZ_CP011509	Archangium gephyra strain DSM 2261 chromosome, complete genome
11. NW_013562494	Exophiala xenobiotica strain CBS 118157 unplaced genomic scaffold supercont1.1, whole genome shotgun sequence
12. NW_015622513	Exophiala spinifera strain CBS 89968 unplaced genomic scaffold supercont1.1, whole genome shotgun sequence
13. NT_165934	Aspergillus terreus NIH2624 scaffold_11 genomic scaffold, whole genome shotgun sequence
14. RRCJ01000004	Pyricularia sp. CBS 133598 strain NI919 Pyricularia_sp_LS_NI919_Scf_4, whole genome shotgun sequence
15. NC_007196	Aspergillus fumigatus Af293 chromosome 3, whole genome shotgun sequence
16. NW_022984632	Aspergillus tanneri strain NIH1004 chromosome Unknown AapTan_5_pilon_pilon_pilon, whole genome shotgun sequence
17. NT_107009	Aspergillus nidulans FGSC A4 chromosome I map unlocalized scaffold_7, whole genome shotgun sequence
18. NC_036442	Aspergillus oryzae RIB40 DNA, chromosome 8
19. NW_017971433	Talaromyces atroroseus strain IBT 11181 chromosome Unknown scaffold_1, whole genome shotgun sequence
20. NW_014574584	Penicillium digitatum Pd1 unplaced genomic scaffold Scaffold_41, whole genome shotgun sequence
21. CM002798	Penicillium chrysogenum strain P2niaD18 chromosome I, whole genome shotgun sequence
22. NT_165981	Chaetomium globosum CBS 148.51 scaffold_6 genomic scaffold, whole genome shotgun sequence
23. NW_003299164	Microsporum canis CBS 113480 supercont1.6 genomic scaffold, whole genome shotgun sequence
24. NC_049562	Talaromyces rugulosus chromosome II, complete sequence
25. NW_003299169	Microsporum canis CBS 113480 supercont1.1 genomic scaffold, whole genome shotgun sequence
26. NW_003315111	Trichophyton benhamiae CBS 112371 chromosome Unknown contig01145.0, whole genome shotgun sequence
27. NW_003456428	Trichophyton rubrum CBS 118892 genomic scaffold supercont2.1, whole genome shotgun sequence
28. NW_003345201	Nannizzia gypsea CBS 118893 supercont1.1 genomic scaffold, whole genome shotgun sequence
29. NC_035793	Pochonia chlamydosporia 170 chromosome 4, whole genome shotgun sequence
30. NZ_CP033464	Brevibacillus laterosporus strain 1821L chromosome, complete genome
31. NZ_CP011509	Archangium gephyra strain DSM 2261 chromosome, complete genome
32. NZ_CP017968	Paenibacillus polymyxa strain YC0573 chromosome, complete genome
33. NZ_CP011512	Paenibacillus peoriae strain HS311 chromosome, complete genome
34. NZ_CP041405	Paenibacillus thiaminolyticus strain NRRL B-4156 chromosome
35. NZ_CP034248	Paenibacillus lentus strain DSM 25539 chromosome, complete genome
36. NZ_KB891762	Streptomyces sp. LaPpAH-165 B070DRAFT_scaffold_0.1, whole genome shotgun sequence
37. NZ_LISW01000001	Streptomyces sp. CB01249 scaffold1, whole genome shotgun sequence
38. NZ_CP032697	Rhizobium jaguaris strain CCGE525 plasmid pRCCGE525a, complete sequence
39. NW_001939244	Pyrenophora tritici-repentis Pt-1C-BFP supercont1.1, whole genome shotgun sequence
40. NC_030961	Colletotrichum higginsianum IMI 349063 chromosome 9, whole genome shotgun sequence
41. NW_023336267	Colletotrichum scovillei strain TJNH1 chromosome Unknown Contig00009, whole genome shotgun sequence
42. NW_011942160	Metarhizium robertsii ARSEF 23 MAA_Scf_2, whole genome shotgun sequence
43. NW_014574716	Metarhizium brunneum ARSEF 3297 chromosome Unknown scaffold_1, whole genome shotgun sequence
44. NW_014013633	Trichoderma atroviride IMI 206040 chromosome Unknown TRIATcontig_23, whole genome shotgun sequence
45. NC_035791	Pochonia chlamydosporia 170 chromosome 2, whole genome shotgun sequence
46. NW_006271973	Cordyceps militaris CM01 unplaced genomic scaffold CCM_S00005, whole genome shotgun sequence
47. NW_008481797	Cyphellophora europaea CBS 101466 unplaced genomic scaffold supercont1.1, whole genome shotgun sequence
48. NW_015622507	Exophiala mesophila strain CBS 40295 unplaced genomic scaffold supercont1.4, whole genome shotgun sequence
49. NW_013550598	Rhinocladiella mackenziei CBS 650.93 unplaced genomic scaffold supercont1.7, whole genome shotgun sequence
50. NZ_GL905390	Paenibacillus polymyxa ATCC 842 PPt02_scaffold1, whole genome shotgun sequence


Details:

>>
1. NW_011371358
Source: Paracoccidioides brasiliensis Pb18 unplaced genomic scaffold supercont2.1, whole genome shotgun sequence
Type: NRPS
Number of proteins with BLAST hits to this cluster: 3
Cumulative BLAST score: 2972

Table of genes, locations, strands and annotations of subject cluster:

Table of Blast hits (query gene, subject gene, %identity, blast score, %coverage, e-value):
input.path1.gene640	PADG_00099	51	443	120.7161125319693	1.8e-122
input.path1.gene641	PADG_00103	50	1108	78.61519607843137	0.0
input.path1.gene642	PADG_00102	49	1421	100.38461538461539	0.0


>>
2. NW_004504310
Source: Coccidioides immitis RS genomic scaffold supercont3.1, whole genome shotgun sequence
Type: NRPS
Number of proteins with BLAST hits to this cluster: 3
Cumulative BLAST score: 2931

Table of genes, locations, strands and annotations of subject cluster:

Table of Blast hits (query gene, subject gene, %identity, blast score, %coverage, e-value):
input.path1.gene640	CIMG_01434	51	436	123.01790281329923	2.8e-120
input.path1.gene641	CIMG_01430	49	1093	78.61519607843137	0.0
input.path1.gene642	CIMG_01429	48	1402	101.08974358974359	0.0


>>
3. NW_003052500
Source: Uncinocarpus reesii 1704 scaffold_1 genomic scaffold, whole genome shotgun sequence
Type: NRPS
Number of proteins with BLAST hits to this cluster: 3
Cumulative BLAST score: 2931

Table of genes, locations, strands and annotations of subject cluster:

Table of Blast hits (query gene, subject gene, %identity, blast score, %coverage, e-value):
input.path1.gene640	UREG_01350	50	430	121.48337595907928	2.6e-118
input.path1.gene641	UREG_01347	49	1083	78.49264705882352	0.0
input.path1.gene642	UREG_01346	49	1418	100.25641025641025	0.0


>>
4. NW_003315979
Source: Coccidioides posadasii C735 delta SOWgp chromosome Unknown gcontig_1105601051781, whole genome shotgun sequence
Type: NRPS
Number of proteins with BLAST hits to this cluster: 3
Cumulative BLAST score: 2923

Table of genes, locations, strands and annotations of subject cluster:

Table of Blast hits (query gene, subject gene, %identity, blast score, %coverage, e-value):
input.path1.gene640	CPC735_050020	48	421	126.59846547314577	9.4e-116
input.path1.gene641	CPC735_050050	49	1094	78.61519607843137	0.0
input.path1.gene642	CPC735_050060	48	1408	100.8974358974359	0.0


>>
5. NW_015971594
Source: Cladophialophora bantiana CBS 173.52 unplaced genomic scaffold supercont1.20, whole genome shotgun sequence
Type: NRPS
Number of proteins with BLAST hits to this cluster: 3
Cumulative BLAST score: 2462

Table of genes, locations, strands and annotations of subject cluster:

Table of Blast hits (query gene, subject gene, %identity, blast score, %coverage, e-value):
input.path1.gene640	Z519_11082	52	441	119.94884910485935	8.7e-122
input.path1.gene641	Z519_11085	48	1094	78.67647058823529	0.0
input.path1.gene642	Z519_11086	35	927	104.42307692307693	2.1e-267


>>
6. NW_013550610
Source: Fonsecaea pedrosoi CBS 271.37 unplaced genomic scaffold supercont1.6, whole genome shotgun sequence
Type: NRPS
Number of proteins with BLAST hits to this cluster: 3
Cumulative BLAST score: 2444

Table of genes, locations, strands and annotations of subject cluster:

Table of Blast hits (query gene, subject gene, %identity, blast score, %coverage, e-value):
input.path1.gene640	Z517_10054	54	446	114.83375959079285	3.5e-123
input.path1.gene641	Z517_10057	48	1080	78.67647058823529	0.0
input.path1.gene642	Z517_10058	35	918	111.85897435897436	7.4e-265


>>
7. NW_022474213
Source: Venustampulla echinocandica strain BP 5553 chromosome Unknown scaffold_8, whole genome shotgun sequence
Type: NRPS
Number of proteins with BLAST hits to this cluster: 3
Cumulative BLAST score: 2415

Table of genes, locations, strands and annotations of subject cluster:

Table of Blast hits (query gene, subject gene, %identity, blast score, %coverage, e-value):
input.path1.gene640	BP5553_08730	48	406	114.57800511508951	3.1e-111
input.path1.gene641	BP5553_08736	47	1054	78.79901960784314	9.1e-306
input.path1.gene642	BP5553_08735	36	955	109.23076923076923	7.2e-276


>>
8. NW_008481827
Source: Cladophialophora carrionii CBS 160.54 unplaced genomic scaffold supercont1.2, whole genome shotgun sequence
Type: NRPS
Number of proteins with BLAST hits to this cluster: 3
Cumulative BLAST score: 2414

Table of genes, locations, strands and annotations of subject cluster:

Table of Blast hits (query gene, subject gene, %identity, blast score, %coverage, e-value):
input.path1.gene640	G647_02317	53	443	115.08951406649616	1.8e-122
input.path1.gene641	G647_02320	47	1050	78.73774509803921	1.7e-304
input.path1.gene642	G647_02321	36	921	104.61538461538463	1.5e-265


>>
9. NW_015971668
Source: Fonsecaea multimorphosa CBS 102226 unplaced genomic scaffold supercont1.34, whole genome shotgun sequence
Type: NRPS
Number of proteins with BLAST hits to this cluster: 3
Cumulative BLAST score: 2413

Table of genes, locations, strands and annotations of subject cluster:

Table of Blast hits (query gene, subject gene, %identity, blast score, %coverage, e-value):
input.path1.gene640	Z520_10931	51	441	119.94884910485935	1.1e-121
input.path1.gene641	Z520_10934	47	1061	78.67647058823529	7.4e-308
input.path1.gene642	Z520_10935	34	911	110.57692307692308	9.1e-263


>>
10. NZ_CP011509
Source: Archangium gephyra strain DSM 2261 chromosome, complete genome
Type: NRPS,T1PKS
Number of proteins with BLAST hits to this cluster: 3
Cumulative BLAST score: 615

Table of genes, locations, strands and annotations of subject cluster:
AA314_RS25190	AA314_RS25190	6399012	6399524	+	translation_initiation_factor_IF-3

Table of Blast hits (query gene, subject gene, %identity, blast score, %coverage, e-value):
input.path1.gene640	AA314_RS25250	31	109	67.26342710997443	1e-21
input.path1.gene641	AA314_RS25235	33	182	26.715686274509803	4e-43
input.path1.gene642	AA314_RS25240	31	170	28.46153846153846	2e-39
input.path1.gene642	AA314_RS25245	32	154	28.653846153846153	1.1e-34


>>
11. NW_013562494
Source: Exophiala xenobiotica strain CBS 118157 unplaced genomic scaffold supercont1.1, whole genome shotgun sequence
Type: NRPS
Number of proteins with BLAST hits to this cluster: 4
Cumulative BLAST score: 3316

Table of genes, locations, strands and annotations of subject cluster:

Table of Blast hits (query gene, subject gene, %identity, blast score, %coverage, e-value):
input.path1.gene640	PV05_00029	52	438	115.6010230179028	7.4e-121
input.path1.gene640	PV05_00029_XP_013320344.1	52	438	115.6010230179028	7.4e-121
input.path1.gene640	PV05_00029_XP_013320345.1	51	433	119.43734015345268	2.4e-119
input.path1.gene641	PV05_00033	45	988	79.47303921568627	1e-285
input.path1.gene642	PV05_00030	36	958	109.23076923076923	6.5e-277
input.path1.gene646	PV05_00035	39	61	24.429223744292237	2.1e-07


>>
12. NW_015622513
Source: Exophiala spinifera strain CBS 89968 unplaced genomic scaffold supercont1.1, whole genome shotgun sequence
Type: NRPS
Number of proteins with BLAST hits to this cluster: 4
Cumulative BLAST score: 2531

Table of genes, locations, strands and annotations of subject cluster:

Table of Blast hits (query gene, subject gene, %identity, blast score, %coverage, e-value):
input.path1.gene640	PV08_02212	52	433	115.08951406649616	2.4e-119
input.path1.gene641	PV08_02216	47	1069	79.8406862745098	2.7e-310
input.path1.gene642	PV08_02213	36	969	110.00000000000001	4.8e-280
input.path1.gene646	PV08_02206	39	60	24.65753424657534	4.7e-07


>>
13. NT_165934
Source: Aspergillus terreus NIH2624 scaffold_11 genomic scaffold, whole genome shotgun sequence
Type: NRPS,T1PKS
Number of proteins with BLAST hits to this cluster: 2
Cumulative BLAST score: 2414

Table of genes, locations, strands and annotations of subject cluster:

Table of Blast hits (query gene, subject gene, %identity, blast score, %coverage, e-value):
input.path1.gene641	ATEG_07489	48	1067	79.22794117647058	1.8e-309
input.path1.gene642	ATEG_07488	48	1347	100.44871794871794	0.0


>>
14. RRCJ01000004
Source: Pyricularia sp. CBS 133598 strain NI919 Pyricularia_sp_LS_NI919_Scf_4, whole genome shotgun sequence
Type: NRPS
Number of proteins with BLAST hits to this cluster: 2
Cumulative BLAST score: 2162

Table of genes, locations, strands and annotations of subject cluster:

Table of Blast hits (query gene, subject gene, %identity, blast score, %coverage, e-value):
input.path1.gene641	PspLS_04326	42	926	82.72058823529412	2.9e-267
input.path1.gene642	PspLS_04325	45	1236	103.33333333333334	0.0


>>
15. NC_007196
Source: Aspergillus fumigatus Af293 chromosome 3, whole genome shotgun sequence
Type: NRPS
Number of proteins with BLAST hits to this cluster: 2
Cumulative BLAST score: 2027

Table of genes, locations, strands and annotations of subject cluster:

Table of Blast hits (query gene, subject gene, %identity, blast score, %coverage, e-value):
input.path1.gene641	AFUA_3G03430	48	1067	78.61519607843137	1.8e-309
input.path1.gene642	AFUA_3G03420	37	960	110.12820512820514	1.7e-277


>>
16. NW_022984632
Source: Aspergillus tanneri strain NIH1004 chromosome Unknown AapTan_5_pilon_pilon_pilon, whole genome shotgun sequence
Type: NRPS
Number of proteins with BLAST hits to this cluster: 2
Cumulative BLAST score: 2012

Table of genes, locations, strands and annotations of subject cluster:

Table of Blast hits (query gene, subject gene, %identity, blast score, %coverage, e-value):
input.path1.gene641	ATNIH1004_010602	47	1065	78.9828431372549	6.7e-309
input.path1.gene642	ATNIH1004_010603	35	947	110.57692307692308	1.5e-273


>>
17. NT_107009
Source: Aspergillus nidulans FGSC A4 chromosome I map unlocalized scaffold_7, whole genome shotgun sequence
Type: NRPS
Number of proteins with BLAST hits to this cluster: 2
Cumulative BLAST score: 2007

Table of genes, locations, strands and annotations of subject cluster:

Table of Blast hits (query gene, subject gene, %identity, blast score, %coverage, e-value):
input.path1.gene641	AN6237.2	48	1068	78.86029411764706	8e-310
input.path1.gene642	AN6236.2	35	939	111.15384615384616	5.3e-271


>>
18. NC_036442
Source: Aspergillus oryzae RIB40 DNA, chromosome 8
Type: NRPS
Number of proteins with BLAST hits to this cluster: 2
Cumulative BLAST score: 1981

Table of genes, locations, strands and annotations of subject cluster:

Table of Blast hits (query gene, subject gene, %identity, blast score, %coverage, e-value):
input.path1.gene641	AO090103000166	47	1068	79.04411764705883	6.1e-310
input.path1.gene642	AO090103000167	35	913	110.83333333333334	3.1e-263


>>
19. NW_017971433
Source: Talaromyces atroroseus strain IBT 11181 chromosome Unknown scaffold_1, whole genome shotgun sequence
Type: NRPS
Number of proteins with BLAST hits to this cluster: 2
Cumulative BLAST score: 1958

Table of genes, locations, strands and annotations of subject cluster:

Table of Blast hits (query gene, subject gene, %identity, blast score, %coverage, e-value):
input.path1.gene641	UA08_01920	47	1059	79.65686274509804	4.8e-307
input.path1.gene642	UA08_01921	35	899	110.44871794871796	6.1e-259


>>
20. NW_014574584
Source: Penicillium digitatum Pd1 unplaced genomic scaffold Scaffold_41, whole genome shotgun sequence
Type: NRPS
Number of proteins with BLAST hits to this cluster: 2
Cumulative BLAST score: 1930

Table of genes, locations, strands and annotations of subject cluster:

Table of Blast hits (query gene, subject gene, %identity, blast score, %coverage, e-value):
input.path1.gene641	PDIP_64370	45	999	79.41176470588235	3.5e-289
input.path1.gene642	PDIP_64360	35	931	110.19230769230771	1.1e-268


>>
21. CM002798
Source: Penicillium chrysogenum strain P2niaD18 chromosome I, whole genome shotgun sequence
Type: NRPS
Number of proteins with BLAST hits to this cluster: 2
Cumulative BLAST score: 1929

Table of genes, locations, strands and annotations of subject cluster:

Table of Blast hits (query gene, subject gene, %identity, blast score, %coverage, e-value):
input.path1.gene641	EN45_007560	45	1017	78.86029411764706	1.2e-294
input.path1.gene642	EN45_007550	35	912	110.7051282051282	7e-263


>>
22. NT_165981
Source: Chaetomium globosum CBS 148.51 scaffold_6 genomic scaffold, whole genome shotgun sequence
Type: NRPS
Number of proteins with BLAST hits to this cluster: 2
Cumulative BLAST score: 1780

Table of genes, locations, strands and annotations of subject cluster:

Table of Blast hits (query gene, subject gene, %identity, blast score, %coverage, e-value):
input.path1.gene641	CHGG_09474	45	990	80.45343137254902	2.1e-286
input.path1.gene642	CHGG_09475	35	790	107.43589743589743	3.1e-226


>>
23. NW_003299164
Source: Microsporum canis CBS 113480 supercont1.6 genomic scaffold, whole genome shotgun sequence
Type: NRPS
Number of proteins with BLAST hits to this cluster: 2
Cumulative BLAST score: 928

Table of genes, locations, strands and annotations of subject cluster:

Table of Blast hits (query gene, subject gene, %identity, blast score, %coverage, e-value):
input.path1.gene641	MCYG_06184	31	524	76.59313725490196	4.3e-146
input.path1.gene642	MCYG_06185	31	404	56.98717948717948	4.7e-110


>>
24. NC_049562
Source: Talaromyces rugulosus chromosome II, complete sequence
Type: NRPS,NRPS-like
Number of proteins with BLAST hits to this cluster: 2
Cumulative BLAST score: 925

Table of genes, locations, strands and annotations of subject cluster:

Table of Blast hits (query gene, subject gene, %identity, blast score, %coverage, e-value):
input.path1.gene641	TRUGW13939_02482	31	467	76.9607843137255	4.7e-129
input.path1.gene642	TRUGW13939_02483	34	458	62.05128205128205	2.1e-126


>>
25. NW_003299169
Source: Microsporum canis CBS 113480 supercont1.1 genomic scaffold, whole genome shotgun sequence
Type: NRPS
Number of proteins with BLAST hits to this cluster: 2
Cumulative BLAST score: 896

Table of genes, locations, strands and annotations of subject cluster:

Table of Blast hits (query gene, subject gene, %identity, blast score, %coverage, e-value):
input.path1.gene641	MCYG_00852	33	591	77.87990196078431	2.2e-166
input.path1.gene642	MCYG_00851	39	305	30.320512820512818	3.9e-80


>>
26. NW_003315111
Source: Trichophyton benhamiae CBS 112371 chromosome Unknown contig01145.0, whole genome shotgun sequence
Type: NRPS
Number of proteins with BLAST hits to this cluster: 2
Cumulative BLAST score: 860

Table of genes, locations, strands and annotations of subject cluster:

Table of Blast hits (query gene, subject gene, %identity, blast score, %coverage, e-value):
input.path1.gene641	ARB_05580	33	566	76.1642156862745	1.3e-158
input.path1.gene642	ARB_05579	38	294	29.03846153846154	5.3e-77


>>
27. NW_003456428
Source: Trichophyton rubrum CBS 118892 genomic scaffold supercont2.1, whole genome shotgun sequence
Type: NRPS
Number of proteins with BLAST hits to this cluster: 2
Cumulative BLAST score: 851

Table of genes, locations, strands and annotations of subject cluster:

Table of Blast hits (query gene, subject gene, %identity, blast score, %coverage, e-value):
input.path1.gene641	TERG_00402	33	557	76.65441176470588	3.5e-156
input.path1.gene642	TERG_08653	38	294	28.974358974358978	9e-77


>>
28. NW_003345201
Source: Nannizzia gypsea CBS 118893 supercont1.1 genomic scaffold, whole genome shotgun sequence
Type: NRPS
Number of proteins with BLAST hits to this cluster: 2
Cumulative BLAST score: 807

Table of genes, locations, strands and annotations of subject cluster:

Table of Blast hits (query gene, subject gene, %identity, blast score, %coverage, e-value):
input.path1.gene641	MGYG_01115	31	510	75.30637254901961	6.4e-142
input.path1.gene642	MGYG_01116	39	297	29.230769230769234	1.1e-77


>>
29. NC_035793
Source: Pochonia chlamydosporia 170 chromosome 4, whole genome shotgun sequence
Type: NRPS-like
Number of proteins with BLAST hits to this cluster: 2
Cumulative BLAST score: 694

Table of genes, locations, strands and annotations of subject cluster:

Table of Blast hits (query gene, subject gene, %identity, blast score, %coverage, e-value):
input.path1.gene641	VFPPC_10104	32	525	76.4093137254902	1.5e-146
input.path1.gene642	VFPPC_10103	32	169	23.974358974358974	2.6e-39


>>
30. NZ_CP033464
Source: Brevibacillus laterosporus strain 1821L chromosome, complete genome
Type: NRPS
Number of proteins with BLAST hits to this cluster: 2
Cumulative BLAST score: 658

Table of genes, locations, strands and annotations of subject cluster:
EEL30_RS19145	EEL30_RS19145	3936482	3937915	+	FAD-binding_oxidoreductase
EEL30_RS19265	EEL30_RS19265	3978083	3979049	+	MFS_transporter

Table of Blast hits (query gene, subject gene, %identity, blast score, %coverage, e-value):
input.path1.gene641	EEL30_RS19240	33	186	29.411764705882355	2.1e-44
input.path1.gene641	EEL30_RS19140	41	168	14.399509803921568	6e-39
input.path1.gene641	EEL30_RS19135	42	155	14.338235294117647	4e-35
input.path1.gene642	EEL30_RS19130	31	149	23.525641025641026	2.1e-33


>>
31. NZ_CP011509
Source: Archangium gephyra strain DSM 2261 chromosome, complete genome
Type: NRPS,T1PKS
Number of proteins with BLAST hits to this cluster: 2
Cumulative BLAST score: 398

Table of genes, locations, strands and annotations of subject cluster:
AA314_RS19910	AA314_RS19910	5007915	5009717	-	SAVED_domain-containing_protein

Table of Blast hits (query gene, subject gene, %identity, blast score, %coverage, e-value):
input.path1.gene641	AA314_RS19835	37	230	25.183823529411764	1.7e-57
input.path1.gene642	AA314_RS19840	32	168	28.910256410256412	7.4e-39


>>
32. NZ_CP017968
Source: Paenibacillus polymyxa strain YC0573 chromosome, complete genome
Type: NRPS
Number of proteins with BLAST hits to this cluster: 2
Cumulative BLAST score: 339

Table of genes, locations, strands and annotations of subject cluster:
PPYC2_RS23330	PPYC2_RS23330	5245607	5245900	-	RNA_polymerase_sigma_factor
PPYC2_RS23390	PPYC2_RS23390	5257028	5257126	-	DNA-binding_response_regulator
PPYC2_RS23465	PPYC2_RS23465	5305140	5305340	-	diaminobutyrate--2-oxoglutarate_transaminase
PPYC2_RS23480	PPYC2_RS23480	5307686	5308032	-	helix-turn-helix_domain-containing_protein
PPYC2_RS23490	PPYC2_RS23490	5309059	5309148	-	DNA-binding_response_regulator
PPYC2_RS23495	PPYC2_RS23495	5309211	5310569	-	sensor_histidine_kinase_KdpD

Table of Blast hits (query gene, subject gene, %identity, blast score, %coverage, e-value):
input.path1.gene641	PPYC2_RS23445	33	181	27.879901960784316	6.8e-43
input.path1.gene642	PPYC2_RS23440	31	158	28.78205128205128	7.7e-36


>>
33. NZ_CP011512
Source: Paenibacillus peoriae strain HS311 chromosome, complete genome
Type: NRPS
Number of proteins with BLAST hits to this cluster: 2
Cumulative BLAST score: 338

Table of genes, locations, strands and annotations of subject cluster:
ABE82_RS27555	ABE82_RS27555	5118770	5119063	-	RNA_polymerase_sigma_factor
ABE82_RS27560	ABE82_RS27560	5129555	5129653	-	winged_helix-turn-helix_domain-containing_protein
ABE82_RS27565	ABE82_RS27565	5177750	5177950	-	diaminobutyrate--2-oxoglutarate_transaminase
ABE82_RS22180	ABE82_RS22180	5180298	5180644	-	helix-turn-helix_transcriptional_regulator
ABE82_RS28055	ABE82_RS28055	5181630	5181719	-	response_regulator_transcription_factor
ABE82_RS27570	ABE82_RS27570	5181782	5183044	-	sensor_histidine_kinase_KdpD

Table of Blast hits (query gene, subject gene, %identity, blast score, %coverage, e-value):
input.path1.gene641	ABE82_RS22145	33	180	28.247549019607842	2e-42
input.path1.gene642	ABE82_RS22140	31	158	28.78205128205128	7.7e-36


>>
34. NZ_CP041405
Source: Paenibacillus thiaminolyticus strain NRRL B-4156 chromosome
Type: NRPS
Number of proteins with BLAST hits to this cluster: 2
Cumulative BLAST score: 337

Table of genes, locations, strands and annotations of subject cluster:
FLT43_RS13415	FLT43_RS13415	2977647	2977772	-	IS3_family_transposase
FLT43_RS13495	FLT43_RS13495	3029396	3029623	-	group_II_intron_reverse_transcriptase/maturase
FLT43_RS13500	FLT43_RS13500	3029793	3030179	-	group_II_intron_reverse_transcriptase/maturase
FLT43_RS13510	FLT43_RS13510	3030875	3031921	-	N-acetyltransferase
FLT43_RS13520	FLT43_RS13520	3032999	3033109	+	DNA_cytosine_methyltransferase
FLT43_RS13530	FLT43_RS13530	3035321	3035929	-	Tn7_transposase_TnsA_N-terminal_domain-containing_protein

Table of Blast hits (query gene, subject gene, %identity, blast score, %coverage, e-value):
input.path1.gene641	FLT43_RS13465	33	182	27.879901960784316	4e-43
input.path1.gene642	FLT43_RS13460	31	155	28.78205128205128	5e-35


>>
35. NZ_CP034248
Source: Paenibacillus lentus strain DSM 25539 chromosome, complete genome
Type: NRPS
Number of proteins with BLAST hits to this cluster: 2
Cumulative BLAST score: 336

Table of genes, locations, strands and annotations of subject cluster:
EIM92_RS07585	EIM92_RS07585	1683176	1698254	+	amino_acid_adenylation_domain-containing_protein
EIM92_RS07620	EIM92_RS07620	1727586	1728827	-	MFS_transporter

Table of Blast hits (query gene, subject gene, %identity, blast score, %coverage, e-value):
input.path1.gene641	EIM92_RS07600	33	180	28.002450980392158	1.2e-42
input.path1.gene642	EIM92_RS07605	31	156	28.846153846153843	2.9e-35


>>
36. NZ_KB891762
Source: Streptomyces sp. LaPpAH-165 B070DRAFT_scaffold_0.1, whole genome shotgun sequence
Type: NRPS
Number of proteins with BLAST hits to this cluster: 2
Cumulative BLAST score: 319

Table of genes, locations, strands and annotations of subject cluster:

Table of Blast hits (query gene, subject gene, %identity, blast score, %coverage, e-value):
input.path1.gene641	B070_RS0100500	31	170	30.208333333333332	2e-39
input.path1.gene642	B070_RS0100495	32	149	27.500000000000004	2.1e-33


>>
37. NZ_LISW01000001
Source: Streptomyces sp. CB01249 scaffold1, whole genome shotgun sequence
Type: NRPS
Number of proteins with BLAST hits to this cluster: 2
Cumulative BLAST score: 317

Table of genes, locations, strands and annotations of subject cluster:

Table of Blast hits (query gene, subject gene, %identity, blast score, %coverage, e-value):
input.path1.gene641	AMK18_RS06025	31	169	30.208333333333332	2.7e-39
input.path1.gene642	AMK18_RS06030	32	148	27.564102564102566	4.7e-33


>>
38. NZ_CP032697
Source: Rhizobium jaguaris strain CCGE525 plasmid pRCCGE525a, complete sequence
Type: LAP,NRPS,NRPS-like
Number of proteins with BLAST hits to this cluster: 2
Cumulative BLAST score: 286

Table of genes, locations, strands and annotations of subject cluster:
CCGE525_RS37050	CCGE525_RS37050	50017	50314	-	helix-turn-helix_domain-containing_protein
CCGE525_RS37060	CCGE525_RS37060	51358	51856	+	MucR_family_transcriptional_regulator
CCGE525_RS37065	CCGE525_RS37065	51828	52519	-	transposase
CCGE525_RS37160	CCGE525_RS37160	70667	70951	-	DDE-type_integrase/transposase/recombinase
CCGE525_RS37290	CCGE525_RS37290	130061	130249	+	hypothetical_protein
CCGE525_RS37365	CCGE525_RS37365	143509	143631	-	DeoR_family_transcriptional_regulator
CCGE525_RS37370	CCGE525_RS37370	144233	145496	+	ISKra4_family_transposase
CCGE525_RS37480	CCGE525_RS37480	165426	165827	+	CYTH_domain-containing_protein
CCGE525_RS37485	CCGE525_RS37485	165841	166064	+	NADP-dependent_oxidoreductase_yncb

Table of Blast hits (query gene, subject gene, %identity, blast score, %coverage, e-value):
input.path1.gene641	CCGE525_RS37195	39	130	12.928921568627452	1.4e-27
input.path1.gene642	CCGE525_RS37190	31	156	28.205128205128204	1.7e-35


>>
39. NW_001939244
Source: Pyrenophora tritici-repentis Pt-1C-BFP supercont1.1, whole genome shotgun sequence
Type: NRPS
Number of proteins with BLAST hits to this cluster: 3
Cumulative BLAST score: 4269

Table of genes, locations, strands and annotations of subject cluster:

Table of Blast hits (query gene, subject gene, %identity, blast score, %coverage, e-value):
input.path1.gene640	PTRG_01678	65	610	123.01790281329923	1.1e-172
input.path1.gene641	PTRG_01681	61	1434	77.75735294117648	0.0
input.path1.gene641	PTRG_01680	58	423	22.54901960784314	7.9e-116
input.path1.gene642	PTRG_01683	60	1802	99.35897435897436	0.0


>>
40. NC_030961
Source: Colletotrichum higginsianum IMI 349063 chromosome 9, whole genome shotgun sequence
Type: NRPS
Number of proteins with BLAST hits to this cluster: 3
Cumulative BLAST score: 2971

Table of genes, locations, strands and annotations of subject cluster:

Table of Blast hits (query gene, subject gene, %identity, blast score, %coverage, e-value):
input.path1.gene640	CH63R_12699	51	438	117.64705882352942	7.4e-121
input.path1.gene641	CH63R_12704	50	1127	79.5343137254902	0.0
input.path1.gene642	CH63R_12697	50	1406	99.67948717948718	0.0


>>
41. NW_023336267
Source: Colletotrichum scovillei strain TJNH1 chromosome Unknown Contig00009, whole genome shotgun sequence
Type: NRPS
Number of proteins with BLAST hits to this cluster: 3
Cumulative BLAST score: 2929

Table of genes, locations, strands and annotations of subject cluster:

Table of Blast hits (query gene, subject gene, %identity, blast score, %coverage, e-value):
input.path1.gene640	HER10_EVM0001035	52	439	114.3222506393862	3.3e-121
input.path1.gene641	HER10_EVM0008190	48	1089	80.26960784313727	0.0
input.path1.gene642	HER10_EVM0002215	49	1401	100.12820512820512	0.0


>>
42. NW_011942160
Source: Metarhizium robertsii ARSEF 23 MAA_Scf_2, whole genome shotgun sequence
Type: NRPS
Number of proteins with BLAST hits to this cluster: 3
Cumulative BLAST score: 2914

Table of genes, locations, strands and annotations of subject cluster:

Table of Blast hits (query gene, subject gene, %identity, blast score, %coverage, e-value):
input.path1.gene640	MAA_05335	52	442	115.34526854219949	3.9e-122
input.path1.gene641	MAA_05338	50	1116	79.35049019607843	0.0
input.path1.gene642	MAA_05334	47	1356	101.6025641025641	0.0


>>
43. NW_014574716
Source: Metarhizium brunneum ARSEF 3297 chromosome Unknown scaffold_1, whole genome shotgun sequence
Type: NRPS
Number of proteins with BLAST hits to this cluster: 3
Cumulative BLAST score: 2911

Table of genes, locations, strands and annotations of subject cluster:

Table of Blast hits (query gene, subject gene, %identity, blast score, %coverage, e-value):
input.path1.gene640	MBR_01851	52	443	115.34526854219949	1.8e-122
input.path1.gene641	MBR_01848	49	1113	79.71813725490196	0.0
input.path1.gene642	MBR_01852	47	1355	101.02564102564102	0.0


>>
44. NW_014013633
Source: Trichoderma atroviride IMI 206040 chromosome Unknown TRIATcontig_23, whole genome shotgun sequence
Type: NRPS
Number of proteins with BLAST hits to this cluster: 3
Cumulative BLAST score: 2901

Table of genes, locations, strands and annotations of subject cluster:

Table of Blast hits (query gene, subject gene, %identity, blast score, %coverage, e-value):
input.path1.gene640	TRIATDRAFT_152707	52	447	115.85677749360615	1.2e-123
input.path1.gene641	TRIATDRAFT_161084	51	1125	77.57352941176471	0.0
input.path1.gene642	TRIATDRAFT_39887	47	1329	101.98717948717947	0.0


>>
45. NC_035791
Source: Pochonia chlamydosporia 170 chromosome 2, whole genome shotgun sequence
Type: NRPS
Number of proteins with BLAST hits to this cluster: 3
Cumulative BLAST score: 2897

Table of genes, locations, strands and annotations of subject cluster:

Table of Blast hits (query gene, subject gene, %identity, blast score, %coverage, e-value):
input.path1.gene640	VFPPC_03466	52	445	116.87979539641944	6e-123
input.path1.gene641	VFPPC_03464	50	1126	78.49264705882352	0.0
input.path1.gene642	VFPPC_03467	47	1326	100.96153846153845	0.0


>>
46. NW_006271973
Source: Cordyceps militaris CM01 unplaced genomic scaffold CCM_S00005, whole genome shotgun sequence
Type: NRPS
Number of proteins with BLAST hits to this cluster: 3
Cumulative BLAST score: 2837

Table of genes, locations, strands and annotations of subject cluster:

Table of Blast hits (query gene, subject gene, %identity, blast score, %coverage, e-value):
input.path1.gene640	CCM_06701	51	435	115.85677749360615	6.3e-120
input.path1.gene641	CCM_06697	47	1067	79.5343137254902	1.8e-309
input.path1.gene642	CCM_06702	46	1335	101.53846153846153	0.0


>>
47. NW_008481797
Source: Cyphellophora europaea CBS 101466 unplaced genomic scaffold supercont1.1, whole genome shotgun sequence
Type: NRPS
Number of proteins with BLAST hits to this cluster: 3
Cumulative BLAST score: 2786

Table of genes, locations, strands and annotations of subject cluster:

Table of Blast hits (query gene, subject gene, %identity, blast score, %coverage, e-value):
input.path1.gene640	HMPREF1541_00534	54	448	114.57800511508951	9.3e-124
input.path1.gene641	HMPREF1541_00532	48	1033	78.49264705882352	2.2e-299
input.path1.gene642	HMPREF1541_00538	46	1305	98.46153846153847	0.0


>>
48. NW_015622507
Source: Exophiala mesophila strain CBS 40295 unplaced genomic scaffold supercont1.4, whole genome shotgun sequence
Type: NRPS
Number of proteins with BLAST hits to this cluster: 3
Cumulative BLAST score: 2723

Table of genes, locations, strands and annotations of subject cluster:

Table of Blast hits (query gene, subject gene, %identity, blast score, %coverage, e-value):
input.path1.gene640	PV10_05613	49	433	121.73913043478262	3.1e-119
input.path1.gene641	PV10_05615	45	965	79.04411764705883	9.5e-279
input.path1.gene642	PV10_05618	47	1325	100.76923076923077	0.0


>>
49. NW_013550598
Source: Rhinocladiella mackenziei CBS 650.93 unplaced genomic scaffold supercont1.7, whole genome shotgun sequence
Type: NRPS
Number of proteins with BLAST hits to this cluster: 3
Cumulative BLAST score: 1401

Table of genes, locations, strands and annotations of subject cluster:

Table of Blast hits (query gene, subject gene, %identity, blast score, %coverage, e-value):
input.path1.gene640	Z518_08931	44	370	114.06649616368287	1.9e-100
input.path1.gene641	Z518_08927	32	532	75.30637254901961	1.2e-148
input.path1.gene642	Z518_08930	35	499	61.6025641025641	1.1e-138


>>
50. NZ_GL905390
Source: Paenibacillus polymyxa ATCC 842 PPt02_scaffold1, whole genome shotgun sequence
Type: NRPS,T1PKS
Number of proteins with BLAST hits to this cluster: 3
Cumulative BLAST score: 560

Table of genes, locations, strands and annotations of subject cluster:
PPT_RS29225	PPT_RS29225	2779056	2779200	+	IS30_family_transposase
PPT_RS28435	PPT_RS28435	2779172	2779719	-	prolyl_oligopeptidase_family_serine_peptidase
PPT_RS27865	PPT_RS27865	2779787	2780116	-	alpha/beta_hydrolase
PPT_RS0112870	PPT_RS0112870	2787658	2788362	-	SDR_family_NAD(P)-dependent_oxidoreductase
PPT_RS0112900	PPT_RS0112900	2825123	2825413	+	short-chain_dehydrogenase
PPT_RS28445	PPT_RS28445	2827400	2827615	-	tyrosine-type_recombinase/integrase
PPT_RS0112910	PPT_RS0112910	2827617	2827937	-	LuxR_family_transcriptional_regulator

Table of Blast hits (query gene, subject gene, %identity, blast score, %coverage, e-value):
input.path1.gene640	PPT_RS27880	32	99	67.51918158567774	1.1e-18
input.path1.gene641	PPT_RS0112730	35	150	16.78921568627451	1.3e-33
input.path1.gene641	PPT_RS0112725	34	149	14.093137254901961	2.9e-33
input.path1.gene642	PPT_RS27870	32	162	24.294871794871796	4.1e-37
