## Supplementary Results for "A draft genome of the ascomycotal fungal species *Pseudopithomyces maydicus* (family *Didymosphaeriaceae*)": contig_12_c3.txt

ClusterBlast scores for contig_12

Table of genes, locations, strands and annotations of query cluster:
input.path1.gene681	2482003	2483606	+		
input.path1.gene682	2485369	2488446	+		
input.path1.gene683	2490879	2491863	+		
input.path1.gene684	2493524	2493843	+		
input.path1.gene685	2496885	2498346	-		
input.path1.gene686	2499384	2503240	-		
input.path1.gene687	2503815	2508545	-		
input.path1.gene688	2509118	2510832	-		
input.path1.gene689	2519212	2520209	+		


Significant hits: 
1. NW_006271975	Cordyceps militaris CM01 unplaced genomic scaffold CCM_S00007, whole genome shotgun sequence


Details:

>>
1. NW_006271975
Source: Cordyceps militaris CM01 unplaced genomic scaffold CCM_S00007, whole genome shotgun sequence
Type: NRPS-like
Number of proteins with BLAST hits to this cluster: 2
Cumulative BLAST score: 282

Table of genes, locations, strands and annotations of subject cluster:

Table of Blast hits (query gene, subject gene, %identity, blast score, %coverage, e-value):
input.path1.gene686	CCM_08331	33	197	37.71486349848331	7.3e-48
input.path1.gene687	CCM_08334	36	85	15.992970123022848	4.6e-14
