## Supplementary Results for "A draft genome of the ascomycotal fungal species *Pseudopithomyces maydicus* (family *Didymosphaeriaceae*)": contig_12_c4.txt

ClusterBlast scores for contig_12

Table of genes, locations, strands and annotations of query cluster:
input.path1.gene794	2926205	2929258	+		
input.path1.gene795	2930227	2932249	-		
input.path1.gene796	2933016	2934698	+		
input.path1.gene797	2936553	2937756	-		
input.path1.gene798	2938052	2939399	+		
input.path1.gene799	2940210	2941036	+		
input.path1.gene800	2941754	2942703	-		
input.path1.gene801	2942795	2943325	+		
input.path1.gene802	2944428	2945553	+		
input.path1.gene803	2947401	2949535	-		
input.path1.gene804	2950659	2956336	+		
input.path1.gene805	2956653	2957742	+		
input.path1.gene806	2957875	2960882	+		
input.path1.gene807	2961299	2962724	+		
input.path1.gene808	2963266	2964148	-		
input.path1.gene809	2965014	2965992	+		
input.path1.gene810	2966576	2968158	-		
input.path1.gene811	2968432	2971157	+		
input.path1.gene812	2972015	2973145	+		
input.path1.gene813	2974911	2975903	+		


Significant hits: 
1. NW_022984630	Aspergillus tanneri strain NIH1004 chromosome Unknown AapTan_3_pilon_pilon_pilon, whole genome shotgun sequence
2. NT_107015	Aspergillus nidulans FGSC A4 chromosome VIII map unlocalized scaffold_1, whole genome shotgun sequence
3. NW_014574694	Metarhizium brunneum ARSEF 3297 chromosome Unknown scaffold_23, whole genome shotgun sequence
4. NW_019154043	Pochonia chlamydosporia 170 chromosome Unknown PCv3seq00008, whole genome shotgun sequence
5. NW_006913046	Cladophialophora yegresii CBS 114405 aczIN-supercont1.2, whole genome shotgun sequence
6. NW_008481833	Cladophialophora carrionii CBS 160.54 unplaced genomic scaffold supercont1.8, whole genome shotgun sequence
7. NW_011942151	Metarhizium robertsii ARSEF 23 MAA_Scf_11, whole genome shotgun sequence
8. NW_023336272	Colletotrichum scovillei strain TJNH1 chromosome Unknown Contig00014, whole genome shotgun sequence
9. NC_016458	Thermothielavioides terrestris NRRL 8126 chromosome 2, complete sequence
10. NW_022984634	Aspergillus tanneri strain NIH1004 chromosome Unknown AapTan_7_pilon_pilon_pilon, whole genome shotgun sequence
11. NW_017971453	Talaromyces atroroseus strain IBT 11181 chromosome Unknown scaffold_21, whole genome shotgun sequence
12. CM002798	Penicillium chrysogenum strain P2niaD18 chromosome I, whole genome shotgun sequence
13. NW_006912905	Capronia epimyces CBS 606.96 aczIQ-supercont1.2, whole genome shotgun sequence
14. NC_049561	Talaromyces rugulosus chromosome I, complete sequence
15. NC_016474	Thermothelomyces thermophilus ATCC 42464 chromosome 3, complete sequence
16. NW_006917098	Pestalotiopsis fici W106-1 unplaced genomic scaffold PFICI_Scf_8, whole genome shotgun sequence
17. NC_007197	Aspergillus fumigatus Af293 chromosome 4, whole genome shotgun sequence
18. NT_165935	Aspergillus terreus NIH2624 scaffold_12 genomic scaffold, whole genome shotgun sequence
19. NW_019716255	Ramularia collo-cygni strain URUG2 genome assembly, contig: RCC_scaffold01, whole genome shotgun sequence
20. NC_049561	Talaromyces rugulosus chromosome I, complete sequence
21. NC_016459	Thermothielavioides terrestris NRRL 8126 chromosome 3, complete sequence
22. CP051141	Peltaster fructicola strain LNHT1506 chromosome 3
23. NW_001914858	Podospora anserina S mat+ genomic DNA chromosome 2, supercontig 2
24. NW_014574708	Metarhizium brunneum ARSEF 3297 chromosome Unknown scaffold_9, whole genome shotgun sequence
25. NW_011942149	Metarhizium robertsii ARSEF 23 MAA_Scf_1, whole genome shotgun sequence
26. NC_036439	Aspergillus oryzae RIB40 DNA, chromosome 5
27. NW_008481832	Cladophialophora carrionii CBS 160.54 unplaced genomic scaffold supercont1.7, whole genome shotgun sequence
28. NW_001820812	Sclerotinia sclerotiorum 1980 UF-70 scaffold_24 genomic scaffold, whole genome shotgun sequence
29. NW_015971651	Fonsecaea multimorphosa CBS 102226 unplaced genomic scaffold supercont1.17, whole genome shotgun sequence
30. NW_013550607	Fonsecaea pedrosoi CBS 271.37 unplaced genomic scaffold supercont1.9, whole genome shotgun sequence
31. NW_017387254	Fonsecaea erecta strain CBS 125763 chromosome Unknown Scaff11, whole genome shotgun sequence
32. NC_007195	Aspergillus fumigatus Af293 chromosome 2, whole genome shotgun sequence
33. NW_017971432	Talaromyces atroroseus strain IBT 11181 chromosome Unknown scaffold_0, whole genome shotgun sequence
34. NC_049564	Talaromyces rugulosus chromosome IV, complete sequence
35. NW_007360988	Glarea lozoyensis ATCC 20868 chromosome Unknown GLAREA14, whole genome shotgun sequence
36. NW_014574713	Metarhizium brunneum ARSEF 3297 chromosome Unknown scaffold_4, whole genome shotgun sequence
37. NW_011942182	Metarhizium robertsii ARSEF 23 MAA_Scf_4, whole genome shotgun sequence
38. NW_006271971	Cordyceps militaris CM01 unplaced genomic scaffold CCM_S00003, whole genome shotgun sequence
39. NW_022984633	Aspergillus tanneri strain NIH1004 chromosome Unknown AapTan_6_pilon_pilon_pilon, whole genome shotgun sequence
40. CM000574	Fusarium graminearum PH-1 chromosome 1, whole genome shotgun sequence
41. NC_026474	Fusarium graminearum PH-1 chromosome 1, whole genome shotgun sequence
42. NW_011942150	Metarhizium robertsii ARSEF 23 MAA_Scf_10, whole genome shotgun sequence
43. NC_038012	Fusarium venenatum strain A3/5 genome assembly, chromosome: I
44. NW_014013635	Trichoderma atroviride IMI 206040 chromosome Unknown TRIATcontig_21, whole genome shotgun sequence
45. NC_035791	Pochonia chlamydosporia 170 chromosome 2, whole genome shotgun sequence
46. NW_023336287	Aspergillus tubingensis WU-2223L DNA, scaffold_13, whole genome shotgun sequence
47. NW_022984628	Aspergillus tanneri strain NIH1004 chromosome Unknown AapTan_1_pilon_pilon_pilon, whole genome shotgun sequence
48. NT_107006	Aspergillus nidulans FGSC A4 chromosome IV map unlocalized scaffold_10, whole genome shotgun sequence
49. NW_006917091	Pestalotiopsis fici W106-1 unplaced genomic scaffold PFICI_Scf_1, whole genome shotgun sequence
50. NW_020167549	Pseudogymnoascus destructans isolate 20631-21 unplaced genomic scaffold scaffold_30, whole genome shotgun sequence


Details:

>>
1. NW_022984630
Source: Aspergillus tanneri strain NIH1004 chromosome Unknown AapTan_3_pilon_pilon_pilon, whole genome shotgun sequence
Type: T1PKS
Number of proteins with BLAST hits to this cluster: 11
Cumulative BLAST score: 4444

Table of genes, locations, strands and annotations of subject cluster:

Table of Blast hits (query gene, subject gene, %identity, blast score, %coverage, e-value):
input.path1.gene797	ATNIH1004_003972	40	295	99.5	6.1e-78
input.path1.gene799	ATNIH1004_003956	71	75	16.853932584269664	8.6e-12
input.path1.gene801	ATNIH1004_003958	73	220	91.25	1.3e-55
input.path1.gene802	ATNIH1004_003959	63	158	59.45945945945946	9.1e-37
input.path1.gene803	ATNIH1004_003967	31	150	62.035541195476576	4.9e-34
input.path1.gene804	ATNIH1004_003961	60	1909	100.28344671201815	0.0
input.path1.gene805	ATNIH1004_003961	49	261	96.74267100977198	9.7e-68
input.path1.gene806	ATNIH1004_003962	45	82	23.098591549295776	7.2e-14
input.path1.gene807	ATNIH1004_003963	45	383	99.77168949771689	2.4e-104
input.path1.gene808	ATNIH1004_003964	62	342	100.0	2.9e-92
input.path1.gene811	ATNIH1004_003960	66	416	56.787762906309744	5.3e-114
input.path1.gene811	ATNIH1004_003965	39	153	38.81453154875717	4.9e-35


>>
2. NT_107015
Source: Aspergillus nidulans FGSC A4 chromosome VIII map unlocalized scaffold_1, whole genome shotgun sequence
Type: T1PKS
Number of proteins with BLAST hits to this cluster: 10
Cumulative BLAST score: 4600

Table of genes, locations, strands and annotations of subject cluster:

Table of Blast hits (query gene, subject gene, %identity, blast score, %coverage, e-value):
input.path1.gene798	AN0145.2	43	80	37.71043771043771	3.9e-13
input.path1.gene799	AN0148.2	70	71	17.228464419475657	1.6e-10
input.path1.gene801	AN0146.2	76	252	98.75	2.4e-65
input.path1.gene802	AN0145.2	66	148	50.27027027027027	7.2e-34
input.path1.gene804	AN0150.2	64	2106	101.24716553287982	0.0
input.path1.gene805	AN0151.2	39	198	92.50814332247556	7.7e-49
input.path1.gene807	AN0152.2	46	373	93.83561643835617	3.3e-101
input.path1.gene808	AN0152.2	52	268	101.50943396226415	6.9e-70
input.path1.gene810	AN0147.2	67	705	105.46558704453442	4.1e-201
input.path1.gene811	AN0149.2	64	399	56.022944550669216	5.1e-109


>>
3. NW_014574694
Source: Metarhizium brunneum ARSEF 3297 chromosome Unknown scaffold_23, whole genome shotgun sequence
Type: NRPS,T1PKS
Number of proteins with BLAST hits to this cluster: 7
Cumulative BLAST score: 3585

Table of genes, locations, strands and annotations of subject cluster:

Table of Blast hits (query gene, subject gene, %identity, blast score, %coverage, e-value):
input.path1.gene799	MBR_09472	65	72	17.97752808988764	9.5e-11
input.path1.gene804	MBR_09479	56	1930	100.51020408163265	0.0
input.path1.gene806	MBR_09474	46	74	22.816901408450704	2.5e-11
input.path1.gene807	MBR_09475	47	396	95.4337899543379	4.7e-108
input.path1.gene808	MBR_09477	70	396	100.0	2.9e-108
input.path1.gene809	MBR_09476	52	336	99.6923076923077	4.3e-90
input.path1.gene811	MBR_09478	62	381	55.25812619502868	1.4e-103


>>
4. NW_019154043
Source: Pochonia chlamydosporia 170 chromosome Unknown PCv3seq00008, whole genome shotgun sequence
Type: T1PKS
Number of proteins with BLAST hits to this cluster: 7
Cumulative BLAST score: 3514

Table of genes, locations, strands and annotations of subject cluster:

Table of Blast hits (query gene, subject gene, %identity, blast score, %coverage, e-value):
input.path1.gene799	VFPPC_06838	52	70	20.973782771535582	3.6e-10
input.path1.gene801	VFPPC_06847	78	254	96.25	8.1e-66
input.path1.gene804	VFPPC_06844	57	1919	100.28344671201815	0.0
input.path1.gene805	VFPPC_14125	61	180	45.27687296416938	2.8e-43
input.path1.gene808	VFPPC_06846	53	285	99.62264150943396	7.1e-75
input.path1.gene810	VFPPC_06848	47	438	97.36842105263158	7.2e-121
input.path1.gene811	VFPPC_06843	59	368	55.64053537284895	9.6e-100


>>
5. NW_006913046
Source: Cladophialophora yegresii CBS 114405 aczIN-supercont1.2, whole genome shotgun sequence
Type: T1PKS
Number of proteins with BLAST hits to this cluster: 7
Cumulative BLAST score: 3939

Table of genes, locations, strands and annotations of subject cluster:

Table of Blast hits (query gene, subject gene, %identity, blast score, %coverage, e-value):
input.path1.gene799	A1O7_02406	55	67	20.973782771535582	1.8e-09
input.path1.gene801	A1O7_02398	79	261	96.25	6.6e-68
input.path1.gene802	A1O7_02397	55	199	85.94594594594595	2.7e-49
input.path1.gene804	A1O7_02402	57	1973	100.85034013605443	0.0
input.path1.gene805	A1O7_02400	42	82	42.34527687296417	6.2e-14
input.path1.gene810	A1O7_02392	49	521	110.32388663967612	8.4e-146
input.path1.gene810	A1O7_02395	48	462	97.77327935222672	7.9e-128
input.path1.gene811	A1O7_02403	61	374	55.64053537284895	1.3e-101


>>
6. NW_008481833
Source: Cladophialophora carrionii CBS 160.54 unplaced genomic scaffold supercont1.8, whole genome shotgun sequence
Type: T1PKS
Number of proteins with BLAST hits to this cluster: 7
Cumulative BLAST score: 3870

Table of genes, locations, strands and annotations of subject cluster:

Table of Blast hits (query gene, subject gene, %identity, blast score, %coverage, e-value):
input.path1.gene799	G647_08918	53	67	19.850187265917604	3.1e-09
input.path1.gene801	G647_08910	79	261	96.25	5.1e-68
input.path1.gene802	G647_08909	56	200	84.32432432432432	2.1e-49
input.path1.gene804	G647_08914	57	1979	100.56689342403628	0.0
input.path1.gene805	G647_08912	36	71	42.34527687296417	1.9e-10
input.path1.gene810	G647_08907	49	460	97.77327935222672	1.8e-127
input.path1.gene810	G647_08904	50	453	92.91497975708502	3.7e-125
input.path1.gene811	G647_08915	62	379	55.64053537284895	4.2e-103


>>
7. NW_011942151
Source: Metarhizium robertsii ARSEF 23 MAA_Scf_11, whole genome shotgun sequence
Type: NRPS,T1PKS,terpene
Number of proteins with BLAST hits to this cluster: 7
Cumulative BLAST score: 3607

Table of genes, locations, strands and annotations of subject cluster:

Table of Blast hits (query gene, subject gene, %identity, blast score, %coverage, e-value):
input.path1.gene799	MAA_06568	59	72	20.224719101123593	7.3e-11
input.path1.gene804	MAA_06575	56	1932	100.51020408163265	0.0
input.path1.gene805	MAA_11604	41	97	42.671009771986974	1.9e-18
input.path1.gene807	MAA_06570	46	398	100.22831050228311	9.5e-109
input.path1.gene808	MAA_06572	69	391	100.0	5.4e-107
input.path1.gene809	MAA_06571	52	334	99.6923076923077	9.6e-90
input.path1.gene811	MAA_06574	63	383	55.25812619502868	4.9e-104


>>
8. NW_023336272
Source: Colletotrichum scovillei strain TJNH1 chromosome Unknown Contig00014, whole genome shotgun sequence
Type: NRPS,T1PKS
Number of proteins with BLAST hits to this cluster: 6
Cumulative BLAST score: 3343

Table of genes, locations, strands and annotations of subject cluster:

Table of Blast hits (query gene, subject gene, %identity, blast score, %coverage, e-value):
input.path1.gene801	HER10_EVM0008728	60	191	102.49999999999999	4.9e-47
input.path1.gene802	HER10_EVM0012982	55	197	87.56756756756758	1.4e-48
input.path1.gene804	HER10_EVM0004288	55	1862	101.98412698412697	0.0
input.path1.gene807	HER10_EVM0009122	50	429	98.85844748858447	5e-118
input.path1.gene808	HER10_EVM0007584	55	294	101.88679245283019	1.2e-77
input.path1.gene811	HER10_EVM0004967	57	370	56.40535372848948	2.5e-100


>>
9. NC_016458
Source: Thermothielavioides terrestris NRRL 8126 chromosome 2, complete sequence
Type: T1PKS
Number of proteins with BLAST hits to this cluster: 6
Cumulative BLAST score: 3007

Table of genes, locations, strands and annotations of subject cluster:

Table of Blast hits (query gene, subject gene, %identity, blast score, %coverage, e-value):
input.path1.gene798	THITE_2114221	33	74	66.32996632996633	2.1e-11
input.path1.gene799	THITE_152256	50	73	30.711610486891384	4.3e-11
input.path1.gene804	THITE_35447	58	1911	101.07709750566893	0.0
input.path1.gene807	THITE_37176	46	359	99.77168949771689	4.9e-97
input.path1.gene808	THITE_2077799	44	241	98.49056603773585	1.2e-61
input.path1.gene811	THITE_2049009	58	349	56.40535372848948	7.9e-94


>>
10. NW_022984634
Source: Aspergillus tanneri strain NIH1004 chromosome Unknown AapTan_7_pilon_pilon_pilon, whole genome shotgun sequence
Type: T1PKS
Number of proteins with BLAST hits to this cluster: 7
Cumulative BLAST score: 3478

Table of genes, locations, strands and annotations of subject cluster:

Table of Blast hits (query gene, subject gene, %identity, blast score, %coverage, e-value):
input.path1.gene798	ATNIH1004_008022	38	102	55.218855218855225	7.3e-20
input.path1.gene799	ATNIH1004_008020	55	64	20.59925093632959	1.5e-08
input.path1.gene804	ATNIH1004_008023	61	2091	100.39682539682539	0.0
input.path1.gene805	ATNIH1004_008015	62	184	42.34527687296417	1.5e-44
input.path1.gene807	ATNIH1004_008013	39	342	101.36986301369863	6.2e-92
input.path1.gene808	ATNIH1004_008018	57	296	98.49056603773585	2.4e-78
input.path1.gene811	ATNIH1004_008024	63	399	56.787762906309744	5.1e-109


>>
11. NW_017971453
Source: Talaromyces atroroseus strain IBT 11181 chromosome Unknown scaffold_21, whole genome shotgun sequence
Type: T1PKS
Number of proteins with BLAST hits to this cluster: 6
Cumulative BLAST score: 3390

Table of genes, locations, strands and annotations of subject cluster:

Table of Blast hits (query gene, subject gene, %identity, blast score, %coverage, e-value):
input.path1.gene799	UA08_09364	67	81	19.475655430711612	1.6e-13
input.path1.gene801	UA08_09372	79	260	96.25	1.1e-67
input.path1.gene802	UA08_09373	56	210	92.97297297297298	1.5e-52
input.path1.gene804	UA08_09374	57	1960	100.28344671201815	0.0
input.path1.gene808	UA08_09371	58	300	99.24528301886792	1.6e-79
input.path1.gene810	UA08_09363	56	579	103.03643724696356	2.6e-163


>>
12. CM002798
Source: Penicillium chrysogenum strain P2niaD18 chromosome I, whole genome shotgun sequence
Type: T1PKS
Number of proteins with BLAST hits to this cluster: 6
Cumulative BLAST score: 3350

Table of genes, locations, strands and annotations of subject cluster:

Table of Blast hits (query gene, subject gene, %identity, blast score, %coverage, e-value):
input.path1.gene798	EN45_038670	33	69	52.18855218855219	6.9e-10
input.path1.gene802	EN45_038650	62	223	88.64864864864866	1.8e-56
input.path1.gene804	EN45_038660	60	1681	84.24036281179139	0.0
input.path1.gene807	EN45_038620	56	492	98.85844748858447	3.7e-137
input.path1.gene808	EN45_038630	60	328	100.0	5.6e-88
input.path1.gene810	EN45_038570	54	557	103.23886639676114	1.1e-156


>>
13. NW_006912905
Source: Capronia epimyces CBS 606.96 aczIQ-supercont1.2, whole genome shotgun sequence
Type: T1PKS
Number of proteins with BLAST hits to this cluster: 6
Cumulative BLAST score: 3171

Table of genes, locations, strands and annotations of subject cluster:

Table of Blast hits (query gene, subject gene, %identity, blast score, %coverage, e-value):
input.path1.gene801	A1O3_02703	70	229	96.25	2.8e-58
input.path1.gene802	A1O3_02702	53	194	84.32432432432432	8.8e-48
input.path1.gene804	A1O3_02705	59	1919	103.11791383219955	0.0
input.path1.gene805	A1O3_02700	44	143	46.579804560260584	5e-32
input.path1.gene808	A1O3_02701	57	311	100.0	7.1e-83
input.path1.gene811	A1O3_02704	62	375	55.64053537284895	1e-101


>>
14. NC_049561
Source: Talaromyces rugulosus chromosome I, complete sequence
Type: T1PKS
Number of proteins with BLAST hits to this cluster: 6
Cumulative BLAST score: 2367

Table of genes, locations, strands and annotations of subject cluster:

Table of Blast hits (query gene, subject gene, %identity, blast score, %coverage, e-value):
input.path1.gene799	TRUGW13939_00040	55	59	16.479400749063668	4.9e-07
input.path1.gene801	TRUGW13939_00030	81	269	98.75	3.2e-70
input.path1.gene802	TRUGW13939_00029	54	200	86.48648648648648	1.6e-49
input.path1.gene804	TRUGW13939_00034	45	1378	100.45351473922904	0.0
input.path1.gene805	TRUGW13939_00031	61	182	45.60260586319218	5.7e-44
input.path1.gene811	TRUGW13939_00039	47	279	54.875717017208416	1e-72


>>
15. NC_016474
Source: Thermothelomyces thermophilus ATCC 42464 chromosome 3, complete sequence
Type: T1PKS
Number of proteins with BLAST hits to this cluster: 6
Cumulative BLAST score: 3545

Table of genes, locations, strands and annotations of subject cluster:

Table of Blast hits (query gene, subject gene, %identity, blast score, %coverage, e-value):
input.path1.gene799	MYCTH_107295	64	66	16.853932584269664	4e-09
input.path1.gene804	MYCTH_101261	61	1953	102.09750566893423	0.0
input.path1.gene805	MYCTH_2061639	63	185	42.671009771986974	8.9e-45
input.path1.gene808	MYCTH_2305631	55	301	98.86792452830188	7.3e-80
input.path1.gene810	MYCTH_50396	59	626	103.44129554655869	3.2e-177
input.path1.gene811	MYCTH_2110682	66	414	56.787762906309744	1.5e-113


>>
16. NW_006917098
Source: Pestalotiopsis fici W106-1 unplaced genomic scaffold PFICI_Scf_8, whole genome shotgun sequence
Type: T1PKS
Number of proteins with BLAST hits to this cluster: 6
Cumulative BLAST score: 3333

Table of genes, locations, strands and annotations of subject cluster:

Table of Blast hits (query gene, subject gene, %identity, blast score, %coverage, e-value):
input.path1.gene799	PFICI_10833	60	77	20.59925093632959	3e-12
input.path1.gene804	PFICI_10824	62	2100	100.45351473922904	0.0
input.path1.gene805	PFICI_10827	57	86	20.521172638436482	7.3e-15
input.path1.gene805	PFICI_10826	33	69	42.34527687296417	5.4e-10
input.path1.gene807	PFICI_10835	44	382	99.08675799086758	5.4e-104
input.path1.gene808	PFICI_10829	54	216	73.58490566037736	4e-54
input.path1.gene811	PFICI_10825	67	403	56.022944550669216	3.5e-110


>>
17. NC_007197
Source: Aspergillus fumigatus Af293 chromosome 4, whole genome shotgun sequence
Type: T1PKS
Number of proteins with BLAST hits to this cluster: 6
Cumulative BLAST score: 3320

Table of genes, locations, strands and annotations of subject cluster:

Table of Blast hits (query gene, subject gene, %identity, blast score, %coverage, e-value):
input.path1.gene799	AFUA_4G14540	51	63	20.59925093632959	3.4e-08
input.path1.gene804	AFUA_4G14560	61	2039	100.68027210884354	0.0
input.path1.gene805	AFUA_4G14470	59	175	43.97394136807817	1.2e-41
input.path1.gene807	AFUA_4G14500	43	356	102.73972602739727	5.4e-96
input.path1.gene808	AFUA_4G14520	59	304	98.49056603773585	8.6e-81
input.path1.gene811	AFUA_4G14570	57	383	60.9942638623327	4.9e-104


>>
18. NT_165935
Source: Aspergillus terreus NIH2624 scaffold_12 genomic scaffold, whole genome shotgun sequence
Type: NRPS,T1PKS
Number of proteins with BLAST hits to this cluster: 6
Cumulative BLAST score: 3230

Table of genes, locations, strands and annotations of subject cluster:

Table of Blast hits (query gene, subject gene, %identity, blast score, %coverage, e-value):
input.path1.gene798	ATEG_08452	32	87	62.62626262626263	2.4e-15
input.path1.gene804	ATEG_08451	61	2053	100.34013605442176	0.0
input.path1.gene805	ATEG_08457	66	167	37.785016286644954	2.5e-39
input.path1.gene807	ATEG_08459	41	347	100.4566210045662	2.5e-93
input.path1.gene808	ATEG_08455	59	178	56.60377358490566	9.3e-43
input.path1.gene811	ATEG_08450	64	398	56.40535372848948	1.1e-108


>>
19. NW_019716255
Source: Ramularia collo-cygni strain URUG2 genome assembly, contig: RCC_scaffold01, whole genome shotgun sequence
Type: T1PKS
Number of proteins with BLAST hits to this cluster: 6
Cumulative BLAST score: 2991

Table of genes, locations, strands and annotations of subject cluster:

Table of Blast hits (query gene, subject gene, %identity, blast score, %coverage, e-value):
input.path1.gene799	RCC_01331	58	59	16.10486891385768	8.3e-07
input.path1.gene801	RCC_01333	77	259	96.25	2.5e-67
input.path1.gene802	RCC_01335	60	210	88.64864864864866	2e-52
input.path1.gene804	RCC_01337	56	1814	100.45351473922904	0.0
input.path1.gene808	RCC_01332	53	287	99.24528301886792	1.1e-75
input.path1.gene811	RCC_01336	57	362	55.64053537284895	9e-98


>>
20. NC_049561
Source: Talaromyces rugulosus chromosome I, complete sequence
Type: T1PKS
Number of proteins with BLAST hits to this cluster: 5
Cumulative BLAST score: 2767

Table of genes, locations, strands and annotations of subject cluster:

Table of Blast hits (query gene, subject gene, %identity, blast score, %coverage, e-value):
input.path1.gene798	TRUGW13939_01158	35	87	56.22895622895623	2.4e-15
input.path1.gene799	TRUGW13939_01157	58	71	18.726591760299627	1.6e-10
input.path1.gene804	TRUGW13939_01165	60	1981	100.2267573696145	0.0
input.path1.gene805	TRUGW13939_01162	60	176	43.32247557003257	4.1e-42
input.path1.gene805	TRUGW13939_01163	36	78	42.99674267100978	1.5e-12
input.path1.gene811	TRUGW13939_01164	61	374	55.64053537284895	1.8e-101


>>
21. NC_016459
Source: Thermothielavioides terrestris NRRL 8126 chromosome 3, complete sequence
Type: T1PKS
Number of proteins with BLAST hits to this cluster: 4
Cumulative BLAST score: 857

Table of genes, locations, strands and annotations of subject cluster:

Table of Blast hits (query gene, subject gene, %identity, blast score, %coverage, e-value):
input.path1.gene797	THITE_2116299	33	153	80.25	6.3e-35
input.path1.gene803	THITE_160963	31	220	98.86914378029078	3.8e-55
input.path1.gene804	THITE_2050784	34	321	37.358276643990926	7.8e-85
input.path1.gene809	THITE_2116303	34	163	100.30769230769229	5e-38


>>
22. CP051141
Source: Peltaster fructicola strain LNHT1506 chromosome 3
Type: T1PKS
Number of proteins with BLAST hits to this cluster: 5
Cumulative BLAST score: 3134

Table of genes, locations, strands and annotations of subject cluster:

Table of Blast hits (query gene, subject gene, %identity, blast score, %coverage, e-value):
input.path1.gene804	AMS68_004773	60	2006	100.39682539682539	0.0
input.path1.gene806	AMS68_004775	45	66	23.661971830985916	5.3e-09
input.path1.gene807	AMS68_004769	47	410	99.77168949771689	1.8e-112
input.path1.gene808	AMS68_004767	53	259	100.37735849056604	5.4e-67
input.path1.gene811	AMS68_004772	64	393	56.022944550669216	2.8e-107


>>
23. NW_001914858
Source: Podospora anserina S mat+ genomic DNA chromosome 2, supercontig 2
Type: T1PKS
Number of proteins with BLAST hits to this cluster: 4
Cumulative BLAST score: 1576

Table of genes, locations, strands and annotations of subject cluster:

Table of Blast hits (query gene, subject gene, %identity, blast score, %coverage, e-value):
input.path1.gene797	PODANS_2_7430	35	149	59.25	9.2e-34
input.path1.gene804	PODANS_2_7410	35	992	101.92743764172336	6e-287
input.path1.gene807	PODANS_2_7450	34	207	86.5296803652968	2.4e-51
input.path1.gene808	PODANS_2_7440	43	228	98.49056603773585	7.9e-58


>>
24. NW_014574708
Source: Metarhizium brunneum ARSEF 3297 chromosome Unknown scaffold_9, whole genome shotgun sequence
Type: NRPS,T1PKS
Number of proteins with BLAST hits to this cluster: 3
Cumulative BLAST score: 1383

Table of genes, locations, strands and annotations of subject cluster:

Table of Blast hits (query gene, subject gene, %identity, blast score, %coverage, e-value):
input.path1.gene797	MBR_06608	32	153	93.5	4.9e-35
input.path1.gene804	MBR_06615	39	1101	100.6235827664399	0.0
input.path1.gene805	MBR_06614	43	129	42.99674267100978	7.5e-28


>>
25. NW_011942149
Source: Metarhizium robertsii ARSEF 23 MAA_Scf_1, whole genome shotgun sequence
Type: NRPS,T1PKS
Number of proteins with BLAST hits to this cluster: 3
Cumulative BLAST score: 1375

Table of genes, locations, strands and annotations of subject cluster:

Table of Blast hits (query gene, subject gene, %identity, blast score, %coverage, e-value):
input.path1.gene797	MAA_08362	32	154	93.5	2.2e-35
input.path1.gene804	MAA_10747	39	1093	100.79365079365078	0.0
input.path1.gene805	MAA_08369	43	128	42.99674267100978	9.8e-28


>>
26. NC_036439
Source: Aspergillus oryzae RIB40 DNA, chromosome 5
Type: T1PKS
Number of proteins with BLAST hits to this cluster: 4
Cumulative BLAST score: 1867

Table of genes, locations, strands and annotations of subject cluster:

Table of Blast hits (query gene, subject gene, %identity, blast score, %coverage, e-value):
input.path1.gene795	AO090701000525	32	212	101.22699386503066	1.4e-52
input.path1.gene797	AO090701000528	32	171	87.25	1.7e-40
input.path1.gene804	AO090701000530	40	1258	105.66893424036282	0.0
input.path1.gene811	AO090701000529	48	226	45.506692160611856	5.9e-57


>>
27. NW_008481832
Source: Cladophialophora carrionii CBS 160.54 unplaced genomic scaffold supercont1.7, whole genome shotgun sequence
Type: T1PKS
Number of proteins with BLAST hits to this cluster: 3
Cumulative BLAST score: 1559

Table of genes, locations, strands and annotations of subject cluster:

Table of Blast hits (query gene, subject gene, %identity, blast score, %coverage, e-value):
input.path1.gene801	G647_08244	52	158	94.375	6e-37
input.path1.gene802	G647_08243	40	140	86.48648648648648	1.5e-31
input.path1.gene804	G647_08240	42	1261	99.20634920634922	0.0


>>
28. NW_001820812
Source: Sclerotinia sclerotiorum 1980 UF-70 scaffold_24 genomic scaffold, whole genome shotgun sequence
Type: T1PKS
Number of proteins with BLAST hits to this cluster: 3
Cumulative BLAST score: 1523

Table of genes, locations, strands and annotations of subject cluster:

Table of Blast hits (query gene, subject gene, %identity, blast score, %coverage, e-value):
input.path1.gene801	SS1G_13315	50	155	95.625	3.9e-36
input.path1.gene802	SS1G_13314	42	162	94.05405405405406	3.7e-38
input.path1.gene804	SS1G_13322	43	1206	99.31972789115646	0.0


>>
29. NW_015971651
Source: Fonsecaea multimorphosa CBS 102226 unplaced genomic scaffold supercont1.17, whole genome shotgun sequence
Type: T1PKS
Number of proteins with BLAST hits to this cluster: 3
Cumulative BLAST score: 1521

Table of genes, locations, strands and annotations of subject cluster:

Table of Blast hits (query gene, subject gene, %identity, blast score, %coverage, e-value):
input.path1.gene801	Z520_08009	54	162	95.625	4.2e-38
input.path1.gene802	Z520_08008	52	104	45.40540540540541	9.2e-21
input.path1.gene804	Z520_08005	43	1255	99.60317460317461	0.0


>>
30. NW_013550607
Source: Fonsecaea pedrosoi CBS 271.37 unplaced genomic scaffold supercont1.9, whole genome shotgun sequence
Type: T1PKS
Number of proteins with BLAST hits to this cluster: 3
Cumulative BLAST score: 1511

Table of genes, locations, strands and annotations of subject cluster:

Table of Blast hits (query gene, subject gene, %identity, blast score, %coverage, e-value):
input.path1.gene801	Z517_12195	51	144	88.125	6.9e-33
input.path1.gene802	Z517_12194	40	143	86.48648648648648	1.8e-32
input.path1.gene804	Z517_12191	42	1224	92.85714285714286	0.0


>>
31. NW_017387254
Source: Fonsecaea erecta strain CBS 125763 chromosome Unknown Scaff11, whole genome shotgun sequence
Type: T1PKS
Number of proteins with BLAST hits to this cluster: 3
Cumulative BLAST score: 1493

Table of genes, locations, strands and annotations of subject cluster:

Table of Blast hits (query gene, subject gene, %identity, blast score, %coverage, e-value):
input.path1.gene801	AYL99_10303	47	128	84.375	5.1e-28
input.path1.gene802	AYL99_10302	47	118	58.91891891891892	6.1e-25
input.path1.gene804	AYL99_10299	42	1247	100.51020408163265	0.0


>>
32. NC_007195
Source: Aspergillus fumigatus Af293 chromosome 2, whole genome shotgun sequence
Type: T1PKS
Number of proteins with BLAST hits to this cluster: 3
Cumulative BLAST score: 1464

Table of genes, locations, strands and annotations of subject cluster:

Table of Blast hits (query gene, subject gene, %identity, blast score, %coverage, e-value):
input.path1.gene801	AFUA_2G17560	48	131	91.25	6.1e-29
input.path1.gene802	AFUA_2G17580	43	159	87.56756756756758	4.1e-37
input.path1.gene804	AFUA_2G17600	40	1174	99.31972789115646	0.0


>>
33. NW_017971432
Source: Talaromyces atroroseus strain IBT 11181 chromosome Unknown scaffold_0, whole genome shotgun sequence
Type: T1PKS
Number of proteins with BLAST hits to this cluster: 3
Cumulative BLAST score: 1452

Table of genes, locations, strands and annotations of subject cluster:

Table of Blast hits (query gene, subject gene, %identity, blast score, %coverage, e-value):
input.path1.gene801	UA08_00429	47	143	94.375	2.6e-32
input.path1.gene802	UA08_00428	46	163	86.48648648648648	2.8e-38
input.path1.gene804	UA08_00425	40	1146	99.82993197278913	0.0


>>
34. NC_049564
Source: Talaromyces rugulosus chromosome IV, complete sequence
Type: T1PKS
Number of proteins with BLAST hits to this cluster: 3
Cumulative BLAST score: 1446

Table of genes, locations, strands and annotations of subject cluster:

Table of Blast hits (query gene, subject gene, %identity, blast score, %coverage, e-value):
input.path1.gene801	TRUGW13939_08435	48	147	94.375	1.4e-33
input.path1.gene802	TRUGW13939_08434	42	159	88.10810810810811	3.1e-37
input.path1.gene804	TRUGW13939_08432	40	1140	99.88662131519274	0.0


>>
35. NW_007360988
Source: Glarea lozoyensis ATCC 20868 chromosome Unknown GLAREA14, whole genome shotgun sequence
Type: T1PKS
Number of proteins with BLAST hits to this cluster: 3
Cumulative BLAST score: 1366

Table of genes, locations, strands and annotations of subject cluster:

Table of Blast hits (query gene, subject gene, %identity, blast score, %coverage, e-value):
input.path1.gene801	GLAREA_06716	51	163	96.25	1.4e-38
input.path1.gene802	GLAREA_06715	40	146	85.4054054054054	3.6e-33
input.path1.gene804	GLAREA_06717	37	1057	100.34013605442176	2e-306


>>
36. NW_014574713
Source: Metarhizium brunneum ARSEF 3297 chromosome Unknown scaffold_4, whole genome shotgun sequence
Type: T1PKS
Number of proteins with BLAST hits to this cluster: 3
Cumulative BLAST score: 1353

Table of genes, locations, strands and annotations of subject cluster:

Table of Blast hits (query gene, subject gene, %identity, blast score, %coverage, e-value):
input.path1.gene801	MBR_03975	49	156	96.25	2.3e-36
input.path1.gene802	MBR_03976	39	129	76.75675675675676	3.5e-28
input.path1.gene804	MBR_03974	39	1068	99.60317460317461	8.6e-310


>>
37. NW_011942182
Source: Metarhizium robertsii ARSEF 23 MAA_Scf_4, whole genome shotgun sequence
Type: T1PKS
Number of proteins with BLAST hits to this cluster: 3
Cumulative BLAST score: 1343

Table of genes, locations, strands and annotations of subject cluster:

Table of Blast hits (query gene, subject gene, %identity, blast score, %coverage, e-value):
input.path1.gene801	MAA_03240	50	156	96.25	1.8e-36
input.path1.gene802	MAA_03241	40	128	76.21621621621621	5.9e-28
input.path1.gene804	MAA_03239	39	1059	99.54648526077098	3.1e-307


>>
38. NW_006271971
Source: Cordyceps militaris CM01 unplaced genomic scaffold CCM_S00003, whole genome shotgun sequence
Type: T1PKS
Number of proteins with BLAST hits to this cluster: 3
Cumulative BLAST score: 1297

Table of genes, locations, strands and annotations of subject cluster:

Table of Blast hits (query gene, subject gene, %identity, blast score, %coverage, e-value):
input.path1.gene801	CCM_03662	47	147	96.25	1.1e-33
input.path1.gene802	CCM_03661	46	161	88.64864864864866	8.3e-38
input.path1.gene804	CCM_03663	37	989	92.57369614512471	5.1e-286


>>
39. NW_022984633
Source: Aspergillus tanneri strain NIH1004 chromosome Unknown AapTan_6_pilon_pilon_pilon, whole genome shotgun sequence
Type: T1PKS
Number of proteins with BLAST hits to this cluster: 2
Cumulative BLAST score: 1273

Table of genes, locations, strands and annotations of subject cluster:

Table of Blast hits (query gene, subject gene, %identity, blast score, %coverage, e-value):
input.path1.gene804	ATNIH1004_008649	40	1126	98.69614512471655	0.0
input.path1.gene805	ATNIH1004_008650	52	147	41.042345276872965	2e-33


>>
40. CM000574
Source: Fusarium graminearum PH-1 chromosome 1, whole genome shotgun sequence
Type: NRPS,T1PKS
Number of proteins with BLAST hits to this cluster: 2
Cumulative BLAST score: 1247

Table of genes, locations, strands and annotations of subject cluster:

Table of Blast hits (query gene, subject gene, %identity, blast score, %coverage, e-value):
input.path1.gene804	FGSG_02324	41	1109	92.17687074829932	0.0
input.path1.gene805	FGSG_02325	47	138	41.36807817589577	1.2e-30


>>
41. NC_026474
Source: Fusarium graminearum PH-1 chromosome 1, whole genome shotgun sequence
Type: NRPS,T1PKS
Number of proteins with BLAST hits to this cluster: 2
Cumulative BLAST score: 1247

Table of genes, locations, strands and annotations of subject cluster:

Table of Blast hits (query gene, subject gene, %identity, blast score, %coverage, e-value):
input.path1.gene804	FGSG_02324	41	1109	92.17687074829932	0.0
input.path1.gene805	FGSG_02325	47	138	41.36807817589577	1.2e-30


>>
42. NW_011942150
Source: Metarhizium robertsii ARSEF 23 MAA_Scf_10, whole genome shotgun sequence
Type: T1PKS
Number of proteins with BLAST hits to this cluster: 2
Cumulative BLAST score: 1229

Table of genes, locations, strands and annotations of subject cluster:

Table of Blast hits (query gene, subject gene, %identity, blast score, %coverage, e-value):
input.path1.gene804	MAA_07745	40	1095	92.3469387755102	0.0
input.path1.gene805	MAA_07746	46	134	41.69381107491857	2.3e-29


>>
43. NC_038012
Source: Fusarium venenatum strain A3/5 genome assembly, chromosome: I
Type: NRPS,T1PKS
Number of proteins with BLAST hits to this cluster: 2
Cumulative BLAST score: 1222

Table of genes, locations, strands and annotations of subject cluster:

Table of Blast hits (query gene, subject gene, %identity, blast score, %coverage, e-value):
input.path1.gene804	FVRRES_02715	40	1082	92.12018140589569	0.0
input.path1.gene805	FVRRES_02716	43	140	43.648208469055376	2.5e-31


>>
44. NW_014013635
Source: Trichoderma atroviride IMI 206040 chromosome Unknown TRIATcontig_21, whole genome shotgun sequence
Type: T1PKS
Number of proteins with BLAST hits to this cluster: 2
Cumulative BLAST score: 1211

Table of genes, locations, strands and annotations of subject cluster:

Table of Blast hits (query gene, subject gene, %identity, blast score, %coverage, e-value):
input.path1.gene804	TRIATDRAFT_217154	40	1076	92.68707482993197	0.0
input.path1.gene805	TRIATDRAFT_154816	43	135	47.23127035830619	8e-30


>>
45. NC_035791
Source: Pochonia chlamydosporia 170 chromosome 2, whole genome shotgun sequence
Type: T1PKS
Number of proteins with BLAST hits to this cluster: 2
Cumulative BLAST score: 1196

Table of genes, locations, strands and annotations of subject cluster:

Table of Blast hits (query gene, subject gene, %identity, blast score, %coverage, e-value):
input.path1.gene804	VFPPC_11725	40	1108	93.14058956916101	0.0
input.path1.gene805	VFPPC_11724	35	88	42.01954397394137	1.1e-15


>>
46. NW_023336287
Source: Aspergillus tubingensis WU-2223L DNA, scaffold_13, whole genome shotgun sequence
Type: T1PKS
Number of proteins with BLAST hits to this cluster: 2
Cumulative BLAST score: 1192

Table of genes, locations, strands and annotations of subject cluster:

Table of Blast hits (query gene, subject gene, %identity, blast score, %coverage, e-value):
input.path1.gene804	AtWU_10461	37	1078	99.03628117913831	0.0
input.path1.gene805	AtWU_10460	43	114	42.01954397394137	1.5e-23


>>
47. NW_022984628
Source: Aspergillus tanneri strain NIH1004 chromosome Unknown AapTan_1_pilon_pilon_pilon, whole genome shotgun sequence
Type: T1PKS
Number of proteins with BLAST hits to this cluster: 2
Cumulative BLAST score: 1148

Table of genes, locations, strands and annotations of subject cluster:

Table of Blast hits (query gene, subject gene, %identity, blast score, %coverage, e-value):
input.path1.gene804	ATNIH1004_003522	37	1053	98.75283446712018	2.2e-305
input.path1.gene805	ATNIH1004_003523	34	95	42.671009771986974	1.2e-17


>>
48. NT_107006
Source: Aspergillus nidulans FGSC A4 chromosome IV map unlocalized scaffold_10, whole genome shotgun sequence
Type: T1PKS
Number of proteins with BLAST hits to this cluster: 3
Cumulative BLAST score: 1115

Table of genes, locations, strands and annotations of subject cluster:

Table of Blast hits (query gene, subject gene, %identity, blast score, %coverage, e-value):
input.path1.gene798	AN7819.2	35	87	50.505050505050505	1.9e-15
input.path1.gene799	AN7820.2	52	71	23.970037453183522	1.2e-10
input.path1.gene804	AN7825.2	34	957	100.45351473922904	2.1e-276


>>
49. NW_006917091
Source: Pestalotiopsis fici W106-1 unplaced genomic scaffold PFICI_Scf_1, whole genome shotgun sequence
Type: NRPS-like,T1PKS,indole
Number of proteins with BLAST hits to this cluster: 3
Cumulative BLAST score: 880

Table of genes, locations, strands and annotations of subject cluster:

Table of Blast hits (query gene, subject gene, %identity, blast score, %coverage, e-value):
input.path1.gene796	PFICI_00369	37	295	110.44776119402985	6.1e-78
input.path1.gene804	PFICI_00366	31	379	52.49433106575964	2.4e-102
input.path1.gene809	PFICI_00371	38	206	99.07692307692308	5.1e-51


>>
50. NW_020167549
Source: Pseudogymnoascus destructans isolate 20631-21 unplaced genomic scaffold scaffold_30, whole genome shotgun sequence
Type: T1PKS
Number of proteins with BLAST hits to this cluster: 3
Cumulative BLAST score: 2388

Table of genes, locations, strands and annotations of subject cluster:

Table of Blast hits (query gene, subject gene, %identity, blast score, %coverage, e-value):
input.path1.gene798	VC83_08898	36	85	47.13804713804714	1.2e-14
input.path1.gene804	VC83_08896	55	1936	100.79365079365078	0.0
input.path1.gene811	VC83_08897	59	367	56.21414913957935	2.8e-99
