## Supplementary Results for "A draft genome of the ascomycotal fungal species *Pseudopithomyces maydicus* (family *Didymosphaeriaceae*)": contig_13_c1.txt

ClusterBlast scores for contig_13

Table of genes, locations, strands and annotations of query cluster:
input.path1.gene217	743998	745304	+		
input.path1.gene218	746583	747448	-		
input.path1.gene219	748440	749102	-		
input.path1.gene220	753378	755774	-		
input.path1.gene221	757765	759723	+		
input.path1.gene222	761089	765032	+		
input.path1.gene223	765028	766926	-		
input.path1.gene224	769493	773444	-		
input.path1.gene225	782932	784069	-		


Significant hits: 
1. NC_049563	Talaromyces rugulosus chromosome III, complete sequence
2. NW_019716264	Ramularia collo-cygni strain URUG2 genome assembly, contig: RCC_scaffold10, whole genome shotgun sequence
3. NW_015971148	Sporothrix schenckii 1099-18 chromosome Unknown Cont47, whole genome shotgun sequence
4. NC_018218	Zymoseptoria tritici IPO323 chromosome 1, whole genome shotgun sequence
5. NZ_VLHW01000001	Pseudomonas syringae pv. dysoxyli strain CFBP 2356 CL_dyso01, whole genome shotgun sequence
6. NC_014718	Paraburkholderia rhizoxinica HKI 454 plasmid pBRH01, complete sequence
7. NZ_WJPS01000001	Pseudomonas syringae strain P73 CL_7301, whole genome shotgun sequence
8. NZ_LT629769	Pseudomonas syringae strain 31R1 chromosome I
9. NZ_FPJB01000001	Pseudomonas sp. NFACC10-1, whole genome shotgun sequence
10. NZ_LT222319	Pseudomonas cerasi isolate Sour cherry (Prunus cerasus) symptomatic leaf chromosome 1
11. NZ_KQ976354	Scytonema hofmannii PCC 7110 Scaffold1, whole genome shotgun sequence
12. NZ_CP006256	Pseudomonas syringae pv. syringae HS191 chromosome, complete genome
13. NZ_CP013732	Burkholderia cepacia JBK9 chromosome 3, complete sequence
14. NZ_FOIB01000007	Myxococcus fulvus strain DSM 16525, whole genome shotgun sequence
15. NZ_CP015529	Rhodococcus sp. WB1 chromosome, complete genome
16. NZ_CP050952	Rhodococcus sp. DMU1 chromosome, complete genome
17. NZ_CP011341	Rhodococcus aetherivorans strain IcdP1 chromosome, complete genome
18. NZ_CP012201	Massilia sp. NR 4-1, complete genome
19. NZ_QEOB01000033	Paraburkholderia unamae strain SCZa-39 Ga0215850_133, whole genome shotgun sequence
20. NZ_CP045351	Vibrio sp. THAF100 plasmid pTHAF100_a, complete sequence
21. NZ_KQ976354	Scytonema hofmannii PCC 7110 Scaffold1, whole genome shotgun sequence
22. NZ_CP034539	Streptomyces cyaneochromogenes strain MK-45 chromosome, complete genome
23. NZ_CM001149	Rhodococcus hoagii ATCC 33707 chromosome, whole genome shotgun sequence
24. NC_014659	Rhodococcus hoagii 103S, complete genome
25. NZ_CP027793	Rhodococcus hoagii strain DSSKP-R-001 chromosome, complete genome
26. NZ_MOBR01000009	Pseudomonas frederiksbergensis strain 38F7 38F7_contig_9, whole genome shotgun sequence
27. NZ_QTTW01000003	Rhodococcus sp. OK269 Ga0189761_103, whole genome shotgun sequence
28. NZ_JMEY01000019	Rhodococcus fascians A3b contig22, whole genome shotgun sequence
29. NZ_KI519398	Rhodococcus sp. JG-3 K414DRAFT_scaffold00001.1, whole genome shotgun sequence
30. NZ_JMEQ01000021	Rhodococcus fascians LMG 3602 contig46, whole genome shotgun sequence
31. NZ_JMEV01000002	Rhodococcus fascians A76 contig2, whole genome shotgun sequence
32. NZ_RAII01000001	Streptomyces sp. E1N211 s-211_S3_L001_R1_001, whole genome shotgun sequence
33. NZ_SMFR01000001	Nocardia alba strain DSM 44684 Ga0244504_101, whole genome shotgun sequence
34. NZ_LMFE01000001	Nocardia sp. Root136 contig_1, whole genome shotgun sequence
35. NZ_KI912151	Fischerella sp. PCC 9605 FIS9605DRAFT_scaffold4.4, whole genome shotgun sequence
36. NZ_CP046172	Nocardia arthritidis strain AUSMDU00012717 chromosome, complete genome
37. NZ_WXEW01000012	Herbidospora sp. NEAU-GS84 Scaffold12, whole genome shotgun sequence
38. NZ_BBXF01000001	Herbidospora daliensis strain NBRC 106372, whole genome shotgun sequence
39. NZ_CP019717	Paenibacillus larvae subsp. larvae strain Eric_V chromosome, complete genome
40. NC_023134	Paenibacillus larvae subsp. larvae DSM 25430, complete sequence
41. NZ_CP019687	Paenibacillus larvae subsp. larvae strain ATCC-9545 chromosome, complete genome
42. NZ_CP041061	Micromonospora sp. HM134 chromosome, complete genome
43. NZ_FNOG01000002	Lysobacter enzymogenes strain ATCC 29487, whole genome shotgun sequence
44. NZ_AP014940	Lysobacter enzymogenes strain M497-1
45. NZ_QXXQ01000017	Gemmobacter lutimaris strain YJ-T1-11 scaffold_17, whole genome shotgun sequence
46. NZ_WWEU01000004	Vibrio sp. OCN044 NODE_4_length_493155_cov_31.134218, whole genome shotgun sequence
47. NZ_AP018288	Nostoc sp. NIES-4103 DNA, nearly complete genome
48. NZ_JABUMT010000016	Myxococcus sp. CA056 NODE_16_length_238008_cov_21.6875, whole genome shotgun sequence
49. NZ_CP028369	Streptomyces sp. P3 chromosome, complete genome
50. NZ_CADFGO010000001	Trinickia caryophylli strain LMG 2155, whole genome shotgun sequence


Details:

>>
1. NC_049563
Source: Talaromyces rugulosus chromosome III, complete sequence
Type: NRPS-like
Number of proteins with BLAST hits to this cluster: 2
Cumulative BLAST score: 2002

Table of genes, locations, strands and annotations of subject cluster:

Table of Blast hits (query gene, subject gene, %identity, blast score, %coverage, e-value):
input.path1.gene222	TRUGW13939_06756	69	1775	100.23328149300154	0.0
input.path1.gene223	TRUGW13939_06755	31	227	94.04761904761905	4.4e-57


>>
2. NW_019716264
Source: Ramularia collo-cygni strain URUG2 genome assembly, contig: RCC_scaffold10, whole genome shotgun sequence
Type: NRPS-like
Number of proteins with BLAST hits to this cluster: 2
Cumulative BLAST score: 1765

Table of genes, locations, strands and annotations of subject cluster:

Table of Blast hits (query gene, subject gene, %identity, blast score, %coverage, e-value):
input.path1.gene221	RCC_06628	35	73	33.33333333333333	5.4e-11
input.path1.gene222	RCC_06627	66	1692	101.5552099533437	0.0


>>
3. NW_015971148
Source: Sporothrix schenckii 1099-18 chromosome Unknown Cont47, whole genome shotgun sequence
Type: NRPS-like
Number of proteins with BLAST hits to this cluster: 2
Cumulative BLAST score: 1386

Table of genes, locations, strands and annotations of subject cluster:

Table of Blast hits (query gene, subject gene, %identity, blast score, %coverage, e-value):
input.path1.gene221	SPSK_03174	37	126	42.00913242009132	5.3e-27
input.path1.gene222	SPSK_02939	50	1260	110.26438569206843	0.0


>>
4. NC_018218
Source: Zymoseptoria tritici IPO323 chromosome 1, whole genome shotgun sequence
Type: NRPS-like
Number of proteins with BLAST hits to this cluster: 2
Cumulative BLAST score: 1791

Table of genes, locations, strands and annotations of subject cluster:

Table of Blast hits (query gene, subject gene, %identity, blast score, %coverage, e-value):
input.path1.gene221	MYCGRDRAFT_34982	34	96	42.00913242009132	7.7e-18
input.path1.gene222	MYCGRDRAFT_107072	65	1695	102.79937791601866	0.0


>>
5. NZ_VLHW01000001
Source: Pseudomonas syringae pv. dysoxyli strain CFBP 2356 CL_dyso01, whole genome shotgun sequence
Type: NRPS
Number of proteins with BLAST hits to this cluster: 2
Cumulative BLAST score: 897

Table of genes, locations, strands and annotations of subject cluster:
PsdCFBP2356_RS05105	PsdCFBP2356_RS05105	1167570	1168320	+	SDR_family_oxidoreductase

Table of Blast hits (query gene, subject gene, %identity, blast score, %coverage, e-value):
input.path1.gene221	PsdCFBP2356_RS05100	38	97	29.22374429223744	4.5e-18
input.path1.gene222	PsdCFBP2356_RS05050	33	283	46.81181959564541	1e-73
input.path1.gene222	PsdCFBP2356_RS05055	33	267	47.82270606531882	7.4e-69
input.path1.gene222	PsdCFBP2356_RS05040	32	250	49.37791601866252	7.2e-64


>>
6. NC_014718
Source: Paraburkholderia rhizoxinica HKI 454 plasmid pBRH01, complete sequence
Type: NRPS,betalactone,phosphonate,terpene
Number of proteins with BLAST hits to this cluster: 2
Cumulative BLAST score: 896

Table of genes, locations, strands and annotations of subject cluster:
RBRH_RS19415	RBRH_RS19415	583389	583604	+	Tn3_family_transposase
RBRH_RS14390	RBRH_RS14390	584953	586309	-	hypothetical_protein
RBRH_RS19420	RBRH_RS19420	586482	586894	+	hypothetical_protein
RBRH_RS14510	RBRH_RS14510	611818	613591	-	cation:proton_antiporter
RBRH_RS14520	RBRH_RS14520	614897	616181	+	dicarboxylate/amino_acid:cation_symporter
RBRH_RS19425	RBRH_RS19425	627509	627736	+	IS5/IS1182_family_transposase
RBRH_RS18125	RBRH_RS18125	638940	639600	-	hypothetical_protein
RBRH_RS14640	RBRH_RS14640	674703	675209	+	transposase

Table of Blast hits (query gene, subject gene, %identity, blast score, %coverage, e-value):
input.path1.gene217	RBRH_RS14365	36	116	42.54807692307692	6.8e-24
input.path1.gene222	RBRH_RS18140	31	269	48.91135303265941	2.6e-69
input.path1.gene222	RBRH_RS16555	31	256	49.14463452566096	1.3e-65
input.path1.gene222	RBRH_RS16810	32	255	46.500777604976676	2.2e-65


>>
7. NZ_WJPS01000001
Source: Pseudomonas syringae strain P73 CL_7301, whole genome shotgun sequence
Type: NRPS
Number of proteins with BLAST hits to this cluster: 2
Cumulative BLAST score: 896

Table of genes, locations, strands and annotations of subject cluster:
PscP73CL_RS02110	PscP73CL_RS02110	455726	456476	-	SDR_family_oxidoreductase

Table of Blast hits (query gene, subject gene, %identity, blast score, %coverage, e-value):
input.path1.gene221	PscP73CL_RS02115	38	97	29.22374429223744	4.5e-18
input.path1.gene222	PscP73CL_RS02165	32	277	46.96734059097978	9.4e-72
input.path1.gene222	PscP73CL_RS02160	32	268	47.82270606531882	4.4e-69
input.path1.gene222	PscP73CL_RS02175	33	254	49.37791601866252	5e-65


>>
8. NZ_LT629769
Source: Pseudomonas syringae strain 31R1 chromosome I
Type: NRPS
Number of proteins with BLAST hits to this cluster: 2
Cumulative BLAST score: 895

Table of genes, locations, strands and annotations of subject cluster:
BLV36_RS04645	BLV36_RS04645	1034358	1035105	+	SDR_family_oxidoreductase

Table of Blast hits (query gene, subject gene, %identity, blast score, %coverage, e-value):
input.path1.gene221	BLV36_RS04635	38	97	29.22374429223744	4.5e-18
input.path1.gene222	BLV36_RS04585	33	281	47.43390357698289	5e-73
input.path1.gene222	BLV36_RS04590	33	267	47.82270606531882	5.7e-69
input.path1.gene222	BLV36_RS04575	32	250	49.37791601866252	7.2e-64


>>
9. NZ_FPJB01000001
Source: Pseudomonas sp. NFACC10-1, whole genome shotgun sequence
Type: NRPS
Number of proteins with BLAST hits to this cluster: 2
Cumulative BLAST score: 891

Table of genes, locations, strands and annotations of subject cluster:
BT119_RS26205	BT119_RS26205	434028	434778	+	SDR_family_oxidoreductase

Table of Blast hits (query gene, subject gene, %identity, blast score, %coverage, e-value):
input.path1.gene221	BT119_RS01745	38	97	29.22374429223744	4.5e-18
input.path1.gene222	BT119_RS01695	32	280	46.81181959564541	6.5e-73
input.path1.gene222	BT119_RS01685	33	267	47.82270606531882	9.7e-69
input.path1.gene222	BT119_RS01680	33	247	49.53343701399689	6.1e-63


>>
10. NZ_LT222319
Source: Pseudomonas cerasi isolate Sour cherry (Prunus cerasus) symptomatic leaf chromosome 1
Type: NRPS
Number of proteins with BLAST hits to this cluster: 2
Cumulative BLAST score: 891

Table of genes, locations, strands and annotations of subject cluster:
PCPL58_RS18750	PCPL58_RS18750	3657003	3657753	-	SDR_family_oxidoreductase

Table of Blast hits (query gene, subject gene, %identity, blast score, %coverage, e-value):
input.path1.gene221	PCPL58_RS18755	38	97	29.22374429223744	4.5e-18
input.path1.gene222	PCPL58_RS18805	32	273	47.43390357698289	1e-70
input.path1.gene222	PCPL58_RS18800	32	266	47.82270606531882	1.7e-68
input.path1.gene222	PCPL58_RS18815	33	255	49.37791601866252	3.8e-65


>>
11. NZ_KQ976354
Source: Scytonema hofmannii PCC 7110 Scaffold1, whole genome shotgun sequence
Type: NRPS,T1PKS
Number of proteins with BLAST hits to this cluster: 2
Cumulative BLAST score: 890

Table of genes, locations, strands and annotations of subject cluster:
WA1_RS54120	WA1_RS54120	10344251	10344350	-	IS5/IS1182_family_transposase

Table of Blast hits (query gene, subject gene, %identity, blast score, %coverage, e-value):
input.path1.gene217	WA1_RS43330	36	108	41.82692307692308	1.4e-21
input.path1.gene222	WA1_RS43550	32	283	47.97822706065319	1e-73
input.path1.gene222	WA1_RS43595	32	256	45.800933125972	1.7e-65
input.path1.gene222	WA1_RS43580	32	243	46.26749611197511	1.5e-61


>>
12. NZ_CP006256
Source: Pseudomonas syringae pv. syringae HS191 chromosome, complete genome
Type: NRPS
Number of proteins with BLAST hits to this cluster: 2
Cumulative BLAST score: 890

Table of genes, locations, strands and annotations of subject cluster:
PSYRH_RS16315	PSYRH_RS16315	3800485	3801235	-	SDR_family_oxidoreductase

Table of Blast hits (query gene, subject gene, %identity, blast score, %coverage, e-value):
input.path1.gene221	PSYRH_RS16320	38	97	29.22374429223744	4.5e-18
input.path1.gene222	PSYRH_RS16380	33	277	46.96734059097978	5.5e-72
input.path1.gene222	PSYRH_RS16375	32	264	47.82270606531882	8.2e-68
input.path1.gene222	PSYRH_RS16390	32	252	48.98911353032659	2.5e-64


>>
13. NZ_CP013732
Source: Burkholderia cepacia JBK9 chromosome 3, complete sequence
Type: NRPS,T1PKS,betalactone
Number of proteins with BLAST hits to this cluster: 2
Cumulative BLAST score: 876

Table of genes, locations, strands and annotations of subject cluster:
P350_RS40155	P350_RS40155	653436	653805	-	EamA_family_transporter
P350_RS39475	P350_RS39475	778659	778883	-	aldehyde_dehydrogenase_family_protein

Table of Blast hits (query gene, subject gene, %identity, blast score, %coverage, e-value):
input.path1.gene221	P350_RS35355	35	105	42.00913242009132	1.3e-20
input.path1.gene222	P350_RS35280	32	267	50.62208398133748	7.4e-69
input.path1.gene222	P350_RS35210	32	259	46.734059097978225	1.5e-66
input.path1.gene222	P350_RS35230	31	245	46.8895800933126	2.3e-62


>>
14. NZ_FOIB01000007
Source: Myxococcus fulvus strain DSM 16525, whole genome shotgun sequence
Type: NRPS,T1PKS
Number of proteins with BLAST hits to this cluster: 2
Cumulative BLAST score: 874

Table of genes, locations, strands and annotations of subject cluster:

Table of Blast hits (query gene, subject gene, %identity, blast score, %coverage, e-value):
input.path1.gene217	BMY20_RS27560	38	120	41.82692307692308	4.7e-25
input.path1.gene222	BMY20_RS27480	31	259	47.045101088646966	2.6e-66
input.path1.gene222	BMY20_RS27475	32	248	48.52255054432349	2.7e-63
input.path1.gene222	BMY20_RS27485	33	247	48.83359253499223	6.1e-63


>>
15. NZ_CP015529
Source: Rhodococcus sp. WB1 chromosome, complete genome
Type: NRPS
Number of proteins with BLAST hits to this cluster: 2
Cumulative BLAST score: 846

Table of genes, locations, strands and annotations of subject cluster:

Table of Blast hits (query gene, subject gene, %identity, blast score, %coverage, e-value):
input.path1.gene221	A4U64_RS06280	32	76	36.757990867579906	6.3e-12
input.path1.gene222	A4U64_RS06405	32	267	48.289269051321924	7.4e-69
input.path1.gene222	A4U64_RS06400	33	253	47.66718506998445	8.5e-65
input.path1.gene222	A4U64_RS06415	33	250	48.600311041990665	7.2e-64


>>
16. NZ_CP050952
Source: Rhodococcus sp. DMU1 chromosome, complete genome
Type: NRPS
Number of proteins with BLAST hits to this cluster: 2
Cumulative BLAST score: 845

Table of genes, locations, strands and annotations of subject cluster:

Table of Blast hits (query gene, subject gene, %identity, blast score, %coverage, e-value):
input.path1.gene221	HFP48_RS18985	31	74	36.3013698630137	3.1e-11
input.path1.gene222	HFP48_RS18860	32	266	48.289269051321924	1.3e-68
input.path1.gene222	HFP48_RS18865	33	253	47.66718506998445	8.5e-65
input.path1.gene222	HFP48_RS18850	33	252	48.600311041990665	3.2e-64


>>
17. NZ_CP011341
Source: Rhodococcus aetherivorans strain IcdP1 chromosome, complete genome
Type: NRPS
Number of proteins with BLAST hits to this cluster: 2
Cumulative BLAST score: 838

Table of genes, locations, strands and annotations of subject cluster:
AAT18_RS18235	AAT18_RS18235	4019169	4032832	-	amino_acid_adenylation_domain-containing_protein
AAT18_RS27685	AAT18_RS27685	4060275	4061741	-	non-ribosomal_peptide_synthetase
AAT18_RS18280	AAT18_RS18280	4112559	4113400	-	CoA_ester_lyase
AAT18_RS18340	AAT18_RS18340	4127281	4128611	-	MHS_family_MFS_transporter
AAT18_RS18355	AAT18_RS18355	4130008	4130771	-	ethanolamine_ammonia-lyase_subunit_EutC

Table of Blast hits (query gene, subject gene, %identity, blast score, %coverage, e-value):
input.path1.gene221	AAT18_RS18385	32	74	36.757990867579906	2.4e-11
input.path1.gene222	AAT18_RS18255	32	266	48.289269051321924	1.3e-68
input.path1.gene222	AAT18_RS18265	33	250	48.13374805598756	9.4e-64
input.path1.gene222	AAT18_RS18245	32	248	48.600311041990665	3.6e-63


>>
18. NZ_CP012201
Source: Massilia sp. NR 4-1, complete genome
Type: NRPS
Number of proteins with BLAST hits to this cluster: 2
Cumulative BLAST score: 748

Table of genes, locations, strands and annotations of subject cluster:

Table of Blast hits (query gene, subject gene, %identity, blast score, %coverage, e-value):
input.path1.gene217	ACZ75_RS03720	55	207	67.3076923076923	3.9e-51
input.path1.gene222	ACZ75_RS03705	35	294	48.83359253499223	5.7e-77
input.path1.gene222	ACZ75_RS03700	32	247	47.12286158631415	6.1e-63


>>
19. NZ_QEOB01000033
Source: Paraburkholderia unamae strain SCZa-39 Ga0215850_133, whole genome shotgun sequence
Type: NRPS
Number of proteins with BLAST hits to this cluster: 2
Cumulative BLAST score: 666

Table of genes, locations, strands and annotations of subject cluster:
C7402_RS40810	C7402_RS40810	44655	46230	-	IS3_family_transposase
C7402_RS40870	C7402_RS40870	71298	72460	+	IS3_family_transposase
C7402_RS40880	C7402_RS40880	74156	75223	-	site-specific_integrase
C7402_RS40885	C7402_RS40885	75327	75449	+	IS5/IS1182_family_transposase
C7402_RS40890	C7402_RS40890	75571	75669	-	integrase

Table of Blast hits (query gene, subject gene, %identity, blast score, %coverage, e-value):
input.path1.gene221	C7402_RS40730	34	108	42.00913242009132	2.6e-21
input.path1.gene222	C7402_RS40820	33	279	47.900466562986004	1.9e-72
input.path1.gene222	C7402_RS47235	33	279	47.900466562986004	2.5e-72


>>
20. NZ_CP045351
Source: Vibrio sp. THAF100 plasmid pTHAF100_a, complete sequence
Type: NRPS
Number of proteins with BLAST hits to this cluster: 2
Cumulative BLAST score: 655

Table of genes, locations, strands and annotations of subject cluster:

Table of Blast hits (query gene, subject gene, %identity, blast score, %coverage, e-value):
input.path1.gene221	FIV01_RS16405	34	111	43.37899543378995	3e-22
input.path1.gene222	FIV01_RS16475	32	282	46.8895800933126	2.9e-73
input.path1.gene222	FIV01_RS16480	32	262	52.56609642301711	1.8e-67


>>
21. NZ_KQ976354
Source: Scytonema hofmannii PCC 7110 Scaffold1, whole genome shotgun sequence
Type: NRPS
Number of proteins with BLAST hits to this cluster: 2
Cumulative BLAST score: 641

Table of genes, locations, strands and annotations of subject cluster:

Table of Blast hits (query gene, subject gene, %identity, blast score, %coverage, e-value):
input.path1.gene221	WA1_RS20760	40	102	28.538812785388128	1.1e-19
input.path1.gene222	WA1_RS20810	32	272	46.96734059097978	2.3e-70
input.path1.gene222	WA1_RS20815	32	267	47.27838258164852	9.7e-69


>>
22. NZ_CP034539
Source: Streptomyces cyaneochromogenes strain MK-45 chromosome, complete genome
Type: NRPS
Number of proteins with BLAST hits to this cluster: 2
Cumulative BLAST score: 611

Table of genes, locations, strands and annotations of subject cluster:
EJ357_RS01060	EJ357_RS01060	302507	302890	+	IS701_family_transposase
EJ357_RS01100	EJ357_RS01100	310815	310934	-	fumarylacetoacetate_hydrolase_family_protein
EJ357_RS01115	EJ357_RS01115	313094	313417	+	hypothetical_protein

Table of Blast hits (query gene, subject gene, %identity, blast score, %coverage, e-value):
input.path1.gene221	EJ357_RS01070	37	121	41.32420091324201	1.7e-25
input.path1.gene222	EJ357_RS00980	32	250	46.8895800933126	1.2e-63
input.path1.gene222	EJ357_RS00975	31	240	46.8895800933126	1.3e-60


>>
23. NZ_CM001149
Source: Rhodococcus hoagii ATCC 33707 chromosome, whole genome shotgun sequence
Type: NRPS
Number of proteins with BLAST hits to this cluster: 2
Cumulative BLAST score: 603

Table of genes, locations, strands and annotations of subject cluster:

Table of Blast hits (query gene, subject gene, %identity, blast score, %coverage, e-value):
input.path1.gene221	HMPREF0724_RS21960	31	82	44.74885844748858	1.2e-13
input.path1.gene222	HMPREF0724_RS22075	32	267	48.21150855365474	5.7e-69
input.path1.gene222	HMPREF0724_RS22080	32	254	48.67807153965785	6.5e-65


>>
24. NC_014659
Source: Rhodococcus hoagii 103S, complete genome
Type: NRPS
Number of proteins with BLAST hits to this cluster: 2
Cumulative BLAST score: 601

Table of genes, locations, strands and annotations of subject cluster:

Table of Blast hits (query gene, subject gene, %identity, blast score, %coverage, e-value):
input.path1.gene221	REQ_RS17745	31	82	44.74885844748858	1.2e-13
input.path1.gene222	REQ_RS17630	32	269	48.52255054432349	2.6e-69
input.path1.gene222	REQ_RS17625	31	250	48.367029548989116	7.2e-64


>>
25. NZ_CP027793
Source: Rhodococcus hoagii strain DSSKP-R-001 chromosome, complete genome
Type: NRPS
Number of proteins with BLAST hits to this cluster: 2
Cumulative BLAST score: 601

Table of genes, locations, strands and annotations of subject cluster:

Table of Blast hits (query gene, subject gene, %identity, blast score, %coverage, e-value):
input.path1.gene221	C7H75_RS18840	31	82	44.74885844748858	1.2e-13
input.path1.gene222	C7H75_RS18725	32	268	48.21150855365474	3.3e-69
input.path1.gene222	C7H75_RS18720	31	251	47.27838258164852	5.5e-64


>>
26. NZ_MOBR01000009
Source: Pseudomonas frederiksbergensis strain 38F7 38F7_contig_9, whole genome shotgun sequence
Type: NRPS,hserlactone
Number of proteins with BLAST hits to this cluster: 2
Cumulative BLAST score: 591

Table of genes, locations, strands and annotations of subject cluster:

Table of Blast hits (query gene, subject gene, %identity, blast score, %coverage, e-value):
input.path1.gene221	BK667_RS14190	36	72	29.908675799086758	1.6e-10
input.path1.gene222	BK667_RS14045	31	263	47.97822706065319	1.4e-67
input.path1.gene222	BK667_RS14270	32	256	47.51166407465008	1.3e-65


>>
27. NZ_QTTW01000003
Source: Rhodococcus sp. OK269 Ga0189761_103, whole genome shotgun sequence
Type: NRPS
Number of proteins with BLAST hits to this cluster: 2
Cumulative BLAST score: 588

Table of genes, locations, strands and annotations of subject cluster:

Table of Blast hits (query gene, subject gene, %identity, blast score, %coverage, e-value):
input.path1.gene221	C8K35_RS07370	34	78	37.44292237442922	2.2e-12
input.path1.gene222	C8K35_RS07295	32	269	48.83359253499223	2.6e-69
input.path1.gene222	C8K35_RS07275	33	241	46.81181959564541	5.7e-61


>>
28. NZ_JMEY01000019
Source: Rhodococcus fascians A3b contig22, whole genome shotgun sequence
Type: NRPS
Number of proteins with BLAST hits to this cluster: 2
Cumulative BLAST score: 575

Table of genes, locations, strands and annotations of subject cluster:
BH94_RS30590	BH94_RS30590	144805	145128	-	hypothetical_protein
BH94_RS28890	BH94_RS28890	164402	164654	-	ethanolamine_permease

Table of Blast hits (query gene, subject gene, %identity, blast score, %coverage, e-value):
input.path1.gene221	BH94_RS0110765	34	80	36.07305936073059	5.7e-13
input.path1.gene222	BH94_RS0110685	31	250	47.82270606531882	1.2e-63
input.path1.gene222	BH94_RS30580	32	245	47.82270606531882	3e-62


>>
29. NZ_KI519398
Source: Rhodococcus sp. JG-3 K414DRAFT_scaffold00001.1, whole genome shotgun sequence
Type: NRPS
Number of proteins with BLAST hits to this cluster: 2
Cumulative BLAST score: 567

Table of genes, locations, strands and annotations of subject cluster:
K414_RS28890	K414_RS28890	2376268	2376363	-	TetR/AcrR_family_transcriptional_regulator
K414_RS28940	K414_RS28940	2398352	2398675	+	hypothetical_protein

Table of Blast hits (query gene, subject gene, %identity, blast score, %coverage, e-value):
input.path1.gene221	K414_RS0112760	33	71	36.07305936073059	2.7e-10
input.path1.gene222	K414_RS0112860	31	249	47.82270606531882	2.1e-63
input.path1.gene222	K414_RS28950	32	247	47.82270606531882	1e-62


>>
30. NZ_JMEQ01000021
Source: Rhodococcus fascians LMG 3602 contig46, whole genome shotgun sequence
Type: NRPS
Number of proteins with BLAST hits to this cluster: 2
Cumulative BLAST score: 567

Table of genes, locations, strands and annotations of subject cluster:
BI02_RS26935	BI02_RS26935	185826	186146	-	hypothetical_protein
BI02_RS25785	BI02_RS25785	211030	211276	-	ethanolamine_permease

Table of Blast hits (query gene, subject gene, %identity, blast score, %coverage, e-value):
input.path1.gene221	BI02_RS0118800	32	70	36.07305936073059	5.9e-10
input.path1.gene222	BI02_RS0118695	31	251	47.82270606531882	5.5e-64
input.path1.gene222	BI02_RS26925	32	246	47.82270606531882	1.4e-62


>>
31. NZ_JMEV01000002
Source: Rhodococcus fascians A76 contig2, whole genome shotgun sequence
Type: NRPS
Number of proteins with BLAST hits to this cluster: 2
Cumulative BLAST score: 564

Table of genes, locations, strands and annotations of subject cluster:
BH97_RS06290	BH97_RS06290	1151685	1154810	+	DUF87_domain-containing_protein

Table of Blast hits (query gene, subject gene, %identity, blast score, %coverage, e-value):
input.path1.gene221	BH97_RS06470	32	79	42.00913242009132	7.5e-13
input.path1.gene222	BH97_RS06400	31	246	46.96734059097978	1.4e-62
input.path1.gene222	BH97_RS06405	31	239	48.67807153965785	1.7e-60


>>
32. NZ_RAII01000001
Source: Streptomyces sp. E1N211 s-211_S3_L001_R1_001, whole genome shotgun sequence
Type: NRPS,PKS-like,betalactone
Number of proteins with BLAST hits to this cluster: 2
Cumulative BLAST score: 555

Table of genes, locations, strands and annotations of subject cluster:
EBH12_RS22790	EBH12_RS22790	4809310	4810044	-	AfsR/SARP_family_transcriptional_regulator
EBH12_RS22895	EBH12_RS22895	4829916	4830236	+	NADP_oxidoreductase
EBH12_RS22930	EBH12_RS22930	4836511	4838167	+	DNA_alkylation_response_protein
EBH12_RS23030	EBH12_RS23030	4867092	4867346	+	2-oxoglutarate_dehydrogenase,_E2_component,_dihydrolipoamide_succinyltransferase
EBH12_RS23035	EBH12_RS23035	4867408	4867980	-	CDP-alcohol_phosphatidyltransferase_family_protein
EBH12_RS23045	EBH12_RS23045	4870108	4871715	+	sugar_phosphotransferase
EBH12_RS23055	EBH12_RS23055	4872530	4872788	-	MFS_transporter

Table of Blast hits (query gene, subject gene, %identity, blast score, %coverage, e-value):
input.path1.gene220	EBH12_RS22920	45	151	43.5960591133005	2.4e-34
input.path1.gene222	EBH12_RS22970	31	404	74.10575427682737	5.1e-110


>>
33. NZ_SMFR01000001
Source: Nocardia alba strain DSM 44684 Ga0244504_101, whole genome shotgun sequence
Type: NRPS
Number of proteins with BLAST hits to this cluster: 2
Cumulative BLAST score: 534

Table of genes, locations, strands and annotations of subject cluster:
DFR71_RS01630	DFR71_RS01630	367994	369223	-	NAD(P)/FAD-dependent_oxidoreductase
DFR71_RS01835	DFR71_RS01835	424841	425526	+	DUF899_family_protein
DFR71_RS01855	DFR71_RS01855	427943	428152	-	transcriptional_regulator

Table of Blast hits (query gene, subject gene, %identity, blast score, %coverage, e-value):
input.path1.gene221	DFR71_RS01595	36	113	41.32420091324201	4.7e-23
input.path1.gene222	DFR71_RS01615	32	421	81.18195956454122	4e-115


>>
34. NZ_LMFE01000001
Source: Nocardia sp. Root136 contig_1, whole genome shotgun sequence
Type: NRPS
Number of proteins with BLAST hits to this cluster: 2
Cumulative BLAST score: 518

Table of genes, locations, strands and annotations of subject cluster:
ASD42_RS08155	ASD42_RS08155	1749990	1751228	-	NAD(P)/FAD-dependent_oxidoreductase

Table of Blast hits (query gene, subject gene, %identity, blast score, %coverage, e-value):
input.path1.gene221	ASD42_RS08065	34	104	41.32420091324201	2.2e-20
input.path1.gene222	ASD42_RS08140	32	414	81.41524105754276	4.9e-113


>>
35. NZ_KI912151
Source: Fischerella sp. PCC 9605 FIS9605DRAFT_scaffold4.4, whole genome shotgun sequence
Type: NRPS
Number of proteins with BLAST hits to this cluster: 2
Cumulative BLAST score: 459

Table of genes, locations, strands and annotations of subject cluster:
FIS9605_RS38290	FIS9605_RS38290	331191	332771	-	choline_dehydrogenase

Table of Blast hits (query gene, subject gene, %identity, blast score, %coverage, e-value):
input.path1.gene221	FIS9605_RS0126420	38	106	32.87671232876712	5.7e-21
input.path1.gene221	FIS9605_RS38290	46	80	21.689497716894977	5.7e-13
input.path1.gene222	FIS9605_RS38275	33	273	45.95645412130638	1.4e-70


>>
36. NZ_CP046172
Source: Nocardia arthritidis strain AUSMDU00012717 chromosome, complete genome
Type: NRPS,T1PKS,T3PKS,lassopeptide
Number of proteins with BLAST hits to this cluster: 2
Cumulative BLAST score: 457

Table of genes, locations, strands and annotations of subject cluster:

Table of Blast hits (query gene, subject gene, %identity, blast score, %coverage, e-value):
input.path1.gene221	F5544_RS31855	32	91	42.00913242009132	2.5e-16
input.path1.gene222	F5544_RS31915	32	366	75.34992223950233	9e-99


>>
37. NZ_WXEW01000012
Source: Herbidospora sp. NEAU-GS84 Scaffold12, whole genome shotgun sequence
Type: NRPS,T3PKS
Number of proteins with BLAST hits to this cluster: 2
Cumulative BLAST score: 426

Table of genes, locations, strands and annotations of subject cluster:

Table of Blast hits (query gene, subject gene, %identity, blast score, %coverage, e-value):
input.path1.gene220	GT755_RS35010	48	163	44.33497536945813	4.8e-38
input.path1.gene222	GT755_RS35125	33	263	46.96734059097978	1.1e-67


>>
38. NZ_BBXF01000001
Source: Herbidospora daliensis strain NBRC 106372, whole genome shotgun sequence
Type: NRPS,T3PKS
Number of proteins with BLAST hits to this cluster: 2
Cumulative BLAST score: 422

Table of genes, locations, strands and annotations of subject cluster:
AW274_RS00670	AW274_RS00670	154585	155970	+	mucin-5B

Table of Blast hits (query gene, subject gene, %identity, blast score, %coverage, e-value):
input.path1.gene220	AW274_RS00815	49	161	44.08866995073892	1.8e-37
input.path1.gene222	AW274_RS00700	31	261	55.7542768273717	4.1e-67


>>
39. NZ_CP019717
Source: Paenibacillus larvae subsp. larvae strain Eric_V chromosome, complete genome
Type: NRPS,phosphonate
Number of proteins with BLAST hits to this cluster: 2
Cumulative BLAST score: 420

Table of genes, locations, strands and annotations of subject cluster:
ERICV_RS12270	ERICV_RS12270	2298627	2299394	+	hypothetical_protein
ERICV_RS12275	ERICV_RS12275	2300031	2300480	+	hypothetical_protein
ERICV_RS12300	ERICV_RS12300	2317838	2318068	+	GntR_family_transcriptional_regulator
ERICV_RS12340	ERICV_RS12340	2326025	2327664	+	sensor_histidine_kinase
ERICV_RS12345	ERICV_RS12345	2327713	2328156	+	response_regulator

Table of Blast hits (query gene, subject gene, %identity, blast score, %coverage, e-value):
input.path1.gene217	ERICV_RS12325	41	146	40.144230769230774	6.2e-33
input.path1.gene222	ERICV_RS12295	33	274	46.96734059097978	7.9e-71


>>
40. NC_023134
Source: Paenibacillus larvae subsp. larvae DSM 25430, complete sequence
Type: NRPS,phosphonate
Number of proteins with BLAST hits to this cluster: 2
Cumulative BLAST score: 419

Table of genes, locations, strands and annotations of subject cluster:
ERIC2_RS21190	ERIC2_RS21190	1987194	1987427	-	HTH_domain-containing_protein
ERIC2_RS21195	ERIC2_RS21195	1987719	1988159	-	response_regulator
ERIC2_RS09415	ERIC2_RS09415	1988208	1989847	-	sensor_histidine_kinase
ERIC2_RS09440	ERIC2_RS09440	1994256	1995444	-	MFS_transporter
ERIC2_RS21205	ERIC2_RS21205	2011007	2011456	-	hypothetical_protein
ERIC2_RS21210	ERIC2_RS21210	2012093	2012860	-	hypothetical_protein

Table of Blast hits (query gene, subject gene, %identity, blast score, %coverage, e-value):
input.path1.gene217	ERIC2_RS09430	41	146	40.144230769230774	6.2e-33
input.path1.gene222	ERIC2_RS09460	33	273	46.96734059097978	1e-70


>>
41. NZ_CP019687
Source: Paenibacillus larvae subsp. larvae strain ATCC-9545 chromosome, complete genome
Type: NRPS,phosphonate
Number of proteins with BLAST hits to this cluster: 2
Cumulative BLAST score: 419

Table of genes, locations, strands and annotations of subject cluster:
BXP28_RS04320	BXP28_RS04320	795493	795965	-	GNAT_family_N-acetyltransferase
BXP28_RS04325	BXP28_RS04325	796274	797301	-	Gfo/Idh/MocA_family_oxidoreductase
BXP28_RS04410	BXP28_RS04410	810092	810859	+	hypothetical_protein
BXP28_RS04415	BXP28_RS04415	811496	811945	+	hypothetical_protein
BXP28_RS04475	BXP28_RS04475	833034	834222	+	MFS_transporter
BXP28_RS04500	BXP28_RS04500	838631	840270	+	sensor_histidine_kinase
BXP28_RS04505	BXP28_RS04505	840319	840762	+	response_regulator
BXP28_RS04520	BXP28_RS04520	842531	843765	+	ATP-grasp_domain-containing_protein
BXP28_RS04585	BXP28_RS04585	860858	862098	+	lipase

Table of Blast hits (query gene, subject gene, %identity, blast score, %coverage, e-value):
input.path1.gene217	BXP28_RS04485	41	146	40.144230769230774	8.1e-33
input.path1.gene222	BXP28_RS04455	33	273	46.96734059097978	1e-70


>>
42. NZ_CP041061
Source: Micromonospora sp. HM134 chromosome, complete genome
Type: NRPS,betalactone
Number of proteins with BLAST hits to this cluster: 2
Cumulative BLAST score: 418

Table of genes, locations, strands and annotations of subject cluster:
FJK98_RS11945	FJK98_RS11945	2701848	2702219	+	MerR_family_transcriptional_regulator
FJK98_RS11980	FJK98_RS11980	2713315	2714214	+	ATP-binding_cassette_domain-containing_protein
FJK98_RS11985	FJK98_RS11985	2714226	2714557	+	kinase

Table of Blast hits (query gene, subject gene, %identity, blast score, %coverage, e-value):
input.path1.gene220	FJK98_RS11990	46	171	47.29064039408867	1.8e-40
input.path1.gene222	FJK98_RS11870	32	247	46.81181959564541	7.9e-63


>>
43. NZ_FNOG01000002
Source: Lysobacter enzymogenes strain ATCC 29487, whole genome shotgun sequence
Type: NRPS,T1PKS
Number of proteins with BLAST hits to this cluster: 2
Cumulative BLAST score: 408

Table of genes, locations, strands and annotations of subject cluster:

Table of Blast hits (query gene, subject gene, %identity, blast score, %coverage, e-value):
input.path1.gene221	BLU84_RS06850	38	132	41.0958904109589	1.3e-28
input.path1.gene222	BLU84_RS06780	33	276	49.06687402799378	1.2e-71


>>
44. NZ_AP014940
Source: Lysobacter enzymogenes strain M497-1
Type: NRPS,T1PKS
Number of proteins with BLAST hits to this cluster: 2
Cumulative BLAST score: 405

Table of genes, locations, strands and annotations of subject cluster:

Table of Blast hits (query gene, subject gene, %identity, blast score, %coverage, e-value):
input.path1.gene221	LEN_RS18945	38	131	41.0958904109589	2.2e-28
input.path1.gene222	LEN_RS18875	32	274	49.22239502332815	6.1e-71


>>
45. NZ_QXXQ01000017
Source: Gemmobacter lutimaris strain YJ-T1-11 scaffold_17, whole genome shotgun sequence
Type: NRPS
Number of proteins with BLAST hits to this cluster: 2
Cumulative BLAST score: 393

Table of genes, locations, strands and annotations of subject cluster:

Table of Blast hits (query gene, subject gene, %identity, blast score, %coverage, e-value):
input.path1.gene221	D2N39_RS19570	41	139	43.15068493150685	6.1e-31
input.path1.gene222	D2N39_RS19525	33	254	46.96734059097978	5e-65


>>
46. NZ_WWEU01000004
Source: Vibrio sp. OCN044 NODE_4_length_493155_cov_31.134218, whole genome shotgun sequence
Type: NRPS
Number of proteins with BLAST hits to this cluster: 2
Cumulative BLAST score: 389

Table of genes, locations, strands and annotations of subject cluster:
GTG28_RS13585	GTG28_RS13585	285314	286172	-	glutathione-dependent_disulfide-bond_oxidoreductase

Table of Blast hits (query gene, subject gene, %identity, blast score, %coverage, e-value):
input.path1.gene221	GTG28_RS13695	34	111	43.37899543378995	3e-22
input.path1.gene222	GTG28_RS13635	32	278	46.8895800933126	4.2e-72


>>
47. NZ_AP018288
Source: Nostoc sp. NIES-4103 DNA, nearly complete genome
Type: NRPS
Number of proteins with BLAST hits to this cluster: 2
Cumulative BLAST score: 388

Table of genes, locations, strands and annotations of subject cluster:
CDC32_RS00530	CDC32_RS00530	143711	144136	+	hypothetical_protein

Table of Blast hits (query gene, subject gene, %identity, blast score, %coverage, e-value):
input.path1.gene221	CDC32_RS00435	40	125	41.55251141552511	1.2e-26
input.path1.gene222	CDC32_RS00460	32	263	47.900466562986004	1.1e-67


>>
48. NZ_JABUMT010000016
Source: Myxococcus sp. CA056 NODE_16_length_238008_cov_21.6875, whole genome shotgun sequence
Type: NRPS,T1PKS
Number of proteins with BLAST hits to this cluster: 2
Cumulative BLAST score: 382

Table of genes, locations, strands and annotations of subject cluster:

Table of Blast hits (query gene, subject gene, %identity, blast score, %coverage, e-value):
input.path1.gene217	HUA76_RS38225	38	125	42.30769230769231	1.5e-26
input.path1.gene222	HUA76_RS38145	31	257	47.97822706065319	1e-65


>>
49. NZ_CP028369
Source: Streptomyces sp. P3 chromosome, complete genome
Type: NRPS
Number of proteins with BLAST hits to this cluster: 2
Cumulative BLAST score: 381

Table of genes, locations, strands and annotations of subject cluster:
C6376_RS42460	C6376_RS42460	9594394	9594624	+	hypothetical_protein

Table of Blast hits (query gene, subject gene, %identity, blast score, %coverage, e-value):
input.path1.gene220	C6376_RS42430	40	141	49.26108374384236	1.5e-31
input.path1.gene222	C6376_RS42400	32	240	46.578538102643854	7.4e-61


>>
50. NZ_CADFGO010000001
Source: Trinickia caryophylli strain LMG 2155, whole genome shotgun sequence
Type: NRPS
Number of proteins with BLAST hits to this cluster: 2
Cumulative BLAST score: 377

Table of genes, locations, strands and annotations of subject cluster:

Table of Blast hits (query gene, subject gene, %identity, blast score, %coverage, e-value):
input.path1.gene221	HWY14_RS03915	40	118	37.44292237442922	2.5e-24
input.path1.gene222	HWY14_RS03965	34	259	48.755832037325035	2e-66
