## Supplementary Results for "A draft genome of the ascomycotal fungal species *Pseudopithomyces maydicus* (family *Didymosphaeriaceae*)": contig_14_c1.txt

ClusterBlast scores for contig_14

Table of genes, locations, strands and annotations of query cluster:
input.path1.gene61	213855	215306	-		
input.path1.gene62	216933	218753	+		
input.path1.gene63	220052	220644	-		
input.path1.gene64	220665	221421	+		
input.path1.gene65	221788	223727	-		
input.path1.gene66	225959	226926	+		
input.path1.gene67	227251	228223	-		
input.path1.gene68	229215	235818	+		
input.path1.gene69	238547	239041	-		
input.path1.gene70	239757	241522	+		
input.path1.gene71	242010	243638	-		
input.path1.gene72	247732	248306	+		
input.path1.gene73	253075	254298	+		


Significant hits: 
1. NT_165977	Chaetomium globosum CBS 148.51 scaffold_2 genomic scaffold, whole genome shotgun sequence
2. NW_003299166	Microsporum canis CBS 113480 supercont1.4 genomic scaffold, whole genome shotgun sequence
3. NW_021167090	Sodiomyces alkalinus F11 unplaced genomic scaffold SODALscaffold_12, whole genome shotgun sequence
4. NW_022474213	Venustampulla echinocandica strain BP 5553 chromosome Unknown scaffold_8, whole genome shotgun sequence
5. NW_003345199	Nannizzia gypsea CBS 118893 supercont1.3 genomic scaffold, whole genome shotgun sequence
6. NC_049565	Talaromyces rugulosus chromosome V, complete sequence
7. NW_003299163	Microsporum canis CBS 113480 supercont1.7 genomic scaffold, whole genome shotgun sequence
8. NW_006271969	Cordyceps militaris CM01 unplaced genomic scaffold CCM_S00001, whole genome shotgun sequence
9. NW_023336279	Aspergillus tubingensis WU-2223L DNA, scaffold_5, whole genome shotgun sequence
10. NT_165933	Aspergillus terreus NIH2624 scaffold_10 genomic scaffold, whole genome shotgun sequence
11. NW_017971444	Talaromyces atroroseus strain IBT 11181 chromosome Unknown scaffold_12, whole genome shotgun sequence
12. NW_023336282	Aspergillus tubingensis WU-2223L DNA, scaffold_8, whole genome shotgun sequence
13. NW_020167523	Pseudogymnoascus destructans isolate 20631-21 unplaced genomic scaffold scaffold_4, whole genome shotgun sequence
14. NW_022194798	Fusarium proliferatum ET1 genome assembly, contig: FPRO_scaffold009, whole genome shotgun sequence
15. NC_030962	Colletotrichum higginsianum IMI 349063 chromosome 10, whole genome shotgun sequence
16. NW_019716276	Ramularia collo-cygni strain URUG2 genome assembly, contig: RCC_scaffold22, whole genome shotgun sequence
17. NC_018213	Zymoseptoria tritici IPO323 chromosome 6, whole genome shotgun sequence
18. NW_023336265	Colletotrichum scovillei strain TJNH1 chromosome Unknown Contig00007, whole genome shotgun sequence
19. NW_022984635	Aspergillus tanneri strain NIH1004 chromosome Unknown AapTan_8_pilon_pilon_pilon, whole genome shotgun sequence
20. NW_013562490	Exophiala xenobiotica strain CBS 118157 unplaced genomic scaffold supercont1.5, whole genome shotgun sequence
21. CM000596	Fusarium oxysporum f. sp. lycopersici 4287 chromosome 8, whole genome shotgun sequence
22. NC_030993	Fusarium oxysporum f. sp. lycopersici 4287 chromosome 8, whole genome shotgun sequence
23. NW_006271969	Cordyceps militaris CM01 unplaced genomic scaffold CCM_S00001, whole genome shotgun sequence
24. NW_015622519	Exophiala spinifera strain CBS 89968 unplaced genomic scaffold supercont1.7, whole genome shotgun sequence
25. NW_006912904	Capronia epimyces CBS 606.96 aczIQ-supercont1.1, whole genome shotgun sequence
26. NW_017971443	Talaromyces atroroseus strain IBT 11181 chromosome Unknown scaffold_11, whole genome shotgun sequence
27. NW_015971652	Fonsecaea multimorphosa CBS 102226 unplaced genomic scaffold supercont1.18, whole genome shotgun sequence
28. NC_036632	Fusarium fujikuroi IMI 58289 draft genome, chromosome FFUJ_chr11
29. NW_022474213	Venustampulla echinocandica strain BP 5553 chromosome Unknown scaffold_8, whole genome shotgun sequence
30. NW_013550604	Rhinocladiella mackenziei CBS 650.93 unplaced genomic scaffold supercont1.1, whole genome shotgun sequence
31. NW_008481799	Cyphellophora europaea CBS 101466 unplaced genomic scaffold supercont1.11, whole genome shotgun sequence
32. NW_023336268	Colletotrichum scovillei strain TJNH1 chromosome Unknown Contig00010, whole genome shotgun sequence
33. NW_007360991	Glarea lozoyensis ATCC 20868 chromosome Unknown GLAREA17, whole genome shotgun sequence
34. NW_023336282	Aspergillus tubingensis WU-2223L DNA, scaffold_8, whole genome shotgun sequence
35. NC_030960	Colletotrichum higginsianum IMI 349063 chromosome 8, whole genome shotgun sequence
36. NZ_CP034352	Streptomyces sp. W1SF4 plasmid p2, complete sequence
37. NZ_CP015163	Amycolatopsis albispora strain WP1 chromosome, complete genome
38. NZ_GG657752	Streptomyces sp. C supercont1.3, whole genome shotgun sequence
39. NZ_JAAWWP010000017	Streptomyces sp. LD120 Scaffold17, whole genome shotgun sequence
40. NW_006912895	Capronia coronata CBS 617.96 acBCG-supercont1.3, whole genome shotgun sequence
41. NW_003052498	Uncinocarpus reesii 1704 scaffold_3 genomic scaffold, whole genome shotgun sequence
42. NC_018210	Zymoseptoria tritici IPO323 chromosome 9, whole genome shotgun sequence
43. NW_023336282	Aspergillus tubingensis WU-2223L DNA, scaffold_8, whole genome shotgun sequence
44. NZ_PVTF01000015	Umezawaea tangerina strain DSM 44720 Ga0180964_115, whole genome shotgun sequence
45. NW_014574615	Penicillium digitatum Pd1 unplaced genomic scaffold Scaffold_10, whole genome shotgun sequence
46. CM000574	Fusarium graminearum PH-1 chromosome 1, whole genome shotgun sequence
47. NC_026474	Fusarium graminearum PH-1 chromosome 1, whole genome shotgun sequence
48. NC_035793	Pochonia chlamydosporia 170 chromosome 4, whole genome shotgun sequence
49. NC_009380	Salinispora tropica CNB-440, complete sequence
50. NZ_CP010519	Streptomyces albus strain DSM 41398 chromosome, complete genome


Details:

>>
1. NT_165977
Source: Chaetomium globosum CBS 148.51 scaffold_2 genomic scaffold, whole genome shotgun sequence
Type: T1PKS
Number of proteins with BLAST hits to this cluster: 4
Cumulative BLAST score: 1636

Table of genes, locations, strands and annotations of subject cluster:

Table of Blast hits (query gene, subject gene, %identity, blast score, %coverage, e-value):
input.path1.gene67	CHGG_03102	35	86	64.56310679611651	4.9e-15
input.path1.gene68	CHGG_03101	34	1233	116.81457262961233	0.0
input.path1.gene69	CHGG_03102	33	79	88.19444444444444	3.2e-13
input.path1.gene70	CHGG_03101	31	238	116.34146341463413	1.2e-60


>>
2. NW_003299166
Source: Microsporum canis CBS 113480 supercont1.4 genomic scaffold, whole genome shotgun sequence
Type: NRPS,T1PKS
Number of proteins with BLAST hits to this cluster: 3
Cumulative BLAST score: 1580

Table of genes, locations, strands and annotations of subject cluster:

Table of Blast hits (query gene, subject gene, %identity, blast score, %coverage, e-value):
input.path1.gene67	MCYG_04544	44	86	53.883495145631066	2.9e-15
input.path1.gene68	MCYG_04545	38	1425	109.71508640822046	0.0
input.path1.gene69	MCYG_04544	35	69	61.111111111111114	4.3e-10


>>
3. NW_021167090
Source: Sodiomyces alkalinus F11 unplaced genomic scaffold SODALscaffold_12, whole genome shotgun sequence
Type: T1PKS
Number of proteins with BLAST hits to this cluster: 3
Cumulative BLAST score: 1564

Table of genes, locations, strands and annotations of subject cluster:

Table of Blast hits (query gene, subject gene, %identity, blast score, %coverage, e-value):
input.path1.gene67	SODALDRAFT_394970	48	59	29.126213592233007	4.9e-07
input.path1.gene68	SODALDRAFT_284490	40	1446	103.03596450256889	0.0
input.path1.gene69	SODALDRAFT_394970	37	59	52.083333333333336	2.6e-07


>>
4. NW_022474213
Source: Venustampulla echinocandica strain BP 5553 chromosome Unknown scaffold_8, whole genome shotgun sequence
Type: T1PKS
Number of proteins with BLAST hits to this cluster: 3
Cumulative BLAST score: 1550

Table of genes, locations, strands and annotations of subject cluster:

Table of Blast hits (query gene, subject gene, %identity, blast score, %coverage, e-value):
input.path1.gene67	BP5553_08501	43	66	35.92233009708738	4e-09
input.path1.gene68	BP5553_08500	37	1381	108.5007006071929	0.0
input.path1.gene69	BP5553_08501	44	103	81.25	1.6e-20


>>
5. NW_003345199
Source: Nannizzia gypsea CBS 118893 supercont1.3 genomic scaffold, whole genome shotgun sequence
Type: NRPS,T1PKS
Number of proteins with BLAST hits to this cluster: 3
Cumulative BLAST score: 1547

Table of genes, locations, strands and annotations of subject cluster:

Table of Blast hits (query gene, subject gene, %identity, blast score, %coverage, e-value):
input.path1.gene67	MGYG_03810	41	62	36.407766990291265	7.6e-08
input.path1.gene68	MGYG_03811	36	1382	108.5007006071929	0.0
input.path1.gene69	MGYG_03810	43	103	81.25	1.6e-20


>>
6. NC_049565
Source: Talaromyces rugulosus chromosome V, complete sequence
Type: T1PKS
Number of proteins with BLAST hits to this cluster: 3
Cumulative BLAST score: 1509

Table of genes, locations, strands and annotations of subject cluster:

Table of Blast hits (query gene, subject gene, %identity, blast score, %coverage, e-value):
input.path1.gene67	TRUGW13939_09153	46	59	30.582524271844658	3.8e-07
input.path1.gene68	TRUGW13939_09154	35	1351	109.8085007006072	0.0
input.path1.gene69	TRUGW13939_09153	45	99	70.13888888888889	3e-19


>>
7. NW_003299163
Source: Microsporum canis CBS 113480 supercont1.7 genomic scaffold, whole genome shotgun sequence
Type: T1PKS,T3PKS
Number of proteins with BLAST hits to this cluster: 3
Cumulative BLAST score: 1488

Table of genes, locations, strands and annotations of subject cluster:

Table of Blast hits (query gene, subject gene, %identity, blast score, %coverage, e-value):
input.path1.gene67	MCYG_07716	38	56	35.92233009708738	3.2e-06
input.path1.gene68	MCYG_07715	36	1348	109.15460065390003	0.0
input.path1.gene69	MCYG_07716	41	84	68.05555555555556	7.6e-15


>>
8. NW_006271969
Source: Cordyceps militaris CM01 unplaced genomic scaffold CCM_S00001, whole genome shotgun sequence
Type: T1PKS
Number of proteins with BLAST hits to this cluster: 3
Cumulative BLAST score: 1459

Table of genes, locations, strands and annotations of subject cluster:

Table of Blast hits (query gene, subject gene, %identity, blast score, %coverage, e-value):
input.path1.gene67	CCM_01517	33	63	57.76699029126213	3.4e-08
input.path1.gene68	CCM_01518	36	1302	108.87435777673984	0.0
input.path1.gene69	CCM_01517	41	94	70.83333333333334	7.4e-18


>>
9. NW_023336279
Source: Aspergillus tubingensis WU-2223L DNA, scaffold_5, whole genome shotgun sequence
Type: NRPS,T1PKS
Number of proteins with BLAST hits to this cluster: 3
Cumulative BLAST score: 1346

Table of genes, locations, strands and annotations of subject cluster:

Table of Blast hits (query gene, subject gene, %identity, blast score, %coverage, e-value):
input.path1.gene67	AtWU_03525	40	74	59.22330097087378	1.5e-11
input.path1.gene68	AtWU_03524	34	1161	108.5007006071929	0.0
input.path1.gene69	AtWU_03525	42	111	90.97222222222221	7.6e-23


>>
10. NT_165933
Source: Aspergillus terreus NIH2624 scaffold_10 genomic scaffold, whole genome shotgun sequence
Type: T1PKS
Number of proteins with BLAST hits to this cluster: 3
Cumulative BLAST score: 1283

Table of genes, locations, strands and annotations of subject cluster:

Table of Blast hits (query gene, subject gene, %identity, blast score, %coverage, e-value):
input.path1.gene67	ATEG_07281	40	57	37.86407766990291	2.4e-06
input.path1.gene68	ATEG_07282	33	1127	110.2288650163475	0.0
input.path1.gene69	ATEG_07281	41	99	87.5	3e-19


>>
11. NW_017971444
Source: Talaromyces atroroseus strain IBT 11181 chromosome Unknown scaffold_12, whole genome shotgun sequence
Type: T1PKS
Number of proteins with BLAST hits to this cluster: 4
Cumulative BLAST score: 2080

Table of genes, locations, strands and annotations of subject cluster:

Table of Blast hits (query gene, subject gene, %identity, blast score, %coverage, e-value):
input.path1.gene67	UA08_07743	45	69	35.43689320388349	4.8e-10
input.path1.gene68	UA08_07744	40	1621	109.38813638486688	0.0
input.path1.gene69	UA08_07742	41	100	88.19444444444444	1.8e-19
input.path1.gene69	UA08_07743	33	84	95.83333333333334	1.3e-14
input.path1.gene70	UA08_07740	35	206	108.53658536585367	5e-51


>>
12. NW_023336282
Source: Aspergillus tubingensis WU-2223L DNA, scaffold_8, whole genome shotgun sequence
Type: NRPS,T1PKS
Number of proteins with BLAST hits to this cluster: 4
Cumulative BLAST score: 1944

Table of genes, locations, strands and annotations of subject cluster:

Table of Blast hits (query gene, subject gene, %identity, blast score, %coverage, e-value):
input.path1.gene67	AtWU_06981	37	66	47.0873786407767	5.3e-09
input.path1.gene68	AtWU_06980	34	1173	110.6492293320878	0.0
input.path1.gene68	AtWU_06965	32	546	59.69173283512378	1.4e-152
input.path1.gene69	AtWU_06981	42	96	86.11111111111111	1.9e-18
input.path1.gene70	AtWU_06964	31	63	29.51219512195122	6.8e-08


>>
13. NW_020167523
Source: Pseudogymnoascus destructans isolate 20631-21 unplaced genomic scaffold scaffold_4, whole genome shotgun sequence
Type: T1PKS
Number of proteins with BLAST hits to this cluster: 3
Cumulative BLAST score: 1332

Table of genes, locations, strands and annotations of subject cluster:

Table of Blast hits (query gene, subject gene, %identity, blast score, %coverage, e-value):
input.path1.gene68	VC83_02202	32	1098	119.66370854740775	0.0
input.path1.gene69	VC83_02203	34	62	71.52777777777779	4.1e-08
input.path1.gene70	VC83_02204	43	172	50.73170731707317	7.9e-41


>>
14. NW_022194798
Source: Fusarium proliferatum ET1 genome assembly, contig: FPRO_scaffold009, whole genome shotgun sequence
Type: T1PKS
Number of proteins with BLAST hits to this cluster: 3
Cumulative BLAST score: 1313

Table of genes, locations, strands and annotations of subject cluster:

Table of Blast hits (query gene, subject gene, %identity, blast score, %coverage, e-value):
input.path1.gene68	FPRO_14907	31	1080	119.75712283979449	0.0
input.path1.gene69	FPRO_14906	34	76	86.11111111111111	3.5e-12
input.path1.gene70	FPRO_14905	42	157	53.90243902439025	3.4e-36


>>
15. NC_030962
Source: Colletotrichum higginsianum IMI 349063 chromosome 10, whole genome shotgun sequence
Type: T1PKS
Number of proteins with BLAST hits to this cluster: 3
Cumulative BLAST score: 2069

Table of genes, locations, strands and annotations of subject cluster:

Table of Blast hits (query gene, subject gene, %identity, blast score, %coverage, e-value):
input.path1.gene67	CH63R_13916	37	69	55.8252427184466	3.6e-10
input.path1.gene68	CH63R_13915	36	1412	109.24801494628677	0.0
input.path1.gene68	CH63R_13918	45	518	29.098552078468003	3.1e-144
input.path1.gene69	CH63R_13916	35	70	92.36111111111111	1.9e-10


>>
16. NW_019716276
Source: Ramularia collo-cygni strain URUG2 genome assembly, contig: RCC_scaffold22, whole genome shotgun sequence
Type: T1PKS
Number of proteins with BLAST hits to this cluster: 3
Cumulative BLAST score: 2045

Table of genes, locations, strands and annotations of subject cluster:

Table of Blast hits (query gene, subject gene, %identity, blast score, %coverage, e-value):
input.path1.gene67	RCC_10364	32	56	64.56310679611651	4.2e-06
input.path1.gene68	RCC_10366	35	1247	108.7809434843531	0.0
input.path1.gene68	RCC_10366_XP_023631365.1	49	649	32.92853806632415	1.2e-183
input.path1.gene69	RCC_10364	36	93	86.80555555555556	1.6e-17


>>
17. NC_018213
Source: Zymoseptoria tritici IPO323 chromosome 6, whole genome shotgun sequence
Type: T1PKS
Number of proteins with BLAST hits to this cluster: 2
Cumulative BLAST score: 2381

Table of genes, locations, strands and annotations of subject cluster:

Table of Blast hits (query gene, subject gene, %identity, blast score, %coverage, e-value):
input.path1.gene68	MYCGRDRAFT_72709	54	2241	100.56048575432041	0.0
input.path1.gene69	MYCGRDRAFT_93616	51	140	88.88888888888889	1.5e-31


>>
18. NW_023336265
Source: Colletotrichum scovillei strain TJNH1 chromosome Unknown Contig00007, whole genome shotgun sequence
Type: T1PKS
Number of proteins with BLAST hits to this cluster: 2
Cumulative BLAST score: 1486

Table of genes, locations, strands and annotations of subject cluster:

Table of Blast hits (query gene, subject gene, %identity, blast score, %coverage, e-value):
input.path1.gene68	HER10_EVM0007585	37	1399	109.48155067725362	0.0
input.path1.gene69	HER10_EVM0002811	37	87	90.97222222222221	1.5e-15


>>
19. NW_022984635
Source: Aspergillus tanneri strain NIH1004 chromosome Unknown AapTan_8_pilon_pilon_pilon, whole genome shotgun sequence
Type: T1PKS
Number of proteins with BLAST hits to this cluster: 2
Cumulative BLAST score: 1446

Table of genes, locations, strands and annotations of subject cluster:

Table of Blast hits (query gene, subject gene, %identity, blast score, %coverage, e-value):
input.path1.gene68	ATNIH1004_011613	36	1346	108.640822045773	0.0
input.path1.gene69	ATNIH1004_011612	40	100	87.5	1.8e-19


>>
20. NW_013562490
Source: Exophiala xenobiotica strain CBS 118157 unplaced genomic scaffold supercont1.5, whole genome shotgun sequence
Type: T1PKS
Number of proteins with BLAST hits to this cluster: 3
Cumulative BLAST score: 1444

Table of genes, locations, strands and annotations of subject cluster:

Table of Blast hits (query gene, subject gene, %identity, blast score, %coverage, e-value):
input.path1.gene68	PV05_09282	32	1180	118.21578701541335	0.0
input.path1.gene69	PV05_09280	35	96	87.5	3.3e-18
input.path1.gene70	PV05_09281	41	168	54.390243902439025	1.1e-39


>>
21. CM000596
Source: Fusarium oxysporum f. sp. lycopersici 4287 chromosome 8, whole genome shotgun sequence
Type: T1PKS
Number of proteins with BLAST hits to this cluster: 3
Cumulative BLAST score: 1393

Table of genes, locations, strands and annotations of subject cluster:

Table of Blast hits (query gene, subject gene, %identity, blast score, %coverage, e-value):
input.path1.gene68	FOXG_02741	32	1163	117.281644091546	0.0
input.path1.gene69	FOXG_02738	41	96	94.44444444444444	3.3e-18
input.path1.gene70	FOXG_02737	35	134	55.1219512195122	2.4e-29


>>
22. NC_030993
Source: Fusarium oxysporum f. sp. lycopersici 4287 chromosome 8, whole genome shotgun sequence
Type: T1PKS
Number of proteins with BLAST hits to this cluster: 3
Cumulative BLAST score: 1393

Table of genes, locations, strands and annotations of subject cluster:

Table of Blast hits (query gene, subject gene, %identity, blast score, %coverage, e-value):
input.path1.gene68	FOXG_02741	32	1163	117.281644091546	0.0
input.path1.gene69	FOXG_02738	41	96	94.44444444444444	3.3e-18
input.path1.gene70	FOXG_02737	35	134	55.1219512195122	2.4e-29


>>
23. NW_006271969
Source: Cordyceps militaris CM01 unplaced genomic scaffold CCM_S00001, whole genome shotgun sequence
Type: T1PKS
Number of proteins with BLAST hits to this cluster: 2
Cumulative BLAST score: 1305

Table of genes, locations, strands and annotations of subject cluster:

Table of Blast hits (query gene, subject gene, %identity, blast score, %coverage, e-value):
input.path1.gene68	CCM_00603	36	1218	104.99766464269034	0.0
input.path1.gene69	CCM_00604	44	87	70.13888888888889	1.5e-15


>>
24. NW_015622519
Source: Exophiala spinifera strain CBS 89968 unplaced genomic scaffold supercont1.7, whole genome shotgun sequence
Type: NRPS-like,T1PKS
Number of proteins with BLAST hits to this cluster: 2
Cumulative BLAST score: 1266

Table of genes, locations, strands and annotations of subject cluster:

Table of Blast hits (query gene, subject gene, %identity, blast score, %coverage, e-value):
input.path1.gene68	PV08_09991	33	1160	110.92947220924802	0.0
input.path1.gene69	PV08_09990	41	106	92.36111111111111	3.2e-21


>>
25. NW_006912904
Source: Capronia epimyces CBS 606.96 aczIQ-supercont1.1, whole genome shotgun sequence
Type: NRPS-like,T1PKS
Number of proteins with BLAST hits to this cluster: 2
Cumulative BLAST score: 1258

Table of genes, locations, strands and annotations of subject cluster:

Table of Blast hits (query gene, subject gene, %identity, blast score, %coverage, e-value):
input.path1.gene68	A1O3_01335	33	1177	114.66604390471741	0.0
input.path1.gene69	A1O3_01336	40	81	56.94444444444444	6.5e-14


>>
26. NW_017971443
Source: Talaromyces atroroseus strain IBT 11181 chromosome Unknown scaffold_11, whole genome shotgun sequence
Type: NRPS-like,T1PKS
Number of proteins with BLAST hits to this cluster: 2
Cumulative BLAST score: 1208

Table of genes, locations, strands and annotations of subject cluster:

Table of Blast hits (query gene, subject gene, %identity, blast score, %coverage, e-value):
input.path1.gene68	UA08_07371	31	1129	117.79542269967305	0.0
input.path1.gene69	UA08_07372	35	79	91.66666666666666	3.2e-13


>>
27. NW_015971652
Source: Fonsecaea multimorphosa CBS 102226 unplaced genomic scaffold supercont1.18, whole genome shotgun sequence
Type: NRPS-like,T1PKS
Number of proteins with BLAST hits to this cluster: 2
Cumulative BLAST score: 1208

Table of genes, locations, strands and annotations of subject cluster:

Table of Blast hits (query gene, subject gene, %identity, blast score, %coverage, e-value):
input.path1.gene68	Z520_08261	32	1115	112.05044371788884	0.0
input.path1.gene69	Z520_08260	38	93	84.02777777777779	1.6e-17


>>
28. NC_036632
Source: Fusarium fujikuroi IMI 58289 draft genome, chromosome FFUJ_chr11
Type: T1PKS
Number of proteins with BLAST hits to this cluster: 2
Cumulative BLAST score: 1148

Table of genes, locations, strands and annotations of subject cluster:

Table of Blast hits (query gene, subject gene, %identity, blast score, %coverage, e-value):
input.path1.gene68	FFUJ_11199	31	1073	119.75712283979449	2.8e-311
input.path1.gene69	FFUJ_11198	34	75	86.11111111111111	4.6e-12


>>
29. NW_022474213
Source: Venustampulla echinocandica strain BP 5553 chromosome Unknown scaffold_8, whole genome shotgun sequence
Type: NRPS,T1PKS
Number of proteins with BLAST hits to this cluster: 3
Cumulative BLAST score: 2306

Table of genes, locations, strands and annotations of subject cluster:

Table of Blast hits (query gene, subject gene, %identity, blast score, %coverage, e-value):
input.path1.gene67	BP5553_08672	40	76	53.398058252427184	5.1e-12
input.path1.gene68	BP5553_08674	35	1391	117.14152265296592	0.0
input.path1.gene68	BP5553_08690	35	593	54.553946753853346	9.8e-167
input.path1.gene70	BP5553_08673	34	246	117.07317073170731	4.3e-63


>>
30. NW_013550604
Source: Rhinocladiella mackenziei CBS 650.93 unplaced genomic scaffold supercont1.1, whole genome shotgun sequence
Type: NRPS-like,T1PKS
Number of proteins with BLAST hits to this cluster: 3
Cumulative BLAST score: 1889

Table of genes, locations, strands and annotations of subject cluster:

Table of Blast hits (query gene, subject gene, %identity, blast score, %coverage, e-value):
input.path1.gene67	Z518_00506	39	80	59.22330097087378	2.7e-13
input.path1.gene67	Z518_00497	33	61	62.62135922330098	1.3e-07
input.path1.gene68	Z518_00509	41	1649	109.52825782344699	0.0
input.path1.gene69	Z518_00497	40	99	87.5	3.9e-19


>>
31. NW_008481799
Source: Cyphellophora europaea CBS 101466 unplaced genomic scaffold supercont1.11, whole genome shotgun sequence
Type: NRPS-like,T1PKS
Number of proteins with BLAST hits to this cluster: 3
Cumulative BLAST score: 1758

Table of genes, locations, strands and annotations of subject cluster:

Table of Blast hits (query gene, subject gene, %identity, blast score, %coverage, e-value):
input.path1.gene67	HMPREF1541_09830	35	77	62.62135922330098	2.3e-12
input.path1.gene68	HMPREF1541_09820	40	1586	110.32227930873424	0.0
input.path1.gene69	HMPREF1541_09824	38	95	87.5	4.3e-18


>>
32. NW_023336268
Source: Colletotrichum scovillei strain TJNH1 chromosome Unknown Contig00010, whole genome shotgun sequence
Type: T1PKS
Number of proteins with BLAST hits to this cluster: 3
Cumulative BLAST score: 1649

Table of genes, locations, strands and annotations of subject cluster:

Table of Blast hits (query gene, subject gene, %identity, blast score, %coverage, e-value):
input.path1.gene67	HER10_EVM0003178	38	71	51.94174757281553	1.3e-10
input.path1.gene68	HER10_EVM0005077	37	1396	110.55581503970106	0.0
input.path1.gene69	HER10_EVM0000089	38	95	86.80555555555556	4.3e-18
input.path1.gene69	HER10_EVM0003178	36	87	86.80555555555556	1.2e-15


>>
33. NW_007360991
Source: Glarea lozoyensis ATCC 20868 chromosome Unknown GLAREA17, whole genome shotgun sequence
Type: T1PKS
Number of proteins with BLAST hits to this cluster: 3
Cumulative BLAST score: 1648

Table of genes, locations, strands and annotations of subject cluster:

Table of Blast hits (query gene, subject gene, %identity, blast score, %coverage, e-value):
input.path1.gene67	GLAREA_11990	42	61	32.5242718446602	1.7e-07
input.path1.gene68	GLAREA_11992	35	1369	116.7211583372256	0.0
input.path1.gene70	GLAREA_11989	36	218	101.95121951219512	1.6e-54


>>
34. NW_023336282
Source: Aspergillus tubingensis WU-2223L DNA, scaffold_8, whole genome shotgun sequence
Type: NRPS-like,T1PKS
Number of proteins with BLAST hits to this cluster: 3
Cumulative BLAST score: 1328

Table of genes, locations, strands and annotations of subject cluster:

Table of Blast hits (query gene, subject gene, %identity, blast score, %coverage, e-value):
input.path1.gene67	AtWU_06731	40	76	53.883495145631066	3e-12
input.path1.gene68	AtWU_06733	34	1156	108.26716487622605	0.0
input.path1.gene69	AtWU_06731	38	96	84.02777777777779	1.9e-18


>>
35. NC_030960
Source: Colletotrichum higginsianum IMI 349063 chromosome 8, whole genome shotgun sequence
Type: T1PKS
Number of proteins with BLAST hits to this cluster: 3
Cumulative BLAST score: 1306

Table of genes, locations, strands and annotations of subject cluster:

Table of Blast hits (query gene, subject gene, %identity, blast score, %coverage, e-value):
input.path1.gene67	CH63R_11490	31	59	49.51456310679612	4.9e-07
input.path1.gene68	CH63R_11486	31	1124	121.2050443717889	0.0
input.path1.gene70	CH63R_11493	46	123	30.48780487804878	4.2e-26


>>
36. NZ_CP034352
Source: Streptomyces sp. W1SF4 plasmid p2, complete sequence
Type: NRPS,NRPS-like,T1PKS,T2PKS,butyrolactone,melanin
Number of proteins with BLAST hits to this cluster: 2
Cumulative BLAST score: 3115

Table of genes, locations, strands and annotations of subject cluster:
D1J60_RS36170	D1J60_RS36170	1038	1672	+	TetR_family_transcriptional_regulator
D1J60_RS36175	D1J60_RS36175	1727	1906	-	hypothetical_protein
D1J60_RS36435	D1J60_RS36435	136120	141220	-	acyltransferase_domain-containing_protein
D1J60_RS37620	D1J60_RS37620	183563	183817	-	ATP-binding_protein
D1J60_RS36485	D1J60_RS36485	184321	185095	-	AfsR/SARP_family_transcriptional_regulator
D1J60_RS36505	D1J60_RS36505	189346	192303	+	acyltransferase_domain-containing_protein
D1J60_RS36675	D1J60_RS36675	227432	229621	+	MMPL_family_transporter
D1J60_RS36725	D1J60_RS36725	243784	245379	-	acyl--CoA_ligase
D1J60_RS36990	D1J60_RS36990	298697	299017	-	TetR_family_transcriptional_regulator

Table of Blast hits (query gene, subject gene, %identity, blast score, %coverage, e-value):
input.path1.gene68	D1J60_RS36290	31	402	47.59458197104157	2.5e-109
input.path1.gene68	D1J60_RS36420	32	384	40.6352171882298	5.3e-104
input.path1.gene68	D1J60_RS36335	32	379	41.61606725829052	1.7e-102
input.path1.gene68	D1J60_RS36415	32	377	41.10228865016347	8.5e-102
input.path1.gene68	D1J60_RS36410	32	375	40.54180289584306	4.2e-101
input.path1.gene68	D1J60_RS36305	32	371	40.58851004203643	6.1e-100
input.path1.gene68	D1J60_RS36300	32	367	41.0555815039701	1.1e-98
input.path1.gene68	D1J60_RS36440	32	363	41.10228865016347	2.2e-97
input.path1.gene70	D1J60_RS36285	34	97	40.97560975609756	4.2e-18


>>
37. NZ_CP015163
Source: Amycolatopsis albispora strain WP1 chromosome, complete genome
Type: NRPS,NRPS-like,T1PKS,terpene
Number of proteins with BLAST hits to this cluster: 2
Cumulative BLAST score: 2757

Table of genes, locations, strands and annotations of subject cluster:
A4R43_RS39030	A4R43_RS39030	8345071	8352591	+	SDR_family_NAD(P)-dependent_oxidoreductase
A4R43_RS39035	A4R43_RS39035	8353480	8356767	+	SDR_family_NAD(P)-dependent_oxidoreductase
A4R43_RS39040	A4R43_RS39040	8357896	8362125	+	SDR_family_NAD(P)-dependent_oxidoreductase
A4R43_RS39045	A4R43_RS39045	8363884	8365188	+	hypothetical_protein
A4R43_RS39050	A4R43_RS39050	8365813	8366907	+	acyltransferase_domain-containing_protein
A4R43_RS39055	A4R43_RS39055	8367028	8368269	+	SDR_family_NAD(P)-dependent_oxidoreductase
A4R43_RS39060	A4R43_RS39060	8368276	8368779	+	hypothetical_protein
A4R43_RS39065	A4R43_RS39065	8368849	8371344	+	type_I_polyketide_synthase
A4R43_RS39070	A4R43_RS39070	8371813	8372652	+	SDR_family_NAD(P)-dependent_oxidoreductase
A4R43_RS39075	A4R43_RS39075	8372662	8373174	+	modular_polyketide_synthase
A4R43_RS39080	A4R43_RS39080	8373274	8373993	+	polyketide_synthase
A4R43_RS39085	A4R43_RS39085	8374315	8374653	+	hypothetical_protein
A4R43_RS39090	A4R43_RS39090	8374807	8375709	+	acyltransferase_domain-containing_protein
A4R43_RS39095	A4R43_RS39095	8376436	8377137	+	SDR_family_NAD(P)-dependent_oxidoreductase

Table of Blast hits (query gene, subject gene, %identity, blast score, %coverage, e-value):
input.path1.gene68	A4R43_RS39030	32	396	46.146660439047174	1.4e-107
input.path1.gene68	A4R43_RS38980	31	392	44.511910322279306	3.3e-106
input.path1.gene68	A4R43_RS39105	31	385	47.314339093881365	4.1e-104
input.path1.gene68	A4R43_RS38975	33	381	35.07706679121906	7.7e-103
input.path1.gene68	A4R43_RS38985	32	374	40.26156001868286	5.5e-101
input.path1.gene68	A4R43_RS39040	33	374	35.03035964502569	9.4e-101
input.path1.gene68	A4R43_RS39065	32	363	40.26156001868286	2.2e-97
input.path1.gene70	A4R43_RS38945	34	92	42.926829268292686	1.4e-16


>>
38. NZ_GG657752
Source: Streptomyces sp. C supercont1.3, whole genome shotgun sequence
Type: T1PKS,melanin
Number of proteins with BLAST hits to this cluster: 2
Cumulative BLAST score: 2686

Table of genes, locations, strands and annotations of subject cluster:
SSNG_RS37050	SSNG_RS37050	179341	181220	-	fatty_acyl-AMP_ligase
SSNG_RS40850	SSNG_RS40850	181397	182231	+	AfsR/SARP_family_transcriptional_regulator
SSNG_RS37090	SSNG_RS37090	190550	191476	+	polyketide_synthase
SSNG_RS40860	SSNG_RS40860	191839	193233	+	acyltransferase_domain-containing_protein
SSNG_RS40865	SSNG_RS40865	193666	194394	+	polyketide_synthase
SSNG_RS40870	SSNG_RS40870	194722	195141	+	hypothetical_protein
SSNG_RS40875	SSNG_RS40875	196063	197166	+	polyketide_synthase
SSNG_RS42825	SSNG_RS42825	224552	225076	+	hypothetical_protein
SSNG_RS37125	SSNG_RS37125	241789	243446	+	polyketide_synthase
SSNG_RS37135	SSNG_RS37135	246155	247028	+	polyketide_synthase
SSNG_RS40930	SSNG_RS40930	291589	292938	+	acyl--CoA_ligase
SSNG_RS40935	SSNG_RS40935	293506	296685	+	acyltransferase_domain-containing_protein
SSNG_RS40940	SSNG_RS40940	296932	300849	+	SDR_family_NAD(P)-dependent_oxidoreductase
SSNG_RS40945	SSNG_RS40945	301975	302829	+	SDR_family_NAD(P)-dependent_oxidoreductase
SSNG_RS42830	SSNG_RS42830	315637	316272	-	hypothetical_protein
SSNG_RS37275	SSNG_RS37275	329857	330916	+	beta-ketoacyl_synthase
SSNG_RS37280	SSNG_RS37280	331754	332761	+	acyltransferase_domain-containing_protein
SSNG_RS37345	SSNG_RS37345	353603	354040	-	beta-ketoacyl-ACP_synthase
SSNG_RS37350	SSNG_RS37350	354337	354840	-	beta-ketoacyl-ACP_synthase

Table of Blast hits (query gene, subject gene, %identity, blast score, %coverage, e-value):
input.path1.gene68	SSNG_RS37145	31	375	41.0555815039701	3.2e-101
input.path1.gene68	SSNG_RS41560	32	375	40.49509574964969	4.2e-101
input.path1.gene68	SSNG_RS37265	31	375	40.44838860345633	4.2e-101
input.path1.gene68	SSNG_RS40935	32	369	40.58851004203643	1.8e-99
input.path1.gene68	SSNG_RS37110	35	368	35.35730966837926	3.9e-99
input.path1.gene68	SSNG_RS40885	32	366	41.242410088743576	2.6e-98
input.path1.gene68	SSNG_RS37255	32	362	40.91546006539	2.8e-97
input.path1.gene70	SSNG_RS37305	34	96	40.97560975609756	5.5e-18


>>
39. NZ_JAAWWP010000017
Source: Streptomyces sp. LD120 Scaffold17, whole genome shotgun sequence
Type: T1PKS
Number of proteins with BLAST hits to this cluster: 2
Cumulative BLAST score: 1979

Table of genes, locations, strands and annotations of subject cluster:
HFV08_RS23905	HFV08_RS23905	20251	20691	-	hypothetical_protein
HFV08_RS23915	HFV08_RS23915	21370	22473	-	ATP-binding_protein
HFV08_RS24010	HFV08_RS24010	78299	80727	-	type_I_polyketide_synthase
HFV08_RS24015	HFV08_RS24015	80785	81453	-	SDR_family_NAD(P)-dependent_oxidoreductase
HFV08_RS24020	HFV08_RS24020	82102	85077	-	acyltransferase_domain-containing_protein
HFV08_RS24025	HFV08_RS24025	86017	87291	-	acyl--CoA_ligase

Table of Blast hits (query gene, subject gene, %identity, blast score, %coverage, e-value):
input.path1.gene68	HFV08_RS23965	32	391	45.25922466137319	5.7e-106
input.path1.gene68	HFV08_RS23960	31	384	48.52872489490892	5.3e-104
input.path1.gene68	HFV08_RS23970	33	384	40.728631480616535	5.3e-104
input.path1.gene68	HFV08_RS24020	32	369	40.96216721158337	2.3e-99
input.path1.gene68	HFV08_RS23975	33	355	37.83278841662774	3.5e-95
input.path1.gene70	HFV08_RS23955	32	96	45.1219512195122	5.5e-18


>>
40. NW_006912895
Source: Capronia coronata CBS 617.96 acBCG-supercont1.3, whole genome shotgun sequence
Type: T1PKS
Number of proteins with BLAST hits to this cluster: 2
Cumulative BLAST score: 1503

Table of genes, locations, strands and annotations of subject cluster:

Table of Blast hits (query gene, subject gene, %identity, blast score, %coverage, e-value):
input.path1.gene67	A1O1_04522	33	62	67.96116504854369	7.6e-08
input.path1.gene68	A1O1_04527	38	1441	110.13545072396076	0.0


>>
41. NW_003052498
Source: Uncinocarpus reesii 1704 scaffold_3 genomic scaffold, whole genome shotgun sequence
Type: T1PKS
Number of proteins with BLAST hits to this cluster: 2
Cumulative BLAST score: 1394

Table of genes, locations, strands and annotations of subject cluster:

Table of Blast hits (query gene, subject gene, %identity, blast score, %coverage, e-value):
input.path1.gene68	UREG_05998	43	925	57.26296123306865	8.4e-267
input.path1.gene68	UREG_05997	33	382	36.01120971508641	3.4e-103
input.path1.gene69	UREG_06000	40	87	81.25	9e-16


>>
42. NC_018210
Source: Zymoseptoria tritici IPO323 chromosome 9, whole genome shotgun sequence
Type: T1PKS
Number of proteins with BLAST hits to this cluster: 2
Cumulative BLAST score: 1379

Table of genes, locations, strands and annotations of subject cluster:

Table of Blast hits (query gene, subject gene, %identity, blast score, %coverage, e-value):
input.path1.gene68	MYCGRDRAFT_47832	32	1119	119.29005137786082	0.0
input.path1.gene70	MYCGRDRAFT_117758	34	260	117.80487804878048	2.9e-67


>>
43. NW_023336282
Source: Aspergillus tubingensis WU-2223L DNA, scaffold_8, whole genome shotgun sequence
Type: T1PKS
Number of proteins with BLAST hits to this cluster: 2
Cumulative BLAST score: 1338

Table of genes, locations, strands and annotations of subject cluster:

Table of Blast hits (query gene, subject gene, %identity, blast score, %coverage, e-value):
input.path1.gene68	AtWU_06660	33	1238	118.68285847734703	0.0
input.path1.gene69	AtWU_06655	39	100	87.5	1.8e-19


>>
44. NZ_PVTF01000015
Source: Umezawaea tangerina strain DSM 44720 Ga0180964_115, whole genome shotgun sequence
Type: T1PKS
Number of proteins with BLAST hits to this cluster: 2
Cumulative BLAST score: 1274

Table of genes, locations, strands and annotations of subject cluster:

Table of Blast hits (query gene, subject gene, %identity, blast score, %coverage, e-value):
input.path1.gene68	CLV43_RS39420	31	421	52.26529659037833	3.9e-115
input.path1.gene68	CLV43_RS39410	31	411	52.91919663708548	6.9e-112
input.path1.gene68	CLV43_RS48190	32	369	40.681924334423165	3e-99
input.path1.gene70	CLV43_RS39465	31	73	53.65853658536586	5e-11


>>
45. NW_014574615
Source: Penicillium digitatum Pd1 unplaced genomic scaffold Scaffold_10, whole genome shotgun sequence
Type: T1PKS
Number of proteins with BLAST hits to this cluster: 2
Cumulative BLAST score: 1236

Table of genes, locations, strands and annotations of subject cluster:

Table of Blast hits (query gene, subject gene, %identity, blast score, %coverage, e-value):
input.path1.gene68	PDIP_18550	32	1134	118.40261560018683	0.0
input.path1.gene70	PDIP_18570	31	102	50.48780487804878	1e-19


>>
46. CM000574
Source: Fusarium graminearum PH-1 chromosome 1, whole genome shotgun sequence
Type: NRPS,T1PKS
Number of proteins with BLAST hits to this cluster: 2
Cumulative BLAST score: 1233

Table of genes, locations, strands and annotations of subject cluster:

Table of Blast hits (query gene, subject gene, %identity, blast score, %coverage, e-value):
input.path1.gene68	FGSG_10548	32	1135	109.85520784680057	0.0
input.path1.gene69	FGSG_10545	37	98	87.5	5.1e-19


>>
47. NC_026474
Source: Fusarium graminearum PH-1 chromosome 1, whole genome shotgun sequence
Type: NRPS,T1PKS
Number of proteins with BLAST hits to this cluster: 2
Cumulative BLAST score: 1233

Table of genes, locations, strands and annotations of subject cluster:

Table of Blast hits (query gene, subject gene, %identity, blast score, %coverage, e-value):
input.path1.gene68	FGSG_10548	32	1135	109.85520784680057	0.0
input.path1.gene69	FGSG_10545	37	98	87.5	5.1e-19


>>
48. NC_035793
Source: Pochonia chlamydosporia 170 chromosome 4, whole genome shotgun sequence
Type: T1PKS
Number of proteins with BLAST hits to this cluster: 2
Cumulative BLAST score: 1224

Table of genes, locations, strands and annotations of subject cluster:

Table of Blast hits (query gene, subject gene, %identity, blast score, %coverage, e-value):
input.path1.gene68	VFPPC_08495	32	1076	108.54740775338627	0.0
input.path1.gene70	VFPPC_08497	37	148	53.65853658536586	1.6e-33


>>
49. NC_009380
Source: Salinispora tropica CNB-440, complete sequence
Type: NRPS,T1PKS,arylpolyene
Number of proteins with BLAST hits to this cluster: 2
Cumulative BLAST score: 1223

Table of genes, locations, strands and annotations of subject cluster:
STROP_RS24860	STROP_RS24860	3099064	3100299	+	IS701_family_transposase
STROP_RS13970	STROP_RS13970	3185638	3186186	-	hypothetical_protein
STROP_RS14275	STROP_RS14275	3261674	3261988	-	IS5/IS1182_family_transposase

Table of Blast hits (query gene, subject gene, %identity, blast score, %coverage, e-value):
input.path1.gene68	STROP_RS13965	31	393	48.248482017748714	1.1e-106
input.path1.gene68	STROP_RS13950	31	384	46.240074731433914	5.3e-104
input.path1.gene68	STROP_RS13960	33	378	40.49509574964969	6.5e-102
input.path1.gene71	STROP_RS14115	35	68	53.333333333333336	1e-09


>>
50. NZ_CP010519
Source: Streptomyces albus strain DSM 41398 chromosome, complete genome
Type: NRPS-like,T1PKS
Number of proteins with BLAST hits to this cluster: 2
Cumulative BLAST score: 1218

Table of genes, locations, strands and annotations of subject cluster:
SLNWT_RS04080	SLNWT_RS04080	1182186	1182626	-	hypothetical_protein
SLNWT_RS04090	SLNWT_RS04090	1183305	1184408	-	ATP-binding_protein

Table of Blast hits (query gene, subject gene, %identity, blast score, %coverage, e-value):
input.path1.gene68	SLNWT_RS04140	32	388	42.830453059318074	3.7e-105
input.path1.gene68	SLNWT_RS04145	33	380	40.728631480616535	1.3e-102
input.path1.gene68	SLNWT_RS33870	32	353	38.25315273236805	1.3e-94
input.path1.gene70	SLNWT_RS04130	32	97	51.463414634146346	3.2e-18
