## Supplementary Results for "A draft genome of the ascomycotal fungal species *Pseudopithomyces maydicus* (family *Didymosphaeriaceae*)": contig_15_c1.txt

ClusterBlast scores for contig_15

Table of genes, locations, strands and annotations of query cluster:
input.path1.gene358	1242961	1245992	+		
input.path1.gene359	1247557	1249435	+		
input.path1.gene360	1249856	1250747	-		
input.path1.gene361	1254245	1255114	+		
input.path1.gene362	1257334	1264006	+		
input.path1.gene363	1266913	1269650	+		
input.path1.gene364	1270669	1276298	-		
input.path1.gene365	1278576	1281682	-		


Significant hits: 
1. NW_007360987	Glarea lozoyensis ATCC 20868 chromosome Unknown GLAREA13, whole genome shotgun sequence
2. NC_035796	Pochonia chlamydosporia 170 chromosome 7, whole genome shotgun sequence
3. NZ_FRCX01000002	Duganella sacchari strain Sac-22, whole genome shotgun sequence
4. NZ_LT607752	Micromonospora rifamycinica strain DSM 44983 chromosome I
5. NZ_LT906483	Mycolicibacterium thermoresistibile strain NCTC10409 chromosome 1
6. NZ_SLYY01000015	Streptomyces sp. BK205 Ga0307705_115, whole genome shotgun sequence
7. NC_017030	Corallococcus coralloides DSM 2259, complete sequence
8. NZ_CP034669	Corallococcus coralloides strain B035 chromosome, complete genome
9. NW_001939246	Pyrenophora tritici-repentis Pt-1C-BFP supercont1.3, whole genome shotgun sequence


Details:

>>
1. NW_007360987
Source: Glarea lozoyensis ATCC 20868 chromosome Unknown GLAREA13, whole genome shotgun sequence
Type: NRPS,T1PKS,betalactone
Number of proteins with BLAST hits to this cluster: 2
Cumulative BLAST score: 1590

Table of genes, locations, strands and annotations of subject cluster:

Table of Blast hits (query gene, subject gene, %identity, blast score, %coverage, e-value):
input.path1.gene361	GLAREA_10041	31	73	77.12177121771218	4.3e-11
input.path1.gene362	GLAREA_10034	42	1517	120.09106431417189	0.0


>>
2. NC_035796
Source: Pochonia chlamydosporia 170 chromosome 7, whole genome shotgun sequence
Type: T1PKS
Number of proteins with BLAST hits to this cluster: 2
Cumulative BLAST score: 612

Table of genes, locations, strands and annotations of subject cluster:

Table of Blast hits (query gene, subject gene, %identity, blast score, %coverage, e-value):
input.path1.gene362	VFPPC_16558	37	278	30.278884462151396	5.7e-72
input.path1.gene363	VFPPC_09580	34	334	95.46783625730994	2.6e-89


>>
3. NZ_FRCX01000002
Source: Duganella sacchari strain Sac-22, whole genome shotgun sequence
Type: NRPS,hserlactone,transAT-PKS
Number of proteins with BLAST hits to this cluster: 2
Cumulative BLAST score: 426

Table of genes, locations, strands and annotations of subject cluster:

Table of Blast hits (query gene, subject gene, %identity, blast score, %coverage, e-value):
input.path1.gene361	BUA36_RS05220	31	99	85.60885608856088	5.6e-19
input.path1.gene362	BUA36_RS05370	43	167	11.952191235059761	1.1e-38
input.path1.gene362	BUA36_RS05365	33	160	19.97723392145703	2.3e-36


>>
4. NZ_LT607752
Source: Micromonospora rifamycinica strain DSM 44983 chromosome I
Type: NRPS,T1PKS,betalactone
Number of proteins with BLAST hits to this cluster: 2
Cumulative BLAST score: 302

Table of genes, locations, strands and annotations of subject cluster:

Table of Blast hits (query gene, subject gene, %identity, blast score, %coverage, e-value):
input.path1.gene361	GA0070623_RS18990	36	131	84.87084870848709	1e-28
input.path1.gene362	GA0070623_RS19000	32	171	20.20489470688674	7.6e-40


>>
5. NZ_LT906483
Source: Mycolicibacterium thermoresistibile strain NCTC10409 chromosome 1
Type: NRPS,T1PKS
Number of proteins with BLAST hits to this cluster: 2
Cumulative BLAST score: 300

Table of genes, locations, strands and annotations of subject cluster:
CKW28_RS10855	CKW28_RS10855	2326870	2327815	+	DUF4129_domain-containing_protein
CKW28_RS11070	CKW28_RS11070	2380820	2381266	+	NUDIX_domain-containing_protein

Table of Blast hits (query gene, subject gene, %identity, blast score, %coverage, e-value):
input.path1.gene361	CKW28_RS10910	36	126	92.25092250922509	5.6e-27
input.path1.gene362	CKW28_RS10985	45	174	12.066021627774616	8.9e-41


>>
6. NZ_SLYY01000015
Source: Streptomyces sp. BK205 Ga0307705_115, whole genome shotgun sequence
Type: NRPS,T1PKS,amglyccycl
Number of proteins with BLAST hits to this cluster: 2
Cumulative BLAST score: 293

Table of genes, locations, strands and annotations of subject cluster:
EV578_RS34965	EV578_RS34965	151897	152964	+	NAD(P)-dependent_alcohol_dehydrogenase
EV578_RS35245	EV578_RS35245	229887	230456	-	ATP-binding_protein

Table of Blast hits (query gene, subject gene, %identity, blast score, %coverage, e-value):
input.path1.gene361	EV578_RS35165	33	119	88.56088560885608	6.9e-25
input.path1.gene362	EV578_RS35200	31	174	27.831531018782012	1.2e-40


>>
7. NC_017030
Source: Corallococcus coralloides DSM 2259, complete sequence
Type: NRPS,T1PKS
Number of proteins with BLAST hits to this cluster: 2
Cumulative BLAST score: 274

Table of genes, locations, strands and annotations of subject cluster:

Table of Blast hits (query gene, subject gene, %identity, blast score, %coverage, e-value):
input.path1.gene361	COCOR_RS18015	35	116	81.54981549815497	5.8e-24
input.path1.gene362	COCOR_RS17995	40	158	12.17985202048947	5.1e-36


>>
8. NZ_CP034669
Source: Corallococcus coralloides strain B035 chromosome, complete genome
Type: NRPS,T1PKS
Number of proteins with BLAST hits to this cluster: 2
Cumulative BLAST score: 274

Table of genes, locations, strands and annotations of subject cluster:
EJ065_RS17545	EJ065_RS17545	4304977	4305375	+	AAA_family_ATPase

Table of Blast hits (query gene, subject gene, %identity, blast score, %coverage, e-value):
input.path1.gene361	EJ065_RS17600	35	116	81.54981549815497	5.8e-24
input.path1.gene362	EJ065_RS17580	40	158	12.17985202048947	6.6e-36


>>
9. NW_001939246
Source: Pyrenophora tritici-repentis Pt-1C-BFP supercont1.3, whole genome shotgun sequence
Type: T1PKS
Number of proteins with BLAST hits to this cluster: 2
Cumulative BLAST score: 1491

Table of genes, locations, strands and annotations of subject cluster:

Table of Blast hits (query gene, subject gene, %identity, blast score, %coverage, e-value):
input.path1.gene358	PTRG_03678	72	1397	100.10438413361169	0.0
input.path1.gene360	PTRG_03681	40	94	74.13793103448276	1.6e-17
