## Supplementary Results for "A draft genome of the ascomycotal fungal species *Pseudopithomyces maydicus* (family *Didymosphaeriaceae*)": contig_15_c2.txt

ClusterBlast scores for contig_15

Table of genes, locations, strands and annotations of query cluster:
input.path1.gene694	2563734	2564084	-		
input.path1.gene695	2565366	2568222	-		
input.path1.gene696	2568582	2569438	+		
input.path1.gene697	2570064	2570503	-		
input.path1.gene698	2574676	2575281	+		
input.path1.gene699	2575288	2579908	-		
input.path1.gene700	2580251	2583454	-		
input.path1.gene701	2588183	2590617	-		
input.path1.gene702	2591254	2591763	+		
input.path1.gene703	2591931	2592980	-		
input.path1.gene704	2593431	2594510	+		
input.path1.gene705	2594999	2596072	-		
input.path1.gene706	2597802	2602222	+		
input.path1.gene707	2602351	2603383	-		


Significant hits: 
1. NC_015312	Pseudonocardia dioxanivorans CB1190, complete sequence
2. NZ_CP019724	Streptomyces pactum strain ACT12 chromosome, complete genome
3. NZ_CP019724	Streptomyces pactum strain ACT12 chromosome, complete genome
4. NZ_FOAZ01000001	Streptacidiphilus jiangxiensis strain CGMCC 4.1857, whole genome shotgun sequence
5. NZ_CP010407	Streptomyces vietnamensis strain GIMV4.0001 chromosome, complete genome
6. NZ_CP034550	Saccharothrix syringae strain NRRL B-16468 chromosome, complete genome
7. NZ_FOPR01000001	Pseudomonas syringae strain BS3829, whole genome shotgun sequence
8. NZ_CP013743	Streptomyces sp. CdTB01 chromosome, complete genome
9. NC_009142	Saccharopolyspora erythraea NRRL 2338, complete genome
10. NZ_GL877878	Saccharopolyspora spinosa NRRL 18395 Scaffold000001, whole genome shotgun sequence
11. NZ_CP015867	Streptomyces parvulus strain 2297 plasmid pSPA1, complete sequence
12. NZ_CP013854	Pseudonocardia sp. HH130630-07, complete genome
13. NZ_LIWA01000004	Streptomyces sp. CB01883 scaffold4, whole genome shotgun sequence
14. NZ_CP040752	Streptomyces rectiverticillatus strain B15-08 chromosome, complete genome
15. NC_015953	Streptomyces sp. SirexAA-E, complete sequence
16. NZ_CP054938	Streptomyces harbinensis strain NA02264 chromosome, complete genome
17. NZ_JTIY01000001	Streptomyces sp. AcH 505 scaffold00001, whole genome shotgun sequence
18. NZ_WOFH01000003	Actinomadura sp. NEAU-AAG5 Scaffold3, whole genome shotgun sequence
19. NZ_CP024894	Amycolatopsis sp. AA4 chromosome, complete genome
20. NZ_CP010519	Streptomyces albus strain DSM 41398 chromosome, complete genome
21. NZ_SDLS01000004	Amycolatopsis sp. CFH S0740 Scaffold4, whole genome shotgun sequence
22. NZ_JAAWWP010000014	Streptomyces sp. LD120 Scaffold14, whole genome shotgun sequence
23. NC_015675	Mesorhizobium opportunistum WSM2075, complete sequence
24. NZ_CP020567	Kitasatospora aureofaciens strain DM-1 chromosome, complete genome
25. NZ_MPLE01000001	Streptomyces viridifaciens strain DSM 40239 plasmid unnamed scaffold1_size, whole genome shotgun sequence
26. NC_015957	Streptomyces violaceusniger Tu 4113, complete sequence
27. NZ_AP019620	Streptomyces antimycoticus strain NBRC 100767
28. NZ_CP003987	Streptomyces sp. 769 chromosome, complete genome
29. NZ_CM002285	Streptomyces roseochromogenus subsp. oscitans DS 12.976 chromosome, whole genome shotgun sequence
30. NZ_KB907214	Actinomadura atramentaria DSM 43919 strain SF2197 G339DRAFT_scaffold00007.7, whole genome shotgun sequence
31. NC_020504	Streptomyces davaonensis JCM 4913, complete genome
32. NZ_CP017157	Streptomyces lydicus strain 103 chromosome, complete genome
33. NZ_LQCI01000007	Amycolatopsis regifaucium strain GY080 contig_15, whole genome shotgun sequence
34. NZ_CP019779	Streptomyces sp. MOE7 chromosome, complete genome
35. NZ_BBYM01000032	Microtetraspora niveoalba strain NBRC 15239, whole genome shotgun sequence
36. NZ_CP054936	Streptomyces seoulensis strain A01 chromosome
37. NZ_MTBP01000002	Actinomadura sp. RB29 Contig_2, whole genome shotgun sequence
38. NZ_CDME01000002	Kibdelosporangium sp. MJ126-NF4, whole genome shotgun sequence
39. NZ_CP019457	Streptomyces lydicus strain GS93 isolate 23 chromosome, complete genome
40. NZ_KQ947992	Streptomyces curacoi strain DSM 40107 PRJNA299217_s009, whole genome shotgun sequence
41. NC_016109	Kitasatospora setae KM-6054, complete genome
42. NZ_BEWA01000010	Streptomyces rishiriensis strain NBRC 13407, whole genome shotgun sequence
43. NZ_CP022961	Plantactinospora sp. KBS50 chromosome, complete genome
44. NZ_LMFE01000008	Nocardia sp. Root136 contig_3, whole genome shotgun sequence
45. NZ_CP015588	Streptomyces alfalfae strain ACCC40021 chromosome, complete genome
46. NZ_CP015588	Streptomyces alfalfae strain ACCC40021 chromosome, complete genome
47. NZ_CP054690	Rhodococcus sp. W8901 chromosome, complete genome
48. NZ_KB913037	Amycolatopsis balhimycina FH 1894 strain DSM 44591 A3CEDRAFT_scaffold1.1, whole genome shotgun sequence
49. NZ_PVZV01000003	Actinocorallia populi strain A251 contig3, whole genome shotgun sequence
50. NZ_QQAT01000012	Rhodococcus sp. AG1013 Ga0222397_112, whole genome shotgun sequence


Details:

>>
1. NC_015312
Source: Pseudonocardia dioxanivorans CB1190, complete sequence
Type: NRPS
Number of proteins with BLAST hits to this cluster: 2
Cumulative BLAST score: 343

Table of genes, locations, strands and annotations of subject cluster:

Table of Blast hits (query gene, subject gene, %identity, blast score, %coverage, e-value):
input.path1.gene699	PSED_RS01275	33	207	65.68627450980392	5.1e-51
input.path1.gene700	PSED_RS01270	33	136	34.07325194228635	1.8e-29


>>
2. NZ_CP019724
Source: Streptomyces pactum strain ACT12 chromosome, complete genome
Type: NRPS,T2PKS,other
Number of proteins with BLAST hits to this cluster: 2
Cumulative BLAST score: 985

Table of genes, locations, strands and annotations of subject cluster:
B1H29_RS00390	B1H29_RS00390	92919	93521	-	alcohol_dehydrogenase_catalytic_domain-containing_protein
B1H29_RS38345	B1H29_RS38345	152049	152147	-	FAD/NAD(P)-binding_protein
B1H29_RS38350	B1H29_RS38350	222457	222567	-	polyhydroxybutyrate_depolymerase

Table of Blast hits (query gene, subject gene, %identity, blast score, %coverage, e-value):
input.path1.gene699	B1H29_RS00825	34	190	57.98319327731093	8.4e-46
input.path1.gene699	B1H29_RS00785	31	168	58.26330532212886	2e-39
input.path1.gene699	B1H29_RS00630	32	160	57.56302521008403	5.4e-37
input.path1.gene699	B1H29_RS00790	32	156	58.40336134453782	7.8e-36
input.path1.gene700	B1H29_RS00475	35	162	34.295227524972255	1.8e-37
input.path1.gene700	B1H29_RS00485	34	149	34.96115427302996	2.1e-33


>>
3. NZ_CP019724
Source: Streptomyces pactum strain ACT12 chromosome, complete genome
Type: NRPS,T2PKS,other
Number of proteins with BLAST hits to this cluster: 2
Cumulative BLAST score: 985

Table of genes, locations, strands and annotations of subject cluster:
B1H29_RS38720	B1H29_RS38720	8328227	8328337	+	polyhydroxybutyrate_depolymerase
B1H29_RS38725	B1H29_RS38725	8398647	8398745	+	FAD/NAD(P)-binding_protein
B1H29_RS36585	B1H29_RS36585	8457273	8457875	+	alcohol_dehydrogenase_catalytic_domain-containing_protein

Table of Blast hits (query gene, subject gene, %identity, blast score, %coverage, e-value):
input.path1.gene699	B1H29_RS36150	34	190	57.98319327731093	8.4e-46
input.path1.gene699	B1H29_RS36190	31	168	58.26330532212886	2e-39
input.path1.gene699	B1H29_RS36345	32	160	57.56302521008403	5.4e-37
input.path1.gene699	B1H29_RS36185	32	156	58.40336134453782	7.8e-36
input.path1.gene700	B1H29_RS36500	35	162	34.295227524972255	1.8e-37
input.path1.gene700	B1H29_RS36490	34	149	34.96115427302996	2.1e-33


>>
4. NZ_FOAZ01000001
Source: Streptacidiphilus jiangxiensis strain CGMCC 4.1857, whole genome shotgun sequence
Type: NRPS
Number of proteins with BLAST hits to this cluster: 2
Cumulative BLAST score: 694

Table of genes, locations, strands and annotations of subject cluster:
BMY61_RS02380	BMY61_RS02380	541536	543329	-	hypothetical_protein
BMY61_RS02440	BMY61_RS02440	559639	560388	-	RNA_polymerase_sigma_factor_SigF

Table of Blast hits (query gene, subject gene, %identity, blast score, %coverage, e-value):
input.path1.gene695	BMY61_RS02300	41	551	82.4188129899216	1.4e-154
input.path1.gene700	BMY61_RS02375	33	143	35.405105438401776	8.7e-32


>>
5. NZ_CP010407
Source: Streptomyces vietnamensis strain GIMV4.0001 chromosome, complete genome
Type: NRPS,T1PKS,T2PKS
Number of proteins with BLAST hits to this cluster: 2
Cumulative BLAST score: 670

Table of genes, locations, strands and annotations of subject cluster:
SVTN_RS28460	SVTN_RS28460	6334809	6335754	-	SDR_family_NAD(P)-dependent_oxidoreductase

Table of Blast hits (query gene, subject gene, %identity, blast score, %coverage, e-value):
input.path1.gene699	SVTN_RS28670	34	195	58.543417366946784	2.6e-47
input.path1.gene699	SVTN_RS28625	33	184	58.543417366946784	3.5e-44
input.path1.gene699	SVTN_RS28620	32	152	58.26330532212886	1.5e-34
input.path1.gene700	SVTN_RS28750	31	139	34.96115427302996	2.1e-30


>>
6. NZ_CP034550
Source: Saccharothrix syringae strain NRRL B-16468 chromosome, complete genome
Type: NRPS,T2PKS
Number of proteins with BLAST hits to this cluster: 2
Cumulative BLAST score: 651

Table of genes, locations, strands and annotations of subject cluster:
EKG83_RS32120	EKG83_RS32120	7609229	7610120	+	TetR_family_transcriptional_regulator
EKG83_RS32320	EKG83_RS32320	7693327	7693569	-	monooxygenase

Table of Blast hits (query gene, subject gene, %identity, blast score, %coverage, e-value):
input.path1.gene699	EKG83_RS32325	33	194	59.66386554621849	4.4e-47
input.path1.gene699	EKG83_RS32270	33	168	56.582633053221286	2e-39
input.path1.gene700	EKG83_RS32175	33	154	35.62708102108768	4.9e-35
input.path1.gene700	EKG83_RS32170	31	135	34.85016648168702	3.1e-29


>>
7. NZ_FOPR01000001
Source: Pseudomonas syringae strain BS3829, whole genome shotgun sequence
Type: NRPS
Number of proteins with BLAST hits to this cluster: 2
Cumulative BLAST score: 649

Table of genes, locations, strands and annotations of subject cluster:
BM038_RS27440	BM038_RS27440	835947	836744	+	hypothetical_protein
BM038_RS27445	BM038_RS27445	836871	837164	+	transposase
BM038_RS03800	BM038_RS03800	869186	871724	+	phosphoketolase_family_protein

Table of Blast hits (query gene, subject gene, %identity, blast score, %coverage, e-value):
input.path1.gene695	BM038_RS03805	36	508	93.72900335946248	1e-141
input.path1.gene700	BM038_RS03755	31	141	36.847946725860155	4.3e-31


>>
8. NZ_CP013743
Source: Streptomyces sp. CdTB01 chromosome, complete genome
Type: NRPS,T2PKS
Number of proteins with BLAST hits to this cluster: 2
Cumulative BLAST score: 640

Table of genes, locations, strands and annotations of subject cluster:
AS200_RS00835	AS200_RS00835	209122	209624	+	CBS_domain-containing_protein
AS200_RS00890	AS200_RS00890	220042	221054	-	cysteine_synthase_family_protein
AS200_RS00920	AS200_RS00920	242103	243089	-	hypothetical_protein
AS200_RS47470	AS200_RS47470	268363	268458	-	ArsR_family_transcriptional_regulator

Table of Blast hits (query gene, subject gene, %identity, blast score, %coverage, e-value):
input.path1.gene699	AS200_RS01235	32	165	58.543417366946784	1.7e-38
input.path1.gene699	AS200_RS01230	32	164	60.08403361344538	3.8e-38
input.path1.gene700	AS200_RS46060	32	156	34.07325194228635	1.3e-35
input.path1.gene700	AS200_RS00910	33	155	35.07214206437292	3.8e-35


>>
9. NC_009142
Source: Saccharopolyspora erythraea NRRL 2338, complete genome
Type: NRPS
Number of proteins with BLAST hits to this cluster: 2
Cumulative BLAST score: 638

Table of genes, locations, strands and annotations of subject cluster:

Table of Blast hits (query gene, subject gene, %identity, blast score, %coverage, e-value):
input.path1.gene695	SACE_RS06275	37	496	92.83314669652856	6.8e-138
input.path1.gene700	SACE_RS06390	32	142	34.73917869034406	1.9e-31


>>
10. NZ_GL877878
Source: Saccharopolyspora spinosa NRRL 18395 Scaffold000001, whole genome shotgun sequence
Type: NRPS
Number of proteins with BLAST hits to this cluster: 2
Cumulative BLAST score: 634

Table of genes, locations, strands and annotations of subject cluster:
SSPN_RS51050	SSPN_RS51050	3081219	3082330	-	hypothetical_protein
SSPN_RS43930	SSPN_RS43930	3088843	3090222	-	sodium/solute_symporter
SSPN_RS54740	SSPN_RS54740	3091032	3091295	-	hypothetical_protein
SSPN_RS0115470	SSPN_RS0115470	3091727	3092092	-	Na+/galactose_cotransporter
SSPN_RS43940	SSPN_RS43940	3094242	3095078	-	transposase
SSPN_RS0115490	SSPN_RS0115490	3095169	3095732	-	transposase
SSPN_RS43950	SSPN_RS43950	3100817	3102107	-	radical_SAM_protein
SSPN_RS51080	SSPN_RS51080	3131571	3131777	-	IS256_family_transposase
SSPN_RS51085	SSPN_RS51085	3132198	3132359	-	beta-ketoacyl-[acyl-carrier-protein]_synthase_family_protein
SSPN_RS53735	SSPN_RS53735	3134437	3134539	-	pyridoxamine_5'-phosphate_oxidase_family_protein

Table of Blast hits (query gene, subject gene, %identity, blast score, %coverage, e-value):
input.path1.gene695	SSPN_RS0115440	42	490	71.33258678611423	2.9e-136
input.path1.gene700	SSPN_RS0115335	32	144	35.294117647058826	3.9e-32


>>
11. NZ_CP015867
Source: Streptomyces parvulus strain 2297 plasmid pSPA1, complete sequence
Type: NRPS,NRPS-like,T1PKS,T2PKS,betalactone,butyrolactone,other
Number of proteins with BLAST hits to this cluster: 2
Cumulative BLAST score: 623

Table of genes, locations, strands and annotations of subject cluster:
Spa2297_RS35105	Spa2297_RS35105	6647	6874	-	DUF2267_domain-containing_protein
Spa2297_RS32790	Spa2297_RS32790	187594	189282	+	acyl-CoA_dehydrogenase
Spa2297_RS33070	Spa2297_RS33070	311892	312449	-	antibiotic_biosynthesis_protein
Spa2297_RS33245	Spa2297_RS33245	384716	385153	+	pentachlorophenol_monooxygenase

Table of Blast hits (query gene, subject gene, %identity, blast score, %coverage, e-value):
input.path1.gene699	Spa2297_RS33155	36	220	57.98319327731093	4.4e-55
input.path1.gene699	Spa2297_RS32645	34	197	56.44257703081232	5.2e-48
input.path1.gene699	Spa2297_RS33245	36	67	16.946778711484594	6.3e-09
input.path1.gene700	Spa2297_RS33130	32	139	35.516093229744726	1.6e-30


>>
12. NZ_CP013854
Source: Pseudonocardia sp. HH130630-07, complete genome
Type: NRPS
Number of proteins with BLAST hits to this cluster: 2
Cumulative BLAST score: 616

Table of genes, locations, strands and annotations of subject cluster:
AFB00_RS13730	AFB00_RS13730	2819788	2820782	+	ABC_transporter_substrate-binding_protein
AFB00_RS13840	AFB00_RS13840	2852804	2854284	+	amidohydrolase_family_protein

Table of Blast hits (query gene, subject gene, %identity, blast score, %coverage, e-value):
input.path1.gene695	AFB00_RS13785	36	475	88.3538633818589	1.2e-131
input.path1.gene700	AFB00_RS13765	33	141	34.85016648168702	3.3e-31


>>
13. NZ_LIWA01000004
Source: Streptomyces sp. CB01883 scaffold4, whole genome shotgun sequence
Type: NRPS
Number of proteins with BLAST hits to this cluster: 2
Cumulative BLAST score: 519

Table of genes, locations, strands and annotations of subject cluster:

Table of Blast hits (query gene, subject gene, %identity, blast score, %coverage, e-value):
input.path1.gene699	AMK32_RS20740	33	191	58.68347338935575	2.9e-46
input.path1.gene700	AMK32_RS20780	33	167	35.18312985571587	7.3e-39
input.path1.gene700	AMK32_RS20775	32	161	36.5149833518313	3.1e-37


>>
14. NZ_CP040752
Source: Streptomyces rectiverticillatus strain B15-08 chromosome, complete genome
Type: NRPS,T1PKS,T2PKS
Number of proteins with BLAST hits to this cluster: 2
Cumulative BLAST score: 496

Table of genes, locations, strands and annotations of subject cluster:
FGW37_RS04365	FGW37_RS04365	1097376	1098584	-	MFS_transporter

Table of Blast hits (query gene, subject gene, %identity, blast score, %coverage, e-value):
input.path1.gene699	FGW37_RS04180	35	186	57.14285714285714	7.1e-45
input.path1.gene699	FGW37_RS04215	31	162	56.86274509803921	1.4e-37
input.path1.gene700	FGW37_RS04355	33	148	36.958934517203105	3.5e-33


>>
15. NC_015953
Source: Streptomyces sp. SirexAA-E, complete sequence
Type: NRPS,T2PKS,butyrolactone
Number of proteins with BLAST hits to this cluster: 2
Cumulative BLAST score: 493

Table of genes, locations, strands and annotations of subject cluster:
SACTE_RS33580	SACTE_RS33580	7226642	7226988	-	xanthine_dehydrogenase_family_protein_molybdopterin-binding_subunit
SACTE_RS31595	SACTE_RS31595	7234031	7234228	+	hypothetical_protein

Table of Blast hits (query gene, subject gene, %identity, blast score, %coverage, e-value):
input.path1.gene699	SACTE_RS31500	33	173	58.123249299719895	1.1e-40
input.path1.gene699	SACTE_RS31465	32	172	57.422969187675065	1.4e-40
input.path1.gene700	SACTE_RS31310	32	148	37.40288568257492	2.7e-33


>>
16. NZ_CP054938
Source: Streptomyces harbinensis strain NA02264 chromosome, complete genome
Type: NRPS,T2PKS,oligosaccharide
Number of proteins with BLAST hits to this cluster: 2
Cumulative BLAST score: 489

Table of genes, locations, strands and annotations of subject cluster:

Table of Blast hits (query gene, subject gene, %identity, blast score, %coverage, e-value):
input.path1.gene699	HUT13_RS24880	34	188	57.56302521008403	3.2e-45
input.path1.gene699	HUT13_RS24915	31	155	57.84313725490197	1.7e-35
input.path1.gene700	HUT13_RS24825	32	146	34.73917869034406	1.7e-32


>>
17. NZ_JTIY01000001
Source: Streptomyces sp. AcH 505 scaffold00001, whole genome shotgun sequence
Type: NRPS,T2PKS,lanthipeptide
Number of proteins with BLAST hits to this cluster: 2
Cumulative BLAST score: 486

Table of genes, locations, strands and annotations of subject cluster:
HY68_RS26535	HY68_RS26535	5924956	5928129	-	type_2_lantipeptide_synthetase_LanM
HY68_RS26550	HY68_RS26550	5930311	5939154	-	amino_acid_adenylation_domain-containing_protein
HY68_RS26590	HY68_RS26590	5947634	5950264	-	amino_acid_adenylation_domain-containing_protein
HY68_RS26620	HY68_RS26620	5967673	5968632	+	chitinase
HY68_RS42250	HY68_RS42250	5969151	5969297	+	ABC_transporter_permease
HY68_RS26740	HY68_RS26740	5992073	5992711	-	5,10-methylene_tetrahydromethanopterin_reductase

Table of Blast hits (query gene, subject gene, %identity, blast score, %coverage, e-value):
input.path1.gene699	HY68_RS26670	34	181	58.26330532212886	3.9e-43
input.path1.gene699	HY68_RS26660	32	162	57.98319327731093	1.9e-37
input.path1.gene700	HY68_RS26550	32	143	35.84905660377358	8.7e-32


>>
18. NZ_WOFH01000003
Source: Actinomadura sp. NEAU-AAG5 Scaffold3, whole genome shotgun sequence
Type: NRPS,NRPS-like,T1PKS,blactam
Number of proteins with BLAST hits to this cluster: 2
Cumulative BLAST score: 479

Table of genes, locations, strands and annotations of subject cluster:
GNZ18_RS40765	GNZ18_RS40765	243139	244056	-	SDR_family_NAD(P)-dependent_oxidoreductase
GNZ18_RS40770	GNZ18_RS40770	244258	245247	-	polyketide_synthase_dehydratase_domain-containing_protein
GNZ18_RS40775	GNZ18_RS40775	246382	247647	-	polyketide_synthase
GNZ18_RS40780	GNZ18_RS40780	247690	248199	-	hypothetical_protein
GNZ18_RS40785	GNZ18_RS40785	248215	249246	-	SDR_family_NAD(P)-dependent_oxidoreductase
GNZ18_RS40790	GNZ18_RS40790	250207	251586	-	acyltransferase_domain-containing_protein
GNZ18_RS40795	GNZ18_RS40795	251701	252762	-	polyketide_synthase

Table of Blast hits (query gene, subject gene, %identity, blast score, %coverage, e-value):
input.path1.gene699	GNZ18_RS09120	32	184	58.123249299719895	4.6e-44
input.path1.gene700	GNZ18_RS09510	33	153	35.62708102108768	1.4e-34
input.path1.gene700	GNZ18_RS09515	31	142	34.295227524972255	1.9e-31


>>
19. NZ_CP024894
Source: Amycolatopsis sp. AA4 chromosome, complete genome
Type: NRPS,other,terpene
Number of proteins with BLAST hits to this cluster: 2
Cumulative BLAST score: 476

Table of genes, locations, strands and annotations of subject cluster:
CU254_RS15930	CU254_RS15930	3349441	3349671	-	GNAT_family_N-acetyltransferase
CU254_RS15950	CU254_RS15950	3353428	3354768	-	amino_acid_permease

Table of Blast hits (query gene, subject gene, %identity, blast score, %coverage, e-value):
input.path1.gene699	CU254_RS15990	32	173	56.86274509803921	8.1e-41
input.path1.gene700	CU254_RS16005	33	160	34.62819089900111	1.2e-36
input.path1.gene700	CU254_RS16090	32	143	33.85127635960045	1.1e-31


>>
20. NZ_CP010519
Source: Streptomyces albus strain DSM 41398 chromosome, complete genome
Type: NRPS
Number of proteins with BLAST hits to this cluster: 2
Cumulative BLAST score: 473

Table of genes, locations, strands and annotations of subject cluster:
SLNWT_RS00810	SLNWT_RS00810	187404	187706	-	transcriptional_regulator
SLNWT_RS34700	SLNWT_RS34700	205139	220561	-	amino_acid_adenylation_domain-containing_protein

Table of Blast hits (query gene, subject gene, %identity, blast score, %coverage, e-value):
input.path1.gene699	SLNWT_RS00920	34	193	62.04481792717087	9.9e-47
input.path1.gene700	SLNWT_RS00870	31	147	35.516093229744726	6e-33
input.path1.gene700	SLNWT_RS00875	31	133	35.405105438401776	9e-29


>>
21. NZ_SDLS01000004
Source: Amycolatopsis sp. CFH S0740 Scaffold4, whole genome shotgun sequence
Type: NRPS,terpene
Number of proteins with BLAST hits to this cluster: 2
Cumulative BLAST score: 471

Table of genes, locations, strands and annotations of subject cluster:
EIZ45_RS09465	EIZ45_RS09465	173754	173888	-	universal_stress_protein
EIZ45_RS46040	EIZ45_RS46040	192995	194128	-	non-ribosomal_peptide_synthetase
EIZ45_RS46045	EIZ45_RS46045	194231	195820	-	amino_acid_adenylation_domain-containing_protein

Table of Blast hits (query gene, subject gene, %identity, blast score, %coverage, e-value):
input.path1.gene699	EIZ45_RS09645	32	171	57.002801120448176	4e-40
input.path1.gene700	EIZ45_RS09630	33	159	34.85016648168702	1.5e-36
input.path1.gene700	EIZ45_RS09550	31	141	33.85127635960045	4.3e-31


>>
22. NZ_JAAWWP010000014
Source: Streptomyces sp. LD120 Scaffold14, whole genome shotgun sequence
Type: NRPS
Number of proteins with BLAST hits to this cluster: 2
Cumulative BLAST score: 461

Table of genes, locations, strands and annotations of subject cluster:

Table of Blast hits (query gene, subject gene, %identity, blast score, %coverage, e-value):
input.path1.gene699	HFV08_RS21760	33	174	60.22408963585434	4.7e-41
input.path1.gene700	HFV08_RS21710	31	151	35.516093229744726	5.4e-34
input.path1.gene700	HFV08_RS21715	31	136	35.405105438401776	1.4e-29


>>
23. NC_015675
Source: Mesorhizobium opportunistum WSM2075, complete sequence
Type: NRPS-like
Number of proteins with BLAST hits to this cluster: 2
Cumulative BLAST score: 455

Table of genes, locations, strands and annotations of subject cluster:

Table of Blast hits (query gene, subject gene, %identity, blast score, %coverage, e-value):
input.path1.gene699	MESOP_RS15940	44	319	57.422969187675065	9.1e-85
input.path1.gene700	MESOP_RS16030	31	136	33.85127635960045	1.8e-29


>>
24. NZ_CP020567
Source: Kitasatospora aureofaciens strain DM-1 chromosome, complete genome
Type: NRPS,NRPS-like,T1PKS,T3PKS
Number of proteins with BLAST hits to this cluster: 2
Cumulative BLAST score: 446

Table of genes, locations, strands and annotations of subject cluster:
B6264_RS24935	B6264_RS24935	5582127	5582882	-	zinc-binding_dehydrogenase
B6264_RS24940	B6264_RS24940	5582888	5583408	+	transposase
B6264_RS24980	B6264_RS24980	5590437	5591219	-	IS982_family_transposase
B6264_RS24985	B6264_RS24985	5591243	5591518	-	transposase
B6264_RS24990	B6264_RS24990	5591624	5591752	+	IS110_family_transposase
B6264_RS24995	B6264_RS24995	5591774	5592477	-	transposase
B6264_RS25010	B6264_RS25010	5595776	5595985	+	helix-turn-helix_domain-containing_protein
B6264_RS33345	B6264_RS33345	5754049	5757381	+	SDR_family_NAD(P)-dependent_oxidoreductase
B6264_RS33350	B6264_RS33350	5757430	5757903	+	hypothetical_protein

Table of Blast hits (query gene, subject gene, %identity, blast score, %coverage, e-value):
input.path1.gene699	B6264_RS25195	31	142	50.700280112044815	2e-31
input.path1.gene700	B6264_RS25055	33	155	34.85016648168702	2.9e-35
input.path1.gene700	B6264_RS25060	32	149	34.73917869034406	2.1e-33


>>
25. NZ_MPLE01000001
Source: Streptomyces viridifaciens strain DSM 40239 plasmid unnamed scaffold1_size, whole genome shotgun sequence
Type: NRPS,T1PKS,T3PKS
Number of proteins with BLAST hits to this cluster: 2
Cumulative BLAST score: 443

Table of genes, locations, strands and annotations of subject cluster:
BOQ63_RS04585	BOQ63_RS04585	1011690	1012466	+	IS110_family_transposase
BOQ63_RS04635	BOQ63_RS04635	1023140	1023349	+	helix-turn-helix_domain-containing_protein
BOQ63_RS46480	BOQ63_RS46480	1078587	1078790	+	SDR_family_NAD(P)-dependent_oxidoreductase
BOQ63_RS46485	BOQ63_RS46485	1078834	1078923	+	SMI1/KNR4_family_protein
BOQ63_RS44640	BOQ63_RS44640	1078969	1079217	-	SMI1/KNR4_family_protein
BOQ63_RS44655	BOQ63_RS44655	1140809	1141462	-	KR_domain-containing_protein
BOQ63_RS44660	BOQ63_RS44660	1143285	1152354	+	SDR_family_NAD(P)-dependent_oxidoreductase
BOQ63_RS46525	BOQ63_RS46525	1190198	1190410	+	long-chain_fatty_acid--CoA_ligase
BOQ63_RS05135	BOQ63_RS05135	1191047	1193159	-	aminodeoxychorismate_synthase_component_I
BOQ63_RS05165	BOQ63_RS05165	1196861	1198255	-	glycosyltransferase_family_1_protein

Table of Blast hits (query gene, subject gene, %identity, blast score, %coverage, e-value):
input.path1.gene699	BOQ63_RS04830	31	141	50.700280112044815	4.5e-31
input.path1.gene700	BOQ63_RS04680	32	153	34.85016648168702	1.1e-34
input.path1.gene700	BOQ63_RS04685	32	149	34.73917869034406	2.1e-33


>>
26. NC_015957
Source: Streptomyces violaceusniger Tu 4113, complete sequence
Type: NRPS
Number of proteins with BLAST hits to this cluster: 2
Cumulative BLAST score: 425

Table of genes, locations, strands and annotations of subject cluster:
STRVI_RS47800	STRVI_RS47800	3233945	3234928	-	hypothetical_protein
STRVI_RS14130	STRVI_RS14130	3283110	3283644	-	TetR/AcrR_family_transcriptional_regulator
STRVI_RS14170	STRVI_RS14170	3294053	3294460	-	IS5_family_transposase
STRVI_RS47805	STRVI_RS47805	3294459	3294761	+	transposase
STRVI_RS47810	STRVI_RS47810	3294772	3295647	+	hypothetical_protein

Table of Blast hits (query gene, subject gene, %identity, blast score, %coverage, e-value):
input.path1.gene699	STRVI_RS14045	39	266	58.96358543417367	7e-69
input.path1.gene700	STRVI_RS46300	33	159	36.736958934517205	1.5e-36


>>
27. NZ_AP019620
Source: Streptomyces antimycoticus strain NBRC 100767
Type: NRPS
Number of proteins with BLAST hits to this cluster: 2
Cumulative BLAST score: 402

Table of genes, locations, strands and annotations of subject cluster:
GYK45_RS09995	GYK45_RS09995	2508708	2509732	+	hypothetical_protein
GYK45_RS10030	GYK45_RS10030	2513850	2515432	+	FAD-dependent_monooxygenase
GYK45_RS10055	GYK45_RS10055	2521637	2522118	+	ABC_transporter_permease

Table of Blast hits (query gene, subject gene, %identity, blast score, %coverage, e-value):
input.path1.gene699	GYK45_RS10030	40	242	53.641456582633054	1.1e-61
input.path1.gene700	GYK45_RS09970	33	160	36.736958934517205	1.2e-36


>>
28. NZ_CP003987
Source: Streptomyces sp. 769 chromosome, complete genome
Type: NRPS,betalactone
Number of proteins with BLAST hits to this cluster: 2
Cumulative BLAST score: 384

Table of genes, locations, strands and annotations of subject cluster:
GZL_RS33540	GZL_RS33540	7839846	7840303	-	ketosteroid_isomerase
GZL_RS47965	GZL_RS47965	7845830	7845901	-	VOC_family_protein
GZL_RS33615	GZL_RS33615	7887256	7889829	+	non-ribosomal_peptide_synthetase
GZL_RS46695	GZL_RS46695	7898694	7898931	+	IS5/IS1182_family_transposase

Table of Blast hits (query gene, subject gene, %identity, blast score, %coverage, e-value):
input.path1.gene699	GZL_RS33565	35	220	59.103641456582636	4.4e-55
input.path1.gene700	GZL_RS33600	32	164	35.516093229744726	6.2e-38


>>
29. NZ_CM002285
Source: Streptomyces roseochromogenus subsp. oscitans DS 12.976 chromosome, whole genome shotgun sequence
Type: NRPS
Number of proteins with BLAST hits to this cluster: 2
Cumulative BLAST score: 374

Table of genes, locations, strands and annotations of subject cluster:
M878_RS65690	M878_RS65690	3697765	3698788	-	DNA-directed_RNA_polymerase_subunit_alpha
M878_RS48000	M878_RS48000	3763028	3764095	+	alpha-hydroxy-acid_oxidizing_protein

Table of Blast hits (query gene, subject gene, %identity, blast score, %coverage, e-value):
input.path1.gene696	M878_RS65685	42	76	42.16417910447761	6.6e-12
input.path1.gene700	M878_RS65910	33	151	35.07214206437292	5.4e-34
input.path1.gene700	M878_RS65915	33	147	34.85016648168702	7.8e-33


>>
30. NZ_KB907214
Source: Actinomadura atramentaria DSM 43919 strain SF2197 G339DRAFT_scaffold00007.7, whole genome shotgun sequence
Type: NRPS
Number of proteins with BLAST hits to this cluster: 2
Cumulative BLAST score: 371

Table of genes, locations, strands and annotations of subject cluster:
G339_RS39480	G339_RS39480	253731	254660	-	hypothetical_protein
G339_RS39485	G339_RS39485	254973	257006	-	amino_acid_adenylation_domain-containing_protein
G339_RS39490	G339_RS39490	258000	259559	-	amino_acid_adenylation_domain-containing_protein
G339_RS39495	G339_RS39495	260835	261983	-	non-ribosomal_peptide_synthetase
G339_RS39500	G339_RS39500	262059	264047	-	amino_acid_adenylation_domain-containing_protein
G339_RS39505	G339_RS39505	264102	265085	-	AMP-binding_protein
G339_RS38970	G339_RS38970	290341	290463	-	MFS_transporter_permease

Table of Blast hits (query gene, subject gene, %identity, blast score, %coverage, e-value):
input.path1.gene696	G339_RS0118560	43	82	41.7910447761194	9.2e-14
input.path1.gene700	G339_RS34350	35	150	35.07214206437292	7.1e-34
input.path1.gene700	G339_RS39490	34	139	35.405105438401776	1.6e-30


>>
31. NC_020504
Source: Streptomyces davaonensis JCM 4913, complete genome
Type: T1PKS,T2PKS
Number of proteins with BLAST hits to this cluster: 2
Cumulative BLAST score: 371

Table of genes, locations, strands and annotations of subject cluster:
BN159_RS12280	BN159_RS12280	2840910	2841297	+	hypothetical_protein
BN159_RS43820	BN159_RS43820	2841294	2841529	+	hypothetical_protein
BN159_RS43825	BN159_RS43825	2842649	2844012	+	hypothetical_protein
BN159_RS12330	BN159_RS12330	2854765	2855409	-	sugar_phosphate_isomerase

Table of Blast hits (query gene, subject gene, %identity, blast score, %coverage, e-value):
input.path1.gene699	BN159_RS12065	34	184	52.10084033613446	4.6e-44
input.path1.gene700	BN159_RS42825	31	187	46.392896781354054	6.8e-45


>>
32. NZ_CP017157
Source: Streptomyces lydicus strain 103 chromosome, complete genome
Type: NRPS,terpene
Number of proteins with BLAST hits to this cluster: 2
Cumulative BLAST score: 369

Table of genes, locations, strands and annotations of subject cluster:
SL103_RS37545	SL103_RS37545	4543353	4543634	+	HIT_domain-containing_protein
SL103_RS36945	SL103_RS36945	4547708	4548094	+	hypothetical_protein

Table of Blast hits (query gene, subject gene, %identity, blast score, %coverage, e-value):
input.path1.gene699	SL103_RS19710	36	223	59.2436974789916	5.2e-56
input.path1.gene700	SL103_RS19635	31	146	36.40399556048835	1e-32


>>
33. NZ_LQCI01000007
Source: Amycolatopsis regifaucium strain GY080 contig_15, whole genome shotgun sequence
Type: NRPS
Number of proteins with BLAST hits to this cluster: 2
Cumulative BLAST score: 367

Table of genes, locations, strands and annotations of subject cluster:
AVL48_RS38930	AVL48_RS38930	36815	36967	-	epoxide_hydrolase
AVL48_RS09110	AVL48_RS09110	89675	89827	+	NADP-dependent_oxidoreductase
AVL48_RS38155	AVL48_RS38155	98311	98444	+	hypothetical_protein

Table of Blast hits (query gene, subject gene, %identity, blast score, %coverage, e-value):
input.path1.gene699	AVL48_RS09050	34	207	58.96358543417367	6.6e-51
input.path1.gene700	AVL48_RS09085	34	160	35.294117647058826	9e-37


>>
34. NZ_CP019779
Source: Streptomyces sp. MOE7 chromosome, complete genome
Type: NRPS,terpene
Number of proteins with BLAST hits to this cluster: 2
Cumulative BLAST score: 365

Table of genes, locations, strands and annotations of subject cluster:
STRMOE7_RS35890	STRMOE7_RS35890	8179775	8180161	-	GNAT_family_N-acetyltransferase
STRMOE7_RS37555	STRMOE7_RS37555	8191032	8191550	-	hypothetical_protein
STRMOE7_RS35995	STRMOE7_RS35995	8205523	8206776	-	MFS_transporter

Table of Blast hits (query gene, subject gene, %identity, blast score, %coverage, e-value):
input.path1.gene699	STRMOE7_RS35900	36	218	58.40336134453782	2.2e-54
input.path1.gene700	STRMOE7_RS35975	32	147	35.294117647058826	6e-33


>>
35. NZ_BBYM01000032
Source: Microtetraspora niveoalba strain NBRC 15239, whole genome shotgun sequence
Type: NRPS
Number of proteins with BLAST hits to this cluster: 2
Cumulative BLAST score: 364

Table of genes, locations, strands and annotations of subject cluster:

Table of Blast hits (query gene, subject gene, %identity, blast score, %coverage, e-value):
input.path1.gene699	AWQ70_RS28295	36	214	59.103641456582636	4.1e-53
input.path1.gene700	AWQ70_RS28320	32	150	35.405105438401776	9.3e-34


>>
36. NZ_CP054936
Source: Streptomyces seoulensis strain A01 chromosome
Type: NRPS
Number of proteins with BLAST hits to this cluster: 2
Cumulative BLAST score: 363

Table of genes, locations, strands and annotations of subject cluster:
HUT11_RS01070	HUT11_RS01070	300248	301929	-	murein_biosynthesis_integral_membrane_protein_MurJ

Table of Blast hits (query gene, subject gene, %identity, blast score, %coverage, e-value):
input.path1.gene699	HUT11_RS01015	34	183	58.123249299719895	6e-44
input.path1.gene700	HUT11_RS01000	36	180	34.85016648168702	8.4e-43


>>
37. NZ_MTBP01000002
Source: Actinomadura sp. RB29 Contig_2, whole genome shotgun sequence
Type: NRPS
Number of proteins with BLAST hits to this cluster: 2
Cumulative BLAST score: 362

Table of genes, locations, strands and annotations of subject cluster:

Table of Blast hits (query gene, subject gene, %identity, blast score, %coverage, e-value):
input.path1.gene696	BTM25_RS20035	41	80	46.26865671641791	2.7e-13
input.path1.gene700	BTM25_RS20090	32	151	34.96115427302996	4.2e-34
input.path1.gene700	BTM25_RS20095	32	131	35.516093229744726	5.8e-28


>>
38. NZ_CDME01000002
Source: Kibdelosporangium sp. MJ126-NF4, whole genome shotgun sequence
Type: NRPS,T1PKS
Number of proteins with BLAST hits to this cluster: 2
Cumulative BLAST score: 360

Table of genes, locations, strands and annotations of subject cluster:
BN1199_RS54435	BN1199_RS54435	1892029	1892228	-	hypothetical_protein

Table of Blast hits (query gene, subject gene, %identity, blast score, %coverage, e-value):
input.path1.gene699	BN1199_RS08335	37	217	57.422969187675065	3.7e-54
input.path1.gene700	BN1199_RS08380	31	143	34.96115427302996	1.5e-31


>>
39. NZ_CP019457
Source: Streptomyces lydicus strain GS93 isolate 23 chromosome, complete genome
Type: NRPS,terpene
Number of proteins with BLAST hits to this cluster: 2
Cumulative BLAST score: 360

Table of genes, locations, strands and annotations of subject cluster:
BXT88_RS37665	BXT88_RS37665	8041815	8042063	-	HIT_domain-containing_protein

Table of Blast hits (query gene, subject gene, %identity, blast score, %coverage, e-value):
input.path1.gene699	BXT88_RS35580	35	212	59.103641456582636	1.6e-52
input.path1.gene700	BXT88_RS35630	32	148	35.294117647058826	4.6e-33


>>
40. NZ_KQ947992
Source: Streptomyces curacoi strain DSM 40107 PRJNA299217_s009, whole genome shotgun sequence
Type: NRPS
Number of proteins with BLAST hits to this cluster: 2
Cumulative BLAST score: 355

Table of genes, locations, strands and annotations of subject cluster:
AQI70_RS38785	AQI70_RS38785	253701	253790	-	hypothetical_protein
AQI70_RS29290	AQI70_RS29290	253803	254539	+	extradiol_ring-cleavage_dioxygenase

Table of Blast hits (query gene, subject gene, %identity, blast score, %coverage, e-value):
input.path1.gene699	AQI70_RS29305	33	202	59.103641456582636	1.6e-49
input.path1.gene700	AQI70_RS29255	33	153	34.406215316315205	1.1e-34


>>
41. NC_016109
Source: Kitasatospora setae KM-6054, complete genome
Type: NRPS,butyrolactone
Number of proteins with BLAST hits to this cluster: 2
Cumulative BLAST score: 348

Table of genes, locations, strands and annotations of subject cluster:
KSE_RS39405	KSE_RS39405	6565204	6565765	+	hypothetical_protein

Table of Blast hits (query gene, subject gene, %identity, blast score, %coverage, e-value):
input.path1.gene699	KSE_RS28895	36	210	57.14285714285714	4.6e-52
input.path1.gene700	KSE_RS28785	32	138	34.184239733629305	3.6e-30


>>
42. NZ_BEWA01000010
Source: Streptomyces rishiriensis strain NBRC 13407, whole genome shotgun sequence
Type: NRPS,PKS-like,T2PKS,oligosaccharide
Number of proteins with BLAST hits to this cluster: 2
Cumulative BLAST score: 345

Table of genes, locations, strands and annotations of subject cluster:
Sri02f_RS18400	Sri02f_RS18400	181920	182087	+	alpha-L-arabinofuranosidase
Sri02f_RS18410	Sri02f_RS18410	182567	182866	-	NUDIX_domain-containing_protein
Sri02f_RS18480	Sri02f_RS18480	192003	192317	+	transposase

Table of Blast hits (query gene, subject gene, %identity, blast score, %coverage, e-value):
input.path1.gene699	Sri02f_RS18065	37	205	58.26330532212886	1.5e-50
input.path1.gene700	Sri02f_RS18245	31	140	37.06992230854606	7.3e-31


>>
43. NZ_CP022961
Source: Plantactinospora sp. KBS50 chromosome, complete genome
Type: NRPS,T1PKS,bacteriocin,blactam,transAT-PKS
Number of proteins with BLAST hits to this cluster: 2
Cumulative BLAST score: 344

Table of genes, locations, strands and annotations of subject cluster:
CIK06_RS17715	CIK06_RS17715	4075398	4076533	+	saccharopine_dehydrogenase
CIK06_RS17785	CIK06_RS17785	4094336	4095480	-	acyl-CoA_dehydrogenase
CIK06_RS17890	CIK06_RS17890	4117280	4118398	+	hypothetical_protein
CIK06_RS29500	CIK06_RS29500	4171593	4171826	+	KR_domain-containing_protein
CIK06_RS29510	CIK06_RS29510	4175027	4175896	+	beta-ketoacyl_synthase
CIK06_RS29515	CIK06_RS29515	4176350	4176997	+	SDR_family_NAD(P)-dependent_oxidoreductase
CIK06_RS29520	CIK06_RS29520	4177010	4179919	+	acyltransferase_domain-containing_protein
CIK06_RS29525	CIK06_RS29525	4180016	4180960	+	polyketide_synthase
CIK06_RS29530	CIK06_RS29530	4181129	4182655	+	SDR_family_NAD(P)-dependent_oxidoreductase
CIK06_RS18045	CIK06_RS18045	4182976	4185877	+	acyltransferase_domain-containing_protein
CIK06_RS18055	CIK06_RS18055	4187388	4188179	+	zinc-binding_dehydrogenase
CIK06_RS18070	CIK06_RS18070	4194942	4201509	+	SDR_family_NAD(P)-dependent_oxidoreductase
CIK06_RS29540	CIK06_RS29540	4219649	4219849	+	AAA_family_ATPase
CIK06_RS18270	CIK06_RS18270	4249541	4250242	+	AMP-binding_protein
CIK06_RS18450	CIK06_RS18450	4299117	4301188	+	dipeptide_ABC_transporter_ATP-binding_protein

Table of Blast hits (query gene, subject gene, %identity, blast score, %coverage, e-value):
input.path1.gene699	CIK06_RS18220	31	191	69.88795518207283	3.8e-46
input.path1.gene700	CIK06_RS18405	31	153	37.29189789123196	1.4e-34


>>
44. NZ_LMFE01000008
Source: Nocardia sp. Root136 contig_3, whole genome shotgun sequence
Type: NRPS
Number of proteins with BLAST hits to this cluster: 2
Cumulative BLAST score: 336

Table of genes, locations, strands and annotations of subject cluster:
ASD42_RS33945	ASD42_RS33945	223397	223867	-	hypothetical_protein

Table of Blast hits (query gene, subject gene, %identity, blast score, %coverage, e-value):
input.path1.gene699	ASD42_RS22265	33	191	59.38375350140056	3.8e-46
input.path1.gene700	ASD42_RS22310	31	145	34.517203107658155	2.3e-32


>>
45. NZ_CP015588
Source: Streptomyces alfalfae strain ACCC40021 chromosome, complete genome
Type: NRPS,PKS-like,T1PKS,butyrolactone
Number of proteins with BLAST hits to this cluster: 2
Cumulative BLAST score: 334

Table of genes, locations, strands and annotations of subject cluster:
A7J05_RS00625	A7J05_RS00625	152239	153054	+	ATP-binding_protein

Table of Blast hits (query gene, subject gene, %identity, blast score, %coverage, e-value):
input.path1.gene699	A7J05_RS00580	31	154	58.40336134453782	3.9e-35
input.path1.gene700	A7J05_RS00380	35	180	36.18201997780244	6.4e-43


>>
46. NZ_CP015588
Source: Streptomyces alfalfae strain ACCC40021 chromosome, complete genome
Type: NRPS,PKS-like,T1PKS,butyrolactone
Number of proteins with BLAST hits to this cluster: 2
Cumulative BLAST score: 334

Table of genes, locations, strands and annotations of subject cluster:
A7J05_RS37265	A7J05_RS37265	8518893	8524543	-	KR_domain-containing_protein

Table of Blast hits (query gene, subject gene, %identity, blast score, %coverage, e-value):
input.path1.gene699	A7J05_RS37095	31	154	58.40336134453782	3.9e-35
input.path1.gene700	A7J05_RS37295	35	180	36.18201997780244	6.4e-43


>>
47. NZ_CP054690
Source: Rhodococcus sp. W8901 chromosome, complete genome
Type: NRPS
Number of proteins with BLAST hits to this cluster: 2
Cumulative BLAST score: 327

Table of genes, locations, strands and annotations of subject cluster:
HUN07_RS13565	HUN07_RS13565	2863628	2863873	-	lipase
HUN07_RS13585	HUN07_RS13585	2868093	2869514	-	rifampin_monooxygenase

Table of Blast hits (query gene, subject gene, %identity, blast score, %coverage, e-value):
input.path1.gene699	HUN07_RS13585	34	193	59.38375350140056	5.8e-47
input.path1.gene700	HUN07_RS13630	31	134	36.625971143174255	6.9e-29


>>
48. NZ_KB913037
Source: Amycolatopsis balhimycina FH 1894 strain DSM 44591 A3CEDRAFT_scaffold1.1, whole genome shotgun sequence
Type: NRPS,T1PKS,T2PKS,oligosaccharide
Number of proteins with BLAST hits to this cluster: 2
Cumulative BLAST score: 325

Table of genes, locations, strands and annotations of subject cluster:
A3CE_RS56255	A3CE_RS56255	1925307	1925426	+	AAA_family_ATPase
A3CE_RS56275	A3CE_RS56275	2006458	2006550	+	IS5/IS1182_family_transposase

Table of Blast hits (query gene, subject gene, %identity, blast score, %coverage, e-value):
input.path1.gene699	A3CE_RS50455	32	179	61.20448179271709	1.5e-42
input.path1.gene700	A3CE_RS0108150	33	146	36.847946725860155	1.7e-32


>>
49. NZ_PVZV01000003
Source: Actinocorallia populi strain A251 contig3, whole genome shotgun sequence
Type: NRPS
Number of proteins with BLAST hits to this cluster: 2
Cumulative BLAST score: 324

Table of genes, locations, strands and annotations of subject cluster:
C2865_RS15445	C2865_RS15445	495473	496291	+	MBL_fold_metallo-hydrolase
C2865_RS15450	C2865_RS15450	496298	496887	+	TetR_family_transcriptional_regulator
C2865_RS15460	C2865_RS15460	497778	498116	+	IS256_family_transposase

Table of Blast hits (query gene, subject gene, %identity, blast score, %coverage, e-value):
input.path1.gene699	C2865_RS15480	33	146	52.801120448179276	1.4e-32
input.path1.gene700	C2865_RS15515	34	178	36.5149833518313	4.1e-42


>>
50. NZ_QQAT01000012
Source: Rhodococcus sp. AG1013 Ga0222397_112, whole genome shotgun sequence
Type: NRPS
Number of proteins with BLAST hits to this cluster: 2
Cumulative BLAST score: 321

Table of genes, locations, strands and annotations of subject cluster:
DEU38_RS15760	DEU38_RS15760	35491	35703	-	lipase
DEU38_RS15780	DEU38_RS15780	37227	38939	-	VWA_domain-containing_protein

Table of Blast hits (query gene, subject gene, %identity, blast score, %coverage, e-value):
input.path1.gene699	DEU38_RS15790	33	187	59.38375350140056	5.4e-45
input.path1.gene700	DEU38_RS15835	31	134	36.625971143174255	6.9e-29
