## Supplementary Results for "A draft genome of the ascomycotal fungal species *Pseudopithomyces maydicus* (family *Didymosphaeriaceae*)": contig_17_c1.txt

ClusterBlast scores for contig_17

Table of genes, locations, strands and annotations of query cluster:
input.path1.gene42	149408	149599	-		
input.path1.gene43	151491	154493	-		
input.path1.gene44	156595	158037	+		
input.path1.gene45	158216	159902	-		
input.path1.gene46	160794	161132	+		
input.path1.gene47	162089	162927	-		
input.path1.gene48	163359	163686	+		
input.path1.gene49	164244	165782	+		


Significant hits: 
1. NW_001939248	Pyrenophora tritici-repentis Pt-1C-BFP supercont1.5, whole genome shotgun sequence
2. NW_001939248	Pyrenophora tritici-repentis Pt-1C-BFP supercont1.5, whole genome shotgun sequence
3. CP042201	Venturia effusa strain albino chromosome 17, complete sequence


Details:

>>
1. NW_001939248
Source: Pyrenophora tritici-repentis Pt-1C-BFP supercont1.5, whole genome shotgun sequence
Type: terpene
Number of proteins with BLAST hits to this cluster: 4
Cumulative BLAST score: 853

Table of genes, locations, strands and annotations of subject cluster:

Table of Blast hits (query gene, subject gene, %identity, blast score, %coverage, e-value):
input.path1.gene45	PTRG_05738	36	183	67.77041942604856	3.8e-44
input.path1.gene47	PTRG_05742	53	143	59.28853754940712	3.2e-32
input.path1.gene48	PTRG_05741	85	147	98.83720930232558	7.5e-34
input.path1.gene49	PTRG_05740	51	380	147.8688524590164	1.4e-103


>>
2. NW_001939248
Source: Pyrenophora tritici-repentis Pt-1C-BFP supercont1.5, whole genome shotgun sequence
Type: terpene
Number of proteins with BLAST hits to this cluster: 2
Cumulative BLAST score: 710

Table of genes, locations, strands and annotations of subject cluster:

Table of Blast hits (query gene, subject gene, %identity, blast score, %coverage, e-value):
input.path1.gene44	PTRG_05472	39	183	87.60806916426513	3.8e-44
input.path1.gene45	PTRG_05471	69	527	89.1832229580574	1.4e-147


>>
3. CP042201
Source: Venturia effusa strain albino chromosome 17, complete sequence
Type: NRPS-like
Number of proteins with BLAST hits to this cluster: 2
Cumulative BLAST score: 437

Table of genes, locations, strands and annotations of subject cluster:

Table of Blast hits (query gene, subject gene, %identity, blast score, %coverage, e-value):
input.path1.gene48	FKW77_006602	65	116	91.86046511627907	1.4e-24
input.path1.gene49	FKW77_006606	45	321	149.18032786885246	1e-85
