## Supplementary Results for "A draft genome of the ascomycotal fungal species *Pseudopithomyces maydicus* (family *Didymosphaeriaceae*)": contig_17_c2.txt

ClusterBlast scores for contig_17

Table of genes, locations, strands and annotations of query cluster:
input.path1.gene911	3156599	3157274	+		
input.path1.gene912	3160347	3161385	+		
input.path1.gene913	3161578	3162784	+		
input.path1.gene914	3163026	3163666	-		
input.path1.gene915	3165231	3167880	+		
input.path1.gene916	3168586	3169140	+		
input.path1.gene917	3174261	3175829	+		
input.path1.gene918	3176184	3182827	-		
input.path1.gene919	3184519	3185567	-		
input.path1.gene920	3187737	3189162	-		
input.path1.gene921	3190504	3191011	+		
input.path1.gene922	3192718	3193765	+		
input.path1.gene923	3195617	3196007	+		
input.path1.gene924	3197131	3199004	-		


Significant hits: 
1. NZ_CP012159	Chondromyces crocatus strain Cm c5 chromosome, complete genome
2. NW_013550603	Rhinocladiella mackenziei CBS 650.93 unplaced genomic scaffold supercont1.2, whole genome shotgun sequence
3. NW_008481832	Cladophialophora carrionii CBS 160.54 unplaced genomic scaffold supercont1.7, whole genome shotgun sequence
4. NW_015971651	Fonsecaea multimorphosa CBS 102226 unplaced genomic scaffold supercont1.17, whole genome shotgun sequence
5. NW_017387254	Fonsecaea erecta strain CBS 125763 chromosome Unknown Scaff11, whole genome shotgun sequence
6. NW_013550607	Fonsecaea pedrosoi CBS 271.37 unplaced genomic scaffold supercont1.9, whole genome shotgun sequence
7. NW_015622518	Exophiala spinifera strain CBS 89968 unplaced genomic scaffold supercont1.6, whole genome shotgun sequence
8. NC_021191	Actinoplanes sp. N902-109, complete genome
9. NC_016582	Streptomyces bingchenggensis BCW-1, complete sequence
10. NC_016459	Thermothielavioides terrestris NRRL 8126 chromosome 3, complete sequence
11. NC_005027	Rhodopirellula baltica SH 1, complete genome
12. NZ_PVTF01000011	Umezawaea tangerina strain DSM 44720 Ga0180964_111, whole genome shotgun sequence
13. NZ_MQUQ01000018	Amycolatopsis coloradensis strain DSM 44225 contig_18, whole genome shotgun sequence
14. NT_165976	Chaetomium globosum CBS 148.51 scaffold_1 genomic scaffold, whole genome shotgun sequence
15. NZ_CP045480	Amycolatopsis sp. YIM 10 chromosome, complete genome


Details:

>>
1. NZ_CP012159
Source: Chondromyces crocatus strain Cm c5 chromosome, complete genome
Type: NRPS,T1PKS
Number of proteins with BLAST hits to this cluster: 2
Cumulative BLAST score: 3639

Table of genes, locations, strands and annotations of subject cluster:

Table of Blast hits (query gene, subject gene, %identity, blast score, %coverage, e-value):
input.path1.gene918	CMC5_RS26985	36	472	44.75766567754698	3.1e-130
input.path1.gene918	CMC5_RS26990	33	471	53.36300692383779	5.3e-130
input.path1.gene918	CMC5_RS26955	35	469	44.95548961424332	2e-129
input.path1.gene918	CMC5_RS26960	35	468	44.80712166172107	3.4e-129
input.path1.gene918	CMC5_RS26980	33	454	44.80712166172107	5.2e-125
input.path1.gene918	CMC5_RS26965	32	444	53.16518298714145	7e-122
input.path1.gene918	CMC5_RS27010	32	391	46.38971315529179	5.4e-106
input.path1.gene918	CMC5_RS27005	33	388	42.82888229475766	3.5e-105
input.path1.gene919	CMC5_RS27025	35	82	52.38095238095239	6.4e-14


>>
2. NW_013550603
Source: Rhinocladiella mackenziei CBS 650.93 unplaced genomic scaffold supercont1.2, whole genome shotgun sequence
Type: T1PKS
Number of proteins with BLAST hits to this cluster: 2
Cumulative BLAST score: 1914

Table of genes, locations, strands and annotations of subject cluster:

Table of Blast hits (query gene, subject gene, %identity, blast score, %coverage, e-value):
input.path1.gene918	Z518_03561	46	1565	92.63105835806132	0.0
input.path1.gene919	Z518_03557	54	349	99.04761904761905	2.8e-94


>>
3. NW_008481832
Source: Cladophialophora carrionii CBS 160.54 unplaced genomic scaffold supercont1.7, whole genome shotgun sequence
Type: T1PKS
Number of proteins with BLAST hits to this cluster: 2
Cumulative BLAST score: 1868

Table of genes, locations, strands and annotations of subject cluster:

Table of Blast hits (query gene, subject gene, %identity, blast score, %coverage, e-value):
input.path1.gene918	G647_08240	45	1507	92.58160237388724	0.0
input.path1.gene919	G647_08245	56	361	99.36507936507937	7.1e-98


>>
4. NW_015971651
Source: Fonsecaea multimorphosa CBS 102226 unplaced genomic scaffold supercont1.17, whole genome shotgun sequence
Type: T1PKS
Number of proteins with BLAST hits to this cluster: 2
Cumulative BLAST score: 1865

Table of genes, locations, strands and annotations of subject cluster:

Table of Blast hits (query gene, subject gene, %identity, blast score, %coverage, e-value):
input.path1.gene918	Z520_08005	41	1499	106.97329376854599	0.0
input.path1.gene919	Z520_08010	58	366	99.36507936507937	2.2e-99


>>
5. NW_017387254
Source: Fonsecaea erecta strain CBS 125763 chromosome Unknown Scaff11, whole genome shotgun sequence
Type: T1PKS
Number of proteins with BLAST hits to this cluster: 2
Cumulative BLAST score: 1861

Table of genes, locations, strands and annotations of subject cluster:

Table of Blast hits (query gene, subject gene, %identity, blast score, %coverage, e-value):
input.path1.gene918	AYL99_10299	41	1496	107.02274975272007	0.0
input.path1.gene919	AYL99_10304	57	365	99.36507936507937	6.4e-99


>>
6. NW_013550607
Source: Fonsecaea pedrosoi CBS 271.37 unplaced genomic scaffold supercont1.9, whole genome shotgun sequence
Type: T1PKS
Number of proteins with BLAST hits to this cluster: 2
Cumulative BLAST score: 1848

Table of genes, locations, strands and annotations of subject cluster:

Table of Blast hits (query gene, subject gene, %identity, blast score, %coverage, e-value):
input.path1.gene918	Z517_12191	41	1487	106.77546983184965	0.0
input.path1.gene919	Z517_12196	57	361	99.68253968253968	9.3e-98


>>
7. NW_015622518
Source: Exophiala spinifera strain CBS 89968 unplaced genomic scaffold supercont1.6, whole genome shotgun sequence
Type: T1PKS
Number of proteins with BLAST hits to this cluster: 2
Cumulative BLAST score: 1621

Table of genes, locations, strands and annotations of subject cluster:

Table of Blast hits (query gene, subject gene, %identity, blast score, %coverage, e-value):
input.path1.gene918	PV08_08641	45	1536	94.31256181998022	0.0
input.path1.gene919	PV08_08638	31	85	50.79365079365079	1.3e-14


>>
8. NC_021191
Source: Actinoplanes sp. N902-109, complete genome
Type: T1PKS
Number of proteins with BLAST hits to this cluster: 2
Cumulative BLAST score: 1281

Table of genes, locations, strands and annotations of subject cluster:
L083_RS43460	L083_RS43460	2651371	2652237	-	SDR_family_NAD(P)-dependent_oxidoreductase
L083_RS43465	L083_RS43465	2652505	2653632	-	beta-ketoacyl_synthase
L083_RS43475	L083_RS43475	2655193	2659158	-	acyltransferase_domain-containing_protein
L083_RS43480	L083_RS43480	2659279	2660865	-	hypothetical_protein
L083_RS12105	L083_RS12105	2691701	2692675	-	alpha/beta_hydrolase
L083_RS12155	L083_RS12155	2704010	2704359	+	glyoxalase

Table of Blast hits (query gene, subject gene, %identity, blast score, %coverage, e-value):
input.path1.gene918	L083_RS43485	31	409	44.90603363006924	2.5e-111
input.path1.gene918	L083_RS12090	31	391	44.01582591493571	4.1e-106
input.path1.gene918	L083_RS40290	31	385	45.79624134520277	2.9e-104
input.path1.gene919	L083_RS12060	31	96	81.58730158730158	7.3e-18


>>
9. NC_016582
Source: Streptomyces bingchenggensis BCW-1, complete sequence
Type: PKS-like,T1PKS
Number of proteins with BLAST hits to this cluster: 2
Cumulative BLAST score: 831

Table of genes, locations, strands and annotations of subject cluster:
SBI_RS07245	SBI_RS07245	2058351	2059574	+	histidine_kinase
SBI_RS07265	SBI_RS07265	2062797	2063115	+	PPOX_class_F420-dependent_oxidoreductase
SBI_RS53795	SBI_RS53795	2101119	2106818	-	acyltransferase_domain-containing_protein
SBI_RS53800	SBI_RS53800	2109876	2109968	-	IS5/IS1182_family_transposase

Table of Blast hits (query gene, subject gene, %identity, blast score, %coverage, e-value):
input.path1.gene918	SBI_RS53790	31	368	45.25222551928783	3.7e-99
input.path1.gene918	SBI_RS53795	31	346	45.40059347181009	2e-92
input.path1.gene919	SBI_RS07255	34	117	73.01587301587301	3e-24


>>
10. NC_016459
Source: Thermothielavioides terrestris NRRL 8126 chromosome 3, complete sequence
Type: T1PKS
Number of proteins with BLAST hits to this cluster: 2
Cumulative BLAST score: 553

Table of genes, locations, strands and annotations of subject cluster:

Table of Blast hits (query gene, subject gene, %identity, blast score, %coverage, e-value):
input.path1.gene918	THITE_2050784	36	380	33.7784371909001	1.2e-102
input.path1.gene919	THITE_2116297	35	173	98.73015873015873	2.7e-41


>>
11. NC_005027
Source: Rhodopirellula baltica SH 1, complete genome
Type: T1PKS
Number of proteins with BLAST hits to this cluster: 2
Cumulative BLAST score: 551

Table of genes, locations, strands and annotations of subject cluster:

Table of Blast hits (query gene, subject gene, %identity, blast score, %coverage, e-value):
input.path1.gene918	RB_RS24870	34	471	44.90603363006924	4.1e-130
input.path1.gene922	RB_RS24900	33	80	61.206896551724135	3.5e-13


>>
12. NZ_PVTF01000011
Source: Umezawaea tangerina strain DSM 44720 Ga0180964_111, whole genome shotgun sequence
Type: NRPS,T1PKS,T3PKS
Number of proteins with BLAST hits to this cluster: 2
Cumulative BLAST score: 512

Table of genes, locations, strands and annotations of subject cluster:

Table of Blast hits (query gene, subject gene, %identity, blast score, %coverage, e-value):
input.path1.gene918	CLV43_RS33265	31	379	45.25222551928783	2.1e-102
input.path1.gene919	CLV43_RS33240	31	133	101.26984126984127	3.1e-29


>>
13. NZ_MQUQ01000018
Source: Amycolatopsis coloradensis strain DSM 44225 contig_18, whole genome shotgun sequence
Type: T1PKS
Number of proteins with BLAST hits to this cluster: 2
Cumulative BLAST score: 434

Table of genes, locations, strands and annotations of subject cluster:
BS329_RS42905	BS329_RS42905	56993	57118	+	GntR_family_transcriptional_regulator
BS329_RS30970	BS329_RS30970	65709	65870	+	endonuclease/exonuclease/phosphatase_family_protein

Table of Blast hits (query gene, subject gene, %identity, blast score, %coverage, e-value):
input.path1.gene918	BS329_RS30895	32	357	42.631058358061324	1.1e-95
input.path1.gene919	BS329_RS30820	33	77	56.507936507936506	2.7e-12


>>
14. NT_165976
Source: Chaetomium globosum CBS 148.51 scaffold_1 genomic scaffold, whole genome shotgun sequence
Type: NRPS-like,T1PKS
Number of proteins with BLAST hits to this cluster: 2
Cumulative BLAST score: 433

Table of genes, locations, strands and annotations of subject cluster:

Table of Blast hits (query gene, subject gene, %identity, blast score, %coverage, e-value):
input.path1.gene918	CHGG_00046	31	353	45.79624134520277	1.6e-94
input.path1.gene919	CHGG_00038	44	80	33.33333333333333	4.1e-13


>>
15. NZ_CP045480
Source: Amycolatopsis sp. YIM 10 chromosome, complete genome
Type: T1PKS
Number of proteins with BLAST hits to this cluster: 2
Cumulative BLAST score: 427

Table of genes, locations, strands and annotations of subject cluster:
YIM_RS33355	YIM_RS33355	7083431	7084368	+	SDR_family_NAD(P)-dependent_oxidoreductase

Table of Blast hits (query gene, subject gene, %identity, blast score, %coverage, e-value):
input.path1.gene914	YIM_RS33480	34	72	73.95833333333334	5.2e-11
input.path1.gene918	YIM_RS33440	31	355	44.362017804154306	4.3e-95
