## Supplementary Results for "A draft genome of the ascomycotal fungal species *Pseudopithomyces maydicus* (family *Didymosphaeriaceae*)": contig_20_c1.txt

ClusterBlast scores for contig_20

Table of genes, locations, strands and annotations of query cluster:
input.path1.gene1	786	10189	-		
input.path1.gene2	11967	23431	-		
input.path1.gene3	25447	26907	+		
input.path1.gene4	27810	28082	+		
input.path1.gene5	30573	30883	+		
input.path1.gene6	31184	32981	-		
input.path1.gene7	37540	39122	-		
input.path1.gene8	40242	40627	+		
input.path1.gene9	42794	43182	-		


Significant hits: 
1. NW_022983863	Arthroderma uncinatum strain CBS 119779 chromosome Unknown scaffold1, whole genome shotgun sequence
2. NW_003299166	Microsporum canis CBS 113480 supercont1.4 genomic scaffold, whole genome shotgun sequence
3. NW_003315112	Trichophyton benhamiae CBS 112371 chromosome Unknown contig01989.5, whole genome shotgun sequence
4. NW_003315027	Verticillium alfalfae VaMs.102 supercont1.12 genomic scaffold, whole genome shotgun sequence
5. NT_165937	Aspergillus terreus NIH2624 scaffold_14 genomic scaffold, whole genome shotgun sequence
6. NT_107012	Aspergillus nidulans FGSC A4 chromosome II map unlocalized scaffold_4, whole genome shotgun sequence
7. NW_003345194	Nannizzia gypsea CBS 118893 supercont1.8 genomic scaffold, whole genome shotgun sequence
8. NW_003456426	Trichophyton rubrum CBS 118892 genomic scaffold supercont2.3, whole genome shotgun sequence
9. NW_022474210	Venustampulla echinocandica strain BP 5553 chromosome Unknown scaffold_5, whole genome shotgun sequence
10. NZ_CP034669	Corallococcus coralloides strain B035 chromosome, complete genome
11. NC_017030	Corallococcus coralloides DSM 2259, complete sequence
12. NC_008095	Myxococcus xanthus DK 1622, complete sequence
13. NC_020126	Myxococcus stipitatus DSM 14675, complete sequence
14. NZ_CP022203	Myxococcus macrosporus DSM 14697 chromosome, complete genome
15. NW_019716257	Ramularia collo-cygni strain URUG2 genome assembly, contig: RCC_scaffold03, whole genome shotgun sequence
16. NZ_CP034669	Corallococcus coralloides strain B035 chromosome, complete genome
17. NW_001820834	Sclerotinia sclerotiorum 1980 UF-70 scaffold_2 genomic scaffold, whole genome shotgun sequence
18. NT_107005	Aspergillus nidulans FGSC A4 chromosome II map unlocalized scaffold_11, whole genome shotgun sequence
19. NT_165935	Aspergillus terreus NIH2624 scaffold_12 genomic scaffold, whole genome shotgun sequence
20. NW_022984631	Aspergillus tanneri strain NIH1004 chromosome Unknown AapTan_4_pilon_pilon_pilon, whole genome shotgun sequence


Details:

>>
1. NW_022983863
Source: Arthroderma uncinatum strain CBS 119779 chromosome Unknown scaffold1, whole genome shotgun sequence
Type: NRPS
Number of proteins with BLAST hits to this cluster: 3
Cumulative BLAST score: 2453

Table of genes, locations, strands and annotations of subject cluster:

Table of Blast hits (query gene, subject gene, %identity, blast score, %coverage, e-value):
input.path1.gene1	GIQ15_01756	33	552	45.86884031122638	3.2e-154
input.path1.gene2	GIQ15_01756	31	1683	101.25101985314116	0.0
input.path1.gene7	GIQ15_01753	38	218	70.11494252873564	1.3e-54


>>
2. NW_003299166
Source: Microsporum canis CBS 113480 supercont1.4 genomic scaffold, whole genome shotgun sequence
Type: NRPS
Number of proteins with BLAST hits to this cluster: 3
Cumulative BLAST score: 2444

Table of genes, locations, strands and annotations of subject cluster:

Table of Blast hits (query gene, subject gene, %identity, blast score, %coverage, e-value):
input.path1.gene1	MCYG_04787	32	535	46.054094108929235	4e-149
input.path1.gene2	MCYG_04787	31	1686	101.60456894207235	0.0
input.path1.gene7	MCYG_04784	39	223	70.11494252873564	4.2e-56


>>
3. NW_003315112
Source: Trichophyton benhamiae CBS 112371 chromosome Unknown contig01989.5, whole genome shotgun sequence
Type: NRPS
Number of proteins with BLAST hits to this cluster: 3
Cumulative BLAST score: 2385

Table of genes, locations, strands and annotations of subject cluster:

Table of Blast hits (query gene, subject gene, %identity, blast score, %coverage, e-value):
input.path1.gene1	ARB_05131	31	509	45.49833271582067	1.8e-141
input.path1.gene2	ARB_05131	31	1674	101.71335327712809	0.0
input.path1.gene7	ARB_05133	35	202	73.33333333333333	7.6e-50


>>
4. NW_003315027
Source: Verticillium alfalfae VaMs.102 supercont1.12 genomic scaffold, whole genome shotgun sequence
Type: NRPS
Number of proteins with BLAST hits to this cluster: 2
Cumulative BLAST score: 3670

Table of genes, locations, strands and annotations of subject cluster:

Table of Blast hits (query gene, subject gene, %identity, blast score, %coverage, e-value):
input.path1.gene1	VDBG_08381	33	582	45.60948499444238	2.2e-163
input.path1.gene1	VDBG_08382	34	575	44.97962208225268	3.5e-161
input.path1.gene1	VDBG_08380	32	542	46.054094108929235	3.3e-151
input.path1.gene2	VDBG_08382	31	1000	64.53630677182485	3.5e-289
input.path1.gene2	VDBG_08381	31	971	61.73511014413925	2.3e-280


>>
5. NT_165937
Source: Aspergillus terreus NIH2624 scaffold_14 genomic scaffold, whole genome shotgun sequence
Type: NRPS
Number of proteins with BLAST hits to this cluster: 2
Cumulative BLAST score: 1478

Table of genes, locations, strands and annotations of subject cluster:

Table of Blast hits (query gene, subject gene, %identity, blast score, %coverage, e-value):
input.path1.gene1	ATEG_09064	32	573	47.6102260096332	1.3e-160
input.path1.gene1	ATEG_09068	31	479	45.905891070766955	2.6e-132
input.path1.gene2	ATEG_09068	41	426	16.37204242589067	3.6e-116


>>
6. NT_107012
Source: Aspergillus nidulans FGSC A4 chromosome II map unlocalized scaffold_4, whole genome shotgun sequence
Type: NRPS
Number of proteins with BLAST hits to this cluster: 2
Cumulative BLAST score: 1388

Table of genes, locations, strands and annotations of subject cluster:

Table of Blast hits (query gene, subject gene, %identity, blast score, %coverage, e-value):
input.path1.gene1	AN3496.2	31	488	45.94294183030752	3.3e-135
input.path1.gene2	AN3496.2	31	471	28.47429970084308	7.4e-130
input.path1.gene2	AN3495.2	31	429	28.610280119662768	3.2e-117


>>
7. NW_003345194
Source: Nannizzia gypsea CBS 118893 supercont1.8 genomic scaffold, whole genome shotgun sequence
Type: NRPS
Number of proteins with BLAST hits to this cluster: 2
Cumulative BLAST score: 1062

Table of genes, locations, strands and annotations of subject cluster:

Table of Blast hits (query gene, subject gene, %identity, blast score, %coverage, e-value):
input.path1.gene2	MGYG_08146	33	847	43.35055751971716	6.5e-243
input.path1.gene7	MGYG_08144	38	215	70.57471264367817	1.5e-53


>>
8. NW_003456426
Source: Trichophyton rubrum CBS 118892 genomic scaffold supercont2.3, whole genome shotgun sequence
Type: NRPS
Number of proteins with BLAST hits to this cluster: 2
Cumulative BLAST score: 701

Table of genes, locations, strands and annotations of subject cluster:

Table of Blast hits (query gene, subject gene, %identity, blast score, %coverage, e-value):
input.path1.gene1	TERG_03194	31	493	45.60948499444238	1e-136
input.path1.gene7	TERG_03192	36	208	70.11494252873564	1.1e-51


>>
9. NW_022474210
Source: Venustampulla echinocandica strain BP 5553 chromosome Unknown scaffold_5, whole genome shotgun sequence
Type: NRPS
Number of proteins with BLAST hits to this cluster: 2
Cumulative BLAST score: 668

Table of genes, locations, strands and annotations of subject cluster:

Table of Blast hits (query gene, subject gene, %identity, blast score, %coverage, e-value):
input.path1.gene1	BP5553_06962	31	510	46.683957021118935	1.1e-141
input.path1.gene7	BP5553_06959	32	158	68.27586206896552	2.1e-36


>>
10. NZ_CP034669
Source: Corallococcus coralloides strain B035 chromosome, complete genome
Type: NRPS
Number of proteins with BLAST hits to this cluster: 2
Cumulative BLAST score: 594

Table of genes, locations, strands and annotations of subject cluster:

Table of Blast hits (query gene, subject gene, %identity, blast score, %coverage, e-value):
input.path1.gene1	EJ065_RS19000	33	491	42.645424231196735	6.6e-136
input.path1.gene7	EJ065_RS18975	31	103	55.172413793103445	6.3e-20


>>
11. NC_017030
Source: Corallococcus coralloides DSM 2259, complete sequence
Type: NRPS
Number of proteins with BLAST hits to this cluster: 2
Cumulative BLAST score: 589

Table of genes, locations, strands and annotations of subject cluster:
COCOR_RS19705	COCOR_RS19705	4919069	4919665	+	hypothetical_protein

Table of Blast hits (query gene, subject gene, %identity, blast score, %coverage, e-value):
input.path1.gene1	COCOR_RS19655	32	488	42.97888106706188	5.6e-135
input.path1.gene7	COCOR_RS19630	31	101	51.49425287356322	1.8e-19


>>
12. NC_008095
Source: Myxococcus xanthus DK 1622, complete sequence
Type: NRPS,T1PKS
Number of proteins with BLAST hits to this cluster: 2
Cumulative BLAST score: 583

Table of genes, locations, strands and annotations of subject cluster:
MXAN_RS37745	MXAN_RS37745	4887695	4889056	-	non-ribosomal_peptide_synthetase

Table of Blast hits (query gene, subject gene, %identity, blast score, %coverage, e-value):
input.path1.gene1	MXAN_RS19445	32	479	42.86772878844016	2e-132
input.path1.gene7	MXAN_RS19535	31	104	55.172413793103445	3.7e-20


>>
13. NC_020126
Source: Myxococcus stipitatus DSM 14675, complete sequence
Type: NRPS,T1PKS
Number of proteins with BLAST hits to this cluster: 2
Cumulative BLAST score: 573

Table of genes, locations, strands and annotations of subject cluster:
MYSTI_RS19300	MYSTI_RS19300	4939987	4940810	+	hypothetical_protein

Table of Blast hits (query gene, subject gene, %identity, blast score, %coverage, e-value):
input.path1.gene1	MYSTI_RS19390	32	476	43.164134864764726	1.7e-131
input.path1.gene7	MYSTI_RS19335	31	97	51.49425287356322	2.6e-18


>>
14. NZ_CP022203
Source: Myxococcus macrosporus DSM 14697 chromosome, complete genome
Type: NRPS,T1PKS
Number of proteins with BLAST hits to this cluster: 2
Cumulative BLAST score: 551

Table of genes, locations, strands and annotations of subject cluster:
MYMAC_RS19735	MYMAC_RS19735	4840739	4841215	-	LysM_peptidoglycan-binding_domain-containing_protein
MYMAC_RS37240	MYMAC_RS37240	4841215	4841625	-	hypothetical_protein

Table of Blast hits (query gene, subject gene, %identity, blast score, %coverage, e-value):
input.path1.gene1	MYMAC_RS19755	31	451	43.38643942200815	4.5e-124
input.path1.gene7	MYMAC_RS19825	31	100	51.49425287356322	4.1e-19


>>
15. NW_019716257
Source: Ramularia collo-cygni strain URUG2 genome assembly, contig: RCC_scaffold03, whole genome shotgun sequence
Type: NRPS
Number of proteins with BLAST hits to this cluster: 2
Cumulative BLAST score: 544

Table of genes, locations, strands and annotations of subject cluster:

Table of Blast hits (query gene, subject gene, %identity, blast score, %coverage, e-value):
input.path1.gene2	RCC_02695	36	388	17.269513190100625	6.3e-105
input.path1.gene7	RCC_02687	32	156	64.82758620689654	4.8e-36


>>
16. NZ_CP034669
Source: Corallococcus coralloides strain B035 chromosome, complete genome
Type: NRPS,T1PKS
Number of proteins with BLAST hits to this cluster: 2
Cumulative BLAST score: 536

Table of genes, locations, strands and annotations of subject cluster:

Table of Blast hits (query gene, subject gene, %identity, blast score, %coverage, e-value):
input.path1.gene1	EJ065_RS09995	35	465	34.19785105594665	5.1e-128
input.path1.gene7	EJ065_RS10085	31	71	50.114942528735625	3.4e-10


>>
17. NW_001820834
Source: Sclerotinia sclerotiorum 1980 UF-70 scaffold_2 genomic scaffold, whole genome shotgun sequence
Type: NRPS
Number of proteins with BLAST hits to this cluster: 2
Cumulative BLAST score: 524

Table of genes, locations, strands and annotations of subject cluster:

Table of Blast hits (query gene, subject gene, %identity, blast score, %coverage, e-value):
input.path1.gene1	SS1G_01265	33	432	34.1237495368655	2.8e-118
input.path1.gene7	SS1G_01266	31	92	48.275862068965516	1.1e-16


>>
18. NT_107005
Source: Aspergillus nidulans FGSC A4 chromosome II map unlocalized scaffold_11, whole genome shotgun sequence
Type: terpene
Number of proteins with BLAST hits to this cluster: 2
Cumulative BLAST score: 754

Table of genes, locations, strands and annotations of subject cluster:

Table of Blast hits (query gene, subject gene, %identity, blast score, %coverage, e-value):
input.path1.gene6	AN8262.2	37	222	101.23456790123457	6.6e-56
input.path1.gene7	AN8261.2	85	532	71.72413793103448	4.2e-149


>>
19. NT_165935
Source: Aspergillus terreus NIH2624 scaffold_12 genomic scaffold, whole genome shotgun sequence
Type: terpene
Number of proteins with BLAST hits to this cluster: 2
Cumulative BLAST score: 750

Table of genes, locations, strands and annotations of subject cluster:

Table of Blast hits (query gene, subject gene, %identity, blast score, %coverage, e-value):
input.path1.gene6	ATEG_08000	37	218	100.24691358024691	1.6e-54
input.path1.gene7	ATEG_08001	74	532	93.10344827586206	4.2e-149


>>
20. NW_022984631
Source: Aspergillus tanneri strain NIH1004 chromosome Unknown AapTan_4_pilon_pilon_pilon, whole genome shotgun sequence
Type: terpene
Number of proteins with BLAST hits to this cluster: 2
Cumulative BLAST score: 697

Table of genes, locations, strands and annotations of subject cluster:

Table of Blast hits (query gene, subject gene, %identity, blast score, %coverage, e-value):
input.path1.gene6	ATNIH1004_006262	35	166	88.39506172839506	7.3e-39
input.path1.gene7	ATNIH1004_006263	73	531	86.66666666666667	7.1e-149
