## Supplementary Results for "A draft genome of the ascomycotal fungal species *Pseudopithomyces maydicus* (family *Didymosphaeriaceae*)": contig_23_c1.txt

ClusterBlast scores for contig_23

Table of genes, locations, strands and annotations of query cluster:
input.path1.gene1	60	4537	-		
input.path1.gene2	6898	8103	+		
input.path1.gene3	14997	16023	-		
input.path1.gene4	18891	21873	-		


Significant hits: 
1. NW_023336279	Aspergillus tubingensis WU-2223L DNA, scaffold_5, whole genome shotgun sequence
2. NW_007361001	Glarea lozoyensis ATCC 20868 chromosome Unknown GLAREA6, whole genome shotgun sequence


Details:

>>
1. NW_023336279
Source: Aspergillus tubingensis WU-2223L DNA, scaffold_5, whole genome shotgun sequence
Type: NRPS
Number of proteins with BLAST hits to this cluster: 2
Cumulative BLAST score: 906

Table of genes, locations, strands and annotations of subject cluster:

Table of Blast hits (query gene, subject gene, %identity, blast score, %coverage, e-value):
input.path1.gene1	AtWU_03451	33	754	102.4932614555256	2.3e-215
input.path1.gene3	AtWU_03450	31	152	97.36070381231671	7.1e-35


>>
2. NW_007361001
Source: Glarea lozoyensis ATCC 20868 chromosome Unknown GLAREA6, whole genome shotgun sequence
Type: NRPS
Number of proteins with BLAST hits to this cluster: 2
Cumulative BLAST score: 591

Table of genes, locations, strands and annotations of subject cluster:

Table of Blast hits (query gene, subject gene, %identity, blast score, %coverage, e-value):
input.path1.gene1	GLAREA_02479	35	436	50.40431266846361	1.4e-119
input.path1.gene3	GLAREA_02481	31	155	96.48093841642229	1.1e-35
