## Supplementary Results for "A draft genome of the ascomycotal fungal species *Pseudopithomyces maydicus* (family *Didymosphaeriaceae*)": contig_26_c1.txt

ClusterBlast scores for contig_26

Table of genes, locations, strands and annotations of query cluster:
input.path1.gene4	23887	26076	-		
input.path1.gene5	26630	27581	+		
input.path1.gene6	28332	30248	-		
input.path1.gene7	32351	33176	-		
input.path1.gene8	34040	35455	-		
input.path1.gene9	38251	41216	-		
input.path1.gene10	42400	49473	+		
input.path1.gene11	50365	51754	-		
input.path1.gene12	55938	57640	+		
input.path1.gene13	62925	63822	+		
input.path1.gene14	64061	65396	-		


Significant hits: 
1. NW_022474205	Venustampulla echinocandica strain BP 5553 chromosome Unknown scaffold_0, whole genome shotgun sequence
2. NW_020194484	Amorphotheca resinae ATCC 22711 unplaced genomic scaffold M430scaffold_14, whole genome shotgun sequence
3. NW_022983863	Arthroderma uncinatum strain CBS 119779 chromosome Unknown scaffold1, whole genome shotgun sequence
4. NW_021167084	Sodiomyces alkalinus F11 unplaced genomic scaffold SODALscaffold_6, whole genome shotgun sequence
5. NC_049565	Talaromyces rugulosus chromosome V, complete sequence
6. NW_006271974	Cordyceps militaris CM01 unplaced genomic scaffold CCM_S00006, whole genome shotgun sequence
7. NW_023336268	Colletotrichum scovillei strain TJNH1 chromosome Unknown Contig00010, whole genome shotgun sequence
8. NW_022984632	Aspergillus tanneri strain NIH1004 chromosome Unknown AapTan_5_pilon_pilon_pilon, whole genome shotgun sequence
9. NT_107007	Aspergillus nidulans FGSC A4 chromosome IV map unlocalized scaffold_9, whole genome shotgun sequence
10. NW_007360999	Glarea lozoyensis ATCC 20868 chromosome Unknown GLAREA4, whole genome shotgun sequence
11. NW_001820833	Sclerotinia sclerotiorum 1980 UF-70 scaffold_3 genomic scaffold, whole genome shotgun sequence
12. NW_017971434	Talaromyces atroroseus strain IBT 11181 chromosome Unknown scaffold_2, whole genome shotgun sequence
13. NT_165929	Aspergillus terreus NIH2624 scaffold_6 genomic scaffold, whole genome shotgun sequence
14. NW_006917100	Pestalotiopsis fici W106-1 unplaced genomic scaffold PFICI_Scf_10, whole genome shotgun sequence
15. NW_006917095	Pestalotiopsis fici W106-1 unplaced genomic scaffold PFICI_Scf_5, whole genome shotgun sequence
16. NW_023336280	Aspergillus tubingensis WU-2223L DNA, scaffold_6, whole genome shotgun sequence
17. NC_016457	Thermothielavioides terrestris NRRL 8126 chromosome 1, complete sequence
18. NW_022474217	Venustampulla echinocandica strain BP 5553 chromosome Unknown scaffold_12, whole genome shotgun sequence
19. NC_007201	Aspergillus fumigatus Af293 chromosome 8, whole genome shotgun sequence
20. RRCJ01000013	Pyricularia sp. CBS 133598 strain NI919 Pyricularia_sp_LS_NI919_Scf_13, whole genome shotgun sequence
21. NW_003613579	Podospora anserina S mat+ genomic DNA chromosome 6, supercontig 4
22. NW_013562492	Exophiala xenobiotica strain CBS 118157 unplaced genomic scaffold supercont1.3, whole genome shotgun sequence
23. NC_018212	Zymoseptoria tritici IPO323 chromosome 7, whole genome shotgun sequence
24. NW_014574689	Metarhizium brunneum ARSEF 3297 chromosome Unknown scaffold_28, whole genome shotgun sequence
25. NW_011942171	Metarhizium robertsii ARSEF 23 MAA_Scf_3, whole genome shotgun sequence
26. NW_006917097	Pestalotiopsis fici W106-1 unplaced genomic scaffold PFICI_Scf_7, whole genome shotgun sequence
27. NW_023336260	Colletotrichum scovillei strain TJNH1 chromosome Unknown Contig00002, whole genome shotgun sequence
28. NC_030954	Colletotrichum higginsianum IMI 349063 chromosome 1, whole genome shotgun sequence
29. NW_019716265	Ramularia collo-cygni strain URUG2 genome assembly, contig: RCC_scaffold11, whole genome shotgun sequence
30. NZ_CP031194	Streptomyces paludis strain GSSD-12 chromosome, complete genome
31. NZ_CP034352	Streptomyces sp. W1SF4 plasmid p2, complete sequence
32. NC_016111	Streptomyces cattleya NRRL 8057 = DSM 46488, complete genome
33. NC_030962	Colletotrichum higginsianum IMI 349063 chromosome 10, whole genome shotgun sequence
34. NW_006912895	Capronia coronata CBS 617.96 acBCG-supercont1.3, whole genome shotgun sequence
35. NZ_KN323065	Burkholderia pseudomallei MSHR1000 scaffold1, whole genome shotgun sequence
36. NZ_CP013395	Burkholderia vietnamiensis strain FL-2-3-30-S1-D0 chromosome 3, complete sequence
37. NC_018217	Zymoseptoria tritici IPO323 chromosome 2, whole genome shotgun sequence
38. NW_019716258	Ramularia collo-cygni strain URUG2 genome assembly, contig: RCC_scaffold04, whole genome shotgun sequence
39. NZ_CP048641	Streptomyces aureoverticillatus strain HN6 chromosome, complete genome
40. NC_016109	Kitasatospora setae KM-6054, complete genome
41. NC_030961	Colletotrichum higginsianum IMI 349063 chromosome 9, whole genome shotgun sequence
42. NZ_LR130759	Mycobacterium basiliense isolate 901379 chromosome I
43. NZ_CP016795	Streptomyces olivaceus strain KLBMP 5084 chromosome, complete genome
44. NZ_CP030771	Streptomyces sp. YIM 121038 chromosome, complete genome
45. NT_165972	Aspergillus terreus NIH2624 scaffold_1 genomic scaffold, whole genome shotgun sequence
46. NZ_CP040916	Streptomyces spectabilis strain NRRL 2792 chromosome, complete genome
47. NZ_CP023690	Streptomyces spectabilis strain ATCC 27465 chromosome, complete genome
48. NC_021191	Actinoplanes sp. N902-109, complete genome
49. NZ_LIVY01000012	Streptomyces sp. CB02460 scaffold12, whole genome shotgun sequence
50. NZ_CP053892	Actinomadura verrucosospora strain NRRL:B18236 chromosome


Details:

>>
1. NW_022474205
Source: Venustampulla echinocandica strain BP 5553 chromosome Unknown scaffold_0, whole genome shotgun sequence
Type: T1PKS
Number of proteins with BLAST hits to this cluster: 6
Cumulative BLAST score: 3661

Table of genes, locations, strands and annotations of subject cluster:

Table of Blast hits (query gene, subject gene, %identity, blast score, %coverage, e-value):
input.path1.gene4	BP5553_02214	34	298	94.5945945945946	1.3e-78
input.path1.gene5	BP5553_02213	47	221	91.4179104477612	1.3e-55
input.path1.gene6	BP5553_02212	48	404	87.1401151631478	2.1e-110
input.path1.gene7	BP5553_02211	54	288	95.98540145985402	6.6e-76
input.path1.gene9	BP5553_02210	51	294	48.888888888888886	2e-77
input.path1.gene10	BP5553_02209	51	2156	111.9700748129676	0.0


>>
2. NW_020194484
Source: Amorphotheca resinae ATCC 22711 unplaced genomic scaffold M430scaffold_14, whole genome shotgun sequence
Type: T1PKS,terpene
Number of proteins with BLAST hits to this cluster: 6
Cumulative BLAST score: 3617

Table of genes, locations, strands and annotations of subject cluster:

Table of Blast hits (query gene, subject gene, %identity, blast score, %coverage, e-value):
input.path1.gene4	M430DRAFT_23063	34	301	92.7927927927928	1.5e-79
input.path1.gene5	M430DRAFT_110425	45	208	90.29850746268657	1.1e-51
input.path1.gene6	M430DRAFT_70069	43	430	94.81765834932821	3.5e-118
input.path1.gene7	M430DRAFT_23060	53	277	99.27007299270073	1.5e-72
input.path1.gene9	M430DRAFT_110432	54	303	48.376068376068375	4.3e-80
input.path1.gene9	M430DRAFT_23063	32	101	31.965811965811962	2.5e-19
input.path1.gene10	M430DRAFT_110834	48	1997	112.36907730673316	0.0


>>
3. NW_022983863
Source: Arthroderma uncinatum strain CBS 119779 chromosome Unknown scaffold1, whole genome shotgun sequence
Type: T1PKS
Number of proteins with BLAST hits to this cluster: 6
Cumulative BLAST score: 3208

Table of genes, locations, strands and annotations of subject cluster:

Table of Blast hits (query gene, subject gene, %identity, blast score, %coverage, e-value):
input.path1.gene4	GIQ15_00206	38	332	94.05405405405406	1.1e-88
input.path1.gene5	GIQ15_00207	40	182	96.64179104477611	5e-44
input.path1.gene6	GIQ15_00208	40	346	96.92898272552783	5.1e-93
input.path1.gene7	GIQ15_00209	47	265	103.2846715328467	7.8e-69
input.path1.gene7	GIQ15_00203	42	211	97.44525547445255	1e-52
input.path1.gene9	GIQ15_00205	48	259	49.743589743589745	7e-67
input.path1.gene9	GIQ15_00206	32	104	32.99145299145299	2.9e-20
input.path1.gene10	GIQ15_00204	40	1509	111.52119700748129	0.0


>>
4. NW_021167084
Source: Sodiomyces alkalinus F11 unplaced genomic scaffold SODALscaffold_6, whole genome shotgun sequence
Type: T1PKS
Number of proteins with BLAST hits to this cluster: 5
Cumulative BLAST score: 2610

Table of genes, locations, strands and annotations of subject cluster:

Table of Blast hits (query gene, subject gene, %identity, blast score, %coverage, e-value):
input.path1.gene4	SODALDRAFT_312698	37	304	91.53153153153153	1.8e-80
input.path1.gene5	SODALDRAFT_333919	47	201	87.68656716417911	8e-50
input.path1.gene6	SODALDRAFT_333920	39	331	95.58541266794626	2.2e-88
input.path1.gene9	SODALDRAFT_333923	46	210	43.24786324786325	4.9e-52
input.path1.gene10	SODALDRAFT_360496	40	1564	113.01745635910225	0.0


>>
5. NC_049565
Source: Talaromyces rugulosus chromosome V, complete sequence
Type: T1PKS
Number of proteins with BLAST hits to this cluster: 6
Cumulative BLAST score: 3174

Table of genes, locations, strands and annotations of subject cluster:

Table of Blast hits (query gene, subject gene, %identity, blast score, %coverage, e-value):
input.path1.gene4	TRUGW13939_09506	37	364	90.45045045045045	1.5e-98
input.path1.gene5	TRUGW13939_09505	43	192	91.4179104477612	6.3e-47
input.path1.gene6	TRUGW13939_09504	45	363	85.79654510556622	4e-98
input.path1.gene7	TRUGW13939_09517	40	211	96.71532846715328	1.3e-52
input.path1.gene9	TRUGW13939_09502	45	262	48.717948717948715	1.4e-67
input.path1.gene9	TRUGW13939_09506	37	166	38.80341880341881	8.1e-39
input.path1.gene10	TRUGW13939_09501	41	1616	116.75810473815463	0.0


>>
6. NW_006271974
Source: Cordyceps militaris CM01 unplaced genomic scaffold CCM_S00006, whole genome shotgun sequence
Type: NRPS,T1PKS
Number of proteins with BLAST hits to this cluster: 6
Cumulative BLAST score: 3786

Table of genes, locations, strands and annotations of subject cluster:

Table of Blast hits (query gene, subject gene, %identity, blast score, %coverage, e-value):
input.path1.gene4	CCM_08012	43	439	90.45045045045045	6.2e-121
input.path1.gene5	CCM_08014	44	212	98.13432835820896	5.9e-53
input.path1.gene6	CCM_08013	48	450	95.39347408829175	3.3e-124
input.path1.gene7	CCM_08011	52	275	98.90510948905109	4.4e-72
input.path1.gene9	CCM_08015	50	295	49.23076923076923	1.2e-77
input.path1.gene9	CCM_08012	34	185	47.521367521367516	1.3e-44
input.path1.gene10	CCM_08016	47	1930	113.76558603491272	0.0


>>
7. NW_023336268
Source: Colletotrichum scovillei strain TJNH1 chromosome Unknown Contig00010, whole genome shotgun sequence
Type: T1PKS
Number of proteins with BLAST hits to this cluster: 5
Cumulative BLAST score: 3574

Table of genes, locations, strands and annotations of subject cluster:

Table of Blast hits (query gene, subject gene, %identity, blast score, %coverage, e-value):
input.path1.gene4	HER10_EVM0006921	35	306	95.67567567567568	4.8e-81
input.path1.gene4	HER10_EVM0007360	34	277	91.8918918918919	2.4e-72
input.path1.gene6	HER10_EVM0000180	44	438	94.81765834932821	9.9e-121
input.path1.gene7	HER10_EVM0005921	49	270	99.27007299270073	2.4e-70
input.path1.gene9	HER10_EVM0001559	57	317	48.376068376068375	2.8e-84
input.path1.gene9	HER10_EVM0007360	33	106	36.06837606837607	1e-20
input.path1.gene9	HER10_EVM0006921	33	106	32.64957264957265	1e-20
input.path1.gene10	HER10_EVM0012096	43	1754	113.11720698254364	0.0


>>
8. NW_022984632
Source: Aspergillus tanneri strain NIH1004 chromosome Unknown AapTan_5_pilon_pilon_pilon, whole genome shotgun sequence
Type: T1PKS
Number of proteins with BLAST hits to this cluster: 6
Cumulative BLAST score: 3090

Table of genes, locations, strands and annotations of subject cluster:

Table of Blast hits (query gene, subject gene, %identity, blast score, %coverage, e-value):
input.path1.gene4	ATNIH1004_009502	36	292	93.69369369369369	7.1e-77
input.path1.gene5	ATNIH1004_009501	42	185	93.28358208955224	7.7e-45
input.path1.gene6	ATNIH1004_009497	39	351	95.20153550863724	2.1e-94
input.path1.gene7	ATNIH1004_009503	51	285	96.71532846715328	7.3e-75
input.path1.gene9	ATNIH1004_009500	50	280	49.23076923076923	5e-73
input.path1.gene9	ATNIH1004_009502	32	106	34.52991452991453	7.6e-21
input.path1.gene10	ATNIH1004_009499	41	1591	113.01745635910225	0.0


>>
9. NT_107007
Source: Aspergillus nidulans FGSC A4 chromosome IV map unlocalized scaffold_9, whole genome shotgun sequence
Type: T1PKS
Number of proteins with BLAST hits to this cluster: 4
Cumulative BLAST score: 2256

Table of genes, locations, strands and annotations of subject cluster:

Table of Blast hits (query gene, subject gene, %identity, blast score, %coverage, e-value):
input.path1.gene4	AN7082.2	35	217	64.14414414414415	5e-54
input.path1.gene6	AN7081.2	44	408	95.20153550863724	8.4e-112
input.path1.gene9	AN7083.2	46	215	43.58974358974359	1.2e-53
input.path1.gene10	AN7084.2	38	1416	114.0149625935162	0.0


>>
10. NW_007360999
Source: Glarea lozoyensis ATCC 20868 chromosome Unknown GLAREA4, whole genome shotgun sequence
Type: T1PKS
Number of proteins with BLAST hits to this cluster: 4
Cumulative BLAST score: 662

Table of genes, locations, strands and annotations of subject cluster:

Table of Blast hits (query gene, subject gene, %identity, blast score, %coverage, e-value):
input.path1.gene4	GLAREA_09954	34	251	86.66666666666667	1.8e-64
input.path1.gene7	GLAREA_09949	34	156	97.08029197080292	3.9e-36
input.path1.gene9	GLAREA_09954	32	97	34.01709401709402	4.6e-18
input.path1.gene10	GLAREA_09953	35	158	13.815461346633418	9.9e-36


>>
11. NW_001820833
Source: Sclerotinia sclerotiorum 1980 UF-70 scaffold_3 genomic scaffold, whole genome shotgun sequence
Type: T1PKS
Number of proteins with BLAST hits to this cluster: 3
Cumulative BLAST score: 2255

Table of genes, locations, strands and annotations of subject cluster:

Table of Blast hits (query gene, subject gene, %identity, blast score, %coverage, e-value):
input.path1.gene6	SS1G_02212	45	423	95.77735124760078	2.5e-116
input.path1.gene9	SS1G_02210	41	205	49.059829059829056	2.1e-50
input.path1.gene10	SS1G_02211	43	1627	106.33416458852867	0.0


>>
12. NW_017971434
Source: Talaromyces atroroseus strain IBT 11181 chromosome Unknown scaffold_2, whole genome shotgun sequence
Type: T1PKS
Number of proteins with BLAST hits to this cluster: 3
Cumulative BLAST score: 2030

Table of genes, locations, strands and annotations of subject cluster:

Table of Blast hits (query gene, subject gene, %identity, blast score, %coverage, e-value):
input.path1.gene4	UA08_02263	34	270	94.23423423423424	3.8e-70
input.path1.gene9	UA08_02260	48	269	49.059829059829056	1.2e-69
input.path1.gene10	UA08_02259	39	1491	115.46134663341645	0.0


>>
13. NT_165929
Source: Aspergillus terreus NIH2624 scaffold_6 genomic scaffold, whole genome shotgun sequence
Type: T1PKS
Number of proteins with BLAST hits to this cluster: 3
Cumulative BLAST score: 1940

Table of genes, locations, strands and annotations of subject cluster:

Table of Blast hits (query gene, subject gene, %identity, blast score, %coverage, e-value):
input.path1.gene7	ATEG_04721	37	150	84.67153284671532	2.8e-34
input.path1.gene9	ATEG_04719	43	220	48.717948717948715	6.2e-55
input.path1.gene10	ATEG_04718	41	1570	113.81546134663341	0.0


>>
14. NW_006917100
Source: Pestalotiopsis fici W106-1 unplaced genomic scaffold PFICI_Scf_10, whole genome shotgun sequence
Type: T1PKS
Number of proteins with BLAST hits to this cluster: 3
Cumulative BLAST score: 1874

Table of genes, locations, strands and annotations of subject cluster:

Table of Blast hits (query gene, subject gene, %identity, blast score, %coverage, e-value):
input.path1.gene8	PFICI_12428	33	167	120.97902097902097	1.8e-39
input.path1.gene9	PFICI_12430	44	203	43.07692307692308	7.8e-50
input.path1.gene10	PFICI_12429	40	1504	113.06733167082295	0.0


>>
15. NW_006917095
Source: Pestalotiopsis fici W106-1 unplaced genomic scaffold PFICI_Scf_5, whole genome shotgun sequence
Type: T1PKS
Number of proteins with BLAST hits to this cluster: 4
Cumulative BLAST score: 3146

Table of genes, locations, strands and annotations of subject cluster:

Table of Blast hits (query gene, subject gene, %identity, blast score, %coverage, e-value):
input.path1.gene4	PFICI_07949	37	371	90.81081081081082	2.1e-100
input.path1.gene6	PFICI_07952	45	396	94.04990403071018	3.3e-108
input.path1.gene9	PFICI_07953	39	236	49.914529914529915	8.3e-60
input.path1.gene9	PFICI_07949	33	141	35.72649572649573	2.1e-31
input.path1.gene10	PFICI_07954	42	1707	112.36907730673316	0.0
input.path1.gene10	PFICI_07957	33	295	26.83291770573566	3e-77


>>
16. NW_023336280
Source: Aspergillus tubingensis WU-2223L DNA, scaffold_6, whole genome shotgun sequence
Type: T1PKS
Number of proteins with BLAST hits to this cluster: 4
Cumulative BLAST score: 3026

Table of genes, locations, strands and annotations of subject cluster:

Table of Blast hits (query gene, subject gene, %identity, blast score, %coverage, e-value):
input.path1.gene4	AtWU_04534	35	290	94.95495495495496	3.5e-76
input.path1.gene4	AtWU_04538	35	266	85.76576576576576	7.1e-69
input.path1.gene6	AtWU_04532	43	385	95.58541266794626	7.6e-105
input.path1.gene9	AtWU_04535	45	233	48.376068376068375	4.1e-59
input.path1.gene9	AtWU_04534	31	104	34.52991452991453	4.9e-20
input.path1.gene9	AtWU_04538	33	104	31.965811965811962	4.9e-20
input.path1.gene10	AtWU_04536	42	1644	114.3640897755611	0.0


>>
17. NC_016457
Source: Thermothielavioides terrestris NRRL 8126 chromosome 1, complete sequence
Type: T1PKS
Number of proteins with BLAST hits to this cluster: 4
Cumulative BLAST score: 2605

Table of genes, locations, strands and annotations of subject cluster:

Table of Blast hits (query gene, subject gene, %identity, blast score, %coverage, e-value):
input.path1.gene4	THITE_2141120	40	311	69.36936936936937	1.5e-82
input.path1.gene6	THITE_2107714	41	375	101.91938579654511	1e-101
input.path1.gene9	THITE_2107710	45	254	48.717948717948715	3e-65
input.path1.gene9	THITE_2141120	41	158	30.94017094017094	2.9e-36
input.path1.gene10	THITE_40673	39	1507	119.8503740648379	0.0


>>
18. NW_022474217
Source: Venustampulla echinocandica strain BP 5553 chromosome Unknown scaffold_12, whole genome shotgun sequence
Type: NRPS,T1PKS
Number of proteins with BLAST hits to this cluster: 2
Cumulative BLAST score: 1616

Table of genes, locations, strands and annotations of subject cluster:

Table of Blast hits (query gene, subject gene, %identity, blast score, %coverage, e-value):
input.path1.gene9	BP5553_09811	42	244	50.256410256410255	3.1e-62
input.path1.gene10	BP5553_09812	38	1372	116.10972568578552	0.0


>>
19. NC_007201
Source: Aspergillus fumigatus Af293 chromosome 8, whole genome shotgun sequence
Type: NRPS,T1PKS,indole
Number of proteins with BLAST hits to this cluster: 2
Cumulative BLAST score: 1610

Table of genes, locations, strands and annotations of subject cluster:

Table of Blast hits (query gene, subject gene, %identity, blast score, %coverage, e-value):
input.path1.gene9	AFUA_8G00380	43	246	50.427350427350426	8.1e-63
input.path1.gene10	AFUA_8G00370	37	1364	117.15710723192021	0.0


>>
20. RRCJ01000013
Source: Pyricularia sp. CBS 133598 strain NI919 Pyricularia_sp_LS_NI919_Scf_13, whole genome shotgun sequence
Type: T1PKS
Number of proteins with BLAST hits to this cluster: 2
Cumulative BLAST score: 1561

Table of genes, locations, strands and annotations of subject cluster:

Table of Blast hits (query gene, subject gene, %identity, blast score, %coverage, e-value):
input.path1.gene9	PspLS_10071	40	212	51.28205128205128	1.3e-52
input.path1.gene10	PspLS_10070	36	1349	122.89276807980049	0.0


>>
21. NW_003613579
Source: Podospora anserina S mat+ genomic DNA chromosome 6, supercontig 4
Type: T1PKS
Number of proteins with BLAST hits to this cluster: 4
Cumulative BLAST score: 1079

Table of genes, locations, strands and annotations of subject cluster:

Table of Blast hits (query gene, subject gene, %identity, blast score, %coverage, e-value):
input.path1.gene4	PODANS_6_10360	33	247	84.50450450450451	4.5e-63
input.path1.gene7	PODANS_6_10350	43	224	100.0	1.2e-56
input.path1.gene9	PODANS_6_10360	36	113	30.256410256410255	6.2e-23
input.path1.gene10	PODANS_6_10330	31	495	64.23940149625935	2.6e-137


>>
22. NW_013562492
Source: Exophiala xenobiotica strain CBS 118157 unplaced genomic scaffold supercont1.3, whole genome shotgun sequence
Type: T1PKS
Number of proteins with BLAST hits to this cluster: 2
Cumulative BLAST score: 379

Table of genes, locations, strands and annotations of subject cluster:

Table of Blast hits (query gene, subject gene, %identity, blast score, %coverage, e-value):
input.path1.gene9	PV05_04655	38	203	53.504273504273506	7.8e-50
input.path1.gene10	PV05_04656	36	176	13.815461346633418	2.1e-41


>>
23. NC_018212
Source: Zymoseptoria tritici IPO323 chromosome 7, whole genome shotgun sequence
Type: T1PKS
Number of proteins with BLAST hits to this cluster: 3
Cumulative BLAST score: 1982

Table of genes, locations, strands and annotations of subject cluster:

Table of Blast hits (query gene, subject gene, %identity, blast score, %coverage, e-value):
input.path1.gene4	MYCGRDRAFT_45271	32	243	95.31531531531532	3.8e-62
input.path1.gene9	MYCGRDRAFT_44980	50	279	48.547008547008545	1.1e-72
input.path1.gene10	MYCGRDRAFT_45348	39	1460	113.91521197007481	0.0


>>
24. NW_014574689
Source: Metarhizium brunneum ARSEF 3297 chromosome Unknown scaffold_28, whole genome shotgun sequence
Type: T1PKS
Number of proteins with BLAST hits to this cluster: 3
Cumulative BLAST score: 1869

Table of genes, locations, strands and annotations of subject cluster:

Table of Blast hits (query gene, subject gene, %identity, blast score, %coverage, e-value):
input.path1.gene7	MBR_09989	39	167	90.14598540145985	2.2e-39
input.path1.gene9	MBR_09987	43	211	50.085470085470085	2.9e-52
input.path1.gene10	MBR_09985	39	1491	110.32418952618454	0.0


>>
25. NW_011942171
Source: Metarhizium robertsii ARSEF 23 MAA_Scf_3, whole genome shotgun sequence
Type: T1PKS
Number of proteins with BLAST hits to this cluster: 3
Cumulative BLAST score: 1852

Table of genes, locations, strands and annotations of subject cluster:

Table of Blast hits (query gene, subject gene, %identity, blast score, %coverage, e-value):
input.path1.gene7	MAA_03278	35	133	90.14598540145985	2.7e-29
input.path1.gene9	MAA_03276	41	215	52.478632478632484	1.2e-53
input.path1.gene10	MAA_03274	39	1504	112.46882793017457	0.0


>>
26. NW_006917097
Source: Pestalotiopsis fici W106-1 unplaced genomic scaffold PFICI_Scf_7, whole genome shotgun sequence
Type: T1PKS
Number of proteins with BLAST hits to this cluster: 3
Cumulative BLAST score: 629

Table of genes, locations, strands and annotations of subject cluster:

Table of Blast hits (query gene, subject gene, %identity, blast score, %coverage, e-value):
input.path1.gene6	PFICI_10087	33	256	96.1612284069098	5.3e-66
input.path1.gene7	PFICI_10090	42	211	97.44525547445255	1.3e-52
input.path1.gene10	PFICI_10088	37	162	13.865336658354115	4e-37


>>
27. NW_023336260
Source: Colletotrichum scovillei strain TJNH1 chromosome Unknown Contig00002, whole genome shotgun sequence
Type: T1PKS
Number of proteins with BLAST hits to this cluster: 3
Cumulative BLAST score: 618

Table of genes, locations, strands and annotations of subject cluster:

Table of Blast hits (query gene, subject gene, %identity, blast score, %coverage, e-value):
input.path1.gene7	HER10_EVM0007722	41	208	98.90510948905109	8.7e-52
input.path1.gene8	HER10_EVM0005941	45	83	31.11888111888112	3.4e-14
input.path1.gene10	HER10_EVM0007815	36	327	29.02743142144638	7.2e-87


>>
28. NC_030954
Source: Colletotrichum higginsianum IMI 349063 chromosome 1, whole genome shotgun sequence
Type: T1PKS
Number of proteins with BLAST hits to this cluster: 3
Cumulative BLAST score: 494

Table of genes, locations, strands and annotations of subject cluster:

Table of Blast hits (query gene, subject gene, %identity, blast score, %coverage, e-value):
input.path1.gene7	CH63R_00155	35	177	115.32846715328466	1.6e-42
input.path1.gene8	CH63R_00146	32	84	59.79020979020979	2.6e-14
input.path1.gene10	CH63R_00145	33	233	27.830423940149622	2.4e-58


>>
29. NW_019716265
Source: Ramularia collo-cygni strain URUG2 genome assembly, contig: RCC_scaffold11, whole genome shotgun sequence
Type: T1PKS
Number of proteins with BLAST hits to this cluster: 2
Cumulative BLAST score: 2071

Table of genes, locations, strands and annotations of subject cluster:

Table of Blast hits (query gene, subject gene, %identity, blast score, %coverage, e-value):
input.path1.gene9	RCC_07518	52	301	50.256410256410255	2.1e-79
input.path1.gene10	RCC_07520	45	1770	112.06982543640898	0.0


>>
30. NZ_CP031194
Source: Streptomyces paludis strain GSSD-12 chromosome, complete genome
Type: NRPS,T1PKS
Number of proteins with BLAST hits to this cluster: 2
Cumulative BLAST score: 647

Table of genes, locations, strands and annotations of subject cluster:
DVK44_RS35535	DVK44_RS35535	8241321	8245454	+	acyltransferase_domain-containing_protein
DVK44_RS36870	DVK44_RS36870	8245455	8245634	+	hypothetical_protein
DVK44_RS35540	DVK44_RS35540	8246097	8246816	+	SDR_family_NAD(P)-dependent_oxidoreductase
DVK44_RS35545	DVK44_RS35545	8246955	8260757	+	SDR_family_NAD(P)-dependent_oxidoreductase
DVK44_RS35550	DVK44_RS35550	8261094	8262218	+	SDR_family_NAD(P)-dependent_oxidoreductase
DVK44_RS35695	DVK44_RS35695	8320606	8320803	+	XRE_family_transcriptional_regulator
DVK44_RS35755	DVK44_RS35755	8340402	8341031	-	SDR_family_NAD(P)-dependent_oxidoreductase
DVK44_RS35760	DVK44_RS35760	8341662	8342183	-	alcohol_dehydrogenase_catalytic_domain-containing_protein
DVK44_RS35765	DVK44_RS35765	8342544	8343608	-	polyketide_synthase_dehydratase_domain-containing_protein
DVK44_RS35770	DVK44_RS35770	8343651	8344448	-	acyltransferase_domain-containing_protein
DVK44_RS35775	DVK44_RS35775	8344593	8345003	-	polyketide_synthase
DVK44_RS35780	DVK44_RS35780	8345331	8346038	-	polyketide_synthase
DVK44_RS35785	DVK44_RS35785	8346555	8347457	-	acyltransferase_domain-containing_protein
DVK44_RS35790	DVK44_RS35790	8347683	8348024	-	hypothetical_protein
DVK44_RS35795	DVK44_RS35795	8348226	8349659	-	hypothetical_protein
DVK44_RS35800	DVK44_RS35800	8349672	8350679	-	SDR_family_NAD(P)-dependent_oxidoreductase
DVK44_RS35805	DVK44_RS35805	8350881	8351954	-	polyketide_synthase_dehydratase_domain-containing_protein
DVK44_RS35810	DVK44_RS35810	8353185	8354519	-	polyketide_synthase_docking_domain-containing_protein
DVK44_RS36600	DVK44_RS36600	8362741	8363079	-	hypothetical_protein
DVK44_RS35845	DVK44_RS35845	8364373	8365641	-	polyketide_synthase
DVK44_RS35890	DVK44_RS35890	8383663	8384239	+	DUF4157_domain-containing_protein

Table of Blast hits (query gene, subject gene, %identity, blast score, %coverage, e-value):
input.path1.gene7	DVK44_RS35915	36	136	88.32116788321169	5.5e-30
input.path1.gene10	DVK44_RS35630	31	258	30.872817955112218	5.4e-66
input.path1.gene10	DVK44_RS35850	35	253	25.835411471321695	1.7e-64


>>
31. NZ_CP034352
Source: Streptomyces sp. W1SF4 plasmid p2, complete sequence
Type: NRPS,NRPS-like,T1PKS,T2PKS,butyrolactone,melanin
Number of proteins with BLAST hits to this cluster: 2
Cumulative BLAST score: 554

Table of genes, locations, strands and annotations of subject cluster:
D1J60_RS36170	D1J60_RS36170	1038	1672	+	TetR_family_transcriptional_regulator
D1J60_RS36175	D1J60_RS36175	1727	1906	-	hypothetical_protein
D1J60_RS36435	D1J60_RS36435	136120	141220	-	acyltransferase_domain-containing_protein
D1J60_RS37620	D1J60_RS37620	183563	183817	-	ATP-binding_protein
D1J60_RS36485	D1J60_RS36485	184321	185095	-	AfsR/SARP_family_transcriptional_regulator
D1J60_RS36505	D1J60_RS36505	189346	192303	+	acyltransferase_domain-containing_protein
D1J60_RS36675	D1J60_RS36675	227432	229621	+	MMPL_family_transporter
D1J60_RS36725	D1J60_RS36725	243784	245379	-	acyl--CoA_ligase
D1J60_RS36990	D1J60_RS36990	298697	299017	-	TetR_family_transcriptional_regulator

Table of Blast hits (query gene, subject gene, %identity, blast score, %coverage, e-value):
input.path1.gene7	D1J60_RS36285	32	82	61.31386861313869	7.2e-14
input.path1.gene10	D1J60_RS36415	33	238	25.187032418952622	7.5e-60
input.path1.gene10	D1J60_RS36450	33	234	26.184538653366584	8.3e-59


>>
32. NC_016111
Source: Streptomyces cattleya NRRL 8057 = DSM 46488, complete genome
Type: CDPS,T1PKS
Number of proteins with BLAST hits to this cluster: 2
Cumulative BLAST score: 552

Table of genes, locations, strands and annotations of subject cluster:
SCAT_RS08835	SCAT_RS08835	198983	199468	+	MBL_fold_metallo-hydrolase
SCAT_RS38670	SCAT_RS38670	209547	209711	+	coenzyme_F420-0:L-glutamate_ligase
SCAT_RS38260	SCAT_RS38260	223578	224482	-	hypothetical_protein
SCAT_RS36850	SCAT_RS36850	229458	229861	-	tautomerase_family_protein
SCAT_RS36855	SCAT_RS36855	230843	231022	-	guanosine_monophosphate_reductase

Table of Blast hits (query gene, subject gene, %identity, blast score, %coverage, e-value):
input.path1.gene9	SCAT_RS08945	36	138	45.81196581196581	2.4e-30
input.path1.gene10	SCAT_RS08820	31	270	35.062344139650875	1.4e-69
input.path1.gene10	SCAT_RS08805	34	144	13.41645885286783	1.1e-31


>>
33. NC_030962
Source: Colletotrichum higginsianum IMI 349063 chromosome 10, whole genome shotgun sequence
Type: T1PKS
Number of proteins with BLAST hits to this cluster: 2
Cumulative BLAST score: 484

Table of genes, locations, strands and annotations of subject cluster:

Table of Blast hits (query gene, subject gene, %identity, blast score, %coverage, e-value):
input.path1.gene7	CH63R_13917	43	225	99.63503649635037	6.9e-57
input.path1.gene10	CH63R_13918	32	259	30.972568578553616	4.1e-66


>>
34. NW_006912895
Source: Capronia coronata CBS 617.96 acBCG-supercont1.3, whole genome shotgun sequence
Type: T1PKS
Number of proteins with BLAST hits to this cluster: 2
Cumulative BLAST score: 432

Table of genes, locations, strands and annotations of subject cluster:

Table of Blast hits (query gene, subject gene, %identity, blast score, %coverage, e-value):
input.path1.gene7	A1O1_04523	53	278	98.54014598540147	9e-73
input.path1.gene10	A1O1_04517	34	154	13.466334164588527	1.1e-34


>>
35. NZ_KN323065
Source: Burkholderia pseudomallei MSHR1000 scaffold1, whole genome shotgun sequence
Type: PKS-like,transAT-PKS
Number of proteins with BLAST hits to this cluster: 2
Cumulative BLAST score: 428

Table of genes, locations, strands and annotations of subject cluster:
Y597_RS21380	Y597_RS21380	4797181	4797649	+	potassium_transporter_TrkA
Y597_RS21385	Y597_RS21385	4797750	4814755	-	SDR_family_NAD(P)-dependent_oxidoreductase
Y597_RS21490	Y597_RS21490	4849225	4850472	-	MFS_transporter

Table of Blast hits (query gene, subject gene, %identity, blast score, %coverage, e-value):
input.path1.gene7	Y597_RS21350	35	129	88.6861313868613	5.1e-28
input.path1.gene10	Y597_RS21390	33	151	13.16708229426434	1.2e-33
input.path1.gene10	Y597_RS21385	33	148	13.017456359102244	1e-32


>>
36. NZ_CP013395
Source: Burkholderia vietnamiensis strain FL-2-3-30-S1-D0 chromosome 3, complete sequence
Type: NRPS,T1PKS
Number of proteins with BLAST hits to this cluster: 2
Cumulative BLAST score: 409

Table of genes, locations, strands and annotations of subject cluster:
WJ02_RS25490	WJ02_RS25490	192706	192963	+	helix-turn-helix_domain-containing_protein
WJ02_RS25505	WJ02_RS25505	195015	196524	-	aminotransferase_class_I/II-fold_pyridoxal_phosphate-dependent_enzyme
WJ02_RS31780	WJ02_RS31780	198697	198864	-	IS256_family_transposase
WJ02_RS25540	WJ02_RS25540	201617	202075	+	LuxR_family_transcriptional_regulator
WJ02_RS31785	WJ02_RS31785	202797	202952	-	EamA/RhaT_family_transporter

Table of Blast hits (query gene, subject gene, %identity, blast score, %coverage, e-value):
input.path1.gene7	WJ02_RS25525	31	74	79.1970802919708	1.5e-11
input.path1.gene10	WJ02_RS25460	38	174	13.96508728179551	1.3e-40
input.path1.gene10	WJ02_RS25450	38	161	13.316708229426432	1.2e-36


>>
37. NC_018217
Source: Zymoseptoria tritici IPO323 chromosome 2, whole genome shotgun sequence
Type: T1PKS
Number of proteins with BLAST hits to this cluster: 2
Cumulative BLAST score: 407

Table of genes, locations, strands and annotations of subject cluster:

Table of Blast hits (query gene, subject gene, %identity, blast score, %coverage, e-value):
input.path1.gene4	MYCGRDRAFT_36644	32	259	94.5945945945946	1.1e-66
input.path1.gene10	MYCGRDRAFT_9788	34	148	13.765586034912719	1e-32


>>
38. NW_019716258
Source: Ramularia collo-cygni strain URUG2 genome assembly, contig: RCC_scaffold04, whole genome shotgun sequence
Type: T1PKS,terpene
Number of proteins with BLAST hits to this cluster: 2
Cumulative BLAST score: 406

Table of genes, locations, strands and annotations of subject cluster:

Table of Blast hits (query gene, subject gene, %identity, blast score, %coverage, e-value):
input.path1.gene4	RCC_03444	32	249	94.77477477477477	6.9e-64
input.path1.gene10	RCC_03448	35	157	14.01496259351621	1.7e-35


>>
39. NZ_CP048641
Source: Streptomyces aureoverticillatus strain HN6 chromosome, complete genome
Type: T1PKS
Number of proteins with BLAST hits to this cluster: 2
Cumulative BLAST score: 404

Table of genes, locations, strands and annotations of subject cluster:
G3H79_RS03770	G3H79_RS03770	1081786	1082475	-	SDR_family_NAD(P)-dependent_oxidoreductase
G3H79_RS03775	G3H79_RS03775	1082893	1084824	-	acyltransferase_domain-containing_protein
G3H79_RS03780	G3H79_RS03780	1085068	1087023	-	hypothetical_protein
G3H79_RS03785	G3H79_RS03785	1087084	1088025	-	SDR_family_NAD(P)-dependent_oxidoreductase
G3H79_RS03790	G3H79_RS03790	1088572	1089480	-	acyltransferase_domain-containing_protein
G3H79_RS03795	G3H79_RS03795	1089847	1096566	-	SDR_family_NAD(P)-dependent_oxidoreductase
G3H79_RS03810	G3H79_RS03810	1121156	1121674	-	polyketide_synthase
G3H79_RS03815	G3H79_RS03815	1121684	1122382	-	SDR_family_NAD(P)-dependent_oxidoreductase
G3H79_RS03820	G3H79_RS03820	1125437	1126663	-	polyketide_synthase
G3H79_RS04075	G3H79_RS04075	1241812	1242810	-	Gfo/Idh/MocA_family_oxidoreductase

Table of Blast hits (query gene, subject gene, %identity, blast score, %coverage, e-value):
input.path1.gene7	G3H79_RS03955	35	127	88.32116788321169	2e-27
input.path1.gene10	G3H79_RS03755	31	277	35.261845386533665	8.6e-72


>>
40. NC_016109
Source: Kitasatospora setae KM-6054, complete genome
Type: NRPS,T1PKS,transAT-PKS
Number of proteins with BLAST hits to this cluster: 2
Cumulative BLAST score: 397

Table of genes, locations, strands and annotations of subject cluster:
KSE_RS42480	KSE_RS42480	7957789	7957881	+	HEAT_repeat_domain-containing_protein
KSE_RS40580	KSE_RS40580	7958636	7958893	-	IS5/IS1182_family_transposase

Table of Blast hits (query gene, subject gene, %identity, blast score, %coverage, e-value):
input.path1.gene7	KSE_RS34885	32	77	90.14598540145985	3e-12
input.path1.gene10	KSE_RS34910	37	169	13.366583541147131	3.3e-39
input.path1.gene10	KSE_RS34920	37	151	13.216957605985039	9.3e-34


>>
41. NC_030961
Source: Colletotrichum higginsianum IMI 349063 chromosome 9, whole genome shotgun sequence
Type: NRPS,T1PKS
Number of proteins with BLAST hits to this cluster: 2
Cumulative BLAST score: 376

Table of genes, locations, strands and annotations of subject cluster:

Table of Blast hits (query gene, subject gene, %identity, blast score, %coverage, e-value):
input.path1.gene6	CH63R_13248	34	230	86.9481765834933	4.1e-58
input.path1.gene10	CH63R_13259	35	146	13.266832917705736	3.9e-32


>>
42. NZ_LR130759
Source: Mycobacterium basiliense isolate 901379 chromosome I
Type: NRPS,T1PKS
Number of proteins with BLAST hits to this cluster: 2
Cumulative BLAST score: 370

Table of genes, locations, strands and annotations of subject cluster:
MB901379_RS13185	MB901379_RS13185	3086759	3087523	+	polyketide_synthase
MB901379_RS13190	MB901379_RS13190	3087779	3088255	+	hypothetical_protein
MB901379_RS13195	MB901379_RS13195	3089243	3090313	+	polyketide_synthase_dehydratase_domain-containing_protein
MB901379_RS13200	MB901379_RS13200	3090401	3091489	+	SDR_family_oxidoreductase
MB901379_RS13205	MB901379_RS13205	3091538	3091996	+	hypothetical_protein
MB901379_RS13210	MB901379_RS13210	3092072	3094228	+	type_I_polyketide_synthase
MB901379_RS23905	MB901379_RS23905	3097911	3098742	+	polyketide_synthase
MB901379_RS24000	MB901379_RS24000	3113476	3115542	+	type_I_polyketide_synthase
MB901379_RS24005	MB901379_RS24005	3115987	3117153	+	polyketide_synthase_dehydratase_domain-containing_protein
MB901379_RS24010	MB901379_RS24010	3117265	3117972	+	alcohol_dehydrogenase_catalytic_domain-containing_protein
MB901379_RS24015	MB901379_RS24015	3118582	3119241	+	SDR_family_NAD(P)-dependent_oxidoreductase
MB901379_RS24020	MB901379_RS24020	3119254	3119775	+	polyketide_synthase
MB901379_RS24025	MB901379_RS24025	3119863	3120588	+	polyketide_synthase
MB901379_RS24030	MB901379_RS24030	3120871	3121326	+	polyketide_synthase
MB901379_RS24035	MB901379_RS24035	3122344	3123444	+	polyketide_synthase_dehydratase_domain-containing_protein
MB901379_RS24040	MB901379_RS24040	3123556	3124263	+	alcohol_dehydrogenase_catalytic_domain-containing_protein
MB901379_RS13345	MB901379_RS13345	3168802	3169369	-	hypothetical_protein

Table of Blast hits (query gene, subject gene, %identity, blast score, %coverage, e-value):
input.path1.gene7	MB901379_RS13335	35	131	100.72992700729928	1e-28
input.path1.gene10	MB901379_RS13260	34	239	25.835411471321695	2.6e-60


>>
43. NZ_CP016795
Source: Streptomyces olivaceus strain KLBMP 5084 chromosome, complete genome
Type: NRPS,NRPS-like,T1PKS
Number of proteins with BLAST hits to this cluster: 2
Cumulative BLAST score: 366

Table of genes, locations, strands and annotations of subject cluster:
BC342_RS36325	BC342_RS36325	7094038	7095701	+	NAD(P)-dependent_oxidoreductase
BC342_RS30915	BC342_RS30915	7108967	7110057	+	isochorismate_synthase
BC342_RS31015	BC342_RS31015	7139451	7140293	+	sugar_ABC_transporter_ATP-binding_protein
BC342_RS31065	BC342_RS31065	7149042	7150358	+	aromatic_acid/H+_symport_family_MFS_transporter

Table of Blast hits (query gene, subject gene, %identity, blast score, %coverage, e-value):
input.path1.gene7	BC342_RS30820	36	132	88.32116788321169	7.9e-29
input.path1.gene10	BC342_RS37210	32	234	25.486284289276806	8.3e-59


>>
44. NZ_CP030771
Source: Streptomyces sp. YIM 121038 chromosome, complete genome
Type: T1PKS
Number of proteins with BLAST hits to this cluster: 2
Cumulative BLAST score: 358

Table of genes, locations, strands and annotations of subject cluster:
C9F11_RS33980	C9F11_RS33980	7945827	7948763	+	type_I_polyketide_synthase
C9F11_RS33985	C9F11_RS33985	7949394	7950674	+	SDR_family_NAD(P)-dependent_oxidoreductase
C9F11_RS33990	C9F11_RS33990	7950699	7951196	+	hypothetical_protein
C9F11_RS47610	C9F11_RS47610	7951290	7953962	+	type_I_polyketide_synthase
C9F11_RS34010	C9F11_RS34010	7954695	7956539	+	SDR_family_NAD(P)-dependent_oxidoreductase

Table of Blast hits (query gene, subject gene, %identity, blast score, %coverage, e-value):
input.path1.gene7	C9F11_RS34040	37	130	88.6861313868613	2.3e-28
input.path1.gene10	C9F11_RS34015	32	228	24.83790523690773	7.8e-57


>>
45. NT_165972
Source: Aspergillus terreus NIH2624 scaffold_1 genomic scaffold, whole genome shotgun sequence
Type: T1PKS
Number of proteins with BLAST hits to this cluster: 2
Cumulative BLAST score: 350

Table of genes, locations, strands and annotations of subject cluster:

Table of Blast hits (query gene, subject gene, %identity, blast score, %coverage, e-value):
input.path1.gene6	ATEG_00914	35	97	30.326295585412666	4.1e-18
input.path1.gene10	ATEG_00913	31	253	32.568578553615964	1.7e-64


>>
46. NZ_CP040916
Source: Streptomyces spectabilis strain NRRL 2792 chromosome, complete genome
Type: T1PKS
Number of proteins with BLAST hits to this cluster: 2
Cumulative BLAST score: 349

Table of genes, locations, strands and annotations of subject cluster:

Table of Blast hits (query gene, subject gene, %identity, blast score, %coverage, e-value):
input.path1.gene7	FH965_RS33320	36	129	88.6861313868613	6.7e-28
input.path1.gene10	FH965_RS33295	33	220	25.386533665835408	2.1e-54


>>
47. NZ_CP023690
Source: Streptomyces spectabilis strain ATCC 27465 chromosome, complete genome
Type: T1PKS
Number of proteins with BLAST hits to this cluster: 2
Cumulative BLAST score: 347

Table of genes, locations, strands and annotations of subject cluster:

Table of Blast hits (query gene, subject gene, %identity, blast score, %coverage, e-value):
input.path1.gene7	CP982_RS34715	37	131	88.6861313868613	1e-28
input.path1.gene10	CP982_RS34690	32	216	25.13715710723192	3.1e-53


>>
48. NC_021191
Source: Actinoplanes sp. N902-109, complete genome
Type: T1PKS
Number of proteins with BLAST hits to this cluster: 2
Cumulative BLAST score: 344

Table of genes, locations, strands and annotations of subject cluster:
L083_RS43460	L083_RS43460	2651371	2652237	-	SDR_family_NAD(P)-dependent_oxidoreductase
L083_RS43465	L083_RS43465	2652505	2653632	-	beta-ketoacyl_synthase
L083_RS43475	L083_RS43475	2655193	2659158	-	acyltransferase_domain-containing_protein
L083_RS43480	L083_RS43480	2659279	2660865	-	hypothetical_protein
L083_RS12105	L083_RS12105	2691701	2692675	-	alpha/beta_hydrolase
L083_RS12155	L083_RS12155	2704010	2704359	+	glyoxalase

Table of Blast hits (query gene, subject gene, %identity, blast score, %coverage, e-value):
input.path1.gene7	L083_RS12065	33	119	91.97080291970804	4.1e-25
input.path1.gene10	L083_RS12090	32	225	25.13715710723192	3.9e-56


>>
49. NZ_LIVY01000012
Source: Streptomyces sp. CB02460 scaffold12, whole genome shotgun sequence
Type: NRPS,PKS-like,T1PKS,butyrolactone,oligosaccharide
Number of proteins with BLAST hits to this cluster: 2
Cumulative BLAST score: 340

Table of genes, locations, strands and annotations of subject cluster:
AMK30_RS30945	AMK30_RS30945	2529	2738	+	IS5/IS1182_family_transposase
AMK30_RS33875	AMK30_RS33875	59682	59795	+	collagen-like_protein
AMK30_RS33880	AMK30_RS33880	60613	61563	-	polyketide_synthase
AMK30_RS33885	AMK30_RS33885	62896	63372	-	hypothetical_protein
AMK30_RS33890	AMK30_RS33890	64123	73215	-	SDR_family_NAD(P)-dependent_oxidoreductase
AMK30_RS31200	AMK30_RS31200	80502	92239	-	SDR_family_NAD(P)-dependent_oxidoreductase
AMK30_RS31435	AMK30_RS31435	164323	165213	-	enediyne_biosynthesis_protein
AMK30_RS33895	AMK30_RS33895	176609	176773	-	HAD-IB_family_hydrolase

Table of Blast hits (query gene, subject gene, %identity, blast score, %coverage, e-value):
input.path1.gene7	AMK30_RS31065	33	111	100.36496350364963	1.9e-22
input.path1.gene10	AMK30_RS31220	32	229	25.6857855361596	2.7e-57


>>
50. NZ_CP053892
Source: Actinomadura verrucosospora strain NRRL:B18236 chromosome
Type: NRPS,T1PKS
Number of proteins with BLAST hits to this cluster: 2
Cumulative BLAST score: 338

Table of genes, locations, strands and annotations of subject cluster:
ACTIVE_RS04015	ACTIVE_RS04015	873488	873766	-	hypothetical_protein
ACTIVE_RS04090	ACTIVE_RS04090	904142	905212	-	SDR_family_NAD(P)-dependent_oxidoreductase
ACTIVE_RS04095	ACTIVE_RS04095	906023	913513	-	SDR_family_NAD(P)-dependent_oxidoreductase
ACTIVE_RS04100	ACTIVE_RS04100	915140	918031	-	type_I_polyketide_synthase
ACTIVE_RS04125	ACTIVE_RS04125	923824	928365	-	KR_domain-containing_protein
ACTIVE_RS04130	ACTIVE_RS04130	928882	936180	-	acyltransferase_domain-containing_protein
ACTIVE_RS04135	ACTIVE_RS04135	936571	937920	-	polyketide_synthase
ACTIVE_RS04150	ACTIVE_RS04150	958665	963170	-	SDR_family_NAD(P)-dependent_oxidoreductase
ACTIVE_RS04155	ACTIVE_RS04155	963444	964958	-	polyketide_synthase_docking_domain-containing_protein

Table of Blast hits (query gene, subject gene, %identity, blast score, %coverage, e-value):
input.path1.gene7	ACTIVE_RS04265	32	68	64.96350364963503	1.4e-09
input.path1.gene10	ACTIVE_RS04060	34	270	27.680798004987533	1.8e-69
