## Supplementary Results for "A draft genome of the ascomycotal fungal species *Pseudopithomyces maydicus* (family *Didymosphaeriaceae*)": contig_26_c2.txt

ClusterBlast scores for contig_26

Table of genes, locations, strands and annotations of query cluster:
input.path1.gene202	739460	741433	+		
input.path1.gene203	746577	747002	+		
input.path1.gene204	747015	751535	-		
input.path1.gene205	752565	753169	+		
input.path1.gene206	753642	754399	-		
input.path1.gene207	756918	763489	-		
input.path1.gene208	775206	778416	+		


Significant hits: 
1. NW_019716270	Ramularia collo-cygni strain URUG2 genome assembly, contig: RCC_scaffold16, whole genome shotgun sequence
2. NW_001939245	Pyrenophora tritici-repentis Pt-1C-BFP supercont1.2, whole genome shotgun sequence
3. NC_018208	Zymoseptoria tritici IPO323 chromosome 11, whole genome shotgun sequence
4. CP051142	Peltaster fructicola strain LNHT1506 chromosome 4
5. CP042190	Venturia effusa strain albino chromosome 6, complete sequence
6. NW_006911258	Baudoinia panamericana UAMH 10762 unplaced genomic scaffold BAUCOscaffold_2, whole genome shotgun sequence
7. NW_006763039	Marssonina brunnea f. sp. 'multigermtubi' MB_m1 unplaced genomic scaffold M6_S00001, whole genome shotgun sequence
8. NW_022474218	Venustampulla echinocandica strain BP 5553 chromosome Unknown scaffold_13, whole genome shotgun sequence
9. NW_020194471	Amorphotheca resinae ATCC 22711 unplaced genomic scaffold M430scaffold_1, whole genome shotgun sequence
10. NW_007360986	Glarea lozoyensis ATCC 20868 chromosome Unknown GLAREA12, whole genome shotgun sequence
11. NW_020167541	Pseudogymnoascus destructans isolate 20631-21 unplaced genomic scaffold scaffold_22, whole genome shotgun sequence
12. NW_015971149	Sporothrix schenckii 1099-18 chromosome Unknown Cont48, whole genome shotgun sequence
13. NT_165976	Chaetomium globosum CBS 148.51 scaffold_1 genomic scaffold, whole genome shotgun sequence
14. NC_016473	Thermothelomyces thermophilus ATCC 42464 chromosome 2, complete sequence
15. NW_003315038	Verticillium alfalfae VaMs.102 supercont1.1 genomic scaffold, whole genome shotgun sequence
16. NZ_CP021983	Halomicronema hongdechloris C2206 chromosome


Details:

>>
1. NW_019716270
Source: Ramularia collo-cygni strain URUG2 genome assembly, contig: RCC_scaffold16, whole genome shotgun sequence
Type: T1PKS
Number of proteins with BLAST hits to this cluster: 3
Cumulative BLAST score: 3705

Table of genes, locations, strands and annotations of subject cluster:

Table of Blast hits (query gene, subject gene, %identity, blast score, %coverage, e-value):
input.path1.gene206	RCC_09027	76	280	91.95979899497488	1.3e-73
input.path1.gene207	RCC_09026	63	2623	104.0360873694207	0.0
input.path1.gene208	RCC_09025	49	802	120.44334975369458	4e-230


>>
2. NW_001939245
Source: Pyrenophora tritici-repentis Pt-1C-BFP supercont1.2, whole genome shotgun sequence
Type: T1PKS
Number of proteins with BLAST hits to this cluster: 3
Cumulative BLAST score: 4525

Table of genes, locations, strands and annotations of subject cluster:

Table of Blast hits (query gene, subject gene, %identity, blast score, %coverage, e-value):
input.path1.gene206	PTRG_03324	85	329	100.0	3.2e-88
input.path1.gene207	PTRG_03323	76	3136	100.47483380816715	0.0
input.path1.gene208	PTRG_03318	64	1060	123.89162561576354	8.2e-308


>>
3. NC_018208
Source: Zymoseptoria tritici IPO323 chromosome 11, whole genome shotgun sequence
Type: T1PKS
Number of proteins with BLAST hits to this cluster: 3
Cumulative BLAST score: 3711

Table of genes, locations, strands and annotations of subject cluster:

Table of Blast hits (query gene, subject gene, %identity, blast score, %coverage, e-value):
input.path1.gene206	MYCGRDRAFT_96591	66	255	97.98994974874373	3.5e-66
input.path1.gene207	MYCGRDRAFT_96592	63	2600	103.79867046533712	0.0
input.path1.gene208	MYCGRDRAFT_87993	51	856	120.6896551724138	2.4e-246


>>
4. CP051142
Source: Peltaster fructicola strain LNHT1506 chromosome 4
Type: T1PKS
Number of proteins with BLAST hits to this cluster: 3
Cumulative BLAST score: 3680

Table of genes, locations, strands and annotations of subject cluster:

Table of Blast hits (query gene, subject gene, %identity, blast score, %coverage, e-value):
input.path1.gene206	AMS68_006177	67	258	92.46231155778895	7e-67
input.path1.gene207	AMS68_006178	65	2707	103.60873694207027	0.0
input.path1.gene208	AMS68_006180	53	715	88.54679802955665	6.5e-204


>>
5. CP042190
Source: Venturia effusa strain albino chromosome 6, complete sequence
Type: T1PKS
Number of proteins with BLAST hits to this cluster: 2
Cumulative BLAST score: 3601

Table of genes, locations, strands and annotations of subject cluster:

Table of Blast hits (query gene, subject gene, %identity, blast score, %coverage, e-value):
input.path1.gene207	FKW77_000779	66	2750	104.7483380816714	0.0
input.path1.gene208	FKW77_000689	52	851	121.42857142857142	7.6e-245


>>
6. NW_006911258
Source: Baudoinia panamericana UAMH 10762 unplaced genomic scaffold BAUCOscaffold_2, whole genome shotgun sequence
Type: T1PKS
Number of proteins with BLAST hits to this cluster: 2
Cumulative BLAST score: 3149

Table of genes, locations, strands and annotations of subject cluster:

Table of Blast hits (query gene, subject gene, %identity, blast score, %coverage, e-value):
input.path1.gene207	BAUCODRAFT_30707	66	2319	89.79107312440647	0.0
input.path1.gene208	BAUCODRAFT_154260	51	830	120.44334975369458	1.4e-238


>>
7. NW_006763039
Source: Marssonina brunnea f. sp. 'multigermtubi' MB_m1 unplaced genomic scaffold M6_S00001, whole genome shotgun sequence
Type: T1PKS
Number of proteins with BLAST hits to this cluster: 2
Cumulative BLAST score: 2367

Table of genes, locations, strands and annotations of subject cluster:

Table of Blast hits (query gene, subject gene, %identity, blast score, %coverage, e-value):
input.path1.gene207	MBM_00260	49	1933	103.51377018043684	0.0
input.path1.gene208	MBM_00257	34	434	107.75862068965519	2.2e-119


>>
8. NW_022474218
Source: Venustampulla echinocandica strain BP 5553 chromosome Unknown scaffold_13, whole genome shotgun sequence
Type: T1PKS
Number of proteins with BLAST hits to this cluster: 2
Cumulative BLAST score: 2359

Table of genes, locations, strands and annotations of subject cluster:

Table of Blast hits (query gene, subject gene, %identity, blast score, %coverage, e-value):
input.path1.gene207	BP5553_10011	48	1908	103.3713200379867	0.0
input.path1.gene208	BP5553_10013	33	451	117.73399014778325	2.3e-124


>>
9. NW_020194471
Source: Amorphotheca resinae ATCC 22711 unplaced genomic scaffold M430scaffold_1, whole genome shotgun sequence
Type: T1PKS
Number of proteins with BLAST hits to this cluster: 2
Cumulative BLAST score: 2334

Table of genes, locations, strands and annotations of subject cluster:

Table of Blast hits (query gene, subject gene, %identity, blast score, %coverage, e-value):
input.path1.gene207	M430DRAFT_93061	48	1897	103.0389363722697	0.0
input.path1.gene208	M430DRAFT_94388	32	437	116.25615763546797	3.4e-120


>>
10. NW_007360986
Source: Glarea lozoyensis ATCC 20868 chromosome Unknown GLAREA12, whole genome shotgun sequence
Type: T1PKS
Number of proteins with BLAST hits to this cluster: 2
Cumulative BLAST score: 2263

Table of genes, locations, strands and annotations of subject cluster:

Table of Blast hits (query gene, subject gene, %identity, blast score, %coverage, e-value):
input.path1.gene207	GLAREA_10585	48	1862	102.84900284900284	0.0
input.path1.gene208	GLAREA_10583	38	401	78.20197044334975	1.6e-109


>>
11. NW_020167541
Source: Pseudogymnoascus destructans isolate 20631-21 unplaced genomic scaffold scaffold_22, whole genome shotgun sequence
Type: T1PKS
Number of proteins with BLAST hits to this cluster: 2
Cumulative BLAST score: 2222

Table of genes, locations, strands and annotations of subject cluster:

Table of Blast hits (query gene, subject gene, %identity, blast score, %coverage, e-value):
input.path1.gene207	VC83_07905	45	1833	104.22602089268756	0.0
input.path1.gene208	VC83_07908	34	389	116.7487684729064	1.1e-105


>>
12. NW_015971149
Source: Sporothrix schenckii 1099-18 chromosome Unknown Cont48, whole genome shotgun sequence
Type: T1PKS
Number of proteins with BLAST hits to this cluster: 2
Cumulative BLAST score: 2198

Table of genes, locations, strands and annotations of subject cluster:

Table of Blast hits (query gene, subject gene, %identity, blast score, %coverage, e-value):
input.path1.gene207	SPSK_00653	46	1788	105.27065527065527	0.0
input.path1.gene208	SPSK_00677	42	410	64.77832512315271	4.5e-112


>>
13. NT_165976
Source: Chaetomium globosum CBS 148.51 scaffold_1 genomic scaffold, whole genome shotgun sequence
Type: T1PKS
Number of proteins with BLAST hits to this cluster: 2
Cumulative BLAST score: 2124

Table of genes, locations, strands and annotations of subject cluster:

Table of Blast hits (query gene, subject gene, %identity, blast score, %coverage, e-value):
input.path1.gene207	CHGG_00542	43	1704	104.41595441595442	0.0
input.path1.gene208	CHGG_00539	36	420	86.33004926108374	4.3e-115


>>
14. NC_016473
Source: Thermothelomyces thermophilus ATCC 42464 chromosome 2, complete sequence
Type: T1PKS
Number of proteins with BLAST hits to this cluster: 2
Cumulative BLAST score: 2073

Table of genes, locations, strands and annotations of subject cluster:

Table of Blast hits (query gene, subject gene, %identity, blast score, %coverage, e-value):
input.path1.gene207	MYCTH_2300170	44	1665	107.0275403608737	0.0
input.path1.gene208	MYCTH_2059558	38	408	78.69458128078819	2.2e-111


>>
15. NW_003315038
Source: Verticillium alfalfae VaMs.102 supercont1.1 genomic scaffold, whole genome shotgun sequence
Type: T1PKS
Number of proteins with BLAST hits to this cluster: 2
Cumulative BLAST score: 1972

Table of genes, locations, strands and annotations of subject cluster:

Table of Blast hits (query gene, subject gene, %identity, blast score, %coverage, e-value):
input.path1.gene207	VDBG_00580	46	1659	98.71794871794873	0.0
input.path1.gene208	VDBG_00583	37	313	74.01477832512316	5.7e-83


>>
16. NZ_CP021983
Source: Halomicronema hongdechloris C2206 chromosome
Type: NRPS-like,T1PKS
Number of proteins with BLAST hits to this cluster: 2
Cumulative BLAST score: 501

Table of genes, locations, strands and annotations of subject cluster:
XM38_RS24135	XM38_RS24135	5285315	5287113	-	ShlB/FhaC/HecB_family_hemolysin_secretion/activation_protein
XM38_RS24215	XM38_RS24215	5311766	5311837	+	DUF3598_family_protein

Table of Blast hits (query gene, subject gene, %identity, blast score, %coverage, e-value):
input.path1.gene205	XM38_RS24240	45	91	75.67567567567568	8.4e-17
input.path1.gene207	XM38_RS24155	32	410	44.396961063627735	8.9e-112
