## Supplementary Results for "A draft genome of the ascomycotal fungal species *Pseudopithomyces maydicus* (family *Didymosphaeriaceae*)": contig_31_c1.txt

ClusterBlast scores for contig_31

Table of genes, locations, strands and annotations of query cluster:
input.path1.gene167	586492	588372	-		
input.path1.gene168	589277	590058	+		
input.path1.gene169	590312	592067	-		
input.path1.gene170	593622	594983	+		
input.path1.gene171	596789	602033	+		
input.path1.gene172	603404	604118	+		
input.path1.gene173	605392	609288	+		
input.path1.gene174	609852	610539	+		
input.path1.gene175	610760	611714	-		
input.path1.gene176	612514	613519	-		
input.path1.gene177	618698	620245	-		
input.path1.gene178	621780	623606	-		


Significant hits: 
1. NW_001914858	Podospora anserina S mat+ genomic DNA chromosome 2, supercontig 2


Details:

>>
1. NW_001914858
Source: Podospora anserina S mat+ genomic DNA chromosome 2, supercontig 2
Type: NRPS
Number of proteins with BLAST hits to this cluster: 2
Cumulative BLAST score: 291

Table of genes, locations, strands and annotations of subject cluster:

Table of Blast hits (query gene, subject gene, %identity, blast score, %coverage, e-value):
input.path1.gene177	PODANS_2_7820	40	190	72.33502538071066	4.6e-46
input.path1.gene178	PODANS_2_7800	33	101	45.98540145985402	3e-19
