## Supplementary Results for "A draft genome of the ascomycotal fungal species *Pseudopithomyces maydicus* (family *Didymosphaeriaceae*)": contig_32_c1.txt

ClusterBlast scores for contig_32

Table of genes, locations, strands and annotations of query cluster:
input.path1.gene171	619649	620129	+		
input.path1.gene172	622952	623971	+		
input.path1.gene173	626033	627314	-		
input.path1.gene174	630452	631010	+		
input.path1.gene175	634733	635246	+		
input.path1.gene176	635480	637571	-		
input.path1.gene177	638089	643360	+		
input.path1.gene178	645232	645777	-		
input.path1.gene179	647508	648473	+		
input.path1.gene180	649745	650600	-		
input.path1.gene181	651140	655720	+		
input.path1.gene182	655746	656661	-		


Significant hits: 
1. NW_001939248	Pyrenophora tritici-repentis Pt-1C-BFP supercont1.5, whole genome shotgun sequence
2. NW_017264203	Xylona heveae TC161 unplaced genomic scaffold L228scaffold_11, whole genome shotgun sequence
3. NW_023336278	Aspergillus tubingensis WU-2223L DNA, scaffold_4, whole genome shotgun sequence
4. NW_020194484	Amorphotheca resinae ATCC 22711 unplaced genomic scaffold M430scaffold_14, whole genome shotgun sequence
5. NC_007197	Aspergillus fumigatus Af293 chromosome 4, whole genome shotgun sequence
6. NW_022474205	Venustampulla echinocandica strain BP 5553 chromosome Unknown scaffold_0, whole genome shotgun sequence
7. NC_036436	Aspergillus oryzae RIB40 DNA, chromosome 2
8. NT_165928	Aspergillus terreus NIH2624 scaffold_5 genomic scaffold, whole genome shotgun sequence
9. NW_022984630	Aspergillus tanneri strain NIH1004 chromosome Unknown AapTan_3_pilon_pilon_pilon, whole genome shotgun sequence
10. NW_006763065	Marssonina brunnea f. sp. 'multigermtubi' MB_m1 unplaced genomic scaffold M6_S00027, whole genome shotgun sequence
11. CM002800	Penicillium chrysogenum strain P2niaD18 chromosome III, whole genome shotgun sequence
12. NW_004504310	Coccidioides immitis RS genomic scaffold supercont3.1, whole genome shotgun sequence
13. NW_003315979	Coccidioides posadasii C735 delta SOWgp chromosome Unknown gcontig_1105601051781, whole genome shotgun sequence
14. NW_003052498	Uncinocarpus reesii 1704 scaffold_3 genomic scaffold, whole genome shotgun sequence
15. NW_014574601	Penicillium digitatum Pd1 unplaced genomic scaffold Scaffold_24, whole genome shotgun sequence
16. NW_007360994	Glarea lozoyensis ATCC 20868 chromosome Unknown GLAREA2, whole genome shotgun sequence
17. NW_017263917	Colletotrichum higginsianum IMI 349063 chromosome 7 map unlocalized, whole genome shotgun sequence
18. NW_006912893	Capronia coronata CBS 617.96 acBCG-supercont1.1, whole genome shotgun sequence
19. NW_008481808	Cyphellophora europaea CBS 101466 unplaced genomic scaffold supercont1.2, whole genome shotgun sequence
20. NW_008751650	Exophiala dermatitidis NIH/UT8656 unplaced genomic scaffold supercont1.5, whole genome shotgun sequence
21. NW_006912905	Capronia epimyces CBS 606.96 aczIQ-supercont1.2, whole genome shotgun sequence
22. NW_015622516	Exophiala spinifera strain CBS 89968 unplaced genomic scaffold supercont1.4, whole genome shotgun sequence
23. NW_015971648	Fonsecaea multimorphosa CBS 102226 unplaced genomic scaffold supercont1.14, whole genome shotgun sequence
24. NW_017387248	Fonsecaea erecta strain CBS 125763 chromosome Unknown Scaff5, whole genome shotgun sequence
25. NW_013550611	Fonsecaea pedrosoi CBS 271.37 unplaced genomic scaffold supercont1.5, whole genome shotgun sequence
26. NW_015971588	Cladophialophora bantiana CBS 173.52 unplaced genomic scaffold supercont1.14, whole genome shotgun sequence
27. NW_015622504	Exophiala mesophila strain CBS 40295 unplaced genomic scaffold supercont1.1, whole genome shotgun sequence
28. NW_001820825	Sclerotinia sclerotiorum 1980 UF-70 scaffold_11 genomic scaffold, whole genome shotgun sequence
29. NT_107010	Aspergillus nidulans FGSC A4 chromosome V map unlocalized scaffold_6, whole genome shotgun sequence
30. NW_006913045	Cladophialophora yegresii CBS 114405 aczIN-supercont1.1, whole genome shotgun sequence
31. NW_008481816	Cladophialophora carrionii CBS 160.54 unplaced genomic scaffold supercont1.1, whole genome shotgun sequence
32. NC_035790	Pochonia chlamydosporia 170 chromosome 1, whole genome shotgun sequence
33. CM000576	Fusarium graminearum PH-1 chromosome 3, whole genome shotgun sequence
34. NC_026476	Fusarium graminearum PH-1 chromosome 3, whole genome shotgun sequence
35. NC_038014	Fusarium venenatum strain A3/5 genome assembly, chromosome: III


Details:

>>
1. NW_001939248
Source: Pyrenophora tritici-repentis Pt-1C-BFP supercont1.5, whole genome shotgun sequence
Type: NRPS-like
Number of proteins with BLAST hits to this cluster: 2
Cumulative BLAST score: 2749

Table of genes, locations, strands and annotations of subject cluster:

Table of Blast hits (query gene, subject gene, %identity, blast score, %coverage, e-value):
input.path1.gene176	PTRG_05642	53	689	100.78003120124805	3e-196
input.path1.gene177	PTRG_05643	86	2060	80.42876901798064	0.0


>>
2. NW_017264203
Source: Xylona heveae TC161 unplaced genomic scaffold L228scaffold_11, whole genome shotgun sequence
Type: NRPS-like
Number of proteins with BLAST hits to this cluster: 2
Cumulative BLAST score: 1919

Table of genes, locations, strands and annotations of subject cluster:

Table of Blast hits (query gene, subject gene, %identity, blast score, %coverage, e-value):
input.path1.gene176	L228DRAFT_250331	38	420	105.30421216848673	2.6e-115
input.path1.gene177	L228DRAFT_223787	61	1499	81.74273858921161	0.0


>>
3. NW_023336278
Source: Aspergillus tubingensis WU-2223L DNA, scaffold_4, whole genome shotgun sequence
Type: NRPS-like
Number of proteins with BLAST hits to this cluster: 2
Cumulative BLAST score: 1854

Table of genes, locations, strands and annotations of subject cluster:

Table of Blast hits (query gene, subject gene, %identity, blast score, %coverage, e-value):
input.path1.gene176	AtWU_02994	39	374	83.61934477379094	1.6e-101
input.path1.gene177	AtWU_02993	62	1480	81.60442600276625	0.0


>>
4. NW_020194484
Source: Amorphotheca resinae ATCC 22711 unplaced genomic scaffold M430scaffold_14, whole genome shotgun sequence
Type: NRPS-like
Number of proteins with BLAST hits to this cluster: 2
Cumulative BLAST score: 1851

Table of genes, locations, strands and annotations of subject cluster:

Table of Blast hits (query gene, subject gene, %identity, blast score, %coverage, e-value):
input.path1.gene176	M430DRAFT_148398	39	416	99.37597503900156	6.4e-114
input.path1.gene177	M430DRAFT_45981	60	1435	81.25864453665284	0.0


>>
5. NC_007197
Source: Aspergillus fumigatus Af293 chromosome 4, whole genome shotgun sequence
Type: NRPS-like
Number of proteins with BLAST hits to this cluster: 2
Cumulative BLAST score: 1847

Table of genes, locations, strands and annotations of subject cluster:

Table of Blast hits (query gene, subject gene, %identity, blast score, %coverage, e-value):
input.path1.gene176	AFUA_4G11230	40	378	84.55538221528862	1.9e-102
input.path1.gene177	AFUA_4G11240	61	1469	81.32780082987551	0.0


>>
6. NW_022474205
Source: Venustampulla echinocandica strain BP 5553 chromosome Unknown scaffold_0, whole genome shotgun sequence
Type: NRPS-like
Number of proteins with BLAST hits to this cluster: 2
Cumulative BLAST score: 1837

Table of genes, locations, strands and annotations of subject cluster:

Table of Blast hits (query gene, subject gene, %identity, blast score, %coverage, e-value):
input.path1.gene176	BP5553_00252	38	409	97.97191887675507	7.9e-112
input.path1.gene177	BP5553_00251	60	1428	81.25864453665284	0.0


>>
7. NC_036436
Source: Aspergillus oryzae RIB40 DNA, chromosome 2
Type: NRPS
Number of proteins with BLAST hits to this cluster: 2
Cumulative BLAST score: 1823

Table of genes, locations, strands and annotations of subject cluster:

Table of Blast hits (query gene, subject gene, %identity, blast score, %coverage, e-value):
input.path1.gene176	AO090003001098	39	365	84.399375975039	1e-98
input.path1.gene177	AO090003001097	61	1458	81.18948824343015	0.0


>>
8. NT_165928
Source: Aspergillus terreus NIH2624 scaffold_5 genomic scaffold, whole genome shotgun sequence
Type: NRPS-like
Number of proteins with BLAST hits to this cluster: 2
Cumulative BLAST score: 1822

Table of genes, locations, strands and annotations of subject cluster:

Table of Blast hits (query gene, subject gene, %identity, blast score, %coverage, e-value):
input.path1.gene176	ATEG_03958	38	346	84.399375975039	4.8e-93
input.path1.gene177	ATEG_03957	61	1476	81.32780082987551	0.0


>>
9. NW_022984630
Source: Aspergillus tanneri strain NIH1004 chromosome Unknown AapTan_3_pilon_pilon_pilon, whole genome shotgun sequence
Type: NRPS-like
Number of proteins with BLAST hits to this cluster: 2
Cumulative BLAST score: 1815

Table of genes, locations, strands and annotations of subject cluster:

Table of Blast hits (query gene, subject gene, %identity, blast score, %coverage, e-value):
input.path1.gene176	ATNIH1004_004605	39	365	83.77535101404057	1e-98
input.path1.gene177	ATNIH1004_004604	60	1450	82.01936376210234	0.0


>>
10. NW_006763065
Source: Marssonina brunnea f. sp. 'multigermtubi' MB_m1 unplaced genomic scaffold M6_S00027, whole genome shotgun sequence
Type: NRPS-like
Number of proteins with BLAST hits to this cluster: 2
Cumulative BLAST score: 1807

Table of genes, locations, strands and annotations of subject cluster:

Table of Blast hits (query gene, subject gene, %identity, blast score, %coverage, e-value):
input.path1.gene176	MBM_08980	37	411	99.37597503900156	2.1e-112
input.path1.gene177	MBM_08979	59	1396	80.9820193637621	0.0


>>
11. CM002800
Source: Penicillium chrysogenum strain P2niaD18 chromosome III, whole genome shotgun sequence
Type: NRPS-like
Number of proteins with BLAST hits to this cluster: 2
Cumulative BLAST score: 1794

Table of genes, locations, strands and annotations of subject cluster:

Table of Blast hits (query gene, subject gene, %identity, blast score, %coverage, e-value):
input.path1.gene176	EN45_099150	38	364	85.49141965678628	1.7e-98
input.path1.gene177	EN45_099160	60	1430	80.77455048409405	0.0


>>
12. NW_004504310
Source: Coccidioides immitis RS genomic scaffold supercont3.1, whole genome shotgun sequence
Type: NRPS-like
Number of proteins with BLAST hits to this cluster: 2
Cumulative BLAST score: 1787

Table of genes, locations, strands and annotations of subject cluster:

Table of Blast hits (query gene, subject gene, %identity, blast score, %coverage, e-value):
input.path1.gene176	CIMG_01494	37	348	85.33541341653667	1.3e-93
input.path1.gene177	CIMG_01491	60	1439	81.12033195020747	0.0


>>
13. NW_003315979
Source: Coccidioides posadasii C735 delta SOWgp chromosome Unknown gcontig_1105601051781, whole genome shotgun sequence
Type: NRPS-like
Number of proteins with BLAST hits to this cluster: 2
Cumulative BLAST score: 1781

Table of genes, locations, strands and annotations of subject cluster:

Table of Blast hits (query gene, subject gene, %identity, blast score, %coverage, e-value):
input.path1.gene176	CPC735_049510	37	346	85.33541341653667	8.2e-93
input.path1.gene177	CPC735_049520	60	1435	81.12033195020747	0.0


>>
14. NW_003052498
Source: Uncinocarpus reesii 1704 scaffold_3 genomic scaffold, whole genome shotgun sequence
Type: NRPS-like
Number of proteins with BLAST hits to this cluster: 2
Cumulative BLAST score: 1768

Table of genes, locations, strands and annotations of subject cluster:

Table of Blast hits (query gene, subject gene, %identity, blast score, %coverage, e-value):
input.path1.gene176	UREG_05105	36	332	79.71918876755069	9.4e-89
input.path1.gene177	UREG_05106	60	1436	81.18948824343015	0.0


>>
15. NW_014574601
Source: Penicillium digitatum Pd1 unplaced genomic scaffold Scaffold_24, whole genome shotgun sequence
Type: NRPS-like
Number of proteins with BLAST hits to this cluster: 2
Cumulative BLAST score: 1763

Table of genes, locations, strands and annotations of subject cluster:

Table of Blast hits (query gene, subject gene, %identity, blast score, %coverage, e-value):
input.path1.gene176	PDIP_33710	37	331	82.21528861154445	2.7e-88
input.path1.gene177	PDIP_33720	60	1432	80.77455048409405	0.0


>>
16. NW_007360994
Source: Glarea lozoyensis ATCC 20868 chromosome Unknown GLAREA2, whole genome shotgun sequence
Type: NRPS-like
Number of proteins with BLAST hits to this cluster: 2
Cumulative BLAST score: 1758

Table of genes, locations, strands and annotations of subject cluster:

Table of Blast hits (query gene, subject gene, %identity, blast score, %coverage, e-value):
input.path1.gene176	GLAREA_03382	36	384	98.28393135725429	2.1e-104
input.path1.gene177	GLAREA_03383	59	1374	79.94467496542185	0.0


>>
17. NW_017263917
Source: Colletotrichum higginsianum IMI 349063 chromosome 7 map unlocalized, whole genome shotgun sequence
Type: NRPS-like,betalactone
Number of proteins with BLAST hits to this cluster: 2
Cumulative BLAST score: 1749

Table of genes, locations, strands and annotations of subject cluster:

Table of Blast hits (query gene, subject gene, %identity, blast score, %coverage, e-value):
input.path1.gene176	CH63R_10367	36	394	101.24804992199688	2.6e-107
input.path1.gene177	CH63R_10366	58	1355	80.77455048409405	0.0


>>
18. NW_006912893
Source: Capronia coronata CBS 617.96 acBCG-supercont1.1, whole genome shotgun sequence
Type: NRPS
Number of proteins with BLAST hits to this cluster: 2
Cumulative BLAST score: 1698

Table of genes, locations, strands and annotations of subject cluster:

Table of Blast hits (query gene, subject gene, %identity, blast score, %coverage, e-value):
input.path1.gene176	A1O1_00419	35	333	90.6396255850234	5.5e-89
input.path1.gene177	A1O1_00420	58	1365	81.67358229598894	0.0


>>
19. NW_008481808
Source: Cyphellophora europaea CBS 101466 unplaced genomic scaffold supercont1.2, whole genome shotgun sequence
Type: NRPS-like
Number of proteins with BLAST hits to this cluster: 2
Cumulative BLAST score: 1695

Table of genes, locations, strands and annotations of subject cluster:

Table of Blast hits (query gene, subject gene, %identity, blast score, %coverage, e-value):
input.path1.gene176	HMPREF1541_02344	35	344	88.9235569422777	3.1e-92
input.path1.gene177	HMPREF1541_02345	58	1351	81.95020746887967	0.0


>>
20. NW_008751650
Source: Exophiala dermatitidis NIH/UT8656 unplaced genomic scaffold supercont1.5, whole genome shotgun sequence
Type: NRPS
Number of proteins with BLAST hits to this cluster: 2
Cumulative BLAST score: 1683

Table of genes, locations, strands and annotations of subject cluster:

Table of Blast hits (query gene, subject gene, %identity, blast score, %coverage, e-value):
input.path1.gene176	HMPREF1120_06044	34	334	90.48361934477379	2.5e-89
input.path1.gene177	HMPREF1120_06043	58	1349	81.53526970954357	0.0


>>
21. NW_006912905
Source: Capronia epimyces CBS 606.96 aczIQ-supercont1.2, whole genome shotgun sequence
Type: NRPS-like
Number of proteins with BLAST hits to this cluster: 2
Cumulative BLAST score: 1682

Table of genes, locations, strands and annotations of subject cluster:

Table of Blast hits (query gene, subject gene, %identity, blast score, %coverage, e-value):
input.path1.gene176	A1O3_02451	34	320	88.14352574102963	4.8e-85
input.path1.gene177	A1O3_02452	58	1362	79.94467496542185	0.0


>>
22. NW_015622516
Source: Exophiala spinifera strain CBS 89968 unplaced genomic scaffold supercont1.4, whole genome shotgun sequence
Type: NRPS-like
Number of proteins with BLAST hits to this cluster: 2
Cumulative BLAST score: 1681

Table of genes, locations, strands and annotations of subject cluster:

Table of Blast hits (query gene, subject gene, %identity, blast score, %coverage, e-value):
input.path1.gene176	PV08_05694	36	332	90.01560062402496	9.4e-89
input.path1.gene177	PV08_05695	59	1349	79.59889349930843	0.0


>>
23. NW_015971648
Source: Fonsecaea multimorphosa CBS 102226 unplaced genomic scaffold supercont1.14, whole genome shotgun sequence
Type: NRPS-like
Number of proteins with BLAST hits to this cluster: 2
Cumulative BLAST score: 1678

Table of genes, locations, strands and annotations of subject cluster:

Table of Blast hits (query gene, subject gene, %identity, blast score, %coverage, e-value):
input.path1.gene176	Z520_07232	36	331	88.45553822152887	2.1e-88
input.path1.gene177	Z520_07233	57	1347	82.78008298755186	0.0


>>
24. NW_017387248
Source: Fonsecaea erecta strain CBS 125763 chromosome Unknown Scaff5, whole genome shotgun sequence
Type: NRPS-like
Number of proteins with BLAST hits to this cluster: 2
Cumulative BLAST score: 1657

Table of genes, locations, strands and annotations of subject cluster:

Table of Blast hits (query gene, subject gene, %identity, blast score, %coverage, e-value):
input.path1.gene176	AYL99_06901	35	325	88.45553822152887	1.5e-86
input.path1.gene177	AYL99_06902	56	1332	82.57261410788381	0.0


>>
25. NW_013550611
Source: Fonsecaea pedrosoi CBS 271.37 unplaced genomic scaffold supercont1.5, whole genome shotgun sequence
Type: NRPS
Number of proteins with BLAST hits to this cluster: 2
Cumulative BLAST score: 1656

Table of genes, locations, strands and annotations of subject cluster:

Table of Blast hits (query gene, subject gene, %identity, blast score, %coverage, e-value):
input.path1.gene176	Z517_07622	35	318	88.45553822152887	1.4e-84
input.path1.gene177	Z517_07621	57	1338	82.71092669432917	0.0


>>
26. NW_015971588
Source: Cladophialophora bantiana CBS 173.52 unplaced genomic scaffold supercont1.14, whole genome shotgun sequence
Type: NRPS-like
Number of proteins with BLAST hits to this cluster: 2
Cumulative BLAST score: 1650

Table of genes, locations, strands and annotations of subject cluster:

Table of Blast hits (query gene, subject gene, %identity, blast score, %coverage, e-value):
input.path1.gene176	Z519_08935	35	321	88.45553822152887	1.7e-85
input.path1.gene177	Z519_08934	55	1329	83.67911479944675	0.0


>>
27. NW_015622504
Source: Exophiala mesophila strain CBS 40295 unplaced genomic scaffold supercont1.1, whole genome shotgun sequence
Type: NRPS-like
Number of proteins with BLAST hits to this cluster: 2
Cumulative BLAST score: 1636

Table of genes, locations, strands and annotations of subject cluster:

Table of Blast hits (query gene, subject gene, %identity, blast score, %coverage, e-value):
input.path1.gene176	PV10_01918	34	327	87.98751950078002	3.9e-87
input.path1.gene177	PV10_01917	57	1309	81.05117565698478	0.0


>>
28. NW_001820825
Source: Sclerotinia sclerotiorum 1980 UF-70 scaffold_11 genomic scaffold, whole genome shotgun sequence
Type: NRPS-like
Number of proteins with BLAST hits to this cluster: 2
Cumulative BLAST score: 1631

Table of genes, locations, strands and annotations of subject cluster:

Table of Blast hits (query gene, subject gene, %identity, blast score, %coverage, e-value):
input.path1.gene176	SS1G_08186	45	235	45.86583463338533	2e-59
input.path1.gene177	SS1G_08185	58	1396	80.91286307053942	0.0


>>
29. NT_107010
Source: Aspergillus nidulans FGSC A4 chromosome V map unlocalized scaffold_6, whole genome shotgun sequence
Type: NRPS-like
Number of proteins with BLAST hits to this cluster: 2
Cumulative BLAST score: 1812

Table of genes, locations, strands and annotations of subject cluster:

Table of Blast hits (query gene, subject gene, %identity, blast score, %coverage, e-value):
input.path1.gene176	AN5612.2	36	340	83.61934477379094	3.4e-91
input.path1.gene177	AN5610.2	62	1472	81.12033195020747	0.0


>>
30. NW_006913045
Source: Cladophialophora yegresii CBS 114405 aczIN-supercont1.1, whole genome shotgun sequence
Type: NRPS-like
Number of proteins with BLAST hits to this cluster: 2
Cumulative BLAST score: 1690

Table of genes, locations, strands and annotations of subject cluster:

Table of Blast hits (query gene, subject gene, %identity, blast score, %coverage, e-value):
input.path1.gene176	A1O7_00933	36	334	88.29953198127926	3.2e-89
input.path1.gene177	A1O7_00939	58	1356	81.74273858921161	0.0


>>
31. NW_008481816
Source: Cladophialophora carrionii CBS 160.54 unplaced genomic scaffold supercont1.1, whole genome shotgun sequence
Type: NRPS
Number of proteins with BLAST hits to this cluster: 2
Cumulative BLAST score: 1684

Table of genes, locations, strands and annotations of subject cluster:

Table of Blast hits (query gene, subject gene, %identity, blast score, %coverage, e-value):
input.path1.gene176	G647_00996	34	329	88.29953198127926	1e-87
input.path1.gene177	G647_01002	58	1355	83.74827109266944	0.0


>>
32. NC_035790
Source: Pochonia chlamydosporia 170 chromosome 1, whole genome shotgun sequence
Type: NRPS-like
Number of proteins with BLAST hits to this cluster: 2
Cumulative BLAST score: 1326

Table of genes, locations, strands and annotations of subject cluster:

Table of Blast hits (query gene, subject gene, %identity, blast score, %coverage, e-value):
input.path1.gene171	VFPPC_00982	55	96	77.35849056603774	1.4e-18
input.path1.gene177	VFPPC_00985	54	1230	80.9820193637621	0.0


>>
33. CM000576
Source: Fusarium graminearum PH-1 chromosome 3, whole genome shotgun sequence
Type: NRPS-like
Number of proteins with BLAST hits to this cluster: 2
Cumulative BLAST score: 1319

Table of genes, locations, strands and annotations of subject cluster:

Table of Blast hits (query gene, subject gene, %identity, blast score, %coverage, e-value):
input.path1.gene171	FGSG_12858	59	100	77.35849056603774	1.3e-19
input.path1.gene177	FGSG_06041	53	1219	81.53526970954357	0.0


>>
34. NC_026476
Source: Fusarium graminearum PH-1 chromosome 3, whole genome shotgun sequence
Type: NRPS-like
Number of proteins with BLAST hits to this cluster: 2
Cumulative BLAST score: 1319

Table of genes, locations, strands and annotations of subject cluster:

Table of Blast hits (query gene, subject gene, %identity, blast score, %coverage, e-value):
input.path1.gene171	FGSG_12858	59	100	77.35849056603774	1.3e-19
input.path1.gene177	FGSG_06041	53	1219	81.53526970954357	0.0


>>
35. NC_038014
Source: Fusarium venenatum strain A3/5 genome assembly, chromosome: III
Type: NRPS-like
Number of proteins with BLAST hits to this cluster: 2
Cumulative BLAST score: 1319

Table of genes, locations, strands and annotations of subject cluster:

Table of Blast hits (query gene, subject gene, %identity, blast score, %coverage, e-value):
input.path1.gene171	FVRRES_09623	57	99	77.35849056603774	1.7e-19
input.path1.gene177	FVRRES_09626	53	1220	81.53526970954357	0.0
