## Supplementary Results for "A draft genome of the ascomycotal fungal species *Pseudopithomyces maydicus* (family *Didymosphaeriaceae*)": contig_32_c2.txt

ClusterBlast scores for contig_32

Table of genes, locations, strands and annotations of query cluster:
input.path1.gene490	1790499	1792745	+		
input.path1.gene491	1794279	1795344	+		
input.path1.gene492	1796187	1796687	-		
input.path1.gene493	1797854	1799036	-		
input.path1.gene494	1801871	1803614	-		
input.path1.gene495	1804625	1805974	-		
input.path1.gene496	1807889	1808228	+		
input.path1.gene497	1808251	1808702	-		
input.path1.gene498	1809457	1809835	-		
input.path1.gene499	1809957	1813058	+		
input.path1.gene500	1815452	1815937	+		
input.path1.gene501	1817535	1820394	-		
input.path1.gene502	1820666	1821429	+		
input.path1.gene503	1821623	1822084	-		
input.path1.gene504	1823485	1827330	-		
input.path1.gene505	1827942	1828495	-		
input.path1.gene506	1829391	1830537	+		
input.path1.gene507	1831455	1832854	-		


Significant hits: 
1. NC_049562	Talaromyces rugulosus chromosome II, complete sequence
2. NC_036442	Aspergillus oryzae RIB40 DNA, chromosome 8
3. NW_011942215	Metarhizium robertsii ARSEF 23 MAA_Scf_7, whole genome shotgun sequence
4. NW_014574711	Metarhizium brunneum ARSEF 3297 chromosome Unknown scaffold_6, whole genome shotgun sequence
5. NC_035793	Pochonia chlamydosporia 170 chromosome 4, whole genome shotgun sequence
6. CM004174	Drechmeria coniospora strain ARSEF 6962 chromosome 01, whole genome shotgun sequence
7. NW_020167549	Pseudogymnoascus destructans isolate 20631-21 unplaced genomic scaffold scaffold_30, whole genome shotgun sequence
8. CM000595	Fusarium oxysporum f. sp. lycopersici 4287 chromosome 7, whole genome shotgun sequence
9. NC_030992	Fusarium oxysporum f. sp. lycopersici 4287 chromosome 7, whole genome shotgun sequence
10. NC_049565	Talaromyces rugulosus chromosome V, complete sequence
11. NZ_LT607409	Micromonospora chokoriensis strain DSM 45160 chromosome I
12. NC_049561	Talaromyces rugulosus chromosome I, complete sequence
13. NW_019154044	Pochonia chlamydosporia 170 chromosome Unknown PCv3seq00009, whole genome shotgun sequence
14. NZ_CP013441	Burkholderia vietnamiensis strain HI2297 chromosome 3, complete sequence
15. NW_006912895	Capronia coronata CBS 617.96 acBCG-supercont1.3, whole genome shotgun sequence
16. NZ_CADFFH010000007	Burkholderia vietnamiensis strain BCC0128, whole genome shotgun sequence
17. NZ_CADFFI010000006	Burkholderia vietnamiensis strain BCC0136, whole genome shotgun sequence
18. NZ_CP013434	Burkholderia vietnamiensis strain AU1233 chromosome 2, complete sequence
19. NZ_CP013456	Burkholderia vietnamiensis strain MSMB608WGS chromosome 3, complete sequence
20. NZ_CADFFL010000002	Burkholderia vietnamiensis strain BCC1409, whole genome shotgun sequence
21. NZ_FMVZ01000016	Burkholderia vietnamiensis strain WPB, whole genome shotgun sequence
22. NZ_CADFER010000002	Burkholderia vietnamiensis strain BCC0046, whole genome shotgun sequence
23. NZ_CP009632	Burkholderia vietnamiensis LMG 10929 chromosome III, complete sequence
24. NZ_CADFFW010000002	Burkholderia vietnamiensis strain BCC1163, whole genome shotgun sequence
25. NC_017922	Burkholderia sp. KJ006 chromosome 3, complete sequence
26. NZ_CP053611	Burkholderia sp. MBR-1 chromosome 3, complete sequence
27. NZ_CADFEV010000013	Burkholderia vietnamiensis strain BCC0151, whole genome shotgun sequence
28. NW_017971432	Talaromyces atroroseus strain IBT 11181 chromosome Unknown scaffold_0, whole genome shotgun sequence
29. CM000574	Fusarium graminearum PH-1 chromosome 1, whole genome shotgun sequence
30. NC_026474	Fusarium graminearum PH-1 chromosome 1, whole genome shotgun sequence
31. NW_017264193	Xylona heveae TC161 unplaced genomic scaffold L228scaffold_1, whole genome shotgun sequence
32. NW_003456426	Trichophyton rubrum CBS 118892 genomic scaffold supercont2.3, whole genome shotgun sequence
33. NC_038012	Fusarium venenatum strain A3/5 genome assembly, chromosome: I
34. NW_003345192	Nannizzia gypsea CBS 118893 supercont1.10 genomic scaffold, whole genome shotgun sequence
35. NW_003315103	Trichophyton benhamiae CBS 112371 chromosome Unknown contig00075.49, whole genome shotgun sequence
36. NW_022984633	Aspergillus tanneri strain NIH1004 chromosome Unknown AapTan_6_pilon_pilon_pilon, whole genome shotgun sequence
37. NW_017971433	Talaromyces atroroseus strain IBT 11181 chromosome Unknown scaffold_1, whole genome shotgun sequence
38. NW_006271969	Cordyceps militaris CM01 unplaced genomic scaffold CCM_S00001, whole genome shotgun sequence
39. NW_001820829	Sclerotinia sclerotiorum 1980 UF-70 scaffold_7 genomic scaffold, whole genome shotgun sequence
40. NW_003299161	Microsporum canis CBS 113480 supercont1.9 genomic scaffold, whole genome shotgun sequence
41. NC_030962	Colletotrichum higginsianum IMI 349063 chromosome 10, whole genome shotgun sequence
42. NT_165927	Aspergillus terreus NIH2624 scaffold_4 genomic scaffold, whole genome shotgun sequence
43. NW_006912904	Capronia epimyces CBS 606.96 aczIQ-supercont1.1, whole genome shotgun sequence
44. NW_017971438	Talaromyces atroroseus strain IBT 11181 chromosome Unknown scaffold_6, whole genome shotgun sequence
45. NZ_FNOG01000002	Lysobacter enzymogenes strain ATCC 29487, whole genome shotgun sequence
46. NZ_JACJIA010000006	Actinomadura namibiensis strain DSM 44197 Ga0415225_06, whole genome shotgun sequence
47. NZ_AP014940	Lysobacter enzymogenes strain M497-1
48. NC_016787	Corynebacterium diphtheriae HC03, complete sequence
49. NZ_MIOB01000012	Corynebacterium diphtheriae strain ST395-KZN-2015-45792 45792_contig_2, whole genome shotgun sequence
50. NZ_CADFFS010000002	Burkholderia vietnamiensis strain BCC1176, whole genome shotgun sequence


Details:

>>
1. NC_049562
Source: Talaromyces rugulosus chromosome II, complete sequence
Type: T1PKS
Number of proteins with BLAST hits to this cluster: 2
Cumulative BLAST score: 690

Table of genes, locations, strands and annotations of subject cluster:

Table of Blast hits (query gene, subject gene, %identity, blast score, %coverage, e-value):
input.path1.gene495	TRUGW13939_04213	37	171	85.79881656804734	1.5e-40
input.path1.gene499	TRUGW13939_04214	50	519	68.46057571964957	8.8e-145


>>
2. NC_036442
Source: Aspergillus oryzae RIB40 DNA, chromosome 8
Type: T1PKS
Number of proteins with BLAST hits to this cluster: 2
Cumulative BLAST score: 628

Table of genes, locations, strands and annotations of subject cluster:

Table of Blast hits (query gene, subject gene, %identity, blast score, %coverage, e-value):
input.path1.gene495	AO090010000047	34	124	77.2189349112426	1.6e-26
input.path1.gene499	AO090010000048	42	504	91.36420525657071	2.2e-140


>>
3. NW_011942215
Source: Metarhizium robertsii ARSEF 23 MAA_Scf_7, whole genome shotgun sequence
Type: T1PKS
Number of proteins with BLAST hits to this cluster: 2
Cumulative BLAST score: 414

Table of genes, locations, strands and annotations of subject cluster:

Table of Blast hits (query gene, subject gene, %identity, blast score, %coverage, e-value):
input.path1.gene495	MAA_00788	33	142	82.54437869822485	7.2e-32
input.path1.gene499	MAA_00789	31	272	72.090112640801	1.9e-70


>>
4. NW_014574711
Source: Metarhizium brunneum ARSEF 3297 chromosome Unknown scaffold_6, whole genome shotgun sequence
Type: T1PKS
Number of proteins with BLAST hits to this cluster: 2
Cumulative BLAST score: 414

Table of genes, locations, strands and annotations of subject cluster:

Table of Blast hits (query gene, subject gene, %identity, blast score, %coverage, e-value):
input.path1.gene495	MBR_05713	32	142	82.54437869822485	7.2e-32
input.path1.gene499	MBR_05714	31	272	72.090112640801	1.9e-70


>>
5. NC_035793
Source: Pochonia chlamydosporia 170 chromosome 4, whole genome shotgun sequence
Type: T1PKS
Number of proteins with BLAST hits to this cluster: 2
Cumulative BLAST score: 408

Table of genes, locations, strands and annotations of subject cluster:

Table of Blast hits (query gene, subject gene, %identity, blast score, %coverage, e-value):
input.path1.gene495	VFPPC_08494	34	148	84.61538461538461	1.3e-33
input.path1.gene499	VFPPC_08495	31	260	72.21526908635795	4.3e-67


>>
6. CM004174
Source: Drechmeria coniospora strain ARSEF 6962 chromosome 01, whole genome shotgun sequence
Type: T1PKS
Number of proteins with BLAST hits to this cluster: 2
Cumulative BLAST score: 406

Table of genes, locations, strands and annotations of subject cluster:

Table of Blast hits (query gene, subject gene, %identity, blast score, %coverage, e-value):
input.path1.gene495	DCS_01788	34	163	88.75739644970415	5.2e-38
input.path1.gene499	DCS_01787	34	243	55.319148936170215	7.1e-62


>>
7. NW_020167549
Source: Pseudogymnoascus destructans isolate 20631-21 unplaced genomic scaffold scaffold_30, whole genome shotgun sequence
Type: T1PKS
Number of proteins with BLAST hits to this cluster: 2
Cumulative BLAST score: 1268

Table of genes, locations, strands and annotations of subject cluster:

Table of Blast hits (query gene, subject gene, %identity, blast score, %coverage, e-value):
input.path1.gene499	VC83_08896	48	508	69.83729662077597	1.6e-141
input.path1.gene504	VC83_08893	50	760	85.37946428571429	2.5e-217


>>
8. CM000595
Source: Fusarium oxysporum f. sp. lycopersici 4287 chromosome 7, whole genome shotgun sequence
Type: T1PKS
Number of proteins with BLAST hits to this cluster: 2
Cumulative BLAST score: 1167

Table of genes, locations, strands and annotations of subject cluster:

Table of Blast hits (query gene, subject gene, %identity, blast score, %coverage, e-value):
input.path1.gene495	FOXG_04755	31	139	85.79881656804734	8e-31
input.path1.gene499	FOXG_04757	51	514	71.33917396745932	2.8e-143
input.path1.gene499	FOXG_04757_KNB01536.1	51	514	71.33917396745932	2.8e-143


>>
9. NC_030992
Source: Fusarium oxysporum f. sp. lycopersici 4287 chromosome 7, whole genome shotgun sequence
Type: T1PKS
Number of proteins with BLAST hits to this cluster: 2
Cumulative BLAST score: 1167

Table of genes, locations, strands and annotations of subject cluster:

Table of Blast hits (query gene, subject gene, %identity, blast score, %coverage, e-value):
input.path1.gene495	FOXG_04755	31	139	85.79881656804734	8e-31
input.path1.gene499	FOXG_04757	51	514	71.33917396745932	2.8e-143
input.path1.gene499	FOXG_04757_XP_018239581.1	51	514	71.33917396745932	2.8e-143


>>
10. NC_049565
Source: Talaromyces rugulosus chromosome V, complete sequence
Type: T1PKS
Number of proteins with BLAST hits to this cluster: 2
Cumulative BLAST score: 1050

Table of genes, locations, strands and annotations of subject cluster:

Table of Blast hits (query gene, subject gene, %identity, blast score, %coverage, e-value):
input.path1.gene494	TRUGW13939_10273	34	292	99.13793103448276	9.7e-77
input.path1.gene499	TRUGW13939_10273	44	501	85.1063829787234	1.9e-139
input.path1.gene499	TRUGW13939_10272	33	257	72.96620775969961	3.7e-66


>>
11. NZ_LT607409
Source: Micromonospora chokoriensis strain DSM 45160 chromosome I
Type: T1PKS
Number of proteins with BLAST hits to this cluster: 2
Cumulative BLAST score: 950

Table of genes, locations, strands and annotations of subject cluster:
GA0070612_RS31865	GA0070612_RS31865	3743792	3746314	+	type_I_polyketide_synthase
GA0070612_RS31870	GA0070612_RS31870	3746426	3747421	+	hypothetical_protein
GA0070612_RS31875	GA0070612_RS31875	3747794	3748534	+	SDR_family_NAD(P)-dependent_oxidoreductase
GA0070612_RS31880	GA0070612_RS31880	3749138	3750511	+	polyketide_synthase
GA0070612_RS31885	GA0070612_RS31885	3750764	3751660	+	acyltransferase_domain-containing_protein

Table of Blast hits (query gene, subject gene, %identity, blast score, %coverage, e-value):
input.path1.gene494	GA0070612_RS17435	31	198	96.55172413793103	1.9e-48
input.path1.gene499	GA0070612_RS31865	33	265	69.71214017521903	2.3e-68
input.path1.gene499	GA0070612_RS31880	36	249	54.31789737171464	1.3e-63
input.path1.gene499	GA0070612_RS17450	32	238	70.08760951188987	2.3e-60


>>
12. NC_049561
Source: Talaromyces rugulosus chromosome I, complete sequence
Type: T1PKS
Number of proteins with BLAST hits to this cluster: 2
Cumulative BLAST score: 891

Table of genes, locations, strands and annotations of subject cluster:

Table of Blast hits (query gene, subject gene, %identity, blast score, %coverage, e-value):
input.path1.gene494	TRUGW13939_01411	39	366	93.96551724137932	5.3e-99
input.path1.gene494	TRUGW13939_01410	32	212	102.75862068965517	1.3e-52
input.path1.gene499	TRUGW13939_01413	41	313	57.446808510638306	5.6e-83


>>
13. NW_019154044
Source: Pochonia chlamydosporia 170 chromosome Unknown PCv3seq00009, whole genome shotgun sequence
Type: NRPS,T1PKS
Number of proteins with BLAST hits to this cluster: 2
Cumulative BLAST score: 839

Table of genes, locations, strands and annotations of subject cluster:

Table of Blast hits (query gene, subject gene, %identity, blast score, %coverage, e-value):
input.path1.gene494	VFPPC_10567	41	373	98.27586206896551	5.7e-101
input.path1.gene499	VFPPC_10579	35	240	54.192740926157704	6e-61
input.path1.gene499	VFPPC_10573	32	226	68.8360450563204	9e-57


>>
14. NZ_CP013441
Source: Burkholderia vietnamiensis strain HI2297 chromosome 3, complete sequence
Type: NRPS,T1PKS
Number of proteins with BLAST hits to this cluster: 2
Cumulative BLAST score: 778

Table of genes, locations, strands and annotations of subject cluster:

Table of Blast hits (query gene, subject gene, %identity, blast score, %coverage, e-value):
input.path1.gene494	WK31_RS26305	31	198	97.24137931034483	1.5e-48
input.path1.gene499	WK31_RS26215	37	302	67.95994993742178	1.3e-79
input.path1.gene499	WK31_RS26205	36	278	66.83354192740926	2.6e-72


>>
15. NW_006912895
Source: Capronia coronata CBS 617.96 acBCG-supercont1.3, whole genome shotgun sequence
Type: T1PKS
Number of proteins with BLAST hits to this cluster: 2
Cumulative BLAST score: 778

Table of genes, locations, strands and annotations of subject cluster:

Table of Blast hits (query gene, subject gene, %identity, blast score, %coverage, e-value):
input.path1.gene495	A1O1_04521	38	203	84.61538461538461	4.5e-50
input.path1.gene499	A1O1_04517	37	324	72.46558197747184	4.2e-86
input.path1.gene499	A1O1_04527	32	251	71.46433041301627	2.6e-64


>>
16. NZ_CADFFH010000007
Source: Burkholderia vietnamiensis strain BCC0128, whole genome shotgun sequence
Type: NRPS,T1PKS
Number of proteins with BLAST hits to this cluster: 2
Cumulative BLAST score: 775

Table of genes, locations, strands and annotations of subject cluster:

Table of Blast hits (query gene, subject gene, %identity, blast score, %coverage, e-value):
input.path1.gene494	HWX54_RS16515	31	196	97.24137931034483	7.3e-48
input.path1.gene499	HWX54_RS16610	36	306	67.70963704630788	6.9e-81
input.path1.gene499	HWX54_RS16620	35	273	68.33541927409262	8.4e-71


>>
17. NZ_CADFFI010000006
Source: Burkholderia vietnamiensis strain BCC0136, whole genome shotgun sequence
Type: NRPS,T1PKS
Number of proteins with BLAST hits to this cluster: 2
Cumulative BLAST score: 773

Table of genes, locations, strands and annotations of subject cluster:

Table of Blast hits (query gene, subject gene, %identity, blast score, %coverage, e-value):
input.path1.gene494	HW535_RS09925	31	196	97.24137931034483	9.5e-48
input.path1.gene499	HW535_RS09830	36	304	67.95994993742178	2.6e-80
input.path1.gene499	HW535_RS09820	35	273	68.33541927409262	6.4e-71


>>
18. NZ_CP013434
Source: Burkholderia vietnamiensis strain AU1233 chromosome 2, complete sequence
Type: NRPS,T1PKS
Number of proteins with BLAST hits to this cluster: 2
Cumulative BLAST score: 772

Table of genes, locations, strands and annotations of subject cluster:

Table of Blast hits (query gene, subject gene, %identity, blast score, %coverage, e-value):
input.path1.gene494	WK23_RS25265	31	195	97.24137931034483	1.2e-47
input.path1.gene499	WK23_RS25355	36	304	67.83479349186483	2.6e-80
input.path1.gene499	WK23_RS25365	35	273	68.33541927409262	8.4e-71


>>
19. NZ_CP013456
Source: Burkholderia vietnamiensis strain MSMB608WGS chromosome 3, complete sequence
Type: NRPS,T1PKS
Number of proteins with BLAST hits to this cluster: 2
Cumulative BLAST score: 772

Table of genes, locations, strands and annotations of subject cluster:

Table of Blast hits (query gene, subject gene, %identity, blast score, %coverage, e-value):
input.path1.gene494	WL96_RS25640	31	195	97.24137931034483	1.2e-47
input.path1.gene499	WL96_RS25730	36	304	67.83479349186483	2.6e-80
input.path1.gene499	WL96_RS25740	35	273	68.33541927409262	8.4e-71


>>
20. NZ_CADFFL010000002
Source: Burkholderia vietnamiensis strain BCC1409, whole genome shotgun sequence
Type: NRPS,T1PKS
Number of proteins with BLAST hits to this cluster: 2
Cumulative BLAST score: 772

Table of genes, locations, strands and annotations of subject cluster:

Table of Blast hits (query gene, subject gene, %identity, blast score, %coverage, e-value):
input.path1.gene494	HWX70_RS07400	31	193	97.24137931034483	4.7e-47
input.path1.gene499	HWX70_RS07500	37	305	67.83479349186483	1.5e-80
input.path1.gene499	HWX70_RS07510	35	274	68.33541927409262	3.8e-71


>>
21. NZ_FMVZ01000016
Source: Burkholderia vietnamiensis strain WPB, whole genome shotgun sequence
Type: NRPS,T1PKS
Number of proteins with BLAST hits to this cluster: 2
Cumulative BLAST score: 772

Table of genes, locations, strands and annotations of subject cluster:

Table of Blast hits (query gene, subject gene, %identity, blast score, %coverage, e-value):
input.path1.gene494	BLQ23_RS22700	31	193	97.24137931034483	4.7e-47
input.path1.gene499	BLQ23_RS22795	37	304	67.95994993742178	3.4e-80
input.path1.gene499	BLQ23_RS22805	36	275	68.33541927409262	1.3e-71


>>
22. NZ_CADFER010000002
Source: Burkholderia vietnamiensis strain BCC0046, whole genome shotgun sequence
Type: NRPS,T1PKS
Number of proteins with BLAST hits to this cluster: 2
Cumulative BLAST score: 771

Table of genes, locations, strands and annotations of subject cluster:
HWX77_RS07575	HWX77_RS07575	526765	527211	-	cupin_domain-containing_protein
HWX77_RS07595	HWX77_RS07595	531811	532026	+	IS256_family_transposase

Table of Blast hits (query gene, subject gene, %identity, blast score, %coverage, e-value):
input.path1.gene494	HWX77_RS07725	31	194	97.24137931034483	2.8e-47
input.path1.gene499	HWX77_RS07630	36	305	67.83479349186483	1.5e-80
input.path1.gene499	HWX77_RS07620	35	272	68.33541927409262	1.4e-70


>>
23. NZ_CP009632
Source: Burkholderia vietnamiensis LMG 10929 chromosome III, complete sequence
Type: NRPS,T1PKS
Number of proteins with BLAST hits to this cluster: 2
Cumulative BLAST score: 770

Table of genes, locations, strands and annotations of subject cluster:

Table of Blast hits (query gene, subject gene, %identity, blast score, %coverage, e-value):
input.path1.gene494	AK36_RS28300	31	195	97.24137931034483	1.2e-47
input.path1.gene499	AK36_RS28210	36	304	67.83479349186483	2.6e-80
input.path1.gene499	AK36_RS28200	35	271	68.33541927409262	2.4e-70


>>
24. NZ_CADFFW010000002
Source: Burkholderia vietnamiensis strain BCC1163, whole genome shotgun sequence
Type: NRPS,T1PKS
Number of proteins with BLAST hits to this cluster: 2
Cumulative BLAST score: 770

Table of genes, locations, strands and annotations of subject cluster:
HW462_RS05050	HW462_RS05050	460993	461652	-	transposase
HW462_RS05055	HW462_RS05055	462022	463400	-	ISNCY_family_transposase

Table of Blast hits (query gene, subject gene, %identity, blast score, %coverage, e-value):
input.path1.gene494	HW462_RS04915	31	194	97.24137931034483	2.8e-47
input.path1.gene499	HW462_RS05015	36	304	67.83479349186483	2.6e-80
input.path1.gene499	HW462_RS05025	35	272	68.33541927409262	1.9e-70


>>
25. NC_017922
Source: Burkholderia sp. KJ006 chromosome 3, complete sequence
Type: NRPS,T1PKS
Number of proteins with BLAST hits to this cluster: 2
Cumulative BLAST score: 770

Table of genes, locations, strands and annotations of subject cluster:

Table of Blast hits (query gene, subject gene, %identity, blast score, %coverage, e-value):
input.path1.gene494	MYA_RS26840	31	193	97.24137931034483	4.7e-47
input.path1.gene499	MYA_RS26745	37	304	67.95994993742178	3.4e-80
input.path1.gene499	MYA_RS26735	35	273	68.33541927409262	6.4e-71


>>
26. NZ_CP053611
Source: Burkholderia sp. MBR-1 chromosome 3, complete sequence
Type: NRPS,T1PKS
Number of proteins with BLAST hits to this cluster: 2
Cumulative BLAST score: 769

Table of genes, locations, strands and annotations of subject cluster:

Table of Blast hits (query gene, subject gene, %identity, blast score, %coverage, e-value):
input.path1.gene494	MBR110_RS27010	31	193	97.24137931034483	4.7e-47
input.path1.gene499	MBR110_RS26915	36	304	67.95994993742178	2.6e-80
input.path1.gene499	MBR110_RS26905	35	272	68.33541927409262	1.9e-70


>>
27. NZ_CADFEV010000013
Source: Burkholderia vietnamiensis strain BCC0151, whole genome shotgun sequence
Type: NRPS,T1PKS
Number of proteins with BLAST hits to this cluster: 2
Cumulative BLAST score: 769

Table of genes, locations, strands and annotations of subject cluster:

Table of Blast hits (query gene, subject gene, %identity, blast score, %coverage, e-value):
input.path1.gene494	HWX57_RS13260	31	193	97.24137931034483	6.1e-47
input.path1.gene499	HWX57_RS13180	36	304	67.70963704630788	2.6e-80
input.path1.gene499	HWX57_RS13170	35	272	68.33541927409262	1.9e-70


>>
28. NW_017971432
Source: Talaromyces atroroseus strain IBT 11181 chromosome Unknown scaffold_0, whole genome shotgun sequence
Type: T1PKS
Number of proteins with BLAST hits to this cluster: 2
Cumulative BLAST score: 733

Table of genes, locations, strands and annotations of subject cluster:

Table of Blast hits (query gene, subject gene, %identity, blast score, %coverage, e-value):
input.path1.gene494	UA08_00992	42	404	94.82758620689656	2.3e-110
input.path1.gene499	UA08_00993	39	329	61.95244055068836	7.6e-88


>>
29. CM000574
Source: Fusarium graminearum PH-1 chromosome 1, whole genome shotgun sequence
Type: NRPS,T1PKS
Number of proteins with BLAST hits to this cluster: 2
Cumulative BLAST score: 670

Table of genes, locations, strands and annotations of subject cluster:

Table of Blast hits (query gene, subject gene, %identity, blast score, %coverage, e-value):
input.path1.gene495	FGSG_02326	32	145	87.27810650887574	8.6e-33
input.path1.gene499	FGSG_02324	51	525	68.96120150187734	9.4e-147


>>
30. NC_026474
Source: Fusarium graminearum PH-1 chromosome 1, whole genome shotgun sequence
Type: NRPS,T1PKS
Number of proteins with BLAST hits to this cluster: 2
Cumulative BLAST score: 670

Table of genes, locations, strands and annotations of subject cluster:

Table of Blast hits (query gene, subject gene, %identity, blast score, %coverage, e-value):
input.path1.gene495	FGSG_02326	32	145	87.27810650887574	8.6e-33
input.path1.gene499	FGSG_02324	51	525	68.96120150187734	9.4e-147


>>
31. NW_017264193
Source: Xylona heveae TC161 unplaced genomic scaffold L228scaffold_1, whole genome shotgun sequence
Type: T1PKS
Number of proteins with BLAST hits to this cluster: 2
Cumulative BLAST score: 660

Table of genes, locations, strands and annotations of subject cluster:

Table of Blast hits (query gene, subject gene, %identity, blast score, %coverage, e-value):
input.path1.gene495	L228DRAFT_235439	33	138	83.4319526627219	1e-30
input.path1.gene499	L228DRAFT_235437	45	522	79.09887359198999	8e-146


>>
32. NW_003456426
Source: Trichophyton rubrum CBS 118892 genomic scaffold supercont2.3, whole genome shotgun sequence
Type: T1PKS
Number of proteins with BLAST hits to this cluster: 2
Cumulative BLAST score: 652

Table of genes, locations, strands and annotations of subject cluster:

Table of Blast hits (query gene, subject gene, %identity, blast score, %coverage, e-value):
input.path1.gene495	TERG_02845	33	146	87.86982248520711	5e-33
input.path1.gene499	TERG_02850	44	506	79.59949937421777	5.9e-141


>>
33. NC_038012
Source: Fusarium venenatum strain A3/5 genome assembly, chromosome: I
Type: NRPS,T1PKS
Number of proteins with BLAST hits to this cluster: 2
Cumulative BLAST score: 652

Table of genes, locations, strands and annotations of subject cluster:

Table of Blast hits (query gene, subject gene, %identity, blast score, %coverage, e-value):
input.path1.gene495	FVRRES_02717	32	141	87.27810650887574	1.2e-31
input.path1.gene499	FVRRES_02715	50	511	68.71088861076345	1.8e-142


>>
34. NW_003345192
Source: Nannizzia gypsea CBS 118893 supercont1.10 genomic scaffold, whole genome shotgun sequence
Type: T1PKS
Number of proteins with BLAST hits to this cluster: 2
Cumulative BLAST score: 651

Table of genes, locations, strands and annotations of subject cluster:

Table of Blast hits (query gene, subject gene, %identity, blast score, %coverage, e-value):
input.path1.gene495	MGYG_08689	33	148	87.86982248520711	1e-33
input.path1.gene499	MGYG_08684	48	503	68.46057571964957	2.9e-140


>>
35. NW_003315103
Source: Trichophyton benhamiae CBS 112371 chromosome Unknown contig00075.49, whole genome shotgun sequence
Type: T1PKS
Number of proteins with BLAST hits to this cluster: 2
Cumulative BLAST score: 649

Table of genes, locations, strands and annotations of subject cluster:

Table of Blast hits (query gene, subject gene, %identity, blast score, %coverage, e-value):
input.path1.gene495	ARB_07989	33	147	87.86982248520711	2.9e-33
input.path1.gene499	ARB_07994	44	502	79.72465581977471	6.5e-140


>>
36. NW_022984633
Source: Aspergillus tanneri strain NIH1004 chromosome Unknown AapTan_6_pilon_pilon_pilon, whole genome shotgun sequence
Type: T1PKS
Number of proteins with BLAST hits to this cluster: 2
Cumulative BLAST score: 646

Table of genes, locations, strands and annotations of subject cluster:

Table of Blast hits (query gene, subject gene, %identity, blast score, %coverage, e-value):
input.path1.gene495	ATNIH1004_008651	33	144	86.09467455621301	1.9e-32
input.path1.gene499	ATNIH1004_008649	45	502	78.72340425531915	1.1e-139


>>
37. NW_017971433
Source: Talaromyces atroroseus strain IBT 11181 chromosome Unknown scaffold_1, whole genome shotgun sequence
Type: T1PKS
Number of proteins with BLAST hits to this cluster: 2
Cumulative BLAST score: 636

Table of genes, locations, strands and annotations of subject cluster:

Table of Blast hits (query gene, subject gene, %identity, blast score, %coverage, e-value):
input.path1.gene495	UA08_01176	34	139	72.48520710059172	6.1e-31
input.path1.gene499	UA08_01172	47	497	71.08886107634544	2.7e-138


>>
38. NW_006271969
Source: Cordyceps militaris CM01 unplaced genomic scaffold CCM_S00001, whole genome shotgun sequence
Type: T1PKS
Number of proteins with BLAST hits to this cluster: 2
Cumulative BLAST score: 626

Table of genes, locations, strands and annotations of subject cluster:

Table of Blast hits (query gene, subject gene, %identity, blast score, %coverage, e-value):
input.path1.gene495	CCM_01923	34	129	79.58579881656804	6.3e-28
input.path1.gene499	CCM_01921	43	497	91.23904881101377	2.7e-138


>>
39. NW_001820829
Source: Sclerotinia sclerotiorum 1980 UF-70 scaffold_7 genomic scaffold, whole genome shotgun sequence
Type: T1PKS
Number of proteins with BLAST hits to this cluster: 2
Cumulative BLAST score: 622

Table of genes, locations, strands and annotations of subject cluster:

Table of Blast hits (query gene, subject gene, %identity, blast score, %coverage, e-value):
input.path1.gene495	SS1G_05678	33	129	68.93491124260355	4.9e-28
input.path1.gene499	SS1G_05681	47	493	70.71339173967459	3e-137


>>
40. NW_003299161
Source: Microsporum canis CBS 113480 supercont1.9 genomic scaffold, whole genome shotgun sequence
Type: T1PKS
Number of proteins with BLAST hits to this cluster: 2
Cumulative BLAST score: 613

Table of genes, locations, strands and annotations of subject cluster:

Table of Blast hits (query gene, subject gene, %identity, blast score, %coverage, e-value):
input.path1.gene495	MCYG_08540	36	119	59.76331360946746	5e-25
input.path1.gene499	MCYG_08535	47	494	68.46057571964957	2.3e-137


>>
41. NC_030962
Source: Colletotrichum higginsianum IMI 349063 chromosome 10, whole genome shotgun sequence
Type: NRPS-like,T1PKS
Number of proteins with BLAST hits to this cluster: 2
Cumulative BLAST score: 609

Table of genes, locations, strands and annotations of subject cluster:

Table of Blast hits (query gene, subject gene, %identity, blast score, %coverage, e-value):
input.path1.gene495	CH63R_14355	31	109	69.23076923076923	8.9e-22
input.path1.gene499	CH63R_14350	37	500	103.12891113892366	4.2e-139


>>
42. NT_165927
Source: Aspergillus terreus NIH2624 scaffold_4 genomic scaffold, whole genome shotgun sequence
Type: T1PKS
Number of proteins with BLAST hits to this cluster: 2
Cumulative BLAST score: 557

Table of genes, locations, strands and annotations of subject cluster:

Table of Blast hits (query gene, subject gene, %identity, blast score, %coverage, e-value):
input.path1.gene495	ATEG_03439	32	114	78.69822485207101	2.1e-23
input.path1.gene499	ATEG_03432	31	225	68.08510638297872	1.5e-56
input.path1.gene499	ATEG_03446	32	218	55.56946182728411	2.5e-54


>>
43. NW_006912904
Source: Capronia epimyces CBS 606.96 aczIQ-supercont1.1, whole genome shotgun sequence
Type: T1PKS
Number of proteins with BLAST hits to this cluster: 2
Cumulative BLAST score: 529

Table of genes, locations, strands and annotations of subject cluster:

Table of Blast hits (query gene, subject gene, %identity, blast score, %coverage, e-value):
input.path1.gene495	A1O3_01769	38	207	85.50295857988166	3.1e-51
input.path1.gene499	A1O3_01765	37	322	69.96245306633291	9.3e-86


>>
44. NW_017971438
Source: Talaromyces atroroseus strain IBT 11181 chromosome Unknown scaffold_6, whole genome shotgun sequence
Type: T1PKS
Number of proteins with BLAST hits to this cluster: 2
Cumulative BLAST score: 499

Table of genes, locations, strands and annotations of subject cluster:

Table of Blast hits (query gene, subject gene, %identity, blast score, %coverage, e-value):
input.path1.gene495	UA08_04988	38	187	86.3905325443787	2.6e-45
input.path1.gene499	UA08_04992	37	312	69.33667083854819	9.6e-83


>>
45. NZ_FNOG01000002
Source: Lysobacter enzymogenes strain ATCC 29487, whole genome shotgun sequence
Type: NRPS,T1PKS
Number of proteins with BLAST hits to this cluster: 2
Cumulative BLAST score: 487

Table of genes, locations, strands and annotations of subject cluster:

Table of Blast hits (query gene, subject gene, %identity, blast score, %coverage, e-value):
input.path1.gene494	BLU84_RS06850	31	197	96.20689655172414	4.3e-48
input.path1.gene499	BLU84_RS06785	35	290	69.33667083854819	6.6e-76


>>
46. NZ_JACJIA010000006
Source: Actinomadura namibiensis strain DSM 44197 Ga0415225_06, whole genome shotgun sequence
Type: T1PKS,other
Number of proteins with BLAST hits to this cluster: 2
Cumulative BLAST score: 483

Table of genes, locations, strands and annotations of subject cluster:

Table of Blast hits (query gene, subject gene, %identity, blast score, %coverage, e-value):
input.path1.gene494	HNR61_RS23165	31	210	94.82758620689656	4.9e-52
input.path1.gene499	HNR61_RS22890	34	273	60.575719649561954	8.4e-71


>>
47. NZ_AP014940
Source: Lysobacter enzymogenes strain M497-1
Type: NRPS,T1PKS
Number of proteins with BLAST hits to this cluster: 2
Cumulative BLAST score: 475

Table of genes, locations, strands and annotations of subject cluster:

Table of Blast hits (query gene, subject gene, %identity, blast score, %coverage, e-value):
input.path1.gene494	LEN_RS18945	31	195	96.20689655172414	2.1e-47
input.path1.gene499	LEN_RS18880	34	280	69.33667083854819	5.3e-73


>>
48. NC_016787
Source: Corynebacterium diphtheriae HC03, complete sequence
Type: NRPS,T1PKS
Number of proteins with BLAST hits to this cluster: 2
Cumulative BLAST score: 459

Table of genes, locations, strands and annotations of subject cluster:
CDHC03_RS12025	CDHC03_RS12025	2200996	2201584	-	DNA_primase
CDHC03_RS10450	CDHC03_RS10450	2255846	2257395	-	acyl-CoA_carboxylase_subunit_beta

Table of Blast hits (query gene, subject gene, %identity, blast score, %coverage, e-value):
input.path1.gene494	CDHC03_RS10525	31	205	96.03448275862068	1.2e-50
input.path1.gene499	CDHC03_RS10460	32	254	80.10012515644556	3.1e-65


>>
49. NZ_MIOB01000012
Source: Corynebacterium diphtheriae strain ST395-KZN-2015-45792 45792_contig_2, whole genome shotgun sequence
Type: T1PKS
Number of proteins with BLAST hits to this cluster: 2
Cumulative BLAST score: 459

Table of genes, locations, strands and annotations of subject cluster:
BHF79_RS03915	BHF79_RS03915	49621	50714	-	hypothetical_protein

Table of Blast hits (query gene, subject gene, %identity, blast score, %coverage, e-value):
input.path1.gene494	BHF79_RS03990	31	205	96.03448275862068	1.2e-50
input.path1.gene499	BHF79_RS03925	32	254	80.10012515644556	3.1e-65


>>
50. NZ_CADFFS010000002
Source: Burkholderia vietnamiensis strain BCC1176, whole genome shotgun sequence
Type: NRPS
Number of proteins with BLAST hits to this cluster: 2
Cumulative BLAST score: 458

Table of genes, locations, strands and annotations of subject cluster:
HW519_RS07450	HW519_RS07450	523933	537397	-	amino_acid_adenylation_domain-containing_protein
HW519_RS07485	HW519_RS07485	550205	550864	-	transposase
HW519_RS07490	HW519_RS07490	551021	551086	-	hypothetical_protein

Table of Blast hits (query gene, subject gene, %identity, blast score, %coverage, e-value):
input.path1.gene494	HW519_RS07360	31	193	97.24137931034483	4.7e-47
input.path1.gene499	HW519_RS07460	35	265	68.33541927409262	1.3e-68
