## Supplementary Results for "A draft genome of the ascomycotal fungal species *Pseudopithomyces maydicus* (family *Didymosphaeriaceae*)": contig_32_c3.txt

ClusterBlast scores for contig_32

Table of genes, locations, strands and annotations of query cluster:
input.path1.gene885	3138049	3139703	-		
input.path1.gene886	3141781	3142924	-		
input.path1.gene887	3145213	3146156	+		
input.path1.gene888	3146892	3147343	-		
input.path1.gene889	3148273	3149705	-		
input.path1.gene890	3150534	3151265	+		
input.path1.gene891	3154000	3154464	-		
input.path1.gene892	3155450	3156116	-		
input.path1.gene893	3157579	3164311	+		
input.path1.gene894	3168713	3169230	-		
input.path1.gene895	3171654	3177360	-		
input.path1.gene896	3178403	3179732	+		
input.path1.gene897	3181316	3182359	+		
input.path1.gene898	3182593	3184390	-		
input.path1.gene899	3185516	3187164	+		
input.path1.gene900	3189780	3193436	-		


Significant hits: 
1. NT_165934	Aspergillus terreus NIH2624 scaffold_11 genomic scaffold, whole genome shotgun sequence
2. NT_107015	Aspergillus nidulans FGSC A4 chromosome VIII map unlocalized scaffold_1, whole genome shotgun sequence
3. CM002799	Penicillium chrysogenum strain P2niaD18 chromosome II, whole genome shotgun sequence
4. NW_014574709	Metarhizium brunneum ARSEF 3297 chromosome Unknown scaffold_8, whole genome shotgun sequence
5. NW_011942171	Metarhizium robertsii ARSEF 23 MAA_Scf_3, whole genome shotgun sequence
6. NT_107014	Aspergillus nidulans FGSC A4 chromosome VII map unlocalized scaffold_2, whole genome shotgun sequence
7. NW_004504311	Coccidioides immitis RS genomic scaffold supercont3.4, whole genome shotgun sequence
8. NW_003315987	Coccidioides posadasii C735 delta SOWgp chromosome Unknown gcontig_1105601051765, whole genome shotgun sequence
9. NW_022983866	Arthroderma uncinatum strain CBS 119779 chromosome Unknown scaffold4, whole genome shotgun sequence
10. NW_003299166	Microsporum canis CBS 113480 supercont1.4 genomic scaffold, whole genome shotgun sequence
11. NW_022194798	Fusarium proliferatum ET1 genome assembly, contig: FPRO_scaffold009, whole genome shotgun sequence
12. NC_036632	Fusarium fujikuroi IMI 58289 draft genome, chromosome FFUJ_chr11
13. NW_022984629	Aspergillus tanneri strain NIH1004 chromosome Unknown AapTan_2_pilon_pilon_pilon, whole genome shotgun sequence
14. NZ_JH109153	Methylobacter tundripaludum SV96 Mettuscaffold_2, whole genome shotgun sequence
15. NT_165980	Chaetomium globosum CBS 148.51 scaffold_5 genomic scaffold, whole genome shotgun sequence
16. NC_036439	Aspergillus oryzae RIB40 DNA, chromosome 5
17. NT_165982	Chaetomium globosum CBS 148.51 scaffold_7 genomic scaffold, whole genome shotgun sequence
18. NZ_RBWU01000008	Actinomadura pelletieri DSM 43383 Ga0170373_108, whole genome shotgun sequence
19. CM004176	Drechmeria coniospora strain ARSEF 6962 chromosome 03, whole genome shotgun sequence
20. NW_022983865	Arthroderma uncinatum strain CBS 119779 chromosome Unknown scaffold3, whole genome shotgun sequence
21. NT_165939	Aspergillus terreus NIH2624 scaffold_16 genomic scaffold, whole genome shotgun sequence
22. NZ_CP017708	Moorea producens JHB chromosome
23. NZ_CP017599	Moorea producens PAL-8-15-08-1 chromosome, complete genome
24. NZ_MKZR01000001	Moorea producens 3L Ga0081465_101, whole genome shotgun sequence
25. CM000574	Fusarium graminearum PH-1 chromosome 1, whole genome shotgun sequence
26. NC_026474	Fusarium graminearum PH-1 chromosome 1, whole genome shotgun sequence
27. NZ_CP012949	Streptomyces ambofaciens strain DSM 40697 chromosome, complete genome
28. NZ_JABBWY010000003	Microbispora sp. H10885 NODE_3, whole genome shotgun sequence
29. NZ_CP034279	Streptomyces ficellus strain NRRL 8067 chromosome, complete genome
30. NZ_WUTW01000001	Actinomadura rayongensis strain DSM 102126 90-285609803_CB02-ArV1_contig_0, whole genome shotgun sequence
31. NZ_FWXV01000001	Kibdelosporangium aridum strain DSM 43828, whole genome shotgun sequence
32. NZ_CP024052	Micromonospora sp. WMMA2032 chromosome
33. NZ_LOJP01000001	Actinoplanes sp. TFC3 Actinoplanes_sp._strain_TFC3_1, whole genome shotgun sequence
34. NC_014623	Stigmatella aurantiaca DW4/3-1, complete sequence
35. NZ_KQ948860	Streptomyces bungoensis strain DSM 41781 PRJNA299233_s010, whole genome shotgun sequence
36. NZ_FNIE01000004	Streptomyces guanduensis strain CGMCC 4.2022, whole genome shotgun sequence
37. NZ_CP029189	Streptomyces venezuelae strain ATCC 21018 chromosome, complete genome
38. NZ_NKYF01000001	Pseudonocardia sp. MH-G8 Scaffold1, whole genome shotgun sequence
39. NZ_JACHJH010000002	Streptomyces olivoverticillatus strain CECT 3266 Ga0436946_02, whole genome shotgun sequence
40. NZ_JACHJG010000002	Streptomyces netropsis strain CECT 3265 Ga0436945_02, whole genome shotgun sequence
41. NZ_CP027297	Streptomyces sp. SGAir0924 chromosome, complete genome
42. NZ_AP019620	Streptomyces antimycoticus strain NBRC 100767
43. NZ_KQ948926	Streptomyces caeruleatus strain NRRL B-24802 PRJNA299234_s003, whole genome shotgun sequence
44. NW_007360999	Glarea lozoyensis ATCC 20868 chromosome Unknown GLAREA4, whole genome shotgun sequence
45. NT_165982	Chaetomium globosum CBS 148.51 scaffold_7 genomic scaffold, whole genome shotgun sequence
46. NW_003316003	Coccidioides posadasii C735 delta SOWgp chromosome Unknown gcontig_1105601051733, whole genome shotgun sequence
47. NW_022984635	Aspergillus tanneri strain NIH1004 chromosome Unknown AapTan_8_pilon_pilon_pilon, whole genome shotgun sequence
48. NW_014574704	Metarhizium brunneum ARSEF 3297 chromosome Unknown scaffold_13, whole genome shotgun sequence
49. NW_011942160	Metarhizium robertsii ARSEF 23 MAA_Scf_2, whole genome shotgun sequence
50. NW_023336265	Colletotrichum scovillei strain TJNH1 chromosome Unknown Contig00007, whole genome shotgun sequence


Details:

>>
1. NT_165934
Source: Aspergillus terreus NIH2624 scaffold_11 genomic scaffold, whole genome shotgun sequence
Type: T1PKS
Number of proteins with BLAST hits to this cluster: 6
Cumulative BLAST score: 4589

Table of genes, locations, strands and annotations of subject cluster:

Table of Blast hits (query gene, subject gene, %identity, blast score, %coverage, e-value):
input.path1.gene891	ATEG_07664	55	78	96.05263157894737	3.8e-13
input.path1.gene892	ATEG_07664	45	162	120.88607594936708	3.2e-38
input.path1.gene893	ATEG_07659	46	1954	115.07936507936508	0.0
input.path1.gene895	ATEG_07661	55	1772	94.59161147902869	0.0
input.path1.gene896	ATEG_07662	48	410	96.51972157772622	2.4e-112
input.path1.gene897	ATEG_07665	51	213	85.71428571428571	2.6e-53


>>
2. NT_107015
Source: Aspergillus nidulans FGSC A4 chromosome VIII map unlocalized scaffold_1, whole genome shotgun sequence
Type: T1PKS
Number of proteins with BLAST hits to this cluster: 6
Cumulative BLAST score: 4361

Table of genes, locations, strands and annotations of subject cluster:

Table of Blast hits (query gene, subject gene, %identity, blast score, %coverage, e-value):
input.path1.gene891	AN1031.2	55	74	96.05263157894737	5.5e-12
input.path1.gene892	AN1031.2	46	154	109.49367088607596	1.1e-35
input.path1.gene893	AN1036.2	44	1805	114.00560224089635	0.0
input.path1.gene895	AN1034.2	54	1740	96.19205298013244	0.0
input.path1.gene896	AN1033.2	43	371	99.07192575406032	1.6e-100
input.path1.gene897	AN1030.2	51	217	87.64478764478764	2.3e-54


>>
3. CM002799
Source: Penicillium chrysogenum strain P2niaD18 chromosome II, whole genome shotgun sequence
Type: T1PKS
Number of proteins with BLAST hits to this cluster: 3
Cumulative BLAST score: 3085

Table of genes, locations, strands and annotations of subject cluster:

Table of Blast hits (query gene, subject gene, %identity, blast score, %coverage, e-value):
input.path1.gene893	EN45_068530	41	1664	113.63211951447245	0.0
input.path1.gene895	EN45_068520	41	1154	94.4260485651214	0.0
input.path1.gene896	EN45_068510	36	267	98.14385150812065	1.9e-69


>>
4. NW_014574709
Source: Metarhizium brunneum ARSEF 3297 chromosome Unknown scaffold_8, whole genome shotgun sequence
Type: T1PKS
Number of proteins with BLAST hits to this cluster: 3
Cumulative BLAST score: 5683

Table of genes, locations, strands and annotations of subject cluster:

Table of Blast hits (query gene, subject gene, %identity, blast score, %coverage, e-value):
input.path1.gene893	MBR_06260	62	2847	113.8655462184874	0.0
input.path1.gene895	MBR_06257	65	2242	101.21412803532009	0.0
input.path1.gene896	MBR_06258	65	594	100.23201856148492	1.2e-167


>>
5. NW_011942171
Source: Metarhizium robertsii ARSEF 23 MAA_Scf_3, whole genome shotgun sequence
Type: NRPS,T1PKS
Number of proteins with BLAST hits to this cluster: 3
Cumulative BLAST score: 5628

Table of genes, locations, strands and annotations of subject cluster:

Table of Blast hits (query gene, subject gene, %identity, blast score, %coverage, e-value):
input.path1.gene893	MAA_09944	61	2756	113.8655462184874	0.0
input.path1.gene895	MAA_09947	66	2272	100.44150110375276	0.0
input.path1.gene896	MAA_09946	66	600	100.69605568445476	1.6e-169


>>
6. NT_107014
Source: Aspergillus nidulans FGSC A4 chromosome VII map unlocalized scaffold_2, whole genome shotgun sequence
Type: T1PKS
Number of proteins with BLAST hits to this cluster: 3
Cumulative BLAST score: 5256

Table of genes, locations, strands and annotations of subject cluster:

Table of Blast hits (query gene, subject gene, %identity, blast score, %coverage, e-value):
input.path1.gene893	AN2035.2	59	2621	111.29785247432307	0.0
input.path1.gene895	AN2032.2	61	2069	98.28918322295806	0.0
input.path1.gene896	AN2033.2	64	566	98.83990719257541	3.4e-159


>>
7. NW_004504311
Source: Coccidioides immitis RS genomic scaffold supercont3.4, whole genome shotgun sequence
Type: T1PKS
Number of proteins with BLAST hits to this cluster: 3
Cumulative BLAST score: 3538

Table of genes, locations, strands and annotations of subject cluster:

Table of Blast hits (query gene, subject gene, %identity, blast score, %coverage, e-value):
input.path1.gene893	CIMG_05569	42	1721	111.76470588235294	0.0
input.path1.gene895	CIMG_05571	46	1446	98.45474613686534	0.0
input.path1.gene896	CIMG_05570	45	371	97.91183294663574	9.4e-101


>>
8. NW_003315987
Source: Coccidioides posadasii C735 delta SOWgp chromosome Unknown gcontig_1105601051765, whole genome shotgun sequence
Type: T1PKS
Number of proteins with BLAST hits to this cluster: 3
Cumulative BLAST score: 3529

Table of genes, locations, strands and annotations of subject cluster:

Table of Blast hits (query gene, subject gene, %identity, blast score, %coverage, e-value):
input.path1.gene893	CPC735_066950	42	1721	110.50420168067228	0.0
input.path1.gene895	CPC735_066970	46	1434	93.92935982339957	0.0
input.path1.gene896	CPC735_066960	46	374	98.14385150812065	1.9e-101


>>
9. NW_022983866
Source: Arthroderma uncinatum strain CBS 119779 chromosome Unknown scaffold4, whole genome shotgun sequence
Type: T1PKS
Number of proteins with BLAST hits to this cluster: 3
Cumulative BLAST score: 3460

Table of genes, locations, strands and annotations of subject cluster:

Table of Blast hits (query gene, subject gene, %identity, blast score, %coverage, e-value):
input.path1.gene893	GIQ15_05885	42	1654	107.00280112044818	0.0
input.path1.gene895	GIQ15_05883	45	1433	98.17880794701986	0.0
input.path1.gene896	GIQ15_05884	46	373	99.53596287703016	4.2e-101


>>
10. NW_003299166
Source: Microsporum canis CBS 113480 supercont1.4 genomic scaffold, whole genome shotgun sequence
Type: T1PKS
Number of proteins with BLAST hits to this cluster: 3
Cumulative BLAST score: 3385

Table of genes, locations, strands and annotations of subject cluster:

Table of Blast hits (query gene, subject gene, %identity, blast score, %coverage, e-value):
input.path1.gene893	MCYG_04404	41	1602	109.99066293183941	0.0
input.path1.gene895	MCYG_04406	45	1400	93.10154525386314	0.0
input.path1.gene896	MCYG_04405	47	383	96.98375870069606	4.1e-104


>>
11. NW_022194798
Source: Fusarium proliferatum ET1 genome assembly, contig: FPRO_scaffold009, whole genome shotgun sequence
Type: T1PKS
Number of proteins with BLAST hits to this cluster: 3
Cumulative BLAST score: 3362

Table of genes, locations, strands and annotations of subject cluster:

Table of Blast hits (query gene, subject gene, %identity, blast score, %coverage, e-value):
input.path1.gene893	FPRO_15779	41	1683	114.47245564892623	0.0
input.path1.gene895	FPRO_15788	50	1314	77.48344370860927	0.0
input.path1.gene896	FPRO_15787	45	365	96.51972157772622	8.8e-99


>>
12. NC_036632
Source: Fusarium fujikuroi IMI 58289 draft genome, chromosome FFUJ_chr11
Type: T1PKS
Number of proteins with BLAST hits to this cluster: 3
Cumulative BLAST score: 3361

Table of genes, locations, strands and annotations of subject cluster:

Table of Blast hits (query gene, subject gene, %identity, blast score, %coverage, e-value):
input.path1.gene893	FFUJ_12066	41	1683	114.47245564892623	0.0
input.path1.gene895	FFUJ_12074	50	1310	77.48344370860927	0.0
input.path1.gene896	FFUJ_12073	45	368	96.51972157772622	7.9e-100


>>
13. NW_022984629
Source: Aspergillus tanneri strain NIH1004 chromosome Unknown AapTan_2_pilon_pilon_pilon, whole genome shotgun sequence
Type: T1PKS,terpene
Number of proteins with BLAST hits to this cluster: 3
Cumulative BLAST score: 1896

Table of genes, locations, strands and annotations of subject cluster:

Table of Blast hits (query gene, subject gene, %identity, blast score, %coverage, e-value):
input.path1.gene893	ATNIH1004_001811	36	493	39.72922502334267	1.1e-136
input.path1.gene895	ATNIH1004_001825	40	1154	92.99116997792495	0.0
input.path1.gene896	ATNIH1004_001824	35	249	99.30394431554525	7e-64


>>
14. NZ_JH109153
Source: Methylobacter tundripaludum SV96 Mettuscaffold_2, whole genome shotgun sequence
Type: NRPS,T1PKS
Number of proteins with BLAST hits to this cluster: 3
Cumulative BLAST score: 744

Table of genes, locations, strands and annotations of subject cluster:
METTU_RS16440	METTU_RS16440	811339	812286	-	NADH-quinone_oxidoreductase_subunit_NuoH

Table of Blast hits (query gene, subject gene, %identity, blast score, %coverage, e-value):
input.path1.gene893	METTU_RS16360	32	310	34.73389355742297	1.3e-81
input.path1.gene895	METTU_RS16365	31	343	46.19205298013245	1.5e-91
input.path1.gene897	METTU_RS16355	34	91	80.6949806949807	1.5e-16


>>
15. NT_165980
Source: Chaetomium globosum CBS 148.51 scaffold_5 genomic scaffold, whole genome shotgun sequence
Type: T1PKS
Number of proteins with BLAST hits to this cluster: 4
Cumulative BLAST score: 3774

Table of genes, locations, strands and annotations of subject cluster:

Table of Blast hits (query gene, subject gene, %identity, blast score, %coverage, e-value):
input.path1.gene892	CHGG_07643	57	85	41.139240506329116	4.9e-15
input.path1.gene893	CHGG_07638	43	1698	110.78431372549021	0.0
input.path1.gene895	CHGG_07645	57	1205	58.94039735099338	0.0
input.path1.gene895	CHGG_07646	45	389	27.81456953642384	1.4e-105
input.path1.gene896	CHGG_07644	47	397	96.75174013921114	1.6e-108


>>
16. NC_036439
Source: Aspergillus oryzae RIB40 DNA, chromosome 5
Type: T1PKS
Number of proteins with BLAST hits to this cluster: 4
Cumulative BLAST score: 3581

Table of genes, locations, strands and annotations of subject cluster:

Table of Blast hits (query gene, subject gene, %identity, blast score, %coverage, e-value):
input.path1.gene893	AO090701000826	42	1708	111.06442577030813	0.0
input.path1.gene895	AO090701000831	45	1445	96.08167770419426	0.0
input.path1.gene896	AO090701000828	48	208	52.90023201856149	1.1e-51
input.path1.gene896	AO090701000827	42	140	37.58700696055685	4.6e-31
input.path1.gene897	AO090701000821	33	80	78.76447876447877	2.6e-13


>>
17. NT_165982
Source: Chaetomium globosum CBS 148.51 scaffold_7 genomic scaffold, whole genome shotgun sequence
Type: T1PKS
Number of proteins with BLAST hits to this cluster: 2
Cumulative BLAST score: 2138

Table of genes, locations, strands and annotations of subject cluster:

Table of Blast hits (query gene, subject gene, %identity, blast score, %coverage, e-value):
input.path1.gene893	CHGG_09585	36	1237	112.41830065359477	0.0
input.path1.gene895	CHGG_09586	35	901	93.4878587196468	1.9e-259


>>
18. NZ_RBWU01000008
Source: Actinomadura pelletieri DSM 43383 Ga0170373_108, whole genome shotgun sequence
Type: PKS-like,T1PKS
Number of proteins with BLAST hits to this cluster: 2
Cumulative BLAST score: 1800

Table of genes, locations, strands and annotations of subject cluster:
BZB76_RS31890	BZB76_RS31890	257852	258844	+	S-methyl-5-thioribose-1-phosphate_isomerase

Table of Blast hits (query gene, subject gene, %identity, blast score, %coverage, e-value):
input.path1.gene893	BZB76_RS35060	33	307	34.12698412698413	1.1e-80
input.path1.gene893	BZB76_RS31800	31	302	38.51540616246499	2.7e-79
input.path1.gene893	BZB76_RS31685	32	284	31.045751633986928	9.8e-74
input.path1.gene893	BZB76_RS31715	31	268	30.11204481792717	7.2e-69
input.path1.gene895	BZB76_RS35060	32	335	47.682119205298015	3.1e-89
input.path1.gene895	BZB76_RS31800	31	304	47.90286975717439	7.7e-80


>>
19. CM004176
Source: Drechmeria coniospora strain ARSEF 6962 chromosome 03, whole genome shotgun sequence
Type: T1PKS
Number of proteins with BLAST hits to this cluster: 3
Cumulative BLAST score: 1635

Table of genes, locations, strands and annotations of subject cluster:

Table of Blast hits (query gene, subject gene, %identity, blast score, %coverage, e-value):
input.path1.gene895	DCS_05930	41	1176	92.82560706401766	0.0
input.path1.gene896	DCS_05929	39	280	95.35962877030161	3.7e-73
input.path1.gene897	DCS_05926	44	179	82.23938223938224	5.4e-43


>>
20. NW_022983865
Source: Arthroderma uncinatum strain CBS 119779 chromosome Unknown scaffold3, whole genome shotgun sequence
Type: T1PKS
Number of proteins with BLAST hits to this cluster: 2
Cumulative BLAST score: 1478

Table of genes, locations, strands and annotations of subject cluster:

Table of Blast hits (query gene, subject gene, %identity, blast score, %coverage, e-value):
input.path1.gene893	GIQ15_03969	58	308	13.071895424836603	4.8e-81
input.path1.gene895	GIQ15_03968	49	1170	74.17218543046357	0.0


>>
21. NT_165939
Source: Aspergillus terreus NIH2624 scaffold_16 genomic scaffold, whole genome shotgun sequence
Type: T1PKS
Number of proteins with BLAST hits to this cluster: 3
Cumulative BLAST score: 1431

Table of genes, locations, strands and annotations of subject cluster:

Table of Blast hits (query gene, subject gene, %identity, blast score, %coverage, e-value):
input.path1.gene892	ATEG_09967	38	124	110.12658227848102	1.2e-26
input.path1.gene893	ATEG_09968	34	1187	111.67133520074697	0.0
input.path1.gene897	ATEG_09963	39	120	72.58687258687259	3e-25


>>
22. NZ_CP017708
Source: Moorea producens JHB chromosome
Type: NRPS,T1PKS
Number of proteins with BLAST hits to this cluster: 2
Cumulative BLAST score: 1403

Table of genes, locations, strands and annotations of subject cluster:
BJP36_RS17885	BJP36_RS17885	4803728	4803964	+	type_II_toxin-antitoxin_system_RelE/ParE_family_toxin
BJP36_RS17890	BJP36_RS17890	4804142	4804725	+	Uma2_family_endonuclease

Table of Blast hits (query gene, subject gene, %identity, blast score, %coverage, e-value):
input.path1.gene893	BJP36_RS17855	31	299	34.64052287581699	2.2e-78
input.path1.gene893	BJP36_RS17865	31	299	34.64052287581699	2.9e-78
input.path1.gene895	BJP36_RS17860	31	403	52.59381898454746	1.2e-109
input.path1.gene895	BJP36_RS17855	31	402	51.04856512141281	2.7e-109


>>
23. NZ_CP017599
Source: Moorea producens PAL-8-15-08-1 chromosome, complete genome
Type: NRPS,PKS-like,T1PKS,T3PKS
Number of proteins with BLAST hits to this cluster: 2
Cumulative BLAST score: 1326

Table of genes, locations, strands and annotations of subject cluster:
BJP34_RS45830	BJP34_RS45830	7892977	7893444	-	PAS_domain_S-box_protein
BJP34_RS41220	BJP34_RS41220	7957288	7961292	-	type_I_polyketide_synthase
BJP34_RS41225	BJP34_RS41225	7961353	7961973	-	zinc-binding_dehydrogenase
BJP34_RS41230	BJP34_RS41230	7962049	7962291	-	alcohol_dehydrogenase_catalytic_domain-containing_protein
BJP34_RS41235	BJP34_RS41235	7962418	7963176	-	enoyl-CoA_hydratase
BJP34_RS29120	BJP34_RS29120	7974663	7975856	+	ISAs1_family_transposase

Table of Blast hits (query gene, subject gene, %identity, blast score, %coverage, e-value):
input.path1.gene893	BJP34_RS29070	31	322	40.14939309056956	3.2e-85
input.path1.gene893	BJP34_RS29080	31	310	37.44164332399627	1.3e-81
input.path1.gene893	BJP34_RS29085	32	284	30.905695611577965	7.5e-74
input.path1.gene895	BJP34_RS29080	32	410	51.82119205298014	1e-111


>>
24. NZ_MKZR01000001
Source: Moorea producens 3L Ga0081465_101, whole genome shotgun sequence
Type: NRPS,PKS-like,T1PKS,T3PKS
Number of proteins with BLAST hits to this cluster: 2
Cumulative BLAST score: 1326

Table of genes, locations, strands and annotations of subject cluster:
BI334_RS38505	BI334_RS38505	6571079	6571216	-	succinate--CoA_ligase_subunit_beta
BI334_RS35730	BI334_RS35730	6637960	6641109	-	type_I_polyketide_synthase
BI334_RS35735	BI334_RS35735	6641170	6641790	-	zinc-binding_dehydrogenase
BI334_RS38520	BI334_RS38520	6641866	6642108	-	alcohol_dehydrogenase_catalytic_domain-containing_protein
BI334_RS35740	BI334_RS35740	6642220	6642993	-	enoyl-CoA_hydratase
BI334_RS35745	BI334_RS35745	6656019	6656960	-	IS630_family_transposase
BI334_RS27460	BI334_RS27460	6657253	6657375	+	transposase
BI334_RS27465	BI334_RS27465	6657961	6659527	+	group_II_intron_reverse_transcriptase/maturase
BI334_RS27475	BI334_RS27475	6659902	6660270	+	transposase
BI334_RS38525	BI334_RS38525	6660912	6662284	+	IS200/IS605_family_element_transposase_accessory_protein_TnpB
BI334_RS27505	BI334_RS27505	6668503	6669138	+	transposase
BI334_RS27510	BI334_RS27510	6669128	6669641	+	transposase
BI334_RS27530	BI334_RS27530	6672828	6673022	-	hypothetical_protein

Table of Blast hits (query gene, subject gene, %identity, blast score, %coverage, e-value):
input.path1.gene893	BI334_RS27395	31	321	39.77591036414566	7.2e-85
input.path1.gene893	BI334_RS27405	31	314	37.488328664799255	6.8e-83
input.path1.gene893	BI334_RS27410	33	284	30.905695611577965	7.5e-74
input.path1.gene895	BI334_RS27405	32	407	52.26269315673289	8.5e-111


>>
25. CM000574
Source: Fusarium graminearum PH-1 chromosome 1, whole genome shotgun sequence
Type: NRPS,T1PKS
Number of proteins with BLAST hits to this cluster: 2
Cumulative BLAST score: 1306

Table of genes, locations, strands and annotations of subject cluster:

Table of Blast hits (query gene, subject gene, %identity, blast score, %coverage, e-value):
input.path1.gene893	FGSG_12126	31	960	106.1624649859944	4e-277
input.path1.gene895	FGSG_02395	31	346	46.24724061810154	2.3e-92


>>
26. NC_026474
Source: Fusarium graminearum PH-1 chromosome 1, whole genome shotgun sequence
Type: NRPS,T1PKS
Number of proteins with BLAST hits to this cluster: 2
Cumulative BLAST score: 1306

Table of genes, locations, strands and annotations of subject cluster:

Table of Blast hits (query gene, subject gene, %identity, blast score, %coverage, e-value):
input.path1.gene893	FGSG_12126	31	960	106.1624649859944	4e-277
input.path1.gene895	FGSG_02395	31	346	46.24724061810154	2.3e-92


>>
27. NZ_CP012949
Source: Streptomyces ambofaciens strain DSM 40697 chromosome, complete genome
Type: T1PKS,transAT-PKS
Number of proteins with BLAST hits to this cluster: 2
Cumulative BLAST score: 1165

Table of genes, locations, strands and annotations of subject cluster:
SAM40697_RS36460	SAM40697_RS36460	7451968	7452164	+	hypothetical_protein
SAM40697_RS37470	SAM40697_RS37470	7523024	7524433	+	hypothetical_protein
SAM40697_RS36550	SAM40697_RS36550	7524776	7525687	+	acyltransferase_domain-containing_protein
SAM40697_RS38195	SAM40697_RS38195	7525868	7527121	+	SDR_family_NAD(P)-dependent_oxidoreductase
SAM40697_RS37475	SAM40697_RS37475	7527182	7529599	+	acyltransferase_domain-containing_protein
SAM40697_RS38020	SAM40697_RS38020	7530164	7530445	+	hypothetical_protein
SAM40697_RS37480	SAM40697_RS37480	7530905	7531600	+	SDR_family_NAD(P)-dependent_oxidoreductase
SAM40697_RS37835	SAM40697_RS37835	7531610	7532107	+	hypothetical_protein
SAM40697_RS37485	SAM40697_RS37485	7532204	7533502	+	polyketide_synthase
SAM40697_RS38025	SAM40697_RS38025	7534574	7534966	+	hypothetical_protein
SAM40697_RS36600	SAM40697_RS36600	7535405	7536151	+	SDR_family_NAD(P)-dependent_oxidoreductase
SAM40697_RS36605	SAM40697_RS36605	7536215	7536670	+	hypothetical_protein
SAM40697_RS36610	SAM40697_RS36610	7536785	7539307	+	type_I_polyketide_synthase
SAM40697_RS37490	SAM40697_RS37490	7540034	7540615	+	SDR_family_NAD(P)-dependent_oxidoreductase
SAM40697_RS37840	SAM40697_RS37840	7575832	7578531	+	type_I_polyketide_synthase
SAM40697_RS37510	SAM40697_RS37510	7578838	7579884	+	SDR_family_NAD(P)-dependent_oxidoreductase
SAM40697_RS37515	SAM40697_RS37515	7579891	7580745	+	polyketide_synthase
SAM40697_RS37520	SAM40697_RS37520	7581049	7581570	+	polyketide_synthase
SAM40697_RS36715	SAM40697_RS36715	7582066	7582977	+	acyltransferase_domain-containing_protein
SAM40697_RS36720	SAM40697_RS36720	7583071	7583919	+	modular_polyketide_synthase
SAM40697_RS36725	SAM40697_RS36725	7584538	7584744	+	alcohol_dehydrogenase_catalytic_domain-containing_protein
SAM40697_RS37525	SAM40697_RS37525	7585321	7585998	+	SDR_family_NAD(P)-dependent_oxidoreductase

Table of Blast hits (query gene, subject gene, %identity, blast score, %coverage, e-value):
input.path1.gene893	SAM40697_RS37495	31	279	34.73389355742297	3.1e-72
input.path1.gene893	SAM40697_RS32685	31	257	30.859010270774977	9.8e-66
input.path1.gene895	SAM40697_RS32680	31	317	46.35761589403973	1.2e-83
input.path1.gene895	SAM40697_RS32675	31	312	46.799116997792495	2.8e-82


>>
28. NZ_JABBWY010000003
Source: Microbispora sp. H10885 NODE_3, whole genome shotgun sequence
Type: NRPS,T1PKS
Number of proteins with BLAST hits to this cluster: 2
Cumulative BLAST score: 948

Table of genes, locations, strands and annotations of subject cluster:
HHX37_RS10380	HHX37_RS10380	412956	413450	+	cellulose_binding_domain-containing_protein
HHX37_RS10410	HHX37_RS10410	421410	422231	-	SDR_family_NAD(P)-dependent_oxidoreductase
HHX37_RS10415	HHX37_RS10415	422520	423827	-	polyketide_synthase_dehydratase_domain-containing_protein
HHX37_RS10420	HHX37_RS10420	424032	424415	-	acyltransferase_domain-containing_protein
HHX37_RS10425	HHX37_RS10425	424623	425060	-	hypothetical_protein
HHX37_RS10430	HHX37_RS10430	425130	426305	-	type_I_polyketide_synthase
HHX37_RS10435	HHX37_RS10435	426726	427484	-	SDR_family_NAD(P)-dependent_oxidoreductase
HHX37_RS10440	HHX37_RS10440	428205	430829	-	type_I_polyketide_synthase
HHX37_RS10445	HHX37_RS10445	430947	431756	-	modular_polyketide_synthase
HHX37_RS10450	HHX37_RS10450	431793	432320	-	acyltransferase_domain-containing_protein
HHX37_RS10455	HHX37_RS10455	433071	433796	-	polyketide_synthase
HHX37_RS10615	HHX37_RS10615	478942	481650	+	type_I_polyketide_synthase
HHX37_RS10620	HHX37_RS10620	482734	483540	+	SDR_family_NAD(P)-dependent_oxidoreductase
HHX37_RS10625	HHX37_RS10625	483550	484089	+	3-ketoacyl-ACP_synthase
HHX37_RS10630	HHX37_RS10630	484189	485686	+	hypothetical_protein
HHX37_RS10635	HHX37_RS10635	486599	487924	+	SDR_family_NAD(P)-dependent_oxidoreductase
HHX37_RS10640	HHX37_RS10640	487949	488425	+	hypothetical_protein
HHX37_RS10645	HHX37_RS10645	488435	490147	+	hypothetical_protein
HHX37_RS10650	HHX37_RS10650	490127	490984	+	acyltransferase_domain-containing_protein
HHX37_RS10655	HHX37_RS10655	491099	491881	+	polyketide_synthase_dehydratase_domain-containing_protein
HHX37_RS10660	HHX37_RS10660	491861	492856	+	SDR_family_NAD(P)-dependent_oxidoreductase
HHX37_RS10665	HHX37_RS10665	492866	493360	+	hypothetical_protein
HHX37_RS10670	HHX37_RS10670	493361	494956	+	hypothetical_protein
HHX37_RS10675	HHX37_RS10675	495098	495958	+	acyltransferase_domain-containing_protein
HHX37_RS10680	HHX37_RS10680	496106	496885	+	polyketide_synthase_dehydratase_domain-containing_protein
HHX37_RS10685	HHX37_RS10685	497315	497998	+	SDR_family_NAD(P)-dependent_oxidoreductase
HHX37_RS10790	HHX37_RS10790	532397	533183	-	IS5_family_transposase
HHX37_RS10890	HHX37_RS10890	598944	599870	+	ACP_S-malonyltransferase

Table of Blast hits (query gene, subject gene, %identity, blast score, %coverage, e-value):
input.path1.gene893	HHX37_RS10910	31	305	40.94304388422035	5.4e-80
input.path1.gene893	HHX37_RS10915	31	302	37.39495798319328	2.7e-79
input.path1.gene895	HHX37_RS10910	31	341	50.55187637969095	7.4e-91


>>
29. NZ_CP034279
Source: Streptomyces ficellus strain NRRL 8067 chromosome, complete genome
Type: NRPS,T1PKS,butyrolactone
Number of proteins with BLAST hits to this cluster: 2
Cumulative BLAST score: 919

Table of genes, locations, strands and annotations of subject cluster:
EIZ62_RS18925	EIZ62_RS18925	4201906	4202106	-	NUDIX_hydrolase

Table of Blast hits (query gene, subject gene, %identity, blast score, %coverage, e-value):
input.path1.gene893	EIZ62_RS19055	34	299	30.532212885154063	3.8e-78
input.path1.gene893	EIZ62_RS19060	33	259	30.018674136321195	2.6e-66
input.path1.gene895	EIZ62_RS19055	32	361	51.21412803532009	4.1e-97


>>
30. NZ_WUTW01000001
Source: Actinomadura rayongensis strain DSM 102126 90-285609803_CB02-ArV1_contig_0, whole genome shotgun sequence
Type: T1PKS
Number of proteins with BLAST hits to this cluster: 2
Cumulative BLAST score: 892

Table of genes, locations, strands and annotations of subject cluster:
GQ466_RS05675	GQ466_RS05675	1200444	1208639	+	SDR_family_NAD(P)-dependent_oxidoreductase

Table of Blast hits (query gene, subject gene, %identity, blast score, %coverage, e-value):
input.path1.gene893	GQ466_RS05680	31	282	34.50046685340803	4.9e-73
input.path1.gene893	GQ466_RS05665	31	268	30.29878618113912	5.6e-69
input.path1.gene895	GQ466_RS05660	32	342	51.21412803532009	3.3e-91


>>
31. NZ_FWXV01000001
Source: Kibdelosporangium aridum strain DSM 43828, whole genome shotgun sequence
Type: PKS-like,T1PKS
Number of proteins with BLAST hits to this cluster: 2
Cumulative BLAST score: 842

Table of genes, locations, strands and annotations of subject cluster:
B9C43_RS00875	B9C43_RS00875	249598	264798	+	SDR_family_NAD(P)-dependent_oxidoreductase

Table of Blast hits (query gene, subject gene, %identity, blast score, %coverage, e-value):
input.path1.gene893	B9C43_RS00870	31	261	30.905695611577965	8.9e-67
input.path1.gene893	B9C43_RS00880	31	255	30.859010270774977	6.4e-65
input.path1.gene895	B9C43_RS00875	31	326	47.95805739514349	2.5e-86


>>
32. NZ_CP024052
Source: Micromonospora sp. WMMA2032 chromosome
Type: NRPS-like,T1PKS
Number of proteins with BLAST hits to this cluster: 2
Cumulative BLAST score: 834

Table of genes, locations, strands and annotations of subject cluster:
CO540_RS05160	CO540_RS05160	1134854	1139826	-	SDR_family_NAD(P)-dependent_oxidoreductase
CO540_RS05235	CO540_RS05235	1280835	1281329	-	zinc-binding_dehydrogenase
CO540_RS05240	CO540_RS05240	1281522	1281941	-	alcohol_dehydrogenase_catalytic_domain-containing_protein
CO540_RS05245	CO540_RS05245	1282242	1294178	-	SDR_family_NAD(P)-dependent_oxidoreductase

Table of Blast hits (query gene, subject gene, %identity, blast score, %coverage, e-value):
input.path1.gene893	CO540_RS05180	32	289	34.31372549019608	2.3e-75
input.path1.gene893	CO540_RS05170	31	285	34.780578898225954	4.4e-74
input.path1.gene895	CO540_RS05165	31	260	38.741721854304636	1.3e-66


>>
33. NZ_LOJP01000001
Source: Actinoplanes sp. TFC3 Actinoplanes_sp._strain_TFC3_1, whole genome shotgun sequence
Type: NRPS-like,T1PKS
Number of proteins with BLAST hits to this cluster: 2
Cumulative BLAST score: 832

Table of genes, locations, strands and annotations of subject cluster:
TFC3_RS37925	TFC3_RS37925	2509064	2509401	-	hypothetical_protein

Table of Blast hits (query gene, subject gene, %identity, blast score, %coverage, e-value):
input.path1.gene893	TFC3_RS11165	31	258	30.718954248366014	5.7e-66
input.path1.gene893	TFC3_RS11160	31	247	30.62558356676004	1e-62
input.path1.gene895	TFC3_RS11155	31	327	46.688741721854306	8.5e-87


>>
34. NC_014623
Source: Stigmatella aurantiaca DW4/3-1, complete sequence
Type: T1PKS
Number of proteins with BLAST hits to this cluster: 2
Cumulative BLAST score: 672

Table of genes, locations, strands and annotations of subject cluster:
STAUR_RS25740	STAUR_RS25740	6478977	6479597	+	IS3_family_transposase
STAUR_RS44425	STAUR_RS44425	6481276	6481739	-	transposase

Table of Blast hits (query gene, subject gene, %identity, blast score, %coverage, e-value):
input.path1.gene893	STAUR_RS25710	31	304	37.068160597572366	7e-80
input.path1.gene895	STAUR_RS25705	31	368	51.49006622516556	3.3e-99


>>
35. NZ_KQ948860
Source: Streptomyces bungoensis strain DSM 41781 PRJNA299233_s010, whole genome shotgun sequence
Type: PKS-like,T1PKS
Number of proteins with BLAST hits to this cluster: 2
Cumulative BLAST score: 633

Table of genes, locations, strands and annotations of subject cluster:

Table of Blast hits (query gene, subject gene, %identity, blast score, %coverage, e-value):
input.path1.gene893	AQJ66_RS21060	31	297	37.488328664799255	1.1e-77
input.path1.gene895	AQJ66_RS21065	31	336	46.13686534216336	1.4e-89


>>
36. NZ_FNIE01000004
Source: Streptomyces guanduensis strain CGMCC 4.2022, whole genome shotgun sequence
Type: NRPS,T1PKS,T3PKS,bottromycin,lanthipeptide,thioamide-NRP
Number of proteins with BLAST hits to this cluster: 2
Cumulative BLAST score: 632

Table of genes, locations, strands and annotations of subject cluster:
BLR33_RS10090	BLR33_RS10090	229593	230129	+	GNAT_family_N-acetyltransferase
BLR33_RS34295	BLR33_RS34295	230502	230597	+	IS5/IS1182_family_transposase

Table of Blast hits (query gene, subject gene, %identity, blast score, %coverage, e-value):
input.path1.gene893	BLR33_RS10475	32	311	33.98692810457516	7.5e-82
input.path1.gene895	BLR33_RS10470	31	321	44.64679911699779	8e-85


>>
37. NZ_CP029189
Source: Streptomyces venezuelae strain ATCC 21018 chromosome, complete genome
Type: T1PKS
Number of proteins with BLAST hits to this cluster: 2
Cumulative BLAST score: 630

Table of genes, locations, strands and annotations of subject cluster:
DEJ51_RS04630	DEJ51_RS04630	1127850	1138787	-	SDR_family_NAD(P)-dependent_oxidoreductase
DEJ51_RS04635	DEJ51_RS04635	1138908	1141028	-	polyketide_synthase
DEJ51_RS04640	DEJ51_RS04640	1144578	1145786	-	polyketide_synthase
DEJ51_RS04665	DEJ51_RS04665	1193217	1193729	-	hypothetical_protein
DEJ51_RS04680	DEJ51_RS04680	1195910	1196026	-	DUF2023_domain-containing_protein

Table of Blast hits (query gene, subject gene, %identity, blast score, %coverage, e-value):
input.path1.gene893	DEJ51_RS04650	31	304	36.22782446311858	7e-80
input.path1.gene895	DEJ51_RS04645	31	326	50.77262693156733	1.5e-86


>>
38. NZ_NKYF01000001
Source: Pseudonocardia sp. MH-G8 Scaffold1, whole genome shotgun sequence
Type: T1PKS
Number of proteins with BLAST hits to this cluster: 2
Cumulative BLAST score: 617

Table of genes, locations, strands and annotations of subject cluster:
CFP66_RS00310	CFP66_RS00310	64103	64402	-	DUF4190_domain-containing_protein
CFP66_RS00430	CFP66_RS00430	101907	103169	+	crotonyl-CoA_carboxylase/reductase

Table of Blast hits (query gene, subject gene, %identity, blast score, %coverage, e-value):
input.path1.gene893	CFP66_RS00345	31	282	32.25957049486461	4.9e-73
input.path1.gene895	CFP66_RS00340	31	335	46.7439293598234	4.1e-89


>>
39. NZ_JACHJH010000002
Source: Streptomyces olivoverticillatus strain CECT 3266 Ga0436946_02, whole genome shotgun sequence
Type: NRPS,NRPS-like,T1PKS,butyrolactone
Number of proteins with BLAST hits to this cluster: 2
Cumulative BLAST score: 585

Table of genes, locations, strands and annotations of subject cluster:
FHS39_RS06710	FHS39_RS06710	316604	317638	-	SDR_family_NAD(P)-dependent_oxidoreductase
FHS39_RS06715	FHS39_RS06715	318503	319534	-	acyltransferase_domain-containing_protein
FHS39_RS06720	FHS39_RS06720	319691	321565	-	beta-ketoacyl_synthase
FHS39_RS06725	FHS39_RS06725	321581	322669	-	SDR_family_NAD(P)-dependent_oxidoreductase
FHS39_RS06730	FHS39_RS06730	323510	324409	-	acyltransferase_domain-containing_protein
FHS39_RS06735	FHS39_RS06735	324719	326374	-	type_I_polyketide_synthase

Table of Blast hits (query gene, subject gene, %identity, blast score, %coverage, e-value):
input.path1.gene893	FHS39_RS06675	31	275	36.507936507936506	5.9e-71
input.path1.gene895	FHS39_RS06670	31	310	47.019867549668874	1.4e-81


>>
40. NZ_JACHJG010000002
Source: Streptomyces netropsis strain CECT 3265 Ga0436945_02, whole genome shotgun sequence
Type: LAP,T1PKS,bacteriocin,terpene,thiopeptide
Number of proteins with BLAST hits to this cluster: 2
Cumulative BLAST score: 584

Table of genes, locations, strands and annotations of subject cluster:
FHS38_RS06700	FHS38_RS06700	423397	424696	+	DUF692_domain-containing_protein
FHS38_RS06895	FHS38_RS06895	561319	562359	-	peptidase_M50

Table of Blast hits (query gene, subject gene, %identity, blast score, %coverage, e-value):
input.path1.gene893	FHS38_RS06735	31	273	34.45378151260504	1.7e-70
input.path1.gene895	FHS38_RS06740	31	311	47.35099337748344	4.8e-82


>>
41. NZ_CP027297
Source: Streptomyces sp. SGAir0924 chromosome, complete genome
Type: T1PKS
Number of proteins with BLAST hits to this cluster: 2
Cumulative BLAST score: 584

Table of genes, locations, strands and annotations of subject cluster:

Table of Blast hits (query gene, subject gene, %identity, blast score, %coverage, e-value):
input.path1.gene893	C1N79_RS31690	32	248	30.485527544351076	5.9e-63
input.path1.gene895	C1N79_RS31695	31	336	53.03532008830022	1.8e-89


>>
42. NZ_AP019620
Source: Streptomyces antimycoticus strain NBRC 100767
Type: T1PKS
Number of proteins with BLAST hits to this cluster: 2
Cumulative BLAST score: 567

Table of genes, locations, strands and annotations of subject cluster:
GYK45_RS29785	GYK45_RS29785	7021302	7031575	+	SDR_family_NAD(P)-dependent_oxidoreductase
GYK45_RS29795	GYK45_RS29795	7057172	7062147	+	SDR_family_NAD(P)-dependent_oxidoreductase
GYK45_RS29800	GYK45_RS29800	7062408	7078045	+	SDR_family_NAD(P)-dependent_oxidoreductase
GYK45_RS29810	GYK45_RS29810	7099735	7109348	+	SDR_family_NAD(P)-dependent_oxidoreductase
GYK45_RS29815	GYK45_RS29815	7109405	7126620	+	SDR_family_NAD(P)-dependent_oxidoreductase
GYK45_RS29820	GYK45_RS29820	7126683	7138486	+	SDR_family_NAD(P)-dependent_oxidoreductase
GYK45_RS29865	GYK45_RS29865	7150963	7151897	+	LysR_family_transcriptional_regulator

Table of Blast hits (query gene, subject gene, %identity, blast score, %coverage, e-value):
input.path1.gene893	GYK45_RS29810	31	238	27.871148459383754	4.7e-60
input.path1.gene895	GYK45_RS29805	31	329	47.40618101545254	1.7e-87


>>
43. NZ_KQ948926
Source: Streptomyces caeruleatus strain NRRL B-24802 PRJNA299234_s003, whole genome shotgun sequence
Type: T1PKS,T2PKS
Number of proteins with BLAST hits to this cluster: 2
Cumulative BLAST score: 566

Table of genes, locations, strands and annotations of subject cluster:
AQJ67_RS47515	AQJ67_RS47515	116953	117051	-	ATP-binding_protein
AQJ67_RS48160	AQJ67_RS48160	117049	117141	+	sugar_ABC_transporter_permease
AQJ67_RS46650	AQJ67_RS46650	117341	117445	+	carbohydrate_ABC_transporter_permease
AQJ67_RS09290	AQJ67_RS09290	119046	119390	+	carbohydrate_ABC_transporter_permease
AQJ67_RS09515	AQJ67_RS09515	176230	177408	-	MFS_transporter
AQJ67_RS09600	AQJ67_RS09600	197182	197709	-	TetR/AcrR_family_transcriptional_regulator
AQJ67_RS44900	AQJ67_RS44900	197812	198565	+	ATP-binding_cassette_domain-containing_protein
AQJ67_RS09610	AQJ67_RS09610	198579	199233	+	ABC_transporter_permease

Table of Blast hits (query gene, subject gene, %identity, blast score, %coverage, e-value):
input.path1.gene893	AQJ67_RS09350	32	307	35.01400560224089	8.3e-81
input.path1.gene895	AQJ67_RS09345	31	259	38.30022075055188	2.2e-66


>>
44. NW_007360999
Source: Glarea lozoyensis ATCC 20868 chromosome Unknown GLAREA4, whole genome shotgun sequence
Type: T1PKS
Number of proteins with BLAST hits to this cluster: 4
Cumulative BLAST score: 1897

Table of genes, locations, strands and annotations of subject cluster:

Table of Blast hits (query gene, subject gene, %identity, blast score, %coverage, e-value):
input.path1.gene892	GLAREA_09805	40	128	121.51898734177216	8.6e-28
input.path1.gene895	GLAREA_09802	43	1300	94.48123620309052	0.0
input.path1.gene896	GLAREA_09799	37	270	99.76798143851508	3.8e-70
input.path1.gene897	GLAREA_09803	47	199	82.62548262548263	3.8e-49


>>
45. NT_165982
Source: Chaetomium globosum CBS 148.51 scaffold_7 genomic scaffold, whole genome shotgun sequence
Type: T1PKS
Number of proteins with BLAST hits to this cluster: 4
Cumulative BLAST score: 1607

Table of genes, locations, strands and annotations of subject cluster:

Table of Blast hits (query gene, subject gene, %identity, blast score, %coverage, e-value):
input.path1.gene892	CHGG_10019	38	114	109.49367088607596	7.6e-24
input.path1.gene895	CHGG_10027	40	1104	93.4878587196468	0.0
input.path1.gene896	CHGG_10021	37	257	91.87935034802784	2e-66
input.path1.gene897	CHGG_10030	40	132	86.10038610038609	5.7e-29


>>
46. NW_003316003
Source: Coccidioides posadasii C735 delta SOWgp chromosome Unknown gcontig_1105601051733, whole genome shotgun sequence
Type: T1PKS
Number of proteins with BLAST hits to this cluster: 2
Cumulative BLAST score: 1517

Table of genes, locations, strands and annotations of subject cluster:

Table of Blast hits (query gene, subject gene, %identity, blast score, %coverage, e-value):
input.path1.gene895	CPC735_034200	42	1236	93.26710816777042	0.0
input.path1.gene896	CPC735_034190	37	281	96.2877030162413	1.7e-73


>>
47. NW_022984635
Source: Aspergillus tanneri strain NIH1004 chromosome Unknown AapTan_8_pilon_pilon_pilon, whole genome shotgun sequence
Type: T1PKS
Number of proteins with BLAST hits to this cluster: 2
Cumulative BLAST score: 1506

Table of genes, locations, strands and annotations of subject cluster:

Table of Blast hits (query gene, subject gene, %identity, blast score, %coverage, e-value):
input.path1.gene895	ATNIH1004_011374	43	1225	93.32229580573951	0.0
input.path1.gene896	ATNIH1004_011375	36	281	101.39211136890951	1.3e-73


>>
48. NW_014574704
Source: Metarhizium brunneum ARSEF 3297 chromosome Unknown scaffold_13, whole genome shotgun sequence
Type: T1PKS
Number of proteins with BLAST hits to this cluster: 2
Cumulative BLAST score: 1491

Table of genes, locations, strands and annotations of subject cluster:

Table of Blast hits (query gene, subject gene, %identity, blast score, %coverage, e-value):
input.path1.gene895	MBR_07911	42	1248	93.87417218543047	0.0
input.path1.gene896	MBR_07910	34	243	91.1832946635731	2.9e-62


>>
49. NW_011942160
Source: Metarhizium robertsii ARSEF 23 MAA_Scf_2, whole genome shotgun sequence
Type: T1PKS
Number of proteins with BLAST hits to this cluster: 2
Cumulative BLAST score: 1484

Table of genes, locations, strands and annotations of subject cluster:

Table of Blast hits (query gene, subject gene, %identity, blast score, %coverage, e-value):
input.path1.gene895	MAA_07925	42	1242	93.65342163355407	0.0
input.path1.gene896	MAA_07924	34	242	91.1832946635731	6.6e-62


>>
50. NW_023336265
Source: Colletotrichum scovillei strain TJNH1 chromosome Unknown Contig00007, whole genome shotgun sequence
Type: T1PKS
Number of proteins with BLAST hits to this cluster: 2
Cumulative BLAST score: 1395

Table of genes, locations, strands and annotations of subject cluster:

Table of Blast hits (query gene, subject gene, %identity, blast score, %coverage, e-value):
input.path1.gene892	HER10_EVM0008012	31	95	117.08860759493672	4.7e-18
input.path1.gene893	HER10_EVM0001067	36	1300	111.95144724556489	0.0
