## Supplementary Results for "A draft genome of the ascomycotal fungal species *Pseudopithomyces maydicus* (family *Didymosphaeriaceae*)": contig_35_c1.txt

ClusterBlast scores for contig_35

Table of genes, locations, strands and annotations of query cluster:
input.path1.gene1	40862	41782	+		
input.path1.gene2	43543	44105	-		
input.path1.gene3	46845	58495	-		
input.path1.gene4	62097	69893	+		
input.path1.gene5	80634	83097	-		
input.path1.gene6	83613	87826	-		


Significant hits: 
1. CM000575	Fusarium graminearum PH-1 chromosome 2, whole genome shotgun sequence
2. NC_026475	Fusarium graminearum PH-1 chromosome 2, whole genome shotgun sequence
3. NW_007360987	Glarea lozoyensis ATCC 20868 chromosome Unknown GLAREA13, whole genome shotgun sequence
4. NW_022474214	Venustampulla echinocandica strain BP 5553 chromosome Unknown scaffold_9, whole genome shotgun sequence
5. NW_022474217	Venustampulla echinocandica strain BP 5553 chromosome Unknown scaffold_12, whole genome shotgun sequence
6. NC_038012	Fusarium venenatum strain A3/5 genome assembly, chromosome: I
7. NT_107014	Aspergillus nidulans FGSC A4 chromosome VII map unlocalized scaffold_2, whole genome shotgun sequence
8. NW_006271969	Cordyceps militaris CM01 unplaced genomic scaffold CCM_S00001, whole genome shotgun sequence
9. NT_165979	Chaetomium globosum CBS 148.51 scaffold_4 genomic scaffold, whole genome shotgun sequence
10. NW_022984629	Aspergillus tanneri strain NIH1004 chromosome Unknown AapTan_2_pilon_pilon_pilon, whole genome shotgun sequence
11. NW_022474208	Venustampulla echinocandica strain BP 5553 chromosome Unknown scaffold_3, whole genome shotgun sequence
12. NC_030957	Colletotrichum higginsianum IMI 349063 chromosome 4, whole genome shotgun sequence


Details:

>>
1. CM000575
Source: Fusarium graminearum PH-1 chromosome 2, whole genome shotgun sequence
Type: NRPS,T1PKS
Number of proteins with BLAST hits to this cluster: 3
Cumulative BLAST score: 7321

Table of genes, locations, strands and annotations of subject cluster:

Table of Blast hits (query gene, subject gene, %identity, blast score, %coverage, e-value):
input.path1.gene2	FGSG_08207	68	226	87.27272727272727	2.4e-57
input.path1.gene3	FGSG_08209	51	3797	111.25714285714285	0.0
input.path1.gene4	FGSG_08208	65	3298	102.78232405891981	0.0


>>
2. NC_026475
Source: Fusarium graminearum PH-1 chromosome 2, whole genome shotgun sequence
Type: NRPS,T1PKS
Number of proteins with BLAST hits to this cluster: 3
Cumulative BLAST score: 7321

Table of genes, locations, strands and annotations of subject cluster:

Table of Blast hits (query gene, subject gene, %identity, blast score, %coverage, e-value):
input.path1.gene2	FGSG_08207	68	226	87.27272727272727	2.4e-57
input.path1.gene3	FGSG_08209	51	3797	111.25714285714285	0.0
input.path1.gene4	FGSG_08208	65	3298	102.78232405891981	0.0


>>
3. NW_007360987
Source: Glarea lozoyensis ATCC 20868 chromosome Unknown GLAREA13, whole genome shotgun sequence
Type: NRPS,T1PKS,betalactone
Number of proteins with BLAST hits to this cluster: 2
Cumulative BLAST score: 3042

Table of genes, locations, strands and annotations of subject cluster:

Table of Blast hits (query gene, subject gene, %identity, blast score, %coverage, e-value):
input.path1.gene3	GLAREA_10035	37	2016	91.91428571428571	0.0
input.path1.gene4	GLAREA_10034	31	1026	107.48772504091653	5.2e-297


>>
4. NW_022474214
Source: Venustampulla echinocandica strain BP 5553 chromosome Unknown scaffold_9, whole genome shotgun sequence
Type: NRPS,T1PKS
Number of proteins with BLAST hits to this cluster: 2
Cumulative BLAST score: 2463

Table of genes, locations, strands and annotations of subject cluster:

Table of Blast hits (query gene, subject gene, %identity, blast score, %coverage, e-value):
input.path1.gene3	BP5553_08847	31	1431	91.14285714285715	0.0
input.path1.gene4	BP5553_08846	31	1032	105.23731587561376	5.5e-299


>>
5. NW_022474217
Source: Venustampulla echinocandica strain BP 5553 chromosome Unknown scaffold_12, whole genome shotgun sequence
Type: NRPS,T1PKS
Number of proteins with BLAST hits to this cluster: 2
Cumulative BLAST score: 2227

Table of genes, locations, strands and annotations of subject cluster:

Table of Blast hits (query gene, subject gene, %identity, blast score, %coverage, e-value):
input.path1.gene3	BP5553_09896	33	912	50.51428571428571	1.2e-262
input.path1.gene4	BP5553_09895	33	1315	105.40098199672667	0.0


>>
6. NC_038012
Source: Fusarium venenatum strain A3/5 genome assembly, chromosome: I
Type: NRPS,T1PKS
Number of proteins with BLAST hits to this cluster: 2
Cumulative BLAST score: 3717

Table of genes, locations, strands and annotations of subject cluster:

Table of Blast hits (query gene, subject gene, %identity, blast score, %coverage, e-value):
input.path1.gene3	FVRRES_02321	38	1894	84.6	0.0
input.path1.gene4	FVRRES_02325	41	1823	105.27823240589198	0.0


>>
7. NT_107014
Source: Aspergillus nidulans FGSC A4 chromosome VII map unlocalized scaffold_2, whole genome shotgun sequence
Type: NRPS,T1PKS
Number of proteins with BLAST hits to this cluster: 2
Cumulative BLAST score: 3690

Table of genes, locations, strands and annotations of subject cluster:

Table of Blast hits (query gene, subject gene, %identity, blast score, %coverage, e-value):
input.path1.gene3	AN2545.2	33	1561	90.74285714285715	0.0
input.path1.gene4	AN2547.2	47	2129	103.23240589198035	0.0


>>
8. NW_006271969
Source: Cordyceps militaris CM01 unplaced genomic scaffold CCM_S00001, whole genome shotgun sequence
Type: NRPS,T1PKS
Number of proteins with BLAST hits to this cluster: 2
Cumulative BLAST score: 3044

Table of genes, locations, strands and annotations of subject cluster:

Table of Blast hits (query gene, subject gene, %identity, blast score, %coverage, e-value):
input.path1.gene3	CCM_01285	34	1029	51.857142857142854	1.1e-297
input.path1.gene4	CCM_01282	45	2015	103.76432078559739	0.0


>>
9. NT_165979
Source: Chaetomium globosum CBS 148.51 scaffold_4 genomic scaffold, whole genome shotgun sequence
Type: NRPS,T1PKS
Number of proteins with BLAST hits to this cluster: 2
Cumulative BLAST score: 2993

Table of genes, locations, strands and annotations of subject cluster:

Table of Blast hits (query gene, subject gene, %identity, blast score, %coverage, e-value):
input.path1.gene3	CHGG_04477	35	1193	67.2	0.0
input.path1.gene4	CHGG_04475	43	1800	100.94108019639934	0.0


>>
10. NW_022984629
Source: Aspergillus tanneri strain NIH1004 chromosome Unknown AapTan_2_pilon_pilon_pilon, whole genome shotgun sequence
Type: NRPS,T1PKS,indole
Number of proteins with BLAST hits to this cluster: 2
Cumulative BLAST score: 2418

Table of genes, locations, strands and annotations of subject cluster:

Table of Blast hits (query gene, subject gene, %identity, blast score, %coverage, e-value):
input.path1.gene3	ATNIH1004_001005	31	1310	89.02857142857142	0.0
input.path1.gene3	ATNIH1004_001002	32	542	32.2	4.2e-151
input.path1.gene4	ATNIH1004_001008	34	566	49.8772504091653	1.1e-158


>>
11. NW_022474208
Source: Venustampulla echinocandica strain BP 5553 chromosome Unknown scaffold_3, whole genome shotgun sequence
Type: NRPS,T1PKS
Number of proteins with BLAST hits to this cluster: 2
Cumulative BLAST score: 1933

Table of genes, locations, strands and annotations of subject cluster:

Table of Blast hits (query gene, subject gene, %identity, blast score, %coverage, e-value):
input.path1.gene2	BP5553_05687	38	76	56.96969696969697	3.1e-12
input.path1.gene3	BP5553_05693	38	1857	89.82857142857142	0.0


>>
12. NC_030957
Source: Colletotrichum higginsianum IMI 349063 chromosome 4, whole genome shotgun sequence
Type: T1PKS,terpene
Number of proteins with BLAST hits to this cluster: 2
Cumulative BLAST score: 1217

Table of genes, locations, strands and annotations of subject cluster:

Table of Blast hits (query gene, subject gene, %identity, blast score, %coverage, e-value):
input.path1.gene2	CH63R_06526	32	81	81.81818181818183	1.3e-13
input.path1.gene4	CH63R_06535	31	1136	107.36497545008183	0.0
