## Supplementary Results for "A draft genome of the ascomycotal fungal species *Pseudopithomyces maydicus* (family *Didymosphaeriaceae*)": contig_39_c1.txt

ClusterBlast scores for contig_39

Table of genes, locations, strands and annotations of query cluster:
input.path1.gene1	34390	35395	-		
input.path1.gene2	37341	38982	-		
input.path1.gene3	42827	44802	-		
input.path1.gene4	45460	46453	+		
input.path1.gene5	48296	54032	-		
input.path1.gene6	56625	57209	+		
input.path1.gene7	59341	60951	-		
input.path1.gene8	61627	62420	+		
input.path1.gene9	64284	64940	+		
input.path1.gene10	67176	69624	-		
input.path1.gene11	71025	71969	+		
input.path1.gene12	72458	73215	-		


Significant hits: 
1. NT_165931	Aspergillus terreus NIH2624 scaffold_8 genomic scaffold, whole genome shotgun sequence
2. NC_036623	Fusarium fujikuroi IMI 58289 draft genome, chromosome FFUJ_chr02
3. NW_022194791	Fusarium proliferatum ET1 genome assembly, contig: FPRO_scaffold002, whole genome shotgun sequence
4. NC_038015	Fusarium venenatum strain A3/5 genome assembly, chromosome: IIII
5. CM000577	Fusarium graminearum PH-1 chromosome 4, whole genome shotgun sequence
6. NC_026477	Fusarium graminearum PH-1 chromosome 4, whole genome shotgun sequence
7. NC_030957	Colletotrichum higginsianum IMI 349063 chromosome 4, whole genome shotgun sequence
8. NW_023336269	Colletotrichum scovillei strain TJNH1 chromosome Unknown Contig00011, whole genome shotgun sequence
9. NC_049562	Talaromyces rugulosus chromosome II, complete sequence
10. NW_023336287	Aspergillus tubingensis WU-2223L DNA, scaffold_13, whole genome shotgun sequence
11. NC_036442	Aspergillus oryzae RIB40 DNA, chromosome 8
12. NW_022984633	Aspergillus tanneri strain NIH1004 chromosome Unknown AapTan_6_pilon_pilon_pilon, whole genome shotgun sequence
13. NW_006917094	Pestalotiopsis fici W106-1 unplaced genomic scaffold PFICI_Scf_4, whole genome shotgun sequence
14. CM000590	Fusarium oxysporum f. sp. lycopersici 4287 chromosome 2, whole genome shotgun sequence
15. NC_030987	Fusarium oxysporum f. sp. lycopersici 4287 chromosome 2, whole genome shotgun sequence
16. CM000595	Fusarium oxysporum f. sp. lycopersici 4287 chromosome 7, whole genome shotgun sequence
17. NC_030992	Fusarium oxysporum f. sp. lycopersici 4287 chromosome 7, whole genome shotgun sequence
18. NW_017264193	Xylona heveae TC161 unplaced genomic scaffold L228scaffold_1, whole genome shotgun sequence
19. NW_003345192	Nannizzia gypsea CBS 118893 supercont1.10 genomic scaffold, whole genome shotgun sequence
20. NW_003315103	Trichophyton benhamiae CBS 112371 chromosome Unknown contig00075.49, whole genome shotgun sequence
21. NW_003456426	Trichophyton rubrum CBS 118892 genomic scaffold supercont2.3, whole genome shotgun sequence
22. NW_003299161	Microsporum canis CBS 113480 supercont1.9 genomic scaffold, whole genome shotgun sequence
23. NW_022984632	Aspergillus tanneri strain NIH1004 chromosome Unknown AapTan_5_pilon_pilon_pilon, whole genome shotgun sequence
24. NW_015971598	Cladophialophora bantiana CBS 173.52 unplaced genomic scaffold supercont1.24, whole genome shotgun sequence
25. NW_006271969	Cordyceps militaris CM01 unplaced genomic scaffold CCM_S00001, whole genome shotgun sequence
26. NT_165935	Aspergillus terreus NIH2624 scaffold_12 genomic scaffold, whole genome shotgun sequence
27. NW_022984631	Aspergillus tanneri strain NIH1004 chromosome Unknown AapTan_4_pilon_pilon_pilon, whole genome shotgun sequence
28. NC_038012	Fusarium venenatum strain A3/5 genome assembly, chromosome: I
29. NW_022984633	Aspergillus tanneri strain NIH1004 chromosome Unknown AapTan_6_pilon_pilon_pilon, whole genome shotgun sequence
30. NW_001820829	Sclerotinia sclerotiorum 1980 UF-70 scaffold_7 genomic scaffold, whole genome shotgun sequence
31. CM000574	Fusarium graminearum PH-1 chromosome 1, whole genome shotgun sequence
32. NC_026474	Fusarium graminearum PH-1 chromosome 1, whole genome shotgun sequence
33. NW_006917098	Pestalotiopsis fici W106-1 unplaced genomic scaffold PFICI_Scf_8, whole genome shotgun sequence
34. NC_035791	Pochonia chlamydosporia 170 chromosome 2, whole genome shotgun sequence
35. NC_036626	Fusarium fujikuroi IMI 58289 draft genome, chromosome FFUJ_chr05
36. NW_022194796	Fusarium proliferatum ET1 genome assembly, contig: FPRO_scaffold007, whole genome shotgun sequence
37. CP051141	Peltaster fructicola strain LNHT1506 chromosome 3
38. NW_014574694	Metarhizium brunneum ARSEF 3297 chromosome Unknown scaffold_23, whole genome shotgun sequence
39. NW_011942151	Metarhizium robertsii ARSEF 23 MAA_Scf_11, whole genome shotgun sequence
40. NW_013550595	Rhinocladiella mackenziei CBS 650.93 unplaced genomic scaffold supercont1.10, whole genome shotgun sequence
41. NW_017971433	Talaromyces atroroseus strain IBT 11181 chromosome Unknown scaffold_1, whole genome shotgun sequence
42. NC_030962	Colletotrichum higginsianum IMI 349063 chromosome 10, whole genome shotgun sequence
43. NW_004504311	Coccidioides immitis RS genomic scaffold supercont3.4, whole genome shotgun sequence
44. NW_003316019	Coccidioides posadasii C735 delta SOWgp chromosome Unknown gcontig_1105601051701, whole genome shotgun sequence
45. NW_001914849	Podospora anserina S mat+ genomic DNA chromosome 5, supercontig 10
46. NW_023336265	Colletotrichum scovillei strain TJNH1 chromosome Unknown Contig00007, whole genome shotgun sequence
47. NC_049565	Talaromyces rugulosus chromosome V, complete sequence
48. NW_003345200	Nannizzia gypsea CBS 118893 supercont1.2 genomic scaffold, whole genome shotgun sequence
49. CM002800	Penicillium chrysogenum strain P2niaD18 chromosome III, whole genome shotgun sequence
50. NW_022984630	Aspergillus tanneri strain NIH1004 chromosome Unknown AapTan_3_pilon_pilon_pilon, whole genome shotgun sequence


Details:

>>
1. NT_165931
Source: Aspergillus terreus NIH2624 scaffold_8 genomic scaffold, whole genome shotgun sequence
Type: T1PKS
Number of proteins with BLAST hits to this cluster: 6
Cumulative BLAST score: 2623

Table of genes, locations, strands and annotations of subject cluster:

Table of Blast hits (query gene, subject gene, %identity, blast score, %coverage, e-value):
input.path1.gene1	ATEG_06208	52	361	103.40557275541795	1.2e-97
input.path1.gene2	ATEG_06204	62	156	31.564245810055862	3.9e-36
input.path1.gene3	ATEG_06210	54	558	100.0	8.9e-157
input.path1.gene4	ATEG_06209	53	356	112.62798634812287	3.6e-96
input.path1.gene5	ATEG_06206	57	1050	55.33453887884268	2.3e-304
input.path1.gene6	ATEG_06203	64	142	67.10526315789474	4.3e-32


>>
2. NC_036623
Source: Fusarium fujikuroi IMI 58289 draft genome, chromosome FFUJ_chr02
Type: T1PKS
Number of proteins with BLAST hits to this cluster: 2
Cumulative BLAST score: 1972

Table of genes, locations, strands and annotations of subject cluster:

Table of Blast hits (query gene, subject gene, %identity, blast score, %coverage, e-value):
input.path1.gene5	FFUJ_03984	53	1894	111.99517781796263	0.0
input.path1.gene6	FFUJ_03985	40	78	66.44736842105263	5.8e-13


>>
3. NW_022194791
Source: Fusarium proliferatum ET1 genome assembly, contig: FPRO_scaffold002, whole genome shotgun sequence
Type: T1PKS
Number of proteins with BLAST hits to this cluster: 2
Cumulative BLAST score: 1972

Table of genes, locations, strands and annotations of subject cluster:

Table of Blast hits (query gene, subject gene, %identity, blast score, %coverage, e-value):
input.path1.gene5	FPRO_03539	53	1894	111.99517781796263	0.0
input.path1.gene6	FPRO_03538	40	78	66.44736842105263	5.8e-13


>>
4. NC_038015
Source: Fusarium venenatum strain A3/5 genome assembly, chromosome: IIII
Type: T1PKS
Number of proteins with BLAST hits to this cluster: 2
Cumulative BLAST score: 1961

Table of genes, locations, strands and annotations of subject cluster:

Table of Blast hits (query gene, subject gene, %identity, blast score, %coverage, e-value):
input.path1.gene5	FVRRES_13609	53	1886	112.41711874623267	0.0
input.path1.gene6	FVRRES_13608	38	75	65.78947368421053	6.4e-12


>>
5. CM000577
Source: Fusarium graminearum PH-1 chromosome 4, whole genome shotgun sequence
Type: T1PKS
Number of proteins with BLAST hits to this cluster: 2
Cumulative BLAST score: 1942

Table of genes, locations, strands and annotations of subject cluster:

Table of Blast hits (query gene, subject gene, %identity, blast score, %coverage, e-value):
input.path1.gene5	FGSG_09182	52	1869	111.99517781796263	0.0
input.path1.gene6	FGSG_09183	35	73	67.10526315789474	2.4e-11


>>
6. NC_026477
Source: Fusarium graminearum PH-1 chromosome 4, whole genome shotgun sequence
Type: T1PKS
Number of proteins with BLAST hits to this cluster: 2
Cumulative BLAST score: 1942

Table of genes, locations, strands and annotations of subject cluster:

Table of Blast hits (query gene, subject gene, %identity, blast score, %coverage, e-value):
input.path1.gene5	FGSG_09182	52	1869	111.99517781796263	0.0
input.path1.gene6	FGSG_09183	35	73	67.10526315789474	2.4e-11


>>
7. NC_030957
Source: Colletotrichum higginsianum IMI 349063 chromosome 4, whole genome shotgun sequence
Type: T1PKS
Number of proteins with BLAST hits to this cluster: 2
Cumulative BLAST score: 1028

Table of genes, locations, strands and annotations of subject cluster:

Table of Blast hits (query gene, subject gene, %identity, blast score, %coverage, e-value):
input.path1.gene5	CH63R_06245	38	864	82.27848101265823	1.4e-248
input.path1.gene6	CH63R_06246	43	87	67.10526315789474	9.5e-16
input.path1.gene6	CH63R_06247	42	77	65.78947368421053	1.7e-12


>>
8. NW_023336269
Source: Colletotrichum scovillei strain TJNH1 chromosome Unknown Contig00011, whole genome shotgun sequence
Type: T1PKS
Number of proteins with BLAST hits to this cluster: 2
Cumulative BLAST score: 1013

Table of genes, locations, strands and annotations of subject cluster:

Table of Blast hits (query gene, subject gene, %identity, blast score, %coverage, e-value):
input.path1.gene5	HER10_EVM0004425	38	852	83.90596745027125	7e-245
input.path1.gene6	HER10_EVM0004116	42	85	67.10526315789474	4.7e-15
input.path1.gene6	HER10_EVM0006957	42	76	65.78947368421053	3.7e-12


>>
9. NC_049562
Source: Talaromyces rugulosus chromosome II, complete sequence
Type: T1PKS
Number of proteins with BLAST hits to this cluster: 2
Cumulative BLAST score: 998

Table of genes, locations, strands and annotations of subject cluster:

Table of Blast hits (query gene, subject gene, %identity, blast score, %coverage, e-value):
input.path1.gene5	TRUGW13939_04214	38	896	87.34177215189874	4.2e-258
input.path1.gene6	TRUGW13939_04213	49	102	65.78947368421053	4.9e-20


>>
10. NW_023336287
Source: Aspergillus tubingensis WU-2223L DNA, scaffold_13, whole genome shotgun sequence
Type: T1PKS
Number of proteins with BLAST hits to this cluster: 2
Cumulative BLAST score: 884

Table of genes, locations, strands and annotations of subject cluster:

Table of Blast hits (query gene, subject gene, %identity, blast score, %coverage, e-value):
input.path1.gene5	AtWU_10461	44	810	58.22784810126582	4e-232
input.path1.gene6	AtWU_10462	41	74	64.47368421052632	8.3e-12


>>
11. NC_036442
Source: Aspergillus oryzae RIB40 DNA, chromosome 8
Type: T1PKS
Number of proteins with BLAST hits to this cluster: 2
Cumulative BLAST score: 872

Table of genes, locations, strands and annotations of subject cluster:

Table of Blast hits (query gene, subject gene, %identity, blast score, %coverage, e-value):
input.path1.gene5	AO090010000048	44	786	57.80590717299579	4.7e-225
input.path1.gene6	AO090010000047	42	86	67.10526315789474	2.1e-15


>>
12. NW_022984633
Source: Aspergillus tanneri strain NIH1004 chromosome Unknown AapTan_6_pilon_pilon_pilon, whole genome shotgun sequence
Type: T1PKS
Number of proteins with BLAST hits to this cluster: 2
Cumulative BLAST score: 2487

Table of genes, locations, strands and annotations of subject cluster:

Table of Blast hits (query gene, subject gene, %identity, blast score, %coverage, e-value):
input.path1.gene2	ATNIH1004_008279	45	431	152.51396648044692	1.1e-118
input.path1.gene5	ATNIH1004_008281	57	2056	112.35684147076552	0.0


>>
13. NW_006917094
Source: Pestalotiopsis fici W106-1 unplaced genomic scaffold PFICI_Scf_4, whole genome shotgun sequence
Type: T1PKS
Number of proteins with BLAST hits to this cluster: 2
Cumulative BLAST score: 2180

Table of genes, locations, strands and annotations of subject cluster:

Table of Blast hits (query gene, subject gene, %identity, blast score, %coverage, e-value):
input.path1.gene2	PFICI_06559	49	277	79.60893854748603	2e-72
input.path1.gene5	PFICI_06561	53	1903	114.34599156118144	0.0


>>
14. CM000590
Source: Fusarium oxysporum f. sp. lycopersici 4287 chromosome 2, whole genome shotgun sequence
Type: T1PKS
Number of proteins with BLAST hits to this cluster: 2
Cumulative BLAST score: 1906

Table of genes, locations, strands and annotations of subject cluster:

Table of Blast hits (query gene, subject gene, %identity, blast score, %coverage, e-value):
input.path1.gene5	FOXG_05816	51	1827	111.99517781796263	0.0
input.path1.gene6	FOXG_05817	40	79	66.44736842105263	4.4e-13


>>
15. NC_030987
Source: Fusarium oxysporum f. sp. lycopersici 4287 chromosome 2, whole genome shotgun sequence
Type: T1PKS
Number of proteins with BLAST hits to this cluster: 2
Cumulative BLAST score: 1906

Table of genes, locations, strands and annotations of subject cluster:

Table of Blast hits (query gene, subject gene, %identity, blast score, %coverage, e-value):
input.path1.gene5	FOXG_05816	51	1827	111.99517781796263	0.0
input.path1.gene6	FOXG_05817	40	79	66.44736842105263	4.4e-13


>>
16. CM000595
Source: Fusarium oxysporum f. sp. lycopersici 4287 chromosome 7, whole genome shotgun sequence
Type: T1PKS
Number of proteins with BLAST hits to this cluster: 2
Cumulative BLAST score: 1642

Table of genes, locations, strands and annotations of subject cluster:

Table of Blast hits (query gene, subject gene, %identity, blast score, %coverage, e-value):
input.path1.gene5	FOXG_04757	44	773	58.22784810126582	5.4e-221
input.path1.gene5	FOXG_04757_KNB01536.1	44	773	58.22784810126582	5.4e-221
input.path1.gene6	FOXG_04755	46	96	64.47368421052632	2.1e-18


>>
17. NC_030992
Source: Fusarium oxysporum f. sp. lycopersici 4287 chromosome 7, whole genome shotgun sequence
Type: T1PKS
Number of proteins with BLAST hits to this cluster: 2
Cumulative BLAST score: 1642

Table of genes, locations, strands and annotations of subject cluster:

Table of Blast hits (query gene, subject gene, %identity, blast score, %coverage, e-value):
input.path1.gene5	FOXG_04757	44	773	58.22784810126582	5.4e-221
input.path1.gene5	FOXG_04757_XP_018239581.1	44	773	58.22784810126582	5.4e-221
input.path1.gene6	FOXG_04755	46	96	64.47368421052632	2.1e-18


>>
18. NW_017264193
Source: Xylona heveae TC161 unplaced genomic scaffold L228scaffold_1, whole genome shotgun sequence
Type: T1PKS
Number of proteins with BLAST hits to this cluster: 2
Cumulative BLAST score: 1314

Table of genes, locations, strands and annotations of subject cluster:

Table of Blast hits (query gene, subject gene, %identity, blast score, %coverage, e-value):
input.path1.gene5	L228DRAFT_235437	47	1213	88.24593128390596	0.0
input.path1.gene6	L228DRAFT_235439	53	101	65.78947368421053	8.3e-20


>>
19. NW_003345192
Source: Nannizzia gypsea CBS 118893 supercont1.10 genomic scaffold, whole genome shotgun sequence
Type: T1PKS
Number of proteins with BLAST hits to this cluster: 2
Cumulative BLAST score: 1249

Table of genes, locations, strands and annotations of subject cluster:

Table of Blast hits (query gene, subject gene, %identity, blast score, %coverage, e-value):
input.path1.gene5	MGYG_08684	45	1143	90.59674502712477	0.0
input.path1.gene6	MGYG_08689	51	106	67.76315789473685	2.6e-21


>>
20. NW_003315103
Source: Trichophyton benhamiae CBS 112371 chromosome Unknown contig00075.49, whole genome shotgun sequence
Type: T1PKS
Number of proteins with BLAST hits to this cluster: 2
Cumulative BLAST score: 1243

Table of genes, locations, strands and annotations of subject cluster:

Table of Blast hits (query gene, subject gene, %identity, blast score, %coverage, e-value):
input.path1.gene5	ARB_07994	44	1137	90.23508137432188	0.0
input.path1.gene6	ARB_07989	51	106	67.76315789473685	3.4e-21


>>
21. NW_003456426
Source: Trichophyton rubrum CBS 118892 genomic scaffold supercont2.3, whole genome shotgun sequence
Type: T1PKS
Number of proteins with BLAST hits to this cluster: 2
Cumulative BLAST score: 1236

Table of genes, locations, strands and annotations of subject cluster:

Table of Blast hits (query gene, subject gene, %identity, blast score, %coverage, e-value):
input.path1.gene5	TERG_02850	44	1130	90.29535864978902	0.0
input.path1.gene6	TERG_02845	51	106	67.76315789473685	3.4e-21


>>
22. NW_003299161
Source: Microsporum canis CBS 113480 supercont1.9 genomic scaffold, whole genome shotgun sequence
Type: T1PKS
Number of proteins with BLAST hits to this cluster: 2
Cumulative BLAST score: 1151

Table of genes, locations, strands and annotations of subject cluster:

Table of Blast hits (query gene, subject gene, %identity, blast score, %coverage, e-value):
input.path1.gene5	MCYG_08535	43	1059	88.84870403857745	3.8e-307
input.path1.gene6	MCYG_08540	47	92	67.10526315789474	5.1e-17


>>
23. NW_022984632
Source: Aspergillus tanneri strain NIH1004 chromosome Unknown AapTan_5_pilon_pilon_pilon, whole genome shotgun sequence
Type: T1PKS
Number of proteins with BLAST hits to this cluster: 2
Cumulative BLAST score: 1112

Table of genes, locations, strands and annotations of subject cluster:

Table of Blast hits (query gene, subject gene, %identity, blast score, %coverage, e-value):
input.path1.gene5	ATNIH1004_009518	40	1026	87.82399035563593	3.5e-297
input.path1.gene6	ATNIH1004_009523	43	86	66.44736842105263	2.1e-15


>>
24. NW_015971598
Source: Cladophialophora bantiana CBS 173.52 unplaced genomic scaffold supercont1.24, whole genome shotgun sequence
Type: T1PKS,indole,terpene
Number of proteins with BLAST hits to this cluster: 2
Cumulative BLAST score: 1075

Table of genes, locations, strands and annotations of subject cluster:

Table of Blast hits (query gene, subject gene, %identity, blast score, %coverage, e-value):
input.path1.gene5	Z519_11905	37	807	88.84870403857745	2.6e-231
input.path1.gene7	Z519_11908	34	268	97.80876494023904	1.7e-69


>>
25. NW_006271969
Source: Cordyceps militaris CM01 unplaced genomic scaffold CCM_S00001, whole genome shotgun sequence
Type: T1PKS
Number of proteins with BLAST hits to this cluster: 2
Cumulative BLAST score: 1054

Table of genes, locations, strands and annotations of subject cluster:

Table of Blast hits (query gene, subject gene, %identity, blast score, %coverage, e-value):
input.path1.gene5	CCM_01921	40	968	86.13622664255576	1.1e-279
input.path1.gene6	CCM_01923	43	86	68.42105263157895	3.6e-15


>>
26. NT_165935
Source: Aspergillus terreus NIH2624 scaffold_12 genomic scaffold, whole genome shotgun sequence
Type: NRPS,T1PKS
Number of proteins with BLAST hits to this cluster: 2
Cumulative BLAST score: 1046

Table of genes, locations, strands and annotations of subject cluster:

Table of Blast hits (query gene, subject gene, %identity, blast score, %coverage, e-value):
input.path1.gene3	ATEG_08445	43	340	84.16206261510129	3.8e-91
input.path1.gene5	ATEG_08451	40	706	58.64978902953587	6.2e-201


>>
27. NW_022984631
Source: Aspergillus tanneri strain NIH1004 chromosome Unknown AapTan_4_pilon_pilon_pilon, whole genome shotgun sequence
Type: T1PKS,indole
Number of proteins with BLAST hits to this cluster: 2
Cumulative BLAST score: 1012

Table of genes, locations, strands and annotations of subject cluster:

Table of Blast hits (query gene, subject gene, %identity, blast score, %coverage, e-value):
input.path1.gene4	ATNIH1004_005536	47	299	112.62798634812287	5.2e-79
input.path1.gene5	ATNIH1004_005531	33	713	81.31404460518384	5.1e-203


>>
28. NC_038012
Source: Fusarium venenatum strain A3/5 genome assembly, chromosome: I
Type: NRPS,T1PKS
Number of proteins with BLAST hits to this cluster: 2
Cumulative BLAST score: 970

Table of genes, locations, strands and annotations of subject cluster:

Table of Blast hits (query gene, subject gene, %identity, blast score, %coverage, e-value):
input.path1.gene5	FVRRES_02715	39	880	82.33875828812538	3.1e-253
input.path1.gene6	FVRRES_02717	39	90	71.05263157894737	1.9e-16


>>
29. NW_022984633
Source: Aspergillus tanneri strain NIH1004 chromosome Unknown AapTan_6_pilon_pilon_pilon, whole genome shotgun sequence
Type: T1PKS
Number of proteins with BLAST hits to this cluster: 2
Cumulative BLAST score: 967

Table of genes, locations, strands and annotations of subject cluster:

Table of Blast hits (query gene, subject gene, %identity, blast score, %coverage, e-value):
input.path1.gene5	ATNIH1004_008649	37	877	89.45147679324894	2.6e-252
input.path1.gene6	ATNIH1004_008651	45	90	67.10526315789474	1.9e-16


>>
30. NW_001820829
Source: Sclerotinia sclerotiorum 1980 UF-70 scaffold_7 genomic scaffold, whole genome shotgun sequence
Type: T1PKS
Number of proteins with BLAST hits to this cluster: 2
Cumulative BLAST score: 953

Table of genes, locations, strands and annotations of subject cluster:

Table of Blast hits (query gene, subject gene, %identity, blast score, %coverage, e-value):
input.path1.gene5	SS1G_05681	40	863	79.86738999397227	3e-248
input.path1.gene6	SS1G_05678	45	90	63.81578947368421	1.9e-16


>>
31. CM000574
Source: Fusarium graminearum PH-1 chromosome 1, whole genome shotgun sequence
Type: NRPS,T1PKS
Number of proteins with BLAST hits to this cluster: 2
Cumulative BLAST score: 925

Table of genes, locations, strands and annotations of subject cluster:

Table of Blast hits (query gene, subject gene, %identity, blast score, %coverage, e-value):
input.path1.gene5	FGSG_02324	40	835	80.6509945750452	1.2e-239
input.path1.gene6	FGSG_02326	41	90	69.73684210526315	1.9e-16


>>
32. NC_026474
Source: Fusarium graminearum PH-1 chromosome 1, whole genome shotgun sequence
Type: NRPS,T1PKS
Number of proteins with BLAST hits to this cluster: 2
Cumulative BLAST score: 925

Table of genes, locations, strands and annotations of subject cluster:

Table of Blast hits (query gene, subject gene, %identity, blast score, %coverage, e-value):
input.path1.gene5	FGSG_02324	40	835	80.6509945750452	1.2e-239
input.path1.gene6	FGSG_02326	41	90	69.73684210526315	1.9e-16


>>
33. NW_006917098
Source: Pestalotiopsis fici W106-1 unplaced genomic scaffold PFICI_Scf_8, whole genome shotgun sequence
Type: T1PKS
Number of proteins with BLAST hits to this cluster: 2
Cumulative BLAST score: 916

Table of genes, locations, strands and annotations of subject cluster:

Table of Blast hits (query gene, subject gene, %identity, blast score, %coverage, e-value):
input.path1.gene1	PFICI_10834	32	176	99.38080495356037	5.7e-42
input.path1.gene5	PFICI_10824	34	740	80.5304400241109	3e-211


>>
34. NC_035791
Source: Pochonia chlamydosporia 170 chromosome 2, whole genome shotgun sequence
Type: T1PKS
Number of proteins with BLAST hits to this cluster: 2
Cumulative BLAST score: 900

Table of genes, locations, strands and annotations of subject cluster:

Table of Blast hits (query gene, subject gene, %identity, blast score, %coverage, e-value):
input.path1.gene2	VFPPC_11721	36	135	61.452513966480446	9.4e-30
input.path1.gene5	VFPPC_11725	42	765	58.71006630500302	8.6e-219


>>
35. NC_036626
Source: Fusarium fujikuroi IMI 58289 draft genome, chromosome FFUJ_chr05
Type: T1PKS
Number of proteins with BLAST hits to this cluster: 2
Cumulative BLAST score: 867

Table of genes, locations, strands and annotations of subject cluster:

Table of Blast hits (query gene, subject gene, %identity, blast score, %coverage, e-value):
input.path1.gene5	FFUJ_06742	44	769	58.16757082579868	1e-219
input.path1.gene6	FFUJ_06744	47	98	64.47368421052632	5.4e-19


>>
36. NW_022194796
Source: Fusarium proliferatum ET1 genome assembly, contig: FPRO_scaffold007, whole genome shotgun sequence
Type: T1PKS
Number of proteins with BLAST hits to this cluster: 2
Cumulative BLAST score: 856

Table of genes, locations, strands and annotations of subject cluster:

Table of Blast hits (query gene, subject gene, %identity, blast score, %coverage, e-value):
input.path1.gene5	FPRO_07275	43	758	58.22784810126582	1.8e-216
input.path1.gene6	FPRO_07277	47	98	64.47368421052632	5.4e-19


>>
37. CP051141
Source: Peltaster fructicola strain LNHT1506 chromosome 3
Type: T1PKS
Number of proteins with BLAST hits to this cluster: 2
Cumulative BLAST score: 848

Table of genes, locations, strands and annotations of subject cluster:

Table of Blast hits (query gene, subject gene, %identity, blast score, %coverage, e-value):
input.path1.gene1	AMS68_004768	32	155	98.14241486068111	7.9e-36
input.path1.gene5	AMS68_004773	39	693	58.529234478601566	7.1e-197


>>
38. NW_014574694
Source: Metarhizium brunneum ARSEF 3297 chromosome Unknown scaffold_23, whole genome shotgun sequence
Type: NRPS,T1PKS
Number of proteins with BLAST hits to this cluster: 2
Cumulative BLAST score: 832

Table of genes, locations, strands and annotations of subject cluster:

Table of Blast hits (query gene, subject gene, %identity, blast score, %coverage, e-value):
input.path1.gene1	MBR_09475	33	178	102.47678018575851	8.7e-43
input.path1.gene5	MBR_09479	41	654	48.583484026522	2.1e-185


>>
39. NW_011942151
Source: Metarhizium robertsii ARSEF 23 MAA_Scf_11, whole genome shotgun sequence
Type: NRPS,T1PKS,terpene
Number of proteins with BLAST hits to this cluster: 2
Cumulative BLAST score: 825

Table of genes, locations, strands and annotations of subject cluster:

Table of Blast hits (query gene, subject gene, %identity, blast score, %coverage, e-value):
input.path1.gene1	MAA_06570	33	178	102.47678018575851	8.7e-43
input.path1.gene5	MAA_06575	41	647	48.583484026522	4.4e-183


>>
40. NW_013550595
Source: Rhinocladiella mackenziei CBS 650.93 unplaced genomic scaffold supercont1.10, whole genome shotgun sequence
Type: T1PKS
Number of proteins with BLAST hits to this cluster: 2
Cumulative BLAST score: 816

Table of genes, locations, strands and annotations of subject cluster:

Table of Blast hits (query gene, subject gene, %identity, blast score, %coverage, e-value):
input.path1.gene5	Z518_10961	32	608	80.40988547317662	2.3e-171
input.path1.gene7	Z518_10954	31	208	88.64541832669323	1.2e-51


>>
41. NW_017971433
Source: Talaromyces atroroseus strain IBT 11181 chromosome Unknown scaffold_1, whole genome shotgun sequence
Type: T1PKS
Number of proteins with BLAST hits to this cluster: 2
Cumulative BLAST score: 723

Table of genes, locations, strands and annotations of subject cluster:

Table of Blast hits (query gene, subject gene, %identity, blast score, %coverage, e-value):
input.path1.gene5	UA08_01172	32	624	81.13321277878241	2.4e-176
input.path1.gene6	UA08_01176	48	99	68.42105263157895	2.4e-19


>>
42. NC_030962
Source: Colletotrichum higginsianum IMI 349063 chromosome 10, whole genome shotgun sequence
Type: NRPS-like,T1PKS
Number of proteins with BLAST hits to this cluster: 2
Cumulative BLAST score: 605

Table of genes, locations, strands and annotations of subject cluster:

Table of Blast hits (query gene, subject gene, %identity, blast score, %coverage, e-value):
input.path1.gene5	CH63R_14350	35	531	57.986738999397225	3.6e-148
input.path1.gene6	CH63R_14355	38	74	65.13157894736842	8.3e-12


>>
43. NW_004504311
Source: Coccidioides immitis RS genomic scaffold supercont3.4, whole genome shotgun sequence
Type: NRPS-like,T1PKS
Number of proteins with BLAST hits to this cluster: 2
Cumulative BLAST score: 335

Table of genes, locations, strands and annotations of subject cluster:

Table of Blast hits (query gene, subject gene, %identity, blast score, %coverage, e-value):
input.path1.gene5	CIMG_04688	41	279	22.78481012658228	2.4e-72
input.path1.gene6	CIMG_04683	37	56	58.55263157894737	2.4e-06


>>
44. NW_003316019
Source: Coccidioides posadasii C735 delta SOWgp chromosome Unknown gcontig_1105601051701, whole genome shotgun sequence
Type: NRPS-like,T1PKS
Number of proteins with BLAST hits to this cluster: 2
Cumulative BLAST score: 334

Table of genes, locations, strands and annotations of subject cluster:

Table of Blast hits (query gene, subject gene, %identity, blast score, %coverage, e-value):
input.path1.gene5	CPC735_071310	40	277	22.78481012658228	7.1e-72
input.path1.gene6	CPC735_071370	37	57	58.55263157894737	1.4e-06


>>
45. NW_001914849
Source: Podospora anserina S mat+ genomic DNA chromosome 5, supercontig 10
Type: NRPS-like,T1PKS
Number of proteins with BLAST hits to this cluster: 2
Cumulative BLAST score: 328

Table of genes, locations, strands and annotations of subject cluster:

Table of Blast hits (query gene, subject gene, %identity, blast score, %coverage, e-value):
input.path1.gene5	PODANS_5_11010	38	268	23.32730560578662	5.6e-69
input.path1.gene6	PODANS_5_11060	37	60	58.55263157894737	2.1e-07


>>
46. NW_023336265
Source: Colletotrichum scovillei strain TJNH1 chromosome Unknown Contig00007, whole genome shotgun sequence
Type: T1PKS
Number of proteins with BLAST hits to this cluster: 2
Cumulative BLAST score: 887

Table of genes, locations, strands and annotations of subject cluster:

Table of Blast hits (query gene, subject gene, %identity, blast score, %coverage, e-value):
input.path1.gene7	HER10_EVM0003064	62	630	103.38645418326693	1.3e-178
input.path1.gene8	HER10_EVM0000803	63	257	102.64550264550265	1.1e-66


>>
47. NC_049565
Source: Talaromyces rugulosus chromosome V, complete sequence
Type: T1PKS
Number of proteins with BLAST hits to this cluster: 2
Cumulative BLAST score: 938

Table of genes, locations, strands and annotations of subject cluster:

Table of Blast hits (query gene, subject gene, %identity, blast score, %coverage, e-value):
input.path1.gene3	TRUGW13939_09155	35	307	99.8158379373849	2.7e-81
input.path1.gene7	TRUGW13939_09157	64	631	101.39442231075697	7.6e-179


>>
48. NW_003345200
Source: Nannizzia gypsea CBS 118893 supercont1.2 genomic scaffold, whole genome shotgun sequence
Type: NRPS
Number of proteins with BLAST hits to this cluster: 2
Cumulative BLAST score: 643

Table of genes, locations, strands and annotations of subject cluster:

Table of Blast hits (query gene, subject gene, %identity, blast score, %coverage, e-value):
input.path1.gene1	MGYG_02583	39	242	104.02476780185759	6.4e-62
input.path1.gene3	MGYG_02588	42	401	98.34254143646409	1.1e-109


>>
49. CM002800
Source: Penicillium chrysogenum strain P2niaD18 chromosome III, whole genome shotgun sequence
Type: T1PKS
Number of proteins with BLAST hits to this cluster: 2
Cumulative BLAST score: 460

Table of genes, locations, strands and annotations of subject cluster:

Table of Blast hits (query gene, subject gene, %identity, blast score, %coverage, e-value):
input.path1.gene2	EN45_097560	35	112	50.5586592178771	1.1e-22
input.path1.gene3	EN45_097450	38	348	98.15837937384899	1.1e-93


>>
50. NW_022984630
Source: Aspergillus tanneri strain NIH1004 chromosome Unknown AapTan_3_pilon_pilon_pilon, whole genome shotgun sequence
Type: T1PKS
Number of proteins with BLAST hits to this cluster: 2
Cumulative BLAST score: 428

Table of genes, locations, strands and annotations of subject cluster:

Table of Blast hits (query gene, subject gene, %identity, blast score, %coverage, e-value):
input.path1.gene2	ATNIH1004_003832	34	127	65.64245810055866	3.3e-27
input.path1.gene3	ATNIH1004_003837	34	301	101.84162062615101	2e-79
