## Supplementary Results for "A draft genome of the ascomycotal fungal species *Pseudopithomyces maydicus* (family *Didymosphaeriaceae*)": contig_39_c3.txt

ClusterBlast scores for contig_39

Table of genes, locations, strands and annotations of query cluster:
input.path1.gene292	1104234	1105474	-		
input.path1.gene293	1107050	1109588	+		
input.path1.gene294	1111179	1113289	+		
input.path1.gene295	1113914	1114964	+		
input.path1.gene296	1117055	1118409	+		
input.path1.gene297	1120064	1122531	+		
input.path1.gene298	1122899	1128493	-		
input.path1.gene299	1129097	1129641	+		
input.path1.gene300	1131510	1134380	-		
input.path1.gene301	1147434	1150632	+		
input.path1.gene302	1152678	1153583	+		


Significant hits: 
1. NC_049565	Talaromyces rugulosus chromosome V, complete sequence
2. NZ_LR733556	Flavobacterium sp. 9AF isolate Flavobacterium sp. 9AF, whole genome shotgun sequence
3. NW_022474219	Venustampulla echinocandica strain BP 5553 chromosome Unknown scaffold_14, whole genome shotgun sequence
4. NT_165977	Chaetomium globosum CBS 148.51 scaffold_2 genomic scaffold, whole genome shotgun sequence
5. NZ_AP018195	Scytonema sp. HK-05 plasmid plasmid1 DNA, nearly complete genome
6. NZ_AP018268	Scytonema sp. NIES-4073 DNA, nearly complete genome
7. NZ_KL662191	[Leptolyngbya] sp. JSC-1 Osccy1DRAFT_CYJSC1_DRAF_scaffold00069.1, whole genome shotgun sequence
8. NZ_CM001979	Dickeya sp. NCPPB 3274 chromosome, whole genome shotgun sequence
9. NZ_KQ976354	Scytonema hofmannii PCC 7110 Scaffold1, whole genome shotgun sequence
10. NC_010628	Nostoc punctiforme PCC 73102, complete sequence
11. NZ_CP023484	Dickeya sp. Secpp 1600 chromosome, complete genome
12. NZ_CP025003	Dickeya fangzhongdai strain DSM 101947 chromosome, complete genome
13. NZ_CP020872	Dickeya fangzhongdai strain PA1 chromosome, complete genome
14. NZ_CM001860	Dickeya solani GBBC 2040 chromosome, whole genome shotgun sequence
15. NC_013592	Dickeya zeae Ech586, complete sequence
16. NZ_QJKB01000008	Undibacterium pigrum strain DSM 19792 Ga0244539_108, whole genome shotgun sequence
17. NZ_CM001984	Dickeya sp. MK7 chromosome, whole genome shotgun sequence
18. NZ_CP054698	Nostoc edaphicum CCNP1411 chromosome, complete genome
19. NC_011726	Rippkaea orientalis PCC 8801, complete sequence
20. NZ_AP018441	Undibacterium sp. YM2
21. NZ_CP031515	Dickeya zeae strain EC2 chromosome, complete genome
22. NZ_CM001977	Dickeya zeae NCPPB 2538 chromosome, whole genome shotgun sequence
23. NZ_CP023467	Dickeya dadantii strain DSM 18020 chromosome, complete genome
24. NZ_CM001980	Dickeya zeae NCPPB 3531 chromosome, whole genome shotgun sequence
25. NZ_CM001985	Dickeya zeae MK19 chromosome, whole genome shotgun sequence
26. NZ_CM001858	Dickeya zeae NCPPB 3532 chromosome, whole genome shotgun sequence
27. NZ_CP033622	Dickeya zeae strain CE1 chromosome, complete genome
28. NZ_CM001982	Dickeya dadantii NCPPB 3537 chromosome, whole genome shotgun sequence
29. NC_013161	Rippkaea orientalis PCC 8802, complete sequence
30. NC_020126	Myxococcus stipitatus DSM 14675, complete sequence
31. NC_008095	Myxococcus xanthus DK 1622, complete sequence
32. NC_026507	Neurospora crassa OR74A linkage group VII, whole genome shotgun sequence
33. NW_007360994	Glarea lozoyensis ATCC 20868 chromosome Unknown GLAREA2, whole genome shotgun sequence
34. NZ_BCMN01000006	Streptomyces turgidiscabies strain T45, whole genome shotgun sequence
35. NZ_CP030864	Streptomyces globosus strain LZH-48 plasmid unnamed2, complete sequence
36. NT_165934	Aspergillus terreus NIH2624 scaffold_11 genomic scaffold, whole genome shotgun sequence
37. NZ_KQ947996	Streptomyces curacoi strain DSM 40107 PRJNA299217_s013, whole genome shotgun sequence
38. NC_013093	Actinosynnema mirum DSM 43827, complete sequence
39. NZ_CP023445	Actinosynnema pretiosum strain X47 chromosome, complete genome
40. NZ_CP009754	Streptomyces sp. CCM_MD2014 chromosome, complete genome
41. NZ_KZ195574	Streptomyces sp. CS113 scaffold00001, whole genome shotgun sequence
42. NZ_CP056041	Streptomyces chartreusis strain NA02069 chromosome, complete genome
43. NZ_CP023689	Streptomyces chartreusis strain ATCC 14922 chromosome, complete genome
44. NZ_RDBL01000014	Streptomyces sp. col6 Scaffold_14, whole genome shotgun sequence
45. NZ_SZNQ01000001	Streptomyces lasalocidi strain X-537 Scaffold1, whole genome shotgun sequence
46. NZ_CP010429	Spirosoma radiotolerans strain DG5A chromosome, complete genome
47. NZ_CP043761	Paenibacillus sp. 37 chromosome, complete genome
48. NZ_KQ948925	Streptomyces caeruleatus strain NRRL B-24802 PRJNA299234_s002, whole genome shotgun sequence
49. NZ_LN831790	Streptomyces leeuwenhoekii strain type strain (C34 = DSM 42122 = NRRL B-24963) chromosome 1
50. NZ_CP022961	Plantactinospora sp. KBS50 chromosome, complete genome


Details:

>>
1. NC_049565
Source: Talaromyces rugulosus chromosome V, complete sequence
Type: NRPS,T1PKS
Number of proteins with BLAST hits to this cluster: 2
Cumulative BLAST score: 797

Table of genes, locations, strands and annotations of subject cluster:

Table of Blast hits (query gene, subject gene, %identity, blast score, %coverage, e-value):
input.path1.gene298	TRUGW13939_10316	31	224	40.65040650406504	6.3e-56
input.path1.gene300	TRUGW13939_10315	39	573	116.32911392405063	3e-161


>>
2. NZ_LR733556
Source: Flavobacterium sp. 9AF isolate Flavobacterium sp. 9AF, whole genome shotgun sequence
Type: NRPS,T1PKS
Number of proteins with BLAST hits to this cluster: 2
Cumulative BLAST score: 325

Table of genes, locations, strands and annotations of subject cluster:
GVY45_RS03070	GVY45_RS03070	640980	642161	+	tyrosine-type_recombinase/integrase
GVY45_RS03145	GVY45_RS03145	653869	654140	+	helix-turn-helix_domain-containing_protein
GVY45_RS03150	GVY45_RS03150	654340	654992	+	ATPase

Table of Blast hits (query gene, subject gene, %identity, blast score, %coverage, e-value):
input.path1.gene298	GVY45_RS03185	31	220	32.38482384823848	1.2e-54
input.path1.gene300	GVY45_RS03180	36	105	20.126582278481013	2.3e-20


>>
3. NW_022474219
Source: Venustampulla echinocandica strain BP 5553 chromosome Unknown scaffold_14, whole genome shotgun sequence
Type: NRPS,T1PKS
Number of proteins with BLAST hits to this cluster: 2
Cumulative BLAST score: 1933

Table of genes, locations, strands and annotations of subject cluster:

Table of Blast hits (query gene, subject gene, %identity, blast score, %coverage, e-value):
input.path1.gene298	BP5553_10299	45	1146	91.19241192411924	0.0
input.path1.gene300	BP5553_10291	48	787	114.68354430379746	1e-225


>>
4. NT_165977
Source: Chaetomium globosum CBS 148.51 scaffold_2 genomic scaffold, whole genome shotgun sequence
Type: NRPS,T1PKS
Number of proteins with BLAST hits to this cluster: 2
Cumulative BLAST score: 1637

Table of genes, locations, strands and annotations of subject cluster:

Table of Blast hits (query gene, subject gene, %identity, blast score, %coverage, e-value):
input.path1.gene298	CHGG_02378	42	402	37.669376693766935	2.2e-109
input.path1.gene298	CHGG_02376	40	322	32.859078590785906	2.9e-85
input.path1.gene300	CHGG_02374	54	913	115.18987341772151	2.1e-263


>>
5. NZ_AP018195
Source: Scytonema sp. HK-05 plasmid plasmid1 DNA, nearly complete genome
Type: NRPS,T1PKS,microviridin
Number of proteins with BLAST hits to this cluster: 2
Cumulative BLAST score: 649

Table of genes, locations, strands and annotations of subject cluster:
CA739_RS37910	CA739_RS37910	326752	326898	-	hypothetical_protein
CA739_RS37925	CA739_RS37925	329902	330363	-	MvdC_family_ATP-grasp_ribosomal_peptide_maturase
CA739_RS37935	CA739_RS37935	331498	331770	-	ABC_transporter
CA739_RS37975	CA739_RS37975	336522	336623	+	transposase

Table of Blast hits (query gene, subject gene, %identity, blast score, %coverage, e-value):
input.path1.gene298	CA739_RS38030	32	223	33.13008130081301	1.1e-55
input.path1.gene300	CA739_RS37900	40	111	22.78481012658228	5.5e-22
input.path1.gene300	CA739_RS37905	41	107	19.367088607594937	7.9e-21
input.path1.gene300	CA739_RS37890	42	104	19.240506329113924	5.1e-20
input.path1.gene300	CA739_RS37795	36	104	19.367088607594937	6.7e-20


>>
6. NZ_AP018268
Source: Scytonema sp. NIES-4073 DNA, nearly complete genome
Type: NRPS,T1PKS
Number of proteins with BLAST hits to this cluster: 2
Cumulative BLAST score: 543

Table of genes, locations, strands and annotations of subject cluster:
CDC39_RS08325	CDC39_RS08325	1541890	1542018	-	TetR_family_transcriptional_regulator
CDC39_RS08395	CDC39_RS08395	1575837	1576052	+	transcriptional_regulator
CDC39_RS08445	CDC39_RS08445	1621469	1621646	-	ISKra4_family_transposase
CDC39_RS08455	CDC39_RS08455	1623095	1624063	-	hypothetical_protein
CDC39_RS08460	CDC39_RS08460	1624060	1624463	-	mechanosensitive_ion_channel_protein_MscS
CDC39_RS08650	CDC39_RS08650	1699915	1700133	+	MbtH_domain_protein
CDC39_RS08655	CDC39_RS08655	1700290	1700400	+	MbtH_domain_protein
CDC39_RS08720	CDC39_RS08720	1712158	1712406	+	transposase

Table of Blast hits (query gene, subject gene, %identity, blast score, %coverage, e-value):
input.path1.gene298	CDC39_RS08350	31	222	33.06233062330624	3.1e-55
input.path1.gene298	CDC39_RS08595	31	215	32.046070460704605	2.9e-53
input.path1.gene300	CDC39_RS08440	41	106	19.367088607594937	1.3e-20


>>
7. NZ_KL662191
Source: [Leptolyngbya] sp. JSC-1 Osccy1DRAFT_CYJSC1_DRAF_scaffold00069.1, whole genome shotgun sequence
Type: NRPS,NRPS-like,T1PKS
Number of proteins with BLAST hits to this cluster: 2
Cumulative BLAST score: 441

Table of genes, locations, strands and annotations of subject cluster:
OSCCY1_RS09320	OSCCY1_RS09320	2281953	2286083	-	peptidase
OSCCY1_RS09395	OSCCY1_RS09395	2342302	2344064	+	polyketide_synthase

Table of Blast hits (query gene, subject gene, %identity, blast score, %coverage, e-value):
input.path1.gene298	OSCCY1_RS09390	33	214	31.70731707317073	6.6e-53
input.path1.gene300	OSCCY1_RS09360	41	118	22.911392405063292	3.4e-24
input.path1.gene300	OSCCY1_RS09355	38	109	22.78481012658228	1.2e-21


>>
8. NZ_CM001979
Source: Dickeya sp. NCPPB 3274 chromosome, whole genome shotgun sequence
Type: NRPS,transAT-PKS
Number of proteins with BLAST hits to this cluster: 2
Cumulative BLAST score: 331

Table of genes, locations, strands and annotations of subject cluster:

Table of Blast hits (query gene, subject gene, %identity, blast score, %coverage, e-value):
input.path1.gene298	DXX3274_RS13125	33	221	31.300813008130078	5.4e-55
input.path1.gene300	DXX3274_RS13155	42	110	20.253164556962027	7.1e-22


>>
9. NZ_KQ976354
Source: Scytonema hofmannii PCC 7110 Scaffold1, whole genome shotgun sequence
Type: NRPS,microviridin
Number of proteins with BLAST hits to this cluster: 2
Cumulative BLAST score: 331

Table of genes, locations, strands and annotations of subject cluster:
WA1_RS15490	WA1_RS15490	3712567	3714927	-	hypothetical_protein
WA1_RS15505	WA1_RS15505	3719729	3741925	+	non-ribosomal_peptide_synthase/polyketide_synthase

Table of Blast hits (query gene, subject gene, %identity, blast score, %coverage, e-value):
input.path1.gene298	WA1_RS15540	32	218	31.571815718157183	4.5e-54
input.path1.gene300	WA1_RS15500	41	113	22.78481012658228	1.4e-22


>>
10. NC_010628
Source: Nostoc punctiforme PCC 73102, complete sequence
Type: NRPS,T1PKS,bacteriocin,lanthidin,transAT-PKS-like
Number of proteins with BLAST hits to this cluster: 2
Cumulative BLAST score: 329

Table of genes, locations, strands and annotations of subject cluster:
NPUN_RS16975	NPUN_RS16975	4170222	4171166	-	abortive_phage_infection_protein
NPUN_RS40115	NPUN_RS40115	4171188	4172353	-	IS630_family_transposase
NPUN_RS17195	NPUN_RS17195	4239986	4240778	+	IS5_family_transposase
NPUN_RS39195	NPUN_RS39195	4241090	4241359	+	hypothetical_protein
NPUN_RS17205	NPUN_RS17205	4242382	4242626	+	tyrosine-type_recombinase/integrase
NPUN_RS17210	NPUN_RS17210	4242933	4243309	+	response_regulator
NPUN_RS41990	NPUN_RS41990	4247654	4248250	-	hypothetical_protein
NPUN_RS40120	NPUN_RS40120	4252166	4253333	+	IS630_family_transposase
NPUN_RS17275	NPUN_RS17275	4253435	4254833	-	transposase
NPUN_RS17280	NPUN_RS17280	4254993	4255893	-	transposase

Table of Blast hits (query gene, subject gene, %identity, blast score, %coverage, e-value):
input.path1.gene298	NPUN_RS17360	32	219	31.910569105691057	2e-54
input.path1.gene300	NPUN_RS17020	39	110	21.139240506329113	7.1e-22


>>
11. NZ_CP023484
Source: Dickeya sp. Secpp 1600 chromosome, complete genome
Type: NRPS,transAT-PKS
Number of proteins with BLAST hits to this cluster: 2
Cumulative BLAST score: 327

Table of genes, locations, strands and annotations of subject cluster:
CPA59_RS13510	CPA59_RS13510	3030461	3031100	-	AEC_family_transporter

Table of Blast hits (query gene, subject gene, %identity, blast score, %coverage, e-value):
input.path1.gene298	CPA59_RS13545	33	218	31.300813008130078	4.5e-54
input.path1.gene300	CPA59_RS13575	41	109	20.253164556962027	1.6e-21


>>
12. NZ_CP025003
Source: Dickeya fangzhongdai strain DSM 101947 chromosome, complete genome
Type: NRPS,transAT-PKS
Number of proteins with BLAST hits to this cluster: 2
Cumulative BLAST score: 327

Table of genes, locations, strands and annotations of subject cluster:
CVE23_RS13335	CVE23_RS13335	2982634	2983273	-	AEC_family_transporter

Table of Blast hits (query gene, subject gene, %identity, blast score, %coverage, e-value):
input.path1.gene298	CVE23_RS13370	33	218	31.300813008130078	4.5e-54
input.path1.gene300	CVE23_RS13400	41	109	20.253164556962027	1.6e-21


>>
13. NZ_CP020872
Source: Dickeya fangzhongdai strain PA1 chromosome, complete genome
Type: NRPS,transAT-PKS
Number of proteins with BLAST hits to this cluster: 2
Cumulative BLAST score: 327

Table of genes, locations, strands and annotations of subject cluster:
B6N31_RS13165	B6N31_RS13165	2970145	2970784	-	AEC_family_transporter

Table of Blast hits (query gene, subject gene, %identity, blast score, %coverage, e-value):
input.path1.gene298	B6N31_RS13200	33	218	31.300813008130078	5.9e-54
input.path1.gene300	B6N31_RS13230	41	109	20.253164556962027	1.6e-21


>>
14. NZ_CM001860
Source: Dickeya solani GBBC 2040 chromosome, whole genome shotgun sequence
Type: NRPS,transAT-PKS
Number of proteins with BLAST hits to this cluster: 2
Cumulative BLAST score: 326

Table of genes, locations, strands and annotations of subject cluster:
DSOGBBC2040_RS22420	DSOGBBC2040_RS22420	3047848	3049133	-	cytochrome_P450
DSOGBBC2040_RS13735	DSOGBBC2040_RS13735	3094576	3094842	-	LysR_family_transcriptional_regulator

Table of Blast hits (query gene, subject gene, %identity, blast score, %coverage, e-value):
input.path1.gene298	DSOGBBC2040_RS13660	33	220	32.046070460704605	9.1e-55
input.path1.gene300	DSOGBBC2040_RS22425	43	106	20.253164556962027	1.3e-20


>>
15. NC_013592
Source: Dickeya zeae Ech586, complete sequence
Type: NRPS,transAT-PKS
Number of proteins with BLAST hits to this cluster: 2
Cumulative BLAST score: 326

Table of genes, locations, strands and annotations of subject cluster:

Table of Blast hits (query gene, subject gene, %identity, blast score, %coverage, e-value):
input.path1.gene298	DD586_RS06835	33	220	31.300813008130078	1.6e-54
input.path1.gene300	DD586_RS06805	39	106	24.556962025316455	1e-20


>>
16. NZ_QJKB01000008
Source: Undibacterium pigrum strain DSM 19792 Ga0244539_108, whole genome shotgun sequence
Type: NRPS,T1PKS
Number of proteins with BLAST hits to this cluster: 2
Cumulative BLAST score: 326

Table of genes, locations, strands and annotations of subject cluster:

Table of Blast hits (query gene, subject gene, %identity, blast score, %coverage, e-value):
input.path1.gene298	DFR42_RS20730	32	218	33.60433604336043	3.5e-54
input.path1.gene300	DFR42_RS20720	39	108	22.911392405063292	3.5e-21


>>
17. NZ_CM001984
Source: Dickeya sp. MK7 chromosome, whole genome shotgun sequence
Type: NRPS,transAT-PKS
Number of proteins with BLAST hits to this cluster: 2
Cumulative BLAST score: 326

Table of genes, locations, strands and annotations of subject cluster:
DXXMK7_RS12715	DXXMK7_RS12715	2925675	2926314	-	AEC_family_transporter
DXXMK7_RS22325	DXXMK7_RS22325	2971572	2971856	+	hypothetical_protein

Table of Blast hits (query gene, subject gene, %identity, blast score, %coverage, e-value):
input.path1.gene298	DXXMK7_RS12750	33	217	31.300813008130078	1e-53
input.path1.gene300	DXXMK7_RS12780	41	109	20.253164556962027	1.6e-21


>>
18. NZ_CP054698
Source: Nostoc edaphicum CCNP1411 chromosome, complete genome
Type: NRPS,NRPS-like,T1PKS
Number of proteins with BLAST hits to this cluster: 2
Cumulative BLAST score: 326

Table of genes, locations, strands and annotations of subject cluster:
HUN01_RS33055	HUN01_RS33055	7232401	7237089	-	polyketide_synthase_dehydratase_domain-containing_protein
HUN01_RS33060	HUN01_RS33060	7237390	7238991	-	polyketide_synthase
HUN01_RS33065	HUN01_RS33065	7239387	7240772	-	SDR_family_NAD(P)-dependent_oxidoreductase
HUN01_RS33070	HUN01_RS33070	7244118	7247324	-	type_I_polyketide_synthase
HUN01_RS33135	HUN01_RS33135	7271922	7273290	-	IS4_family_transposase
HUN01_RS33175	HUN01_RS33175	7298691	7304903	+	amino_acid_adenylation_domain-containing_protein
HUN01_RS33195	HUN01_RS33195	7318503	7318964	+	non-ribosomal_peptide_synthetase
HUN01_RS33200	HUN01_RS33200	7319073	7322186	+	amino_acid_adenylation_domain-containing_protein
HUN01_RS33235	HUN01_RS33235	7329399	7329935	+	siderophore_biosynthesis_protein
HUN01_RS33250	HUN01_RS33250	7336861	7337235	+	DUF3102_domain-containing_protein

Table of Blast hits (query gene, subject gene, %identity, blast score, %coverage, e-value):
input.path1.gene298	HUN01_RS33160	31	213	36.24661246612466	1.5e-52
input.path1.gene300	HUN01_RS33070	42	113	21.012658227848103	1.4e-22


>>
19. NC_011726
Source: Rippkaea orientalis PCC 8801, complete sequence
Type: NRPS,T1PKS
Number of proteins with BLAST hits to this cluster: 2
Cumulative BLAST score: 325

Table of genes, locations, strands and annotations of subject cluster:
PCC8801_RS22705	PCC8801_RS22705	3118347	3118511	-	IS200/IS605_family_transposase
PCC8801_RS22710	PCC8801_RS22710	3120389	3120548	+	serine_protease

Table of Blast hits (query gene, subject gene, %identity, blast score, %coverage, e-value):
input.path1.gene298	PCC8801_RS14805	32	218	33.87533875338754	5.9e-54
input.path1.gene300	PCC8801_RS14770	41	107	19.240506329113924	7.9e-21


>>
20. NZ_AP018441
Source: Undibacterium sp. YM2
Type: NRPS,T1PKS,siderophore
Number of proteins with BLAST hits to this cluster: 2
Cumulative BLAST score: 325

Table of genes, locations, strands and annotations of subject cluster:

Table of Blast hits (query gene, subject gene, %identity, blast score, %coverage, e-value):
input.path1.gene298	UNDYM_RS18035	32	215	33.33333333333333	3.8e-53
input.path1.gene300	UNDYM_RS18045	36	110	26.962025316455694	9.3e-22


>>
21. NZ_CP031515
Source: Dickeya zeae strain EC2 chromosome, complete genome
Type: NRPS,transAT-PKS
Number of proteins with BLAST hits to this cluster: 2
Cumulative BLAST score: 324

Table of genes, locations, strands and annotations of subject cluster:
DWV07_RS06805	DWV07_RS06805	1530562	1531175	-	IS3_family_transposase
DWV07_RS06825	DWV07_RS06825	1533696	1534260	+	TetR/AcrR_family_transcriptional_regulator
DWV07_RS06855	DWV07_RS06855	1568535	1569247	+	IS5_family_transposase
DWV07_RS06860	DWV07_RS06860	1569259	1569393	-	helix-turn-helix_domain-containing_protein

Table of Blast hits (query gene, subject gene, %identity, blast score, %coverage, e-value):
input.path1.gene298	DWV07_RS06760	33	221	31.300813008130078	7e-55
input.path1.gene300	DWV07_RS06730	38	103	20.506329113924053	1.1e-19


>>
22. NZ_CM001977
Source: Dickeya zeae NCPPB 2538 chromosome, whole genome shotgun sequence
Type: NRPS,transAT-PKS
Number of proteins with BLAST hits to this cluster: 2
Cumulative BLAST score: 323

Table of genes, locations, strands and annotations of subject cluster:

Table of Blast hits (query gene, subject gene, %identity, blast score, %coverage, e-value):
input.path1.gene298	DZE2538_RS06655	33	219	31.300813008130078	2e-54
input.path1.gene300	DZE2538_RS06625	38	104	20.506329113924053	5.1e-20


>>
23. NZ_CP023467
Source: Dickeya dadantii strain DSM 18020 chromosome, complete genome
Type: NRPS,transAT-PKS
Number of proteins with BLAST hits to this cluster: 2
Cumulative BLAST score: 323

Table of genes, locations, strands and annotations of subject cluster:
CO076_RS14585	CO076_RS14585	3242623	3242952	+	hypothetical_protein
CO076_RS14625	CO076_RS14625	3249737	3250000	-	LysR_family_transcriptional_regulator

Table of Blast hits (query gene, subject gene, %identity, blast score, %coverage, e-value):
input.path1.gene298	CO076_RS14545	32	218	32.046070460704605	4.5e-54
input.path1.gene300	CO076_RS14575	43	105	20.253164556962027	3e-20


>>
24. NZ_CM001980
Source: Dickeya zeae NCPPB 3531 chromosome, whole genome shotgun sequence
Type: NRPS,transAT-PKS
Number of proteins with BLAST hits to this cluster: 2
Cumulative BLAST score: 322

Table of genes, locations, strands and annotations of subject cluster:
DZE3531_RS21065	DZE3531_RS21065	1565296	1565898	-	IS3_family_transposase
DZE3531_RS21070	DZE3531_RS21070	1575281	1575370	-	VOC_family_protein
DZE3531_RS21075	DZE3531_RS21075	1593343	1593417	-	N-acetyltransferase

Table of Blast hits (query gene, subject gene, %identity, blast score, %coverage, e-value):
input.path1.gene298	DZE3531_RS06845	33	219	31.300813008130078	2e-54
input.path1.gene300	DZE3531_RS06815	38	103	20.506329113924053	1.1e-19


>>
25. NZ_CM001985
Source: Dickeya zeae MK19 chromosome, whole genome shotgun sequence
Type: NRPS,transAT-PKS
Number of proteins with BLAST hits to this cluster: 2
Cumulative BLAST score: 322

Table of genes, locations, strands and annotations of subject cluster:
DZEMK19_RS21075	DZEMK19_RS21075	1589362	1589448	-	VOC_family_protein

Table of Blast hits (query gene, subject gene, %identity, blast score, %coverage, e-value):
input.path1.gene298	DZEMK19_RS06970	33	219	31.300813008130078	2e-54
input.path1.gene300	DZEMK19_RS06940	36	103	23.29113924050633	1.1e-19


>>
26. NZ_CM001858
Source: Dickeya zeae NCPPB 3532 chromosome, whole genome shotgun sequence
Type: NRPS,transAT-PKS
Number of proteins with BLAST hits to this cluster: 2
Cumulative BLAST score: 321

Table of genes, locations, strands and annotations of subject cluster:
DZE3532_RS22195	DZE3532_RS22195	1442556	1442830	+	IS630_family_transposase

Table of Blast hits (query gene, subject gene, %identity, blast score, %coverage, e-value):
input.path1.gene298	DZE3532_RS06715	33	218	31.300813008130078	3.5e-54
input.path1.gene300	DZE3532_RS06685	36	103	23.29113924050633	1.1e-19


>>
27. NZ_CP033622
Source: Dickeya zeae strain CE1 chromosome, complete genome
Type: NRPS,transAT-PKS
Number of proteins with BLAST hits to this cluster: 2
Cumulative BLAST score: 321

Table of genes, locations, strands and annotations of subject cluster:
DWG24_RS13920	DWG24_RS13920	3146122	3146396	-	IS630_family_transposase

Table of Blast hits (query gene, subject gene, %identity, blast score, %coverage, e-value):
input.path1.gene298	DWG24_RS13810	33	218	31.300813008130078	3.5e-54
input.path1.gene300	DWG24_RS13840	38	103	20.506329113924053	1.1e-19


>>
28. NZ_CM001982
Source: Dickeya dadantii NCPPB 3537 chromosome, whole genome shotgun sequence
Type: NRPS,transAT-PKS
Number of proteins with BLAST hits to this cluster: 2
Cumulative BLAST score: 321

Table of genes, locations, strands and annotations of subject cluster:
DDA3537_RS13605	DDA3537_RS13605	3092512	3093655	+	iron-containing_alcohol_dehydrogenase
DDA3537_RS21470	DDA3537_RS21470	3149653	3149982	+	hypothetical_protein
DDA3537_RS13780	DDA3537_RS13780	3156619	3156885	-	LysR_family_transcriptional_regulator

Table of Blast hits (query gene, subject gene, %identity, blast score, %coverage, e-value):
input.path1.gene298	DDA3537_RS13705	33	217	31.368563685636857	1.3e-53
input.path1.gene300	DDA3537_RS13735	43	104	20.253164556962027	6.7e-20


>>
29. NC_013161
Source: Rippkaea orientalis PCC 8802, complete sequence
Type: NRPS,T1PKS
Number of proteins with BLAST hits to this cluster: 2
Cumulative BLAST score: 321

Table of genes, locations, strands and annotations of subject cluster:
CYAN8802_RS23255	CYAN8802_RS23255	3105275	3105826	+	transposase
CYAN8802_RS14870	CYAN8802_RS14870	3169612	3172874	-	acyltransferase_domain-containing_protein
CYAN8802_RS22920	CYAN8802_RS22920	3177985	3178144	-	serine_protease

Table of Blast hits (query gene, subject gene, %identity, blast score, %coverage, e-value):
input.path1.gene298	CYAN8802_RS14825	31	215	33.87533875338754	5e-53
input.path1.gene300	CYAN8802_RS14860	41	106	19.240506329113924	1e-20


>>
30. NC_020126
Source: Myxococcus stipitatus DSM 14675, complete sequence
Type: NRPS,T1PKS
Number of proteins with BLAST hits to this cluster: 2
Cumulative BLAST score: 319

Table of genes, locations, strands and annotations of subject cluster:
MYSTI_RS10200	MYSTI_RS10200	2574926	2584558	-	amino_acid_adenylation_domain-containing_protein
MYSTI_RS10280	MYSTI_RS10280	2608393	2610033	+	hypothetical_protein

Table of Blast hits (query gene, subject gene, %identity, blast score, %coverage, e-value):
input.path1.gene298	MYSTI_RS10120	33	214	31.910569105691057	8.6e-53
input.path1.gene300	MYSTI_RS10210	40	105	23.037974683544306	2.3e-20


>>
31. NC_008095
Source: Myxococcus xanthus DK 1622, complete sequence
Type: NRPS,T1PKS
Number of proteins with BLAST hits to this cluster: 2
Cumulative BLAST score: 317

Table of genes, locations, strands and annotations of subject cluster:

Table of Blast hits (query gene, subject gene, %identity, blast score, %coverage, e-value):
input.path1.gene298	MXAN_RS21385	32	214	34.552845528455286	6.6e-53
input.path1.gene300	MXAN_RS21375	41	103	19.367088607594937	8.7e-20


>>
32. NC_026507
Source: Neurospora crassa OR74A linkage group VII, whole genome shotgun sequence
Type: T1PKS
Number of proteins with BLAST hits to this cluster: 2
Cumulative BLAST score: 1472

Table of genes, locations, strands and annotations of subject cluster:

Table of Blast hits (query gene, subject gene, %identity, blast score, %coverage, e-value):
input.path1.gene297	NCU08397	34	432	103.49462365591397	7.7e-119
input.path1.gene300	NCU08399_XP_011395280.1	35	520	117.59493670886076	3e-145
input.path1.gene300	NCU08399	35	520	117.59493670886076	3e-145


>>
33. NW_007360994
Source: Glarea lozoyensis ATCC 20868 chromosome Unknown GLAREA2, whole genome shotgun sequence
Type: NRPS,T1PKS
Number of proteins with BLAST hits to this cluster: 2
Cumulative BLAST score: 738

Table of genes, locations, strands and annotations of subject cluster:

Table of Blast hits (query gene, subject gene, %identity, blast score, %coverage, e-value):
input.path1.gene296	GLAREA_03906	36	183	83.16062176165802	4.2e-44
input.path1.gene300	GLAREA_03913	39	555	116.20253164556962	8.4e-156


>>
34. NZ_BCMN01000006
Source: Streptomyces turgidiscabies strain T45, whole genome shotgun sequence
Type: T1PKS,hglE-KS,terpene
Number of proteins with BLAST hits to this cluster: 2
Cumulative BLAST score: 565

Table of genes, locations, strands and annotations of subject cluster:
AV140_RS47855	AV140_RS47855	273327	273461	-	IS110_family_transposase

Table of Blast hits (query gene, subject gene, %identity, blast score, %coverage, e-value):
input.path1.gene293	AV140_RS15160	38	450	104.31654676258992	2.6e-124
input.path1.gene300	AV140_RS15195	36	115	34.43037974683544	2.2e-23


>>
35. NZ_CP030864
Source: Streptomyces globosus strain LZH-48 plasmid unnamed2, complete sequence
Type: NRPS,NRPS-like,T2PKS,butyrolactone,ectoine,hglE-KS,linaridin,transAT-PKS
Number of proteins with BLAST hits to this cluster: 2
Cumulative BLAST score: 484

Table of genes, locations, strands and annotations of subject cluster:
C0216_RS31350	C0216_RS31350	25068	25503	+	transposase
C0216_RS33995	C0216_RS33995	31513	32988	-	hypothetical_protein
C0216_RS34000	C0216_RS34000	33058	33558	-	hypothetical_protein
C0216_RS34005	C0216_RS34005	33571	34563	-	SDR_family_NAD(P)-dependent_oxidoreductase
C0216_RS34010	C0216_RS34010	34585	35604	-	polyketide_synthase_dehydratase_domain-containing_protein
C0216_RS34015	C0216_RS34015	36697	38229	-	polyketide_synthase_docking_domain-containing_protein
C0216_RS31715	C0216_RS31715	175226	175600	-	hypothetical_protein
C0216_RS31790	C0216_RS31790	191392	191821	-	DUF1772_domain-containing_protein
C0216_RS34025	C0216_RS34025	296568	297596	+	NAD-dependent_epimerase/dehydratase_family_protein
C0216_RS32170	C0216_RS32170	303897	303992	+	short-chain_dehydrogenase

Table of Blast hits (query gene, subject gene, %identity, blast score, %coverage, e-value):
input.path1.gene296	C0216_RS31500	47	267	77.97927461139896	2.9e-69
input.path1.gene300	C0216_RS31395	44	111	19.62025316455696	5.5e-22
input.path1.gene300	C0216_RS31375	38	106	21.645569620253163	1e-20


>>
36. NT_165934
Source: Aspergillus terreus NIH2624 scaffold_11 genomic scaffold, whole genome shotgun sequence
Type: T1PKS
Number of proteins with BLAST hits to this cluster: 2
Cumulative BLAST score: 472

Table of genes, locations, strands and annotations of subject cluster:

Table of Blast hits (query gene, subject gene, %identity, blast score, %coverage, e-value):
input.path1.gene300	ATEG_07659	31	305	112.65822784810126	1.2e-80
input.path1.gene302	ATEG_07669	33	167	90.45936395759718	1.8e-39


>>
37. NZ_KQ947996
Source: Streptomyces curacoi strain DSM 40107 PRJNA299217_s013, whole genome shotgun sequence
Type: NRPS,T1PKS
Number of proteins with BLAST hits to this cluster: 2
Cumulative BLAST score: 453

Table of genes, locations, strands and annotations of subject cluster:
AQI70_RS33640	AQI70_RS33640	16296	17110	+	oxidoreductase
AQI70_RS33735	AQI70_RS33735	47069	48596	-	metallophosphoesterase
AQI70_RS33795	AQI70_RS33795	59739	61079	-	ATP-binding_protein

Table of Blast hits (query gene, subject gene, %identity, blast score, %coverage, e-value):
input.path1.gene296	AQI70_RS33835	46	244	78.49740932642487	1.5e-62
input.path1.gene300	AQI70_RS33815	38	105	22.40506329113924	3e-20
input.path1.gene300	AQI70_RS33730	38	104	21.645569620253163	6.7e-20


>>
38. NC_013093
Source: Actinosynnema mirum DSM 43827, complete sequence
Type: T1PKS
Number of proteins with BLAST hits to this cluster: 2
Cumulative BLAST score: 381

Table of genes, locations, strands and annotations of subject cluster:
AMIR_RS23350	AMIR_RS23350	5586137	5587270	+	acyl-CoA/acyl-ACP_dehydrogenase
AMIR_RS40545	AMIR_RS40545	5595337	5597907	-	type_I_polyketide_synthase
AMIR_RS38395	AMIR_RS38395	5598529	5599728	-	KR_domain-containing_protein
AMIR_RS39510	AMIR_RS39510	5600362	5603115	-	type_I_polyketide_synthase
AMIR_RS38415	AMIR_RS38415	5603629	5604543	-	acyltransferase_domain-containing_protein
AMIR_RS39515	AMIR_RS39515	5605204	5606481	-	polyketide_synthase

Table of Blast hits (query gene, subject gene, %identity, blast score, %coverage, e-value):
input.path1.gene293	AMIR_RS23430	33	264	82.87769784172662	2.6e-68
input.path1.gene300	AMIR_RS39515	42	117	19.746835443037973	5.8e-24


>>
39. NZ_CP023445
Source: Actinosynnema pretiosum strain X47 chromosome, complete genome
Type: T1PKS
Number of proteins with BLAST hits to this cluster: 2
Cumulative BLAST score: 372

Table of genes, locations, strands and annotations of subject cluster:
CNX65_RS23355	CNX65_RS23355	5513325	5513885	+	response_regulator
CNX65_RS23360	CNX65_RS23360	5513886	5515016	-	serine_hydrolase
CNX65_RS35645	CNX65_RS35645	5531187	5533763	-	acyltransferase_domain-containing_protein
CNX65_RS35650	CNX65_RS35650	5533857	5534267	-	hypothetical_protein
CNX65_RS35655	CNX65_RS35655	5534388	5535587	-	KR_domain-containing_protein
CNX65_RS35660	CNX65_RS35660	5536227	5539181	-	acyltransferase_domain-containing_protein
CNX65_RS35665	CNX65_RS35665	5539827	5540408	-	acyltransferase_domain-containing_protein
CNX65_RS35670	CNX65_RS35670	5540997	5542496	-	hypothetical_protein

Table of Blast hits (query gene, subject gene, %identity, blast score, %coverage, e-value):
input.path1.gene293	CNX65_RS23455	34	256	73.66906474820144	7.1e-66
input.path1.gene300	CNX65_RS35670	42	116	19.746835443037973	1.3e-23


>>
40. NZ_CP009754
Source: Streptomyces sp. CCM_MD2014 chromosome, complete genome
Type: NRPS-like,T1PKS
Number of proteins with BLAST hits to this cluster: 2
Cumulative BLAST score: 364

Table of genes, locations, strands and annotations of subject cluster:
NI25_RS37470	NI25_RS37470	2260860	2261006	-	toxin
NI25_RS38425	NI25_RS38425	2293418	2294812	-	polyketide_synthase
NI25_RS38430	NI25_RS38430	2295164	2296444	-	polyketide_synthase

Table of Blast hits (query gene, subject gene, %identity, blast score, %coverage, e-value):
input.path1.gene296	NI25_RS09765	48	259	76.94300518134715	4.6e-67
input.path1.gene300	NI25_RS09835	40	105	19.746835443037973	3e-20


>>
41. NZ_KZ195574
Source: Streptomyces sp. CS113 scaffold00001, whole genome shotgun sequence
Type: NRPS-like,T1PKS
Number of proteins with BLAST hits to this cluster: 2
Cumulative BLAST score: 361

Table of genes, locations, strands and annotations of subject cluster:

Table of Blast hits (query gene, subject gene, %identity, blast score, %coverage, e-value):
input.path1.gene296	B9W62_RS09415	47	257	76.94300518134715	2.3e-66
input.path1.gene300	B9W62_RS09490	37	104	22.78481012658228	5.1e-20


>>
42. NZ_CP056041
Source: Streptomyces chartreusis strain NA02069 chromosome, complete genome
Type: NRPS,T1PKS
Number of proteins with BLAST hits to this cluster: 2
Cumulative BLAST score: 354

Table of genes, locations, strands and annotations of subject cluster:

Table of Blast hits (query gene, subject gene, %identity, blast score, %coverage, e-value):
input.path1.gene296	HUT05_RS44275	47	249	78.49740932642487	4.8e-64
input.path1.gene300	HUT05_RS44150	39	105	22.40506329113924	2.3e-20


>>
43. NZ_CP023689
Source: Streptomyces chartreusis strain ATCC 14922 chromosome, complete genome
Type: NRPS,T1PKS
Number of proteins with BLAST hits to this cluster: 2
Cumulative BLAST score: 352

Table of genes, locations, strands and annotations of subject cluster:
CP983_RS41525	CP983_RS41525	9370410	9371200	+	maleylpyruvate_isomerase_family_mycothiol-dependent_enzyme

Table of Blast hits (query gene, subject gene, %identity, blast score, %coverage, e-value):
input.path1.gene296	CP983_RS41515	47	247	78.49740932642487	3.1e-63
input.path1.gene300	CP983_RS41390	39	105	22.40506329113924	2.3e-20


>>
44. NZ_RDBL01000014
Source: Streptomyces sp. col6 Scaffold_14, whole genome shotgun sequence
Type: T1PKS
Number of proteins with BLAST hits to this cluster: 2
Cumulative BLAST score: 340

Table of genes, locations, strands and annotations of subject cluster:

Table of Blast hits (query gene, subject gene, %identity, blast score, %coverage, e-value):
input.path1.gene296	EAO73_RS24705	40	217	79.53367875647669	2e-54
input.path1.gene300	EAO73_RS24760	46	123	19.746835443037973	8.1e-26


>>
45. NZ_SZNQ01000001
Source: Streptomyces lasalocidi strain X-537 Scaffold1, whole genome shotgun sequence
Type: T1PKS
Number of proteins with BLAST hits to this cluster: 2
Cumulative BLAST score: 312

Table of genes, locations, strands and annotations of subject cluster:
E4U91_RS38520	E4U91_RS38520	268563	270044	+	polyketide_synthase_docking_domain-containing_protein
E4U91_RS00945	E4U91_RS00945	270294	271187	+	acyltransferase_domain-containing_protein
E4U91_RS00950	E4U91_RS00950	271599	278027	+	SDR_family_NAD(P)-dependent_oxidoreductase
E4U91_RS00965	E4U91_RS00965	288393	290420	+	type_I_polyketide_synthase
E4U91_RS38525	E4U91_RS38525	290970	292001	+	polyketide_synthase_dehydratase_domain-containing_protein
E4U91_RS00975	E4U91_RS00975	292038	292946	+	alcohol_dehydrogenase_catalytic_domain-containing_protein
E4U91_RS38395	E4U91_RS38395	293565	297362	+	SDR_family_NAD(P)-dependent_oxidoreductase
E4U91_RS38530	E4U91_RS38530	298035	298661	+	KR_domain-containing_protein
E4U91_RS01015	E4U91_RS01015	312195	313353	+	beta-lactamase_family_protein

Table of Blast hits (query gene, subject gene, %identity, blast score, %coverage, e-value):
input.path1.gene296	E4U91_RS00880	36	197	80.82901554404145	2.2e-48
input.path1.gene300	E4U91_RS00935	42	115	23.924050632911392	2.9e-23


>>
46. NZ_CP010429
Source: Spirosoma radiotolerans strain DG5A chromosome, complete genome
Type: NRPS,T1PKS
Number of proteins with BLAST hits to this cluster: 2
Cumulative BLAST score: 280

Table of genes, locations, strands and annotations of subject cluster:

Table of Blast hits (query gene, subject gene, %identity, blast score, %coverage, e-value):
input.path1.gene296	SD10_RS02220	34	171	80.05181347150258	2.2e-40
input.path1.gene300	SD10_RS02295	39	109	19.367088607594937	1.2e-21


>>
47. NZ_CP043761
Source: Paenibacillus sp. 37 chromosome, complete genome
Type: NRPS,NRPS-like,ladderane,transAT-PKS
Number of proteins with BLAST hits to this cluster: 2
Cumulative BLAST score: 258

Table of genes, locations, strands and annotations of subject cluster:
F0220_RS22660	F0220_RS22660	5276334	5277784	+	IS1182_family_transposase
F0220_RS22760	F0220_RS22760	5298072	5298708	+	IS30_family_transposase
F0220_RS22765	F0220_RS22765	5298982	5300263	+	IS3_family_transposase
F0220_RS22865	F0220_RS22865	5325096	5325185	+	short-chain_dehydrogenase
F0220_RS23030	F0220_RS23030	5389286	5389471	-	GNAT_family_N-acetyltransferase

Table of Blast hits (query gene, subject gene, %identity, blast score, %coverage, e-value):
input.path1.gene296	F0220_RS23010	31	149	80.31088082901555	8.8e-34
input.path1.gene300	F0220_RS22925	39	109	22.40506329113924	1.2e-21


>>
48. NZ_KQ948925
Source: Streptomyces caeruleatus strain NRRL B-24802 PRJNA299234_s002, whole genome shotgun sequence
Type: NRPS,T1PKS
Number of proteins with BLAST hits to this cluster: 2
Cumulative BLAST score: 175

Table of genes, locations, strands and annotations of subject cluster:
AQJ67_RS05625	AQJ67_RS05625	150567	151055	-	1,4-beta-xylanase
AQJ67_RS46590	AQJ67_RS46590	153203	153772	-	polyketide_synthase
AQJ67_RS44435	AQJ67_RS44435	153902	155956	-	NAD-dependent_epimerase/dehydratase_family_protein
AQJ67_RS44760	AQJ67_RS44760	209239	209952	-	ATP-binding_protein

Table of Blast hits (query gene, subject gene, %identity, blast score, %coverage, e-value):
input.path1.gene296	AQJ67_RS05625	61	71	13.21243523316062	1.8e-10
input.path1.gene300	AQJ67_RS05770	39	104	21.645569620253163	6.7e-20


>>
49. NZ_LN831790
Source: Streptomyces leeuwenhoekii strain type strain (C34 = DSM 42122 = NRRL B-24963) chromosome 1
Type: NRPS-like
Number of proteins with BLAST hits to this cluster: 2
Cumulative BLAST score: 486

Table of genes, locations, strands and annotations of subject cluster:

Table of Blast hits (query gene, subject gene, %identity, blast score, %coverage, e-value):
input.path1.gene293	BN2145_RS33550	32	229	69.64028776978417	9.3e-58
input.path1.gene296	BN2145_RS33555	48	257	76.94300518134715	1.8e-66


>>
50. NZ_CP022961
Source: Plantactinospora sp. KBS50 chromosome, complete genome
Type: NRPS
Number of proteins with BLAST hits to this cluster: 2
Cumulative BLAST score: 477

Table of genes, locations, strands and annotations of subject cluster:
CIK06_RS15845	CIK06_RS15845	3562771	3564646	-	CBS_domain-containing_protein
CIK06_RS15860	CIK06_RS15860	3566944	3568086	-	sodium:proton_antiporter
CIK06_RS15890	CIK06_RS15890	3571537	3572076	+	anti-sigma_factor_antagonist
CIK06_RS15900	CIK06_RS15900	3572928	3573698	+	hypothetical_protein
CIK06_RS15935	CIK06_RS15935	3583685	3584623	+	hypothetical_protein
CIK06_RS16015	CIK06_RS16015	3615099	3616026	-	ParA_family_protein

Table of Blast hits (query gene, subject gene, %identity, blast score, %coverage, e-value):
input.path1.gene293	CIK06_RS15975	33	223	63.30935251798561	8.7e-56
input.path1.gene296	CIK06_RS15920	45	254	75.12953367875647	1.5e-65
