## Supplementary Results for "A draft genome of the ascomycotal fungal species *Pseudopithomyces maydicus* (family *Didymosphaeriaceae*)": input.path1.gene387_searchgtr.html

Search Query


### **SEARCHGTr input**

**Sequence name:**    
FASTA-formatted sequence:   
>input.path1.gene387
MEYHGVTFNFAFNYVDHELGQASVILSVAYEIAIQSDWDVTIASFASLQSQVDALNRTFETTMNWYAIPGKSMKDCLAANGLDFLPIHAPGVKGAVKAYRECLPYVIAPWTPEEYDRHCAGSFKDHIVNEQPYQGVVWKYSVNALSNGLSSIFSNFDPAIPYLVPSTPSIDFPLTIPRNVHGCGPILLPLPPTSSSSPTSTPPTILFNLGSHTKYTSVETNAIISALRILLASHPTLHVTWKYQAADAYAEEHNGRSIAALPPRISSRITQIPWLSHTPLTLLSQPSTVLTVHHGGANSFHEALAAGVPQVICPRWLDTYEFARRAEWLGVGVMGNERAAPGFEDRELAEAMGKVLGGEAEEEKKSKGKSYRDIPEPPGGSFIYGHFRYMISMGNGEAEMKWQGVQEKNNDPHTPSLSPFPHHAVFEVGAWAARVALDIITLTAMGRDFGAIQNADSQLAVVYSRVVEPTLGHMLIAILRIYLPSCVVEALPIKSNHDQAAAMHTIRGLCRELLHEKKETRSDGKDILSTTSRRNQNQGSATGKARGGNAVGRDAGAGTGVCRGSSEILSPDFIDHARAV  
  
