## Supplementary Results for "A draft genome of the ascomycotal fungal species *Pseudopithomyces maydicus* (family *Didymosphaeriaceae*)": index.html

assembly - 39 region(s) - antiSMASH results


antiSMASH version 6.0.1

Download

- Download all results
- Download GenBank summary file
- Download log file

About

Help

Contact

Select genomic region:

Overview

1.1

2.1

2.2

2.3

4.1

4.2

4.3

4.4

5.1

6.1

7.1

7.2

7.3

9.1

9.2

12.1

14.1

17.1

17.2

19.1

21.1

22.1

22.2

23.1

23.2

23.3

26.1

29.1

29.2

29.3

30.1

30.2

30.3

30.4

34.1

34.2

34.3

36.1

36.2

### Identified secondary metabolite regions using strictness 'relaxed'

**contig\_1**

| Region | Type | From | To | Most similar known cluster | | Similarity |
| --- | --- | --- | --- | --- | --- | --- |
| Region&nbsp1.1 | T1PKS | 156,319 | 204,648 |  | | |

**contig\_10**

| Region | Type | From | To | Most similar known cluster | | Similarity |
| --- | --- | --- | --- | --- | --- | --- |
| Region&nbsp2.1 | T1PKS | 133,010 | 174,203 |  | | |
| Region&nbsp2.2 | T1PKS | 637,823 | 682,907 | alternariol | Terpene | 100% |
| Region&nbsp2.3 | NRPS-like | 864,201 | 909,865 |  | | |

**contig\_12**

| Region | Type | From | To | Most similar known cluster | | Similarity |
| --- | --- | --- | --- | --- | --- | --- |
| Region&nbsp4.1 | T1PKS,NRPS | 1,263,496 | 1,311,834 | burnettramic acid A | Alkaloid + Polyketide:Iterative type I | 33% |
| Region&nbsp4.2 | NRPS | 2,325,183 | 2,369,957 | dimethylcoprogen | NRP | 100% |
| Region&nbsp4.3 | NRPS-like | 2,479,385 | 2,523,240 |  | | |
| Region&nbsp4.4 | indole,T1PKS | 2,923,017 | 2,976,336 | secalonic acids | Polyketide | 50% |

**contig\_13**

| Region | Type | From | To | Most similar known cluster | | Similarity |
| --- | --- | --- | --- | --- | --- | --- |
| Region&nbsp5.1 | NRPS-like | 741,090 | 785,032 |  | | |

**contig\_14**

| Region | Type | From | To | Most similar known cluster | | Similarity |
| --- | --- | --- | --- | --- | --- | --- |
| Region&nbsp6.1 | T1PKS | 209,216 | 255,818 |  | | |

**contig\_15**

| Region | Type | From | To | Most similar known cluster | | Similarity |
| --- | --- | --- | --- | --- | --- | --- |
| Region&nbsp7.1 | T1PKS | 1,237,335 | 1,284,006 |  | | |
| Region&nbsp7.2 | NRPS-like | 2,560,252 | 2,603,454 |  | | |
| Region&nbsp7.3 | terpene | 2,691,940 | 2,713,034 |  | | |

**contig\_17**

| Region | Type | From | To | Most similar known cluster | | Similarity |
| --- | --- | --- | --- | --- | --- | --- |
| Region&nbsp9.1 | terpene | 148,217 | 169,902 |  | | |
| Region&nbsp9.2 | T1PKS | 3,156,185 | 3,202,827 | melanin | Polyketide | 100% |

**contig\_20**

| Region | Type | From | To | Most similar known cluster | | Similarity |
| --- | --- | --- | --- | --- | --- | --- |
| Region&nbsp12.1 | NRPS | 1 | 43,431 |  | | |

**contig\_23**

| Region | Type | From | To | Most similar known cluster | | Similarity |
| --- | --- | --- | --- | --- | --- | --- |
| Region&nbsp14.1 | NRPS | 1 | 24,537 |  | | |

**contig\_26**

| Region | Type | From | To | Most similar known cluster | | Similarity |
| --- | --- | --- | --- | --- | --- | --- |
| Region&nbsp17.1 | T1PKS | 23,888 | 65,396 |  | | |
| Region&nbsp17.2 | T1PKS | 736,919 | 783,489 | melanin | Polyketide | 100% |

**contig\_28**

| Region | Type | From | To | Most similar known cluster | | Similarity |
| --- | --- | --- | --- | --- | --- | --- |
| Region&nbsp19.1 | T1PKS | 5,917 | 54,518 |  | | |

**contig\_3**

| Region | Type | From | To | Most similar known cluster | | Similarity |
| --- | --- | --- | --- | --- | --- | --- |
| Region&nbsp21.1 | T1PKS | 486,327 | 531,102 |  | | |

**contig\_31**

| Region | Type | From | To | Most similar known cluster | | Similarity |
| --- | --- | --- | --- | --- | --- | --- |
| Region&nbsp22.1 | NRPS | 585,393 | 629,288 |  | | |
| Region&nbsp22.2 | T1PKS | 1,112,429 | 1,160,321 |  | | |

**contig\_32**

| Region | Type | From | To | Most similar known cluster | | Similarity |
| --- | --- | --- | --- | --- | --- | --- |
| Region&nbsp23.1 | NRPS-like | 618,090 | 663,360 |  | | |
| Region&nbsp23.2 | T1PKS | 1,790,500 | 1,832,854 | 1,3,6,8-tetrahydroxynaphthalene | Polyketide | 100% |
| Region&nbsp23.3 | T1PKS | 3,137,580 | 3,197,360 | chaetoviridin E / 11-epichaetomugilin A | Polyketide | 16% |

**contig\_35**

| Region | Type | From | To | Most similar known cluster | | Similarity |
| --- | --- | --- | --- | --- | --- | --- |
| Region&nbsp26.1 | NRPS,T1PKS | 26,846 | 89,893 |  | | |

**contig\_39**

| Region | Type | From | To | Most similar known cluster | | Similarity |
| --- | --- | --- | --- | --- | --- | --- |
| Region&nbsp29.1 | T1PKS | 34,391 | 73,215 |  | | |
| Region&nbsp29.2 | T1PKS | 1,044,010 | 1,083,080 |  | | |
| Region&nbsp29.3 | NRPS,T1PKS | 1,102,900 | 1,154,380 |  | | |

**contig\_4**

| Region | Type | From | To | Most similar known cluster | | Similarity |
| --- | --- | --- | --- | --- | --- | --- |
| Region&nbsp30.1 | NRPS | 354,414 | 392,004 | aspirochlorine | NRP | 13% |
| Region&nbsp30.2 | T1PKS | 1,279,350 | 1,324,830 | (-)-Mellein | Polyketide | 100% |
| Region&nbsp30.3 | NRPS | 1,830,601 | 1,896,090 |  | | |
| Region&nbsp30.4 | indole | 2,306,177 | 2,327,439 |  | | |

**contig\_6**

| Region | Type | From | To | Most similar known cluster | | Similarity |
| --- | --- | --- | --- | --- | --- | --- |
| Region&nbsp34.1 | T1PKS | 87,743 | 140,198 | radicicol | Polyketide | 60% |
| Region&nbsp34.2 | NRPS | 575,471 | 624,218 |  | | |
| Region&nbsp34.3 | T1PKS | 1,814,903 | 1,861,987 |  | | |

**contig\_9**

| Region | Type | From | To | Most similar known cluster | | Similarity |
| --- | --- | --- | --- | --- | --- | --- |
| Region&nbsp36.1 | terpene,NRPS | 1,341,734 | 1,419,185 |  | | |
| Region&nbsp36.2 | terpene | 1,785,179 | 1,806,398 | squalestatin S1 | Terpene | 40% |

| Region | Type | From | To | Most similar known cluster | | Similarity |
| --- | --- | --- | --- | --- | --- | --- |
| Region&nbsp1.1 | T1PKS | 156,319 | 204,648 |  | | |
| Region&nbsp2.1 | T1PKS | 133,010 | 174,203 |  | | |
| Region&nbsp2.2 | T1PKS | 637,823 | 682,907 | alternariol | Terpene | 100% |
| Region&nbsp2.3 | NRPS-like | 864,201 | 909,865 |  | | |
| Region&nbsp4.1 | T1PKS,NRPS | 1,263,496 | 1,311,834 | burnettramic acid A | Alkaloid + Polyketide:Iterative type I | 33% |
| Region&nbsp4.2 | NRPS | 2,325,183 | 2,369,957 | dimethylcoprogen | NRP | 100% |
| Region&nbsp4.3 | NRPS-like | 2,479,385 | 2,523,240 |  | | |
| Region&nbsp4.4 | indole,T1PKS | 2,923,017 | 2,976,336 | secalonic acids | Polyketide | 50% |
| Region&nbsp5.1 | NRPS-like | 741,090 | 785,032 |  | | |
| Region&nbsp6.1 | T1PKS | 209,216 | 255,818 |  | | |
| Region&nbsp7.1 | T1PKS | 1,237,335 | 1,284,006 |  | | |
| Region&nbsp7.2 | NRPS-like | 2,560,252 | 2,603,454 |  | | |
| Region&nbsp7.3 | terpene | 2,691,940 | 2,713,034 |  | | |
| Region&nbsp9.1 | terpene | 148,217 | 169,902 |  | | |
| Region&nbsp9.2 | T1PKS | 3,156,185 | 3,202,827 | melanin | Polyketide | 100% |
| Region&nbsp12.1 | NRPS | 1 | 43,431 |  | | |
| Region&nbsp14.1 | NRPS | 1 | 24,537 |  | | |
| Region&nbsp17.1 | T1PKS | 23,888 | 65,396 |  | | |
| Region&nbsp17.2 | T1PKS | 736,919 | 783,489 | melanin | Polyketide | 100% |
| Region&nbsp19.1 | T1PKS | 5,917 | 54,518 |  | | |
| Region&nbsp21.1 | T1PKS | 486,327 | 531,102 |  | | |
| Region&nbsp22.1 | NRPS | 585,393 | 629,288 |  | | |
| Region&nbsp22.2 | T1PKS | 1,112,429 | 1,160,321 |  | | |
| Region&nbsp23.1 | NRPS-like | 618,090 | 663,360 |  | | |
| Region&nbsp23.2 | T1PKS | 1,790,500 | 1,832,854 | 1,3,6,8-tetrahydroxynaphthalene | Polyketide | 100% |
| Region&nbsp23.3 | T1PKS | 3,137,580 | 3,197,360 | chaetoviridin E / 11-epichaetomugilin A | Polyketide | 16% |
| Region&nbsp26.1 | NRPS,T1PKS | 26,846 | 89,893 |  | | |
| Region&nbsp29.1 | T1PKS | 34,391 | 73,215 |  | | |
| Region&nbsp29.2 | T1PKS | 1,044,010 | 1,083,080 |  | | |
| Region&nbsp29.3 | NRPS,T1PKS | 1,102,900 | 1,154,380 |  | | |
| Region&nbsp30.1 | NRPS | 354,414 | 392,004 | aspirochlorine | NRP | 13% |
| Region&nbsp30.2 | T1PKS | 1,279,350 | 1,324,830 | (-)-Mellein | Polyketide | 100% |
| Region&nbsp30.3 | NRPS | 1,830,601 | 1,896,090 |  | | |
| Region&nbsp30.4 | indole | 2,306,177 | 2,327,439 |  | | |
| Region&nbsp34.1 | T1PKS | 87,743 | 140,198 | radicicol | Polyketide | 60% |
| Region&nbsp34.2 | NRPS | 575,471 | 624,218 |  | | |
| Region&nbsp34.3 | T1PKS | 1,814,903 | 1,861,987 |  | | |
| Region&nbsp36.1 | terpene,NRPS | 1,341,734 | 1,419,185 |  | | |
| Region&nbsp36.2 | terpene | 1,785,179 | 1,806,398 | squalestatin S1 | Terpene | 40% |

No secondary metabolite regions were found in these records:
:   **contig\_11**
:   **contig\_16**
:   **contig\_18**
:   **contig\_19**
:   **contig\_21**
:   **contig\_24**
:   **contig\_25**
:   **contig\_27**
:   **contig\_29**
:   **contig\_33**
:   **contig\_34**
:   **contig\_37**
:   **contig\_38**
:   **contig\_40**
:   **contig\_41**
:   **contig\_5**
:   **contig\_7**

Compact view

contig\_1 - Region 1 - T1PKS

Shows the layout of the region, marking coding sequences and areas of interest. Clicking a gene will select it and show any relevant details. Clicking an area feature (e.g. a candidate cluster) will select all coding sequences within that area. Double clicking an area feature will zoom to that area. Multiple genes and area features can be selected by clicking them while holding the Ctrl key.  
More detailed help is available here.

Download region GenBank file

Download region SVG

Location: 156,319 - 204,648 nt. (total: 48,330 nt)
Show pHMM detection rules used

T1PKS: cds(PKS\_AT and (PKS\_KS or ene\_KS or mod\_KS or hyb\_KS or itr\_KS or tra\_KS))

#### Legend:

core biosynthetic genes

additional biosynthetic genes

transport-related genes

regulatory genes

other genes

resistance

reset view

zoom to selection

Gene details

Shows details of the most recently selected gene, including names, products, location, and other annotations.

Select a gene to view the details available for it

NRPS/PKS domains

ClusterBlast

KnownClusterBlast

SubClusterBlast

MIBiG comparison

Pfam domains

Detailed domain annotation

Shows NRPS- and PKS-related domains for each feature that contains them. Click on each domain for more information about the domain's location, consensus monomer prediction, and other details.  
A glossary is available here.

Selected features only

Show module domains

Similar gene clusters

Shows clusters from the antiSMASH database and other clusters of interest that are similar to the current region. Genes marked with the same colour are interrelated. White genes have no relationship.  
Click on reference genes to show details of similarities to genes within the current region.  
Click on an accession to open that entry in the antiSMASH database (if applicable).

All hits

NC\_035795 (191603-237741): Pochonia chlamydosporia 170 chromosome 6, whole ge... (16% of genes show similarity), T1PKS

NW\_007360999 (1402989-1446170): Glarea lozoyensis ATCC 20868 chromosome Unkno... (11% of genes show similarity), T1PKS

NZ\_KL647090 (198809-315466): Streptomyces sp. NRRL S-623 Doro1 scaffold1, who... (3% of genes show similarity), NRPS,PKS-like,T1PKS,transAT-PKS

NZ\_CP054926 (233409-350060): Streptomyces fulvissimus strain NA06532 chromoso... (3% of genes show similarity), NRPS,PKS-like,T1PKS,transAT-PKS

NZ\_WWGV01000024 (105328-220093): Streptomyces sp. SID8356 SID8356.c24, whole ... (4% of genes show similarity), NRPS,PKS-like,T1PKS,transAT-PKS

NZ\_WWGS01000014 (126482-241157): Streptomyces sp. SID8359 SID8359.c14, whole ... (4% of genes show similarity), NRPS,PKS-like,T1PKS,transAT-PKS

NZ\_CP030930 (108461-226163): Streptomyces cavourensis strain TJ430 chromosome... (4% of genes show similarity), NRPS,PKS-like,T1PKS,transAT-PKS

NZ\_CP025018 (1027611-1151572): Streptomyces sp. M56 (3% of genes show similarity), NRPS,T1PKS,T3PKS,betalactone

NZ\_CP048289 (5996076-6064491): Streptacidiphilus sp. P02-A3a chromosome, comp... (5% of genes show similarity), T1PKS

NZ\_LJSN01000003 (3474570-3580413): Streptomyces noursei strain JCM 4701 scaff... (15% of genes show similarity), T1PKS
Download graphic

Similar known gene clusters

Shows clusters from the MiBIG database that are similar to the current region. Genes marked with the same colour are interrelated. White genes have no relationship.  
Click on reference genes to show details of similarities to genes within the current region.  
Click on an accession to open that entry in the MiBIG database.

No matches found.

Similar subclusters

Shows sub-cluster units that are similar to the current region. Genes marked with the same colour are interrelated. White genes have no relationship.  
Click on reference genes to show details of similarities to genes within the current region.

No matches found.

Similar gene clusters

Shows careas that are similar to the current region to a reference database.  
Mouseover a score cell in the table to get a breakdown of how the score was calculated.The MIBiG database.  
  
Click on an accession to open that entry in the MIBiG database.

Analysis type:

Protocluster to Region
Region to Region

| Reference | T1PKS | Similarity score | Type | Compound(s) | Organism |
| --- | --- | --- | --- | --- | --- |
| BGC0001858.1 |  | 0.32 | Polyketide | alternapyrone B, alternapyrone C, alternapyrone D, alternapyrone E, alternapyrone F | Parastagonospora nodorum SN15 |
| BGC0000688.1 |  | 0.30 | Terpene | copalyl diphosphate | Diaporthe amygdali |
| BGC0001280.1 |  | 0.27 | Polyketide | betaenone C, betaenone A | Phoma betae |
| BGC0001264.1 |  | 0.27 | Polyketide | betaenone A, betaenone B, betaenone C | Phoma betae |
| BGC0000685.1 |  | 0.26 | Terpene | brassicicene C | Alternaria brassicicola ATCC 96836 |
| BGC0001141.1 |  | 0.25 | Polyketide | 4-epi-15-epi-brefeldin A | Penicillium brefeldianum |
| BGC0000003.1 |  | 0.24 | Polyketide | AF-toxin | Alternaria alternata |
| BGC0001995.1 |  | 0.24 | Terpene | heptelidic acid | Aspergillus oryzae RIB40 |
| BGC0001909.1 |  | 0.23 | Polyketide | strobilurin | Strobilurus tenacellus |
| BGC0000811.1 |  | 0.22 | Alkaloid | fumigaclavine C | Aspergillus fumigatus Af293 |

| Reference | Aggregated | Similarity score | Type | Compound(s) | Organism |
| --- | --- | --- | --- | --- | --- |
| BGC0001858.1 |  | 0.65 | Polyketide | alternapyrone B, alternapyrone C, alternapyrone D, alternapyrone E, alternapyrone F | Parastagonospora nodorum SN15 |
| BGC0000688.1 |  | 0.63 | Terpene | copalyl diphosphate | Diaporthe amygdali |
| BGC0001280.1 |  | 0.61 | Polyketide | betaenone C, betaenone A | Phoma betae |
| BGC0001264.1 |  | 0.61 | Polyketide | betaenone A, betaenone B, betaenone C | Phoma betae |
| BGC0000685.1 |  | 0.60 | Terpene | brassicicene C | Alternaria brassicicola ATCC 96836 |
| BGC0001141.1 |  | 0.59 | Polyketide | 4-epi-15-epi-brefeldin A | Penicillium brefeldianum |
| BGC0000003.1 |  | 0.57 | Polyketide | AF-toxin | Alternaria alternata |
| BGC0001995.1 |  | 0.57 | Terpene | heptelidic acid | Aspergillus oryzae RIB40 |
| BGC0001909.1 |  | 0.56 | Polyketide | strobilurin | Strobilurus tenacellus |
| BGC0000811.1 |  | 0.55 | Alkaloid | fumigaclavine C | Aspergillus fumigatus Af293 |

Detailed Pfam domain annotation

Shows Pfam domains found in each gene within the region.
Click on each domain for more information about the domain's
accession, location, description, and any relevant Gene Ontology.
Domains with a bold border have Gene Ontology information.

Selected features only

NRPS/PKS products

NRPS/PKS monomers

Predicted core structure(s)

Shows estimated product structure and polymer for each candidate cluster in the region. To show the product, click on the expander or the candidate cluster feature drawn in the overview.

For candidate cluster 1, location 156318 - 204648:

Rough prediction of core scaffold based on assumed PKS/NRPS colinearity; tailoring reactions not taken into account

**Polymer prediction:**
:   (pk)

  
Direct lookup in NORINE database:
strict
or
relaxed

Link to NORINE database query form

NRPS/PKS monomer predictions

Shows the predicted monomers for each adynelation domain and acyltransferase within genes. Each gene prediction can be expanded to view detailed predictions of each domain. Each prediction can be expanded to view the predictions by tool (and, for some tools, further expanded for extra details).

**input.path1.gene39**: pk

:   **PKS\_AT (497..793)**: pk

    ATSignature: Malonyl-CoA

    Top 3 matches:
    :   Malonyl-CoA: 79.2%
    :   Methylmalonyl-CoA: 62.5%
    :   2-Methylbutyryl-CoA: 58.3%

      
    minowa: Methylmalonyl-CoA

    Prediction, score:
    :   Methylmalonyl-CoA: 84.5


        Methoxymalonyl-CoA: 78.1


        Malonyl-CoA: 51.3


        Ethylmalonyl-CoA: 46.0


        Isobutyryl-CoA: 38.1


        Propionyl-CoA: 35.7


        Benzoyl-CoA: 34.1


        2-Methylbutyryl-CoA: 26.5


        fatty\_acid: 25.5


        inactive: 21.8


        trans-1,2-CPDA: 17.8


        CHC-CoA: 13.5


        3-Methylbutyryl-CoA: 11.8


        Acetyl-CoA: 0.0

contig\_10 - Region 1 - T1PKS

Shows the layout of the region, marking coding sequences and areas of interest. Clicking a gene will select it and show any relevant details. Clicking an area feature (e.g. a candidate cluster) will select all coding sequences within that area. Double clicking an area feature will zoom to that area. Multiple genes and area features can be selected by clicking them while holding the Ctrl key.  
More detailed help is available here.

Download region GenBank file

Download region SVG

Location: 133,010 - 174,203 nt. (total: 41,194 nt)
Show pHMM detection rules used

T1PKS: cds(PKS\_AT and (PKS\_KS or ene\_KS or mod\_KS or hyb\_KS or itr\_KS or tra\_KS))

#### Legend:

core biosynthetic genes

additional biosynthetic genes

transport-related genes

regulatory genes

other genes

resistance

reset view

zoom to selection

Gene details

Shows details of the most recently selected gene, including names, products, location, and other annotations.

Select a gene to view the details available for it

NRPS/PKS domains

ClusterBlast

KnownClusterBlast

SubClusterBlast

MIBiG comparison

Pfam domains

Detailed domain annotation

Shows NRPS- and PKS-related domains for each feature that contains them. Click on each domain for more information about the domain's location, consensus monomer prediction, and other details.  
A glossary is available here.

Selected features only

Show module domains

Similar gene clusters

Shows clusters from the antiSMASH database and other clusters of interest that are similar to the current region. Genes marked with the same colour are interrelated. White genes have no relationship.  
Click on reference genes to show details of similarities to genes within the current region.  
Click on an accession to open that entry in the antiSMASH database (if applicable).

All hits

NW\_001820829 (166655-214112): Sclerotinia sclerotiorum 1980 UF-70 scaffold 7 ... (40% of genes show similarity), T1PKS

NW\_009276941 (357077-408070): Verticillium dahliae VdLs.17 supercont1.26 geno... (21% of genes show similarity), NRPS-like,T1PKS

NW\_022984633 (567506-613385): Aspergillus tanneri strain NIH1004 chromosome U... (18% of genes show similarity), T1PKS

NC\_030962 (3003402-3051438): Colletotrichum higginsianum IMI 349063 chromosom... (20% of genes show similarity), T1PKS,indole

CM000595 (183841-220794): Fusarium oxysporum f. sp. lycopersici 4287 chromoso... (18% of genes show similarity), T1PKS

NC\_030992 (183841-220794): Fusarium oxysporum f. sp. lycopersici 4287 chromos... (18% of genes show similarity), T1PKS

NZ\_JAAALO010000001 (810907-919842): Micromonospora sp. NEAU-HG-1 Scaffold1, w... (11% of genes show similarity), NRPS,T1PKS,bacteriocin,bottromycin,transAT-PKS,transAT-PKS-like

NW\_003315103 (395199-442024): Trichophyton benhamiae CBS 112371 chromosome Un... (10% of genes show similarity), T1PKS

NW\_003456426 (342748-389572): Trichophyton rubrum CBS 118892 genomic scaffold... (9% of genes show similarity), T1PKS

NW\_003345192 (200075-244874): Nannizzia gypsea CBS 118893 supercont1.10 genom... (10% of genes show similarity), T1PKS
Download graphic

Similar known gene clusters

Shows clusters from the MiBIG database that are similar to the current region. Genes marked with the same colour are interrelated. White genes have no relationship.  
Click on reference genes to show details of similarities to genes within the current region.  
Click on an accession to open that entry in the MiBIG database.

No matches found.

Similar subclusters

Shows sub-cluster units that are similar to the current region. Genes marked with the same colour are interrelated. White genes have no relationship.  
Click on reference genes to show details of similarities to genes within the current region.

No matches found.

Similar gene clusters

Shows careas that are similar to the current region to a reference database.  
Mouseover a score cell in the table to get a breakdown of how the score was calculated.The MIBiG database.  
  
Click on an accession to open that entry in the MIBiG database.

Analysis type:

Protocluster to Region
Region to Region

| Reference | T1PKS | Similarity score | Type | Compound(s) | Organism |
| --- | --- | --- | --- | --- | --- |
| BGC0001068.1 |  | 0.25 | Terpene, Polyketide | pyripyropene A | unidentified unclassified sequences. |
| BGC0000056.1 |  | 0.23 | Polyketide | esperamicin | Actinomadura verrucosospora |
| BGC0001858.1 |  | 0.22 | Polyketide | alternapyrone B, alternapyrone C, alternapyrone D, alternapyrone E, alternapyrone F | Parastagonospora nodorum SN15 |
| BGC0000046.1 |  | 0.19 | Polyketide | depudecin | Alternaria brassicicola |
| BGC0000107.1 |  | 0.19 | Polyketide | naphthopyrone | Aspergillus nidulans FGSC A4 |
| BGC0001265.1 |  | 0.19 | Polyketide | melanin | Bipolaris oryzae |
| BGC0001304.1 |  | 0.19 | Polyketide | aflavarin | Aspergillus flavus NRRL3357 |
| BGC0001284.1 |  | 0.18 | Terpene | alternariol | Parastagonospora nodorum SN15 |
| BGC0001257.1 |  | 0.18 | Polyketide | 1,3,6,8-tetrahydroxynaphthalene | Nodulisporium sp. ATCC74245 |
| BGC0000156.1 |  | 0.18 | Polyketide | TAN-1612, 1-(2,3,5,10-tetrahydroxy-7-methoxy-4-oxo-1,2,3,4-tetrahydroanthracen-2-yl)pentane-2,4-dione, desmethyl TAN-1612 | Aspergillus niger |

| Reference | Aggregated | Similarity score | Type | Compound(s) | Organism |
| --- | --- | --- | --- | --- | --- |
| BGC0001068.1 |  | 0.58 | Terpene, Polyketide | pyripyropene A | unidentified unclassified sequences. |
| BGC0000056.1 |  | 0.57 | Polyketide | esperamicin | Actinomadura verrucosospora |
| BGC0001858.1 |  | 0.55 | Polyketide | alternapyrone B, alternapyrone C, alternapyrone D, alternapyrone E, alternapyrone F | Parastagonospora nodorum SN15 |
| BGC0000046.1 |  | 0.51 | Polyketide | depudecin | Alternaria brassicicola |
| BGC0000107.1 |  | 0.51 | Polyketide | naphthopyrone | Aspergillus nidulans FGSC A4 |
| BGC0001265.1 |  | 0.50 | Polyketide | melanin | Bipolaris oryzae |
| BGC0001304.1 |  | 0.50 | Polyketide | aflavarin | Aspergillus flavus NRRL3357 |
| BGC0001284.1 |  | 0.50 | Terpene | alternariol | Parastagonospora nodorum SN15 |
| BGC0001257.1 |  | 0.50 | Polyketide | 1,3,6,8-tetrahydroxynaphthalene | Nodulisporium sp. ATCC74245 |
| BGC0000156.1 |  | 0.50 | Polyketide | TAN-1612, 1-(2,3,5,10-tetrahydroxy-7-methoxy-4-oxo-1,2,3,4-tetrahydroanthracen-2-yl)pentane-2,4-dione, desmethyl TAN-1612 | Aspergillus niger |

Detailed Pfam domain annotation

Shows Pfam domains found in each gene within the region.
Click on each domain for more information about the domain's
accession, location, description, and any relevant Gene Ontology.
Domains with a bold border have Gene Ontology information.

Selected features only

contig\_10 - Region 2 - T1PKS

Shows the layout of the region, marking coding sequences and areas of interest. Clicking a gene will select it and show any relevant details. Clicking an area feature (e.g. a candidate cluster) will select all coding sequences within that area. Double clicking an area feature will zoom to that area. Multiple genes and area features can be selected by clicking them while holding the Ctrl key.  
More detailed help is available here.

Download region GenBank file

Download region SVG

Location: 637,823 - 682,907 nt. (total: 45,085 nt)
Show pHMM detection rules used

T1PKS: cds(PKS\_AT and (PKS\_KS or ene\_KS or mod\_KS or hyb\_KS or itr\_KS or tra\_KS))

#### Legend:

core biosynthetic genes

additional biosynthetic genes

transport-related genes

regulatory genes

other genes

resistance

reset view

zoom to selection

Gene details

Shows details of the most recently selected gene, including names, products, location, and other annotations.

Select a gene to view the details available for it

NRPS/PKS domains

ClusterBlast

KnownClusterBlast

SubClusterBlast

MIBiG comparison

Pfam domains

Detailed domain annotation

Shows NRPS- and PKS-related domains for each feature that contains them. Click on each domain for more information about the domain's location, consensus monomer prediction, and other details.  
A glossary is available here.

Selected features only

Show module domains

Similar gene clusters

Shows clusters from the antiSMASH database and other clusters of interest that are similar to the current region. Genes marked with the same colour are interrelated. White genes have no relationship.  
Click on reference genes to show details of similarities to genes within the current region.  
Click on an accession to open that entry in the antiSMASH database (if applicable).

All hits

NW\_022984631 (964303-1009823): Aspergillus tanneri strain NIH1004 chromosome ... (18% of genes show similarity), T1PKS,indole

NC\_007200 (16016-61589): Aspergillus fumigatus Af293 chromosome 7, whole geno... (20% of genes show similarity), T1PKS,indole

NW\_003299166 (1162703-1201813): Microsporum canis CBS 113480 supercont1.4 gen... (11% of genes show similarity), T1PKS

CM000595 (183841-220794): Fusarium oxysporum f. sp. lycopersici 4287 chromoso... (18% of genes show similarity), T1PKS

NC\_030992 (183841-220794): Fusarium oxysporum f. sp. lycopersici 4287 chromos... (18% of genes show similarity), T1PKS

NZ\_CP030864 (11545-320938): Streptomyces globosus strain LZH-48 plasmid unnam... (2% of genes show similarity), NRPS,NRPS-like,T2PKS,butyrolactone,ectoine,hglE-KS,linaridin,transAT-PKS

NW\_017971433 (261493-301685): Talaromyces atroroseus strain IBT 11181 chromos... (16% of genes show similarity), T1PKS

NW\_022984634 (4255890-4301503): Aspergillus tanneri strain NIH1004 chromosome... (10% of genes show similarity), T1PKS

NC\_007197 (3816930-3862461): Aspergillus fumigatus Af293 chromosome 4, whole ... (9% of genes show similarity), T1PKS

NW\_019154043 (172137-217646): Pochonia chlamydosporia 170 chromosome Unknown ... (12% of genes show similarity), T1PKS
Download graphic

Similar known gene clusters

Shows clusters from the MiBIG database that are similar to the current region. Genes marked with the same colour are interrelated. White genes have no relationship.  
Click on reference genes to show details of similarities to genes within the current region.  
Click on an accession to open that entry in the MiBIG database.

All hits

alternariol
Download graphic

Similar subclusters

Shows sub-cluster units that are similar to the current region. Genes marked with the same colour are interrelated. White genes have no relationship.  
Click on reference genes to show details of similarities to genes within the current region.

No matches found.

Similar gene clusters

Shows careas that are similar to the current region to a reference database.  
Mouseover a score cell in the table to get a breakdown of how the score was calculated.The MIBiG database.  
  
Click on an accession to open that entry in the MIBiG database.

Analysis type:

Protocluster to Region
Region to Region

| Reference | T1PKS | Similarity score | Type | Compound(s) | Organism |
| --- | --- | --- | --- | --- | --- |
| BGC0000013.1 |  | 0.34 | Polyketide | alternariol | Aspergillus nidulans FGSC A4 |
| BGC0000161.1 |  | 0.31 | Polyketide | isoterrein | Aspergillus terreus NIH2624 |
| BGC0001144.1 |  | 0.27 | Polyketide | neosartoricin B | Trichophyton tonsurans CBS 112818 |
| BGC0001541.1 |  | 0.27 | Polyketide | cercosporin | Cercospora beticola |
| BGC0000048.1 |  | 0.25 | Polyketide | dothistromin | Dothistroma septosporum |
| BGC0001284.1 |  | 0.24 | Terpene | alternariol | Parastagonospora nodorum SN15 |
| BGC0001542.1 |  | 0.23 | Polyketide | cercosporin | Cercospora zeina |
| BGC0001583.1 |  | 0.22 | Polyketide | emodin | Escovopsis weberi |
| BGC0001258.1 |  | 0.22 | Polyketide | 1,3,6,8-tetrahydroxynaphthalene | Glarea lozoyensis |
| BGC0001265.1 |  | 0.22 | Polyketide | melanin | Bipolaris oryzae |

| Reference | Aggregated | Similarity score | Type | Compound(s) | Organism |
| --- | --- | --- | --- | --- | --- |
| BGC0000013.1 |  | 0.67 | Polyketide | alternariol | Aspergillus nidulans FGSC A4 |
| BGC0000161.1 |  | 0.64 | Polyketide | isoterrein | Aspergillus terreus NIH2624 |
| BGC0001144.1 |  | 0.61 | Polyketide | neosartoricin B | Trichophyton tonsurans CBS 112818 |
| BGC0001541.1 |  | 0.61 | Polyketide | cercosporin | Cercospora beticola |
| BGC0000048.1 |  | 0.59 | Polyketide | dothistromin | Dothistroma septosporum |
| BGC0001284.1 |  | 0.58 | Terpene | alternariol | Parastagonospora nodorum SN15 |
| BGC0001542.1 |  | 0.56 | Polyketide | cercosporin | Cercospora zeina |
| BGC0001583.1 |  | 0.55 | Polyketide | emodin | Escovopsis weberi |
| BGC0001258.1 |  | 0.55 | Polyketide | 1,3,6,8-tetrahydroxynaphthalene | Glarea lozoyensis |
| BGC0001265.1 |  | 0.55 | Polyketide | melanin | Bipolaris oryzae |

Detailed Pfam domain annotation

Shows Pfam domains found in each gene within the region.
Click on each domain for more information about the domain's
accession, location, description, and any relevant Gene Ontology.
Domains with a bold border have Gene Ontology information.

Selected features only

NRPS/PKS products

NRPS/PKS monomers

Predicted core structure(s)

Shows estimated product structure and polymer for each candidate cluster in the region. To show the product, click on the expander or the candidate cluster feature drawn in the overview.

For candidate cluster 2, location 637822 - 682907:

Rough prediction of core scaffold based on assumed PKS/NRPS colinearity; tailoring reactions not taken into account

**Polymer prediction:**
:   (mal)

  
Direct lookup in NORINE database:
strict
or
relaxed

Link to NORINE database query form

NRPS/PKS monomer predictions

Shows the predicted monomers for each adynelation domain and acyltransferase within genes. Each gene prediction can be expanded to view detailed predictions of each domain. Each prediction can be expanded to view the predictions by tool (and, for some tools, further expanded for extra details).

**input.path1.gene178**: mal

:   **PKS\_AT (517..782)**: mal

    ATSignature: Malonyl-CoA

    Top 3 matches:
    :   Malonyl-CoA: 58.3%
    :   inactive: 54.2%

      
    minowa: inactive

    Prediction, score:
    :   inactive: 51.4


        Malonyl-CoA: 48.4


        Methoxymalonyl-CoA: 28.3


        Benzoyl-CoA: 18.4


        Methylmalonyl-CoA: 16.5


        Ethylmalonyl-CoA: 10.0


        Propionyl-CoA: 9.3


        2-Methylbutyryl-CoA: 7.1


        trans-1,2-CPDA: 0.0


        fatty\_acid: 0.0


        Isobutyryl-CoA: 0.0


        CHC-CoA: 0.0


        Acetyl-CoA: 0.0


        3-Methylbutyryl-CoA: 0.0

contig\_10 - Region 3 - NRPS-like

Shows the layout of the region, marking coding sequences and areas of interest. Clicking a gene will select it and show any relevant details. Clicking an area feature (e.g. a candidate cluster) will select all coding sequences within that area. Double clicking an area feature will zoom to that area. Multiple genes and area features can be selected by clicking them while holding the Ctrl key.  
More detailed help is available here.

Download region GenBank file

Download region SVG

Location: 864,201 - 909,865 nt. (total: 45,665 nt)
Show pHMM detection rules used

NRPS-like: cds((PP-binding or NAD\_binding\_4) and (AMP-binding or A-OX))

#### Legend:

core biosynthetic genes

additional biosynthetic genes

transport-related genes

regulatory genes

other genes

resistance

reset view

zoom to selection

Gene details

Shows details of the most recently selected gene, including names, products, location, and other annotations.

Select a gene to view the details available for it

NRPS/PKS domains

ClusterBlast

KnownClusterBlast

SubClusterBlast

MIBiG comparison

Pfam domains

Detailed domain annotation

Shows NRPS- and PKS-related domains for each feature that contains them. Click on each domain for more information about the domain's location, consensus monomer prediction, and other details.  
A glossary is available here.

Selected features only

Show module domains

Similar gene clusters

Shows clusters from the antiSMASH database and other clusters of interest that are similar to the current region. Genes marked with the same colour are interrelated. White genes have no relationship.  
Click on reference genes to show details of similarities to genes within the current region.  
Click on an accession to open that entry in the antiSMASH database (if applicable).

All hits

NW\_017264204 (354553-412140): Xylona heveae TC161 unplaced genomic scaffold L... (28% of genes show similarity), NRPS

CP051139 (1539386-1594036): Peltaster fructicola strain LNHT1506 chromosome 1 (25% of genes show similarity), NRPS

NW\_022474218 (481115-535740): Venustampulla echinocandica strain BP 5553 chro... (21% of genes show similarity), NRPS

CM000590 (2613642-2668705): Fusarium oxysporum f. sp. lycopersici 4287 chromo... (33% of genes show similarity), NRPS

NC\_030987 (2613642-2668705): Fusarium oxysporum f. sp. lycopersici 4287 chrom... (33% of genes show similarity), NRPS

NW\_001939252 (607119-653701): Pyrenophora tritici-repentis Pt-1C-BFP supercon... (30% of genes show similarity), NRPS

NW\_019716256 (1521806-1576138): Ramularia collo-cygni strain URUG2 genome ass... (23% of genes show similarity), NRPS

NW\_020194477 (737075-791773): Amorphotheca resinae ATCC 22711 unplaced genomi... (25% of genes show similarity), NRPS

NZ\_CP046621 (2054103-2144309): Pseudomonas alkylphenolica strain Neo chromoso... (12% of genes show similarity), NRPS

NZ\_VNJK01000001 (2614574-2704845): Paenibacillus sp. N4 contig1, whole genome... (8% of genes show similarity), NRPS
Download graphic

Similar known gene clusters

Shows clusters from the MiBIG database that are similar to the current region. Genes marked with the same colour are interrelated. White genes have no relationship.  
Click on reference genes to show details of similarities to genes within the current region.  
Click on an accession to open that entry in the MiBIG database.

No matches found.

Similar subclusters

Shows sub-cluster units that are similar to the current region. Genes marked with the same colour are interrelated. White genes have no relationship.  
Click on reference genes to show details of similarities to genes within the current region.

No matches found.

Similar gene clusters

Shows careas that are similar to the current region to a reference database.  
Mouseover a score cell in the table to get a breakdown of how the score was calculated.The MIBiG database.  
  
Click on an accession to open that entry in the MIBiG database.

Analysis type:

Protocluster to Region
Region to Region

| Reference | NRPS-like | NRPS-like | Similarity score | Type | Compound(s) | Organism |
| --- | --- | --- | --- | --- | --- | --- |
| BGC0000426.1 |  |  | 0.63 | NRP | sevadicin | Paenibacillus larvae |
| BGC0000375.1 |  |  | 0.47 | NRP | indigoidine | Streptomyces chromofuscus |
| BGC0001806.1 |  |  | 0.45 | NRP | tolaasin A | Pseudomonas tolaasii |
| BGC0001641.1 |  |  | 0.45 | NRP | kolossin | Photorhabdus laumondii subsp. laumondii TTO1 |
| BGC0001132.1 |  |  | 0.44 | NRP | xenotetrapeptide | Xenorhabdus nematophila ATCC 19061 |
| BGC0001825.1 |  |  | 0.44 | NRP | xenematide | Xenorhabdus nematophila AN6/1 |
| BGC0002075.1 |  |  | 0.43 | NRP, Alkaloid | Pyreudione A, Pyreudione B, Pyreudione C, Pyreudione D, Pyreudione E | Pseudomonas fluorescens |
| BGC0001128.1 |  |  | 0.43 | NRP | luminmide | Photorhabdus laumondii subsp. laumondii TTO1 |
| BGC0001527.1 |  |  | 0.43 | Other | basidioferrin | Gelatoporia subvermispora |
| BGC0001636.1 |  |  | 0.42 | NRP | KK-1 | Curvularia clavata |

| Reference | Aggregated | Similarity score | Type | Compound(s) | Organism |
| --- | --- | --- | --- | --- | --- |
| BGC0000426.1 |  | 0.65 | NRP | sevadicin | Paenibacillus larvae |
| BGC0000375.1 |  | 0.57 | NRP | indigoidine | Streptomyces chromofuscus |
| BGC0001806.1 |  | 0.56 | NRP | tolaasin A | Pseudomonas tolaasii |
| BGC0001641.1 |  | 0.55 | NRP | kolossin | Photorhabdus laumondii subsp. laumondii TTO1 |
| BGC0001132.1 |  | 0.55 | NRP | xenotetrapeptide | Xenorhabdus nematophila ATCC 19061 |
| BGC0001825.1 |  | 0.55 | NRP | xenematide | Xenorhabdus nematophila AN6/1 |
| BGC0002075.1 |  | 0.54 | NRP, Alkaloid | Pyreudione A, Pyreudione B, Pyreudione C, Pyreudione D, Pyreudione E | Pseudomonas fluorescens |
| BGC0001128.1 |  | 0.54 | NRP | luminmide | Photorhabdus laumondii subsp. laumondii TTO1 |
| BGC0001527.1 |  | 0.54 | Other | basidioferrin | Gelatoporia subvermispora |
| BGC0001636.1 |  | 0.54 | NRP | KK-1 | Curvularia clavata |

Detailed Pfam domain annotation

Shows Pfam domains found in each gene within the region.
Click on each domain for more information about the domain's
accession, location, description, and any relevant Gene Ontology.
Domains with a bold border have Gene Ontology information.

Selected features only

contig\_12 - Region 1 - NRPS,T1PKS

Shows the layout of the region, marking coding sequences and areas of interest. Clicking a gene will select it and show any relevant details. Clicking an area feature (e.g. a candidate cluster) will select all coding sequences within that area. Double clicking an area feature will zoom to that area. Multiple genes and area features can be selected by clicking them while holding the Ctrl key.  
More detailed help is available here.

Download region GenBank file

Download region SVG

Location: 1,263,496 - 1,311,834 nt. (total: 48,339 nt)
Show pHMM detection rules used

T1PKS: cds(PKS\_AT and (PKS\_KS or ene\_KS or mod\_KS or hyb\_KS or itr\_KS or tra\_KS))  
NRPS: cds(Condensation and (AMP-binding or A-OX))

#### Legend:

core biosynthetic genes

additional biosynthetic genes

transport-related genes

regulatory genes

other genes

resistance

reset view

zoom to selection

Gene details

Shows details of the most recently selected gene, including names, products, location, and other annotations.

Select a gene to view the details available for it

NRPS/PKS domains

ClusterBlast

KnownClusterBlast

SubClusterBlast

MIBiG comparison

Pfam domains

Detailed domain annotation

Shows NRPS- and PKS-related domains for each feature that contains them. Click on each domain for more information about the domain's location, consensus monomer prediction, and other details.  
A glossary is available here.

Selected features only

Show module domains

Similar gene clusters

Shows clusters from the antiSMASH database and other clusters of interest that are similar to the current region. Genes marked with the same colour are interrelated. White genes have no relationship.  
Click on reference genes to show details of similarities to genes within the current region.  
Click on an accession to open that entry in the antiSMASH database (if applicable).

All hits

NW\_022984629 (4004137-4052179): Aspergillus tanneri strain NIH1004 chromosome... (21% of genes show similarity), NRPS,T1PKS

NW\_022474207 (2348451-2400505): Venustampulla echinocandica strain BP 5553 ch... (20% of genes show similarity), NRPS,T1PKS

NW\_007360993 (574614-627444): Glarea lozoyensis ATCC 20868 chromosome Unknown... (20% of genes show similarity), NRPS,T1PKS

NW\_022983866 (1121166-1165427): Arthroderma uncinatum strain CBS 119779 chrom... (21% of genes show similarity), T1PKS

NW\_006917091 (912491-957103): Pestalotiopsis fici W106-1 unplaced genomic sca... (21% of genes show similarity), T1PKS

NW\_014574692 (312996-360813): Metarhizium brunneum ARSEF 3297 chromosome Unkn... (18% of genes show similarity), T1PKS

NW\_011942153 (260108-307925): Metarhizium robertsii ARSEF 23 MAA Scf 13, whol... (18% of genes show similarity), T1PKS

NC\_036436 (621182-751686): Aspergillus oryzae RIB40 DNA, chromosome 2 (7% of genes show similarity), NRPS,T1PKS

NW\_006917095 (2028347-2112797): Pestalotiopsis fici W106-1 unplaced genomic s... (12% of genes show similarity), T1PKS

NW\_022474213 (921678-1013392): Venustampulla echinocandica strain BP 5553 chr... (10% of genes show similarity), NRPS,T1PKS
Download graphic

Similar known gene clusters

Shows clusters from the MiBIG database that are similar to the current region. Genes marked with the same colour are interrelated. White genes have no relationship.  
Click on reference genes to show details of similarities to genes within the current region.  
Click on an accession to open that entry in the MiBIG database.

All hits

burnettramic acid A

burnettramic acid
Download graphic

Similar subclusters

Shows sub-cluster units that are similar to the current region. Genes marked with the same colour are interrelated. White genes have no relationship.  
Click on reference genes to show details of similarities to genes within the current region.

No matches found.

Similar gene clusters

Shows careas that are similar to the current region to a reference database.  
Mouseover a score cell in the table to get a breakdown of how the score was calculated.The MIBiG database.  
  
Click on an accession to open that entry in the MIBiG database.

Analysis type:

Protocluster to Region
Region to Region

| Reference | NRPS | T1PKS | Similarity score | Type | Compound(s) | Organism |
| --- | --- | --- | --- | --- | --- | --- |
| BGC0001068.1 |  |  | 0.60 | Terpene, Polyketide | pyripyropene A | unidentified unclassified sequences. |
| BGC0001858.1 |  |  | 0.48 | Polyketide | alternapyrone B, alternapyrone C, alternapyrone D, alternapyrone E, alternapyrone F | Parastagonospora nodorum SN15 |
| BGC0001280.1 |  |  | 0.46 | Polyketide | betaenone C, betaenone A | Phoma betae |
| BGC0001264.1 |  |  | 0.46 | Polyketide | betaenone A, betaenone B, betaenone C | Phoma betae |
| BGC0000003.1 |  |  | 0.40 | Polyketide | AF-toxin | Alternaria alternata |
| BGC0000156.1 |  |  | 0.40 | Polyketide | TAN-1612, 1-(2,3,5,10-tetrahydroxy-7-methoxy-4-oxo-1,2,3,4-tetrahydroanthracen-2-yl)pentane-2,4-dione, desmethyl TAN-1612 | Aspergillus niger |
| BGC0001254.1 |  |  | 0.39 | Polyketide | ACT-Toxin II | Alternaria alternata |
| BGC0000064.1 |  |  | 0.38 | Polyketide | fusarin | Fusarium verticillioides |
| BGC0001268.1 |  |  | 0.38 | NRP, Polyketide | fusarin | Fusarium fujikuroi |
| BGC0001124.1 |  |  | 0.38 | Polyketide | pyranonigrin E | Aspergillus niger ATCC 1015 |

| Reference | Aggregated | Similarity score | Type | Compound(s) | Organism |
| --- | --- | --- | --- | --- | --- |
| BGC0001068.1 |  | 0.63 | Terpene, Polyketide | pyripyropene A | unidentified unclassified sequences. |
| BGC0001858.1 |  | 0.58 | Polyketide | alternapyrone B, alternapyrone C, alternapyrone D, alternapyrone E, alternapyrone F | Parastagonospora nodorum SN15 |
| BGC0001280.1 |  | 0.56 | Polyketide | betaenone C, betaenone A | Phoma betae |
| BGC0001264.1 |  | 0.56 | Polyketide | betaenone A, betaenone B, betaenone C | Phoma betae |
| BGC0000003.1 |  | 0.53 | Polyketide | AF-toxin | Alternaria alternata |
| BGC0000156.1 |  | 0.52 | Polyketide | TAN-1612, 1-(2,3,5,10-tetrahydroxy-7-methoxy-4-oxo-1,2,3,4-tetrahydroanthracen-2-yl)pentane-2,4-dione, desmethyl TAN-1612 | Aspergillus niger |
| BGC0001254.1 |  | 0.52 | Polyketide | ACT-Toxin II | Alternaria alternata |
| BGC0000064.1 |  | 0.51 | Polyketide | fusarin | Fusarium verticillioides |
| BGC0001268.1 |  | 0.51 | NRP, Polyketide | fusarin | Fusarium fujikuroi |
| BGC0001124.1 |  | 0.51 | Polyketide | pyranonigrin E | Aspergillus niger ATCC 1015 |

Detailed Pfam domain annotation

Shows Pfam domains found in each gene within the region.
Click on each domain for more information about the domain's
accession, location, description, and any relevant Gene Ontology.
Domains with a bold border have Gene Ontology information.

Selected features only

NRPS/PKS products

NRPS/PKS monomers

Predicted core structure(s)

Shows estimated product structure and polymer for each candidate cluster in the region. To show the product, click on the expander or the candidate cluster feature drawn in the overview.

For candidate cluster 1, location 1263495 - 1311834:

Rough prediction of core scaffold based on assumed PKS/NRPS colinearity; tailoring reactions not taken into account

**Polymer prediction:**
:   (pk)

  
Direct lookup in NORINE database:
strict
or
relaxed

Link to NORINE database query form

NRPS/PKS monomer predictions

Shows the predicted monomers for each adynelation domain and acyltransferase within genes. Each gene prediction can be expanded to view detailed predictions of each domain. Each prediction can be expanded to view the predictions by tool (and, for some tools, further expanded for extra details).

**input.path1.gene392**: pk

:   **PKS\_AT (366..684)**: pk

    ATSignature: Malonyl-CoA

    Top 3 matches:
    :   Malonyl-CoA: 79.2%
    :   Ethylmalonyl-CoA: 62.5%
    :   Methylmalonyl-CoA: 62.5%

      
    minowa: Methylmalonyl-CoA

    Prediction, score:
    :   Methylmalonyl-CoA: 83.3


        Methoxymalonyl-CoA: 67.1


        Ethylmalonyl-CoA: 49.6


        Isobutyryl-CoA: 40.6


        Malonyl-CoA: 37.8


        Propionyl-CoA: 30.3


        2-Methylbutyryl-CoA: 28.2


        trans-1,2-CPDA: 23.7


        Benzoyl-CoA: 21.7


        fatty\_acid: 20.3


        3-Methylbutyryl-CoA: 14.7


        CHC-CoA: 12.3


        Acetyl-CoA: 10.1


        inactive: 9.0

contig\_12 - Region 2 - NRPS

Shows the layout of the region, marking coding sequences and areas of interest. Clicking a gene will select it and show any relevant details. Clicking an area feature (e.g. a candidate cluster) will select all coding sequences within that area. Double clicking an area feature will zoom to that area. Multiple genes and area features can be selected by clicking them while holding the Ctrl key.  
More detailed help is available here.

Download region GenBank file

Download region SVG

Location: 2,325,183 - 2,369,957 nt. (total: 44,775 nt)
Show pHMM detection rules used

NRPS: cds(Condensation and (AMP-binding or A-OX))

#### Legend:

core biosynthetic genes

additional biosynthetic genes

transport-related genes

regulatory genes

other genes

resistance

reset view

zoom to selection

Gene details

Shows details of the most recently selected gene, including names, products, location, and other annotations.

Select a gene to view the details available for it

NRPS/PKS domains

ClusterBlast

KnownClusterBlast

SubClusterBlast

MIBiG comparison

Pfam domains

Detailed domain annotation

Shows NRPS- and PKS-related domains for each feature that contains them. Click on each domain for more information about the domain's location, consensus monomer prediction, and other details.  
A glossary is available here.

Selected features only

Show module domains

Similar gene clusters

Shows clusters from the antiSMASH database and other clusters of interest that are similar to the current region. Genes marked with the same colour are interrelated. White genes have no relationship.  
Click on reference genes to show details of similarities to genes within the current region.  
Click on an accession to open that entry in the antiSMASH database (if applicable).

All hits

NW\_011371358 (308375-361830): Paracoccidioides brasiliensis Pb18 unplaced gen... (23% of genes show similarity), NRPS

NW\_004504310 (4681983-4736885): Coccidioides immitis RS genomic scaffold supe... (17% of genes show similarity), NRPS

NW\_003052500 (3331052-3385198): Uncinocarpus reesii 1704 scaffold 1 genomic s... (16% of genes show similarity), NRPS

NW\_003315979 (1767537-1821260): Coccidioides posadasii C735 delta SOWgp chrom... (18% of genes show similarity), NRPS

NW\_015971594 (673293-731762): Cladophialophora bantiana CBS 173.52 unplaced g... (16% of genes show similarity), NRPS

NW\_013550610 (3320149-3386717): Fonsecaea pedrosoi CBS 271.37 unplaced genomi... (15% of genes show similarity), NRPS

NW\_022474213 (1109585-1168941): Venustampulla echinocandica strain BP 5553 ch... (18% of genes show similarity), NRPS

NW\_008481827 (678973-734548): Cladophialophora carrionii CBS 160.54 unplaced ... (18% of genes show similarity), NRPS

NW\_015971668 (45805-117806): Fonsecaea multimorphosa CBS 102226 unplaced geno... (13% of genes show similarity), NRPS

NZ\_CP011509 (6389853-6491908): Archangium gephyra strain DSM 2261 chromosome,... (9% of genes show similarity), NRPS,T1PKS
Download graphic

Similar known gene clusters

Shows clusters from the MiBIG database that are similar to the current region. Genes marked with the same colour are interrelated. White genes have no relationship.  
Click on reference genes to show details of similarities to genes within the current region.  
Click on an accession to open that entry in the MiBIG database.

All hits

dimethylcoprogen
Download graphic

Similar subclusters

Shows sub-cluster units that are similar to the current region. Genes marked with the same colour are interrelated. White genes have no relationship.  
Click on reference genes to show details of similarities to genes within the current region.

No matches found.

Similar gene clusters

Shows careas that are similar to the current region to a reference database.  
Mouseover a score cell in the table to get a breakdown of how the score was calculated.The MIBiG database.  
  
Click on an accession to open that entry in the MIBiG database.

Analysis type:

Protocluster to Region
Region to Region

| Reference | NRPS | Similarity score | Type | Compound(s) | Organism |
| --- | --- | --- | --- | --- | --- |
| BGC0001249.1 |  | 0.43 | NRP | dimethylcoprogen | Alternaria alternata |
| BGC0000900.1 |  | 0.33 | Other | ferrichrome | Aspergillus oryzae |
| BGC0000348.1 |  | 0.33 | NRP | ergovaline | Epichloe festucae var. lolii |
| BGC0001261.1 |  | 0.26 | NRP | AM-toxin | Alternaria alternata |
| BGC0000357.1 |  | 0.25 | NRP | cyclo-(D-Phe-L-Phe-D-Val-L-Val), cyclo-(D-Tyr-L-Phe-D-Val-L-Val), cyclo-(D-Tyr-L-Trp-D-Val-L-Val), cyclo-(D-Phe-L-Trp-D-Val-L-Val), cyclo-(D-Phe-L-Phe-D-Val-L-Ile), cyclo-(D-Phe-L-Phe-D-Ile-L-Val), cyclo-(D-Tyr-L-Trp-D-Val-L-Ile), cyclo-(D-Tyr-L-Trp-D-Ile-L-Val), cyclo-(D-Tyr-L-Phe-D-Val-L-Ile), cyclo-(D-Tyr-L-Phe-D-Ile-L-Val) | Penicillium rubens Wisconsin 54-1255 |
| BGC0001517.1 |  | 0.25 | NRP | asperphenamate | Aspergillus terreus NIH2624 |
| BGC0001717.1 |  | 0.24 | Alkaloid | okaramine B | Penicillium simplicissimum |
| BGC0001718.1 |  | 0.23 | NRP | okaramine D | Aspergillus aculeatus ATCC 16872 |
| BGC0001132.1 |  | 0.22 | NRP | xenotetrapeptide | Xenorhabdus nematophila ATCC 19061 |
| BGC0001545.1 |  | 0.22 | NRP | chrysogine | Fusarium graminearum PH-1 |

| Reference | Aggregated | Similarity score | Type | Compound(s) | Organism |
| --- | --- | --- | --- | --- | --- |
| BGC0001249.1 |  | 0.74 | NRP | dimethylcoprogen | Alternaria alternata |
| BGC0000900.1 |  | 0.67 | Other | ferrichrome | Aspergillus oryzae |
| BGC0000348.1 |  | 0.66 | NRP | ergovaline | Epichloe festucae var. lolii |
| BGC0001261.1 |  | 0.59 | NRP | AM-toxin | Alternaria alternata |
| BGC0000357.1 |  | 0.58 | NRP | cyclo-(D-Phe-L-Phe-D-Val-L-Val), cyclo-(D-Tyr-L-Phe-D-Val-L-Val), cyclo-(D-Tyr-L-Trp-D-Val-L-Val), cyclo-(D-Phe-L-Trp-D-Val-L-Val), cyclo-(D-Phe-L-Phe-D-Val-L-Ile), cyclo-(D-Phe-L-Phe-D-Ile-L-Val), cyclo-(D-Tyr-L-Trp-D-Val-L-Ile), cyclo-(D-Tyr-L-Trp-D-Ile-L-Val), cyclo-(D-Tyr-L-Phe-D-Val-L-Ile), cyclo-(D-Tyr-L-Phe-D-Ile-L-Val) | Penicillium rubens Wisconsin 54-1255 |
| BGC0001517.1 |  | 0.58 | NRP | asperphenamate | Aspergillus terreus NIH2624 |
| BGC0001717.1 |  | 0.57 | Alkaloid | okaramine B | Penicillium simplicissimum |
| BGC0001718.1 |  | 0.57 | NRP | okaramine D | Aspergillus aculeatus ATCC 16872 |
| BGC0001132.1 |  | 0.56 | NRP | xenotetrapeptide | Xenorhabdus nematophila ATCC 19061 |
| BGC0001545.1 |  | 0.56 | NRP | chrysogine | Fusarium graminearum PH-1 |

Detailed Pfam domain annotation

Shows Pfam domains found in each gene within the region.
Click on each domain for more information about the domain's
accession, location, description, and any relevant Gene Ontology.
Domains with a bold border have Gene Ontology information.

Selected features only

NRPS/PKS products

NRPS/PKS monomers

Predicted core structure(s)

Shows estimated product structure and polymer for each candidate cluster in the region. To show the product, click on the expander or the candidate cluster feature drawn in the overview.

For candidate cluster 2, location 2325182 - 2369957:

Rough prediction of core scaffold based on assumed PKS/NRPS colinearity; tailoring reactions not taken into account

**Polymer prediction:**
:   (X)

  
Direct lookup in NORINE database:
strict
or
relaxed

Link to NORINE database query form

NRPS/PKS monomer predictions

Shows the predicted monomers for each adynelation domain and acyltransferase within genes. Each gene prediction can be expanded to view detailed predictions of each domain. Each prediction can be expanded to view the predictions by tool (and, for some tools, further expanded for extra details).

**input.path1.gene640**: X

:   Search NORINE for peptide:
    strict
    or
    relaxed
  
:   **AMP-binding (64..385)**: X

    NRPSPredictor2: (unknown)

    SVM prediction details:
    :   Predicted physicochemical class:
        :   N/A

        Large clusters prediction:
        :   N/A

        Small clusters prediction:
        :   N/A

        Single AA prediction:
        :   N/A

    Stachelhaus prediction details:
    :   Stachelhaus sequence:
        :   gllhiasptk

        Nearest Stachelhaus code:
        :   N, A

        Stachelhaus code match:
        :   0% (weak)

  
**input.path1.gene642**: X

:   Search NORINE for peptide:
    strict
    or
    relaxed
  
:   **AMP-binding (0..245)**: X

    NRPSPredictor2: hydrophobic-aromatic

    SVM prediction details:
    :   Predicted physicochemical class:
        :   hydrophobic-aromatic

        Large clusters prediction:
        :   N/A

        Small clusters prediction:
        :   N/A

        Single AA prediction:
        :   N/A

    Stachelhaus prediction details:
    :   Stachelhaus sequence:
        :   dvdgagcvgk

        Nearest Stachelhaus code:
        :   N, A

        Stachelhaus code match:
        :   0% (weak)

contig\_12 - Region 3 - NRPS-like

Shows the layout of the region, marking coding sequences and areas of interest. Clicking a gene will select it and show any relevant details. Clicking an area feature (e.g. a candidate cluster) will select all coding sequences within that area. Double clicking an area feature will zoom to that area. Multiple genes and area features can be selected by clicking them while holding the Ctrl key.  
More detailed help is available here.

Download region GenBank file

Download region SVG

Location: 2,479,385 - 2,523,240 nt. (total: 43,856 nt)
Show pHMM detection rules used

NRPS-like: cds((PP-binding or NAD\_binding\_4) and (AMP-binding or A-OX))

#### Legend:

core biosynthetic genes

additional biosynthetic genes

transport-related genes

regulatory genes

other genes

resistance

reset view

zoom to selection

Gene details

Shows details of the most recently selected gene, including names, products, location, and other annotations.

Select a gene to view the details available for it

NRPS/PKS domains

ClusterBlast

KnownClusterBlast

SubClusterBlast

MIBiG comparison

Pfam domains

Detailed domain annotation

Shows NRPS- and PKS-related domains for each feature that contains them. Click on each domain for more information about the domain's location, consensus monomer prediction, and other details.  
A glossary is available here.

Selected features only

Show module domains

Similar gene clusters

Shows clusters from the antiSMASH database and other clusters of interest that are similar to the current region. Genes marked with the same colour are interrelated. White genes have no relationship.  
Click on reference genes to show details of similarities to genes within the current region.  
Click on an accession to open that entry in the antiSMASH database (if applicable).

All hits

NW\_006271975 (324992-369230): Cordyceps militaris CM01 unplaced genomic scaff... (14% of genes show similarity), NRPS-like
Download graphic

Similar known gene clusters

Shows clusters from the MiBIG database that are similar to the current region. Genes marked with the same colour are interrelated. White genes have no relationship.  
Click on reference genes to show details of similarities to genes within the current region.  
Click on an accession to open that entry in the MiBIG database.

No matches found.

Similar subclusters

Shows sub-cluster units that are similar to the current region. Genes marked with the same colour are interrelated. White genes have no relationship.  
Click on reference genes to show details of similarities to genes within the current region.

No matches found.

Similar gene clusters

Shows careas that are similar to the current region to a reference database.  
Mouseover a score cell in the table to get a breakdown of how the score was calculated.The MIBiG database.  
  
Click on an accession to open that entry in the MIBiG database.

Analysis type:

Protocluster to Region
Region to Region

| Reference | NRPS-like | Similarity score | Type | Compound(s) | Organism |
| --- | --- | --- | --- | --- | --- |
| BGC0000099.1 |  | 0.16 | Polyketide | monascorubrin | Talaromyces marneffei |
| BGC0001374.1 |  | 0.14 | Other | thienodolin | Streptomyces albogriseolus |
| BGC0001896.1 |  | 0.13 | Other | carbazomycin B | Streptomyces luteoverticillatus |
| BGC0001390.1 |  | 0.12 | NRP, Polyketide | LL-Z1272beta | Stachybotrys bisbyi |
| BGC0001600.1 |  | 0.10 | Polyketide | fusarielin H | Fusarium graminearum |
| BGC0001722.1 |  | 0.09 | Polyketide | aspernidine A | Aspergillus nidulans FGSC A4 |
| BGC0001182.1 |  | 0.09 | NRP, Polyketide | chaetoglobosins | Chaetomium globosum CBS 148.51 |
| BGC0001404.1 |  | 0.09 | Polyketide | sorbicillin | Penicillium rubens Wisconsin 54-1255 |
| BGC0000876.1 |  | 0.09 | Other | neopolyoxin C | Streptomyces tendae |
| BGC0001626.1 |  | 0.08 | Polyketide | isoindolinone | Stachybotrys chlorohalonata IBT 40285 |

| Reference | Aggregated | Similarity score | Type | Compound(s) | Organism |
| --- | --- | --- | --- | --- | --- |
| BGC0000099.1 |  | 0.46 | Polyketide | monascorubrin | Talaromyces marneffei |
| BGC0001374.1 |  | 0.43 | Other | thienodolin | Streptomyces albogriseolus |
| BGC0001896.1 |  | 0.42 | Other | carbazomycin B | Streptomyces luteoverticillatus |
| BGC0001390.1 |  | 0.40 | NRP, Polyketide | LL-Z1272beta | Stachybotrys bisbyi |
| BGC0001600.1 |  | 0.36 | Polyketide | fusarielin H | Fusarium graminearum |
| BGC0001722.1 |  | 0.34 | Polyketide | aspernidine A | Aspergillus nidulans FGSC A4 |
| BGC0001182.1 |  | 0.34 | NRP, Polyketide | chaetoglobosins | Chaetomium globosum CBS 148.51 |
| BGC0001404.1 |  | 0.33 | Polyketide | sorbicillin | Penicillium rubens Wisconsin 54-1255 |
| BGC0000876.1 |  | 0.32 | Other | neopolyoxin C | Streptomyces tendae |
| BGC0001626.1 |  | 0.32 | Polyketide | isoindolinone | Stachybotrys chlorohalonata IBT 40285 |

Detailed Pfam domain annotation

Shows Pfam domains found in each gene within the region.
Click on each domain for more information about the domain's
accession, location, description, and any relevant Gene Ontology.
Domains with a bold border have Gene Ontology information.

Selected features only

NRPS/PKS products

NRPS/PKS monomers

Predicted core structure(s)

Shows estimated product structure and polymer for each candidate cluster in the region. To show the product, click on the expander or the candidate cluster feature drawn in the overview.

For candidate cluster 3, location 2479384 - 2523240:

Rough prediction of core scaffold based on assumed PKS/NRPS colinearity; tailoring reactions not taken into account

**Polymer prediction:**
:   (X)

  
Direct lookup in NORINE database:
strict
or
relaxed

Link to NORINE database query form

NRPS/PKS monomer predictions

Shows the predicted monomers for each adynelation domain and acyltransferase within genes. Each gene prediction can be expanded to view detailed predictions of each domain. Each prediction can be expanded to view the predictions by tool (and, for some tools, further expanded for extra details).

**input.path1.gene686**: X

:   Search NORINE for peptide:
    strict
    or
    relaxed
  
:   **AMP-binding (56..282)**: X

    NRPSPredictor2: (unknown)

    SVM prediction details:
    :   Predicted physicochemical class:
        :   N/A

        Large clusters prediction:
        :   N/A

        Small clusters prediction:
        :   N/A

        Single AA prediction:
        :   N/A

    Stachelhaus prediction details:
    :   Stachelhaus sequence:
        :   --t------k

        Nearest Stachelhaus code:
        :   N, A

        Stachelhaus code match:
        :   0% (weak)

contig\_12 - Region 4 - T1PKS,indole

Shows the layout of the region, marking coding sequences and areas of interest. Clicking a gene will select it and show any relevant details. Clicking an area feature (e.g. a candidate cluster) will select all coding sequences within that area. Double clicking an area feature will zoom to that area. Multiple genes and area features can be selected by clicking them while holding the Ctrl key.  
More detailed help is available here.

Download region GenBank file

Download region SVG

Location: 2,923,017 - 2,976,336 nt. (total: 53,320 nt)
Show pHMM detection rules used

indole: (indsynth or dmat or indole\_PTase)  
T1PKS: cds(PKS\_AT and (PKS\_KS or ene\_KS or mod\_KS or hyb\_KS or itr\_KS or tra\_KS))

#### Legend:

core biosynthetic genes

additional biosynthetic genes

transport-related genes

regulatory genes

other genes

resistance

reset view

zoom to selection

Gene details

Shows details of the most recently selected gene, including names, products, location, and other annotations.

Select a gene to view the details available for it

NRPS/PKS domains

ClusterBlast

KnownClusterBlast

SubClusterBlast

MIBiG comparison

Pfam domains

Detailed domain annotation

Shows NRPS- and PKS-related domains for each feature that contains them. Click on each domain for more information about the domain's location, consensus monomer prediction, and other details.  
A glossary is available here.

Selected features only

Show module domains

Similar gene clusters

Shows clusters from the antiSMASH database and other clusters of interest that are similar to the current region. Genes marked with the same colour are interrelated. White genes have no relationship.  
Click on reference genes to show details of similarities to genes within the current region.  
Click on an accession to open that entry in the antiSMASH database (if applicable).

All hits

NW\_022984630 (495088-539069): Aspergillus tanneri strain NIH1004 chromosome U... (55% of genes show similarity), T1PKS

NT\_107015 (458256-500846): Aspergillus nidulans FGSC A4 chromosome VIII map u... (61% of genes show similarity), T1PKS

NW\_014574694 (86125-153989): Metarhizium brunneum ARSEF 3297 chromosome Unkno... (26% of genes show similarity), NRPS,T1PKS

NW\_019154043 (172137-217646): Pochonia chlamydosporia 170 chromosome Unknown ... (43% of genes show similarity), T1PKS

NW\_006913046 (1285411-1330974): Cladophialophora yegresii CBS 114405 aczIN-su... (47% of genes show similarity), T1PKS

NW\_008481833 (674692-720244): Cladophialophora carrionii CBS 160.54 unplaced ... (47% of genes show similarity), T1PKS

NW\_011942151 (90937-172848): Metarhizium robertsii ARSEF 23 MAA Scf 11, whole... (25% of genes show similarity), NRPS,T1PKS,terpene

NW\_023336272 (555100-616490): Colletotrichum scovillei strain TJNH1 chromosom... (50% of genes show similarity), NRPS,T1PKS

NC\_016458 (5151387-5191687): Thermothielavioides terrestris NRRL 8126 chromos... (37% of genes show similarity), T1PKS

NW\_022984634 (4255890-4301503): Aspergillus tanneri strain NIH1004 chromosome... (36% of genes show similarity), T1PKS
Download graphic

Similar known gene clusters

Shows clusters from the MiBIG database that are similar to the current region. Genes marked with the same colour are interrelated. White genes have no relationship.  
Click on reference genes to show details of similarities to genes within the current region.  
Click on an accession to open that entry in the MiBIG database.

All hits

secalonic acids

neosartorin

trypacidin

RES-1214-2

endocrocin

emodin

TAN-1612 / 1-(2,3,5,10-tetrahydroxy-7-methoxy-4-oxo-1,2,3,4-tetrahydroanthracen-2-yl)pentane-2,4-dione / desmethyl TAN-1612

chrysoxanthone A / chrysoxanthone B / chrysoxanthone C

notoamide A
Download graphic

Similar subclusters

Shows sub-cluster units that are similar to the current region. Genes marked with the same colour are interrelated. White genes have no relationship.  
Click on reference genes to show details of similarities to genes within the current region.

No matches found.

Similar gene clusters

Shows careas that are similar to the current region to a reference database.  
Mouseover a score cell in the table to get a breakdown of how the score was calculated.The MIBiG database.  
  
Click on an accession to open that entry in the MIBiG database.

Analysis type:

Protocluster to Region
Region to Region

| Reference | indole | T1PKS | Similarity score | Type | Compound(s) | Organism |
| --- | --- | --- | --- | --- | --- | --- |
| BGC0001542.1 |  |  | 0.40 | Polyketide | cercosporin | Cercospora zeina |
| BGC0000013.1 |  |  | 0.38 | Polyketide | alternariol | Aspergillus nidulans FGSC A4 |
| BGC0001541.1 |  |  | 0.38 | Polyketide | cercosporin | Cercospora beticola |
| BGC0001260.1 |  |  | 0.33 | Terpene | terpendole E | Tolypocladium album |
| BGC0000685.1 |  |  | 0.31 | Terpene | brassicicene C | Alternaria brassicicola ATCC 96836 |
| BGC0000161.1 |  |  | 0.31 | Polyketide | isoterrein | Aspergillus terreus NIH2624 |
| BGC0001374.1 |  |  | 0.31 | Other | thienodolin | Streptomyces albogriseolus |
| BGC0001906.1 |  |  | 0.28 | Polyketide | naphthalene | Daldinia eschscholzii IFB-TL01 |
| BGC0001068.1 |  |  | 0.28 | Terpene, Polyketide | pyripyropene A | unidentified unclassified sequences. |
| BGC0001304.1 |  |  | 0.28 | Polyketide | aflavarin | Aspergillus flavus NRRL3357 |

| Reference | Aggregated | Similarity score | Type | Compound(s) | Organism |
| --- | --- | --- | --- | --- | --- |
| BGC0000013.1 |  | 0.71 | Polyketide | alternariol | Aspergillus nidulans FGSC A4 |
| BGC0000161.1 |  | 0.64 | Polyketide | isoterrein | Aspergillus terreus NIH2624 |
| BGC0001068.1 |  | 0.62 | Terpene, Polyketide | pyripyropene A | unidentified unclassified sequences. |
| BGC0001338.1 |  | 0.59 | Polyketide | citrinin | Monascus ruber |
| BGC0001858.1 |  | 0.58 | Polyketide | alternapyrone B, alternapyrone C, alternapyrone D, alternapyrone E, alternapyrone F | Parastagonospora nodorum SN15 |
| BGC0001284.1 |  | 0.57 | Terpene | alternariol | Parastagonospora nodorum SN15 |
| BGC0001265.1 |  | 0.57 | Polyketide | melanin | Bipolaris oryzae |
| BGC0001258.1 |  | 0.57 | Polyketide | 1,3,6,8-tetrahydroxynaphthalene | Glarea lozoyensis |
| BGC0001257.1 |  | 0.56 | Polyketide | 1,3,6,8-tetrahydroxynaphthalene | Nodulisporium sp. ATCC74245 |
| BGC0001144.1 |  | 0.56 | Polyketide | neosartoricin B | Trichophyton tonsurans CBS 112818 |

Detailed Pfam domain annotation

Shows Pfam domains found in each gene within the region.
Click on each domain for more information about the domain's
accession, location, description, and any relevant Gene Ontology.
Domains with a bold border have Gene Ontology information.

Selected features only

NRPS/PKS products

NRPS/PKS monomers

Predicted core structure(s)

Shows estimated product structure and polymer for each candidate cluster in the region. To show the product, click on the expander or the candidate cluster feature drawn in the overview.

For candidate cluster 4, location 2923016 - 2976336:

Rough prediction of core scaffold based on assumed PKS/NRPS colinearity; tailoring reactions not taken into account

**Polymer prediction:**
:   (mal)

  
Direct lookup in NORINE database:
strict
or
relaxed

---

For candidate cluster 6, location 2930659 - 2976336:

Rough prediction of core scaffold based on assumed PKS/NRPS colinearity; tailoring reactions not taken into account

**Polymer prediction:**
:   (mal)

  
Direct lookup in NORINE database:
strict
or
relaxed

Link to NORINE database query form

NRPS/PKS monomer predictions

Shows the predicted monomers for each adynelation domain and acyltransferase within genes. Each gene prediction can be expanded to view detailed predictions of each domain. Each prediction can be expanded to view the predictions by tool (and, for some tools, further expanded for extra details).

**input.path1.gene804**: mal

:   **PKS\_AT (925..1228)**: mal

    ATSignature: Malonyl-CoA

    Top 3 matches:
    :   Malonyl-CoA: 75.0%
    :   inactive: 62.5%
    :   Methylmalonyl-CoA: 54.2%

      
    minowa: Malonyl-CoA

    Prediction, score:
    :   Malonyl-CoA: 88.4


        inactive: 74.1


        Methylmalonyl-CoA: 53.6


        Methoxymalonyl-CoA: 47.8


        Propionyl-CoA: 42.5


        Isobutyryl-CoA: 23.9


        2-Methylbutyryl-CoA: 22.2


        Benzoyl-CoA: 21.9


        Ethylmalonyl-CoA: 19.2


        fatty\_acid: 18.2


        CHC-CoA: 14.8


        trans-1,2-CPDA: 14.2


        3-Methylbutyryl-CoA: 12.1


        Acetyl-CoA: 11.6

contig\_13 - Region 1 - NRPS-like

Shows the layout of the region, marking coding sequences and areas of interest. Clicking a gene will select it and show any relevant details. Clicking an area feature (e.g. a candidate cluster) will select all coding sequences within that area. Double clicking an area feature will zoom to that area. Multiple genes and area features can be selected by clicking them while holding the Ctrl key.  
More detailed help is available here.

Download region GenBank file

Download region SVG

Location: 741,090 - 785,032 nt. (total: 43,943 nt)
Show pHMM detection rules used

NRPS-like: cds((PP-binding or NAD\_binding\_4) and (AMP-binding or A-OX))

#### Legend:

core biosynthetic genes

additional biosynthetic genes

transport-related genes

regulatory genes

other genes

resistance

reset view

zoom to selection

Gene details

Shows details of the most recently selected gene, including names, products, location, and other annotations.

Select a gene to view the details available for it

NRPS/PKS domains

ClusterBlast

KnownClusterBlast

SubClusterBlast

MIBiG comparison

Pfam domains

Detailed domain annotation

Shows NRPS- and PKS-related domains for each feature that contains them. Click on each domain for more information about the domain's location, consensus monomer prediction, and other details.  
A glossary is available here.

Selected features only

Show module domains

Similar gene clusters

Shows clusters from the antiSMASH database and other clusters of interest that are similar to the current region. Genes marked with the same colour are interrelated. White genes have no relationship.  
Click on reference genes to show details of similarities to genes within the current region.  
Click on an accession to open that entry in the antiSMASH database (if applicable).

All hits

NC\_049563 (5665253-5709096): Talaromyces rugulosus chromosome III, complete s... (14% of genes show similarity), NRPS-like

NW\_019716264 (12526-43637): Ramularia collo-cygni strain URUG2 genome assembl... (15% of genes show similarity), NRPS-like

NW\_015971148 (336061-377773): Sporothrix schenckii 1099-18 chromosome Unknown... (10% of genes show similarity), NRPS-like

NC\_018218 (2171437-2215442): Zymoseptoria tritici IPO323 chromosome 1, whole ... (15% of genes show similarity), NRPS-like

NZ\_VLHW01000001 (1081672-1171208): Pseudomonas syringae pv. dysoxyli strain C... (8% of genes show similarity), NRPS

NC\_014718 (578820-691899): Paraburkholderia rhizoxinica HKI 454 plasmid pBRH0... (5% of genes show similarity), NRPS,betalactone,phosphonate,terpene

NZ\_WJPS01000001 (454313-541031): Pseudomonas syringae strain P73 CL 7301, who... (8% of genes show similarity), NRPS

NZ\_LT629769 (949574-1036517): Pseudomonas syringae strain 31R1 chromosome I (8% of genes show similarity), NRPS

NZ\_FPJB01000001 (348284-438096): Pseudomonas sp. NFACC10-1, whole genome shot... (8% of genes show similarity), NRPS

NZ\_LT222319 (3655590-3742143): Pseudomonas cerasi isolate Sour cherry (Prunus... (8% of genes show similarity), NRPS
Download graphic

Similar known gene clusters

Shows clusters from the MiBIG database that are similar to the current region. Genes marked with the same colour are interrelated. White genes have no relationship.  
Click on reference genes to show details of similarities to genes within the current region.  
Click on an accession to open that entry in the MiBIG database.

No matches found.

Similar subclusters

Shows sub-cluster units that are similar to the current region. Genes marked with the same colour are interrelated. White genes have no relationship.  
Click on reference genes to show details of similarities to genes within the current region.

No matches found.

Similar gene clusters

Shows careas that are similar to the current region to a reference database.  
Mouseover a score cell in the table to get a breakdown of how the score was calculated.The MIBiG database.  
  
Click on an accession to open that entry in the MIBiG database.

Analysis type:

Protocluster to Region
Region to Region

| Reference | NRPS-like | Similarity score | Type | Compound(s) | Organism |
| --- | --- | --- | --- | --- | --- |
| BGC0001399.1 |  | 0.25 | NRP | fellutamide B | Aspergillus nidulans FGSC A4 |
| BGC0001168.1 |  | 0.21 | NRP | livipeptin | Streptomyces lividans 1326 |
| BGC0001900.1 |  | 0.21 | NRP | fragin | Burkholderia cenocepacia H111 |
| BGC0001641.1 |  | 0.18 | NRP | kolossin | Photorhabdus laumondii subsp. laumondii TTO1 |
| BGC0001128.1 |  | 0.18 | NRP | luminmide | Photorhabdus laumondii subsp. laumondii TTO1 |
| BGC0001833.1 |  | 0.18 | NRP | icosalide A, icosalide B | Burkholderia gladioli |
| BGC0001135.1 |  | 0.18 | NRP | bicornutin A1, bicornutin A2 | Xenorhabdus budapestensis |
| BGC0001132.1 |  | 0.18 | NRP | xenotetrapeptide | Xenorhabdus nematophila ATCC 19061 |
| BGC0001479.1 |  | 0.18 | NRP | anabaenopeptin NZ857, nostamide A | Nostoc punctiforme PCC 73102 |
| BGC0001844.1 |  | 0.18 | NRP | holrhizin | Paraburkholderia rhizoxinica HKI 454 |

| Reference | Aggregated | Similarity score | Type | Compound(s) | Organism |
| --- | --- | --- | --- | --- | --- |
| BGC0001399.1 |  | 0.59 | NRP | fellutamide B | Aspergillus nidulans FGSC A4 |
| BGC0001168.1 |  | 0.54 | NRP | livipeptin | Streptomyces lividans 1326 |
| BGC0001900.1 |  | 0.53 | NRP | fragin | Burkholderia cenocepacia H111 |
| BGC0001641.1 |  | 0.50 | NRP | kolossin | Photorhabdus laumondii subsp. laumondii TTO1 |
| BGC0001128.1 |  | 0.50 | NRP | luminmide | Photorhabdus laumondii subsp. laumondii TTO1 |
| BGC0001833.1 |  | 0.49 | NRP | icosalide A, icosalide B | Burkholderia gladioli |
| BGC0001135.1 |  | 0.49 | NRP | bicornutin A1, bicornutin A2 | Xenorhabdus budapestensis |
| BGC0001132.1 |  | 0.49 | NRP | xenotetrapeptide | Xenorhabdus nematophila ATCC 19061 |
| BGC0001479.1 |  | 0.49 | NRP | anabaenopeptin NZ857, nostamide A | Nostoc punctiforme PCC 73102 |
| BGC0001844.1 |  | 0.49 | NRP | holrhizin | Paraburkholderia rhizoxinica HKI 454 |

Detailed Pfam domain annotation

Shows Pfam domains found in each gene within the region.
Click on each domain for more information about the domain's
accession, location, description, and any relevant Gene Ontology.
Domains with a bold border have Gene Ontology information.

Selected features only

NRPS/PKS products

NRPS/PKS monomers

Predicted core structure(s)

Shows estimated product structure and polymer for each candidate cluster in the region. To show the product, click on the expander or the candidate cluster feature drawn in the overview.

For candidate cluster 1, location 741089 - 785032:

Rough prediction of core scaffold based on assumed PKS/NRPS colinearity; tailoring reactions not taken into account

**Polymer prediction:**
:   (ala)

  
Direct lookup in NORINE database:
strict
or
relaxed

Link to NORINE database query form

NRPS/PKS monomer predictions

Shows the predicted monomers for each adynelation domain and acyltransferase within genes. Each gene prediction can be expanded to view detailed predictions of each domain. Each prediction can be expanded to view the predictions by tool (and, for some tools, further expanded for extra details).

**input.path1.gene222**: ala

:   Search NORINE for peptide:
    strict
    or
    relaxed
  
:   **AMP-binding (13..419)**: ala

    NRPSPredictor2: ala

    SVM prediction details:
    :   Predicted physicochemical class:
        :   hydrophobic-aliphatic

        Large clusters prediction:
        :   N/A

        Small clusters prediction:
        :   N/A

        Single AA prediction:
        :   ala

    Stachelhaus prediction details:
    :   Stachelhaus sequence:
        :   nvwlwnvevk

        Nearest Stachelhaus code:
        :   N, A

        Stachelhaus code match:
        :   0% (weak)

contig\_14 - Region 1 - T1PKS

Shows the layout of the region, marking coding sequences and areas of interest. Clicking a gene will select it and show any relevant details. Clicking an area feature (e.g. a candidate cluster) will select all coding sequences within that area. Double clicking an area feature will zoom to that area. Multiple genes and area features can be selected by clicking them while holding the Ctrl key.  
More detailed help is available here.

Download region GenBank file

Download region SVG

Location: 209,216 - 255,818 nt. (total: 46,603 nt)
Show pHMM detection rules used

T1PKS: cds(PKS\_AT and (PKS\_KS or ene\_KS or mod\_KS or hyb\_KS or itr\_KS or tra\_KS))

#### Legend:

core biosynthetic genes

additional biosynthetic genes

transport-related genes

regulatory genes

other genes

resistance

reset view

zoom to selection

Gene details

Shows details of the most recently selected gene, including names, products, location, and other annotations.

Select a gene to view the details available for it

NRPS/PKS domains

ClusterBlast

KnownClusterBlast

SubClusterBlast

MIBiG comparison

Pfam domains

Detailed domain annotation

Shows NRPS- and PKS-related domains for each feature that contains them. Click on each domain for more information about the domain's location, consensus monomer prediction, and other details.  
A glossary is available here.

Selected features only

Show module domains

Similar gene clusters

Shows clusters from the antiSMASH database and other clusters of interest that are similar to the current region. Genes marked with the same colour are interrelated. White genes have no relationship.  
Click on reference genes to show details of similarities to genes within the current region.  
Click on an accession to open that entry in the antiSMASH database (if applicable).

All hits

NT\_165977 (3041183-3093199): Chaetomium globosum CBS 148.51 scaffold 2 genomi... (16% of genes show similarity), T1PKS

NW\_003299166 (1419813-1463169): Microsporum canis CBS 113480 supercont1.4 gen... (13% of genes show similarity), NRPS,T1PKS

NW\_021167090 (680283-725597): Sodiomyces alkalinus F11 unplaced genomic scaff... (12% of genes show similarity), T1PKS

NW\_022474213 (391991-435464): Venustampulla echinocandica strain BP 5553 chro... (13% of genes show similarity), T1PKS

NW\_003345199 (1003727-1098367): Nannizzia gypsea CBS 118893 supercont1.3 geno... (7% of genes show similarity), NRPS,T1PKS

NC\_049565 (1728455-1770196): Talaromyces rugulosus chromosome V, complete seq... (14% of genes show similarity), T1PKS

NW\_003299163 (1364680-1417022): Microsporum canis CBS 113480 supercont1.7 gen... (11% of genes show similarity), T1PKS,T3PKS

NW\_006271969 (4877275-4917236): Cordyceps militaris CM01 unplaced genomic sca... (15% of genes show similarity), T1PKS

NW\_023336279 (547097-649987): Aspergillus tubingensis WU-2223L DNA, scaffold ... (5% of genes show similarity), NRPS,T1PKS

NT\_165933 (1206996-1265490): Aspergillus terreus NIH2624 scaffold 10 genomic ... (10% of genes show similarity), T1PKS
Download graphic

Similar known gene clusters

Shows clusters from the MiBIG database that are similar to the current region. Genes marked with the same colour are interrelated. White genes have no relationship.  
Click on reference genes to show details of similarities to genes within the current region.  
Click on an accession to open that entry in the MiBIG database.

No matches found.

Similar subclusters

Shows sub-cluster units that are similar to the current region. Genes marked with the same colour are interrelated. White genes have no relationship.  
Click on reference genes to show details of similarities to genes within the current region.

No matches found.

Similar gene clusters

Shows careas that are similar to the current region to a reference database.  
Mouseover a score cell in the table to get a breakdown of how the score was calculated.The MIBiG database.  
  
Click on an accession to open that entry in the MIBiG database.

Analysis type:

Protocluster to Region
Region to Region

| Reference | T1PKS | Similarity score | Type | Compound(s) | Organism |
| --- | --- | --- | --- | --- | --- |
| BGC0001909.1 |  | 0.25 | Polyketide | strobilurin | Strobilurus tenacellus |
| BGC0000056.1 |  | 0.23 | Polyketide | esperamicin | Actinomadura verrucosospora |
| BGC0001998.1 |  | 0.21 | Polyketide | aspernidgulene A1, aspernidgulene A2, aspernidgulene B1 | Aspergillus nidulans FGSC A4 |
| BGC0001400.1 |  | 0.21 | Polyketide | citreoviridin | Aspergillus terreus NIH2624 |
| BGC0001276.1 |  | 0.21 | Polyketide | 6-methylsalicyclic acid | Aspergillus terreus |
| BGC0001275.1 |  | 0.21 | Polyketide | 6-methylsalicyclic acid | Glarea lozoyensis |
| BGC0001244.1 |  | 0.21 | Polyketide | (-)-Mellein | Parastagonospora nodorum |
| BGC0001273.1 |  | 0.21 | Polyketide | asperlactone | Aspergillus ochraceus |
| BGC0001068.1 |  | 0.20 | Terpene, Polyketide | pyripyropene A | unidentified unclassified sequences. |
| BGC0001858.1 |  | 0.20 | Polyketide | alternapyrone B, alternapyrone C, alternapyrone D, alternapyrone E, alternapyrone F | Parastagonospora nodorum SN15 |

| Reference | Aggregated | Similarity score | Type | Compound(s) | Organism |
| --- | --- | --- | --- | --- | --- |
| BGC0001909.1 |  | 0.59 | Polyketide | strobilurin | Strobilurus tenacellus |
| BGC0000056.1 |  | 0.56 | Polyketide | esperamicin | Actinomadura verrucosospora |
| BGC0001998.1 |  | 0.54 | Polyketide | aspernidgulene A1, aspernidgulene A2, aspernidgulene B1 | Aspergillus nidulans FGSC A4 |
| BGC0001400.1 |  | 0.54 | Polyketide | citreoviridin | Aspergillus terreus NIH2624 |
| BGC0001276.1 |  | 0.54 | Polyketide | 6-methylsalicyclic acid | Aspergillus terreus |
| BGC0001275.1 |  | 0.54 | Polyketide | 6-methylsalicyclic acid | Glarea lozoyensis |
| BGC0001244.1 |  | 0.53 | Polyketide | (-)-Mellein | Parastagonospora nodorum |
| BGC0001273.1 |  | 0.53 | Polyketide | asperlactone | Aspergillus ochraceus |
| BGC0001068.1 |  | 0.53 | Terpene, Polyketide | pyripyropene A | unidentified unclassified sequences. |
| BGC0001858.1 |  | 0.52 | Polyketide | alternapyrone B, alternapyrone C, alternapyrone D, alternapyrone E, alternapyrone F | Parastagonospora nodorum SN15 |

Detailed Pfam domain annotation

Shows Pfam domains found in each gene within the region.
Click on each domain for more information about the domain's
accession, location, description, and any relevant Gene Ontology.
Domains with a bold border have Gene Ontology information.

Selected features only

NRPS/PKS products

NRPS/PKS monomers

Predicted core structure(s)

Shows estimated product structure and polymer for each candidate cluster in the region. To show the product, click on the expander or the candidate cluster feature drawn in the overview.

For candidate cluster 1, location 209215 - 255818:

Rough prediction of core scaffold based on assumed PKS/NRPS colinearity; tailoring reactions not taken into account

**Polymer prediction:**
:   (pk)

  
Direct lookup in NORINE database:
strict
or
relaxed

Link to NORINE database query form

NRPS/PKS monomer predictions

Shows the predicted monomers for each adynelation domain and acyltransferase within genes. Each gene prediction can be expanded to view detailed predictions of each domain. Each prediction can be expanded to view the predictions by tool (and, for some tools, further expanded for extra details).

**input.path1.gene68**: pk

:   **PKS\_AT (480..801)**: pk

    ATSignature: Malonyl-CoA

    Top 3 matches:
    :   Malonyl-CoA: 75.0%
    :   Methylmalonyl-CoA: 62.5%
    :   Propionyl-CoA: 58.3%

      
    minowa: Methoxymalonyl-CoA

    Prediction, score:
    :   Methoxymalonyl-CoA: 74.8


        Methylmalonyl-CoA: 74.3


        Malonyl-CoA: 57.3


        Isobutyryl-CoA: 48.4


        Propionyl-CoA: 32.1


        Benzoyl-CoA: 29.3


        trans-1,2-CPDA: 19.0


        inactive: 19.0


        fatty\_acid: 19.0


        2-Methylbutyryl-CoA: 17.6


        3-Methylbutyryl-CoA: 15.4


        CHC-CoA: 0.0


        Acetyl-CoA: -3.6


        Ethylmalonyl-CoA: -4.3

contig\_15 - Region 1 - T1PKS

Shows the layout of the region, marking coding sequences and areas of interest. Clicking a gene will select it and show any relevant details. Clicking an area feature (e.g. a candidate cluster) will select all coding sequences within that area. Double clicking an area feature will zoom to that area. Multiple genes and area features can be selected by clicking them while holding the Ctrl key.  
More detailed help is available here.

Download region GenBank file

Download region SVG

Location: 1,237,335 - 1,284,006 nt. (total: 46,672 nt)
Show pHMM detection rules used

T1PKS: cds(PKS\_AT and (PKS\_KS or ene\_KS or mod\_KS or hyb\_KS or itr\_KS or tra\_KS))

#### Legend:

core biosynthetic genes

additional biosynthetic genes

transport-related genes

regulatory genes

other genes

resistance

reset view

zoom to selection

Gene details

Shows details of the most recently selected gene, including names, products, location, and other annotations.

Select a gene to view the details available for it

NRPS/PKS domains

ClusterBlast

KnownClusterBlast

SubClusterBlast

MIBiG comparison

Pfam domains

Detailed domain annotation

Shows NRPS- and PKS-related domains for each feature that contains them. Click on each domain for more information about the domain's location, consensus monomer prediction, and other details.  
A glossary is available here.

Selected features only

Show module domains

Similar gene clusters

Shows clusters from the antiSMASH database and other clusters of interest that are similar to the current region. Genes marked with the same colour are interrelated. White genes have no relationship.  
Click on reference genes to show details of similarities to genes within the current region.  
Click on an accession to open that entry in the antiSMASH database (if applicable).

All hits

NW\_007360987 (200861-291572): Glarea lozoyensis ATCC 20868 chromosome Unknown... (8% of genes show similarity), NRPS,T1PKS,betalactone

NC\_035796 (1939470-1976606): Pochonia chlamydosporia 170 chromosome 7, whole ... (16% of genes show similarity), T1PKS

NZ\_FRCX01000002 (270749-377659): Duganella sacchari strain Sac-22, whole geno... (5% of genes show similarity), NRPS,hserlactone,transAT-PKS

NZ\_LT607752 (4505012-4584315): Micromonospora rifamycinica strain DSM 44983 c... (3% of genes show similarity), NRPS,T1PKS,betalactone

NZ\_LT906483 (2326869-2393668): Mycolicibacterium thermoresistibile strain NCT... (3% of genes show similarity), NRPS,T1PKS

NZ\_SLYY01000015 (135638-242033): Streptomyces sp. BK205 Ga0307705 115, whole ... (2% of genes show similarity), NRPS,T1PKS,amglyccycl

NC\_017030 (4412962-4485151): Corallococcus coralloides DSM 2259, complete seq... (5% of genes show similarity), NRPS,T1PKS

NZ\_CP034669 (4290899-4361767): Corallococcus coralloides strain B035 chromoso... (4% of genes show similarity), NRPS,T1PKS

NW\_001939246 (883554-927495): Pyrenophora tritici-repentis Pt-1C-BFP supercon... (14% of genes show similarity), T1PKS
Download graphic

Similar known gene clusters

Shows clusters from the MiBIG database that are similar to the current region. Genes marked with the same colour are interrelated. White genes have no relationship.  
Click on reference genes to show details of similarities to genes within the current region.  
Click on an accession to open that entry in the MiBIG database.

No matches found.

Similar subclusters

Shows sub-cluster units that are similar to the current region. Genes marked with the same colour are interrelated. White genes have no relationship.  
Click on reference genes to show details of similarities to genes within the current region.

No matches found.

Similar gene clusters

Shows careas that are similar to the current region to a reference database.  
Mouseover a score cell in the table to get a breakdown of how the score was calculated.The MIBiG database.  
  
Click on an accession to open that entry in the MIBiG database.

Analysis type:

Protocluster to Region
Region to Region

| Reference | T1PKS | Similarity score | Type | Compound(s) | Organism |
| --- | --- | --- | --- | --- | --- |
| BGC0000046.1 |  | 0.31 | Polyketide | depudecin | Alternaria brassicicola |
| BGC0001068.1 |  | 0.25 | Terpene, Polyketide | pyripyropene A | unidentified unclassified sequences. |
| BGC0001858.1 |  | 0.19 | Polyketide | alternapyrone B, alternapyrone C, alternapyrone D, alternapyrone E, alternapyrone F | Parastagonospora nodorum SN15 |
| BGC0001252.1 |  | 0.19 | Polyketide | UNII-YC2Q1O94PT | Alternaria alternata |
| BGC0001606.1 |  | 0.18 | Polyketide | gibepyrone-A | Fusarium fujikuroi IMI 58289 |
| BGC0001124.1 |  | 0.18 | Polyketide | pyranonigrin E | Aspergillus niger ATCC 1015 |
| BGC0001254.1 |  | 0.18 | Polyketide | ACT-Toxin II | Alternaria alternata |
| BGC0001400.1 |  | 0.16 | Polyketide | citreoviridin | Aspergillus terreus NIH2624 |
| BGC0001998.1 |  | 0.16 | Polyketide | aspernidgulene A1, aspernidgulene A2, aspernidgulene B1 | Aspergillus nidulans FGSC A4 |
| BGC0001118.1 |  | 0.15 | Polyketide | endocrocin | Aspergillus fumigatus Af293 |

| Reference | Aggregated | Similarity score | Type | Compound(s) | Organism |
| --- | --- | --- | --- | --- | --- |
| BGC0000046.1 |  | 0.64 | Polyketide | depudecin | Alternaria brassicicola |
| BGC0001068.1 |  | 0.58 | Terpene, Polyketide | pyripyropene A | unidentified unclassified sequences. |
| BGC0001858.1 |  | 0.51 | Polyketide | alternapyrone B, alternapyrone C, alternapyrone D, alternapyrone E, alternapyrone F | Parastagonospora nodorum SN15 |
| BGC0001252.1 |  | 0.51 | Polyketide | UNII-YC2Q1O94PT | Alternaria alternata |
| BGC0001606.1 |  | 0.49 | Polyketide | gibepyrone-A | Fusarium fujikuroi IMI 58289 |
| BGC0001124.1 |  | 0.49 | Polyketide | pyranonigrin E | Aspergillus niger ATCC 1015 |
| BGC0001254.1 |  | 0.49 | Polyketide | ACT-Toxin II | Alternaria alternata |
| BGC0001400.1 |  | 0.47 | Polyketide | citreoviridin | Aspergillus terreus NIH2624 |
| BGC0001998.1 |  | 0.47 | Polyketide | aspernidgulene A1, aspernidgulene A2, aspernidgulene B1 | Aspergillus nidulans FGSC A4 |
| BGC0001118.1 |  | 0.46 | Polyketide | endocrocin | Aspergillus fumigatus Af293 |

Detailed Pfam domain annotation

Shows Pfam domains found in each gene within the region.
Click on each domain for more information about the domain's
accession, location, description, and any relevant Gene Ontology.
Domains with a bold border have Gene Ontology information.

Selected features only

contig\_15 - Region 2 - NRPS-like

Shows the layout of the region, marking coding sequences and areas of interest. Clicking a gene will select it and show any relevant details. Clicking an area feature (e.g. a candidate cluster) will select all coding sequences within that area. Double clicking an area feature will zoom to that area. Multiple genes and area features can be selected by clicking them while holding the Ctrl key.  
More detailed help is available here.

Download region GenBank file

Download region SVG

Location: 2,560,252 - 2,603,454 nt. (total: 43,203 nt)
Show pHMM detection rules used

NRPS-like: cds((PP-binding or NAD\_binding\_4) and (AMP-binding or A-OX))

#### Legend:

core biosynthetic genes

additional biosynthetic genes

transport-related genes

regulatory genes

other genes

resistance

reset view

zoom to selection

Gene details

Shows details of the most recently selected gene, including names, products, location, and other annotations.

Select a gene to view the details available for it

NRPS/PKS domains

ClusterBlast

KnownClusterBlast

SubClusterBlast

MIBiG comparison

Pfam domains

Detailed domain annotation

Shows NRPS- and PKS-related domains for each feature that contains them. Click on each domain for more information about the domain's location, consensus monomer prediction, and other details.  
A glossary is available here.

Selected features only

Show module domains

Similar gene clusters

Shows clusters from the antiSMASH database and other clusters of interest that are similar to the current region. Genes marked with the same colour are interrelated. White genes have no relationship.  
Click on reference genes to show details of similarities to genes within the current region.  
Click on an accession to open that entry in the antiSMASH database (if applicable).

All hits

NC\_015312 (234323-317006): Pseudonocardia dioxanivorans CB1190, complete sequ... (3% of genes show similarity), NRPS

NZ\_CP019724 (<92918->222567): Streptomyces pactum strain ACT12 chromosome, co... (6% of genes show similarity), NRPS,T2PKS,other

NZ\_CP019724 (8328160-8459066): Streptomyces pactum strain ACT12 chromosome, c... (6% of genes show similarity), NRPS,T2PKS,other

NZ\_FOAZ01000001 (501540-577120): Streptacidiphilus jiangxiensis strain CGMCC ... (3% of genes show similarity), NRPS

NZ\_CP010407 (6327232-6440759): Streptomyces vietnamensis strain GIMV4.0001 ch... (3% of genes show similarity), NRPS,T1PKS,T2PKS

NZ\_CP034550 (7603482-7727935): Saccharothrix syringae strain NRRL B-16468 chr... (5% of genes show similarity), NRPS,T2PKS

NZ\_FOPR01000001 (828149-881769): Pseudomonas syringae strain BS3829, whole ge... (6% of genes show similarity), NRPS

NZ\_CP013743 (204849-316559): Streptomyces sp. CdTB01 chromosome, complete genome (4% of genes show similarity), NRPS,T2PKS

NC\_009142 (1408066-1461769): Saccharopolyspora erythraea NRRL 2338, complete ... (4% of genes show similarity), NRPS

NZ\_GL877878 (3038168-3144077): Saccharopolyspora spinosa NRRL 18395 Scaffold0... (2% of genes show similarity), NRPS
Download graphic

Similar known gene clusters

Shows clusters from the MiBIG database that are similar to the current region. Genes marked with the same colour are interrelated. White genes have no relationship.  
Click on reference genes to show details of similarities to genes within the current region.  
Click on an accession to open that entry in the MiBIG database.

No matches found.

Similar subclusters

Shows sub-cluster units that are similar to the current region. Genes marked with the same colour are interrelated. White genes have no relationship.  
Click on reference genes to show details of similarities to genes within the current region.

No matches found.

Similar gene clusters

Shows careas that are similar to the current region to a reference database.  
Mouseover a score cell in the table to get a breakdown of how the score was calculated.The MIBiG database.  
  
Click on an accession to open that entry in the MIBiG database.

Analysis type:

Protocluster to Region
Region to Region

| Reference | NRPS-like | Similarity score | Type | Compound(s) | Organism |
| --- | --- | --- | --- | --- | --- |
| BGC0001168.1 |  | 0.21 | NRP | livipeptin | Streptomyces lividans 1326 |
| BGC0001900.1 |  | 0.21 | NRP | fragin | Burkholderia cenocepacia H111 |
| BGC0001132.1 |  | 0.18 | NRP | xenotetrapeptide | Xenorhabdus nematophila ATCC 19061 |
| BGC0001641.1 |  | 0.18 | NRP | kolossin | Photorhabdus laumondii subsp. laumondii TTO1 |
| BGC0001825.1 |  | 0.18 | NRP | xenematide | Xenorhabdus nematophila AN6/1 |
| BGC0001844.1 |  | 0.17 | NRP | holrhizin | Paraburkholderia rhizoxinica HKI 454 |
| BGC0001833.1 |  | 0.17 | NRP | icosalide A, icosalide B | Burkholderia gladioli |
| BGC0001135.1 |  | 0.17 | NRP | bicornutin A1, bicornutin A2 | Xenorhabdus budapestensis |
| BGC0001220.1 |  | 0.17 | NRP | aculeacin A | Aspergillus japonicus |
| BGC0001873.1 |  | 0.17 | NRP | pyrrolizixenamide A | Xenorhabdus szentirmaii DSM 16338 |

| Reference | Aggregated | Similarity score | Type | Compound(s) | Organism |
| --- | --- | --- | --- | --- | --- |
| BGC0001168.1 |  | 0.54 | NRP | livipeptin | Streptomyces lividans 1326 |
| BGC0001900.1 |  | 0.53 | NRP | fragin | Burkholderia cenocepacia H111 |
| BGC0001132.1 |  | 0.49 | NRP | xenotetrapeptide | Xenorhabdus nematophila ATCC 19061 |
| BGC0001641.1 |  | 0.49 | NRP | kolossin | Photorhabdus laumondii subsp. laumondii TTO1 |
| BGC0001825.1 |  | 0.49 | NRP | xenematide | Xenorhabdus nematophila AN6/1 |
| BGC0001844.1 |  | 0.49 | NRP | holrhizin | Paraburkholderia rhizoxinica HKI 454 |
| BGC0001833.1 |  | 0.49 | NRP | icosalide A, icosalide B | Burkholderia gladioli |
| BGC0001135.1 |  | 0.49 | NRP | bicornutin A1, bicornutin A2 | Xenorhabdus budapestensis |
| BGC0001220.1 |  | 0.49 | NRP | aculeacin A | Aspergillus japonicus |
| BGC0001873.1 |  | 0.48 | NRP | pyrrolizixenamide A | Xenorhabdus szentirmaii DSM 16338 |

Detailed Pfam domain annotation

Shows Pfam domains found in each gene within the region.
Click on each domain for more information about the domain's
accession, location, description, and any relevant Gene Ontology.
Domains with a bold border have Gene Ontology information.

Selected features only

contig\_15 - Region 3 - terpene

Shows the layout of the region, marking coding sequences and areas of interest. Clicking a gene will select it and show any relevant details. Clicking an area feature (e.g. a candidate cluster) will select all coding sequences within that area. Double clicking an area feature will zoom to that area. Multiple genes and area features can be selected by clicking them while holding the Ctrl key.  
More detailed help is available here.

Download region GenBank file

Download region SVG

Location: 2,691,940 - 2,713,034 nt. (total: 21,095 nt)
Show pHMM detection rules used

terpene: (Terpene\_synth or Terpene\_synth\_C or phytoene\_synt or Lycopene\_cycl or terpene\_cyclase or NapT7 or fung\_ggpps or fung\_ggpps2 or trichodiene\_synth or TRI5)

#### Legend:

core biosynthetic genes

additional biosynthetic genes

transport-related genes

regulatory genes

other genes

resistance

reset view

zoom to selection

Gene details

Shows details of the most recently selected gene, including names, products, location, and other annotations.

Select a gene to view the details available for it

ClusterBlast

KnownClusterBlast

SubClusterBlast

MIBiG comparison

Pfam domains

Similar gene clusters

Shows clusters from the antiSMASH database and other clusters of interest that are similar to the current region. Genes marked with the same colour are interrelated. White genes have no relationship.  
Click on reference genes to show details of similarities to genes within the current region.  
Click on an accession to open that entry in the antiSMASH database (if applicable).

No significant ClusterBlast hits found.

Similar known gene clusters

Shows clusters from the MiBIG database that are similar to the current region. Genes marked with the same colour are interrelated. White genes have no relationship.  
Click on reference genes to show details of similarities to genes within the current region.  
Click on an accession to open that entry in the MiBIG database.

No matches found.

Similar subclusters

Shows sub-cluster units that are similar to the current region. Genes marked with the same colour are interrelated. White genes have no relationship.  
Click on reference genes to show details of similarities to genes within the current region.

No matches found.

Similar gene clusters

Shows careas that are similar to the current region to a reference database.  
Mouseover a score cell in the table to get a breakdown of how the score was calculated.The MIBiG database.  
  
Click on an accession to open that entry in the MIBiG database.

Analysis type:

Protocluster to Region
Region to Region

| Reference | terpene | Similarity score | Type | Compound(s) | Organism |
| --- | --- | --- | --- | --- | --- |
| BGC0000686.1 |  | 0.14 | Terpene | helvolic acid | Aspergillus fumigatus Af293 |
| BGC0001996.1 |  | 0.13 | Other | oryzine A, oryzine B | Aspergillus oryzae RIB40 |
| BGC0000010.1 |  | 0.13 | Polyketide | aflatoxin | Aspergillus flavus |
| BGC0002013.1 |  | 0.12 | RiPP | curacozole | Streptomyces curacoi |
| BGC0001991.1 |  | 0.12 | Polyketide | toblerol A, toblerol B, toblerol C, toblerol D, toblerol E, toblerol F, toblerol G, toblerol H | Methylorubrum extorquens AM1 |
| BGC0000134.1 |  | 0.08 | Polyketide | radicicol | Pochonia chlamydosporia |
| BGC0000819.1 |  | 0.07 | NRP, Alkaloid | paraherquamide | Penicillium fellutanum |
| BGC0000065.1 |  | 0.05 | Polyketide | rustmicin | Streptomyces galbus |
| BGC0000009.1 |  | 0.05 | Polyketide | aflatoxin | Aspergillus nomius |
| BGC0000007.1 |  | 0.05 | Polyketide | aflatoxin | Aspergillus flavus |

| Reference | Aggregated | Similarity score | Type | Compound(s) | Organism |
| --- | --- | --- | --- | --- | --- |
| BGC0000686.1 |  | 0.43 | Terpene | helvolic acid | Aspergillus fumigatus Af293 |
| BGC0001996.1 |  | 0.41 | Other | oryzine A, oryzine B | Aspergillus oryzae RIB40 |
| BGC0000010.1 |  | 0.41 | Polyketide | aflatoxin | Aspergillus flavus |
| BGC0002013.1 |  | 0.40 | RiPP | curacozole | Streptomyces curacoi |
| BGC0001991.1 |  | 0.40 | Polyketide | toblerol A, toblerol B, toblerol C, toblerol D, toblerol E, toblerol F, toblerol G, toblerol H | Methylorubrum extorquens AM1 |
| BGC0000134.1 |  | 0.32 | Polyketide | radicicol | Pochonia chlamydosporia |
| BGC0000819.1 |  | 0.28 | NRP, Alkaloid | paraherquamide | Penicillium fellutanum |
| BGC0000065.1 |  | 0.22 | Polyketide | rustmicin | Streptomyces galbus |
| BGC0000009.1 |  | 0.21 | Polyketide | aflatoxin | Aspergillus nomius |
| BGC0000007.1 |  | 0.21 | Polyketide | aflatoxin | Aspergillus flavus |

Detailed Pfam domain annotation

Shows Pfam domains found in each gene within the region.
Click on each domain for more information about the domain's
accession, location, description, and any relevant Gene Ontology.
Domains with a bold border have Gene Ontology information.

Selected features only

contig\_17 - Region 1 - terpene

Shows the layout of the region, marking coding sequences and areas of interest. Clicking a gene will select it and show any relevant details. Clicking an area feature (e.g. a candidate cluster) will select all coding sequences within that area. Double clicking an area feature will zoom to that area. Multiple genes and area features can be selected by clicking them while holding the Ctrl key.  
More detailed help is available here.

Download region GenBank file

Download region SVG

Location: 148,217 - 169,902 nt. (total: 21,686 nt)
Show pHMM detection rules used

terpene: (Terpene\_synth or Terpene\_synth\_C or phytoene\_synt or Lycopene\_cycl or terpene\_cyclase or NapT7 or fung\_ggpps or fung\_ggpps2 or trichodiene\_synth or TRI5)

#### Legend:

core biosynthetic genes

additional biosynthetic genes

transport-related genes

regulatory genes

other genes

resistance

reset view

zoom to selection

Gene details

Shows details of the most recently selected gene, including names, products, location, and other annotations.

Select a gene to view the details available for it

ClusterBlast

KnownClusterBlast

SubClusterBlast

MIBiG comparison

Pfam domains

Similar gene clusters

Shows clusters from the antiSMASH database and other clusters of interest that are similar to the current region. Genes marked with the same colour are interrelated. White genes have no relationship.  
Click on reference genes to show details of similarities to genes within the current region.  
Click on an accession to open that entry in the antiSMASH database (if applicable).

All hits

NW\_001939248 (1292694-1314983): Pyrenophora tritici-repentis Pt-1C-BFP superc... (44% of genes show similarity), terpene

NW\_001939248 (494344-513275): Pyrenophora tritici-repentis Pt-1C-BFP supercon... (22% of genes show similarity), terpene

CP042201 (1414941-1458802): Venturia effusa strain albino chromosome 17, comp... (13% of genes show similarity), NRPS-like
Download graphic

Similar known gene clusters

Shows clusters from the MiBIG database that are similar to the current region. Genes marked with the same colour are interrelated. White genes have no relationship.  
Click on reference genes to show details of similarities to genes within the current region.  
Click on an accession to open that entry in the MiBIG database.

No matches found.

Similar subclusters

Shows sub-cluster units that are similar to the current region. Genes marked with the same colour are interrelated. White genes have no relationship.  
Click on reference genes to show details of similarities to genes within the current region.

No matches found.

Similar gene clusters

Shows careas that are similar to the current region to a reference database.  
Mouseover a score cell in the table to get a breakdown of how the score was calculated.The MIBiG database.  
  
Click on an accession to open that entry in the MIBiG database.

Analysis type:

Protocluster to Region
Region to Region

| Reference | terpene | Similarity score | Type | Compound(s) | Organism |
| --- | --- | --- | --- | --- | --- |
| BGC0000673.1 |  | 0.17 | Terpene | pimara-8(14),15-diene | Aspergillus nidulans FGSC A4 |
| BGC0001659.1 |  | 0.15 | Terpene | mangicol A | Fusarium equiseti |
| BGC0000676.1 |  | 0.14 | Terpene | aphidicolin, aphidicolan-16β-ol, 3-deoxyaphidicolin, 17-deoxyaphidicolin | Phoma betae |
| BGC0000688.1 |  | 0.11 | Terpene | copalyl diphosphate | Diaporthe amygdali |
| BGC0001082.1 |  | 0.08 | Terpene | paxilline, paspaline, 13-dehydroxypaxilline, paspaline B | Penicillium paxilli |
| BGC0001969.1 |  | 0.07 | Terpene | asperterpenoid A | Talaromyces wortmannii |
| BGC0001776.1 |  | 0.06 | Terpene | shearinine D | Penicillium janthinellum |
| BGC0000646.1 |  | 0.04 | Terpene | β-carotein | uncultured bacterium |
| BGC0001604.1 |  | 0.04 | Terpene | gibberellin | Fusarium fujikuroi |

| Reference | Aggregated | Similarity score | Type | Compound(s) | Organism |
| --- | --- | --- | --- | --- | --- |
| BGC0000673.1 |  | 0.49 | Terpene | pimara-8(14),15-diene | Aspergillus nidulans FGSC A4 |
| BGC0001659.1 |  | 0.46 | Terpene | mangicol A | Fusarium equiseti |
| BGC0000676.1 |  | 0.43 | Terpene | aphidicolin, aphidicolan-16β-ol, 3-deoxyaphidicolin, 17-deoxyaphidicolin | Phoma betae |
| BGC0000688.1 |  | 0.38 | Terpene | copalyl diphosphate | Diaporthe amygdali |
| BGC0001082.1 |  | 0.30 | Terpene | paxilline, paspaline, 13-dehydroxypaxilline, paspaline B | Penicillium paxilli |
| BGC0001969.1 |  | 0.28 | Terpene | asperterpenoid A | Talaromyces wortmannii |
| BGC0001776.1 |  | 0.26 | Terpene | shearinine D | Penicillium janthinellum |
| BGC0000646.1 |  | 0.20 | Terpene | β-carotein | uncultured bacterium |
| BGC0001604.1 |  | 0.19 | Terpene | gibberellin | Fusarium fujikuroi |

Detailed Pfam domain annotation

Shows Pfam domains found in each gene within the region.
Click on each domain for more information about the domain's
accession, location, description, and any relevant Gene Ontology.
Domains with a bold border have Gene Ontology information.

Selected features only

contig\_17 - Region 2 - T1PKS

Shows the layout of the region, marking coding sequences and areas of interest. Clicking a gene will select it and show any relevant details. Clicking an area feature (e.g. a candidate cluster) will select all coding sequences within that area. Double clicking an area feature will zoom to that area. Multiple genes and area features can be selected by clicking them while holding the Ctrl key.  
More detailed help is available here.

Download region GenBank file

Download region SVG

Location: 3,156,185 - 3,202,827 nt. (total: 46,643 nt)
Show pHMM detection rules used

T1PKS: cds(PKS\_AT and (PKS\_KS or ene\_KS or mod\_KS or hyb\_KS or itr\_KS or tra\_KS))

#### Legend:

core biosynthetic genes

additional biosynthetic genes

transport-related genes

regulatory genes

other genes

resistance

reset view

zoom to selection

Gene details

Shows details of the most recently selected gene, including names, products, location, and other annotations.

Select a gene to view the details available for it

NRPS/PKS domains

ClusterBlast

KnownClusterBlast

SubClusterBlast

MIBiG comparison

Pfam domains

Detailed domain annotation

Shows NRPS- and PKS-related domains for each feature that contains them. Click on each domain for more information about the domain's location, consensus monomer prediction, and other details.  
A glossary is available here.

Selected features only

Show module domains

Similar gene clusters

Shows clusters from the antiSMASH database and other clusters of interest that are similar to the current region. Genes marked with the same colour are interrelated. White genes have no relationship.  
Click on reference genes to show details of similarities to genes within the current region.  
Click on an accession to open that entry in the antiSMASH database (if applicable).

All hits

NZ\_CP012159 (7479336-7584891): Chondromyces crocatus strain Cm c5 chromosome,... (32% of genes show similarity), NRPS,T1PKS

NW\_013550603 (4984406-5031188): Rhinocladiella mackenziei CBS 650.93 unplaced... (13% of genes show similarity), T1PKS

NW\_008481832 (391284-438019): Cladophialophora carrionii CBS 160.54 unplaced ... (15% of genes show similarity), T1PKS

NW\_015971651 (252180-298965): Fonsecaea multimorphosa CBS 102226 unplaced gen... (18% of genes show similarity), T1PKS

NW\_017387254 (270821-317619): Fonsecaea erecta strain CBS 125763 chromosome U... (16% of genes show similarity), T1PKS

NW\_013550607 (222843-269664): Fonsecaea pedrosoi CBS 271.37 unplaced genomic ... (14% of genes show similarity), T1PKS

NW\_015622518 (1932984-1979646): Exophiala spinifera strain CBS 89968 unplaced... (13% of genes show similarity), T1PKS

NC\_021191 (2626192-2710257): Actinoplanes sp. N902-109, complete genome (9% of genes show similarity), T1PKS

NC\_016582 (2058350-2126205): Streptomyces bingchenggensis BCW-1, complete seq... (6% of genes show similarity), PKS-like,T1PKS

NC\_016459 (4360-72261): Thermothielavioides terrestris NRRL 8126 chromosome 3... (9% of genes show similarity), T1PKS
Download graphic

Similar known gene clusters

Shows clusters from the MiBIG database that are similar to the current region. Genes marked with the same colour are interrelated. White genes have no relationship.  
Click on reference genes to show details of similarities to genes within the current region.  
Click on an accession to open that entry in the MiBIG database.

All hits

melanin

1,3,6,8-tetrahydroxynaphthalene
Download graphic

Similar subclusters

Shows sub-cluster units that are similar to the current region. Genes marked with the same colour are interrelated. White genes have no relationship.  
Click on reference genes to show details of similarities to genes within the current region.

No matches found.

Similar gene clusters

Shows careas that are similar to the current region to a reference database.  
Mouseover a score cell in the table to get a breakdown of how the score was calculated.The MIBiG database.  
  
Click on an accession to open that entry in the MIBiG database.

Analysis type:

Protocluster to Region
Region to Region

| Reference | T1PKS | Similarity score | Type | Compound(s) | Organism |
| --- | --- | --- | --- | --- | --- |
| BGC0001258.1 |  | 0.38 | Polyketide | 1,3,6,8-tetrahydroxynaphthalene | Glarea lozoyensis |
| BGC0001265.1 |  | 0.38 | Polyketide | melanin | Bipolaris oryzae |
| BGC0001257.1 |  | 0.37 | Polyketide | 1,3,6,8-tetrahydroxynaphthalene | Nodulisporium sp. ATCC74245 |
| BGC0001284.1 |  | 0.36 | Terpene | alternariol | Parastagonospora nodorum SN15 |
| BGC0000107.1 |  | 0.36 | Polyketide | naphthopyrone | Aspergillus nidulans FGSC A4 |
| BGC0001906.1 |  | 0.36 | Polyketide | naphthalene | Daldinia eschscholzii IFB-TL01 |
| BGC0000156.1 |  | 0.36 | Polyketide | TAN-1612, 1-(2,3,5,10-tetrahydroxy-7-methoxy-4-oxo-1,2,3,4-tetrahydroanthracen-2-yl)pentane-2,4-dione, desmethyl TAN-1612 | Aspergillus niger |
| BGC0000057.1 |  | 0.33 | Polyketide | F9775A, F9775B, orsellinic acid | Aspergillus nidulans FGSC A4 |
| BGC0000684.1 |  | 0.32 | Terpene | asperthecin | Aspergillus nidulans FGSC A4 |
| BGC0001266.1 |  | 0.32 | Polyketide | grayanic acid | Cladonia grayi |

| Reference | Aggregated | Similarity score | Type | Compound(s) | Organism |
| --- | --- | --- | --- | --- | --- |
| BGC0001258.1 |  | 0.70 | Polyketide | 1,3,6,8-tetrahydroxynaphthalene | Glarea lozoyensis |
| BGC0001265.1 |  | 0.70 | Polyketide | melanin | Bipolaris oryzae |
| BGC0001257.1 |  | 0.70 | Polyketide | 1,3,6,8-tetrahydroxynaphthalene | Nodulisporium sp. ATCC74245 |
| BGC0001284.1 |  | 0.69 | Terpene | alternariol | Parastagonospora nodorum SN15 |
| BGC0000107.1 |  | 0.69 | Polyketide | naphthopyrone | Aspergillus nidulans FGSC A4 |
| BGC0001906.1 |  | 0.69 | Polyketide | naphthalene | Daldinia eschscholzii IFB-TL01 |
| BGC0000156.1 |  | 0.68 | Polyketide | TAN-1612, 1-(2,3,5,10-tetrahydroxy-7-methoxy-4-oxo-1,2,3,4-tetrahydroanthracen-2-yl)pentane-2,4-dione, desmethyl TAN-1612 | Aspergillus niger |
| BGC0000057.1 |  | 0.66 | Polyketide | F9775A, F9775B, orsellinic acid | Aspergillus nidulans FGSC A4 |
| BGC0000684.1 |  | 0.66 | Terpene | asperthecin | Aspergillus nidulans FGSC A4 |
| BGC0001266.1 |  | 0.65 | Polyketide | grayanic acid | Cladonia grayi |

Detailed Pfam domain annotation

Shows Pfam domains found in each gene within the region.
Click on each domain for more information about the domain's
accession, location, description, and any relevant Gene Ontology.
Domains with a bold border have Gene Ontology information.

Selected features only

NRPS/PKS products

NRPS/PKS monomers

Predicted core structure(s)

Shows estimated product structure and polymer for each candidate cluster in the region. To show the product, click on the expander or the candidate cluster feature drawn in the overview.

For candidate cluster 2, location 3156184 - 3202827:

Rough prediction of core scaffold based on assumed PKS/NRPS colinearity; tailoring reactions not taken into account

**Polymer prediction:**
:   (mal)

  
Direct lookup in NORINE database:
strict
or
relaxed

Link to NORINE database query form

NRPS/PKS monomer predictions

Shows the predicted monomers for each adynelation domain and acyltransferase within genes. Each gene prediction can be expanded to view detailed predictions of each domain. Each prediction can be expanded to view the predictions by tool (and, for some tools, further expanded for extra details).

**input.path1.gene918**: mal

:   **PKS\_AT (786..1082)**: mal

    ATSignature: Malonyl-CoA

    Top 3 matches:
    :   Malonyl-CoA: 70.8%
    :   inactive: 66.7%
    :   Methylmalonyl-CoA: 54.2%

      
    minowa: Malonyl-CoA

    Prediction, score:
    :   Malonyl-CoA: 113.4


        inactive: 70.0


        Methylmalonyl-CoA: 60.2


        Methoxymalonyl-CoA: 56.7


        Propionyl-CoA: 42.7


        Isobutyryl-CoA: 29.9


        fatty\_acid: 27.5


        Ethylmalonyl-CoA: 26.9


        Acetyl-CoA: 25.8


        2-Methylbutyryl-CoA: 25.6


        Benzoyl-CoA: 22.0


        CHC-CoA: 19.9


        trans-1,2-CPDA: 15.9


        3-Methylbutyryl-CoA: 15.2

contig\_20 - Region 1 - NRPS

Shows the layout of the region, marking coding sequences and areas of interest. Clicking a gene will select it and show any relevant details. Clicking an area feature (e.g. a candidate cluster) will select all coding sequences within that area. Double clicking an area feature will zoom to that area. Multiple genes and area features can be selected by clicking them while holding the Ctrl key.  
More detailed help is available here.

Download region GenBank file

Download region SVG

Location: 1 - 43,431 nt. (total: 43,431 nt)
Show pHMM detection rules used

Region on contig edge.

NRPS: cds(Condensation and (AMP-binding or A-OX))

#### Legend:

core biosynthetic genes

additional biosynthetic genes

transport-related genes

regulatory genes

other genes

resistance

reset view

zoom to selection

Gene details

Shows details of the most recently selected gene, including names, products, location, and other annotations.

Select a gene to view the details available for it

NRPS/PKS domains

ClusterBlast

KnownClusterBlast

SubClusterBlast

MIBiG comparison

Pfam domains

Detailed domain annotation

Shows NRPS- and PKS-related domains for each feature that contains them. Click on each domain for more information about the domain's location, consensus monomer prediction, and other details.  
A glossary is available here.

Selected features only

Show module domains

Similar gene clusters

Shows clusters from the antiSMASH database and other clusters of interest that are similar to the current region. Genes marked with the same colour are interrelated. White genes have no relationship.  
Click on reference genes to show details of similarities to genes within the current region.  
Click on an accession to open that entry in the antiSMASH database (if applicable).

All hits

NW\_022983863 (5871271-5921179): Arthroderma uncinatum strain CBS 119779 chrom... (15% of genes show similarity), NRPS

NW\_003299166 (2130053-2177991): Microsporum canis CBS 113480 supercont1.4 gen... (13% of genes show similarity), NRPS

NW\_003315112 (1173041-1224848): Trichophyton benhamiae CBS 112371 chromosome ... (13% of genes show similarity), NRPS

NW\_003315027 (453020-520877): Verticillium alfalfae VaMs.102 supercont1.12 ge... (20% of genes show similarity), NRPS

NT\_165937 (139863-204269): Aspergillus terreus NIH2624 scaffold 14 genomic sc... (9% of genes show similarity), NRPS

NT\_107012 (44177-106681): Aspergillus nidulans FGSC A4 chromosome II map unlo... (11% of genes show similarity), NRPS

NW\_003345194 (767990-812854): Nannizzia gypsea CBS 118893 supercont1.8 genomi... (13% of genes show similarity), NRPS

NW\_003456426 (1219119-1268117): Trichophyton rubrum CBS 118892 genomic scaffo... (11% of genes show similarity), NRPS

NW\_022474210 (169476-217015): Venustampulla echinocandica strain BP 5553 chro... (16% of genes show similarity), NRPS

NZ\_CP034669 (4665507-4751192): Corallococcus coralloides strain B035 chromoso... (5% of genes show similarity), NRPS
Download graphic

Similar known gene clusters

Shows clusters from the MiBIG database that are similar to the current region. Genes marked with the same colour are interrelated. White genes have no relationship.  
Click on reference genes to show details of similarities to genes within the current region.  
Click on an accession to open that entry in the MiBIG database.

No matches found.

Similar subclusters

Shows sub-cluster units that are similar to the current region. Genes marked with the same colour are interrelated. White genes have no relationship.  
Click on reference genes to show details of similarities to genes within the current region.

No matches found.

Similar gene clusters

Shows careas that are similar to the current region to a reference database.  
Mouseover a score cell in the table to get a breakdown of how the score was calculated.The MIBiG database.  
  
Click on an accession to open that entry in the MIBiG database.

Analysis type:

Protocluster to Region
Region to Region

| Reference | NRPS | Similarity score | Type | Compound(s) | Organism |
| --- | --- | --- | --- | --- | --- |
| BGC0000342.1 |  | 0.35 | NRP | enniatin | Fusarium equiseti |
| BGC0000357.1 |  | 0.30 | NRP | cyclo-(D-Phe-L-Phe-D-Val-L-Val), cyclo-(D-Tyr-L-Phe-D-Val-L-Val), cyclo-(D-Tyr-L-Trp-D-Val-L-Val), cyclo-(D-Phe-L-Trp-D-Val-L-Val), cyclo-(D-Phe-L-Phe-D-Val-L-Ile), cyclo-(D-Phe-L-Phe-D-Ile-L-Val), cyclo-(D-Tyr-L-Trp-D-Val-L-Ile), cyclo-(D-Tyr-L-Trp-D-Ile-L-Val), cyclo-(D-Tyr-L-Phe-D-Val-L-Ile), cyclo-(D-Tyr-L-Phe-D-Ile-L-Val) | Penicillium rubens Wisconsin 54-1255 |
| BGC0000313.1 |  | 0.29 | NRP | beauvericin | Beauveria bassiana |
| BGC0001825.1 |  | 0.29 | NRP | xenematide | Xenorhabdus nematophila AN6/1 |
| BGC0000396.1 |  | 0.29 | NRP | nodularin | Nostoc sp. 73.1 |
| BGC0000901.1 |  | 0.28 | Other | ferrichrome | Aspergillus niger |
| BGC0001261.1 |  | 0.27 | NRP | AM-toxin | Alternaria alternata |
| BGC0000426.1 |  | 0.27 | NRP | sevadicin | Paenibacillus larvae |
| BGC0000307.1 |  | 0.27 | NRP | AbT1 | Aureobasidium pullulans |
| BGC0001166.1 |  | 0.27 | NRP | HC-toxin | Alternaria jesenskae |

| Reference | Aggregated | Similarity score | Type | Compound(s) | Organism |
| --- | --- | --- | --- | --- | --- |
| BGC0000342.1 |  | 0.68 | NRP | enniatin | Fusarium equiseti |
| BGC0000357.1 |  | 0.63 | NRP | cyclo-(D-Phe-L-Phe-D-Val-L-Val), cyclo-(D-Tyr-L-Phe-D-Val-L-Val), cyclo-(D-Tyr-L-Trp-D-Val-L-Val), cyclo-(D-Phe-L-Trp-D-Val-L-Val), cyclo-(D-Phe-L-Phe-D-Val-L-Ile), cyclo-(D-Phe-L-Phe-D-Ile-L-Val), cyclo-(D-Tyr-L-Trp-D-Val-L-Ile), cyclo-(D-Tyr-L-Trp-D-Ile-L-Val), cyclo-(D-Tyr-L-Phe-D-Val-L-Ile), cyclo-(D-Tyr-L-Phe-D-Ile-L-Val) | Penicillium rubens Wisconsin 54-1255 |
| BGC0000313.1 |  | 0.63 | NRP | beauvericin | Beauveria bassiana |
| BGC0001825.1 |  | 0.62 | NRP | xenematide | Xenorhabdus nematophila AN6/1 |
| BGC0000396.1 |  | 0.62 | NRP | nodularin | Nostoc sp. 73.1 |
| BGC0000901.1 |  | 0.62 | Other | ferrichrome | Aspergillus niger |
| BGC0001261.1 |  | 0.61 | NRP | AM-toxin | Alternaria alternata |
| BGC0000426.1 |  | 0.61 | NRP | sevadicin | Paenibacillus larvae |
| BGC0000307.1 |  | 0.61 | NRP | AbT1 | Aureobasidium pullulans |
| BGC0001166.1 |  | 0.60 | NRP | HC-toxin | Alternaria jesenskae |

Detailed Pfam domain annotation

Shows Pfam domains found in each gene within the region.
Click on each domain for more information about the domain's
accession, location, description, and any relevant Gene Ontology.
Domains with a bold border have Gene Ontology information.

Selected features only

NRPS/PKS products

NRPS/PKS monomers

Predicted core structure(s)

Shows estimated product structure and polymer for each candidate cluster in the region. To show the product, click on the expander or the candidate cluster feature drawn in the overview.

For candidate cluster 1, location 0 - 43431:

Rough prediction of core scaffold based on assumed PKS/NRPS colinearity; tailoring reactions not taken into account

**Polymer prediction:**
:   (D-val - D-X - X) + (phe - X)

  
Direct lookup in NORINE database:
strict
or
relaxed

Link to NORINE database query form

NRPS/PKS monomer predictions

Shows the predicted monomers for each adynelation domain and acyltransferase within genes. Each gene prediction can be expanded to view detailed predictions of each domain. Each prediction can be expanded to view the predictions by tool (and, for some tools, further expanded for extra details).

**input.path1.gene1**: phe - X

:   Search NORINE for peptide:
    strict
    or
    relaxed
  
:   **AMP-binding (718..1123)**: phe

    NRPSPredictor2: phe

    SVM prediction details:
    :   Predicted physicochemical class:
        :   N/A

        Large clusters prediction:
        :   phe, trp, phg, tyr, bht

        Small clusters prediction:
        :   phe, trp

        Single AA prediction:
        :   phe

    Stachelhaus prediction details:
    :   Stachelhaus sequence:
        :   dawlcgcvck

        Nearest Stachelhaus code:
        :   N, A

        Stachelhaus code match:
        :   0% (weak)
:   **AMP-binding (2171..2426)**: X

    NRPSPredictor2: gly, ala, val, leu, ile, abu, iva

    SVM prediction details:
    :   Predicted physicochemical class:
        :   hydrophobic-aliphatic

        Large clusters prediction:
        :   gly, ala, val, leu, ile, abu, iva

        Small clusters prediction:
        :   N/A

        Single AA prediction:
        :   N/A

    Stachelhaus prediction details:
    :   Stachelhaus sequence:
        :   ey-w--tgvk

        Nearest Stachelhaus code:
        :   N, A

        Stachelhaus code match:
        :   0% (weak)

  
**input.path1.gene2**: val - X - X

:   Search NORINE for peptide:
    strict
    or
    relaxed
  
:   **AMP-binding (55..448)**: val

    NRPSPredictor2: val

    SVM prediction details:
    :   Predicted physicochemical class:
        :   hydrophobic-aliphatic

        Large clusters prediction:
        :   gly, ala, val, leu, ile, abu, iva

        Small clusters prediction:
        :   val, leu, ile, abu, iva

        Single AA prediction:
        :   val

    Stachelhaus prediction details:
    :   Stachelhaus sequence:
        :   daafvggvfk

        Nearest Stachelhaus code:
        :   N, A

        Stachelhaus code match:
        :   0% (weak)
:   **AMP-binding (1589..1981)**: X

    NRPSPredictor2: hydrophobic-aliphatic

    SVM prediction details:
    :   Predicted physicochemical class:
        :   hydrophobic-aliphatic

        Large clusters prediction:
        :   N/A

        Small clusters prediction:
        :   N/A

        Single AA prediction:
        :   N/A

    Stachelhaus prediction details:
    :   Stachelhaus sequence:
        :   datlvgavvk

        Nearest Stachelhaus code:
        :   N, A

        Stachelhaus code match:
        :   0% (weak)
:   **AMP-binding (3082..3478)**: X

    NRPSPredictor2: val, leu, ile, abu, iva

    SVM prediction details:
    :   Predicted physicochemical class:
        :   hydrophobic-aliphatic

        Large clusters prediction:
        :   gly, ala, val, leu, ile, abu, iva

        Small clusters prediction:
        :   val, leu, ile, abu, iva

        Single AA prediction:
        :   N/A

    Stachelhaus prediction details:
    :   Stachelhaus sequence:
        :   agafcgtgfk

        Nearest Stachelhaus code:
        :   N, A

        Stachelhaus code match:
        :   0% (weak)

contig\_23 - Region 1 - NRPS

Shows the layout of the region, marking coding sequences and areas of interest. Clicking a gene will select it and show any relevant details. Clicking an area feature (e.g. a candidate cluster) will select all coding sequences within that area. Double clicking an area feature will zoom to that area. Multiple genes and area features can be selected by clicking them while holding the Ctrl key.  
More detailed help is available here.

Download region GenBank file

Download region SVG

Location: 1 - 24,537 nt. (total: 24,537 nt)
Show pHMM detection rules used

Region on contig edge.

NRPS: cds(Condensation and (AMP-binding or A-OX))

#### Legend:

core biosynthetic genes

additional biosynthetic genes

transport-related genes

regulatory genes

other genes

resistance

reset view

zoom to selection

Gene details

Shows details of the most recently selected gene, including names, products, location, and other annotations.

Select a gene to view the details available for it

NRPS/PKS domains

ClusterBlast

KnownClusterBlast

SubClusterBlast

MIBiG comparison

Pfam domains

Detailed domain annotation

Shows NRPS- and PKS-related domains for each feature that contains them. Click on each domain for more information about the domain's location, consensus monomer prediction, and other details.  
A glossary is available here.

Selected features only

Show module domains

Similar gene clusters

Shows clusters from the antiSMASH database and other clusters of interest that are similar to the current region. Genes marked with the same colour are interrelated. White genes have no relationship.  
Click on reference genes to show details of similarities to genes within the current region.  
Click on an accession to open that entry in the antiSMASH database (if applicable).

All hits

NW\_023336279 (253632-306603): Aspergillus tubingensis WU-2223L DNA, scaffold ... (13% of genes show similarity), NRPS

NW\_007361001 (524266-566887): Glarea lozoyensis ATCC 20868 chromosome Unknown... (11% of genes show similarity), NRPS
Download graphic

Similar known gene clusters

Shows clusters from the MiBIG database that are similar to the current region. Genes marked with the same colour are interrelated. White genes have no relationship.  
Click on reference genes to show details of similarities to genes within the current region.  
Click on an accession to open that entry in the MiBIG database.

No matches found.

Similar subclusters

Shows sub-cluster units that are similar to the current region. Genes marked with the same colour are interrelated. White genes have no relationship.  
Click on reference genes to show details of similarities to genes within the current region.

No matches found.

Similar gene clusters

Shows careas that are similar to the current region to a reference database.  
Mouseover a score cell in the table to get a breakdown of how the score was calculated.The MIBiG database.  
  
Click on an accession to open that entry in the MIBiG database.

Analysis type:

Protocluster to Region
Region to Region

| Reference | NRPS | Similarity score | Type | Compound(s) | Organism |
| --- | --- | --- | --- | --- | --- |
| BGC0000357.1 |  | 0.26 | NRP | cyclo-(D-Phe-L-Phe-D-Val-L-Val), cyclo-(D-Tyr-L-Phe-D-Val-L-Val), cyclo-(D-Tyr-L-Trp-D-Val-L-Val), cyclo-(D-Phe-L-Trp-D-Val-L-Val), cyclo-(D-Phe-L-Phe-D-Val-L-Ile), cyclo-(D-Phe-L-Phe-D-Ile-L-Val), cyclo-(D-Tyr-L-Trp-D-Val-L-Ile), cyclo-(D-Tyr-L-Trp-D-Ile-L-Val), cyclo-(D-Tyr-L-Phe-D-Val-L-Ile), cyclo-(D-Tyr-L-Phe-D-Ile-L-Val) | Penicillium rubens Wisconsin 54-1255 |
| BGC0001220.1 |  | 0.22 | NRP | aculeacin A | Aspergillus japonicus |
| BGC0001249.1 |  | 0.22 | NRP | dimethylcoprogen | Alternaria alternata |
| BGC0001261.1 |  | 0.22 | NRP | AM-toxin | Alternaria alternata |
| BGC0000900.1 |  | 0.22 | Other | ferrichrome | Aspergillus oryzae |
| BGC0000342.1 |  | 0.22 | NRP | enniatin | Fusarium equiseti |
| BGC0001240.1 |  | 0.22 | NRP | serinocyclin A, serinocyclin B | Metarhizium robertsii |
| BGC0000313.1 |  | 0.22 | NRP | beauvericin | Beauveria bassiana |
| BGC0000307.1 |  | 0.21 | NRP | AbT1 | Aureobasidium pullulans |
| BGC0001166.1 |  | 0.21 | NRP | HC-toxin | Alternaria jesenskae |

| Reference | Aggregated | Similarity score | Type | Compound(s) | Organism |
| --- | --- | --- | --- | --- | --- |
| BGC0000357.1 |  | 0.59 | NRP | cyclo-(D-Phe-L-Phe-D-Val-L-Val), cyclo-(D-Tyr-L-Phe-D-Val-L-Val), cyclo-(D-Tyr-L-Trp-D-Val-L-Val), cyclo-(D-Phe-L-Trp-D-Val-L-Val), cyclo-(D-Phe-L-Phe-D-Val-L-Ile), cyclo-(D-Phe-L-Phe-D-Ile-L-Val), cyclo-(D-Tyr-L-Trp-D-Val-L-Ile), cyclo-(D-Tyr-L-Trp-D-Ile-L-Val), cyclo-(D-Tyr-L-Phe-D-Val-L-Ile), cyclo-(D-Tyr-L-Phe-D-Ile-L-Val) | Penicillium rubens Wisconsin 54-1255 |
| BGC0001220.1 |  | 0.55 | NRP | aculeacin A | Aspergillus japonicus |
| BGC0001249.1 |  | 0.55 | NRP | dimethylcoprogen | Alternaria alternata |
| BGC0001261.1 |  | 0.55 | NRP | AM-toxin | Alternaria alternata |
| BGC0000900.1 |  | 0.55 | Other | ferrichrome | Aspergillus oryzae |
| BGC0000342.1 |  | 0.55 | NRP | enniatin | Fusarium equiseti |
| BGC0001240.1 |  | 0.55 | NRP | serinocyclin A, serinocyclin B | Metarhizium robertsii |
| BGC0000313.1 |  | 0.55 | NRP | beauvericin | Beauveria bassiana |
| BGC0000307.1 |  | 0.54 | NRP | AbT1 | Aureobasidium pullulans |
| BGC0001166.1 |  | 0.54 | NRP | HC-toxin | Alternaria jesenskae |

Detailed Pfam domain annotation

Shows Pfam domains found in each gene within the region.
Click on each domain for more information about the domain's
accession, location, description, and any relevant Gene Ontology.
Domains with a bold border have Gene Ontology information.

Selected features only

NRPS/PKS products

NRPS/PKS monomers

Predicted core structure(s)

Shows estimated product structure and polymer for each candidate cluster in the region. To show the product, click on the expander or the candidate cluster feature drawn in the overview.

For candidate cluster 1, location 0 - 24537:

Rough prediction of core scaffold based on assumed PKS/NRPS colinearity; tailoring reactions not taken into account

**Polymer prediction:**
:   (D-val)

  
Direct lookup in NORINE database:
strict
or
relaxed

Link to NORINE database query form

NRPS/PKS monomer predictions

Shows the predicted monomers for each adynelation domain and acyltransferase within genes. Each gene prediction can be expanded to view detailed predictions of each domain. Each prediction can be expanded to view the predictions by tool (and, for some tools, further expanded for extra details).

**input.path1.gene1**: val

:   Search NORINE for peptide:
    strict
    or
    relaxed
  
:   **AMP-binding (41..436)**: val

    NRPSPredictor2: val

    SVM prediction details:
    :   Predicted physicochemical class:
        :   hydrophobic-aliphatic

        Large clusters prediction:
        :   gly, ala, val, leu, ile, abu, iva

        Small clusters prediction:
        :   val, leu, ile, abu, iva

        Single AA prediction:
        :   val

    Stachelhaus prediction details:
    :   Stachelhaus sequence:
        :   daafvggvfk

        Nearest Stachelhaus code:
        :   N, A

        Stachelhaus code match:
        :   0% (weak)

contig\_26 - Region 1 - T1PKS

Shows the layout of the region, marking coding sequences and areas of interest. Clicking a gene will select it and show any relevant details. Clicking an area feature (e.g. a candidate cluster) will select all coding sequences within that area. Double clicking an area feature will zoom to that area. Multiple genes and area features can be selected by clicking them while holding the Ctrl key.  
More detailed help is available here.

Download region GenBank file

Download region SVG

Location: 23,888 - 65,396 nt. (total: 41,509 nt)
Show pHMM detection rules used

T1PKS: cds(PKS\_AT and (PKS\_KS or ene\_KS or mod\_KS or hyb\_KS or itr\_KS or tra\_KS))

#### Legend:

core biosynthetic genes

additional biosynthetic genes

transport-related genes

regulatory genes

other genes

resistance

reset view

zoom to selection

Gene details

Shows details of the most recently selected gene, including names, products, location, and other annotations.

Select a gene to view the details available for it

NRPS/PKS domains

ClusterBlast

KnownClusterBlast

SubClusterBlast

MIBiG comparison

Pfam domains

Detailed domain annotation

Shows NRPS- and PKS-related domains for each feature that contains them. Click on each domain for more information about the domain's location, consensus monomer prediction, and other details.  
A glossary is available here.

Selected features only

Show module domains

Similar gene clusters

Shows clusters from the antiSMASH database and other clusters of interest that are similar to the current region. Genes marked with the same colour are interrelated. White genes have no relationship.  
Click on reference genes to show details of similarities to genes within the current region.  
Click on an accession to open that entry in the antiSMASH database (if applicable).

All hits

NW\_022474205 (7029281-7077036): Venustampulla echinocandica strain BP 5553 ch... (42% of genes show similarity), T1PKS

NW\_020194484 (12340-73776): Amorphotheca resinae ATCC 22711 unplaced genomic ... (50% of genes show similarity), T1PKS,terpene

NW\_022983863 (624230-690958): Arthroderma uncinatum strain CBS 119779 chromos... (28% of genes show similarity), T1PKS

NW\_021167084 (1847567-1897553): Sodiomyces alkalinus F11 unplaced genomic sca... (31% of genes show similarity), T1PKS

NC\_049565 (2708272-2802515): Talaromyces rugulosus chromosome V, complete seq... (22% of genes show similarity), T1PKS

NW\_006271974 (2331315-2395136): Cordyceps militaris CM01 unplaced genomic sca... (40% of genes show similarity), NRPS,T1PKS

NW\_023336268 (2893694-2960350): Colletotrichum scovillei strain TJNH1 chromos... (33% of genes show similarity), T1PKS

NW\_022984632 (251759-297619): Aspergillus tanneri strain NIH1004 chromosome U... (42% of genes show similarity), T1PKS

NT\_107007 (292411-381895): Aspergillus nidulans FGSC A4 chromosome IV map unl... (15% of genes show similarity), T1PKS

NW\_007360999 (1771796-1819725): Glarea lozoyensis ATCC 20868 chromosome Unkno... (18% of genes show similarity), T1PKS
Download graphic

Similar known gene clusters

Shows clusters from the MiBIG database that are similar to the current region. Genes marked with the same colour are interrelated. White genes have no relationship.  
Click on reference genes to show details of similarities to genes within the current region.  
Click on an accession to open that entry in the MiBIG database.

No matches found.

Similar subclusters

Shows sub-cluster units that are similar to the current region. Genes marked with the same colour are interrelated. White genes have no relationship.  
Click on reference genes to show details of similarities to genes within the current region.

No matches found.

Similar gene clusters

Shows careas that are similar to the current region to a reference database.  
Mouseover a score cell in the table to get a breakdown of how the score was calculated.The MIBiG database.  
  
Click on an accession to open that entry in the MIBiG database.

Analysis type:

Protocluster to Region
Region to Region

| Reference | T1PKS | Similarity score | Type | Compound(s) | Organism |
| --- | --- | --- | --- | --- | --- |
| BGC0000688.1 |  | 0.29 | Terpene | copalyl diphosphate | Diaporthe amygdali |
| BGC0000631.1 |  | 0.29 | Terpene | botrydial | Botrytis cinerea B05.10 |
| BGC0001969.1 |  | 0.28 | Terpene | asperterpenoid A | Talaromyces wortmannii |
| BGC0001858.1 |  | 0.28 | Polyketide | alternapyrone B, alternapyrone C, alternapyrone D, alternapyrone E, alternapyrone F | Parastagonospora nodorum SN15 |
| BGC0001475.1 |  | 0.27 | Polyketide | aspterric acid | Aspergillus terreus NIH2624 |
| BGC0001811.1 |  | 0.27 | Terpene | trichodiene-11-one | Fusarium asiaticum |
| BGC0001277.1 |  | 0.27 | Terpene | nivalenol, vomitoxin, 3-acetyldeoxynivalenol, 15-acetyldeoxynivalenol, neosolaniol, calonectrin, apotrichodiol, isotrichotriol, 15-deacetylcalonectrin, T-2 toxin, 3-acetyl T-2 toxin, trichodiene | Fusarium graminearum |
| BGC0000676.1 |  | 0.26 | Terpene | aphidicolin, aphidicolan-16β-ol, 3-deoxyaphidicolin, 17-deoxyaphidicolin | Phoma betae |
| BGC0000003.1 |  | 0.24 | Polyketide | AF-toxin | Alternaria alternata |
| BGC0000046.1 |  | 0.23 | Polyketide | depudecin | Alternaria brassicicola |

| Reference | Aggregated | Similarity score | Type | Compound(s) | Organism |
| --- | --- | --- | --- | --- | --- |
| BGC0000688.1 |  | 0.63 | Terpene | copalyl diphosphate | Diaporthe amygdali |
| BGC0000631.1 |  | 0.62 | Terpene | botrydial | Botrytis cinerea B05.10 |
| BGC0001969.1 |  | 0.62 | Terpene | asperterpenoid A | Talaromyces wortmannii |
| BGC0001858.1 |  | 0.62 | Polyketide | alternapyrone B, alternapyrone C, alternapyrone D, alternapyrone E, alternapyrone F | Parastagonospora nodorum SN15 |
| BGC0001475.1 |  | 0.61 | Polyketide | aspterric acid | Aspergillus terreus NIH2624 |
| BGC0001811.1 |  | 0.61 | Terpene | trichodiene-11-one | Fusarium asiaticum |
| BGC0001277.1 |  | 0.60 | Terpene | nivalenol, vomitoxin, 3-acetyldeoxynivalenol, 15-acetyldeoxynivalenol, neosolaniol, calonectrin, apotrichodiol, isotrichotriol, 15-deacetylcalonectrin, T-2 toxin, 3-acetyl T-2 toxin, trichodiene | Fusarium graminearum |
| BGC0000676.1 |  | 0.60 | Terpene | aphidicolin, aphidicolan-16β-ol, 3-deoxyaphidicolin, 17-deoxyaphidicolin | Phoma betae |
| BGC0000003.1 |  | 0.58 | Polyketide | AF-toxin | Alternaria alternata |
| BGC0000046.1 |  | 0.56 | Polyketide | depudecin | Alternaria brassicicola |

Detailed Pfam domain annotation

Shows Pfam domains found in each gene within the region.
Click on each domain for more information about the domain's
accession, location, description, and any relevant Gene Ontology.
Domains with a bold border have Gene Ontology information.

Selected features only

contig\_26 - Region 2 - T1PKS

Shows the layout of the region, marking coding sequences and areas of interest. Clicking a gene will select it and show any relevant details. Clicking an area feature (e.g. a candidate cluster) will select all coding sequences within that area. Double clicking an area feature will zoom to that area. Multiple genes and area features can be selected by clicking them while holding the Ctrl key.  
More detailed help is available here.

Download region GenBank file

Download region SVG

Location: 736,919 - 783,489 nt. (total: 46,571 nt)
Show pHMM detection rules used

T1PKS: cds(PKS\_AT and (PKS\_KS or ene\_KS or mod\_KS or hyb\_KS or itr\_KS or tra\_KS))

#### Legend:

core biosynthetic genes

additional biosynthetic genes

transport-related genes

regulatory genes

other genes

resistance

reset view

zoom to selection

Gene details

Shows details of the most recently selected gene, including names, products, location, and other annotations.

Select a gene to view the details available for it

NRPS/PKS domains

ClusterBlast

KnownClusterBlast

SubClusterBlast

MIBiG comparison

Pfam domains

Detailed domain annotation

Shows NRPS- and PKS-related domains for each feature that contains them. Click on each domain for more information about the domain's location, consensus monomer prediction, and other details.  
A glossary is available here.

Selected features only

Show module domains

Similar gene clusters

Shows clusters from the antiSMASH database and other clusters of interest that are similar to the current region. Genes marked with the same colour are interrelated. White genes have no relationship.  
Click on reference genes to show details of similarities to genes within the current region.  
Click on an accession to open that entry in the antiSMASH database (if applicable).

All hits

NW\_019716270 (370523-417118): Ramularia collo-cygni strain URUG2 genome assem... (17% of genes show similarity), T1PKS

NW\_001939245 (3019281-3065857): Pyrenophora tritici-repentis Pt-1C-BFP superc... (18% of genes show similarity), T1PKS

NC\_018208 (566455-613034): Zymoseptoria tritici IPO323 chromosome 11, whole g... (21% of genes show similarity), T1PKS

CP051142 (1836611-1883130): Peltaster fructicola strain LNHT1506 chromosome 4 (21% of genes show similarity), T1PKS

CP042190 (213425-260153): Venturia effusa strain albino chromosome 6, complet... (25% of genes show similarity), T1PKS

NW\_006911258 (1739237-1784893): Baudoinia panamericana UAMH 10762 unplaced ge... (8% of genes show similarity), T1PKS

NW\_006763039 (2048727-2099747): Marssonina brunnea f. sp. 'multigermtubi' MB ... (18% of genes show similarity), T1PKS

NW\_022474218 (39162-85806): Venustampulla echinocandica strain BP 5553 chromo... (16% of genes show similarity), T1PKS

NW\_020194471 (1753062-1797008): Amorphotheca resinae ATCC 22711 unplaced geno... (12% of genes show similarity), T1PKS

NW\_007360986 (378551-425838): Glarea lozoyensis ATCC 20868 chromosome Unknown... (22% of genes show similarity), T1PKS
Download graphic

Similar known gene clusters

Shows clusters from the MiBIG database that are similar to the current region. Genes marked with the same colour are interrelated. White genes have no relationship.  
Click on reference genes to show details of similarities to genes within the current region.  
Click on an accession to open that entry in the MiBIG database.

All hits

melanin

1,3,6,8-tetrahydroxynaphthalene

1,3,6,8-tetrahydroxynaphthalene
Download graphic

Similar subclusters

Shows sub-cluster units that are similar to the current region. Genes marked with the same colour are interrelated. White genes have no relationship.  
Click on reference genes to show details of similarities to genes within the current region.

No matches found.

Similar gene clusters

Shows careas that are similar to the current region to a reference database.  
Mouseover a score cell in the table to get a breakdown of how the score was calculated.The MIBiG database.  
  
Click on an accession to open that entry in the MIBiG database.

Analysis type:

Protocluster to Region
Region to Region

| Reference | T1PKS | Similarity score | Type | Compound(s) | Organism |
| --- | --- | --- | --- | --- | --- |
| BGC0001265.1 |  | 0.48 | Polyketide | melanin | Bipolaris oryzae |
| BGC0001258.1 |  | 0.38 | Polyketide | 1,3,6,8-tetrahydroxynaphthalene | Glarea lozoyensis |
| BGC0001257.1 |  | 0.38 | Polyketide | 1,3,6,8-tetrahydroxynaphthalene | Nodulisporium sp. ATCC74245 |
| BGC0000107.1 |  | 0.37 | Polyketide | naphthopyrone | Aspergillus nidulans FGSC A4 |
| BGC0001284.1 |  | 0.35 | Terpene | alternariol | Parastagonospora nodorum SN15 |
| BGC0001906.1 |  | 0.30 | Polyketide | naphthalene | Daldinia eschscholzii IFB-TL01 |
| BGC0000156.1 |  | 0.29 | Polyketide | TAN-1612, 1-(2,3,5,10-tetrahydroxy-7-methoxy-4-oxo-1,2,3,4-tetrahydroanthracen-2-yl)pentane-2,4-dione, desmethyl TAN-1612 | Aspergillus niger |
| BGC0001583.1 |  | 0.29 | Polyketide | emodin | Escovopsis weberi |
| BGC0000121.1 |  | 0.26 | Polyketide | RES-1214-2 | Pestalotiopsis fici |
| BGC0000057.1 |  | 0.25 | Polyketide | F9775A, F9775B, orsellinic acid | Aspergillus nidulans FGSC A4 |

| Reference | Aggregated | Similarity score | Type | Compound(s) | Organism |
| --- | --- | --- | --- | --- | --- |
| BGC0001265.1 |  | 0.77 | Polyketide | melanin | Bipolaris oryzae |
| BGC0001258.1 |  | 0.70 | Polyketide | 1,3,6,8-tetrahydroxynaphthalene | Glarea lozoyensis |
| BGC0001257.1 |  | 0.70 | Polyketide | 1,3,6,8-tetrahydroxynaphthalene | Nodulisporium sp. ATCC74245 |
| BGC0000107.1 |  | 0.69 | Polyketide | naphthopyrone | Aspergillus nidulans FGSC A4 |
| BGC0001284.1 |  | 0.68 | Terpene | alternariol | Parastagonospora nodorum SN15 |
| BGC0001906.1 |  | 0.64 | Polyketide | naphthalene | Daldinia eschscholzii IFB-TL01 |
| BGC0000156.1 |  | 0.62 | Polyketide | TAN-1612, 1-(2,3,5,10-tetrahydroxy-7-methoxy-4-oxo-1,2,3,4-tetrahydroanthracen-2-yl)pentane-2,4-dione, desmethyl TAN-1612 | Aspergillus niger |
| BGC0001583.1 |  | 0.62 | Polyketide | emodin | Escovopsis weberi |
| BGC0000121.1 |  | 0.59 | Polyketide | RES-1214-2 | Pestalotiopsis fici |
| BGC0000057.1 |  | 0.58 | Polyketide | F9775A, F9775B, orsellinic acid | Aspergillus nidulans FGSC A4 |

Detailed Pfam domain annotation

Shows Pfam domains found in each gene within the region.
Click on each domain for more information about the domain's
accession, location, description, and any relevant Gene Ontology.
Domains with a bold border have Gene Ontology information.

Selected features only

NRPS/PKS products

NRPS/PKS monomers

Predicted core structure(s)

Shows estimated product structure and polymer for each candidate cluster in the region. To show the product, click on the expander or the candidate cluster feature drawn in the overview.

For candidate cluster 2, location 736918 - 783489:

Rough prediction of core scaffold based on assumed PKS/NRPS colinearity; tailoring reactions not taken into account

**Polymer prediction:**
:   (pk)

  
Direct lookup in NORINE database:
strict
or
relaxed

Link to NORINE database query form

NRPS/PKS monomer predictions

Shows the predicted monomers for each adynelation domain and acyltransferase within genes. Each gene prediction can be expanded to view detailed predictions of each domain. Each prediction can be expanded to view the predictions by tool (and, for some tools, further expanded for extra details).

**input.path1.gene207**: pk

:   **PKS\_AT (900..1148)**: pk

    ATSignature: Malonyl-CoA

    Top 3 matches:
    :   Malonyl-CoA: 66.7%

      
    minowa: Methoxymalonyl-CoA

    Prediction, score:
    :   Methoxymalonyl-CoA: 39.6


        Methylmalonyl-CoA: 38.0


        Propionyl-CoA: 14.2


        fatty\_acid: 12.5


        Ethylmalonyl-CoA: 8.7


        trans-1,2-CPDA: 0.0


        Isobutyryl-CoA: 0.0


        CHC-CoA: 0.0


        Benzoyl-CoA: 0.0


        Acetyl-CoA: 0.0


        3-Methylbutyryl-CoA: 0.0


        2-Methylbutyryl-CoA: 0.0


        Malonyl-CoA: -3.0


        inactive: -4.3

contig\_28 - Region 1 - T1PKS

Shows the layout of the region, marking coding sequences and areas of interest. Clicking a gene will select it and show any relevant details. Clicking an area feature (e.g. a candidate cluster) will select all coding sequences within that area. Double clicking an area feature will zoom to that area. Multiple genes and area features can be selected by clicking them while holding the Ctrl key.  
More detailed help is available here.

Download region GenBank file

Download region SVG

Location: 5,917 - 54,518 nt. (total: 48,602 nt)
Show pHMM detection rules used

T1PKS: cds(PKS\_AT and (PKS\_KS or ene\_KS or mod\_KS or hyb\_KS or itr\_KS or tra\_KS))

#### Legend:

core biosynthetic genes

additional biosynthetic genes

transport-related genes

regulatory genes

other genes

resistance

reset view

zoom to selection

Gene details

Shows details of the most recently selected gene, including names, products, location, and other annotations.

Select a gene to view the details available for it

NRPS/PKS domains

ClusterBlast

KnownClusterBlast

SubClusterBlast

MIBiG comparison

Pfam domains

Detailed domain annotation

Shows NRPS- and PKS-related domains for each feature that contains them. Click on each domain for more information about the domain's location, consensus monomer prediction, and other details.  
A glossary is available here.

Selected features only

Show module domains

Similar gene clusters

Shows clusters from the antiSMASH database and other clusters of interest that are similar to the current region. Genes marked with the same colour are interrelated. White genes have no relationship.  
Click on reference genes to show details of similarities to genes within the current region.  
Click on an accession to open that entry in the antiSMASH database (if applicable).

All hits

NW\_001939250 (1616542-1663778): Pyrenophora tritici-repentis Pt-1C-BFP superc... (33% of genes show similarity), T1PKS

NZ\_JHXQ01000006 (44122-99285): Rhizobium undicola ORS 992 = ATCC 700741 T424D... (5% of genes show similarity), T1PKS

NZ\_JACHJK010000016 (95643-150850): Streptomyces echinatus strain CECT 3313 Ga... (6% of genes show similarity), NRPS,T1PKS

NW\_022474205 (107314-149544): Venustampulla echinocandica strain BP 5553 chro... (21% of genes show similarity), T1PKS

NZ\_LR134356 (3157977-3224691): Mycolicibacterium aurum strain NCTC10437 chrom... (10% of genes show similarity), NRPS,T1PKS

NZ\_CM002271 (6545259-6848841): Streptomyces sp. GBA 94-10 chromosome, whole g... (5% of genes show similarity), NRPS,NRPS-like,T1PKS,T3PKS,lanthipeptide

NZ\_AP022574 (3690047-3755967): Mycolicibacterium psychrotolerans strain JCM 1... (10% of genes show similarity), NRPS,T1PKS

NZ\_CP045480 (2297107-2388505): Amycolatopsis sp. YIM 10 chromosome, complete ... (8% of genes show similarity), T1PKS

NZ\_CP054932 (6291872-6432263): Actinomadura sp. NAK00032 chromosome, complete... (5% of genes show similarity), NRPS,T1PKS,other

NZ\_CP012672 (10824439-10880437): Sorangium cellulosum strain So ce836 chromos... (7% of genes show similarity), T1PKS
Download graphic

Similar known gene clusters

Shows clusters from the MiBIG database that are similar to the current region. Genes marked with the same colour are interrelated. White genes have no relationship.  
Click on reference genes to show details of similarities to genes within the current region.  
Click on an accession to open that entry in the MiBIG database.

No matches found.

Similar subclusters

Shows sub-cluster units that are similar to the current region. Genes marked with the same colour are interrelated. White genes have no relationship.  
Click on reference genes to show details of similarities to genes within the current region.

No matches found.

Similar gene clusters

Shows careas that are similar to the current region to a reference database.  
Mouseover a score cell in the table to get a breakdown of how the score was calculated.The MIBiG database.  
  
Click on an accession to open that entry in the MIBiG database.

Analysis type:

Protocluster to Region
Region to Region

| Reference | T1PKS | Similarity score | Type | Compound(s) | Organism |
| --- | --- | --- | --- | --- | --- |
| BGC0001858.1 |  | 0.33 | Polyketide | alternapyrone B, alternapyrone C, alternapyrone D, alternapyrone E, alternapyrone F | Parastagonospora nodorum SN15 |
| BGC0001909.1 |  | 0.32 | Polyketide | strobilurin | Strobilurus tenacellus |
| BGC0000037.1 |  | 0.30 | Polyketide | cichorine | Aspergillus nidulans FGSC A4 |
| BGC0001068.1 |  | 0.28 | Terpene, Polyketide | pyripyropene A | unidentified unclassified sequences. |
| BGC0001969.1 |  | 0.27 | Terpene | asperterpenoid A | Talaromyces wortmannii |
| BGC0001141.1 |  | 0.26 | Polyketide | 4-epi-15-epi-brefeldin A | Penicillium brefeldianum |
| BGC0001923.1 |  | 0.26 | Terpene, Polyketide | ascochlorin | Acremonium egyptiacum |
| BGC0000003.1 |  | 0.26 | Polyketide | AF-toxin | Alternaria alternata |
| BGC0001277.1 |  | 0.25 | Terpene | nivalenol, vomitoxin, 3-acetyldeoxynivalenol, 15-acetyldeoxynivalenol, neosolaniol, calonectrin, apotrichodiol, isotrichotriol, 15-deacetylcalonectrin, T-2 toxin, 3-acetyl T-2 toxin, trichodiene | Fusarium graminearum |
| BGC0001104.1 |  | 0.24 | NRP, Polyketide | myxovirescin A1 | Myxococcus xanthus DK 1622 |

| Reference | Aggregated | Similarity score | Type | Compound(s) | Organism |
| --- | --- | --- | --- | --- | --- |
| BGC0001858.1 |  | 0.67 | Polyketide | alternapyrone B, alternapyrone C, alternapyrone D, alternapyrone E, alternapyrone F | Parastagonospora nodorum SN15 |
| BGC0001909.1 |  | 0.65 | Polyketide | strobilurin | Strobilurus tenacellus |
| BGC0000037.1 |  | 0.63 | Polyketide | cichorine | Aspergillus nidulans FGSC A4 |
| BGC0001068.1 |  | 0.61 | Terpene, Polyketide | pyripyropene A | unidentified unclassified sequences. |
| BGC0001969.1 |  | 0.60 | Terpene | asperterpenoid A | Talaromyces wortmannii |
| BGC0001141.1 |  | 0.60 | Polyketide | 4-epi-15-epi-brefeldin A | Penicillium brefeldianum |
| BGC0001923.1 |  | 0.59 | Terpene, Polyketide | ascochlorin | Acremonium egyptiacum |
| BGC0000003.1 |  | 0.59 | Polyketide | AF-toxin | Alternaria alternata |
| BGC0001277.1 |  | 0.59 | Terpene | nivalenol, vomitoxin, 3-acetyldeoxynivalenol, 15-acetyldeoxynivalenol, neosolaniol, calonectrin, apotrichodiol, isotrichotriol, 15-deacetylcalonectrin, T-2 toxin, 3-acetyl T-2 toxin, trichodiene | Fusarium graminearum |
| BGC0001104.1 |  | 0.58 | NRP, Polyketide | myxovirescin A1 | Myxococcus xanthus DK 1622 |

Detailed Pfam domain annotation

Shows Pfam domains found in each gene within the region.
Click on each domain for more information about the domain's
accession, location, description, and any relevant Gene Ontology.
Domains with a bold border have Gene Ontology information.

Selected features only

NRPS/PKS products

NRPS/PKS monomers

Predicted core structure(s)

Shows estimated product structure and polymer for each candidate cluster in the region. To show the product, click on the expander or the candidate cluster feature drawn in the overview.

For candidate cluster 1, location 5916 - 54518:

Rough prediction of core scaffold based on assumed PKS/NRPS colinearity; tailoring reactions not taken into account

**Polymer prediction:**
:   (mal)

  
Direct lookup in NORINE database:
strict
or
relaxed

Link to NORINE database query form

NRPS/PKS monomer predictions

Shows the predicted monomers for each adynelation domain and acyltransferase within genes. Each gene prediction can be expanded to view detailed predictions of each domain. Each prediction can be expanded to view the predictions by tool (and, for some tools, further expanded for extra details).

**input.path1.gene5**: mal

:   **PKS\_AT (703..991)**: mal

    ATSignature: Malonyl-CoA

    Top 3 matches:
    :   Malonyl-CoA: 66.7%
    :   inactive: 58.3%
    :   Methylmalonyl-CoA: 58.3%

      
    minowa: Malonyl-CoA

    Prediction, score:
    :   Malonyl-CoA: 73.9


        Methoxymalonyl-CoA: 53.2


        Methylmalonyl-CoA: 50.1


        inactive: 41.3


        Ethylmalonyl-CoA: 34.8


        2-Methylbutyryl-CoA: 27.6


        Propionyl-CoA: 24.5


        fatty\_acid: 22.7


        CHC-CoA: 22.3


        trans-1,2-CPDA: 20.8


        Acetyl-CoA: 13.7


        Benzoyl-CoA: 12.4


        Isobutyryl-CoA: 12.2


        3-Methylbutyryl-CoA: 8.2

  
**input.path1.gene7**: mal

:   **PKS\_AT (6..267)**: mal

    ATSignature: Malonyl-CoA

    Top 3 matches:
    :   Malonyl-CoA: 62.5%
    :   2-Methylbutyryl-CoA: 58.3%
    :   Methylmalonyl-CoA: 58.3%

      
    minowa: Malonyl-CoA

    Prediction, score:
    :   Malonyl-CoA: 58.5


        Methylmalonyl-CoA: 55.4


        Methoxymalonyl-CoA: 49.0


        Propionyl-CoA: 25.8


        Ethylmalonyl-CoA: 23.6


        Isobutyryl-CoA: 15.7


        inactive: 12.9


        fatty\_acid: 12.2


        trans-1,2-CPDA: 0.0


        CHC-CoA: 0.0


        Benzoyl-CoA: 0.0


        Acetyl-CoA: 0.0


        3-Methylbutyryl-CoA: 0.0


        2-Methylbutyryl-CoA: 0.0

contig\_3 - Region 1 - T1PKS

Shows the layout of the region, marking coding sequences and areas of interest. Clicking a gene will select it and show any relevant details. Clicking an area feature (e.g. a candidate cluster) will select all coding sequences within that area. Double clicking an area feature will zoom to that area. Multiple genes and area features can be selected by clicking them while holding the Ctrl key.  
More detailed help is available here.

Download region GenBank file

Download region SVG

Location: 486,327 - 531,102 nt. (total: 44,776 nt)
Show pHMM detection rules used

T1PKS: cds(PKS\_AT and (PKS\_KS or ene\_KS or mod\_KS or hyb\_KS or itr\_KS or tra\_KS))

#### Legend:

core biosynthetic genes

additional biosynthetic genes

transport-related genes

regulatory genes

other genes

resistance

reset view

zoom to selection

Gene details

Shows details of the most recently selected gene, including names, products, location, and other annotations.

Select a gene to view the details available for it

NRPS/PKS domains

ClusterBlast

KnownClusterBlast

SubClusterBlast

MIBiG comparison

Pfam domains

Detailed domain annotation

Shows NRPS- and PKS-related domains for each feature that contains them. Click on each domain for more information about the domain's location, consensus monomer prediction, and other details.  
A glossary is available here.

Selected features only

Show module domains

Similar gene clusters

Shows clusters from the antiSMASH database and other clusters of interest that are similar to the current region. Genes marked with the same colour are interrelated. White genes have no relationship.  
Click on reference genes to show details of similarities to genes within the current region.  
Click on an accession to open that entry in the antiSMASH database (if applicable).

All hits

NW\_006271975 (2592207-2639723): Cordyceps militaris CM01 unplaced genomic sca... (20% of genes show similarity), T1PKS

NW\_006917091 (912491-957103): Pestalotiopsis fici W106-1 unplaced genomic sca... (14% of genes show similarity), T1PKS

NW\_003052497 (1093806-1145882): Uncinocarpus reesii 1704 scaffold 4 genomic s... (13% of genes show similarity), NRPS,T1PKS
Download graphic

Similar known gene clusters

Shows clusters from the MiBIG database that are similar to the current region. Genes marked with the same colour are interrelated. White genes have no relationship.  
Click on reference genes to show details of similarities to genes within the current region.  
Click on an accession to open that entry in the MiBIG database.

No matches found.

Similar subclusters

Shows sub-cluster units that are similar to the current region. Genes marked with the same colour are interrelated. White genes have no relationship.  
Click on reference genes to show details of similarities to genes within the current region.

No matches found.

Similar gene clusters

Shows careas that are similar to the current region to a reference database.  
Mouseover a score cell in the table to get a breakdown of how the score was calculated.The MIBiG database.  
  
Click on an accession to open that entry in the MIBiG database.

Analysis type:

Protocluster to Region
Region to Region

| Reference | T1PKS | Similarity score | Type | Compound(s) | Organism |
| --- | --- | --- | --- | --- | --- |
| BGC0000046.1 |  | 0.18 | Polyketide | depudecin | Alternaria brassicicola |
| BGC0000107.1 |  | 0.17 | Polyketide | naphthopyrone | Aspergillus nidulans FGSC A4 |
| BGC0001254.1 |  | 0.16 | Polyketide | ACT-Toxin II | Alternaria alternata |
| BGC0001068.1 |  | 0.16 | Terpene, Polyketide | pyripyropene A | unidentified unclassified sequences. |
| BGC0001124.1 |  | 0.16 | Polyketide | pyranonigrin E | Aspergillus niger ATCC 1015 |
| BGC0000037.1 |  | 0.16 | Polyketide | cichorine | Aspergillus nidulans FGSC A4 |
| BGC0001252.1 |  | 0.15 | Polyketide | UNII-YC2Q1O94PT | Alternaria alternata |
| BGC0001268.1 |  | 0.14 | NRP, Polyketide | fusarin | Fusarium fujikuroi |
| BGC0000064.1 |  | 0.14 | Polyketide | fusarin | Fusarium verticillioides |
| BGC0001400.1 |  | 0.14 | Polyketide | citreoviridin | Aspergillus terreus NIH2624 |

| Reference | Aggregated | Similarity score | Type | Compound(s) | Organism |
| --- | --- | --- | --- | --- | --- |
| BGC0000046.1 |  | 0.50 | Polyketide | depudecin | Alternaria brassicicola |
| BGC0000107.1 |  | 0.48 | Polyketide | naphthopyrone | Aspergillus nidulans FGSC A4 |
| BGC0001254.1 |  | 0.47 | Polyketide | ACT-Toxin II | Alternaria alternata |
| BGC0001068.1 |  | 0.47 | Terpene, Polyketide | pyripyropene A | unidentified unclassified sequences. |
| BGC0001124.1 |  | 0.46 | Polyketide | pyranonigrin E | Aspergillus niger ATCC 1015 |
| BGC0000037.1 |  | 0.46 | Polyketide | cichorine | Aspergillus nidulans FGSC A4 |
| BGC0001252.1 |  | 0.45 | Polyketide | UNII-YC2Q1O94PT | Alternaria alternata |
| BGC0001268.1 |  | 0.44 | NRP, Polyketide | fusarin | Fusarium fujikuroi |
| BGC0000064.1 |  | 0.44 | Polyketide | fusarin | Fusarium verticillioides |
| BGC0001400.1 |  | 0.43 | Polyketide | citreoviridin | Aspergillus terreus NIH2624 |

Detailed Pfam domain annotation

Shows Pfam domains found in each gene within the region.
Click on each domain for more information about the domain's
accession, location, description, and any relevant Gene Ontology.
Domains with a bold border have Gene Ontology information.

Selected features only

contig\_31 - Region 1 - NRPS

Shows the layout of the region, marking coding sequences and areas of interest. Clicking a gene will select it and show any relevant details. Clicking an area feature (e.g. a candidate cluster) will select all coding sequences within that area. Double clicking an area feature will zoom to that area. Multiple genes and area features can be selected by clicking them while holding the Ctrl key.  
More detailed help is available here.

Download region GenBank file

Download region SVG

Location: 585,393 - 629,288 nt. (total: 43,896 nt)
Show pHMM detection rules used

NRPS: cds(Condensation and (AMP-binding or A-OX))

#### Legend:

core biosynthetic genes

additional biosynthetic genes

transport-related genes

regulatory genes

other genes

resistance

reset view

zoom to selection

Gene details

Shows details of the most recently selected gene, including names, products, location, and other annotations.

Select a gene to view the details available for it

NRPS/PKS domains

ClusterBlast

KnownClusterBlast

SubClusterBlast

MIBiG comparison

Pfam domains

Detailed domain annotation

Shows NRPS- and PKS-related domains for each feature that contains them. Click on each domain for more information about the domain's location, consensus monomer prediction, and other details.  
A glossary is available here.

Selected features only

Show module domains

Similar gene clusters

Shows clusters from the antiSMASH database and other clusters of interest that are similar to the current region. Genes marked with the same colour are interrelated. White genes have no relationship.  
Click on reference genes to show details of similarities to genes within the current region.  
Click on an accession to open that entry in the antiSMASH database (if applicable).

All hits

NW\_001914858 (2760303-2808125): Podospora anserina S mat+ genomic DNA chromos... (11% of genes show similarity), NRPS
Download graphic

Similar known gene clusters

Shows clusters from the MiBIG database that are similar to the current region. Genes marked with the same colour are interrelated. White genes have no relationship.  
Click on reference genes to show details of similarities to genes within the current region.  
Click on an accession to open that entry in the MiBIG database.

No matches found.

Similar subclusters

Shows sub-cluster units that are similar to the current region. Genes marked with the same colour are interrelated. White genes have no relationship.  
Click on reference genes to show details of similarities to genes within the current region.

No matches found.

Similar gene clusters

Shows careas that are similar to the current region to a reference database.  
Mouseover a score cell in the table to get a breakdown of how the score was calculated.The MIBiG database.  
  
Click on an accession to open that entry in the MIBiG database.

Analysis type:

Protocluster to Region
Region to Region

| Reference | NRPS | Similarity score | Type | Compound(s) | Organism |
| --- | --- | --- | --- | --- | --- |
| BGC0001313.1 |  | 0.13 | Terpene | arabidiol-baruol | Arabidopsis thaliana |
| BGC0000353.1 |  | 0.10 | NRP | tacrolimus | Streptomyces sp. MA6548 |
| BGC0001544.1 |  | 0.07 | NRP, Polyketide | chlorflavonin | Aspergillus campestris IBT 28561 |
| BGC0002019.1 |  | 0.06 | Terpene | tiancilactone | Streptomyces sp. CB03234 |
| BGC0000220.1 |  | 0.06 | Polyketide | enterocin | Streptomyces maritimus |
| BGC0001502.1 |  | 0.06 | NRP | amonabactin P 750 | Aeromonas hydrophila subsp. hydrophila ATCC 7966 |
| BGC0000168.1 |  | 0.06 | Polyketide | viridicatumtoxin, previridicatumtoxin, 5-hydroxyanthrotainin, 8-O-desmethylanthrotainin | Penicillium aethiopicum |
| BGC0001007.1 |  | 0.06 | Polyketide, NRP | lymphostin, neolymphostinol B, lymphostinol, neolymphostin b | Salinispora arenicola CNS-205 |
| BGC0001006.1 |  | 0.05 | NRP, Polyketide | lymphostin, neolymphostinol B, lymphostinol, neolymphostin B | Salinispora tropica CNB-440 |
| BGC0001298.1 |  | 0.05 | Polyketide | 4-Z-annimycin | Streptomyces calvus |

| Reference | Aggregated | Similarity score | Type | Compound(s) | Organism |
| --- | --- | --- | --- | --- | --- |
| BGC0001313.1 |  | 0.41 | Terpene | arabidiol-baruol | Arabidopsis thaliana |
| BGC0000353.1 |  | 0.35 | NRP | tacrolimus | Streptomyces sp. MA6548 |
| BGC0001544.1 |  | 0.29 | NRP, Polyketide | chlorflavonin | Aspergillus campestris IBT 28561 |
| BGC0002019.1 |  | 0.26 | Terpene | tiancilactone | Streptomyces sp. CB03234 |
| BGC0000220.1 |  | 0.25 | Polyketide | enterocin | Streptomyces maritimus |
| BGC0001502.1 |  | 0.25 | NRP | amonabactin P 750 | Aeromonas hydrophila subsp. hydrophila ATCC 7966 |
| BGC0000168.1 |  | 0.24 | Polyketide | viridicatumtoxin, previridicatumtoxin, 5-hydroxyanthrotainin, 8-O-desmethylanthrotainin | Penicillium aethiopicum |
| BGC0001007.1 |  | 0.24 | Polyketide, NRP | lymphostin, neolymphostinol B, lymphostinol, neolymphostin b | Salinispora arenicola CNS-205 |
| BGC0001006.1 |  | 0.23 | NRP, Polyketide | lymphostin, neolymphostinol B, lymphostinol, neolymphostin B | Salinispora tropica CNB-440 |
| BGC0001298.1 |  | 0.21 | Polyketide | 4-Z-annimycin | Streptomyces calvus |

Detailed Pfam domain annotation

Shows Pfam domains found in each gene within the region.
Click on each domain for more information about the domain's
accession, location, description, and any relevant Gene Ontology.
Domains with a bold border have Gene Ontology information.

Selected features only

contig\_31 - Region 2 - T1PKS

Shows the layout of the region, marking coding sequences and areas of interest. Clicking a gene will select it and show any relevant details. Clicking an area feature (e.g. a candidate cluster) will select all coding sequences within that area. Double clicking an area feature will zoom to that area. Multiple genes and area features can be selected by clicking them while holding the Ctrl key.  
More detailed help is available here.

Download region GenBank file

Download region SVG

Location: 1,112,429 - 1,160,321 nt. (total: 47,893 nt)
Show pHMM detection rules used

T1PKS: cds(PKS\_AT and (PKS\_KS or ene\_KS or mod\_KS or hyb\_KS or itr\_KS or tra\_KS))

#### Legend:

core biosynthetic genes

additional biosynthetic genes

transport-related genes

regulatory genes

other genes

resistance

reset view

zoom to selection

Gene details

Shows details of the most recently selected gene, including names, products, location, and other annotations.

Select a gene to view the details available for it

NRPS/PKS domains

ClusterBlast

KnownClusterBlast

SubClusterBlast

MIBiG comparison

Pfam domains

Detailed domain annotation

Shows NRPS- and PKS-related domains for each feature that contains them. Click on each domain for more information about the domain's location, consensus monomer prediction, and other details.  
A glossary is available here.

Selected features only

Show module domains

Similar gene clusters

Shows clusters from the antiSMASH database and other clusters of interest that are similar to the current region. Genes marked with the same colour are interrelated. White genes have no relationship.  
Click on reference genes to show details of similarities to genes within the current region.  
Click on an accession to open that entry in the antiSMASH database (if applicable).

All hits

NW\_007360985 (1400169-1447192): Glarea lozoyensis ATCC 20868 chromosome Unkno... (23% of genes show similarity), T1PKS

NW\_015971142 (1762145-1804164): Sporothrix schenckii 1099-18 chromosome Unkno... (33% of genes show similarity), T1PKS

NC\_016457 (8723684-8767633): Thermothielavioides terrestris NRRL 8126 chromos... (27% of genes show similarity), T1PKS

NW\_006917098 (949178-996160): Pestalotiopsis fici W106-1 unplaced genomic sca... (23% of genes show similarity), T1PKS

RRCJ01000010 (1439907-1555741): Pyricularia sp. CBS 133598 strain NI919 Pyric... (9% of genes show similarity), T1PKS

NW\_009276967 (457874-504921): Verticillium dahliae VdLs.17 supercont1.20 geno... (30% of genes show similarity), T1PKS

NW\_003315038 (479797-514119): Verticillium alfalfae VaMs.102 supercont1.1 gen... (27% of genes show similarity), T1PKS

NW\_006763060 (505883-553571): Marssonina brunnea f. sp. 'multigermtubi' MB m1... (30% of genes show similarity), T1PKS

NC\_049562 (4122890-4164474): Talaromyces rugulosus chromosome II, complete se... (25% of genes show similarity), T1PKS

NW\_023336259 (2343266-2388301): Colletotrichum scovillei strain TJNH1 chromos... (33% of genes show similarity), T1PKS
Download graphic

Similar known gene clusters

Shows clusters from the MiBIG database that are similar to the current region. Genes marked with the same colour are interrelated. White genes have no relationship.  
Click on reference genes to show details of similarities to genes within the current region.  
Click on an accession to open that entry in the MiBIG database.

No matches found.

Similar subclusters

Shows sub-cluster units that are similar to the current region. Genes marked with the same colour are interrelated. White genes have no relationship.  
Click on reference genes to show details of similarities to genes within the current region.

No matches found.

Similar gene clusters

Shows careas that are similar to the current region to a reference database.  
Mouseover a score cell in the table to get a breakdown of how the score was calculated.The MIBiG database.  
  
Click on an accession to open that entry in the MIBiG database.

Analysis type:

Protocluster to Region
Region to Region

| Reference | T1PKS | Similarity score | Type | Compound(s) | Organism |
| --- | --- | --- | --- | --- | --- |
| BGC0000046.1 |  | 0.32 | Polyketide | depudecin | Alternaria brassicicola |
| BGC0001254.1 |  | 0.22 | Polyketide | ACT-Toxin II | Alternaria alternata |
| BGC0001400.1 |  | 0.21 | Polyketide | citreoviridin | Aspergillus terreus NIH2624 |
| BGC0001124.1 |  | 0.21 | Polyketide | pyranonigrin E | Aspergillus niger ATCC 1015 |
| BGC0001606.1 |  | 0.21 | Polyketide | gibepyrone-A | Fusarium fujikuroi IMI 58289 |
| BGC0000107.1 |  | 0.21 | Polyketide | naphthopyrone | Aspergillus nidulans FGSC A4 |
| BGC0001909.1 |  | 0.20 | Polyketide | strobilurin | Strobilurus tenacellus |
| BGC0001273.1 |  | 0.20 | Polyketide | asperlactone | Aspergillus ochraceus |
| BGC0001252.1 |  | 0.20 | Polyketide | UNII-YC2Q1O94PT | Alternaria alternata |
| BGC0001281.1 |  | 0.18 | Polyketide | ustilagic acid | Ustilago maydis 521 |

| Reference | Aggregated | Similarity score | Type | Compound(s) | Organism |
| --- | --- | --- | --- | --- | --- |
| BGC0000046.1 |  | 0.65 | Polyketide | depudecin | Alternaria brassicicola |
| BGC0001254.1 |  | 0.55 | Polyketide | ACT-Toxin II | Alternaria alternata |
| BGC0001400.1 |  | 0.54 | Polyketide | citreoviridin | Aspergillus terreus NIH2624 |
| BGC0001124.1 |  | 0.54 | Polyketide | pyranonigrin E | Aspergillus niger ATCC 1015 |
| BGC0001606.1 |  | 0.54 | Polyketide | gibepyrone-A | Fusarium fujikuroi IMI 58289 |
| BGC0000107.1 |  | 0.53 | Polyketide | naphthopyrone | Aspergillus nidulans FGSC A4 |
| BGC0001909.1 |  | 0.53 | Polyketide | strobilurin | Strobilurus tenacellus |
| BGC0001273.1 |  | 0.53 | Polyketide | asperlactone | Aspergillus ochraceus |
| BGC0001252.1 |  | 0.52 | Polyketide | UNII-YC2Q1O94PT | Alternaria alternata |
| BGC0001281.1 |  | 0.49 | Polyketide | ustilagic acid | Ustilago maydis 521 |

Detailed Pfam domain annotation

Shows Pfam domains found in each gene within the region.
Click on each domain for more information about the domain's
accession, location, description, and any relevant Gene Ontology.
Domains with a bold border have Gene Ontology information.

Selected features only

NRPS/PKS products

NRPS/PKS monomers

Predicted core structure(s)

Shows estimated product structure and polymer for each candidate cluster in the region. To show the product, click on the expander or the candidate cluster feature drawn in the overview.

For candidate cluster 2, location 1112428 - 1160321:

Rough prediction of core scaffold based on assumed PKS/NRPS colinearity; tailoring reactions not taken into account

**Polymer prediction:**
:   (pk)

  
Direct lookup in NORINE database:
strict
or
relaxed

Link to NORINE database query form

NRPS/PKS monomer predictions

Shows the predicted monomers for each adynelation domain and acyltransferase within genes. Each gene prediction can be expanded to view detailed predictions of each domain. Each prediction can be expanded to view the predictions by tool (and, for some tools, further expanded for extra details).

**input.path1.gene300**: pk

:   **PKS\_AT (567..884)**: pk

    ATSignature: Malonyl-CoA

    Top 3 matches:
    :   Malonyl-CoA: 79.2%
    :   Methylmalonyl-CoA: 66.7%
    :   Ethylmalonyl-CoA: 58.3%

      
    minowa: Methoxymalonyl-CoA

    Prediction, score:
    :   Methoxymalonyl-CoA: 84.8


        Methylmalonyl-CoA: 82.8


        Malonyl-CoA: 73.7


        Isobutyryl-CoA: 61.3


        Ethylmalonyl-CoA: 55.3


        trans-1,2-CPDA: 48.2


        Propionyl-CoA: 36.0


        2-Methylbutyryl-CoA: 34.4


        Benzoyl-CoA: 31.2


        inactive: 21.2


        fatty\_acid: 17.2


        3-Methylbutyryl-CoA: 13.5


        CHC-CoA: 9.2


        Acetyl-CoA: 9.1

contig\_32 - Region 1 - NRPS-like

Shows the layout of the region, marking coding sequences and areas of interest. Clicking a gene will select it and show any relevant details. Clicking an area feature (e.g. a candidate cluster) will select all coding sequences within that area. Double clicking an area feature will zoom to that area. Multiple genes and area features can be selected by clicking them while holding the Ctrl key.  
More detailed help is available here.

Download region GenBank file

Download region SVG

Location: 618,090 - 663,360 nt. (total: 45,271 nt)
Show pHMM detection rules used

NRPS-like: cds((PP-binding or NAD\_binding\_4) and (AMP-binding or A-OX))

#### Legend:

core biosynthetic genes

additional biosynthetic genes

transport-related genes

regulatory genes

other genes

resistance

reset view

zoom to selection

Gene details

Shows details of the most recently selected gene, including names, products, location, and other annotations.

Select a gene to view the details available for it

NRPS/PKS domains

ClusterBlast

KnownClusterBlast

SubClusterBlast

MIBiG comparison

Pfam domains

Detailed domain annotation

Shows NRPS- and PKS-related domains for each feature that contains them. Click on each domain for more information about the domain's location, consensus monomer prediction, and other details.  
A glossary is available here.

Selected features only

Show module domains

Similar gene clusters

Shows clusters from the antiSMASH database and other clusters of interest that are similar to the current region. Genes marked with the same colour are interrelated. White genes have no relationship.  
Click on reference genes to show details of similarities to genes within the current region.  
Click on an accession to open that entry in the antiSMASH database (if applicable).

All hits

NW\_001939248 (971466-1014527): Pyrenophora tritici-repentis Pt-1C-BFP superco... (11% of genes show similarity), NRPS-like

NW\_017264203 (284168-327921): Xylona heveae TC161 unplaced genomic scaffold L... (15% of genes show similarity), NRPS-like

NW\_023336278 (1339198-1376990): Aspergillus tubingensis WU-2223L DNA, scaffol... (18% of genes show similarity), NRPS-like

NW\_020194484 (860721-904328): Amorphotheca resinae ATCC 22711 unplaced genomi... (12% of genes show similarity), NRPS-like

NC\_007197 (2914762-2959218): Aspergillus fumigatus Af293 chromosome 4, whole ... (16% of genes show similarity), NRPS-like

NW\_022474205 (807987-876555): Venustampulla echinocandica strain BP 5553 chro... (7% of genes show similarity), NRPS-like

NC\_036436 (5056198-5100650): Aspergillus oryzae RIB40 DNA, chromosome 2 (15% of genes show similarity), NRPS

NT\_165928 (1095178-1139640): Aspergillus terreus NIH2624 scaffold 5 genomic s... (16% of genes show similarity), NRPS-like

NW\_022984630 (2409364-2453077): Aspergillus tanneri strain NIH1004 chromosome... (14% of genes show similarity), NRPS-like

NW\_006763065 (184681-228240): Marssonina brunnea f. sp. 'multigermtubi' MB m1... (16% of genes show similarity), NRPS-like
Download graphic

Similar known gene clusters

Shows clusters from the MiBIG database that are similar to the current region. Genes marked with the same colour are interrelated. White genes have no relationship.  
Click on reference genes to show details of similarities to genes within the current region.  
Click on an accession to open that entry in the MiBIG database.

No matches found.

Similar subclusters

Shows sub-cluster units that are similar to the current region. Genes marked with the same colour are interrelated. White genes have no relationship.  
Click on reference genes to show details of similarities to genes within the current region.

No matches found.

Similar gene clusters

Shows careas that are similar to the current region to a reference database.  
Mouseover a score cell in the table to get a breakdown of how the score was calculated.The MIBiG database.  
  
Click on an accession to open that entry in the MIBiG database.

Analysis type:

Protocluster to Region
Region to Region

| Reference | NRPS-like | Similarity score | Type | Compound(s) | Organism |
| --- | --- | --- | --- | --- | --- |
| BGC0000308.1 |  | 0.21 | NRP | aureusimine | Staphylococcus aureus subsp. aureus str. JKD6008 |
| BGC0001900.1 |  | 0.20 | NRP | fragin | Burkholderia cenocepacia H111 |
| BGC0001168.1 |  | 0.20 | NRP | livipeptin | Streptomyces lividans 1326 |
| BGC0001261.1 |  | 0.20 | NRP | AM-toxin | Alternaria alternata |
| BGC0001132.1 |  | 0.20 | NRP | xenotetrapeptide | Xenorhabdus nematophila ATCC 19061 |
| BGC0001135.1 |  | 0.19 | NRP | bicornutin A1, bicornutin A2 | Xenorhabdus budapestensis |
| BGC0001128.1 |  | 0.19 | NRP | luminmide | Photorhabdus laumondii subsp. laumondii TTO1 |
| BGC0001561.1 |  | 0.19 | NRP | curacomycin | Streptomyces curacoi |
| BGC0000334.1 |  | 0.18 | NRP | cyclosporine | Tolypocladium inflatum NRRL8044 |
| BGC0000876.1 |  | 0.18 | Other | neopolyoxin C | Streptomyces tendae |

| Reference | Aggregated | Similarity score | Type | Compound(s) | Organism |
| --- | --- | --- | --- | --- | --- |
| BGC0000308.1 |  | 0.54 | NRP | aureusimine | Staphylococcus aureus subsp. aureus str. JKD6008 |
| BGC0001900.1 |  | 0.53 | NRP | fragin | Burkholderia cenocepacia H111 |
| BGC0001168.1 |  | 0.53 | NRP | livipeptin | Streptomyces lividans 1326 |
| BGC0001261.1 |  | 0.53 | NRP | AM-toxin | Alternaria alternata |
| BGC0001132.1 |  | 0.52 | NRP | xenotetrapeptide | Xenorhabdus nematophila ATCC 19061 |
| BGC0001135.1 |  | 0.51 | NRP | bicornutin A1, bicornutin A2 | Xenorhabdus budapestensis |
| BGC0001128.1 |  | 0.51 | NRP | luminmide | Photorhabdus laumondii subsp. laumondii TTO1 |
| BGC0001561.1 |  | 0.51 | NRP | curacomycin | Streptomyces curacoi |
| BGC0000334.1 |  | 0.50 | NRP | cyclosporine | Tolypocladium inflatum NRRL8044 |
| BGC0000876.1 |  | 0.49 | Other | neopolyoxin C | Streptomyces tendae |

Detailed Pfam domain annotation

Shows Pfam domains found in each gene within the region.
Click on each domain for more information about the domain's
accession, location, description, and any relevant Gene Ontology.
Domains with a bold border have Gene Ontology information.

Selected features only

NRPS/PKS products

NRPS/PKS monomers

Predicted core structure(s)

Shows estimated product structure and polymer for each candidate cluster in the region. To show the product, click on the expander or the candidate cluster feature drawn in the overview.

For candidate cluster 1, location 618089 - 663360:

Rough prediction of core scaffold based on assumed PKS/NRPS colinearity; tailoring reactions not taken into account

**Polymer prediction:**
:   (X)

  
Direct lookup in NORINE database:
strict
or
relaxed

Link to NORINE database query form

NRPS/PKS monomer predictions

Shows the predicted monomers for each adynelation domain and acyltransferase within genes. Each gene prediction can be expanded to view detailed predictions of each domain. Each prediction can be expanded to view the predictions by tool (and, for some tools, further expanded for extra details).

**input.path1.gene177**: X

:   Search NORINE for peptide:
    strict
    or
    relaxed
  
:   **AMP-binding (76..541)**: X

    NRPSPredictor2: asp, asn, glu, gln, aad

    SVM prediction details:
    :   Predicted physicochemical class:
        :   hydrophobic-aliphatic

        Large clusters prediction:
        :   asp, asn, glu, gln, aad

        Small clusters prediction:
        :   N/A

        Single AA prediction:
        :   N/A

    Stachelhaus prediction details:
    :   Stachelhaus sequence:
        :   dprhfvmrak

        Nearest Stachelhaus code:
        :   N, A

        Stachelhaus code match:
        :   0% (weak)

contig\_32 - Region 2 - T1PKS

Shows the layout of the region, marking coding sequences and areas of interest. Clicking a gene will select it and show any relevant details. Clicking an area feature (e.g. a candidate cluster) will select all coding sequences within that area. Double clicking an area feature will zoom to that area. Multiple genes and area features can be selected by clicking them while holding the Ctrl key.  
More detailed help is available here.

Download region GenBank file

Download region SVG

Location: 1,790,500 - 1,832,854 nt. (total: 42,355 nt)
Show pHMM detection rules used

T1PKS: cds(PKS\_AT and (PKS\_KS or ene\_KS or mod\_KS or hyb\_KS or itr\_KS or tra\_KS))

#### Legend:

core biosynthetic genes

additional biosynthetic genes

transport-related genes

regulatory genes

other genes

resistance

reset view

zoom to selection

Gene details

Shows details of the most recently selected gene, including names, products, location, and other annotations.

Select a gene to view the details available for it

NRPS/PKS domains

ClusterBlast

KnownClusterBlast

SubClusterBlast

MIBiG comparison

Pfam domains

Detailed domain annotation

Shows NRPS- and PKS-related domains for each feature that contains them. Click on each domain for more information about the domain's location, consensus monomer prediction, and other details.  
A glossary is available here.

Selected features only

Show module domains

Similar gene clusters

Shows clusters from the antiSMASH database and other clusters of interest that are similar to the current region. Genes marked with the same colour are interrelated. White genes have no relationship.  
Click on reference genes to show details of similarities to genes within the current region.  
Click on an accession to open that entry in the antiSMASH database (if applicable).

All hits

NC\_049562 (5064149-5105472): Talaromyces rugulosus chromosome II, complete se... (14% of genes show similarity), T1PKS

NC\_036442 (3256010-3302476): Aspergillus oryzae RIB40 DNA, chromosome 8 (16% of genes show similarity), T1PKS

NW\_011942215 (524193-566139): Metarhizium robertsii ARSEF 23 MAA Scf 7, whole... (16% of genes show similarity), T1PKS

NW\_014574711 (1529497-1577032): Metarhizium brunneum ARSEF 3297 chromosome Un... (18% of genes show similarity), T1PKS

NC\_035793 (3639294-3686908): Pochonia chlamydosporia 170 chromosome 4, whole ... (15% of genes show similarity), T1PKS

CM004174 (6841316-6883283): Drechmeria coniospora strain ARSEF 6962 chromosom... (25% of genes show similarity), T1PKS

NW\_020167549 (97131-142659): Pseudogymnoascus destructans isolate 20631-21 un... (22% of genes show similarity), T1PKS

CM000595 (183841-220794): Fusarium oxysporum f. sp. lycopersici 4287 chromoso... (18% of genes show similarity), T1PKS

NC\_030992 (183841-220794): Fusarium oxysporum f. sp. lycopersici 4287 chromos... (18% of genes show similarity), T1PKS

NC\_049565 (5043304-5100528): Talaromyces rugulosus chromosome V, complete seq... (18% of genes show similarity), T1PKS
Download graphic

Similar known gene clusters

Shows clusters from the MiBIG database that are similar to the current region. Genes marked with the same colour are interrelated. White genes have no relationship.  
Click on reference genes to show details of similarities to genes within the current region.  
Click on an accession to open that entry in the MiBIG database.

All hits

1,3,6,8-tetrahydroxynaphthalene

melanin

1,3,6,8-tetrahydroxynaphthalene

alternariol
Download graphic

Similar subclusters

Shows sub-cluster units that are similar to the current region. Genes marked with the same colour are interrelated. White genes have no relationship.  
Click on reference genes to show details of similarities to genes within the current region.

No matches found.

Similar gene clusters

Shows careas that are similar to the current region to a reference database.  
Mouseover a score cell in the table to get a breakdown of how the score was calculated.The MIBiG database.  
  
Click on an accession to open that entry in the MIBiG database.

Analysis type:

Protocluster to Region
Region to Region

| Reference | T1PKS | Similarity score | Type | Compound(s) | Organism |
| --- | --- | --- | --- | --- | --- |
| BGC0000056.1 |  | 0.22 | Polyketide | esperamicin | Actinomadura verrucosospora |
| BGC0001265.1 |  | 0.20 | Polyketide | melanin | Bipolaris oryzae |
| BGC0001284.1 |  | 0.20 | Terpene | alternariol | Parastagonospora nodorum SN15 |
| BGC0000156.1 |  | 0.20 | Polyketide | TAN-1612, 1-(2,3,5,10-tetrahydroxy-7-methoxy-4-oxo-1,2,3,4-tetrahydroanthracen-2-yl)pentane-2,4-dione, desmethyl TAN-1612 | Aspergillus niger |
| BGC0001258.1 |  | 0.20 | Polyketide | 1,3,6,8-tetrahydroxynaphthalene | Glarea lozoyensis |
| BGC0001257.1 |  | 0.20 | Polyketide | 1,3,6,8-tetrahydroxynaphthalene | Nodulisporium sp. ATCC74245 |
| BGC0001906.1 |  | 0.20 | Polyketide | naphthalene | Daldinia eschscholzii IFB-TL01 |
| BGC0000013.1 |  | 0.20 | Polyketide | alternariol | Aspergillus nidulans FGSC A4 |
| BGC0000107.1 |  | 0.19 | Polyketide | naphthopyrone | Aspergillus nidulans FGSC A4 |
| BGC0000046.1 |  | 0.19 | Polyketide | depudecin | Alternaria brassicicola |

| Reference | Aggregated | Similarity score | Type | Compound(s) | Organism |
| --- | --- | --- | --- | --- | --- |
| BGC0000056.1 |  | 0.55 | Polyketide | esperamicin | Actinomadura verrucosospora |
| BGC0001265.1 |  | 0.53 | Polyketide | melanin | Bipolaris oryzae |
| BGC0001284.1 |  | 0.53 | Terpene | alternariol | Parastagonospora nodorum SN15 |
| BGC0000156.1 |  | 0.53 | Polyketide | TAN-1612, 1-(2,3,5,10-tetrahydroxy-7-methoxy-4-oxo-1,2,3,4-tetrahydroanthracen-2-yl)pentane-2,4-dione, desmethyl TAN-1612 | Aspergillus niger |
| BGC0001258.1 |  | 0.53 | Polyketide | 1,3,6,8-tetrahydroxynaphthalene | Glarea lozoyensis |
| BGC0001257.1 |  | 0.52 | Polyketide | 1,3,6,8-tetrahydroxynaphthalene | Nodulisporium sp. ATCC74245 |
| BGC0001906.1 |  | 0.52 | Polyketide | naphthalene | Daldinia eschscholzii IFB-TL01 |
| BGC0000013.1 |  | 0.52 | Polyketide | alternariol | Aspergillus nidulans FGSC A4 |
| BGC0000107.1 |  | 0.51 | Polyketide | naphthopyrone | Aspergillus nidulans FGSC A4 |
| BGC0000046.1 |  | 0.51 | Polyketide | depudecin | Alternaria brassicicola |

Detailed Pfam domain annotation

Shows Pfam domains found in each gene within the region.
Click on each domain for more information about the domain's
accession, location, description, and any relevant Gene Ontology.
Domains with a bold border have Gene Ontology information.

Selected features only

contig\_32 - Region 3 - T1PKS

Shows the layout of the region, marking coding sequences and areas of interest. Clicking a gene will select it and show any relevant details. Clicking an area feature (e.g. a candidate cluster) will select all coding sequences within that area. Double clicking an area feature will zoom to that area. Multiple genes and area features can be selected by clicking them while holding the Ctrl key.  
More detailed help is available here.

Download region GenBank file

Download region SVG

Location: 3,137,580 - 3,197,360 nt. (total: 59,781 nt)
Show pHMM detection rules used

T1PKS: cds(PKS\_AT and (PKS\_KS or ene\_KS or mod\_KS or hyb\_KS or itr\_KS or tra\_KS))

#### Legend:

core biosynthetic genes

additional biosynthetic genes

transport-related genes

regulatory genes

other genes

resistance

reset view

zoom to selection

Gene details

Shows details of the most recently selected gene, including names, products, location, and other annotations.

Select a gene to view the details available for it

NRPS/PKS domains

ClusterBlast

KnownClusterBlast

SubClusterBlast

MIBiG comparison

Pfam domains

Detailed domain annotation

Shows NRPS- and PKS-related domains for each feature that contains them. Click on each domain for more information about the domain's location, consensus monomer prediction, and other details.  
A glossary is available here.

Selected features only

Show module domains

Similar gene clusters

Shows clusters from the antiSMASH database and other clusters of interest that are similar to the current region. Genes marked with the same colour are interrelated. White genes have no relationship.  
Click on reference genes to show details of similarities to genes within the current region.  
Click on an accession to open that entry in the antiSMASH database (if applicable).

All hits

NT\_165934 (634984-694556): Aspergillus terreus NIH2624 scaffold 11 genomic sc... (27% of genes show similarity), T1PKS

NT\_107015 (3162503-3223371): Aspergillus nidulans FGSC A4 chromosome VIII map... (33% of genes show similarity), T1PKS

CM002799 (5899416-5951896): Penicillium chrysogenum strain P2niaD18 chromosom... (23% of genes show similarity), T1PKS

NW\_014574709 (<912->101750): Metarhizium brunneum ARSEF 3297 chromosome Unkno... (9% of genes show similarity), T1PKS

NW\_011942171 (5254002-5365378): Metarhizium robertsii ARSEF 23 MAA Scf 3, who... (14% of genes show similarity), NRPS,T1PKS

NT\_107014 (1931488-1984178): Aspergillus nidulans FGSC A4 chromosome VII map ... (21% of genes show similarity), T1PKS

NW\_004504311 (3651688-3709946): Coccidioides immitis RS genomic scaffold supe... (15% of genes show similarity), T1PKS

NW\_003315987 (867603-925836): Coccidioides posadasii C735 delta SOWgp chromos... (20% of genes show similarity), T1PKS

NW\_022983866 (1749407-1792896): Arthroderma uncinatum strain CBS 119779 chrom... (23% of genes show similarity), T1PKS

NW\_003299166 (1066347-1122248): Microsporum canis CBS 113480 supercont1.4 gen... (13% of genes show similarity), T1PKS
Download graphic

Similar known gene clusters

Shows clusters from the MiBIG database that are similar to the current region. Genes marked with the same colour are interrelated. White genes have no relationship.  
Click on reference genes to show details of similarities to genes within the current region.  
Click on an accession to open that entry in the MiBIG database.

All hits

chaetoviridin E / 11-epichaetomugilin A

asperfuranone
Download graphic

Similar subclusters

Shows sub-cluster units that are similar to the current region. Genes marked with the same colour are interrelated. White genes have no relationship.  
Click on reference genes to show details of similarities to genes within the current region.

No matches found.

Similar gene clusters

Shows careas that are similar to the current region to a reference database.  
Mouseover a score cell in the table to get a breakdown of how the score was calculated.The MIBiG database.  
  
Click on an accession to open that entry in the MIBiG database.

Analysis type:

Protocluster to Region
Region to Region

| Reference | T1PKS | Similarity score | Type | Compound(s) | Organism |
| --- | --- | --- | --- | --- | --- |
| BGC0000099.1 |  | 0.33 | Polyketide | monascorubrin | Talaromyces marneffei |
| BGC0001909.1 |  | 0.30 | Polyketide | strobilurin | Strobilurus tenacellus |
| BGC0001858.1 |  | 0.29 | Polyketide | alternapyrone B, alternapyrone C, alternapyrone D, alternapyrone E, alternapyrone F | Parastagonospora nodorum SN15 |
| BGC0001542.1 |  | 0.26 | Polyketide | cercosporin | Cercospora zeina |
| BGC0000161.1 |  | 0.26 | Polyketide | isoterrein | Aspergillus terreus NIH2624 |
| BGC0001541.1 |  | 0.25 | Polyketide | cercosporin | Cercospora beticola |
| BGC0001068.1 |  | 0.23 | Terpene, Polyketide | pyripyropene A | unidentified unclassified sequences. |
| BGC0001338.1 |  | 0.23 | Polyketide | citrinin | Monascus ruber |
| BGC0000046.1 |  | 0.23 | Polyketide | depudecin | Alternaria brassicicola |
| BGC0000861.1 |  | 0.23 | Other | eicosapentaenoic acid | Shewanella sp. BR-2 |

| Reference | Aggregated | Similarity score | Type | Compound(s) | Organism |
| --- | --- | --- | --- | --- | --- |
| BGC0000099.1 |  | 0.66 | Polyketide | monascorubrin | Talaromyces marneffei |
| BGC0001909.1 |  | 0.64 | Polyketide | strobilurin | Strobilurus tenacellus |
| BGC0001858.1 |  | 0.62 | Polyketide | alternapyrone B, alternapyrone C, alternapyrone D, alternapyrone E, alternapyrone F | Parastagonospora nodorum SN15 |
| BGC0001542.1 |  | 0.60 | Polyketide | cercosporin | Cercospora zeina |
| BGC0000161.1 |  | 0.60 | Polyketide | isoterrein | Aspergillus terreus NIH2624 |
| BGC0001541.1 |  | 0.58 | Polyketide | cercosporin | Cercospora beticola |
| BGC0001068.1 |  | 0.57 | Terpene, Polyketide | pyripyropene A | unidentified unclassified sequences. |
| BGC0001338.1 |  | 0.56 | Polyketide | citrinin | Monascus ruber |
| BGC0000046.1 |  | 0.56 | Polyketide | depudecin | Alternaria brassicicola |
| BGC0000861.1 |  | 0.56 | Other | eicosapentaenoic acid | Shewanella sp. BR-2 |

Detailed Pfam domain annotation

Shows Pfam domains found in each gene within the region.
Click on each domain for more information about the domain's
accession, location, description, and any relevant Gene Ontology.
Domains with a bold border have Gene Ontology information.

Selected features only

NRPS/PKS products

NRPS/PKS monomers

Predicted core structure(s)

Shows estimated product structure and polymer for each candidate cluster in the region. To show the product, click on the expander or the candidate cluster feature drawn in the overview.

For candidate cluster 3, location 3137579 - 3197360:

Rough prediction of core scaffold based on assumed PKS/NRPS colinearity; tailoring reactions not taken into account

**Polymer prediction:**
:   (pk) + (pk)

  
Direct lookup in NORINE database:
strict
or
relaxed

Link to NORINE database query form

NRPS/PKS monomer predictions

Shows the predicted monomers for each adynelation domain and acyltransferase within genes. Each gene prediction can be expanded to view detailed predictions of each domain. Each prediction can be expanded to view the predictions by tool (and, for some tools, further expanded for extra details).

**input.path1.gene893**: pk

:   **PKS\_AT (326..655)**: pk

    ATSignature: Malonyl-CoA

    Top 3 matches:
    :   Malonyl-CoA: 62.5%
    :   Methylmalonyl-CoA: 54.2%

      
    minowa: Methoxymalonyl-CoA

    Prediction, score:
    :   Methoxymalonyl-CoA: 55.6


        Methylmalonyl-CoA: 51.0


        Malonyl-CoA: 38.0


        Isobutyryl-CoA: 35.4


        Ethylmalonyl-CoA: 20.0


        Benzoyl-CoA: 17.2


        Propionyl-CoA: 15.9


        inactive: 11.7


        3-Methylbutyryl-CoA: 10.8


        CHC-CoA: 7.7


        2-Methylbutyryl-CoA: 3.1


        trans-1,2-CPDA: 0.0


        fatty\_acid: 0.0


        Acetyl-CoA: 0.0

  
**input.path1.gene895**: pk

:   **PKS\_AT (878..1168)**: pk

    ATSignature: Malonyl-CoA

    Top 3 matches:
    :   Malonyl-CoA: 62.5%
    :   Ethylmalonyl-CoA: 58.3%
    :   Methylmalonyl-CoA: 54.2%

      
    minowa: Methoxymalonyl-CoA

    Prediction, score:
    :   Methoxymalonyl-CoA: 60.7


        inactive: 58.4


        Malonyl-CoA: 57.6


        Methylmalonyl-CoA: 55.1


        Ethylmalonyl-CoA: 36.7


        CHC-CoA: 29.0


        Propionyl-CoA: 28.0


        Acetyl-CoA: 23.0


        trans-1,2-CPDA: 22.7


        2-Methylbutyryl-CoA: 22.6


        Isobutyryl-CoA: 21.3


        Benzoyl-CoA: 17.7


        fatty\_acid: 13.8


        3-Methylbutyryl-CoA: 9.6

contig\_35 - Region 1 - NRPS,T1PKS

Shows the layout of the region, marking coding sequences and areas of interest. Clicking a gene will select it and show any relevant details. Clicking an area feature (e.g. a candidate cluster) will select all coding sequences within that area. Double clicking an area feature will zoom to that area. Multiple genes and area features can be selected by clicking them while holding the Ctrl key.  
More detailed help is available here.

Download region GenBank file

Download region SVG

Location: 26,846 - 89,893 nt. (total: 63,048 nt)
Show pHMM detection rules used

NRPS: cds(Condensation and (AMP-binding or A-OX))  
T1PKS: cds(PKS\_AT and (PKS\_KS or ene\_KS or mod\_KS or hyb\_KS or itr\_KS or tra\_KS))

#### Legend:

core biosynthetic genes

additional biosynthetic genes

transport-related genes

regulatory genes

other genes

resistance

reset view

zoom to selection

Gene details

Shows details of the most recently selected gene, including names, products, location, and other annotations.

Select a gene to view the details available for it

NRPS/PKS domains

ClusterBlast

KnownClusterBlast

SubClusterBlast

MIBiG comparison

Pfam domains

Detailed domain annotation

Shows NRPS- and PKS-related domains for each feature that contains them. Click on each domain for more information about the domain's location, consensus monomer prediction, and other details.  
A glossary is available here.

Selected features only

Show module domains

Similar gene clusters

Shows clusters from the antiSMASH database and other clusters of interest that are similar to the current region. Genes marked with the same colour are interrelated. White genes have no relationship.  
Click on reference genes to show details of similarities to genes within the current region.  
Click on an accession to open that entry in the antiSMASH database (if applicable).

All hits

CM000575 (2561027-2621917): Fusarium graminearum PH-1 chromosome 2, whole gen... (17% of genes show similarity), NRPS,T1PKS

NC\_026475 (2561027-2621917): Fusarium graminearum PH-1 chromosome 2, whole ge... (17% of genes show similarity), NRPS,T1PKS

NW\_007360987 (200861-291572): Glarea lozoyensis ATCC 20868 chromosome Unknown... (8% of genes show similarity), NRPS,T1PKS,betalactone

NW\_022474214 (256395-313016): Venustampulla echinocandica strain BP 5553 chro... (14% of genes show similarity), NRPS,T1PKS

NW\_022474217 (345692-404245): Venustampulla echinocandica strain BP 5553 chro... (15% of genes show similarity), NRPS,T1PKS

NC\_038012 (6515913-6598624): Fusarium venenatum strain A3/5 genome assembly, ... (11% of genes show similarity), NRPS,T1PKS

NT\_107014 (3567979-3645403): Aspergillus nidulans FGSC A4 chromosome VII map ... (12% of genes show similarity), NRPS,T1PKS

NW\_006271969 (4113130-4163983): Cordyceps militaris CM01 unplaced genomic sca... (16% of genes show similarity), NRPS,T1PKS

NT\_165979 (2279535-2344621): Chaetomium globosum CBS 148.51 scaffold 4 genomi... (11% of genes show similarity), NRPS,T1PKS

NW\_022984629 (3081705-3186571): Aspergillus tanneri strain NIH1004 chromosome... (15% of genes show similarity), NRPS,T1PKS,indole
Download graphic

Similar known gene clusters

Shows clusters from the MiBIG database that are similar to the current region. Genes marked with the same colour are interrelated. White genes have no relationship.  
Click on reference genes to show details of similarities to genes within the current region.  
Click on an accession to open that entry in the MiBIG database.

No matches found.

Similar subclusters

Shows sub-cluster units that are similar to the current region. Genes marked with the same colour are interrelated. White genes have no relationship.  
Click on reference genes to show details of similarities to genes within the current region.

No matches found.

Similar gene clusters

Shows careas that are similar to the current region to a reference database.  
Mouseover a score cell in the table to get a breakdown of how the score was calculated.The MIBiG database.  
  
Click on an accession to open that entry in the MIBiG database.

Analysis type:

Protocluster to Region
Region to Region

| Reference | NRPS | T1PKS | Similarity score | Type | Compound(s) | Organism |
| --- | --- | --- | --- | --- | --- | --- |
| BGC0001261.1 |  |  | 0.66 | NRP | AM-toxin | Alternaria alternata |
| BGC0001249.1 |  |  | 0.65 | NRP | dimethylcoprogen | Alternaria alternata |
| BGC0000900.1 |  |  | 0.62 | Other | ferrichrome | Aspergillus oryzae |
| BGC0001718.1 |  |  | 0.61 | NRP | okaramine D | Aspergillus aculeatus ATCC 16872 |
| BGC0000348.1 |  |  | 0.58 | NRP | ergovaline | Epichloe festucae var. lolii |
| BGC0001132.1 |  |  | 0.56 | NRP | xenotetrapeptide | Xenorhabdus nematophila ATCC 19061 |
| BGC0001858.1 |  |  | 0.54 | Polyketide | alternapyrone B, alternapyrone C, alternapyrone D, alternapyrone E, alternapyrone F | Parastagonospora nodorum SN15 |
| BGC0001135.1 |  |  | 0.53 | NRP | bicornutin A1, bicornutin A2 | Xenorhabdus budapestensis |
| BGC0001128.1 |  |  | 0.53 | NRP | luminmide | Photorhabdus laumondii subsp. laumondii TTO1 |
| BGC0001166.1 |  |  | 0.53 | NRP | HC-toxin | Alternaria jesenskae |

| Reference | Aggregated | Similarity score | Type | Compound(s) | Organism |
| --- | --- | --- | --- | --- | --- |
| BGC0001261.1 |  | 0.66 | NRP | AM-toxin | Alternaria alternata |
| BGC0001249.1 |  | 0.66 | NRP | dimethylcoprogen | Alternaria alternata |
| BGC0000900.1 |  | 0.64 | Other | ferrichrome | Aspergillus oryzae |
| BGC0001718.1 |  | 0.64 | NRP | okaramine D | Aspergillus aculeatus ATCC 16872 |
| BGC0000348.1 |  | 0.63 | NRP | ergovaline | Epichloe festucae var. lolii |
| BGC0001132.1 |  | 0.62 | NRP | xenotetrapeptide | Xenorhabdus nematophila ATCC 19061 |
| BGC0001858.1 |  | 0.61 | Polyketide | alternapyrone B, alternapyrone C, alternapyrone D, alternapyrone E, alternapyrone F | Parastagonospora nodorum SN15 |
| BGC0001135.1 |  | 0.60 | NRP | bicornutin A1, bicornutin A2 | Xenorhabdus budapestensis |
| BGC0001128.1 |  | 0.60 | NRP | luminmide | Photorhabdus laumondii subsp. laumondii TTO1 |
| BGC0001166.1 |  | 0.60 | NRP | HC-toxin | Alternaria jesenskae |

Detailed Pfam domain annotation

Shows Pfam domains found in each gene within the region.
Click on each domain for more information about the domain's
accession, location, description, and any relevant Gene Ontology.
Domains with a bold border have Gene Ontology information.

Selected features only

NRPS/PKS products

NRPS/PKS monomers

Predicted core structure(s)

Shows estimated product structure and polymer for each candidate cluster in the region. To show the product, click on the expander or the candidate cluster feature drawn in the overview.

For candidate cluster 1, location 26845 - 89893:

Rough prediction of core scaffold based on assumed PKS/NRPS colinearity; tailoring reactions not taken into account

**Polymer prediction:**
:   (ala - X - X) + (pk)

  
Direct lookup in NORINE database:
strict
or
relaxed

---

For candidate cluster 2, location 26845 - 78495:

Rough prediction of core scaffold based on assumed PKS/NRPS colinearity; tailoring reactions not taken into account

**Polymer prediction:**
:   (ala - X - X) + (pk)

  
Direct lookup in NORINE database:
strict
or
relaxed

---

For candidate cluster 3, location 42097 - 89893:

Rough prediction of core scaffold based on assumed PKS/NRPS colinearity; tailoring reactions not taken into account

**Polymer prediction:**
:   (ala - X - X) + (pk)

  
Direct lookup in NORINE database:
strict
or
relaxed

Link to NORINE database query form

NRPS/PKS monomer predictions

Shows the predicted monomers for each adynelation domain and acyltransferase within genes. Each gene prediction can be expanded to view detailed predictions of each domain. Each prediction can be expanded to view the predictions by tool (and, for some tools, further expanded for extra details).

**input.path1.gene3**: ala - X - X

:   Search NORINE for peptide:
    strict
    or
    relaxed
  
:   **AMP-binding (267..665)**: ala

    NRPSPredictor2: ala

    SVM prediction details:
    :   Predicted physicochemical class:
        :   hydrophobic-aliphatic

        Large clusters prediction:
        :   N/A

        Small clusters prediction:
        :   gly, ala

        Single AA prediction:
        :   ala

    Stachelhaus prediction details:
    :   Stachelhaus sequence:
        :   dvysvfaifk

        Nearest Stachelhaus code:
        :   N, A

        Stachelhaus code match:
        :   0% (weak)
:   **AMP-binding (1362..1761)**: X

    NRPSPredictor2: hydrophobic-aliphatic

    SVM prediction details:
    :   Predicted physicochemical class:
        :   hydrophobic-aliphatic

        Large clusters prediction:
        :   N/A

        Small clusters prediction:
        :   N/A

        Single AA prediction:
        :   N/A

    Stachelhaus prediction details:
    :   Stachelhaus sequence:
        :   dpqvqvivyk

        Nearest Stachelhaus code:
        :   N, A

        Stachelhaus code match:
        :   0% (weak)
:   **AMP-binding (2464..2862)**: X

    NRPSPredictor2: hydrophobic-aliphatic

    SVM prediction details:
    :   Predicted physicochemical class:
        :   hydrophobic-aliphatic

        Large clusters prediction:
        :   N/A

        Small clusters prediction:
        :   N/A

        Single AA prediction:
        :   N/A

    Stachelhaus prediction details:
    :   Stachelhaus sequence:
        :   dvsqlmsiyk

        Nearest Stachelhaus code:
        :   N, A

        Stachelhaus code match:
        :   0% (weak)

  
**input.path1.gene4**: pk

:   **PKS\_AT (490..808)**: pk

    ATSignature: Malonyl-CoA

    Top 3 matches:
    :   Malonyl-CoA: 66.7%
    :   Methylmalonyl-CoA: 54.2%
    :   Ethylmalonyl-CoA: 54.2%

      
    minowa: Methoxymalonyl-CoA

    Prediction, score:
    :   Methoxymalonyl-CoA: 65.6


        Methylmalonyl-CoA: 64.1


        Malonyl-CoA: 45.5


        Ethylmalonyl-CoA: 38.4


        Propionyl-CoA: 33.8


        trans-1,2-CPDA: 32.4


        2-Methylbutyryl-CoA: 22.1


        fatty\_acid: 18.3


        Isobutyryl-CoA: 18.0


        Benzoyl-CoA: 17.8


        Acetyl-CoA: 15.6


        inactive: 15.4


        CHC-CoA: 15.4


        3-Methylbutyryl-CoA: 0.0

contig\_39 - Region 1 - T1PKS

Shows the layout of the region, marking coding sequences and areas of interest. Clicking a gene will select it and show any relevant details. Clicking an area feature (e.g. a candidate cluster) will select all coding sequences within that area. Double clicking an area feature will zoom to that area. Multiple genes and area features can be selected by clicking them while holding the Ctrl key.  
More detailed help is available here.

Download region GenBank file

Download region SVG

Location: 34,391 - 73,215 nt. (total: 38,825 nt)
Show pHMM detection rules used

T1PKS: cds(PKS\_AT and (PKS\_KS or ene\_KS or mod\_KS or hyb\_KS or itr\_KS or tra\_KS))

#### Legend:

core biosynthetic genes

additional biosynthetic genes

transport-related genes

regulatory genes

other genes

resistance

reset view

zoom to selection

Gene details

Shows details of the most recently selected gene, including names, products, location, and other annotations.

Select a gene to view the details available for it

NRPS/PKS domains

ClusterBlast

KnownClusterBlast

SubClusterBlast

MIBiG comparison

Pfam domains

Detailed domain annotation

Shows NRPS- and PKS-related domains for each feature that contains them. Click on each domain for more information about the domain's location, consensus monomer prediction, and other details.  
A glossary is available here.

Selected features only

Show module domains

Similar gene clusters

Shows clusters from the antiSMASH database and other clusters of interest that are similar to the current region. Genes marked with the same colour are interrelated. White genes have no relationship.  
Click on reference genes to show details of similarities to genes within the current region.  
Click on an accession to open that entry in the antiSMASH database (if applicable).

All hits

NT\_165931 (1584470-1627478): Aspergillus terreus NIH2624 scaffold 8 genomic s... (35% of genes show similarity), T1PKS

NC\_036623 (4276540-4323612): Fusarium fujikuroi IMI 58289 draft genome, chrom... (10% of genes show similarity), T1PKS

NW\_022194791 (779496-826088): Fusarium proliferatum ET1 genome assembly, cont... (10% of genes show similarity), T1PKS

NC\_038015 (7239050-7286189): Fusarium venenatum strain A3/5 genome assembly, ... (11% of genes show similarity), T1PKS

CM000577 (7275310-7321978): Fusarium graminearum PH-1 chromosome 4, whole gen... (10% of genes show similarity), T1PKS

NC\_026477 (7275310-7321978): Fusarium graminearum PH-1 chromosome 4, whole ge... (10% of genes show similarity), T1PKS

NC\_030957 (3106295-3150324): Colletotrichum higginsianum IMI 349063 chromosom... (17% of genes show similarity), T1PKS

NW\_023336269 (1206036-1252444): Colletotrichum scovillei strain TJNH1 chromos... (18% of genes show similarity), T1PKS

NC\_049562 (5064149-5105472): Talaromyces rugulosus chromosome II, complete se... (14% of genes show similarity), T1PKS

NW\_023336287 (462128-503095): Aspergillus tubingensis WU-2223L DNA, scaffold ... (13% of genes show similarity), T1PKS
Download graphic

Similar known gene clusters

Shows clusters from the MiBIG database that are similar to the current region. Genes marked with the same colour are interrelated. White genes have no relationship.  
Click on reference genes to show details of similarities to genes within the current region.  
Click on an accession to open that entry in the MiBIG database.

No matches found.

Similar subclusters

Shows sub-cluster units that are similar to the current region. Genes marked with the same colour are interrelated. White genes have no relationship.  
Click on reference genes to show details of similarities to genes within the current region.

No matches found.

Similar gene clusters

Shows careas that are similar to the current region to a reference database.  
Mouseover a score cell in the table to get a breakdown of how the score was calculated.The MIBiG database.  
  
Click on an accession to open that entry in the MIBiG database.

Analysis type:

Protocluster to Region
Region to Region

| Reference | T1PKS | Similarity score | Type | Compound(s) | Organism |
| --- | --- | --- | --- | --- | --- |
| BGC0000156.1 |  | 0.35 | Polyketide | TAN-1612, 1-(2,3,5,10-tetrahydroxy-7-methoxy-4-oxo-1,2,3,4-tetrahydroanthracen-2-yl)pentane-2,4-dione, desmethyl TAN-1612 | Aspergillus niger |
| BGC0000107.1 |  | 0.35 | Polyketide | naphthopyrone | Aspergillus nidulans FGSC A4 |
| BGC0001906.1 |  | 0.34 | Polyketide | naphthalene | Daldinia eschscholzii IFB-TL01 |
| BGC0001258.1 |  | 0.34 | Polyketide | 1,3,6,8-tetrahydroxynaphthalene | Glarea lozoyensis |
| BGC0001257.1 |  | 0.34 | Polyketide | 1,3,6,8-tetrahydroxynaphthalene | Nodulisporium sp. ATCC74245 |
| BGC0001265.1 |  | 0.34 | Polyketide | melanin | Bipolaris oryzae |
| BGC0000057.1 |  | 0.33 | Polyketide | F9775A, F9775B, orsellinic acid | Aspergillus nidulans FGSC A4 |
| BGC0001284.1 |  | 0.33 | Terpene | alternariol | Parastagonospora nodorum SN15 |
| BGC0001304.1 |  | 0.31 | Polyketide | aflavarin | Aspergillus flavus NRRL3357 |
| BGC0001242.1 |  | 0.30 | Polyketide | oxyjavanicin | Fusarium fujikuroi |

| Reference | Aggregated | Similarity score | Type | Compound(s) | Organism |
| --- | --- | --- | --- | --- | --- |
| BGC0000156.1 |  | 0.68 | Polyketide | TAN-1612, 1-(2,3,5,10-tetrahydroxy-7-methoxy-4-oxo-1,2,3,4-tetrahydroanthracen-2-yl)pentane-2,4-dione, desmethyl TAN-1612 | Aspergillus niger |
| BGC0000107.1 |  | 0.68 | Polyketide | naphthopyrone | Aspergillus nidulans FGSC A4 |
| BGC0001906.1 |  | 0.68 | Polyketide | naphthalene | Daldinia eschscholzii IFB-TL01 |
| BGC0001258.1 |  | 0.67 | Polyketide | 1,3,6,8-tetrahydroxynaphthalene | Glarea lozoyensis |
| BGC0001257.1 |  | 0.67 | Polyketide | 1,3,6,8-tetrahydroxynaphthalene | Nodulisporium sp. ATCC74245 |
| BGC0001265.1 |  | 0.67 | Polyketide | melanin | Bipolaris oryzae |
| BGC0000057.1 |  | 0.66 | Polyketide | F9775A, F9775B, orsellinic acid | Aspergillus nidulans FGSC A4 |
| BGC0001284.1 |  | 0.66 | Terpene | alternariol | Parastagonospora nodorum SN15 |
| BGC0001304.1 |  | 0.65 | Polyketide | aflavarin | Aspergillus flavus NRRL3357 |
| BGC0001242.1 |  | 0.64 | Polyketide | oxyjavanicin | Fusarium fujikuroi |

Detailed Pfam domain annotation

Shows Pfam domains found in each gene within the region.
Click on each domain for more information about the domain's
accession, location, description, and any relevant Gene Ontology.
Domains with a bold border have Gene Ontology information.

Selected features only

NRPS/PKS products

NRPS/PKS monomers

Predicted core structure(s)

Shows estimated product structure and polymer for each candidate cluster in the region. To show the product, click on the expander or the candidate cluster feature drawn in the overview.

For candidate cluster 1, location 34390 - 73215:

Rough prediction of core scaffold based on assumed PKS/NRPS colinearity; tailoring reactions not taken into account

**Polymer prediction:**
:   (mal)

  
Direct lookup in NORINE database:
strict
or
relaxed

Link to NORINE database query form

NRPS/PKS monomer predictions

Shows the predicted monomers for each adynelation domain and acyltransferase within genes. Each gene prediction can be expanded to view detailed predictions of each domain. Each prediction can be expanded to view the predictions by tool (and, for some tools, further expanded for extra details).

**input.path1.gene5**: mal

:   **PKS\_AT (440..737)**: mal

    ATSignature: Malonyl-CoA

    Top 3 matches:
    :   Malonyl-CoA: 79.2%
    :   inactive: 58.3%
    :   Methylmalonyl-CoA: 54.2%

      
    minowa: Malonyl-CoA

    Prediction, score:
    :   Malonyl-CoA: 106.1


        inactive: 68.1


        Methoxymalonyl-CoA: 56.7


        Methylmalonyl-CoA: 53.8


        Ethylmalonyl-CoA: 28.8


        Propionyl-CoA: 21.3


        Isobutyryl-CoA: 18.4


        fatty\_acid: 18.2


        2-Methylbutyryl-CoA: 17.8


        CHC-CoA: 15.4


        Benzoyl-CoA: 14.2


        3-Methylbutyryl-CoA: 13.9


        Acetyl-CoA: 12.0


        trans-1,2-CPDA: 9.1

contig\_39 - Region 2 - T1PKS

Shows the layout of the region, marking coding sequences and areas of interest. Clicking a gene will select it and show any relevant details. Clicking an area feature (e.g. a candidate cluster) will select all coding sequences within that area. Double clicking an area feature will zoom to that area. Multiple genes and area features can be selected by clicking them while holding the Ctrl key.  
More detailed help is available here.

Download region GenBank file

Download region SVG

Location: 1,044,010 - 1,083,080 nt. (total: 39,071 nt)
Show pHMM detection rules used

T1PKS: cds(PKS\_AT and (PKS\_KS or ene\_KS or mod\_KS or hyb\_KS or itr\_KS or tra\_KS))

#### Legend:

core biosynthetic genes

additional biosynthetic genes

transport-related genes

regulatory genes

other genes

resistance

reset view

zoom to selection

Gene details

Shows details of the most recently selected gene, including names, products, location, and other annotations.

Select a gene to view the details available for it

NRPS/PKS domains

ClusterBlast

KnownClusterBlast

SubClusterBlast

MIBiG comparison

Pfam domains

Detailed domain annotation

Shows NRPS- and PKS-related domains for each feature that contains them. Click on each domain for more information about the domain's location, consensus monomer prediction, and other details.  
A glossary is available here.

Selected features only

Show module domains

Similar gene clusters

Shows clusters from the antiSMASH database and other clusters of interest that are similar to the current region. Genes marked with the same colour are interrelated. White genes have no relationship.  
Click on reference genes to show details of similarities to genes within the current region.  
Click on an accession to open that entry in the antiSMASH database (if applicable).

All hits

NZ\_CP022438 (1767719-1834144): Streptomyces peucetius subsp. caesius ATCC 279... (12% of genes show similarity), T1PKS

NW\_007360985 (1145957-1199332): Glarea lozoyensis ATCC 20868 chromosome Unkno... (13% of genes show similarity), T1PKS

NZ\_CP023690 (3216937-3334738): Streptomyces spectabilis strain ATCC 27465 chr... (4% of genes show similarity), NRPS,T1PKS

NZ\_CP040916 (2991833-3110679): Streptomyces spectabilis strain NRRL 2792 chro... (2% of genes show similarity), NRPS,T1PKS
Download graphic

Similar known gene clusters

Shows clusters from the MiBIG database that are similar to the current region. Genes marked with the same colour are interrelated. White genes have no relationship.  
Click on reference genes to show details of similarities to genes within the current region.  
Click on an accession to open that entry in the MiBIG database.

No matches found.

Similar subclusters

Shows sub-cluster units that are similar to the current region. Genes marked with the same colour are interrelated. White genes have no relationship.  
Click on reference genes to show details of similarities to genes within the current region.

No matches found.

Similar gene clusters

Shows careas that are similar to the current region to a reference database.  
Mouseover a score cell in the table to get a breakdown of how the score was calculated.The MIBiG database.  
  
Click on an accession to open that entry in the MIBiG database.

Analysis type:

Protocluster to Region
Region to Region

| Reference | T1PKS | Similarity score | Type | Compound(s) | Organism |
| --- | --- | --- | --- | --- | --- |
| BGC0001265.1 |  | 0.33 | Polyketide | melanin | Bipolaris oryzae |
| BGC0001257.1 |  | 0.33 | Polyketide | 1,3,6,8-tetrahydroxynaphthalene | Nodulisporium sp. ATCC74245 |
| BGC0001258.1 |  | 0.32 | Polyketide | 1,3,6,8-tetrahydroxynaphthalene | Glarea lozoyensis |
| BGC0001284.1 |  | 0.32 | Terpene | alternariol | Parastagonospora nodorum SN15 |
| BGC0000107.1 |  | 0.32 | Polyketide | naphthopyrone | Aspergillus nidulans FGSC A4 |
| BGC0000057.1 |  | 0.31 | Polyketide | F9775A, F9775B, orsellinic acid | Aspergillus nidulans FGSC A4 |
| BGC0001583.1 |  | 0.27 | Polyketide | emodin | Escovopsis weberi |
| BGC0000156.1 |  | 0.27 | Polyketide | TAN-1612, 1-(2,3,5,10-tetrahydroxy-7-methoxy-4-oxo-1,2,3,4-tetrahydroanthracen-2-yl)pentane-2,4-dione, desmethyl TAN-1612 | Aspergillus niger |
| BGC0001906.1 |  | 0.27 | Polyketide | naphthalene | Daldinia eschscholzii IFB-TL01 |
| BGC0000121.1 |  | 0.23 | Polyketide | RES-1214-2 | Pestalotiopsis fici |

| Reference | Aggregated | Similarity score | Type | Compound(s) | Organism |
| --- | --- | --- | --- | --- | --- |
| BGC0001265.1 |  | 0.66 | Polyketide | melanin | Bipolaris oryzae |
| BGC0001257.1 |  | 0.66 | Polyketide | 1,3,6,8-tetrahydroxynaphthalene | Nodulisporium sp. ATCC74245 |
| BGC0001258.1 |  | 0.66 | Polyketide | 1,3,6,8-tetrahydroxynaphthalene | Glarea lozoyensis |
| BGC0001284.1 |  | 0.66 | Terpene | alternariol | Parastagonospora nodorum SN15 |
| BGC0000107.1 |  | 0.65 | Polyketide | naphthopyrone | Aspergillus nidulans FGSC A4 |
| BGC0000057.1 |  | 0.65 | Polyketide | F9775A, F9775B, orsellinic acid | Aspergillus nidulans FGSC A4 |
| BGC0001583.1 |  | 0.61 | Polyketide | emodin | Escovopsis weberi |
| BGC0000156.1 |  | 0.60 | Polyketide | TAN-1612, 1-(2,3,5,10-tetrahydroxy-7-methoxy-4-oxo-1,2,3,4-tetrahydroanthracen-2-yl)pentane-2,4-dione, desmethyl TAN-1612 | Aspergillus niger |
| BGC0001906.1 |  | 0.60 | Polyketide | naphthalene | Daldinia eschscholzii IFB-TL01 |
| BGC0000121.1 |  | 0.57 | Polyketide | RES-1214-2 | Pestalotiopsis fici |

Detailed Pfam domain annotation

Shows Pfam domains found in each gene within the region.
Click on each domain for more information about the domain's
accession, location, description, and any relevant Gene Ontology.
Domains with a bold border have Gene Ontology information.

Selected features only

NRPS/PKS products

NRPS/PKS monomers

Predicted core structure(s)

Shows estimated product structure and polymer for each candidate cluster in the region. To show the product, click on the expander or the candidate cluster feature drawn in the overview.

For candidate cluster 2, location 1044009 - 1083080:

Rough prediction of core scaffold based on assumed PKS/NRPS colinearity; tailoring reactions not taken into account

**Polymer prediction:**
:   (mal)

  
Direct lookup in NORINE database:
strict
or
relaxed

Link to NORINE database query form

NRPS/PKS monomer predictions

Shows the predicted monomers for each adynelation domain and acyltransferase within genes. Each gene prediction can be expanded to view detailed predictions of each domain. Each prediction can be expanded to view the predictions by tool (and, for some tools, further expanded for extra details).

**input.path1.gene283**: mal

:   **PKS\_AT (888..1188)**: mal

    ATSignature: Malonyl-CoA

    Top 3 matches:
    :   Malonyl-CoA: 75.0%
    :   inactive: 66.7%
    :   Methylmalonyl-CoA: 62.5%

      
    minowa: Malonyl-CoA

    Prediction, score:
    :   Malonyl-CoA: 135.1


        inactive: 84.2


        Methoxymalonyl-CoA: 67.9


        Methylmalonyl-CoA: 67.3


        Ethylmalonyl-CoA: 38.7


        Isobutyryl-CoA: 27.0


        Propionyl-CoA: 25.2


        Benzoyl-CoA: 23.3


        2-Methylbutyryl-CoA: 17.4


        trans-1,2-CPDA: 14.5


        Acetyl-CoA: 13.8


        CHC-CoA: 11.4


        3-Methylbutyryl-CoA: 9.5


        fatty\_acid: 0.0

contig\_39 - Region 3 - NRPS,T1PKS

Shows the layout of the region, marking coding sequences and areas of interest. Clicking a gene will select it and show any relevant details. Clicking an area feature (e.g. a candidate cluster) will select all coding sequences within that area. Double clicking an area feature will zoom to that area. Multiple genes and area features can be selected by clicking them while holding the Ctrl key.  
More detailed help is available here.

Download region GenBank file

Download region SVG

Location: 1,102,900 - 1,154,380 nt. (total: 51,481 nt)
Show pHMM detection rules used

NRPS: cds(Condensation and (AMP-binding or A-OX))  
T1PKS: cds(PKS\_AT and (PKS\_KS or ene\_KS or mod\_KS or hyb\_KS or itr\_KS or tra\_KS))

#### Legend:

core biosynthetic genes

additional biosynthetic genes

transport-related genes

regulatory genes

other genes

resistance

reset view

zoom to selection

Gene details

Shows details of the most recently selected gene, including names, products, location, and other annotations.

Select a gene to view the details available for it

NRPS/PKS domains

ClusterBlast

KnownClusterBlast

SubClusterBlast

MIBiG comparison

Pfam domains

Detailed domain annotation

Shows NRPS- and PKS-related domains for each feature that contains them. Click on each domain for more information about the domain's location, consensus monomer prediction, and other details.  
A glossary is available here.

Selected features only

Show module domains

Similar gene clusters

Shows clusters from the antiSMASH database and other clusters of interest that are similar to the current region. Genes marked with the same colour are interrelated. White genes have no relationship.  
Click on reference genes to show details of similarities to genes within the current region.  
Click on an accession to open that entry in the antiSMASH database (if applicable).

All hits

NC\_049565 (5170731-5222831): Talaromyces rugulosus chromosome V, complete seq... (9% of genes show similarity), NRPS,T1PKS

NZ\_LR733556 (639454-714742): Flavobacterium sp. 9AF isolate Flavobacterium sp... (4% of genes show similarity), NRPS,T1PKS

NW\_022474219 (274912-345725): Venustampulla echinocandica strain BP 5553 chro... (10% of genes show similarity), NRPS,T1PKS

NT\_165977 (760296-807762): Chaetomium globosum CBS 148.51 scaffold 2 genomic ... (17% of genes show similarity), NRPS,T1PKS

NZ\_AP018195 (207519-380160): Scytonema sp. HK-05 plasmid plasmid1 DNA, nearly... (6% of genes show similarity), NRPS,T1PKS,microviridin

NZ\_AP018268 (1523285-1713097): Scytonema sp. NIES-4073 DNA, nearly complete g... (2% of genes show similarity), NRPS,T1PKS

NZ\_KL662191 (<2281952-2385351): [Leptolyngbya] sp. JSC-1 Osccy1DRAFT CYJSC1 D... (8% of genes show similarity), NRPS,NRPS-like,T1PKS

NZ\_CM001979 (2880117-2956585): Dickeya sp. NCPPB 3274 chromosome, whole genom... (4% of genes show similarity), NRPS,transAT-PKS

NZ\_KQ976354 (3589179-3832343): Scytonema hofmannii PCC 7110 Scaffold1, whole ... (2% of genes show similarity), NRPS,microviridin

NC\_010628 (4167586-4349893): Nostoc punctiforme PCC 73102, complete sequence (1% of genes show similarity), NRPS,T1PKS,bacteriocin,lanthidin,transAT-PKS-like
Download graphic

Similar known gene clusters

Shows clusters from the MiBIG database that are similar to the current region. Genes marked with the same colour are interrelated. White genes have no relationship.  
Click on reference genes to show details of similarities to genes within the current region.  
Click on an accession to open that entry in the MiBIG database.

No matches found.

Similar subclusters

Shows sub-cluster units that are similar to the current region. Genes marked with the same colour are interrelated. White genes have no relationship.  
Click on reference genes to show details of similarities to genes within the current region.

No matches found.

Similar gene clusters

Shows careas that are similar to the current region to a reference database.  
Mouseover a score cell in the table to get a breakdown of how the score was calculated.The MIBiG database.  
  
Click on an accession to open that entry in the MIBiG database.

Analysis type:

Protocluster to Region
Region to Region

| Reference | NRPS | T1PKS | Similarity score | Type | Compound(s) | Organism |
| --- | --- | --- | --- | --- | --- | --- |
| BGC0001874.1 |  |  | 0.52 | NRP, Polyketide | cyclopiazonic acid | Aspergillus flavus |
| BGC0000064.1 |  |  | 0.42 | Polyketide | fusarin | Fusarium verticillioides |
| BGC0001268.1 |  |  | 0.42 | NRP, Polyketide | fusarin | Fusarium fujikuroi |
| BGC0000900.1 |  |  | 0.41 | Other | ferrichrome | Aspergillus oryzae |
| BGC0000357.1 |  |  | 0.41 | NRP | cyclo-(D-Phe-L-Phe-D-Val-L-Val), cyclo-(D-Tyr-L-Phe-D-Val-L-Val), cyclo-(D-Tyr-L-Trp-D-Val-L-Val), cyclo-(D-Phe-L-Trp-D-Val-L-Val), cyclo-(D-Phe-L-Phe-D-Val-L-Ile), cyclo-(D-Phe-L-Phe-D-Ile-L-Val), cyclo-(D-Tyr-L-Trp-D-Val-L-Ile), cyclo-(D-Tyr-L-Trp-D-Ile-L-Val), cyclo-(D-Tyr-L-Phe-D-Val-L-Ile), cyclo-(D-Tyr-L-Phe-D-Ile-L-Val) | Penicillium rubens Wisconsin 54-1255 |
| BGC0001240.1 |  |  | 0.41 | NRP | serinocyclin A, serinocyclin B | Metarhizium robertsii |
| BGC0001068.1 |  |  | 0.41 | Terpene, Polyketide | pyripyropene A | unidentified unclassified sequences. |
| BGC0001724.1 |  |  | 0.41 | NRP, Polyketide | oxaleimide C | Penicillium oxalicum 114-2 |
| BGC0001261.1 |  |  | 0.41 | NRP | AM-toxin | Alternaria alternata |
| BGC0001220.1 |  |  | 0.41 | NRP | aculeacin A | Aspergillus japonicus |

| Reference | Aggregated | Similarity score | Type | Compound(s) | Organism |
| --- | --- | --- | --- | --- | --- |
| BGC0001874.1 |  | 0.59 | NRP, Polyketide | cyclopiazonic acid | Aspergillus flavus |
| BGC0000064.1 |  | 0.54 | Polyketide | fusarin | Fusarium verticillioides |
| BGC0001268.1 |  | 0.54 | NRP, Polyketide | fusarin | Fusarium fujikuroi |
| BGC0000900.1 |  | 0.53 | Other | ferrichrome | Aspergillus oryzae |
| BGC0000357.1 |  | 0.53 | NRP | cyclo-(D-Phe-L-Phe-D-Val-L-Val), cyclo-(D-Tyr-L-Phe-D-Val-L-Val), cyclo-(D-Tyr-L-Trp-D-Val-L-Val), cyclo-(D-Phe-L-Trp-D-Val-L-Val), cyclo-(D-Phe-L-Phe-D-Val-L-Ile), cyclo-(D-Phe-L-Phe-D-Ile-L-Val), cyclo-(D-Tyr-L-Trp-D-Val-L-Ile), cyclo-(D-Tyr-L-Trp-D-Ile-L-Val), cyclo-(D-Tyr-L-Phe-D-Val-L-Ile), cyclo-(D-Tyr-L-Phe-D-Ile-L-Val) | Penicillium rubens Wisconsin 54-1255 |
| BGC0001240.1 |  | 0.53 | NRP | serinocyclin A, serinocyclin B | Metarhizium robertsii |
| BGC0001068.1 |  | 0.53 | Terpene, Polyketide | pyripyropene A | unidentified unclassified sequences. |
| BGC0001724.1 |  | 0.53 | NRP, Polyketide | oxaleimide C | Penicillium oxalicum 114-2 |
| BGC0001261.1 |  | 0.53 | NRP | AM-toxin | Alternaria alternata |
| BGC0001220.1 |  | 0.53 | NRP | aculeacin A | Aspergillus japonicus |

Detailed Pfam domain annotation

Shows Pfam domains found in each gene within the region.
Click on each domain for more information about the domain's
accession, location, description, and any relevant Gene Ontology.
Domains with a bold border have Gene Ontology information.

Selected features only

NRPS/PKS products

NRPS/PKS monomers

Predicted core structure(s)

Shows estimated product structure and polymer for each candidate cluster in the region. To show the product, click on the expander or the candidate cluster feature drawn in the overview.

For candidate cluster 3, location 1102899 - 1154380:

Rough prediction of core scaffold based on assumed PKS/NRPS colinearity; tailoring reactions not taken into account

**Polymer prediction:**
:   (X)

  
Direct lookup in NORINE database:
strict
or
relaxed

---

For candidate cluster 4, location 1102899 - 1148493:

Rough prediction of core scaffold based on assumed PKS/NRPS colinearity; tailoring reactions not taken into account

**Polymer prediction:**
:   (X)

  
Direct lookup in NORINE database:
strict
or
relaxed

---

For candidate cluster 5, location 1111510 - 1154380:

Rough prediction of core scaffold based on assumed PKS/NRPS colinearity; tailoring reactions not taken into account

**Polymer prediction:**
:   (X)

  
Direct lookup in NORINE database:
strict
or
relaxed

Link to NORINE database query form

NRPS/PKS monomer predictions

Shows the predicted monomers for each adynelation domain and acyltransferase within genes. Each gene prediction can be expanded to view detailed predictions of each domain. Each prediction can be expanded to view the predictions by tool (and, for some tools, further expanded for extra details).

**input.path1.gene298**: X

:   Search NORINE for peptide:
    strict
    or
    relaxed
  
:   **AMP-binding (726..1009)**: X

    NRPSPredictor2: val, leu, ile, abu, iva

    SVM prediction details:
    :   Predicted physicochemical class:
        :   hydrophilic

        Large clusters prediction:
        :   N/A

        Small clusters prediction:
        :   val, leu, ile, abu, iva

        Single AA prediction:
        :   N/A

    Stachelhaus prediction details:
    :   Stachelhaus sequence:
        :   daimwgsisk

        Nearest Stachelhaus code:
        :   N, A

        Stachelhaus code match:
        :   0% (weak)

  
**input.path1.gene300**: pk

:   **PKS\_AT (213..531)**: pk

    ATSignature: Malonyl-CoA

    Top 3 matches:
    :   Malonyl-CoA: 75.0%
    :   Methylmalonyl-CoA: 70.8%
    :   Ethylmalonyl-CoA: 58.3%

      
    minowa: Methylmalonyl-CoA

    Prediction, score:
    :   Methylmalonyl-CoA: 72.2


        Methoxymalonyl-CoA: 53.3


        Isobutyryl-CoA: 44.9


        Ethylmalonyl-CoA: 40.9


        Malonyl-CoA: 39.1


        Benzoyl-CoA: 28.8


        Acetyl-CoA: 18.9


        trans-1,2-CPDA: 17.9


        Propionyl-CoA: 15.6


        fatty\_acid: 14.5


        CHC-CoA: 11.4


        2-Methylbutyryl-CoA: 9.6


        3-Methylbutyryl-CoA: 8.1


        inactive: 0.0

contig\_4 - Region 1 - NRPS

Shows the layout of the region, marking coding sequences and areas of interest. Clicking a gene will select it and show any relevant details. Clicking an area feature (e.g. a candidate cluster) will select all coding sequences within that area. Double clicking an area feature will zoom to that area. Multiple genes and area features can be selected by clicking them while holding the Ctrl key.  
More detailed help is available here.

Download region GenBank file

Download region SVG

Location: 354,414 - 392,004 nt. (total: 37,591 nt)
Show pHMM detection rules used

NRPS: cds(Condensation and (AMP-binding or A-OX))

#### Legend:

core biosynthetic genes

additional biosynthetic genes

transport-related genes

regulatory genes

other genes

resistance

reset view

zoom to selection

Gene details

Shows details of the most recently selected gene, including names, products, location, and other annotations.

Select a gene to view the details available for it

NRPS/PKS domains

ClusterBlast

KnownClusterBlast

SubClusterBlast

MIBiG comparison

Pfam domains

Detailed domain annotation

Shows NRPS- and PKS-related domains for each feature that contains them. Click on each domain for more information about the domain's location, consensus monomer prediction, and other details.  
A glossary is available here.

Selected features only

Show module domains

Similar gene clusters

Shows clusters from the antiSMASH database and other clusters of interest that are similar to the current region. Genes marked with the same colour are interrelated. White genes have no relationship.  
Click on reference genes to show details of similarities to genes within the current region.  
Click on an accession to open that entry in the antiSMASH database (if applicable).

All hits

NC\_007199 (2323509-2373250): Aspergillus fumigatus Af293 chromosome 6, whole ... (13% of genes show similarity), NRPS

NW\_022984634 (297326-341201): Aspergillus tanneri strain NIH1004 chromosome U... (16% of genes show similarity), NRPS

NZ\_FWXV01000013 (88093-154130): Kibdelosporangium aridum strain DSM 43828, wh... (7% of genes show similarity), NRPS

NZ\_BBXF01000001 (471923-526170): Herbidospora daliensis strain NBRC 106372, w... (5% of genes show similarity), NRPS

NZ\_JXCA02000019 (428514-508874): Scytonema tolypothrichoides VB-61278 scaffol... (5% of genes show similarity), NRPS,T1PKS

NZ\_FNQS01000007 (107273-167999): Lonsdalea quercina strain ATCC 29281, whole ... (6% of genes show similarity), NRPS

NZ\_CP023009 (1742714-1805023): Lonsdalea britannica strain 477 chromosome, co... (5% of genes show similarity), NRPS

NZ\_FO818637 (3625327-3741808): Xenorhabdus bovienii strain CS03 (4% of genes show similarity), NRPS,T1PKS

NZ\_CP036282 (3281882-3325923): Rhodoferax sediminis strain Gr-4 chromosome, c... (4% of genes show similarity), NRPS-like

NZ\_CP015698 (4082749-4126763): Curvibacter sp. AEP1-3 genome (4% of genes show similarity), NRPS-like
Download graphic

Similar known gene clusters

Shows clusters from the MiBIG database that are similar to the current region. Genes marked with the same colour are interrelated. White genes have no relationship.  
Click on reference genes to show details of similarities to genes within the current region.  
Click on an accession to open that entry in the MiBIG database.

All hits

aspirochlorine
Download graphic

Similar subclusters

Shows sub-cluster units that are similar to the current region. Genes marked with the same colour are interrelated. White genes have no relationship.  
Click on reference genes to show details of similarities to genes within the current region.

No matches found.

Similar gene clusters

Shows careas that are similar to the current region to a reference database.  
Mouseover a score cell in the table to get a breakdown of how the score was calculated.The MIBiG database.  
  
Click on an accession to open that entry in the MIBiG database.

Analysis type:

Protocluster to Region
Region to Region

| Reference | NRPS | Similarity score | Type | Compound(s) | Organism |
| --- | --- | --- | --- | --- | --- |
| BGC0001261.1 |  | 0.17 | NRP | AM-toxin | Alternaria alternata |
| BGC0000357.1 |  | 0.17 | NRP | cyclo-(D-Phe-L-Phe-D-Val-L-Val), cyclo-(D-Tyr-L-Phe-D-Val-L-Val), cyclo-(D-Tyr-L-Trp-D-Val-L-Val), cyclo-(D-Phe-L-Trp-D-Val-L-Val), cyclo-(D-Phe-L-Phe-D-Val-L-Ile), cyclo-(D-Phe-L-Phe-D-Ile-L-Val), cyclo-(D-Tyr-L-Trp-D-Val-L-Ile), cyclo-(D-Tyr-L-Trp-D-Ile-L-Val), cyclo-(D-Tyr-L-Phe-D-Val-L-Ile), cyclo-(D-Tyr-L-Phe-D-Ile-L-Val) | Penicillium rubens Wisconsin 54-1255 |
| BGC0000260.1 |  | 0.14 | Polyketide | prodigiosin | Hahella chejuensis KCTC 2396 |
| BGC0000417.1 |  | 0.14 | NRP | rhodochelin | Rhodococcus jostii RHA1 |
| BGC0001888.1 |  | 0.14 | Other | mannosylerythritol lipid A, mannosylerythritol lipid B, mannosylerythritol lipid C | Ustilago maydis 521 |
| BGC0001516.1 |  | 0.14 | NRP | aspergillic acid | Aspergillus flavus NRRL3357 |
| BGC0001026.1 |  | 0.14 | NRP, Polyketide | NG-391 | Metarhizium anisopliae |
| BGC0001791.1 |  | 0.13 | NRP | sunshinamide | Gynuella sunshinyii YC6258 |
| BGC0000306.1 |  | 0.13 | NRP | arylomycin | Streptomyces filamentosus NRRL 11379 |
| BGC0001954.1 |  | 0.13 | NRP, Polyketide | wortmanamide A, wortmanamide B | Talaromyces wortmannii |

| Reference | Aggregated | Similarity score | Type | Compound(s) | Organism |
| --- | --- | --- | --- | --- | --- |
| BGC0001261.1 |  | 0.48 | NRP | AM-toxin | Alternaria alternata |
| BGC0000357.1 |  | 0.48 | NRP | cyclo-(D-Phe-L-Phe-D-Val-L-Val), cyclo-(D-Tyr-L-Phe-D-Val-L-Val), cyclo-(D-Tyr-L-Trp-D-Val-L-Val), cyclo-(D-Phe-L-Trp-D-Val-L-Val), cyclo-(D-Phe-L-Phe-D-Val-L-Ile), cyclo-(D-Phe-L-Phe-D-Ile-L-Val), cyclo-(D-Tyr-L-Trp-D-Val-L-Ile), cyclo-(D-Tyr-L-Trp-D-Ile-L-Val), cyclo-(D-Tyr-L-Phe-D-Val-L-Ile), cyclo-(D-Tyr-L-Phe-D-Ile-L-Val) | Penicillium rubens Wisconsin 54-1255 |
| BGC0000260.1 |  | 0.44 | Polyketide | prodigiosin | Hahella chejuensis KCTC 2396 |
| BGC0000417.1 |  | 0.44 | NRP | rhodochelin | Rhodococcus jostii RHA1 |
| BGC0001888.1 |  | 0.43 | Other | mannosylerythritol lipid A, mannosylerythritol lipid B, mannosylerythritol lipid C | Ustilago maydis 521 |
| BGC0001516.1 |  | 0.43 | NRP | aspergillic acid | Aspergillus flavus NRRL3357 |
| BGC0001026.1 |  | 0.43 | NRP, Polyketide | NG-391 | Metarhizium anisopliae |
| BGC0001791.1 |  | 0.42 | NRP | sunshinamide | Gynuella sunshinyii YC6258 |
| BGC0000306.1 |  | 0.42 | NRP | arylomycin | Streptomyces filamentosus NRRL 11379 |
| BGC0001954.1 |  | 0.42 | NRP, Polyketide | wortmanamide A, wortmanamide B | Talaromyces wortmannii |

Detailed Pfam domain annotation

Shows Pfam domains found in each gene within the region.
Click on each domain for more information about the domain's
accession, location, description, and any relevant Gene Ontology.
Domains with a bold border have Gene Ontology information.

Selected features only

contig\_4 - Region 2 - T1PKS

Shows the layout of the region, marking coding sequences and areas of interest. Clicking a gene will select it and show any relevant details. Clicking an area feature (e.g. a candidate cluster) will select all coding sequences within that area. Double clicking an area feature will zoom to that area. Multiple genes and area features can be selected by clicking them while holding the Ctrl key.  
More detailed help is available here.

Download region GenBank file

Download region SVG

Location: 1,279,350 - 1,324,830 nt. (total: 45,481 nt)
Show pHMM detection rules used

T1PKS: cds(PKS\_AT and (PKS\_KS or ene\_KS or mod\_KS or hyb\_KS or itr\_KS or tra\_KS))

#### Legend:

core biosynthetic genes

additional biosynthetic genes

transport-related genes

regulatory genes

other genes

resistance

reset view

zoom to selection

Gene details

Shows details of the most recently selected gene, including names, products, location, and other annotations.

Select a gene to view the details available for it

NRPS/PKS domains

ClusterBlast

KnownClusterBlast

SubClusterBlast

MIBiG comparison

Pfam domains

Detailed domain annotation

Shows NRPS- and PKS-related domains for each feature that contains them. Click on each domain for more information about the domain's location, consensus monomer prediction, and other details.  
A glossary is available here.

Selected features only

Show module domains

Similar gene clusters

Shows clusters from the antiSMASH database and other clusters of interest that are similar to the current region. Genes marked with the same colour are interrelated. White genes have no relationship.  
Click on reference genes to show details of similarities to genes within the current region.  
Click on an accession to open that entry in the antiSMASH database (if applicable).

All hits

NZ\_CP034352 (1037-312401): Streptomyces sp. W1SF4 plasmid p2, complete sequence (3% of genes show similarity), NRPS,NRPS-like,T1PKS,T2PKS,butyrolactone,melanin

NZ\_CP016824 (54347-384189): Streptomyces sampsonii strain KJ40 chromosome, co... (2% of genes show similarity), NRPS,NRPS-like,T1PKS,T2PKS,butyrolactone,ectoine,lanthipeptide,transAT-PKS

NZ\_CP056773 (6686330-6958067): Streptomyces violascens strain YIM 100212 chro... (3% of genes show similarity), NRPS,NRPS-like,T1PKS,lanthipeptide

NZ\_CM002271 (6545259-6848841): Streptomyces sp. GBA 94-10 chromosome, whole g... (2% of genes show similarity), NRPS,NRPS-like,T1PKS,T3PKS,lanthipeptide

NC\_020990 (6567851-6838639): Streptomyces albidoflavus, complete sequence (3% of genes show similarity), NRPS,NRPS-like,T1PKS,lanthipeptide

NZ\_CP047147 (63289-333437): Streptomyces sp. GF20 chromosome, complete genome (3% of genes show similarity), NRPS,NRPS-like,T1PKS,lanthipeptide

NZ\_CP031742 (6968727-7175511): Streptomyces koyangensis strain VK-A60T chromo... (5% of genes show similarity), NRPS,NRPS-like,T1PKS

NZ\_AMPN02000002 (6662429-6928181): Streptomyces sp. SM8 Scaffold 2, whole gen... (3% of genes show similarity), NRPS,NRPS-like,T1PKS,lanthipeptide,transAT-PKS

NZ\_CP024052 (1100414-1313530): Micromonospora sp. WMMA2032 chromosome (7% of genes show similarity), NRPS-like,T1PKS

NZ\_SMBL01000011 (165355-242085): Streptomyces sp. BK215 Ga0307669 111, whole ... (7% of genes show similarity), T1PKS,lanthipeptide
Download graphic

Similar known gene clusters

Shows clusters from the MiBIG database that are similar to the current region. Genes marked with the same colour are interrelated. White genes have no relationship.  
Click on reference genes to show details of similarities to genes within the current region.  
Click on an accession to open that entry in the MiBIG database.

All hits

(-)-Mellein

6-methylsalicyclic acid

6-methylsalicyclic acid

asperlactone
Download graphic

Similar subclusters

Shows sub-cluster units that are similar to the current region. Genes marked with the same colour are interrelated. White genes have no relationship.  
Click on reference genes to show details of similarities to genes within the current region.

No matches found.

Similar gene clusters

Shows careas that are similar to the current region to a reference database.  
Mouseover a score cell in the table to get a breakdown of how the score was calculated.The MIBiG database.  
  
Click on an accession to open that entry in the MIBiG database.

Analysis type:

Protocluster to Region
Region to Region

| Reference | T1PKS | Similarity score | Type | Compound(s) | Organism |
| --- | --- | --- | --- | --- | --- |
| BGC0001244.1 |  | 0.40 | Polyketide | (-)-Mellein | Parastagonospora nodorum |
| BGC0001276.1 |  | 0.39 | Polyketide | 6-methylsalicyclic acid | Aspergillus terreus |
| BGC0001275.1 |  | 0.38 | Polyketide | 6-methylsalicyclic acid | Glarea lozoyensis |
| BGC0001273.1 |  | 0.37 | Polyketide | asperlactone | Aspergillus ochraceus |
| BGC0000160.1 |  | 0.28 | Polyketide | terreic acid | Aspergillus terreus NIH2624 |
| BGC0000170.1 |  | 0.28 | Polyketide | yanuthone D | Aspergillus niger ATCC 1015 |
| BGC0000120.1 |  | 0.26 | Polyketide | patulin | Penicillium expansum |
| BGC0001858.1 |  | 0.24 | Polyketide | alternapyrone B, alternapyrone C, alternapyrone D, alternapyrone E, alternapyrone F | Parastagonospora nodorum SN15 |
| BGC0000056.1 |  | 0.23 | Polyketide | esperamicin | Actinomadura verrucosospora |
| BGC0001998.1 |  | 0.21 | Polyketide | aspernidgulene A1, aspernidgulene A2, aspernidgulene B1 | Aspergillus nidulans FGSC A4 |

| Reference | Aggregated | Similarity score | Type | Compound(s) | Organism |
| --- | --- | --- | --- | --- | --- |
| BGC0001244.1 |  | 0.72 | Polyketide | (-)-Mellein | Parastagonospora nodorum |
| BGC0001276.1 |  | 0.71 | Polyketide | 6-methylsalicyclic acid | Aspergillus terreus |
| BGC0001275.1 |  | 0.70 | Polyketide | 6-methylsalicyclic acid | Glarea lozoyensis |
| BGC0001273.1 |  | 0.70 | Polyketide | asperlactone | Aspergillus ochraceus |
| BGC0000160.1 |  | 0.62 | Polyketide | terreic acid | Aspergillus terreus NIH2624 |
| BGC0000170.1 |  | 0.61 | Polyketide | yanuthone D | Aspergillus niger ATCC 1015 |
| BGC0000120.1 |  | 0.59 | Polyketide | patulin | Penicillium expansum |
| BGC0001858.1 |  | 0.58 | Polyketide | alternapyrone B, alternapyrone C, alternapyrone D, alternapyrone E, alternapyrone F | Parastagonospora nodorum SN15 |
| BGC0000056.1 |  | 0.56 | Polyketide | esperamicin | Actinomadura verrucosospora |
| BGC0001998.1 |  | 0.54 | Polyketide | aspernidgulene A1, aspernidgulene A2, aspernidgulene B1 | Aspergillus nidulans FGSC A4 |

Detailed Pfam domain annotation

Shows Pfam domains found in each gene within the region.
Click on each domain for more information about the domain's
accession, location, description, and any relevant Gene Ontology.
Domains with a bold border have Gene Ontology information.

Selected features only

NRPS/PKS products

NRPS/PKS monomers

Predicted core structure(s)

Shows estimated product structure and polymer for each candidate cluster in the region. To show the product, click on the expander or the candidate cluster feature drawn in the overview.

For candidate cluster 2, location 1279349 - 1324830:

Rough prediction of core scaffold based on assumed PKS/NRPS colinearity; tailoring reactions not taken into account

**Polymer prediction:**
:   (pk)

  
Direct lookup in NORINE database:
strict
or
relaxed

Link to NORINE database query form

NRPS/PKS monomer predictions

Shows the predicted monomers for each adynelation domain and acyltransferase within genes. Each gene prediction can be expanded to view detailed predictions of each domain. Each prediction can be expanded to view the predictions by tool (and, for some tools, further expanded for extra details).

**input.path1.gene345**: pk

:   **PKS\_AT (579..871)**: pk

    ATSignature: Malonyl-CoA

    Top 3 matches:
    :   Malonyl-CoA: 66.7%
    :   Methoxymalonyl-CoA: 62.5%
    :   Methylmalonyl-CoA: 62.5%

      
    minowa: Methylmalonyl-CoA

    Prediction, score:
    :   Methylmalonyl-CoA: 112.7


        Methoxymalonyl-CoA: 91.5


        Ethylmalonyl-CoA: 79.0


        Isobutyryl-CoA: 65.0


        Malonyl-CoA: 61.7


        trans-1,2-CPDA: 47.8


        2-Methylbutyryl-CoA: 39.5


        Benzoyl-CoA: 39.0


        Propionyl-CoA: 36.0


        Acetyl-CoA: 32.5


        CHC-CoA: 30.1


        3-Methylbutyryl-CoA: 27.1


        fatty\_acid: 24.8


        inactive: 21.4

contig\_4 - Region 3 - NRPS

Shows the layout of the region, marking coding sequences and areas of interest. Clicking a gene will select it and show any relevant details. Clicking an area feature (e.g. a candidate cluster) will select all coding sequences within that area. Double clicking an area feature will zoom to that area. Multiple genes and area features can be selected by clicking them while holding the Ctrl key.  
More detailed help is available here.

Download region GenBank file

Download region SVG

Location: 1,830,601 - 1,896,090 nt. (total: 65,490 nt)
Show pHMM detection rules used

NRPS: cds(Condensation and (AMP-binding or A-OX))

#### Legend:

core biosynthetic genes

additional biosynthetic genes

transport-related genes

regulatory genes

other genes

resistance

reset view

zoom to selection

Gene details

Shows details of the most recently selected gene, including names, products, location, and other annotations.

Select a gene to view the details available for it

NRPS/PKS domains

ClusterBlast

KnownClusterBlast

SubClusterBlast

MIBiG comparison

Pfam domains

Detailed domain annotation

Shows NRPS- and PKS-related domains for each feature that contains them. Click on each domain for more information about the domain's location, consensus monomer prediction, and other details.  
A glossary is available here.

Selected features only

Show module domains

Similar gene clusters

Shows clusters from the antiSMASH database and other clusters of interest that are similar to the current region. Genes marked with the same colour are interrelated. White genes have no relationship.  
Click on reference genes to show details of similarities to genes within the current region.  
Click on an accession to open that entry in the antiSMASH database (if applicable).

All hits

NC\_036442 (328729-377356): Aspergillus oryzae RIB40 DNA, chromosome 8 (25% of genes show similarity), NRPS

NW\_001939244 (5159157-5221187): Pyrenophora tritici-repentis Pt-1C-BFP superc... (16% of genes show similarity), NRPS

NW\_003456428 (3794045-3858539): Trichophyton rubrum CBS 118892 genomic scaffo... (16% of genes show similarity), NRPS

NW\_003299167 (206715-267084): Microsporum canis CBS 113480 supercont1.3 genom... (18% of genes show similarity), NRPS

NW\_022983863 (1077260-1137112): Arthroderma uncinatum strain CBS 119779 chrom... (20% of genes show similarity), NRPS

NW\_003316003 (190578-250482): Coccidioides posadasii C735 delta SOWgp chromos... (22% of genes show similarity), NRPS

NW\_017971444 (292150-370888): Talaromyces atroroseus strain IBT 11181 chromos... (13% of genes show similarity), NRPS

NC\_016458 (118795-184185): Thermothielavioides terrestris NRRL 8126 chromosom... (18% of genes show similarity), NRPS

NC\_016477 (4027230-4092286): Thermothelomyces thermophilus ATCC 42464 chromos... (25% of genes show similarity), NRPS

CM000574 (7452834-7544494): Fusarium graminearum PH-1 chromosome 1, whole gen... (7% of genes show similarity), NRPS,T1PKS
Download graphic

Similar known gene clusters

Shows clusters from the MiBIG database that are similar to the current region. Genes marked with the same colour are interrelated. White genes have no relationship.  
Click on reference genes to show details of similarities to genes within the current region.  
Click on an accession to open that entry in the MiBIG database.

No matches found.

Similar subclusters

Shows sub-cluster units that are similar to the current region. Genes marked with the same colour are interrelated. White genes have no relationship.  
Click on reference genes to show details of similarities to genes within the current region.

No matches found.

Similar gene clusters

Shows careas that are similar to the current region to a reference database.  
Mouseover a score cell in the table to get a breakdown of how the score was calculated.The MIBiG database.  
  
Click on an accession to open that entry in the MIBiG database.

Analysis type:

Protocluster to Region
Region to Region

| Reference | NRPS | Similarity score | Type | Compound(s) | Organism |
| --- | --- | --- | --- | --- | --- |
| BGC0001166.1 |  | 0.28 | NRP | HC-toxin | Alternaria jesenskae |
| BGC0000357.1 |  | 0.28 | NRP | cyclo-(D-Phe-L-Phe-D-Val-L-Val), cyclo-(D-Tyr-L-Phe-D-Val-L-Val), cyclo-(D-Tyr-L-Trp-D-Val-L-Val), cyclo-(D-Phe-L-Trp-D-Val-L-Val), cyclo-(D-Phe-L-Phe-D-Val-L-Ile), cyclo-(D-Phe-L-Phe-D-Ile-L-Val), cyclo-(D-Tyr-L-Trp-D-Val-L-Ile), cyclo-(D-Tyr-L-Trp-D-Ile-L-Val), cyclo-(D-Tyr-L-Phe-D-Val-L-Ile), cyclo-(D-Tyr-L-Phe-D-Ile-L-Val) | Penicillium rubens Wisconsin 54-1255 |
| BGC0001240.1 |  | 0.27 | NRP | serinocyclin A, serinocyclin B | Metarhizium robertsii |
| BGC0001128.1 |  | 0.26 | NRP | luminmide | Photorhabdus laumondii subsp. laumondii TTO1 |
| BGC0001479.1 |  | 0.25 | NRP | anabaenopeptin NZ857, nostamide A | Nostoc punctiforme PCC 73102 |
| BGC0001132.1 |  | 0.25 | NRP | xenotetrapeptide | Xenorhabdus nematophila ATCC 19061 |
| BGC0001767.1 |  | 0.23 | NRP | salinichelins | Salinispora pacifica CNY331 |
| BGC0000325.1 |  | 0.21 | NRP | coelichelin | Streptomyces coelicolor A3(2) |
| BGC0000304.1 |  | 0.21 | NRP | apicidin | Fusarium incarnatum |
| BGC0001768.1 |  | 0.19 | NRP | sansalvamide | [Nectria] haematococca mpVI 77-13-4 |

| Reference | Aggregated | Similarity score | Type | Compound(s) | Organism |
| --- | --- | --- | --- | --- | --- |
| BGC0001166.1 |  | 0.62 | NRP | HC-toxin | Alternaria jesenskae |
| BGC0000357.1 |  | 0.62 | NRP | cyclo-(D-Phe-L-Phe-D-Val-L-Val), cyclo-(D-Tyr-L-Phe-D-Val-L-Val), cyclo-(D-Tyr-L-Trp-D-Val-L-Val), cyclo-(D-Phe-L-Trp-D-Val-L-Val), cyclo-(D-Phe-L-Phe-D-Val-L-Ile), cyclo-(D-Phe-L-Phe-D-Ile-L-Val), cyclo-(D-Tyr-L-Trp-D-Val-L-Ile), cyclo-(D-Tyr-L-Trp-D-Ile-L-Val), cyclo-(D-Tyr-L-Phe-D-Val-L-Ile), cyclo-(D-Tyr-L-Phe-D-Ile-L-Val) | Penicillium rubens Wisconsin 54-1255 |
| BGC0001240.1 |  | 0.60 | NRP | serinocyclin A, serinocyclin B | Metarhizium robertsii |
| BGC0001128.1 |  | 0.60 | NRP | luminmide | Photorhabdus laumondii subsp. laumondii TTO1 |
| BGC0001479.1 |  | 0.59 | NRP | anabaenopeptin NZ857, nostamide A | Nostoc punctiforme PCC 73102 |
| BGC0001132.1 |  | 0.59 | NRP | xenotetrapeptide | Xenorhabdus nematophila ATCC 19061 |
| BGC0001767.1 |  | 0.56 | NRP | salinichelins | Salinispora pacifica CNY331 |
| BGC0000325.1 |  | 0.54 | NRP | coelichelin | Streptomyces coelicolor A3(2) |
| BGC0000304.1 |  | 0.53 | NRP | apicidin | Fusarium incarnatum |
| BGC0001768.1 |  | 0.51 | NRP | sansalvamide | [Nectria] haematococca mpVI 77-13-4 |

Detailed Pfam domain annotation

Shows Pfam domains found in each gene within the region.
Click on each domain for more information about the domain's
accession, location, description, and any relevant Gene Ontology.
Domains with a bold border have Gene Ontology information.

Selected features only

NRPS/PKS products

NRPS/PKS monomers

Predicted core structure(s)

Shows estimated product structure and polymer for each candidate cluster in the region. To show the product, click on the expander or the candidate cluster feature drawn in the overview.

For candidate cluster 3, location 1830600 - 1896090:

Rough prediction of core scaffold based on assumed PKS/NRPS colinearity; tailoring reactions not taken into account

**Polymer prediction:**
:   (D-X - X - D-X - X - X)

  
Direct lookup in NORINE database:
strict
or
relaxed

Link to NORINE database query form

NRPS/PKS monomer predictions

Shows the predicted monomers for each adynelation domain and acyltransferase within genes. Each gene prediction can be expanded to view detailed predictions of each domain. Each prediction can be expanded to view the predictions by tool (and, for some tools, further expanded for extra details).

**input.path1.gene501**: X - X - X - X - X

:   Search NORINE for peptide:
    strict
    or
    relaxed
  
:   **AMP-binding (254..635)**: X

    NRPSPredictor2: (unknown)

    SVM prediction details:
    :   Predicted physicochemical class:
        :   N/A

        Large clusters prediction:
        :   N/A

        Small clusters prediction:
        :   N/A

        Single AA prediction:
        :   N/A

    Stachelhaus prediction details:
    :   Stachelhaus sequence:
        :   dslsimgi-k

        Nearest Stachelhaus code:
        :   N, A

        Stachelhaus code match:
        :   0% (weak)
:   **AMP-binding (1634..2004)**: X

    NRPSPredictor2: glu, gln

    SVM prediction details:
    :   Predicted physicochemical class:
        :   hydrophilic

        Large clusters prediction:
        :   asp, asn, glu, gln, aad

        Small clusters prediction:
        :   glu, gln

        Single AA prediction:
        :   N/A

    Stachelhaus prediction details:
    :   Stachelhaus sequence:
        :   dnedagqvnk

        Nearest Stachelhaus code:
        :   N, A

        Stachelhaus code match:
        :   0% (weak)
:   **AMP-binding (2699..3090)**: X

    NRPSPredictor2: hydrophobic-aliphatic

    SVM prediction details:
    :   Predicted physicochemical class:
        :   hydrophobic-aliphatic

        Large clusters prediction:
        :   N/A

        Small clusters prediction:
        :   N/A

        Single AA prediction:
        :   N/A

    Stachelhaus prediction details:
    :   Stachelhaus sequence:
        :   daiclvgaik

        Nearest Stachelhaus code:
        :   N, A

        Stachelhaus code match:
        :   0% (weak)
:   **AMP-binding (4238..4511)**: X

    NRPSPredictor2: (unknown)

    SVM prediction details:
    :   Predicted physicochemical class:
        :   N/A

        Large clusters prediction:
        :   N/A

        Small clusters prediction:
        :   N/A

        Single AA prediction:
        :   N/A

    Stachelhaus prediction details:
    :   Stachelhaus sequence:
        :   ----vgavvk

        Nearest Stachelhaus code:
        :   N, A

        Stachelhaus code match:
        :   0% (weak)
:   **AMP-binding (5203..5565)**: X

    NRPSPredictor2: gly, ala, val, leu, ile, abu, iva

    SVM prediction details:
    :   Predicted physicochemical class:
        :   hydrophobic-aliphatic

        Large clusters prediction:
        :   gly, ala, val, leu, ile, abu, iva

        Small clusters prediction:
        :   N/A

        Single AA prediction:
        :   N/A

    Stachelhaus prediction details:
    :   Stachelhaus sequence:
        :   ---caaavik

        Nearest Stachelhaus code:
        :   N, A

        Stachelhaus code match:
        :   0% (weak)

contig\_4 - Region 4 - indole

Shows the layout of the region, marking coding sequences and areas of interest. Clicking a gene will select it and show any relevant details. Clicking an area feature (e.g. a candidate cluster) will select all coding sequences within that area. Double clicking an area feature will zoom to that area. Multiple genes and area features can be selected by clicking them while holding the Ctrl key.  
More detailed help is available here.

Download region GenBank file

Download region SVG

Location: 2,306,177 - 2,327,439 nt. (total: 21,263 nt)
Show pHMM detection rules used

indole: (indsynth or dmat or indole\_PTase)

#### Legend:

core biosynthetic genes

additional biosynthetic genes

transport-related genes

regulatory genes

other genes

resistance

reset view

zoom to selection

Gene details

Shows details of the most recently selected gene, including names, products, location, and other annotations.

Select a gene to view the details available for it

ClusterBlast

KnownClusterBlast

SubClusterBlast

MIBiG comparison

Pfam domains

Similar gene clusters

Shows clusters from the antiSMASH database and other clusters of interest that are similar to the current region. Genes marked with the same colour are interrelated. White genes have no relationship.  
Click on reference genes to show details of similarities to genes within the current region.  
Click on an accession to open that entry in the antiSMASH database (if applicable).

No significant ClusterBlast hits found.

Similar known gene clusters

Shows clusters from the MiBIG database that are similar to the current region. Genes marked with the same colour are interrelated. White genes have no relationship.  
Click on reference genes to show details of similarities to genes within the current region.  
Click on an accession to open that entry in the MiBIG database.

No matches found.

Similar subclusters

Shows sub-cluster units that are similar to the current region. Genes marked with the same colour are interrelated. White genes have no relationship.  
Click on reference genes to show details of similarities to genes within the current region.

No matches found.

Similar gene clusters

Shows careas that are similar to the current region to a reference database.  
Mouseover a score cell in the table to get a breakdown of how the score was calculated.The MIBiG database.  
  
Click on an accession to open that entry in the MIBiG database.

Analysis type:

Protocluster to Region
Region to Region

| Reference | indole | Similarity score | Type | Compound(s) | Organism |
| --- | --- | --- | --- | --- | --- |
| BGC0001534.1 |  | 0.17 | Other | branched-chain fatty acids | Streptomyces filamentosus |
| BGC0001535.1 |  | 0.17 | Other | branched-chain fatty acids | Streptomyces filamentosus |
| BGC0001628.1 |  | 0.06 | NRP | JBIR-78, JBIR-95 | Kibdelosporangium sp. AK-AA56 |
| BGC0001372.1 |  | 0.05 | Terpene | penigequinolone A | Penicillium thymicola |
| BGC0001567.1 |  | 0.05 | NRP | cysteoamide | Streptomyces lincolnensis |
| BGC0001737.1 |  | 0.04 | NRP, Polyketide | phenalamide | Corallococcus coralloides |
| BGC0001394.1 |  | 0.04 | NRP, Polyketide | phenalamide A2 | Myxococcus stipitatus DSM 14675 |
| BGC0000111.1 |  | 0.04 | Polyketide | neocarzilin A, neocarzilin B | Streptomyces carzinostaticus |

| Reference | Aggregated | Similarity score | Type | Compound(s) | Organism |
| --- | --- | --- | --- | --- | --- |
| BGC0001534.1 |  | 0.48 | Other | branched-chain fatty acids | Streptomyces filamentosus |
| BGC0001535.1 |  | 0.48 | Other | branched-chain fatty acids | Streptomyces filamentosus |
| BGC0001628.1 |  | 0.25 | NRP | JBIR-78, JBIR-95 | Kibdelosporangium sp. AK-AA56 |
| BGC0001372.1 |  | 0.23 | Terpene | penigequinolone A | Penicillium thymicola |
| BGC0001567.1 |  | 0.22 | NRP | cysteoamide | Streptomyces lincolnensis |
| BGC0001737.1 |  | 0.19 | NRP, Polyketide | phenalamide | Corallococcus coralloides |
| BGC0001394.1 |  | 0.19 | NRP, Polyketide | phenalamide A2 | Myxococcus stipitatus DSM 14675 |
| BGC0000111.1 |  | 0.18 | Polyketide | neocarzilin A, neocarzilin B | Streptomyces carzinostaticus |

Detailed Pfam domain annotation

Shows Pfam domains found in each gene within the region.
Click on each domain for more information about the domain's
accession, location, description, and any relevant Gene Ontology.
Domains with a bold border have Gene Ontology information.

Selected features only

contig\_6 - Region 1 - T1PKS

Shows the layout of the region, marking coding sequences and areas of interest. Clicking a gene will select it and show any relevant details. Clicking an area feature (e.g. a candidate cluster) will select all coding sequences within that area. Double clicking an area feature will zoom to that area. Multiple genes and area features can be selected by clicking them while holding the Ctrl key.  
More detailed help is available here.

Download region GenBank file

Download region SVG

Location: 87,743 - 140,198 nt. (total: 52,456 nt)
Show pHMM detection rules used

T1PKS: cds(PKS\_AT and (PKS\_KS or ene\_KS or mod\_KS or hyb\_KS or itr\_KS or tra\_KS))

#### Legend:

core biosynthetic genes

additional biosynthetic genes

transport-related genes

regulatory genes

other genes

resistance

reset view

zoom to selection

Gene details

Shows details of the most recently selected gene, including names, products, location, and other annotations.

Select a gene to view the details available for it

NRPS/PKS domains

ClusterBlast

KnownClusterBlast

SubClusterBlast

MIBiG comparison

Pfam domains

Detailed domain annotation

Shows NRPS- and PKS-related domains for each feature that contains them. Click on each domain for more information about the domain's location, consensus monomer prediction, and other details.  
A glossary is available here.

Selected features only

Show module domains

Similar gene clusters

Shows clusters from the antiSMASH database and other clusters of interest that are similar to the current region. Genes marked with the same colour are interrelated. White genes have no relationship.  
Click on reference genes to show details of similarities to genes within the current region.  
Click on an accession to open that entry in the antiSMASH database (if applicable).

All hits

NT\_165980 (3959334-4034642): Chaetomium globosum CBS 148.51 scaffold 5 genomi... (21% of genes show similarity), T1PKS

NC\_035795 (2016121-2083159): Pochonia chlamydosporia 170 chromosome 6, whole ... (17% of genes show similarity), T1PKS

NW\_015971638 (789307-847606): Fonsecaea multimorphosa CBS 102226 unplaced gen... (16% of genes show similarity), T1PKS

NW\_015622519 (1364690-1433320): Exophiala spinifera strain CBS 89968 unplaced... (13% of genes show similarity), T1PKS

NW\_013562494 (2311114-2377735): Exophiala xenobiotica strain CBS 118157 unpla... (12% of genes show similarity), T1PKS

NW\_023336282 (1773961-1890170): Aspergillus tubingensis WU-2223L DNA, scaffol... (11% of genes show similarity), NRPS,T1PKS

NZ\_CP023445 (3511456-3591164): Actinosynnema pretiosum strain X47 chromosome,... (15% of genes show similarity), NRPS,T1PKS

NW\_003345196 (1322-36166): Nannizzia gypsea CBS 118893 supercont1.6 genomic s... (14% of genes show similarity), T1PKS,indole

NZ\_CP006850 (5145309-5194926): Nocardia nova SH22a chromosome, complete genome (4% of genes show similarity), NRPS,T1PKS

NC\_021191 (2626192-2710257): Actinoplanes sp. N902-109, complete genome (4% of genes show similarity), T1PKS
Download graphic

Similar known gene clusters

Shows clusters from the MiBIG database that are similar to the current region. Genes marked with the same colour are interrelated. White genes have no relationship.  
Click on reference genes to show details of similarities to genes within the current region.  
Click on an accession to open that entry in the MiBIG database.

All hits

radicicol

hypothemycin

hypothemycin

trans-resorcylide

zearalenone

lasiodiplodin

dehydrocurvularin
Download graphic

Similar subclusters

Shows sub-cluster units that are similar to the current region. Genes marked with the same colour are interrelated. White genes have no relationship.  
Click on reference genes to show details of similarities to genes within the current region.

No matches found.

Similar gene clusters

Shows careas that are similar to the current region to a reference database.  
Mouseover a score cell in the table to get a breakdown of how the score was calculated.The MIBiG database.  
  
Click on an accession to open that entry in the MIBiG database.

Analysis type:

Protocluster to Region
Region to Region

| Reference | T1PKS | Similarity score | Type | Compound(s) | Organism |
| --- | --- | --- | --- | --- | --- |
| BGC0000046.1 |  | 0.32 | Polyketide | depudecin | Alternaria brassicicola |
| BGC0000030.1 |  | 0.30 | Polyketide | bikaverin | Fusarium fujikuroi IMI 58289 |
| BGC0001909.1 |  | 0.26 | Polyketide | strobilurin | Strobilurus tenacellus |
| BGC0001254.1 |  | 0.23 | Polyketide | ACT-Toxin II | Alternaria alternata |
| BGC0001858.1 |  | 0.23 | Polyketide | alternapyrone B, alternapyrone C, alternapyrone D, alternapyrone E, alternapyrone F | Parastagonospora nodorum SN15 |
| BGC0001998.1 |  | 0.22 | Polyketide | aspernidgulene A1, aspernidgulene A2, aspernidgulene B1 | Aspergillus nidulans FGSC A4 |
| BGC0001446.1 |  | 0.22 | Polyketide | asparasone A | Aspergillus flavus NRRL3357 |
| BGC0001906.1 |  | 0.22 | Polyketide | naphthalene | Daldinia eschscholzii IFB-TL01 |
| BGC0001258.1 |  | 0.22 | Polyketide | 1,3,6,8-tetrahydroxynaphthalene | Glarea lozoyensis |
| BGC0001246.1 |  | 0.22 | Polyketide | trans-resorcylide | Sarocladium zeae |

| Reference | Aggregated | Similarity score | Type | Compound(s) | Organism |
| --- | --- | --- | --- | --- | --- |
| BGC0000046.1 |  | 0.65 | Polyketide | depudecin | Alternaria brassicicola |
| BGC0000030.1 |  | 0.64 | Polyketide | bikaverin | Fusarium fujikuroi IMI 58289 |
| BGC0001909.1 |  | 0.60 | Polyketide | strobilurin | Strobilurus tenacellus |
| BGC0001254.1 |  | 0.56 | Polyketide | ACT-Toxin II | Alternaria alternata |
| BGC0001858.1 |  | 0.56 | Polyketide | alternapyrone B, alternapyrone C, alternapyrone D, alternapyrone E, alternapyrone F | Parastagonospora nodorum SN15 |
| BGC0001998.1 |  | 0.55 | Polyketide | aspernidgulene A1, aspernidgulene A2, aspernidgulene B1 | Aspergillus nidulans FGSC A4 |
| BGC0001446.1 |  | 0.55 | Polyketide | asparasone A | Aspergillus flavus NRRL3357 |
| BGC0001906.1 |  | 0.55 | Polyketide | naphthalene | Daldinia eschscholzii IFB-TL01 |
| BGC0001258.1 |  | 0.55 | Polyketide | 1,3,6,8-tetrahydroxynaphthalene | Glarea lozoyensis |
| BGC0001246.1 |  | 0.55 | Polyketide | trans-resorcylide | Sarocladium zeae |

Detailed Pfam domain annotation

Shows Pfam domains found in each gene within the region.
Click on each domain for more information about the domain's
accession, location, description, and any relevant Gene Ontology.
Domains with a bold border have Gene Ontology information.

Selected features only

NRPS/PKS products

NRPS/PKS monomers

Predicted core structure(s)

Shows estimated product structure and polymer for each candidate cluster in the region. To show the product, click on the expander or the candidate cluster feature drawn in the overview.

For candidate cluster 1, location 87742 - 140198:

Rough prediction of core scaffold based on assumed PKS/NRPS colinearity; tailoring reactions not taken into account

**Polymer prediction:**
:   (pk) + (mal)

  
Direct lookup in NORINE database:
strict
or
relaxed

Link to NORINE database query form

NRPS/PKS monomer predictions

Shows the predicted monomers for each adynelation domain and acyltransferase within genes. Each gene prediction can be expanded to view detailed predictions of each domain. Each prediction can be expanded to view the predictions by tool (and, for some tools, further expanded for extra details).

**input.path1.gene24**: mal

:   **PKS\_AT (635..932)**: mal

    ATSignature: Malonyl-CoA

    Top 3 matches:
    :   Malonyl-CoA: 66.7%
    :   Methylmalonyl-CoA: 58.3%
    :   inactive: 58.3%

      
    minowa: Malonyl-CoA

    Prediction, score:
    :   Malonyl-CoA: 105.4


        Methoxymalonyl-CoA: 67.0


        inactive: 56.0


        Methylmalonyl-CoA: 55.7


        Isobutyryl-CoA: 45.6


        Benzoyl-CoA: 33.0


        Acetyl-CoA: 27.1


        Propionyl-CoA: 26.7


        Ethylmalonyl-CoA: 23.9


        2-Methylbutyryl-CoA: 19.6


        CHC-CoA: 18.1


        fatty\_acid: 17.5


        3-Methylbutyryl-CoA: 16.4


        trans-1,2-CPDA: 14.7

  
**input.path1.gene29**: pk

:   **PKS\_AT (577..761)**: pk

    ATSignature: Malonyl-CoA

    Top 3 matches:
    :   Malonyl-CoA: 54.2%

      
    minowa: Methylmalonyl-CoA

    Prediction, score:
    :   Methylmalonyl-CoA: 42.0


        Methoxymalonyl-CoA: 40.4


        Malonyl-CoA: 35.7


        Isobutyryl-CoA: 18.9


        trans-1,2-CPDA: 18.7


        Propionyl-CoA: 18.7


        CHC-CoA: 17.8


        2-Methylbutyryl-CoA: 17.5


        Benzoyl-CoA: 16.4


        Ethylmalonyl-CoA: 15.2


        fatty\_acid: 10.5


        inactive: 0.0


        Acetyl-CoA: 0.0


        3-Methylbutyryl-CoA: 0.0

contig\_6 - Region 2 - NRPS

Shows the layout of the region, marking coding sequences and areas of interest. Clicking a gene will select it and show any relevant details. Clicking an area feature (e.g. a candidate cluster) will select all coding sequences within that area. Double clicking an area feature will zoom to that area. Multiple genes and area features can be selected by clicking them while holding the Ctrl key.  
More detailed help is available here.

Download region GenBank file

Download region SVG

Location: 575,471 - 624,218 nt. (total: 48,748 nt)
Show pHMM detection rules used

NRPS: cds(Condensation and (AMP-binding or A-OX))

#### Legend:

core biosynthetic genes

additional biosynthetic genes

transport-related genes

regulatory genes

other genes

resistance

reset view

zoom to selection

Gene details

Shows details of the most recently selected gene, including names, products, location, and other annotations.

Select a gene to view the details available for it

NRPS/PKS domains

ClusterBlast

KnownClusterBlast

SubClusterBlast

MIBiG comparison

Pfam domains

Detailed domain annotation

Shows NRPS- and PKS-related domains for each feature that contains them. Click on each domain for more information about the domain's location, consensus monomer prediction, and other details.  
A glossary is available here.

Selected features only

Show module domains

Similar gene clusters

Shows clusters from the antiSMASH database and other clusters of interest that are similar to the current region. Genes marked with the same colour are interrelated. White genes have no relationship.  
Click on reference genes to show details of similarities to genes within the current region.  
Click on an accession to open that entry in the antiSMASH database (if applicable).

All hits

NC\_049562 (6635017-6678323): Talaromyces rugulosus chromosome II, complete se... (12% of genes show similarity), NRPS

NZ\_KQ976354 (3589179-3832343): Scytonema hofmannii PCC 7110 Scaffold1, whole ... (9% of genes show similarity), NRPS,microviridin

NZ\_FYDI01000001 (301554-398938): Pseudomonas sp. Irchel 3A18 isolate Pseudomo... (10% of genes show similarity), NRPS

NZ\_LN847264 (2342050-2456415): Pseudomonas sp. CCOS 191 chromosome I (5% of genes show similarity), NRPS

NZ\_QPDJ01000011 (55430-167518): Pseudomonas coronafaciens pv. garcae strain 4... (5% of genes show similarity), NRPS,acyl\_amino\_acids,hserlactone

NZ\_CP026386 (6103849-6217077): Pseudomonas sp. PONIH3 chromosome, complete ge... (6% of genes show similarity), NRPS

NZ\_SLXS01000001 (724943-860633): Tumebacillus sp. BK434 Ga0307714 101, whole ... (8% of genes show similarity), NRPS,T1PKS

NZ\_CP007039 (3694917-3791847): Pseudomonas cichorii JBC1 chromosome, complete... (9% of genes show similarity), NRPS

NZ\_LT962481 (3809539-3909878): Pseudomonas syringae pv. syringae isolate CFBP... (7% of genes show similarity), NRPS,hserlactone

NZ\_CP005969 (2096619-2195299): Pseudomonas syringae pv. syringae B301D chromo... (7% of genes show similarity), NRPS,hserlactone
Download graphic

Similar known gene clusters

Shows clusters from the MiBIG database that are similar to the current region. Genes marked with the same colour are interrelated. White genes have no relationship.  
Click on reference genes to show details of similarities to genes within the current region.  
Click on an accession to open that entry in the MiBIG database.

No matches found.

Similar subclusters

Shows sub-cluster units that are similar to the current region. Genes marked with the same colour are interrelated. White genes have no relationship.  
Click on reference genes to show details of similarities to genes within the current region.

No matches found.

Similar gene clusters

Shows careas that are similar to the current region to a reference database.  
Mouseover a score cell in the table to get a breakdown of how the score was calculated.The MIBiG database.  
  
Click on an accession to open that entry in the MIBiG database.

Analysis type:

Protocluster to Region
Region to Region

| Reference | NRPS | Similarity score | Type | Compound(s) | Organism |
| --- | --- | --- | --- | --- | --- |
| BGC0001399.1 |  | 0.20 | NRP | fellutamide B | Aspergillus nidulans FGSC A4 |
| BGC0000357.1 |  | 0.19 | NRP | cyclo-(D-Phe-L-Phe-D-Val-L-Val), cyclo-(D-Tyr-L-Phe-D-Val-L-Val), cyclo-(D-Tyr-L-Trp-D-Val-L-Val), cyclo-(D-Phe-L-Trp-D-Val-L-Val), cyclo-(D-Phe-L-Phe-D-Val-L-Ile), cyclo-(D-Phe-L-Phe-D-Ile-L-Val), cyclo-(D-Tyr-L-Trp-D-Val-L-Ile), cyclo-(D-Tyr-L-Trp-D-Ile-L-Val), cyclo-(D-Tyr-L-Phe-D-Val-L-Ile), cyclo-(D-Tyr-L-Phe-D-Ile-L-Val) | Penicillium rubens Wisconsin 54-1255 |
| BGC0000901.1 |  | 0.19 | Other | ferrichrome | Aspergillus niger |
| BGC0001679.1 |  | 0.19 | NRP | N-Acetyltryptophan | Aspergillus nidulans FGSC A4 |
| BGC0000900.1 |  | 0.19 | Other | ferrichrome | Aspergillus oryzae |
| BGC0000313.1 |  | 0.19 | NRP | beauvericin | Beauveria bassiana |
| BGC0000342.1 |  | 0.18 | NRP | enniatin | Fusarium equiseti |
| BGC0001240.1 |  | 0.18 | NRP | serinocyclin A, serinocyclin B | Metarhizium robertsii |
| BGC0001220.1 |  | 0.18 | NRP | aculeacin A | Aspergillus japonicus |
| BGC0000307.1 |  | 0.18 | NRP | AbT1 | Aureobasidium pullulans |

| Reference | Aggregated | Similarity score | Type | Compound(s) | Organism |
| --- | --- | --- | --- | --- | --- |
| BGC0001399.1 |  | 0.52 | NRP | fellutamide B | Aspergillus nidulans FGSC A4 |
| BGC0000357.1 |  | 0.51 | NRP | cyclo-(D-Phe-L-Phe-D-Val-L-Val), cyclo-(D-Tyr-L-Phe-D-Val-L-Val), cyclo-(D-Tyr-L-Trp-D-Val-L-Val), cyclo-(D-Phe-L-Trp-D-Val-L-Val), cyclo-(D-Phe-L-Phe-D-Val-L-Ile), cyclo-(D-Phe-L-Phe-D-Ile-L-Val), cyclo-(D-Tyr-L-Trp-D-Val-L-Ile), cyclo-(D-Tyr-L-Trp-D-Ile-L-Val), cyclo-(D-Tyr-L-Phe-D-Val-L-Ile), cyclo-(D-Tyr-L-Phe-D-Ile-L-Val) | Penicillium rubens Wisconsin 54-1255 |
| BGC0000901.1 |  | 0.51 | Other | ferrichrome | Aspergillus niger |
| BGC0001679.1 |  | 0.51 | NRP | N-Acetyltryptophan | Aspergillus nidulans FGSC A4 |
| BGC0000900.1 |  | 0.51 | Other | ferrichrome | Aspergillus oryzae |
| BGC0000313.1 |  | 0.50 | NRP | beauvericin | Beauveria bassiana |
| BGC0000342.1 |  | 0.50 | NRP | enniatin | Fusarium equiseti |
| BGC0001240.1 |  | 0.50 | NRP | serinocyclin A, serinocyclin B | Metarhizium robertsii |
| BGC0001220.1 |  | 0.50 | NRP | aculeacin A | Aspergillus japonicus |
| BGC0000307.1 |  | 0.50 | NRP | AbT1 | Aureobasidium pullulans |

Detailed Pfam domain annotation

Shows Pfam domains found in each gene within the region.
Click on each domain for more information about the domain's
accession, location, description, and any relevant Gene Ontology.
Domains with a bold border have Gene Ontology information.

Selected features only

contig\_6 - Region 3 - T1PKS

Shows the layout of the region, marking coding sequences and areas of interest. Clicking a gene will select it and show any relevant details. Clicking an area feature (e.g. a candidate cluster) will select all coding sequences within that area. Double clicking an area feature will zoom to that area. Multiple genes and area features can be selected by clicking them while holding the Ctrl key.  
More detailed help is available here.

Download region GenBank file

Download region SVG

Location: 1,814,903 - 1,861,987 nt. (total: 47,085 nt)
Show pHMM detection rules used

T1PKS: cds(PKS\_AT and (PKS\_KS or ene\_KS or mod\_KS or hyb\_KS or itr\_KS or tra\_KS))

#### Legend:

core biosynthetic genes

additional biosynthetic genes

transport-related genes

regulatory genes

other genes

resistance

reset view

zoom to selection

Gene details

Shows details of the most recently selected gene, including names, products, location, and other annotations.

Select a gene to view the details available for it

NRPS/PKS domains

ClusterBlast

KnownClusterBlast

SubClusterBlast

MIBiG comparison

Pfam domains

Detailed domain annotation

Shows NRPS- and PKS-related domains for each feature that contains them. Click on each domain for more information about the domain's location, consensus monomer prediction, and other details.  
A glossary is available here.

Selected features only

Show module domains

Similar gene clusters

Shows clusters from the antiSMASH database and other clusters of interest that are similar to the current region. Genes marked with the same colour are interrelated. White genes have no relationship.  
Click on reference genes to show details of similarities to genes within the current region.  
Click on an accession to open that entry in the antiSMASH database (if applicable).

All hits

NW\_011942164 (4001-59385): Metarhizium robertsii ARSEF 23 MAA Scf 23, whole g... (20% of genes show similarity), NRPS-like,T1PKS

NW\_014574700 (5068-60461): Metarhizium brunneum ARSEF 3297 chromosome Unknown... (21% of genes show similarity), NRPS-like,T1PKS

NC\_016459 (3622511-3676948): Thermothielavioides terrestris NRRL 8126 chromos... (18% of genes show similarity), NRPS-like,T1PKS

NW\_008481799 (398930-462807): Cyphellophora europaea CBS 101466 unplaced geno... (9% of genes show similarity), NRPS-like,T1PKS

NW\_013550604 (1410002-1468918): Rhinocladiella mackenziei CBS 650.93 unplaced... (10% of genes show similarity), NRPS-like,T1PKS

CM000593 (155300-223684): Fusarium oxysporum f. sp. lycopersici 4287 chromoso... (4% of genes show similarity), NRPS-like,T1PKS

NC\_030990 (155300-223684): Fusarium oxysporum f. sp. lycopersici 4287 chromos... (4% of genes show similarity), NRPS-like,T1PKS

NC\_036624 (49805-115416): Fusarium fujikuroi IMI 58289 draft genome, chromoso... (8% of genes show similarity), NRPS-like,T1PKS

NW\_022983866 (1121166-1165427): Arthroderma uncinatum strain CBS 119779 chrom... (14% of genes show similarity), T1PKS

NW\_003345200 (2075136-2126392): Nannizzia gypsea CBS 118893 supercont1.2 geno... (10% of genes show similarity), NRPS-like,T1PKS
Download graphic

Similar known gene clusters

Shows clusters from the MiBIG database that are similar to the current region. Genes marked with the same colour are interrelated. White genes have no relationship.  
Click on reference genes to show details of similarities to genes within the current region.  
Click on an accession to open that entry in the MiBIG database.

No matches found.

Similar subclusters

Shows sub-cluster units that are similar to the current region. Genes marked with the same colour are interrelated. White genes have no relationship.  
Click on reference genes to show details of similarities to genes within the current region.

No matches found.

Similar gene clusters

Shows careas that are similar to the current region to a reference database.  
Mouseover a score cell in the table to get a breakdown of how the score was calculated.The MIBiG database.  
  
Click on an accession to open that entry in the MIBiG database.

Analysis type:

Protocluster to Region
Region to Region

| Reference | T1PKS | Similarity score | Type | Compound(s) | Organism |
| --- | --- | --- | --- | --- | --- |
| BGC0001068.1 |  | 0.24 | Terpene, Polyketide | pyripyropene A | unidentified unclassified sequences. |
| BGC0001858.1 |  | 0.19 | Polyketide | alternapyrone B, alternapyrone C, alternapyrone D, alternapyrone E, alternapyrone F | Parastagonospora nodorum SN15 |
| BGC0001254.1 |  | 0.19 | Polyketide | ACT-Toxin II | Alternaria alternata |
| BGC0000046.1 |  | 0.19 | Polyketide | depudecin | Alternaria brassicicola |
| BGC0001252.1 |  | 0.18 | Polyketide | UNII-YC2Q1O94PT | Alternaria alternata |
| BGC0001400.1 |  | 0.18 | Polyketide | citreoviridin | Aspergillus terreus NIH2624 |
| BGC0001998.1 |  | 0.18 | Polyketide | aspernidgulene A1, aspernidgulene A2, aspernidgulene B1 | Aspergillus nidulans FGSC A4 |
| BGC0001124.1 |  | 0.18 | Polyketide | pyranonigrin E | Aspergillus niger ATCC 1015 |
| BGC0001268.1 |  | 0.17 | NRP, Polyketide | fusarin | Fusarium fujikuroi |
| BGC0000064.1 |  | 0.17 | Polyketide | fusarin | Fusarium verticillioides |

| Reference | Aggregated | Similarity score | Type | Compound(s) | Organism |
| --- | --- | --- | --- | --- | --- |
| BGC0001068.1 |  | 0.58 | Terpene, Polyketide | pyripyropene A | unidentified unclassified sequences. |
| BGC0001858.1 |  | 0.52 | Polyketide | alternapyrone B, alternapyrone C, alternapyrone D, alternapyrone E, alternapyrone F | Parastagonospora nodorum SN15 |
| BGC0001254.1 |  | 0.51 | Polyketide | ACT-Toxin II | Alternaria alternata |
| BGC0000046.1 |  | 0.51 | Polyketide | depudecin | Alternaria brassicicola |
| BGC0001252.1 |  | 0.49 | Polyketide | UNII-YC2Q1O94PT | Alternaria alternata |
| BGC0001400.1 |  | 0.49 | Polyketide | citreoviridin | Aspergillus terreus NIH2624 |
| BGC0001998.1 |  | 0.49 | Polyketide | aspernidgulene A1, aspernidgulene A2, aspernidgulene B1 | Aspergillus nidulans FGSC A4 |
| BGC0001124.1 |  | 0.49 | Polyketide | pyranonigrin E | Aspergillus niger ATCC 1015 |
| BGC0001268.1 |  | 0.49 | NRP, Polyketide | fusarin | Fusarium fujikuroi |
| BGC0000064.1 |  | 0.49 | Polyketide | fusarin | Fusarium verticillioides |

Detailed Pfam domain annotation

Shows Pfam domains found in each gene within the region.
Click on each domain for more information about the domain's
accession, location, description, and any relevant Gene Ontology.
Domains with a bold border have Gene Ontology information.

Selected features only

contig\_9 - Region 1 - NRPS,terpene

Shows the layout of the region, marking coding sequences and areas of interest. Clicking a gene will select it and show any relevant details. Clicking an area feature (e.g. a candidate cluster) will select all coding sequences within that area. Double clicking an area feature will zoom to that area. Multiple genes and area features can be selected by clicking them while holding the Ctrl key.  
More detailed help is available here.

Download region GenBank file

Download region SVG

Location: 1,341,734 - 1,419,185 nt. (total: 77,452 nt)
Show pHMM detection rules used

terpene: (Terpene\_synth or Terpene\_synth\_C or phytoene\_synt or Lycopene\_cycl or terpene\_cyclase or NapT7 or fung\_ggpps or fung\_ggpps2 or trichodiene\_synth or TRI5)  
NRPS: cds(Condensation and (AMP-binding or A-OX))

#### Legend:

core biosynthetic genes

additional biosynthetic genes

transport-related genes

regulatory genes

other genes

resistance

reset view

zoom to selection

Gene details

Shows details of the most recently selected gene, including names, products, location, and other annotations.

Select a gene to view the details available for it

NRPS/PKS domains

ClusterBlast

KnownClusterBlast

SubClusterBlast

MIBiG comparison

Pfam domains

Detailed domain annotation

Shows NRPS- and PKS-related domains for each feature that contains them. Click on each domain for more information about the domain's location, consensus monomer prediction, and other details.  
A glossary is available here.

Selected features only

Show module domains

Similar gene clusters

Shows clusters from the antiSMASH database and other clusters of interest that are similar to the current region. Genes marked with the same colour are interrelated. White genes have no relationship.  
Click on reference genes to show details of similarities to genes within the current region.  
Click on an accession to open that entry in the antiSMASH database (if applicable).

All hits

NW\_017971440 (1299138-1363215): Talaromyces atroroseus strain IBT 11181 chrom... (10% of genes show similarity), NRPS

CM004174 (10735158-10801765): Drechmeria coniospora strain ARSEF 6962 chromos... (17% of genes show similarity), NRPS

NW\_003315027 (453020-520877): Verticillium alfalfae VaMs.102 supercont1.12 ge... (26% of genes show similarity), NRPS

NT\_165937 (139863-204269): Aspergillus terreus NIH2624 scaffold 14 genomic sc... (9% of genes show similarity), NRPS

NT\_165929 (178497-230302): Aspergillus terreus NIH2624 scaffold 6 genomic sca... (11% of genes show similarity), NRPS

NC\_035790 (391322-515501): Pochonia chlamydosporia 170 chromosome 1, whole ge... (15% of genes show similarity), NRPS

NC\_038012 (79994-172014): Fusarium venenatum strain A3/5 genome assembly, chr... (11% of genes show similarity), NRPS,T1PKS

NW\_017387250 (1103710-1123306): Fonsecaea erecta strain CBS 125763 chromosome... (33% of genes show similarity), terpene

NW\_013550602 (1647725-1671658): Rhinocladiella mackenziei CBS 650.93 unplaced... (18% of genes show similarity), terpene

NW\_022984629 (3081705-3186571): Aspergillus tanneri strain NIH1004 chromosome... (25% of genes show similarity), NRPS,T1PKS,indole
Download graphic

Similar known gene clusters

Shows clusters from the MiBIG database that are similar to the current region. Genes marked with the same colour are interrelated. White genes have no relationship.  
Click on reference genes to show details of similarities to genes within the current region.  
Click on an accession to open that entry in the MiBIG database.

No matches found.

Similar subclusters

Shows sub-cluster units that are similar to the current region. Genes marked with the same colour are interrelated. White genes have no relationship.  
Click on reference genes to show details of similarities to genes within the current region.

No matches found.

Similar gene clusters

Shows careas that are similar to the current region to a reference database.  
Mouseover a score cell in the table to get a breakdown of how the score was calculated.The MIBiG database.  
  
Click on an accession to open that entry in the MIBiG database.

Analysis type:

Protocluster to Region
Region to Region

| Reference | terpene | NRPS | Similarity score | Type | Compound(s) | Organism |
| --- | --- | --- | --- | --- | --- | --- |
| BGC0000465.1 |  |  | 0.32 | NRP | xenortide A, xenortide B, xenortide C, xenortide D | Xenorhabdus nematophila ATCC 19061 |
| BGC0002071.1 |  |  | 0.32 | NRP | Virginiafactin | Pseudomonas sp. QS1027 |
| BGC0001679.1 |  |  | 0.30 | NRP | N-Acetyltryptophan | Aspergillus nidulans FGSC A4 |
| BGC0000901.1 |  |  | 0.28 | Other | ferrichrome | Aspergillus niger |
| BGC0000457.1 |  |  | 0.28 | NRP | vicibactin | Rhizobium etli CFN 42 |
| BGC0001434.1 |  |  | 0.27 | NRP | nematophin | Xenorhabdus budapestensis |
| BGC0001698.1 |  |  | 0.27 | NRP | nevaltophin A, nevaltophin B, nevaltophin C, nevaltophin D | Xenorhabdus budapestensis |
| BGC0001399.1 |  |  | 0.27 | NRP | fellutamide B | Aspergillus nidulans FGSC A4 |
| BGC0001166.1 |  |  | 0.27 | NRP | HC-toxin | Alternaria jesenskae |
| BGC0000313.1 |  |  | 0.26 | NRP | beauvericin | Beauveria bassiana |

| Reference | Aggregated | Similarity score | Type | Compound(s) | Organism |
| --- | --- | --- | --- | --- | --- |
| BGC0000465.1 |  | 0.66 | NRP | xenortide A, xenortide B, xenortide C, xenortide D | Xenorhabdus nematophila ATCC 19061 |
| BGC0002071.1 |  | 0.65 | NRP | Virginiafactin | Pseudomonas sp. QS1027 |
| BGC0001679.1 |  | 0.64 | NRP | N-Acetyltryptophan | Aspergillus nidulans FGSC A4 |
| BGC0000457.1 |  | 0.63 | NRP | vicibactin | Rhizobium etli CFN 42 |
| BGC0000460.1 |  | 0.62 | NRP | vulnibactin | Vibrio vulnificus CMCP6 |
| BGC0000901.1 |  | 0.62 | Other | ferrichrome | Aspergillus niger |
| BGC0001434.1 |  | 0.61 | NRP | nematophin | Xenorhabdus budapestensis |
| BGC0001698.1 |  | 0.61 | NRP | nevaltophin A, nevaltophin B, nevaltophin C, nevaltophin D | Xenorhabdus budapestensis |
| BGC0001399.1 |  | 0.61 | NRP | fellutamide B | Aspergillus nidulans FGSC A4 |
| BGC0001166.1 |  | 0.60 | NRP | HC-toxin | Alternaria jesenskae |

Detailed Pfam domain annotation

Shows Pfam domains found in each gene within the region.
Click on each domain for more information about the domain's
accession, location, description, and any relevant Gene Ontology.
Domains with a bold border have Gene Ontology information.

Selected features only

NRPS/PKS products

NRPS/PKS monomers

Predicted core structure(s)

Shows estimated product structure and polymer for each candidate cluster in the region. To show the product, click on the expander or the candidate cluster feature drawn in the overview.

For candidate cluster 1, location 1341733 - 1419185:

Rough prediction of core scaffold based on assumed PKS/NRPS colinearity; tailoring reactions not taken into account

**Polymer prediction:**
:   (D-X) + (X)

  
Direct lookup in NORINE database:
strict
or
relaxed

---

For candidate cluster 3, location 1358137 - 1419185:

Rough prediction of core scaffold based on assumed PKS/NRPS colinearity; tailoring reactions not taken into account

**Polymer prediction:**
:   (D-X) + (X)

  
Direct lookup in NORINE database:
strict
or
relaxed

Link to NORINE database query form

NRPS/PKS monomer predictions

Shows the predicted monomers for each adynelation domain and acyltransferase within genes. Each gene prediction can be expanded to view detailed predictions of each domain. Each prediction can be expanded to view the predictions by tool (and, for some tools, further expanded for extra details).

**input.path1.gene394**: X

:   Search NORINE for peptide:
    strict
    or
    relaxed
  
:   **AMP-binding (6..225)**: X

    NRPSPredictor2: hydrophobic-aliphatic

    SVM prediction details:
    :   Predicted physicochemical class:
        :   hydrophobic-aliphatic

        Large clusters prediction:
        :   N/A

        Small clusters prediction:
        :   N/A

        Single AA prediction:
        :   N/A

    Stachelhaus prediction details:
    :   Stachelhaus sequence:
        :   en-d--viyk

        Nearest Stachelhaus code:
        :   N, A

        Stachelhaus code match:
        :   0% (weak)

  
**input.path1.gene398**: X

:   Search NORINE for peptide:
    strict
    or
    relaxed
  
:   **AMP-binding (390..759)**: X

    NRPSPredictor2: hydrophobic-aliphatic

    SVM prediction details:
    :   Predicted physicochemical class:
        :   hydrophobic-aliphatic

        Large clusters prediction:
        :   N/A

        Small clusters prediction:
        :   N/A

        Single AA prediction:
        :   N/A

    Stachelhaus prediction details:
    :   Stachelhaus sequence:
        :   dv-lvgavlk

        Nearest Stachelhaus code:
        :   N, A

        Stachelhaus code match:
        :   0% (weak)

  
**input.path1.gene399**: X - X

:   Search NORINE for peptide:
    strict
    or
    relaxed
  
:   **AMP-binding (886..1282)**: X

    NRPSPredictor2: val, leu, ile, abu, iva

    SVM prediction details:
    :   Predicted physicochemical class:
        :   hydrophobic-aliphatic

        Large clusters prediction:
        :   gly, ala, val, leu, ile, abu, iva

        Small clusters prediction:
        :   val, leu, ile, abu, iva

        Single AA prediction:
        :   N/A

    Stachelhaus prediction details:
    :   Stachelhaus sequence:
        :   dslfvggvfk

        Nearest Stachelhaus code:
        :   N, A

        Stachelhaus code match:
        :   0% (weak)
:   **AMP-binding (1774..2091)**: X

    NRPSPredictor2: val, leu, ile, abu, iva

    SVM prediction details:
    :   Predicted physicochemical class:
        :   hydrophobic-aliphatic

        Large clusters prediction:
        :   gly, ala, val, leu, ile, abu, iva

        Small clusters prediction:
        :   val, leu, ile, abu, iva

        Single AA prediction:
        :   N/A

    Stachelhaus prediction details:
    :   Stachelhaus sequence:
        :   dvgfvgsiwk

        Nearest Stachelhaus code:
        :   N, A

        Stachelhaus code match:
        :   0% (weak)

  
**input.path1.gene401**: X

:   Search NORINE for peptide:
    strict
    or
    relaxed
  
:   **AMP-binding (26..283)**: X

    NRPSPredictor2: gly, ala, val, leu, ile, abu, iva

    SVM prediction details:
    :   Predicted physicochemical class:
        :   hydrophobic-aliphatic

        Large clusters prediction:
        :   gly, ala, val, leu, ile, abu, iva

        Small clusters prediction:
        :   N/A

        Single AA prediction:
        :   N/A

    Stachelhaus prediction details:
    :   Stachelhaus sequence:
        :   igfqdgvgyk

        Nearest Stachelhaus code:
        :   N, A

        Stachelhaus code match:
        :   0% (weak)

contig\_9 - Region 2 - terpene

Shows the layout of the region, marking coding sequences and areas of interest. Clicking a gene will select it and show any relevant details. Clicking an area feature (e.g. a candidate cluster) will select all coding sequences within that area. Double clicking an area feature will zoom to that area. Multiple genes and area features can be selected by clicking them while holding the Ctrl key.  
More detailed help is available here.

Download region GenBank file

Download region SVG

Location: 1,785,179 - 1,806,398 nt. (total: 21,220 nt)
Show pHMM detection rules used

terpene: (Terpene\_synth or Terpene\_synth\_C or phytoene\_synt or Lycopene\_cycl or terpene\_cyclase or NapT7 or fung\_ggpps or fung\_ggpps2 or trichodiene\_synth or TRI5)

#### Legend:

core biosynthetic genes

additional biosynthetic genes

transport-related genes

regulatory genes

other genes

resistance

reset view

zoom to selection

Gene details

Shows details of the most recently selected gene, including names, products, location, and other annotations.

Select a gene to view the details available for it

ClusterBlast

KnownClusterBlast

SubClusterBlast

MIBiG comparison

Pfam domains

Similar gene clusters

Shows clusters from the antiSMASH database and other clusters of interest that are similar to the current region. Genes marked with the same colour are interrelated. White genes have no relationship.  
Click on reference genes to show details of similarities to genes within the current region.  
Click on an accession to open that entry in the antiSMASH database (if applicable).

All hits

NW\_001939250 (1746352-1767983): Pyrenophora tritici-repentis Pt-1C-BFP superc... (42% of genes show similarity), terpene

CM000590 (1383523-1405073): Fusarium oxysporum f. sp. lycopersici 4287 chromo... (50% of genes show similarity), terpene

NC\_030987 (1383523-1405073): Fusarium oxysporum f. sp. lycopersici 4287 chrom... (50% of genes show similarity), terpene

NC\_026507 (1953450-1975103): Neurospora crassa OR74A linkage group VII, whole... (42% of genes show similarity), terpene

NW\_020194478 (1246381-1267991): Amorphotheca resinae ATCC 22711 unplaced geno... (22% of genes show similarity), terpene

NW\_013562494 (4881346-4902884): Exophiala xenobiotica strain CBS 118157 unpla... (22% of genes show similarity), terpene

NW\_013550612 (3623167-3644725): Fonsecaea pedrosoi CBS 271.37 unplaced genomi... (28% of genes show similarity), terpene

NW\_015971664 (214925-236492): Fonsecaea multimorphosa CBS 102226 unplaced gen... (25% of genes show similarity), terpene

NW\_015971575 (1296644-1318223): Cladophialophora bantiana CBS 173.52 unplaced... (22% of genes show similarity), terpene

NW\_006763066 (356300-377896): Marssonina brunnea f. sp. 'multigermtubi' MB m1... (25% of genes show similarity), terpene
Download graphic

Similar known gene clusters

Shows clusters from the MiBIG database that are similar to the current region. Genes marked with the same colour are interrelated. White genes have no relationship.  
Click on reference genes to show details of similarities to genes within the current region.  
Click on an accession to open that entry in the MiBIG database.

All hits

squalestatin S1
Download graphic

Similar subclusters

Shows sub-cluster units that are similar to the current region. Genes marked with the same colour are interrelated. White genes have no relationship.  
Click on reference genes to show details of similarities to genes within the current region.

No matches found.

Similar gene clusters

Shows careas that are similar to the current region to a reference database.  
Mouseover a score cell in the table to get a breakdown of how the score was calculated.The MIBiG database.  
  
Click on an accession to open that entry in the MIBiG database.

Analysis type:

Protocluster to Region
Region to Region

| Reference | terpene | Similarity score | Type | Compound(s) | Organism |
| --- | --- | --- | --- | --- | --- |
| BGC0001839.1 |  | 0.25 | Terpene | squalestatin S1 | Aspergillus sp. Z5 |
| BGC0000774.1 |  | 0.14 | Saccharide | lipopolysaccharide | Xanthomonas campestris pv. campestris |
| BGC0001422.1 |  | 0.04 | NRP, Polyketide | myxochromide A | Myxococcus fulvus HW-1 |
| BGC0001339.1 |  | 0.04 | Polyketide | squalestatin S1 | Phoma sp. MF5453 |

| Reference | Aggregated | Similarity score | Type | Compound(s) | Organism |
| --- | --- | --- | --- | --- | --- |
| BGC0001839.1 |  | 0.59 | Terpene | squalestatin S1 | Aspergillus sp. Z5 |
| BGC0000774.1 |  | 0.43 | Saccharide | lipopolysaccharide | Xanthomonas campestris pv. campestris |
| BGC0001422.1 |  | 0.17 | NRP, Polyketide | myxochromide A | Myxococcus fulvus HW-1 |
| BGC0001339.1 |  | 0.17 | Polyketide | squalestatin S1 | Phoma sp. MF5453 |

Detailed Pfam domain annotation

Shows Pfam domains found in each gene within the region.
Click on each domain for more information about the domain's
accession, location, description, and any relevant Gene Ontology.
Domains with a bold border have Gene Ontology information.

Selected features only

If you have found antiSMASH useful, please cite us.
