## Supplementary Results for "A draft genome of the ascomycotal fungal species *Pseudopithomyces maydicus* (family *Didymosphaeriaceae*)": antismash.js

```
/*! antismash-js, version: 0.8.2 */
var viewer=function(t){var e={};function n(r){if(e[r])return e[r].exports;var i=e[r]={i:r,l:!1,exports:{}};return t[r].call(i.exports,i,i.exports,n),i.l=!0,i.exports}return n.m=t,n.c=e,n.d=function(t,e,r){n.o(t,e)||Object.defineProperty(t,e,{enumerable:!0,get:r})},n.r=function(t){"undefined"!=typeof Symbol&&Symbol.toStringTag&&Object.defineProperty(t,Symbol.toStringTag,{value:"Module"}),Object.defineProperty(t,"__esModule",{value:!0})},n.t=function(t,e){if(1&e&&(t=n(t)),8&e)return t;if(4&e&&"object"==typeof t&&t&&t.__esModule)return t;var r=Object.create(null);if(n.r(r),Object.defineProperty(r,"default",{enumerable:!0,value:t}),2&e&&"string"!=typeof t)for(var i in t)n.d(r,i,function(e){return t[e]}.bind(null,i));return r},n.n=function(t){var e=t&&t.__esModule?function(){return t.default}:function(){return t};return n.d(e,"a",e),e},n.o=function(t,e){return Object.prototype.hasOwnProperty.call(t,e)},n.p="",n(n.s=19)}([function(t,e,n){"use strict";n.r(e),n.d(e,"create",(function(){return yt})),n.d(e,"creator",(function(){return l})),n.d(e,"local",(function(){return xt})),n.d(e,"matcher",(function(){return v})),n.d(e,"mouse",(function(){return _t})),n.d(e,"namespace",(function(){return o})),n.d(e,"namespaces",(function(){return i})),n.d(e,"clientPoint",(function(){return Ct})),n.d(e,"select",(function(){return mt})),n.d(e,"selectAll",(function(){return St})),n.d(e,"selection",(function(){return vt})),n.d(e,"selector",(function(){return c})),n.d(e,"selectorAll",(function(){return d})),n.d(e,"style",(function(){return P})),n.d(e,"touch",(function(){return Mt})),n.d(e,"touches",(function(){return Tt})),n.d(e,"window",(function(){return k})),n.d(e,"event",(function(){return rt})),n.d(e,"customEvent",(function(){return ut}));var r="http://www.w3.org/1999/xhtml",i={svg:"http://www.w3.org/2000/svg",xhtml:r,xlink:"http://www.w3.org/1999/xlink",xml:"http://www.w3.org/XML/1998/namespace",xmlns:"http://www.w3.org/2000/xmlns/"},o=function(t){var e=t+="",n=e.indexOf(":");return n>=0&&"xmlns"!==(e=t.slice(0,n))&&(t=t.slice(n+1)),i.hasOwnProperty(e)?{space:i[e],local:t}:t};function a(t){return function(){var e=this.ownerDocument,n=this.namespaceURI;return n===r&&e.documentElement.namespaceURI===r?e.createElement(t):e.createElementNS(n,t)}}function s(t){return function(){return this.ownerDocument.createElementNS(t.space,t.local)}}var l=function(t){var e=o(t);return(e.local?s:a)(e)};function u(){}var c=function(t){return null==t?u:function(){return this.querySelector(t)}};function h(){return[]}var d=function(t){return null==t?h:function(){return this.querySelectorAll(t)}},g=function(t){return function(){return this.matches(t)}};if("undefined"!=typeof document){var f=document.documentElement;if(!f.matches){var p=f.webkitMatchesSelector||f.msMatchesSelector||f.mozMatchesSelector||f.oMatchesSelector;g=function(t){return function(){return p.call(this,t)}}}}var v=g,m=function(t){return new Array(t.length)};function y(t,e){this.ownerDocument=t.ownerDocument,this.namespaceURI=t.namespaceURI,this._next=null,this._parent=t,this.__data__=e}y.prototype={constructor:y,appendChild:function(t){return this._parent.insertBefore(t,this._next)},insertBefore:function(t,e){return this._parent.insertBefore(t,e)},querySelector:function(t){return this._parent.querySelector(t)},querySelectorAll:function(t){return this._parent.querySelectorAll(t)}};function b(t,e,n,r,i,o){for(var a,s=0,l=e.length,u=o.length;s<u;++s)(a=e[s])?(a.__data__=o[s],r[s]=a):n[s]=new y(t,o[s]);for(;s<l;++s)(a=e[s])&&(i[s]=a)}function x(t,e,n,r,i,o,a){var s,l,u,c={},h=e.length,d=o.length,g=new Array(h);for(s=0;s<h;++s)(l=e[s])&&(g[s]=u="$"+a.call(l,l.__data__,s,e),u in c?i[s]=l:c[u]=l);for(s=0;s<d;++s)(l=c[u="$"+a.call(t,o[s],s,o)])?(r[s]=l,l.__data__=o[s],c[u]=null):n[s]=new y(t,o[s]);for(s=0;s<h;++s)(l=e[s])&&c[g[s]]===l&&(i[s]=l)}function w(t,e){return t<e?-1:t>e?1:t>=e?0:NaN}function A(t){return function(){this.removeAttribute(t)}}function C(t){return function(){this.removeAttributeNS(t.space,t.local)}}function _(t,e){return function(){this.setAttribute(t,e)}}function S(t,e){return function(){this.setAttributeNS(t.space,t.local,e)}}function M(t,e){return function(){var n=e.apply(this,arguments);null==n?this.removeAttribute(t):this.setAttribute(t,n)}}function T(t,e){return function(){var n=e.apply(this,arguments);null==n?this.removeAttributeNS(t.space,t.local):this.setAttributeNS(t.space,t.local,n)}}var k=function(t){return t.ownerDocument&&t.ownerDocument.defaultView||t.document&&t||t.defaultView};function R(t){return function(){this.style.removeProperty(t)}}function N(t,e,n){return function(){this.style.setProperty(t,e,n)}}function $(t,e,n){return function(){var r=e.apply(this,arguments);null==r?this.style.removeProperty(t):this.style.setProperty(t,r,n)}}function P(t,e){return t.style.getPropertyValue(e)||k(t).getComputedStyle(t,null).getPropertyValue(e)}function L(t){return function(){delete this[t]}}function I(t,e){return function(){this[t]=e}}function D(t,e){return function(){var n=e.apply(this,arguments);null==n?delete this[t]:this[t]=n}}function O(t){return t.trim().split(/^|\s+/)}function E(t){return t.classList||new B(t)}function B(t){this._node=t,this._names=O(t.getAttribute("class")||"")}function j(t,e){for(var n=E(t),r=-1,i=e.length;++r<i;)n.add(e[r])}function F(t,e){for(var n=E(t),r=-1,i=e.length;++r<i;)n.remove(e[r])}function z(t){return function(){j(this,t)}}function H(t){return function(){F(this,t)}}function U(t,e){return function(){(e.apply(this,arguments)?j:F)(this,t)}}B.prototype={add:function(t){this._names.indexOf(t)<0&&(this._names.push(t),this._node.setAttribute("class",this._names.join(" ")))},remove:function(t){var e=this._names.indexOf(t);e>=0&&(this._names.splice(e,1),this._node.setAttribute("class",this._names.join(" ")))},contains:function(t){return this._names.indexOf(t)>=0}};function V(){this.textContent=""}function q(t){return function(){this.textContent=t}}function W(t){return function(){var e=t.apply(this,arguments);this.textContent=null==e?"":e}}function Y(){this.innerHTML=""}function X(t){return function(){this.innerHTML=t}}function Z(t){return function(){var e=t.apply(this,arguments);this.innerHTML=null==e?"":e}}function G(){this.nextSibling&&this.parentNode.appendChild(this)}function K(){this.previousSibling&&this.parentNode.insertBefore(this,this.parentNode.firstChild)}function Q(){return null}function J(){var t=this.parentNode;t&&t.removeChild(this)}function tt(){return this.parentNode.insertBefore(this.cloneNode(!1),this.nextSibling)}function et(){return this.parentNode.insertBefore(this.cloneNode(!0),this.nextSibling)}var nt={},rt=null;"undefined"!=typeof document&&("onmouseenter"in document.documentElement||(nt={mouseenter:"mouseover",mouseleave:"mouseout"}));function it(t,e,n){return t=ot(t,e,n),function(e){var n=e.relatedTarget;n&&(n===this||8&n.compareDocumentPosition(this))||t.call(this,e)}}function ot(t,e,n){return function(r){var i=rt;rt=r;try{t.call(this,this.__data__,e,n)}finally{rt=i}}}function at(t){return t.trim().split(/^|\s+/).map((function(t){var e="",n=t.indexOf(".");return n>=0&&(e=t.slice(n+1),t=t.slice(0,n)),{type:t,name:e}}))}function st(t){return function(){var e=this.__on;if(e){for(var n,r=0,i=-1,o=e.length;r<o;++r)n=e[r],t.type&&n.type!==t.type||n.name!==t.name?e[++i]=n:this.removeEventListener(n.type,n.listener,n.capture);++i?e.length=i:delete this.__on}}}function lt(t,e,n){var r=nt.hasOwnProperty(t.type)?it:ot;return function(i,o,a){var s,l=this.__on,u=r(e,o,a);if(l)for(var c=0,h=l.length;c<h;++c)if((s=l[c]).type===t.type&&s.name===t.name)return this.removeEventListener(s.type,s.listener,s.capture),this.addEventListener(s.type,s.listener=u,s.capture=n),void(s.value=e);this.addEventListener(t.type,u,n),s={type:t.type,name:t.name,value:e,listener:u,capture:n},l?l.push(s):this.__on=[s]}}function ut(t,e,n,r){var i=rt;t.sourceEvent=rt,rt=t;try{return e.apply(n,r)}finally{rt=i}}function ct(t,e,n){var r=k(t),i=r.CustomEvent;"function"==typeof i?i=new i(e,n):(i=r.document.createEvent("Event"),n?(i.initEvent(e,n.bubbles,n.cancelable),i.detail=n.detail):i.initEvent(e,!1,!1)),t.dispatchEvent(i)}function ht(t,e){return function(){return ct(this,t,e)}}function dt(t,e){return function(){return ct(this,t,e.apply(this,arguments))}}var gt=[null];function ft(t,e){this._groups=t,this._parents=e}function pt(){return new ft([[document.documentElement]],gt)}ft.prototype=pt.prototype={constructor:ft,select:function(t){"function"!=typeof t&&(t=c(t));for(var e=this._groups,n=e.length,r=new Array(n),i=0;i<n;++i)for(var o,a,s=e[i],l=s.length,u=r[i]=new Array(l),h=0;h<l;++h)(o=s[h])&&(a=t.call(o,o.__data__,h,s))&&("__data__"in o&&(a.__data__=o.__data__),u[h]=a);return new ft(r,this._parents)},selectAll:function(t){"function"!=typeof t&&(t=d(t));for(var e=this._groups,n=e.length,r=[],i=[],o=0;o<n;++o)for(var a,s=e[o],l=s.length,u=0;u<l;++u)(a=s[u])&&(r.push(t.call(a,a.__data__,u,s)),i.push(a));return new ft(r,i)},filter:function(t){"function"!=typeof t&&(t=v(t));for(var e=this._groups,n=e.length,r=new Array(n),i=0;i<n;++i)for(var o,a=e[i],s=a.length,l=r[i]=[],u=0;u<s;++u)(o=a[u])&&t.call(o,o.__data__,u,a)&&l.push(o);return new ft(r,this._parents)},data:function(t,e){if(!t)return f=new Array(this.size()),c=-1,this.each((function(t){f[++c]=t})),f;var n,r=e?x:b,i=this._parents,o=this._groups;"function"!=typeof t&&(n=t,t=function(){return n});for(var a=o.length,s=new Array(a),l=new Array(a),u=new Array(a),c=0;c<a;++c){var h=i[c],d=o[c],g=d.length,f=t.call(h,h&&h.__data__,c,i),p=f.length,v=l[c]=new Array(p),m=s[c]=new Array(p);r(h,d,v,m,u[c]=new Array(g),f,e);for(var y,w,A=0,C=0;A<p;++A)if(y=v[A]){for(A>=C&&(C=A+1);!(w=m[C])&&++C<p;);y._next=w||null}}return(s=new ft(s,i))._enter=l,s._exit=u,s},enter:function(){return new ft(this._enter||this._groups.map(m),this._parents)},exit:function(){return new ft(this._exit||this._groups.map(m),this._parents)},merge:function(t){for(var e=this._groups,n=t._groups,r=e.length,i=n.length,o=Math.min(r,i),a=new Array(r),s=0;s<o;++s)for(var l,u=e[s],c=n[s],h=u.length,d=a[s]=new Array(h),g=0;g<h;++g)(l=u[g]||c[g])&&(d[g]=l);for(;s<r;++s)a[s]=e[s];return new ft(a,this._parents)},order:function(){for(var t=this._groups,e=-1,n=t.length;++e<n;)for(var r,i=t[e],o=i.length-1,a=i[o];--o>=0;)(r=i[o])&&(a&&a!==r.nextSibling&&a.parentNode.insertBefore(r,a),a=r);return this},sort:function(t){function e(e,n){return e&&n?t(e.__data__,n.__data__):!e-!n}t||(t=w);for(var n=this._groups,r=n.length,i=new Array(r),o=0;o<r;++o){for(var a,s=n[o],l=s.length,u=i[o]=new Array(l),c=0;c<l;++c)(a=s[c])&&(u[c]=a);u.sort(e)}return new ft(i,this._parents).order()},call:function(){var t=arguments[0];return arguments[0]=this,t.apply(null,arguments),this},nodes:function(){var t=new Array(this.size()),e=-1;return this.each((function(){t[++e]=this})),t},node:function(){for(var t=this._groups,e=0,n=t.length;e<n;++e)for(var r=t[e],i=0,o=r.length;i<o;++i){var a=r[i];if(a)return a}return null},size:function(){var t=0;return this.each((function(){++t})),t},empty:function(){return!this.node()},each:function(t){for(var e=this._groups,n=0,r=e.length;n<r;++n)for(var i,o=e[n],a=0,s=o.length;a<s;++a)(i=o[a])&&t.call(i,i.__data__,a,o);return this},attr:function(t,e){var n=o(t);if(arguments.length<2){var r=this.node();return n.local?r.getAttributeNS(n.space,n.local):r.getAttribute(n)}return this.each((null==e?n.local?C:A:"function"==typeof e?n.local?T:M:n.local?S:_)(n,e))},style:function(t,e,n){return arguments.length>1?this.each((null==e?R:"function"==typeof e?$:N)(t,e,null==n?"":n)):P(this.node(),t)},property:function(t,e){return arguments.length>1?this.each((null==e?L:"function"==typeof e?D:I)(t,e)):this.node()[t]},classed:function(t,e){var n=O(t+"");if(arguments.length<2){for(var r=E(this.node()),i=-1,o=n.length;++i<o;)if(!r.contains(n[i]))return!1;return!0}return this.each(("function"==typeof e?U:e?z:H)(n,e))},text:function(t){return arguments.length?this.each(null==t?V:("function"==typeof t?W:q)(t)):this.node().textContent},html:function(t){return arguments.length?this.each(null==t?Y:("function"==typeof t?Z:X)(t)):this.node().innerHTML},raise:function(){return this.each(G)},lower:function(){return this.each(K)},append:function(t){var e="function"==typeof t?t:l(t);return this.select((function(){return this.appendChild(e.apply(this,arguments))}))},insert:function(t,e){var n="function"==typeof t?t:l(t),r=null==e?Q:"function"==typeof e?e:c(e);return this.select((function(){return this.insertBefore(n.apply(this,arguments),r.apply(this,arguments)||null)}))},remove:function(){return this.each(J)},clone:function(t){return this.select(t?et:tt)},datum:function(t){return arguments.length?this.property("__data__",t):this.node().__data__},on:function(t,e,n){var r,i,o=at(t+""),a=o.length;if(!(arguments.length<2)){for(s=e?lt:st,null==n&&(n=!1),r=0;r<a;++r)this.each(s(o[r],e,n));return this}var s=this.node().__on;if(s)for(var l,u=0,c=s.length;u<c;++u)for(r=0,l=s[u];r<a;++r)if((i=o[r]).type===l.type&&i.name===l.name)return l.value},dispatch:function(t,e){return this.each(("function"==typeof e?dt:ht)(t,e))}};var vt=pt,mt=function(t){return"string"==typeof t?new ft([[document.querySelector(t)]],[document.documentElement]):new ft([[t]],gt)},yt=function(t){return mt(l(t).call(document.documentElement))},bt=0;function xt(){return new wt}function wt(){this._="@"+(++bt).toString(36)}wt.prototype=xt.prototype={constructor:wt,get:function(t){for(var e=this._;!(e in t);)if(!(t=t.parentNode))return;return t[e]},set:function(t,e){return t[this._]=e},remove:function(t){return this._ in t&&delete t[this._]},toString:function(){return this._}};var At=function(){for(var t,e=rt;t=e.sourceEvent;)e=t;return e},Ct=function(t,e){var n=t.ownerSVGElement||t;if(n.createSVGPoint){var r=n.createSVGPoint();return r.x=e.clientX,r.y=e.clientY,[(r=r.matrixTransform(t.getScreenCTM().inverse())).x,r.y]}var i=t.getBoundingClientRect();return[e.clientX-i.left-t.clientLeft,e.clientY-i.top-t.clientTop]},_t=function(t){var e=At();return e.changedTouches&&(e=e.changedTouches[0]),Ct(t,e)},St=function(t){return"string"==typeof t?new ft([document.querySelectorAll(t)],[document.documentElement]):new ft([null==t?[]:t],gt)},Mt=function(t,e,n){arguments.length<3&&(n=e,e=At().changedTouches);for(var r,i=0,o=e?e.length:0;i<o;++i)if((r=e[i]).identifier===n)return Ct(t,r);return null},Tt=function(t,e){null==e&&(e=At().touches);for(var n=0,r=e?e.length:0,i=new Array(r);n<r;++n)i[n]=Ct(t,e[n]);return i}},function(t,e,n){"use strict";n.d(e,"a",(function(){return y})),n.d(e,"f",(function(){return A})),n.d(e,"d",(function(){return T})),n.d(e,"e",(function(){return D})),n.d(e,"c",(function(){return H})),n.d(e,"b",(function(){return J}));var r=function(t,e,n){t.prototype=e.prototype=n,n.constructor=t};function i(t,e){var n=Object.create(t.prototype);for(var r in e)n[r]=e[r];return n}function o(){}var a="\\s*([+-]?\\d+)\\s*",s="\\s*([+-]?\\d*\\.?\\d+(?:[eE][+-]?\\d+)?)\\s*",l="\\s*([+-]?\\d*\\.?\\d+(?:[eE][+-]?\\d+)?)%\\s*",u=/^#([0-9a-f]{3})$/,c=/^#([0-9a-f]{6})$/,h=new RegExp("^rgb\\("+[a,a,a]+"\\)$"),d=new RegExp("^rgb\\("+[l,l,l]+"\\)$"),g=new RegExp("^rgba\\("+[a,a,a,s]+"\\)$"),f=new RegExp("^rgba\\("+[l,l,l,s]+"\\)$"),p=new RegExp("^hsl\\("+[s,l,l]+"\\)$"),v=new RegExp("^hsla\\("+[s,l,l,s]+"\\)$"),m={aliceblue:15792383,antiquewhite:16444375,aqua:65535,aquamarine:8388564,azure:15794175,beige:16119260,bisque:16770244,black:0,blanchedalmond:16772045,blue:255,blueviolet:9055202,brown:10824234,burlywood:14596231,cadetblue:6266528,chartreuse:8388352,chocolate:13789470,coral:16744272,cornflowerblue:6591981,cornsilk:16775388,crimson:14423100,cyan:65535,darkblue:139,darkcyan:35723,darkgoldenrod:12092939,darkgray:11119017,darkgreen:25600,darkgrey:11119017,darkkhaki:12433259,darkmagenta:9109643,darkolivegreen:5597999,darkorange:16747520,darkorchid:10040012,darkred:9109504,darksalmon:15308410,darkseagreen:9419919,darkslateblue:4734347,darkslategray:3100495,darkslategrey:3100495,darkturquoise:52945,darkviolet:9699539,deeppink:16716947,deepskyblue:49151,dimgray:6908265,dimgrey:6908265,dodgerblue:2003199,firebrick:11674146,floralwhite:16775920,forestgreen:2263842,fuchsia:16711935,gainsboro:14474460,ghostwhite:16316671,gold:16766720,goldenrod:14329120,gray:8421504,green:32768,greenyellow:11403055,grey:8421504,honeydew:15794160,hotpink:16738740,indianred:13458524,indigo:4915330,ivory:16777200,khaki:15787660,lavender:15132410,lavenderblush:16773365,lawngreen:8190976,lemonchiffon:16775885,lightblue:11393254,lightcoral:15761536,lightcyan:14745599,lightgoldenrodyellow:16448210,lightgray:13882323,lightgreen:9498256,lightgrey:13882323,lightpink:16758465,lightsalmon:16752762,lightseagreen:2142890,lightskyblue:8900346,lightslategray:7833753,lightslategrey:7833753,lightsteelblue:11584734,lightyellow:16777184,lime:65280,limegreen:3329330,linen:16445670,magenta:16711935,maroon:8388608,mediumaquamarine:6737322,mediumblue:205,mediumorchid:12211667,mediumpurple:9662683,mediumseagreen:3978097,mediumslateblue:8087790,mediumspringgreen:64154,mediumturquoise:4772300,mediumvioletred:13047173,midnightblue:1644912,mintcream:16121850,mistyrose:16770273,moccasin:16770229,navajowhite:16768685,navy:128,oldlace:16643558,olive:8421376,olivedrab:7048739,orange:16753920,orangered:16729344,orchid:14315734,palegoldenrod:15657130,palegreen:10025880,paleturquoise:11529966,palevioletred:14381203,papayawhip:16773077,peachpuff:16767673,peru:13468991,pink:16761035,plum:14524637,powderblue:11591910,purple:8388736,rebeccapurple:6697881,red:16711680,rosybrown:12357519,royalblue:4286945,saddlebrown:9127187,salmon:16416882,sandybrown:16032864,seagreen:3050327,seashell:16774638,sienna:10506797,silver:12632256,skyblue:8900331,slateblue:6970061,slategray:7372944,slategrey:7372944,snow:16775930,springgreen:65407,steelblue:4620980,tan:13808780,teal:32896,thistle:14204888,tomato:16737095,turquoise:4251856,violet:15631086,wheat:16113331,white:16777215,whitesmoke:16119285,yellow:16776960,yellowgreen:10145074};function y(t){var e;return t=(t+"").trim().toLowerCase(),(e=u.exec(t))?new C((e=parseInt(e[1],16))>>8&15|e>>4&240,e>>4&15|240&e,(15&e)<<4|15&e,1):(e=c.exec(t))?b(parseInt(e[1],16)):(e=h.exec(t))?new C(e[1],e[2],e[3],1):(e=d.exec(t))?new C(255*e[1]/100,255*e[2]/100,255*e[3]/100,1):(e=g.exec(t))?x(e[1],e[2],e[3],e[4]):(e=f.exec(t))?x(255*e[1]/100,255*e[2]/100,255*e[3]/100,e[4]):(e=p.exec(t))?S(e[1],e[2]/100,e[3]/100,1):(e=v.exec(t))?S(e[1],e[2]/100,e[3]/100,e[4]):m.hasOwnProperty(t)?b(m[t]):"transparent"===t?new C(NaN,NaN,NaN,0):null}function b(t){return new C(t>>16&255,t>>8&255,255&t,1)}function x(t,e,n,r){return r<=0&&(t=e=n=NaN),new C(t,e,n,r)}function w(t){return t instanceof o||(t=y(t)),t?new C((t=t.rgb()).r,t.g,t.b,t.opacity):new C}function A(t,e,n,r){return 1===arguments.length?w(t):new C(t,e,n,null==r?1:r)}function C(t,e,n,r){this.r=+t,this.g=+e,this.b=+n,this.opacity=+r}function _(t){return((t=Math.max(0,Math.min(255,Math.round(t)||0)))<16?"0":"")+t.toString(16)}function S(t,e,n,r){return r<=0?t=e=n=NaN:n<=0||n>=1?t=e=NaN:e<=0&&(t=NaN),new k(t,e,n,r)}function M(t){if(t instanceof k)return new k(t.h,t.s,t.l,t.opacity);if(t instanceof o||(t=y(t)),!t)return new k;if(t instanceof k)return t;var e=(t=t.rgb()).r/255,n=t.g/255,r=t.b/255,i=Math.min(e,n,r),a=Math.max(e,n,r),s=NaN,l=a-i,u=(a+i)/2;return l?(s=e===a?(n-r)/l+6*(n<r):n===a?(r-e)/l+2:(e-n)/l+4,l/=u<.5?a+i:2-a-i,s*=60):l=u>0&&u<1?0:s,new k(s,l,u,t.opacity)}function T(t,e,n,r){return 1===arguments.length?M(t):new k(t,e,n,null==r?1:r)}function k(t,e,n,r){this.h=+t,this.s=+e,this.l=+n,this.opacity=+r}function R(t,e,n){return 255*(t<60?e+(n-e)*t/60:t<180?n:t<240?e+(n-e)*(240-t)/60:e)}r(o,y,{displayable:function(){return this.rgb().displayable()},hex:function(){return this.rgb().hex()},toString:function(){return this.rgb()+""}}),r(C,A,i(o,{brighter:function(t){return t=null==t?1/.7:Math.pow(1/.7,t),new C(this.r*t,this.g*t,this.b*t,this.opacity)},darker:function(t){return t=null==t?.7:Math.pow(.7,t),new C(this.r*t,this.g*t,this.b*t,this.opacity)},rgb:function(){return this},displayable:function(){return 0<=this.r&&this.r<=255&&0<=this.g&&this.g<=255&&0<=this.b&&this.b<=255&&0<=this.opacity&&this.opacity<=1},hex:function(){return"#"+_(this.r)+_(this.g)+_(this.b)},toString:function(){var t=this.opacity;return(1===(t=isNaN(t)?1:Math.max(0,Math.min(1,t)))?"rgb(":"rgba(")+Math.max(0,Math.min(255,Math.round(this.r)||0))+", "+Math.max(0,Math.min(255,Math.round(this.g)||0))+", "+Math.max(0,Math.min(255,Math.round(this.b)||0))+(1===t?")":", "+t+")")}})),r(k,T,i(o,{brighter:function(t){return t=null==t?1/.7:Math.pow(1/.7,t),new k(this.h,this.s,this.l*t,this.opacity)},darker:function(t){return t=null==t?.7:Math.pow(.7,t),new k(this.h,this.s,this.l*t,this.opacity)},rgb:function(){var t=this.h%360+360*(this.h<0),e=isNaN(t)||isNaN(this.s)?0:this.s,n=this.l,r=n+(n<.5?n:1-n)*e,i=2*n-r;return new C(R(t>=240?t-240:t+120,i,r),R(t,i,r),R(t<120?t+240:t-120,i,r),this.opacity)},displayable:function(){return(0<=this.s&&this.s<=1||isNaN(this.s))&&0<=this.l&&this.l<=1&&0<=this.opacity&&this.opacity<=1}}));var N=Math.PI/180,$=180/Math.PI,P=6/29,L=3*P*P;function I(t){if(t instanceof O)return new O(t.l,t.a,t.b,t.opacity);if(t instanceof U){if(isNaN(t.h))return new O(t.l,0,0,t.opacity);var e=t.h*N;return new O(t.l,Math.cos(e)*t.c,Math.sin(e)*t.c,t.opacity)}t instanceof C||(t=w(t));var n,r,i=F(t.r),o=F(t.g),a=F(t.b),s=E((.2225045*i+.7168786*o+.0606169*a)/1);return i===o&&o===a?n=r=s:(n=E((.4360747*i+.3850649*o+.1430804*a)/.96422),r=E((.0139322*i+.0971045*o+.7141733*a)/.82521)),new O(116*s-16,500*(n-s),200*(s-r),t.opacity)}function D(t,e,n,r){return 1===arguments.length?I(t):new O(t,e,n,null==r?1:r)}function O(t,e,n,r){this.l=+t,this.a=+e,this.b=+n,this.opacity=+r}function E(t){return t>.008856451679035631?Math.pow(t,1/3):t/L+4/29}function B(t){return t>P?t*t*t:L*(t-4/29)}function j(t){return 255*(t<=.0031308?12.92*t:1.055*Math.pow(t,1/2.4)-.055)}function F(t){return(t/=255)<=.04045?t/12.92:Math.pow((t+.055)/1.055,2.4)}function z(t){if(t instanceof U)return new U(t.h,t.c,t.l,t.opacity);if(t instanceof O||(t=I(t)),0===t.a&&0===t.b)return new U(NaN,0,t.l,t.opacity);var e=Math.atan2(t.b,t.a)*$;return new U(e<0?e+360:e,Math.sqrt(t.a*t.a+t.b*t.b),t.l,t.opacity)}function H(t,e,n,r){return 1===arguments.length?z(t):new U(t,e,n,null==r?1:r)}function U(t,e,n,r){this.h=+t,this.c=+e,this.l=+n,this.opacity=+r}r(O,D,i(o,{brighter:function(t){return new O(this.l+18*(null==t?1:t),this.a,this.b,this.opacity)},darker:function(t){return new O(this.l-18*(null==t?1:t),this.a,this.b,this.opacity)},rgb:function(){var t=(this.l+16)/116,e=isNaN(this.a)?t:t+this.a/500,n=isNaN(this.b)?t:t-this.b/200;return new C(j(3.1338561*(e=.96422*B(e))-1.6168667*(t=1*B(t))-.4906146*(n=.82521*B(n))),j(-.9787684*e+1.9161415*t+.033454*n),j(.0719453*e-.2289914*t+1.4052427*n),this.opacity)}})),r(U,H,i(o,{brighter:function(t){return new U(this.h,this.c,this.l+18*(null==t?1:t),this.opacity)},darker:function(t){return new U(this.h,this.c,this.l-18*(null==t?1:t),this.opacity)},rgb:function(){return I(this).rgb()}}));var V=-.14861,q=1.78277,W=-.29227,Y=-.90649,X=1.97294,Z=X*Y,G=X*q,K=q*W-Y*V;function Q(t){if(t instanceof tt)return new tt(t.h,t.s,t.l,t.opacity);t instanceof C||(t=w(t));var e=t.r/255,n=t.g/255,r=t.b/255,i=(K*r+Z*e-G*n)/(K+Z-G),o=r-i,a=(X*(n-i)-W*o)/Y,s=Math.sqrt(a*a+o*o)/(X*i*(1-i)),l=s?Math.atan2(a,o)*$-120:NaN;return new tt(l<0?l+360:l,s,i,t.opacity)}function J(t,e,n,r){return 1===arguments.length?Q(t):new tt(t,e,n,null==r?1:r)}function tt(t,e,n,r){this.h=+t,this.s=+e,this.l=+n,this.opacity=+r}r(tt,J,i(o,{brighter:function(t){return t=null==t?1/.7:Math.pow(1/.7,t),new tt(this.h,this.s,this.l*t,this.opacity)},darker:function(t){return t=null==t?.7:Math.pow(.7,t),new tt(this.h,this.s,this.l*t,this.opacity)},rgb:function(){var t=isNaN(this.h)?0:(this.h+120)*N,e=+this.l,n=isNaN(this.s)?0:this.s*e*(1-e),r=Math.cos(t),i=Math.sin(t);return new C(255*(e+n*(V*r+q*i)),255*(e+n*(W*r+Y*i)),255*(e+n*(X*r)),this.opacity)}}))},function(t,e,n){"use strict";n.d(e,"a",(function(){return C})),n.d(e,"b",(function(){return f})),n.d(e,"d",(function(){return _})),n.d(e,"e",(function(){return A})),n.d(e,"f",(function(){return R})),n.d(e,"g",(function(){return N})),n.d(e,"c",(function(){return c}));var r=n(1);function i(t,e,n,r,i){var o=t*t,a=o*t;return((1-3*t+3*o-a)*e+(4-6*o+3*a)*n+(1+3*t+3*o-3*a)*r+a*i)/6}var o=function(t){return function(){return t}};function a(t,e){return function(n){return t+n*e}}function s(t,e){var n=e-t;return n?a(t,n>180||n<-180?n-360*Math.round(n/360):n):o(isNaN(t)?e:t)}function l(t){return 1==(t=+t)?u:function(e,n){return n-e?function(t,e,n){return t=Math.pow(t,n),e=Math.pow(e,n)-t,n=1/n,function(r){return Math.pow(t+r*e,n)}}(e,n,t):o(isNaN(e)?n:e)}}function u(t,e){var n=e-t;return n?a(t,n):o(isNaN(t)?e:t)}var c=function t(e){var n=l(e);function i(t,e){var i=n((t=Object(r.f)(t)).r,(e=Object(r.f)(e)).r),o=n(t.g,e.g),a=n(t.b,e.b),s=u(t.opacity,e.opacity);return function(e){return t.r=i(e),t.g=o(e),t.b=a(e),t.opacity=s(e),t+""}}return i.gamma=t,i}(1);function h(t){return function(e){var n,i,o=e.length,a=new Array(o),s=new Array(o),l=new Array(o);for(n=0;n<o;++n)i=Object(r.f)(e[n]),a[n]=i.r||0,s[n]=i.g||0,l[n]=i.b||0;return a=t(a),s=t(s),l=t(l),i.opacity=1,function(t){return i.r=a(t),i.g=s(t),i.b=l(t),i+""}}}h((function(t){var e=t.length-1;return function(n){var r=n<=0?n=0:n>=1?(n=1,e-1):Math.floor(n*e),o=t[r],a=t[r+1],s=r>0?t[r-1]:2*o-a,l=r<e-1?t[r+2]:2*a-o;return i((n-r/e)*e,s,o,a,l)}})),h((function(t){var e=t.length;return function(n){var r=Math.floor(((n%=1)<0?++n:n)*e),o=t[(r+e-1)%e],a=t[r%e],s=t[(r+1)%e],l=t[(r+2)%e];return i((n-r/e)*e,o,a,s,l)}}));var d=function(t,e){var n,r=e?e.length:0,i=t?Math.min(r,t.length):0,o=new Array(i),a=new Array(r);for(n=0;n<i;++n)o[n]=C(t[n],e[n]);for(;n<r;++n)a[n]=e[n];return function(t){for(n=0;n<i;++n)a[n]=o[n](t);return a}},g=function(t,e){var n=new Date;return e-=t=+t,function(r){return n.setTime(t+e*r),n}},f=function(t,e){return e-=t=+t,function(n){return t+e*n}},p=function(t,e){var n,r={},i={};for(n in null!==t&&"object"==typeof t||(t={}),null!==e&&"object"==typeof e||(e={}),e)n in t?r[n]=C(t[n],e[n]):i[n]=e[n];return function(t){for(n in r)i[n]=r[n](t);return i}},v=/[-+]?(?:\d+\.?\d*|\.?\d+)(?:[eE][-+]?\d+)?/g,m=new RegExp(v.source,"g");var y,b,x,w,A=function(t,e){var n,r,i,o=v.lastIndex=m.lastIndex=0,a=-1,s=[],l=[];for(t+="",e+="";(n=v.exec(t))&&(r=m.exec(e));)(i=r.index)>o&&(i=e.slice(o,i),s[a]?s[a]+=i:s[++a]=i),(n=n[0])===(r=r[0])?s[a]?s[a]+=r:s[++a]=r:(s[++a]=null,l.push({i:a,x:f(n,r)})),o=m.lastIndex;return o<e.length&&(i=e.slice(o),s[a]?s[a]+=i:s[++a]=i),s.length<2?l[0]?function(t){return function(e){return t(e)+""}}(l[0].x):function(t){return function(){return t}}(e):(e=l.length,function(t){for(var n,r=0;r<e;++r)s[(n=l[r]).i]=n.x(t);return s.join("")})},C=function(t,e){var n,i=typeof e;return null==e||"boolean"===i?o(e):("number"===i?f:"string"===i?(n=Object(r.a)(e))?(e=n,c):A:e instanceof r.a?c:e instanceof Date?g:Array.isArray(e)?d:"function"!=typeof e.valueOf&&"function"!=typeof e.toString||isNaN(e)?p:f)(t,e)},_=function(t,e){return e-=t=+t,function(n){return Math.round(t+e*n)}},S=180/Math.PI,M={translateX:0,translateY:0,rotate:0,skewX:0,scaleX:1,scaleY:1},T=function(t,e,n,r,i,o){var a,s,l;return(a=Math.sqrt(t*t+e*e))&&(t/=a,e/=a),(l=t*n+e*r)&&(n-=t*l,r-=e*l),(s=Math.sqrt(n*n+r*r))&&(n/=s,r/=s,l/=s),t*r<e*n&&(t=-t,e=-e,l=-l,a=-a),{translateX:i,translateY:o,rotate:Math.atan2(e,t)*S,skewX:Math.atan(l)*S,scaleX:a,scaleY:s}};function k(t,e,n,r){function i(t){return t.length?t.pop()+" ":""}return function(o,a){var s=[],l=[];return o=t(o),a=t(a),function(t,r,i,o,a,s){if(t!==i||r!==o){var l=a.push("translate(",null,e,null,n);s.push({i:l-4,x:f(t,i)},{i:l-2,x:f(r,o)})}else(i||o)&&a.push("translate("+i+e+o+n)}(o.translateX,o.translateY,a.translateX,a.translateY,s,l),function(t,e,n,o){t!==e?(t-e>180?e+=360:e-t>180&&(t+=360),o.push({i:n.push(i(n)+"rotate(",null,r)-2,x:f(t,e)})):e&&n.push(i(n)+"rotate("+e+r)}(o.rotate,a.rotate,s,l),function(t,e,n,o){t!==e?o.push({i:n.push(i(n)+"skewX(",null,r)-2,x:f(t,e)}):e&&n.push(i(n)+"skewX("+e+r)}(o.skewX,a.skewX,s,l),function(t,e,n,r,o,a){if(t!==n||e!==r){var s=o.push(i(o)+"scale(",null,",",null,")");a.push({i:s-4,x:f(t,n)},{i:s-2,x:f(e,r)})}else 1===n&&1===r||o.push(i(o)+"scale("+n+","+r+")")}(o.scaleX,o.scaleY,a.scaleX,a.scaleY,s,l),o=a=null,function(t){for(var e,n=-1,r=l.length;++n<r;)s[(e=l[n]).i]=e.x(t);return s.join("")}}}var R=k((function(t){return"none"===t?M:(y||(y=document.createElement("DIV"),b=document.documentElement,x=document.defaultView),y.style.transform=t,t=x.getComputedStyle(b.appendChild(y),null).getPropertyValue("transform"),b.removeChild(y),t=t.slice(7,-1).split(","),T(+t[0],+t[1],+t[2],+t[3],+t[4],+t[5]))}),"px, ","px)","deg)"),N=k((function(t){return null==t?M:(w||(w=document.createElementNS("http://www.w3.org/2000/svg","g")),w.setAttribute("transform",t),(t=w.transform.baseVal.consolidate())?(t=t.matrix,T(t.a,t.b,t.c,t.d,t.e,t.f)):M)}),", ",")",")");Math.SQRT2;function $(t){return function(e,n){var i=t((e=Object(r.d)(e)).h,(n=Object(r.d)(n)).h),o=u(e.s,n.s),a=u(e.l,n.l),s=u(e.opacity,n.opacity);return function(t){return e.h=i(t),e.s=o(t),e.l=a(t),e.opacity=s(t),e+""}}}$(s),$(u);function P(t){return function(e,n){var i=t((e=Object(r.c)(e)).h,(n=Object(r.c)(n)).h),o=u(e.c,n.c),a=u(e.l,n.l),s=u(e.opacity,n.opacity);return function(t){return e.h=i(t),e.c=o(t),e.l=a(t),e.opacity=s(t),e+""}}}P(s),P(u);function L(t){return function e(n){function i(e,i){var o=t((e=Object(r.b)(e)).h,(i=Object(r.b)(i)).h),a=u(e.s,i.s),s=u(e.l,i.l),l=u(e.opacity,i.opacity);return function(t){return e.h=o(t),e.s=a(t),e.l=s(Math.pow(t,n)),e.opacity=l(t),e+""}}return n=+n,i.gamma=e,i}(1)}L(s),L(u)},function(t,e){class n{constructor(t,e){0==arguments.length?(this.x=0,this.y=0):1==arguments.length?(this.x=t.x,this.y=t.y):(this.x=t,this.y=e)}clone(){return new n(this.x,this.y)}toString(){return"("+this.x+","+this.y+")"}add(t){return this.x+=t.x,this.y+=t.y,this}subtract(t){return this.x-=t.x,this.y-=t.y,this}divide(t){return this.x/=t,this.y/=t,this}multiply(t){return this.x*=t.x,this.y*=t.y,this}multiplyScalar(t){return this.x*=t,this.y*=t,this}invert(){return this.x=-this.x,this.y=-this.y,this}angle(){return Math.atan2(this.y,this.x)}distance(t){return Math.sqrt((t.x-this.x)*(t.x-this.x)+(t.y-this.y)*(t.y-this.y))}distanceSq(t){return(t.x-this.x)*(t.x-this.x)+(t.y-this.y)*(t.y-this.y)}clockwise(t){let e=this.y*t.x,n=this.x*t.y;return e>n?-1:e===n?0:1}relativeClockwise(t,e){let n=(this.y-t.y)*(e.x-t.x),r=(this.x-t.x)*(e.y-t.y);return n>r?-1:n===r?0:1}rotate(t){let e=new n(0,0),r=Math.cos(t),i=Math.sin(t);return e.x=this.x*r-this.y*i,e.y=this.x*i+this.y*r,this.x=e.x,this.y=e.y,this}rotateAround(t,e){let n=Math.sin(t),r=Math.cos(t);this.x-=e.x,this.y-=e.y;let i=this.x*r-this.y*n,o=this.x*n+this.y*r;return this.x=i+e.x,this.y=o+e.y,this}rotateTo(t,e,r=0){this.x+=.001,this.y-=.001;let i=n.subtract(this,e),o=n.subtract(t,e),a=n.angle(o,i);return this.rotateAround(a+r,e),this}rotateAwayFrom(t,e,n){this.rotateAround(n,e);let r=this.distanceSq(t);this.rotateAround(-2*n,e),this.distanceSq(t)<r&&this.rotateAround(2*n,e)}getRotateAwayFromAngle(t,e,n){let r=this.clone();r.rotateAround(n,e);let i=r.distanceSq(t);return r.rotateAround(-2*n,e),r.distanceSq(t)<i?n:-n}getRotateTowardsAngle(t,e,n){let r=this.clone();r.rotateAround(n,e);let i=r.distanceSq(t);return r.rotateAround(-2*n,e),r.distanceSq(t)>i?n:-n}getRotateToAngle(t,e){let r=n.subtract(this,e),i=n.subtract(t,e),o=n.angle(i,r);return Number.isNaN(o)?0:o}isInPolygon(t){let e=!1;for(let n=0,r=t.length-1;n<t.length;r=n++)t[n].y>this.y!=t[r].y>this.y&&this.x<(t[r].x-t[n].x)*(this.y-t[n].y)/(t[r].y-t[n].y)+t[n].x&&(e=!e);return e}length(){return Math.sqrt(this.x*this.x+this.y*this.y)}lengthSq(){return this.x*this.x+this.y*this.y}normalize(){return this.divide(this.length()),this}normalized(){return n.divideScalar(this,this.length())}whichSide(t,e){return(this.x-t.x)*(e.y-t.y)-(this.y-t.y)*(e.x-t.x)}sameSideAs(t,e,n){let r=this.whichSide(t,e),i=n.whichSide(t,e);return r<0&&i<0||0==r&&0==i||r>0&&i>0}static add(t,e){return new n(t.x+e.x,t.y+e.y)}static subtract(t,e){return new n(t.x-e.x,t.y-e.y)}static multiply(t,e){return new n(t.x*e.x,t.y*e.y)}static multiplyScalar(t,e){return new n(t.x,t.y).multiplyScalar(e)}static midpoint(t,e){return new n((t.x+e.x)/2,(t.y+e.y)/2)}static normals(t,e){let r=n.subtract(e,t);return[new n(-r.y,r.x),new n(r.y,-r.x)]}static units(t,e){let r=n.subtract(e,t);return[new n(-r.y,r.x).normalize(),new n(r.y,-r.x).normalize()]}static divide(t,e){return new n(t.x/e.x,t.y/e.y)}static divideScalar(t,e){return new n(t.x/e,t.y/e)}static dot(t,e){return t.x*e.x+t.y*e.y}static angle(t,e){let r=n.dot(t,e);return Math.acos(r/(t.length()*e.length()))}static threePointangle(t,e,r){let i=n.subtract(e,t),o=n.subtract(r,e),a=t.distance(e),s=e.distance(r);return Math.acos(n.dot(i,o)/(a*s))}static scalarProjection(t,e){let r=e.normalized();return n.dot(t,r)}static averageDirection(t){let e=new n(0,0);for(var r=0;r<t.length;r++){let n=t[r];e.add(n)}return e.normalize()}}t.exports=n},function(t,e,n){const r=n(9),i=n(10),o=n(3);n(12);class a{constructor(t,e=0,n=0){this.id=null,this.value=t,this.position=new o(e||0,n||0),this.previousPosition=new o(0,0),this.parentVertexId=null,this.children=Array(),this.spanningTreeChildren=Array(),this.edges=Array(),this.positioned=!1,this.angle=null,this.dir=1,this.neighbourCount=0,this.neighbours=Array(),this.neighbouringElements=Array(),this.forcePositioned=!1}setPosition(t,e){this.position.x=t,this.position.y=e}setPositionFromVector(t){this.position.x=t.x,this.position.y=t.y}addChild(t){this.children.push(t),this.neighbours.push(t),this.neighbourCount++}addRingbondChild(t,e){if(this.children.push(t),this.value.bracket){let n=1;0===this.id&&0===this.value.bracket.hcount&&(n=0),1===this.value.bracket.hcount&&0===e&&(n=2),1===this.value.bracket.hcount&&1===e&&(n=this.neighbours.length<3?2:3),null===this.value.bracket.hcount&&0===e&&(n=1),null===this.value.bracket.hcount&&1===e&&(n=this.neighbours.length<3?1:2),this.neighbours.splice(n,0,t)}else this.neighbours.push(t);this.neighbourCount++}setParentVertexId(t){this.neighbourCount++,this.parentVertexId=t,this.neighbours.push(t)}isTerminal(){return!!this.value.hasAttachedPseudoElements||(null===this.parentVertexId&&this.children.length<2||0===this.children.length)}clone(){let t=new a(this.value,this.position.x,this.position.y);return t.id=this.id,t.previousPosition=new o(this.previousPosition.x,this.previousPosition.y),t.parentVertexId=this.parentVertexId,t.children=i.clone(this.children),t.spanningTreeChildren=i.clone(this.spanningTreeChildren),t.edges=i.clone(this.edges),t.positioned=this.positioned,t.angle=this.angle,t.forcePositioned=this.forcePositioned,t}equals(t){return this.id===t.id}getAngle(t=null,e=!1){let n=null;return n=t?o.subtract(this.position,t):o.subtract(this.position,this.previousPosition),e?r.toDeg(n.angle()):n.angle()}getTextDirection(t){let e=this.getDrawnNeighbours(t),n=Array();for(let r=0;r<e.length;r++)n.push(this.getAngle(t[e[r]].position));let i=r.meanAngle(n),o=Math.PI/2;return i=Math.round(Math.round(i/o)*o),2===i?"down":-2===i?"up":0===i||-0===i?"right":3===i||-3===i?"left":"down"}getNeighbours(t=null){if(null===t)return this.neighbours.slice();let e=Array();for(let n=0;n<this.neighbours.length;n++)this.neighbours[n]!==t&&e.push(this.neighbours[n]);return e}getDrawnNeighbours(t){let e=Array();for(let n=0;n<this.neighbours.length;n++)t[this.neighbours[n]].value.isDrawn&&e.push(this.neighbours[n]);return e}getNeighbourCount(){return this.neighbourCount}getSpanningTreeNeighbours(t=null){let e=Array();for(let n=0;n<this.spanningTreeChildren.length;n++)void 0!==t&&t==this.spanningTreeChildren[n]||e.push(this.spanningTreeChildren[n]);return null!=this.parentVertexId&&(void 0!==t&&t==this.parentVertexId||e.push(this.parentVertexId)),e}getNextInRing(t,e,n){let r=this.getNeighbours();for(let o=0;o<r.length;o++)if(i.contains(t[r[o]].value.rings,{value:e})&&r[o]!=n)return r[o];return null}}t.exports=a},function(t,e,n){const r=n(10),i=n(3),o=(n(4),n(16));class a{constructor(t){this.id=null,this.members=t,this.edges=[],this.insiders=[],this.neighbours=[],this.positioned=!1,this.center=new i(0,0),this.rings=[],this.isBridged=!1,this.isPartOfBridged=!1,this.isSpiro=!1,this.isFused=!1,this.centralAngle=0,this.canFlip=!0}clone(){let t=new a(this.members);return t.id=this.id,t.insiders=r.clone(this.insiders),t.neighbours=r.clone(this.neighbours),t.positioned=this.positioned,t.center=this.center.clone(),t.rings=r.clone(this.rings),t.isBridged=this.isBridged,t.isPartOfBridged=this.isPartOfBridged,t.isSpiro=this.isSpiro,t.isFused=this.isFused,t.centralAngle=this.centralAngle,t.canFlip=this.canFlip,t}getSize(){return this.members.length}getPolygon(t){let e=[];for(let n=0;n<this.members.length;n++)e.push(t[this.members[n]].position);return e}getAngle(){return Math.PI-this.centralAngle}eachMember(t,e,n,r){let i=n=n||0===n?n:this.members[0],o=0;for(;null!=i&&o<100;){let a=i;e(a),i=t[i].getNextInRing(t,this.id,r),r=a,i==n&&(i=null),o++}}getOrderedNeighbours(t){let e=Array(this.neighbours.length);for(let n=0;n<this.neighbours.length;n++){let r=o.getVertices(t,this.id,this.neighbours[n]);e[n]={n:r.length,neighbour:this.neighbours[n]}}return e.sort((function(t,e){return e.n-t.n})),e}isBenzeneLike(t){let e=this.getDoubleBondCount(t),n=this.members.length;return 3===e&&6===n||2===e&&5===n}getDoubleBondCount(t){let e=0;for(let n=0;n<this.members.length;n++){let r=t[this.members[n]].value;"="!==r.bondType&&"="!==r.branchBond||e++}return e}contains(t){for(let e=0;e<this.members.length;e++)if(this.members[e]==t)return!0;return!1}}t.exports=a},function(t,e,n){"use strict";n.r(e),n.d(e,"scaleBand",(function(){return R})),n.d(e,"scalePoint",(function(){return N})),n.d(e,"scaleIdentity",(function(){return nt})),n.d(e,"scaleLinear",(function(){return et})),n.d(e,"scaleLog",(function(){return ct})),n.d(e,"scaleOrdinal",(function(){return k})),n.d(e,"scaleImplicit",(function(){return T})),n.d(e,"scalePow",(function(){return dt})),n.d(e,"scaleSqrt",(function(){return gt})),n.d(e,"scaleQuantile",(function(){return ft})),n.d(e,"scaleQuantize",(function(){return pt})),n.d(e,"scaleThreshold",(function(){return vt})),n.d(e,"scaleTime",(function(){return Tn})),n.d(e,"scaleUtc",(function(){return kn})),n.d(e,"scaleSequential",(function(){return Rn})),n.d(e,"scaleDiverging",(function(){return Nn}));var r=function(t,e){return t<e?-1:t>e?1:t>=e?0:NaN},i=function(t){var e;return 1===t.length&&(e=t,t=function(t,n){return r(e(t),n)}),{left:function(e,n,r,i){for(null==r&&(r=0),null==i&&(i=e.length);r<i;){var o=r+i>>>1;t(e[o],n)<0?r=o+1:i=o}return r},right:function(e,n,r,i){for(null==r&&(r=0),null==i&&(i=e.length);r<i;){var o=r+i>>>1;t(e[o],n)>0?i=o:r=o+1}return r}}};var o=i(r),a=o.right,s=(o.left,a);var l=function(t){return null===t?NaN:+t},u=Array.prototype,c=(u.slice,u.map,function(t,e,n){t=+t,e=+e,n=(i=arguments.length)<2?(e=t,t=0,1):i<3?1:+n;for(var r=-1,i=0|Math.max(0,Math.ceil((e-t)/n)),o=new Array(i);++r<i;)o[r]=t+r*n;return o}),h=Math.sqrt(50),d=Math.sqrt(10),g=Math.sqrt(2),f=function(t,e,n){var r,i,o,a,s=-1;if(n=+n,(t=+t)===(e=+e)&&n>0)return[t];if((r=e<t)&&(i=t,t=e,e=i),0===(a=p(t,e,n))||!isFinite(a))return[];if(a>0)for(t=Math.ceil(t/a),e=Math.floor(e/a),o=new Array(i=Math.ceil(e-t+1));++s<i;)o[s]=(t+s)*a;else for(t=Math.floor(t*a),e=Math.ceil(e*a),o=new Array(i=Math.ceil(t-e+1));++s<i;)o[s]=(t-s)/a;return r&&o.reverse(),o};function p(t,e,n){var r=(e-t)/Math.max(0,n),i=Math.floor(Math.log(r)/Math.LN10),o=r/Math.pow(10,i);return i>=0?(o>=h?10:o>=d?5:o>=g?2:1)*Math.pow(10,i):-Math.pow(10,-i)/(o>=h?10:o>=d?5:o>=g?2:1)}function v(t,e,n){var r=Math.abs(e-t)/Math.max(0,n),i=Math.pow(10,Math.floor(Math.log(r)/Math.LN10)),o=r/i;return o>=h?i*=10:o>=d?i*=5:o>=g&&(i*=2),e<t?-i:i}var m=function(t,e,n){if(null==n&&(n=l),r=t.length){if((e=+e)<=0||r<2)return+n(t[0],0,t);if(e>=1)return+n(t[r-1],r-1,t);var r,i=(r-1)*e,o=Math.floor(i),a=+n(t[o],o,t);return a+(+n(t[o+1],o+1,t)-a)*(i-o)}};function y(){}function b(t,e){var n=new y;if(t instanceof y)t.each((function(t,e){n.set(e,t)}));else if(Array.isArray(t)){var r,i=-1,o=t.length;if(null==e)for(;++i<o;)n.set(i,t[i]);else for(;++i<o;)n.set(e(r=t[i],i,t),r)}else if(t)for(var a in t)n.set(a,t[a]);return n}y.prototype=b.prototype={constructor:y,has:function(t){return"$"+t in this},get:function(t){return this["$"+t]},set:function(t,e){return this["$"+t]=e,this},remove:function(t){var e="$"+t;return e in this&&delete this[e]},clear:function(){for(var t in this)"$"===t[0]&&delete this[t]},keys:function(){var t=[];for(var e in this)"$"===e[0]&&t.push(e.slice(1));return t},values:function(){var t=[];for(var e in this)"$"===e[0]&&t.push(this[e]);return t},entries:function(){var t=[];for(var e in this)"$"===e[0]&&t.push({key:e.slice(1),value:this[e]});return t},size:function(){var t=0;for(var e in this)"$"===e[0]&&++t;return t},empty:function(){for(var t in this)if("$"===t[0])return!1;return!0},each:function(t){for(var e in this)"$"===e[0]&&t(this[e],e.slice(1),this)}};var x=b;function w(){}var A=x.prototype;function C(t,e){var n=new w;if(t instanceof w)t.each((function(t){n.add(t)}));else if(t){var r=-1,i=t.length;if(null==e)for(;++r<i;)n.add(t[r]);else for(;++r<i;)n.add(e(t[r],r,t))}return n}w.prototype=C.prototype={constructor:w,has:A.has,add:function(t){return this["$"+(t+="")]=t,this},remove:A.remove,clear:A.clear,values:A.keys,size:A.size,empty:A.empty,each:A.each};var _=Array.prototype,S=_.map,M=_.slice,T={name:"implicit"};function k(t){var e=x(),n=[],r=T;function i(i){var o=i+"",a=e.get(o);if(!a){if(r!==T)return r;e.set(o,a=n.push(i))}return t[(a-1)%t.length]}return t=null==t?[]:M.call(t),i.domain=function(t){if(!arguments.length)return n.slice();n=[],e=x();for(var r,o,a=-1,s=t.length;++a<s;)e.has(o=(r=t[a])+"")||e.set(o,n.push(r));return i},i.range=function(e){return arguments.length?(t=M.call(e),i):t.slice()},i.unknown=function(t){return arguments.length?(r=t,i):r},i.copy=function(){return k().domain(n).range(t).unknown(r)},i}function R(){var t,e,n=k().unknown(void 0),r=n.domain,i=n.range,o=[0,1],a=!1,s=0,l=0,u=.5;function h(){var n=r().length,h=o[1]<o[0],d=o[h-0],g=o[1-h];t=(g-d)/Math.max(1,n-s+2*l),a&&(t=Math.floor(t)),d+=(g-d-t*(n-s))*u,e=t*(1-s),a&&(d=Math.round(d),e=Math.round(e));var f=c(n).map((function(e){return d+t*e}));return i(h?f.reverse():f)}return delete n.unknown,n.domain=function(t){return arguments.length?(r(t),h()):r()},n.range=function(t){return arguments.length?(o=[+t[0],+t[1]],h()):o.slice()},n.rangeRound=function(t){return o=[+t[0],+t[1]],a=!0,h()},n.bandwidth=function(){return e},n.step=function(){return t},n.round=function(t){return arguments.length?(a=!!t,h()):a},n.padding=function(t){return arguments.length?(s=l=Math.max(0,Math.min(1,t)),h()):s},n.paddingInner=function(t){return arguments.length?(s=Math.max(0,Math.min(1,t)),h()):s},n.paddingOuter=function(t){return arguments.length?(l=Math.max(0,Math.min(1,t)),h()):l},n.align=function(t){return arguments.length?(u=Math.max(0,Math.min(1,t)),h()):u},n.copy=function(){return R().domain(r()).range(o).round(a).paddingInner(s).paddingOuter(l).align(u)},h()}function N(){return function t(e){var n=e.copy;return e.padding=e.paddingOuter,delete e.paddingInner,delete e.paddingOuter,e.copy=function(){return t(n())},e}(R().paddingInner(1))}var $=n(2),P=function(t){return function(){return t}},L=function(t){return+t},I=[0,1];function D(t,e){return(e-=t=+t)?function(n){return(n-t)/e}:P(e)}function O(t,e,n,r){var i=t[0],o=t[1],a=e[0],s=e[1];return o<i?(i=n(o,i),a=r(s,a)):(i=n(i,o),a=r(a,s)),function(t){return a(i(t))}}function E(t,e,n,r){var i=Math.min(t.length,e.length)-1,o=new Array(i),a=new Array(i),l=-1;for(t[i]<t[0]&&(t=t.slice().reverse(),e=e.slice().reverse());++l<i;)o[l]=n(t[l],t[l+1]),a[l]=r(e[l],e[l+1]);return function(e){var n=s(t,e,1,i)-1;return a[n](o[n](e))}}function B(t,e){return e.domain(t.domain()).range(t.range()).interpolate(t.interpolate()).clamp(t.clamp())}function j(t,e){var n,r,i,o=I,a=I,s=$.a,l=!1;function u(){return n=Math.min(o.length,a.length)>2?E:O,r=i=null,c}function c(e){return(r||(r=n(o,a,l?function(t){return function(e,n){var r=t(e=+e,n=+n);return function(t){return t<=e?0:t>=n?1:r(t)}}}(t):t,s)))(+e)}return c.invert=function(t){return(i||(i=n(a,o,D,l?function(t){return function(e,n){var r=t(e=+e,n=+n);return function(t){return t<=0?e:t>=1?n:r(t)}}}(e):e)))(+t)},c.domain=function(t){return arguments.length?(o=S.call(t,L),u()):o.slice()},c.range=function(t){return arguments.length?(a=M.call(t),u()):a.slice()},c.rangeRound=function(t){return a=M.call(t),s=$.d,u()},c.clamp=function(t){return arguments.length?(l=!!t,u()):l},c.interpolate=function(t){return arguments.length?(s=t,u()):s},u()}var F=function(t,e){if((n=(t=e?t.toExponential(e-1):t.toExponential()).indexOf("e"))<0)return null;var n,r=t.slice(0,n);return[r.length>1?r[0]+r.slice(2):r,+t.slice(n+1)]},z=function(t){return(t=F(Math.abs(t)))?t[1]:NaN},H=/^(?:(.)?([<>=^]))?([+\-( ])?([$#])?(0)?(\d+)?(,)?(\.\d+)?(~)?([a-z%])?$/i;function U(t){return new V(t)}function V(t){if(!(e=H.exec(t)))throw new Error("invalid format: "+t);var e;this.fill=e[1]||" ",this.align=e[2]||">",this.sign=e[3]||"-",this.symbol=e[4]||"",this.zero=!!e[5],this.width=e[6]&&+e[6],this.comma=!!e[7],this.precision=e[8]&&+e[8].slice(1),this.trim=!!e[9],this.type=e[10]||""}U.prototype=V.prototype,V.prototype.toString=function(){return this.fill+this.align+this.sign+this.symbol+(this.zero?"0":"")+(null==this.width?"":Math.max(1,0|this.width))+(this.comma?",":"")+(null==this.precision?"":"."+Math.max(0,0|this.precision))+(this.trim?"~":"")+this.type};var q,W,Y,X,Z=function(t,e){var n=F(t,e);if(!n)return t+"";var r=n[0],i=n[1];return i<0?"0."+new Array(-i).join("0")+r:r.length>i+1?r.slice(0,i+1)+"."+r.slice(i+1):r+new Array(i-r.length+2).join("0")},G={"%":function(t,e){return(100*t).toFixed(e)},b:function(t){return Math.round(t).toString(2)},c:function(t){return t+""},d:function(t){return Math.round(t).toString(10)},e:function(t,e){return t.toExponential(e)},f:function(t,e){return t.toFixed(e)},g:function(t,e){return t.toPrecision(e)},o:function(t){return Math.round(t).toString(8)},p:function(t,e){return Z(100*t,e)},r:Z,s:function(t,e){var n=F(t,e);if(!n)return t+"";var r=n[0],i=n[1],o=i-(q=3*Math.max(-8,Math.min(8,Math.floor(i/3))))+1,a=r.length;return o===a?r:o>a?r+new Array(o-a+1).join("0"):o>0?r.slice(0,o)+"."+r.slice(o):"0."+new Array(1-o).join("0")+F(t,Math.max(0,e+o-1))[0]},X:function(t){return Math.round(t).toString(16).toUpperCase()},x:function(t){return Math.round(t).toString(16)}},K=function(t){return t},Q=["y","z","a","f","p","n","µ","m","","k","M","G","T","P","E","Z","Y"];W=function(t){var e,n,r=t.grouping&&t.thousands?(e=t.grouping,n=t.thousands,function(t,r){for(var i=t.length,o=[],a=0,s=e[0],l=0;i>0&&s>0&&(l+s+1>r&&(s=Math.max(1,r-l)),o.push(t.substring(i-=s,i+s)),!((l+=s+1)>r));)s=e[a=(a+1)%e.length];return o.reverse().join(n)}):K,i=t.currency,o=t.decimal,a=t.numerals?function(t){return function(e){return e.replace(/[0-9]/g,(function(e){return t[+e]}))}}(t.numerals):K,s=t.percent||"%";function l(t){var e=(t=U(t)).fill,n=t.align,l=t.sign,u=t.symbol,c=t.zero,h=t.width,d=t.comma,g=t.precision,f=t.trim,p=t.type;"n"===p?(d=!0,p="g"):G[p]||(null==g&&(g=12),f=!0,p="g"),(c||"0"===e&&"="===n)&&(c=!0,e="0",n="=");var v="$"===u?i[0]:"#"===u&&/[boxX]/.test(p)?"0"+p.toLowerCase():"",m="$"===u?i[1]:/[%p]/.test(p)?s:"",y=G[p],b=/[defgprs%]/.test(p);function x(t){var i,s,u,x=v,w=m;if("c"===p)w=y(t)+w,t="";else{var A=(t=+t)<0;if(t=y(Math.abs(t),g),f&&(t=function(t){t:for(var e,n=t.length,r=1,i=-1;r<n;++r)switch(t[r]){case".":i=e=r;break;case"0":0===i&&(i=r),e=r;break;default:if(i>0){if(!+t[r])break t;i=0}}return i>0?t.slice(0,i)+t.slice(e+1):t}(t)),A&&0==+t&&(A=!1),x=(A?"("===l?l:"-":"-"===l||"("===l?"":l)+x,w=("s"===p?Q[8+q/3]:"")+w+(A&&"("===l?")":""),b)for(i=-1,s=t.length;++i<s;)if(48>(u=t.charCodeAt(i))||u>57){w=(46===u?o+t.slice(i+1):t.slice(i))+w,t=t.slice(0,i);break}}d&&!c&&(t=r(t,1/0));var C=x.length+t.length+w.length,_=C<h?new Array(h-C+1).join(e):"";switch(d&&c&&(t=r(_+t,_.length?h-w.length:1/0),_=""),n){case"<":t=x+t+w+_;break;case"=":t=x+_+t+w;break;case"^":t=_.slice(0,C=_.length>>1)+x+t+w+_.slice(C);break;default:t=_+x+t+w}return a(t)}return g=null==g?6:/[gprs]/.test(p)?Math.max(1,Math.min(21,g)):Math.max(0,Math.min(20,g)),x.toString=function(){return t+""},x}return{format:l,formatPrefix:function(t,e){var n=l(((t=U(t)).type="f",t)),r=3*Math.max(-8,Math.min(8,Math.floor(z(e)/3))),i=Math.pow(10,-r),o=Q[8+r/3];return function(t){return n(i*t)+o}}}}({decimal:".",thousands:",",grouping:[3],currency:["$",""]}),Y=W.format,X=W.formatPrefix;var J=function(t,e,n){var r,i=t[0],o=t[t.length-1],a=v(i,o,null==e?10:e);switch((n=U(null==n?",f":n)).type){case"s":var s=Math.max(Math.abs(i),Math.abs(o));return null!=n.precision||isNaN(r=function(t,e){return Math.max(0,3*Math.max(-8,Math.min(8,Math.floor(z(e)/3)))-z(Math.abs(t)))}(a,s))||(n.precision=r),X(n,s);case"":case"e":case"g":case"p":case"r":null!=n.precision||isNaN(r=function(t,e){return t=Math.abs(t),e=Math.abs(e)-t,Math.max(0,z(e)-z(t))+1}(a,Math.max(Math.abs(i),Math.abs(o))))||(n.precision=r-("e"===n.type));break;case"f":case"%":null!=n.precision||isNaN(r=function(t){return Math.max(0,-z(Math.abs(t)))}(a))||(n.precision=r-2*("%"===n.type))}return Y(n)};function tt(t){var e=t.domain;return t.ticks=function(t){var n=e();return f(n[0],n[n.length-1],null==t?10:t)},t.tickFormat=function(t,n){return J(e(),t,n)},t.nice=function(n){null==n&&(n=10);var r,i=e(),o=0,a=i.length-1,s=i[o],l=i[a];return l<s&&(r=s,s=l,l=r,r=o,o=a,a=r),(r=p(s,l,n))>0?r=p(s=Math.floor(s/r)*r,l=Math.ceil(l/r)*r,n):r<0&&(r=p(s=Math.ceil(s*r)/r,l=Math.floor(l*r)/r,n)),r>0?(i[o]=Math.floor(s/r)*r,i[a]=Math.ceil(l/r)*r,e(i)):r<0&&(i[o]=Math.ceil(s*r)/r,i[a]=Math.floor(l*r)/r,e(i)),t},t}function et(){var t=j(D,$.b);return t.copy=function(){return B(t,et())},tt(t)}function nt(){var t=[0,1];function e(t){return+t}return e.invert=e,e.domain=e.range=function(n){return arguments.length?(t=S.call(n,L),e):t.slice()},e.copy=function(){return nt().domain(t)},tt(e)}var rt=function(t,e){var n,r=0,i=(t=t.slice()).length-1,o=t[r],a=t[i];return a<o&&(n=r,r=i,i=n,n=o,o=a,a=n),t[r]=e.floor(o),t[i]=e.ceil(a),t};function it(t,e){return(e=Math.log(e/t))?function(n){return Math.log(n/t)/e}:P(e)}function ot(t,e){return t<0?function(n){return-Math.pow(-e,n)*Math.pow(-t,1-n)}:function(n){return Math.pow(e,n)*Math.pow(t,1-n)}}function at(t){return isFinite(t)?+("1e"+t):t<0?0:t}function st(t){return 10===t?at:t===Math.E?Math.exp:function(e){return Math.pow(t,e)}}function lt(t){return t===Math.E?Math.log:10===t&&Math.log10||2===t&&Math.log2||(t=Math.log(t),function(e){return Math.log(e)/t})}function ut(t){return function(e){return-t(-e)}}function ct(){var t=j(it,ot).domain([1,10]),e=t.domain,n=10,r=lt(10),i=st(10);function o(){return r=lt(n),i=st(n),e()[0]<0&&(r=ut(r),i=ut(i)),t}return t.base=function(t){return arguments.length?(n=+t,o()):n},t.domain=function(t){return arguments.length?(e(t),o()):e()},t.ticks=function(t){var o,a=e(),s=a[0],l=a[a.length-1];(o=l<s)&&(d=s,s=l,l=d);var u,c,h,d=r(s),g=r(l),p=null==t?10:+t,v=[];if(!(n%1)&&g-d<p){if(d=Math.round(d)-1,g=Math.round(g)+1,s>0){for(;d<g;++d)for(c=1,u=i(d);c<n;++c)if(!((h=u*c)<s)){if(h>l)break;v.push(h)}}else for(;d<g;++d)for(c=n-1,u=i(d);c>=1;--c)if(!((h=u*c)<s)){if(h>l)break;v.push(h)}}else v=f(d,g,Math.min(g-d,p)).map(i);return o?v.reverse():v},t.tickFormat=function(e,o){if(null==o&&(o=10===n?".0e":","),"function"!=typeof o&&(o=Y(o)),e===1/0)return o;null==e&&(e=10);var a=Math.max(1,n*e/t.ticks().length);return function(t){var e=t/i(Math.round(r(t)));return e*n<n-.5&&(e*=n),e<=a?o(t):""}},t.nice=function(){return e(rt(e(),{floor:function(t){return i(Math.floor(r(t)))},ceil:function(t){return i(Math.ceil(r(t)))}}))},t.copy=function(){return B(t,ct().base(n))},t}function ht(t,e){return t<0?-Math.pow(-t,e):Math.pow(t,e)}function dt(){var t=1,e=j((function(e,n){return(n=ht(n,t)-(e=ht(e,t)))?function(r){return(ht(r,t)-e)/n}:P(n)}),(function(e,n){return n=ht(n,t)-(e=ht(e,t)),function(r){return ht(e+n*r,1/t)}})),n=e.domain;return e.exponent=function(e){return arguments.length?(t=+e,n(n())):t},e.copy=function(){return B(e,dt().exponent(t))},tt(e)}function gt(){return dt().exponent(.5)}function ft(){var t=[],e=[],n=[];function i(){var r=0,i=Math.max(1,e.length);for(n=new Array(i-1);++r<i;)n[r-1]=m(t,r/i);return o}function o(t){if(!isNaN(t=+t))return e[s(n,t)]}return o.invertExtent=function(r){var i=e.indexOf(r);return i<0?[NaN,NaN]:[i>0?n[i-1]:t[0],i<n.length?n[i]:t[t.length-1]]},o.domain=function(e){if(!arguments.length)return t.slice();t=[];for(var n,o=0,a=e.length;o<a;++o)null==(n=e[o])||isNaN(n=+n)||t.push(n);return t.sort(r),i()},o.range=function(t){return arguments.length?(e=M.call(t),i()):e.slice()},o.quantiles=function(){return n.slice()},o.copy=function(){return ft().domain(t).range(e)},o}function pt(){var t=0,e=1,n=1,r=[.5],i=[0,1];function o(t){if(t<=t)return i[s(r,t,0,n)]}function a(){var i=-1;for(r=new Array(n);++i<n;)r[i]=((i+1)*e-(i-n)*t)/(n+1);return o}return o.domain=function(n){return arguments.length?(t=+n[0],e=+n[1],a()):[t,e]},o.range=function(t){return arguments.length?(n=(i=M.call(t)).length-1,a()):i.slice()},o.invertExtent=function(o){var a=i.indexOf(o);return a<0?[NaN,NaN]:a<1?[t,r[0]]:a>=n?[r[n-1],e]:[r[a-1],r[a]]},o.copy=function(){return pt().domain([t,e]).range(i)},tt(o)}function vt(){var t=[.5],e=[0,1],n=1;function r(r){if(r<=r)return e[s(t,r,0,n)]}return r.domain=function(i){return arguments.length?(t=M.call(i),n=Math.min(t.length,e.length-1),r):t.slice()},r.range=function(i){return arguments.length?(e=M.call(i),n=Math.min(t.length,e.length-1),r):e.slice()},r.invertExtent=function(n){var r=e.indexOf(n);return[t[r-1],t[r]]},r.copy=function(){return vt().domain(t).range(e)},r}var mt=new Date,yt=new Date;function bt(t,e,n,r){function i(e){return t(e=new Date(+e)),e}return i.floor=i,i.ceil=function(n){return t(n=new Date(n-1)),e(n,1),t(n),n},i.round=function(t){var e=i(t),n=i.ceil(t);return t-e<n-t?e:n},i.offset=function(t,n){return e(t=new Date(+t),null==n?1:Math.floor(n)),t},i.range=function(n,r,o){var a,s=[];if(n=i.ceil(n),o=null==o?1:Math.floor(o),!(n<r&&o>0))return s;do{s.push(a=new Date(+n)),e(n,o),t(n)}while(a<n&&n<r);return s},i.filter=function(n){return bt((function(e){if(e>=e)for(;t(e),!n(e);)e.setTime(e-1)}),(function(t,r){if(t>=t)if(r<0)for(;++r<=0;)for(;e(t,-1),!n(t););else for(;--r>=0;)for(;e(t,1),!n(t););}))},n&&(i.count=function(e,r){return mt.setTime(+e),yt.setTime(+r),t(mt),t(yt),Math.floor(n(mt,yt))},i.every=function(t){return t=Math.floor(t),isFinite(t)&&t>0?t>1?i.filter(r?function(e){return r(e)%t==0}:function(e){return i.count(0,e)%t==0}):i:null}),i}var xt=bt((function(){}),(function(t,e){t.setTime(+t+e)}),(function(t,e){return e-t}));xt.every=function(t){return t=Math.floor(t),isFinite(t)&&t>0?t>1?bt((function(e){e.setTime(Math.floor(e/t)*t)}),(function(e,n){e.setTime(+e+n*t)}),(function(e,n){return(n-e)/t})):xt:null};var wt=xt,At=(xt.range,bt((function(t){t.setTime(1e3*Math.floor(t/1e3))}),(function(t,e){t.setTime(+t+1e3*e)}),(function(t,e){return(e-t)/1e3}),(function(t){return t.getUTCSeconds()}))),Ct=At,_t=(At.range,bt((function(t){t.setTime(6e4*Math.floor(t/6e4))}),(function(t,e){t.setTime(+t+6e4*e)}),(function(t,e){return(e-t)/6e4}),(function(t){return t.getMinutes()}))),St=_t,Mt=(_t.range,bt((function(t){var e=6e4*t.getTimezoneOffset()%36e5;e<0&&(e+=36e5),t.setTime(36e5*Math.floor((+t-e)/36e5)+e)}),(function(t,e){t.setTime(+t+36e5*e)}),(function(t,e){return(e-t)/36e5}),(function(t){return t.getHours()}))),Tt=Mt,kt=(Mt.range,bt((function(t){t.setHours(0,0,0,0)}),(function(t,e){t.setDate(t.getDate()+e)}),(function(t,e){return(e-t-6e4*(e.getTimezoneOffset()-t.getTimezoneOffset()))/864e5}),(function(t){return t.getDate()-1}))),Rt=kt;kt.range;function Nt(t){return bt((function(e){e.setDate(e.getDate()-(e.getDay()+7-t)%7),e.setHours(0,0,0,0)}),(function(t,e){t.setDate(t.getDate()+7*e)}),(function(t,e){return(e-t-6e4*(e.getTimezoneOffset()-t.getTimezoneOffset()))/6048e5}))}var $t=Nt(0),Pt=Nt(1),Lt=Nt(2),It=Nt(3),Dt=Nt(4),Ot=Nt(5),Et=Nt(6),Bt=($t.range,Pt.range,Lt.range,It.range,Dt.range,Ot.range,Et.range,bt((function(t){t.setDate(1),t.setHours(0,0,0,0)}),(function(t,e){t.setMonth(t.getMonth()+e)}),(function(t,e){return e.getMonth()-t.getMonth()+12*(e.getFullYear()-t.getFullYear())}),(function(t){return t.getMonth()}))),jt=Bt,Ft=(Bt.range,bt((function(t){t.setMonth(0,1),t.setHours(0,0,0,0)}),(function(t,e){t.setFullYear(t.getFullYear()+e)}),(function(t,e){return e.getFullYear()-t.getFullYear()}),(function(t){return t.getFullYear()})));Ft.every=function(t){return isFinite(t=Math.floor(t))&&t>0?bt((function(e){e.setFullYear(Math.floor(e.getFullYear()/t)*t),e.setMonth(0,1),e.setHours(0,0,0,0)}),(function(e,n){e.setFullYear(e.getFullYear()+n*t)})):null};var zt=Ft,Ht=(Ft.range,bt((function(t){t.setUTCSeconds(0,0)}),(function(t,e){t.setTime(+t+6e4*e)}),(function(t,e){return(e-t)/6e4}),(function(t){return t.getUTCMinutes()}))),Ut=Ht,Vt=(Ht.range,bt((function(t){t.setUTCMinutes(0,0,0)}),(function(t,e){t.setTime(+t+36e5*e)}),(function(t,e){return(e-t)/36e5}),(function(t){return t.getUTCHours()}))),qt=Vt,Wt=(Vt.range,bt((function(t){t.setUTCHours(0,0,0,0)}),(function(t,e){t.setUTCDate(t.getUTCDate()+e)}),(function(t,e){return(e-t)/864e5}),(function(t){return t.getUTCDate()-1}))),Yt=Wt;Wt.range;function Xt(t){return bt((function(e){e.setUTCDate(e.getUTCDate()-(e.getUTCDay()+7-t)%7),e.setUTCHours(0,0,0,0)}),(function(t,e){t.setUTCDate(t.getUTCDate()+7*e)}),(function(t,e){return(e-t)/6048e5}))}var Zt=Xt(0),Gt=Xt(1),Kt=Xt(2),Qt=Xt(3),Jt=Xt(4),te=Xt(5),ee=Xt(6),ne=(Zt.range,Gt.range,Kt.range,Qt.range,Jt.range,te.range,ee.range,bt((function(t){t.setUTCDate(1),t.setUTCHours(0,0,0,0)}),(function(t,e){t.setUTCMonth(t.getUTCMonth()+e)}),(function(t,e){return e.getUTCMonth()-t.getUTCMonth()+12*(e.getUTCFullYear()-t.getUTCFullYear())}),(function(t){return t.getUTCMonth()}))),re=ne,ie=(ne.range,bt((function(t){t.setUTCMonth(0,1),t.setUTCHours(0,0,0,0)}),(function(t,e){t.setUTCFullYear(t.getUTCFullYear()+e)}),(function(t,e){return e.getUTCFullYear()-t.getUTCFullYear()}),(function(t){return t.getUTCFullYear()})));ie.every=function(t){return isFinite(t=Math.floor(t))&&t>0?bt((function(e){e.setUTCFullYear(Math.floor(e.getUTCFullYear()/t)*t),e.setUTCMonth(0,1),e.setUTCHours(0,0,0,0)}),(function(e,n){e.setUTCFullYear(e.getUTCFullYear()+n*t)})):null};var oe=ie;ie.range;function ae(t){if(0<=t.y&&t.y<100){var e=new Date(-1,t.m,t.d,t.H,t.M,t.S,t.L);return e.setFullYear(t.y),e}return new Date(t.y,t.m,t.d,t.H,t.M,t.S,t.L)}function se(t){if(0<=t.y&&t.y<100){var e=new Date(Date.UTC(-1,t.m,t.d,t.H,t.M,t.S,t.L));return e.setUTCFullYear(t.y),e}return new Date(Date.UTC(t.y,t.m,t.d,t.H,t.M,t.S,t.L))}function le(t){return{y:t,m:0,d:1,H:0,M:0,S:0,L:0}}var ue,ce,he,de,ge={"-":"",_:" ",0:"0"},fe=/^\s*\d+/,pe=/^%/,ve=/[\\^$*+?|[\]().{}]/g;function me(t,e,n){var r=t<0?"-":"",i=(r?-t:t)+"",o=i.length;return r+(o<n?new Array(n-o+1).join(e)+i:i)}function ye(t){return t.replace(ve,"\\$&")}function be(t){return new RegExp("^(?:"+t.map(ye).join("|")+")","i")}function xe(t){for(var e={},n=-1,r=t.length;++n<r;)e[t[n].toLowerCase()]=n;return e}function we(t,e,n){var r=fe.exec(e.slice(n,n+1));return r?(t.w=+r[0],n+r[0].length):-1}function Ae(t,e,n){var r=fe.exec(e.slice(n,n+1));return r?(t.u=+r[0],n+r[0].length):-1}function Ce(t,e,n){var r=fe.exec(e.slice(n,n+2));return r?(t.U=+r[0],n+r[0].length):-1}function _e(t,e,n){var r=fe.exec(e.slice(n,n+2));return r?(t.V=+r[0],n+r[0].length):-1}function Se(t,e,n){var r=fe.exec(e.slice(n,n+2));return r?(t.W=+r[0],n+r[0].length):-1}function Me(t,e,n){var r=fe.exec(e.slice(n,n+4));return r?(t.y=+r[0],n+r[0].length):-1}function Te(t,e,n){var r=fe.exec(e.slice(n,n+2));return r?(t.y=+r[0]+(+r[0]>68?1900:2e3),n+r[0].length):-1}function ke(t,e,n){var r=/^(Z)|([+-]\d\d)(?::?(\d\d))?/.exec(e.slice(n,n+6));return r?(t.Z=r[1]?0:-(r[2]+(r[3]||"00")),n+r[0].length):-1}function Re(t,e,n){var r=fe.exec(e.slice(n,n+2));return r?(t.m=r[0]-1,n+r[0].length):-1}function Ne(t,e,n){var r=fe.exec(e.slice(n,n+2));return r?(t.d=+r[0],n+r[0].length):-1}function $e(t,e,n){var r=fe.exec(e.slice(n,n+3));return r?(t.m=0,t.d=+r[0],n+r[0].length):-1}function Pe(t,e,n){var r=fe.exec(e.slice(n,n+2));return r?(t.H=+r[0],n+r[0].length):-1}function Le(t,e,n){var r=fe.exec(e.slice(n,n+2));return r?(t.M=+r[0],n+r[0].length):-1}function Ie(t,e,n){var r=fe.exec(e.slice(n,n+2));return r?(t.S=+r[0],n+r[0].length):-1}function De(t,e,n){var r=fe.exec(e.slice(n,n+3));return r?(t.L=+r[0],n+r[0].length):-1}function Oe(t,e,n){var r=fe.exec(e.slice(n,n+6));return r?(t.L=Math.floor(r[0]/1e3),n+r[0].length):-1}function Ee(t,e,n){var r=pe.exec(e.slice(n,n+1));return r?n+r[0].length:-1}function Be(t,e,n){var r=fe.exec(e.slice(n));return r?(t.Q=+r[0],n+r[0].length):-1}function je(t,e,n){var r=fe.exec(e.slice(n));return r?(t.Q=1e3*+r[0],n+r[0].length):-1}function Fe(t,e){return me(t.getDate(),e,2)}function ze(t,e){return me(t.getHours(),e,2)}function He(t,e){return me(t.getHours()%12||12,e,2)}function Ue(t,e){return me(1+Rt.count(zt(t),t),e,3)}function Ve(t,e){return me(t.getMilliseconds(),e,3)}function qe(t,e){return Ve(t,e)+"000"}function We(t,e){return me(t.getMonth()+1,e,2)}function Ye(t,e){return me(t.getMinutes(),e,2)}function Xe(t,e){return me(t.getSeconds(),e,2)}function Ze(t){var e=t.getDay();return 0===e?7:e}function Ge(t,e){return me($t.count(zt(t),t),e,2)}function Ke(t,e){var n=t.getDay();return t=n>=4||0===n?Dt(t):Dt.ceil(t),me(Dt.count(zt(t),t)+(4===zt(t).getDay()),e,2)}function Qe(t){return t.getDay()}function Je(t,e){return me(Pt.count(zt(t),t),e,2)}function tn(t,e){return me(t.getFullYear()%100,e,2)}function en(t,e){return me(t.getFullYear()%1e4,e,4)}function nn(t){var e=t.getTimezoneOffset();return(e>0?"-":(e*=-1,"+"))+me(e/60|0,"0",2)+me(e%60,"0",2)}function rn(t,e){return me(t.getUTCDate(),e,2)}function on(t,e){return me(t.getUTCHours(),e,2)}function an(t,e){return me(t.getUTCHours()%12||12,e,2)}function sn(t,e){return me(1+Yt.count(oe(t),t),e,3)}function ln(t,e){return me(t.getUTCMilliseconds(),e,3)}function un(t,e){return ln(t,e)+"000"}function cn(t,e){return me(t.getUTCMonth()+1,e,2)}function hn(t,e){return me(t.getUTCMinutes(),e,2)}function dn(t,e){return me(t.getUTCSeconds(),e,2)}function gn(t){var e=t.getUTCDay();return 0===e?7:e}function fn(t,e){return me(Zt.count(oe(t),t),e,2)}function pn(t,e){var n=t.getUTCDay();return t=n>=4||0===n?Jt(t):Jt.ceil(t),me(Jt.count(oe(t),t)+(4===oe(t).getUTCDay()),e,2)}function vn(t){return t.getUTCDay()}function mn(t,e){return me(Gt.count(oe(t),t),e,2)}function yn(t,e){return me(t.getUTCFullYear()%100,e,2)}function bn(t,e){return me(t.getUTCFullYear()%1e4,e,4)}function xn(){return"+0000"}function wn(){return"%"}function An(t){return+t}function Cn(t){return Math.floor(+t/1e3)}!function(t){ue=function(t){var e=t.dateTime,n=t.date,r=t.time,i=t.periods,o=t.days,a=t.shortDays,s=t.months,l=t.shortMonths,u=be(i),c=xe(i),h=be(o),d=xe(o),g=be(a),f=xe(a),p=be(s),v=xe(s),m=be(l),y=xe(l),b={a:function(t){return a[t.getDay()]},A:function(t){return o[t.getDay()]},b:function(t){return l[t.getMonth()]},B:function(t){return s[t.getMonth()]},c:null,d:Fe,e:Fe,f:qe,H:ze,I:He,j:Ue,L:Ve,m:We,M:Ye,p:function(t){return i[+(t.getHours()>=12)]},Q:An,s:Cn,S:Xe,u:Ze,U:Ge,V:Ke,w:Qe,W:Je,x:null,X:null,y:tn,Y:en,Z:nn,"%":wn},x={a:function(t){return a[t.getUTCDay()]},A:function(t){return o[t.getUTCDay()]},b:function(t){return l[t.getUTCMonth()]},B:function(t){return s[t.getUTCMonth()]},c:null,d:rn,e:rn,f:un,H:on,I:an,j:sn,L:ln,m:cn,M:hn,p:function(t){return i[+(t.getUTCHours()>=12)]},Q:An,s:Cn,S:dn,u:gn,U:fn,V:pn,w:vn,W:mn,x:null,X:null,y:yn,Y:bn,Z:xn,"%":wn},w={a:function(t,e,n){var r=g.exec(e.slice(n));return r?(t.w=f[r[0].toLowerCase()],n+r[0].length):-1},A:function(t,e,n){var r=h.exec(e.slice(n));return r?(t.w=d[r[0].toLowerCase()],n+r[0].length):-1},b:function(t,e,n){var r=m.exec(e.slice(n));return r?(t.m=y[r[0].toLowerCase()],n+r[0].length):-1},B:function(t,e,n){var r=p.exec(e.slice(n));return r?(t.m=v[r[0].toLowerCase()],n+r[0].length):-1},c:function(t,n,r){return _(t,e,n,r)},d:Ne,e:Ne,f:Oe,H:Pe,I:Pe,j:$e,L:De,m:Re,M:Le,p:function(t,e,n){var r=u.exec(e.slice(n));return r?(t.p=c[r[0].toLowerCase()],n+r[0].length):-1},Q:Be,s:je,S:Ie,u:Ae,U:Ce,V:_e,w:we,W:Se,x:function(t,e,r){return _(t,n,e,r)},X:function(t,e,n){return _(t,r,e,n)},y:Te,Y:Me,Z:ke,"%":Ee};function A(t,e){return function(n){var r,i,o,a=[],s=-1,l=0,u=t.length;for(n instanceof Date||(n=new Date(+n));++s<u;)37===t.charCodeAt(s)&&(a.push(t.slice(l,s)),null!=(i=ge[r=t.charAt(++s)])?r=t.charAt(++s):i="e"===r?" ":"0",(o=e[r])&&(r=o(n,i)),a.push(r),l=s+1);return a.push(t.slice(l,s)),a.join("")}}function C(t,e){return function(n){var r,i,o=le(1900);if(_(o,t,n+="",0)!=n.length)return null;if("Q"in o)return new Date(o.Q);if("p"in o&&(o.H=o.H%12+12*o.p),"V"in o){if(o.V<1||o.V>53)return null;"w"in o||(o.w=1),"Z"in o?(i=(r=se(le(o.y))).getUTCDay(),r=i>4||0===i?Gt.ceil(r):Gt(r),r=Yt.offset(r,7*(o.V-1)),o.y=r.getUTCFullYear(),o.m=r.getUTCMonth(),o.d=r.getUTCDate()+(o.w+6)%7):(i=(r=e(le(o.y))).getDay(),r=i>4||0===i?Pt.ceil(r):Pt(r),r=Rt.offset(r,7*(o.V-1)),o.y=r.getFullYear(),o.m=r.getMonth(),o.d=r.getDate()+(o.w+6)%7)}else("W"in o||"U"in o)&&("w"in o||(o.w="u"in o?o.u%7:"W"in o?1:0),i="Z"in o?se(le(o.y)).getUTCDay():e(le(o.y)).getDay(),o.m=0,o.d="W"in o?(o.w+6)%7+7*o.W-(i+5)%7:o.w+7*o.U-(i+6)%7);return"Z"in o?(o.H+=o.Z/100|0,o.M+=o.Z%100,se(o)):e(o)}}function _(t,e,n,r){for(var i,o,a=0,s=e.length,l=n.length;a<s;){if(r>=l)return-1;if(37===(i=e.charCodeAt(a++))){if(i=e.charAt(a++),!(o=w[i in ge?e.charAt(a++):i])||(r=o(t,n,r))<0)return-1}else if(i!=n.charCodeAt(r++))return-1}return r}return b.x=A(n,b),b.X=A(r,b),b.c=A(e,b),x.x=A(n,x),x.X=A(r,x),x.c=A(e,x),{format:function(t){var e=A(t+="",b);return e.toString=function(){return t},e},parse:function(t){var e=C(t+="",ae);return e.toString=function(){return t},e},utcFormat:function(t){var e=A(t+="",x);return e.toString=function(){return t},e},utcParse:function(t){var e=C(t,se);return e.toString=function(){return t},e}}}(t),ce=ue.format,ue.parse,he=ue.utcFormat,de=ue.utcParse}({dateTime:"%x, %X",date:"%-m/%-d/%Y",time:"%-I:%M:%S %p",periods:["AM","PM"],days:["Sunday","Monday","Tuesday","Wednesday","Thursday","Friday","Saturday"],shortDays:["Sun","Mon","Tue","Wed","Thu","Fri","Sat"],months:["January","February","March","April","May","June","July","August","September","October","November","December"],shortMonths:["Jan","Feb","Mar","Apr","May","Jun","Jul","Aug","Sep","Oct","Nov","Dec"]});Date.prototype.toISOString||he("%Y-%m-%dT%H:%M:%S.%LZ");+new Date("2000-01-01T00:00:00.000Z")||de("%Y-%m-%dT%H:%M:%S.%LZ");function _n(t){return new Date(t)}function Sn(t){return t instanceof Date?+t:+new Date(+t)}function Mn(t,e,n,r,o,a,s,l,u){var c=j(D,$.b),h=c.invert,d=c.domain,g=u(".%L"),f=u(":%S"),p=u("%I:%M"),m=u("%I %p"),y=u("%a %d"),b=u("%b %d"),x=u("%B"),w=u("%Y"),A=[[s,1,1e3],[s,5,5e3],[s,15,15e3],[s,30,3e4],[a,1,6e4],[a,5,3e5],[a,15,9e5],[a,30,18e5],[o,1,36e5],[o,3,108e5],[o,6,216e5],[o,12,432e5],[r,1,864e5],[r,2,1728e5],[n,1,6048e5],[e,1,2592e6],[e,3,7776e6],[t,1,31536e6]];function C(i){return(s(i)<i?g:a(i)<i?f:o(i)<i?p:r(i)<i?m:e(i)<i?n(i)<i?y:b:t(i)<i?x:w)(i)}function _(e,n,r,o){if(null==e&&(e=10),"number"==typeof e){var a=Math.abs(r-n)/e,s=i((function(t){return t[2]})).right(A,a);s===A.length?(o=v(n/31536e6,r/31536e6,e),e=t):s?(o=(s=A[a/A[s-1][2]<A[s][2]/a?s-1:s])[1],e=s[0]):(o=Math.max(v(n,r,e),1),e=l)}return null==o?e:e.every(o)}return c.invert=function(t){return new Date(h(t))},c.domain=function(t){return arguments.length?d(S.call(t,Sn)):d().map(_n)},c.ticks=function(t,e){var n,r=d(),i=r[0],o=r[r.length-1],a=o<i;return a&&(n=i,i=o,o=n),n=(n=_(t,i,o,e))?n.range(i,o+1):[],a?n.reverse():n},c.tickFormat=function(t,e){return null==e?C:u(e)},c.nice=function(t,e){var n=d();return(t=_(t,n[0],n[n.length-1],e))?d(rt(n,t)):c},c.copy=function(){return B(c,Mn(t,e,n,r,o,a,s,l,u))},c}var Tn=function(){return Mn(zt,jt,$t,Rt,Tt,St,Ct,wt,ce).domain([new Date(2e3,0,1),new Date(2e3,0,2)])},kn=function(){return Mn(oe,re,Zt,Yt,qt,Ut,Ct,wt,he).domain([Date.UTC(2e3,0,1),Date.UTC(2e3,0,2)])};function Rn(t){var e=0,n=1,r=1,i=!1;function o(n){var o=(n-e)*r;return t(i?Math.max(0,Math.min(1,o)):o)}return o.domain=function(t){return arguments.length?(e=+t[0],n=+t[1],r=e===n?0:1/(n-e),o):[e,n]},o.clamp=function(t){return arguments.length?(i=!!t,o):i},o.interpolator=function(e){return arguments.length?(t=e,o):t},o.copy=function(){return Rn(t).domain([e,n]).clamp(i)},tt(o)}function Nn(t){var e=0,n=.5,r=1,i=1,o=1,a=!1;function s(e){var r=.5+((e=+e)-n)*(e<n?i:o);return t(a?Math.max(0,Math.min(1,r)):r)}return s.domain=function(t){return arguments.length?(e=+t[0],n=+t[1],r=+t[2],i=e===n?0:.5/(n-e),o=n===r?0:.5/(r-n),s):[e,n,r]},s.clamp=function(t){return arguments.length?(a=!!t,s):a},s.interpolator=function(e){return arguments.length?(t=e,s):t},s.copy=function(){return Nn(t).domain([e,n,r]).clamp(a)},tt(s)}},function(t,e,n){"use strict";Object.defineProperty(e,"__esModule",{value:!0}),e.clipboardCopyConstruct=e.copyToClipboard=void 0,e.copyToClipboard=function(){var t=document.createElement("textarea");t.style.position="fixed",t.style.top="0",t.style.left="0",t.style.width="1px",t.style.height="1px",t.value=$(this).attr("data-seq")||"",document.body.appendChild(t),t.select();try{document.execCommand("copy"),alert("Sequence copied to clipboard")}finally{document.body.removeChild(t)}},e.clipboardCopyConstruct=function(t){return'<span class="clipboard-copy" data-seq="'+t+'">Copy to clipboard</span>'}},function(t,e,n){"use strict";Object.defineProperty(e,"__esModule",{value:!0}),e.tag_to_id=e.drawRegion=e.locusToFullId=e.selectOrfsByLoci=void 0;var r=n(34),i=n(33),o=n(6),a=n(0);n(32);var s=n(7),l=n(13),u=100;function c(t){return I?"u"+(O-1)+"-region"+I.idx+"-"+p(t):""+p(t)}function h(t){var e=L+14+P,n=L+14+u-P,r=L+14+u/2;if(1===t.strand){var i=D(t.start),o=Math.max(D(t.end)-2*P,i);return i+","+e+" "+o+","+e+" "+D(t.end)+","+r+" "+o+","+n+" "+i+","+n}if(-1===t.strand){var a=D(t.start),s=D(t.end),l=Math.min(D(t.start)+2*P,s);return a+","+r+" "+l+","+e+" "+s+","+e+" "+s+","+n+" "+l+","+n}return t.start+","+e+" "+t.end+","+e+" "+t.end+","+n+" "+t.start+","+n}function d(t,e,n,r){var i=n+14+e,o=n+28+e-r,a=Math.floor(D((t.start+t.end)/2));return a+","+i+" "+(a-5)+","+o+" "+(a+5)+","+o}function g(t,e,n){return Math.max(t,Math.min(e,n))}function f(t,e){return"biosynthetic"!==t.type&&"biosynthetic"!==e.type||"biosynthetic"===t.type&&"biosynthetic"===e.type?t.start-e.start:"biosynthetic"===t.type?1:-1}function p(t){return t.replace(/(:|\.)/g,"-").replace(/-svgeneorf/g,"_orf")}function v(t){t[0].classList.contains("svgene-selected-orf")?(_(t),0===$(".svgene-selected-orf").length&&C()):(0===$(".svgene-selected-orf").length&&_(),C(t)),x(t,"cds")}function m(t){$(".legend-selected").removeClass("legend-selected"),t.ctrlKey||t.metaKey?v($(this)):y($(this))}function y(t){if(I){var e=a.selectAll(t.toArray()),n=e.datum(),r=$(".focus-panel-content-"+I.anchor);r.html(n.description).find(".collapser").click(l.toggleCollapserHandler),$(".clipboard-copy",r).off("click").click(s.copyToClipboard),e.classed("svgene-selected-orf")&&1===$(".svgene-orf.svgene-selected-orf").length?C():(_(),C(t)),x(t,"cds")}}function b(t){l.toggleCollapser($(".collapser-level-candidate.expanded"));var e=t.product.split(":")[0].split(" ")[1];l.toggleCollapser($(".collapser-level-candidate.collapser-target-CC"+e))}function x(t,e){var n=a.selectAll(t.toArray()).datum();l.toggleCollapser($(".collapser-target-"+p(n.locus_tag)+".collapser-level-"+e))}function w(t,e){$(".legend-selected").removeClass("legend-selected"),$(".svgene-orf").each((function(n){var r=a.selectAll($(this).toArray()).datum();t<=r.start&&r.end<=e&&(C($(this)),x($(this),"candidate"),x($(this),"protocluster"))}))}function A(t,e){var n=e?"1":"0.5";a.selectAll(t.toArray()).attr("opacity",n).classed("svgene-selected-orf",e).each((function(t){a.selectAll("#svgene-minimap-orf-"+p(t.locus_tag)).attr("opacity",n).classed("svgene-selected-orf",e);var r=c(t.locus_tag);e?($("#"+r+"-domains").show(),$("#"+r+"-pfam-domains").show()):($("#"+r+"-domains").hide(),$("#"+r+"-pfam-domains").hide())}))}function C(t){void 0===t&&(t=$(".svgene-orf, .svgene-minimap-orf")),A(t,!0)}function _(t){void 0===t&&(t=$(".svgene-selected-orf")),A(t,!1)}function S(t){var e=$(this).attr("data-id");if(void 0!==e){var n=".contains-tta-codon";if(n="legend-tta-codon"===e?".contains-tta-codon":"legend-resistance"===e?".svgene-resistance-orf":"."+e.replace("legend-","svgene-"),$(this).hasClass("legend-selected"))return $(this).removeClass("legend-selected"),_($(n)),void(0===$(".legend-selected").length&&C());(t.ctrlKey||t.metaKey)&&0!==$(".legend-selected").length||($(".legend-selected").removeClass("legend-selected"),_()),$(this).addClass("legend-selected"),C($(n))}}function M(t){var e=-1,n=-1;$(".svgene-selected-orf").each((function(t){var r=a.selectAll($(this).toArray()).datum();e=-1===e?Math.min(r.start,r.end):Math.min(e,r.start,r.end),n=-1===n?Math.max(r.start,r.end):Math.max(n,r.start,r.end)})),-1!==e&&-1!==n&&T(e,n)}function T(t,e,n){if(I){var r=0;void 0===n&&(n=!1,r=500),t-=1,e+=1,D.domain([t,e]);var i=t+(e-t)/2;if(a.selectAll(".svgene-orf,.svgene-orf-bg").transition().duration(r).attr("points",(function(t){return h(t)})),a.selectAll(".svgene-locustag").transition().duration(r).attr("x",(function(t){return t.start<i?D(t.start):D(t.end)})).attr("text-anchor",(function(t){return t.start<i?"start":"end"})),a.selectAll(".cluster-background").transition().duration(r).attr("width",(function(t){return D(t.neighbouring_end)-D(t.neighbouring_start)})).attr("x",(function(t){return D(t.neighbouring_start)})),a.selectAll(".cluster-core").transition().duration(r).attr("width",(function(t){return D(t.end)-D(t.start)})).attr("x",(function(t){return D(t.start)})),a.selectAll(".cluster-line").transition().duration(r).attr("x1",(function(t){return D(t.neighbouring_start)})).attr("x2",(function(t){return D(t.neighbouring_end)})),a.selectAll(".clusterlabel").transition().duration(r).attr("x",(function(n){return D((Math.max(n.start,t+1)+Math.min(n.end,e-1))/2)})),a.selectAll(".svgene-resistance").filter((function(t){return t&&!0===t.resistance})).attr("x",(function(t){return D(t.start)})).attr("width",(function(t){return D(t.end)-D(t.start)})),a.selectAll(".svgene-tta-codon").filter((function(t){return void 0!==t})).transition().duration(r).attr("points",(function(t){return d(t,u,L,P)})),null!==R&&a.select(".svgene-axis").transition().duration(r).call(R),!n){0===$(".svgene-selected-orf").length&&C();var o=N(t),s=N(e);a.select("#svgene-minimap-grabber-a-"+I.anchor).transition().duration(r).attr("x",o),a.select("#svgene-minimap-grabber-b-"+I.anchor).transition().duration(r).attr("x",s),a.select("#svgene-minimap-window-"+I.anchor).transition().duration(r).attr("x",o).attr("width",s-o)}}}function k(t){null!==I&&($(".legend-selected").removeClass("legend-selected"),C(),T(I.start,I.end),l.toggleCollapser($(".collapser-level-cds.expanded")))}e.selectOrfsByLoci=function(t,e){if(I&&!(t.length<1)){for(var n=$("#"+c(t[0])+"-svgeneorf"),r=0,i=t.slice(1);r<i.length;r++){var o=i[r];n=n.add("#"+c(o)+"-svgeneorf")}e?v(n):y(n)}},e.locusToFullId=c,e.drawRegion=function(t,e,n,s,l){I&&(a.select("#"+I.anchor+"-svg").selectAll("*").remove(),$(".legend-selected").removeClass("legend-selected"));var c=e;if(null!==(I=e)){void 0===s&&(s=c.start),void 0===l&&(l=c.end);for(var v=!1,y=0,x=c.orfs;y<x.length;y++){if(x[y].resistance){v=!0;break}}var A=n+28+12*Math.max.apply(Math,c.clusters.map((function(t){return t.height})))+40+20+(v||c.ttaCodons.length>0?Math.floor(2*n/3):0),C=(c.idx,a.select("#"+t)),E=$("#"+t).parent().width()||700;C.selectAll("svg").remove(),C.selectAll("div").remove();var B=C.append("svg").attr("height",A).attr("width","100%").attr("viewbox","-1 0 "+E+" "+A),j=[],F=[],z=[];j.push.apply(j,c.orfs.sort(f)),F.push.apply(F,c.clusters?c.clusters:[]),z.push.apply(z,c.ttaCodons?c.ttaCodons:[]);var H=O++;P=n/10,D=o.scaleLinear().domain([s,l]).range([0,E]),function(t,e,n,r,i,o,s,l,c,g){if(null!==I){var f=I.idx;L=12*Math.max.apply(Math,n.map((function(t){return t.height})))+14,u=o,t.append("line").attr("x1",0).attr("y1",L+14+o/2).attr("x2",s).attr("y2",L+14+o/2).attr("class","centerline");var v=t.selectAll("g.cluster-bar-group").data(n).enter().append("g").attr("class",(function(t){return"candidatecluster"===t.kind?"candidate-"+t.product.split(" ")[2].replace("chemical_",""):"subregion"===t.kind&&0===t.prefix.length?"svgene-border-"+t.tool:""})).on("click",(function(t){$(".svgene-selected-orf").length!==e.length&&(a.event.ctrlKey||a.event.metaKey)||_(),w(t.neighbouring_start,t.neighbouring_end),b(t)})).on("dblclick",(function(t){w(t.neighbouring_start,t.neighbouring_end),b(t),T(t.neighbouring_start,t.neighbouring_end)}));v.append("rect").attr("width",(function(t){return D(t.neighbouring_end)-D(t.neighbouring_start)})).attr("height",10).attr("x",(function(t){return D(t.neighbouring_start)})).attr("y",(function(t){return 12*t.height+l})).attr("opacity","0.5").attr("class",(function(t){return"subregion"===t.kind?"cluster-background":"cluster-background "+t.product})).style("stroke-width","0"),v.append("line").attr("x1",(function(t){return D(t.neighbouring_start)})).attr("y1",(function(t){return 12*t.height+l+5})).attr("x2",(function(t){return D(t.neighbouring_end)})).attr("y2",(function(t){return 12*t.height+l+5})).attr("class","cluster-line"),v.append("rect").attr("width",(function(t){return D(t.end)-D(t.start)})).attr("height",10).attr("x",(function(t){return D(t.start)})).attr("y",(function(t){return 12*t.height+l})).attr("class",(function(t){return"subregion"===t.kind?t.prefix?"cluster-core":"cluster-core svgene-border-"+t.tool:"cluster-core "+t.product})).style("stroke","black"),v.append("text").attr("x",(function(t){return D((t.start+t.end)/2)})).attr("y",(function(t){return"protocluster"===t.kind?12*(t.height-1)-2+10+l:12*t.height-2+10+l})).style("font-size","xx-small").attr("class","clusterlabel").attr("text-anchor","middle").style("pointer-events","none").text((function(t){return t.prefix+t.product.replace("_"," ")}));var m=t.selectAll("g.orf-group").data(e).enter().append("g").attr("class","orf-group");m.append("polygon").attr("points",(function(t){return h(t)})).attr("class","svgene-orf-bg").attr("id",(function(t){return"u"+i+"-region"+f+"-"+p(t.locus_tag)+"-svgeneorf-bg"})).style("fill","white"),m.append("polygon").attr("points",(function(t){return h(t)})).attr("class",(function(t){return"svgene-type-"+t.type+" svgene-orf svgene-selected-orf"})).attr("id",(function(t){return"u"+i+"-region"+f+"-"+p(t.locus_tag)+"-svgeneorf"})).attr("opacity","1");for(var y=L+14+o+2,x=(t.selectAll("rect.svgene-resist").data(e.filter((function(t){return t.resistance}))).enter().append("rect").attr("class","svgene-resistance").attr("width",(function(t){return D(t.end)-D(t.start)})).attr("height",7).attr("x",(function(t){return D(t.start)})).attr("y",y),0),A=e.filter((function(t){return t.resistance}));x<A.length;x++){var C=A[x],S="#u"+i+"-region"+f+"-"+p(C.locus_tag)+"-svgeneorf";a.select(S).classed("svgene-resistance-orf",!0)}t.selectAll("polyline.svgene-tta-codon").data(r).enter().append("polyline").attr("points",(function(t){return d(t,o,L,l)})).attr("class","svgene-tta-codon"),t.selectAll("text.svgene-locustag").data(e).enter().append("text").attr("x",(function(t){return D(t.start)<s/2?D(t.start):D(t.end)})).attr("text-anchor",(function(t){return D(t.start)<s/2?"start":"end"})).attr("y",L+14).attr("class","svgene-locustag").attr("id",(function(t){return"u"+i+"-region"+f+"-"+p(t.locus_tag)+"-label"})).text((function(t){return t.locus_tag}));for(var M=0,k=r;M<k.length;M++)for(var R=0,N=k[M].containedBy;R<N.length;R++){var P=N[R];S="#u"+i+"-region"+f+"-"+p(P)+"-svgeneorf";(C=a.select(S)).classed("contains-tta-codon",!0)}}}(B,j,F,z,H,n,E,P),void 0!==c.label&&B.append("text").text(c.label).attr("class","svgene-regionlabel").attr("x",(function(){var t=this.getComputedTextLength();return E-t-5})).attr("y",14).attr("font-size",14),R=r.axisBottom(D),B.append("g").attr("class","svgene-axis").attr("transform","translate(0,"+(A-40-20)+")").call(R),function(t,e,n,r,s,l,u){if(!I)return;N=o.scaleLinear().domain([e,n]).range([Math.floor(.25*u),Math.floor(.75*u)]).nice();var c=s/4;t.append("g").attr("class","svgene-minimap"),t.append("line").attr("x1",N(e)).attr("y1",r).attr("x2",N(n)).attr("y2",r).attr("class","centerline");t.selectAll("rect.svgene-minimap-orf").data(l).enter().append("rect").attr("width",(function(t){return N(t.end)-N(t.start)})).attr("height",c).attr("x",(function(t){return N(t.start)})).attr("y",r-c/2).attr("class",(function(t){return"svgene-type-"+t.type+" svgene-selected-orf svgene-minimap-orf"})).attr("id",(function(t){return"svgene-minimap-orf-"+p(t.locus_tag)}));var h=N(e),d=N(n),f=N(Math.min(n,e+1e3)),v=N(Math.max(e,n-1e3));t.append("rect").attr("x",N(e)).attr("y",r-s/2).attr("width",N(n)-N(e)).attr("height",s).style("cursor","grab").attr("id","svgene-minimap-window-"+I.anchor).attr("opacity","0.2").style("fill","blue").call(i.drag().on("drag",(function(){if(null!==I){Math.max(0,a.event.x-parseInt(a.select(this).attr("x"),10));var t=parseInt(a.select(this).attr("width"),10),e=parseInt(a.select(this).attr("x"),10)+a.event.dx,n=g(h,e,v);e<h?t=Math.max(f-h,t-(h-e)):e+t>d&&(n=Math.min(e,v),t=d-n),n=g(h,n,v);var r=Math.max(f,n+t);a.select(this).attr("x",n).attr("width",t),a.select("#svgene-minimap-grabber-a-"+I.anchor).attr("x",n),a.select("#svgene-minimap-grabber-b-"+I.anchor).attr("x",r),T(N.invert(n),N.invert(r),!0)}}))),t.append("rect").attr("x",N(e)).attr("y",r-s/2).attr("id","svgene-minimap-grabber-a-"+I.anchor).attr("width",2).attr("height",s).style("cursor","ew-resize").call(i.drag().on("drag",(function(t,e){if(I){var n=parseInt(a.select("#svgene-minimap-grabber-b-"+I.anchor).attr("x"),10),r=Math.min(N(N.invert(n)-1e3),d,Math.max(a.event.x,h));a.select(this).transition().duration(10).attr("x",r),a.select("#svgene-minimap-window-"+I.anchor).transition().duration(10).attr("x",r).attr("width",n-r),T(N.invert(r),N.invert(n),!0)}}))),t.append("rect").attr("x",N(n)).attr("y",r-s/2).attr("id","svgene-minimap-grabber-b-"+I.anchor).attr("width",2).attr("height",s).style("cursor","ew-resize").call(i.drag().on("drag",(function(t,e){if(I){var n=parseInt(a.select("#svgene-minimap-grabber-a-"+I.anchor).attr("x"),10),r=Math.max(N(N.invert(n)+1e3),f,Math.min(a.event.x,d));a.select(this).transition().duration(10).attr("x",r),a.select("#svgene-minimap-window-"+I.anchor).transition().duration(10).attr("width",r-n),T(N.invert(n),N.invert(r),!0)}})))}(B,c.start,c.end,A-20,40,j,E),$(".svgene-orf").mouseover((function(t){var e=$(this).attr("id");void 0!==e&&(e=e.replace("-svgeneorf","-label"),$("#"+e).show())})).mouseout((function(t){var e=$(this).attr("id");void 0!==e&&(e=e.replace("-svgeneorf","-label"),$("#"+e).hide())})).click(m),$(".svgene-textarea").click((function(t){return t.stopPropagation()})),$(".legend-selector").unbind("click").click(S),$(".zoom-in").unbind("click").click(M),$(".zoom-reset").unbind("click").click(k),1===O&&$(document).keyup((function(t){var e=t.keyCode;82===e?k(t):90===e&&M(t)}))}},e.tag_to_id=p;var R=null,N=o.scaleLinear().domain([0,100]).range([0,100]),P=0,L=0,I=null,D=o.scaleLinear().domain([0,100]).range([0,100]),O=0},function(t,e){class n{static round(t,e){return e=e||1,Number(Math.round(t+"e"+e)+"e-"+e)}static meanAngle(t){let e=0,n=0;for(var r=0;r<t.length;r++)e+=Math.sin(t[r]),n+=Math.cos(t[r]);return Math.atan2(e/t.length,n/t.length)}static innerAngle(t){return n.toRad(180*(t-2)/t)}static polyCircumradius(t,e){return t/(2*Math.sin(Math.PI/e))}static apothem(t,e){return t*Math.cos(Math.PI/e)}static apothemFromSideLength(t,e){let r=n.polyCircumradius(t,e);return n.apothem(r,e)}static centralAngle(t){return n.toRad(360/t)}static toDeg(t){return t*n.degFactor}static toRad(t){return t*n.radFactor}static parityOfPermutation(t){let e=new Uint8Array(t.length),n=0,r=function(n,i=0){return 1===e[n]?i:(i++,e[n]=1,r(t[n],i))};for(var i=0;i<t.length;i++){if(1===e[i])continue;n+=1-r(i)%2}return n%2?-1:1}static get radFactor(){return Math.PI/180}static get degFactor(){return 180/Math.PI}static get twoPI(){return 2*Math.PI}}t.exports=n},function(t,e){class n{static clone(t){let e=Array.isArray(t)?Array():{};for(let r in t){let i=t[r];"function"==typeof i.clone?e[r]=i.clone():e[r]="object"==typeof i?n.clone(i):i}return e}static equals(t,e){if(t.length!==e.length)return!1;let n=t.slice().sort(),r=e.slice().sort();for(var i=0;i<n.length;i++)if(n[i]!==r[i])return!1;return!0}static print(t){if(0==t.length)return"";let e="(";for(let n=0;n<t.length;n++)e+=t[n].id?t[n].id+", ":t[n]+", ";return e=e.substring(0,e.length-2),e+")"}static each(t,e){for(let n=0;n<t.length;n++)e(t[n])}static get(t,e,n){for(let r=0;r<t.length;r++)if(t[r][e]==n)return t[r]}static contains(t,e){if(e.property||e.func){if(e.func){for(let n=0;n<t.length;n++)if(e.func(t[n]))return!0}else for(let n=0;n<t.length;n++)if(t[n][e.property]==e.value)return!0}else for(let n=0;n<t.length;n++)if(t[n]==e.value)return!0;return!1}static intersection(t,e){let n=new Array;for(let r=0;r<t.length;r++)for(let i=0;i<e.length;i++)t[r]===e[i]&&n.push(t[r]);return n}static unique(t){let e={};return t.filter((function(t){return void 0===e[t]&&(e[t]=!0)}))}static count(t,e){let n=0;for(let r=0;r<t.length;r++)t[r]===e&&n++;return n}static toggle(t,e){let n=Array(),r=!1;for(let i=0;i<t.length;i++)t[i]!==e?n.push(t[i]):r=!0;return r||n.push(e),n}static remove(t,e){let n=Array();for(let r=0;r<t.length;r++)t[r]!==e&&n.push(t[r]);return n}static removeUnique(t,e){let n=t.indexOf(e);return n>-1&&t.splice(n,1),t}static removeAll(t,e){return t.filter((function(t){return-1===e.indexOf(t)}))}static merge(t,e){let n=new Array(t.length+e.length);for(let e=0;e<t.length;e++)n[e]=t[e];for(let r=0;r<e.length;r++)n[t.length+r]=e[r];return n}static containsAll(t,e){let n=0;for(let r=0;r<t.length;r++)for(let i=0;i<e.length;i++)t[r]===e[i]&&n++;return n===e.length}static sortByAtomicNumberDesc(t){let e=t.map((function(t,e){return{index:e,value:t.atomicNumber.split(".").map(Number)}}));return e.sort((function(t,e){let n=Math.min(e.value.length,t.value.length),r=0;for(;r<n&&e.value[r]===t.value[r];)r++;return r===n?e.value.length-t.value.length:e.value[r]-t.value[r]})),e.map((function(e){return t[e.index]}))}static deepCopy(t){let e=Array();for(let r=0;r<t.length;r++){let i=t[r];e[r]=i instanceof Array?n.deepCopy(i):i}return e}}t.exports=n},function(t,e,n){"use strict";n.d(e,"a",(function(){return u}));var r={value:function(){}};function i(){for(var t,e=0,n=arguments.length,r={};e<n;++e){if(!(t=arguments[e]+"")||t in r)throw new Error("illegal type: "+t);r[t]=[]}return new o(r)}function o(t){this._=t}function a(t,e){return t.trim().split(/^|\s+/).map((function(t){var n="",r=t.indexOf(".");if(r>=0&&(n=t.slice(r+1),t=t.slice(0,r)),t&&!e.hasOwnProperty(t))throw new Error("unknown type: "+t);return{type:t,name:n}}))}function s(t,e){for(var n,r=0,i=t.length;r<i;++r)if((n=t[r]).name===e)return n.value}function l(t,e,n){for(var i=0,o=t.length;i<o;++i)if(t[i].name===e){t[i]=r,t=t.slice(0,i).concat(t.slice(i+1));break}return null!=n&&t.push({name:e,value:n}),t}o.prototype=i.prototype={constructor:o,on:function(t,e){var n,r=this._,i=a(t+"",r),o=-1,u=i.length;if(!(arguments.length<2)){if(null!=e&&"function"!=typeof e)throw new Error("invalid callback: "+e);for(;++o<u;)if(n=(t=i[o]).type)r[n]=l(r[n],t.name,e);else if(null==e)for(n in r)r[n]=l(r[n],t.name,null);return this}for(;++o<u;)if((n=(t=i[o]).type)&&(n=s(r[n],t.name)))return n},copy:function(){var t={},e=this._;for(var n in e)t[n]=e[n].slice();return new o(t)},call:function(t,e){if((n=arguments.length-2)>0)for(var n,r,i=new Array(n),o=0;o<n;++o)i[o]=arguments[o+2];if(!this._.hasOwnProperty(t))throw new Error("unknown type: "+t);for(o=0,n=(r=this._[t]).length;o<n;++o)r[o].value.apply(e,i)},apply:function(t,e,n){if(!this._.hasOwnProperty(t))throw new Error("unknown type: "+t);for(var r=this._[t],i=0,o=r.length;i<o;++i)r[i].value.apply(e,n)}};var u=i},function(t,e,n){const r=n(10);n(4),n(5);class i{constructor(t,e="-"){this.element=1===t.length?t.toUpperCase():t,this.drawExplicit=!1,this.ringbonds=Array(),this.rings=Array(),this.bondType=e,this.branchBond=null,this.isBridge=!1,this.isBridgeNode=!1,this.originalRings=Array(),this.bridgedRing=null,this.anchoredRings=Array(),this.bracket=null,this.plane=0,this.attachedPseudoElements={},this.hasAttachedPseudoElements=!1,this.isDrawn=!0,this.isConnectedToRing=!1,this.neighbouringElements=Array(),this.isPartOfAromaticRing=t!==this.element,this.bondCount=0,this.chirality="",this.isStereoCenter=!1,this.priority=0,this.mainChain=!1,this.hydrogenDirection="down",this.subtreeDepth=1,this.hasHydrogen=!1}addNeighbouringElement(t){this.neighbouringElements.push(t)}attachPseudoElement(t,e,n=0,r=0){null===n&&(n=0),null===r&&(r=0);let i=n+t+r;this.attachedPseudoElements[i]?this.attachedPseudoElements[i].count+=1:this.attachedPseudoElements[i]={element:t,count:1,hydrogenCount:n,previousElement:e,charge:r},this.hasAttachedPseudoElements=!0}getAttachedPseudoElements(){let t={},e=this;return Object.keys(this.attachedPseudoElements).sort().forEach((function(n){t[n]=e.attachedPseudoElements[n]})),t}getAttachedPseudoElementsCount(){return Object.keys(this.attachedPseudoElements).length}isHeteroAtom(){return"C"!==this.element&&"H"!==this.element}addAnchoredRing(t){r.contains(this.anchoredRings,{value:t})||this.anchoredRings.push(t)}getRingbondCount(){return this.ringbonds.length}backupRings(){this.originalRings=Array(this.rings.length);for(let t=0;t<this.rings.length;t++)this.originalRings[t]=this.rings[t]}restoreRings(){this.rings=Array(this.originalRings.length);for(let t=0;t<this.originalRings.length;t++)this.rings[t]=this.originalRings[t]}haveCommonRingbond(t,e){for(let n=0;n<t.ringbonds.length;n++)for(let r=0;r<e.ringbonds.length;r++)if(t.ringbonds[n].id==e.ringbonds[r].id)return!0;return!1}neighbouringElementsEqual(t){if(t.length!==this.neighbouringElements.length)return!1;t.sort(),this.neighbouringElements.sort();for(var e=0;e<this.neighbouringElements.length;e++)if(t[e]!==this.neighbouringElements[e])return!1;return!0}getAtomicNumber(){return i.atomicNumbers[this.element]}getMaxBonds(){return i.maxBonds[this.element]}static get maxBonds(){return{H:1,C:4,N:3,O:2,P:3,S:2,B:3,F:1,I:1,Cl:1,Br:1}}static get atomicNumbers(){return{H:1,He:2,Li:3,Be:4,B:5,b:5,C:6,c:6,N:7,n:7,O:8,o:8,F:9,Ne:10,Na:11,Mg:12,Al:13,Si:14,P:15,p:15,S:16,s:16,Cl:17,Ar:18,K:19,Ca:20,Sc:21,Ti:22,V:23,Cr:24,Mn:25,Fe:26,Co:27,Ni:28,Cu:29,Zn:30,Ga:31,Ge:32,As:33,Se:34,Br:35,Kr:36,Rb:37,Sr:38,Y:39,Zr:40,Nb:41,Mo:42,Tc:43,Ru:44,Rh:45,Pd:46,Ag:47,Cd:48,In:49,Sn:50,Sb:51,Te:52,I:53,Xe:54,Cs:55,Ba:56,La:57,Ce:58,Pr:59,Nd:60,Pm:61,Sm:62,Eu:63,Gd:64,Tb:65,Dy:66,Ho:67,Er:68,Tm:69,Yb:70,Lu:71,Hf:72,Ta:73,W:74,Re:75,Os:76,Ir:77,Pt:78,Au:79,Hg:80,Tl:81,Pb:82,Bi:83,Po:84,At:85,Rn:86,Fr:87,Ra:88,Ac:89,Th:90,Pa:91,U:92,Np:93,Pu:94,Am:95,Cm:96,Bk:97,Cf:98,Es:99,Fm:100,Md:101,No:102,Lr:103,Rf:104,Db:105,Sg:106,Bh:107,Hs:108,Mt:109,Ds:110,Rg:111,Cn:112,Uut:113,Uuq:114,Uup:115,Uuh:116,Uus:117,Uuo:118}}static get mass(){return{H:1,He:2,Li:3,Be:4,B:5,b:5,C:6,c:6,N:7,n:7,O:8,o:8,F:9,Ne:10,Na:11,Mg:12,Al:13,Si:14,P:15,p:15,S:16,s:16,Cl:17,Ar:18,K:19,Ca:20,Sc:21,Ti:22,V:23,Cr:24,Mn:25,Fe:26,Co:27,Ni:28,Cu:29,Zn:30,Ga:31,Ge:32,As:33,Se:34,Br:35,Kr:36,Rb:37,Sr:38,Y:39,Zr:40,Nb:41,Mo:42,Tc:43,Ru:44,Rh:45,Pd:46,Ag:47,Cd:48,In:49,Sn:50,Sb:51,Te:52,I:53,Xe:54,Cs:55,Ba:56,La:57,Ce:58,Pr:59,Nd:60,Pm:61,Sm:62,Eu:63,Gd:64,Tb:65,Dy:66,Ho:67,Er:68,Tm:69,Yb:70,Lu:71,Hf:72,Ta:73,W:74,Re:75,Os:76,Ir:77,Pt:78,Au:79,Hg:80,Tl:81,Pb:82,Bi:83,Po:84,At:85,Rn:86,Fr:87,Ra:88,Ac:89,Th:90,Pa:91,U:92,Np:93,Pu:94,Am:95,Cm:96,Bk:97,Cf:98,Es:99,Fm:100,Md:101,No:102,Lr:103,Rf:104,Db:105,Sg:106,Bh:107,Hs:108,Mt:109,Ds:110,Rg:111,Cn:112,Uut:113,Uuq:114,Uup:115,Uuh:116,Uus:117,Uuo:118}}}t.exports=i},function(t,e,n){"use strict";function r(t){t.toggleClass("expanded"),t.hasClass("expanded")?t.next().css("display","block"):t.next().removeAttr("style")}Object.defineProperty(e,"__esModule",{value:!0}),e.toggleCollapserHandler=e.toggleCollapser=void 0,e.toggleCollapser=r,e.toggleCollapserHandler=function(t){r($(this))}},function(t,e,n){"use strict";Object.defineProperty(e,"__esModule",{value:!0}),e.downloadSvg=e.initDownloadButtons=void 0;var r="http://www.w3.org/2000/svg",i="http://www.w3.org/1999/xlink",o="http://www.w3.org/2000/xmlns/";function a(t){if(!t.target)return!0;var n=t.target,r=n.getAttribute("data-tag"),i=n.getAttribute("data-filename");return!(!r||!i)&&(e.downloadSvg(r,i),!1)}function s(t,e){for(var n=getComputedStyle(t),r="",i=0,o=n.length;i<o;i++){var a=n[i],s=n.getPropertyValue(a);s!==e.getPropertyValue(a)&&(r+=a+":"+s+";")}t.setAttribute("style",r)}e.initDownloadButtons=function(){document.querySelectorAll(".download-svg").forEach((function(t){t.removeEventListener("click",a,!1),t.addEventListener("click",a,!1)}))},e.downloadSvg=function(t,e){var n,a=document.querySelector("#"+t);if(a&&((n="svg"===a.nodeName?a:a.children.item(0))&&"svg"===n.nodeName)){var l=n.cloneNode(!0);!function(t){t.hasAttribute("version")||t.setAttribute("version","1.1");t.hasAttributeNS(o,"xmlns")||t.setAttributeNS(o,"xmlns",r);t.hasAttributeNS(o,"xmlns:xlink")||t.setAttributeNS(o,"xmlns:xlink",i);document.body.appendChild(t),function(t){var e=document.createElementNS(r,"svg");document.body.appendChild(e);var n=getComputedStyle(e);document.body.removeChild(e);var i=function(t){var e=[];return e.push(t),function t(e,n){if(!(null==e?void 0:e.hasChildNodes()))return;var r=e.firstChild;for(;r;)r.nodeType===Node.ELEMENT_NODE&&"SCRIPT"!==r.nodeName&&(n.push(r),t(r,n)),r=r.nextSibling}(t,e),e}(t),o=i.length;for(;o--;)s(i[o],n)}(t),document.body.removeChild(t)}(l);var u=(new XMLSerializer).serializeToString(l),c="data:image/svg+xml;base64,"+btoa(u),h=document.createElement("a");h.href=c,h.download=e;var d=function(t){setTimeout((function(){var e;null===(e=null==t?void 0:t.target)||void 0===e||e.removeEventListener("click",d)}),200)};h.addEventListener("click",d,!1),h.click()}}},function(t,e,n){const r=n(3);class i{constructor(t=new r(0,0),e=new r(0,0),n=null,i=null,o=!1,a=!1){this.from=t,this.to=e,this.elementFrom=n,this.elementTo=i,this.chiralFrom=o,this.chiralTo=a}clone(){return new i(this.from.clone(),this.to.clone(),this.elementFrom,this.elementTo)}getLength(){return Math.sqrt(Math.pow(this.to.x-this.from.x,2)+Math.pow(this.to.y-this.from.y,2))}getAngle(){return r.subtract(this.getRightVector(),this.getLeftVector()).angle()}getRightVector(){return this.from.x<this.to.x?this.to:this.from}getLeftVector(){return this.from.x<this.to.x?this.from:this.to}getRightElement(){return this.from.x<this.to.x?this.elementTo:this.elementFrom}getLeftElement(){return this.from.x<this.to.x?this.elementFrom:this.elementTo}getRightChiral(){return this.from.x<this.to.x?this.chiralTo:this.chiralFrom}getLeftChiral(){return this.from.x<this.to.x?this.chiralFrom:this.chiralTo}setRightVector(t,e){return this.from.x<this.to.x?(this.to.x=t,this.to.y=e):(this.from.x=t,this.from.y=e),this}setLeftVector(t,e){return this.from.x<this.to.x?(this.from.x=t,this.from.y=e):(this.to.x=t,this.to.y=e),this}rotateToXAxis(){let t=this.getLeftVector();return this.setRightVector(t.x+this.getLength(),t.y),this}rotate(t){let e=this.getLeftVector(),n=this.getRightVector(),r=Math.sin(t),i=Math.cos(t),o=i*(n.x-e.x)-r*(n.y-e.y)+e.x,a=r*(n.x-e.x)-i*(n.y-e.y)+e.y;return this.setRightVector(o,a),this}shortenFrom(t){let e=r.subtract(this.to,this.from);return e.normalize(),e.multiplyScalar(t),this.from.add(e),this}shortenTo(t){let e=r.subtract(this.from,this.to);return e.normalize(),e.multiplyScalar(t),this.to.add(e),this}shortenRight(t){return this.from.x<this.to.x?this.shortenTo(t):this.shortenFrom(t),this}shortenLeft(t){return this.from.x<this.to.x?this.shortenFrom(t):this.shortenTo(t),this}shorten(t){let e=r.subtract(this.from,this.to);return e.normalize(),e.multiplyScalar(t/2),this.to.add(e),this.from.subtract(e),this}}t.exports=i},function(t,e,n){n(4),n(5);t.exports=class{constructor(t,e){this.id=null,this.firstRingId=t.id,this.secondRingId=e.id,this.vertices=new Set;for(var n=0;n<t.members.length;n++){let r=t.members[n];for(let t=0;t<e.members.length;t++){r===e.members[t]&&this.addVertex(r)}}}addVertex(t){this.vertices.add(t)}updateOther(t,e){this.firstRingId===e?this.secondRingId=t:this.firstRingId=t}containsRing(t){return this.firstRingId===t||this.secondRingId===t}isBridge(t){if(this.vertices.size>2)return!0;for(let e of this.vertices)if(t[e].value.rings.length>2)return!0;return!1}static isBridge(t,e,n,r){let i=null;for(let o=0;o<t.length;o++)if(i=t[o],i.firstRingId===n&&i.secondRingId===r||i.firstRingId===r&&i.secondRingId===n)return i.isBridge(e);return!1}static getNeighbours(t,e){let n=[];for(let r=0;r<t.length;r++){let i=t[r];i.firstRingId===e?n.push(i.secondRingId):i.secondRingId===e&&n.push(i.firstRingId)}return n}static getVertices(t,e,n){for(let r=0;r<t.length;r++){let i=t[r];if(i.firstRingId===e&&i.secondRingId===n||i.firstRingId===n&&i.secondRingId===e)return[...i.vertices]}}}},function(t,e){class n{constructor(t,e,n=1){this.id=null,this.sourceId=t,this.targetId=e,this.weight=n,this.bondType="-",this.isPartOfAromaticRing=!1,this.center=!1,this.wedge=""}setBondType(t){this.bondType=t,this.weight=n.bonds[t]}static get bonds(){return{"-":1,"/":1,"\\":1,"=":2,"#":3,$:4}}}t.exports=n},function(t,e,n){const r=n(9),i=(n(3),n(4)),o=n(17),a=(n(5),n(12));class s{constructor(t,e=!1){this.vertices=Array(),this.edges=Array(),this.vertexIdsToEdgeId={},this.isomeric=e,this._time=0,this._init(t)}_init(t,e=0,n=null,r=!1){let s=new a(t.atom.element?t.atom.element:t.atom,t.bond);s.branchBond=t.branchBond,s.ringbonds=t.ringbonds,s.bracket=t.atom.element?t.atom:null;let l=new i(s),u=this.vertices[n];if(this.addVertex(l),null!==n){l.setParentVertexId(n),l.value.addNeighbouringElement(u.value.element),u.addChild(l.id),u.value.addNeighbouringElement(s.element),u.spanningTreeChildren.push(l.id);let t=new o(n,l.id,1),e=null;r?(t.setBondType(l.value.branchBond||"-"),e=l.id,t.setBondType(l.value.branchBond||"-"),e=l.id):(t.setBondType(u.value.bondType||"-"),e=u.id);this.addEdge(t)}let c=t.ringbondCount+1;s.bracket&&(c+=s.bracket.hcount);let h=0;if(s.bracket&&s.bracket.chirality){s.isStereoCenter=!0,h=s.bracket.hcount;for(var d=0;d<h;d++)this._init({atom:"H",isBracket:"false",branches:Array(),branchCount:0,ringbonds:Array(),ringbondCount:!1,next:null,hasNext:!1,bond:"-"},d,l.id,!0)}for(d=0;d<t.branchCount;d++)this._init(t.branches[d],d+c,l.id,!0);t.hasNext&&this._init(t.next,t.branchCount+c,l.id)}clear(){this.vertices=Array(),this.edges=Array(),this.vertexIdsToEdgeId={}}addVertex(t){return t.id=this.vertices.length,this.vertices.push(t),t.id}addEdge(t){let e=this.vertices[t.sourceId],n=this.vertices[t.targetId];return t.id=this.edges.length,this.edges.push(t),this.vertexIdsToEdgeId[t.sourceId+"_"+t.targetId]=t.id,this.vertexIdsToEdgeId[t.targetId+"_"+t.sourceId]=t.id,t.isPartOfAromaticRing=e.value.isPartOfAromaticRing&&n.value.isPartOfAromaticRing,e.value.bondCount+=t.weight,n.value.bondCount+=t.weight,e.edges.push(t.id),n.edges.push(t.id),t.id}getEdge(t,e){let n=this.vertexIdsToEdgeId[t+"_"+e];return void 0===n?null:this.edges[n]}getEdges(t){let e=Array(),n=this.vertices[t];for(var r=0;r<n.neighbours.length;r++)e.push(this.vertexIdsToEdgeId[t+"_"+n.neighbours[r]]);return e}hasEdge(t,e){return void 0!==this.vertexIdsToEdgeId[t+"_"+e]}getVertexList(){let t=[this.vertices.length];for(var e=0;e<this.vertices.length;e++)t[e]=this.vertices[e].id;return t}getEdgeList(){let t=Array(this.edges.length);for(var e=0;e<this.edges.length;e++)t[e]=[this.edges[e].sourceId,this.edges[e].targetId];return t}getAdjacencyMatrix(){let t=this.vertices.length,e=Array(t);for(var n=0;n<t;n++)e[n]=new Array(t),e[n].fill(0);for(n=0;n<this.edges.length;n++){let t=this.edges[n];e[t.sourceId][t.targetId]=1,e[t.targetId][t.sourceId]=1}return e}getComponentsAdjacencyMatrix(){let t=this.vertices.length,e=Array(t),n=this.getBridges();for(var r=0;r<t;r++)e[r]=new Array(t),e[r].fill(0);for(r=0;r<this.edges.length;r++){let t=this.edges[r];e[t.sourceId][t.targetId]=1,e[t.targetId][t.sourceId]=1}for(r=0;r<n.length;r++)e[n[r][0]][n[r][1]]=0,e[n[r][1]][n[r][0]]=0;return e}getSubgraphAdjacencyMatrix(t){let e=t.length,n=Array(e);for(var r=0;r<e;r++){n[r]=new Array(e),n[r].fill(0);for(var i=0;i<e;i++)r!==i&&this.hasEdge(t[r],t[i])&&(n[r][i]=1)}return n}getDistanceMatrix(){let t=this.vertices.length,e=this.getAdjacencyMatrix(),n=Array(t);for(var r=0;r<t;r++)n[r]=Array(t),n[r].fill(1/0);for(r=0;r<t;r++)for(var i=0;i<t;i++)1===e[r][i]&&(n[r][i]=1);for(var o=0;o<t;o++)for(r=0;r<t;r++)for(i=0;i<t;i++)n[r][i]>n[r][o]+n[o][i]&&(n[r][i]=n[r][o]+n[o][i]);return n}getSubgraphDistanceMatrix(t){let e=t.length,n=this.getSubgraphAdjacencyMatrix(t),r=Array(e);for(var i=0;i<e;i++)r[i]=Array(e),r[i].fill(1/0);for(i=0;i<e;i++)for(var o=0;o<e;o++)1===n[i][o]&&(r[i][o]=1);for(var a=0;a<e;a++)for(i=0;i<e;i++)for(o=0;o<e;o++)r[i][o]>r[i][a]+r[a][o]&&(r[i][o]=r[i][a]+r[a][o]);return r}getAdjacencyList(){let t=this.vertices.length,e=Array(t);for(var n=0;n<t;n++){e[n]=[];for(var r=0;r<t;r++)n!==r&&this.hasEdge(this.vertices[n].id,this.vertices[r].id)&&e[n].push(r)}return e}getSubgraphAdjacencyList(t){let e=t.length,n=Array(e);for(var r=0;r<e;r++){n[r]=Array();for(var i=0;i<e;i++)r!==i&&this.hasEdge(t[r],t[i])&&n[r].push(i)}return n}getBridges(){let t=this.vertices.length,e=new Array(t),n=new Array(t),r=new Array(t),i=new Array(t),o=this.getAdjacencyList(),a=Array();e.fill(!1),i.fill(null),this._time=0;for(var s=0;s<t;s++)e[s]||this._bridgeDfs(s,e,n,r,i,o,a);return a}traverseBF(t,e){let n=this.vertices.length,r=new Array(n);r.fill(!1);for(var i=[t];i.length>0;){let t=i.shift(),n=this.vertices[t];e(n);for(var o=0;o<n.neighbours.length;o++){let t=n.neighbours[o];r[t]||(r[t]=!0,i.push(t))}}}getTreeDepth(t,e){if(null===t||null===e)return 0;let n=this.vertices[t].getSpanningTreeNeighbours(e),r=0;for(var i=0;i<n.length;i++){let e=n[i],o=this.getTreeDepth(e,t);o>r&&(r=o)}return r+1}traverseTree(t,e,n,r=Number.MAX_SAFE_INTEGER,i=!1,o=1,a=null){if(null===a&&(a=new Uint8Array(this.vertices.length)),o>r+1||1===a[t])return;a[t]=1;let s=this.vertices[t],l=s.getNeighbours(e);(!i||o>1)&&n(s);for(var u=0;u<l.length;u++)this.traverseTree(l[u],t,n,r,i,o+1,a)}kkLayout(t,e,n,i,o){let a=o;for(var s=t.length;s--;){var l=this.vertices[t[s]].neighbours.length}let u=this.getSubgraphDistanceMatrix(t),c=t.length,h=r.polyCircumradius(500,c),d=r.centralAngle(c),g=0,f=new Float32Array(c),p=new Float32Array(c),v=Array(c);for(s=c;s--;){let n=this.vertices[t[s]];n.positioned?(f[s]=n.position.x,p[s]=n.position.y):(f[s]=e.x+Math.cos(g)*h,p[s]=e.y+Math.sin(g)*h),v[s]=n.positioned,g+=d}let m=Array(c);for(s=c;s--;){m[s]=new Array(c);for(l=c;l--;)m[s][l]=o*u[s][l]}let y=Array(c);for(s=c;s--;){y[s]=Array(c);for(l=c;l--;)y[s][l]=a*Math.pow(u[s][l],-2)}let b,x,w,A,C,_,S,M=Array(c),T=new Float32Array(c),k=new Float32Array(c);for(s=c;s--;)M[s]=Array(c);for(s=c;s--;){b=f[s],x=p[s],w=0,A=0;let t=c;for(;t--;)s!==t&&(C=f[t],_=p[t],S=1/Math.sqrt((b-C)*(b-C)+(x-_)*(x-_)),M[s][t]=[y[s][t]*(b-C-m[s][t]*(b-C)*S),y[s][t]*(x-_-m[s][t]*(x-_)*S)],M[t][s]=M[s][t],w+=M[s][t][0],A+=M[s][t][1]);T[s]=w,k[s]=A}let R=function(t){return[T[t]*T[t]+k[t]*k[t],T[t],k[t]]},N=function(){let t=0,e=0,n=0,r=0;for(s=c;s--;){let[i,o,a]=R(s);i>t&&!1===v[s]&&(t=i,e=s,n=o,r=a)}return[e,t,n,r]},$=function(t,e,n){let r=0,i=0,o=0,a=f[t],l=p[t],u=m[t],h=y[t];for(s=c;s--;){if(s===t)continue;let e=f[s],n=p[s],c=u[s],d=h[s],g=(a-e)*(a-e),v=1/Math.pow(g+(l-n)*(l-n),1.5);r+=d*(1-c*(l-n)*(l-n)*v),i+=d*(1-c*g*v),o+=d*(c*(a-e)*(l-n)*v)}0===r&&(r=.1),0===i&&(i=.1),0===o&&(o=.1);let d=e/r+n/o;d/=o/r-i/o;let g=-(o*d+e)/r;f[t]+=g,p[t]+=d;let v,b,x,w,A,C=M[t];for(e=0,n=0,a=f[t],l=p[t],s=c;s--;)t!==s&&(v=f[s],b=p[s],x=C[s][0],w=C[s][1],A=1/Math.sqrt((a-v)*(a-v)+(l-b)*(l-b)),g=h[s]*(a-v-u[s]*(a-v)*A),d=h[s]*(l-b-u[s]*(l-b)*A),C[s]=[g,d],e+=g,n+=d,T[s]+=g-x,k[s]+=d-w);T[t]=e,k[t]=n},P=1e9,L=0,I=0,D=0,O=0,E=0,B=0;for(;P>.1&&2e3>E;)for(E++,[L,P,I,D]=N(),O=P,B=0;O>.1&&50>B;)B++,$(L,I,D),[O,I,D]=R(L);for(s=c;s--;){let e=t[s],n=this.vertices[e];n.position.x=f[s],n.position.y=p[s],n.positioned=!0,n.forcePositioned=!0}}_bridgeDfs(t,e,n,r,i,o,a){e[t]=!0,n[t]=r[t]=++this._time;for(var s=0;s<o[t].length;s++){let l=o[t][s];e[l]?l!==i[t]&&(r[t]=Math.min(r[t],n[l])):(i[l]=t,this._bridgeDfs(l,e,n,r,i,o,a),r[t]=Math.min(r[t],r[l]),r[l]>n[t]&&a.push([t,l]))}}static getConnectedComponents(t){let e=t.length,n=new Array(e),r=new Array;n.fill(!1);for(var i=0;i<e;i++)if(!n[i]){let e=Array();n[i]=!0,e.push(i),s._ccGetDfs(i,n,t,e),e.length>1&&r.push(e)}return r}static getConnectedComponentCount(t){let e=t.length,n=new Array(e),r=0;n.fill(!1);for(var i=0;i<e;i++)n[i]||(n[i]=!0,r++,s._ccCountDfs(i,n,t));return r}static _ccCountDfs(t,e,n){for(var r=0;r<n[t].length;r++){n[t][r]&&!e[r]&&t!==r&&(e[r]=!0,s._ccCountDfs(r,e,n))}}static _ccGetDfs(t,e,n,r){for(var i=0;i<n[t].length;i++){n[t][i]&&!e[i]&&t!==i&&(e[i]=!0,r.push(i),s._ccGetDfs(i,e,n,r))}}}t.exports=s},function(t,e,n){"use strict";Object.defineProperty(e,"__esModule",{value:!0}),e.start=e.getAnchor=void 0;var r=n(7),i=n(20),o=n(13),a=n(21),s=n(22),l=n(14),u=n(23),c=n(24),h=n(25),d=n(26),g=n(8),f=n(14);Object.defineProperty(e,"downloadSvg",{enumerable:!0,get:function(){return f.downloadSvg}});var p=null,v=null,m=null;function y(t){t.preventDefault(),$("#downloadmenu").fadeToggle("fast","linear")}function b(){var t=window.location.hash.substring(1);return t||"overview"}function x(){setTimeout((function(){$(".page").hide(),$(".regbutton").removeClass("active");var t=b();$("#"+t).show(),"overview"!==t&&($(".regbutton."+t).addClass("active"),void 0!==p[t]&&g.drawRegion(t+"-svg",p[t],20),$("#"+t+"-details-svg").length>0&&u.drawDomains(t+"-details-svg",v.nrpspks[t],25),$("#"+t+"-pfam-details-svg").length>0&&c.drawPfamDomains(""+t,v.pfam[t],25),$("#"+t+" .clusterblast-selector").change(),t in m&&"antismash.modules.cluster_compare"in m[t]&&a.setComparisonData(t,m[t]["antismash.modules.cluster_compare"],p[t]),$("#"+t+" .comparison-selector").change(),$("#"+t+" * .body-details-header-active").first().click())}),1)}function w(){var t=p.order,e=b(),n="overview";if("overview"===e)n=t[0];else{var r=t.indexOf(e);r!==t.length-1&&(n=t[r+1])}window.location.href="#"+n,x()}function A(){var t=p.order,e=b(),n="";if("overview"===e)n=t[t.length-1];else{var r=t.indexOf(e);0!==r&&(n=t[r-1])}window.location.href="#"+n,x()}function C(t,e){var n=t+"-active",r=b(),i=$("#"+r).find("."+t),o=$("#"+r).find("."+n);i&&o&&o&&(e>0?o.is(i.last())?i.first().trigger("click"):o.next().trigger("click"):o.is(i.first())?i.last().trigger("click"):o.prev().trigger("click"))}function _(t){var e=t.keyCode;37===e||87===e?A():39!==e&&69!==e||w(),"overview"!==b()&&(65===e?C("body-details-header",-1):83===e?C("body-details-header",1):68===e?C("sidepanel-details-header",-1):70===e?C("sidepanel-details-header",1):77===e&&$("input.show-module-domains").first().click())}function S(t){t.preventDefault();var e=($(this).attr("id")||"").replace(/-header/,"");if(e){var n=$("#"+e);"none"===n.css("display")?$(this).text("Hide pHMM detection rules used"):$(this).text("Show pHMM detection rules used"),n.fadeToggle("fast","linear")}}e.getAnchor=b,e.start=function(t,e,n,a){h.createRecordOverviews(a),p=t,v=e,m=n,document.addEventListener("keyup",_,!1),$("#download").click(y),$("#next-region").click(w),$("#prev-region").click(A),$(".regbutton").click((function(){var t=$(this).children().first().attr("href");void 0!==t&&(window.location.href=t,x())})).mouseover((function(){var t=($(this).attr("class")||"").split(" ");if(!(t.length<2)&&"separator"!==t[1]){var e=function(t){switch(t){case"nrps":return"NRPS";case"t1pks":return"Type I PKS";case"t2pks":return"Type II PKS";case"t3pks":return"Type III PKS";case"t4pks":return"Type IV PKS";default:return t}}(t[1]),n=$("#region-type");n.data("orig_text",n.text()),n.text(e+":")}})).mouseout((function(){var t=$("#region-type");t.text(t.data("orig_text"))})),$(".clusterblast-selector").change((function(){var t=($(this).attr("id")||"nonexistant").replace("-select",""),e=""+$(this).val();e&&$.get(e,(function(e){$("#"+t+"-svg").html(e),i.init(t+"-svg")}),"html"),$("#"+t+"-download").off("click"),$("#"+t+"-download").click((function(){return window.open(""+$("#"+t+"-select").val(),"_blank")}))})),$(".comparison-selector").change((function(){var t=($(this).attr("id")||"nonexistant").replace("-selector","");$("#"+t).siblings().removeClass("comparison-container-active");var e=$(this).attr("data-tag");$("#"+t+"-"+$(this).val()).addClass("comparison-container-active").find(".heat-row-"+e).first().click()})),$(".cluster-rules-header").click(S),x(),$("input.overview-switch-compact").change((function(){$(this).prop("checked")?($("#single-record-tables").hide(),$("#compact-record-table").show()):($("#single-record-tables").show(),$("#compact-record-table").hide())})).trigger("change"),$(".linked-row").off("click").click((function(){var t=$(this).attr("data-anchor");t&&(window.location.href=t,x())})),$("input.domains-selected-only").change((function(){$("input.domains-selected-only").prop("checked",$(this).prop("checked")),u.redrawDomains(),c.redrawPfamDomains()})),u.createModuleHandlers(),d.drawStructures(),s.setupDetails(t.order),$(".help-icon").off("click").click((function(){$(this).toggleClass("active"),$(this).next().toggle()})),$(".help-tooltip").off("click").click((function(){$(this).hide(),$(this).prev().removeClass("active")})),$(".clipboard-copy").off("click").click(r.copyToClipboard),$(".collapser").click(o.toggleCollapserHandler),l.initDownloadButtons()}},function(t,e,n){"use strict";Object.defineProperty(e,"__esModule",{value:!0}),e.init=void 0;var r=null;function i(t){var e=$("#"+$(this).attr("id")+"-tooltip");if(r&&(clearTimeout(e.data("timeout")),r.hide()),r=e,"none"===e.css("display")){var n=$(this).offset();void 0===n&&(n={left:0,top:0});var i=setTimeout((function(){return e.slideUp("fast")}),5e3);e.css("top",n.top+10).css("left",n.left+5).show().click((function(){$(this).hide()})).data("timeout",i).mouseover((function(){return clearTimeout(e.data("timeout"))})).mouseout((function(){clearTimeout(e.data("timeout")),i=setTimeout((function(){return e.slideUp("fast")}),5e3),e.data("timeout",i)}))}else e.hide()}e.init=function(t){$("#"+t+" .clusterblast-orf").each((function(){var e=$(this);!function(t,e){var n=t.attr("id"),r=$("<div>");r.addClass("clusterblast-locustag"),r.attr("id",n+"-label");var i=t.attr("locus_tag");i?r.text(i):r.text("unknown");$("#"+e).append(r),t.mouseover((function(e){var i=t.offset();void 0===i&&(i={left:0,top:32}),r.css("top",i.top-32),r.css("left",i.left),$("#"+n+"-label").show()})).mouseout((function(t){$("#"+n+"-label").hide()}))}(e,t),function(t,e){var n=t.attr("id"),r=$("<div>");r.addClass("clusterblast-tooltip"),r.attr("id",n+"-tooltip"),r.html((t.attr("description")||"").replace("[br]","<br>")),$("#"+e).append(r),t.click(i)}(e,t)}))}},function(t,e,n){"use strict";Object.defineProperty(e,"__esModule",{value:!0}),e.setComparisonData=void 0;var r=n(6),i=n(0),o=n(8),a=null,s=null;function l(t,e,n,r,i){var o=n+r/2,a=n-r/2,s=n;if(i){var l,u;if(1===t.strand)return(l=Math.floor(e(t.start)))+","+o+" "+(u=Math.floor(Math.min(e(t.end)+r/2,l)))+","+o+" "+Math.floor(e(t.end))+","+s+" "+u+","+a+" "+l+","+a;if(-1===t.strand){var c=Math.floor(e(t.start)),h=Math.floor(e(t.end));return c+","+s+" "+(d=Math.floor(Math.max(e(t.start)-r/2,h)))+","+o+" "+h+","+o+" "+h+","+a+" "+d+","+a}return t.start+","+o+" "+t.end+","+o+" "+t.end+","+a+" "+t.start+","+a}if(1===t.strand)return(l=Math.floor(e(t.start)))+","+o+" "+(u=Math.floor(Math.max(e(t.end)-r/2,l)))+","+o+" "+Math.floor(e(t.end))+","+s+" "+u+","+a+" "+l+","+a;if(-1===t.strand){var d;c=Math.floor(e(t.start)),h=Math.floor(e(t.end));return c+","+s+" "+(d=Math.floor(Math.min(e(t.start)+r/2,h)))+","+o+" "+h+","+o+" "+h+","+a+" "+d+","+a}return t.start+","+o+" "+e(t.end)+","+o+" "+e(t.end)+","+a+" "+e(t.start)+","+a}function u(t,e){var n=window.location.hash.substring(1),u=document.getElementById("comparison-"+e+"-"+n+"-selector");if(u&&u.value){var c=$("#"+t+" * .heat-row-"+e);c.off("click").click((function(){$(".heat-row-"+e).css("border",""),$(this).css("border","1px solid black");var t=$(this),c="comparison-"+e+"-"+n+"-svg";$("#"+c).remove();var h=i.select("#comparison-"+e+"-"+n).append("svg").attr("width","100%").attr("id",c),d=a[e][u.value],g=t.attr("data-accession");g&&(!function(t,e,n,i){if(null!==i&&null!==s){var a=s.end-s.start,u=s.idx,c=r.scaleLinear().domain([s.start,s.end]).range([0,800]),h=r.scaleLinear().domain([i.start,i.end]).range([0,800]);i.reverse&&(h=r.scaleLinear().domain([i.end,i.start]).range([0,800])),t.attr("viewport","0 164 0 "+a).attr("height",164).attr("width",800),t.append("text").attr("x",0).attr("y",20).attr("text-anchor","start").text("Query"),t.append("text").attr("x",0).attr("y",155).attr("text-anchor","start").text("Reference: "+(e.indexOf("-mibig-")>-1?n.split(":")[0]:n)),t.append("line").attr("x1",0).attr("y1",54.5).attr("x2",a).attr("y2",54.5).attr("class","centerline"),t.append("line").attr("x1",0).attr("y1",109.5).attr("x2",a).attr("y2",109.5).attr("class","centerline");t.selectAll("line.link").data(i.links).enter().append("line").attr("x1",(function(t){return c(t.query_loc)})).attr("y1",59.5).attr("x2",(function(t){return h(t.subject_loc)})).attr("y2",104.5).attr("class","link").attr("class","centerline");var d=t.selectAll("g.cc-svg-orf-group").data(s.orfs).enter().append("g").attr("class","cc-svg-orf-group");d.append("polygon").attr("points",(function(t){return l(t,c,54.5,10)})).attr("class","cc-svg-orf-bg").attr("id",(function(t){return e+"-"+o.tag_to_id(t.locus_tag)+"-bg"})).style("fill","white"),d.append("polygon").attr("points",(function(t){return l(t,c,54.5,10)})).attr("class",(function(t){return"cc-svg-orf svgene-type-"+t.type})).attr("id",(function(t){return e+"-"+o.tag_to_id(t.locus_tag)})).attr("opacity","1").attr("data-locus",(function(t){return t.locus_tag}));t.selectAll("text.svgene-locustag").data(s.orfs).enter().append("text").attr("x",(function(t){return c(t.start)<a/2?c(t.start):c(t.end)})).attr("text-anchor",(function(t){return c(t.start)<a/2?"start":"end"})).attr("y",36.5).attr("class","cc-svg-locustag").attr("id",(function(t){return e+"-"+o.tag_to_id(t.locus_tag)+"-tag"})).text((function(t){return t.locus_tag}));var g=t.selectAll("g.cc-svg-reforf-group").data(i.genes).enter().append("g").attr("class","cc-svg-reforf-group");g.append("polygon").attr("points",(function(t){return l(t,h,109.5,10,i.reverse)})).attr("class","cc-svg-reforf-bg").attr("id",(function(t){return e+"-ref"+o.tag_to_id(t.locus_tag)+"-bg"})).style("fill","white"),g.append("polygon").attr("points",(function(t){return l(t,h,109.5,10,i.reverse)})).attr("class",(function(t){return"cc-svg-reforf svgene-type-"+t.function})).attr("id",(function(t){return e+"-ref"+o.tag_to_id(t.locus_tag)})).attr("data-locus",(function(t){return t.locus_tag})).style("opacity",(function(t){return t.linked[""+u]?"1":"0.5"}));t.selectAll("text.svgene-locustag").data(i.genes).enter().append("text").attr("x",(function(t){return h(t.start)<a/2?h(t.start):h(t.end)})).attr("text-anchor",(function(t){return h(t.end)<a/2?"start":"end"})).attr("y",127.5).attr("class","cc-svg-locustag").attr("id",(function(t){return e+"-ref"+o.tag_to_id(t.locus_tag)+"-tag"})).text((function(t){return t.locus_tag}))}}(h,c,g,d.reference_clusters[g]),$(".cc-svg-orf").mouseover((function(t){var e=$(this).attr("id");void 0!==e&&$("#"+e+"-tag").show()})).mouseout((function(t){var e=$(this).attr("id");void 0!==e&&$("#"+e+"-tag").hide()})).click((function(t){var e=$(this).attr("data-locus");e&&o.selectOrfsByLoci([e],t.ctrlKey||t.metaKey)})),$(".cc-svg-reforf").mouseover((function(t){var e=$(this).attr("id");void 0!==e&&$("#"+e+"-tag").show()})).mouseout((function(t){var e=$(this).attr("id");void 0!==e&&$("#"+e+"-tag").hide()})))})),c.first().click()}}e.setComparisonData=function(t,e,n){for(var r in a=e,s=n,e)e.hasOwnProperty(r)&&u(t,r)}},function(t,e,n){"use strict";function r(t,e){i($(this),t);var n=$("#"+t+" * .sidepanel-details-header."+$(this).attr("data-name"));n.length>0&&a(n,t)}function i(t,e){$("#"+e+" * .body-details-section").hide(),$("#"+e+" * .body-details-header-active").toggleClass("body-details-header-active"),t.addClass("body-details-header-active"),$("#"+e+" * .body-details-section."+t.attr("data-name")).show()}function o(t,e){a($(this),t);var n=$("#"+t+" * .body-details-header."+$(this).attr("data-name"));n.length>0&&i(n,t)}function a(t,e){$("#"+e+" * .sidepanel-details-section").hide(),$("#"+e+" * .sidepanel-details-header-active").toggleClass("sidepanel-details-header-active"),t.addClass("sidepanel-details-header-active"),$("#"+e+" * .sidepanel-details-section."+t.attr("data-name")).show()}Object.defineProperty(e,"__esModule",{value:!0}),e.setupDetails=void 0,e.setupDetails=function(t){$(".body-details-header").off("click"),$(".body-details-section").hide();for(var e=0,n=t;e<n.length;e++){var i=n[e];$("#"+i+" * .body-details-header").click($.proxy(r,null,i)).first().trigger("click")}$(".sidepanel-details-header").off("click"),$(".sidepanel-details-section").hide();for(var a=0,s=t;a<s.length;a++){i=s[a];$("#"+i+" * .sidepanel-details-header").click($.proxy(o,null,i)).first().trigger("click")}}},function(t,e,n){"use strict";Object.defineProperty(e,"__esModule",{value:!0}),e.createModuleHandlers=e.redrawDomains=e.drawDomains=void 0;var r,i=n(35),o=n(6),a=n(0),s=n(7),l=n(8),u={text_height:14,unique_id:0,version:"0.0.1"};function c(t,e,n,r,o){var a=i.path(),s=e(t.start),l=e(t.end),u=n+r;return"head"===t.multi_cds?(a.moveTo(l+o,n),a.lineTo(l,n+r/3),a.lineTo(l+o,n+2*r/3),a.lineTo(l,u),a.lineTo(s+o,u),a.arcTo(s,u,s,n,o),a.lineTo(s,n+o),a.arcTo(s,n,l,n,o)):(a.moveTo(l-o,n),a.arcTo(l,n,l,u,o),a.lineTo(l,u-o),a.arcTo(l,u,s,u,o),a.lineTo(s-o,u),a.lineTo(s,n+2*r/3),a.lineTo(s-o,n+r/3),a.lineTo(s,n)),a.closePath(),a.toString()}function h(t,e,n,r,o,a,s,l){var h=(a+o)*n+4,d=t.append("g").attr("class","domain-group");d.append("text").text(e.id).attr("x",0).attr("y",h+.7*a).attr("class","jsdomain-orflabel").attr("data-locus",e.id),d.selectAll("rect.jsdomain-module").data(e.modules.filter((function(t){return!t.multi_cds}))).enter().append("rect").attr("x",(function(t){return l(t.start)})).attr("y",h-3).attr("width",(function(t){return l(t.end)-l(t.start)})).attr("height",a+6).attr("rx",5).attr("class",(function(t){return t.complete?"jsdomain-module":"jsdomain-module jsdomain-incomplete-module"})),d.selectAll("text.jsdomain-match").data(e.modules.filter((function(t){return t.multi_cds}))).enter().append("text").text((function(t){return t.match_id})).attr("x",(function(t){return l("head"===t.multi_cds?t.end:t.start)+("head"===t.multi_cds?7:-5)})).attr("y",(function(t){return h+("head"===t.multi_cds?2:4)*a/4-2})).attr("text-anchor","middle").attr("class","jsdomain-match"),d.selectAll("path.jsdomain-module").data(e.modules.filter((function(t){return t.multi_cds}))).enter().append("path").attr("d",(function(t){return c(t,l,h-3,a+6,5)})).attr("class","jsdomain-module"),d.append("line").attr("y1",h+a/2).attr("y2",h+a/2).attr("x1",l(0)).attr("x2",l(e.sequence.length)).attr("class","jsdomain-line"),d.append("line").attr("x1",l(0)).attr("x2",l(0)).attr("y1",h+a/4).attr("y2",h+3*a/4).attr("class","jsdomain-line"),d.append("line").attr("x1",l(e.sequence.length)).attr("x2",l(e.sequence.length)).attr("y1",h+a/4).attr("y2",h+3*a/4).attr("class","jsdomain-line"),d.selectAll("rect.jsdomain-domain").data(e.domains).enter().append("rect").attr("x",(function(t){return l(t.start)})).attr("y",h).attr("rx",17).attr("ry",17).attr("width",(function(t){return l(t.end)-l(t.start)})).attr("height",a).attr("data-id",(function(t,e){return"details-orf-"+r+"-"+e+"-tooltip"})).attr("class",(function(t){return"jsdomain-domain "+t.html_class})).attr("stroke-width",1),d.selectAll("text.jsdomain-text").data(e.domains).enter().append("text").text((function(t){return t.abbreviation})).attr("x",(function(t){return l((t.start+t.end)/2)})).attr("text-anchor","middle").attr("y",h+.7*a).attr("class","jsdomain-text").attr("font-size",u.text_height).attr("font-weight","bold");var g=d.selectAll("g.jsdomain-module-lid").data(e.modules.filter((function(t){return t.complete&&!t.multi_cds}))).enter().append("g").attr("class","jsdomain-module-lid");g.append("rect").attr("x",(function(t){return l(t.start)})).attr("y",h-3).attr("width",(function(t){return l(t.end)-l(t.start)})).attr("height",a+6).attr("rx",5).attr("class","jsdomain-module-lid-body"),g.append("text").text((function(t){return t.monomer||"no prediction"})).attr("x",(function(t){return l((t.start+t.end)/2)})).attr("y",(function(t){return h+a*(t.iterative?.5:.7)})).attr("text-anchor","middle").attr("class","jsdomain-text");var f=d.selectAll("g.jsdomain-module-lid-fragment").data(e.modules.filter((function(t){return t.multi_cds}))).enter().append("g").attr("class","jsdomain-module-lid jsdomain-module-lid-fragment");f.append("path").attr("d",(function(t){return c(t,l,h-3,a+6,5)})).attr("class","jsdomain-module-lid-body"),f.append("text").text((function(t){return t.monomer||"no prediction"})).attr("x",(function(t){return l((t.start+t.end)/2)})).attr("y",(function(t){return h+a*(t.iterative?.5:.7)})).attr("text-anchor","middle").attr("class","jsdomain-text");var p=i.path();p.moveTo(50,50),p.arc(0,50,50,0,Math.PI/180*45,!0),p.moveTo(50,50),p.lineTo(10,40);var v=p.toString();g.append("path").attr("d",v).attr("stroke",(function(t){return t.iterative?"black":"none"})).attr("stroke-width","5px").attr("fill","none").attr("stroke-linecap","round").attr("transform",(function(t){return"translate("+l((t.start+t.end)/2)+", "+(h+.6*a)+") scale(0.1, 0.1)"}))}function d(t){var e=$(this).attr("data-id");if(void 0!==e){var n=$("#"+e);if(r&&(clearTimeout(n.data("timeout")),r.hide()),r=n,"none"===n.css("display")){var i=$(this).offset();void 0===i&&(i={left:0,top:0});var o=setTimeout((function(){return n.slideUp("fast")}),5e3);n.css("top",i.top+30).css("left",i.left+10).show().click((function(){$(this).hide()})).data("timeout",o).mouseover((function(){return clearTimeout(n.data("timeout"))})).mouseout((function(){clearTimeout(n.data("timeout")),o=setTimeout((function(){return n.slideUp("fast")}),5e3),n.data("timeout",o)}))}else n.hide()}}function g(t){if($(this).hide(),t.clientX&&t.clientY){var e=document.elementFromPoint(t.clientX,t.clientY);e&&$(e).click()}$(this).show()}function f(){$("input.domains-selected-only").prop("checked")?($(".jsdomain-svg").hide(),$(".jsdomain-svg-singles").show()):($(".jsdomain-svg").show(),$(".jsdomain-svg-singles").hide()),$("input.show-module-domains").prop("checked")?$(".jsdomain-module-lid").hide():$(".jsdomain-module-lid").show()}e.drawDomains=function(t,e,n){var r=a.select("#"+t),i=n,c=$("#"+t).width()||700;r.selectAll("svg.jsdomain-svg").remove(),r.selectAll("svg.jsdomain-svg-single").remove();for(var p=(i+10)*e.orfs.length,v=r.append("svg").attr("height",p).attr("width","100%").attr("viewbox","-1 -1 "+c+" "+p).attr("class","jsdomain-svg"),m=0,y="",b=0,x=e.orfs;b<x.length;b++){var w=x[b];m=Math.max(m,w.sequence.length),y.length<w.id.length&&(y=w.id)}var A=v.append("g").append("text").text(y).attr("x",0).attr("y",0).attr("id","dummy-label"),C=A.node().getComputedTextLength()||15*y.length;A.remove();for(var _=o.scaleLinear().domain([1,1.02*m]).range([C+10,c]),S=r.append("div").attr("class","jsdomain-svg-singles"),M=i+8,T=function(t){var n=e.orfs[t],o=u.unique_id++;h(S.append("svg").attr("height",M).attr("width","100%").attr("viewbox","-1 -1 "+c+" "+M).attr("class","jsdomain-svg-single").attr("id",l.locusToFullId(n.id)+"-domains"),n,0,o,10,i,0,_),h(v,n,t,o,10,i,0,_),r.append("div").attr("id","details-orf-"+o).selectAll("div.jsdomain-tooltip").data(n.domains).enter().append("div").attr("class","jsdomain-tooltip").attr("id",(function(t,e){return"details-orf-"+o+"-"+e+"-tooltip"})).html((function(t){return function(t,e){var n=t.type+"<br>Location: "+t.start+"-"+t.end+" AA<br>";t.napdoslink.length>0&&(n+='<a href="'+t.napdoslink+'" target="_blank">Analyze with NaPDoS</a><br>');if(n+='<a href="'+t.blastlink+'" target="_blank">Run BlastP on this domain</a><br>',t.predictions.length>0){n+="<dl><dt>Substrate predictions:</dt>";for(var r=0,i=t.predictions;r<i.length;r++){var o=i[r];n+="<dd>-"+o[0]+": "+o[1]+"</dd>"}n+="</dl>"}n+="AA sequence: "+s.clipboardCopyConstruct(t.sequence)+"<br>",t.dna_sequence&&(n+="Nucleotide sequence: "+s.clipboardCopyConstruct(t.dna_sequence)+"<br>");return n}(t)})),$(".jsdomain-tooltip .clipboard-copy").off("click").click(s.copyToClipboard)},k=0;k<e.orfs.length;k++)T(k);$(".jsdomain-orflabel").off("click").click((function(t){$("#"+l.locusToFullId($(this).attr("data-locus")||"none")+"-svgeneorf").trigger(t)})),a.selectAll("g.domain-group").data(e.orfs),$(".jsdomain-domain").click(d),$(".jsdomain-module-lid").click(g),f()},e.redrawDomains=f,e.createModuleHandlers=function(){$("input.show-module-domains").change((function(){$("input.show-module-domains").prop("checked",$(this).prop("checked")),f()}))}},function(t,e,n){"use strict";Object.defineProperty(e,"__esModule",{value:!0}),e.redrawPfamDomains=e.drawPfamDomains=void 0;var r,i=n(6),o=n(0),a=n(7),s=n(8),l={text_height:14,unique_id:0,version:"0.0.1"};function u(t,e,n,r,i,o,a,s){var u=(o+i)*n+4,c=t.append("g").attr("class","domain-group");c.append("text").text(e.id).attr("x",0).attr("y",u+.7*o).attr("class","pfam-domain-orflabel").attr("data-locus",e.id),c.append("line").attr("y1",u+o/2).attr("y2",u+o/2).attr("x1",s(0)).attr("x2",s(e.seqLength)).attr("class","pfam-domain-line"),c.append("line").attr("x1",s(0)).attr("x2",s(0)).attr("y1",u+o/4).attr("y2",u+3*o/4).attr("class","pfam-domain-line"),c.append("line").attr("x1",s(e.seqLength)).attr("x2",s(e.seqLength)).attr("y1",u+o/4).attr("y2",u+3*o/4).attr("class","pfam-domain-line"),c.selectAll("rect.pfam-domain-domain").data(e.pfams).enter().append("rect").attr("x",(function(t){return s(t.start)})).attr("y",u).attr("rx",17).attr("ry",17).attr("width",(function(t){return s(t.end)-s(t.start)})).attr("height",o).attr("data-id",(function(t,e){return"pfam-details-orf-"+r+"-"+e+"-tooltip"})).attr("class",(function(t){return"pfam-domain-domain "+t.html_class})).attr("stroke-width",(function(t){return t.go_terms.length>0?3:1})),c.selectAll("text.pfam-domain-text").data(e.pfams).enter().append("text").text((function(t){var e=s(t.end)-s(t.start),n=(t.name.match(/[a-z]/g)||[]).length,r=(t.name.match(/[A-Z]/g)||[]).length;return e>16*r+18*(t.name.length-r-n)+8*n?t.name:"..."})).attr("x",(function(t){return s((t.start+t.end)/2)})).attr("text-anchor","middle").attr("y",u+.7*o).attr("class",(function(t){return"pfam-domain-text "+t.html_class})).attr("font-size",l.text_height).attr("font-weight","bold")}function c(t){var e=$(this).attr("data-id");if(void 0!==e){var n=$("#"+e);if(r&&(clearTimeout(n.data("timeout")),r.hide()),r=n,"none"===n.css("display")){var i=$(this).offset();void 0===i&&(i={left:0,top:0});var o=setTimeout((function(){return n.slideUp("fast")}),5e3);n.css("top",i.top+30).css("left",i.left+10).show().click((function(){$(this).hide()})).data("timeout",o).mouseover((function(){return clearTimeout(n.data("timeout"))})).mouseout((function(){clearTimeout(n.data("timeout")),o=setTimeout((function(){return n.slideUp("fast")}),5e3),n.data("timeout",o)}))}else n.hide()}}function h(){$("input.domains-selected-only").prop("checked")?($(".pfam-domain-svg").hide(),$(".pfam-domain-svg-singles").show()):($(".pfam-domain-svg").show(),$(".pfam-domain-svg-singles").hide())}e.drawPfamDomains=function(t,e,n){$("."+t+"-pfam-details").off(".firstClick").one("click.firstClick",(function(){!function(t,e,n){if($("#"+t).find("svg.pfam-domain-svg").length>0)return;for(var r=o.select("#"+t),d=n,g=$("#"+t).width()||1200,f=(d+10)*e.pfamOrfs.length,p=r.append("svg").attr("height",f).attr("width",g).attr("viewbox","-1 -1 "+g+" "+f).attr("class","pfam-domain-svg"),v=0,m="",y=0,b=e.pfamOrfs;y<b.length;y++){var x=b[y];v=Math.max(v,x.seqLength||0),m.length<x.id.length&&(m=x.id)}var w=p.append("g").append("text").text(m).attr("x",0).attr("y",0).attr("id","dummy-label"),A=w.node().getComputedTextLength()||10*m.length;w.remove();for(var C=i.scaleLinear().domain([1,1.02*v]).range([A+10,g]),_=r.append("div").attr("class","pfam-domain-svg-singles"),S=d+8,M=function(t){var n=e.pfamOrfs[t],i=l.unique_id++;u(_.append("svg").attr("height",S).attr("width","100%").attr("viewbox","-1 -1 "+g+" "+S).attr("class","pfam-domain-svg-single").attr("id",s.locusToFullId(n.id)+"-pfam-domains"),n,0,i,10,d,0,C),u(p,n,t,i,10,d,0,C),$("#"+s.locusToFullId(n.id)+"-svgeneorf")[0].classList.contains("svgene-selected-orf")||$("#"+s.locusToFullId(n.id)+"-pfam-domains").hide(),r.append("div").attr("id","pfam-details-orf-"+i).selectAll("div.pfam-domain-tooltip").data(n.pfams).enter().append("div").attr("class","pfam-domain-tooltip").attr("id",(function(t,e){return"pfam-details-orf-"+i+"-"+e+"-tooltip"})).html((function(t){return function(t,e){var n="<a class='external-link' target='_blank' href=\"https://pfam.xfam.org/family/"+t.accession+'">'+t.accession+"</a> - "+t.name+"<br>";if(n+="Location: "+t.start+"-"+t.end+" AA<br>",n+="Score: "+t.score+", E-Value: "+t.evalue+"<br>",n+=t.description+"<br><br>",t.go_terms.length>0){n+="Gene Ontologies:<br>";for(var r=0,i=t.go_terms;r<i.length;r++){var o=i[r];n+="&nbsp;"+o+"<br>"}n+="<br>"}return n+="AA sequence: "+a.clipboardCopyConstruct(t.translation)+"<br>"}(t)})),$(".pfam-domain-tooltip .clipboard-copy").off("click").click(a.copyToClipboard)},T=0;T<e.pfamOrfs.length;T++)M(T);$(".pfam-domain-orflabel").off("click").click((function(t){$("#"+s.locusToFullId($(this).attr("data-locus")||"none")+"-svgeneorf").trigger(t)})),o.selectAll("g.domain-group").data(e.pfamOrfs),$(".pfam-domain-domain").click(c),h()}(t+"-pfam-details-svg",e,n)}))},e.redrawPfamDomains=h},function(t,e,n){"use strict";Object.defineProperty(e,"__esModule",{value:!0}),e.createRecordOverviews=void 0;var r=n(6),i=n(0);function o(t,e,n){if(0!==e.length){var o=i.select("#record-minimap-"+t);o.attr("width",810).attr("height",40);var a=r.scaleLinear().domain([0,n]).range([0,800]);o.append("line").attr("x1",0).attr("y1",20).attr("x2",800).attr("y2",20).attr("class","minimap-record-end centerline"),o.append("line").attr("x1",0).attr("y1",0).attr("x2",0).attr("y2",40).attr("class","minimap-record-end centerline"),o.append("line").attr("x1",800).attr("y1",0).attr("x2",800).attr("y2",40).attr("class","centerline"),o.selectAll("line.minimap-region-label-line").data(e).enter().append("line").attr("class","minimap-region-label-line centerline").attr("x1",(function(t){return a((t.start+t.end)/2)})).attr("y1",(function(t){return parseInt(t.anchor.split("c")[1],10)%2==0?25:15})).attr("x2",(function(t){return a((t.start+t.end)/2)})).attr("y2",(function(t){return parseInt(t.anchor.split("c")[1],10)%2==0?30:10})),o.selectAll("rect.minimap-region").data(e).enter().append("rect").attr("class",(function(t){return"minimap-region "+(t.products.length>1?"hybrid":t.products[0])})).attr("y",15).attr("x",(function(t){return a(t.start)})).attr("width",(function(t){return a(t.end-t.start)})).attr("height",10).style("stroke","black"),o.selectAll("text.minimap-region-label").data(e).enter().append("text").attr("x",(function(t){return a((t.start+t.end)/2)})).attr("y",20).attr("font-size","xx-small").attr("class","minimap-region-label").attr("text-anchor","middle").attr("dy",(function(t){return(parseInt(t.anchor.split("c")[1],10)%2==0?"2":"-1.2")+"em"})).text((function(t){return t.anchor.split("c")[1]}))}}e.createRecordOverviews=function(t){for(var e=0,n=0,r=t;n<r.length;n++){var i=r[n];e+=1,0!==i.regions.length&&o(e,i.regions,i.length)}}},function(t,e,n){"use strict";var r=this&&this.__importDefault||function(t){return t&&t.__esModule?t:{default:t}};Object.defineProperty(e,"__esModule",{value:!0}),e.drawStructures=e.drawStructure=void 0;var i=r(n(27));function o(){var t=this,e={height:200,padding:30,width:($(t).parent().width()||270)-20},n=$(t).parent(),r=$(t).is(":visible");r||(n.detach(t.id),$("body").append(t));var o=new i.default.Drawer(e);i.default.parse(t.getAttribute("data-smiles"),(function(e){return o.draw(e,t,"light",!1)})),r||n.prepend(t),$(this).click((function(t){$("#"+$(this).attr("id")+"-modal").show();var e=$("#"+$(this).attr("id")+"-modal").children(".modal-content").children(".smiles-canvas-modal").first();if(void 0!==e){var n={height:e.height()||500,padding:30,width:(e.width()||700)+80},r=e.parent().parent().hasClass("expanded");r||e.parent().css("display","block");var o=new i.default.Drawer(n),a=e[0];i.default.parse(a.getAttribute("data-smiles"),(function(t){return o.draw(t,a,"light",!1)})),r||$(e).parent().removeAttr("style"),$("#"+$(this).attr("id")+"-modal").children(".modal-content").css("top",$(this).position().top-n.height/2+"px")}}))}e.drawStructure=o,e.drawStructures=function(){$(".smiles-canvas").each(o),$(".modal-container").off("click").click((function(){return $(".modal-container").hide()}))}},function(t,e,n){const r=n(28),i=n(31);var o=!("undefined"==typeof window||!window.document||!window.document.createElement),a={Version:"1.0.0"};a.Drawer=r,a.Parser=i,a.clean=function(t){return t.replace(/[^A-Za-z0-9@\.\+\-\?!\(\)\[\]\{\}/\\=#\$:\*]/g,"")},a.apply=function(t,e="canvas[data-smiles]",n="light",i=null){let o=new r(t),s=document.querySelectorAll(e);for(var l=0;l<s.length;l++){let t=s[l];a.parse(t.getAttribute("data-smiles"),(function(e){o.draw(e,t,n,!1)}),(function(t){i&&i(t)}))}},a.parse=function(t,e,n){try{e&&e(i.parse(t))}catch(t){n&&n(t)}},o&&(window.SmilesDrawer=a),Array.prototype.fill||Object.defineProperty(Array.prototype,"fill",{value:function(t){if(null==this)throw new TypeError("this is null or not defined");for(var e=Object(this),n=e.length>>>0,r=arguments[1],i=r>>0,o=i<0?Math.max(n+i,0):Math.min(i,n),a=arguments[2],s=void 0===a?n:a>>0,l=s<0?Math.max(n+s,0):Math.min(s,n);o<l;)e[o]=t,o++;return e}}),t.exports=a},function(t,e,n){const r=n(9),i=n(10),o=n(3),a=n(15),s=(n(4),n(17)),l=n(12),u=n(5),c=n(16),h=n(29),d=n(18),g=n(30);t.exports=class{constructor(t){this.graph=null,this.doubleBondConfigCount=0,this.doubleBondConfig=null,this.ringIdCounter=0,this.ringConnectionIdCounter=0,this.canvasWrapper=null,this.totalOverlapScore=0,this.defaultOptions={width:500,height:500,bondThickness:.6,bondLength:15,shortBondLength:.85,bondSpacing:.18*15,atomVisualization:"default",isomeric:!0,debug:!1,terminalCarbons:!1,explicitHydrogens:!1,overlapSensitivity:.42,overlapResolutionIterations:1,compactDrawing:!0,fontSizeLarge:5,fontSizeSmall:3,padding:20,themes:{dark:{C:"#fff",O:"#e74c3c",N:"#3498db",F:"#27ae60",CL:"#16a085",BR:"#d35400",I:"#8e44ad",P:"#d35400",S:"#f1c40f",B:"#e67e22",SI:"#e67e22",H:"#fff",BACKGROUND:"#141414"},light:{C:"#222",O:"#e74c3c",N:"#3498db",F:"#27ae60",CL:"#16a085",BR:"#d35400",I:"#8e44ad",P:"#d35400",S:"#f1c40f",B:"#e67e22",SI:"#e67e22",H:"#222",BACKGROUND:"#fff"}}},this.opts=this.extend(!0,this.defaultOptions,t),this.opts.halfBondSpacing=this.opts.bondSpacing/2,this.opts.bondLengthSq=this.opts.bondLength*this.opts.bondLength,this.opts.halfFontSizeLarge=this.opts.fontSizeLarge/2,this.opts.quarterFontSizeLarge=this.opts.fontSizeLarge/4,this.opts.fifthFontSizeSmall=this.opts.fontSizeSmall/5,this.theme=this.opts.themes.dark}extend(){let t=this,e={},n=!1,r=0,i=arguments.length;"[object Boolean]"===Object.prototype.toString.call(arguments[0])&&(n=arguments[0],r++);let o=function(r){for(var i in r)Object.prototype.hasOwnProperty.call(r,i)&&(n&&"[object Object]"===Object.prototype.toString.call(r[i])?e[i]=t.extend(!0,e[i],r[i]):e[i]=r[i])};for(;r<i;r++){o(arguments[r])}return e}draw(t,e,n="light",i=!1){if(this.data=t,this.infoOnly=i,this.infoOnly||(this.canvasWrapper=new h(e,this.opts.themes[n],this.opts)),this.ringIdCounter=0,this.ringConnectionIdCounter=0,this.graph=new d(t,this.opts.isomeric),this.rings=Array(),this.ringConnections=Array(),this.originalRings=Array(),this.originalRingConnections=Array(),this.bridgedRing=!1,this.doubleBondConfigCount=null,this.doubleBondConfig=null,this.initRings(),this.initHydrogens(),!this.infoOnly){this.position(),this.restoreRingInformation(),this.resolvePrimaryOverlaps();let t=this.getOverlapScore();this.totalOverlapScore=this.getOverlapScore().total;for(var o=0;o<this.opts.overlapResolutionIterations;o++)for(var a=0;a<this.graph.edges.length;a++){let e=this.graph.edges[a];if(this.isEdgeRotatable(e)){let n=this.graph.getTreeDepth(e.sourceId,e.targetId),i=this.graph.getTreeDepth(e.targetId,e.sourceId),o=e.targetId,a=e.sourceId;if(n>i&&(o=e.sourceId,a=e.targetId),this.getSubtreeOverlapScore(a,o,t.vertexScores).value>this.opts.overlapSensitivity){let e=this.graph.vertices[o],n=this.graph.vertices[a],i=n.getNeighbours(o);if(1===i.length){let t=this.graph.vertices[i[0]],o=t.position.getRotateAwayFromAngle(e.position,n.position,r.toRad(120));this.rotateSubtree(t.id,n.id,o,n.position);let a=this.getOverlapScore().total;a>this.totalOverlapScore?this.rotateSubtree(t.id,n.id,-o,n.position):this.totalOverlapScore=a}else if(2===i.length){if(0!==n.value.rings.length&&0!==e.value.rings.length)continue;let t=this.graph.vertices[i[0]],o=this.graph.vertices[i[1]];if(1===t.value.rings.length&&1===o.value.rings.length){if(t.value.rings[0]!==o.value.rings[0])continue}else{if(0!==t.value.rings.length||0!==o.value.rings.length)continue;{let i=t.position.getRotateAwayFromAngle(e.position,n.position,r.toRad(120)),a=o.position.getRotateAwayFromAngle(e.position,n.position,r.toRad(120));this.rotateSubtree(t.id,n.id,i,n.position),this.rotateSubtree(o.id,n.id,a,n.position);let s=this.getOverlapScore().total;s>this.totalOverlapScore?(this.rotateSubtree(t.id,n.id,-i,n.position),this.rotateSubtree(o.id,n.id,-a,n.position)):this.totalOverlapScore=s}}}t=this.getOverlapScore()}}}this.resolveSecondaryOverlaps(t.scores),this.opts.isomeric&&this.annotateStereochemistry(),this.opts.compactDrawing&&"default"===this.opts.atomVisualization&&this.initPseudoElements(),this.rotateDrawing(),this.canvasWrapper.scale(this.graph.vertices),this.drawEdges(this.opts.debug),this.drawVertices(this.opts.debug),this.canvasWrapper.reset()}}edgeRingCount(t){let e=this.graph.edges[t],n=this.graph.vertices[e.sourceId],r=this.graph.vertices[e.targetId];return Math.min(n.value.rings.length,r.value.rings.length)}getBridgedRings(){let t=Array();for(var e=0;e<this.rings.length;e++)this.rings[e].isBridged&&t.push(this.rings[e]);return t}getFusedRings(){let t=Array();for(var e=0;e<this.rings.length;e++)this.rings[e].isFused&&t.push(this.rings[e]);return t}getSpiros(){let t=Array();for(var e=0;e<this.rings.length;e++)this.rings[e].isSpiro&&t.push(this.rings[e]);return t}printRingInfo(){let t="";for(var e=0;e<this.rings.length;e++){const n=this.rings[e];t+=n.id+";",t+=n.members.length+";",t+=n.neighbours.length+";",t+=n.isSpiro?"true;":"false;",t+=n.isFused?"true;":"false;",t+=n.isBridged?"true;":"false;",t+=n.rings.length+";",t+="\n"}return t}rotateDrawing(){let t=0,e=0,n=0;for(var r=0;r<this.graph.vertices.length;r++){let o=this.graph.vertices[r];if(o.value.isDrawn)for(var i=r+1;i<this.graph.vertices.length;i++){let a=this.graph.vertices[i];if(!a.value.isDrawn)continue;let s=o.position.distanceSq(a.position);s>n&&(n=s,t=r,e=i)}}let a=-o.subtract(this.graph.vertices[t].position,this.graph.vertices[e].position).angle();if(!isNaN(a)){let t=a%.523599;t<.2617995?a-=t:a+=.523599-t;for(r=0;r<this.graph.vertices.length;r++)r!==e&&this.graph.vertices[r].position.rotateAround(a,this.graph.vertices[e].position);for(r=0;r<this.rings.length;r++)this.rings[r].center.rotateAround(a,this.graph.vertices[e].position)}}getTotalOverlapScore(){return this.totalOverlapScore}getRingCount(){return this.rings.length}hasBridgedRing(){return this.bridgedRing}getHeavyAtomCount(){let t=0;for(var e=0;e<this.graph.vertices.length;e++)"H"!==this.graph.vertices[e].value.element&&t++;return t}getMolecularFormula(){let t="",e=new Map;for(var n=0;n<this.graph.vertices.length;n++){let t=this.graph.vertices[n].value;if(e.has(t.element)?e.set(t.element,e.get(t.element)+1):e.set(t.element,1),t.bracket&&!t.bracket.chirality&&(e.has("H")?e.set("H",e.get("H")+t.bracket.hcount):e.set("H",t.bracket.hcount)),!t.bracket){let n=l.maxBonds[t.element]-t.bondCount;t.isPartOfAromaticRing&&n--,e.has("H")?e.set("H",e.get("H")+n):e.set("H",n)}}if(e.has("C")){let n=e.get("C");t+="C"+(n>1?n:""),e.delete("C")}if(e.has("H")){let n=e.get("H");t+="H"+(n>1?n:""),e.delete("H")}return Object.keys(l.atomicNumbers).sort().map(n=>{if(e.has(n)){let r=e.get(n);t+=n+(r>1?r:"")}}),t}getRingbondType(t,e){if(t.value.getRingbondCount()<1||e.value.getRingbondCount()<1)return null;for(var n=0;n<t.value.ringbonds.length;n++)for(var r=0;r<e.value.ringbonds.length;r++)if(t.value.ringbonds[n].id===e.value.ringbonds[r].id)return"-"===t.value.ringbonds[n].bondType?e.value.ringbonds[r].bond:t.value.ringbonds[n].bond;return null}initRings(){let t=new Map;for(var e=this.graph.vertices.length-1;e>=0;e--){let r=this.graph.vertices[e];if(0!==r.value.ringbonds.length)for(var n=0;n<r.value.ringbonds.length;n++){let e=r.value.ringbonds[n].id,i=r.value.ringbonds[n].bond;if(t.has(e)){let o=r.id,a=t.get(e)[0],l=t.get(e)[1],u=new s(o,a,1);u.setBondType(l||i||"-");let c=this.graph.addEdge(u),h=this.graph.vertices[a];r.addRingbondChild(a,n),r.value.addNeighbouringElement(h.value.element),h.addRingbondChild(o,n),h.value.addNeighbouringElement(r.value.element),r.edges.push(c),h.edges.push(c),t.delete(e)}else t.set(e,[r.id,i])}}let r=g.getRings(this.graph);if(null!==r){for(e=0;e<r.length;e++){let t=[...r[e]],i=this.addRing(new u(t));for(n=0;n<t.length;n++)this.graph.vertices[t[n]].value.rings.push(i)}for(e=0;e<this.rings.length-1;e++)for(n=e+1;n<this.rings.length;n++){let t=this.rings[e],r=this.rings[n],i=new c(t,r);i.vertices.size>0&&this.addRingConnection(i)}for(e=0;e<this.rings.length;e++){let t=this.rings[e];t.neighbours=c.getNeighbours(this.ringConnections,t.id)}for(e=0;e<this.rings.length;e++){let t=this.rings[e];this.graph.vertices[t.members[0]].value.addAnchoredRing(t.id)}for(this.backupRingInformation();this.rings.length>0;){let t=-1;for(e=0;e<this.rings.length;e++){let n=this.rings[e];this.isPartOfBridgedRing(n.id)&&!n.isBridged&&(t=n.id)}if(-1===t)break;let n=this.getRing(t),r=this.getBridgedRingRings(n.id);this.bridgedRing=!0,this.createBridgedRing(r,n.members[0]);for(e=0;e<r.length;e++)this.removeRing(r[e])}}}initHydrogens(){if(!this.opts.explicitHydrogens)for(var t=0;t<this.graph.vertices.length;t++){let e=this.graph.vertices[t];if("H"!==e.value.element)continue;let n=this.graph.vertices[e.neighbours[0]];n.value.hasHydrogen=!0,(!n.value.isStereoCenter||n.value.rings.length<2&&!n.value.bridgedRing||n.value.bridgedRing&&n.value.originalRings.length<2)&&(e.value.isDrawn=!1)}}getBridgedRingRings(t){let e=Array(),n=this,r=function(t){let i=n.getRing(t);e.push(t);for(var o=0;o<i.neighbours.length;o++){let a=i.neighbours[o];-1===e.indexOf(a)&&a!==t&&c.isBridge(n.ringConnections,n.graph.vertices,t,a)&&r(a)}};return r(t),i.unique(e)}isPartOfBridgedRing(t){for(var e=0;e<this.ringConnections.length;e++)if(this.ringConnections[e].containsRing(t)&&this.ringConnections[e].isBridge(this.graph.vertices))return!0;return!1}createBridgedRing(t,e){let n=new Set,r=new Set,o=new Set;for(var a=0;a<t.length;a++){let e=this.getRing(t[a]);e.isPartOfBridged=!0;for(var s=0;s<e.members.length;s++)r.add(e.members[s]);for(s=0;s<e.neighbours.length;s++){let n=e.neighbours[s];-1===t.indexOf(n)&&o.add(e.neighbours[s])}}let l=new Set;for(let e of r){let r=this.graph.vertices[e],o=i.intersection(t,r.value.rings);1===r.value.rings.length||1===o.length?n.add(r.id):l.add(r.id)}Array();let c=Array();for(let t of l){let e=this.graph.vertices[t],r=!1;for(let t=0;t<e.edges.length;t++)1===this.edgeRingCount(e.edges[t])&&(r=!0);r?(e.value.isBridgeNode=!0,n.add(e.id)):(e.value.isBridge=!0,n.add(e.id))}let h=new u([...n]);h.isBridged=!0,h.neighbours=[...o];for(a=0;a<t.length;a++)h.rings.push(this.getRing(t[a]).clone());this.addRing(h);for(a=0;a<h.members.length;a++)this.graph.vertices[h.members[a]].value.bridgedRing=h.id;for(a=0;a<c.length;a++){this.graph.vertices[c[a]].value.rings=Array()}for(let e of n){let n=this.graph.vertices[e];n.value.rings=i.removeAll(n.value.rings,t),n.value.rings.push(h.id)}for(a=0;a<t.length;a++)for(s=a+1;s<t.length;s++)this.removeRingConnectionsBetween(t[a],t[s]);for(let e of o){let n=this.getRingConnections(e,t);for(s=0;s<n.length;s++)this.getRingConnection(n[s]).updateOther(h.id,e);this.getRing(e).neighbours.push(h.id)}return h}areVerticesInSameRing(t,e){for(var n=0;n<t.value.rings.length;n++)for(var r=0;r<e.value.rings.length;r++)if(t.value.rings[n]===e.value.rings[r])return!0;return!1}getCommonRings(t,e){let n=Array();for(var r=0;r<t.value.rings.length;r++)for(var i=0;i<e.value.rings.length;i++)t.value.rings[r]==e.value.rings[i]&&n.push(t.value.rings[r]);return n}getLargestOrAromaticCommonRing(t,e){let n=this.getCommonRings(t,e),r=0,i=null;for(var o=0;o<n.length;o++){let t=this.getRing(n[o]),e=t.getSize();if(t.isBenzeneLike(this.graph.vertices))return t;e>r&&(r=e,i=t)}return i}getVerticesAt(t,e,n){let r=Array();for(var i=0;i<this.graph.vertices.length;i++){let o=this.graph.vertices[i];o.id!==n&&o.positioned&&(t.distanceSq(o.position)<=e*e&&r.push(o.id))}return r}getClosestVertex(t){let e=99999,n=null;for(var r=0;r<this.graph.vertices.length;r++){let i=this.graph.vertices[r];if(i.id===t.id)continue;let o=t.position.distanceSq(i.position);o<e&&(e=o,n=i)}return n}addRing(t){return t.id=this.ringIdCounter++,this.rings.push(t),t.id}removeRing(t){this.rings=this.rings.filter((function(e){return e.id!==t})),this.ringConnections=this.ringConnections.filter((function(e){return e.firstRingId!==t&&e.secondRingId!==t}));for(var e=0;e<this.rings.length;e++){let n=this.rings[e];n.neighbours=n.neighbours.filter((function(e){return e!==t}))}}getRing(t){for(var e=0;e<this.rings.length;e++)if(this.rings[e].id==t)return this.rings[e]}addRingConnection(t){return t.id=this.ringConnectionIdCounter++,this.ringConnections.push(t),t.id}removeRingConnection(t){this.ringConnections=this.ringConnections.filter((function(e){return e.id!==t}))}removeRingConnectionsBetween(t,e){let n=Array();for(var r=0;r<this.ringConnections.length;r++){let i=this.ringConnections[r];(i.firstRingId===t&&i.secondRingId===e||i.firstRingId===e&&i.secondRingId===t)&&n.push(i.id)}for(r=0;r<n.length;r++)this.removeRingConnection(n[r])}getRingConnection(t){for(var e=0;e<this.ringConnections.length;e++)if(this.ringConnections[e].id==t)return this.ringConnections[e]}getRingConnections(t,e){let n=Array();for(var r=0;r<this.ringConnections.length;r++){let o=this.ringConnections[r];for(var i=0;i<e.length;i++){let r=e[i];(o.firstRingId===t&&o.secondRingId===r||o.firstRingId===r&&o.secondRingId===t)&&n.push(o.id)}}return n}getOverlapScore(){let t=0,e=new Float32Array(this.graph.vertices.length);for(var n=0;n<this.graph.vertices.length;n++)e[n]=0;for(n=0;n<this.graph.vertices.length;n++)for(var r=this.graph.vertices.length;--r>n;){let i=this.graph.vertices[n],a=this.graph.vertices[r];if(!i.value.isDrawn||!a.value.isDrawn)continue;let s=o.subtract(i.position,a.position).lengthSq();if(s<this.opts.bondLengthSq){let i=(this.opts.bondLength-Math.sqrt(s))/this.opts.bondLength;t+=i,e[n]+=i,e[r]+=i}}let i=Array();for(n=0;n<this.graph.vertices.length;n++)i.push({id:n,score:e[n]});return i.sort((function(t,e){return e.score-t.score})),{total:t,scores:i,vertexScores:e}}chooseSide(t,e,n){let r=t.getNeighbours(e.id),o=e.getNeighbours(t.id),a=r.length,s=o.length,l=i.merge(r,o),u=[0,0];for(var c=0;c<l.length;c++){this.graph.vertices[l[c]].position.sameSideAs(t.position,e.position,n[0])?u[0]++:u[1]++}let h=[0,0];for(c=0;c<this.graph.vertices.length;c++){this.graph.vertices[c].position.sameSideAs(t.position,e.position,n[0])?h[0]++:h[1]++}return{totalSideCount:h,totalPosition:h[0]>h[1]?0:1,sideCount:u,position:u[0]>u[1]?0:1,anCount:a,bnCount:s}}setRingCenter(t){let e=t.getSize(),n=new o(0,0);for(var r=0;r<e;r++)n.add(this.graph.vertices[t.members[r]].position);t.center=n.divide(e)}getSubringCenter(t,e){let n=e.value.originalRings,r=t.center,i=Number.MAX_VALUE;for(var o=0;o<n.length;o++)for(var a=0;a<t.rings.length;a++)n[o]===t.rings[a].id&&t.rings[a].getSize()<i&&(r=t.rings[a].center,i=t.rings[a].getSize());return r}drawEdges(t){let e=this,n=Array(this.graph.edges.length);if(n.fill(!1),this.graph.traverseBF(0,(function(r){let i=e.graph.getEdges(r.id);for(var o=0;o<i.length;o++){let r=i[o];n[r]||(n[r]=!0,e.drawEdge(r,t))}})),!this.bridgedRing)for(var r=0;r<this.rings.length;r++){let t=this.rings[r];this.isRingAromatic(t)&&this.canvasWrapper.drawAromaticityRing(t)}}drawEdge(t,e){let n=this,r=this.graph.edges[t],s=this.graph.vertices[r.sourceId],l=this.graph.vertices[r.targetId],u=s.value.element,c=l.value.element;if(!(s.value.isDrawn&&l.value.isDrawn||"default"!==this.opts.atomVisualization))return;let h=s.position,d=l.position,g=this.getEdgeNormals(r),f=i.clone(g);if(f[0].multiplyScalar(10).add(h),f[1].multiplyScalar(10).add(h),"="===r.bondType||"="===this.getRingbondType(s,l)||r.isPartOfAromaticRing&&this.bridgedRing){let t=this.areVerticesInSameRing(s,l),e=this.chooseSide(s,l,f);if(t){let t=this.getLargestOrAromaticCommonRing(s,l).center;g[0].multiplyScalar(n.opts.bondSpacing),g[1].multiplyScalar(n.opts.bondSpacing);let e=null;e=t.sameSideAs(s.position,l.position,o.add(h,g[0]))?new a(o.add(h,g[0]),o.add(d,g[0]),u,c):new a(o.add(h,g[1]),o.add(d,g[1]),u,c),e.shorten(this.opts.bondLength-this.opts.shortBondLength*this.opts.bondLength),r.isPartOfAromaticRing?this.canvasWrapper.drawLine(e,!0):this.canvasWrapper.drawLine(e),this.canvasWrapper.drawLine(new a(h,d,u,c))}else if(r.center||s.isTerminal()&&l.isTerminal()){g[0].multiplyScalar(n.opts.halfBondSpacing),g[1].multiplyScalar(n.opts.halfBondSpacing);let t=new a(o.add(h,g[0]),o.add(d,g[0]),u,c),e=new a(o.add(h,g[1]),o.add(d,g[1]),u,c);this.canvasWrapper.drawLine(t),this.canvasWrapper.drawLine(e)}else if(0==e.anCount&&e.bnCount>1||0==e.bnCount&&e.anCount>1){g[0].multiplyScalar(n.opts.halfBondSpacing),g[1].multiplyScalar(n.opts.halfBondSpacing);let t=new a(o.add(h,g[0]),o.add(d,g[0]),u,c),e=new a(o.add(h,g[1]),o.add(d,g[1]),u,c);this.canvasWrapper.drawLine(t),this.canvasWrapper.drawLine(e)}else if(e.sideCount[0]>e.sideCount[1]){g[0].multiplyScalar(n.opts.bondSpacing),g[1].multiplyScalar(n.opts.bondSpacing);let t=new a(o.add(h,g[0]),o.add(d,g[0]),u,c);t.shorten(this.opts.bondLength-this.opts.shortBondLength*this.opts.bondLength),this.canvasWrapper.drawLine(t),this.canvasWrapper.drawLine(new a(h,d,u,c))}else if(e.sideCount[0]<e.sideCount[1]){g[0].multiplyScalar(n.opts.bondSpacing),g[1].multiplyScalar(n.opts.bondSpacing);let t=new a(o.add(h,g[1]),o.add(d,g[1]),u,c);t.shorten(this.opts.bondLength-this.opts.shortBondLength*this.opts.bondLength),this.canvasWrapper.drawLine(t),this.canvasWrapper.drawLine(new a(h,d,u,c))}else if(e.totalSideCount[0]>e.totalSideCount[1]){g[0].multiplyScalar(n.opts.bondSpacing),g[1].multiplyScalar(n.opts.bondSpacing);let t=new a(o.add(h,g[0]),o.add(d,g[0]),u,c);t.shorten(this.opts.bondLength-this.opts.shortBondLength*this.opts.bondLength),this.canvasWrapper.drawLine(t),this.canvasWrapper.drawLine(new a(h,d,u,c))}else if(e.totalSideCount[0]<=e.totalSideCount[1]){g[0].multiplyScalar(n.opts.bondSpacing),g[1].multiplyScalar(n.opts.bondSpacing);let t=new a(o.add(h,g[1]),o.add(d,g[1]),u,c);t.shorten(this.opts.bondLength-this.opts.shortBondLength*this.opts.bondLength),this.canvasWrapper.drawLine(t),this.canvasWrapper.drawLine(new a(h,d,u,c))}}else if("#"===r.bondType){g[0].multiplyScalar(n.opts.bondSpacing/1.5),g[1].multiplyScalar(n.opts.bondSpacing/1.5);let t=new a(o.add(h,g[0]),o.add(d,g[0]),u,c),e=new a(o.add(h,g[1]),o.add(d,g[1]),u,c);this.canvasWrapper.drawLine(t),this.canvasWrapper.drawLine(e),this.canvasWrapper.drawLine(new a(h,d,u,c))}else if("."===r.bondType);else{let t=s.value.isStereoCenter,e=l.value.isStereoCenter;"up"===r.wedge?this.canvasWrapper.drawWedge(new a(h,d,u,c,t,e)):"down"===r.wedge?this.canvasWrapper.drawDashedWedge(new a(h,d,u,c,t,e)):this.canvasWrapper.drawLine(new a(h,d,u,c,t,e))}if(e){let e=o.midpoint(h,d);this.canvasWrapper.drawDebugText(e.x,e.y,"e: "+t)}}drawVertices(t){var e=this.graph.vertices.length;for(e=0;e<this.graph.vertices.length;e++){let n=this.graph.vertices[e],r=n.value,a=0,s=0,u=n.value.bondCount,c=r.element,h=l.maxBonds[c]-u,d=n.getTextDirection(this.graph.vertices),g=!(!this.opts.terminalCarbons&&"C"===c&&!r.hasAttachedPseudoElements)&&n.isTerminal(),f="C"===r.element;if("N"===r.element&&r.isPartOfAromaticRing&&(h=0),r.bracket&&(h=r.bracket.hcount,a=r.bracket.charge,s=r.bracket.isotope),"allballs"===this.opts.atomVisualization)this.canvasWrapper.drawBall(n.position.x,n.position.y,c);else if(r.isDrawn&&(!f||r.drawExplicit||g||r.hasAttachedPseudoElements)||1===this.graph.vertices.length)"default"===this.opts.atomVisualization?this.canvasWrapper.drawText(n.position.x,n.position.y,c,h,d,g,a,s,r.getAttachedPseudoElements()):"balls"===this.opts.atomVisualization&&this.canvasWrapper.drawBall(n.position.x,n.position.y,c);else if(2===n.getNeighbourCount()&&1==n.forcePositioned){let t=this.graph.vertices[n.neighbours[0]].position,e=this.graph.vertices[n.neighbours[1]].position,r=o.threePointangle(n.position,t,e);Math.abs(Math.PI-r)<.1&&this.canvasWrapper.drawPoint(n.position.x,n.position.y,c)}if(t){let t="v: "+n.id+" "+i.print(r.ringbonds);this.canvasWrapper.drawDebugText(n.position.x,n.position.y,t)}}if(this.opts.debug)for(e=0;e<this.rings.length;e++){let t=this.rings[e].center;this.canvasWrapper.drawDebugPoint(t.x,t.y,"r: "+this.rings[e].id)}}position(){let t=null;for(var e=0;e<this.graph.vertices.length;e++)if(null!==this.graph.vertices[e].value.bridgedRing){t=this.graph.vertices[e];break}for(e=0;e<this.rings.length;e++)this.rings[e].isBridged&&(t=this.graph.vertices[this.rings[e].members[0]]);this.rings.length>0&&null===t&&(t=this.graph.vertices[this.rings[0].members[0]]),null===t&&(t=this.graph.vertices[0]),this.createNextBond(t,null,0)}backupRingInformation(){this.originalRings=Array(),this.originalRingConnections=Array();for(var t=0;t<this.rings.length;t++)this.originalRings.push(this.rings[t]);for(t=0;t<this.ringConnections.length;t++)this.originalRingConnections.push(this.ringConnections[t]);for(t=0;t<this.graph.vertices.length;t++)this.graph.vertices[t].value.backupRings()}restoreRingInformation(){let t=this.getBridgedRings();this.rings=Array(),this.ringConnections=Array();for(var e=0;e<t.length;e++){let r=t[e];for(var n=0;n<r.rings.length;n++){let t=r.rings[n];this.originalRings[t.id].center=t.center}}for(e=0;e<this.originalRings.length;e++)this.rings.push(this.originalRings[e]);for(e=0;e<this.originalRingConnections.length;e++)this.ringConnections.push(this.originalRingConnections[e]);for(e=0;e<this.graph.vertices.length;e++)this.graph.vertices[e].value.restoreRings()}createRing(t,e=null,n=null,i=null){if(t.positioned)return;e=e||new o(0,0);let a=t.getOrderedNeighbours(this.ringConnections),s=n?o.subtract(n.position,e).angle():0,l=r.polyCircumradius(this.opts.bondLength,t.getSize()),u=r.centralAngle(t.getSize());t.centralAngle=u;let h=s,d=this,g=n?n.id:null;if(-1===t.members.indexOf(g)&&(n&&(n.positioned=!1),g=t.members[0]),t.isBridged){this.graph.kkLayout(t.members.slice(),e,n.id,t,this.opts.bondLength),t.positioned=!0,this.setRingCenter(t),e=t.center;for(var f=0;f<t.rings.length;f++)this.setRingCenter(t.rings[f])}else t.eachMember(this.graph.vertices,(function(n){let r=d.graph.vertices[n];r.positioned||r.setPosition(e.x+Math.cos(h)*l,e.y+Math.sin(h)*l),h+=u,(!t.isBridged||t.rings.length<3)&&(r.angle=h,r.positioned=!0)}),g,i?i.id:null);t.positioned=!0,t.center=e;for(f=0;f<a.length;f++){let n=this.getRing(a[f].neighbour);if(n.positioned)continue;let i=c.getVertices(this.ringConnections,t.id,n.id);if(2===i.length){t.isFused=!0,n.isFused=!0;let a=this.graph.vertices[i[0]],s=this.graph.vertices[i[1]],l=o.midpoint(a.position,s.position),u=o.normals(a.position,s.position);u[0].normalize(),u[1].normalize();let c=r.polyCircumradius(this.opts.bondLength,n.getSize()),h=r.apothem(c,n.getSize());u[0].multiplyScalar(h).add(l),u[1].multiplyScalar(h).add(l);let d=u[0];o.subtract(e,u[1]).lengthSq()>o.subtract(e,u[0]).lengthSq()&&(d=u[1]);let g=o.subtract(a.position,d),f=o.subtract(s.position,d);-1===g.clockwise(f)?n.positioned||this.createRing(n,d,a,s):n.positioned||this.createRing(n,d,s,a)}else if(1===i.length){t.isSpiro=!0,n.isSpiro=!0;let a=this.graph.vertices[i[0]],s=o.subtract(e,a.position);s.invert(),s.normalize();let l=r.polyCircumradius(this.opts.bondLength,n.getSize());s.multiplyScalar(l),s.add(a.position),n.positioned||this.createRing(n,s,a)}}for(f=0;f<t.members.length;f++){let e=this.graph.vertices[t.members[f]],n=e.neighbours;for(var p=0;p<n.length;p++){let t=this.graph.vertices[n[p]];t.positioned||(t.value.isConnectedToRing=!0,this.createNextBond(t,e,0))}}}rotateSubtree(t,e,n,r){let i=this;this.graph.traverseTree(t,e,(function(t){t.position.rotateAround(n,r);for(var e=0;e<t.value.anchoredRings.length;e++){let o=i.rings[t.value.anchoredRings[e]];o&&o.center.rotateAround(n,r)}}))}getSubtreeOverlapScore(t,e,n){let r=this,i=0,a=new o(0,0),s=0;return this.graph.traverseTree(t,e,(function(t){if(!t.value.isDrawn)return;let e=n[t.id];e>r.opts.overlapSensitivity&&(i+=e,s++);let o=r.graph.vertices[t.id].position.clone();o.multiplyScalar(e),a.add(o)})),a.divide(i),{value:i/s,center:a}}getCurrentCenterOfMass(){let t=new o(0,0),e=0;for(var n=0;n<this.graph.vertices.length;n++){let r=this.graph.vertices[n];r.positioned&&(t.add(r.position),e++)}return t.divide(e)}getCurrentCenterOfMassInNeigbourhood(t,e=2*this.opts.bondLength){let n=new o(0,0),r=0,i=e*e;for(var a=0;a<this.graph.vertices.length;a++){let e=this.graph.vertices[a];e.positioned&&t.distanceSq(e.position)<i&&(n.add(e.position),r++)}return n.divide(r)}resolvePrimaryOverlaps(){let t=Array(),e=Array(this.graph.vertices.length);for(var n=0;n<this.rings.length;n++){let o=this.rings[n];for(var r=0;r<o.members.length;r++){let n=this.graph.vertices[o.members[r]];if(e[n.id])continue;e[n.id]=!0;let a=this.getNonRingNeighbours(n.id);if(a.length>1){let e=Array();for(var i=0;i<n.value.rings.length;i++)e.push(n.value.rings[i]);t.push({common:n,rings:e,vertices:a})}else if(1===a.length&&2===n.value.rings.length){let e=Array();for(i=0;i<n.value.rings.length;i++)e.push(n.value.rings[i]);t.push({common:n,rings:e,vertices:a})}}}for(n=0;n<t.length;n++){let e=t[n];if(2===e.vertices.length){let t=e.vertices[0],n=e.vertices[1];if(!t.value.isDrawn||!n.value.isDrawn)continue;let r=(2*Math.PI-this.getRing(e.rings[0]).getAngle())/6;this.rotateSubtree(t.id,e.common.id,r,e.common.position),this.rotateSubtree(n.id,e.common.id,-r,e.common.position);let i=this.getOverlapScore(),o=this.getSubtreeOverlapScore(t.id,e.common.id,i.vertexScores),a=this.getSubtreeOverlapScore(n.id,e.common.id,i.vertexScores),s=o.value+a.value;this.rotateSubtree(t.id,e.common.id,-2*r,e.common.position),this.rotateSubtree(n.id,e.common.id,2*r,e.common.position),i=this.getOverlapScore(),o=this.getSubtreeOverlapScore(t.id,e.common.id,i.vertexScores),a=this.getSubtreeOverlapScore(n.id,e.common.id,i.vertexScores),o.value+a.value>s&&(this.rotateSubtree(t.id,e.common.id,2*r,e.common.position),this.rotateSubtree(n.id,e.common.id,-2*r,e.common.position))}else 1===e.vertices.length&&e.rings.length}}resolveSecondaryOverlaps(t){for(var e=0;e<t.length;e++)if(t[e].score>this.opts.overlapSensitivity){let n=this.graph.vertices[t[e].id];if(n.isTerminal()){let t=this.getClosestVertex(n);if(t){let e=null;e=t.isTerminal()?0===t.id?this.graph.vertices[1].position:t.previousPosition:0===t.id?this.graph.vertices[1].position:t.position;let i=0===n.id?this.graph.vertices[1].position:n.previousPosition;n.position.rotateAwayFrom(e,i,r.toRad(20))}}}}getLastVertexWithAngle(t){let e=0,n=null;for(;!e&&t;)n=this.graph.vertices[t],e=n.angle,t=n.parentVertexId;return n}createNextBond(t,e=null,n=0,a=!1,s=!1){if(t.positioned&&!s)return;let l=!1;if(e){let n=this.graph.getEdge(t.id,e.id);"/"!==n.bondType&&"\\"!==n.bondType||++this.doubleBondConfigCount%2!=1||null===this.doubleBondConfig&&(this.doubleBondConfig=n.bondType,l=!0,null===e.parentVertexId&&t.value.branchBond&&("/"===this.doubleBondConfig?this.doubleBondConfig="\\":"\\"===this.doubleBondConfig&&(this.doubleBondConfig="/")))}if(!s)if(e)if(e.value.rings.length>0){let n=e.neighbours,r=null,a=new o(0,0);if(null===e.value.bridgedRing&&e.value.rings.length>1)for(var u=0;u<n.length;u++){let t=this.graph.vertices[n[u]];if(i.containsAll(t.value.rings,e.value.rings)){r=t;break}}if(null===r){for(u=0;u<n.length;u++){let t=this.graph.vertices[n[u]];t.positioned&&this.areVerticesInSameRing(t,e)&&a.add(o.subtract(t.position,e.position))}a.invert().normalize().multiplyScalar(this.opts.bondLength).add(e.position)}else a=r.position.clone().rotateAround(Math.PI,e.position);t.previousPosition=e.position,t.setPositionFromVector(a),t.positioned=!0}else{let r=new o(this.opts.bondLength,0);r.rotate(n),r.add(e.position),t.setPositionFromVector(r),t.previousPosition=e.position,t.positioned=!0}else{let e=new o(this.opts.bondLength,0);e.rotate(r.toRad(-60)),t.previousPosition=e,t.setPosition(this.opts.bondLength,0),t.angle=r.toRad(-60),null===t.value.bridgedRing&&(t.positioned=!0)}if(null!==t.value.bridgedRing){let e=this.getRing(t.value.bridgedRing);if(!e.positioned){let n=o.subtract(t.previousPosition,t.position);n.invert(),n.normalize();let i=r.polyCircumradius(this.opts.bondLength,e.members.length);n.multiplyScalar(i),n.add(t.position),this.createRing(e,n,t)}}else if(t.value.rings.length>0){let e=this.getRing(t.value.rings[0]);if(!e.positioned){let n=o.subtract(t.previousPosition,t.position);n.invert(),n.normalize();let i=r.polyCircumradius(this.opts.bondLength,e.getSize());n.multiplyScalar(i),n.add(t.position),this.createRing(e,n,t)}}else{t.value.isStereoCenter;let n=t.getNeighbours(),s=Array();for(u=0;u<n.length;u++)this.graph.vertices[n[u]].value.isDrawn&&s.push(n[u]);e&&(s=i.remove(s,e.id));let c=t.getAngle();if(1===s.length){let n=this.graph.vertices[s[0]];if("#"===t.value.bondType||e&&"#"===e.value.bondType||"="===t.value.bondType&&e&&0===e.value.rings.length&&"="===e.value.bondType&&"-"!==t.value.branchBond){if(t.value.drawExplicit=!1,e){this.graph.getEdge(t.id,e.id).center=!0}this.graph.getEdge(t.id,n.id).center=!0,("#"===t.value.bondType||e&&"#"===e.value.bondType)&&(n.angle=0),n.drawExplicit=!0,this.createNextBond(n,t,c+n.angle)}else if(e&&e.value.rings.length>0){let e=r.toRad(60),i=-e,a=new o(this.opts.bondLength,0),s=new o(this.opts.bondLength,0);a.rotate(e).add(t.position),s.rotate(i).add(t.position);let l=this.getCurrentCenterOfMass(),u=a.distanceSq(l),h=s.distanceSq(l);n.angle=u<h?i:e,this.createNextBond(n,t,c+n.angle)}else{let r=t.angle;if(e&&e.neighbours.length>3)r=r>0?Math.min(1.0472,r):r<0?Math.max(-1.0472,r):1.0472;else if(!r){r=this.getLastVertexWithAngle(t.id).angle,r||(r=1.0472)}if(e&&!l){let e=this.graph.getEdge(t.id,n.id).bondType;"/"===e?("/"===this.doubleBondConfig||"\\"===this.doubleBondConfig&&(r=-r),this.doubleBondConfig=null):"\\"===e&&("/"===this.doubleBondConfig?r=-r:this.doubleBondConfig,this.doubleBondConfig=null)}n.angle=a?r:-r,this.createNextBond(n,t,c+n.angle)}}else if(2===s.length){let n=t.angle;n||(n=1.0472);let r=this.graph.getTreeDepth(s[0],t.id),i=this.graph.getTreeDepth(s[1],t.id),o=this.graph.vertices[s[0]],a=this.graph.vertices[s[1]];o.value.subtreeDepth=r,a.value.subtreeDepth=i;let l=this.graph.getTreeDepth(e?e.id:null,t.id);e&&(e.value.subtreeDepth=l);let u=0,h=1;"C"===a.value.element&&"C"!==o.value.element&&i>1&&r<5?(u=1,h=0):"C"!==a.value.element&&"C"===o.value.element&&r>1&&i<5?(u=0,h=1):i>r&&(u=1,h=0);let d=this.graph.vertices[s[u]],g=this.graph.vertices[s[h]],f=(this.graph.getEdge(t.id,d.id),this.graph.getEdge(t.id,g.id),!1);l<r&&l<i&&(f=!0),g.angle=n,d.angle=-n,"\\"===this.doubleBondConfig?"\\"===g.value.branchBond&&(g.angle=-n,d.angle=n):"/"===this.doubleBondConfig&&"/"===g.value.branchBond&&(g.angle=-n,d.angle=n),this.createNextBond(g,t,c+g.angle,f),this.createNextBond(d,t,c+d.angle,f)}else if(3===s.length){let n=this.graph.getTreeDepth(s[0],t.id),i=this.graph.getTreeDepth(s[1],t.id),o=this.graph.getTreeDepth(s[2],t.id),a=this.graph.vertices[s[0]],l=this.graph.vertices[s[1]],u=this.graph.vertices[s[2]];a.value.subtreeDepth=n,l.value.subtreeDepth=i,u.value.subtreeDepth=o,i>n&&i>o?(a=this.graph.vertices[s[1]],l=this.graph.vertices[s[0]],u=this.graph.vertices[s[2]]):o>n&&o>i&&(a=this.graph.vertices[s[2]],l=this.graph.vertices[s[0]],u=this.graph.vertices[s[1]]),e&&e.value.rings.length<1&&a.value.rings.length<1&&l.value.rings.length<1&&u.value.rings.length<1&&1===this.graph.getTreeDepth(l.id,t.id)&&1===this.graph.getTreeDepth(u.id,t.id)&&this.graph.getTreeDepth(a.id,t.id)>1?(a.angle=-t.angle,t.angle>=0?(l.angle=r.toRad(30),u.angle=r.toRad(90)):(l.angle=-r.toRad(30),u.angle=-r.toRad(90)),this.createNextBond(a,t,c+a.angle),this.createNextBond(l,t,c+l.angle),this.createNextBond(u,t,c+u.angle)):(a.angle=0,l.angle=r.toRad(90),u.angle=-r.toRad(90),this.createNextBond(a,t,c+a.angle),this.createNextBond(l,t,c+l.angle),this.createNextBond(u,t,c+u.angle))}else if(4===s.length){let e=this.graph.getTreeDepth(s[0],t.id),n=this.graph.getTreeDepth(s[1],t.id),i=this.graph.getTreeDepth(s[2],t.id),o=this.graph.getTreeDepth(s[3],t.id),a=this.graph.vertices[s[0]],l=this.graph.vertices[s[1]],u=this.graph.vertices[s[2]],h=this.graph.vertices[s[3]];a.value.subtreeDepth=e,l.value.subtreeDepth=n,u.value.subtreeDepth=i,h.value.subtreeDepth=o,n>e&&n>i&&n>o?(a=this.graph.vertices[s[1]],l=this.graph.vertices[s[0]],u=this.graph.vertices[s[2]],h=this.graph.vertices[s[3]]):i>e&&i>n&&i>o?(a=this.graph.vertices[s[2]],l=this.graph.vertices[s[0]],u=this.graph.vertices[s[1]],h=this.graph.vertices[s[3]]):o>e&&o>n&&o>i&&(a=this.graph.vertices[s[3]],l=this.graph.vertices[s[0]],u=this.graph.vertices[s[1]],h=this.graph.vertices[s[2]]),a.angle=-r.toRad(36),l.angle=r.toRad(36),u.angle=-r.toRad(108),h.angle=r.toRad(108),this.createNextBond(a,t,c+a.angle),this.createNextBond(l,t,c+l.angle),this.createNextBond(u,t,c+u.angle),this.createNextBond(h,t,c+h.angle)}}}getCommonRingbondNeighbour(t){let e=t.neighbours;for(var n=0;n<e.length;n++){let r=this.graph.vertices[e[n]];if(i.containsAll(r.value.rings,t.value.rings))return r}return null}isPointInRing(t){for(var e=0;e<this.rings.length;e++){let n=this.rings[e];if(!n.positioned)continue;let i=r.polyCircumradius(this.opts.bondLength,n.getSize()),o=i*i;if(t.distanceSq(n.center)<o)return!0}return!1}isEdgeInRing(t){let e=this.graph.vertices[t.sourceId],n=this.graph.vertices[t.targetId];return this.areVerticesInSameRing(e,n)}isEdgeRotatable(t){let e=this.graph.vertices[t.sourceId],n=this.graph.vertices[t.targetId];return"-"===t.bondType&&(!e.isTerminal()&&!n.isTerminal()&&!(e.value.rings.length>0&&n.value.rings.length>0&&this.areVerticesInSameRing(e,n)))}isRingAromatic(t){for(var e=0;e<t.members.length;e++){if(!this.graph.vertices[t.members[e]].value.isPartOfAromaticRing)return!1}return!0}getEdgeNormals(t){let e=this.graph.vertices[t.sourceId].position,n=this.graph.vertices[t.targetId].position;return o.units(e,n)}getNonRingNeighbours(t){let e=Array(),n=this.graph.vertices[t],r=n.neighbours;for(var o=0;o<r.length;o++){let t=this.graph.vertices[r[o]];0===i.intersection(n.value.rings,t.value.rings).length&&0==t.value.isBridge&&e.push(t)}return e}annotateStereochemistry(){for(var t=0;t<this.graph.vertices.length;t++){let o=this.graph.vertices[t];if(!o.value.isStereoCenter)continue;let a=o.getNeighbours(),s=a.length,l=Array(s);for(var e=0;e<s;e++){let t=new Uint8Array(this.graph.vertices.length),r=Array(Array());t[o.id]=1,this.visitStereochemistry(a[e],o.id,t,r,10,0);for(var n=0;n<r.length;n++)r[n].sort((function(t,e){return e-t}));l[e]=[e,r]}let u=0,c=0;for(e=0;e<l.length;e++){l[e][1].length>u&&(u=l[e][1].length);for(n=0;n<l[e][1].length;n++)l[e][1][n].length>c&&(c=l[e][1][n].length)}for(e=0;e<l.length;e++){let t=u-l[e][1].length;for(n=0;n<t;n++)l[e][1].push([]);l[e][1].push([a[e]]);for(n=0;n<l[e][1].length;n++){let t=c-l[e][1][n].length;for(var i=0;i<t;i++)l[e][1][n].push(0)}}l.sort((function(t,e){for(var n=0;n<t[1].length;n++)for(var r=0;r<t[1][n].length;r++){if(t[1][n][r]>e[1][n][r])return-1;if(t[1][n][r]<e[1][n][r])return 1}return 0}));let h=new Uint8Array(s);for(e=0;e<s;e++)h[e]=l[e][0],o.value.priority=e;let d=this.graph.vertices[a[h[0]]].position,g=this.graph.vertices[a[h[1]]].position,f=this.graph.vertices[a[h[2]]].position,p=d.relativeClockwise(g,o.position),v=(d.relativeClockwise(f,o.position),-1===p),m="@"===o.value.bracket.chirality?-1:1,y=r.parityOfPermutation(h)*m==1?"R":"S",b="down",x="up";(v&&"R"!==y||!v&&"S"!==y)&&(o.value.hydrogenDirection="up",b="up",x="down"),o.value.hasHydrogen&&(this.graph.getEdge(o.id,a[h[h.length-1]]).wedge=b);let w=new Array(a.length-1),A=o.value.rings.length>1&&o.value.hasHydrogen,C=o.value.hasHydrogen?1:0;for(e=0;e<h.length-C;e++){w[e]=new Uint32Array(2);let t=this.graph.vertices[a[h[e]]];w[e][0]+=t.value.isStereoCenter?0:1e5,w[e][0]+=this.areVerticesInSameRing(t,o)?0:1e4,w[e][0]+=t.value.isHeteroAtom()?1e3:0,w[e][0]-=0===t.value.subtreeDepth?1e3:0,w[e][0]+=1e3-t.value.subtreeDepth,w[e][1]=a[h[e]]}if(w.sort((function(t,e){return t[0]>e[0]?-1:t[0]<e[0]?1:0})),!A){let t=w[0][1];if(o.value.hasHydrogen)this.graph.getEdge(o.id,t).wedge=x;else{let n=x;for(e=h.length-1;e>=0&&(n=n===b?x:b,a[h[e]]!==t);e--);this.graph.getEdge(o.id,t).wedge=n}}o.value.chirality=y}}visitStereochemistry(t,e,n,r,i,o,a=0){n[t]=1;let s=this.graph.vertices[t],l=s.value.getAtomicNumber();r.length<=o&&r.push(Array());for(var u=0;u<this.graph.getEdge(t,e).weight;u++)r[o].push(1e3*a+l);let c=this.graph.vertices[t].neighbours;for(u=0;u<c.length;u++)1!==n[c[u]]&&o<i-1&&this.visitStereochemistry(c[u],t,n.slice(),r,i,o+1,l);if(o<i-1){let e=0;for(u=0;u<c.length;u++)e+=this.graph.getEdge(t,c[u]).weight;for(u=0;u<s.value.getMaxBonds()-e;u++)r.length<=o+1&&r.push(Array()),r[o+1].push(1e3*l+1)}}initPseudoElements(){for(var t=0;t<this.graph.vertices.length;t++){const n=this.graph.vertices[t],r=n.neighbours;let i=Array(r.length);for(var e=0;e<r.length;e++)i[e]=this.graph.vertices[r[e]];if(n.getNeighbourCount()<3||n.value.rings.length>0)continue;if("P"===n.value.element)continue;if("C"===n.value.element&&3===i.length&&"N"===i[0].value.element&&"N"===i[1].value.element&&"N"===i[2].value.element)continue;let o=0,a=0;for(e=0;e<i.length;e++){let t=i[e],n=t.value.element,r=t.getNeighbourCount();"C"!==n&&"H"!==n&&1===r&&o++,r>1&&a++}if(a>1||o<2)continue;let s=null;for(e=0;e<i.length;e++){let t=i[e];t.getNeighbourCount()>1&&(s=t)}for(e=0;e<i.length;e++){let t=i[e];if(t.getNeighbourCount()>1)continue;t.value.isDrawn=!1;let r=l.maxBonds[t.value.element]-t.value.bondCount,o="";t.value.bracket&&(r=t.value.bracket.hcount,o=t.value.bracket.charge||0),n.value.attachPseudoElement(t.value.element,s?s.value.element:null,r,o)}}for(t=0;t<this.graph.vertices.length;t++){const n=this.graph.vertices[t],r=n.value,i=r.element;if("C"===i||"H"===i||!r.isDrawn)continue;const o=n.neighbours;let a=Array(o.length);for(e=0;e<o.length;e++)a[e]=this.graph.vertices[o[e]];for(e=0;e<a.length;e++){let t=a[e].value;if(!t.hasAttachedPseudoElements||2!==t.getAttachedPseudoElementsCount())continue;const r=t.getAttachedPseudoElements();r.hasOwnProperty("0O")&&r.hasOwnProperty("3C")&&(t.isDrawn=!1,n.value.attachPseudoElement("Ac","",0))}}}}},function(t,e,n){const r=n(9),i=n(3);n(15),n(4),n(5);t.exports=class{constructor(t,e,n){this.canvas="string"==typeof t||t instanceof String?document.getElementById(t):t,this.ctx=this.canvas.getContext("2d"),this.colors=e,this.opts=n,this.drawingWidth=0,this.drawingHeight=0,this.offsetX=0,this.offsetY=0,this.fontLarge=this.opts.fontSizeLarge+"pt Helvetica, Arial, sans-serif",this.fontSmall=this.opts.fontSizeSmall+"pt Helvetica, Arial, sans-serif",this.updateSize(this.opts.width,this.opts.height),this.ctx.font=this.fontLarge,this.hydrogenWidth=this.ctx.measureText("H").width,this.halfHydrogenWidth=this.hydrogenWidth/2,this.halfBondThickness=this.opts.bondThickness/2}updateSize(t,e){this.devicePixelRatio=window.devicePixelRatio||1,this.backingStoreRatio=this.ctx.webkitBackingStorePixelRatio||this.ctx.mozBackingStorePixelRatio||this.ctx.msBackingStorePixelRatio||this.ctx.oBackingStorePixelRatio||this.ctx.backingStorePixelRatio||1,this.ratio=this.devicePixelRatio/this.backingStoreRatio,1!==this.ratio?(this.canvas.width=t*this.ratio,this.canvas.height=e*this.ratio,this.canvas.style.width=t+"px",this.canvas.style.height=e+"px",this.ctx.setTransform(this.ratio,0,0,this.ratio,0,0)):(this.canvas.width=t*this.ratio,this.canvas.height=e*this.ratio)}setTheme(t){this.colors=t}scale(t){let e=-Number.MAX_VALUE,n=-Number.MAX_VALUE,r=Number.MAX_VALUE,i=Number.MAX_VALUE;for(var o=0;o<t.length;o++){if(!t[o].value.isDrawn)continue;let a=t[o].position;e<a.x&&(e=a.x),n<a.y&&(n=a.y),r>a.x&&(r=a.x),i>a.y&&(i=a.y)}var a=this.opts.padding;e+=a,n+=a,r-=a,i-=a,this.drawingWidth=e-r,this.drawingHeight=n-i;var s=this.canvas.offsetWidth/this.drawingWidth,l=this.canvas.offsetHeight/this.drawingHeight,u=s<l?s:l;this.ctx.scale(u,u),this.offsetX=-r,this.offsetY=-i,s<l?this.offsetY+=this.canvas.offsetHeight/(2*u)-this.drawingHeight/2:this.offsetX+=this.canvas.offsetWidth/(2*u)-this.drawingWidth/2}reset(){this.ctx.setTransform(1,0,0,1,0,0)}getColor(t){return(t=t.toUpperCase())in this.colors?this.colors[t]:this.colors.C}drawCircle(t,e,n,i,o=!0,a=!1,s=""){let l=this.ctx,u=this.offsetX,c=this.offsetY;l.save(),l.lineWidth=1.5,l.beginPath(),l.arc(t+u,e+c,n,0,r.twoPI,!0),l.closePath(),a?(o?(l.fillStyle="#f00",l.fill()):(l.strokeStyle="#f00",l.stroke()),this.drawDebugText(t,e,s)):o?(l.fillStyle=i,l.fill()):(l.strokeStyle=i,l.stroke()),l.restore()}drawLine(t,e=!1,n=1){let r=this.ctx,i=this.offsetX,o=this.offsetY,a=t.clone().shorten(4),s=a.getLeftVector().clone(),l=a.getRightVector().clone();s.x+=i,s.y+=o,l.x+=i,l.y+=o,e||(r.save(),r.globalCompositeOperation="destination-out",r.beginPath(),r.moveTo(s.x,s.y),r.lineTo(l.x,l.y),r.lineCap="round",r.lineWidth=this.opts.bondThickness+1.2,r.strokeStyle=this.getColor("BACKGROUND"),r.stroke(),r.globalCompositeOperation="source-over",r.restore()),s=t.getLeftVector().clone(),l=t.getRightVector().clone(),s.x+=i,s.y+=o,l.x+=i,l.y+=o,r.save(),r.beginPath(),r.moveTo(s.x,s.y),r.lineTo(l.x,l.y),r.lineCap="round",r.lineWidth=this.opts.bondThickness;let u=this.ctx.createLinearGradient(s.x,s.y,l.x,l.y);u.addColorStop(.4,this.getColor(t.getLeftElement())||this.getColor("C")),u.addColorStop(.6,this.getColor(t.getRightElement())||this.getColor("C")),e&&(r.setLineDash([1,1.5]),r.lineWidth=this.opts.bondThickness/1.5),n<1&&(r.globalAlpha=n),r.strokeStyle=u,r.stroke(),r.restore()}drawWedge(t,e=1){if(isNaN(t.from.x)||isNaN(t.from.y)||isNaN(t.to.x)||isNaN(t.to.y))return;let n=this.ctx,r=this.offsetX,o=this.offsetY,a=t.clone().shorten(5),s=a.getLeftVector().clone(),l=a.getRightVector().clone();s.x+=r,s.y+=o,l.x+=r,l.y+=o,s=t.getLeftVector().clone(),l=t.getRightVector().clone(),s.x+=r,s.y+=o,l.x+=r,l.y+=o,n.save();let u=i.normals(s,l);u[0].normalize(),u[1].normalize();let c=s,h=l;t.getRightChiral()&&(c=l,h=s);let d=i.add(c,i.multiplyScalar(u[0],this.halfBondThickness)),g=i.add(h,i.multiplyScalar(u[0],1.5+this.halfBondThickness)),f=i.add(h,i.multiplyScalar(u[1],1.5+this.halfBondThickness)),p=i.add(c,i.multiplyScalar(u[1],this.halfBondThickness));n.beginPath(),n.moveTo(d.x,d.y),n.lineTo(g.x,g.y),n.lineTo(f.x,f.y),n.lineTo(p.x,p.y);let v=this.ctx.createRadialGradient(l.x,l.y,this.opts.bondLength,l.x,l.y,0);v.addColorStop(.4,this.getColor(t.getLeftElement())||this.getColor("C")),v.addColorStop(.6,this.getColor(t.getRightElement())||this.getColor("C")),n.fillStyle=v,n.fill(),n.restore()}drawDashedWedge(t){if(isNaN(t.from.x)||isNaN(t.from.y)||isNaN(t.to.x)||isNaN(t.to.y))return;let e=this.ctx,n=this.offsetX,r=this.offsetY,o=t.getLeftVector().clone(),a=t.getRightVector().clone();o.x+=n,o.y+=r,a.x+=n,a.y+=r,e.save();let s=i.normals(o,a);s[0].normalize(),s[1].normalize();let l,u,c,h,d=t.getRightChiral(),g=t.clone();d?(l=a,u=o,g.shortenRight(1),c=g.getRightVector().clone(),h=g.getLeftVector().clone()):(l=o,u=a,g.shortenLeft(1),c=g.getLeftVector().clone(),h=g.getRightVector().clone()),c.x+=n,c.y+=r,h.x+=n,h.y+=r;let f=i.subtract(u,l).normalize();e.strokeStyle=this.getColor("C"),e.lineCap="round",e.lineWidth=this.opts.bondThickness,e.beginPath();let p=t.getLength(),v=1.25/(p/(3*this.opts.bondThickness)),m=!1;for(var y=0;y<1;y+=v){let n=i.multiplyScalar(f,y*p),r=i.add(l,n),o=1.5*y,a=i.multiplyScalar(s[0],o);!m&&y>.5&&(e.stroke(),e.beginPath(),e.strokeStyle=this.getColor(t.getRightElement())||this.getColor("C"),m=!0),r.subtract(a),e.moveTo(r.x,r.y),r.add(i.multiplyScalar(a,2)),e.lineTo(r.x,r.y)}e.stroke(),e.restore()}drawDebugText(t,e,n){let r=this.ctx;r.save(),r.font="5px Droid Sans, sans-serif",r.textAlign="start",r.textBaseline="top",r.fillStyle="#ff0000",r.fillText(n,t+this.offsetX,e+this.offsetY),r.restore()}drawBall(t,e,n){let i=this.ctx;i.save(),i.beginPath(),i.arc(t+this.offsetX,e+this.offsetY,this.opts.bondLength/4.5,0,r.twoPI,!1),i.fillStyle=this.getColor(n),i.fill(),i.restore()}drawPoint(t,e,n){let i=this.ctx,o=this.offsetX,a=this.offsetY;i.save(),i.globalCompositeOperation="destination-out",i.beginPath(),i.arc(t+o,e+a,1.5,0,r.twoPI,!0),i.closePath(),i.fill(),i.globalCompositeOperation="source-over",i.beginPath(),i.arc(t+this.offsetX,e+this.offsetY,.75,0,r.twoPI,!1),i.fillStyle=this.getColor(n),i.fill(),i.restore()}drawText(t,e,n,i,o,a,s,l,u={}){let c=this.ctx,h=this.offsetX,d=this.offsetY;c.save(),c.textAlign="start",c.textBaseline="alphabetic";let g="",f=0;s&&(g=this.getChargeText(s),c.font=this.fontSmall,f=c.measureText(g).width);let p="0",v=0;l>0&&(p=l.toString(),c.font=this.fontSmall,v=c.measureText(p).width),1===s&&"N"===n&&u.hasOwnProperty("0O")&&u.hasOwnProperty("0O-1")&&(u={"0O":{element:"O",count:2,hydrogenCount:0,previousElement:"C",charge:""}},s=0),c.font=this.fontLarge,c.fillStyle=this.getColor("BACKGROUND");let m=c.measureText(n);m.totalWidth=m.width+f,m.height=parseInt(this.fontLarge,10);let y=m.width>this.opts.fontSizeLarge?m.width:this.opts.fontSizeLarge;y/=1.5,c.globalCompositeOperation="destination-out",c.beginPath(),c.arc(t+h,e+d,y,0,r.twoPI,!0),c.closePath(),c.fill(),c.globalCompositeOperation="source-over";let b=-m.width/2,x=-m.width/2;c.fillStyle=this.getColor(n),c.fillText(n,t+h+b,e+this.opts.halfFontSizeLarge+d),b+=m.width,s&&(c.font=this.fontSmall,c.fillText(g,t+h+b,e-this.opts.fifthFontSizeSmall+d),b+=f),l>0&&(c.font=this.fontSmall,c.fillText(p,t+h+x-v,e-this.opts.fifthFontSizeSmall+d),x-=v),c.font=this.fontLarge;let w=0,A=0;if(1===i){let n=t+h,r=e+d+this.opts.halfFontSizeLarge;w=this.hydrogenWidth,x-=w,"left"===o?n+=x:"right"===o||"up"===o&&a||"down"===o&&a?n+=b:"up"!==o||a?"down"!==o||a||(r+=this.opts.fontSizeLarge+this.opts.quarterFontSizeLarge,n-=this.halfHydrogenWidth):(r-=this.opts.fontSizeLarge+this.opts.quarterFontSizeLarge,n-=this.halfHydrogenWidth),c.fillText("H",n,r),b+=w}else if(i>1){let n=t+h,r=e+d+this.opts.halfFontSizeLarge;w=this.hydrogenWidth,c.font=this.fontSmall,A=c.measureText(i).width,x-=w+A,"left"===o?n+=x:"right"===o||"up"===o&&a||"down"===o&&a?n+=b:"up"!==o||a?"down"!==o||a||(r+=this.opts.fontSizeLarge+this.opts.quarterFontSizeLarge,n-=this.halfHydrogenWidth):(r-=this.opts.fontSizeLarge+this.opts.quarterFontSizeLarge,n-=this.halfHydrogenWidth),c.font=this.fontLarge,c.fillText("H",n,r),c.font=this.fontSmall,c.fillText(i,n+this.halfHydrogenWidth+A,r+this.opts.fifthFontSizeSmall),b+=w+this.halfHydrogenWidth+A}for(let n in u){if(!u.hasOwnProperty(n))continue;let r=0,i=0,a=u[n].element,s=u[n].count,l=u[n].hydrogenCount,g=u[n].charge;c.font=this.fontLarge,s>1&&l>0&&(r=c.measureText("(").width,i=c.measureText(")").width);let f=c.measureText(a).width,p=0,v="",m=0;w=0,l>0&&(w=this.hydrogenWidth),c.font=this.fontSmall,s>1&&(p=c.measureText(s).width),0!==g&&(v=this.getChargeText(g),m=c.measureText(v).width),A=0,l>1&&(A=c.measureText(l).width),c.font=this.fontLarge;let y=t+h,C=e+d+this.opts.halfFontSizeLarge;c.fillStyle=this.getColor(a),s>0&&(x-=p),s>1&&l>0&&("left"===o?(x-=i,c.fillText(")",y+x,C)):(c.fillText("(",y+b,C),b+=r)),"left"===o?(x-=f,c.fillText(a,y+x,C)):(c.fillText(a,y+b,C),b+=f),l>0&&("left"===o?(x-=w+A,c.fillText("H",y+x,C),l>1&&(c.font=this.fontSmall,c.fillText(l,y+x+w,C+this.opts.fifthFontSizeSmall))):(c.fillText("H",y+b,C),b+=w,l>1&&(c.font=this.fontSmall,c.fillText(l,y+b,C+this.opts.fifthFontSizeSmall),b+=A))),c.font=this.fontLarge,s>1&&l>0&&("left"===o?(x-=r,c.fillText("(",y+x,C)):(c.fillText(")",y+b,C),b+=i)),c.font=this.fontSmall,s>1&&("left"===o?c.fillText(s,y+x+r+i+w+A+f,C+this.opts.fifthFontSizeSmall):(c.fillText(s,y+b,C+this.opts.fifthFontSizeSmall),b+=p)),0!==g&&("left"===o?c.fillText(v,y+x+r+i+w+A+f,e-this.opts.fifthFontSizeSmall+d):(c.fillText(v,y+b,e-this.opts.fifthFontSizeSmall+d),b+=m))}c.restore()}getChargeText(t){return 1===t?"+":2===t?"2+":-1===t?"-":-2===t?"2-":""}drawDebugPoint(t,e,n="",r="#f00"){this.drawCircle(t,e,2,r,!0,!0,n)}drawAromaticityRing(t){let e=this.ctx,n=r.apothemFromSideLength(this.opts.bondLength,t.getSize());e.save(),e.strokeStyle=this.getColor("C"),e.lineWidth=this.opts.bondThickness,e.beginPath(),e.arc(t.center.x+this.offsetX,t.center.y+this.offsetY,n-this.opts.bondSpacing,0,2*Math.PI,!0),e.closePath(),e.stroke(),e.restore()}clear(){this.ctx.clearRect(0,0,this.canvas.offsetWidth,this.canvas.offsetHeight)}}},function(t,e,n){const r=n(18);class i{static getRings(t){let e=t.getComponentsAdjacencyMatrix();if(0===e.length)return null;let n=r.getConnectedComponents(e),o=Array();for(var a=0;a<n.length;a++){let e=n[a],r=t.getSubgraphAdjacencyMatrix([...e]),u=new Uint16Array(r.length),c=new Uint16Array(r.length);for(var s=0;s<r.length;s++){c[s]=0,u[s]=0;for(var l=0;l<r[s].length;l++)u[s]+=r[s][l]}let h=0;for(s=0;s<r.length;s++)for(l=s+1;l<r.length;l++)h+=r[s][l];let d=h-r.length+1,g=!0;for(s=0;s<u.length;s++)3!==u[s]&&(g=!1);if(g&&(d=2+h-r.length),1===d){o.push([...e]);continue}let{d:f,pe:p,pe_prime:v}=i.getPathIncludedDistanceMatrices(r),m=i.getRingCandidates(f,p,v),y=i.getSSSR(m,f,r,p,v,u,c,d);for(s=0;s<y.length;s++){let t=Array(y[s].size),n=0;for(let r of y[s])t[n++]=e[r];o.push(t)}}return o}static matrixToString(t){let e="";for(var n=0;n<t.length;n++){for(var r=0;r<t[n].length;r++)e+=t[n][r]+" ";e+="\n"}return e}static getPathIncludedDistanceMatrices(t){let e=t.length,n=Array(e),r=Array(e),i=Array(e);for(var o=0,a=0,s=0,l=e;l--;){n[l]=Array(e),r[l]=Array(e),i[l]=Array(e);for(var u=e;u--;)n[l][u]=l===u||1===t[l][u]?t[l][u]:Number.POSITIVE_INFINITY,1===n[l][u]?r[l][u]=[[[l,u]]]:r[l][u]=Array(),i[l][u]=Array()}for(var c=e;c--;)for(l=e;l--;)for(u=e;u--;){const t=n[l][u],e=n[l][c]+n[c][u];if(t>e){if(t===e+1)for(i[l][u]=[r[l][u].length],o=r[l][u].length;o--;)for(i[l][u][o]=[r[l][u][o].length],a=r[l][u][o].length;a--;)for(i[l][u][o][a]=[r[l][u][o][a].length],s=r[l][u][o][a].length;s--;)i[l][u][o][a][s]=[r[l][u][o][a][0],r[l][u][o][a][1]];else i[l][u]=Array();for(n[l][u]=e,r[l][u]=[[]],o=r[l][c][0].length;o--;)r[l][u][0].push(r[l][c][0][o]);for(o=r[c][u][0].length;o--;)r[l][u][0].push(r[c][u][0][o])}else if(t===e){if(r[l][c].length&&r[c][u].length)if(r[l][u].length){let t=Array();for(o=r[l][c][0].length;o--;)t.push(r[l][c][0][o]);for(o=r[c][u][0].length;o--;)t.push(r[c][u][0][o]);r[l][u].push(t)}else{let t=Array();for(o=r[l][c][0].length;o--;)t.push(r[l][c][0][o]);for(o=r[c][u][0].length;o--;)t.push(r[c][u][0][o]);r[l][u][0]=t}}else if(t===e-1){if(i[l][u].length){let t=Array();for(o=r[l][c][0].length;o--;)t.push(r[l][c][0][o]);for(o=r[c][u][0].length;o--;)t.push(r[c][u][0][o]);i[l][u].push(t)}else{let t=Array();for(o=r[l][c][0].length;o--;)t.push(r[l][c][0][o]);for(o=r[c][u][0].length;o--;)t.push(r[c][u][0][o]);i[l][u][0]=t}}}return{d:n,pe:r,pe_prime:i}}static getRingCandidates(t,e,n){let r=t.length,i=Array(),o=0;for(let a=0;a<r;a++)for(let s=0;s<r;s++)0===t[a][s]||1===e[a][s].length&&0===n[a][s]||(o=0!==n[a][s].length?2*(t[a][s]+.5):2*t[a][s],o!==1/0&&i.push([o,e[a][s],n[a][s]]));return i.sort((function(t,e){return t[0]-e[0]})),i}static getSSSR(t,e,n,r,o,a,s,l){let u=Array(),c=Array();for(let e=0;e<t.length;e++)if(t[e][0]%2!=0)for(let r=0;r<t[e][2].length;r++){let o=t[e][1][0].concat(t[e][2][r]);for(var h=0;h<o.length;h++)o[h][0].constructor===Array&&(o[h]=o[h][0]);let d=i.bondsToAtoms(o);if(i.getBondCount(d,n)!==d.size||i.pathSetsContain(u,d,o,c,a,s)||(u.push(d),c=c.concat(o)),u.length>l)return u}else for(let r=0;r<t[e][1].length-1;r++){let o=t[e][1][r].concat(t[e][1][r+1]);for(h=0;h<o.length;h++)o[h][0].constructor===Array&&(o[h]=o[h][0]);let d=i.bondsToAtoms(o);if(i.getBondCount(d,n)!==d.size||i.pathSetsContain(u,d,o,c,a,s)||(u.push(d),c=c.concat(o)),u.length>l)return u}return u}static getEdgeCount(t){let e=0,n=t.length;for(var r=n-1;r--;)for(var i=n;i--;)1===t[r][i]&&e++;return e}static getEdgeList(t){let e=t.length,n=Array();for(var r=e-1;r--;)for(var i=e;i--;)1===t[r][i]&&n.push([r,i]);return n}static bondsToAtoms(t){let e=new Set;for(var n=t.length;n--;)e.add(t[n][0]),e.add(t[n][1]);return e}static getBondCount(t,e){let n=0;for(let r of t)for(let i of t)r!==i&&(n+=e[r][i]);return n/2}static pathSetsContain(t,e,n,r,o,a){for(var s=t.length;s--;){if(i.isSupersetOf(e,t[s]))return!0;if(t[s].size===e.size&&i.areSetsEqual(t[s],e))return!0}let l=0,u=!1;for(s=n.length;s--;)for(var c=r.length;c--;)(n[s][0]===r[c][0]&&n[s][1]===r[c][1]||n[s][1]===r[c][0]&&n[s][0]===r[c][1])&&l++,l===n.length&&(u=!0);let h=!1;if(u)for(let t of e)if(a[t]<o[t]){h=!0;break}if(u&&!h)return!0;for(let t of e)a[t]++;return!1}static areSetsEqual(t,e){if(t.size!==e.size)return!1;for(let n of t)if(!e.has(n))return!1;return!0}static isSupersetOf(t,e){for(var n of e)if(!t.has(n))return!1;return!0}}t.exports=i},function(t,e){t.exports=function(){"use strict";function t(e,n,r,i){this.message=e,this.expected=n,this.found=r,this.location=i,this.name="SyntaxError","function"==typeof Error.captureStackTrace&&Error.captureStackTrace(this,t)}return function(t,e){function n(){this.constructor=t}n.prototype=e.prototype,t.prototype=new n}(t,Error),t.buildMessage=function(t,e){var n={literal:function(t){return'"'+i(t.text)+'"'},class:function(t){var e,n="";for(e=0;e<t.parts.length;e++)n+=t.parts[e]instanceof Array?o(t.parts[e][0])+"-"+o(t.parts[e][1]):o(t.parts[e]);return"["+(t.inverted?"^":"")+n+"]"},any:function(t){return"any character"},end:function(t){return"end of input"},other:function(t){return t.description}};function r(t){return t.charCodeAt(0).toString(16).toUpperCase()}function i(t){return t.replace(/\\/g,"\\\\").replace(/"/g,'\\"').replace(/\0/g,"\\0").replace(/\t/g,"\\t").replace(/\n/g,"\\n").replace(/\r/g,"\\r").replace(/[\x00-\x0F]/g,(function(t){return"\\x0"+r(t)})).replace(/[\x10-\x1F\x7F-\x9F]/g,(function(t){return"\\x"+r(t)}))}function o(t){return t.replace(/\\/g,"\\\\").replace(/\]/g,"\\]").replace(/\^/g,"\\^").replace(/-/g,"\\-").replace(/\0/g,"\\0").replace(/\t/g,"\\t").replace(/\n/g,"\\n").replace(/\r/g,"\\r").replace(/[\x00-\x0F]/g,(function(t){return"\\x0"+r(t)})).replace(/[\x10-\x1F\x7F-\x9F]/g,(function(t){return"\\x"+r(t)}))}return"Expected "+function(t){var e,r,i,o=new Array(t.length);for(e=0;e<t.length;e++)o[e]=(i=t[e],n[i.type](i));if(o.sort(),o.length>0){for(e=1,r=1;e<o.length;e++)o[e-1]!==o[e]&&(o[r]=o[e],r++);o.length=r}switch(o.length){case 1:return o[0];case 2:return o[0]+" or "+o[1];default:return o.slice(0,-1).join(", ")+", or "+o[o.length-1]}}(t)+" but "+function(t){return t?'"'+i(t)+'"':"end of input"}(e)+" found."},{SyntaxError:t,parse:function(e,n){if(n=void 0!==n?n:{},e.split("(").length-1!==e.split(")").length-1)throw it("The number of opening parentheses does not match the number of closing parentheses.",0);var r,i={},o={chain:at},a=at,s=J("(",!1),l=J(")",!1),u=/^[\-=#$:\/\\.]/,c=tt(["-","=","#","$",":","/","\\","."],!1,!1),h=J("[",!1),d=J("se",!1),g=J("as",!1),f=J("]",!1),p=J("B",!1),v=J("r",!1),m=J("C",!1),y=J("l",!1),b=/^[NOPSFI]/,x=tt(["N","O","P","S","F","I"],!1,!1),w=/^[bcnops]/,A=tt(["b","c","n","o","p","s"],!1,!1),C=J("*",!1),_=/^[A-Z]/,S=tt([["A","Z"]],!1,!1),M=/^[a-z]/,T=tt([["a","z"]],!1,!1),k=J("%",!1),R=/^[1-9]/,N=tt([["1","9"]],!1,!1),$=/^[0-9]/,P=tt([["0","9"]],!1,!1),L=J("@",!1),I=J("TH",!1),D=/^[12]/,O=tt(["1","2"],!1,!1),E=J("AL",!1),B=J("SP",!1),j=/^[1-3]/,F=tt([["1","3"]],!1,!1),z=J("TB",!1),H=J("OH",!1),U=J("+",!1),V=J("-",!1),q=J("H",!1),W=J(":",!1),Y=/^[0]/,X=tt(["0"],!1,!1),Z=0,G=[{line:1,column:1}],K=0,Q=[];if("startRule"in n){if(!(n.startRule in o))throw new Error("Can't start parsing from rule \""+n.startRule+'".');a=o[n.startRule]}function J(t,e){return{type:"literal",text:t,ignoreCase:e}}function tt(t,e,n){return{type:"class",parts:t,inverted:e,ignoreCase:n}}function et(t){var n,r=G[t];if(r)return r;for(n=t-1;!G[n];)n--;for(r={line:(r=G[n]).line,column:r.column};n<t;)10===e.charCodeAt(n)?(r.line++,r.column=1):r.column++,n++;return G[t]=r,r}function nt(t,e){var n=et(t),r=et(e);return{start:{offset:t,line:n.line,column:n.column},end:{offset:e,line:r.line,column:r.column}}}function rt(t){Z<K||(Z>K&&(K=Z,Q=[]),Q.push(t))}function it(e,n){return new t(e,null,null,n)}function ot(e,n,r){return new t(t.buildMessage(e,n),e,n,r)}function at(){var t,n,r,o,a,s,l,u,c,w;if(t=Z,n=Z,(r=function(){var t,n;t=Z,(n=function(){var t,n,r,o;t=Z,n=Z,66===e.charCodeAt(Z)?(r="B",Z++):(r=i,rt(p));r!==i?(114===e.charCodeAt(Z)?(o="r",Z++):(o=i,rt(v)),o===i&&(o=null),o!==i?n=r=[r,o]:(Z=n,n=i)):(Z=n,n=i);n===i&&(n=Z,67===e.charCodeAt(Z)?(r="C",Z++):(r=i,rt(m)),r!==i?(108===e.charCodeAt(Z)?(o="l",Z++):(o=i,rt(y)),o===i&&(o=null),o!==i?n=r=[r,o]:(Z=n,n=i)):(Z=n,n=i),n===i&&(b.test(e.charAt(Z))?(n=e.charAt(Z),Z++):(n=i,rt(x))));n!==i&&(t,n=(a=n).length>1?a.join(""):a);var a;return t=n}())===i&&(n=ut())===i&&(n=function(){var t,n,r,o,a,s,l,u,c,p;t=Z,n=Z,91===e.charCodeAt(Z)?(r="[",Z++):(r=i,rt(h));r!==i?((o=function(){var t,n,r,o,a;t=Z,n=Z,R.test(e.charAt(Z))?(r=e.charAt(Z),Z++):(r=i,rt(N));r!==i?($.test(e.charAt(Z))?(o=e.charAt(Z),Z++):(o=i,rt(P)),o===i&&(o=null),o!==i?($.test(e.charAt(Z))?(a=e.charAt(Z),Z++):(a=i,rt(P)),a===i&&(a=null),a!==i?n=r=[r,o,a]:(Z=n,n=i)):(Z=n,n=i)):(Z=n,n=i);n!==i&&(t,n=Number(n.join("")));return t=n}())===i&&(o=null),o!==i?("se"===e.substr(Z,2)?(a="se",Z+=2):(a=i,rt(d)),a===i&&("as"===e.substr(Z,2)?(a="as",Z+=2):(a=i,rt(g)),a===i&&(a=ut())===i&&(a=function(){var t,n,r,o;t=Z,n=Z,_.test(e.charAt(Z))?(r=e.charAt(Z),Z++):(r=i,rt(S));r!==i?(M.test(e.charAt(Z))?(o=e.charAt(Z),Z++):(o=i,rt(T)),o===i&&(o=null),o!==i?n=r=[r,o]:(Z=n,n=i)):(Z=n,n=i);n!==i&&(t,n=n.join(""));return t=n}())===i&&(a=ct())),a!==i?((s=function(){var t,n,r,o,a,s,l;t=Z,n=Z,64===e.charCodeAt(Z)?(r="@",Z++):(r=i,rt(L));r!==i?(64===e.charCodeAt(Z)?(o="@",Z++):(o=i,rt(L)),o===i&&(o=Z,"TH"===e.substr(Z,2)?(a="TH",Z+=2):(a=i,rt(I)),a!==i?(D.test(e.charAt(Z))?(s=e.charAt(Z),Z++):(s=i,rt(O)),s!==i?o=a=[a,s]:(Z=o,o=i)):(Z=o,o=i),o===i&&(o=Z,"AL"===e.substr(Z,2)?(a="AL",Z+=2):(a=i,rt(E)),a!==i?(D.test(e.charAt(Z))?(s=e.charAt(Z),Z++):(s=i,rt(O)),s!==i?o=a=[a,s]:(Z=o,o=i)):(Z=o,o=i),o===i&&(o=Z,"SP"===e.substr(Z,2)?(a="SP",Z+=2):(a=i,rt(B)),a!==i?(j.test(e.charAt(Z))?(s=e.charAt(Z),Z++):(s=i,rt(F)),s!==i?o=a=[a,s]:(Z=o,o=i)):(Z=o,o=i),o===i&&(o=Z,"TB"===e.substr(Z,2)?(a="TB",Z+=2):(a=i,rt(z)),a!==i?(R.test(e.charAt(Z))?(s=e.charAt(Z),Z++):(s=i,rt(N)),s!==i?($.test(e.charAt(Z))?(l=e.charAt(Z),Z++):(l=i,rt(P)),l===i&&(l=null),l!==i?o=a=[a,s,l]:(Z=o,o=i)):(Z=o,o=i)):(Z=o,o=i),o===i&&(o=Z,"OH"===e.substr(Z,2)?(a="OH",Z+=2):(a=i,rt(H)),a!==i?(R.test(e.charAt(Z))?(s=e.charAt(Z),Z++):(s=i,rt(N)),s!==i?($.test(e.charAt(Z))?(l=e.charAt(Z),Z++):(l=i,rt(P)),l===i&&(l=null),l!==i?o=a=[a,s,l]:(Z=o,o=i)):(Z=o,o=i)):(Z=o,o=i)))))),o===i&&(o=null),o!==i?n=r=[r,o]:(Z=n,n=i)):(Z=n,n=i);n!==i&&(t,n=(u=n)[1]?"@"==u[1]?"@@":u[1].join("").replace(",",""):"@");var u;return t=n}())===i&&(s=null),s!==i?((l=function(){var t,n,r,o;t=Z,n=Z,72===e.charCodeAt(Z)?(r="H",Z++):(r=i,rt(q));r!==i?($.test(e.charAt(Z))?(o=e.charAt(Z),Z++):(o=i,rt(P)),o===i&&(o=null),o!==i?n=r=[r,o]:(Z=n,n=i)):(Z=n,n=i);n!==i&&(t,n=(a=n)[1]?Number(a[1]):1);var a;return t=n}())===i&&(l=null),l!==i?((u=function(){var t,n;t=Z,(n=function(){var t,n,r,o,a,s;t=Z,n=Z,43===e.charCodeAt(Z)?(r="+",Z++):(r=i,rt(U));r!==i?(43===e.charCodeAt(Z)?(o="+",Z++):(o=i,rt(U)),o===i&&(o=Z,R.test(e.charAt(Z))?(a=e.charAt(Z),Z++):(a=i,rt(N)),a!==i?($.test(e.charAt(Z))?(s=e.charAt(Z),Z++):(s=i,rt(P)),s===i&&(s=null),s!==i?o=a=[a,s]:(Z=o,o=i)):(Z=o,o=i)),o===i&&(o=null),o!==i?n=r=[r,o]:(Z=n,n=i)):(Z=n,n=i);n!==i&&(t,n=(l=n)[1]?"+"!=l[1]?Number(l[1].join("")):2:1);var l;return t=n}())===i&&(n=function(){var t,n,r,o,a,s;t=Z,n=Z,45===e.charCodeAt(Z)?(r="-",Z++):(r=i,rt(V));r!==i?(45===e.charCodeAt(Z)?(o="-",Z++):(o=i,rt(V)),o===i&&(o=Z,R.test(e.charAt(Z))?(a=e.charAt(Z),Z++):(a=i,rt(N)),a!==i?($.test(e.charAt(Z))?(s=e.charAt(Z),Z++):(s=i,rt(P)),s===i&&(s=null),s!==i?o=a=[a,s]:(Z=o,o=i)):(Z=o,o=i)),o===i&&(o=null),o!==i?n=r=[r,o]:(Z=n,n=i)):(Z=n,n=i);n!==i&&(t,n=(l=n)[1]?"-"!=l[1]?-Number(l[1].join("")):-2:-1);var l;return t=n}());n!==i&&(t,n=n);return t=n}())===i&&(u=null),u!==i?((c=function(){var t,n,r,o,a,s,l;t=Z,n=Z,58===e.charCodeAt(Z)?(r=":",Z++):(r=i,rt(W));if(r!==i){if(o=Z,R.test(e.charAt(Z))?(a=e.charAt(Z),Z++):(a=i,rt(N)),a!==i){for(s=[],$.test(e.charAt(Z))?(l=e.charAt(Z),Z++):(l=i,rt(P));l!==i;)s.push(l),$.test(e.charAt(Z))?(l=e.charAt(Z),Z++):(l=i,rt(P));s!==i?o=a=[a,s]:(Z=o,o=i)}else Z=o,o=i;o===i&&(Y.test(e.charAt(Z))?(o=e.charAt(Z),Z++):(o=i,rt(X))),o!==i?n=r=[r,o]:(Z=n,n=i)}else Z=n,n=i;n!==i&&(t,u=n,n=Number(u[1][0]+u[1][1].join("")));var u;return t=n}())===i&&(c=null),c!==i?(93===e.charCodeAt(Z)?(p="]",Z++):(p=i,rt(f)),p!==i?n=r=[r,o,a,s,l,u,c,p]:(Z=n,n=i)):(Z=n,n=i)):(Z=n,n=i)):(Z=n,n=i)):(Z=n,n=i)):(Z=n,n=i)):(Z=n,n=i)):(Z=n,n=i);n!==i&&(t,n={isotope:(v=n)[1],element:v[2],chirality:v[3],hcount:v[4],charge:v[5],class:v[6]});var v;return t=n}())===i&&(n=ct());n!==i&&(t,n=n);return t=n}())!==i){for(o=[],a=st();a!==i;)o.push(a),a=st();if(o!==i){for(a=[],s=Z,(l=lt())===i&&(l=null),l!==i&&(u=ht())!==i?s=l=[l,u]:(Z=s,s=i);s!==i;)a.push(s),s=Z,(l=lt())===i&&(l=null),l!==i&&(u=ht())!==i?s=l=[l,u]:(Z=s,s=i);if(a!==i){for(s=[],l=st();l!==i;)s.push(l),l=st();if(s!==i)if((l=lt())===i&&(l=null),l!==i)if((u=at())===i&&(u=null),u!==i){for(c=[],w=st();w!==i;)c.push(w),w=st();c!==i?n=r=[r,o,a,s,l,u,c]:(Z=n,n=i)}else Z=n,n=i;else Z=n,n=i;else Z=n,n=i}else Z=n,n=i}else Z=n,n=i}else Z=n,n=i;return n!==i&&(t,n=function(t){for(var e=[],n=[],r=0;r<t[1].length;r++)e.push(t[1][r]);for(r=0;r<t[2].length;r++){var i=t[2][r][0]?t[2][r][0]:"-";n.push({bond:i,id:t[2][r][1]})}for(r=0;r<t[3].length;r++)e.push(t[3][r]);for(r=0;r<t[6].length;r++)e.push(t[6][r]);return{atom:t[0],isBracket:!!t[0].element,branches:e,branchCount:e.length,ringbonds:n,ringbondCount:n.length,bond:t[4]?t[4]:"-",next:t[5],hasNext:!!t[5]}}(n)),t=n}function st(){var t,n,r,o,a,u,c,h;return t=Z,n=Z,40===e.charCodeAt(Z)?(r="(",Z++):(r=i,rt(s)),r!==i?((o=lt())===i&&(o=null),o!==i&&(a=at())!==i?(41===e.charCodeAt(Z)?(u=")",Z++):(u=i,rt(l)),u!==i?n=r=[r,o,a,u]:(Z=n,n=i)):(Z=n,n=i)):(Z=n,n=i),n!==i&&(t,h=(c=n)[1]?c[1]:"-",c[2].branchBond=h,n=c[2]),t=n}function lt(){var t,n;return t=Z,u.test(e.charAt(Z))?(n=e.charAt(Z),Z++):(n=i,rt(c)),n!==i&&(t,n=n),t=n}function ut(){var t,n;return t=Z,w.test(e.charAt(Z))?(n=e.charAt(Z),Z++):(n=i,rt(A)),n!==i&&(t,n=n),t=n}function ct(){var t,n;return t=Z,42===e.charCodeAt(Z)?(n="*",Z++):(n=i,rt(C)),n!==i&&(t,n=n),t=n}function ht(){var t,n,r,o,a,s;return t=Z,n=Z,37===e.charCodeAt(Z)?(r="%",Z++):(r=i,rt(k)),r!==i?(R.test(e.charAt(Z))?(o=e.charAt(Z),Z++):(o=i,rt(N)),o!==i?($.test(e.charAt(Z))?(a=e.charAt(Z),Z++):(a=i,rt(P)),a!==i?n=r=[r,o,a]:(Z=n,n=i)):(Z=n,n=i)):(Z=n,n=i),n===i&&($.test(e.charAt(Z))?(n=e.charAt(Z),Z++):(n=i,rt(P))),n!==i&&(t,n=1==(s=n).length?Number(s):Number(s.join("").replace("%",""))),t=n}if((r=a())!==i&&Z===e.length)return r;throw r!==i&&Z<e.length&&rt({type:"end"}),ot(Q,K<e.length?e.charAt(K):null,K<e.length?nt(K,K+1):nt(K,K))}}}()},function(t,e,n){"use strict";n.r(e),n.d(e,"transition",(function(){return tt})),n.d(e,"active",(function(){return st})),n.d(e,"interrupt",(function(){return R}));var r,i,o=n(0),a=n(11),s=0,l=0,u=0,c=0,h=0,d=0,g="object"==typeof performance&&performance.now?performance:Date,f="object"==typeof window&&window.requestAnimationFrame?window.requestAnimationFrame.bind(window):function(t){setTimeout(t,17)};function p(){return h||(f(v),h=g.now()+d)}function v(){h=0}function m(){this._call=this._time=this._next=null}function y(t,e,n){var r=new m;return r.restart(t,e,n),r}function b(){h=(c=g.now())+d,s=l=0;try{!function(){p(),++s;for(var t,e=r;e;)(t=h-e._time)>=0&&e._call.call(null,t),e=e._next;--s}()}finally{s=0,function(){var t,e,n=r,o=1/0;for(;n;)n._call?(o>n._time&&(o=n._time),t=n,n=n._next):(e=n._next,n._next=null,n=t?t._next=e:r=e);i=t,w(o)}(),h=0}}function x(){var t=g.now(),e=t-c;e>1e3&&(d-=e,c=t)}function w(t){s||(l&&(l=clearTimeout(l)),t-h>24?(t<1/0&&(l=setTimeout(b,t-g.now()-d)),u&&(u=clearInterval(u))):(u||(c=g.now(),u=setInterval(x,1e3)),s=1,f(b)))}m.prototype=y.prototype={constructor:m,restart:function(t,e,n){if("function"!=typeof t)throw new TypeError("callback is not a function");n=(null==n?p():+n)+(null==e?0:+e),this._next||i===this||(i?i._next=this:r=this,i=this),this._call=t,this._time=n,w()},stop:function(){this._call&&(this._call=null,this._time=1/0,w())}};var A=function(t,e,n){var r=new m;return e=null==e?0:+e,r.restart((function(n){r.stop(),t(n+e)}),e,n),r},C=Object(a.a)("start","end","interrupt"),_=[],S=function(t,e,n,r,i,o){var a=t.__transition;if(a){if(n in a)return}else t.__transition={};!function(t,e,n){var r,i=t.__transition;function o(l){var u,c,h,d;if(1!==n.state)return s();for(u in i)if((d=i[u]).name===n.name){if(3===d.state)return A(o);4===d.state?(d.state=6,d.timer.stop(),d.on.call("interrupt",t,t.__data__,d.index,d.group),delete i[u]):+u<e&&(d.state=6,d.timer.stop(),delete i[u])}if(A((function(){3===n.state&&(n.state=4,n.timer.restart(a,n.delay,n.time),a(l))})),n.state=2,n.on.call("start",t,t.__data__,n.index,n.group),2===n.state){for(n.state=3,r=new Array(h=n.tween.length),u=0,c=-1;u<h;++u)(d=n.tween[u].value.call(t,t.__data__,n.index,n.group))&&(r[++c]=d);r.length=c+1}}function a(e){for(var i=e<n.duration?n.ease.call(null,e/n.duration):(n.timer.restart(s),n.state=5,1),o=-1,a=r.length;++o<a;)r[o].call(null,i);5===n.state&&(n.on.call("end",t,t.__data__,n.index,n.group),s())}function s(){for(var r in n.state=6,n.timer.stop(),delete i[e],i)return;delete t.__transition}i[e]=n,n.timer=y((function(t){n.state=1,n.timer.restart(o,n.delay,n.time),n.delay<=t&&o(t-n.delay)}),0,n.time)}(t,n,{name:e,index:r,group:i,on:C,tween:_,time:o.time,delay:o.delay,duration:o.duration,ease:o.ease,timer:null,state:0})};function M(t,e){var n=k(t,e);if(n.state>0)throw new Error("too late; already scheduled");return n}function T(t,e){var n=k(t,e);if(n.state>2)throw new Error("too late; already started");return n}function k(t,e){var n=t.__transition;if(!n||!(n=n[e]))throw new Error("transition not found");return n}var R=function(t,e){var n,r,i,o=t.__transition,a=!0;if(o){for(i in e=null==e?null:e+"",o)(n=o[i]).name===e?(r=n.state>2&&n.state<5,n.state=6,n.timer.stop(),r&&n.on.call("interrupt",t,t.__data__,n.index,n.group),delete o[i]):a=!1;a&&delete t.__transition}},N=n(2);function $(t,e){var n,r;return function(){var i=T(this,t),o=i.tween;if(o!==n)for(var a=0,s=(r=n=o).length;a<s;++a)if(r[a].name===e){(r=r.slice()).splice(a,1);break}i.tween=r}}function P(t,e,n){var r,i;if("function"!=typeof n)throw new Error;return function(){var o=T(this,t),a=o.tween;if(a!==r){i=(r=a).slice();for(var s={name:e,value:n},l=0,u=i.length;l<u;++l)if(i[l].name===e){i[l]=s;break}l===u&&i.push(s)}o.tween=i}}function L(t,e,n){var r=t._id;return t.each((function(){var t=T(this,r);(t.value||(t.value={}))[e]=n.apply(this,arguments)})),function(t){return k(t,r).value[e]}}var I=n(1),D=function(t,e){var n;return("number"==typeof e?N.b:e instanceof I.a?N.c:(n=Object(I.a)(e))?(e=n,N.c):N.e)(t,e)};function O(t){return function(){this.removeAttribute(t)}}function E(t){return function(){this.removeAttributeNS(t.space,t.local)}}function B(t,e,n){var r,i;return function(){var o=this.getAttribute(t);return o===n?null:o===r?i:i=e(r=o,n)}}function j(t,e,n){var r,i;return function(){var o=this.getAttributeNS(t.space,t.local);return o===n?null:o===r?i:i=e(r=o,n)}}function F(t,e,n){var r,i,o;return function(){var a,s=n(this);if(null!=s)return(a=this.getAttribute(t))===s?null:a===r&&s===i?o:o=e(r=a,i=s);this.removeAttribute(t)}}function z(t,e,n){var r,i,o;return function(){var a,s=n(this);if(null!=s)return(a=this.getAttributeNS(t.space,t.local))===s?null:a===r&&s===i?o:o=e(r=a,i=s);this.removeAttributeNS(t.space,t.local)}}function H(t,e){function n(){var n=this,r=e.apply(n,arguments);return r&&function(e){n.setAttributeNS(t.space,t.local,r(e))}}return n._value=e,n}function U(t,e){function n(){var n=this,r=e.apply(n,arguments);return r&&function(e){n.setAttribute(t,r(e))}}return n._value=e,n}function V(t,e){return function(){M(this,t).delay=+e.apply(this,arguments)}}function q(t,e){return e=+e,function(){M(this,t).delay=e}}function W(t,e){return function(){T(this,t).duration=+e.apply(this,arguments)}}function Y(t,e){return e=+e,function(){T(this,t).duration=e}}function X(t,e){if("function"!=typeof e)throw new Error;return function(){T(this,t).ease=e}}function Z(t,e,n){var r,i,o=function(t){return(t+"").trim().split(/^|\s+/).every((function(t){var e=t.indexOf(".");return e>=0&&(t=t.slice(0,e)),!t||"start"===t}))}(e)?M:T;return function(){var a=o(this,t),s=a.on;s!==r&&(i=(r=s).copy()).on(e,n),a.on=i}}var G=o.selection.prototype.constructor;function K(t,e,n){function r(){var r=this,i=e.apply(r,arguments);return i&&function(e){r.style.setProperty(t,i(e),n)}}return r._value=e,r}var Q=0;function J(t,e,n,r){this._groups=t,this._parents=e,this._name=n,this._id=r}function tt(t){return Object(o.selection)().transition(t)}function et(){return++Q}var nt=o.selection.prototype;J.prototype=tt.prototype={constructor:J,select:function(t){var e=this._name,n=this._id;"function"!=typeof t&&(t=Object(o.selector)(t));for(var r=this._groups,i=r.length,a=new Array(i),s=0;s<i;++s)for(var l,u,c=r[s],h=c.length,d=a[s]=new Array(h),g=0;g<h;++g)(l=c[g])&&(u=t.call(l,l.__data__,g,c))&&("__data__"in l&&(u.__data__=l.__data__),d[g]=u,S(d[g],e,n,g,d,k(l,n)));return new J(a,this._parents,e,n)},selectAll:function(t){var e=this._name,n=this._id;"function"!=typeof t&&(t=Object(o.selectorAll)(t));for(var r=this._groups,i=r.length,a=[],s=[],l=0;l<i;++l)for(var u,c=r[l],h=c.length,d=0;d<h;++d)if(u=c[d]){for(var g,f=t.call(u,u.__data__,d,c),p=k(u,n),v=0,m=f.length;v<m;++v)(g=f[v])&&S(g,e,n,v,f,p);a.push(f),s.push(u)}return new J(a,s,e,n)},filter:function(t){"function"!=typeof t&&(t=Object(o.matcher)(t));for(var e=this._groups,n=e.length,r=new Array(n),i=0;i<n;++i)for(var a,s=e[i],l=s.length,u=r[i]=[],c=0;c<l;++c)(a=s[c])&&t.call(a,a.__data__,c,s)&&u.push(a);return new J(r,this._parents,this._name,this._id)},merge:function(t){if(t._id!==this._id)throw new Error;for(var e=this._groups,n=t._groups,r=e.length,i=n.length,o=Math.min(r,i),a=new Array(r),s=0;s<o;++s)for(var l,u=e[s],c=n[s],h=u.length,d=a[s]=new Array(h),g=0;g<h;++g)(l=u[g]||c[g])&&(d[g]=l);for(;s<r;++s)a[s]=e[s];return new J(a,this._parents,this._name,this._id)},selection:function(){return new G(this._groups,this._parents)},transition:function(){for(var t=this._name,e=this._id,n=et(),r=this._groups,i=r.length,o=0;o<i;++o)for(var a,s=r[o],l=s.length,u=0;u<l;++u)if(a=s[u]){var c=k(a,e);S(a,t,n,u,s,{time:c.time+c.delay+c.duration,delay:0,duration:c.duration,ease:c.ease})}return new J(r,this._parents,t,n)},call:nt.call,nodes:nt.nodes,node:nt.node,size:nt.size,empty:nt.empty,each:nt.each,on:function(t,e){var n=this._id;return arguments.length<2?k(this.node(),n).on.on(t):this.each(Z(n,t,e))},attr:function(t,e){var n=Object(o.namespace)(t),r="transform"===n?N.g:D;return this.attrTween(t,"function"==typeof e?(n.local?z:F)(n,r,L(this,"attr."+t,e)):null==e?(n.local?E:O)(n):(n.local?j:B)(n,r,e+""))},attrTween:function(t,e){var n="attr."+t;if(arguments.length<2)return(n=this.tween(n))&&n._value;if(null==e)return this.tween(n,null);if("function"!=typeof e)throw new Error;var r=Object(o.namespace)(t);return this.tween(n,(r.local?H:U)(r,e))},style:function(t,e,n){var r="transform"==(t+="")?N.f:D;return null==e?this.styleTween(t,function(t,e){var n,r,i;return function(){var a=Object(o.style)(this,t),s=(this.style.removeProperty(t),Object(o.style)(this,t));return a===s?null:a===n&&s===r?i:i=e(n=a,r=s)}}(t,r)).on("end.style."+t,function(t){return function(){this.style.removeProperty(t)}}(t)):this.styleTween(t,"function"==typeof e?function(t,e,n){var r,i,a;return function(){var s=Object(o.style)(this,t),l=n(this);return null==l&&(this.style.removeProperty(t),l=Object(o.style)(this,t)),s===l?null:s===r&&l===i?a:a=e(r=s,i=l)}}(t,r,L(this,"style."+t,e)):function(t,e,n){var r,i;return function(){var a=Object(o.style)(this,t);return a===n?null:a===r?i:i=e(r=a,n)}}(t,r,e+""),n)},styleTween:function(t,e,n){var r="style."+(t+="");if(arguments.length<2)return(r=this.tween(r))&&r._value;if(null==e)return this.tween(r,null);if("function"!=typeof e)throw new Error;return this.tween(r,K(t,e,null==n?"":n))},text:function(t){return this.tween("text","function"==typeof t?function(t){return function(){var e=t(this);this.textContent=null==e?"":e}}(L(this,"text",t)):function(t){return function(){this.textContent=t}}(null==t?"":t+""))},remove:function(){return this.on("end.remove",(t=this._id,function(){var e=this.parentNode;for(var n in this.__transition)if(+n!==t)return;e&&e.removeChild(this)}));var t},tween:function(t,e){var n=this._id;if(t+="",arguments.length<2){for(var r,i=k(this.node(),n).tween,o=0,a=i.length;o<a;++o)if((r=i[o]).name===t)return r.value;return null}return this.each((null==e?$:P)(n,t,e))},delay:function(t){var e=this._id;return arguments.length?this.each(("function"==typeof t?V:q)(e,t)):k(this.node(),e).delay},duration:function(t){var e=this._id;return arguments.length?this.each(("function"==typeof t?W:Y)(e,t)):k(this.node(),e).duration},ease:function(t){var e=this._id;return arguments.length?this.each(X(e,t)):k(this.node(),e).ease}};(function t(e){function n(t){return Math.pow(t,e)}return e=+e,n.exponent=t,n})(3),function t(e){function n(t){return 1-Math.pow(1-t,e)}return e=+e,n.exponent=t,n}(3),function t(e){function n(t){return((t*=2)<=1?Math.pow(t,e):2-Math.pow(2-t,e))/2}return e=+e,n.exponent=t,n}(3),Math.PI;(function t(e){function n(t){return t*t*((e+1)*t-e)}return e=+e,n.overshoot=t,n})(1.70158),function t(e){function n(t){return--t*t*((e+1)*t+e)+1}return e=+e,n.overshoot=t,n}(1.70158),function t(e){function n(t){return((t*=2)<1?t*t*((e+1)*t-e):(t-=2)*t*((e+1)*t+e)+2)/2}return e=+e,n.overshoot=t,n}(1.70158);var rt=2*Math.PI,it=(function t(e,n){var r=Math.asin(1/(e=Math.max(1,e)))*(n/=rt);function i(t){return e*Math.pow(2,10*--t)*Math.sin((r-t)/n)}return i.amplitude=function(e){return t(e,n*rt)},i.period=function(n){return t(e,n)},i}(1,.3),function t(e,n){var r=Math.asin(1/(e=Math.max(1,e)))*(n/=rt);function i(t){return 1-e*Math.pow(2,-10*(t=+t))*Math.sin((t+r)/n)}return i.amplitude=function(e){return t(e,n*rt)},i.period=function(n){return t(e,n)},i}(1,.3),function t(e,n){var r=Math.asin(1/(e=Math.max(1,e)))*(n/=rt);function i(t){return((t=2*t-1)<0?e*Math.pow(2,10*t)*Math.sin((r-t)/n):2-e*Math.pow(2,-10*t)*Math.sin((r+t)/n))/2}return i.amplitude=function(e){return t(e,n*rt)},i.period=function(n){return t(e,n)},i}(1,.3),{time:null,delay:0,duration:250,ease:function(t){return((t*=2)<=1?t*t*t:(t-=2)*t*t+2)/2}});function ot(t,e){for(var n;!(n=t.__transition)||!(n=n[e]);)if(!(t=t.parentNode))return it.time=p(),it;return n}o.selection.prototype.interrupt=function(t){return this.each((function(){R(this,t)}))},o.selection.prototype.transition=function(t){var e,n;t instanceof J?(e=t._id,t=t._name):(e=et(),(n=it).time=p(),t=null==t?null:t+"");for(var r=this._groups,i=r.length,o=0;o<i;++o)for(var a,s=r[o],l=s.length,u=0;u<l;++u)(a=s[u])&&S(a,t,e,u,s,n||ot(a,e));return new J(r,this._parents,t,e)};var at=[null],st=function(t,e){var n,r,i=t.__transition;if(i)for(r in e=null==e?null:e+"",i)if((n=i[r]).state>1&&n.name===e)return new J([[t]],at,e,+r);return null}},function(t,e,n){"use strict";n.r(e),n.d(e,"drag",(function(){return p})),n.d(e,"dragDisable",(function(){return s})),n.d(e,"dragEnable",(function(){return l}));var r=n(11),i=n(0);function o(){i.event.stopImmediatePropagation()}var a=function(){i.event.preventDefault(),i.event.stopImmediatePropagation()},s=function(t){var e=t.document.documentElement,n=Object(i.select)(t).on("dragstart.drag",a,!0);"onselectstart"in e?n.on("selectstart.drag",a,!0):(e.__noselect=e.style.MozUserSelect,e.style.MozUserSelect="none")};function l(t,e){var n=t.document.documentElement,r=Object(i.select)(t).on("dragstart.drag",null);e&&(r.on("click.drag",a,!0),setTimeout((function(){r.on("click.drag",null)}),0)),"onselectstart"in n?r.on("selectstart.drag",null):(n.style.MozUserSelect=n.__noselect,delete n.__noselect)}var u=function(t){return function(){return t}};function c(t,e,n,r,i,o,a,s,l,u){this.target=t,this.type=e,this.subject=n,this.identifier=r,this.active=i,this.x=o,this.y=a,this.dx=s,this.dy=l,this._=u}function h(){return!i.event.button}function d(){return this.parentNode}function g(t){return null==t?{x:i.event.x,y:i.event.y}:t}function f(){return"ontouchstart"in this}c.prototype.on=function(){var t=this._.on.apply(this._,arguments);return t===this._?this:t};var p=function(){var t,e,n,p,v=h,m=d,y=g,b=f,x={},w=Object(r.a)("start","drag","end"),A=0,C=0;function _(t){t.on("mousedown.drag",S).filter(b).on("touchstart.drag",k).on("touchmove.drag",R).on("touchend.drag touchcancel.drag",N).style("touch-action","none").style("-webkit-tap-highlight-color","rgba(0,0,0,0)")}function S(){if(!p&&v.apply(this,arguments)){var r=$("mouse",m.apply(this,arguments),i.mouse,this,arguments);r&&(Object(i.select)(i.event.view).on("mousemove.drag",M,!0).on("mouseup.drag",T,!0),s(i.event.view),o(),n=!1,t=i.event.clientX,e=i.event.clientY,r("start"))}}function M(){if(a(),!n){var r=i.event.clientX-t,o=i.event.clientY-e;n=r*r+o*o>C}x.mouse("drag")}function T(){Object(i.select)(i.event.view).on("mousemove.drag mouseup.drag",null),l(i.event.view,n),a(),x.mouse("end")}function k(){if(v.apply(this,arguments)){var t,e,n=i.event.changedTouches,r=m.apply(this,arguments),a=n.length;for(t=0;t<a;++t)(e=$(n[t].identifier,r,i.touch,this,arguments))&&(o(),e("start"))}}function R(){var t,e,n=i.event.changedTouches,r=n.length;for(t=0;t<r;++t)(e=x[n[t].identifier])&&(a(),e("drag"))}function N(){var t,e,n=i.event.changedTouches,r=n.length;for(p&&clearTimeout(p),p=setTimeout((function(){p=null}),500),t=0;t<r;++t)(e=x[n[t].identifier])&&(o(),e("end"))}function $(t,e,n,r,o){var a,s,l,u=n(e,t),h=w.copy();if(Object(i.customEvent)(new c(_,"beforestart",a,t,A,u[0],u[1],0,0,h),(function(){return null!=(i.event.subject=a=y.apply(r,o))&&(s=a.x-u[0]||0,l=a.y-u[1]||0,!0)})))return function d(g){var f,p=u;switch(g){case"start":x[t]=d,f=A++;break;case"end":delete x[t],--A;case"drag":u=n(e,t),f=A}Object(i.customEvent)(new c(_,g,a,t,f,u[0]+s,u[1]+l,u[0]-p[0],u[1]-p[1],h),h.apply,h,[g,r,o])}}return _.filter=function(t){return arguments.length?(v="function"==typeof t?t:u(!!t),_):v},_.container=function(t){return arguments.length?(m="function"==typeof t?t:u(t),_):m},_.subject=function(t){return arguments.length?(y="function"==typeof t?t:u(t),_):y},_.touchable=function(t){return arguments.length?(b="function"==typeof t?t:u(!!t),_):b},_.on=function(){var t=w.on.apply(w,arguments);return t===w?_:t},_.clickDistance=function(t){return arguments.length?(C=(t=+t)*t,_):Math.sqrt(C)},_}},function(t,e,n){"use strict";n.r(e),n.d(e,"axisTop",(function(){return h})),n.d(e,"axisRight",(function(){return d})),n.d(e,"axisBottom",(function(){return g})),n.d(e,"axisLeft",(function(){return f}));var r=Array.prototype.slice,i=function(t){return t};function o(t){return"translate("+(t+.5)+",0)"}function a(t){return"translate(0,"+(t+.5)+")"}function s(t){return function(e){return+t(e)}}function l(t){var e=Math.max(0,t.bandwidth()-1)/2;return t.round()&&(e=Math.round(e)),function(n){return+t(n)+e}}function u(){return!this.__axis}function c(t,e){var n=[],c=null,h=null,d=6,g=6,f=3,p=1===t||4===t?-1:1,v=4===t||2===t?"x":"y",m=1===t||3===t?o:a;function y(r){var o=null==c?e.ticks?e.ticks.apply(e,n):e.domain():c,a=null==h?e.tickFormat?e.tickFormat.apply(e,n):i:h,y=Math.max(d,0)+f,b=e.range(),x=+b[0]+.5,w=+b[b.length-1]+.5,A=(e.bandwidth?l:s)(e.copy()),C=r.selection?r.selection():r,_=C.selectAll(".domain").data([null]),S=C.selectAll(".tick").data(o,e).order(),M=S.exit(),T=S.enter().append("g").attr("class","tick"),k=S.select("line"),R=S.select("text");_=_.merge(_.enter().insert("path",".tick").attr("class","domain").attr("stroke","currentColor")),S=S.merge(T),k=k.merge(T.append("line").attr("stroke","currentColor").attr(v+"2",p*d)),R=R.merge(T.append("text").attr("fill","currentColor").attr(v,p*y).attr("dy",1===t?"0em":3===t?"0.71em":"0.32em")),r!==C&&(_=_.transition(r),S=S.transition(r),k=k.transition(r),R=R.transition(r),M=M.transition(r).attr("opacity",1e-6).attr("transform",(function(t){return isFinite(t=A(t))?m(t):this.getAttribute("transform")})),T.attr("opacity",1e-6).attr("transform",(function(t){var e=this.parentNode.__axis;return m(e&&isFinite(e=e(t))?e:A(t))}))),M.remove(),_.attr("d",4===t||2==t?g?"M"+p*g+","+x+"H0.5V"+w+"H"+p*g:"M0.5,"+x+"V"+w:g?"M"+x+","+p*g+"V0.5H"+w+"V"+p*g:"M"+x+",0.5H"+w),S.attr("opacity",1).attr("transform",(function(t){return m(A(t))})),k.attr(v+"2",p*d),R.attr(v,p*y).text(a),C.filter(u).attr("fill","none").attr("font-size",10).attr("font-family","sans-serif").attr("text-anchor",2===t?"start":4===t?"end":"middle"),C.each((function(){this.__axis=A}))}return y.scale=function(t){return arguments.length?(e=t,y):e},y.ticks=function(){return n=r.call(arguments),y},y.tickArguments=function(t){return arguments.length?(n=null==t?[]:r.call(t),y):n.slice()},y.tickValues=function(t){return arguments.length?(c=null==t?null:r.call(t),y):c&&c.slice()},y.tickFormat=function(t){return arguments.length?(h=t,y):h},y.tickSize=function(t){return arguments.length?(d=g=+t,y):d},y.tickSizeInner=function(t){return arguments.length?(d=+t,y):d},y.tickSizeOuter=function(t){return arguments.length?(g=+t,y):g},y.tickPadding=function(t){return arguments.length?(f=+t,y):f},y}function h(t){return c(1,t)}function d(t){return c(2,t)}function g(t){return c(3,t)}function f(t){return c(4,t)}},function(t,e,n){"use strict";n.r(e),n.d(e,"path",(function(){return l}));var r=Math.PI,i=2*r,o=i-1e-6;function a(){this._x0=this._y0=this._x1=this._y1=null,this._=""}function s(){return new a}a.prototype=s.prototype={constructor:a,moveTo:function(t,e){this._+="M"+(this._x0=this._x1=+t)+","+(this._y0=this._y1=+e)},closePath:function(){null!==this._x1&&(this._x1=this._x0,this._y1=this._y0,this._+="Z")},lineTo:function(t,e){this._+="L"+(this._x1=+t)+","+(this._y1=+e)},quadraticCurveTo:function(t,e,n,r){this._+="Q"+ +t+","+ +e+","+(this._x1=+n)+","+(this._y1=+r)},bezierCurveTo:function(t,e,n,r,i,o){this._+="C"+ +t+","+ +e+","+ +n+","+ +r+","+(this._x1=+i)+","+(this._y1=+o)},arcTo:function(t,e,n,i,o){t=+t,e=+e,n=+n,i=+i,o=+o;var a=this._x1,s=this._y1,l=n-t,u=i-e,c=a-t,h=s-e,d=c*c+h*h;if(o<0)throw new Error("negative radius: "+o);if(null===this._x1)this._+="M"+(this._x1=t)+","+(this._y1=e);else if(d>1e-6)if(Math.abs(h*l-u*c)>1e-6&&o){var g=n-a,f=i-s,p=l*l+u*u,v=g*g+f*f,m=Math.sqrt(p),y=Math.sqrt(d),b=o*Math.tan((r-Math.acos((p+d-v)/(2*m*y)))/2),x=b/y,w=b/m;Math.abs(x-1)>1e-6&&(this._+="L"+(t+x*c)+","+(e+x*h)),this._+="A"+o+","+o+",0,0,"+ +(h*g>c*f)+","+(this._x1=t+w*l)+","+(this._y1=e+w*u)}else this._+="L"+(this._x1=t)+","+(this._y1=e);else;},arc:function(t,e,n,a,s,l){t=+t,e=+e;var u=(n=+n)*Math.cos(a),c=n*Math.sin(a),h=t+u,d=e+c,g=1^l,f=l?a-s:s-a;if(n<0)throw new Error("negative radius: "+n);null===this._x1?this._+="M"+h+","+d:(Math.abs(this._x1-h)>1e-6||Math.abs(this._y1-d)>1e-6)&&(this._+="L"+h+","+d),n&&(f<0&&(f=f%i+i),f>o?this._+="A"+n+","+n+",0,1,"+g+","+(t-u)+","+(e-c)+"A"+n+","+n+",0,1,"+g+","+(this._x1=h)+","+(this._y1=d):f>1e-6&&(this._+="A"+n+","+n+",0,"+ +(f>=r)+","+g+","+(this._x1=t+n*Math.cos(s))+","+(this._y1=e+n*Math.sin(s))))},rect:function(t,e,n,r){this._+="M"+(this._x0=this._x1=+t)+","+(this._y0=this._y1=+e)+"h"+ +n+"v"+ +r+"h"+-n+"Z"},toString:function(){return this._}};var l=s}]);
```
