## Supplementary Results for "A draft genome of the ascomycotal fungal species *Pseudopithomyces maydicus* (family *Didymosphaeriaceae*)": jquery.js

```
/*! jQuery v1.9.1 | (c) 2005, 2012 jQuery Foundation, Inc. | jquery.org/license
//@ sourceMappingURL=jquery.min.map
*/(function(e,t){var n,r,i=typeof t,o=e.document,a=e.location,s=e.jQuery,u=e.$,l={},c=[],p="1.9.1",f=c.concat,d=c.push,h=c.slice,g=c.indexOf,m=l.toString,y=l.hasOwnProperty,v=p.trim,b=function(e,t){return new b.fn.init(e,t,r)},x=/[+-]?(?:\d*\.|)\d+(?:[eE][+-]?\d+|)/.source,w=/\S+/g,T=/^[\s\uFEFF\xA0]+|[\s\uFEFF\xA0]+$/g,N=/^(?:(<[\w\W]+>)[^>]*|#([\w-]*))$/,C=/^<(\w+)\s*\/?>(?:<\/\1>|)$/,k=/^[\],:{}\s]*$/,E=/(?:^|:|,)(?:\s*\[)+/g,S=/\\(?:["\\\/bfnrt]|u[\da-fA-F]{4})/g,A=/"[^"\\\r\n]*"|true|false|null|-?(?:\d+\.|)\d+(?:[eE][+-]?\d+|)/g,j=/^-ms-/,D=/-([\da-z])/gi,L=function(e,t){return t.toUpperCase()},H=function(e){(o.addEventListener||"load"===e.type||"complete"===o.readyState)&&(q(),b.ready())},q=function(){o.addEventListener?(o.removeEventListener("DOMContentLoaded",H,!1),e.removeEventListener("load",H,!1)):(o.detachEvent("onreadystatechange",H),e.detachEvent("onload",H))};b.fn=b.prototype={jquery:p,constructor:b,init:function(e,n,r){var i,a;if(!e)return this;if("string"==typeof e){if(i="<"===e.charAt(0)&&">"===e.charAt(e.length-1)&&e.length>=3?[null,e,null]:N.exec(e),!i||!i[1]&&n)return!n||n.jquery?(n||r).find(e):this.constructor(n).find(e);if(i[1]){if(n=n instanceof b?n[0]:n,b.merge(this,b.parseHTML(i[1],n&&n.nodeType?n.ownerDocument||n:o,!0)),C.test(i[1])&&b.isPlainObject(n))for(i in n)b.isFunction(this[i])?this[i](n[i]):this.attr(i,n[i]);return this}if(a=o.getElementById(i[2]),a&&a.parentNode){if(a.id!==i[2])return r.find(e);this.length=1,this[0]=a}return this.context=o,this.selector=e,this}return e.nodeType?(this.context=this[0]=e,this.length=1,this):b.isFunction(e)?r.ready(e):(e.selector!==t&&(this.selector=e.selector,this.context=e.context),b.makeArray(e,this))},selector:"",length:0,size:function(){return this.length},toArray:function(){return h.call(this)},get:function(e){return null==e?this.toArray():0>e?this[this.length+e]:this[e]},pushStack:function(e){var t=b.merge(this.constructor(),e);return t.prevObject=this,t.context=this.context,t},each:function(e,t){return b.each(this,e,t)},ready:function(e){return b.ready.promise().done(e),this},slice:function(){return this.pushStack(h.apply(this,arguments))},first:function(){return this.eq(0)},last:function(){return this.eq(-1)},eq:function(e){var t=this.length,n=+e+(0>e?t:0);return this.pushStack(n>=0&&t>n?[this[n]]:[])},map:function(e){return this.pushStack(b.map(this,function(t,n){return e.call(t,n,t)}))},end:function(){return this.prevObject||this.constructor(null)},push:d,sort:[].sort,splice:[].splice},b.fn.init.prototype=b.fn,b.extend=b.fn.extend=function(){var e,n,r,i,o,a,s=arguments[0]||{},u=1,l=arguments.length,c=!1;for("boolean"==typeof s&&(c=s,s=arguments[1]||{},u=2),"object"==typeof s||b.isFunction(s)||(s={}),l===u&&(s=this,--u);l>u;u++)if(null!=(o=arguments[u]))for(i in o)e=s[i],r=o[i],s!==r&&(c&&r&&(b.isPlainObject(r)||(n=b.isArray(r)))?(n?(n=!1,a=e&&b.isArray(e)?e:[]):a=e&&b.isPlainObject(e)?e:{},s[i]=b.extend(c,a,r)):r!==t&&(s[i]=r));return s},b.extend({noConflict:function(t){return e.$===b&&(e.$=u),t&&e.jQuery===b&&(e.jQuery=s),b},isReady:!1,readyWait:1,holdReady:function(e){e?b.readyWait++:b.ready(!0)},ready:function(e){if(e===!0?!--b.readyWait:!b.isReady){if(!o.body)return setTimeout(b.ready);b.isReady=!0,e!==!0&&--b.readyWait>0||(n.resolveWith(o,[b]),b.fn.trigger&&b(o).trigger("ready").off("ready"))}},isFunction:function(e){return"function"===b.type(e)},isArray:Array.isArray||function(e){return"array"===b.type(e)},isWindow:function(e){return null!=e&&e==e.window},isNumeric:function(e){return!isNaN(parseFloat(e))&&isFinite(e)},type:function(e){return null==e?e+"":"object"==typeof e||"function"==typeof e?l[m.call(e)]||"object":typeof e},isPlainObject:function(e){if(!e||"object"!==b.type(e)||e.nodeType||b.isWindow(e))return!1;try{if(e.constructor&&!y.call(e,"constructor")&&!y.call(e.constructor.prototype,"isPrototypeOf"))return!1}catch(n){return!1}var r;for(r in e);return r===t||y.call(e,r)},isEmptyObject:function(e){var t;for(t in e)return!1;return!0},error:function(e){throw Error(e)},parseHTML:function(e,t,n){if(!e||"string"!=typeof e)return null;"boolean"==typeof t&&(n=t,t=!1),t=t||o;var r=C.exec(e),i=!n&&[];return r?[t.createElement(r[1])]:(r=b.buildFragment([e],t,i),i&&b(i).remove(),b.merge([],r.childNodes))},parseJSON:function(n){return e.JSON&&e.JSON.parse?e.JSON.parse(n):null===n?n:"string"==typeof n&&(n=b.trim(n),n&&k.test(n.replace(S,"@").replace(A,"]").replace(E,"")))?Function("return "+n)():(b.error("Invalid JSON: "+n),t)},parseXML:function(n){var r,i;if(!n||"string"!=typeof n)return null;try{e.DOMParser?(i=new DOMParser,r=i.parseFromString(n,"text/xml")):(r=new ActiveXObject("Microsoft.XMLDOM"),r.async="false",r.loadXML(n))}catch(o){r=t}return r&&r.documentElement&&!r.getElementsByTagName("parsererror").length||b.error("Invalid XML: "+n),r},noop:function(){},globalEval:function(t){t&&b.trim(t)&&(e.execScript||function(t){e.eval.call(e,t)})(t)},camelCase:function(e){return e.replace(j,"ms-").replace(D,L)},nodeName:function(e,t){return e.nodeName&&e.nodeName.toLowerCase()===t.toLowerCase()},each:function(e,t,n){var r,i=0,o=e.length,a=M(e);if(n){if(a){for(;o>i;i++)if(r=t.apply(e[i],n),r===!1)break}else for(i in e)if(r=t.apply(e[i],n),r===!1)break}else if(a){for(;o>i;i++)if(r=t.call(e[i],i,e[i]),r===!1)break}else for(i in e)if(r=t.call(e[i],i,e[i]),r===!1)break;return e},trim:v&&!v.call("\ufeff\u00a0")?function(e){return null==e?"":v.call(e)}:function(e){return null==e?"":(e+"").replace(T,"")},makeArray:function(e,t){var n=t||[];return null!=e&&(M(Object(e))?b.merge(n,"string"==typeof e?[e]:e):d.call(n,e)),n},inArray:function(e,t,n){var r;if(t){if(g)return g.call(t,e,n);for(r=t.length,n=n?0>n?Math.max(0,r+n):n:0;r>n;n++)if(n in t&&t[n]===e)return n}return-1},merge:function(e,n){var r=n.length,i=e.length,o=0;if("number"==typeof r)for(;r>o;o++)e[i++]=n[o];else while(n[o]!==t)e[i++]=n[o++];return e.length=i,e},grep:function(e,t,n){var r,i=[],o=0,a=e.length;for(n=!!n;a>o;o++)r=!!t(e[o],o),n!==r&&i.push(e[o]);return i},map:function(e,t,n){var r,i=0,o=e.length,a=M(e),s=[];if(a)for(;o>i;i++)r=t(e[i],i,n),null!=r&&(s[s.length]=r);else for(i in e)r=t(e[i],i,n),null!=r&&(s[s.length]=r);return f.apply([],s)},guid:1,proxy:function(e,n){var r,i,o;return"string"==typeof n&&(o=e[n],n=e,e=o),b.isFunction(e)?(r=h.call(arguments,2),i=function(){return e.apply(n||this,r.concat(h.call(arguments)))},i.guid=e.guid=e.guid||b.guid++,i):t},access:function(e,n,r,i,o,a,s){var u=0,l=e.length,c=null==r;if("object"===b.type(r)){o=!0;for(u in r)b.access(e,n,u,r[u],!0,a,s)}else if(i!==t&&(o=!0,b.isFunction(i)||(s=!0),c&&(s?(n.call(e,i),n=null):(c=n,n=function(e,t,n){return c.call(b(e),n)})),n))for(;l>u;u++)n(e[u],r,s?i:i.call(e[u],u,n(e[u],r)));return o?e:c?n.call(e):l?n(e[0],r):a},now:function(){return(new Date).getTime()}}),b.ready.promise=function(t){if(!n)if(n=b.Deferred(),"complete"===o.readyState)setTimeout(b.ready);else if(o.addEventListener)o.addEventListener("DOMContentLoaded",H,!1),e.addEventListener("load",H,!1);else{o.attachEvent("onreadystatechange",H),e.attachEvent("onload",H);var r=!1;try{r=null==e.frameElement&&o.documentElement}catch(i){}r&&r.doScroll&&function a(){if(!b.isReady){try{r.doScroll("left")}catch(e){return setTimeout(a,50)}q(),b.ready()}}()}return n.promise(t)},b.each("Boolean Number String Function Array Date RegExp Object Error".split(" "),function(e,t){l["[object "+t+"]"]=t.toLowerCase()});function M(e){var t=e.length,n=b.type(e);return b.isWindow(e)?!1:1===e.nodeType&&t?!0:"array"===n||"function"!==n&&(0===t||"number"==typeof t&&t>0&&t-1 in e)}r=b(o);var _={};function F(e){var t=_[e]={};return b.each(e.match(w)||[],function(e,n){t[n]=!0}),t}b.Callbacks=function(e){e="string"==typeof e?_[e]||F(e):b.extend({},e);var n,r,i,o,a,s,u=[],l=!e.once&&[],c=function(t){for(r=e.memory&&t,i=!0,a=s||0,s=0,o=u.length,n=!0;u&&o>a;a++)if(u[a].apply(t[0],t[1])===!1&&e.stopOnFalse){r=!1;break}n=!1,u&&(l?l.length&&c(l.shift()):r?u=[]:p.disable())},p={add:function(){if(u){var t=u.length;(function i(t){b.each(t,function(t,n){var r=b.type(n);"function"===r?e.unique&&p.has(n)||u.push(n):n&&n.length&&"string"!==r&&i(n)})})(arguments),n?o=u.length:r&&(s=t,c(r))}return this},remove:function(){return u&&b.each(arguments,function(e,t){var r;while((r=b.inArray(t,u,r))>-1)u.splice(r,1),n&&(o>=r&&o--,a>=r&&a--)}),this},has:function(e){return e?b.inArray(e,u)>-1:!(!u||!u.length)},empty:function(){return u=[],this},disable:function(){return u=l=r=t,this},disabled:function(){return!u},lock:function(){return l=t,r||p.disable(),this},locked:function(){return!l},fireWith:function(e,t){return t=t||[],t=[e,t.slice?t.slice():t],!u||i&&!l||(n?l.push(t):c(t)),this},fire:function(){return p.fireWith(this,arguments),this},fired:function(){return!!i}};return p},b.extend({Deferred:function(e){var t=[["resolve","done",b.Callbacks("once memory"),"resolved"],["reject","fail",b.Callbacks("once memory"),"rejected"],["notify","progress",b.Callbacks("memory")]],n="pending",r={state:function(){return n},always:function(){return i.done(arguments).fail(arguments),this},then:function(){var e=arguments;return b.Deferred(function(n){b.each(t,function(t,o){var a=o[0],s=b.isFunction(e[t])&&e[t];i[o[1]](function(){var e=s&&s.apply(this,arguments);e&&b.isFunction(e.promise)?e.promise().done(n.resolve).fail(n.reject).progress(n.notify):n[a+"With"](this===r?n.promise():this,s?[e]:arguments)})}),e=null}).promise()},promise:function(e){return null!=e?b.extend(e,r):r}},i={};return r.pipe=r.then,b.each(t,function(e,o){var a=o[2],s=o[3];r[o[1]]=a.add,s&&a.add(function(){n=s},t[1^e][2].disable,t[2][2].lock),i[o[0]]=function(){return i[o[0]+"With"](this===i?r:this,arguments),this},i[o[0]+"With"]=a.fireWith}),r.promise(i),e&&e.call(i,i),i},when:function(e){var t=0,n=h.call(arguments),r=n.length,i=1!==r||e&&b.isFunction(e.promise)?r:0,o=1===i?e:b.Deferred(),a=function(e,t,n){return function(r){t[e]=this,n[e]=arguments.length>1?h.call(arguments):r,n===s?o.notifyWith(t,n):--i||o.resolveWith(t,n)}},s,u,l;if(r>1)for(s=Array(r),u=Array(r),l=Array(r);r>t;t++)n[t]&&b.isFunction(n[t].promise)?n[t].promise().done(a(t,l,n)).fail(o.reject).progress(a(t,u,s)):--i;return i||o.resolveWith(l,n),o.promise()}}),b.support=function(){var t,n,r,a,s,u,l,c,p,f,d=o.createElement("div");if(d.setAttribute("className","t"),d.innerHTML="  <link/><table></table><a href='/a'>a</a><input type='checkbox'/>",n=d.getElementsByTagName("*"),r=d.getElementsByTagName("a")[0],!n||!r||!n.length)return{};s=o.createElement("select"),l=s.appendChild(o.createElement("option")),a=d.getElementsByTagName("input")[0],r.style.cssText="top:1px;float:left;opacity:.5",t={getSetAttribute:"t"!==d.className,leadingWhitespace:3===d.firstChild.nodeType,tbody:!d.getElementsByTagName("tbody").length,htmlSerialize:!!d.getElementsByTagName("link").length,style:/top/.test(r.getAttribute("style")),hrefNormalized:"/a"===r.getAttribute("href"),opacity:/^0.5/.test(r.style.opacity),cssFloat:!!r.style.cssFloat,checkOn:!!a.value,optSelected:l.selected,enctype:!!o.createElement("form").enctype,html5Clone:"<:nav></:nav>"!==o.createElement("nav").cloneNode(!0).outerHTML,boxModel:"CSS1Compat"===o.compatMode,deleteExpando:!0,noCloneEvent:!0,inlineBlockNeedsLayout:!1,shrinkWrapBlocks:!1,reliableMarginRight:!0,boxSizingReliable:!0,pixelPosition:!1},a.checked=!0,t.noCloneChecked=a.cloneNode(!0).checked,s.disabled=!0,t.optDisabled=!l.disabled;try{delete d.test}catch(h){t.deleteExpando=!1}a=o.createElement("input"),a.setAttribute("value",""),t.input=""===a.getAttribute("value"),a.value="t",a.setAttribute("type","radio"),t.radioValue="t"===a.value,a.setAttribute("checked","t"),a.setAttribute("name","t"),u=o.createDocumentFragment(),u.appendChild(a),t.appendChecked=a.checked,t.checkClone=u.cloneNode(!0).cloneNode(!0).lastChild.checked,d.attachEvent&&(d.attachEvent("onclick",function(){t.noCloneEvent=!1}),d.cloneNode(!0).click());for(f in{submit:!0,change:!0,focusin:!0})d.setAttribute(c="on"+f,"t"),t[f+"Bubbles"]=c in e||d.attributes[c].expando===!1;return d.style.backgroundClip="content-box",d.cloneNode(!0).style.backgroundClip="",t.clearCloneStyle="content-box"===d.style.backgroundClip,b(function(){var n,r,a,s="padding:0;margin:0;border:0;display:block;box-sizing:content-box;-moz-box-sizing:content-box;-webkit-box-sizing:content-box;",u=o.getElementsByTagName("body")[0];u&&(n=o.createElement("div"),n.style.cssText="border:0;width:0;height:0;position:absolute;top:0;left:-9999px;margin-top:1px",u.appendChild(n).appendChild(d),d.innerHTML="<table><tr><td></td><td>t</td></tr></table>",a=d.getElementsByTagName("td"),a[0].style.cssText="padding:0;margin:0;border:0;display:none",p=0===a[0].offsetHeight,a[0].style.display="",a[1].style.display="none",t.reliableHiddenOffsets=p&&0===a[0].offsetHeight,d.innerHTML="",d.style.cssText="box-sizing:border-box;-moz-box-sizing:border-box;-webkit-box-sizing:border-box;padding:1px;border:1px;display:block;width:4px;margin-top:1%;position:absolute;top:1%;",t.boxSizing=4===d.offsetWidth,t.doesNotIncludeMarginInBodyOffset=1!==u.offsetTop,e.getComputedStyle&&(t.pixelPosition="1%"!==(e.getComputedStyle(d,null)||{}).top,t.boxSizingReliable="4px"===(e.getComputedStyle(d,null)||{width:"4px"}).width,r=d.appendChild(o.createElement("div")),r.style.cssText=d.style.cssText=s,r.style.marginRight=r.style.width="0",d.style.width="1px",t.reliableMarginRight=!parseFloat((e.getComputedStyle(r,null)||{}).marginRight)),typeof d.style.zoom!==i&&(d.innerHTML="",d.style.cssText=s+"width:1px;padding:1px;display:inline;zoom:1",t.inlineBlockNeedsLayout=3===d.offsetWidth,d.style.display="block",d.innerHTML="<div></div>",d.firstChild.style.width="5px",t.shrinkWrapBlocks=3!==d.offsetWidth,t.inlineBlockNeedsLayout&&(u.style.zoom=1)),u.removeChild(n),n=d=a=r=null)}),n=s=u=l=r=a=null,t}();var O=/(?:\{[\s\S]*\}|\[[\s\S]*\])$/,B=/([A-Z])/g;function P(e,n,r,i){if(b.acceptData(e)){var o,a,s=b.expando,u="string"==typeof n,l=e.nodeType,p=l?b.cache:e,f=l?e[s]:e[s]&&s;if(f&&p[f]&&(i||p[f].data)||!u||r!==t)return f||(l?e[s]=f=c.pop()||b.guid++:f=s),p[f]||(p[f]={},l||(p[f].toJSON=b.noop)),("object"==typeof n||"function"==typeof n)&&(i?p[f]=b.extend(p[f],n):p[f].data=b.extend(p[f].data,n)),o=p[f],i||(o.data||(o.data={}),o=o.data),r!==t&&(o[b.camelCase(n)]=r),u?(a=o[n],null==a&&(a=o[b.camelCase(n)])):a=o,a}}function R(e,t,n){if(b.acceptData(e)){var r,i,o,a=e.nodeType,s=a?b.cache:e,u=a?e[b.expando]:b.expando;if(s[u]){if(t&&(o=n?s[u]:s[u].data)){b.isArray(t)?t=t.concat(b.map(t,b.camelCase)):t in o?t=[t]:(t=b.camelCase(t),t=t in o?[t]:t.split(" "));for(r=0,i=t.length;i>r;r++)delete o[t[r]];if(!(n?$:b.isEmptyObject)(o))return}(n||(delete s[u].data,$(s[u])))&&(a?b.cleanData([e],!0):b.support.deleteExpando||s!=s.window?delete s[u]:s[u]=null)}}}b.extend({cache:{},expando:"jQuery"+(p+Math.random()).replace(/\D/g,""),noData:{embed:!0,object:"clsid:D27CDB6E-AE6D-11cf-96B8-444553540000",applet:!0},hasData:function(e){return e=e.nodeType?b.cache[e[b.expando]]:e[b.expando],!!e&&!$(e)},data:function(e,t,n){return P(e,t,n)},removeData:function(e,t){return R(e,t)},_data:function(e,t,n){return P(e,t,n,!0)},_removeData:function(e,t){return R(e,t,!0)},acceptData:function(e){if(e.nodeType&&1!==e.nodeType&&9!==e.nodeType)return!1;var t=e.nodeName&&b.noData[e.nodeName.toLowerCase()];return!t||t!==!0&&e.getAttribute("classid")===t}}),b.fn.extend({data:function(e,n){var r,i,o=this[0],a=0,s=null;if(e===t){if(this.length&&(s=b.data(o),1===o.nodeType&&!b._data(o,"parsedAttrs"))){for(r=o.attributes;r.length>a;a++)i=r[a].name,i.indexOf("data-")||(i=b.camelCase(i.slice(5)),W(o,i,s[i]));b._data(o,"parsedAttrs",!0)}return s}return"object"==typeof e?this.each(function(){b.data(this,e)}):b.access(this,function(n){return n===t?o?W(o,e,b.data(o,e)):null:(this.each(function(){b.data(this,e,n)}),t)},null,n,arguments.length>1,null,!0)},removeData:function(e){return this.each(function(){b.removeData(this,e)})}});function W(e,n,r){if(r===t&&1===e.nodeType){var i="data-"+n.replace(B,"-$1").toLowerCase();if(r=e.getAttribute(i),"string"==typeof r){try{r="true"===r?!0:"false"===r?!1:"null"===r?null:+r+""===r?+r:O.test(r)?b.parseJSON(r):r}catch(o){}b.data(e,n,r)}else r=t}return r}function $(e){var t;for(t in e)if(("data"!==t||!b.isEmptyObject(e[t]))&&"toJSON"!==t)return!1;return!0}b.extend({queue:function(e,n,r){var i;return e?(n=(n||"fx")+"queue",i=b._data(e,n),r&&(!i||b.isArray(r)?i=b._data(e,n,b.makeArray(r)):i.push(r)),i||[]):t},dequeue:function(e,t){t=t||"fx";var n=b.queue(e,t),r=n.length,i=n.shift(),o=b._queueHooks(e,t),a=function(){b.dequeue(e,t)};"inprogress"===i&&(i=n.shift(),r--),o.cur=i,i&&("fx"===t&&n.unshift("inprogress"),delete o.stop,i.call(e,a,o)),!r&&o&&o.empty.fire()},_queueHooks:function(e,t){var n=t+"queueHooks";return b._data(e,n)||b._data(e,n,{empty:b.Callbacks("once memory").add(function(){b._removeData(e,t+"queue"),b._removeData(e,n)})})}}),b.fn.extend({queue:function(e,n){var r=2;return"string"!=typeof e&&(n=e,e="fx",r--),r>arguments.length?b.queue(this[0],e):n===t?this:this.each(function(){var t=b.queue(this,e,n);b._queueHooks(this,e),"fx"===e&&"inprogress"!==t[0]&&b.dequeue(this,e)})},dequeue:function(e){return this.each(function(){b.dequeue(this,e)})},delay:function(e,t){return e=b.fx?b.fx.speeds[e]||e:e,t=t||"fx",this.queue(t,function(t,n){var r=setTimeout(t,e);n.stop=function(){clearTimeout(r)}})},clearQueue:function(e){return this.queue(e||"fx",[])},promise:function(e,n){var r,i=1,o=b.Deferred(),a=this,s=this.length,u=function(){--i||o.resolveWith(a,[a])};"string"!=typeof e&&(n=e,e=t),e=e||"fx";while(s--)r=b._data(a[s],e+"queueHooks"),r&&r.empty&&(i++,r.empty.add(u));return u(),o.promise(n)}});var I,z,X=/[\t\r\n]/g,U=/\r/g,V=/^(?:input|select|textarea|button|object)$/i,Y=/^(?:a|area)$/i,J=/^(?:checked|selected|autofocus|autoplay|async|controls|defer|disabled|hidden|loop|multiple|open|readonly|required|scoped)$/i,G=/^(?:checked|selected)$/i,Q=b.support.getSetAttribute,K=b.support.input;b.fn.extend({attr:function(e,t){return b.access(this,b.attr,e,t,arguments.length>1)},removeAttr:function(e){return this.each(function(){b.removeAttr(this,e)})},prop:function(e,t){return b.access(this,b.prop,e,t,arguments.length>1)},removeProp:function(e){return e=b.propFix[e]||e,this.each(function(){try{this[e]=t,delete this[e]}catch(n){}})},addClass:function(e){var t,n,r,i,o,a=0,s=this.length,u="string"==typeof e&&e;if(b.isFunction(e))return this.each(function(t){b(this).addClass(e.call(this,t,this.className))});if(u)for(t=(e||"").match(w)||[];s>a;a++)if(n=this[a],r=1===n.nodeType&&(n.className?(" "+n.className+" ").replace(X," "):" ")){o=0;while(i=t[o++])0>r.indexOf(" "+i+" ")&&(r+=i+" ");n.className=b.trim(r)}return this},removeClass:function(e){var t,n,r,i,o,a=0,s=this.length,u=0===arguments.length||"string"==typeof e&&e;if(b.isFunction(e))return this.each(function(t){b(this).removeClass(e.call(this,t,this.className))});if(u)for(t=(e||"").match(w)||[];s>a;a++)if(n=this[a],r=1===n.nodeType&&(n.className?(" "+n.className+" ").replace(X," "):"")){o=0;while(i=t[o++])while(r.indexOf(" "+i+" ")>=0)r=r.replace(" "+i+" "," ");n.className=e?b.trim(r):""}return this},toggleClass:function(e,t){var n=typeof e,r="boolean"==typeof t;return b.isFunction(e)?this.each(function(n){b(this).toggleClass(e.call(this,n,this.className,t),t)}):this.each(function(){if("string"===n){var o,a=0,s=b(this),u=t,l=e.match(w)||[];while(o=l[a++])u=r?u:!s.hasClass(o),s[u?"addClass":"removeClass"](o)}else(n===i||"boolean"===n)&&(this.className&&b._data(this,"__className__",this.className),this.className=this.className||e===!1?"":b._data(this,"__className__")||"")})},hasClass:function(e){var t=" "+e+" ",n=0,r=this.length;for(;r>n;n++)if(1===this[n].nodeType&&(" "+this[n].className+" ").replace(X," ").indexOf(t)>=0)return!0;return!1},val:function(e){var n,r,i,o=this[0];{if(arguments.length)return i=b.isFunction(e),this.each(function(n){var o,a=b(this);1===this.nodeType&&(o=i?e.call(this,n,a.val()):e,null==o?o="":"number"==typeof o?o+="":b.isArray(o)&&(o=b.map(o,function(e){return null==e?"":e+""})),r=b.valHooks[this.type]||b.valHooks[this.nodeName.toLowerCase()],r&&"set"in r&&r.set(this,o,"value")!==t||(this.value=o))});if(o)return r=b.valHooks[o.type]||b.valHooks[o.nodeName.toLowerCase()],r&&"get"in r&&(n=r.get(o,"value"))!==t?n:(n=o.value,"string"==typeof n?n.replace(U,""):null==n?"":n)}}}),b.extend({valHooks:{option:{get:function(e){var t=e.attributes.value;return!t||t.specified?e.value:e.text}},select:{get:function(e){var t,n,r=e.options,i=e.selectedIndex,o="select-one"===e.type||0>i,a=o?null:[],s=o?i+1:r.length,u=0>i?s:o?i:0;for(;s>u;u++)if(n=r[u],!(!n.selected&&u!==i||(b.support.optDisabled?n.disabled:null!==n.getAttribute("disabled"))||n.parentNode.disabled&&b.nodeName(n.parentNode,"optgroup"))){if(t=b(n).val(),o)return t;a.push(t)}return a},set:function(e,t){var n=b.makeArray(t);return b(e).find("option").each(function(){this.selected=b.inArray(b(this).val(),n)>=0}),n.length||(e.selectedIndex=-1),n}}},attr:function(e,n,r){var o,a,s,u=e.nodeType;if(e&&3!==u&&8!==u&&2!==u)return typeof e.getAttribute===i?b.prop(e,n,r):(a=1!==u||!b.isXMLDoc(e),a&&(n=n.toLowerCase(),o=b.attrHooks[n]||(J.test(n)?z:I)),r===t?o&&a&&"get"in o&&null!==(s=o.get(e,n))?s:(typeof e.getAttribute!==i&&(s=e.getAttribute(n)),null==s?t:s):null!==r?o&&a&&"set"in o&&(s=o.set(e,r,n))!==t?s:(e.setAttribute(n,r+""),r):(b.removeAttr(e,n),t))},removeAttr:function(e,t){var n,r,i=0,o=t&&t.match(w);if(o&&1===e.nodeType)while(n=o[i++])r=b.propFix[n]||n,J.test(n)?!Q&&G.test(n)?e[b.camelCase("default-"+n)]=e[r]=!1:e[r]=!1:b.attr(e,n,""),e.removeAttribute(Q?n:r)},attrHooks:{type:{set:function(e,t){if(!b.support.radioValue&&"radio"===t&&b.nodeName(e,"input")){var n=e.value;return e.setAttribute("type",t),n&&(e.value=n),t}}}},propFix:{tabindex:"tabIndex",readonly:"readOnly","for":"htmlFor","class":"className",maxlength:"maxLength",cellspacing:"cellSpacing",cellpadding:"cellPadding",rowspan:"rowSpan",colspan:"colSpan",usemap:"useMap",frameborder:"frameBorder",contenteditable:"contentEditable"},prop:function(e,n,r){var i,o,a,s=e.nodeType;if(e&&3!==s&&8!==s&&2!==s)return a=1!==s||!b.isXMLDoc(e),a&&(n=b.propFix[n]||n,o=b.propHooks[n]),r!==t?o&&"set"in o&&(i=o.set(e,r,n))!==t?i:e[n]=r:o&&"get"in o&&null!==(i=o.get(e,n))?i:e[n]},propHooks:{tabIndex:{get:function(e){var n=e.getAttributeNode("tabindex");return n&&n.specified?parseInt(n.value,10):V.test(e.nodeName)||Y.test(e.nodeName)&&e.href?0:t}}}}),z={get:function(e,n){var r=b.prop(e,n),i="boolean"==typeof r&&e.getAttribute(n),o="boolean"==typeof r?K&&Q?null!=i:G.test(n)?e[b.camelCase("default-"+n)]:!!i:e.getAttributeNode(n);return o&&o.value!==!1?n.toLowerCase():t},set:function(e,t,n){return t===!1?b.removeAttr(e,n):K&&Q||!G.test(n)?e.setAttribute(!Q&&b.propFix[n]||n,n):e[b.camelCase("default-"+n)]=e[n]=!0,n}},K&&Q||(b.attrHooks.value={get:function(e,n){var r=e.getAttributeNode(n);return b.nodeName(e,"input")?e.defaultValue:r&&r.specified?r.value:t},set:function(e,n,r){return b.nodeName(e,"input")?(e.defaultValue=n,t):I&&I.set(e,n,r)}}),Q||(I=b.valHooks.button={get:function(e,n){var r=e.getAttributeNode(n);return r&&("id"===n||"name"===n||"coords"===n?""!==r.value:r.specified)?r.value:t},set:function(e,n,r){var i=e.getAttributeNode(r);return i||e.setAttributeNode(i=e.ownerDocument.createAttribute(r)),i.value=n+="","value"===r||n===e.getAttribute(r)?n:t}},b.attrHooks.contenteditable={get:I.get,set:function(e,t,n){I.set(e,""===t?!1:t,n)}},b.each(["width","height"],function(e,n){b.attrHooks[n]=b.extend(b.attrHooks[n],{set:function(e,r){return""===r?(e.setAttribute(n,"auto"),r):t}})})),b.support.hrefNormalized||(b.each(["href","src","width","height"],function(e,n){b.attrHooks[n]=b.extend(b.attrHooks[n],{get:function(e){var r=e.getAttribute(n,2);return null==r?t:r}})}),b.each(["href","src"],function(e,t){b.propHooks[t]={get:function(e){return e.getAttribute(t,4)}}})),b.support.style||(b.attrHooks.style={get:function(e){return e.style.cssText||t},set:function(e,t){return e.style.cssText=t+""}}),b.support.optSelected||(b.propHooks.selected=b.extend(b.propHooks.selected,{get:function(e){var t=e.parentNode;return t&&(t.selectedIndex,t.parentNode&&t.parentNode.selectedIndex),null}})),b.support.enctype||(b.propFix.enctype="encoding"),b.support.checkOn||b.each(["radio","checkbox"],function(){b.valHooks[this]={get:function(e){return null===e.getAttribute("value")?"on":e.value}}}),b.each(["radio","checkbox"],function(){b.valHooks[this]=b.extend(b.valHooks[this],{set:function(e,n){return b.isArray(n)?e.checked=b.inArray(b(e).val(),n)>=0:t}})});var Z=/^(?:input|select|textarea)$/i,et=/^key/,tt=/^(?:mouse|contextmenu)|click/,nt=/^(?:focusinfocus|focusoutblur)$/,rt=/^([^.]*)(?:\.(.+)|)$/;function it(){return!0}function ot(){return!1}b.event={global:{},add:function(e,n,r,o,a){var s,u,l,c,p,f,d,h,g,m,y,v=b._data(e);if(v){r.handler&&(c=r,r=c.handler,a=c.selector),r.guid||(r.guid=b.guid++),(u=v.events)||(u=v.events={}),(f=v.handle)||(f=v.handle=function(e){return typeof b===i||e&&b.event.triggered===e.type?t:b.event.dispatch.apply(f.elem,arguments)},f.elem=e),n=(n||"").match(w)||[""],l=n.length;while(l--)s=rt.exec(n[l])||[],g=y=s[1],m=(s[2]||"").split(".").sort(),p=b.event.special[g]||{},g=(a?p.delegateType:p.bindType)||g,p=b.event.special[g]||{},d=b.extend({type:g,origType:y,data:o,handler:r,guid:r.guid,selector:a,needsContext:a&&b.expr.match.needsContext.test(a),namespace:m.join(".")},c),(h=u[g])||(h=u[g]=[],h.delegateCount=0,p.setup&&p.setup.call(e,o,m,f)!==!1||(e.addEventListener?e.addEventListener(g,f,!1):e.attachEvent&&e.attachEvent("on"+g,f))),p.add&&(p.add.call(e,d),d.handler.guid||(d.handler.guid=r.guid)),a?h.splice(h.delegateCount++,0,d):h.push(d),b.event.global[g]=!0;e=null}},remove:function(e,t,n,r,i){var o,a,s,u,l,c,p,f,d,h,g,m=b.hasData(e)&&b._data(e);if(m&&(c=m.events)){t=(t||"").match(w)||[""],l=t.length;while(l--)if(s=rt.exec(t[l])||[],d=g=s[1],h=(s[2]||"").split(".").sort(),d){p=b.event.special[d]||{},d=(r?p.delegateType:p.bindType)||d,f=c[d]||[],s=s[2]&&RegExp("(^|\\.)"+h.join("\\.(?:.*\\.|)")+"(\\.|$)"),u=o=f.length;while(o--)a=f[o],!i&&g!==a.origType||n&&n.guid!==a.guid||s&&!s.test(a.namespace)||r&&r!==a.selector&&("**"!==r||!a.selector)||(f.splice(o,1),a.selector&&f.delegateCount--,p.remove&&p.remove.call(e,a));u&&!f.length&&(p.teardown&&p.teardown.call(e,h,m.handle)!==!1||b.removeEvent(e,d,m.handle),delete c[d])}else for(d in c)b.event.remove(e,d+t[l],n,r,!0);b.isEmptyObject(c)&&(delete m.handle,b._removeData(e,"events"))}},trigger:function(n,r,i,a){var s,u,l,c,p,f,d,h=[i||o],g=y.call(n,"type")?n.type:n,m=y.call(n,"namespace")?n.namespace.split("."):[];if(l=f=i=i||o,3!==i.nodeType&&8!==i.nodeType&&!nt.test(g+b.event.triggered)&&(g.indexOf(".")>=0&&(m=g.split("."),g=m.shift(),m.sort()),u=0>g.indexOf(":")&&"on"+g,n=n[b.expando]?n:new b.Event(g,"object"==typeof n&&n),n.isTrigger=!0,n.namespace=m.join("."),n.namespace_re=n.namespace?RegExp("(^|\\.)"+m.join("\\.(?:.*\\.|)")+"(\\.|$)"):null,n.result=t,n.target||(n.target=i),r=null==r?[n]:b.makeArray(r,[n]),p=b.event.special[g]||{},a||!p.trigger||p.trigger.apply(i,r)!==!1)){if(!a&&!p.noBubble&&!b.isWindow(i)){for(c=p.delegateType||g,nt.test(c+g)||(l=l.parentNode);l;l=l.parentNode)h.push(l),f=l;f===(i.ownerDocument||o)&&h.push(f.defaultView||f.parentWindow||e)}d=0;while((l=h[d++])&&!n.isPropagationStopped())n.type=d>1?c:p.bindType||g,s=(b._data(l,"events")||{})[n.type]&&b._data(l,"handle"),s&&s.apply(l,r),s=u&&l[u],s&&b.acceptData(l)&&s.apply&&s.apply(l,r)===!1&&n.preventDefault();if(n.type=g,!(a||n.isDefaultPrevented()||p._default&&p._default.apply(i.ownerDocument,r)!==!1||"click"===g&&b.nodeName(i,"a")||!b.acceptData(i)||!u||!i[g]||b.isWindow(i))){f=i[u],f&&(i[u]=null),b.event.triggered=g;try{i[g]()}catch(v){}b.event.triggered=t,f&&(i[u]=f)}return n.result}},dispatch:function(e){e=b.event.fix(e);var n,r,i,o,a,s=[],u=h.call(arguments),l=(b._data(this,"events")||{})[e.type]||[],c=b.event.special[e.type]||{};if(u[0]=e,e.delegateTarget=this,!c.preDispatch||c.preDispatch.call(this,e)!==!1){s=b.event.handlers.call(this,e,l),n=0;while((o=s[n++])&&!e.isPropagationStopped()){e.currentTarget=o.elem,a=0;while((i=o.handlers[a++])&&!e.isImmediatePropagationStopped())(!e.namespace_re||e.namespace_re.test(i.namespace))&&(e.handleObj=i,e.data=i.data,r=((b.event.special[i.origType]||{}).handle||i.handler).apply(o.elem,u),r!==t&&(e.result=r)===!1&&(e.preventDefault(),e.stopPropagation()))}return c.postDispatch&&c.postDispatch.call(this,e),e.result}},handlers:function(e,n){var r,i,o,a,s=[],u=n.delegateCount,l=e.target;if(u&&l.nodeType&&(!e.button||"click"!==e.type))for(;l!=this;l=l.parentNode||this)if(1===l.nodeType&&(l.disabled!==!0||"click"!==e.type)){for(o=[],a=0;u>a;a++)i=n[a],r=i.selector+" ",o[r]===t&&(o[r]=i.needsContext?b(r,this).index(l)>=0:b.find(r,this,null,[l]).length),o[r]&&o.push(i);o.length&&s.push({elem:l,handlers:o})}return n.length>u&&s.push({elem:this,handlers:n.slice(u)}),s},fix:function(e){if(e[b.expando])return e;var t,n,r,i=e.type,a=e,s=this.fixHooks[i];s||(this.fixHooks[i]=s=tt.test(i)?this.mouseHooks:et.test(i)?this.keyHooks:{}),r=s.props?this.props.concat(s.props):this.props,e=new b.Event(a),t=r.length;while(t--)n=r[t],e[n]=a[n];return e.target||(e.target=a.srcElement||o),3===e.target.nodeType&&(e.target=e.target.parentNode),e.metaKey=!!e.metaKey,s.filter?s.filter(e,a):e},props:"altKey bubbles cancelable ctrlKey currentTarget eventPhase metaKey relatedTarget shiftKey target timeStamp view which".split(" "),fixHooks:{},keyHooks:{props:"char charCode key keyCode".split(" "),filter:function(e,t){return null==e.which&&(e.which=null!=t.charCode?t.charCode:t.keyCode),e}},mouseHooks:{props:"button buttons clientX clientY fromElement offsetX offsetY pageX pageY screenX screenY toElement".split(" "),filter:function(e,n){var r,i,a,s=n.button,u=n.fromElement;return null==e.pageX&&null!=n.clientX&&(i=e.target.ownerDocument||o,a=i.documentElement,r=i.body,e.pageX=n.clientX+(a&&a.scrollLeft||r&&r.scrollLeft||0)-(a&&a.clientLeft||r&&r.clientLeft||0),e.pageY=n.clientY+(a&&a.scrollTop||r&&r.scrollTop||0)-(a&&a.clientTop||r&&r.clientTop||0)),!e.relatedTarget&&u&&(e.relatedTarget=u===e.target?n.toElement:u),e.which||s===t||(e.which=1&s?1:2&s?3:4&s?2:0),e}},special:{load:{noBubble:!0},click:{trigger:function(){return b.nodeName(this,"input")&&"checkbox"===this.type&&this.click?(this.click(),!1):t}},focus:{trigger:function(){if(this!==o.activeElement&&this.focus)try{return this.focus(),!1}catch(e){}},delegateType:"focusin"},blur:{trigger:function(){return this===o.activeElement&&this.blur?(this.blur(),!1):t},delegateType:"focusout"},beforeunload:{postDispatch:function(e){e.result!==t&&(e.originalEvent.returnValue=e.result)}}},simulate:function(e,t,n,r){var i=b.extend(new b.Event,n,{type:e,isSimulated:!0,originalEvent:{}});r?b.event.trigger(i,null,t):b.event.dispatch.call(t,i),i.isDefaultPrevented()&&n.preventDefault()}},b.removeEvent=o.removeEventListener?function(e,t,n){e.removeEventListener&&e.removeEventListener(t,n,!1)}:function(e,t,n){var r="on"+t;e.detachEvent&&(typeof e[r]===i&&(e[r]=null),e.detachEvent(r,n))},b.Event=function(e,n){return this instanceof b.Event?(e&&e.type?(this.originalEvent=e,this.type=e.type,this.isDefaultPrevented=e.defaultPrevented||e.returnValue===!1||e.getPreventDefault&&e.getPreventDefault()?it:ot):this.type=e,n&&b.extend(this,n),this.timeStamp=e&&e.timeStamp||b.now(),this[b.expando]=!0,t):new b.Event(e,n)},b.Event.prototype={isDefaultPrevented:ot,isPropagationStopped:ot,isImmediatePropagationStopped:ot,preventDefault:function(){var e=this.originalEvent;this.isDefaultPrevented=it,e&&(e.preventDefault?e.preventDefault():e.returnValue=!1)},stopPropagation:function(){var e=this.originalEvent;this.isPropagationStopped=it,e&&(e.stopPropagation&&e.stopPropagation(),e.cancelBubble=!0)},stopImmediatePropagation:function(){this.isImmediatePropagationStopped=it,this.stopPropagation()}},b.each({mouseenter:"mouseover",mouseleave:"mouseout"},function(e,t){b.event.special[e]={delegateType:t,bindType:t,handle:function(e){var n,r=this,i=e.relatedTarget,o=e.handleObj;
return(!i||i!==r&&!b.contains(r,i))&&(e.type=o.origType,n=o.handler.apply(this,arguments),e.type=t),n}}}),b.support.submitBubbles||(b.event.special.submit={setup:function(){return b.nodeName(this,"form")?!1:(b.event.add(this,"click._submit keypress._submit",function(e){var n=e.target,r=b.nodeName(n,"input")||b.nodeName(n,"button")?n.form:t;r&&!b._data(r,"submitBubbles")&&(b.event.add(r,"submit._submit",function(e){e._submit_bubble=!0}),b._data(r,"submitBubbles",!0))}),t)},postDispatch:function(e){e._submit_bubble&&(delete e._submit_bubble,this.parentNode&&!e.isTrigger&&b.event.simulate("submit",this.parentNode,e,!0))},teardown:function(){return b.nodeName(this,"form")?!1:(b.event.remove(this,"._submit"),t)}}),b.support.changeBubbles||(b.event.special.change={setup:function(){return Z.test(this.nodeName)?(("checkbox"===this.type||"radio"===this.type)&&(b.event.add(this,"propertychange._change",function(e){"checked"===e.originalEvent.propertyName&&(this._just_changed=!0)}),b.event.add(this,"click._change",function(e){this._just_changed&&!e.isTrigger&&(this._just_changed=!1),b.event.simulate("change",this,e,!0)})),!1):(b.event.add(this,"beforeactivate._change",function(e){var t=e.target;Z.test(t.nodeName)&&!b._data(t,"changeBubbles")&&(b.event.add(t,"change._change",function(e){!this.parentNode||e.isSimulated||e.isTrigger||b.event.simulate("change",this.parentNode,e,!0)}),b._data(t,"changeBubbles",!0))}),t)},handle:function(e){var n=e.target;return this!==n||e.isSimulated||e.isTrigger||"radio"!==n.type&&"checkbox"!==n.type?e.handleObj.handler.apply(this,arguments):t},teardown:function(){return b.event.remove(this,"._change"),!Z.test(this.nodeName)}}),b.support.focusinBubbles||b.each({focus:"focusin",blur:"focusout"},function(e,t){var n=0,r=function(e){b.event.simulate(t,e.target,b.event.fix(e),!0)};b.event.special[t]={setup:function(){0===n++&&o.addEventListener(e,r,!0)},teardown:function(){0===--n&&o.removeEventListener(e,r,!0)}}}),b.fn.extend({on:function(e,n,r,i,o){var a,s;if("object"==typeof e){"string"!=typeof n&&(r=r||n,n=t);for(a in e)this.on(a,n,r,e[a],o);return this}if(null==r&&null==i?(i=n,r=n=t):null==i&&("string"==typeof n?(i=r,r=t):(i=r,r=n,n=t)),i===!1)i=ot;else if(!i)return this;return 1===o&&(s=i,i=function(e){return b().off(e),s.apply(this,arguments)},i.guid=s.guid||(s.guid=b.guid++)),this.each(function(){b.event.add(this,e,i,r,n)})},one:function(e,t,n,r){return this.on(e,t,n,r,1)},off:function(e,n,r){var i,o;if(e&&e.preventDefault&&e.handleObj)return i=e.handleObj,b(e.delegateTarget).off(i.namespace?i.origType+"."+i.namespace:i.origType,i.selector,i.handler),this;if("object"==typeof e){for(o in e)this.off(o,n,e[o]);return this}return(n===!1||"function"==typeof n)&&(r=n,n=t),r===!1&&(r=ot),this.each(function(){b.event.remove(this,e,r,n)})},bind:function(e,t,n){return this.on(e,null,t,n)},unbind:function(e,t){return this.off(e,null,t)},delegate:function(e,t,n,r){return this.on(t,e,n,r)},undelegate:function(e,t,n){return 1===arguments.length?this.off(e,"**"):this.off(t,e||"**",n)},trigger:function(e,t){return this.each(function(){b.event.trigger(e,t,this)})},triggerHandler:function(e,n){var r=this[0];return r?b.event.trigger(e,n,r,!0):t}}),function(e,t){var n,r,i,o,a,s,u,l,c,p,f,d,h,g,m,y,v,x="sizzle"+-new Date,w=e.document,T={},N=0,C=0,k=it(),E=it(),S=it(),A=typeof t,j=1<<31,D=[],L=D.pop,H=D.push,q=D.slice,M=D.indexOf||function(e){var t=0,n=this.length;for(;n>t;t++)if(this[t]===e)return t;return-1},_="[\\x20\\t\\r\\n\\f]",F="(?:\\\\.|[\\w-]|[^\\x00-\\xa0])+",O=F.replace("w","w#"),B="([*^$|!~]?=)",P="\\["+_+"*("+F+")"+_+"*(?:"+B+_+"*(?:(['\"])((?:\\\\.|[^\\\\])*?)\\3|("+O+")|)|)"+_+"*\\]",R=":("+F+")(?:\\(((['\"])((?:\\\\.|[^\\\\])*?)\\3|((?:\\\\.|[^\\\\()[\\]]|"+P.replace(3,8)+")*)|.*)\\)|)",W=RegExp("^"+_+"+|((?:^|[^\\\\])(?:\\\\.)*)"+_+"+$","g"),$=RegExp("^"+_+"*,"+_+"*"),I=RegExp("^"+_+"*([\\x20\\t\\r\\n\\f>+~])"+_+"*"),z=RegExp(R),X=RegExp("^"+O+"$"),U={ID:RegExp("^#("+F+")"),CLASS:RegExp("^\\.("+F+")"),NAME:RegExp("^\\[name=['\"]?("+F+")['\"]?\\]"),TAG:RegExp("^("+F.replace("w","w*")+")"),ATTR:RegExp("^"+P),PSEUDO:RegExp("^"+R),CHILD:RegExp("^:(only|first|last|nth|nth-last)-(child|of-type)(?:\\("+_+"*(even|odd|(([+-]|)(\\d*)n|)"+_+"*(?:([+-]|)"+_+"*(\\d+)|))"+_+"*\\)|)","i"),needsContext:RegExp("^"+_+"*[>+~]|:(even|odd|eq|gt|lt|nth|first|last)(?:\\("+_+"*((?:-\\d)?\\d*)"+_+"*\\)|)(?=[^-]|$)","i")},V=/[\x20\t\r\n\f]*[+~]/,Y=/^[^{]+\{\s*\[native code/,J=/^(?:#([\w-]+)|(\w+)|\.([\w-]+))$/,G=/^(?:input|select|textarea|button)$/i,Q=/^h\d$/i,K=/'|\\/g,Z=/\=[\x20\t\r\n\f]*([^'"\]]*)[\x20\t\r\n\f]*\]/g,et=/\\([\da-fA-F]{1,6}[\x20\t\r\n\f]?|.)/g,tt=function(e,t){var n="0x"+t-65536;return n!==n?t:0>n?String.fromCharCode(n+65536):String.fromCharCode(55296|n>>10,56320|1023&n)};try{q.call(w.documentElement.childNodes,0)[0].nodeType}catch(nt){q=function(e){var t,n=[];while(t=this[e++])n.push(t);return n}}function rt(e){return Y.test(e+"")}function it(){var e,t=[];return e=function(n,r){return t.push(n+=" ")>i.cacheLength&&delete e[t.shift()],e[n]=r}}function ot(e){return e[x]=!0,e}function at(e){var t=p.createElement("div");try{return e(t)}catch(n){return!1}finally{t=null}}function st(e,t,n,r){var i,o,a,s,u,l,f,g,m,v;if((t?t.ownerDocument||t:w)!==p&&c(t),t=t||p,n=n||[],!e||"string"!=typeof e)return n;if(1!==(s=t.nodeType)&&9!==s)return[];if(!d&&!r){if(i=J.exec(e))if(a=i[1]){if(9===s){if(o=t.getElementById(a),!o||!o.parentNode)return n;if(o.id===a)return n.push(o),n}else if(t.ownerDocument&&(o=t.ownerDocument.getElementById(a))&&y(t,o)&&o.id===a)return n.push(o),n}else{if(i[2])return H.apply(n,q.call(t.getElementsByTagName(e),0)),n;if((a=i[3])&&T.getByClassName&&t.getElementsByClassName)return H.apply(n,q.call(t.getElementsByClassName(a),0)),n}if(T.qsa&&!h.test(e)){if(f=!0,g=x,m=t,v=9===s&&e,1===s&&"object"!==t.nodeName.toLowerCase()){l=ft(e),(f=t.getAttribute("id"))?g=f.replace(K,"\\$&"):t.setAttribute("id",g),g="[id='"+g+"'] ",u=l.length;while(u--)l[u]=g+dt(l[u]);m=V.test(e)&&t.parentNode||t,v=l.join(",")}if(v)try{return H.apply(n,q.call(m.querySelectorAll(v),0)),n}catch(b){}finally{f||t.removeAttribute("id")}}}return wt(e.replace(W,"$1"),t,n,r)}a=st.isXML=function(e){var t=e&&(e.ownerDocument||e).documentElement;return t?"HTML"!==t.nodeName:!1},c=st.setDocument=function(e){var n=e?e.ownerDocument||e:w;return n!==p&&9===n.nodeType&&n.documentElement?(p=n,f=n.documentElement,d=a(n),T.tagNameNoComments=at(function(e){return e.appendChild(n.createComment("")),!e.getElementsByTagName("*").length}),T.attributes=at(function(e){e.innerHTML="<select></select>";var t=typeof e.lastChild.getAttribute("multiple");return"boolean"!==t&&"string"!==t}),T.getByClassName=at(function(e){return e.innerHTML="<div class='hidden e'></div><div class='hidden'></div>",e.getElementsByClassName&&e.getElementsByClassName("e").length?(e.lastChild.className="e",2===e.getElementsByClassName("e").length):!1}),T.getByName=at(function(e){e.id=x+0,e.innerHTML="<a name='"+x+"'></a><div name='"+x+"'></div>",f.insertBefore(e,f.firstChild);var t=n.getElementsByName&&n.getElementsByName(x).length===2+n.getElementsByName(x+0).length;return T.getIdNotName=!n.getElementById(x),f.removeChild(e),t}),i.attrHandle=at(function(e){return e.innerHTML="<a href='#'></a>",e.firstChild&&typeof e.firstChild.getAttribute!==A&&"#"===e.firstChild.getAttribute("href")})?{}:{href:function(e){return e.getAttribute("href",2)},type:function(e){return e.getAttribute("type")}},T.getIdNotName?(i.find.ID=function(e,t){if(typeof t.getElementById!==A&&!d){var n=t.getElementById(e);return n&&n.parentNode?[n]:[]}},i.filter.ID=function(e){var t=e.replace(et,tt);return function(e){return e.getAttribute("id")===t}}):(i.find.ID=function(e,n){if(typeof n.getElementById!==A&&!d){var r=n.getElementById(e);return r?r.id===e||typeof r.getAttributeNode!==A&&r.getAttributeNode("id").value===e?[r]:t:[]}},i.filter.ID=function(e){var t=e.replace(et,tt);return function(e){var n=typeof e.getAttributeNode!==A&&e.getAttributeNode("id");return n&&n.value===t}}),i.find.TAG=T.tagNameNoComments?function(e,n){return typeof n.getElementsByTagName!==A?n.getElementsByTagName(e):t}:function(e,t){var n,r=[],i=0,o=t.getElementsByTagName(e);if("*"===e){while(n=o[i++])1===n.nodeType&&r.push(n);return r}return o},i.find.NAME=T.getByName&&function(e,n){return typeof n.getElementsByName!==A?n.getElementsByName(name):t},i.find.CLASS=T.getByClassName&&function(e,n){return typeof n.getElementsByClassName===A||d?t:n.getElementsByClassName(e)},g=[],h=[":focus"],(T.qsa=rt(n.querySelectorAll))&&(at(function(e){e.innerHTML="<select><option selected=''></option></select>",e.querySelectorAll("[selected]").length||h.push("\\["+_+"*(?:checked|disabled|ismap|multiple|readonly|selected|value)"),e.querySelectorAll(":checked").length||h.push(":checked")}),at(function(e){e.innerHTML="<input type='hidden' i=''/>",e.querySelectorAll("[i^='']").length&&h.push("[*^$]="+_+"*(?:\"\"|'')"),e.querySelectorAll(":enabled").length||h.push(":enabled",":disabled"),e.querySelectorAll("*,:x"),h.push(",.*:")})),(T.matchesSelector=rt(m=f.matchesSelector||f.mozMatchesSelector||f.webkitMatchesSelector||f.oMatchesSelector||f.msMatchesSelector))&&at(function(e){T.disconnectedMatch=m.call(e,"div"),m.call(e,"[s!='']:x"),g.push("!=",R)}),h=RegExp(h.join("|")),g=RegExp(g.join("|")),y=rt(f.contains)||f.compareDocumentPosition?function(e,t){var n=9===e.nodeType?e.documentElement:e,r=t&&t.parentNode;return e===r||!(!r||1!==r.nodeType||!(n.contains?n.contains(r):e.compareDocumentPosition&&16&e.compareDocumentPosition(r)))}:function(e,t){if(t)while(t=t.parentNode)if(t===e)return!0;return!1},v=f.compareDocumentPosition?function(e,t){var r;return e===t?(u=!0,0):(r=t.compareDocumentPosition&&e.compareDocumentPosition&&e.compareDocumentPosition(t))?1&r||e.parentNode&&11===e.parentNode.nodeType?e===n||y(w,e)?-1:t===n||y(w,t)?1:0:4&r?-1:1:e.compareDocumentPosition?-1:1}:function(e,t){var r,i=0,o=e.parentNode,a=t.parentNode,s=[e],l=[t];if(e===t)return u=!0,0;if(!o||!a)return e===n?-1:t===n?1:o?-1:a?1:0;if(o===a)return ut(e,t);r=e;while(r=r.parentNode)s.unshift(r);r=t;while(r=r.parentNode)l.unshift(r);while(s[i]===l[i])i++;return i?ut(s[i],l[i]):s[i]===w?-1:l[i]===w?1:0},u=!1,[0,0].sort(v),T.detectDuplicates=u,p):p},st.matches=function(e,t){return st(e,null,null,t)},st.matchesSelector=function(e,t){if((e.ownerDocument||e)!==p&&c(e),t=t.replace(Z,"='$1']"),!(!T.matchesSelector||d||g&&g.test(t)||h.test(t)))try{var n=m.call(e,t);if(n||T.disconnectedMatch||e.document&&11!==e.document.nodeType)return n}catch(r){}return st(t,p,null,[e]).length>0},st.contains=function(e,t){return(e.ownerDocument||e)!==p&&c(e),y(e,t)},st.attr=function(e,t){var n;return(e.ownerDocument||e)!==p&&c(e),d||(t=t.toLowerCase()),(n=i.attrHandle[t])?n(e):d||T.attributes?e.getAttribute(t):((n=e.getAttributeNode(t))||e.getAttribute(t))&&e[t]===!0?t:n&&n.specified?n.value:null},st.error=function(e){throw Error("Syntax error, unrecognized expression: "+e)},st.uniqueSort=function(e){var t,n=[],r=1,i=0;if(u=!T.detectDuplicates,e.sort(v),u){for(;t=e[r];r++)t===e[r-1]&&(i=n.push(r));while(i--)e.splice(n[i],1)}return e};function ut(e,t){var n=t&&e,r=n&&(~t.sourceIndex||j)-(~e.sourceIndex||j);if(r)return r;if(n)while(n=n.nextSibling)if(n===t)return-1;return e?1:-1}function lt(e){return function(t){var n=t.nodeName.toLowerCase();return"input"===n&&t.type===e}}function ct(e){return function(t){var n=t.nodeName.toLowerCase();return("input"===n||"button"===n)&&t.type===e}}function pt(e){return ot(function(t){return t=+t,ot(function(n,r){var i,o=e([],n.length,t),a=o.length;while(a--)n[i=o[a]]&&(n[i]=!(r[i]=n[i]))})})}o=st.getText=function(e){var t,n="",r=0,i=e.nodeType;if(i){if(1===i||9===i||11===i){if("string"==typeof e.textContent)return e.textContent;for(e=e.firstChild;e;e=e.nextSibling)n+=o(e)}else if(3===i||4===i)return e.nodeValue}else for(;t=e[r];r++)n+=o(t);return n},i=st.selectors={cacheLength:50,createPseudo:ot,match:U,find:{},relative:{">":{dir:"parentNode",first:!0}," ":{dir:"parentNode"},"+":{dir:"previousSibling",first:!0},"~":{dir:"previousSibling"}},preFilter:{ATTR:function(e){return e[1]=e[1].replace(et,tt),e[3]=(e[4]||e[5]||"").replace(et,tt),"~="===e[2]&&(e[3]=" "+e[3]+" "),e.slice(0,4)},CHILD:function(e){return e[1]=e[1].toLowerCase(),"nth"===e[1].slice(0,3)?(e[3]||st.error(e[0]),e[4]=+(e[4]?e[5]+(e[6]||1):2*("even"===e[3]||"odd"===e[3])),e[5]=+(e[7]+e[8]||"odd"===e[3])):e[3]&&st.error(e[0]),e},PSEUDO:function(e){var t,n=!e[5]&&e[2];return U.CHILD.test(e[0])?null:(e[4]?e[2]=e[4]:n&&z.test(n)&&(t=ft(n,!0))&&(t=n.indexOf(")",n.length-t)-n.length)&&(e[0]=e[0].slice(0,t),e[2]=n.slice(0,t)),e.slice(0,3))}},filter:{TAG:function(e){return"*"===e?function(){return!0}:(e=e.replace(et,tt).toLowerCase(),function(t){return t.nodeName&&t.nodeName.toLowerCase()===e})},CLASS:function(e){var t=k[e+" "];return t||(t=RegExp("(^|"+_+")"+e+"("+_+"|$)"))&&k(e,function(e){return t.test(e.className||typeof e.getAttribute!==A&&e.getAttribute("class")||"")})},ATTR:function(e,t,n){return function(r){var i=st.attr(r,e);return null==i?"!="===t:t?(i+="","="===t?i===n:"!="===t?i!==n:"^="===t?n&&0===i.indexOf(n):"*="===t?n&&i.indexOf(n)>-1:"$="===t?n&&i.slice(-n.length)===n:"~="===t?(" "+i+" ").indexOf(n)>-1:"|="===t?i===n||i.slice(0,n.length+1)===n+"-":!1):!0}},CHILD:function(e,t,n,r,i){var o="nth"!==e.slice(0,3),a="last"!==e.slice(-4),s="of-type"===t;return 1===r&&0===i?function(e){return!!e.parentNode}:function(t,n,u){var l,c,p,f,d,h,g=o!==a?"nextSibling":"previousSibling",m=t.parentNode,y=s&&t.nodeName.toLowerCase(),v=!u&&!s;if(m){if(o){while(g){p=t;while(p=p[g])if(s?p.nodeName.toLowerCase()===y:1===p.nodeType)return!1;h=g="only"===e&&!h&&"nextSibling"}return!0}if(h=[a?m.firstChild:m.lastChild],a&&v){c=m[x]||(m[x]={}),l=c[e]||[],d=l[0]===N&&l[1],f=l[0]===N&&l[2],p=d&&m.childNodes[d];while(p=++d&&p&&p[g]||(f=d=0)||h.pop())if(1===p.nodeType&&++f&&p===t){c[e]=[N,d,f];break}}else if(v&&(l=(t[x]||(t[x]={}))[e])&&l[0]===N)f=l[1];else while(p=++d&&p&&p[g]||(f=d=0)||h.pop())if((s?p.nodeName.toLowerCase()===y:1===p.nodeType)&&++f&&(v&&((p[x]||(p[x]={}))[e]=[N,f]),p===t))break;return f-=i,f===r||0===f%r&&f/r>=0}}},PSEUDO:function(e,t){var n,r=i.pseudos[e]||i.setFilters[e.toLowerCase()]||st.error("unsupported pseudo: "+e);return r[x]?r(t):r.length>1?(n=[e,e,"",t],i.setFilters.hasOwnProperty(e.toLowerCase())?ot(function(e,n){var i,o=r(e,t),a=o.length;while(a--)i=M.call(e,o[a]),e[i]=!(n[i]=o[a])}):function(e){return r(e,0,n)}):r}},pseudos:{not:ot(function(e){var t=[],n=[],r=s(e.replace(W,"$1"));return r[x]?ot(function(e,t,n,i){var o,a=r(e,null,i,[]),s=e.length;while(s--)(o=a[s])&&(e[s]=!(t[s]=o))}):function(e,i,o){return t[0]=e,r(t,null,o,n),!n.pop()}}),has:ot(function(e){return function(t){return st(e,t).length>0}}),contains:ot(function(e){return function(t){return(t.textContent||t.innerText||o(t)).indexOf(e)>-1}}),lang:ot(function(e){return X.test(e||"")||st.error("unsupported lang: "+e),e=e.replace(et,tt).toLowerCase(),function(t){var n;do if(n=d?t.getAttribute("xml:lang")||t.getAttribute("lang"):t.lang)return n=n.toLowerCase(),n===e||0===n.indexOf(e+"-");while((t=t.parentNode)&&1===t.nodeType);return!1}}),target:function(t){var n=e.location&&e.location.hash;return n&&n.slice(1)===t.id},root:function(e){return e===f},focus:function(e){return e===p.activeElement&&(!p.hasFocus||p.hasFocus())&&!!(e.type||e.href||~e.tabIndex)},enabled:function(e){return e.disabled===!1},disabled:function(e){return e.disabled===!0},checked:function(e){var t=e.nodeName.toLowerCase();return"input"===t&&!!e.checked||"option"===t&&!!e.selected},selected:function(e){return e.parentNode&&e.parentNode.selectedIndex,e.selected===!0},empty:function(e){for(e=e.firstChild;e;e=e.nextSibling)if(e.nodeName>"@"||3===e.nodeType||4===e.nodeType)return!1;return!0},parent:function(e){return!i.pseudos.empty(e)},header:function(e){return Q.test(e.nodeName)},input:function(e){return G.test(e.nodeName)},button:function(e){var t=e.nodeName.toLowerCase();return"input"===t&&"button"===e.type||"button"===t},text:function(e){var t;return"input"===e.nodeName.toLowerCase()&&"text"===e.type&&(null==(t=e.getAttribute("type"))||t.toLowerCase()===e.type)},first:pt(function(){return[0]}),last:pt(function(e,t){return[t-1]}),eq:pt(function(e,t,n){return[0>n?n+t:n]}),even:pt(function(e,t){var n=0;for(;t>n;n+=2)e.push(n);return e}),odd:pt(function(e,t){var n=1;for(;t>n;n+=2)e.push(n);return e}),lt:pt(function(e,t,n){var r=0>n?n+t:n;for(;--r>=0;)e.push(r);return e}),gt:pt(function(e,t,n){var r=0>n?n+t:n;for(;t>++r;)e.push(r);return e})}};for(n in{radio:!0,checkbox:!0,file:!0,password:!0,image:!0})i.pseudos[n]=lt(n);for(n in{submit:!0,reset:!0})i.pseudos[n]=ct(n);function ft(e,t){var n,r,o,a,s,u,l,c=E[e+" "];if(c)return t?0:c.slice(0);s=e,u=[],l=i.preFilter;while(s){(!n||(r=$.exec(s)))&&(r&&(s=s.slice(r[0].length)||s),u.push(o=[])),n=!1,(r=I.exec(s))&&(n=r.shift(),o.push({value:n,type:r[0].replace(W," ")}),s=s.slice(n.length));for(a in i.filter)!(r=U[a].exec(s))||l[a]&&!(r=l[a](r))||(n=r.shift(),o.push({value:n,type:a,matches:r}),s=s.slice(n.length));if(!n)break}return t?s.length:s?st.error(e):E(e,u).slice(0)}function dt(e){var t=0,n=e.length,r="";for(;n>t;t++)r+=e[t].value;return r}function ht(e,t,n){var i=t.dir,o=n&&"parentNode"===i,a=C++;return t.first?function(t,n,r){while(t=t[i])if(1===t.nodeType||o)return e(t,n,r)}:function(t,n,s){var u,l,c,p=N+" "+a;if(s){while(t=t[i])if((1===t.nodeType||o)&&e(t,n,s))return!0}else while(t=t[i])if(1===t.nodeType||o)if(c=t[x]||(t[x]={}),(l=c[i])&&l[0]===p){if((u=l[1])===!0||u===r)return u===!0}else if(l=c[i]=[p],l[1]=e(t,n,s)||r,l[1]===!0)return!0}}function gt(e){return e.length>1?function(t,n,r){var i=e.length;while(i--)if(!e[i](t,n,r))return!1;return!0}:e[0]}function mt(e,t,n,r,i){var o,a=[],s=0,u=e.length,l=null!=t;for(;u>s;s++)(o=e[s])&&(!n||n(o,r,i))&&(a.push(o),l&&t.push(s));return a}function yt(e,t,n,r,i,o){return r&&!r[x]&&(r=yt(r)),i&&!i[x]&&(i=yt(i,o)),ot(function(o,a,s,u){var l,c,p,f=[],d=[],h=a.length,g=o||xt(t||"*",s.nodeType?[s]:s,[]),m=!e||!o&&t?g:mt(g,f,e,s,u),y=n?i||(o?e:h||r)?[]:a:m;if(n&&n(m,y,s,u),r){l=mt(y,d),r(l,[],s,u),c=l.length;while(c--)(p=l[c])&&(y[d[c]]=!(m[d[c]]=p))}if(o){if(i||e){if(i){l=[],c=y.length;while(c--)(p=y[c])&&l.push(m[c]=p);i(null,y=[],l,u)}c=y.length;while(c--)(p=y[c])&&(l=i?M.call(o,p):f[c])>-1&&(o[l]=!(a[l]=p))}}else y=mt(y===a?y.splice(h,y.length):y),i?i(null,a,y,u):H.apply(a,y)})}function vt(e){var t,n,r,o=e.length,a=i.relative[e[0].type],s=a||i.relative[" "],u=a?1:0,c=ht(function(e){return e===t},s,!0),p=ht(function(e){return M.call(t,e)>-1},s,!0),f=[function(e,n,r){return!a&&(r||n!==l)||((t=n).nodeType?c(e,n,r):p(e,n,r))}];for(;o>u;u++)if(n=i.relative[e[u].type])f=[ht(gt(f),n)];else{if(n=i.filter[e[u].type].apply(null,e[u].matches),n[x]){for(r=++u;o>r;r++)if(i.relative[e[r].type])break;return yt(u>1&&gt(f),u>1&&dt(e.slice(0,u-1)).replace(W,"$1"),n,r>u&&vt(e.slice(u,r)),o>r&&vt(e=e.slice(r)),o>r&&dt(e))}f.push(n)}return gt(f)}function bt(e,t){var n=0,o=t.length>0,a=e.length>0,s=function(s,u,c,f,d){var h,g,m,y=[],v=0,b="0",x=s&&[],w=null!=d,T=l,C=s||a&&i.find.TAG("*",d&&u.parentNode||u),k=N+=null==T?1:Math.random()||.1;for(w&&(l=u!==p&&u,r=n);null!=(h=C[b]);b++){if(a&&h){g=0;while(m=e[g++])if(m(h,u,c)){f.push(h);break}w&&(N=k,r=++n)}o&&((h=!m&&h)&&v--,s&&x.push(h))}if(v+=b,o&&b!==v){g=0;while(m=t[g++])m(x,y,u,c);if(s){if(v>0)while(b--)x[b]||y[b]||(y[b]=L.call(f));y=mt(y)}H.apply(f,y),w&&!s&&y.length>0&&v+t.length>1&&st.uniqueSort(f)}return w&&(N=k,l=T),x};return o?ot(s):s}s=st.compile=function(e,t){var n,r=[],i=[],o=S[e+" "];if(!o){t||(t=ft(e)),n=t.length;while(n--)o=vt(t[n]),o[x]?r.push(o):i.push(o);o=S(e,bt(i,r))}return o};function xt(e,t,n){var r=0,i=t.length;for(;i>r;r++)st(e,t[r],n);return n}function wt(e,t,n,r){var o,a,u,l,c,p=ft(e);if(!r&&1===p.length){if(a=p[0]=p[0].slice(0),a.length>2&&"ID"===(u=a[0]).type&&9===t.nodeType&&!d&&i.relative[a[1].type]){if(t=i.find.ID(u.matches[0].replace(et,tt),t)[0],!t)return n;e=e.slice(a.shift().value.length)}o=U.needsContext.test(e)?0:a.length;while(o--){if(u=a[o],i.relative[l=u.type])break;if((c=i.find[l])&&(r=c(u.matches[0].replace(et,tt),V.test(a[0].type)&&t.parentNode||t))){if(a.splice(o,1),e=r.length&&dt(a),!e)return H.apply(n,q.call(r,0)),n;break}}}return s(e,p)(r,t,d,n,V.test(e)),n}i.pseudos.nth=i.pseudos.eq;function Tt(){}i.filters=Tt.prototype=i.pseudos,i.setFilters=new Tt,c(),st.attr=b.attr,b.find=st,b.expr=st.selectors,b.expr[":"]=b.expr.pseudos,b.unique=st.uniqueSort,b.text=st.getText,b.isXMLDoc=st.isXML,b.contains=st.contains}(e);var at=/Until$/,st=/^(?:parents|prev(?:Until|All))/,ut=/^.[^:#\[\.,]*$/,lt=b.expr.match.needsContext,ct={children:!0,contents:!0,next:!0,prev:!0};b.fn.extend({find:function(e){var t,n,r,i=this.length;if("string"!=typeof e)return r=this,this.pushStack(b(e).filter(function(){for(t=0;i>t;t++)if(b.contains(r[t],this))return!0}));for(n=[],t=0;i>t;t++)b.find(e,this[t],n);return n=this.pushStack(i>1?b.unique(n):n),n.selector=(this.selector?this.selector+" ":"")+e,n},has:function(e){var t,n=b(e,this),r=n.length;return this.filter(function(){for(t=0;r>t;t++)if(b.contains(this,n[t]))return!0})},not:function(e){return this.pushStack(ft(this,e,!1))},filter:function(e){return this.pushStack(ft(this,e,!0))},is:function(e){return!!e&&("string"==typeof e?lt.test(e)?b(e,this.context).index(this[0])>=0:b.filter(e,this).length>0:this.filter(e).length>0)},closest:function(e,t){var n,r=0,i=this.length,o=[],a=lt.test(e)||"string"!=typeof e?b(e,t||this.context):0;for(;i>r;r++){n=this[r];while(n&&n.ownerDocument&&n!==t&&11!==n.nodeType){if(a?a.index(n)>-1:b.find.matchesSelector(n,e)){o.push(n);break}n=n.parentNode}}return this.pushStack(o.length>1?b.unique(o):o)},index:function(e){return e?"string"==typeof e?b.inArray(this[0],b(e)):b.inArray(e.jquery?e[0]:e,this):this[0]&&this[0].parentNode?this.first().prevAll().length:-1},add:function(e,t){var n="string"==typeof e?b(e,t):b.makeArray(e&&e.nodeType?[e]:e),r=b.merge(this.get(),n);return this.pushStack(b.unique(r))},addBack:function(e){return this.add(null==e?this.prevObject:this.prevObject.filter(e))}}),b.fn.andSelf=b.fn.addBack;function pt(e,t){do e=e[t];while(e&&1!==e.nodeType);return e}b.each({parent:function(e){var t=e.parentNode;return t&&11!==t.nodeType?t:null},parents:function(e){return b.dir(e,"parentNode")},parentsUntil:function(e,t,n){return b.dir(e,"parentNode",n)},next:function(e){return pt(e,"nextSibling")},prev:function(e){return pt(e,"previousSibling")},nextAll:function(e){return b.dir(e,"nextSibling")},prevAll:function(e){return b.dir(e,"previousSibling")},nextUntil:function(e,t,n){return b.dir(e,"nextSibling",n)},prevUntil:function(e,t,n){return b.dir(e,"previousSibling",n)},siblings:function(e){return b.sibling((e.parentNode||{}).firstChild,e)},children:function(e){return b.sibling(e.firstChild)},contents:function(e){return b.nodeName(e,"iframe")?e.contentDocument||e.contentWindow.document:b.merge([],e.childNodes)}},function(e,t){b.fn[e]=function(n,r){var i=b.map(this,t,n);return at.test(e)||(r=n),r&&"string"==typeof r&&(i=b.filter(r,i)),i=this.length>1&&!ct[e]?b.unique(i):i,this.length>1&&st.test(e)&&(i=i.reverse()),this.pushStack(i)}}),b.extend({filter:function(e,t,n){return n&&(e=":not("+e+")"),1===t.length?b.find.matchesSelector(t[0],e)?[t[0]]:[]:b.find.matches(e,t)},dir:function(e,n,r){var i=[],o=e[n];while(o&&9!==o.nodeType&&(r===t||1!==o.nodeType||!b(o).is(r)))1===o.nodeType&&i.push(o),o=o[n];return i},sibling:function(e,t){var n=[];for(;e;e=e.nextSibling)1===e.nodeType&&e!==t&&n.push(e);return n}});function ft(e,t,n){if(t=t||0,b.isFunction(t))return b.grep(e,function(e,r){var i=!!t.call(e,r,e);return i===n});if(t.nodeType)return b.grep(e,function(e){return e===t===n});if("string"==typeof t){var r=b.grep(e,function(e){return 1===e.nodeType});if(ut.test(t))return b.filter(t,r,!n);t=b.filter(t,r)}return b.grep(e,function(e){return b.inArray(e,t)>=0===n})}function dt(e){var t=ht.split("|"),n=e.createDocumentFragment();if(n.createElement)while(t.length)n.createElement(t.pop());return n}var ht="abbr|article|aside|audio|bdi|canvas|data|datalist|details|figcaption|figure|footer|header|hgroup|mark|meter|nav|output|progress|section|summary|time|video",gt=/ jQuery\d+="(?:null|\d+)"/g,mt=RegExp("<(?:"+ht+")[\\s/>]","i"),yt=/^\s+/,vt=/<(?!area|br|col|embed|hr|img|input|link|meta|param)(([\w:]+)[^>]*)\/>/gi,bt=/<([\w:]+)/,xt=/<tbody/i,wt=/<|&#?\w+;/,Tt=/<(?:script|style|link)/i,Nt=/^(?:checkbox|radio)$/i,Ct=/checked\s*(?:[^=]|=\s*.checked.)/i,kt=/^$|\/(?:java|ecma)script/i,Et=/^true\/(.*)/,St=/^\s*<!(?:\[CDATA\[|--)|(?:\]\]|--)>\s*$/g,At={option:[1,"<select multiple='multiple'>","</select>"],legend:[1,"<fieldset>","</fieldset>"],area:[1,"<map>","</map>"],param:[1,"<object>","</object>"],thead:[1,"<table>","</table>"],tr:[2,"<table><tbody>","</tbody></table>"],col:[2,"<table><tbody></tbody><colgroup>","</colgroup></table>"],td:[3,"<table><tbody><tr>","</tr></tbody></table>"],_default:b.support.htmlSerialize?[0,"",""]:[1,"X<div>","</div>"]},jt=dt(o),Dt=jt.appendChild(o.createElement("div"));At.optgroup=At.option,At.tbody=At.tfoot=At.colgroup=At.caption=At.thead,At.th=At.td,b.fn.extend({text:function(e){return b.access(this,function(e){return e===t?b.text(this):this.empty().append((this[0]&&this[0].ownerDocument||o).createTextNode(e))},null,e,arguments.length)},wrapAll:function(e){if(b.isFunction(e))return this.each(function(t){b(this).wrapAll(e.call(this,t))});if(this[0]){var t=b(e,this[0].ownerDocument).eq(0).clone(!0);this[0].parentNode&&t.insertBefore(this[0]),t.map(function(){var e=this;while(e.firstChild&&1===e.firstChild.nodeType)e=e.firstChild;return e}).append(this)}return this},wrapInner:function(e){return b.isFunction(e)?this.each(function(t){b(this).wrapInner(e.call(this,t))}):this.each(function(){var t=b(this),n=t.contents();n.length?n.wrapAll(e):t.append(e)})},wrap:function(e){var t=b.isFunction(e);return this.each(function(n){b(this).wrapAll(t?e.call(this,n):e)})},unwrap:function(){return this.parent().each(function(){b.nodeName(this,"body")||b(this).replaceWith(this.childNodes)}).end()},append:function(){return this.domManip(arguments,!0,function(e){(1===this.nodeType||11===this.nodeType||9===this.nodeType)&&this.appendChild(e)})},prepend:function(){return this.domManip(arguments,!0,function(e){(1===this.nodeType||11===this.nodeType||9===this.nodeType)&&this.insertBefore(e,this.firstChild)})},before:function(){return this.domManip(arguments,!1,function(e){this.parentNode&&this.parentNode.insertBefore(e,this)})},after:function(){return this.domManip(arguments,!1,function(e){this.parentNode&&this.parentNode.insertBefore(e,this.nextSibling)})},remove:function(e,t){var n,r=0;for(;null!=(n=this[r]);r++)(!e||b.filter(e,[n]).length>0)&&(t||1!==n.nodeType||b.cleanData(Ot(n)),n.parentNode&&(t&&b.contains(n.ownerDocument,n)&&Mt(Ot(n,"script")),n.parentNode.removeChild(n)));return this},empty:function(){var e,t=0;for(;null!=(e=this[t]);t++){1===e.nodeType&&b.cleanData(Ot(e,!1));while(e.firstChild)e.removeChild(e.firstChild);e.options&&b.nodeName(e,"select")&&(e.options.length=0)}return this},clone:function(e,t){return e=null==e?!1:e,t=null==t?e:t,this.map(function(){return b.clone(this,e,t)})},html:function(e){return b.access(this,function(e){var n=this[0]||{},r=0,i=this.length;if(e===t)return 1===n.nodeType?n.innerHTML.replace(gt,""):t;if(!("string"!=typeof e||Tt.test(e)||!b.support.htmlSerialize&&mt.test(e)||!b.support.leadingWhitespace&&yt.test(e)||At[(bt.exec(e)||["",""])[1].toLowerCase()])){e=e.replace(vt,"<$1></$2>");try{for(;i>r;r++)n=this[r]||{},1===n.nodeType&&(b.cleanData(Ot(n,!1)),n.innerHTML=e);n=0}catch(o){}}n&&this.empty().append(e)},null,e,arguments.length)},replaceWith:function(e){var t=b.isFunction(e);return t||"string"==typeof e||(e=b(e).not(this).detach()),this.domManip([e],!0,function(e){var t=this.nextSibling,n=this.parentNode;n&&(b(this).remove(),n.insertBefore(e,t))})},detach:function(e){return this.remove(e,!0)},domManip:function(e,n,r){e=f.apply([],e);var i,o,a,s,u,l,c=0,p=this.length,d=this,h=p-1,g=e[0],m=b.isFunction(g);if(m||!(1>=p||"string"!=typeof g||b.support.checkClone)&&Ct.test(g))return this.each(function(i){var o=d.eq(i);m&&(e[0]=g.call(this,i,n?o.html():t)),o.domManip(e,n,r)});if(p&&(l=b.buildFragment(e,this[0].ownerDocument,!1,this),i=l.firstChild,1===l.childNodes.length&&(l=i),i)){for(n=n&&b.nodeName(i,"tr"),s=b.map(Ot(l,"script"),Ht),a=s.length;p>c;c++)o=l,c!==h&&(o=b.clone(o,!0,!0),a&&b.merge(s,Ot(o,"script"))),r.call(n&&b.nodeName(this[c],"table")?Lt(this[c],"tbody"):this[c],o,c);if(a)for(u=s[s.length-1].ownerDocument,b.map(s,qt),c=0;a>c;c++)o=s[c],kt.test(o.type||"")&&!b._data(o,"globalEval")&&b.contains(u,o)&&(o.src?b.ajax({url:o.src,type:"GET",dataType:"script",async:!1,global:!1,"throws":!0}):b.globalEval((o.text||o.textContent||o.innerHTML||"").replace(St,"")));l=i=null}return this}});function Lt(e,t){return e.getElementsByTagName(t)[0]||e.appendChild(e.ownerDocument.createElement(t))}function Ht(e){var t=e.getAttributeNode("type");return e.type=(t&&t.specified)+"/"+e.type,e}function qt(e){var t=Et.exec(e.type);return t?e.type=t[1]:e.removeAttribute("type"),e}function Mt(e,t){var n,r=0;for(;null!=(n=e[r]);r++)b._data(n,"globalEval",!t||b._data(t[r],"globalEval"))}function _t(e,t){if(1===t.nodeType&&b.hasData(e)){var n,r,i,o=b._data(e),a=b._data(t,o),s=o.events;if(s){delete a.handle,a.events={};for(n in s)for(r=0,i=s[n].length;i>r;r++)b.event.add(t,n,s[n][r])}a.data&&(a.data=b.extend({},a.data))}}function Ft(e,t){var n,r,i;if(1===t.nodeType){if(n=t.nodeName.toLowerCase(),!b.support.noCloneEvent&&t[b.expando]){i=b._data(t);for(r in i.events)b.removeEvent(t,r,i.handle);t.removeAttribute(b.expando)}"script"===n&&t.text!==e.text?(Ht(t).text=e.text,qt(t)):"object"===n?(t.parentNode&&(t.outerHTML=e.outerHTML),b.support.html5Clone&&e.innerHTML&&!b.trim(t.innerHTML)&&(t.innerHTML=e.innerHTML)):"input"===n&&Nt.test(e.type)?(t.defaultChecked=t.checked=e.checked,t.value!==e.value&&(t.value=e.value)):"option"===n?t.defaultSelected=t.selected=e.defaultSelected:("input"===n||"textarea"===n)&&(t.defaultValue=e.defaultValue)}}b.each({appendTo:"append",prependTo:"prepend",insertBefore:"before",insertAfter:"after",replaceAll:"replaceWith"},function(e,t){b.fn[e]=function(e){var n,r=0,i=[],o=b(e),a=o.length-1;for(;a>=r;r++)n=r===a?this:this.clone(!0),b(o[r])[t](n),d.apply(i,n.get());return this.pushStack(i)}});function Ot(e,n){var r,o,a=0,s=typeof e.getElementsByTagName!==i?e.getElementsByTagName(n||"*"):typeof e.querySelectorAll!==i?e.querySelectorAll(n||"*"):t;if(!s)for(s=[],r=e.childNodes||e;null!=(o=r[a]);a++)!n||b.nodeName(o,n)?s.push(o):b.merge(s,Ot(o,n));return n===t||n&&b.nodeName(e,n)?b.merge([e],s):s}function Bt(e){Nt.test(e.type)&&(e.defaultChecked=e.checked)}b.extend({clone:function(e,t,n){var r,i,o,a,s,u=b.contains(e.ownerDocument,e);if(b.support.html5Clone||b.isXMLDoc(e)||!mt.test("<"+e.nodeName+">")?o=e.cloneNode(!0):(Dt.innerHTML=e.outerHTML,Dt.removeChild(o=Dt.firstChild)),!(b.support.noCloneEvent&&b.support.noCloneChecked||1!==e.nodeType&&11!==e.nodeType||b.isXMLDoc(e)))for(r=Ot(o),s=Ot(e),a=0;null!=(i=s[a]);++a)r[a]&&Ft(i,r[a]);if(t)if(n)for(s=s||Ot(e),r=r||Ot(o),a=0;null!=(i=s[a]);a++)_t(i,r[a]);else _t(e,o);return r=Ot(o,"script"),r.length>0&&Mt(r,!u&&Ot(e,"script")),r=s=i=null,o},buildFragment:function(e,t,n,r){var i,o,a,s,u,l,c,p=e.length,f=dt(t),d=[],h=0;for(;p>h;h++)if(o=e[h],o||0===o)if("object"===b.type(o))b.merge(d,o.nodeType?[o]:o);else if(wt.test(o)){s=s||f.appendChild(t.createElement("div")),u=(bt.exec(o)||["",""])[1].toLowerCase(),c=At[u]||At._default,s.innerHTML=c[1]+o.replace(vt,"<$1></$2>")+c[2],i=c[0];while(i--)s=s.lastChild;if(!b.support.leadingWhitespace&&yt.test(o)&&d.push(t.createTextNode(yt.exec(o)[0])),!b.support.tbody){o="table"!==u||xt.test(o)?"<table>"!==c[1]||xt.test(o)?0:s:s.firstChild,i=o&&o.childNodes.length;while(i--)b.nodeName(l=o.childNodes[i],"tbody")&&!l.childNodes.length&&o.removeChild(l)
}b.merge(d,s.childNodes),s.textContent="";while(s.firstChild)s.removeChild(s.firstChild);s=f.lastChild}else d.push(t.createTextNode(o));s&&f.removeChild(s),b.support.appendChecked||b.grep(Ot(d,"input"),Bt),h=0;while(o=d[h++])if((!r||-1===b.inArray(o,r))&&(a=b.contains(o.ownerDocument,o),s=Ot(f.appendChild(o),"script"),a&&Mt(s),n)){i=0;while(o=s[i++])kt.test(o.type||"")&&n.push(o)}return s=null,f},cleanData:function(e,t){var n,r,o,a,s=0,u=b.expando,l=b.cache,p=b.support.deleteExpando,f=b.event.special;for(;null!=(n=e[s]);s++)if((t||b.acceptData(n))&&(o=n[u],a=o&&l[o])){if(a.events)for(r in a.events)f[r]?b.event.remove(n,r):b.removeEvent(n,r,a.handle);l[o]&&(delete l[o],p?delete n[u]:typeof n.removeAttribute!==i?n.removeAttribute(u):n[u]=null,c.push(o))}}});var Pt,Rt,Wt,$t=/alpha\([^)]*\)/i,It=/opacity\s*=\s*([^)]*)/,zt=/^(top|right|bottom|left)$/,Xt=/^(none|table(?!-c[ea]).+)/,Ut=/^margin/,Vt=RegExp("^("+x+")(.*)$","i"),Yt=RegExp("^("+x+")(?!px)[a-z%]+$","i"),Jt=RegExp("^([+-])=("+x+")","i"),Gt={BODY:"block"},Qt={position:"absolute",visibility:"hidden",display:"block"},Kt={letterSpacing:0,fontWeight:400},Zt=["Top","Right","Bottom","Left"],en=["Webkit","O","Moz","ms"];function tn(e,t){if(t in e)return t;var n=t.charAt(0).toUpperCase()+t.slice(1),r=t,i=en.length;while(i--)if(t=en[i]+n,t in e)return t;return r}function nn(e,t){return e=t||e,"none"===b.css(e,"display")||!b.contains(e.ownerDocument,e)}function rn(e,t){var n,r,i,o=[],a=0,s=e.length;for(;s>a;a++)r=e[a],r.style&&(o[a]=b._data(r,"olddisplay"),n=r.style.display,t?(o[a]||"none"!==n||(r.style.display=""),""===r.style.display&&nn(r)&&(o[a]=b._data(r,"olddisplay",un(r.nodeName)))):o[a]||(i=nn(r),(n&&"none"!==n||!i)&&b._data(r,"olddisplay",i?n:b.css(r,"display"))));for(a=0;s>a;a++)r=e[a],r.style&&(t&&"none"!==r.style.display&&""!==r.style.display||(r.style.display=t?o[a]||"":"none"));return e}b.fn.extend({css:function(e,n){return b.access(this,function(e,n,r){var i,o,a={},s=0;if(b.isArray(n)){for(o=Rt(e),i=n.length;i>s;s++)a[n[s]]=b.css(e,n[s],!1,o);return a}return r!==t?b.style(e,n,r):b.css(e,n)},e,n,arguments.length>1)},show:function(){return rn(this,!0)},hide:function(){return rn(this)},toggle:function(e){var t="boolean"==typeof e;return this.each(function(){(t?e:nn(this))?b(this).show():b(this).hide()})}}),b.extend({cssHooks:{opacity:{get:function(e,t){if(t){var n=Wt(e,"opacity");return""===n?"1":n}}}},cssNumber:{columnCount:!0,fillOpacity:!0,fontWeight:!0,lineHeight:!0,opacity:!0,orphans:!0,widows:!0,zIndex:!0,zoom:!0},cssProps:{"float":b.support.cssFloat?"cssFloat":"styleFloat"},style:function(e,n,r,i){if(e&&3!==e.nodeType&&8!==e.nodeType&&e.style){var o,a,s,u=b.camelCase(n),l=e.style;if(n=b.cssProps[u]||(b.cssProps[u]=tn(l,u)),s=b.cssHooks[n]||b.cssHooks[u],r===t)return s&&"get"in s&&(o=s.get(e,!1,i))!==t?o:l[n];if(a=typeof r,"string"===a&&(o=Jt.exec(r))&&(r=(o[1]+1)*o[2]+parseFloat(b.css(e,n)),a="number"),!(null==r||"number"===a&&isNaN(r)||("number"!==a||b.cssNumber[u]||(r+="px"),b.support.clearCloneStyle||""!==r||0!==n.indexOf("background")||(l[n]="inherit"),s&&"set"in s&&(r=s.set(e,r,i))===t)))try{l[n]=r}catch(c){}}},css:function(e,n,r,i){var o,a,s,u=b.camelCase(n);return n=b.cssProps[u]||(b.cssProps[u]=tn(e.style,u)),s=b.cssHooks[n]||b.cssHooks[u],s&&"get"in s&&(a=s.get(e,!0,r)),a===t&&(a=Wt(e,n,i)),"normal"===a&&n in Kt&&(a=Kt[n]),""===r||r?(o=parseFloat(a),r===!0||b.isNumeric(o)?o||0:a):a},swap:function(e,t,n,r){var i,o,a={};for(o in t)a[o]=e.style[o],e.style[o]=t[o];i=n.apply(e,r||[]);for(o in t)e.style[o]=a[o];return i}}),e.getComputedStyle?(Rt=function(t){return e.getComputedStyle(t,null)},Wt=function(e,n,r){var i,o,a,s=r||Rt(e),u=s?s.getPropertyValue(n)||s[n]:t,l=e.style;return s&&(""!==u||b.contains(e.ownerDocument,e)||(u=b.style(e,n)),Yt.test(u)&&Ut.test(n)&&(i=l.width,o=l.minWidth,a=l.maxWidth,l.minWidth=l.maxWidth=l.width=u,u=s.width,l.width=i,l.minWidth=o,l.maxWidth=a)),u}):o.documentElement.currentStyle&&(Rt=function(e){return e.currentStyle},Wt=function(e,n,r){var i,o,a,s=r||Rt(e),u=s?s[n]:t,l=e.style;return null==u&&l&&l[n]&&(u=l[n]),Yt.test(u)&&!zt.test(n)&&(i=l.left,o=e.runtimeStyle,a=o&&o.left,a&&(o.left=e.currentStyle.left),l.left="fontSize"===n?"1em":u,u=l.pixelLeft+"px",l.left=i,a&&(o.left=a)),""===u?"auto":u});function on(e,t,n){var r=Vt.exec(t);return r?Math.max(0,r[1]-(n||0))+(r[2]||"px"):t}function an(e,t,n,r,i){var o=n===(r?"border":"content")?4:"width"===t?1:0,a=0;for(;4>o;o+=2)"margin"===n&&(a+=b.css(e,n+Zt[o],!0,i)),r?("content"===n&&(a-=b.css(e,"padding"+Zt[o],!0,i)),"margin"!==n&&(a-=b.css(e,"border"+Zt[o]+"Width",!0,i))):(a+=b.css(e,"padding"+Zt[o],!0,i),"padding"!==n&&(a+=b.css(e,"border"+Zt[o]+"Width",!0,i)));return a}function sn(e,t,n){var r=!0,i="width"===t?e.offsetWidth:e.offsetHeight,o=Rt(e),a=b.support.boxSizing&&"border-box"===b.css(e,"boxSizing",!1,o);if(0>=i||null==i){if(i=Wt(e,t,o),(0>i||null==i)&&(i=e.style[t]),Yt.test(i))return i;r=a&&(b.support.boxSizingReliable||i===e.style[t]),i=parseFloat(i)||0}return i+an(e,t,n||(a?"border":"content"),r,o)+"px"}function un(e){var t=o,n=Gt[e];return n||(n=ln(e,t),"none"!==n&&n||(Pt=(Pt||b("<iframe frameborder='0' width='0' height='0'/>").css("cssText","display:block !important")).appendTo(t.documentElement),t=(Pt[0].contentWindow||Pt[0].contentDocument).document,t.write("<!doctype html><html><body>"),t.close(),n=ln(e,t),Pt.detach()),Gt[e]=n),n}function ln(e,t){var n=b(t.createElement(e)).appendTo(t.body),r=b.css(n[0],"display");return n.remove(),r}b.each(["height","width"],function(e,n){b.cssHooks[n]={get:function(e,r,i){return r?0===e.offsetWidth&&Xt.test(b.css(e,"display"))?b.swap(e,Qt,function(){return sn(e,n,i)}):sn(e,n,i):t},set:function(e,t,r){var i=r&&Rt(e);return on(e,t,r?an(e,n,r,b.support.boxSizing&&"border-box"===b.css(e,"boxSizing",!1,i),i):0)}}}),b.support.opacity||(b.cssHooks.opacity={get:function(e,t){return It.test((t&&e.currentStyle?e.currentStyle.filter:e.style.filter)||"")?.01*parseFloat(RegExp.$1)+"":t?"1":""},set:function(e,t){var n=e.style,r=e.currentStyle,i=b.isNumeric(t)?"alpha(opacity="+100*t+")":"",o=r&&r.filter||n.filter||"";n.zoom=1,(t>=1||""===t)&&""===b.trim(o.replace($t,""))&&n.removeAttribute&&(n.removeAttribute("filter"),""===t||r&&!r.filter)||(n.filter=$t.test(o)?o.replace($t,i):o+" "+i)}}),b(function(){b.support.reliableMarginRight||(b.cssHooks.marginRight={get:function(e,n){return n?b.swap(e,{display:"inline-block"},Wt,[e,"marginRight"]):t}}),!b.support.pixelPosition&&b.fn.position&&b.each(["top","left"],function(e,n){b.cssHooks[n]={get:function(e,r){return r?(r=Wt(e,n),Yt.test(r)?b(e).position()[n]+"px":r):t}}})}),b.expr&&b.expr.filters&&(b.expr.filters.hidden=function(e){return 0>=e.offsetWidth&&0>=e.offsetHeight||!b.support.reliableHiddenOffsets&&"none"===(e.style&&e.style.display||b.css(e,"display"))},b.expr.filters.visible=function(e){return!b.expr.filters.hidden(e)}),b.each({margin:"",padding:"",border:"Width"},function(e,t){b.cssHooks[e+t]={expand:function(n){var r=0,i={},o="string"==typeof n?n.split(" "):[n];for(;4>r;r++)i[e+Zt[r]+t]=o[r]||o[r-2]||o[0];return i}},Ut.test(e)||(b.cssHooks[e+t].set=on)});var cn=/%20/g,pn=/\[\]$/,fn=/\r?\n/g,dn=/^(?:submit|button|image|reset|file)$/i,hn=/^(?:input|select|textarea|keygen)/i;b.fn.extend({serialize:function(){return b.param(this.serializeArray())},serializeArray:function(){return this.map(function(){var e=b.prop(this,"elements");return e?b.makeArray(e):this}).filter(function(){var e=this.type;return this.name&&!b(this).is(":disabled")&&hn.test(this.nodeName)&&!dn.test(e)&&(this.checked||!Nt.test(e))}).map(function(e,t){var n=b(this).val();return null==n?null:b.isArray(n)?b.map(n,function(e){return{name:t.name,value:e.replace(fn,"\r\n")}}):{name:t.name,value:n.replace(fn,"\r\n")}}).get()}}),b.param=function(e,n){var r,i=[],o=function(e,t){t=b.isFunction(t)?t():null==t?"":t,i[i.length]=encodeURIComponent(e)+"="+encodeURIComponent(t)};if(n===t&&(n=b.ajaxSettings&&b.ajaxSettings.traditional),b.isArray(e)||e.jquery&&!b.isPlainObject(e))b.each(e,function(){o(this.name,this.value)});else for(r in e)gn(r,e[r],n,o);return i.join("&").replace(cn,"+")};function gn(e,t,n,r){var i;if(b.isArray(t))b.each(t,function(t,i){n||pn.test(e)?r(e,i):gn(e+"["+("object"==typeof i?t:"")+"]",i,n,r)});else if(n||"object"!==b.type(t))r(e,t);else for(i in t)gn(e+"["+i+"]",t[i],n,r)}b.each("blur focus focusin focusout load resize scroll unload click dblclick mousedown mouseup mousemove mouseover mouseout mouseenter mouseleave change select submit keydown keypress keyup error contextmenu".split(" "),function(e,t){b.fn[t]=function(e,n){return arguments.length>0?this.on(t,null,e,n):this.trigger(t)}}),b.fn.hover=function(e,t){return this.mouseenter(e).mouseleave(t||e)};var mn,yn,vn=b.now(),bn=/\?/,xn=/#.*$/,wn=/([?&])_=[^&]*/,Tn=/^(.*?):[ \t]*([^\r\n]*)\r?$/gm,Nn=/^(?:about|app|app-storage|.+-extension|file|res|widget):$/,Cn=/^(?:GET|HEAD)$/,kn=/^\/\//,En=/^([\w.+-]+:)(?:\/\/([^\/?#:]*)(?::(\d+)|)|)/,Sn=b.fn.load,An={},jn={},Dn="*/".concat("*");try{yn=a.href}catch(Ln){yn=o.createElement("a"),yn.href="",yn=yn.href}mn=En.exec(yn.toLowerCase())||[];function Hn(e){return function(t,n){"string"!=typeof t&&(n=t,t="*");var r,i=0,o=t.toLowerCase().match(w)||[];if(b.isFunction(n))while(r=o[i++])"+"===r[0]?(r=r.slice(1)||"*",(e[r]=e[r]||[]).unshift(n)):(e[r]=e[r]||[]).push(n)}}function qn(e,n,r,i){var o={},a=e===jn;function s(u){var l;return o[u]=!0,b.each(e[u]||[],function(e,u){var c=u(n,r,i);return"string"!=typeof c||a||o[c]?a?!(l=c):t:(n.dataTypes.unshift(c),s(c),!1)}),l}return s(n.dataTypes[0])||!o["*"]&&s("*")}function Mn(e,n){var r,i,o=b.ajaxSettings.flatOptions||{};for(i in n)n[i]!==t&&((o[i]?e:r||(r={}))[i]=n[i]);return r&&b.extend(!0,e,r),e}b.fn.load=function(e,n,r){if("string"!=typeof e&&Sn)return Sn.apply(this,arguments);var i,o,a,s=this,u=e.indexOf(" ");return u>=0&&(i=e.slice(u,e.length),e=e.slice(0,u)),b.isFunction(n)?(r=n,n=t):n&&"object"==typeof n&&(a="POST"),s.length>0&&b.ajax({url:e,type:a,dataType:"html",data:n}).done(function(e){o=arguments,s.html(i?b("<div>").append(b.parseHTML(e)).find(i):e)}).complete(r&&function(e,t){s.each(r,o||[e.responseText,t,e])}),this},b.each(["ajaxStart","ajaxStop","ajaxComplete","ajaxError","ajaxSuccess","ajaxSend"],function(e,t){b.fn[t]=function(e){return this.on(t,e)}}),b.each(["get","post"],function(e,n){b[n]=function(e,r,i,o){return b.isFunction(r)&&(o=o||i,i=r,r=t),b.ajax({url:e,type:n,dataType:o,data:r,success:i})}}),b.extend({active:0,lastModified:{},etag:{},ajaxSettings:{url:yn,type:"GET",isLocal:Nn.test(mn[1]),global:!0,processData:!0,async:!0,contentType:"application/x-www-form-urlencoded; charset=UTF-8",accepts:{"*":Dn,text:"text/plain",html:"text/html",xml:"application/xml, text/xml",json:"application/json, text/javascript"},contents:{xml:/xml/,html:/html/,json:/json/},responseFields:{xml:"responseXML",text:"responseText"},converters:{"* text":e.String,"text html":!0,"text json":b.parseJSON,"text xml":b.parseXML},flatOptions:{url:!0,context:!0}},ajaxSetup:function(e,t){return t?Mn(Mn(e,b.ajaxSettings),t):Mn(b.ajaxSettings,e)},ajaxPrefilter:Hn(An),ajaxTransport:Hn(jn),ajax:function(e,n){"object"==typeof e&&(n=e,e=t),n=n||{};var r,i,o,a,s,u,l,c,p=b.ajaxSetup({},n),f=p.context||p,d=p.context&&(f.nodeType||f.jquery)?b(f):b.event,h=b.Deferred(),g=b.Callbacks("once memory"),m=p.statusCode||{},y={},v={},x=0,T="canceled",N={readyState:0,getResponseHeader:function(e){var t;if(2===x){if(!c){c={};while(t=Tn.exec(a))c[t[1].toLowerCase()]=t[2]}t=c[e.toLowerCase()]}return null==t?null:t},getAllResponseHeaders:function(){return 2===x?a:null},setRequestHeader:function(e,t){var n=e.toLowerCase();return x||(e=v[n]=v[n]||e,y[e]=t),this},overrideMimeType:function(e){return x||(p.mimeType=e),this},statusCode:function(e){var t;if(e)if(2>x)for(t in e)m[t]=[m[t],e[t]];else N.always(e[N.status]);return this},abort:function(e){var t=e||T;return l&&l.abort(t),k(0,t),this}};if(h.promise(N).complete=g.add,N.success=N.done,N.error=N.fail,p.url=((e||p.url||yn)+"").replace(xn,"").replace(kn,mn[1]+"//"),p.type=n.method||n.type||p.method||p.type,p.dataTypes=b.trim(p.dataType||"*").toLowerCase().match(w)||[""],null==p.crossDomain&&(r=En.exec(p.url.toLowerCase()),p.crossDomain=!(!r||r[1]===mn[1]&&r[2]===mn[2]&&(r[3]||("http:"===r[1]?80:443))==(mn[3]||("http:"===mn[1]?80:443)))),p.data&&p.processData&&"string"!=typeof p.data&&(p.data=b.param(p.data,p.traditional)),qn(An,p,n,N),2===x)return N;u=p.global,u&&0===b.active++&&b.event.trigger("ajaxStart"),p.type=p.type.toUpperCase(),p.hasContent=!Cn.test(p.type),o=p.url,p.hasContent||(p.data&&(o=p.url+=(bn.test(o)?"&":"?")+p.data,delete p.data),p.cache===!1&&(p.url=wn.test(o)?o.replace(wn,"$1_="+vn++):o+(bn.test(o)?"&":"?")+"_="+vn++)),p.ifModified&&(b.lastModified[o]&&N.setRequestHeader("If-Modified-Since",b.lastModified[o]),b.etag[o]&&N.setRequestHeader("If-None-Match",b.etag[o])),(p.data&&p.hasContent&&p.contentType!==!1||n.contentType)&&N.setRequestHeader("Content-Type",p.contentType),N.setRequestHeader("Accept",p.dataTypes[0]&&p.accepts[p.dataTypes[0]]?p.accepts[p.dataTypes[0]]+("*"!==p.dataTypes[0]?", "+Dn+"; q=0.01":""):p.accepts["*"]);for(i in p.headers)N.setRequestHeader(i,p.headers[i]);if(p.beforeSend&&(p.beforeSend.call(f,N,p)===!1||2===x))return N.abort();T="abort";for(i in{success:1,error:1,complete:1})N[i](p[i]);if(l=qn(jn,p,n,N)){N.readyState=1,u&&d.trigger("ajaxSend",[N,p]),p.async&&p.timeout>0&&(s=setTimeout(function(){N.abort("timeout")},p.timeout));try{x=1,l.send(y,k)}catch(C){if(!(2>x))throw C;k(-1,C)}}else k(-1,"No Transport");function k(e,n,r,i){var c,y,v,w,T,C=n;2!==x&&(x=2,s&&clearTimeout(s),l=t,a=i||"",N.readyState=e>0?4:0,r&&(w=_n(p,N,r)),e>=200&&300>e||304===e?(p.ifModified&&(T=N.getResponseHeader("Last-Modified"),T&&(b.lastModified[o]=T),T=N.getResponseHeader("etag"),T&&(b.etag[o]=T)),204===e?(c=!0,C="nocontent"):304===e?(c=!0,C="notmodified"):(c=Fn(p,w),C=c.state,y=c.data,v=c.error,c=!v)):(v=C,(e||!C)&&(C="error",0>e&&(e=0))),N.status=e,N.statusText=(n||C)+"",c?h.resolveWith(f,[y,C,N]):h.rejectWith(f,[N,C,v]),N.statusCode(m),m=t,u&&d.trigger(c?"ajaxSuccess":"ajaxError",[N,p,c?y:v]),g.fireWith(f,[N,C]),u&&(d.trigger("ajaxComplete",[N,p]),--b.active||b.event.trigger("ajaxStop")))}return N},getScript:function(e,n){return b.get(e,t,n,"script")},getJSON:function(e,t,n){return b.get(e,t,n,"json")}});function _n(e,n,r){var i,o,a,s,u=e.contents,l=e.dataTypes,c=e.responseFields;for(s in c)s in r&&(n[c[s]]=r[s]);while("*"===l[0])l.shift(),o===t&&(o=e.mimeType||n.getResponseHeader("Content-Type"));if(o)for(s in u)if(u[s]&&u[s].test(o)){l.unshift(s);break}if(l[0]in r)a=l[0];else{for(s in r){if(!l[0]||e.converters[s+" "+l[0]]){a=s;break}i||(i=s)}a=a||i}return a?(a!==l[0]&&l.unshift(a),r[a]):t}function Fn(e,t){var n,r,i,o,a={},s=0,u=e.dataTypes.slice(),l=u[0];if(e.dataFilter&&(t=e.dataFilter(t,e.dataType)),u[1])for(i in e.converters)a[i.toLowerCase()]=e.converters[i];for(;r=u[++s];)if("*"!==r){if("*"!==l&&l!==r){if(i=a[l+" "+r]||a["* "+r],!i)for(n in a)if(o=n.split(" "),o[1]===r&&(i=a[l+" "+o[0]]||a["* "+o[0]])){i===!0?i=a[n]:a[n]!==!0&&(r=o[0],u.splice(s--,0,r));break}if(i!==!0)if(i&&e["throws"])t=i(t);else try{t=i(t)}catch(c){return{state:"parsererror",error:i?c:"No conversion from "+l+" to "+r}}}l=r}return{state:"success",data:t}}b.ajaxSetup({accepts:{script:"text/javascript, application/javascript, application/ecmascript, application/x-ecmascript"},contents:{script:/(?:java|ecma)script/},converters:{"text script":function(e){return b.globalEval(e),e}}}),b.ajaxPrefilter("script",function(e){e.cache===t&&(e.cache=!1),e.crossDomain&&(e.type="GET",e.global=!1)}),b.ajaxTransport("script",function(e){if(e.crossDomain){var n,r=o.head||b("head")[0]||o.documentElement;return{send:function(t,i){n=o.createElement("script"),n.async=!0,e.scriptCharset&&(n.charset=e.scriptCharset),n.src=e.url,n.onload=n.onreadystatechange=function(e,t){(t||!n.readyState||/loaded|complete/.test(n.readyState))&&(n.onload=n.onreadystatechange=null,n.parentNode&&n.parentNode.removeChild(n),n=null,t||i(200,"success"))},r.insertBefore(n,r.firstChild)},abort:function(){n&&n.onload(t,!0)}}}});var On=[],Bn=/(=)\?(?=&|$)|\?\?/;b.ajaxSetup({jsonp:"callback",jsonpCallback:function(){var e=On.pop()||b.expando+"_"+vn++;return this[e]=!0,e}}),b.ajaxPrefilter("json jsonp",function(n,r,i){var o,a,s,u=n.jsonp!==!1&&(Bn.test(n.url)?"url":"string"==typeof n.data&&!(n.contentType||"").indexOf("application/x-www-form-urlencoded")&&Bn.test(n.data)&&"data");return u||"jsonp"===n.dataTypes[0]?(o=n.jsonpCallback=b.isFunction(n.jsonpCallback)?n.jsonpCallback():n.jsonpCallback,u?n[u]=n[u].replace(Bn,"$1"+o):n.jsonp!==!1&&(n.url+=(bn.test(n.url)?"&":"?")+n.jsonp+"="+o),n.converters["script json"]=function(){return s||b.error(o+" was not called"),s[0]},n.dataTypes[0]="json",a=e[o],e[o]=function(){s=arguments},i.always(function(){e[o]=a,n[o]&&(n.jsonpCallback=r.jsonpCallback,On.push(o)),s&&b.isFunction(a)&&a(s[0]),s=a=t}),"script"):t});var Pn,Rn,Wn=0,$n=e.ActiveXObject&&function(){var e;for(e in Pn)Pn[e](t,!0)};function In(){try{return new e.XMLHttpRequest}catch(t){}}function zn(){try{return new e.ActiveXObject("Microsoft.XMLHTTP")}catch(t){}}b.ajaxSettings.xhr=e.ActiveXObject?function(){return!this.isLocal&&In()||zn()}:In,Rn=b.ajaxSettings.xhr(),b.support.cors=!!Rn&&"withCredentials"in Rn,Rn=b.support.ajax=!!Rn,Rn&&b.ajaxTransport(function(n){if(!n.crossDomain||b.support.cors){var r;return{send:function(i,o){var a,s,u=n.xhr();if(n.username?u.open(n.type,n.url,n.async,n.username,n.password):u.open(n.type,n.url,n.async),n.xhrFields)for(s in n.xhrFields)u[s]=n.xhrFields[s];n.mimeType&&u.overrideMimeType&&u.overrideMimeType(n.mimeType),n.crossDomain||i["X-Requested-With"]||(i["X-Requested-With"]="XMLHttpRequest");try{for(s in i)u.setRequestHeader(s,i[s])}catch(l){}u.send(n.hasContent&&n.data||null),r=function(e,i){var s,l,c,p;try{if(r&&(i||4===u.readyState))if(r=t,a&&(u.onreadystatechange=b.noop,$n&&delete Pn[a]),i)4!==u.readyState&&u.abort();else{p={},s=u.status,l=u.getAllResponseHeaders(),"string"==typeof u.responseText&&(p.text=u.responseText);try{c=u.statusText}catch(f){c=""}s||!n.isLocal||n.crossDomain?1223===s&&(s=204):s=p.text?200:404}}catch(d){i||o(-1,d)}p&&o(s,c,p,l)},n.async?4===u.readyState?setTimeout(r):(a=++Wn,$n&&(Pn||(Pn={},b(e).unload($n)),Pn[a]=r),u.onreadystatechange=r):r()},abort:function(){r&&r(t,!0)}}}});var Xn,Un,Vn=/^(?:toggle|show|hide)$/,Yn=RegExp("^(?:([+-])=|)("+x+")([a-z%]*)$","i"),Jn=/queueHooks$/,Gn=[nr],Qn={"*":[function(e,t){var n,r,i=this.createTween(e,t),o=Yn.exec(t),a=i.cur(),s=+a||0,u=1,l=20;if(o){if(n=+o[2],r=o[3]||(b.cssNumber[e]?"":"px"),"px"!==r&&s){s=b.css(i.elem,e,!0)||n||1;do u=u||".5",s/=u,b.style(i.elem,e,s+r);while(u!==(u=i.cur()/a)&&1!==u&&--l)}i.unit=r,i.start=s,i.end=o[1]?s+(o[1]+1)*n:n}return i}]};function Kn(){return setTimeout(function(){Xn=t}),Xn=b.now()}function Zn(e,t){b.each(t,function(t,n){var r=(Qn[t]||[]).concat(Qn["*"]),i=0,o=r.length;for(;o>i;i++)if(r[i].call(e,t,n))return})}function er(e,t,n){var r,i,o=0,a=Gn.length,s=b.Deferred().always(function(){delete u.elem}),u=function(){if(i)return!1;var t=Xn||Kn(),n=Math.max(0,l.startTime+l.duration-t),r=n/l.duration||0,o=1-r,a=0,u=l.tweens.length;for(;u>a;a++)l.tweens[a].run(o);return s.notifyWith(e,[l,o,n]),1>o&&u?n:(s.resolveWith(e,[l]),!1)},l=s.promise({elem:e,props:b.extend({},t),opts:b.extend(!0,{specialEasing:{}},n),originalProperties:t,originalOptions:n,startTime:Xn||Kn(),duration:n.duration,tweens:[],createTween:function(t,n){var r=b.Tween(e,l.opts,t,n,l.opts.specialEasing[t]||l.opts.easing);return l.tweens.push(r),r},stop:function(t){var n=0,r=t?l.tweens.length:0;if(i)return this;for(i=!0;r>n;n++)l.tweens[n].run(1);return t?s.resolveWith(e,[l,t]):s.rejectWith(e,[l,t]),this}}),c=l.props;for(tr(c,l.opts.specialEasing);a>o;o++)if(r=Gn[o].call(l,e,c,l.opts))return r;return Zn(l,c),b.isFunction(l.opts.start)&&l.opts.start.call(e,l),b.fx.timer(b.extend(u,{elem:e,anim:l,queue:l.opts.queue})),l.progress(l.opts.progress).done(l.opts.done,l.opts.complete).fail(l.opts.fail).always(l.opts.always)}function tr(e,t){var n,r,i,o,a;for(i in e)if(r=b.camelCase(i),o=t[r],n=e[i],b.isArray(n)&&(o=n[1],n=e[i]=n[0]),i!==r&&(e[r]=n,delete e[i]),a=b.cssHooks[r],a&&"expand"in a){n=a.expand(n),delete e[r];for(i in n)i in e||(e[i]=n[i],t[i]=o)}else t[r]=o}b.Animation=b.extend(er,{tweener:function(e,t){b.isFunction(e)?(t=e,e=["*"]):e=e.split(" ");var n,r=0,i=e.length;for(;i>r;r++)n=e[r],Qn[n]=Qn[n]||[],Qn[n].unshift(t)},prefilter:function(e,t){t?Gn.unshift(e):Gn.push(e)}});function nr(e,t,n){var r,i,o,a,s,u,l,c,p,f=this,d=e.style,h={},g=[],m=e.nodeType&&nn(e);n.queue||(c=b._queueHooks(e,"fx"),null==c.unqueued&&(c.unqueued=0,p=c.empty.fire,c.empty.fire=function(){c.unqueued||p()}),c.unqueued++,f.always(function(){f.always(function(){c.unqueued--,b.queue(e,"fx").length||c.empty.fire()})})),1===e.nodeType&&("height"in t||"width"in t)&&(n.overflow=[d.overflow,d.overflowX,d.overflowY],"inline"===b.css(e,"display")&&"none"===b.css(e,"float")&&(b.support.inlineBlockNeedsLayout&&"inline"!==un(e.nodeName)?d.zoom=1:d.display="inline-block")),n.overflow&&(d.overflow="hidden",b.support.shrinkWrapBlocks||f.always(function(){d.overflow=n.overflow[0],d.overflowX=n.overflow[1],d.overflowY=n.overflow[2]}));for(i in t)if(a=t[i],Vn.exec(a)){if(delete t[i],u=u||"toggle"===a,a===(m?"hide":"show"))continue;g.push(i)}if(o=g.length){s=b._data(e,"fxshow")||b._data(e,"fxshow",{}),"hidden"in s&&(m=s.hidden),u&&(s.hidden=!m),m?b(e).show():f.done(function(){b(e).hide()}),f.done(function(){var t;b._removeData(e,"fxshow");for(t in h)b.style(e,t,h[t])});for(i=0;o>i;i++)r=g[i],l=f.createTween(r,m?s[r]:0),h[r]=s[r]||b.style(e,r),r in s||(s[r]=l.start,m&&(l.end=l.start,l.start="width"===r||"height"===r?1:0))}}function rr(e,t,n,r,i){return new rr.prototype.init(e,t,n,r,i)}b.Tween=rr,rr.prototype={constructor:rr,init:function(e,t,n,r,i,o){this.elem=e,this.prop=n,this.easing=i||"swing",this.options=t,this.start=this.now=this.cur(),this.end=r,this.unit=o||(b.cssNumber[n]?"":"px")},cur:function(){var e=rr.propHooks[this.prop];return e&&e.get?e.get(this):rr.propHooks._default.get(this)},run:function(e){var t,n=rr.propHooks[this.prop];return this.pos=t=this.options.duration?b.easing[this.easing](e,this.options.duration*e,0,1,this.options.duration):e,this.now=(this.end-this.start)*t+this.start,this.options.step&&this.options.step.call(this.elem,this.now,this),n&&n.set?n.set(this):rr.propHooks._default.set(this),this}},rr.prototype.init.prototype=rr.prototype,rr.propHooks={_default:{get:function(e){var t;return null==e.elem[e.prop]||e.elem.style&&null!=e.elem.style[e.prop]?(t=b.css(e.elem,e.prop,""),t&&"auto"!==t?t:0):e.elem[e.prop]},set:function(e){b.fx.step[e.prop]?b.fx.step[e.prop](e):e.elem.style&&(null!=e.elem.style[b.cssProps[e.prop]]||b.cssHooks[e.prop])?b.style(e.elem,e.prop,e.now+e.unit):e.elem[e.prop]=e.now}}},rr.propHooks.scrollTop=rr.propHooks.scrollLeft={set:function(e){e.elem.nodeType&&e.elem.parentNode&&(e.elem[e.prop]=e.now)}},b.each(["toggle","show","hide"],function(e,t){var n=b.fn[t];b.fn[t]=function(e,r,i){return null==e||"boolean"==typeof e?n.apply(this,arguments):this.animate(ir(t,!0),e,r,i)}}),b.fn.extend({fadeTo:function(e,t,n,r){return this.filter(nn).css("opacity",0).show().end().animate({opacity:t},e,n,r)},animate:function(e,t,n,r){var i=b.isEmptyObject(e),o=b.speed(t,n,r),a=function(){var t=er(this,b.extend({},e),o);a.finish=function(){t.stop(!0)},(i||b._data(this,"finish"))&&t.stop(!0)};return a.finish=a,i||o.queue===!1?this.each(a):this.queue(o.queue,a)},stop:function(e,n,r){var i=function(e){var t=e.stop;delete e.stop,t(r)};return"string"!=typeof e&&(r=n,n=e,e=t),n&&e!==!1&&this.queue(e||"fx",[]),this.each(function(){var t=!0,n=null!=e&&e+"queueHooks",o=b.timers,a=b._data(this);if(n)a[n]&&a[n].stop&&i(a[n]);else for(n in a)a[n]&&a[n].stop&&Jn.test(n)&&i(a[n]);for(n=o.length;n--;)o[n].elem!==this||null!=e&&o[n].queue!==e||(o[n].anim.stop(r),t=!1,o.splice(n,1));(t||!r)&&b.dequeue(this,e)})},finish:function(e){return e!==!1&&(e=e||"fx"),this.each(function(){var t,n=b._data(this),r=n[e+"queue"],i=n[e+"queueHooks"],o=b.timers,a=r?r.length:0;for(n.finish=!0,b.queue(this,e,[]),i&&i.cur&&i.cur.finish&&i.cur.finish.call(this),t=o.length;t--;)o[t].elem===this&&o[t].queue===e&&(o[t].anim.stop(!0),o.splice(t,1));for(t=0;a>t;t++)r[t]&&r[t].finish&&r[t].finish.call(this);delete n.finish})}});function ir(e,t){var n,r={height:e},i=0;for(t=t?1:0;4>i;i+=2-t)n=Zt[i],r["margin"+n]=r["padding"+n]=e;return t&&(r.opacity=r.width=e),r}b.each({slideDown:ir("show"),slideUp:ir("hide"),slideToggle:ir("toggle"),fadeIn:{opacity:"show"},fadeOut:{opacity:"hide"},fadeToggle:{opacity:"toggle"}},function(e,t){b.fn[e]=function(e,n,r){return this.animate(t,e,n,r)}}),b.speed=function(e,t,n){var r=e&&"object"==typeof e?b.extend({},e):{complete:n||!n&&t||b.isFunction(e)&&e,duration:e,easing:n&&t||t&&!b.isFunction(t)&&t};return r.duration=b.fx.off?0:"number"==typeof r.duration?r.duration:r.duration in b.fx.speeds?b.fx.speeds[r.duration]:b.fx.speeds._default,(null==r.queue||r.queue===!0)&&(r.queue="fx"),r.old=r.complete,r.complete=function(){b.isFunction(r.old)&&r.old.call(this),r.queue&&b.dequeue(this,r.queue)},r},b.easing={linear:function(e){return e},swing:function(e){return.5-Math.cos(e*Math.PI)/2}},b.timers=[],b.fx=rr.prototype.init,b.fx.tick=function(){var e,n=b.timers,r=0;for(Xn=b.now();n.length>r;r++)e=n[r],e()||n[r]!==e||n.splice(r--,1);n.length||b.fx.stop(),Xn=t},b.fx.timer=function(e){e()&&b.timers.push(e)&&b.fx.start()},b.fx.interval=13,b.fx.start=function(){Un||(Un=setInterval(b.fx.tick,b.fx.interval))},b.fx.stop=function(){clearInterval(Un),Un=null},b.fx.speeds={slow:600,fast:200,_default:400},b.fx.step={},b.expr&&b.expr.filters&&(b.expr.filters.animated=function(e){return b.grep(b.timers,function(t){return e===t.elem}).length}),b.fn.offset=function(e){if(arguments.length)return e===t?this:this.each(function(t){b.offset.setOffset(this,e,t)});var n,r,o={top:0,left:0},a=this[0],s=a&&a.ownerDocument;if(s)return n=s.documentElement,b.contains(n,a)?(typeof a.getBoundingClientRect!==i&&(o=a.getBoundingClientRect()),r=or(s),{top:o.top+(r.pageYOffset||n.scrollTop)-(n.clientTop||0),left:o.left+(r.pageXOffset||n.scrollLeft)-(n.clientLeft||0)}):o},b.offset={setOffset:function(e,t,n){var r=b.css(e,"position");"static"===r&&(e.style.position="relative");var i=b(e),o=i.offset(),a=b.css(e,"top"),s=b.css(e,"left"),u=("absolute"===r||"fixed"===r)&&b.inArray("auto",[a,s])>-1,l={},c={},p,f;u?(c=i.position(),p=c.top,f=c.left):(p=parseFloat(a)||0,f=parseFloat(s)||0),b.isFunction(t)&&(t=t.call(e,n,o)),null!=t.top&&(l.top=t.top-o.top+p),null!=t.left&&(l.left=t.left-o.left+f),"using"in t?t.using.call(e,l):i.css(l)}},b.fn.extend({position:function(){if(this[0]){var e,t,n={top:0,left:0},r=this[0];return"fixed"===b.css(r,"position")?t=r.getBoundingClientRect():(e=this.offsetParent(),t=this.offset(),b.nodeName(e[0],"html")||(n=e.offset()),n.top+=b.css(e[0],"borderTopWidth",!0),n.left+=b.css(e[0],"borderLeftWidth",!0)),{top:t.top-n.top-b.css(r,"marginTop",!0),left:t.left-n.left-b.css(r,"marginLeft",!0)}}},offsetParent:function(){return this.map(function(){var e=this.offsetParent||o.documentElement;while(e&&!b.nodeName(e,"html")&&"static"===b.css(e,"position"))e=e.offsetParent;return e||o.documentElement})}}),b.each({scrollLeft:"pageXOffset",scrollTop:"pageYOffset"},function(e,n){var r=/Y/.test(n);b.fn[e]=function(i){return b.access(this,function(e,i,o){var a=or(e);return o===t?a?n in a?a[n]:a.document.documentElement[i]:e[i]:(a?a.scrollTo(r?b(a).scrollLeft():o,r?o:b(a).scrollTop()):e[i]=o,t)},e,i,arguments.length,null)}});function or(e){return b.isWindow(e)?e:9===e.nodeType?e.defaultView||e.parentWindow:!1}b.each({Height:"height",Width:"width"},function(e,n){b.each({padding:"inner"+e,content:n,"":"outer"+e},function(r,i){b.fn[i]=function(i,o){var a=arguments.length&&(r||"boolean"!=typeof i),s=r||(i===!0||o===!0?"margin":"border");return b.access(this,function(n,r,i){var o;return b.isWindow(n)?n.document.documentElement["client"+e]:9===n.nodeType?(o=n.documentElement,Math.max(n.body["scroll"+e],o["scroll"+e],n.body["offset"+e],o["offset"+e],o["client"+e])):i===t?b.css(n,r,s):b.style(n,r,i,s)},n,a?i:t,a,null)}})}),e.jQuery=e.$=b,"function"==typeof define&&define.amd&&define.amd.jQuery&&define("jquery",[],function(){return b})})(window);
```
