## Supplementary Results for "A draft genome of the ascomycotal fungal species *Pseudopithomyces maydicus* (family *Didymosphaeriaceae*)": jquery.tablesorter.min.js

```
(function($){$.extend({tablesorter:new
function(){var parsers=[],widgets=[];this.defaults={cssHeader:"header",cssAsc:"headerSortUp",cssDesc:"headerSortDown",cssChildRow:"expand-child",sortInitialOrder:"asc",sortMultiSortKey:"shiftKey",sortForce:null,sortAppend:null,sortLocaleCompare:true,textExtraction:"simple",parsers:{},widgets:[],widgetZebra:{css:["even","odd"]},headers:{},widthFixed:false,cancelSelection:true,sortList:[],headerList:[],dateFormat:"us",decimal:'/\.|\,/g',onRenderHeader:null,selectorHeaders:'thead th',debug:false};function benchmark(s,d){log(s+","+(new Date().getTime()-d.getTime())+"ms");}this.benchmark=benchmark;function log(s){if(typeof console!="undefined"&&typeof console.debug!="undefined"){console.log(s);}else{alert(s);}}function buildParserCache(table,$headers){if(table.config.debug){var parsersDebug="";}if(table.tBodies.length==0)return;var rows=table.tBodies[0].rows;if(rows[0]){var list=[],cells=rows[0].cells,l=cells.length;for(var i=0;i<l;i++){var p=false;if($.metadata&&($($headers[i]).metadata()&&$($headers[i]).metadata().sorter)){p=getParserById($($headers[i]).metadata().sorter);}else if((table.config.headers[i]&&table.config.headers[i].sorter)){p=getParserById(table.config.headers[i].sorter);}if(!p){p=detectParserForColumn(table,rows,-1,i);}if(table.config.debug){parsersDebug+="column:"+i+" parser:"+p.id+"\n";}list.push(p);}}if(table.config.debug){log(parsersDebug);}return list;};function detectParserForColumn(table,rows,rowIndex,cellIndex){var l=parsers.length,node=false,nodeValue=false,keepLooking=true;while(nodeValue==''&&keepLooking){rowIndex++;if(rows[rowIndex]){node=getNodeFromRowAndCellIndex(rows,rowIndex,cellIndex);nodeValue=trimAndGetNodeText(table.config,node);if(table.config.debug){log('Checking if value was empty on row:'+rowIndex);}}else{keepLooking=false;}}for(var i=1;i<l;i++){if(parsers[i].is(nodeValue,table,node)){return parsers[i];}}return parsers[0];}function getNodeFromRowAndCellIndex(rows,rowIndex,cellIndex){return rows[rowIndex].cells[cellIndex];}function trimAndGetNodeText(config,node){return $.trim(getElementText(config,node));}function getParserById(name){var l=parsers.length;for(var i=0;i<l;i++){if(parsers[i].id.toLowerCase()==name.toLowerCase()){return parsers[i];}}return false;}function buildCache(table){if(table.config.debug){var cacheTime=new Date();}var totalRows=(table.tBodies[0]&&table.tBodies[0].rows.length)||0,totalCells=(table.tBodies[0].rows[0]&&table.tBodies[0].rows[0].cells.length)||0,parsers=table.config.parsers,cache={row:[],normalized:[]};for(var i=0;i<totalRows;++i){var c=$(table.tBodies[0].rows[i]),cols=[];if(c.hasClass(table.config.cssChildRow)){cache.row[cache.row.length-1]=cache.row[cache.row.length-1].add(c);continue;}cache.row.push(c);for(var j=0;j<totalCells;++j){cols.push(parsers[j].format(getElementText(table.config,c[0].cells[j]),table,c[0].cells[j]));}cols.push(cache.normalized.length);cache.normalized.push(cols);cols=null;};if(table.config.debug){benchmark("Building cache for "+totalRows+" rows:",cacheTime);}return cache;};function getElementText(config,node){var text="";if(!node)return"";if(!config.supportsTextContent)config.supportsTextContent=node.textContent||false;if(config.textExtraction=="simple"){if(config.supportsTextContent){text=node.textContent;}else{if(node.childNodes[0]&&node.childNodes[0].hasChildNodes()){text=node.childNodes[0].innerHTML;}else{text=node.innerHTML;}}}else{if(typeof(config.textExtraction)=="function"){text=config.textExtraction(node);}else{text=$(node).text();}}return text;}function appendToTable(table,cache){if(table.config.debug){var appendTime=new Date()}var c=cache,r=c.row,n=c.normalized,totalRows=n.length,checkCell=(n[0].length-1),tableBody=$(table.tBodies[0]),rows=[];for(var i=0;i<totalRows;i++){var pos=n[i][checkCell];rows.push(r[pos]);if(!table.config.appender){var l=r[pos].length;for(var j=0;j<l;j++){tableBody[0].appendChild(r[pos][j]);}}}if(table.config.appender){table.config.appender(table,rows);}rows=null;if(table.config.debug){benchmark("Rebuilt table:",appendTime);}applyWidget(table);setTimeout(function(){$(table).trigger("sortEnd");},0);};function buildHeaders(table){if(table.config.debug){var time=new Date();}var meta=($.metadata)?true:false;var header_index=computeTableHeaderCellIndexes(table);$tableHeaders=$(table.config.selectorHeaders,table).each(function(index){this.column=header_index[this.parentNode.rowIndex+"-"+this.cellIndex];this.order=formatSortingOrder(table.config.sortInitialOrder);this.count=this.order;if(checkHeaderMetadata(this)||checkHeaderOptions(table,index))this.sortDisabled=true;if(checkHeaderOptionsSortingLocked(table,index))this.order=this.lockedOrder=checkHeaderOptionsSortingLocked(table,index);if(!this.sortDisabled){var $th=$(this).addClass(table.config.cssHeader);if(table.config.onRenderHeader)table.config.onRenderHeader.apply($th);}table.config.headerList[index]=this;});if(table.config.debug){benchmark("Built headers:",time);log($tableHeaders);}return $tableHeaders;};function computeTableHeaderCellIndexes(t){var matrix=[];var lookup={};var thead=t.getElementsByTagName('THEAD')[0];var trs=thead.getElementsByTagName('TR');for(var i=0;i<trs.length;i++){var cells=trs[i].cells;for(var j=0;j<cells.length;j++){var c=cells[j];var rowIndex=c.parentNode.rowIndex;var cellId=rowIndex+"-"+c.cellIndex;var rowSpan=c.rowSpan||1;var colSpan=c.colSpan||1
var firstAvailCol;if(typeof(matrix[rowIndex])=="undefined"){matrix[rowIndex]=[];}for(var k=0;k<matrix[rowIndex].length+1;k++){if(typeof(matrix[rowIndex][k])=="undefined"){firstAvailCol=k;break;}}lookup[cellId]=firstAvailCol;for(var k=rowIndex;k<rowIndex+rowSpan;k++){if(typeof(matrix[k])=="undefined"){matrix[k]=[];}var matrixrow=matrix[k];for(var l=firstAvailCol;l<firstAvailCol+colSpan;l++){matrixrow[l]="x";}}}}return lookup;}function checkCellColSpan(table,rows,row){var arr=[],r=table.tHead.rows,c=r[row].cells;for(var i=0;i<c.length;i++){var cell=c[i];if(cell.colSpan>1){arr=arr.concat(checkCellColSpan(table,headerArr,row++));}else{if(table.tHead.length==1||(cell.rowSpan>1||!r[row+1])){arr.push(cell);}}}return arr;};function checkHeaderMetadata(cell){if(($.metadata)&&($(cell).metadata().sorter===false)){return true;};return false;}function checkHeaderOptions(table,i){if((table.config.headers[i])&&(table.config.headers[i].sorter===false)){return true;};return false;}function checkHeaderOptionsSortingLocked(table,i){if((table.config.headers[i])&&(table.config.headers[i].lockedOrder))return table.config.headers[i].lockedOrder;return false;}function applyWidget(table){var c=table.config.widgets;var l=c.length;for(var i=0;i<l;i++){getWidgetById(c[i]).format(table);}}function getWidgetById(name){var l=widgets.length;for(var i=0;i<l;i++){if(widgets[i].id.toLowerCase()==name.toLowerCase()){return widgets[i];}}};function formatSortingOrder(v){if(typeof(v)!="Number"){return(v.toLowerCase()=="desc")?1:0;}else{return(v==1)?1:0;}}function isValueInArray(v,a){var l=a.length;for(var i=0;i<l;i++){if(a[i][0]==v){return true;}}return false;}function setHeadersCss(table,$headers,list,css){$headers.removeClass(css[0]).removeClass(css[1]);var h=[];$headers.each(function(offset){if(!this.sortDisabled){h[this.column]=$(this);}});var l=list.length;for(var i=0;i<l;i++){h[list[i][0]].addClass(css[list[i][1]]);}}function fixColumnWidth(table,$headers){var c=table.config;if(c.widthFixed){var colgroup=$('<colgroup>');$("tr:first td",table.tBodies[0]).each(function(){colgroup.append($('<col>').css('width',$(this).width()));});$(table).prepend(colgroup);};}function updateHeaderSortCount(table,sortList){var c=table.config,l=sortList.length;for(var i=0;i<l;i++){var s=sortList[i],o=c.headerList[s[0]];o.count=s[1];o.count++;}}function multisort(table,sortList,cache){if(table.config.debug){var sortTime=new Date();}var dynamicExp="var sortWrapper = function(a,b) {",l=sortList.length;for(var i=0;i<l;i++){var c=sortList[i][0];var order=sortList[i][1];var s=(table.config.parsers[c].type=="text")?((order==0)?makeSortFunction("text","asc",c):makeSortFunction("text","desc",c)):((order==0)?makeSortFunction("numeric","asc",c):makeSortFunction("numeric","desc",c));var e="e"+i;dynamicExp+="var "+e+" = "+s;dynamicExp+="if("+e+") { return "+e+"; } ";dynamicExp+="else { ";}var orgOrderCol=cache.normalized[0].length-1;dynamicExp+="return a["+orgOrderCol+"]-b["+orgOrderCol+"];";for(var i=0;i<l;i++){dynamicExp+="}; ";}dynamicExp+="return 0; ";dynamicExp+="}; ";if(table.config.debug){benchmark("Evaling expression:"+dynamicExp,new Date());}eval(dynamicExp);cache.normalized.sort(sortWrapper);if(table.config.debug){benchmark("Sorting on "+sortList.toString()+" and dir "+order+" time:",sortTime);}return cache;};function makeSortFunction(type,direction,index){var a="a["+index+"]",b="b["+index+"]";if(type=='text'&&direction=='asc'){return"("+a+" == "+b+" ? 0 : ("+a+" === null ? Number.POSITIVE_INFINITY : ("+b+" === null ? Number.NEGATIVE_INFINITY : ("+a+" < "+b+") ? -1 : 1 )));";}else if(type=='text'&&direction=='desc'){return"("+a+" == "+b+" ? 0 : ("+a+" === null ? Number.POSITIVE_INFINITY : ("+b+" === null ? Number.NEGATIVE_INFINITY : ("+b+" < "+a+") ? -1 : 1 )));";}else if(type=='numeric'&&direction=='asc'){return"("+a+" === null && "+b+" === null) ? 0 :("+a+" === null ? Number.POSITIVE_INFINITY : ("+b+" === null ? Number.NEGATIVE_INFINITY : "+a+" - "+b+"));";}else if(type=='numeric'&&direction=='desc'){return"("+a+" === null && "+b+" === null) ? 0 :("+a+" === null ? Number.POSITIVE_INFINITY : ("+b+" === null ? Number.NEGATIVE_INFINITY : "+b+" - "+a+"));";}};function makeSortText(i){return"((a["+i+"] < b["+i+"]) ? -1 : ((a["+i+"] > b["+i+"]) ? 1 : 0));";};function makeSortTextDesc(i){return"((b["+i+"] < a["+i+"]) ? -1 : ((b["+i+"] > a["+i+"]) ? 1 : 0));";};function makeSortNumeric(i){return"a["+i+"]-b["+i+"];";};function makeSortNumericDesc(i){return"b["+i+"]-a["+i+"];";};function sortText(a,b){if(table.config.sortLocaleCompare)return a.localeCompare(b);return((a<b)?-1:((a>b)?1:0));};function sortTextDesc(a,b){if(table.config.sortLocaleCompare)return b.localeCompare(a);return((b<a)?-1:((b>a)?1:0));};function sortNumeric(a,b){return a-b;};function sortNumericDesc(a,b){return b-a;};function getCachedSortType(parsers,i){return parsers[i].type;};this.construct=function(settings){return this.each(function(){if(!this.tHead||!this.tBodies)return;var $this,$document,$headers,cache,config,shiftDown=0,sortOrder;this.config={};config=$.extend(this.config,$.tablesorter.defaults,settings);$this=$(this);$.data(this,"tablesorter",config);$headers=buildHeaders(this);this.config.parsers=buildParserCache(this,$headers);cache=buildCache(this);var sortCSS=[config.cssDesc,config.cssAsc];fixColumnWidth(this);$headers.click(function(e){var totalRows=($this[0].tBodies[0]&&$this[0].tBodies[0].rows.length)||0;if(!this.sortDisabled&&totalRows>0){$this.trigger("sortStart");var $cell=$(this);var i=this.column;this.order=this.count++%2;if(this.lockedOrder)this.order=this.lockedOrder;if(!e[config.sortMultiSortKey]){config.sortList=[];if(config.sortForce!=null){var a=config.sortForce;for(var j=0;j<a.length;j++){if(a[j][0]!=i){config.sortList.push(a[j]);}}}config.sortList.push([i,this.order]);}else{if(isValueInArray(i,config.sortList)){for(var j=0;j<config.sortList.length;j++){var s=config.sortList[j],o=config.headerList[s[0]];if(s[0]==i){o.count=s[1];o.count++;s[1]=o.count%2;}}}else{config.sortList.push([i,this.order]);}};setTimeout(function(){setHeadersCss($this[0],$headers,config.sortList,sortCSS);appendToTable($this[0],multisort($this[0],config.sortList,cache));},1);return false;}}).mousedown(function(){if(config.cancelSelection){this.onselectstart=function(){return false};return false;}});$this.bind("update",function(){var me=this;setTimeout(function(){me.config.parsers=buildParserCache(me,$headers);cache=buildCache(me);},1);}).bind("updateCell",function(e,cell){var config=this.config;var pos=[(cell.parentNode.rowIndex-1),cell.cellIndex];cache.normalized[pos[0]][pos[1]]=config.parsers[pos[1]].format(getElementText(config,cell),cell);}).bind("sorton",function(e,list){$(this).trigger("sortStart");config.sortList=list;var sortList=config.sortList;updateHeaderSortCount(this,sortList);setHeadersCss(this,$headers,sortList,sortCSS);appendToTable(this,multisort(this,sortList,cache));}).bind("appendCache",function(){appendToTable(this,cache);}).bind("applyWidgetId",function(e,id){getWidgetById(id).format(this);}).bind("applyWidgets",function(){applyWidget(this);});if($.metadata&&($(this).metadata()&&$(this).metadata().sortlist)){config.sortList=$(this).metadata().sortlist;}if(config.sortList.length>0){$this.trigger("sorton",[config.sortList]);}applyWidget(this);});};this.addParser=function(parser){var l=parsers.length,a=true;for(var i=0;i<l;i++){if(parsers[i].id.toLowerCase()==parser.id.toLowerCase()){a=false;}}if(a){parsers.push(parser);};};this.addWidget=function(widget){widgets.push(widget);};this.formatFloat=function(s){var i=parseFloat(s);return(isNaN(i))?0:i;};this.formatInt=function(s){var i=parseInt(s);return(isNaN(i))?0:i;};this.isDigit=function(s,config){return/^[-+]?\d*$/.test($.trim(s.replace(/[,.']/g,'')));};this.clearTableBody=function(table){if($.browser.msie){function empty(){while(this.firstChild)this.removeChild(this.firstChild);}empty.apply(table.tBodies[0]);}else{table.tBodies[0].innerHTML="";}};}});$.fn.extend({tablesorter:$.tablesorter.construct});var ts=$.tablesorter;ts.addParser({id:"text",is:function(s){return true;},format:function(s){return $.trim(s.toLocaleLowerCase());},type:"text"});ts.addParser({id:"digit",is:function(s,table){var c=table.config;return $.tablesorter.isDigit(s,c);},format:function(s){return $.tablesorter.formatFloat(s);},type:"numeric"});ts.addParser({id:"currency",is:function(s){return/^[£$€?.]/.test(s);},format:function(s){return $.tablesorter.formatFloat(s.replace(new RegExp(/[£$€]/g),""));},type:"numeric"});ts.addParser({id:"ipAddress",is:function(s){return/^\d{2,3}[\.]\d{2,3}[\.]\d{2,3}[\.]\d{2,3}$/.test(s);},format:function(s){var a=s.split("."),r="",l=a.length;for(var i=0;i<l;i++){var item=a[i];if(item.length==2){r+="0"+item;}else{r+=item;}}return $.tablesorter.formatFloat(r);},type:"numeric"});ts.addParser({id:"url",is:function(s){return/^(https?|ftp|file):\/\/$/.test(s);},format:function(s){return jQuery.trim(s.replace(new RegExp(/(https?|ftp|file):\/\//),''));},type:"text"});ts.addParser({id:"isoDate",is:function(s){return/^\d{4}[\/-]\d{1,2}[\/-]\d{1,2}$/.test(s);},format:function(s){return $.tablesorter.formatFloat((s!="")?new Date(s.replace(new RegExp(/-/g),"/")).getTime():"0");},type:"numeric"});ts.addParser({id:"percent",is:function(s){return/\%$/.test($.trim(s));},format:function(s){return $.tablesorter.formatFloat(s.replace(new RegExp(/%/g),""));},type:"numeric"});ts.addParser({id:"usLongDate",is:function(s){return s.match(new RegExp(/^[A-Za-z]{3,10}\.? [0-9]{1,2}, ([0-9]{4}|'?[0-9]{2}) (([0-2]?[0-9]:[0-5][0-9])|([0-1]?[0-9]:[0-5][0-9]\s(AM|PM)))$/));},format:function(s){return $.tablesorter.formatFloat(new Date(s).getTime());},type:"numeric"});ts.addParser({id:"shortDate",is:function(s){return/\d{1,2}[\/\-]\d{1,2}[\/\-]\d{2,4}/.test(s);},format:function(s,table){var c=table.config;s=s.replace(/\-/g,"/");if(c.dateFormat=="us"){s=s.replace(/(\d{1,2})[\/\-](\d{1,2})[\/\-](\d{4})/,"$3/$1/$2");}else if(c.dateFormat=="uk"){s=s.replace(/(\d{1,2})[\/\-](\d{1,2})[\/\-](\d{4})/,"$3/$2/$1");}else if(c.dateFormat=="dd/mm/yy"||c.dateFormat=="dd-mm-yy"){s=s.replace(/(\d{1,2})[\/\-](\d{1,2})[\/\-](\d{2})/,"$1/$2/$3");}return $.tablesorter.formatFloat(new Date(s).getTime());},type:"numeric"});ts.addParser({id:"time",is:function(s){return/^(([0-2]?[0-9]:[0-5][0-9])|([0-1]?[0-9]:[0-5][0-9]\s(am|pm)))$/.test(s);},format:function(s){return $.tablesorter.formatFloat(new Date("2000/01/01 "+s).getTime());},type:"numeric"});ts.addParser({id:"metadata",is:function(s){return false;},format:function(s,table,cell){var c=table.config,p=(!c.parserMetadataName)?'sortValue':c.parserMetadataName;return $(cell).metadata()[p];},type:"numeric"});ts.addWidget({id:"zebra",format:function(table){if(table.config.debug){var time=new Date();}var $tr,row=-1,odd;$("tr:visible",table.tBodies[0]).each(function(i){$tr=$(this);if(!$tr.hasClass(table.config.cssChildRow))row++;odd=(row%2==0);$tr.removeClass(table.config.widgetZebra.css[odd?0:1]).addClass(table.config.widgetZebra.css[odd?1:0])});if(table.config.debug){$.tablesorter.benchmark("Applying Zebra widget",time);}}});})(jQuery);
```
