## Supplementary Results for "A draft genome of the ascomycotal fungal species *Pseudopithomyces maydicus* (family *Didymosphaeriaceae*)": contig_4_c1.txt

ClusterBlast scores for contig_4

Table of genes, locations, strands and annotations of query cluster:
input.path1.gene103	354413	355557	-		
input.path1.gene104	360322	361219	-		
input.path1.gene105	361306	362766	+		
input.path1.gene106	363258	364317	+		
input.path1.gene107	364516	365570	-		
input.path1.gene108	365875	366276	+		
input.path1.gene109	371338	373741	+		
input.path1.gene110	376034	377180	+		
input.path1.gene111	377444	379239	-		
input.path1.gene112	381789	382134	+		
input.path1.gene113	383003	384665	-		
input.path1.gene114	388890	389490	+		
input.path1.gene115	390223	392004	+		


Significant hits: 
1. BGC0001123	aspirochlorine


Details:

>>
1. BGC0001123
Source: aspirochlorine
Type: NRP
Number of proteins with BLAST hits to this cluster: 3
Cumulative BLAST score: 1037

Table of genes, locations, strands and annotations of subject cluster:

Table of Blast hits (query gene, subject gene, %identity, blast score, %coverage, e-value):
input.path1.gene106	AO090001000036	60	187	49.82332155477032	3.3e-47
input.path1.gene111	AO090001000031	48	508	90.65255731922399	1.7e-143
input.path1.gene113	AO090001000039	48	342	71.04166666666667	8e-94
