## Supplementary Results for "A draft genome of the ascomycotal fungal species *Pseudopithomyces maydicus* (family *Didymosphaeriaceae*)": contig_4_c2.txt

ClusterBlast scores for contig_4

Table of genes, locations, strands and annotations of query cluster:
input.path1.gene340	1284316	1284607	+		
input.path1.gene341	1286354	1288678	+		
input.path1.gene342	1288948	1289596	-		
input.path1.gene343	1294518	1295517	+		
input.path1.gene344	1295611	1297093	-		
input.path1.gene345	1299349	1304830	-		
input.path1.gene346	1307660	1309419	-		
input.path1.gene347	1309571	1310220	+		
input.path1.gene348	1311642	1312949	+		
input.path1.gene349	1313508	1315245	-		
input.path1.gene350	1315986	1317843	+		
input.path1.gene351	1320309	1321467	-		


Significant hits: 
1. BGC0001244	(-)-Mellein
2. BGC0001276	6-methylsalicyclic acid
3. BGC0001275	6-methylsalicyclic acid
4. BGC0001273	asperlactone


Details:

>>
1. BGC0001244
Source: (-)-Mellein
Type: Polyketide
Number of proteins with BLAST hits to this cluster: 1
Cumulative BLAST score: 1889

Table of genes, locations, strands and annotations of subject cluster:

Table of Blast hits (query gene, subject gene, %identity, blast score, %coverage, e-value):
input.path1.gene345	SNOG_00477	54	1889	100.11160714285714	0.0


>>
2. BGC0001276
Source: 6-methylsalicyclic acid
Type: Polyketide
Number of proteins with BLAST hits to this cluster: 1
Cumulative BLAST score: 1810

Table of genes, locations, strands and annotations of subject cluster:
BAA20102.2	BAA20102.2	1102	6575	+	6-methylsalicylic_acid_synthase

Table of Blast hits (query gene, subject gene, %identity, blast score, %coverage, e-value):
input.path1.gene345	BAA20102.2	52	1810	98.88392857142857	0.0


>>
3. BGC0001275
Source: 6-methylsalicyclic acid
Type: Polyketide
Number of proteins with BLAST hits to this cluster: 1
Cumulative BLAST score: 1636

Table of genes, locations, strands and annotations of subject cluster:
AAX35547.1	AAX35547.1	1064	6614	+	polyketide_syntase_2

Table of Blast hits (query gene, subject gene, %identity, blast score, %coverage, e-value):
input.path1.gene345	AAX35547.1	48	1636	100.16741071428572	0.0


>>
4. BGC0001273
Source: asperlactone
Type: Polyketide
Number of proteins with BLAST hits to this cluster: 1
Cumulative BLAST score: 1602

Table of genes, locations, strands and annotations of subject cluster:
AAS98200.1	AAS98200.1	1	5301	+	MSAS-type_polyketide_synthase

Table of Blast hits (query gene, subject gene, %identity, blast score, %coverage, e-value):
input.path1.gene345	AAS98200.1	47	1602	97.76785714285714	0.0
