## Supplementary Results for "A draft genome of the ascomycotal fungal species *Pseudopithomyces maydicus* (family *Didymosphaeriaceae*)": contig_9_c2.txt

ClusterBlast scores for contig_9

Table of genes, locations, strands and annotations of query cluster:
input.path1.gene511	1787953	1788622	+		
input.path1.gene512	1790829	1792452	-		
input.path1.gene513	1793072	1793939	-		
input.path1.gene514	1795178	1796398	-		
input.path1.gene515	1803508	1805696	+		


Significant hits: 
1. BGC0001839	squalestatin S1


Details:

>>
1. BGC0001839
Source: squalestatin S1
Type: Terpene
Number of proteins with BLAST hits to this cluster: 2
Cumulative BLAST score: 1065

Table of genes, locations, strands and annotations of subject cluster:
gene1	gene1	153	639	-	
gene2	gene2	1396	3141	-	
gene3	gene3	3426	15590	-	
gene4	gene4	16469	17614	+	
gene5	gene5	17760	19345	-	

Table of Blast hits (query gene, subject gene, %identity, blast score, %coverage, e-value):
input.path1.gene512	gene3	65	533	96.34615384615385	3.5e-151
input.path1.gene514	gene2	66	532	97.41602067183463	5.7e-151
