## Supplementary Results for "A draft genome of the ascomycotal fungal species *Pseudopithomyces maydicus* (family *Didymosphaeriaceae*)": contig_10_c2.txt

ClusterBlast scores for contig_10

Table of genes, locations, strands and annotations of query cluster:
input.path1.gene173	638688	639604	-		
input.path1.gene174	640219	642139	+		
input.path1.gene175	644124	648112	-		
input.path1.gene176	651038	652466	+		
input.path1.gene177	653296	656734	-		
input.path1.gene178	657822	662907	+		
input.path1.gene179	663094	664224	-		
input.path1.gene180	666162	668145	+		
input.path1.gene181	668454	668627	+		
input.path1.gene182	669026	671092	-		
input.path1.gene183	672052	673122	-		
input.path1.gene184	674274	676506	+		
input.path1.gene185	678483	681267	+		
input.path1.gene186	681672	682178	+		


Significant hits: 
1. BGC0001284	alternariol


Details:

>>
1. BGC0001284
Source: alternariol
Type: Terpene
Number of proteins with BLAST hits to this cluster: 1
Cumulative BLAST score: 1437

Table of genes, locations, strands and annotations of subject cluster:
AKN45693.1	AKN45693.1	1	5292	+	polyketide_synthase

Table of Blast hits (query gene, subject gene, %identity, blast score, %coverage, e-value):
input.path1.gene178	AKN45693.1	50	1437	133.4987593052109	0.0
