## Supplementary Results for "A draft genome of the ascomycotal fungal species *Pseudopithomyces maydicus* (family *Didymosphaeriaceae*)": contig_12_c2.txt

ClusterBlast scores for contig_12

Table of genes, locations, strands and annotations of query cluster:
input.path1.gene638	2325386	2326715	-		
input.path1.gene639	2329982	2331603	+		
input.path1.gene640	2331896	2333866	-		
input.path1.gene641	2336738	2343315	+		
input.path1.gene642	2345182	2349957	+		
input.path1.gene643	2350125	2351567	-		
input.path1.gene644	2352390	2353742	-		
input.path1.gene645	2363625	2365011	+		
input.path1.gene646	2365115	2367616	-		
input.path1.gene647	2367807	2369320	+		


Significant hits: 
1. BGC0001249	dimethylcoprogen


Details:

>>
1. BGC0001249
Source: dimethylcoprogen
Type: NRP
Number of proteins with BLAST hits to this cluster: 1
Cumulative BLAST score: 1948

Table of genes, locations, strands and annotations of subject cluster:
AFN69082.1	AFN69082.1	664	5994	+	nonribosomal_peptide_synthetase_6

Table of Blast hits (query gene, subject gene, %identity, blast score, %coverage, e-value):
input.path1.gene642	AFN69082.1	64	1948	98.97435897435898	0.0
