## Supplementary Results for "A draft genome of the ascomycotal fungal species *Pseudopithomyces maydicus* (family *Didymosphaeriaceae*)": contig_17_c2.txt

ClusterBlast scores for contig_17

Table of genes, locations, strands and annotations of query cluster:
input.path1.gene911	3156599	3157274	+		
input.path1.gene912	3160347	3161385	+		
input.path1.gene913	3161578	3162784	+		
input.path1.gene914	3163026	3163666	-		
input.path1.gene915	3165231	3167880	+		
input.path1.gene916	3168586	3169140	+		
input.path1.gene917	3174261	3175829	+		
input.path1.gene918	3176184	3182827	-		
input.path1.gene919	3184519	3185567	-		
input.path1.gene920	3187737	3189162	-		
input.path1.gene921	3190504	3191011	+		
input.path1.gene922	3192718	3193765	+		
input.path1.gene923	3195617	3196007	+		
input.path1.gene924	3197131	3199004	-		


Significant hits: 
1. BGC0001265	melanin
2. BGC0001258	1,3,6,8-tetrahydroxynaphthalene


Details:

>>
1. BGC0001265
Source: melanin
Type: Polyketide
Number of proteins with BLAST hits to this cluster: 1
Cumulative BLAST score: 1699

Table of genes, locations, strands and annotations of subject cluster:
BAD22832.1	BAD22832.1	668	7238	+	polyketide_synthase

Table of Blast hits (query gene, subject gene, %identity, blast score, %coverage, e-value):
input.path1.gene918	BAD22832.1	47	1699	91.69139465875371	0.0


>>
2. BGC0001258
Source: 1,3,6,8-tetrahydroxynaphthalene
Type: Polyketide
Number of proteins with BLAST hits to this cluster: 1
Cumulative BLAST score: 1562

Table of genes, locations, strands and annotations of subject cluster:
AAN59953.1	AAN59953.1	44	7265	+	polyketide_synthase_1

Table of Blast hits (query gene, subject gene, %identity, blast score, %coverage, e-value):
input.path1.gene918	AAN59953.1	47	1562	87.784371909001	0.0
