## Supplementary Results for "A draft genome of the ascomycotal fungal species *Pseudopithomyces maydicus* (family *Didymosphaeriaceae*)": contig_26_c2.txt

ClusterBlast scores for contig_26

Table of genes, locations, strands and annotations of query cluster:
input.path1.gene202	739460	741433	+		
input.path1.gene203	746577	747002	+		
input.path1.gene204	747015	751535	-		
input.path1.gene205	752565	753169	+		
input.path1.gene206	753642	754399	-		
input.path1.gene207	756918	763489	-		
input.path1.gene208	775206	778416	+		


Significant hits: 
1. BGC0001265	melanin
2. BGC0001258	1,3,6,8-tetrahydroxynaphthalene
3. BGC0001257	1,3,6,8-tetrahydroxynaphthalene


Details:

>>
1. BGC0001265
Source: melanin
Type: Polyketide
Number of proteins with BLAST hits to this cluster: 1
Cumulative BLAST score: 3216

Table of genes, locations, strands and annotations of subject cluster:
BAD22832.1	BAD22832.1	668	7238	+	polyketide_synthase

Table of Blast hits (query gene, subject gene, %identity, blast score, %coverage, e-value):
input.path1.gene207	BAD22832.1	76	3216	102.80151946818614	0.0


>>
2. BGC0001258
Source: 1,3,6,8-tetrahydroxynaphthalene
Type: Polyketide
Number of proteins with BLAST hits to this cluster: 1
Cumulative BLAST score: 1858

Table of genes, locations, strands and annotations of subject cluster:
AAN59953.1	AAN59953.1	44	7265	+	polyketide_synthase_1

Table of Blast hits (query gene, subject gene, %identity, blast score, %coverage, e-value):
input.path1.gene207	AAN59953.1	48	1858	102.37416904083571	0.0


>>
3. BGC0001257
Source: 1,3,6,8-tetrahydroxynaphthalene
Type: Polyketide
Number of proteins with BLAST hits to this cluster: 1
Cumulative BLAST score: 1839

Table of genes, locations, strands and annotations of subject cluster:
AAD38786.1	AAD38786.1	400	7066	+	polyketide_synthase

Table of Blast hits (query gene, subject gene, %identity, blast score, %coverage, e-value):
input.path1.gene207	AAD38786.1	47	1839	103.94112060778728	0.0
