## Supplementary Results for "A draft genome of the ascomycotal fungal species *Pseudopithomyces maydicus* (family *Didymosphaeriaceae*)": contig_32_c3.txt

ClusterBlast scores for contig_32

Table of genes, locations, strands and annotations of query cluster:
input.path1.gene885	3138049	3139703	-		
input.path1.gene886	3141781	3142924	-		
input.path1.gene887	3145213	3146156	+		
input.path1.gene888	3146892	3147343	-		
input.path1.gene889	3148273	3149705	-		
input.path1.gene890	3150534	3151265	+		
input.path1.gene891	3154000	3154464	-		
input.path1.gene892	3155450	3156116	-		
input.path1.gene893	3157579	3164311	+		
input.path1.gene894	3168713	3169230	-		
input.path1.gene895	3171654	3177360	-		
input.path1.gene896	3178403	3179732	+		
input.path1.gene897	3181316	3182359	+		
input.path1.gene898	3182593	3184390	-		
input.path1.gene899	3185516	3187164	+		
input.path1.gene900	3189780	3193436	-		


Significant hits: 
1. BGC0001405	chaetoviridin E / 11-epichaetomugilin A
2. BGC0000022	asperfuranone


Details:

>>
1. BGC0001405
Source: chaetoviridin E / 11-epichaetomugilin A
Type: Polyketide
Number of proteins with BLAST hits to this cluster: 4
Cumulative BLAST score: 1767

Table of genes, locations, strands and annotations of subject cluster:

Table of Blast hits (query gene, subject gene, %identity, blast score, %coverage, e-value):
input.path1.gene891	CHGG_07643	55	80	96.05263157894737	1.2e-15
input.path1.gene892	CHGG_07643	57	85	41.139240506329116	7.6e-17
input.path1.gene895	CHGG_07645	57	1205	58.94039735099338	0.0
input.path1.gene896	CHGG_07644	47	397	96.75174013921114	2.5e-110


>>
2. BGC0000022
Source: asperfuranone
Type: Polyketide
Number of proteins with BLAST hits to this cluster: 3
Cumulative BLAST score: 2111

Table of genes, locations, strands and annotations of subject cluster:

Table of Blast hits (query gene, subject gene, %identity, blast score, %coverage, e-value):
input.path1.gene892	AN1031.2	46	154	109.49367088607596	1.7e-37
input.path1.gene895	AN1034.2	54	1740	96.19205298013244	0.0
input.path1.gene897	AN1030.2	51	217	87.64478764478764	3.6e-56
