## Supplementary Results for "A draft genome of the ascomycotal fungal species *Pseudopithomyces maydicus* (family *Didymosphaeriaceae*)": input.path1.gene1_mibig_hits.html

| MIBiG Protein | Description | MIBiG Cluster | MiBiG Product | % ID | % Coverage | BLAST Score | E-value |
| --- | --- | --- | --- | --- | --- | --- | --- |
| AGO59038.1 | PtaI | BGC0000121 | Polyketide | 32.0 | 99.4 | 176.0 | 8.7e-44 |
| PKY07883.1 | S-adenosyl-L-methionine-dependent\_methyltransferase | BGC0001544 | NRP + Polyketide | 34.0 | 99.7 | 170.0 | 3.7e-42 |
