## Supplementary Results for "A draft genome of the ascomycotal fungal species *Pseudopithomyces maydicus* (family *Didymosphaeriaceae*)": input.path1.gene2_mibig_hits.html

| MIBiG Protein | Description | MIBiG Cluster | MiBiG Product | % ID | % Coverage | BLAST Score | E-value |
| --- | --- | --- | --- | --- | --- | --- | --- |
| XP\_007301851.1 | cytochrome\_P450 | BGC0001617 | Terpene | 28.0 | 65.9 | 101.0 | 3e-21 |
| AFW60201.1 | benzoxazinone\_synthesis4 | BGC0000810 | Alkaloid | 28.0 | 59.2 | 81.0 | 2.5e-15 |
| AFW60200.1 | benzoxazinone\_synthesis3 | BGC0000810 | Alkaloid | 28.0 | 58.9 | 77.0 | 3.6e-14 |
| AFW60202.1 | benzoxazinone\_synthesis5 | BGC0000810 | Alkaloid | 27.0 | 49.4 | 74.0 | 4e-13 |
| MAA\_10044 | benzoate\_4-monooxygenase\_cytochrome\_P450 | BGC0000337 | NRP | 28.0 | 50.8 | 70.0 | 5.7e-12 |
| chr3.CM0292.110.r2.m |  | BGC0001317 | Terpene | 30.0 | 47.2 | 70.0 | 7.5e-12 |
| EAL89312.1 | cytochrome\_P450\_monooxygenase,\_putative | BGC0000686 | Terpene | 29.0 | 47.8 | 68.0 | 2.8e-11 |
| NP\_199073.1 | cytochrome\_P450\_71A16 | BGC0000669 | Terpene | 27.0 | 45.0 | 66.0 | 1.1e-10 |
| ATZ56106.1 | Bcbot1 | BGC0000631 | Terpene | 30.0 | 40.5 | 64.0 | 3.1e-10 |
| EAL89314.2 | cytochrome\_P450\_monooxygenase,\_putative | BGC0000686 | Terpene | 35.0 | 33.0 | 64.0 | 4.1e-10 |
| AFW60212.1 | benzoxazinone\_synthesis2 | BGC0000810 | Alkaloid | 27.0 | 50.0 | 62.0 | 1.6e-09 |
| AFW60213.1 | benzoxazinone\_synthesis2 | BGC0000810 | Alkaloid | 27.0 | 50.0 | 62.0 | 1.6e-09 |
| BAD83680.1 | cytochrome\_P-450 | BGC0000012 | Polyketide | 25.0 | 53.9 | 54.0 | 5.5e-07 |
| BBB04330.1 | cytochrome\_P450 | BGC0001717 | Alkaloid | 33.0 | 28.2 | 52.0 | 2.1e-06 |
