## Supplementary Results for "A draft genome of the ascomycotal fungal species *Pseudopithomyces maydicus* (family *Didymosphaeriaceae*)": input.path1.gene3_mibig_hits.html

| MIBiG Protein | Description | MIBiG Cluster | MiBiG Product | % ID | % Coverage | BLAST Score | E-value |
| --- | --- | --- | --- | --- | --- | --- | --- |
| ACZ57546.1 | predicted\_MFS\_transporter | BGC0000046 | Polyketide:Iterative type I | 36.0 | 93.7 | 321.0 | 2.2e-87 |
| AIG62136.1 | MFS\_transporter | BGC0000120 | Polyketide:Iterative type I | 37.0 | 98.3 | 308.0 | 1.9e-83 |
| BAZ95819.1 | cpaN1\_MFS\_transporter | BGC0001563 | NRP + Polyketide | 34.0 | 97.4 | 300.0 | 3.9e-81 |
| CAP93755.1 |  | BGC0001882 | Polyketide | 26.0 | 92.4 | 196.0 | 1e-49 |
| BAE60006.1 |  | BGC0001518 | Terpene | 28.0 | 93.7 | 193.0 | 1.2e-48 |
| DAB41650.1 | MFS\_transporter | BGC0001583 | Polyketide | 29.0 | 80.7 | 192.0 | 1.5e-48 |
| ARP51716.1 | toxin\_efflux\_transporter\_MFS | BGC0001741 | NRP + Polyketide | 28.0 | 100.4 | 187.0 | 6.3e-47 |
| ACD39756.1 | major\_facilitator\_superfamily\_transporter | BGC0000076 | Polyketide | 28.0 | 98.3 | 186.0 | 1.1e-46 |
| ACD39765.1 | major\_facilitator\_superfamily\_transporter | BGC0000077 | Polyketide | 28.0 | 98.3 | 186.0 | 1.1e-46 |
| BAD29973.1 | transporter\_protein | BGC0000676 | Terpene | 28.0 | 81.0 | 185.0 | 1.8e-46 |
| BAZ95831.1 | MFS\_transporter\_cpaI | BGC0001563 | NRP + Polyketide | 26.0 | 100.0 | 184.0 | 4.1e-46 |
| CAP96439.1 | Transporter | BGC0000420 | NRP | 28.0 | 97.6 | 183.0 | 7e-46 |
| AVY05519.1 | major\_facilitator\_superfamily\_transporter | BGC0001571 | Terpene | 28.0 | 100.6 | 183.0 | 9.2e-46 |
| RWQ92172.1 | putative\_MFS\_transporter | BGC0002030 | Polyketide | 27.0 | 93.0 | 183.0 | 1.2e-45 |
| BBG28481.1 | putative\_MFS\_toxin\_efflux\_pump\_CdmB | BGC0001926 | Polyketide | 28.0 | 99.4 | 182.0 | 1.6e-45 |
| CCE28984.1 | probable\_DHA14-like\_major\_facilitator;\_ABC\_transporter | BGC0001365 | NRP | 27.0 | 95.0 | 169.0 | 1.4e-41 |
| BAV69308.1 | PrhG | BGC0001729 | Polyketide + Terpene | 27.0 | 94.5 | 169.0 | 1.8e-41 |
| EAL88822.1 | MFS\_gliotoxin\_efflux\_transporter\_GliA | BGC0000361 | NRP | 27.0 | 97.2 | 168.0 | 2.3e-41 |
| AAD34558.1 | unknown | BGC0000088 | Polyketide | 27.0 | 97.1 | 160.0 | 6.4e-39 |
| BAE71313.1 | putative\_ABC\_transporter | BGC0000004 | Polyketide | 26.0 | 88.4 | 160.0 | 8.3e-39 |
| ABA02247.1 | efflux\_pump | BGC0000098 | Polyketide | 28.0 | 96.3 | 160.0 | 8.3e-39 |
| AAS90046.1 | AflT | BGC0000009 | Polyketide | 26.0 | 90.8 | 159.0 | 1.9e-38 |
| EAU36749.1 | predicted\_protein | BGC0000292 | NRP | 31.0 | 74.8 | 158.0 | 3.2e-38 |
| AAS89998.1 | AflT | BGC0000007 | Polyketide | 26.0 | 90.8 | 158.0 | 4.1e-38 |
| XP\_001798920.1 | MFS\_transporter | BGC0001865 | Polyketide:Iterative type I | 28.0 | 83.1 | 158.0 | 4.1e-38 |
| AAS90069.1 | AflT | BGC0000010 | Polyketide | 25.0 | 96.3 | 154.0 | 6e-37 |
| AAS90021.1 | AflT | BGC0000008 | Polyketide | 26.0 | 97.1 | 150.0 | 6.6e-36 |
| ACZ66257.1 | APS11 | BGC0000304 | NRP | 25.0 | 100.4 | 149.0 | 1.5e-35 |
| AAS90092.1 | AflT | BGC0000006 | Polyketide | 25.0 | 97.1 | 148.0 | 3.3e-35 |
| PIB01159.1 | putative\_HC-toxin\_efflux\_carrier\_TOXA | BGC0001541 | Polyketide | 27.0 | 88.8 | 148.0 | 3.3e-35 |
| BAC20568.1 | efflux\_pump | BGC0000039 | Polyketide | 27.0 | 98.2 | 147.0 | 5.6e-35 |
| ACS68556.1 | major\_facilitator\_superfamily\_protein | BGC0001026 | NRP + Polyketide | 24.0 | 92.3 | 133.0 | 1.4e-30 |
| EAA65599.1 | hypothetical\_protein | BGC0000022 | Polyketide | 26.0 | 101.7 | 129.0 | 1.2e-29 |
| AEO57490.1 | general\_substrate\_transporter | BGC0001449 | NRP + Alkaloid + Polyketide:Iterative type I | 24.0 | 97.1 | 126.0 | 1e-28 |
| AAM33663.1 | putative\_efflux\_protein | BGC0000230 | Polyketide:Type II | 24.0 | 91.7 | 118.0 | 4.7e-26 |
| AAM94765.1 | CalT1 | BGC0000033 | Polyketide | 24.0 | 81.8 | 102.0 | 2.1e-21 |
| QCS37513.1 | PyiT | BGC0001982 | NRP + Polyketide | 21.0 | 99.1 | 93.0 | 1.2e-18 |
| AQP25575.1 | MFS\_family\_transporter | BGC0001590 | Polyketide | 25.0 | 81.8 | 67.0 | 9.6e-11 |
| CAP12594.1 | elloramycin\_permease | BGC0000219 | Polyketide:Type II + Saccharide:Hybrid/tailoring | 24.0 | 90.4 | 65.0 | 2.8e-10 |
