## Supplementary Results for "A draft genome of the ascomycotal fungal species *Pseudopithomyces maydicus* (family *Didymosphaeriaceae*)": input.path1.gene4_mibig_hits.html

| MIBiG Protein | Description | MIBiG Cluster | MiBiG Product | % ID | % Coverage | BLAST Score | E-value |
| --- | --- | --- | --- | --- | --- | --- | --- |
| BAK64647.1 | putative\_quinone\_oxidoreductase | BGC0000135 | Polyketide | 28.0 | 64.8 | 76.0 | 8.5e-14 |
| ALV82389.1 | alcohol\_dehydrogenase | BGC0001370 | NRP | 35.0 | 34.8 | 57.0 | 5.3e-08 |
