## Supplementary Results for "A draft genome of the ascomycotal fungal species *Pseudopithomyces maydicus* (family *Didymosphaeriaceae*)": input.path1.gene5_mibig_hits.html

| MIBiG Protein | Description | MIBiG Cluster | MiBiG Product | % ID | % Coverage | BLAST Score | E-value |
| --- | --- | --- | --- | --- | --- | --- | --- |
| CCE67070.1 | polyketide\_synthase | BGC0001242 | Polyketide | 53.0 | 112.0 | 1894.0 | 0.0 |
| RWQ92175.1 | putative\_polyketide\_synthase | BGC0002030 | Polyketide | 43.0 | 86.4 | 1091.0 | 0.0 |
| XP\_001798923.1 | polyketide\_synthase | BGC0001865 | Polyketide:Iterative type I | 41.0 | 86.3 | 1004.0 | 1.7e-292 |
| AAS89999.1 | PksA | BGC0000007 | Polyketide | 41.0 | 84.6 | 963.0 | 4.3e-280 |
| AAS90093.1 | PksA | BGC0000006 | Polyketide | 41.0 | 84.6 | 963.0 | 5.6e-280 |
| BAE71314.1 | polyketide\_synthase | BGC0000004 | Polyketide | 41.0 | 84.6 | 960.0 | 4.8e-279 |
| AAS90022.1 | PksA | BGC0000008 | Polyketide | 40.0 | 84.6 | 957.0 | 2.4e-278 |
| EED57518.1 | polyketide\_synthase,\_putative | BGC0001446 | Polyketide:Iterative type I | 42.0 | 78.9 | 948.0 | 1.9e-275 |
| CBF74114.1 | Conidial\_yellow\_pigment\_biosynthesis\_polyketide\_synthase\_(PKS)(EC\_2.3.1.-)\_[Source:UniProtKB/Swiss-Prot;Acc:Q03149] | BGC0000107 | Polyketide | 39.0 | 85.7 | 943.0 | 4.6e-274 |
| AAS90047.1 | PksA | BGC0000009 | Polyketide | 41.0 | 83.7 | 941.0 | 1.3e-273 |
| AAZ95017.1 | polyketide\_synthase | BGC0000048 | Polyketide | 38.0 | 93.5 | 939.0 | 6.7e-273 |
| ACH72912.1 | AflC | BGC0000011 | Polyketide | 39.0 | 90.5 | 930.0 | 3.1e-270 |
| PIB02405.1 | CTB1 | BGC0001541 | Polyketide | 39.0 | 87.4 | 916.0 | 4.6e-266 |
| ARU80380.1 | polyketide\_synthase | BGC0001542 | Polyketide | 38.0 | 87.0 | 916.0 | 4.6e-266 |
| AAC49191.1 | putative\_polyketide\_synthase | BGC0000152 | Polyketide | 38.0 | 90.9 | 910.0 | 4.3e-264 |
| AAN59953.1 | polyketide\_synthase\_1 | BGC0001258 | Polyketide | 37.0 | 86.7 | 856.0 | 5.7e-248 |
| CCT67991.1 | bikaverin\_cluster-polyketide\_synthase | BGC0000030 | Polyketide | 38.0 | 79.9 | 843.0 | 6.5e-244 |
| BAD22832.1 | polyketide\_synthase | BGC0001265 | Polyketide | 35.0 | 91.7 | 842.0 | 1.1e-243 |
| EAU38791.1 | hypothetical\_protein | BGC0000161 | Polyketide:Iterative type I | 37.0 | 86.4 | 839.0 | 7.2e-243 |
| gene6 |  | BGC0001906 | Polyketide | 37.0 | 80.7 | 788.0 | 2.5e-227 |
| EED21099.1 | polyketide\_synthase,\_putative | BGC0001578 | Polyketide | 43.0 | 58.3 | 775.0 | 1.3e-223 |
| AAD38786.1 | polyketide\_synthase | BGC0001257 | Polyketide | 36.0 | 81.3 | 770.0 | 5.4e-222 |
| EAL89339.1 | polyketide\_synthase,\_putative | BGC0001403 | Polyketide | 34.0 | 79.9 | 756.0 | 8e-218 |
| AGO59040.1 | PtaA | BGC0000121 | Polyketide | 34.0 | 80.5 | 740.0 | 4.5e-213 |
| AUW31184.1 | putative\_type\_I\_PKS | BGC0001489 | Polyketide | 41.0 | 57.5 | 731.0 | 3.6e-210 |
| EGD99348.1 | polyketide\_synthase | BGC0001144 | Polyketide | 34.0 | 82.3 | 723.0 | 9.8e-208 |
| ADM79459.1 | PKS16\_protein | BGC0001266 | Polyketide | 35.0 | 81.1 | 719.0 | 1.4e-206 |
| PKX92308.1 | putative\_polyketide\_synthase | BGC0001988 | Polyketide | 33.0 | 81.1 | 712.0 | 1.7e-204 |
| CCE31584.1 | polyketide\_synthase\_that\_catalyse\_the\_condensation\_of\_one\_acetyl-CoA\_and\_six\_malonyl-CoA\_resulting\_in\_formation\_of\_nor-rubrofusarin | BGC0001886 | Polyketide | 33.0 | 82.2 | 698.0 | 2e-200 |
| EAL84397.1 | polyketide\_synthase | BGC0001118 | Polyketide:Iterative type I | 40.0 | 58.0 | 696.0 | 7.5e-200 |
| ADI24926.1 | VrtA | BGC0000168 | Polyketide:Iterative type I | 39.0 | 60.7 | 695.0 | 2.2e-199 |
| AEN83889.1 | AdaA | BGC0000156 | Polyketide:Iterative type I | 39.0 | 58.3 | 684.0 | 5.1e-196 |
| CBF70387.1 | polyketide\_synthase,\_putative\_(JCVI) | BGC0000684 | Terpene | 32.0 | 81.7 | 681.0 | 2.5e-195 |
| AKN45693.1 | polyketide\_synthase | BGC0001284 | Terpene | 33.0 | 80.2 | 670.0 | 7.6e-192 |
| EED53479.1 | polyketide\_synthase,\_putative | BGC0001304 | Polyketide | 38.0 | 57.6 | 666.0 | 1.4e-190 |
| DAB41653.1 | polyketide\_synthase | BGC0001583 | Polyketide | 34.0 | 73.9 | 645.0 | 2e-184 |
| ADI24953.1 | GsfA | BGC0000070 | Polyketide:Iterative type I | 38.0 | 55.5 | 638.0 | 4.2e-182 |
| KKP00966.1 | RADS2\_nonreducing\_polyketide\_synthase | BGC0001901 | Polyketide | 32.0 | 80.3 | 577.0 | 8.7e-164 |
| CBF79143.1 | polyketide\_synthase,\_putative\_(JCVI) | BGC0000013 | Polyketide | 36.0 | 58.6 | 571.0 | 3.7e-162 |
| AGC95321.1 | CurS2 | BGC0000045 | Polyketide | 37.0 | 56.2 | 550.0 | 6.7e-156 |
| ACD39753.1 | non-reducing\_polyketide\_synthase | BGC0000076 | Polyketide | 29.0 | 85.6 | 547.0 | 5.7e-155 |
| ACD39762.1 | non-reducing\_polyketide\_synthase | BGC0000077 | Polyketide | 28.0 | 85.6 | 546.0 | 1.6e-154 |
| ALI92655.1 | CitS\_citrinin\_polyketide\_synthase | BGC0001338 | Polyketide:Iterative type I | 24.0 | 132.3 | 537.0 | 7.6e-152 |
| AHV78247.1 | LasS2 | BGC0001245 | Polyketide | 30.0 | 80.2 | 535.0 | 3.8e-151 |
| AHV78253.1 | ResS2 | BGC0001246 | Polyketide | 38.0 | 49.1 | 533.0 | 1.4e-150 |
| CBF83139.1 | polyketide\_synthase,\_putative\_(JCVI) | BGC0001722 | Polyketide | 27.0 | 113.2 | 530.0 | 9.3e-150 |
| EAA59563.1 | polyketide\_synthase | BGC0000057 | Polyketide:Iterative type I | 33.0 | 59.1 | 522.0 | 1.9e-147 |
| EHA28237.1 | hypothetical\_protein | BGC0001143 | Polyketide | 26.0 | 127.7 | 520.0 | 1.3e-146 |
| ABB90282.1 | polyketide\_synthase | BGC0001057 | NRP + Polyketide | 39.0 | 46.0 | 514.0 | 6.9e-145 |
| QCO93110.1 | polyketide\_synthase | BGC0001977 | Other | 24.0 | 137.7 | 476.0 | 1.6e-133 |
| ACD39770.1 | non-reducing\_polyketide\_synthase | BGC0000134 | Polyketide | 36.0 | 46.5 | 475.0 | 4.7e-133 |
| EED18001.1 | NR-PKS | BGC0000154 | Polyketide:Iterative type I | 24.0 | 137.9 | 465.0 | 2.8e-130 |
| AGN71604.1 | conidial\_yellow\_pigment\_biosynthesis\_polyketide\_synthase | BGC0000027 | Polyketide:Iterative type I | 52.0 | 25.9 | 449.0 | 2.1e-125 |
| ADH01663.1 | putative\_polyketide\_synthase\_PKS3 | BGC0000099 | Polyketide | 50.0 | 25.6 | 420.0 | 1.8e-116 |
| CAP95404.1 |  | BGC0001404 | Polyketide | 23.0 | 125.4 | 414.0 | 9.8e-115 |
| CAQ18828.1 | polyketide\_synthase | BGC0000954 | NRP + Polyketide:Modular type I | 31.0 | 51.8 | 377.0 | 1e-103 |
| AEU11005.1 | NpnA | BGC0001029 | NRP + Polyketide | 31.0 | 50.8 | 373.0 | 1.5e-102 |
| AGC45620.1 | polyketide\_synthase | BGC0001394 | NRP + Polyketide | 31.0 | 51.5 | 373.0 | 1.9e-102 |
| XP\_011392701.1 | hypothetical\_protein | BGC0001281 | Polyketide | 30.0 | 51.8 | 370.0 | 1.2e-101 |
| AQW44888.1 | polyketide\_synthase | BGC0001737 | NRP + Polyketide | 31.0 | 51.4 | 370.0 | 1.2e-101 |
| CAQ34920.1 | polyketide\_synthase | BGC0000986 | NRP + Polyketide | 30.0 | 57.1 | 369.0 | 3.6e-101 |
| CBD77748.1 | polyketide\_synthase | BGC0000974 | NRP + Polyketide | 30.0 | 56.0 | 366.0 | 3e-100 |
| AIR74917.1 | polyketide\_synthase | BGC0001559 | RiPP | 30.0 | 56.0 | 366.0 | 3e-100 |
| EAA65602.1 | hypothetical\_protein | BGC0000022 | Polyketide | 45.0 | 26.3 | 365.0 | 4e-100 |
| CAQ18834.1 | polyketide\_synthase | BGC0000954 | NRP + Polyketide:Modular type I | 32.0 | 50.7 | 362.0 | 3.4e-99 |
| AQA28562.1 | type\_I\_polyketide\_synthase | BGC0001663 | Polyketide | 29.0 | 57.9 | 362.0 | 3.4e-99 |
| ACR33078.1 | polyketide\_synthase | BGC0000017 | Alkaloid + Polyketide:Modular type I | 29.0 | 57.8 | 361.0 | 9.8e-99 |
| CAD19087.1 | StiC\_protein | BGC0000153 | NRP + Polyketide:Modular type I | 30.0 | 56.8 | 359.0 | 2.2e-98 |
| AHB82053.1 | polyketide\_synthase | BGC0001019 | NRP + Polyketide:Modular type I | 31.0 | 51.4 | 358.0 | 4.9e-98 |
| EJP62792.1 | polyketide\_synthase | BGC0001720 | Polyketide | 33.0 | 48.9 | 358.0 | 6.4e-98 |
| ADN13832.1 | Polyketide\_Synthase | BGC0001164 | Polyketide:Modular type I | 30.0 | 51.5 | 357.0 | 1.1e-97 |
| CBD77736.1 | polyketide\_synthase | BGC0000974 | NRP + Polyketide | 30.0 | 56.5 | 357.0 | 1.4e-97 |
| AIR74912.1 | polyketide\_synthase | BGC0001559 | RiPP | 30.0 | 56.5 | 357.0 | 1.4e-97 |
| AWM95789.1 | non-reduciing\_polyketide\_synthase\_methylorcinaldehyde\_synthase | BGC0001827 | Polyketide | 44.0 | 25.6 | 356.0 | 1.8e-97 |
| ABM21570.1 | crpB | BGC0000975 | NRP + Polyketide | 32.0 | 43.7 | 356.0 | 2.4e-97 |
| ADF88276.1 | polyketide\_synthase | BGC0000981 | NRP + Polyketide | 28.0 | 56.5 | 354.0 | 7e-97 |
| WP\_026247674.1 | type\_I\_polyketide\_synthase | BGC0001332 | NRP + Polyketide | 30.0 | 50.8 | 352.0 | 3.5e-96 |
| AHB82064.1 | polyketide\_synthase | BGC0001231 | NRP + Polyketide:Modular type I | 30.0 | 51.1 | 351.0 | 7.8e-96 |
| AVI26388.1 | polyketide\_synthase | BGC0001800 | NRP + Polyketide | 32.0 | 41.8 | 351.0 | 7.8e-96 |
| ABX60152.1 | polyketide\_synthase | BGC0000978 | NRP + Alkaloid + Polyketide:Modular type I | 28.0 | 56.5 | 351.0 | 1e-95 |
| ACV42478.1 | polyketide\_synthase | BGC0000043 | Polyketide | 30.0 | 54.6 | 349.0 | 2.3e-95 |
| AEE88277.1 | CurM | BGC0000976 | NRP + Polyketide:Modular type I | 30.0 | 54.6 | 349.0 | 2.3e-95 |
| AAT70108.1 | CurM | BGC0001165 | NRP + Polyketide:Modular type I | 30.0 | 54.6 | 349.0 | 2.3e-95 |
| AEE88282.1 | CurH | BGC0000976 | NRP + Polyketide:Modular type I | 29.0 | 56.2 | 349.0 | 3.9e-95 |
| CAD89777.1 | MelF\_protein | BGC0001010 | NRP + Polyketide:Modular type I | 32.0 | 42.6 | 349.0 | 3.9e-95 |
| AAT70103.1 | CurH | BGC0001165 | NRP + Polyketide:Modular type I | 29.0 | 56.2 | 349.0 | 3.9e-95 |
| ABX60153.1 | polyketide\_synthase | BGC0000978 | NRP + Alkaloid + Polyketide:Modular type I | 29.0 | 54.0 | 348.0 | 6.6e-95 |
| ADF88275.1 | polyketide\_synthase | BGC0000981 | NRP + Polyketide | 29.0 | 54.0 | 347.0 | 1.5e-94 |
| AMB48442.1 | polyketide\_synthase | BGC0001357 | Polyketide | 29.0 | 64.0 | 346.0 | 3.3e-94 |
| QDA77058.1 | polyketide\_synthase | BGC0002026 | NRP | 29.0 | 56.2 | 345.0 | 5.6e-94 |
| CAD19093.1 | StiJ\_protein | BGC0000153 | NRP + Polyketide:Modular type I | 30.0 | 51.1 | 343.0 | 1.6e-93 |
| AFU82617.1 | polyketide\_synthase | BGC0000998 | NRP + Polyketide | 30.0 | 51.8 | 343.0 | 1.6e-93 |
| AXN93613.1 | PuwE | BGC0001953 | NRP | 29.0 | 53.9 | 341.0 | 6.2e-93 |
| AHA12079.1 | polyketide\_synthase\_type\_1 | BGC0001172 | NRP + Polyketide:Modular type I | 31.0 | 51.0 | 341.0 | 8e-93 |
| AZH23819.1 | MgiR | BGC0001971 | NRP + Polyketide | 31.0 | 44.0 | 341.0 | 8e-93 |
| AVI26389.1 | polyketide\_synthase | BGC0001800 | NRP + Polyketide | 31.0 | 51.1 | 341.0 | 1e-92 |
| AIW82282.1 | PuwE | BGC0001125 | NRP + Polyketide | 32.0 | 44.1 | 340.0 | 1.4e-92 |
| AXN93601.1 | PuwE | BGC0001952 | NRP | 29.0 | 54.1 | 339.0 | 2.3e-92 |
| WP\_035121546.1 | type\_I\_polyketide\_synthase | BGC0001467 | NRP:Cyclic depsipeptide + Polyketide:Modular type I | 31.0 | 47.7 | 339.0 | 4e-92 |
| ATP76242.1 | NdaC | BGC0001705 | NRP + Polyketide | 32.0 | 44.2 | 337.0 | 8.9e-92 |
| CAD19086.1 | StiB\_protein | BGC0000153 | NRP + Polyketide:Modular type I | 28.0 | 53.4 | 337.0 | 1.5e-91 |
| TGZ15168.1 | hypothetical\_protein | BGC0002032 | Polyketide | 29.0 | 52.4 | 336.0 | 3.4e-91 |
| DAB41916.1 | ArzN\_-\_PKS\_(KS,\_AT,\_OMT,\_KR,\_ACP) | BGC0001884 | NRP + Polyketide | 32.0 | 41.8 | 335.0 | 5.8e-91 |
| CAQ34919.1 | polyketide\_synthase | BGC0000986 | NRP + Polyketide | 30.0 | 51.2 | 334.0 | 7.5e-91 |
| AQA28563.1 | type\_I\_polyketide\_synthase | BGC0001663 | Polyketide | 33.0 | 41.9 | 334.0 | 1.3e-90 |
| CAD19090.1 | StiF\_protein | BGC0000153 | NRP + Polyketide:Modular type I | 31.0 | 50.4 | 333.0 | 2.2e-90 |
| ctg1\_orf16 |  | BGC0001457 | NRP | 29.0 | 51.8 | 333.0 | 2.2e-90 |
| AAW03329.1 | CtaF | BGC0000982 | NRP + Polyketide | 28.0 | 50.9 | 332.0 | 2.9e-90 |
| AEE88289.1 | CurA | BGC0000976 | NRP + Polyketide:Modular type I | 32.0 | 44.1 | 332.0 | 3.7e-90 |
| AAT70096.1 | CurA | BGC0001165 | NRP + Polyketide:Modular type I | 32.0 | 44.1 | 332.0 | 3.7e-90 |
| AQH32483.1 | hybrid\_peptide\_synthetase/polyketide\_synthase | BGC0001667 | NRP + Polyketide | 31.0 | 44.5 | 332.0 | 3.7e-90 |
| ACN69988.1 | polyketide\_synthase | BGC0000079 | Polyketide | 30.0 | 51.5 | 332.0 | 4.9e-90 |
| AZH23817.1 | MgiQ | BGC0001971 | NRP + Polyketide | 30.0 | 48.9 | 332.0 | 4.9e-90 |
| AAS98782.1 | polyketide\_synthase | BGC0001001 | NRP + Polyketide | 30.0 | 44.7 | 331.0 | 6.4e-90 |
| QDA77059.1 | polyketide\_synthase/nonribosomal\_peptide\_synthetase | BGC0002026 | NRP | 30.0 | 51.6 | 331.0 | 6.4e-90 |
| AZH23791.1 | MgcH | BGC0001970 | NRP + Polyketide | 28.0 | 53.7 | 331.0 | 1.1e-89 |
| CAD19092.1 | StiH\_protein | BGC0000153 | NRP + Polyketide:Modular type I | 30.0 | 51.2 | 330.0 | 1.4e-89 |
| AZF85946.1 | type\_I\_polyketide\_synthase | BGC0001963 | NRP + Polyketide | 30.0 | 52.4 | 330.0 | 1.4e-89 |
| ACR50791.1 | putative\_polyketide\_synthase | BGC0000163 | Polyketide | 28.0 | 51.2 | 330.0 | 1.9e-89 |
| AAK57187.1 | MxaC | BGC0001022 | NRP + Polyketide | 30.0 | 53.2 | 329.0 | 2.4e-89 |
| AEE88283.1 | CurG | BGC0000976 | NRP + Polyketide:Modular type I | 29.0 | 52.8 | 329.0 | 3.2e-89 |
| AAT70102.1 | CurG | BGC0001165 | NRP + Polyketide:Modular type I | 29.0 | 52.8 | 329.0 | 3.2e-89 |
| AZH23823.1 | MgiK | BGC0001971 | NRP + Polyketide | 32.0 | 41.8 | 329.0 | 3.2e-89 |
| AAF19814.1 | MtaF | BGC0001024 | NRP + Polyketide:Modular type I | 32.0 | 40.9 | 329.0 | 4.1e-89 |
| DAB41918.1 | ArzP\_-\_PKS\_(KS,\_AT,\_OMT,\_ACP,\_TE) | BGC0001884 | NRP + Polyketide | 30.0 | 52.2 | 328.0 | 5.4e-89 |
| CBD77738.1 | polyketide\_synthase | BGC0000974 | NRP + Polyketide | 30.0 | 51.8 | 328.0 | 7e-89 |
| AAF26923.1 | polyketide\_synthase | BGC0000988 | NRP + Polyketide | 28.0 | 58.8 | 328.0 | 7e-89 |
| AIR74913.1 | polyketide\_synthase | BGC0001559 | RiPP | 30.0 | 51.8 | 328.0 | 7e-89 |
| BBF25315.1 | polyketide\_synthase | BGC0001923 | Terpene + Polyketide | 29.0 | 56.3 | 328.0 | 7e-89 |
| AID65222.1 | putative\_aspartate\_racemase | BGC0000335 | NRP | 34.0 | 40.7 | 327.0 | 1.2e-88 |
| ADZ24998.1 | polyketide\_synthase | BGC0000380 | NRP + Polyketide:Modular type I | 29.0 | 51.8 | 326.0 | 2.7e-88 |
| AEU11006.1 | NpnB | BGC0001029 | NRP + Polyketide | 32.0 | 41.2 | 326.0 | 3.5e-88 |
| AIW82279.1 | PuwB | BGC0001125 | NRP + Polyketide | 31.0 | 48.9 | 326.0 | 3.5e-88 |
| ACB46197.1 | polyketide\_synthase | BGC0000989 | NRP + Polyketide | 28.0 | 58.8 | 325.0 | 6e-88 |
| ADB12493.1 | EpoF | BGC0000990 | NRP + Polyketide | 28.0 | 58.8 | 325.0 | 6e-88 |
| CAL58685.1 | polyketide\_synthase | BGC0000149 | Polyketide:Modular type I | 29.0 | 50.7 | 323.0 | 2.3e-87 |
| AZH23793.1 | MgcK | BGC0001970 | NRP + Polyketide | 29.0 | 46.6 | 323.0 | 2.3e-87 |
| ABX60161.1 | mixed\_NRPS/PKS | BGC0000978 | NRP + Alkaloid + Polyketide:Modular type I | 30.0 | 48.9 | 322.0 | 3e-87 |
| AQW44892.1 | polyketide\_synthase | BGC0001737 | NRP + Polyketide | 31.0 | 44.5 | 322.0 | 3e-87 |
| AAF62885.1 | EpoF | BGC0000991 | NRP + Polyketide | 28.0 | 58.8 | 322.0 | 5e-87 |
| AAO62585.1 | peptide\_sythetase\_polyketide\_synthase\_fusion\_protein | BGC0001016 | NRP + Polyketide | 32.0 | 44.3 | 321.0 | 8.6e-87 |
| ABO15860.1 | polyketide\_synthase | BGC0000130 | Polyketide | 30.0 | 51.8 | 320.0 | 1.5e-86 |
| AZH23821.1 | MgiH | BGC0001971 | NRP + Polyketide | 28.0 | 53.7 | 320.0 | 1.5e-86 |
| ADF88279.1 | mixed\_NRPS/PKS | BGC0000981 | NRP + Polyketide | 30.0 | 48.9 | 320.0 | 1.9e-86 |
| CAD89775.1 | MelD\_protein | BGC0001010 | NRP + Polyketide:Modular type I | 29.0 | 48.0 | 320.0 | 1.9e-86 |
| AAF00958.1 | mcyE | BGC0001017 | NRP + Polyketide:Modular type I | 29.0 | 51.4 | 319.0 | 2.5e-86 |
| AAW03328.1 | CtaE | BGC0000982 | NRP + Polyketide | 32.0 | 41.6 | 319.0 | 4.3e-86 |
| ATY12793.1 | type\_I\_polyketide\_synthase | BGC0001504 | Polyketide | 29.0 | 51.7 | 319.0 | 4.3e-86 |
| AXN93580.1 | PuwE | BGC0001950 | NRP | 30.0 | 48.1 | 318.0 | 7.3e-86 |
| AXN93589.1 | PuwE | BGC0001951 | NRP | 30.0 | 48.1 | 318.0 | 7.3e-86 |
| CAO98879.1 | polyketide\_synthase\_AufD | BGC0000023 | Polyketide:Modular type I | 28.0 | 53.0 | 317.0 | 9.5e-86 |
| ACR33079.1 | polyketide\_synthase | BGC0000017 | Alkaloid + Polyketide:Modular type I | 30.0 | 42.7 | 317.0 | 1.2e-85 |
| AHA38199.1 | GphF | BGC0000069 | Polyketide | 28.0 | 58.4 | 317.0 | 1.2e-85 |
| AAQ90174.1 | polyketide\_synthase\_type\_I | BGC0000128 | Polyketide | 29.0 | 50.6 | 317.0 | 1.2e-85 |
| ACY13415.1 | KR\_domain\_protein | BGC0001367 | NRP + Polyketide | 28.0 | 50.7 | 317.0 | 1.2e-85 |
| AHH99920.1 | PKS\_I | BGC0000002 | Polyketide | 29.0 | 51.6 | 317.0 | 1.6e-85 |
| CAL58686.1 | polyketide\_synthase | BGC0000149 | Polyketide:Modular type I | 28.0 | 55.9 | 317.0 | 1.6e-85 |
| CAQ18833.1 | polyketide\_synthase | BGC0000954 | NRP + Polyketide:Modular type I | 31.0 | 41.2 | 316.0 | 2.1e-85 |
| ABP55493.1 | thioester\_reductase\_domain | BGC0001006 | NRP + Polyketide | 28.0 | 52.0 | 316.0 | 2.1e-85 |
| AAF00959.1 | mcyD | BGC0001017 | NRP + Polyketide:Modular type I | 28.0 | 57.9 | 316.0 | 2.1e-85 |
| AWS21279.1 | type\_I\_polyketide\_synthase | BGC0001934 | Polyketide | 29.0 | 51.2 | 316.0 | 2.1e-85 |
| AZY91989.1 | polyketide\_synthase | BGC0002022 | Polyketide | 29.0 | 51.2 | 316.0 | 2.1e-85 |
| ADY00130.1 | polyketide\_synthase | BGC0000104 | Terpene + Polyketide:Iterative type I | 27.0 | 60.4 | 316.0 | 2.8e-85 |
| AVI57433.1 | AbmB1 | BGC0001694 | Polyketide | 31.0 | 41.3 | 315.0 | 6.2e-85 |
| AVV61984.1 | type\_I\_modular\_polyketide\_synthase | BGC0001477 | NRP + Polyketide:Modular type I | 28.0 | 52.0 | 314.0 | 8.1e-85 |
| AGC45621.1 | polyketide\_synthase | BGC0001394 | NRP + Polyketide | 28.0 | 51.1 | 314.0 | 1.1e-84 |
| EHK80167.1 | modular\_polyketide\_synthase | BGC0001447 | Polyketide | 28.0 | 52.1 | 314.0 | 1.4e-84 |
| AZF85932.1 | type\_I\_polyketide\_synthase | BGC0001963 | NRP + Polyketide | 31.0 | 44.9 | 313.0 | 2.3e-84 |
| AMB48441.1 | polyketide\_synthase | BGC0001357 | Polyketide | 29.0 | 45.1 | 312.0 | 3.1e-84 |
| AXN93597.1 | PuwB | BGC0001952 | NRP | 29.0 | 49.1 | 312.0 | 3.1e-84 |
| AXI91552.1 | FunP1 | BGC0001944 | Polyketide | 28.0 | 53.6 | 312.0 | 4e-84 |
| ACO94484.1 | polyketide\_synthase\_type\_I | BGC0000097 | Polyketide:Modular type I | 28.0 | 52.4 | 312.0 | 5.2e-84 |
| AHA12078.1 | polyketide\_synthase\_type\_1 | BGC0001172 | NRP + Polyketide:Modular type I | 32.0 | 41.5 | 312.0 | 5.2e-84 |
| AHH34186.1 | polyketide\_synthase | BGC0001161 | Polyketide:Modular type I | 30.0 | 42.5 | 311.0 | 6.8e-84 |
| AAF19813.1 | MtaE | BGC0001024 | NRP + Polyketide:Modular type I | 33.0 | 41.8 | 311.0 | 8.9e-84 |
| AAZ77693.1 | ChlA1 | BGC0000036 | Polyketide:Modular type I + Polyketide:Iterative type I + Saccharide:Oligosaccharide | 27.0 | 52.9 | 311.0 | 1.2e-83 |
| AEE88284.1 | CurF | BGC0000976 | NRP + Polyketide:Modular type I | 29.0 | 49.5 | 310.0 | 1.5e-83 |
| AFI57007.1 | QmnA3 | BGC0000133 | Polyketide | 29.0 | 51.5 | 310.0 | 2e-83 |
| CDM36726.1 | Beta-ketoacyl\_synthase | BGC0001360 | Polyketide | 28.0 | 58.3 | 310.0 | 2e-83 |
| AZH23787.1 | MgcQ | BGC0001970 | NRP + Polyketide | 30.0 | 43.9 | 309.0 | 3.4e-83 |
| AGC45623.1 | polyketide\_synthase | BGC0001394 | NRP + Polyketide | 30.0 | 46.5 | 309.0 | 4.4e-83 |
| AAT70101.1 | CurF | BGC0001165 | NRP + Polyketide:Modular type I | 29.0 | 49.5 | 308.0 | 5.8e-83 |
| AAS98781.1 | polyketide\_synthase | BGC0001001 | NRP + Polyketide | 29.0 | 50.0 | 307.0 | 9.8e-83 |
| AVX51098.1 | nysI | BGC0001709 | Polyketide | 28.0 | 54.7 | 307.0 | 9.8e-83 |
| AXG22406.1 | type\_I\_polyketide\_synthase | BGC0002024 | Polyketide | 28.0 | 52.0 | 307.0 | 9.8e-83 |
| CAL58687.1 | polyketide\_synthase | BGC0000149 | Polyketide:Modular type I | 28.0 | 52.9 | 307.0 | 1.3e-82 |
| ABB88521.1 | polyketide\_synthase\_type\_I | BGC0000050 | Polyketide | 29.0 | 51.6 | 307.0 | 1.7e-82 |
| CAD29793.1 | polyketide\_synthase\_type\_I | BGC0001015 | NRP + Polyketide | 27.0 | 58.6 | 307.0 | 1.7e-82 |
| CAD29794.1 | peptide\_synthetase | BGC0001015 | NRP + Polyketide | 30.0 | 50.9 | 307.0 | 1.7e-82 |
| AHH34189.1 | polyketide\_synthase | BGC0001162 | Polyketide:Modular type I | 30.0 | 43.5 | 307.0 | 1.7e-82 |
| ctg1\_orf27 |  | BGC0000096 | Polyketide | 30.0 | 52.7 | 306.0 | 2.9e-82 |
| ACA99172.1 | polyketide\_synthase | BGC0001160 | Polyketide:Modular type I | 29.0 | 50.5 | 306.0 | 2.9e-82 |
| ADC79639.1 | TamAIII | BGC0001052 | NRP + Polyketide:Modular type I | 30.0 | 46.7 | 305.0 | 3.7e-82 |
| AEH42490.1 | polyketide\_synthase | BGC0000032 | Polyketide | 28.0 | 51.3 | 305.0 | 4.9e-82 |
| AAO62582.1 | polyketide\_synthase\_peptide\_sythetase\_fusion\_protein | BGC0001016 | NRP + Polyketide | 30.0 | 54.5 | 305.0 | 6.4e-82 |
| ADH04639.1 | TgaA | BGC0001051 | NRP + Polyketide:Modular type I | 31.0 | 42.2 | 305.0 | 6.4e-82 |
| AFV96142.1 | polyketide\_synthase | BGC0001064 | Polyketide:Modular type I + Polyketide:Type III | 29.0 | 41.5 | 305.0 | 6.4e-82 |
| ARU81122.1 | CylH | BGC0001566 | Polyketide | 29.0 | 41.5 | 305.0 | 6.4e-82 |
| AEH42491.1 | polyketide\_synthase | BGC0000032 | Polyketide | 28.0 | 50.9 | 304.0 | 8.3e-82 |
| AFI57005.1 | QmnA1 | BGC0000133 | Polyketide | 29.0 | 50.3 | 304.0 | 8.3e-82 |
| BAC68129.1 | modular\_polyketide\_synthase | BGC0000059 | Polyketide | 28.0 | 51.7 | 304.0 | 1.1e-81 |
| CAQ34928.1 | polyketide\_synthase | BGC0000986 | NRP + Polyketide | 30.0 | 46.9 | 304.0 | 1.1e-81 |
| ADA69241.1 | cis-AT\_polyketide\_synthase | BGC0001071 | NRP + Polyketide:Modular type I + Polyketide:Trans-AT type I | 29.0 | 51.4 | 304.0 | 1.1e-81 |
| BAC68126.1 | modular\_polyketide\_synthase | BGC0000059 | Polyketide | 27.0 | 56.1 | 304.0 | 1.4e-81 |
| AFL48528.1 | laidlomycin\_polyketide\_synthase\_(module\_7\_and\_module\_8) | BGC0000084 | Polyketide | 29.0 | 51.2 | 304.0 | 1.4e-81 |
| ADM46358.1 | polyketide\_synthase | BGC0000106 | Polyketide | 28.0 | 51.1 | 304.0 | 1.4e-81 |
| ANI24099.1 | polyketide\_synthase | BGC0001235 | NRP + Polyketide | 28.0 | 51.4 | 304.0 | 1.4e-81 |
| EHK80166.1 | beta-ketoacyl\_synthase | BGC0001447 | Polyketide | 28.0 | 51.8 | 304.0 | 1.4e-81 |
| AVV61985.1 | beta-ketoacyl\_synthase | BGC0001477 | NRP + Polyketide:Modular type I | 29.0 | 51.4 | 304.0 | 1.4e-81 |
| ANH11409.1 | SceN | BGC0001908 | Polyketide | 28.0 | 52.8 | 304.0 | 1.4e-81 |
| ACC40922.1 | polyketide\_synthase,\_Pks8 | BGC0001665 | Polyketide | 27.0 | 55.7 | 303.0 | 1.9e-81 |
| AAS98784.1 | polyketide\_synthase | BGC0001001 | NRP + Polyketide | 30.0 | 44.7 | 302.0 | 3.2e-81 |
| ATP76239.1 | NdaF | BGC0001705 | NRP + Polyketide | 29.0 | 53.5 | 302.0 | 3.2e-81 |
| CBD77746.1 | non-ribosomal\_peptide\_synthetase/polyketide\_synthase | BGC0000974 | NRP + Polyketide | 31.0 | 41.2 | 302.0 | 4.1e-81 |
| AIR74926.1 | polyketide\_synthase | BGC0001559 | RiPP | 31.0 | 41.2 | 302.0 | 4.1e-81 |
| BAO66529.1 | type\_I\_polyketide\_synthase | BGC0000042 | Polyketide | 27.0 | 51.2 | 302.0 | 5.4e-81 |
| ABB88519.1 | polyketide\_synthase\_type\_I | BGC0000050 | Polyketide | 29.0 | 52.7 | 302.0 | 5.4e-81 |
| sipP3 | Type\_I\_Modular\_PKS | BGC0001452 | Polyketide | 27.0 | 57.0 | 302.0 | 5.4e-81 |
| AEU17899.1 | putative\_type\_I\_PKS | BGC0001072 | Saccharide + Polyketide:Modular type I + Polyketide:Type II + Other:Aminocoumarin | 29.0 | 51.5 | 301.0 | 9.2e-81 |
| BAP34733.1 | type\_I\_polyketide\_synthase | BGC0000078 | Polyketide | 27.0 | 56.2 | 300.0 | 1.2e-80 |
| AAG23263.1 | polyketide\_synthase\_extender\_modules\_5-7 | BGC0000148 | Polyketide | 28.0 | 53.4 | 300.0 | 1.6e-80 |
| AXG22405.1 | type\_I\_polyketide\_synthase | BGC0002024 | Polyketide | 28.0 | 51.4 | 300.0 | 1.6e-80 |
| AAW03327.1 | CtaD | BGC0000982 | NRP + Polyketide | 28.0 | 47.6 | 300.0 | 2.1e-80 |
| CAD89776.1 | MelE\_protein | BGC0001010 | NRP + Polyketide:Modular type I | 31.0 | 41.7 | 300.0 | 2.1e-80 |
| BBA66512.1 | type\_I\_polyketide\_synthase | BGC0001495 | Polyketide | 28.0 | 52.3 | 300.0 | 2.1e-80 |
| AZF85945.1 | type\_I\_polyketide\_synthase | BGC0001963 | NRP + Polyketide | 30.0 | 47.1 | 300.0 | 2.1e-80 |
| ADU85988.1 | putative\_iterative\_type\_I\_polyketide\_synthase | BGC0000165 | Polyketide:Modular type I | 29.0 | 51.1 | 299.0 | 2.7e-80 |
| CAD15508.1 | polyketide\_synthase/non-ribosomal\_peptide\_synthetase | BGC0001014 | NRP:NRP siderophore + Polyketide:Modular type I + Polyketide:Iterative type I | 29.0 | 51.8 | 299.0 | 2.7e-80 |
| ALP32042.1 | CycB | BGC0001293 | Polyketide | 27.0 | 53.3 | 299.0 | 2.7e-80 |
| CAD29795.1 | peptide\_synthetase | BGC0001015 | NRP + Polyketide | 30.0 | 45.4 | 299.0 | 3.5e-80 |
| BAJ16467.1 | polyketide\_synthase | BGC0000058 | Polyketide | 28.0 | 52.9 | 298.0 | 7.8e-80 |
| EGJ35088.1 | Polyketide\_synthase | BGC0001163 | Polyketide:Modular type I | 30.0 | 43.9 | 297.0 | 1e-79 |
| ctg1\_orf7 |  | BGC0000053 | Polyketide | 31.0 | 40.8 | 297.0 | 1.3e-79 |
| AEZ64504.1 | Herc | BGC0001065 | Polyketide | 31.0 | 41.7 | 297.0 | 1.3e-79 |
| AAD03048.1 | type\_I\_polyketide\_synthase | BGC0000041 | Polyketide | 29.0 | 53.9 | 297.0 | 1.7e-79 |
| AAX98189.1 | polyketide\_synthase\_type\_I | BGC0000052 | Polyketide | 28.0 | 51.6 | 297.0 | 1.7e-79 |
| BAO66543.1 | type\_I\_polyketide\_synthase | BGC0000042 | Polyketide | 27.0 | 52.3 | 296.0 | 2.3e-79 |
| BAD08360.1 | polyketide\_synthase\_modules\_7-8 | BGC0000167 | Polyketide | 30.0 | 41.5 | 296.0 | 2.3e-79 |
| AXN93586.1 | PuwB | BGC0001951 | NRP | 28.0 | 52.1 | 296.0 | 2.3e-79 |
| AXN93577.1 | PuwB | BGC0001950 | NRP | 28.0 | 52.1 | 296.0 | 3e-79 |
| AAF26920.1 | polyketide\_synthase | BGC0000988 | NRP + Polyketide | 28.0 | 48.9 | 295.0 | 5.1e-79 |
| ADB12491.1 | EpoD | BGC0000990 | NRP + Polyketide | 29.0 | 47.7 | 295.0 | 6.6e-79 |
| AAF62883.1 | epoD | BGC0000991 | NRP + Polyketide | 29.0 | 47.7 | 295.0 | 6.6e-79 |
| BAB69194.1 | modular\_polyketide\_synthase | BGC0000117 | Polyketide | 28.0 | 51.5 | 294.0 | 8.6e-79 |
| AWR88399.1 | putative\_beta-ketoacyl\_synthase | BGC0001522 | Polyketide | 26.0 | 56.8 | 294.0 | 1.1e-78 |
| ACB46195.1 | polyketide\_synthase | BGC0000989 | NRP + Polyketide | 29.0 | 47.7 | 294.0 | 1.5e-78 |
| WP\_055480220.1 | type\_I\_polyketide\_synthase | BGC0001653 | Polyketide | 28.0 | 52.1 | 294.0 | 1.5e-78 |
| AAF62882.1 | EpoC | BGC0000991 | NRP + Polyketide | 28.0 | 48.9 | 293.0 | 1.9e-78 |
| ADC79637.1 | TamAI | BGC0001052 | NRP + Polyketide:Modular type I | 29.0 | 49.2 | 293.0 | 1.9e-78 |
| AUO16401.1 | polyketide\_synthase | BGC0001700 | Polyketide | 30.0 | 41.9 | 293.0 | 1.9e-78 |
| AXG22407.1 | type\_I\_polyketide\_synthase | BGC0002024 | Polyketide | 27.0 | 55.5 | 293.0 | 1.9e-78 |
| BAO66539.1 | type\_I\_polyketide\_synthase | BGC0000042 | Polyketide | 28.0 | 55.0 | 293.0 | 2.5e-78 |
| AHB82070.1 | polyketide\_synthase | BGC0001231 | NRP + Polyketide:Modular type I | 28.0 | 50.8 | 293.0 | 2.5e-78 |
| AAP42855.1 | NanA1 | BGC0000105 | Polyketide | 30.0 | 43.2 | 292.0 | 3.3e-78 |
| TGZ15167.1 | polyketide\_synthase | BGC0002032 | Polyketide | 31.0 | 42.1 | 292.0 | 3.3e-78 |
| AHH99921.1 | PKS\_I | BGC0000002 | Polyketide | 26.0 | 56.9 | 292.0 | 4.3e-78 |
| ABM21569.1 | crpA | BGC0000975 | NRP + Polyketide | 30.0 | 41.8 | 292.0 | 5.6e-78 |
| ADB12490.1 | EpoC | BGC0000990 | NRP + Polyketide | 28.0 | 48.9 | 292.0 | 5.6e-78 |
| AAP42857.1 | NanA3 | BGC0000105 | Polyketide | 28.0 | 48.5 | 291.0 | 7.3e-78 |
| CAQ34918.1 | nonribosomal\_peptide\_synthetase/\_polyketide\_synthase | BGC0000986 | NRP + Polyketide | 28.0 | 46.4 | 291.0 | 7.3e-78 |
| AAF26921.1 | polyketide\_synthase | BGC0000988 | NRP + Polyketide | 29.0 | 47.7 | 291.0 | 7.3e-78 |
| AFU82616.1 | polyketide\_synthase | BGC0000998 | NRP + Polyketide | 27.0 | 51.5 | 291.0 | 7.3e-78 |
| CAA16183.1 | polyketide\_synthase | BGC0001063 | NRP + Polyketide | 28.0 | 57.4 | 291.0 | 7.3e-78 |
| AQH32482.1 | type\_1\_polyketide\_synthase | BGC0001667 | NRP + Polyketide | 27.0 | 58.3 | 291.0 | 7.3e-78 |
| ABK32257.1 | AmbC | BGC0000014 | Polyketide | 28.0 | 53.0 | 290.0 | 1.2e-77 |
| AXI91546.1 | FunP7 | BGC0001944 | Polyketide | 28.0 | 50.7 | 290.0 | 1.6e-77 |
| AAC38076.1 | polyketide\_synthase\_type\_I | BGC0000127 | Polyketide | 29.0 | 41.2 | 290.0 | 2.1e-77 |
| ABJ97437.1 | MerA | BGC0001012 | NRP + Polyketide | 29.0 | 44.4 | 289.0 | 2.8e-77 |
| ABV83221.1 | CppI | BGC0000116 | Polyketide | 28.0 | 51.6 | 289.0 | 3.6e-77 |
| BAP34734.1 | type\_I\_polyketide\_synthase | BGC0000078 | Polyketide | 27.0 | 56.4 | 289.0 | 4.7e-77 |
| ABK32289.1 | JerC | BGC0000080 | Polyketide | 28.0 | 51.8 | 289.0 | 4.7e-77 |
| CAD19085.1 | StiA\_protein | BGC0000153 | NRP + Polyketide:Modular type I | 29.0 | 50.6 | 289.0 | 4.7e-77 |
| AVI26390.1 | polyketide\_synthase\_/\_nonribosomal\_peptide\_synthase\_hybrid | BGC0001800 | NRP + Polyketide | 27.0 | 51.3 | 289.0 | 4.7e-77 |
| ABO15888.1 | polyketide\_synthase | BGC0000132 | Polyketide | 31.0 | 42.3 | 288.0 | 6.2e-77 |
| ABI94379.1 | tautomycetin\_biosynthetic\_PKS | BGC0000157 | Polyketide | 28.0 | 53.7 | 288.0 | 6.2e-77 |
| WP\_106731933.1 | type\_I\_polyketide\_synthase | BGC0001332 | NRP + Polyketide | 29.0 | 52.2 | 288.0 | 6.2e-77 |
| AQW44891.1 | polyketide\_synthase | BGC0001737 | NRP + Polyketide | 30.0 | 41.1 | 288.0 | 6.2e-77 |
| ABP55220.1 | beta-ketoacyl\_synthase | BGC0000142 | Polyketide | 27.0 | 57.0 | 288.0 | 8.1e-77 |
| ABC84456.1 | NigAI | BGC0000114 | Polyketide:Modular type I | 30.0 | 41.9 | 287.0 | 1.1e-76 |
| CAO98850.1 | polyketide\_synthase\_AufG | BGC0000023 | Polyketide:Modular type I | 31.0 | 42.0 | 287.0 | 1.4e-76 |
| AAQ82567.1 | FscE | BGC0000061 | Polyketide | 28.0 | 51.7 | 287.0 | 1.4e-76 |
| ABV91286.1 | type\_I\_modular\_polyketide\_synthase | BGC0000158 | Polyketide:Modular type I | 28.0 | 53.5 | 287.0 | 1.4e-76 |
| ALA09371.1 | type\_I\_modular\_PKS | BGC0001303 | Polyketide | 28.0 | 52.7 | 287.0 | 1.4e-76 |
| CAL58684.1 | polyketide\_synthase | BGC0000149 | Polyketide:Modular type I | 27.0 | 51.2 | 287.0 | 1.8e-76 |
| ctg1\_orf15 |  | BGC0001457 | NRP | 29.0 | 50.5 | 287.0 | 1.8e-76 |
| OJF16266.1 | AceP4 | BGC0001491 | Polyketide | 29.0 | 50.6 | 287.0 | 1.8e-76 |
| ACO94472.1 | polyketide\_synthase\_type\_I | BGC0000029 | Polyketide:Modular type I | 27.0 | 50.9 | 286.0 | 3.1e-76 |
| APZ78832.1 | polyketide\_synthase | BGC0001430 | NRP:Cyclic depsipeptide + Polyketide:Iterative type I | 29.0 | 46.3 | 286.0 | 3.1e-76 |
| OAP25821.1 | Phenolphthiocerol\_synthesis\_polyketide\_synthase\_type\_I\_Pks15/1 | BGC0001658 | Polyketide | 27.0 | 50.6 | 286.0 | 3.1e-76 |
| BAQ21939.1 | putative\_type\_I\_polyketide\_synthase | BGC0001204 | Polyketide | 30.0 | 45.4 | 285.0 | 4e-76 |
| AWR88393.1 | putative\_beta-ketoacyl\_synthase | BGC0001522 | Polyketide | 26.0 | 55.4 | 285.0 | 4e-76 |
| AAZ94386.1 | modular\_polyketide\_synthase | BGC0000040 | Polyketide | 31.0 | 41.6 | 285.0 | 5.2e-76 |
| ABC87509.1 | polyketide\_synthase | BGC0001011 | NRP + Polyketide | 29.0 | 46.2 | 285.0 | 5.2e-76 |
| ctg1\_orf20 |  | BGC0001013 | NRP + Polyketide | 29.0 | 46.2 | 285.0 | 5.2e-76 |
| BAT51065.1 | type\_I\_polyketide\_synthase | BGC0001296 | Polyketide | 31.0 | 41.5 | 285.0 | 5.2e-76 |
| AFL48525.1 | laidlomycin\_polyketide\_synthase\_(loading\_module\_and\_module\_1) | BGC0000084 | Polyketide | 28.0 | 51.8 | 284.0 | 8.9e-76 |
| BAG85026.1 | putative\_polyketide\_synthase | BGC0000086 | Polyketide | 27.0 | 52.6 | 284.0 | 8.9e-76 |
| CAQ64686.1 | lasalocid\_modular\_polyketide\_synthase | BGC0000087 | Polyketide | 27.0 | 52.6 | 284.0 | 8.9e-76 |
| sipP2 | Type\_I\_Modular\_PKS | BGC0001452 | Polyketide | 27.0 | 52.9 | 284.0 | 8.9e-76 |
| ctg1\_15 |  | BGC0001931 | Polyketide | 28.0 | 51.7 | 284.0 | 8.9e-76 |
| AAK57188.1 | MxaD | BGC0001022 | NRP + Polyketide | 29.0 | 41.2 | 284.0 | 1.2e-75 |
| AIT55261.1 | polyketide\_synthase | BGC0000072 | Polyketide:Modular type I | 29.0 | 50.9 | 284.0 | 1.5e-75 |
| AGY62753.1 | EbeA | BGC0000051 | Polyketide | 28.0 | 44.8 | 283.0 | 2e-75 |
| SCN11949.1 | ebeA-type\_I\_polyketide\_synthase\_KSQ-ATa-ACP | BGC0001580 | Polyketide | 28.0 | 44.8 | 283.0 | 2e-75 |
| WP\_083502114.1 | type\_I\_polyketide\_synthase | BGC0001653 | Polyketide | 28.0 | 51.1 | 283.0 | 2e-75 |
| AWR88405.1 | putative\_phosphopantetheine-binding\_domain-\_containing\_prot\_ein | BGC0001522 | Polyketide | 27.0 | 52.6 | 283.0 | 2.6e-75 |
| CAE46843.1 | Type\_I\_modular\_polyketide\_synthase | BGC0000103 | Polyketide | 30.0 | 42.6 | 282.0 | 3.4e-75 |
| CAE46851.1 | Type\_I\_modular\_polyketide\_synthase | BGC0000103 | Polyketide | 30.0 | 42.6 | 282.0 | 3.4e-75 |
| BAR73020.1 | putative\_PKS\_(KS-AT-DH-KR-ACP-KS-AT-DH-KR-ACP-KS-AT-DH-KR-ACP) | BGC0001194 | Polyketide | 30.0 | 40.9 | 282.0 | 3.4e-75 |
| sipP1 | Type\_I\_Modular\_PKS | BGC0001452 | Polyketide | 27.0 | 52.9 | 282.0 | 3.4e-75 |
| ACC40923.1 | polyketide\_synthase\_Pks9 | BGC0001665 | Polyketide | 29.0 | 43.2 | 282.0 | 3.4e-75 |
| AAG23262.1 | polyketide\_synthase\_extender\_modules\_8-10 | BGC0000148 | Polyketide | 29.0 | 41.5 | 282.0 | 4.4e-75 |
| AIT55263.1 | polyketide\_synthase | BGC0000072 | Polyketide:Modular type I | 30.0 | 42.0 | 282.0 | 5.8e-75 |
| ADZ24995.1 | non-ribosomal\_peptide\_synthase/polyketide\_synthase | BGC0000380 | NRP + Polyketide:Modular type I | 28.0 | 49.6 | 282.0 | 5.8e-75 |
| AKD43765.1 | HerE | BGC0001349 | NRP + Polyketide | 27.0 | 51.1 | 282.0 | 5.8e-75 |
| CAJ88175.1 | putative\_polyketide\_synthase\_B | BGC0000151 | Polyketide:Modular type I + Saccharide:Hybrid/tailoring | 27.0 | 51.5 | 280.0 | 1.7e-74 |
| AFU82615.1 | polyketide\_synthase | BGC0000998 | NRP + Polyketide | 27.0 | 51.1 | 280.0 | 1.7e-74 |
| APZ78742.1 | polyketide\_synthase | BGC0001422 | NRP:Cyclic depsipeptide + Polyketide:Iterative type I | 29.0 | 41.7 | 280.0 | 1.7e-74 |
| AWR88404.1 | putative\_beta-ketoacyl\_synthase | BGC0001522 | Polyketide | 27.0 | 52.8 | 280.0 | 2.2e-74 |
| BAR73007.1 | putative\_PKS\_(ACP-KS-AT-DH-KR-ACP-KS-AT-DH-ER-KR-ACP) | BGC0001194 | Polyketide | 28.0 | 51.9 | 279.0 | 2.9e-74 |
| ALA09355.1 | type\_I\_modular\_PKS | BGC0001303 | Polyketide | 28.0 | 51.3 | 279.0 | 2.9e-74 |
| WP\_035122279.1 | type\_I\_polyketide\_synthase | BGC0001467 | NRP:Cyclic depsipeptide + Polyketide:Modular type I | 27.0 | 50.9 | 279.0 | 2.9e-74 |
| ABK32288.1 | JerB | BGC0000080 | Polyketide | 30.0 | 41.5 | 279.0 | 3.7e-74 |
| AZF85947.1 | type\_I\_polyketide\_synthase | BGC0001963 | NRP + Polyketide | 30.0 | 42.4 | 279.0 | 3.7e-74 |
| AAQ84145.1 | Plm5 | BGC0000123 | Polyketide | 26.0 | 57.7 | 279.0 | 4.9e-74 |
| QBF51757.1 | type\_I\_polyketide\_synthase | BGC0001856 | Polyketide:Modular type I | 30.0 | 41.6 | 278.0 | 6.4e-74 |
| ABK32255.1 | AmbA | BGC0000014 | Polyketide | 27.0 | 57.4 | 278.0 | 8.4e-74 |
| ACO94468.1 | polyketide\_synthase\_type\_I | BGC0000029 | Polyketide:Modular type I | 26.0 | 50.9 | 278.0 | 8.4e-74 |
| AAG23265.1 | polyketide\_synthase\_extender\_module\_2 | BGC0000148 | Polyketide | 28.0 | 41.7 | 278.0 | 8.4e-74 |
| AFD30954.1 | CrmA | BGC0000966 | NRP + Polyketide | 31.0 | 41.7 | 278.0 | 8.4e-74 |
| AKD43761.1 | HerD | BGC0001349 | NRP + Polyketide | 27.0 | 51.0 | 278.0 | 8.4e-74 |
| ACB46194.1 | polyketide\_synthase | BGC0000989 | NRP + Polyketide | 27.0 | 48.9 | 277.0 | 1.1e-73 |
| BAV32159.1 | polyketide\_synthase | BGC0001373 | Polyketide | 28.0 | 52.7 | 277.0 | 1.1e-73 |
| AAU04878.1 | polyketide\_synthase | BGC0000365 | NRP | 28.0 | 41.7 | 277.0 | 1.4e-73 |
| AAG13918.1 | megalomicin\_6-deoxyerythronolide\_B\_synthase\_2 | BGC0000092 | Polyketide | 26.0 | 52.3 | 277.0 | 1.9e-73 |
| ACO94483.1 | polyketide\_synthase\_type\_I | BGC0000097 | Polyketide:Modular type I | 28.0 | 42.0 | 277.0 | 1.9e-73 |
| KFA69335.1 | hypothetical\_protein | BGC0001626 | Polyketide | 27.0 | 48.1 | 277.0 | 1.9e-73 |
| CAQ18832.1 | polyketide\_synthase | BGC0000954 | NRP + Polyketide:Modular type I | 29.0 | 47.1 | 276.0 | 2.4e-73 |
| AAS98787.1 | polyketide\_synthase/thioesterase | BGC0001001 | NRP + Polyketide | 30.0 | 41.7 | 276.0 | 2.4e-73 |
| WP\_030498975.1 | type\_I\_polyketide\_synthase | BGC0001327 | NRP:Cyclic depsipeptide + Polyketide:Modular type I | 30.0 | 41.5 | 276.0 | 2.4e-73 |
| APZ78844.1 | polyketide\_synthase | BGC0001431 | NRP:Cyclic depsipeptide + Polyketide:Iterative type I | 31.0 | 42.3 | 276.0 | 2.4e-73 |
| BAG85030.1 | putative\_polyketide\_synthase | BGC0000086 | Polyketide | 27.0 | 52.7 | 276.0 | 3.2e-73 |
| CAQ64690.1 | lasalocid\_modular\_polyketide\_synthase | BGC0000087 | Polyketide | 27.0 | 52.7 | 276.0 | 3.2e-73 |
| BAF85844.1 | modular\_polyketide\_synthase | BGC0000109 | Polyketide | 28.0 | 47.0 | 276.0 | 3.2e-73 |
| AFL48527.1 | laidlomycin\_polyketide\_synthase\_(module\_3\_and\_module\_4) | BGC0000084 | Polyketide | 27.0 | 53.0 | 275.0 | 4.1e-73 |
| AHB82059.1 | non\_ribosomal\_peptide\_synthetase/polyketide\_synthase | BGC0001019 | NRP + Polyketide:Modular type I | 30.0 | 41.8 | 275.0 | 4.1e-73 |
| CAQ52624.1 | type\_I\_polyketide\_synthase,\_modules\_7-8 | BGC0001066 | Polyketide:Modular type I | 28.0 | 43.9 | 275.0 | 4.1e-73 |
| AAS98200.1 | MSAS-type\_polyketide\_synthase | BGC0001273 | Polyketide | 27.0 | 55.0 | 275.0 | 4.1e-73 |
| SCO70308.1 | Type\_I\_polyketide\_synthase | BGC0001433 | Polyketide:Modular type I | 29.0 | 41.6 | 275.0 | 4.1e-73 |
| ABK32287.1 | JerA | BGC0000080 | Polyketide | 27.0 | 59.4 | 275.0 | 5.4e-73 |
| AHB82051.1 | polyketide\_synthase | BGC0001019 | NRP + Polyketide:Modular type I | 30.0 | 41.0 | 275.0 | 7.1e-73 |
| CAQ52626.1 | type\_I\_polyketide\_synthase,\_loading\_module\_and\_modules\_1-3 | BGC0001066 | Polyketide:Modular type I | 30.0 | 43.5 | 275.0 | 7.1e-73 |
| BBA66513.1 | type\_I\_polyketide\_synthase | BGC0001495 | Polyketide | 27.0 | 51.2 | 275.0 | 7.1e-73 |
| ARS01473.1 | NcmAI | BGC0001702 | NRP + Polyketide | 31.0 | 44.5 | 275.0 | 7.1e-73 |
| ABK32256.1 | AmbB | BGC0000014 | Polyketide | 30.0 | 42.0 | 274.0 | 9.2e-73 |
| ACY06287.1 | type\_I\_polyketide\_synthase | BGC0001042 | NRP + Polyketide | 27.0 | 51.8 | 274.0 | 9.2e-73 |
| ADH04659.1 | TugC | BGC0001342 | NRP + Polyketide | 26.0 | 58.2 | 274.0 | 9.2e-73 |
| ABC84458.1 | NigAIII | BGC0000114 | Polyketide:Modular type I | 29.0 | 45.9 | 274.0 | 1.2e-72 |
| CAG28678.1 | polyketide\_synthase | BGC0001023 | NRP + Polyketide:Modular type I | 30.0 | 42.2 | 274.0 | 1.2e-72 |
| AKD43769.1 | HerA2 | BGC0001349 | NRP + Polyketide | 29.0 | 41.3 | 274.0 | 1.2e-72 |
| APZ78754.1 | polyketide\_synthase | BGC0001423 | NRP:Cyclic depsipeptide + Polyketide:Iterative type I | 29.0 | 41.9 | 274.0 | 1.2e-72 |
| AQM37582.1 | polyketide\_synthase | BGC0001424 | NRP:Cyclic depsipeptide + Polyketide:Iterative type I | 30.0 | 41.7 | 274.0 | 1.2e-72 |
| APZ78820.1 | polyketide\_synthase | BGC0001429 | NRP:Cyclic depsipeptide + Polyketide:Iterative type I | 30.0 | 42.2 | 274.0 | 1.2e-72 |
| QBF51759.1 | type\_I\_polyketide\_synthase | BGC0001856 | Polyketide:Modular type I | 28.0 | 43.0 | 274.0 | 1.2e-72 |
| AAG23264.1 | polyketide\_synthase\_loading\_and\_extender\_module\_1 | BGC0000148 | Polyketide | 28.0 | 44.1 | 274.0 | 1.6e-72 |
| CAQ43078.1 | polyketide\_synthase | BGC0000970 | NRP + Polyketide:Modular type I | 28.0 | 47.0 | 274.0 | 1.6e-72 |
| AGY30676.1 | Ann4 | BGC0001298 | Polyketide | 27.0 | 51.4 | 273.0 | 2.1e-72 |
| APZ78780.1 | polyketide\_synthase | BGC0001426 | NRP:Cyclic depsipeptide + Polyketide:Iterative type I | 28.0 | 46.0 | 273.0 | 2.1e-72 |
| AAQ82568.1 | FscD | BGC0000061 | Polyketide | 29.0 | 42.4 | 273.0 | 2.7e-72 |
| CAQ64692.1 | lasalocid\_modular\_polyketide\_synthase | BGC0000087 | Polyketide | 27.0 | 52.4 | 273.0 | 2.7e-72 |
| CCA29203.1 | non-ribosomal\_peptide\_synthetase/polyketide\_synthase | BGC0000955 | NRP + Polyketide:Modular type I | 27.0 | 51.7 | 273.0 | 2.7e-72 |
| APZ78714.1 | polyketide\_synthase | BGC0001420 | NRP:Cyclic depsipeptide + Polyketide:Iterative type I | 30.0 | 41.7 | 273.0 | 2.7e-72 |
| APZ78727.1 | polyketide\_synthase | BGC0001421 | NRP:Cyclic depsipeptide + Polyketide:Iterative type I | 29.0 | 41.7 | 273.0 | 2.7e-72 |
| WP\_039806854.1 | type\_I\_polyketide\_synthase | BGC0002001 | NRP + Polyketide | 31.0 | 41.4 | 273.0 | 2.7e-72 |
| BAO98805.1 | putative\_polyketide\_synthase | BGC0001002 | NRP + Polyketide | 29.0 | 41.8 | 272.0 | 3.5e-72 |
| AGI99497.1 | type\_I\_polyketide\_synthase | BGC0001004 | Polyketide:Modular type I | 28.0 | 53.0 | 272.0 | 3.5e-72 |
| BAF02921.1 | type\_I\_polyketide\_synthase | BGC0000073 | Polyketide | 29.0 | 41.3 | 272.0 | 4.6e-72 |
| AFU82614.1 | mixed\_NRPS\_PKS | BGC0000998 | NRP + Polyketide | 30.0 | 40.9 | 272.0 | 4.6e-72 |
| APZ78690.1 | polyketide\_synthase | BGC0001418 | NRP:Cyclic depsipeptide + Polyketide:Iterative type I | 30.0 | 41.5 | 272.0 | 6e-72 |
| ATG32077.1 | polyketide\_synthase | BGC0001750 | NRP + Polyketide | 30.0 | 45.5 | 272.0 | 6e-72 |
| AGC09484.1 | LobS1 | BGC0001183 | Polyketide | 28.0 | 53.2 | 271.0 | 7.8e-72 |
| APZ78702.1 | polyketide\_synthase | BGC0001419 | NRP:Cyclic depsipeptide + Polyketide:Iterative type I | 30.0 | 41.5 | 271.0 | 7.8e-72 |
| APZ78793.1 | polyketide\_synthase | BGC0001427 | NRP:Cyclic depsipeptide + Polyketide:Iterative type I | 29.0 | 41.7 | 271.0 | 7.8e-72 |
| AAM70355.1 | CalO5 | BGC0000033 | Polyketide | 26.0 | 56.4 | 271.0 | 1e-71 |
| ABP55222.1 | beta-ketoacyl\_synthase | BGC0000142 | Polyketide | 26.0 | 51.2 | 271.0 | 1e-71 |
| CAJ46689.1 | polyketide\_synthase | BGC0000969 | NRP:Cyclic depsipeptide + Polyketide:Modular type I | 29.0 | 42.3 | 271.0 | 1e-71 |
| CAM00064.1 | EryAII\_Erythromycin\_polyketide\_synthase\_modules\_3\_and\_4 | BGC0000055 | Polyketide:Modular type I + Saccharide:Hybrid/tailoring | 27.0 | 53.1 | 270.0 | 1.3e-71 |
| AAF86393.1 | FkbB | BGC0000994 | NRP + Polyketide | 28.0 | 55.2 | 270.0 | 1.3e-71 |
| ABW96541.1 | type\_I\_modular\_polyketide\_synthase | BGC0000159 | Polyketide:Modular type I | 29.0 | 41.2 | 270.0 | 1.7e-71 |
| CBD77732.1 | polyketide\_synthase | BGC0000974 | NRP + Polyketide | 31.0 | 41.8 | 270.0 | 1.7e-71 |
| CAF05651.1 | TubF\_protein | BGC0001053 | NRP + Polyketide | 28.0 | 52.1 | 270.0 | 1.7e-71 |
| AIR74910.1 | polyketide\_synthase | BGC0001559 | RiPP | 31.0 | 41.8 | 270.0 | 1.7e-71 |
| ACO94496.1 | polyketide\_synthase\_type\_I | BGC0000097 | Polyketide:Modular type I | 26.0 | 51.1 | 270.0 | 2.3e-71 |
| BAE93722.1 | type\_I\_polyketide\_synthase | BGC0000164 | Polyketide | 25.0 | 54.2 | 270.0 | 2.3e-71 |
| AGC09487.1 | LobS5 | BGC0001183 | Polyketide | 28.0 | 45.6 | 270.0 | 2.3e-71 |
| APZ78807.1 | polyketide\_synthase | BGC0001428 | NRP:Cyclic depsipeptide + Polyketide:Iterative type I | 30.0 | 41.8 | 270.0 | 2.3e-71 |
| APZ78854.1 | polyketide\_synthase | BGC0001432 | NRP:Cyclic depsipeptide + Polyketide:Iterative type I | 30.0 | 41.4 | 270.0 | 2.3e-71 |
| AVV61979.1 | beta-ketoacyl\_synthase | BGC0001477 | NRP + Polyketide:Modular type I | 29.0 | 41.7 | 270.0 | 2.3e-71 |
| AVX51099.1 | NysJ | BGC0001709 | Polyketide | 27.0 | 50.8 | 270.0 | 2.3e-71 |
| AIT55260.1 | polyketide\_synthase | BGC0000072 | Polyketide:Modular type I | 29.0 | 41.2 | 269.0 | 3e-71 |
| BAG85032.1 | putative\_polyketide\_synthase | BGC0000086 | Polyketide | 27.0 | 51.4 | 269.0 | 3e-71 |
| AEZ53945.1 | polyketide\_synthase | BGC0000144 | Polyketide:Modular type I | 30.0 | 41.5 | 269.0 | 3e-71 |
| CBA11584.1 | polyketide\_synthase\_type\_I | BGC0001046 | NRP + Polyketide:Modular type I + Saccharide:Hybrid/tailoring | 26.0 | 52.3 | 269.0 | 3e-71 |
| AWS21278.1 | type\_I\_polyketide\_synthase | BGC0001934 | Polyketide | 28.0 | 52.1 | 269.0 | 3e-71 |
| AZY91987.1 | polyketide\_synthase | BGC0002022 | Polyketide | 28.0 | 52.1 | 269.0 | 3e-71 |
| AAP85335.1 | type\_I\_PKS | BGC0000233 | Polyketide | 25.0 | 56.9 | 269.0 | 3.9e-71 |
| AAA79984.2 | soraphen\_polyketide\_synthase\_B | BGC0000147 | Polyketide:Modular type I | 25.0 | 59.5 | 269.0 | 5.1e-71 |
| AIA58899.1 | HRPKS | BGC0001141 | Polyketide:Iterative type I | 30.0 | 45.9 | 269.0 | 5.1e-71 |
| ctg1\_orf521 |  | BGC0001199 | Polyketide | 27.0 | 52.3 | 268.0 | 6.6e-71 |
| AJW65409.1 | type\_I\_modular\_polyketide\_synthase | BGC0001195 | NRP + Polyketide | 30.0 | 42.2 | 268.0 | 8.6e-71 |
| AEK75502.1 | type\_1\_polyketide\_synthase | BGC0000001 | Polyketide:Modular type I | 29.0 | 44.7 | 267.0 | 1.1e-70 |
| AAC38075.1 | polyketide\_synthase\_type\_I | BGC0000127 | Polyketide | 27.0 | 51.2 | 267.0 | 1.5e-70 |
| ABI91470.1 | beta-ketoacyl\_synthase | BGC0001094 | NRP + Polyketide | 31.0 | 41.2 | 267.0 | 1.9e-70 |
| AAZ94387.1 | modular\_polyketide\_synthase | BGC0000040 | Polyketide | 29.0 | 43.2 | 266.0 | 3.3e-70 |
| CAM00062.1 | EryAI\_Erythromycin\_polyketide\_synthase\_modules\_1\_and\_2 | BGC0000055 | Polyketide:Modular type I + Saccharide:Hybrid/tailoring | 26.0 | 51.2 | 266.0 | 3.3e-70 |
| CAE02605.1 | polyketide\_synthase\_type\_I | BGC0000024 | Polyketide:Modular type I | 29.0 | 42.1 | 265.0 | 4.3e-70 |
| ACO94456.1 | polyketide\_synthase\_type\_I | BGC0000029 | Polyketide:Modular type I | 28.0 | 41.6 | 265.0 | 4.3e-70 |
| ARS01477.1 | NcmAV | BGC0001702 | NRP + Polyketide | 27.0 | 48.8 | 265.0 | 4.3e-70 |
| QDA77044.1 | polyketide\_synthase | BGC0002025 | NRP | 28.0 | 50.6 | 265.0 | 4.3e-70 |
| BAQ25511.1 | type\_I\_polyketide\_synthase | BGC0001288 | Polyketide | 30.0 | 43.9 | 265.0 | 5.6e-70 |
| AWC08662.1 | polyketide\_synthase\_type\_I | BGC0001932 | Polyketide | 27.0 | 41.6 | 265.0 | 5.6e-70 |
| CAD19089.1 | StiE\_protein | BGC0000153 | NRP + Polyketide:Modular type I | 29.0 | 41.2 | 264.0 | 9.6e-70 |
| ALP32046.1 | CycF | BGC0001293 | Polyketide | 28.0 | 40.9 | 264.0 | 9.6e-70 |
| ART41209.1 | AdrD | BGC0001508 | Polyketide | 27.0 | 50.0 | 264.0 | 1.2e-69 |
| AAO65798.1 | monensin\_polyketide\_synthase\_modules\_3\_and\_4 | BGC0000100 | Polyketide | 28.0 | 45.0 | 264.0 | 1.6e-69 |
| CAE46850.1 | Type\_I\_modular\_polyketide\_synthase | BGC0000103 | Polyketide | 28.0 | 43.1 | 264.0 | 1.6e-69 |
| AHD05619.1 | putative\_polyketide\_synthase\_subunit | BGC0001033 | NRP + Polyketide | 28.0 | 41.5 | 264.0 | 1.6e-69 |
| CAA60462.1 | polyketide\_synthase | BGC0001040 | NRP + Polyketide | 26.0 | 52.8 | 264.0 | 1.6e-69 |
| AJW65407.1 | type\_I\_modular\_polyketide\_synthase | BGC0001195 | NRP + Polyketide | 29.0 | 42.1 | 264.0 | 1.6e-69 |
| ANZ52461.1 | MonAIII | BGC0001670 | Polyketide | 28.0 | 45.0 | 264.0 | 1.6e-69 |
| ACB46487.1 | polyketide\_synthase | BGC0000082 | Polyketide | 26.0 | 58.5 | 263.0 | 2.1e-69 |
| ASK38717.1 | polyketide\_synthase | BGC0001557 | Polyketide | 28.0 | 46.3 | 263.0 | 2.1e-69 |
| ABK32259.1 | AmbE | BGC0000014 | Polyketide | 29.0 | 41.5 | 263.0 | 2.8e-69 |
| ABV97151.1 | AMP-dependent\_synthetase\_and\_ligase | BGC0000137 | Polyketide | 27.0 | 41.0 | 263.0 | 2.8e-69 |
| BAV69313.1 | PrhL | BGC0001729 | Polyketide + Terpene | 28.0 | 49.4 | 263.0 | 2.8e-69 |
| TXD00025.1 | SDR\_family\_NAD(P)-dependent\_oxidoreductase | BGC0001877 | Polyketide | 27.0 | 42.2 | 263.0 | 2.8e-69 |
| ABY21538.1 | AngAI | BGC0000018 | Polyketide | 26.0 | 52.7 | 262.0 | 3.6e-69 |
| AIG62146.1 | 6-methylsalicylic\_acid\_synthase | BGC0000120 | Polyketide:Iterative type I | 27.0 | 51.1 | 262.0 | 3.6e-69 |
| AHD05614.1 | putative\_non-ribosomal\_peptide\_ligase/\_polyketide\_synthase\_hybrid | BGC0001033 | NRP + Polyketide | 27.0 | 51.1 | 262.0 | 4.7e-69 |
| AGY30677.1 | Ann5 | BGC0001298 | Polyketide | 28.0 | 42.3 | 262.0 | 4.7e-69 |
| AKD43768.1 | HerA1 | BGC0001349 | NRP + Polyketide | 27.0 | 52.0 | 262.0 | 4.7e-69 |
| APZ78678.1 | polyketide\_synthase | BGC0001417 | NRP:Cyclic depsipeptide + Polyketide:Iterative type I | 29.0 | 41.5 | 262.0 | 4.7e-69 |
| BAD97694.1 | Aft9-1 | BGC0000003 | Polyketide | 30.0 | 42.1 | 262.0 | 6.2e-69 |
| CAD55506.1 | CpkA;\_Polyketide\_synthase\_loading\_module,\_and\_modules\_1\_and\_2 | BGC0000038 | Polyketide:Modular type I | 29.0 | 44.7 | 262.0 | 6.2e-69 |
| EHA28244.1 | hypothetical\_protein | BGC0001143 | Polyketide | 27.0 | 52.8 | 262.0 | 6.2e-69 |
| ANY10600.1 | polyketide\_synthase | BGC0001773 | Polyketide | 25.0 | 52.6 | 261.0 | 8.1e-69 |
| AWH12936.1 | StmA | BGC0001939 | Polyketide | 27.0 | 52.0 | 261.0 | 8.1e-69 |
| BAN19720.1 | polyketide\_synthase | BGC0001252 | Polyketide | 29.0 | 43.7 | 261.0 | 1.1e-68 |
| AAQ90173.1 | polyketide\_synthase\_type\_I | BGC0000128 | Polyketide | 30.0 | 41.0 | 260.0 | 1.4e-68 |
| ABP55223.1 | beta-ketoacyl\_synthase | BGC0000142 | Polyketide | 26.0 | 52.0 | 260.0 | 1.8e-68 |
| AAK19883.1 | soraphen\_polyketide\_synthase\_A | BGC0000147 | Polyketide:Modular type I | 30.0 | 41.2 | 260.0 | 1.8e-68 |
| BAE93729.1 | type\_I\_polyketide\_synthase | BGC0000164 | Polyketide | 28.0 | 42.0 | 260.0 | 1.8e-68 |
| BBD17742.1 | polyketide\_synthase | BGC0001918 | NRP + Polyketide | 30.0 | 42.4 | 260.0 | 1.8e-68 |
| AAC68815.1 | FK506\_polyketide\_synthase | BGC0000353 | NRP | 27.0 | 50.6 | 260.0 | 2.4e-68 |
| ABI91466.1 | beta-ketoacyl\_synthase | BGC0001094 | NRP + Polyketide | 30.0 | 42.4 | 260.0 | 2.4e-68 |
| BAC20566.1 | polyketide\_synthase | BGC0000039 | Polyketide | 27.0 | 53.6 | 259.0 | 4e-68 |
| AAC69329.1 | type\_I\_polyketide\_synthase\_PikAI | BGC0000094 | Polyketide:Modular type I + Saccharide:Hybrid/tailoring | 27.0 | 52.1 | 259.0 | 4e-68 |
| ACB37740.1 | putative\_type\_I\_polyketide\_synthase | BGC0000162 | Polyketide | 27.0 | 51.1 | 259.0 | 5.2e-68 |
| CAJ76298.1 | putative\_hybrid\_polyketide-non-ribosomal\_peptide\_synthetase | BGC0000972 | NRP + Polyketide:Modular type I + Polyketide:Trans-AT type I | 29.0 | 40.8 | 259.0 | 5.2e-68 |
| ARS01474.1 | NcmAII | BGC0001702 | NRP + Polyketide | 27.0 | 56.1 | 259.0 | 5.2e-68 |
| BBG28498.1 | putative\_polyketide\_synthase | BGC0001913 | Polyketide | 28.0 | 45.1 | 259.0 | 5.2e-68 |
| BAE93728.1 | type\_I\_polyketide\_synthase | BGC0000164 | Polyketide | 29.0 | 41.4 | 258.0 | 6.8e-68 |
| ABL74938.1 | PKS | BGC0001048 | NRP:Glycopeptide + Polyketide:Modular type I + Saccharide:Hybrid/tailoring | 26.0 | 54.8 | 258.0 | 6.8e-68 |
| WP\_019032754.1 | type\_I\_polyketide\_synthase | BGC0001331 | NRP:Cyclic depsipeptide + Polyketide:Modular type I | 28.0 | 51.1 | 258.0 | 6.8e-68 |
| CAA60459.1 | polyketide\_synthase | BGC0001040 | NRP + Polyketide | 26.0 | 52.9 | 258.0 | 8.9e-68 |
| ASZ00148.1 | polyketide\_synthase | BGC0001785 | Polyketide | 29.0 | 41.5 | 258.0 | 8.9e-68 |
| BAD83684.1 | PKSN\_polyketide\_synthase\_for\_alternapyrone\_biosynthesis | BGC0000012 | Polyketide | 28.0 | 44.5 | 257.0 | 2e-67 |
| WP\_053065267.1 | type\_I\_polyketide\_synthase | BGC0001330 | NRP:Cyclic depsipeptide + Polyketide:Modular type I | 27.0 | 51.1 | 257.0 | 2e-67 |
| CQR60497.1 | Polyketide\_synthase,\_type\_I,\_modules:\_loading,\_1,\_2\_and\_3 | BGC0001287 | Polyketide | 27.0 | 42.6 | 256.0 | 2.6e-67 |
| WP\_053138504.1 | type\_I\_polyketide\_synthase | BGC0002033 | Polyketide | 27.0 | 52.4 | 256.0 | 3.4e-67 |
| AGZ15475.1 | putative\_type\_1\_modular\_polyketide\_synthase | BGC0001036 | NRP + Polyketide | 28.0 | 41.3 | 255.0 | 4.4e-67 |
| WP\_051137606.1 | type\_I\_polyketide\_synthase | BGC0002011 | Polyketide | 26.0 | 50.5 | 255.0 | 7.6e-67 |
| ACZ65476.1 | type\_I\_modular\_polyketide\_synthase | BGC0000140 | Polyketide | 28.0 | 47.4 | 254.0 | 9.9e-67 |
| CAA60460.1 | polyketide\_synthase | BGC0001040 | NRP + Polyketide | 26.0 | 53.0 | 254.0 | 9.9e-67 |
| AIT55264.1 | polyketide\_synthase | BGC0000072 | Polyketide:Modular type I | 29.0 | 50.2 | 254.0 | 1.3e-66 |
| BAG84248.1 | putative\_polyketide\_synthase | BGC0000257 | Polyketide | 27.0 | 42.4 | 254.0 | 1.3e-66 |
| ADU86002.1 | putative\_modular\_polyketide\_synthase | BGC0000165 | Polyketide:Modular type I | 27.0 | 50.2 | 253.0 | 1.7e-66 |
| AGI99496.1 | Type\_I\_polyketide\_synthase | BGC0001004 | Polyketide:Modular type I | 26.0 | 50.0 | 253.0 | 1.7e-66 |
| AKA59088.1 | type-I\_PKS | BGC0001619 | Polyketide | 29.0 | 45.1 | 253.0 | 1.7e-66 |
| CAL58682.1 | polyketide\_synthase | BGC0000149 | Polyketide:Modular type I | 26.0 | 54.4 | 253.0 | 2.2e-66 |
| BAD08359.1 | polyketide\_synthase\_modules\_5-6 | BGC0000167 | Polyketide | 26.0 | 51.8 | 253.0 | 2.2e-66 |
| AQW44873.1 | polyketide\_synthase | BGC0001761 | Polyketide | 29.0 | 41.2 | 253.0 | 2.2e-66 |
| ADU86004.1 | putative\_modular\_polyketide\_synthase | BGC0000165 | Polyketide:Modular type I | 27.0 | 53.6 | 253.0 | 2.9e-66 |
| BAG17643.1 | putative\_NRPS-type-I\_PKS\_fusion\_protein | BGC0001043 | NRP + Polyketide | 30.0 | 39.8 | 253.0 | 2.9e-66 |
| WP\_016638481.1 | type\_I\_polyketide\_synthase | BGC0001519 | NRP + Polyketide | 28.0 | 41.5 | 252.0 | 3.8e-66 |
| CCC55921.1 | non-ribosomal\_peptide\_synthetase/polyketide\_synthase\_hybrid\_protein | BGC0000973 | NRP + Polyketide:Modular type I | 30.0 | 41.6 | 252.0 | 6.4e-66 |
| AHB82063.1 | polyketide\_synthase | BGC0001231 | NRP + Polyketide:Modular type I | 35.0 | 26.9 | 252.0 | 6.4e-66 |
| ABB88522.1 | polyketide\_synthase\_type\_I | BGC0000050 | Polyketide | 29.0 | 44.2 | 251.0 | 8.4e-66 |
| AAG02357.1 | polyketide\_synthase | BGC0000963 | NRP:Glycopeptide + Polyketide:Modular type I + Saccharide:Hybrid/tailoring | 27.0 | 52.1 | 251.0 | 8.4e-66 |
| WP\_019032757.1 | type\_I\_polyketide\_synthase | BGC0001331 | NRP:Cyclic depsipeptide + Polyketide:Modular type I | 26.0 | 59.6 | 251.0 | 8.4e-66 |
| AJW65408.1 | type\_I\_modular\_polyketide\_synthase | BGC0001195 | NRP + Polyketide | 29.0 | 41.9 | 251.0 | 1.1e-65 |
| AAO65796.1 | monensin\_polyketide\_synthase\_loading\_module\_and\_module\_1 | BGC0000100 | Polyketide | 29.0 | 43.5 | 250.0 | 1.4e-65 |
| ANZ52459.1 | MonAI | BGC0001670 | Polyketide | 29.0 | 43.5 | 250.0 | 1.4e-65 |
| BAH02268.1 | polyketide\_synthase | BGC0000126 | Polyketide | 27.0 | 52.9 | 250.0 | 2.4e-65 |
| AHA38203.1 | GphJ | BGC0000069 | Polyketide | 29.0 | 42.3 | 249.0 | 3.2e-65 |
| AMY15068.1 | hexaketide\_synthase\_MF-SQHKS | BGC0001339 | Polyketide:Iterative type I | 26.0 | 54.1 | 249.0 | 3.2e-65 |
| AAD43562.2 | Fum1p | BGC0000062 | Polyketide | 28.0 | 48.2 | 249.0 | 4.2e-65 |
| ACB46488.1 | polyketide\_synthase | BGC0000082 | Polyketide | 28.0 | 45.1 | 248.0 | 5.4e-65 |
| AEF33079.1 | polyketide\_synthase | BGC0001039 | NRP + Polyketide | 29.0 | 42.0 | 248.0 | 5.4e-65 |
| AHB82057.1 | polyketide\_synthase | BGC0001019 | NRP + Polyketide:Modular type I | 29.0 | 41.5 | 248.0 | 7.1e-65 |
| CAF05649.1 | TubD\_protein | BGC0001053 | NRP + Polyketide | 29.0 | 41.5 | 248.0 | 7.1e-65 |
| EAA65604.1 | hypothetical\_protein | BGC0000022 | Polyketide | 29.0 | 44.4 | 248.0 | 9.3e-65 |
| ACZ57548.1 | polyketide\_synthase | BGC0000046 | Polyketide:Iterative type I | 25.0 | 59.6 | 248.0 | 9.3e-65 |
| AEC13072.1 | fosF | BGC0000060 | Polyketide | 26.0 | 47.8 | 248.0 | 9.3e-65 |
| AAK57190.1 | MxaF | BGC0001022 | NRP + Polyketide | 29.0 | 39.5 | 248.0 | 9.3e-65 |
| ACY13414.1 | amino\_acid\_adenylation\_domain\_protein | BGC0001367 | NRP + Polyketide | 28.0 | 41.5 | 248.0 | 9.3e-65 |
| BAB69198.1 | modular\_polyketide\_synthase | BGC0000117 | Polyketide | 27.0 | 47.0 | 247.0 | 1.2e-64 |
| ABP55221.1 | acyl\_transferase\_domain\_protein | BGC0000142 | Polyketide | 26.0 | 51.8 | 247.0 | 1.2e-64 |
| ADU85981.1 | putative\_modular\_polyketide\_synthase | BGC0000165 | Polyketide:Modular type I | 26.0 | 52.1 | 247.0 | 1.2e-64 |
| AGC24271.1 | prlQ | BGC0001038 | NRP + Polyketide:Modular type I | 26.0 | 53.0 | 247.0 | 1.2e-64 |
| ARS01475.1 | NcmAIII | BGC0001702 | NRP + Polyketide | 26.0 | 52.3 | 247.0 | 1.2e-64 |
| AEH42473.1 | polyketide\_synthase | BGC0000032 | Polyketide | 28.0 | 41.2 | 247.0 | 1.6e-64 |
| CCE88377.1 | non-ribosomal\_peptide\_synthetase/polyketide\_synthase | BGC0001034 | NRP + Polyketide:Modular type I | 28.0 | 50.9 | 247.0 | 1.6e-64 |
| AKG06376.1 | polyketide\_synthase\_type\_1 | BGC0001830 | Polyketide | 29.0 | 42.1 | 247.0 | 1.6e-64 |
| ABY21540.1 | AngAIII | BGC0000018 | Polyketide | 25.0 | 53.5 | 246.0 | 2.7e-64 |
| BAC76491.1 | lankamycin\_synthase\_LkmAIII | BGC0000085 | Polyketide | 28.0 | 42.0 | 246.0 | 2.7e-64 |
| AJO72736.1 | Type\_I\_modular\_polyketide\_synthase | BGC0001381 | Polyketide | 27.0 | 41.0 | 246.0 | 2.7e-64 |
| AGC09485.1 | LobS2 | BGC0001183 | Polyketide | 26.0 | 58.6 | 246.0 | 3.5e-64 |
| PHM26614.1 | Phthiocerol\_synthesis\_polyketide\_synthase\_type\_I\_PpsE | BGC0001130 | NRP + Polyketide | 29.0 | 43.4 | 245.0 | 4.6e-64 |
| ADH04657.1 | TugA | BGC0001342 | NRP + Polyketide | 29.0 | 39.6 | 245.0 | 4.6e-64 |
| ADH04680.1 | hybrid\_polyketide\_synthase/non-ribosomal\_peptide\_synthetase | BGC0001344 | NRP + Polyketide | 28.0 | 41.2 | 245.0 | 4.6e-64 |
| EAA36364.1 | hypothetical\_protein | BGC0001697 | Polyketide | 27.0 | 50.3 | 245.0 | 4.6e-64 |
| ACY06289.1 | type\_I\_polyketide\_synthase | BGC0001042 | NRP + Polyketide | 26.0 | 53.0 | 245.0 | 6e-64 |
| AAO65807.1 | monensin\_polyketide\_synthase\_module\_10 | BGC0000100 | Polyketide | 25.0 | 43.0 | 245.0 | 7.8e-64 |
| ANZ52470.1 | MonAVII | BGC0001670 | Polyketide | 25.0 | 43.0 | 245.0 | 7.8e-64 |
| AFL48533.1 | laidlomycin\_polyketide\_synthase\_(module\_10) | BGC0000084 | Polyketide | 27.0 | 41.1 | 244.0 | 1e-63 |
| WP\_047890614.1 | type\_I\_polyketide\_synthase | BGC0001330 | NRP:Cyclic depsipeptide + Polyketide:Modular type I | 27.0 | 51.5 | 244.0 | 1e-63 |
| ADH04641.1 | TgaC | BGC0001051 | NRP + Polyketide:Modular type I | 26.0 | 52.1 | 244.0 | 1.3e-63 |
| PKX88487.1 | polyketide\_synthase | BGC0001708 | Polyketide + Terpene | 31.0 | 37.9 | 243.0 | 1.7e-63 |
| ACB12550.1 | Fum1 | BGC0000063 | Polyketide | 28.0 | 46.7 | 243.0 | 2.3e-63 |
| ABA02240.1 | polyketide\_synthase | BGC0000098 | Polyketide | 27.0 | 53.6 | 243.0 | 2.3e-63 |
| AEU17898.1 | putative\_type\_I\_PKS | BGC0001072 | Saccharide + Polyketide:Modular type I + Polyketide:Type II + Other:Aminocoumarin | 27.0 | 41.9 | 243.0 | 2.3e-63 |
| AIW00670.1 | mellein\_synthase | BGC0001244 | Polyketide | 26.0 | 56.1 | 243.0 | 2.3e-63 |
| TXD00033.1 | SDR\_family\_NAD(P)-dependent\_oxidoreductase | BGC0001877 | Polyketide | 26.0 | 42.9 | 243.0 | 2.3e-63 |
| BAO66542.1 | type\_I\_polyketide\_synthase | BGC0000042 | Polyketide | 27.0 | 41.7 | 243.0 | 3e-63 |
| BAC57032.1 | protomycinolide\_IV\_synthase\_5 | BGC0000102 | Polyketide | 28.0 | 43.5 | 243.0 | 3e-63 |
| ADM46360.1 | polyketide\_synthase | BGC0000106 | Polyketide | 26.0 | 42.5 | 243.0 | 3e-63 |
| KGO40478.1 | Acyl\_transferase/acyl\_hydrolase/lysophospholipase | BGC0001205 | Polyketide | 26.0 | 52.8 | 243.0 | 3e-63 |
| AHF22854.1 | MarL | BGC0000091 | Polyketide | 29.0 | 41.7 | 242.0 | 3.9e-63 |
| EAL89230.2 | LovB-like\_polyketide\_synthase,\_putative | BGC0000129 | Polyketide | 29.0 | 42.7 | 242.0 | 3.9e-63 |
| CAL58681.1 | polyketide\_synthase | BGC0000149 | Polyketide:Modular type I | 27.0 | 49.8 | 242.0 | 3.9e-63 |
| orf3 | polyketide\_synthase | BGC0001432 | NRP:Cyclic depsipeptide + Polyketide:Iterative type I | 30.0 | 39.6 | 242.0 | 3.9e-63 |
| APZ78858.1 | polyketide\_synthase | BGC0001432 | NRP:Cyclic depsipeptide + Polyketide:Iterative type I | 30.0 | 39.6 | 242.0 | 3.9e-63 |
| EAU32819.1 | 6-methylsalicylic\_acid\_synthase | BGC0000160 | Polyketide | 28.0 | 41.3 | 242.0 | 5.1e-63 |
| ACN64831.1 | PokM1 | BGC0001061 | Polyketide:Iterative type I + Polyketide:Type II + Saccharide:Hybrid/tailoring | 30.0 | 42.5 | 242.0 | 6.6e-63 |
| CAO98852.1 | polyketide\_synthase\_AufI | BGC0000023 | Polyketide:Modular type I | 27.0 | 50.1 | 241.0 | 8.7e-63 |
| AHB82052.1 | polyketide\_synthase | BGC0001019 | NRP + Polyketide:Modular type I | 35.0 | 26.8 | 241.0 | 8.7e-63 |
| ctg1\_orf28 |  | BGC0000096 | Polyketide | 29.0 | 42.4 | 241.0 | 1.1e-62 |
| ACB37741.1 | putative\_type\_I\_polyketide\_synthase | BGC0000162 | Polyketide | 26.0 | 49.2 | 241.0 | 1.1e-62 |
| CBJ89766.1 | Polyketide\_synthase\_involved\_in\_xenocoumacin\_synthesis | BGC0001054 | NRP + Polyketide:Modular type I | 27.0 | 43.2 | 241.0 | 1.1e-62 |
| CAE02602.1 | polyketide\_synthase\_type\_I | BGC0000024 | Polyketide:Modular type I | 27.0 | 41.5 | 240.0 | 1.9e-62 |
| ACR50785.1 | polyketide\_synthase | BGC0000163 | Polyketide | 27.0 | 42.8 | 240.0 | 2.5e-62 |
| ACY06288.1 | type\_I\_polyketide\_synthase | BGC0001042 | NRP + Polyketide | 26.0 | 52.7 | 240.0 | 2.5e-62 |
| ADH04682.1 | polyketide\_synthase | BGC0001344 | NRP + Polyketide | 28.0 | 43.4 | 240.0 | 2.5e-62 |
| CRI73798.1 | CongD\_protein | BGC0001215 | NRP | 26.0 | 46.9 | 239.0 | 3.3e-62 |
| BAA20102.2 | 6-methylsalicylic\_acid\_synthase | BGC0001276 | Polyketide | 28.0 | 41.3 | 239.0 | 4.3e-62 |
| ANF07288.1 | hrPKS | BGC0001340 | Polyketide:Iterative type I | 27.0 | 46.8 | 239.0 | 4.3e-62 |
| AHH99923.1 | PKS\_I | BGC0000002 | Polyketide | 25.0 | 51.2 | 238.0 | 5.6e-62 |
| ARE67853.1 | AbsB1 | BGC0001492 | Polyketide | 29.0 | 41.2 | 238.0 | 5.6e-62 |
| ABC87510.1 | polyketide\_synthase | BGC0001011 | NRP + Polyketide | 27.0 | 45.6 | 238.0 | 7.3e-62 |
| ctg1\_orf21 |  | BGC0001013 | NRP + Polyketide | 27.0 | 45.6 | 238.0 | 7.3e-62 |
| CAI94713.1 | putative\_polyketide\_synthase | BGC0000141 | Polyketide | 27.0 | 41.4 | 238.0 | 9.6e-62 |
| AEZ64505.1 | Herb | BGC0001065 | Polyketide | 28.0 | 42.7 | 238.0 | 9.6e-62 |
| BAC76492.1 | lankamycin\_synthase\_LkmAII | BGC0000085 | Polyketide | 25.0 | 52.3 | 237.0 | 1.6e-61 |
| CAP95405.1 |  | BGC0001404 | Polyketide | 30.0 | 39.6 | 237.0 | 1.6e-61 |
| ABI93779.1 | GdmPKS | BGC0000068 | Polyketide | 26.0 | 46.7 | 237.0 | 2.1e-61 |
| CAI94682.1 | putative\_polyketide\_synthase | BGC0000141 | Polyketide | 27.0 | 43.8 | 237.0 | 2.1e-61 |
| CAL80821.1 | sylD-like\_NRPS/PKS | BGC0000997 | NRP + Polyketide | 27.0 | 41.2 | 237.0 | 2.1e-61 |
| APZ78767.1 | polyketide\_synthase | BGC0001425 | NRP:Cyclic depsipeptide + Polyketide:Iterative type I | 27.0 | 46.1 | 237.0 | 2.1e-61 |
| AHA38202.1 | GphI | BGC0000069 | Polyketide | 29.0 | 40.3 | 236.0 | 2.8e-61 |
| EHA22196.1 | polyketide\_synthase | BGC0000170 | Polyketide | 26.0 | 54.1 | 236.0 | 3.6e-61 |
| ADF88262.1 | mixed\_nonribosomal\_peptide\_synthetase/\_polyketide\_synthase | BGC0000979 | NRP + Polyketide | 32.0 | 31.0 | 236.0 | 3.6e-61 |
| ADF88265.1 | mixed\_nonribosomal\_peptide\_synthetase/\_polyketide\_synthase | BGC0000980 | NRP + Polyketide | 32.0 | 31.0 | 236.0 | 3.6e-61 |
| CAD70195.1 | non-ribosomal\_peptide\_synthetase | BGC0001047 | NRP + Polyketide | 26.0 | 50.8 | 236.0 | 3.6e-61 |
| ABX37384.1 | Beta-ketoacyl\_synthase | BGC0000984 | NRP + Polyketide | 29.0 | 42.7 | 235.0 | 4.8e-61 |
| BAB69196.1 | modular\_polyketide\_synthase | BGC0000117 | Polyketide | 28.0 | 42.3 | 235.0 | 6.2e-61 |
| CCM44338.1 | Polyketide\_synthase | BGC0001056 | NRP + Polyketide:Modular type I + Polyketide:PUFA synthase or related | 28.0 | 41.3 | 235.0 | 6.2e-61 |
| ABC87511.1 | polyketide\_synthase | BGC0001011 | NRP + Polyketide | 25.0 | 55.6 | 234.0 | 1.4e-60 |
| ctg1\_orf22 |  | BGC0001013 | NRP + Polyketide | 25.0 | 55.6 | 234.0 | 1.4e-60 |
| ASZ00147.1 | polyketide\_synthase | BGC0001785 | Polyketide | 27.0 | 43.0 | 234.0 | 1.4e-60 |
| AHA38201.1 | GphH | BGC0000069 | Polyketide | 30.0 | 35.4 | 233.0 | 2.4e-60 |
| AAF86396.1 | FkbA | BGC0000994 | NRP + Polyketide | 27.0 | 50.4 | 233.0 | 3.1e-60 |
| BBA66511.1 | type\_I\_polyketide\_synthase | BGC0001495 | Polyketide | 27.0 | 44.8 | 233.0 | 3.1e-60 |
| CAL69597.1 | PKS-NRPS | BGC0001049 | NRP + Polyketide:Iterative type I | 27.0 | 52.1 | 232.0 | 5.3e-60 |
| antaD | Type\_I\_PKS | BGC0001455 | NRP + Polyketide | 28.0 | 42.4 | 232.0 | 6.9e-60 |
| AAT28740.1 | FUSS | BGC0000064 | Polyketide | 26.0 | 49.7 | 231.0 | 9e-60 |
| BAC57028.1 | protomycinolide\_IV\_synthase\_1 | BGC0000102 | Polyketide | 27.0 | 41.2 | 231.0 | 9e-60 |
| EAQ86385.1 | hypothetical\_protein | BGC0001405 | Polyketide | 28.0 | 41.4 | 231.0 | 9e-60 |
| ABY21541.1 | AngAIV | BGC0000018 | Polyketide | 25.0 | 51.2 | 231.0 | 1.2e-59 |
| WP\_019634550.1 | type\_I\_polyketide\_synthase | BGC0001443 | NRP + Polyketide | 29.0 | 40.3 | 230.0 | 1.5e-59 |
| ctg1\_orf0002 |  | BGC0001068 | Terpene + Polyketide | 29.0 | 43.0 | 230.0 | 2e-59 |
| API82664.1 | putative\_polyketide\_synthase | BGC0001677 | Polyketide | 26.0 | 53.9 | 230.0 | 2e-59 |
| AHV78252.1 | ResS1 | BGC0001246 | Polyketide | 28.0 | 43.5 | 230.0 | 2.6e-59 |
| AAY28227.1 | HbmAIII | BGC0000074 | Polyketide | 27.0 | 42.0 | 229.0 | 3.4e-59 |
| CAJ76291.1 | putative\_polyketide\_synthase | BGC0000972 | NRP + Polyketide:Modular type I + Polyketide:Trans-AT type I | 27.0 | 41.4 | 229.0 | 3.4e-59 |
| BAK64649.1 | polyketide\_synthase | BGC0000135 | Polyketide | 29.0 | 38.1 | 229.0 | 4.5e-59 |
| QBF51754.1 | type\_I\_polyketide\_synthase | BGC0001856 | Polyketide:Modular type I | 27.0 | 42.6 | 229.0 | 4.5e-59 |
| BAB69195.1 | modular\_polyketide\_synthase | BGC0000117 | Polyketide | 26.0 | 42.3 | 228.0 | 5.8e-59 |
| ADN43685.1 | DmbS | BGC0001136 | NRP + Polyketide:Iterative type I | 27.0 | 51.7 | 228.0 | 7.6e-59 |
| B073\_RS40860 | type\_I\_polyketide\_synthase | BGC0001332 | NRP + Polyketide | 27.0 | 44.2 | 228.0 | 7.6e-59 |
| AWX24483.1 | type\_I\_polyketide\_synthase | BGC0001695 | NRP | 28.0 | 40.6 | 228.0 | 7.6e-59 |
| BAK64650.1 | polyketide\_synthase | BGC0000135 | Polyketide | 25.0 | 47.0 | 228.0 | 9.9e-59 |
| AMY15057.1 | tetraketide\_synthase\_MF-SQTKS | BGC0001339 | Polyketide:Iterative type I | 27.0 | 45.9 | 228.0 | 9.9e-59 |
| ALA09356.1 | type\_I\_modular\_PKS | BGC0001303 | Polyketide | 26.0 | 52.1 | 227.0 | 1.3e-58 |
| EAQ86392.1 | hypothetical\_protein | BGC0001405 | Polyketide | 30.0 | 35.9 | 227.0 | 1.3e-58 |
| EPE34340.1 | polyketide\_synthase | BGC0001035 | Polyketide + NRP + Other:Aminocoumarin | 25.0 | 47.6 | 227.0 | 1.7e-58 |
| AFP73394.1 | FusA | BGC0001268 | NRP + Polyketide | 26.0 | 49.7 | 227.0 | 1.7e-58 |
| AAD34559.1 | polyketide\_synthase | BGC0000088 | Polyketide | 25.0 | 53.2 | 227.0 | 2.2e-58 |
| BBD17760.1 | polyketide\_synthase | BGC0001919 | NRP + Polyketide | 28.0 | 40.6 | 226.0 | 3.8e-58 |
| AHD05615.1 | putative\_non-ribosomal\_peptide\_ligase/\_polyketide\_synthase\_hybrid | BGC0001033 | NRP + Polyketide | 26.0 | 40.9 | 225.0 | 4.9e-58 |
| AKA54627.1 | PKS | BGC0001216 | NRP + Polyketide | 28.0 | 41.3 | 225.0 | 6.4e-58 |
| CCT75967.1 | polyketide\_synthase | BGC0001606 | Polyketide | 28.0 | 45.3 | 225.0 | 6.4e-58 |
| ARP51711.1 | PKS-NRPS\_hybrid\_protein | BGC0001741 | NRP + Polyketide | 28.0 | 50.6 | 225.0 | 6.4e-58 |
| EAU29529.1 | hypothetical\_protein | BGC0000682 | Terpene | 26.0 | 49.0 | 225.0 | 8.4e-58 |
| ANZ22995.1 | ZinA | BGC0001828 | Polyketide | 27.0 | 40.8 | 224.0 | 1.1e-57 |
| EHA52508.1 | mycocerosic\_acid\_synthase | BGC0001749 | Polyketide | 30.0 | 39.8 | 224.0 | 1.4e-57 |
| TXD00034.1 | SDR\_family\_NAD(P)-dependent\_oxidoreductase | BGC0001877 | Polyketide | 26.0 | 41.7 | 224.0 | 1.4e-57 |
| EAU29808.1 | hypothetical\_protein | BGC0001400 | Polyketide | 27.0 | 51.6 | 223.0 | 1.9e-57 |
| AAX35547.1 | polyketide\_syntase\_2 | BGC0001275 | Polyketide | 27.0 | 45.9 | 223.0 | 2.4e-57 |
| AUD08663.1 | iPKS-NRPS | BGC0001553 | NRP + Polyketide | 27.0 | 45.8 | 223.0 | 2.4e-57 |
| KFA69336.1 | hypothetical\_protein | BGC0001626 | Polyketide | 30.0 | 30.0 | 222.0 | 7.1e-57 |
| BBG28484.1 | polyketide\_synthase\_CdmE | BGC0001926 | Polyketide | 27.0 | 39.8 | 221.0 | 9.3e-57 |
| gene3 |  | BGC0002035 | NRP + Polyketide | 25.0 | 50.4 | 221.0 | 9.3e-57 |
| AWW87422.1 | type\_I\_polyketide\_synthase | BGC0001755 | Polyketide | 27.0 | 41.7 | 220.0 | 1.6e-56 |
| ARE67851.1 | AbsB3 | BGC0001492 | Polyketide | 26.0 | 42.4 | 220.0 | 2.1e-56 |
| AAC69330.1 | type\_I\_polyketide\_synthase\_PikAII | BGC0000094 | Polyketide:Modular type I + Saccharide:Hybrid/tailoring | 27.0 | 41.0 | 218.0 | 7.9e-56 |
| EYT83439.1 | beta-ketoacyl\_synthase | BGC0001213 | Polyketide | 28.0 | 41.0 | 218.0 | 7.9e-56 |
| ATZ45185.1 | Bcboa9 | BGC0001892 | Polyketide | 30.0 | 39.3 | 218.0 | 7.9e-56 |
| AXM42948.1 | type\_1\_polyketide\_synthase | BGC0001941 | NRP + Polyketide | 26.0 | 48.7 | 218.0 | 1e-55 |
| AHB38498.1 | polyketide\_synthase | BGC0000346 | NRP + Polyketide:Modular type I | 27.0 | 39.7 | 217.0 | 1.8e-55 |
| AAF19810.1 | MtaB | BGC0001024 | NRP + Polyketide:Modular type I | 25.0 | 49.5 | 217.0 | 2.3e-55 |
| AP234\_RS14720 | hypothetical\_protein | BGC0001653 | Polyketide | 25.0 | 51.0 | 216.0 | 3e-55 |
| WP\_106967775.1 | type\_I\_polyketide\_synthase | BGC0001519 | NRP + Polyketide | 26.0 | 42.8 | 216.0 | 3.9e-55 |
| ACS68554.1 | hybrid\_PKS-NRPS\_protein | BGC0001026 | NRP + Polyketide | 26.0 | 51.2 | 215.0 | 6.7e-55 |
| ASX95227.1 | IlaE | BGC0001620 | Polyketide | 25.0 | 43.8 | 215.0 | 6.7e-55 |
| AFA26384.1 | polyketide\_synthase\_A | BGC0001874 | NRP + Polyketide | 26.0 | 48.2 | 214.0 | 1.5e-54 |
| QCX41945.1 | Amc8 | BGC0001958 | Other | 28.0 | 40.3 | 213.0 | 2.5e-54 |
| AHN85651.1 | Phn2 | BGC0000122 | Polyketide:Modular type I | 26.0 | 48.9 | 212.0 | 5.6e-54 |
| AAS79459.1 | polyketide\_synthase\_subunit | BGC0000035 | Polyketide | 26.0 | 41.7 | 211.0 | 9.6e-54 |
| KFL51883.1 | amino\_acid\_adenylation\_protein | BGC0001711 | NRP + Polyketide | 28.0 | 36.3 | 211.0 | 1.3e-53 |
| AUS29495.1 | polyketide\_synthase | BGC0001030 | NRP + Polyketide | 26.0 | 40.7 | 208.0 | 1.1e-52 |
| AKC54422.1 | fumosorinone\_biosynthesis\_polyketide\_synthase | BGC0001218 | NRP + Polyketide | 25.0 | 52.6 | 207.0 | 1.4e-52 |
| ATQ39432.1 | PKS | BGC0001565 | NRP | 26.0 | 49.9 | 207.0 | 1.4e-52 |
| CAJ88177.1 | putative\_type\_I\_polyketide\_synthase | BGC0000151 | Polyketide:Modular type I + Saccharide:Hybrid/tailoring | 27.0 | 41.8 | 207.0 | 2.4e-52 |
| ABO15861.1 | polyketide\_synthase | BGC0000130 | Polyketide | 28.0 | 38.2 | 206.0 | 3.1e-52 |
| BAC20564.1 | polyketide\_synthase | BGC0000039 | Polyketide | 25.0 | 51.5 | 205.0 | 9e-52 |
| BAF92601.1 | iterative\_type\_I\_PKS | BGC0000118 | Polyketide | 28.0 | 44.3 | 205.0 | 9e-52 |
| ACJ24875.1 | 6-methylsalicylic\_acid\_synthase | BGC0000119 | Polyketide:Iterative type I + Saccharide:Hybrid/tailoring | 28.0 | 44.3 | 205.0 | 9e-52 |
| AGJ76601.1 | HglE | BGC0000869 | Other | 28.0 | 40.1 | 205.0 | 9e-52 |
| BAV19380.1 | NRPS-like\_enzyme | BGC0001390 | NRP + Polyketide | 28.0 | 30.3 | 205.0 | 9e-52 |
| ATL73033.1 | type\_I\_modular\_polyketide\_synthase | BGC0001807 | NRP + Polyketide | 27.0 | 39.6 | 205.0 | 9e-52 |
| QCX41916.1 | Mhr10 | BGC0001956 | Polyketide | 28.0 | 40.8 | 204.0 | 1.2e-51 |
| BAK26562.1 | PKS-NRPS\_hybrid | BGC0000977 | NRP + Polyketide | 26.0 | 48.0 | 204.0 | 1.5e-51 |
| WP\_010639241.1 | type\_I\_polyketide\_synthase | BGC0000958 | NRP:Cyclic depsipeptide + Polyketide:Modular type I | 27.0 | 41.2 | 202.0 | 5.8e-51 |
| BBC43184.1 | PKS-NRPS\_hybrid | BGC0001738 | NRP + Polyketide | 26.0 | 51.4 | 201.0 | 7.6e-51 |
| CAO91861.1 | PKS-NRPS\_hybrid | BGC0000968 | NRP + Polyketide:Iterative type I | 26.0 | 48.1 | 201.0 | 9.9e-51 |
| AAV66110.2 | fusaridione\_A\_synthetase | BGC0000992 | NRP + Polyketide | 25.0 | 50.8 | 201.0 | 1.3e-50 |
| AEO57481.1 | PKS-NRPSs | BGC0001449 | NRP + Alkaloid + Polyketide:Iterative type I | 25.0 | 53.4 | 201.0 | 1.3e-50 |
| ATZ45182.1 | Bcboa6 | BGC0001892 | Polyketide | 25.0 | 50.3 | 200.0 | 2.9e-50 |
| XP\_659388.1 | hypothetical\_protein | BGC0001998 | Polyketide | 27.0 | 43.2 | 199.0 | 3.8e-50 |
| AFV30248.1 | polyketide\_synthase | BGC0000075 | Polyketide | 24.0 | 45.3 | 199.0 | 4.9e-50 |
| EAU38971.1 | hypothetical\_protein | BGC0001122 | NRP + Polyketide:Iterative type I | 24.0 | 60.0 | 199.0 | 4.9e-50 |
| ABJ97439.1 | MerC | BGC0001012 | NRP + Polyketide | 25.0 | 50.0 | 198.0 | 6.4e-50 |
| AKA59437.1 | polyketide\_synthase | BGC0001202 | NRP + Polyketide | 27.0 | 36.6 | 194.0 | 1.6e-48 |
| CAQ34917.1 | polyketide\_synthase | BGC0000986 | NRP + Polyketide | 32.0 | 26.6 | 192.0 | 4.6e-48 |
| ACO79122.1 | Type\_I\_fatty\_acid\_synthase\_ArsA | BGC0000284 | Polyketide | 26.0 | 38.5 | 184.0 | 1.6e-45 |
| EAL85113.2 | hybrid\_PKS-NRPS\_enzyme | BGC0001037 | NRP + Polyketide:Iterative type I | 24.0 | 51.2 | 182.0 | 6.2e-45 |
| XP\_001220460.1 | hypothetical\_protein | BGC0001182 | NRP + Polyketide:Iterative type I | 25.0 | 51.1 | 181.0 | 8.1e-45 |
| KFL51881.1 | beta-ketoacyl\_synthase | BGC0001711 | NRP + Polyketide | 30.0 | 25.3 | 180.0 | 3.1e-44 |
| AEC13079.1 | fosA | BGC0000060 | Polyketide | 31.0 | 28.1 | 177.0 | 1.5e-43 |
| AEZ53953.1 | polyketide\_synthase | BGC0000144 | Polyketide:Modular type I | 26.0 | 34.4 | 177.0 | 1.5e-43 |
| AKQ22698.1 | malonyl\_CoA-acyl\_carrier\_protein\_transacylase | BGC0001186 | Polyketide | 28.0 | 27.8 | 175.0 | 7.6e-43 |
| ATX68127.1 | malonyl\_CoA-acyl\_carrier\_protein\_transacylase | BGC0001795 | Polyketide | 28.0 | 28.5 | 175.0 | 1e-42 |
| AJY78093.1 | polyketide\_synthase | BGC0001902 | NRP + Polyketide | 32.0 | 26.7 | 174.0 | 1.3e-42 |
| AXA20093.1 | trans-AT\_PKS\_LgaD | BGC0001946 | NRP + Polyketide | 29.0 | 31.1 | 174.0 | 1.7e-42 |
| BAQ25512.1 | type\_I\_polyketide\_synthase | BGC0001288 | Polyketide | 27.0 | 28.3 | 172.0 | 4.9e-42 |
| CAJ57411.1 | polyketide\_synthase\_type\_I | BGC0000176 | Polyketide + NRP | 28.0 | 31.0 | 170.0 | 1.9e-41 |
| CAL69894.1 | RhiF\_protein | BGC0001112 | NRP + Polyketide:Trans-AT type I | 29.0 | 29.4 | 170.0 | 2.5e-41 |
| AZF85941.1 | type\_I\_polyketide\_synthase | BGC0001963 | NRP + Polyketide | 28.0 | 26.6 | 170.0 | 3.2e-41 |
| EWM62997.1 | non-ribosomal\_peptide\_synthetase | BGC0001328 | NRP:Cyclic depsipeptide + Polyketide:Modular type I | 32.0 | 23.5 | 168.0 | 7.1e-41 |
| ADA69239.2 | trans-AT\_hybrid\_polyketide\_synthase-NRPS | BGC0001071 | NRP + Polyketide:Modular type I + Polyketide:Trans-AT type I | 27.0 | 27.3 | 168.0 | 9.3e-41 |
| AKQ22696.1 | malonyl\_CoA-acyl\_carrier\_protein\_transacylase | BGC0001186 | Polyketide | 27.0 | 30.1 | 167.0 | 2.1e-40 |
| AAM12909.2 | MmpA | BGC0000182 | Polyketide:Iterative type I + Polyketide:Trans-AT type I | 28.0 | 26.7 | 166.0 | 2.7e-40 |
| AKQ22669.1 | malonyl\_CoA-acyl\_carrier\_protein\_transacylase | BGC0001656 | Polyketide | 27.0 | 28.5 | 166.0 | 4.6e-40 |
| AKQ22682.1 | malonyl\_CoA-acyl\_carrier\_protein\_transacylase | BGC0001656 | Polyketide | 27.0 | 27.5 | 163.0 | 2.3e-39 |
| ADD82941.1 | Bat3 | BGC0001099 | NRP + Polyketide:Modular type I + Polyketide:Trans-AT type I | 27.0 | 30.3 | 163.0 | 3e-39 |
| PHM26606.1 | malonyl\_CoA-acyl\_carrier\_protein\_transacylase | BGC0001130 | NRP + Polyketide | 26.0 | 39.1 | 163.0 | 3e-39 |
| AJQ95708.1 | polyketide\_synthase\_modules-related\_protein | BGC0001644 | Polyketide | 27.0 | 28.6 | 163.0 | 3e-39 |
| CCC21123.1 | type-I\_polyketide\_synthases | BGC0000171 | Polyketide:Modular type I | 30.0 | 20.1 | 162.0 | 5.1e-39 |
| ABI91469.1 | beta-ketoacyl\_synthase | BGC0001094 | NRP + Polyketide | 30.0 | 26.6 | 162.0 | 6.7e-39 |
| OEI73466.1 | hypothetical\_protein | BGC0001520 | Polyketide | 29.0 | 26.9 | 161.0 | 1.1e-38 |
| ABC34599.1 | polyketide\_synthase,\_putative | BGC0000186 | NRP + Polyketide:Modular type I | 28.0 | 27.5 | 161.0 | 1.5e-38 |
| WP\_078625924.1 | type\_I\_polyketide\_synthase | BGC0002010 | NRP + Polyketide | 24.0 | 46.5 | 160.0 | 1.9e-38 |
| CTQ34882.1 | AtcE;\_polyketide\_synthase,\_modules\_5-7 | BGC0001301 | Polyketide | 29.0 | 29.1 | 160.0 | 3.3e-38 |
| CAG23959.2 | polyketide\_synthase\_of\_type\_I | BGC0001089 | Polyketide + NRP | 26.0 | 28.8 | 159.0 | 4.3e-38 |
| AVR48535.1 | CusC | BGC0001564 | NRP + Polyketide | 25.0 | 27.7 | 159.0 | 4.3e-38 |
| CBJ82077.1 | hypothetical\_protein | BGC0001872 | Polyketide | 26.0 | 38.8 | 158.0 | 7.4e-38 |
| ABS90472.1 | PKS | BGC0001106 | NRP + Polyketide | 32.0 | 20.1 | 158.0 | 1.3e-37 |
| CAQ18838.1 | polyketide\_synthase | BGC0000954 | NRP + Polyketide:Modular type I | 28.0 | 27.1 | 157.0 | 2.1e-37 |
| ABC35796.1 | polyketide\_synthase,\_putative | BGC0001102 | NRP:Beta-lactam + Polyketide:Modular type I | 30.0 | 23.9 | 156.0 | 2.8e-37 |
| WP\_018960020.1 | type\_I\_polyketide\_synthase | BGC0002010 | NRP + Polyketide | 25.0 | 39.3 | 156.0 | 3.7e-37 |
| ctg1\_orf5 |  | BGC0001329 | Polyketide + NRP:Cyclic depsipeptide | 30.0 | 21.4 | 155.0 | 6.3e-37 |
| AXA20091.1 | hybrid\_trans-AT\_PKS/NRPS\_LgaB | BGC0001946 | NRP + Polyketide | 30.0 | 26.8 | 155.0 | 6.3e-37 |
| RAT98517.1 | trans-acyltransferase\_polyketide\_synthase | BGC0001470 | Polyketide:Trans-AT type I | 29.0 | 24.3 | 155.0 | 1.1e-36 |
| AMH40443.1 | PKS | BGC0001350 | Polyketide | 27.0 | 29.5 | 154.0 | 1.4e-36 |
| ATX68109.1 | malonyl\_CoA-acyl\_carrier\_protein\_transacylase | BGC0001772 | Polyketide | 28.0 | 28.3 | 154.0 | 1.4e-36 |
| CAG23964.1 | polyketide\_synthase\_type\_I | BGC0000181 | Polyketide | 24.0 | 36.3 | 153.0 | 2.4e-36 |
| AIJ04681.1 | polyketide\_synthase | BGC0001383 | Polyketide | 24.0 | 36.2 | 153.0 | 3.1e-36 |
| ASA76632.1 | polyketide\_synthase\_non-ribosomal\_peptide\_synthetase\_hybrid | BGC0001751 | NRP + Polyketide | 31.0 | 20.0 | 152.0 | 5.3e-36 |
| ALD83687.1 | tAT\_polyketide\_synthase | BGC0001300 | Polyketide | 27.0 | 29.0 | 152.0 | 6.9e-36 |
| AAM94794.1 | CalE8 | BGC0000033 | Polyketide | 25.0 | 40.1 | 151.0 | 1.5e-35 |
| AAY89049.1 | polyketide\_synthase | BGC0001069 | NRP + Polyketide:Trans-AT type I | 29.0 | 20.2 | 149.0 | 5.9e-35 |
| CAE51182.1 | RemE\_protein | BGC0000264 | Polyketide:Type II | 29.0 | 22.5 | 147.0 | 1.7e-34 |
| CCM44330.1 | Polyketide\_synthase | BGC0001056 | NRP + Polyketide:Modular type I + Polyketide:PUFA synthase or related | 24.0 | 38.3 | 147.0 | 2.2e-34 |
| ABI91465.1 | beta-ketoacyl\_synthase | BGC0001094 | NRP + Polyketide | 31.0 | 20.9 | 146.0 | 2.9e-34 |
| CCA89326.1 | mixed\_trans-AT\_type\_I\_polyketide\_synthase/nonribosomal\_peptide\_synthetase | BGC0001111 | NRP + Polyketide:Trans-AT type I | 29.0 | 20.6 | 146.0 | 3.8e-34 |
| AMK92560.1 | enediyne\_polyketide\_synthase | BGC0001815 | Polyketide | 26.0 | 37.4 | 146.0 | 3.8e-34 |
| ATX68111.1 | malonyl\_CoA-acyl\_carrier\_protein\_transacylase | BGC0001772 | Polyketide | 28.0 | 20.2 | 146.0 | 3.8e-34 |
| ACY06292.1 | modular\_polyketide\_synthase | BGC0001042 | NRP + Polyketide | 26.0 | 39.7 | 146.0 | 5e-34 |
| AYJ71721.1 | non-ribosomal\_peptide\_synthetase | BGC0001942 | NRP + Polyketide | 28.0 | 22.7 | 146.0 | 5e-34 |
| AAQ17110.2 | enediyne\_polyketide\_synthase | BGC0001008 | Polyketide:Iterative type I + Polyketide:Enediyne type I | 24.0 | 39.7 | 145.0 | 6.5e-34 |
| ALD83703.1 | tAT\_polyketide\_synthase | BGC0001299 | Polyketide | 31.0 | 21.2 | 144.0 | 1.1e-33 |
| AXA20092.1 | trans-AT\_PKS\_LgaC | BGC0001946 | NRP + Polyketide | 29.0 | 23.6 | 144.0 | 1.1e-33 |
| AFX60318.1 | polyketide\_synthase | BGC0001031 | NRP + Polyketide | 27.0 | 24.8 | 144.0 | 1.9e-33 |
| AKQ22681.1 | malonyl\_CoA-acyl\_carrier\_protein\_transacylase | BGC0001656 | Polyketide | 26.0 | 29.2 | 143.0 | 2.5e-33 |
| ATV95639.1 | type\_I\_PKS | BGC0001503 | Polyketide | 23.0 | 39.8 | 143.0 | 3.2e-33 |
| RAT98529.1 | trans-acyltransferase\_polyketide\_synthase | BGC0001470 | Polyketide:Trans-AT type I | 30.0 | 20.0 | 143.0 | 4.2e-33 |
| BBA20952.1 | type\_I\_polyketide\_synthase | BGC0001763 | NRP + Polyketide | 27.0 | 28.5 | 143.0 | 4.2e-33 |
| ALD83704.1 | tAT\_polyketide\_synthase | BGC0001299 | Polyketide | 30.0 | 20.0 | 141.0 | 1.2e-32 |
| ATX68125.1 | malonyl\_CoA-acyl\_carrier\_protein\_transacylase | BGC0001795 | Polyketide | 30.0 | 20.5 | 141.0 | 1.2e-32 |
| ABP73645.1 | SalA | BGC0000145 | Polyketide | 24.0 | 28.8 | 141.0 | 1.6e-32 |
| ACR50796.1 | putative\_polyketide\_synthase | BGC0000163 | Polyketide | 31.0 | 21.0 | 140.0 | 2.1e-32 |
| ABP53498.1 | PKS\_(ACP-AT-AT-KS-ACP-C) | BGC0001041 | NRP + Polyketide | 24.0 | 30.5 | 140.0 | 2.7e-32 |
| BAP05594.1 | calF | BGC0000967 | NRP + Polyketide:Trans-AT type I | 28.0 | 26.9 | 139.0 | 3.6e-32 |
| AGN11883.1 | tstI | BGC0001114 | NRP + Polyketide | 28.0 | 27.2 | 139.0 | 3.6e-32 |
| ABP55141.1 | beta-ketoacyl\_synthase | BGC0000150 | NRP + Polyketide:Enediyne type I | 24.0 | 39.7 | 139.0 | 4.6e-32 |
| ACM79805.1 | ZmaA | BGC0001059 | NRP + Polyketide | 30.0 | 20.6 | 137.0 | 1.8e-31 |
| ADH01490.1 | type\_I\_polyketide\_synthase | BGC0001096 | NRP + Polyketide | 28.0 | 27.7 | 136.0 | 3e-31 |
| AIC32695.1 | FR9I | BGC0001113 | NRP + Polyketide | 28.0 | 27.7 | 136.0 | 3e-31 |
| AAS47564.1 | mixed\_type\_I\_polyketide\_synthase/nonribosomal\_peptide\_synthetase | BGC0001108 | Polyketide:Trans-AT type I | 27.0 | 27.2 | 136.0 | 5.1e-31 |
| ctg1\_orf6 |  | BGC0001109 | NRP + Polyketide | 27.0 | 27.2 | 136.0 | 5.1e-31 |
| AAM78012.1 | warhead-forming\_iterative\_polyketide\_synthase | BGC0000112 | Polyketide:Iterative type I + Polyketide:Enediyne type I | 25.0 | 40.3 | 133.0 | 3.3e-30 |
| WP\_055469549.1 | type\_I\_polyketide\_synthase | BGC0001537 | Polyketide | 27.0 | 26.1 | 132.0 | 5.7e-30 |
| AEC04364.1 | polyketide\_synthase | BGC0000178 | Polyketide:Trans-AT type I | 25.0 | 28.0 | 131.0 | 1.3e-29 |
| AFX60341.1 | polyketide\_synthase | BGC0001032 | NRP + Polyketide | 26.0 | 24.8 | 130.0 | 2.2e-29 |
| AFV52145.1 | polyketide\_synthase | BGC0000081 | Polyketide:Iterative type I + Polyketide:Enediyne type I | 24.0 | 38.1 | 130.0 | 2.8e-29 |
| ANY94470.1 | enediyne\_polyketide\_synthase | BGC0001584 | Polyketide | 23.0 | 38.6 | 129.0 | 3.7e-29 |
| CAN89634.1 | putative\_polyketide\_synthase | BGC0001070 | NRP + Polyketide:Modular type I + Polyketide:Trans-AT type I | 28.0 | 20.4 | 129.0 | 4.8e-29 |
| AAL06699.1 | polyketide\_synthase | BGC0000965 | Polyketide:Iterative type I + Polyketide:Enediyne type I | 23.0 | 38.8 | 129.0 | 6.3e-29 |
| ALU98461.1 | erythronolide\_synthase | BGC0001397 | NRP + Polyketide | 23.0 | 38.8 | 129.0 | 6.3e-29 |
| AJQ95706.1 | polyketide\_synthase\_modules-related\_protein | BGC0001644 | Polyketide | 26.0 | 20.1 | 128.0 | 8.2e-29 |
| AAP92148.1 | EspE | BGC0000056 | Polyketide | 25.0 | 39.9 | 126.0 | 4.1e-28 |
| OAQ83765.1 | KR\_domain-containing\_protein | BGC0001358 | Polyketide | 29.0 | 20.1 | 125.0 | 9.1e-28 |
| EYT83433.1 | hypothetical\_protein | BGC0001213 | Polyketide | 24.0 | 38.3 | 124.0 | 2e-27 |
| ALD82521.1 | polyketide\_synthase | BGC0001212 | NRP + Polyketide | 24.0 | 53.2 | 121.0 | 1.7e-26 |
| AXA20090.1 | hybrid\_trans-AT\_PKS/NRPS\_LgaA | BGC0001946 | NRP + Polyketide | 26.0 | 27.3 | 120.0 | 2.2e-26 |
| AAG31130.1 | MxcG | BGC0001345 | NRP | 26.0 | 27.7 | 120.0 | 2.9e-26 |
| CBK62729.1 |  | BGC0001115 | NRP + Polyketide | 24.0 | 28.3 | 113.0 | 2.7e-24 |
| ALE27513.1 | putative\_polyketide\_synthase | BGC0001292 | Other | 24.0 | 40.4 | 110.0 | 3e-23 |
| AJY78091.1 | polyketide\_synthase | BGC0001902 | NRP + Polyketide | 24.0 | 43.5 | 105.0 | 7.4e-22 |
| ABC87512.1 | polyketide\_synthase | BGC0001011 | NRP + Polyketide | 27.0 | 21.3 | 97.0 | 2e-19 |
| ABI22133.1 | putative\_non-ribosomal\_peptide\_synthetase | BGC0000422 | NRP | 25.0 | 20.1 | 91.0 | 1.4e-17 |
| ABW71853.1 | nonribosomal\_peptide\_synthetase | BGC0000303 | NRP | 25.0 | 23.7 | 83.0 | 3e-15 |
| ATP76246.1 | SpuB | BGC0001748 | NRP + Polyketide | 22.0 | 25.7 | 83.0 | 3.9e-15 |
| AEC14349.1 | nonribosomal\_peptide\_synthetase | BGC0000377 | NRP | 24.0 | 20.3 | 77.0 | 2.8e-13 |
