## Supplementary Results for "A draft genome of the ascomycotal fungal species *Pseudopithomyces maydicus* (family *Didymosphaeriaceae*)": input.path1.gene6_mibig_hits.html

| MIBiG Protein | Description | MIBiG Cluster | MiBiG Product | % ID | % Coverage | BLAST Score | E-value |
| --- | --- | --- | --- | --- | --- | --- | --- |
| ADI24954.1 | GsfB | BGC0000070 | Polyketide:Iterative type I | 39.0 | 65.8 | 69.0 | 4.1e-12 |
| ADM79461.1 | O-methyltransferase | BGC0001266 | Polyketide | 44.0 | 65.1 | 69.0 | 7.1e-12 |
