## Supplementary Results for "A draft genome of the ascomycotal fungal species *Pseudopithomyces maydicus* (family *Didymosphaeriaceae*)": input.path1.gene7_mibig_hits.html

| MIBiG Protein | Description | MIBiG Cluster | MiBiG Product | % ID | % Coverage | BLAST Score | E-value |
| --- | --- | --- | --- | --- | --- | --- | --- |
| PLB46274.1 | putative\_allantoate\_permease | BGC0001712 | Other | 38.0 | 89.0 | 275.0 | 2.1e-73 |
| KGO40481.1 | Major\_facilitator\_superfamily\_domain,\_general\_substrate\_transporter | BGC0001205 | Polyketide | 29.0 | 97.0 | 196.0 | 9.7e-50 |
| MAA\_10050 | vitamin\_H\_transporter,\_putative | BGC0000337 | NRP | 32.0 | 81.3 | 177.0 | 6.1e-44 |
| KGO40484.1 | Major\_facilitator\_superfamily\_domain,\_general\_substrate\_transporter | BGC0001205 | Polyketide | 23.0 | 84.5 | 128.0 | 2.5e-29 |
