## Supplementary Results for "A draft genome of the ascomycotal fungal species *Pseudopithomyces maydicus* (family *Didymosphaeriaceae*)": input.path1.gene9_mibig_hits.html

| MIBiG Protein | Description | MIBiG Cluster | MiBiG Product | % ID | % Coverage | BLAST Score | E-value |
| --- | --- | --- | --- | --- | --- | --- | --- |
| AIA58903.1 | thiohydrolase | BGC0001141 | Polyketide:Iterative type I | 55.0 | 50.1 | 317.0 | 3.4e-86 |
| EAL85128.1 | acyltransferase | BGC0001067 | Terpene + Polyketide:Iterative type I | 43.0 | 50.4 | 246.0 | 1.2e-64 |
| CCT72379.1 | related\_to\_DltD\_N-terminal\_domain\_protein | BGC0001305 | Polyketide | 38.0 | 52.8 | 219.0 | 1.2e-56 |
| CBF77085.1 | cytochrome\_P450,\_putative\_(Eurofung) | BGC0001679 | NRP | 30.0 | 34.4 | 125.0 | 3.2e-28 |
| AAK33073.1 | cytochrome\_P450 | BGC0001278 | Terpene | 33.0 | 33.8 | 121.0 | 3.5e-27 |
| XP\_001213595.1 | hypothetical\_protein | BGC0001475 | Polyketide | 31.0 | 36.9 | 121.0 | 3.5e-27 |
| BAX01960.1 | trichodiene\_oxygenase | BGC0001811 | Terpene | 32.0 | 34.0 | 121.0 | 6e-27 |
| AAK53577.1 | trichodiene\_oxygenase | BGC0000930 | Other | 32.0 | 34.0 | 120.0 | 7.9e-27 |
| AAK33083.1 | putative\_cytochrome\_P450 | BGC0001277 | Terpene | 32.0 | 34.0 | 119.0 | 2.3e-26 |
| PLB46285.1 | cytochrome\_P450 | BGC0001712 | Other | 30.0 | 36.1 | 115.0 | 2.5e-25 |
| AZQ56743.1 | cytochrome\_P450 | BGC0001969 | Terpene | 31.0 | 33.8 | 108.0 | 5.3e-23 |
| ADM79460.1 | P450\_protein | BGC0001266 | Polyketide | 36.0 | 27.9 | 106.0 | 1.5e-22 |
| XP\_001826052.1 |  | BGC0001995 | Terpene | 27.0 | 35.0 | 103.0 | 9.9e-22 |
| BAV32149.1 | cytochrome\_P450\_monooxygenase | BGC0001373 | Polyketide | 27.0 | 34.2 | 97.0 | 9.3e-20 |
| AIA58896.1 | putative\_cytochrome\_P450 | BGC0001141 | Polyketide:Iterative type I | 28.0 | 33.5 | 92.0 | 1.8e-18 |
| WP\_020636849.1 | alpha/beta\_fold\_hydrolase | BGC0002011 | Polyketide | 29.0 | 43.9 | 91.0 | 5.1e-18 |
| BAH23996.1 | cytochrome\_P450 | BGC0000356 | NRP + Alkaloid | 29.0 | 24.4 | 61.0 | 7.4e-09 |
| AGC83578.1 | P450\_monooxygenase | BGC0000818 | NRP | 28.0 | 23.6 | 56.0 | 1.4e-07 |
| ADM34140.1 | P450 | BGC0001084 | NRP + Terpene + Alkaloid | 28.0 | 23.6 | 56.0 | 1.8e-07 |
| BAD83682.1 | cytochrome\_P-450 | BGC0000012 | Polyketide | 30.0 | 26.7 | 56.0 | 2.4e-07 |
| BAE56591.1 |  | BGC0001123 | NRP | 31.0 | 25.0 | 55.0 | 3.1e-07 |
