## Supplementary Results for "A draft genome of the ascomycotal fungal species *Pseudopithomyces maydicus* (family *Didymosphaeriaceae*)": input.path1.gene10_mibig_hits.html

| MIBiG Protein | Description | MIBiG Cluster | MiBiG Product | % ID | % Coverage | BLAST Score | E-value |
| --- | --- | --- | --- | --- | --- | --- | --- |
| AIA58899.1 | HRPKS | BGC0001141 | Polyketide:Iterative type I | 42.0 | 112.3 | 1675.0 | 0.0 |
| EAL85129.1 | polyketide\_synthase | BGC0001067 | Terpene + Polyketide:Iterative type I | 37.0 | 117.2 | 1364.0 | 0.0 |
| ATZ45185.1 | Bcboa9 | BGC0001892 | Polyketide | 30.0 | 112.4 | 860.0 | 4.8e-249 |
| AAD43562.2 | Fum1p | BGC0000062 | Polyketide | 30.0 | 117.9 | 856.0 | 9e-248 |
| EPE34340.1 | polyketide\_synthase | BGC0001035 | Polyketide + NRP + Other:Aminocoumarin | 29.0 | 120.6 | 849.0 | 1.1e-245 |
| ACB12550.1 | Fum1 | BGC0000063 | Polyketide | 29.0 | 122.9 | 846.0 | 7.1e-245 |
| EHA19289.1 | hypothetical\_protein | BGC0001124 | Polyketide | 30.0 | 111.6 | 820.0 | 7.1e-237 |
| BBG28498.1 | putative\_polyketide\_synthase | BGC0001913 | Polyketide | 28.0 | 122.7 | 816.0 | 1e-235 |
| EAA36364.1 | hypothetical\_protein | BGC0001697 | Polyketide | 28.0 | 112.0 | 803.0 | 6.9e-232 |
| OAQ83765.1 | KR\_domain-containing\_protein | BGC0001358 | Polyketide | 29.0 | 114.4 | 795.0 | 2.5e-229 |
| ABA02240.1 | polyketide\_synthase | BGC0000098 | Polyketide | 28.0 | 123.3 | 783.0 | 9.7e-226 |
| AGC95324.1 | CurS1 | BGC0000045 | Polyketide | 29.0 | 113.9 | 781.0 | 2.8e-225 |
| BAC20566.1 | polyketide\_synthase | BGC0000039 | Polyketide | 28.0 | 120.2 | 777.0 | 6.9e-224 |
| ACD39774.1 | reducing\_polyketide\_synthase | BGC0000134 | Polyketide | 30.0 | 112.5 | 773.0 | 7.7e-223 |
| EHA28244.1 | hypothetical\_protein | BGC0001143 | Polyketide | 28.0 | 122.7 | 773.0 | 1e-222 |
| AKL78824.1 | GLPKS3 | BGC0001187 | NRP:Lipopeptide + Polyketide:Iterative type I | 29.0 | 113.0 | 772.0 | 2.2e-222 |
| AHV78245.1 | LasS1 | BGC0001245 | Polyketide | 29.0 | 114.4 | 772.0 | 2.2e-222 |
| AHV78252.1 | ResS1 | BGC0001246 | Polyketide | 29.0 | 115.1 | 770.0 | 5e-222 |
| ATQ39432.1 | PKS | BGC0001565 | NRP | 28.0 | 121.1 | 766.0 | 1.2e-220 |
| ABB90283.1 | polyketide\_synthase | BGC0001057 | NRP + Polyketide | 29.0 | 113.7 | 765.0 | 1.6e-220 |
| ACD39758.1 | reducing\_polyketide\_synthase | BGC0000076 | Polyketide | 28.0 | 114.5 | 761.0 | 3.9e-219 |
| ACD39767.1 | reducing\_polyketide\_synthase | BGC0000077 | Polyketide | 28.0 | 114.5 | 761.0 | 3.9e-219 |
| AAD34559.1 | polyketide\_synthase | BGC0000088 | Polyketide | 28.0 | 121.6 | 760.0 | 8.8e-219 |
| ASK38717.1 | polyketide\_synthase | BGC0001557 | Polyketide | 29.0 | 116.5 | 759.0 | 1.5e-218 |
| AMY15068.1 | hexaketide\_synthase\_MF-SQHKS | BGC0001339 | Polyketide:Iterative type I | 28.0 | 117.8 | 743.0 | 1.1e-213 |
| EWG54266.1 | hypothetical\_protein | BGC0001190 | Polyketide | 28.0 | 112.4 | 742.0 | 2.5e-213 |
| BAD83684.1 | PKSN\_polyketide\_synthase\_for\_alternapyrone\_biosynthesis | BGC0000012 | Polyketide | 28.0 | 123.1 | 738.0 | 2.1e-212 |
| OAQ83760.1 | polyketide\_synthase | BGC0001358 | Polyketide | 27.0 | 123.4 | 737.0 | 4.7e-212 |
| ACZ57548.1 | polyketide\_synthase | BGC0000046 | Polyketide:Iterative type I | 28.0 | 110.9 | 736.0 | 1e-211 |
| BAJ09789.1 | polyketide\_synthase | BGC0000146 | Polyketide | 27.0 | 124.0 | 734.0 | 3.9e-211 |
| CBF87072.1 | polyketide\_synthase,\_putative\_(Eurofung) | BGC0001290 | NRP | 28.0 | 119.1 | 730.0 | 5.7e-210 |
| CCT75967.1 | polyketide\_synthase | BGC0001606 | Polyketide | 27.0 | 121.1 | 725.0 | 2.4e-208 |
| CBX99534.1 | similar\_to\_polyketide\_synthase | BGC0001899 | Polyketide | 29.0 | 120.9 | 720.0 | 1e-206 |
| ANF07288.1 | hrPKS | BGC0001340 | Polyketide:Iterative type I | 27.0 | 124.5 | 699.0 | 1.4e-200 |
| CAP95405.1 |  | BGC0001404 | Polyketide | 27.0 | 123.7 | 691.0 | 3.8e-198 |
| AMY15057.1 | tetraketide\_synthase\_MF-SQTKS | BGC0001339 | Polyketide:Iterative type I | 30.0 | 97.9 | 681.0 | 3e-195 |
| EAA65604.1 | hypothetical\_protein | BGC0000022 | Polyketide | 27.0 | 122.9 | 666.0 | 1e-190 |
| EHA52508.1 | mycocerosic\_acid\_synthase | BGC0001749 | Polyketide | 26.0 | 116.7 | 653.0 | 1.2e-186 |
| EAQ86385.1 | hypothetical\_protein | BGC0001405 | Polyketide | 26.0 | 117.5 | 612.0 | 2.9e-174 |
| KKP00963.1 | fatty\_acid\_synthase\_S-acetyltransferase | BGC0001901 | Polyketide | 27.0 | 95.4 | 604.0 | 4.7e-172 |
| RWQ92174.1 | KR\_domain-containing\_protein | BGC0002030 | Polyketide | 25.0 | 110.0 | 567.0 | 1.1e-160 |
| AQM58285.1 | polyketide\_synthase | BGC0001816 | NRP + Polyketide | 27.0 | 100.8 | 555.0 | 2.5e-157 |
| BAV32159.1 | polyketide\_synthase | BGC0001373 | Polyketide | 27.0 | 98.2 | 554.0 | 7.3e-157 |
| QCC63000.1 | BII-rafflesfungin\_polyketide\_synthase | BGC0001966 | NRP | 28.0 | 95.5 | 549.0 | 3.1e-155 |
| CAQ18830.1 | polyketide\_synthase | BGC0000954 | NRP + Polyketide:Modular type I | 26.0 | 114.1 | 536.0 | 1.6e-151 |
| AGC45624.1 | polyketide\_synthase | BGC0001394 | NRP + Polyketide | 25.0 | 106.8 | 527.0 | 1.2e-148 |
| AQW44889.1 | polyketide\_synthase | BGC0001737 | NRP + Polyketide | 26.0 | 113.0 | 517.0 | 9.9e-146 |
| gene4 |  | BGC0001907 | Polyketide | 26.0 | 106.3 | 514.0 | 1.1e-144 |
| BAN19720.1 | polyketide\_synthase | BGC0001252 | Polyketide | 27.0 | 95.0 | 505.0 | 3.9e-142 |
| CAJ46690.1 | polyketide\_synthase | BGC0000969 | NRP:Cyclic depsipeptide + Polyketide:Modular type I | 25.0 | 107.1 | 496.0 | 1.8e-139 |
| CAQ18832.1 | polyketide\_synthase | BGC0000954 | NRP + Polyketide:Modular type I | 26.0 | 107.7 | 495.0 | 4e-139 |
| AAF62883.1 | epoD | BGC0000991 | NRP + Polyketide | 26.0 | 108.4 | 483.0 | 1.6e-135 |
| ADB12491.1 | EpoD | BGC0000990 | NRP + Polyketide | 26.0 | 108.4 | 480.0 | 1.7e-134 |
| AAF26921.1 | polyketide\_synthase | BGC0000988 | NRP + Polyketide | 25.0 | 108.4 | 479.0 | 3e-134 |
| QDA77058.1 | polyketide\_synthase | BGC0002026 | NRP | 26.0 | 107.5 | 478.0 | 6.6e-134 |
| BAZ95823.1 | PKS-NRPS\_hybrid\_cpaA | BGC0001563 | NRP + Polyketide | 25.0 | 112.1 | 477.0 | 8.7e-134 |
| ACB46195.1 | polyketide\_synthase | BGC0000989 | NRP + Polyketide | 25.0 | 108.3 | 476.0 | 2.5e-133 |
| AAT28740.1 | FUSS | BGC0000064 | Polyketide | 25.0 | 110.4 | 471.0 | 8.1e-132 |
| AFP73394.1 | FusA | BGC0001268 | NRP + Polyketide | 25.0 | 110.4 | 463.0 | 1.3e-129 |
| AAV66110.2 | fusaridione\_A\_synthetase | BGC0000992 | NRP + Polyketide | 24.0 | 120.0 | 462.0 | 3.8e-129 |
| KGO40478.1 | Acyl\_transferase/acyl\_hydrolase/lysophospholipase | BGC0001205 | Polyketide | 27.0 | 93.3 | 453.0 | 1.3e-126 |
| gene3 |  | BGC0002035 | NRP + Polyketide | 24.0 | 117.2 | 449.0 | 2.5e-125 |
| ACS68554.1 | hybrid\_PKS-NRPS\_protein | BGC0001026 | NRP + Polyketide | 24.0 | 115.3 | 446.0 | 2.8e-124 |
| AGO86662.1 | equisetin\_synthetase | BGC0001255 | NRP + Polyketide | 24.0 | 118.9 | 445.0 | 4.8e-124 |
| ADN43685.1 | DmbS | BGC0001136 | NRP + Polyketide:Iterative type I | 25.0 | 118.7 | 443.0 | 1.8e-123 |
| ATZ45182.1 | Bcboa6 | BGC0001892 | Polyketide | 23.0 | 117.5 | 443.0 | 1.8e-123 |
| AQA28562.1 | type\_I\_polyketide\_synthase | BGC0001663 | Polyketide | 24.0 | 119.2 | 443.0 | 2.4e-123 |
| QBC19710.1 | TwmB | BGC0001954 | NRP + Polyketide | 24.0 | 116.5 | 440.0 | 1.5e-122 |
| CAQ18828.1 | polyketide\_synthase | BGC0000954 | NRP + Polyketide:Modular type I | 25.0 | 106.5 | 438.0 | 4.5e-122 |
| CAL58681.1 | polyketide\_synthase | BGC0000149 | Polyketide:Modular type I | 25.0 | 107.7 | 436.0 | 2.9e-121 |
| CCT72377.1 | probable\_polyketide\_synthase | BGC0001305 | Polyketide | 24.0 | 117.6 | 435.0 | 3.8e-121 |
| AUS29495.1 | polyketide\_synthase | BGC0001030 | NRP + Polyketide | 26.0 | 98.8 | 433.0 | 1.9e-120 |
| CAD19090.1 | StiF\_protein | BGC0000153 | NRP + Polyketide:Modular type I | 24.0 | 109.9 | 431.0 | 9.3e-120 |
| CAO91861.1 | PKS-NRPS\_hybrid | BGC0000968 | NRP + Polyketide:Iterative type I | 24.0 | 116.0 | 429.0 | 2.7e-119 |
| BAJ14522.1 | polyketide\_synthase | BGC0001254 | Polyketide | 24.0 | 119.8 | 429.0 | 2.7e-119 |
| EED49862.1 | hybrid\_PKS/NRPS\_enzyme,\_putative | BGC0001445 | NRP + Polyketide:Iterative type I | 23.0 | 117.1 | 428.0 | 7.9e-119 |
| CAL58684.1 | polyketide\_synthase | BGC0000149 | Polyketide:Modular type I | 25.0 | 107.1 | 427.0 | 1e-118 |
| AZH23788.1 | MgcR | BGC0001970 | NRP + Polyketide | 23.0 | 108.2 | 426.0 | 2.3e-118 |
| BAD97694.1 | Aft9-1 | BGC0000003 | Polyketide | 29.0 | 55.5 | 416.0 | 2.4e-115 |
| EAW09117.1 | hybrid\_NRPS/PKS\_enzyme,\_putative | BGC0000983 | NRP + Polyketide:Iterative type I | 28.0 | 60.3 | 409.0 | 2.9e-113 |
| BAQ25466.1 | polyketide\_synthase | BGC0001280 | Polyketide | 25.0 | 116.3 | 404.0 | 7.2e-112 |
| AAQ82567.1 | FscE | BGC0000061 | Polyketide | 24.0 | 105.7 | 398.0 | 8.7e-110 |
| ADC45538.1 | modular\_polyketide\_synthase | BGC0000093 | Polyketide | 24.0 | 111.6 | 397.0 | 1.5e-109 |
| EPS29069.1 | hypothetical\_protein | BGC0001724 | NRP + Polyketide | 29.0 | 60.7 | 397.0 | 1.5e-109 |
| AEO57481.1 | PKS-NRPSs | BGC0001449 | NRP + Alkaloid + Polyketide:Iterative type I | 29.0 | 57.9 | 386.0 | 2.6e-106 |
| QCS37521.1 | PyiS | BGC0001982 | NRP + Polyketide | 28.0 | 61.4 | 383.0 | 1.7e-105 |
| CAL69597.1 | PKS-NRPS | BGC0001049 | NRP + Polyketide:Iterative type I | 28.0 | 63.4 | 378.0 | 5.5e-104 |
| CBF80487.1 | hybrid\_PKS-NRPS\_(Eurofung) | BGC0000959 | NRP + Polyketide:Iterative type I | 27.0 | 61.3 | 374.0 | 1.4e-102 |
| ARP51711.1 | PKS-NRPS\_hybrid\_protein | BGC0001741 | NRP + Polyketide | 28.0 | 57.4 | 372.0 | 5.1e-102 |
| ANR02555.1 | LodN | BGC0001648 | Polyketide | 24.0 | 106.0 | 370.0 | 2e-101 |
| ACF35445.1 | mbcAI | BGC0000090 | Polyketide | 24.0 | 107.0 | 368.0 | 9.7e-101 |
| QBE85649.1 | BuaA | BGC0001978 | NRP + Polyketide | 27.0 | 60.5 | 366.0 | 3.7e-100 |
| AAD03047.1 | type\_I\_polyketide\_synthase | BGC0000041 | Polyketide | 23.0 | 105.7 | 361.0 | 1.2e-98 |
| EAU38971.1 | hypothetical\_protein | BGC0001122 | NRP + Polyketide:Iterative type I | 26.0 | 60.7 | 358.0 | 1e-97 |
| EAL85113.2 | hybrid\_PKS-NRPS\_enzyme | BGC0001037 | NRP + Polyketide:Iterative type I | 28.0 | 58.6 | 357.0 | 1.7e-97 |
| AAZ94389.1 | modular\_polyketide\_synthase | BGC0000040 | Polyketide | 25.0 | 108.9 | 356.0 | 3.8e-97 |
| BAF85843.1 | modular\_polyketide\_synthase | BGC0000109 | Polyketide | 23.0 | 109.1 | 353.0 | 2.5e-96 |
| AFA26384.1 | polyketide\_synthase\_A | BGC0001874 | NRP + Polyketide | 28.0 | 61.1 | 352.0 | 3.2e-96 |
| XP\_659388.1 | hypothetical\_protein | BGC0001998 | Polyketide | 29.0 | 57.6 | 351.0 | 9.4e-96 |
| BAC20564.1 | polyketide\_synthase | BGC0000039 | Polyketide | 28.0 | 57.8 | 350.0 | 2.1e-95 |
| AJW65409.1 | type\_I\_modular\_polyketide\_synthase | BGC0001195 | NRP + Polyketide | 24.0 | 106.8 | 347.0 | 1e-94 |
| ADB23403.1 | polyketide\_synthase\_type\_I | BGC0001062 | Polyketide | 24.0 | 107.3 | 347.0 | 1.8e-94 |
| BBC43184.1 | PKS-NRPS\_hybrid | BGC0001738 | NRP + Polyketide | 27.0 | 59.0 | 345.0 | 5.1e-94 |
| ABA02239.1 | polyketide\_synthase | BGC0000098 | Polyketide | 27.0 | 59.6 | 340.0 | 1.7e-92 |
| ADB12492.1 | EpoE | BGC0000990 | NRP + Polyketide | 23.0 | 109.1 | 334.0 | 1.6e-90 |
| BAK26562.1 | PKS-NRPS\_hybrid | BGC0000977 | NRP + Polyketide | 28.0 | 56.3 | 333.0 | 2.6e-90 |
| ACB46196.1 | polyketide\_synthase | BGC0000989 | NRP + Polyketide | 23.0 | 108.2 | 332.0 | 4.5e-90 |
| AAF62884.1 | EpoE | BGC0000991 | NRP + Polyketide | 23.0 | 108.8 | 331.0 | 1.3e-89 |
| AAF26922.1 | polyketide\_synthase | BGC0000988 | NRP + Polyketide | 23.0 | 108.9 | 329.0 | 2.9e-89 |
| AEH42474.1 | polyketide\_synthase | BGC0000032 | Polyketide | 23.0 | 105.6 | 325.0 | 5.5e-88 |
| MAA\_10033 | polyketide\_synthase,\_putative | BGC0000337 | NRP | 28.0 | 51.2 | 322.0 | 6.1e-87 |
| CCE88378.1 | polyketide\_synthase | BGC0001034 | NRP + Polyketide:Modular type I | 23.0 | 105.9 | 319.0 | 3e-86 |
| AQW44891.1 | polyketide\_synthase | BGC0001737 | NRP + Polyketide | 26.0 | 58.2 | 314.0 | 1.3e-84 |
| AEE88279.1 | CurK | BGC0000976 | NRP + Polyketide:Modular type I | 23.0 | 103.6 | 313.0 | 2.8e-84 |
| AAT70106.1 | CurK | BGC0001165 | NRP + Polyketide:Modular type I | 23.0 | 103.6 | 313.0 | 2.8e-84 |
| AAK57189.1 | MxaE | BGC0001022 | NRP + Polyketide | 26.0 | 57.0 | 304.0 | 1.3e-81 |
| CCE88376.1 | polyketide\_synthase | BGC0001034 | NRP + Polyketide:Modular type I | 24.0 | 104.2 | 304.0 | 1.3e-81 |
| AGC45620.1 | polyketide\_synthase | BGC0001394 | NRP + Polyketide | 26.0 | 55.7 | 304.0 | 1.7e-81 |
| AQW44893.1 | polyketide\_synthase | BGC0001737 | NRP + Polyketide | 27.0 | 54.3 | 303.0 | 2.2e-81 |
| CAD19091.1 | StiG\_protein | BGC0000153 | NRP + Polyketide:Modular type I | 26.0 | 56.2 | 301.0 | 8.5e-81 |
| CAJ46689.1 | polyketide\_synthase | BGC0000969 | NRP:Cyclic depsipeptide + Polyketide:Modular type I | 27.0 | 55.4 | 301.0 | 8.5e-81 |
| CCE88380.1 | polyketide\_synthase | BGC0001034 | NRP + Polyketide:Modular type I | 26.0 | 54.3 | 301.0 | 8.5e-81 |
| AQW44888.1 | polyketide\_synthase | BGC0001737 | NRP + Polyketide | 26.0 | 55.7 | 301.0 | 1.1e-80 |
| AGC45621.1 | polyketide\_synthase | BGC0001394 | NRP + Polyketide | 26.0 | 56.3 | 299.0 | 4.2e-80 |
| AQW44892.1 | polyketide\_synthase | BGC0001737 | NRP + Polyketide | 27.0 | 54.0 | 298.0 | 9.4e-80 |
| XP\_001220460.1 | hypothetical\_protein | BGC0001182 | NRP + Polyketide:Iterative type I | 25.0 | 58.0 | 295.0 | 4.7e-79 |
| ACR33077.1 | polyketide\_synthase | BGC0000017 | Alkaloid + Polyketide:Modular type I | 23.0 | 102.6 | 294.0 | 1.4e-78 |
| ACB46197.1 | polyketide\_synthase | BGC0000989 | NRP + Polyketide | 23.0 | 105.3 | 289.0 | 4.4e-77 |
| ABL86391.1 | hybrid\_polyketide\_synthase\_and\_nonribosomal\_peptide\_synthetase | BGC0000999 | NRP + Polyketide | 26.0 | 54.9 | 289.0 | 4.4e-77 |
| AAS98783.1 | polyketide\_synthase/nonribosomal\_peptide\_synthase\_hybrid | BGC0001001 | NRP + Polyketide | 23.0 | 103.6 | 288.0 | 7.5e-77 |
| ADB12493.1 | EpoF | BGC0000990 | NRP + Polyketide | 23.0 | 105.7 | 287.0 | 2.2e-76 |
| AAF62885.1 | EpoF | BGC0000991 | NRP + Polyketide | 23.0 | 105.7 | 284.0 | 1.4e-75 |
| AAF26923.1 | polyketide\_synthase | BGC0000988 | NRP + Polyketide | 23.0 | 105.1 | 282.0 | 4.1e-75 |
| AAK19883.1 | soraphen\_polyketide\_synthase\_A | BGC0000147 | Polyketide:Modular type I | 34.0 | 32.0 | 280.0 | 2.7e-74 |
| AZH23818.1 | MgiI | BGC0001971 | NRP + Polyketide | 23.0 | 102.6 | 277.0 | 1.3e-73 |
| BBG28484.1 | polyketide\_synthase\_CdmE | BGC0001926 | Polyketide | 29.0 | 39.6 | 277.0 | 1.7e-73 |
| CBD77734.1 | polyketide\_synthase | BGC0000974 | NRP + Polyketide | 28.0 | 42.4 | 275.0 | 6.5e-73 |
| AIR74911.1 | polyketide\_synthase | BGC0001559 | RiPP | 28.0 | 42.4 | 275.0 | 6.5e-73 |
| AZH23789.1 | MgcI | BGC0001970 | NRP + Polyketide | 23.0 | 103.4 | 275.0 | 8.5e-73 |
| AAM81586.2 | putative\_type\_I\_polyketide\_synthase | BGC0000047 | Polyketide | 23.0 | 105.2 | 272.0 | 4.2e-72 |
| ABK32263.1 | AmbH | BGC0000014 | Polyketide | 27.0 | 43.4 | 271.0 | 1.2e-71 |
| ARM20280.1 | polyketide\_synthase | BGC0001523 | Polyketide | 24.0 | 104.8 | 270.0 | 1.6e-71 |
| CAQ43075.1 | polyketide\_synthase | BGC0000970 | NRP + Polyketide:Modular type I | 24.0 | 108.5 | 270.0 | 2.1e-71 |
| AAS79461.1 | polyketide\_synthase\_subunit | BGC0000035 | Polyketide | 23.0 | 102.1 | 269.0 | 4.7e-71 |
| AAU04878.1 | polyketide\_synthase | BGC0000365 | NRP | 24.0 | 100.9 | 269.0 | 4.7e-71 |
| ABC84457.1 | NigAII | BGC0000114 | Polyketide:Modular type I | 24.0 | 103.0 | 268.0 | 8e-71 |
| BAO98805.1 | putative\_polyketide\_synthase | BGC0001002 | NRP + Polyketide | 25.0 | 56.0 | 266.0 | 4e-70 |
| CAA60460.1 | polyketide\_synthase | BGC0001040 | NRP + Polyketide | 35.0 | 26.0 | 266.0 | 4e-70 |
| WP\_042799407.1 | type\_I\_polyketide\_synthase | BGC0001283 | Polyketide | 22.0 | 105.1 | 266.0 | 4e-70 |
| AVI57434.1 | AbmB2 | BGC0001694 | Polyketide | 23.0 | 106.8 | 266.0 | 4e-70 |
| AVV61983.1 | type\_I\_modular\_polyketide\_synthase | BGC0001477 | NRP + Polyketide:Modular type I | 34.0 | 28.1 | 265.0 | 6.8e-70 |
| ADC45586.1 | modular\_polyketide\_synthase | BGC0000093 | Polyketide | 24.0 | 100.8 | 264.0 | 1.5e-69 |
| ALD82522.1 | polyketide\_synthase | BGC0001212 | NRP + Polyketide | 26.0 | 54.2 | 264.0 | 1.5e-69 |
| AAZ77694.1 | ChlA2 | BGC0000036 | Polyketide:Modular type I + Polyketide:Iterative type I + Saccharide:Oligosaccharide | 31.0 | 31.3 | 263.0 | 2.6e-69 |
| AAF19813.1 | MtaE | BGC0001024 | NRP + Polyketide:Modular type I | 27.0 | 43.0 | 261.0 | 9.8e-69 |
| AUD08663.1 | iPKS-NRPS | BGC0001553 | NRP + Polyketide | 25.0 | 58.7 | 261.0 | 9.8e-69 |
| CEF75886.1 |  | BGC0001600 | Polyketide | 28.0 | 42.6 | 261.0 | 1.3e-68 |
| CAA60462.1 | polyketide\_synthase | BGC0001040 | NRP + Polyketide | 35.0 | 25.5 | 260.0 | 2.2e-68 |
| ANC94964.1 | AlmHIII | BGC0001396 | Polyketide | 24.0 | 105.1 | 260.0 | 2.2e-68 |
| ACB37755.1 | putative\_type\_I\_polyketide\_synthase | BGC0000162 | Polyketide | 24.0 | 101.4 | 260.0 | 2.8e-68 |
| EAU29808.1 | hypothetical\_protein | BGC0001400 | Polyketide | 24.0 | 99.6 | 258.0 | 8.3e-68 |
| AVX51108.1 | nysC | BGC0001709 | Polyketide | 33.0 | 25.8 | 258.0 | 8.3e-68 |
| AXN93610.1 | PuwB | BGC0001953 | NRP | 32.0 | 27.7 | 258.0 | 8.3e-68 |
| AXN93577.1 | PuwB | BGC0001950 | NRP | 27.0 | 49.0 | 258.0 | 1.1e-67 |
| AXN93586.1 | PuwB | BGC0001951 | NRP | 27.0 | 49.0 | 258.0 | 1.1e-67 |
| AXN93597.1 | PuwB | BGC0001952 | NRP | 32.0 | 27.7 | 258.0 | 1.1e-67 |
| ACR50791.1 | putative\_polyketide\_synthase | BGC0000163 | Polyketide | 24.0 | 103.1 | 257.0 | 2.4e-67 |
| AGY62754.1 | EbeB | BGC0000051 | Polyketide | 35.0 | 26.1 | 256.0 | 3.1e-67 |
| SCN11950.1 | EbeB-type\_I\_polyketide\_synthase | BGC0001580 | Polyketide | 35.0 | 26.1 | 256.0 | 3.1e-67 |
| ctg1\_orf521 |  | BGC0001199 | Polyketide | 34.0 | 26.3 | 256.0 | 4.1e-67 |
| ADH04657.1 | TugA | BGC0001342 | NRP + Polyketide | 29.0 | 37.2 | 256.0 | 4.1e-67 |
| ACR50775.1 | polyketide\_synthase | BGC0000163 | Polyketide | 34.0 | 25.0 | 255.0 | 7e-67 |
| ABW96540.1 | type\_I\_modular\_polyketide\_synthase | BGC0000159 | Polyketide:Modular type I | 32.0 | 30.4 | 254.0 | 1.2e-66 |
| AEU17897.1 | putative\_type\_I\_PKS | BGC0001072 | Saccharide + Polyketide:Modular type I + Polyketide:Type II + Other:Aminocoumarin | 33.0 | 26.3 | 254.0 | 1.2e-66 |
| ctg1\_orf255 |  | BGC0001200 | Polyketide | 33.0 | 26.4 | 254.0 | 1.2e-66 |
| AJO72736.1 | Type\_I\_modular\_polyketide\_synthase | BGC0001381 | Polyketide | 32.0 | 30.6 | 254.0 | 1.2e-66 |
| AFV30250.1 | polyketide\_synthase | BGC0000075 | Polyketide | 32.0 | 31.7 | 254.0 | 1.6e-66 |
| BAH02268.1 | polyketide\_synthase | BGC0000126 | Polyketide | 34.0 | 26.2 | 254.0 | 1.6e-66 |
| SCN11952.1 | ebeD-type\_I\_polyketide\_synthase | BGC0001580 | Polyketide | 34.0 | 25.1 | 254.0 | 1.6e-66 |
| CAA60459.1 | polyketide\_synthase | BGC0001040 | NRP + Polyketide | 35.0 | 25.7 | 253.0 | 2e-66 |
| IF55\_RS32375 | beta-ketoacyl\_synthase | BGC0001348 | Polyketide:Modular type I | 32.0 | 31.4 | 253.0 | 2e-66 |
| AJO72742.1 | Type\_I\_modular\_polyketide\_synthase | BGC0001381 | Polyketide | 33.0 | 25.2 | 253.0 | 2e-66 |
| ARM20284.1 | polyketide\_synthase | BGC0001523 | Polyketide | 33.0 | 25.4 | 253.0 | 2e-66 |
| ctg1\_orf253 |  | BGC0001200 | Polyketide | 32.0 | 27.6 | 253.0 | 3.5e-66 |
| AAP42857.1 | NanA3 | BGC0000105 | Polyketide | 31.0 | 32.5 | 252.0 | 4.5e-66 |
| CAJ88175.1 | putative\_polyketide\_synthase\_B | BGC0000151 | Polyketide:Modular type I + Saccharide:Hybrid/tailoring | 33.0 | 25.1 | 252.0 | 4.5e-66 |
| AAB66506.1 | tylactone\_synthase\_modules\_4\_&\_5 | BGC0000166 | Polyketide | 31.0 | 31.1 | 252.0 | 4.5e-66 |
| AEE88282.1 | CurH | BGC0000976 | NRP + Polyketide:Modular type I | 23.0 | 106.4 | 252.0 | 4.5e-66 |
| AAT70103.1 | CurH | BGC0001165 | NRP + Polyketide:Modular type I | 23.0 | 106.4 | 252.0 | 4.5e-66 |
| ctg1\_orf254 |  | BGC0001200 | Polyketide | 32.0 | 26.3 | 252.0 | 4.5e-66 |
| BAO66529.1 | type\_I\_polyketide\_synthase | BGC0000042 | Polyketide | 31.0 | 30.5 | 252.0 | 5.9e-66 |
| DAB41915.1 | ArzM\_-\_PKS\_(KS,\_AT,\_DH,\_MT,\_ER,\_KR,\_ACP) | BGC0001884 | NRP + Polyketide | 25.0 | 56.5 | 252.0 | 5.9e-66 |
| ACN69991.1 | polyketide\_synthase | BGC0000079 | Polyketide | 33.0 | 25.8 | 251.0 | 1e-65 |
| AAF71767.1 | nysJ | BGC0000115 | Polyketide:Modular type I + Saccharide:Hybrid/tailoring | 34.0 | 25.3 | 251.0 | 1.3e-65 |
| ABC84460.1 | NigAV | BGC0000114 | Polyketide:Modular type I | 24.0 | 105.9 | 250.0 | 2.9e-65 |
| AEZ53946.1 | polyketide\_synthase | BGC0000144 | Polyketide:Modular type I | 33.0 | 25.6 | 250.0 | 2.9e-65 |
| AEZ64504.1 | Herc | BGC0001065 | Polyketide | 35.0 | 24.7 | 250.0 | 2.9e-65 |
| CAD89776.1 | MelE\_protein | BGC0001010 | NRP + Polyketide:Modular type I | 26.0 | 42.9 | 249.0 | 3.8e-65 |
| EHK80166.1 | beta-ketoacyl\_synthase | BGC0001447 | Polyketide | 31.0 | 29.7 | 249.0 | 3.8e-65 |
| EHK80170.1 | acyl\_transferase | BGC0001447 | Polyketide | 34.0 | 25.9 | 249.0 | 3.8e-65 |
| ANR02553.1 | LodL | BGC0001648 | Polyketide | 34.0 | 25.6 | 249.0 | 3.8e-65 |
| AEZ54377.1 | PieA4 | BGC0000124 | Polyketide | 34.0 | 26.7 | 249.0 | 5e-65 |
| BAD08358.1 | polyketide\_synthase\_modules\_4 | BGC0000167 | Polyketide | 32.0 | 28.2 | 248.0 | 6.6e-65 |
| AIW82279.1 | PuwB | BGC0001125 | NRP + Polyketide | 26.0 | 49.4 | 248.0 | 6.6e-65 |
| WP\_055469549.1 | type\_I\_polyketide\_synthase | BGC0001537 | Polyketide | 34.0 | 27.8 | 248.0 | 6.6e-65 |
| AHH99926.1 | PKS\_I | BGC0000002 | Polyketide | 32.0 | 25.6 | 248.0 | 8.6e-65 |
| ABW96542.1 | type\_I\_modular\_polyketide\_synthase | BGC0000159 | Polyketide:Modular type I | 32.0 | 28.1 | 248.0 | 8.6e-65 |
| AAZ77698.1 | ChlA5 | BGC0000036 | Polyketide:Modular type I + Polyketide:Iterative type I + Saccharide:Oligosaccharide | 33.0 | 25.2 | 248.0 | 1.1e-64 |
| CAM00064.1 | EryAII\_Erythromycin\_polyketide\_synthase\_modules\_3\_and\_4 | BGC0000055 | Polyketide:Modular type I + Saccharide:Hybrid/tailoring | 34.0 | 24.6 | 248.0 | 1.1e-64 |
| ABC84459.1 | NigAIV | BGC0000114 | Polyketide:Modular type I | 23.0 | 101.4 | 248.0 | 1.1e-64 |
| ABS90471.1 | PKS\_type\_I | BGC0001106 | NRP + Polyketide | 34.0 | 25.4 | 248.0 | 1.1e-64 |
| CAO98848.1 | polyketide\_synthase\_AufE | BGC0000023 | Polyketide:Modular type I | 31.0 | 32.1 | 247.0 | 1.5e-64 |
| AJW65408.1 | type\_I\_modular\_polyketide\_synthase | BGC0001195 | NRP + Polyketide | 34.0 | 25.9 | 247.0 | 1.5e-64 |
| AAO65800.1 | monensin\_polyketide\_synthase\_modules\_7\_and\_8 | BGC0000100 | Polyketide | 24.0 | 103.2 | 247.0 | 1.9e-64 |
| AAF71776.1 | nysC | BGC0000115 | Polyketide:Modular type I + Saccharide:Hybrid/tailoring | 32.0 | 25.2 | 247.0 | 1.9e-64 |
| ANZ52463.1 | MonAV | BGC0001670 | Polyketide | 24.0 | 103.2 | 247.0 | 1.9e-64 |
| AUO16423.1 | polyketide\_synthase | BGC0001700 | Polyketide | 33.0 | 25.4 | 247.0 | 1.9e-64 |
| ABY21540.1 | AngAIII | BGC0000018 | Polyketide | 23.0 | 104.0 | 247.0 | 2.5e-64 |
| BAK64637.1 | polyketide\_synthase | BGC0000135 | Polyketide | 30.0 | 30.9 | 246.0 | 4.3e-64 |
| AAW03328.1 | CtaE | BGC0000982 | NRP + Polyketide | 25.0 | 42.5 | 246.0 | 4.3e-64 |
| AHH99925.1 | PKS\_I | BGC0000002 | Polyketide | 30.0 | 32.7 | 245.0 | 5.6e-64 |
| QBF51756.1 | type\_I\_polyketide\_synthase | BGC0001856 | Polyketide:Modular type I | 31.0 | 27.4 | 245.0 | 5.6e-64 |
| AJW65407.1 | type\_I\_modular\_polyketide\_synthase | BGC0001195 | NRP + Polyketide | 34.0 | 25.3 | 245.0 | 7.3e-64 |
| AUO16397.1 | polyketide\_synthase | BGC0001700 | Polyketide | 33.0 | 26.1 | 245.0 | 7.3e-64 |
| AEZ64505.1 | Herb | BGC0001065 | Polyketide | 32.0 | 26.5 | 245.0 | 9.5e-64 |
| ANR02554.1 | LodM | BGC0001648 | Polyketide | 33.0 | 28.7 | 245.0 | 9.5e-64 |
| AIT55261.1 | polyketide\_synthase | BGC0000072 | Polyketide:Modular type I | 27.0 | 41.7 | 244.0 | 1.2e-63 |
| BAF85838.1 | modular\_polyketide\_synthase | BGC0000109 | Polyketide | 34.0 | 25.4 | 244.0 | 1.2e-63 |
| BAK64638.1 | polyketide\_synthase | BGC0000135 | Polyketide | 32.0 | 24.8 | 244.0 | 1.2e-63 |
| BAC76492.1 | lankamycin\_synthase\_LkmAII | BGC0000085 | Polyketide | 29.0 | 30.7 | 244.0 | 1.6e-63 |
| AAG23265.1 | polyketide\_synthase\_extender\_module\_2 | BGC0000148 | Polyketide | 33.0 | 25.2 | 244.0 | 1.6e-63 |
| BAR73007.1 | putative\_PKS\_(ACP-KS-AT-DH-KR-ACP-KS-AT-DH-ER-KR-ACP) | BGC0001194 | Polyketide | 32.0 | 27.9 | 244.0 | 1.6e-63 |
| ACR50785.1 | polyketide\_synthase | BGC0000163 | Polyketide | 33.0 | 26.0 | 243.0 | 2.1e-63 |
| CQR60496.1 | Polyketide\_synthase,\_type\_I,\_modules:\_4,\_5\_and\_6 | BGC0001287 | Polyketide | 25.0 | 55.9 | 243.0 | 2.1e-63 |
| EHK80169.1 | acyl\_transferase | BGC0001447 | Polyketide | 32.0 | 26.9 | 243.0 | 2.8e-63 |
| AEZ53952.1 | polyketide\_synthase | BGC0000144 | Polyketide:Modular type I | 33.0 | 25.4 | 243.0 | 3.6e-63 |
| AAQ82565.1 | FscB | BGC0000061 | Polyketide | 33.0 | 24.3 | 242.0 | 4.7e-63 |
| EWM62998.1 | mycocerosic\_acid\_synthase | BGC0001328 | NRP:Cyclic depsipeptide + Polyketide:Modular type I | 34.0 | 25.5 | 242.0 | 4.7e-63 |
| ATY46587.1 | polyketide\_synthase | BGC0001666 | Polyketide | 32.0 | 28.9 | 242.0 | 4.7e-63 |
| ASZ00148.1 | polyketide\_synthase | BGC0001785 | Polyketide | 25.0 | 56.4 | 242.0 | 8e-63 |
| AHH99921.1 | PKS\_I | BGC0000002 | Polyketide | 32.0 | 24.8 | 241.0 | 1e-62 |
| CAE46850.1 | Type\_I\_modular\_polyketide\_synthase | BGC0000103 | Polyketide | 33.0 | 27.0 | 241.0 | 1e-62 |
| AAP42873.1 | NanA11 | BGC0000105 | Polyketide | 31.0 | 32.1 | 241.0 | 1e-62 |
| ABJ97438.1 | MerB | BGC0001012 | NRP + Polyketide | 32.0 | 25.4 | 241.0 | 1e-62 |
| CAO98849.1 | polyketide\_synthase\_AufF | BGC0000023 | Polyketide:Modular type I | 26.0 | 45.7 | 241.0 | 1.4e-62 |
| ABC84458.1 | NigAIII | BGC0000114 | Polyketide:Modular type I | 34.0 | 24.9 | 241.0 | 1.4e-62 |
| CAL58682.1 | polyketide\_synthase | BGC0000149 | Polyketide:Modular type I | 25.0 | 55.5 | 241.0 | 1.4e-62 |
| AUO16401.1 | polyketide\_synthase | BGC0001700 | Polyketide | 32.0 | 28.9 | 241.0 | 1.4e-62 |
| BAO66519.1 | type\_I\_polyketide\_synthase | BGC0000042 | Polyketide | 31.0 | 29.7 | 240.0 | 1.8e-62 |
| AHE80994.1 | PieA4 | BGC0001169 | Polyketide:Modular type I | 32.0 | 26.0 | 240.0 | 2.3e-62 |
| CAE46851.1 | Type\_I\_modular\_polyketide\_synthase | BGC0000103 | Polyketide | 33.0 | 27.0 | 240.0 | 3e-62 |
| ABC84470.1 | NIGAVIII | BGC0000114 | Polyketide:Modular type I | 30.0 | 32.7 | 239.0 | 4e-62 |
| CBD77738.1 | polyketide\_synthase | BGC0000974 | NRP + Polyketide | 26.0 | 43.4 | 239.0 | 4e-62 |
| ABC87510.1 | polyketide\_synthase | BGC0001011 | NRP + Polyketide | 33.0 | 25.3 | 239.0 | 4e-62 |
| ctg1\_orf21 |  | BGC0001013 | NRP + Polyketide | 33.0 | 25.3 | 239.0 | 4e-62 |
| AIR74913.1 | polyketide\_synthase | BGC0001559 | RiPP | 26.0 | 43.4 | 239.0 | 4e-62 |
| CCE88377.1 | non-ribosomal\_peptide\_synthetase/polyketide\_synthase | BGC0001034 | NRP + Polyketide:Modular type I | 28.0 | 38.5 | 239.0 | 5.2e-62 |
| AWC08662.1 | polyketide\_synthase\_type\_I | BGC0001932 | Polyketide | 33.0 | 25.8 | 238.0 | 6.8e-62 |
| CAE02605.1 | polyketide\_synthase\_type\_I | BGC0000024 | Polyketide:Modular type I | 33.0 | 26.5 | 238.0 | 1.2e-61 |
| AAP42859.1 | NanA5 | BGC0000105 | Polyketide | 34.0 | 25.0 | 238.0 | 1.2e-61 |
| AVV61984.1 | type\_I\_modular\_polyketide\_synthase | BGC0001477 | NRP + Polyketide:Modular type I | 33.0 | 25.5 | 238.0 | 1.2e-61 |
| AFL48527.1 | laidlomycin\_polyketide\_synthase\_(module\_3\_and\_module\_4) | BGC0000084 | Polyketide | 32.0 | 25.7 | 237.0 | 1.5e-61 |
| AKA59091.1 | type-I\_PKS | BGC0001619 | Polyketide | 32.0 | 27.6 | 237.0 | 1.5e-61 |
| BAG85030.1 | putative\_polyketide\_synthase | BGC0000086 | Polyketide | 34.0 | 25.2 | 237.0 | 2e-61 |
| CAQ64690.1 | lasalocid\_modular\_polyketide\_synthase | BGC0000087 | Polyketide | 34.0 | 25.2 | 237.0 | 2e-61 |
| ACR50774.1 | polyketide\_synthase | BGC0000163 | Polyketide | 33.0 | 25.8 | 237.0 | 2e-61 |
| BAV56012.1 | PKS\_(KS-AT-DH-ER-KR-ACP-TE) | BGC0001597 | Polyketide | 32.0 | 26.4 | 237.0 | 2e-61 |
| AAC01711.1 | RifB | BGC0000136 | Polyketide | 25.0 | 55.6 | 237.0 | 2.6e-61 |
| AEZ53949.1 | polyketide\_synthase | BGC0000144 | Polyketide:Modular type I | 33.0 | 26.0 | 236.0 | 3.4e-61 |
| CAD19089.1 | StiE\_protein | BGC0000153 | NRP + Polyketide:Modular type I | 26.0 | 42.2 | 236.0 | 3.4e-61 |
| AGZ15474.1 | putative\_type\_I\_polyketide\_synthase | BGC0001036 | NRP + Polyketide | 32.0 | 27.6 | 236.0 | 3.4e-61 |
| QDA77044.1 | polyketide\_synthase | BGC0002025 | NRP | 32.0 | 26.1 | 236.0 | 3.4e-61 |
| ABV91286.1 | type\_I\_modular\_polyketide\_synthase | BGC0000158 | Polyketide:Modular type I | 32.0 | 25.7 | 236.0 | 4.4e-61 |
| WP\_055469545.1 | type\_I\_polyketide\_synthase | BGC0001537 | Polyketide | 34.0 | 24.9 | 236.0 | 4.4e-61 |
| TXD00266.1 | SDR\_family\_NAD(P)-dependent\_oxidoreductase | BGC0001877 | Polyketide | 29.0 | 30.7 | 236.0 | 4.4e-61 |
| ABC87512.1 | polyketide\_synthase | BGC0001011 | NRP + Polyketide | 34.0 | 24.5 | 235.0 | 5.7e-61 |
| ctg1\_orf23 |  | BGC0001013 | NRP + Polyketide | 34.0 | 24.5 | 235.0 | 5.7e-61 |
| WP\_016638480.1 | type\_I\_polyketide\_synthase | BGC0001519 | NRP + Polyketide | 33.0 | 25.5 | 235.0 | 5.7e-61 |
| CAO85897.1 | modular\_polyketide\_synthase\_NorB | BGC0000110 | Polyketide:Modular type I | 34.0 | 26.1 | 235.0 | 7.5e-61 |
| ABI94379.1 | tautomycetin\_biosynthetic\_PKS | BGC0000157 | Polyketide | 33.0 | 25.6 | 235.0 | 7.5e-61 |
| ARV85763.1 | PieA4\_type\_I\_PKS | BGC0001742 | Polyketide | 31.0 | 28.7 | 235.0 | 7.5e-61 |
| ARW71485.1 | type\_I\_PKS\_module\_4,\_module\_5 | BGC0001812 | Polyketide | 31.0 | 25.7 | 235.0 | 7.5e-61 |
| CAJ88176.1 | putative\_polyketide\_synthase\_B | BGC0000151 | Polyketide:Modular type I + Saccharide:Hybrid/tailoring | 33.0 | 24.9 | 235.0 | 9.8e-61 |
| BAV56011.1 | PKS\_(KS-AT-DH-ER-KR-ACP-KS-AT-DH-ER-KR-ACP) | BGC0001597 | Polyketide | 30.0 | 30.9 | 235.0 | 9.8e-61 |
| ANZ22986.1 | ZinC | BGC0001828 | Polyketide | 29.0 | 31.4 | 235.0 | 9.8e-61 |
| AKG06377.1 | polyketide\_synthase\_type\_1 | BGC0001830 | Polyketide | 31.0 | 25.3 | 235.0 | 9.8e-61 |
| ACN69988.1 | polyketide\_synthase | BGC0000079 | Polyketide | 33.0 | 25.0 | 234.0 | 1.3e-60 |
| ATL73034.1 | type\_I\_modular\_polyketide\_synthase | BGC0001807 | NRP + Polyketide | 31.0 | 26.0 | 234.0 | 1.3e-60 |
| CCP20050.1 | divL3\_protein | BGC0001119 | Polyketide:Modular type I | 32.0 | 25.5 | 234.0 | 1.7e-60 |
| AKD43522.1 | Type\_I\_polyketide\_synthase | BGC0001409 | Polyketide | 22.0 | 104.4 | 234.0 | 1.7e-60 |
| ATL73033.1 | type\_I\_modular\_polyketide\_synthase | BGC0001807 | NRP + Polyketide | 29.0 | 31.9 | 234.0 | 1.7e-60 |
| CAQ52624.1 | type\_I\_polyketide\_synthase,\_modules\_7-8 | BGC0001066 | Polyketide:Modular type I | 32.0 | 26.4 | 233.0 | 2.2e-60 |
| BAQ25513.1 | type\_I\_polyketide\_synthase | BGC0001288 | Polyketide | 32.0 | 25.6 | 233.0 | 2.2e-60 |
| ABB86408.1 | GelA | BGC0000067 | Polyketide | 32.0 | 28.0 | 233.0 | 2.9e-60 |
| BAC57032.1 | protomycinolide\_IV\_synthase\_5 | BGC0000102 | Polyketide | 25.0 | 54.4 | 233.0 | 2.9e-60 |
| CAQ18838.1 | polyketide\_synthase | BGC0000954 | NRP + Polyketide:Modular type I | 27.0 | 39.6 | 233.0 | 2.9e-60 |
| DAB41916.1 | ArzN\_-\_PKS\_(KS,\_AT,\_OMT,\_KR,\_ACP) | BGC0001884 | NRP + Polyketide | 27.0 | 37.6 | 233.0 | 2.9e-60 |
| ABK32291.1 | JerE | BGC0000080 | Polyketide | 27.0 | 42.4 | 233.0 | 3.7e-60 |
| AAO65798.1 | monensin\_polyketide\_synthase\_modules\_3\_and\_4 | BGC0000100 | Polyketide | 33.0 | 25.7 | 233.0 | 3.7e-60 |
| ANZ52461.1 | MonAIII | BGC0001670 | Polyketide | 33.0 | 25.7 | 233.0 | 3.7e-60 |
| KFL51883.1 | amino\_acid\_adenylation\_protein | BGC0001711 | NRP + Polyketide | 25.0 | 42.2 | 233.0 | 3.7e-60 |
| AAG13918.1 | megalomicin\_6-deoxyerythronolide\_B\_synthase\_2 | BGC0000092 | Polyketide | 32.0 | 25.3 | 232.0 | 4.9e-60 |
| ACY13415.1 | KR\_domain\_protein | BGC0001367 | NRP + Polyketide | 32.0 | 25.2 | 232.0 | 4.9e-60 |
| AAO06916.1 | GdmAI | BGC0000066 | Polyketide | 32.0 | 28.0 | 232.0 | 6.4e-60 |
| ABV97152.1 | Beta-ketoacyl\_synthase | BGC0000137 | Polyketide | 24.0 | 55.3 | 232.0 | 8.3e-60 |
| BAD08373.1 | polyketide\_synthase\_modules\_1-3 | BGC0000167 | Polyketide | 31.0 | 25.3 | 232.0 | 8.3e-60 |
| AAS98777.1 | polyketide\_synthetase | BGC0001001 | NRP + Polyketide | 26.0 | 42.6 | 232.0 | 8.3e-60 |
| CAF05649.1 | TubD\_protein | BGC0001053 | NRP + Polyketide | 22.0 | 100.1 | 232.0 | 8.3e-60 |
| AAC69330.1 | type\_I\_polyketide\_synthase\_PikAII | BGC0000094 | Polyketide:Modular type I + Saccharide:Hybrid/tailoring | 31.0 | 26.3 | 231.0 | 1.1e-59 |
| AAY28225.1 | HbmAI | BGC0000074 | Polyketide | 32.0 | 26.3 | 231.0 | 1.4e-59 |
| ABW96541.1 | type\_I\_modular\_polyketide\_synthase | BGC0000159 | Polyketide:Modular type I | 33.0 | 25.3 | 231.0 | 1.4e-59 |
| BAV56006.1 | PKS\_(ACP-KS-AT-DH-ER-KR-ACP-KS-AT-KR-ACP) | BGC0001597 | Polyketide | 32.0 | 26.4 | 231.0 | 1.4e-59 |
| AXI91546.1 | FunP7 | BGC0001944 | Polyketide | 33.0 | 25.1 | 230.0 | 1.8e-59 |
| ADX66459.1 | ScnS4 | BGC0000108 | Polyketide | 26.0 | 45.3 | 230.0 | 2.4e-59 |
| AEH42491.1 | polyketide\_synthase | BGC0000032 | Polyketide | 31.0 | 27.5 | 230.0 | 3.2e-59 |
| ACB46471.1 | polyketide\_synthase | BGC0000082 | Polyketide | 32.0 | 25.0 | 230.0 | 3.2e-59 |
| AAO65797.1 | monensin\_polyketide\_synthase\_module\_2 | BGC0000100 | Polyketide | 30.0 | 26.8 | 230.0 | 3.2e-59 |
| AFU82616.1 | polyketide\_synthase | BGC0000998 | NRP + Polyketide | 32.0 | 26.6 | 230.0 | 3.2e-59 |
| ANZ52460.1 | MonAII | BGC0001670 | Polyketide | 30.0 | 26.8 | 230.0 | 3.2e-59 |
| CAQ18834.1 | polyketide\_synthase | BGC0000954 | NRP + Polyketide:Modular type I | 27.0 | 44.7 | 229.0 | 4.1e-59 |
| ADH04680.1 | hybrid\_polyketide\_synthase/non-ribosomal\_peptide\_synthetase | BGC0001344 | NRP + Polyketide | 23.0 | 99.6 | 229.0 | 5.4e-59 |
| ACC40921.1 | polyketide\_synthase\_Pks7 | BGC0001665 | Polyketide | 32.0 | 26.2 | 229.0 | 5.4e-59 |
| AWH12669.1 | RmpB | BGC0001759 | Polyketide | 24.0 | 55.7 | 229.0 | 5.4e-59 |
| AGI99482.1 | Type\_I\_polyketide\_synthase | BGC0001004 | Polyketide:Modular type I | 32.0 | 26.4 | 228.0 | 7e-59 |
| ADC79618.1 | BafAIII | BGC0000028 | Polyketide:Modular type I | 30.0 | 30.0 | 228.0 | 9.2e-59 |
| AAZ77696.1 | ChlA3 | BGC0000036 | Polyketide:Modular type I + Polyketide:Iterative type I + Saccharide:Oligosaccharide | 32.0 | 25.6 | 228.0 | 9.2e-59 |
| BBA66512.1 | type\_I\_polyketide\_synthase | BGC0001495 | Polyketide | 34.0 | 26.2 | 228.0 | 9.2e-59 |
| QBF51758.1 | type\_I\_polyketide\_synthase | BGC0001856 | Polyketide:Modular type I | 31.0 | 28.9 | 228.0 | 9.2e-59 |
| AHA38200.1 | GphG | BGC0000069 | Polyketide | 31.0 | 28.0 | 227.0 | 1.6e-58 |
| AAO65799.1 | monensin\_polyketide\_synthase\_modules\_5\_and\_6 | BGC0000100 | Polyketide | 30.0 | 30.5 | 227.0 | 1.6e-58 |
| ABJ97439.1 | MerC | BGC0001012 | NRP + Polyketide | 34.0 | 24.5 | 227.0 | 1.6e-58 |
| ANZ52462.1 | MonAIV | BGC0001670 | Polyketide | 30.0 | 30.5 | 227.0 | 1.6e-58 |
| AZH23819.1 | MgiR | BGC0001971 | NRP + Polyketide | 26.0 | 43.2 | 227.0 | 2e-58 |
| CAC20920.1 | PimS3\_protein | BGC0000125 | Polyketide | 25.0 | 53.3 | 227.0 | 2.7e-58 |
| AQT01394.1 | SgnS3 | BGC0001690 | Polyketide | 25.0 | 53.3 | 227.0 | 2.7e-58 |
| QBF51757.1 | type\_I\_polyketide\_synthase | BGC0001856 | Polyketide:Modular type I | 31.0 | 24.7 | 227.0 | 2.7e-58 |
| AFL48528.1 | laidlomycin\_polyketide\_synthase\_(module\_7\_and\_module\_8) | BGC0000084 | Polyketide | 30.0 | 28.0 | 226.0 | 3.5e-58 |
| CAE45671.1 | borrelidin\_polyketide\_synthase,\_type\_I | BGC0000031 | Polyketide:Modular type I | 31.0 | 26.1 | 225.0 | 5.9e-58 |
| CAC20919.1 | PimS4\_protein | BGC0000125 | Polyketide | 26.0 | 45.2 | 225.0 | 5.9e-58 |
| AFU82617.1 | polyketide\_synthase | BGC0000998 | NRP + Polyketide | 26.0 | 41.1 | 225.0 | 5.9e-58 |
| AQT01395.1 | SgnS4 | BGC0001690 | Polyketide | 26.0 | 45.2 | 225.0 | 5.9e-58 |
| ALV82345.1 | borrelidin\_type\_I\_polyketide\_synthase | BGC0001533 | Polyketide | 31.0 | 26.1 | 225.0 | 7.8e-58 |
| TXD00265.1 | SDR\_family\_NAD(P)-dependent\_oxidoreductase | BGC0001877 | Polyketide | 30.0 | 27.5 | 225.0 | 7.8e-58 |
| BAB69194.1 | modular\_polyketide\_synthase | BGC0000117 | Polyketide | 31.0 | 26.3 | 225.0 | 1e-57 |
| CAQ18833.1 | polyketide\_synthase | BGC0000954 | NRP + Polyketide:Modular type I | 26.0 | 40.5 | 224.0 | 1.3e-57 |
| AQW44871.1 | polyketide\_synthase | BGC0001761 | Polyketide | 31.0 | 25.9 | 224.0 | 1.7e-57 |
| CAQ18829.1 | polyketide\_synthase | BGC0000954 | NRP + Polyketide:Modular type I | 25.0 | 42.5 | 223.0 | 2.3e-57 |
| AGC09499.1 | LobS4 | BGC0001183 | Polyketide | 31.0 | 26.4 | 223.0 | 2.3e-57 |
| OJF16269.1 | AceP2 | BGC0001491 | Polyketide | 30.0 | 32.1 | 223.0 | 2.3e-57 |
| AAP42858.1 | NanA4 | BGC0000105 | Polyketide | 32.0 | 24.4 | 223.0 | 3.9e-57 |
| CAQ52622.1 | type\_I\_polyketide\_synthase,\_modules\_4-5 | BGC0001066 | Polyketide:Modular type I | 31.0 | 26.3 | 223.0 | 3.9e-57 |
| BAQ25512.1 | type\_I\_polyketide\_synthase | BGC0001288 | Polyketide | 33.0 | 25.1 | 223.0 | 3.9e-57 |
| AFV96142.1 | polyketide\_synthase | BGC0001064 | Polyketide:Modular type I + Polyketide:Type III | 25.0 | 42.7 | 222.0 | 8.6e-57 |
| ARU81122.1 | CylH | BGC0001566 | Polyketide | 25.0 | 42.7 | 222.0 | 8.6e-57 |
| ASX95227.1 | IlaE | BGC0001620 | Polyketide | 32.0 | 28.6 | 222.0 | 8.6e-57 |
| AFL48529.1 | laidlomycin\_polyketide\_synthase\_(module\_5\_and\_module\_6) | BGC0000084 | Polyketide | 31.0 | 26.5 | 221.0 | 1.5e-56 |
| AAA79984.2 | soraphen\_polyketide\_synthase\_B | BGC0000147 | Polyketide:Modular type I | 25.0 | 42.6 | 220.0 | 1.9e-56 |
| AAU93806.2 | polyketide\_synthase\_modules\_3\_and\_4 | BGC0000054 | Polyketide | 31.0 | 23.6 | 219.0 | 4.3e-56 |
| CRI73799.1 | CongC\_protein | BGC0001215 | NRP | 32.0 | 26.6 | 219.0 | 4.3e-56 |
| BBA20952.1 | type\_I\_polyketide\_synthase | BGC0001763 | NRP + Polyketide | 31.0 | 28.3 | 219.0 | 4.3e-56 |
| AZF85941.1 | type\_I\_polyketide\_synthase | BGC0001963 | NRP + Polyketide | 30.0 | 31.1 | 218.0 | 9.5e-56 |
| AAC46026.1 | polyketide\_synthase\_modules\_4\_and\_5 | BGC0000113 | Polyketide | 31.0 | 25.4 | 218.0 | 1.2e-55 |
| ARM20277.1 | polyketide\_synthase | BGC0001523 | Polyketide | 26.0 | 42.7 | 218.0 | 1.2e-55 |
| WP\_107408739.1 | type\_I\_polyketide\_synthase | BGC0002033 | Polyketide | 31.0 | 25.8 | 217.0 | 1.6e-55 |
| CCP20048.1 | divL1\_protein | BGC0001119 | Polyketide:Modular type I | 30.0 | 31.2 | 217.0 | 2.1e-55 |
| AKA59093.1 | type-I\_PKS | BGC0001619 | Polyketide | 24.0 | 54.5 | 217.0 | 2.1e-55 |
| ANZ22995.1 | ZinA | BGC0001828 | Polyketide | 32.0 | 26.6 | 217.0 | 2.8e-55 |
| WP\_083502114.1 | type\_I\_polyketide\_synthase | BGC0001653 | Polyketide | 28.0 | 34.7 | 216.0 | 3.6e-55 |
| AUO16399.1 | polyketide\_synthase | BGC0001700 | Polyketide | 27.0 | 42.2 | 216.0 | 3.6e-55 |
| ADC45515.1 | modular\_polyketide\_synthase | BGC0000093 | Polyketide | 24.0 | 55.0 | 216.0 | 4.7e-55 |
| ACC40922.1 | polyketide\_synthase,\_Pks8 | BGC0001665 | Polyketide | 32.0 | 24.7 | 215.0 | 6.2e-55 |
| AAF86396.1 | FkbA | BGC0000994 | NRP + Polyketide | 33.0 | 23.5 | 215.0 | 1e-54 |
| AGY30676.1 | Ann4 | BGC0001298 | Polyketide | 27.0 | 43.3 | 214.0 | 1.8e-54 |
| ARE67852.1 | AbsB2 | BGC0001492 | Polyketide | 31.0 | 25.5 | 214.0 | 1.8e-54 |
| ctg1\_13 |  | BGC0001931 | Polyketide | 32.0 | 26.1 | 214.0 | 1.8e-54 |
| BAF02922.1 | type\_I\_polyketide\_synthase | BGC0000073 | Polyketide | 32.0 | 24.4 | 213.0 | 3.1e-54 |
| BAF02925.1 | type\_I\_polyketide\_synthase | BGC0000073 | Polyketide | 33.0 | 24.8 | 213.0 | 3.1e-54 |
| ACF35447.1 | mbcAIII | BGC0000090 | Polyketide | 30.0 | 32.1 | 213.0 | 3.1e-54 |
| ALP32045.1 | CycE | BGC0001293 | Polyketide | 25.0 | 54.8 | 213.0 | 3.1e-54 |
| AFL48526.1 | laidlomycin\_polyketide\_synthase\_(module\_2) | BGC0000084 | Polyketide | 28.0 | 31.0 | 213.0 | 4e-54 |
| orf3 | polyketide\_synthase | BGC0001432 | NRP:Cyclic depsipeptide + Polyketide:Iterative type I | 26.0 | 38.6 | 212.0 | 5.2e-54 |
| APZ78858.1 | polyketide\_synthase | BGC0001432 | NRP:Cyclic depsipeptide + Polyketide:Iterative type I | 26.0 | 38.6 | 212.0 | 5.2e-54 |
| AAK57190.1 | MxaF | BGC0001022 | NRP + Polyketide | 29.0 | 34.1 | 212.0 | 6.8e-54 |
| ADH04641.1 | TgaC | BGC0001051 | NRP + Polyketide:Modular type I | 25.0 | 40.8 | 211.0 | 1.2e-53 |
| AAF19810.1 | MtaB | BGC0001024 | NRP + Polyketide:Modular type I | 28.0 | 34.2 | 210.0 | 2e-53 |
| ADH04660.1 | TugD | BGC0001342 | NRP + Polyketide | 26.0 | 42.5 | 210.0 | 2e-53 |
| BAF02924.1 | type\_I\_polyketide\_synthase | BGC0000073 | Polyketide | 32.0 | 24.7 | 210.0 | 3.4e-53 |
| ANZ22989.1 | ZinF | BGC0001828 | Polyketide | 30.0 | 26.1 | 210.0 | 3.4e-53 |
| ANR02552.1 | LodK | BGC0001648 | Polyketide | 25.0 | 42.1 | 209.0 | 4.4e-53 |
| ctg1\_orf29 |  | BGC0000096 | Polyketide | 31.0 | 25.5 | 208.0 | 9.8e-53 |
| AAF86392.1 | FkbC | BGC0000994 | NRP + Polyketide | 33.0 | 24.5 | 207.0 | 1.7e-52 |
| BAE93722.1 | type\_I\_polyketide\_synthase | BGC0000164 | Polyketide | 29.0 | 30.2 | 207.0 | 2.2e-52 |
| WP\_055480219.1 | type\_I\_polyketide\_synthase | BGC0001653 | Polyketide | 32.0 | 25.2 | 207.0 | 2.9e-52 |
| WP\_053138504.1 | type\_I\_polyketide\_synthase | BGC0002033 | Polyketide | 27.0 | 40.0 | 205.0 | 8.3e-52 |
| WP\_035121546.1 | type\_I\_polyketide\_synthase | BGC0001467 | NRP:Cyclic depsipeptide + Polyketide:Modular type I | 25.0 | 43.8 | 205.0 | 1.1e-51 |
| BAJ16467.1 | polyketide\_synthase | BGC0000058 | Polyketide | 26.0 | 39.1 | 203.0 | 2.4e-51 |
| AAY89049.1 | polyketide\_synthase | BGC0001069 | NRP + Polyketide:Trans-AT type I | 25.0 | 40.5 | 203.0 | 2.4e-51 |
| ABI94380.1 | tautomycetin\_biosynthetic\_PKS | BGC0000157 | Polyketide | 26.0 | 42.6 | 203.0 | 4.1e-51 |
| CAQ34918.1 | nonribosomal\_peptide\_synthetase/\_polyketide\_synthase | BGC0000986 | NRP + Polyketide | 24.0 | 55.7 | 202.0 | 7e-51 |
| BAQ21940.1 | putative\_Type\_I\_polyketide\_synthase | BGC0001204 | Polyketide | 28.0 | 33.6 | 202.0 | 7e-51 |
| CAJ76298.1 | putative\_hybrid\_polyketide-non-ribosomal\_peptide\_synthetase | BGC0000972 | NRP + Polyketide:Modular type I + Polyketide:Trans-AT type I | 23.0 | 77.9 | 201.0 | 9.2e-51 |
| ALD82524.1 | polyketide\_synthase | BGC0001212 | NRP + Polyketide | 29.0 | 33.4 | 201.0 | 1.6e-50 |
| ABV91287.1 | type\_I\_modular\_polyketide\_synthase | BGC0000158 | Polyketide:Modular type I | 26.0 | 42.5 | 198.0 | 7.8e-50 |
| CCE88381.1 | polyketide\_synthase | BGC0001034 | NRP + Polyketide:Modular type I | 25.0 | 38.5 | 198.0 | 1e-49 |
| AUO16422.1 | polyketide\_synthase | BGC0001700 | Polyketide | 25.0 | 42.8 | 198.0 | 1e-49 |
| AKG06378.1 | polyketide\_synthase\_type\_1 | BGC0001830 | Polyketide | 26.0 | 42.1 | 197.0 | 2.3e-49 |
| BAQ21939.1 | putative\_type\_I\_polyketide\_synthase | BGC0001204 | Polyketide | 30.0 | 25.8 | 196.0 | 5e-49 |
| ARM20282.1 | polyketide\_synthase | BGC0001523 | Polyketide | 25.0 | 42.9 | 196.0 | 5e-49 |
| ABI91470.1 | beta-ketoacyl\_synthase | BGC0001094 | NRP + Polyketide | 23.0 | 59.2 | 193.0 | 2.5e-48 |
| AQZ37113.1 | polyketide\_synthase | BGC0001511 | Polyketide | 31.0 | 24.6 | 193.0 | 2.5e-48 |
| ALP32043.1 | CycC | BGC0001293 | Polyketide | 27.0 | 34.0 | 193.0 | 3.3e-48 |
| CAO98852.1 | polyketide\_synthase\_AufI | BGC0000023 | Polyketide:Modular type I | 26.0 | 35.2 | 190.0 | 2.8e-47 |
| WP\_020636845.1 | type\_I\_polyketide\_synthase | BGC0002011 | Polyketide | 31.0 | 24.1 | 190.0 | 2.8e-47 |
| CBF74114.1 | Conidial\_yellow\_pigment\_biosynthesis\_polyketide\_synthase\_(PKS)(EC\_2.3.1.-)\_[Source:UniProtKB/Swiss-Prot;Acc:Q03149] | BGC0000107 | Polyketide | 25.0 | 42.6 | 182.0 | 5.8e-45 |
| ADZ24998.1 | polyketide\_synthase | BGC0000380 | NRP + Polyketide:Modular type I | 25.0 | 36.5 | 177.0 | 1.9e-43 |
| ABB05104.1 | LipPks3 | BGC0001003 | NRP:Lipopeptide + Polyketide:Modular type I + Saccharide:Hybrid/tailoring | 26.0 | 37.3 | 176.0 | 3.2e-43 |
| QBF51760.1 | type\_I\_polyketide\_synthase | BGC0001856 | Polyketide:Modular type I | 26.0 | 33.6 | 176.0 | 3.2e-43 |
| sipP1 | Type\_I\_Modular\_PKS | BGC0001452 | Polyketide | 27.0 | 34.4 | 175.0 | 7.1e-43 |
| ARM20278.1 | polyketide\_synthase | BGC0001523 | Polyketide | 27.0 | 33.9 | 173.0 | 3.5e-42 |
| ARM20283.1 | polyketide\_synthase | BGC0001523 | Polyketide | 26.0 | 34.2 | 172.0 | 6e-42 |
| APZ78854.1 | polyketide\_synthase | BGC0001432 | NRP:Cyclic depsipeptide + Polyketide:Iterative type I | 22.0 | 102.7 | 172.0 | 7.8e-42 |
| ATG32077.1 | polyketide\_synthase | BGC0001750 | NRP + Polyketide | 28.0 | 26.4 | 169.0 | 5.1e-41 |
| CBD77736.1 | polyketide\_synthase | BGC0000974 | NRP + Polyketide | 26.0 | 36.5 | 169.0 | 6.6e-41 |
| ADH04639.1 | TgaA | BGC0001051 | NRP + Polyketide:Modular type I | 25.0 | 33.9 | 169.0 | 6.6e-41 |
| AIR74912.1 | polyketide\_synthase | BGC0001559 | RiPP | 26.0 | 36.5 | 169.0 | 6.6e-41 |
| CCT67991.1 | bikaverin\_cluster-polyketide\_synthase | BGC0000030 | Polyketide | 24.0 | 42.9 | 168.0 | 8.6e-41 |
| AQH32483.1 | hybrid\_peptide\_synthetase/polyketide\_synthase | BGC0001667 | NRP + Polyketide | 23.0 | 41.3 | 168.0 | 1.1e-40 |
| AUO16403.1 | polyketide\_synthase | BGC0001700 | Polyketide | 26.0 | 34.1 | 167.0 | 1.9e-40 |
| OAP25820.1 | Erythronolide\_synthase,\_modules\_1\_and\_2 | BGC0001658 | Polyketide | 30.0 | 23.2 | 166.0 | 3.3e-40 |
| BAF85839.1 | modular\_polyketide\_synthase | BGC0000109 | Polyketide | 26.0 | 36.9 | 166.0 | 5.6e-40 |
| QDA77059.1 | polyketide\_synthase/nonribosomal\_peptide\_synthetase | BGC0002026 | NRP | 30.0 | 21.8 | 166.0 | 5.6e-40 |
| BAD38873.1 | polyketide\_synthase | BGC0000111 | Polyketide | 27.0 | 26.1 | 165.0 | 7.3e-40 |
| TXD00261.1 | AMP-binding\_protein | BGC0001877 | Polyketide | 26.0 | 33.4 | 163.0 | 3.6e-39 |
| sipP2 | Type\_I\_Modular\_PKS | BGC0001452 | Polyketide | 28.0 | 34.7 | 163.0 | 4.7e-39 |
| CAD55506.1 | CpkA;\_Polyketide\_synthase\_loading\_module,\_and\_modules\_1\_and\_2 | BGC0000038 | Polyketide:Modular type I | 29.0 | 24.9 | 161.0 | 1.1e-38 |
| CAO98850.1 | polyketide\_synthase\_AufG | BGC0000023 | Polyketide:Modular type I | 26.0 | 36.3 | 161.0 | 1.4e-38 |
| AXM42950.1 | polyketide\_synthase | BGC0001941 | NRP + Polyketide | 29.0 | 21.6 | 161.0 | 1.4e-38 |
| AAM54075.1 | polyketide\_synthase | BGC0000020 | Polyketide | 28.0 | 24.7 | 161.0 | 1.8e-38 |
| CAD29795.1 | peptide\_synthetase | BGC0001015 | NRP + Polyketide | 23.0 | 41.2 | 160.0 | 2.3e-38 |
| AEC13079.1 | fosA | BGC0000060 | Polyketide | 23.0 | 43.7 | 160.0 | 3.1e-38 |
| AAP85335.1 | type\_I\_PKS | BGC0000233 | Polyketide | 24.0 | 30.8 | 160.0 | 4e-38 |
| ANR02556.1 | LodO | BGC0001648 | Polyketide | 25.0 | 33.7 | 160.0 | 4e-38 |
| CBA11582.1 | polyketide\_synthase\_type\_I | BGC0001046 | NRP + Polyketide:Modular type I + Saccharide:Hybrid/tailoring | 26.0 | 32.0 | 158.0 | 8.9e-38 |
| ANH11414.1 | SceS | BGC0001908 | Polyketide | 26.0 | 33.4 | 158.0 | 8.9e-38 |
| AMYAL\_RS48925 | polyketide\_synthase | BGC0002011 | Polyketide | 27.0 | 38.1 | 158.0 | 8.9e-38 |
| ABB88521.1 | polyketide\_synthase\_type\_I | BGC0000050 | Polyketide | 26.0 | 31.1 | 158.0 | 1.2e-37 |
| ACB37740.1 | putative\_type\_I\_polyketide\_synthase | BGC0000162 | Polyketide | 27.0 | 32.0 | 156.0 | 4.4e-37 |
| CQR60494.1 | Polyketide\_synthase,\_type\_I,\_module\_8 | BGC0001287 | Polyketide | 29.0 | 23.6 | 156.0 | 4.4e-37 |
| AWW87422.1 | type\_I\_polyketide\_synthase | BGC0001755 | Polyketide | 26.0 | 33.9 | 155.0 | 7.6e-37 |
| BAJ16471.1 | polyketide\_synthase | BGC0000058 | Polyketide | 26.0 | 33.5 | 155.0 | 9.9e-37 |
| DAB41484.1 | nonribosomal\_peptide\_synthetase/polyketide\_synthase\_type\_I | BGC0001766 | NRP | 30.0 | 22.9 | 154.0 | 1.7e-36 |
| AGC09484.1 | LobS1 | BGC0001183 | Polyketide | 27.0 | 33.2 | 152.0 | 8.4e-36 |
| AAC38075.1 | polyketide\_synthase\_type\_I | BGC0000127 | Polyketide | 29.0 | 20.7 | 150.0 | 2.4e-35 |
| CAD19093.1 | StiJ\_protein | BGC0000153 | NRP + Polyketide:Modular type I | 28.0 | 22.3 | 150.0 | 2.4e-35 |
| ABB88522.1 | polyketide\_synthase\_type\_I | BGC0000050 | Polyketide | 26.0 | 32.4 | 150.0 | 3.2e-35 |
| AHN85651.1 | Phn2 | BGC0000122 | Polyketide:Modular type I | 24.0 | 35.8 | 149.0 | 7.1e-35 |
| CAQ18835.1 | polyketide\_synthase | BGC0000954 | NRP + Polyketide:Modular type I | 25.0 | 34.2 | 149.0 | 7.1e-35 |
| ASZ00147.1 | polyketide\_synthase | BGC0001785 | Polyketide | 31.0 | 20.4 | 148.0 | 9.2e-35 |
| ABP55210.1 | beta-ketoacyl\_synthase | BGC0000142 | Polyketide | 26.0 | 33.5 | 148.0 | 1.2e-34 |
| AGI99497.1 | type\_I\_polyketide\_synthase | BGC0001004 | Polyketide:Modular type I | 27.0 | 33.9 | 148.0 | 1.2e-34 |
| OAP25815.1 | Phenolphthiocerol\_synthesis\_polyketide\_synthase\_type\_I\_Pks15/1 | BGC0001658 | Polyketide | 23.0 | 50.3 | 146.0 | 6e-34 |
| TXD00024.1 | SDR\_family\_NAD(P)-dependent\_oxidoreductase | BGC0001877 | Polyketide | 29.0 | 23.4 | 144.0 | 1.3e-33 |
| APZ78832.1 | polyketide\_synthase | BGC0001430 | NRP:Cyclic depsipeptide + Polyketide:Iterative type I | 25.0 | 25.6 | 144.0 | 1.7e-33 |
| BAB69199.1 | modular\_polyketide\_synthase | BGC0000117 | Polyketide | 25.0 | 30.9 | 144.0 | 2.3e-33 |
| WP\_051137606.1 | type\_I\_polyketide\_synthase | BGC0002011 | Polyketide | 27.0 | 35.1 | 144.0 | 2.3e-33 |
| ATG32075.1 | polyketide\_synthase | BGC0001750 | NRP + Polyketide | 28.0 | 24.2 | 143.0 | 5.1e-33 |
| ANR02551.1 | LodJ | BGC0001648 | Polyketide | 29.0 | 22.0 | 141.0 | 1.1e-32 |
| ARM20281.1 | polyketide\_synthase | BGC0001523 | Polyketide | 29.0 | 21.4 | 141.0 | 1.5e-32 |
| BAE93730.1 | type\_I\_polyketide\_synthase | BGC0000164 | Polyketide | 26.0 | 33.8 | 139.0 | 4.3e-32 |
| ADH04640.1 | TgaB | BGC0001051 | NRP + Polyketide:Modular type I | 29.0 | 23.6 | 139.0 | 4.3e-32 |
| BAE93731.1 | type\_I\_polyketide\_synthase | BGC0000164 | Polyketide | 25.0 | 31.8 | 139.0 | 7.3e-32 |
| ARW71486.1 | type\_I\_PKS\_module\_6 | BGC0001812 | Polyketide | 29.0 | 22.4 | 139.0 | 7.3e-32 |
| ctg1\_orf522 |  | BGC0001199 | Polyketide | 26.0 | 31.2 | 138.0 | 9.6e-32 |
| AQZ37095.1 | polyketide\_synthase | BGC0001511 | Polyketide | 26.0 | 31.0 | 138.0 | 1.6e-31 |
| AHH99924.1 | PKS\_I | BGC0000002 | Polyketide | 26.0 | 36.3 | 137.0 | 2.8e-31 |
| BAC76491.1 | lankamycin\_synthase\_LkmAIII | BGC0000085 | Polyketide | 30.0 | 21.9 | 137.0 | 2.8e-31 |
| AAZ94390.1 | modular\_polyketide\_synthase | BGC0000040 | Polyketide | 29.0 | 21.8 | 136.0 | 6.2e-31 |
| AAM54078.1 | polyketide\_synthase | BGC0000020 | Polyketide | 26.0 | 34.2 | 135.0 | 1.1e-30 |
| ABV83229.1 | CppB | BGC0000116 | Polyketide | 29.0 | 21.7 | 133.0 | 4e-30 |
| CAJ88187.2 | Type\_I\_modular\_polyketide\_synthase | BGC0000151 | Polyketide:Modular type I + Saccharide:Hybrid/tailoring | 27.0 | 23.3 | 133.0 | 5.2e-30 |
| AAS90093.1 | PksA | BGC0000006 | Polyketide | 26.0 | 21.8 | 132.0 | 6.9e-30 |
| BAE71314.1 | polyketide\_synthase | BGC0000004 | Polyketide | 27.0 | 21.8 | 132.0 | 8.9e-30 |
| AAS89999.1 | PksA | BGC0000007 | Polyketide | 26.0 | 21.8 | 131.0 | 1.5e-29 |
| AAS90022.1 | PksA | BGC0000008 | Polyketide | 26.0 | 21.8 | 130.0 | 2.6e-29 |
| ANY10591.1 | polyketide\_synthase | BGC0001773 | Polyketide | 25.0 | 41.6 | 130.0 | 3.4e-29 |
| WP\_020636817.1 | type\_I\_polyketide\_synthase | BGC0002011 | Polyketide | 30.0 | 24.1 | 129.0 | 5.8e-29 |
| AWS21279.1 | type\_I\_polyketide\_synthase | BGC0001934 | Polyketide | 29.0 | 21.4 | 128.0 | 1.3e-28 |
| AZY91989.1 | polyketide\_synthase | BGC0002022 | Polyketide | 29.0 | 21.4 | 128.0 | 1.3e-28 |
| CBD77746.1 | non-ribosomal\_peptide\_synthetase/polyketide\_synthase | BGC0000974 | NRP + Polyketide | 26.0 | 27.2 | 126.0 | 3.8e-28 |
| AIR74926.1 | polyketide\_synthase | BGC0001559 | RiPP | 26.0 | 27.2 | 126.0 | 3.8e-28 |
| AAS90047.1 | PksA | BGC0000009 | Polyketide | 26.0 | 21.9 | 126.0 | 4.9e-28 |
| EHA22196.1 | polyketide\_synthase | BGC0000170 | Polyketide | 25.0 | 27.5 | 126.0 | 4.9e-28 |
| ctg1\_orf523 |  | BGC0001199 | Polyketide | 29.0 | 21.7 | 124.0 | 2.4e-27 |
| CAD19092.1 | StiH\_protein | BGC0000153 | NRP + Polyketide:Modular type I | 25.0 | 22.4 | 123.0 | 3.2e-27 |
| BAB69193.1 |  | BGC0000117 | Polyketide | 27.0 | 21.6 | 122.0 | 7.1e-27 |
| CQR60497.1 | Polyketide\_synthase,\_type\_I,\_modules:\_loading,\_1,\_2\_and\_3 | BGC0001287 | Polyketide | 27.0 | 22.2 | 121.0 | 1.6e-26 |
| ctg1\_orf30 |  | BGC0000096 | Polyketide | 27.0 | 22.7 | 120.0 | 2.7e-26 |
| EAU32819.1 | 6-methylsalicylic\_acid\_synthase | BGC0000160 | Polyketide | 24.0 | 25.5 | 116.0 | 3.9e-25 |
| BAA20102.2 | 6-methylsalicylic\_acid\_synthase | BGC0001276 | Polyketide | 24.0 | 25.5 | 116.0 | 6.6e-25 |
| AAM54077.1 | polyketide\_synthase | BGC0000020 | Polyketide | 31.0 | 21.5 | 115.0 | 1.1e-24 |
| AJO72735.1 | Type\_I\_modular\_polyketide\_synthase | BGC0001381 | Polyketide | 26.0 | 20.8 | 113.0 | 4.3e-24 |
| sipP4 | Type\_I\_Modular\_PKS | BGC0001452 | Polyketide | 26.0 | 21.9 | 112.0 | 9.6e-24 |
| ADM46356.1 | polyketide\_synthase | BGC0000106 | Polyketide | 27.0 | 22.3 | 110.0 | 3.6e-23 |
| AWR88398.1 | putative\_beta-ketoacyl\_synthase | BGC0001522 | Polyketide | 25.0 | 29.8 | 108.0 | 1.1e-22 |
| CAL58683.1 | polyketide\_synthase | BGC0000149 | Polyketide:Modular type I | 29.0 | 20.2 | 107.0 | 3.1e-22 |
| ABF87031.1 | non-ribosomal\_peptide\_synthetase/polyketide\_synthase | BGC0000393 | NRP + Polyketide:Modular type I | 27.0 | 20.7 | 106.0 | 4e-22 |
