## Supplementary Results for "A draft genome of the ascomycotal fungal species *Pseudopithomyces maydicus* (family *Didymosphaeriaceae*)": input.path1.gene22_mibig_hits.html

| MIBiG Protein | Description | MIBiG Cluster | MiBiG Product | % ID | % Coverage | BLAST Score | E-value |
| --- | --- | --- | --- | --- | --- | --- | --- |
| EAU38977.1 | predicted\_protein | BGC0001122 | NRP + Polyketide:Iterative type I | 37.0 | 88.5 | 277.0 | 3.7e-74 |
| CAP93741.1 |  | BGC0001882 | Polyketide | 34.0 | 83.8 | 252.0 | 1.3e-66 |
| ATY69599.1 | antibiotic\_efflux\_protein | BGC0001823 | NRP + Polyketide | 28.0 | 81.8 | 161.0 | 3.9e-39 |
| AAF00219.1 | transporter | BGC0000277 | Polyketide | 27.0 | 82.0 | 146.0 | 9.9e-35 |
| AAM94765.1 | CalT1 | BGC0000033 | Polyketide | 24.0 | 82.5 | 143.0 | 1.1e-33 |
| ATY69558.1 | antibiotic\_efflux\_protein | BGC0001611 | NRP + Polyketide | 26.0 | 81.7 | 134.0 | 6.7e-31 |
| AAQ08935.1 | putative\_membrane\_transporter | BGC0000224 | Polyketide:Type II | 27.0 | 82.0 | 133.0 | 1.1e-30 |
| EAQ86390.1 | hypothetical\_protein | BGC0001405 | Polyketide | 31.0 | 52.2 | 130.0 | 9.6e-30 |
| ATU31811.1 | MFS\_transporter | BGC0001814 | NRP | 26.0 | 83.2 | 129.0 | 1.6e-29 |
| ABV91295.1 | putative\_multidrug\_transporter | BGC0000158 | Polyketide:Modular type I | 25.0 | 86.6 | 117.0 | 6.4e-26 |
| CAE51185.1 | RemN\_protein | BGC0000264 | Polyketide:Type II | 29.0 | 84.3 | 113.0 | 1.2e-24 |
| AGO50605.1 | transporter | BGC0000229 | Polyketide:Type II + Saccharide:Hybrid/tailoring | 25.0 | 77.6 | 106.0 | 1.9e-22 |
| CAH10123.1 | putative\_transporter | BGC0000268 | Polyketide | 25.0 | 80.6 | 106.0 | 1.9e-22 |
| AAL15595.1 | Sim17 | BGC0000270 | Polyketide | 24.0 | 82.2 | 104.0 | 4.3e-22 |
| AAK06799.1 | simocyclinone-specific\_efflux\_pump | BGC0001072 | Saccharide + Polyketide:Modular type I + Polyketide:Type II + Other:Aminocoumarin | 23.0 | 82.2 | 102.0 | 2.1e-21 |
| CCE31569.1 | probable\_aflatoxin\_efflux\_pump\_AFLT | BGC0001886 | Polyketide | 26.0 | 50.6 | 101.0 | 6.2e-21 |
| AEI98661.1 | CtcR | BGC0000209 | Polyketide | 44.0 | 22.2 | 94.0 | 4.5e-19 |
| AEO57490.1 | general\_substrate\_transporter | BGC0001449 | NRP + Alkaloid + Polyketide:Iterative type I | 32.0 | 28.4 | 86.0 | 2.1e-16 |
| BAZ95831.1 | MFS\_transporter\_cpaI | BGC0001563 | NRP + Polyketide | 29.0 | 39.2 | 81.0 | 3.9e-15 |
| BAE56594.1 |  | BGC0001123 | NRP | 29.0 | 28.2 | 79.0 | 1.5e-14 |
| ACB37757.1 | putative\_multidrug\_export\_protein | BGC0000162 | Polyketide | 26.0 | 79.2 | 75.0 | 3.7e-13 |
| ABC87523.1 | putative\_drug\_efflux\_transporter | BGC0001011 | NRP + Polyketide | 35.0 | 22.0 | 74.0 | 4.8e-13 |
| AMY15059.1 | MFS\_transporter | BGC0001339 | Polyketide:Iterative type I | 35.0 | 20.6 | 70.0 | 1.2e-11 |
| CAA09636.1 | putative\_transmembrane\_protein | BGC0000227 | Polyketide:Type II | 34.0 | 22.6 | 69.0 | 2e-11 |
| ADE34501.1 | ssfR | BGC0000269 | Polyketide:Type II + Saccharide:Hybrid/tailoring | 34.0 | 21.2 | 69.0 | 2e-11 |
| AIL50170.1 | putative\_transport\_protein | BGC0000213 | Polyketide:Type II | 24.0 | 70.0 | 67.0 | 5.9e-11 |
| ACB46467.1 | efflux\_permease | BGC0000082 | Polyketide | 25.0 | 66.7 | 66.0 | 1.3e-10 |
| ALS30797.1 | itaconate\_transport\_protein\_1 | BGC0001286 | Other | 30.0 | 23.1 | 59.0 | 1.6e-08 |
| AGI99475.1 | efflux\_permease | BGC0001004 | Polyketide:Modular type I | 25.0 | 71.6 | 59.0 | 2.7e-08 |
| AWM72923.1 | drug\_resistance\_transporter | BGC0001719 | Polyketide | 21.0 | 66.7 | 59.0 | 2.7e-08 |
| ATL73060.1 | efflux\_permease | BGC0001807 | NRP + Polyketide | 25.0 | 71.6 | 59.0 | 2.7e-08 |
| AGC09506.1 | LobA4 | BGC0001183 | Polyketide | 25.0 | 70.5 | 58.0 | 4.6e-08 |
| ctg1\_orf263 |  | BGC0001200 | Polyketide | 22.0 | 50.8 | 50.0 | 7.4e-06 |
