## Supplementary Results for "A draft genome of the ascomycotal fungal species *Pseudopithomyces maydicus* (family *Didymosphaeriaceae*)": input.path1.gene24_mibig_hits.html

| MIBiG Protein | Description | MIBiG Cluster | MiBiG Product | % ID | % Coverage | BLAST Score | E-value |
| --- | --- | --- | --- | --- | --- | --- | --- |
| ACD39770.1 | non-reducing\_polyketide\_synthase | BGC0000134 | Polyketide | 62.0 | 107.0 | 2240.0 | 0.0 |
| AHV78253.1 | ResS2 | BGC0001246 | Polyketide | 56.0 | 113.3 | 2047.0 | 0.0 |
| ACD39753.1 | non-reducing\_polyketide\_synthase | BGC0000076 | Polyketide | 56.0 | 107.5 | 1981.0 | 0.0 |
| ACD39762.1 | non-reducing\_polyketide\_synthase | BGC0000077 | Polyketide | 56.0 | 107.5 | 1979.0 | 0.0 |
| ABB90282.1 | polyketide\_synthase | BGC0001057 | NRP + Polyketide | 56.0 | 106.7 | 1940.0 | 0.0 |
| AHV78247.1 | LasS2 | BGC0001245 | Polyketide | 57.0 | 108.9 | 1921.0 | 0.0 |
| AGC95321.1 | CurS2 | BGC0000045 | Polyketide | 54.0 | 108.6 | 1810.0 | 0.0 |
| ADM79459.1 | PKS16\_protein | BGC0001266 | Polyketide | 36.0 | 85.3 | 870.0 | 5.1e-252 |
| EAA59563.1 | polyketide\_synthase | BGC0000057 | Polyketide:Iterative type I | 37.0 | 79.0 | 785.0 | 1.3e-226 |
| AAN59953.1 | polyketide\_synthase\_1 | BGC0001258 | Polyketide | 36.0 | 78.9 | 782.0 | 1.8e-225 |
| KKP00966.1 | RADS2\_nonreducing\_polyketide\_synthase | BGC0001901 | Polyketide | 35.0 | 79.1 | 761.0 | 2.5e-219 |
| BAD22832.1 | polyketide\_synthase | BGC0001265 | Polyketide | 36.0 | 79.5 | 753.0 | 6.9e-217 |
| AUW31184.1 | putative\_type\_I\_PKS | BGC0001489 | Polyketide | 35.0 | 80.0 | 748.0 | 2.9e-215 |
| EAU38791.1 | hypothetical\_protein | BGC0000161 | Polyketide:Iterative type I | 35.0 | 76.9 | 743.0 | 7.2e-214 |
| gene6 |  | BGC0001906 | Polyketide | 37.0 | 73.3 | 741.0 | 2.7e-213 |
| CBF74114.1 | Conidial\_yellow\_pigment\_biosynthesis\_polyketide\_synthase\_(PKS)(EC\_2.3.1.-)\_[Source:UniProtKB/Swiss-Prot;Acc:Q03149] | BGC0000107 | Polyketide | 35.0 | 74.1 | 741.0 | 3.6e-213 |
| AAD38786.1 | polyketide\_synthase | BGC0001257 | Polyketide | 36.0 | 78.6 | 731.0 | 3.7e-210 |
| RWQ92175.1 | putative\_polyketide\_synthase | BGC0002030 | Polyketide | 34.0 | 79.0 | 723.0 | 1e-207 |
| AGO59040.1 | PtaA | BGC0000121 | Polyketide | 34.0 | 79.6 | 719.0 | 1.1e-206 |
| EED21099.1 | polyketide\_synthase,\_putative | BGC0001578 | Polyketide | 35.0 | 74.0 | 718.0 | 1.9e-206 |
| EAL89339.1 | polyketide\_synthase,\_putative | BGC0001403 | Polyketide | 34.0 | 80.1 | 713.0 | 1e-204 |
| AAS89999.1 | PksA | BGC0000007 | Polyketide | 35.0 | 77.7 | 711.0 | 3.9e-204 |
| AAS90093.1 | PksA | BGC0000006 | Polyketide | 35.0 | 76.4 | 707.0 | 4.4e-203 |
| BAE71314.1 | polyketide\_synthase | BGC0000004 | Polyketide | 35.0 | 76.4 | 705.0 | 2.2e-202 |
| AAS90022.1 | PksA | BGC0000008 | Polyketide | 35.0 | 76.4 | 705.0 | 2.2e-202 |
| EAL84397.1 | polyketide\_synthase | BGC0001118 | Polyketide:Iterative type I | 35.0 | 74.4 | 700.0 | 5.3e-201 |
| PKX92308.1 | putative\_polyketide\_synthase | BGC0001988 | Polyketide | 34.0 | 79.6 | 698.0 | 2.6e-200 |
| EGD99348.1 | polyketide\_synthase | BGC0001144 | Polyketide | 35.0 | 73.8 | 695.0 | 2.2e-199 |
| AAS90047.1 | PksA | BGC0000009 | Polyketide | 33.0 | 83.1 | 694.0 | 3.8e-199 |
| CCE67070.1 | polyketide\_synthase | BGC0001242 | Polyketide | 34.0 | 78.7 | 693.0 | 1.1e-198 |
| CCE31584.1 | polyketide\_synthase\_that\_catalyse\_the\_condensation\_of\_one\_acetyl-CoA\_and\_six\_malonyl-CoA\_resulting\_in\_formation\_of\_nor-rubrofusarin | BGC0001886 | Polyketide | 33.0 | 80.4 | 691.0 | 4.2e-198 |
| CCT67991.1 | bikaverin\_cluster-polyketide\_synthase | BGC0000030 | Polyketide | 33.0 | 79.3 | 684.0 | 4e-196 |
| ADI24953.1 | GsfA | BGC0000070 | Polyketide:Iterative type I | 33.0 | 77.5 | 679.0 | 1.3e-194 |
| EED57518.1 | polyketide\_synthase,\_putative | BGC0001446 | Polyketide:Iterative type I | 32.0 | 85.6 | 670.0 | 1e-191 |
| AKN45693.1 | polyketide\_synthase | BGC0001284 | Terpene | 32.0 | 78.6 | 669.0 | 1.3e-191 |
| ACH72912.1 | AflC | BGC0000011 | Polyketide | 34.0 | 79.0 | 665.0 | 1.9e-190 |
| ADI24926.1 | VrtA | BGC0000168 | Polyketide:Iterative type I | 33.0 | 79.0 | 664.0 | 4.2e-190 |
| DAB41653.1 | polyketide\_synthase | BGC0001583 | Polyketide | 33.0 | 83.2 | 657.0 | 5.2e-188 |
| AAC49191.1 | putative\_polyketide\_synthase | BGC0000152 | Polyketide | 35.0 | 74.5 | 656.0 | 1.2e-187 |
| EED53479.1 | polyketide\_synthase,\_putative | BGC0001304 | Polyketide | 32.0 | 78.6 | 654.0 | 4.4e-187 |
| AEN83889.1 | AdaA | BGC0000156 | Polyketide:Iterative type I | 32.0 | 79.7 | 640.0 | 8.6e-183 |
| CBF70387.1 | polyketide\_synthase,\_putative\_(JCVI) | BGC0000684 | Terpene | 33.0 | 77.5 | 636.0 | 1.2e-181 |
| AAZ95017.1 | polyketide\_synthase | BGC0000048 | Polyketide | 32.0 | 80.5 | 634.0 | 4.7e-181 |
| CBF79143.1 | polyketide\_synthase,\_putative\_(JCVI) | BGC0000013 | Polyketide | 32.0 | 83.2 | 621.0 | 3.2e-177 |
| XP\_001798923.1 | polyketide\_synthase | BGC0001865 | Polyketide:Iterative type I | 32.0 | 75.3 | 614.0 | 3.9e-175 |
| ARU80380.1 | polyketide\_synthase | BGC0001542 | Polyketide | 32.0 | 74.4 | 612.0 | 2.5e-174 |
| PIB02405.1 | CTB1 | BGC0001541 | Polyketide | 31.0 | 74.5 | 608.0 | 2.8e-173 |
| EJP62792.1 | polyketide\_synthase | BGC0001720 | Polyketide | 30.0 | 71.2 | 480.0 | 8.7e-135 |
| AQW44888.1 | polyketide\_synthase | BGC0001737 | NRP + Polyketide | 33.0 | 58.5 | 480.0 | 1.1e-134 |
| AGC45620.1 | polyketide\_synthase | BGC0001394 | NRP + Polyketide | 32.0 | 60.0 | 474.0 | 8.1e-133 |
| AIW82282.1 | PuwE | BGC0001125 | NRP + Polyketide | 34.0 | 55.2 | 472.0 | 3.1e-132 |
| AZH23818.1 | MgiI | BGC0001971 | NRP + Polyketide | 33.0 | 60.1 | 468.0 | 5.8e-131 |
| ACR33078.1 | polyketide\_synthase | BGC0000017 | Alkaloid + Polyketide:Modular type I | 32.0 | 58.3 | 465.0 | 4.9e-130 |
| AZH23819.1 | MgiR | BGC0001971 | NRP + Polyketide | 33.0 | 54.9 | 465.0 | 4.9e-130 |
| AID65222.1 | putative\_aspartate\_racemase | BGC0000335 | NRP | 34.0 | 53.2 | 462.0 | 2.4e-129 |
| AMB48442.1 | polyketide\_synthase | BGC0001357 | Polyketide | 33.0 | 54.4 | 462.0 | 4.2e-129 |
| AEE88289.1 | CurA | BGC0000976 | NRP + Polyketide:Modular type I | 32.0 | 55.1 | 459.0 | 2.1e-128 |
| AAT70096.1 | CurA | BGC0001165 | NRP + Polyketide:Modular type I | 32.0 | 55.1 | 459.0 | 2.1e-128 |
| ADZ24998.1 | polyketide\_synthase | BGC0000380 | NRP + Polyketide:Modular type I | 32.0 | 59.2 | 459.0 | 2.7e-128 |
| AQA28563.1 | type\_I\_polyketide\_synthase | BGC0001663 | Polyketide | 33.0 | 54.4 | 458.0 | 3.5e-128 |
| XP\_011392701.1 | hypothetical\_protein | BGC0001281 | Polyketide | 27.0 | 91.2 | 456.0 | 2.3e-127 |
| AZH23788.1 | MgcR | BGC0001970 | NRP + Polyketide | 32.0 | 56.0 | 455.0 | 3e-127 |
| AXN93601.1 | PuwE | BGC0001952 | NRP | 32.0 | 57.2 | 452.0 | 3.3e-126 |
| CAD19086.1 | StiB\_protein | BGC0000153 | NRP + Polyketide:Modular type I | 34.0 | 54.0 | 451.0 | 5.6e-126 |
| AEE88280.1 | CurJ | BGC0000976 | NRP + Polyketide:Modular type I | 33.0 | 54.9 | 451.0 | 7.4e-126 |
| AAT70105.1 | CurJ | BGC0001165 | NRP + Polyketide:Modular type I | 33.0 | 54.9 | 451.0 | 7.4e-126 |
| AXN93613.1 | PuwE | BGC0001953 | NRP | 32.0 | 56.7 | 451.0 | 7.4e-126 |
| CAD19093.1 | StiJ\_protein | BGC0000153 | NRP + Polyketide:Modular type I | 34.0 | 53.5 | 450.0 | 1.3e-125 |
| AXN93597.1 | PuwB | BGC0001952 | NRP | 32.0 | 58.4 | 450.0 | 1.6e-125 |
| AXN93580.1 | PuwE | BGC0001950 | NRP | 32.0 | 58.8 | 448.0 | 3.7e-125 |
| AXN93589.1 | PuwE | BGC0001951 | NRP | 32.0 | 58.8 | 448.0 | 3.7e-125 |
| AIW82279.1 | PuwB | BGC0001125 | NRP + Polyketide | 31.0 | 59.0 | 446.0 | 1.4e-124 |
| AQW44891.1 | polyketide\_synthase | BGC0001737 | NRP + Polyketide | 32.0 | 59.1 | 446.0 | 1.4e-124 |
| AAS98777.1 | polyketide\_synthetase | BGC0001001 | NRP + Polyketide | 32.0 | 54.1 | 445.0 | 4e-124 |
| ABM21570.1 | crpB | BGC0000975 | NRP + Polyketide | 32.0 | 54.4 | 443.0 | 1.2e-123 |
| DAB41915.1 | ArzM\_-\_PKS\_(KS,\_AT,\_DH,\_MT,\_ER,\_KR,\_ACP) | BGC0001884 | NRP + Polyketide | 31.0 | 59.7 | 443.0 | 1.5e-123 |
| AZH23791.1 | MgcH | BGC0001970 | NRP + Polyketide | 32.0 | 56.2 | 443.0 | 1.5e-123 |
| AGC45621.1 | polyketide\_synthase | BGC0001394 | NRP + Polyketide | 31.0 | 59.2 | 443.0 | 2e-123 |
| AHB82053.1 | polyketide\_synthase | BGC0001019 | NRP + Polyketide:Modular type I | 32.0 | 60.6 | 441.0 | 5.8e-123 |
| AAF19814.1 | MtaF | BGC0001024 | NRP + Polyketide:Modular type I | 33.0 | 53.9 | 441.0 | 7.6e-123 |
| CAD89777.1 | MelF\_protein | BGC0001010 | NRP + Polyketide:Modular type I | 32.0 | 54.0 | 438.0 | 3.8e-122 |
| ATX68116.1 | malonyl\_CoA-acyl\_carrier\_protein\_transacylase | BGC0001772 | Polyketide | 32.0 | 57.0 | 438.0 | 3.8e-122 |
| AZH23821.1 | MgiH | BGC0001971 | NRP + Polyketide | 32.0 | 56.0 | 438.0 | 3.8e-122 |
| AAS98782.1 | polyketide\_synthase | BGC0001001 | NRP + Polyketide | 31.0 | 56.8 | 438.0 | 4.9e-122 |
| DAB41916.1 | ArzN\_-\_PKS\_(KS,\_AT,\_OMT,\_KR,\_ACP) | BGC0001884 | NRP + Polyketide | 32.0 | 55.2 | 437.0 | 1.1e-121 |
| AAW03328.1 | CtaE | BGC0000982 | NRP + Polyketide | 34.0 | 54.6 | 436.0 | 1.4e-121 |
| BBF25315.1 | polyketide\_synthase | BGC0001923 | Terpene + Polyketide | 27.0 | 79.9 | 436.0 | 1.4e-121 |
| ACR33079.1 | polyketide\_synthase | BGC0000017 | Alkaloid + Polyketide:Modular type I | 33.0 | 53.7 | 436.0 | 1.9e-121 |
| AQA28562.1 | type\_I\_polyketide\_synthase | BGC0001663 | Polyketide | 31.0 | 59.2 | 436.0 | 2.5e-121 |
| AVI26390.1 | polyketide\_synthase\_/\_nonribosomal\_peptide\_synthase\_hybrid | BGC0001800 | NRP + Polyketide | 33.0 | 54.9 | 436.0 | 2.5e-121 |
| AEE88278.1 | CurL | BGC0000976 | NRP + Polyketide:Modular type I | 32.0 | 54.9 | 435.0 | 5.5e-121 |
| AAT70107.1 | CurL | BGC0001165 | NRP + Polyketide:Modular type I | 32.0 | 54.9 | 435.0 | 5.5e-121 |
| ADF88276.1 | polyketide\_synthase | BGC0000981 | NRP + Polyketide | 33.0 | 53.6 | 434.0 | 7.1e-121 |
| AEE88282.1 | CurH | BGC0000976 | NRP + Polyketide:Modular type I | 32.0 | 53.9 | 433.0 | 1.2e-120 |
| AAT70103.1 | CurH | BGC0001165 | NRP + Polyketide:Modular type I | 32.0 | 53.9 | 433.0 | 1.2e-120 |
| AAF19813.1 | MtaE | BGC0001024 | NRP + Polyketide:Modular type I | 34.0 | 54.5 | 433.0 | 1.6e-120 |
| AXN93610.1 | PuwB | BGC0001953 | NRP | 32.0 | 59.1 | 433.0 | 1.6e-120 |
| CAQ34920.1 | polyketide\_synthase | BGC0000986 | NRP + Polyketide | 33.0 | 58.5 | 433.0 | 2.1e-120 |
| ADY00130.1 | polyketide\_synthase | BGC0000104 | Terpene + Polyketide:Iterative type I | 27.0 | 75.7 | 432.0 | 2.7e-120 |
| AEE88279.1 | CurK | BGC0000976 | NRP + Polyketide:Modular type I | 31.0 | 58.2 | 432.0 | 2.7e-120 |
| AAT70106.1 | CurK | BGC0001165 | NRP + Polyketide:Modular type I | 31.0 | 58.2 | 432.0 | 2.7e-120 |
| ctg1\_orf16 |  | BGC0001457 | NRP | 31.0 | 59.4 | 432.0 | 2.7e-120 |
| ADF88280.1 | polyketide\_synthase | BGC0000981 | NRP + Polyketide | 32.0 | 53.2 | 432.0 | 3.5e-120 |
| ABX60162.1 | polyketide\_synthase | BGC0000978 | NRP + Alkaloid + Polyketide:Modular type I | 32.0 | 53.2 | 431.0 | 4.6e-120 |
| CAQ18829.1 | polyketide\_synthase | BGC0000954 | NRP + Polyketide:Modular type I | 32.0 | 59.1 | 431.0 | 6e-120 |
| CAQ18835.1 | polyketide\_synthase | BGC0000954 | NRP + Polyketide:Modular type I | 32.0 | 59.7 | 431.0 | 7.9e-120 |
| CAQ18834.1 | polyketide\_synthase | BGC0000954 | NRP + Polyketide:Modular type I | 33.0 | 53.6 | 429.0 | 3e-119 |
| AEE88281.1 | CurI | BGC0000976 | NRP + Polyketide:Modular type I | 32.0 | 54.0 | 429.0 | 3e-119 |
| AAT70104.1 | CurI | BGC0001165 | NRP + Polyketide:Modular type I | 32.0 | 54.0 | 429.0 | 3e-119 |
| ABX60152.1 | polyketide\_synthase | BGC0000978 | NRP + Alkaloid + Polyketide:Modular type I | 32.0 | 53.6 | 428.0 | 3.9e-119 |
| AHA12078.1 | polyketide\_synthase\_type\_1 | BGC0001172 | NRP + Polyketide:Modular type I | 33.0 | 55.6 | 428.0 | 5.1e-119 |
| AXN93586.1 | PuwB | BGC0001951 | NRP | 30.0 | 62.4 | 428.0 | 6.7e-119 |
| AMB48441.1 | polyketide\_synthase | BGC0001357 | Polyketide | 31.0 | 55.5 | 427.0 | 1.1e-118 |
| AZH23820.1 | MgiG | BGC0001971 | NRP + Polyketide | 31.0 | 54.5 | 427.0 | 1.1e-118 |
| ABX60163.1 | polyketide\_synthase | BGC0000978 | NRP + Alkaloid + Polyketide:Modular type I | 31.0 | 55.1 | 426.0 | 1.5e-118 |
| ADN13832.1 | Polyketide\_Synthase | BGC0001164 | Polyketide:Modular type I | 32.0 | 53.5 | 426.0 | 1.5e-118 |
| AZH23790.1 | MgcG | BGC0001970 | NRP + Polyketide | 31.0 | 54.5 | 426.0 | 1.5e-118 |
| CAD19087.1 | StiC\_protein | BGC0000153 | NRP + Polyketide:Modular type I | 32.0 | 58.2 | 426.0 | 2.5e-118 |
| CAD19091.1 | StiG\_protein | BGC0000153 | NRP + Polyketide:Modular type I | 32.0 | 59.6 | 425.0 | 4.3e-118 |
| ATX68115.1 | malonyl\_CoA-acyl\_carrier\_protein\_transacylase | BGC0001772 | Polyketide | 32.0 | 54.8 | 425.0 | 4.3e-118 |
| AXN93577.1 | PuwB | BGC0001950 | NRP | 30.0 | 62.6 | 425.0 | 5.7e-118 |
| ADF88277.1 | polyketide\_synthase | BGC0000981 | NRP + Polyketide | 31.0 | 54.0 | 424.0 | 7.4e-118 |
| KFA69335.1 | hypothetical\_protein | BGC0001626 | Polyketide | 28.0 | 74.7 | 424.0 | 7.4e-118 |
| CBD77736.1 | polyketide\_synthase | BGC0000974 | NRP + Polyketide | 32.0 | 59.0 | 424.0 | 9.6e-118 |
| AIR74912.1 | polyketide\_synthase | BGC0001559 | RiPP | 32.0 | 59.0 | 424.0 | 9.6e-118 |
| KFL51883.1 | amino\_acid\_adenylation\_protein | BGC0001711 | NRP + Polyketide | 31.0 | 55.2 | 424.0 | 9.6e-118 |
| AVI26388.1 | polyketide\_synthase | BGC0001800 | NRP + Polyketide | 31.0 | 56.3 | 423.0 | 1.3e-117 |
| AGC45624.1 | polyketide\_synthase | BGC0001394 | NRP + Polyketide | 31.0 | 59.7 | 421.0 | 4.8e-117 |
| AAK57188.1 | MxaD | BGC0001022 | NRP + Polyketide | 31.0 | 58.9 | 421.0 | 6.2e-117 |
| AVI26389.1 | polyketide\_synthase | BGC0001800 | NRP + Polyketide | 32.0 | 53.9 | 421.0 | 6.2e-117 |
| AQW44889.1 | polyketide\_synthase | BGC0001737 | NRP + Polyketide | 31.0 | 59.3 | 421.0 | 8.2e-117 |
| AAU04878.1 | polyketide\_synthase | BGC0000365 | NRP | 30.0 | 62.5 | 420.0 | 1.4e-116 |
| AGC45619.1 | polyketide\_synthase | BGC0001394 | NRP + Polyketide | 32.0 | 58.8 | 420.0 | 1.4e-116 |
| AAK57189.1 | MxaE | BGC0001022 | NRP + Polyketide | 31.0 | 58.9 | 420.0 | 1.8e-116 |
| AHB82064.1 | polyketide\_synthase | BGC0001231 | NRP + Polyketide:Modular type I | 32.0 | 53.7 | 419.0 | 2.4e-116 |
| WP\_035121546.1 | type\_I\_polyketide\_synthase | BGC0001467 | NRP:Cyclic depsipeptide + Polyketide:Modular type I | 32.0 | 55.3 | 419.0 | 3.1e-116 |
| AXM42950.1 | polyketide\_synthase | BGC0001941 | NRP + Polyketide | 32.0 | 53.5 | 419.0 | 3.1e-116 |
| ABX60153.1 | polyketide\_synthase | BGC0000978 | NRP + Alkaloid + Polyketide:Modular type I | 32.0 | 53.2 | 418.0 | 5.3e-116 |
| CAD19090.1 | StiF\_protein | BGC0000153 | NRP + Polyketide:Modular type I | 28.0 | 73.6 | 417.0 | 9e-116 |
| ADF88275.1 | polyketide\_synthase | BGC0000981 | NRP + Polyketide | 31.0 | 53.2 | 417.0 | 1.2e-115 |
| AHH34186.1 | polyketide\_synthase | BGC0001161 | Polyketide:Modular type I | 31.0 | 56.1 | 416.0 | 1.5e-115 |
| CAO98850.1 | polyketide\_synthase\_AufG | BGC0000023 | Polyketide:Modular type I | 33.0 | 53.8 | 416.0 | 2e-115 |
| CDM36726.1 | Beta-ketoacyl\_synthase | BGC0001360 | Polyketide | 27.0 | 76.0 | 415.0 | 4.5e-115 |
| CAD19088.1 | StiD\_protein | BGC0000153 | NRP + Polyketide:Modular type I | 32.0 | 55.1 | 415.0 | 5.8e-115 |
| BAV19379.1 | polyketide\_synthase | BGC0001390 | NRP + Polyketide | 27.0 | 75.7 | 415.0 | 5.8e-115 |
| AHA38203.1 | GphJ | BGC0000069 | Polyketide | 33.0 | 54.3 | 414.0 | 7.6e-115 |
| AQW44890.1 | polyketide\_synthase | BGC0001737 | NRP + Polyketide | 32.0 | 59.0 | 414.0 | 7.6e-115 |
| AHH34189.1 | polyketide\_synthase | BGC0001162 | Polyketide:Modular type I | 31.0 | 57.3 | 413.0 | 1.3e-114 |
| CAO98879.1 | polyketide\_synthase\_AufD | BGC0000023 | Polyketide:Modular type I | 31.0 | 61.3 | 413.0 | 1.7e-114 |
| AZH23817.1 | MgiQ | BGC0001971 | NRP + Polyketide | 30.0 | 56.6 | 412.0 | 2.9e-114 |
| CAQ18828.1 | polyketide\_synthase | BGC0000954 | NRP + Polyketide:Modular type I | 33.0 | 53.3 | 411.0 | 5e-114 |
| AAF00959.1 | mcyD | BGC0001017 | NRP + Polyketide:Modular type I | 31.0 | 55.3 | 411.0 | 5e-114 |
| EGJ35088.1 | Polyketide\_synthase | BGC0001163 | Polyketide:Modular type I | 31.0 | 57.6 | 410.0 | 1.4e-113 |
| AZH23787.1 | MgcQ | BGC0001970 | NRP + Polyketide | 31.0 | 56.5 | 410.0 | 1.4e-113 |
| AFU82617.1 | polyketide\_synthase | BGC0000998 | NRP + Polyketide | 31.0 | 53.9 | 408.0 | 4.2e-113 |
| ADX66472.1 | ScnS1 | BGC0000108 | Polyketide | 30.0 | 59.3 | 408.0 | 7.2e-113 |
| AAK57187.1 | MxaC | BGC0001022 | NRP + Polyketide | 32.0 | 54.8 | 407.0 | 9.3e-113 |
| CAG28678.1 | polyketide\_synthase | BGC0001023 | NRP + Polyketide:Modular type I | 33.0 | 54.4 | 407.0 | 9.3e-113 |
| APZ78820.1 | polyketide\_synthase | BGC0001429 | NRP:Cyclic depsipeptide + Polyketide:Iterative type I | 33.0 | 54.4 | 407.0 | 9.3e-113 |
| ABK32288.1 | JerB | BGC0000080 | Polyketide | 32.0 | 53.2 | 407.0 | 1.2e-112 |
| AAW03329.1 | CtaF | BGC0000982 | NRP + Polyketide | 32.0 | 53.5 | 407.0 | 1.2e-112 |
| CAO98849.1 | polyketide\_synthase\_AufF | BGC0000023 | Polyketide:Modular type I | 31.0 | 58.9 | 406.0 | 1.6e-112 |
| AQW44893.1 | polyketide\_synthase | BGC0001737 | NRP + Polyketide | 31.0 | 59.0 | 406.0 | 2.1e-112 |
| CAD19092.1 | StiH\_protein | BGC0000153 | NRP + Polyketide:Modular type I | 31.0 | 54.3 | 405.0 | 3.5e-112 |
| AAF26921.1 | polyketide\_synthase | BGC0000988 | NRP + Polyketide | 30.0 | 58.7 | 405.0 | 3.5e-112 |
| APZ78844.1 | polyketide\_synthase | BGC0001431 | NRP:Cyclic depsipeptide + Polyketide:Iterative type I | 33.0 | 54.5 | 405.0 | 3.5e-112 |
| ACB46195.1 | polyketide\_synthase | BGC0000989 | NRP + Polyketide | 30.0 | 58.6 | 404.0 | 6.1e-112 |
| ADB12491.1 | EpoD | BGC0000990 | NRP + Polyketide | 30.0 | 58.6 | 404.0 | 7.9e-112 |
| AAF62883.1 | epoD | BGC0000991 | NRP + Polyketide | 30.0 | 58.6 | 404.0 | 7.9e-112 |
| APZ78854.1 | polyketide\_synthase | BGC0001432 | NRP:Cyclic depsipeptide + Polyketide:Iterative type I | 31.0 | 58.5 | 404.0 | 7.9e-112 |
| ABK32256.1 | AmbB | BGC0000014 | Polyketide | 32.0 | 53.6 | 403.0 | 1.3e-111 |
| CAD89776.1 | MelE\_protein | BGC0001010 | NRP + Polyketide:Modular type I | 32.0 | 54.4 | 403.0 | 1.3e-111 |
| AGC45622.1 | polyketide\_synthase | BGC0001394 | NRP + Polyketide | 30.0 | 59.0 | 403.0 | 1.8e-111 |
| ASZ00149.1 | polyketide\_synthase | BGC0001785 | Polyketide | 31.0 | 58.1 | 403.0 | 1.8e-111 |
| AWS21279.1 | type\_I\_polyketide\_synthase | BGC0001934 | Polyketide | 32.0 | 51.8 | 402.0 | 3e-111 |
| AZY91989.1 | polyketide\_synthase | BGC0002022 | Polyketide | 32.0 | 51.8 | 402.0 | 3e-111 |
| ADH04641.1 | TgaC | BGC0001051 | NRP + Polyketide:Modular type I | 31.0 | 59.9 | 402.0 | 3.9e-111 |
| ATP76241.1 | NdaD | BGC0001705 | NRP + Polyketide | 30.0 | 54.0 | 402.0 | 3.9e-111 |
| AQH32482.1 | type\_1\_polyketide\_synthase | BGC0001667 | NRP + Polyketide | 30.0 | 55.4 | 401.0 | 5.1e-111 |
| CAQ18833.1 | polyketide\_synthase | BGC0000954 | NRP + Polyketide:Modular type I | 31.0 | 54.5 | 401.0 | 6.7e-111 |
| ABK32289.1 | JerC | BGC0000080 | Polyketide | 30.0 | 58.9 | 400.0 | 1.1e-110 |
| WP\_051137606.1 | type\_I\_polyketide\_synthase | BGC0002011 | Polyketide | 31.0 | 54.0 | 400.0 | 1.1e-110 |
| APZ78714.1 | polyketide\_synthase | BGC0001420 | NRP:Cyclic depsipeptide + Polyketide:Iterative type I | 32.0 | 55.6 | 399.0 | 2.5e-110 |
| AVI57433.1 | AbmB1 | BGC0001694 | Polyketide | 31.0 | 53.6 | 399.0 | 2.5e-110 |
| ABK32263.1 | AmbH | BGC0000014 | Polyketide | 31.0 | 56.3 | 399.0 | 3.3e-110 |
| APZ78754.1 | polyketide\_synthase | BGC0001423 | NRP:Cyclic depsipeptide + Polyketide:Iterative type I | 32.0 | 54.4 | 399.0 | 3.3e-110 |
| CAO98847.1 | polyketide\_synthase\_AufC | BGC0000023 | Polyketide:Modular type I | 32.0 | 54.0 | 398.0 | 4.3e-110 |
| AEE88284.1 | CurF | BGC0000976 | NRP + Polyketide:Modular type I | 30.0 | 55.5 | 398.0 | 4.3e-110 |
| APZ78690.1 | polyketide\_synthase | BGC0001418 | NRP:Cyclic depsipeptide + Polyketide:Iterative type I | 32.0 | 55.0 | 398.0 | 4.3e-110 |
| ART41209.1 | AdrD | BGC0001508 | Polyketide | 26.0 | 74.4 | 398.0 | 4.3e-110 |
| AQW44892.1 | polyketide\_synthase | BGC0001737 | NRP + Polyketide | 30.0 | 58.9 | 398.0 | 4.3e-110 |
| AGC45623.1 | polyketide\_synthase | BGC0001394 | NRP + Polyketide | 29.0 | 58.7 | 398.0 | 5.7e-110 |
| AAO62584.1 | polyketide\_synthase\_type\_1 | BGC0001016 | NRP + Polyketide | 30.0 | 53.2 | 398.0 | 7.4e-110 |
| ANI24099.1 | polyketide\_synthase | BGC0001235 | NRP + Polyketide | 32.0 | 59.0 | 397.0 | 1.3e-109 |
| AAT70101.1 | CurF | BGC0001165 | NRP + Polyketide:Modular type I | 30.0 | 55.5 | 396.0 | 1.6e-109 |
| APZ78727.1 | polyketide\_synthase | BGC0001421 | NRP:Cyclic depsipeptide + Polyketide:Iterative type I | 32.0 | 54.4 | 396.0 | 1.6e-109 |
| APZ78832.1 | polyketide\_synthase | BGC0001430 | NRP:Cyclic depsipeptide + Polyketide:Iterative type I | 32.0 | 55.2 | 396.0 | 1.6e-109 |
| AAF26922.1 | polyketide\_synthase | BGC0000988 | NRP + Polyketide | 31.0 | 58.4 | 396.0 | 2.2e-109 |
| AHN85651.1 | Phn2 | BGC0000122 | Polyketide:Modular type I | 31.0 | 53.2 | 396.0 | 2.8e-109 |
| ATY46587.1 | polyketide\_synthase | BGC0001666 | Polyketide | 31.0 | 60.2 | 396.0 | 2.8e-109 |
| ATY12793.1 | type\_I\_polyketide\_synthase | BGC0001504 | Polyketide | 30.0 | 59.4 | 394.0 | 6.3e-109 |
| WP\_020636845.1 | type\_I\_polyketide\_synthase | BGC0002011 | Polyketide | 32.0 | 55.1 | 394.0 | 6.3e-109 |
| AIT55259.1 | polyketide\_synthase | BGC0000072 | Polyketide:Modular type I | 32.0 | 53.8 | 394.0 | 8.2e-109 |
| APZ78702.1 | polyketide\_synthase | BGC0001419 | NRP:Cyclic depsipeptide + Polyketide:Iterative type I | 32.0 | 55.0 | 394.0 | 8.2e-109 |
| AQT01382.1 | SgnS1 | BGC0001690 | Polyketide | 30.0 | 59.8 | 394.0 | 8.2e-109 |
| CAC20931.1 | PimS1\_protein | BGC0000125 | Polyketide | 30.0 | 59.8 | 393.0 | 2.4e-108 |
| ANH11412.1 | SceQ | BGC0001908 | Polyketide | 31.0 | 59.3 | 393.0 | 2.4e-108 |
| ASZ00151.1 | polyketide\_synthase | BGC0001785 | Polyketide | 32.0 | 59.9 | 392.0 | 4.1e-108 |
| AIT55260.1 | polyketide\_synthase | BGC0000072 | Polyketide:Modular type I | 31.0 | 59.1 | 391.0 | 5.3e-108 |
| ADZ24996.1 | polyketide\_synthase | BGC0000380 | NRP + Polyketide:Modular type I | 30.0 | 59.8 | 391.0 | 5.3e-108 |
| ADB12492.1 | EpoE | BGC0000990 | NRP + Polyketide | 31.0 | 58.4 | 391.0 | 5.3e-108 |
| AAF62884.1 | EpoE | BGC0000991 | NRP + Polyketide | 31.0 | 58.4 | 391.0 | 6.9e-108 |
| AAO62585.1 | peptide\_sythetase\_polyketide\_synthase\_fusion\_protein | BGC0001016 | NRP + Polyketide | 31.0 | 54.2 | 391.0 | 9e-108 |
| CBD77738.1 | polyketide\_synthase | BGC0000974 | NRP + Polyketide | 32.0 | 53.7 | 390.0 | 1.2e-107 |
| ALD82522.1 | polyketide\_synthase | BGC0001212 | NRP + Polyketide | 30.0 | 59.2 | 390.0 | 1.2e-107 |
| AIR74913.1 | polyketide\_synthase | BGC0001559 | RiPP | 32.0 | 53.7 | 390.0 | 1.2e-107 |
| ABK32258.1 | AmbD | BGC0000014 | Polyketide | 30.0 | 59.1 | 390.0 | 1.5e-107 |
| APZ78678.1 | polyketide\_synthase | BGC0001417 | NRP:Cyclic depsipeptide + Polyketide:Iterative type I | 32.0 | 55.3 | 390.0 | 1.5e-107 |
| ACB46196.1 | polyketide\_synthase | BGC0000989 | NRP + Polyketide | 30.0 | 58.4 | 389.0 | 2.6e-107 |
| AZH23793.1 | MgcK | BGC0001970 | NRP + Polyketide | 30.0 | 55.4 | 389.0 | 2.6e-107 |
| AQH32481.1 | hybrid\_polyketide\_synthase/peptide\_synthetase | BGC0001667 | NRP + Polyketide | 31.0 | 55.0 | 388.0 | 4.5e-107 |
| AVI57434.1 | AbmB2 | BGC0001694 | Polyketide | 30.0 | 58.9 | 388.0 | 4.5e-107 |
| ABK32290.1 | JerD | BGC0000080 | Polyketide | 30.0 | 58.9 | 388.0 | 5.9e-107 |
| WP\_042799407.1 | type\_I\_polyketide\_synthase | BGC0001283 | Polyketide | 31.0 | 53.4 | 388.0 | 5.9e-107 |
| ATP76239.1 | NdaF | BGC0001705 | NRP + Polyketide | 31.0 | 55.6 | 388.0 | 5.9e-107 |
| ATP76242.1 | NdaC | BGC0001705 | NRP + Polyketide | 31.0 | 54.6 | 388.0 | 5.9e-107 |
| CAQ18830.1 | polyketide\_synthase | BGC0000954 | NRP + Polyketide:Modular type I | 31.0 | 59.3 | 388.0 | 7.7e-107 |
| AAK57186.1 | MxaB2 | BGC0001022 | NRP + Polyketide | 31.0 | 52.2 | 388.0 | 7.7e-107 |
| APZ78780.1 | polyketide\_synthase | BGC0001426 | NRP:Cyclic depsipeptide + Polyketide:Iterative type I | 31.0 | 54.4 | 387.0 | 1e-106 |
| EAA65602.1 | hypothetical\_protein | BGC0000022 | Polyketide | 28.0 | 78.4 | 387.0 | 1.3e-106 |
| AAS98784.1 | polyketide\_synthase | BGC0001001 | NRP + Polyketide | 30.0 | 55.2 | 387.0 | 1.3e-106 |
| CAD29793.1 | polyketide\_synthase\_type\_I | BGC0001015 | NRP + Polyketide | 29.0 | 55.5 | 387.0 | 1.3e-106 |
| APZ78793.1 | polyketide\_synthase | BGC0001427 | NRP:Cyclic depsipeptide + Polyketide:Iterative type I | 31.0 | 54.4 | 386.0 | 2.2e-106 |
| BAV69313.1 | PrhL | BGC0001729 | Polyketide + Terpene | 27.0 | 74.2 | 386.0 | 2.9e-106 |
| AGY30677.1 | Ann5 | BGC0001298 | Polyketide | 30.0 | 59.9 | 385.0 | 5e-106 |
| AAZ77696.1 | ChlA3 | BGC0000036 | Polyketide:Modular type I + Polyketide:Iterative type I + Saccharide:Oligosaccharide | 31.0 | 58.6 | 384.0 | 8.5e-106 |
| CAD19089.1 | StiE\_protein | BGC0000153 | NRP + Polyketide:Modular type I | 30.0 | 57.3 | 384.0 | 1.1e-105 |
| DAB41918.1 | ArzP\_-\_PKS\_(KS,\_AT,\_OMT,\_ACP,\_TE) | BGC0001884 | NRP + Polyketide | 30.0 | 54.6 | 384.0 | 1.1e-105 |
| CAJ46689.1 | polyketide\_synthase | BGC0000969 | NRP:Cyclic depsipeptide + Polyketide:Modular type I | 30.0 | 59.3 | 383.0 | 1.4e-105 |
| AWC08662.1 | polyketide\_synthase\_type\_I | BGC0001932 | Polyketide | 29.0 | 53.8 | 383.0 | 2.5e-105 |
| BAF85844.1 | modular\_polyketide\_synthase | BGC0000109 | Polyketide | 30.0 | 58.9 | 382.0 | 3.2e-105 |
| AAG23263.1 | polyketide\_synthase\_extender\_modules\_5-7 | BGC0000148 | Polyketide | 30.0 | 58.2 | 382.0 | 4.2e-105 |
| AAF00958.1 | mcyE | BGC0001017 | NRP + Polyketide:Modular type I | 30.0 | 54.5 | 382.0 | 4.2e-105 |
| CBD77734.1 | polyketide\_synthase | BGC0000974 | NRP + Polyketide | 31.0 | 53.9 | 381.0 | 7.2e-105 |
| AIR74911.1 | polyketide\_synthase | BGC0001559 | RiPP | 31.0 | 53.9 | 381.0 | 7.2e-105 |
| CAQ43077.1 | polyketide\_synthase | BGC0000970 | NRP + Polyketide:Modular type I | 30.0 | 59.1 | 381.0 | 9.4e-105 |
| APZ78742.1 | polyketide\_synthase | BGC0001422 | NRP:Cyclic depsipeptide + Polyketide:Iterative type I | 31.0 | 54.4 | 381.0 | 9.4e-105 |
| ASZ00150.1 | polyketide\_synthase | BGC0001785 | Polyketide | 30.0 | 58.6 | 381.0 | 9.4e-105 |
| ctg1\_orf253 |  | BGC0001200 | Polyketide | 29.0 | 58.6 | 380.0 | 1.2e-104 |
| AQH32483.1 | hybrid\_peptide\_synthetase/polyketide\_synthase | BGC0001667 | NRP + Polyketide | 30.0 | 57.6 | 380.0 | 1.2e-104 |
| BAO66529.1 | type\_I\_polyketide\_synthase | BGC0000042 | Polyketide | 30.0 | 58.9 | 380.0 | 1.6e-104 |
| APZ78807.1 | polyketide\_synthase | BGC0001428 | NRP:Cyclic depsipeptide + Polyketide:Iterative type I | 31.0 | 54.4 | 380.0 | 1.6e-104 |
| CAD19085.1 | StiA\_protein | BGC0000153 | NRP + Polyketide:Modular type I | 31.0 | 52.6 | 379.0 | 2.7e-104 |
| ctg1\_orf256 |  | BGC0001200 | Polyketide | 30.0 | 58.5 | 379.0 | 2.7e-104 |
| ACY13415.1 | KR\_domain\_protein | BGC0001367 | NRP + Polyketide | 31.0 | 53.4 | 379.0 | 2.7e-104 |
| ALP32046.1 | CycF | BGC0001293 | Polyketide | 30.0 | 59.0 | 379.0 | 3.6e-104 |
| AQM37582.1 | polyketide\_synthase | BGC0001424 | NRP:Cyclic depsipeptide + Polyketide:Iterative type I | 31.0 | 54.4 | 379.0 | 3.6e-104 |
| AAG23262.1 | polyketide\_synthase\_extender\_modules\_8-10 | BGC0000148 | Polyketide | 29.0 | 58.8 | 378.0 | 4.6e-104 |
| CAQ43075.1 | polyketide\_synthase | BGC0000970 | NRP + Polyketide:Modular type I | 29.0 | 65.3 | 378.0 | 6.1e-104 |
| TGZ15168.1 | hypothetical\_protein | BGC0002032 | Polyketide | 30.0 | 58.5 | 378.0 | 7.9e-104 |
| AEU17899.1 | putative\_type\_I\_PKS | BGC0001072 | Saccharide + Polyketide:Modular type I + Polyketide:Type II + Other:Aminocoumarin | 32.0 | 53.9 | 377.0 | 1e-103 |
| ABK32257.1 | AmbC | BGC0000014 | Polyketide | 30.0 | 58.5 | 377.0 | 1.4e-103 |
| BAC68129.1 | modular\_polyketide\_synthase | BGC0000059 | Polyketide | 31.0 | 54.1 | 377.0 | 1.4e-103 |
| BAD08360.1 | polyketide\_synthase\_modules\_7-8 | BGC0000167 | Polyketide | 30.0 | 58.2 | 377.0 | 1.4e-103 |
| AAG23264.1 | polyketide\_synthase\_loading\_and\_extender\_module\_1 | BGC0000148 | Polyketide | 30.0 | 54.9 | 376.0 | 1.8e-103 |
| ADH04639.1 | TgaA | BGC0001051 | NRP + Polyketide:Modular type I | 31.0 | 55.2 | 376.0 | 1.8e-103 |
| OJF16266.1 | AceP4 | BGC0001491 | Polyketide | 31.0 | 55.0 | 376.0 | 1.8e-103 |
| ADH04660.1 | TugD | BGC0001342 | NRP + Polyketide | 30.0 | 54.3 | 376.0 | 2.3e-103 |
| WP\_055480220.1 | type\_I\_polyketide\_synthase | BGC0001653 | Polyketide | 30.0 | 58.6 | 376.0 | 2.3e-103 |
| AAW03325.1 | CtaB | BGC0000982 | NRP + Polyketide | 31.0 | 54.2 | 376.0 | 3e-103 |
| ADH04659.1 | TugC | BGC0001342 | NRP + Polyketide | 30.0 | 55.0 | 376.0 | 3e-103 |
| ctg1\_orf254 |  | BGC0001200 | Polyketide | 29.0 | 58.2 | 375.0 | 3.9e-103 |
| QDA77044.1 | polyketide\_synthase | BGC0002025 | NRP | 29.0 | 63.9 | 375.0 | 5.1e-103 |
| CAQ43079.1 | polyketide\_synthase | BGC0000970 | NRP + Polyketide:Modular type I | 30.0 | 56.7 | 374.0 | 6.7e-103 |
| AMYAL\_RS48925 | polyketide\_synthase | BGC0002011 | Polyketide | 30.0 | 58.3 | 374.0 | 6.7e-103 |
| CAD15508.1 | polyketide\_synthase/non-ribosomal\_peptide\_synthetase | BGC0001014 | NRP:NRP siderophore + Polyketide:Modular type I + Polyketide:Iterative type I | 30.0 | 60.6 | 374.0 | 8.8e-103 |
| AWR88393.1 | putative\_beta-ketoacyl\_synthase | BGC0001522 | Polyketide | 29.0 | 60.4 | 374.0 | 8.8e-103 |
| CAD55506.1 | CpkA;\_Polyketide\_synthase\_loading\_module,\_and\_modules\_1\_and\_2 | BGC0000038 | Polyketide:Modular type I | 31.0 | 54.3 | 374.0 | 1.1e-102 |
| BAH02268.1 | polyketide\_synthase | BGC0000126 | Polyketide | 31.0 | 54.6 | 374.0 | 1.1e-102 |
| CAD29794.1 | peptide\_synthetase | BGC0001015 | NRP + Polyketide | 31.0 | 54.3 | 373.0 | 1.5e-102 |
| sipP5 | Type\_I\_Modular\_PKS | BGC0001452 | Polyketide | 29.0 | 60.4 | 373.0 | 2e-102 |
| AAX98189.1 | polyketide\_synthase\_type\_I | BGC0000052 | Polyketide | 30.0 | 59.2 | 372.0 | 3.3e-102 |
| AEC13069.1 | fosC | BGC0000060 | Polyketide | 31.0 | 54.0 | 372.0 | 3.3e-102 |
| AQW44873.1 | polyketide\_synthase | BGC0001761 | Polyketide | 31.0 | 59.1 | 372.0 | 3.3e-102 |
| AAQ82567.1 | FscE | BGC0000061 | Polyketide | 30.0 | 54.8 | 372.0 | 4.3e-102 |
| ABC84458.1 | NigAIII | BGC0000114 | Polyketide:Modular type I | 30.0 | 60.8 | 372.0 | 4.3e-102 |
| ACA99172.1 | polyketide\_synthase | BGC0001160 | Polyketide:Modular type I | 30.0 | 54.4 | 371.0 | 5.7e-102 |
| ctg1\_orf255 |  | BGC0001200 | Polyketide | 30.0 | 59.5 | 371.0 | 5.7e-102 |
| ALA09371.1 | type\_I\_modular\_PKS | BGC0001303 | Polyketide | 30.0 | 58.9 | 371.0 | 5.7e-102 |
| PKX88487.1 | polyketide\_synthase | BGC0001708 | Polyketide + Terpene | 26.0 | 76.4 | 371.0 | 5.7e-102 |
| BAD08359.1 | polyketide\_synthase\_modules\_5-6 | BGC0000167 | Polyketide | 30.0 | 58.4 | 371.0 | 7.4e-102 |
| ctg1\_orf7 |  | BGC0000053 | Polyketide | 30.0 | 54.5 | 371.0 | 9.7e-102 |
| AAN32979.1 | BarE | BGC0000962 | NRP + Polyketide:Modular type I | 30.0 | 56.1 | 371.0 | 9.7e-102 |
| EHA28237.1 | hypothetical\_protein | BGC0001143 | Polyketide | 25.0 | 77.7 | 371.0 | 9.7e-102 |
| EHK80167.1 | modular\_polyketide\_synthase | BGC0001447 | Polyketide | 30.0 | 54.6 | 371.0 | 9.7e-102 |
| AAG23265.1 | polyketide\_synthase\_extender\_module\_2 | BGC0000148 | Polyketide | 29.0 | 58.7 | 370.0 | 1.7e-101 |
| CAE46843.1 | Type\_I\_modular\_polyketide\_synthase | BGC0000103 | Polyketide | 30.0 | 56.4 | 369.0 | 2.2e-101 |
| CAE46851.1 | Type\_I\_modular\_polyketide\_synthase | BGC0000103 | Polyketide | 30.0 | 56.4 | 369.0 | 2.2e-101 |
| AFI57005.1 | QmnA1 | BGC0000133 | Polyketide | 31.0 | 53.9 | 369.0 | 2.2e-101 |
| AZH23823.1 | MgiK | BGC0001971 | NRP + Polyketide | 29.0 | 56.2 | 368.0 | 4.8e-101 |
| CBD77746.1 | non-ribosomal\_peptide\_synthetase/polyketide\_synthase | BGC0000974 | NRP + Polyketide | 30.0 | 55.4 | 367.0 | 1.1e-100 |
| AIR74926.1 | polyketide\_synthase | BGC0001559 | RiPP | 30.0 | 55.4 | 367.0 | 1.1e-100 |
| ASZ00148.1 | polyketide\_synthase | BGC0001785 | Polyketide | 30.0 | 58.5 | 367.0 | 1.1e-100 |
| BAO66539.1 | type\_I\_polyketide\_synthase | BGC0000042 | Polyketide | 30.0 | 55.6 | 367.0 | 1.4e-100 |
| AAO62582.1 | polyketide\_synthase\_peptide\_sythetase\_fusion\_protein | BGC0001016 | NRP + Polyketide | 31.0 | 53.7 | 367.0 | 1.4e-100 |
| CAI94682.1 | putative\_polyketide\_synthase | BGC0000141 | Polyketide | 30.0 | 55.9 | 366.0 | 1.8e-100 |
| OAP25821.1 | Phenolphthiocerol\_synthesis\_polyketide\_synthase\_type\_I\_Pks15/1 | BGC0001658 | Polyketide | 29.0 | 54.3 | 366.0 | 2.4e-100 |
| ACB46485.1 | polyketide\_synthase | BGC0000082 | Polyketide | 30.0 | 53.8 | 366.0 | 3.1e-100 |
| ACB37755.1 | putative\_type\_I\_polyketide\_synthase | BGC0000162 | Polyketide | 29.0 | 59.6 | 365.0 | 4.1e-100 |
| QBF51759.1 | type\_I\_polyketide\_synthase | BGC0001856 | Polyketide:Modular type I | 30.0 | 54.0 | 365.0 | 4.1e-100 |
| CAD89775.1 | MelD\_protein | BGC0001010 | NRP + Polyketide:Modular type I | 28.0 | 60.0 | 365.0 | 5.3e-100 |
| ACF35445.1 | mbcAI | BGC0000090 | Polyketide | 30.0 | 54.7 | 364.0 | 6.9e-100 |
| BAO66528.1 | type\_I\_polyketide\_synthase | BGC0000042 | Polyketide | 30.0 | 58.7 | 364.0 | 9.1e-100 |
| AFV30249.1 | polyketide\_synthase | BGC0000075 | Polyketide | 28.0 | 58.3 | 364.0 | 9.1e-100 |
| ABV97152.1 | Beta-ketoacyl\_synthase | BGC0000137 | Polyketide | 30.0 | 58.2 | 364.0 | 9.1e-100 |
| AAW03327.1 | CtaD | BGC0000982 | NRP + Polyketide | 29.0 | 59.7 | 364.0 | 9.1e-100 |
| AWH12669.1 | RmpB | BGC0001759 | Polyketide | 30.0 | 59.7 | 364.0 | 1.2e-99 |
| TXD00034.1 | SDR\_family\_NAD(P)-dependent\_oxidoreductase | BGC0001877 | Polyketide | 30.0 | 56.8 | 364.0 | 1.2e-99 |
| AXM42951.1 | polyketide\_synthase | BGC0001941 | NRP + Polyketide | 30.0 | 53.9 | 364.0 | 1.2e-99 |
| ACR50774.1 | polyketide\_synthase | BGC0000163 | Polyketide | 29.0 | 60.0 | 363.0 | 1.5e-99 |
| CCP20047.1 | divK\_protein | BGC0001119 | Polyketide:Modular type I | 31.0 | 54.9 | 363.0 | 1.5e-99 |
| CAQ52624.1 | type\_I\_polyketide\_synthase,\_modules\_7-8 | BGC0001066 | Polyketide:Modular type I | 29.0 | 54.8 | 363.0 | 2e-99 |
| BAQ25511.1 | type\_I\_polyketide\_synthase | BGC0001288 | Polyketide | 30.0 | 56.2 | 363.0 | 2e-99 |
| ANR02549.1 | LodH | BGC0001648 | Polyketide | 31.0 | 55.2 | 363.0 | 2e-99 |
| TXD00025.1 | SDR\_family\_NAD(P)-dependent\_oxidoreductase | BGC0001877 | Polyketide | 30.0 | 54.0 | 363.0 | 2e-99 |
| AGI99496.1 | Type\_I\_polyketide\_synthase | BGC0001004 | Polyketide:Modular type I | 30.0 | 61.8 | 363.0 | 2.6e-99 |
| AFU82616.1 | polyketide\_synthase | BGC0000998 | NRP + Polyketide | 29.0 | 58.9 | 362.0 | 3.4e-99 |
| BAQ21939.1 | putative\_type\_I\_polyketide\_synthase | BGC0001204 | Polyketide | 30.0 | 54.3 | 362.0 | 3.4e-99 |
| AAP42857.1 | NanA3 | BGC0000105 | Polyketide | 28.0 | 58.5 | 362.0 | 4.5e-99 |
| BAO66541.1 | type\_I\_polyketide\_synthase | BGC0000042 | Polyketide | 30.0 | 56.8 | 361.0 | 5.9e-99 |
| AAF71766.1 | nysI | BGC0000115 | Polyketide:Modular type I + Saccharide:Hybrid/tailoring | 31.0 | 53.0 | 361.0 | 5.9e-99 |
| CAO98852.1 | polyketide\_synthase\_AufI | BGC0000023 | Polyketide:Modular type I | 32.0 | 51.8 | 361.0 | 7.7e-99 |
| CAF05651.1 | TubF\_protein | BGC0001053 | NRP + Polyketide | 30.0 | 59.4 | 361.0 | 7.7e-99 |
| WP\_020636817.1 | type\_I\_polyketide\_synthase | BGC0002011 | Polyketide | 29.0 | 53.0 | 361.0 | 7.7e-99 |
| AAQ82568.1 | FscD | BGC0000061 | Polyketide | 31.0 | 54.2 | 361.0 | 1e-98 |
| AAG13917.1 | megalomicin\_6-deoxyerythronolide\_B\_synthase\_1 | BGC0000092 | Polyketide | 32.0 | 53.9 | 361.0 | 1e-98 |
| AAC69329.1 | type\_I\_polyketide\_synthase\_PikAI | BGC0000094 | Polyketide:Modular type I + Saccharide:Hybrid/tailoring | 30.0 | 54.5 | 361.0 | 1e-98 |
| CAL58683.1 | polyketide\_synthase | BGC0000149 | Polyketide:Modular type I | 29.0 | 54.8 | 361.0 | 1e-98 |
| CAD17792.1 | probable\_non\_ribosomal\_peptide\_synthetase\_protein | BGC0001754 | NRP + Polyketide | 29.0 | 56.0 | 361.0 | 1e-98 |
| SCO70308.1 | Type\_I\_polyketide\_synthase | BGC0001433 | Polyketide:Modular type I | 30.0 | 54.3 | 361.0 | 1e-98 |
| ANY10600.1 | polyketide\_synthase | BGC0001773 | Polyketide | 30.0 | 53.3 | 361.0 | 1e-98 |
| ABL86391.1 | hybrid\_polyketide\_synthase\_and\_nonribosomal\_peptide\_synthetase | BGC0000999 | NRP + Polyketide | 29.0 | 53.7 | 360.0 | 1.3e-98 |
| BAJ16467.1 | polyketide\_synthase | BGC0000058 | Polyketide | 31.0 | 54.3 | 360.0 | 1.7e-98 |
| ABV97155.1 | Acyl\_transferase | BGC0000137 | Polyketide | 30.0 | 54.9 | 359.0 | 2.2e-98 |
| AEE88283.1 | CurG | BGC0000976 | NRP + Polyketide:Modular type I | 29.0 | 54.9 | 359.0 | 2.2e-98 |
| AAT70102.1 | CurG | BGC0001165 | NRP + Polyketide:Modular type I | 29.0 | 54.9 | 359.0 | 2.2e-98 |
| BAC68126.1 | modular\_polyketide\_synthase | BGC0000059 | Polyketide | 29.0 | 53.7 | 359.0 | 2.9e-98 |
| AAP42873.1 | NanA11 | BGC0000105 | Polyketide | 29.0 | 58.5 | 359.0 | 2.9e-98 |
| BAK64638.1 | polyketide\_synthase | BGC0000135 | Polyketide | 29.0 | 60.5 | 359.0 | 2.9e-98 |
| QCO93110.1 | polyketide\_synthase | BGC0001977 | Other | 25.0 | 82.8 | 359.0 | 2.9e-98 |
| ACR50785.1 | polyketide\_synthase | BGC0000163 | Polyketide | 30.0 | 54.6 | 359.0 | 3.8e-98 |
| AZF85932.1 | type\_I\_polyketide\_synthase | BGC0001963 | NRP + Polyketide | 29.0 | 54.0 | 359.0 | 3.8e-98 |
| ACN69989.1 | polyketide\_synthase | BGC0000079 | Polyketide | 29.0 | 60.9 | 358.0 | 5e-98 |
| CAO85897.1 | modular\_polyketide\_synthase\_NorB | BGC0000110 | Polyketide:Modular type I | 29.0 | 59.0 | 358.0 | 5e-98 |
| CAD29795.1 | peptide\_synthetase | BGC0001015 | NRP + Polyketide | 31.0 | 54.4 | 358.0 | 5e-98 |
| ADC79637.1 | TamAI | BGC0001052 | NRP + Polyketide:Modular type I | 31.0 | 55.0 | 358.0 | 6.5e-98 |
| BBA66513.1 | type\_I\_polyketide\_synthase | BGC0001495 | Polyketide | 30.0 | 53.6 | 358.0 | 6.5e-98 |
| AAZ77693.1 | ChlA1 | BGC0000036 | Polyketide:Modular type I + Polyketide:Iterative type I + Saccharide:Oligosaccharide | 31.0 | 56.5 | 358.0 | 8.5e-98 |
| BAP34763.1 | type\_I\_polyketide\_synthase | BGC0000078 | Polyketide | 29.0 | 60.0 | 358.0 | 8.5e-98 |
| AAQ90173.1 | polyketide\_synthase\_type\_I | BGC0000128 | Polyketide | 31.0 | 53.7 | 358.0 | 8.5e-98 |
| ANH11409.1 | SceN | BGC0001908 | Polyketide | 29.0 | 58.2 | 357.0 | 1.1e-97 |
| EAU29529.1 | hypothetical\_protein | BGC0000682 | Terpene | 30.0 | 53.5 | 357.0 | 1.4e-97 |
| CAQ34929.1 | putative\_polyketide\_synthase | BGC0000986 | NRP + Polyketide | 34.0 | 46.3 | 357.0 | 1.4e-97 |
| BAR73007.1 | putative\_PKS\_(ACP-KS-AT-DH-KR-ACP-KS-AT-DH-ER-KR-ACP) | BGC0001194 | Polyketide | 29.0 | 60.1 | 357.0 | 1.4e-97 |
| AAC01711.1 | RifB | BGC0000136 | Polyketide | 29.0 | 59.6 | 356.0 | 1.9e-97 |
| CAJ88186.1 | putative\_modular\_polyketide\_synthase | BGC0000151 | Polyketide:Modular type I + Saccharide:Hybrid/tailoring | 30.0 | 55.3 | 356.0 | 1.9e-97 |
| AGC09487.1 | LobS5 | BGC0001183 | Polyketide | 29.0 | 55.3 | 356.0 | 1.9e-97 |
| CQR60496.1 | Polyketide\_synthase,\_type\_I,\_modules:\_4,\_5\_and\_6 | BGC0001287 | Polyketide | 29.0 | 58.3 | 356.0 | 1.9e-97 |
| ADH01663.1 | putative\_polyketide\_synthase\_PKS3 | BGC0000099 | Polyketide | 25.0 | 74.7 | 356.0 | 2.5e-97 |
| BAH02271.1 | polyketide\_synthase | BGC0000126 | Polyketide | 30.0 | 53.4 | 356.0 | 2.5e-97 |
| BAT51065.1 | type\_I\_polyketide\_synthase | BGC0001296 | Polyketide | 31.0 | 53.4 | 356.0 | 3.2e-97 |
| AKA59093.1 | type-I\_PKS | BGC0001619 | Polyketide | 29.0 | 56.7 | 356.0 | 3.2e-97 |
| ABB52544.1 | putative\_type\_I\_polyketide\_synthase | BGC0000047 | Polyketide | 30.0 | 53.3 | 355.0 | 4.2e-97 |
| ARW71486.1 | type\_I\_PKS\_module\_6 | BGC0001812 | Polyketide | 29.0 | 53.7 | 355.0 | 4.2e-97 |
| AGN71604.1 | conidial\_yellow\_pigment\_biosynthesis\_polyketide\_synthase | BGC0000027 | Polyketide:Iterative type I | 25.0 | 74.9 | 355.0 | 5.5e-97 |
| AAF71767.1 | nysJ | BGC0000115 | Polyketide:Modular type I + Saccharide:Hybrid/tailoring | 29.0 | 58.6 | 355.0 | 5.5e-97 |
| ABV83222.1 | CppJ | BGC0000116 | Polyketide | 30.0 | 60.7 | 355.0 | 5.5e-97 |
| ABC84469.1 | NigAIX | BGC0000114 | Polyketide:Modular type I | 29.0 | 59.4 | 354.0 | 7.2e-97 |
| AAB66505.1 | tylactone\_synthase\_module\_3 | BGC0000166 | Polyketide | 29.0 | 60.9 | 354.0 | 7.2e-97 |
| BAG17643.1 | putative\_NRPS-type-I\_PKS\_fusion\_protein | BGC0001043 | NRP + Polyketide | 31.0 | 53.8 | 354.0 | 7.2e-97 |
| AVV61983.1 | type\_I\_modular\_polyketide\_synthase | BGC0001477 | NRP + Polyketide:Modular type I | 29.0 | 60.0 | 354.0 | 7.2e-97 |
| ADB23403.1 | polyketide\_synthase\_type\_I | BGC0001062 | Polyketide | 31.0 | 50.3 | 354.0 | 9.4e-97 |
| ABV97151.1 | AMP-dependent\_synthetase\_and\_ligase | BGC0000137 | Polyketide | 30.0 | 54.1 | 354.0 | 1.2e-96 |
| AHB82059.1 | non\_ribosomal\_peptide\_synthetase/polyketide\_synthase | BGC0001019 | NRP + Polyketide:Modular type I | 31.0 | 54.7 | 353.0 | 1.6e-96 |
| ACR50775.1 | polyketide\_synthase | BGC0000163 | Polyketide | 29.0 | 56.7 | 353.0 | 2.1e-96 |
| AGI99494.1 | Type\_I\_polyketide\_synthase | BGC0001004 | Polyketide:Modular type I | 29.0 | 55.5 | 353.0 | 2.1e-96 |
| CAA16183.1 | polyketide\_synthase | BGC0001063 | NRP + Polyketide | 30.0 | 52.2 | 353.0 | 2.1e-96 |
| APZ78767.1 | polyketide\_synthase | BGC0001425 | NRP:Cyclic depsipeptide + Polyketide:Iterative type I | 30.0 | 54.4 | 353.0 | 2.1e-96 |
| AAM81584.2 | putative\_type\_I\_polyketide\_synthase | BGC0000047 | Polyketide | 30.0 | 54.9 | 352.0 | 2.7e-96 |
| BAE93722.1 | type\_I\_polyketide\_synthase | BGC0000164 | Polyketide | 28.0 | 59.7 | 352.0 | 3.6e-96 |
| ACN69990.1 | polyketide\_synthase | BGC0000079 | Polyketide | 29.0 | 58.8 | 352.0 | 4.7e-96 |
| ACB37741.1 | putative\_type\_I\_polyketide\_synthase | BGC0000162 | Polyketide | 29.0 | 59.3 | 352.0 | 4.7e-96 |
| AAK57190.1 | MxaF | BGC0001022 | NRP + Polyketide | 29.0 | 52.0 | 352.0 | 4.7e-96 |
| AAD03047.1 | type\_I\_polyketide\_synthase | BGC0000041 | Polyketide | 30.0 | 59.9 | 351.0 | 6.1e-96 |
| AAZ94387.1 | modular\_polyketide\_synthase | BGC0000040 | Polyketide | 31.0 | 56.6 | 351.0 | 7.9e-96 |
| ADM46359.1 | polyketide\_synthase | BGC0000106 | Polyketide | 29.0 | 58.0 | 351.0 | 7.9e-96 |
| ACR50773.1 | polyketide\_synthase | BGC0000163 | Polyketide | 27.0 | 70.7 | 351.0 | 7.9e-96 |
| CAQ34918.1 | nonribosomal\_peptide\_synthetase/\_polyketide\_synthase | BGC0000986 | NRP + Polyketide | 26.0 | 77.3 | 351.0 | 7.9e-96 |
| AWW87422.1 | type\_I\_polyketide\_synthase | BGC0001755 | Polyketide | 31.0 | 54.9 | 351.0 | 7.9e-96 |
| CAE02605.1 | polyketide\_synthase\_type\_I | BGC0000024 | Polyketide:Modular type I | 28.0 | 58.9 | 351.0 | 1e-95 |
| AAZ94389.1 | modular\_polyketide\_synthase | BGC0000040 | Polyketide | 29.0 | 59.6 | 350.0 | 1.4e-95 |
| ACB37743.1 | putative\_type\_I\_polyketide\_synthase | BGC0000162 | Polyketide | 30.0 | 53.7 | 350.0 | 1.4e-95 |
| AAP42859.1 | NanA5 | BGC0000105 | Polyketide | 29.0 | 61.5 | 350.0 | 1.8e-95 |
| ABC84456.1 | NigAI | BGC0000114 | Polyketide:Modular type I | 29.0 | 55.6 | 350.0 | 1.8e-95 |
| AFD30954.1 | CrmA | BGC0000966 | NRP + Polyketide | 29.0 | 56.7 | 350.0 | 1.8e-95 |
| ABX60161.1 | mixed\_NRPS/PKS | BGC0000978 | NRP + Alkaloid + Polyketide:Modular type I | 28.0 | 55.0 | 350.0 | 1.8e-95 |
| ABJ97437.1 | MerA | BGC0001012 | NRP + Polyketide | 30.0 | 55.3 | 350.0 | 1.8e-95 |
| ALA09358.1 | type\_I\_modular\_PKS | BGC0001303 | Polyketide | 29.0 | 59.4 | 350.0 | 1.8e-95 |
| CAQ64686.1 | lasalocid\_modular\_polyketide\_synthase | BGC0000087 | Polyketide | 31.0 | 53.9 | 349.0 | 2.3e-95 |
| AAG13919.1 | megalomicin\_6-deoxyerythronolide\_B\_synthase\_3 | BGC0000092 | Polyketide | 32.0 | 54.4 | 349.0 | 2.3e-95 |
| CCE88377.1 | non-ribosomal\_peptide\_synthetase/polyketide\_synthase | BGC0001034 | NRP + Polyketide:Modular type I | 30.0 | 52.9 | 349.0 | 2.3e-95 |
| ABC87509.1 | polyketide\_synthase | BGC0001011 | NRP + Polyketide | 30.0 | 55.7 | 349.0 | 3e-95 |
| ctg1\_orf20 |  | BGC0001013 | NRP + Polyketide | 30.0 | 55.7 | 349.0 | 3e-95 |
| ctg1\_orf521 |  | BGC0001199 | Polyketide | 30.0 | 58.3 | 349.0 | 3e-95 |
| ADH04682.1 | polyketide\_synthase | BGC0001344 | NRP + Polyketide | 30.0 | 53.9 | 349.0 | 3e-95 |
| ACN69991.1 | polyketide\_synthase | BGC0000079 | Polyketide | 29.0 | 58.2 | 348.0 | 5.1e-95 |
| BAH02269.1 | polyketide\_synthase | BGC0000126 | Polyketide | 29.0 | 58.9 | 348.0 | 5.1e-95 |
| AAC01713.1 | RifD | BGC0000136 | Polyketide | 30.0 | 53.7 | 348.0 | 6.7e-95 |
| AAK19883.1 | soraphen\_polyketide\_synthase\_A | BGC0000147 | Polyketide:Modular type I | 30.0 | 53.2 | 348.0 | 6.7e-95 |
| CAQ52626.1 | type\_I\_polyketide\_synthase,\_loading\_module\_and\_modules\_1-3 | BGC0001066 | Polyketide:Modular type I | 29.0 | 55.7 | 348.0 | 6.7e-95 |
| AAS79462.1 | polyketide\_synthase\_subunit | BGC0000035 | Polyketide | 31.0 | 53.7 | 347.0 | 8.8e-95 |
| AWO77084.1 | hybrid\_non-ribosomal\_peptide\_synthetase/type\_I\_polyketide\_synthase | BGC0001556 | NRP + Polyketide | 29.0 | 57.5 | 347.0 | 1.1e-94 |
| AWM95789.1 | non-reduciing\_polyketide\_synthase\_methylorcinaldehyde\_synthase | BGC0001827 | Polyketide | 26.0 | 76.8 | 347.0 | 1.1e-94 |
| AAM81586.2 | putative\_type\_I\_polyketide\_synthase | BGC0000047 | Polyketide | 31.0 | 53.6 | 347.0 | 1.5e-94 |
| ALD82524.1 | polyketide\_synthase | BGC0001212 | NRP + Polyketide | 29.0 | 62.1 | 347.0 | 1.5e-94 |
| ADF88279.1 | mixed\_NRPS/PKS | BGC0000981 | NRP + Polyketide | 28.0 | 55.0 | 346.0 | 2e-94 |
| AVX51099.1 | NysJ | BGC0001709 | Polyketide | 29.0 | 58.6 | 346.0 | 2.6e-94 |
| ANH11414.1 | SceS | BGC0001908 | Polyketide | 29.0 | 58.6 | 346.0 | 2.6e-94 |
| QBF51754.1 | type\_I\_polyketide\_synthase | BGC0001856 | Polyketide:Modular type I | 29.0 | 58.3 | 346.0 | 2.6e-94 |
| ABV83221.1 | CppI | BGC0000116 | Polyketide | 29.0 | 56.2 | 346.0 | 3.3e-94 |
| AWC08658.1 | polyketide\_synthase\_type\_I | BGC0001932 | Polyketide | 30.0 | 53.7 | 346.0 | 3.3e-94 |
| ANZ22991.1 | ZinG | BGC0001828 | Polyketide | 29.0 | 54.4 | 346.0 | 3.3e-94 |
| AEZ53945.1 | polyketide\_synthase | BGC0000144 | Polyketide:Modular type I | 30.0 | 55.2 | 345.0 | 4.4e-94 |
| TGZ15165.1 | hypothetical\_protein | BGC0002032 | Polyketide | 29.0 | 53.7 | 345.0 | 4.4e-94 |
| AAU93806.2 | polyketide\_synthase\_modules\_3\_and\_4 | BGC0000054 | Polyketide | 28.0 | 58.3 | 345.0 | 5.7e-94 |
| ACZ65476.1 | type\_I\_modular\_polyketide\_synthase | BGC0000140 | Polyketide | 31.0 | 48.8 | 345.0 | 5.7e-94 |
| AHB82062.1 | polyketide\_synthase | BGC0001231 | NRP + Polyketide:Modular type I | 29.0 | 54.9 | 345.0 | 5.7e-94 |
| AAP42855.1 | NanA1 | BGC0000105 | Polyketide | 30.0 | 54.7 | 344.0 | 7.4e-94 |
| AAF71776.1 | nysC | BGC0000115 | Polyketide:Modular type I + Saccharide:Hybrid/tailoring | 29.0 | 59.6 | 344.0 | 9.7e-94 |
| AHA38200.1 | GphG | BGC0000069 | Polyketide | 31.0 | 48.9 | 343.0 | 1.7e-93 |
| orf3 | polyketide\_synthase | BGC0001432 | NRP:Cyclic depsipeptide + Polyketide:Iterative type I | 30.0 | 52.2 | 343.0 | 1.7e-93 |
| APZ78858.1 | polyketide\_synthase | BGC0001432 | NRP:Cyclic depsipeptide + Polyketide:Iterative type I | 30.0 | 52.2 | 343.0 | 1.7e-93 |
| ARM20279.1 | polyketide\_synthase | BGC0001523 | Polyketide | 29.0 | 56.1 | 343.0 | 1.7e-93 |
| ADC79638.1 | TamAII | BGC0001052 | NRP + Polyketide:Modular type I | 29.0 | 53.9 | 343.0 | 2.2e-93 |
| WP\_053065268.1 | type\_I\_polyketide\_synthase | BGC0001330 | NRP:Cyclic depsipeptide + Polyketide:Modular type I | 35.0 | 39.1 | 343.0 | 2.2e-93 |
| AAZ94386.1 | modular\_polyketide\_synthase | BGC0000040 | Polyketide | 30.0 | 54.1 | 342.0 | 2.8e-93 |
| ABO15888.1 | polyketide\_synthase | BGC0000132 | Polyketide | 30.0 | 47.1 | 342.0 | 2.8e-93 |
| AGC09484.1 | LobS1 | BGC0001183 | Polyketide | 30.0 | 56.1 | 342.0 | 2.8e-93 |
| ABP55210.1 | beta-ketoacyl\_synthase | BGC0000142 | Polyketide | 28.0 | 60.4 | 342.0 | 3.7e-93 |
| AAK83194.1 | polyketide\_synthase | BGC0000026 | Saccharide:Oligosaccharide | 31.0 | 54.6 | 342.0 | 4.8e-93 |
| AEU11005.1 | NpnA | BGC0001029 | NRP + Polyketide | 29.0 | 55.5 | 342.0 | 4.8e-93 |
| AWH12668.1 | RmpC | BGC0001759 | Polyketide | 29.0 | 59.5 | 342.0 | 4.8e-93 |
| CAM00064.1 | EryAII\_Erythromycin\_polyketide\_synthase\_modules\_3\_and\_4 | BGC0000055 | Polyketide:Modular type I + Saccharide:Hybrid/tailoring | 28.0 | 58.6 | 341.0 | 6.3e-93 |
| AFL48525.1 | laidlomycin\_polyketide\_synthase\_(loading\_module\_and\_module\_1) | BGC0000084 | Polyketide | 29.0 | 58.5 | 341.0 | 6.3e-93 |
| AAC01712.2 | RifC | BGC0000136 | Polyketide | 28.0 | 61.6 | 341.0 | 8.2e-93 |
| AFU82614.1 | mixed\_NRPS\_PKS | BGC0000998 | NRP + Polyketide | 28.0 | 59.8 | 341.0 | 8.2e-93 |
| ADH04657.1 | TugA | BGC0001342 | NRP + Polyketide | 31.0 | 52.3 | 341.0 | 8.2e-93 |
| AXI91545.1 | FunP8 | BGC0001944 | Polyketide | 28.0 | 59.3 | 341.0 | 1.1e-92 |
| ACB46471.1 | polyketide\_synthase | BGC0000082 | Polyketide | 28.0 | 58.7 | 340.0 | 1.4e-92 |
| SCN11949.1 | ebeA-type\_I\_polyketide\_synthase\_KSQ-ATa-ACP | BGC0001580 | Polyketide | 29.0 | 57.0 | 340.0 | 1.4e-92 |
| BAC76491.1 | lankamycin\_synthase\_LkmAIII | BGC0000085 | Polyketide | 29.0 | 53.7 | 340.0 | 1.8e-92 |
| AAO65796.1 | monensin\_polyketide\_synthase\_loading\_module\_and\_module\_1 | BGC0000100 | Polyketide | 29.0 | 55.6 | 340.0 | 1.8e-92 |
| ANC94964.1 | AlmHIII | BGC0001396 | Polyketide | 29.0 | 59.3 | 340.0 | 1.8e-92 |
| ANZ52459.1 | MonAI | BGC0001670 | Polyketide | 29.0 | 55.6 | 340.0 | 1.8e-92 |
| AHH99921.1 | PKS\_I | BGC0000002 | Polyketide | 28.0 | 58.0 | 339.0 | 2.4e-92 |
| BAD08373.1 | polyketide\_synthase\_modules\_1-3 | BGC0000167 | Polyketide | 28.0 | 58.5 | 339.0 | 2.4e-92 |
| CCP20049.1 | divL2\_protein | BGC0001119 | Polyketide:Modular type I | 29.0 | 55.8 | 339.0 | 2.4e-92 |
| ABW96541.1 | type\_I\_modular\_polyketide\_synthase | BGC0000159 | Polyketide:Modular type I | 27.0 | 61.7 | 339.0 | 3.1e-92 |
| BAO98805.1 | putative\_polyketide\_synthase | BGC0001002 | NRP + Polyketide | 26.0 | 68.9 | 339.0 | 3.1e-92 |
| AGC09499.1 | LobS4 | BGC0001183 | Polyketide | 28.0 | 59.4 | 339.0 | 3.1e-92 |
| WP\_039806854.1 | type\_I\_polyketide\_synthase | BGC0002001 | NRP + Polyketide | 30.0 | 55.9 | 339.0 | 3.1e-92 |
| BAG84248.1 | putative\_polyketide\_synthase | BGC0000257 | Polyketide | 29.0 | 55.2 | 339.0 | 4.1e-92 |
| AHB82052.1 | polyketide\_synthase | BGC0001019 | NRP + Polyketide:Modular type I | 34.0 | 38.9 | 339.0 | 4.1e-92 |
| WP\_019032756.1 | type\_I\_polyketide\_synthase | BGC0001331 | NRP:Cyclic depsipeptide + Polyketide:Modular type I | 35.0 | 39.2 | 339.0 | 4.1e-92 |
| AVX51098.1 | nysI | BGC0001709 | Polyketide | 29.0 | 55.8 | 339.0 | 4.1e-92 |
| AEP40940.1 | polyketide\_synthase\_type\_I | BGC0000021 | Polyketide | 30.0 | 53.8 | 338.0 | 7e-92 |
| AAD03048.1 | type\_I\_polyketide\_synthase | BGC0000041 | Polyketide | 29.0 | 54.2 | 338.0 | 7e-92 |
| CAQ18839.1 | hybrid\_polyketide\_synthase/nonribosomal\_polypetide\_synthetase | BGC0000954 | NRP + Polyketide:Modular type I | 29.0 | 54.4 | 338.0 | 7e-92 |
| AGI99497.1 | type\_I\_polyketide\_synthase | BGC0001004 | Polyketide:Modular type I | 29.0 | 56.1 | 338.0 | 7e-92 |
| ABI91470.1 | beta-ketoacyl\_synthase | BGC0001094 | NRP + Polyketide | 29.0 | 53.9 | 337.0 | 9.1e-92 |
| ALA09357.1 | type\_I\_modular\_PKS | BGC0001303 | Polyketide | 29.0 | 58.7 | 337.0 | 1.6e-91 |
| AAO65798.1 | monensin\_polyketide\_synthase\_modules\_3\_and\_4 | BGC0000100 | Polyketide | 29.0 | 56.4 | 336.0 | 2e-91 |
| ACB37740.1 | putative\_type\_I\_polyketide\_synthase | BGC0000162 | Polyketide | 28.0 | 55.6 | 336.0 | 2e-91 |
| ARE67853.1 | AbsB1 | BGC0001492 | Polyketide | 29.0 | 54.0 | 336.0 | 2e-91 |
| WP\_083502114.1 | type\_I\_polyketide\_synthase | BGC0001653 | Polyketide | 29.0 | 55.2 | 336.0 | 2e-91 |
| ANZ52461.1 | MonAIII | BGC0001670 | Polyketide | 29.0 | 56.4 | 336.0 | 2e-91 |
| AXI91549.1 | FunP4 | BGC0001944 | Polyketide | 29.0 | 58.8 | 336.0 | 2e-91 |
| ABB88523.1 | polyketide\_synthase\_type\_I | BGC0000050 | Polyketide | 29.0 | 62.8 | 335.0 | 4.5e-91 |
| AEZ64505.1 | Herb | BGC0001065 | Polyketide | 30.0 | 52.0 | 335.0 | 4.5e-91 |
| AKG06378.1 | polyketide\_synthase\_type\_1 | BGC0001830 | Polyketide | 29.0 | 60.0 | 335.0 | 4.5e-91 |
| AAQ84144.1 | Plm4 | BGC0000123 | Polyketide | 29.0 | 51.4 | 335.0 | 5.9e-91 |
| AWH12936.1 | StmA | BGC0001939 | Polyketide | 29.0 | 54.5 | 335.0 | 5.9e-91 |
| ABV91286.1 | type\_I\_modular\_polyketide\_synthase | BGC0000158 | Polyketide:Modular type I | 30.0 | 53.3 | 334.0 | 1e-90 |
| B073\_RS40860 | type\_I\_polyketide\_synthase | BGC0001332 | NRP + Polyketide | 30.0 | 55.4 | 334.0 | 1e-90 |
| BAC57028.1 | protomycinolide\_IV\_synthase\_1 | BGC0000102 | Polyketide | 29.0 | 55.5 | 333.0 | 1.7e-90 |
| AHF22854.1 | MarL | BGC0000091 | Polyketide | 28.0 | 53.4 | 333.0 | 2.2e-90 |
| AAZ94390.1 | modular\_polyketide\_synthase | BGC0000040 | Polyketide | 30.0 | 55.3 | 332.0 | 2.9e-90 |
| ABI94379.1 | tautomycetin\_biosynthetic\_PKS | BGC0000157 | Polyketide | 30.0 | 53.3 | 332.0 | 2.9e-90 |
| AHD05619.1 | putative\_polyketide\_synthase\_subunit | BGC0001033 | NRP + Polyketide | 29.0 | 54.7 | 332.0 | 2.9e-90 |
| AAZ94388.1 | nodular\_polyketide\_synthase | BGC0000040 | Polyketide | 30.0 | 53.9 | 332.0 | 3.8e-90 |
| CBA11583.1 | polyketide\_synthase\_type\_I | BGC0001046 | NRP + Polyketide:Modular type I + Saccharide:Hybrid/tailoring | 29.0 | 55.5 | 332.0 | 3.8e-90 |
| ACB46488.1 | polyketide\_synthase | BGC0000082 | Polyketide | 30.0 | 53.4 | 332.0 | 5e-90 |
| ABV83223.1 | CppK | BGC0000116 | Polyketide | 29.0 | 55.1 | 332.0 | 5e-90 |
| ALV82320.1 | borrelidin\_type\_I\_polyketide\_synthase | BGC0001533 | Polyketide | 28.0 | 53.9 | 331.0 | 6.5e-90 |
| ABY21538.1 | AngAI | BGC0000018 | Polyketide | 30.0 | 54.2 | 331.0 | 8.5e-90 |
| EAQ86392.1 | hypothetical\_protein | BGC0001405 | Polyketide | 30.0 | 47.9 | 331.0 | 1.1e-89 |
| AAG13918.1 | megalomicin\_6-deoxyerythronolide\_B\_synthase\_2 | BGC0000092 | Polyketide | 28.0 | 59.9 | 330.0 | 1.9e-89 |
| AHD05614.1 | putative\_non-ribosomal\_peptide\_ligase/\_polyketide\_synthase\_hybrid | BGC0001033 | NRP + Polyketide | 29.0 | 54.4 | 330.0 | 1.9e-89 |
| AAX35547.1 | polyketide\_syntase\_2 | BGC0001275 | Polyketide | 30.0 | 55.1 | 330.0 | 1.9e-89 |
| AEK75504.1 | type\_1\_polyketide\_synthase | BGC0000001 | Polyketide:Modular type I | 29.0 | 53.3 | 329.0 | 2.5e-89 |
| AAX98190.1 | polyketide\_synthase\_type\_I | BGC0000052 | Polyketide | 29.0 | 54.0 | 329.0 | 2.5e-89 |
| CAO98848.1 | polyketide\_synthase\_AufE | BGC0000023 | Polyketide:Modular type I | 31.0 | 43.5 | 329.0 | 4.2e-89 |
| AEK75502.1 | type\_1\_polyketide\_synthase | BGC0000001 | Polyketide:Modular type I | 29.0 | 54.3 | 328.0 | 5.5e-89 |
| ABB88519.1 | polyketide\_synthase\_type\_I | BGC0000050 | Polyketide | 30.0 | 56.0 | 328.0 | 5.5e-89 |
| ABL74938.1 | PKS | BGC0001048 | NRP:Glycopeptide + Polyketide:Modular type I + Saccharide:Hybrid/tailoring | 30.0 | 55.0 | 328.0 | 5.5e-89 |
| CAE45669.1 | borrelidin\_polyketide\_synthase,\_type\_I | BGC0000031 | Polyketide:Modular type I | 28.0 | 53.9 | 328.0 | 7.2e-89 |
| BAW32334.1 | hybrid\_cis-AT\_polyketide\_synthase\_-\_nonribosomal\_peptide\_synthetase | BGC0001631 | NRP + Polyketide | 30.0 | 52.6 | 328.0 | 7.2e-89 |
| CAJ88177.1 | putative\_type\_I\_polyketide\_synthase | BGC0000151 | Polyketide:Modular type I + Saccharide:Hybrid/tailoring | 30.0 | 49.7 | 327.0 | 1.2e-88 |
| AHH99922.1 | PKS\_I | BGC0000002 | Polyketide | 30.0 | 53.8 | 326.0 | 2.7e-88 |
| AHH99923.1 | PKS\_I | BGC0000002 | Polyketide | 29.0 | 53.3 | 326.0 | 2.7e-88 |
| AGC09485.1 | LobS2 | BGC0001183 | Polyketide | 29.0 | 61.8 | 326.0 | 2.7e-88 |
| ABC84460.1 | NigAV | BGC0000114 | Polyketide:Modular type I | 29.0 | 59.2 | 325.0 | 6.1e-88 |
| WP\_055469548.1 | type\_I\_polyketide\_synthase | BGC0001537 | Polyketide | 29.0 | 54.0 | 325.0 | 6.1e-88 |
| WP\_107408739.1 | type\_I\_polyketide\_synthase | BGC0002033 | Polyketide | 29.0 | 54.9 | 325.0 | 6.1e-88 |
| AAC46026.1 | polyketide\_synthase\_modules\_4\_and\_5 | BGC0000113 | Polyketide | 27.0 | 67.1 | 324.0 | 8e-88 |
| ALV82341.1 | borrelidin\_type\_I\_polyketide\_synthase | BGC0001533 | Polyketide | 28.0 | 53.7 | 324.0 | 8e-88 |
| CAE45670.1 | borrelidin\_polyketide\_synthase,\_type\_I | BGC0000031 | Polyketide:Modular type I | 28.0 | 53.4 | 324.0 | 1e-87 |
| ADU86004.1 | putative\_modular\_polyketide\_synthase | BGC0000165 | Polyketide:Modular type I | 29.0 | 54.6 | 324.0 | 1e-87 |
| AAP85335.1 | type\_I\_PKS | BGC0000233 | Polyketide | 27.0 | 58.5 | 324.0 | 1e-87 |
| AWS21278.1 | type\_I\_polyketide\_synthase | BGC0001934 | Polyketide | 28.0 | 54.3 | 324.0 | 1.4e-87 |
| AZY91987.1 | polyketide\_synthase | BGC0002022 | Polyketide | 28.0 | 54.3 | 324.0 | 1.4e-87 |
| ACF35447.1 | mbcAIII | BGC0000090 | Polyketide | 29.0 | 59.1 | 323.0 | 2.3e-87 |
| AAV66110.2 | fusaridione\_A\_synthetase | BGC0000992 | NRP + Polyketide | 29.0 | 50.6 | 322.0 | 4e-87 |
| AJO72737.1 | Type\_I\_modular\_polyketide\_synthase | BGC0001381 | Polyketide | 28.0 | 53.7 | 322.0 | 5.2e-87 |
| AAP42858.1 | NanA4 | BGC0000105 | Polyketide | 29.0 | 60.9 | 321.0 | 8.8e-87 |
| BAK64649.1 | polyketide\_synthase | BGC0000135 | Polyketide | 31.0 | 50.2 | 321.0 | 1.1e-86 |
| CCC55921.1 | non-ribosomal\_peptide\_synthetase/polyketide\_synthase\_hybrid\_protein | BGC0000973 | NRP + Polyketide:Modular type I | 28.0 | 56.3 | 321.0 | 1.1e-86 |
| ACO94472.1 | polyketide\_synthase\_type\_I | BGC0000029 | Polyketide:Modular type I | 29.0 | 58.6 | 320.0 | 1.5e-86 |
| AIA58899.1 | HRPKS | BGC0001141 | Polyketide:Iterative type I | 30.0 | 56.5 | 319.0 | 3.3e-86 |
| CAI94713.1 | putative\_polyketide\_synthase | BGC0000141 | Polyketide | 29.0 | 59.4 | 319.0 | 4.4e-86 |
| AGC24271.1 | prlQ | BGC0001038 | NRP + Polyketide:Modular type I | 28.0 | 54.2 | 319.0 | 4.4e-86 |
| BAQ25512.1 | type\_I\_polyketide\_synthase | BGC0001288 | Polyketide | 26.0 | 64.4 | 319.0 | 4.4e-86 |
| ADU86002.1 | putative\_modular\_polyketide\_synthase | BGC0000165 | Polyketide:Modular type I | 29.0 | 53.6 | 317.0 | 9.7e-86 |
| CAD70195.1 | non-ribosomal\_peptide\_synthetase | BGC0001047 | NRP + Polyketide | 28.0 | 53.9 | 317.0 | 1.3e-85 |
| WP\_079080698.1 | type\_I\_polyketide\_synthase | BGC0001537 | Polyketide | 29.0 | 53.3 | 316.0 | 2.8e-85 |
| AAC68815.1 | FK506\_polyketide\_synthase | BGC0000353 | NRP | 29.0 | 56.2 | 315.0 | 4.8e-85 |
| ALA09354.1 | type\_I\_modular\_PKS | BGC0001303 | Polyketide | 29.0 | 49.4 | 315.0 | 4.8e-85 |
| AAR87760.2 | ZmaK | BGC0001059 | NRP + Polyketide | 29.0 | 53.9 | 315.0 | 6.3e-85 |
| CDG12864.1 | non-ribosomal\_peptide\_synthetase | BGC0001415 | NRP | 29.0 | 56.0 | 314.0 | 8.2e-85 |
| ASX95227.1 | IlaE | BGC0001620 | Polyketide | 28.0 | 56.6 | 314.0 | 1.1e-84 |
| CAJ76291.1 | putative\_polyketide\_synthase | BGC0000972 | NRP + Polyketide:Modular type I + Polyketide:Trans-AT type I | 28.0 | 59.4 | 313.0 | 2.4e-84 |
| CBA11584.1 | polyketide\_synthase\_type\_I | BGC0001046 | NRP + Polyketide:Modular type I + Saccharide:Hybrid/tailoring | 28.0 | 53.6 | 313.0 | 2.4e-84 |
| AGY30676.1 | Ann4 | BGC0001298 | Polyketide | 29.0 | 55.5 | 312.0 | 4.1e-84 |
| AAY89049.1 | polyketide\_synthase | BGC0001069 | NRP + Polyketide:Trans-AT type I | 28.0 | 52.9 | 311.0 | 7e-84 |
| ARP51711.1 | PKS-NRPS\_hybrid\_protein | BGC0001741 | NRP + Polyketide | 30.0 | 50.3 | 309.0 | 2.7e-83 |
| ACC40923.1 | polyketide\_synthase\_Pks9 | BGC0001665 | Polyketide | 28.0 | 53.5 | 309.0 | 4.5e-83 |
| AKA59088.1 | type-I\_PKS | BGC0001619 | Polyketide | 29.0 | 54.9 | 308.0 | 7.7e-83 |
| AAG02357.1 | polyketide\_synthase | BGC0000963 | NRP:Glycopeptide + Polyketide:Modular type I + Saccharide:Hybrid/tailoring | 29.0 | 54.3 | 307.0 | 1.3e-82 |
| AXM42948.1 | type\_1\_polyketide\_synthase | BGC0001941 | NRP + Polyketide | 33.0 | 39.7 | 307.0 | 1.3e-82 |
| AEZ64504.1 | Herc | BGC0001065 | Polyketide | 28.0 | 58.0 | 306.0 | 2.2e-82 |
| AWX24483.1 | type\_I\_polyketide\_synthase | BGC0001695 | NRP | 28.0 | 54.4 | 306.0 | 2.2e-82 |
| ADU85981.1 | putative\_modular\_polyketide\_synthase | BGC0000165 | Polyketide:Modular type I | 28.0 | 62.0 | 305.0 | 3.8e-82 |
| CBJ89760.1 | Polyketide\_synthase\_involved\_in\_xenocoumacin\_synthesis | BGC0001054 | NRP + Polyketide:Modular type I | 27.0 | 53.6 | 305.0 | 6.5e-82 |
| AGY62753.1 | EbeA | BGC0000051 | Polyketide | 28.0 | 53.9 | 304.0 | 8.5e-82 |
| BAG85026.1 | putative\_polyketide\_synthase | BGC0000086 | Polyketide | 31.0 | 48.9 | 303.0 | 2.5e-81 |
| CBW54671.1 | polyketide\_synthase/non\_ribosomal\_peptide\_synthetase | BGC0000971 | NRP + Polyketide:Modular type I | 29.0 | 49.6 | 303.0 | 2.5e-81 |
| BBA66511.1 | type\_I\_polyketide\_synthase | BGC0001495 | Polyketide | 29.0 | 53.5 | 302.0 | 3.2e-81 |
| AFP87524.1 | type\_I\_polyketide\_synthase | BGC0001159 | NRP + Polyketide:Modular type I | 33.0 | 39.7 | 302.0 | 4.2e-81 |
| AKA54627.1 | PKS | BGC0001216 | NRP + Polyketide | 27.0 | 54.2 | 302.0 | 4.2e-81 |
| ctg1\_orf27 |  | BGC0000096 | Polyketide | 29.0 | 49.6 | 301.0 | 7.2e-81 |
| AGO86662.1 | equisetin\_synthetase | BGC0001255 | NRP + Polyketide | 28.0 | 49.6 | 301.0 | 7.2e-81 |
| ARE67851.1 | AbsB3 | BGC0001492 | Polyketide | 28.0 | 51.9 | 301.0 | 9.4e-81 |
| BAD97694.1 | Aft9-1 | BGC0000003 | Polyketide | 29.0 | 49.6 | 300.0 | 1.2e-80 |
| BAP34740.1 | type\_I\_polyketide\_synthase | BGC0000078 | Polyketide | 29.0 | 56.5 | 300.0 | 1.2e-80 |
| CAQ34917.1 | polyketide\_synthase | BGC0000986 | NRP + Polyketide | 33.0 | 41.0 | 300.0 | 2.1e-80 |
| WP\_106967775.1 | type\_I\_polyketide\_synthase | BGC0001519 | NRP + Polyketide | 27.0 | 60.7 | 299.0 | 4.7e-80 |
| CAJ76298.1 | putative\_hybrid\_polyketide-non-ribosomal\_peptide\_synthetase | BGC0000972 | NRP + Polyketide:Modular type I + Polyketide:Trans-AT type I | 28.0 | 53.9 | 298.0 | 8e-80 |
| AKG06375.1 | polyketide\_synthase\_type\_1 | BGC0001830 | Polyketide | 27.0 | 54.2 | 296.0 | 3e-79 |
| BBG28484.1 | polyketide\_synthase\_CdmE | BGC0001926 | Polyketide | 28.0 | 56.3 | 295.0 | 4e-79 |
| EAQ86385.1 | hypothetical\_protein | BGC0001405 | Polyketide | 29.0 | 45.7 | 295.0 | 5.2e-79 |
| KGO40478.1 | Acyl\_transferase/acyl\_hydrolase/lysophospholipase | BGC0001205 | Polyketide | 30.0 | 48.6 | 295.0 | 6.8e-79 |
| WP\_078620910.1 | acyltransferase\_domain-containing\_protein | BGC0001519 | NRP + Polyketide | 28.0 | 54.0 | 295.0 | 6.8e-79 |
| CAN93347.1 | Polyketide\_synthase | BGC0000179 | Polyketide:Trans-AT type I | 35.0 | 31.9 | 294.0 | 8.8e-79 |
| ANZ22995.1 | ZinA | BGC0001828 | Polyketide | 28.0 | 53.6 | 294.0 | 8.8e-79 |
| WP\_053138504.1 | type\_I\_polyketide\_synthase | BGC0002033 | Polyketide | 27.0 | 55.3 | 294.0 | 8.8e-79 |
| ANC94966.1 | AlmHI | BGC0001396 | Polyketide | 28.0 | 54.0 | 293.0 | 2e-78 |
| BBD17742.1 | polyketide\_synthase | BGC0001918 | NRP + Polyketide | 28.0 | 54.6 | 293.0 | 2.6e-78 |
| antaD | Type\_I\_PKS | BGC0001455 | NRP + Polyketide | 29.0 | 55.0 | 292.0 | 3.4e-78 |
| ctg1\_15 |  | BGC0001931 | Polyketide | 28.0 | 49.7 | 291.0 | 7.5e-78 |
| ADF88262.1 | mixed\_nonribosomal\_peptide\_synthetase/\_polyketide\_synthase | BGC0000979 | NRP + Polyketide | 29.0 | 44.3 | 291.0 | 9.8e-78 |
| ADF88265.1 | mixed\_nonribosomal\_peptide\_synthetase/\_polyketide\_synthase | BGC0000980 | NRP + Polyketide | 29.0 | 44.3 | 291.0 | 9.8e-78 |
| AGZ15472.1 | putative\_modular\_polyketide\_synthase | BGC0001036 | NRP + Polyketide | 28.0 | 54.7 | 290.0 | 1.7e-77 |
| AQZ37113.1 | polyketide\_synthase | BGC0001511 | Polyketide | 28.0 | 57.2 | 290.0 | 1.7e-77 |
| ARS01473.1 | NcmAI | BGC0001702 | NRP + Polyketide | 30.0 | 48.5 | 290.0 | 1.7e-77 |
| ASA76631.1 | polyketide\_synthase | BGC0001751 | NRP + Polyketide | 34.0 | 35.1 | 289.0 | 2.8e-77 |
| XP\_659388.1 | hypothetical\_protein | BGC0001998 | Polyketide | 25.0 | 77.1 | 289.0 | 2.8e-77 |
| ACR50795.1 | putative\_polyketide\_synthase | BGC0000163 | Polyketide | 31.0 | 38.1 | 289.0 | 4.8e-77 |
| AAS79459.1 | polyketide\_synthase\_subunit | BGC0000035 | Polyketide | 29.0 | 54.0 | 287.0 | 1.4e-76 |
| AJQ95706.1 | polyketide\_synthase\_modules-related\_protein | BGC0001644 | Polyketide | 34.0 | 32.3 | 287.0 | 1.4e-76 |
| BAJ16468.1 | polyketide\_synthase | BGC0000058 | Polyketide | 27.0 | 54.0 | 287.0 | 1.8e-76 |
| ABO15861.1 | polyketide\_synthase | BGC0000130 | Polyketide | 33.0 | 40.0 | 285.0 | 4.1e-76 |
| ACZ57548.1 | polyketide\_synthase | BGC0000046 | Polyketide:Iterative type I | 27.0 | 57.2 | 284.0 | 9.1e-76 |
| AAY32964.1 | DszA | BGC0001093 | NRP + Polyketide | 35.0 | 31.6 | 284.0 | 9.1e-76 |
| AMY15057.1 | tetraketide\_synthase\_MF-SQTKS | BGC0001339 | Polyketide:Iterative type I | 28.0 | 49.0 | 284.0 | 1.2e-75 |
| BBD17760.1 | polyketide\_synthase | BGC0001919 | NRP + Polyketide | 27.0 | 53.7 | 284.0 | 1.6e-75 |
| AAM54075.1 | polyketide\_synthase | BGC0000020 | Polyketide | 27.0 | 57.3 | 282.0 | 3.5e-75 |
| BBC43184.1 | PKS-NRPS\_hybrid | BGC0001738 | NRP + Polyketide | 28.0 | 51.3 | 281.0 | 7.7e-75 |
| ASA76643.1 | polyketide\_synthase | BGC0001751 | NRP + Polyketide | 33.0 | 32.5 | 281.0 | 7.7e-75 |
| CBK62724.1 |  | BGC0001115 | NRP + Polyketide | 33.0 | 32.6 | 279.0 | 2.9e-74 |
| CAE52339.1 | Polyketide\_non-ribosomal\_peptide\_synthase | BGC0001088 | NRP + Polyketide | 32.0 | 36.3 | 279.0 | 5e-74 |
| CAP95405.1 |  | BGC0001404 | Polyketide | 27.0 | 50.3 | 278.0 | 6.5e-74 |
| CBJ89766.1 | Polyketide\_synthase\_involved\_in\_xenocoumacin\_synthesis | BGC0001054 | NRP + Polyketide:Modular type I | 27.0 | 52.2 | 278.0 | 8.5e-74 |
| AHV78245.1 | LasS1 | BGC0001245 | Polyketide | 28.0 | 55.6 | 278.0 | 8.5e-74 |
| AJO72743.1 | Type\_I\_modular\_polyketide\_synthase | BGC0001381 | Polyketide | 28.0 | 53.4 | 278.0 | 8.5e-74 |
| ARR97036.1 | SphC | BGC0001780 | NRP | 35.0 | 30.3 | 277.0 | 1.1e-73 |
| AEC13079.1 | fosA | BGC0000060 | Polyketide | 31.0 | 41.2 | 275.0 | 7.2e-73 |
| AKQ22696.1 | malonyl\_CoA-acyl\_carrier\_protein\_transacylase | BGC0001186 | Polyketide | 32.0 | 32.5 | 275.0 | 7.2e-73 |
| ACY01401.1 | AT-less\_polyketide\_synthase | BGC0000083 | Polyketide:Modular type I + Polyketide:Trans-AT type I | 35.0 | 31.2 | 274.0 | 9.4e-73 |
| AEC04361.1 | polyketide\_synthase | BGC0000178 | Polyketide:Trans-AT type I | 33.0 | 31.2 | 274.0 | 9.4e-73 |
| ADI59533.1 | CorK | BGC0001091 | NRP + Polyketide | 30.0 | 39.2 | 274.0 | 1.2e-72 |
| AFX60309.1 | polyketide\_synthase | BGC0001031 | NRP + Polyketide | 33.0 | 31.2 | 274.0 | 1.6e-72 |
| CBF87072.1 | polyketide\_synthase,\_putative\_(Eurofung) | BGC0001290 | NRP | 28.0 | 51.6 | 274.0 | 1.6e-72 |
| AEC04357.1 | polyketide\_synthase | BGC0000178 | Polyketide:Trans-AT type I | 32.0 | 36.5 | 273.0 | 2.7e-72 |
| ATX68125.1 | malonyl\_CoA-acyl\_carrier\_protein\_transacylase | BGC0001795 | Polyketide | 33.0 | 31.2 | 272.0 | 4.7e-72 |
| AAS47564.1 | mixed\_type\_I\_polyketide\_synthase/nonribosomal\_peptide\_synthetase | BGC0001108 | Polyketide:Trans-AT type I | 33.0 | 32.2 | 272.0 | 6.1e-72 |
| RAT98525.1 | trans-acyltransferase\_polyketide\_synthase | BGC0001470 | Polyketide:Trans-AT type I | 32.0 | 32.5 | 272.0 | 6.1e-72 |
| AKQ22669.1 | malonyl\_CoA-acyl\_carrier\_protein\_transacylase | BGC0001656 | Polyketide | 32.0 | 31.7 | 272.0 | 6.1e-72 |
| ctg1\_orf6 |  | BGC0001109 | NRP + Polyketide | 33.0 | 32.2 | 271.0 | 1e-71 |
| AMYAL\_RS48910 | polyketide\_synthase | BGC0002011 | Polyketide | 33.0 | 34.1 | 271.0 | 1e-71 |
| CAG23968.1 | polyketide\_synthase\_type\_I | BGC0000181 | Polyketide | 31.0 | 33.5 | 270.0 | 1.4e-71 |
| ADN68476.1 | sorA | BGC0000184 | Polyketide:Trans-AT type I | 33.0 | 31.2 | 270.0 | 1.4e-71 |
| AAV97870.1 | OnnB | BGC0001105 | NRP + Polyketide:Trans-AT type I | 33.0 | 31.5 | 270.0 | 1.4e-71 |
| AIJ04686.1 | polyketide\_synthase | BGC0001383 | Polyketide | 31.0 | 33.9 | 270.0 | 1.4e-71 |
| ADD82941.1 | Bat3 | BGC0001099 | NRP + Polyketide:Modular type I + Polyketide:Trans-AT type I | 33.0 | 31.9 | 270.0 | 1.8e-71 |
| AMH40443.1 | PKS | BGC0001350 | Polyketide | 31.0 | 36.3 | 270.0 | 1.8e-71 |
| AAV97877.1 | OnnI | BGC0001105 | NRP + Polyketide:Trans-AT type I | 32.0 | 33.1 | 269.0 | 4e-71 |
| CAN93348.1 | polyketide\_synthase | BGC0000179 | Polyketide:Trans-AT type I | 31.0 | 32.6 | 269.0 | 5.2e-71 |
| AKQ22680.1 | malonyl\_CoA-acyl\_carrier\_protein\_transacylase | BGC0001656 | Polyketide | 31.0 | 34.3 | 269.0 | 5.2e-71 |
| WP\_010639241.1 | type\_I\_polyketide\_synthase | BGC0000958 | NRP:Cyclic depsipeptide + Polyketide:Modular type I | 27.0 | 53.7 | 268.0 | 6.8e-71 |
| OAP25815.1 | Phenolphthiocerol\_synthesis\_polyketide\_synthase\_type\_I\_Pks15/1 | BGC0001658 | Polyketide | 30.0 | 38.8 | 268.0 | 6.8e-71 |
| ATX68111.1 | malonyl\_CoA-acyl\_carrier\_protein\_transacylase | BGC0001772 | Polyketide | 31.0 | 31.9 | 268.0 | 6.8e-71 |
| AFX60311.1 | polyketide\_synthase | BGC0001031 | NRP + Polyketide | 33.0 | 32.0 | 268.0 | 8.8e-71 |
| AAS47562.1 | mixed\_type\_I\_polyketide\_synthase\_-\_peptide\_synthetase | BGC0001108 | Polyketide:Trans-AT type I | 33.0 | 28.8 | 267.0 | 1.5e-70 |
| ctg1\_orf8 |  | BGC0001109 | NRP + Polyketide | 33.0 | 28.8 | 267.0 | 1.5e-70 |
| ATX68127.1 | malonyl\_CoA-acyl\_carrier\_protein\_transacylase | BGC0001795 | Polyketide | 32.0 | 32.5 | 267.0 | 2e-70 |
| AFN27480.1 | pks\_BonA | BGC0000173 | Polyketide:Modular type I | 32.0 | 32.7 | 264.0 | 1.7e-69 |
| CAJ57409.1 | polyketide\_synthase\_type\_I | BGC0000176 | Polyketide + NRP | 33.0 | 31.3 | 264.0 | 1.7e-69 |
| ATX68124.1 | malonyl\_CoA-acyl\_carrier\_protein\_transacylase | BGC0001795 | Polyketide | 31.0 | 31.9 | 264.0 | 1.7e-69 |
| EAL85129.1 | polyketide\_synthase | BGC0001067 | Terpene + Polyketide:Iterative type I | 27.0 | 57.5 | 263.0 | 2.2e-69 |
| ASA76644.1 | polyketide\_synthase | BGC0001751 | NRP + Polyketide | 32.0 | 31.3 | 263.0 | 2.2e-69 |
| AFX60332.1 | polyketide\_synthase | BGC0001032 | NRP + Polyketide | 32.0 | 31.3 | 263.0 | 2.8e-69 |
| ALD83703.1 | tAT\_polyketide\_synthase | BGC0001299 | Polyketide | 32.0 | 31.6 | 263.0 | 2.8e-69 |
| RAT98517.1 | trans-acyltransferase\_polyketide\_synthase | BGC0001470 | Polyketide:Trans-AT type I | 30.0 | 31.8 | 263.0 | 2.8e-69 |
| CAG23964.1 | polyketide\_synthase\_type\_I | BGC0000181 | Polyketide | 37.0 | 26.3 | 262.0 | 3.7e-69 |
| ABC36687.1 | polyketide\_synthase | BGC0000964 | NRP:Cyclic depsipeptide + Polyketide:Trans-AT type I | 32.0 | 31.6 | 262.0 | 3.7e-69 |
| CBK62733.1 |  | BGC0001115 | NRP + Polyketide | 34.0 | 30.5 | 262.0 | 3.7e-69 |
| ATY69589.1 | type\_I\_polyketide\_synthase | BGC0001823 | NRP + Polyketide | 33.0 | 31.7 | 262.0 | 4.9e-69 |
| AIJ04681.1 | polyketide\_synthase | BGC0001383 | Polyketide | 36.0 | 26.3 | 261.0 | 8.3e-69 |
| AMH40421.1 | PKS | BGC0001350 | Polyketide | 32.0 | 33.9 | 261.0 | 1.1e-68 |
| ATX68112.1 | malonyl\_CoA-acyl\_carrier\_protein\_transacylase | BGC0001772 | Polyketide | 31.0 | 32.4 | 260.0 | 1.4e-68 |
| AIU36104.1 | LglE | BGC0000180 | Polyketide:Trans-AT type I | 33.0 | 31.5 | 260.0 | 1.8e-68 |
| ADH01487.1 | polyketide\_synthase | BGC0001096 | NRP + Polyketide | 33.0 | 31.5 | 260.0 | 1.8e-68 |
| CCA89326.1 | mixed\_trans-AT\_type\_I\_polyketide\_synthase/nonribosomal\_peptide\_synthetase | BGC0001111 | NRP + Polyketide:Trans-AT type I | 33.0 | 31.3 | 260.0 | 1.8e-68 |
| CAL69894.1 | RhiF\_protein | BGC0001112 | NRP + Polyketide:Trans-AT type I | 32.0 | 32.5 | 260.0 | 1.8e-68 |
| AIC32693.1 | FR9DEF | BGC0001113 | NRP + Polyketide | 33.0 | 31.5 | 260.0 | 1.8e-68 |
| ABP57747.1 | DepC | BGC0000993 | NRP:Cyclic depsipeptide + Polyketide:Modular type I | 33.0 | 30.6 | 260.0 | 2.4e-68 |
| ERM18798.1 | polyketide\_synthase | BGC0000172 | Polyketide | 32.0 | 31.9 | 259.0 | 5.4e-68 |
| AAM12909.2 | MmpA | BGC0000182 | Polyketide:Iterative type I + Polyketide:Trans-AT type I | 32.0 | 31.1 | 259.0 | 5.4e-68 |
| AFX60334.1 | polyketide\_synthase | BGC0001032 | NRP + Polyketide | 33.0 | 31.2 | 259.0 | 5.4e-68 |
| AEC04363.1 | polyketide\_synthase | BGC0000178 | Polyketide:Trans-AT type I | 31.0 | 33.1 | 258.0 | 7e-68 |
| ADH01489.1 | type\_I\_polyketide\_synthase | BGC0001096 | NRP + Polyketide | 34.0 | 31.2 | 258.0 | 7e-68 |
| AIC32694.1 | FR9GH | BGC0001113 | NRP + Polyketide | 34.0 | 31.2 | 258.0 | 7e-68 |
| AKQ22698.1 | malonyl\_CoA-acyl\_carrier\_protein\_transacylase | BGC0001186 | Polyketide | 32.0 | 31.2 | 258.0 | 7e-68 |
| EWM62997.1 | non-ribosomal\_peptide\_synthetase | BGC0001328 | NRP:Cyclic depsipeptide + Polyketide:Modular type I | 32.0 | 36.5 | 258.0 | 9.2e-68 |
| ACY01391.1 | AT-less\_polyketide\_synthase | BGC0000177 | Polyketide:Modular type I + Polyketide:Trans-AT type I | 34.0 | 31.3 | 257.0 | 1.2e-67 |
| CAN93352.1 | polyketide\_synthase | BGC0000179 | Polyketide:Trans-AT type I | 33.0 | 31.8 | 257.0 | 1.2e-67 |
| OEI73462.1 | hypothetical\_protein | BGC0001520 | Polyketide | 33.0 | 32.0 | 257.0 | 1.2e-67 |
| ANY10591.1 | polyketide\_synthase | BGC0001773 | Polyketide | 31.0 | 39.7 | 257.0 | 1.2e-67 |
| ADA69239.2 | trans-AT\_hybrid\_polyketide\_synthase-NRPS | BGC0001071 | NRP + Polyketide:Modular type I + Polyketide:Trans-AT type I | 31.0 | 32.6 | 257.0 | 1.6e-67 |
| CTQ34881.1 | AtcD;\_polyketide\_synthase,\_modules\_1-4 | BGC0001301 | Polyketide | 35.0 | 28.2 | 257.0 | 2e-67 |
| AJQ95705.1 | polyketide\_synthase\_modules-related\_protein | BGC0001644 | Polyketide | 32.0 | 31.8 | 257.0 | 2e-67 |
| CUX96955.1 | TmcH | BGC0001829 | NRP + Polyketide | 27.0 | 53.3 | 257.0 | 2e-67 |
| AKQ22699.1 | malonyl\_CoA-acyl\_carrier\_protein\_transacylase | BGC0001186 | Polyketide | 32.0 | 31.2 | 256.0 | 3.5e-67 |
| AIJ04680.1 | polyketide\_synthase | BGC0001383 | Polyketide | 31.0 | 31.4 | 256.0 | 3.5e-67 |
| AKQ22681.1 | malonyl\_CoA-acyl\_carrier\_protein\_transacylase | BGC0001656 | Polyketide | 29.0 | 35.7 | 256.0 | 3.5e-67 |
| BAF50727.1 | hybrid\_polyketide\_synthase-non\_ribosomal\_peptide\_synthetase | BGC0001116 | NRP + Polyketide | 33.0 | 32.4 | 254.0 | 1e-66 |
| ATG32078.1 | polyketide\_synthase | BGC0001750 | NRP + Polyketide | 34.0 | 27.6 | 254.0 | 1e-66 |
| CAG23965.1 | polyketide\_synthase\_type\_I | BGC0000181 | Polyketide | 31.0 | 31.4 | 254.0 | 1.3e-66 |
| CAG23966.1 | polyketide\_synthase\_type\_I | BGC0000181 | Polyketide | 33.0 | 28.2 | 254.0 | 1.3e-66 |
| CBJ89764.1 | Polyketide\_synthase\_involved\_in\_xenocoumacin\_synthesis | BGC0001054 | NRP + Polyketide:Modular type I | 25.0 | 53.5 | 254.0 | 1.3e-66 |
| RAT98518.1 | trans-acyltransferase\_polyketide\_synthase | BGC0001470 | Polyketide:Trans-AT type I | 31.0 | 32.8 | 254.0 | 1.3e-66 |
| QCX41945.1 | Amc8 | BGC0001958 | Other | 28.0 | 53.9 | 254.0 | 1.3e-66 |
| CAL69890.1 | RhiC\_protein | BGC0001112 | NRP + Polyketide:Trans-AT type I | 31.0 | 31.4 | 253.0 | 1.7e-66 |
| QCX41916.1 | Mhr10 | BGC0001956 | Polyketide | 28.0 | 53.6 | 253.0 | 1.7e-66 |
| AIJ04683.1 | polyketide\_synthase | BGC0001383 | Polyketide | 33.0 | 28.2 | 253.0 | 2.3e-66 |
| OEI73461.1 | hypothetical\_protein | BGC0001520 | Polyketide | 31.0 | 31.3 | 253.0 | 2.3e-66 |
| AAF19810.1 | MtaB | BGC0001024 | NRP + Polyketide:Modular type I | 35.0 | 27.2 | 252.0 | 3.8e-66 |
| ABF85931.1 | non-ribosomal\_peptide\_synthase/polyketide\_synthase\_Ta1 | BGC0001025 | NRP + Polyketide:Trans-AT type I | 31.0 | 31.0 | 252.0 | 5e-66 |
| CCA89329.1 | trans-AT\_type\_I\_polyketide\_synthase | BGC0001111 | NRP + Polyketide:Trans-AT type I | 32.0 | 31.1 | 252.0 | 5e-66 |
| ALD83688.1 | tAT\_polyketide\_synthase | BGC0001300 | Polyketide | 31.0 | 31.2 | 252.0 | 6.6e-66 |
| CTQ34882.1 | AtcE;\_polyketide\_synthase,\_modules\_5-7 | BGC0001301 | Polyketide | 32.0 | 30.9 | 252.0 | 6.6e-66 |
| AIJ04685.1 | polyketide\_synthase | BGC0001383 | Polyketide | 32.0 | 28.1 | 251.0 | 8.6e-66 |
| CAG23977.1 | polyketide\_synthase\_type\_I | BGC0000176 | Polyketide + NRP | 31.0 | 31.4 | 251.0 | 1.1e-65 |
| AJQ95708.1 | polyketide\_synthase\_modules-related\_protein | BGC0001644 | Polyketide | 30.0 | 34.6 | 250.0 | 1.5e-65 |
| CAG23969.1 | polyketide\_synthase\_type\_I | BGC0000181 | Polyketide | 32.0 | 28.1 | 250.0 | 1.9e-65 |
| ADN68477.1 | SorB | BGC0000184 | Polyketide:Trans-AT type I | 33.0 | 31.8 | 250.0 | 1.9e-65 |
| CAN89632.1 | putative\_polyketide\_synthase | BGC0001070 | NRP + Polyketide:Modular type I + Polyketide:Trans-AT type I | 31.0 | 34.5 | 250.0 | 1.9e-65 |
| ABI91469.1 | beta-ketoacyl\_synthase | BGC0001094 | NRP + Polyketide | 33.0 | 30.9 | 250.0 | 1.9e-65 |
| ALD83687.1 | tAT\_polyketide\_synthase | BGC0001300 | Polyketide | 31.0 | 30.9 | 250.0 | 1.9e-65 |
| ATQ39432.1 | PKS | BGC0001565 | NRP | 25.0 | 58.0 | 250.0 | 1.9e-65 |
| AGN11882.1 | tstGH | BGC0001114 | NRP + Polyketide | 34.0 | 30.9 | 250.0 | 2.5e-65 |
| ADA82585.1 | hybrid\_trans-AT\_polyketide\_synthase\_-\_nonribosomal\_peptide\_synthetase | BGC0001110 | NRP + Polyketide:Trans-AT type I | 33.0 | 32.3 | 249.0 | 3.3e-65 |
| ACR12418.1 | modular\_polyketide\_synthase,\_type\_I\_PKS | BGC0000185 | Polyketide | 32.0 | 31.3 | 248.0 | 5.6e-65 |
| CAG23957.2 | hybrid\_NRPS/PKS\_protein | BGC0001089 | Polyketide + NRP | 32.0 | 31.8 | 248.0 | 5.6e-65 |
| AJQ95707.1 | polyketide\_synthase\_modules-related\_protein | BGC0001644 | Polyketide | 30.0 | 31.1 | 248.0 | 7.3e-65 |
| BAE93740.1 | type\_I\_polyketide\_synthase-related\_protein | BGC0000164 | Polyketide | 32.0 | 33.2 | 248.0 | 9.5e-65 |
| ALD83704.1 | tAT\_polyketide\_synthase | BGC0001299 | Polyketide | 32.0 | 32.5 | 248.0 | 9.5e-65 |
| CAG23958.2 | polyketide\_synthase\_of\_type\_I | BGC0001089 | Polyketide + NRP | 31.0 | 31.3 | 247.0 | 1.2e-64 |
| AGN11881.1 | tstDEF | BGC0001114 | NRP + Polyketide | 33.0 | 31.5 | 247.0 | 1.2e-64 |
| ABI91467.1 | beta-ketoacyl\_synthase | BGC0001094 | NRP + Polyketide | 31.0 | 30.8 | 247.0 | 1.6e-64 |
| AHA38199.1 | GphF | BGC0000069 | Polyketide | 34.0 | 30.9 | 247.0 | 2.1e-64 |
| ADD82940.1 | Bat2 | BGC0001099 | NRP + Polyketide:Modular type I + Polyketide:Trans-AT type I | 30.0 | 35.7 | 247.0 | 2.1e-64 |
| CCG06113.1 | type\_I\_polyketide\_synthase | BGC0001543 | Polyketide | 31.0 | 32.4 | 247.0 | 2.1e-64 |
| CAL69893.1 | RhiE\_protein | BGC0001112 | NRP + Polyketide:Trans-AT type I | 32.0 | 30.8 | 246.0 | 2.8e-64 |
| ATY69569.1 | type\_I\_polyketide\_synthase | BGC0001611 | NRP + Polyketide | 32.0 | 34.8 | 246.0 | 2.8e-64 |
| AVR48533.1 | CusA | BGC0001564 | NRP + Polyketide | 31.0 | 31.3 | 245.0 | 4.7e-64 |
| ATX68126.1 | malonyl\_CoA-acyl\_carrier\_protein\_transacylase | BGC0001795 | Polyketide | 30.0 | 32.4 | 245.0 | 4.7e-64 |
| ABM63527.1 | BryB | BGC0000174 | Polyketide | 31.0 | 30.9 | 244.0 | 1e-63 |
| ABM63528.1 | BryC | BGC0000174 | Polyketide | 31.0 | 31.9 | 244.0 | 1e-63 |
| AMH40423.1 | PKS | BGC0001350 | Polyketide | 32.0 | 32.0 | 244.0 | 1e-63 |
| AGN74894.1 | nonribosomal\_peptide\_synthetase/polyketide\_synthase\_hybrid\_protein | BGC0000459 | NRP:Cyclic depsipeptide + Polyketide:Trans-AT type I | 33.0 | 32.6 | 244.0 | 1.4e-63 |
| CAG23959.2 | polyketide\_synthase\_of\_type\_I | BGC0001089 | Polyketide + NRP | 32.0 | 31.0 | 244.0 | 1.4e-63 |
| AAY32965.1 | DszB | BGC0001093 | NRP + Polyketide | 31.0 | 31.9 | 243.0 | 2.3e-63 |
| DAC80077.1 | PKS | BGC0001836 | Polyketide:Trans-AT type I | 32.0 | 31.5 | 243.0 | 2.3e-63 |
| AAP42872.1 | NanA9 | BGC0000105 | Polyketide | 29.0 | 39.1 | 243.0 | 3e-63 |
| CAG23960.2 | hybrid\_NRPS/PKS\_protein | BGC0001089 | Polyketide + NRP | 31.0 | 31.1 | 242.0 | 5.2e-63 |
| WP\_106980515.1 | type\_I\_polyketide\_synthase | BGC0001348 | Polyketide:Modular type I | 29.0 | 36.4 | 242.0 | 5.2e-63 |
| CTQ34883.1 | AtcF;\_polyketide\_synthase,\_modules\_8-10 | BGC0001301 | Polyketide | 31.0 | 31.7 | 242.0 | 6.8e-63 |
| ATX68109.1 | malonyl\_CoA-acyl\_carrier\_protein\_transacylase | BGC0001772 | Polyketide | 30.0 | 32.2 | 241.0 | 8.9e-63 |
| ALD83686.1 | tAT\_polyketide\_synthase | BGC0001300 | Polyketide | 31.0 | 31.3 | 241.0 | 1.2e-62 |
| BAP05595.1 | calG | BGC0000967 | NRP + Polyketide:Trans-AT type I | 33.0 | 31.1 | 240.0 | 1.5e-62 |
| ADI59531.1 | CorI | BGC0001091 | NRP + Polyketide | 32.0 | 31.6 | 240.0 | 1.5e-62 |
| RAT98530.1 | trans-acyltransferase\_polyketide\_synthase | BGC0001470 | Polyketide:Trans-AT type I | 30.0 | 31.4 | 240.0 | 2e-62 |
| CCG06109.1 | type\_I\_polyketide\_synthase | BGC0001543 | Polyketide | 32.0 | 32.2 | 240.0 | 2e-62 |
| AEC04356.1 | polyketide\_synthase | BGC0000178 | Polyketide:Trans-AT type I | 32.0 | 31.2 | 239.0 | 4.4e-62 |
| ADA69237.1 | trans-AT\_polyketide\_synthase | BGC0001071 | NRP + Polyketide:Modular type I + Polyketide:Trans-AT type I | 30.0 | 31.2 | 239.0 | 4.4e-62 |
| AKQ22697.1 | malonyl\_CoA-acyl\_carrier\_protein\_transacylase | BGC0001186 | Polyketide | 32.0 | 31.2 | 239.0 | 4.4e-62 |
| ATX68110.1 | malonyl\_CoA-acyl\_carrier\_protein\_transacylase | BGC0001772 | Polyketide | 31.0 | 31.2 | 239.0 | 4.4e-62 |
| ADN68478.1 | SorC | BGC0000184 | Polyketide:Trans-AT type I | 31.0 | 31.2 | 238.0 | 5.7e-62 |
| ADN68479.1 | SorD | BGC0000184 | Polyketide:Trans-AT type I | 32.0 | 30.9 | 238.0 | 5.7e-62 |
| RAT98527.1 | trans-acyltransferase\_polyketide\_synthase | BGC0001470 | Polyketide:Trans-AT type I | 30.0 | 31.0 | 238.0 | 5.7e-62 |
| AKA59448.1 | polyketide\_synthase | BGC0001203 | NRP + Polyketide | 30.0 | 39.6 | 238.0 | 9.8e-62 |
| BBA21072.1 | putative\_modular\_polyketide\_synthase | BGC0001740 | NRP + Polyketide | 28.0 | 39.5 | 238.0 | 9.8e-62 |
| AXA20092.1 | trans-AT\_PKS\_LgaC | BGC0001946 | NRP + Polyketide | 31.0 | 33.4 | 237.0 | 1.7e-61 |
| DAC80073.1 | PKS | BGC0001836 | Polyketide:Trans-AT type I | 30.0 | 31.7 | 237.0 | 1.7e-61 |
| AJY78091.1 | polyketide\_synthase | BGC0001902 | NRP + Polyketide | 31.0 | 30.2 | 237.0 | 1.7e-61 |
| ERM18797.1 | polyketide\_synthase | BGC0000172 | Polyketide | 29.0 | 32.9 | 236.0 | 3.7e-61 |
| AAY32966.1 | DszC | BGC0001093 | NRP + Polyketide | 32.0 | 32.4 | 235.0 | 4.9e-61 |
| AKQ22682.1 | malonyl\_CoA-acyl\_carrier\_protein\_transacylase | BGC0001656 | Polyketide | 31.0 | 31.2 | 235.0 | 4.9e-61 |
| CAN89634.1 | putative\_polyketide\_synthase | BGC0001070 | NRP + Polyketide:Modular type I + Polyketide:Trans-AT type I | 34.0 | 26.2 | 234.0 | 1.1e-60 |
| AEH42473.1 | polyketide\_synthase | BGC0000032 | Polyketide | 28.0 | 39.7 | 234.0 | 1.4e-60 |
| CAN93349.1 | polyketide\_synthase | BGC0000179 | Polyketide:Trans-AT type I | 30.0 | 32.7 | 234.0 | 1.4e-60 |
| BAP05597.1 | calI | BGC0000967 | NRP + Polyketide:Trans-AT type I | 32.0 | 31.7 | 234.0 | 1.4e-60 |
| BAD38875.1 | polyketide\_synthase | BGC0000111 | Polyketide | 30.0 | 34.4 | 233.0 | 1.8e-60 |
| DAC80074.1 | PKS | BGC0001836 | Polyketide:Trans-AT type I | 30.0 | 32.1 | 233.0 | 1.8e-60 |
| AMH40422.1 | PKS | BGC0001350 | Polyketide | 32.0 | 31.9 | 233.0 | 2.4e-60 |
| ACY13414.1 | amino\_acid\_adenylation\_domain\_protein | BGC0001367 | NRP + Polyketide | 32.0 | 26.2 | 233.0 | 2.4e-60 |
| CAL69889.1 | RhiB\_protein | BGC0001112 | NRP + Polyketide:Trans-AT type I | 31.0 | 32.6 | 232.0 | 5.4e-60 |
| WP\_055469550.1 | type\_I\_polyketide\_synthase | BGC0001537 | Polyketide | 26.0 | 56.2 | 232.0 | 5.4e-60 |
| ACR13997.1 | modular\_polyketide\_synthase,\_type\_I\_PKS | BGC0000185 | Polyketide | 29.0 | 33.5 | 232.0 | 7e-60 |
| ABM63537.1 | BryA | BGC0000174 | Polyketide | 28.0 | 32.6 | 231.0 | 9.2e-60 |
| AAY89050.1 | polyketide\_synthase | BGC0001069 | NRP + Polyketide:Trans-AT type I | 31.0 | 30.3 | 231.0 | 1.2e-59 |
| BAC76474.1 | type\_I\_polyketide\_synthase\_LkcC | BGC0001100 | NRP + Polyketide | 31.0 | 31.2 | 231.0 | 1.2e-59 |
| AWS21290.1 | type\_I\_polyketide\_synthase | BGC0001934 | Polyketide | 31.0 | 39.6 | 231.0 | 1.2e-59 |
| AZY91988.1 | polyketide\_synthase | BGC0002022 | Polyketide | 31.0 | 39.6 | 231.0 | 1.2e-59 |
| AGN74892.1 | nonribosomal\_peptide\_synthetase/polyketide\_synthase\_hybrid\_protein | BGC0000459 | NRP:Cyclic depsipeptide + Polyketide:Trans-AT type I | 32.0 | 31.9 | 230.0 | 1.6e-59 |
| BAC76471.1 | type\_I\_polyketide\_synthase\_LkcF | BGC0001100 | NRP + Polyketide | 33.0 | 31.0 | 230.0 | 1.6e-59 |
| ABF89568.1 | polyketide\_synthase | BGC0001025 | NRP + Polyketide:Trans-AT type I | 30.0 | 31.8 | 230.0 | 2e-59 |
| AAN85522.1 | hybrid\_nonribosomal\_peptide\_synthetase\_/\_polyketide\_synthase | BGC0001101 | NRP + Polyketide:Modular type I + Polyketide:Trans-AT type I | 32.0 | 30.2 | 230.0 | 2e-59 |
| ASA76642.1 | polyketide\_synthase | BGC0001751 | NRP + Polyketide | 33.0 | 26.8 | 230.0 | 2e-59 |
| XP\_001220460.1 | hypothetical\_protein | BGC0001182 | NRP + Polyketide:Iterative type I | 28.0 | 41.1 | 229.0 | 3.5e-59 |
| CBK62731.1 |  | BGC0001115 | NRP + Polyketide | 29.0 | 33.5 | 229.0 | 4.6e-59 |
| AKQ22670.1 | malonyl\_CoA-acyl\_carrier\_protein\_transacylase | BGC0001656 | Polyketide | 30.0 | 30.9 | 228.0 | 5.9e-59 |
| ABS90472.1 | PKS | BGC0001106 | NRP + Polyketide | 31.0 | 31.8 | 228.0 | 7.8e-59 |
| BAD38874.1 | polyketide\_synthase | BGC0000111 | Polyketide | 32.0 | 31.8 | 228.0 | 1e-58 |
| AAC38075.1 | polyketide\_synthase\_type\_I | BGC0000127 | Polyketide | 31.0 | 33.3 | 228.0 | 1e-58 |
| ABI91466.1 | beta-ketoacyl\_synthase | BGC0001094 | NRP + Polyketide | 31.0 | 31.5 | 227.0 | 1.3e-58 |
| BAP05594.1 | calF | BGC0000967 | NRP + Polyketide:Trans-AT type I | 30.0 | 30.8 | 226.0 | 3e-58 |
| WP\_003598535.1 | SDR\_family\_NAD(P)-dependent\_oxidoreductase | BGC0001991 | Polyketide | 30.0 | 31.2 | 226.0 | 3e-58 |
| ADN68480.1 | SorE | BGC0000184 | Polyketide:Trans-AT type I | 32.0 | 32.1 | 226.0 | 3.9e-58 |
| BBA84070.1 | type\_I\_polyketide\_synthase | BGC0001916 | Polyketide | 33.0 | 31.3 | 225.0 | 5e-58 |
| QCC63000.1 | BII-rafflesfungin\_polyketide\_synthase | BGC0001966 | NRP | 35.0 | 26.4 | 225.0 | 6.6e-58 |
| ADB23402.1 | polyketide\_synthase\_type\_I | BGC0001062 | Polyketide | 35.0 | 23.4 | 224.0 | 1.1e-57 |
| AJO72734.1 | Type\_I\_modular\_polyketide\_synthase | BGC0001381 | Polyketide | 34.0 | 25.5 | 223.0 | 2.5e-57 |
| AAM54076.1 | polyketide\_synthase | BGC0000020 | Polyketide | 34.0 | 24.8 | 222.0 | 4.3e-57 |
| DAC80076.1 | PKS | BGC0001836 | Polyketide:Trans-AT type I | 31.0 | 28.9 | 222.0 | 7.3e-57 |
| AQZ37114.1 | polyketide\_synthase | BGC0001511 | Polyketide | 34.0 | 24.8 | 221.0 | 9.5e-57 |
| AAN85523.1 | polyketide\_synthase | BGC0001101 | NRP + Polyketide:Modular type I + Polyketide:Trans-AT type I | 34.0 | 25.8 | 221.0 | 1.2e-56 |
| AWH12667.1 | RmpD1 | BGC0001759 | Polyketide | 32.0 | 26.2 | 219.0 | 3.6e-56 |
| DAC76733.1 | type\_I\_polyketide\_synthase | BGC0001885 | Polyketide | 29.0 | 39.3 | 219.0 | 3.6e-56 |
| BAJ09789.1 | polyketide\_synthase | BGC0000146 | Polyketide | 32.0 | 27.3 | 219.0 | 4.7e-56 |
| OEI73463.1 | hypothetical\_protein | BGC0001520 | Polyketide | 30.0 | 31.2 | 219.0 | 4.7e-56 |
| AQW44870.1 | polyketide\_synthase | BGC0001761 | Polyketide | 32.0 | 25.7 | 219.0 | 4.7e-56 |
| BBA20952.1 | type\_I\_polyketide\_synthase | BGC0001763 | NRP + Polyketide | 27.0 | 41.2 | 219.0 | 4.7e-56 |
| ABC34675.1 | polyketide\_synthase,\_putative | BGC0000186 | NRP + Polyketide:Modular type I | 30.0 | 30.1 | 217.0 | 1.4e-55 |
| ADH01484.1 | putative\_type-I\_PKS | BGC0001096 | NRP + Polyketide | 31.0 | 31.3 | 214.0 | 1.5e-54 |
| AIC32692.1 | FR9C | BGC0001113 | NRP + Polyketide | 31.0 | 31.3 | 214.0 | 1.5e-54 |
| ADA82581.1 | trans-AT\_polyketide\_synthase | BGC0001110 | NRP + Polyketide:Trans-AT type I | 32.0 | 26.3 | 213.0 | 2.6e-54 |
| BAL90255.1 | putative\_beta-ketoacyl\_synthase | BGC0002021 | Polyketide | 32.0 | 25.5 | 212.0 | 5.8e-54 |
| CCG06115.1 | hybrid\_NRPS/PKS | BGC0001543 | Polyketide | 30.0 | 32.8 | 210.0 | 1.7e-53 |
| CAL69891.1 | RhiD\_protein | BGC0001112 | NRP + Polyketide:Trans-AT type I | 28.0 | 31.1 | 209.0 | 3.7e-53 |
| AHD05679.1 | putative\_non-ribosomal\_peptide\_ligase/\_polyketide\_synthase\_hybrid | BGC0000402 | NRP | 28.0 | 31.5 | 209.0 | 4.9e-53 |
| ABC35796.1 | polyketide\_synthase,\_putative | BGC0001102 | NRP:Beta-lactam + Polyketide:Modular type I | 31.0 | 30.5 | 208.0 | 6.4e-53 |
| AGN11880.1 | tstC | BGC0001114 | NRP + Polyketide | 30.0 | 31.1 | 208.0 | 8.3e-53 |
| ATY69557.1 | type\_I\_polyketide\_synthase | BGC0001611 | NRP + Polyketide | 33.0 | 22.7 | 208.0 | 8.3e-53 |
| ABS90478.1 | PKS | BGC0001106 | NRP + Polyketide | 30.0 | 34.6 | 207.0 | 1.4e-52 |
| ASA76633.1 | polyketide\_synthase | BGC0001751 | NRP + Polyketide | 30.0 | 31.7 | 203.0 | 2e-51 |
| ALJ49922.1 | TtmG | BGC0001236 | Polyketide | 32.0 | 23.3 | 201.0 | 7.8e-51 |
| BAP05593.1 | calE | BGC0000967 | NRP + Polyketide:Trans-AT type I | 30.0 | 26.3 | 200.0 | 3e-50 |
| OAQ83760.1 | polyketide\_synthase | BGC0001358 | Polyketide | 27.0 | 38.7 | 195.0 | 7.3e-49 |
| ABP57746.1 | DepB | BGC0000993 | NRP:Cyclic depsipeptide + Polyketide:Modular type I | 30.0 | 33.5 | 192.0 | 6.2e-48 |
| ABC34832.1 | polyketide\_synthase | BGC0000186 | NRP + Polyketide:Modular type I | 30.0 | 31.3 | 191.0 | 8.1e-48 |
| CCG06108.1 | type\_I\_polyketide\_synthase | BGC0001543 | Polyketide | 30.0 | 22.4 | 190.0 | 1.8e-47 |
| AFR69332.1 | polyketide\_synthase\_SpiB | BGC0001045 | NRP:Cyclic depsipeptide + Polyketide:Modular type I | 30.0 | 33.8 | 190.0 | 2.3e-47 |
| ATY69600.1 | type\_I\_polyketide\_synthase | BGC0001823 | NRP + Polyketide | 32.0 | 22.9 | 188.0 | 1.2e-46 |
| ABC38737.1 | polyketide\_synthase | BGC0000964 | NRP:Cyclic depsipeptide + Polyketide:Trans-AT type I | 30.0 | 33.4 | 184.0 | 1.3e-45 |
| AAM12934.1 | MmpF | BGC0000182 | Polyketide:Iterative type I + Polyketide:Trans-AT type I | 28.0 | 30.4 | 183.0 | 2.9e-45 |
| ABC35027.1 | JamP | BGC0000961 | NRP + Polyketide | 27.0 | 33.4 | 181.0 | 1.4e-44 |
| BAP05596.1 | calH | BGC0000967 | NRP + Polyketide:Trans-AT type I | 31.0 | 26.3 | 181.0 | 1.4e-44 |
| CAJ76289.1 | putative\_hybrid\_non-ribosomal\_peptide-polyketide\_synthetase | BGC0000972 | NRP + Polyketide:Modular type I + Polyketide:Trans-AT type I | 28.0 | 30.9 | 179.0 | 4.1e-44 |
| BAD55609.1 | putative\_polyketide\_synthase | BGC0001027 | NRP + Polyketide | 31.0 | 21.6 | 179.0 | 5.4e-44 |
| AXA20091.1 | hybrid\_trans-AT\_PKS/NRPS\_LgaB | BGC0001946 | NRP + Polyketide | 28.0 | 33.5 | 178.0 | 1.2e-43 |
| AFL68054.1 | beta-ketoacyl\_synthase\_family\_protein,phosphopantetheine-containing\_protein | BGC0001524 | NRP + Polyketide | 25.0 | 30.9 | 177.0 | 2.1e-43 |
| ALJ49911.1 | TlmG | BGC0001237 | Polyketide | 32.0 | 22.1 | 176.0 | 4.6e-43 |
| ATY69551.1 | hybrid\_nonribosomal\_peptide\_synthetase/type\_I\_polyketide\_synthase | BGC0001611 | NRP + Polyketide | 28.0 | 26.1 | 165.0 | 8.1e-40 |
| WP\_030185025.1 | NAD-dependent\_epimerase/dehydratase\_family\_protein | BGC0001813 | NRP | 29.0 | 26.2 | 164.0 | 1.4e-39 |
| ACB47048.1 | DynE8 | BGC0001060 | Polyketide | 25.0 | 51.9 | 158.0 | 9.9e-38 |
| ABB69082.1 | putative\_pyrrolyl-deta-ketoacyl\_ACP\_synthase | BGC0000260 | Polyketide | 29.0 | 29.0 | 141.0 | 9.6e-33 |
| ctg1\_13 |  | BGC0001931 | Polyketide | 28.0 | 26.1 | 135.0 | 8.9e-31 |
| AFX60321.1 | acyl\_transferase | BGC0001031 | NRP + Polyketide | 28.0 | 21.4 | 120.0 | 2.3e-26 |
| AFX60344.1 | acyl\_transferase | BGC0001032 | NRP + Polyketide | 28.0 | 20.9 | 118.0 | 1.1e-25 |
| AAF86396.1 | FkbA | BGC0000994 | NRP + Polyketide | 27.0 | 26.0 | 113.0 | 3.6e-24 |
| BAQ25481.1 | type\_I\_polyketide\_synthase | BGC0001288 | Polyketide | 28.0 | 21.4 | 113.0 | 4.8e-24 |
| ADB23391.1 | ketosynthase | BGC0001062 | Polyketide | 23.0 | 23.5 | 112.0 | 6.2e-24 |
| ABP73645.1 | SalA | BGC0000145 | Polyketide | 26.0 | 28.9 | 111.0 | 1.1e-23 |
| ABP53498.1 | PKS\_(ACP-AT-AT-KS-ACP-C) | BGC0001041 | NRP + Polyketide | 25.0 | 28.9 | 109.0 | 4e-23 |
| BAJ52681.1 | putative\_ketoacyl\_synthase | BGC0000222 | Polyketide | 24.0 | 24.1 | 104.0 | 1.3e-21 |
| AHA81977.1 | KSα | BGC0000199 | Polyketide:Type II + Saccharide:Hybrid/tailoring | 24.0 | 23.9 | 99.0 | 4.2e-20 |
| ARD70901.1 | beta-ketoacyl\_synthase | BGC0001693 | Polyketide | 23.0 | 23.8 | 98.0 | 9.3e-20 |
| ALJ99855.1 | FlsA | BGC0001904 | Polyketide | 24.0 | 23.5 | 97.0 | 1.6e-19 |
| QDG00826.1 | beta-ketoacyl\_synthase | BGC0002028 | Polyketide | 24.0 | 23.7 | 97.0 | 1.6e-19 |
| CAJ42320.1 | ketoacyl\_synthase | BGC0000273 | Polyketide:Type II + Saccharide:Hybrid/tailoring | 24.0 | 23.8 | 97.0 | 2.7e-19 |
| CAH10117.1 | putative\_ketoacyl\_synthase | BGC0000268 | Polyketide | 23.0 | 23.6 | 96.0 | 6e-19 |
| AAK06784.1 | putative\_ketosynthase\_SimA1 | BGC0001072 | Saccharide + Polyketide:Modular type I + Polyketide:Type II + Other:Aminocoumarin | 23.0 | 23.5 | 95.0 | 7.9e-19 |
| CAE17527.1 | ketosynthase | BGC0000210 | Polyketide:Type II + Saccharide:Oligosaccharide | 23.0 | 23.8 | 94.0 | 1.3e-18 |
| BAJ07842.1 | putative\_ketosynthase | BGC0000232 | Polyketide | 24.0 | 23.8 | 94.0 | 1.8e-18 |
| ACN64834.1 | PokP1 | BGC0001061 | Polyketide:Iterative type I + Polyketide:Type II + Saccharide:Hybrid/tailoring | 23.0 | 23.4 | 91.0 | 1.5e-17 |
| AEI98666.1 | CtcW | BGC0000209 | Polyketide | 23.0 | 23.9 | 90.0 | 3.3e-17 |
| EHM27505.1 | ketoacyl\_synthase | BGC0000235 | Polyketide | 22.0 | 23.9 | 83.0 | 4e-15 |
| AFJ52674.1 | ketoacyl\_synthase\_alpha-subunit | BGC0001073 | NRP + Polyketide | 24.0 | 24.6 | 82.0 | 5.3e-15 |
