## Supplementary Results for "A draft genome of the ascomycotal fungal species *Pseudopithomyces maydicus* (family *Didymosphaeriaceae*)": input.path1.gene26_mibig_hits.html

| MIBiG Protein | Description | MIBiG Cluster | MiBiG Product | % ID | % Coverage | BLAST Score | E-value |
| --- | --- | --- | --- | --- | --- | --- | --- |
| EWG54280.1 | hypothetical\_protein | BGC0001190 | Polyketide | 42.0 | 98.6 | 259.0 | 6.8e-69 |
| EWG54279.1 | hypothetical\_protein | BGC0001190 | Polyketide | 42.0 | 98.6 | 259.0 | 6.8e-69 |
| CAP95407.1 |  | BGC0001404 | Polyketide | 42.0 | 95.4 | 252.0 | 1.1e-66 |
| ALI92656.1 | MRR1\_Major\_Facilitator\_Superfamily\_(MFS)\_protein | BGC0001338 | Polyketide:Iterative type I | 34.0 | 94.6 | 188.0 | 1.9e-47 |
| KFH48706.1 | putative\_MFS-type\_transporter-like\_protein | BGC0000317 | NRP | 32.0 | 99.2 | 175.0 | 1.3e-43 |
| XP\_001827195.1 |  | BGC0001996 | Other | 32.0 | 99.7 | 175.0 | 2.2e-43 |
| AGO86666.1 | putative\_MFS\_transporter | BGC0001255 | NRP + Polyketide | 33.0 | 87.7 | 162.0 | 1.1e-39 |
| XP\_023093494.1 |  | BGC0001995 | Terpene | 30.0 | 98.6 | 155.0 | 2.4e-37 |
| EED18004.1 | conserved\_hypothetical\_protein | BGC0000154 | Polyketide:Iterative type I | 28.0 | 98.6 | 150.0 | 4.4e-36 |
| CBF87867.1 | MFS\_multidrug\_transporter,\_putative\_(AFU\_orthologue;\_AFUA\_1G10370) | BGC0001699 | Polyketide | 29.0 | 94.0 | 150.0 | 5.8e-36 |
| PIB02403.1 | putative\_transporter | BGC0001541 | Polyketide | 29.0 | 97.5 | 147.0 | 4.9e-35 |
| BAK26557.1 | putative\_MFS\_multidrug\_transporter | BGC0000977 | NRP + Polyketide | 28.0 | 95.4 | 143.0 | 7.1e-34 |
| ARU80382.1 | MFS\_transporter | BGC0001542 | Polyketide | 29.0 | 98.1 | 140.0 | 4.6e-33 |
| EAQ86381.1 | hypothetical\_protein | BGC0001405 | Polyketide | 26.0 | 90.7 | 120.0 | 6.4e-27 |
| ASK38705.1 | major\_facilitator\_superfamily\_transporter | BGC0001557 | Polyketide | 26.0 | 99.2 | 114.0 | 3.5e-25 |
| EED57520.1 | efflux\_pump\_antibiotic\_resistance\_protein,\_putative | BGC0001446 | Polyketide:Iterative type I | 33.0 | 67.0 | 109.0 | 1.1e-23 |
| XP\_001827204.1 |  | BGC0001996 | Other | 26.0 | 77.9 | 96.0 | 1e-19 |
