## Supplementary Results for "A draft genome of the ascomycotal fungal species *Pseudopithomyces maydicus* (family *Didymosphaeriaceae*)": input.path1.gene29_mibig_hits.html

| MIBiG Protein | Description | MIBiG Cluster | MiBiG Product | % ID | % Coverage | BLAST Score | E-value |
| --- | --- | --- | --- | --- | --- | --- | --- |
| ACD39774.1 | reducing\_polyketide\_synthase | BGC0000134 | Polyketide | 60.0 | 105.6 | 2767.0 | 0.0 |
| ACD39758.1 | reducing\_polyketide\_synthase | BGC0000076 | Polyketide | 57.0 | 105.5 | 2590.0 | 0.0 |
| ACD39767.1 | reducing\_polyketide\_synthase | BGC0000077 | Polyketide | 57.0 | 105.5 | 2589.0 | 0.0 |
| ABB90283.1 | polyketide\_synthase | BGC0001057 | NRP + Polyketide | 57.0 | 104.9 | 2555.0 | 0.0 |
| AHV78252.1 | ResS1 | BGC0001246 | Polyketide | 54.0 | 105.6 | 2466.0 | 0.0 |
| AHV78245.1 | LasS1 | BGC0001245 | Polyketide | 55.0 | 107.2 | 2460.0 | 0.0 |
| AGC95324.1 | CurS1 | BGC0000045 | Polyketide | 54.0 | 106.1 | 2382.0 | 0.0 |
| KKP00963.1 | fatty\_acid\_synthase\_S-acetyltransferase | BGC0001901 | Polyketide | 34.0 | 107.8 | 1233.0 | 0.0 |
| EWG54266.1 | hypothetical\_protein | BGC0001190 | Polyketide | 33.0 | 106.8 | 1145.0 | 0.0 |
| BAD83684.1 | PKSN\_polyketide\_synthase\_for\_alternapyrone\_biosynthesis | BGC0000012 | Polyketide | 31.0 | 114.5 | 1095.0 | 0.0 |
| BBG28498.1 | putative\_polyketide\_synthase | BGC0001913 | Polyketide | 32.0 | 114.6 | 1094.0 | 0.0 |
| ASK38717.1 | polyketide\_synthase | BGC0001557 | Polyketide | 32.0 | 108.7 | 1056.0 | 5.2e-308 |
| ANF07288.1 | hrPKS | BGC0001340 | Polyketide:Iterative type I | 30.0 | 115.3 | 1053.0 | 4.4e-307 |
| EHA28244.1 | hypothetical\_protein | BGC0001143 | Polyketide | 31.0 | 113.7 | 1051.0 | 1.7e-306 |
| AKL78824.1 | GLPKS3 | BGC0001187 | NRP:Lipopeptide + Polyketide:Iterative type I | 32.0 | 104.9 | 1037.0 | 3.2e-302 |
| EHA19289.1 | hypothetical\_protein | BGC0001124 | Polyketide | 32.0 | 107.4 | 1035.0 | 1.6e-301 |
| ABA02240.1 | polyketide\_synthase | BGC0000098 | Polyketide | 31.0 | 115.1 | 1026.0 | 7.5e-299 |
| AMY15057.1 | tetraketide\_synthase\_MF-SQTKS | BGC0001339 | Polyketide:Iterative type I | 29.0 | 115.2 | 1006.0 | 8e-293 |
| BAC20566.1 | polyketide\_synthase | BGC0000039 | Polyketide | 29.0 | 114.6 | 984.0 | 2.5e-286 |
| AAD34559.1 | polyketide\_synthase | BGC0000088 | Polyketide | 30.0 | 114.5 | 983.0 | 7.2e-286 |
| EAQ86385.1 | hypothetical\_protein | BGC0001405 | Polyketide | 32.0 | 107.5 | 977.0 | 4e-284 |
| CAP95405.1 |  | BGC0001404 | Polyketide | 29.0 | 115.7 | 958.0 | 1.5e-278 |
| EAA65604.1 | hypothetical\_protein | BGC0000022 | Polyketide | 30.0 | 114.2 | 941.0 | 2.4e-273 |
| CBX99534.1 | similar\_to\_polyketide\_synthase | BGC0001899 | Polyketide | 30.0 | 114.2 | 941.0 | 2.4e-273 |
| CBF87072.1 | polyketide\_synthase,\_putative\_(Eurofung) | BGC0001290 | NRP | 30.0 | 111.8 | 939.0 | 9.2e-273 |
| QCC63000.1 | BII-rafflesfungin\_polyketide\_synthase | BGC0001966 | NRP | 28.0 | 116.9 | 899.0 | 1.4e-260 |
| BBG28484.1 | polyketide\_synthase\_CdmE | BGC0001926 | Polyketide | 28.0 | 114.9 | 888.0 | 2.4e-257 |
| EPE34340.1 | polyketide\_synthase | BGC0001035 | Polyketide + NRP + Other:Aminocoumarin | 28.0 | 113.9 | 867.0 | 4.4e-251 |
| ACB12550.1 | Fum1 | BGC0000063 | Polyketide | 28.0 | 114.3 | 858.0 | 2.7e-248 |
| AAD43562.2 | Fum1p | BGC0000062 | Polyketide | 28.0 | 114.7 | 855.0 | 2.3e-247 |
| AIA58899.1 | HRPKS | BGC0001141 | Polyketide:Iterative type I | 29.0 | 108.5 | 851.0 | 2.5e-246 |
| ATZ45185.1 | Bcboa9 | BGC0001892 | Polyketide | 29.0 | 107.2 | 849.0 | 1.6e-245 |
| OAQ83760.1 | polyketide\_synthase | BGC0001358 | Polyketide | 29.0 | 114.5 | 834.0 | 3.2e-241 |
| EAL85129.1 | polyketide\_synthase | BGC0001067 | Terpene + Polyketide:Iterative type I | 29.0 | 110.6 | 832.0 | 1.6e-240 |
| KGO40478.1 | Acyl\_transferase/acyl\_hydrolase/lysophospholipase | BGC0001205 | Polyketide | 29.0 | 108.1 | 830.0 | 7.8e-240 |
| ATQ39432.1 | PKS | BGC0001565 | NRP | 28.0 | 114.5 | 829.0 | 1.7e-239 |
| BAV32159.1 | polyketide\_synthase | BGC0001373 | Polyketide | 28.0 | 109.5 | 800.0 | 6.7e-231 |
| BAJ09789.1 | polyketide\_synthase | BGC0000146 | Polyketide | 28.0 | 118.0 | 799.0 | 1.9e-230 |
| AQM58285.1 | polyketide\_synthase | BGC0001816 | NRP + Polyketide | 27.0 | 113.3 | 797.0 | 5.6e-230 |
| BAN19720.1 | polyketide\_synthase | BGC0001252 | Polyketide | 27.0 | 113.6 | 794.0 | 6.2e-229 |
| EAA36364.1 | hypothetical\_protein | BGC0001697 | Polyketide | 29.0 | 108.2 | 792.0 | 2.4e-228 |
| ACZ57548.1 | polyketide\_synthase | BGC0000046 | Polyketide:Iterative type I | 29.0 | 106.6 | 787.0 | 5.8e-227 |
| CAQ18830.1 | polyketide\_synthase | BGC0000954 | NRP + Polyketide:Modular type I | 30.0 | 104.3 | 780.0 | 7.1e-225 |
| AMY15068.1 | hexaketide\_synthase\_MF-SQHKS | BGC0001339 | Polyketide:Iterative type I | 27.0 | 112.7 | 774.0 | 6.7e-223 |
| CAJ46690.1 | polyketide\_synthase | BGC0000969 | NRP:Cyclic depsipeptide + Polyketide:Modular type I | 29.0 | 103.8 | 772.0 | 1.5e-222 |
| OAQ83765.1 | KR\_domain-containing\_protein | BGC0001358 | Polyketide | 28.0 | 109.0 | 769.0 | 1.6e-221 |
| AGC45624.1 | polyketide\_synthase | BGC0001394 | NRP + Polyketide | 29.0 | 102.7 | 764.0 | 6.9e-220 |
| AUS29495.1 | polyketide\_synthase | BGC0001030 | NRP + Polyketide | 27.0 | 114.5 | 760.0 | 7.6e-219 |
| AQW44889.1 | polyketide\_synthase | BGC0001737 | NRP + Polyketide | 29.0 | 102.1 | 757.0 | 4.9e-218 |
| CCT75967.1 | polyketide\_synthase | BGC0001606 | Polyketide | 26.0 | 116.0 | 754.0 | 4.2e-217 |
| CCE88376.1 | polyketide\_synthase | BGC0001034 | NRP + Polyketide:Modular type I | 29.0 | 102.5 | 731.0 | 5e-210 |
| QDA77058.1 | polyketide\_synthase | BGC0002026 | NRP | 29.0 | 102.8 | 730.0 | 8.5e-210 |
| CCE88378.1 | polyketide\_synthase | BGC0001034 | NRP + Polyketide:Modular type I | 28.0 | 102.5 | 728.0 | 3.2e-209 |
| CAQ43075.1 | polyketide\_synthase | BGC0000970 | NRP + Polyketide:Modular type I | 28.0 | 105.7 | 715.0 | 3.7e-205 |
| CAQ18828.1 | polyketide\_synthase | BGC0000954 | NRP + Polyketide:Modular type I | 29.0 | 103.5 | 709.0 | 1.5e-203 |
| RWQ92174.1 | KR\_domain-containing\_protein | BGC0002030 | Polyketide | 27.0 | 105.2 | 703.0 | 1.1e-201 |
| ACR33077.1 | polyketide\_synthase | BGC0000017 | Alkaloid + Polyketide:Modular type I | 27.0 | 103.2 | 694.0 | 5.1e-199 |
| AAF26923.1 | polyketide\_synthase | BGC0000988 | NRP + Polyketide | 28.0 | 103.3 | 694.0 | 5.1e-199 |
| ADB12493.1 | EpoF | BGC0000990 | NRP + Polyketide | 28.0 | 103.2 | 689.0 | 1.7e-197 |
| AAF62885.1 | EpoF | BGC0000991 | NRP + Polyketide | 28.0 | 103.2 | 685.0 | 3.1e-196 |
| ACB46197.1 | polyketide\_synthase | BGC0000989 | NRP + Polyketide | 28.0 | 103.5 | 680.0 | 1e-194 |
| AZH23789.1 | MgcI | BGC0001970 | NRP + Polyketide | 28.0 | 103.3 | 680.0 | 1.3e-194 |
| AZH23818.1 | MgiI | BGC0001971 | NRP + Polyketide | 27.0 | 103.3 | 676.0 | 1.1e-193 |
| EHA52508.1 | mycocerosic\_acid\_synthase | BGC0001749 | Polyketide | 28.0 | 112.4 | 676.0 | 1.9e-193 |
| CAQ18832.1 | polyketide\_synthase | BGC0000954 | NRP + Polyketide:Modular type I | 29.0 | 96.5 | 664.0 | 7.4e-190 |
| AQA28562.1 | type\_I\_polyketide\_synthase | BGC0001663 | Polyketide | 26.0 | 108.2 | 656.0 | 1.6e-187 |
| WP\_042799407.1 | type\_I\_polyketide\_synthase | BGC0001283 | Polyketide | 28.0 | 105.3 | 654.0 | 4.5e-187 |
| CAD19090.1 | StiF\_protein | BGC0000153 | NRP + Polyketide:Modular type I | 28.0 | 96.4 | 652.0 | 2.2e-186 |
| AAS98783.1 | polyketide\_synthase/nonribosomal\_peptide\_synthase\_hybrid | BGC0001001 | NRP + Polyketide | 27.0 | 102.5 | 648.0 | 3.2e-185 |
| DAB41915.1 | ArzM\_-\_PKS\_(KS,\_AT,\_DH,\_MT,\_ER,\_KR,\_ACP) | BGC0001884 | NRP + Polyketide | 26.0 | 114.4 | 639.0 | 2.6e-182 |
| AIW82279.1 | PuwB | BGC0001125 | NRP + Polyketide | 26.0 | 107.4 | 631.0 | 4.1e-180 |
| AEE88282.1 | CurH | BGC0000976 | NRP + Polyketide:Modular type I | 27.0 | 96.3 | 629.0 | 2.7e-179 |
| AAT70103.1 | CurH | BGC0001165 | NRP + Polyketide:Modular type I | 27.0 | 96.3 | 629.0 | 2.7e-179 |
| BAD97694.1 | Aft9-1 | BGC0000003 | Polyketide | 26.0 | 108.4 | 628.0 | 3.5e-179 |
| ACB37755.1 | putative\_type\_I\_polyketide\_synthase | BGC0000162 | Polyketide | 28.0 | 99.4 | 626.0 | 1.7e-178 |
| ABI94379.1 | tautomycetin\_biosynthetic\_PKS | BGC0000157 | Polyketide | 27.0 | 94.6 | 622.0 | 3.2e-177 |
| ABV91286.1 | type\_I\_modular\_polyketide\_synthase | BGC0000158 | Polyketide:Modular type I | 27.0 | 94.6 | 621.0 | 5.5e-177 |
| AXN93577.1 | PuwB | BGC0001950 | NRP | 25.0 | 115.1 | 617.0 | 1e-175 |
| AAT28740.1 | FUSS | BGC0000064 | Polyketide | 27.0 | 106.1 | 614.0 | 5.2e-175 |
| AAZ77696.1 | ChlA3 | BGC0000036 | Polyketide:Modular type I + Polyketide:Iterative type I + Saccharide:Oligosaccharide | 27.0 | 102.3 | 614.0 | 6.8e-175 |
| AXN93586.1 | PuwB | BGC0001951 | NRP | 25.0 | 115.1 | 614.0 | 6.8e-175 |
| AZH23788.1 | MgcR | BGC0001970 | NRP + Polyketide | 27.0 | 95.8 | 614.0 | 8.8e-175 |
| AFP73394.1 | FusA | BGC0001268 | NRP + Polyketide | 27.0 | 106.9 | 612.0 | 3.4e-174 |
| BBA66513.1 | type\_I\_polyketide\_synthase | BGC0001495 | Polyketide | 28.0 | 101.1 | 612.0 | 3.4e-174 |
| AEE88279.1 | CurK | BGC0000976 | NRP + Polyketide:Modular type I | 27.0 | 97.6 | 611.0 | 7.5e-174 |
| AAT70106.1 | CurK | BGC0001165 | NRP + Polyketide:Modular type I | 27.0 | 97.6 | 611.0 | 7.5e-174 |
| AKD43522.1 | Type\_I\_polyketide\_synthase | BGC0001409 | Polyketide | 27.0 | 104.9 | 610.0 | 1.3e-173 |
| ABW96541.1 | type\_I\_modular\_polyketide\_synthase | BGC0000159 | Polyketide:Modular type I | 27.0 | 95.4 | 609.0 | 2.8e-173 |
| TXD00025.1 | SDR\_family\_NAD(P)-dependent\_oxidoreductase | BGC0001877 | Polyketide | 28.0 | 99.0 | 602.0 | 2e-171 |
| ANC94964.1 | AlmHIII | BGC0001396 | Polyketide | 27.0 | 104.6 | 600.0 | 1.3e-170 |
| AHH99921.1 | PKS\_I | BGC0000002 | Polyketide | 26.0 | 101.9 | 599.0 | 2.9e-170 |
| AAG23265.1 | polyketide\_synthase\_extender\_module\_2 | BGC0000148 | Polyketide | 28.0 | 92.1 | 597.0 | 1.1e-169 |
| BAG85026.1 | putative\_polyketide\_synthase | BGC0000086 | Polyketide | 26.0 | 102.2 | 596.0 | 1.9e-169 |
| CAQ64686.1 | lasalocid\_modular\_polyketide\_synthase | BGC0000087 | Polyketide | 26.0 | 102.2 | 596.0 | 1.9e-169 |
| ABI94380.1 | tautomycetin\_biosynthetic\_PKS | BGC0000157 | Polyketide | 27.0 | 94.9 | 596.0 | 1.9e-169 |
| ABA02239.1 | polyketide\_synthase | BGC0000098 | Polyketide | 26.0 | 113.4 | 594.0 | 7.2e-169 |
| BAC20564.1 | polyketide\_synthase | BGC0000039 | Polyketide | 25.0 | 114.3 | 591.0 | 6.1e-168 |
| ACB46471.1 | polyketide\_synthase | BGC0000082 | Polyketide | 27.0 | 99.5 | 590.0 | 1.4e-167 |
| ABV91287.1 | type\_I\_modular\_polyketide\_synthase | BGC0000158 | Polyketide:Modular type I | 27.0 | 94.9 | 590.0 | 1.4e-167 |
| AJW65409.1 | type\_I\_modular\_polyketide\_synthase | BGC0001195 | NRP + Polyketide | 28.0 | 101.1 | 589.0 | 1.8e-167 |
| CAA60462.1 | polyketide\_synthase | BGC0001040 | NRP + Polyketide | 28.0 | 94.4 | 589.0 | 2.3e-167 |
| BAF85843.1 | modular\_polyketide\_synthase | BGC0000109 | Polyketide | 27.0 | 101.8 | 584.0 | 9.8e-166 |
| AVV61984.1 | type\_I\_modular\_polyketide\_synthase | BGC0001477 | NRP + Polyketide:Modular type I | 28.0 | 99.7 | 582.0 | 3.7e-165 |
| QDA77044.1 | polyketide\_synthase | BGC0002025 | NRP | 26.0 | 101.1 | 582.0 | 3.7e-165 |
| AQW44890.1 | polyketide\_synthase | BGC0001737 | NRP + Polyketide | 27.0 | 102.1 | 581.0 | 6.3e-165 |
| AAG13918.1 | megalomicin\_6-deoxyerythronolide\_B\_synthase\_2 | BGC0000092 | Polyketide | 27.0 | 95.7 | 580.0 | 1.1e-164 |
| CAO91861.1 | PKS-NRPS\_hybrid | BGC0000968 | NRP + Polyketide:Iterative type I | 26.0 | 112.4 | 579.0 | 1.8e-164 |
| AAS98781.1 | polyketide\_synthase | BGC0001001 | NRP + Polyketide | 25.0 | 109.5 | 579.0 | 2.4e-164 |
| CAA60460.1 | polyketide\_synthase | BGC0001040 | NRP + Polyketide | 27.0 | 101.6 | 579.0 | 2.4e-164 |
| ABY21540.1 | AngAIII | BGC0000018 | Polyketide | 28.0 | 103.0 | 578.0 | 5.4e-164 |
| BAO66519.1 | type\_I\_polyketide\_synthase | BGC0000042 | Polyketide | 26.0 | 102.7 | 578.0 | 5.4e-164 |
| AWW87422.1 | type\_I\_polyketide\_synthase | BGC0001755 | Polyketide | 27.0 | 95.6 | 574.0 | 7.8e-163 |
| AGC45619.1 | polyketide\_synthase | BGC0001394 | NRP + Polyketide | 27.0 | 102.6 | 572.0 | 3.9e-162 |
| CAG28678.1 | polyketide\_synthase | BGC0001023 | NRP + Polyketide:Modular type I | 27.0 | 102.8 | 570.0 | 1.1e-161 |
| APZ78820.1 | polyketide\_synthase | BGC0001429 | NRP:Cyclic depsipeptide + Polyketide:Iterative type I | 27.0 | 102.8 | 570.0 | 1.1e-161 |
| BAQ25512.1 | type\_I\_polyketide\_synthase | BGC0001288 | Polyketide | 27.0 | 101.1 | 569.0 | 1.9e-161 |
| QBF51757.1 | type\_I\_polyketide\_synthase | BGC0001856 | Polyketide:Modular type I | 26.0 | 101.5 | 569.0 | 2.5e-161 |
| APZ78844.1 | polyketide\_synthase | BGC0001431 | NRP:Cyclic depsipeptide + Polyketide:Iterative type I | 26.0 | 103.6 | 563.0 | 1.4e-159 |
| BAB69194.1 | modular\_polyketide\_synthase | BGC0000117 | Polyketide | 28.0 | 92.3 | 557.0 | 9.8e-158 |
| AAQ82567.1 | FscE | BGC0000061 | Polyketide | 27.0 | 101.9 | 552.0 | 3.2e-156 |
| CAJ88186.1 | putative\_modular\_polyketide\_synthase | BGC0000151 | Polyketide:Modular type I + Saccharide:Hybrid/tailoring | 27.0 | 93.5 | 551.0 | 5.4e-156 |
| ACC40921.1 | polyketide\_synthase\_Pks7 | BGC0001665 | Polyketide | 26.0 | 101.6 | 547.0 | 7.8e-155 |
| AEC13079.1 | fosA | BGC0000060 | Polyketide | 27.0 | 102.3 | 545.0 | 3.9e-154 |
| CAJ88187.2 | Type\_I\_modular\_polyketide\_synthase | BGC0000151 | Polyketide:Modular type I + Saccharide:Hybrid/tailoring | 26.0 | 103.0 | 545.0 | 3.9e-154 |
| ACF35445.1 | mbcAI | BGC0000090 | Polyketide | 28.0 | 99.5 | 539.0 | 3.6e-152 |
| ADC45586.1 | modular\_polyketide\_synthase | BGC0000093 | Polyketide | 27.0 | 101.5 | 537.0 | 1.1e-151 |
| AHH99926.1 | PKS\_I | BGC0000002 | Polyketide | 26.0 | 94.2 | 537.0 | 1.4e-151 |
| CAJ88176.1 | putative\_polyketide\_synthase\_B | BGC0000151 | Polyketide:Modular type I + Saccharide:Hybrid/tailoring | 28.0 | 93.2 | 531.0 | 7.5e-150 |
| ATY12793.1 | type\_I\_polyketide\_synthase | BGC0001504 | Polyketide | 27.0 | 100.0 | 531.0 | 7.5e-150 |
| ACS68554.1 | hybrid\_PKS-NRPS\_protein | BGC0001026 | NRP + Polyketide | 42.0 | 33.2 | 528.0 | 6.4e-149 |
| AFU82616.1 | polyketide\_synthase | BGC0000998 | NRP + Polyketide | 26.0 | 95.1 | 527.0 | 1.4e-148 |
| ACY06286.1 | polyketide\_synthase | BGC0001042 | NRP + Polyketide | 26.0 | 101.4 | 527.0 | 1.4e-148 |
| BAQ21939.1 | putative\_type\_I\_polyketide\_synthase | BGC0001204 | Polyketide | 26.0 | 92.2 | 523.0 | 1.6e-147 |
| ctg1\_orf255 |  | BGC0001200 | Polyketide | 27.0 | 101.0 | 522.0 | 3.5e-147 |
| QBC19710.1 | TwmB | BGC0001954 | NRP + Polyketide | 33.0 | 52.5 | 514.0 | 9.5e-145 |
| AFA26384.1 | polyketide\_synthase\_A | BGC0001874 | NRP + Polyketide | 42.0 | 33.5 | 512.0 | 4.7e-144 |
| gene4 |  | BGC0001907 | Polyketide | 25.0 | 103.8 | 512.0 | 4.7e-144 |
| ACF35447.1 | mbcAIII | BGC0000090 | Polyketide | 27.0 | 94.0 | 510.0 | 1.4e-143 |
| BAK26562.1 | PKS-NRPS\_hybrid | BGC0000977 | NRP + Polyketide | 42.0 | 33.8 | 499.0 | 3.2e-140 |
| BAZ95823.1 | PKS-NRPS\_hybrid\_cpaA | BGC0001563 | NRP + Polyketide | 38.0 | 38.1 | 495.0 | 3.5e-139 |
| AIT55263.1 | polyketide\_synthase | BGC0000072 | Polyketide:Modular type I | 26.0 | 102.8 | 494.0 | 7.8e-139 |
| ctg1\_orf254 |  | BGC0001200 | Polyketide | 26.0 | 100.0 | 493.0 | 1.3e-138 |
| AEO57481.1 | PKS-NRPSs | BGC0001449 | NRP + Alkaloid + Polyketide:Iterative type I | 41.0 | 33.1 | 493.0 | 2.3e-138 |
| AEE88280.1 | CurJ | BGC0000976 | NRP + Polyketide:Modular type I | 25.0 | 97.4 | 492.0 | 3e-138 |
| AAT70105.1 | CurJ | BGC0001165 | NRP + Polyketide:Modular type I | 25.0 | 97.4 | 492.0 | 3e-138 |
| EPS29069.1 | hypothetical\_protein | BGC0001724 | NRP + Polyketide | 40.0 | 33.4 | 491.0 | 6.6e-138 |
| EAU38971.1 | hypothetical\_protein | BGC0001122 | NRP + Polyketide:Iterative type I | 41.0 | 33.4 | 489.0 | 2.5e-137 |
| AGO86662.1 | equisetin\_synthetase | BGC0001255 | NRP + Polyketide | 42.0 | 33.2 | 485.0 | 6.2e-136 |
| BBC43184.1 | PKS-NRPS\_hybrid | BGC0001738 | NRP + Polyketide | 41.0 | 33.2 | 485.0 | 6.2e-136 |
| AAV66110.2 | fusaridione\_A\_synthetase | BGC0000992 | NRP + Polyketide | 41.0 | 33.6 | 483.0 | 1.8e-135 |
| ARP51711.1 | PKS-NRPS\_hybrid\_protein | BGC0001741 | NRP + Polyketide | 41.0 | 34.0 | 483.0 | 2.4e-135 |
| CBF80487.1 | hybrid\_PKS-NRPS\_(Eurofung) | BGC0000959 | NRP + Polyketide:Iterative type I | 41.0 | 33.6 | 482.0 | 3.1e-135 |
| EAW09117.1 | hybrid\_NRPS/PKS\_enzyme,\_putative | BGC0000983 | NRP + Polyketide:Iterative type I | 41.0 | 32.8 | 480.0 | 2e-134 |
| CEF75886.1 |  | BGC0001600 | Polyketide | 39.0 | 33.9 | 480.0 | 2e-134 |
| AHA38200.1 | GphG | BGC0000069 | Polyketide | 27.0 | 105.9 | 479.0 | 2.6e-134 |
| BAJ14522.1 | polyketide\_synthase | BGC0001254 | Polyketide | 41.0 | 33.7 | 478.0 | 5.8e-134 |
| EED49862.1 | hybrid\_PKS/NRPS\_enzyme,\_putative | BGC0001445 | NRP + Polyketide:Iterative type I | 41.0 | 32.9 | 478.0 | 7.6e-134 |
| CAO98852.1 | polyketide\_synthase\_AufI | BGC0000023 | Polyketide:Modular type I | 24.0 | 116.2 | 477.0 | 9.9e-134 |
| ACR50791.1 | putative\_polyketide\_synthase | BGC0000163 | Polyketide | 26.0 | 100.7 | 475.0 | 6.4e-133 |
| AQW44888.1 | polyketide\_synthase | BGC0001737 | NRP + Polyketide | 33.0 | 50.8 | 474.0 | 1.1e-132 |
| CCT72377.1 | probable\_polyketide\_synthase | BGC0001305 | Polyketide | 40.0 | 33.8 | 473.0 | 1.9e-132 |
| ATZ45182.1 | Bcboa6 | BGC0001892 | Polyketide | 40.0 | 33.8 | 473.0 | 1.9e-132 |
| orf3 | polyketide\_synthase | BGC0001432 | NRP:Cyclic depsipeptide + Polyketide:Iterative type I | 26.0 | 103.6 | 470.0 | 2.1e-131 |
| APZ78858.1 | polyketide\_synthase | BGC0001432 | NRP:Cyclic depsipeptide + Polyketide:Iterative type I | 26.0 | 103.6 | 470.0 | 2.1e-131 |
| QBE85649.1 | BuaA | BGC0001978 | NRP + Polyketide | 40.0 | 33.3 | 469.0 | 3.5e-131 |
| ADN43685.1 | DmbS | BGC0001136 | NRP + Polyketide:Iterative type I | 40.0 | 33.9 | 468.0 | 7.8e-131 |
| CAL69597.1 | PKS-NRPS | BGC0001049 | NRP + Polyketide:Iterative type I | 40.0 | 33.6 | 467.0 | 1e-130 |
| AGC45622.1 | polyketide\_synthase | BGC0001394 | NRP + Polyketide | 30.0 | 52.2 | 467.0 | 1.3e-130 |
| EFL02193.1 | amino\_acid\_adenylation\_domain-containing\_protein | BGC0000996 | NRP + Polyketide:Iterative type I | 31.0 | 54.9 | 466.0 | 2.3e-130 |
| gene3 |  | BGC0002035 | NRP + Polyketide | 39.0 | 33.1 | 466.0 | 3e-130 |
| XP\_659388.1 | hypothetical\_protein | BGC0001998 | Polyketide | 39.0 | 34.2 | 462.0 | 5.6e-129 |
| QCS37521.1 | PyiS | BGC0001982 | NRP + Polyketide | 39.0 | 33.0 | 460.0 | 1.3e-128 |
| AKC54422.1 | fumosorinone\_biosynthesis\_polyketide\_synthase | BGC0001218 | NRP + Polyketide | 39.0 | 34.3 | 459.0 | 2.8e-128 |
| BAQ25466.1 | polyketide\_synthase | BGC0001280 | Polyketide | 39.0 | 33.6 | 459.0 | 3.6e-128 |
| AQW44893.1 | polyketide\_synthase | BGC0001737 | NRP + Polyketide | 30.0 | 51.0 | 457.0 | 1.1e-127 |
| CAD19085.1 | StiA\_protein | BGC0000153 | NRP + Polyketide:Modular type I | 25.0 | 105.7 | 456.0 | 3.1e-127 |
| AAF26922.1 | polyketide\_synthase | BGC0000988 | NRP + Polyketide | 25.0 | 103.4 | 453.0 | 1.5e-126 |
| ctg1\_orf15 |  | BGC0001457 | NRP | 39.0 | 31.8 | 452.0 | 4.4e-126 |
| AGC45620.1 | polyketide\_synthase | BGC0001394 | NRP + Polyketide | 38.0 | 33.3 | 451.0 | 7.6e-126 |
| AAK57189.1 | MxaE | BGC0001022 | NRP + Polyketide | 32.0 | 45.7 | 449.0 | 2.9e-125 |
| AAW03328.1 | CtaE | BGC0000982 | NRP + Polyketide | 37.0 | 32.0 | 448.0 | 8.4e-125 |
| AAF19813.1 | MtaE | BGC0001024 | NRP + Polyketide:Modular type I | 37.0 | 32.2 | 446.0 | 3.2e-124 |
| BAG17643.1 | putative\_NRPS-type-I\_PKS\_fusion\_protein | BGC0001043 | NRP + Polyketide | 32.0 | 50.0 | 445.0 | 5.4e-124 |
| AAF19814.1 | MtaF | BGC0001024 | NRP + Polyketide:Modular type I | 39.0 | 32.9 | 445.0 | 7.1e-124 |
| CAD19086.1 | StiB\_protein | BGC0000153 | NRP + Polyketide:Modular type I | 38.0 | 32.6 | 443.0 | 1.6e-123 |
| APD26279.1 | PtmA | BGC0001726 | NRP + Polyketide | 32.0 | 51.9 | 443.0 | 1.6e-123 |
| ADA69241.1 | cis-AT\_polyketide\_synthase | BGC0001071 | NRP + Polyketide:Modular type I + Polyketide:Trans-AT type I | 25.0 | 84.1 | 443.0 | 2.1e-123 |
| AQW44873.1 | polyketide\_synthase | BGC0001761 | Polyketide | 30.0 | 54.2 | 443.0 | 2.1e-123 |
| ADZ24998.1 | polyketide\_synthase | BGC0000380 | NRP + Polyketide:Modular type I | 25.0 | 104.2 | 443.0 | 2.7e-123 |
| AAF19810.1 | MtaB | BGC0001024 | NRP + Polyketide:Modular type I | 26.0 | 101.2 | 442.0 | 4.6e-123 |
| AAK57187.1 | MxaC | BGC0001022 | NRP + Polyketide | 39.0 | 32.2 | 436.0 | 1.9e-121 |
| CAO98850.1 | polyketide\_synthase\_AufG | BGC0000023 | Polyketide:Modular type I | 37.0 | 32.9 | 434.0 | 1.3e-120 |
| CAD89776.1 | MelE\_protein | BGC0001010 | NRP + Polyketide:Modular type I | 36.0 | 32.0 | 432.0 | 3.6e-120 |
| AHA12078.1 | polyketide\_synthase\_type\_1 | BGC0001172 | NRP + Polyketide:Modular type I | 38.0 | 32.8 | 432.0 | 4.8e-120 |
| CAQ18829.1 | polyketide\_synthase | BGC0000954 | NRP + Polyketide:Modular type I | 37.0 | 31.9 | 431.0 | 6.2e-120 |
| APZ78832.1 | polyketide\_synthase | BGC0001430 | NRP:Cyclic depsipeptide + Polyketide:Iterative type I | 38.0 | 32.9 | 429.0 | 3.1e-119 |
| AJD77023.1 | IkaA | BGC0001435 | NRP + Polyketide:Iterative type I | 32.0 | 49.1 | 429.0 | 4e-119 |
| BAV56012.1 | PKS\_(KS-AT-DH-ER-KR-ACP-TE) | BGC0001597 | Polyketide | 25.0 | 104.7 | 429.0 | 4e-119 |
| AGC45623.1 | polyketide\_synthase | BGC0001394 | NRP + Polyketide | 30.0 | 52.4 | 428.0 | 5.3e-119 |
| CAO98847.1 | polyketide\_synthase\_AufC | BGC0000023 | Polyketide:Modular type I | 37.0 | 32.7 | 428.0 | 6.9e-119 |
| CBD77732.1 | polyketide\_synthase | BGC0000974 | NRP + Polyketide | 38.0 | 32.2 | 428.0 | 9e-119 |
| ABM21570.1 | crpB | BGC0000975 | NRP + Polyketide | 36.0 | 32.2 | 428.0 | 9e-119 |
| AEU11005.1 | NpnA | BGC0001029 | NRP + Polyketide | 37.0 | 31.4 | 428.0 | 9e-119 |
| AIR74910.1 | polyketide\_synthase | BGC0001559 | RiPP | 38.0 | 32.2 | 428.0 | 9e-119 |
| APZ78702.1 | polyketide\_synthase | BGC0001419 | NRP:Cyclic depsipeptide + Polyketide:Iterative type I | 38.0 | 33.0 | 427.0 | 1.2e-118 |
| AQM37582.1 | polyketide\_synthase | BGC0001424 | NRP:Cyclic depsipeptide + Polyketide:Iterative type I | 37.0 | 32.7 | 427.0 | 1.2e-118 |
| CCE88377.1 | non-ribosomal\_peptide\_synthetase/polyketide\_synthase | BGC0001034 | NRP + Polyketide:Modular type I | 25.0 | 107.1 | 426.0 | 2e-118 |
| APZ78754.1 | polyketide\_synthase | BGC0001423 | NRP:Cyclic depsipeptide + Polyketide:Iterative type I | 36.0 | 32.7 | 426.0 | 3.4e-118 |
| CAF05649.1 | TubD\_protein | BGC0001053 | NRP + Polyketide | 25.0 | 91.4 | 425.0 | 5.8e-118 |
| APZ78714.1 | polyketide\_synthase | BGC0001420 | NRP:Cyclic depsipeptide + Polyketide:Iterative type I | 37.0 | 33.0 | 425.0 | 5.8e-118 |
| APZ78854.1 | polyketide\_synthase | BGC0001432 | NRP:Cyclic depsipeptide + Polyketide:Iterative type I | 38.0 | 33.0 | 425.0 | 5.8e-118 |
| DAB41916.1 | ArzN\_-\_PKS\_(KS,\_AT,\_OMT,\_KR,\_ACP) | BGC0001884 | NRP + Polyketide | 37.0 | 32.5 | 425.0 | 5.8e-118 |
| CAQ18838.1 | polyketide\_synthase | BGC0000954 | NRP + Polyketide:Modular type I | 39.0 | 32.5 | 425.0 | 7.6e-118 |
| CAQ18833.1 | polyketide\_synthase | BGC0000954 | NRP + Polyketide:Modular type I | 37.0 | 32.5 | 424.0 | 9.9e-118 |
| APZ78678.1 | polyketide\_synthase | BGC0001417 | NRP:Cyclic depsipeptide + Polyketide:Iterative type I | 37.0 | 33.1 | 424.0 | 9.9e-118 |
| APZ78690.1 | polyketide\_synthase | BGC0001418 | NRP:Cyclic depsipeptide + Polyketide:Iterative type I | 37.0 | 33.0 | 424.0 | 9.9e-118 |
| CAD19089.1 | StiE\_protein | BGC0000153 | NRP + Polyketide:Modular type I | 36.0 | 32.1 | 424.0 | 1.3e-117 |
| CBD77748.1 | polyketide\_synthase | BGC0000974 | NRP + Polyketide | 38.0 | 32.2 | 424.0 | 1.3e-117 |
| APZ78727.1 | polyketide\_synthase | BGC0001421 | NRP:Cyclic depsipeptide + Polyketide:Iterative type I | 36.0 | 32.7 | 424.0 | 1.3e-117 |
| AIR74917.1 | polyketide\_synthase | BGC0001559 | RiPP | 38.0 | 32.2 | 424.0 | 1.3e-117 |
| BAK64649.1 | polyketide\_synthase | BGC0000135 | Polyketide | 30.0 | 50.9 | 423.0 | 2.2e-117 |
| APZ78793.1 | polyketide\_synthase | BGC0001427 | NRP:Cyclic depsipeptide + Polyketide:Iterative type I | 37.0 | 32.5 | 423.0 | 2.2e-117 |
| CQR60495.1 | Polyketide\_synthase,\_type\_I,\_module\_7 | BGC0001287 | Polyketide | 30.0 | 50.8 | 423.0 | 2.9e-117 |
| AHB82064.1 | polyketide\_synthase | BGC0001231 | NRP + Polyketide:Modular type I | 36.0 | 32.4 | 422.0 | 3.8e-117 |
| AAZ77673.1 | ChlB1 | BGC0000036 | Polyketide:Modular type I + Polyketide:Iterative type I + Saccharide:Oligosaccharide | 38.0 | 32.7 | 422.0 | 4.9e-117 |
| AIT55259.1 | polyketide\_synthase | BGC0000072 | Polyketide:Modular type I | 38.0 | 32.5 | 421.0 | 6.4e-117 |
| AAK57186.1 | MxaB2 | BGC0001022 | NRP + Polyketide | 36.0 | 32.4 | 421.0 | 8.4e-117 |
| AEU11006.1 | NpnB | BGC0001029 | NRP + Polyketide | 37.0 | 32.5 | 421.0 | 8.4e-117 |
| APZ78780.1 | polyketide\_synthase | BGC0001426 | NRP:Cyclic depsipeptide + Polyketide:Iterative type I | 37.0 | 32.5 | 421.0 | 8.4e-117 |
| APZ78807.1 | polyketide\_synthase | BGC0001428 | NRP:Cyclic depsipeptide + Polyketide:Iterative type I | 37.0 | 32.5 | 421.0 | 8.4e-117 |
| AHH34186.1 | polyketide\_synthase | BGC0001161 | Polyketide:Modular type I | 37.0 | 33.2 | 421.0 | 1.1e-116 |
| ADN13832.1 | Polyketide\_Synthase | BGC0001164 | Polyketide:Modular type I | 36.0 | 31.3 | 421.0 | 1.1e-116 |
| CAJ88177.1 | putative\_type\_I\_polyketide\_synthase | BGC0000151 | Polyketide:Modular type I + Saccharide:Hybrid/tailoring | 31.0 | 49.8 | 420.0 | 1.9e-116 |
| AZH23817.1 | MgiQ | BGC0001971 | NRP + Polyketide | 24.0 | 97.4 | 420.0 | 1.9e-116 |
| AIT55262.1 | polyketide\_synthase | BGC0000072 | Polyketide:Modular type I | 37.0 | 32.1 | 420.0 | 2.4e-116 |
| AID65222.1 | putative\_aspartate\_racemase | BGC0000335 | NRP | 38.0 | 32.3 | 420.0 | 2.4e-116 |
| ACY13414.1 | amino\_acid\_adenylation\_domain\_protein | BGC0001367 | NRP + Polyketide | 29.0 | 52.0 | 419.0 | 3.2e-116 |
| CAD19087.1 | StiC\_protein | BGC0000153 | NRP + Polyketide:Modular type I | 38.0 | 32.5 | 419.0 | 4.2e-116 |
| AZH23821.1 | MgiH | BGC0001971 | NRP + Polyketide | 36.0 | 32.9 | 419.0 | 4.2e-116 |
| AAQ90173.1 | polyketide\_synthase\_type\_I | BGC0000128 | Polyketide | 37.0 | 32.2 | 418.0 | 5.4e-116 |
| ABY66019.1 | 6-methylsalicylic\_acid\_synthase | BGC0001008 | Polyketide:Iterative type I + Polyketide:Enediyne type I | 37.0 | 32.5 | 418.0 | 5.4e-116 |
| ABX60163.1 | polyketide\_synthase | BGC0000978 | NRP + Alkaloid + Polyketide:Modular type I | 35.0 | 35.2 | 418.0 | 7.1e-116 |
| AFV96138.1 | polyketide\_synthase | BGC0001064 | Polyketide:Modular type I + Polyketide:Type III | 32.0 | 39.0 | 418.0 | 7.1e-116 |
| AHA12079.1 | polyketide\_synthase\_type\_1 | BGC0001172 | NRP + Polyketide:Modular type I | 36.0 | 32.0 | 418.0 | 7.1e-116 |
| ARU81118.1 | CylD | BGC0001566 | Polyketide | 32.0 | 39.0 | 418.0 | 7.1e-116 |
| AIT55264.1 | polyketide\_synthase | BGC0000072 | Polyketide:Modular type I | 37.0 | 32.5 | 418.0 | 9.3e-116 |
| AAK19883.1 | soraphen\_polyketide\_synthase\_A | BGC0000147 | Polyketide:Modular type I | 36.0 | 33.0 | 418.0 | 9.3e-116 |
| APZ78742.1 | polyketide\_synthase | BGC0001422 | NRP:Cyclic depsipeptide + Polyketide:Iterative type I | 37.0 | 32.8 | 418.0 | 9.3e-116 |
| ADF88277.1 | polyketide\_synthase | BGC0000981 | NRP + Polyketide | 35.0 | 35.5 | 417.0 | 1.2e-115 |
| BAV56006.1 | PKS\_(ACP-KS-AT-DH-ER-KR-ACP-KS-AT-KR-ACP) | BGC0001597 | Polyketide | 24.0 | 103.8 | 417.0 | 1.2e-115 |
| CAQ43076.1 | polyketide\_synthase | BGC0000970 | NRP + Polyketide:Modular type I | 38.0 | 32.3 | 417.0 | 1.6e-115 |
| BAQ25507.1 | type\_I\_polyketide\_synthase | BGC0001288 | Polyketide | 31.0 | 45.4 | 416.0 | 2.1e-115 |
| ADC79638.1 | TamAII | BGC0001052 | NRP + Polyketide:Modular type I | 30.0 | 50.9 | 416.0 | 2.7e-115 |
| QDA77059.1 | polyketide\_synthase/nonribosomal\_peptide\_synthetase | BGC0002026 | NRP | 36.0 | 34.8 | 416.0 | 2.7e-115 |
| CAQ34928.1 | polyketide\_synthase | BGC0000986 | NRP + Polyketide | 37.0 | 32.0 | 416.0 | 3.5e-115 |
| ABK32287.1 | JerA | BGC0000080 | Polyketide | 36.0 | 33.0 | 415.0 | 4.6e-115 |
| ABK32255.1 | AmbA | BGC0000014 | Polyketide | 36.0 | 33.0 | 415.0 | 6e-115 |
| AHH34189.1 | polyketide\_synthase | BGC0001162 | Polyketide:Modular type I | 36.0 | 33.9 | 415.0 | 6e-115 |
| ABO15860.1 | polyketide\_synthase | BGC0000130 | Polyketide | 36.0 | 33.4 | 415.0 | 7.9e-115 |
| CAD19091.1 | StiG\_protein | BGC0000153 | NRP + Polyketide:Modular type I | 36.0 | 32.5 | 415.0 | 7.9e-115 |
| ADB12492.1 | EpoE | BGC0000990 | NRP + Polyketide | 36.0 | 32.4 | 414.0 | 1e-114 |
| AAF62884.1 | EpoE | BGC0000991 | NRP + Polyketide | 36.0 | 32.4 | 414.0 | 1e-114 |
| ATV95616.1 | 6-methylsalicylic\_acid\_synthase | BGC0001503 | Polyketide | 35.0 | 32.5 | 414.0 | 1e-114 |
| CAQ18835.1 | polyketide\_synthase | BGC0000954 | NRP + Polyketide:Modular type I | 36.0 | 33.3 | 413.0 | 1.8e-114 |
| AEE88278.1 | CurL | BGC0000976 | NRP + Polyketide:Modular type I | 36.0 | 32.7 | 413.0 | 1.8e-114 |
| AAS98777.1 | polyketide\_synthetase | BGC0001001 | NRP + Polyketide | 35.0 | 32.4 | 413.0 | 1.8e-114 |
| AAT70107.1 | CurL | BGC0001165 | NRP + Polyketide:Modular type I | 36.0 | 32.7 | 413.0 | 1.8e-114 |
| ADH04680.1 | hybrid\_polyketide\_synthase/non-ribosomal\_peptide\_synthetase | BGC0001344 | NRP + Polyketide | 26.0 | 91.2 | 413.0 | 3e-114 |
| ACR33078.1 | polyketide\_synthase | BGC0000017 | Alkaloid + Polyketide:Modular type I | 36.0 | 32.2 | 412.0 | 3.9e-114 |
| CAD89777.1 | MelF\_protein | BGC0001010 | NRP + Polyketide:Modular type I | 36.0 | 32.6 | 412.0 | 3.9e-114 |
| AOE23578.1 | FoxBII | BGC0001598 | NRP + Polyketide | 36.0 | 32.5 | 412.0 | 5.1e-114 |
| ctg1\_orf16 |  | BGC0001457 | NRP | 37.0 | 32.2 | 411.0 | 6.7e-114 |
| AZH23791.1 | MgcH | BGC0001970 | NRP + Polyketide | 36.0 | 32.8 | 411.0 | 6.7e-114 |
| EGJ35088.1 | Polyketide\_synthase | BGC0001163 | Polyketide:Modular type I | 35.0 | 34.3 | 411.0 | 8.7e-114 |
| AWS21279.1 | type\_I\_polyketide\_synthase | BGC0001934 | Polyketide | 36.0 | 31.9 | 411.0 | 8.7e-114 |
| AZY91989.1 | polyketide\_synthase | BGC0002022 | Polyketide | 36.0 | 31.9 | 411.0 | 8.7e-114 |
| ACB46196.1 | polyketide\_synthase | BGC0000989 | NRP + Polyketide | 36.0 | 32.4 | 411.0 | 1.1e-113 |
| CAO98879.1 | polyketide\_synthase\_AufD | BGC0000023 | Polyketide:Modular type I | 36.0 | 34.0 | 410.0 | 1.5e-113 |
| AAQ82568.1 | FscD | BGC0000061 | Polyketide | 36.0 | 32.3 | 410.0 | 1.5e-113 |
| AMB48442.1 | polyketide\_synthase | BGC0001357 | Polyketide | 36.0 | 32.1 | 410.0 | 1.5e-113 |
| AXM42950.1 | polyketide\_synthase | BGC0001941 | NRP + Polyketide | 36.0 | 32.0 | 410.0 | 1.5e-113 |
| AQX77694.1 | NocP | BGC0001704 | Other | 36.0 | 31.5 | 410.0 | 1.9e-113 |
| CAL58681.1 | polyketide\_synthase | BGC0000149 | Polyketide:Modular type I | 37.0 | 31.6 | 409.0 | 3.3e-113 |
| AKL71649.1 | NocP | BGC0001703 | Other | 37.0 | 31.9 | 409.0 | 3.3e-113 |
| CAD19092.1 | StiH\_protein | BGC0000153 | NRP + Polyketide:Modular type I | 36.0 | 32.5 | 409.0 | 4.3e-113 |
| AAC38075.1 | polyketide\_synthase\_type\_I | BGC0000127 | Polyketide | 36.0 | 31.7 | 408.0 | 5.6e-113 |
| CAJ46689.1 | polyketide\_synthase | BGC0000969 | NRP:Cyclic depsipeptide + Polyketide:Modular type I | 36.0 | 32.9 | 408.0 | 7.4e-113 |
| AIW82282.1 | PuwE | BGC0001125 | NRP + Polyketide | 36.0 | 32.3 | 407.0 | 1.3e-112 |
| CAQ34919.1 | polyketide\_synthase | BGC0000986 | NRP + Polyketide | 36.0 | 32.1 | 407.0 | 1.6e-112 |
| AHB82053.1 | polyketide\_synthase | BGC0001019 | NRP + Polyketide:Modular type I | 35.0 | 32.6 | 407.0 | 1.6e-112 |
| CCE88380.1 | polyketide\_synthase | BGC0001034 | NRP + Polyketide:Modular type I | 36.0 | 32.1 | 406.0 | 2.1e-112 |
| WP\_053065267.1 | type\_I\_polyketide\_synthase | BGC0001330 | NRP:Cyclic depsipeptide + Polyketide:Modular type I | 36.0 | 32.1 | 406.0 | 2.8e-112 |
| ACV42478.1 | polyketide\_synthase | BGC0000043 | Polyketide | 36.0 | 32.5 | 406.0 | 3.7e-112 |
| AEE88277.1 | CurM | BGC0000976 | NRP + Polyketide:Modular type I | 36.0 | 32.5 | 406.0 | 3.7e-112 |
| CCE88381.1 | polyketide\_synthase | BGC0001034 | NRP + Polyketide:Modular type I | 36.0 | 31.8 | 406.0 | 3.7e-112 |
| AAT70108.1 | CurM | BGC0001165 | NRP + Polyketide:Modular type I | 36.0 | 32.5 | 406.0 | 3.7e-112 |
| EHK80163.1 | acyl\_transferase | BGC0001447 | Polyketide | 35.0 | 32.6 | 406.0 | 3.7e-112 |
| TXD00034.1 | SDR\_family\_NAD(P)-dependent\_oxidoreductase | BGC0001877 | Polyketide | 36.0 | 32.0 | 405.0 | 6.2e-112 |
| AAG23264.1 | polyketide\_synthase\_loading\_and\_extender\_module\_1 | BGC0000148 | Polyketide | 35.0 | 32.2 | 404.0 | 1.1e-111 |
| AQW44892.1 | polyketide\_synthase | BGC0001737 | NRP + Polyketide | 36.0 | 32.2 | 404.0 | 1.1e-111 |
| AEP40940.1 | polyketide\_synthase\_type\_I | BGC0000021 | Polyketide | 36.0 | 33.6 | 404.0 | 1.4e-111 |
| AAS98782.1 | polyketide\_synthase | BGC0001001 | NRP + Polyketide | 35.0 | 33.2 | 404.0 | 1.4e-111 |
| APZ78767.1 | polyketide\_synthase | BGC0001425 | NRP:Cyclic depsipeptide + Polyketide:Iterative type I | 36.0 | 32.2 | 404.0 | 1.4e-111 |
| ATP76239.1 | NdaF | BGC0001705 | NRP + Polyketide | 35.0 | 32.0 | 404.0 | 1.4e-111 |
| AFP87523.1 | type\_I\_polyketide\_synthase | BGC0001159 | NRP + Polyketide:Modular type I | 36.0 | 32.5 | 403.0 | 2.4e-111 |
| BAQ25513.1 | type\_I\_polyketide\_synthase | BGC0001288 | Polyketide | 30.0 | 50.4 | 403.0 | 3.1e-111 |
| ADH04657.1 | TugA | BGC0001342 | NRP + Polyketide | 36.0 | 33.2 | 403.0 | 3.1e-111 |
| AEK75502.1 | type\_1\_polyketide\_synthase | BGC0000001 | Polyketide:Modular type I | 37.0 | 31.9 | 402.0 | 4e-111 |
| AAW03329.1 | CtaF | BGC0000982 | NRP + Polyketide | 37.0 | 32.3 | 402.0 | 4e-111 |
| ADH04660.1 | TugD | BGC0001342 | NRP + Polyketide | 35.0 | 32.8 | 402.0 | 4e-111 |
| AFI57005.1 | QmnA1 | BGC0000133 | Polyketide | 38.0 | 31.6 | 402.0 | 5.3e-111 |
| CBD77734.1 | polyketide\_synthase | BGC0000974 | NRP + Polyketide | 34.0 | 32.7 | 402.0 | 5.3e-111 |
| AAO62582.1 | polyketide\_synthase\_peptide\_sythetase\_fusion\_protein | BGC0001016 | NRP + Polyketide | 35.0 | 32.0 | 402.0 | 5.3e-111 |
| WP\_019032757.1 | type\_I\_polyketide\_synthase | BGC0001331 | NRP:Cyclic depsipeptide + Polyketide:Modular type I | 35.0 | 32.0 | 402.0 | 5.3e-111 |
| AIR74911.1 | polyketide\_synthase | BGC0001559 | RiPP | 34.0 | 32.7 | 402.0 | 5.3e-111 |
| ACN64831.1 | PokM1 | BGC0001061 | Polyketide:Iterative type I + Polyketide:Type II + Saccharide:Hybrid/tailoring | 36.0 | 32.8 | 401.0 | 6.9e-111 |
| ctg1\_orf3 |  | BGC0001329 | Polyketide + NRP:Cyclic depsipeptide | 35.0 | 32.5 | 401.0 | 6.9e-111 |
| AWO77084.1 | hybrid\_non-ribosomal\_peptide\_synthetase/type\_I\_polyketide\_synthase | BGC0001556 | NRP + Polyketide | 35.0 | 34.3 | 401.0 | 9e-111 |
| AAG13917.1 | megalomicin\_6-deoxyerythronolide\_B\_synthase\_1 | BGC0000092 | Polyketide | 35.0 | 32.7 | 400.0 | 1.5e-110 |
| CBD77738.1 | polyketide\_synthase | BGC0000974 | NRP + Polyketide | 36.0 | 31.9 | 399.0 | 2.6e-110 |
| AIR74913.1 | polyketide\_synthase | BGC0001559 | RiPP | 36.0 | 31.9 | 399.0 | 2.6e-110 |
| AQH32481.1 | hybrid\_polyketide\_synthase/peptide\_synthetase | BGC0001667 | NRP + Polyketide | 35.0 | 32.1 | 399.0 | 2.6e-110 |
| BAC57028.1 | protomycinolide\_IV\_synthase\_1 | BGC0000102 | Polyketide | 37.0 | 31.8 | 399.0 | 3.4e-110 |
| AAC01712.2 | RifC | BGC0000136 | Polyketide | 30.0 | 51.0 | 399.0 | 3.4e-110 |
| AAY42396.1 | Polyketide\_synthase | BGC0001000 | NRP:Lipopeptide + Polyketide:Modular type I | 33.0 | 39.0 | 399.0 | 4.5e-110 |
| BAC68129.1 | modular\_polyketide\_synthase | BGC0000059 | Polyketide | 34.0 | 33.1 | 398.0 | 5.8e-110 |
| CAC20921.1 | PimS2\_protein | BGC0000125 | Polyketide | 36.0 | 32.4 | 398.0 | 5.8e-110 |
| ABX60162.1 | polyketide\_synthase | BGC0000978 | NRP + Alkaloid + Polyketide:Modular type I | 36.0 | 31.7 | 398.0 | 5.8e-110 |
| AQT01393.1 | SgnS2 | BGC0001690 | Polyketide | 36.0 | 32.4 | 398.0 | 5.8e-110 |
| ACB46488.1 | polyketide\_synthase | BGC0000082 | Polyketide | 36.0 | 33.0 | 398.0 | 7.6e-110 |
| CAQ43078.1 | polyketide\_synthase | BGC0000970 | NRP + Polyketide:Modular type I | 36.0 | 32.9 | 398.0 | 7.6e-110 |
| ADF88280.1 | polyketide\_synthase | BGC0000981 | NRP + Polyketide | 36.0 | 31.8 | 398.0 | 7.6e-110 |
| WP\_026247674.1 | type\_I\_polyketide\_synthase | BGC0001332 | NRP + Polyketide | 35.0 | 32.5 | 398.0 | 7.6e-110 |
| AIT55261.1 | polyketide\_synthase | BGC0000072 | Polyketide:Modular type I | 36.0 | 33.0 | 398.0 | 1e-109 |
| ATX68116.1 | malonyl\_CoA-acyl\_carrier\_protein\_transacylase | BGC0001772 | Polyketide | 34.0 | 33.3 | 398.0 | 1e-109 |
| AVI26389.1 | polyketide\_synthase | BGC0001800 | NRP + Polyketide | 36.0 | 33.1 | 398.0 | 1e-109 |
| FS847\_01985 | type\_I\_polyketide\_synthase | BGC0001877 | Polyketide | 33.0 | 32.4 | 398.0 | 1e-109 |
| AXN93601.1 | PuwE | BGC0001952 | NRP | 35.0 | 32.5 | 398.0 | 1e-109 |
| AMYAL\_RS48925 | polyketide\_synthase | BGC0002011 | Polyketide | 31.0 | 42.5 | 398.0 | 1e-109 |
| AEZ64503.1 | Herd | BGC0001065 | Polyketide | 29.0 | 44.4 | 397.0 | 1.3e-109 |
| AAW03325.1 | CtaB | BGC0000982 | NRP + Polyketide | 35.0 | 32.5 | 397.0 | 1.7e-109 |
| AWC08657.1 | polyketide\_synthase\_type\_I | BGC0001932 | Polyketide | 35.0 | 32.2 | 397.0 | 1.7e-109 |
| AWS21278.1 | type\_I\_polyketide\_synthase | BGC0001934 | Polyketide | 37.0 | 32.1 | 397.0 | 1.7e-109 |
| AZY91987.1 | polyketide\_synthase | BGC0002022 | Polyketide | 37.0 | 32.1 | 397.0 | 1.7e-109 |
| ABV97152.1 | Beta-ketoacyl\_synthase | BGC0000137 | Polyketide | 31.0 | 44.2 | 396.0 | 2.2e-109 |
| AHH99925.1 | PKS\_I | BGC0000002 | Polyketide | 28.0 | 50.2 | 396.0 | 2.9e-109 |
| AAX98184.1 | polyketide\_synthase\_type\_I | BGC0000052 | Polyketide | 34.0 | 33.1 | 396.0 | 2.9e-109 |
| AEE88281.1 | CurI | BGC0000976 | NRP + Polyketide:Modular type I | 34.0 | 33.3 | 396.0 | 3.8e-109 |
| AAT70104.1 | CurI | BGC0001165 | NRP + Polyketide:Modular type I | 34.0 | 33.3 | 396.0 | 3.8e-109 |
| AHB82065.1 | polyketide\_synthase | BGC0001231 | NRP + Polyketide:Modular type I | 36.0 | 31.0 | 395.0 | 4.9e-109 |
| TXD00033.1 | SDR\_family\_NAD(P)-dependent\_oxidoreductase | BGC0001877 | Polyketide | 28.0 | 52.1 | 395.0 | 4.9e-109 |
| AHH99922.1 | PKS\_I | BGC0000002 | Polyketide | 35.0 | 31.6 | 395.0 | 6.5e-109 |
| AAP42855.1 | NanA1 | BGC0000105 | Polyketide | 35.0 | 33.6 | 395.0 | 6.5e-109 |
| BAC57032.1 | protomycinolide\_IV\_synthase\_5 | BGC0000102 | Polyketide | 29.0 | 51.4 | 394.0 | 8.4e-109 |
| AMB48441.1 | polyketide\_synthase | BGC0001357 | Polyketide | 34.0 | 32.8 | 394.0 | 8.4e-109 |
| ACO94471.1 | polyketide\_synthase\_type\_I | BGC0000029 | Polyketide:Modular type I | 28.0 | 51.3 | 394.0 | 1.1e-108 |
| ABB05103.1 | LipPks2 | BGC0001003 | NRP:Lipopeptide + Polyketide:Modular type I + Saccharide:Hybrid/tailoring | 34.0 | 32.2 | 394.0 | 1.1e-108 |
| AHB82054.1 | polyketide\_synthase | BGC0001019 | NRP + Polyketide:Modular type I | 36.0 | 30.7 | 394.0 | 1.1e-108 |
| AKA59090.1 | type-I\_PKS | BGC0001619 | Polyketide | 36.0 | 32.2 | 394.0 | 1.4e-108 |
| AEE88289.1 | CurA | BGC0000976 | NRP + Polyketide:Modular type I | 35.0 | 32.4 | 393.0 | 1.9e-108 |
| CAD89773.1 | MelB\_protein | BGC0001010 | NRP + Polyketide:Modular type I | 35.0 | 33.2 | 393.0 | 1.9e-108 |
| AAT70096.1 | CurA | BGC0001165 | NRP + Polyketide:Modular type I | 35.0 | 32.4 | 393.0 | 1.9e-108 |
| AWR88404.1 | putative\_beta-ketoacyl\_synthase | BGC0001522 | Polyketide | 35.0 | 33.5 | 393.0 | 1.9e-108 |
| ADX66461.1 | ScnS2 | BGC0000108 | Polyketide | 36.0 | 32.0 | 393.0 | 2.5e-108 |
| AAC01711.1 | RifB | BGC0000136 | Polyketide | 29.0 | 52.9 | 393.0 | 2.5e-108 |
| CAD19093.1 | StiJ\_protein | BGC0000153 | NRP + Polyketide:Modular type I | 36.0 | 31.8 | 393.0 | 2.5e-108 |
| AAF26921.1 | polyketide\_synthase | BGC0000988 | NRP + Polyketide | 36.0 | 33.2 | 393.0 | 2.5e-108 |
| AWH12668.1 | RmpC | BGC0001759 | Polyketide | 29.0 | 51.0 | 393.0 | 2.5e-108 |
| ACR33079.1 | polyketide\_synthase | BGC0000017 | Alkaloid + Polyketide:Modular type I | 36.0 | 32.3 | 393.0 | 3.2e-108 |
| ANI24099.1 | polyketide\_synthase | BGC0001235 | NRP + Polyketide | 36.0 | 33.0 | 393.0 | 3.2e-108 |
| AXN93613.1 | PuwE | BGC0001953 | NRP | 35.0 | 32.3 | 393.0 | 3.2e-108 |
| AZF85946.1 | type\_I\_polyketide\_synthase | BGC0001963 | NRP + Polyketide | 35.0 | 32.4 | 393.0 | 3.2e-108 |
| ABB05102.1 | LipPks1 | BGC0001003 | NRP:Lipopeptide + Polyketide:Modular type I + Saccharide:Hybrid/tailoring | 35.0 | 32.5 | 392.0 | 4.2e-108 |
| AAF00958.1 | mcyE | BGC0001017 | NRP + Polyketide:Modular type I | 34.0 | 31.8 | 392.0 | 4.2e-108 |
| ALD82521.1 | polyketide\_synthase | BGC0001212 | NRP + Polyketide | 25.0 | 104.1 | 392.0 | 4.2e-108 |
| AAQ82561.1 | FscA | BGC0000061 | Polyketide | 36.0 | 32.6 | 392.0 | 5.5e-108 |
| AAF71775.1 | nysB | BGC0000115 | Polyketide:Modular type I + Saccharide:Hybrid/tailoring | 35.0 | 31.9 | 392.0 | 5.5e-108 |
| CAQ18834.1 | polyketide\_synthase | BGC0000954 | NRP + Polyketide:Modular type I | 33.0 | 39.6 | 392.0 | 5.5e-108 |
| ABV99085.1 | thioester\_reductase\_domain | BGC0001007 | Polyketide + NRP | 35.0 | 33.1 | 392.0 | 5.5e-108 |
| ADH04640.1 | TgaB | BGC0001051 | NRP + Polyketide:Modular type I | 36.0 | 31.9 | 392.0 | 5.5e-108 |
| sipP2 | Type\_I\_Modular\_PKS | BGC0001452 | Polyketide | 35.0 | 33.5 | 392.0 | 5.5e-108 |
| ASZ00149.1 | polyketide\_synthase | BGC0001785 | Polyketide | 35.0 | 32.8 | 392.0 | 5.5e-108 |
| ABO15861.1 | polyketide\_synthase | BGC0000130 | Polyketide | 35.0 | 33.2 | 391.0 | 7.1e-108 |
| BAC76493.1 | lankamycin\_synthase\_LkmAI | BGC0000085 | Polyketide | 34.0 | 33.9 | 391.0 | 9.3e-108 |
| ADF88275.1 | polyketide\_synthase | BGC0000981 | NRP + Polyketide | 36.0 | 31.7 | 391.0 | 9.3e-108 |
| ADZ24997.1 | polyketide\_synthase | BGC0000380 | NRP + Polyketide:Modular type I | 35.0 | 32.8 | 391.0 | 1.2e-107 |
| AGC24270.1 | prlP | BGC0001038 | NRP + Polyketide:Modular type I | 36.0 | 32.9 | 391.0 | 1.2e-107 |
| AAO65796.1 | monensin\_polyketide\_synthase\_loading\_module\_and\_module\_1 | BGC0000100 | Polyketide | 36.0 | 33.3 | 390.0 | 1.6e-107 |
| AFV96142.1 | polyketide\_synthase | BGC0001064 | Polyketide:Modular type I + Polyketide:Type III | 35.0 | 32.3 | 390.0 | 1.6e-107 |
| ARU81122.1 | CylH | BGC0001566 | Polyketide | 35.0 | 32.3 | 390.0 | 1.6e-107 |
| ANZ52459.1 | MonAI | BGC0001670 | Polyketide | 36.0 | 33.3 | 390.0 | 1.6e-107 |
| AJY78092.1 | polyketide\_synthase | BGC0001902 | NRP + Polyketide | 35.0 | 33.3 | 390.0 | 1.6e-107 |
| BAJ16467.1 | polyketide\_synthase | BGC0000058 | Polyketide | 34.0 | 33.2 | 390.0 | 2.1e-107 |
| AHB82063.1 | polyketide\_synthase | BGC0001231 | NRP + Polyketide:Modular type I | 36.0 | 30.0 | 390.0 | 2.1e-107 |
| CAQ43077.1 | polyketide\_synthase | BGC0000970 | NRP + Polyketide:Modular type I | 37.0 | 32.7 | 389.0 | 2.7e-107 |
| CBD77736.1 | polyketide\_synthase | BGC0000974 | NRP + Polyketide | 35.0 | 32.4 | 389.0 | 2.7e-107 |
| AIR74912.1 | polyketide\_synthase | BGC0001559 | RiPP | 35.0 | 32.4 | 389.0 | 2.7e-107 |
| AZF85932.1 | type\_I\_polyketide\_synthase | BGC0001963 | NRP + Polyketide | 35.0 | 32.7 | 389.0 | 2.7e-107 |
| AZF85947.1 | type\_I\_polyketide\_synthase | BGC0001963 | NRP + Polyketide | 34.0 | 32.9 | 389.0 | 2.7e-107 |
| BAF85838.1 | modular\_polyketide\_synthase | BGC0000109 | Polyketide | 35.0 | 32.9 | 389.0 | 3.5e-107 |
| ADH04641.1 | TgaC | BGC0001051 | NRP + Polyketide:Modular type I | 35.0 | 31.7 | 389.0 | 3.5e-107 |
| AVI57433.1 | AbmB1 | BGC0001694 | Polyketide | 36.0 | 30.3 | 389.0 | 3.5e-107 |
| CAD29794.1 | peptide\_synthetase | BGC0001015 | NRP + Polyketide | 33.0 | 32.0 | 389.0 | 4.6e-107 |
| AVX51107.1 | nysB | BGC0001709 | Polyketide | 35.0 | 32.6 | 389.0 | 4.6e-107 |
| ACN69988.1 | polyketide\_synthase | BGC0000079 | Polyketide | 35.0 | 32.9 | 388.0 | 6e-107 |
| ABX60153.1 | polyketide\_synthase | BGC0000978 | NRP + Alkaloid + Polyketide:Modular type I | 36.0 | 31.7 | 388.0 | 6e-107 |
| ANR02551.1 | LodJ | BGC0001648 | Polyketide | 34.0 | 32.2 | 388.0 | 6e-107 |
| AVI26390.1 | polyketide\_synthase\_/\_nonribosomal\_peptide\_synthase\_hybrid | BGC0001800 | NRP + Polyketide | 35.0 | 32.0 | 388.0 | 6e-107 |
| AZH23819.1 | MgiR | BGC0001971 | NRP + Polyketide | 34.0 | 32.7 | 388.0 | 6e-107 |
| AEC13080.1 | fosB | BGC0000060 | Polyketide | 35.0 | 33.1 | 388.0 | 7.9e-107 |
| BAK64637.1 | polyketide\_synthase | BGC0000135 | Polyketide | 35.0 | 32.3 | 388.0 | 7.9e-107 |
| QBF51755.1 | type\_I\_polyketide\_synthase | BGC0001856 | Polyketide:Modular type I | 28.0 | 51.7 | 388.0 | 7.9e-107 |
| AAF71766.1 | nysI | BGC0000115 | Polyketide:Modular type I + Saccharide:Hybrid/tailoring | 34.0 | 32.2 | 388.0 | 1e-106 |
| AUD08663.1 | iPKS-NRPS | BGC0001553 | NRP + Polyketide | 35.0 | 34.3 | 388.0 | 1e-106 |
| ABJ97439.1 | MerC | BGC0001012 | NRP + Polyketide | 35.0 | 32.5 | 387.0 | 1.3e-106 |
| ATX68115.1 | malonyl\_CoA-acyl\_carrier\_protein\_transacylase | BGC0001772 | Polyketide | 35.0 | 33.1 | 387.0 | 1.3e-106 |
| AXM42948.1 | type\_1\_polyketide\_synthase | BGC0001941 | NRP + Polyketide | 35.0 | 29.6 | 387.0 | 1.3e-106 |
| AAF62883.1 | epoD | BGC0000991 | NRP + Polyketide | 36.0 | 33.2 | 387.0 | 1.8e-106 |
| ACC40922.1 | polyketide\_synthase,\_Pks8 | BGC0001665 | Polyketide | 31.0 | 44.2 | 387.0 | 1.8e-106 |
| ARW71486.1 | type\_I\_PKS\_module\_6 | BGC0001812 | Polyketide | 35.0 | 32.3 | 387.0 | 1.8e-106 |
| AHN85651.1 | Phn2 | BGC0000122 | Polyketide:Modular type I | 34.0 | 32.9 | 386.0 | 2.3e-106 |
| AWC08663.1 | polyketide\_synthase\_type\_I | BGC0001932 | Polyketide | 35.0 | 32.4 | 386.0 | 2.3e-106 |
| AKG06378.1 | polyketide\_synthase\_type\_1 | BGC0001830 | Polyketide | 35.0 | 32.4 | 386.0 | 2.3e-106 |
| AAM70355.1 | CalO5 | BGC0000033 | Polyketide | 36.0 | 32.2 | 386.0 | 3.9e-106 |
| BAO66529.1 | type\_I\_polyketide\_synthase | BGC0000042 | Polyketide | 35.0 | 33.0 | 386.0 | 3.9e-106 |
| CQR60496.1 | Polyketide\_synthase,\_type\_I,\_modules:\_4,\_5\_and\_6 | BGC0001287 | Polyketide | 35.0 | 32.3 | 386.0 | 3.9e-106 |
| ADB12491.1 | EpoD | BGC0000990 | NRP + Polyketide | 36.0 | 33.2 | 385.0 | 5.1e-106 |
| CCE88379.1 | polyketide\_synthase | BGC0001034 | NRP + Polyketide:Modular type I | 36.0 | 32.0 | 385.0 | 5.1e-106 |
| WP\_052165465.1 | type\_I\_polyketide\_synthase | BGC0001327 | NRP:Cyclic depsipeptide + Polyketide:Modular type I | 34.0 | 32.3 | 385.0 | 5.1e-106 |
| TXD00024.1 | SDR\_family\_NAD(P)-dependent\_oxidoreductase | BGC0001877 | Polyketide | 35.0 | 32.4 | 385.0 | 5.1e-106 |
| AXN93580.1 | PuwE | BGC0001950 | NRP | 34.0 | 33.7 | 385.0 | 5.1e-106 |
| AXN93589.1 | PuwE | BGC0001951 | NRP | 34.0 | 33.7 | 385.0 | 5.1e-106 |
| AQV04230.1 | SwnK | BGC0001794 | Polyketide | 33.0 | 31.8 | 385.0 | 6.7e-106 |
| ABK32263.1 | AmbH | BGC0000014 | Polyketide | 34.0 | 32.7 | 384.0 | 8.7e-106 |
| ALD82523.1 | polyketide\_synthase | BGC0001212 | NRP + Polyketide | 34.0 | 34.3 | 384.0 | 8.7e-106 |
| ALD82524.1 | polyketide\_synthase | BGC0001212 | NRP + Polyketide | 34.0 | 34.5 | 384.0 | 8.7e-106 |
| AVX51098.1 | nysI | BGC0001709 | Polyketide | 34.0 | 32.2 | 384.0 | 8.7e-106 |
| AAK83194.1 | polyketide\_synthase | BGC0000026 | Saccharide:Oligosaccharide | 33.0 | 37.5 | 384.0 | 1.1e-105 |
| AEZ53952.1 | polyketide\_synthase | BGC0000144 | Polyketide:Modular type I | 34.0 | 32.9 | 384.0 | 1.1e-105 |
| ACB46195.1 | polyketide\_synthase | BGC0000989 | NRP + Polyketide | 36.0 | 33.2 | 384.0 | 1.1e-105 |
| AJO72742.1 | Type\_I\_modular\_polyketide\_synthase | BGC0001381 | Polyketide | 25.0 | 97.8 | 384.0 | 1.1e-105 |
| EWM63000.1 | non-ribosomal\_peptide\_synthetase | BGC0001328 | NRP:Cyclic depsipeptide + Polyketide:Modular type I | 34.0 | 32.3 | 384.0 | 1.5e-105 |
| B073\_RS40860 | type\_I\_polyketide\_synthase | BGC0001332 | NRP + Polyketide | 35.0 | 33.2 | 384.0 | 1.5e-105 |
| AWH12669.1 | RmpB | BGC0001759 | Polyketide | 29.0 | 50.7 | 384.0 | 1.5e-105 |
| MAA\_10033 | polyketide\_synthase,\_putative | BGC0000337 | NRP | 33.0 | 35.0 | 383.0 | 1.9e-105 |
| CAD17792.1 | probable\_non\_ribosomal\_peptide\_synthetase\_protein | BGC0001754 | NRP + Polyketide | 35.0 | 33.2 | 383.0 | 1.9e-105 |
| ANY10600.1 | polyketide\_synthase | BGC0001773 | Polyketide | 34.0 | 32.1 | 383.0 | 1.9e-105 |
| CAO98849.1 | polyketide\_synthase\_AufF | BGC0000023 | Polyketide:Modular type I | 36.0 | 32.5 | 383.0 | 2.5e-105 |
| BAF92601.1 | iterative\_type\_I\_PKS | BGC0000118 | Polyketide | 35.0 | 33.5 | 383.0 | 2.5e-105 |
| ACJ24875.1 | 6-methylsalicylic\_acid\_synthase | BGC0000119 | Polyketide:Iterative type I + Saccharide:Hybrid/tailoring | 35.0 | 33.5 | 383.0 | 2.5e-105 |
| BAE93722.1 | type\_I\_polyketide\_synthase | BGC0000164 | Polyketide | 32.0 | 32.8 | 383.0 | 2.5e-105 |
| CAE45669.1 | borrelidin\_polyketide\_synthase,\_type\_I | BGC0000031 | Polyketide:Modular type I | 31.0 | 42.3 | 383.0 | 3.3e-105 |
| ACO94484.1 | polyketide\_synthase\_type\_I | BGC0000097 | Polyketide:Modular type I | 34.0 | 32.1 | 383.0 | 3.3e-105 |
| ABC87512.1 | polyketide\_synthase | BGC0001011 | NRP + Polyketide | 34.0 | 32.5 | 382.0 | 4.3e-105 |
| ctg1\_orf23 |  | BGC0001013 | NRP + Polyketide | 34.0 | 32.5 | 382.0 | 4.3e-105 |
| AHH99919.1 | PKS\_I | BGC0000002 | Polyketide | 29.0 | 50.7 | 382.0 | 5.7e-105 |
| AAX98187.1 | polyketide\_synthase\_type\_I | BGC0000052 | Polyketide | 34.0 | 32.7 | 382.0 | 5.7e-105 |
| CAJ88184.1 | putative\_modular\_polyketide\_synthase | BGC0000151 | Polyketide:Modular type I + Saccharide:Hybrid/tailoring | 35.0 | 32.6 | 382.0 | 5.7e-105 |
| ADH04658.1 | TugB | BGC0001342 | NRP + Polyketide | 35.0 | 32.5 | 382.0 | 5.7e-105 |
| OJF16266.1 | AceP4 | BGC0001491 | Polyketide | 35.0 | 32.5 | 382.0 | 5.7e-105 |
| ALV82320.1 | borrelidin\_type\_I\_polyketide\_synthase | BGC0001533 | Polyketide | 31.0 | 42.3 | 382.0 | 5.7e-105 |
| QBF51760.1 | type\_I\_polyketide\_synthase | BGC0001856 | Polyketide:Modular type I | 34.0 | 32.3 | 382.0 | 5.7e-105 |
| AZH23787.1 | MgcQ | BGC0001970 | NRP + Polyketide | 34.0 | 32.4 | 382.0 | 5.7e-105 |
| ACB37740.1 | putative\_type\_I\_polyketide\_synthase | BGC0000162 | Polyketide | 34.0 | 32.5 | 381.0 | 7.4e-105 |
| ABY83164.1 | Azi26 | BGC0000960 | NRP + Polyketide | 34.0 | 33.0 | 381.0 | 7.4e-105 |
| CAQ43079.1 | polyketide\_synthase | BGC0000970 | NRP + Polyketide:Modular type I | 34.0 | 33.4 | 381.0 | 7.4e-105 |
| AGI99497.1 | type\_I\_polyketide\_synthase | BGC0001004 | Polyketide:Modular type I | 37.0 | 32.7 | 381.0 | 7.4e-105 |
| AHH99920.1 | PKS\_I | BGC0000002 | Polyketide | 35.0 | 31.9 | 381.0 | 9.6e-105 |
| AAC69332.1 | type\_I\_polyketide\_synthase\_PikAIV | BGC0000094 | Polyketide:Modular type I + Saccharide:Hybrid/tailoring | 33.0 | 32.2 | 381.0 | 9.6e-105 |
| AEZ54378.1 | PieA5 | BGC0000124 | Polyketide | 35.0 | 32.0 | 381.0 | 9.6e-105 |
| ADZ24996.1 | polyketide\_synthase | BGC0000380 | NRP + Polyketide:Modular type I | 35.0 | 32.7 | 381.0 | 9.6e-105 |
| AWC08655.1 | polyketide\_synthase\_type\_I | BGC0001932 | Polyketide | 34.0 | 32.3 | 381.0 | 9.6e-105 |
| QBF51758.1 | type\_I\_polyketide\_synthase | BGC0001856 | Polyketide:Modular type I | 34.0 | 32.5 | 381.0 | 9.6e-105 |
| AXI91545.1 | FunP8 | BGC0001944 | Polyketide | 28.0 | 50.8 | 381.0 | 9.6e-105 |
| ANR02555.1 | LodN | BGC0001648 | Polyketide | 36.0 | 32.0 | 381.0 | 1.3e-104 |
| AQV04224.1 | SwnK | BGC0001793 | Polyketide | 34.0 | 32.4 | 381.0 | 1.3e-104 |
| ABK32259.1 | AmbE | BGC0000014 | Polyketide | 35.0 | 32.8 | 380.0 | 1.6e-104 |
| AEZ53950.1 | polyketide\_synthase | BGC0000144 | Polyketide:Modular type I | 34.0 | 33.0 | 380.0 | 1.6e-104 |
| ACR50785.1 | polyketide\_synthase | BGC0000163 | Polyketide | 33.0 | 33.9 | 380.0 | 1.6e-104 |
| ARV85765.1 | PieA6\_type\_I\_PKS | BGC0001742 | Polyketide | 35.0 | 33.4 | 380.0 | 1.6e-104 |
| CAM00065.1 | EryAIII\_Erythromycin\_polyketide\_synthase\_modules\_5\_and\_6 | BGC0000055 | Polyketide:Modular type I + Saccharide:Hybrid/tailoring | 34.0 | 32.1 | 380.0 | 2.1e-104 |
| ABV83229.1 | CppB | BGC0000116 | Polyketide | 35.0 | 32.3 | 380.0 | 2.1e-104 |
| AEU17898.1 | putative\_type\_I\_PKS | BGC0001072 | Saccharide + Polyketide:Modular type I + Polyketide:Type II + Other:Aminocoumarin | 34.0 | 31.8 | 379.0 | 2.8e-104 |
| BAC68127.1 | modular\_polyketide\_synthase | BGC0000059 | Polyketide | 35.0 | 32.4 | 379.0 | 3.7e-104 |
| BAH02269.1 | polyketide\_synthase | BGC0000126 | Polyketide | 33.0 | 32.2 | 379.0 | 3.7e-104 |
| AFU82617.1 | polyketide\_synthase | BGC0000998 | NRP + Polyketide | 35.0 | 32.6 | 379.0 | 3.7e-104 |
| AAX98186.1 | polyketide\_synthase\_type\_I | BGC0000052 | Polyketide | 34.0 | 32.1 | 379.0 | 4.8e-104 |
| ACN69989.1 | polyketide\_synthase | BGC0000079 | Polyketide | 27.0 | 51.8 | 379.0 | 4.8e-104 |
| BAQ25483.1 | type\_I\_polyketide\_synthase | BGC0001288 | Polyketide | 34.0 | 32.2 | 379.0 | 4.8e-104 |
| TXD00026.1 | SDR\_family\_NAD(P)-dependent\_oxidoreductase | BGC0001877 | Polyketide | 34.0 | 32.7 | 379.0 | 4.8e-104 |
| ABB88520.1 | polyketide\_synthase\_type\_I | BGC0000050 | Polyketide | 35.0 | 32.3 | 378.0 | 6.2e-104 |
| AGC09484.1 | LobS1 | BGC0001183 | Polyketide | 36.0 | 32.7 | 378.0 | 8.2e-104 |
| BAF85844.1 | modular\_polyketide\_synthase | BGC0000109 | Polyketide | 34.0 | 32.5 | 378.0 | 1.1e-103 |
| CAQ34920.1 | polyketide\_synthase | BGC0000986 | NRP + Polyketide | 34.0 | 32.2 | 378.0 | 1.1e-103 |
| AAX98189.1 | polyketide\_synthase\_type\_I | BGC0000052 | Polyketide | 34.0 | 32.5 | 377.0 | 1.4e-103 |
| CAO85893.1 | modular\_polyketide\_synthase\_NorA | BGC0000110 | Polyketide:Modular type I | 33.0 | 32.3 | 377.0 | 1.4e-103 |
| AEZ53945.1 | polyketide\_synthase | BGC0000144 | Polyketide:Modular type I | 36.0 | 32.5 | 377.0 | 1.4e-103 |
| ARS01476.1 | NcmAIV | BGC0001702 | NRP + Polyketide | 35.0 | 32.1 | 377.0 | 1.4e-103 |
| AAS79460.1 | polyketide\_synthase\_subunit | BGC0000035 | Polyketide | 33.0 | 33.0 | 377.0 | 1.8e-103 |
| CBZ41585.1 | Type\_I\_modular\_polyketide\_synthase | BGC0000151 | Polyketide:Modular type I + Saccharide:Hybrid/tailoring | 29.0 | 50.9 | 376.0 | 2.4e-103 |
| BAO66539.1 | type\_I\_polyketide\_synthase | BGC0000042 | Polyketide | 35.0 | 32.7 | 376.0 | 3.1e-103 |
| AAM77986.1 | iterative\_type\_I\_polyketide\_synthase | BGC0000112 | Polyketide:Iterative type I + Polyketide:Enediyne type I | 35.0 | 31.3 | 376.0 | 3.1e-103 |
| AAQ90174.1 | polyketide\_synthase\_type\_I | BGC0000128 | Polyketide | 33.0 | 32.7 | 376.0 | 3.1e-103 |
| IF55\_RS32375 | beta-ketoacyl\_synthase | BGC0001348 | Polyketide:Modular type I | 33.0 | 33.1 | 376.0 | 3.1e-103 |
| AXM42951.1 | polyketide\_synthase | BGC0001941 | NRP + Polyketide | 34.0 | 33.5 | 376.0 | 3.1e-103 |
| AAX98190.1 | polyketide\_synthase\_type\_I | BGC0000052 | Polyketide | 33.0 | 34.2 | 376.0 | 4.1e-103 |
| ALP32045.1 | CycE | BGC0001293 | Polyketide | 34.0 | 33.2 | 376.0 | 4.1e-103 |
| TXD00265.1 | SDR\_family\_NAD(P)-dependent\_oxidoreductase | BGC0001877 | Polyketide | 27.0 | 53.3 | 376.0 | 4.1e-103 |
| AAZ94389.1 | modular\_polyketide\_synthase | BGC0000040 | Polyketide | 34.0 | 32.1 | 375.0 | 5.3e-103 |
| BAE93729.1 | type\_I\_polyketide\_synthase | BGC0000164 | Polyketide | 34.0 | 32.2 | 375.0 | 5.3e-103 |
| ALA09356.1 | type\_I\_modular\_PKS | BGC0001303 | Polyketide | 35.0 | 32.2 | 375.0 | 5.3e-103 |
| OAP25821.1 | Phenolphthiocerol\_synthesis\_polyketide\_synthase\_type\_I\_Pks15/1 | BGC0001658 | Polyketide | 27.0 | 50.4 | 375.0 | 5.3e-103 |
| AKG06379.1 | polyketide\_synthase\_type\_1 | BGC0001830 | Polyketide | 35.0 | 32.3 | 375.0 | 5.3e-103 |
| AZF85917.1 | type\_I\_polyketide\_synthase | BGC0001963 | NRP + Polyketide | 34.0 | 32.0 | 375.0 | 5.3e-103 |
| BAP34734.1 | type\_I\_polyketide\_synthase | BGC0000078 | Polyketide | 34.0 | 32.9 | 375.0 | 6.9e-103 |
| ANC94966.1 | AlmHI | BGC0001396 | Polyketide | 36.0 | 30.4 | 375.0 | 6.9e-103 |
| AWC08662.1 | polyketide\_synthase\_type\_I | BGC0001932 | Polyketide | 34.0 | 32.1 | 375.0 | 6.9e-103 |
| AAP42867.1 | NanA7 | BGC0000105 | Polyketide | 34.0 | 32.7 | 374.0 | 9e-103 |
| AKD43769.1 | HerA2 | BGC0001349 | NRP + Polyketide | 34.0 | 32.5 | 374.0 | 9e-103 |
| BAO66542.1 | type\_I\_polyketide\_synthase | BGC0000042 | Polyketide | 33.0 | 33.2 | 374.0 | 1.2e-102 |
| BAB69195.1 | modular\_polyketide\_synthase | BGC0000117 | Polyketide | 32.0 | 32.4 | 374.0 | 1.2e-102 |
| AAC01713.1 | RifD | BGC0000136 | Polyketide | 35.0 | 32.1 | 374.0 | 1.2e-102 |
| BAD38874.1 | polyketide\_synthase | BGC0000111 | Polyketide | 35.0 | 32.5 | 374.0 | 1.5e-102 |
| CBA11584.1 | polyketide\_synthase\_type\_I | BGC0001046 | NRP + Polyketide:Modular type I + Saccharide:Hybrid/tailoring | 36.0 | 29.5 | 374.0 | 1.5e-102 |
| AWC08661.1 | polyketide\_synthase\_type\_I | BGC0001932 | Polyketide | 34.0 | 32.0 | 374.0 | 1.5e-102 |
| ABV97151.1 | AMP-dependent\_synthetase\_and\_ligase | BGC0000137 | Polyketide | 34.0 | 32.0 | 373.0 | 2e-102 |
| ctg1\_orf10 |  | BGC0000053 | Polyketide | 34.0 | 31.4 | 373.0 | 2.6e-102 |
| BAB69198.1 | modular\_polyketide\_synthase | BGC0000117 | Polyketide | 32.0 | 33.0 | 373.0 | 2.6e-102 |
| AAM81584.2 | putative\_type\_I\_polyketide\_synthase | BGC0000047 | Polyketide | 35.0 | 30.4 | 373.0 | 3.4e-102 |
| AAX98188.1 | polyketide\_synthase\_type\_I | BGC0000052 | Polyketide | 33.0 | 33.2 | 373.0 | 3.4e-102 |
| ABP73645.1 | SalA | BGC0000145 | Polyketide | 36.0 | 32.3 | 373.0 | 3.4e-102 |
| AWC08659.1 | polyketide\_synthase\_type\_I | BGC0001932 | Polyketide | 34.0 | 32.2 | 373.0 | 3.4e-102 |
| AAS79463.1 | polyketide\_synthase\_subunit | BGC0000035 | Polyketide | 34.0 | 33.1 | 372.0 | 4.5e-102 |
| CAI94682.1 | putative\_polyketide\_synthase | BGC0000141 | Polyketide | 34.0 | 32.1 | 372.0 | 4.5e-102 |
| AEP40939.1 | polyketide\_synthase\_type\_I | BGC0000021 | Polyketide | 35.0 | 32.0 | 372.0 | 5.8e-102 |
| AAM81585.1 | putative\_type\_I\_polyketide\_synthase | BGC0000047 | Polyketide | 34.0 | 32.9 | 372.0 | 5.8e-102 |
| BAR73017.1 | putative\_PKS\_(KS-AT-KR-ACP-KS-AT-DH-KR-ACP) | BGC0001194 | Polyketide | 35.0 | 33.0 | 372.0 | 5.8e-102 |
| AWC08660.1 | polyketide\_synthase\_type\_I | BGC0001932 | Polyketide | 34.0 | 32.2 | 372.0 | 5.8e-102 |
| AAF26920.1 | polyketide\_synthase | BGC0000988 | NRP + Polyketide | 36.0 | 32.5 | 371.0 | 7.6e-102 |
| AAF19812.1 | MtaD | BGC0001024 | NRP + Polyketide:Modular type I | 35.0 | 33.8 | 371.0 | 7.6e-102 |
| ADH04639.1 | TgaA | BGC0001051 | NRP + Polyketide:Modular type I | 33.0 | 33.3 | 371.0 | 7.6e-102 |
| BAF02926.1 | type\_I\_polyketide\_synthase | BGC0000073 | Polyketide | 34.0 | 32.7 | 371.0 | 1e-101 |
| ACB46486.1 | polyketide\_synthase | BGC0000082 | Polyketide | 33.0 | 33.4 | 371.0 | 1e-101 |
| BAQ21947.1 | putative\_type\_I\_polyketide\_synthase | BGC0001204 | Polyketide | 34.0 | 33.2 | 371.0 | 1e-101 |
| AWC08656.1 | polyketide\_synthase\_type\_I | BGC0001932 | Polyketide | 34.0 | 32.2 | 371.0 | 1e-101 |
| ADC45534.1 | modular\_polyketide\_synthase | BGC0000093 | Polyketide | 34.0 | 33.8 | 371.0 | 1.3e-101 |
| AAG23266.1 | polyketide\_synthase\_extender\_modules\_3-4 | BGC0000148 | Polyketide | 33.0 | 31.8 | 371.0 | 1.3e-101 |
| ADU85988.1 | putative\_iterative\_type\_I\_polyketide\_synthase | BGC0000165 | Polyketide:Modular type I | 35.0 | 32.7 | 371.0 | 1.3e-101 |
| AAF26919.1 | polyketide\_synthase | BGC0000988 | NRP + Polyketide | 34.0 | 32.4 | 371.0 | 1.3e-101 |
| BBA66511.1 | type\_I\_polyketide\_synthase | BGC0001495 | Polyketide | 37.0 | 31.7 | 371.0 | 1.3e-101 |
| ARS01477.1 | NcmAV | BGC0001702 | NRP + Polyketide | 36.0 | 30.3 | 371.0 | 1.3e-101 |
| AXI91549.1 | FunP4 | BGC0001944 | Polyketide | 34.0 | 32.0 | 371.0 | 1.3e-101 |
| AXG22406.1 | type\_I\_polyketide\_synthase | BGC0002024 | Polyketide | 34.0 | 32.0 | 371.0 | 1.3e-101 |
| ADX66470.1 | ScnS0 | BGC0000108 | Polyketide | 34.0 | 32.1 | 370.0 | 1.7e-101 |
| ABV97155.1 | Acyl\_transferase | BGC0000137 | Polyketide | 34.0 | 32.4 | 370.0 | 1.7e-101 |
| CAL58683.1 | polyketide\_synthase | BGC0000149 | Polyketide:Modular type I | 34.0 | 32.3 | 370.0 | 1.7e-101 |
| ABX60152.1 | polyketide\_synthase | BGC0000978 | NRP + Alkaloid + Polyketide:Modular type I | 34.0 | 32.4 | 370.0 | 1.7e-101 |
| ADF88276.1 | polyketide\_synthase | BGC0000981 | NRP + Polyketide | 34.0 | 32.4 | 370.0 | 1.7e-101 |
| ACB46192.1 | polyketide\_synthase | BGC0000989 | NRP + Polyketide | 35.0 | 32.4 | 370.0 | 1.7e-101 |
| ABP53498.1 | PKS\_(ACP-AT-AT-KS-ACP-C) | BGC0001041 | NRP + Polyketide | 36.0 | 32.6 | 370.0 | 1.7e-101 |
| ARV85761.1 | PieA2\_type\_I\_PKS | BGC0001742 | Polyketide | 34.0 | 32.2 | 370.0 | 1.7e-101 |
| ABV83222.1 | CppJ | BGC0000116 | Polyketide | 34.0 | 31.9 | 370.0 | 2.2e-101 |
| AAF62880.1 | EpoA | BGC0000991 | NRP + Polyketide | 35.0 | 32.4 | 370.0 | 2.2e-101 |
| AHB82052.1 | polyketide\_synthase | BGC0001019 | NRP + Polyketide:Modular type I | 35.0 | 29.3 | 370.0 | 2.2e-101 |
| ANC94965.1 | AlmHII | BGC0001396 | Polyketide | 34.0 | 32.9 | 370.0 | 2.2e-101 |
| ANZ22987.1 | ZinD | BGC0001828 | Polyketide | 35.0 | 32.0 | 370.0 | 2.2e-101 |
| AXG22407.1 | type\_I\_polyketide\_synthase | BGC0002024 | Polyketide | 35.0 | 31.8 | 370.0 | 2.2e-101 |
| ACA99172.1 | polyketide\_synthase | BGC0001160 | Polyketide:Modular type I | 35.0 | 31.6 | 369.0 | 2.9e-101 |
| XP\_001220460.1 | hypothetical\_protein | BGC0001182 | NRP + Polyketide:Iterative type I | 41.0 | 26.2 | 369.0 | 2.9e-101 |
| ASZ00151.1 | polyketide\_synthase | BGC0001785 | Polyketide | 34.0 | 32.8 | 369.0 | 2.9e-101 |
| AVI26388.1 | polyketide\_synthase | BGC0001800 | NRP + Polyketide | 34.0 | 33.2 | 369.0 | 2.9e-101 |
| AAS79462.1 | polyketide\_synthase\_subunit | BGC0000035 | Polyketide | 34.0 | 32.3 | 369.0 | 3.8e-101 |
| ABP55221.1 | acyl\_transferase\_domain\_protein | BGC0000142 | Polyketide | 34.0 | 32.3 | 369.0 | 3.8e-101 |
| EYT83439.1 | beta-ketoacyl\_synthase | BGC0001213 | Polyketide | 33.0 | 39.5 | 369.0 | 3.8e-101 |
| ADB12490.1 | EpoC | BGC0000990 | NRP + Polyketide | 36.0 | 32.5 | 369.0 | 5e-101 |
| sipP4 | Type\_I\_Modular\_PKS | BGC0001452 | Polyketide | 35.0 | 32.3 | 369.0 | 5e-101 |
| AAB66508.1 | tylactone\_synthase\_module\_7 | BGC0000166 | Polyketide | 33.0 | 32.5 | 368.0 | 6.5e-101 |
| ANZ22991.1 | ZinG | BGC0001828 | Polyketide | 33.0 | 32.2 | 368.0 | 1.1e-100 |
| AAQ82566.1 | FscF | BGC0000061 | Polyketide | 34.0 | 32.4 | 367.0 | 1.4e-100 |
| BAF02923.1 | type\_I\_polyketide\_synthase | BGC0000073 | Polyketide | 34.0 | 32.6 | 367.0 | 1.4e-100 |
| ACO94500.1 | polyketide\_synthase\_type\_I | BGC0000097 | Polyketide:Modular type I | 30.0 | 42.2 | 367.0 | 1.4e-100 |
| ANZ22985.1 | ZinB | BGC0001828 | Polyketide | 34.0 | 32.6 | 367.0 | 1.4e-100 |
| AZF85945.1 | type\_I\_polyketide\_synthase | BGC0001963 | NRP + Polyketide | 33.0 | 33.7 | 367.0 | 1.4e-100 |
| AAU93805.2 | polyketide\_synthase\_modules\_5\_and\_6 | BGC0000054 | Polyketide | 34.0 | 32.7 | 367.0 | 1.9e-100 |
| ctg1\_orf7 |  | BGC0000053 | Polyketide | 35.0 | 31.7 | 366.0 | 2.5e-100 |
| ADB12488.1 | EpoA | BGC0000990 | NRP + Polyketide | 34.0 | 32.4 | 366.0 | 2.5e-100 |
| AAF62882.1 | EpoC | BGC0000991 | NRP + Polyketide | 35.0 | 32.5 | 366.0 | 2.5e-100 |
| CAD29793.1 | polyketide\_synthase\_type\_I | BGC0001015 | NRP + Polyketide | 30.0 | 48.1 | 366.0 | 2.5e-100 |
| QDA77045.1 | polyketide\_synthase/nonribosomal\_peptide\_synthetase | BGC0002025 | NRP | 33.0 | 32.4 | 366.0 | 2.5e-100 |
| BAD08359.1 | polyketide\_synthase\_modules\_5-6 | BGC0000167 | Polyketide | 32.0 | 32.3 | 366.0 | 3.2e-100 |
| AAF86392.1 | FkbC | BGC0000994 | NRP + Polyketide | 31.0 | 42.1 | 366.0 | 3.2e-100 |
| CQR60493.1 | Polyketide\_synthase,\_type\_I,\_modules:\_9\_and\_10 | BGC0001287 | Polyketide | 34.0 | 32.7 | 366.0 | 3.2e-100 |
| ACO94460.1 | polyketide\_synthase\_type\_I | BGC0000029 | Polyketide:Modular type I | 34.0 | 34.0 | 366.0 | 4.2e-100 |
| AAM81586.2 | putative\_type\_I\_polyketide\_synthase | BGC0000047 | Polyketide | 34.0 | 32.7 | 366.0 | 4.2e-100 |
| QBF51759.1 | type\_I\_polyketide\_synthase | BGC0001856 | Polyketide:Modular type I | 33.0 | 32.6 | 366.0 | 4.2e-100 |
| AAG23263.1 | polyketide\_synthase\_extender\_modules\_5-7 | BGC0000148 | Polyketide | 34.0 | 32.2 | 365.0 | 5.5e-100 |
| ACR50775.1 | polyketide\_synthase | BGC0000163 | Polyketide | 34.0 | 33.1 | 365.0 | 5.5e-100 |
| AEZ64505.1 | Herb | BGC0001065 | Polyketide | 36.0 | 30.5 | 365.0 | 5.5e-100 |
| ALJ49910.1 | TlmH | BGC0001237 | Polyketide | 35.0 | 32.4 | 365.0 | 7.2e-100 |
| ABY21538.1 | AngAI | BGC0000018 | Polyketide | 35.0 | 32.3 | 364.0 | 1.2e-99 |
| CAE45670.1 | borrelidin\_polyketide\_synthase,\_type\_I | BGC0000031 | Polyketide:Modular type I | 34.0 | 32.4 | 364.0 | 1.2e-99 |
| CAJ88175.1 | putative\_polyketide\_synthase\_B | BGC0000151 | Polyketide:Modular type I + Saccharide:Hybrid/tailoring | 34.0 | 32.1 | 364.0 | 1.2e-99 |
| BAE93728.1 | type\_I\_polyketide\_synthase | BGC0000164 | Polyketide | 34.0 | 32.6 | 364.0 | 1.6e-99 |
| EHA22196.1 | polyketide\_synthase | BGC0000170 | Polyketide | 35.0 | 32.9 | 364.0 | 1.6e-99 |
| ARW71483.1 | type\_I\_PKS\_loading\_module,\_module\_1,\_module\_2 | BGC0001812 | Polyketide | 35.0 | 32.2 | 364.0 | 1.6e-99 |
| ARW71487.1 | type\_I\_PKS\_module\_7 | BGC0001812 | Polyketide | 34.0 | 32.2 | 364.0 | 1.6e-99 |
| ACO94456.1 | polyketide\_synthase\_type\_I | BGC0000029 | Polyketide:Modular type I | 33.0 | 32.6 | 363.0 | 2.1e-99 |
| AAZ94390.1 | modular\_polyketide\_synthase | BGC0000040 | Polyketide | 34.0 | 32.5 | 363.0 | 2.1e-99 |
| CAM00062.1 | EryAI\_Erythromycin\_polyketide\_synthase\_modules\_1\_and\_2 | BGC0000055 | Polyketide:Modular type I + Saccharide:Hybrid/tailoring | 33.0 | 31.9 | 363.0 | 2.1e-99 |
| AFL48532.1 | laidlomycin\_polyketide\_synthase\_(module\_11\_and\_module\_12) | BGC0000084 | Polyketide | 34.0 | 32.5 | 363.0 | 2.1e-99 |
| BAB69193.1 |  | BGC0000117 | Polyketide | 34.0 | 32.6 | 363.0 | 2.1e-99 |
| ARE67853.1 | AbsB1 | BGC0001492 | Polyketide | 36.0 | 32.3 | 363.0 | 2.1e-99 |
| CAJ88185.2 | Type\_I\_modular\_polyketide\_synthase | BGC0000151 | Polyketide:Modular type I + Saccharide:Hybrid/tailoring | 34.0 | 33.1 | 363.0 | 2.7e-99 |
| WP\_053065268.1 | type\_I\_polyketide\_synthase | BGC0001330 | NRP:Cyclic depsipeptide + Polyketide:Modular type I | 36.0 | 30.1 | 363.0 | 2.7e-99 |
| AHA38203.1 | GphJ | BGC0000069 | Polyketide | 34.0 | 33.6 | 363.0 | 3.5e-99 |
| ABC84469.1 | NigAIX | BGC0000114 | Polyketide:Modular type I | 34.0 | 32.3 | 363.0 | 3.5e-99 |
| AEU17899.1 | putative\_type\_I\_PKS | BGC0001072 | Saccharide + Polyketide:Modular type I + Polyketide:Type II + Other:Aminocoumarin | 34.0 | 31.7 | 363.0 | 3.5e-99 |
| AHE80996.1 | PieA6 | BGC0001169 | Polyketide:Modular type I | 34.0 | 32.8 | 363.0 | 3.5e-99 |
| EHK80167.1 | modular\_polyketide\_synthase | BGC0001447 | Polyketide | 33.0 | 33.4 | 363.0 | 3.5e-99 |
| AAU04878.1 | polyketide\_synthase | BGC0000365 | NRP | 33.0 | 32.6 | 362.0 | 4.6e-99 |
| AJO72736.1 | Type\_I\_modular\_polyketide\_synthase | BGC0001381 | Polyketide | 33.0 | 32.7 | 362.0 | 4.6e-99 |
| ABV83221.1 | CppI | BGC0000116 | Polyketide | 33.0 | 32.5 | 362.0 | 6.1e-99 |
| CAC20930.1 | PimS0\_protein | BGC0000125 | Polyketide | 33.0 | 33.9 | 362.0 | 6.1e-99 |
| AAF00959.1 | mcyD | BGC0001017 | NRP + Polyketide:Modular type I | 29.0 | 49.3 | 362.0 | 6.1e-99 |
| AQT01384.1 | SgnS0 | BGC0001690 | Polyketide | 33.0 | 33.9 | 362.0 | 6.1e-99 |
| AAZ94386.1 | modular\_polyketide\_synthase | BGC0000040 | Polyketide | 34.0 | 32.9 | 361.0 | 1e-98 |
| ACR50773.1 | polyketide\_synthase | BGC0000163 | Polyketide | 32.0 | 36.5 | 361.0 | 1e-98 |
| CAQ18839.1 | hybrid\_polyketide\_synthase/nonribosomal\_polypetide\_synthetase | BGC0000954 | NRP + Polyketide:Modular type I | 35.0 | 33.6 | 361.0 | 1e-98 |
| ACO94483.1 | polyketide\_synthase\_type\_I | BGC0000097 | Polyketide:Modular type I | 33.0 | 32.6 | 361.0 | 1.3e-98 |
| AEZ54379.1 | PieA6 | BGC0000124 | Polyketide | 33.0 | 33.3 | 361.0 | 1.3e-98 |
| CAQ34929.1 | putative\_polyketide\_synthase | BGC0000986 | NRP + Polyketide | 36.0 | 29.0 | 361.0 | 1.3e-98 |
| AKA59088.1 | type-I\_PKS | BGC0001619 | Polyketide | 35.0 | 32.1 | 361.0 | 1.3e-98 |
| AAX98191.1 | polyketide\_synthase\_type\_I | BGC0000052 | Polyketide | 32.0 | 32.9 | 360.0 | 1.8e-98 |
| BAK64650.1 | polyketide\_synthase | BGC0000135 | Polyketide | 33.0 | 33.5 | 360.0 | 1.8e-98 |
| ACB46194.1 | polyketide\_synthase | BGC0000989 | NRP + Polyketide | 35.0 | 32.5 | 360.0 | 1.8e-98 |
| WP\_019032756.1 | type\_I\_polyketide\_synthase | BGC0001331 | NRP:Cyclic depsipeptide + Polyketide:Modular type I | 35.0 | 30.2 | 360.0 | 1.8e-98 |
| AAS79461.1 | polyketide\_synthase\_subunit | BGC0000035 | Polyketide | 34.0 | 32.0 | 360.0 | 2.3e-98 |
| AGC24271.1 | prlQ | BGC0001038 | NRP + Polyketide:Modular type I | 34.0 | 31.8 | 360.0 | 2.3e-98 |
| ONK09689.1 | Beta-ketoacyl-acyl-carrier-protein\_synthase\_I | BGC0001647 | Polyketide | 33.0 | 32.2 | 360.0 | 2.3e-98 |
| BAQ25511.1 | type\_I\_polyketide\_synthase | BGC0001288 | Polyketide | 33.0 | 35.4 | 359.0 | 3e-98 |
| BAT51067.1 | type\_I\_polyketide\_synthase | BGC0001296 | Polyketide | 32.0 | 33.5 | 359.0 | 3e-98 |
| AFU82615.1 | polyketide\_synthase | BGC0000998 | NRP + Polyketide | 36.0 | 30.2 | 359.0 | 3.9e-98 |
| AGY30676.1 | Ann4 | BGC0001298 | Polyketide | 33.0 | 32.1 | 359.0 | 3.9e-98 |
| AWR88398.1 | putative\_beta-ketoacyl\_synthase | BGC0001522 | Polyketide | 34.0 | 32.3 | 359.0 | 3.9e-98 |
| CAL58686.1 | polyketide\_synthase | BGC0000149 | Polyketide:Modular type I | 33.0 | 32.8 | 359.0 | 5.1e-98 |
| ACY06290.1 | type\_I\_polyketide\_synthase | BGC0001042 | NRP + Polyketide | 33.0 | 33.9 | 359.0 | 5.1e-98 |
| CAO98848.1 | polyketide\_synthase\_AufE | BGC0000023 | Polyketide:Modular type I | 36.0 | 29.5 | 358.0 | 6.7e-98 |
| ABC84471.1 | NigAVII | BGC0000114 | Polyketide:Modular type I | 32.0 | 32.6 | 358.0 | 6.7e-98 |
| BAK64638.1 | polyketide\_synthase | BGC0000135 | Polyketide | 34.0 | 32.7 | 358.0 | 6.7e-98 |
| ALA09357.1 | type\_I\_modular\_PKS | BGC0001303 | Polyketide | 28.0 | 42.9 | 358.0 | 6.7e-98 |
| AWH12937.1 | StmB | BGC0001939 | Polyketide | 33.0 | 31.8 | 358.0 | 6.7e-98 |
| AHA38201.1 | GphH | BGC0000069 | Polyketide | 35.0 | 32.9 | 358.0 | 8.7e-98 |
| AAO65806.1 | monensin\_polyketide\_synthase\_modules\_11\_and\_12 | BGC0000100 | Polyketide | 33.0 | 32.5 | 358.0 | 8.7e-98 |
| AGC09499.1 | LobS4 | BGC0001183 | Polyketide | 33.0 | 33.2 | 358.0 | 8.7e-98 |
| AAX35547.1 | polyketide\_syntase\_2 | BGC0001275 | Polyketide | 33.0 | 33.2 | 358.0 | 8.7e-98 |
| ALV82341.1 | borrelidin\_type\_I\_polyketide\_synthase | BGC0001533 | Polyketide | 34.0 | 32.4 | 358.0 | 8.7e-98 |
| ANZ52469.1 | MonAVIII | BGC0001670 | Polyketide | 33.0 | 32.5 | 358.0 | 8.7e-98 |
| AXI91552.1 | FunP1 | BGC0001944 | Polyketide | 33.0 | 32.4 | 358.0 | 8.7e-98 |
| AAC69329.1 | type\_I\_polyketide\_synthase\_PikAI | BGC0000094 | Polyketide:Modular type I + Saccharide:Hybrid/tailoring | 33.0 | 33.7 | 358.0 | 1.1e-97 |
| CAL58687.1 | polyketide\_synthase | BGC0000149 | Polyketide:Modular type I | 32.0 | 32.2 | 358.0 | 1.1e-97 |
| AGI99482.1 | Type\_I\_polyketide\_synthase | BGC0001004 | Polyketide:Modular type I | 33.0 | 33.2 | 358.0 | 1.1e-97 |
| AAS98200.1 | MSAS-type\_polyketide\_synthase | BGC0001273 | Polyketide | 33.0 | 33.6 | 358.0 | 1.1e-97 |
| ADC79618.1 | BafAIII | BGC0000028 | Polyketide:Modular type I | 33.0 | 33.3 | 357.0 | 1.5e-97 |
| AAP42857.1 | NanA3 | BGC0000105 | Polyketide | 33.0 | 33.3 | 357.0 | 1.5e-97 |
| AKD43764.1 | HerF | BGC0001349 | NRP + Polyketide | 33.0 | 32.3 | 357.0 | 1.5e-97 |
| ATG32075.1 | polyketide\_synthase | BGC0001750 | NRP + Polyketide | 34.0 | 32.5 | 357.0 | 1.5e-97 |
| ADC45535.1 | modular\_polyketide\_synthase | BGC0000093 | Polyketide | 32.0 | 34.3 | 357.0 | 1.9e-97 |
| ACY06289.1 | type\_I\_polyketide\_synthase | BGC0001042 | NRP + Polyketide | 36.0 | 32.2 | 357.0 | 1.9e-97 |
| AEP40935.1 | polyketide\_synthase\_type\_I | BGC0000021 | Polyketide | 33.0 | 31.5 | 356.0 | 2.5e-97 |
| AGI99496.1 | Type\_I\_polyketide\_synthase | BGC0001004 | Polyketide:Modular type I | 32.0 | 32.9 | 356.0 | 2.5e-97 |
| AGZ15472.1 | putative\_modular\_polyketide\_synthase | BGC0001036 | NRP + Polyketide | 34.0 | 32.2 | 356.0 | 2.5e-97 |
| BAQ25481.1 | type\_I\_polyketide\_synthase | BGC0001288 | Polyketide | 31.0 | 36.6 | 356.0 | 2.5e-97 |
| ANY10590.1 | polyketide\_synthase | BGC0001773 | Polyketide | 34.0 | 32.9 | 356.0 | 2.5e-97 |
| ACO94499.1 | polyketide\_synthase\_type\_I | BGC0000097 | Polyketide:Modular type I | 33.0 | 32.3 | 356.0 | 3.3e-97 |
| BAC57029.1 | protomycinolide\_IV\_synthase\_2 | BGC0000102 | Polyketide | 33.0 | 33.1 | 356.0 | 3.3e-97 |
| TXD00266.1 | SDR\_family\_NAD(P)-dependent\_oxidoreductase | BGC0001877 | Polyketide | 31.0 | 33.0 | 356.0 | 3.3e-97 |
| ADX66472.1 | ScnS1 | BGC0000108 | Polyketide | 32.0 | 32.8 | 356.0 | 4.3e-97 |
| ABG02264.1 | SalB | BGC0000143 | Polyketide | 34.0 | 32.4 | 355.0 | 5.7e-97 |
| sipP3 | Type\_I\_Modular\_PKS | BGC0001452 | Polyketide | 33.0 | 32.8 | 355.0 | 5.7e-97 |
| BAW32323.1 | hybrid\_cis-AT\_polyketide\_synthase\_-\_nonribosomal\_peptide\_synthetase | BGC0001630 | NRP + Polyketide | 34.0 | 32.5 | 355.0 | 5.7e-97 |
| AAP42856.1 | NanA2 | BGC0000105 | Polyketide | 33.0 | 32.7 | 355.0 | 7.4e-97 |
| AWR88399.1 | putative\_beta-ketoacyl\_synthase | BGC0001522 | Polyketide | 33.0 | 33.2 | 355.0 | 7.4e-97 |
| AEE88284.1 | CurF | BGC0000976 | NRP + Polyketide:Modular type I | 32.0 | 33.4 | 354.0 | 9.7e-97 |
| BAQ25482.1 | type\_I\_polyketide\_synthase | BGC0001288 | Polyketide | 35.0 | 30.8 | 354.0 | 9.7e-97 |
| ctg1\_orf256 |  | BGC0001200 | Polyketide | 32.0 | 33.6 | 354.0 | 1.3e-96 |
| CAO85896.1 | protein\_modular\_polyketide\_synthase\_NorA' | BGC0000110 | Polyketide:Modular type I | 32.0 | 31.9 | 354.0 | 1.6e-96 |
| ALP32042.1 | CycB | BGC0001293 | Polyketide | 34.0 | 32.3 | 353.0 | 2.2e-96 |
| AQH32482.1 | type\_1\_polyketide\_synthase | BGC0001667 | NRP + Polyketide | 28.0 | 48.2 | 353.0 | 2.2e-96 |
| AAT70101.1 | CurF | BGC0001165 | NRP + Polyketide:Modular type I | 32.0 | 33.4 | 353.0 | 2.8e-96 |
| ADC79620.1 | BafAV | BGC0000028 | Polyketide:Modular type I | 33.0 | 32.7 | 352.0 | 3.7e-96 |
| BAO66543.1 | type\_I\_polyketide\_synthase | BGC0000042 | Polyketide | 33.0 | 32.3 | 352.0 | 3.7e-96 |
| CAQ34917.1 | polyketide\_synthase | BGC0000986 | NRP + Polyketide | 36.0 | 29.2 | 352.0 | 3.7e-96 |
| AAO62584.1 | polyketide\_synthase\_type\_1 | BGC0001016 | NRP + Polyketide | 32.0 | 39.9 | 352.0 | 3.7e-96 |
| WP\_048832936.1 | polyketide\_synthase | BGC0001348 | Polyketide:Modular type I | 34.0 | 31.0 | 352.0 | 3.7e-96 |
| ADC45515.1 | modular\_polyketide\_synthase | BGC0000093 | Polyketide | 34.0 | 31.5 | 352.0 | 4.8e-96 |
| ABP55223.1 | beta-ketoacyl\_synthase | BGC0000142 | Polyketide | 34.0 | 32.3 | 352.0 | 4.8e-96 |
| KFL51881.1 | beta-ketoacyl\_synthase | BGC0001711 | NRP + Polyketide | 40.0 | 23.2 | 352.0 | 4.8e-96 |
| AKG06375.1 | polyketide\_synthase\_type\_1 | BGC0001830 | Polyketide | 34.0 | 32.4 | 352.0 | 4.8e-96 |
| BAQ21940.1 | putative\_Type\_I\_polyketide\_synthase | BGC0001204 | Polyketide | 32.0 | 33.3 | 352.0 | 6.3e-96 |
| SCO70310.1 | Type\_I\_polyketide\_synthase | BGC0001433 | Polyketide:Modular type I | 33.0 | 32.2 | 352.0 | 6.3e-96 |
| AXI91550.1 | FunP3 | BGC0001944 | Polyketide | 33.0 | 33.6 | 352.0 | 6.3e-96 |
| BAH02268.1 | polyketide\_synthase | BGC0000126 | Polyketide | 32.0 | 32.8 | 351.0 | 8.2e-96 |
| AAC38076.1 | polyketide\_synthase\_type\_I | BGC0000127 | Polyketide | 32.0 | 32.3 | 351.0 | 1.1e-95 |
| QBF51756.1 | type\_I\_polyketide\_synthase | BGC0001856 | Polyketide:Modular type I | 33.0 | 29.6 | 351.0 | 1.4e-95 |
| AEC13071.1 | fosE | BGC0000060 | Polyketide | 31.0 | 33.9 | 350.0 | 1.8e-95 |
| AEH42473.1 | polyketide\_synthase | BGC0000032 | Polyketide | 33.0 | 32.2 | 349.0 | 3.1e-95 |
| AAZ77693.1 | ChlA1 | BGC0000036 | Polyketide:Modular type I + Polyketide:Iterative type I + Saccharide:Oligosaccharide | 33.0 | 33.3 | 349.0 | 3.1e-95 |
| AAR16521.1 | RimA | BGC0000138 | Polyketide | 34.0 | 32.0 | 349.0 | 3.1e-95 |
| CAA60459.1 | polyketide\_synthase | BGC0001040 | NRP + Polyketide | 34.0 | 31.7 | 349.0 | 3.1e-95 |
| BAP34740.1 | type\_I\_polyketide\_synthase | BGC0000078 | Polyketide | 32.0 | 33.7 | 349.0 | 5.3e-95 |
| AAB66504.1 | tylactone\_synthase\_starter\_module\_and\_modules\_1\_&\_2 | BGC0000166 | Polyketide | 32.0 | 35.2 | 349.0 | 5.3e-95 |
| AJO72734.1 | Type\_I\_modular\_polyketide\_synthase | BGC0001381 | Polyketide | 34.0 | 32.2 | 349.0 | 5.3e-95 |
| CAJ76298.1 | putative\_hybrid\_polyketide-non-ribosomal\_peptide\_synthetase | BGC0000972 | NRP + Polyketide:Modular type I + Polyketide:Trans-AT type I | 25.0 | 89.2 | 348.0 | 6.9e-95 |
| AAD03048.1 | type\_I\_polyketide\_synthase | BGC0000041 | Polyketide | 33.0 | 32.6 | 348.0 | 9e-95 |
| ABC84458.1 | NigAIII | BGC0000114 | Polyketide:Modular type I | 32.0 | 33.6 | 348.0 | 9e-95 |
| AIG62146.1 | 6-methylsalicylic\_acid\_synthase | BGC0000120 | Polyketide:Iterative type I | 33.0 | 32.2 | 348.0 | 9e-95 |
| AIW00670.1 | mellein\_synthase | BGC0001244 | Polyketide | 32.0 | 31.6 | 348.0 | 9e-95 |
| AXI91547.1 | FunP6 | BGC0001944 | Polyketide | 32.0 | 32.9 | 348.0 | 9e-95 |
| AAS79459.1 | polyketide\_synthase\_subunit | BGC0000035 | Polyketide | 34.0 | 30.2 | 347.0 | 1.2e-94 |
| AFL48527.1 | laidlomycin\_polyketide\_synthase\_(module\_3\_and\_module\_4) | BGC0000084 | Polyketide | 32.0 | 33.2 | 347.0 | 1.2e-94 |
| ACB37742.1 | putative\_type\_I\_polyketide\_synthase | BGC0000162 | Polyketide | 32.0 | 32.3 | 347.0 | 1.2e-94 |
| AAP85336.1 | type\_I\_PKS | BGC0000233 | Polyketide | 34.0 | 32.4 | 347.0 | 1.2e-94 |
| CAD15508.1 | polyketide\_synthase/non-ribosomal\_peptide\_synthetase | BGC0001014 | NRP:NRP siderophore + Polyketide:Modular type I + Polyketide:Iterative type I | 34.0 | 33.0 | 347.0 | 2e-94 |
| AHF22854.1 | MarL | BGC0000091 | Polyketide | 33.0 | 32.0 | 346.0 | 3.4e-94 |
| AEC13072.1 | fosF | BGC0000060 | Polyketide | 32.0 | 33.3 | 346.0 | 4.5e-94 |
| CAO85898.1 | modular\_polyketide\_synthase\_NorC | BGC0000110 | Polyketide:Modular type I | 34.0 | 29.1 | 346.0 | 4.5e-94 |
| ABP55220.1 | beta-ketoacyl\_synthase | BGC0000142 | Polyketide | 34.0 | 31.7 | 346.0 | 4.5e-94 |
| CAQ34918.1 | nonribosomal\_peptide\_synthetase/\_polyketide\_synthase | BGC0000986 | NRP + Polyketide | 31.0 | 39.4 | 346.0 | 4.5e-94 |
| ADH04682.1 | polyketide\_synthase | BGC0001344 | NRP + Polyketide | 35.0 | 32.8 | 346.0 | 4.5e-94 |
| AKD43768.1 | HerA1 | BGC0001349 | NRP + Polyketide | 33.0 | 33.0 | 346.0 | 4.5e-94 |
| ARV85764.1 | PieA5\_type\_I\_PKS | BGC0001742 | Polyketide | 32.0 | 33.6 | 345.0 | 5.9e-94 |
| KFL51883.1 | amino\_acid\_adenylation\_protein | BGC0001711 | NRP + Polyketide | 35.0 | 29.3 | 345.0 | 7.7e-94 |
| ANH11413.1 | SceR | BGC0001908 | Polyketide | 31.0 | 32.6 | 345.0 | 7.7e-94 |
| ANZ22989.1 | ZinF | BGC0001828 | Polyketide | 33.0 | 33.9 | 345.0 | 7.7e-94 |
| AZH23790.1 | MgcG | BGC0001970 | NRP + Polyketide | 32.0 | 32.2 | 345.0 | 7.7e-94 |
| AZH23820.1 | MgiG | BGC0001971 | NRP + Polyketide | 32.0 | 32.2 | 345.0 | 7.7e-94 |
| BAF02921.1 | type\_I\_polyketide\_synthase | BGC0000073 | Polyketide | 34.0 | 32.8 | 344.0 | 1e-93 |
| BAP34733.1 | type\_I\_polyketide\_synthase | BGC0000078 | Polyketide | 33.0 | 32.6 | 344.0 | 1e-93 |
| WP\_055469545.1 | type\_I\_polyketide\_synthase | BGC0001537 | Polyketide | 33.0 | 31.7 | 344.0 | 1e-93 |
| ACY13415.1 | KR\_domain\_protein | BGC0001367 | NRP + Polyketide | 34.0 | 32.4 | 344.0 | 1.3e-93 |
| ATP76241.1 | NdaD | BGC0001705 | NRP + Polyketide | 30.0 | 41.0 | 344.0 | 1.3e-93 |
| ACZ65476.1 | type\_I\_modular\_polyketide\_synthase | BGC0000140 | Polyketide | 33.0 | 33.0 | 343.0 | 2.9e-93 |
| AFU82614.1 | mixed\_NRPS\_PKS | BGC0000998 | NRP + Polyketide | 34.0 | 32.8 | 343.0 | 2.9e-93 |
| ABX60161.1 | mixed\_NRPS/PKS | BGC0000978 | NRP + Alkaloid + Polyketide:Modular type I | 34.0 | 32.6 | 341.0 | 8.5e-93 |
| BBA84067.1 | type\_I\_polyketide\_synthase | BGC0001916 | Polyketide | 33.0 | 32.0 | 341.0 | 8.5e-93 |
| WP\_083502114.1 | type\_I\_polyketide\_synthase | BGC0001653 | Polyketide | 33.0 | 32.2 | 341.0 | 1.4e-92 |
| ABI91466.1 | beta-ketoacyl\_synthase | BGC0001094 | NRP + Polyketide | 33.0 | 32.0 | 340.0 | 2.5e-92 |
| CAF05651.1 | TubF\_protein | BGC0001053 | NRP + Polyketide | 33.0 | 33.7 | 339.0 | 3.2e-92 |
| CAE46843.1 | Type\_I\_modular\_polyketide\_synthase | BGC0000103 | Polyketide | 33.0 | 32.7 | 339.0 | 4.2e-92 |
| CAE46851.1 | Type\_I\_modular\_polyketide\_synthase | BGC0000103 | Polyketide | 33.0 | 32.7 | 339.0 | 4.2e-92 |
| AVX51099.1 | NysJ | BGC0001709 | Polyketide | 33.0 | 32.2 | 339.0 | 4.2e-92 |
| AEP40934.1 | polyketide\_synthase\_type\_I | BGC0000021 | Polyketide | 36.0 | 26.8 | 339.0 | 5.5e-92 |
| ADM46356.1 | polyketide\_synthase | BGC0000106 | Polyketide | 32.0 | 32.6 | 339.0 | 5.5e-92 |
| DAB41918.1 | ArzP\_-\_PKS\_(KS,\_AT,\_OMT,\_ACP,\_TE) | BGC0001884 | NRP + Polyketide | 34.0 | 32.4 | 339.0 | 5.5e-92 |
| ABB88519.1 | polyketide\_synthase\_type\_I | BGC0000050 | Polyketide | 34.0 | 34.1 | 338.0 | 7.2e-92 |
| ADF88262.1 | mixed\_nonribosomal\_peptide\_synthetase/\_polyketide\_synthase | BGC0000979 | NRP + Polyketide | 33.0 | 32.6 | 338.0 | 7.2e-92 |
| ADF88265.1 | mixed\_nonribosomal\_peptide\_synthetase/\_polyketide\_synthase | BGC0000980 | NRP + Polyketide | 33.0 | 32.6 | 338.0 | 7.2e-92 |
| ADF88279.1 | mixed\_NRPS/PKS | BGC0000981 | NRP + Polyketide | 33.0 | 32.6 | 338.0 | 7.2e-92 |
| ctg1\_orf253 |  | BGC0001200 | Polyketide | 34.0 | 32.3 | 338.0 | 7.2e-92 |
| AZH23793.1 | MgcK | BGC0001970 | NRP + Polyketide | 32.0 | 32.8 | 338.0 | 7.2e-92 |
| AHD05619.1 | putative\_polyketide\_synthase\_subunit | BGC0001033 | NRP + Polyketide | 33.0 | 32.7 | 338.0 | 9.4e-92 |
| ATG32077.1 | polyketide\_synthase | BGC0001750 | NRP + Polyketide | 33.0 | 34.9 | 338.0 | 9.4e-92 |
| AFR69333.1 | polyketide\_synthase\_SpiC1 | BGC0001045 | NRP:Cyclic depsipeptide + Polyketide:Modular type I | 39.0 | 23.7 | 337.0 | 1.6e-91 |
| AAS98784.1 | polyketide\_synthase | BGC0001001 | NRP + Polyketide | 33.0 | 32.8 | 337.0 | 2.1e-91 |
| AEZ64504.1 | Herc | BGC0001065 | Polyketide | 33.0 | 30.7 | 336.0 | 4.6e-91 |
| AHB82070.1 | polyketide\_synthase | BGC0001231 | NRP + Polyketide:Modular type I | 34.0 | 32.4 | 336.0 | 4.6e-91 |
| EJK79843.1 | amino\_acid\_adenylation\_enzyme/thioester\_reductase\_family\_protein | BGC0000436 | NRP | 34.0 | 32.4 | 335.0 | 6.1e-91 |
| AHE80993.1 | PieA3 | BGC0001169 | Polyketide:Modular type I | 34.0 | 30.1 | 335.0 | 6.1e-91 |
| AEZ54376.1 | PieA3 | BGC0000124 | Polyketide | 33.0 | 29.6 | 335.0 | 7.9e-91 |
| AHA38199.1 | GphF | BGC0000069 | Polyketide | 35.0 | 29.3 | 334.0 | 1.4e-90 |
| BAC76492.1 | lankamycin\_synthase\_LkmAII | BGC0000085 | Polyketide | 32.0 | 32.6 | 334.0 | 1.8e-90 |
| EWM62997.1 | non-ribosomal\_peptide\_synthetase | BGC0001328 | NRP:Cyclic depsipeptide + Polyketide:Modular type I | 36.0 | 25.1 | 334.0 | 1.8e-90 |
| WP\_039806854.1 | type\_I\_polyketide\_synthase | BGC0002001 | NRP + Polyketide | 34.0 | 32.3 | 334.0 | 1.8e-90 |
| ABI91470.1 | beta-ketoacyl\_synthase | BGC0001094 | NRP + Polyketide | 32.0 | 32.2 | 333.0 | 3e-90 |
| WP\_035122279.1 | type\_I\_polyketide\_synthase | BGC0001467 | NRP:Cyclic depsipeptide + Polyketide:Modular type I | 33.0 | 33.3 | 332.0 | 3.9e-90 |
| ARS01473.1 | NcmAI | BGC0001702 | NRP + Polyketide | 36.0 | 28.5 | 332.0 | 3.9e-90 |
| SCN11949.1 | ebeA-type\_I\_polyketide\_synthase\_KSQ-ATa-ACP | BGC0001580 | Polyketide | 33.0 | 30.2 | 332.0 | 6.7e-90 |
| AHD05614.1 | putative\_non-ribosomal\_peptide\_ligase/\_polyketide\_synthase\_hybrid | BGC0001033 | NRP + Polyketide | 34.0 | 32.2 | 331.0 | 1.1e-89 |
| ADZ24995.1 | non-ribosomal\_peptide\_synthase/polyketide\_synthase | BGC0000380 | NRP + Polyketide:Modular type I | 35.0 | 33.2 | 331.0 | 1.5e-89 |
| ABM21569.1 | crpA | BGC0000975 | NRP + Polyketide | 23.0 | 98.2 | 329.0 | 3.3e-89 |
| ACR50795.1 | putative\_polyketide\_synthase | BGC0000163 | Polyketide | 33.0 | 30.5 | 328.0 | 7.4e-89 |
| ARO38317.1 | nonribosomal\_peptide\_synthetase | BGC0001560 | NRP + Polyketide | 32.0 | 32.8 | 327.0 | 2.2e-88 |
| AZH23823.1 | MgiK | BGC0001971 | NRP + Polyketide | 31.0 | 32.8 | 327.0 | 2.2e-88 |
| ABY21542.1 | AngAV | BGC0000018 | Polyketide | 31.0 | 33.7 | 326.0 | 2.8e-88 |
| AXA20096.1 | trans-AT\_PKS\_LgaG | BGC0001946 | NRP + Polyketide | 37.0 | 23.3 | 326.0 | 3.7e-88 |
| CAC22145.1 | CpkB;\_Polyketide\_synthase\_modules\_3\_and\_4 | BGC0000038 | Polyketide:Modular type I | 31.0 | 32.9 | 326.0 | 4.8e-88 |
| AAN32979.1 | BarE | BGC0000962 | NRP + Polyketide:Modular type I | 33.0 | 33.6 | 325.0 | 6.3e-88 |
| OAP25811.1 | Phenolphthiocerol\_synthesis\_polyketide\_synthase\_type\_I\_Pks15/1 | BGC0001658 | Polyketide | 33.0 | 30.3 | 324.0 | 1.1e-87 |
| DAB41653.1 | polyketide\_synthase | BGC0001583 | Polyketide | 32.0 | 34.5 | 324.0 | 1.4e-87 |
| AEK75504.1 | type\_1\_polyketide\_synthase | BGC0000001 | Polyketide:Modular type I | 33.0 | 31.8 | 324.0 | 1.8e-87 |
| CAE02606.1 | polyketide\_synthase\_type\_I | BGC0000024 | Polyketide:Modular type I | 33.0 | 29.1 | 323.0 | 2.4e-87 |
| AAC46024.1 | polyketide\_synthase\_modules\_1\_and\_2 | BGC0000113 | Polyketide | 32.0 | 31.7 | 323.0 | 2.4e-87 |
| AFP87524.1 | type\_I\_polyketide\_synthase | BGC0001159 | NRP + Polyketide:Modular type I | 34.0 | 29.8 | 323.0 | 2.4e-87 |
| AAR87760.2 | ZmaK | BGC0001059 | NRP + Polyketide | 33.0 | 32.4 | 322.0 | 4.1e-87 |
| EAU29808.1 | hypothetical\_protein | BGC0001400 | Polyketide | 37.0 | 26.8 | 322.0 | 4.1e-87 |
| AHB82057.1 | polyketide\_synthase | BGC0001019 | NRP + Polyketide:Modular type I | 33.0 | 32.6 | 322.0 | 5.3e-87 |
| AEP40932.1 | polyketide\_synthase\_type\_I | BGC0000021 | Polyketide | 28.0 | 43.5 | 321.0 | 9.1e-87 |
| WP\_106731933.1 | type\_I\_polyketide\_synthase | BGC0001332 | NRP + Polyketide | 33.0 | 32.8 | 321.0 | 1.2e-86 |
| CAE14173.1 | hypothetical\_protein | BGC0000383 | NRP + Polyketide:Modular type I | 33.0 | 32.7 | 321.0 | 1.5e-86 |
| CCA29203.1 | non-ribosomal\_peptide\_synthetase/polyketide\_synthase | BGC0000955 | NRP + Polyketide:Modular type I | 34.0 | 32.8 | 321.0 | 1.5e-86 |
| AAS98787.1 | polyketide\_synthase/thioesterase | BGC0001001 | NRP + Polyketide | 33.0 | 33.0 | 321.0 | 1.5e-86 |
| sipP5 | Type\_I\_Modular\_PKS | BGC0001452 | Polyketide | 31.0 | 32.9 | 320.0 | 2e-86 |
| AHE80992.1 | PieA2 | BGC0001169 | Polyketide:Modular type I | 31.0 | 31.8 | 319.0 | 4.5e-86 |
| CBD77746.1 | non-ribosomal\_peptide\_synthetase/polyketide\_synthase | BGC0000974 | NRP + Polyketide | 34.0 | 32.1 | 318.0 | 1e-85 |
| AIR74926.1 | polyketide\_synthase | BGC0001559 | RiPP | 34.0 | 32.1 | 318.0 | 1e-85 |
| EED21099.1 | polyketide\_synthase,\_putative | BGC0001578 | Polyketide | 30.0 | 39.0 | 317.0 | 1.3e-85 |
| OAP25815.1 | Phenolphthiocerol\_synthesis\_polyketide\_synthase\_type\_I\_Pks15/1 | BGC0001658 | Polyketide | 32.0 | 32.3 | 317.0 | 1.3e-85 |
| WP\_019032754.1 | type\_I\_polyketide\_synthase | BGC0001331 | NRP:Cyclic depsipeptide + Polyketide:Modular type I | 33.0 | 32.2 | 317.0 | 1.7e-85 |
| CAN89634.1 | putative\_polyketide\_synthase | BGC0001070 | NRP + Polyketide:Modular type I + Polyketide:Trans-AT type I | 38.0 | 23.4 | 316.0 | 2.9e-85 |
| AKN45693.1 | polyketide\_synthase | BGC0001284 | Terpene | 32.0 | 32.4 | 316.0 | 2.9e-85 |
| ADI24926.1 | VrtA | BGC0000168 | Polyketide:Iterative type I | 32.0 | 33.1 | 316.0 | 3.8e-85 |
| WP\_047890614.1 | type\_I\_polyketide\_synthase | BGC0001330 | NRP:Cyclic depsipeptide + Polyketide:Modular type I | 33.0 | 32.3 | 316.0 | 5e-85 |
| EJP62792.1 | polyketide\_synthase | BGC0001720 | Polyketide | 32.0 | 32.6 | 316.0 | 5e-85 |
| EGD99348.1 | polyketide\_synthase | BGC0001144 | Polyketide | 32.0 | 33.2 | 315.0 | 6.5e-85 |
| AGN74892.1 | nonribosomal\_peptide\_synthetase/polyketide\_synthase\_hybrid\_protein | BGC0000459 | NRP:Cyclic depsipeptide + Polyketide:Trans-AT type I | 37.0 | 24.5 | 314.0 | 1.4e-84 |
| CCE31584.1 | polyketide\_synthase\_that\_catalyse\_the\_condensation\_of\_one\_acetyl-CoA\_and\_six\_malonyl-CoA\_resulting\_in\_formation\_of\_nor-rubrofusarin | BGC0001886 | Polyketide | 32.0 | 32.7 | 313.0 | 2.5e-84 |
| AXN93610.1 | PuwB | BGC0001953 | NRP | 34.0 | 28.1 | 313.0 | 2.5e-84 |
| AAD03047.1 | type\_I\_polyketide\_synthase | BGC0000041 | Polyketide | 34.0 | 30.8 | 313.0 | 3.2e-84 |
| ERM18799.1 | polyketide\_synthase | BGC0000172 | Polyketide | 35.0 | 24.6 | 313.0 | 3.2e-84 |
| ABF87031.1 | non-ribosomal\_peptide\_synthetase/polyketide\_synthase | BGC0000393 | NRP + Polyketide:Modular type I | 32.0 | 32.0 | 313.0 | 3.2e-84 |
| CCM44338.1 | Polyketide\_synthase | BGC0001056 | NRP + Polyketide:Modular type I + Polyketide:PUFA synthase or related | 33.0 | 32.3 | 313.0 | 3.2e-84 |
| ABX37384.1 | Beta-ketoacyl\_synthase | BGC0000984 | NRP + Polyketide | 32.0 | 33.2 | 312.0 | 5.5e-84 |
| ABP57747.1 | DepC | BGC0000993 | NRP:Cyclic depsipeptide + Polyketide:Modular type I | 37.0 | 23.1 | 312.0 | 5.5e-84 |
| ABS90471.1 | PKS\_type\_I | BGC0001106 | NRP + Polyketide | 33.0 | 32.5 | 312.0 | 5.5e-84 |
| ALD83688.1 | tAT\_polyketide\_synthase | BGC0001300 | Polyketide | 37.0 | 23.9 | 311.0 | 9.4e-84 |
| WP\_030498975.1 | type\_I\_polyketide\_synthase | BGC0001327 | NRP:Cyclic depsipeptide + Polyketide:Modular type I | 33.0 | 32.3 | 311.0 | 9.4e-84 |
| ARR97039.1 | SphF | BGC0001780 | NRP | 37.0 | 24.3 | 311.0 | 9.4e-84 |
| AAF15892.2 | nosB | BGC0001028 | Polyketide + NRP:Cyclic depsipeptide | 32.0 | 32.8 | 311.0 | 1.2e-83 |
| ACC40923.1 | polyketide\_synthase\_Pks9 | BGC0001665 | Polyketide | 32.0 | 32.1 | 310.0 | 2.1e-83 |
| AMH40422.1 | PKS | BGC0001350 | Polyketide | 36.0 | 23.4 | 310.0 | 2.7e-83 |
| ACY01400.1 | AT-less\_polyketide\_synthase | BGC0000083 | Polyketide:Modular type I + Polyketide:Trans-AT type I | 36.0 | 24.1 | 309.0 | 3.6e-83 |
| ASA76631.1 | polyketide\_synthase | BGC0001751 | NRP + Polyketide | 40.0 | 21.2 | 309.0 | 3.6e-83 |
| ARE67851.1 | AbsB3 | BGC0001492 | Polyketide | 31.0 | 32.9 | 309.0 | 4.7e-83 |
| AWR88393.1 | putative\_beta-ketoacyl\_synthase | BGC0001522 | Polyketide | 31.0 | 33.2 | 309.0 | 4.7e-83 |
| AZF85934.1 | type\_I\_polyketide\_synthase | BGC0001963 | NRP + Polyketide | 33.0 | 29.4 | 309.0 | 4.7e-83 |
| ACN69991.1 | polyketide\_synthase | BGC0000079 | Polyketide | 35.0 | 31.1 | 309.0 | 6.1e-83 |
| AKQ22681.1 | malonyl\_CoA-acyl\_carrier\_protein\_transacylase | BGC0001656 | Polyketide | 37.0 | 23.2 | 309.0 | 6.1e-83 |
| AFV52200.1 | polyketide\_synthase\_module | BGC0000081 | Polyketide:Iterative type I + Polyketide:Enediyne type I | 33.0 | 33.5 | 308.0 | 7.9e-83 |
| CAL69894.1 | RhiF\_protein | BGC0001112 | NRP + Polyketide:Trans-AT type I | 36.0 | 24.5 | 308.0 | 7.9e-83 |
| AKQ22698.1 | malonyl\_CoA-acyl\_carrier\_protein\_transacylase | BGC0001186 | Polyketide | 37.0 | 23.2 | 308.0 | 7.9e-83 |
| CCA89329.1 | trans-AT\_type\_I\_polyketide\_synthase | BGC0001111 | NRP + Polyketide:Trans-AT type I | 37.0 | 23.0 | 308.0 | 1e-82 |
| ATX68111.1 | malonyl\_CoA-acyl\_carrier\_protein\_transacylase | BGC0001772 | Polyketide | 34.0 | 25.0 | 307.0 | 1.8e-82 |
| AJQ95706.1 | polyketide\_synthase\_modules-related\_protein | BGC0001644 | Polyketide | 36.0 | 25.3 | 307.0 | 2.3e-82 |
| AAY89053.1 | polyketide\_synthase | BGC0001069 | NRP + Polyketide:Trans-AT type I | 40.0 | 20.5 | 306.0 | 3e-82 |
| AFN27483.1 | pks\_BonD | BGC0000173 | Polyketide:Modular type I | 37.0 | 23.3 | 305.0 | 5.1e-82 |
| CAD29795.1 | peptide\_synthetase | BGC0001015 | NRP + Polyketide | 30.0 | 33.2 | 305.0 | 6.7e-82 |
| BBA21069.1 | putative\_modular\_polyketide\_synthase | BGC0001740 | NRP + Polyketide | 35.0 | 28.3 | 305.0 | 6.7e-82 |
| AKG06377.1 | polyketide\_synthase\_type\_1 | BGC0001830 | Polyketide | 38.0 | 23.7 | 305.0 | 6.7e-82 |
| ABO15888.1 | polyketide\_synthase | BGC0000132 | Polyketide | 35.0 | 26.5 | 305.0 | 8.8e-82 |
| AIU36104.1 | LglE | BGC0000180 | Polyketide:Trans-AT type I | 36.0 | 23.2 | 305.0 | 8.8e-82 |
| ALD82522.1 | polyketide\_synthase | BGC0001212 | NRP + Polyketide | 40.0 | 20.2 | 305.0 | 8.8e-82 |
| AXN93597.1 | PuwB | BGC0001952 | NRP | 35.0 | 25.8 | 305.0 | 8.8e-82 |
| ASA76632.1 | polyketide\_synthase\_non-ribosomal\_peptide\_synthetase\_hybrid | BGC0001751 | NRP + Polyketide | 37.0 | 23.0 | 304.0 | 1.5e-81 |
| ATL73033.1 | type\_I\_modular\_polyketide\_synthase | BGC0001807 | NRP + Polyketide | 32.0 | 31.0 | 304.0 | 2e-81 |
| ABL74938.1 | PKS | BGC0001048 | NRP:Glycopeptide + Polyketide:Modular type I + Saccharide:Hybrid/tailoring | 33.0 | 31.6 | 303.0 | 3.3e-81 |
| CCC55921.1 | non-ribosomal\_peptide\_synthetase/polyketide\_synthase\_hybrid\_protein | BGC0000973 | NRP + Polyketide:Modular type I | 32.0 | 31.7 | 302.0 | 4.4e-81 |
| AAQ82565.1 | FscB | BGC0000061 | Polyketide | 33.0 | 31.1 | 302.0 | 7.4e-81 |
| ABI91467.1 | beta-ketoacyl\_synthase | BGC0001094 | NRP + Polyketide | 36.0 | 23.6 | 302.0 | 7.4e-81 |
| AGN11881.1 | tstDEF | BGC0001114 | NRP + Polyketide | 39.0 | 20.2 | 302.0 | 7.4e-81 |
| AAM12913.2 | MmpD | BGC0000182 | Polyketide:Iterative type I + Polyketide:Trans-AT type I | 37.0 | 23.0 | 301.0 | 9.7e-81 |
| ADI59532.1 | CorJ | BGC0001091 | NRP + Polyketide | 34.0 | 25.4 | 301.0 | 9.7e-81 |
| ABI91469.1 | beta-ketoacyl\_synthase | BGC0001094 | NRP + Polyketide | 38.0 | 23.9 | 301.0 | 9.7e-81 |
| ALD83704.1 | tAT\_polyketide\_synthase | BGC0001299 | Polyketide | 36.0 | 23.8 | 301.0 | 9.7e-81 |
| CAN93349.1 | polyketide\_synthase | BGC0000179 | Polyketide:Trans-AT type I | 37.0 | 23.0 | 301.0 | 1.3e-80 |
| AIU36103.1 | LglD | BGC0000180 | Polyketide:Trans-AT type I | 37.0 | 23.3 | 301.0 | 1.3e-80 |
| ABS90470.1 | NRPS/PKS | BGC0001106 | NRP + Polyketide | 35.0 | 27.6 | 301.0 | 1.3e-80 |
| ALD83702.1 | tAT\_polyketide\_synthase | BGC0001299 | Polyketide | 37.0 | 22.8 | 301.0 | 1.3e-80 |
| CDG12864.1 | non-ribosomal\_peptide\_synthetase | BGC0001415 | NRP | 32.0 | 32.9 | 301.0 | 1.3e-80 |
| ARR97037.1 | SphD | BGC0001780 | NRP | 36.0 | 24.0 | 300.0 | 1.7e-80 |
| AAZ77698.1 | ChlA5 | BGC0000036 | Polyketide:Modular type I + Polyketide:Iterative type I + Saccharide:Oligosaccharide | 31.0 | 31.9 | 300.0 | 2.2e-80 |
| ANR02553.1 | LodL | BGC0001648 | Polyketide | 33.0 | 31.6 | 300.0 | 2.2e-80 |
| ARW71485.1 | type\_I\_PKS\_module\_4,\_module\_5 | BGC0001812 | Polyketide | 34.0 | 30.6 | 300.0 | 2.2e-80 |
| AAY32965.1 | DszB | BGC0001093 | NRP + Polyketide | 35.0 | 26.5 | 300.0 | 2.8e-80 |
| AAP42873.1 | NanA11 | BGC0000105 | Polyketide | 33.0 | 31.0 | 299.0 | 3.7e-80 |
| AQH32483.1 | hybrid\_peptide\_synthetase/polyketide\_synthase | BGC0001667 | NRP + Polyketide | 29.0 | 32.9 | 299.0 | 3.7e-80 |
| AGY62759.1 | EbeG | BGC0000051 | Polyketide | 36.0 | 23.2 | 299.0 | 4.8e-80 |
| WP\_019634550.1 | type\_I\_polyketide\_synthase | BGC0001443 | NRP + Polyketide | 33.0 | 31.7 | 299.0 | 4.8e-80 |
| SCN11953.1 | ebeE-type\_I\_polyketide\_synthase | BGC0001580 | Polyketide | 36.0 | 23.2 | 299.0 | 4.8e-80 |
| AGO59040.1 | PtaA | BGC0000121 | Polyketide | 30.0 | 32.8 | 299.0 | 6.3e-80 |
| AAF00957.1 | mcyG | BGC0001017 | NRP + Polyketide:Modular type I | 30.0 | 34.0 | 299.0 | 6.3e-80 |
| ABF92489.1 | mixed\_type\_I\_polyketide\_synthase\_-\_peptide\_synthetase | BGC0001025 | NRP + Polyketide:Trans-AT type I | 36.0 | 24.8 | 299.0 | 6.3e-80 |
| ASA76642.1 | polyketide\_synthase | BGC0001751 | NRP + Polyketide | 37.0 | 23.4 | 299.0 | 6.3e-80 |
| CBF74114.1 | Conidial\_yellow\_pigment\_biosynthesis\_polyketide\_synthase\_(PKS)(EC\_2.3.1.-)\_[Source:UniProtKB/Swiss-Prot;Acc:Q03149] | BGC0000107 | Polyketide | 31.0 | 32.6 | 298.0 | 8.2e-80 |
| CAG23957.2 | hybrid\_NRPS/PKS\_protein | BGC0001089 | Polyketide + NRP | 35.0 | 25.1 | 298.0 | 8.2e-80 |
| DAC76734.1 | type\_I\_polyketide\_synthase/non-ribosomal\_peptide\_synthetase | BGC0001885 | Polyketide | 39.0 | 21.9 | 298.0 | 8.2e-80 |
| AEN83889.1 | AdaA | BGC0000156 | Polyketide:Iterative type I | 31.0 | 32.9 | 298.0 | 1.1e-79 |
| AHD05615.1 | putative\_non-ribosomal\_peptide\_ligase/\_polyketide\_synthase\_hybrid | BGC0001033 | NRP + Polyketide | 31.0 | 32.1 | 298.0 | 1.1e-79 |
| CAG23977.1 | polyketide\_synthase\_type\_I | BGC0000176 | Polyketide + NRP | 33.0 | 26.1 | 297.0 | 1.8e-79 |
| ADH01487.1 | polyketide\_synthase | BGC0001096 | NRP + Polyketide | 34.0 | 25.7 | 297.0 | 1.8e-79 |
| AIC32693.1 | FR9DEF | BGC0001113 | NRP + Polyketide | 34.0 | 25.7 | 297.0 | 1.8e-79 |
| ADB23403.1 | polyketide\_synthase\_type\_I | BGC0001062 | Polyketide | 37.0 | 23.8 | 297.0 | 2.4e-79 |
| AJY78093.1 | polyketide\_synthase | BGC0001902 | NRP + Polyketide | 34.0 | 30.4 | 296.0 | 4.1e-79 |
| AFX60311.1 | polyketide\_synthase | BGC0001031 | NRP + Polyketide | 37.0 | 24.1 | 295.0 | 5.3e-79 |
| AAP42872.1 | NanA9 | BGC0000105 | Polyketide | 32.0 | 29.2 | 295.0 | 7e-79 |
| ABC34832.1 | polyketide\_synthase | BGC0000186 | NRP + Polyketide:Modular type I | 38.0 | 20.4 | 295.0 | 7e-79 |
| DAC80074.1 | PKS | BGC0001836 | Polyketide:Trans-AT type I | 35.0 | 23.5 | 295.0 | 7e-79 |
| ADY00130.1 | polyketide\_synthase | BGC0000104 | Terpene + Polyketide:Iterative type I | 31.0 | 32.7 | 295.0 | 9.1e-79 |
| ABC87510.1 | polyketide\_synthase | BGC0001011 | NRP + Polyketide | 32.0 | 33.9 | 295.0 | 9.1e-79 |
| ctg1\_orf21 |  | BGC0001013 | NRP + Polyketide | 32.0 | 33.9 | 295.0 | 9.1e-79 |
| AAF71767.1 | nysJ | BGC0000115 | Polyketide:Modular type I + Saccharide:Hybrid/tailoring | 34.0 | 30.0 | 294.0 | 1.2e-78 |
| BAB69192.1 | modular\_polyketide\_synthase | BGC0000117 | Polyketide | 34.0 | 30.0 | 294.0 | 1.2e-78 |
| ACY01391.1 | AT-less\_polyketide\_synthase | BGC0000177 | Polyketide:Modular type I + Polyketide:Trans-AT type I | 36.0 | 23.6 | 294.0 | 1.2e-78 |
| ADD82941.1 | Bat3 | BGC0001099 | NRP + Polyketide:Modular type I + Polyketide:Trans-AT type I | 35.0 | 23.5 | 294.0 | 1.2e-78 |
| AJW65407.1 | type\_I\_modular\_polyketide\_synthase | BGC0001195 | NRP + Polyketide | 32.0 | 31.1 | 294.0 | 1.2e-78 |
| ADN68477.1 | SorB | BGC0000184 | Polyketide:Trans-AT type I | 37.0 | 23.7 | 294.0 | 1.6e-78 |
| AKA54627.1 | PKS | BGC0001216 | NRP + Polyketide | 32.0 | 32.8 | 294.0 | 1.6e-78 |
| AMH40443.1 | PKS | BGC0001350 | Polyketide | 37.0 | 24.3 | 294.0 | 1.6e-78 |
| DAC80073.1 | PKS | BGC0001836 | Polyketide:Trans-AT type I | 35.0 | 22.9 | 294.0 | 1.6e-78 |
| AFN27480.1 | pks\_BonA | BGC0000173 | Polyketide:Modular type I | 33.0 | 31.6 | 294.0 | 2e-78 |
| AFD30954.1 | CrmA | BGC0000966 | NRP + Polyketide | 32.0 | 32.7 | 294.0 | 2e-78 |
| AFX60336.1 | polyketide\_synthase | BGC0001032 | NRP + Polyketide | 34.0 | 24.7 | 294.0 | 2e-78 |
| AVX51108.1 | nysC | BGC0001709 | Polyketide | 35.0 | 30.0 | 294.0 | 2e-78 |
| ABW96542.1 | type\_I\_modular\_polyketide\_synthase | BGC0000159 | Polyketide:Modular type I | 33.0 | 31.0 | 293.0 | 3.5e-78 |
| AAV97870.1 | OnnB | BGC0001105 | NRP + Polyketide:Trans-AT type I | 35.0 | 24.1 | 293.0 | 3.5e-78 |
| AHE80994.1 | PieA4 | BGC0001169 | Polyketide:Modular type I | 36.0 | 24.0 | 293.0 | 3.5e-78 |
| EAL89339.1 | polyketide\_synthase,\_putative | BGC0001403 | Polyketide | 29.0 | 32.7 | 293.0 | 3.5e-78 |
| CAN89632.1 | putative\_polyketide\_synthase | BGC0001070 | NRP + Polyketide:Modular type I + Polyketide:Trans-AT type I | 35.0 | 24.7 | 292.0 | 4.5e-78 |
| ACY01401.1 | AT-less\_polyketide\_synthase | BGC0000083 | Polyketide:Modular type I + Polyketide:Trans-AT type I | 34.0 | 24.3 | 292.0 | 5.9e-78 |
| AAF71776.1 | nysC | BGC0000115 | Polyketide:Modular type I + Saccharide:Hybrid/tailoring | 35.0 | 29.9 | 292.0 | 5.9e-78 |
| AEZ54377.1 | PieA4 | BGC0000124 | Polyketide | 36.0 | 24.3 | 292.0 | 5.9e-78 |
| ERM18798.1 | polyketide\_synthase | BGC0000172 | Polyketide | 38.0 | 20.4 | 292.0 | 5.9e-78 |
| ABC34675.1 | polyketide\_synthase,\_putative | BGC0000186 | NRP + Polyketide:Modular type I | 37.0 | 26.5 | 292.0 | 5.9e-78 |
| RAT98529.1 | trans-acyltransferase\_polyketide\_synthase | BGC0001470 | Polyketide:Trans-AT type I | 35.0 | 24.7 | 292.0 | 5.9e-78 |
| AAY89050.1 | polyketide\_synthase | BGC0001069 | NRP + Polyketide:Trans-AT type I | 37.0 | 23.7 | 292.0 | 7.7e-78 |
| AJW65408.1 | type\_I\_modular\_polyketide\_synthase | BGC0001195 | NRP + Polyketide | 36.0 | 23.5 | 291.0 | 1e-77 |
| RAT98527.1 | trans-acyltransferase\_polyketide\_synthase | BGC0001470 | Polyketide:Trans-AT type I | 36.0 | 23.0 | 291.0 | 1e-77 |
| AWS21290.1 | type\_I\_polyketide\_synthase | BGC0001934 | Polyketide | 32.0 | 31.3 | 291.0 | 1e-77 |
| RAT98517.1 | trans-acyltransferase\_polyketide\_synthase | BGC0001470 | Polyketide:Trans-AT type I | 35.0 | 23.1 | 291.0 | 1.3e-77 |
| ATX68112.1 | malonyl\_CoA-acyl\_carrier\_protein\_transacylase | BGC0001772 | Polyketide | 33.0 | 23.9 | 291.0 | 1.3e-77 |
| CCT67991.1 | bikaverin\_cluster-polyketide\_synthase | BGC0000030 | Polyketide | 31.0 | 32.8 | 290.0 | 1.7e-77 |
| AFL48528.1 | laidlomycin\_polyketide\_synthase\_(module\_7\_and\_module\_8) | BGC0000084 | Polyketide | 37.0 | 23.2 | 290.0 | 1.7e-77 |
| AKA59091.1 | type-I\_PKS | BGC0001619 | Polyketide | 32.0 | 31.1 | 290.0 | 1.7e-77 |
| ATY46587.1 | polyketide\_synthase | BGC0001666 | Polyketide | 35.0 | 29.8 | 290.0 | 1.7e-77 |
| ATG32074.1 | putative\_nonfunctional\_polyketide\_synthase\_module | BGC0001750 | NRP + Polyketide | 35.0 | 23.8 | 290.0 | 1.7e-77 |
| EAL84397.1 | polyketide\_synthase | BGC0001118 | Polyketide:Iterative type I | 30.0 | 32.5 | 290.0 | 2.2e-77 |
| PHM26614.1 | Phthiocerol\_synthesis\_polyketide\_synthase\_type\_I\_PpsE | BGC0001130 | NRP + Polyketide | 31.0 | 33.0 | 290.0 | 2.2e-77 |
| AAZ77694.1 | ChlA2 | BGC0000036 | Polyketide:Modular type I + Polyketide:Iterative type I + Saccharide:Oligosaccharide | 32.0 | 31.1 | 290.0 | 2.9e-77 |
| ctg1\_orf5 |  | BGC0001329 | Polyketide + NRP:Cyclic depsipeptide | 35.0 | 25.6 | 290.0 | 2.9e-77 |
| AVV61983.1 | type\_I\_modular\_polyketide\_synthase | BGC0001477 | NRP + Polyketide:Modular type I | 32.0 | 30.9 | 290.0 | 2.9e-77 |
| AEH59109.1 | keto-hydroxyglutarate-aldolase/polyketide\_synthase | BGC0000385 | NRP | 35.0 | 23.7 | 289.0 | 3.8e-77 |
| ABF88102.1 | polyketide\_synthase | BGC0001025 | NRP + Polyketide:Trans-AT type I | 36.0 | 23.3 | 289.0 | 3.8e-77 |
| BBA21072.1 | putative\_modular\_polyketide\_synthase | BGC0001740 | NRP + Polyketide | 37.0 | 23.0 | 289.0 | 3.8e-77 |
| AZY91988.1 | polyketide\_synthase | BGC0002022 | Polyketide | 32.0 | 29.8 | 289.0 | 3.8e-77 |
| ABJ97438.1 | MerB | BGC0001012 | NRP + Polyketide | 33.0 | 31.1 | 289.0 | 5e-77 |
| BAF50722.1 | polyketide\_synthase | BGC0001116 | NRP + Polyketide | 37.0 | 20.6 | 289.0 | 5e-77 |
| ADM79459.1 | PKS16\_protein | BGC0001266 | Polyketide | 30.0 | 32.9 | 289.0 | 5e-77 |
| ASX95227.1 | IlaE | BGC0001620 | Polyketide | 28.0 | 50.2 | 289.0 | 5e-77 |
| AJQ95708.1 | polyketide\_synthase\_modules-related\_protein | BGC0001644 | Polyketide | 32.0 | 23.7 | 289.0 | 5e-77 |
| ATY69589.1 | type\_I\_polyketide\_synthase | BGC0001823 | NRP + Polyketide | 35.0 | 24.4 | 289.0 | 5e-77 |
| gene6 |  | BGC0001906 | Polyketide | 31.0 | 32.6 | 289.0 | 5e-77 |
| CAN93347.1 | Polyketide\_synthase | BGC0000179 | Polyketide:Trans-AT type I | 36.0 | 25.4 | 289.0 | 6.5e-77 |
| CBF70387.1 | polyketide\_synthase,\_putative\_(JCVI) | BGC0000684 | Terpene | 30.0 | 32.9 | 289.0 | 6.5e-77 |
| BBA21068.1 | putative\_non-ribosomal\_peptide\_synthetase | BGC0001740 | NRP + Polyketide | 35.0 | 24.1 | 289.0 | 6.5e-77 |
| AAA79984.2 | soraphen\_polyketide\_synthase\_B | BGC0000147 | Polyketide:Modular type I | 32.0 | 30.7 | 288.0 | 8.5e-77 |
| AEU17897.1 | putative\_type\_I\_PKS | BGC0001072 | Saccharide + Polyketide:Modular type I + Polyketide:Type II + Other:Aminocoumarin | 33.0 | 31.1 | 288.0 | 8.5e-77 |
| RAT98518.1 | trans-acyltransferase\_polyketide\_synthase | BGC0001470 | Polyketide:Trans-AT type I | 36.0 | 23.5 | 288.0 | 8.5e-77 |
| CAL58684.1 | polyketide\_synthase | BGC0000149 | Polyketide:Modular type I | 32.0 | 31.0 | 288.0 | 1.1e-76 |
| ADN68484.1 | sorI | BGC0000184 | Polyketide:Trans-AT type I | 35.0 | 24.3 | 287.0 | 1.5e-76 |
| CCA89327.1 | trans-AT\_type\_I\_polyketide\_synthase | BGC0001111 | NRP + Polyketide:Trans-AT type I | 35.0 | 23.1 | 287.0 | 1.5e-76 |
| ALJ49922.1 | TtmG | BGC0001236 | Polyketide | 37.0 | 21.2 | 287.0 | 1.5e-76 |
| BBA20952.1 | type\_I\_polyketide\_synthase | BGC0001763 | NRP + Polyketide | 28.0 | 50.2 | 287.0 | 1.5e-76 |
| ABK32288.1 | JerB | BGC0000080 | Polyketide | 33.0 | 32.2 | 287.0 | 1.9e-76 |
| AEZ53946.1 | polyketide\_synthase | BGC0000144 | Polyketide:Modular type I | 36.0 | 22.5 | 287.0 | 1.9e-76 |
| CAJ76291.1 | putative\_polyketide\_synthase | BGC0000972 | NRP + Polyketide:Modular type I + Polyketide:Trans-AT type I | 32.0 | 32.0 | 287.0 | 1.9e-76 |
| BAJ16468.1 | polyketide\_synthase | BGC0000058 | Polyketide | 33.0 | 30.9 | 287.0 | 2.5e-76 |
| ADN68483.1 | sorH | BGC0000184 | Polyketide:Trans-AT type I | 35.0 | 23.9 | 287.0 | 2.5e-76 |
| AEC04357.1 | polyketide\_synthase | BGC0000178 | Polyketide:Trans-AT type I | 35.0 | 23.0 | 286.0 | 3.2e-76 |
| EHK80169.1 | acyl\_transferase | BGC0001447 | Polyketide | 33.0 | 31.0 | 286.0 | 3.2e-76 |
| ART41209.1 | AdrD | BGC0001508 | Polyketide | 31.0 | 32.9 | 286.0 | 3.2e-76 |
| CAQ52624.1 | type\_I\_polyketide\_synthase,\_modules\_7-8 | BGC0001066 | Polyketide:Modular type I | 33.0 | 30.8 | 286.0 | 4.2e-76 |
| EHK80170.1 | acyl\_transferase | BGC0001447 | Polyketide | 33.0 | 30.9 | 286.0 | 4.2e-76 |
| RAT98530.1 | trans-acyltransferase\_polyketide\_synthase | BGC0001470 | Polyketide:Trans-AT type I | 35.0 | 23.6 | 286.0 | 4.2e-76 |
| ACR50796.1 | putative\_polyketide\_synthase | BGC0000163 | Polyketide | 35.0 | 23.4 | 285.0 | 5.5e-76 |
| OEI73460.1 | hypothetical\_protein | BGC0001520 | Polyketide | 35.0 | 24.2 | 285.0 | 5.5e-76 |
| AFX60332.1 | polyketide\_synthase | BGC0001032 | NRP + Polyketide | 34.0 | 23.6 | 285.0 | 7.2e-76 |
| AKA59437.1 | polyketide\_synthase | BGC0001202 | NRP + Polyketide | 31.0 | 32.6 | 285.0 | 9.4e-76 |
| ANR02554.1 | LodM | BGC0001648 | Polyketide | 33.0 | 30.7 | 285.0 | 9.4e-76 |
| ADI24953.1 | GsfA | BGC0000070 | Polyketide:Iterative type I | 30.0 | 32.3 | 284.0 | 1.2e-75 |
| AFX60309.1 | polyketide\_synthase | BGC0001031 | NRP + Polyketide | 34.0 | 25.2 | 284.0 | 1.2e-75 |
| AAY89052.1 | polyketide\_synthase | BGC0001069 | NRP + Polyketide:Trans-AT type I | 37.0 | 24.3 | 284.0 | 1.2e-75 |
| OEI73466.1 | hypothetical\_protein | BGC0001520 | Polyketide | 35.0 | 23.5 | 284.0 | 1.2e-75 |
| CTQ34882.1 | AtcE;\_polyketide\_synthase,\_modules\_5-7 | BGC0001301 | Polyketide | 34.0 | 23.0 | 284.0 | 1.6e-75 |
| CCG06113.1 | type\_I\_polyketide\_synthase | BGC0001543 | Polyketide | 34.0 | 23.8 | 284.0 | 1.6e-75 |
| CAN93351.1 | polyketide\_synthase | BGC0000179 | Polyketide:Trans-AT type I | 35.0 | 23.0 | 284.0 | 2.1e-75 |
| CAQ34916.1 | polyketide\_synthase | BGC0000986 | NRP + Polyketide | 37.0 | 20.1 | 284.0 | 2.1e-75 |
| AAS47562.1 | mixed\_type\_I\_polyketide\_synthase\_-\_peptide\_synthetase | BGC0001108 | Polyketide:Trans-AT type I | 35.0 | 22.4 | 284.0 | 2.1e-75 |
| ctg1\_orf8 |  | BGC0001109 | NRP + Polyketide | 35.0 | 22.4 | 284.0 | 2.1e-75 |
| AAD38786.1 | polyketide\_synthase | BGC0001257 | Polyketide | 30.0 | 32.8 | 284.0 | 2.1e-75 |
| BAF50727.1 | hybrid\_polyketide\_synthase-non\_ribosomal\_peptide\_synthetase | BGC0001116 | NRP + Polyketide | 35.0 | 24.7 | 283.0 | 2.7e-75 |
| BBF25315.1 | polyketide\_synthase | BGC0001923 | Terpene + Polyketide | 31.0 | 33.4 | 283.0 | 2.7e-75 |
| ABB88521.1 | polyketide\_synthase\_type\_I | BGC0000050 | Polyketide | 38.0 | 23.5 | 283.0 | 3.6e-75 |
| BAD08358.1 | polyketide\_synthase\_modules\_4 | BGC0000167 | Polyketide | 33.0 | 31.4 | 283.0 | 3.6e-75 |
| CAJ57409.1 | polyketide\_synthase\_type\_I | BGC0000176 | Polyketide + NRP | 35.0 | 25.1 | 283.0 | 3.6e-75 |
| ACY01392.1 | AT-less\_polyketide\_synthase | BGC0000177 | Polyketide:Modular type I + Polyketide:Trans-AT type I | 35.0 | 23.7 | 283.0 | 3.6e-75 |
| CAM00064.1 | EryAII\_Erythromycin\_polyketide\_synthase\_modules\_3\_and\_4 | BGC0000055 | Polyketide:Modular type I + Saccharide:Hybrid/tailoring | 39.0 | 23.1 | 282.0 | 4.7e-75 |
| AEZ53949.1 | polyketide\_synthase | BGC0000144 | Polyketide:Modular type I | 38.0 | 23.2 | 282.0 | 4.7e-75 |
| ABW96540.1 | type\_I\_modular\_polyketide\_synthase | BGC0000159 | Polyketide:Modular type I | 31.0 | 31.5 | 282.0 | 4.7e-75 |
| ADI59531.1 | CorI | BGC0001091 | NRP + Polyketide | 33.0 | 23.5 | 282.0 | 6.1e-75 |
| ABK32256.1 | AmbB | BGC0000014 | Polyketide | 33.0 | 32.4 | 282.0 | 8e-75 |
| CAJ87591.1 | putative\_peptide/polyketide\_synthetase | BGC0001055 | NRP + Polyketide | 31.0 | 32.5 | 281.0 | 1e-74 |
| CRI73799.1 | CongC\_protein | BGC0001215 | NRP | 33.0 | 32.1 | 281.0 | 1e-74 |
| ABM63528.1 | BryC | BGC0000174 | Polyketide | 34.0 | 24.5 | 281.0 | 1.4e-74 |
| AFO59866.1 | ChxE | BGC0000175 | Polyketide:Trans-AT type I | 35.0 | 23.3 | 280.0 | 1.8e-74 |
| CCA89326.1 | mixed\_trans-AT\_type\_I\_polyketide\_synthase/nonribosomal\_peptide\_synthetase | BGC0001111 | NRP + Polyketide:Trans-AT type I | 32.0 | 26.6 | 280.0 | 1.8e-74 |
| CCP20048.1 | divL1\_protein | BGC0001119 | Polyketide:Modular type I | 32.0 | 31.1 | 280.0 | 1.8e-74 |
| CDM36726.1 | Beta-ketoacyl\_synthase | BGC0001360 | Polyketide | 30.0 | 32.8 | 280.0 | 1.8e-74 |
| ADN68476.1 | sorA | BGC0000184 | Polyketide:Trans-AT type I | 34.0 | 24.0 | 280.0 | 2.3e-74 |
| AKQ22680.1 | malonyl\_CoA-acyl\_carrier\_protein\_transacylase | BGC0001656 | Polyketide | 33.0 | 23.9 | 280.0 | 2.3e-74 |
| ATL73034.1 | type\_I\_modular\_polyketide\_synthase | BGC0001807 | NRP + Polyketide | 34.0 | 29.9 | 280.0 | 2.3e-74 |
| BAG85027.1 | putative\_polyketide\_synthase | BGC0000086 | Polyketide | 32.0 | 31.8 | 280.0 | 3e-74 |
| CAQ64687.1 | lasalocid\_modular\_polyketide\_synthase | BGC0000087 | Polyketide | 32.0 | 31.8 | 280.0 | 3e-74 |
| ctg1\_orf521 |  | BGC0001199 | Polyketide | 35.0 | 23.4 | 280.0 | 3e-74 |
| AAB66506.1 | tylactone\_synthase\_modules\_4\_&\_5 | BGC0000166 | Polyketide | 36.0 | 23.3 | 279.0 | 4e-74 |
| AEC04361.1 | polyketide\_synthase | BGC0000178 | Polyketide:Trans-AT type I | 34.0 | 23.7 | 279.0 | 4e-74 |
| AEC04362.1 | polyketide\_synthase | BGC0000178 | Polyketide:Trans-AT type I | 33.0 | 24.8 | 279.0 | 4e-74 |
| AFX60340.1 | polyketide\_synthase | BGC0001032 | NRP + Polyketide | 32.0 | 30.4 | 279.0 | 4e-74 |
| ABS90478.1 | PKS | BGC0001106 | NRP + Polyketide | 36.0 | 25.8 | 279.0 | 4e-74 |
| CTQ34883.1 | AtcF;\_polyketide\_synthase,\_modules\_8-10 | BGC0001301 | Polyketide | 35.0 | 23.5 | 279.0 | 4e-74 |
| EWM62998.1 | mycocerosic\_acid\_synthase | BGC0001328 | NRP:Cyclic depsipeptide + Polyketide:Modular type I | 36.0 | 22.3 | 279.0 | 4e-74 |
| RAT98525.1 | trans-acyltransferase\_polyketide\_synthase | BGC0001470 | Polyketide:Trans-AT type I | 34.0 | 24.0 | 279.0 | 4e-74 |
| ARM20280.1 | polyketide\_synthase | BGC0001523 | Polyketide | 32.0 | 30.0 | 279.0 | 4e-74 |
| ARV85763.1 | PieA4\_type\_I\_PKS | BGC0001742 | Polyketide | 32.0 | 31.1 | 279.0 | 4e-74 |
| PKX92308.1 | putative\_polyketide\_synthase | BGC0001988 | Polyketide | 29.0 | 33.9 | 279.0 | 4e-74 |
| CAG23964.1 | polyketide\_synthase\_type\_I | BGC0000181 | Polyketide | 33.0 | 23.7 | 279.0 | 5.2e-74 |
| ABF89568.1 | polyketide\_synthase | BGC0001025 | NRP + Polyketide:Trans-AT type I | 35.0 | 23.1 | 279.0 | 5.2e-74 |
| SCN11952.1 | ebeD-type\_I\_polyketide\_synthase | BGC0001580 | Polyketide | 38.0 | 22.2 | 279.0 | 5.2e-74 |
| EAA59563.1 | polyketide\_synthase | BGC0000057 | Polyketide:Iterative type I | 31.0 | 33.6 | 279.0 | 6.7e-74 |
| ABC34154.1 | thiotemplate\_mechanism\_natural\_product\_synthetase | BGC0000961 | NRP + Polyketide | 34.0 | 25.8 | 279.0 | 6.7e-74 |
| ACY01402.1 | AT-less\_polyketide\_synthase | BGC0000083 | Polyketide:Modular type I + Polyketide:Trans-AT type I | 34.0 | 24.5 | 278.0 | 8.8e-74 |
| AGN74894.1 | nonribosomal\_peptide\_synthetase/polyketide\_synthase\_hybrid\_protein | BGC0000459 | NRP:Cyclic depsipeptide + Polyketide:Trans-AT type I | 35.0 | 23.1 | 278.0 | 8.8e-74 |
| AJQ95704.1 | polyketide\_synthase\_modules-related\_protein | BGC0001644 | Polyketide | 32.0 | 24.5 | 278.0 | 8.8e-74 |
| CAG23965.1 | polyketide\_synthase\_type\_I | BGC0000181 | Polyketide | 33.0 | 25.2 | 278.0 | 1.1e-73 |
| ADC45538.1 | modular\_polyketide\_synthase | BGC0000093 | Polyketide | 34.0 | 29.9 | 277.0 | 1.5e-73 |
| CAA73127.1 | HMWP1\_protein | BGC0000467 | NRP | 31.0 | 32.7 | 277.0 | 1.5e-73 |
| OEI73462.1 | hypothetical\_protein | BGC0001520 | Polyketide | 35.0 | 24.0 | 277.0 | 1.5e-73 |
| AEC04363.1 | polyketide\_synthase | BGC0000178 | Polyketide:Trans-AT type I | 35.0 | 24.9 | 277.0 | 2e-73 |
| WP\_010639241.1 | type\_I\_polyketide\_synthase | BGC0000958 | NRP:Cyclic depsipeptide + Polyketide:Modular type I | 32.0 | 29.6 | 277.0 | 2e-73 |
| CAL69889.1 | RhiB\_protein | BGC0001112 | NRP + Polyketide:Trans-AT type I | 33.0 | 24.9 | 277.0 | 2e-73 |
| ALD83687.1 | tAT\_polyketide\_synthase | BGC0001300 | Polyketide | 37.0 | 21.2 | 277.0 | 2.6e-73 |
| AIJ04680.1 | polyketide\_synthase | BGC0001383 | Polyketide | 33.0 | 24.6 | 277.0 | 2.6e-73 |
| OEI73463.1 | hypothetical\_protein | BGC0001520 | Polyketide | 34.0 | 23.7 | 277.0 | 2.6e-73 |
| CAQ52622.1 | type\_I\_polyketide\_synthase,\_modules\_4-5 | BGC0001066 | Polyketide:Modular type I | 36.0 | 23.3 | 276.0 | 3.3e-73 |
| ADI59533.1 | CorK | BGC0001091 | NRP + Polyketide | 32.0 | 25.6 | 276.0 | 3.3e-73 |
| AAY32964.1 | DszA | BGC0001093 | NRP + Polyketide | 36.0 | 23.5 | 276.0 | 3.3e-73 |
| AWX24483.1 | type\_I\_polyketide\_synthase | BGC0001695 | NRP | 32.0 | 32.5 | 276.0 | 3.3e-73 |
| ASA76643.1 | polyketide\_synthase | BGC0001751 | NRP + Polyketide | 35.0 | 24.0 | 276.0 | 3.3e-73 |
| CAE45671.1 | borrelidin\_polyketide\_synthase,\_type\_I | BGC0000031 | Polyketide:Modular type I | 32.0 | 30.5 | 276.0 | 4.4e-73 |
| ABM63527.1 | BryB | BGC0000174 | Polyketide | 35.0 | 23.8 | 275.0 | 5.7e-73 |
| AIJ04681.1 | polyketide\_synthase | BGC0001383 | Polyketide | 33.0 | 23.7 | 275.0 | 5.7e-73 |
| ALV82345.1 | borrelidin\_type\_I\_polyketide\_synthase | BGC0001533 | Polyketide | 32.0 | 30.5 | 275.0 | 5.7e-73 |
| DAC80077.1 | PKS | BGC0001836 | Polyketide:Trans-AT type I | 34.0 | 24.0 | 275.0 | 5.7e-73 |
| AAC69330.1 | type\_I\_polyketide\_synthase\_PikAII | BGC0000094 | Polyketide:Modular type I + Saccharide:Hybrid/tailoring | 32.0 | 31.9 | 275.0 | 7.5e-73 |
| ABC38101.1 | polyketide\_synthase | BGC0000964 | NRP:Cyclic depsipeptide + Polyketide:Trans-AT type I | 36.0 | 23.3 | 275.0 | 7.5e-73 |
| BAP05593.1 | calE | BGC0000967 | NRP + Polyketide:Trans-AT type I | 34.0 | 23.4 | 275.0 | 7.5e-73 |
| DAC76730.1 | type\_I\_polyketide\_synthase | BGC0001885 | Polyketide | 35.0 | 24.7 | 275.0 | 7.5e-73 |
| AVI57434.1 | AbmB2 | BGC0001694 | Polyketide | 33.0 | 31.3 | 274.0 | 1.3e-72 |
| AMH40421.1 | PKS | BGC0001350 | Polyketide | 34.0 | 22.9 | 274.0 | 1.7e-72 |
| AMH40423.1 | PKS | BGC0001350 | Polyketide | 37.0 | 23.5 | 274.0 | 1.7e-72 |
| EHK80166.1 | beta-ketoacyl\_synthase | BGC0001447 | Polyketide | 36.0 | 23.0 | 274.0 | 1.7e-72 |
| BAP05589.1 | calA | BGC0000967 | NRP + Polyketide:Trans-AT type I | 34.0 | 25.6 | 274.0 | 2.2e-72 |
| AFR69335.1 | polyketide\_synthase\_SpiC2 | BGC0001045 | NRP:Cyclic depsipeptide + Polyketide:Modular type I | 36.0 | 24.0 | 274.0 | 2.2e-72 |
| BAG85030.1 | putative\_polyketide\_synthase | BGC0000086 | Polyketide | 31.0 | 31.0 | 273.0 | 2.8e-72 |
| CAQ64690.1 | lasalocid\_modular\_polyketide\_synthase | BGC0000087 | Polyketide | 31.0 | 31.0 | 273.0 | 2.8e-72 |
| BAC76471.1 | type\_I\_polyketide\_synthase\_LkcF | BGC0001100 | NRP + Polyketide | 34.0 | 23.1 | 273.0 | 2.8e-72 |
| WP\_003598535.1 | SDR\_family\_NAD(P)-dependent\_oxidoreductase | BGC0001991 | Polyketide | 32.0 | 26.5 | 273.0 | 2.8e-72 |
| CAN89635.1 | putative\_polyketide\_synthase | BGC0001070 | NRP + Polyketide:Modular type I + Polyketide:Trans-AT type I | 37.0 | 21.9 | 273.0 | 3.7e-72 |
| OJF16269.1 | AceP2 | BGC0001491 | Polyketide | 32.0 | 31.4 | 273.0 | 3.7e-72 |
| ASA76633.1 | polyketide\_synthase | BGC0001751 | NRP + Polyketide | 35.0 | 24.8 | 273.0 | 3.7e-72 |
| ANZ22986.1 | ZinC | BGC0001828 | Polyketide | 32.0 | 31.1 | 273.0 | 3.7e-72 |
| BAE93740.1 | type\_I\_polyketide\_synthase-related\_protein | BGC0000164 | Polyketide | 34.0 | 24.4 | 272.0 | 4.8e-72 |
| CAD15512.1 | polyketide\_synthase | BGC0001014 | NRP:NRP siderophore + Polyketide:Modular type I + Polyketide:Iterative type I | 35.0 | 24.1 | 272.0 | 4.8e-72 |
| ABC84459.1 | NigAIV | BGC0000114 | Polyketide:Modular type I | 33.0 | 29.8 | 272.0 | 6.3e-72 |
| CAG23958.2 | polyketide\_synthase\_of\_type\_I | BGC0001089 | Polyketide + NRP | 32.0 | 24.2 | 272.0 | 6.3e-72 |
| BAD38875.1 | polyketide\_synthase | BGC0000111 | Polyketide | 33.0 | 24.7 | 272.0 | 8.2e-72 |
| BAP05595.1 | calG | BGC0000967 | NRP + Polyketide:Trans-AT type I | 35.0 | 23.4 | 272.0 | 8.2e-72 |
| AFX60317.1 | polyketide\_synthase | BGC0001031 | NRP + Polyketide | 34.0 | 24.2 | 272.0 | 8.2e-72 |
| CCP20050.1 | divL3\_protein | BGC0001119 | Polyketide:Modular type I | 32.0 | 31.0 | 272.0 | 8.2e-72 |
| AAC46026.1 | polyketide\_synthase\_modules\_4\_and\_5 | BGC0000113 | Polyketide | 36.0 | 23.3 | 271.0 | 1.1e-71 |
| BAV19379.1 | polyketide\_synthase | BGC0001390 | NRP + Polyketide | 30.0 | 32.2 | 271.0 | 1.1e-71 |
| AFV30250.1 | polyketide\_synthase | BGC0000075 | Polyketide | 34.0 | 24.2 | 271.0 | 1.4e-71 |
| AJQ95705.1 | polyketide\_synthase\_modules-related\_protein | BGC0001644 | Polyketide | 35.0 | 25.8 | 271.0 | 1.4e-71 |
| ARR97040.1 | SphG | BGC0001780 | NRP | 35.0 | 22.9 | 271.0 | 1.4e-71 |
| WP\_012753526.1 | polyketide\_synthase | BGC0001991 | Polyketide | 33.0 | 24.2 | 271.0 | 1.4e-71 |
| AFV30248.1 | polyketide\_synthase | BGC0000075 | Polyketide | 31.0 | 31.6 | 270.0 | 2.4e-71 |
| CAO85897.1 | modular\_polyketide\_synthase\_NorB | BGC0000110 | Polyketide:Modular type I | 36.0 | 22.7 | 270.0 | 2.4e-71 |
| AAM12911.1 | MmpB | BGC0000182 | Polyketide:Iterative type I + Polyketide:Trans-AT type I | 34.0 | 25.7 | 270.0 | 2.4e-71 |
| BAC76476.1 | multifunctional\_polyketide-peptide\_synthase\_LkcA | BGC0001100 | NRP + Polyketide | 36.0 | 23.3 | 270.0 | 2.4e-71 |
| ADD82940.1 | Bat2 | BGC0001099 | NRP + Polyketide:Modular type I + Polyketide:Trans-AT type I | 35.0 | 22.8 | 270.0 | 3.1e-71 |
| ABS90475.1 | PKS | BGC0001106 | NRP + Polyketide | 34.0 | 26.5 | 269.0 | 4.1e-71 |
| ATV82110.1 | PKS | BGC0001909 | Polyketide | 30.0 | 33.4 | 269.0 | 4.1e-71 |
| WP\_016638480.1 | type\_I\_polyketide\_synthase | BGC0001519 | NRP + Polyketide | 32.0 | 31.9 | 269.0 | 5.3e-71 |
| ATX68127.1 | malonyl\_CoA-acyl\_carrier\_protein\_transacylase | BGC0001795 | Polyketide | 34.0 | 23.4 | 269.0 | 5.3e-71 |
| BAE93731.1 | type\_I\_polyketide\_synthase | BGC0000164 | Polyketide | 32.0 | 30.8 | 269.0 | 7e-71 |
| AXA20093.1 | trans-AT\_PKS\_LgaD | BGC0001946 | NRP + Polyketide | 35.0 | 25.4 | 269.0 | 7e-71 |
| CAQ52623.1 | type\_I\_polyketide\_synthase,\_module\_6 | BGC0001066 | Polyketide:Modular type I | 32.0 | 30.1 | 268.0 | 9.1e-71 |
| CCC21123.1 | type-I\_polyketide\_synthases | BGC0000171 | Polyketide:Modular type I | 33.0 | 24.4 | 268.0 | 1.2e-70 |
| ADN68479.1 | SorD | BGC0000184 | Polyketide:Trans-AT type I | 36.0 | 21.8 | 268.0 | 1.2e-70 |
| AFL48529.1 | laidlomycin\_polyketide\_synthase\_(module\_5\_and\_module\_6) | BGC0000084 | Polyketide | 32.0 | 30.0 | 267.0 | 1.6e-70 |
| BAP05590.1 | calB | BGC0000967 | NRP + Polyketide:Trans-AT type I | 35.0 | 23.9 | 267.0 | 1.6e-70 |
| AAS47564.1 | mixed\_type\_I\_polyketide\_synthase/nonribosomal\_peptide\_synthetase | BGC0001108 | Polyketide:Trans-AT type I | 34.0 | 23.1 | 267.0 | 1.6e-70 |
| ctg1\_orf6 |  | BGC0001109 | NRP + Polyketide | 34.0 | 23.1 | 267.0 | 1.6e-70 |
| DAC76729.1 | type\_I\_polyketide\_synthase | BGC0001885 | Polyketide | 33.0 | 22.7 | 267.0 | 1.6e-70 |
| AAO06918.1 | GdmAIII | BGC0000066 | Polyketide | 30.0 | 30.9 | 267.0 | 2e-70 |
| AAY28225.1 | HbmAI | BGC0000074 | Polyketide | 35.0 | 24.1 | 267.0 | 2e-70 |
| AFN27482.1 | pks\_BonC | BGC0000173 | Polyketide:Modular type I | 35.0 | 24.1 | 267.0 | 2e-70 |
| BAP05594.1 | calF | BGC0000967 | NRP + Polyketide:Trans-AT type I | 34.0 | 23.1 | 267.0 | 2e-70 |
| ARM20284.1 | polyketide\_synthase | BGC0001523 | Polyketide | 31.0 | 31.0 | 267.0 | 2e-70 |
| ABF85931.1 | non-ribosomal\_peptide\_synthase/polyketide\_synthase\_Ta1 | BGC0001025 | NRP + Polyketide:Trans-AT type I | 35.0 | 23.5 | 267.0 | 2.7e-70 |
| ALD83686.1 | tAT\_polyketide\_synthase | BGC0001300 | Polyketide | 30.0 | 29.0 | 267.0 | 2.7e-70 |
| ALI92655.1 | CitS\_citrinin\_polyketide\_synthase | BGC0001338 | Polyketide:Iterative type I | 31.0 | 33.1 | 267.0 | 2.7e-70 |
| CBK62724.1 |  | BGC0001115 | NRP + Polyketide | 33.0 | 24.2 | 266.0 | 3.5e-70 |
| AEK75503.1 | type\_1\_polyketide\_synthase | BGC0000001 | Polyketide:Modular type I | 32.0 | 30.7 | 266.0 | 4.5e-70 |
| ABC84460.1 | NigAV | BGC0000114 | Polyketide:Modular type I | 31.0 | 31.3 | 266.0 | 4.5e-70 |
| BAC76474.1 | type\_I\_polyketide\_synthase\_LkcC | BGC0001100 | NRP + Polyketide | 32.0 | 25.3 | 266.0 | 4.5e-70 |
| EHK80165.1 | beta-ketoacyl\_synthase | BGC0001447 | Polyketide | 37.0 | 23.2 | 266.0 | 4.5e-70 |
| KKP00966.1 | RADS2\_nonreducing\_polyketide\_synthase | BGC0001901 | Polyketide | 28.0 | 32.7 | 266.0 | 4.5e-70 |
| antaD | Type\_I\_PKS | BGC0001455 | NRP + Polyketide | 31.0 | 33.6 | 265.0 | 5.9e-70 |
| AUO16401.1 | polyketide\_synthase | BGC0001700 | Polyketide | 36.0 | 22.6 | 265.0 | 5.9e-70 |
| BBA21074.1 | putative\_modular\_polyketide\_synthase | BGC0001740 | NRP + Polyketide | 34.0 | 24.0 | 265.0 | 7.7e-70 |
| ATX68109.1 | malonyl\_CoA-acyl\_carrier\_protein\_transacylase | BGC0001772 | Polyketide | 34.0 | 24.0 | 265.0 | 7.7e-70 |
| AUO16397.1 | polyketide\_synthase | BGC0001700 | Polyketide | 32.0 | 32.3 | 265.0 | 1e-69 |
| AWM95789.1 | non-reduciing\_polyketide\_synthase\_methylorcinaldehyde\_synthase | BGC0001827 | Polyketide | 29.0 | 34.0 | 265.0 | 1e-69 |
| ABC35796.1 | polyketide\_synthase,\_putative | BGC0001102 | NRP:Beta-lactam + Polyketide:Modular type I | 33.0 | 25.7 | 264.0 | 1.3e-69 |
| AXI91546.1 | FunP7 | BGC0001944 | Polyketide | 32.0 | 30.4 | 264.0 | 1.3e-69 |
| AGZ15474.1 | putative\_type\_I\_polyketide\_synthase | BGC0001036 | NRP + Polyketide | 32.0 | 31.4 | 264.0 | 1.7e-69 |
| AAM12909.2 | MmpA | BGC0000182 | Polyketide:Iterative type I + Polyketide:Trans-AT type I | 32.0 | 27.6 | 264.0 | 2.2e-69 |
| ABC35522.1 | thiotemplate\_mechanism\_natural\_product\_synthetase | BGC0000186 | NRP + Polyketide:Modular type I | 33.0 | 23.7 | 264.0 | 2.2e-69 |
| ADN68480.1 | SorE | BGC0000184 | Polyketide:Trans-AT type I | 33.0 | 24.7 | 263.0 | 2.9e-69 |
| AAN59953.1 | polyketide\_synthase\_1 | BGC0001258 | Polyketide | 29.0 | 32.5 | 263.0 | 2.9e-69 |
| ANY10599.1 | polyketide\_synthase | BGC0001773 | Polyketide | 32.0 | 29.9 | 263.0 | 2.9e-69 |
| AFX60334.1 | polyketide\_synthase | BGC0001032 | NRP + Polyketide | 34.0 | 24.4 | 263.0 | 3.8e-69 |
| AQW44874.1 | polyketide\_synthase | BGC0001761 | Polyketide | 33.0 | 25.2 | 263.0 | 3.8e-69 |
| ABB86408.1 | GelA | BGC0000067 | Polyketide | 34.0 | 24.0 | 262.0 | 5e-69 |
| ABB86410.1 | GelC | BGC0000067 | Polyketide | 30.0 | 30.9 | 262.0 | 5e-69 |
| AFX60313.1 | polyketide\_synthase | BGC0001031 | NRP + Polyketide | 32.0 | 27.7 | 262.0 | 5e-69 |
| CBK62733.1 |  | BGC0001115 | NRP + Polyketide | 32.0 | 27.6 | 262.0 | 5e-69 |
| ctg1\_13 |  | BGC0001931 | Polyketide | 32.0 | 30.7 | 262.0 | 5e-69 |
| AAO06916.1 | GdmAI | BGC0000066 | Polyketide | 34.0 | 24.0 | 262.0 | 6.5e-69 |
| CAG23968.1 | polyketide\_synthase\_type\_I | BGC0000181 | Polyketide | 33.0 | 23.7 | 262.0 | 6.5e-69 |
| CAL69890.1 | RhiC\_protein | BGC0001112 | NRP + Polyketide:Trans-AT type I | 33.0 | 24.0 | 262.0 | 6.5e-69 |
| WP\_079080698.1 | type\_I\_polyketide\_synthase | BGC0001537 | Polyketide | 36.0 | 24.0 | 262.0 | 8.5e-69 |
| AAY28227.1 | HbmAIII | BGC0000074 | Polyketide | 30.0 | 30.9 | 261.0 | 1.1e-68 |
| AAO65797.1 | monensin\_polyketide\_synthase\_module\_2 | BGC0000100 | Polyketide | 34.0 | 23.2 | 261.0 | 1.1e-68 |
| ACR13997.1 | modular\_polyketide\_synthase,\_type\_I\_PKS | BGC0000185 | Polyketide | 32.0 | 25.5 | 261.0 | 1.1e-68 |
| AIJ04686.1 | polyketide\_synthase | BGC0001383 | Polyketide | 33.0 | 23.7 | 261.0 | 1.1e-68 |
| AKQ22669.1 | malonyl\_CoA-acyl\_carrier\_protein\_transacylase | BGC0001656 | Polyketide | 33.0 | 24.1 | 261.0 | 1.1e-68 |
| ANZ52460.1 | MonAII | BGC0001670 | Polyketide | 34.0 | 23.2 | 261.0 | 1.1e-68 |
| WP\_055469549.1 | type\_I\_polyketide\_synthase | BGC0001537 | Polyketide | 33.0 | 31.3 | 261.0 | 1.5e-68 |
| CAE02605.1 | polyketide\_synthase\_type\_I | BGC0000024 | Polyketide:Modular type I | 31.0 | 29.6 | 260.0 | 1.9e-68 |
| AAY32966.1 | DszC | BGC0001093 | NRP + Polyketide | 34.0 | 23.7 | 260.0 | 1.9e-68 |
| AVR48533.1 | CusA | BGC0001564 | NRP + Polyketide | 34.0 | 23.8 | 260.0 | 1.9e-68 |
| AAP42859.1 | NanA5 | BGC0000105 | Polyketide | 31.0 | 30.2 | 260.0 | 2.5e-68 |
| AKQ22697.1 | malonyl\_CoA-acyl\_carrier\_protein\_transacylase | BGC0001186 | Polyketide | 33.0 | 23.2 | 260.0 | 2.5e-68 |
| ctg1\_orf29 |  | BGC0000096 | Polyketide | 32.0 | 29.8 | 260.0 | 3.2e-68 |
| BAD08373.1 | polyketide\_synthase\_modules\_1-3 | BGC0000167 | Polyketide | 34.0 | 24.2 | 260.0 | 3.2e-68 |
| CAJ57411.1 | polyketide\_synthase\_type\_I | BGC0000176 | Polyketide + NRP | 33.0 | 25.3 | 260.0 | 3.2e-68 |
| CBJ89764.1 | Polyketide\_synthase\_involved\_in\_xenocoumacin\_synthesis | BGC0001054 | NRP + Polyketide:Modular type I | 28.0 | 31.9 | 260.0 | 3.2e-68 |
| AKQ22696.1 | malonyl\_CoA-acyl\_carrier\_protein\_transacylase | BGC0001186 | Polyketide | 34.0 | 23.5 | 260.0 | 3.2e-68 |
| CCG06109.1 | type\_I\_polyketide\_synthase | BGC0001543 | Polyketide | 33.0 | 23.4 | 260.0 | 3.2e-68 |
| BAC57030.1 | protomycinolide\_IV\_synthase\_3 | BGC0000102 | Polyketide | 31.0 | 31.1 | 259.0 | 5.5e-68 |
| BAP05591.1 | calC | BGC0000967 | NRP + Polyketide:Trans-AT type I | 35.0 | 21.0 | 259.0 | 5.5e-68 |
| AAF86396.1 | FkbA | BGC0000994 | NRP + Polyketide | 35.0 | 23.1 | 259.0 | 7.2e-68 |
| AAO65800.1 | monensin\_polyketide\_synthase\_modules\_7\_and\_8 | BGC0000100 | Polyketide | 35.0 | 22.1 | 258.0 | 9.4e-68 |
| ANZ52463.1 | MonAV | BGC0001670 | Polyketide | 35.0 | 22.1 | 258.0 | 9.4e-68 |
| AZF85941.1 | type\_I\_polyketide\_synthase | BGC0001963 | NRP + Polyketide | 31.0 | 31.1 | 258.0 | 9.4e-68 |
| AME18003.1 | enediyne\_polyketie\_synthase | BGC0001804 | Polyketide | 28.0 | 34.8 | 257.0 | 1.6e-67 |
| ACD39770.1 | non-reducing\_polyketide\_synthase | BGC0000134 | Polyketide | 30.0 | 32.1 | 257.0 | 2.1e-67 |
| AAV97877.1 | OnnI | BGC0001105 | NRP + Polyketide:Trans-AT type I | 33.0 | 24.7 | 257.0 | 2.1e-67 |
| BAE93730.1 | type\_I\_polyketide\_synthase | BGC0000164 | Polyketide | 30.0 | 29.9 | 257.0 | 2.7e-67 |
| ABB90282.1 | polyketide\_synthase | BGC0001057 | NRP + Polyketide | 30.0 | 32.9 | 257.0 | 2.7e-67 |
| AGN71604.1 | conidial\_yellow\_pigment\_biosynthesis\_polyketide\_synthase | BGC0000027 | Polyketide:Iterative type I | 29.0 | 32.2 | 256.0 | 4.7e-67 |
| AFX60341.1 | polyketide\_synthase | BGC0001032 | NRP + Polyketide | 32.0 | 24.3 | 255.0 | 6.1e-67 |
| ADN68478.1 | SorC | BGC0000184 | Polyketide:Trans-AT type I | 34.0 | 25.0 | 255.0 | 8e-67 |
| WP\_055480219.1 | type\_I\_polyketide\_synthase | BGC0001653 | Polyketide | 36.0 | 24.0 | 255.0 | 1e-66 |
| AAO65799.1 | monensin\_polyketide\_synthase\_modules\_5\_and\_6 | BGC0000100 | Polyketide | 30.0 | 32.1 | 254.0 | 1.4e-66 |
| ANZ52462.1 | MonAIV | BGC0001670 | Polyketide | 30.0 | 32.1 | 254.0 | 1.4e-66 |
| ABC84470.1 | NIGAVIII | BGC0000114 | Polyketide:Modular type I | 31.0 | 30.8 | 254.0 | 1.8e-66 |
| AEH42474.1 | polyketide\_synthase | BGC0000032 | Polyketide | 31.0 | 30.8 | 253.0 | 3e-66 |
| CTQ34881.1 | AtcD;\_polyketide\_synthase,\_modules\_1-4 | BGC0001301 | Polyketide | 33.0 | 23.9 | 253.0 | 3e-66 |
| AFN27481.1 | pks\_BonB | BGC0000173 | Polyketide:Modular type I | 33.0 | 23.6 | 253.0 | 4e-66 |
| ABM63537.1 | BryA | BGC0000174 | Polyketide | 32.0 | 25.1 | 253.0 | 4e-66 |
| WP\_107408739.1 | type\_I\_polyketide\_synthase | BGC0002033 | Polyketide | 31.0 | 30.7 | 253.0 | 4e-66 |
| CAN93352.1 | polyketide\_synthase | BGC0000179 | Polyketide:Trans-AT type I | 32.0 | 26.8 | 252.0 | 5.2e-66 |
| ABI93779.1 | GdmPKS | BGC0000068 | Polyketide | 33.0 | 23.3 | 252.0 | 8.8e-66 |
| AFX60318.1 | polyketide\_synthase | BGC0001031 | NRP + Polyketide | 31.0 | 26.3 | 252.0 | 8.8e-66 |
| CAE52339.1 | Polyketide\_non-ribosomal\_peptide\_synthase | BGC0001088 | NRP + Polyketide | 35.0 | 23.4 | 251.0 | 1.2e-65 |
| EED18001.1 | NR-PKS | BGC0000154 | Polyketide:Iterative type I | 29.0 | 33.6 | 250.0 | 2.6e-65 |
| ABM63530.1 | BryD | BGC0000174 | Polyketide | 33.0 | 23.5 | 250.0 | 2.6e-65 |
| ABC84457.1 | NigAII | BGC0000114 | Polyketide:Modular type I | 31.0 | 31.0 | 250.0 | 3.4e-65 |
| BAR73007.1 | putative\_PKS\_(ACP-KS-AT-DH-KR-ACP-KS-AT-DH-ER-KR-ACP) | BGC0001194 | Polyketide | 34.0 | 22.3 | 250.0 | 3.4e-65 |
| AXA20090.1 | hybrid\_trans-AT\_PKS/NRPS\_LgaA | BGC0001946 | NRP + Polyketide | 31.0 | 23.6 | 249.0 | 5.7e-65 |
| ACY06288.1 | type\_I\_polyketide\_synthase | BGC0001042 | NRP + Polyketide | 35.0 | 22.9 | 248.0 | 9.8e-65 |
| AEH42491.1 | polyketide\_synthase | BGC0000032 | Polyketide | 30.0 | 30.8 | 248.0 | 1.3e-64 |
| ACD39753.1 | non-reducing\_polyketide\_synthase | BGC0000076 | Polyketide | 29.0 | 32.5 | 248.0 | 1.3e-64 |
| ACD39762.1 | non-reducing\_polyketide\_synthase | BGC0000077 | Polyketide | 29.0 | 32.5 | 248.0 | 1.3e-64 |
| CBK62731.1 |  | BGC0001115 | NRP + Polyketide | 32.0 | 24.4 | 248.0 | 1.3e-64 |
| AAP42858.1 | NanA4 | BGC0000105 | Polyketide | 35.0 | 23.7 | 247.0 | 2.2e-64 |
| CBJ89760.1 | Polyketide\_synthase\_involved\_in\_xenocoumacin\_synthesis | BGC0001054 | NRP + Polyketide:Modular type I | 29.0 | 32.0 | 246.0 | 4.8e-64 |
| AAU93806.2 | polyketide\_synthase\_modules\_3\_and\_4 | BGC0000054 | Polyketide | 34.0 | 24.1 | 245.0 | 6.3e-64 |
| ACM79805.1 | ZmaA | BGC0001059 | NRP + Polyketide | 34.0 | 23.0 | 245.0 | 6.3e-64 |
| DAC80076.1 | PKS | BGC0001836 | Polyketide:Trans-AT type I | 33.0 | 23.5 | 245.0 | 6.3e-64 |
| ABC33986.1 | polyketide\_synthase,\_putative | BGC0000186 | NRP + Polyketide:Modular type I | 33.0 | 23.2 | 245.0 | 8.3e-64 |
| BAF50721.1 | hybrid\_non\_ribosomal\_peptide\_synthetase-polyketide\_synthase | BGC0001116 | NRP + Polyketide | 33.0 | 23.7 | 244.0 | 1.4e-63 |
| AFL48526.1 | laidlomycin\_polyketide\_synthase\_(module\_2) | BGC0000084 | Polyketide | 34.0 | 23.7 | 244.0 | 1.8e-63 |
| BAB69199.1 | modular\_polyketide\_synthase | BGC0000117 | Polyketide | 34.0 | 22.1 | 244.0 | 1.8e-63 |
| BAV56011.1 | PKS\_(KS-AT-DH-ER-KR-ACP-KS-AT-DH-ER-KR-ACP) | BGC0001597 | Polyketide | 33.0 | 24.1 | 240.0 | 2e-62 |
| WP\_020636845.1 | type\_I\_polyketide\_synthase | BGC0002011 | Polyketide | 34.0 | 22.2 | 240.0 | 2e-62 |
| CAL69893.1 | RhiE\_protein | BGC0001112 | NRP + Polyketide:Trans-AT type I | 32.0 | 23.5 | 240.0 | 3.5e-62 |
| BAH02270.1 | polyketide\_synthase | BGC0000126 | Polyketide | 34.0 | 24.0 | 239.0 | 5.9e-62 |
| ANZ22995.1 | ZinA | BGC0001828 | Polyketide | 31.0 | 32.2 | 239.0 | 5.9e-62 |
| CAL69888.1 | RhiA\_protein | BGC0001112 | NRP + Polyketide:Trans-AT type I | 32.0 | 23.9 | 238.0 | 7.7e-62 |
| ABK32260.1 | AmbF | BGC0000014 | Polyketide | 31.0 | 23.6 | 238.0 | 1e-61 |
| XP\_011392701.1 | hypothetical\_protein | BGC0001281 | Polyketide | 28.0 | 33.6 | 238.0 | 1.3e-61 |
| AAL06699.1 | polyketide\_synthase | BGC0000965 | Polyketide:Iterative type I + Polyketide:Enediyne type I | 29.0 | 35.0 | 237.0 | 2.2e-61 |
| ALU98461.1 | erythronolide\_synthase | BGC0001397 | NRP + Polyketide | 29.0 | 35.0 | 237.0 | 2.2e-61 |
| AGY62758.1 | EbeF | BGC0000051 | Polyketide | 29.0 | 33.0 | 236.0 | 3.8e-61 |
| AKQ22670.1 | malonyl\_CoA-acyl\_carrier\_protein\_transacylase | BGC0001656 | Polyketide | 32.0 | 20.4 | 236.0 | 3.8e-61 |
| ADA69239.2 | trans-AT\_hybrid\_polyketide\_synthase-NRPS | BGC0001071 | NRP + Polyketide:Modular type I + Polyketide:Trans-AT type I | 34.0 | 23.9 | 235.0 | 6.5e-61 |
| AHV78247.1 | LasS2 | BGC0001245 | Polyketide | 29.0 | 32.2 | 234.0 | 1.5e-60 |
| WP\_055469551.1 | SDR\_family\_NAD(P)-dependent\_oxidoreductase | BGC0001537 | Polyketide | 34.0 | 23.4 | 233.0 | 3.2e-60 |
| ABM63529.1 | BryX | BGC0000174 | Polyketide | 32.0 | 23.7 | 233.0 | 4.2e-60 |
| CAJ76289.1 | putative\_hybrid\_non-ribosomal\_peptide-polyketide\_synthetase | BGC0000972 | NRP + Polyketide:Modular type I + Polyketide:Trans-AT type I | 31.0 | 23.3 | 233.0 | 4.2e-60 |
| CBK62718.1 |  | BGC0001115 | NRP + Polyketide | 33.0 | 23.3 | 232.0 | 5.5e-60 |
| BAF02922.1 | type\_I\_polyketide\_synthase | BGC0000073 | Polyketide | 34.0 | 22.1 | 230.0 | 2.7e-59 |
| AEH59110.1 | polyketide\_synthase\_of\_type\_I | BGC0000385 | NRP | 32.0 | 24.0 | 230.0 | 2.7e-59 |
| BAF02925.1 | type\_I\_polyketide\_synthase | BGC0000073 | Polyketide | 34.0 | 22.4 | 230.0 | 3.6e-59 |
| ABC34108.1 | JamP | BGC0000961 | NRP + Polyketide | 28.0 | 32.2 | 230.0 | 3.6e-59 |
| ANY94470.1 | enediyne\_polyketide\_synthase | BGC0001584 | Polyketide | 27.0 | 35.3 | 229.0 | 4.7e-59 |
| BAF02924.1 | type\_I\_polyketide\_synthase | BGC0000073 | Polyketide | 34.0 | 22.4 | 227.0 | 1.8e-58 |
| orf1 | polyketide\_synthase | BGC0001432 | NRP:Cyclic depsipeptide + Polyketide:Iterative type I | 27.0 | 39.1 | 227.0 | 3e-58 |
| APZ78852.1 | polyketide\_synthase | BGC0001432 | NRP:Cyclic depsipeptide + Polyketide:Iterative type I | 27.0 | 39.1 | 227.0 | 3e-58 |
| AGN11883.1 | tstI | BGC0001114 | NRP + Polyketide | 33.0 | 23.0 | 226.0 | 5.2e-58 |
| ADH01485.1 | putative\_mixed\_polyketide\_synthase/non-ribosomal\_peptide\_synthetase | BGC0001096 | NRP + Polyketide | 36.0 | 24.6 | 223.0 | 2.6e-57 |
| AAM12934.1 | MmpF | BGC0000182 | Polyketide:Iterative type I + Polyketide:Trans-AT type I | 34.0 | 20.8 | 222.0 | 5.7e-57 |
| AVR48535.1 | CusC | BGC0001564 | NRP + Polyketide | 32.0 | 24.6 | 221.0 | 1.7e-56 |
| AGN11880.1 | tstC | BGC0001114 | NRP + Polyketide | 32.0 | 23.5 | 220.0 | 2.2e-56 |
| ADH01484.1 | putative\_type-I\_PKS | BGC0001096 | NRP + Polyketide | 30.0 | 24.4 | 219.0 | 4.9e-56 |
| AIC32692.1 | FR9C | BGC0001113 | NRP + Polyketide | 30.0 | 24.4 | 219.0 | 4.9e-56 |
| EYT83433.1 | hypothetical\_protein | BGC0001213 | Polyketide | 28.0 | 33.9 | 219.0 | 4.9e-56 |
| CBF83139.1 | polyketide\_synthase,\_putative\_(JCVI) | BGC0001722 | Polyketide | 28.0 | 32.7 | 219.0 | 6.3e-56 |
| EAQ86392.1 | hypothetical\_protein | BGC0001405 | Polyketide | 28.0 | 33.4 | 217.0 | 3.1e-55 |
| ADO69378.1 | beta-ketoacyl\_synthase-like\_protein,\_Acyl\_transferase\_subunit | BGC0000044 | Polyketide | 26.0 | 38.2 | 213.0 | 2.7e-54 |
| CAG23961.2 | polyketide\_synthase\_of\_type\_I | BGC0001089 | Polyketide + NRP | 29.0 | 24.4 | 213.0 | 2.7e-54 |
| AAK57185.1 | MxaB1 | BGC0001022 | NRP + Polyketide | 28.0 | 28.3 | 210.0 | 3.8e-53 |
| CBK62729.1 |  | BGC0001115 | NRP + Polyketide | 29.0 | 23.8 | 208.0 | 1.5e-52 |
| EAA65602.1 | hypothetical\_protein | BGC0000022 | Polyketide | 27.0 | 34.6 | 206.0 | 4.3e-52 |
| AHH25595.1 | PKS | BGC0000957 | NRP + Polyketide | 31.0 | 20.5 | 203.0 | 4.7e-51 |
| ADO69381.1 | Beta-ketoacyl\_synthase | BGC0000044 | Polyketide | 31.0 | 20.9 | 200.0 | 3e-50 |
| BAD38873.1 | polyketide\_synthase | BGC0000111 | Polyketide | 28.0 | 28.0 | 195.0 | 9.8e-49 |
| AAL01062.1 | omega-3\_polyunsaturated\_fatty\_acid\_synthase\_PfaC | BGC0000865 | Other | 30.0 | 20.2 | 190.0 | 4.1e-47 |
| ACI12950.1 | PfaC | BGC0000861 | Other | 30.0 | 21.2 | 188.0 | 1.2e-46 |
| XP\_011392698.1 | hypothetical\_protein | BGC0001281 | Polyketide | 27.0 | 24.0 | 176.0 | 3.6e-43 |
| ABF00128.1 | polyunsaturated\_fatty\_acid\_synthase | BGC0000863 | Other | 28.0 | 20.8 | 165.0 | 1.4e-39 |
| BAF50720.1 | hybrid\_non\_ribosomal\_peptide\_synthetase-polyketide\_synthase | BGC0001116 | NRP + Polyketide | 28.0 | 24.9 | 163.0 | 3.2e-39 |
