## Supplementary Results for "A draft genome of the ascomycotal fungal species *Pseudopithomyces maydicus* (family *Didymosphaeriaceae*)": input.path1.gene32_mibig_hits.html

| MIBiG Protein | Description | MIBiG Cluster | MiBiG Product | % ID | % Coverage | BLAST Score | E-value |
| --- | --- | --- | --- | --- | --- | --- | --- |
| AIG62134.1 | patulin\_synthase | BGC0000120 | Polyketide:Iterative type I | 45.0 | 108.6 | 247.0 | 2.5e-65 |
| EAU32818.1 | predicted\_protein | BGC0000160 | Polyketide | 37.0 | 108.6 | 175.0 | 1.6e-43 |
| AAS90106.1 | VBS | BGC0000006 | Polyketide | 35.0 | 111.7 | 161.0 | 2.4e-39 |
| BAE71331.1 | versicolorin\_B\_synthase | BGC0000004 | Polyketide | 35.0 | 111.7 | 160.0 | 3.1e-39 |
| AAS90066.1 | VBS | BGC0000009 | Polyketide | 35.0 | 111.3 | 160.0 | 3.1e-39 |
| AAS90088.1 | VBS | BGC0000010 | Polyketide | 35.0 | 111.7 | 160.0 | 3.1e-39 |
| AAS90019.1 | VBS | BGC0000007 | Polyketide | 35.0 | 111.7 | 160.0 | 5.3e-39 |
| AAS90042.1 | VBS | BGC0000008 | Polyketide | 35.0 | 111.3 | 160.0 | 5.3e-39 |
| ACH72898.1 | AflK | BGC0000011 | Polyketide | 35.0 | 112.8 | 155.0 | 1e-37 |
| BAQ25461.1 | putative\_dehydrogenase | BGC0001280 | Polyketide | 22.0 | 120.7 | 61.0 | 3.4e-09 |
