## Supplementary Results for "A draft genome of the ascomycotal fungal species *Pseudopithomyces maydicus* (family *Didymosphaeriaceae*)": input.path1.gene33_mibig_hits.html

| MIBiG Protein | Description | MIBiG Cluster | MiBiG Product | % ID | % Coverage | BLAST Score | E-value |
| --- | --- | --- | --- | --- | --- | --- | --- |
| RWQ92168.1 | O-methyltransferase-domain-containing\_protein | BGC0002030 | Polyketide | 42.0 | 93.8 | 250.0 | 4.3e-66 |
| CCT67993.1 | bikaverin\_cluster-O-methyltransferase | BGC0000030 | Polyketide | 37.0 | 92.7 | 206.0 | 7.2e-53 |
| ADM79461.1 | O-methyltransferase | BGC0001266 | Polyketide | 34.0 | 77.3 | 139.0 | 1.1e-32 |
| BAJ09788.1 | O-methyltransferase | BGC0000146 | Polyketide | 29.0 | 103.1 | 111.0 | 2.4e-24 |
| OAQ83753.1 | sterigmatocystin\_8-O-methyltransferase | BGC0001358 | Polyketide | 25.0 | 97.7 | 90.0 | 5.7e-18 |
| ANY57881.1 | PenC | BGC0001372 | Terpene | 33.0 | 37.5 | 66.0 | 1.2e-10 |
| AHZ61883.1 | O-methyltransferase | BGC0000240 | Polyketide:Type II + Saccharide:Hybrid/tailoring | 32.0 | 33.1 | 59.0 | 1.8e-08 |
| ARD70890.1 | O-methyltransferase | BGC0001693 | Polyketide | 25.0 | 70.6 | 57.0 | 7e-08 |
| ABP54665.1 | O-methyltransferase,\_family\_2 | BGC0000241 | Polyketide:Type II + Saccharide:Hybrid/tailoring | 31.0 | 33.1 | 56.0 | 1.2e-07 |
