## Supplementary Results for "A draft genome of the ascomycotal fungal species *Pseudopithomyces maydicus* (family *Didymosphaeriaceae*)": input.path1.gene34_mibig_hits.html

| MIBiG Protein | Description | MIBiG Cluster | MiBiG Product | % ID | % Coverage | BLAST Score | E-value |
| --- | --- | --- | --- | --- | --- | --- | --- |
| EHA52497.1 | hypothetical\_protein | BGC0001749 | Polyketide | 30.0 | 80.4 | 171.0 | 3.6e-42 |
| EAA36372.1 | hypothetical\_protein | BGC0001697 | Polyketide | 29.0 | 80.1 | 162.0 | 2.2e-39 |
| AVY05517.1 | 6-hydroxy-D-nicotine\_oxidase | BGC0001571 | Terpene | 29.0 | 83.6 | 154.0 | 4.6e-37 |
| EAA36373.1 | hypothetical\_protein | BGC0001697 | Polyketide | 29.0 | 83.6 | 151.0 | 3.9e-36 |
| EHA28243.1 | hypothetical\_protein | BGC0001143 | Polyketide | 26.0 | 83.8 | 149.0 | 1.5e-35 |
| EAL89048.1 | FAD-dependent\_oxidoreductase | BGC0000355 | NRP | 28.0 | 84.7 | 138.0 | 4.4e-32 |
| XP\_011325834.1 | hypothetical\_protein | BGC0001545 | NRP | 27.0 | 89.9 | 129.0 | 2e-29 |
| ADY16695.1 | TqaG | BGC0001142 | NRP | 26.0 | 79.5 | 126.0 | 1e-28 |
| EHA28234.1 | FAD/FMN-containing\_dehydrogenase | BGC0001143 | Polyketide | 24.0 | 76.9 | 123.0 | 1.1e-27 |
| AAF81732.1 | putative\_FAD-dependent\_oxygenase\_EncM | BGC0000220 | Polyketide:Type II | 29.0 | 42.4 | 95.0 | 3.3e-19 |
| BBE36455.1 | oxidase | BGC0001922 | Polyketide | 35.0 | 33.6 | 94.0 | 7.3e-19 |
| ACB46474.1 | FAD-dependent\_oxidoreductase | BGC0000082 | Polyketide | 33.0 | 39.7 | 91.0 | 4.7e-18 |
| AGC09496.1 | LobB3 | BGC0001183 | Polyketide | 32.0 | 38.2 | 91.0 | 4.7e-18 |
| WP\_026290905.1 | FAD-binding\_oxidoreductase | BGC0002010 | NRP + Polyketide | 28.0 | 49.4 | 91.0 | 4.7e-18 |
| AGI99485.1 | FAD-dependent\_oxidoreductase | BGC0001004 | Polyketide:Modular type I | 32.0 | 38.2 | 89.0 | 1.4e-17 |
| BAD83683.1 | FAD/FMN-dependent\_oxygenase/oxidase | BGC0000012 | Polyketide | 29.0 | 46.7 | 83.0 | 9.9e-16 |
| ABP55175.1 | FAD\_linked\_oxidase\_domain\_protein | BGC0000150 | NRP + Polyketide:Enediyne type I | 27.0 | 48.2 | 83.0 | 1.3e-15 |
| BAJ09785.1 | oxidase/Diels-Alderase | BGC0000146 | Polyketide | 32.0 | 35.8 | 81.0 | 4.9e-15 |
| AGA37275.1 | oxidoreductase | BGC0000819 | NRP + Alkaloid | 32.0 | 39.1 | 81.0 | 6.4e-15 |
| EAQ86396.1 | hypothetical\_protein | BGC0001405 | Polyketide | 32.0 | 33.2 | 77.0 | 9.2e-14 |
| AGZ20488.1 | putative\_FAD-binding\_oxidoreductase | BGC0001776 | Terpene | 27.0 | 36.5 | 68.0 | 3.3e-11 |
| KGO40470.1 | FAD\_linked\_oxidase,\_N-terminal | BGC0001205 | Polyketide | 35.0 | 21.4 | 67.0 | 9.6e-11 |
| BAU61564.1 | putative\_FAD\_binding\_domain\_protein | BGC0001375 | Terpene | 27.0 | 37.6 | 63.0 | 1.1e-09 |
| ACD39759.1 | alcohol\_oxidase | BGC0000076 | Polyketide | 32.0 | 20.5 | 53.0 | 1.4e-06 |
| ACD39768.1 | alcohol\_oxidase | BGC0000077 | Polyketide | 32.0 | 20.5 | 53.0 | 1.4e-06 |
