## Supplementary Results for "A draft genome of the ascomycotal fungal species *Pseudopithomyces maydicus* (family *Didymosphaeriaceae*)": input.path1.gene37_mibig_hits.html

| MIBiG Protein | Description | MIBiG Cluster | MiBiG Product | % ID | % Coverage | BLAST Score | E-value |
| --- | --- | --- | --- | --- | --- | --- | --- |
| BAV32173.1 | putative\_MFS\_sugar\_transporter | BGC0001373 | Polyketide | 31.0 | 96.1 | 246.0 | 8.4e-65 |
| XP\_023094066.1 |  | BGC0001996 | Other | 26.0 | 88.3 | 141.0 | 3.8e-33 |
| AEZ53939.1 | putative\_glucose-6-phosphate\_1-dehydrogenase | BGC0000144 | Polyketide:Modular type I | 27.0 | 80.0 | 103.0 | 1.1e-21 |
| ADC45555.1 | sugar\_transporter | BGC0000093 | Polyketide | 26.0 | 75.5 | 75.0 | 2.6e-13 |
| CDG12868.1 | putative\_drug\_efflux\_protein | BGC0001415 | NRP | 27.0 | 28.2 | 50.0 | 8.8e-06 |
