## Supplementary Results for "A draft genome of the ascomycotal fungal species *Pseudopithomyces maydicus* (family *Didymosphaeriaceae*)": input.path1.gene38_mibig_hits.html

| MIBiG Protein | Description | MIBiG Cluster | MiBiG Product | % ID | % Coverage | BLAST Score | E-value |
| --- | --- | --- | --- | --- | --- | --- | --- |
| ABB52530.1 | beta\_glucosidase | BGC0000047 | Polyketide | 32.0 | 97.0 | 331.0 | 4.4e-90 |
| ANC94980.1 | AlmRII | BGC0001396 | Polyketide | 31.0 | 97.0 | 328.0 | 2.8e-89 |
| BAC76488.1 | putative\_beta-glycosidase | BGC0000085 | Polyketide | 31.0 | 99.4 | 326.0 | 1.8e-88 |
| AAS79445.1 | putative\_beta-glucosidase | BGC0000035 | Polyketide | 30.0 | 97.0 | 316.0 | 1.1e-85 |
| AAM88355.1 | NbmF | BGC0000899 | Other | 30.0 | 98.3 | 311.0 | 6.1e-84 |
| CAM00073.1 | beta-D-glucosidase | BGC0000055 | Polyketide:Modular type I + Saccharide:Hybrid/tailoring | 29.0 | 97.6 | 309.0 | 2.3e-83 |
| AAC68679.1 | beta-glucosidase | BGC0000898 | Saccharide:Hybrid/tailoring | 31.0 | 93.8 | 302.0 | 1.7e-81 |
| AGZ20486.1 | putative\_beta-glucosidase | BGC0001776 | Terpene | 26.0 | 102.1 | 210.0 | 1.5e-53 |
| AAY32974.1 | glycosyl\_hydrolase | BGC0001093 | NRP + Polyketide | 26.0 | 102.6 | 182.0 | 3.3e-45 |
| ADN68487.1 | SorL | BGC0000184 | Polyketide:Trans-AT type I | 33.0 | 42.4 | 161.0 | 4.6e-39 |
| AGC95327.1 | putative\_glycosyl\_hydrolase | BGC0000045 | Polyketide | 31.0 | 48.4 | 160.0 | 1.7e-38 |
| AEH59053.1 | glycosyl\_hydrolase\_family\_3/N\_terminal\_domain\_protein | BGC0000385 | NRP | 21.0 | 90.4 | 142.0 | 3.7e-33 |
| BAV56273.1 | hypothetical\_protein | BGC0001657 | NRP | 22.0 | 90.6 | 138.0 | 5.4e-32 |
| EAA59569.1 | glucosidase,\_putative | BGC0000057 | Polyketide:Iterative type I | 23.0 | 103.6 | 137.0 | 9.2e-32 |
