## Supplementary Results for "A draft genome of the ascomycotal fungal species *Pseudopithomyces maydicus* (family *Didymosphaeriaceae*)": input.path1.gene40_mibig_hits.html

| MIBiG Protein | Description | MIBiG Cluster | MiBiG Product | % ID | % Coverage | BLAST Score | E-value |
| --- | --- | --- | --- | --- | --- | --- | --- |
| BBF25319.1 | cytochrome\_P450 | BGC0001923 | Terpene + Polyketide | 34.0 | 100.0 | 128.0 | 1.4e-29 |
| ANV81301.1 | C13\_oxidase | BGC0001604 | Terpene | 37.0 | 94.5 | 121.0 | 1.7e-27 |
| XP\_001826049.1 |  | BGC0001995 | Terpene | 30.0 | 94.5 | 94.0 | 2.2e-19 |
| ATZ45183.1 | Bcboa7 | BGC0001892 | Polyketide | 31.0 | 96.8 | 94.0 | 2.9e-19 |
| EAU36745.1 | hypothetical\_protein | BGC0000292 | NRP | 29.0 | 95.4 | 91.0 | 2.5e-18 |
| BAV32161.1 | cytochrome\_P450\_monooxygenase | BGC0001373 | Polyketide | 30.0 | 95.4 | 84.0 | 1.8e-16 |
| ASK38704.1 | cytochrome\_P450 | BGC0001557 | Polyketide | 28.0 | 97.7 | 84.0 | 3e-16 |
| AGO86661.1 | putative\_cytochrome\_p450 | BGC0001255 | NRP + Polyketide | 28.0 | 94.5 | 82.0 | 6.7e-16 |
| ANV81298.1 | C20\_oxidase | BGC0001604 | Terpene | 28.0 | 95.9 | 81.0 | 2e-15 |
| BAC20565.1 | cytochrome\_P450 | BGC0000039 | Polyketide | 28.0 | 95.4 | 79.0 | 5.7e-15 |
| ANV81296.1 | kaurenoxidase | BGC0001604 | Terpene | 26.0 | 95.9 | 77.0 | 2.2e-14 |
| ANV81297.1 | GA14\_synthase | BGC0001604 | Terpene | 26.0 | 95.9 | 76.0 | 4.8e-14 |
| CAP96442.1 | P450\_monooxygenase | BGC0000420 | NRP | 27.0 | 97.7 | 71.0 | 1.6e-12 |
| BBG28489.1 | cytochrome\_P450\_monooxygenase\_CdmJ | BGC0001926 | Polyketide | 28.0 | 96.3 | 70.0 | 3.5e-12 |
| ctg1\_orf0002 |  | BGC0000685 | Terpene | 27.0 | 89.4 | 66.0 | 5e-11 |
| BAH24001.1 | cytochrome\_P450 | BGC0000356 | NRP + Alkaloid | 24.0 | 94.9 | 63.0 | 4.2e-10 |
| AGZ20487.1 | cytochrome\_P450\_monooxygenase | BGC0001776 | Terpene | 26.0 | 97.2 | 61.0 | 2.7e-09 |
| BAM84045.1 | cytochrome\_P450\_monooxygenase | BGC0001260 | Terpene | 24.0 | 95.4 | 58.0 | 1.4e-08 |
| QCS37514.1 | PyiG | BGC0001982 | NRP + Polyketide | 26.0 | 99.5 | 57.0 | 2.3e-08 |
| AAK11527.1 | PaxQ | BGC0001082 | Terpene | 28.0 | 67.3 | 55.0 | 1.5e-07 |
| AGA37282.1 | P450\_monooxygenase | BGC0000819 | NRP + Alkaloid | 25.0 | 90.3 | 52.0 | 7.5e-07 |
| BBD84642.1 | putative\_cytochrome\_P450 | BGC0001775 | Terpene | 29.0 | 75.6 | 51.0 | 2.2e-06 |
| BBD84646.1 | putative\_cytochrome\_P450 | BGC0001775 | Terpene | 28.0 | 94.9 | 50.0 | 4.8e-06 |
