## Supplementary Results for "A draft genome of the ascomycotal fungal species *Pseudopithomyces maydicus* (family *Didymosphaeriaceae*)": input.path1.gene41_mibig_hits.html

| MIBiG Protein | Description | MIBiG Cluster | MiBiG Product | % ID | % Coverage | BLAST Score | E-value |
| --- | --- | --- | --- | --- | --- | --- | --- |
| BAC20568.1 | efflux\_pump | BGC0000039 | Polyketide | 35.0 | 132.9 | 286.0 | 5.5e-77 |
| AAD34558.1 | unknown | BGC0000088 | Polyketide | 33.0 | 130.6 | 263.0 | 5e-70 |
| ABA02247.1 | efflux\_pump | BGC0000098 | Polyketide | 33.0 | 129.8 | 252.0 | 1.5e-66 |
| EAA65599.1 | hypothetical\_protein | BGC0000022 | Polyketide | 35.0 | 124.6 | 245.0 | 1.1e-64 |
| AGN71625.1 | putative\_HC-toxin\_efflux\_carrier\_TOXA | BGC0000027 | Polyketide:Iterative type I | 36.0 | 103.9 | 223.0 | 5.7e-58 |
| BAE60006.1 |  | BGC0001518 | Terpene | 30.0 | 129.8 | 220.0 | 4.8e-57 |
| ARP51716.1 | toxin\_efflux\_transporter\_MFS | BGC0001741 | NRP + Polyketide | 30.0 | 130.1 | 213.0 | 5.9e-55 |
| ADY16699.1 | TqaJ | BGC0001142 | NRP | 30.0 | 129.0 | 212.0 | 1.3e-54 |
| CAP96439.1 | Transporter | BGC0000420 | NRP | 28.0 | 129.5 | 210.0 | 3.8e-54 |
| RWQ92172.1 | putative\_MFS\_transporter | BGC0002030 | Polyketide | 30.0 | 129.8 | 207.0 | 3.2e-53 |
| CCE28984.1 | probable\_DHA14-like\_major\_facilitator;\_ABC\_transporter | BGC0001365 | NRP | 27.0 | 132.4 | 191.0 | 3.1e-48 |
| EAL88822.1 | MFS\_gliotoxin\_efflux\_transporter\_GliA | BGC0000361 | NRP | 26.0 | 131.6 | 174.0 | 4e-43 |
| AVY05519.1 | major\_facilitator\_superfamily\_transporter | BGC0001571 | Terpene | 27.0 | 141.5 | 170.0 | 4.4e-42 |
| CAP93755.1 |  | BGC0001882 | Polyketide | 26.0 | 128.8 | 170.0 | 4.4e-42 |
| EPS29061.1 | hypothetical\_protein | BGC0001724 | NRP + Polyketide | 28.0 | 131.9 | 168.0 | 1.7e-41 |
| PKX92296.1 | MFS\_general\_substrate\_transporter | BGC0001988 | Polyketide | 27.0 | 125.1 | 165.0 | 1.4e-40 |
| DAB41650.1 | MFS\_transporter | BGC0001583 | Polyketide | 26.0 | 131.9 | 165.0 | 1.8e-40 |
| AAS89998.1 | AflT | BGC0000007 | Polyketide | 49.0 | 46.1 | 160.0 | 7.7e-39 |
| AAS90046.1 | AflT | BGC0000009 | Polyketide | 51.0 | 46.1 | 160.0 | 7.7e-39 |
| BAE71313.1 | putative\_ABC\_transporter | BGC0000004 | Polyketide | 49.0 | 46.1 | 157.0 | 5e-38 |
| AAS90092.1 | AflT | BGC0000006 | Polyketide | 49.0 | 46.1 | 156.0 | 8.5e-38 |
| AAS90069.1 | AflT | BGC0000010 | Polyketide | 49.0 | 46.1 | 156.0 | 8.5e-38 |
| AAS90021.1 | AflT | BGC0000008 | Polyketide | 48.0 | 46.1 | 155.0 | 2.5e-37 |
| CCE31569.1 | probable\_aflatoxin\_efflux\_pump\_AFLT | BGC0001886 | Polyketide | 25.0 | 130.6 | 154.0 | 3.2e-37 |
| EAU36749.1 | predicted\_protein | BGC0000292 | NRP | 29.0 | 114.5 | 145.0 | 2e-34 |
| BBG28481.1 | putative\_MFS\_toxin\_efflux\_pump\_CdmB | BGC0001926 | Polyketide | 44.0 | 43.0 | 145.0 | 2e-34 |
| BAD29973.1 | transporter\_protein | BGC0000676 | Terpene | 43.0 | 47.2 | 140.0 | 4.8e-33 |
| ctg1\_orf0002 |  | BGC0000688 | Terpene | 43.0 | 43.0 | 128.0 | 1.9e-29 |
| ADM34144.1 | efflux\_pump | BGC0001084 | NRP + Terpene + Alkaloid | 42.0 | 48.7 | 125.0 | 1.6e-28 |
| XP\_001798920.1 | MFS\_transporter | BGC0001865 | Polyketide:Iterative type I | 23.0 | 114.8 | 124.0 | 2.8e-28 |
| AGC95323.1 | CurE | BGC0000045 | Polyketide | 39.0 | 43.0 | 123.0 | 8e-28 |
| BAV69308.1 | PrhG | BGC0001729 | Polyketide + Terpene | 43.0 | 41.5 | 123.0 | 1e-27 |
| AHV78251.1 | ResE | BGC0001246 | Polyketide | 36.0 | 49.0 | 118.0 | 2.6e-26 |
| CBF76046.1 | conserved\_hypothetical\_protein | BGC0001399 | NRP | 40.0 | 37.3 | 107.0 | 5.9e-23 |
| BAZ95819.1 | cpaN1\_MFS\_transporter | BGC0001563 | NRP + Polyketide | 35.0 | 48.2 | 106.0 | 1.3e-22 |
| ACD39756.1 | major\_facilitator\_superfamily\_transporter | BGC0000076 | Polyketide | 34.0 | 45.3 | 105.0 | 1.7e-22 |
| ACD39765.1 | major\_facilitator\_superfamily\_transporter | BGC0000077 | Polyketide | 34.0 | 45.3 | 105.0 | 1.7e-22 |
| ACZ57546.1 | predicted\_MFS\_transporter | BGC0000046 | Polyketide:Iterative type I | 38.0 | 42.5 | 103.0 | 6.6e-22 |
| AEO57490.1 | general\_substrate\_transporter | BGC0001449 | NRP + Alkaloid + Polyketide:Iterative type I | 37.0 | 42.5 | 99.0 | 9.5e-21 |
| PIB01159.1 | putative\_HC-toxin\_efflux\_carrier\_TOXA | BGC0001541 | Polyketide | 31.0 | 56.2 | 99.0 | 1.6e-20 |
| ACZ66257.1 | APS11 | BGC0000304 | NRP | 31.0 | 47.2 | 89.0 | 1.7e-17 |
| AAL15595.1 | Sim17 | BGC0000270 | Polyketide | 33.0 | 47.9 | 87.0 | 4.9e-17 |
| AAK06799.1 | simocyclinone-specific\_efflux\_pump | BGC0001072 | Saccharide + Polyketide:Modular type I + Polyketide:Type II + Other:Aminocoumarin | 32.0 | 47.9 | 85.0 | 2.4e-16 |
| ADI24949.1 | GsfJ | BGC0000070 | Polyketide:Iterative type I | 29.0 | 69.7 | 83.0 | 7e-16 |
| BAZ95831.1 | MFS\_transporter\_cpaI | BGC0001563 | NRP + Polyketide | 31.0 | 44.6 | 81.0 | 4.6e-15 |
| CAE51185.1 | RemN\_protein | BGC0000264 | Polyketide:Type II | 32.0 | 46.9 | 78.0 | 3e-14 |
| BBE36450.1 | transporter | BGC0001922 | Polyketide | 33.0 | 40.4 | 78.0 | 3e-14 |
| AMK92567.1 | tetracenomycin\_c\_resistance\_and\_export\_protein | BGC0001815 | Polyketide | 29.0 | 44.0 | 77.0 | 6.6e-14 |
| KDQ70104.1 | multidrug\_MFS\_transporter | BGC0001538 | NRP + Polyketide | 32.0 | 40.4 | 77.0 | 6.6e-14 |
| AME18021.1 | transporter | BGC0001804 | Polyketide | 26.0 | 85.8 | 76.0 | 1.5e-13 |
| ABC87523.1 | putative\_drug\_efflux\_transporter | BGC0001011 | NRP + Polyketide | 32.0 | 41.7 | 73.0 | 9.5e-13 |
| ARO49560.1 | MFS\_transporter | BGC0001440 | Other | 29.0 | 51.6 | 68.0 | 3.1e-11 |
| ABX71155.1 | Lcz38 | BGC0000237 | Polyketide | 28.0 | 47.4 | 66.0 | 1.2e-10 |
| AGC24258.1 | prlD | BGC0001038 | NRP + Polyketide:Modular type I | 29.0 | 44.3 | 65.0 | 2.6e-10 |
| AQW35028.1 | Multidrug\_MFS\_transporter | BGC0001675 | Polyketide | 33.0 | 39.6 | 63.0 | 9.8e-10 |
| CBA63674.1 | MFS\_griseobactin\_exporter | BGC0000368 | NRP | 33.0 | 37.3 | 62.0 | 1.3e-09 |
| WP\_019634551.1 | MFS\_transporter | BGC0001443 | NRP + Polyketide | 28.0 | 46.6 | 60.0 | 8.3e-09 |
| CAF34033.1 | putative\_transmembrane\_efflux\_protein | BGC0000689 | Saccharide | 30.0 | 47.2 | 59.0 | 1.1e-08 |
| CAF31444.1 | putative\_gentamicin\_exporter | BGC0000696 | Saccharide | 30.0 | 47.2 | 59.0 | 1.1e-08 |
| ARV75721.1 | hypothetical\_protein | BGC0001603 | Saccharide | 30.0 | 47.2 | 59.0 | 1.1e-08 |
| EDY47104.1 | cephamycin\_export\_protein\_cmcT | BGC0000319 | NRP:Beta-lactam | 30.0 | 47.4 | 59.0 | 1.4e-08 |
| WP\_051025816.1 | MFS\_transporter | BGC0001819 | Polyketide | 26.0 | 56.2 | 58.0 | 2.4e-08 |
| ABX71121.1 | Lct38 | BGC0000238 | Polyketide | 28.0 | 40.2 | 57.0 | 4.1e-08 |
| AYV61423.1 | MFS\_transporter | BGC0001965 | Other | 30.0 | 47.7 | 57.0 | 7e-08 |
| ACN38360.1 | putative\_sisomicin\_exporter | BGC0000714 | Saccharide | 27.0 | 55.7 | 54.0 | 6e-07 |
| BAJ52670.1 | putative\_transporter | BGC0000222 | Polyketide | 32.0 | 37.0 | 53.0 | 1e-06 |
