## Supplementary Results for "A draft genome of the ascomycotal fungal species *Pseudopithomyces maydicus* (family *Didymosphaeriaceae*)": input.path1.gene42_mibig_hits.html

| MIBiG Protein | Description | MIBiG Cluster | MiBiG Product | % ID | % Coverage | BLAST Score | E-value |
| --- | --- | --- | --- | --- | --- | --- | --- |
| BAD29974.1 | P450\_monooxygenase\_2 | BGC0000676 | Terpene | 31.0 | 95.8 | 197.0 | 4.3e-50 |
| ASK38704.1 | cytochrome\_P450 | BGC0001557 | Polyketide | 33.0 | 56.9 | 176.0 | 1e-43 |
| XP\_001826049.1 |  | BGC0001995 | Terpene | 28.0 | 102.4 | 168.0 | 2.1e-41 |
| AAK11527.1 | PaxQ | BGC0001082 | Terpene | 33.0 | 60.3 | 165.0 | 3.1e-40 |
| ART41207.1 | AdrA | BGC0001508 | Polyketide | 28.0 | 101.2 | 158.0 | 2.2e-38 |
| EAW09122.1 | cytochrome\_P450 | BGC0000983 | NRP + Polyketide:Iterative type I | 28.0 | 102.6 | 157.0 | 6.4e-38 |
| EAU29532.1 | conserved\_hypothetical\_protein | BGC0000682 | Terpene | 35.0 | 64.7 | 154.0 | 5.4e-37 |
| AGZ20487.1 | cytochrome\_P450\_monooxygenase | BGC0001776 | Terpene | 32.0 | 59.1 | 153.0 | 1.2e-36 |
| ANV81297.1 | GA14\_synthase | BGC0001604 | Terpene | 31.0 | 64.9 | 151.0 | 3.5e-36 |
| BAW27598.1 | citreohybridonol\_synthase | BGC0001547 | Terpene | 26.0 | 103.6 | 150.0 | 6e-36 |
| BAW27601.1 | putative\_cytochrome\_P450\_monooxygenase | BGC0001547 | Terpene | 26.0 | 100.4 | 150.0 | 7.9e-36 |
| EBA27371.1 | cytochrome\_P450\_monooxygenase,\_putative | BGC0000129 | Polyketide | 28.0 | 92.3 | 149.0 | 1.3e-35 |
| EAL94100.1 | cytochrome\_P450\_monooxygenase,\_putative | BGC0000811 | Alkaloid | 34.0 | 60.1 | 148.0 | 2.3e-35 |
| BAV69305.1 | PrhD | BGC0001729 | Polyketide + Terpene | 34.0 | 64.1 | 146.0 | 8.7e-35 |
| QCS37514.1 | PyiG | BGC0001982 | NRP + Polyketide | 27.0 | 85.9 | 140.0 | 6.2e-33 |
| BAV32161.1 | cytochrome\_P450\_monooxygenase | BGC0001373 | Polyketide | 32.0 | 55.2 | 139.0 | 1.1e-32 |
| CAP96442.1 | P450\_monooxygenase | BGC0000420 | NRP | 27.0 | 87.5 | 139.0 | 1.8e-32 |
| AVY05525.1 | cytochrome\_P450\_monooxygenase | BGC0001571 | Terpene | 26.0 | 99.0 | 138.0 | 4e-32 |
| BAE56598.1 |  | BGC0001123 | NRP | 32.0 | 54.4 | 137.0 | 5.3e-32 |
| BAM84048.1 | cytochrome\_P450\_monooxygenase | BGC0001260 | Terpene | 33.0 | 58.3 | 137.0 | 5.3e-32 |
| ATZ45183.1 | Bcboa7 | BGC0001892 | Polyketide | 32.0 | 57.1 | 137.0 | 5.3e-32 |
| BBD84642.1 | putative\_cytochrome\_P450 | BGC0001775 | Terpene | 34.0 | 58.5 | 136.0 | 9e-32 |
| AAS92544.1 | SirB | BGC0001044 | NRP + Polyketide | 34.0 | 54.0 | 136.0 | 1.5e-31 |
| AGO86661.1 | putative\_cytochrome\_p450 | BGC0001255 | NRP + Polyketide | 25.0 | 95.8 | 134.0 | 4.5e-31 |
| BAH24001.1 | cytochrome\_P450 | BGC0000356 | NRP + Alkaloid | 28.0 | 88.9 | 127.0 | 7.1e-29 |
| ANV81301.1 | C13\_oxidase | BGC0001604 | Terpene | 27.0 | 63.3 | 126.0 | 9.3e-29 |
| DAB41646.1 | cytochrome\_P450\_monooxygenase | BGC0001777 | Terpene | 31.0 | 52.8 | 125.0 | 2.1e-28 |
| AGC83579.1 | P450\_monooxygenase | BGC0000818 | NRP | 33.0 | 51.4 | 124.0 | 3.5e-28 |
| EPE34336.1 | Cytochrome\_P450 | BGC0001035 | Polyketide + NRP + Other:Aminocoumarin | 31.0 | 59.1 | 124.0 | 6e-28 |
| AAD34552.1 | cytochrome\_P450\_monooxygenase | BGC0000088 | Polyketide | 30.0 | 59.1 | 123.0 | 7.9e-28 |
| ctg1\_orf0002 |  | BGC0000685 | Terpene | 30.0 | 59.9 | 120.0 | 6.7e-27 |
| EAW09118.1 | cytochrome\_P450\_oxidoreductase\_GliF | BGC0000983 | NRP + Polyketide:Iterative type I | 30.0 | 68.8 | 120.0 | 6.7e-27 |
| ctg1\_orf7 |  | BGC0000685 | Terpene | 25.0 | 86.5 | 118.0 | 2.5e-26 |
| EAU36745.1 | hypothetical\_protein | BGC0000292 | NRP | 28.0 | 59.1 | 113.0 | 1.1e-24 |
| AMM63168.1 | AniF2 | BGC0001371 | NRP | 28.0 | 56.5 | 91.0 | 4.3e-18 |
| ctg1\_orf003 |  | BGC0001068 | Terpene + Polyketide | 36.0 | 27.8 | 82.0 | 1.5e-15 |
| CAL69595.1 | hypothetical\_protein | BGC0001049 | NRP + Polyketide:Iterative type I | 25.0 | 56.0 | 81.0 | 5.9e-15 |
| ADN43683.1 | DmbB | BGC0001136 | NRP + Polyketide:Iterative type I | 25.0 | 56.0 | 81.0 | 5.9e-15 |
| AKC54421.1 | cytochrome\_P450\_monooxygenase | BGC0001218 | NRP + Polyketide | 25.0 | 56.9 | 77.0 | 8.5e-14 |
| BAE56589.1 |  | BGC0001123 | NRP | 39.0 | 21.8 | 76.0 | 1.4e-13 |
| ctg1\_orf00001 |  | BGC0000685 | Terpene | 39.0 | 21.2 | 75.0 | 3.2e-13 |
| EED53482.1 | cytochrome\_P450,\_putative | BGC0001304 | Polyketide | 29.0 | 32.9 | 71.0 | 3.6e-12 |
| AZH23827.1 | MgiT | BGC0001971 | NRP + Polyketide | 26.0 | 49.0 | 59.0 | 1.4e-08 |
| AUTOORF\_00003 |  | BGC0001322 | Terpene | 24.0 | 54.8 | 57.0 | 5.3e-08 |
| pseudo106205\_112773 |  | BGC0001322 | Terpene | 24.0 | 54.8 | 57.0 | 5.3e-08 |
| ADO85579.1 | PntI | BGC0000653 | Terpene | 28.0 | 41.3 | 56.0 | 1.2e-07 |
| CAJ42333.1 | cytochrome\_P450 | BGC0000273 | Polyketide:Type II + Saccharide:Hybrid/tailoring | 29.0 | 37.7 | 52.0 | 2.2e-06 |
