## Supplementary Results for "A draft genome of the ascomycotal fungal species *Pseudopithomyces maydicus* (family *Didymosphaeriaceae*)": input.path1.gene45_mibig_hits.html

| MIBiG Protein | Description | MIBiG Cluster | MiBiG Product | % ID | % Coverage | BLAST Score | E-value |
| --- | --- | --- | --- | --- | --- | --- | --- |
| BAD29970.1 | geranylgeranyldiphosphate\_synthase | BGC0000676 | Terpene | 51.0 | 67.1 | 317.0 | 4.4e-86 |
| ctg1\_orf003 |  | BGC0000688 | Terpene | 49.0 | 75.1 | 312.0 | 1.1e-84 |
| AGZ20472.2 | geranylgeranyl\_diphosphate\_synthase | BGC0001776 | Terpene | 47.0 | 65.8 | 278.0 | 2.3e-74 |
| AAK11531.1 | PaxG | BGC0001082 | Terpene | 46.0 | 65.8 | 273.0 | 5.6e-73 |
| CBF85177.1 | conserved\_hypothetical\_protein | BGC0000673 | Terpene | 47.0 | 65.8 | 269.0 | 1.4e-71 |
| AQM58281.1 | isoprenoid\_synthase | BGC0001816 | NRP + Polyketide | 40.0 | 66.4 | 219.0 | 1.3e-56 |
| AZQ56744.1 | sesterterpene\_synthase | BGC0001969 | Terpene | 37.0 | 68.2 | 200.0 | 6e-51 |
| APY21859.1 | FmsB | BGC0001659 | Terpene | 38.0 | 68.2 | 194.0 | 3.3e-49 |
| ANV81299.1 | GGPP\_synthase | BGC0001604 | Terpene | 32.0 | 73.1 | 175.0 | 1.6e-43 |
| ACI04497.1 | CrtE | BGC0000646 | Terpene | 28.0 | 56.7 | 65.0 | 2.3e-10 |
