## Supplementary Results for "A draft genome of the ascomycotal fungal species *Pseudopithomyces maydicus* (family *Didymosphaeriaceae*)": input.path1.gene67_mibig_hits.html

| MIBiG Protein | Description | MIBiG Cluster | MiBiG Product | % ID | % Coverage | BLAST Score | E-value |
| --- | --- | --- | --- | --- | --- | --- | --- |
| KGO40477.1 | Serine\_hydrolase\_FSH | BGC0001205 | Polyketide | 35.0 | 42.2 | 60.0 | 4.4e-09 |
| EWG54261.1 | hypothetical\_protein | BGC0001190 | Polyketide | 36.0 | 35.0 | 50.0 | 3.5e-06 |
