## Supplementary Results for "A draft genome of the ascomycotal fungal species *Pseudopithomyces maydicus* (family *Didymosphaeriaceae*)": input.path1.gene69_mibig_hits.html

| MIBiG Protein | Description | MIBiG Cluster | MiBiG Product | % ID | % Coverage | BLAST Score | E-value |
| --- | --- | --- | --- | --- | --- | --- | --- |
| ANF07287.1 | hydrolase\_341 | BGC0001340 | Polyketide:Iterative type I | 39.0 | 91.0 | 99.0 | 3.5e-21 |
| KGO40477.1 | Serine\_hydrolase\_FSH | BGC0001205 | Polyketide | 34.0 | 57.6 | 58.0 | 1.2e-08 |
