## Supplementary Results for "A draft genome of the ascomycotal fungal species *Pseudopithomyces maydicus* (family *Didymosphaeriaceae*)": input.path1.gene70_mibig_hits.html

| MIBiG Protein | Description | MIBiG Cluster | MiBiG Product | % ID | % Coverage | BLAST Score | E-value |
| --- | --- | --- | --- | --- | --- | --- | --- |
| AFU65888.1 | DacR2 | BGC0000216 | Polyketide | 38.0 | 42.9 | 111.0 | 2.6e-24 |
| ADG86335.1 | transporter | BGC0000190 | Polyketide | 37.0 | 43.4 | 111.0 | 4.4e-24 |
| AAQ08935.1 | putative\_membrane\_transporter | BGC0000224 | Polyketide:Type II | 38.0 | 45.1 | 110.0 | 5.7e-24 |
| BAV16993.1 | putative\_transporter | BGC0001384 | Polyketide | 30.0 | 82.9 | 109.0 | 9.7e-24 |
| KIS69144.1 | Major\_Facilitator\_invovled\_in\_MEL\_transport | BGC0001888 | Other | 33.0 | 52.2 | 106.0 | 8.2e-23 |
| AAF00219.1 | transporter | BGC0000277 | Polyketide | 28.0 | 96.6 | 106.0 | 1.1e-22 |
| ACP19369.1 | SaqJ1 | BGC0000267 | Polyketide:Type II + Saccharide:Oligosaccharide | 25.0 | 113.2 | 106.0 | 1.4e-22 |
| ARO44650.1 | transporter | BGC0001769 | Polyketide | 32.0 | 61.7 | 105.0 | 1.8e-22 |
| CAH10123.1 | putative\_transporter | BGC0000268 | Polyketide | 32.0 | 52.0 | 104.0 | 5.3e-22 |
| AAL15595.1 | Sim17 | BGC0000270 | Polyketide | 29.0 | 70.7 | 102.0 | 2e-21 |
| CAF60521.1 | putative\_efflux\_protein | BGC0000704 | Saccharide | 35.0 | 47.1 | 102.0 | 2e-21 |
| CAF31575.1 | putative\_kanamycin\_efflux\_protein | BGC0000705 | Saccharide | 35.0 | 47.1 | 102.0 | 2e-21 |
| AHA12095.1 | transporter | BGC0001172 | NRP + Polyketide:Modular type I | 26.0 | 110.0 | 102.0 | 2e-21 |
| AHW57792.1 | PgaJ3 | BGC0000262 | Polyketide:Type II + Saccharide:Hybrid/tailoring | 26.0 | 110.0 | 101.0 | 2.6e-21 |
| AGO50605.1 | transporter | BGC0000229 | Polyketide:Type II + Saccharide:Hybrid/tailoring | 33.0 | 51.0 | 100.0 | 7.7e-21 |
| AAK06799.1 | simocyclinone-specific\_efflux\_pump | BGC0001072 | Saccharide + Polyketide:Modular type I + Polyketide:Type II + Other:Aminocoumarin | 29.0 | 70.7 | 99.0 | 1e-20 |
| ACB46467.1 | efflux\_permease | BGC0000082 | Polyketide | 34.0 | 51.7 | 98.0 | 2.9e-20 |
| AEO57490.1 | general\_substrate\_transporter | BGC0001449 | NRP + Alkaloid + Polyketide:Iterative type I | 30.0 | 52.9 | 96.0 | 1.5e-19 |
| BAE56594.1 |  | BGC0001123 | NRP | 28.0 | 52.9 | 92.0 | 1.2e-18 |
| OWA25250.1 | MFS\_transporter | BGC0001438 | Polyketide + Saccharide:Hybrid/tailoring | 28.0 | 73.2 | 91.0 | 3.6e-18 |
| EYT83437.1 | multidrug\_MFS\_transporter | BGC0001213 | Polyketide | 31.0 | 44.9 | 90.0 | 6.1e-18 |
| AMY15059.1 | MFS\_transporter | BGC0001339 | Polyketide:Iterative type I | 24.0 | 112.7 | 90.0 | 6.1e-18 |
| CAP93741.1 |  | BGC0001882 | Polyketide | 23.0 | 104.6 | 89.0 | 1e-17 |
| AQW35077.1 | MFS\_transporter | BGC0001675 | Polyketide | 28.0 | 80.2 | 89.0 | 1.4e-17 |
| BAQ25491.1 | multidrug\_MFS\_(major\_facilitator\_superfamily)\_transporter | BGC0001288 | Polyketide | 34.0 | 45.9 | 88.0 | 3e-17 |
| ABX71121.1 | Lct38 | BGC0000238 | Polyketide | 30.0 | 47.3 | 86.0 | 1.5e-16 |
| AMY15055.1 | MFS\_transporter | BGC0001339 | Polyketide:Iterative type I | 28.0 | 51.0 | 84.0 | 5.7e-16 |
| AIL50170.1 | putative\_transport\_protein | BGC0000213 | Polyketide:Type II | 30.0 | 54.4 | 83.0 | 7.5e-16 |
| ACN38355.1 | putative\_transmembrane\_efflux\_protein | BGC0000714 | Saccharide | 32.0 | 51.5 | 83.0 | 9.7e-16 |
| ACS68556.1 | major\_facilitator\_superfamily\_protein | BGC0001026 | NRP + Polyketide | 28.0 | 53.4 | 83.0 | 9.7e-16 |
| PIB01159.1 | putative\_HC-toxin\_efflux\_carrier\_TOXA | BGC0001541 | Polyketide | 28.0 | 54.6 | 82.0 | 1.7e-15 |
| CAF34033.1 | putative\_transmembrane\_efflux\_protein | BGC0000689 | Saccharide | 31.0 | 51.5 | 80.0 | 8.3e-15 |
| CAF31444.1 | putative\_gentamicin\_exporter | BGC0000696 | Saccharide | 31.0 | 51.5 | 80.0 | 8.3e-15 |
| ARV75721.1 | hypothetical\_protein | BGC0001603 | Saccharide | 31.0 | 51.5 | 80.0 | 8.3e-15 |
| CAE51185.1 | RemN\_protein | BGC0000264 | Polyketide:Type II | 25.0 | 109.8 | 79.0 | 1.1e-14 |
| AFW04557.1 | drug\_resistance\_transporter | BGC0001783 | Other | 31.0 | 44.6 | 79.0 | 1.1e-14 |
| AFW04581.1 | drug\_resistance\_transporter | BGC0001783 | Other | 30.0 | 41.2 | 78.0 | 2.4e-14 |
| AAO65328.1 | putative\_transmembrane\_efflux\_protein | BGC0000236 | Polyketide | 32.0 | 53.4 | 77.0 | 5.3e-14 |
| XP\_007301853.1 | MFS\_general\_substrate\_transporter | BGC0001617 | Terpene | 28.0 | 53.2 | 77.0 | 5.3e-14 |
| CBF76046.1 | conserved\_hypothetical\_protein | BGC0001399 | NRP | 27.0 | 55.4 | 76.0 | 1.2e-13 |
| AXL88822.1 | MFS\_transporter | BGC0001895 | Polyketide | 28.0 | 68.5 | 76.0 | 1.6e-13 |
| AAQ08939.1 | putative\_membrane\_transporter | BGC0000224 | Polyketide:Type II | 30.0 | 51.0 | 73.0 | 1e-12 |
| QCE20600.1 | AsmE | BGC0001961 | NRP | 27.0 | 53.4 | 72.0 | 1.3e-12 |
| AFL48523.1 | laidlomycin\_efflux\_protein | BGC0000084 | Polyketide | 29.0 | 52.4 | 72.0 | 1.7e-12 |
| ARG41907.1 | IstD | BGC0001622 | Polyketide | 30.0 | 52.7 | 72.0 | 1.7e-12 |
| ACN38360.1 | putative\_sisomicin\_exporter | BGC0000714 | Saccharide | 22.0 | 95.9 | 72.0 | 2.2e-12 |
| AAO65793.1 | putative\_monensin\_resistance\_protein | BGC0000100 | Polyketide | 30.0 | 52.4 | 70.0 | 6.5e-12 |
| ANZ52456.1 | MonT | BGC0001670 | Polyketide | 30.0 | 52.4 | 70.0 | 6.5e-12 |
| ACU62762.1 | drug\_resistance\_transporter,\_EmrB/QacA\_subfamily | BGC0001392 | RiPP | 33.0 | 33.7 | 69.0 | 1.5e-11 |
| ATJ34010.1 | MFS\_transporter | BGC0001497 | Polyketide | 29.0 | 45.1 | 69.0 | 1.5e-11 |
| AKC91607.1 | transporter | BGC0001302 | Other | 32.0 | 35.4 | 69.0 | 1.9e-11 |
| ACZ57546.1 | predicted\_MFS\_transporter | BGC0000046 | Polyketide:Iterative type I | 29.0 | 43.9 | 68.0 | 3.2e-11 |
| ASK38705.1 | major\_facilitator\_superfamily\_transporter | BGC0001557 | Polyketide | 28.0 | 35.6 | 63.0 | 1e-09 |
| BAZ95819.1 | cpaN1\_MFS\_transporter | BGC0001563 | NRP + Polyketide | 26.0 | 46.6 | 62.0 | 1.4e-09 |
| AAP69589.1 | putative\_transmembrane\_efflux\_protein | BGC0000226 | Polyketide | 30.0 | 42.2 | 62.0 | 2.3e-09 |
| WP\_015507660.1 | MFS\_transporter | BGC0001792 | NRP | 27.0 | 45.1 | 61.0 | 3e-09 |
| AAD32747.1 | Mct | BGC0000915 | Other:Aminocoumarin | 27.0 | 52.4 | 59.0 | 1.2e-08 |
| BAN59741.1 | MFS\_transporter | BGC0001075 | Terpene + Polyketide | 30.0 | 32.2 | 59.0 | 1.5e-08 |
| CTQ34878.1 | AtcA;\_major\_facilitator\_superfamily\_(MFS)\_transporter;\_12\_transmembrane\_helices | BGC0001301 | Polyketide | 23.0 | 81.2 | 59.0 | 2e-08 |
| ADI24938.1 | VrtL | BGC0000168 | Polyketide:Iterative type I | 36.0 | 29.5 | 58.0 | 3.4e-08 |
| XP\_023093494.1 |  | BGC0001995 | Terpene | 29.0 | 25.9 | 56.0 | 9.8e-08 |
| ATY72541.1 | MFS\_transporter | BGC0001574 | NRP | 28.0 | 32.2 | 56.0 | 1.7e-07 |
| AGN74873.1 | major\_facilitator\_transporter | BGC0000459 | NRP:Cyclic depsipeptide + Polyketide:Trans-AT type I | 26.0 | 82.2 | 54.0 | 4.8e-07 |
