## Supplementary Results for "A draft genome of the ascomycotal fungal species *Pseudopithomyces maydicus* (family *Didymosphaeriaceae*)": input.path1.gene103_mibig_hits.html

| MIBiG Protein | Description | MIBiG Cluster | MiBiG Product | % ID | % Coverage | BLAST Score | E-value |
| --- | --- | --- | --- | --- | --- | --- | --- |
| BAH23997.1 | O-methyltransferase | BGC0000356 | NRP + Alkaloid | 25.0 | 62.9 | 55.0 | 1.9e-07 |
