## Supplementary Results for "A draft genome of the ascomycotal fungal species *Pseudopithomyces maydicus* (family *Didymosphaeriaceae*)": input.path1.gene106_mibig_hits.html

| MIBiG Protein | Description | MIBiG Cluster | MiBiG Product | % ID | % Coverage | BLAST Score | E-value |
| --- | --- | --- | --- | --- | --- | --- | --- |
| BAE56599.1 |  | BGC0001123 | NRP | 60.0 | 49.8 | 187.0 | 3.3e-47 |
| PKX88479.1 | methyltransferase | BGC0001708 | Polyketide + Terpene | 39.0 | 88.0 | 178.0 | 2e-44 |
| XP\_002373812.1 | GA4\_desaturase\_family\_protein | BGC0001516 | NRP | 33.0 | 69.6 | 83.0 | 6.7e-16 |
