## Supplementary Results for "A draft genome of the ascomycotal fungal species *Pseudopithomyces maydicus* (family *Didymosphaeriaceae*)": input.path1.gene107_mibig_hits.html

| MIBiG Protein | Description | MIBiG Cluster | MiBiG Product | % ID | % Coverage | BLAST Score | E-value |
| --- | --- | --- | --- | --- | --- | --- | --- |
| BAE56600.1 |  | BGC0001123 | NRP | 35.0 | 100.4 | 156.0 | 5.5e-38 |
| AAS92555.1 | SirT | BGC0001044 | NRP + Polyketide | 27.0 | 123.8 | 133.0 | 5e-31 |
| CCE28990.1 | uncharacterized\_protein | BGC0001365 | NRP | 29.0 | 121.0 | 128.0 | 1.6e-29 |
| WP\_044620356.1 | NAD(P)/FAD-dependent\_oxidoreductase | BGC0001791 | NRP | 27.0 | 98.8 | 81.0 | 2.2e-15 |
| EFG10348.1 | Thioredoxin\_reductase | BGC0000373 | NRP | 29.0 | 56.5 | 69.0 | 6.7e-12 |
| AFV30261.1 | FAD-dependent\_pyridine\_nucleotide-disulfide\_oxidoreductase | BGC0000075 | Polyketide | 31.0 | 42.3 | 62.0 | 8.2e-10 |
| AJI44183.1 | thioredoxin\_reductase | BGC0001193 | NRP | 29.0 | 59.7 | 60.0 | 5.3e-09 |
