## Supplementary Results for "A draft genome of the ascomycotal fungal species *Pseudopithomyces maydicus* (family *Didymosphaeriaceae*)": input.path1.gene109_mibig_hits.html

| MIBiG Protein | Description | MIBiG Cluster | MiBiG Product | % ID | % Coverage | BLAST Score | E-value |
| --- | --- | --- | --- | --- | --- | --- | --- |
| EHK22005.1 | putative\_non-ribosomal\_peptide\_synthetase\_GliP | BGC0001609 | NRP | 38.0 | 43.1 | 182.0 | 1.6e-45 |
| AAS92545.1 | SirP | BGC0001044 | NRP + Polyketide | 39.0 | 44.3 | 173.0 | 1.3e-42 |
| EAL92291.2 | nonribosomal\_peptide\_synthtease | BGC0000372 | NRP | 39.0 | 43.6 | 171.0 | 2.9e-42 |
| CCE28989.1 | non-ribosomal\_peptide\_synthetase | BGC0001365 | NRP | 38.0 | 38.1 | 150.0 | 8.9e-36 |
| MAA\_10036 | nonribosomal\_peptide\_synthase\_GliP-like,\_putative | BGC0000337 | NRP | 35.0 | 47.7 | 143.0 | 1.4e-33 |
| DAB41661.1 | nonribosomal\_peptide\_synthetase | BGC0001585 | Alkaloid | 39.0 | 33.3 | 131.0 | 5.6e-30 |
| ctg1\_orf10 |  | BGC0000321 | NRP | 36.0 | 35.9 | 123.0 | 1.5e-27 |
| BAE56606.1 |  | BGC0001123 | NRP | 25.0 | 47.0 | 106.0 | 1.1e-22 |
| APZ78729.1 | nonribosomal\_peptide\_synthetase | BGC0001421 | NRP:Cyclic depsipeptide + Polyketide:Iterative type I | 34.0 | 45.4 | 105.0 | 2.5e-22 |
| BBA20967.1 | nonribosomal\_peptide\_synthetase | BGC0001763 | NRP + Polyketide | 34.0 | 43.4 | 103.0 | 9.6e-22 |
| ASX95241.1 | IlaS | BGC0001620 | Polyketide | 35.0 | 42.2 | 102.0 | 1.6e-21 |
| CAJ34381.1 | NRPS\_protein | BGC0000445 | NRP:Cyclic depsipeptide | 31.0 | 53.6 | 101.0 | 4.7e-21 |
| ALV86867.1 | Tlo21 | BGC0001406 | NRP | 36.0 | 34.5 | 98.0 | 4e-20 |
| AAS47562.1 | mixed\_type\_I\_polyketide\_synthase\_-\_peptide\_synthetase | BGC0001108 | Polyketide:Trans-AT type I | 40.0 | 30.6 | 95.0 | 2.6e-19 |
| ctg1\_orf8 |  | BGC0001109 | NRP + Polyketide | 40.0 | 30.6 | 95.0 | 2.6e-19 |
| ABG94125.1 | non-ribosomal\_peptide\_synthetase | BGC0000417 | NRP | 32.0 | 41.8 | 90.0 | 8.4e-18 |
| AAN32978.1 | BarD | BGC0000962 | NRP + Polyketide:Modular type I | 30.0 | 32.4 | 90.0 | 8.4e-18 |
| CCM44336.1 | Nonribosomal\_peptide\_synthetase | BGC0001056 | NRP + Polyketide:Modular type I + Polyketide:PUFA synthase or related | 34.0 | 34.2 | 90.0 | 8.4e-18 |
| AAU39360.1 | lichenysin\_synthase\_LchAB | BGC0000381 | NRP | 34.0 | 32.2 | 88.0 | 3.2e-17 |
| APZ78809.1 | nonribosomal\_peptide\_synthetase | BGC0001428 | NRP:Cyclic depsipeptide + Polyketide:Iterative type I | 27.0 | 50.9 | 88.0 | 4.2e-17 |
| BAH43871.1 | truncated\_linear\_pentadecapeptide\_gramicidin\_synthetase\_LgrC | BGC0000367 | NRP | 26.0 | 53.6 | 87.0 | 7.1e-17 |
| APZ78756.1 | nonribosomal\_peptide\_synthetase | BGC0001423 | NRP:Cyclic depsipeptide + Polyketide:Iterative type I | 26.0 | 50.9 | 87.0 | 9.3e-17 |
| CAM59606.1 | non-ribosomal\_peptide\_synthetase | BGC0000297 | NRP:Glycopeptide + Polyketide:Other + Saccharide:Hybrid/tailoring | 28.0 | 47.3 | 86.0 | 1.2e-16 |
| ctg1\_orf000000 |  | BGC0000901 | Other | 25.0 | 54.3 | 86.0 | 1.6e-16 |
| APZ78782.1 | nonribosomal\_peptide\_synthetase | BGC0001426 | NRP:Cyclic depsipeptide + Polyketide:Iterative type I | 26.0 | 50.9 | 86.0 | 1.6e-16 |
| CAP93139.1 | cyclic\_hydrophobic\_tetrapeptide | BGC0000357 | NRP:Cyclic depsipeptide | 26.0 | 47.5 | 86.0 | 2.1e-16 |
| APZ78795.1 | nonribosomal\_peptide\_synthetase | BGC0001427 | NRP:Cyclic depsipeptide + Polyketide:Iterative type I | 26.0 | 50.9 | 86.0 | 2.1e-16 |
| APZ78680.1 | nonribosomal\_peptide\_synthetase | BGC0001417 | NRP:Cyclic depsipeptide + Polyketide:Iterative type I | 25.0 | 51.2 | 85.0 | 2.7e-16 |
| APZ78716.1 | nonribosomal\_peptide\_synthetase | BGC0001420 | NRP:Cyclic depsipeptide + Polyketide:Iterative type I | 25.0 | 51.8 | 85.0 | 2.7e-16 |
| APZ78692.1 | nonribosomal\_peptide\_synthetase | BGC0001418 | NRP:Cyclic depsipeptide + Polyketide:Iterative type I | 25.0 | 51.8 | 84.0 | 4.6e-16 |
| APZ78704.1 | nonribosomal\_peptide\_synthetase | BGC0001419 | NRP:Cyclic depsipeptide + Polyketide:Iterative type I | 25.0 | 51.8 | 84.0 | 4.6e-16 |
| APZ78769.1 | nonribosomal\_peptide\_synthetase | BGC0001425 | NRP:Cyclic depsipeptide + Polyketide:Iterative type I | 26.0 | 50.9 | 84.0 | 4.6e-16 |
| APZ78744.1 | nonribosomal\_peptide\_synthetase | BGC0001422 | NRP:Cyclic depsipeptide + Polyketide:Iterative type I | 25.0 | 51.2 | 84.0 | 6e-16 |
| BAE98162.1 | putative\_non-ribosomal\_peptide\_synthetase | BGC0000339 | NRP | 37.0 | 27.0 | 84.0 | 7.8e-16 |
| AAF19815.1 | mtaG | BGC0001024 | NRP + Polyketide:Modular type I | 34.0 | 35.8 | 84.0 | 7.8e-16 |
| ABD65958.1 | nonribosomal\_peptide\_synthetase | BGC0000341 | NRP | 26.0 | 59.3 | 83.0 | 1e-15 |
| AQZ69228.1 | hypothetical\_protein | BGC0001635 | NRP + Polyketide | 27.0 | 46.4 | 83.0 | 1.3e-15 |
| CAJ45639.1 | vanchrobactin\_non\_ribosomal\_peptide\_synthetase | BGC0000454 | NRP | 39.0 | 24.2 | 82.0 | 1.7e-15 |
| OAQ83772.1 | nonribosomal\_peptide\_synthase | BGC0001358 | Polyketide | 25.0 | 47.5 | 82.0 | 1.7e-15 |
| OAL11435.1 | non-ribosomal\_peptide\_synthetase | BGC0001570 | NRP | 35.0 | 28.8 | 82.0 | 1.7e-15 |
| ctg1\_orf20 |  | BGC0001767 | NRP | 29.0 | 40.4 | 82.0 | 1.7e-15 |
| EME52990.1 | amino\_acid\_adenylation\_protein | BGC0001460 | NRP:Glycopeptide | 31.0 | 35.9 | 82.0 | 2.3e-15 |
| ctg1\_orf17 |  | BGC0001457 | NRP | 34.0 | 35.9 | 82.0 | 3e-15 |
| CAE02631.1 | surfactin\_synthetase\_B\_ | BGC0000433 | NRP:Lipopeptide | 28.0 | 31.9 | 81.0 | 3.9e-15 |
| AEI58865.1 | peptide\_synthetase | BGC0000455 | NRP | 30.0 | 35.8 | 81.0 | 3.9e-15 |
| APZ78856.1 | nonribosomal\_peptide\_synthetase | BGC0001432 | NRP:Cyclic depsipeptide + Polyketide:Iterative type I | 25.0 | 51.4 | 81.0 | 3.9e-15 |
| BAH43870.1 | putative\_linear\_pentadecapeptide\_gramicidin\_synthetase\_LgrB | BGC0000367 | NRP | 24.0 | 53.6 | 81.0 | 5.1e-15 |
| WP\_013310343.1 | non-ribosomal\_peptide\_synthetase | BGC0001993 | NRP | 30.0 | 35.2 | 81.0 | 6.6e-15 |
| BAW32324.1 | nonribosomal\_peptide\_synthetase | BGC0001630 | NRP + Polyketide | 24.0 | 59.8 | 80.0 | 8.7e-15 |
| ABB90279.1 | non-ribosomal\_peptide\_synthetase | BGC0001057 | NRP + Polyketide | 27.0 | 47.2 | 80.0 | 1.1e-14 |
| AAF01762.1 | AM-toxin\_synthetase | BGC0001261 | NRP | 26.0 | 55.3 | 80.0 | 1.1e-14 |
| AQZ42163.1 | putative\_nonribosomal\_peptide\_synthase | BGC0001820 | NRP | 33.0 | 37.5 | 80.0 | 1.1e-14 |
| AFR69334.1 | nonribosomal\_peptide\_synthetase\_SpiDE1 | BGC0001045 | NRP:Cyclic depsipeptide + Polyketide:Modular type I | 25.0 | 49.5 | 79.0 | 1.5e-14 |
| CAA16182.1 | putative\_peptide\_synthase | BGC0001063 | NRP + Polyketide | 30.0 | 45.4 | 79.0 | 1.5e-14 |
| EWS95122.1 | hypothetical\_protein | BGC0000306 | NRP:Lipopeptide | 29.0 | 34.9 | 79.0 | 1.9e-14 |
| AAY37654.1 | Amino\_acid\_adenylation | BGC0000437 | NRP | 30.0 | 42.3 | 79.0 | 2.5e-14 |
| AGS77309.1 | NRPS\_modules\_4-6 | BGC0001178 | NRP:Glycopeptide | 25.0 | 47.5 | 79.0 | 2.5e-14 |
| OKA09423.1 | non-ribosomal\_peptide\_synthetase | BGC0001459 | NRP:Glycopeptide | 30.0 | 35.9 | 79.0 | 2.5e-14 |
| BAV56270.1 |  | BGC0001657 | NRP | 30.0 | 35.4 | 79.0 | 2.5e-14 |
| OLZ52457.1 | non-ribosomal\_peptide\_synthetase | BGC0001462 | NRP:Glycopeptide | 26.0 | 49.3 | 78.0 | 3.3e-14 |
| AEW31015.1 | plipastatin\_synthetase | BGC0000407 | NRP | 29.0 | 31.3 | 77.0 | 5.6e-14 |
| AIE77059.1 | peptide\_synthetase | BGC0000418 | NRP | 26.0 | 48.2 | 77.0 | 7.3e-14 |
| BAB69699.1 | iturin\_A\_synthetase\_B | BGC0001098 | NRP + Polyketide | 29.0 | 32.2 | 77.0 | 7.3e-14 |
| ALV82356.1 | CDA\_peptide\_synthetase\_I | BGC0001370 | NRP | 26.0 | 46.3 | 77.0 | 7.3e-14 |
| ALV82388.1 | CDA\_peptide\_synthetase\_III | BGC0001370 | NRP | 28.0 | 34.7 | 77.0 | 7.3e-14 |
| EFL06867.1 | predicted\_protein | BGC0000300 | NRP | 29.0 | 44.0 | 77.0 | 9.6e-14 |
| CAD91221.1 | putative\_non-ribosomal\_peptide\_synthetase,\_module\_3 | BGC0000289 | NRP:Glycopeptide + Saccharide:Hybrid/tailoring | 25.0 | 46.4 | 76.0 | 1.3e-13 |
| CDG12864.1 | non-ribosomal\_peptide\_synthetase | BGC0001415 | NRP | 32.0 | 38.3 | 76.0 | 1.3e-13 |
| EME52989.1 | amino\_acid\_adenylation\_protein | BGC0001460 | NRP:Glycopeptide | 26.0 | 46.8 | 76.0 | 1.3e-13 |
| ABS74180.1 | bacillomycin\_D\_synthetase\_B | BGC0001090 | Polyketide + NRP:Lipopeptide | 28.0 | 31.9 | 76.0 | 1.6e-13 |
| ALV82384.1 | CDA\_peptide\_synthetase\_II | BGC0001370 | NRP | 28.0 | 32.6 | 76.0 | 1.6e-13 |
| ATY37608.1 | BreC | BGC0001536 | NRP | 28.0 | 37.5 | 76.0 | 1.6e-13 |
| KPN90369.1 | NunE | BGC0001416 | NRP | 28.0 | 30.8 | 76.0 | 2.1e-13 |
| BAW32334.1 | hybrid\_cis-AT\_polyketide\_synthase\_-\_nonribosomal\_peptide\_synthetase | BGC0001631 | NRP + Polyketide | 24.0 | 47.7 | 76.0 | 2.1e-13 |
| ctg1\_orf003 |  | BGC0000334 | NRP | 23.0 | 49.1 | 75.0 | 2.8e-13 |
| AAC06348.1 | bacitracin\_synthetase\_3 | BGC0000310 | NRP | 28.0 | 30.8 | 75.0 | 3.6e-13 |
| ABS74179.1 | bacillomycin\_D\_synthetase\_C | BGC0001090 | Polyketide + NRP:Lipopeptide | 30.0 | 33.3 | 74.0 | 4.8e-13 |
| AAM80537.1 | StaC | BGC0000290 | NRP:Glycopeptide | 26.0 | 46.6 | 74.0 | 6.2e-13 |
| ABS74208.1 | fengycin\_synthetase\_B | BGC0001095 | NRP | 31.0 | 38.8 | 74.0 | 6.2e-13 |
| BAO66533.1 | nonribosomal\_peptide\_synthase | BGC0000042 | Polyketide | 33.0 | 34.9 | 74.0 | 8.1e-13 |
| AGA37269.1 | NRPS | BGC0000819 | NRP + Alkaloid | 25.0 | 47.0 | 74.0 | 8.1e-13 |
| CAN89663.1 | putative\_non-ribosomal\_peptide\_synthetase | BGC0001070 | NRP + Polyketide:Modular type I + Polyketide:Trans-AT type I | 29.0 | 34.9 | 74.0 | 8.1e-13 |
| PHM26612.1 | pvdj | BGC0001130 | NRP + Polyketide | 26.0 | 47.3 | 74.0 | 8.1e-13 |
| CAD91212.1 | putative\_non-ribosomal\_peptide\_synthetase,\_modules\_4-6 | BGC0000289 | NRP:Glycopeptide + Saccharide:Hybrid/tailoring | 25.0 | 48.2 | 73.0 | 1.1e-12 |
| AIG79241.1 | Hypothetical\_protein | BGC0000419 | Saccharide + NRP:Glycopeptide | 25.0 | 46.6 | 73.0 | 1.1e-12 |
| AJV88375.1 | MfnC | BGC0001214 | NRP | 30.0 | 40.4 | 73.0 | 1.1e-12 |
| ALK27914.1 | non-ribosomal\_peptide\_synthase | BGC0001233 | NRP | 31.0 | 31.0 | 73.0 | 1.1e-12 |
| ABA73954.1 | putative\_non-ribosomal\_peptide\_synthetase | BGC0001842 | NRP:Lipopeptide | 27.0 | 33.3 | 73.0 | 1.1e-12 |
| CAJ76292.1 | putative\_non-ribosomal\_peptide\_synthase | BGC0000972 | NRP + Polyketide:Modular type I + Polyketide:Trans-AT type I | 34.0 | 35.9 | 73.0 | 1.4e-12 |
| AOC89000.1 | putative\_nonribosomal\_peptide\_synthetase | BGC0001652 | NRP | 34.0 | 31.1 | 73.0 | 1.4e-12 |
| AEH41794.1 | HrmP | BGC0000374 | NRP:Cyclic depsipeptide | 25.0 | 50.2 | 72.0 | 1.8e-12 |
| CZT62785.1 | Non-ribosomal\_peptide\_synthase,\_involved\_in\_Hassallidin\_biosynthesis | BGC0001614 | NRP | 25.0 | 49.6 | 72.0 | 1.8e-12 |
| QBG38782.1 | Atr21 | BGC0001975 | NRP | 30.0 | 33.3 | 72.0 | 1.8e-12 |
| AAX31557.1 | peptide\_synthetase\_1 | BGC0000336 | NRP | 29.0 | 32.9 | 72.0 | 2.4e-12 |
| CAB38517.1 | CDA\_peptide\_synthetase\_II\_(CdaPs2) | BGC0000315 | NRP:Ca+-dependent lipopeptide | 28.0 | 32.9 | 72.0 | 3.1e-12 |
| CAD55498.1 | CDA\_peptide\_synthetase\_III\_(CdaPs3) | BGC0000315 | NRP:Ca+-dependent lipopeptide | 29.0 | 36.1 | 72.0 | 3.1e-12 |
| DAB41477.1 | nonribosomal\_peptide\_synthetase | BGC0001766 | NRP | 31.0 | 29.2 | 72.0 | 3.1e-12 |
| AAK81826.1 | peptide\_synthetase | BGC0000326 | NRP | 23.0 | 49.3 | 71.0 | 4e-12 |
| CAG15011.1 | peptide\_synthetase,\_module\_4-6 | BGC0000441 | NRP | 24.0 | 46.6 | 71.0 | 4e-12 |
| DAB41476.1 | nonribosomal\_peptide\_synthetase | BGC0001766 | NRP | 26.0 | 30.6 | 71.0 | 4e-12 |
| ATU31794.1 | NRPS | BGC0001814 | NRP | 29.0 | 33.6 | 71.0 | 4e-12 |
| QED55422.1 | nonribosomal\_peptide\_synthetase | BGC0001984 | NRP | 28.0 | 32.4 | 71.0 | 4e-12 |
| AAQ93484.1 | CmaA | BGC0000328 | NRP | 31.0 | 40.4 | 71.0 | 5.3e-12 |
| AGA37267.1 | NRPS | BGC0000816 | NRP + Alkaloid | 25.0 | 45.6 | 71.0 | 5.3e-12 |
| BAV56271.1 |  | BGC0001657 | NRP | 28.0 | 32.2 | 71.0 | 5.3e-12 |
| ABB69081.1 | putative\_L-prolyl-AMP\_ligase | BGC0000260 | Polyketide | 32.0 | 36.1 | 71.0 | 6.9e-12 |
| CAB38518.1 | CDA\_peptide\_synthetase\_I\_(CdaPs1) | BGC0000315 | NRP:Ca+-dependent lipopeptide | 26.0 | 47.2 | 71.0 | 6.9e-12 |
| AEH41793.1 | HrmO | BGC0000374 | NRP:Cyclic depsipeptide | 25.0 | 57.3 | 71.0 | 6.9e-12 |
| RAT94090.1 | NRPS | BGC0001469 | NRP | 27.0 | 33.8 | 71.0 | 6.9e-12 |
| AAM80538.1 | StaB | BGC0000290 | NRP:Glycopeptide | 22.0 | 46.4 | 70.0 | 9e-12 |
| AEA30274.1 | peptide\_synthetase | BGC0000429 | Polyketide + NRP:Cyclic depsipeptide | 28.0 | 32.6 | 70.0 | 9e-12 |
| AED90003.1 | non-ribosomal\_peptide\_synthetase\_ThaB | BGC0000443 | NRP:Beta-lactam | 25.0 | 33.1 | 70.0 | 9e-12 |
| CCP45167.1 | Peptide\_synthetase\_MbtF\_(peptide\_synthase) | BGC0001021 | NRP + Polyketide | 25.0 | 44.8 | 70.0 | 9e-12 |
| CRG85572.1 | nonribosomal\_peptide\_synthase,\_putative | BGC0001402 | NRP | 24.0 | 48.4 | 70.0 | 9e-12 |
| ACR33075.1 | Proline\_adenylation\_protein | BGC0000017 | Alkaloid + Polyketide:Modular type I | 28.0 | 39.5 | 70.0 | 1.2e-11 |
| AHZ20774.1 | non-ribosomal\_peptide\_synthase | BGC0000369 | NRP + Saccharide:Hybrid/tailoring | 24.0 | 37.5 | 70.0 | 1.2e-11 |
| AOA33122.1 | Nonribosomal\_peptide\_synthetase | BGC0001346 | NRP:Cyclic depsipeptide | 26.0 | 35.4 | 70.0 | 1.2e-11 |
| BBA21073.1 | putative\_non-ribosomal\_peptide\_synthetase | BGC0001740 | NRP + Polyketide | 30.0 | 33.1 | 70.0 | 1.2e-11 |
| OKA09424.1 | non-ribosomal\_peptide\_synthetase | BGC0001459 | NRP:Glycopeptide | 24.0 | 46.3 | 69.0 | 1.5e-11 |
| CAC48360.1 | peptide\_synthetase | BGC0000311 | NRP | 27.0 | 32.6 | 69.0 | 2e-11 |
| CAE53351.1 | non-ribosomal\_peptide\_synthetase | BGC0000440 | NRP:Glycopeptide | 25.0 | 47.0 | 69.0 | 2e-11 |
| CAG15010.1 | peptide\_synthetase,\_module\_3 | BGC0000441 | NRP | 25.0 | 47.0 | 69.0 | 2e-11 |
| CAE53352.1 | non-ribosomal\_peptide\_synthetase | BGC0000440 | NRP:Glycopeptide | 24.0 | 47.0 | 68.0 | 3.4e-11 |
| AEI58866.1 | peptide\_synthetase | BGC0000455 | NRP | 23.0 | 48.9 | 68.0 | 3.4e-11 |
| AEP18656.1 | WAPS1 | BGC0000461 | NRP | 28.0 | 34.0 | 68.0 | 3.4e-11 |
| CCJ67647.1 | JagC | BGC0001127 | NRP | 26.0 | 33.3 | 68.0 | 3.4e-11 |
| AJW76710.1 | DsaH | BGC0001196 | NRP | 28.0 | 35.9 | 68.0 | 3.4e-11 |
| ALG65317.1 | Cal19 | BGC0001297 | NRP | 29.0 | 41.1 | 68.0 | 3.4e-11 |
| AKC91857.1 | nonribosomal\_peptide\_synthetase | BGC0001414 | NRP | 26.0 | 38.1 | 68.0 | 3.4e-11 |
| KFL51886.1 | amino\_acid\_adenylation\_protein | BGC0001711 | NRP + Polyketide | 25.0 | 47.5 | 68.0 | 3.4e-11 |
| AAY37655.1 | Amino\_acid\_adenylation | BGC0000437 | NRP | 27.0 | 29.5 | 67.0 | 7.6e-11 |
| AAO72425.1 | syringopeptin\_synthetase\_C | BGC0000438 | NRP | 27.0 | 29.5 | 67.0 | 7.6e-11 |
| AAF08796.1 | MycB | BGC0001103 | NRP + Polyketide | 26.0 | 33.1 | 67.0 | 7.6e-11 |
| ctg1\_orf1265 |  | BGC0001752 | NRP | 27.0 | 32.0 | 67.0 | 7.6e-11 |
| XP\_003044554.1 | hypothetical\_protein | BGC0001768 | NRP | 23.0 | 53.0 | 67.0 | 7.6e-11 |
| ATO51563.1 | non-ribosomal\_peptide\_synthetase | BGC0001796 | NRP | 25.0 | 33.3 | 67.0 | 9.9e-11 |
| AIG79242.1 | Non-ribosomal\_peptide\_synthetase | BGC0000419 | Saccharide + NRP:Glycopeptide | 23.0 | 46.6 | 66.0 | 1.3e-10 |
| CBF73453.1 | nonribosomal\_peptide\_synthase,\_putative\_(JCVI) | BGC0001515 | NRP | 26.0 | 45.9 | 66.0 | 1.7e-10 |
| QBC75022.1 | non-ribosomal\_peptide\_synthetase | BGC0001968 | NRP | 29.0 | 34.2 | 66.0 | 1.7e-10 |
| AHD05678.1 | nonribosomal\_peptide\_ligase\_subunit | BGC0000402 | NRP | 25.0 | 33.8 | 66.0 | 2.2e-10 |
| ALK27915.1 | non-ribosomal\_peptide\_synthase | BGC0001233 | NRP | 27.0 | 29.9 | 66.0 | 2.2e-10 |
| CZT62784.1 | Non-ribosomal\_peptide\_synthase,\_involved\_in\_Hassallidin\_biosynthesis | BGC0001614 | NRP | 27.0 | 29.9 | 66.0 | 2.2e-10 |
| BAX64247.1 | NRPS | BGC0001623 | NRP + Polyketide | 26.0 | 34.2 | 66.0 | 2.2e-10 |
| QED55423.1 | nonribosomal\_peptide\_synthetase | BGC0001984 | NRP | 28.0 | 30.2 | 65.0 | 2.9e-10 |
| AAT09804.1 | NocA | BGC0000395 | NRP | 28.0 | 39.1 | 65.0 | 3.8e-10 |
| AIE77058.1 | peptide\_synthetase\_module\_3 | BGC0000418 | NRP | 23.0 | 46.6 | 65.0 | 3.8e-10 |
| AAY37647.1 | Amino\_acid\_adenylation | BGC0000437 | NRP | 24.0 | 29.9 | 65.0 | 3.8e-10 |
| AGS77308.1 | NRPS\_module\_3 | BGC0001178 | NRP:Glycopeptide | 27.0 | 30.2 | 65.0 | 3.8e-10 |
| BAX64246.1 | NRPS | BGC0001623 | NRP + Polyketide | 25.0 | 41.3 | 65.0 | 3.8e-10 |
| AEA30273.1 | peptide\_synthetase | BGC0000429 | Polyketide + NRP:Cyclic depsipeptide | 26.0 | 38.1 | 64.0 | 4.9e-10 |
| ALG65318.1 | Cal18 | BGC0001297 | NRP | 26.0 | 35.4 | 64.0 | 4.9e-10 |
| AQM58286.1 | non-ribosomal\_peptide\_synthase | BGC0001816 | NRP + Polyketide | 26.0 | 46.3 | 64.0 | 4.9e-10 |
| QBG38783.1 | Atr22 | BGC0001975 | NRP | 25.0 | 32.6 | 64.0 | 4.9e-10 |
| AID65222.1 | putative\_aspartate\_racemase | BGC0000335 | NRP | 28.0 | 32.0 | 64.0 | 6.4e-10 |
| AWI62626.1 | nonribosomal\_peptide\_synthetase | BGC0001943 | NRP | 26.0 | 31.7 | 64.0 | 6.4e-10 |
| CDG17987.1 | Putative\_Ornithine\_racemase\_(fragment) | BGC0000464 | NRP:Cyclic depsipeptide | 23.0 | 31.7 | 64.0 | 8.4e-10 |
| BAD55613.1 | putative\_non-ribosomal\_peptide\_synthetase | BGC0001027 | NRP + Polyketide | 26.0 | 40.7 | 64.0 | 8.4e-10 |
| ARU08075.1 | mlcM | BGC0001448 | NRP:Ca+-dependent lipopeptide | 27.0 | 33.1 | 64.0 | 8.4e-10 |
| QBC75021.1 | non-ribosomal\_peptide\_synthetase | BGC0001968 | NRP | 26.0 | 33.5 | 64.0 | 8.4e-10 |
| AEG64698.1 | LpmD | BGC0000379 | NRP | 28.0 | 32.9 | 63.0 | 1.1e-09 |
| AAT09805.1 | NocB | BGC0000395 | NRP | 23.0 | 47.3 | 63.0 | 1.1e-09 |
| WP\_013184463.1 | non-ribosomal\_peptide\_synthetase | BGC0001716 | NRP | 27.0 | 29.2 | 63.0 | 1.1e-09 |
| AYA22334.1 | KerC | BGC0001955 | Other | 23.0 | 51.8 | 63.0 | 1.1e-09 |
| AAO39110.1 | AdmP | BGC0000956 | NRP:Beta-lactam + Polyketide:Type II | 33.0 | 32.0 | 63.0 | 1.4e-09 |
| AQZ69227.1 | hypothetical\_protein | BGC0001635 | NRP + Polyketide | 25.0 | 34.2 | 63.0 | 1.4e-09 |
| BAP34707.1 | AMP-dependent\_synthetase\_and\_ligase | BGC0000078 | Polyketide | 32.0 | 28.1 | 62.0 | 1.9e-09 |
| CBF87069.1 | nonribosomal\_peptide\_synthase,\_putative\_(Eurofung) | BGC0001290 | NRP | 23.0 | 49.1 | 62.0 | 1.9e-09 |
| OLZ50885.1 | non-ribosomal\_peptide\_synthetase | BGC0001461 | NRP:Glycopeptide | 23.0 | 52.0 | 62.0 | 1.9e-09 |
| KUM80514.1 | hypothetical\_protein | BGC0001562 | NRP | 27.0 | 32.9 | 62.0 | 1.9e-09 |
| AQX14497.1 | monobactam\_NRPS\_scaffold\_4 | BGC0001672 | NRP | 24.0 | 34.9 | 62.0 | 2.4e-09 |
| AAX31558.1 | peptide\_synthetase\_2 | BGC0000336 | NRP | 25.0 | 32.6 | 61.0 | 5.4e-09 |
| CAC48361.1 | peptide\_synthetase | BGC0000311 | NRP | 27.0 | 33.6 | 61.0 | 7.1e-09 |
| ACM79812.1 | ZmaQ | BGC0001059 | NRP + Polyketide | 26.0 | 32.6 | 61.0 | 7.1e-09 |
| AAF08795.1 | MycA | BGC0001103 | NRP + Polyketide | 25.0 | 50.7 | 61.0 | 7.1e-09 |
| QBC75023.1 | non-ribosomal\_peptide\_synthetase | BGC0001968 | NRP | 26.0 | 34.5 | 60.0 | 1.2e-08 |
| ACZ55944.1 | non-ribosomal\_peptide\_synthetase | BGC0000302 | NRP | 26.0 | 33.5 | 59.0 | 1.6e-08 |
| AFK57215.1 | DidD | BGC0000985 | Polyketide + NRP:Cyclic depsipeptide | 25.0 | 30.1 | 59.0 | 1.6e-08 |
| EXU96269.1 | nonribosomal\_peptide\_synthetase,\_serinocyclin\_synthetase\_NPS1 | BGC0001240 | NRP | 30.0 | 29.5 | 59.0 | 1.6e-08 |
| CAM56771.1 |  | BGC0000354 | NRP | 27.0 | 30.1 | 59.0 | 2.1e-08 |
| AAZ23078.1 | peptide\_synthetase | BGC0000291 | NRP | 27.0 | 31.5 | 59.0 | 2.7e-08 |
| ABC36785.1 | peptide\_synthetase,\_putative | BGC0000964 | NRP:Cyclic depsipeptide + Polyketide:Trans-AT type I | 23.0 | 48.4 | 58.0 | 4.6e-08 |
| BBB04327.1 | nonribosomal\_peptide\_synthetase | BGC0001717 | Alkaloid | 24.0 | 60.0 | 58.0 | 4.6e-08 |
| AVI26390.1 | polyketide\_synthase\_/\_nonribosomal\_peptide\_synthase\_hybrid | BGC0001800 | NRP + Polyketide | 25.0 | 33.5 | 57.0 | 1e-07 |
| BAH43766.1 | tyrocidine\_synthetase\_III | BGC0000452 | NRP | 25.0 | 45.0 | 56.0 | 1.3e-07 |
| AHB82072.1 | non\_ribosomal\_peptide\_synthetase/polyketide\_synthase | BGC0001231 | NRP + Polyketide:Modular type I | 23.0 | 35.9 | 56.0 | 1.7e-07 |
| ACA97576.1 | PmxA | BGC0000408 | NRP | 23.0 | 39.9 | 56.0 | 2.3e-07 |
| AEZ51516.1 | pmxA | BGC0001153 | NRP:Lipopeptide | 24.0 | 40.2 | 55.0 | 3e-07 |
| AAC06347.1 | bacitracin\_synthetase\_2 | BGC0000310 | NRP | 23.0 | 31.1 | 55.0 | 3.9e-07 |
| AAF17280.1 | nosC | BGC0001028 | Polyketide + NRP:Cyclic depsipeptide | 25.0 | 30.8 | 55.0 | 3.9e-07 |
| ABC34483.1 | nonribosomal\_peptide\_synthetase,\_putative | BGC0000961 | NRP + Polyketide | 29.0 | 34.0 | 54.0 | 5.1e-07 |
| AEC14347.1 | nonribosomal\_peptide\_synthetase | BGC0000377 | NRP | 25.0 | 32.0 | 54.0 | 8.7e-07 |
| WP\_013310342.1 | non-ribosomal\_peptide\_synthetase | BGC0001993 | NRP | 22.0 | 31.3 | 54.0 | 8.7e-07 |
| AAG29789.1 | acyl-CoA\_synthetase | BGC0000833 | Saccharide:Hybrid/tailoring + Other:Aminocoumarin | 34.0 | 23.7 | 53.0 | 1.5e-06 |
| ERM18795.1 | surfactin\_synthase\_subunit\_1 | BGC0000172 | Polyketide | 23.0 | 35.2 | 52.0 | 1.9e-06 |
| WP\_047197778.1 | non-ribosomal\_peptide\_synthetase | BGC0002003 | NRP | 25.0 | 32.6 | 52.0 | 1.9e-06 |
| AHI59108.1 | locillomycin\_synthase\_A | BGC0001005 | NRP + Polyketide | 22.0 | 51.8 | 52.0 | 2.5e-06 |
| ATP76243.1 | NdaA | BGC0001705 | NRP + Polyketide | 25.0 | 35.6 | 51.0 | 4.3e-06 |
