## Supplementary Results for "A draft genome of the ascomycotal fungal species *Pseudopithomyces maydicus* (family *Didymosphaeriaceae*)": input.path1.gene110_mibig_hits.html

| MIBiG Protein | Description | MIBiG Cluster | MiBiG Product | % ID | % Coverage | BLAST Score | E-value |
| --- | --- | --- | --- | --- | --- | --- | --- |
| BAE56606.1 |  | BGC0001123 | NRP | 31.0 | 84.0 | 136.0 | 9e-32 |
| EAU36744.1 | predicted\_protein | BGC0000292 | NRP | 32.0 | 92.4 | 129.0 | 8.4e-30 |
