## Supplementary Results for "A draft genome of the ascomycotal fungal species *Pseudopithomyces maydicus* (family *Didymosphaeriaceae*)": input.path1.gene111_mibig_hits.html

| MIBiG Protein | Description | MIBiG Cluster | MiBiG Product | % ID | % Coverage | BLAST Score | E-value |
| --- | --- | --- | --- | --- | --- | --- | --- |
| BAE56594.1 |  | BGC0001123 | NRP | 48.0 | 90.7 | 508.0 | 1.7e-143 |
| AEO57490.1 | general\_substrate\_transporter | BGC0001449 | NRP + Alkaloid + Polyketide:Iterative type I | 40.0 | 92.9 | 404.0 | 2e-112 |
| ACS68556.1 | major\_facilitator\_superfamily\_protein | BGC0001026 | NRP + Polyketide | 39.0 | 95.9 | 401.0 | 2.2e-111 |
| QBC19708.1 | TwmF | BGC0001954 | NRP + Polyketide | 37.0 | 91.0 | 376.0 | 5.9e-104 |
| QCS37513.1 | PyiT | BGC0001982 | NRP + Polyketide | 37.0 | 93.5 | 367.0 | 4.7e-101 |
| AMY15055.1 | MFS\_transporter | BGC0001339 | Polyketide:Iterative type I | 33.0 | 92.4 | 296.0 | 7.8e-80 |
| KIS69144.1 | Major\_Facilitator\_invovled\_in\_MEL\_transport | BGC0001888 | Other | 31.0 | 92.9 | 253.0 | 7.5e-67 |
| DAB41650.1 | MFS\_transporter | BGC0001583 | Polyketide | 28.0 | 93.5 | 200.0 | 9.9e-51 |
| AAF00219.1 | transporter | BGC0000277 | Polyketide | 30.0 | 81.8 | 192.0 | 2.1e-48 |
| CAF60521.1 | putative\_efflux\_protein | BGC0000704 | Saccharide | 31.0 | 70.4 | 190.0 | 1e-47 |
| CAF31575.1 | putative\_kanamycin\_efflux\_protein | BGC0000705 | Saccharide | 31.0 | 70.4 | 190.0 | 1e-47 |
| AAM94765.1 | CalT1 | BGC0000033 | Polyketide | 29.0 | 77.2 | 184.0 | 5.6e-46 |
| ABA02247.1 | efflux\_pump | BGC0000098 | Polyketide | 27.0 | 88.5 | 181.0 | 2.8e-45 |
| AAD34558.1 | unknown | BGC0000088 | Polyketide | 27.0 | 88.7 | 180.0 | 6.2e-45 |
| BAC20568.1 | efflux\_pump | BGC0000039 | Polyketide | 28.0 | 77.4 | 179.0 | 1.4e-44 |
| ATY69599.1 | antibiotic\_efflux\_protein | BGC0001823 | NRP + Polyketide | 28.0 | 83.6 | 179.0 | 1.4e-44 |
| BAE71313.1 | putative\_ABC\_transporter | BGC0000004 | Polyketide | 30.0 | 73.2 | 177.0 | 5.3e-44 |
| ADY16699.1 | TqaJ | BGC0001142 | NRP | 29.0 | 79.9 | 175.0 | 2.6e-43 |
| AAS90069.1 | AflT | BGC0000010 | Polyketide | 28.0 | 86.2 | 175.0 | 3.4e-43 |
| PKX92296.1 | MFS\_general\_substrate\_transporter | BGC0001988 | Polyketide | 28.0 | 76.4 | 173.0 | 7.6e-43 |
| BBG28481.1 | putative\_MFS\_toxin\_efflux\_pump\_CdmB | BGC0001926 | Polyketide | 27.0 | 86.9 | 171.0 | 4.9e-42 |
| AGC95323.1 | CurE | BGC0000045 | Polyketide | 26.0 | 94.2 | 169.0 | 1.9e-41 |
| BAV69308.1 | PrhG | BGC0001729 | Polyketide + Terpene | 27.0 | 87.7 | 168.0 | 3.2e-41 |
| AHW57792.1 | PgaJ3 | BGC0000262 | Polyketide:Type II + Saccharide:Hybrid/tailoring | 27.0 | 89.8 | 168.0 | 4.2e-41 |
| BAV16993.1 | putative\_transporter | BGC0001384 | Polyketide | 26.0 | 81.7 | 165.0 | 3.5e-40 |
| ATU31811.1 | MFS\_transporter | BGC0001814 | NRP | 25.0 | 91.4 | 165.0 | 3.5e-40 |
| BAZ95831.1 | MFS\_transporter\_cpaI | BGC0001563 | NRP + Polyketide | 27.0 | 93.3 | 164.0 | 4.6e-40 |
| CCE31569.1 | probable\_aflatoxin\_efflux\_pump\_AFLT | BGC0001886 | Polyketide | 26.0 | 87.7 | 164.0 | 4.6e-40 |
| RWQ92172.1 | putative\_MFS\_transporter | BGC0002030 | Polyketide | 27.0 | 75.5 | 162.0 | 2.3e-39 |
| EAU38977.1 | predicted\_protein | BGC0001122 | NRP + Polyketide:Iterative type I | 26.0 | 98.4 | 161.0 | 3.9e-39 |
| AFU65888.1 | DacR2 | BGC0000216 | Polyketide | 25.0 | 88.7 | 160.0 | 1.1e-38 |
| ACP19369.1 | SaqJ1 | BGC0000267 | Polyketide:Type II + Saccharide:Oligosaccharide | 28.0 | 77.6 | 151.0 | 3.1e-36 |
| AHA12095.1 | transporter | BGC0001172 | NRP + Polyketide:Modular type I | 29.0 | 68.4 | 151.0 | 4e-36 |
| ADM34144.1 | efflux\_pump | BGC0001084 | NRP + Terpene + Alkaloid | 26.0 | 91.4 | 150.0 | 9e-36 |
| ARO44650.1 | transporter | BGC0001769 | Polyketide | 26.0 | 83.2 | 149.0 | 2e-35 |
| AGO50605.1 | transporter | BGC0000229 | Polyketide:Type II + Saccharide:Hybrid/tailoring | 26.0 | 76.2 | 142.0 | 2.4e-33 |
| ABC87523.1 | putative\_drug\_efflux\_transporter | BGC0001011 | NRP + Polyketide | 29.0 | 73.5 | 138.0 | 4.6e-32 |
| OWA25250.1 | MFS\_transporter | BGC0001438 | Polyketide + Saccharide:Hybrid/tailoring | 30.0 | 63.0 | 134.0 | 5.1e-31 |
| ACN38355.1 | putative\_transmembrane\_efflux\_protein | BGC0000714 | Saccharide | 26.0 | 71.8 | 131.0 | 4.3e-30 |
| KDQ70104.1 | multidrug\_MFS\_transporter | BGC0001538 | NRP + Polyketide | 24.0 | 83.8 | 129.0 | 1.6e-29 |
| CAF34033.1 | putative\_transmembrane\_efflux\_protein | BGC0000689 | Saccharide | 26.0 | 76.7 | 128.0 | 4.8e-29 |
| CAF31444.1 | putative\_gentamicin\_exporter | BGC0000696 | Saccharide | 26.0 | 76.7 | 128.0 | 4.8e-29 |
| ARV75721.1 | hypothetical\_protein | BGC0001603 | Saccharide | 26.0 | 76.7 | 128.0 | 4.8e-29 |
| ADI24949.1 | GsfJ | BGC0000070 | Polyketide:Iterative type I | 25.0 | 88.7 | 126.0 | 1.4e-28 |
| CBF76046.1 | conserved\_hypothetical\_protein | BGC0001399 | NRP | 25.0 | 65.4 | 126.0 | 1.4e-28 |
| EAU36749.1 | predicted\_protein | BGC0000292 | NRP | 25.0 | 82.4 | 126.0 | 1.8e-28 |
| AIG62136.1 | MFS\_transporter | BGC0000120 | Polyketide:Iterative type I | 25.0 | 81.8 | 122.0 | 2.6e-27 |
| BAQ25491.1 | multidrug\_MFS\_(major\_facilitator\_superfamily)\_transporter | BGC0001288 | Polyketide | 26.0 | 69.5 | 122.0 | 2.6e-27 |
| AAZ77686.1 | ChlG | BGC0000036 | Polyketide:Modular type I + Polyketide:Iterative type I + Saccharide:Oligosaccharide | 26.0 | 70.7 | 119.0 | 1.7e-26 |
| BAG85020.1 | putative\_resistance\_protein | BGC0000086 | Polyketide | 24.0 | 80.4 | 115.0 | 2.5e-25 |
| CAQ64681.1 | putative\_resistance\_protein,\_transporter | BGC0000087 | Polyketide | 24.0 | 80.4 | 115.0 | 2.5e-25 |
| AQW35077.1 | MFS\_transporter | BGC0001675 | Polyketide | 24.0 | 85.9 | 110.0 | 1e-23 |
| ANR02546.1 | LodE | BGC0001648 | Polyketide | 25.0 | 75.7 | 108.0 | 3e-23 |
| CAC44196.1 | putative\_actinorhodin\_transporter | BGC0000194 | Polyketide:Type II | 25.0 | 77.4 | 102.0 | 2.1e-21 |
| BAC10683.1 | putative\_translocase\_protein | BGC0000821 | Other:Aminocoumarin | 25.0 | 71.1 | 101.0 | 6.2e-21 |
| CAC93723.1 | putative\_integral\_membrane\_transporter | BGC0000822 | Alkaloid | 25.0 | 71.1 | 101.0 | 6.2e-21 |
| EHM27501.1 | putative\_export\_protein | BGC0000235 | Polyketide | 25.0 | 73.5 | 97.0 | 6.9e-20 |
| AGN71625.1 | putative\_HC-toxin\_efflux\_carrier\_TOXA | BGC0000027 | Polyketide:Iterative type I | 25.0 | 67.5 | 92.0 | 1.7e-18 |
| PPQ57485.1 | MFS\_transporter | BGC0002016 | Polyketide | 25.0 | 72.3 | 89.0 | 1.9e-17 |
| ADZ13560.1 | YtkR6 | BGC0000466 | NRP | 24.0 | 82.7 | 87.0 | 7.1e-17 |
| RLV71200.1 | Export\_protein | BGC0001846 | NRP + Saccharide:Hybrid/tailoring | 25.0 | 73.2 | 87.0 | 7.1e-17 |
| ADC79648.1 | TamJ | BGC0001052 | NRP + Polyketide:Modular type I | 23.0 | 72.7 | 84.0 | 4.6e-16 |
| AGN74899.1 | major\_facilitator\_transporter | BGC0000459 | NRP:Cyclic depsipeptide + Polyketide:Trans-AT type I | 25.0 | 67.9 | 76.0 | 1.3e-13 |
| ABP55162.1 | major\_facilitator\_superfamily\_MFS\_1 | BGC0000150 | NRP + Polyketide:Enediyne type I | 23.0 | 75.0 | 75.0 | 3.7e-13 |
| ATE50865.1 | transporter\_CxnT | BGC0001485 | Other | 24.0 | 71.8 | 62.0 | 2.5e-09 |
| AVL27077.1 | MFS\_transporter | BGC0001546 | Alkaloid | 24.0 | 71.8 | 62.0 | 2.5e-09 |
| CCP20037.1 | divA\_protein | BGC0001119 | Polyketide:Modular type I | 27.0 | 32.6 | 52.0 | 3.3e-06 |
