## Supplementary Results for "A draft genome of the ascomycotal fungal species *Pseudopithomyces maydicus* (family *Didymosphaeriaceae*)": input.path1.gene113_mibig_hits.html

| MIBiG Protein | Description | MIBiG Cluster | MiBiG Product | % ID | % Coverage | BLAST Score | E-value |
| --- | --- | --- | --- | --- | --- | --- | --- |
| ctg1\_orf0002 |  | BGC0000321 | NRP | 63.0 | 70.2 | 436.0 | 4.1e-122 |
| AQZ42159.1 | putative\_O-methyltransferase | BGC0001820 | NRP | 61.0 | 70.2 | 424.0 | 2.7e-118 |
| DAB41656.1 | methyltransferase | BGC0001585 | Alkaloid | 49.0 | 93.3 | 374.0 | 3.2e-103 |
| BAE56602.1 |  | BGC0001123 | NRP | 48.0 | 71.0 | 342.0 | 8e-94 |
| EHK21998.1 | hypothetical\_protein | BGC0001609 | NRP | 48.0 | 70.6 | 334.0 | 3.7e-91 |
| EAL88819.2 | O-methyltransferase\_GliM | BGC0000361 | NRP | 46.0 | 70.0 | 316.0 | 8e-86 |
| AAS92548.1 | SirM | BGC0001044 | NRP + Polyketide | 40.0 | 70.6 | 277.0 | 3.2e-74 |
| BAE56609.1 |  | BGC0001123 | NRP | 42.0 | 70.6 | 276.0 | 7e-74 |
| EAU36746.1 | conserved\_hypothetical\_protein | BGC0000292 | NRP | 34.0 | 89.4 | 221.0 | 2.7e-57 |
| EAL92292.1 | O-methyltransferase | BGC0000372 | NRP | 34.0 | 70.6 | 218.0 | 3e-56 |
| AUO15565.1 | methyltransferase | BGC0001513 | Polyketide | 28.0 | 67.9 | 100.0 | 9e-21 |
| CAM34360.1 | putative\_O-methyltransferase | BGC0000242 | Polyketide | 28.0 | 64.4 | 95.0 | 2.9e-19 |
| ctg1\_orf14 |  | BGC0001366 | Polyketide | 26.0 | 68.5 | 89.0 | 1.6e-17 |
| AUI41033.1 | O-methyltransferase | BGC0001512 | Polyketide | 26.0 | 71.9 | 86.0 | 1e-16 |
| EAL89337.1 | O-methyltransferase,\_putative | BGC0001403 | Polyketide | 23.0 | 81.9 | 84.0 | 3.9e-16 |
| CAX48665.1 | phenazine\_N-methyltransferase | BGC0001080 | Other:Phenazine | 26.0 | 70.4 | 84.0 | 6.7e-16 |
| AEI98654.1 | CtcK | BGC0000209 | Polyketide | 29.0 | 63.3 | 83.0 | 1.1e-15 |
| ACN64843.1 | PokMT2 | BGC0001061 | Polyketide:Iterative type I + Polyketide:Type II + Saccharide:Hybrid/tailoring | 28.0 | 71.9 | 78.0 | 3.7e-14 |
| AKD43506.1 | Methtyltransferase | BGC0001409 | Polyketide | 28.0 | 69.0 | 74.0 | 5.3e-13 |
| AQP25568.1 | O-methyltransferase | BGC0001590 | Polyketide | 28.0 | 62.9 | 74.0 | 5.3e-13 |
| CAM58795.1 | C-dimethyltransferse | BGC0000204 | Polyketide:Type II | 27.0 | 70.0 | 73.0 | 1.2e-12 |
| AAZ78330.1 | OxyF | BGC0000254 | Polyketide | 26.0 | 65.2 | 72.0 | 2e-12 |
| ATN39909.1 | MstO | BGC0001664 | Terpene | 25.0 | 34.8 | 59.0 | 1.8e-08 |
