## Supplementary Results for "A draft genome of the ascomycotal fungal species *Pseudopithomyces maydicus* (family *Didymosphaeriaceae*)": input.path1.gene137_mibig_hits.html

| MIBiG Protein | Description | MIBiG Cluster | MiBiG Product | % ID | % Coverage | BLAST Score | E-value |
| --- | --- | --- | --- | --- | --- | --- | --- |
| EAW09117.1 | hybrid\_NRPS/PKS\_enzyme,\_putative | BGC0000983 | NRP + Polyketide:Iterative type I | 34.0 | 96.5 | 666.0 | 1.1e-190 |
| AAV66110.2 | fusaridione\_A\_synthetase | BGC0000992 | NRP + Polyketide | 33.0 | 99.0 | 658.0 | 2.3e-188 |
| EPS29069.1 | hypothetical\_protein | BGC0001724 | NRP + Polyketide | 33.0 | 97.2 | 639.0 | 1.5e-182 |
| AGO86662.1 | equisetin\_synthetase | BGC0001255 | NRP + Polyketide | 32.0 | 96.5 | 638.0 | 3.3e-182 |
| CEF75886.1 |  | BGC0001600 | Polyketide | 33.0 | 100.5 | 632.0 | 1.4e-180 |
| QBC19710.1 | TwmB | BGC0001954 | NRP + Polyketide | 34.0 | 95.8 | 630.0 | 5.2e-180 |
| BBC43184.1 | PKS-NRPS\_hybrid | BGC0001738 | NRP + Polyketide | 33.0 | 96.6 | 629.0 | 1.2e-179 |
| QBE85649.1 | BuaA | BGC0001978 | NRP + Polyketide | 33.0 | 96.5 | 629.0 | 1.2e-179 |
| BAZ95823.1 | PKS-NRPS\_hybrid\_cpaA | BGC0001563 | NRP + Polyketide | 33.0 | 97.1 | 615.0 | 1.7e-175 |
| ARP51711.1 | PKS-NRPS\_hybrid\_protein | BGC0001741 | NRP + Polyketide | 33.0 | 99.5 | 609.0 | 1.2e-173 |
| gene3 |  | BGC0002035 | NRP + Polyketide | 32.0 | 99.1 | 599.0 | 9.9e-171 |
| ABA02239.1 | polyketide\_synthase | BGC0000098 | Polyketide | 33.0 | 98.6 | 599.0 | 1.7e-170 |
| AEO57481.1 | PKS-NRPSs | BGC0001449 | NRP + Alkaloid + Polyketide:Iterative type I | 33.0 | 98.0 | 593.0 | 7.1e-169 |
| BAJ14522.1 | polyketide\_synthase | BGC0001254 | Polyketide | 32.0 | 97.0 | 591.0 | 2.7e-168 |
| BAQ25466.1 | polyketide\_synthase | BGC0001280 | Polyketide | 31.0 | 98.5 | 590.0 | 7.8e-168 |
| ctg1\_orf0002 |  | BGC0001068 | Terpene + Polyketide | 32.0 | 96.5 | 588.0 | 2.3e-167 |
| QCS37521.1 | PyiS | BGC0001982 | NRP + Polyketide | 31.0 | 98.3 | 588.0 | 3e-167 |
| EED49862.1 | hybrid\_PKS/NRPS\_enzyme,\_putative | BGC0001445 | NRP + Polyketide:Iterative type I | 31.0 | 96.9 | 579.0 | 1.1e-164 |
| ADN43685.1 | DmbS | BGC0001136 | NRP + Polyketide:Iterative type I | 31.0 | 100.5 | 577.0 | 4e-164 |
| CBF80487.1 | hybrid\_PKS-NRPS\_(Eurofung) | BGC0000959 | NRP + Polyketide:Iterative type I | 32.0 | 97.5 | 576.0 | 8.9e-164 |
| CAO91861.1 | PKS-NRPS\_hybrid | BGC0000968 | NRP + Polyketide:Iterative type I | 31.0 | 100.5 | 573.0 | 9.9e-163 |
| EAL85113.2 | hybrid\_PKS-NRPS\_enzyme | BGC0001037 | NRP + Polyketide:Iterative type I | 31.0 | 98.8 | 573.0 | 9.9e-163 |
| CAL69597.1 | PKS-NRPS | BGC0001049 | NRP + Polyketide:Iterative type I | 31.0 | 100.7 | 571.0 | 4.9e-162 |
| EAU38971.1 | hypothetical\_protein | BGC0001122 | NRP + Polyketide:Iterative type I | 32.0 | 97.5 | 569.0 | 1.9e-161 |
| EAL89230.2 | LovB-like\_polyketide\_synthase,\_putative | BGC0000129 | Polyketide | 31.0 | 91.2 | 563.0 | 7.8e-160 |
| ACS68554.1 | hybrid\_PKS-NRPS\_protein | BGC0001026 | NRP + Polyketide | 31.0 | 96.1 | 561.0 | 3.9e-159 |
| AKC54422.1 | fumosorinone\_biosynthesis\_polyketide\_synthase | BGC0001218 | NRP + Polyketide | 30.0 | 100.6 | 550.0 | 5.2e-156 |
| XP\_001220460.1 | hypothetical\_protein | BGC0001182 | NRP + Polyketide:Iterative type I | 32.0 | 83.9 | 518.0 | 3.8e-146 |
| EHA19289.1 | hypothetical\_protein | BGC0001124 | Polyketide | 29.0 | 93.6 | 411.0 | 3.8e-114 |
| BAC20564.1 | polyketide\_synthase | BGC0000039 | Polyketide | 38.0 | 49.3 | 396.0 | 2.2e-109 |
| CCT72377.1 | probable\_polyketide\_synthase | BGC0001305 | Polyketide | 37.0 | 49.5 | 394.0 | 6.3e-109 |
| AKL78824.1 | GLPKS3 | BGC0001187 | NRP:Lipopeptide + Polyketide:Iterative type I | 28.0 | 91.3 | 386.0 | 2.2e-106 |
| ATZ45182.1 | Bcboa6 | BGC0001892 | Polyketide | 37.0 | 48.2 | 385.0 | 3.8e-106 |
| EAA36364.1 | hypothetical\_protein | BGC0001697 | Polyketide | 27.0 | 100.0 | 381.0 | 7.2e-105 |
| XP\_659388.1 | hypothetical\_protein | BGC0001998 | Polyketide | 37.0 | 50.9 | 380.0 | 9.4e-105 |
| KKP00963.1 | fatty\_acid\_synthase\_S-acetyltransferase | BGC0001901 | Polyketide | 28.0 | 93.8 | 369.0 | 1.7e-101 |
| BAK26562.1 | PKS-NRPS\_hybrid | BGC0000977 | NRP + Polyketide | 34.0 | 56.3 | 367.0 | 1.1e-100 |
| CBX99534.1 | similar\_to\_polyketide\_synthase | BGC0001899 | Polyketide | 27.0 | 94.2 | 366.0 | 2.4e-100 |
| AFA26384.1 | polyketide\_synthase\_A | BGC0001874 | NRP + Polyketide | 34.0 | 56.3 | 364.0 | 7e-100 |
| ANF07288.1 | hrPKS | BGC0001340 | Polyketide:Iterative type I | 25.0 | 97.8 | 361.0 | 7.7e-99 |
| KGO40478.1 | Acyl\_transferase/acyl\_hydrolase/lysophospholipase | BGC0001205 | Polyketide | 28.0 | 80.8 | 359.0 | 1.7e-98 |
| AAT28740.1 | FUSS | BGC0000064 | Polyketide | 36.0 | 49.2 | 358.0 | 5e-98 |
| AFP73394.1 | FusA | BGC0001268 | NRP + Polyketide | 37.0 | 48.3 | 358.0 | 5e-98 |
| ACB12550.1 | Fum1 | BGC0000063 | Polyketide | 26.0 | 100.1 | 355.0 | 4.2e-97 |
| EHA52508.1 | mycocerosic\_acid\_synthase | BGC0001749 | Polyketide | 27.0 | 94.6 | 354.0 | 7.2e-97 |
| AHV78252.1 | ResS1 | BGC0001246 | Polyketide | 28.0 | 93.4 | 354.0 | 9.4e-97 |
| AAD43562.2 | Fum1p | BGC0000062 | Polyketide | 26.0 | 100.2 | 352.0 | 2.7e-96 |
| AGC95324.1 | CurS1 | BGC0000045 | Polyketide | 26.0 | 99.2 | 348.0 | 4e-95 |
| ACD39774.1 | reducing\_polyketide\_synthase | BGC0000134 | Polyketide | 26.0 | 96.1 | 347.0 | 1.2e-94 |
| AQW44889.1 | polyketide\_synthase | BGC0001737 | NRP + Polyketide | 27.0 | 93.9 | 342.0 | 2.8e-93 |
| AKG06377.1 | polyketide\_synthase\_type\_1 | BGC0001830 | Polyketide | 27.0 | 89.0 | 333.0 | 1.7e-90 |
| EWG54266.1 | hypothetical\_protein | BGC0001190 | Polyketide | 26.0 | 92.9 | 322.0 | 3e-87 |
| AGC45624.1 | polyketide\_synthase | BGC0001394 | NRP + Polyketide | 27.0 | 90.6 | 319.0 | 2e-86 |
| EFL02193.1 | amino\_acid\_adenylation\_domain-containing\_protein | BGC0000996 | NRP + Polyketide:Iterative type I | 29.0 | 75.4 | 319.0 | 2.6e-86 |
| AJD77023.1 | IkaA | BGC0001435 | NRP + Polyketide:Iterative type I | 29.0 | 75.4 | 317.0 | 7.5e-86 |
| BAJ09789.1 | polyketide\_synthase | BGC0000146 | Polyketide | 26.0 | 96.6 | 317.0 | 9.8e-86 |
| ABL86391.1 | hybrid\_polyketide\_synthase\_and\_nonribosomal\_peptide\_synthetase | BGC0000999 | NRP + Polyketide | 27.0 | 88.8 | 316.0 | 1.7e-85 |
| APD26279.1 | PtmA | BGC0001726 | NRP + Polyketide | 27.0 | 89.9 | 316.0 | 1.7e-85 |
| QDA77058.1 | polyketide\_synthase | BGC0002026 | NRP | 28.0 | 89.6 | 316.0 | 1.7e-85 |
| ACB46195.1 | polyketide\_synthase | BGC0000989 | NRP + Polyketide | 26.0 | 94.1 | 313.0 | 1.4e-84 |
| ADB12491.1 | EpoD | BGC0000990 | NRP + Polyketide | 26.0 | 100.3 | 313.0 | 1.4e-84 |
| ABB90283.1 | polyketide\_synthase | BGC0001057 | NRP + Polyketide | 25.0 | 91.0 | 313.0 | 1.8e-84 |
| AAF62883.1 | epoD | BGC0000991 | NRP + Polyketide | 26.0 | 100.3 | 311.0 | 5.4e-84 |
| AAF26921.1 | polyketide\_synthase | BGC0000988 | NRP + Polyketide | 26.0 | 100.4 | 311.0 | 7e-84 |
| DAB41915.1 | ArzM\_-\_PKS\_(KS,\_AT,\_DH,\_MT,\_ER,\_KR,\_ACP) | BGC0001884 | NRP + Polyketide | 27.0 | 90.1 | 310.0 | 1.2e-83 |
| CCE88376.1 | polyketide\_synthase | BGC0001034 | NRP + Polyketide:Modular type I | 27.0 | 90.4 | 310.0 | 1.6e-83 |
| CAJ46690.1 | polyketide\_synthase | BGC0000969 | NRP:Cyclic depsipeptide + Polyketide:Modular type I | 27.0 | 87.6 | 309.0 | 3.5e-83 |
| EAU29808.1 | hypothetical\_protein | BGC0001400 | Polyketide | 35.0 | 44.2 | 309.0 | 3.5e-83 |
| ADZ24998.1 | polyketide\_synthase | BGC0000380 | NRP + Polyketide:Modular type I | 27.0 | 88.5 | 305.0 | 3.8e-82 |
| AAK57187.1 | MxaC | BGC0001022 | NRP + Polyketide | 26.0 | 90.2 | 304.0 | 6.6e-82 |
| ADH04641.1 | TgaC | BGC0001051 | NRP + Polyketide:Modular type I | 26.0 | 90.3 | 300.0 | 1.2e-80 |
| WP\_042799407.1 | type\_I\_polyketide\_synthase | BGC0001283 | Polyketide | 26.0 | 90.5 | 296.0 | 1.8e-79 |
| AUO16397.1 | polyketide\_synthase | BGC0001700 | Polyketide | 25.0 | 99.4 | 294.0 | 1.2e-78 |
| ADZ24996.1 | polyketide\_synthase | BGC0000380 | NRP + Polyketide:Modular type I | 27.0 | 89.6 | 292.0 | 3.4e-78 |
| CAQ18830.1 | polyketide\_synthase | BGC0000954 | NRP + Polyketide:Modular type I | 27.0 | 88.8 | 292.0 | 3.4e-78 |
| APZ78780.1 | polyketide\_synthase | BGC0001426 | NRP:Cyclic depsipeptide + Polyketide:Iterative type I | 27.0 | 89.0 | 292.0 | 3.4e-78 |
| ARW71485.1 | type\_I\_PKS\_module\_4,\_module\_5 | BGC0001812 | Polyketide | 26.0 | 88.6 | 290.0 | 9.8e-78 |
| APZ78793.1 | polyketide\_synthase | BGC0001427 | NRP:Cyclic depsipeptide + Polyketide:Iterative type I | 27.0 | 89.2 | 289.0 | 3.7e-77 |
| APZ78807.1 | polyketide\_synthase | BGC0001428 | NRP:Cyclic depsipeptide + Polyketide:Iterative type I | 27.0 | 89.3 | 288.0 | 4.9e-77 |
| ctg1\_orf29 |  | BGC0000096 | Polyketide | 26.0 | 91.3 | 288.0 | 6.3e-77 |
| AWO77084.1 | hybrid\_non-ribosomal\_peptide\_synthetase/type\_I\_polyketide\_synthase | BGC0001556 | NRP + Polyketide | 27.0 | 82.8 | 288.0 | 6.3e-77 |
| CCE88378.1 | polyketide\_synthase | BGC0001034 | NRP + Polyketide:Modular type I | 25.0 | 97.2 | 287.0 | 8.3e-77 |
| CAO98847.1 | polyketide\_synthase\_AufC | BGC0000023 | Polyketide:Modular type I | 27.0 | 92.0 | 287.0 | 1.4e-76 |
| SCO70308.1 | Type\_I\_polyketide\_synthase | BGC0001433 | Polyketide:Modular type I | 26.0 | 90.4 | 287.0 | 1.4e-76 |
| APZ78742.1 | polyketide\_synthase | BGC0001422 | NRP:Cyclic depsipeptide + Polyketide:Iterative type I | 27.0 | 87.9 | 286.0 | 1.8e-76 |
| APZ78702.1 | polyketide\_synthase | BGC0001419 | NRP:Cyclic depsipeptide + Polyketide:Iterative type I | 27.0 | 89.4 | 286.0 | 2.4e-76 |
| OAP25821.1 | Phenolphthiocerol\_synthesis\_polyketide\_synthase\_type\_I\_Pks15/1 | BGC0001658 | Polyketide | 26.0 | 87.1 | 285.0 | 5.4e-76 |
| AQM37582.1 | polyketide\_synthase | BGC0001424 | NRP:Cyclic depsipeptide + Polyketide:Iterative type I | 27.0 | 88.8 | 284.0 | 7e-76 |
| AXN93613.1 | PuwE | BGC0001953 | NRP | 35.0 | 45.8 | 284.0 | 7e-76 |
| AAB66506.1 | tylactone\_synthase\_modules\_4\_&\_5 | BGC0000166 | Polyketide | 26.0 | 89.4 | 284.0 | 9.2e-76 |
| AXN93601.1 | PuwE | BGC0001952 | NRP | 34.0 | 48.3 | 283.0 | 2e-75 |
| APZ78714.1 | polyketide\_synthase | BGC0001420 | NRP:Cyclic depsipeptide + Polyketide:Iterative type I | 27.0 | 89.4 | 282.0 | 3.5e-75 |
| APZ78690.1 | polyketide\_synthase | BGC0001418 | NRP:Cyclic depsipeptide + Polyketide:Iterative type I | 27.0 | 89.4 | 282.0 | 4.5e-75 |
| AAP42873.1 | NanA11 | BGC0000105 | Polyketide | 26.0 | 92.9 | 281.0 | 5.9e-75 |
| ADH04660.1 | TugD | BGC0001342 | NRP + Polyketide | 26.0 | 89.3 | 280.0 | 1e-74 |
| ANY10588.1 | polyketide\_synthase | BGC0001773 | Polyketide | 27.0 | 86.5 | 280.0 | 1.3e-74 |
| CAG28678.1 | polyketide\_synthase | BGC0001023 | NRP + Polyketide:Modular type I | 27.0 | 89.3 | 280.0 | 1.7e-74 |
| APZ78820.1 | polyketide\_synthase | BGC0001429 | NRP:Cyclic depsipeptide + Polyketide:Iterative type I | 27.0 | 89.3 | 280.0 | 1.7e-74 |
| AAZ77697.1 | ChlA4 | BGC0000036 | Polyketide:Modular type I + Polyketide:Iterative type I + Saccharide:Oligosaccharide | 26.0 | 88.8 | 279.0 | 2.3e-74 |
| APZ78844.1 | polyketide\_synthase | BGC0001431 | NRP:Cyclic depsipeptide + Polyketide:Iterative type I | 27.0 | 90.2 | 279.0 | 2.3e-74 |
| CCE88381.1 | polyketide\_synthase | BGC0001034 | NRP + Polyketide:Modular type I | 24.0 | 96.9 | 278.0 | 5e-74 |
| AAD34559.1 | polyketide\_synthase | BGC0000088 | Polyketide | 32.0 | 51.5 | 278.0 | 6.6e-74 |
| BBA66512.1 | type\_I\_polyketide\_synthase | BGC0001495 | Polyketide | 26.0 | 87.3 | 277.0 | 1.1e-73 |
| AIW82282.1 | PuwE | BGC0001125 | NRP + Polyketide | 32.0 | 54.9 | 276.0 | 1.9e-73 |
| ACO94498.1 | polyketide\_synthase\_type\_I | BGC0000097 | Polyketide:Modular type I | 26.0 | 86.1 | 276.0 | 2.5e-73 |
| BAN19720.1 | polyketide\_synthase | BGC0001252 | Polyketide | 25.0 | 81.1 | 275.0 | 3.3e-73 |
| AHH99919.1 | PKS\_I | BGC0000002 | Polyketide | 26.0 | 88.2 | 275.0 | 4.3e-73 |
| AAF19810.1 | MtaB | BGC0001024 | NRP + Polyketide:Modular type I | 27.0 | 77.7 | 274.0 | 7.3e-73 |
| AQM58285.1 | polyketide\_synthase | BGC0001816 | NRP + Polyketide | 24.0 | 92.7 | 274.0 | 1.2e-72 |
| CAP95405.1 |  | BGC0001404 | Polyketide | 31.0 | 55.4 | 273.0 | 1.6e-72 |
| AIT55259.1 | polyketide\_synthase | BGC0000072 | Polyketide:Modular type I | 28.0 | 70.1 | 273.0 | 2.1e-72 |
| CAQ34919.1 | polyketide\_synthase | BGC0000986 | NRP + Polyketide | 34.0 | 47.5 | 273.0 | 2.1e-72 |
| APZ78678.1 | polyketide\_synthase | BGC0001417 | NRP:Cyclic depsipeptide + Polyketide:Iterative type I | 26.0 | 88.9 | 273.0 | 2.1e-72 |
| EHK80165.1 | beta-ketoacyl\_synthase | BGC0001447 | Polyketide | 25.0 | 86.6 | 273.0 | 2.1e-72 |
| ABB88521.1 | polyketide\_synthase\_type\_I | BGC0000050 | Polyketide | 25.0 | 94.0 | 272.0 | 2.8e-72 |
| AEE88280.1 | CurJ | BGC0000976 | NRP + Polyketide:Modular type I | 32.0 | 48.8 | 272.0 | 3.6e-72 |
| ABB05104.1 | LipPks3 | BGC0001003 | NRP:Lipopeptide + Polyketide:Modular type I + Saccharide:Hybrid/tailoring | 25.0 | 88.9 | 272.0 | 3.6e-72 |
| AAT70105.1 | CurJ | BGC0001165 | NRP + Polyketide:Modular type I | 32.0 | 48.8 | 272.0 | 3.6e-72 |
| AUD08663.1 | iPKS-NRPS | BGC0001553 | NRP + Polyketide | 25.0 | 94.3 | 272.0 | 3.6e-72 |
| APZ78767.1 | polyketide\_synthase | BGC0001425 | NRP:Cyclic depsipeptide + Polyketide:Iterative type I | 26.0 | 88.9 | 271.0 | 6.1e-72 |
| AAZ77673.1 | ChlB1 | BGC0000036 | Polyketide:Modular type I + Polyketide:Iterative type I + Saccharide:Oligosaccharide | 27.0 | 79.4 | 271.0 | 8e-72 |
| ABA02240.1 | polyketide\_synthase | BGC0000098 | Polyketide | 31.0 | 50.9 | 270.0 | 1e-71 |
| AAC46025.1 | polyketide\_synthase\_module\_3 | BGC0000113 | Polyketide | 26.0 | 88.2 | 270.0 | 1e-71 |
| AEE88281.1 | CurI | BGC0000976 | NRP + Polyketide:Modular type I | 28.0 | 70.6 | 270.0 | 1e-71 |
| ABB05105.1 | LipPks4 | BGC0001003 | NRP:Lipopeptide + Polyketide:Modular type I + Saccharide:Hybrid/tailoring | 25.0 | 90.3 | 270.0 | 1e-71 |
| AAT70104.1 | CurI | BGC0001165 | NRP + Polyketide:Modular type I | 28.0 | 70.6 | 270.0 | 1e-71 |
| WP\_055480219.1 | type\_I\_polyketide\_synthase | BGC0001653 | Polyketide | 25.0 | 93.7 | 270.0 | 1e-71 |
| CAJ88186.1 | putative\_modular\_polyketide\_synthase | BGC0000151 | Polyketide:Modular type I + Saccharide:Hybrid/tailoring | 25.0 | 88.0 | 269.0 | 2.3e-71 |
| AGC45621.1 | polyketide\_synthase | BGC0001394 | NRP + Polyketide | 32.0 | 47.9 | 269.0 | 2.3e-71 |
| CAD19089.1 | StiE\_protein | BGC0000153 | NRP + Polyketide:Modular type I | 31.0 | 48.0 | 269.0 | 3.1e-71 |
| AAK57189.1 | MxaE | BGC0001022 | NRP + Polyketide | 32.0 | 47.9 | 269.0 | 3.1e-71 |
| AXI91549.1 | FunP4 | BGC0001944 | Polyketide | 26.0 | 86.2 | 269.0 | 3.1e-71 |
| AWH12664.1 | RmpE2 | BGC0001759 | Polyketide | 25.0 | 95.6 | 269.0 | 4e-71 |
| QDA77059.1 | polyketide\_synthase/nonribosomal\_peptide\_synthetase | BGC0002026 | NRP | 28.0 | 70.3 | 269.0 | 4e-71 |
| BAC20566.1 | polyketide\_synthase | BGC0000039 | Polyketide | 32.0 | 49.2 | 268.0 | 5.2e-71 |
| ABK32257.1 | AmbC | BGC0000014 | Polyketide | 26.0 | 88.8 | 267.0 | 8.9e-71 |
| AIT55263.1 | polyketide\_synthase | BGC0000072 | Polyketide:Modular type I | 32.0 | 48.7 | 267.0 | 8.9e-71 |
| EHA28244.1 | hypothetical\_protein | BGC0001143 | Polyketide | 31.0 | 48.5 | 267.0 | 8.9e-71 |
| ABW96540.1 | type\_I\_modular\_polyketide\_synthase | BGC0000159 | Polyketide:Modular type I | 25.0 | 88.7 | 267.0 | 1.2e-70 |
| AAF86393.1 | FkbB | BGC0000994 | NRP + Polyketide | 24.0 | 95.0 | 267.0 | 1.2e-70 |
| AAF00958.1 | mcyE | BGC0001017 | NRP + Polyketide:Modular type I | 28.0 | 66.3 | 267.0 | 1.2e-70 |
| ADX66459.1 | ScnS4 | BGC0000108 | Polyketide | 25.0 | 92.2 | 267.0 | 1.5e-70 |
| ANI24099.1 | polyketide\_synthase | BGC0001235 | NRP + Polyketide | 26.0 | 89.0 | 267.0 | 1.5e-70 |
| ASK38717.1 | polyketide\_synthase | BGC0001557 | Polyketide | 30.0 | 50.2 | 267.0 | 1.5e-70 |
| ATY12793.1 | type\_I\_polyketide\_synthase | BGC0001504 | Polyketide | 26.0 | 85.8 | 267.0 | 1.5e-70 |
| AXI91545.1 | FunP8 | BGC0001944 | Polyketide | 27.0 | 87.6 | 267.0 | 1.5e-70 |
| AXN93597.1 | PuwB | BGC0001952 | NRP | 32.0 | 48.3 | 267.0 | 1.5e-70 |
| ATP76239.1 | NdaF | BGC0001705 | NRP + Polyketide | 27.0 | 65.9 | 266.0 | 2e-70 |
| CAC20920.1 | PimS3\_protein | BGC0000125 | Polyketide | 25.0 | 97.9 | 266.0 | 2.6e-70 |
| AQT01394.1 | SgnS3 | BGC0001690 | Polyketide | 25.0 | 97.9 | 266.0 | 2.6e-70 |
| AQW44891.1 | polyketide\_synthase | BGC0001737 | NRP + Polyketide | 32.0 | 48.2 | 266.0 | 2.6e-70 |
| AGY62758.1 | EbeF | BGC0000051 | Polyketide | 25.0 | 88.7 | 265.0 | 3.4e-70 |
| CBF87072.1 | polyketide\_synthase,\_putative\_(Eurofung) | BGC0001290 | NRP | 33.0 | 49.2 | 264.0 | 9.8e-70 |
| EAQ86385.1 | hypothetical\_protein | BGC0001405 | Polyketide | 33.0 | 48.8 | 264.0 | 9.8e-70 |
| BBA66513.1 | type\_I\_polyketide\_synthase | BGC0001495 | Polyketide | 26.0 | 86.4 | 264.0 | 9.8e-70 |
| AEU11006.1 | NpnB | BGC0001029 | NRP + Polyketide | 32.0 | 47.7 | 264.0 | 1.3e-69 |
| WP\_055469548.1 | type\_I\_polyketide\_synthase | BGC0001537 | Polyketide | 25.0 | 92.2 | 263.0 | 2.2e-69 |
| BAP34763.1 | type\_I\_polyketide\_synthase | BGC0000078 | Polyketide | 25.0 | 89.7 | 262.0 | 2.9e-69 |
| CAL58682.1 | polyketide\_synthase | BGC0000149 | Polyketide:Modular type I | 26.0 | 90.0 | 262.0 | 2.9e-69 |
| CAQ43075.1 | polyketide\_synthase | BGC0000970 | NRP + Polyketide:Modular type I | 26.0 | 90.0 | 262.0 | 2.9e-69 |
| AJO72736.1 | Type\_I\_modular\_polyketide\_synthase | BGC0001381 | Polyketide | 26.0 | 88.0 | 262.0 | 2.9e-69 |
| AWH12669.1 | RmpB | BGC0001759 | Polyketide | 24.0 | 89.2 | 262.0 | 2.9e-69 |
| ACD39758.1 | reducing\_polyketide\_synthase | BGC0000076 | Polyketide | 31.0 | 49.4 | 262.0 | 3.7e-69 |
| ACD39767.1 | reducing\_polyketide\_synthase | BGC0000077 | Polyketide | 31.0 | 49.4 | 262.0 | 3.7e-69 |
| AAK19883.1 | soraphen\_polyketide\_synthase\_A | BGC0000147 | Polyketide:Modular type I | 25.0 | 88.7 | 262.0 | 3.7e-69 |
| AAS98783.1 | polyketide\_synthase/nonribosomal\_peptide\_synthase\_hybrid | BGC0001001 | NRP + Polyketide | 31.0 | 48.7 | 262.0 | 3.7e-69 |
| CAA60460.1 | polyketide\_synthase | BGC0001040 | NRP + Polyketide | 25.0 | 85.0 | 262.0 | 3.7e-69 |
| ACV42478.1 | polyketide\_synthase | BGC0000043 | Polyketide | 33.0 | 47.7 | 262.0 | 4.9e-69 |
| AEE88277.1 | CurM | BGC0000976 | NRP + Polyketide:Modular type I | 33.0 | 47.7 | 262.0 | 4.9e-69 |
| AAT70108.1 | CurM | BGC0001165 | NRP + Polyketide:Modular type I | 33.0 | 47.7 | 262.0 | 4.9e-69 |
| AIW82279.1 | PuwB | BGC0001125 | NRP + Polyketide | 34.0 | 48.8 | 261.0 | 6.4e-69 |
| AZH23818.1 | MgiI | BGC0001971 | NRP + Polyketide | 31.0 | 48.5 | 261.0 | 6.4e-69 |
| AAS98782.1 | polyketide\_synthase | BGC0001001 | NRP + Polyketide | 28.0 | 64.9 | 261.0 | 8.3e-69 |
| AXN93610.1 | PuwB | BGC0001953 | NRP | 32.0 | 49.5 | 261.0 | 8.3e-69 |
| CAC20919.1 | PimS4\_protein | BGC0000125 | Polyketide | 25.0 | 96.9 | 260.0 | 1.1e-68 |
| AQT01395.1 | SgnS4 | BGC0001690 | Polyketide | 25.0 | 96.9 | 260.0 | 1.1e-68 |
| AZH23791.1 | MgcH | BGC0001970 | NRP + Polyketide | 28.0 | 66.3 | 260.0 | 1.9e-68 |
| AAF86392.1 | FkbC | BGC0000994 | NRP + Polyketide | 26.0 | 86.0 | 259.0 | 2.4e-68 |
| AAO62582.1 | polyketide\_synthase\_peptide\_sythetase\_fusion\_protein | BGC0001016 | NRP + Polyketide | 27.0 | 66.0 | 259.0 | 2.4e-68 |
| AHE80994.1 | PieA4 | BGC0001169 | Polyketide:Modular type I | 25.0 | 88.5 | 259.0 | 2.4e-68 |
| WP\_026247674.1 | type\_I\_polyketide\_synthase | BGC0001332 | NRP + Polyketide | 32.0 | 48.0 | 259.0 | 3.2e-68 |
| AQH32481.1 | hybrid\_polyketide\_synthase/peptide\_synthetase | BGC0001667 | NRP + Polyketide | 31.0 | 47.2 | 258.0 | 5.4e-68 |
| AQW44888.1 | polyketide\_synthase | BGC0001737 | NRP + Polyketide | 32.0 | 48.3 | 258.0 | 5.4e-68 |
| ABK32289.1 | JerC | BGC0000080 | Polyketide | 26.0 | 88.1 | 258.0 | 7e-68 |
| AZH23789.1 | MgcI | BGC0001970 | NRP + Polyketide | 31.0 | 48.5 | 258.0 | 7e-68 |
| ABK32290.1 | JerD | BGC0000080 | Polyketide | 26.0 | 89.0 | 257.0 | 1.2e-67 |
| AHA12079.1 | polyketide\_synthase\_type\_1 | BGC0001172 | NRP + Polyketide:Modular type I | 33.0 | 48.4 | 257.0 | 1.6e-67 |
| ctg1\_orf15 |  | BGC0001457 | NRP | 31.0 | 46.8 | 257.0 | 1.6e-67 |
| CAQ43076.1 | polyketide\_synthase | BGC0000970 | NRP + Polyketide:Modular type I | 31.0 | 48.4 | 256.0 | 2e-67 |
| AGC45620.1 | polyketide\_synthase | BGC0001394 | NRP + Polyketide | 32.0 | 48.3 | 256.0 | 2e-67 |
| RWQ92174.1 | KR\_domain-containing\_protein | BGC0002030 | Polyketide | 30.0 | 48.3 | 256.0 | 2e-67 |
| AXI91546.1 | FunP7 | BGC0001944 | Polyketide | 26.0 | 89.3 | 256.0 | 2e-67 |
| AAC68815.1 | FK506\_polyketide\_synthase | BGC0000353 | NRP | 25.0 | 85.3 | 256.0 | 2.7e-67 |
| CAQ18834.1 | polyketide\_synthase | BGC0000954 | NRP + Polyketide:Modular type I | 27.0 | 69.4 | 256.0 | 2.7e-67 |
| AAK57188.1 | MxaD | BGC0001022 | NRP + Polyketide | 30.0 | 47.9 | 256.0 | 2.7e-67 |
| AZH23821.1 | MgiH | BGC0001971 | NRP + Polyketide | 27.0 | 66.1 | 256.0 | 2.7e-67 |
| AAZ77696.1 | ChlA3 | BGC0000036 | Polyketide:Modular type I + Polyketide:Iterative type I + Saccharide:Oligosaccharide | 24.0 | 88.5 | 255.0 | 3.5e-67 |
| BAD83684.1 | PKSN\_polyketide\_synthase\_for\_alternapyrone\_biosynthesis | BGC0000012 | Polyketide | 30.0 | 51.2 | 255.0 | 4.6e-67 |
| AAF86396.1 | FkbA | BGC0000994 | NRP + Polyketide | 25.0 | 86.3 | 255.0 | 4.6e-67 |
| AAF26920.1 | polyketide\_synthase | BGC0000988 | NRP + Polyketide | 26.0 | 89.3 | 255.0 | 6e-67 |
| AAF62882.1 | EpoC | BGC0000991 | NRP + Polyketide | 26.0 | 101.2 | 255.0 | 6e-67 |
| AAF19814.1 | MtaF | BGC0001024 | NRP + Polyketide:Modular type I | 31.0 | 44.9 | 255.0 | 6e-67 |
| gene4 |  | BGC0001907 | Polyketide | 31.0 | 50.6 | 255.0 | 6e-67 |
| AAC01711.1 | RifB | BGC0000136 | Polyketide | 24.0 | 88.3 | 254.0 | 1e-66 |
| AAC01713.1 | RifD | BGC0000136 | Polyketide | 26.0 | 78.8 | 254.0 | 1e-66 |
| WP\_079080698.1 | type\_I\_polyketide\_synthase | BGC0001537 | Polyketide | 25.0 | 86.3 | 253.0 | 1.3e-66 |
| AWH12936.1 | StmA | BGC0001939 | Polyketide | 31.0 | 47.9 | 253.0 | 1.3e-66 |
| AIT55262.1 | polyketide\_synthase | BGC0000072 | Polyketide:Modular type I | 32.0 | 48.1 | 253.0 | 1.7e-66 |
| AHH99924.1 | PKS\_I | BGC0000002 | Polyketide | 25.0 | 85.2 | 253.0 | 2.3e-66 |
| ACB46194.1 | polyketide\_synthase | BGC0000989 | NRP + Polyketide | 26.0 | 89.4 | 253.0 | 2.3e-66 |
| AZH23788.1 | MgcR | BGC0001970 | NRP + Polyketide | 31.0 | 48.8 | 253.0 | 2.3e-66 |
| ABM21570.1 | crpB | BGC0000975 | NRP + Polyketide | 31.0 | 48.2 | 252.0 | 3e-66 |
| CAD29794.1 | peptide\_synthetase | BGC0001015 | NRP + Polyketide | 31.0 | 47.2 | 252.0 | 3e-66 |
| AQA28563.1 | type\_I\_polyketide\_synthase | BGC0001663 | Polyketide | 32.0 | 48.5 | 252.0 | 3e-66 |
| ADB12490.1 | EpoC | BGC0000990 | NRP + Polyketide | 26.0 | 100.9 | 252.0 | 3.9e-66 |
| AKD43764.1 | HerF | BGC0001349 | NRP + Polyketide | 26.0 | 83.7 | 252.0 | 3.9e-66 |
| ATZ45185.1 | Bcboa9 | BGC0001892 | Polyketide | 29.0 | 49.0 | 252.0 | 3.9e-66 |
| CAQ18833.1 | polyketide\_synthase | BGC0000954 | NRP + Polyketide:Modular type I | 32.0 | 48.0 | 252.0 | 5e-66 |
| AEE88289.1 | CurA | BGC0000976 | NRP + Polyketide:Modular type I | 31.0 | 48.5 | 252.0 | 5e-66 |
| AAS98777.1 | polyketide\_synthetase | BGC0001001 | NRP + Polyketide | 32.0 | 49.0 | 252.0 | 5e-66 |
| AEU11005.1 | NpnA | BGC0001029 | NRP + Polyketide | 31.0 | 47.8 | 252.0 | 5e-66 |
| AAT70096.1 | CurA | BGC0001165 | NRP + Polyketide:Modular type I | 31.0 | 48.5 | 252.0 | 5e-66 |
| ABI94380.1 | tautomycetin\_biosynthetic\_PKS | BGC0000157 | Polyketide | 24.0 | 87.4 | 251.0 | 6.6e-66 |
| AMY15057.1 | tetraketide\_synthase\_MF-SQTKS | BGC0001339 | Polyketide:Iterative type I | 28.0 | 54.8 | 251.0 | 6.6e-66 |
| ADZ24997.1 | polyketide\_synthase | BGC0000380 | NRP + Polyketide:Modular type I | 31.0 | 48.6 | 251.0 | 8.6e-66 |
| AEC13070.1 | fosD | BGC0000060 | Polyketide | 27.0 | 71.0 | 250.0 | 1.9e-65 |
| AID65222.1 | putative\_aspartate\_racemase | BGC0000335 | NRP | 32.0 | 48.1 | 249.0 | 2.5e-65 |
| AAF19813.1 | MtaE | BGC0001024 | NRP + Polyketide:Modular type I | 30.0 | 48.6 | 249.0 | 2.5e-65 |
| ABV91287.1 | type\_I\_modular\_polyketide\_synthase | BGC0000158 | Polyketide:Modular type I | 24.0 | 87.1 | 249.0 | 3.3e-65 |
| AMY15068.1 | hexaketide\_synthase\_MF-SQHKS | BGC0001339 | Polyketide:Iterative type I | 31.0 | 47.9 | 249.0 | 3.3e-65 |
| AWS21279.1 | type\_I\_polyketide\_synthase | BGC0001934 | Polyketide | 32.0 | 48.7 | 249.0 | 3.3e-65 |
| AZY91989.1 | polyketide\_synthase | BGC0002022 | Polyketide | 32.0 | 48.7 | 249.0 | 3.3e-65 |
| CAQ18835.1 | polyketide\_synthase | BGC0000954 | NRP + Polyketide:Modular type I | 26.0 | 76.5 | 248.0 | 4.3e-65 |
| ADF88276.1 | polyketide\_synthase | BGC0000981 | NRP + Polyketide | 30.0 | 48.2 | 248.0 | 4.3e-65 |
| AMB48442.1 | polyketide\_synthase | BGC0001357 | Polyketide | 31.0 | 47.7 | 248.0 | 4.3e-65 |
| DAB41916.1 | ArzN\_-\_PKS\_(KS,\_AT,\_OMT,\_KR,\_ACP) | BGC0001884 | NRP + Polyketide | 31.0 | 47.9 | 248.0 | 4.3e-65 |
| AIT55264.1 | polyketide\_synthase | BGC0000072 | Polyketide:Modular type I | 32.0 | 43.6 | 248.0 | 5.6e-65 |
| AAK57186.1 | MxaB2 | BGC0001022 | NRP + Polyketide | 32.0 | 47.8 | 248.0 | 5.6e-65 |
| ABK32258.1 | AmbD | BGC0000014 | Polyketide | 26.0 | 88.9 | 248.0 | 7.3e-65 |
| CAD19087.1 | StiC\_protein | BGC0000153 | NRP + Polyketide:Modular type I | 32.0 | 48.8 | 248.0 | 7.3e-65 |
| CAQ18829.1 | polyketide\_synthase | BGC0000954 | NRP + Polyketide:Modular type I | 32.0 | 47.9 | 248.0 | 7.3e-65 |
| AEE88282.1 | CurH | BGC0000976 | NRP + Polyketide:Modular type I | 31.0 | 48.1 | 248.0 | 7.3e-65 |
| AAT70103.1 | CurH | BGC0001165 | NRP + Polyketide:Modular type I | 31.0 | 48.1 | 248.0 | 7.3e-65 |
| ACR33078.1 | polyketide\_synthase | BGC0000017 | Alkaloid + Polyketide:Modular type I | 31.0 | 47.8 | 247.0 | 9.5e-65 |
| ADX66460.1 | ScnS3 | BGC0000108 | Polyketide | 25.0 | 93.6 | 247.0 | 9.5e-65 |
| AAW03329.1 | CtaF | BGC0000982 | NRP + Polyketide | 31.0 | 44.9 | 247.0 | 9.5e-65 |
| CAD89777.1 | MelF\_protein | BGC0001010 | NRP + Polyketide:Modular type I | 30.0 | 48.5 | 247.0 | 9.5e-65 |
| AAF00959.1 | mcyD | BGC0001017 | NRP + Polyketide:Modular type I | 32.0 | 47.5 | 247.0 | 9.5e-65 |
| AHB82065.1 | polyketide\_synthase | BGC0001231 | NRP + Polyketide:Modular type I | 32.0 | 48.0 | 247.0 | 1.2e-64 |
| BAQ25507.1 | type\_I\_polyketide\_synthase | BGC0001288 | Polyketide | 32.0 | 44.9 | 247.0 | 1.2e-64 |
| ADC79618.1 | BafAIII | BGC0000028 | Polyketide:Modular type I | 24.0 | 88.8 | 247.0 | 1.6e-64 |
| AAZ77694.1 | ChlA2 | BGC0000036 | Polyketide:Modular type I + Polyketide:Iterative type I + Saccharide:Oligosaccharide | 25.0 | 89.3 | 247.0 | 1.6e-64 |
| ABX60162.1 | polyketide\_synthase | BGC0000978 | NRP + Alkaloid + Polyketide:Modular type I | 30.0 | 47.2 | 247.0 | 1.6e-64 |
| ctg1\_orf3 |  | BGC0001329 | Polyketide + NRP:Cyclic depsipeptide | 32.0 | 47.3 | 247.0 | 1.6e-64 |
| CAQ52626.1 | type\_I\_polyketide\_synthase,\_loading\_module\_and\_modules\_1-3 | BGC0001066 | Polyketide:Modular type I | 25.0 | 89.3 | 246.0 | 2.8e-64 |
| ADN13832.1 | Polyketide\_Synthase | BGC0001164 | Polyketide:Modular type I | 30.0 | 48.9 | 246.0 | 2.8e-64 |
| ABB88519.1 | polyketide\_synthase\_type\_I | BGC0000050 | Polyketide | 27.0 | 72.2 | 245.0 | 3.6e-64 |
| CAD19092.1 | StiH\_protein | BGC0000153 | NRP + Polyketide:Modular type I | 31.0 | 47.7 | 245.0 | 3.6e-64 |
| AIW00670.1 | mellein\_synthase | BGC0001244 | Polyketide | 24.0 | 80.1 | 245.0 | 3.6e-64 |
| ABK32255.1 | AmbA | BGC0000014 | Polyketide | 32.0 | 47.4 | 245.0 | 4.7e-64 |
| ACO94472.1 | polyketide\_synthase\_type\_I | BGC0000029 | Polyketide:Modular type I | 27.0 | 73.2 | 245.0 | 4.7e-64 |
| CAQ18828.1 | polyketide\_synthase | BGC0000954 | NRP + Polyketide:Modular type I | 32.0 | 48.7 | 245.0 | 4.7e-64 |
| ABX60152.1 | polyketide\_synthase | BGC0000978 | NRP + Alkaloid + Polyketide:Modular type I | 30.0 | 48.2 | 245.0 | 4.7e-64 |
| BAG17643.1 | putative\_NRPS-type-I\_PKS\_fusion\_protein | BGC0001043 | NRP + Polyketide | 27.0 | 75.4 | 245.0 | 4.7e-64 |
| AFV96138.1 | polyketide\_synthase | BGC0001064 | Polyketide:Modular type I + Polyketide:Type III | 31.0 | 48.6 | 245.0 | 4.7e-64 |
| ARU81118.1 | CylD | BGC0001566 | Polyketide | 31.0 | 48.6 | 245.0 | 4.7e-64 |
| AQW44893.1 | polyketide\_synthase | BGC0001737 | NRP + Polyketide | 31.0 | 47.9 | 245.0 | 4.7e-64 |
| ABY21541.1 | AngAIV | BGC0000018 | Polyketide | 26.0 | 74.0 | 245.0 | 6.2e-64 |
| CAD19090.1 | StiF\_protein | BGC0000153 | NRP + Polyketide:Modular type I | 31.0 | 47.4 | 245.0 | 6.2e-64 |
| ADF88280.1 | polyketide\_synthase | BGC0000981 | NRP + Polyketide | 31.0 | 47.2 | 245.0 | 6.2e-64 |
| AGC45622.1 | polyketide\_synthase | BGC0001394 | NRP + Polyketide | 31.0 | 48.0 | 245.0 | 6.2e-64 |
| AQA28562.1 | type\_I\_polyketide\_synthase | BGC0001663 | Polyketide | 31.0 | 47.8 | 245.0 | 6.2e-64 |
| IF55\_RS36525 | polyketide\_synthase | BGC0001348 | Polyketide:Modular type I | 26.0 | 70.3 | 244.0 | 8e-64 |
| WP\_037966908.1 | type\_I\_polyketide\_synthase | BGC0001348 | Polyketide:Modular type I | 26.0 | 70.3 | 244.0 | 8e-64 |
| ATX68116.1 | malonyl\_CoA-acyl\_carrier\_protein\_transacylase | BGC0001772 | Polyketide | 31.0 | 48.8 | 244.0 | 8e-64 |
| ACO94484.1 | polyketide\_synthase\_type\_I | BGC0000097 | Polyketide:Modular type I | 26.0 | 82.7 | 244.0 | 1.1e-63 |
| AHB82054.1 | polyketide\_synthase | BGC0001019 | NRP + Polyketide:Modular type I | 32.0 | 44.9 | 244.0 | 1.1e-63 |
| CQR60494.1 | Polyketide\_synthase,\_type\_I,\_module\_8 | BGC0001287 | Polyketide | 25.0 | 79.6 | 244.0 | 1.1e-63 |
| CQR60497.1 | Polyketide\_synthase,\_type\_I,\_modules:\_loading,\_1,\_2\_and\_3 | BGC0001287 | Polyketide | 26.0 | 70.8 | 244.0 | 1.1e-63 |
| AQX77694.1 | NocP | BGC0001704 | Other | 27.0 | 64.6 | 244.0 | 1.1e-63 |
| ACA99172.1 | polyketide\_synthase | BGC0001160 | Polyketide:Modular type I | 32.0 | 46.6 | 243.0 | 1.4e-63 |
| OJF16270.1 | AceP3 | BGC0001491 | Polyketide | 25.0 | 94.6 | 243.0 | 1.4e-63 |
| CAD19088.1 | StiD\_protein | BGC0000153 | NRP + Polyketide:Modular type I | 30.0 | 48.2 | 243.0 | 1.8e-63 |
| AGC24271.1 | prlQ | BGC0001038 | NRP + Polyketide:Modular type I | 27.0 | 69.8 | 243.0 | 1.8e-63 |
| AKD43761.1 | HerD | BGC0001349 | NRP + Polyketide | 25.0 | 84.0 | 243.0 | 1.8e-63 |
| BAV32159.1 | polyketide\_synthase | BGC0001373 | Polyketide | 31.0 | 49.3 | 243.0 | 1.8e-63 |
| BAF85838.1 | modular\_polyketide\_synthase | BGC0000109 | Polyketide | 25.0 | 89.9 | 243.0 | 2.3e-63 |
| CAD19091.1 | StiG\_protein | BGC0000153 | NRP + Polyketide:Modular type I | 32.0 | 45.4 | 243.0 | 2.3e-63 |
| AQW44890.1 | polyketide\_synthase | BGC0001737 | NRP + Polyketide | 31.0 | 48.5 | 243.0 | 2.3e-63 |
| ABY66019.1 | 6-methylsalicylic\_acid\_synthase | BGC0001008 | Polyketide:Iterative type I + Polyketide:Enediyne type I | 31.0 | 49.0 | 242.0 | 3.1e-63 |
| AKD43768.1 | HerA1 | BGC0001349 | NRP + Polyketide | 31.0 | 48.1 | 242.0 | 3.1e-63 |
| AUO16402.1 | polyketide\_synthase | BGC0001700 | Polyketide | 26.0 | 71.2 | 242.0 | 3.1e-63 |
| ACR33079.1 | polyketide\_synthase | BGC0000017 | Alkaloid + Polyketide:Modular type I | 31.0 | 48.5 | 242.0 | 4e-63 |
| ADB12492.1 | EpoE | BGC0000990 | NRP + Polyketide | 30.0 | 47.9 | 242.0 | 4e-63 |
| AAF62884.1 | EpoE | BGC0000991 | NRP + Polyketide | 30.0 | 47.9 | 242.0 | 4e-63 |
| ACZ57548.1 | polyketide\_synthase | BGC0000046 | Polyketide:Iterative type I | 30.0 | 49.5 | 242.0 | 5.2e-63 |
| ABK32287.1 | JerA | BGC0000080 | Polyketide | 30.0 | 47.5 | 242.0 | 5.2e-63 |
| CAD19086.1 | StiB\_protein | BGC0000153 | NRP + Polyketide:Modular type I | 29.0 | 47.9 | 242.0 | 5.2e-63 |
| ADN68477.1 | SorB | BGC0000184 | Polyketide:Trans-AT type I | 37.0 | 32.7 | 242.0 | 5.2e-63 |
| AHA12078.1 | polyketide\_synthase\_type\_1 | BGC0001172 | NRP + Polyketide:Modular type I | 32.0 | 48.8 | 242.0 | 5.2e-63 |
| AXN93580.1 | PuwE | BGC0001950 | NRP | 38.0 | 32.1 | 242.0 | 5.2e-63 |
| AXN93589.1 | PuwE | BGC0001951 | NRP | 38.0 | 32.1 | 242.0 | 5.2e-63 |
| QCC63000.1 | BII-rafflesfungin\_polyketide\_synthase | BGC0001966 | NRP | 37.0 | 32.4 | 242.0 | 5.2e-63 |
| ACR33077.1 | polyketide\_synthase | BGC0000017 | Alkaloid + Polyketide:Modular type I | 29.0 | 48.8 | 241.0 | 6.8e-63 |
| CAD19085.1 | StiA\_protein | BGC0000153 | NRP + Polyketide:Modular type I | 31.0 | 47.1 | 241.0 | 6.8e-63 |
| CBD77732.1 | polyketide\_synthase | BGC0000974 | NRP + Polyketide | 32.0 | 48.5 | 241.0 | 6.8e-63 |
| AIR74910.1 | polyketide\_synthase | BGC0001559 | RiPP | 32.0 | 48.5 | 241.0 | 6.8e-63 |
| AVI26389.1 | polyketide\_synthase | BGC0001800 | NRP + Polyketide | 31.0 | 44.8 | 241.0 | 6.8e-63 |
| AAK83194.1 | polyketide\_synthase | BGC0000026 | Saccharide:Oligosaccharide | 32.0 | 45.2 | 241.0 | 8.9e-63 |
| ACB46196.1 | polyketide\_synthase | BGC0000989 | NRP + Polyketide | 30.0 | 47.9 | 241.0 | 8.9e-63 |
| AIT55260.1 | polyketide\_synthase | BGC0000072 | Polyketide:Modular type I | 29.0 | 48.5 | 240.0 | 1.5e-62 |
| CAQ34928.1 | polyketide\_synthase | BGC0000986 | NRP + Polyketide | 32.0 | 48.2 | 240.0 | 1.5e-62 |
| AAF26922.1 | polyketide\_synthase | BGC0000988 | NRP + Polyketide | 30.0 | 47.9 | 240.0 | 1.5e-62 |
| AHH99925.1 | PKS\_I | BGC0000002 | Polyketide | 27.0 | 67.6 | 240.0 | 2e-62 |
| ACO94500.1 | polyketide\_synthase\_type\_I | BGC0000097 | Polyketide:Modular type I | 27.0 | 72.4 | 240.0 | 2e-62 |
| AAC46024.1 | polyketide\_synthase\_modules\_1\_and\_2 | BGC0000113 | Polyketide | 30.0 | 48.3 | 240.0 | 2e-62 |
| AAU04878.1 | polyketide\_synthase | BGC0000365 | NRP | 30.0 | 49.7 | 240.0 | 2e-62 |
| ACN64831.1 | PokM1 | BGC0001061 | Polyketide:Iterative type I + Polyketide:Type II + Saccharide:Hybrid/tailoring | 32.0 | 40.7 | 240.0 | 2e-62 |
| AHB82064.1 | polyketide\_synthase | BGC0001231 | NRP + Polyketide:Modular type I | 28.0 | 62.3 | 240.0 | 2e-62 |
| ATV95616.1 | 6-methylsalicylic\_acid\_synthase | BGC0001503 | Polyketide | 29.0 | 48.8 | 240.0 | 2e-62 |
| BAT51066.1 | type\_I\_polyketide\_synthase | BGC0001296 | Polyketide | 30.0 | 48.4 | 239.0 | 2.6e-62 |
| AGC45619.1 | polyketide\_synthase | BGC0001394 | NRP + Polyketide | 31.0 | 48.1 | 239.0 | 2.6e-62 |
| CAJ88175.1 | putative\_polyketide\_synthase\_B | BGC0000151 | Polyketide:Modular type I + Saccharide:Hybrid/tailoring | 27.0 | 68.8 | 239.0 | 3.4e-62 |
| CBD77738.1 | polyketide\_synthase | BGC0000974 | NRP + Polyketide | 32.0 | 45.0 | 239.0 | 3.4e-62 |
| AIR74913.1 | polyketide\_synthase | BGC0001559 | RiPP | 32.0 | 45.0 | 239.0 | 3.4e-62 |
| AXM42950.1 | polyketide\_synthase | BGC0001941 | NRP + Polyketide | 32.0 | 45.3 | 239.0 | 3.4e-62 |
| CAL58681.1 | polyketide\_synthase | BGC0000149 | Polyketide:Modular type I | 31.0 | 48.0 | 238.0 | 4.4e-62 |
| AJO72734.1 | Type\_I\_modular\_polyketide\_synthase | BGC0001381 | Polyketide | 40.0 | 32.3 | 238.0 | 4.4e-62 |
| CCT75967.1 | polyketide\_synthase | BGC0001606 | Polyketide | 28.0 | 49.8 | 238.0 | 4.4e-62 |
| BAJ16469.1 | polyketide\_synthase | BGC0000058 | Polyketide | 25.0 | 78.5 | 238.0 | 5.8e-62 |
| AEE88278.1 | CurL | BGC0000976 | NRP + Polyketide:Modular type I | 30.0 | 49.2 | 238.0 | 5.8e-62 |
| AAT70107.1 | CurL | BGC0001165 | NRP + Polyketide:Modular type I | 30.0 | 49.2 | 238.0 | 5.8e-62 |
| KFL51883.1 | amino\_acid\_adenylation\_protein | BGC0001711 | NRP + Polyketide | 30.0 | 47.9 | 238.0 | 5.8e-62 |
| BAJ16468.1 | polyketide\_synthase | BGC0000058 | Polyketide | 29.0 | 49.5 | 238.0 | 7.5e-62 |
| BAF85843.1 | modular\_polyketide\_synthase | BGC0000109 | Polyketide | 24.0 | 90.2 | 238.0 | 7.5e-62 |
| CAD19093.1 | StiJ\_protein | BGC0000153 | NRP + Polyketide:Modular type I | 30.0 | 47.9 | 238.0 | 7.5e-62 |
| AGC24270.1 | prlP | BGC0001038 | NRP + Polyketide:Modular type I | 31.0 | 48.2 | 238.0 | 7.5e-62 |
| BBG28484.1 | polyketide\_synthase\_CdmE | BGC0001926 | Polyketide | 30.0 | 50.6 | 238.0 | 7.5e-62 |
| ACO94456.1 | polyketide\_synthase\_type\_I | BGC0000029 | Polyketide:Modular type I | 28.0 | 58.4 | 237.0 | 9.8e-62 |
| ABC84456.1 | NigAI | BGC0000114 | Polyketide:Modular type I | 30.0 | 45.0 | 237.0 | 9.8e-62 |
| BAB69199.1 | modular\_polyketide\_synthase | BGC0000117 | Polyketide | 30.0 | 48.2 | 237.0 | 9.8e-62 |
| ABC87511.1 | polyketide\_synthase | BGC0001011 | NRP + Polyketide | 30.0 | 47.9 | 237.0 | 9.8e-62 |
| ctg1\_orf22 |  | BGC0001013 | NRP + Polyketide | 30.0 | 47.9 | 237.0 | 9.8e-62 |
| AEU17897.1 | putative\_type\_I\_PKS | BGC0001072 | Saccharide + Polyketide:Modular type I + Polyketide:Type II + Other:Aminocoumarin | 26.0 | 91.0 | 237.0 | 9.8e-62 |
| BAB69198.1 | modular\_polyketide\_synthase | BGC0000117 | Polyketide | 30.0 | 49.8 | 237.0 | 1.3e-61 |
| AAY42396.1 | Polyketide\_synthase | BGC0001000 | NRP:Lipopeptide + Polyketide:Modular type I | 31.0 | 48.3 | 237.0 | 1.3e-61 |
| AKD43765.1 | HerE | BGC0001349 | NRP + Polyketide | 25.0 | 74.6 | 237.0 | 1.3e-61 |
| TXD00024.1 | SDR\_family\_NAD(P)-dependent\_oxidoreductase | BGC0001877 | Polyketide | 30.0 | 48.0 | 237.0 | 1.7e-61 |
| CAJ46689.1 | polyketide\_synthase | BGC0000969 | NRP:Cyclic depsipeptide + Polyketide:Modular type I | 32.0 | 48.5 | 236.0 | 2.2e-61 |
| CAQ43077.1 | polyketide\_synthase | BGC0000970 | NRP + Polyketide:Modular type I | 26.0 | 91.5 | 236.0 | 2.2e-61 |
| ABO15861.1 | polyketide\_synthase | BGC0000130 | Polyketide | 34.0 | 39.7 | 236.0 | 2.9e-61 |
| ADU85988.1 | putative\_iterative\_type\_I\_polyketide\_synthase | BGC0000165 | Polyketide:Modular type I | 30.0 | 44.4 | 236.0 | 2.9e-61 |
| OAQ83760.1 | polyketide\_synthase | BGC0001358 | Polyketide | 30.0 | 51.4 | 236.0 | 2.9e-61 |
| AUO16398.1 | polyketide\_synthase | BGC0001700 | Polyketide | 25.0 | 71.5 | 236.0 | 2.9e-61 |
| AUO16400.1 | polyketide\_synthase | BGC0001700 | Polyketide | 29.0 | 48.5 | 236.0 | 2.9e-61 |
| BAK64649.1 | polyketide\_synthase | BGC0000135 | Polyketide | 31.0 | 48.9 | 235.0 | 3.7e-61 |
| EPE34340.1 | polyketide\_synthase | BGC0001035 | Polyketide + NRP + Other:Aminocoumarin | 27.0 | 62.0 | 235.0 | 3.7e-61 |
| OAP25811.1 | Phenolphthiocerol\_synthesis\_polyketide\_synthase\_type\_I\_Pks15/1 | BGC0001658 | Polyketide | 32.0 | 43.8 | 235.0 | 3.7e-61 |
| ACB46485.1 | polyketide\_synthase | BGC0000082 | Polyketide | 26.0 | 70.1 | 235.0 | 4.9e-61 |
| SCO70309.1 | Type\_I\_polyketide\_synthase | BGC0001433 | Polyketide:Modular type I | 25.0 | 75.9 | 235.0 | 4.9e-61 |
| TXD00025.1 | SDR\_family\_NAD(P)-dependent\_oxidoreductase | BGC0001877 | Polyketide | 24.0 | 91.7 | 235.0 | 4.9e-61 |
| EAA65604.1 | hypothetical\_protein | BGC0000022 | Polyketide | 29.0 | 50.8 | 235.0 | 6.4e-61 |
| ADN68480.1 | SorE | BGC0000184 | Polyketide:Trans-AT type I | 36.0 | 32.6 | 235.0 | 6.4e-61 |
| ADF88275.1 | polyketide\_synthase | BGC0000981 | NRP + Polyketide | 31.0 | 47.7 | 235.0 | 6.4e-61 |
| ABC87510.1 | polyketide\_synthase | BGC0001011 | NRP + Polyketide | 30.0 | 46.1 | 235.0 | 6.4e-61 |
| ctg1\_orf21 |  | BGC0001013 | NRP + Polyketide | 30.0 | 46.1 | 235.0 | 6.4e-61 |
| APZ78727.1 | polyketide\_synthase | BGC0001421 | NRP:Cyclic depsipeptide + Polyketide:Iterative type I | 31.0 | 48.1 | 235.0 | 6.4e-61 |
| ANR02553.1 | LodL | BGC0001648 | Polyketide | 30.0 | 46.9 | 235.0 | 6.4e-61 |
| ATP76241.1 | NdaD | BGC0001705 | NRP + Polyketide | 25.0 | 82.4 | 235.0 | 6.4e-61 |
| AWH12670.1 | RmpA2 | BGC0001759 | Polyketide | 27.0 | 69.9 | 235.0 | 6.4e-61 |
| BAR73007.1 | putative\_PKS\_(ACP-KS-AT-DH-KR-ACP-KS-AT-DH-ER-KR-ACP) | BGC0001194 | Polyketide | 31.0 | 44.5 | 234.0 | 8.3e-61 |
| APZ78754.1 | polyketide\_synthase | BGC0001423 | NRP:Cyclic depsipeptide + Polyketide:Iterative type I | 31.0 | 48.1 | 234.0 | 8.3e-61 |
| AVV61984.1 | type\_I\_modular\_polyketide\_synthase | BGC0001477 | NRP + Polyketide:Modular type I | 32.0 | 44.3 | 234.0 | 8.3e-61 |
| ACO94483.1 | polyketide\_synthase\_type\_I | BGC0000097 | Polyketide:Modular type I | 30.0 | 47.9 | 234.0 | 1.1e-60 |
| CBZ41585.1 | Type\_I\_modular\_polyketide\_synthase | BGC0000151 | Polyketide:Modular type I + Saccharide:Hybrid/tailoring | 25.0 | 88.5 | 234.0 | 1.1e-60 |
| AAK57190.1 | MxaF | BGC0001022 | NRP + Polyketide | 32.0 | 44.6 | 234.0 | 1.1e-60 |
| AHV78245.1 | LasS1 | BGC0001245 | Polyketide | 29.0 | 50.3 | 234.0 | 1.1e-60 |
| AEH42491.1 | polyketide\_synthase | BGC0000032 | Polyketide | 25.0 | 77.5 | 233.0 | 1.4e-60 |
| WP\_035121546.1 | type\_I\_polyketide\_synthase | BGC0001467 | NRP:Cyclic depsipeptide + Polyketide:Modular type I | 31.0 | 50.0 | 233.0 | 1.4e-60 |
| AUO16401.1 | polyketide\_synthase | BGC0001700 | Polyketide | 29.0 | 48.3 | 233.0 | 1.4e-60 |
| AXG22407.1 | type\_I\_polyketide\_synthase | BGC0002024 | Polyketide | 30.0 | 47.4 | 233.0 | 1.4e-60 |
| CAD55506.1 | CpkA;\_Polyketide\_synthase\_loading\_module,\_and\_modules\_1\_and\_2 | BGC0000038 | Polyketide:Modular type I | 27.0 | 72.3 | 233.0 | 1.9e-60 |
| AFU82617.1 | polyketide\_synthase | BGC0000998 | NRP + Polyketide | 32.0 | 45.9 | 233.0 | 1.9e-60 |
| BAQ21939.1 | putative\_type\_I\_polyketide\_synthase | BGC0001204 | Polyketide | 32.0 | 48.8 | 233.0 | 1.9e-60 |
| ARM20282.1 | polyketide\_synthase | BGC0001523 | Polyketide | 29.0 | 48.2 | 233.0 | 1.9e-60 |
| BAC57029.1 | protomycinolide\_IV\_synthase\_2 | BGC0000102 | Polyketide | 31.0 | 45.4 | 233.0 | 2.4e-60 |
| ABO15860.1 | polyketide\_synthase | BGC0000130 | Polyketide | 31.0 | 47.8 | 233.0 | 2.4e-60 |
| CAQ34920.1 | polyketide\_synthase | BGC0000986 | NRP + Polyketide | 30.0 | 47.6 | 233.0 | 2.4e-60 |
| ADC79637.1 | TamAI | BGC0001052 | NRP + Polyketide:Modular type I | 30.0 | 48.0 | 233.0 | 2.4e-60 |
| AJW65407.1 | type\_I\_modular\_polyketide\_synthase | BGC0001195 | NRP + Polyketide | 27.0 | 63.2 | 233.0 | 2.4e-60 |
| ADH04657.1 | TugA | BGC0001342 | NRP + Polyketide | 31.0 | 47.1 | 232.0 | 3.2e-60 |
| AEP40935.1 | polyketide\_synthase\_type\_I | BGC0000021 | Polyketide | 29.0 | 47.2 | 232.0 | 4.1e-60 |
| CAO98879.1 | polyketide\_synthase\_AufD | BGC0000023 | Polyketide:Modular type I | 30.0 | 48.2 | 232.0 | 4.1e-60 |
| CAD29793.1 | polyketide\_synthase\_type\_I | BGC0001015 | NRP + Polyketide | 24.0 | 82.6 | 232.0 | 4.1e-60 |
| ATX68109.1 | malonyl\_CoA-acyl\_carrier\_protein\_transacylase | BGC0001772 | Polyketide | 31.0 | 42.3 | 232.0 | 4.1e-60 |
| AXG22406.1 | type\_I\_polyketide\_synthase | BGC0002024 | Polyketide | 29.0 | 47.9 | 232.0 | 4.1e-60 |
| AHH99922.1 | PKS\_I | BGC0000002 | Polyketide | 26.0 | 68.4 | 232.0 | 5.4e-60 |
| BAH02271.1 | polyketide\_synthase | BGC0000126 | Polyketide | 26.0 | 70.6 | 232.0 | 5.4e-60 |
| CAL58685.1 | polyketide\_synthase | BGC0000149 | Polyketide:Modular type I | 31.0 | 48.1 | 232.0 | 5.4e-60 |
| AKQ22696.1 | malonyl\_CoA-acyl\_carrier\_protein\_transacylase | BGC0001186 | Polyketide | 32.0 | 41.0 | 232.0 | 5.4e-60 |
| ALD82522.1 | polyketide\_synthase | BGC0001212 | NRP + Polyketide | 38.0 | 31.8 | 232.0 | 5.4e-60 |
| EHK80167.1 | modular\_polyketide\_synthase | BGC0001447 | Polyketide | 29.0 | 47.7 | 232.0 | 5.4e-60 |
| ALJ49921.1 | TtmH | BGC0001236 | Polyketide | 24.0 | 90.7 | 231.0 | 7e-60 |
| AKD43522.1 | Type\_I\_polyketide\_synthase | BGC0001409 | Polyketide | 29.0 | 48.5 | 231.0 | 7e-60 |
| ARS01477.1 | NcmAV | BGC0001702 | NRP + Polyketide | 30.0 | 47.9 | 231.0 | 7e-60 |
| BAK64637.1 | polyketide\_synthase | BGC0000135 | Polyketide | 30.0 | 47.1 | 231.0 | 9.2e-60 |
| BAP05593.1 | calE | BGC0000967 | NRP + Polyketide:Trans-AT type I | 31.0 | 40.1 | 231.0 | 9.2e-60 |
| AVV61981.1 | type\_I\_modular\_PKS | BGC0001477 | NRP + Polyketide:Modular type I | 29.0 | 48.8 | 231.0 | 9.2e-60 |
| OJF16272.1 | AceP1 | BGC0001491 | Polyketide | 27.0 | 66.6 | 231.0 | 9.2e-60 |
| AXM42951.1 | polyketide\_synthase | BGC0001941 | NRP + Polyketide | 32.0 | 44.1 | 231.0 | 9.2e-60 |
| EHA22196.1 | polyketide\_synthase | BGC0000170 | Polyketide | 31.0 | 45.6 | 230.0 | 1.2e-59 |
| ABX60153.1 | polyketide\_synthase | BGC0000978 | NRP + Alkaloid + Polyketide:Modular type I | 31.0 | 47.8 | 230.0 | 1.6e-59 |
| AHB82063.1 | polyketide\_synthase | BGC0001231 | NRP + Polyketide:Modular type I | 29.0 | 48.0 | 230.0 | 1.6e-59 |
| EWM62997.1 | non-ribosomal\_peptide\_synthetase | BGC0001328 | NRP:Cyclic depsipeptide + Polyketide:Modular type I | 36.0 | 32.1 | 230.0 | 1.6e-59 |
| AAM70355.1 | CalO5 | BGC0000033 | Polyketide | 31.0 | 45.5 | 230.0 | 2.1e-59 |
| AAG13917.1 | megalomicin\_6-deoxyerythronolide\_B\_synthase\_1 | BGC0000092 | Polyketide | 30.0 | 45.2 | 230.0 | 2.1e-59 |
| AAC38076.1 | polyketide\_synthase\_type\_I | BGC0000127 | Polyketide | 30.0 | 48.4 | 230.0 | 2.1e-59 |
| WP\_053065268.1 | type\_I\_polyketide\_synthase | BGC0001330 | NRP:Cyclic depsipeptide + Polyketide:Modular type I | 32.0 | 43.4 | 230.0 | 2.1e-59 |
| WP\_020636845.1 | type\_I\_polyketide\_synthase | BGC0002011 | Polyketide | 31.0 | 46.5 | 230.0 | 2.1e-59 |
| ACO94499.1 | polyketide\_synthase\_type\_I | BGC0000097 | Polyketide:Modular type I | 30.0 | 47.2 | 229.0 | 2.7e-59 |
| ACY06287.1 | type\_I\_polyketide\_synthase | BGC0001042 | NRP + Polyketide | 30.0 | 47.2 | 229.0 | 2.7e-59 |
| AUO16399.1 | polyketide\_synthase | BGC0001700 | Polyketide | 29.0 | 48.2 | 229.0 | 2.7e-59 |
| ASZ00147.1 | polyketide\_synthase | BGC0001785 | Polyketide | 26.0 | 70.7 | 229.0 | 2.7e-59 |
| AAU93806.2 | polyketide\_synthase\_modules\_3\_and\_4 | BGC0000054 | Polyketide | 30.0 | 44.6 | 229.0 | 3.5e-59 |
| AAG13918.1 | megalomicin\_6-deoxyerythronolide\_B\_synthase\_2 | BGC0000092 | Polyketide | 29.0 | 48.2 | 229.0 | 3.5e-59 |
| AKL71649.1 | NocP | BGC0001703 | Other | 29.0 | 48.2 | 229.0 | 3.5e-59 |
| AVI26390.1 | polyketide\_synthase\_/\_nonribosomal\_peptide\_synthase\_hybrid | BGC0001800 | NRP + Polyketide | 31.0 | 48.4 | 229.0 | 3.5e-59 |
| ctg1\_13 |  | BGC0001931 | Polyketide | 29.0 | 49.3 | 229.0 | 3.5e-59 |
| AXN93577.1 | PuwB | BGC0001950 | NRP | 39.0 | 31.3 | 229.0 | 3.5e-59 |
| AXN93586.1 | PuwB | BGC0001951 | NRP | 39.0 | 31.3 | 229.0 | 3.5e-59 |
| BAC68128.1 | modular\_polyketide\_synthase | BGC0000059 | Polyketide | 29.0 | 49.2 | 228.0 | 4.6e-59 |
| AAC69330.1 | type\_I\_polyketide\_synthase\_PikAII | BGC0000094 | Polyketide:Modular type I + Saccharide:Hybrid/tailoring | 29.0 | 49.2 | 228.0 | 4.6e-59 |
| ALD82521.1 | polyketide\_synthase | BGC0001212 | NRP + Polyketide | 37.0 | 32.9 | 228.0 | 4.6e-59 |
| ACB37741.1 | putative\_type\_I\_polyketide\_synthase | BGC0000162 | Polyketide | 29.0 | 48.9 | 228.0 | 6e-59 |
| SCO70310.1 | Type\_I\_polyketide\_synthase | BGC0001433 | Polyketide:Modular type I | 30.0 | 48.1 | 228.0 | 6e-59 |
| ARM20280.1 | polyketide\_synthase | BGC0001523 | Polyketide | 29.0 | 47.4 | 228.0 | 6e-59 |
| AOE23578.1 | FoxBII | BGC0001598 | NRP + Polyketide | 25.0 | 81.2 | 228.0 | 6e-59 |
| AJY78092.1 | polyketide\_synthase | BGC0001902 | NRP + Polyketide | 31.0 | 44.2 | 228.0 | 6e-59 |
| AEP40934.1 | polyketide\_synthase\_type\_I | BGC0000021 | Polyketide | 29.0 | 49.1 | 228.0 | 7.8e-59 |
| ABC84457.1 | NigAII | BGC0000114 | Polyketide:Modular type I | 29.0 | 48.2 | 228.0 | 7.8e-59 |
| AFU82616.1 | polyketide\_synthase | BGC0000998 | NRP + Polyketide | 31.0 | 44.5 | 228.0 | 7.8e-59 |
| ALD83704.1 | tAT\_polyketide\_synthase | BGC0001299 | Polyketide | 33.0 | 41.0 | 228.0 | 7.8e-59 |
| APZ78832.1 | polyketide\_synthase | BGC0001430 | NRP:Cyclic depsipeptide + Polyketide:Iterative type I | 30.0 | 48.5 | 228.0 | 7.8e-59 |
| AAC69331.1 | type\_I\_polyketide\_synthase\_PikAIII | BGC0000094 | Polyketide:Modular type I + Saccharide:Hybrid/tailoring | 28.0 | 48.1 | 227.0 | 1e-58 |
| BAH02268.1 | polyketide\_synthase | BGC0000126 | Polyketide | 28.0 | 48.8 | 227.0 | 1e-58 |
| AAF62880.1 | EpoA | BGC0000991 | NRP + Polyketide | 32.0 | 45.3 | 227.0 | 1e-58 |
| ADH04639.1 | TgaA | BGC0001051 | NRP + Polyketide:Modular type I | 31.0 | 45.1 | 227.0 | 1e-58 |
| ANH11415.1 | SceT | BGC0001908 | Polyketide | 28.0 | 61.6 | 227.0 | 1e-58 |
| CAC20921.1 | PimS2\_protein | BGC0000125 | Polyketide | 28.0 | 47.5 | 227.0 | 1.3e-58 |
| ADB12488.1 | EpoA | BGC0000990 | NRP + Polyketide | 32.0 | 45.3 | 227.0 | 1.3e-58 |
| ABC87509.1 | polyketide\_synthase | BGC0001011 | NRP + Polyketide | 29.0 | 48.9 | 227.0 | 1.3e-58 |
| ctg1\_orf20 |  | BGC0001013 | NRP + Polyketide | 29.0 | 48.9 | 227.0 | 1.3e-58 |
| CAD17792.1 | probable\_non\_ribosomal\_peptide\_synthetase\_protein | BGC0001754 | NRP + Polyketide | 31.0 | 48.2 | 227.0 | 1.3e-58 |
| AQT01393.1 | SgnS2 | BGC0001690 | Polyketide | 28.0 | 47.5 | 227.0 | 1.3e-58 |
| AQW44873.1 | polyketide\_synthase | BGC0001761 | Polyketide | 30.0 | 48.3 | 227.0 | 1.3e-58 |
| CAO98850.1 | polyketide\_synthase\_AufG | BGC0000023 | Polyketide:Modular type I | 31.0 | 45.4 | 227.0 | 1.7e-58 |
| CAM00062.1 | EryAI\_Erythromycin\_polyketide\_synthase\_modules\_1\_and\_2 | BGC0000055 | Polyketide:Modular type I + Saccharide:Hybrid/tailoring | 29.0 | 44.9 | 227.0 | 1.7e-58 |
| CAM00065.1 | EryAIII\_Erythromycin\_polyketide\_synthase\_modules\_5\_and\_6 | BGC0000055 | Polyketide:Modular type I + Saccharide:Hybrid/tailoring | 27.0 | 70.2 | 227.0 | 1.7e-58 |
| ADU86003.1 | putative\_modular\_polyketide\_synthase | BGC0000165 | Polyketide:Modular type I | 29.0 | 48.4 | 227.0 | 1.7e-58 |
| AEP40940.1 | polyketide\_synthase\_type\_I | BGC0000021 | Polyketide | 33.0 | 43.6 | 226.0 | 2.3e-58 |
| BAP34739.1 | type\_I\_polyketide\_synthase | BGC0000078 | Polyketide | 31.0 | 47.5 | 226.0 | 2.3e-58 |
| BAH02269.1 | polyketide\_synthase | BGC0000126 | Polyketide | 28.0 | 48.8 | 226.0 | 2.3e-58 |
| ATX68114.1 | malonyl\_CoA-acyl\_carrier\_protein\_transacylase | BGC0001772 | Polyketide | 25.0 | 65.5 | 226.0 | 2.3e-58 |
| AHH99923.1 | PKS\_I | BGC0000002 | Polyketide | 29.0 | 47.5 | 226.0 | 3e-58 |
| ctg1\_orf7 |  | BGC0000053 | Polyketide | 32.0 | 44.3 | 226.0 | 3e-58 |
| ACB37740.1 | putative\_type\_I\_polyketide\_synthase | BGC0000162 | Polyketide | 30.0 | 48.6 | 226.0 | 3e-58 |
| ACR50785.1 | polyketide\_synthase | BGC0000163 | Polyketide | 28.0 | 48.5 | 226.0 | 3e-58 |
| CAG23977.1 | polyketide\_synthase\_type\_I | BGC0000176 | Polyketide + NRP | 35.0 | 33.1 | 226.0 | 3e-58 |
| ACB46192.1 | polyketide\_synthase | BGC0000989 | NRP + Polyketide | 32.0 | 45.3 | 226.0 | 3e-58 |
| ADB12493.1 | EpoF | BGC0000990 | NRP + Polyketide | 32.0 | 44.4 | 226.0 | 3e-58 |
| ALD82523.1 | polyketide\_synthase | BGC0001212 | NRP + Polyketide | 36.0 | 32.0 | 226.0 | 3e-58 |
| BAO66542.1 | type\_I\_polyketide\_synthase | BGC0000042 | Polyketide | 30.0 | 46.9 | 225.0 | 3.9e-58 |
| AAC46027.1 | polyketide\_synthase\_module\_6 | BGC0000113 | Polyketide | 30.0 | 49.4 | 225.0 | 3.9e-58 |
| AAA79984.2 | soraphen\_polyketide\_synthase\_B | BGC0000147 | Polyketide:Modular type I | 29.0 | 45.8 | 225.0 | 3.9e-58 |
| AFP87523.1 | type\_I\_polyketide\_synthase | BGC0001159 | NRP + Polyketide:Modular type I | 30.0 | 48.0 | 225.0 | 3.9e-58 |
| ARM20279.1 | polyketide\_synthase | BGC0001523 | Polyketide | 28.0 | 49.5 | 225.0 | 3.9e-58 |
| KFL51881.1 | beta-ketoacyl\_synthase | BGC0001711 | NRP + Polyketide | 38.0 | 31.9 | 225.0 | 3.9e-58 |
| AAQ82565.1 | FscB | BGC0000061 | Polyketide | 29.0 | 51.8 | 225.0 | 5.1e-58 |
| BAC76493.1 | lankamycin\_synthase\_LkmAI | BGC0000085 | Polyketide | 30.0 | 45.6 | 225.0 | 5.1e-58 |
| ABV97151.1 | AMP-dependent\_synthetase\_and\_ligase | BGC0000137 | Polyketide | 25.0 | 70.2 | 225.0 | 5.1e-58 |
| CCE88377.1 | non-ribosomal\_peptide\_synthetase/polyketide\_synthase | BGC0001034 | NRP + Polyketide:Modular type I | 36.0 | 31.1 | 225.0 | 5.1e-58 |
| BAQ21940.1 | putative\_Type\_I\_polyketide\_synthase | BGC0001204 | Polyketide | 30.0 | 44.8 | 225.0 | 5.1e-58 |
| APZ78854.1 | polyketide\_synthase | BGC0001432 | NRP:Cyclic depsipeptide + Polyketide:Iterative type I | 29.0 | 48.5 | 225.0 | 5.1e-58 |
| QBF51758.1 | type\_I\_polyketide\_synthase | BGC0001856 | Polyketide:Modular type I | 29.0 | 48.8 | 225.0 | 5.1e-58 |
| FS847\_01985 | type\_I\_polyketide\_synthase | BGC0001877 | Polyketide | 29.0 | 48.8 | 225.0 | 5.1e-58 |
| ADU86004.1 | putative\_modular\_polyketide\_synthase | BGC0000165 | Polyketide:Modular type I | 26.0 | 85.8 | 225.0 | 6.6e-58 |
| CAQ18838.1 | polyketide\_synthase | BGC0000954 | NRP + Polyketide:Modular type I | 31.0 | 42.4 | 225.0 | 6.6e-58 |
| CCE88380.1 | polyketide\_synthase | BGC0001034 | NRP + Polyketide:Modular type I | 29.0 | 47.5 | 225.0 | 6.6e-58 |
| AAS98200.1 | MSAS-type\_polyketide\_synthase | BGC0001273 | Polyketide | 34.0 | 32.4 | 225.0 | 6.6e-58 |
| WP\_052165465.1 | type\_I\_polyketide\_synthase | BGC0001327 | NRP:Cyclic depsipeptide + Polyketide:Modular type I | 29.0 | 48.0 | 225.0 | 6.6e-58 |
| orf3 | polyketide\_synthase | BGC0001432 | NRP:Cyclic depsipeptide + Polyketide:Iterative type I | 29.0 | 47.6 | 225.0 | 6.6e-58 |
| APZ78858.1 | polyketide\_synthase | BGC0001432 | NRP:Cyclic depsipeptide + Polyketide:Iterative type I | 29.0 | 47.6 | 225.0 | 6.6e-58 |
| AIG62146.1 | 6-methylsalicylic\_acid\_synthase | BGC0000120 | Polyketide:Iterative type I | 30.0 | 45.0 | 224.0 | 8.6e-58 |
| CBD77736.1 | polyketide\_synthase | BGC0000974 | NRP + Polyketide | 31.0 | 44.4 | 224.0 | 8.6e-58 |
| AAF26923.1 | polyketide\_synthase | BGC0000988 | NRP + Polyketide | 31.0 | 48.5 | 224.0 | 8.6e-58 |
| AGC45623.1 | polyketide\_synthase | BGC0001394 | NRP + Polyketide | 29.0 | 46.9 | 224.0 | 8.6e-58 |
| AIR74912.1 | polyketide\_synthase | BGC0001559 | RiPP | 31.0 | 44.4 | 224.0 | 8.6e-58 |
| AKA59090.1 | type-I\_PKS | BGC0001619 | Polyketide | 36.0 | 32.4 | 224.0 | 8.6e-58 |
| AQH32482.1 | type\_1\_polyketide\_synthase | BGC0001667 | NRP + Polyketide | 25.0 | 83.2 | 224.0 | 8.6e-58 |
| AQW44892.1 | polyketide\_synthase | BGC0001737 | NRP + Polyketide | 30.0 | 47.5 | 224.0 | 8.6e-58 |
| QBF51760.1 | type\_I\_polyketide\_synthase | BGC0001856 | Polyketide:Modular type I | 30.0 | 48.4 | 224.0 | 8.6e-58 |
| TXD00034.1 | SDR\_family\_NAD(P)-dependent\_oxidoreductase | BGC0001877 | Polyketide | 26.0 | 70.4 | 224.0 | 8.6e-58 |
| AHA38201.1 | GphH | BGC0000069 | Polyketide | 35.0 | 33.6 | 224.0 | 1.1e-57 |
| AHA38203.1 | GphJ | BGC0000069 | Polyketide | 31.0 | 47.9 | 224.0 | 1.1e-57 |
| BAF92601.1 | iterative\_type\_I\_PKS | BGC0000118 | Polyketide | 38.0 | 32.4 | 224.0 | 1.1e-57 |
| ACJ24875.1 | 6-methylsalicylic\_acid\_synthase | BGC0000119 | Polyketide:Iterative type I + Saccharide:Hybrid/tailoring | 38.0 | 32.4 | 224.0 | 1.1e-57 |
| BAD08359.1 | polyketide\_synthase\_modules\_5-6 | BGC0000167 | Polyketide | 28.0 | 47.2 | 224.0 | 1.1e-57 |
| BAG84248.1 | putative\_polyketide\_synthase | BGC0000257 | Polyketide | 31.0 | 49.1 | 224.0 | 1.1e-57 |
| ACY06288.1 | type\_I\_polyketide\_synthase | BGC0001042 | NRP + Polyketide | 29.0 | 48.4 | 224.0 | 1.1e-57 |
| EHK80168.1 | acyl\_transferase | BGC0001447 | Polyketide | 30.0 | 47.1 | 224.0 | 1.1e-57 |
| WP\_106967775.1 | type\_I\_polyketide\_synthase | BGC0001519 | NRP + Polyketide | 36.0 | 32.4 | 224.0 | 1.1e-57 |
| ATG32075.1 | polyketide\_synthase | BGC0001750 | NRP + Polyketide | 23.0 | 94.2 | 224.0 | 1.1e-57 |
| AVI26388.1 | polyketide\_synthase | BGC0001800 | NRP + Polyketide | 30.0 | 49.5 | 224.0 | 1.1e-57 |
| WP\_051137606.1 | type\_I\_polyketide\_synthase | BGC0002011 | Polyketide | 30.0 | 44.5 | 224.0 | 1.1e-57 |
| CAO98849.1 | polyketide\_synthase\_AufF | BGC0000023 | Polyketide:Modular type I | 30.0 | 48.9 | 223.0 | 1.5e-57 |
| AEH42490.1 | polyketide\_synthase | BGC0000032 | Polyketide | 25.0 | 70.5 | 223.0 | 1.5e-57 |
| BAF85839.1 | modular\_polyketide\_synthase | BGC0000109 | Polyketide | 27.0 | 64.2 | 223.0 | 1.5e-57 |
| ABY83164.1 | Azi26 | BGC0000960 | NRP + Polyketide | 29.0 | 51.3 | 223.0 | 1.5e-57 |
| ABX60163.1 | polyketide\_synthase | BGC0000978 | NRP + Alkaloid + Polyketide:Modular type I | 29.0 | 48.1 | 223.0 | 1.5e-57 |
| OAQ83765.1 | KR\_domain-containing\_protein | BGC0001358 | Polyketide | 26.0 | 60.0 | 223.0 | 1.5e-57 |
| WP\_083502114.1 | type\_I\_polyketide\_synthase | BGC0001653 | Polyketide | 24.0 | 89.2 | 223.0 | 1.5e-57 |
| ACC40921.1 | polyketide\_synthase\_Pks7 | BGC0001665 | Polyketide | 28.0 | 47.7 | 223.0 | 1.5e-57 |
| AWH12671.1 | RmpA1 | BGC0001759 | Polyketide | 30.0 | 47.5 | 223.0 | 1.5e-57 |
| AEP40936.1 | polyketide\_synthase\_type\_I | BGC0000021 | Polyketide | 29.0 | 48.5 | 223.0 | 1.9e-57 |
| CBW54671.1 | polyketide\_synthase/non\_ribosomal\_peptide\_synthetase | BGC0000971 | NRP + Polyketide:Modular type I | 30.0 | 48.6 | 223.0 | 1.9e-57 |
| AHB82052.1 | polyketide\_synthase | BGC0001019 | NRP + Polyketide:Modular type I | 27.0 | 64.9 | 223.0 | 1.9e-57 |
| EWM63000.1 | non-ribosomal\_peptide\_synthetase | BGC0001328 | NRP:Cyclic depsipeptide + Polyketide:Modular type I | 29.0 | 47.9 | 223.0 | 1.9e-57 |
| AQW44871.1 | polyketide\_synthase | BGC0001761 | Polyketide | 36.0 | 32.4 | 223.0 | 1.9e-57 |
| ATX68115.1 | malonyl\_CoA-acyl\_carrier\_protein\_transacylase | BGC0001772 | Polyketide | 29.0 | 49.2 | 223.0 | 1.9e-57 |
| AAZ94390.1 | modular\_polyketide\_synthase | BGC0000040 | Polyketide | 29.0 | 47.9 | 223.0 | 2.5e-57 |
| BAF02921.1 | type\_I\_polyketide\_synthase | BGC0000073 | Polyketide | 28.0 | 47.9 | 223.0 | 2.5e-57 |
| ABM63528.1 | BryC | BGC0000174 | Polyketide | 36.0 | 33.3 | 223.0 | 2.5e-57 |
| AGI99496.1 | Type\_I\_polyketide\_synthase | BGC0001004 | Polyketide:Modular type I | 30.0 | 48.1 | 223.0 | 2.5e-57 |
| ABV99085.1 | thioester\_reductase\_domain | BGC0001007 | Polyketide + NRP | 26.0 | 70.8 | 223.0 | 2.5e-57 |
| ABJ97438.1 | MerB | BGC0001012 | NRP + Polyketide | 31.0 | 43.8 | 223.0 | 2.5e-57 |
| ARM20277.1 | polyketide\_synthase | BGC0001523 | Polyketide | 28.0 | 49.1 | 223.0 | 2.5e-57 |
| AJY78091.1 | polyketide\_synthase | BGC0001902 | NRP + Polyketide | 29.0 | 58.7 | 223.0 | 2.5e-57 |
| AZF85946.1 | type\_I\_polyketide\_synthase | BGC0001963 | NRP + Polyketide | 34.0 | 32.7 | 223.0 | 2.5e-57 |
| CAE46843.1 | Type\_I\_modular\_polyketide\_synthase | BGC0000103 | Polyketide | 29.0 | 49.5 | 222.0 | 3.3e-57 |
| BAB69195.1 | modular\_polyketide\_synthase | BGC0000117 | Polyketide | 29.0 | 48.8 | 222.0 | 3.3e-57 |
| AGZ15474.1 | putative\_type\_I\_polyketide\_synthase | BGC0001036 | NRP + Polyketide | 29.0 | 49.2 | 222.0 | 3.3e-57 |
| BAR73020.1 | putative\_PKS\_(KS-AT-DH-KR-ACP-KS-AT-DH-KR-ACP-KS-AT-DH-KR-ACP) | BGC0001194 | Polyketide | 29.0 | 46.9 | 222.0 | 3.3e-57 |
| sipP3 | Type\_I\_Modular\_PKS | BGC0001452 | Polyketide | 29.0 | 44.8 | 222.0 | 3.3e-57 |
| AKQ22669.1 | malonyl\_CoA-acyl\_carrier\_protein\_transacylase | BGC0001656 | Polyketide | 34.0 | 32.8 | 222.0 | 3.3e-57 |
| AEK75502.1 | type\_1\_polyketide\_synthase | BGC0000001 | Polyketide:Modular type I | 31.0 | 47.5 | 222.0 | 4.3e-57 |
| AEZ54375.1 | PieA2 | BGC0000124 | Polyketide | 29.0 | 47.7 | 222.0 | 4.3e-57 |
| CAI94682.1 | putative\_polyketide\_synthase | BGC0000141 | Polyketide | 29.0 | 49.2 | 222.0 | 4.3e-57 |
| CAA16183.1 | polyketide\_synthase | BGC0001063 | NRP + Polyketide | 31.0 | 48.5 | 222.0 | 4.3e-57 |
| EHK80163.1 | acyl\_transferase | BGC0001447 | Polyketide | 29.0 | 46.9 | 222.0 | 4.3e-57 |
| TXD00026.1 | SDR\_family\_NAD(P)-dependent\_oxidoreductase | BGC0001877 | Polyketide | 31.0 | 44.2 | 222.0 | 4.3e-57 |
| AXM42948.1 | type\_1\_polyketide\_synthase | BGC0001941 | NRP + Polyketide | 27.0 | 60.8 | 222.0 | 4.3e-57 |
| ABK32263.1 | AmbH | BGC0000014 | Polyketide | 29.0 | 48.5 | 222.0 | 5.6e-57 |
| ADM46359.1 | polyketide\_synthase | BGC0000106 | Polyketide | 29.0 | 44.6 | 222.0 | 5.6e-57 |
| ADX66461.1 | ScnS2 | BGC0000108 | Polyketide | 28.0 | 47.5 | 222.0 | 5.6e-57 |
| AAC46026.1 | polyketide\_synthase\_modules\_4\_and\_5 | BGC0000113 | Polyketide | 29.0 | 47.3 | 222.0 | 5.6e-57 |
| BAB69194.1 | modular\_polyketide\_synthase | BGC0000117 | Polyketide | 28.0 | 49.0 | 222.0 | 5.6e-57 |
| ADF88277.1 | polyketide\_synthase | BGC0000981 | NRP + Polyketide | 29.0 | 48.1 | 222.0 | 5.6e-57 |
| AAF62885.1 | EpoF | BGC0000991 | NRP + Polyketide | 31.0 | 48.5 | 222.0 | 5.6e-57 |
| AGI99482.1 | Type\_I\_polyketide\_synthase | BGC0001004 | Polyketide:Modular type I | 29.0 | 48.8 | 222.0 | 5.6e-57 |
| ADI59532.1 | CorJ | BGC0001091 | NRP + Polyketide | 36.0 | 32.9 | 222.0 | 5.6e-57 |
| WP\_019032757.1 | type\_I\_polyketide\_synthase | BGC0001331 | NRP:Cyclic depsipeptide + Polyketide:Modular type I | 31.0 | 47.5 | 222.0 | 5.6e-57 |
| EHK80169.1 | acyl\_transferase | BGC0001447 | Polyketide | 36.0 | 32.4 | 222.0 | 5.6e-57 |
| WP\_016638480.1 | type\_I\_polyketide\_synthase | BGC0001519 | NRP + Polyketide | 27.0 | 60.6 | 222.0 | 5.6e-57 |
| ANR02552.1 | LodK | BGC0001648 | Polyketide | 30.0 | 48.1 | 222.0 | 5.6e-57 |
| AHH99920.1 | PKS\_I | BGC0000002 | Polyketide | 26.0 | 67.5 | 221.0 | 7.3e-57 |
| BAJ16470.1 | polyketide\_synthase | BGC0000058 | Polyketide | 29.0 | 48.2 | 221.0 | 7.3e-57 |
| ABV83223.1 | CppK | BGC0000116 | Polyketide | 25.0 | 77.4 | 221.0 | 7.3e-57 |
| ABW96541.1 | type\_I\_modular\_polyketide\_synthase | BGC0000159 | Polyketide:Modular type I | 29.0 | 48.4 | 221.0 | 7.3e-57 |
| BAE93729.1 | type\_I\_polyketide\_synthase | BGC0000164 | Polyketide | 29.0 | 45.6 | 221.0 | 7.3e-57 |
| ACY13414.1 | amino\_acid\_adenylation\_domain\_protein | BGC0001367 | NRP + Polyketide | 29.0 | 47.6 | 221.0 | 7.3e-57 |
| AXA20091.1 | hybrid\_trans-AT\_PKS/NRPS\_LgaB | BGC0001946 | NRP + Polyketide | 34.0 | 34.2 | 221.0 | 7.3e-57 |
| ANR02555.1 | LodN | BGC0001648 | Polyketide | 30.0 | 47.6 | 221.0 | 7.3e-57 |
| AUO16422.1 | polyketide\_synthase | BGC0001700 | Polyketide | 28.0 | 48.4 | 221.0 | 7.3e-57 |
| ABY21540.1 | AngAIII | BGC0000018 | Polyketide | 29.0 | 47.9 | 221.0 | 9.5e-57 |
| BAF02926.1 | type\_I\_polyketide\_synthase | BGC0000073 | Polyketide | 29.0 | 48.7 | 221.0 | 9.5e-57 |
| ACB46197.1 | polyketide\_synthase | BGC0000989 | NRP + Polyketide | 31.0 | 48.5 | 221.0 | 9.5e-57 |
| ALP32046.1 | CycF | BGC0001293 | Polyketide | 29.0 | 48.2 | 221.0 | 9.5e-57 |
| EHK80166.1 | beta-ketoacyl\_synthase | BGC0001447 | Polyketide | 29.0 | 48.3 | 221.0 | 9.5e-57 |
| AWR88399.1 | putative\_beta-ketoacyl\_synthase | BGC0001522 | Polyketide | 29.0 | 44.8 | 221.0 | 9.5e-57 |
| AWW87422.1 | type\_I\_polyketide\_synthase | BGC0001755 | Polyketide | 29.0 | 63.1 | 221.0 | 9.5e-57 |
| ANZ22991.1 | ZinG | BGC0001828 | Polyketide | 29.0 | 47.2 | 221.0 | 9.5e-57 |
| WP\_020636817.1 | type\_I\_polyketide\_synthase | BGC0002011 | Polyketide | 30.0 | 48.5 | 221.0 | 9.5e-57 |
| AAX98186.1 | polyketide\_synthase\_type\_I | BGC0000052 | Polyketide | 29.0 | 48.6 | 220.0 | 1.2e-56 |
| BAG85027.1 | putative\_polyketide\_synthase | BGC0000086 | Polyketide | 28.0 | 47.5 | 220.0 | 1.2e-56 |
| BAE93730.1 | type\_I\_polyketide\_synthase | BGC0000164 | Polyketide | 34.0 | 32.3 | 220.0 | 1.2e-56 |
| AAO62584.1 | polyketide\_synthase\_type\_1 | BGC0001016 | NRP + Polyketide | 24.0 | 81.9 | 220.0 | 1.2e-56 |
| CBA11582.1 | polyketide\_synthase\_type\_I | BGC0001046 | NRP + Polyketide:Modular type I + Saccharide:Hybrid/tailoring | 28.0 | 48.3 | 220.0 | 1.2e-56 |
| AWC08657.1 | polyketide\_synthase\_type\_I | BGC0001932 | Polyketide | 29.0 | 55.6 | 220.0 | 1.2e-56 |
| QDA77044.1 | polyketide\_synthase | BGC0002025 | NRP | 37.0 | 32.2 | 220.0 | 1.2e-56 |
| AAW03325.1 | CtaB | BGC0000982 | NRP + Polyketide | 30.0 | 48.2 | 220.0 | 1.6e-56 |
| CAN89634.1 | putative\_polyketide\_synthase | BGC0001070 | NRP + Polyketide:Modular type I + Polyketide:Trans-AT type I | 37.0 | 32.7 | 220.0 | 1.6e-56 |
| ADI59531.1 | CorI | BGC0001091 | NRP + Polyketide | 36.0 | 33.1 | 220.0 | 1.6e-56 |
| ADH04659.1 | TugC | BGC0001342 | NRP + Polyketide | 29.0 | 50.1 | 220.0 | 1.6e-56 |
| ARM20278.1 | polyketide\_synthase | BGC0001523 | Polyketide | 28.0 | 47.9 | 220.0 | 1.6e-56 |
| ASA76643.1 | polyketide\_synthase | BGC0001751 | NRP + Polyketide | 33.0 | 39.3 | 220.0 | 1.6e-56 |
| AWH12937.1 | StmB | BGC0001939 | Polyketide | 29.0 | 47.5 | 220.0 | 1.6e-56 |
| ACF35445.1 | mbcAI | BGC0000090 | Polyketide | 29.0 | 48.6 | 220.0 | 2.1e-56 |
| AEZ53945.1 | polyketide\_synthase | BGC0000144 | Polyketide:Modular type I | 30.0 | 47.9 | 220.0 | 2.1e-56 |
| ADN68483.1 | sorH | BGC0000184 | Polyketide:Trans-AT type I | 34.0 | 32.9 | 220.0 | 2.1e-56 |
| WP\_019032756.1 | type\_I\_polyketide\_synthase | BGC0001331 | NRP:Cyclic depsipeptide + Polyketide:Modular type I | 36.0 | 32.4 | 220.0 | 2.1e-56 |
| BBA84068.1 | type\_I\_polyketide\_synthase | BGC0001916 | Polyketide | 30.0 | 45.5 | 220.0 | 2.1e-56 |
| TXD00265.1 | SDR\_family\_NAD(P)-dependent\_oxidoreductase | BGC0001877 | Polyketide | 28.0 | 50.6 | 220.0 | 2.1e-56 |
| BAJ16467.1 | polyketide\_synthase | BGC0000058 | Polyketide | 29.0 | 64.9 | 219.0 | 2.8e-56 |
| ACB46471.1 | polyketide\_synthase | BGC0000082 | Polyketide | 30.0 | 44.2 | 219.0 | 2.8e-56 |
| AHN85651.1 | Phn2 | BGC0000122 | Polyketide:Modular type I | 34.0 | 32.3 | 219.0 | 2.8e-56 |
| AAF26919.1 | polyketide\_synthase | BGC0000988 | NRP + Polyketide | 32.0 | 45.7 | 219.0 | 2.8e-56 |
| AGZ15473.1 | putative\_type\_I\_polyketide\_synthase | BGC0001036 | NRP + Polyketide | 36.0 | 32.4 | 219.0 | 2.8e-56 |
| ABP53498.1 | PKS\_(ACP-AT-AT-KS-ACP-C) | BGC0001041 | NRP + Polyketide | 30.0 | 48.2 | 219.0 | 2.8e-56 |
| IF55\_RS32375 | beta-ketoacyl\_synthase | BGC0001348 | Polyketide:Modular type I | 28.0 | 48.2 | 219.0 | 2.8e-56 |
| AWW87423.1 | type\_I\_polyketide\_synthase | BGC0001755 | Polyketide | 30.0 | 45.6 | 219.0 | 2.8e-56 |
| ATX68127.1 | malonyl\_CoA-acyl\_carrier\_protein\_transacylase | BGC0001795 | Polyketide | 35.0 | 32.9 | 219.0 | 2.8e-56 |
| AKG06376.1 | polyketide\_synthase\_type\_1 | BGC0001830 | Polyketide | 29.0 | 48.8 | 219.0 | 2.8e-56 |
| AEP40938.1 | polyketide\_synthase\_type\_I | BGC0000021 | Polyketide | 34.0 | 32.8 | 219.0 | 3.6e-56 |
| ACO94471.1 | polyketide\_synthase\_type\_I | BGC0000029 | Polyketide:Modular type I | 29.0 | 47.5 | 219.0 | 3.6e-56 |
| AAU93807.2 | polyketide\_synthase\_modules\_1\_and\_2 | BGC0000054 | Polyketide | 29.0 | 44.0 | 219.0 | 3.6e-56 |
| AAY28226.1 | HbmAII | BGC0000074 | Polyketide | 27.0 | 48.7 | 219.0 | 3.6e-56 |
| AFL48529.1 | laidlomycin\_polyketide\_synthase\_(module\_5\_and\_module\_6) | BGC0000084 | Polyketide | 29.0 | 44.6 | 219.0 | 3.6e-56 |
| AAQ90173.1 | polyketide\_synthase\_type\_I | BGC0000128 | Polyketide | 30.0 | 48.1 | 219.0 | 3.6e-56 |
| CCA89328.1 | mixed\_trans-AT\_type\_I\_polyketide\_synthase/nonribosomal\_peptide\_synthetase | BGC0001111 | NRP + Polyketide:Trans-AT type I | 36.0 | 32.2 | 219.0 | 3.6e-56 |
| AFP87524.1 | type\_I\_polyketide\_synthase | BGC0001159 | NRP + Polyketide:Modular type I | 35.0 | 32.0 | 219.0 | 3.6e-56 |
| AWC08661.1 | polyketide\_synthase\_type\_I | BGC0001932 | Polyketide | 29.0 | 44.6 | 219.0 | 3.6e-56 |
| AVX51107.1 | nysB | BGC0001709 | Polyketide | 25.0 | 70.8 | 219.0 | 3.6e-56 |
| ABC84469.1 | NigAIX | BGC0000114 | Polyketide:Modular type I | 29.0 | 48.5 | 218.0 | 4.7e-56 |
| CAN89636.1 | putative\_polyketide\_synthase | BGC0001070 | NRP + Polyketide:Modular type I + Polyketide:Trans-AT type I | 29.0 | 44.9 | 218.0 | 4.7e-56 |
| AJW65409.1 | type\_I\_modular\_polyketide\_synthase | BGC0001195 | NRP + Polyketide | 29.0 | 48.8 | 218.0 | 4.7e-56 |
| AGY30676.1 | Ann4 | BGC0001298 | Polyketide | 29.0 | 48.1 | 218.0 | 4.7e-56 |
| WP\_107408739.1 | type\_I\_polyketide\_synthase | BGC0002033 | Polyketide | 29.0 | 49.4 | 218.0 | 4.7e-56 |
| BAF02925.1 | type\_I\_polyketide\_synthase | BGC0000073 | Polyketide | 30.0 | 48.1 | 218.0 | 6.2e-56 |
| AFL48532.1 | laidlomycin\_polyketide\_synthase\_(module\_11\_and\_module\_12) | BGC0000084 | Polyketide | 29.0 | 47.8 | 218.0 | 6.2e-56 |
| ABC84460.1 | NigAV | BGC0000114 | Polyketide:Modular type I | 28.0 | 49.0 | 218.0 | 6.2e-56 |
| AAQ84147.1 | Plm7 | BGC0000123 | Polyketide | 29.0 | 48.2 | 218.0 | 6.2e-56 |
| AEZ53952.1 | polyketide\_synthase | BGC0000144 | Polyketide:Modular type I | 26.0 | 51.7 | 218.0 | 6.2e-56 |
| ABP73645.1 | SalA | BGC0000145 | Polyketide | 30.0 | 48.2 | 218.0 | 6.2e-56 |
| CAL58684.1 | polyketide\_synthase | BGC0000149 | Polyketide:Modular type I | 29.0 | 48.8 | 218.0 | 6.2e-56 |
| AHB82053.1 | polyketide\_synthase | BGC0001019 | NRP + Polyketide:Modular type I | 30.0 | 47.9 | 218.0 | 6.2e-56 |
| ALA09371.1 | type\_I\_modular\_PKS | BGC0001303 | Polyketide | 29.0 | 48.2 | 218.0 | 6.2e-56 |
| AKA59089.1 | type-I\_PKS | BGC0001619 | Polyketide | 29.0 | 47.9 | 218.0 | 6.2e-56 |
| ANR02551.1 | LodJ | BGC0001648 | Polyketide | 28.0 | 47.9 | 218.0 | 6.2e-56 |
| CAE02602.1 | polyketide\_synthase\_type\_I | BGC0000024 | Polyketide:Modular type I | 29.0 | 48.2 | 218.0 | 8.1e-56 |
| AAX98184.1 | polyketide\_synthase\_type\_I | BGC0000052 | Polyketide | 29.0 | 49.2 | 218.0 | 8.1e-56 |
| AEC13079.1 | fosA | BGC0000060 | Polyketide | 38.0 | 32.5 | 218.0 | 8.1e-56 |
| ABP55221.1 | acyl\_transferase\_domain\_protein | BGC0000142 | Polyketide | 30.0 | 41.9 | 218.0 | 8.1e-56 |
| ABJ97439.1 | MerC | BGC0001012 | NRP + Polyketide | 29.0 | 48.9 | 218.0 | 8.1e-56 |
| AJO72735.1 | Type\_I\_modular\_polyketide\_synthase | BGC0001381 | Polyketide | 24.0 | 89.0 | 218.0 | 8.1e-56 |
| AJO72743.1 | Type\_I\_modular\_polyketide\_synthase | BGC0001381 | Polyketide | 27.0 | 69.6 | 218.0 | 8.1e-56 |
| OAP25819.1 | Phenolphthiocerol\_synthesis\_polyketide\_synthase\_type\_I\_Pks15/1 | BGC0001658 | Polyketide | 30.0 | 46.9 | 218.0 | 8.1e-56 |
| AXI91548.1 | FunP5 | BGC0001944 | Polyketide | 29.0 | 49.1 | 218.0 | 8.1e-56 |
| QDA77045.1 | polyketide\_synthase/nonribosomal\_peptide\_synthetase | BGC0002025 | NRP | 29.0 | 45.3 | 218.0 | 8.1e-56 |
| CAM00064.1 | EryAII\_Erythromycin\_polyketide\_synthase\_modules\_3\_and\_4 | BGC0000055 | Polyketide:Modular type I + Saccharide:Hybrid/tailoring | 28.0 | 48.0 | 217.0 | 1.1e-55 |
| AHA38199.1 | GphF | BGC0000069 | Polyketide | 26.0 | 82.1 | 217.0 | 1.1e-55 |
| ctg1\_orf28 |  | BGC0000096 | Polyketide | 29.0 | 49.3 | 217.0 | 1.1e-55 |
| ABW96542.1 | type\_I\_modular\_polyketide\_synthase | BGC0000159 | Polyketide:Modular type I | 28.0 | 48.6 | 217.0 | 1.1e-55 |
| AMH40423.1 | PKS | BGC0001350 | Polyketide | 35.0 | 33.1 | 217.0 | 1.1e-55 |
| WP\_016638481.1 | type\_I\_polyketide\_synthase | BGC0001519 | NRP + Polyketide | 29.0 | 44.4 | 217.0 | 1.1e-55 |
| AWW87424.1 | type\_I\_polyketide\_synthase | BGC0001755 | Polyketide | 30.0 | 45.1 | 217.0 | 1.1e-55 |
| QBF51754.1 | type\_I\_polyketide\_synthase | BGC0001856 | Polyketide:Modular type I | 29.0 | 48.8 | 217.0 | 1.1e-55 |
| ACB46487.1 | polyketide\_synthase | BGC0000082 | Polyketide | 29.0 | 47.5 | 217.0 | 1.4e-55 |
| BAG85028.1 | putative\_polyketide\_synthase | BGC0000086 | Polyketide | 28.0 | 48.0 | 217.0 | 1.4e-55 |
| CAQ64688.1 | lasalocid\_modular\_polyketide\_synthase | BGC0000087 | Polyketide | 28.0 | 48.0 | 217.0 | 1.4e-55 |
| AAG23264.1 | polyketide\_synthase\_loading\_and\_extender\_module\_1 | BGC0000148 | Polyketide | 30.0 | 44.6 | 217.0 | 1.4e-55 |
| ACR50795.1 | putative\_polyketide\_synthase | BGC0000163 | Polyketide | 34.0 | 33.0 | 217.0 | 1.4e-55 |
| AGI99495.1 | Type\_I\_polyketide\_synthase | BGC0001004 | Polyketide:Modular type I | 29.0 | 48.2 | 217.0 | 1.4e-55 |
| ABC87512.1 | polyketide\_synthase | BGC0001011 | NRP + Polyketide | 29.0 | 47.5 | 217.0 | 1.4e-55 |
| ctg1\_orf23 |  | BGC0001013 | NRP + Polyketide | 29.0 | 47.5 | 217.0 | 1.4e-55 |
| AWS21278.1 | type\_I\_polyketide\_synthase | BGC0001934 | Polyketide | 35.0 | 31.9 | 217.0 | 1.4e-55 |
| AZF85932.1 | type\_I\_polyketide\_synthase | BGC0001963 | NRP + Polyketide | 29.0 | 49.1 | 217.0 | 1.4e-55 |
| AZY91987.1 | polyketide\_synthase | BGC0002022 | Polyketide | 35.0 | 31.9 | 217.0 | 1.4e-55 |
| AAM54077.1 | polyketide\_synthase | BGC0000020 | Polyketide | 30.0 | 47.9 | 217.0 | 1.8e-55 |
| CAO98848.1 | polyketide\_synthase\_AufE | BGC0000023 | Polyketide:Modular type I | 26.0 | 64.6 | 217.0 | 1.8e-55 |
| AAX98191.1 | polyketide\_synthase\_type\_I | BGC0000052 | Polyketide | 34.0 | 32.4 | 217.0 | 1.8e-55 |
| ABK32288.1 | JerB | BGC0000080 | Polyketide | 31.0 | 45.5 | 217.0 | 1.8e-55 |
| ADM46356.1 | polyketide\_synthase | BGC0000106 | Polyketide | 28.0 | 47.9 | 217.0 | 1.8e-55 |
| BAH02270.1 | polyketide\_synthase | BGC0000126 | Polyketide | 29.0 | 48.5 | 217.0 | 1.8e-55 |
| ADC79639.1 | TamAIII | BGC0001052 | NRP + Polyketide:Modular type I | 30.0 | 45.6 | 217.0 | 1.8e-55 |
| AIA58899.1 | HRPKS | BGC0001141 | Polyketide:Iterative type I | 31.0 | 48.3 | 217.0 | 1.8e-55 |
| AGY30677.1 | Ann5 | BGC0001298 | Polyketide | 30.0 | 45.4 | 217.0 | 1.8e-55 |
| SCN11952.1 | ebeD-type\_I\_polyketide\_synthase | BGC0001580 | Polyketide | 29.0 | 47.8 | 217.0 | 1.8e-55 |
| AWC08660.1 | polyketide\_synthase\_type\_I | BGC0001932 | Polyketide | 29.0 | 49.5 | 217.0 | 1.8e-55 |
| ARW71486.1 | type\_I\_PKS\_module\_6 | BGC0001812 | Polyketide | 28.0 | 48.5 | 217.0 | 1.8e-55 |
| AKG06378.1 | polyketide\_synthase\_type\_1 | BGC0001830 | Polyketide | 28.0 | 47.4 | 217.0 | 1.8e-55 |
| AEP40939.1 | polyketide\_synthase\_type\_I | BGC0000021 | Polyketide | 29.0 | 48.1 | 216.0 | 2.3e-55 |
| AAS79463.1 | polyketide\_synthase\_subunit | BGC0000035 | Polyketide | 29.0 | 45.2 | 216.0 | 2.3e-55 |
| AAD03047.1 | type\_I\_polyketide\_synthase | BGC0000041 | Polyketide | 29.0 | 48.7 | 216.0 | 2.3e-55 |
| BAC76492.1 | lankamycin\_synthase\_LkmAII | BGC0000085 | Polyketide | 34.0 | 33.3 | 216.0 | 2.3e-55 |
| BAG85030.1 | putative\_polyketide\_synthase | BGC0000086 | Polyketide | 28.0 | 47.9 | 216.0 | 2.3e-55 |
| CAQ64687.1 | lasalocid\_modular\_polyketide\_synthase | BGC0000087 | Polyketide | 28.0 | 48.0 | 216.0 | 2.3e-55 |
| CAQ64690.1 | lasalocid\_modular\_polyketide\_synthase | BGC0000087 | Polyketide | 28.0 | 47.9 | 216.0 | 2.3e-55 |
| CAJ88184.1 | putative\_modular\_polyketide\_synthase | BGC0000151 | Polyketide:Modular type I + Saccharide:Hybrid/tailoring | 29.0 | 44.2 | 216.0 | 2.3e-55 |
| EAU32819.1 | 6-methylsalicylic\_acid\_synthase | BGC0000160 | Polyketide | 29.0 | 44.0 | 216.0 | 2.3e-55 |
| AAB66508.1 | tylactone\_synthase\_module\_7 | BGC0000166 | Polyketide | 29.0 | 48.8 | 216.0 | 2.3e-55 |
| ABB05103.1 | LipPks2 | BGC0001003 | NRP:Lipopeptide + Polyketide:Modular type I + Saccharide:Hybrid/tailoring | 28.0 | 49.1 | 216.0 | 2.3e-55 |
| AVV61979.1 | beta-ketoacyl\_synthase | BGC0001477 | NRP + Polyketide:Modular type I | 29.0 | 45.1 | 216.0 | 2.3e-55 |
| ARM20283.1 | polyketide\_synthase | BGC0001523 | Polyketide | 27.0 | 49.3 | 216.0 | 2.3e-55 |
| AEZ54379.1 | PieA6 | BGC0000124 | Polyketide | 29.0 | 48.2 | 216.0 | 3.1e-55 |
| AAG23265.1 | polyketide\_synthase\_extender\_module\_2 | BGC0000148 | Polyketide | 29.0 | 44.7 | 216.0 | 3.1e-55 |
| CAQ34917.1 | polyketide\_synthase | BGC0000986 | NRP + Polyketide | 33.0 | 39.2 | 216.0 | 3.1e-55 |
| ADC79638.1 | TamAII | BGC0001052 | NRP + Polyketide:Modular type I | 29.0 | 48.5 | 216.0 | 3.1e-55 |
| AVV61989.1 | beta-ketoacyl\_synthase | BGC0001477 | NRP + Polyketide:Modular type I | 30.0 | 43.7 | 216.0 | 3.1e-55 |
| ALV82341.1 | borrelidin\_type\_I\_polyketide\_synthase | BGC0001533 | Polyketide | 28.0 | 43.6 | 216.0 | 3.1e-55 |
| ANH11414.1 | SceS | BGC0001908 | Polyketide | 28.0 | 47.5 | 216.0 | 3.1e-55 |
| ACR50774.1 | polyketide\_synthase | BGC0000163 | Polyketide | 28.0 | 48.2 | 215.0 | 4e-55 |
| AGZ15475.1 | putative\_type\_1\_modular\_polyketide\_synthase | BGC0001036 | NRP + Polyketide | 29.0 | 45.0 | 215.0 | 4e-55 |
| AGC09484.1 | LobS1 | BGC0001183 | Polyketide | 28.0 | 47.4 | 215.0 | 4e-55 |
| AVX51098.1 | nysI | BGC0001709 | Polyketide | 28.0 | 49.1 | 215.0 | 4e-55 |
| ADC45535.1 | modular\_polyketide\_synthase | BGC0000093 | Polyketide | 30.0 | 46.0 | 215.0 | 5.2e-55 |
| AAC69332.1 | type\_I\_polyketide\_synthase\_PikAIV | BGC0000094 | Polyketide:Modular type I + Saccharide:Hybrid/tailoring | 28.0 | 48.5 | 215.0 | 5.2e-55 |
| ADM46358.1 | polyketide\_synthase | BGC0000106 | Polyketide | 27.0 | 47.9 | 215.0 | 5.2e-55 |
| AAC01710.1 | RifA | BGC0000136 | Polyketide | 29.0 | 47.6 | 215.0 | 5.2e-55 |
| CAA60462.1 | polyketide\_synthase | BGC0001040 | NRP + Polyketide | 30.0 | 45.6 | 215.0 | 5.2e-55 |
| BAQ25511.1 | type\_I\_polyketide\_synthase | BGC0001288 | Polyketide | 28.0 | 47.6 | 215.0 | 5.2e-55 |
| AKD43753.1 | HerB | BGC0001349 | NRP + Polyketide | 29.0 | 45.7 | 215.0 | 5.2e-55 |
| AKD43769.1 | HerA2 | BGC0001349 | NRP + Polyketide | 29.0 | 49.0 | 215.0 | 5.2e-55 |
| ASZ00150.1 | polyketide\_synthase | BGC0001785 | Polyketide | 28.0 | 48.0 | 215.0 | 5.2e-55 |
| AAQ82567.1 | FscE | BGC0000061 | Polyketide | 28.0 | 48.4 | 215.0 | 6.8e-55 |
| ABB52545.1 | putative\_type\_I\_polyketide\_synthase | BGC0000047 | Polyketide | 30.0 | 45.9 | 215.0 | 6.8e-55 |
| AAX98189.1 | polyketide\_synthase\_type\_I | BGC0000052 | Polyketide | 28.0 | 48.5 | 215.0 | 6.8e-55 |
| BAB69193.1 |  | BGC0000117 | Polyketide | 30.0 | 45.9 | 215.0 | 6.8e-55 |
| AGI99497.1 | type\_I\_polyketide\_synthase | BGC0001004 | Polyketide:Modular type I | 27.0 | 48.0 | 215.0 | 6.8e-55 |
| AEU17899.1 | putative\_type\_I\_PKS | BGC0001072 | Saccharide + Polyketide:Modular type I + Polyketide:Type II + Other:Aminocoumarin | 28.0 | 48.8 | 215.0 | 6.8e-55 |
| ANC94962.1 | AlmHV | BGC0001396 | Polyketide | 30.0 | 45.9 | 215.0 | 6.8e-55 |
| AUO16403.1 | polyketide\_synthase | BGC0001700 | Polyketide | 27.0 | 48.9 | 215.0 | 6.8e-55 |
| AMYAL\_RS48925 | polyketide\_synthase | BGC0002011 | Polyketide | 29.0 | 48.0 | 215.0 | 6.8e-55 |
| ABK32256.1 | AmbB | BGC0000014 | Polyketide | 36.0 | 33.0 | 214.0 | 8.9e-55 |
| ADN68476.1 | sorA | BGC0000184 | Polyketide:Trans-AT type I | 33.0 | 40.4 | 214.0 | 8.9e-55 |
| BAR73017.1 | putative\_PKS\_(KS-AT-KR-ACP-KS-AT-DH-KR-ACP) | BGC0001194 | Polyketide | 29.0 | 47.5 | 214.0 | 8.9e-55 |
| CRI73799.1 | CongC\_protein | BGC0001215 | NRP | 29.0 | 48.9 | 214.0 | 8.9e-55 |
| AKA59088.1 | type-I\_PKS | BGC0001619 | Polyketide | 29.0 | 45.1 | 214.0 | 8.9e-55 |
| ARV85765.1 | PieA6\_type\_I\_PKS | BGC0001742 | Polyketide | 29.0 | 48.1 | 214.0 | 8.9e-55 |
| BBG28498.1 | putative\_polyketide\_synthase | BGC0001913 | Polyketide | 27.0 | 52.5 | 214.0 | 8.9e-55 |
| AAQ82568.1 | FscD | BGC0000061 | Polyketide | 33.0 | 31.7 | 214.0 | 1.2e-54 |
| ADA82585.1 | hybrid\_trans-AT\_polyketide\_synthase\_-\_nonribosomal\_peptide\_synthetase | BGC0001110 | NRP + Polyketide:Trans-AT type I | 33.0 | 31.8 | 214.0 | 1.2e-54 |
| BAA20102.2 | 6-methylsalicylic\_acid\_synthase | BGC0001276 | Polyketide | 29.0 | 44.0 | 214.0 | 1.2e-54 |
| BAQ25513.1 | type\_I\_polyketide\_synthase | BGC0001288 | Polyketide | 28.0 | 46.5 | 214.0 | 1.2e-54 |
| OAP25820.1 | Erythronolide\_synthase,\_modules\_1\_and\_2 | BGC0001658 | Polyketide | 28.0 | 47.9 | 214.0 | 1.2e-54 |
| AWC08659.1 | polyketide\_synthase\_type\_I | BGC0001932 | Polyketide | 29.0 | 48.9 | 214.0 | 1.2e-54 |
| ATY46594.1 | polyketide\_synthase | BGC0001666 | Polyketide | 28.0 | 48.2 | 214.0 | 1.2e-54 |
| BBA21068.1 | putative\_non-ribosomal\_peptide\_synthetase | BGC0001740 | NRP + Polyketide | 36.0 | 32.7 | 214.0 | 1.2e-54 |
| AWW87425.1 | polyketide\_synthase | BGC0001755 | Polyketide | 35.0 | 32.4 | 214.0 | 1.2e-54 |
| AAZ94391.1 | modular\_polyketide\_synthase | BGC0000040 | Polyketide | 29.0 | 47.5 | 213.0 | 1.5e-54 |
| ABV83222.1 | CppJ | BGC0000116 | Polyketide | 28.0 | 44.9 | 213.0 | 1.5e-54 |
| AFI57007.1 | QmnA3 | BGC0000133 | Polyketide | 28.0 | 50.5 | 213.0 | 1.5e-54 |
| ERM18799.1 | polyketide\_synthase | BGC0000172 | Polyketide | 35.0 | 32.6 | 213.0 | 1.5e-54 |
| BAQ25483.1 | type\_I\_polyketide\_synthase | BGC0001288 | Polyketide | 29.0 | 51.1 | 213.0 | 1.5e-54 |
| ALD83688.1 | tAT\_polyketide\_synthase | BGC0001300 | Polyketide | 34.0 | 33.7 | 213.0 | 1.5e-54 |
| ATY46595.1 | polyketide\_synthase | BGC0001666 | Polyketide | 28.0 | 48.3 | 213.0 | 1.5e-54 |
| ANH11409.1 | SceN | BGC0001908 | Polyketide | 29.0 | 48.6 | 213.0 | 1.5e-54 |
| TGZ15165.1 | hypothetical\_protein | BGC0002032 | Polyketide | 29.0 | 48.2 | 213.0 | 1.5e-54 |
| ctg1\_orf11 |  | BGC0000053 | Polyketide | 34.0 | 32.7 | 213.0 | 2e-54 |
| AIT55261.1 | polyketide\_synthase | BGC0000072 | Polyketide:Modular type I | 35.0 | 32.5 | 213.0 | 2e-54 |
| BAG85029.1 | putative\_polyketide\_synthase | BGC0000086 | Polyketide | 30.0 | 48.1 | 213.0 | 2e-54 |
| CAQ64689.1 | lasalocid\_modular\_polyketide\_synthase | BGC0000087 | Polyketide | 30.0 | 48.1 | 213.0 | 2e-54 |
| AAG13919.1 | megalomicin\_6-deoxyerythronolide\_B\_synthase\_3 | BGC0000092 | Polyketide | 29.0 | 49.2 | 213.0 | 2e-54 |
| AEZ54376.1 | PieA3 | BGC0000124 | Polyketide | 25.0 | 78.2 | 213.0 | 2e-54 |
| BAD08358.1 | polyketide\_synthase\_modules\_4 | BGC0000167 | Polyketide | 33.0 | 32.8 | 213.0 | 2e-54 |
| ctg1\_orf523 |  | BGC0001199 | Polyketide | 31.0 | 39.4 | 213.0 | 2e-54 |
| BAQ25482.1 | type\_I\_polyketide\_synthase | BGC0001288 | Polyketide | 29.0 | 47.2 | 213.0 | 2e-54 |
| AQZ37094.1 | polyketide\_synthase | BGC0001511 | Polyketide | 31.0 | 43.2 | 213.0 | 2e-54 |
| AKQ22680.1 | malonyl\_CoA-acyl\_carrier\_protein\_transacylase | BGC0001656 | Polyketide | 35.0 | 33.0 | 213.0 | 2e-54 |
| ANY10589.1 | polyketide\_synthase | BGC0001773 | Polyketide | 29.0 | 48.2 | 213.0 | 2e-54 |
| ANZ22987.1 | ZinD | BGC0001828 | Polyketide | 27.0 | 48.6 | 213.0 | 2e-54 |
| QBF51759.1 | type\_I\_polyketide\_synthase | BGC0001856 | Polyketide:Modular type I | 29.0 | 48.8 | 213.0 | 2e-54 |
| AFV30247.1 | polyketide\_synthase | BGC0000075 | Polyketide | 30.0 | 44.3 | 213.0 | 2.6e-54 |
| AEZ53946.1 | polyketide\_synthase | BGC0000144 | Polyketide:Modular type I | 30.0 | 48.2 | 213.0 | 2.6e-54 |
| AAG23263.1 | polyketide\_synthase\_extender\_modules\_5-7 | BGC0000148 | Polyketide | 34.0 | 32.7 | 213.0 | 2.6e-54 |
| AEH59109.1 | keto-hydroxyglutarate-aldolase/polyketide\_synthase | BGC0000385 | NRP | 31.0 | 40.3 | 213.0 | 2.6e-54 |
| AEZ64503.1 | Herd | BGC0001065 | Polyketide | 29.0 | 48.2 | 213.0 | 2.6e-54 |
| BAQ21948.1 | putative\_type\_I\_polyketide\_synthase | BGC0001204 | Polyketide | 28.0 | 48.5 | 213.0 | 2.6e-54 |
| CQR60493.1 | Polyketide\_synthase,\_type\_I,\_modules:\_9\_and\_10 | BGC0001287 | Polyketide | 30.0 | 44.9 | 213.0 | 2.6e-54 |
| RAT98530.1 | trans-acyltransferase\_polyketide\_synthase | BGC0001470 | Polyketide:Trans-AT type I | 31.0 | 39.6 | 213.0 | 2.6e-54 |
| AAQ90174.1 | polyketide\_synthase\_type\_I | BGC0000128 | Polyketide | 30.0 | 44.2 | 212.0 | 3.4e-54 |
| ABP55222.1 | beta-ketoacyl\_synthase | BGC0000142 | Polyketide | 35.0 | 31.4 | 212.0 | 3.4e-54 |
| AAG23262.1 | polyketide\_synthase\_extender\_modules\_8-10 | BGC0000148 | Polyketide | 29.0 | 45.3 | 212.0 | 3.4e-54 |
| BAP05589.1 | calA | BGC0000967 | NRP + Polyketide:Trans-AT type I | 34.0 | 32.6 | 212.0 | 3.4e-54 |
| AHH34186.1 | polyketide\_synthase | BGC0001161 | Polyketide:Modular type I | 34.0 | 34.0 | 212.0 | 3.4e-54 |
| AWR88398.1 | putative\_beta-ketoacyl\_synthase | BGC0001522 | Polyketide | 35.0 | 31.2 | 212.0 | 3.4e-54 |
| AP234\_RS14720 | hypothetical\_protein | BGC0001653 | Polyketide | 31.0 | 42.6 | 212.0 | 3.4e-54 |
| ANY10590.1 | polyketide\_synthase | BGC0001773 | Polyketide | 28.0 | 53.9 | 212.0 | 3.4e-54 |
| BAO66529.1 | type\_I\_polyketide\_synthase | BGC0000042 | Polyketide | 30.0 | 50.6 | 212.0 | 4.4e-54 |
| ACN69988.1 | polyketide\_synthase | BGC0000079 | Polyketide | 28.0 | 48.4 | 212.0 | 4.4e-54 |
| ABK32291.1 | JerE | BGC0000080 | Polyketide | 34.0 | 33.1 | 212.0 | 4.4e-54 |
| ctg1\_orf30 |  | BGC0000096 | Polyketide | 35.0 | 32.3 | 212.0 | 4.4e-54 |
| AAO65796.1 | monensin\_polyketide\_synthase\_loading\_module\_and\_module\_1 | BGC0000100 | Polyketide | 28.0 | 47.7 | 212.0 | 4.4e-54 |
| CAC20931.1 | PimS1\_protein | BGC0000125 | Polyketide | 30.0 | 45.9 | 212.0 | 4.4e-54 |
| AAG23266.1 | polyketide\_synthase\_extender\_modules\_3-4 | BGC0000148 | Polyketide | 29.0 | 47.4 | 212.0 | 4.4e-54 |
| CAJ88187.2 | Type\_I\_modular\_polyketide\_synthase | BGC0000151 | Polyketide:Modular type I + Saccharide:Hybrid/tailoring | 25.0 | 63.5 | 212.0 | 4.4e-54 |
| ADD82939.1 | Bat1 | BGC0001099 | NRP + Polyketide:Modular type I + Polyketide:Trans-AT type I | 30.0 | 43.3 | 212.0 | 4.4e-54 |
| ALA09354.1 | type\_I\_modular\_PKS | BGC0001303 | Polyketide | 34.0 | 32.0 | 212.0 | 4.4e-54 |
| WP\_053065267.1 | type\_I\_polyketide\_synthase | BGC0001330 | NRP:Cyclic depsipeptide + Polyketide:Modular type I | 35.0 | 32.4 | 212.0 | 4.4e-54 |
| WP\_063764078.1 | polyketide\_synthase | BGC0001348 | Polyketide:Modular type I | 29.0 | 51.1 | 212.0 | 4.4e-54 |
| ANC94965.1 | AlmHII | BGC0001396 | Polyketide | 28.0 | 48.2 | 212.0 | 4.4e-54 |
| ANZ52459.1 | MonAI | BGC0001670 | Polyketide | 28.0 | 47.7 | 212.0 | 4.4e-54 |
| QBF51756.1 | type\_I\_polyketide\_synthase | BGC0001856 | Polyketide:Modular type I | 30.0 | 48.6 | 212.0 | 4.4e-54 |
| QBF51757.1 | type\_I\_polyketide\_synthase | BGC0001856 | Polyketide:Modular type I | 28.0 | 48.0 | 212.0 | 4.4e-54 |
| BAD97694.1 | Aft9-1 | BGC0000003 | Polyketide | 30.0 | 40.8 | 212.0 | 5.8e-54 |
| ADC79619.1 | BafAIV | BGC0000028 | Polyketide:Modular type I | 35.0 | 32.2 | 212.0 | 5.8e-54 |
| ADC79620.1 | BafAV | BGC0000028 | Polyketide:Modular type I | 29.0 | 47.9 | 212.0 | 5.8e-54 |
| CAQ64691.1 | lasalocid\_modular\_polyketide\_synthase | BGC0000087 | Polyketide | 27.0 | 47.8 | 212.0 | 5.8e-54 |
| AAC69329.1 | type\_I\_polyketide\_synthase\_PikAI | BGC0000094 | Polyketide:Modular type I + Saccharide:Hybrid/tailoring | 35.0 | 32.4 | 212.0 | 5.8e-54 |
| AAO65798.1 | monensin\_polyketide\_synthase\_modules\_3\_and\_4 | BGC0000100 | Polyketide | 29.0 | 47.9 | 212.0 | 5.8e-54 |
| AAP42859.1 | NanA5 | BGC0000105 | Polyketide | 28.0 | 49.5 | 212.0 | 5.8e-54 |
| ABC84458.1 | NigAIII | BGC0000114 | Polyketide:Modular type I | 28.0 | 49.5 | 212.0 | 5.8e-54 |
| ABI94379.1 | tautomycetin\_biosynthetic\_PKS | BGC0000157 | Polyketide | 30.0 | 45.8 | 212.0 | 5.8e-54 |
| ABV91286.1 | type\_I\_modular\_polyketide\_synthase | BGC0000158 | Polyketide:Modular type I | 30.0 | 45.8 | 212.0 | 5.8e-54 |
| ADB23403.1 | polyketide\_synthase\_type\_I | BGC0001062 | Polyketide | 29.0 | 49.0 | 212.0 | 5.8e-54 |
| AWC08658.1 | polyketide\_synthase\_type\_I | BGC0001932 | Polyketide | 29.0 | 44.6 | 212.0 | 5.8e-54 |
| ANZ52461.1 | MonAIII | BGC0001670 | Polyketide | 29.0 | 47.9 | 212.0 | 5.8e-54 |
| ASA76632.1 | polyketide\_synthase\_non-ribosomal\_peptide\_synthetase\_hybrid | BGC0001751 | NRP + Polyketide | 35.0 | 32.7 | 212.0 | 5.8e-54 |
| CAE45670.1 | borrelidin\_polyketide\_synthase,\_type\_I | BGC0000031 | Polyketide:Modular type I | 28.0 | 43.5 | 211.0 | 7.5e-54 |
| AAZ94388.1 | nodular\_polyketide\_synthase | BGC0000040 | Polyketide | 28.0 | 47.7 | 211.0 | 7.5e-54 |
| BAJ16471.1 | polyketide\_synthase | BGC0000058 | Polyketide | 28.0 | 49.2 | 211.0 | 7.5e-54 |
| AAP42856.1 | NanA2 | BGC0000105 | Polyketide | 28.0 | 48.6 | 211.0 | 7.5e-54 |
| CBD77734.1 | polyketide\_synthase | BGC0000974 | NRP + Polyketide | 36.0 | 31.9 | 211.0 | 7.5e-54 |
| CAQ34929.1 | putative\_polyketide\_synthase | BGC0000986 | NRP + Polyketide | 29.0 | 44.2 | 211.0 | 7.5e-54 |
| AEZ64505.1 | Herb | BGC0001065 | Polyketide | 30.0 | 46.2 | 211.0 | 7.5e-54 |
| AEU17898.1 | putative\_type\_I\_PKS | BGC0001072 | Saccharide + Polyketide:Modular type I + Polyketide:Type II + Other:Aminocoumarin | 28.0 | 49.1 | 211.0 | 7.5e-54 |
| AHE80992.1 | PieA2 | BGC0001169 | Polyketide:Modular type I | 32.0 | 32.1 | 211.0 | 7.5e-54 |
| ARM20281.1 | polyketide\_synthase | BGC0001523 | Polyketide | 28.0 | 48.7 | 211.0 | 7.5e-54 |
| AIR74911.1 | polyketide\_synthase | BGC0001559 | RiPP | 36.0 | 31.9 | 211.0 | 7.5e-54 |
| AVX51108.1 | nysC | BGC0001709 | Polyketide | 28.0 | 47.9 | 211.0 | 7.5e-54 |
| ARV85760.1 | PieA1\_type\_I\_PKS | BGC0001742 | Polyketide | 28.0 | 48.7 | 211.0 | 7.5e-54 |
| ATX68112.1 | malonyl\_CoA-acyl\_carrier\_protein\_transacylase | BGC0001772 | Polyketide | 34.0 | 33.1 | 211.0 | 7.5e-54 |
| CAQ64692.1 | lasalocid\_modular\_polyketide\_synthase | BGC0000087 | Polyketide | 28.0 | 47.9 | 211.0 | 9.9e-54 |
| BAC57032.1 | protomycinolide\_IV\_synthase\_5 | BGC0000102 | Polyketide | 29.0 | 48.7 | 211.0 | 9.9e-54 |
| AFI57005.1 | QmnA1 | BGC0000133 | Polyketide | 29.0 | 46.5 | 211.0 | 9.9e-54 |
| CAQ43079.1 | polyketide\_synthase | BGC0000970 | NRP + Polyketide:Modular type I | 30.0 | 49.8 | 211.0 | 9.9e-54 |
| BAR73021.1 | putative\_PKS\_(KS-AT-KR-ACP) | BGC0001194 | Polyketide | 29.0 | 43.6 | 211.0 | 9.9e-54 |
| ALA09356.1 | type\_I\_modular\_PKS | BGC0001303 | Polyketide | 30.0 | 42.1 | 211.0 | 9.9e-54 |
| WP\_048832936.1 | polyketide\_synthase | BGC0001348 | Polyketide:Modular type I | 28.0 | 49.8 | 211.0 | 9.9e-54 |
| ARW71483.1 | type\_I\_PKS\_loading\_module,\_module\_1,\_module\_2 | BGC0001812 | Polyketide | 31.0 | 44.5 | 211.0 | 9.9e-54 |
| AEK75503.1 | type\_1\_polyketide\_synthase | BGC0000001 | Polyketide:Modular type I | 34.0 | 31.9 | 210.0 | 1.3e-53 |
| BAC68126.1 | modular\_polyketide\_synthase | BGC0000059 | Polyketide | 29.0 | 44.6 | 210.0 | 1.3e-53 |
| BAF02924.1 | type\_I\_polyketide\_synthase | BGC0000073 | Polyketide | 26.0 | 47.2 | 210.0 | 1.3e-53 |
| AFV30250.1 | polyketide\_synthase | BGC0000075 | Polyketide | 27.0 | 48.4 | 210.0 | 1.3e-53 |
| ctg1\_orf16 |  | BGC0001457 | NRP | 34.0 | 32.3 | 210.0 | 1.3e-53 |
| AVV61980.1 | type\_I\_modular\_PKS | BGC0001477 | NRP + Polyketide:Modular type I | 34.0 | 32.3 | 210.0 | 1.3e-53 |
| ATL73034.1 | type\_I\_modular\_polyketide\_synthase | BGC0001807 | NRP + Polyketide | 30.0 | 48.7 | 210.0 | 1.3e-53 |
| DAC80076.1 | PKS | BGC0001836 | Polyketide:Trans-AT type I | 32.0 | 40.5 | 210.0 | 1.3e-53 |
| ACO94470.1 | polyketide\_synthase\_type\_I | BGC0000029 | Polyketide:Modular type I | 29.0 | 47.1 | 210.0 | 1.7e-53 |
| CAE45668.1 | borrelidin\_polyketide\_synthase,\_type\_I | BGC0000031 | Polyketide:Modular type I | 29.0 | 43.8 | 210.0 | 1.7e-53 |
| AAC01712.2 | RifC | BGC0000136 | Polyketide | 29.0 | 47.6 | 210.0 | 1.7e-53 |
| CAL58686.1 | polyketide\_synthase | BGC0000149 | Polyketide:Modular type I | 29.0 | 47.0 | 210.0 | 1.7e-53 |
| AVV61982.1 | type\_I\_modular\_PKS | BGC0001477 | NRP + Polyketide:Modular type I | 29.0 | 48.9 | 210.0 | 1.7e-53 |
| ASA76631.1 | polyketide\_synthase | BGC0001751 | NRP + Polyketide | 32.0 | 39.4 | 210.0 | 1.7e-53 |
| AXI91550.1 | FunP3 | BGC0001944 | Polyketide | 28.0 | 56.8 | 210.0 | 1.7e-53 |
| AHH99926.1 | PKS\_I | BGC0000002 | Polyketide | 25.0 | 67.1 | 210.0 | 2.2e-53 |
| BAF02923.1 | type\_I\_polyketide\_synthase | BGC0000073 | Polyketide | 33.0 | 32.8 | 210.0 | 2.2e-53 |
| BAF02927.1 | type\_I\_polyketide\_synthase | BGC0000073 | Polyketide | 28.0 | 48.0 | 210.0 | 2.2e-53 |
| ACB46486.1 | polyketide\_synthase | BGC0000082 | Polyketide | 29.0 | 47.4 | 210.0 | 2.2e-53 |
| AFL48526.1 | laidlomycin\_polyketide\_synthase\_(module\_2) | BGC0000084 | Polyketide | 27.0 | 47.9 | 210.0 | 2.2e-53 |
| CAJ88185.2 | Type\_I\_modular\_polyketide\_synthase | BGC0000151 | Polyketide:Modular type I + Saccharide:Hybrid/tailoring | 28.0 | 49.2 | 210.0 | 2.2e-53 |
| ACR50773.1 | polyketide\_synthase | BGC0000163 | Polyketide | 28.0 | 48.8 | 210.0 | 2.2e-53 |
| AFU82615.1 | polyketide\_synthase | BGC0000998 | NRP + Polyketide | 31.0 | 43.8 | 210.0 | 2.2e-53 |
| ADB23402.1 | polyketide\_synthase\_type\_I | BGC0001062 | Polyketide | 37.0 | 32.4 | 210.0 | 2.2e-53 |
| ABI91470.1 | beta-ketoacyl\_synthase | BGC0001094 | NRP + Polyketide | 24.0 | 88.3 | 210.0 | 2.2e-53 |
| CCP20051.1 | divM\_protein | BGC0001119 | Polyketide:Modular type I | 33.0 | 33.5 | 210.0 | 2.2e-53 |
| CRI73798.1 | CongD\_protein | BGC0001215 | NRP | 33.0 | 32.0 | 210.0 | 2.2e-53 |
| CTQ34883.1 | AtcF;\_polyketide\_synthase,\_modules\_8-10 | BGC0001301 | Polyketide | 34.0 | 33.4 | 210.0 | 2.2e-53 |
| ALA09357.1 | type\_I\_modular\_PKS | BGC0001303 | Polyketide | 35.0 | 31.1 | 210.0 | 2.2e-53 |
| ANH11410.1 | SceO | BGC0001908 | Polyketide | 29.0 | 49.2 | 210.0 | 2.2e-53 |
| ctg1\_12 |  | BGC0001931 | Polyketide | 35.0 | 32.2 | 210.0 | 2.2e-53 |
| ABC84461.1 | NigAVI | BGC0000114 | Polyketide:Modular type I | 28.0 | 50.8 | 209.0 | 2.9e-53 |
| AAF71766.1 | nysI | BGC0000115 | Polyketide:Modular type I + Saccharide:Hybrid/tailoring | 28.0 | 49.2 | 209.0 | 2.9e-53 |
| AFI57006.1 | QmnA2 | BGC0000133 | Polyketide | 28.0 | 48.5 | 209.0 | 2.9e-53 |
| CCP20047.1 | divK\_protein | BGC0001119 | Polyketide:Modular type I | 27.0 | 48.9 | 209.0 | 2.9e-53 |
| CQR60496.1 | Polyketide\_synthase,\_type\_I,\_modules:\_4,\_5\_and\_6 | BGC0001287 | Polyketide | 28.0 | 48.0 | 209.0 | 2.9e-53 |
| OJF16266.1 | AceP4 | BGC0001491 | Polyketide | 27.0 | 47.5 | 209.0 | 2.9e-53 |
| ANY10600.1 | polyketide\_synthase | BGC0001773 | Polyketide | 28.0 | 49.4 | 209.0 | 2.9e-53 |
| AQV04230.1 | SwnK | BGC0001794 | Polyketide | 34.0 | 31.7 | 209.0 | 2.9e-53 |
| BAF02922.1 | type\_I\_polyketide\_synthase | BGC0000073 | Polyketide | 27.0 | 47.9 | 209.0 | 3.7e-53 |
| AFL48527.1 | laidlomycin\_polyketide\_synthase\_(module\_3\_and\_module\_4) | BGC0000084 | Polyketide | 29.0 | 48.3 | 209.0 | 3.7e-53 |
| BAD38874.1 | polyketide\_synthase | BGC0000111 | Polyketide | 35.0 | 32.4 | 209.0 | 3.7e-53 |
| ACY01392.1 | AT-less\_polyketide\_synthase | BGC0000177 | Polyketide:Modular type I + Polyketide:Trans-AT type I | 32.0 | 33.2 | 209.0 | 3.7e-53 |
| ABC36687.1 | polyketide\_synthase | BGC0000964 | NRP:Cyclic depsipeptide + Polyketide:Trans-AT type I | 36.0 | 28.8 | 209.0 | 3.7e-53 |
| CAD15508.1 | polyketide\_synthase/non-ribosomal\_peptide\_synthetase | BGC0001014 | NRP:NRP siderophore + Polyketide:Modular type I + Polyketide:Iterative type I | 31.0 | 45.1 | 209.0 | 3.7e-53 |
| ABF89568.1 | polyketide\_synthase | BGC0001025 | NRP + Polyketide:Trans-AT type I | 31.0 | 43.2 | 209.0 | 3.7e-53 |
| ABI91466.1 | beta-ketoacyl\_synthase | BGC0001094 | NRP + Polyketide | 35.0 | 32.1 | 209.0 | 3.7e-53 |
| ABC35796.1 | polyketide\_synthase,\_putative | BGC0001102 | NRP:Beta-lactam + Polyketide:Modular type I | 36.0 | 32.4 | 209.0 | 3.7e-53 |
| ctg1\_orf524 |  | BGC0001199 | Polyketide | 30.0 | 40.1 | 209.0 | 3.7e-53 |
| ALA09358.1 | type\_I\_modular\_PKS | BGC0001303 | Polyketide | 29.0 | 46.6 | 209.0 | 3.7e-53 |
| sipP4 | Type\_I\_Modular\_PKS | BGC0001452 | Polyketide | 35.0 | 31.2 | 209.0 | 3.7e-53 |
| AVI57433.1 | AbmB1 | BGC0001694 | Polyketide | 29.0 | 45.2 | 209.0 | 3.7e-53 |
| ABY21539.1 | AngAII | BGC0000018 | Polyketide | 33.0 | 32.4 | 208.0 | 4.9e-53 |
| AAO65797.1 | monensin\_polyketide\_synthase\_module\_2 | BGC0000100 | Polyketide | 28.0 | 48.2 | 208.0 | 4.9e-53 |
| AAB66505.1 | tylactone\_synthase\_module\_3 | BGC0000166 | Polyketide | 28.0 | 49.5 | 208.0 | 4.9e-53 |
| CAD89773.1 | MelB\_protein | BGC0001010 | NRP + Polyketide:Modular type I | 29.0 | 48.8 | 208.0 | 4.9e-53 |
| ADH04640.1 | TgaB | BGC0001051 | NRP + Polyketide:Modular type I | 30.0 | 44.4 | 208.0 | 4.9e-53 |
| AHE80996.1 | PieA6 | BGC0001169 | Polyketide:Modular type I | 33.0 | 32.0 | 208.0 | 4.9e-53 |
| AGC09486.1 | LobS3 | BGC0001183 | Polyketide | 30.0 | 44.8 | 208.0 | 4.9e-53 |
| AJW65410.1 | type\_I\_modular\_polyketide\_synthase | BGC0001195 | NRP + Polyketide | 34.0 | 32.2 | 208.0 | 4.9e-53 |
| BAQ25481.1 | type\_I\_polyketide\_synthase | BGC0001288 | Polyketide | 33.0 | 32.1 | 208.0 | 4.9e-53 |
| ALV82345.1 | borrelidin\_type\_I\_polyketide\_synthase | BGC0001533 | Polyketide | 29.0 | 43.3 | 208.0 | 4.9e-53 |
| ANZ52460.1 | MonAII | BGC0001670 | Polyketide | 28.0 | 48.2 | 208.0 | 4.9e-53 |
| CAE02606.1 | polyketide\_synthase\_type\_I | BGC0000024 | Polyketide:Modular type I | 25.0 | 64.1 | 208.0 | 6.4e-53 |
| AAZ94387.1 | modular\_polyketide\_synthase | BGC0000040 | Polyketide | 28.0 | 49.1 | 208.0 | 6.4e-53 |
| ACO94488.1 | polyketide\_synthase\_type\_I | BGC0000097 | Polyketide:Modular type I | 29.0 | 44.5 | 208.0 | 6.4e-53 |
| CBZ41586.1 | Type\_I\_modular\_polyketide\_synthase | BGC0000151 | Polyketide:Modular type I + Saccharide:Hybrid/tailoring | 25.0 | 68.5 | 208.0 | 6.4e-53 |
| AGI99494.1 | Type\_I\_polyketide\_synthase | BGC0001004 | Polyketide:Modular type I | 26.0 | 66.3 | 208.0 | 6.4e-53 |
| AGC09487.1 | LobS5 | BGC0001183 | Polyketide | 26.0 | 66.5 | 208.0 | 6.4e-53 |
| TGZ15167.1 | polyketide\_synthase | BGC0002032 | Polyketide | 28.0 | 49.3 | 208.0 | 6.4e-53 |
| AAS79460.1 | polyketide\_synthase\_subunit | BGC0000035 | Polyketide | 28.0 | 48.2 | 208.0 | 8.3e-53 |
| ABB52544.1 | putative\_type\_I\_polyketide\_synthase | BGC0000047 | Polyketide | 33.0 | 32.4 | 208.0 | 8.3e-53 |
| AGY62754.1 | EbeB | BGC0000051 | Polyketide | 34.0 | 32.1 | 208.0 | 8.3e-53 |
| BAG85026.1 | putative\_polyketide\_synthase | BGC0000086 | Polyketide | 28.0 | 48.1 | 208.0 | 8.3e-53 |
| CAQ64686.1 | lasalocid\_modular\_polyketide\_synthase | BGC0000087 | Polyketide | 28.0 | 48.1 | 208.0 | 8.3e-53 |
| AAP42858.1 | NanA4 | BGC0000105 | Polyketide | 28.0 | 50.5 | 208.0 | 8.3e-53 |
| AAQ84145.1 | Plm5 | BGC0000123 | Polyketide | 28.0 | 47.3 | 208.0 | 8.3e-53 |
| AAM12913.2 | MmpD | BGC0000182 | Polyketide:Iterative type I + Polyketide:Trans-AT type I | 35.0 | 33.0 | 208.0 | 8.3e-53 |
| AHE80993.1 | PieA3 | BGC0001169 | Polyketide:Modular type I | 24.0 | 82.6 | 208.0 | 8.3e-53 |
| AJW65408.1 | type\_I\_modular\_polyketide\_synthase | BGC0001195 | NRP + Polyketide | 28.0 | 49.1 | 208.0 | 8.3e-53 |
| BAQ25512.1 | type\_I\_polyketide\_synthase | BGC0001288 | Polyketide | 32.0 | 32.0 | 208.0 | 8.3e-53 |
| ALV82346.1 | borrelidin\_type\_I\_polyketide\_synthase | BGC0001533 | Polyketide | 29.0 | 43.7 | 208.0 | 8.3e-53 |
| SCN11950.1 | EbeB-type\_I\_polyketide\_synthase | BGC0001580 | Polyketide | 34.0 | 32.1 | 208.0 | 8.3e-53 |
| AVI57435.1 | AbmB3 | BGC0001694 | Polyketide | 28.0 | 44.9 | 208.0 | 8.3e-53 |
| ARW71487.1 | type\_I\_PKS\_module\_7 | BGC0001812 | Polyketide | 29.0 | 44.6 | 208.0 | 8.3e-53 |
| AAC46028.1 | polyketide\_synthase\_module\_7 | BGC0000113 | Polyketide | 30.0 | 46.7 | 207.0 | 1.1e-52 |
| ABC84470.1 | NIGAVIII | BGC0000114 | Polyketide:Modular type I | 28.0 | 49.0 | 207.0 | 1.1e-52 |
| ABV83229.1 | CppB | BGC0000116 | Polyketide | 26.0 | 70.3 | 207.0 | 1.1e-52 |
| AEZ54377.1 | PieA4 | BGC0000124 | Polyketide | 28.0 | 47.6 | 207.0 | 1.1e-52 |
| ACB37742.1 | putative\_type\_I\_polyketide\_synthase | BGC0000162 | Polyketide | 29.0 | 48.2 | 207.0 | 1.1e-52 |
| ACB37755.1 | putative\_type\_I\_polyketide\_synthase | BGC0000162 | Polyketide | 28.0 | 45.0 | 207.0 | 1.1e-52 |
| BAE93740.1 | type\_I\_polyketide\_synthase-related\_protein | BGC0000164 | Polyketide | 35.0 | 33.1 | 207.0 | 1.1e-52 |
| CAQ52622.1 | type\_I\_polyketide\_synthase,\_modules\_4-5 | BGC0001066 | Polyketide:Modular type I | 28.0 | 45.7 | 207.0 | 1.1e-52 |
| AGN11881.1 | tstDEF | BGC0001114 | NRP + Polyketide | 34.0 | 32.7 | 207.0 | 1.1e-52 |
| AZF85917.1 | type\_I\_polyketide\_synthase | BGC0001963 | NRP + Polyketide | 27.0 | 48.7 | 207.0 | 1.1e-52 |
| AGY62756.1 | EbeD | BGC0000051 | Polyketide | 33.0 | 32.1 | 207.0 | 1.4e-52 |
| BAC68129.1 | modular\_polyketide\_synthase | BGC0000059 | Polyketide | 29.0 | 48.2 | 207.0 | 1.4e-52 |
| ABB86408.1 | GelA | BGC0000067 | Polyketide | 28.0 | 49.0 | 207.0 | 1.4e-52 |
| ADC45586.1 | modular\_polyketide\_synthase | BGC0000093 | Polyketide | 28.0 | 48.7 | 207.0 | 1.4e-52 |
| ABV97152.1 | Beta-ketoacyl\_synthase | BGC0000137 | Polyketide | 29.0 | 44.2 | 207.0 | 1.4e-52 |
| CAG23966.1 | polyketide\_synthase\_type\_I | BGC0000181 | Polyketide | 31.0 | 34.3 | 207.0 | 1.4e-52 |
| ADH04658.1 | TugB | BGC0001342 | NRP + Polyketide | 29.0 | 47.9 | 207.0 | 1.4e-52 |
| AWC08655.1 | polyketide\_synthase\_type\_I | BGC0001932 | Polyketide | 33.0 | 32.0 | 207.0 | 1.4e-52 |
| ARS01473.1 | NcmAI | BGC0001702 | NRP + Polyketide | 33.0 | 32.5 | 207.0 | 1.4e-52 |
| WP\_012753526.1 | polyketide\_synthase | BGC0001991 | Polyketide | 34.0 | 33.1 | 207.0 | 1.4e-52 |
| BAO66543.1 | type\_I\_polyketide\_synthase | BGC0000042 | Polyketide | 29.0 | 48.3 | 207.0 | 1.9e-52 |
| AAM81585.1 | putative\_type\_I\_polyketide\_synthase | BGC0000047 | Polyketide | 33.0 | 31.9 | 207.0 | 1.9e-52 |
| ACB46488.1 | polyketide\_synthase | BGC0000082 | Polyketide | 33.0 | 32.4 | 207.0 | 1.9e-52 |
| BAG85032.1 | putative\_polyketide\_synthase | BGC0000086 | Polyketide | 28.0 | 47.2 | 207.0 | 1.9e-52 |
| AAO65806.1 | monensin\_polyketide\_synthase\_modules\_11\_and\_12 | BGC0000100 | Polyketide | 28.0 | 48.0 | 207.0 | 1.9e-52 |
| AAP42874.1 | NanA8 | BGC0000105 | Polyketide | 28.0 | 48.5 | 207.0 | 1.9e-52 |
| AEZ54378.1 | PieA5 | BGC0000124 | Polyketide | 28.0 | 49.3 | 207.0 | 1.9e-52 |
| CAI94713.1 | putative\_polyketide\_synthase | BGC0000141 | Polyketide | 29.0 | 46.3 | 207.0 | 1.9e-52 |
| CAN93350.1 | polyketide\_synthase | BGC0000179 | Polyketide:Trans-AT type I | 34.0 | 32.4 | 207.0 | 1.9e-52 |
| AFX60336.1 | polyketide\_synthase | BGC0001032 | NRP + Polyketide | 33.0 | 33.0 | 207.0 | 1.9e-52 |
| CAQ52623.1 | type\_I\_polyketide\_synthase,\_module\_6 | BGC0001066 | Polyketide:Modular type I | 33.0 | 32.1 | 207.0 | 1.9e-52 |
| ctg1\_orf522 |  | BGC0001199 | Polyketide | 31.0 | 39.6 | 207.0 | 1.9e-52 |
| BAQ21946.1 | putative\_type\_I\_polyketide\_synthase | BGC0001204 | Polyketide | 33.0 | 32.3 | 207.0 | 1.9e-52 |
| ANC94963.1 | AlmHIV | BGC0001396 | Polyketide | 33.0 | 32.4 | 207.0 | 1.9e-52 |
| AQZ37095.1 | polyketide\_synthase | BGC0001511 | Polyketide | 35.0 | 32.2 | 207.0 | 1.9e-52 |
| ATY69569.1 | type\_I\_polyketide\_synthase | BGC0001611 | NRP + Polyketide | 36.0 | 32.0 | 207.0 | 1.9e-52 |
| BBA84067.1 | type\_I\_polyketide\_synthase | BGC0001916 | Polyketide | 29.0 | 47.9 | 207.0 | 1.9e-52 |
| ANZ52469.1 | MonAVIII | BGC0001670 | Polyketide | 28.0 | 48.0 | 207.0 | 1.9e-52 |
| ARR97037.1 | SphD | BGC0001780 | NRP | 30.0 | 42.4 | 207.0 | 1.9e-52 |
| AAX98187.1 | polyketide\_synthase\_type\_I | BGC0000052 | Polyketide | 34.0 | 32.4 | 206.0 | 2.4e-52 |
| ACN69990.1 | polyketide\_synthase | BGC0000079 | Polyketide | 34.0 | 32.1 | 206.0 | 2.4e-52 |
| ABG02264.1 | SalB | BGC0000143 | Polyketide | 34.0 | 32.3 | 206.0 | 2.4e-52 |
| ADU86002.1 | putative\_modular\_polyketide\_synthase | BGC0000165 | Polyketide:Modular type I | 34.0 | 32.1 | 206.0 | 2.4e-52 |
| ALP32043.1 | CycC | BGC0001293 | Polyketide | 29.0 | 47.8 | 206.0 | 2.4e-52 |
| AQT01382.1 | SgnS1 | BGC0001690 | Polyketide | 32.0 | 32.0 | 206.0 | 2.4e-52 |
| ARW71484.1 | type\_I\_PKS\_module\_3 | BGC0001812 | Polyketide | 33.0 | 33.6 | 206.0 | 2.4e-52 |
| AXI91551.1 | FunP2 | BGC0001944 | Polyketide | 34.0 | 32.0 | 206.0 | 2.4e-52 |
| CAE45671.1 | borrelidin\_polyketide\_synthase,\_type\_I | BGC0000031 | Polyketide:Modular type I | 28.0 | 43.3 | 206.0 | 3.2e-52 |
| ACY01402.1 | AT-less\_polyketide\_synthase | BGC0000083 | Polyketide:Modular type I + Polyketide:Trans-AT type I | 32.0 | 32.7 | 206.0 | 3.2e-52 |
| ADC45515.1 | modular\_polyketide\_synthase | BGC0000093 | Polyketide | 30.0 | 46.6 | 206.0 | 3.2e-52 |
| ADC45534.1 | modular\_polyketide\_synthase | BGC0000093 | Polyketide | 27.0 | 48.7 | 206.0 | 3.2e-52 |
| CAO85897.1 | modular\_polyketide\_synthase\_NorB | BGC0000110 | Polyketide:Modular type I | 33.0 | 32.0 | 206.0 | 3.2e-52 |
| ABB05102.1 | LipPks1 | BGC0001003 | NRP:Lipopeptide + Polyketide:Modular type I + Saccharide:Hybrid/tailoring | 28.0 | 49.6 | 206.0 | 3.2e-52 |
| AFR69333.1 | polyketide\_synthase\_SpiC1 | BGC0001045 | NRP:Cyclic depsipeptide + Polyketide:Modular type I | 36.0 | 31.9 | 206.0 | 3.2e-52 |
| B073\_RS40860 | type\_I\_polyketide\_synthase | BGC0001332 | NRP + Polyketide | 29.0 | 48.4 | 206.0 | 3.2e-52 |
| OJF16268.1 | AceP6 | BGC0001491 | Polyketide | 33.0 | 31.9 | 206.0 | 3.2e-52 |
| AQZ37114.1 | polyketide\_synthase | BGC0001511 | Polyketide | 23.0 | 109.4 | 206.0 | 3.2e-52 |
| AZF85941.1 | type\_I\_polyketide\_synthase | BGC0001963 | NRP + Polyketide | 29.0 | 47.3 | 206.0 | 3.2e-52 |
| AHA38202.1 | GphI | BGC0000069 | Polyketide | 29.0 | 44.9 | 205.0 | 4.1e-52 |
| AFV30251.1 | polyketide\_synthase | BGC0000075 | Polyketide | 34.0 | 32.0 | 205.0 | 4.1e-52 |
| BAG85031.1 | putative\_polyketide\_synthase | BGC0000086 | Polyketide | 26.0 | 48.2 | 205.0 | 4.1e-52 |
| ALD82524.1 | polyketide\_synthase | BGC0001212 | NRP + Polyketide | 35.0 | 32.3 | 205.0 | 4.1e-52 |
| AIJ04683.1 | polyketide\_synthase | BGC0001383 | Polyketide | 31.0 | 34.3 | 205.0 | 4.1e-52 |
| ARE67852.1 | AbsB2 | BGC0001492 | Polyketide | 27.0 | 53.3 | 205.0 | 4.1e-52 |
| AWH12665.1 | RmpE1 | BGC0001759 | Polyketide | 33.0 | 32.1 | 205.0 | 4.1e-52 |
| AMYAL\_RS48910 | polyketide\_synthase | BGC0002011 | Polyketide | 31.0 | 42.6 | 205.0 | 4.1e-52 |
| AAM81586.2 | putative\_type\_I\_polyketide\_synthase | BGC0000047 | Polyketide | 33.0 | 32.3 | 205.0 | 5.4e-52 |
| AGY62755.1 | EbeC | BGC0000051 | Polyketide | 27.0 | 47.9 | 205.0 | 5.4e-52 |
| AAO06916.1 | GdmAI | BGC0000066 | Polyketide | 28.0 | 49.0 | 205.0 | 5.4e-52 |
| ABI93779.1 | GdmPKS | BGC0000068 | Polyketide | 27.0 | 48.8 | 205.0 | 5.4e-52 |
| CAE46851.1 | Type\_I\_modular\_polyketide\_synthase | BGC0000103 | Polyketide | 34.0 | 32.4 | 205.0 | 5.4e-52 |
| ABV97155.1 | Acyl\_transferase | BGC0000137 | Polyketide | 33.0 | 32.1 | 205.0 | 5.4e-52 |
| CAL58683.1 | polyketide\_synthase | BGC0000149 | Polyketide:Modular type I | 29.0 | 48.5 | 205.0 | 5.4e-52 |
| BAE93739.1 | type\_I\_polyketide\_synthase-related\_protein | BGC0000164 | Polyketide | 34.0 | 32.8 | 205.0 | 5.4e-52 |
| AFX60317.1 | polyketide\_synthase | BGC0001031 | NRP + Polyketide | 33.0 | 33.1 | 205.0 | 5.4e-52 |
| ADI59533.1 | CorK | BGC0001091 | NRP + Polyketide | 34.0 | 32.7 | 205.0 | 5.4e-52 |
| ctg1\_orf255 |  | BGC0001200 | Polyketide | 27.0 | 49.2 | 205.0 | 5.4e-52 |
| ALP32042.1 | CycB | BGC0001293 | Polyketide | 28.0 | 47.9 | 205.0 | 5.4e-52 |
| SCN11951.1 | ebeC-type\_I\_polyketide\_synthase | BGC0001580 | Polyketide | 27.0 | 47.9 | 205.0 | 5.4e-52 |
| AWC08656.1 | polyketide\_synthase\_type\_I | BGC0001932 | Polyketide | 28.0 | 48.8 | 205.0 | 5.4e-52 |
| ACO94460.1 | polyketide\_synthase\_type\_I | BGC0000029 | Polyketide:Modular type I | 29.0 | 44.6 | 205.0 | 7.1e-52 |
| ADC45516.1 | modular\_polyketide\_synthase | BGC0000093 | Polyketide | 30.0 | 45.6 | 205.0 | 7.1e-52 |
| AAQ84146.1 | Plm6 | BGC0000123 | Polyketide | 27.0 | 49.3 | 205.0 | 7.1e-52 |
| ABV97154.1 | Beta-ketoacyl\_synthase | BGC0000137 | Polyketide | 28.0 | 43.6 | 205.0 | 7.1e-52 |
| ABF92489.1 | mixed\_type\_I\_polyketide\_synthase\_-\_peptide\_synthetase | BGC0001025 | NRP + Polyketide:Trans-AT type I | 32.0 | 32.8 | 205.0 | 7.1e-52 |
| AUS29495.1 | polyketide\_synthase | BGC0001030 | NRP + Polyketide | 34.0 | 32.7 | 205.0 | 7.1e-52 |
| CAA60459.1 | polyketide\_synthase | BGC0001040 | NRP + Polyketide | 34.0 | 31.8 | 205.0 | 7.1e-52 |
| AHH34189.1 | polyketide\_synthase | BGC0001162 | Polyketide:Modular type I | 33.0 | 35.0 | 205.0 | 7.1e-52 |
| AGC09485.1 | LobS2 | BGC0001183 | Polyketide | 34.0 | 32.3 | 205.0 | 7.1e-52 |
| ANC94966.1 | AlmHI | BGC0001396 | Polyketide | 34.0 | 32.4 | 205.0 | 7.1e-52 |
| sipP1 | Type\_I\_Modular\_PKS | BGC0001452 | Polyketide | 33.0 | 32.3 | 205.0 | 7.1e-52 |
| AKG06375.1 | polyketide\_synthase\_type\_1 | BGC0001830 | Polyketide | 35.0 | 32.1 | 205.0 | 7.1e-52 |
| WP\_079030450.1 | type\_I\_polyketide\_synthase | BGC0002033 | Polyketide | 34.0 | 32.3 | 205.0 | 7.1e-52 |
| AAM81584.2 | putative\_type\_I\_polyketide\_synthase | BGC0000047 | Polyketide | 33.0 | 32.4 | 204.0 | 9.2e-52 |
| BAC68127.1 | modular\_polyketide\_synthase | BGC0000059 | Polyketide | 33.0 | 32.6 | 204.0 | 9.2e-52 |
| AFL48528.1 | laidlomycin\_polyketide\_synthase\_(module\_7\_and\_module\_8) | BGC0000084 | Polyketide | 27.0 | 44.0 | 204.0 | 9.2e-52 |
| ctg1\_orf27 |  | BGC0000096 | Polyketide | 28.0 | 48.7 | 204.0 | 9.2e-52 |
| AAO65801.1 | monensin\_polyketide\_synthase\_module\_9 | BGC0000100 | Polyketide | 27.0 | 45.0 | 204.0 | 9.2e-52 |
| ADX66472.1 | ScnS1 | BGC0000108 | Polyketide | 27.0 | 49.8 | 204.0 | 9.2e-52 |
| ABC84459.1 | NigAIV | BGC0000114 | Polyketide:Modular type I | 29.0 | 46.0 | 204.0 | 9.2e-52 |
| BAK64638.1 | polyketide\_synthase | BGC0000135 | Polyketide | 33.0 | 32.4 | 204.0 | 9.2e-52 |
| ABP55223.1 | beta-ketoacyl\_synthase | BGC0000142 | Polyketide | 33.0 | 31.2 | 204.0 | 9.2e-52 |
| EGJ35088.1 | Polyketide\_synthase | BGC0001163 | Polyketide:Modular type I | 33.0 | 35.5 | 204.0 | 9.2e-52 |
| ALD83703.1 | tAT\_polyketide\_synthase | BGC0001299 | Polyketide | 35.0 | 32.7 | 204.0 | 9.2e-52 |
| ANR02549.1 | LodH | BGC0001648 | Polyketide | 32.0 | 32.3 | 204.0 | 9.2e-52 |
| AWC08662.1 | polyketide\_synthase\_type\_I | BGC0001932 | Polyketide | 32.0 | 32.4 | 204.0 | 9.2e-52 |
| ANZ52464.1 | MonAVI | BGC0001670 | Polyketide | 27.0 | 45.0 | 204.0 | 9.2e-52 |
| BBA20952.1 | type\_I\_polyketide\_synthase | BGC0001763 | NRP + Polyketide | 36.0 | 31.9 | 204.0 | 9.2e-52 |
| TGZ15168.1 | hypothetical\_protein | BGC0002032 | Polyketide | 33.0 | 32.4 | 204.0 | 9.2e-52 |
| ABK32259.1 | AmbE | BGC0000014 | Polyketide | 33.0 | 32.2 | 204.0 | 1.2e-51 |
| AAM54078.1 | polyketide\_synthase | BGC0000020 | Polyketide | 35.0 | 32.2 | 204.0 | 1.2e-51 |
| BAF85837.1 | modular\_polyketide\_synthase | BGC0000109 | Polyketide | 27.0 | 48.2 | 204.0 | 1.2e-51 |
| CAO85893.1 | modular\_polyketide\_synthase\_NorA | BGC0000110 | Polyketide:Modular type I | 27.0 | 48.1 | 204.0 | 1.2e-51 |
| ABP55220.1 | beta-ketoacyl\_synthase | BGC0000142 | Polyketide | 33.0 | 31.2 | 204.0 | 1.2e-51 |
| AEZ53951.1 | polyketide\_synthase | BGC0000144 | Polyketide:Modular type I | 27.0 | 51.8 | 204.0 | 1.2e-51 |
| CAN93348.1 | polyketide\_synthase | BGC0000179 | Polyketide:Trans-AT type I | 32.0 | 44.0 | 204.0 | 1.2e-51 |
| AAY89051.1 | hybrid\_nonribosomal\_peptide\_synthetase/polyketide\_synthase | BGC0001069 | NRP + Polyketide:Trans-AT type I | 32.0 | 39.9 | 204.0 | 1.2e-51 |
| BAT51067.1 | type\_I\_polyketide\_synthase | BGC0001296 | Polyketide | 28.0 | 50.5 | 204.0 | 1.2e-51 |
| AXI91552.1 | FunP1 | BGC0001944 | Polyketide | 33.0 | 32.0 | 204.0 | 1.2e-51 |
| AFL48533.1 | laidlomycin\_polyketide\_synthase\_(module\_10) | BGC0000084 | Polyketide | 28.0 | 47.6 | 203.0 | 1.6e-51 |
| ACR50782.1 | polyketide\_synthase | BGC0000163 | Polyketide | 28.0 | 48.8 | 203.0 | 1.6e-51 |
| CAL69889.1 | RhiB\_protein | BGC0001112 | NRP + Polyketide:Trans-AT type I | 34.0 | 33.4 | 203.0 | 1.6e-51 |
| ALA09355.1 | type\_I\_modular\_PKS | BGC0001303 | Polyketide | 29.0 | 46.9 | 203.0 | 1.6e-51 |
| AWR88404.1 | putative\_beta-ketoacyl\_synthase | BGC0001522 | Polyketide | 33.0 | 32.3 | 203.0 | 1.6e-51 |
| AXG22405.1 | type\_I\_polyketide\_synthase | BGC0002024 | Polyketide | 29.0 | 45.0 | 203.0 | 1.6e-51 |
| ADC79616.1 | BafAI | BGC0000028 | Polyketide:Modular type I | 32.0 | 32.3 | 203.0 | 2.1e-51 |
| ABB88520.1 | polyketide\_synthase\_type\_I | BGC0000050 | Polyketide | 34.0 | 32.7 | 203.0 | 2.1e-51 |
| AFL48520.1 | laidlomycin\_polyketide\_synthase\_(module\_9) | BGC0000084 | Polyketide | 26.0 | 44.6 | 203.0 | 2.1e-51 |
| ADM46357.1 | polyketide\_synthase | BGC0000106 | Polyketide | 27.0 | 48.2 | 203.0 | 2.1e-51 |
| CAJ57409.1 | polyketide\_synthase\_type\_I | BGC0000176 | Polyketide + NRP | 30.0 | 42.9 | 203.0 | 2.1e-51 |
| ACY01391.1 | AT-less\_polyketide\_synthase | BGC0000177 | Polyketide:Modular type I + Polyketide:Trans-AT type I | 33.0 | 31.1 | 203.0 | 2.1e-51 |
| ACY06286.1 | polyketide\_synthase | BGC0001042 | NRP + Polyketide | 32.0 | 32.4 | 203.0 | 2.1e-51 |
| AVV61985.1 | beta-ketoacyl\_synthase | BGC0001477 | NRP + Polyketide:Modular type I | 27.0 | 48.7 | 203.0 | 2.1e-51 |
| ASX95227.1 | IlaE | BGC0001620 | Polyketide | 36.0 | 31.9 | 203.0 | 2.1e-51 |
| AWC08663.1 | polyketide\_synthase\_type\_I | BGC0001932 | Polyketide | 28.0 | 48.5 | 203.0 | 2.1e-51 |
| AVX51100.1 | nysK | BGC0001709 | Polyketide | 33.0 | 32.2 | 203.0 | 2.1e-51 |
| AWH12668.1 | RmpC | BGC0001759 | Polyketide | 28.0 | 46.9 | 203.0 | 2.1e-51 |
| TXD00261.1 | AMP-binding\_protein | BGC0001877 | Polyketide | 34.0 | 33.1 | 203.0 | 2.1e-51 |
| FS847\_01975 | type\_I\_polyketide\_synthase | BGC0001877 | Polyketide | 33.0 | 32.8 | 203.0 | 2.1e-51 |
| AAZ77699.1 | ChlA6 | BGC0000036 | Polyketide:Modular type I + Polyketide:Iterative type I + Saccharide:Oligosaccharide | 28.0 | 48.3 | 203.0 | 2.7e-51 |
| BAO66519.1 | type\_I\_polyketide\_synthase | BGC0000042 | Polyketide | 34.0 | 32.9 | 203.0 | 2.7e-51 |
| BAP34734.1 | type\_I\_polyketide\_synthase | BGC0000078 | Polyketide | 33.0 | 32.7 | 203.0 | 2.7e-51 |
| ctg1\_orf31 |  | BGC0000096 | Polyketide | 35.0 | 32.4 | 203.0 | 2.7e-51 |
| CAE46850.1 | Type\_I\_modular\_polyketide\_synthase | BGC0000103 | Polyketide | 34.0 | 31.8 | 203.0 | 2.7e-51 |
| AAC01714.1 | RifE | BGC0000136 | Polyketide | 33.0 | 32.1 | 203.0 | 2.7e-51 |
| BAO98805.1 | putative\_polyketide\_synthase | BGC0001002 | NRP + Polyketide | 28.0 | 48.5 | 203.0 | 2.7e-51 |
| AJQ95708.1 | polyketide\_synthase\_modules-related\_protein | BGC0001644 | Polyketide | 31.0 | 33.5 | 203.0 | 2.7e-51 |
| ANR02556.1 | LodO | BGC0001648 | Polyketide | 26.0 | 48.6 | 203.0 | 2.7e-51 |
| AXI91547.1 | FunP6 | BGC0001944 | Polyketide | 28.0 | 58.5 | 203.0 | 2.7e-51 |
| AZF85945.1 | type\_I\_polyketide\_synthase | BGC0001963 | NRP + Polyketide | 33.0 | 32.4 | 203.0 | 2.7e-51 |
| CAE02605.1 | polyketide\_synthase\_type\_I | BGC0000024 | Polyketide:Modular type I | 33.0 | 31.9 | 202.0 | 3.5e-51 |
| AAS79461.1 | polyketide\_synthase\_subunit | BGC0000035 | Polyketide | 33.0 | 32.1 | 202.0 | 3.5e-51 |
| ABB88522.1 | polyketide\_synthase\_type\_I | BGC0000050 | Polyketide | 33.0 | 32.3 | 202.0 | 3.5e-51 |
| BAC76491.1 | lankamycin\_synthase\_LkmAIII | BGC0000085 | Polyketide | 28.0 | 48.7 | 202.0 | 3.5e-51 |
| AAF71768.1 | nysK | BGC0000115 | Polyketide:Modular type I + Saccharide:Hybrid/tailoring | 34.0 | 31.9 | 202.0 | 3.5e-51 |
| AAF71774.1 | nysA | BGC0000115 | Polyketide:Modular type I + Saccharide:Hybrid/tailoring | 33.0 | 32.4 | 202.0 | 3.5e-51 |
| AAQ84156.1 | Plm1 | BGC0000123 | Polyketide | 33.0 | 32.6 | 202.0 | 3.5e-51 |
| AEE88279.1 | CurK | BGC0000976 | NRP + Polyketide:Modular type I | 33.0 | 31.9 | 202.0 | 3.5e-51 |
| AFX60340.1 | polyketide\_synthase | BGC0001032 | NRP + Polyketide | 32.0 | 33.1 | 202.0 | 3.5e-51 |
| AAT70106.1 | CurK | BGC0001165 | NRP + Polyketide:Modular type I | 33.0 | 31.9 | 202.0 | 3.5e-51 |
| AWR88405.1 | putative\_phosphopantetheine-binding\_domain-\_containing\_prot\_ein | BGC0001522 | Polyketide | 33.0 | 32.3 | 202.0 | 3.5e-51 |
| WP\_055480220.1 | type\_I\_polyketide\_synthase | BGC0001653 | Polyketide | 30.0 | 44.2 | 202.0 | 3.5e-51 |
| AAS79462.1 | polyketide\_synthase\_subunit | BGC0000035 | Polyketide | 28.0 | 48.3 | 202.0 | 4.6e-51 |
| AAO65807.1 | monensin\_polyketide\_synthase\_module\_10 | BGC0000100 | Polyketide | 27.0 | 48.0 | 202.0 | 4.6e-51 |
| ABC84471.1 | NigAVII | BGC0000114 | Polyketide:Modular type I | 28.0 | 44.4 | 202.0 | 4.6e-51 |
| ABP57747.1 | DepC | BGC0000993 | NRP:Cyclic depsipeptide + Polyketide:Modular type I | 35.0 | 29.0 | 202.0 | 4.6e-51 |
| CCE88379.1 | polyketide\_synthase | BGC0001034 | NRP + Polyketide:Modular type I | 27.0 | 47.8 | 202.0 | 4.6e-51 |
| BBA66511.1 | type\_I\_polyketide\_synthase | BGC0001495 | Polyketide | 32.0 | 43.3 | 202.0 | 4.6e-51 |
| OAP25815.1 | Phenolphthiocerol\_synthesis\_polyketide\_synthase\_type\_I\_Pks15/1 | BGC0001658 | Polyketide | 23.0 | 81.2 | 202.0 | 4.6e-51 |
| ANZ52470.1 | MonAVII | BGC0001670 | Polyketide | 27.0 | 48.0 | 202.0 | 4.6e-51 |
| AAZ77698.1 | ChlA5 | BGC0000036 | Polyketide:Modular type I + Polyketide:Iterative type I + Saccharide:Oligosaccharide | 27.0 | 48.4 | 201.0 | 6e-51 |
| AAU93805.2 | polyketide\_synthase\_modules\_5\_and\_6 | BGC0000054 | Polyketide | 34.0 | 32.2 | 201.0 | 6e-51 |
| ABB86409.1 | GelB | BGC0000067 | Polyketide | 27.0 | 48.8 | 201.0 | 6e-51 |
| AAY28225.1 | HbmAI | BGC0000074 | Polyketide | 28.0 | 48.8 | 201.0 | 6e-51 |
| AAY28227.1 | HbmAIII | BGC0000074 | Polyketide | 27.0 | 48.7 | 201.0 | 6e-51 |
| ACY01400.1 | AT-less\_polyketide\_synthase | BGC0000083 | Polyketide:Modular type I + Polyketide:Trans-AT type I | 35.0 | 33.8 | 201.0 | 6e-51 |
| AAB66507.1 | tylactone\_synthase\_module\_6 | BGC0000166 | Polyketide | 28.0 | 50.0 | 201.0 | 6e-51 |
| AAS47562.1 | mixed\_type\_I\_polyketide\_synthase\_-\_peptide\_synthetase | BGC0001108 | Polyketide:Trans-AT type I | 33.0 | 32.8 | 201.0 | 6e-51 |
| ctg1\_orf8 |  | BGC0000053 | Polyketide | 28.0 | 45.4 | 198.0 | 8.6e-50 |
| ALD83686.1 | tAT\_polyketide\_synthase | BGC0001300 | Polyketide | 36.0 | 32.2 | 201.0 | 6e-51 |
| AKD43763.1 | HerG | BGC0001349 | NRP + Polyketide | 28.0 | 43.6 | 201.0 | 6e-51 |
| BAO66528.1 | type\_I\_polyketide\_synthase | BGC0000042 | Polyketide | 27.0 | 48.8 | 201.0 | 7.8e-51 |
| ABB88523.1 | polyketide\_synthase\_type\_I | BGC0000050 | Polyketide | 33.0 | 32.0 | 201.0 | 7.8e-51 |
| AAO06918.1 | GdmAIII | BGC0000066 | Polyketide | 26.0 | 48.9 | 201.0 | 7.8e-51 |
| AAF71775.1 | nysB | BGC0000115 | Polyketide:Modular type I + Saccharide:Hybrid/tailoring | 28.0 | 47.3 | 201.0 | 7.8e-51 |
| ABV83221.1 | CppI | BGC0000116 | Polyketide | 25.0 | 68.0 | 201.0 | 7.8e-51 |
| ABV83228.1 | CppA | BGC0000116 | Polyketide | 28.0 | 48.2 | 201.0 | 7.8e-51 |
| AEZ54374.1 | PieA1 | BGC0000124 | Polyketide | 27.0 | 47.5 | 201.0 | 7.8e-51 |
| ADR01054.1 | Noc-11 | BGC0000609 | RiPP:Thiopeptide | 33.0 | 32.4 | 201.0 | 7.8e-51 |
| AFX60313.1 | polyketide\_synthase | BGC0001031 | NRP + Polyketide | 32.0 | 33.1 | 201.0 | 7.8e-51 |
| CCA89326.1 | mixed\_trans-AT\_type\_I\_polyketide\_synthase/nonribosomal\_peptide\_synthetase | BGC0001111 | NRP + Polyketide:Trans-AT type I | 32.0 | 39.3 | 201.0 | 7.8e-51 |
| EGD99348.1 | polyketide\_synthase | BGC0001144 | Polyketide | 28.0 | 48.8 | 201.0 | 7.8e-51 |
| BAV19379.1 | polyketide\_synthase | BGC0001390 | NRP + Polyketide | 29.0 | 45.6 | 201.0 | 7.8e-51 |
| ASZ00151.1 | polyketide\_synthase | BGC0001785 | Polyketide | 32.0 | 32.7 | 201.0 | 7.8e-51 |
| ACY01401.1 | AT-less\_polyketide\_synthase | BGC0000083 | Polyketide:Modular type I + Polyketide:Trans-AT type I | 31.0 | 32.8 | 201.0 | 1e-50 |
| CAO85896.1 | protein\_modular\_polyketide\_synthase\_NorA' | BGC0000110 | Polyketide:Modular type I | 28.0 | 47.9 | 201.0 | 1e-50 |
| CAQ43078.1 | polyketide\_synthase | BGC0000970 | NRP + Polyketide:Modular type I | 33.0 | 31.7 | 201.0 | 1e-50 |
| CBA11584.1 | polyketide\_synthase\_type\_I | BGC0001046 | NRP + Polyketide:Modular type I + Saccharide:Hybrid/tailoring | 33.0 | 32.0 | 201.0 | 1e-50 |
| ctg1\_orf254 |  | BGC0001200 | Polyketide | 32.0 | 32.1 | 201.0 | 1e-50 |
| ARV85761.1 | PieA2\_type\_I\_PKS | BGC0001742 | Polyketide | 27.0 | 49.1 | 201.0 | 1e-50 |
| ANH11413.1 | SceR | BGC0001908 | Polyketide | 33.0 | 31.0 | 201.0 | 1e-50 |
| ATY69589.1 | type\_I\_polyketide\_synthase | BGC0001823 | NRP + Polyketide | 34.0 | 32.7 | 201.0 | 1e-50 |
| AAO06917.1 | GdmAII | BGC0000066 | Polyketide | 27.0 | 48.8 | 200.0 | 1.3e-50 |
| BAQ21947.1 | putative\_type\_I\_polyketide\_synthase | BGC0001204 | Polyketide | 31.0 | 32.0 | 200.0 | 1.3e-50 |
| AKA59093.1 | type-I\_PKS | BGC0001619 | Polyketide | 29.0 | 47.0 | 200.0 | 1.3e-50 |
| BBA84070.1 | type\_I\_polyketide\_synthase | BGC0001916 | Polyketide | 29.0 | 46.6 | 200.0 | 1.3e-50 |
| DAC76734.1 | type\_I\_polyketide\_synthase/non-ribosomal\_peptide\_synthetase | BGC0001885 | Polyketide | 34.0 | 33.3 | 200.0 | 1.3e-50 |
| TGZ15166.1 | hypothetical\_protein | BGC0002032 | Polyketide | 33.0 | 32.9 | 200.0 | 1.3e-50 |
| AHH99921.1 | PKS\_I | BGC0000002 | Polyketide | 27.0 | 47.9 | 200.0 | 1.7e-50 |
| AEC13071.1 | fosE | BGC0000060 | Polyketide | 32.0 | 33.3 | 200.0 | 1.7e-50 |
| BAC57031.1 | protomycinolide\_IV\_synthase\_4 | BGC0000102 | Polyketide | 32.0 | 32.0 | 200.0 | 1.7e-50 |
| ACY06289.1 | type\_I\_polyketide\_synthase | BGC0001042 | NRP + Polyketide | 33.0 | 31.8 | 200.0 | 1.7e-50 |
| ARV85764.1 | PieA5\_type\_I\_PKS | BGC0001742 | Polyketide | 33.0 | 32.3 | 200.0 | 1.7e-50 |
| ASZ00148.1 | polyketide\_synthase | BGC0001785 | Polyketide | 27.0 | 47.8 | 200.0 | 1.7e-50 |
| AQV04224.1 | SwnK | BGC0001793 | Polyketide | 32.0 | 31.7 | 200.0 | 1.7e-50 |
| AAQ82566.1 | FscF | BGC0000061 | Polyketide | 28.0 | 42.7 | 200.0 | 2.3e-50 |
| CAC22144.1 | CpkC;\_Polyketide\_synthase\_module\_5 | BGC0000038 | Polyketide:Modular type I | 32.0 | 31.9 | 200.0 | 2.3e-50 |
| AAF71776.1 | nysC | BGC0000115 | Polyketide:Modular type I + Saccharide:Hybrid/tailoring | 33.0 | 31.9 | 200.0 | 2.3e-50 |
| BAB69196.1 | modular\_polyketide\_synthase | BGC0000117 | Polyketide | 27.0 | 48.1 | 200.0 | 2.3e-50 |
| ABI91469.1 | beta-ketoacyl\_synthase | BGC0001094 | NRP + Polyketide | 31.0 | 40.4 | 200.0 | 2.3e-50 |
| CCA89329.1 | trans-AT\_type\_I\_polyketide\_synthase | BGC0001111 | NRP + Polyketide:Trans-AT type I | 32.0 | 39.5 | 200.0 | 2.3e-50 |
| AKA59092.1 | type-I\_PKS | BGC0001619 | Polyketide | 32.0 | 32.0 | 200.0 | 2.3e-50 |
| CAC22145.1 | CpkB;\_Polyketide\_synthase\_modules\_3\_and\_4 | BGC0000038 | Polyketide:Modular type I | 27.0 | 45.8 | 199.0 | 3e-50 |
| ABB86410.1 | GelC | BGC0000067 | Polyketide | 26.0 | 48.9 | 199.0 | 3e-50 |
| ACN69992.1 | polyketide\_synthase | BGC0000079 | Polyketide | 28.0 | 45.1 | 199.0 | 3e-50 |
| ABV83230.1 | CppC | BGC0000116 | Polyketide | 28.0 | 48.6 | 199.0 | 3e-50 |
| ABP55210.1 | beta-ketoacyl\_synthase | BGC0000142 | Polyketide | 34.0 | 32.5 | 199.0 | 3e-50 |
| ABJ97437.1 | MerA | BGC0001012 | NRP + Polyketide | 32.0 | 32.0 | 199.0 | 3e-50 |
| CQR60495.1 | Polyketide\_synthase,\_type\_I,\_module\_7 | BGC0001287 | Polyketide | 32.0 | 32.1 | 199.0 | 3e-50 |
| AJO72737.1 | Type\_I\_modular\_polyketide\_synthase | BGC0001381 | Polyketide | 33.0 | 31.9 | 199.0 | 3e-50 |
| AAQ82561.1 | FscA | BGC0000061 | Polyketide | 29.0 | 48.4 | 199.0 | 3.9e-50 |
| BAC57030.1 | protomycinolide\_IV\_synthase\_3 | BGC0000102 | Polyketide | 33.0 | 32.1 | 199.0 | 3.9e-50 |
| AAQ84157.1 | Plm2-3 | BGC0000123 | Polyketide | 27.0 | 50.4 | 198.0 | 5.1e-50 |
| ABG02263.1 | SalA | BGC0000143 | Polyketide | 34.0 | 32.1 | 198.0 | 5.1e-50 |
| AAY89052.1 | polyketide\_synthase | BGC0001069 | NRP + Polyketide:Trans-AT type I | 31.0 | 41.8 | 198.0 | 5.1e-50 |
| AVI57434.1 | AbmB2 | BGC0001694 | Polyketide | 33.0 | 32.5 | 198.0 | 5.1e-50 |
| AEZ53953.1 | polyketide\_synthase | BGC0000144 | Polyketide:Modular type I | 29.0 | 42.3 | 198.0 | 6.6e-50 |
| BAE93722.1 | type\_I\_polyketide\_synthase | BGC0000164 | Polyketide | 31.0 | 32.2 | 198.0 | 6.6e-50 |
| CCC21123.1 | type-I\_polyketide\_synthases | BGC0000171 | Polyketide:Modular type I | 31.0 | 39.6 | 198.0 | 6.6e-50 |
| AQZ37113.1 | polyketide\_synthase | BGC0001511 | Polyketide | 34.0 | 32.2 | 198.0 | 6.6e-50 |
| AAX98190.1 | polyketide\_synthase\_type\_I | BGC0000052 | Polyketide | 33.0 | 32.0 | 198.0 | 8.6e-50 |
| AEZ53948.1 | polyketide\_synthase | BGC0000144 | Polyketide:Modular type I | 27.0 | 51.8 | 198.0 | 8.6e-50 |
| ADU85981.1 | putative\_modular\_polyketide\_synthase | BGC0000165 | Polyketide:Modular type I | 33.0 | 32.4 | 198.0 | 8.6e-50 |
| CAN89632.1 | putative\_polyketide\_synthase | BGC0001070 | NRP + Polyketide:Modular type I + Polyketide:Trans-AT type I | 34.0 | 32.0 | 198.0 | 8.6e-50 |
| CAL69894.1 | RhiF\_protein | BGC0001112 | NRP + Polyketide:Trans-AT type I | 33.0 | 32.0 | 198.0 | 8.6e-50 |
| RAT98518.1 | trans-acyltransferase\_polyketide\_synthase | BGC0001470 | Polyketide:Trans-AT type I | 36.0 | 28.2 | 198.0 | 8.6e-50 |
| ARE67853.1 | AbsB1 | BGC0001492 | Polyketide | 38.0 | 31.7 | 198.0 | 8.6e-50 |
| TXD00033.1 | SDR\_family\_NAD(P)-dependent\_oxidoreductase | BGC0001877 | Polyketide | 32.0 | 32.0 | 198.0 | 8.6e-50 |
| AHH25595.1 | PKS | BGC0000957 | NRP + Polyketide | 29.0 | 46.2 | 197.0 | 1.1e-49 |
| ARS01474.1 | NcmAII | BGC0001702 | NRP + Polyketide | 29.0 | 42.3 | 197.0 | 1.1e-49 |
| ASZ00149.1 | polyketide\_synthase | BGC0001785 | Polyketide | 28.0 | 46.5 | 197.0 | 1.1e-49 |
| AZH23819.1 | MgiR | BGC0001971 | NRP + Polyketide | 31.0 | 40.1 | 197.0 | 1.1e-49 |
| AAM54075.1 | polyketide\_synthase | BGC0000020 | Polyketide | 34.0 | 32.2 | 197.0 | 1.5e-49 |
| CAE45669.1 | borrelidin\_polyketide\_synthase,\_type\_I | BGC0000031 | Polyketide:Modular type I | 28.0 | 43.7 | 197.0 | 1.5e-49 |
| AAW03328.1 | CtaE | BGC0000982 | NRP + Polyketide | 34.0 | 32.3 | 197.0 | 1.5e-49 |
| ALV82320.1 | borrelidin\_type\_I\_polyketide\_synthase | BGC0001533 | Polyketide | 28.0 | 43.7 | 197.0 | 1.5e-49 |
| QBF51755.1 | type\_I\_polyketide\_synthase | BGC0001856 | Polyketide:Modular type I | 32.0 | 32.3 | 197.0 | 1.5e-49 |
| QBF51769.1 | type\_I\_polyketide\_synthase | BGC0001856 | Polyketide:Modular type I | 33.0 | 32.9 | 197.0 | 1.5e-49 |
| TXD00266.1 | SDR\_family\_NAD(P)-dependent\_oxidoreductase | BGC0001877 | Polyketide | 30.0 | 32.5 | 197.0 | 1.5e-49 |
| ACO94468.1 | polyketide\_synthase\_type\_I | BGC0000029 | Polyketide:Modular type I | 33.0 | 31.7 | 196.0 | 1.9e-49 |
| ABC84465.1 | NigAX | BGC0000114 | Polyketide:Modular type I | 29.0 | 36.2 | 196.0 | 1.9e-49 |
| BAB69192.1 | modular\_polyketide\_synthase | BGC0000117 | Polyketide | 32.0 | 32.9 | 196.0 | 1.9e-49 |
| ACZ65476.1 | type\_I\_modular\_polyketide\_synthase | BGC0000140 | Polyketide | 29.0 | 50.0 | 196.0 | 1.9e-49 |
| AAS98784.1 | polyketide\_synthase | BGC0001001 | NRP + Polyketide | 28.0 | 48.5 | 196.0 | 1.9e-49 |
| CTQ34881.1 | AtcD;\_polyketide\_synthase,\_modules\_1-4 | BGC0001301 | Polyketide | 35.0 | 32.2 | 196.0 | 1.9e-49 |
| ANZ22995.1 | ZinA | BGC0001828 | Polyketide | 32.0 | 32.2 | 196.0 | 1.9e-49 |
| AAZ94386.1 | modular\_polyketide\_synthase | BGC0000040 | Polyketide | 33.0 | 32.3 | 196.0 | 2.5e-49 |
| AFV30248.1 | polyketide\_synthase | BGC0000075 | Polyketide | 32.0 | 32.0 | 196.0 | 2.5e-49 |
| AFL48525.1 | laidlomycin\_polyketide\_synthase\_(loading\_module\_and\_module\_1) | BGC0000084 | Polyketide | 30.0 | 48.4 | 196.0 | 2.5e-49 |
| ACR50775.1 | polyketide\_synthase | BGC0000163 | Polyketide | 27.0 | 50.3 | 196.0 | 2.5e-49 |
| OJF16267.1 | AceP5 | BGC0001491 | Polyketide | 26.0 | 68.3 | 196.0 | 2.5e-49 |
| WP\_015031692.1 | type\_I\_polyketide\_synthase | BGC0001819 | Polyketide | 29.0 | 45.5 | 196.0 | 2.5e-49 |
| ANZ22986.1 | ZinC | BGC0001828 | Polyketide | 32.0 | 32.1 | 196.0 | 2.5e-49 |
| AAO65800.1 | monensin\_polyketide\_synthase\_modules\_7\_and\_8 | BGC0000100 | Polyketide | 27.0 | 49.4 | 196.0 | 3.3e-49 |
| AAF71767.1 | nysJ | BGC0000115 | Polyketide:Modular type I + Saccharide:Hybrid/tailoring | 31.0 | 32.4 | 196.0 | 3.3e-49 |
| CAJ88176.1 | putative\_polyketide\_synthase\_B | BGC0000151 | Polyketide:Modular type I + Saccharide:Hybrid/tailoring | 28.0 | 47.7 | 196.0 | 3.3e-49 |
| ACY01390.1 | AT-less\_polyketide\_synthase | BGC0000177 | Polyketide:Modular type I + Polyketide:Trans-AT type I | 35.0 | 32.5 | 196.0 | 3.3e-49 |
| CCP20048.1 | divL1\_protein | BGC0001119 | Polyketide:Modular type I | 32.0 | 32.3 | 196.0 | 3.3e-49 |
| WP\_055469549.1 | type\_I\_polyketide\_synthase | BGC0001537 | Polyketide | 33.0 | 32.4 | 196.0 | 3.3e-49 |
| ANZ52463.1 | MonAV | BGC0001670 | Polyketide | 27.0 | 49.4 | 196.0 | 3.3e-49 |
| AWH12938.1 | StmC | BGC0001939 | Polyketide | 33.0 | 32.1 | 196.0 | 3.3e-49 |
| AZF85934.1 | type\_I\_polyketide\_synthase | BGC0001963 | NRP + Polyketide | 34.0 | 32.8 | 196.0 | 3.3e-49 |
| AZF85947.1 | type\_I\_polyketide\_synthase | BGC0001963 | NRP + Polyketide | 27.0 | 43.9 | 196.0 | 3.3e-49 |
| ADC79617.1 | BafAII | BGC0000028 | Polyketide:Modular type I | 33.0 | 32.2 | 195.0 | 4.3e-49 |
| BAP34733.1 | type\_I\_polyketide\_synthase | BGC0000078 | Polyketide | 33.0 | 32.5 | 195.0 | 4.3e-49 |
| BAD08360.1 | polyketide\_synthase\_modules\_7-8 | BGC0000167 | Polyketide | 25.0 | 65.2 | 195.0 | 4.3e-49 |
| ALD83687.1 | tAT\_polyketide\_synthase | BGC0001300 | Polyketide | 33.0 | 32.1 | 195.0 | 4.3e-49 |
| AJQ95706.1 | polyketide\_synthase\_modules-related\_protein | BGC0001644 | Polyketide | 30.0 | 41.1 | 195.0 | 4.3e-49 |
| ARS01475.1 | NcmAIII | BGC0001702 | NRP + Polyketide | 29.0 | 47.6 | 195.0 | 4.3e-49 |
| AEC13069.1 | fosC | BGC0000060 | Polyketide | 33.0 | 32.5 | 195.0 | 5.6e-49 |
| BAP34740.1 | type\_I\_polyketide\_synthase | BGC0000078 | Polyketide | 33.0 | 31.3 | 195.0 | 5.6e-49 |
| CBA11583.1 | polyketide\_synthase\_type\_I | BGC0001046 | NRP + Polyketide:Modular type I + Saccharide:Hybrid/tailoring | 33.0 | 32.3 | 195.0 | 5.6e-49 |
| ctg1\_orf521 |  | BGC0001199 | Polyketide | 35.0 | 32.4 | 195.0 | 5.6e-49 |
| BAT51065.1 | type\_I\_polyketide\_synthase | BGC0001296 | Polyketide | 29.0 | 48.5 | 195.0 | 5.6e-49 |
| ARM20284.1 | polyketide\_synthase | BGC0001523 | Polyketide | 27.0 | 44.4 | 195.0 | 5.6e-49 |
| BAV56006.1 | PKS\_(ACP-KS-AT-DH-ER-KR-ACP-KS-AT-KR-ACP) | BGC0001597 | Polyketide | 32.0 | 32.4 | 195.0 | 5.6e-49 |
| ANR02557.1 | LodP | BGC0001648 | Polyketide | 32.0 | 32.5 | 195.0 | 5.6e-49 |
| ABY21538.1 | AngAI | BGC0000018 | Polyketide | 30.0 | 42.7 | 195.0 | 7.3e-49 |
| CAE45672.1 | Borrelidin\_polyketide\_synthase,\_type\_I | BGC0000031 | Polyketide:Modular type I | 28.0 | 44.5 | 195.0 | 7.3e-49 |
| AAX98185.1 | polyketide\_synthase\_type\_I | BGC0000052 | Polyketide | 33.0 | 32.3 | 195.0 | 7.3e-49 |
| ABV97153.1 | Beta-ketoacyl\_synthase | BGC0000137 | Polyketide | 28.0 | 43.4 | 195.0 | 7.3e-49 |
| BAD08373.1 | polyketide\_synthase\_modules\_1-3 | BGC0000167 | Polyketide | 27.0 | 48.3 | 195.0 | 7.3e-49 |
| CAQ52624.1 | type\_I\_polyketide\_synthase,\_modules\_7-8 | BGC0001066 | Polyketide:Modular type I | 32.0 | 32.8 | 195.0 | 7.3e-49 |
| AMH40421.1 | PKS | BGC0001350 | Polyketide | 31.0 | 38.1 | 195.0 | 7.3e-49 |
| ALV82335.1 | borrelidin\_type\_I\_polyketide\_synthase | BGC0001533 | Polyketide | 28.0 | 44.5 | 195.0 | 7.3e-49 |
| AUO16423.1 | polyketide\_synthase | BGC0001700 | Polyketide | 27.0 | 44.4 | 195.0 | 7.3e-49 |
| ANH11412.1 | SceQ | BGC0001908 | Polyketide | 32.0 | 32.4 | 195.0 | 7.3e-49 |
| WP\_015031691.1 | type\_I\_polyketide\_synthase | BGC0001819 | Polyketide | 31.0 | 31.8 | 195.0 | 7.3e-49 |
| WP\_053138504.1 | type\_I\_polyketide\_synthase | BGC0002033 | Polyketide | 33.0 | 32.3 | 195.0 | 7.3e-49 |
| AAX98192.1 | polyketide\_synthase\_type\_I | BGC0000052 | Polyketide | 32.0 | 32.4 | 194.0 | 9.5e-49 |
| ACB37743.1 | putative\_type\_I\_polyketide\_synthase | BGC0000162 | Polyketide | 28.0 | 48.2 | 194.0 | 9.5e-49 |
| CAN89631.1 | putative\_polyketide\_synthase | BGC0001070 | NRP + Polyketide:Modular type I + Polyketide:Trans-AT type I | 36.0 | 29.8 | 194.0 | 9.5e-49 |
| CAE52339.1 | Polyketide\_non-ribosomal\_peptide\_synthase | BGC0001088 | NRP + Polyketide | 33.0 | 32.8 | 194.0 | 9.5e-49 |
| CAG23957.2 | hybrid\_NRPS/PKS\_protein | BGC0001089 | Polyketide + NRP | 28.0 | 41.6 | 194.0 | 9.5e-49 |
| ANR02554.1 | LodM | BGC0001648 | Polyketide | 28.0 | 47.5 | 194.0 | 9.5e-49 |
| AWH12667.1 | RmpD1 | BGC0001759 | Polyketide | 32.0 | 32.3 | 194.0 | 9.5e-49 |
| ATX68111.1 | malonyl\_CoA-acyl\_carrier\_protein\_transacylase | BGC0001772 | Polyketide | 30.0 | 39.2 | 194.0 | 9.5e-49 |
| ACN69991.1 | polyketide\_synthase | BGC0000079 | Polyketide | 27.0 | 45.0 | 194.0 | 1.2e-48 |
| CAD89776.1 | MelE\_protein | BGC0001010 | NRP + Polyketide:Modular type I | 34.0 | 32.3 | 194.0 | 1.2e-48 |
| AHE80995.1 | PieA5 | BGC0001169 | Polyketide:Modular type I | 33.0 | 32.0 | 194.0 | 1.2e-48 |
| BAR73019.1 | putative\_PKS\_(KS-AT-DH-KR-ACP-TE) | BGC0001194 | Polyketide | 34.0 | 31.3 | 194.0 | 1.2e-48 |
| ARE67851.1 | AbsB3 | BGC0001492 | Polyketide | 25.0 | 71.4 | 194.0 | 1.2e-48 |
| OEI73463.1 | hypothetical\_protein | BGC0001520 | Polyketide | 33.0 | 33.1 | 194.0 | 1.2e-48 |
| ANY10599.1 | polyketide\_synthase | BGC0001773 | Polyketide | 33.0 | 32.9 | 194.0 | 1.2e-48 |
| CAG23959.2 | polyketide\_synthase\_of\_type\_I | BGC0001089 | Polyketide + NRP | 33.0 | 33.1 | 193.0 | 1.6e-48 |
| AAZ94389.1 | modular\_polyketide\_synthase | BGC0000040 | Polyketide | 33.0 | 31.9 | 193.0 | 2.1e-48 |
| AAO65799.1 | monensin\_polyketide\_synthase\_modules\_5\_and\_6 | BGC0000100 | Polyketide | 26.0 | 48.3 | 193.0 | 2.1e-48 |
| ACY06290.1 | type\_I\_polyketide\_synthase | BGC0001042 | NRP + Polyketide | 32.0 | 32.1 | 193.0 | 2.1e-48 |
| AEZ64504.1 | Herc | BGC0001065 | Polyketide | 28.0 | 50.8 | 193.0 | 2.1e-48 |
| ABS90470.1 | NRPS/PKS | BGC0001106 | NRP + Polyketide | 33.0 | 32.7 | 193.0 | 2.1e-48 |
| CTQ34882.1 | AtcE;\_polyketide\_synthase,\_modules\_5-7 | BGC0001301 | Polyketide | 34.0 | 32.7 | 193.0 | 2.1e-48 |
| WP\_106980515.1 | type\_I\_polyketide\_synthase | BGC0001348 | Polyketide:Modular type I | 31.0 | 37.3 | 193.0 | 2.1e-48 |
| WP\_063764077.1 | hypothetical\_protein | BGC0001348 | Polyketide:Modular type I | 31.0 | 32.2 | 193.0 | 2.1e-48 |
| ANZ52462.1 | MonAIV | BGC0001670 | Polyketide | 26.0 | 48.3 | 193.0 | 2.1e-48 |
| DAC80077.1 | PKS | BGC0001836 | Polyketide:Trans-AT type I | 31.0 | 32.0 | 193.0 | 2.1e-48 |
| ACO94496.1 | polyketide\_synthase\_type\_I | BGC0000097 | Polyketide:Modular type I | 28.0 | 47.7 | 193.0 | 2.8e-48 |
| CAL58687.1 | polyketide\_synthase | BGC0000149 | Polyketide:Modular type I | 29.0 | 48.2 | 193.0 | 2.8e-48 |
| AAP85335.1 | type\_I\_PKS | BGC0000233 | Polyketide | 31.0 | 33.5 | 193.0 | 2.8e-48 |
| AAP85336.1 | type\_I\_PKS | BGC0000233 | Polyketide | 34.0 | 32.6 | 193.0 | 2.8e-48 |
| EAU29529.1 | hypothetical\_protein | BGC0000682 | Terpene | 24.0 | 91.2 | 193.0 | 2.8e-48 |
| CAG23960.2 | hybrid\_NRPS/PKS\_protein | BGC0001089 | Polyketide + NRP | 29.0 | 39.8 | 193.0 | 2.8e-48 |
| CCP20050.1 | divL3\_protein | BGC0001119 | Polyketide:Modular type I | 32.0 | 32.3 | 193.0 | 2.8e-48 |
| ARR97036.1 | SphC | BGC0001780 | NRP | 32.0 | 37.7 | 193.0 | 2.8e-48 |
| ANZ22988.1 | ZinE | BGC0001828 | Polyketide | 32.0 | 32.7 | 193.0 | 2.8e-48 |
| ctg1\_15 |  | BGC0001931 | Polyketide | 32.0 | 41.6 | 193.0 | 2.8e-48 |
| ARV85762.1 | PieA3\_type\_I\_PKS | BGC0001742 | Polyketide | 32.0 | 32.3 | 192.0 | 3.6e-48 |
| WP\_053138522.1 | type\_I\_polyketide\_synthase | BGC0002033 | Polyketide | 34.0 | 32.3 | 192.0 | 3.6e-48 |
| BAE93728.1 | type\_I\_polyketide\_synthase | BGC0000164 | Polyketide | 27.0 | 47.3 | 192.0 | 4.7e-48 |
| ADH01489.1 | type\_I\_polyketide\_synthase | BGC0001096 | NRP + Polyketide | 31.0 | 40.0 | 192.0 | 4.7e-48 |
| AWR88393.1 | putative\_beta-ketoacyl\_synthase | BGC0001522 | Polyketide | 32.0 | 31.2 | 192.0 | 4.7e-48 |
| ONK09689.1 | Beta-ketoacyl-acyl-carrier-protein\_synthase\_I | BGC0001647 | Polyketide | 27.0 | 48.9 | 192.0 | 4.7e-48 |
| AKG06379.1 | polyketide\_synthase\_type\_1 | BGC0001830 | Polyketide | 28.0 | 47.5 | 192.0 | 4.7e-48 |
| ctg1\_14 |  | BGC0001931 | Polyketide | 33.0 | 32.3 | 192.0 | 4.7e-48 |
| AAP42855.1 | NanA1 | BGC0000105 | Polyketide | 29.0 | 50.0 | 191.0 | 6.2e-48 |
| CAJ88177.1 | putative\_type\_I\_polyketide\_synthase | BGC0000151 | Polyketide:Modular type I + Saccharide:Hybrid/tailoring | 34.0 | 32.2 | 191.0 | 6.2e-48 |
| ctg1\_orf256 |  | BGC0001200 | Polyketide | 33.0 | 32.7 | 191.0 | 6.2e-48 |
| WP\_055480218.1 | type\_I\_polyketide\_synthase | BGC0001653 | Polyketide | 33.0 | 32.3 | 191.0 | 6.2e-48 |
| AAS90047.1 | PksA | BGC0000009 | Polyketide | 33.0 | 32.7 | 191.0 | 8.1e-48 |
| BAC57028.1 | protomycinolide\_IV\_synthase\_1 | BGC0000102 | Polyketide | 32.0 | 32.1 | 191.0 | 8.1e-48 |
| AAP42860.1 | NanA6 | BGC0000105 | Polyketide | 31.0 | 32.0 | 191.0 | 8.1e-48 |
| ADM46360.1 | polyketide\_synthase | BGC0000106 | Polyketide | 32.0 | 32.2 | 191.0 | 8.1e-48 |
| CAO85898.1 | modular\_polyketide\_synthase\_NorC | BGC0000110 | Polyketide:Modular type I | 26.0 | 47.1 | 191.0 | 8.1e-48 |
| AEZ53947.1 | polyketide\_synthase | BGC0000144 | Polyketide:Modular type I | 27.0 | 49.0 | 191.0 | 8.1e-48 |
| AKA59091.1 | type-I\_PKS | BGC0001619 | Polyketide | 32.0 | 33.7 | 191.0 | 8.1e-48 |
| AVX51106.1 | nysA | BGC0001709 | Polyketide | 33.0 | 32.4 | 191.0 | 8.1e-48 |
| AQW44874.1 | polyketide\_synthase | BGC0001761 | Polyketide | 30.0 | 39.3 | 191.0 | 8.1e-48 |
| ANZ22985.1 | ZinB | BGC0001828 | Polyketide | 31.0 | 32.4 | 191.0 | 8.1e-48 |
| WP\_003598535.1 | SDR\_family\_NAD(P)-dependent\_oxidoreductase | BGC0001991 | Polyketide | 34.0 | 32.7 | 191.0 | 8.1e-48 |
| ACR50791.1 | putative\_polyketide\_synthase | BGC0000163 | Polyketide | 33.0 | 32.0 | 191.0 | 1.1e-47 |
| ABF85931.1 | non-ribosomal\_peptide\_synthase/polyketide\_synthase\_Ta1 | BGC0001025 | NRP + Polyketide:Trans-AT type I | 33.0 | 33.3 | 191.0 | 1.1e-47 |
| AHE80991.1 | PieA1 | BGC0001169 | Polyketide:Modular type I | 32.0 | 32.3 | 191.0 | 1.1e-47 |
| EYT83439.1 | beta-ketoacyl\_synthase | BGC0001213 | Polyketide | 38.0 | 26.1 | 191.0 | 1.1e-47 |
| AAX35547.1 | polyketide\_syntase\_2 | BGC0001275 | Polyketide | 31.0 | 31.9 | 191.0 | 1.1e-47 |
| ATY46587.1 | polyketide\_synthase | BGC0001666 | Polyketide | 33.0 | 32.5 | 191.0 | 1.1e-47 |
| BAO66541.1 | type\_I\_polyketide\_synthase | BGC0000042 | Polyketide | 33.0 | 31.0 | 190.0 | 1.4e-47 |
| ACF35447.1 | mbcAIII | BGC0000090 | Polyketide | 32.0 | 32.4 | 190.0 | 1.4e-47 |
| BAF85844.1 | modular\_polyketide\_synthase | BGC0000109 | Polyketide | 33.0 | 32.3 | 190.0 | 1.4e-47 |
| WP\_055469545.1 | type\_I\_polyketide\_synthase | BGC0001537 | Polyketide | 33.0 | 32.1 | 190.0 | 1.4e-47 |
| ATL73033.1 | type\_I\_modular\_polyketide\_synthase | BGC0001807 | NRP + Polyketide | 33.0 | 32.3 | 190.0 | 1.4e-47 |
| BAO66539.1 | type\_I\_polyketide\_synthase | BGC0000042 | Polyketide | 29.0 | 44.2 | 190.0 | 1.8e-47 |
| AEC13080.1 | fosB | BGC0000060 | Polyketide | 28.0 | 44.6 | 190.0 | 1.8e-47 |
| AFN27483.1 | pks\_BonD | BGC0000173 | Polyketide:Modular type I | 33.0 | 35.8 | 190.0 | 1.8e-47 |
| CAN93347.1 | Polyketide\_synthase | BGC0000179 | Polyketide:Trans-AT type I | 32.0 | 32.9 | 190.0 | 1.8e-47 |
| EHK80171.1 | modular\_polyketide\_synthase | BGC0001447 | Polyketide | 32.0 | 32.3 | 190.0 | 1.8e-47 |
| sipP5 | Type\_I\_Modular\_PKS | BGC0001452 | Polyketide | 32.0 | 31.2 | 190.0 | 1.8e-47 |
| SCN11949.1 | ebeA-type\_I\_polyketide\_synthase\_KSQ-ATa-ACP | BGC0001580 | Polyketide | 37.0 | 27.4 | 190.0 | 1.8e-47 |
| BAV56012.1 | PKS\_(KS-AT-DH-ER-KR-ACP-TE) | BGC0001597 | Polyketide | 32.0 | 32.7 | 190.0 | 1.8e-47 |
| CAO98852.1 | polyketide\_synthase\_AufI | BGC0000023 | Polyketide:Modular type I | 33.0 | 32.5 | 190.0 | 2.4e-47 |
| AAQ82564.1 | FscC | BGC0000061 | Polyketide | 32.0 | 31.9 | 190.0 | 2.4e-47 |
| BAE71314.1 | polyketide\_synthase | BGC0000004 | Polyketide | 33.0 | 32.7 | 189.0 | 3.1e-47 |
| AAS90093.1 | PksA | BGC0000006 | Polyketide | 33.0 | 32.7 | 189.0 | 3.1e-47 |
| AAS90022.1 | PksA | BGC0000008 | Polyketide | 33.0 | 32.7 | 189.0 | 3.1e-47 |
| AFV30249.1 | polyketide\_synthase | BGC0000075 | Polyketide | 32.0 | 32.1 | 189.0 | 3.1e-47 |
| ACN69989.1 | polyketide\_synthase | BGC0000079 | Polyketide | 32.0 | 32.0 | 189.0 | 3.1e-47 |
| ABP55493.1 | thioester\_reductase\_domain | BGC0001006 | NRP + Polyketide | 26.0 | 49.8 | 189.0 | 3.1e-47 |
| CAL69891.1 | RhiD\_protein | BGC0001112 | NRP + Polyketide:Trans-AT type I | 31.0 | 33.0 | 189.0 | 3.1e-47 |
| AXA20093.1 | trans-AT\_PKS\_LgaD | BGC0001946 | NRP + Polyketide | 34.0 | 31.1 | 189.0 | 3.1e-47 |
| AAZ77693.1 | ChlA1 | BGC0000036 | Polyketide:Modular type I + Polyketide:Iterative type I + Saccharide:Oligosaccharide | 30.0 | 46.5 | 189.0 | 4e-47 |
| AEC13072.1 | fosF | BGC0000060 | Polyketide | 31.0 | 32.2 | 189.0 | 4e-47 |
| AAN85523.1 | polyketide\_synthase | BGC0001101 | NRP + Polyketide:Modular type I + Polyketide:Trans-AT type I | 31.0 | 39.0 | 189.0 | 4e-47 |
| AAS47564.1 | mixed\_type\_I\_polyketide\_synthase/nonribosomal\_peptide\_synthetase | BGC0001108 | Polyketide:Trans-AT type I | 33.0 | 33.1 | 189.0 | 4e-47 |
| sipP2 | Type\_I\_Modular\_PKS | BGC0001452 | Polyketide | 32.0 | 31.9 | 189.0 | 4e-47 |
| ACC40922.1 | polyketide\_synthase,\_Pks8 | BGC0001665 | Polyketide | 32.0 | 32.3 | 189.0 | 4e-47 |
| ARS01476.1 | NcmAIV | BGC0001702 | NRP + Polyketide | 32.0 | 32.1 | 189.0 | 4e-47 |
| AAX98188.1 | polyketide\_synthase\_type\_I | BGC0000052 | Polyketide | 32.0 | 32.2 | 188.0 | 5.2e-47 |
| ACR12418.1 | modular\_polyketide\_synthase,\_type\_I\_PKS | BGC0000185 | Polyketide | 31.0 | 32.2 | 188.0 | 5.2e-47 |
| CAN89635.1 | putative\_polyketide\_synthase | BGC0001070 | NRP + Polyketide:Modular type I + Polyketide:Trans-AT type I | 32.0 | 31.5 | 188.0 | 5.2e-47 |
| ATX68125.1 | malonyl\_CoA-acyl\_carrier\_protein\_transacylase | BGC0001795 | Polyketide | 29.0 | 40.2 | 188.0 | 5.2e-47 |
| AAP42867.1 | NanA7 | BGC0000105 | Polyketide | 27.0 | 47.9 | 188.0 | 6.8e-47 |
| AFN27480.1 | pks\_BonA | BGC0000173 | Polyketide:Modular type I | 32.0 | 31.5 | 188.0 | 6.8e-47 |
| ctg1\_orf6 |  | BGC0001109 | NRP + Polyketide | 33.0 | 33.1 | 188.0 | 6.8e-47 |
| AZH23793.1 | MgcK | BGC0001970 | NRP + Polyketide | 28.0 | 48.7 | 188.0 | 6.8e-47 |
| BAK64650.1 | polyketide\_synthase | BGC0000135 | Polyketide | 31.0 | 32.4 | 188.0 | 8.9e-47 |
| AEC04356.1 | polyketide\_synthase | BGC0000178 | Polyketide:Trans-AT type I | 34.0 | 28.1 | 188.0 | 8.9e-47 |
| ABI91464.1 | beta-ketoacyl\_synthase | BGC0001094 | NRP + Polyketide | 32.0 | 32.1 | 188.0 | 8.9e-47 |
| AQW44870.1 | polyketide\_synthase | BGC0001761 | Polyketide | 30.0 | 40.0 | 188.0 | 8.9e-47 |
| AAM54076.1 | polyketide\_synthase | BGC0000020 | Polyketide | 33.0 | 29.9 | 187.0 | 1.2e-46 |
| ADH04682.1 | polyketide\_synthase | BGC0001344 | NRP + Polyketide | 27.0 | 48.0 | 187.0 | 1.2e-46 |
| ART41209.1 | AdrD | BGC0001508 | Polyketide | 29.0 | 45.7 | 187.0 | 1.2e-46 |
| AVX51099.1 | NysJ | BGC0001709 | Polyketide | 30.0 | 35.3 | 187.0 | 1.2e-46 |
| ATX68110.1 | malonyl\_CoA-acyl\_carrier\_protein\_transacylase | BGC0001772 | Polyketide | 29.0 | 39.5 | 187.0 | 1.2e-46 |
| ctg1\_orf9 |  | BGC0000053 | Polyketide | 31.0 | 32.2 | 187.0 | 1.5e-46 |
| BAD38875.1 | polyketide\_synthase | BGC0000111 | Polyketide | 32.0 | 32.6 | 187.0 | 1.5e-46 |
| AEC04357.1 | polyketide\_synthase | BGC0000178 | Polyketide:Trans-AT type I | 33.0 | 32.0 | 187.0 | 1.5e-46 |
| AFX60334.1 | polyketide\_synthase | BGC0001032 | NRP + Polyketide | 31.0 | 35.2 | 187.0 | 1.5e-46 |
| RAT98527.1 | trans-acyltransferase\_polyketide\_synthase | BGC0001470 | Polyketide:Trans-AT type I | 33.0 | 32.3 | 187.0 | 1.5e-46 |
| ctg1\_11 |  | BGC0001931 | Polyketide | 33.0 | 32.4 | 187.0 | 1.5e-46 |
| ADC45538.1 | modular\_polyketide\_synthase | BGC0000093 | Polyketide | 27.0 | 48.9 | 186.0 | 2e-46 |
| CBD77748.1 | polyketide\_synthase | BGC0000974 | NRP + Polyketide | 33.0 | 32.1 | 186.0 | 2e-46 |
| AVV61983.1 | type\_I\_modular\_polyketide\_synthase | BGC0001477 | NRP + Polyketide:Modular type I | 31.0 | 32.0 | 186.0 | 2e-46 |
| AIR74917.1 | polyketide\_synthase | BGC0001559 | RiPP | 33.0 | 32.1 | 186.0 | 2e-46 |
| AJQ95704.1 | polyketide\_synthase\_modules-related\_protein | BGC0001644 | Polyketide | 29.0 | 45.5 | 186.0 | 2e-46 |
| AAM77986.1 | iterative\_type\_I\_polyketide\_synthase | BGC0000112 | Polyketide:Iterative type I + Polyketide:Enediyne type I | 27.0 | 47.2 | 186.0 | 2.6e-46 |
| AAB66504.1 | tylactone\_synthase\_starter\_module\_and\_modules\_1\_&\_2 | BGC0000166 | Polyketide | 34.0 | 32.3 | 186.0 | 3.4e-46 |
| AKQ22699.1 | malonyl\_CoA-acyl\_carrier\_protein\_transacylase | BGC0001186 | Polyketide | 29.0 | 42.9 | 186.0 | 3.4e-46 |
| BBA21074.1 | putative\_modular\_polyketide\_synthase | BGC0001740 | NRP + Polyketide | 29.0 | 45.2 | 186.0 | 3.4e-46 |
| AAS79459.1 | polyketide\_synthase\_subunit | BGC0000035 | Polyketide | 33.0 | 32.4 | 185.0 | 4.4e-46 |
| CAL69893.1 | RhiE\_protein | BGC0001112 | NRP + Polyketide:Trans-AT type I | 32.0 | 32.1 | 185.0 | 4.4e-46 |
| AZH23823.1 | MgiK | BGC0001971 | NRP + Polyketide | 29.0 | 49.2 | 185.0 | 4.4e-46 |
| AHF22854.1 | MarL | BGC0000091 | Polyketide | 29.0 | 48.1 | 185.0 | 5.8e-46 |
| AEC04362.1 | polyketide\_synthase | BGC0000178 | Polyketide:Trans-AT type I | 33.0 | 33.1 | 185.0 | 5.8e-46 |
| CAQ18839.1 | hybrid\_polyketide\_synthase/nonribosomal\_polypetide\_synthetase | BGC0000954 | NRP + Polyketide:Modular type I | 28.0 | 45.0 | 185.0 | 5.8e-46 |
| AEE88283.1 | CurG | BGC0000976 | NRP + Polyketide:Modular type I | 27.0 | 48.5 | 185.0 | 5.8e-46 |
| AAT70102.1 | CurG | BGC0001165 | NRP + Polyketide:Modular type I | 27.0 | 48.5 | 185.0 | 5.8e-46 |
| AAS89999.1 | PksA | BGC0000007 | Polyketide | 33.0 | 32.7 | 185.0 | 7.6e-46 |
| ADN68482.1 | sorG | BGC0000184 | Polyketide:Trans-AT type I | 34.0 | 32.1 | 185.0 | 7.6e-46 |
| CAN89633.1 | putative\_hybrid\_non-ribosomal\_peptide\_synthetase/polyketide\_synthase | BGC0001070 | NRP + Polyketide:Modular type I + Polyketide:Trans-AT type I | 33.0 | 32.6 | 185.0 | 7.6e-46 |
| AAY32964.1 | DszA | BGC0001093 | NRP + Polyketide | 32.0 | 31.7 | 185.0 | 7.6e-46 |
| DAC76730.1 | type\_I\_polyketide\_synthase | BGC0001885 | Polyketide | 34.0 | 33.0 | 185.0 | 7.6e-46 |
| WP\_053138519.1 | type\_I\_polyketide\_synthase | BGC0002033 | Polyketide | 33.0 | 32.2 | 185.0 | 7.6e-46 |
| AEK75504.1 | type\_1\_polyketide\_synthase | BGC0000001 | Polyketide:Modular type I | 32.0 | 32.1 | 184.0 | 9.9e-46 |
| RAT98529.1 | trans-acyltransferase\_polyketide\_synthase | BGC0001470 | Polyketide:Trans-AT type I | 30.0 | 39.0 | 184.0 | 9.9e-46 |
| AXA20092.1 | trans-AT\_PKS\_LgaC | BGC0001946 | NRP + Polyketide | 33.0 | 31.6 | 184.0 | 9.9e-46 |
| AEN83889.1 | AdaA | BGC0000156 | Polyketide:Iterative type I | 28.0 | 44.8 | 184.0 | 1.3e-45 |
| ACR13997.1 | modular\_polyketide\_synthase,\_type\_I\_PKS | BGC0000185 | Polyketide | 33.0 | 32.1 | 184.0 | 1.3e-45 |
| ABC34832.1 | polyketide\_synthase | BGC0000186 | NRP + Polyketide:Modular type I | 34.0 | 29.4 | 184.0 | 1.3e-45 |
| AUW31184.1 | putative\_type\_I\_PKS | BGC0001489 | Polyketide | 27.0 | 45.2 | 184.0 | 1.3e-45 |
| BAV56011.1 | PKS\_(KS-AT-DH-ER-KR-ACP-KS-AT-DH-ER-KR-ACP) | BGC0001597 | Polyketide | 31.0 | 32.7 | 184.0 | 1.3e-45 |
| AKQ22682.1 | malonyl\_CoA-acyl\_carrier\_protein\_transacylase | BGC0001656 | Polyketide | 28.0 | 47.9 | 184.0 | 1.3e-45 |
| AJQ95707.1 | polyketide\_synthase\_modules-related\_protein | BGC0001644 | Polyketide | 28.0 | 42.5 | 183.0 | 1.7e-45 |
| PKX88487.1 | polyketide\_synthase | BGC0001708 | Polyketide + Terpene | 29.0 | 46.2 | 183.0 | 1.7e-45 |
| BAV69313.1 | PrhL | BGC0001729 | Polyketide + Terpene | 29.0 | 45.8 | 183.0 | 1.7e-45 |
| ADY00130.1 | polyketide\_synthase | BGC0000104 | Terpene + Polyketide:Iterative type I | 29.0 | 45.2 | 183.0 | 2.2e-45 |
| BBA21072.1 | putative\_modular\_polyketide\_synthase | BGC0001740 | NRP + Polyketide | 32.0 | 32.1 | 183.0 | 2.2e-45 |
| AGZ15472.1 | putative\_modular\_polyketide\_synthase | BGC0001036 | NRP + Polyketide | 27.0 | 48.5 | 183.0 | 2.9e-45 |
| AAY32965.1 | DszB | BGC0001093 | NRP + Polyketide | 29.0 | 41.2 | 183.0 | 2.9e-45 |
| AAP42857.1 | NanA3 | BGC0000105 | Polyketide | 31.0 | 31.8 | 182.0 | 3.8e-45 |
| AFO59866.1 | ChxE | BGC0000175 | Polyketide:Trans-AT type I | 30.0 | 39.4 | 182.0 | 3.8e-45 |
| AKQ22698.1 | malonyl\_CoA-acyl\_carrier\_protein\_transacylase | BGC0001186 | Polyketide | 31.0 | 32.2 | 182.0 | 3.8e-45 |
| KFA69335.1 | hypothetical\_protein | BGC0001626 | Polyketide | 29.0 | 50.1 | 182.0 | 3.8e-45 |
| AJQ95705.1 | polyketide\_synthase\_modules-related\_protein | BGC0001644 | Polyketide | 30.0 | 43.3 | 182.0 | 3.8e-45 |
| DAC80073.1 | PKS | BGC0001836 | Polyketide:Trans-AT type I | 29.0 | 44.4 | 182.0 | 3.8e-45 |
| OEI73460.1 | hypothetical\_protein | BGC0001520 | Polyketide | 29.0 | 40.0 | 182.0 | 4.9e-45 |
| ANZ22989.1 | ZinF | BGC0001828 | Polyketide | 32.0 | 32.0 | 182.0 | 4.9e-45 |
| ACR50796.1 | putative\_polyketide\_synthase | BGC0000163 | Polyketide | 29.0 | 47.0 | 181.0 | 6.4e-45 |
| AEC04363.1 | polyketide\_synthase | BGC0000178 | Polyketide:Trans-AT type I | 32.0 | 31.1 | 181.0 | 6.4e-45 |
| ADD82940.1 | Bat2 | BGC0001099 | NRP + Polyketide:Modular type I + Polyketide:Trans-AT type I | 29.0 | 39.6 | 181.0 | 6.4e-45 |
| ERM18798.1 | polyketide\_synthase | BGC0000172 | Polyketide | 30.0 | 39.3 | 181.0 | 8.4e-45 |
| RAT98517.1 | trans-acyltransferase\_polyketide\_synthase | BGC0001470 | Polyketide:Trans-AT type I | 33.0 | 31.7 | 181.0 | 8.4e-45 |
| RAT98525.1 | trans-acyltransferase\_polyketide\_synthase | BGC0001470 | Polyketide:Trans-AT type I | 33.0 | 32.0 | 181.0 | 8.4e-45 |
| OJF16269.1 | AceP2 | BGC0001491 | Polyketide | 31.0 | 32.9 | 181.0 | 8.4e-45 |
| OEI73461.1 | hypothetical\_protein | BGC0001520 | Polyketide | 32.0 | 32.0 | 181.0 | 8.4e-45 |
| AAM12911.1 | MmpB | BGC0000182 | Polyketide:Iterative type I + Polyketide:Trans-AT type I | 34.0 | 32.1 | 181.0 | 1.1e-44 |
| AAY89050.1 | polyketide\_synthase | BGC0001069 | NRP + Polyketide:Trans-AT type I | 32.0 | 31.2 | 181.0 | 1.1e-44 |
| RAT98528.1 | trans-acyltransferase\_polyketide\_synthase | BGC0001470 | Polyketide:Trans-AT type I | 32.0 | 32.1 | 181.0 | 1.1e-44 |
| ADN68478.1 | SorC | BGC0000184 | Polyketide:Trans-AT type I | 30.0 | 43.8 | 180.0 | 1.4e-44 |
| ACF35446.1 | mbcAII | BGC0000090 | Polyketide | 26.0 | 46.7 | 180.0 | 1.9e-44 |
| AAN85522.1 | hybrid\_nonribosomal\_peptide\_synthetase\_/\_polyketide\_synthase | BGC0001101 | NRP + Polyketide:Modular type I + Polyketide:Trans-AT type I | 32.0 | 33.3 | 180.0 | 1.9e-44 |
| AAM12909.2 | MmpA | BGC0000182 | Polyketide:Iterative type I + Polyketide:Trans-AT type I | 29.0 | 38.7 | 180.0 | 2.4e-44 |
| BAP05596.1 | calH | BGC0000967 | NRP + Polyketide:Trans-AT type I | 27.0 | 39.7 | 180.0 | 2.4e-44 |
| AFX60311.1 | polyketide\_synthase | BGC0001031 | NRP + Polyketide | 32.0 | 33.1 | 180.0 | 2.4e-44 |
| CAG23961.2 | polyketide\_synthase\_of\_type\_I | BGC0001089 | Polyketide + NRP | 30.0 | 31.4 | 180.0 | 2.4e-44 |
| AXA20096.1 | trans-AT\_PKS\_LgaG | BGC0001946 | NRP + Polyketide | 33.0 | 32.3 | 180.0 | 2.4e-44 |
| ATX68124.1 | malonyl\_CoA-acyl\_carrier\_protein\_transacylase | BGC0001795 | Polyketide | 32.0 | 33.1 | 180.0 | 2.4e-44 |
| AFN27481.1 | pks\_BonB | BGC0000173 | Polyketide:Modular type I | 32.0 | 32.7 | 179.0 | 3.2e-44 |
| CAF05649.1 | TubD\_protein | BGC0001053 | NRP + Polyketide | 27.0 | 47.5 | 179.0 | 3.2e-44 |
| BAC76474.1 | type\_I\_polyketide\_synthase\_LkcC | BGC0001100 | NRP + Polyketide | 30.0 | 39.4 | 179.0 | 3.2e-44 |
| AGC09499.1 | LobS4 | BGC0001183 | Polyketide | 26.0 | 48.5 | 179.0 | 3.2e-44 |
| AMH40443.1 | PKS | BGC0001350 | Polyketide | 31.0 | 32.0 | 179.0 | 3.2e-44 |
| AKQ22681.1 | malonyl\_CoA-acyl\_carrier\_protein\_transacylase | BGC0001656 | Polyketide | 32.0 | 32.3 | 179.0 | 3.2e-44 |
| RAT98519.1 | trans-acyltransferase\_polyketide\_synthase | BGC0001470 | Polyketide:Trans-AT type I | 27.0 | 46.5 | 178.0 | 5.4e-44 |
| OEI73466.1 | hypothetical\_protein | BGC0001520 | Polyketide | 30.0 | 38.2 | 178.0 | 5.4e-44 |
| AAD03048.1 | type\_I\_polyketide\_synthase | BGC0000041 | Polyketide | 27.0 | 45.2 | 178.0 | 7.1e-44 |
| CAG23964.1 | polyketide\_synthase\_type\_I | BGC0000181 | Polyketide | 29.0 | 40.0 | 178.0 | 7.1e-44 |
| ADN68479.1 | SorD | BGC0000184 | Polyketide:Trans-AT type I | 30.0 | 39.8 | 178.0 | 7.1e-44 |
| CAL69890.1 | RhiC\_protein | BGC0001112 | NRP + Polyketide:Trans-AT type I | 31.0 | 40.5 | 178.0 | 7.1e-44 |
| DAC80074.1 | PKS | BGC0001836 | Polyketide:Trans-AT type I | 32.0 | 31.7 | 178.0 | 7.1e-44 |
| AFX60309.1 | polyketide\_synthase | BGC0001031 | NRP + Polyketide | 31.0 | 32.1 | 178.0 | 9.2e-44 |
| CAD70195.1 | non-ribosomal\_peptide\_synthetase | BGC0001047 | NRP + Polyketide | 28.0 | 44.9 | 178.0 | 9.2e-44 |
| BAC76471.1 | type\_I\_polyketide\_synthase\_LkcF | BGC0001100 | NRP + Polyketide | 33.0 | 32.1 | 178.0 | 9.2e-44 |
| ACY13415.1 | KR\_domain\_protein | BGC0001367 | NRP + Polyketide | 33.0 | 32.6 | 178.0 | 9.2e-44 |
| OEI73464.1 | hypothetical\_protein | BGC0001520 | Polyketide | 28.0 | 46.4 | 178.0 | 9.2e-44 |
| CAN93349.1 | polyketide\_synthase | BGC0000179 | Polyketide:Trans-AT type I | 28.0 | 44.9 | 177.0 | 1.2e-43 |
| AIU36103.1 | LglD | BGC0000180 | Polyketide:Trans-AT type I | 31.0 | 32.4 | 177.0 | 1.2e-43 |
| AAY89049.1 | polyketide\_synthase | BGC0001069 | NRP + Polyketide:Trans-AT type I | 30.0 | 39.4 | 177.0 | 1.2e-43 |
| ABI91467.1 | beta-ketoacyl\_synthase | BGC0001094 | NRP + Polyketide | 33.0 | 31.9 | 177.0 | 1.2e-43 |
| EJK79843.1 | amino\_acid\_adenylation\_enzyme/thioester\_reductase\_family\_protein | BGC0000436 | NRP | 28.0 | 43.0 | 177.0 | 1.6e-43 |
| ctg1\_orf253 |  | BGC0001200 | Polyketide | 34.0 | 31.9 | 177.0 | 1.6e-43 |
| AJO72742.1 | Type\_I\_modular\_polyketide\_synthase | BGC0001381 | Polyketide | 32.0 | 32.2 | 177.0 | 1.6e-43 |
| CAQ34918.1 | nonribosomal\_peptide\_synthetase/\_polyketide\_synthase | BGC0000986 | NRP + Polyketide | 29.0 | 45.9 | 176.0 | 2.1e-43 |
| BBF25315.1 | polyketide\_synthase | BGC0001923 | Terpene + Polyketide | 30.0 | 40.4 | 176.0 | 2.1e-43 |
| ABC34176.1 | polyketide\_synthase,\_putative | BGC0000186 | NRP + Polyketide:Modular type I | 32.0 | 30.5 | 176.0 | 2.7e-43 |
| CAQ18832.1 | polyketide\_synthase | BGC0000954 | NRP + Polyketide:Modular type I | 28.0 | 48.1 | 176.0 | 2.7e-43 |
| AHD05614.1 | putative\_non-ribosomal\_peptide\_ligase/\_polyketide\_synthase\_hybrid | BGC0001033 | NRP + Polyketide | 28.0 | 48.8 | 176.0 | 2.7e-43 |
| CAG23967.1 | polyketide\_synthase\_type\_I | BGC0000181 | Polyketide | 29.0 | 38.9 | 176.0 | 3.5e-43 |
| CCA89327.1 | trans-AT\_type\_I\_polyketide\_synthase | BGC0001111 | NRP + Polyketide:Trans-AT type I | 30.0 | 32.2 | 176.0 | 3.5e-43 |
| WP\_047890614.1 | type\_I\_polyketide\_synthase | BGC0001330 | NRP:Cyclic depsipeptide + Polyketide:Modular type I | 28.0 | 48.3 | 176.0 | 3.5e-43 |
| ABF88102.1 | polyketide\_synthase | BGC0001025 | NRP + Polyketide:Trans-AT type I | 31.0 | 39.6 | 175.0 | 4.6e-43 |
| CBK62733.1 |  | BGC0001115 | NRP + Polyketide | 31.0 | 31.9 | 175.0 | 4.6e-43 |
| gene5 |  | BGC0001907 | Polyketide | 31.0 | 31.8 | 175.0 | 4.6e-43 |
| CAG23965.1 | polyketide\_synthase\_type\_I | BGC0000181 | Polyketide | 27.0 | 40.3 | 175.0 | 6e-43 |
| AAV97870.1 | OnnB | BGC0001105 | NRP + Polyketide:Trans-AT type I | 29.0 | 43.5 | 175.0 | 6e-43 |
| CDM36726.1 | Beta-ketoacyl\_synthase | BGC0001360 | Polyketide | 28.0 | 45.5 | 175.0 | 6e-43 |
| AIJ04680.1 | polyketide\_synthase | BGC0001383 | Polyketide | 27.0 | 40.3 | 175.0 | 6e-43 |
| AIJ04684.1 | polyketide\_synthase | BGC0001383 | Polyketide | 31.0 | 32.3 | 175.0 | 6e-43 |
| AAY89053.1 | polyketide\_synthase | BGC0001069 | NRP + Polyketide:Trans-AT type I | 32.0 | 32.6 | 174.0 | 1e-42 |
| EAQ84779.1 | hypothetical\_protein | BGC0001219 | Polyketide | 32.0 | 29.3 | 174.0 | 1e-42 |
| OEI73462.1 | hypothetical\_protein | BGC0001520 | Polyketide | 31.0 | 33.4 | 174.0 | 1e-42 |
| CAC20930.1 | PimS0\_protein | BGC0000125 | Polyketide | 31.0 | 31.9 | 173.0 | 1.7e-42 |
| AQT01384.1 | SgnS0 | BGC0001690 | Polyketide | 31.0 | 31.9 | 173.0 | 1.7e-42 |
| BAF50727.1 | hybrid\_polyketide\_synthase-non\_ribosomal\_peptide\_synthetase | BGC0001116 | NRP + Polyketide | 32.0 | 33.1 | 173.0 | 2.3e-42 |
| AWS21290.1 | type\_I\_polyketide\_synthase | BGC0001934 | Polyketide | 32.0 | 31.3 | 173.0 | 2.3e-42 |
| AZY91988.1 | polyketide\_synthase | BGC0002022 | Polyketide | 32.0 | 31.3 | 173.0 | 2.3e-42 |
| ABX37384.1 | Beta-ketoacyl\_synthase | BGC0000984 | NRP + Polyketide | 27.0 | 48.4 | 173.0 | 3e-42 |
| ALJ49911.1 | TlmG | BGC0001237 | Polyketide | 35.0 | 28.9 | 173.0 | 3e-42 |
| AAC38075.1 | polyketide\_synthase\_type\_I | BGC0000127 | Polyketide | 32.0 | 32.4 | 172.0 | 3.9e-42 |
| ADA69237.1 | trans-AT\_polyketide\_synthase | BGC0001071 | NRP + Polyketide:Modular type I + Polyketide:Trans-AT type I | 31.0 | 32.1 | 172.0 | 3.9e-42 |
| CAN93351.1 | polyketide\_synthase | BGC0000179 | Polyketide:Trans-AT type I | 29.0 | 33.4 | 172.0 | 5.1e-42 |
| PKY07881.1 | hypothetical\_protein | BGC0001544 | NRP + Polyketide | 29.0 | 33.0 | 172.0 | 5.1e-42 |
| ARR97040.1 | SphG | BGC0001780 | NRP | 30.0 | 34.0 | 172.0 | 5.1e-42 |
| CAJ57411.1 | polyketide\_synthase\_type\_I | BGC0000176 | Polyketide + NRP | 29.0 | 42.7 | 171.0 | 6.6e-42 |
| AFX60332.1 | polyketide\_synthase | BGC0001032 | NRP + Polyketide | 32.0 | 32.6 | 171.0 | 6.6e-42 |
| ASA76644.1 | polyketide\_synthase | BGC0001751 | NRP + Polyketide | 30.0 | 39.8 | 171.0 | 6.6e-42 |
| DAC76729.1 | type\_I\_polyketide\_synthase | BGC0001885 | Polyketide | 28.0 | 39.6 | 171.0 | 6.6e-42 |
| CAG23983.1 | polyketide\_synthase\_type\_I | BGC0000176 | Polyketide + NRP | 32.0 | 32.0 | 171.0 | 8.7e-42 |
| ADN68484.1 | sorI | BGC0000184 | Polyketide:Trans-AT type I | 33.0 | 29.2 | 171.0 | 8.7e-42 |
| BAP05591.1 | calC | BGC0000967 | NRP + Polyketide:Trans-AT type I | 30.0 | 32.1 | 171.0 | 8.7e-42 |
| BAP05594.1 | calF | BGC0000967 | NRP + Polyketide:Trans-AT type I | 32.0 | 32.5 | 171.0 | 8.7e-42 |
| AIJ04681.1 | polyketide\_synthase | BGC0001383 | Polyketide | 30.0 | 32.6 | 171.0 | 1.1e-41 |
| CCG06113.1 | type\_I\_polyketide\_synthase | BGC0001543 | Polyketide | 28.0 | 40.6 | 171.0 | 1.1e-41 |
| ANY10591.1 | polyketide\_synthase | BGC0001773 | Polyketide | 26.0 | 70.0 | 171.0 | 1.1e-41 |
| ADX66470.1 | ScnS0 | BGC0000108 | Polyketide | 30.0 | 31.9 | 170.0 | 2.5e-41 |
| AEC04361.1 | polyketide\_synthase | BGC0000178 | Polyketide:Trans-AT type I | 29.0 | 39.0 | 170.0 | 2.5e-41 |
| WP\_010639241.1 | type\_I\_polyketide\_synthase | BGC0000958 | NRP:Cyclic depsipeptide + Polyketide:Modular type I | 31.0 | 32.9 | 170.0 | 2.5e-41 |
| ADH01487.1 | polyketide\_synthase | BGC0001096 | NRP + Polyketide | 33.0 | 27.8 | 170.0 | 2.5e-41 |
| AIC32693.1 | FR9DEF | BGC0001113 | NRP + Polyketide | 33.0 | 27.8 | 170.0 | 2.5e-41 |
| ATG32078.1 | polyketide\_synthase | BGC0001750 | NRP + Polyketide | 33.0 | 31.9 | 170.0 | 2.5e-41 |
| ABC38101.1 | polyketide\_synthase | BGC0000964 | NRP:Cyclic depsipeptide + Polyketide:Trans-AT type I | 29.0 | 39.9 | 169.0 | 3.3e-41 |
| ANC94964.1 | AlmHIII | BGC0001396 | Polyketide | 33.0 | 27.8 | 169.0 | 3.3e-41 |
| RAT98526.1 | trans-acyltransferase\_polyketide\_synthase | BGC0001470 | Polyketide:Trans-AT type I | 30.0 | 33.1 | 169.0 | 3.3e-41 |
| CAN93352.1 | polyketide\_synthase | BGC0000179 | Polyketide:Trans-AT type I | 32.0 | 28.6 | 169.0 | 4.3e-41 |
| ABC35522.1 | thiotemplate\_mechanism\_natural\_product\_synthetase | BGC0000186 | NRP + Polyketide:Modular type I | 32.0 | 33.3 | 169.0 | 4.3e-41 |
| CCA29203.1 | non-ribosomal\_peptide\_synthetase/polyketide\_synthase | BGC0000955 | NRP + Polyketide:Modular type I | 27.0 | 48.4 | 169.0 | 4.3e-41 |
| ADA82581.1 | trans-AT\_polyketide\_synthase | BGC0001110 | NRP + Polyketide:Trans-AT type I | 33.0 | 32.3 | 169.0 | 4.3e-41 |
| BAW32323.1 | hybrid\_cis-AT\_polyketide\_synthase\_-\_nonribosomal\_peptide\_synthetase | BGC0001630 | NRP + Polyketide | 29.0 | 44.1 | 169.0 | 4.3e-41 |
| ABI91465.1 | beta-ketoacyl\_synthase | BGC0001094 | NRP + Polyketide | 31.0 | 32.1 | 168.0 | 5.6e-41 |
| AKA54627.1 | PKS | BGC0001216 | NRP + Polyketide | 24.0 | 71.1 | 168.0 | 5.6e-41 |
| AVR48533.1 | CusA | BGC0001564 | NRP + Polyketide | 28.0 | 41.0 | 168.0 | 5.6e-41 |
| AHA38200.1 | GphG | BGC0000069 | Polyketide | 35.0 | 26.9 | 168.0 | 7.3e-41 |
| AAN32979.1 | BarE | BGC0000962 | NRP + Polyketide:Modular type I | 25.0 | 56.0 | 168.0 | 7.3e-41 |
| ALD83702.1 | tAT\_polyketide\_synthase | BGC0001299 | Polyketide | 30.0 | 33.3 | 168.0 | 7.3e-41 |
| ATP76242.1 | NdaC | BGC0001705 | NRP + Polyketide | 26.0 | 46.2 | 168.0 | 7.3e-41 |
| ERM18797.1 | polyketide\_synthase | BGC0000172 | Polyketide | 32.0 | 29.5 | 168.0 | 9.6e-41 |
| AAO62585.1 | peptide\_sythetase\_polyketide\_synthase\_fusion\_protein | BGC0001016 | NRP + Polyketide | 26.0 | 49.3 | 168.0 | 9.6e-41 |
| AJY78093.1 | polyketide\_synthase | BGC0001902 | NRP + Polyketide | 29.0 | 44.3 | 168.0 | 9.6e-41 |
| CBF79143.1 | polyketide\_synthase,\_putative\_(JCVI) | BGC0000013 | Polyketide | 29.0 | 32.7 | 166.0 | 2.1e-40 |
| ADD82941.1 | Bat3 | BGC0001099 | NRP + Polyketide:Modular type I + Polyketide:Trans-AT type I | 31.0 | 31.8 | 166.0 | 2.1e-40 |
| CCG06109.1 | type\_I\_polyketide\_synthase | BGC0001543 | Polyketide | 29.0 | 36.5 | 166.0 | 2.1e-40 |
| AIJ04685.1 | polyketide\_synthase | BGC0001383 | Polyketide | 31.0 | 33.2 | 166.0 | 2.8e-40 |
| BAP05590.1 | calB | BGC0000967 | NRP + Polyketide:Trans-AT type I | 30.0 | 39.4 | 166.0 | 3.6e-40 |
| ABL74938.1 | PKS | BGC0001048 | NRP:Glycopeptide + Polyketide:Modular type I + Saccharide:Hybrid/tailoring | 27.0 | 48.8 | 165.0 | 4.7e-40 |
| AMH40422.1 | PKS | BGC0001350 | Polyketide | 29.0 | 32.5 | 165.0 | 4.7e-40 |
| AFN27482.1 | pks\_BonC | BGC0000173 | Polyketide:Modular type I | 31.0 | 31.8 | 165.0 | 8.1e-40 |
| ACR13065.1 | modular\_polyketide\_synthase,\_type\_I\_PKS | BGC0000185 | Polyketide | 31.0 | 28.8 | 165.0 | 8.1e-40 |
| ABC33986.1 | polyketide\_synthase,\_putative | BGC0000186 | NRP + Polyketide:Modular type I | 32.0 | 31.3 | 165.0 | 8.1e-40 |
| ABM21569.1 | crpA | BGC0000975 | NRP + Polyketide | 26.0 | 49.7 | 165.0 | 8.1e-40 |
| ABY21542.1 | AngAV | BGC0000018 | Polyketide | 25.0 | 50.0 | 164.0 | 1.1e-39 |
| CAG23969.1 | polyketide\_synthase\_type\_I | BGC0000181 | Polyketide | 31.0 | 33.9 | 164.0 | 1.1e-39 |
| AKQ22670.1 | malonyl\_CoA-acyl\_carrier\_protein\_transacylase | BGC0001656 | Polyketide | 29.0 | 41.2 | 164.0 | 1.1e-39 |
| AAO65790.1 | ketosynthase-like\_protein | BGC0000100 | Polyketide | 27.0 | 42.7 | 164.0 | 1.4e-39 |
| CAF05651.1 | TubF\_protein | BGC0001053 | NRP + Polyketide | 26.0 | 48.6 | 164.0 | 1.4e-39 |
| AAF00957.1 | mcyG | BGC0001017 | NRP + Polyketide:Modular type I | 26.0 | 49.5 | 163.0 | 1.8e-39 |
| ADH04680.1 | hybrid\_polyketide\_synthase/non-ribosomal\_peptide\_synthetase | BGC0001344 | NRP + Polyketide | 26.0 | 47.6 | 163.0 | 1.8e-39 |
| AFR69335.1 | polyketide\_synthase\_SpiC2 | BGC0001045 | NRP:Cyclic depsipeptide + Polyketide:Modular type I | 32.0 | 32.3 | 163.0 | 2.4e-39 |
| ctg1\_orf5 |  | BGC0001329 | Polyketide + NRP:Cyclic depsipeptide | 27.0 | 48.2 | 163.0 | 2.4e-39 |
| AQH32483.1 | hybrid\_peptide\_synthetase/polyketide\_synthase | BGC0001667 | NRP + Polyketide | 25.0 | 46.3 | 163.0 | 2.4e-39 |
| ATY69600.1 | type\_I\_polyketide\_synthase | BGC0001823 | NRP + Polyketide | 31.0 | 31.5 | 163.0 | 2.4e-39 |
| AAP42872.1 | NanA9 | BGC0000105 | Polyketide | 28.0 | 32.5 | 163.0 | 3.1e-39 |
| ABC34675.1 | polyketide\_synthase,\_putative | BGC0000186 | NRP + Polyketide:Modular type I | 30.0 | 33.0 | 163.0 | 3.1e-39 |
| ADA69241.1 | cis-AT\_polyketide\_synthase | BGC0001071 | NRP + Polyketide:Modular type I + Polyketide:Trans-AT type I | 27.0 | 43.3 | 162.0 | 4e-39 |
| ATG32074.1 | putative\_nonfunctional\_polyketide\_synthase\_module | BGC0001750 | NRP + Polyketide | 31.0 | 33.2 | 162.0 | 4e-39 |
| CAJ57408.1 | polyketide\_synthase\_type\_I | BGC0000176 | Polyketide + NRP | 32.0 | 33.4 | 162.0 | 5.3e-39 |
| ABO15888.1 | polyketide\_synthase | BGC0000132 | Polyketide | 31.0 | 34.7 | 161.0 | 6.9e-39 |
| AIU36104.1 | LglE | BGC0000180 | Polyketide:Trans-AT type I | 30.0 | 31.3 | 161.0 | 6.9e-39 |
| AHB82059.1 | non\_ribosomal\_peptide\_synthetase/polyketide\_synthase | BGC0001019 | NRP + Polyketide:Modular type I | 28.0 | 45.9 | 161.0 | 6.9e-39 |
| PHM26614.1 | Phthiocerol\_synthesis\_polyketide\_synthase\_type\_I\_PpsE | BGC0001130 | NRP + Polyketide | 26.0 | 49.4 | 161.0 | 6.9e-39 |
| AHB82072.1 | non\_ribosomal\_peptide\_synthetase/polyketide\_synthase | BGC0001231 | NRP + Polyketide:Modular type I | 28.0 | 45.4 | 161.0 | 6.9e-39 |
| CAE14173.1 | hypothetical\_protein | BGC0000383 | NRP + Polyketide:Modular type I | 28.0 | 47.9 | 161.0 | 9e-39 |
| ADI59534.1 | CorL | BGC0001091 | NRP + Polyketide | 31.0 | 32.7 | 161.0 | 9e-39 |
| AAR16521.1 | RimA | BGC0000138 | Polyketide | 29.0 | 31.8 | 161.0 | 1.2e-38 |
| CAG23978.1 | polyketide\_synthase\_type\_I | BGC0000176 | Polyketide + NRP | 28.0 | 43.6 | 161.0 | 1.2e-38 |
| AAW03327.1 | CtaD | BGC0000982 | NRP + Polyketide | 26.0 | 46.7 | 161.0 | 1.2e-38 |
| ALI92655.1 | CitS\_citrinin\_polyketide\_synthase | BGC0001338 | Polyketide:Iterative type I | 29.0 | 32.1 | 161.0 | 1.2e-38 |
| ABM63527.1 | BryB | BGC0000174 | Polyketide | 29.0 | 38.7 | 160.0 | 1.5e-38 |
| ADH01488.1 | polyketide\_synthase | BGC0001096 | NRP + Polyketide | 28.0 | 34.7 | 160.0 | 1.5e-38 |
| AIC32694.1 | FR9GH | BGC0001113 | NRP + Polyketide | 28.0 | 34.7 | 160.0 | 1.5e-38 |
| EAA59563.1 | polyketide\_synthase | BGC0000057 | Polyketide:Iterative type I | 27.0 | 49.1 | 160.0 | 2.6e-38 |
| CAG23968.1 | polyketide\_synthase\_type\_I | BGC0000181 | Polyketide | 30.0 | 28.8 | 160.0 | 2.6e-38 |
| AHV78247.1 | LasS2 | BGC0001245 | Polyketide | 26.0 | 44.9 | 160.0 | 2.6e-38 |
| DAB41918.1 | ArzP\_-\_PKS\_(KS,\_AT,\_OMT,\_ACP,\_TE) | BGC0001884 | NRP + Polyketide | 27.0 | 48.5 | 160.0 | 2.6e-38 |
| AEP40932.1 | polyketide\_synthase\_type\_I | BGC0000021 | Polyketide | 28.0 | 35.5 | 159.0 | 3.4e-38 |
| AFL48521.1 | beta-ketosynthase | BGC0000084 | Polyketide | 26.0 | 41.6 | 159.0 | 3.4e-38 |
| CBD77746.1 | non-ribosomal\_peptide\_synthetase/polyketide\_synthase | BGC0000974 | NRP + Polyketide | 29.0 | 43.8 | 159.0 | 3.4e-38 |
| AIR74926.1 | polyketide\_synthase | BGC0001559 | RiPP | 29.0 | 43.8 | 159.0 | 3.4e-38 |
| AMB48441.1 | polyketide\_synthase | BGC0001357 | Polyketide | 29.0 | 34.5 | 159.0 | 4.4e-38 |
| CAJ76298.1 | putative\_hybrid\_polyketide-non-ribosomal\_peptide\_synthetase | BGC0000972 | NRP + Polyketide:Modular type I + Polyketide:Trans-AT type I | 27.0 | 47.5 | 158.0 | 5.8e-38 |
| AIJ04686.1 | polyketide\_synthase | BGC0001383 | Polyketide | 30.0 | 28.8 | 158.0 | 5.8e-38 |
| AGN74894.1 | nonribosomal\_peptide\_synthetase/polyketide\_synthase\_hybrid\_protein | BGC0000459 | NRP:Cyclic depsipeptide + Polyketide:Trans-AT type I | 31.0 | 34.4 | 158.0 | 7.6e-38 |
| ATY69557.1 | type\_I\_polyketide\_synthase | BGC0001611 | NRP + Polyketide | 30.0 | 38.2 | 158.0 | 7.6e-38 |
| AGY62753.1 | EbeA | BGC0000051 | Polyketide | 41.0 | 20.1 | 158.0 | 9.9e-38 |
| BAP05595.1 | calG | BGC0000967 | NRP + Polyketide:Trans-AT type I | 31.0 | 32.7 | 158.0 | 9.9e-38 |
| ABM63537.1 | BryA | BGC0000174 | Polyketide | 31.0 | 30.8 | 157.0 | 1.3e-37 |
| CCP20049.1 | divL2\_protein | BGC0001119 | Polyketide:Modular type I | 28.0 | 32.0 | 157.0 | 1.7e-37 |
| ATX68126.1 | malonyl\_CoA-acyl\_carrier\_protein\_transacylase | BGC0001795 | Polyketide | 32.0 | 29.4 | 157.0 | 1.7e-37 |
| BAD38873.1 | polyketide\_synthase | BGC0000111 | Polyketide | 28.0 | 45.4 | 156.0 | 2.9e-37 |
| AGN11882.1 | tstGH | BGC0001114 | NRP + Polyketide | 29.0 | 32.6 | 156.0 | 2.9e-37 |
| AFL68054.1 | beta-ketoacyl\_synthase\_family\_protein,phosphopantetheine-containing\_protein | BGC0001524 | NRP + Polyketide | 29.0 | 30.5 | 156.0 | 2.9e-37 |
| ABM63529.1 | BryX | BGC0000174 | Polyketide | 30.0 | 31.9 | 156.0 | 3.8e-37 |
| CBK62731.1 |  | BGC0001115 | NRP + Polyketide | 29.0 | 31.9 | 156.0 | 3.8e-37 |
| WP\_039806854.1 | type\_I\_polyketide\_synthase | BGC0002001 | NRP + Polyketide | 27.0 | 46.9 | 155.0 | 6.4e-37 |
| ABM63530.1 | BryD | BGC0000174 | Polyketide | 28.0 | 39.3 | 155.0 | 8.4e-37 |
| ALJ49922.1 | TtmG | BGC0001236 | Polyketide | 32.0 | 30.3 | 155.0 | 8.4e-37 |
| BAE93725.1 | type\_I\_polyketide\_synthase | BGC0000164 | Polyketide | 30.0 | 32.1 | 154.0 | 1.1e-36 |
| AWM95789.1 | non-reduciing\_polyketide\_synthase\_methylorcinaldehyde\_synthase | BGC0001827 | Polyketide | 29.0 | 32.6 | 154.0 | 1.1e-36 |
| AEZ53950.1 | polyketide\_synthase | BGC0000144 | Polyketide:Modular type I | 29.0 | 32.1 | 154.0 | 1.4e-36 |
| AHB38498.1 | polyketide\_synthase | BGC0000346 | NRP + Polyketide:Modular type I | 28.0 | 43.9 | 154.0 | 1.4e-36 |
| AEH59110.1 | polyketide\_synthase\_of\_type\_I | BGC0000385 | NRP | 30.0 | 29.7 | 154.0 | 1.4e-36 |
| ACC40923.1 | polyketide\_synthase\_Pks9 | BGC0001665 | Polyketide | 30.0 | 36.3 | 154.0 | 1.4e-36 |
| EED18001.1 | NR-PKS | BGC0000154 | Polyketide:Iterative type I | 31.0 | 29.4 | 153.0 | 1.9e-36 |
| BAC68125.1 | modular\_polyketide\_synthase | BGC0000059 | Polyketide | 29.0 | 31.1 | 153.0 | 2.4e-36 |
| BAE93731.1 | type\_I\_polyketide\_synthase | BGC0000164 | Polyketide | 29.0 | 32.1 | 152.0 | 4.2e-36 |
| CCG06115.1 | hybrid\_NRPS/PKS | BGC0001543 | Polyketide | 27.0 | 39.4 | 152.0 | 4.2e-36 |
| AQW44872.1 | polyketide\_synthase | BGC0001761 | Polyketide | 29.0 | 32.2 | 152.0 | 4.2e-36 |
| ATG32077.1 | polyketide\_synthase | BGC0001750 | NRP + Polyketide | 31.0 | 33.4 | 152.0 | 5.4e-36 |
| WP\_004571777.1 | non-ribosomal\_peptide\_synthase | BGC0001760 | NRP | 30.0 | 32.9 | 151.0 | 7.1e-36 |
| AFV96142.1 | polyketide\_synthase | BGC0001064 | Polyketide:Modular type I + Polyketide:Type III | 28.0 | 35.0 | 151.0 | 1.2e-35 |
| BAF50721.1 | hybrid\_non\_ribosomal\_peptide\_synthetase-polyketide\_synthase | BGC0001116 | NRP + Polyketide | 29.0 | 32.6 | 151.0 | 1.2e-35 |
| ARU81122.1 | CylH | BGC0001566 | Polyketide | 28.0 | 35.0 | 151.0 | 1.2e-35 |
| CAJ76291.1 | putative\_polyketide\_synthase | BGC0000972 | NRP + Polyketide:Modular type I + Polyketide:Trans-AT type I | 30.0 | 34.9 | 150.0 | 1.6e-35 |
| AHD05619.1 | putative\_polyketide\_synthase\_subunit | BGC0001033 | NRP + Polyketide | 29.0 | 34.5 | 150.0 | 1.6e-35 |
| ABS90472.1 | PKS | BGC0001106 | NRP + Polyketide | 28.0 | 39.3 | 150.0 | 1.6e-35 |
| PPE\_RS11475 | aminotransferase\_class\_III-fold\_pyridoxal\_phosphate-dependent\_enzyme | BGC0001993 | NRP | 29.0 | 32.6 | 150.0 | 1.6e-35 |
| ABX60161.1 | mixed\_NRPS/PKS | BGC0000978 | NRP + Alkaloid + Polyketide:Modular type I | 31.0 | 31.8 | 150.0 | 2.1e-35 |
| CAQ34916.1 | polyketide\_synthase | BGC0000986 | NRP + Polyketide | 27.0 | 39.4 | 150.0 | 2.1e-35 |
| CAL80821.1 | sylD-like\_NRPS/PKS | BGC0000997 | NRP + Polyketide | 31.0 | 32.9 | 150.0 | 2.1e-35 |
| CAD29795.1 | peptide\_synthetase | BGC0001015 | NRP + Polyketide | 25.0 | 49.5 | 150.0 | 2.1e-35 |
| CBJ89764.1 | Polyketide\_synthase\_involved\_in\_xenocoumacin\_synthesis | BGC0001054 | NRP + Polyketide:Modular type I | 25.0 | 44.1 | 149.0 | 3.5e-35 |
| DAC76735.1 | type\_I\_polyketide\_synthase | BGC0001885 | Polyketide | 31.0 | 32.7 | 149.0 | 3.5e-35 |
| BBD17742.1 | polyketide\_synthase | BGC0001918 | NRP + Polyketide | 26.0 | 44.7 | 149.0 | 3.5e-35 |
| CBF69451.1 | polyketide\_synthase,\_putative\_(JCVI) | BGC0000037 | Polyketide | 28.0 | 43.7 | 149.0 | 4.6e-35 |
| ADF88262.1 | mixed\_nonribosomal\_peptide\_synthetase/\_polyketide\_synthase | BGC0000979 | NRP + Polyketide | 31.0 | 31.8 | 148.0 | 6e-35 |
| ADF88265.1 | mixed\_nonribosomal\_peptide\_synthetase/\_polyketide\_synthase | BGC0000980 | NRP + Polyketide | 31.0 | 31.8 | 148.0 | 6e-35 |
| ADF88279.1 | mixed\_NRPS/PKS | BGC0000981 | NRP + Polyketide | 31.0 | 31.8 | 148.0 | 6e-35 |
| ADZ24995.1 | non-ribosomal\_peptide\_synthase/polyketide\_synthase | BGC0000380 | NRP + Polyketide:Modular type I | 26.0 | 44.1 | 148.0 | 7.8e-35 |
| CCC55921.1 | non-ribosomal\_peptide\_synthetase/polyketide\_synthase\_hybrid\_protein | BGC0000973 | NRP + Polyketide:Modular type I | 27.0 | 45.9 | 148.0 | 7.8e-35 |
| AEH42474.1 | polyketide\_synthase | BGC0000032 | Polyketide | 32.0 | 25.8 | 148.0 | 1e-34 |
| AFD30954.1 | CrmA | BGC0000966 | NRP + Polyketide | 27.0 | 48.5 | 147.0 | 1.3e-34 |
| AHD05679.1 | putative\_non-ribosomal\_peptide\_ligase/\_polyketide\_synthase\_hybrid | BGC0000402 | NRP | 30.0 | 29.6 | 147.0 | 1.7e-34 |
| AAR87760.2 | ZmaK | BGC0001059 | NRP + Polyketide | 30.0 | 35.5 | 146.0 | 2.3e-34 |
| CAP95404.1 |  | BGC0001404 | Polyketide | 29.0 | 31.0 | 146.0 | 2.3e-34 |
| ADH01490.1 | type\_I\_polyketide\_synthase | BGC0001096 | NRP + Polyketide | 30.0 | 31.8 | 146.0 | 3e-34 |
| AAF19812.1 | MtaD | BGC0001024 | NRP + Polyketide:Modular type I | 29.0 | 32.7 | 146.0 | 3e-34 |
| AIC32695.1 | FR9I | BGC0001113 | NRP + Polyketide | 30.0 | 31.8 | 146.0 | 3e-34 |
| AAS98781.1 | polyketide\_synthase | BGC0001001 | NRP + Polyketide | 27.0 | 36.3 | 146.0 | 3.9e-34 |
| AEF33079.1 | polyketide\_synthase | BGC0001039 | NRP + Polyketide | 27.0 | 45.2 | 146.0 | 3.9e-34 |
| CDG12864.1 | non-ribosomal\_peptide\_synthetase | BGC0001415 | NRP | 27.0 | 39.6 | 146.0 | 3.9e-34 |
| AFK57216.1 | DidE | BGC0000985 | Polyketide + NRP:Cyclic depsipeptide | 31.0 | 33.0 | 145.0 | 6.6e-34 |
| AAG02357.1 | polyketide\_synthase | BGC0000963 | NRP:Glycopeptide + Polyketide:Modular type I + Saccharide:Hybrid/tailoring | 29.0 | 32.6 | 144.0 | 1.1e-33 |
| BAC76476.1 | multifunctional\_polyketide-peptide\_synthase\_LkcA | BGC0001100 | NRP + Polyketide | 29.0 | 28.9 | 143.0 | 1.9e-33 |
| CBF83139.1 | polyketide\_synthase,\_putative\_(JCVI) | BGC0001722 | Polyketide | 31.0 | 27.5 | 143.0 | 2.5e-33 |
| TGZ15164.1 | hypothetical\_protein | BGC0002032 | Polyketide | 28.0 | 30.6 | 143.0 | 2.5e-33 |
| AAQ84144.1 | Plm4 | BGC0000123 | Polyketide | 30.0 | 26.5 | 142.0 | 4.3e-33 |
| CBF70387.1 | polyketide\_synthase,\_putative\_(JCVI) | BGC0000684 | Terpene | 30.0 | 31.4 | 142.0 | 4.3e-33 |
| BAD55609.1 | putative\_polyketide\_synthase | BGC0001027 | NRP + Polyketide | 28.0 | 32.3 | 142.0 | 4.3e-33 |
| QCO93110.1 | polyketide\_synthase | BGC0001977 | Other | 29.0 | 32.1 | 142.0 | 4.3e-33 |
| AAY32966.1 | DszC | BGC0001093 | NRP + Polyketide | 30.0 | 32.7 | 142.0 | 5.6e-33 |
| WP\_055469550.1 | type\_I\_polyketide\_synthase | BGC0001537 | Polyketide | 27.0 | 40.6 | 141.0 | 1.3e-32 |
| AGN11883.1 | tstI | BGC0001114 | NRP + Polyketide | 29.0 | 32.3 | 140.0 | 2.1e-32 |
| ALJ49910.1 | TlmH | BGC0001237 | Polyketide | 27.0 | 31.5 | 139.0 | 3.6e-32 |
| EJP62792.1 | polyketide\_synthase | BGC0001720 | Polyketide | 28.0 | 31.7 | 139.0 | 3.6e-32 |
| ATQ39432.1 | PKS | BGC0001565 | NRP | 24.0 | 53.2 | 139.0 | 4.8e-32 |
| AXA20090.1 | hybrid\_trans-AT\_PKS/NRPS\_LgaA | BGC0001946 | NRP + Polyketide | 30.0 | 32.7 | 139.0 | 4.8e-32 |
| DAB41653.1 | polyketide\_synthase | BGC0001583 | Polyketide | 29.0 | 33.1 | 138.0 | 6.2e-32 |
| AKQ22697.1 | malonyl\_CoA-acyl\_carrier\_protein\_transacylase | BGC0001186 | Polyketide | 29.0 | 30.5 | 138.0 | 8.1e-32 |
| AHB82051.1 | polyketide\_synthase | BGC0001019 | NRP + Polyketide:Modular type I | 30.0 | 31.4 | 138.0 | 1.1e-31 |
| ATY12792.1 | polyketide\_synthase | BGC0001504 | Polyketide | 30.0 | 32.4 | 137.0 | 1.4e-31 |
| ctg1\_orf10 |  | BGC0000053 | Polyketide | 26.0 | 34.3 | 137.0 | 1.8e-31 |
| AKA59448.1 | polyketide\_synthase | BGC0001203 | NRP + Polyketide | 27.0 | 42.0 | 136.0 | 3.1e-31 |
| ABI91468.1 | beta-ketoacyl\_synthase | BGC0001094 | NRP + Polyketide | 29.0 | 31.2 | 136.0 | 4e-31 |
| EAL84397.1 | polyketide\_synthase | BGC0001118 | Polyketide:Iterative type I | 29.0 | 30.3 | 135.0 | 5.3e-31 |
| AEC04364.1 | polyketide\_synthase | BGC0000178 | Polyketide:Trans-AT type I | 30.0 | 30.1 | 134.0 | 1.2e-30 |
| ADH01484.1 | putative\_type-I\_PKS | BGC0001096 | NRP + Polyketide | 28.0 | 31.4 | 134.0 | 1.2e-30 |
| CAJ87591.1 | putative\_peptide/polyketide\_synthetase | BGC0001055 | NRP + Polyketide | 27.0 | 43.2 | 134.0 | 1.2e-30 |
| AIC32692.1 | FR9C | BGC0001113 | NRP + Polyketide | 28.0 | 31.4 | 134.0 | 1.2e-30 |
| ABB90282.1 | polyketide\_synthase | BGC0001057 | NRP + Polyketide | 27.0 | 30.6 | 134.0 | 1.5e-30 |
| ADF88260.1 | polyketide\_synthase | BGC0000979 | NRP + Polyketide | 33.0 | 21.2 | 133.0 | 2e-30 |
| ADF88263.1 | polyketide\_synthase | BGC0000980 | NRP + Polyketide | 33.0 | 21.2 | 133.0 | 2e-30 |
| CAA73127.1 | HMWP1\_protein | BGC0000467 | NRP | 32.0 | 27.6 | 133.0 | 2.6e-30 |
| CCE67070.1 | polyketide\_synthase | BGC0001242 | Polyketide | 27.0 | 31.9 | 133.0 | 2.6e-30 |
| WP\_035122279.1 | type\_I\_polyketide\_synthase | BGC0001467 | NRP:Cyclic depsipeptide + Polyketide:Modular type I | 28.0 | 32.7 | 133.0 | 2.6e-30 |
| AHB82057.1 | polyketide\_synthase | BGC0001019 | NRP + Polyketide:Modular type I | 28.0 | 32.4 | 132.0 | 4.5e-30 |
| WP\_019634550.1 | type\_I\_polyketide\_synthase | BGC0001443 | NRP + Polyketide | 29.0 | 37.3 | 132.0 | 4.5e-30 |
| ARR97039.1 | SphF | BGC0001780 | NRP | 34.0 | 20.1 | 131.0 | 7.6e-30 |
| ALP32045.1 | CycE | BGC0001293 | Polyketide | 27.0 | 34.3 | 131.0 | 9.9e-30 |
| AAB81123.1 | unknown | BGC0000862 | Other | 26.0 | 34.4 | 131.0 | 1.3e-29 |
| CBK62718.1 |  | BGC0001115 | NRP + Polyketide | 27.0 | 32.3 | 131.0 | 1.3e-29 |
| BAL90255.1 | putative\_beta-ketoacyl\_synthase | BGC0002021 | Polyketide | 29.0 | 32.7 | 131.0 | 1.3e-29 |
| AAL01060.1 | omega-3\_polyunsaturated\_fatty\_acid\_synthase\_PfaA | BGC0000865 | Other | 26.0 | 35.0 | 130.0 | 1.7e-29 |
| ABS90471.1 | PKS\_type\_I | BGC0001106 | NRP + Polyketide | 29.0 | 32.1 | 130.0 | 1.7e-29 |
| AEZ53949.1 | polyketide\_synthase | BGC0000144 | Polyketide:Modular type I | 26.0 | 35.1 | 130.0 | 2.2e-29 |
| AHB82062.1 | polyketide\_synthase | BGC0001231 | NRP + Polyketide:Modular type I | 27.0 | 30.8 | 130.0 | 2.2e-29 |
| AGN74892.1 | nonribosomal\_peptide\_synthetase/polyketide\_synthase\_hybrid\_protein | BGC0000459 | NRP:Cyclic depsipeptide + Polyketide:Trans-AT type I | 37.0 | 20.5 | 129.0 | 2.9e-29 |
| CCP45170.1 | Polyketide\_synthetase\_MbtC\_(polyketide\_synthase) | BGC0001021 | NRP + Polyketide | 26.0 | 32.0 | 129.0 | 2.9e-29 |
| CAG23958.2 | polyketide\_synthase\_of\_type\_I | BGC0001089 | Polyketide + NRP | 32.0 | 25.8 | 129.0 | 2.9e-29 |
| QCX41945.1 | Amc8 | BGC0001958 | Other | 27.0 | 43.6 | 129.0 | 2.9e-29 |
| AFX60341.1 | polyketide\_synthase | BGC0001032 | NRP + Polyketide | 31.0 | 26.2 | 129.0 | 3.8e-29 |
| ACD39770.1 | non-reducing\_polyketide\_synthase | BGC0000134 | Polyketide | 27.0 | 31.1 | 129.0 | 4.9e-29 |
| CAD89775.1 | MelD\_protein | BGC0001010 | NRP + Polyketide:Modular type I | 27.0 | 35.9 | 128.0 | 1.1e-28 |
| AHV78253.1 | ResS2 | BGC0001246 | Polyketide | 27.0 | 32.3 | 126.0 | 2.4e-28 |
| CBF74114.1 | Conidial\_yellow\_pigment\_biosynthesis\_polyketide\_synthase\_(PKS)(EC\_2.3.1.-)\_[Source:UniProtKB/Swiss-Prot;Acc:Q03149] | BGC0000107 | Polyketide | 26.0 | 31.6 | 126.0 | 4.2e-28 |
| AGN11880.1 | tstC | BGC0001114 | NRP + Polyketide | 28.0 | 34.4 | 126.0 | 4.2e-28 |
| WP\_030498975.1 | type\_I\_polyketide\_synthase | BGC0001327 | NRP:Cyclic depsipeptide + Polyketide:Modular type I | 29.0 | 32.4 | 126.0 | 4.2e-28 |
| EWM63003.1 | polyketide\_synthase\_type\_I | BGC0001328 | NRP:Cyclic depsipeptide + Polyketide:Modular type I | 29.0 | 32.4 | 125.0 | 5.4e-28 |
| ABF87031.1 | non-ribosomal\_peptide\_synthetase/polyketide\_synthase | BGC0000393 | NRP + Polyketide:Modular type I | 27.0 | 33.0 | 125.0 | 7.1e-28 |
| CBJ89766.1 | Polyketide\_synthase\_involved\_in\_xenocoumacin\_synthesis | BGC0001054 | NRP + Polyketide:Modular type I | 29.0 | 31.2 | 124.0 | 1.6e-27 |
| ABC35027.1 | JamP | BGC0000961 | NRP + Polyketide | 30.0 | 27.8 | 123.0 | 2.1e-27 |
| ACD39753.1 | non-reducing\_polyketide\_synthase | BGC0000076 | Polyketide | 28.0 | 30.6 | 123.0 | 2.7e-27 |
| ACD39762.1 | non-reducing\_polyketide\_synthase | BGC0000077 | Polyketide | 28.0 | 30.6 | 123.0 | 2.7e-27 |
| BAW32334.1 | hybrid\_cis-AT\_polyketide\_synthase\_-\_nonribosomal\_peptide\_synthetase | BGC0001631 | NRP + Polyketide | 26.0 | 35.2 | 123.0 | 2.7e-27 |
| antaD | Type\_I\_PKS | BGC0001455 | NRP + Polyketide | 27.0 | 32.3 | 123.0 | 3.5e-27 |
| CBJ82077.1 | hypothetical\_protein | BGC0001872 | Polyketide | 27.0 | 31.7 | 122.0 | 4.6e-27 |
| AFB35635.1 | KS-AT-ACP | BGC0000935 | Polyketide + Other:Aminocoumarin | 31.0 | 27.0 | 122.0 | 6e-27 |
| AGC65516.1 | NRPS/PKS\_hybrid | BGC0001050 | NRP:Lipopeptide + Polyketide:Trans-AT type I | 28.0 | 33.1 | 121.0 | 1e-26 |
| AXM42949.1 | hybrid\_type\_1\_PKS/NRPS | BGC0001941 | NRP + Polyketide | 28.0 | 32.7 | 121.0 | 1e-26 |
| ACG60781.1 | PKS(KS/AT/MT/KR/ACP) | BGC0001058 | NRP:Glycopeptide + Polyketide:Modular type I + Saccharide:Hybrid/tailoring | 32.0 | 20.2 | 121.0 | 1.3e-26 |
| WP\_019032754.1 | type\_I\_polyketide\_synthase | BGC0001331 | NRP:Cyclic depsipeptide + Polyketide:Modular type I | 28.0 | 32.4 | 121.0 | 1.3e-26 |
| ARO38317.1 | nonribosomal\_peptide\_synthetase | BGC0001560 | NRP + Polyketide | 26.0 | 35.2 | 120.0 | 2.3e-26 |
| AGC95321.1 | CurS2 | BGC0000045 | Polyketide | 27.0 | 33.4 | 119.0 | 3e-26 |
| WP\_106731933.1 | type\_I\_polyketide\_synthase | BGC0001332 | NRP + Polyketide | 31.0 | 26.9 | 119.0 | 5.1e-26 |
| AGO59040.1 | PtaA | BGC0000121 | Polyketide | 27.0 | 31.7 | 118.0 | 6.7e-26 |
| EED21099.1 | polyketide\_synthase,\_putative | BGC0001578 | Polyketide | 27.0 | 31.6 | 118.0 | 6.7e-26 |
| AEH42473.1 | polyketide\_synthase | BGC0000032 | Polyketide | 28.0 | 30.4 | 118.0 | 8.7e-26 |
| CCT67991.1 | bikaverin\_cluster-polyketide\_synthase | BGC0000030 | Polyketide | 28.0 | 32.3 | 118.0 | 1.1e-25 |
| EYT83433.1 | hypothetical\_protein | BGC0001213 | Polyketide | 27.0 | 33.6 | 117.0 | 1.9e-25 |
| ADH01485.1 | putative\_mixed\_polyketide\_synthase/non-ribosomal\_peptide\_synthetase | BGC0001096 | NRP + Polyketide | 31.0 | 22.5 | 115.0 | 5.6e-25 |
| AAF15892.2 | nosB | BGC0001028 | Polyketide + NRP:Cyclic depsipeptide | 28.0 | 32.8 | 115.0 | 7.4e-25 |
| PHM26606.1 | malonyl\_CoA-acyl\_carrier\_protein\_transacylase | BGC0001130 | NRP + Polyketide | 24.0 | 32.1 | 115.0 | 7.4e-25 |
| ASA76633.1 | polyketide\_synthase | BGC0001751 | NRP + Polyketide | 30.0 | 26.8 | 115.0 | 7.4e-25 |
| AAM12925.2 | MmpE | BGC0000182 | Polyketide:Iterative type I + Polyketide:Trans-AT type I | 26.0 | 28.7 | 114.0 | 9.6e-25 |
| AFR69332.1 | polyketide\_synthase\_SpiB | BGC0001045 | NRP:Cyclic depsipeptide + Polyketide:Modular type I | 28.0 | 32.7 | 114.0 | 1.3e-24 |
| CCM44330.1 | Polyketide\_synthase | BGC0001056 | NRP + Polyketide:Modular type I + Polyketide:PUFA synthase or related | 25.0 | 35.2 | 114.0 | 1.6e-24 |
| CBK62729.1 |  | BGC0001115 | NRP + Polyketide | 25.0 | 32.1 | 114.0 | 1.6e-24 |
| AEP40925.1 | polyketide\_synthase\_type\_I | BGC0000021 | Polyketide | 27.0 | 40.3 | 113.0 | 2.1e-24 |
| PKX92308.1 | putative\_polyketide\_synthase | BGC0001988 | Polyketide | 29.0 | 30.0 | 113.0 | 2.8e-24 |
| CCM44338.1 | Polyketide\_synthase | BGC0001056 | NRP + Polyketide:Modular type I + Polyketide:PUFA synthase or related | 25.0 | 33.0 | 113.0 | 3.7e-24 |
| CCG06108.1 | type\_I\_polyketide\_synthase | BGC0001543 | Polyketide | 27.0 | 25.8 | 113.0 | 3.7e-24 |
| ATY69584.1 | hybrid\_nonribosomal\_peptide\_synthetase/type\_I\_polyketide\_synthase | BGC0001823 | NRP + Polyketide | 28.0 | 33.1 | 113.0 | 3.7e-24 |
| AGJ76601.1 | HglE | BGC0000869 | Other | 25.0 | 32.0 | 112.0 | 6.2e-24 |
| CAL69888.1 | RhiA\_protein | BGC0001112 | NRP + Polyketide:Trans-AT type I | 26.0 | 31.7 | 112.0 | 6.2e-24 |
| AEH42487.1 | polyketide\_synthase | BGC0000032 | Polyketide | 27.0 | 24.3 | 111.0 | 1.4e-23 |
| AAS98787.1 | polyketide\_synthase/thioesterase | BGC0001001 | NRP + Polyketide | 26.0 | 33.2 | 111.0 | 1.4e-23 |
| ABP57746.1 | DepB | BGC0000993 | NRP:Cyclic depsipeptide + Polyketide:Modular type I | 27.0 | 32.7 | 110.0 | 1.8e-23 |
| ABC34108.1 | JamP | BGC0000961 | NRP + Polyketide | 26.0 | 34.0 | 110.0 | 2.4e-23 |
| AHD05615.1 | putative\_non-ribosomal\_peptide\_ligase/\_polyketide\_synthase\_hybrid | BGC0001033 | NRP + Polyketide | 27.0 | 31.9 | 110.0 | 2.4e-23 |
| KKP00966.1 | RADS2\_nonreducing\_polyketide\_synthase | BGC0001901 | Polyketide | 26.0 | 29.1 | 109.0 | 3.1e-23 |
| AHD05620.1 | putative\_polyketide\_synthase\_subunit | BGC0001033 | NRP + Polyketide | 30.0 | 25.7 | 109.0 | 5.3e-23 |
| ABK32260.1 | AmbF | BGC0000014 | Polyketide | 27.0 | 28.5 | 106.0 | 2.6e-22 |
| API82664.1 | putative\_polyketide\_synthase | BGC0001677 | Polyketide | 25.0 | 40.0 | 104.0 | 1.7e-21 |
| BAC76470.1 | type\_I\_polyketide\_synthase\_LkcG | BGC0001100 | NRP + Polyketide | 27.0 | 25.4 | 102.0 | 6.4e-21 |
| BBD17760.1 | polyketide\_synthase | BGC0001919 | NRP + Polyketide | 27.0 | 32.5 | 102.0 | 6.4e-21 |
| ADO69378.1 | beta-ketoacyl\_synthase-like\_protein,\_Acyl\_transferase\_subunit | BGC0000044 | Polyketide | 27.0 | 24.8 | 101.0 | 8.4e-21 |
| AWX24483.1 | type\_I\_polyketide\_synthase | BGC0001695 | NRP | 27.0 | 31.3 | 101.0 | 8.4e-21 |
| CUX96955.1 | TmcH | BGC0001829 | NRP + Polyketide | 32.0 | 20.2 | 101.0 | 1.4e-20 |
| CAJ76285.1 | putative\_polyketide\_synthase | BGC0000972 | NRP + Polyketide:Modular type I + Polyketide:Trans-AT type I | 26.0 | 32.6 | 100.0 | 1.9e-20 |
| ABC38737.1 | polyketide\_synthase | BGC0000964 | NRP:Cyclic depsipeptide + Polyketide:Trans-AT type I | 26.0 | 32.7 | 98.0 | 7.1e-20 |
| AEE88284.1 | CurF | BGC0000976 | NRP + Polyketide:Modular type I | 24.0 | 37.3 | 98.0 | 7.1e-20 |
| AAT70101.1 | CurF | BGC0001165 | NRP + Polyketide:Modular type I | 24.0 | 37.3 | 98.0 | 7.1e-20 |
| CAJ76289.1 | putative\_hybrid\_non-ribosomal\_peptide-polyketide\_synthetase | BGC0000972 | NRP + Polyketide:Modular type I + Polyketide:Trans-AT type I | 27.0 | 28.3 | 96.0 | 4.6e-19 |
| EAL89339.1 | polyketide\_synthase,\_putative | BGC0001403 | Polyketide | 29.0 | 20.6 | 94.0 | 1e-18 |
| BAF50720.1 | hybrid\_non\_ribosomal\_peptide\_synthetase-polyketide\_synthase | BGC0001116 | NRP + Polyketide | 25.0 | 33.6 | 91.0 | 1.1e-17 |
| WP\_030185025.1 | NAD-dependent\_epimerase/dehydratase\_family\_protein | BGC0001813 | NRP | 27.0 | 20.1 | 81.0 | 1.2e-14 |
| DAB41484.1 | nonribosomal\_peptide\_synthetase/polyketide\_synthase\_type\_I | BGC0001766 | NRP | 25.0 | 22.0 | 70.0 | 2.1e-11 |
| AAO39107.1 | AdmM | BGC0000956 | NRP:Beta-lactam + Polyketide:Type II | 27.0 | 22.1 | 69.0 | 4.6e-11 |
| AWH12666.1 | RmpD2 | BGC0001759 | Polyketide | 26.0 | 22.6 | 68.0 | 7.9e-11 |
| WP\_078625924.1 | type\_I\_polyketide\_synthase | BGC0002010 | NRP + Polyketide | 26.0 | 32.7 | 67.0 | 1.8e-10 |
| AZH23787.1 | MgcQ | BGC0001970 | NRP + Polyketide | 22.0 | 35.3 | 64.0 | 1.5e-09 |
