## Supplementary Results for "A draft genome of the ascomycotal fungal species *Pseudopithomyces maydicus* (family *Didymosphaeriaceae*)": input.path1.gene173_mibig_hits.html

| MIBiG Protein | Description | MIBiG Cluster | MiBiG Product | % ID | % Coverage | BLAST Score | E-value |
| --- | --- | --- | --- | --- | --- | --- | --- |
| TXD00261.1 | AMP-binding\_protein | BGC0001877 | Polyketide | 28.0 | 48.8 | 139.0 | 2.4e-32 |
| AAC68815.1 | FK506\_polyketide\_synthase | BGC0000353 | NRP | 29.0 | 45.0 | 139.0 | 3.1e-32 |
| AAF86393.1 | FkbB | BGC0000994 | NRP + Polyketide | 29.0 | 44.8 | 138.0 | 8.9e-32 |
| AGY30675.1 | Ann3 | BGC0001298 | Polyketide | 29.0 | 49.7 | 137.0 | 1.2e-31 |
| QBF51769.1 | type\_I\_polyketide\_synthase | BGC0001856 | Polyketide:Modular type I | 27.0 | 49.9 | 134.0 | 1.3e-30 |
| ABV97151.1 | AMP-dependent\_synthetase\_and\_ligase | BGC0000137 | Polyketide | 28.0 | 46.7 | 127.0 | 1.2e-28 |
| ABV99085.1 | thioester\_reductase\_domain | BGC0001007 | Polyketide + NRP | 29.0 | 45.3 | 126.0 | 2.7e-28 |
| CAA60460.1 | polyketide\_synthase | BGC0001040 | NRP + Polyketide | 27.0 | 44.8 | 126.0 | 3.5e-28 |
| ABP55493.1 | thioester\_reductase\_domain | BGC0001006 | NRP + Polyketide | 28.0 | 45.4 | 125.0 | 4.6e-28 |
| PKY07881.1 | hypothetical\_protein | BGC0001544 | NRP + Polyketide | 30.0 | 35.0 | 107.0 | 1.7e-22 |
| CAI94705.1 | putative\_acid\_AMP\_ligase | BGC0000141 | Polyketide | 27.0 | 47.6 | 106.0 | 3.8e-22 |
| AAF81723.1 | putative\_acyl-CoA\_ligase\_EncH | BGC0000220 | Polyketide:Type II | 25.0 | 44.4 | 87.0 | 1.4e-16 |
| YP\_856992.1 | 2,3-dihydroxybenzoate-AMP\_ligase | BGC0001502 | NRP | 25.0 | 46.0 | 78.0 | 6.4e-14 |
| AGN74895.1 | nonribosomal\_peptide\_synthetase/polyketide\_synthase\_hybrid\_protein | BGC0000459 | NRP:Cyclic depsipeptide + Polyketide:Trans-AT type I | 24.0 | 44.8 | 76.0 | 4.2e-13 |
| AQX14441.1 | EM5400\_NRPS\_scaffold | BGC0001671 | NRP | 24.0 | 43.8 | 74.0 | 1.6e-12 |
| CDG12864.1 | non-ribosomal\_peptide\_synthetase | BGC0001415 | NRP | 24.0 | 51.5 | 71.0 | 7.9e-12 |
| AAM12937.1 | MupU | BGC0000182 | Polyketide:Iterative type I + Polyketide:Trans-AT type I | 26.0 | 46.8 | 69.0 | 5.1e-11 |
| AAY37647.1 | Amino\_acid\_adenylation | BGC0000437 | NRP | 23.0 | 44.5 | 67.0 | 1.1e-10 |
| AAF99707.2 | syringopeptin\_synthetase | BGC0000438 | NRP | 22.0 | 43.6 | 62.0 | 3.7e-09 |
| AAY37655.1 | Amino\_acid\_adenylation | BGC0000437 | NRP | 21.0 | 44.4 | 59.0 | 5.3e-08 |
