## Supplementary Results for "A draft genome of the ascomycotal fungal species *Pseudopithomyces maydicus* (family *Didymosphaeriaceae*)": input.path1.gene177_mibig_hits.html

| MIBiG Protein | Description | MIBiG Cluster | MiBiG Product | % ID | % Coverage | BLAST Score | E-value |
| --- | --- | --- | --- | --- | --- | --- | --- |
| AAT12283.1 | LtxA | BGC0000384 | NRP | 29.0 | 73.3 | 379.0 | 3e-104 |
| AAO23334.1 | NcpB | BGC0000397 | NRP | 31.0 | 60.0 | 378.0 | 5.2e-104 |
| EOY45602.1 | non-ribosomal\_peptide\_synthetase | BGC0001168 | NRP | 28.0 | 72.8 | 331.0 | 7.2e-90 |
| ATD51280.1 | nonribosomal\_peptide\_synthase | BGC0001650 | NRP | 28.0 | 73.9 | 320.0 | 1.7e-86 |
| AQW44894.1 | non-ribosomal\_peptide\_synthetase | BGC0001737 | NRP + Polyketide | 32.0 | 48.1 | 308.0 | 6.6e-83 |
| ABI22133.1 | putative\_non-ribosomal\_peptide\_synthetase | BGC0000422 | NRP | 27.0 | 66.5 | 304.0 | 7.3e-82 |
| AAK57184.1 | MxaA | BGC0001022 | NRP + Polyketide | 32.0 | 47.9 | 303.0 | 1.6e-81 |
| AGC45618.1 | non-ribosomal\_peptide\_synthetase | BGC0001394 | NRP + Polyketide | 31.0 | 47.8 | 294.0 | 1.3e-78 |
| OAQ83772.1 | nonribosomal\_peptide\_synthase | BGC0001358 | Polyketide | 25.0 | 79.7 | 270.0 | 1.2e-71 |
| AIW82283.1 | PuwF | BGC0001125 | NRP + Polyketide | 30.0 | 47.9 | 269.0 | 2.6e-71 |
| BAO84866.1 | putative\_non-ribosomal\_peptide\_synthetase | BGC0000414 | NRP | 30.0 | 48.4 | 266.0 | 2.9e-70 |
| AIW82284.1 | PuwG | BGC0001125 | NRP + Polyketide | 29.0 | 48.3 | 264.0 | 1.1e-69 |
| CBF73453.1 | nonribosomal\_peptide\_synthase,\_putative\_(JCVI) | BGC0001515 | NRP | 26.0 | 66.0 | 263.0 | 2.4e-69 |
| ACN39015.1 | putative\_nonribosomal\_peptide\_synthetase\_TomB | BGC0000448 | NRP | 32.0 | 43.7 | 262.0 | 5.4e-69 |
| AGD80618.1 | non-ribosomal\_peptide\_synthetase | BGC0000394 | NRP | 29.0 | 48.5 | 261.0 | 7.1e-69 |
| ATD51278.1 | nonribosomal\_peptide\_synthase | BGC0001650 | NRP | 31.0 | 43.3 | 257.0 | 1e-67 |
| WP\_047197779.1 | non-ribosomal\_peptide\_synthetase | BGC0002003 | NRP | 29.0 | 49.7 | 257.0 | 1.3e-67 |
| BAP27942.1 | nonribosomal\_peptide\_synthetase | BGC0001085 | NRP + Terpene | 30.0 | 48.6 | 256.0 | 2.3e-67 |
| AOC89001.1 | putative\_nonribosomal\_peptide\_synthetase | BGC0001652 | NRP | 32.0 | 43.0 | 255.0 | 5.1e-67 |
| CAB38518.1 | CDA\_peptide\_synthetase\_I\_(CdaPs1) | BGC0000315 | NRP:Ca+-dependent lipopeptide | 31.0 | 48.3 | 253.0 | 1.5e-66 |
| AFO85453.1 | non-ribosomal\_peptide\_synthetase | BGC0000391 | NRP | 31.0 | 50.1 | 253.0 | 1.9e-66 |
| WP\_003981346.1 | non-ribosomal\_peptide\_synthetase | BGC0001813 | NRP | 29.0 | 47.8 | 251.0 | 9.5e-66 |
| AEH59100.1 | amino\_acid\_adenylation\_domain-containing\_protein/NRPS | BGC0000385 | NRP | 29.0 | 47.6 | 250.0 | 1.2e-65 |
| AAU34203.1 | mannopeptimycin\_peptide\_synthetase\_MppB | BGC0000388 | NRP | 28.0 | 48.6 | 249.0 | 2.8e-65 |
| ABY83163.1 | Azi25 | BGC0000960 | NRP + Polyketide | 29.0 | 56.9 | 248.0 | 8.1e-65 |
| QDA77059.1 | polyketide\_synthase/nonribosomal\_peptide\_synthetase | BGC0002026 | NRP | 30.0 | 48.7 | 246.0 | 2.3e-64 |
| ALV82356.1 | CDA\_peptide\_synthetase\_I | BGC0001370 | NRP | 30.0 | 48.3 | 246.0 | 3.1e-64 |
| CDG76959.1 | non-ribosomal\_peptide\_synthetase,\_terminal\_component | BGC0001805 | NRP | 30.0 | 43.5 | 243.0 | 2e-63 |
| AAO23333.1 | NcpA | BGC0000397 | NRP | 32.0 | 37.2 | 242.0 | 4.4e-63 |
| ADG27358.1 | peptide\_synthetase | BGC0000296 | NRP | 29.0 | 49.9 | 242.0 | 5.8e-63 |
| ABF87031.1 | non-ribosomal\_peptide\_synthetase/polyketide\_synthase | BGC0000393 | NRP + Polyketide:Modular type I | 28.0 | 48.3 | 241.0 | 9.9e-63 |
| AXN93614.1 | PuwF | BGC0001953 | NRP | 33.0 | 35.0 | 240.0 | 1.3e-62 |
| ABB90279.1 | non-ribosomal\_peptide\_synthetase | BGC0001057 | NRP + Polyketide | 27.0 | 58.1 | 240.0 | 1.7e-62 |
| AFH75322.1 | nonribosomal\_peptide\_synthetase | BGC0000425 | NRP:Cyclic depsipeptide | 29.0 | 48.0 | 239.0 | 2.9e-62 |
| AGM14934.1 | xantholysin\_synthetase\_C | BGC0000463 | NRP:Lipopeptide | 29.0 | 48.3 | 239.0 | 2.9e-62 |
| ALG65319.1 | Cal17 | BGC0001297 | NRP | 30.0 | 47.8 | 238.0 | 4.9e-62 |
| ANZ15840.1 | non-ribosomal\_peptide\_synthase/amino\_acid\_adenylation\_enzyme | BGC0001569 | NRP | 29.0 | 49.1 | 238.0 | 6.4e-62 |
| CCJ67640.1 | TaaE | BGC0000447 | NRP:Lipopeptide | 29.0 | 49.7 | 237.0 | 1.4e-61 |
| KUM80514.1 | hypothetical\_protein | BGC0001562 | NRP | 29.0 | 47.6 | 237.0 | 1.4e-61 |
| AXN93616.1 | PuwH | BGC0001953 | NRP | 29.0 | 52.3 | 237.0 | 1.4e-61 |
| CUX79061.1 | Octapeptin\_synthase\_subunit\_B | BGC0001715 | NRP | 31.0 | 36.4 | 237.0 | 1.9e-61 |
| AXN93603.1 | PuwH | BGC0001952 | NRP | 28.0 | 52.3 | 237.0 | 1.9e-61 |
| CAQ34921.1 | nonribosomal\_peptide\_synthetase | BGC0000986 | NRP + Polyketide | 31.0 | 40.1 | 235.0 | 5.4e-61 |
| WP\_013428324.1 | non-ribosomal\_peptide\_synthetase | BGC0001758 | NRP | 28.0 | 47.9 | 235.0 | 7.1e-61 |
| ATJ04411.1 | NRPS,\_TomB\_binding | BGC0001637 | NRP | 29.0 | 43.5 | 234.0 | 9.2e-61 |
| ABW17376.1 | PsoB | BGC0000411 | NRP | 29.0 | 50.6 | 233.0 | 1.6e-60 |
| AAD44234.1 | PstB | BGC0000362 | NRP | 30.0 | 46.7 | 232.0 | 4.6e-60 |
| CAY48788.1 | putative\_non-ribosomal\_peptide\_synthetase | BGC0001312 | NRP | 29.0 | 47.6 | 231.0 | 7.8e-60 |
| AGZ15458.1 | putative\_non-ribosomal\_peptide\_synthetase | BGC0001036 | NRP + Polyketide | 28.0 | 48.8 | 230.0 | 1.3e-59 |
| WP\_041826607.1 | non-ribosomal\_peptide\_synthetase | BGC0001792 | NRP | 28.0 | 52.1 | 230.0 | 1.3e-59 |
| KPN90376.1 | NupC | BGC0001416 | NRP | 28.0 | 47.8 | 230.0 | 1.7e-59 |
| AHB82072.1 | non\_ribosomal\_peptide\_synthetase/polyketide\_synthase | BGC0001231 | NRP + Polyketide:Modular type I | 28.0 | 49.0 | 230.0 | 2.3e-59 |
| ADA69239.2 | trans-AT\_hybrid\_polyketide\_synthase-NRPS | BGC0001071 | NRP + Polyketide:Modular type I + Polyketide:Trans-AT type I | 29.0 | 48.0 | 229.0 | 3e-59 |
| EWS95124.1 | hypothetical\_protein | BGC0000306 | NRP:Lipopeptide | 30.0 | 47.2 | 228.0 | 5.1e-59 |
| CAK15814.1 | putative\_non-ribosomal\_peptide\_synthetase,\_terminal\_component | BGC0000344 | NRP | 28.0 | 48.6 | 228.0 | 5.1e-59 |
| WP\_011146892.1 | non-ribosomal\_peptide\_synthetase | BGC0001641 | NRP | 28.0 | 48.6 | 228.0 | 5.1e-59 |
| AJV88375.1 | MfnC | BGC0001214 | NRP | 28.0 | 47.0 | 228.0 | 6.6e-59 |
| AHZ34238.1 | CipA | BGC0001389 | NRP | 30.0 | 48.5 | 228.0 | 6.6e-59 |
| AFJ23826.1 | WLIP\_synthetase\_C | BGC0001838 | NRP | 29.0 | 49.2 | 228.0 | 8.6e-59 |
| AAO56328.1 | non-ribosomal\_peptide\_synthetase\_SyfA | BGC0000435 | NRP | 28.0 | 47.6 | 227.0 | 1.9e-58 |
| AEA30272.1 | peptide\_synthetase | BGC0000429 | Polyketide + NRP:Cyclic depsipeptide | 29.0 | 48.1 | 226.0 | 2.5e-58 |
| ATU31794.1 | NRPS | BGC0001814 | NRP | 28.0 | 47.9 | 226.0 | 3.3e-58 |
| PHM49485.1 | Amino\_acid\_adenylation | BGC0001131 | NRP | 30.0 | 39.5 | 225.0 | 4.3e-58 |
| AXN93590.1 | PuwF-G | BGC0001951 | NRP | 28.0 | 48.7 | 225.0 | 7.3e-58 |
| APU91751.1 | Non-Ribosomal\_Peptide\_Synthetase | BGC0001806 | NRP | 28.0 | 48.3 | 224.0 | 9.6e-58 |
| WP\_047197778.1 | non-ribosomal\_peptide\_synthetase | BGC0002003 | NRP | 30.0 | 41.7 | 224.0 | 1.2e-57 |
| WP\_047195042.1 | non-ribosomal\_peptide\_synthetase | BGC0002003 | NRP | 27.0 | 53.5 | 223.0 | 1.6e-57 |
| AXN93581.1 | PuwF-G | BGC0001950 | NRP | 28.0 | 49.0 | 223.0 | 1.6e-57 |
| QBG38782.1 | Atr21 | BGC0001975 | NRP | 28.0 | 48.1 | 223.0 | 1.6e-57 |
| CCA89328.1 | mixed\_trans-AT\_type\_I\_polyketide\_synthase/nonribosomal\_peptide\_synthetase | BGC0001111 | NRP + Polyketide:Trans-AT type I | 27.0 | 52.7 | 223.0 | 2.1e-57 |
| AJF34463.1 | Txo1 | BGC0001207 | NRP | 29.0 | 47.7 | 223.0 | 2.8e-57 |
| AWI62628.1 | nonribosomal\_peptide\_synthetase | BGC0001943 | NRP | 33.0 | 36.0 | 223.0 | 2.8e-57 |
| WP\_010639240.1 | non-ribosomal\_peptide\_synthetase | BGC0000958 | NRP:Cyclic depsipeptide + Polyketide:Modular type I | 29.0 | 48.4 | 222.0 | 3.6e-57 |
| ARU08075.1 | mlcM | BGC0001448 | NRP:Ca+-dependent lipopeptide | 29.0 | 49.4 | 222.0 | 4.7e-57 |
| AGM16414.1 | paenibacterin\_synthetase\_C | BGC0000400 | NRP | 28.0 | 47.6 | 222.0 | 6.2e-57 |
| AAL33758.1 | putative\_non-ribosomal\_peptide\_synthetase | BGC0000421 | NRP | 29.0 | 47.6 | 222.0 | 6.2e-57 |
| BAH22764.1 | nonribosomal\_peptide\_synthetase | BGC0001018 | NRP + Polyketide | 30.0 | 38.3 | 222.0 | 6.2e-57 |
| BAW32323.1 | hybrid\_cis-AT\_polyketide\_synthase\_-\_nonribosomal\_peptide\_synthetase | BGC0001630 | NRP + Polyketide | 30.0 | 37.9 | 222.0 | 6.2e-57 |
| AYJ71720.1 | non-ribosomal\_peptide\_synthetase | BGC0001942 | NRP + Polyketide | 28.0 | 48.1 | 222.0 | 6.2e-57 |
| AAZ23078.1 | peptide\_synthetase | BGC0000291 | NRP | 29.0 | 47.9 | 221.0 | 1.1e-56 |
| WP\_039806856.1 | non-ribosomal\_peptide\_synthetase | BGC0002001 | NRP + Polyketide | 31.0 | 37.1 | 221.0 | 1.1e-56 |
| AAT01807.1 | non-ribosomal\_peptide\_synthetase | BGC0000365 | NRP | 31.0 | 45.9 | 220.0 | 1.8e-56 |
| XP\_002373813.1 | NRPS-like\_enzyme,\_putative | BGC0001516 | NRP | 30.0 | 42.1 | 220.0 | 2.4e-56 |
| AJW76710.1 | DsaH | BGC0001196 | NRP | 31.0 | 36.1 | 219.0 | 3.1e-56 |
| BAH43765.1 | tyrocidine\_synthetase\_II | BGC0000452 | NRP | 26.0 | 47.7 | 218.0 | 5.2e-56 |
| AAZ03552.1 | McnC | BGC0000332 | NRP | 29.0 | 42.9 | 218.0 | 6.8e-56 |
| CDG17985.1 | Putative\_Ornithine\_racemase\_(fragment) | BGC0000464 | NRP:Cyclic depsipeptide | 27.0 | 47.6 | 218.0 | 8.9e-56 |
| CBF76038.1 | nonribosomal\_peptide\_synthase,\_putative\_(Eurofung) | BGC0001399 | NRP | 27.0 | 63.2 | 218.0 | 8.9e-56 |
| AFJ23825.1 | WLIP\_synthetase\_B | BGC0001838 | NRP | 28.0 | 48.1 | 217.0 | 1.2e-55 |
| CAF05648.1 | TubC\_protein | BGC0001053 | NRP + Polyketide | 28.0 | 47.7 | 217.0 | 1.5e-55 |
| AFP87549.1 | NrpS | BGC0001135 | NRP | 29.0 | 42.9 | 217.0 | 1.5e-55 |
| EAU38971.1 | hypothetical\_protein | BGC0001122 | NRP + Polyketide:Iterative type I | 26.0 | 68.5 | 217.0 | 2e-55 |
| CDG17981.1 | Non-ribosomal\_peptide\_synthetase | BGC0000464 | NRP:Cyclic depsipeptide | 27.0 | 48.1 | 216.0 | 3.4e-55 |
| QED55423.1 | nonribosomal\_peptide\_synthetase | BGC0001984 | NRP | 29.0 | 46.9 | 216.0 | 3.4e-55 |
| CDG17982.1 | Non-ribosomal\_peptide\_synthetase | BGC0000464 | NRP:Cyclic depsipeptide | 28.0 | 48.3 | 215.0 | 4.4e-55 |
| CDG17986.1 | Non-ribosomal\_peptide\_synthetase | BGC0000464 | NRP:Cyclic depsipeptide | 28.0 | 46.6 | 215.0 | 7.6e-55 |
| ALK27914.1 | non-ribosomal\_peptide\_synthase | BGC0001233 | NRP | 30.0 | 45.4 | 215.0 | 7.6e-55 |
| AXN93602.1 | PuwF-G | BGC0001952 | NRP | 30.0 | 36.6 | 215.0 | 7.6e-55 |
| AAG31130.1 | MxcG | BGC0001345 | NRP | 30.0 | 44.1 | 214.0 | 9.9e-55 |
| ABA73955.1 | putative\_non-ribosomal\_peptide\_synthetase | BGC0001842 | NRP:Lipopeptide | 30.0 | 36.5 | 214.0 | 9.9e-55 |
| AHF21228.1 | TriD | BGC0000449 | NRP | 29.0 | 49.9 | 214.0 | 1.3e-54 |
| WP\_006051170.1 | non-ribosomal\_peptide\_synthetase | BGC0001999 | NRP | 28.0 | 55.5 | 213.0 | 1.7e-54 |
| KPN90369.1 | NunE | BGC0001416 | NRP | 28.0 | 50.3 | 213.0 | 2.2e-54 |
| ABH06368.2 | MassB | BGC0000389 | NRP:Cyclic depsipeptide | 27.0 | 48.2 | 213.0 | 2.9e-54 |
| EFE73312.1 | nonribosomal\_peptide\_synthetase | BGC0000431 | NRP:Cyclic depsipeptide | 29.0 | 48.5 | 213.0 | 2.9e-54 |
| CBJ90082.1 | Non\_Ribosomal\_peptide\_synthetase\_(-succinylbenzoate--CoA\_ligase) | BGC0001132 | NRP | 30.0 | 38.7 | 213.0 | 2.9e-54 |
| WP\_019032755.1 | non-ribosomal\_peptide\_synthetase | BGC0001331 | NRP:Cyclic depsipeptide + Polyketide:Modular type I | 31.0 | 36.0 | 212.0 | 3.8e-54 |
| AAX31558.1 | peptide\_synthetase\_2 | BGC0000336 | NRP | 27.0 | 47.5 | 212.0 | 4.9e-54 |
| BAF50711.1 | non\_ribosomal\_peptide\_synthetase\_for\_virginiamycin\_S | BGC0001116 | NRP + Polyketide | 29.0 | 47.9 | 212.0 | 4.9e-54 |
| AKA54626.1 | NRPS | BGC0001216 | NRP + Polyketide | 29.0 | 47.6 | 212.0 | 4.9e-54 |
| CDN62030.1 | Peptide\_synthetase | BGC0001900 | NRP | 28.0 | 48.5 | 212.0 | 4.9e-54 |
| ABW17375.1 | PsoA | BGC0000411 | NRP | 28.0 | 48.4 | 212.0 | 6.4e-54 |
| ABJ97436.1 | MerP | BGC0001012 | NRP + Polyketide | 27.0 | 48.8 | 211.0 | 8.4e-54 |
| WP\_079164394.1 | non-ribosomal\_peptide\_synthetase | BGC0001567 | NRP | 30.0 | 36.9 | 211.0 | 1.1e-53 |
| QED88055.1 | nonribosomal\_peptide\_synthetase | BGC0001967 | NRP | 27.0 | 48.7 | 211.0 | 1.1e-53 |
| AGC09528.1 | NRPS | BGC0001183 | Polyketide | 27.0 | 47.4 | 210.0 | 1.4e-53 |
| ALV86866.1 | Tlo20 | BGC0001406 | NRP | 29.0 | 49.8 | 210.0 | 1.4e-53 |
| BAV56271.1 |  | BGC0001657 | NRP | 31.0 | 34.9 | 210.0 | 1.4e-53 |
| AAX31557.1 | peptide\_synthetase\_1 | BGC0000336 | NRP | 28.0 | 49.3 | 210.0 | 1.9e-53 |
| antaC | NRPS | BGC0001455 | NRP + Polyketide | 28.0 | 49.4 | 210.0 | 1.9e-53 |
| ABR67744.1 | CmnA | BGC0000316 | NRP | 27.0 | 48.6 | 210.0 | 2.4e-53 |
| CBZ42146.1 | putative\_non-ribosomal\_peptide\_synthetase | BGC0001117 | NRP | 31.0 | 36.0 | 210.0 | 2.4e-53 |
| AXN93615.1 | PuwG | BGC0001953 | NRP | 30.0 | 36.6 | 210.0 | 2.4e-53 |
| CAJ77696.1 | MPS2\_protein | BGC0000363 | NRP | 30.0 | 44.3 | 209.0 | 3.2e-53 |
| ATO51563.1 | non-ribosomal\_peptide\_synthetase | BGC0001796 | NRP | 29.0 | 40.0 | 209.0 | 3.2e-53 |
| ABD65958.1 | nonribosomal\_peptide\_synthetase | BGC0000341 | NRP | 29.0 | 36.1 | 209.0 | 4.2e-53 |
| BAH43871.1 | truncated\_linear\_pentadecapeptide\_gramicidin\_synthetase\_LgrC | BGC0000367 | NRP | 31.0 | 34.0 | 209.0 | 4.2e-53 |
| QBG38783.1 | Atr22 | BGC0001975 | NRP | 28.0 | 49.2 | 209.0 | 4.2e-53 |
| AJF34464.1 | Txo2 | BGC0001207 | NRP | 28.0 | 48.4 | 208.0 | 5.4e-53 |
| AQM37584.1 | nonribosomal\_peptide\_synthetase | BGC0001424 | NRP:Cyclic depsipeptide + Polyketide:Iterative type I | 29.0 | 50.0 | 208.0 | 5.4e-53 |
| ABU70377.1 | hypothetical\_protein | BGC0001890 | NRP | 30.0 | 36.9 | 208.0 | 7.1e-53 |
| BAI63288.1 | putative\_non-ribosomal\_peptide\_synthetase | BGC0000434 | NRP | 30.0 | 38.2 | 207.0 | 1.6e-52 |
| CDG17987.1 | Putative\_Ornithine\_racemase\_(fragment) | BGC0000464 | NRP:Cyclic depsipeptide | 28.0 | 45.3 | 207.0 | 1.6e-52 |
| AEG64697.1 | LpmC | BGC0000379 | NRP | 30.0 | 35.4 | 207.0 | 2.1e-52 |
| ABC94347.1 | vicibactin\_biosynthesis\_non-ribosomal\_peptide\_synthase\_protein | BGC0000457 | NRP | 30.0 | 36.0 | 207.0 | 2.1e-52 |
| ABQ96384.2 | fusaricidin\_synthetase | BGC0001152 | Polyketide + NRP:Lipopeptide | 29.0 | 36.2 | 206.0 | 3.5e-52 |
| ABV79986.1 | ApnB | BGC0000301 | NRP | 28.0 | 37.6 | 205.0 | 4.6e-52 |
| BBA21071.1 | putative\_non-ribosomal\_peptide\_synthetase | BGC0001740 | NRP + Polyketide | 28.0 | 47.7 | 205.0 | 4.6e-52 |
| CAJ77716.1 | Mps2\_protein | BGC0000364 | NRP | 30.0 | 44.3 | 205.0 | 6e-52 |
| ATU31795.1 | NRPS | BGC0001814 | NRP | 28.0 | 49.7 | 205.0 | 7.8e-52 |
| AXN93613.1 | PuwE | BGC0001953 | NRP | 28.0 | 38.2 | 204.0 | 1e-51 |
| CAN89656.1 | putative\_non-ribosomal\_peptide\_synthetase | BGC0001070 | NRP + Polyketide:Modular type I + Polyketide:Trans-AT type I | 31.0 | 35.8 | 204.0 | 1.3e-51 |
| APZ78756.1 | nonribosomal\_peptide\_synthetase | BGC0001423 | NRP:Cyclic depsipeptide + Polyketide:Iterative type I | 29.0 | 49.6 | 204.0 | 1.3e-51 |
| BBD17759.1 | non-ribosomal\_peptide\_synthetase | BGC0001919 | NRP + Polyketide | 28.0 | 48.9 | 204.0 | 1.3e-51 |
| AJW76709.1 | DsaG | BGC0001196 | NRP | 28.0 | 50.7 | 203.0 | 1.7e-51 |
| ctg1\_orf003 |  | BGC0000334 | NRP | 26.0 | 58.8 | 203.0 | 2.3e-51 |
| AAY91420.2 | non-ribosomal\_peptide\_synthetase\_OfaB | BGC0000399 | NRP:Cyclic depsipeptide | 30.0 | 37.5 | 203.0 | 2.3e-51 |
| AXN93601.1 | PuwE | BGC0001952 | NRP | 27.0 | 38.2 | 203.0 | 2.3e-51 |
| AAS98786.1 | nonribosomal\_peptide\_synthetase | BGC0001001 | NRP + Polyketide | 30.0 | 36.4 | 202.0 | 3.9e-51 |
| ANZ15839.1 | peptide\_synthetase\_ScpsB | BGC0001569 | NRP | 27.0 | 48.1 | 202.0 | 5.1e-51 |
| QBC75022.1 | non-ribosomal\_peptide\_synthetase | BGC0001968 | NRP | 31.0 | 39.9 | 202.0 | 5.1e-51 |
| CAD91212.1 | putative\_non-ribosomal\_peptide\_synthetase,\_modules\_4-6 | BGC0000289 | NRP:Glycopeptide + Saccharide:Hybrid/tailoring | 31.0 | 34.5 | 201.0 | 6.6e-51 |
| ANI24100.1 | nonribosomal\_peptide\_synthetase | BGC0001235 | NRP + Polyketide | 32.0 | 35.7 | 201.0 | 6.6e-51 |
| WP\_053065269.1 | non-ribosomal\_peptide\_synthetase | BGC0001330 | NRP:Cyclic depsipeptide + Polyketide:Modular type I | 30.0 | 36.7 | 201.0 | 6.6e-51 |
| Xekj\_RS17945 | non-ribosomal\_peptide\_synthetase | BGC0001826 | NRP | 26.0 | 48.5 | 201.0 | 6.6e-51 |
| AAU34202.1 | mannopeptimycin\_peptide\_synthetase\_MppA | BGC0000388 | NRP | 31.0 | 35.8 | 201.0 | 8.7e-51 |
| BAI63283.1 | putative\_non-ribosomal\_peptide\_synthetase | BGC0000434 | NRP | 28.0 | 48.7 | 201.0 | 8.7e-51 |
| QBC75021.1 | non-ribosomal\_peptide\_synthetase | BGC0001968 | NRP | 32.0 | 35.5 | 201.0 | 8.7e-51 |
| QBG38784.1 | Atr23 | BGC0001975 | NRP | 27.0 | 49.0 | 201.0 | 8.7e-51 |
| CAM56770.1 |  | BGC0000354 | NRP | 30.0 | 37.8 | 200.0 | 1.5e-50 |
| CBJ79915.1 | putative\_Phenylalanine\_racemase\_(ATP-hydrolyzing) | BGC0001133 | NRP | 26.0 | 48.0 | 200.0 | 1.5e-50 |
| KFL51886.1 | amino\_acid\_adenylation\_protein | BGC0001711 | NRP + Polyketide | 27.0 | 49.2 | 200.0 | 1.5e-50 |
| AXN93575.1 | PuwA | BGC0001950 | NRP | 29.0 | 36.3 | 200.0 | 1.5e-50 |
| AXN93584.1 | PuwA | BGC0001951 | NRP | 29.0 | 36.3 | 200.0 | 1.5e-50 |
| BAD55611.1 | putative\_non-ribosomal\_peptide\_synthetase | BGC0001027 | NRP + Polyketide | 31.0 | 35.2 | 200.0 | 1.9e-50 |
| AAG02355.1 | peptide\_synthetase\_NRPS9-8 | BGC0000963 | NRP:Glycopeptide + Polyketide:Modular type I + Saccharide:Hybrid/tailoring | 26.0 | 47.9 | 200.0 | 2.5e-50 |
| BAF50720.1 | hybrid\_non\_ribosomal\_peptide\_synthetase-polyketide\_synthase | BGC0001116 | NRP + Polyketide | 29.0 | 40.7 | 200.0 | 2.5e-50 |
| AEH41793.1 | HrmO | BGC0000374 | NRP:Cyclic depsipeptide | 31.0 | 34.6 | 199.0 | 3.3e-50 |
| CZT62785.1 | Non-ribosomal\_peptide\_synthase,\_involved\_in\_Hassallidin\_biosynthesis | BGC0001614 | NRP | 30.0 | 36.1 | 199.0 | 3.3e-50 |
| APZ78744.1 | nonribosomal\_peptide\_synthetase | BGC0001422 | NRP:Cyclic depsipeptide + Polyketide:Iterative type I | 28.0 | 49.3 | 198.0 | 5.6e-50 |
| AGM14933.1 | xantholysin\_synthetase\_B | BGC0000463 | NRP:Lipopeptide | 30.0 | 42.8 | 198.0 | 7.3e-50 |
| WP\_051700105.1 | non-ribosomal\_peptide\_synthetase | BGC0001368 | NRP | 27.0 | 47.4 | 198.0 | 7.3e-50 |
| DAB41479.1 | nonribosomal\_peptide\_synthetase | BGC0001766 | NRP | 30.0 | 37.5 | 198.0 | 9.6e-50 |
| WP\_067426775.1 | non-ribosomal\_peptide\_synthetase | BGC0001567 | NRP | 32.0 | 36.7 | 198.0 | 9.6e-50 |
| BAV56270.1 |  | BGC0001657 | NRP | 29.0 | 47.1 | 197.0 | 1.3e-49 |
| DAC76731.1 | methionyl-tRNA\_formyltransferase | BGC0001885 | Polyketide | 27.0 | 47.5 | 197.0 | 1.3e-49 |
| AAY93445.1 | non-ribosomal\_peptide\_synthetase\_PvdL | BGC0000413 | NRP | 30.0 | 35.8 | 196.0 | 2.8e-49 |
| APZ78692.1 | nonribosomal\_peptide\_synthetase | BGC0001418 | NRP:Cyclic depsipeptide + Polyketide:Iterative type I | 30.0 | 35.1 | 196.0 | 2.8e-49 |
| APZ78716.1 | nonribosomal\_peptide\_synthetase | BGC0001420 | NRP:Cyclic depsipeptide + Polyketide:Iterative type I | 30.0 | 35.1 | 196.0 | 2.8e-49 |
| OKA09425.1 | non-ribosomal\_peptide\_synthetase | BGC0001459 | NRP:Glycopeptide | 28.0 | 43.2 | 196.0 | 2.8e-49 |
| AEU11003.1 | NpnC | BGC0001029 | NRP + Polyketide | 30.0 | 37.2 | 196.0 | 3.6e-49 |
| AIW82277.1 | PuwA | BGC0001125 | NRP + Polyketide | 29.0 | 36.3 | 196.0 | 3.6e-49 |
| BAX89998.1 | Non-ribosomal\_peptide\_synthetase | BGC0001628 | NRP | 27.0 | 46.3 | 196.0 | 3.6e-49 |
| CAC17499.1 | putative\_non-ribosomal\_peptide\_synthase | BGC0000324 | NRP | 27.0 | 49.7 | 195.0 | 4.8e-49 |
| AEG64696.1 | LpmB | BGC0000379 | NRP | 28.0 | 48.6 | 195.0 | 6.2e-49 |
| APU91750.1 | Non-Ribosomal\_Peptide\_Synthetase | BGC0001806 | NRP | 30.0 | 35.2 | 195.0 | 6.2e-49 |
| AIE77060.1 | peptide\_synthetase\_module\_7 | BGC0000418 | NRP | 28.0 | 36.3 | 195.0 | 8.1e-49 |
| CAD29797.1 | peptide\_synthetase | BGC0001015 | NRP + Polyketide | 30.0 | 36.2 | 195.0 | 8.1e-49 |
| AJD77023.1 | IkaA | BGC0001435 | NRP + Polyketide:Iterative type I | 28.0 | 48.6 | 195.0 | 8.1e-49 |
| AAF00961.1 | mcyB | BGC0001017 | NRP + Polyketide:Modular type I | 27.0 | 47.2 | 194.0 | 1.1e-48 |
| ABD14711.1 | cesA | BGC0000320 | NRP:Cyclic depsipeptide | 29.0 | 39.8 | 193.0 | 1.8e-48 |
| WP\_016638470.1 | non-ribosomal\_peptide\_synthetase | BGC0001519 | NRP + Polyketide | 30.0 | 39.1 | 193.0 | 2.4e-48 |
| AYJ71721.1 | non-ribosomal\_peptide\_synthetase | BGC0001942 | NRP + Polyketide | 27.0 | 36.7 | 193.0 | 2.4e-48 |
| APZ78856.1 | nonribosomal\_peptide\_synthetase | BGC0001432 | NRP:Cyclic depsipeptide + Polyketide:Iterative type I | 31.0 | 34.6 | 192.0 | 4e-48 |
| KUM80513.1 | hypothetical\_protein | BGC0001562 | NRP | 27.0 | 51.7 | 192.0 | 4e-48 |
| AQZ69228.1 | hypothetical\_protein | BGC0001635 | NRP + Polyketide | 30.0 | 36.3 | 192.0 | 4e-48 |
| CAJ18237.2 | non-ribosomal\_peptide\_synthetase\_B | BGC0000354 | NRP | 28.0 | 45.8 | 191.0 | 6.9e-48 |
| APZ78704.1 | nonribosomal\_peptide\_synthetase | BGC0001419 | NRP:Cyclic depsipeptide + Polyketide:Iterative type I | 30.0 | 35.1 | 191.0 | 6.9e-48 |
| CDG17980.1 | Putative\_Ornithine\_racemase\_(fragment) | BGC0000464 | NRP:Cyclic depsipeptide | 30.0 | 35.1 | 191.0 | 9e-48 |
| CAG23957.2 | hybrid\_NRPS/PKS\_protein | BGC0001089 | Polyketide + NRP | 25.0 | 47.7 | 191.0 | 1.2e-47 |
| ABL86391.1 | hybrid\_polyketide\_synthase\_and\_nonribosomal\_peptide\_synthetase | BGC0000999 | NRP + Polyketide | 26.0 | 48.3 | 190.0 | 1.5e-47 |
| AEH41794.1 | HrmP | BGC0000374 | NRP:Cyclic depsipeptide | 31.0 | 35.3 | 190.0 | 2e-47 |
| AIG79240.1 | Hypothetical\_protein | BGC0000419 | Saccharide + NRP:Glycopeptide | 29.0 | 36.0 | 190.0 | 2e-47 |
| CCP42826.1 | Probable\_peptide\_synthetase\_Nrp\_(peptide\_synthase) | BGC0001627 | NRP | 28.0 | 45.1 | 190.0 | 2e-47 |
| ABW00331.1 | amino\_acid\_adenylation\_domain | BGC0000333 | NRP | 29.0 | 38.7 | 190.0 | 2.6e-47 |
| CAF32362.1 | putative\_non-ribosomal\_peptide\_synthetase | BGC0000712 | Saccharide | 27.0 | 49.4 | 189.0 | 3.4e-47 |
| ATP76243.1 | NdaA | BGC0001705 | NRP + Polyketide | 26.0 | 50.8 | 189.0 | 3.4e-47 |
| CBL93730.1 | NRPS | BGC0000360 | NRP | 30.0 | 36.2 | 189.0 | 4.4e-47 |
| AET98916.1 | putative\_non-ribosomal\_peptide\_synthetase | BGC0000415 | NRP | 27.0 | 46.7 | 189.0 | 4.4e-47 |
| CAC11137.1 | NikP1\_protein | BGC0000876 | Other | 28.0 | 36.2 | 189.0 | 4.4e-47 |
| AGZ15459.1 | putative\_non-ribosomal\_peptide\_synthetase | BGC0001036 | NRP + Polyketide | 26.0 | 47.6 | 189.0 | 4.4e-47 |
| WP\_015507662.1 | non-ribosomal\_peptide\_synthetase | BGC0001792 | NRP | 29.0 | 48.8 | 189.0 | 4.4e-47 |
| BAH04173.1 | putative\_non-ribosomal\_peptide\_synthetase | BGC0000450 | NRP | 27.0 | 46.7 | 188.0 | 5.8e-47 |
| CCJ67648.1 | JagD | BGC0001127 | NRP | 30.0 | 35.7 | 188.0 | 5.8e-47 |
| AFR69334.1 | nonribosomal\_peptide\_synthetase\_SpiDE1 | BGC0001045 | NRP:Cyclic depsipeptide + Polyketide:Modular type I | 29.0 | 34.8 | 188.0 | 7.6e-47 |
| AWI62627.1 | nonribosomal\_peptide\_synthetase | BGC0001943 | NRP | 29.0 | 36.8 | 188.0 | 7.6e-47 |
| KFL51887.1 | amino\_acid\_adenylation\_protein | BGC0001711 | NRP + Polyketide | 29.0 | 37.6 | 188.0 | 9.9e-47 |
| AEF16059.1 | non-ribosomal\_peptide\_synthetase | BGC0000953 | Saccharide:Aminoglycoside | 26.0 | 48.3 | 187.0 | 1.3e-46 |
| CUX96954.1 | TmcG | BGC0001829 | NRP + Polyketide | 29.0 | 36.6 | 187.0 | 1.3e-46 |
| AEH59099.1 | amino\_acid\_adenylation\_domain-containing\_protein/NRPS | BGC0000385 | NRP | 29.0 | 36.0 | 187.0 | 1.7e-46 |
| ABO15860.1 | polyketide\_synthase | BGC0000130 | Polyketide | 28.0 | 37.2 | 186.0 | 2.2e-46 |
| ALG65313.1 | Cal23 | BGC0001297 | NRP | 29.0 | 47.0 | 186.0 | 2.2e-46 |
| APZ78680.1 | nonribosomal\_peptide\_synthetase | BGC0001417 | NRP:Cyclic depsipeptide + Polyketide:Iterative type I | 30.0 | 35.1 | 186.0 | 2.2e-46 |
| AHB82059.1 | non\_ribosomal\_peptide\_synthetase/polyketide\_synthase | BGC0001019 | NRP + Polyketide:Modular type I | 30.0 | 34.6 | 186.0 | 2.9e-46 |
| CAJ76290.1 | putative\_non-ribosomal\_peptide\_synthase | BGC0000972 | NRP + Polyketide:Modular type I + Polyketide:Trans-AT type I | 26.0 | 50.1 | 186.0 | 3.8e-46 |
| EAL85113.2 | hybrid\_PKS-NRPS\_enzyme | BGC0001037 | NRP + Polyketide:Iterative type I | 26.0 | 58.9 | 186.0 | 3.8e-46 |
| ATL73036.1 | amino\_acid\_adenylation\_protein | BGC0001807 | NRP + Polyketide | 27.0 | 50.5 | 186.0 | 3.8e-46 |
| CAL80821.1 | sylD-like\_NRPS/PKS | BGC0000997 | NRP + Polyketide | 26.0 | 47.9 | 185.0 | 4.9e-46 |
| WP\_015507663.1 | non-ribosomal\_peptide\_synthase | BGC0001792 | NRP | 30.0 | 45.1 | 185.0 | 4.9e-46 |
| WP\_004571777.1 | non-ribosomal\_peptide\_synthase | BGC0001760 | NRP | 27.0 | 48.2 | 185.0 | 6.4e-46 |
| AAP92491.1 | nonribosomal\_peptide\_synthetase | BGC0000458 | NRP | 26.0 | 47.1 | 184.0 | 1.1e-45 |
| ABD65957.1 | nonribosomal\_peptide\_synthetase | BGC0000341 | NRP | 29.0 | 38.6 | 184.0 | 1.4e-45 |
| CCP45168.1 | Peptide\_synthetase\_MbtE\_(peptide\_synthase) | BGC0001021 | NRP + Polyketide | 29.0 | 35.8 | 183.0 | 1.9e-45 |
| ADZ24999.1 | non-ribosomal\_peptide\_synthase | BGC0000380 | NRP + Polyketide:Modular type I | 29.0 | 36.7 | 183.0 | 3.2e-45 |
| AGD80623.1 | non-ribosomal\_peptide\_synthetase | BGC0000394 | NRP | 29.0 | 35.1 | 182.0 | 4.2e-45 |
| ALK27916.1 | non-ribosomal\_peptide\_synthase | BGC0001233 | NRP | 30.0 | 36.5 | 182.0 | 4.2e-45 |
| APZ78769.1 | nonribosomal\_peptide\_synthetase | BGC0001425 | NRP:Cyclic depsipeptide + Polyketide:Iterative type I | 28.0 | 49.5 | 182.0 | 4.2e-45 |
| APZ78834.1 | nonribosomal\_peptide\_synthetase | BGC0001430 | NRP:Cyclic depsipeptide + Polyketide:Iterative type I | 29.0 | 35.1 | 182.0 | 4.2e-45 |
| AQI70\_32580 |  | BGC0001561 | NRP | 30.0 | 36.2 | 182.0 | 4.2e-45 |
| AEO14743.1 | NdaA | BGC0000396 | NRP | 26.0 | 51.9 | 182.0 | 5.4e-45 |
| AGN74876.1 | nonribosomal\_peptide\_synthetase | BGC0000459 | NRP:Cyclic depsipeptide + Polyketide:Trans-AT type I | 30.0 | 33.1 | 181.0 | 7.1e-45 |
| APZ78846.1 | nonribosomal\_peptide\_synthetase | BGC0001431 | NRP:Cyclic depsipeptide + Polyketide:Iterative type I | 28.0 | 36.9 | 181.0 | 9.3e-45 |
| AAF86395.1 | FkbP | BGC0000994 | NRP + Polyketide | 30.0 | 36.4 | 181.0 | 1.2e-44 |
| ABV79988.1 | ApnD | BGC0000301 | NRP | 28.0 | 36.4 | 180.0 | 1.6e-44 |
| AHZ34232.1 | CifA | BGC0000323 | NRP:Lipopeptide | 28.0 | 39.8 | 180.0 | 1.6e-44 |
| CAE15637.1 |  | BGC0001128 | NRP | 27.0 | 48.2 | 180.0 | 1.6e-44 |
| AQM37583.1 | nonribosomal\_peptide\_synthetase | BGC0001424 | NRP:Cyclic depsipeptide + Polyketide:Iterative type I | 29.0 | 36.0 | 180.0 | 1.6e-44 |
| CAG29032.1 | nonribosomal\_peptide\_synthetase\_(modules\_3\_to\_6) | BGC0001023 | NRP + Polyketide:Modular type I | 29.0 | 35.1 | 180.0 | 2.1e-44 |
| AJI44167.1 | long-chain-fatty-acid-CoA\_ligase | BGC0001193 | NRP | 26.0 | 47.3 | 180.0 | 2.1e-44 |
| APZ78822.1 | nonribosomal\_peptide\_synthetase | BGC0001429 | NRP:Cyclic depsipeptide + Polyketide:Iterative type I | 29.0 | 35.1 | 180.0 | 2.1e-44 |
| CBJ79916.1 | putative\_Ornithine\_racemase | BGC0001133 | NRP | 29.0 | 35.7 | 180.0 | 2.7e-44 |
| APZ78782.1 | nonribosomal\_peptide\_synthetase | BGC0001426 | NRP:Cyclic depsipeptide + Polyketide:Iterative type I | 28.0 | 49.4 | 179.0 | 3.5e-44 |
| ABO15888.1 | polyketide\_synthase | BGC0000132 | Polyketide | 28.0 | 37.6 | 178.0 | 6e-44 |
| CAM02313.1 | putative\_non-ribosomal\_peptide\_synthetase | BGC0000349 | NRP | 29.0 | 38.2 | 178.0 | 6e-44 |
| APZ78728.1 | nonribosomal\_peptide\_synthetase | BGC0001421 | NRP:Cyclic depsipeptide + Polyketide:Iterative type I | 30.0 | 34.2 | 178.0 | 6e-44 |
| CAJ34367.1 | NRPS\_protein | BGC0000445 | NRP:Cyclic depsipeptide | 26.0 | 47.9 | 178.0 | 1e-43 |
| AKP45395.1 | CysG | BGC0001413 | NRP | 27.0 | 38.9 | 178.0 | 1e-43 |
| APZ78755.1 | nonribosomal\_peptide\_synthetase | BGC0001423 | NRP:Cyclic depsipeptide + Polyketide:Iterative type I | 29.0 | 34.4 | 177.0 | 1.3e-43 |
| ALV86867.1 | Tlo21 | BGC0001406 | NRP | 30.0 | 34.6 | 177.0 | 1.7e-43 |
| ABV56588.1 | KtzH | BGC0000378 | NRP | 28.0 | 45.6 | 176.0 | 3e-43 |
| AGJ76605.1 | HglB | BGC0000869 | Other | 34.0 | 21.2 | 176.0 | 3.9e-43 |
| AFK57215.1 | DidD | BGC0000985 | Polyketide + NRP:Cyclic depsipeptide | 28.0 | 47.4 | 176.0 | 3.9e-43 |
| BAE98162.1 | putative\_non-ribosomal\_peptide\_synthetase | BGC0000339 | NRP | 26.0 | 49.2 | 175.0 | 5.1e-43 |
| AGE11899.1 | nonribosomal\_peptide\_synthetase | BGC0000366 | NRP | 31.0 | 36.6 | 175.0 | 5.1e-43 |
| ABP57748.1 | DepD | BGC0000993 | NRP:Cyclic depsipeptide + Polyketide:Modular type I | 28.0 | 38.7 | 175.0 | 5.1e-43 |
| ctg1\_orf20 |  | BGC0001767 | NRP | 29.0 | 35.4 | 175.0 | 5.1e-43 |
| CAJ96472.1 | non-ribosomal\_peptide\_synthetase | BGC0000330 | NRP:NRP siderophore | 27.0 | 47.6 | 174.0 | 1.1e-42 |
| AFH75321.1 | nonribosomal\_peptide\_synthetase | BGC0000425 | NRP:Cyclic depsipeptide | 28.0 | 34.8 | 174.0 | 1.1e-42 |
| AFK57212.1 | DidA | BGC0000985 | Polyketide + NRP:Cyclic depsipeptide | 29.0 | 46.3 | 174.0 | 1.1e-42 |
| AHH25592.1 | NRPS | BGC0000957 | NRP + Polyketide | 28.0 | 35.3 | 174.0 | 1.5e-42 |
| DAC76736.1 | non-ribosomal\_peptide\_synthetase | BGC0001885 | Polyketide | 29.0 | 36.5 | 174.0 | 1.5e-42 |
| CBZ42140.1 | non-ribosomal\_peptide\_synthetase | BGC0001117 | NRP | 29.0 | 48.1 | 173.0 | 1.9e-42 |
| ACY06285.1 | non-ribosomal\_peptide\_synthetase | BGC0001042 | NRP + Polyketide | 26.0 | 41.8 | 173.0 | 2.5e-42 |
| AHB82058.1 | non\_ribosomal\_peptide\_synthetase | BGC0001019 | NRP + Polyketide:Modular type I | 29.0 | 35.6 | 173.0 | 3.3e-42 |
| WP\_107406787.1 | non-ribosomal\_peptide\_synthetase | BGC0001567 | NRP | 26.0 | 45.8 | 172.0 | 4.3e-42 |
| AAO72425.1 | syringopeptin\_synthetase\_C | BGC0000438 | NRP | 27.0 | 36.7 | 172.0 | 5.6e-42 |
| WP\_030498980.1 | hypothetical\_protein | BGC0001327 | NRP:Cyclic depsipeptide + Polyketide:Modular type I | 29.0 | 41.0 | 171.0 | 9.6e-42 |
| APZ78795.1 | nonribosomal\_peptide\_synthetase | BGC0001427 | NRP:Cyclic depsipeptide + Polyketide:Iterative type I | 28.0 | 48.1 | 171.0 | 9.6e-42 |
| APZ78809.1 | nonribosomal\_peptide\_synthetase | BGC0001428 | NRP:Cyclic depsipeptide + Polyketide:Iterative type I | 28.0 | 48.1 | 171.0 | 9.6e-42 |
| CAD29798.1 | peptide\_synthetase | BGC0001015 | NRP + Polyketide | 25.0 | 47.4 | 171.0 | 1.3e-41 |
| AAF01762.1 | AM-toxin\_synthetase | BGC0001261 | NRP | 26.0 | 47.4 | 171.0 | 1.3e-41 |
| AQH32485.1 | peptide\_synthetase | BGC0001667 | NRP + Polyketide | 26.0 | 37.1 | 171.0 | 1.3e-41 |
| BBA21068.1 | putative\_non-ribosomal\_peptide\_synthetase | BGC0001740 | NRP + Polyketide | 26.0 | 48.5 | 171.0 | 1.3e-41 |
| AAY37655.1 | Amino\_acid\_adenylation | BGC0000437 | NRP | 27.0 | 36.7 | 170.0 | 1.6e-41 |
| QCC62999.1 | BII-rafflesfungin\_nonribosomal\_protein\_synthetase | BGC0001966 | NRP | 26.0 | 48.8 | 170.0 | 1.6e-41 |
| AFH75320.1 | nonribosomal\_peptide\_synthetase | BGC0000425 | NRP:Cyclic depsipeptide | 28.0 | 34.7 | 170.0 | 2.8e-41 |
| AAT01806.1 | non-ribosomal\_peptide\_synthetase | BGC0000365 | NRP | 30.0 | 36.4 | 169.0 | 4.8e-41 |
| ABI22132.1 | putative\_non-ribosomal\_peptide\_synthetase | BGC0000422 | NRP | 29.0 | 38.2 | 169.0 | 4.8e-41 |
| CAJ34381.1 | NRPS\_protein | BGC0000445 | NRP:Cyclic depsipeptide | 29.0 | 38.1 | 168.0 | 1.1e-40 |
| AEP18656.1 | WAPS1 | BGC0000461 | NRP | 29.0 | 37.0 | 168.0 | 1.1e-40 |
| EWM63010.1 | non-ribosomal\_peptide\_synthetase | BGC0001328 | NRP:Cyclic depsipeptide + Polyketide:Modular type I | 29.0 | 41.1 | 168.0 | 1.1e-40 |
| SCO70310.1 | Type\_I\_polyketide\_synthase | BGC0001433 | Polyketide:Modular type I | 29.0 | 34.6 | 168.0 | 1.1e-40 |
| AFP87523.1 | type\_I\_polyketide\_synthase | BGC0001159 | NRP + Polyketide:Modular type I | 27.0 | 36.5 | 167.0 | 1.8e-40 |
| ABC87508.1 | NRPS\_for\_pipecolate\_incorporation | BGC0001011 | NRP + Polyketide | 28.0 | 36.8 | 166.0 | 2.4e-40 |
| ctg1\_orf19 |  | BGC0001013 | NRP + Polyketide | 28.0 | 36.8 | 166.0 | 2.4e-40 |
| APZ78729.1 | nonribosomal\_peptide\_synthetase | BGC0001421 | NRP:Cyclic depsipeptide + Polyketide:Iterative type I | 28.0 | 36.9 | 166.0 | 2.4e-40 |
| AKC91857.1 | nonribosomal\_peptide\_synthetase | BGC0001414 | NRP | 28.0 | 36.5 | 166.0 | 3.1e-40 |
| AQZ71347.1 | hypothetical\_protein | BGC0001635 | NRP + Polyketide | 28.0 | 34.2 | 166.0 | 3.1e-40 |
| AWS21279.1 | type\_I\_polyketide\_synthase | BGC0001934 | Polyketide | 31.0 | 27.1 | 166.0 | 3.1e-40 |
| AZY91989.1 | polyketide\_synthase | BGC0002022 | Polyketide | 31.0 | 27.1 | 166.0 | 3.1e-40 |
| EWS95122.1 | hypothetical\_protein | BGC0000306 | NRP:Lipopeptide | 29.0 | 35.2 | 165.0 | 5.3e-40 |
| AAF19811.1 | mtaC | BGC0001024 | NRP + Polyketide:Modular type I | 25.0 | 43.5 | 165.0 | 6.9e-40 |
| CRI73800.1 | loading\_module\_of\_NRPS-PKS | BGC0001215 | NRP | 28.0 | 36.0 | 165.0 | 6.9e-40 |
| BAT51067.1 | type\_I\_polyketide\_synthase | BGC0001296 | Polyketide | 29.0 | 34.1 | 165.0 | 6.9e-40 |
| CBA63680.1 | nonribosomal\_peptide\_synthetase\_NRPS | BGC0000368 | NRP | 29.0 | 35.9 | 165.0 | 9e-40 |
| BAO84868.1 | putative\_non-ribosomal\_peptide\_synthetase | BGC0000414 | NRP | 28.0 | 36.7 | 164.0 | 1.2e-39 |
| AHB38497.1 | non-ribosomal\_peptide\_synthetase | BGC0000346 | NRP + Polyketide:Modular type I | 30.0 | 35.2 | 164.0 | 1.5e-39 |
| AID65222.1 | putative\_aspartate\_racemase | BGC0000335 | NRP | 28.0 | 34.1 | 163.0 | 2.6e-39 |
| AEI58879.1 | peptide\_synthetase | BGC0000455 | NRP | 28.0 | 35.8 | 163.0 | 2.6e-39 |
| ABV56582.1 | KtzB | BGC0000378 | NRP | 29.0 | 30.4 | 163.0 | 3.4e-39 |
| AHH53506.1 | non-ribosomal\_peptide\_synthetase | BGC0000439 | NRP:Ca+-dependent lipopeptide | 29.0 | 36.7 | 163.0 | 3.4e-39 |
| ATY37608.1 | BreC | BGC0001536 | NRP | 26.0 | 35.8 | 163.0 | 3.4e-39 |
| CAC22144.1 | CpkC;\_Polyketide\_synthase\_module\_5 | BGC0000038 | Polyketide:Modular type I | 29.0 | 36.0 | 162.0 | 4.5e-39 |
| AKC91856.1 | nonribosomal\_peptide\_synthetase | BGC0001414 | NRP | 29.0 | 34.3 | 162.0 | 5.8e-39 |
| AEC14348.1 | nonribosomal\_peptide\_synthetase | BGC0000377 | NRP | 26.0 | 35.5 | 161.0 | 7.6e-39 |
| AAF19815.1 | mtaG | BGC0001024 | NRP + Polyketide:Modular type I | 29.0 | 30.4 | 161.0 | 9.9e-39 |
| CAE52334.1 | non-ribosomal\_peptide\_synthase | BGC0001088 | NRP + Polyketide | 25.0 | 38.6 | 161.0 | 9.9e-39 |
| AFU82614.1 | mixed\_NRPS\_PKS | BGC0000998 | NRP + Polyketide | 26.0 | 48.2 | 161.0 | 1.3e-38 |
| EME52974.1 | non-ribosomal\_peptide\_synthetase | BGC0001460 | NRP:Glycopeptide | 28.0 | 35.8 | 160.0 | 2.2e-38 |
| ARS01470.1 | NcmB | BGC0001702 | NRP + Polyketide | 28.0 | 36.9 | 160.0 | 2.2e-38 |
| ABP55493.1 | thioester\_reductase\_domain | BGC0001006 | NRP + Polyketide | 27.0 | 38.2 | 159.0 | 4.9e-38 |
| OKA09664.1 | non-ribosomal\_peptide\_synthetase | BGC0001459 | NRP:Glycopeptide | 28.0 | 35.9 | 158.0 | 6.4e-38 |
| AKC91849.1 | nonribosomal\_peptide\_synthetase | BGC0001414 | NRP | 29.0 | 34.3 | 158.0 | 1.1e-37 |
| ctg1\_orf1265 |  | BGC0001752 | NRP | 30.0 | 35.0 | 158.0 | 1.1e-37 |
| ADJ63842.1 | Serobactin\_synthetase | BGC0000424 | NRP:NRP siderophore | 27.0 | 35.9 | 157.0 | 1.9e-37 |
| EYT83459.1 | hypothetical\_protein | BGC0001213 | Polyketide | 30.0 | 28.8 | 157.0 | 1.9e-37 |
| ADG27359.1 | peptide\_synthetase | BGC0000296 | NRP | 28.0 | 36.0 | 156.0 | 2.4e-37 |
| AZH23788.1 | MgcR | BGC0001970 | NRP + Polyketide | 29.0 | 26.2 | 156.0 | 2.4e-37 |
| ABV99085.1 | thioester\_reductase\_domain | BGC0001007 | Polyketide + NRP | 28.0 | 33.6 | 156.0 | 3.2e-37 |
| ABL74940.1 | NRPS | BGC0001048 | NRP:Glycopeptide + Polyketide:Modular type I + Saccharide:Hybrid/tailoring | 26.0 | 45.4 | 156.0 | 3.2e-37 |
| CCM44336.1 | Nonribosomal\_peptide\_synthetase | BGC0001056 | NRP + Polyketide:Modular type I + Polyketide:PUFA synthase or related | 30.0 | 26.5 | 156.0 | 4.2e-37 |
| AAF19812.1 | MtaD | BGC0001024 | NRP + Polyketide:Modular type I | 27.0 | 36.2 | 155.0 | 9.3e-37 |
| WP\_013310343.1 | non-ribosomal\_peptide\_synthetase | BGC0001993 | NRP | 27.0 | 35.9 | 155.0 | 9.3e-37 |
| CAJ88610.1 | putative\_non-ribosomal\_peptide\_synthetase | BGC0000327 | NRP | 27.0 | 38.2 | 154.0 | 1.6e-36 |
| BBA84067.1 | type\_I\_polyketide\_synthase | BGC0001916 | Polyketide | 31.0 | 27.1 | 153.0 | 2.1e-36 |
| ABP55169.1 | amino\_acid\_adenylation\_domain | BGC0000150 | NRP + Polyketide:Enediyne type I | 26.0 | 35.9 | 153.0 | 2.7e-36 |
| ABV56604.1 | adenylation\_domain\_protein | BGC0000378 | NRP | 27.0 | 34.4 | 152.0 | 4.6e-36 |
| CAQ71827.1 | non\_ribosomal\_peptide\_synthase,\_antibiotic\_synthesis;\_contains\_1\_condensation\_domain,\_1\_AMP-acid\_ligases\_II\_domain | BGC0001189 | NRP | 28.0 | 34.2 | 152.0 | 4.6e-36 |
| KFH48607.1 | N-like\_protein | BGC0000317 | NRP | 27.0 | 36.6 | 152.0 | 6e-36 |
| ABP57749.1 | DepE | BGC0000993 | NRP:Cyclic depsipeptide + Polyketide:Modular type I | 29.0 | 34.0 | 152.0 | 6e-36 |
| AGN74885.1 | nonribosomal\_peptide\_synthetase | BGC0000459 | NRP:Cyclic depsipeptide + Polyketide:Trans-AT type I | 28.0 | 38.7 | 151.0 | 1e-35 |
| WP\_054234617.1 | non-ribosomal\_peptide\_synthase | BGC0002014 | NRP + Polyketide | 27.0 | 38.2 | 150.0 | 1.8e-35 |
| CAA79245.2 | enniatin\_synthetase | BGC0000342 | NRP | 24.0 | 36.3 | 150.0 | 2.3e-35 |
| AAL15600.1 | SimH | BGC0000270 | Polyketide | 32.0 | 26.3 | 149.0 | 3.9e-35 |
| AAG34184.1 | SimH | BGC0000835 | Polyketide | 32.0 | 26.3 | 149.0 | 3.9e-35 |
| AAK06804.1 | Tyroxyl-AMP-forming\_enzyme | BGC0001072 | Saccharide + Polyketide:Modular type I + Polyketide:Type II + Other:Aminocoumarin | 32.0 | 26.3 | 149.0 | 3.9e-35 |
| EAW16180.1 | nonribosomal\_peptide\_synthase,\_putative | BGC0000293 | NRP | 25.0 | 36.4 | 149.0 | 5.1e-35 |
| AXA20091.1 | hybrid\_trans-AT\_PKS/NRPS\_LgaB | BGC0001946 | NRP + Polyketide | 25.0 | 38.4 | 148.0 | 8.7e-35 |
| AAZ23075.1 | peptide\_synthetase | BGC0000291 | NRP | 29.0 | 36.7 | 146.0 | 3.3e-34 |
| ACO94461.1 | putative\_AMP-dependent\_acyl-CoA\_synthetase/ligase | BGC0000029 | Polyketide:Modular type I | 25.0 | 40.8 | 145.0 | 7.4e-34 |
| WP\_084702182.1 | non-ribosomal\_peptide\_synthetase | BGC0001211 | NRP | 27.0 | 38.0 | 145.0 | 7.4e-34 |
| WP\_050383084.1 | non-ribosomal\_peptide\_synthetase | BGC0001451 | NRP | 27.0 | 36.0 | 144.0 | 9.6e-34 |
| BAW32333.1 | nonribosomal\_peptide\_synthetase | BGC0001631 | NRP + Polyketide | 26.0 | 37.7 | 144.0 | 1.6e-33 |
| AXM43052.1 | non-ribosomal\_peptide\_synthetase | BGC0001945 | NRP | 27.0 | 36.1 | 144.0 | 1.6e-33 |
| AAC32048.1 | FxbC | BGC0000351 | NRP | 26.0 | 50.2 | 143.0 | 3.7e-33 |
| QED88054.1 | nonribosomal\_peptide\_synthetase | BGC0001967 | NRP | 26.0 | 36.4 | 143.0 | 3.7e-33 |
| AZH23819.1 | MgiR | BGC0001971 | NRP + Polyketide | 30.0 | 24.7 | 143.0 | 3.7e-33 |
| ALP32042.1 | CycB | BGC0001293 | Polyketide | 28.0 | 34.4 | 142.0 | 6.2e-33 |
| ADZ45339.1 | non-ribosomal\_peptide\_synthetase | BGC0001020 | NRP + Polyketide | 25.0 | 42.6 | 141.0 | 8.1e-33 |
| AFP73394.1 | FusA | BGC0001268 | NRP + Polyketide | 26.0 | 43.1 | 141.0 | 1.1e-32 |
| AVI26390.1 | polyketide\_synthase\_/\_nonribosomal\_peptide\_synthase\_hybrid | BGC0001800 | NRP + Polyketide | 27.0 | 35.9 | 141.0 | 1.4e-32 |
| AGC65516.1 | NRPS/PKS\_hybrid | BGC0001050 | NRP:Lipopeptide + Polyketide:Trans-AT type I | 26.0 | 38.4 | 138.0 | 1.2e-31 |
| ACG60776.1 | NRPS(AL/ACP/C/A/PCP/C/A) | BGC0001058 | NRP:Glycopeptide + Polyketide:Modular type I + Saccharide:Hybrid/tailoring | 28.0 | 29.4 | 138.0 | 1.2e-31 |
| ABC35796.1 | polyketide\_synthase,\_putative | BGC0001102 | NRP:Beta-lactam + Polyketide:Modular type I | 32.0 | 24.6 | 137.0 | 2e-31 |
| ADL64235.1 | aureusimine\_non-ribosomal\_peptide\_synthetase | BGC0000308 | NRP | 26.0 | 35.8 | 135.0 | 7.6e-31 |
| ACN39728.1 | SibE | BGC0000428 | NRP | 27.0 | 33.0 | 135.0 | 7.6e-31 |
| AAS92545.1 | SirP | BGC0001044 | NRP + Polyketide | 26.0 | 35.8 | 134.0 | 1e-30 |
| AAQ90177.1 | putative\_acyl-CoA\_synthetase | BGC0000128 | Polyketide | 23.0 | 40.9 | 134.0 | 1.3e-30 |
| CAD89775.1 | MelD\_protein | BGC0001010 | NRP + Polyketide:Modular type I | 26.0 | 37.6 | 134.0 | 1.7e-30 |
| AAD24881.1 | putative\_acyl-CoA\_synthetase | BGC0000127 | Polyketide | 25.0 | 41.1 | 131.0 | 8.4e-30 |
| AAT28740.1 | FUSS | BGC0000064 | Polyketide | 26.0 | 42.7 | 131.0 | 1.4e-29 |
| AEP18655.1 | WAPS2 | BGC0000461 | NRP | 26.0 | 36.9 | 130.0 | 2.5e-29 |
| AAK81824.1 | peptide\_synthetase | BGC0000326 | NRP | 33.0 | 20.6 | 129.0 | 3.2e-29 |
| AAZ23077.1 | peptide\_synthetase | BGC0000291 | NRP | 31.0 | 24.7 | 129.0 | 4.2e-29 |
| WP\_064118560.1 | non-ribosomal\_peptide\_synthetase | BGC0001509 | NRP | 28.0 | 28.8 | 128.0 | 1.2e-28 |
| ADY76684.1 | non-ribosomal\_peptide\_synthetase | BGC0000950 | NRP:Uridylpeptide + Other:Nucleoside | 24.0 | 39.2 | 127.0 | 2.1e-28 |
| AXA20090.1 | hybrid\_trans-AT\_PKS/NRPS\_LgaA | BGC0001946 | NRP + Polyketide | 27.0 | 24.7 | 127.0 | 2.1e-28 |
| sipL1 | AMP-dependent\_synthetase\_and\_ligase | BGC0001452 | Polyketide | 24.0 | 43.1 | 126.0 | 3.5e-28 |
| ALG65314.1 | Cal22 | BGC0001297 | NRP | 28.0 | 34.0 | 125.0 | 7.9e-28 |
| WP\_016638469.1 | non-ribosomal\_peptide\_synthetase | BGC0001519 | NRP + Polyketide | 27.0 | 28.0 | 124.0 | 1.3e-27 |
| ACN39014.1 | putative\_nonribosomal\_peptide\_synthetase\_TomA | BGC0000448 | NRP | 27.0 | 32.8 | 123.0 | 2.3e-27 |
| WP\_016638466.1 | amino\_acid\_adenylation\_domain-containing\_protein | BGC0001519 | NRP + Polyketide | 23.0 | 36.2 | 123.0 | 3.9e-27 |
| QDF82255.1 | non-ribosomal\_peptide\_synthetase | BGC0001980 | NRP | 26.0 | 28.8 | 122.0 | 5.1e-27 |
| AIE77059.1 | peptide\_synthetase | BGC0000418 | NRP | 26.0 | 30.1 | 121.0 | 8.7e-27 |
| OTA20325.1 | peptide\_synthase | BGC0001824 | NRP | 26.0 | 35.3 | 121.0 | 1.1e-26 |
| AQV04224.1 | SwnK | BGC0001793 | Polyketide | 27.0 | 28.5 | 121.0 | 1.5e-26 |
| AWR88409.1 | putative\_AMP-dependent\_synthetase\_and\_ligase | BGC0001522 | Polyketide | 24.0 | 43.3 | 120.0 | 1.9e-26 |
| AUS29494.1 | non-ribosomal\_peptide\_synthetase | BGC0001030 | NRP + Polyketide | 24.0 | 47.0 | 118.0 | 9.6e-26 |
| AOA33121.1 | Nonribosomal\_peptide\_synthetase | BGC0001346 | NRP:Cyclic depsipeptide | 26.0 | 29.7 | 118.0 | 1.3e-25 |
| BAC67535.1 | arthrofactin\_synthetase\_B | BGC0000305 | NRP:Lipopeptide | 26.0 | 28.8 | 117.0 | 1.6e-25 |
| AFP87519.1 | proline\_adenyltransferase | BGC0001159 | NRP + Polyketide:Modular type I | 23.0 | 43.1 | 116.0 | 3.7e-25 |
| ABA73954.1 | putative\_non-ribosomal\_peptide\_synthetase | BGC0001842 | NRP:Lipopeptide | 26.0 | 30.0 | 115.0 | 6.3e-25 |
| BAE98155.1 | putative\_non-ribosomal\_peptide\_synthetase | BGC0000339 | NRP | 26.0 | 32.6 | 114.0 | 1.1e-24 |
| CBJ82073.1 | hypothetical\_protein | BGC0001872 | Polyketide | 27.0 | 28.2 | 113.0 | 3.1e-24 |
| AXG22420.1 | proline\_adenyltransferase | BGC0002024 | Polyketide | 24.0 | 29.0 | 109.0 | 3.4e-23 |
| ADN26252.1 | peptide\_synthetase | BGC0000951 | NRP | 23.0 | 35.7 | 109.0 | 4.5e-23 |
| BBC83957.1 | nonribosomal\_peptide\_synthetase | BGC0001636 | NRP | 22.0 | 35.4 | 109.0 | 5.9e-23 |
| AFK57219.1 | DidH | BGC0000985 | Polyketide + NRP:Cyclic depsipeptide | 32.0 | 21.0 | 107.0 | 2.2e-22 |
| CBL93718.1 | NRPS\_didomain\_PCP-C | BGC0000360 | NRP | 25.0 | 33.7 | 106.0 | 3.8e-22 |
| AAY37653.1 | Amino\_acid\_adenylation | BGC0000437 | NRP | 24.0 | 28.7 | 106.0 | 3.8e-22 |
| BAH04162.1 | trsJ | BGC0000450 | NRP | 27.0 | 28.1 | 106.0 | 5e-22 |
| BAD55613.1 | putative\_non-ribosomal\_peptide\_synthetase | BGC0001027 | NRP + Polyketide | 26.0 | 29.4 | 106.0 | 5e-22 |
| WP\_030185025.1 | NAD-dependent\_epimerase/dehydratase\_family\_protein | BGC0001813 | NRP | 27.0 | 27.6 | 106.0 | 5e-22 |
| AGZ15460.1 | putative\_non-ribosomal\_peptide\_synthetase | BGC0001036 | NRP + Polyketide | 25.0 | 27.8 | 104.0 | 1.9e-21 |
| AET98906.1 | putative\_non-ribosomal\_peptide\_synthetase | BGC0000415 | NRP | 26.0 | 29.0 | 103.0 | 2.5e-21 |
| AHA12086.1 | amino\_acid\_adenyltransferase | BGC0001172 | NRP + Polyketide:Modular type I | 25.0 | 22.8 | 98.0 | 7.9e-20 |
| ABL74937.1 | NRPS | BGC0001048 | NRP:Glycopeptide + Polyketide:Modular type I + Saccharide:Hybrid/tailoring | 25.0 | 28.4 | 95.0 | 8.7e-19 |
| OAP25804.1 | Surfactin\_synthase\_subunit\_1 | BGC0001658 | Polyketide | 25.0 | 22.1 | 94.0 | 1.5e-18 |
| AYJ71713.1 | non-ribosomal\_peptide\_synthetase | BGC0001942 | NRP + Polyketide | 24.0 | 33.1 | 94.0 | 1.9e-18 |
| QCE20608.1 | AsmM | BGC0001961 | NRP | 25.0 | 21.9 | 93.0 | 2.5e-18 |
| CAJ87590.1 | putative\_peptide\_synthase | BGC0001055 | NRP + Polyketide | 23.0 | 29.3 | 76.0 | 3.2e-13 |
| AAX98208.1 | amide\_synthetase | BGC0000052 | Polyketide | 22.0 | 36.1 | 72.0 | 6.1e-12 |
| AGY30674.1 | Ann1 | BGC0001298 | Polyketide | 22.0 | 29.9 | 66.0 | 4.3e-10 |
