## Supplementary Results for "A draft genome of the ascomycotal fungal species *Pseudopithomyces maydicus* (family *Didymosphaeriaceae*)": input.path1.gene222_mibig_hits.html

| MIBiG Protein | Description | MIBiG Cluster | MiBiG Product | % ID | % Coverage | BLAST Score | E-value |
| --- | --- | --- | --- | --- | --- | --- | --- |
| AGC45618.1 | non-ribosomal\_peptide\_synthetase | BGC0001394 | NRP + Polyketide | 35.0 | 81.7 | 487.0 | 5.4e-137 |
| AQW44894.1 | non-ribosomal\_peptide\_synthetase | BGC0001737 | NRP + Polyketide | 34.0 | 80.7 | 477.0 | 9.5e-134 |
| AAK57184.1 | MxaA | BGC0001022 | NRP + Polyketide | 34.0 | 81.0 | 475.0 | 3.6e-133 |
| AAO23334.1 | NcpB | BGC0000397 | NRP | 32.0 | 81.3 | 454.0 | 6.6e-127 |
| ATP76246.1 | SpuB | BGC0001748 | NRP + Polyketide | 32.0 | 80.0 | 451.0 | 4.3e-126 |
| BAP27942.1 | nonribosomal\_peptide\_synthetase | BGC0001085 | NRP + Terpene | 33.0 | 81.7 | 447.0 | 6.2e-125 |
| AHJ31215.1 | Long-chain-fatty-acid--CoA\_ligase | BGC0000430 | NRP + Polyketide:Modular type I | 32.0 | 81.8 | 436.0 | 1.1e-121 |
| AOC89001.1 | putative\_nonribosomal\_peptide\_synthetase | BGC0001652 | NRP | 33.0 | 76.6 | 434.0 | 5.4e-121 |
| AFO85453.1 | non-ribosomal\_peptide\_synthetase | BGC0000391 | NRP | 33.0 | 80.9 | 425.0 | 2.5e-118 |
| ACN39015.1 | putative\_nonribosomal\_peptide\_synthetase\_TomB | BGC0000448 | NRP | 32.0 | 80.3 | 416.0 | 1.5e-115 |
| AAT12283.1 | LtxA | BGC0000384 | NRP | 31.0 | 80.7 | 414.0 | 5.8e-115 |
| AGD80618.1 | non-ribosomal\_peptide\_synthetase | BGC0000394 | NRP | 31.0 | 74.1 | 403.0 | 1.3e-111 |
| ACN39727.1 | SibD | BGC0000428 | NRP | 31.0 | 78.3 | 389.0 | 2.6e-107 |
| ATD51280.1 | nonribosomal\_peptide\_synthase | BGC0001650 | NRP | 34.0 | 76.1 | 388.0 | 5.8e-107 |
| EOY45602.1 | non-ribosomal\_peptide\_synthetase | BGC0001168 | NRP | 32.0 | 73.6 | 385.0 | 3.8e-106 |
| BAO84866.1 | putative\_non-ribosomal\_peptide\_synthetase | BGC0000414 | NRP | 30.0 | 74.7 | 383.0 | 1.4e-105 |
| ABI22133.1 | putative\_non-ribosomal\_peptide\_synthetase | BGC0000422 | NRP | 31.0 | 74.8 | 382.0 | 2.4e-105 |
| CAF32362.1 | putative\_non-ribosomal\_peptide\_synthetase | BGC0000712 | Saccharide | 30.0 | 82.5 | 379.0 | 2.1e-104 |
| AEA29644.1 | putative\_nonribosomal\_peptide\_synthetase\_and\_kinurenine\_monooxygenase | BGC0000409 | NRP | 32.0 | 76.7 | 376.0 | 2.3e-103 |
| AAL33758.1 | putative\_non-ribosomal\_peptide\_synthetase | BGC0000421 | NRP | 31.0 | 73.1 | 372.0 | 2.5e-102 |
| AAG31130.1 | MxcG | BGC0001345 | NRP | 30.0 | 82.3 | 369.0 | 2.8e-101 |
| ATD51278.1 | nonribosomal\_peptide\_synthase | BGC0001650 | NRP | 32.0 | 75.0 | 360.0 | 1.3e-98 |
| ATY72525.1 | non-ribosomal\_peptide\_synthetase | BGC0001574 | NRP | 32.0 | 81.7 | 359.0 | 1.7e-98 |
| ABW71853.1 | nonribosomal\_peptide\_synthetase | BGC0000303 | NRP | 31.0 | 76.0 | 356.0 | 1.9e-97 |
| ABY83163.1 | Azi25 | BGC0000960 | NRP + Polyketide | 30.0 | 75.7 | 349.0 | 3e-95 |
| CDG76959.1 | non-ribosomal\_peptide\_synthetase,\_terminal\_component | BGC0001805 | NRP | 28.0 | 74.7 | 338.0 | 5.3e-92 |
| AEC14349.1 | nonribosomal\_peptide\_synthetase | BGC0000377 | NRP | 26.0 | 79.6 | 336.0 | 2e-91 |
| ATJ04411.1 | NRPS,\_TomB\_binding | BGC0001637 | NRP | 28.0 | 74.7 | 336.0 | 2e-91 |
| ADQ55475.1 | NRPS | BGC0000350 | NRP:Beta-lactam | 27.0 | 73.7 | 334.0 | 5.8e-91 |
| ATY37592.1 | BogE | BGC0001532 | NRP | 28.0 | 74.4 | 333.0 | 1.3e-90 |
| AAD44234.1 | PstB | BGC0000362 | NRP | 30.0 | 76.9 | 326.0 | 1.6e-88 |
| CDN62030.1 | Peptide\_synthetase | BGC0001900 | NRP | 28.0 | 81.7 | 325.0 | 3.5e-88 |
| ATJ34002.1 | non-ribosomal\_peptide\_synthetase | BGC0001497 | Polyketide | 29.0 | 85.5 | 318.0 | 4.3e-86 |
| CAJ77716.1 | Mps2\_protein | BGC0000364 | NRP | 30.0 | 78.5 | 317.0 | 7.4e-86 |
| CAJ77696.1 | MPS2\_protein | BGC0000363 | NRP | 30.0 | 74.2 | 315.0 | 3.7e-85 |
| AAT01807.1 | non-ribosomal\_peptide\_synthetase | BGC0000365 | NRP | 28.0 | 77.8 | 295.0 | 5.1e-79 |
| CCP42826.1 | Probable\_peptide\_synthetase\_Nrp\_(peptide\_synthase) | BGC0001627 | NRP | 29.0 | 77.9 | 295.0 | 5.1e-79 |
| AIW82277.1 | PuwA | BGC0001125 | NRP + Polyketide | 34.0 | 48.4 | 292.0 | 2.5e-78 |
| AXN93575.1 | PuwA | BGC0001950 | NRP | 33.0 | 48.3 | 292.0 | 3.3e-78 |
| AXN93584.1 | PuwA | BGC0001951 | NRP | 33.0 | 48.3 | 292.0 | 3.3e-78 |
| QDA77059.1 | polyketide\_synthase/nonribosomal\_peptide\_synthetase | BGC0002026 | NRP | 33.0 | 47.0 | 292.0 | 4.3e-78 |
| CAD17792.1 | probable\_non\_ribosomal\_peptide\_synthetase\_protein | BGC0001754 | NRP + Polyketide | 32.0 | 49.4 | 285.0 | 5.3e-76 |
| AGM16413.1 | paenibacterin\_synthetase\_B | BGC0000400 | NRP | 32.0 | 47.7 | 284.0 | 9e-76 |
| AFP73394.1 | FusA | BGC0001268 | NRP + Polyketide | 29.0 | 71.5 | 283.0 | 1.5e-75 |
| AAU34202.1 | mannopeptimycin\_peptide\_synthetase\_MppA | BGC0000388 | NRP | 33.0 | 47.2 | 282.0 | 2.6e-75 |
| CDG17981.1 | Non-ribosomal\_peptide\_synthetase | BGC0000464 | NRP:Cyclic depsipeptide | 34.0 | 46.3 | 282.0 | 3.4e-75 |
| AEH59100.1 | amino\_acid\_adenylation\_domain-containing\_protein/NRPS | BGC0000385 | NRP | 33.0 | 48.5 | 279.0 | 2.2e-74 |
| WP\_041826607.1 | non-ribosomal\_peptide\_synthetase | BGC0001792 | NRP | 32.0 | 47.8 | 279.0 | 2.2e-74 |
| CDG17986.1 | Non-ribosomal\_peptide\_synthetase | BGC0000464 | NRP:Cyclic depsipeptide | 34.0 | 46.2 | 275.0 | 4.2e-73 |
| AGM16412.1 | paenibacterin\_synthetase\_A | BGC0000400 | NRP | 33.0 | 46.3 | 275.0 | 5.5e-73 |
| CAE15637.1 |  | BGC0001128 | NRP | 33.0 | 46.9 | 274.0 | 7.2e-73 |
| AAT28740.1 | FUSS | BGC0000064 | Polyketide | 28.0 | 72.7 | 274.0 | 9.4e-73 |
| WP\_011146892.1 | non-ribosomal\_peptide\_synthetase | BGC0001641 | NRP | 33.0 | 48.1 | 274.0 | 9.4e-73 |
| EAU38971.1 | hypothetical\_protein | BGC0001122 | NRP + Polyketide:Iterative type I | 28.0 | 79.6 | 274.0 | 1.2e-72 |
| AGZ03651.1 | sevB | BGC0000426 | NRP | 33.0 | 47.0 | 273.0 | 1.6e-72 |
| AEU11006.1 | NpnB | BGC0001029 | NRP + Polyketide | 32.0 | 46.7 | 273.0 | 1.6e-72 |
| AAC06348.1 | bacitracin\_synthetase\_3 | BGC0000310 | NRP | 30.0 | 48.5 | 273.0 | 2.1e-72 |
| CDG17980.1 | Putative\_Ornithine\_racemase\_(fragment) | BGC0000464 | NRP:Cyclic depsipeptide | 33.0 | 49.1 | 272.0 | 4.6e-72 |
| CCJ67637.1 | TaaB | BGC0000447 | NRP:Lipopeptide | 33.0 | 48.0 | 271.0 | 6.1e-72 |
| ATO51563.1 | non-ribosomal\_peptide\_synthetase | BGC0001796 | NRP | 30.0 | 56.0 | 271.0 | 7.9e-72 |
| WP\_019032755.1 | non-ribosomal\_peptide\_synthetase | BGC0001331 | NRP:Cyclic depsipeptide + Polyketide:Modular type I | 34.0 | 48.8 | 270.0 | 1e-71 |
| WP\_013428324.1 | non-ribosomal\_peptide\_synthetase | BGC0001758 | NRP | 31.0 | 46.6 | 270.0 | 1e-71 |
| ABM21572.1 | crpD | BGC0000975 | NRP + Polyketide | 30.0 | 53.4 | 270.0 | 1.8e-71 |
| APU91750.1 | Non-Ribosomal\_Peptide\_Synthetase | BGC0001806 | NRP | 33.0 | 45.9 | 270.0 | 1.8e-71 |
| AXN93592.1 | PuwA | BGC0001952 | NRP | 32.0 | 48.8 | 268.0 | 5.1e-71 |
| WP\_053065269.1 | non-ribosomal\_peptide\_synthetase | BGC0001330 | NRP:Cyclic depsipeptide + Polyketide:Modular type I | 33.0 | 49.2 | 268.0 | 6.7e-71 |
| AFH75329.1 | nonribosomal\_peptide\_synthetase | BGC0000398 | NRP:Cyclic depsipeptide | 32.0 | 47.0 | 267.0 | 8.8e-71 |
| WP\_054234643.1 | non-ribosomal\_peptide\_synthetase | BGC0002014 | NRP + Polyketide | 32.0 | 47.6 | 267.0 | 8.8e-71 |
| BAC67536.1 | arthrofactin\_synthetase\_C | BGC0000305 | NRP:Lipopeptide | 32.0 | 48.8 | 267.0 | 1.5e-70 |
| ABQ96384.2 | fusaricidin\_synthetase | BGC0001152 | Polyketide + NRP:Lipopeptide | 31.0 | 49.3 | 266.0 | 1.9e-70 |
| BAV56270.1 |  | BGC0001657 | NRP | 31.0 | 47.7 | 265.0 | 4.3e-70 |
| CAM59606.1 | non-ribosomal\_peptide\_synthetase | BGC0000297 | NRP:Glycopeptide + Polyketide:Other + Saccharide:Hybrid/tailoring | 32.0 | 49.1 | 265.0 | 5.7e-70 |
| AHZ34238.1 | CipA | BGC0001389 | NRP | 32.0 | 49.2 | 265.0 | 5.7e-70 |
| AXN93605.1 | PuwA | BGC0001953 | NRP | 32.0 | 48.7 | 265.0 | 5.7e-70 |
| ABH06368.2 | MassB | BGC0000389 | NRP:Cyclic depsipeptide | 32.0 | 47.6 | 264.0 | 1.3e-69 |
| ABA23460.1 | Amino\_acid\_adenylation | BGC0000427 | NRP | 32.0 | 46.8 | 264.0 | 1.3e-69 |
| ATY37590.1 | BogC | BGC0001532 | NRP | 30.0 | 47.6 | 264.0 | 1.3e-69 |
| ABW17376.1 | PsoB | BGC0000411 | NRP | 32.0 | 47.6 | 263.0 | 1.7e-69 |
| AFP87549.1 | NrpS | BGC0001135 | NRP | 32.0 | 50.9 | 263.0 | 1.7e-69 |
| QBG38783.1 | Atr22 | BGC0001975 | NRP | 34.0 | 47.4 | 263.0 | 1.7e-69 |
| CAY48788.1 | putative\_non-ribosomal\_peptide\_synthetase | BGC0001312 | NRP | 32.0 | 48.0 | 263.0 | 2.2e-69 |
| BAH22763.1 | nonribosomal\_peptide\_synthetase | BGC0001018 | NRP + Polyketide | 31.0 | 49.5 | 262.0 | 2.8e-69 |
| KFL51887.1 | amino\_acid\_adenylation\_protein | BGC0001711 | NRP + Polyketide | 32.0 | 47.1 | 262.0 | 2.8e-69 |
| WP\_010639240.1 | non-ribosomal\_peptide\_synthetase | BGC0000958 | NRP:Cyclic depsipeptide + Polyketide:Modular type I | 32.0 | 48.3 | 262.0 | 3.7e-69 |
| AFH75320.1 | nonribosomal\_peptide\_synthetase | BGC0000425 | NRP:Cyclic depsipeptide | 32.0 | 45.9 | 262.0 | 4.8e-69 |
| ABL74945.1 | NRPS | BGC0001048 | NRP:Glycopeptide + Polyketide:Modular type I + Saccharide:Hybrid/tailoring | 33.0 | 47.5 | 262.0 | 4.8e-69 |
| ALG65317.1 | Cal19 | BGC0001297 | NRP | 31.0 | 47.7 | 262.0 | 4.8e-69 |
| BBA21073.1 | putative\_non-ribosomal\_peptide\_synthetase | BGC0001740 | NRP + Polyketide | 32.0 | 48.2 | 262.0 | 4.8e-69 |
| QBC75021.1 | non-ribosomal\_peptide\_synthetase | BGC0001968 | NRP | 33.0 | 47.7 | 262.0 | 4.8e-69 |
| EFE73312.1 | nonribosomal\_peptide\_synthetase | BGC0000431 | NRP:Cyclic depsipeptide | 32.0 | 46.9 | 261.0 | 6.3e-69 |
| EED49862.1 | hybrid\_PKS/NRPS\_enzyme,\_putative | BGC0001445 | NRP + Polyketide:Iterative type I | 27.0 | 71.5 | 261.0 | 6.3e-69 |
| WP\_004571779.1 | non-ribosomal\_peptide\_synthetase | BGC0001760 | NRP | 33.0 | 47.7 | 261.0 | 6.3e-69 |
| AVI26390.1 | polyketide\_synthase\_/\_nonribosomal\_peptide\_synthase\_hybrid | BGC0001800 | NRP + Polyketide | 31.0 | 46.4 | 261.0 | 6.3e-69 |
| ACC81021.1 | non-ribosomal\_peptide\_synthetase | BGC0001479 | NRP | 32.0 | 45.7 | 261.0 | 8.2e-69 |
| WP\_013428774.1 | non-ribosomal\_peptide\_synthetase | BGC0001844 | NRP:Lipopeptide | 32.0 | 46.7 | 261.0 | 8.2e-69 |
| AFJ23825.1 | WLIP\_synthetase\_B | BGC0001838 | NRP | 32.0 | 47.4 | 260.0 | 1.1e-68 |
| ACS68554.1 | hybrid\_PKS-NRPS\_protein | BGC0001026 | NRP + Polyketide | 28.0 | 72.6 | 260.0 | 1.4e-68 |
| AAF17281.1 | nosD | BGC0001028 | Polyketide + NRP:Cyclic depsipeptide | 32.0 | 48.1 | 260.0 | 1.4e-68 |
| AQH32486.1 | peptide\_synthetase | BGC0001667 | NRP + Polyketide | 31.0 | 47.0 | 260.0 | 1.4e-68 |
| AYA44686.1 | icosalide\_NRPS | BGC0001833 | NRP:Lipopeptide | 33.0 | 48.5 | 260.0 | 1.4e-68 |
| ACM68684.1 | AerB | BGC0000298 | NRP | 33.0 | 46.7 | 259.0 | 2.4e-68 |
| ABI26077.1 | OciA | BGC0000331 | NRP | 31.0 | 50.7 | 259.0 | 2.4e-68 |
| WP\_047197778.1 | non-ribosomal\_peptide\_synthetase | BGC0002003 | NRP | 33.0 | 46.9 | 259.0 | 2.4e-68 |
| AGM16414.1 | paenibacterin\_synthetase\_C | BGC0000400 | NRP | 31.0 | 46.5 | 259.0 | 3.1e-68 |
| ABC36203.1 | thiotemplate\_mechanism\_natural\_product\_synthetase | BGC0001102 | NRP:Beta-lactam + Polyketide:Modular type I | 30.0 | 49.7 | 259.0 | 3.1e-68 |
| AKP45395.1 | CysG | BGC0001413 | NRP | 31.0 | 47.9 | 259.0 | 3.1e-68 |
| ATL73036.1 | amino\_acid\_adenylation\_protein | BGC0001807 | NRP + Polyketide | 32.0 | 49.4 | 259.0 | 3.1e-68 |
| AAC06346.1 | bacitracin\_synthetase\_1 | BGC0000310 | NRP | 31.0 | 47.6 | 259.0 | 4.1e-68 |
| CAD29799.1 | microcystin\_synthetase | BGC0001015 | NRP + Polyketide | 31.0 | 47.3 | 259.0 | 4.1e-68 |
| AIW82284.1 | PuwG | BGC0001125 | NRP + Polyketide | 32.0 | 47.1 | 259.0 | 4.1e-68 |
| AXN93581.1 | PuwF-G | BGC0001950 | NRP | 32.0 | 47.1 | 259.0 | 4.1e-68 |
| AXN93590.1 | PuwF-G | BGC0001951 | NRP | 32.0 | 47.1 | 259.0 | 4.1e-68 |
| CBA63680.1 | nonribosomal\_peptide\_synthetase\_NRPS | BGC0000368 | NRP | 32.0 | 48.3 | 258.0 | 5.3e-68 |
| AIW82285.1 | PuwH | BGC0001125 | NRP + Polyketide | 32.0 | 46.1 | 258.0 | 6.9e-68 |
| QBG38784.1 | Atr23 | BGC0001975 | NRP | 34.0 | 48.3 | 258.0 | 6.9e-68 |
| AEO14744.1 | NdaB | BGC0000396 | NRP | 30.0 | 47.3 | 257.0 | 9.1e-68 |
| PHM49485.1 | Amino\_acid\_adenylation | BGC0001131 | NRP | 32.0 | 47.9 | 257.0 | 9.1e-68 |
| AKA54626.1 | NRPS | BGC0001216 | NRP + Polyketide | 31.0 | 48.3 | 257.0 | 9.1e-68 |
| KPN93065.1 | NunD | BGC0001416 | NRP | 31.0 | 48.0 | 257.0 | 9.1e-68 |
| WP\_016638470.1 | non-ribosomal\_peptide\_synthetase | BGC0001519 | NRP + Polyketide | 33.0 | 47.3 | 257.0 | 9.1e-68 |
| BAW32323.1 | hybrid\_cis-AT\_polyketide\_synthase\_-\_nonribosomal\_peptide\_synthetase | BGC0001630 | NRP + Polyketide | 31.0 | 49.5 | 257.0 | 9.1e-68 |
| CAJ14037.1 | peptide\_synthetase | BGC0000406 | NRP | 31.0 | 50.4 | 257.0 | 1.2e-67 |
| AEZ51516.1 | pmxA | BGC0001153 | NRP:Lipopeptide | 30.0 | 46.2 | 257.0 | 1.2e-67 |
| CAD17793.1 | probable\_non\_ribosomal\_peptide\_synthetase\_protein | BGC0001754 | NRP + Polyketide | 32.0 | 49.1 | 257.0 | 1.2e-67 |
| CDG17982.1 | Non-ribosomal\_peptide\_synthetase | BGC0000464 | NRP:Cyclic depsipeptide | 32.0 | 47.3 | 257.0 | 1.5e-67 |
| AIW82282.1 | PuwE | BGC0001125 | NRP + Polyketide | 31.0 | 46.8 | 257.0 | 1.5e-67 |
| CAY48789.1 | putative\_non-ribosomal\_peptide\_synthetase | BGC0001312 | NRP | 33.0 | 46.9 | 257.0 | 1.5e-67 |
| AAZ03551.1 | McnB | BGC0000332 | NRP | 30.0 | 49.6 | 256.0 | 2e-67 |
| AHB38515.1 | non-ribosomal\_peptide\_synthetase | BGC0000345 | NRP + Polyketide:Modular type I | 33.0 | 46.7 | 256.0 | 2e-67 |
| ctg1\_orf17 |  | BGC0001457 | NRP | 32.0 | 47.0 | 256.0 | 2e-67 |
| ABI26079.1 | OciC | BGC0000331 | NRP | 31.0 | 48.2 | 256.0 | 2.6e-67 |
| AJV88375.1 | MfnC | BGC0001214 | NRP | 32.0 | 48.1 | 256.0 | 2.6e-67 |
| BAV57451.1 | NRPS\_(A-PCP) | BGC0001912 | NRP | 33.0 | 47.0 | 256.0 | 2.6e-67 |
| CCJ67636.1 | TaaA | BGC0000447 | NRP:Lipopeptide | 31.0 | 48.1 | 255.0 | 3.4e-67 |
| AAF17280.1 | nosC | BGC0001028 | Polyketide + NRP:Cyclic depsipeptide | 31.0 | 51.5 | 255.0 | 3.4e-67 |
| QED55422.1 | nonribosomal\_peptide\_synthetase | BGC0001984 | NRP | 32.0 | 50.6 | 255.0 | 3.4e-67 |
| antaC | NRPS | BGC0001455 | NRP + Polyketide | 31.0 | 47.3 | 255.0 | 4.5e-67 |
| AAZ03552.1 | McnC | BGC0000332 | NRP | 31.0 | 52.6 | 255.0 | 5.9e-67 |
| AJF34464.1 | Txo2 | BGC0001207 | NRP | 32.0 | 46.7 | 255.0 | 5.9e-67 |
| AHZ34240.1 | CipC | BGC0001389 | NRP | 31.0 | 46.9 | 255.0 | 5.9e-67 |
| AAU34203.1 | mannopeptimycin\_peptide\_synthetase\_MppB | BGC0000388 | NRP | 33.0 | 49.5 | 254.0 | 7.7e-67 |
| EFE73313.1 | nonribosomal\_peptide\_synthetase | BGC0000431 | NRP:Cyclic depsipeptide | 31.0 | 50.9 | 254.0 | 7.7e-67 |
| AHF21228.1 | TriD | BGC0000449 | NRP | 31.0 | 50.5 | 254.0 | 7.7e-67 |
| CAQ34921.1 | nonribosomal\_peptide\_synthetase | BGC0000986 | NRP + Polyketide | 32.0 | 49.7 | 254.0 | 7.7e-67 |
| AVR48535.1 | CusC | BGC0001564 | NRP + Polyketide | 31.0 | 46.6 | 254.0 | 7.7e-67 |
| ATP76244.1 | NdaB | BGC0001705 | NRP + Polyketide | 32.0 | 47.1 | 254.0 | 7.7e-67 |
| ABA73956.1 | putative\_non-ribosomal\_peptide\_synthetase | BGC0001842 | NRP:Lipopeptide | 32.0 | 49.2 | 254.0 | 1e-66 |
| KFL51883.1 | amino\_acid\_adenylation\_protein | BGC0001711 | NRP + Polyketide | 31.0 | 47.4 | 253.0 | 1.3e-66 |
| QED88054.1 | nonribosomal\_peptide\_synthetase | BGC0001967 | NRP | 32.0 | 48.3 | 253.0 | 1.3e-66 |
| AAY91420.2 | non-ribosomal\_peptide\_synthetase\_OfaB | BGC0000399 | NRP:Cyclic depsipeptide | 32.0 | 47.3 | 253.0 | 1.7e-66 |
| BAH22765.1 | nonribosomal\_peptide\_synthetase | BGC0001018 | NRP + Polyketide | 31.0 | 48.6 | 253.0 | 1.7e-66 |
| ANI24100.1 | nonribosomal\_peptide\_synthetase | BGC0001235 | NRP + Polyketide | 32.0 | 46.6 | 253.0 | 1.7e-66 |
| AEW31021.1 | plipastatin\_synthetase | BGC0000407 | NRP | 31.0 | 48.8 | 253.0 | 2.2e-66 |
| CDG17985.1 | Putative\_Ornithine\_racemase\_(fragment) | BGC0000464 | NRP:Cyclic depsipeptide | 32.0 | 48.1 | 253.0 | 2.2e-66 |
| ATY37591.1 | BogD | BGC0001532 | NRP | 31.0 | 46.5 | 252.0 | 2.9e-66 |
| AXN93580.1 | PuwE | BGC0001950 | NRP | 32.0 | 46.9 | 252.0 | 2.9e-66 |
| AXN93589.1 | PuwE | BGC0001951 | NRP | 32.0 | 46.9 | 252.0 | 2.9e-66 |
| CBJ90082.1 | Non\_Ribosomal\_peptide\_synthetase\_(-succinylbenzoate--CoA\_ligase) | BGC0001132 | NRP | 32.0 | 46.5 | 252.0 | 3.8e-66 |
| AJF34463.1 | Txo1 | BGC0001207 | NRP | 32.0 | 46.7 | 252.0 | 3.8e-66 |
| AHB82071.1 | non\_ribosomal\_peptide\_synthetase | BGC0001231 | NRP + Polyketide:Modular type I | 32.0 | 49.0 | 252.0 | 3.8e-66 |
| WP\_067426775.1 | non-ribosomal\_peptide\_synthetase | BGC0001567 | NRP | 32.0 | 47.4 | 252.0 | 3.8e-66 |
| AWI62626.1 | nonribosomal\_peptide\_synthetase | BGC0001943 | NRP | 33.0 | 46.2 | 252.0 | 3.8e-66 |
| AAO23333.1 | NcpA | BGC0000397 | NRP | 32.0 | 47.5 | 252.0 | 5e-66 |
| BAH43766.1 | tyrocidine\_synthetase\_III | BGC0000452 | NRP | 31.0 | 47.8 | 252.0 | 5e-66 |
| BAP05597.1 | calI | BGC0000967 | NRP + Polyketide:Trans-AT type I | 30.0 | 48.9 | 252.0 | 5e-66 |
| AAY89051.1 | hybrid\_nonribosomal\_peptide\_synthetase/polyketide\_synthase | BGC0001069 | NRP + Polyketide:Trans-AT type I | 32.0 | 47.1 | 252.0 | 5e-66 |
| CCJ67648.1 | JagD | BGC0001127 | NRP | 32.0 | 47.3 | 252.0 | 5e-66 |
| WP\_079164394.1 | non-ribosomal\_peptide\_synthetase | BGC0001567 | NRP | 32.0 | 47.7 | 252.0 | 5e-66 |
| ANY58984.1 | non-ribosomal\_synthetase | BGC0001615 | NRP | 32.0 | 47.2 | 252.0 | 5e-66 |
| AWX24482.1 | non-ribosomal\_peptide\_synthetase | BGC0001695 | NRP | 32.0 | 47.0 | 252.0 | 5e-66 |
| CUX96954.1 | TmcG | BGC0001829 | NRP + Polyketide | 33.0 | 48.4 | 252.0 | 5e-66 |
| AEU11003.1 | NpnC | BGC0001029 | NRP + Polyketide | 30.0 | 48.6 | 251.0 | 6.5e-66 |
| APZ78782.1 | nonribosomal\_peptide\_synthetase | BGC0001426 | NRP:Cyclic depsipeptide + Polyketide:Iterative type I | 30.0 | 52.5 | 251.0 | 6.5e-66 |
| AXN93582.1 | PuwH | BGC0001950 | NRP | 32.0 | 46.4 | 251.0 | 6.5e-66 |
| AXN93591.1 | PuwH | BGC0001951 | NRP | 32.0 | 46.4 | 251.0 | 6.5e-66 |
| CAK15814.1 | putative\_non-ribosomal\_peptide\_synthetase,\_terminal\_component | BGC0000344 | NRP | 31.0 | 49.1 | 251.0 | 8.5e-66 |
| AGN74876.1 | nonribosomal\_peptide\_synthetase | BGC0000459 | NRP:Cyclic depsipeptide + Polyketide:Trans-AT type I | 33.0 | 47.7 | 251.0 | 8.5e-66 |
| AOA33121.1 | Nonribosomal\_peptide\_synthetase | BGC0001346 | NRP:Cyclic depsipeptide | 32.0 | 47.0 | 251.0 | 8.5e-66 |
| CAK15815.1 | putative\_non\_ribosomal\_peptide\_synthetase | BGC0000344 | NRP | 31.0 | 48.1 | 250.0 | 1.1e-65 |
| AEW31019.1 | plipastatin\_synthetase | BGC0000407 | NRP | 31.0 | 48.8 | 250.0 | 1.1e-65 |
| ADJ63842.1 | Serobactin\_synthetase | BGC0000424 | NRP:NRP siderophore | 32.0 | 46.1 | 250.0 | 1.1e-65 |
| AAY37655.1 | Amino\_acid\_adenylation | BGC0000437 | NRP | 32.0 | 45.6 | 250.0 | 1.1e-65 |
| AHH53507.1 | non-ribosomal\_peptide\_synthetase | BGC0000439 | NRP:Ca+-dependent lipopeptide | 33.0 | 47.5 | 250.0 | 1.1e-65 |
| AGM14933.1 | xantholysin\_synthetase\_B | BGC0000463 | NRP:Lipopeptide | 30.0 | 46.2 | 250.0 | 1.1e-65 |
| CUX79061.1 | Octapeptin\_synthase\_subunit\_B | BGC0001715 | NRP | 30.0 | 49.3 | 250.0 | 1.1e-65 |
| AHD05677.1 | nonribosomal\_peptide\_ligase\_subunit | BGC0000402 | NRP | 31.0 | 46.6 | 250.0 | 1.4e-65 |
| ABS74208.1 | fengycin\_synthetase\_B | BGC0001095 | NRP | 30.0 | 47.7 | 250.0 | 1.4e-65 |
| AHZ34241.1 | CipD | BGC0001389 | NRP | 31.0 | 46.2 | 250.0 | 1.4e-65 |
| APZ78729.1 | nonribosomal\_peptide\_synthetase | BGC0001421 | NRP:Cyclic depsipeptide + Polyketide:Iterative type I | 30.0 | 52.6 | 250.0 | 1.4e-65 |
| CAB15186.3 | siderophore\_2,3-dihydroxybenzoate-glycine-threonine\_trimeric\_ester\_bacillibactin\_synthetase | BGC0000309 | NRP | 31.0 | 47.9 | 250.0 | 1.9e-65 |
| ABI26078.1 | OciB | BGC0000331 | NRP | 29.0 | 49.2 | 250.0 | 1.9e-65 |
| AEP18655.1 | WAPS2 | BGC0000461 | NRP | 31.0 | 49.1 | 250.0 | 1.9e-65 |
| AAO62587.1 | peptide\_sythetase | BGC0001016 | NRP + Polyketide | 32.0 | 47.1 | 250.0 | 1.9e-65 |
| ADA69239.2 | trans-AT\_hybrid\_polyketide\_synthase-NRPS | BGC0001071 | NRP + Polyketide:Modular type I + Polyketide:Trans-AT type I | 33.0 | 46.3 | 250.0 | 1.9e-65 |
| BAF50711.1 | non\_ribosomal\_peptide\_synthetase\_for\_virginiamycin\_S | BGC0001116 | NRP + Polyketide | 32.0 | 50.1 | 250.0 | 1.9e-65 |
| ATY37609.1 | BreD | BGC0001536 | NRP | 30.0 | 48.6 | 250.0 | 1.9e-65 |
| CUX79062.1 | Octapeptin\_synthase\_subunit\_C | BGC0001715 | NRP | 31.0 | 46.3 | 250.0 | 1.9e-65 |
| AFJ14794.1 | PlpE | BGC0000403 | NRP | 31.0 | 46.1 | 249.0 | 2.5e-65 |
| AAN85512.1 | nonribosomal\_peptide\_synthetase | BGC0001101 | NRP + Polyketide:Modular type I + Polyketide:Trans-AT type I | 32.0 | 46.4 | 249.0 | 2.5e-65 |
| QDF82259.1 | non-ribosomal\_peptide\_synthetase | BGC0001980 | NRP | 31.0 | 50.0 | 249.0 | 2.5e-65 |
| AAF15891.2 | nosA | BGC0001028 | Polyketide + NRP:Cyclic depsipeptide | 31.0 | 50.7 | 249.0 | 3.2e-65 |
| AEW31022.1 | plipastatin\_synthetase | BGC0000407 | NRP | 29.0 | 48.6 | 248.0 | 4.2e-65 |
| ADY76664.1 | non-ribosomal\_peptide\_synthetase | BGC0000950 | NRP:Uridylpeptide + Other:Nucleoside | 33.0 | 46.4 | 248.0 | 5.5e-65 |
| AAG02364.1 | peptide\_synthetase\_NRPS2-1 | BGC0000963 | NRP:Glycopeptide + Polyketide:Modular type I + Saccharide:Hybrid/tailoring | 33.0 | 45.1 | 248.0 | 5.5e-65 |
| AAZ03550.1 | McnA | BGC0000332 | NRP | 31.0 | 46.7 | 248.0 | 7.2e-65 |
| AAY91421.3 | non-ribosomal\_peptide\_synthetase\_OfaC | BGC0000399 | NRP:Cyclic depsipeptide | 31.0 | 47.0 | 248.0 | 7.2e-65 |
| ACA97576.1 | PmxA | BGC0000408 | NRP | 30.0 | 46.7 | 248.0 | 7.2e-65 |
| CCA89328.1 | mixed\_trans-AT\_type\_I\_polyketide\_synthase/nonribosomal\_peptide\_synthetase | BGC0001111 | NRP + Polyketide:Trans-AT type I | 32.0 | 47.4 | 248.0 | 7.2e-65 |
| AJM89735.1 | PmxA | BGC0001192 | NRP | 30.0 | 46.2 | 248.0 | 7.2e-65 |
| APZ78769.1 | nonribosomal\_peptide\_synthetase | BGC0001425 | NRP:Cyclic depsipeptide + Polyketide:Iterative type I | 30.0 | 52.6 | 248.0 | 7.2e-65 |
| BAC67534.2 | arthrofactin\_synthetase\_A | BGC0000305 | NRP:Lipopeptide | 32.0 | 47.2 | 247.0 | 9.4e-65 |
| ABV56585.1 | KtzE | BGC0000378 | NRP | 31.0 | 47.7 | 247.0 | 9.4e-65 |
| ABH06369.2 | MassC | BGC0000389 | NRP:Cyclic depsipeptide | 32.0 | 47.3 | 247.0 | 9.4e-65 |
| AAY93355.1 | non-ribosomal\_peptide\_synthetase\_PvdJ | BGC0000413 | NRP | 31.0 | 49.9 | 247.0 | 9.4e-65 |
| CAQ43084.1 | non\_ribosomal\_polypeptide\_synthetase | BGC0000970 | NRP + Polyketide:Modular type I | 30.0 | 47.4 | 247.0 | 9.4e-65 |
| AEZ51520.1 | pmxE | BGC0001153 | NRP:Lipopeptide | 29.0 | 49.9 | 247.0 | 9.4e-65 |
| EWM63002.1 | non-ribosomal\_peptide\_synthetase | BGC0001328 | NRP:Cyclic depsipeptide + Polyketide:Modular type I | 32.0 | 49.2 | 247.0 | 9.4e-65 |
| APZ78744.1 | nonribosomal\_peptide\_synthetase | BGC0001422 | NRP:Cyclic depsipeptide + Polyketide:Iterative type I | 29.0 | 52.7 | 247.0 | 9.4e-65 |
| ANG60379.1 | nonribosomal\_peptide\_synthetase\_BudA | BGC0001698 | NRP | 31.0 | 48.1 | 247.0 | 9.4e-65 |
| CAJ14039.1 | peptide\_synthetase | BGC0000406 | NRP | 30.0 | 47.7 | 247.0 | 1.2e-64 |
| AEA30273.1 | peptide\_synthetase | BGC0000429 | Polyketide + NRP:Cyclic depsipeptide | 31.0 | 46.9 | 247.0 | 1.2e-64 |
| EAL85113.2 | hybrid\_PKS-NRPS\_enzyme | BGC0001037 | NRP + Polyketide:Iterative type I | 26.0 | 77.4 | 247.0 | 1.2e-64 |
| AIW82283.1 | PuwF | BGC0001125 | NRP + Polyketide | 31.0 | 47.0 | 247.0 | 1.2e-64 |
| AJW65406.1 | nonribosomal\_peptide\_synthetase | BGC0001195 | NRP + Polyketide | 31.0 | 47.1 | 247.0 | 1.2e-64 |
| ABA73955.1 | putative\_non-ribosomal\_peptide\_synthetase | BGC0001842 | NRP:Lipopeptide | 32.0 | 46.8 | 247.0 | 1.2e-64 |
| ACA97580.1 | PmxE | BGC0000408 | NRP | 29.0 | 49.9 | 247.0 | 1.6e-64 |
| AET98906.1 | putative\_non-ribosomal\_peptide\_synthetase | BGC0000415 | NRP | 33.0 | 47.3 | 247.0 | 1.6e-64 |
| APZ78795.1 | nonribosomal\_peptide\_synthetase | BGC0001427 | NRP:Cyclic depsipeptide + Polyketide:Iterative type I | 30.0 | 52.6 | 247.0 | 1.6e-64 |
| AAN32981.1 | BarG | BGC0000962 | NRP + Polyketide:Modular type I | 31.0 | 48.1 | 246.0 | 2.1e-64 |
| AAO62588.1 | peptide\_sythetase | BGC0001016 | NRP + Polyketide | 31.0 | 47.4 | 246.0 | 2.1e-64 |
| ctg1\_orf1265 |  | BGC0001752 | NRP | 31.0 | 49.2 | 246.0 | 2.1e-64 |
| AFJ23826.1 | WLIP\_synthetase\_C | BGC0001838 | NRP | 30.0 | 49.5 | 246.0 | 2.1e-64 |
| ABA73954.1 | putative\_non-ribosomal\_peptide\_synthetase | BGC0001842 | NRP:Lipopeptide | 31.0 | 46.7 | 246.0 | 2.1e-64 |
| BBD17759.1 | non-ribosomal\_peptide\_synthetase | BGC0001919 | NRP + Polyketide | 31.0 | 49.9 | 246.0 | 2.1e-64 |
| AGE11898.1 | nonribosomal\_peptide\_synthetase | BGC0000366 | NRP | 32.0 | 48.0 | 246.0 | 2.7e-64 |
| CDG17987.1 | Putative\_Ornithine\_racemase\_(fragment) | BGC0000464 | NRP:Cyclic depsipeptide | 30.0 | 51.9 | 246.0 | 2.7e-64 |
| AJM89738.1 | PmxE | BGC0001192 | NRP | 30.0 | 48.1 | 246.0 | 2.7e-64 |
| AQM37584.1 | nonribosomal\_peptide\_synthetase | BGC0001424 | NRP:Cyclic depsipeptide + Polyketide:Iterative type I | 32.0 | 49.1 | 246.0 | 2.7e-64 |
| CUX79060.1 | Octapeptin\_synthase\_subunit\_A | BGC0001715 | NRP | 30.0 | 47.7 | 246.0 | 2.7e-64 |
| WP\_039806852.1 | non-ribosomal\_peptide\_synthetase | BGC0002001 | NRP + Polyketide | 32.0 | 46.8 | 246.0 | 2.7e-64 |
| AAZ03554.1 | McnE | BGC0000332 | NRP | 30.0 | 48.3 | 245.0 | 3.6e-64 |
| ABW00331.1 | amino\_acid\_adenylation\_domain | BGC0000333 | NRP | 32.0 | 46.3 | 245.0 | 3.6e-64 |
| AHB82058.1 | non\_ribosomal\_peptide\_synthetase | BGC0001019 | NRP + Polyketide:Modular type I | 33.0 | 47.6 | 245.0 | 3.6e-64 |
| AWI62628.1 | nonribosomal\_peptide\_synthetase | BGC0001943 | NRP | 32.0 | 46.0 | 245.0 | 3.6e-64 |
| QBG38782.1 | Atr21 | BGC0001975 | NRP | 33.0 | 47.4 | 245.0 | 3.6e-64 |
| APZ78756.1 | nonribosomal\_peptide\_synthetase | BGC0001423 | NRP:Cyclic depsipeptide + Polyketide:Iterative type I | 32.0 | 47.9 | 245.0 | 4.7e-64 |
| APZ78809.1 | nonribosomal\_peptide\_synthetase | BGC0001428 | NRP:Cyclic depsipeptide + Polyketide:Iterative type I | 31.0 | 46.9 | 245.0 | 4.7e-64 |
| AWI62627.1 | nonribosomal\_peptide\_synthetase | BGC0001943 | NRP | 31.0 | 48.8 | 245.0 | 4.7e-64 |
| WP\_018540607.1 | non-ribosomal\_peptide\_synthetase | BGC0001332 | NRP + Polyketide | 32.0 | 46.6 | 245.0 | 6.1e-64 |
| AXN93603.1 | PuwH | BGC0001952 | NRP | 33.0 | 49.3 | 245.0 | 6.1e-64 |
| CZT62784.1 | Non-ribosomal\_peptide\_synthase,\_involved\_in\_Hassallidin\_biosynthesis | BGC0001614 | NRP | 31.0 | 48.4 | 244.0 | 7.9e-64 |
| CAQ71828.1 | non\_ribosomal\_peptide\_synthase,\_antibiotic\_synthesis;\_contains\_3\_condensation\_domains,\_2\_AMP-acid\_ligases\_II\_domains,\_2\_PP-binding,\_Phosphopantetheine\_attachment\_site | BGC0001189 | NRP | 33.0 | 47.2 | 244.0 | 1e-63 |
| AFJ14795.1 | PlpF | BGC0000403 | NRP | 29.0 | 49.7 | 243.0 | 1.4e-63 |
| AEW31020.1 | plipastatin\_synthetase | BGC0000407 | NRP | 28.0 | 47.9 | 243.0 | 1.4e-63 |
| AGZ15459.1 | putative\_non-ribosomal\_peptide\_synthetase | BGC0001036 | NRP + Polyketide | 32.0 | 48.4 | 243.0 | 1.4e-63 |
| AEF33078.1 | dimodular\_nonribosomal\_peptide\_synthetase | BGC0001039 | NRP + Polyketide | 31.0 | 48.1 | 243.0 | 1.4e-63 |
| WP\_030498974.1 | tyrocidine\_synthase\_3 | BGC0001327 | NRP:Cyclic depsipeptide + Polyketide:Modular type I | 32.0 | 49.7 | 243.0 | 1.4e-63 |
| WP\_047197779.1 | non-ribosomal\_peptide\_synthetase | BGC0002003 | NRP | 31.0 | 48.1 | 243.0 | 1.4e-63 |
| ASA76632.1 | polyketide\_synthase\_non-ribosomal\_peptide\_synthetase\_hybrid | BGC0001751 | NRP + Polyketide | 32.0 | 49.1 | 243.0 | 1.4e-63 |
| AFH75322.1 | nonribosomal\_peptide\_synthetase | BGC0000425 | NRP:Cyclic depsipeptide | 31.0 | 46.7 | 243.0 | 1.8e-63 |
| DAB41476.1 | nonribosomal\_peptide\_synthetase | BGC0001766 | NRP | 31.0 | 49.5 | 243.0 | 1.8e-63 |
| WP\_006051170.1 | non-ribosomal\_peptide\_synthetase | BGC0001999 | NRP | 32.0 | 47.0 | 243.0 | 1.8e-63 |
| AHI59109.1 | locillomycin\_synthase\_B | BGC0001005 | NRP + Polyketide | 29.0 | 47.7 | 243.0 | 2.3e-63 |
| AJM89734.1 | PmxB | BGC0001192 | NRP | 30.0 | 46.8 | 243.0 | 2.3e-63 |
| ATY37588.1 | BogA | BGC0001532 | NRP | 29.0 | 46.7 | 243.0 | 2.3e-63 |
| CZT62794.1 | Non-ribosomal\_peptide\_synthase\_involved\_in\_Hassallidin\_biosynthesis | BGC0001614 | NRP | 32.0 | 47.0 | 243.0 | 2.3e-63 |
| BAV56271.1 |  | BGC0001657 | NRP | 30.0 | 46.8 | 243.0 | 2.3e-63 |
| AXN93616.1 | PuwH | BGC0001953 | NRP | 33.0 | 49.3 | 243.0 | 2.3e-63 |
| AEG64697.1 | LpmC | BGC0000379 | NRP | 32.0 | 49.1 | 242.0 | 3e-63 |
| ACA97577.1 | PmxB | BGC0000408 | NRP | 30.0 | 46.8 | 242.0 | 3e-63 |
| ABW17377.1 | PsoC | BGC0000411 | NRP | 31.0 | 46.7 | 242.0 | 3e-63 |
| AAF00962.1 | mcyC | BGC0001017 | NRP + Polyketide:Modular type I | 29.0 | 46.9 | 242.0 | 3e-63 |
| BAH22764.1 | nonribosomal\_peptide\_synthetase | BGC0001018 | NRP + Polyketide | 30.0 | 48.5 | 242.0 | 3e-63 |
| AAR87760.2 | ZmaK | BGC0001059 | NRP + Polyketide | 28.0 | 47.7 | 242.0 | 3e-63 |
| ARU08074.1 | mlcL | BGC0001448 | NRP:Ca+-dependent lipopeptide | 32.0 | 46.7 | 242.0 | 3e-63 |
| AHZ20784.1 | non-ribosomal\_peptide\_synthase | BGC0000369 | NRP + Saccharide:Hybrid/tailoring | 30.0 | 47.9 | 242.0 | 3.9e-63 |
| AHI59110.1 | locillomycin\_synthase\_C | BGC0001005 | NRP + Polyketide | 29.0 | 51.1 | 242.0 | 3.9e-63 |
| ABS74209.1 | fengycin\_synthetase\_A | BGC0001095 | NRP | 30.0 | 48.5 | 242.0 | 3.9e-63 |
| CDF96614.1 | NRPS | BGC0001149 | NRP:Lipopeptide + Saccharide:Hybrid/tailoring | 30.0 | 47.8 | 242.0 | 3.9e-63 |
| ALK27915.1 | non-ribosomal\_peptide\_synthase | BGC0001233 | NRP | 32.0 | 45.6 | 242.0 | 3.9e-63 |
| AKC91856.1 | nonribosomal\_peptide\_synthetase | BGC0001414 | NRP | 31.0 | 47.5 | 242.0 | 3.9e-63 |
| WP\_047195042.1 | non-ribosomal\_peptide\_synthetase | BGC0002003 | NRP | 31.0 | 47.7 | 242.0 | 3.9e-63 |
| AED90002.1 | non-ribosomal\_peptide\_synthetase\_ThaA | BGC0000443 | NRP:Beta-lactam | 30.0 | 49.0 | 242.0 | 5.1e-63 |
| BAX64247.1 | NRPS | BGC0001623 | NRP + Polyketide | 31.0 | 47.2 | 242.0 | 5.1e-63 |
| AEG64698.1 | LpmD | BGC0000379 | NRP | 30.0 | 48.3 | 241.0 | 6.7e-63 |
| CAG29032.1 | nonribosomal\_peptide\_synthetase\_(modules\_3\_to\_6) | BGC0001023 | NRP + Polyketide:Modular type I | 29.0 | 50.5 | 241.0 | 6.7e-63 |
| CAF05648.1 | TubC\_protein | BGC0001053 | NRP + Polyketide | 32.0 | 46.5 | 241.0 | 6.7e-63 |
| ACM79810.1 | ZmaO | BGC0001059 | NRP + Polyketide | 31.0 | 46.5 | 241.0 | 6.7e-63 |
| ABS74207.1 | fengycin\_synthetase\_C | BGC0001095 | NRP | 30.0 | 48.5 | 241.0 | 6.7e-63 |
| ALV86867.1 | Tlo21 | BGC0001406 | NRP | 32.0 | 48.9 | 241.0 | 6.7e-63 |
| AKC91849.1 | nonribosomal\_peptide\_synthetase | BGC0001414 | NRP | 31.0 | 48.1 | 241.0 | 6.7e-63 |
| APZ78822.1 | nonribosomal\_peptide\_synthetase | BGC0001429 | NRP:Cyclic depsipeptide + Polyketide:Iterative type I | 29.0 | 50.5 | 241.0 | 6.7e-63 |
| BAX90000.1 | Non-ribosomal\_peptide\_synthetase | BGC0001628 | NRP | 33.0 | 45.9 | 241.0 | 6.7e-63 |
| AXN93615.1 | PuwG | BGC0001953 | NRP | 31.0 | 46.9 | 241.0 | 6.7e-63 |
| CAM56771.1 |  | BGC0000354 | NRP | 33.0 | 47.2 | 241.0 | 8.8e-63 |
| APZ78834.1 | nonribosomal\_peptide\_synthetase | BGC0001430 | NRP:Cyclic depsipeptide + Polyketide:Iterative type I | 31.0 | 48.8 | 241.0 | 8.8e-63 |
| WP\_099111429.1 | non-ribosomal\_peptide\_synthetase | BGC0001826 | NRP | 31.0 | 47.2 | 241.0 | 8.8e-63 |
| AXN93602.1 | PuwF-G | BGC0001952 | NRP | 30.0 | 48.1 | 241.0 | 8.8e-63 |
| QDA77045.1 | polyketide\_synthase/nonribosomal\_peptide\_synthetase | BGC0002025 | NRP | 31.0 | 48.1 | 241.0 | 8.8e-63 |
| AAC06347.1 | bacitracin\_synthetase\_2 | BGC0000310 | NRP | 30.0 | 46.5 | 240.0 | 1.1e-62 |
| BBD17741.1 | non-ribosomal\_peptide\_synthetase | BGC0001918 | NRP + Polyketide | 30.0 | 50.6 | 240.0 | 1.1e-62 |
| ACZ55943.1 | non-ribosomal\_peptide\_synthetase | BGC0000302 | NRP | 30.0 | 47.2 | 240.0 | 1.5e-62 |
| CBL93730.1 | NRPS | BGC0000360 | NRP | 32.0 | 50.2 | 240.0 | 1.5e-62 |
| CAE14171.1 |  | BGC0000383 | NRP + Polyketide:Modular type I | 30.0 | 47.4 | 240.0 | 1.5e-62 |
| BAP05589.1 | calA | BGC0000967 | NRP + Polyketide:Trans-AT type I | 34.0 | 37.9 | 240.0 | 1.5e-62 |
| ABS75232.1 | DhbF | BGC0001185 | NRP:NRP siderophore | 31.0 | 48.4 | 240.0 | 1.5e-62 |
| AHB82072.1 | non\_ribosomal\_peptide\_synthetase/polyketide\_synthase | BGC0001231 | NRP + Polyketide:Modular type I | 30.0 | 48.1 | 240.0 | 1.5e-62 |
| AAZ55899.1 | amino\_acid\_adenylation | BGC0000359 | NRP | 32.0 | 45.8 | 240.0 | 2e-62 |
| AAO72425.1 | syringopeptin\_synthetase\_C | BGC0000438 | NRP | 32.0 | 45.3 | 240.0 | 2e-62 |
| AYJ71720.1 | non-ribosomal\_peptide\_synthetase | BGC0001942 | NRP + Polyketide | 31.0 | 49.8 | 240.0 | 2e-62 |
| AAO72424.1 | syringopeptin\_synthetase\_B | BGC0000438 | NRP | 32.0 | 45.6 | 239.0 | 2.6e-62 |
| AAP92491.1 | nonribosomal\_peptide\_synthetase | BGC0000458 | NRP | 31.0 | 47.1 | 239.0 | 2.6e-62 |
| AGO86662.1 | equisetin\_synthetase | BGC0001255 | NRP + Polyketide | 27.0 | 70.2 | 239.0 | 2.6e-62 |
| AQH32485.1 | peptide\_synthetase | BGC0001667 | NRP + Polyketide | 30.0 | 47.7 | 239.0 | 2.6e-62 |
| APU91751.1 | Non-Ribosomal\_Peptide\_Synthetase | BGC0001806 | NRP | 31.0 | 46.7 | 239.0 | 2.6e-62 |
| AWS21279.1 | type\_I\_polyketide\_synthase | BGC0001934 | Polyketide | 33.0 | 38.4 | 239.0 | 2.6e-62 |
| AZY91989.1 | polyketide\_synthase | BGC0002022 | Polyketide | 33.0 | 38.4 | 239.0 | 2.6e-62 |
| ABV79985.1 | ApnA | BGC0000301 | NRP | 30.0 | 48.0 | 239.0 | 3.3e-62 |
| ABF87031.1 | non-ribosomal\_peptide\_synthetase/polyketide\_synthase | BGC0000393 | NRP + Polyketide:Modular type I | 30.0 | 49.1 | 239.0 | 3.3e-62 |
| AEA30272.1 | peptide\_synthetase | BGC0000429 | Polyketide + NRP:Cyclic depsipeptide | 30.0 | 49.8 | 239.0 | 3.3e-62 |
| AHD05615.1 | putative\_non-ribosomal\_peptide\_ligase/\_polyketide\_synthase\_hybrid | BGC0001033 | NRP + Polyketide | 29.0 | 47.4 | 239.0 | 3.3e-62 |
| AAN85501.1 | nonribosomal\_peptide\_synthetase | BGC0001101 | NRP + Polyketide:Modular type I + Polyketide:Trans-AT type I | 30.0 | 50.6 | 239.0 | 3.3e-62 |
| KPN90376.1 | NupC | BGC0001416 | NRP | 31.0 | 46.6 | 239.0 | 3.3e-62 |
| ctg1\_orf1264 |  | BGC0001752 | NRP | 30.0 | 55.5 | 239.0 | 3.3e-62 |
| QDF82254.1 | non-ribosomal\_peptide\_synthetase | BGC0001980 | NRP | 32.0 | 47.2 | 239.0 | 3.3e-62 |
| AEH59099.1 | amino\_acid\_adenylation\_domain-containing\_protein/NRPS | BGC0000385 | NRP | 30.0 | 46.3 | 238.0 | 4.4e-62 |
| CAE02631.1 | surfactin\_synthetase\_B\_ | BGC0000433 | NRP:Lipopeptide | 30.0 | 46.9 | 238.0 | 4.4e-62 |
| APZ78846.1 | nonribosomal\_peptide\_synthetase | BGC0001431 | NRP:Cyclic depsipeptide + Polyketide:Iterative type I | 29.0 | 50.4 | 238.0 | 4.4e-62 |
| AXN93614.1 | PuwF | BGC0001953 | NRP | 30.0 | 48.1 | 238.0 | 4.4e-62 |
| CCJ67640.1 | TaaE | BGC0000447 | NRP:Lipopeptide | 30.0 | 48.7 | 238.0 | 5.7e-62 |
| WP\_043882190.1 | non-ribosomal\_peptide\_synthetase | BGC0001993 | NRP | 31.0 | 45.6 | 238.0 | 5.7e-62 |
| ABV79986.1 | ApnB | BGC0000301 | NRP | 30.0 | 47.8 | 238.0 | 7.4e-62 |
| ACZ55945.1 | non-ribosomal\_peptide\_synthetase | BGC0000302 | NRP | 31.0 | 47.4 | 238.0 | 7.4e-62 |
| BAH43871.1 | truncated\_linear\_pentadecapeptide\_gramicidin\_synthetase\_LgrC | BGC0000367 | NRP | 29.0 | 50.2 | 238.0 | 7.4e-62 |
| CAG23960.2 | hybrid\_NRPS/PKS\_protein | BGC0001089 | Polyketide + NRP | 29.0 | 47.4 | 238.0 | 7.4e-62 |
| AUD11994.1 | OrbI | BGC0001721 | NRP | 31.0 | 47.8 | 238.0 | 7.4e-62 |
| CAL80824.1 | NRPS\_module\_protein | BGC0000997 | NRP + Polyketide | 32.0 | 47.9 | 237.0 | 9.7e-62 |
| ABS74206.1 | fengycin\_synthetase\_D | BGC0001095 | NRP | 31.0 | 46.2 | 237.0 | 9.7e-62 |
| WP\_018540603.1 | non-ribosomal\_peptide\_synthetase | BGC0001332 | NRP + Polyketide | 32.0 | 45.1 | 237.0 | 9.7e-62 |
| AKP45399.1 | CysK | BGC0001413 | NRP | 29.0 | 49.5 | 237.0 | 9.7e-62 |
| KPN93063.1 | NupA | BGC0001416 | NRP | 32.0 | 46.7 | 237.0 | 9.7e-62 |
| CZT62785.1 | Non-ribosomal\_peptide\_synthase,\_involved\_in\_Hassallidin\_biosynthesis | BGC0001614 | NRP | 31.0 | 45.1 | 237.0 | 9.7e-62 |
| CAJ34374.1 | NRPS\_protein | BGC0000445 | NRP:Cyclic depsipeptide | 33.0 | 46.2 | 237.0 | 1.3e-61 |
| BAE98155.1 | putative\_non-ribosomal\_peptide\_synthetase | BGC0000339 | NRP | 32.0 | 49.6 | 237.0 | 1.7e-61 |
| AAY37654.1 | Amino\_acid\_adenylation | BGC0000437 | NRP | 32.0 | 45.6 | 237.0 | 1.7e-61 |
| ABC36785.1 | peptide\_synthetase,\_putative | BGC0000964 | NRP:Cyclic depsipeptide + Polyketide:Trans-AT type I | 33.0 | 45.3 | 237.0 | 1.7e-61 |
| ABC87508.1 | NRPS\_for\_pipecolate\_incorporation | BGC0001011 | NRP + Polyketide | 30.0 | 47.4 | 236.0 | 2.2e-61 |
| ctg1\_orf19 |  | BGC0001013 | NRP + Polyketide | 30.0 | 47.4 | 236.0 | 2.2e-61 |
| DAB41484.1 | nonribosomal\_peptide\_synthetase/polyketide\_synthase\_type\_I | BGC0001766 | NRP | 31.0 | 48.1 | 236.0 | 2.2e-61 |
| ATV95610.1 | NRPS\_A-PCP\_didomain\_protein | BGC0001503 | Polyketide | 31.0 | 47.4 | 236.0 | 2.2e-61 |
| AXM43052.1 | non-ribosomal\_peptide\_synthetase | BGC0001945 | NRP | 32.0 | 47.9 | 236.0 | 2.2e-61 |
| BAH22762.1 | nonribosomal\_peptide\_synthetase | BGC0001018 | NRP + Polyketide | 30.0 | 47.4 | 236.0 | 2.8e-61 |
| AHZ34233.1 | CifB | BGC0000323 | NRP:Lipopeptide | 30.0 | 46.5 | 235.0 | 3.7e-61 |
| BAH04162.1 | trsJ | BGC0000450 | NRP | 32.0 | 47.3 | 235.0 | 3.7e-61 |
| AEZ51517.1 | pmxB | BGC0001153 | NRP:Lipopeptide | 30.0 | 46.8 | 235.0 | 3.7e-61 |
| DAB41479.1 | nonribosomal\_peptide\_synthetase | BGC0001766 | NRP | 33.0 | 47.6 | 235.0 | 3.7e-61 |
| ALG65319.1 | Cal17 | BGC0001297 | NRP | 31.0 | 49.0 | 235.0 | 3.7e-61 |
| BBA20967.1 | nonribosomal\_peptide\_synthetase | BGC0001763 | NRP + Polyketide | 31.0 | 46.8 | 235.0 | 3.7e-61 |
| ABU70377.1 | hypothetical\_protein | BGC0001890 | NRP | 30.0 | 47.0 | 235.0 | 3.7e-61 |
| CAJ18237.2 | non-ribosomal\_peptide\_synthetase\_B | BGC0000354 | NRP | 32.0 | 48.1 | 235.0 | 4.8e-61 |
| AEG64689.1 | LpmA | BGC0000379 | NRP | 28.0 | 52.6 | 235.0 | 4.8e-61 |
| ABL74939.1 | NRPS | BGC0001048 | NRP:Glycopeptide + Polyketide:Modular type I + Saccharide:Hybrid/tailoring | 32.0 | 49.8 | 235.0 | 4.8e-61 |
| ASX95241.1 | IlaS | BGC0001620 | Polyketide | 31.0 | 46.8 | 235.0 | 4.8e-61 |
| AAZ23075.1 | peptide\_synthetase | BGC0000291 | NRP | 32.0 | 47.7 | 235.0 | 6.3e-61 |
| CAB38518.1 | CDA\_peptide\_synthetase\_I\_(CdaPs1) | BGC0000315 | NRP:Ca+-dependent lipopeptide | 31.0 | 47.4 | 235.0 | 6.3e-61 |
| AHZ20774.1 | non-ribosomal\_peptide\_synthase | BGC0000369 | NRP + Saccharide:Hybrid/tailoring | 30.0 | 48.1 | 235.0 | 6.3e-61 |
| AEI70245.1 | nonribosomal\_peptide\_synthetase\_NRPS | BGC0000401 | NRP | 31.0 | 48.4 | 235.0 | 6.3e-61 |
| AAO56329.1 | non-ribosomal\_peptide\_synthetase\_SyfB | BGC0000435 | NRP | 30.0 | 50.2 | 235.0 | 6.3e-61 |
| BAW32324.1 | nonribosomal\_peptide\_synthetase | BGC0001630 | NRP + Polyketide | 31.0 | 45.9 | 235.0 | 6.3e-61 |
| WP\_144411596.1 | amino\_acid\_adenylation\_domain-containing\_protein | BGC0002001 | NRP + Polyketide | 31.0 | 45.7 | 235.0 | 6.3e-61 |
| WP\_039806856.1 | non-ribosomal\_peptide\_synthetase | BGC0002001 | NRP + Polyketide | 31.0 | 45.8 | 235.0 | 6.3e-61 |
| ADL64235.1 | aureusimine\_non-ribosomal\_peptide\_synthetase | BGC0000308 | NRP | 26.0 | 75.0 | 234.0 | 8.2e-61 |
| CCA29203.1 | non-ribosomal\_peptide\_synthetase/polyketide\_synthase | BGC0000955 | NRP + Polyketide:Modular type I | 30.0 | 49.3 | 234.0 | 8.2e-61 |
| CBJ79916.1 | putative\_Ornithine\_racemase | BGC0001133 | NRP | 30.0 | 48.4 | 234.0 | 8.2e-61 |
| BAH43870.1 | putative\_linear\_pentadecapeptide\_gramicidin\_synthetase\_LgrB | BGC0000367 | NRP | 30.0 | 47.0 | 234.0 | 1.1e-60 |
| AAO56328.1 | non-ribosomal\_peptide\_synthetase\_SyfA | BGC0000435 | NRP | 29.0 | 47.9 | 234.0 | 1.1e-60 |
| CAJ45639.1 | vanchrobactin\_non\_ribosomal\_peptide\_synthetase | BGC0000454 | NRP | 29.0 | 47.7 | 234.0 | 1.1e-60 |
| AHZ34239.1 | CipB | BGC0001389 | NRP | 32.0 | 47.5 | 234.0 | 1.1e-60 |
| WP\_064118559.1 | non-ribosomal\_peptide\_synthetase | BGC0001509 | NRP | 30.0 | 49.0 | 234.0 | 1.1e-60 |
| CAB38517.1 | CDA\_peptide\_synthetase\_II\_(CdaPs2) | BGC0000315 | NRP:Ca+-dependent lipopeptide | 30.0 | 48.6 | 233.0 | 1.4e-60 |
| ABS74179.1 | bacillomycin\_D\_synthetase\_C | BGC0001090 | Polyketide + NRP:Lipopeptide | 28.0 | 47.7 | 233.0 | 1.4e-60 |
| CAQ71829.1 | non\_ribosomal\_peptide\_synthase,\_antibiotic\_synthesis;\_contains\_4\_condensation\_domains,\_3\_AMP-acid\_ligases\_II\_domains,\_3\_PP-binding,\_Phosphopantetheine\_attachment\_site\_and\_a\_putative\_thioesterase\_domain | BGC0001189 | NRP | 32.0 | 45.4 | 233.0 | 1.4e-60 |
| ADH04679.1 | non-ribosomal\_peptide\_synthetase | BGC0001344 | NRP + Polyketide | 30.0 | 47.7 | 233.0 | 1.4e-60 |
| AXN93613.1 | PuwE | BGC0001953 | NRP | 28.0 | 47.9 | 233.0 | 1.4e-60 |
| AAF63833.1 | PstD | BGC0000362 | NRP | 32.0 | 48.7 | 233.0 | 1.8e-60 |
| AFJ14793.1 | PlpD | BGC0000403 | NRP | 29.0 | 49.7 | 233.0 | 1.8e-60 |
| ADF88262.1 | mixed\_nonribosomal\_peptide\_synthetase/\_polyketide\_synthase | BGC0000979 | NRP + Polyketide | 31.0 | 49.6 | 233.0 | 1.8e-60 |
| ADF88279.1 | mixed\_NRPS/PKS | BGC0000981 | NRP + Polyketide | 31.0 | 49.6 | 233.0 | 1.8e-60 |
| CAD29798.1 | peptide\_synthetase | BGC0001015 | NRP + Polyketide | 29.0 | 46.8 | 233.0 | 1.8e-60 |
| AHB82056.1 | non\_ribosomal\_peptide\_synthetase | BGC0001019 | NRP + Polyketide:Modular type I | 31.0 | 49.2 | 233.0 | 1.8e-60 |
| AFR69334.1 | nonribosomal\_peptide\_synthetase\_SpiDE1 | BGC0001045 | NRP:Cyclic depsipeptide + Polyketide:Modular type I | 32.0 | 46.4 | 233.0 | 1.8e-60 |
| APZ78856.1 | nonribosomal\_peptide\_synthetase | BGC0001432 | NRP:Cyclic depsipeptide + Polyketide:Iterative type I | 30.0 | 50.7 | 233.0 | 1.8e-60 |
| ADN26248.1 | peptide\_synthetase | BGC0000951 | NRP | 31.0 | 46.4 | 233.0 | 2.4e-60 |
| AHH25592.1 | NRPS | BGC0000957 | NRP + Polyketide | 31.0 | 47.4 | 233.0 | 2.4e-60 |
| CBF80487.1 | hybrid\_PKS-NRPS\_(Eurofung) | BGC0000959 | NRP + Polyketide:Iterative type I | 27.0 | 69.8 | 233.0 | 2.4e-60 |
| AAY32966.1 | DszC | BGC0001093 | NRP + Polyketide | 31.0 | 51.2 | 233.0 | 2.4e-60 |
| AHB82069.1 | non\_ribosomal\_peptide\_synthetase | BGC0001231 | NRP + Polyketide:Modular type I | 31.0 | 47.3 | 233.0 | 2.4e-60 |
| WP\_013310343.1 | non-ribosomal\_peptide\_synthetase | BGC0001993 | NRP | 30.0 | 47.0 | 233.0 | 2.4e-60 |
| QBC75022.1 | non-ribosomal\_peptide\_synthetase | BGC0001968 | NRP | 33.0 | 48.3 | 233.0 | 2.4e-60 |
| ADF88265.1 | mixed\_nonribosomal\_peptide\_synthetase/\_polyketide\_synthase | BGC0000980 | NRP + Polyketide | 31.0 | 49.6 | 232.0 | 3.1e-60 |
| ACM79806.1 | ZmaB | BGC0001059 | NRP + Polyketide | 28.0 | 45.8 | 232.0 | 3.1e-60 |
| WP\_080679150.1 | non-ribosomal\_peptide\_synthetase | BGC0001228 | NRP:Cyclic depsipeptide | 32.0 | 47.6 | 232.0 | 3.1e-60 |
| ATY37608.1 | BreC | BGC0001536 | NRP | 30.0 | 49.8 | 232.0 | 3.1e-60 |
| WP\_013184596.1 | non-ribosomal\_peptide\_synthetase | BGC0001825 | NRP | 31.0 | 47.4 | 232.0 | 3.1e-60 |
| CAE52339.1 | Polyketide\_non-ribosomal\_peptide\_synthase | BGC0001088 | NRP + Polyketide | 28.0 | 48.1 | 232.0 | 4.1e-60 |
| CAM56770.1 |  | BGC0000354 | NRP | 31.0 | 49.5 | 232.0 | 5.3e-60 |
| CAJ46691.1 | non-ribosomal\_peptide\_synthase | BGC0000969 | NRP:Cyclic depsipeptide + Polyketide:Modular type I | 30.0 | 49.5 | 232.0 | 5.3e-60 |
| CBD77746.1 | non-ribosomal\_peptide\_synthetase/polyketide\_synthase | BGC0000974 | NRP + Polyketide | 32.0 | 48.2 | 232.0 | 5.3e-60 |
| ABX60161.1 | mixed\_NRPS/PKS | BGC0000978 | NRP + Alkaloid + Polyketide:Modular type I | 31.0 | 49.6 | 232.0 | 5.3e-60 |
| ACG60782.1 | NRPS(C/A/PCP/C/A/PCP) | BGC0001058 | NRP:Glycopeptide + Polyketide:Modular type I + Saccharide:Hybrid/tailoring | 30.0 | 48.1 | 232.0 | 5.3e-60 |
| AGC09528.1 | NRPS | BGC0001183 | Polyketide | 33.0 | 42.7 | 232.0 | 5.3e-60 |
| AOA33123.1 | Nonribosomal\_peptide\_synthetase | BGC0001346 | NRP:Cyclic depsipeptide | 30.0 | 48.9 | 232.0 | 5.3e-60 |
| AIR74926.1 | polyketide\_synthase | BGC0001559 | RiPP | 32.0 | 48.2 | 232.0 | 5.3e-60 |
| AXN93601.1 | PuwE | BGC0001952 | NRP | 28.0 | 48.8 | 232.0 | 5.3e-60 |
| AFH75330.1 | nonribosomal\_peptide\_synthetase | BGC0000398 | NRP:Cyclic depsipeptide | 30.0 | 49.0 | 231.0 | 7e-60 |
| CAG23957.2 | hybrid\_NRPS/PKS\_protein | BGC0001089 | Polyketide + NRP | 31.0 | 46.7 | 231.0 | 7e-60 |
| ALV86866.1 | Tlo20 | BGC0001406 | NRP | 32.0 | 49.1 | 231.0 | 7e-60 |
| KPN93064.1 | NupB | BGC0001416 | NRP | 28.0 | 51.7 | 231.0 | 7e-60 |
| AAU39361.1 | lichenysin\_synthase\_LchAC | BGC0000381 | NRP | 30.0 | 47.0 | 231.0 | 9.1e-60 |
| PHM26612.1 | pvdj | BGC0001130 | NRP + Polyketide | 29.0 | 45.9 | 231.0 | 9.1e-60 |
| BAB69700.1 | iturin\_A\_synthetase\_C | BGC0001098 | NRP + Polyketide | 29.0 | 47.7 | 230.0 | 1.2e-59 |
| AFH75328.1 | nonribosomal\_peptide\_synthetase | BGC0000398 | NRP:Cyclic depsipeptide | 30.0 | 49.5 | 230.0 | 1.5e-59 |
| CBW54672.1 | non\_ribosomal\_peptide\_synthetase | BGC0000971 | NRP + Polyketide:Modular type I | 31.0 | 48.1 | 230.0 | 1.5e-59 |
| AAF00961.1 | mcyB | BGC0001017 | NRP + Polyketide:Modular type I | 30.0 | 47.0 | 230.0 | 1.5e-59 |
| AHB38497.1 | non-ribosomal\_peptide\_synthetase | BGC0000346 | NRP + Polyketide:Modular type I | 30.0 | 46.4 | 230.0 | 2e-59 |
| AAY91419.3 | non-ribosomal\_peptide\_synthetase\_OfaA | BGC0000399 | NRP:Cyclic depsipeptide | 32.0 | 46.6 | 230.0 | 2e-59 |
| AAY37647.1 | Amino\_acid\_adenylation | BGC0000437 | NRP | 28.0 | 48.5 | 230.0 | 2e-59 |
| CRI73800.1 | loading\_module\_of\_NRPS-PKS | BGC0001215 | NRP | 31.0 | 49.1 | 230.0 | 2e-59 |
| BAW27693.1 | NRPS(C-A-T-TE) | BGC0001764 | NRP | 32.0 | 48.8 | 230.0 | 2e-59 |
| BAI63288.1 | putative\_non-ribosomal\_peptide\_synthetase | BGC0000434 | NRP | 31.0 | 45.5 | 229.0 | 2.6e-59 |
| WP\_064118561.1 | non-ribosomal\_peptide\_synthetase | BGC0001509 | NRP | 30.0 | 46.9 | 229.0 | 2.6e-59 |
| CCA29202.1 | non-ribosomal\_peptide\_synthetase | BGC0000955 | NRP + Polyketide:Modular type I | 30.0 | 48.1 | 229.0 | 3.4e-59 |
| AJW76711.1 | DsaI | BGC0001196 | NRP | 32.0 | 45.3 | 229.0 | 3.4e-59 |
| WP\_084702182.1 | non-ribosomal\_peptide\_synthetase | BGC0001211 | NRP | 30.0 | 49.1 | 229.0 | 3.4e-59 |
| AJV88377.1 | MfnE | BGC0001214 | NRP | 32.0 | 48.1 | 229.0 | 3.4e-59 |
| ACM68690.1 | AerG1 | BGC0000298 | NRP | 31.0 | 47.0 | 228.0 | 4.5e-59 |
| AAY37653.1 | Amino\_acid\_adenylation | BGC0000437 | NRP | 31.0 | 47.0 | 228.0 | 4.5e-59 |
| AVI26393.1 | nonribosomal\_peptide\_synthase | BGC0001800 | NRP + Polyketide | 30.0 | 48.4 | 228.0 | 4.5e-59 |
| ABV56588.1 | KtzH | BGC0000378 | NRP | 30.0 | 48.5 | 228.0 | 5.9e-59 |
| CAJ34382.1 | NRPS\_protein | BGC0000445 | NRP:Cyclic depsipeptide | 31.0 | 45.8 | 228.0 | 5.9e-59 |
| CAO91861.1 | PKS-NRPS\_hybrid | BGC0000968 | NRP + Polyketide:Iterative type I | 25.0 | 73.7 | 228.0 | 5.9e-59 |
| AAF86395.1 | FkbP | BGC0000994 | NRP + Polyketide | 30.0 | 48.0 | 228.0 | 5.9e-59 |
| OLZ50885.1 | non-ribosomal\_peptide\_synthetase | BGC0001461 | NRP:Glycopeptide | 31.0 | 46.1 | 228.0 | 5.9e-59 |
| RAT94091.1 | NRPS | BGC0001469 | NRP | 30.0 | 48.1 | 228.0 | 5.9e-59 |
| BBA21068.1 | putative\_non-ribosomal\_peptide\_synthetase | BGC0001740 | NRP + Polyketide | 29.0 | 47.9 | 228.0 | 5.9e-59 |
| AAT09805.1 | NocB | BGC0000395 | NRP | 32.0 | 47.1 | 228.0 | 7.7e-59 |
| AEH41793.1 | HrmO | BGC0000374 | NRP:Cyclic depsipeptide | 30.0 | 47.7 | 227.0 | 1e-58 |
| AHD05678.1 | nonribosomal\_peptide\_ligase\_subunit | BGC0000402 | NRP | 30.0 | 46.3 | 227.0 | 1e-58 |
| CAE02633.1 | surfactin\_synthetase\_C\_ | BGC0000433 | NRP:Lipopeptide | 29.0 | 47.5 | 227.0 | 1e-58 |
| AHH53508.1 | non-ribosomal\_peptide\_synthetase | BGC0000439 | NRP:Ca+-dependent lipopeptide | 32.0 | 49.8 | 227.0 | 1e-58 |
| AOA33122.1 | Nonribosomal\_peptide\_synthetase | BGC0001346 | NRP:Cyclic depsipeptide | 30.0 | 48.3 | 227.0 | 1e-58 |
| AKC91857.1 | nonribosomal\_peptide\_synthetase | BGC0001414 | NRP | 30.0 | 48.1 | 227.0 | 1e-58 |
| KPN90369.1 | NunE | BGC0001416 | NRP | 29.0 | 47.2 | 227.0 | 1e-58 |
| APZ78680.1 | nonribosomal\_peptide\_synthetase | BGC0001417 | NRP:Cyclic depsipeptide + Polyketide:Iterative type I | 30.0 | 46.9 | 227.0 | 1e-58 |
| WP\_013184464.1 | non-ribosomal\_peptide\_synthetase | BGC0001716 | NRP | 31.0 | 47.6 | 227.0 | 1e-58 |
| WP\_013310342.1 | non-ribosomal\_peptide\_synthetase | BGC0001993 | NRP | 30.0 | 49.0 | 227.0 | 1e-58 |
| AEH41794.1 | HrmP | BGC0000374 | NRP:Cyclic depsipeptide | 29.0 | 49.4 | 227.0 | 1.3e-58 |
| AEW31015.1 | plipastatin\_synthetase | BGC0000407 | NRP | 29.0 | 47.0 | 227.0 | 1.3e-58 |
| EME52989.1 | amino\_acid\_adenylation\_protein | BGC0001460 | NRP:Glycopeptide | 31.0 | 47.4 | 227.0 | 1.3e-58 |
| ATY37589.1 | BogB | BGC0001532 | NRP | 28.0 | 46.3 | 227.0 | 1.3e-58 |
| WP\_078586793.1 | non-ribosomal\_peptide\_synthetase | BGC0001760 | NRP | 30.0 | 46.3 | 227.0 | 1.3e-58 |
| ctg1\_orf20 |  | BGC0001767 | NRP | 30.0 | 50.4 | 227.0 | 1.3e-58 |
| BAV57443.1 | NRPS\_(C-A-PCP-TE) | BGC0001912 | NRP | 31.0 | 48.2 | 227.0 | 1.3e-58 |
| AYA22334.1 | KerC | BGC0001955 | Other | 31.0 | 46.1 | 227.0 | 1.3e-58 |
| BAC67535.1 | arthrofactin\_synthetase\_B | BGC0000305 | NRP:Lipopeptide | 32.0 | 48.6 | 227.0 | 1.7e-58 |
| EWS95122.1 | hypothetical\_protein | BGC0000306 | NRP:Lipopeptide | 32.0 | 47.7 | 227.0 | 1.7e-58 |
| AAY93356.2 | non-ribosomal\_peptide\_synthetase\_PvdI | BGC0000413 | NRP | 31.0 | 47.0 | 227.0 | 1.7e-58 |
| BAH43765.1 | tyrocidine\_synthetase\_II | BGC0000452 | NRP | 29.0 | 51.6 | 227.0 | 1.7e-58 |
| BAP05591.1 | calC | BGC0000967 | NRP + Polyketide:Trans-AT type I | 30.0 | 48.0 | 227.0 | 1.7e-58 |
| AAS98786.1 | nonribosomal\_peptide\_synthetase | BGC0001001 | NRP + Polyketide | 31.0 | 46.8 | 227.0 | 1.7e-58 |
| EFL06867.1 | predicted\_protein | BGC0000300 | NRP | 32.0 | 48.8 | 226.0 | 2.2e-58 |
| AAC68816.1 | FK506\_peptide\_synthetase | BGC0000353 | NRP | 31.0 | 48.2 | 226.0 | 2.2e-58 |
| AAU39359.1 | lichenysin\_synthase\_LchAA | BGC0000381 | NRP | 29.0 | 48.2 | 226.0 | 2.2e-58 |
| AHD05618.1 | putative\_non-ribosomal\_peptide\_ligase\_domain\_protein | BGC0001033 | NRP + Polyketide | 30.0 | 49.3 | 226.0 | 2.2e-58 |
| BAB69699.1 | iturin\_A\_synthetase\_B | BGC0001098 | NRP + Polyketide | 29.0 | 47.4 | 226.0 | 2.2e-58 |
| EWS95124.1 | hypothetical\_protein | BGC0000306 | NRP:Lipopeptide | 29.0 | 47.4 | 226.0 | 2.9e-58 |
| AAF99707.2 | syringopeptin\_synthetase | BGC0000438 | NRP | 30.0 | 47.6 | 226.0 | 2.9e-58 |
| ABX37382.1 | amino\_acid\_adenylation\_domain\_protein | BGC0000984 | NRP + Polyketide | 31.0 | 46.4 | 226.0 | 2.9e-58 |
| ACM79812.1 | ZmaQ | BGC0001059 | NRP + Polyketide | 31.0 | 47.7 | 226.0 | 2.9e-58 |
| OAQ83772.1 | nonribosomal\_peptide\_synthase | BGC0001358 | Polyketide | 26.0 | 72.7 | 226.0 | 2.9e-58 |
| WP\_050383088.1 | non-ribosomal\_peptide\_synthetase | BGC0001451 | NRP | 31.0 | 46.7 | 226.0 | 2.9e-58 |
| BAX89999.1 | Non-ribosomal\_peptide\_synthetase | BGC0001628 | NRP | 31.0 | 47.5 | 226.0 | 2.9e-58 |
| ABV56604.1 | adenylation\_domain\_protein | BGC0000378 | NRP | 32.0 | 45.3 | 225.0 | 3.8e-58 |
| ADZ24999.1 | non-ribosomal\_peptide\_synthase | BGC0000380 | NRP + Polyketide:Modular type I | 30.0 | 47.7 | 225.0 | 3.8e-58 |
| AAU39360.1 | lichenysin\_synthase\_LchAB | BGC0000381 | NRP | 28.0 | 47.6 | 225.0 | 3.8e-58 |
| ACY06285.1 | non-ribosomal\_peptide\_synthetase | BGC0001042 | NRP + Polyketide | 30.0 | 49.1 | 225.0 | 3.8e-58 |
| CCM44336.1 | Nonribosomal\_peptide\_synthetase | BGC0001056 | NRP + Polyketide:Modular type I + Polyketide:PUFA synthase or related | 29.0 | 47.0 | 225.0 | 3.8e-58 |
| AAF08797.1 | MycC | BGC0001103 | NRP + Polyketide | 28.0 | 46.2 | 225.0 | 3.8e-58 |
| WP\_050383094.1 | non-ribosomal\_peptide\_synthetase | BGC0001451 | NRP | 31.0 | 47.3 | 225.0 | 3.8e-58 |
| ADG27358.1 | peptide\_synthetase | BGC0000296 | NRP | 31.0 | 47.7 | 225.0 | 5e-58 |
| AID65224.1 | nonribosomal\_peptide\_synthetase | BGC0000335 | NRP | 31.0 | 47.5 | 225.0 | 5e-58 |
| AAX31559.1 | peptide\_synthetase\_3 | BGC0000336 | NRP | 32.0 | 47.3 | 225.0 | 5e-58 |
| AHF21229.1 | TriE | BGC0000449 | NRP | 29.0 | 46.8 | 225.0 | 5e-58 |
| BAD55611.1 | putative\_non-ribosomal\_peptide\_synthetase | BGC0001027 | NRP + Polyketide | 30.0 | 47.2 | 225.0 | 5e-58 |
| AHZ34243.1 | CipF | BGC0001389 | NRP | 30.0 | 46.7 | 225.0 | 5e-58 |
| APZ78716.1 | nonribosomal\_peptide\_synthetase | BGC0001420 | NRP:Cyclic depsipeptide + Polyketide:Iterative type I | 30.0 | 45.6 | 225.0 | 5e-58 |
| XP\_001217690.1 | hypothetical\_protein | BGC0001517 | NRP | 25.0 | 75.0 | 225.0 | 5e-58 |
| ABD65958.1 | nonribosomal\_peptide\_synthetase | BGC0000341 | NRP | 32.0 | 47.8 | 225.0 | 6.5e-58 |
| AAF08796.1 | MycB | BGC0001103 | NRP + Polyketide | 29.0 | 46.7 | 225.0 | 6.5e-58 |
| AAV97877.1 | OnnI | BGC0001105 | NRP + Polyketide:Trans-AT type I | 31.0 | 47.8 | 225.0 | 6.5e-58 |
| APZ78692.1 | nonribosomal\_peptide\_synthetase | BGC0001418 | NRP:Cyclic depsipeptide + Polyketide:Iterative type I | 29.0 | 45.6 | 225.0 | 6.5e-58 |
| gene3 |  | BGC0002035 | NRP + Polyketide | 26.0 | 73.4 | 225.0 | 6.5e-58 |
| CAJ34381.1 | NRPS\_protein | BGC0000445 | NRP:Cyclic depsipeptide | 30.0 | 45.3 | 224.0 | 8.5e-58 |
| ABX37385.1 | amino\_acid\_adenylation\_domain\_protein | BGC0000984 | NRP + Polyketide | 30.0 | 48.4 | 224.0 | 8.5e-58 |
| BAB69698.1 | iturin\_A\_synthetase\_A | BGC0001098 | NRP + Polyketide | 28.0 | 48.5 | 224.0 | 8.5e-58 |
| DAB41478.1 | nonribosomal\_peptide\_synthetase | BGC0001766 | NRP | 31.0 | 48.2 | 224.0 | 8.5e-58 |
| APZ78704.1 | nonribosomal\_peptide\_synthetase | BGC0001419 | NRP:Cyclic depsipeptide + Polyketide:Iterative type I | 29.0 | 45.6 | 224.0 | 8.5e-58 |
| ACZ55942.1 | non-ribosomal\_peptide\_synthetase | BGC0000302 | NRP | 30.0 | 46.6 | 224.0 | 1.1e-57 |
| ABR67744.1 | CmnA | BGC0000316 | NRP | 32.0 | 48.1 | 224.0 | 1.1e-57 |
| AGJ76605.1 | HglB | BGC0000869 | Other | 39.0 | 26.4 | 224.0 | 1.1e-57 |
| AHD05621.1 | non-ribosomal\_peptide\_ligase\_domain\_protein | BGC0001033 | NRP + Polyketide | 32.0 | 37.2 | 224.0 | 1.1e-57 |
| ALK27916.1 | non-ribosomal\_peptide\_synthase | BGC0001233 | NRP | 31.0 | 45.7 | 224.0 | 1.1e-57 |
| AHZ34242.1 | CipE | BGC0001389 | NRP | 31.0 | 46.5 | 224.0 | 1.1e-57 |
| ABD14711.1 | cesA | BGC0000320 | NRP:Cyclic depsipeptide | 31.0 | 46.1 | 223.0 | 1.4e-57 |
| AAT01806.1 | non-ribosomal\_peptide\_synthetase | BGC0000365 | NRP | 31.0 | 45.6 | 223.0 | 1.4e-57 |
| BAH43764.1 | tyrocidine\_synthetase\_I | BGC0000452 | NRP | 29.0 | 45.6 | 223.0 | 1.4e-57 |
| AGZ15458.1 | putative\_non-ribosomal\_peptide\_synthetase | BGC0001036 | NRP + Polyketide | 30.0 | 46.3 | 223.0 | 1.4e-57 |
| ABS74181.1 | bacillomycin\_D\_synthetase\_A\_ | BGC0001090 | Polyketide + NRP:Lipopeptide | 28.0 | 48.5 | 223.0 | 1.4e-57 |
| ABS74205.1 | fengycin\_synthetase\_E | BGC0001095 | NRP | 29.0 | 47.1 | 223.0 | 1.4e-57 |
| ABS90473.1 | NRPS | BGC0001106 | NRP + Polyketide | 34.0 | 36.1 | 223.0 | 1.4e-57 |
| ALD82526.1 | non-ribosomal\_peptide\_synthase | BGC0001212 | NRP + Polyketide | 32.0 | 48.1 | 223.0 | 1.4e-57 |
| AKC54422.1 | fumosorinone\_biosynthesis\_polyketide\_synthase | BGC0001218 | NRP + Polyketide | 27.0 | 70.5 | 223.0 | 1.4e-57 |
| BAT51067.1 | type\_I\_polyketide\_synthase | BGC0001296 | Polyketide | 37.0 | 32.0 | 223.0 | 1.4e-57 |
| APZ78728.1 | nonribosomal\_peptide\_synthetase | BGC0001421 | NRP:Cyclic depsipeptide + Polyketide:Iterative type I | 30.0 | 47.6 | 223.0 | 1.4e-57 |
| APZ78794.1 | nonribosomal\_peptide\_synthetase | BGC0001427 | NRP:Cyclic depsipeptide + Polyketide:Iterative type I | 33.0 | 36.9 | 223.0 | 1.4e-57 |
| EFL06865.1 | hypothetical\_protein | BGC0000300 | NRP | 31.0 | 45.4 | 223.0 | 1.9e-57 |
| CAB53322.1 | putative\_peptide\_synthetase | BGC0000325 | NRP | 29.0 | 48.6 | 223.0 | 1.9e-57 |
| ABX37383.1 | amino\_acid\_adenylation\_domain\_protein | BGC0000984 | NRP + Polyketide | 31.0 | 49.0 | 223.0 | 1.9e-57 |
| ABJ97436.1 | MerP | BGC0001012 | NRP + Polyketide | 29.0 | 49.4 | 223.0 | 1.9e-57 |
| CAN89663.1 | putative\_non-ribosomal\_peptide\_synthetase | BGC0001070 | NRP + Polyketide:Modular type I + Polyketide:Trans-AT type I | 31.0 | 47.8 | 223.0 | 1.9e-57 |
| CAE52334.1 | non-ribosomal\_peptide\_synthase | BGC0001088 | NRP + Polyketide | 28.0 | 48.3 | 223.0 | 1.9e-57 |
| APZ78679.1 | nonribosomal\_peptide\_synthetase | BGC0001417 | NRP:Cyclic depsipeptide + Polyketide:Iterative type I | 31.0 | 46.4 | 223.0 | 1.9e-57 |
| APZ78768.1 | nonribosomal\_peptide\_synthetase | BGC0001425 | NRP:Cyclic depsipeptide + Polyketide:Iterative type I | 33.0 | 36.9 | 223.0 | 1.9e-57 |
| APZ78808.1 | nonribosomal\_peptide\_synthetase | BGC0001428 | NRP:Cyclic depsipeptide + Polyketide:Iterative type I | 31.0 | 46.7 | 223.0 | 1.9e-57 |
| APZ78855.1 | nonribosomal\_peptide\_synthetase | BGC0001432 | NRP:Cyclic depsipeptide + Polyketide:Iterative type I | 31.0 | 46.4 | 223.0 | 1.9e-57 |
| EME52990.1 | amino\_acid\_adenylation\_protein | BGC0001460 | NRP:Glycopeptide | 31.0 | 46.7 | 223.0 | 1.9e-57 |
| QDF82255.1 | non-ribosomal\_peptide\_synthetase | BGC0001980 | NRP | 31.0 | 47.8 | 223.0 | 1.9e-57 |
| ABO15888.1 | polyketide\_synthase | BGC0000132 | Polyketide | 32.0 | 39.0 | 223.0 | 2.5e-57 |
| CAJ96468.1 | non-ribosomal\_peptide\_synthetase | BGC0000330 | NRP:NRP siderophore | 31.0 | 47.3 | 223.0 | 2.5e-57 |
| ABD65960.1 | nonribosomal\_peptide\_synthetase | BGC0000341 | NRP | 31.0 | 46.8 | 223.0 | 2.5e-57 |
| AAG02358.1 | peptide\_synthetase\_NRPS6 | BGC0000963 | NRP:Glycopeptide + Polyketide:Modular type I + Saccharide:Hybrid/tailoring | 31.0 | 49.3 | 223.0 | 2.5e-57 |
| WP\_082191961.1 | non-ribosomal\_peptide\_synthetase | BGC0001451 | NRP | 32.0 | 47.0 | 223.0 | 2.5e-57 |
| OKA09423.1 | non-ribosomal\_peptide\_synthetase | BGC0001459 | NRP:Glycopeptide | 31.0 | 46.4 | 223.0 | 2.5e-57 |
| AQZ69229.1 | hypothetical\_protein | BGC0001635 | NRP + Polyketide | 31.0 | 47.0 | 223.0 | 2.5e-57 |
| ADG27359.1 | peptide\_synthetase | BGC0000296 | NRP | 34.0 | 37.4 | 222.0 | 3.2e-57 |
| CAC48360.1 | peptide\_synthetase | BGC0000311 | NRP | 30.0 | 50.1 | 222.0 | 3.2e-57 |
| AEG64696.1 | LpmB | BGC0000379 | NRP | 30.0 | 48.7 | 222.0 | 4.2e-57 |
| APZ78715.1 | nonribosomal\_peptide\_synthetase | BGC0001420 | NRP:Cyclic depsipeptide + Polyketide:Iterative type I | 30.0 | 46.9 | 222.0 | 4.2e-57 |
| APZ78755.1 | nonribosomal\_peptide\_synthetase | BGC0001423 | NRP:Cyclic depsipeptide + Polyketide:Iterative type I | 32.0 | 37.2 | 222.0 | 4.2e-57 |
| OKA09424.1 | non-ribosomal\_peptide\_synthetase | BGC0001459 | NRP:Glycopeptide | 31.0 | 47.3 | 222.0 | 4.2e-57 |
| EME52974.1 | non-ribosomal\_peptide\_synthetase | BGC0001460 | NRP:Glycopeptide | 31.0 | 47.2 | 222.0 | 4.2e-57 |
| WP\_013184322.1 | non-ribosomal\_peptide\_synthetase | BGC0001692 | NRP | 30.0 | 49.0 | 222.0 | 4.2e-57 |
| BAO84861.1 | putative\_non-ribosomal\_peptide\_synthetase | BGC0000414 | NRP | 30.0 | 47.2 | 222.0 | 5.5e-57 |
| BAP05590.1 | calB | BGC0000967 | NRP + Polyketide:Trans-AT type I | 30.0 | 48.3 | 222.0 | 5.5e-57 |
| ABS90470.1 | NRPS/PKS | BGC0001106 | NRP + Polyketide | 31.0 | 47.1 | 222.0 | 5.5e-57 |
| APZ78781.1 | nonribosomal\_peptide\_synthetase | BGC0001426 | NRP:Cyclic depsipeptide + Polyketide:Iterative type I | 33.0 | 36.9 | 222.0 | 5.5e-57 |
| ATD51279.1 | nonribosomal\_peptide\_synthase | BGC0001650 | NRP | 30.0 | 55.1 | 222.0 | 5.5e-57 |
| AJY78094.1 | nonribosomal\_peptide\_synthetase | BGC0001902 | NRP + Polyketide | 31.0 | 48.3 | 222.0 | 5.5e-57 |
| AAY93354.1 | non-ribosomal\_peptide\_synthetase\_PvdD | BGC0000413 | NRP | 30.0 | 47.4 | 221.0 | 7.2e-57 |
| AIE77076.1 | peptide\_synthetase | BGC0000418 | NRP | 30.0 | 47.0 | 221.0 | 7.2e-57 |
| AED90003.1 | non-ribosomal\_peptide\_synthetase\_ThaB | BGC0000443 | NRP:Beta-lactam | 29.0 | 48.8 | 221.0 | 7.2e-57 |
| CBD77749.1 | non-ribosomal\_peptide\_synthetase | BGC0000974 | NRP + Polyketide | 32.0 | 47.6 | 221.0 | 7.2e-57 |
| ABV99085.1 | thioester\_reductase\_domain | BGC0001007 | Polyketide + NRP | 31.0 | 40.4 | 221.0 | 7.2e-57 |
| APZ78743.1 | nonribosomal\_peptide\_synthetase | BGC0001422 | NRP:Cyclic depsipeptide + Polyketide:Iterative type I | 32.0 | 36.4 | 221.0 | 7.2e-57 |
| AIR74925.1 | non-ribosomal\_peptide\_synthetase | BGC0001559 | RiPP | 32.0 | 47.6 | 221.0 | 7.2e-57 |
| ABP55169.1 | amino\_acid\_adenylation\_domain | BGC0000150 | NRP + Polyketide:Enediyne type I | 31.0 | 48.2 | 221.0 | 9.4e-57 |
| AHH53506.1 | non-ribosomal\_peptide\_synthetase | BGC0000439 | NRP:Ca+-dependent lipopeptide | 32.0 | 47.7 | 221.0 | 9.4e-57 |
| CAD70195.1 | non-ribosomal\_peptide\_synthetase | BGC0001047 | NRP + Polyketide | 28.0 | 50.3 | 221.0 | 9.4e-57 |
| APZ78703.1 | nonribosomal\_peptide\_synthetase | BGC0001419 | NRP:Cyclic depsipeptide + Polyketide:Iterative type I | 30.0 | 47.1 | 221.0 | 9.4e-57 |
| WP\_050383084.1 | non-ribosomal\_peptide\_synthetase | BGC0001451 | NRP | 31.0 | 47.3 | 221.0 | 9.4e-57 |
| AKQ22680.1 | malonyl\_CoA-acyl\_carrier\_protein\_transacylase | BGC0001656 | Polyketide | 29.0 | 46.7 | 221.0 | 9.4e-57 |
| BBC43184.1 | PKS-NRPS\_hybrid | BGC0001738 | NRP + Polyketide | 25.0 | 71.2 | 221.0 | 9.4e-57 |
| QED88055.1 | nonribosomal\_peptide\_synthetase | BGC0001967 | NRP | 31.0 | 48.1 | 221.0 | 9.4e-57 |
| ABV79987.1 | ApnC | BGC0000301 | NRP | 29.0 | 47.6 | 220.0 | 1.2e-56 |
| BAE98162.1 | putative\_non-ribosomal\_peptide\_synthetase | BGC0000339 | NRP | 30.0 | 47.7 | 220.0 | 1.2e-56 |
| CAE53352.1 | non-ribosomal\_peptide\_synthetase | BGC0000440 | NRP:Glycopeptide | 31.0 | 47.6 | 220.0 | 1.2e-56 |
| ALK27914.1 | non-ribosomal\_peptide\_synthase | BGC0001233 | NRP | 30.0 | 48.4 | 220.0 | 1.2e-56 |
| AQM37583.1 | nonribosomal\_peptide\_synthetase | BGC0001424 | NRP:Cyclic depsipeptide + Polyketide:Iterative type I | 32.0 | 37.2 | 220.0 | 1.2e-56 |
| CAJ21198.2 | non-ribosomal\_peptide\_synthetase | BGC0000297 | NRP:Glycopeptide + Polyketide:Other + Saccharide:Hybrid/tailoring | 31.0 | 47.7 | 220.0 | 1.6e-56 |
| ABA70582.1 | alpha-aminoadypil-cysteinyl-valine\_synthetase | BGC0000404 | NRP | 30.0 | 48.8 | 220.0 | 1.6e-56 |
| ABR12615.1 | ACV\_synthetase | BGC0000405 | NRP:Beta-lactam | 30.0 | 48.8 | 220.0 | 1.6e-56 |
| CAJ46692.1 | non-ribosomal\_peptide\_synthase | BGC0000969 | NRP:Cyclic depsipeptide + Polyketide:Modular type I | 31.0 | 46.7 | 220.0 | 1.6e-56 |
| APZ78691.1 | nonribosomal\_peptide\_synthetase | BGC0001418 | NRP:Cyclic depsipeptide + Polyketide:Iterative type I | 30.0 | 46.9 | 220.0 | 1.6e-56 |
| AJD77023.1 | IkaA | BGC0001435 | NRP + Polyketide:Iterative type I | 29.0 | 48.4 | 220.0 | 1.6e-56 |
| WP\_013184461.1 | non-ribosomal\_peptide\_synthetase | BGC0001716 | NRP | 28.0 | 47.6 | 220.0 | 1.6e-56 |
| AHZ34232.1 | CifA | BGC0000323 | NRP:Lipopeptide | 31.0 | 48.3 | 220.0 | 2.1e-56 |
| CAJ88610.1 | putative\_non-ribosomal\_peptide\_synthetase | BGC0000327 | NRP | 31.0 | 50.2 | 220.0 | 2.1e-56 |
| AHI59108.1 | locillomycin\_synthase\_A | BGC0001005 | NRP + Polyketide | 28.0 | 47.6 | 220.0 | 2.1e-56 |
| AJW76710.1 | DsaH | BGC0001196 | NRP | 29.0 | 47.3 | 220.0 | 2.1e-56 |
| AZF85944.1 | hypothetical\_protein | BGC0001963 | NRP + Polyketide | 30.0 | 45.8 | 220.0 | 2.1e-56 |
| CAJ77695.1 | MPS1\_protein | BGC0000363 | NRP | 29.0 | 48.8 | 219.0 | 2.7e-56 |
| ABP55493.1 | thioester\_reductase\_domain | BGC0001006 | NRP + Polyketide | 34.0 | 39.7 | 219.0 | 2.7e-56 |
| AAZ55900.1 | non-ribosomal\_peptide\_synthase:Amino\_acid\_adenylation | BGC0000359 | NRP | 31.0 | 50.6 | 219.0 | 3.6e-56 |
| AAO62586.1 | peptide\_sythetase | BGC0001016 | NRP + Polyketide | 29.0 | 47.3 | 219.0 | 3.6e-56 |
| BAX64244.1 | NRPS | BGC0001623 | NRP + Polyketide | 31.0 | 51.3 | 219.0 | 3.6e-56 |
| AAG05693.1 | AmbB | BGC0000287 | NRP | 31.0 | 48.8 | 218.0 | 4.7e-56 |
| CAJ96470.1 | non-ribosomal\_peptide\_synthetase | BGC0000330 | NRP:NRP siderophore | 29.0 | 45.1 | 218.0 | 4.7e-56 |
| AAX31557.1 | peptide\_synthetase\_1 | BGC0000336 | NRP | 30.0 | 49.7 | 218.0 | 4.7e-56 |
| AEI58879.1 | peptide\_synthetase | BGC0000455 | NRP | 30.0 | 46.3 | 218.0 | 4.7e-56 |
| AKA59436.1 | non-ribosomal\_peptide\_synthetase | BGC0001202 | NRP + Polyketide | 31.0 | 49.2 | 218.0 | 4.7e-56 |
| AJV88376.1 | MfnD | BGC0001214 | NRP | 32.0 | 47.9 | 218.0 | 4.7e-56 |
| BBA84067.1 | type\_I\_polyketide\_synthase | BGC0001916 | Polyketide | 34.0 | 36.9 | 218.0 | 4.7e-56 |
| QED55421.1 | nonribosomal\_peptide\_synthetase | BGC0001984 | NRP | 31.0 | 46.9 | 218.0 | 4.7e-56 |
| AAZ23077.1 | peptide\_synthetase | BGC0000291 | NRP | 32.0 | 48.2 | 218.0 | 6.1e-56 |
| ACZ55944.1 | non-ribosomal\_peptide\_synthetase | BGC0000302 | NRP | 28.0 | 47.0 | 218.0 | 6.1e-56 |
| ACZ55946.1 | non-ribosomal\_peptide\_synthetase | BGC0000302 | NRP | 30.0 | 47.1 | 218.0 | 6.1e-56 |
| OLZ52457.1 | non-ribosomal\_peptide\_synthetase | BGC0001462 | NRP:Glycopeptide | 31.0 | 47.7 | 218.0 | 6.1e-56 |
| WP\_015507663.1 | non-ribosomal\_peptide\_synthase | BGC0001792 | NRP | 30.0 | 48.1 | 218.0 | 6.1e-56 |
| ABO15860.1 | polyketide\_synthase | BGC0000130 | Polyketide | 31.0 | 39.3 | 218.0 | 8e-56 |
| WP\_064118560.1 | non-ribosomal\_peptide\_synthetase | BGC0001509 | NRP | 32.0 | 46.6 | 218.0 | 8e-56 |
| AVI26392.1 | putative\_nonribosomal\_peptide\_synthase | BGC0001800 | NRP + Polyketide | 29.0 | 49.1 | 218.0 | 8e-56 |
| Xekj\_RS17945 | non-ribosomal\_peptide\_synthetase | BGC0001826 | NRP | 30.0 | 48.8 | 218.0 | 8e-56 |
| CCC55927.1 | hypothetical\_protein | BGC0000973 | NRP + Polyketide:Modular type I | 33.0 | 38.2 | 217.0 | 1e-55 |
| CAN89656.1 | putative\_non-ribosomal\_peptide\_synthetase | BGC0001070 | NRP + Polyketide:Modular type I + Polyketide:Trans-AT type I | 27.0 | 53.2 | 217.0 | 1e-55 |
| CAL69889.1 | RhiB\_protein | BGC0001112 | NRP + Polyketide:Trans-AT type I | 30.0 | 42.1 | 217.0 | 1e-55 |
| AAM80537.1 | StaC | BGC0000290 | NRP:Glycopeptide | 31.0 | 44.8 | 217.0 | 1.4e-55 |
| AAZ23078.1 | peptide\_synthetase | BGC0000291 | NRP | 30.0 | 47.7 | 217.0 | 1.4e-55 |
| AAZ55898.1 | amino\_acid\_adenylation | BGC0000359 | NRP | 32.0 | 49.2 | 217.0 | 1.4e-55 |
| AIG79224.1 | Non-ribosomal\_peptide\_synthetase/andenylation\_domain | BGC0000419 | Saccharide + NRP:Glycopeptide | 30.0 | 47.0 | 217.0 | 1.4e-55 |
| AEI58866.1 | peptide\_synthetase | BGC0000455 | NRP | 31.0 | 46.7 | 217.0 | 1.4e-55 |
| ALV82356.1 | CDA\_peptide\_synthetase\_I | BGC0001370 | NRP | 32.0 | 46.5 | 217.0 | 1.4e-55 |
| ARF06222.1 | non-ribosomal\_peptide\_synthetase | BGC0001593 | NRP | 29.0 | 49.8 | 217.0 | 1.4e-55 |
| AEE88284.1 | CurF | BGC0000976 | NRP + Polyketide:Modular type I | 29.0 | 49.2 | 217.0 | 1.8e-55 |
| ABL86391.1 | hybrid\_polyketide\_synthase\_and\_nonribosomal\_peptide\_synthetase | BGC0000999 | NRP + Polyketide | 28.0 | 48.9 | 217.0 | 1.8e-55 |
| ABF85931.1 | non-ribosomal\_peptide\_synthase/polyketide\_synthase\_Ta1 | BGC0001025 | NRP + Polyketide:Trans-AT type I | 29.0 | 49.8 | 217.0 | 1.8e-55 |
| AAT70101.1 | CurF | BGC0001165 | NRP + Polyketide:Modular type I | 29.0 | 49.2 | 217.0 | 1.8e-55 |
| AUD11993.1 | OrbJ | BGC0001721 | NRP | 29.0 | 49.7 | 217.0 | 1.8e-55 |
| CAD91212.1 | putative\_non-ribosomal\_peptide\_synthetase,\_modules\_4-6 | BGC0000289 | NRP:Glycopeptide + Saccharide:Hybrid/tailoring | 33.0 | 47.7 | 216.0 | 2.3e-55 |
| ANZ15840.1 | non-ribosomal\_peptide\_synthase/amino\_acid\_adenylation\_enzyme | BGC0001569 | NRP | 29.0 | 47.3 | 216.0 | 2.3e-55 |
| AGE11899.1 | nonribosomal\_peptide\_synthetase | BGC0000366 | NRP | 30.0 | 49.1 | 216.0 | 3e-55 |
| AEA30274.1 | peptide\_synthetase | BGC0000429 | Polyketide + NRP:Cyclic depsipeptide | 30.0 | 48.8 | 216.0 | 3e-55 |
| AAF19811.1 | mtaC | BGC0001024 | NRP + Polyketide:Modular type I | 29.0 | 46.8 | 216.0 | 3e-55 |
| ALV82384.1 | CDA\_peptide\_synthetase\_II | BGC0001370 | NRP | 29.0 | 48.4 | 216.0 | 3e-55 |
| WP\_015507662.1 | non-ribosomal\_peptide\_synthetase | BGC0001792 | NRP | 30.0 | 47.3 | 216.0 | 3e-55 |
| WP\_003981346.1 | non-ribosomal\_peptide\_synthetase | BGC0001813 | NRP | 29.0 | 49.1 | 216.0 | 3e-55 |
| ATU31795.1 | NRPS | BGC0001814 | NRP | 32.0 | 47.7 | 216.0 | 3e-55 |
| WP\_039806850.1 | non-ribosomal\_peptide\_synthetase | BGC0002001 | NRP + Polyketide | 29.0 | 49.1 | 216.0 | 3e-55 |
| AAD44233.1 | PstA | BGC0000362 | NRP | 30.0 | 44.9 | 215.0 | 3.9e-55 |
| AIE77059.1 | peptide\_synthetase | BGC0000418 | NRP | 29.0 | 49.5 | 215.0 | 3.9e-55 |
| ARU08075.1 | mlcM | BGC0001448 | NRP:Ca+-dependent lipopeptide | 30.0 | 48.3 | 215.0 | 3.9e-55 |
| ATQ39428.1 | cyclosporin\_C\_synthetase | BGC0001565 | NRP | 30.0 | 50.5 | 215.0 | 3.9e-55 |
| ABI22131.1 | putative\_non-ribosomal\_peptide\_synthetase | BGC0000422 | NRP | 31.0 | 48.5 | 215.0 | 5.2e-55 |
| WP\_078857610.1 | non-ribosomal\_peptide\_synthetase | BGC0001368 | NRP | 31.0 | 45.6 | 215.0 | 5.2e-55 |
| KFL51886.1 | amino\_acid\_adenylation\_protein | BGC0001711 | NRP + Polyketide | 29.0 | 46.5 | 215.0 | 5.2e-55 |
| CAC48361.1 | peptide\_synthetase | BGC0000311 | NRP | 31.0 | 49.8 | 215.0 | 6.7e-55 |
| BAO84868.1 | putative\_non-ribosomal\_peptide\_synthetase | BGC0000414 | NRP | 30.0 | 43.0 | 215.0 | 6.7e-55 |
| CAE02630.1 | surfactin\_synthetase\_A | BGC0000433 | NRP:Lipopeptide | 28.0 | 50.8 | 215.0 | 6.7e-55 |
| CAG15011.1 | peptide\_synthetase,\_module\_4-6 | BGC0000441 | NRP | 31.0 | 47.6 | 215.0 | 6.7e-55 |
| CAC11137.1 | NikP1\_protein | BGC0000876 | Other | 30.0 | 50.6 | 214.0 | 8.8e-55 |
| AHZ20773.1 | non-ribosomal\_peptide\_synthase | BGC0000369 | NRP + Saccharide:Hybrid/tailoring | 30.0 | 45.3 | 214.0 | 1.1e-54 |
| AHD05679.1 | putative\_non-ribosomal\_peptide\_ligase/\_polyketide\_synthase\_hybrid | BGC0000402 | NRP | 29.0 | 46.5 | 214.0 | 1.1e-54 |
| AEI58865.1 | peptide\_synthetase | BGC0000455 | NRP | 30.0 | 46.7 | 214.0 | 1.1e-54 |
| AIW58892.1 | non-ribosomal\_peptide\_synthetase | BGC0001582 | NRP | 30.0 | 48.0 | 214.0 | 1.1e-54 |
| ATX68112.1 | malonyl\_CoA-acyl\_carrier\_protein\_transacylase | BGC0001772 | Polyketide | 29.0 | 45.9 | 214.0 | 1.1e-54 |
| AAC32047.1 | FxbB | BGC0000351 | NRP | 30.0 | 46.4 | 213.0 | 1.5e-54 |
| ABS74180.1 | bacillomycin\_D\_synthetase\_B | BGC0001090 | Polyketide + NRP:Lipopeptide | 29.0 | 48.8 | 213.0 | 1.5e-54 |
| AGN11881.1 | tstDEF | BGC0001114 | NRP + Polyketide | 30.0 | 49.9 | 213.0 | 1.5e-54 |
| AAX31558.1 | peptide\_synthetase\_2 | BGC0000336 | NRP | 30.0 | 47.2 | 213.0 | 2e-54 |
| AIG79243.1 | Non-ribosomal\_peptide\_synthetase | BGC0000419 | Saccharide + NRP:Glycopeptide | 30.0 | 46.7 | 213.0 | 2e-54 |
| AAF00960.1 | mcyA | BGC0001017 | NRP + Polyketide:Modular type I | 32.0 | 37.4 | 213.0 | 2e-54 |
| CZT62792.1 | non-ribosomal\_peptide\_synthase\_involved\_in\_Hassallidin\_biosynthesis | BGC0001614 | NRP | 29.0 | 47.1 | 213.0 | 2e-54 |
| BAW27670.1 | NRPS(A-T) | BGC0001764 | NRP | 29.0 | 49.9 | 213.0 | 2e-54 |
| AZH23822.1 | MgiJ | BGC0001971 | NRP + Polyketide | 32.0 | 36.6 | 213.0 | 2e-54 |
| AQH32484.1 | peptide\_synthetase | BGC0001667 | NRP + Polyketide | 31.0 | 36.9 | 213.0 | 2.6e-54 |
| QBC75023.1 | non-ribosomal\_peptide\_synthetase | BGC0001968 | NRP | 35.0 | 38.0 | 213.0 | 2.6e-54 |
| AAL15600.1 | SimH | BGC0000270 | Polyketide | 29.0 | 44.9 | 212.0 | 3.3e-54 |
| AET98905.1 | putative\_non-ribosomal\_peptide\_synthetase | BGC0000415 | NRP | 32.0 | 37.3 | 212.0 | 3.3e-54 |
| AAG34184.1 | SimH | BGC0000835 | Polyketide | 29.0 | 44.9 | 212.0 | 3.3e-54 |
| BAP05596.1 | calH | BGC0000967 | NRP + Polyketide:Trans-AT type I | 28.0 | 47.7 | 212.0 | 3.3e-54 |
| ADH01485.1 | putative\_mixed\_polyketide\_synthase/non-ribosomal\_peptide\_synthetase | BGC0001096 | NRP + Polyketide | 30.0 | 50.5 | 212.0 | 3.3e-54 |
| AAK06804.1 | Tyroxyl-AMP-forming\_enzyme | BGC0001072 | Saccharide + Polyketide:Modular type I + Polyketide:Type II + Other:Aminocoumarin | 29.0 | 44.9 | 212.0 | 3.3e-54 |
| AAN85493.1 | nonribosomal\_peptide\_synthetase | BGC0001101 | NRP + Polyketide:Modular type I + Polyketide:Trans-AT type I | 29.0 | 49.1 | 212.0 | 3.3e-54 |
| AIC32693.1 | FR9DEF | BGC0001113 | NRP + Polyketide | 30.0 | 50.5 | 212.0 | 3.3e-54 |
| APZ78845.1 | nonribosomal\_peptide\_synthetase | BGC0001431 | NRP:Cyclic depsipeptide + Polyketide:Iterative type I | 33.0 | 36.9 | 212.0 | 3.3e-54 |
| WP\_019634550.1 | type\_I\_polyketide\_synthase | BGC0001443 | NRP + Polyketide | 30.0 | 47.4 | 212.0 | 3.3e-54 |
| ARU08073.1 | mlcK | BGC0001448 | NRP:Ca+-dependent lipopeptide | 27.0 | 77.6 | 212.0 | 3.3e-54 |
| AYA22318.1 | KerE | BGC0001955 | Other | 30.0 | 47.0 | 212.0 | 3.3e-54 |
| AAK81827.1 | peptide\_synthetase | BGC0000326 | NRP | 29.0 | 46.9 | 212.0 | 4.4e-54 |
| AGN74895.1 | nonribosomal\_peptide\_synthetase/polyketide\_synthase\_hybrid\_protein | BGC0000459 | NRP:Cyclic depsipeptide + Polyketide:Trans-AT type I | 30.0 | 46.7 | 212.0 | 4.4e-54 |
| AGM14934.1 | xantholysin\_synthetase\_C | BGC0000463 | NRP:Lipopeptide | 29.0 | 46.7 | 212.0 | 4.4e-54 |
| AFU82614.1 | mixed\_NRPS\_PKS | BGC0000998 | NRP + Polyketide | 30.0 | 49.8 | 212.0 | 4.4e-54 |
| AHD05616.1 | putative\_non-ribosomal\_peptide\_ligase\_domain\_protein | BGC0001033 | NRP + Polyketide | 31.0 | 36.4 | 212.0 | 4.4e-54 |
| CBA11570.1 | non-ribosomal\_peptide\_synthetase | BGC0001046 | NRP + Polyketide:Modular type I + Saccharide:Hybrid/tailoring | 32.0 | 40.4 | 212.0 | 4.4e-54 |
| ALV86868.1 | Tlo22 | BGC0001406 | NRP | 29.0 | 49.5 | 212.0 | 4.4e-54 |
| OLZ50899.1 | non-ribosomal\_peptide\_synthetase | BGC0001461 | NRP:Glycopeptide | 30.0 | 47.0 | 212.0 | 4.4e-54 |
| ALG65314.1 | Cal22 | BGC0001297 | NRP | 29.0 | 47.6 | 212.0 | 5.7e-54 |
| AFH75321.1 | nonribosomal\_peptide\_synthetase | BGC0000425 | NRP:Cyclic depsipeptide | 29.0 | 47.5 | 211.0 | 7.4e-54 |
| AGN74886.1 | nonribosomal\_peptide\_synthetase | BGC0000459 | NRP:Cyclic depsipeptide + Polyketide:Trans-AT type I | 30.0 | 46.8 | 211.0 | 7.4e-54 |
| CCE88377.1 | non-ribosomal\_peptide\_synthetase/polyketide\_synthase | BGC0001034 | NRP + Polyketide:Modular type I | 33.0 | 42.1 | 211.0 | 7.4e-54 |
| AAS47564.1 | mixed\_type\_I\_polyketide\_synthase/nonribosomal\_peptide\_synthetase | BGC0001108 | Polyketide:Trans-AT type I | 28.0 | 49.9 | 211.0 | 7.4e-54 |
| ctg1\_orf6 |  | BGC0001109 | NRP + Polyketide | 28.0 | 49.9 | 211.0 | 7.4e-54 |
| ADH04680.1 | hybrid\_polyketide\_synthase/non-ribosomal\_peptide\_synthetase | BGC0001344 | NRP + Polyketide | 29.0 | 49.1 | 211.0 | 7.4e-54 |
| OLZ52442.1 | non-ribosomal\_peptide\_synthetase | BGC0001462 | NRP:Glycopeptide | 30.0 | 46.9 | 211.0 | 7.4e-54 |
| CBF76038.1 | nonribosomal\_peptide\_synthase,\_putative\_(Eurofung) | BGC0001399 | NRP | 26.0 | 70.3 | 211.0 | 9.7e-54 |
| CCC55922.1 | putative\_non-ribosomal\_peptide\_synthetase | BGC0000973 | NRP + Polyketide:Modular type I | 29.0 | 48.9 | 210.0 | 1.3e-53 |
| AHB82059.1 | non\_ribosomal\_peptide\_synthetase/polyketide\_synthase | BGC0001019 | NRP + Polyketide:Modular type I | 29.0 | 45.3 | 210.0 | 1.3e-53 |
| KUM80514.1 | hypothetical\_protein | BGC0001562 | NRP | 29.0 | 47.6 | 210.0 | 1.3e-53 |
| DAC76734.1 | type\_I\_polyketide\_synthase/non-ribosomal\_peptide\_synthetase | BGC0001885 | Polyketide | 30.0 | 46.7 | 210.0 | 1.3e-53 |
| CAC48369.1 | peptide\_synthetase | BGC0000311 | NRP | 30.0 | 46.9 | 210.0 | 1.7e-53 |
| ABP57749.1 | DepE | BGC0000993 | NRP:Cyclic depsipeptide + Polyketide:Modular type I | 31.0 | 45.3 | 210.0 | 1.7e-53 |
| ALG65318.1 | Cal18 | BGC0001297 | NRP | 29.0 | 49.4 | 210.0 | 1.7e-53 |
| EWM63005.1 | linear\_gramicidin\_synthetase\_LgrC | BGC0001328 | NRP:Cyclic depsipeptide + Polyketide:Modular type I | 34.0 | 35.8 | 210.0 | 1.7e-53 |
| AAK81824.1 | peptide\_synthetase | BGC0000326 | NRP | 30.0 | 49.6 | 210.0 | 2.2e-53 |
| AET98916.1 | putative\_non-ribosomal\_peptide\_synthetase | BGC0000415 | NRP | 30.0 | 47.8 | 210.0 | 2.2e-53 |
| ABG94125.1 | non-ribosomal\_peptide\_synthetase | BGC0000417 | NRP | 30.0 | 48.7 | 210.0 | 2.2e-53 |
| AAG02349.1 | peptide\_synthetase\_NRPS11-10 | BGC0000963 | NRP:Glycopeptide + Polyketide:Modular type I + Saccharide:Hybrid/tailoring | 33.0 | 35.2 | 210.0 | 2.2e-53 |
| CAD29797.1 | peptide\_synthetase | BGC0001015 | NRP + Polyketide | 31.0 | 46.8 | 210.0 | 2.2e-53 |
| BAG17643.1 | putative\_NRPS-type-I\_PKS\_fusion\_protein | BGC0001043 | NRP + Polyketide | 29.0 | 50.5 | 210.0 | 2.2e-53 |
| AAF08795.1 | MycA | BGC0001103 | NRP + Polyketide | 29.0 | 46.2 | 210.0 | 2.2e-53 |
| CBZ42146.1 | putative\_non-ribosomal\_peptide\_synthetase | BGC0001117 | NRP | 28.0 | 47.2 | 210.0 | 2.2e-53 |
| ADN43685.1 | DmbS | BGC0001136 | NRP + Polyketide:Iterative type I | 25.0 | 71.8 | 210.0 | 2.2e-53 |
| AKA59447.1 | non-ribosomal\_peptide\_synthetase | BGC0001203 | NRP + Polyketide | 33.0 | 36.0 | 210.0 | 2.2e-53 |
| AYA22335.1 | KerB | BGC0001955 | Other | 30.0 | 47.0 | 210.0 | 2.2e-53 |
| AAW03326.1 | CtaC | BGC0000982 | NRP + Polyketide | 29.0 | 47.7 | 209.0 | 2.8e-53 |
| CAG29031.1 | nonribosomal\_peptide\_synthetase\_(modules\_1\_and\_2) | BGC0001023 | NRP + Polyketide:Modular type I | 32.0 | 36.9 | 209.0 | 2.8e-53 |
| CBJ79915.1 | putative\_Phenylalanine\_racemase\_(ATP-hydrolyzing) | BGC0001133 | NRP | 29.0 | 48.3 | 209.0 | 2.8e-53 |
| APZ78821.1 | nonribosomal\_peptide\_synthetase | BGC0001429 | NRP:Cyclic depsipeptide + Polyketide:Iterative type I | 32.0 | 36.9 | 209.0 | 2.8e-53 |
| AYJ71712.1 | non-ribosomal\_peptide\_synthetase | BGC0001942 | NRP + Polyketide | 29.0 | 46.7 | 209.0 | 2.8e-53 |
| ALP32042.1 | CycB | BGC0001293 | Polyketide | 33.0 | 36.7 | 209.0 | 3.7e-53 |
| ADZ24995.1 | non-ribosomal\_peptide\_synthase/polyketide\_synthase | BGC0000380 | NRP + Polyketide:Modular type I | 27.0 | 46.5 | 208.0 | 4.8e-53 |
| CAE53353.1 | non-ribosomal\_peptide\_synthetase | BGC0000440 | NRP:Glycopeptide | 31.0 | 47.2 | 208.0 | 4.8e-53 |
| BAH04161.1 | putative\_non-ribosomal\_peptide\_synthetase | BGC0000450 | NRP | 31.0 | 37.0 | 208.0 | 4.8e-53 |
| WP\_016638471.1 | non-ribosomal\_peptide\_synthetase | BGC0001519 | NRP + Polyketide | 29.0 | 48.0 | 208.0 | 4.8e-53 |
| ATY69551.1 | hybrid\_nonribosomal\_peptide\_synthetase/type\_I\_polyketide\_synthase | BGC0001611 | NRP + Polyketide | 28.0 | 50.3 | 208.0 | 4.8e-53 |
| ABP73646.1 | SalB | BGC0000145 | Polyketide | 31.0 | 46.7 | 208.0 | 6.3e-53 |
| AAM80538.1 | StaB | BGC0000290 | NRP:Glycopeptide | 30.0 | 48.8 | 208.0 | 6.3e-53 |
| AAK81825.1 | peptide\_synthetase | BGC0000326 | NRP | 29.0 | 50.0 | 208.0 | 6.3e-53 |
| ABD65957.1 | nonribosomal\_peptide\_synthetase | BGC0000341 | NRP | 30.0 | 48.6 | 208.0 | 6.3e-53 |
| ABL74936.1 | NRPS | BGC0001048 | NRP:Glycopeptide + Polyketide:Modular type I + Saccharide:Hybrid/tailoring | 30.0 | 49.9 | 208.0 | 6.3e-53 |
| ALV82388.1 | CDA\_peptide\_synthetase\_III | BGC0001370 | NRP | 30.0 | 49.1 | 208.0 | 6.3e-53 |
| ARU08063.1 | mlcA | BGC0001448 | NRP:Ca+-dependent lipopeptide | 28.0 | 45.6 | 208.0 | 6.3e-53 |
| WP\_008704344.1 | non-ribosomal\_peptide\_synthetase | BGC0001575 | NRP | 27.0 | 48.0 | 208.0 | 6.3e-53 |
| CAM02313.1 | putative\_non-ribosomal\_peptide\_synthetase | BGC0000349 | NRP | 29.0 | 45.3 | 208.0 | 8.2e-53 |
| BAH43869.1 | linear\_pentadecapeptide\_gramicidin\_synthetase\_LgrA | BGC0000367 | NRP | 29.0 | 47.4 | 208.0 | 8.2e-53 |
| ABP57748.1 | DepD | BGC0000993 | NRP:Cyclic depsipeptide + Polyketide:Modular type I | 30.0 | 46.1 | 208.0 | 8.2e-53 |
| WP\_052165466.1 | non-ribosomal\_peptide\_synthetase | BGC0001327 | NRP:Cyclic depsipeptide + Polyketide:Modular type I | 35.0 | 35.9 | 208.0 | 8.2e-53 |
| OKA09425.1 | non-ribosomal\_peptide\_synthetase | BGC0001459 | NRP:Glycopeptide | 31.0 | 46.8 | 208.0 | 8.2e-53 |
| WP\_078620912.1 | non-ribosomal\_peptide\_synthetase | BGC0001519 | NRP + Polyketide | 29.0 | 47.4 | 208.0 | 8.2e-53 |
| QED55423.1 | nonribosomal\_peptide\_synthetase | BGC0001984 | NRP | 30.0 | 47.5 | 208.0 | 8.2e-53 |
| CAG15012.1 | peptide\_synthetase,\_module\_7 | BGC0000441 | NRP | 31.0 | 47.2 | 207.0 | 1.1e-52 |
| CCJ67638.1 | TaaC | BGC0000447 | NRP:Lipopeptide | 30.0 | 46.5 | 207.0 | 1.1e-52 |
| ABP53497.1 | NRPS\_(ACP-P) | BGC0001041 | NRP + Polyketide | 31.0 | 46.1 | 207.0 | 1.1e-52 |
| WP\_051206727.1 | non-ribosomal\_peptide\_synthetase | BGC0001467 | NRP:Cyclic depsipeptide + Polyketide:Modular type I | 33.0 | 36.5 | 207.0 | 1.4e-52 |
| CBL93723.1 | NRPS\_didomain\_A-PCP | BGC0000360 | NRP | 29.0 | 48.3 | 207.0 | 1.8e-52 |
| CDE97356.1 | plipastatin\_synthase\_subunit\_C | BGC0001686 | NRP | 26.0 | 47.6 | 207.0 | 1.8e-52 |
| CAD91211.1 | putative\_non-ribosomal\_peptide\_synthetase,\_module\_7 | BGC0000289 | NRP:Glycopeptide + Saccharide:Hybrid/tailoring | 31.0 | 47.1 | 206.0 | 2.4e-52 |
| ABV79988.1 | ApnD | BGC0000301 | NRP | 28.0 | 47.4 | 206.0 | 2.4e-52 |
| ABW17375.1 | PsoA | BGC0000411 | NRP | 29.0 | 48.2 | 206.0 | 2.4e-52 |
| AAY93445.1 | non-ribosomal\_peptide\_synthetase\_PvdL | BGC0000413 | NRP | 28.0 | 48.1 | 206.0 | 2.4e-52 |
| CAA60461.1 | pipecolate\_incorporating\_enzyme | BGC0001040 | NRP + Polyketide | 28.0 | 47.0 | 206.0 | 2.4e-52 |
| CAF05649.1 | TubD\_protein | BGC0001053 | NRP + Polyketide | 29.0 | 49.5 | 206.0 | 2.4e-52 |
| AGS77308.1 | NRPS\_module\_3 | BGC0001178 | NRP:Glycopeptide | 30.0 | 45.6 | 206.0 | 2.4e-52 |
| OKA09664.1 | non-ribosomal\_peptide\_synthetase | BGC0001459 | NRP:Glycopeptide | 29.0 | 46.8 | 206.0 | 2.4e-52 |
| DAC76736.1 | non-ribosomal\_peptide\_synthetase | BGC0001885 | Polyketide | 32.0 | 35.1 | 206.0 | 2.4e-52 |
| AZH23792.1 | MgcJ | BGC0001970 | NRP + Polyketide | 32.0 | 36.1 | 206.0 | 2.4e-52 |
| ABI22132.1 | putative\_non-ribosomal\_peptide\_synthetase | BGC0000422 | NRP | 30.0 | 45.5 | 206.0 | 3.1e-52 |
| CBG75492.1 | putative\_NRPS/siderophore\_biosynthesis\_protein | BGC0000423 | NRP | 32.0 | 37.1 | 206.0 | 3.1e-52 |
| BAI63283.1 | putative\_non-ribosomal\_peptide\_synthetase | BGC0000434 | NRP | 29.0 | 48.1 | 206.0 | 3.1e-52 |
| CAN89638.1 | putative\_non-ribosomal\_peptide\_synthetase | BGC0001070 | NRP + Polyketide:Modular type I + Polyketide:Trans-AT type I | 30.0 | 48.0 | 206.0 | 3.1e-52 |
| AQV04224.1 | SwnK | BGC0001793 | Polyketide | 28.0 | 48.0 | 206.0 | 3.1e-52 |
| CAC22144.1 | CpkC;\_Polyketide\_synthase\_module\_5 | BGC0000038 | Polyketide:Modular type I | 34.0 | 33.7 | 205.0 | 4.1e-52 |
| ABC94347.1 | vicibactin\_biosynthesis\_non-ribosomal\_peptide\_synthase\_protein | BGC0000457 | NRP | 29.0 | 46.3 | 205.0 | 4.1e-52 |
| WP\_051700105.1 | non-ribosomal\_peptide\_synthetase | BGC0001368 | NRP | 29.0 | 49.3 | 205.0 | 4.1e-52 |
| ATU31794.1 | NRPS | BGC0001814 | NRP | 28.0 | 47.8 | 205.0 | 4.1e-52 |
| CAD91221.1 | putative\_non-ribosomal\_peptide\_synthetase,\_module\_3 | BGC0000289 | NRP:Glycopeptide + Saccharide:Hybrid/tailoring | 30.0 | 48.4 | 205.0 | 5.3e-52 |
| BAP16689.1 | nonribosomal\_peptide\_synthetase | BGC0000376 | NRP | 29.0 | 47.1 | 205.0 | 5.3e-52 |
| ALG65313.1 | Cal23 | BGC0001297 | NRP | 29.0 | 47.7 | 205.0 | 5.3e-52 |
| BAH04173.1 | putative\_non-ribosomal\_peptide\_synthetase | BGC0000450 | NRP | 30.0 | 47.8 | 205.0 | 7e-52 |
| WP\_013184463.1 | non-ribosomal\_peptide\_synthetase | BGC0001716 | NRP | 28.0 | 46.9 | 205.0 | 7e-52 |
| CAJ96472.1 | non-ribosomal\_peptide\_synthetase | BGC0000330 | NRP:NRP siderophore | 29.0 | 44.8 | 204.0 | 9.1e-52 |
| CAJ77715.1 | Mps1\_protein | BGC0000364 | NRP | 32.0 | 45.0 | 204.0 | 9.1e-52 |
| AAL33756.1 | putative\_non-ribosomal\_peptide\_synthetase | BGC0000421 | NRP | 28.0 | 48.0 | 204.0 | 9.1e-52 |
| AQI70\_32580 |  | BGC0001561 | NRP | 29.0 | 49.5 | 204.0 | 9.1e-52 |
| BAE98156.1 | putative\_non-ribosomal\_peptide\_synthetase | BGC0000339 | NRP | 32.0 | 35.6 | 204.0 | 1.2e-51 |
| AIE77060.1 | peptide\_synthetase\_module\_7 | BGC0000418 | NRP | 31.0 | 47.1 | 204.0 | 1.2e-51 |
| CAE53350.1 | non-ribosomal\_peptide\_synthetase | BGC0000440 | NRP:Glycopeptide | 31.0 | 46.9 | 204.0 | 1.2e-51 |
| CAD89774.1 | MelC\_protein | BGC0001010 | NRP + Polyketide:Modular type I | 28.0 | 46.3 | 204.0 | 1.2e-51 |
| AHD05614.1 | putative\_non-ribosomal\_peptide\_ligase/\_polyketide\_synthase\_hybrid | BGC0001033 | NRP + Polyketide | 28.0 | 46.9 | 203.0 | 2e-51 |
| AGZ03650.1 | sevA | BGC0000426 | NRP | 29.0 | 46.4 | 203.0 | 2.6e-51 |
| CAF05647.1 | TubB\_protein | BGC0001053 | NRP + Polyketide | 32.0 | 36.4 | 203.0 | 2.6e-51 |
| WP\_028678148.1 | non-ribosomal\_peptide\_synthetase | BGC0001228 | NRP:Cyclic depsipeptide | 32.0 | 36.2 | 203.0 | 2.6e-51 |
| WP\_015507661.1 | non-ribosomal\_peptide\_synthetase | BGC0001792 | NRP | 29.0 | 46.9 | 203.0 | 2.6e-51 |
| AYJ71721.1 | non-ribosomal\_peptide\_synthetase | BGC0001942 | NRP + Polyketide | 29.0 | 48.6 | 203.0 | 2.6e-51 |
| AEO14743.1 | NdaA | BGC0000396 | NRP | 32.0 | 37.4 | 202.0 | 3.5e-51 |
| CCJ67639.1 | TaaD | BGC0000447 | NRP:Lipopeptide | 30.0 | 46.8 | 202.0 | 3.5e-51 |
| AHD05617.1 | putative\_non-ribosomal\_peptide\_ligase\_domain\_protein | BGC0001033 | NRP + Polyketide | 28.0 | 47.6 | 202.0 | 3.5e-51 |
| AAR87759.1 | ZmaJ | BGC0001059 | NRP + Polyketide | 28.0 | 41.4 | 202.0 | 3.5e-51 |
| AAN85506.1 | nonribosomal\_peptide\_synthetase | BGC0001101 | NRP + Polyketide:Modular type I + Polyketide:Trans-AT type I | 31.0 | 48.1 | 202.0 | 3.5e-51 |
| AID65222.1 | putative\_aspartate\_racemase | BGC0000335 | NRP | 30.0 | 46.2 | 202.0 | 4.5e-51 |
| AAG29780.1 | peptide\_synthetase-like\_protein | BGC0000833 | Saccharide:Hybrid/tailoring + Other:Aminocoumarin | 29.0 | 46.2 | 202.0 | 4.5e-51 |
| ADA82585.1 | hybrid\_trans-AT\_polyketide\_synthase\_-\_nonribosomal\_peptide\_synthetase | BGC0001110 | NRP + Polyketide:Trans-AT type I | 28.0 | 50.4 | 202.0 | 4.5e-51 |
| CAG15009.1 | peptide\_synthetase,\_module\_1-2 | BGC0000441 | NRP | 30.0 | 46.6 | 201.0 | 5.9e-51 |
| EFL02193.1 | amino\_acid\_adenylation\_domain-containing\_protein | BGC0000996 | NRP + Polyketide:Iterative type I | 32.0 | 40.9 | 201.0 | 5.9e-51 |
| CCP45168.1 | Peptide\_synthetase\_MbtE\_(peptide\_synthase) | BGC0001021 | NRP + Polyketide | 29.0 | 48.1 | 201.0 | 5.9e-51 |
| ACG60773.1 | NRPS(C/A/PCP) | BGC0001058 | NRP:Glycopeptide + Polyketide:Modular type I + Saccharide:Hybrid/tailoring | 30.0 | 46.2 | 201.0 | 5.9e-51 |
| ACG60778.1 | NRPS(A/PCP) | BGC0001058 | NRP:Glycopeptide + Polyketide:Modular type I + Saccharide:Hybrid/tailoring | 32.0 | 47.2 | 201.0 | 5.9e-51 |
| BAF50720.1 | hybrid\_non\_ribosomal\_peptide\_synthetase-polyketide\_synthase | BGC0001116 | NRP + Polyketide | 26.0 | 50.7 | 201.0 | 5.9e-51 |
| KUM80513.1 | hypothetical\_protein | BGC0001562 | NRP | 31.0 | 45.2 | 201.0 | 5.9e-51 |
| CAD32904.2 | non-ribosomal\_peptide\_synthetase\_A | BGC0000354 | NRP | 28.0 | 48.1 | 201.0 | 7.7e-51 |
| AIG79241.1 | Hypothetical\_protein | BGC0000419 | Saccharide + NRP:Glycopeptide | 30.0 | 45.3 | 201.0 | 7.7e-51 |
| CAJ34367.1 | NRPS\_protein | BGC0000445 | NRP:Cyclic depsipeptide | 29.0 | 47.7 | 201.0 | 7.7e-51 |
| PHM26613.1 | pyoverdine\_synthetase\_D | BGC0001130 | NRP + Polyketide | 29.0 | 49.0 | 201.0 | 7.7e-51 |
| ATP76243.1 | NdaA | BGC0001705 | NRP + Polyketide | 31.0 | 36.0 | 201.0 | 7.7e-51 |
| AAM80539.1 | StaA | BGC0000290 | NRP:Glycopeptide | 31.0 | 46.5 | 201.0 | 1e-50 |
| CAC48362.1 | peptide\_synthetase | BGC0000311 | NRP | 31.0 | 47.1 | 201.0 | 1e-50 |
| ACU36654.1 | amino\_acid\_adenylation\_domain\_protein | BGC0000392 | NRP | 29.0 | 48.7 | 201.0 | 1e-50 |
| AAT09804.1 | NocA | BGC0000395 | NRP | 30.0 | 48.2 | 201.0 | 1e-50 |
| AIG79240.1 | Hypothetical\_protein | BGC0000419 | Saccharide + NRP:Glycopeptide | 31.0 | 47.1 | 201.0 | 1e-50 |
| ACU71638.1 | peptide\_synthetase-like\_protein | BGC0001154 | Other | 29.0 | 48.8 | 201.0 | 1e-50 |
| APD26279.1 | PtmA | BGC0001726 | NRP + Polyketide | 29.0 | 48.9 | 201.0 | 1e-50 |
| ARR97036.1 | SphC | BGC0001780 | NRP | 27.0 | 47.1 | 201.0 | 1e-50 |
| AQX14497.1 | monobactam\_NRPS\_scaffold\_4 | BGC0001672 | NRP | 25.0 | 47.4 | 200.0 | 1.7e-50 |
| AHY86403.1 | non\_ribosomal\_peptide\_synthetase | BGC0000329 | NRP | 30.0 | 40.5 | 200.0 | 2.2e-50 |
| AGS77307.1 | NRPS\_modules\_1-2 | BGC0001178 | NRP:Glycopeptide | 31.0 | 46.8 | 200.0 | 2.2e-50 |
| OTA20325.1 | peptide\_synthase | BGC0001824 | NRP | 29.0 | 46.3 | 200.0 | 2.2e-50 |
| AFA26384.1 | polyketide\_synthase\_A | BGC0001874 | NRP + Polyketide | 24.0 | 75.7 | 200.0 | 2.2e-50 |
| DAC76731.1 | methionyl-tRNA\_formyltransferase | BGC0001885 | Polyketide | 30.0 | 45.1 | 200.0 | 2.2e-50 |
| AIE77058.1 | peptide\_synthetase\_module\_3 | BGC0000418 | NRP | 30.0 | 46.1 | 199.0 | 3.8e-50 |
| ADC79642.1 | TamD | BGC0001052 | NRP + Polyketide:Modular type I | 30.0 | 47.8 | 199.0 | 3.8e-50 |
| CAE53351.1 | non-ribosomal\_peptide\_synthetase | BGC0000440 | NRP:Glycopeptide | 29.0 | 45.0 | 198.0 | 5e-50 |
| AHD05627.1 | putative\_non-ribosomal\_peptide\_ligase\_domain\_protein | BGC0001033 | NRP + Polyketide | 29.0 | 46.9 | 198.0 | 5e-50 |
| AGS77309.1 | NRPS\_modules\_4-6 | BGC0001178 | NRP:Glycopeptide | 30.0 | 47.0 | 198.0 | 5e-50 |
| EME52988.1 | amino\_acid\_adenylation\_protein | BGC0001460 | NRP:Glycopeptide | 31.0 | 47.1 | 198.0 | 5e-50 |
| AVV61987.1 | putative\_non-ribosomal\_peptide\_synthetase | BGC0001477 | NRP + Polyketide:Modular type I | 30.0 | 45.9 | 198.0 | 5e-50 |
| EJK79842.1 | amino\_acid\_adenylation\_enzyme/thioester\_reductase\_family\_protein | BGC0000436 | NRP | 28.0 | 47.9 | 198.0 | 6.5e-50 |
| AGZ15460.1 | putative\_non-ribosomal\_peptide\_synthetase | BGC0001036 | NRP + Polyketide | 31.0 | 36.7 | 198.0 | 6.5e-50 |
| CCM44337.1 | Nonribosomal\_peptide\_synthetase | BGC0001056 | NRP + Polyketide:Modular type I + Polyketide:PUFA synthase or related | 30.0 | 46.4 | 198.0 | 6.5e-50 |
| APZ78833.1 | nonribosomal\_peptide\_synthetase | BGC0001430 | NRP:Cyclic depsipeptide + Polyketide:Iterative type I | 29.0 | 46.6 | 198.0 | 6.5e-50 |
| BAX89998.1 | Non-ribosomal\_peptide\_synthetase | BGC0001628 | NRP | 29.0 | 49.2 | 198.0 | 6.5e-50 |
| ABC35522.1 | thiotemplate\_mechanism\_natural\_product\_synthetase | BGC0000186 | NRP + Polyketide:Modular type I | 29.0 | 47.7 | 198.0 | 8.5e-50 |
| AIE77057.1 | peptide\_synthetase | BGC0000418 | NRP | 30.0 | 46.9 | 198.0 | 8.5e-50 |
| AIG79242.1 | Non-ribosomal\_peptide\_synthetase | BGC0000419 | Saccharide + NRP:Glycopeptide | 30.0 | 45.1 | 198.0 | 8.5e-50 |
| CAD91220.1 | putative\_non-ribosomal\_peptide\_synthetase,\_modules\_1-2 | BGC0000289 | NRP:Glycopeptide + Saccharide:Hybrid/tailoring | 30.0 | 45.8 | 197.0 | 1.1e-49 |
| CAG15010.1 | peptide\_synthetase,\_module\_3 | BGC0000441 | NRP | 29.0 | 45.0 | 197.0 | 1.1e-49 |
| AEP18656.1 | WAPS1 | BGC0000461 | NRP | 28.0 | 48.1 | 197.0 | 1.1e-49 |
| BS329\_14150 | non-ribosomal\_peptide\_synthetase | BGC0001462 | NRP:Glycopeptide | 28.0 | 49.0 | 197.0 | 1.1e-49 |
| AGD80623.1 | non-ribosomal\_peptide\_synthetase | BGC0000394 | NRP | 31.0 | 47.1 | 197.0 | 1.5e-49 |
| CBZ42140.1 | non-ribosomal\_peptide\_synthetase | BGC0001117 | NRP | 28.0 | 50.2 | 197.0 | 1.5e-49 |
| AXA20090.1 | hybrid\_trans-AT\_PKS/NRPS\_LgaA | BGC0001946 | NRP + Polyketide | 28.0 | 47.0 | 197.0 | 1.5e-49 |
| AQX14499.1 | monobactam\_NRPS\_scaffold\_1 | BGC0001672 | NRP | 26.0 | 46.9 | 197.0 | 1.5e-49 |
| SCO70310.1 | Type\_I\_polyketide\_synthase | BGC0001433 | Polyketide:Modular type I | 29.0 | 41.1 | 196.0 | 1.9e-49 |
| ARO38317.1 | nonribosomal\_peptide\_synthetase | BGC0001560 | NRP + Polyketide | 29.0 | 47.4 | 196.0 | 1.9e-49 |
| OAL11475.1 | non-ribosomal\_peptide\_synthetase\_module | BGC0001570 | NRP | 29.0 | 49.2 | 196.0 | 1.9e-49 |
| AAZ23076.1 | peptide\_synthetase | BGC0000291 | NRP | 30.0 | 49.3 | 196.0 | 2.5e-49 |
| OLZ50886.1 | non-ribosomal\_peptide\_synthetase | BGC0001461 | NRP:Glycopeptide | 30.0 | 49.5 | 196.0 | 2.5e-49 |
| AWI62629.1 | nonribosomal\_peptide\_synthetase | BGC0001943 | NRP | 32.0 | 36.7 | 196.0 | 2.5e-49 |
| AFK57215.1 | DidD | BGC0000985 | Polyketide + NRP:Cyclic depsipeptide | 29.0 | 48.1 | 196.0 | 3.2e-49 |
| AAN85522.1 | hybrid\_nonribosomal\_peptide\_synthetase\_/\_polyketide\_synthase | BGC0001101 | NRP + Polyketide:Modular type I + Polyketide:Trans-AT type I | 27.0 | 49.1 | 196.0 | 3.2e-49 |
| WP\_078857611.1 | non-ribosomal\_peptide\_synthetase | BGC0001368 | NRP | 30.0 | 48.7 | 196.0 | 3.2e-49 |
| ANG60380.1 | nonribosomal\_peptide\_synthetase\_BudB | BGC0001698 | NRP | 32.0 | 36.2 | 196.0 | 3.2e-49 |
| BBC83957.1 | nonribosomal\_peptide\_synthetase | BGC0001636 | NRP | 29.0 | 48.5 | 196.0 | 3.2e-49 |
| AYA22333.1 | KerD | BGC0001955 | Other | 30.0 | 47.3 | 196.0 | 3.2e-49 |
| EFL06866.1 | predicted\_protein | BGC0000300 | NRP | 28.0 | 45.6 | 195.0 | 4.2e-49 |
| BAK26562.1 | PKS-NRPS\_hybrid | BGC0000977 | NRP + Polyketide | 24.0 | 68.7 | 195.0 | 4.2e-49 |
| CAN89662.1 | putative\_non-ribosomal\_peptide\_synthetase | BGC0001070 | NRP + Polyketide:Modular type I + Polyketide:Trans-AT type I | 30.0 | 48.1 | 195.0 | 4.2e-49 |
| ADH04678.1 | non-ribosomal\_peptide\_synthetase | BGC0001344 | NRP + Polyketide | 32.0 | 36.9 | 195.0 | 4.2e-49 |
| AAG02343.1 | peptide\_synthetase\_NRPS12 | BGC0000963 | NRP:Glycopeptide + Polyketide:Modular type I + Saccharide:Hybrid/tailoring | 29.0 | 47.7 | 195.0 | 5.5e-49 |
| BAH33409.1 | putative\_non-ribosomal\_peptide\_synthetase | BGC0000371 | NRP | 28.0 | 49.2 | 195.0 | 7.2e-49 |
| AJW76709.1 | DsaG | BGC0001196 | NRP | 29.0 | 51.6 | 195.0 | 7.2e-49 |
| AQZ26587.1 | obafluorin\_dimodular\_nonribosomal\_peptide\_synthetase | BGC0001710 | NRP | 29.0 | 48.5 | 195.0 | 7.2e-49 |
| OLZ52456.1 | non-ribosomal\_peptide\_synthetase | BGC0001462 | NRP:Glycopeptide | 30.0 | 47.5 | 195.0 | 7.2e-49 |
| BBA21071.1 | putative\_non-ribosomal\_peptide\_synthetase | BGC0001740 | NRP + Polyketide | 30.0 | 48.6 | 195.0 | 7.2e-49 |
| AQV04230.1 | SwnK | BGC0001794 | Polyketide | 27.0 | 47.5 | 195.0 | 7.2e-49 |
| ATY69584.1 | hybrid\_nonribosomal\_peptide\_synthetase/type\_I\_polyketide\_synthase | BGC0001823 | NRP + Polyketide | 29.0 | 47.7 | 195.0 | 7.2e-49 |
| TXD00259.1 | amino\_acid\_adenylation\_domain-containing\_protein | BGC0001877 | Polyketide | 31.0 | 34.9 | 195.0 | 7.2e-49 |
| BAP16699.1 | nonribosomal\_peptide\_synthetase | BGC0000376 | NRP | 28.0 | 47.4 | 194.0 | 9.4e-49 |
| AAO62582.1 | polyketide\_synthase\_peptide\_sythetase\_fusion\_protein | BGC0001016 | NRP + Polyketide | 27.0 | 49.5 | 194.0 | 9.4e-49 |
| CAD70194.1 | non-ribosomal\_peptide\_synthetase | BGC0001047 | NRP + Polyketide | 29.0 | 46.5 | 194.0 | 9.4e-49 |
| AQZ69228.1 | hypothetical\_protein | BGC0001635 | NRP + Polyketide | 31.0 | 46.0 | 194.0 | 9.4e-49 |
| AFV27434.1 | indigoidine\_synthase | BGC0000375 | NRP | 29.0 | 38.9 | 193.0 | 1.6e-48 |
| CAQ18839.1 | hybrid\_polyketide\_synthase/nonribosomal\_polypetide\_synthetase | BGC0000954 | NRP + Polyketide:Modular type I | 32.0 | 37.7 | 193.0 | 1.6e-48 |
| ERG36782.1 | Ornithine\_racemase | BGC0000299 | NRP | 27.0 | 48.1 | 193.0 | 2.1e-48 |
| WP\_010369425.1 | non-ribosomal\_peptide\_synthetase | BGC0000314 | Polyketide + NRP:Cyclic depsipeptide + Other:Aminocoumarin | 27.0 | 48.1 | 193.0 | 2.1e-48 |
| CBJ90358.1 | putative\_Peptide\_synthetase | BGC0000465 | NRP | 31.0 | 35.3 | 193.0 | 2.1e-48 |
| AAG02355.1 | peptide\_synthetase\_NRPS9-8 | BGC0000963 | NRP:Glycopeptide + Polyketide:Modular type I + Saccharide:Hybrid/tailoring | 29.0 | 49.6 | 193.0 | 2.1e-48 |
| ctg1\_orf13 |  | BGC0001329 | Polyketide + NRP:Cyclic depsipeptide | 29.0 | 50.3 | 193.0 | 2.1e-48 |
| BAW32333.1 | nonribosomal\_peptide\_synthetase | BGC0001631 | NRP + Polyketide | 29.0 | 48.0 | 193.0 | 2.1e-48 |
| CAD55498.1 | CDA\_peptide\_synthetase\_III\_(CdaPs3) | BGC0000315 | NRP:Ca+-dependent lipopeptide | 30.0 | 47.7 | 193.0 | 2.7e-48 |
| CUX96957.1 | TmcJ | BGC0001829 | NRP + Polyketide | 28.0 | 47.9 | 193.0 | 2.7e-48 |
| AEI58867.1 | peptide\_synthetase | BGC0000455 | NRP | 30.0 | 47.1 | 192.0 | 3.6e-48 |
| AEC14346.1 | nonribosomal\_peptide\_synthetase | BGC0000377 | NRP | 26.0 | 45.1 | 192.0 | 4.7e-48 |
| BAD55612.1 | non-ribosomal\_peptide\_synthetase | BGC0001027 | NRP + Polyketide | 30.0 | 47.5 | 192.0 | 4.7e-48 |
| AAK81826.1 | peptide\_synthetase | BGC0000326 | NRP | 32.0 | 35.8 | 191.0 | 6.1e-48 |
| ABD65956.1 | nonribosomal\_peptide\_synthetase | BGC0000341 | NRP | 28.0 | 47.7 | 191.0 | 6.1e-48 |
| BAP16697.1 | nonribosomal\_peptide\_synthetase | BGC0000376 | NRP | 29.0 | 47.1 | 191.0 | 6.1e-48 |
| AAW49319.1 | thaxtomin\_synthetase\_B | BGC0000444 | NRP | 32.0 | 37.9 | 191.0 | 6.1e-48 |
| AFD30953.1 | CrmB | BGC0000966 | NRP + Polyketide | 29.0 | 47.7 | 191.0 | 6.1e-48 |
| CAD89775.1 | MelD\_protein | BGC0001010 | NRP + Polyketide:Modular type I | 31.0 | 36.6 | 191.0 | 6.1e-48 |
| KFL51885.1 | amino\_acid\_adenylation\_protein | BGC0001711 | NRP + Polyketide | 27.0 | 53.4 | 191.0 | 6.1e-48 |
| AJO72717.1 | adenylation\_domain-containing\_protein | BGC0001381 | Polyketide | 29.0 | 47.0 | 191.0 | 8e-48 |
| QBF51786.1 | Nrps | BGC0001856 | Polyketide:Modular type I | 29.0 | 46.9 | 191.0 | 8e-48 |
| TXC99989.1 | non-ribosomal\_peptide\_synthetase | BGC0001877 | Polyketide | 29.0 | 46.8 | 191.0 | 8e-48 |
| AAN65224.1 | peptide\_synthetase-like\_protein | BGC0000832 | Saccharide:Hybrid/tailoring + Other:Aminocoumarin | 28.0 | 47.7 | 191.0 | 1e-47 |
| AQZ69227.1 | hypothetical\_protein | BGC0001635 | NRP + Polyketide | 30.0 | 46.0 | 191.0 | 1e-47 |
| ARR97039.1 | SphF | BGC0001780 | NRP | 27.0 | 47.2 | 191.0 | 1e-47 |
| ABM21571.1 | crpC | BGC0000975 | NRP + Polyketide | 30.0 | 37.1 | 190.0 | 1.4e-47 |
| EDY47118.1 | N-(5-amino-5-carboxypentanoyl)-L-cysteinyl-D-\_valine\_synthase | BGC0000319 | NRP:Beta-lactam | 27.0 | 49.5 | 190.0 | 1.8e-47 |
| BAI63289.1 | putative\_non-ribosomal\_peptide\_synthetase | BGC0000434 | NRP | 31.0 | 36.4 | 190.0 | 1.8e-47 |
| EJK79843.1 | amino\_acid\_adenylation\_enzyme/thioester\_reductase\_family\_protein | BGC0000436 | NRP | 29.0 | 49.1 | 190.0 | 1.8e-47 |
| CAL80821.1 | sylD-like\_NRPS/PKS | BGC0000997 | NRP + Polyketide | 28.0 | 50.9 | 190.0 | 1.8e-47 |
| CAE14173.1 | hypothetical\_protein | BGC0000383 | NRP + Polyketide:Modular type I | 27.0 | 52.8 | 190.0 | 2.3e-47 |
| ABY66004.1 | type\_II\_beta-Tyr\_adenylation\_domain\_protein | BGC0001008 | Polyketide:Iterative type I + Polyketide:Enediyne type I | 29.0 | 47.4 | 190.0 | 2.3e-47 |
| QCC62999.1 | BII-rafflesfungin\_nonribosomal\_protein\_synthetase | BGC0001966 | NRP | 29.0 | 48.6 | 190.0 | 2.3e-47 |
| AAM80536.1 | StaD | BGC0000290 | NRP:Glycopeptide | 31.0 | 47.0 | 189.0 | 3e-47 |
| ABA59547.1 | NRPS | BGC0000453 | NRP:Cyclic depsipeptide | 28.0 | 45.6 | 189.0 | 4e-47 |
| EYT83459.1 | hypothetical\_protein | BGC0001213 | Polyketide | 33.0 | 31.1 | 189.0 | 4e-47 |
| ANZ15839.1 | peptide\_synthetase\_ScpsB | BGC0001569 | NRP | 31.0 | 50.4 | 189.0 | 4e-47 |
| AGN74885.1 | nonribosomal\_peptide\_synthetase | BGC0000459 | NRP:Cyclic depsipeptide + Polyketide:Trans-AT type I | 30.0 | 39.0 | 188.0 | 5.2e-47 |
| CAJ76290.1 | putative\_non-ribosomal\_peptide\_synthase | BGC0000972 | NRP + Polyketide:Modular type I + Polyketide:Trans-AT type I | 28.0 | 48.8 | 188.0 | 5.2e-47 |
| AFK57212.1 | DidA | BGC0000985 | Polyketide + NRP:Cyclic depsipeptide | 29.0 | 48.9 | 188.0 | 5.2e-47 |
| CCJ67647.1 | JagC | BGC0001127 | NRP | 28.0 | 40.3 | 188.0 | 5.2e-47 |
| RLV71193.1 | non-ribosomal\_peptide\_synthetase | BGC0001846 | NRP + Saccharide:Hybrid/tailoring | 29.0 | 45.4 | 188.0 | 5.2e-47 |
| CAI94718.1 | putative\_CoA\_ligase | BGC0000141 | Polyketide | 31.0 | 45.3 | 188.0 | 6.7e-47 |
| ALD82525.1 | non-ribosomal\_peptide\_synthase | BGC0001212 | NRP + Polyketide | 29.0 | 48.3 | 188.0 | 6.7e-47 |
| CBJ90287.1 | peptide\_synthetase | BGC0000416 | NRP | 31.0 | 35.9 | 188.0 | 8.8e-47 |
| CAD29794.1 | peptide\_synthetase | BGC0001015 | NRP + Polyketide | 27.0 | 47.1 | 188.0 | 8.8e-47 |
| WP\_013184317.1 | non-ribosomal\_peptide\_synthetase | BGC0001692 | NRP | 31.0 | 35.9 | 188.0 | 8.8e-47 |
| WP\_010369428.1 | non-ribosomal\_peptide\_synthase | BGC0000314 | Polyketide + NRP:Cyclic depsipeptide + Other:Aminocoumarin | 28.0 | 48.6 | 187.0 | 1.2e-46 |
| CBL93718.1 | NRPS\_didomain\_PCP-C | BGC0000360 | NRP | 29.0 | 47.7 | 187.0 | 1.2e-46 |
| AAW03327.1 | CtaD | BGC0000982 | NRP + Polyketide | 32.0 | 36.4 | 187.0 | 1.2e-46 |
| AZF85940.1 | non-ribosomal\_peptide\_synthase | BGC0001963 | NRP + Polyketide | 32.0 | 37.4 | 187.0 | 1.2e-46 |
| AKA59440.1 | non-ribosomal\_peptide\_synthetase | BGC0001202 | NRP + Polyketide | 28.0 | 47.4 | 187.0 | 1.5e-46 |
| WP\_054234617.1 | non-ribosomal\_peptide\_synthase | BGC0002014 | NRP + Polyketide | 31.0 | 38.1 | 187.0 | 1.5e-46 |
| AEF16059.1 | non-ribosomal\_peptide\_synthetase | BGC0000953 | Saccharide:Aminoglycoside | 28.0 | 46.0 | 186.0 | 2e-46 |
| ADH04660.1 | TugD | BGC0001342 | NRP + Polyketide | 30.0 | 38.0 | 186.0 | 2e-46 |
| AXA20091.1 | hybrid\_trans-AT\_PKS/NRPS\_LgaB | BGC0001946 | NRP + Polyketide | 28.0 | 50.3 | 186.0 | 2e-46 |
| ABL74940.1 | NRPS | BGC0001048 | NRP:Glycopeptide + Polyketide:Modular type I + Saccharide:Hybrid/tailoring | 33.0 | 40.5 | 186.0 | 2.6e-46 |
| AAS47562.1 | mixed\_type\_I\_polyketide\_synthase\_-\_peptide\_synthetase | BGC0001108 | Polyketide:Trans-AT type I | 28.0 | 48.2 | 186.0 | 3.3e-46 |
| ctg1\_orf8 |  | BGC0001109 | NRP + Polyketide | 28.0 | 48.2 | 186.0 | 3.3e-46 |
| AFP87523.1 | type\_I\_polyketide\_synthase | BGC0001159 | NRP + Polyketide:Modular type I | 30.0 | 39.5 | 186.0 | 3.3e-46 |
| DAB41477.1 | nonribosomal\_peptide\_synthetase | BGC0001766 | NRP | 31.0 | 37.4 | 186.0 | 3.3e-46 |
| CAC17499.1 | putative\_non-ribosomal\_peptide\_synthase | BGC0000324 | NRP | 29.0 | 48.1 | 185.0 | 4.4e-46 |
| CBJ90359.1 | putative\_Peptide\_synthetase | BGC0000465 | NRP | 30.0 | 35.4 | 185.0 | 4.4e-46 |
| AAG02356.1 | peptide\_synthetase\_NRPS7 | BGC0000963 | NRP:Glycopeptide + Polyketide:Modular type I + Saccharide:Hybrid/tailoring | 30.0 | 49.6 | 185.0 | 4.4e-46 |
| ABL74937.1 | NRPS | BGC0001048 | NRP:Glycopeptide + Polyketide:Modular type I + Saccharide:Hybrid/tailoring | 29.0 | 45.5 | 185.0 | 4.4e-46 |
| AGD80616.1 | amino\_acid\_adenylation\_domain-containing\_protein | BGC0000394 | NRP | 31.0 | 41.6 | 185.0 | 5.7e-46 |
| WP\_050383082.1 | non-ribosomal\_peptide\_synthetase | BGC0001451 | NRP | 30.0 | 38.3 | 185.0 | 5.7e-46 |
| ABR67750.1 | CmnG | BGC0000316 | NRP | 28.0 | 51.1 | 185.0 | 7.5e-46 |
| CAJ96471.1 | non-ribosomal\_peptide\_synthetase | BGC0000330 | NRP:NRP siderophore | 28.0 | 50.2 | 185.0 | 7.5e-46 |
| AFD30954.1 | CrmA | BGC0000966 | NRP + Polyketide | 28.0 | 46.7 | 185.0 | 7.5e-46 |
| ADZ45339.1 | non-ribosomal\_peptide\_synthetase | BGC0001020 | NRP + Polyketide | 31.0 | 39.3 | 185.0 | 7.5e-46 |
| WP\_053065270.1 | non-ribosomal\_peptide\_synthetase | BGC0001330 | NRP:Cyclic depsipeptide + Polyketide:Modular type I | 31.0 | 35.2 | 185.0 | 7.5e-46 |
| CBL93720.1 | NRPS\_didomain\_PCP-A | BGC0000360 | NRP | 28.0 | 47.9 | 184.0 | 9.7e-46 |
| ATP76239.1 | NdaF | BGC0001705 | NRP + Polyketide | 26.0 | 49.5 | 184.0 | 9.7e-46 |
| AAF00958.1 | mcyE | BGC0001017 | NRP + Polyketide:Modular type I | 27.0 | 46.9 | 183.0 | 1.7e-45 |
| AAF19812.1 | MtaD | BGC0001024 | NRP + Polyketide:Modular type I | 30.0 | 35.9 | 183.0 | 1.7e-45 |
| BS330\_28385 | non-ribosomal\_peptide\_synthetase | BGC0001461 | NRP:Glycopeptide | 28.0 | 46.7 | 183.0 | 1.7e-45 |
| AQZ71347.1 | hypothetical\_protein | BGC0001635 | NRP + Polyketide | 30.0 | 50.0 | 183.0 | 1.7e-45 |
| AZH23819.1 | MgiR | BGC0001971 | NRP + Polyketide | 31.0 | 31.9 | 183.0 | 1.7e-45 |
| AGS77310.1 | NRPS\_module\_7 | BGC0001178 | NRP:Glycopeptide | 29.0 | 46.5 | 183.0 | 2.2e-45 |
| AYA22336.1 | KerA | BGC0001955 | Other | 27.0 | 46.4 | 183.0 | 2.2e-45 |
| BAF50727.1 | hybrid\_polyketide\_synthase-non\_ribosomal\_peptide\_synthetase | BGC0001116 | NRP + Polyketide | 28.0 | 48.1 | 183.0 | 2.8e-45 |
| QCX41944.1 | Amc7 | BGC0001958 | Other | 29.0 | 48.8 | 183.0 | 2.8e-45 |
| ACJ04424.1 | aureobasidin\_A1\_biosynthesis\_complex | BGC0000307 | NRP | 28.0 | 50.0 | 182.0 | 3.7e-45 |
| ADY76684.1 | non-ribosomal\_peptide\_synthetase | BGC0000950 | NRP:Uridylpeptide + Other:Nucleoside | 30.0 | 41.6 | 182.0 | 3.7e-45 |
| WP\_078857609.1 | non-ribosomal\_peptide\_synthetase | BGC0001368 | NRP | 30.0 | 48.5 | 182.0 | 3.7e-45 |
| BAW32334.1 | hybrid\_cis-AT\_polyketide\_synthase\_-\_nonribosomal\_peptide\_synthetase | BGC0001631 | NRP + Polyketide | 28.0 | 48.0 | 182.0 | 3.7e-45 |
| AED90004.1 | non-ribosomal\_peptide\_synthetase\_ThaC1 | BGC0000443 | NRP:Beta-lactam | 28.0 | 48.5 | 182.0 | 4.8e-45 |
| CCC55921.1 | non-ribosomal\_peptide\_synthetase/polyketide\_synthase\_hybrid\_protein | BGC0000973 | NRP + Polyketide:Modular type I | 27.0 | 47.5 | 182.0 | 4.8e-45 |
| WP\_107406787.1 | non-ribosomal\_peptide\_synthetase | BGC0001567 | NRP | 30.0 | 46.7 | 182.0 | 4.8e-45 |
| AAW49318.1 | thaxtomin\_synthetase\_A | BGC0000444 | NRP | 30.0 | 37.7 | 181.0 | 6.3e-45 |
| AAF67501.2 | peptide\_synthetase-like\_protein | BGC0000834 | Saccharide:Hybrid/tailoring + Other:Aminocoumarin | 28.0 | 48.9 | 181.0 | 6.3e-45 |
| AJI44167.1 | long-chain-fatty-acid-CoA\_ligase | BGC0001193 | NRP | 27.0 | 51.0 | 181.0 | 6.3e-45 |
| CDG12865.1 | non-ribosomal\_peptide\_synthetase | BGC0001415 | NRP | 27.0 | 46.1 | 181.0 | 6.3e-45 |
| ACY13414.1 | amino\_acid\_adenylation\_domain\_protein | BGC0001367 | NRP + Polyketide | 28.0 | 48.7 | 181.0 | 8.2e-45 |
| AZH23788.1 | MgcR | BGC0001970 | NRP + Polyketide | 30.0 | 33.4 | 181.0 | 8.2e-45 |
| CBJ90289.1 | peptide\_synthetase | BGC0000416 | NRP | 30.0 | 35.5 | 181.0 | 1.1e-44 |
| ACN39014.1 | putative\_nonribosomal\_peptide\_synthetase\_TomA | BGC0000448 | NRP | 29.0 | 47.1 | 181.0 | 1.1e-44 |
| WP\_030498977.1 | hypothetical\_protein | BGC0001327 | NRP:Cyclic depsipeptide + Polyketide:Modular type I | 28.0 | 49.8 | 181.0 | 1.1e-44 |
| AQX14495.1 | monobactam\_NRPS\_scaffold\_3 | BGC0001672 | NRP | 27.0 | 47.9 | 181.0 | 1.1e-44 |
| WP\_013184319.1 | non-ribosomal\_peptide\_synthetase | BGC0001692 | NRP | 30.0 | 35.5 | 181.0 | 1.1e-44 |
| ABD14712.1 | cesB | BGC0000320 | NRP:Cyclic depsipeptide | 28.0 | 44.6 | 180.0 | 1.4e-44 |
| AEC14347.1 | nonribosomal\_peptide\_synthetase | BGC0000377 | NRP | 27.0 | 44.6 | 180.0 | 1.4e-44 |
| CCA89326.1 | mixed\_trans-AT\_type\_I\_polyketide\_synthase/nonribosomal\_peptide\_synthetase | BGC0001111 | NRP + Polyketide:Trans-AT type I | 27.0 | 61.0 | 180.0 | 1.4e-44 |
| ORC16618.1 | hypothetical\_protein | BGC0001341 | NRP | 27.0 | 45.6 | 180.0 | 1.4e-44 |
| AJO72702.1 | Acyl-CoA\_ligase/oxidoreductase | BGC0001381 | Polyketide | 26.0 | 64.8 | 180.0 | 1.4e-44 |
| ATJ34005.1 | adenylation\_domain-containing\_protein | BGC0001497 | Polyketide | 30.0 | 40.7 | 180.0 | 1.4e-44 |
| QCX41915.1 | Mhr9 | BGC0001956 | Polyketide | 29.0 | 47.5 | 180.0 | 1.8e-44 |
| BGRAMDRAFT\_RS22640 | non-ribosomal\_peptide\_synthetase | BGC0001999 | NRP | 26.0 | 54.6 | 180.0 | 1.8e-44 |
| WP\_010369430.1 | non-ribosomal\_peptide\_synthetase | BGC0000314 | Polyketide + NRP:Cyclic depsipeptide + Other:Aminocoumarin | 28.0 | 49.7 | 180.0 | 2.4e-44 |
| CBK62746.1 |  | BGC0001115 | NRP + Polyketide | 26.0 | 46.7 | 179.0 | 3.1e-44 |
| AZF85929.1 | non-ribosomal\_peptide\_synthase | BGC0001963 | NRP + Polyketide | 28.0 | 47.3 | 179.0 | 3.1e-44 |
| CBL93716.1 | NRPS\_A-domain | BGC0000360 | NRP | 29.0 | 41.2 | 179.0 | 4.1e-44 |
| WP\_081656241.1 | non-ribosomal\_peptide\_synthetase | BGC0001467 | NRP:Cyclic depsipeptide + Polyketide:Modular type I | 31.0 | 40.0 | 179.0 | 4.1e-44 |
| WP\_004571777.1 | non-ribosomal\_peptide\_synthase | BGC0001760 | NRP | 30.0 | 41.1 | 179.0 | 4.1e-44 |
| CBJ90288.1 | Peptide\_synthetase | BGC0000416 | NRP | 29.0 | 36.2 | 178.0 | 5.3e-44 |
| WP\_013184318.1 | non-ribosomal\_peptide\_synthetase | BGC0001692 | NRP | 29.0 | 36.2 | 178.0 | 5.3e-44 |
| KFH48607.1 | N-like\_protein | BGC0000317 | NRP | 27.0 | 48.1 | 178.0 | 7e-44 |
| AEC14348.1 | nonribosomal\_peptide\_synthetase | BGC0000377 | NRP | 28.0 | 45.3 | 178.0 | 7e-44 |
| AAL33757.1 | putative\_non-ribosomal\_peptide\_synthetase | BGC0000421 | NRP | 27.0 | 45.6 | 178.0 | 7e-44 |
| ABC34137.1 | peptide\_synthetase,\_putative | BGC0000961 | NRP + Polyketide | 29.0 | 41.0 | 178.0 | 9.1e-44 |
| CAJ34375.1 | NRPS | BGC0000445 | NRP:Cyclic depsipeptide | 28.0 | 36.4 | 177.0 | 1.2e-43 |
| ANG60381.1 | nonribosomal\_peptide\_synthetase\_BudC | BGC0001698 | NRP | 31.0 | 36.8 | 177.0 | 1.6e-43 |
| AXM42949.1 | hybrid\_type\_1\_PKS/NRPS | BGC0001941 | NRP + Polyketide | 29.0 | 35.9 | 177.0 | 1.6e-43 |
| ABR67749.1 | CmnF | BGC0000316 | NRP | 30.0 | 46.3 | 176.0 | 2e-43 |
| ADI59531.1 | CorI | BGC0001091 | NRP + Polyketide | 27.0 | 50.2 | 176.0 | 2e-43 |
| ADH04641.1 | TgaC | BGC0001051 | NRP + Polyketide:Modular type I | 31.0 | 36.4 | 176.0 | 2.7e-43 |
| ACG60776.1 | NRPS(AL/ACP/C/A/PCP/C/A) | BGC0001058 | NRP:Glycopeptide + Polyketide:Modular type I + Saccharide:Hybrid/tailoring | 31.0 | 40.3 | 176.0 | 2.7e-43 |
| RAT94090.1 | NRPS | BGC0001469 | NRP | 25.0 | 48.4 | 176.0 | 2.7e-43 |
| CAD15508.1 | polyketide\_synthase/non-ribosomal\_peptide\_synthetase | BGC0001014 | NRP:NRP siderophore + Polyketide:Modular type I + Polyketide:Iterative type I | 30.0 | 37.1 | 176.0 | 3.5e-43 |
| AWO77084.1 | hybrid\_non-ribosomal\_peptide\_synthetase/type\_I\_polyketide\_synthase | BGC0001556 | NRP + Polyketide | 26.0 | 51.1 | 176.0 | 3.5e-43 |
| WP\_019032753.1 | non-ribosomal\_peptide\_synthetase | BGC0001331 | NRP:Cyclic depsipeptide + Polyketide:Modular type I | 31.0 | 35.2 | 175.0 | 5.9e-43 |
| CDG12864.1 | non-ribosomal\_peptide\_synthetase | BGC0001415 | NRP | 27.0 | 49.0 | 175.0 | 5.9e-43 |
| WP\_016638469.1 | non-ribosomal\_peptide\_synthetase | BGC0001519 | NRP + Polyketide | 29.0 | 35.5 | 175.0 | 5.9e-43 |
| ACK77757.1 | putative\_indigoidine\_synthase\_IndC | BGC0000727 | Saccharide | 29.0 | 36.3 | 174.0 | 1e-42 |
| CAN89633.1 | putative\_hybrid\_non-ribosomal\_peptide\_synthetase/polyketide\_synthase | BGC0001070 | NRP + Polyketide:Modular type I + Polyketide:Trans-AT type I | 29.0 | 48.1 | 174.0 | 1e-42 |
| CCJ67646.1 | JagB | BGC0001127 | NRP | 28.0 | 47.1 | 174.0 | 1.3e-42 |
| AGN74892.1 | nonribosomal\_peptide\_synthetase/polyketide\_synthase\_hybrid\_protein | BGC0000459 | NRP:Cyclic depsipeptide + Polyketide:Trans-AT type I | 29.0 | 49.7 | 173.0 | 1.7e-42 |
| CAL69597.1 | PKS-NRPS | BGC0001049 | NRP + Polyketide:Iterative type I | 25.0 | 64.5 | 173.0 | 1.7e-42 |
| ARS01470.1 | NcmB | BGC0001702 | NRP + Polyketide | 26.0 | 52.6 | 173.0 | 1.7e-42 |
| BAO98812.1 | putative\_nonribosomal\_peptide\_synthetase | BGC0001002 | NRP + Polyketide | 27.0 | 49.8 | 173.0 | 2.2e-42 |
| ABP57745.1 | DepA | BGC0000993 | NRP:Cyclic depsipeptide + Polyketide:Modular type I | 28.0 | 46.1 | 172.0 | 3.8e-42 |
| DAB41917.1 | ArzO\_-\_NRPS\_(Cy,\_A,\_Ox,\_PCP) | BGC0001884 | NRP + Polyketide | 29.0 | 36.7 | 172.0 | 3.8e-42 |
| AXM42952.1 | non-ribosomal\_peptide\_synthetase | BGC0001941 | NRP + Polyketide | 32.0 | 33.6 | 172.0 | 5e-42 |
| EAL89046.1 | nonribosomal\_peptide\_synthetase | BGC0000355 | NRP | 27.0 | 50.6 | 171.0 | 6.5e-42 |
| AAC83657.1 | pyochelin\_synthetase | BGC0000412 | NRP | 28.0 | 37.4 | 171.0 | 6.5e-42 |
| WP\_013184462.1 | non-ribosomal\_peptide\_synthetase | BGC0001716 | NRP | 26.0 | 47.9 | 171.0 | 6.5e-42 |
| CAD15513.1 | non-ribosomal\_peptide\_synthetase | BGC0001014 | NRP:NRP siderophore + Polyketide:Modular type I + Polyketide:Iterative type I | 30.0 | 36.5 | 171.0 | 8.5e-42 |
| ACB46193.1 | nonribosomal\_peptide\_synthetase | BGC0000989 | NRP + Polyketide | 29.0 | 36.0 | 171.0 | 1.1e-41 |
| AID65225.1 | nonribosomal\_peptide\_synthetase | BGC0000335 | NRP | 27.0 | 48.1 | 170.0 | 1.5e-41 |
| ADB12489.1 | EpoB | BGC0000990 | NRP + Polyketide | 29.0 | 36.0 | 170.0 | 1.5e-41 |
| BAX64246.1 | NRPS | BGC0001623 | NRP + Polyketide | 27.0 | 44.8 | 170.0 | 1.5e-41 |
| WP\_030498980.1 | hypothetical\_protein | BGC0001327 | NRP:Cyclic depsipeptide + Polyketide:Modular type I | 29.0 | 41.1 | 170.0 | 1.9e-41 |
| CAA79245.2 | enniatin\_synthetase | BGC0000342 | NRP | 29.0 | 36.7 | 170.0 | 2.5e-41 |
| AAF26925.1 | nonribosomal\_peptide\_synthetase | BGC0000988 | NRP + Polyketide | 29.0 | 36.0 | 170.0 | 2.5e-41 |
| CCP45167.1 | Peptide\_synthetase\_MbtF\_(peptide\_synthase) | BGC0001021 | NRP + Polyketide | 31.0 | 41.6 | 170.0 | 2.5e-41 |
| AJI44176.1 | nonribosomal\_peptide\_synthetase | BGC0001193 | NRP | 27.0 | 47.9 | 170.0 | 2.5e-41 |
| ATG32076.1 | nonribosomal\_peptide\_synthetase | BGC0001750 | NRP + Polyketide | 30.0 | 36.2 | 170.0 | 2.5e-41 |
| AAF62881.1 | EpoB | BGC0000991 | NRP + Polyketide | 29.0 | 36.0 | 169.0 | 3.2e-41 |
| AFR69331.1 | nonribosomal\_peptide\_synthetase\_SpiA | BGC0001045 | NRP:Cyclic depsipeptide + Polyketide:Modular type I | 26.0 | 48.4 | 169.0 | 3.2e-41 |
| AOZ21320.1 | SulM | BGC0001790 | NRP | 27.0 | 54.9 | 169.0 | 3.2e-41 |
| AFO59871.1 | three-domain\_carboxylic\_acid\_reductase | BGC0000175 | Polyketide:Trans-AT type I | 25.0 | 79.6 | 168.0 | 5.5e-41 |
| CAP96445.1 | NRPS | BGC0000420 | NRP | 27.0 | 51.3 | 168.0 | 5.5e-41 |
| CBJ89761.1 | Non-ribosomal\_peptide\_synthase\_involved\_in\_xenocoumacin\_synthesis | BGC0001054 | NRP + Polyketide:Modular type I | 25.0 | 50.6 | 168.0 | 5.5e-41 |
| ALJ49909.1 | TlmI | BGC0001237 | Polyketide | 27.0 | 51.1 | 168.0 | 5.5e-41 |
| EWM63010.1 | non-ribosomal\_peptide\_synthetase | BGC0001328 | NRP:Cyclic depsipeptide + Polyketide:Modular type I | 29.0 | 41.0 | 168.0 | 5.5e-41 |
| QCE20609.1 | AsmN | BGC0001961 | NRP | 29.0 | 48.1 | 168.0 | 5.5e-41 |
| AFK57221.1 | DidJ | BGC0000985 | Polyketide + NRP:Cyclic depsipeptide | 30.0 | 33.1 | 168.0 | 7.2e-41 |
| AAN65233.1 | acyl-CoA\_synthetase | BGC0000832 | Saccharide:Hybrid/tailoring + Other:Aminocoumarin | 28.0 | 41.7 | 168.0 | 9.4e-41 |
| ABY83142.1 | Azi3 | BGC0000960 | NRP + Polyketide | 27.0 | 48.3 | 167.0 | 1.2e-40 |
| ACN39728.1 | SibE | BGC0000428 | NRP | 29.0 | 48.8 | 166.0 | 2.1e-40 |
| AFL68053.1 | amino\_acid\_adenylation\_enzyme/thioester\_reductase\_family\_protein | BGC0001524 | NRP + Polyketide | 26.0 | 48.1 | 165.0 | 4.7e-40 |
| WP\_044618979.1 | non-ribosomal\_peptide\_synthetase | BGC0001791 | NRP | 24.0 | 47.7 | 165.0 | 4.7e-40 |
| BAN91483.1 | putative\_Aculeacin\_A\_nonribosomal\_peptide\_synthase | BGC0001220 | NRP | 25.0 | 48.4 | 164.0 | 1e-39 |
| YP\_856988.1 | nonribosomal\_peptide\_synthetase | BGC0001502 | NRP | 29.0 | 45.1 | 164.0 | 1e-39 |
| CAJ34370.1 | NRPS\_protein | BGC0000445 | NRP:Cyclic depsipeptide | 30.0 | 31.3 | 164.0 | 1.4e-39 |
| CBF77087.1 | conserved\_hypothetical\_protein | BGC0001679 | NRP | 27.0 | 47.7 | 164.0 | 1.4e-39 |
| CAJ76292.1 | putative\_non-ribosomal\_peptide\_synthase | BGC0000972 | NRP + Polyketide:Modular type I + Polyketide:Trans-AT type I | 26.0 | 47.2 | 163.0 | 1.8e-39 |
| AAF19815.1 | mtaG | BGC0001024 | NRP + Polyketide:Modular type I | 29.0 | 30.7 | 163.0 | 1.8e-39 |
| ADD82940.1 | Bat2 | BGC0001099 | NRP + Polyketide:Modular type I + Polyketide:Trans-AT type I | 27.0 | 47.1 | 163.0 | 2.3e-39 |
| AQX14441.1 | EM5400\_NRPS\_scaffold | BGC0001671 | NRP | 26.0 | 47.4 | 163.0 | 2.3e-39 |
| EFG10344.1 | Non-ribosomal\_peptide\_synthetase | BGC0000373 | NRP | 27.0 | 47.0 | 163.0 | 3e-39 |
| ABC39418.1 | dihydroaeruginoic\_acid\_synthetase | BGC0000964 | NRP:Cyclic depsipeptide + Polyketide:Trans-AT type I | 27.0 | 46.7 | 163.0 | 3e-39 |
| AFK57219.1 | DidH | BGC0000985 | Polyketide + NRP:Cyclic depsipeptide | 29.0 | 45.5 | 163.0 | 3e-39 |
| AQM58286.1 | non-ribosomal\_peptide\_synthase | BGC0001816 | NRP + Polyketide | 26.0 | 52.3 | 162.0 | 5.2e-39 |
| AQH32481.1 | hybrid\_polyketide\_synthase/peptide\_synthetase | BGC0001667 | NRP + Polyketide | 24.0 | 51.2 | 161.0 | 8.8e-39 |
| EPE34341.1 | non-ribosomal\_peptide\_synthetase | BGC0001035 | Polyketide + NRP + Other:Aminocoumarin | 28.0 | 48.0 | 161.0 | 1.2e-38 |
| CCA53804.1 | pyochelin\_synthetase\_F | BGC0001801 | NRP | 29.0 | 37.2 | 161.0 | 1.2e-38 |
| ADN26251.1 | ATP-dependent\_adenylase | BGC0000951 | NRP | 29.0 | 41.8 | 160.0 | 1.5e-38 |
| CBF87069.1 | nonribosomal\_peptide\_synthase,\_putative\_(Eurofung) | BGC0001290 | NRP | 24.0 | 50.3 | 160.0 | 2e-38 |
| AMM63162.1 | AniA | BGC0001371 | NRP | 27.0 | 49.2 | 160.0 | 2e-38 |
| BBB04327.1 | nonribosomal\_peptide\_synthetase | BGC0001717 | Alkaloid | 26.0 | 48.7 | 159.0 | 4.4e-38 |
| AGE11891.1 | nonribosomal\_peptide\_synthetase | BGC0000366 | NRP | 27.0 | 45.6 | 158.0 | 7.5e-38 |
| AAS92545.1 | SirP | BGC0001044 | NRP + Polyketide | 27.0 | 46.6 | 158.0 | 7.5e-38 |
| CAD89778.1 | MelG\_protein | BGC0001010 | NRP + Polyketide:Modular type I | 31.0 | 29.6 | 158.0 | 9.8e-38 |
| CAC17500.1 | putative\_non-ribosomal\_peptide\_synthase | BGC0000324 | NRP | 28.0 | 37.6 | 157.0 | 1.3e-37 |
| AGC65516.1 | NRPS/PKS\_hybrid | BGC0001050 | NRP:Lipopeptide + Polyketide:Trans-AT type I | 26.0 | 46.7 | 157.0 | 1.3e-37 |
| QCE20608.1 | AsmM | BGC0001961 | NRP | 28.0 | 48.7 | 157.0 | 1.3e-37 |
| ATG32071.1 | proline\_specific\_adenylation\_domain-containing\_protein | BGC0001750 | NRP + Polyketide | 27.0 | 35.8 | 157.0 | 1.7e-37 |
| AFK57214.1 | DidC | BGC0000985 | Polyketide + NRP:Cyclic depsipeptide | 26.0 | 47.1 | 156.0 | 2.2e-37 |
| ADY16689.1 | TqaB | BGC0001142 | NRP | 27.0 | 46.7 | 156.0 | 2.2e-37 |
| AOC89000.1 | putative\_nonribosomal\_peptide\_synthetase | BGC0001652 | NRP | 27.0 | 48.1 | 156.0 | 2.2e-37 |
| AGN74898.1 | nonribosomal\_peptide\_synthetase | BGC0000459 | NRP:Cyclic depsipeptide + Polyketide:Trans-AT type I | 28.0 | 40.3 | 156.0 | 3.7e-37 |
| ABC34483.1 | nonribosomal\_peptide\_synthetase,\_putative | BGC0000961 | NRP + Polyketide | 26.0 | 50.3 | 156.0 | 3.7e-37 |
| AAY42398.1 | Nonribosomal\_peptide\_synthetase | BGC0001000 | NRP:Lipopeptide + Polyketide:Modular type I | 27.0 | 36.0 | 155.0 | 4.8e-37 |
| AQX14493.1 | monobactam\_NRPS\_scaffold\_2 | BGC0001672 | NRP | 23.0 | 47.0 | 155.0 | 4.8e-37 |
| XP\_003044554.1 | hypothetical\_protein | BGC0001768 | NRP | 26.0 | 53.8 | 155.0 | 6.3e-37 |
| XP\_020057667.1 | hypothetical\_protein | BGC0001718 | NRP | 26.0 | 47.7 | 155.0 | 8.3e-37 |
| AGA37269.1 | NRPS | BGC0000819 | NRP + Alkaloid | 22.0 | 75.6 | 154.0 | 1.1e-36 |
| AKP45396.1 | CysH | BGC0001413 | NRP | 30.0 | 34.4 | 154.0 | 1.1e-36 |
| BAH23995.1 | nonribosomal\_peptide\_synthetase | BGC0000356 | NRP + Alkaloid | 26.0 | 52.8 | 154.0 | 1.4e-36 |
| AQZ42163.1 | putative\_nonribosomal\_peptide\_synthase | BGC0001820 | NRP | 26.0 | 56.2 | 154.0 | 1.4e-36 |
| AAP92497.1 | nonribosomal\_peptide\_synthetase | BGC0000458 | NRP | 26.0 | 48.1 | 153.0 | 3.1e-36 |
| OAL11435.1 | non-ribosomal\_peptide\_synthetase | BGC0001570 | NRP | 28.0 | 46.7 | 152.0 | 4.1e-36 |
| AAW03330.1 | CtaG | BGC0000982 | NRP + Polyketide | 29.0 | 30.8 | 151.0 | 7e-36 |
| AYJ71713.1 | non-ribosomal\_peptide\_synthetase | BGC0001942 | NRP + Polyketide | 26.0 | 49.0 | 151.0 | 7e-36 |
| EAU36744.1 | predicted\_protein | BGC0000292 | NRP | 27.0 | 47.5 | 151.0 | 9.1e-36 |
| XP\_002379984.1 | nonribosomal\_peptide\_synthase,\_putative | BGC0001621 | NRP | 26.0 | 50.4 | 151.0 | 9.1e-36 |
| BAW27672.1 | NRPS(A-T-C) | BGC0001764 | NRP | 27.0 | 45.6 | 151.0 | 9.1e-36 |
| EAL92291.2 | nonribosomal\_peptide\_synthtease | BGC0000372 | NRP | 26.0 | 46.4 | 151.0 | 1.2e-35 |
| AGA37267.1 | NRPS | BGC0000816 | NRP + Alkaloid | 25.0 | 49.7 | 151.0 | 1.2e-35 |
| CAJ76298.1 | putative\_hybrid\_polyketide-non-ribosomal\_peptide\_synthetase | BGC0000972 | NRP + Polyketide:Modular type I + Polyketide:Trans-AT type I | 25.0 | 50.7 | 150.0 | 1.6e-35 |
| ACR78148.1 | BSLS | BGC0000312 | NRP | 30.0 | 37.5 | 149.0 | 2.7e-35 |
| CAE52342.1 | non-ribosomal\_peptide\_synthase | BGC0001088 | NRP + Polyketide | 27.0 | 37.3 | 149.0 | 2.7e-35 |
| AAY42397.1 | Nonribosomal\_peptide\_synthetase | BGC0001000 | NRP:Lipopeptide + Polyketide:Modular type I | 28.0 | 36.9 | 149.0 | 3.5e-35 |
| CCG06115.1 | hybrid\_NRPS/PKS | BGC0001543 | Polyketide | 28.0 | 33.8 | 149.0 | 3.5e-35 |
| CAJ76289.1 | putative\_hybrid\_non-ribosomal\_peptide-polyketide\_synthetase | BGC0000972 | NRP + Polyketide:Modular type I + Polyketide:Trans-AT type I | 26.0 | 35.1 | 149.0 | 4.5e-35 |
| AGZ15455.1 | non-ribosomal\_peptide\_synthetase | BGC0001036 | NRP + Polyketide | 28.0 | 40.1 | 149.0 | 4.5e-35 |
| WP\_016638466.1 | amino\_acid\_adenylation\_domain-containing\_protein | BGC0001519 | NRP + Polyketide | 29.0 | 40.8 | 149.0 | 4.5e-35 |
| EAL89049.1 | nonribosomal\_peptide\_synthetase | BGC0000355 | NRP | 25.0 | 52.2 | 148.0 | 7.7e-35 |
| YP\_856987.1 | dimodular\_nonribosomal\_peptide\_synthetase | BGC0001502 | NRP | 29.0 | 39.4 | 147.0 | 1.7e-34 |
| EWM63007.1 | linear\_gramicidin\_synthetase\_LgrC | BGC0001328 | NRP:Cyclic depsipeptide + Polyketide:Modular type I | 29.0 | 36.9 | 146.0 | 2.3e-34 |
| ACI30655.1 | BEAS\_beauvericin\_nonribosomal\_cyclodepsipeptide\_synthetase | BGC0000313 | NRP | 29.0 | 37.4 | 146.0 | 3.8e-34 |
| BAR73008.1 | putative\_ATP-dependent\_b-aminoacyl-ACP\_synthetase | BGC0001194 | Polyketide | 27.0 | 41.4 | 146.0 | 3.8e-34 |
| CRG85572.1 | nonribosomal\_peptide\_synthase,\_putative | BGC0001402 | NRP | 24.0 | 52.4 | 145.0 | 5e-34 |
| AMK92578.1 | oxidoreductase | BGC0001815 | Polyketide | 38.0 | 20.3 | 145.0 | 6.6e-34 |
| ABX00600.1 | LmbC | BGC0000907 | Other | 28.0 | 40.5 | 144.0 | 1.1e-33 |
| CBF73453.1 | nonribosomal\_peptide\_synthase,\_putative\_(JCVI) | BGC0001515 | NRP | 26.0 | 49.1 | 143.0 | 1.9e-33 |
| ACO94461.1 | putative\_AMP-dependent\_acyl-CoA\_synthetase/ligase | BGC0000029 | Polyketide:Modular type I | 26.0 | 41.0 | 143.0 | 2.5e-33 |
| AFV52191.1 | NRPS\_adenylation\_domain | BGC0000081 | Polyketide:Iterative type I + Polyketide:Enediyne type I | 28.0 | 35.6 | 143.0 | 2.5e-33 |
| BAO84864.1 | putative\_non-ribosomal\_peptide\_synthetase | BGC0000414 | NRP | 25.0 | 49.1 | 143.0 | 2.5e-33 |
| ctg1\_orf003 |  | BGC0000334 | NRP | 25.0 | 48.7 | 143.0 | 3.3e-33 |
| EAL88817.1 | nonribosomal\_peptide\_synthase\_GliP | BGC0000361 | NRP | 27.0 | 48.2 | 143.0 | 3.3e-33 |
| ADQ74618.1 | amino\_acid\_adenylation\_protein | BGC0000921 | Polyketide + NRP + Other:Shikimate-derived | 25.0 | 47.4 | 143.0 | 3.3e-33 |
| ACZ66258.1 | APS1 | BGC0000304 | NRP | 24.0 | 50.4 | 142.0 | 4.2e-33 |
| CAN89664.1 | putative\_amino\_acid\_adenylating\_enzyme | BGC0001070 | NRP + Polyketide:Modular type I + Polyketide:Trans-AT type I | 29.0 | 40.8 | 141.0 | 7.2e-33 |
| BAE56606.1 |  | BGC0001123 | NRP | 26.0 | 46.6 | 141.0 | 7.2e-33 |
| ANY94451.1 | A-domain\_type\_II\_peptide\_synthetase | BGC0001584 | Polyketide | 29.0 | 41.2 | 141.0 | 7.2e-33 |
| DAB41661.1 | nonribosomal\_peptide\_synthetase | BGC0001585 | Alkaloid | 26.0 | 47.9 | 141.0 | 7.2e-33 |
| AGQ43600.1 | HC-toxin\_synthetase | BGC0001166 | NRP | 25.0 | 49.5 | 141.0 | 9.5e-33 |
| AAS98783.1 | polyketide\_synthase/nonribosomal\_peptide\_synthase\_hybrid | BGC0001001 | NRP + Polyketide | 33.0 | 21.9 | 141.0 | 1.2e-32 |
| AAF01762.1 | AM-toxin\_synthetase | BGC0001261 | NRP | 25.0 | 52.4 | 141.0 | 1.2e-32 |
| AAC83656.1 | dihydroaeruginoic\_acid\_synthetase | BGC0000412 | NRP | 28.0 | 37.1 | 139.0 | 2.8e-32 |
| ALU98443.1 | hypothetical\_protein | BGC0001397 | NRP + Polyketide | 27.0 | 41.7 | 139.0 | 2.8e-32 |
| XP\_011325838.1 | hypothetical\_protein | BGC0001545 | NRP | 26.0 | 49.7 | 139.0 | 2.8e-32 |
| TXD00261.1 | AMP-binding\_protein | BGC0001877 | Polyketide | 26.0 | 50.1 | 139.0 | 2.8e-32 |
| AGC83576.1 | NRPS | BGC0000818 | NRP | 23.0 | 52.3 | 139.0 | 4.7e-32 |
| WP\_051462298.1 | non-ribosomal\_peptide\_synthetase | BGC0001873 | NRP:Lipopeptide | 26.0 | 50.0 | 138.0 | 6.1e-32 |
| AAL06681.1 | A-domain\_type\_II\_peptide\_synthetase | BGC0000965 | Polyketide:Iterative type I + Polyketide:Enediyne type I | 27.0 | 41.8 | 138.0 | 8e-32 |
| BAU50944.1 | nonribosomal\_peptide\_synthetase | BGC0001379 | NRP | 27.0 | 45.7 | 137.0 | 1.4e-31 |
| CCE28989.1 | non-ribosomal\_peptide\_synthetase | BGC0001365 | NRP | 26.0 | 46.9 | 137.0 | 1.8e-31 |
| AFK57220.1 | DidI | BGC0000985 | Polyketide + NRP:Cyclic depsipeptide | 25.0 | 48.8 | 136.0 | 3e-31 |
| ctg1\_orf4 |  | BGC0001329 | Polyketide + NRP:Cyclic depsipeptide | 35.0 | 21.6 | 136.0 | 3e-31 |
| AWR88409.1 | putative\_AMP-dependent\_synthetase\_and\_ligase | BGC0001522 | Polyketide | 26.0 | 41.0 | 136.0 | 3e-31 |
| ADM34138.1 | non-ribosomal\_peptide\_synthetase | BGC0001084 | NRP + Terpene + Alkaloid | 23.0 | 47.8 | 136.0 | 4e-31 |
| BAO66533.1 | nonribosomal\_peptide\_synthase | BGC0000042 | Polyketide | 26.0 | 42.3 | 135.0 | 5.2e-31 |
| ctg1\_orf000000 |  | BGC0000901 | Other | 26.0 | 50.2 | 135.0 | 5.2e-31 |
| CBF87869.1 | nonribosomal\_peptide\_synthase,\_putative\_(Eurofung) | BGC0001699 | Polyketide | 26.0 | 47.5 | 135.0 | 6.8e-31 |
| AQM58287.1 | non-ribosomal\_peptide\_synthase | BGC0001816 | NRP + Polyketide | 24.0 | 47.8 | 135.0 | 6.8e-31 |
| sipL1 | AMP-dependent\_synthetase\_and\_ligase | BGC0001452 | Polyketide | 26.0 | 40.9 | 134.0 | 1.5e-30 |
| MAA\_10043 | non-ribosomal\_peptide\_synthetase | BGC0000337 | NRP | 26.0 | 37.9 | 133.0 | 2e-30 |
| ctg1\_orf18 |  | BGC0001329 | Polyketide + NRP:Cyclic depsipeptide | 32.0 | 32.1 | 133.0 | 2e-30 |
| WP\_032920652.1 | amino\_acid\_adenylation\_domain-containing\_protein | BGC0001813 | NRP | 27.0 | 42.4 | 133.0 | 2e-30 |
| ACO94489.1 | putative\_AMP-dependent\_acyl-CoA\_synthetase/ligase | BGC0000097 | Polyketide:Modular type I | 25.0 | 41.0 | 133.0 | 2.6e-30 |
| OAL11436.1 | hypothetical\_protein | BGC0001570 | NRP | 28.0 | 41.7 | 133.0 | 2.6e-30 |
| ADM46356.1 | polyketide\_synthase | BGC0000106 | Polyketide | 27.0 | 37.6 | 132.0 | 5.7e-30 |
| BAP16684.1 | nonribosomal\_peptide\_synthetase | BGC0000376 | NRP | 24.0 | 47.8 | 132.0 | 5.7e-30 |
| ABV56582.1 | KtzB | BGC0000378 | NRP | 27.0 | 40.2 | 132.0 | 5.7e-30 |
| ABV97151.1 | AMP-dependent\_synthetase\_and\_ligase | BGC0000137 | Polyketide | 25.0 | 49.7 | 131.0 | 7.5e-30 |
| CQR60497.1 | Polyketide\_synthase,\_type\_I,\_modules:\_loading,\_1,\_2\_and\_3 | BGC0001287 | Polyketide | 27.0 | 40.8 | 131.0 | 9.8e-30 |
| QBF51769.1 | type\_I\_polyketide\_synthase | BGC0001856 | Polyketide:Modular type I | 25.0 | 52.0 | 131.0 | 9.8e-30 |
| ACO94492.1 | NRPS\_adenylation\_domain\_protein | BGC0000097 | Polyketide:Modular type I | 26.0 | 40.1 | 131.0 | 1.3e-29 |
| ALA09365.1 | AMP-dependent\_synthetase\_and\_ligase | BGC0001303 | Polyketide | 26.0 | 40.9 | 130.0 | 1.7e-29 |
| AKD43754.1 | HerJ | BGC0001349 | NRP + Polyketide | 26.0 | 39.3 | 129.0 | 2.8e-29 |
| AKD43757.1 | HerL | BGC0001349 | NRP + Polyketide | 26.0 | 40.0 | 129.0 | 3.7e-29 |
| WP\_063738219.1 | amino\_acid\_adenylation\_domain-containing\_protein | BGC0002010 | NRP + Polyketide | 26.0 | 41.7 | 129.0 | 3.7e-29 |
| CCB53264.1 | non-ribosomal\_peptide\_synthetase | BGC0001393 | NRP | 23.0 | 45.6 | 129.0 | 4.9e-29 |
| ERG36783.1 | phenylalanine\_racemase | BGC0000299 | NRP | 29.0 | 32.0 | 128.0 | 6.3e-29 |
| ABB90279.1 | non-ribosomal\_peptide\_synthetase | BGC0001057 | NRP + Polyketide | 25.0 | 47.0 | 128.0 | 6.3e-29 |
| EHK22005.1 | putative\_non-ribosomal\_peptide\_synthetase\_GliP | BGC0001609 | NRP | 25.0 | 48.5 | 127.0 | 1.4e-28 |
| ABP55217.1 | AMP-dependent\_synthetase\_and\_ligase | BGC0000142 | Polyketide | 25.0 | 40.9 | 127.0 | 1.8e-28 |
| ctg1\_orf10 |  | BGC0000321 | NRP | 24.0 | 52.3 | 125.0 | 7e-28 |
| BAP34718.1 | AMP-dependent\_synthetase\_and\_ligase | BGC0000078 | Polyketide | 30.0 | 36.8 | 124.0 | 9.2e-28 |
| ANY57892.1 | PenN | BGC0001372 | Terpene | 24.0 | 50.8 | 124.0 | 9.2e-28 |
| ATY46587.1 | polyketide\_synthase | BGC0001666 | Polyketide | 25.0 | 42.9 | 124.0 | 1.6e-27 |
| AQM58288.1 | non-ribosomal\_peptide\_synthase | BGC0001816 | NRP + Polyketide | 24.0 | 55.4 | 121.0 | 1e-26 |
| ABA59548.1 | NRPS | BGC0000453 | NRP:Cyclic depsipeptide | 25.0 | 47.3 | 119.0 | 3.8e-26 |
| BAP34707.1 | AMP-dependent\_synthetase\_and\_ligase | BGC0000078 | Polyketide | 25.0 | 41.9 | 119.0 | 5e-26 |
| AAC01710.1 | RifA | BGC0000136 | Polyketide | 25.0 | 50.9 | 119.0 | 5e-26 |
| AAC32048.1 | FxbC | BGC0000351 | NRP | 28.0 | 25.2 | 118.0 | 8.6e-26 |
| ARB50207.1 | lysergyl\_peptide\_synthetase\_21 | BGC0001573 | Alkaloid | 25.0 | 47.4 | 118.0 | 1.1e-25 |
| AUS29494.1 | non-ribosomal\_peptide\_synthetase | BGC0001030 | NRP + Polyketide | 26.0 | 41.3 | 117.0 | 1.5e-25 |
| AAN85530.1 | nonribosomal\_peptide\_synthetase,\_discrete\_adenylation\_domain | BGC0001101 | NRP + Polyketide:Modular type I + Polyketide:Trans-AT type I | 27.0 | 40.0 | 117.0 | 1.5e-25 |
| ADY16697.1 | TqaA | BGC0001142 | NRP | 24.0 | 49.2 | 117.0 | 1.5e-25 |
| AWH12671.1 | RmpA1 | BGC0001759 | Polyketide | 25.0 | 51.3 | 117.0 | 1.5e-25 |
| AHH25595.1 | PKS | BGC0000957 | NRP + Polyketide | 27.0 | 34.4 | 115.0 | 5.6e-25 |
| sipL7 | AMP-dependent\_synthetase\_and\_ligase | BGC0001452 | Polyketide | 27.0 | 36.9 | 115.0 | 5.6e-25 |
| AET13879.1 | epichloenin\_A\_synthetase | BGC0001251 | NRP | 28.0 | 42.7 | 115.0 | 7.3e-25 |
| CBA63665.1 | 2,3-dihydro-2,3-dihydroxybenzoate\_dehydrogenase | BGC0000368 | NRP | 30.0 | 20.5 | 114.0 | 1.2e-24 |
| AET13875.1 | epichloenin\_A\_synthetase | BGC0001250 | NRP | 28.0 | 42.6 | 114.0 | 1.2e-24 |
| CAP93139.1 | cyclic\_hydrophobic\_tetrapeptide | BGC0000357 | NRP:Cyclic depsipeptide | 23.0 | 53.6 | 113.0 | 2.1e-24 |
| CCP20047.1 | divK\_protein | BGC0001119 | Polyketide:Modular type I | 25.0 | 41.5 | 113.0 | 2.8e-24 |
| ABM91454.1 | lysergyl\_peptide\_synthetase\_LpsB | BGC0000348 | NRP | 23.0 | 53.5 | 113.0 | 3.6e-24 |
| EXU96269.1 | nonribosomal\_peptide\_synthetase,\_serinocyclin\_synthetase\_NPS1 | BGC0001240 | NRP | 23.0 | 49.5 | 111.0 | 8e-24 |
| AWR88407.1 | putative\_AMP-dependent\_synthetase\_and\_ligase | BGC0001522 | Polyketide | 27.0 | 36.8 | 109.0 | 3.1e-23 |
| ACO94464.1 | NRPS\_adenylation\_domain\_protein | BGC0000029 | Polyketide:Modular type I | 25.0 | 38.8 | 109.0 | 4e-23 |
| ALE27507.1 | 2,3-dihydro-3-hydroxyanthranilic\_acid\_dehydrogenase | BGC0001292 | Other | 30.0 | 20.1 | 109.0 | 5.2e-23 |
| ANH11404.1 | SceI | BGC0001908 | Polyketide | 24.0 | 42.7 | 109.0 | 5.2e-23 |
| ABP55216.1 | AMP-dependent\_synthetase\_and\_ligase | BGC0000142 | Polyketide | 26.0 | 37.6 | 108.0 | 6.8e-23 |
| ADC79613.1 | BafX | BGC0000028 | Polyketide:Modular type I | 24.0 | 38.0 | 106.0 | 2.6e-22 |
| BAO66530.1 | nonribosomal\_peptide\_synthase | BGC0000042 | Polyketide | 25.0 | 36.2 | 105.0 | 7.5e-22 |
| CAH10130.1 | feruloyl-CoA\_synthetase | BGC0000268 | Polyketide | 25.0 | 40.4 | 105.0 | 7.5e-22 |
| ADY76674.1 | non-ribosomal\_peptide\_synthetase | BGC0000950 | NRP:Uridylpeptide + Other:Nucleoside | 24.0 | 41.5 | 104.0 | 9.8e-22 |
| BAD08370.1 | non-ribosomal\_peptide\_synthetase | BGC0000167 | Polyketide | 23.0 | 41.8 | 103.0 | 2.2e-21 |
| OAP25804.1 | Surfactin\_synthase\_subunit\_1 | BGC0001658 | Polyketide | 24.0 | 41.6 | 103.0 | 2.9e-21 |
| CAJ45636.1 | 2,3-dihydroxybenzoate-AMP\_ligase | BGC0000454 | NRP | 24.0 | 41.8 | 102.0 | 3.7e-21 |
| CEF75881.1 |  | BGC0001600 | Polyketide | 26.0 | 26.5 | 102.0 | 3.7e-21 |
| AAX98210.1 | acyl\_CoA\_ligase | BGC0000052 | Polyketide | 25.0 | 37.5 | 101.0 | 8.3e-21 |
| ALA09366.1 | AMP-dependent\_synthetase\_and\_ligase | BGC0001303 | Polyketide | 25.0 | 37.2 | 101.0 | 8.3e-21 |
| ADN26257.1 | ATP-dependent\_adenylase | BGC0000951 | NRP | 24.0 | 37.0 | 100.0 | 2.4e-20 |
| AFV52187.1 | acyl-CoA\_synthetase | BGC0000081 | Polyketide:Iterative type I + Polyketide:Enediyne type I | 25.0 | 42.8 | 99.0 | 4.1e-20 |
| AAF86393.1 | FkbB | BGC0000994 | NRP + Polyketide | 24.0 | 41.7 | 99.0 | 5.4e-20 |
| ORC16617.1 | hypothetical\_protein | BGC0001341 | NRP | 26.0 | 30.6 | 98.0 | 9.2e-20 |
| BAR73011.1 | putative\_ATP-dependent\_aminoacyl-ACP\_synthetase | BGC0001194 | Polyketide | 24.0 | 36.8 | 97.0 | 1.6e-19 |
| ADI58647.1 | 5-Aminolevulinate\_CoA\_ligase | BGC0000187 | Polyketide:Type II | 24.0 | 38.4 | 96.0 | 4.6e-19 |
| AAO39104.1 | AdmJ | BGC0000956 | NRP:Beta-lactam + Polyketide:Type II | 23.0 | 33.8 | 96.0 | 4.6e-19 |
| ABJ90146.1 | MoeA4 | BGC0000805 | Saccharide | 24.0 | 38.1 | 93.0 | 3e-18 |
| CAP20363.1 | equibactin\_siderophore\_biosynthetic\_protein | BGC0000347 | NRP | 21.0 | 36.6 | 92.0 | 3.9e-18 |
| AAC68815.1 | FK506\_polyketide\_synthase | BGC0000353 | NRP | 24.0 | 41.5 | 92.0 | 6.6e-18 |
| AAF81733.1 | putative\_aryl-CoA\_ligase\_EncN | BGC0000220 | Polyketide:Type II | 24.0 | 39.7 | 91.0 | 1.1e-17 |
| ATY12791.1 | long-chain\_fatty\_acid--CoA\_ligase | BGC0001504 | Polyketide | 23.0 | 41.4 | 91.0 | 1.1e-17 |
| ACF35445.1 | mbcAI | BGC0000090 | Polyketide | 25.0 | 35.7 | 89.0 | 4.3e-17 |
| ADH04681.1 | non-ribosomal\_peptide\_synthetase | BGC0001344 | NRP + Polyketide | 25.0 | 42.6 | 88.0 | 7.3e-17 |
| AIL50191.1 | putative\_amide\_synthetase | BGC0000213 | Polyketide:Type II | 23.0 | 39.7 | 85.0 | 6.2e-16 |
| CCA65703.1 | anthranilate-CoA\_ligase | BGC0001343 | Polyketide | 23.0 | 35.8 | 85.0 | 8e-16 |
| BBA21082.1 | putative\_CoA\_ligase | BGC0001740 | NRP + Polyketide | 26.0 | 34.5 | 85.0 | 8e-16 |
| orf5 | benzoate-CoA\_ligase\_family\_protein | BGC0001420 | NRP:Cyclic depsipeptide + Polyketide:Iterative type I | 25.0 | 34.8 | 84.0 | 1.1e-15 |
| APZ78718.1 | benzoate-CoA\_ligase\_family\_protein | BGC0001420 | NRP:Cyclic depsipeptide + Polyketide:Iterative type I | 25.0 | 34.8 | 84.0 | 1.1e-15 |
| DAC74137.1 | AMP-dependent\_synthetase | BGC0002019 | Terpene | 24.0 | 39.6 | 84.0 | 1.8e-15 |
| ACA34358.1 | acyl-CoA\_ligase | BGC0001152 | Polyketide + NRP:Lipopeptide | 23.0 | 40.4 | 82.0 | 4e-15 |
| ASZ00147.1 | polyketide\_synthase | BGC0001785 | Polyketide | 27.0 | 25.3 | 82.0 | 4e-15 |
| CAF05650.1 | TubE\_protein | BGC0001053 | NRP + Polyketide | 23.0 | 49.9 | 81.0 | 1.5e-14 |
| AGY30674.1 | Ann1 | BGC0001298 | Polyketide | 24.0 | 37.6 | 80.0 | 2e-14 |
| OJF16272.1 | AceP1 | BGC0001491 | Polyketide | 23.0 | 47.8 | 80.0 | 2.6e-14 |
| AAN32979.1 | BarE | BGC0000962 | NRP + Polyketide:Modular type I | 23.0 | 29.8 | 79.0 | 3.4e-14 |
| ctg1\_orf00001 |  | BGC0000901 | Other | 21.0 | 36.7 | 78.0 | 9.8e-14 |
| ADI58645.1 | Amide\_synthase | BGC0000187 | Polyketide:Type II | 24.0 | 41.8 | 76.0 | 3.7e-13 |
| AHH34188.1 | fatty\_acyl\_ACP\_ligase | BGC0001162 | Polyketide:Modular type I | 23.0 | 37.0 | 73.0 | 3.2e-12 |
| AHH34187.1 | fatty\_acyl\_ACP\_ligase | BGC0001161 | Polyketide:Modular type I | 24.0 | 37.2 | 72.0 | 5.4e-12 |
| AGD80620.1 | non-ribosomal\_peptide\_synthetase | BGC0000394 | NRP | 27.0 | 22.9 | 72.0 | 7e-12 |
| EGJ35087.1 | Fatty-acyl-ACP\_ligase | BGC0001163 | Polyketide:Modular type I | 23.0 | 36.5 | 72.0 | 7e-12 |
| CAD29795.1 | peptide\_synthetase | BGC0001015 | NRP + Polyketide | 22.0 | 38.5 | 70.0 | 2.1e-11 |
| CBZ42143.1 | putative\_non-ribosomal\_peptide\_synthetase | BGC0001117 | NRP | 27.0 | 21.6 | 69.0 | 4.6e-11 |
| MAA\_10036 | nonribosomal\_peptide\_synthase\_GliP-like,\_putative | BGC0000337 | NRP | 25.0 | 21.6 | 69.0 | 6e-11 |
| AQT01384.1 | SgnS0 | BGC0001690 | Polyketide | 23.0 | 50.7 | 68.0 | 7.8e-11 |
| AXN93598.1 | PuwC | BGC0001952 | NRP | 21.0 | 39.0 | 67.0 | 2.3e-10 |
| AQH32483.1 | hybrid\_peptide\_synthetase/polyketide\_synthase | BGC0001667 | NRP + Polyketide | 23.0 | 40.6 | 66.0 | 5.1e-10 |
| CAE02600.1 | p-nitrobenzoate\_CoA\_ligase | BGC0000024 | Polyketide:Modular type I | 24.0 | 36.9 | 64.0 | 1.1e-09 |
| ADX66470.1 | ScnS0 | BGC0000108 | Polyketide | 23.0 | 50.5 | 63.0 | 2.5e-09 |
| ABJ90147.1 | MoeB4 | BGC0000805 | Saccharide | 22.0 | 23.4 | 63.0 | 3.3e-09 |
| AIL50174.1 | putative\_3,4-AHBA\_carboxyl\_group\_adenylase | BGC0000213 | Polyketide:Type II | 23.0 | 34.5 | 62.0 | 5.6e-09 |
| OAL11476.1 | non-ribosomal\_peptide\_synthetase/polyketide\_synthase | BGC0001570 | NRP | 26.0 | 23.1 | 59.0 | 4.7e-08 |
| ABB05101.1 | LipNrps | BGC0001003 | NRP:Lipopeptide + Polyketide:Modular type I + Saccharide:Hybrid/tailoring | 26.0 | 22.2 | 58.0 | 1.1e-07 |
| AAG13917.1 | megalomicin\_6-deoxyerythronolide\_B\_synthase\_1 | BGC0000092 | Polyketide | 25.0 | 20.6 | 55.0 | 6.8e-07 |
