## Supplementary Results for "A draft genome of the ascomycotal fungal species *Pseudopithomyces maydicus* (family *Didymosphaeriaceae*)": input.path1.gene223_mibig_hits.html

| MIBiG Protein | Description | MIBiG Cluster | MiBiG Product | % ID | % Coverage | BLAST Score | E-value |
| --- | --- | --- | --- | --- | --- | --- | --- |
| KGO40484.1 | Major\_facilitator\_superfamily\_domain,\_general\_substrate\_transporter | BGC0001205 | Polyketide | 30.0 | 94.4 | 191.0 | 3.1e-48 |
