## Supplementary Results for "A draft genome of the ascomycotal fungal species *Pseudopithomyces maydicus* (family *Didymosphaeriaceae*)": input.path1.gene224_mibig_hits.html

| MIBiG Protein | Description | MIBiG Cluster | MiBiG Product | % ID | % Coverage | BLAST Score | E-value |
| --- | --- | --- | --- | --- | --- | --- | --- |
| AGN71623.1 | hydroxylase | BGC0000027 | Polyketide:Iterative type I | 40.0 | 42.2 | 183.0 | 1.3e-45 |
| EHA28235.1 | hypothetical\_protein | BGC0001143 | Polyketide | 39.0 | 42.9 | 171.0 | 3.9e-42 |
| EAQ86391.1 | hypothetical\_protein | BGC0001405 | Polyketide | 36.0 | 41.7 | 161.0 | 5.3e-39 |
| AWM95796.1 | salicylate\_hydroxylase | BGC0001827 | Polyketide | 37.0 | 43.7 | 154.0 | 6.4e-37 |
| QCO93109.1 | monooxygenase | BGC0001977 | Other | 35.0 | 43.2 | 153.0 | 1.4e-36 |
| EAU32816.1 | conserved\_hypothetical\_protein | BGC0000160 | Polyketide | 38.0 | 42.7 | 152.0 | 1.9e-36 |
| EED18000.1 | FAD\_oxygenase | BGC0000154 | Polyketide:Iterative type I | 37.0 | 42.2 | 148.0 | 4.6e-35 |
| CAP95403.1 |  | BGC0001404 | Polyketide | 35.0 | 41.0 | 141.0 | 4.3e-33 |
| CBF83145.1 | conserved\_hypothetical\_protein | BGC0001722 | Polyketide | 33.0 | 43.5 | 139.0 | 1.3e-32 |
| EAA65601.1 | hypothetical\_protein | BGC0000022 | Polyketide | 31.0 | 42.0 | 132.0 | 2.6e-30 |
| EWM63055.1 | monooxygenase | BGC0000679 | Terpene | 28.0 | 41.3 | 104.0 | 5.9e-22 |
| CAB38889.1 | Hexenoyl-S-ACP\_Monooxygenase\_(hcmO) | BGC0000315 | NRP:Ca+-dependent lipopeptide | 33.0 | 41.8 | 99.0 | 1.4e-20 |
| ACO31289.1 | PtmB3 | BGC0001898 | Terpene | 29.0 | 42.2 | 99.0 | 2.5e-20 |
| ADD82995.1 | PtnB3 | BGC0001156 | Terpene | 28.0 | 42.2 | 97.0 | 9.4e-20 |
| AXO35181.1 | putative\_n-hydroxybenzoate\_hydroxylase | BGC0001848 | Other | 28.0 | 40.5 | 96.0 | 2.1e-19 |
| AWF83809.1 | Tropone\_2-monooxygenase | BGC0001935 | Other | 28.0 | 41.3 | 90.0 | 1.1e-17 |
| ALV82350.1 | salicylate\_hydroxylase | BGC0001370 | NRP | 30.0 | 41.0 | 86.0 | 1.7e-16 |
| ATV82119.1 | hydroxylase | BGC0001909 | Polyketide | 27.0 | 48.0 | 82.0 | 3.1e-15 |
| RZB16712.1 | FAD-binding\_protein | BGC0001850 | Other:Shikimate-derived | 25.0 | 41.2 | 70.0 | 1.2e-11 |
| CDF96615.1 | FAD-dependent\_Baeyer—Villiger\_monooxygenase | BGC0001149 | NRP:Lipopeptide + Saccharide:Hybrid/tailoring | 26.0 | 35.4 | 63.0 | 1.1e-09 |
| AEE65479.1 | oxidoreductase | BGC0000223 | Polyketide:Type II | 29.0 | 31.3 | 57.0 | 6.3e-08 |
| ATL73025.1 | oxidoreductase | BGC0001807 | NRP + Polyketide | 26.0 | 41.3 | 57.0 | 1.1e-07 |
| ARE67860.1 | AbsH3 | BGC0001492 | Polyketide | 25.0 | 39.8 | 56.0 | 1.8e-07 |
| BAJ09785.1 | oxidase/Diels-Alderase | BGC0000146 | Polyketide | 28.0 | 24.1 | 53.0 | 1.2e-06 |
