## Supplementary Results for "A draft genome of the ascomycotal fungal species *Pseudopithomyces maydicus* (family *Didymosphaeriaceae*)": input.path1.gene362_mibig_hits.html

| MIBiG Protein | Description | MIBiG Cluster | MiBiG Product | % ID | % Coverage | BLAST Score | E-value |
| --- | --- | --- | --- | --- | --- | --- | --- |
| EPE34340.1 | polyketide\_synthase | BGC0001035 | Polyketide + NRP + Other:Aminocoumarin | 42.0 | 120.1 | 1517.0 | 0.0 |
| ACB12550.1 | Fum1 | BGC0000063 | Polyketide | 38.0 | 122.1 | 1357.0 | 0.0 |
| AAD43562.2 | Fum1p | BGC0000062 | Polyketide | 38.0 | 123.0 | 1336.0 | 0.0 |
| OAQ83765.1 | KR\_domain-containing\_protein | BGC0001358 | Polyketide | 40.0 | 112.7 | 1314.0 | 0.0 |
| OAQ83760.1 | polyketide\_synthase | BGC0001358 | Polyketide | 37.0 | 121.9 | 1274.0 | 0.0 |
| ATQ39432.1 | PKS | BGC0001565 | NRP | 35.0 | 122.1 | 1164.0 | 0.0 |
| AMY15068.1 | hexaketide\_synthase\_MF-SQHKS | BGC0001339 | Polyketide:Iterative type I | 34.0 | 118.6 | 1135.0 | 0.0 |
| gene4 |  | BGC0001907 | Polyketide | 34.0 | 107.9 | 1046.0 | 5.4e-305 |
| RWQ92174.1 | KR\_domain-containing\_protein | BGC0002030 | Polyketide | 34.0 | 112.1 | 1007.0 | 2.1e-293 |
| ABA02240.1 | polyketide\_synthase | BGC0000098 | Polyketide | 28.0 | 122.7 | 749.0 | 1e-215 |
| BAC20566.1 | polyketide\_synthase | BGC0000039 | Polyketide | 28.0 | 123.3 | 742.0 | 2.2e-213 |
| BBG28498.1 | putative\_polyketide\_synthase | BGC0001913 | Polyketide | 29.0 | 125.6 | 737.0 | 4.1e-212 |
| AAD34559.1 | polyketide\_synthase | BGC0000088 | Polyketide | 28.0 | 123.1 | 732.0 | 1.3e-210 |
| EHA28244.1 | hypothetical\_protein | BGC0001143 | Polyketide | 28.0 | 123.2 | 722.0 | 1.8e-207 |
| CCT75967.1 | polyketide\_synthase | BGC0001606 | Polyketide | 27.0 | 123.2 | 706.0 | 1.3e-202 |
| BAD83684.1 | PKSN\_polyketide\_synthase\_for\_alternapyrone\_biosynthesis | BGC0000012 | Polyketide | 27.0 | 124.6 | 698.0 | 2.7e-200 |
| EAA65604.1 | hypothetical\_protein | BGC0000022 | Polyketide | 27.0 | 124.6 | 673.0 | 7.2e-193 |
| CBF87072.1 | polyketide\_synthase,\_putative\_(Eurofung) | BGC0001290 | NRP | 28.0 | 123.4 | 658.0 | 2.4e-188 |
| ANF07288.1 | hrPKS | BGC0001340 | Polyketide:Iterative type I | 28.0 | 126.5 | 643.0 | 1.4e-183 |
| AIA58899.1 | HRPKS | BGC0001141 | Polyketide:Iterative type I | 27.0 | 121.2 | 635.0 | 2.9e-181 |
| CAP95405.1 |  | BGC0001404 | Polyketide | 26.0 | 125.0 | 634.0 | 3.7e-181 |
| BAJ09789.1 | polyketide\_synthase | BGC0000146 | Polyketide | 29.0 | 103.2 | 631.0 | 5.4e-180 |
| ASK38717.1 | polyketide\_synthase | BGC0001557 | Polyketide | 26.0 | 125.1 | 627.0 | 5.9e-179 |
| ABB90283.1 | polyketide\_synthase | BGC0001057 | NRP + Polyketide | 27.0 | 120.6 | 626.0 | 1.3e-178 |
| EAL85129.1 | polyketide\_synthase | BGC0001067 | Terpene + Polyketide:Iterative type I | 26.0 | 125.1 | 618.0 | 3.6e-176 |
| EAA36364.1 | hypothetical\_protein | BGC0001697 | Polyketide | 27.0 | 120.7 | 609.0 | 1.7e-173 |
| EHA19289.1 | hypothetical\_protein | BGC0001124 | Polyketide | 27.0 | 122.9 | 595.0 | 2.5e-169 |
| ACD39758.1 | reducing\_polyketide\_synthase | BGC0000076 | Polyketide | 26.0 | 121.3 | 591.0 | 4.7e-168 |
| ACD39767.1 | reducing\_polyketide\_synthase | BGC0000077 | Polyketide | 26.0 | 121.3 | 591.0 | 4.7e-168 |
| AHV78245.1 | LasS1 | BGC0001245 | Polyketide | 27.0 | 122.2 | 586.0 | 1.5e-166 |
| AMY15057.1 | tetraketide\_synthase\_MF-SQTKS | BGC0001339 | Polyketide:Iterative type I | 26.0 | 108.3 | 585.0 | 3.4e-166 |
| EWG54266.1 | hypothetical\_protein | BGC0001190 | Polyketide | 26.0 | 123.1 | 581.0 | 4.9e-165 |
| AHV78252.1 | ResS1 | BGC0001246 | Polyketide | 26.0 | 122.9 | 580.0 | 8.3e-165 |
| ATZ45185.1 | Bcboa9 | BGC0001892 | Polyketide | 26.0 | 121.0 | 580.0 | 8.3e-165 |
| EAQ86385.1 | hypothetical\_protein | BGC0001405 | Polyketide | 26.0 | 118.8 | 578.0 | 3.2e-164 |
| ACD39774.1 | reducing\_polyketide\_synthase | BGC0000134 | Polyketide | 26.0 | 121.5 | 568.0 | 4.3e-161 |
| ACS68554.1 | hybrid\_PKS-NRPS\_protein | BGC0001026 | NRP + Polyketide | 26.0 | 119.6 | 565.0 | 2.8e-160 |
| AKL78824.1 | GLPKS3 | BGC0001187 | NRP:Lipopeptide + Polyketide:Iterative type I | 26.0 | 120.0 | 563.0 | 1.1e-159 |
| EED49862.1 | hybrid\_PKS/NRPS\_enzyme,\_putative | BGC0001445 | NRP + Polyketide:Iterative type I | 27.0 | 119.1 | 562.0 | 1.8e-159 |
| AGC95324.1 | CurS1 | BGC0000045 | Polyketide | 26.0 | 122.1 | 551.0 | 4.1e-156 |
| MAA\_10033 | polyketide\_synthase,\_putative | BGC0000337 | NRP | 33.0 | 63.5 | 551.0 | 4.1e-156 |
| BAD97694.1 | Aft9-1 | BGC0000003 | Polyketide | 25.0 | 121.3 | 548.0 | 4.6e-155 |
| AQM58285.1 | polyketide\_synthase | BGC0001816 | NRP + Polyketide | 26.0 | 125.1 | 548.0 | 4.6e-155 |
| ACZ57548.1 | polyketide\_synthase | BGC0000046 | Polyketide:Iterative type I | 26.0 | 119.8 | 545.0 | 2.3e-154 |
| ARP51711.1 | PKS-NRPS\_hybrid\_protein | BGC0001741 | NRP + Polyketide | 27.0 | 123.1 | 542.0 | 3.3e-153 |
| CCT72377.1 | probable\_polyketide\_synthase | BGC0001305 | Polyketide | 26.0 | 119.7 | 526.0 | 1.9e-148 |
| EHA52508.1 | mycocerosic\_acid\_synthase | BGC0001749 | Polyketide | 27.0 | 122.3 | 521.0 | 4.6e-147 |
| KKP00963.1 | fatty\_acid\_synthase\_S-acetyltransferase | BGC0001901 | Polyketide | 25.0 | 122.4 | 515.0 | 3.3e-145 |
| QBE85649.1 | BuaA | BGC0001978 | NRP + Polyketide | 26.0 | 120.3 | 508.0 | 4e-143 |
| CBF80487.1 | hybrid\_PKS-NRPS\_(Eurofung) | BGC0000959 | NRP + Polyketide:Iterative type I | 28.0 | 89.2 | 503.0 | 1.3e-141 |
| EAW09117.1 | hybrid\_NRPS/PKS\_enzyme,\_putative | BGC0000983 | NRP + Polyketide:Iterative type I | 27.0 | 87.6 | 463.0 | 1.5e-129 |
| KGO40478.1 | Acyl\_transferase/acyl\_hydrolase/lysophospholipase | BGC0001205 | Polyketide | 25.0 | 122.4 | 460.0 | 9.6e-129 |
| QCC63000.1 | BII-rafflesfungin\_polyketide\_synthase | BGC0001966 | NRP | 25.0 | 105.1 | 456.0 | 1.8e-127 |
| QBC19710.1 | TwmB | BGC0001954 | NRP + Polyketide | 26.0 | 120.1 | 453.0 | 2e-126 |
| AEE88280.1 | CurJ | BGC0000976 | NRP + Polyketide:Modular type I | 24.0 | 113.7 | 443.0 | 1.2e-123 |
| AAT70105.1 | CurJ | BGC0001165 | NRP + Polyketide:Modular type I | 24.0 | 113.7 | 443.0 | 1.2e-123 |
| ABA02239.1 | polyketide\_synthase | BGC0000098 | Polyketide | 28.0 | 88.7 | 439.0 | 3e-122 |
| BAZ95823.1 | PKS-NRPS\_hybrid\_cpaA | BGC0001563 | NRP + Polyketide | 25.0 | 120.6 | 439.0 | 3e-122 |
| QCS37521.1 | PyiS | BGC0001982 | NRP + Polyketide | 27.0 | 89.4 | 436.0 | 1.5e-121 |
| EPS29069.1 | hypothetical\_protein | BGC0001724 | NRP + Polyketide | 28.0 | 78.9 | 431.0 | 6.3e-120 |
| CAL69597.1 | PKS-NRPS | BGC0001049 | NRP + Polyketide:Iterative type I | 26.0 | 92.0 | 427.0 | 1.2e-118 |
| ADN43685.1 | DmbS | BGC0001136 | NRP + Polyketide:Iterative type I | 26.0 | 91.2 | 425.0 | 4.5e-118 |
| ACB46196.1 | polyketide\_synthase | BGC0000989 | NRP + Polyketide | 24.0 | 112.9 | 411.0 | 6.7e-114 |
| ADB12492.1 | EpoE | BGC0000990 | NRP + Polyketide | 24.0 | 113.2 | 410.0 | 1.5e-113 |
| AAF62884.1 | EpoE | BGC0000991 | NRP + Polyketide | 24.0 | 113.4 | 408.0 | 4.3e-113 |
| AAF26922.1 | polyketide\_synthase | BGC0000988 | NRP + Polyketide | 24.0 | 113.4 | 407.0 | 9.7e-113 |
| AQW44889.1 | polyketide\_synthase | BGC0001737 | NRP + Polyketide | 25.0 | 110.6 | 404.0 | 8.2e-112 |
| BAQ25466.1 | polyketide\_synthase | BGC0001280 | Polyketide | 25.0 | 90.8 | 394.0 | 6.5e-109 |
| AHA38200.1 | GphG | BGC0000069 | Polyketide | 25.0 | 112.7 | 394.0 | 8.5e-109 |
| BAJ14522.1 | polyketide\_synthase | BGC0001254 | Polyketide | 26.0 | 96.5 | 391.0 | 7.2e-108 |
| BAV32159.1 | polyketide\_synthase | BGC0001373 | Polyketide | 29.0 | 58.8 | 390.0 | 1.2e-107 |
| AFA26384.1 | polyketide\_synthase\_A | BGC0001874 | NRP + Polyketide | 27.0 | 79.5 | 389.0 | 2.7e-107 |
| AGC45624.1 | polyketide\_synthase | BGC0001394 | NRP + Polyketide | 24.0 | 111.7 | 371.0 | 5.9e-102 |
| ABM21569.1 | crpA | BGC0000975 | NRP + Polyketide | 23.0 | 114.7 | 368.0 | 8.5e-101 |
| APD26279.1 | PtmA | BGC0001726 | NRP + Polyketide | 29.0 | 54.0 | 346.0 | 2.6e-94 |
| BAG17643.1 | putative\_NRPS-type-I\_PKS\_fusion\_protein | BGC0001043 | NRP + Polyketide | 29.0 | 51.5 | 343.0 | 2.2e-93 |
| ACR33077.1 | polyketide\_synthase | BGC0000017 | Alkaloid + Polyketide:Modular type I | 22.0 | 112.6 | 334.0 | 1.4e-90 |
| WP\_035121546.1 | type\_I\_polyketide\_synthase | BGC0001467 | NRP:Cyclic depsipeptide + Polyketide:Modular type I | 24.0 | 106.6 | 330.0 | 2e-89 |
| ATZ45182.1 | Bcboa6 | BGC0001892 | Polyketide | 26.0 | 73.5 | 329.0 | 3.3e-89 |
| gene3 |  | BGC0002035 | NRP + Polyketide | 26.0 | 70.9 | 326.0 | 3.7e-88 |
| EFL02193.1 | amino\_acid\_adenylation\_domain-containing\_protein | BGC0000996 | NRP + Polyketide:Iterative type I | 28.0 | 53.2 | 319.0 | 2.7e-86 |
| ABL86391.1 | hybrid\_polyketide\_synthase\_and\_nonribosomal\_peptide\_synthetase | BGC0000999 | NRP + Polyketide | 28.0 | 51.6 | 319.0 | 3.5e-86 |
| AEE88282.1 | CurH | BGC0000976 | NRP + Polyketide:Modular type I | 22.0 | 118.4 | 316.0 | 3.8e-85 |
| AAT70103.1 | CurH | BGC0001165 | NRP + Polyketide:Modular type I | 22.0 | 118.4 | 316.0 | 3.8e-85 |
| CCE88380.1 | polyketide\_synthase | BGC0001034 | NRP + Polyketide:Modular type I | 29.0 | 51.7 | 314.0 | 1.5e-84 |
| CAJ46689.1 | polyketide\_synthase | BGC0000969 | NRP:Cyclic depsipeptide + Polyketide:Modular type I | 27.0 | 53.6 | 305.0 | 5.2e-82 |
| CBX99534.1 | similar\_to\_polyketide\_synthase | BGC0001899 | Polyketide | 27.0 | 53.4 | 305.0 | 5.2e-82 |
| APZ78832.1 | polyketide\_synthase | BGC0001430 | NRP:Cyclic depsipeptide + Polyketide:Iterative type I | 24.0 | 109.2 | 305.0 | 6.8e-82 |
| CAQ18828.1 | polyketide\_synthase | BGC0000954 | NRP + Polyketide:Modular type I | 23.0 | 114.7 | 300.0 | 1.3e-80 |
| XP\_001220460.1 | hypothetical\_protein | BGC0001182 | NRP + Polyketide:Iterative type I | 25.0 | 76.3 | 299.0 | 2.8e-80 |
| CAQ18830.1 | polyketide\_synthase | BGC0000954 | NRP + Polyketide:Modular type I | 23.0 | 114.4 | 299.0 | 3.7e-80 |
| AQW44873.1 | polyketide\_synthase | BGC0001761 | Polyketide | 28.0 | 51.8 | 291.0 | 1e-77 |
| CAJ46690.1 | polyketide\_synthase | BGC0000969 | NRP:Cyclic depsipeptide + Polyketide:Modular type I | 24.0 | 107.7 | 284.0 | 1.6e-75 |
| AZH23789.1 | MgcI | BGC0001970 | NRP + Polyketide | 23.0 | 110.4 | 282.0 | 3.6e-75 |
| AZH23818.1 | MgiI | BGC0001971 | NRP + Polyketide | 23.0 | 110.4 | 280.0 | 2.3e-74 |
| AEO57481.1 | PKS-NRPSs | BGC0001449 | NRP + Alkaloid + Polyketide:Iterative type I | 38.0 | 30.1 | 279.0 | 3e-74 |
| AXN93577.1 | PuwB | BGC0001950 | NRP | 27.0 | 53.2 | 278.0 | 6.8e-74 |
| AXN93586.1 | PuwB | BGC0001951 | NRP | 27.0 | 53.2 | 278.0 | 6.8e-74 |
| AIW82279.1 | PuwB | BGC0001125 | NRP + Polyketide | 27.0 | 53.3 | 276.0 | 2.6e-73 |
| AAS98783.1 | polyketide\_synthase/nonribosomal\_peptide\_synthase\_hybrid | BGC0001001 | NRP + Polyketide | 22.0 | 110.5 | 275.0 | 5.7e-73 |
| CBD77732.1 | polyketide\_synthase | BGC0000974 | NRP + Polyketide | 35.0 | 30.9 | 264.0 | 1e-69 |
| AIR74910.1 | polyketide\_synthase | BGC0001559 | RiPP | 35.0 | 30.9 | 264.0 | 1e-69 |
| CEF75886.1 |  | BGC0001600 | Polyketide | 25.0 | 68.4 | 264.0 | 1e-69 |
| CAD29794.1 | peptide\_synthetase | BGC0001015 | NRP + Polyketide | 23.0 | 74.3 | 263.0 | 2.3e-69 |
| QDA77058.1 | polyketide\_synthase | BGC0002026 | NRP | 24.0 | 109.2 | 262.0 | 5e-69 |
| AAV66110.2 | fusaridione\_A\_synthetase | BGC0000992 | NRP + Polyketide | 37.0 | 30.6 | 260.0 | 1.5e-68 |
| AEE88279.1 | CurK | BGC0000976 | NRP + Polyketide:Modular type I | 22.0 | 111.1 | 255.0 | 4.7e-67 |
| AAT70106.1 | CurK | BGC0001165 | NRP + Polyketide:Modular type I | 22.0 | 111.1 | 255.0 | 4.7e-67 |
| AXM42948.1 | type\_1\_polyketide\_synthase | BGC0001941 | NRP + Polyketide | 24.0 | 107.5 | 255.0 | 4.7e-67 |
| AAK57189.1 | MxaE | BGC0001022 | NRP + Polyketide | 34.0 | 30.4 | 255.0 | 6.1e-67 |
| BAC20564.1 | polyketide\_synthase | BGC0000039 | Polyketide | 35.0 | 30.9 | 254.0 | 1.4e-66 |
| CAO91861.1 | PKS-NRPS\_hybrid | BGC0000968 | NRP + Polyketide:Iterative type I | 34.0 | 30.3 | 253.0 | 2.3e-66 |
| AAF00958.1 | mcyE | BGC0001017 | NRP + Polyketide:Modular type I | 23.0 | 73.6 | 252.0 | 4e-66 |
| ATP76239.1 | NdaF | BGC0001705 | NRP + Polyketide | 22.0 | 74.2 | 252.0 | 5.2e-66 |
| BBG28484.1 | polyketide\_synthase\_CdmE | BGC0001926 | Polyketide | 34.0 | 31.6 | 252.0 | 5.2e-66 |
| AAO62582.1 | polyketide\_synthase\_peptide\_sythetase\_fusion\_protein | BGC0001016 | NRP + Polyketide | 24.0 | 73.6 | 251.0 | 8.9e-66 |
| AAT28740.1 | FUSS | BGC0000064 | Polyketide | 27.0 | 49.8 | 248.0 | 7.5e-65 |
| WP\_042799407.1 | type\_I\_polyketide\_synthase | BGC0001283 | Polyketide | 23.0 | 114.7 | 247.0 | 1.3e-64 |
| ADZ24998.1 | polyketide\_synthase | BGC0000380 | NRP + Polyketide:Modular type I | 23.0 | 109.3 | 243.0 | 1.8e-63 |
| CCE88378.1 | polyketide\_synthase | BGC0001034 | NRP + Polyketide:Modular type I | 22.0 | 107.0 | 242.0 | 5.4e-63 |
| AAF00959.1 | mcyD | BGC0001017 | NRP + Polyketide:Modular type I | 26.0 | 51.0 | 241.0 | 1.2e-62 |
| XP\_659388.1 | hypothetical\_protein | BGC0001998 | Polyketide | 34.0 | 32.0 | 240.0 | 2.7e-62 |
| AFP73394.1 | FusA | BGC0001268 | NRP + Polyketide | 27.0 | 49.3 | 238.0 | 7.8e-62 |
| AZH23788.1 | MgcR | BGC0001970 | NRP + Polyketide | 23.0 | 107.3 | 237.0 | 1.7e-61 |
| AUS29495.1 | polyketide\_synthase | BGC0001030 | NRP + Polyketide | 33.0 | 32.5 | 236.0 | 3e-61 |
| EAU29808.1 | hypothetical\_protein | BGC0001400 | Polyketide | 35.0 | 30.3 | 235.0 | 5e-61 |
| AFU82617.1 | polyketide\_synthase | BGC0000998 | NRP + Polyketide | 33.0 | 29.8 | 234.0 | 1.1e-60 |
| ABX60162.1 | polyketide\_synthase | BGC0000978 | NRP + Alkaloid + Polyketide:Modular type I | 30.0 | 35.3 | 234.0 | 1.5e-60 |
| CAQ34919.1 | polyketide\_synthase | BGC0000986 | NRP + Polyketide | 32.0 | 29.2 | 233.0 | 2.5e-60 |
| ADF88276.1 | polyketide\_synthase | BGC0000981 | NRP + Polyketide | 24.0 | 51.5 | 233.0 | 3.3e-60 |
| ctg1\_orf0002 |  | BGC0001068 | Terpene + Polyketide | 35.0 | 30.5 | 233.0 | 3.3e-60 |
| BAK26562.1 | PKS-NRPS\_hybrid | BGC0000977 | NRP + Polyketide | 24.0 | 67.9 | 232.0 | 5.6e-60 |
| ADF88280.1 | polyketide\_synthase | BGC0000981 | NRP + Polyketide | 29.0 | 35.3 | 231.0 | 9.5e-60 |
| ABX60152.1 | polyketide\_synthase | BGC0000978 | NRP + Alkaloid + Polyketide:Modular type I | 25.0 | 51.6 | 231.0 | 1.2e-59 |
| AXN93601.1 | PuwE | BGC0001952 | NRP | 30.0 | 30.1 | 230.0 | 1.6e-59 |
| AXN93613.1 | PuwE | BGC0001953 | NRP | 30.0 | 30.0 | 230.0 | 2.1e-59 |
| AWO77084.1 | hybrid\_non-ribosomal\_peptide\_synthetase/type\_I\_polyketide\_synthase | BGC0001556 | NRP + Polyketide | 30.0 | 33.4 | 229.0 | 3.6e-59 |
| EAU38971.1 | hypothetical\_protein | BGC0001122 | NRP + Polyketide:Iterative type I | 24.0 | 71.3 | 228.0 | 6.2e-59 |
| AAF26923.1 | polyketide\_synthase | BGC0000988 | NRP + Polyketide | 22.0 | 107.4 | 227.0 | 1.4e-58 |
| AAF62885.1 | EpoF | BGC0000991 | NRP + Polyketide | 22.0 | 107.9 | 226.0 | 3.1e-58 |
| AAC38075.1 | polyketide\_synthase\_type\_I | BGC0000127 | Polyketide | 31.0 | 29.3 | 225.0 | 6.8e-58 |
| ALD82521.1 | polyketide\_synthase | BGC0001212 | NRP + Polyketide | 35.0 | 27.0 | 225.0 | 6.8e-58 |
| BAN19720.1 | polyketide\_synthase | BGC0001252 | Polyketide | 32.0 | 30.8 | 224.0 | 1.2e-57 |
| ADB12493.1 | EpoF | BGC0000990 | NRP + Polyketide | 22.0 | 107.9 | 222.0 | 4.4e-57 |
| AAF19814.1 | MtaF | BGC0001024 | NRP + Polyketide:Modular type I | 32.0 | 29.1 | 219.0 | 3.7e-56 |
| AWS21279.1 | type\_I\_polyketide\_synthase | BGC0001934 | Polyketide | 30.0 | 29.8 | 219.0 | 3.7e-56 |
| AZY91989.1 | polyketide\_synthase | BGC0002022 | Polyketide | 30.0 | 29.8 | 219.0 | 3.7e-56 |
| ABO15888.1 | polyketide\_synthase | BGC0000132 | Polyketide | 33.0 | 29.9 | 219.0 | 4.9e-56 |
| ABM21570.1 | crpB | BGC0000975 | NRP + Polyketide | 30.0 | 29.5 | 218.0 | 8.3e-56 |
| EAL89230.2 | LovB-like\_polyketide\_synthase,\_putative | BGC0000129 | Polyketide | 33.0 | 30.2 | 218.0 | 1.1e-55 |
| QDA77059.1 | polyketide\_synthase/nonribosomal\_peptide\_synthetase | BGC0002026 | NRP | 31.0 | 29.0 | 217.0 | 1.4e-55 |
| ACB46197.1 | polyketide\_synthase | BGC0000989 | NRP + Polyketide | 22.0 | 107.6 | 215.0 | 5.4e-55 |
| ctg1\_orf27 |  | BGC0000096 | Polyketide | 32.0 | 30.7 | 214.0 | 1.2e-54 |
| CAD89777.1 | MelF\_protein | BGC0001010 | NRP + Polyketide:Modular type I | 31.0 | 29.1 | 212.0 | 4.6e-54 |
| AUD08663.1 | iPKS-NRPS | BGC0001553 | NRP + Polyketide | 30.0 | 33.2 | 211.0 | 1.3e-53 |
| CAQ18834.1 | polyketide\_synthase | BGC0000954 | NRP + Polyketide:Modular type I | 31.0 | 32.3 | 210.0 | 2.3e-53 |
| ABC87510.1 | polyketide\_synthase | BGC0001011 | NRP + Polyketide | 31.0 | 29.5 | 210.0 | 3e-53 |
| ctg1\_orf21 |  | BGC0001013 | NRP + Polyketide | 31.0 | 29.5 | 210.0 | 3e-53 |
| BAH02268.1 | polyketide\_synthase | BGC0000126 | Polyketide | 31.0 | 30.1 | 209.0 | 3.9e-53 |
| CAD19090.1 | StiF\_protein | BGC0000153 | NRP + Polyketide:Modular type I | 22.0 | 108.5 | 209.0 | 3.9e-53 |
| AFU82615.1 | polyketide\_synthase | BGC0000998 | NRP + Polyketide | 33.0 | 25.7 | 209.0 | 5e-53 |
| EWM63000.1 | non-ribosomal\_peptide\_synthetase | BGC0001328 | NRP:Cyclic depsipeptide + Polyketide:Modular type I | 37.0 | 20.3 | 209.0 | 5e-53 |
| ATY12793.1 | type\_I\_polyketide\_synthase | BGC0001504 | Polyketide | 31.0 | 28.9 | 209.0 | 5e-53 |
| ABB05102.1 | LipPks1 | BGC0001003 | NRP:Lipopeptide + Polyketide:Modular type I + Saccharide:Hybrid/tailoring | 30.0 | 31.4 | 208.0 | 8.6e-53 |
| WP\_052165465.1 | type\_I\_polyketide\_synthase | BGC0001327 | NRP:Cyclic depsipeptide + Polyketide:Modular type I | 36.0 | 20.3 | 208.0 | 8.6e-53 |
| AQH32481.1 | hybrid\_polyketide\_synthase/peptide\_synthetase | BGC0001667 | NRP + Polyketide | 33.0 | 21.9 | 208.0 | 8.6e-53 |
| ABY83164.1 | Azi26 | BGC0000960 | NRP + Polyketide | 31.0 | 29.5 | 208.0 | 1.1e-52 |
| CBW54671.1 | polyketide\_synthase/non\_ribosomal\_peptide\_synthetase | BGC0000971 | NRP + Polyketide:Modular type I | 31.0 | 32.3 | 208.0 | 1.1e-52 |
| ARM20281.1 | polyketide\_synthase | BGC0001523 | Polyketide | 29.0 | 31.1 | 208.0 | 1.1e-52 |
| CAD19092.1 | StiH\_protein | BGC0000153 | NRP + Polyketide:Modular type I | 31.0 | 29.8 | 207.0 | 2.5e-52 |
| AEP40940.1 | polyketide\_synthase\_type\_I | BGC0000021 | Polyketide | 31.0 | 33.2 | 206.0 | 4.3e-52 |
| AAF19812.1 | MtaD | BGC0001024 | NRP + Polyketide:Modular type I | 30.0 | 29.4 | 204.0 | 1.2e-51 |
| WP\_051137606.1 | type\_I\_polyketide\_synthase | BGC0002011 | Polyketide | 31.0 | 28.7 | 204.0 | 1.2e-51 |
| CCE88379.1 | polyketide\_synthase | BGC0001034 | NRP + Polyketide:Modular type I | 35.0 | 20.3 | 204.0 | 1.6e-51 |
| AGC09484.1 | LobS1 | BGC0001183 | Polyketide | 32.0 | 29.8 | 203.0 | 2.1e-51 |
| ALA09371.1 | type\_I\_modular\_PKS | BGC0001303 | Polyketide | 31.0 | 28.7 | 203.0 | 2.1e-51 |
| WP\_053138504.1 | type\_I\_polyketide\_synthase | BGC0002033 | Polyketide | 32.0 | 31.3 | 203.0 | 2.8e-51 |
| CAQ18839.1 | hybrid\_polyketide\_synthase/nonribosomal\_polypetide\_synthetase | BGC0000954 | NRP + Polyketide:Modular type I | 35.0 | 20.9 | 203.0 | 3.6e-51 |
| DAB41918.1 | ArzP\_-\_PKS\_(KS,\_AT,\_OMT,\_ACP,\_TE) | BGC0001884 | NRP + Polyketide | 29.0 | 32.4 | 203.0 | 3.6e-51 |
| CAQ34928.1 | polyketide\_synthase | BGC0000986 | NRP + Polyketide | 30.0 | 31.5 | 202.0 | 4.7e-51 |
| AGI99497.1 | type\_I\_polyketide\_synthase | BGC0001004 | Polyketide:Modular type I | 32.0 | 29.1 | 202.0 | 4.7e-51 |
| ANI24099.1 | polyketide\_synthase | BGC0001235 | NRP + Polyketide | 35.0 | 20.8 | 202.0 | 4.7e-51 |
| CAQ34918.1 | nonribosomal\_peptide\_synthetase/\_polyketide\_synthase | BGC0000986 | NRP + Polyketide | 29.0 | 29.3 | 202.0 | 6.2e-51 |
| AFV96138.1 | polyketide\_synthase | BGC0001064 | Polyketide:Modular type I + Polyketide:Type III | 34.0 | 20.3 | 201.0 | 8.1e-51 |
| ARU81118.1 | CylD | BGC0001566 | Polyketide | 34.0 | 20.3 | 201.0 | 8.1e-51 |
| ACC40921.1 | polyketide\_synthase\_Pks7 | BGC0001665 | Polyketide | 30.0 | 29.4 | 201.0 | 1.1e-50 |
| AQW44893.1 | polyketide\_synthase | BGC0001737 | NRP + Polyketide | 33.0 | 25.7 | 201.0 | 1.1e-50 |
| AVI26388.1 | polyketide\_synthase | BGC0001800 | NRP + Polyketide | 22.0 | 111.9 | 201.0 | 1.1e-50 |
| CAO98850.1 | polyketide\_synthase\_AufG | BGC0000023 | Polyketide:Modular type I | 32.0 | 27.9 | 200.0 | 1.8e-50 |
| CAQ18832.1 | polyketide\_synthase | BGC0000954 | NRP + Polyketide:Modular type I | 23.0 | 108.2 | 200.0 | 1.8e-50 |
| AAK57186.1 | MxaB2 | BGC0001022 | NRP + Polyketide | 31.0 | 28.1 | 200.0 | 3.1e-50 |
| CAQ18829.1 | polyketide\_synthase | BGC0000954 | NRP + Polyketide:Modular type I | 35.0 | 20.5 | 199.0 | 5.2e-50 |
| CAD89776.1 | MelE\_protein | BGC0001010 | NRP + Polyketide:Modular type I | 36.0 | 20.6 | 198.0 | 6.8e-50 |
| AMB48442.1 | polyketide\_synthase | BGC0001357 | Polyketide | 34.0 | 20.3 | 198.0 | 6.8e-50 |
| AAC46026.1 | polyketide\_synthase\_modules\_4\_and\_5 | BGC0000113 | Polyketide | 31.0 | 29.9 | 198.0 | 8.9e-50 |
| AWC08663.1 | polyketide\_synthase\_type\_I | BGC0001932 | Polyketide | 29.0 | 31.4 | 198.0 | 8.9e-50 |
| AAG13917.1 | megalomicin\_6-deoxyerythronolide\_B\_synthase\_1 | BGC0000092 | Polyketide | 30.0 | 29.0 | 198.0 | 1.2e-49 |
| ARM20280.1 | polyketide\_synthase | BGC0001523 | Polyketide | 29.0 | 31.2 | 197.0 | 2e-49 |
| ADZ24997.1 | polyketide\_synthase | BGC0000380 | NRP + Polyketide:Modular type I | 33.0 | 22.1 | 196.0 | 2.6e-49 |
| AUO16402.1 | polyketide\_synthase | BGC0001700 | Polyketide | 29.0 | 31.1 | 196.0 | 2.6e-49 |
| AQW44890.1 | polyketide\_synthase | BGC0001737 | NRP + Polyketide | 30.0 | 28.1 | 196.0 | 2.6e-49 |
| AFI57005.1 | QmnA1 | BGC0000133 | Polyketide | 30.0 | 28.7 | 196.0 | 3.4e-49 |
| ABB05105.1 | LipPks4 | BGC0001003 | NRP:Lipopeptide + Polyketide:Modular type I + Saccharide:Hybrid/tailoring | 29.0 | 29.1 | 196.0 | 3.4e-49 |
| AAK57188.1 | MxaD | BGC0001022 | NRP + Polyketide | 32.0 | 27.0 | 196.0 | 3.4e-49 |
| AGC45619.1 | polyketide\_synthase | BGC0001394 | NRP + Polyketide | 31.0 | 28.3 | 196.0 | 3.4e-49 |
| AIT55259.1 | polyketide\_synthase | BGC0000072 | Polyketide:Modular type I | 33.0 | 24.8 | 196.0 | 4.4e-49 |
| ABC84460.1 | NigAV | BGC0000114 | Polyketide:Modular type I | 29.0 | 28.5 | 196.0 | 4.4e-49 |
| AIW82282.1 | PuwE | BGC0001125 | NRP + Polyketide | 33.0 | 20.7 | 195.0 | 5.8e-49 |
| AHE80994.1 | PieA4 | BGC0001169 | Polyketide:Modular type I | 30.0 | 29.0 | 195.0 | 5.8e-49 |
| WP\_026247674.1 | type\_I\_polyketide\_synthase | BGC0001332 | NRP + Polyketide | 34.0 | 20.5 | 195.0 | 5.8e-49 |
| AVI57433.1 | AbmB1 | BGC0001694 | Polyketide | 33.0 | 24.7 | 195.0 | 5.8e-49 |
| CAA60462.1 | polyketide\_synthase | BGC0001040 | NRP + Polyketide | 21.0 | 112.1 | 195.0 | 7.5e-49 |
| WP\_055480219.1 | type\_I\_polyketide\_synthase | BGC0001653 | Polyketide | 30.0 | 29.8 | 195.0 | 7.5e-49 |
| CBD77736.1 | polyketide\_synthase | BGC0000974 | NRP + Polyketide | 30.0 | 29.8 | 195.0 | 9.9e-49 |
| AAW03328.1 | CtaE | BGC0000982 | NRP + Polyketide | 35.0 | 20.6 | 195.0 | 9.9e-49 |
| AIR74912.1 | polyketide\_synthase | BGC0001559 | RiPP | 30.0 | 29.8 | 195.0 | 9.9e-49 |
| AUO16401.1 | polyketide\_synthase | BGC0001700 | Polyketide | 29.0 | 29.5 | 195.0 | 9.9e-49 |
| DAB41915.1 | ArzM\_-\_PKS\_(KS,\_AT,\_DH,\_MT,\_ER,\_KR,\_ACP) | BGC0001884 | NRP + Polyketide | 35.0 | 20.3 | 195.0 | 9.9e-49 |
| DAB41916.1 | ArzN\_-\_PKS\_(KS,\_AT,\_OMT,\_KR,\_ACP) | BGC0001884 | NRP + Polyketide | 34.0 | 20.4 | 195.0 | 9.9e-49 |
| AEP40936.1 | polyketide\_synthase\_type\_I | BGC0000021 | Polyketide | 28.0 | 30.6 | 194.0 | 1.3e-48 |
| BAC57028.1 | protomycinolide\_IV\_synthase\_1 | BGC0000102 | Polyketide | 33.0 | 24.6 | 194.0 | 1.3e-48 |
| AAF86393.1 | FkbB | BGC0000994 | NRP + Polyketide | 32.0 | 27.9 | 194.0 | 1.3e-48 |
| AAF19810.1 | MtaB | BGC0001024 | NRP + Polyketide:Modular type I | 33.0 | 24.8 | 194.0 | 1.3e-48 |
| AGC45622.1 | polyketide\_synthase | BGC0001394 | NRP + Polyketide | 31.0 | 25.0 | 194.0 | 1.3e-48 |
| AWW87422.1 | type\_I\_polyketide\_synthase | BGC0001755 | Polyketide | 30.0 | 29.4 | 194.0 | 1.3e-48 |
| ALD82522.1 | polyketide\_synthase | BGC0001212 | NRP + Polyketide | 34.0 | 20.4 | 194.0 | 1.7e-48 |
| AAG23266.1 | polyketide\_synthase\_extender\_modules\_3-4 | BGC0000148 | Polyketide | 29.0 | 29.4 | 193.0 | 2.2e-48 |
| ctg1\_orf3 |  | BGC0001329 | Polyketide + NRP:Cyclic depsipeptide | 33.0 | 24.2 | 193.0 | 2.2e-48 |
| AEK75503.1 | type\_1\_polyketide\_synthase | BGC0000001 | Polyketide:Modular type I | 28.0 | 32.6 | 193.0 | 2.9e-48 |
| ADB23403.1 | polyketide\_synthase\_type\_I | BGC0001062 | Polyketide | 29.0 | 29.5 | 193.0 | 2.9e-48 |
| AXI91546.1 | FunP7 | BGC0001944 | Polyketide | 31.0 | 28.5 | 193.0 | 2.9e-48 |
| ACR33079.1 | polyketide\_synthase | BGC0000017 | Alkaloid + Polyketide:Modular type I | 34.0 | 20.2 | 193.0 | 3.7e-48 |
| AEP40935.1 | polyketide\_synthase\_type\_I | BGC0000021 | Polyketide | 28.0 | 30.8 | 193.0 | 3.7e-48 |
| AAU93807.2 | polyketide\_synthase\_modules\_1\_and\_2 | BGC0000054 | Polyketide | 30.0 | 29.3 | 193.0 | 3.7e-48 |
| AAF86392.1 | FkbC | BGC0000994 | NRP + Polyketide | 32.0 | 28.7 | 193.0 | 3.7e-48 |
| AAP42857.1 | NanA3 | BGC0000105 | Polyketide | 29.0 | 29.9 | 192.0 | 6.4e-48 |
| AVI26389.1 | polyketide\_synthase | BGC0001800 | NRP + Polyketide | 35.0 | 20.1 | 192.0 | 6.4e-48 |
| CAD89775.1 | MelD\_protein | BGC0001010 | NRP + Polyketide:Modular type I | 30.0 | 29.4 | 191.0 | 8.3e-48 |
| AVI26390.1 | polyketide\_synthase\_/\_nonribosomal\_peptide\_synthase\_hybrid | BGC0001800 | NRP + Polyketide | 28.0 | 32.5 | 191.0 | 8.3e-48 |
| AEP40932.1 | polyketide\_synthase\_type\_I | BGC0000021 | Polyketide | 28.0 | 33.4 | 191.0 | 1.1e-47 |
| ACB46486.1 | polyketide\_synthase | BGC0000082 | Polyketide | 29.0 | 30.8 | 191.0 | 1.1e-47 |
| AAP42874.1 | NanA8 | BGC0000105 | Polyketide | 29.0 | 29.5 | 191.0 | 1.1e-47 |
| XP\_001798923.1 | polyketide\_synthase | BGC0001865 | Polyketide:Iterative type I | 29.0 | 31.9 | 191.0 | 1.1e-47 |
| CAJ88175.1 | putative\_polyketide\_synthase\_B | BGC0000151 | Polyketide:Modular type I + Saccharide:Hybrid/tailoring | 30.0 | 29.1 | 191.0 | 1.4e-47 |
| ADZ24995.1 | non-ribosomal\_peptide\_synthase/polyketide\_synthase | BGC0000380 | NRP + Polyketide:Modular type I | 30.0 | 30.7 | 191.0 | 1.4e-47 |
| EYT83439.1 | beta-ketoacyl\_synthase | BGC0001213 | Polyketide | 32.0 | 30.8 | 191.0 | 1.4e-47 |
| AGY30676.1 | Ann4 | BGC0001298 | Polyketide | 28.0 | 33.2 | 191.0 | 1.4e-47 |
| ADH04657.1 | TugA | BGC0001342 | NRP + Polyketide | 34.0 | 24.6 | 191.0 | 1.4e-47 |
| AUO16422.1 | polyketide\_synthase | BGC0001700 | Polyketide | 28.0 | 31.6 | 191.0 | 1.4e-47 |
| AHH99924.1 | PKS\_I | BGC0000002 | Polyketide | 30.0 | 28.0 | 190.0 | 1.9e-47 |
| CQR60494.1 | Polyketide\_synthase,\_type\_I,\_module\_8 | BGC0001287 | Polyketide | 30.0 | 31.3 | 190.0 | 1.9e-47 |
| CAD17792.1 | probable\_non\_ribosomal\_peptide\_synthetase\_protein | BGC0001754 | NRP + Polyketide | 34.0 | 22.0 | 190.0 | 1.9e-47 |
| ANZ22991.1 | ZinG | BGC0001828 | Polyketide | 29.0 | 29.1 | 190.0 | 1.9e-47 |
| ACB46488.1 | polyketide\_synthase | BGC0000082 | Polyketide | 31.0 | 27.7 | 190.0 | 2.4e-47 |
| AEZ53945.1 | polyketide\_synthase | BGC0000144 | Polyketide:Modular type I | 30.0 | 31.1 | 190.0 | 2.4e-47 |
| ARM20282.1 | polyketide\_synthase | BGC0001523 | Polyketide | 28.0 | 31.2 | 190.0 | 2.4e-47 |
| BAJ16470.1 | polyketide\_synthase | BGC0000058 | Polyketide | 29.0 | 31.9 | 190.0 | 3.2e-47 |
| ACB46487.1 | polyketide\_synthase | BGC0000082 | Polyketide | 30.0 | 27.8 | 190.0 | 3.2e-47 |
| ABJ97438.1 | MerB | BGC0001012 | NRP + Polyketide | 30.0 | 27.8 | 190.0 | 3.2e-47 |
| AHA12079.1 | polyketide\_synthase\_type\_1 | BGC0001172 | NRP + Polyketide:Modular type I | 34.0 | 21.7 | 190.0 | 3.2e-47 |
| AGC45623.1 | polyketide\_synthase | BGC0001394 | NRP + Polyketide | 34.0 | 20.8 | 190.0 | 3.2e-47 |
| ACN69990.1 | polyketide\_synthase | BGC0000079 | Polyketide | 29.0 | 31.2 | 189.0 | 4.1e-47 |
| AAO65797.1 | monensin\_polyketide\_synthase\_module\_2 | BGC0000100 | Polyketide | 30.0 | 27.9 | 189.0 | 4.1e-47 |
| ABX60163.1 | polyketide\_synthase | BGC0000978 | NRP + Alkaloid + Polyketide:Modular type I | 32.0 | 20.5 | 189.0 | 4.1e-47 |
| AQA28562.1 | type\_I\_polyketide\_synthase | BGC0001663 | Polyketide | 33.0 | 20.3 | 189.0 | 4.1e-47 |
| ANZ52460.1 | MonAII | BGC0001670 | Polyketide | 30.0 | 27.9 | 189.0 | 4.1e-47 |
| ANY10590.1 | polyketide\_synthase | BGC0001773 | Polyketide | 29.0 | 29.1 | 189.0 | 4.1e-47 |
| AEP40934.1 | polyketide\_synthase\_type\_I | BGC0000021 | Polyketide | 28.0 | 32.0 | 189.0 | 5.4e-47 |
| AIT55262.1 | polyketide\_synthase | BGC0000072 | Polyketide:Modular type I | 32.0 | 21.2 | 189.0 | 5.4e-47 |
| ABP55210.1 | beta-ketoacyl\_synthase | BGC0000142 | Polyketide | 29.0 | 30.7 | 189.0 | 5.4e-47 |
| ADF88277.1 | polyketide\_synthase | BGC0000981 | NRP + Polyketide | 32.0 | 20.5 | 189.0 | 5.4e-47 |
| AAW03327.1 | CtaD | BGC0000982 | NRP + Polyketide | 29.0 | 29.4 | 189.0 | 5.4e-47 |
| BBA66512.1 | type\_I\_polyketide\_synthase | BGC0001495 | Polyketide | 29.0 | 30.4 | 189.0 | 5.4e-47 |
| AXI91549.1 | FunP4 | BGC0001944 | Polyketide | 31.0 | 28.0 | 189.0 | 5.4e-47 |
| ACO94460.1 | polyketide\_synthase\_type\_I | BGC0000029 | Polyketide:Modular type I | 30.0 | 29.4 | 188.0 | 9.2e-47 |
| BAQ25507.1 | type\_I\_polyketide\_synthase | BGC0001288 | Polyketide | 33.0 | 25.3 | 188.0 | 9.2e-47 |
| ABV83221.1 | CppI | BGC0000116 | Polyketide | 29.0 | 28.4 | 188.0 | 1.2e-46 |
| ABP55222.1 | beta-ketoacyl\_synthase | BGC0000142 | Polyketide | 29.0 | 29.1 | 188.0 | 1.2e-46 |
| AAW03329.1 | CtaF | BGC0000982 | NRP + Polyketide | 29.0 | 29.1 | 188.0 | 1.2e-46 |
| AUO16397.1 | polyketide\_synthase | BGC0001700 | Polyketide | 28.0 | 34.5 | 188.0 | 1.2e-46 |
| AAP42858.1 | NanA4 | BGC0000105 | Polyketide | 28.0 | 32.0 | 187.0 | 1.6e-46 |
| CBZ41585.1 | Type\_I\_modular\_polyketide\_synthase | BGC0000151 | Polyketide:Modular type I + Saccharide:Hybrid/tailoring | 29.0 | 29.3 | 187.0 | 1.6e-46 |
| WP\_055480220.1 | type\_I\_polyketide\_synthase | BGC0001653 | Polyketide | 29.0 | 31.4 | 187.0 | 1.6e-46 |
| ACN69988.1 | polyketide\_synthase | BGC0000079 | Polyketide | 30.0 | 29.1 | 187.0 | 2.1e-46 |
| AEZ53953.1 | polyketide\_synthase | BGC0000144 | Polyketide:Modular type I | 28.0 | 31.2 | 187.0 | 2.1e-46 |
| AQX77694.1 | NocP | BGC0001704 | Other | 35.0 | 20.0 | 187.0 | 2.1e-46 |
| ADC45586.1 | modular\_polyketide\_synthase | BGC0000093 | Polyketide | 29.0 | 29.7 | 186.0 | 2.7e-46 |
| AEU11006.1 | NpnB | BGC0001029 | NRP + Polyketide | 32.0 | 20.4 | 186.0 | 2.7e-46 |
| BAH02270.1 | polyketide\_synthase | BGC0000126 | Polyketide | 29.0 | 29.5 | 186.0 | 3.5e-46 |
| AGC09499.1 | LobS4 | BGC0001183 | Polyketide | 29.0 | 31.9 | 186.0 | 3.5e-46 |
| ctg1\_orf253 |  | BGC0001200 | Polyketide | 30.0 | 29.1 | 186.0 | 3.5e-46 |
| AKA59090.1 | type-I\_PKS | BGC0001619 | Polyketide | 29.0 | 29.0 | 186.0 | 3.5e-46 |
| OAP25821.1 | Phenolphthiocerol\_synthesis\_polyketide\_synthase\_type\_I\_Pks15/1 | BGC0001658 | Polyketide | 27.0 | 28.6 | 186.0 | 3.5e-46 |
| ATX68116.1 | malonyl\_CoA-acyl\_carrier\_protein\_transacylase | BGC0001772 | Polyketide | 32.0 | 20.2 | 186.0 | 3.5e-46 |
| AGY62755.1 | EbeC | BGC0000051 | Polyketide | 29.0 | 29.7 | 185.0 | 6e-46 |
| ACY06286.1 | polyketide\_synthase | BGC0001042 | NRP + Polyketide | 30.0 | 27.9 | 185.0 | 6e-46 |
| EHK80167.1 | modular\_polyketide\_synthase | BGC0001447 | Polyketide | 28.0 | 29.0 | 185.0 | 6e-46 |
| OJF16266.1 | AceP4 | BGC0001491 | Polyketide | 29.0 | 30.0 | 185.0 | 6e-46 |
| SCN11951.1 | ebeC-type\_I\_polyketide\_synthase | BGC0001580 | Polyketide | 29.0 | 29.7 | 185.0 | 6e-46 |
| AZH23819.1 | MgiR | BGC0001971 | NRP + Polyketide | 32.0 | 20.9 | 185.0 | 6e-46 |
| ACD39770.1 | non-reducing\_polyketide\_synthase | BGC0000134 | Polyketide | 27.0 | 32.7 | 185.0 | 7.8e-46 |
| CCP20051.1 | divM\_protein | BGC0001119 | Polyketide:Modular type I | 29.0 | 30.8 | 185.0 | 7.8e-46 |
| AKD43753.1 | HerB | BGC0001349 | NRP + Polyketide | 30.0 | 29.4 | 185.0 | 7.8e-46 |
| ARM20278.1 | polyketide\_synthase | BGC0001523 | Polyketide | 28.0 | 30.9 | 185.0 | 7.8e-46 |
| AQW44892.1 | polyketide\_synthase | BGC0001737 | NRP + Polyketide | 31.0 | 24.8 | 185.0 | 7.8e-46 |
| ATX68115.1 | malonyl\_CoA-acyl\_carrier\_protein\_transacylase | BGC0001772 | Polyketide | 32.0 | 21.0 | 185.0 | 7.8e-46 |
| AAX98187.1 | polyketide\_synthase\_type\_I | BGC0000052 | Polyketide | 28.0 | 29.8 | 185.0 | 1e-45 |
| ACO94488.1 | polyketide\_synthase\_type\_I | BGC0000097 | Polyketide:Modular type I | 30.0 | 29.4 | 185.0 | 1e-45 |
| AAF86396.1 | FkbA | BGC0000994 | NRP + Polyketide | 31.0 | 27.8 | 185.0 | 1e-45 |
| AQW44891.1 | polyketide\_synthase | BGC0001737 | NRP + Polyketide | 31.0 | 25.4 | 185.0 | 1e-45 |
| ATL73034.1 | type\_I\_modular\_polyketide\_synthase | BGC0001807 | NRP + Polyketide | 28.0 | 29.9 | 185.0 | 1e-45 |
| AZH23817.1 | MgiQ | BGC0001971 | NRP + Polyketide | 23.0 | 54.2 | 185.0 | 1e-45 |
| ABW96542.1 | type\_I\_modular\_polyketide\_synthase | BGC0000159 | Polyketide:Modular type I | 29.0 | 29.4 | 184.0 | 1.7e-45 |
| ACY06287.1 | type\_I\_polyketide\_synthase | BGC0001042 | NRP + Polyketide | 28.0 | 29.9 | 183.0 | 2.3e-45 |
| AJO72735.1 | Type\_I\_modular\_polyketide\_synthase | BGC0001381 | Polyketide | 30.0 | 28.5 | 183.0 | 2.3e-45 |
| AGC45620.1 | polyketide\_synthase | BGC0001394 | NRP + Polyketide | 31.0 | 25.7 | 183.0 | 2.3e-45 |
| ctg1\_12 |  | BGC0001931 | Polyketide | 29.0 | 28.1 | 183.0 | 2.3e-45 |
| AAK57187.1 | MxaC | BGC0001022 | NRP + Polyketide | 31.0 | 25.1 | 183.0 | 3e-45 |
| ABK32288.1 | JerB | BGC0000080 | Polyketide | 33.0 | 21.1 | 183.0 | 3.9e-45 |
| BAC57032.1 | protomycinolide\_IV\_synthase\_5 | BGC0000102 | Polyketide | 29.0 | 29.7 | 183.0 | 3.9e-45 |
| ABP55223.1 | beta-ketoacyl\_synthase | BGC0000142 | Polyketide | 29.0 | 29.1 | 182.0 | 5.1e-45 |
| AUO16399.1 | polyketide\_synthase | BGC0001700 | Polyketide | 28.0 | 30.1 | 182.0 | 5.1e-45 |
| AHB82054.1 | polyketide\_synthase | BGC0001019 | NRP + Polyketide:Modular type I | 31.0 | 21.7 | 181.0 | 8.6e-45 |
| OJF16269.1 | AceP2 | BGC0001491 | Polyketide | 28.0 | 28.4 | 181.0 | 8.6e-45 |
| AAX98190.1 | polyketide\_synthase\_type\_I | BGC0000052 | Polyketide | 27.0 | 29.6 | 181.0 | 1.1e-44 |
| ADB12490.1 | EpoC | BGC0000990 | NRP + Polyketide | 30.0 | 26.1 | 181.0 | 1.5e-44 |
| ACO94483.1 | polyketide\_synthase\_type\_I | BGC0000097 | Polyketide:Modular type I | 30.0 | 29.8 | 180.0 | 1.9e-44 |
| AAF62882.1 | EpoC | BGC0000991 | NRP + Polyketide | 30.0 | 26.1 | 180.0 | 1.9e-44 |
| AIT55261.1 | polyketide\_synthase | BGC0000072 | Polyketide:Modular type I | 34.0 | 20.4 | 180.0 | 2.5e-44 |
| CAO98848.1 | polyketide\_synthase\_AufE | BGC0000023 | Polyketide:Modular type I | 31.0 | 21.5 | 180.0 | 3.3e-44 |
| CBD77734.1 | polyketide\_synthase | BGC0000974 | NRP + Polyketide | 31.0 | 20.8 | 180.0 | 3.3e-44 |
| AAF26920.1 | polyketide\_synthase | BGC0000988 | NRP + Polyketide | 30.0 | 26.1 | 180.0 | 3.3e-44 |
| AIR74911.1 | polyketide\_synthase | BGC0001559 | RiPP | 31.0 | 20.8 | 180.0 | 3.3e-44 |
| CAM00062.1 | EryAI\_Erythromycin\_polyketide\_synthase\_modules\_1\_and\_2 | BGC0000055 | Polyketide:Modular type I + Saccharide:Hybrid/tailoring | 29.0 | 29.2 | 179.0 | 4.3e-44 |
| AAC49191.1 | putative\_polyketide\_synthase | BGC0000152 | Polyketide | 29.0 | 26.9 | 179.0 | 4.3e-44 |
| AXM42951.1 | polyketide\_synthase | BGC0001941 | NRP + Polyketide | 32.0 | 24.6 | 179.0 | 4.3e-44 |
| AAZ94388.1 | nodular\_polyketide\_synthase | BGC0000040 | Polyketide | 29.0 | 28.0 | 179.0 | 5.6e-44 |
| AGC45621.1 | polyketide\_synthase | BGC0001394 | NRP + Polyketide | 30.0 | 25.3 | 179.0 | 5.6e-44 |
| WP\_055469545.1 | type\_I\_polyketide\_synthase | BGC0001537 | Polyketide | 29.0 | 30.6 | 179.0 | 5.6e-44 |
| AAZ94389.1 | modular\_polyketide\_synthase | BGC0000040 | Polyketide | 27.0 | 29.7 | 178.0 | 7.3e-44 |
| ACF35445.1 | mbcAI | BGC0000090 | Polyketide | 29.0 | 30.2 | 178.0 | 7.3e-44 |
| AEZ53950.1 | polyketide\_synthase | BGC0000144 | Polyketide:Modular type I | 29.0 | 30.3 | 178.0 | 7.3e-44 |
| ABB90282.1 | polyketide\_synthase | BGC0001057 | NRP + Polyketide | 28.0 | 30.0 | 178.0 | 7.3e-44 |
| ctg1\_orf9 |  | BGC0000053 | Polyketide | 28.0 | 29.6 | 178.0 | 9.5e-44 |
| AEF33079.1 | polyketide\_synthase | BGC0001039 | NRP + Polyketide | 31.0 | 23.1 | 178.0 | 9.5e-44 |
| AKA59088.1 | type-I\_PKS | BGC0001619 | Polyketide | 36.0 | 20.4 | 178.0 | 9.5e-44 |
| ANH11414.1 | SceS | BGC0001908 | Polyketide | 29.0 | 29.1 | 178.0 | 9.5e-44 |
| AGY62754.1 | EbeB | BGC0000051 | Polyketide | 29.0 | 29.8 | 178.0 | 1.2e-43 |
| ctg1\_orf521 |  | BGC0001199 | Polyketide | 30.0 | 29.2 | 178.0 | 1.2e-43 |
| SCN11950.1 | EbeB-type\_I\_polyketide\_synthase | BGC0001580 | Polyketide | 29.0 | 29.8 | 178.0 | 1.2e-43 |
| ABJ97437.1 | MerA | BGC0001012 | NRP + Polyketide | 27.0 | 28.7 | 177.0 | 2.1e-43 |
| AGC24270.1 | prlP | BGC0001038 | NRP + Polyketide:Modular type I | 33.0 | 20.4 | 177.0 | 2.1e-43 |
| TXD00025.1 | SDR\_family\_NAD(P)-dependent\_oxidoreductase | BGC0001877 | Polyketide | 29.0 | 29.9 | 177.0 | 2.1e-43 |
| CAD55506.1 | CpkA;\_Polyketide\_synthase\_loading\_module,\_and\_modules\_1\_and\_2 | BGC0000038 | Polyketide:Modular type I | 28.0 | 30.1 | 176.0 | 2.8e-43 |
| AAQ84144.1 | Plm4 | BGC0000123 | Polyketide | 29.0 | 28.6 | 176.0 | 2.8e-43 |
| AAS98784.1 | polyketide\_synthase | BGC0001001 | NRP + Polyketide | 33.0 | 20.4 | 176.0 | 2.8e-43 |
| AAX35547.1 | polyketide\_syntase\_2 | BGC0001275 | Polyketide | 26.0 | 33.9 | 176.0 | 2.8e-43 |
| ADU86003.1 | putative\_modular\_polyketide\_synthase | BGC0000165 | Polyketide:Modular type I | 29.0 | 30.2 | 176.0 | 3.6e-43 |
| ASZ00148.1 | polyketide\_synthase | BGC0001785 | Polyketide | 29.0 | 29.8 | 176.0 | 3.6e-43 |
| ABW96540.1 | type\_I\_modular\_polyketide\_synthase | BGC0000159 | Polyketide:Modular type I | 28.0 | 31.6 | 176.0 | 4.7e-43 |
| ABC87511.1 | polyketide\_synthase | BGC0001011 | NRP + Polyketide | 28.0 | 29.0 | 176.0 | 4.7e-43 |
| ctg1\_orf22 |  | BGC0001013 | NRP + Polyketide | 28.0 | 29.0 | 176.0 | 4.7e-43 |
| OAP25815.1 | Phenolphthiocerol\_synthesis\_polyketide\_synthase\_type\_I\_Pks15/1 | BGC0001658 | Polyketide | 23.0 | 50.3 | 176.0 | 4.7e-43 |
| AAC38076.1 | polyketide\_synthase\_type\_I | BGC0000127 | Polyketide | 31.0 | 20.0 | 175.0 | 6.2e-43 |
| CAD15508.1 | polyketide\_synthase/non-ribosomal\_peptide\_synthetase | BGC0001014 | NRP:NRP siderophore + Polyketide:Modular type I + Polyketide:Iterative type I | 31.0 | 24.3 | 175.0 | 8.1e-43 |
| AFV30250.1 | polyketide\_synthase | BGC0000075 | Polyketide | 27.0 | 31.1 | 175.0 | 1.1e-42 |
| AEZ53951.1 | polyketide\_synthase | BGC0000144 | Polyketide:Modular type I | 28.0 | 29.2 | 175.0 | 1.1e-42 |
| ctg1\_orf256 |  | BGC0001200 | Polyketide | 28.0 | 29.3 | 175.0 | 1.1e-42 |
| AVI57434.1 | AbmB2 | BGC0001694 | Polyketide | 30.0 | 27.0 | 175.0 | 1.1e-42 |
| AAQ90174.1 | polyketide\_synthase\_type\_I | BGC0000128 | Polyketide | 31.0 | 20.0 | 174.0 | 1.4e-42 |
| AAY89052.1 | polyketide\_synthase | BGC0001069 | NRP + Polyketide:Trans-AT type I | 33.0 | 21.9 | 174.0 | 1.8e-42 |
| ACA99172.1 | polyketide\_synthase | BGC0001160 | Polyketide:Modular type I | 28.0 | 29.9 | 174.0 | 1.8e-42 |
| AKA59089.1 | type-I\_PKS | BGC0001619 | Polyketide | 29.0 | 28.9 | 174.0 | 1.8e-42 |
| OAP25811.1 | Phenolphthiocerol\_synthesis\_polyketide\_synthase\_type\_I\_Pks15/1 | BGC0001658 | Polyketide | 31.0 | 24.7 | 174.0 | 1.8e-42 |
| ABP55493.1 | thioester\_reductase\_domain | BGC0001006 | NRP + Polyketide | 30.0 | 29.7 | 173.0 | 2.3e-42 |
| WP\_020636817.1 | type\_I\_polyketide\_synthase | BGC0002011 | Polyketide | 30.0 | 26.0 | 173.0 | 3.1e-42 |
| ACB46194.1 | polyketide\_synthase | BGC0000989 | NRP + Polyketide | 30.0 | 26.1 | 173.0 | 4e-42 |
| EHK80163.1 | acyl\_transferase | BGC0001447 | Polyketide | 29.0 | 29.4 | 173.0 | 4e-42 |
| AAM77986.1 | iterative\_type\_I\_polyketide\_synthase | BGC0000112 | Polyketide:Iterative type I + Polyketide:Enediyne type I | 30.0 | 26.5 | 172.0 | 5.2e-42 |
| AAZ94386.1 | modular\_polyketide\_synthase | BGC0000040 | Polyketide | 32.0 | 20.7 | 172.0 | 6.8e-42 |
| ABP55220.1 | beta-ketoacyl\_synthase | BGC0000142 | Polyketide | 28.0 | 29.2 | 172.0 | 6.8e-42 |
| CAD19093.1 | StiJ\_protein | BGC0000153 | NRP + Polyketide:Modular type I | 33.0 | 22.6 | 172.0 | 6.8e-42 |
| AHA38202.1 | GphI | BGC0000069 | Polyketide | 29.0 | 29.2 | 171.0 | 8.9e-42 |
| BAE93722.1 | type\_I\_polyketide\_synthase | BGC0000164 | Polyketide | 28.0 | 29.3 | 171.0 | 8.9e-42 |
| BAD08358.1 | polyketide\_synthase\_modules\_4 | BGC0000167 | Polyketide | 21.0 | 112.5 | 171.0 | 1.2e-41 |
| BBA84068.1 | type\_I\_polyketide\_synthase | BGC0001916 | Polyketide | 28.0 | 29.1 | 171.0 | 1.2e-41 |
| ABI93779.1 | GdmPKS | BGC0000068 | Polyketide | 28.0 | 31.9 | 171.0 | 1.5e-41 |
| CBD77746.1 | non-ribosomal\_peptide\_synthetase/polyketide\_synthase | BGC0000974 | NRP + Polyketide | 33.0 | 20.5 | 171.0 | 1.5e-41 |
| AIR74926.1 | polyketide\_synthase | BGC0001559 | RiPP | 33.0 | 20.5 | 171.0 | 1.5e-41 |
| AWH12671.1 | RmpA1 | BGC0001759 | Polyketide | 29.0 | 23.8 | 171.0 | 1.5e-41 |
| CCP20049.1 | divL2\_protein | BGC0001119 | Polyketide:Modular type I | 30.0 | 27.4 | 170.0 | 2e-41 |
| AHB82063.1 | polyketide\_synthase | BGC0001231 | NRP + Polyketide:Modular type I | 31.0 | 23.7 | 170.0 | 2e-41 |
| AMYAL\_RS48925 | polyketide\_synthase | BGC0002011 | Polyketide | 31.0 | 20.3 | 170.0 | 2e-41 |
| ctg1\_orf10 |  | BGC0000053 | Polyketide | 28.0 | 29.6 | 170.0 | 2.6e-41 |
| ACO94456.1 | polyketide\_synthase\_type\_I | BGC0000029 | Polyketide:Modular type I | 28.0 | 29.7 | 170.0 | 3.4e-41 |
| AFL48529.1 | laidlomycin\_polyketide\_synthase\_(module\_5\_and\_module\_6) | BGC0000084 | Polyketide | 29.0 | 25.0 | 169.0 | 4.4e-41 |
| CAI94682.1 | putative\_polyketide\_synthase | BGC0000141 | Polyketide | 27.0 | 29.7 | 169.0 | 5.8e-41 |
| AFV30251.1 | polyketide\_synthase | BGC0000075 | Polyketide | 27.0 | 30.7 | 168.0 | 7.6e-41 |
| BAK64638.1 | polyketide\_synthase | BGC0000135 | Polyketide | 21.0 | 110.6 | 168.0 | 7.6e-41 |
| ctg1\_orf522 |  | BGC0001199 | Polyketide | 28.0 | 29.1 | 168.0 | 7.6e-41 |
| AHH99919.1 | PKS\_I | BGC0000002 | Polyketide | 27.0 | 28.2 | 168.0 | 9.9e-41 |
| ABI91465.1 | beta-ketoacyl\_synthase | BGC0001094 | NRP + Polyketide | 26.0 | 35.6 | 168.0 | 1.3e-40 |
| ctg1\_orf254 |  | BGC0001200 | Polyketide | 27.0 | 29.6 | 168.0 | 1.3e-40 |
| AFV30247.1 | polyketide\_synthase | BGC0000075 | Polyketide | 28.0 | 30.4 | 167.0 | 1.7e-40 |
| ctg1\_orf30 |  | BGC0000096 | Polyketide | 30.0 | 25.2 | 167.0 | 1.7e-40 |
| AAK19883.1 | soraphen\_polyketide\_synthase\_A | BGC0000147 | Polyketide:Modular type I | 30.0 | 21.2 | 167.0 | 1.7e-40 |
| ARO38317.1 | nonribosomal\_peptide\_synthetase | BGC0001560 | NRP + Polyketide | 29.0 | 29.7 | 167.0 | 1.7e-40 |
| RWQ92175.1 | putative\_polyketide\_synthase | BGC0002030 | Polyketide | 26.0 | 34.4 | 167.0 | 1.7e-40 |
| BAG85027.1 | putative\_polyketide\_synthase | BGC0000086 | Polyketide | 29.0 | 20.9 | 167.0 | 2.2e-40 |
| CAQ64687.1 | lasalocid\_modular\_polyketide\_synthase | BGC0000087 | Polyketide | 29.0 | 20.9 | 167.0 | 2.2e-40 |
| CBF74114.1 | Conidial\_yellow\_pigment\_biosynthesis\_polyketide\_synthase\_(PKS)(EC\_2.3.1.-)\_[Source:UniProtKB/Swiss-Prot;Acc:Q03149] | BGC0000107 | Polyketide | 29.0 | 20.3 | 166.0 | 2.9e-40 |
| AEZ53952.1 | polyketide\_synthase | BGC0000144 | Polyketide:Modular type I | 21.0 | 110.4 | 166.0 | 2.9e-40 |
| ABC87509.1 | polyketide\_synthase | BGC0001011 | NRP + Polyketide | 27.0 | 29.4 | 166.0 | 2.9e-40 |
| ctg1\_orf20 |  | BGC0001013 | NRP + Polyketide | 27.0 | 29.4 | 166.0 | 2.9e-40 |
| DAB41653.1 | polyketide\_synthase | BGC0001583 | Polyketide | 27.0 | 29.3 | 166.0 | 2.9e-40 |
| ADA69241.1 | cis-AT\_polyketide\_synthase | BGC0001071 | NRP + Polyketide:Modular type I + Polyketide:Trans-AT type I | 31.0 | 24.4 | 166.0 | 4.9e-40 |
| BAG85028.1 | putative\_polyketide\_synthase | BGC0000086 | Polyketide | 29.0 | 21.2 | 165.0 | 6.4e-40 |
| CAQ64688.1 | lasalocid\_modular\_polyketide\_synthase | BGC0000087 | Polyketide | 29.0 | 21.2 | 165.0 | 6.4e-40 |
| ABV83229.1 | CppB | BGC0000116 | Polyketide | 28.0 | 31.2 | 165.0 | 6.4e-40 |
| AFD30954.1 | CrmA | BGC0000966 | NRP + Polyketide | 33.0 | 22.3 | 165.0 | 6.4e-40 |
| EED21099.1 | polyketide\_synthase,\_putative | BGC0001578 | Polyketide | 27.0 | 29.7 | 165.0 | 6.4e-40 |
| AGC95321.1 | CurS2 | BGC0000045 | Polyketide | 32.0 | 20.1 | 165.0 | 8.4e-40 |
| BAE93729.1 | type\_I\_polyketide\_synthase | BGC0000164 | Polyketide | 32.0 | 22.4 | 165.0 | 8.4e-40 |
| ACB37755.1 | putative\_type\_I\_polyketide\_synthase | BGC0000162 | Polyketide | 28.0 | 27.0 | 165.0 | 1.1e-39 |
| CAC22144.1 | CpkC;\_Polyketide\_synthase\_module\_5 | BGC0000038 | Polyketide:Modular type I | 26.0 | 32.1 | 164.0 | 1.4e-39 |
| CCE67070.1 | polyketide\_synthase | BGC0001242 | Polyketide | 28.0 | 29.5 | 164.0 | 1.4e-39 |
| AEH42474.1 | polyketide\_synthase | BGC0000032 | Polyketide | 27.0 | 27.0 | 164.0 | 1.9e-39 |
| AAZ77696.1 | ChlA3 | BGC0000036 | Polyketide:Modular type I + Polyketide:Iterative type I + Saccharide:Oligosaccharide | 30.0 | 20.4 | 163.0 | 2.4e-39 |
| CQR60496.1 | Polyketide\_synthase,\_type\_I,\_modules:\_4,\_5\_and\_6 | BGC0001287 | Polyketide | 30.0 | 21.5 | 163.0 | 2.4e-39 |
| AEH42490.1 | polyketide\_synthase | BGC0000032 | Polyketide | 30.0 | 21.7 | 163.0 | 4.1e-39 |
| CAO85897.1 | modular\_polyketide\_synthase\_NorB | BGC0000110 | Polyketide:Modular type I | 31.0 | 20.7 | 162.0 | 5.4e-39 |
| WP\_030498975.1 | type\_I\_polyketide\_synthase | BGC0001327 | NRP:Cyclic depsipeptide + Polyketide:Modular type I | 32.0 | 22.3 | 162.0 | 7.1e-39 |
| APZ78807.1 | polyketide\_synthase | BGC0001428 | NRP:Cyclic depsipeptide + Polyketide:Iterative type I | 25.0 | 36.7 | 162.0 | 7.1e-39 |
| CAE02605.1 | polyketide\_synthase\_type\_I | BGC0000024 | Polyketide:Modular type I | 29.0 | 22.4 | 161.0 | 1.2e-38 |
| AAC46027.1 | polyketide\_synthase\_module\_6 | BGC0000113 | Polyketide | 29.0 | 25.0 | 161.0 | 1.2e-38 |
| BAK64637.1 | polyketide\_synthase | BGC0000135 | Polyketide | 28.0 | 27.7 | 161.0 | 1.6e-38 |
| ALV82341.1 | borrelidin\_type\_I\_polyketide\_synthase | BGC0001533 | Polyketide | 30.0 | 20.9 | 160.0 | 2.1e-38 |
| AAQ82567.1 | FscE | BGC0000061 | Polyketide | 27.0 | 28.2 | 160.0 | 2.7e-38 |
| AGI99482.1 | Type\_I\_polyketide\_synthase | BGC0001004 | Polyketide:Modular type I | 29.0 | 25.3 | 159.0 | 6e-38 |
| ALP32043.1 | CycC | BGC0001293 | Polyketide | 31.0 | 20.4 | 159.0 | 6e-38 |
| AKD43522.1 | Type\_I\_polyketide\_synthase | BGC0001409 | Polyketide | 30.0 | 20.9 | 157.0 | 1.7e-37 |
| AAZ95017.1 | polyketide\_synthase | BGC0000048 | Polyketide | 29.0 | 24.6 | 156.0 | 3e-37 |
| ABC84459.1 | NigAIV | BGC0000114 | Polyketide:Modular type I | 28.0 | 24.9 | 156.0 | 3e-37 |
| BAA20102.2 | 6-methylsalicylic\_acid\_synthase | BGC0001276 | Polyketide | 29.0 | 22.4 | 156.0 | 3e-37 |
| AWC08658.1 | polyketide\_synthase\_type\_I | BGC0001932 | Polyketide | 27.0 | 26.0 | 156.0 | 3e-37 |
| CAE45670.1 | borrelidin\_polyketide\_synthase,\_type\_I | BGC0000031 | Polyketide:Modular type I | 30.0 | 20.9 | 156.0 | 3.9e-37 |
| AJO72734.1 | Type\_I\_modular\_polyketide\_synthase | BGC0001381 | Polyketide | 28.0 | 25.9 | 156.0 | 3.9e-37 |
| EAA59563.1 | polyketide\_synthase | BGC0000057 | Polyketide:Iterative type I | 31.0 | 20.7 | 156.0 | 5.1e-37 |
| AUO16400.1 | polyketide\_synthase | BGC0001700 | Polyketide | 28.0 | 22.9 | 156.0 | 5.1e-37 |
| ACB46471.1 | polyketide\_synthase | BGC0000082 | Polyketide | 28.0 | 25.2 | 155.0 | 6.6e-37 |
| ABS90471.1 | PKS\_type\_I | BGC0001106 | NRP + Polyketide | 27.0 | 29.5 | 155.0 | 6.6e-37 |
| QDA77044.1 | polyketide\_synthase | BGC0002025 | NRP | 27.0 | 25.2 | 155.0 | 6.6e-37 |
| ACB46485.1 | polyketide\_synthase | BGC0000082 | Polyketide | 29.0 | 23.3 | 155.0 | 1.1e-36 |
| AJW65408.1 | type\_I\_modular\_polyketide\_synthase | BGC0001195 | NRP + Polyketide | 24.0 | 36.8 | 154.0 | 1.5e-36 |
| AEU17897.1 | putative\_type\_I\_PKS | BGC0001072 | Saccharide + Polyketide:Modular type I + Polyketide:Type II + Other:Aminocoumarin | 26.0 | 30.7 | 154.0 | 1.9e-36 |
| CAA60459.1 | polyketide\_synthase | BGC0001040 | NRP + Polyketide | 26.0 | 32.7 | 153.0 | 2.5e-36 |
| AXM42949.1 | hybrid\_type\_1\_PKS/NRPS | BGC0001941 | NRP + Polyketide | 31.0 | 24.9 | 153.0 | 2.5e-36 |
| BAH02271.1 | polyketide\_synthase | BGC0000126 | Polyketide | 28.0 | 22.5 | 153.0 | 3.3e-36 |
| CAQ52623.1 | type\_I\_polyketide\_synthase,\_module\_6 | BGC0001066 | Polyketide:Modular type I | 26.0 | 25.0 | 153.0 | 3.3e-36 |
| ADM79459.1 | PKS16\_protein | BGC0001266 | Polyketide | 27.0 | 27.1 | 153.0 | 3.3e-36 |
| EED53479.1 | polyketide\_synthase,\_putative | BGC0001304 | Polyketide | 28.0 | 29.2 | 153.0 | 3.3e-36 |
| ANH11409.1 | SceN | BGC0001908 | Polyketide | 29.0 | 20.1 | 153.0 | 3.3e-36 |
| ANZ22986.1 | ZinC | BGC0001828 | Polyketide | 25.0 | 31.2 | 153.0 | 3.3e-36 |
| ABP57746.1 | DepB | BGC0000993 | NRP:Cyclic depsipeptide + Polyketide:Modular type I | 24.0 | 45.3 | 153.0 | 4.3e-36 |
| ASZ00149.1 | polyketide\_synthase | BGC0001785 | Polyketide | 30.0 | 21.1 | 153.0 | 4.3e-36 |
| ADX66461.1 | ScnS2 | BGC0000108 | Polyketide | 28.0 | 20.3 | 152.0 | 5.6e-36 |
| AAC01712.2 | RifC | BGC0000136 | Polyketide | 29.0 | 20.5 | 152.0 | 5.6e-36 |
| AGO59040.1 | PtaA | BGC0000121 | Polyketide | 28.0 | 29.2 | 152.0 | 7.3e-36 |
| AFR69332.1 | polyketide\_synthase\_SpiB | BGC0001045 | NRP:Cyclic depsipeptide + Polyketide:Modular type I | 24.0 | 42.5 | 151.0 | 9.6e-36 |
| AFL48528.1 | laidlomycin\_polyketide\_synthase\_(module\_7\_and\_module\_8) | BGC0000084 | Polyketide | 24.0 | 39.6 | 151.0 | 1.6e-35 |
| AEZ53946.1 | polyketide\_synthase | BGC0000144 | Polyketide:Modular type I | 26.0 | 27.3 | 150.0 | 2.1e-35 |
| CAC20921.1 | PimS2\_protein | BGC0000125 | Polyketide | 29.0 | 20.5 | 150.0 | 2.8e-35 |
| ALV82320.1 | borrelidin\_type\_I\_polyketide\_synthase | BGC0001533 | Polyketide | 30.0 | 20.9 | 150.0 | 2.8e-35 |
| AQT01393.1 | SgnS2 | BGC0001690 | Polyketide | 29.0 | 20.5 | 150.0 | 2.8e-35 |
| ANZ22989.1 | ZinF | BGC0001828 | Polyketide | 25.0 | 28.5 | 149.0 | 3.6e-35 |
| AFU82616.1 | polyketide\_synthase | BGC0000998 | NRP + Polyketide | 26.0 | 30.9 | 149.0 | 4.7e-35 |
| CAA60460.1 | polyketide\_synthase | BGC0001040 | NRP + Polyketide | 25.0 | 34.0 | 149.0 | 4.7e-35 |
| CAJ88177.1 | putative\_type\_I\_polyketide\_synthase | BGC0000151 | Polyketide:Modular type I + Saccharide:Hybrid/tailoring | 29.0 | 24.8 | 148.0 | 8.1e-35 |
| AAO65799.1 | monensin\_polyketide\_synthase\_modules\_5\_and\_6 | BGC0000100 | Polyketide | 27.0 | 24.9 | 148.0 | 1.1e-34 |
| ANZ52462.1 | MonAIV | BGC0001670 | Polyketide | 27.0 | 24.9 | 148.0 | 1.1e-34 |
| CAE45669.1 | borrelidin\_polyketide\_synthase,\_type\_I | BGC0000031 | Polyketide:Modular type I | 30.0 | 20.9 | 148.0 | 1.4e-34 |
| ACB37743.1 | putative\_type\_I\_polyketide\_synthase | BGC0000162 | Polyketide | 28.0 | 22.8 | 147.0 | 1.8e-34 |
| AJW65409.1 | type\_I\_modular\_polyketide\_synthase | BGC0001195 | NRP + Polyketide | 25.0 | 29.8 | 147.0 | 1.8e-34 |
| ADU86002.1 | putative\_modular\_polyketide\_synthase | BGC0000165 | Polyketide:Modular type I | 27.0 | 24.0 | 146.0 | 3.1e-34 |
| EED57518.1 | polyketide\_synthase,\_putative | BGC0001446 | Polyketide:Iterative type I | 29.0 | 20.2 | 146.0 | 3.1e-34 |
| ASA76643.1 | polyketide\_synthase | BGC0001751 | NRP + Polyketide | 23.0 | 42.7 | 146.0 | 3.1e-34 |
| AAZ77694.1 | ChlA2 | BGC0000036 | Polyketide:Modular type I + Polyketide:Iterative type I + Saccharide:Oligosaccharide | 25.0 | 28.6 | 146.0 | 4e-34 |
| CAM00064.1 | EryAII\_Erythromycin\_polyketide\_synthase\_modules\_3\_and\_4 | BGC0000055 | Polyketide:Modular type I + Saccharide:Hybrid/tailoring | 25.0 | 28.5 | 146.0 | 4e-34 |
| EAU38791.1 | hypothetical\_protein | BGC0000161 | Polyketide:Iterative type I | 27.0 | 26.5 | 146.0 | 5.2e-34 |
| AJW65407.1 | type\_I\_modular\_polyketide\_synthase | BGC0001195 | NRP + Polyketide | 25.0 | 27.9 | 146.0 | 5.2e-34 |
| AKA59448.1 | polyketide\_synthase | BGC0001203 | NRP + Polyketide | 22.0 | 74.6 | 146.0 | 5.2e-34 |
| KFA69335.1 | hypothetical\_protein | BGC0001626 | Polyketide | 26.0 | 27.9 | 145.0 | 9e-34 |
| BAP34739.1 | type\_I\_polyketide\_synthase | BGC0000078 | Polyketide | 29.0 | 22.3 | 144.0 | 1.2e-33 |
| ACY06289.1 | type\_I\_polyketide\_synthase | BGC0001042 | NRP + Polyketide | 27.0 | 28.7 | 144.0 | 2e-33 |
| BAV19379.1 | polyketide\_synthase | BGC0001390 | NRP + Polyketide | 30.0 | 20.4 | 144.0 | 2e-33 |
| ACM79805.1 | ZmaA | BGC0001059 | NRP + Polyketide | 26.0 | 30.4 | 143.0 | 2.6e-33 |
| ASZ00150.1 | polyketide\_synthase | BGC0001785 | Polyketide | 30.0 | 22.0 | 143.0 | 2.6e-33 |
| CCT67991.1 | bikaverin\_cluster-polyketide\_synthase | BGC0000030 | Polyketide | 29.0 | 20.7 | 143.0 | 3.4e-33 |
| WP\_055469549.1 | type\_I\_polyketide\_synthase | BGC0001537 | Polyketide | 26.0 | 27.5 | 141.0 | 1.3e-32 |
| ATY46587.1 | polyketide\_synthase | BGC0001666 | Polyketide | 26.0 | 27.8 | 141.0 | 1.3e-32 |
| EAA65602.1 | hypothetical\_protein | BGC0000022 | Polyketide | 27.0 | 30.3 | 140.0 | 2.9e-32 |
| gene6 |  | BGC0001906 | Polyketide | 30.0 | 22.1 | 139.0 | 4.9e-32 |
| ANR02554.1 | LodM | BGC0001648 | Polyketide | 24.0 | 37.6 | 137.0 | 1.9e-31 |
| BAG85026.1 | putative\_polyketide\_synthase | BGC0000086 | Polyketide | 25.0 | 26.9 | 136.0 | 3.2e-31 |
| CAQ64686.1 | lasalocid\_modular\_polyketide\_synthase | BGC0000087 | Polyketide | 25.0 | 26.9 | 136.0 | 3.2e-31 |
| AAD38786.1 | polyketide\_synthase | BGC0001257 | Polyketide | 30.0 | 20.1 | 136.0 | 4.2e-31 |
| BAD08373.1 | polyketide\_synthase\_modules\_1-3 | BGC0000167 | Polyketide | 24.0 | 31.9 | 135.0 | 7.1e-31 |
| AEZ64504.1 | Herc | BGC0001065 | Polyketide | 25.0 | 27.7 | 134.0 | 1.2e-30 |
| ABV97152.1 | Beta-ketoacyl\_synthase | BGC0000137 | Polyketide | 27.0 | 30.8 | 133.0 | 2.7e-30 |
| ACF35446.1 | mbcAII | BGC0000090 | Polyketide | 26.0 | 20.6 | 130.0 | 3e-29 |
| AAG13918.1 | megalomicin\_6-deoxyerythronolide\_B\_synthase\_2 | BGC0000092 | Polyketide | 24.0 | 29.0 | 125.0 | 9.6e-28 |
| AFL48526.1 | laidlomycin\_polyketide\_synthase\_(module\_2) | BGC0000084 | Polyketide | 23.0 | 28.6 | 114.0 | 1.7e-24 |
| WP\_083502114.1 | type\_I\_polyketide\_synthase | BGC0001653 | Polyketide | 23.0 | 28.3 | 109.0 | 5.4e-23 |
| AKA59093.1 | type-I\_PKS | BGC0001619 | Polyketide | 24.0 | 27.7 | 106.0 | 6e-22 |
| APZ78754.1 | polyketide\_synthase | BGC0001423 | NRP:Cyclic depsipeptide + Polyketide:Iterative type I | 27.0 | 25.7 | 102.0 | 5.1e-21 |
| CAG28678.1 | polyketide\_synthase | BGC0001023 | NRP + Polyketide:Modular type I | 30.0 | 20.6 | 102.0 | 8.7e-21 |
| APZ78727.1 | polyketide\_synthase | BGC0001421 | NRP:Cyclic depsipeptide + Polyketide:Iterative type I | 27.0 | 25.7 | 102.0 | 8.7e-21 |
| APZ78820.1 | polyketide\_synthase | BGC0001429 | NRP:Cyclic depsipeptide + Polyketide:Iterative type I | 30.0 | 20.6 | 102.0 | 8.7e-21 |
| APZ78767.1 | polyketide\_synthase | BGC0001425 | NRP:Cyclic depsipeptide + Polyketide:Iterative type I | 29.0 | 20.3 | 96.0 | 4.8e-19 |
