## Supplementary Results for "A draft genome of the ascomycotal fungal species *Pseudopithomyces maydicus* (family *Didymosphaeriaceae*)": input.path1.gene388_mibig_hits.html

| MIBiG Protein | Description | MIBiG Cluster | MiBiG Product | % ID | % Coverage | BLAST Score | E-value |
| --- | --- | --- | --- | --- | --- | --- | --- |
| QBE85642.1 | BuaC | BGC0001978 | NRP + Polyketide | 59.0 | 100.0 | 398.0 | 8.7e-111 |
| AGO86659.1 | equisetin\_enoylreductase | BGC0001255 | NRP + Polyketide | 40.0 | 100.0 | 259.0 | 6.3e-69 |
| BBC43187.1 | trans-enoyl\_reductase | BGC0001738 | NRP + Polyketide | 39.0 | 100.3 | 246.0 | 7.3e-65 |
| CEF75882.1 |  | BGC0001600 | Polyketide | 36.0 | 100.6 | 241.0 | 2.3e-63 |
| AAD34554.1 | enoyl\_reductase | BGC0000088 | Polyketide | 39.0 | 100.6 | 237.0 | 4.4e-62 |
| BAC20562.1 | enoyl\_reductase | BGC0000039 | Polyketide | 40.0 | 94.7 | 234.0 | 2.9e-61 |
| CBF80481.1 | enoylreductase | BGC0000959 | NRP + Polyketide:Iterative type I | 39.0 | 92.4 | 233.0 | 3.7e-61 |
| XP\_001220461.1 | hypothetical\_protein | BGC0001182 | NRP + Polyketide:Iterative type I | 38.0 | 101.8 | 227.0 | 2.7e-59 |
| ARP51714.1 | NADP-dependent\_dehydrogenase\_/enoyl-reductase | BGC0001741 | NRP + Polyketide | 36.0 | 103.5 | 227.0 | 3.5e-59 |
| ABA02243.1 | dehydrogenase | BGC0000098 | Polyketide | 39.0 | 95.9 | 225.0 | 1e-58 |
| EAW09121.1 | zinc-binding\_dehydrogenase\_family\_oxidoreductase,\_putative | BGC0000983 | NRP + Polyketide:Iterative type I | 34.0 | 100.3 | 213.0 | 4e-55 |
| CCT72378.1 | related\_to\_C.carbonum\_toxD\_protein | BGC0001305 | Polyketide | 33.0 | 99.7 | 185.0 | 2e-46 |
| EPS29079.1 | hypothetical\_protein | BGC0001724 | NRP + Polyketide | 31.0 | 98.2 | 176.0 | 7.1e-44 |
| AEO57491.1 | enoylreductase | BGC0001449 | NRP + Alkaloid + Polyketide:Iterative type I | 33.0 | 105.3 | 175.0 | 1.2e-43 |
| QBC19713.1 | TwmE | BGC0001954 | NRP + Polyketide | 32.0 | 99.1 | 154.0 | 2.9e-37 |
| EHA28239.1 | hypothetical\_protein | BGC0001143 | Polyketide | 28.0 | 105.3 | 101.0 | 2.9e-21 |
| AEE88282.1 | CurH | BGC0000976 | NRP + Polyketide:Modular type I | 26.0 | 46.2 | 57.0 | 6.2e-08 |
| AAT70103.1 | CurH | BGC0001165 | NRP + Polyketide:Modular type I | 26.0 | 46.2 | 57.0 | 6.2e-08 |
| CCT75967.1 | polyketide\_synthase | BGC0001606 | Polyketide | 26.0 | 57.3 | 51.0 | 2.6e-06 |
| AEZ53946.1 | polyketide\_synthase | BGC0000144 | Polyketide:Modular type I | 36.0 | 30.4 | 50.0 | 7.6e-06 |
