## Supplementary Results for "A draft genome of the ascomycotal fungal species *Pseudopithomyces maydicus* (family *Didymosphaeriaceae*)": input.path1.gene389_mibig_hits.html

| MIBiG Protein | Description | MIBiG Cluster | MiBiG Product | % ID | % Coverage | BLAST Score | E-value |
| --- | --- | --- | --- | --- | --- | --- | --- |
| AAY28421.1 | phytoene\_synthase | BGC0000630 | Terpene | 29.0 | 63.1 | 94.0 | 4.2e-19 |
| BAE47469.1 | phytoene\_synthase | BGC0000635 | Terpene | 29.0 | 63.1 | 92.0 | 1.2e-18 |
| AAZ73137.1 | phytoene\_synthase | BGC0000640 | Terpene | 28.0 | 64.5 | 92.0 | 2.1e-18 |
