## Supplementary Results for "A draft genome of the ascomycotal fungal species *Pseudopithomyces maydicus* (family *Didymosphaeriaceae*)": input.path1.gene392_mibig_hits.html

| MIBiG Protein | Description | MIBiG Cluster | MiBiG Product | % ID | % Coverage | BLAST Score | E-value |
| --- | --- | --- | --- | --- | --- | --- | --- |
| QBE85649.1 | BuaA | BGC0001978 | NRP + Polyketide | 43.0 | 144.9 | 2799.0 | 0.0 |
| EAW09117.1 | hybrid\_NRPS/PKS\_enzyme,\_putative | BGC0000983 | NRP + Polyketide:Iterative type I | 27.0 | 147.6 | 1253.0 | 0.0 |
| AGO86662.1 | equisetin\_synthetase | BGC0001255 | NRP + Polyketide | 27.0 | 144.9 | 1206.0 | 0.0 |
| ARP51711.1 | PKS-NRPS\_hybrid\_protein | BGC0001741 | NRP + Polyketide | 27.0 | 142.4 | 1203.0 | 0.0 |
| BAZ95823.1 | PKS-NRPS\_hybrid\_cpaA | BGC0001563 | NRP + Polyketide | 27.0 | 137.1 | 1151.0 | 0.0 |
| AEO57481.1 | PKS-NRPSs | BGC0001449 | NRP + Alkaloid + Polyketide:Iterative type I | 26.0 | 147.1 | 1108.0 | 0.0 |
| EPS29069.1 | hypothetical\_protein | BGC0001724 | NRP + Polyketide | 39.0 | 63.6 | 1086.0 | 0.0 |
| EED49862.1 | hybrid\_PKS/NRPS\_enzyme,\_putative | BGC0001445 | NRP + Polyketide:Iterative type I | 26.0 | 139.5 | 1086.0 | 0.0 |
| BBC43184.1 | PKS-NRPS\_hybrid | BGC0001738 | NRP + Polyketide | 37.0 | 62.3 | 1032.0 | 1.6e-300 |
| XP\_001220460.1 | hypothetical\_protein | BGC0001182 | NRP + Polyketide:Iterative type I | 39.0 | 61.2 | 1009.0 | 1.1e-293 |
| QCS37521.1 | PyiS | BGC0001982 | NRP + Polyketide | 38.0 | 61.0 | 1004.0 | 3.5e-292 |
| QBC19710.1 | TwmB | BGC0001954 | NRP + Polyketide | 38.0 | 60.5 | 986.0 | 9.8e-287 |
| BAJ14522.1 | polyketide\_synthase | BGC0001254 | Polyketide | 38.0 | 60.7 | 984.0 | 3.7e-286 |
| CCT72377.1 | probable\_polyketide\_synthase | BGC0001305 | Polyketide | 37.0 | 60.6 | 982.0 | 1.4e-285 |
| AAT28740.1 | FUSS | BGC0000064 | Polyketide | 29.0 | 103.5 | 965.0 | 1.4e-280 |
| ATZ45182.1 | Bcboa6 | BGC0001892 | Polyketide | 35.0 | 66.5 | 962.0 | 1.2e-279 |
| AFP73394.1 | FusA | BGC0001268 | NRP + Polyketide | 29.0 | 103.4 | 961.0 | 2.6e-279 |
| CEF75886.1 |  | BGC0001600 | Polyketide | 38.0 | 60.8 | 961.0 | 2.6e-279 |
| ABA02239.1 | polyketide\_synthase | BGC0000098 | Polyketide | 37.0 | 59.9 | 951.0 | 2.7e-276 |
| CBF80487.1 | hybrid\_PKS-NRPS\_(Eurofung) | BGC0000959 | NRP + Polyketide:Iterative type I | 37.0 | 60.6 | 948.0 | 3e-275 |
| gene3 |  | BGC0002035 | NRP + Polyketide | 36.0 | 61.2 | 945.0 | 1.5e-274 |
| BAC20564.1 | polyketide\_synthase | BGC0000039 | Polyketide | 36.0 | 64.2 | 940.0 | 6.2e-273 |
| ACS68554.1 | hybrid\_PKS-NRPS\_protein | BGC0001026 | NRP + Polyketide | 35.0 | 66.0 | 937.0 | 5.2e-272 |
| BAQ25466.1 | polyketide\_synthase | BGC0001280 | Polyketide | 35.0 | 64.0 | 933.0 | 5.8e-271 |
| EAU29808.1 | hypothetical\_protein | BGC0001400 | Polyketide | 37.0 | 59.2 | 926.0 | 7.1e-269 |
| XP\_659388.1 | hypothetical\_protein | BGC0001998 | Polyketide | 36.0 | 64.8 | 911.0 | 4e-264 |
| CAO91861.1 | PKS-NRPS\_hybrid | BGC0000968 | NRP + Polyketide:Iterative type I | 34.0 | 63.9 | 896.0 | 1.3e-259 |
| ADN43685.1 | DmbS | BGC0001136 | NRP + Polyketide:Iterative type I | 36.0 | 61.6 | 884.0 | 3.1e-256 |
| CAL69597.1 | PKS-NRPS | BGC0001049 | NRP + Polyketide:Iterative type I | 35.0 | 61.9 | 868.0 | 3e-251 |
| AAV66110.2 | fusaridione\_A\_synthetase | BGC0000992 | NRP + Polyketide | 40.0 | 47.7 | 832.0 | 1.4e-240 |
| EAL85113.2 | hybrid\_PKS-NRPS\_enzyme | BGC0001037 | NRP + Polyketide:Iterative type I | 35.0 | 62.3 | 821.0 | 4.2e-237 |
| EAU38971.1 | hypothetical\_protein | BGC0001122 | NRP + Polyketide:Iterative type I | 34.0 | 64.2 | 819.0 | 1.2e-236 |
| BAK26562.1 | PKS-NRPS\_hybrid | BGC0000977 | NRP + Polyketide | 33.0 | 61.3 | 784.0 | 4.3e-226 |
| AFA26384.1 | polyketide\_synthase\_A | BGC0001874 | NRP + Polyketide | 33.0 | 61.4 | 767.0 | 5.5e-221 |
| AKC54422.1 | fumosorinone\_biosynthesis\_polyketide\_synthase | BGC0001218 | NRP + Polyketide | 38.0 | 45.6 | 739.0 | 2.1e-212 |
| EAL89230.2 | LovB-like\_polyketide\_synthase,\_putative | BGC0000129 | Polyketide | 39.0 | 43.3 | 720.0 | 7.7e-207 |
| ctg1\_orf0002 |  | BGC0001068 | Terpene + Polyketide | 38.0 | 44.1 | 713.0 | 1.2e-204 |
| BAD83684.1 | PKSN\_polyketide\_synthase\_for\_alternapyrone\_biosynthesis | BGC0000012 | Polyketide | 29.0 | 62.8 | 521.0 | 6.9e-147 |
| CBF87072.1 | polyketide\_synthase,\_putative\_(Eurofung) | BGC0001290 | NRP | 30.0 | 57.7 | 500.0 | 1.6e-140 |
| AMY15057.1 | tetraketide\_synthase\_MF-SQTKS | BGC0001339 | Polyketide:Iterative type I | 27.0 | 66.8 | 489.0 | 3.8e-137 |
| AAD43562.2 | Fum1p | BGC0000062 | Polyketide | 27.0 | 65.5 | 484.0 | 1.2e-135 |
| EHA19289.1 | hypothetical\_protein | BGC0001124 | Polyketide | 34.0 | 42.5 | 482.0 | 4.6e-135 |
| EPE34340.1 | polyketide\_synthase | BGC0001035 | Polyketide + NRP + Other:Aminocoumarin | 28.0 | 61.6 | 481.0 | 1e-134 |
| ACB12550.1 | Fum1 | BGC0000063 | Polyketide | 27.0 | 65.2 | 477.0 | 1.5e-133 |
| gene4 |  | BGC0001907 | Polyketide | 28.0 | 60.0 | 477.0 | 1.5e-133 |
| AKL78824.1 | GLPKS3 | BGC0001187 | NRP:Lipopeptide + Polyketide:Iterative type I | 28.0 | 61.4 | 474.0 | 1.3e-132 |
| EAA65604.1 | hypothetical\_protein | BGC0000022 | Polyketide | 32.0 | 41.8 | 469.0 | 3.1e-131 |
| AAD34559.1 | polyketide\_synthase | BGC0000088 | Polyketide | 27.0 | 62.6 | 465.0 | 5.8e-130 |
| KGO40478.1 | Acyl\_transferase/acyl\_hydrolase/lysophospholipase | BGC0001205 | Polyketide | 38.0 | 31.8 | 464.0 | 1e-129 |
| OAQ83760.1 | polyketide\_synthase | BGC0001358 | Polyketide | 26.0 | 66.9 | 456.0 | 2.7e-127 |
| CBX99534.1 | similar\_to\_polyketide\_synthase | BGC0001899 | Polyketide | 26.0 | 61.9 | 452.0 | 3.9e-126 |
| KKP00963.1 | fatty\_acid\_synthase\_S-acetyltransferase | BGC0001901 | Polyketide | 28.0 | 58.1 | 444.0 | 1.1e-123 |
| AMY15068.1 | hexaketide\_synthase\_MF-SQHKS | BGC0001339 | Polyketide:Iterative type I | 29.0 | 55.6 | 440.0 | 2.6e-122 |
| RWQ92174.1 | KR\_domain-containing\_protein | BGC0002030 | Polyketide | 29.0 | 50.3 | 438.0 | 7.6e-122 |
| EHA28244.1 | hypothetical\_protein | BGC0001143 | Polyketide | 29.0 | 49.3 | 436.0 | 3.8e-121 |
| BAD97694.1 | Aft9-1 | BGC0000003 | Polyketide | 31.0 | 44.3 | 434.0 | 1.4e-120 |
| ASK38717.1 | polyketide\_synthase | BGC0001557 | Polyketide | 27.0 | 59.9 | 434.0 | 1.4e-120 |
| AGC95324.1 | CurS1 | BGC0000045 | Polyketide | 32.0 | 41.9 | 431.0 | 1.2e-119 |
| CAP95405.1 |  | BGC0001404 | Polyketide | 31.0 | 41.4 | 430.0 | 2.7e-119 |
| ABB90283.1 | polyketide\_synthase | BGC0001057 | NRP + Polyketide | 33.0 | 38.5 | 426.0 | 3.9e-118 |
| ACD39758.1 | reducing\_polyketide\_synthase | BGC0000076 | Polyketide | 32.0 | 38.6 | 425.0 | 5.1e-118 |
| ACD39767.1 | reducing\_polyketide\_synthase | BGC0000077 | Polyketide | 32.0 | 38.6 | 425.0 | 5.1e-118 |
| BAV32159.1 | polyketide\_synthase | BGC0001373 | Polyketide | 31.0 | 40.7 | 425.0 | 6.7e-118 |
| ABA02240.1 | polyketide\_synthase | BGC0000098 | Polyketide | 27.0 | 49.4 | 425.0 | 8.7e-118 |
| ACD39774.1 | reducing\_polyketide\_synthase | BGC0000134 | Polyketide | 27.0 | 60.5 | 425.0 | 8.7e-118 |
| AHV78245.1 | LasS1 | BGC0001245 | Polyketide | 34.0 | 38.7 | 418.0 | 8.2e-116 |
| EAA36364.1 | hypothetical\_protein | BGC0001697 | Polyketide | 31.0 | 40.4 | 418.0 | 1.1e-115 |
| ANF07288.1 | hrPKS | BGC0001340 | Polyketide:Iterative type I | 31.0 | 40.3 | 416.0 | 2.4e-115 |
| ATQ39432.1 | PKS | BGC0001565 | NRP | 26.0 | 64.1 | 412.0 | 4.5e-114 |
| BAC20566.1 | polyketide\_synthase | BGC0000039 | Polyketide | 28.0 | 46.8 | 407.0 | 1.4e-112 |
| AQM58285.1 | polyketide\_synthase | BGC0001816 | NRP + Polyketide | 26.0 | 63.7 | 407.0 | 1.9e-112 |
| BBG28498.1 | putative\_polyketide\_synthase | BGC0001913 | Polyketide | 31.0 | 40.5 | 405.0 | 5.5e-112 |
| EWG54266.1 | hypothetical\_protein | BGC0001190 | Polyketide | 32.0 | 39.9 | 401.0 | 7.9e-111 |
| AHV78252.1 | ResS1 | BGC0001246 | Polyketide | 33.0 | 31.8 | 400.0 | 1.8e-110 |
| EHA52508.1 | mycocerosic\_acid\_synthase | BGC0001749 | Polyketide | 32.0 | 36.3 | 400.0 | 1.8e-110 |
| AIA58899.1 | HRPKS | BGC0001141 | Polyketide:Iterative type I | 31.0 | 37.7 | 399.0 | 3.9e-110 |
| EAL85129.1 | polyketide\_synthase | BGC0001067 | Terpene + Polyketide:Iterative type I | 27.0 | 57.9 | 398.0 | 8.8e-110 |
| EAQ86385.1 | hypothetical\_protein | BGC0001405 | Polyketide | 27.0 | 61.5 | 398.0 | 1.1e-109 |
| QCC63000.1 | BII-rafflesfungin\_polyketide\_synthase | BGC0001966 | NRP | 26.0 | 64.0 | 397.0 | 1.5e-109 |
| BAJ09789.1 | polyketide\_synthase | BGC0000146 | Polyketide | 29.0 | 40.4 | 386.0 | 3.4e-106 |
| AAF62884.1 | EpoE | BGC0000991 | NRP + Polyketide | 28.0 | 52.2 | 385.0 | 5.9e-106 |
| ACB46196.1 | polyketide\_synthase | BGC0000989 | NRP + Polyketide | 28.0 | 52.2 | 383.0 | 2.9e-105 |
| AAF26922.1 | polyketide\_synthase | BGC0000988 | NRP + Polyketide | 28.0 | 52.2 | 382.0 | 5e-105 |
| BBG28484.1 | polyketide\_synthase\_CdmE | BGC0001926 | Polyketide | 32.0 | 34.8 | 382.0 | 5e-105 |
| APD26279.1 | PtmA | BGC0001726 | NRP + Polyketide | 31.0 | 39.0 | 382.0 | 6.5e-105 |
| ADB12492.1 | EpoE | BGC0000990 | NRP + Polyketide | 28.0 | 52.3 | 378.0 | 1.2e-103 |
| EFL02193.1 | amino\_acid\_adenylation\_domain-containing\_protein | BGC0000996 | NRP + Polyketide:Iterative type I | 30.0 | 41.1 | 371.0 | 1.5e-101 |
| AQW44891.1 | polyketide\_synthase | BGC0001737 | NRP + Polyketide | 31.0 | 40.1 | 369.0 | 5.7e-101 |
| AUS29495.1 | polyketide\_synthase | BGC0001030 | NRP + Polyketide | 30.0 | 37.7 | 364.0 | 1.4e-99 |
| CAJ46690.1 | polyketide\_synthase | BGC0000969 | NRP:Cyclic depsipeptide + Polyketide:Modular type I | 31.0 | 36.7 | 361.0 | 9.1e-99 |
| ACZ57548.1 | polyketide\_synthase | BGC0000046 | Polyketide:Iterative type I | 28.0 | 43.2 | 360.0 | 2.6e-98 |
| BAG17643.1 | putative\_NRPS-type-I\_PKS\_fusion\_protein | BGC0001043 | NRP + Polyketide | 31.0 | 38.0 | 358.0 | 7.7e-98 |
| ABL86391.1 | hybrid\_polyketide\_synthase\_and\_nonribosomal\_peptide\_synthetase | BGC0000999 | NRP + Polyketide | 32.0 | 36.3 | 356.0 | 3.8e-97 |
| AAK57187.1 | MxaC | BGC0001022 | NRP + Polyketide | 31.0 | 37.2 | 353.0 | 3.2e-96 |
| AGC45621.1 | polyketide\_synthase | BGC0001394 | NRP + Polyketide | 31.0 | 38.8 | 351.0 | 1.6e-95 |
| AQW44873.1 | polyketide\_synthase | BGC0001761 | Polyketide | 31.0 | 39.8 | 349.0 | 6.1e-95 |
| AQW44889.1 | polyketide\_synthase | BGC0001737 | NRP + Polyketide | 30.0 | 37.6 | 345.0 | 6.7e-94 |
| ADB12491.1 | EpoD | BGC0000990 | NRP + Polyketide | 29.0 | 37.6 | 345.0 | 8.8e-94 |
| AJD77023.1 | IkaA | BGC0001435 | NRP + Polyketide:Iterative type I | 31.0 | 38.4 | 344.0 | 1.5e-93 |
| ACB46195.1 | polyketide\_synthase | BGC0000989 | NRP + Polyketide | 29.0 | 37.9 | 344.0 | 2e-93 |
| AAF62883.1 | epoD | BGC0000991 | NRP + Polyketide | 30.0 | 37.5 | 343.0 | 3.3e-93 |
| AAF26921.1 | polyketide\_synthase | BGC0000988 | NRP + Polyketide | 29.0 | 37.9 | 342.0 | 5.7e-93 |
| AUD08663.1 | iPKS-NRPS | BGC0001553 | NRP + Polyketide | 30.0 | 43.0 | 338.0 | 1.1e-91 |
| AIT55263.1 | polyketide\_synthase | BGC0000072 | Polyketide:Modular type I | 30.0 | 38.2 | 335.0 | 7e-91 |
| AGC45624.1 | polyketide\_synthase | BGC0001394 | NRP + Polyketide | 31.0 | 37.7 | 332.0 | 7.7e-90 |
| ATZ45185.1 | Bcboa9 | BGC0001892 | Polyketide | 31.0 | 32.9 | 332.0 | 7.7e-90 |
| APZ78678.1 | polyketide\_synthase | BGC0001417 | NRP:Cyclic depsipeptide + Polyketide:Iterative type I | 29.0 | 41.3 | 331.0 | 1.3e-89 |
| CAQ43075.1 | polyketide\_synthase | BGC0000970 | NRP + Polyketide:Modular type I | 30.0 | 38.0 | 331.0 | 1.7e-89 |
| CCE88378.1 | polyketide\_synthase | BGC0001034 | NRP + Polyketide:Modular type I | 32.0 | 30.3 | 325.0 | 7.2e-88 |
| QDA77058.1 | polyketide\_synthase | BGC0002026 | NRP | 33.0 | 30.2 | 325.0 | 9.4e-88 |
| APZ78844.1 | polyketide\_synthase | BGC0001431 | NRP:Cyclic depsipeptide + Polyketide:Iterative type I | 29.0 | 41.1 | 324.0 | 1.6e-87 |
| APZ78702.1 | polyketide\_synthase | BGC0001419 | NRP:Cyclic depsipeptide + Polyketide:Iterative type I | 29.0 | 40.5 | 323.0 | 3.6e-87 |
| AGC45620.1 | polyketide\_synthase | BGC0001394 | NRP + Polyketide | 29.0 | 37.6 | 322.0 | 6.1e-87 |
| AAK57189.1 | MxaE | BGC0001022 | NRP + Polyketide | 29.0 | 36.9 | 321.0 | 1e-86 |
| APZ78690.1 | polyketide\_synthase | BGC0001418 | NRP:Cyclic depsipeptide + Polyketide:Iterative type I | 29.0 | 40.5 | 321.0 | 1.4e-86 |
| AZH23788.1 | MgcR | BGC0001970 | NRP + Polyketide | 29.0 | 36.1 | 321.0 | 1.4e-86 |
| APZ78714.1 | polyketide\_synthase | BGC0001420 | NRP:Cyclic depsipeptide + Polyketide:Iterative type I | 29.0 | 40.5 | 321.0 | 1.8e-86 |
| APZ78854.1 | polyketide\_synthase | BGC0001432 | NRP:Cyclic depsipeptide + Polyketide:Iterative type I | 28.0 | 41.3 | 321.0 | 1.8e-86 |
| CAD19091.1 | StiG\_protein | BGC0000153 | NRP + Polyketide:Modular type I | 29.0 | 41.8 | 320.0 | 2.3e-86 |
| AQW44892.1 | polyketide\_synthase | BGC0001737 | NRP + Polyketide | 31.0 | 31.6 | 319.0 | 4e-86 |
| CAQ18830.1 | polyketide\_synthase | BGC0000954 | NRP + Polyketide:Modular type I | 30.0 | 39.2 | 319.0 | 5.2e-86 |
| QDA77044.1 | polyketide\_synthase | BGC0002025 | NRP | 31.0 | 32.0 | 319.0 | 5.2e-86 |
| APZ78754.1 | polyketide\_synthase | BGC0001423 | NRP:Cyclic depsipeptide + Polyketide:Iterative type I | 29.0 | 40.4 | 319.0 | 6.8e-86 |
| ADB12493.1 | EpoF | BGC0000990 | NRP + Polyketide | 27.0 | 37.9 | 318.0 | 1.2e-85 |
| AAF26923.1 | polyketide\_synthase | BGC0000988 | NRP + Polyketide | 28.0 | 38.1 | 317.0 | 2e-85 |
| AWO77084.1 | hybrid\_non-ribosomal\_peptide\_synthetase/type\_I\_polyketide\_synthase | BGC0001556 | NRP + Polyketide | 30.0 | 39.4 | 317.0 | 2e-85 |
| APZ78780.1 | polyketide\_synthase | BGC0001426 | NRP:Cyclic depsipeptide + Polyketide:Iterative type I | 28.0 | 40.9 | 317.0 | 2.6e-85 |
| CAD19089.1 | StiE\_protein | BGC0000153 | NRP + Polyketide:Modular type I | 35.0 | 25.6 | 316.0 | 3.4e-85 |
| CAQ18832.1 | polyketide\_synthase | BGC0000954 | NRP + Polyketide:Modular type I | 29.0 | 37.3 | 316.0 | 3.4e-85 |
| AAF19813.1 | MtaE | BGC0001024 | NRP + Polyketide:Modular type I | 33.0 | 28.6 | 316.0 | 3.4e-85 |
| ACB46197.1 | polyketide\_synthase | BGC0000989 | NRP + Polyketide | 28.0 | 38.3 | 315.0 | 7.5e-85 |
| CCE88381.1 | polyketide\_synthase | BGC0001034 | NRP + Polyketide:Modular type I | 30.0 | 38.5 | 315.0 | 7.5e-85 |
| AQW44893.1 | polyketide\_synthase | BGC0001737 | NRP + Polyketide | 29.0 | 40.8 | 314.0 | 1.3e-84 |
| APZ78727.1 | polyketide\_synthase | BGC0001421 | NRP:Cyclic depsipeptide + Polyketide:Iterative type I | 29.0 | 36.1 | 314.0 | 1.7e-84 |
| AAF62885.1 | EpoF | BGC0000991 | NRP + Polyketide | 27.0 | 37.9 | 314.0 | 2.2e-84 |
| CAG28678.1 | polyketide\_synthase | BGC0001023 | NRP + Polyketide:Modular type I | 28.0 | 41.9 | 312.0 | 4.8e-84 |
| APZ78820.1 | polyketide\_synthase | BGC0001429 | NRP:Cyclic depsipeptide + Polyketide:Iterative type I | 28.0 | 41.9 | 312.0 | 4.8e-84 |
| APZ78742.1 | polyketide\_synthase | BGC0001422 | NRP:Cyclic depsipeptide + Polyketide:Iterative type I | 30.0 | 36.1 | 311.0 | 1.1e-83 |
| APZ78793.1 | polyketide\_synthase | BGC0001427 | NRP:Cyclic depsipeptide + Polyketide:Iterative type I | 28.0 | 40.9 | 311.0 | 1.1e-83 |
| ALD82522.1 | polyketide\_synthase | BGC0001212 | NRP + Polyketide | 27.0 | 39.6 | 311.0 | 1.8e-83 |
| AAC01711.1 | RifB | BGC0000136 | Polyketide | 30.0 | 36.5 | 310.0 | 2.4e-83 |
| AAA79984.2 | soraphen\_polyketide\_synthase\_B | BGC0000147 | Polyketide:Modular type I | 29.0 | 37.8 | 309.0 | 7e-83 |
| AQM37582.1 | polyketide\_synthase | BGC0001424 | NRP:Cyclic depsipeptide + Polyketide:Iterative type I | 30.0 | 36.1 | 307.0 | 1.6e-82 |
| APZ78807.1 | polyketide\_synthase | BGC0001428 | NRP:Cyclic depsipeptide + Polyketide:Iterative type I | 30.0 | 35.9 | 307.0 | 1.6e-82 |
| AQW44888.1 | polyketide\_synthase | BGC0001737 | NRP + Polyketide | 29.0 | 37.0 | 306.0 | 4.5e-82 |
| AWH12669.1 | RmpB | BGC0001759 | Polyketide | 29.0 | 36.3 | 306.0 | 4.5e-82 |
| APZ78832.1 | polyketide\_synthase | BGC0001430 | NRP:Cyclic depsipeptide + Polyketide:Iterative type I | 28.0 | 39.4 | 305.0 | 7.7e-82 |
| AIW82279.1 | PuwB | BGC0001125 | NRP + Polyketide | 28.0 | 39.3 | 304.0 | 1.3e-81 |
| CBD77734.1 | polyketide\_synthase | BGC0000974 | NRP + Polyketide | 33.0 | 27.6 | 304.0 | 2.2e-81 |
| AIR74911.1 | polyketide\_synthase | BGC0001559 | RiPP | 33.0 | 27.6 | 304.0 | 2.2e-81 |
| OAQ83765.1 | KR\_domain-containing\_protein | BGC0001358 | Polyketide | 27.0 | 50.8 | 303.0 | 2.9e-81 |
| CAQ18838.1 | polyketide\_synthase | BGC0000954 | NRP + Polyketide:Modular type I | 35.0 | 25.3 | 300.0 | 3.2e-80 |
| ARM20283.1 | polyketide\_synthase | BGC0001523 | Polyketide | 30.0 | 36.9 | 299.0 | 4.2e-80 |
| ABV97152.1 | Beta-ketoacyl\_synthase | BGC0000137 | Polyketide | 30.0 | 36.2 | 299.0 | 7.2e-80 |
| AUO16401.1 | polyketide\_synthase | BGC0001700 | Polyketide | 32.0 | 28.1 | 298.0 | 9.4e-80 |
| APZ78767.1 | polyketide\_synthase | BGC0001425 | NRP:Cyclic depsipeptide + Polyketide:Iterative type I | 28.0 | 40.6 | 298.0 | 1.2e-79 |
| AQA28562.1 | type\_I\_polyketide\_synthase | BGC0001663 | Polyketide | 26.0 | 42.1 | 297.0 | 2.1e-79 |
| ACN69990.1 | polyketide\_synthase | BGC0000079 | Polyketide | 28.0 | 37.2 | 296.0 | 3.6e-79 |
| ACB37740.1 | putative\_type\_I\_polyketide\_synthase | BGC0000162 | Polyketide | 28.0 | 38.9 | 295.0 | 6.1e-79 |
| AAF19810.1 | MtaB | BGC0001024 | NRP + Polyketide:Modular type I | 28.0 | 37.1 | 295.0 | 6.1e-79 |
| ADH04641.1 | TgaC | BGC0001051 | NRP + Polyketide:Modular type I | 31.0 | 31.6 | 295.0 | 6.1e-79 |
| AAW03328.1 | CtaE | BGC0000982 | NRP + Polyketide | 31.0 | 27.8 | 295.0 | 1e-78 |
| CAD89776.1 | MelE\_protein | BGC0001010 | NRP + Polyketide:Modular type I | 32.0 | 27.3 | 294.0 | 1.4e-78 |
| ALD82524.1 | polyketide\_synthase | BGC0001212 | NRP + Polyketide | 28.0 | 39.8 | 294.0 | 1.4e-78 |
| TXD00034.1 | SDR\_family\_NAD(P)-dependent\_oxidoreductase | BGC0001877 | Polyketide | 36.0 | 25.2 | 294.0 | 1.4e-78 |
| CAL58682.1 | polyketide\_synthase | BGC0000149 | Polyketide:Modular type I | 32.0 | 31.7 | 294.0 | 1.8e-78 |
| AIT55260.1 | polyketide\_synthase | BGC0000072 | Polyketide:Modular type I | 27.0 | 44.0 | 294.0 | 2.3e-78 |
| ASZ00148.1 | polyketide\_synthase | BGC0001785 | Polyketide | 29.0 | 37.0 | 293.0 | 3e-78 |
| CAL58684.1 | polyketide\_synthase | BGC0000149 | Polyketide:Modular type I | 31.0 | 30.3 | 293.0 | 4e-78 |
| ctg1\_orf522 |  | BGC0001199 | Polyketide | 30.0 | 36.4 | 293.0 | 4e-78 |
| CBD77732.1 | polyketide\_synthase | BGC0000974 | NRP + Polyketide | 34.0 | 24.9 | 292.0 | 6.8e-78 |
| AAK57186.1 | MxaB2 | BGC0001022 | NRP + Polyketide | 35.0 | 23.9 | 292.0 | 6.8e-78 |
| AIR74910.1 | polyketide\_synthase | BGC0001559 | RiPP | 34.0 | 24.9 | 292.0 | 6.8e-78 |
| AUO16397.1 | polyketide\_synthase | BGC0001700 | Polyketide | 29.0 | 36.2 | 292.0 | 8.8e-78 |
| CAL58681.1 | polyketide\_synthase | BGC0000149 | Polyketide:Modular type I | 31.0 | 30.3 | 291.0 | 1.2e-77 |
| AUO16403.1 | polyketide\_synthase | BGC0001700 | Polyketide | 31.0 | 30.5 | 291.0 | 1.2e-77 |
| AGC09484.1 | LobS1 | BGC0001183 | Polyketide | 28.0 | 36.8 | 290.0 | 2e-77 |
| AGI99497.1 | type\_I\_polyketide\_synthase | BGC0001004 | Polyketide:Modular type I | 27.0 | 40.6 | 290.0 | 2.6e-77 |
| ARM20280.1 | polyketide\_synthase | BGC0001523 | Polyketide | 32.0 | 28.1 | 290.0 | 2.6e-77 |
| ABM21570.1 | crpB | BGC0000975 | NRP + Polyketide | 33.0 | 24.4 | 289.0 | 5.7e-77 |
| QBF51754.1 | type\_I\_polyketide\_synthase | BGC0001856 | Polyketide:Modular type I | 34.0 | 26.0 | 289.0 | 7.5e-77 |
| AEE88289.1 | CurA | BGC0000976 | NRP + Polyketide:Modular type I | 31.0 | 27.6 | 287.0 | 2.8e-76 |
| AAT70096.1 | CurA | BGC0001165 | NRP + Polyketide:Modular type I | 31.0 | 27.6 | 287.0 | 2.8e-76 |
| BAJ16471.1 | polyketide\_synthase | BGC0000058 | Polyketide | 28.0 | 38.4 | 286.0 | 3.7e-76 |
| ctg1\_orf521 |  | BGC0001199 | Polyketide | 29.0 | 36.8 | 286.0 | 3.7e-76 |
| AZH23789.1 | MgcI | BGC0001970 | NRP + Polyketide | 28.0 | 37.7 | 286.0 | 3.7e-76 |
| CQR60496.1 | Polyketide\_synthase,\_type\_I,\_modules:\_4,\_5\_and\_6 | BGC0001287 | Polyketide | 29.0 | 36.4 | 286.0 | 4.9e-76 |
| ADZ24998.1 | polyketide\_synthase | BGC0000380 | NRP + Polyketide:Modular type I | 26.0 | 43.6 | 285.0 | 6.3e-76 |
| ALV82345.1 | borrelidin\_type\_I\_polyketide\_synthase | BGC0001533 | Polyketide | 29.0 | 36.4 | 285.0 | 8.3e-76 |
| CAE45671.1 | borrelidin\_polyketide\_synthase,\_type\_I | BGC0000031 | Polyketide:Modular type I | 29.0 | 36.4 | 284.0 | 1.4e-75 |
| CAI94713.1 | putative\_polyketide\_synthase | BGC0000141 | Polyketide | 31.0 | 29.9 | 284.0 | 2.4e-75 |
| AIT55259.1 | polyketide\_synthase | BGC0000072 | Polyketide:Modular type I | 32.0 | 27.3 | 283.0 | 3.1e-75 |
| QDA77059.1 | polyketide\_synthase/nonribosomal\_peptide\_synthetase | BGC0002026 | NRP | 34.0 | 25.8 | 283.0 | 3.1e-75 |
| AAK19883.1 | soraphen\_polyketide\_synthase\_A | BGC0000147 | Polyketide:Modular type I | 31.0 | 30.3 | 283.0 | 4.1e-75 |
| CAJ46689.1 | polyketide\_synthase | BGC0000969 | NRP:Cyclic depsipeptide + Polyketide:Modular type I | 32.0 | 26.5 | 283.0 | 4.1e-75 |
| WP\_042799407.1 | type\_I\_polyketide\_synthase | BGC0001283 | Polyketide | 27.0 | 36.8 | 283.0 | 4.1e-75 |
| BAB69198.1 | modular\_polyketide\_synthase | BGC0000117 | Polyketide | 30.0 | 32.1 | 282.0 | 5.4e-75 |
| AXN93597.1 | PuwB | BGC0001952 | NRP | 27.0 | 39.7 | 282.0 | 5.4e-75 |
| ACN69988.1 | polyketide\_synthase | BGC0000079 | Polyketide | 27.0 | 37.2 | 281.0 | 1.2e-74 |
| ARM20281.1 | polyketide\_synthase | BGC0001523 | Polyketide | 32.0 | 27.9 | 281.0 | 1.2e-74 |
| AXN93586.1 | PuwB | BGC0001951 | NRP | 26.0 | 41.1 | 281.0 | 1.2e-74 |
| CAQ34918.1 | nonribosomal\_peptide\_synthetase/\_polyketide\_synthase | BGC0000986 | NRP + Polyketide | 28.0 | 39.4 | 281.0 | 1.6e-74 |
| CAQ34920.1 | polyketide\_synthase | BGC0000986 | NRP + Polyketide | 29.0 | 36.0 | 281.0 | 1.6e-74 |
| AUO16422.1 | polyketide\_synthase | BGC0001700 | Polyketide | 32.0 | 25.9 | 281.0 | 1.6e-74 |
| AXN93577.1 | PuwB | BGC0001950 | NRP | 26.0 | 41.1 | 281.0 | 1.6e-74 |
| ABK32263.1 | AmbH | BGC0000014 | Polyketide | 31.0 | 28.3 | 280.0 | 2e-74 |
| SCO70309.1 | Type\_I\_polyketide\_synthase | BGC0001433 | Polyketide:Modular type I | 28.0 | 40.0 | 280.0 | 2.7e-74 |
| AXM42950.1 | polyketide\_synthase | BGC0001941 | NRP + Polyketide | 32.0 | 26.3 | 280.0 | 2.7e-74 |
| ABK32291.1 | JerE | BGC0000080 | Polyketide | 32.0 | 27.5 | 280.0 | 3.5e-74 |
| AEZ53945.1 | polyketide\_synthase | BGC0000144 | Polyketide:Modular type I | 34.0 | 23.8 | 280.0 | 3.5e-74 |
| ARM20278.1 | polyketide\_synthase | BGC0001523 | Polyketide | 30.0 | 31.2 | 280.0 | 3.5e-74 |
| ACC40921.1 | polyketide\_synthase\_Pks7 | BGC0001665 | Polyketide | 29.0 | 30.3 | 279.0 | 4.5e-74 |
| sipP2 | Type\_I\_Modular\_PKS | BGC0001452 | Polyketide | 31.0 | 31.3 | 278.0 | 1e-73 |
| AMYAL\_RS48925 | polyketide\_synthase | BGC0002011 | Polyketide | 29.0 | 37.4 | 278.0 | 1e-73 |
| TXD00261.1 | AMP-binding\_protein | BGC0001877 | Polyketide | 30.0 | 29.7 | 278.0 | 1.3e-73 |
| ANZ22989.1 | ZinF | BGC0001828 | Polyketide | 27.0 | 41.7 | 277.0 | 1.7e-73 |
| BAB69195.1 | modular\_polyketide\_synthase | BGC0000117 | Polyketide | 32.0 | 25.3 | 277.0 | 2.3e-73 |
| DAB41916.1 | ArzN\_-\_PKS\_(KS,\_AT,\_OMT,\_KR,\_ACP) | BGC0001884 | NRP + Polyketide | 35.0 | 24.0 | 277.0 | 2.3e-73 |
| ADF88279.1 | mixed\_NRPS/PKS | BGC0000981 | NRP + Polyketide | 27.0 | 39.6 | 277.0 | 2.9e-73 |
| ctg1\_orf256 |  | BGC0001200 | Polyketide | 28.0 | 36.5 | 277.0 | 2.9e-73 |
| ANI24099.1 | polyketide\_synthase | BGC0001235 | NRP + Polyketide | 34.0 | 23.7 | 277.0 | 2.9e-73 |
| AWH12936.1 | StmA | BGC0001939 | Polyketide | 28.0 | 37.0 | 277.0 | 2.9e-73 |
| ABX60161.1 | mixed\_NRPS/PKS | BGC0000978 | NRP + Alkaloid + Polyketide:Modular type I | 27.0 | 39.6 | 275.0 | 6.6e-73 |
| AFV96138.1 | polyketide\_synthase | BGC0001064 | Polyketide:Modular type I + Polyketide:Type III | 34.0 | 24.0 | 275.0 | 6.6e-73 |
| AGY30676.1 | Ann4 | BGC0001298 | Polyketide | 33.0 | 25.3 | 275.0 | 6.6e-73 |
| ARU81118.1 | CylD | BGC0001566 | Polyketide | 34.0 | 24.0 | 275.0 | 6.6e-73 |
| CAL58686.1 | polyketide\_synthase | BGC0000149 | Polyketide:Modular type I | 29.0 | 32.2 | 275.0 | 1.1e-72 |
| ACR33079.1 | polyketide\_synthase | BGC0000017 | Alkaloid + Polyketide:Modular type I | 31.0 | 28.0 | 274.0 | 1.5e-72 |
| ABC84460.1 | NigAV | BGC0000114 | Polyketide:Modular type I | 30.0 | 31.0 | 274.0 | 1.5e-72 |
| sipP1 | Type\_I\_Modular\_PKS | BGC0001452 | Polyketide | 30.0 | 31.9 | 274.0 | 1.5e-72 |
| AAS98784.1 | polyketide\_synthase | BGC0001001 | NRP + Polyketide | 30.0 | 28.2 | 273.0 | 3.3e-72 |
| ctg1\_orf15 |  | BGC0001457 | NRP | 34.0 | 25.3 | 273.0 | 4.2e-72 |
| AMB48442.1 | polyketide\_synthase | BGC0001357 | Polyketide | 32.0 | 24.9 | 272.0 | 5.5e-72 |
| ctg1\_15 |  | BGC0001931 | Polyketide | 32.0 | 27.6 | 272.0 | 5.5e-72 |
| CAL58685.1 | polyketide\_synthase | BGC0000149 | Polyketide:Modular type I | 31.0 | 27.5 | 272.0 | 7.2e-72 |
| ADH04660.1 | TugD | BGC0001342 | NRP + Polyketide | 32.0 | 26.1 | 272.0 | 7.2e-72 |
| AAK83194.1 | polyketide\_synthase | BGC0000026 | Saccharide:Oligosaccharide | 30.0 | 30.1 | 271.0 | 1.2e-71 |
| AUO16398.1 | polyketide\_synthase | BGC0001700 | Polyketide | 27.0 | 38.6 | 271.0 | 1.6e-71 |
| ACB46488.1 | polyketide\_synthase | BGC0000082 | Polyketide | 30.0 | 29.6 | 270.0 | 2.1e-71 |
| BAF85838.1 | modular\_polyketide\_synthase | BGC0000109 | Polyketide | 32.0 | 24.1 | 269.0 | 4.7e-71 |
| ABI94380.1 | tautomycetin\_biosynthetic\_PKS | BGC0000157 | Polyketide | 27.0 | 38.3 | 269.0 | 4.7e-71 |
| ABV91287.1 | type\_I\_modular\_polyketide\_synthase | BGC0000158 | Polyketide:Modular type I | 27.0 | 38.3 | 269.0 | 4.7e-71 |
| AUO16399.1 | polyketide\_synthase | BGC0001700 | Polyketide | 33.0 | 25.5 | 269.0 | 4.7e-71 |
| ctg1\_orf27 |  | BGC0000096 | Polyketide | 32.0 | 27.0 | 269.0 | 6.1e-71 |
| ARM20282.1 | polyketide\_synthase | BGC0001523 | Polyketide | 31.0 | 26.0 | 268.0 | 1e-70 |
| AEP40940.1 | polyketide\_synthase\_type\_I | BGC0000021 | Polyketide | 32.0 | 25.2 | 267.0 | 1.8e-70 |
| CAJ88187.2 | Type\_I\_modular\_polyketide\_synthase | BGC0000151 | Polyketide:Modular type I + Saccharide:Hybrid/tailoring | 32.0 | 26.5 | 267.0 | 2.3e-70 |
| BAJ16467.1 | polyketide\_synthase | BGC0000058 | Polyketide | 31.0 | 27.1 | 267.0 | 3e-70 |
| ADC45586.1 | modular\_polyketide\_synthase | BGC0000093 | Polyketide | 30.0 | 25.7 | 267.0 | 3e-70 |
| AFI57005.1 | QmnA1 | BGC0000133 | Polyketide | 28.0 | 38.2 | 267.0 | 3e-70 |
| CAQ34919.1 | polyketide\_synthase | BGC0000986 | NRP + Polyketide | 28.0 | 37.6 | 267.0 | 3e-70 |
| AAZ77673.1 | ChlB1 | BGC0000036 | Polyketide:Modular type I + Polyketide:Iterative type I + Saccharide:Oligosaccharide | 31.0 | 29.9 | 266.0 | 4e-70 |
| CAM00065.1 | EryAIII\_Erythromycin\_polyketide\_synthase\_modules\_5\_and\_6 | BGC0000055 | Polyketide:Modular type I + Saccharide:Hybrid/tailoring | 33.0 | 24.2 | 266.0 | 4e-70 |
| SCN11952.1 | ebeD-type\_I\_polyketide\_synthase | BGC0001580 | Polyketide | 28.0 | 36.6 | 266.0 | 4e-70 |
| ABK32256.1 | AmbB | BGC0000014 | Polyketide | 32.0 | 26.8 | 266.0 | 5.2e-70 |
| ADC45515.1 | modular\_polyketide\_synthase | BGC0000093 | Polyketide | 33.0 | 23.9 | 266.0 | 5.2e-70 |
| ARE67853.1 | AbsB1 | BGC0001492 | Polyketide | 28.0 | 37.7 | 266.0 | 5.2e-70 |
| AJW65407.1 | type\_I\_modular\_polyketide\_synthase | BGC0001195 | NRP + Polyketide | 32.0 | 27.3 | 265.0 | 6.8e-70 |
| AZH23819.1 | MgiR | BGC0001971 | NRP + Polyketide | 31.0 | 23.9 | 265.0 | 6.8e-70 |
| AKD43522.1 | Type\_I\_polyketide\_synthase | BGC0001409 | Polyketide | 27.0 | 41.6 | 265.0 | 8.9e-70 |
| AOE23578.1 | FoxBII | BGC0001598 | NRP + Polyketide | 30.0 | 31.6 | 265.0 | 8.9e-70 |
| AWC08658.1 | polyketide\_synthase\_type\_I | BGC0001932 | Polyketide | 30.0 | 28.6 | 265.0 | 8.9e-70 |
| ATP76241.1 | NdaD | BGC0001705 | NRP + Polyketide | 27.0 | 36.9 | 265.0 | 8.9e-70 |
| BAF85843.1 | modular\_polyketide\_synthase | BGC0000109 | Polyketide | 28.0 | 37.5 | 265.0 | 1.2e-69 |
| WP\_020636845.1 | type\_I\_polyketide\_synthase | BGC0002011 | Polyketide | 29.0 | 31.9 | 264.0 | 2e-69 |
| BAJ16470.1 | polyketide\_synthase | BGC0000058 | Polyketide | 27.0 | 37.4 | 264.0 | 2.6e-69 |
| AEE88278.1 | CurL | BGC0000976 | NRP + Polyketide:Modular type I | 32.0 | 24.6 | 264.0 | 2.6e-69 |
| AAT70107.1 | CurL | BGC0001165 | NRP + Polyketide:Modular type I | 32.0 | 24.6 | 264.0 | 2.6e-69 |
| AVI57433.1 | AbmB1 | BGC0001694 | Polyketide | 27.0 | 37.0 | 264.0 | 2.6e-69 |
| WP\_051137606.1 | type\_I\_polyketide\_synthase | BGC0002011 | Polyketide | 31.0 | 30.3 | 264.0 | 2.6e-69 |
| ABC84459.1 | NigAIV | BGC0000114 | Polyketide:Modular type I | 31.0 | 27.6 | 263.0 | 3.4e-69 |
| AQA28563.1 | type\_I\_polyketide\_synthase | BGC0001663 | Polyketide | 30.0 | 28.9 | 263.0 | 3.4e-69 |
| BAF85839.1 | modular\_polyketide\_synthase | BGC0000109 | Polyketide | 29.0 | 30.3 | 263.0 | 4.4e-69 |
| CAD19090.1 | StiF\_protein | BGC0000153 | NRP + Polyketide:Modular type I | 26.0 | 39.5 | 263.0 | 4.4e-69 |
| AGY62753.1 | EbeA | BGC0000051 | Polyketide | 32.0 | 25.5 | 262.0 | 5.7e-69 |
| CAM00062.1 | EryAI\_Erythromycin\_polyketide\_synthase\_modules\_1\_and\_2 | BGC0000055 | Polyketide:Modular type I + Saccharide:Hybrid/tailoring | 31.0 | 24.6 | 262.0 | 5.7e-69 |
| SCN11949.1 | ebeA-type\_I\_polyketide\_synthase\_KSQ-ATa-ACP | BGC0001580 | Polyketide | 32.0 | 25.5 | 262.0 | 5.7e-69 |
| EHK80163.1 | acyl\_transferase | BGC0001447 | Polyketide | 27.0 | 37.1 | 262.0 | 9.8e-69 |
| AZH23790.1 | MgcG | BGC0001970 | NRP + Polyketide | 31.0 | 24.8 | 262.0 | 9.8e-69 |
| AZH23820.1 | MgiG | BGC0001971 | NRP + Polyketide | 31.0 | 24.8 | 262.0 | 9.8e-69 |
| CAC22145.1 | CpkB;\_Polyketide\_synthase\_modules\_3\_and\_4 | BGC0000038 | Polyketide:Modular type I | 26.0 | 36.8 | 261.0 | 1.3e-68 |
| BAO66529.1 | type\_I\_polyketide\_synthase | BGC0000042 | Polyketide | 31.0 | 29.4 | 261.0 | 1.3e-68 |
| AAQ84145.1 | Plm5 | BGC0000123 | Polyketide | 31.0 | 25.3 | 261.0 | 1.7e-68 |
| AFL48526.1 | laidlomycin\_polyketide\_synthase\_(module\_2) | BGC0000084 | Polyketide | 27.0 | 35.9 | 260.0 | 2.2e-68 |
| ABJ97437.1 | MerA | BGC0001012 | NRP + Polyketide | 28.0 | 38.2 | 260.0 | 2.2e-68 |
| AAF00957.1 | mcyG | BGC0001017 | NRP + Polyketide:Modular type I | 24.0 | 50.1 | 260.0 | 2.9e-68 |
| CAO98849.1 | polyketide\_synthase\_AufF | BGC0000023 | Polyketide:Modular type I | 30.0 | 29.2 | 260.0 | 3.7e-68 |
| ABP55221.1 | acyl\_transferase\_domain\_protein | BGC0000142 | Polyketide | 29.0 | 31.3 | 260.0 | 3.7e-68 |
| AAS98782.1 | polyketide\_synthase | BGC0001001 | NRP + Polyketide | 32.0 | 24.7 | 260.0 | 3.7e-68 |
| CAD55506.1 | CpkA;\_Polyketide\_synthase\_loading\_module,\_and\_modules\_1\_and\_2 | BGC0000038 | Polyketide:Modular type I | 32.0 | 24.8 | 259.0 | 4.9e-68 |
| AFL48529.1 | laidlomycin\_polyketide\_synthase\_(module\_5\_and\_module\_6) | BGC0000084 | Polyketide | 29.0 | 32.0 | 259.0 | 4.9e-68 |
| AAF26919.1 | polyketide\_synthase | BGC0000988 | NRP + Polyketide | 33.0 | 24.2 | 259.0 | 4.9e-68 |
| ASZ00147.1 | polyketide\_synthase | BGC0001785 | Polyketide | 33.0 | 23.8 | 259.0 | 4.9e-68 |
| AIT55264.1 | polyketide\_synthase | BGC0000072 | Polyketide:Modular type I | 34.0 | 23.9 | 259.0 | 6.3e-68 |
| AHH99921.1 | PKS\_I | BGC0000002 | Polyketide | 33.0 | 23.3 | 259.0 | 8.3e-68 |
| ADU85988.1 | putative\_iterative\_type\_I\_polyketide\_synthase | BGC0000165 | Polyketide:Modular type I | 30.0 | 29.4 | 259.0 | 8.3e-68 |
| CAN89636.1 | putative\_polyketide\_synthase | BGC0001070 | NRP + Polyketide:Modular type I + Polyketide:Trans-AT type I | 29.0 | 27.2 | 259.0 | 8.3e-68 |
| TXD00265.1 | SDR\_family\_NAD(P)-dependent\_oxidoreductase | BGC0001877 | Polyketide | 33.0 | 23.9 | 259.0 | 8.3e-68 |
| AZH23793.1 | MgcK | BGC0001970 | NRP + Polyketide | 33.0 | 24.0 | 259.0 | 8.3e-68 |
| CAQ18834.1 | polyketide\_synthase | BGC0000954 | NRP + Polyketide:Modular type I | 34.0 | 24.3 | 258.0 | 1.1e-67 |
| ALA09356.1 | type\_I\_modular\_PKS | BGC0001303 | Polyketide | 30.0 | 27.2 | 258.0 | 1.1e-67 |
| AEC13070.1 | fosD | BGC0000060 | Polyketide | 32.0 | 24.6 | 258.0 | 1.4e-67 |
| BAT51066.1 | type\_I\_polyketide\_synthase | BGC0001296 | Polyketide | 26.0 | 40.8 | 258.0 | 1.4e-67 |
| AAO65799.1 | monensin\_polyketide\_synthase\_modules\_5\_and\_6 | BGC0000100 | Polyketide | 29.0 | 29.8 | 257.0 | 3.2e-67 |
| CBD77738.1 | polyketide\_synthase | BGC0000974 | NRP + Polyketide | 32.0 | 27.1 | 257.0 | 3.2e-67 |
| AIR74913.1 | polyketide\_synthase | BGC0001559 | RiPP | 32.0 | 27.1 | 257.0 | 3.2e-67 |
| ANZ52462.1 | MonAIV | BGC0001670 | Polyketide | 29.0 | 29.8 | 257.0 | 3.2e-67 |
| QBF51758.1 | type\_I\_polyketide\_synthase | BGC0001856 | Polyketide:Modular type I | 33.0 | 23.9 | 257.0 | 3.2e-67 |
| BAC76493.1 | lankamycin\_synthase\_LkmAI | BGC0000085 | Polyketide | 31.0 | 27.4 | 256.0 | 4.1e-67 |
| AZF85917.1 | type\_I\_polyketide\_synthase | BGC0001963 | NRP + Polyketide | 31.0 | 25.8 | 256.0 | 4.1e-67 |
| BAC76492.1 | lankamycin\_synthase\_LkmAII | BGC0000085 | Polyketide | 27.0 | 38.5 | 256.0 | 5.4e-67 |
| AEZ53946.1 | polyketide\_synthase | BGC0000144 | Polyketide:Modular type I | 29.0 | 31.6 | 255.0 | 7e-67 |
| WP\_020636817.1 | type\_I\_polyketide\_synthase | BGC0002011 | Polyketide | 32.0 | 25.7 | 255.0 | 7e-67 |
| AAZ77698.1 | ChlA5 | BGC0000036 | Polyketide:Modular type I + Polyketide:Iterative type I + Saccharide:Oligosaccharide | 29.0 | 29.7 | 255.0 | 9.2e-67 |
| ACB46192.1 | polyketide\_synthase | BGC0000989 | NRP + Polyketide | 33.0 | 24.0 | 255.0 | 9.2e-67 |
| AAF62880.1 | EpoA | BGC0000991 | NRP + Polyketide | 33.0 | 24.2 | 255.0 | 9.2e-67 |
| AAM70355.1 | CalO5 | BGC0000033 | Polyketide | 30.0 | 30.3 | 255.0 | 1.2e-66 |
| AAZ94386.1 | modular\_polyketide\_synthase | BGC0000040 | Polyketide | 34.0 | 23.8 | 255.0 | 1.2e-66 |
| BAG85026.1 | putative\_polyketide\_synthase | BGC0000086 | Polyketide | 32.0 | 25.3 | 254.0 | 1.6e-66 |
| CAQ64686.1 | lasalocid\_modular\_polyketide\_synthase | BGC0000087 | Polyketide | 32.0 | 25.3 | 254.0 | 1.6e-66 |
| AAZ94390.1 | modular\_polyketide\_synthase | BGC0000040 | Polyketide | 29.0 | 28.9 | 254.0 | 2e-66 |
| CAI94682.1 | putative\_polyketide\_synthase | BGC0000141 | Polyketide | 30.0 | 29.4 | 254.0 | 2e-66 |
| CAQ18835.1 | polyketide\_synthase | BGC0000954 | NRP + Polyketide:Modular type I | 30.0 | 30.9 | 253.0 | 2.7e-66 |
| ADB12488.1 | EpoA | BGC0000990 | NRP + Polyketide | 33.0 | 24.2 | 253.0 | 2.7e-66 |
| ADN13832.1 | Polyketide\_Synthase | BGC0001164 | Polyketide:Modular type I | 32.0 | 24.1 | 253.0 | 2.7e-66 |
| AZH23823.1 | MgiK | BGC0001971 | NRP + Polyketide | 31.0 | 23.9 | 253.0 | 2.7e-66 |
| ANR02549.1 | LodH | BGC0001648 | Polyketide | 31.0 | 23.9 | 253.0 | 3.5e-66 |
| AAG13917.1 | megalomicin\_6-deoxyerythronolide\_B\_synthase\_1 | BGC0000092 | Polyketide | 30.0 | 28.1 | 253.0 | 4.6e-66 |
| CAJ88177.1 | putative\_type\_I\_polyketide\_synthase | BGC0000151 | Polyketide:Modular type I + Saccharide:Hybrid/tailoring | 33.0 | 24.3 | 253.0 | 4.6e-66 |
| AAO62584.1 | polyketide\_synthase\_type\_1 | BGC0001016 | NRP + Polyketide | 26.0 | 36.7 | 253.0 | 4.6e-66 |
| AVX51107.1 | nysB | BGC0001709 | Polyketide | 31.0 | 24.0 | 253.0 | 4.6e-66 |
| ABV83229.1 | CppB | BGC0000116 | Polyketide | 29.0 | 26.8 | 252.0 | 5.9e-66 |
| AAZ94389.1 | modular\_polyketide\_synthase | BGC0000040 | Polyketide | 31.0 | 24.3 | 252.0 | 7.8e-66 |
| ADF88276.1 | polyketide\_synthase | BGC0000981 | NRP + Polyketide | 29.0 | 30.9 | 252.0 | 7.8e-66 |
| WP\_035121546.1 | type\_I\_polyketide\_synthase | BGC0001467 | NRP:Cyclic depsipeptide + Polyketide:Modular type I | 29.0 | 28.6 | 252.0 | 7.8e-66 |
| ctg1\_orf7 |  | BGC0000053 | Polyketide | 29.0 | 30.2 | 252.0 | 1e-65 |
| AAO65797.1 | monensin\_polyketide\_synthase\_module\_2 | BGC0000100 | Polyketide | 29.0 | 30.2 | 252.0 | 1e-65 |
| CAQ18839.1 | hybrid\_polyketide\_synthase/nonribosomal\_polypetide\_synthetase | BGC0000954 | NRP + Polyketide:Modular type I | 33.0 | 24.2 | 252.0 | 1e-65 |
| ANZ52460.1 | MonAII | BGC0001670 | Polyketide | 29.0 | 30.2 | 252.0 | 1e-65 |
| AXM42948.1 | type\_1\_polyketide\_synthase | BGC0001941 | NRP + Polyketide | 30.0 | 30.9 | 252.0 | 1e-65 |
| ATX68116.1 | malonyl\_CoA-acyl\_carrier\_protein\_transacylase | BGC0001772 | Polyketide | 31.0 | 24.0 | 251.0 | 1.3e-65 |
| ANR02556.1 | LodO | BGC0001648 | Polyketide | 26.0 | 36.8 | 251.0 | 1.7e-65 |
| AXI91546.1 | FunP7 | BGC0001944 | Polyketide | 27.0 | 38.3 | 251.0 | 1.7e-65 |
| ACF35445.1 | mbcAI | BGC0000090 | Polyketide | 32.0 | 24.1 | 250.0 | 2.3e-65 |
| CAA16183.1 | polyketide\_synthase | BGC0001063 | NRP + Polyketide | 32.0 | 24.2 | 250.0 | 2.3e-65 |
| AAU93807.2 | polyketide\_synthase\_modules\_1\_and\_2 | BGC0000054 | Polyketide | 32.0 | 25.2 | 250.0 | 2.9e-65 |
| CAQ18833.1 | polyketide\_synthase | BGC0000954 | NRP + Polyketide:Modular type I | 32.0 | 24.2 | 250.0 | 2.9e-65 |
| AQH32482.1 | type\_1\_polyketide\_synthase | BGC0001667 | NRP + Polyketide | 26.0 | 36.4 | 250.0 | 2.9e-65 |
| ATX68115.1 | malonyl\_CoA-acyl\_carrier\_protein\_transacylase | BGC0001772 | Polyketide | 30.0 | 27.9 | 250.0 | 2.9e-65 |
| ARM20277.1 | polyketide\_synthase | BGC0001523 | Polyketide | 29.0 | 31.4 | 250.0 | 3.9e-65 |
| AWW87422.1 | type\_I\_polyketide\_synthase | BGC0001755 | Polyketide | 31.0 | 26.2 | 249.0 | 5e-65 |
| AAZ94387.1 | modular\_polyketide\_synthase | BGC0000040 | Polyketide | 32.0 | 25.0 | 249.0 | 6.6e-65 |
| AAZ94388.1 | nodular\_polyketide\_synthase | BGC0000040 | Polyketide | 30.0 | 27.0 | 248.0 | 8.6e-65 |
| AAX98190.1 | polyketide\_synthase\_type\_I | BGC0000052 | Polyketide | 29.0 | 30.0 | 248.0 | 8.6e-65 |
| AAU04878.1 | polyketide\_synthase | BGC0000365 | NRP | 31.0 | 29.0 | 248.0 | 8.6e-65 |
| AAF00958.1 | mcyE | BGC0001017 | NRP + Polyketide:Modular type I | 25.0 | 49.3 | 248.0 | 8.6e-65 |
| AAC69329.1 | type\_I\_polyketide\_synthase\_PikAI | BGC0000094 | Polyketide:Modular type I + Saccharide:Hybrid/tailoring | 34.0 | 24.6 | 248.0 | 1.1e-64 |
| BAB69194.1 | modular\_polyketide\_synthase | BGC0000117 | Polyketide | 26.0 | 39.6 | 248.0 | 1.1e-64 |
| AEE88283.1 | CurG | BGC0000976 | NRP + Polyketide:Modular type I | 32.0 | 24.3 | 248.0 | 1.1e-64 |
| ABX60152.1 | polyketide\_synthase | BGC0000978 | NRP + Alkaloid + Polyketide:Modular type I | 29.0 | 30.9 | 248.0 | 1.1e-64 |
| AAT70102.1 | CurG | BGC0001165 | NRP + Polyketide:Modular type I | 32.0 | 24.3 | 248.0 | 1.1e-64 |
| ACN69992.1 | polyketide\_synthase | BGC0000079 | Polyketide | 32.0 | 24.6 | 248.0 | 1.5e-64 |
| AFD30954.1 | CrmA | BGC0000966 | NRP + Polyketide | 31.0 | 27.1 | 248.0 | 1.5e-64 |
| EJK79843.1 | amino\_acid\_adenylation\_enzyme/thioester\_reductase\_family\_protein | BGC0000436 | NRP | 29.0 | 32.1 | 247.0 | 1.9e-64 |
| ARW71486.1 | type\_I\_PKS\_module\_6 | BGC0001812 | Polyketide | 30.0 | 26.2 | 247.0 | 1.9e-64 |
| ARS01473.1 | NcmAI | BGC0001702 | NRP + Polyketide | 32.0 | 23.9 | 247.0 | 2.5e-64 |
| AAQ82564.1 | FscC | BGC0000061 | Polyketide | 26.0 | 41.2 | 247.0 | 3.3e-64 |
| BAC76491.1 | lankamycin\_synthase\_LkmAIII | BGC0000085 | Polyketide | 31.0 | 28.9 | 247.0 | 3.3e-64 |
| CAO85897.1 | modular\_polyketide\_synthase\_NorB | BGC0000110 | Polyketide:Modular type I | 29.0 | 31.3 | 247.0 | 3.3e-64 |
| AAW03325.1 | CtaB | BGC0000982 | NRP + Polyketide | 29.0 | 27.6 | 247.0 | 3.3e-64 |
| AKG06378.1 | polyketide\_synthase\_type\_1 | BGC0001830 | Polyketide | 29.0 | 25.9 | 247.0 | 3.3e-64 |
| AWS21279.1 | type\_I\_polyketide\_synthase | BGC0001934 | Polyketide | 32.0 | 24.5 | 247.0 | 3.3e-64 |
| AZY91989.1 | polyketide\_synthase | BGC0002022 | Polyketide | 32.0 | 24.5 | 247.0 | 3.3e-64 |
| AHH99925.1 | PKS\_I | BGC0000002 | Polyketide | 28.0 | 31.3 | 246.0 | 4.3e-64 |
| AFV30249.1 | polyketide\_synthase | BGC0000075 | Polyketide | 27.0 | 36.1 | 246.0 | 4.3e-64 |
| ADC45534.1 | modular\_polyketide\_synthase | BGC0000093 | Polyketide | 27.0 | 30.3 | 246.0 | 4.3e-64 |
| BAB69199.1 | modular\_polyketide\_synthase | BGC0000117 | Polyketide | 27.0 | 36.4 | 246.0 | 4.3e-64 |
| AAQ84156.1 | Plm1 | BGC0000123 | Polyketide | 28.0 | 30.3 | 246.0 | 4.3e-64 |
| ABV91286.1 | type\_I\_modular\_polyketide\_synthase | BGC0000158 | Polyketide:Modular type I | 31.0 | 27.1 | 246.0 | 4.3e-64 |
| AAF00959.1 | mcyD | BGC0001017 | NRP + Polyketide:Modular type I | 26.0 | 36.4 | 246.0 | 4.3e-64 |
| SCO70308.1 | Type\_I\_polyketide\_synthase | BGC0001433 | Polyketide:Modular type I | 31.0 | 25.4 | 246.0 | 4.3e-64 |
| AFL48527.1 | laidlomycin\_polyketide\_synthase\_(module\_3\_and\_module\_4) | BGC0000084 | Polyketide | 27.0 | 38.5 | 246.0 | 5.6e-64 |
| ABI94379.1 | tautomycetin\_biosynthetic\_PKS | BGC0000157 | Polyketide | 31.0 | 27.1 | 246.0 | 5.6e-64 |
| ANH11412.1 | SceQ | BGC0001908 | Polyketide | 29.0 | 30.8 | 246.0 | 5.6e-64 |
| ADH04640.1 | TgaB | BGC0001051 | NRP + Polyketide:Modular type I | 27.0 | 31.9 | 245.0 | 7.3e-64 |
| ctg1\_orf254 |  | BGC0001200 | Polyketide | 27.0 | 37.8 | 245.0 | 7.3e-64 |
| TXD00026.1 | SDR\_family\_NAD(P)-dependent\_oxidoreductase | BGC0001877 | Polyketide | 28.0 | 30.8 | 245.0 | 7.3e-64 |
| AAM54077.1 | polyketide\_synthase | BGC0000020 | Polyketide | 32.0 | 25.2 | 245.0 | 9.5e-64 |
| ABP55210.1 | beta-ketoacyl\_synthase | BGC0000142 | Polyketide | 27.0 | 37.2 | 245.0 | 9.5e-64 |
| AHH34189.1 | polyketide\_synthase | BGC0001162 | Polyketide:Modular type I | 28.0 | 29.6 | 245.0 | 9.5e-64 |
| AVI26389.1 | polyketide\_synthase | BGC0001800 | NRP + Polyketide | 31.0 | 24.5 | 245.0 | 9.5e-64 |
| CAO98879.1 | polyketide\_synthase\_AufD | BGC0000023 | Polyketide:Modular type I | 31.0 | 24.0 | 245.0 | 1.2e-63 |
| ABW96540.1 | type\_I\_modular\_polyketide\_synthase | BGC0000159 | Polyketide:Modular type I | 28.0 | 32.8 | 245.0 | 1.2e-63 |
| BAO98805.1 | putative\_polyketide\_synthase | BGC0001002 | NRP + Polyketide | 30.0 | 28.4 | 245.0 | 1.2e-63 |
| ADC45538.1 | modular\_polyketide\_synthase | BGC0000093 | Polyketide | 27.0 | 30.7 | 244.0 | 2.1e-63 |
| ABV83230.1 | CppC | BGC0000116 | Polyketide | 26.0 | 38.7 | 244.0 | 2.1e-63 |
| WP\_053138504.1 | type\_I\_polyketide\_synthase | BGC0002033 | Polyketide | 31.0 | 26.4 | 244.0 | 2.1e-63 |
| WP\_055469549.1 | type\_I\_polyketide\_synthase | BGC0001537 | Polyketide | 29.0 | 30.9 | 243.0 | 2.8e-63 |
| ANR02555.1 | LodN | BGC0001648 | Polyketide | 26.0 | 37.6 | 243.0 | 4.7e-63 |
| AXG22405.1 | type\_I\_polyketide\_synthase | BGC0002024 | Polyketide | 26.0 | 36.4 | 243.0 | 4.7e-63 |
| AAC38076.1 | polyketide\_synthase\_type\_I | BGC0000127 | Polyketide | 33.0 | 24.0 | 242.0 | 6.1e-63 |
| AGC24271.1 | prlQ | BGC0001038 | NRP + Polyketide:Modular type I | 32.0 | 24.4 | 242.0 | 6.1e-63 |
| ALA09371.1 | type\_I\_modular\_PKS | BGC0001303 | Polyketide | 30.0 | 31.1 | 242.0 | 6.1e-63 |
| AFP87523.1 | type\_I\_polyketide\_synthase | BGC0001159 | NRP + Polyketide:Modular type I | 32.0 | 23.6 | 242.0 | 1e-62 |
| BAE93730.1 | type\_I\_polyketide\_synthase | BGC0000164 | Polyketide | 26.0 | 36.7 | 241.0 | 1.8e-62 |
| CAE02605.1 | polyketide\_synthase\_type\_I | BGC0000024 | Polyketide:Modular type I | 26.0 | 36.4 | 240.0 | 2.3e-62 |
| ACZ65476.1 | type\_I\_modular\_polyketide\_synthase | BGC0000140 | Polyketide | 32.0 | 23.8 | 240.0 | 2.3e-62 |
| CAD29793.1 | polyketide\_synthase\_type\_I | BGC0001015 | NRP + Polyketide | 26.0 | 36.9 | 240.0 | 3.1e-62 |
| ctg1\_orf253 |  | BGC0001200 | Polyketide | 30.0 | 27.9 | 240.0 | 3.1e-62 |
| AEZ53948.1 | polyketide\_synthase | BGC0000144 | Polyketide:Modular type I | 29.0 | 27.7 | 240.0 | 4e-62 |
| AHH34186.1 | polyketide\_synthase | BGC0001161 | Polyketide:Modular type I | 28.0 | 29.2 | 240.0 | 4e-62 |
| AHB82070.1 | polyketide\_synthase | BGC0001231 | NRP + Polyketide:Modular type I | 33.0 | 23.5 | 240.0 | 4e-62 |
| sipP5 | Type\_I\_Modular\_PKS | BGC0001452 | Polyketide | 30.0 | 30.2 | 240.0 | 4e-62 |
| QBF51760.1 | type\_I\_polyketide\_synthase | BGC0001856 | Polyketide:Modular type I | 31.0 | 24.2 | 240.0 | 4e-62 |
| ADB23403.1 | polyketide\_synthase\_type\_I | BGC0001062 | Polyketide | 30.0 | 24.2 | 239.0 | 5.2e-62 |
| AZF85932.1 | type\_I\_polyketide\_synthase | BGC0001963 | NRP + Polyketide | 32.0 | 24.2 | 239.0 | 5.2e-62 |
| BAD08360.1 | polyketide\_synthase\_modules\_7-8 | BGC0000167 | Polyketide | 28.0 | 30.3 | 239.0 | 6.8e-62 |
| ANH11409.1 | SceN | BGC0001908 | Polyketide | 29.0 | 30.0 | 239.0 | 6.8e-62 |
| AAP42857.1 | NanA3 | BGC0000105 | Polyketide | 29.0 | 31.5 | 238.0 | 8.9e-62 |
| AAM77986.1 | iterative\_type\_I\_polyketide\_synthase | BGC0000112 | Polyketide:Iterative type I + Polyketide:Enediyne type I | 28.0 | 29.5 | 238.0 | 8.9e-62 |
| AAF71766.1 | nysI | BGC0000115 | Polyketide:Modular type I + Saccharide:Hybrid/tailoring | 31.0 | 25.5 | 238.0 | 1.2e-61 |
| ABV83221.1 | CppI | BGC0000116 | Polyketide | 30.0 | 25.8 | 238.0 | 1.2e-61 |
| ABC84471.1 | NigAVII | BGC0000114 | Polyketide:Modular type I | 28.0 | 28.5 | 238.0 | 1.5e-61 |
| CAD89773.1 | MelB\_protein | BGC0001010 | NRP + Polyketide:Modular type I | 29.0 | 27.8 | 238.0 | 1.5e-61 |
| TXD00024.1 | SDR\_family\_NAD(P)-dependent\_oxidoreductase | BGC0001877 | Polyketide | 31.0 | 24.1 | 238.0 | 1.5e-61 |
| AJO72737.1 | Type\_I\_modular\_polyketide\_synthase | BGC0001381 | Polyketide | 27.0 | 30.0 | 237.0 | 2e-61 |
| AWR88393.1 | putative\_beta-ketoacyl\_synthase | BGC0001522 | Polyketide | 29.0 | 30.2 | 237.0 | 2.6e-61 |
| AHN85651.1 | Phn2 | BGC0000122 | Polyketide:Modular type I | 28.0 | 31.6 | 237.0 | 3.4e-61 |
| AVX51098.1 | nysI | BGC0001709 | Polyketide | 31.0 | 25.1 | 236.0 | 4.4e-61 |
| AHA38203.1 | GphJ | BGC0000069 | Polyketide | 32.0 | 25.4 | 236.0 | 5.8e-61 |
| AEH42474.1 | polyketide\_synthase | BGC0000032 | Polyketide | 29.0 | 31.9 | 235.0 | 7.5e-61 |
| EHK80167.1 | modular\_polyketide\_synthase | BGC0001447 | Polyketide | 30.0 | 24.2 | 235.0 | 7.5e-61 |
| AVV61985.1 | beta-ketoacyl\_synthase | BGC0001477 | NRP + Polyketide:Modular type I | 32.0 | 25.1 | 235.0 | 9.8e-61 |
| AQX77694.1 | NocP | BGC0001704 | Other | 31.0 | 22.6 | 235.0 | 9.8e-61 |
| BAR73017.1 | putative\_PKS\_(KS-AT-KR-ACP-KS-AT-DH-KR-ACP) | BGC0001194 | Polyketide | 31.0 | 24.0 | 235.0 | 1.3e-60 |
| EGJ35088.1 | Polyketide\_synthase | BGC0001163 | Polyketide:Modular type I | 30.0 | 26.1 | 234.0 | 1.7e-60 |
| AKA59091.1 | type-I\_PKS | BGC0001619 | Polyketide | 31.0 | 28.1 | 234.0 | 1.7e-60 |
| AJY78092.1 | polyketide\_synthase | BGC0001902 | NRP + Polyketide | 30.0 | 26.5 | 234.0 | 2.2e-60 |
| AAN32979.1 | BarE | BGC0000962 | NRP + Polyketide:Modular type I | 30.0 | 24.6 | 233.0 | 2.9e-60 |
| AHB82064.1 | polyketide\_synthase | BGC0001231 | NRP + Polyketide:Modular type I | 32.0 | 24.0 | 233.0 | 3.7e-60 |
| ACF35447.1 | mbcAIII | BGC0000090 | Polyketide | 26.0 | 38.5 | 233.0 | 4.9e-60 |
| AVI57434.1 | AbmB2 | BGC0001694 | Polyketide | 29.0 | 30.3 | 233.0 | 4.9e-60 |
| BAD38873.1 | polyketide\_synthase | BGC0000111 | Polyketide | 31.0 | 24.2 | 232.0 | 8.3e-60 |
| ABC84458.1 | NigAIII | BGC0000114 | Polyketide:Modular type I | 26.0 | 39.5 | 232.0 | 8.3e-60 |
| ACY06290.1 | type\_I\_polyketide\_synthase | BGC0001042 | NRP + Polyketide | 29.0 | 30.0 | 232.0 | 8.3e-60 |
| AAP42858.1 | NanA4 | BGC0000105 | Polyketide | 30.0 | 24.9 | 232.0 | 1.1e-59 |
| ADC79637.1 | TamAI | BGC0001052 | NRP + Polyketide:Modular type I | 27.0 | 27.6 | 232.0 | 1.1e-59 |
| AQZ37094.1 | polyketide\_synthase | BGC0001511 | Polyketide | 31.0 | 25.3 | 231.0 | 1.4e-59 |
| ATY46587.1 | polyketide\_synthase | BGC0001666 | Polyketide | 28.0 | 30.1 | 231.0 | 1.4e-59 |
| AKD43761.1 | HerD | BGC0001349 | NRP + Polyketide | 27.0 | 34.0 | 231.0 | 1.9e-59 |
| ANR02551.1 | LodJ | BGC0001648 | Polyketide | 29.0 | 23.7 | 230.0 | 2.4e-59 |
| ANY10589.1 | polyketide\_synthase | BGC0001773 | Polyketide | 28.0 | 31.3 | 230.0 | 2.4e-59 |
| AAF71774.1 | nysA | BGC0000115 | Polyketide:Modular type I + Saccharide:Hybrid/tailoring | 26.0 | 35.8 | 230.0 | 3.2e-59 |
| AQZ37113.1 | polyketide\_synthase | BGC0001511 | Polyketide | 28.0 | 31.3 | 230.0 | 4.1e-59 |
| BAE93728.1 | type\_I\_polyketide\_synthase | BGC0000164 | Polyketide | 31.0 | 24.2 | 229.0 | 5.4e-59 |
| BAW32323.1 | hybrid\_cis-AT\_polyketide\_synthase\_-\_nonribosomal\_peptide\_synthetase | BGC0001630 | NRP + Polyketide | 30.0 | 24.5 | 229.0 | 5.4e-59 |
| BAE93731.1 | type\_I\_polyketide\_synthase | BGC0000164 | Polyketide | 26.0 | 36.9 | 229.0 | 7e-59 |
| AGY62758.1 | EbeF | BGC0000051 | Polyketide | 28.0 | 30.5 | 228.0 | 9.2e-59 |
| BAC68125.1 | modular\_polyketide\_synthase | BGC0000059 | Polyketide | 30.0 | 26.2 | 228.0 | 1.2e-58 |
| BAK64650.1 | polyketide\_synthase | BGC0000135 | Polyketide | 26.0 | 38.5 | 228.0 | 1.2e-58 |
| AAP85336.1 | type\_I\_PKS | BGC0000233 | Polyketide | 32.0 | 24.1 | 228.0 | 1.2e-58 |
| AHA38202.1 | GphI | BGC0000069 | Polyketide | 28.0 | 32.1 | 228.0 | 1.6e-58 |
| CAD19085.1 | StiA\_protein | BGC0000153 | NRP + Polyketide:Modular type I | 33.0 | 22.1 | 228.0 | 1.6e-58 |
| AGI99496.1 | Type\_I\_polyketide\_synthase | BGC0001004 | Polyketide:Modular type I | 31.0 | 23.6 | 228.0 | 1.6e-58 |
| BAT51065.1 | type\_I\_polyketide\_synthase | BGC0001296 | Polyketide | 31.0 | 25.0 | 228.0 | 1.6e-58 |
| BBA84070.1 | type\_I\_polyketide\_synthase | BGC0001916 | Polyketide | 26.0 | 37.5 | 228.0 | 1.6e-58 |
| ADH04639.1 | TgaA | BGC0001051 | NRP + Polyketide:Modular type I | 30.0 | 24.2 | 227.0 | 2e-58 |
| WP\_039806854.1 | type\_I\_polyketide\_synthase | BGC0002001 | NRP + Polyketide | 33.0 | 23.7 | 227.0 | 2e-58 |
| QDA77045.1 | polyketide\_synthase/nonribosomal\_peptide\_synthetase | BGC0002025 | NRP | 31.0 | 24.9 | 227.0 | 2e-58 |
| AHB82059.1 | non\_ribosomal\_peptide\_synthetase/polyketide\_synthase | BGC0001019 | NRP + Polyketide:Modular type I | 32.0 | 24.0 | 227.0 | 2.7e-58 |
| WP\_055469545.1 | type\_I\_polyketide\_synthase | BGC0001537 | Polyketide | 27.0 | 36.7 | 227.0 | 2.7e-58 |
| ANR02552.1 | LodK | BGC0001648 | Polyketide | 28.0 | 27.4 | 227.0 | 2.7e-58 |
| AWC08656.1 | polyketide\_synthase\_type\_I | BGC0001932 | Polyketide | 29.0 | 25.6 | 226.0 | 4.6e-58 |
| BAP34740.1 | type\_I\_polyketide\_synthase | BGC0000078 | Polyketide | 30.0 | 24.6 | 226.0 | 6e-58 |
| AAF86396.1 | FkbA | BGC0000994 | NRP + Polyketide | 31.0 | 23.1 | 226.0 | 6e-58 |
| EAU32819.1 | 6-methylsalicylic\_acid\_synthase | BGC0000160 | Polyketide | 29.0 | 28.9 | 225.0 | 7.8e-58 |
| CAD70195.1 | non-ribosomal\_peptide\_synthetase | BGC0001047 | NRP + Polyketide | 27.0 | 32.0 | 225.0 | 7.8e-58 |
| BAA20102.2 | 6-methylsalicylic\_acid\_synthase | BGC0001276 | Polyketide | 29.0 | 28.9 | 225.0 | 1.3e-57 |
| AKD43765.1 | HerE | BGC0001349 | NRP + Polyketide | 27.0 | 34.1 | 224.0 | 1.7e-57 |
| ACB46485.1 | polyketide\_synthase | BGC0000082 | Polyketide | 29.0 | 25.5 | 224.0 | 2.3e-57 |
| AAF15892.2 | nosB | BGC0001028 | Polyketide + NRP:Cyclic depsipeptide | 30.0 | 23.9 | 224.0 | 2.3e-57 |
| AWC08657.1 | polyketide\_synthase\_type\_I | BGC0001932 | Polyketide | 31.0 | 23.9 | 224.0 | 2.3e-57 |
| AXI91548.1 | FunP5 | BGC0001944 | Polyketide | 27.0 | 31.5 | 224.0 | 2.3e-57 |
| ABV83228.1 | CppA | BGC0000116 | Polyketide | 30.0 | 25.2 | 223.0 | 3.9e-57 |
| BBA66513.1 | type\_I\_polyketide\_synthase | BGC0001495 | Polyketide | 31.0 | 24.2 | 223.0 | 3.9e-57 |
| antaD | Type\_I\_PKS | BGC0001455 | NRP + Polyketide | 29.0 | 26.2 | 223.0 | 5e-57 |
| AAQ82568.1 | FscD | BGC0000061 | Polyketide | 31.0 | 23.9 | 222.0 | 6.6e-57 |
| AFU82616.1 | polyketide\_synthase | BGC0000998 | NRP + Polyketide | 25.0 | 38.9 | 222.0 | 6.6e-57 |
| AJO72734.1 | Type\_I\_modular\_polyketide\_synthase | BGC0001381 | Polyketide | 31.0 | 25.5 | 222.0 | 8.6e-57 |
| CAL80821.1 | sylD-like\_NRPS/PKS | BGC0000997 | NRP + Polyketide | 29.0 | 28.4 | 222.0 | 1.1e-56 |
| OJF16270.1 | AceP3 | BGC0001491 | Polyketide | 27.0 | 30.0 | 222.0 | 1.1e-56 |
| OAP25815.1 | Phenolphthiocerol\_synthesis\_polyketide\_synthase\_type\_I\_Pks15/1 | BGC0001658 | Polyketide | 28.0 | 31.1 | 222.0 | 1.1e-56 |
| AKA54627.1 | PKS | BGC0001216 | NRP + Polyketide | 29.0 | 26.6 | 221.0 | 1.5e-56 |
| ABL74938.1 | PKS | BGC0001048 | NRP:Glycopeptide + Polyketide:Modular type I + Saccharide:Hybrid/tailoring | 30.0 | 23.8 | 220.0 | 2.5e-56 |
| AAM54075.1 | polyketide\_synthase | BGC0000020 | Polyketide | 28.0 | 31.3 | 220.0 | 3.3e-56 |
| AAY42396.1 | Polyketide\_synthase | BGC0001000 | NRP:Lipopeptide + Polyketide:Modular type I | 28.0 | 27.3 | 220.0 | 3.3e-56 |
| ACB37743.1 | putative\_type\_I\_polyketide\_synthase | BGC0000162 | Polyketide | 30.0 | 23.9 | 219.0 | 5.6e-56 |
| orf3 | polyketide\_synthase | BGC0001432 | NRP:Cyclic depsipeptide + Polyketide:Iterative type I | 31.0 | 20.7 | 219.0 | 7.3e-56 |
| APZ78858.1 | polyketide\_synthase | BGC0001432 | NRP:Cyclic depsipeptide + Polyketide:Iterative type I | 31.0 | 20.7 | 219.0 | 7.3e-56 |
| TGZ15164.1 | hypothetical\_protein | BGC0002032 | Polyketide | 29.0 | 25.2 | 218.0 | 9.5e-56 |
| AEF33079.1 | polyketide\_synthase | BGC0001039 | NRP + Polyketide | 31.0 | 23.9 | 218.0 | 1.2e-55 |
| AJO72743.1 | Type\_I\_modular\_polyketide\_synthase | BGC0001381 | Polyketide | 29.0 | 26.3 | 217.0 | 2.1e-55 |
| AAX98186.1 | polyketide\_synthase\_type\_I | BGC0000052 | Polyketide | 32.0 | 21.4 | 216.0 | 4.7e-55 |
| AAX98189.1 | polyketide\_synthase\_type\_I | BGC0000052 | Polyketide | 26.0 | 36.9 | 216.0 | 4.7e-55 |
| AIW00670.1 | mellein\_synthase | BGC0001244 | Polyketide | 26.0 | 32.1 | 216.0 | 4.7e-55 |
| AEH42491.1 | polyketide\_synthase | BGC0000032 | Polyketide | 30.0 | 26.4 | 215.0 | 1.1e-54 |
| ABF87031.1 | non-ribosomal\_peptide\_synthetase/polyketide\_synthase | BGC0000393 | NRP + Polyketide:Modular type I | 30.0 | 27.3 | 215.0 | 1.1e-54 |
| WP\_063764078.1 | polyketide\_synthase | BGC0001348 | Polyketide:Modular type I | 28.0 | 30.0 | 215.0 | 1.4e-54 |
| WP\_030498975.1 | type\_I\_polyketide\_synthase | BGC0001327 | NRP:Cyclic depsipeptide + Polyketide:Modular type I | 31.0 | 23.8 | 214.0 | 1.8e-54 |
| ABW96542.1 | type\_I\_modular\_polyketide\_synthase | BGC0000159 | Polyketide:Modular type I | 27.0 | 32.6 | 214.0 | 2.3e-54 |
| CAJ76291.1 | putative\_polyketide\_synthase | BGC0000972 | NRP + Polyketide:Modular type I + Polyketide:Trans-AT type I | 29.0 | 27.3 | 213.0 | 5.2e-54 |
| AVV61980.1 | type\_I\_modular\_PKS | BGC0001477 | NRP + Polyketide:Modular type I | 25.0 | 32.8 | 213.0 | 5.2e-54 |
| AAD03048.1 | type\_I\_polyketide\_synthase | BGC0000041 | Polyketide | 29.0 | 28.1 | 212.0 | 6.8e-54 |
| AAM54078.1 | polyketide\_synthase | BGC0000020 | Polyketide | 28.0 | 31.3 | 211.0 | 1.5e-53 |
| ACF35446.1 | mbcAII | BGC0000090 | Polyketide | 28.0 | 27.1 | 211.0 | 1.5e-53 |
| CBJ89760.1 | Polyketide\_synthase\_involved\_in\_xenocoumacin\_synthesis | BGC0001054 | NRP + Polyketide:Modular type I | 28.0 | 25.4 | 211.0 | 1.5e-53 |
| WP\_015031691.1 | type\_I\_polyketide\_synthase | BGC0001819 | Polyketide | 28.0 | 25.1 | 211.0 | 1.5e-53 |
| BAD38874.1 | polyketide\_synthase | BGC0000111 | Polyketide | 29.0 | 25.0 | 210.0 | 2.6e-53 |
| BAF92601.1 | iterative\_type\_I\_PKS | BGC0000118 | Polyketide | 31.0 | 24.7 | 210.0 | 2.6e-53 |
| ACJ24875.1 | 6-methylsalicylic\_acid\_synthase | BGC0000119 | Polyketide:Iterative type I + Saccharide:Hybrid/tailoring | 31.0 | 24.7 | 210.0 | 2.6e-53 |
| ACR33077.1 | polyketide\_synthase | BGC0000017 | Alkaloid + Polyketide:Modular type I | 27.0 | 32.8 | 210.0 | 3.4e-53 |
| BAP34763.1 | type\_I\_polyketide\_synthase | BGC0000078 | Polyketide | 28.0 | 30.9 | 210.0 | 3.4e-53 |
| AAX35547.1 | polyketide\_syntase\_2 | BGC0001275 | Polyketide | 26.0 | 30.0 | 210.0 | 3.4e-53 |
| ANY10600.1 | polyketide\_synthase | BGC0001773 | Polyketide | 32.0 | 20.7 | 210.0 | 3.4e-53 |
| ABC87509.1 | polyketide\_synthase | BGC0001011 | NRP + Polyketide | 28.0 | 28.7 | 209.0 | 5.8e-53 |
| ctg1\_orf20 |  | BGC0001013 | NRP + Polyketide | 28.0 | 28.7 | 209.0 | 5.8e-53 |
| BAB69192.1 | modular\_polyketide\_synthase | BGC0000117 | Polyketide | 31.0 | 20.4 | 209.0 | 7.5e-53 |
| ABW96541.1 | type\_I\_modular\_polyketide\_synthase | BGC0000159 | Polyketide:Modular type I | 26.0 | 33.7 | 208.0 | 1.3e-52 |
| ALD82523.1 | polyketide\_synthase | BGC0001212 | NRP + Polyketide | 29.0 | 26.5 | 208.0 | 1.3e-52 |
| CAO85896.1 | protein\_modular\_polyketide\_synthase\_NorA' | BGC0000110 | Polyketide:Modular type I | 29.0 | 26.3 | 208.0 | 1.7e-52 |
| AAL01060.1 | omega-3\_polyunsaturated\_fatty\_acid\_synthase\_PfaA | BGC0000865 | Other | 28.0 | 26.6 | 208.0 | 1.7e-52 |
| AGC09487.1 | LobS5 | BGC0001183 | Polyketide | 28.0 | 26.3 | 207.0 | 2.2e-52 |
| AAW03329.1 | CtaF | BGC0000982 | NRP + Polyketide | 26.0 | 31.4 | 207.0 | 2.9e-52 |
| AFU82615.1 | polyketide\_synthase | BGC0000998 | NRP + Polyketide | 31.0 | 24.3 | 207.0 | 2.9e-52 |
| AHB82072.1 | non\_ribosomal\_peptide\_synthetase/polyketide\_synthase | BGC0001231 | NRP + Polyketide:Modular type I | 29.0 | 27.4 | 207.0 | 3.7e-52 |
| AGI99494.1 | Type\_I\_polyketide\_synthase | BGC0001004 | Polyketide:Modular type I | 27.0 | 26.6 | 206.0 | 4.9e-52 |
| AAS98200.1 | MSAS-type\_polyketide\_synthase | BGC0001273 | Polyketide | 26.0 | 33.4 | 206.0 | 6.4e-52 |
| AVX51106.1 | nysA | BGC0001709 | Polyketide | 26.0 | 36.9 | 206.0 | 6.4e-52 |
| AWH12938.1 | StmC | BGC0001939 | Polyketide | 29.0 | 24.9 | 205.0 | 8.3e-52 |
| PHM26606.1 | malonyl\_CoA-acyl\_carrier\_protein\_transacylase | BGC0001130 | NRP + Polyketide | 29.0 | 24.8 | 204.0 | 1.9e-51 |
| ADH04680.1 | hybrid\_polyketide\_synthase/non-ribosomal\_peptide\_synthetase | BGC0001344 | NRP + Polyketide | 29.0 | 27.6 | 204.0 | 1.9e-51 |
| ACO79122.1 | Type\_I\_fatty\_acid\_synthase\_ArsA | BGC0000284 | Polyketide | 29.0 | 25.7 | 203.0 | 5.4e-51 |
| AQZ37095.1 | polyketide\_synthase | BGC0001511 | Polyketide | 28.0 | 31.3 | 203.0 | 5.4e-51 |
| CCA29203.1 | non-ribosomal\_peptide\_synthetase/polyketide\_synthase | BGC0000955 | NRP + Polyketide:Modular type I | 29.0 | 23.8 | 202.0 | 9.2e-51 |
| CCM44330.1 | Polyketide\_synthase | BGC0001056 | NRP + Polyketide:Modular type I + Polyketide:PUFA synthase or related | 29.0 | 24.9 | 202.0 | 9.2e-51 |
| CBJ82077.1 | hypothetical\_protein | BGC0001872 | Polyketide | 28.0 | 25.1 | 201.0 | 1.6e-50 |
| CAJ76298.1 | putative\_hybrid\_polyketide-non-ribosomal\_peptide\_synthetase | BGC0000972 | NRP + Polyketide:Modular type I + Polyketide:Trans-AT type I | 29.0 | 24.5 | 200.0 | 2.7e-50 |
| ABG02264.1 | SalB | BGC0000143 | Polyketide | 28.0 | 24.4 | 200.0 | 3.5e-50 |
| ADM46359.1 | polyketide\_synthase | BGC0000106 | Polyketide | 26.0 | 32.6 | 200.0 | 4.6e-50 |
| CAE14173.1 | hypothetical\_protein | BGC0000383 | NRP + Polyketide:Modular type I | 29.0 | 25.6 | 200.0 | 4.6e-50 |
| ABJ97438.1 | MerB | BGC0001012 | NRP + Polyketide | 28.0 | 26.3 | 199.0 | 6e-50 |
| AJO72736.1 | Type\_I\_modular\_polyketide\_synthase | BGC0001381 | Polyketide | 27.0 | 33.3 | 199.0 | 6e-50 |
| AAO65798.1 | monensin\_polyketide\_synthase\_modules\_3\_and\_4 | BGC0000100 | Polyketide | 28.0 | 25.5 | 198.0 | 1e-49 |
| ANZ52461.1 | MonAIII | BGC0001670 | Polyketide | 28.0 | 25.5 | 198.0 | 1e-49 |
| ACO94483.1 | polyketide\_synthase\_type\_I | BGC0000097 | Polyketide:Modular type I | 27.0 | 33.9 | 197.0 | 3e-49 |
| AWH12937.1 | StmB | BGC0001939 | Polyketide | 28.0 | 25.1 | 195.0 | 1.5e-48 |
| BAV69313.1 | PrhL | BGC0001729 | Polyketide + Terpene | 27.0 | 28.4 | 194.0 | 1.9e-48 |
| WP\_055480220.1 | type\_I\_polyketide\_synthase | BGC0001653 | Polyketide | 26.0 | 42.0 | 194.0 | 2.5e-48 |
| CCP20049.1 | divL2\_protein | BGC0001119 | Polyketide:Modular type I | 30.0 | 23.2 | 193.0 | 3.3e-48 |
| WP\_048832936.1 | polyketide\_synthase | BGC0001348 | Polyketide:Modular type I | 28.0 | 26.6 | 193.0 | 3.3e-48 |
| CBJ89764.1 | Polyketide\_synthase\_involved\_in\_xenocoumacin\_synthesis | BGC0001054 | NRP + Polyketide:Modular type I | 26.0 | 27.0 | 193.0 | 4.3e-48 |
| AEN83889.1 | AdaA | BGC0000156 | Polyketide:Iterative type I | 26.0 | 32.2 | 191.0 | 1.6e-47 |
| DAB41484.1 | nonribosomal\_peptide\_synthetase/polyketide\_synthase\_type\_I | BGC0001766 | NRP | 31.0 | 23.7 | 189.0 | 8.1e-47 |
| CAQ43077.1 | polyketide\_synthase | BGC0000970 | NRP + Polyketide:Modular type I | 27.0 | 33.2 | 188.0 | 1.4e-46 |
| OJF16272.1 | AceP1 | BGC0001491 | Polyketide | 30.0 | 21.2 | 188.0 | 1.4e-46 |
| ABC87510.1 | polyketide\_synthase | BGC0001011 | NRP + Polyketide | 27.0 | 26.6 | 188.0 | 1.8e-46 |
| ctg1\_orf21 |  | BGC0001013 | NRP + Polyketide | 27.0 | 26.6 | 188.0 | 1.8e-46 |
| AGJ76601.1 | HglE | BGC0000869 | Other | 27.0 | 22.4 | 186.0 | 5.2e-46 |
| BAV19379.1 | polyketide\_synthase | BGC0001390 | NRP + Polyketide | 29.0 | 22.5 | 186.0 | 6.8e-46 |
| BBF25315.1 | polyketide\_synthase | BGC0001923 | Terpene + Polyketide | 25.0 | 28.0 | 185.0 | 8.9e-46 |
| AAB81123.1 | unknown | BGC0000862 | Other | 27.0 | 26.5 | 185.0 | 1.2e-45 |
| CAP95404.1 |  | BGC0001404 | Polyketide | 27.0 | 24.6 | 184.0 | 2.6e-45 |
| AEH42490.1 | polyketide\_synthase | BGC0000032 | Polyketide | 29.0 | 23.8 | 183.0 | 3.4e-45 |
| CCE88377.1 | non-ribosomal\_peptide\_synthetase/polyketide\_synthase | BGC0001034 | NRP + Polyketide:Modular type I | 26.0 | 30.7 | 182.0 | 7.6e-45 |
| AAK57190.1 | MxaF | BGC0001022 | NRP + Polyketide | 26.0 | 25.4 | 181.0 | 1.3e-44 |
| AP234\_RS14720 | hypothetical\_protein | BGC0001653 | Polyketide | 25.0 | 41.6 | 181.0 | 1.3e-44 |
| ALA09355.1 | type\_I\_modular\_PKS | BGC0001303 | Polyketide | 27.0 | 31.6 | 179.0 | 8.4e-44 |
| AID65222.1 | putative\_aspartate\_racemase | BGC0000335 | NRP | 30.0 | 21.0 | 176.0 | 4.1e-43 |
| ABF00127.1 | polyunsaturated\_fatty\_acid\_synthase | BGC0000863 | Other | 27.0 | 24.4 | 176.0 | 7.1e-43 |
| ACY06292.1 | modular\_polyketide\_synthase | BGC0001042 | NRP + Polyketide | 30.0 | 22.6 | 176.0 | 7.1e-43 |
| AXM42951.1 | polyketide\_synthase | BGC0001941 | NRP + Polyketide | 29.0 | 21.2 | 176.0 | 7.1e-43 |
| EAU29529.1 | hypothetical\_protein | BGC0000682 | Terpene | 28.0 | 20.5 | 175.0 | 1.2e-42 |
| AME18003.1 | enediyne\_polyketie\_synthase | BGC0001804 | Polyketide | 29.0 | 24.6 | 173.0 | 6e-42 |
| PKX88487.1 | polyketide\_synthase | BGC0001708 | Polyketide + Terpene | 25.0 | 32.3 | 173.0 | 6e-42 |
| BAQ25507.1 | type\_I\_polyketide\_synthase | BGC0001288 | Polyketide | 27.0 | 27.1 | 172.0 | 7.8e-42 |
| ALE27513.1 | putative\_polyketide\_synthase | BGC0001292 | Other | 27.0 | 26.3 | 172.0 | 7.8e-42 |
| ATV95639.1 | type\_I\_PKS | BGC0001503 | Polyketide | 28.0 | 24.7 | 172.0 | 7.8e-42 |
| AAM78012.1 | warhead-forming\_iterative\_polyketide\_synthase | BGC0000112 | Polyketide:Iterative type I + Polyketide:Enediyne type I | 28.0 | 23.7 | 170.0 | 5.1e-41 |
| AAQ17110.2 | enediyne\_polyketide\_synthase | BGC0001008 | Polyketide:Iterative type I + Polyketide:Enediyne type I | 28.0 | 24.1 | 170.0 | 5.1e-41 |
| EYT83433.1 | hypothetical\_protein | BGC0001213 | Polyketide | 29.0 | 22.1 | 169.0 | 6.6e-41 |
| ALP32046.1 | CycF | BGC0001293 | Polyketide | 29.0 | 21.7 | 167.0 | 2.5e-40 |
| AFV52145.1 | polyketide\_synthase | BGC0000081 | Polyketide:Iterative type I + Polyketide:Enediyne type I | 27.0 | 23.8 | 164.0 | 2.1e-39 |
| BAA89382.2 |  | BGC0000864 | Other | 27.0 | 20.0 | 164.0 | 2.8e-39 |
| ALJ49921.1 | TtmH | BGC0001236 | Polyketide | 24.0 | 32.7 | 163.0 | 4.7e-39 |
| AAL06699.1 | polyketide\_synthase | BGC0000965 | Polyketide:Iterative type I + Polyketide:Enediyne type I | 24.0 | 25.6 | 158.0 | 1.2e-37 |
| ALU98461.1 | erythronolide\_synthase | BGC0001397 | NRP + Polyketide | 24.0 | 25.6 | 158.0 | 1.2e-37 |
| ANY94470.1 | enediyne\_polyketide\_synthase | BGC0001584 | Polyketide | 26.0 | 23.9 | 158.0 | 2e-37 |
| ABP55141.1 | beta-ketoacyl\_synthase | BGC0000150 | NRP + Polyketide:Enediyne type I | 25.0 | 24.3 | 155.0 | 1.3e-36 |
| AKA59448.1 | polyketide\_synthase | BGC0001203 | NRP + Polyketide | 27.0 | 26.1 | 153.0 | 4.9e-36 |
| AAM94794.1 | CalE8 | BGC0000033 | Polyketide | 26.0 | 23.8 | 148.0 | 1.2e-34 |
| ANY10599.1 | polyketide\_synthase | BGC0001773 | Polyketide | 23.0 | 44.1 | 148.0 | 1.6e-34 |
| ANY10588.1 | polyketide\_synthase | BGC0001773 | Polyketide | 23.0 | 40.4 | 146.0 | 7.8e-34 |
| EHK80165.1 | beta-ketoacyl\_synthase | BGC0001447 | Polyketide | 26.0 | 28.7 | 139.0 | 9.6e-32 |
| ACB47048.1 | DynE8 | BGC0001060 | Polyketide | 27.0 | 24.0 | 123.0 | 4.2e-27 |
| AAD44234.1 | PstB | BGC0000362 | NRP | 26.0 | 21.4 | 122.0 | 1.2e-26 |
| BBA66512.1 | type\_I\_polyketide\_synthase | BGC0001495 | Polyketide | 25.0 | 24.0 | 120.0 | 4.6e-26 |
| AAF71776.1 | nysC | BGC0000115 | Polyketide:Modular type I + Saccharide:Hybrid/tailoring | 22.0 | 40.8 | 104.0 | 3.4e-21 |
| AVX51108.1 | nysC | BGC0001709 | Polyketide | 24.0 | 23.3 | 89.0 | 1.1e-16 |
