## Supplementary Results for "A draft genome of the ascomycotal fungal species *Pseudopithomyces maydicus* (family *Didymosphaeriaceae*)": input.path1.gene393_mibig_hits.html

| MIBiG Protein | Description | MIBiG Cluster | MiBiG Product | % ID | % Coverage | BLAST Score | E-value |
| --- | --- | --- | --- | --- | --- | --- | --- |
| AAK33081.1 | trichothecene\_efflux\_pump | BGC0001277 | Terpene | 38.0 | 39.4 | 124.0 | 5.9e-28 |
| AAK53581.1 | trichothecene\_efflux\_pump | BGC0000930 | Other | 37.0 | 39.6 | 120.0 | 8.5e-27 |
| BAX01966.1 | trichothecene\_efflux\_pump | BGC0001811 | Terpene | 37.0 | 39.6 | 120.0 | 8.5e-27 |
| AWH12934.1 | StmW | BGC0001939 | Polyketide | 30.0 | 34.4 | 60.0 | 1e-08 |
| ACB37757.1 | putative\_multidrug\_export\_protein | BGC0000162 | Polyketide | 28.0 | 37.5 | 55.0 | 3.4e-07 |
