## Supplementary Results for "A draft genome of the ascomycotal fungal species *Pseudopithomyces maydicus* (family *Didymosphaeriaceae*)": input.path1.gene394_mibig_hits.html

| MIBiG Protein | Description | MIBiG Cluster | MiBiG Product | % ID | % Coverage | BLAST Score | E-value |
| --- | --- | --- | --- | --- | --- | --- | --- |
| EPE34349.1 | acyl-AMP\_ligase | BGC0001035 | Polyketide + NRP + Other:Aminocoumarin | 45.0 | 44.9 | 95.0 | 1.5e-19 |
| OAQ83766.1 | phenylacetyl-\_ligase\_protein | BGC0001358 | Polyketide | 46.0 | 44.9 | 94.0 | 3.3e-19 |
| CCA65703.1 | anthranilate-CoA\_ligase | BGC0001343 | Polyketide | 42.0 | 43.3 | 91.0 | 1.7e-18 |
| EHA28233.1 | hypothetical\_protein | BGC0001143 | Polyketide | 39.0 | 49.0 | 86.0 | 6.9e-17 |
| AMM63166.1 | AniI | BGC0001371 | NRP | 41.0 | 44.9 | 83.0 | 4.5e-16 |
| AHH25595.1 | PKS | BGC0000957 | NRP + Polyketide | 38.0 | 55.9 | 81.0 | 1.7e-15 |
| AFU65902.1 | DacH | BGC0000216 | Polyketide | 41.0 | 42.9 | 81.0 | 2.2e-15 |
| ABC34346.1 | long-chain-fatty-acid--CoA\_ligase,\_putative | BGC0001102 | NRP:Beta-lactam + Polyketide:Modular type I | 39.0 | 43.3 | 78.0 | 1.9e-14 |
| CAL48957.1 | anthranilate-CoA-ACP\_transferase | BGC0001343 | Polyketide | 33.0 | 56.3 | 77.0 | 4.2e-14 |
| CBF87076.1 | conserved\_hypothetical\_protein | BGC0001290 | NRP | 33.0 | 68.8 | 75.0 | 1.2e-13 |
| AAG31128.1 | MxcE | BGC0001345 | NRP | 42.0 | 39.7 | 74.0 | 2.1e-13 |
| AXL88825.1 | hypothetical\_protein | BGC0001895 | Polyketide | 37.0 | 44.1 | 72.0 | 7.9e-13 |
| QCC62990.1 | AMP\_dependent\_ligase | BGC0001966 | NRP | 39.0 | 45.7 | 70.0 | 3.9e-12 |
| AEI70243.1 | 2,3-dihydroxybenzoate-AMP\_ligase | BGC0000401 | NRP | 38.0 | 39.3 | 70.0 | 5.1e-12 |
| AGY30675.1 | Ann3 | BGC0001298 | Polyketide | 38.0 | 43.7 | 69.0 | 8.8e-12 |
| AAQ90177.1 | putative\_acyl-CoA\_synthetase | BGC0000128 | Polyketide | 33.0 | 41.7 | 68.0 | 1.5e-11 |
| AEZ64574.1 | fatty-acid-CoA\_ligase | BGC0001065 | Polyketide | 38.0 | 43.3 | 67.0 | 2.6e-11 |
| CAQ52626.1 | type\_I\_polyketide\_synthase,\_loading\_module\_and\_modules\_1-3 | BGC0001066 | Polyketide:Modular type I | 39.0 | 38.5 | 67.0 | 4.4e-11 |
| AGE11892.1 | 2,3-dihydroxybenzoate-AMP\_ligase | BGC0000366 | NRP | 42.0 | 38.9 | 66.0 | 7.4e-11 |
| ATV95617.1 | CoA\_ligase | BGC0001503 | Polyketide | 36.0 | 39.3 | 66.0 | 7.4e-11 |
| CBA63660.1 | 2,3-dihydroxybenzoate-AMP\_ligase | BGC0000368 | NRP | 34.0 | 39.3 | 64.0 | 2.8e-10 |
| AQM37583.1 | nonribosomal\_peptide\_synthetase | BGC0001424 | NRP:Cyclic depsipeptide + Polyketide:Iterative type I | 35.0 | 45.7 | 64.0 | 2.8e-10 |
| ATG32071.1 | proline\_specific\_adenylation\_domain-containing\_protein | BGC0001750 | NRP + Polyketide | 33.0 | 41.7 | 63.0 | 6.3e-10 |
| APZ78808.1 | nonribosomal\_peptide\_synthetase | BGC0001428 | NRP:Cyclic depsipeptide + Polyketide:Iterative type I | 33.0 | 44.9 | 62.0 | 8.2e-10 |
| AAO07763.1 | 2,3-dihydroxybenzoate-AMP\_ligase | BGC0000460 | NRP | 40.0 | 38.5 | 62.0 | 1.1e-09 |
| AAD24881.1 | putative\_acyl-CoA\_synthetase | BGC0000127 | Polyketide | 30.0 | 41.7 | 62.0 | 1.4e-09 |
| ACN69986.1 | proline\_adenyltransferase | BGC0000079 | Polyketide | 33.0 | 41.7 | 61.0 | 2.4e-09 |
| AXG22420.1 | proline\_adenyltransferase | BGC0002024 | Polyketide | 31.0 | 41.7 | 61.0 | 3.1e-09 |
| AAN65233.1 | acyl-CoA\_synthetase | BGC0000832 | Saccharide:Hybrid/tailoring + Other:Aminocoumarin | 30.0 | 41.7 | 59.0 | 1.2e-08 |
| AEH42484.1 | adenylation\_for\_L-proline | BGC0000032 | Polyketide | 30.0 | 41.7 | 57.0 | 2.6e-08 |
| ACB12561.1 | Fum16 | BGC0000063 | Polyketide | 46.0 | 24.7 | 57.0 | 2.6e-08 |
| ACU36660.1 | AMP-dependent\_synthetase\_and\_ligase | BGC0000392 | NRP | 39.0 | 41.3 | 57.0 | 4.5e-08 |
| AEH41793.1 | HrmO | BGC0000374 | NRP:Cyclic depsipeptide | 28.0 | 49.8 | 56.0 | 7.7e-08 |
| AAY37650.1 | Amino\_acid\_adenylation | BGC0000437 | NRP | 30.0 | 43.3 | 56.0 | 7.7e-08 |
| AAG29789.1 | acyl-CoA\_synthetase | BGC0000833 | Saccharide:Hybrid/tailoring + Other:Aminocoumarin | 29.0 | 41.7 | 55.0 | 1.3e-07 |
| AAN74819.2 | Fum16p | BGC0000062 | Polyketide | 42.0 | 24.3 | 55.0 | 1.7e-07 |
| ABI22132.1 | putative\_non-ribosomal\_peptide\_synthetase | BGC0000422 | NRP | 31.0 | 44.9 | 54.0 | 2.2e-07 |
| ACA34725.1 | CtnI | BGC0000894 | Other | 31.0 | 39.7 | 54.0 | 2.2e-07 |
| ALI92663.1 | MRR8\_AMP-binding\_enzyme | BGC0001338 | Polyketide:Iterative type I | 31.0 | 39.7 | 54.0 | 2.2e-07 |
| BBA21082.1 | putative\_CoA\_ligase | BGC0001740 | NRP + Polyketide | 26.0 | 44.9 | 54.0 | 2.2e-07 |
| ABB69752.1 | PlaP4 | BGC0000654 | Terpene + Saccharide:Hybrid/tailoring | 29.0 | 41.7 | 53.0 | 5e-07 |
| CAJ87594.1 | 2,3-dihydroxybenzoate-AMP\_ligase | BGC0001055 | NRP + Polyketide | 36.0 | 36.0 | 53.0 | 5e-07 |
| BAC87906.1 | probable\_acinetobactin\_biosynthesis\_protein | BGC0000294 | NRP | 31.0 | 38.5 | 51.0 | 1.9e-06 |
| ABL74940.1 | NRPS | BGC0001048 | NRP:Glycopeptide + Polyketide:Modular type I + Saccharide:Hybrid/tailoring | 32.0 | 44.5 | 51.0 | 2.5e-06 |
| KFH48713.1 | Isopenicillin\_N\_epimerase\_component-like\_protein | BGC0000317 | NRP | 38.0 | 35.2 | 50.0 | 3.2e-06 |
