## Supplementary Results for "A draft genome of the ascomycotal fungal species *Pseudopithomyces maydicus* (family *Didymosphaeriaceae*)": input.path1.gene395_mibig_hits.html

| MIBiG Protein | Description | MIBiG Cluster | MiBiG Product | % ID | % Coverage | BLAST Score | E-value |
| --- | --- | --- | --- | --- | --- | --- | --- |
| CAD19094.1 | methyl\_transferase | BGC0000153 | NRP + Polyketide:Modular type I | 27.0 | 30.7 | 69.0 | 2.7e-11 |
