## Supplementary Results for "A draft genome of the ascomycotal fungal species *Pseudopithomyces maydicus* (family *Didymosphaeriaceae*)": input.path1.gene397_mibig_hits.html

| MIBiG Protein | Description | MIBiG Cluster | MiBiG Product | % ID | % Coverage | BLAST Score | E-value |
| --- | --- | --- | --- | --- | --- | --- | --- |
| CBF73453.1 | nonribosomal\_peptide\_synthase,\_putative\_(JCVI) | BGC0001515 | NRP | 32.0 | 103.2 | 402.0 | 1.8e-111 |
| CBF87069.1 | nonribosomal\_peptide\_synthase,\_putative\_(Eurofung) | BGC0001290 | NRP | 30.0 | 103.1 | 346.0 | 9e-95 |
| ctg1\_orf003 |  | BGC0000334 | NRP | 31.0 | 100.1 | 346.0 | 1.2e-94 |
| EXU96269.1 | nonribosomal\_peptide\_synthetase,\_serinocyclin\_synthetase\_NPS1 | BGC0001240 | NRP | 31.0 | 100.8 | 344.0 | 5.8e-94 |
| EPE34341.1 | non-ribosomal\_peptide\_synthetase | BGC0001035 | Polyketide + NRP + Other:Aminocoumarin | 31.0 | 99.9 | 339.0 | 1.1e-92 |
| XP\_001217690.1 | hypothetical\_protein | BGC0001517 | NRP | 32.0 | 97.1 | 337.0 | 4.2e-92 |
| QCC62999.1 | BII-rafflesfungin\_nonribosomal\_protein\_synthetase | BGC0001966 | NRP | 31.0 | 101.2 | 335.0 | 2.1e-91 |
| MAA\_10043 | non-ribosomal\_peptide\_synthetase | BGC0000337 | NRP | 29.0 | 102.8 | 317.0 | 4.5e-86 |
| AUS29494.1 | non-ribosomal\_peptide\_synthetase | BGC0001030 | NRP + Polyketide | 29.0 | 104.4 | 316.0 | 1e-85 |
| BAN91483.1 | putative\_Aculeacin\_A\_nonribosomal\_peptide\_synthase | BGC0001220 | NRP | 31.0 | 101.5 | 312.0 | 1.9e-84 |
| EAL89049.1 | nonribosomal\_peptide\_synthetase | BGC0000355 | NRP | 29.0 | 101.2 | 305.0 | 2.3e-82 |
| ctg1\_orf00001 |  | BGC0000901 | Other | 30.0 | 96.3 | 304.0 | 5.1e-82 |
| OAQ83772.1 | nonribosomal\_peptide\_synthase | BGC0001358 | Polyketide | 28.0 | 103.5 | 302.0 | 1.5e-81 |
| ACR78148.1 | BSLS | BGC0000312 | NRP | 28.0 | 104.7 | 297.0 | 6.3e-80 |
| BAH23995.1 | nonribosomal\_peptide\_synthetase | BGC0000356 | NRP + Alkaloid | 28.0 | 104.5 | 293.0 | 9e-79 |
| CBF87869.1 | nonribosomal\_peptide\_synthase,\_putative\_(Eurofung) | BGC0001699 | Polyketide | 30.0 | 90.8 | 292.0 | 2.6e-78 |
| AGA37267.1 | NRPS | BGC0000816 | NRP + Alkaloid | 29.0 | 102.3 | 290.0 | 5.9e-78 |
| AAF01762.1 | AM-toxin\_synthetase | BGC0001261 | NRP | 28.0 | 101.3 | 290.0 | 1e-77 |
| XP\_003044554.1 | hypothetical\_protein | BGC0001768 | NRP | 28.0 | 105.6 | 289.0 | 2.2e-77 |
| CAP93139.1 | cyclic\_hydrophobic\_tetrapeptide | BGC0000357 | NRP:Cyclic depsipeptide | 28.0 | 91.9 | 287.0 | 6.5e-77 |
| CBF76038.1 | nonribosomal\_peptide\_synthase,\_putative\_(Eurofung) | BGC0001399 | NRP | 28.0 | 99.6 | 284.0 | 7.2e-76 |
| XP\_011325838.1 | hypothetical\_protein | BGC0001545 | NRP | 28.0 | 101.0 | 281.0 | 3.6e-75 |
| ACI30655.1 | BEAS\_beauvericin\_nonribosomal\_cyclodepsipeptide\_synthetase | BGC0000313 | NRP | 27.0 | 106.3 | 278.0 | 3e-74 |
| BBC83957.1 | nonribosomal\_peptide\_synthetase | BGC0001636 | NRP | 26.0 | 99.5 | 275.0 | 2.5e-73 |
| ctg1\_orf000000 |  | BGC0000900 | Other | 33.0 | 20.9 | 99.0 | 2.5e-20 |
| AQM58286.1 | non-ribosomal\_peptide\_synthase | BGC0001816 | NRP + Polyketide | 29.0 | 103.6 | 271.0 | 4.8e-72 |
| CAA79245.2 | enniatin\_synthetase | BGC0000342 | NRP | 26.0 | 101.3 | 266.0 | 1.2e-70 |
| AGQ43600.1 | HC-toxin\_synthetase | BGC0001166 | NRP | 28.0 | 103.2 | 257.0 | 5.5e-68 |
| ABM91454.1 | lysergyl\_peptide\_synthetase\_LpsB | BGC0000348 | NRP | 28.0 | 89.5 | 254.0 | 6.1e-67 |
| CRG85572.1 | nonribosomal\_peptide\_synthase,\_putative | BGC0001402 | NRP | 27.0 | 94.9 | 254.0 | 6.1e-67 |
| ABB90279.1 | non-ribosomal\_peptide\_synthetase | BGC0001057 | NRP + Polyketide | 28.0 | 88.7 | 248.0 | 2.6e-65 |
| AMM63162.1 | AniA | BGC0001371 | NRP | 29.0 | 89.9 | 248.0 | 4.4e-65 |
| ACZ66258.1 | APS1 | BGC0000304 | NRP | 28.0 | 103.6 | 238.0 | 4.5e-62 |
| BBB04327.1 | nonribosomal\_peptide\_synthetase | BGC0001717 | Alkaloid | 28.0 | 80.9 | 235.0 | 2.9e-61 |
| QDA77059.1 | polyketide\_synthase/nonribosomal\_peptide\_synthetase | BGC0002026 | NRP | 29.0 | 84.0 | 234.0 | 5e-61 |
| ACJ04424.1 | aureobasidin\_A1\_biosynthesis\_complex | BGC0000307 | NRP | 27.0 | 94.8 | 233.0 | 8.5e-61 |
| AGA37269.1 | NRPS | BGC0000819 | NRP + Alkaloid | 26.0 | 101.4 | 233.0 | 1.5e-60 |
| ANY57892.1 | PenN | BGC0001372 | Terpene | 27.0 | 85.1 | 228.0 | 2.7e-59 |
| XP\_002379984.1 | nonribosomal\_peptide\_synthase,\_putative | BGC0001621 | NRP | 26.0 | 89.1 | 225.0 | 2.3e-58 |
| CAP96445.1 | NRPS | BGC0000420 | NRP | 33.0 | 51.2 | 211.0 | 5.9e-54 |
| CAJ21198.2 | non-ribosomal\_peptide\_synthetase | BGC0000297 | NRP:Glycopeptide + Polyketide:Other + Saccharide:Hybrid/tailoring | 27.0 | 102.0 | 209.0 | 1.7e-53 |
| AXN93603.1 | PuwH | BGC0001952 | NRP | 26.0 | 99.2 | 207.0 | 6.5e-53 |
| AIW82285.1 | PuwH | BGC0001125 | NRP + Polyketide | 26.0 | 100.0 | 205.0 | 4.2e-52 |
| QDA77045.1 | polyketide\_synthase/nonribosomal\_peptide\_synthetase | BGC0002025 | NRP | 25.0 | 96.0 | 203.0 | 9.4e-52 |
| AXN93582.1 | PuwH | BGC0001950 | NRP | 26.0 | 104.7 | 203.0 | 1.6e-51 |
| AXN93591.1 | PuwH | BGC0001951 | NRP | 26.0 | 104.7 | 203.0 | 1.6e-51 |
| AXN93616.1 | PuwH | BGC0001953 | NRP | 25.0 | 99.2 | 202.0 | 2.7e-51 |
| APZ78703.1 | nonribosomal\_peptide\_synthetase | BGC0001419 | NRP:Cyclic depsipeptide + Polyketide:Iterative type I | 26.0 | 84.6 | 201.0 | 6.1e-51 |
| AXN93615.1 | PuwG | BGC0001953 | NRP | 27.0 | 84.3 | 201.0 | 6.1e-51 |
| ABI26078.1 | OciB | BGC0000331 | NRP | 26.0 | 100.4 | 200.0 | 8e-51 |
| ADY16689.1 | TqaB | BGC0001142 | NRP | 27.0 | 77.3 | 200.0 | 1.4e-50 |
| APZ78715.1 | nonribosomal\_peptide\_synthetase | BGC0001420 | NRP:Cyclic depsipeptide + Polyketide:Iterative type I | 27.0 | 84.6 | 200.0 | 1.4e-50 |
| APZ78691.1 | nonribosomal\_peptide\_synthetase | BGC0001418 | NRP:Cyclic depsipeptide + Polyketide:Iterative type I | 27.0 | 84.6 | 199.0 | 1.8e-50 |
| ARB50206.1 | lysergyl\_peptide\_synthetase\_2 | BGC0001573 | Alkaloid | 25.0 | 90.0 | 199.0 | 1.8e-50 |
| AXN93614.1 | PuwF | BGC0001953 | NRP | 27.0 | 83.9 | 199.0 | 1.8e-50 |
| APZ78728.1 | nonribosomal\_peptide\_synthetase | BGC0001421 | NRP:Cyclic depsipeptide + Polyketide:Iterative type I | 26.0 | 84.6 | 198.0 | 5.2e-50 |
| ACZ55944.1 | non-ribosomal\_peptide\_synthetase | BGC0000302 | NRP | 26.0 | 98.5 | 195.0 | 2.6e-49 |
| ABI26077.1 | OciA | BGC0000331 | NRP | 25.0 | 98.2 | 192.0 | 2.2e-48 |
| AEG64697.1 | LpmC | BGC0000379 | NRP | 25.0 | 99.6 | 191.0 | 4.8e-48 |
| EAL89046.1 | nonribosomal\_peptide\_synthetase | BGC0000355 | NRP | 27.0 | 77.7 | 187.0 | 7e-47 |
| AEO14744.1 | NdaB | BGC0000396 | NRP | 26.0 | 96.7 | 186.0 | 1.2e-46 |
| ALD82526.1 | non-ribosomal\_peptide\_synthase | BGC0001212 | NRP + Polyketide | 26.0 | 95.9 | 186.0 | 1.2e-46 |
| AEG64696.1 | LpmB | BGC0000379 | NRP | 27.0 | 85.1 | 186.0 | 1.6e-46 |
| EAW16180.1 | nonribosomal\_peptide\_synthase,\_putative | BGC0000293 | NRP | 24.0 | 89.1 | 185.0 | 3.5e-46 |
| ATP76243.1 | NdaA | BGC0001705 | NRP + Polyketide | 27.0 | 78.7 | 183.0 | 1.3e-45 |
| BAE60013.1 |  | BGC0001518 | Terpene | 26.0 | 79.6 | 181.0 | 5e-45 |
| AAK06804.1 | Tyroxyl-AMP-forming\_enzyme | BGC0001072 | Saccharide + Polyketide:Modular type I + Polyketide:Type II + Other:Aminocoumarin | 24.0 | 85.0 | 181.0 | 6.5e-45 |
| AAL15600.1 | SimH | BGC0000270 | Polyketide | 24.0 | 85.0 | 180.0 | 1.1e-44 |
| CAE53351.1 | non-ribosomal\_peptide\_synthetase | BGC0000440 | NRP:Glycopeptide | 25.0 | 98.2 | 180.0 | 1.1e-44 |
| CAG15010.1 | peptide\_synthetase,\_module\_3 | BGC0000441 | NRP | 25.0 | 98.2 | 180.0 | 1.1e-44 |
| AAG34184.1 | SimH | BGC0000835 | Polyketide | 24.0 | 85.0 | 180.0 | 1.1e-44 |
| WP\_010369428.1 | non-ribosomal\_peptide\_synthase | BGC0000314 | Polyketide + NRP:Cyclic depsipeptide + Other:Aminocoumarin | 24.0 | 102.6 | 180.0 | 1.5e-44 |
| ARU08074.1 | mlcL | BGC0001448 | NRP:Ca+-dependent lipopeptide | 26.0 | 84.9 | 178.0 | 3.2e-44 |
| AAU39359.1 | lichenysin\_synthase\_LchAA | BGC0000381 | NRP | 26.0 | 98.6 | 175.0 | 2.7e-43 |
| AAO56328.1 | non-ribosomal\_peptide\_synthetase\_SyfA | BGC0000435 | NRP | 25.0 | 99.9 | 174.0 | 8e-43 |
| AHD05618.1 | putative\_non-ribosomal\_peptide\_ligase\_domain\_protein | BGC0001033 | NRP + Polyketide | 25.0 | 99.6 | 174.0 | 8e-43 |
| OTA20325.1 | peptide\_synthase | BGC0001824 | NRP | 24.0 | 97.7 | 173.0 | 1e-42 |
| CCJ67637.1 | TaaB | BGC0000447 | NRP:Lipopeptide | 27.0 | 88.7 | 173.0 | 1.4e-42 |
| AGS77308.1 | NRPS\_module\_3 | BGC0001178 | NRP:Glycopeptide | 25.0 | 99.2 | 172.0 | 2.3e-42 |
| AEZ51516.1 | pmxA | BGC0001153 | NRP:Lipopeptide | 25.0 | 103.7 | 172.0 | 3e-42 |
| ADJ63842.1 | Serobactin\_synthetase | BGC0000424 | NRP:NRP siderophore | 25.0 | 97.8 | 171.0 | 5.2e-42 |
| CAE02631.1 | surfactin\_synthetase\_B\_ | BGC0000433 | NRP:Lipopeptide | 23.0 | 101.3 | 171.0 | 5.2e-42 |
| AAM80538.1 | StaB | BGC0000290 | NRP:Glycopeptide | 23.0 | 100.8 | 169.0 | 2.6e-41 |
| WP\_010369430.1 | non-ribosomal\_peptide\_synthetase | BGC0000314 | Polyketide + NRP:Cyclic depsipeptide + Other:Aminocoumarin | 24.0 | 99.2 | 168.0 | 3.4e-41 |
| AHZ34232.1 | CifA | BGC0000323 | NRP:Lipopeptide | 25.0 | 89.2 | 168.0 | 3.4e-41 |
| ACM79810.1 | ZmaO | BGC0001059 | NRP + Polyketide | 24.0 | 99.7 | 168.0 | 3.4e-41 |
| AAU39360.1 | lichenysin\_synthase\_LchAB | BGC0000381 | NRP | 28.0 | 80.0 | 167.0 | 7.5e-41 |
| CUX79061.1 | Octapeptin\_synthase\_subunit\_B | BGC0001715 | NRP | 25.0 | 102.3 | 167.0 | 7.5e-41 |
| BS330\_28385 | non-ribosomal\_peptide\_synthetase | BGC0001461 | NRP:Glycopeptide | 25.0 | 96.9 | 166.0 | 1.7e-40 |
| AYA22336.1 | KerA | BGC0001955 | Other | 25.0 | 96.9 | 165.0 | 4.9e-40 |
| ABD65957.1 | nonribosomal\_peptide\_synthetase | BGC0000341 | NRP | 26.0 | 101.7 | 164.0 | 8.3e-40 |
| CUX79060.1 | Octapeptin\_synthase\_subunit\_A | BGC0001715 | NRP | 24.0 | 99.7 | 163.0 | 1.4e-39 |
| AFH75329.1 | nonribosomal\_peptide\_synthetase | BGC0000398 | NRP:Cyclic depsipeptide | 27.0 | 76.2 | 162.0 | 2.4e-39 |
| CCA29203.1 | non-ribosomal\_peptide\_synthetase/polyketide\_synthase | BGC0000955 | NRP + Polyketide:Modular type I | 24.0 | 95.9 | 162.0 | 3.1e-39 |
| AAY91419.3 | non-ribosomal\_peptide\_synthetase\_OfaA | BGC0000399 | NRP:Cyclic depsipeptide | 26.0 | 83.5 | 161.0 | 5.4e-39 |
| AJM89735.1 | PmxA | BGC0001192 | NRP | 23.0 | 106.1 | 161.0 | 5.4e-39 |
| ARB50207.1 | lysergyl\_peptide\_synthetase\_21 | BGC0001573 | Alkaloid | 25.0 | 64.7 | 160.0 | 9.2e-39 |
| WP\_051462298.1 | non-ribosomal\_peptide\_synthetase | BGC0001873 | NRP:Lipopeptide | 23.0 | 105.4 | 160.0 | 1.2e-38 |
| ACA97576.1 | PmxA | BGC0000408 | NRP | 24.0 | 101.4 | 159.0 | 2.7e-38 |
| APZ78692.1 | nonribosomal\_peptide\_synthetase | BGC0001418 | NRP:Cyclic depsipeptide + Polyketide:Iterative type I | 22.0 | 98.1 | 158.0 | 5.9e-38 |
| ABP57749.1 | DepE | BGC0000993 | NRP:Cyclic depsipeptide + Polyketide:Modular type I | 24.0 | 99.4 | 156.0 | 1.7e-37 |
| ALD82525.1 | non-ribosomal\_peptide\_synthase | BGC0001212 | NRP + Polyketide | 27.0 | 67.9 | 156.0 | 1.7e-37 |
| APZ78704.1 | nonribosomal\_peptide\_synthetase | BGC0001419 | NRP:Cyclic depsipeptide + Polyketide:Iterative type I | 22.0 | 98.0 | 156.0 | 1.7e-37 |
| APZ78716.1 | nonribosomal\_peptide\_synthetase | BGC0001420 | NRP:Cyclic depsipeptide + Polyketide:Iterative type I | 22.0 | 98.1 | 156.0 | 1.7e-37 |
| APZ78856.1 | nonribosomal\_peptide\_synthetase | BGC0001432 | NRP:Cyclic depsipeptide + Polyketide:Iterative type I | 22.0 | 98.1 | 156.0 | 2.3e-37 |
| ABD65958.1 | nonribosomal\_peptide\_synthetase | BGC0000341 | NRP | 24.0 | 83.2 | 155.0 | 3.8e-37 |
| AEZ51520.1 | pmxE | BGC0001153 | NRP:Lipopeptide | 24.0 | 102.9 | 155.0 | 3.8e-37 |
| ATD51279.1 | nonribosomal\_peptide\_synthase | BGC0001650 | NRP | 25.0 | 76.8 | 154.0 | 6.6e-37 |
| AGM14934.1 | xantholysin\_synthetase\_C | BGC0000463 | NRP:Lipopeptide | 25.0 | 81.6 | 154.0 | 8.6e-37 |
| ADY16697.1 | TqaA | BGC0001142 | NRP | 27.0 | 48.8 | 153.0 | 1.5e-36 |
| AJM89738.1 | PmxE | BGC0001192 | NRP | 24.0 | 106.0 | 153.0 | 1.5e-36 |
| AFJ14793.1 | PlpD | BGC0000403 | NRP | 25.0 | 88.0 | 151.0 | 5.6e-36 |
| WP\_043882190.1 | non-ribosomal\_peptide\_synthetase | BGC0001993 | NRP | 27.0 | 62.5 | 150.0 | 9.5e-36 |
| AYJ71720.1 | non-ribosomal\_peptide\_synthetase | BGC0001942 | NRP + Polyketide | 23.0 | 96.8 | 149.0 | 1.6e-35 |
| BAH22764.1 | nonribosomal\_peptide\_synthetase | BGC0001018 | NRP + Polyketide | 28.0 | 65.7 | 149.0 | 2.1e-35 |
| AAR87759.1 | ZmaJ | BGC0001059 | NRP + Polyketide | 26.0 | 67.3 | 149.0 | 2.8e-35 |
| WP\_041826607.1 | non-ribosomal\_peptide\_synthetase | BGC0001792 | NRP | 30.0 | 44.3 | 149.0 | 2.8e-35 |
| CAK15814.1 | putative\_non-ribosomal\_peptide\_synthetase,\_terminal\_component | BGC0000344 | NRP | 26.0 | 85.3 | 148.0 | 6.1e-35 |
| AAY37653.1 | Amino\_acid\_adenylation | BGC0000437 | NRP | 24.0 | 83.9 | 146.0 | 2.3e-34 |
| ADG27359.1 | peptide\_synthetase | BGC0000296 | NRP | 25.0 | 82.6 | 145.0 | 3e-34 |
| BAH43765.1 | tyrocidine\_synthetase\_II | BGC0000452 | NRP | 27.0 | 61.5 | 145.0 | 3e-34 |
| CDG17981.1 | Non-ribosomal\_peptide\_synthetase | BGC0000464 | NRP:Cyclic depsipeptide | 23.0 | 103.5 | 145.0 | 4e-34 |
| APZ78834.1 | nonribosomal\_peptide\_synthetase | BGC0001430 | NRP:Cyclic depsipeptide + Polyketide:Iterative type I | 21.0 | 98.2 | 144.0 | 6.8e-34 |
| AXN93581.1 | PuwF-G | BGC0001950 | NRP | 30.0 | 40.7 | 143.0 | 1.2e-33 |
| AXN93590.1 | PuwF-G | BGC0001951 | NRP | 30.0 | 40.7 | 143.0 | 1.2e-33 |
| ABM21571.1 | crpC | BGC0000975 | NRP + Polyketide | 30.0 | 47.8 | 143.0 | 2e-33 |
| BAH43870.1 | putative\_linear\_pentadecapeptide\_gramicidin\_synthetase\_LgrB | BGC0000367 | NRP | 29.0 | 45.6 | 142.0 | 2.6e-33 |
| AEG64698.1 | LpmD | BGC0000379 | NRP | 24.0 | 100.1 | 142.0 | 2.6e-33 |
| ABW17376.1 | PsoB | BGC0000411 | NRP | 26.0 | 83.9 | 142.0 | 2.6e-33 |
| AHF21228.1 | TriD | BGC0000449 | NRP | 31.0 | 47.4 | 141.0 | 4.4e-33 |
| ADM34138.1 | non-ribosomal\_peptide\_synthetase | BGC0001084 | NRP + Terpene + Alkaloid | 36.0 | 30.6 | 141.0 | 5.7e-33 |
| ABQ96384.2 | fusaricidin\_synthetase | BGC0001152 | Polyketide + NRP:Lipopeptide | 26.0 | 61.1 | 140.0 | 1.3e-32 |
| AFN69082.1 | nonribosomal\_peptide\_synthetase\_6 | BGC0001249 | NRP | 27.0 | 62.2 | 138.0 | 6.4e-32 |
| AHF21229.1 | TriE | BGC0000449 | NRP | 22.0 | 109.2 | 136.0 | 2.4e-31 |
| ALG65319.1 | Cal17 | BGC0001297 | NRP | 25.0 | 79.5 | 136.0 | 2.4e-31 |
| KFH48607.1 | N-like\_protein | BGC0000317 | NRP | 21.0 | 100.8 | 135.0 | 3.2e-31 |
| CAJ45639.1 | vanchrobactin\_non\_ribosomal\_peptide\_synthetase | BGC0000454 | NRP | 22.0 | 96.8 | 134.0 | 9.2e-31 |
| CDG17985.1 | Putative\_Ornithine\_racemase\_(fragment) | BGC0000464 | NRP:Cyclic depsipeptide | 26.0 | 61.6 | 133.0 | 1.6e-30 |
| AIW82284.1 | PuwG | BGC0001125 | NRP + Polyketide | 28.0 | 44.2 | 133.0 | 1.6e-30 |
| AGC83576.1 | NRPS | BGC0000818 | NRP | 28.0 | 50.2 | 132.0 | 2.7e-30 |
| AIW82283.1 | PuwF | BGC0001125 | NRP + Polyketide | 31.0 | 41.1 | 132.0 | 2.7e-30 |
| ASA76632.1 | polyketide\_synthase\_non-ribosomal\_peptide\_synthetase\_hybrid | BGC0001751 | NRP + Polyketide | 29.0 | 45.6 | 132.0 | 2.7e-30 |
| ACC81021.1 | non-ribosomal\_peptide\_synthetase | BGC0001479 | NRP | 35.0 | 31.4 | 132.0 | 3.5e-30 |
| AVR48535.1 | CusC | BGC0001564 | NRP + Polyketide | 29.0 | 46.2 | 132.0 | 3.5e-30 |
| AEW31021.1 | plipastatin\_synthetase | BGC0000407 | NRP | 29.0 | 44.9 | 131.0 | 5.9e-30 |
| APZ78680.1 | nonribosomal\_peptide\_synthetase | BGC0001417 | NRP:Cyclic depsipeptide + Polyketide:Iterative type I | 26.0 | 46.7 | 131.0 | 5.9e-30 |
| ATY37608.1 | BreC | BGC0001536 | NRP | 28.0 | 46.0 | 131.0 | 7.8e-30 |
| WP\_015507662.1 | non-ribosomal\_peptide\_synthetase | BGC0001792 | NRP | 30.0 | 47.2 | 131.0 | 7.8e-30 |
| AVI26393.1 | nonribosomal\_peptide\_synthase | BGC0001800 | NRP + Polyketide | 30.0 | 45.3 | 131.0 | 7.8e-30 |
| AKP45399.1 | CysK | BGC0001413 | NRP | 22.0 | 100.3 | 129.0 | 1.7e-29 |
| APZ78846.1 | nonribosomal\_peptide\_synthetase | BGC0001431 | NRP:Cyclic depsipeptide + Polyketide:Iterative type I | 25.0 | 46.5 | 129.0 | 2.3e-29 |
| CZT62794.1 | Non-ribosomal\_peptide\_synthase\_involved\_in\_Hassallidin\_biosynthesis | BGC0001614 | NRP | 21.0 | 91.2 | 129.0 | 2.3e-29 |
| AXN93602.1 | PuwF-G | BGC0001952 | NRP | 33.0 | 33.3 | 129.0 | 2.3e-29 |
| AQM37584.1 | nonribosomal\_peptide\_synthetase | BGC0001424 | NRP:Cyclic depsipeptide + Polyketide:Iterative type I | 27.0 | 44.4 | 128.0 | 5e-29 |
| EFE73313.1 | nonribosomal\_peptide\_synthetase | BGC0000431 | NRP:Cyclic depsipeptide | 26.0 | 60.9 | 128.0 | 6.6e-29 |
| CCJ67646.1 | JagB | BGC0001127 | NRP | 23.0 | 105.6 | 127.0 | 8.6e-29 |
| CAK15815.1 | putative\_non\_ribosomal\_peptide\_synthetase | BGC0000344 | NRP | 24.0 | 75.8 | 127.0 | 1.1e-28 |
| CDG17980.1 | Putative\_Ornithine\_racemase\_(fragment) | BGC0000464 | NRP:Cyclic depsipeptide | 28.0 | 45.2 | 126.0 | 1.9e-28 |
| ANZ15840.1 | non-ribosomal\_peptide\_synthase/amino\_acid\_adenylation\_enzyme | BGC0001569 | NRP | 29.0 | 46.6 | 126.0 | 1.9e-28 |
| ACZ55942.1 | non-ribosomal\_peptide\_synthetase | BGC0000302 | NRP | 28.0 | 42.6 | 126.0 | 2.5e-28 |
| APZ78756.1 | nonribosomal\_peptide\_synthetase | BGC0001423 | NRP:Cyclic depsipeptide + Polyketide:Iterative type I | 26.0 | 44.8 | 125.0 | 3.3e-28 |
| AAG02355.1 | peptide\_synthetase\_NRPS9-8 | BGC0000963 | NRP:Glycopeptide + Polyketide:Modular type I + Saccharide:Hybrid/tailoring | 34.0 | 31.0 | 125.0 | 4.3e-28 |
| WP\_064118560.1 | non-ribosomal\_peptide\_synthetase | BGC0001509 | NRP | 29.0 | 46.5 | 125.0 | 4.3e-28 |
| AGM16413.1 | paenibacterin\_synthetase\_B | BGC0000400 | NRP | 29.0 | 46.4 | 124.0 | 5.6e-28 |
| ABS74180.1 | bacillomycin\_D\_synthetase\_B | BGC0001090 | Polyketide + NRP:Lipopeptide | 28.0 | 44.8 | 124.0 | 7.3e-28 |
| AJF34464.1 | Txo2 | BGC0001207 | NRP | 28.0 | 52.5 | 124.0 | 7.3e-28 |
| ERG36783.1 | phenylalanine\_racemase | BGC0000299 | NRP | 25.0 | 61.1 | 123.0 | 1.6e-27 |
| CAD29799.1 | microcystin\_synthetase | BGC0001015 | NRP + Polyketide | 33.0 | 35.5 | 123.0 | 2.1e-27 |
| WP\_015507663.1 | non-ribosomal\_peptide\_synthase | BGC0001792 | NRP | 29.0 | 46.9 | 123.0 | 2.1e-27 |
| CCJ67645.1 | JagA | BGC0001127 | NRP | 26.0 | 75.2 | 121.0 | 6.2e-27 |
| ATU31794.1 | NRPS | BGC0001814 | NRP | 27.0 | 48.3 | 121.0 | 6.2e-27 |
| ACZ55945.1 | non-ribosomal\_peptide\_synthetase | BGC0000302 | NRP | 35.0 | 29.7 | 121.0 | 8e-27 |
| AEA30274.1 | peptide\_synthetase | BGC0000429 | Polyketide + NRP:Cyclic depsipeptide | 26.0 | 63.5 | 121.0 | 8e-27 |
| BAP05596.1 | calH | BGC0000967 | NRP + Polyketide:Trans-AT type I | 38.0 | 24.7 | 121.0 | 8e-27 |
| CZT62785.1 | Non-ribosomal\_peptide\_synthase,\_involved\_in\_Hassallidin\_biosynthesis | BGC0001614 | NRP | 27.0 | 46.7 | 121.0 | 8e-27 |
| ACG60782.1 | NRPS(C/A/PCP/C/A/PCP) | BGC0001058 | NRP:Glycopeptide + Polyketide:Modular type I + Saccharide:Hybrid/tailoring | 33.0 | 30.5 | 120.0 | 1e-26 |
| APZ78769.1 | nonribosomal\_peptide\_synthetase | BGC0001425 | NRP:Cyclic depsipeptide + Polyketide:Iterative type I | 24.0 | 52.0 | 120.0 | 1e-26 |
| AHB82058.1 | non\_ribosomal\_peptide\_synthetase | BGC0001019 | NRP + Polyketide:Modular type I | 29.0 | 45.1 | 120.0 | 1.4e-26 |
| APZ78795.1 | nonribosomal\_peptide\_synthetase | BGC0001427 | NRP:Cyclic depsipeptide + Polyketide:Iterative type I | 24.0 | 52.0 | 120.0 | 1.4e-26 |
| WP\_013428774.1 | non-ribosomal\_peptide\_synthetase | BGC0001844 | NRP:Lipopeptide | 36.0 | 27.4 | 120.0 | 1.4e-26 |
| ABS74179.1 | bacillomycin\_D\_synthetase\_C | BGC0001090 | Polyketide + NRP:Lipopeptide | 29.0 | 44.7 | 119.0 | 1.8e-26 |
| APZ78744.1 | nonribosomal\_peptide\_synthetase | BGC0001422 | NRP:Cyclic depsipeptide + Polyketide:Iterative type I | 25.0 | 47.2 | 119.0 | 2.3e-26 |
| CAJ77696.1 | MPS2\_protein | BGC0000363 | NRP | 29.0 | 47.8 | 118.0 | 5.2e-26 |
| CAJ77716.1 | Mps2\_protein | BGC0000364 | NRP | 29.0 | 45.8 | 118.0 | 5.2e-26 |
| AEW31019.1 | plipastatin\_synthetase | BGC0000407 | NRP | 27.0 | 44.8 | 118.0 | 5.2e-26 |
| APZ78782.1 | nonribosomal\_peptide\_synthetase | BGC0001426 | NRP:Cyclic depsipeptide + Polyketide:Iterative type I | 24.0 | 52.0 | 118.0 | 5.2e-26 |
| ABI26079.1 | OciC | BGC0000331 | NRP | 36.0 | 28.2 | 118.0 | 6.8e-26 |
| CDG17987.1 | Putative\_Ornithine\_racemase\_(fragment) | BGC0000464 | NRP:Cyclic depsipeptide | 27.0 | 45.5 | 118.0 | 6.8e-26 |
| AGS77309.1 | NRPS\_modules\_4-6 | BGC0001178 | NRP:Glycopeptide | 27.0 | 61.1 | 118.0 | 6.8e-26 |
| CAY48788.1 | putative\_non-ribosomal\_peptide\_synthetase | BGC0001312 | NRP | 27.0 | 47.5 | 118.0 | 6.8e-26 |
| AHZ34233.1 | CifB | BGC0000323 | NRP:Lipopeptide | 29.0 | 47.1 | 117.0 | 8.9e-26 |
| BAH43764.1 | tyrocidine\_synthetase\_I | BGC0000452 | NRP | 29.0 | 38.3 | 117.0 | 8.9e-26 |
| WP\_051700122.1 | non-ribosomal\_peptide\_synthetase | BGC0001368 | NRP | 22.0 | 86.4 | 117.0 | 8.9e-26 |
| ADG27358.1 | peptide\_synthetase | BGC0000296 | NRP | 29.0 | 47.0 | 117.0 | 1.2e-25 |
| AFH75330.1 | nonribosomal\_peptide\_synthetase | BGC0000398 | NRP:Cyclic depsipeptide | 37.0 | 28.4 | 116.0 | 1.5e-25 |
| ABS74181.1 | bacillomycin\_D\_synthetase\_A\_ | BGC0001090 | Polyketide + NRP:Lipopeptide | 27.0 | 44.9 | 116.0 | 2e-25 |
| APZ78743.1 | nonribosomal\_peptide\_synthetase | BGC0001422 | NRP:Cyclic depsipeptide + Polyketide:Iterative type I | 26.0 | 46.1 | 116.0 | 2e-25 |
| AAY37647.1 | Amino\_acid\_adenylation | BGC0000437 | NRP | 27.0 | 44.9 | 116.0 | 2.6e-25 |
| ABA73955.1 | putative\_non-ribosomal\_peptide\_synthetase | BGC0001842 | NRP:Lipopeptide | 27.0 | 45.5 | 116.0 | 2.6e-25 |
| AHZ20773.1 | non-ribosomal\_peptide\_synthase | BGC0000369 | NRP + Saccharide:Hybrid/tailoring | 26.0 | 47.2 | 115.0 | 3.4e-25 |
| ACA97580.1 | PmxE | BGC0000408 | NRP | 28.0 | 49.0 | 115.0 | 3.4e-25 |
| APZ78809.1 | nonribosomal\_peptide\_synthetase | BGC0001428 | NRP:Cyclic depsipeptide + Polyketide:Iterative type I | 26.0 | 45.7 | 115.0 | 3.4e-25 |
| XP\_020057667.1 | hypothetical\_protein | BGC0001718 | NRP | 26.0 | 49.9 | 115.0 | 3.4e-25 |
| BAB69698.1 | iturin\_A\_synthetase\_A | BGC0001098 | NRP + Polyketide | 27.0 | 44.9 | 115.0 | 4.4e-25 |
| CAD17792.1 | probable\_non\_ribosomal\_peptide\_synthetase\_protein | BGC0001754 | NRP + Polyketide | 28.0 | 46.9 | 115.0 | 4.4e-25 |
| AQM58287.1 | non-ribosomal\_peptide\_synthase | BGC0001816 | NRP + Polyketide | 25.0 | 52.5 | 115.0 | 4.4e-25 |
| CAM56771.1 |  | BGC0000354 | NRP | 27.0 | 44.9 | 114.0 | 5.8e-25 |
| AHB82071.1 | non\_ribosomal\_peptide\_synthetase | BGC0001231 | NRP + Polyketide:Modular type I | 28.0 | 43.8 | 114.0 | 5.8e-25 |
| ABH06368.2 | MassB | BGC0000389 | NRP:Cyclic depsipeptide | 28.0 | 47.4 | 114.0 | 9.8e-25 |
| CDG17986.1 | Non-ribosomal\_peptide\_synthetase | BGC0000464 | NRP:Cyclic depsipeptide | 28.0 | 46.1 | 114.0 | 9.8e-25 |
| AAF08795.1 | MycA | BGC0001103 | NRP + Polyketide | 27.0 | 45.3 | 114.0 | 9.8e-25 |
| ACM79812.1 | ZmaQ | BGC0001059 | NRP + Polyketide | 26.0 | 45.3 | 113.0 | 1.3e-24 |
| ABV79987.1 | ApnC | BGC0000301 | NRP | 23.0 | 62.2 | 113.0 | 1.7e-24 |
| ALK27914.1 | non-ribosomal\_peptide\_synthase | BGC0001233 | NRP | 28.0 | 47.0 | 113.0 | 2.2e-24 |
| AJF34463.1 | Txo1 | BGC0001207 | NRP | 30.0 | 38.9 | 112.0 | 2.9e-24 |
| AQM58288.1 | non-ribosomal\_peptide\_synthase | BGC0001816 | NRP + Polyketide | 26.0 | 50.8 | 112.0 | 2.9e-24 |
| CAJ34381.1 | NRPS\_protein | BGC0000445 | NRP:Cyclic depsipeptide | 27.0 | 43.0 | 112.0 | 3.7e-24 |
| ABC36785.1 | peptide\_synthetase,\_putative | BGC0000964 | NRP:Cyclic depsipeptide + Polyketide:Trans-AT type I | 27.0 | 49.6 | 112.0 | 3.7e-24 |
| AQM37583.1 | nonribosomal\_peptide\_synthetase | BGC0001424 | NRP:Cyclic depsipeptide + Polyketide:Iterative type I | 26.0 | 46.1 | 112.0 | 3.7e-24 |
| AGM16414.1 | paenibacterin\_synthetase\_C | BGC0000400 | NRP | 27.0 | 45.3 | 111.0 | 6.4e-24 |
| ABW17377.1 | PsoC | BGC0000411 | NRP | 27.0 | 46.5 | 111.0 | 6.4e-24 |
| AEH59100.1 | amino\_acid\_adenylation\_domain-containing\_protein/NRPS | BGC0000385 | NRP | 37.0 | 22.0 | 111.0 | 8.3e-24 |
| QED88054.1 | nonribosomal\_peptide\_synthetase | BGC0001967 | NRP | 28.0 | 47.0 | 111.0 | 8.3e-24 |
| AAY37655.1 | Amino\_acid\_adenylation | BGC0000437 | NRP | 38.0 | 22.0 | 110.0 | 1.1e-23 |
| AAO72425.1 | syringopeptin\_synthetase\_C | BGC0000438 | NRP | 38.0 | 22.0 | 110.0 | 1.1e-23 |
| OKA09424.1 | non-ribosomal\_peptide\_synthetase | BGC0001459 | NRP:Glycopeptide | 27.0 | 45.3 | 110.0 | 1.1e-23 |
| AGZ03650.1 | sevA | BGC0000426 | NRP | 29.0 | 38.0 | 110.0 | 1.4e-23 |
| AAF08797.1 | MycC | BGC0001103 | NRP + Polyketide | 30.0 | 33.8 | 110.0 | 1.4e-23 |
| APZ78755.1 | nonribosomal\_peptide\_synthetase | BGC0001423 | NRP:Cyclic depsipeptide + Polyketide:Iterative type I | 24.0 | 49.8 | 110.0 | 1.4e-23 |
| CAB38517.1 | CDA\_peptide\_synthetase\_II\_(CdaPs2) | BGC0000315 | NRP:Ca+-dependent lipopeptide | 32.0 | 32.4 | 109.0 | 1.9e-23 |
| ABL74936.1 | NRPS | BGC0001048 | NRP:Glycopeptide + Polyketide:Modular type I + Saccharide:Hybrid/tailoring | 32.0 | 31.9 | 109.0 | 1.9e-23 |
| CCJ67648.1 | JagD | BGC0001127 | NRP | 33.0 | 28.2 | 109.0 | 1.9e-23 |
| AED90003.1 | non-ribosomal\_peptide\_synthetase\_ThaB | BGC0000443 | NRP:Beta-lactam | 26.0 | 45.6 | 109.0 | 2.4e-23 |
| OKA09423.1 | non-ribosomal\_peptide\_synthetase | BGC0001459 | NRP:Glycopeptide | 27.0 | 47.9 | 109.0 | 2.4e-23 |
| AAG05690.1 | AmbE | BGC0000287 | NRP | 23.0 | 74.3 | 109.0 | 3.2e-23 |
| APZ78808.1 | nonribosomal\_peptide\_synthetase | BGC0001428 | NRP:Cyclic depsipeptide + Polyketide:Iterative type I | 25.0 | 46.1 | 109.0 | 3.2e-23 |
| AHI59109.1 | locillomycin\_synthase\_B | BGC0001005 | NRP + Polyketide | 25.0 | 49.6 | 108.0 | 4.1e-23 |
| CAE53352.1 | non-ribosomal\_peptide\_synthetase | BGC0000440 | NRP:Glycopeptide | 26.0 | 44.7 | 108.0 | 5.4e-23 |
| CAG15011.1 | peptide\_synthetase,\_module\_4-6 | BGC0000441 | NRP | 26.0 | 44.7 | 108.0 | 5.4e-23 |
| QED55423.1 | nonribosomal\_peptide\_synthetase | BGC0001984 | NRP | 30.0 | 41.1 | 108.0 | 5.4e-23 |
| AFJ14794.1 | PlpE | BGC0000403 | NRP | 28.0 | 39.3 | 108.0 | 7e-23 |
| QDF82254.1 | non-ribosomal\_peptide\_synthetase | BGC0001980 | NRP | 34.0 | 27.5 | 108.0 | 7e-23 |
| WP\_047195042.1 | non-ribosomal\_peptide\_synthetase | BGC0002003 | NRP | 35.0 | 27.3 | 107.0 | 9.2e-23 |
| CAJ77715.1 | Mps1\_protein | BGC0000364 | NRP | 36.0 | 30.3 | 107.0 | 1.2e-22 |
| AAF17281.1 | nosD | BGC0001028 | Polyketide + NRP:Cyclic depsipeptide | 26.0 | 45.1 | 107.0 | 1.2e-22 |
| AOA33122.1 | Nonribosomal\_peptide\_synthetase | BGC0001346 | NRP:Cyclic depsipeptide | 28.0 | 45.8 | 107.0 | 1.2e-22 |
| BAC67534.2 | arthrofactin\_synthetase\_A | BGC0000305 | NRP:Lipopeptide | 34.0 | 27.5 | 106.0 | 1.6e-22 |
| BAC67535.1 | arthrofactin\_synthetase\_B | BGC0000305 | NRP:Lipopeptide | 27.0 | 45.6 | 106.0 | 1.6e-22 |
| WP\_013184462.1 | non-ribosomal\_peptide\_synthetase | BGC0001716 | NRP | 25.0 | 47.0 | 106.0 | 1.6e-22 |
| ABA73954.1 | putative\_non-ribosomal\_peptide\_synthetase | BGC0001842 | NRP:Lipopeptide | 32.0 | 35.2 | 106.0 | 1.6e-22 |
| APZ78729.1 | nonribosomal\_peptide\_synthetase | BGC0001421 | NRP:Cyclic depsipeptide + Polyketide:Iterative type I | 23.0 | 46.5 | 106.0 | 2e-22 |
| AJV88375.1 | MfnC | BGC0001214 | NRP | 27.0 | 49.8 | 106.0 | 2.7e-22 |
| CAG29032.1 | nonribosomal\_peptide\_synthetase\_(modules\_3\_to\_6) | BGC0001023 | NRP + Polyketide:Modular type I | 23.0 | 64.7 | 105.0 | 3.5e-22 |
| KPN90369.1 | NunE | BGC0001416 | NRP | 33.0 | 23.4 | 105.0 | 3.5e-22 |
| APZ78822.1 | nonribosomal\_peptide\_synthetase | BGC0001429 | NRP:Cyclic depsipeptide + Polyketide:Iterative type I | 23.0 | 64.7 | 105.0 | 3.5e-22 |
| APU91751.1 | Non-Ribosomal\_Peptide\_Synthetase | BGC0001806 | NRP | 39.0 | 22.3 | 105.0 | 3.5e-22 |
| CAJ77695.1 | MPS1\_protein | BGC0000363 | NRP | 35.0 | 30.5 | 105.0 | 4.6e-22 |
| AGM16412.1 | paenibacterin\_synthetase\_A | BGC0000400 | NRP | 28.0 | 34.7 | 105.0 | 4.6e-22 |
| WP\_003981346.1 | non-ribosomal\_peptide\_synthetase | BGC0001813 | NRP | 27.0 | 47.0 | 105.0 | 4.6e-22 |
| ABV56585.1 | KtzE | BGC0000378 | NRP | 35.0 | 27.0 | 104.0 | 6e-22 |
| AED90002.1 | non-ribosomal\_peptide\_synthetase\_ThaA | BGC0000443 | NRP:Beta-lactam | 32.0 | 33.5 | 104.0 | 6e-22 |
| ADF88262.1 | mixed\_nonribosomal\_peptide\_synthetase/\_polyketide\_synthase | BGC0000979 | NRP + Polyketide | 30.0 | 32.8 | 104.0 | 6e-22 |
| ADF88265.1 | mixed\_nonribosomal\_peptide\_synthetase/\_polyketide\_synthase | BGC0000980 | NRP + Polyketide | 30.0 | 32.8 | 104.0 | 6e-22 |
| ADF88279.1 | mixed\_NRPS/PKS | BGC0000981 | NRP + Polyketide | 30.0 | 32.8 | 104.0 | 6e-22 |
| AJY78094.1 | nonribosomal\_peptide\_synthetase | BGC0001902 | NRP + Polyketide | 27.0 | 53.3 | 104.0 | 6e-22 |
| QBC75021.1 | non-ribosomal\_peptide\_synthetase | BGC0001968 | NRP | 26.0 | 45.3 | 104.0 | 6e-22 |
| AAC06347.1 | bacitracin\_synthetase\_2 | BGC0000310 | NRP | 30.0 | 33.5 | 104.0 | 7.8e-22 |
| AAD44234.1 | PstB | BGC0000362 | NRP | 28.0 | 46.0 | 104.0 | 7.8e-22 |
| ABX60161.1 | mixed\_NRPS/PKS | BGC0000978 | NRP + Alkaloid + Polyketide:Modular type I | 30.0 | 32.8 | 104.0 | 7.8e-22 |
| ALV82356.1 | CDA\_peptide\_synthetase\_I | BGC0001370 | NRP | 33.0 | 29.3 | 104.0 | 7.8e-22 |
| APZ78679.1 | nonribosomal\_peptide\_synthetase | BGC0001417 | NRP:Cyclic depsipeptide + Polyketide:Iterative type I | 23.0 | 49.8 | 104.0 | 7.8e-22 |
| AAY91421.3 | non-ribosomal\_peptide\_synthetase\_OfaC | BGC0000399 | NRP:Cyclic depsipeptide | 38.0 | 24.1 | 104.0 | 1e-21 |
| AFH75322.1 | nonribosomal\_peptide\_synthetase | BGC0000425 | NRP:Cyclic depsipeptide | 39.0 | 22.3 | 104.0 | 1e-21 |
| AGM14933.1 | xantholysin\_synthetase\_B | BGC0000463 | NRP:Lipopeptide | 27.0 | 44.9 | 104.0 | 1e-21 |
| AHZ34243.1 | CipF | BGC0001389 | NRP | 26.0 | 46.6 | 104.0 | 1e-21 |
| ATY37590.1 | BogC | BGC0001532 | NRP | 22.0 | 65.7 | 104.0 | 1e-21 |
| AEI58865.1 | peptide\_synthetase | BGC0000455 | NRP | 26.0 | 47.9 | 103.0 | 1.3e-21 |
| AEH59099.1 | amino\_acid\_adenylation\_domain-containing\_protein/NRPS | BGC0000385 | NRP | 37.0 | 22.0 | 103.0 | 1.7e-21 |
| ABW17375.1 | PsoA | BGC0000411 | NRP | 30.0 | 32.1 | 102.0 | 2.3e-21 |
| ACG60772.1 | NRPS(C/A/PCP/Cy/A/PCP/Cy) | BGC0001058 | NRP:Glycopeptide + Polyketide:Modular type I + Saccharide:Hybrid/tailoring | 26.0 | 45.1 | 102.0 | 3e-21 |
| AAT01807.1 | non-ribosomal\_peptide\_synthetase | BGC0000365 | NRP | 28.0 | 44.0 | 102.0 | 3.9e-21 |
| AAO56329.1 | non-ribosomal\_peptide\_synthetase\_SyfB | BGC0000435 | NRP | 25.0 | 47.4 | 102.0 | 3.9e-21 |
| AOA33121.1 | Nonribosomal\_peptide\_synthetase | BGC0001346 | NRP:Cyclic depsipeptide | 34.0 | 27.1 | 102.0 | 3.9e-21 |
| ALV82384.1 | CDA\_peptide\_synthetase\_II | BGC0001370 | NRP | 33.0 | 27.1 | 101.0 | 5e-21 |
| BAW32324.1 | nonribosomal\_peptide\_synthetase | BGC0001630 | NRP + Polyketide | 32.0 | 29.7 | 101.0 | 5e-21 |
| WP\_013184464.1 | non-ribosomal\_peptide\_synthetase | BGC0001716 | NRP | 29.0 | 35.9 | 101.0 | 5e-21 |
| BAH43871.1 | truncated\_linear\_pentadecapeptide\_gramicidin\_synthetase\_LgrC | BGC0000367 | NRP | 26.0 | 47.9 | 101.0 | 6.6e-21 |
| ALK27916.1 | non-ribosomal\_peptide\_synthase | BGC0001233 | NRP | 34.0 | 28.7 | 101.0 | 6.6e-21 |
| ALV86866.1 | Tlo20 | BGC0001406 | NRP | 27.0 | 47.1 | 101.0 | 6.6e-21 |
| OLZ52457.1 | non-ribosomal\_peptide\_synthetase | BGC0001462 | NRP:Glycopeptide | 25.0 | 44.7 | 101.0 | 6.6e-21 |
| AHJ31215.1 | Long-chain-fatty-acid--CoA\_ligase | BGC0000430 | NRP + Polyketide:Modular type I | 32.0 | 26.4 | 101.0 | 8.6e-21 |
| CBJ79915.1 | putative\_Phenylalanine\_racemase\_(ATP-hydrolyzing) | BGC0001133 | NRP | 24.0 | 45.5 | 101.0 | 8.6e-21 |
| AGC09528.1 | NRPS | BGC0001183 | Polyketide | 29.0 | 36.2 | 101.0 | 8.6e-21 |
| ATL73036.1 | amino\_acid\_adenylation\_protein | BGC0001807 | NRP + Polyketide | 29.0 | 36.2 | 101.0 | 8.6e-21 |
| QBG38783.1 | Atr22 | BGC0001975 | NRP | 26.0 | 56.5 | 101.0 | 8.6e-21 |
| AAK81824.1 | peptide\_synthetase | BGC0000326 | NRP | 31.0 | 28.2 | 100.0 | 1.1e-20 |
| AAC32047.1 | FxbB | BGC0000351 | NRP | 28.0 | 44.7 | 100.0 | 1.1e-20 |
| AHI59110.1 | locillomycin\_synthase\_C | BGC0001005 | NRP + Polyketide | 31.0 | 31.4 | 100.0 | 1.1e-20 |
| AAT01806.1 | non-ribosomal\_peptide\_synthetase | BGC0000365 | NRP | 36.0 | 27.1 | 100.0 | 1.5e-20 |
| AEW31022.1 | plipastatin\_synthetase | BGC0000407 | NRP | 30.0 | 35.2 | 100.0 | 1.5e-20 |
| AIG79241.1 | Hypothetical\_protein | BGC0000419 | Saccharide + NRP:Glycopeptide | 29.0 | 36.6 | 100.0 | 1.5e-20 |
| ALK27915.1 | non-ribosomal\_peptide\_synthase | BGC0001233 | NRP | 26.0 | 46.0 | 100.0 | 1.5e-20 |
| EWS95122.1 | hypothetical\_protein | BGC0000306 | NRP:Lipopeptide | 34.0 | 27.1 | 99.0 | 2.5e-20 |
| AQZ71347.1 | hypothetical\_protein | BGC0001635 | NRP + Polyketide | 25.0 | 52.6 | 99.0 | 3.3e-20 |
| AVI26392.1 | putative\_nonribosomal\_peptide\_synthase | BGC0001800 | NRP + Polyketide | 27.0 | 41.6 | 99.0 | 3.3e-20 |
| CAM56770.1 |  | BGC0000354 | NRP | 25.0 | 50.2 | 98.0 | 4.3e-20 |
| AIE77059.1 | peptide\_synthetase | BGC0000418 | NRP | 30.0 | 30.7 | 98.0 | 4.3e-20 |
| ABX37382.1 | amino\_acid\_adenylation\_domain\_protein | BGC0000984 | NRP + Polyketide | 35.0 | 26.2 | 98.0 | 4.3e-20 |
| ARR97036.1 | SphC | BGC0001780 | NRP | 24.0 | 74.6 | 98.0 | 4.3e-20 |
| ATO51563.1 | non-ribosomal\_peptide\_synthetase | BGC0001796 | NRP | 23.0 | 61.6 | 98.0 | 4.3e-20 |
| AAZ03551.1 | McnB | BGC0000332 | NRP | 31.0 | 28.9 | 98.0 | 5.6e-20 |
| CBL93723.1 | NRPS\_didomain\_A-PCP | BGC0000360 | NRP | 38.0 | 23.6 | 98.0 | 5.6e-20 |
| CAJ46692.1 | non-ribosomal\_peptide\_synthase | BGC0000969 | NRP:Cyclic depsipeptide + Polyketide:Modular type I | 24.0 | 46.9 | 98.0 | 5.6e-20 |
| ABV79985.1 | ApnA | BGC0000301 | NRP | 32.0 | 28.8 | 97.0 | 7.3e-20 |
| PHM26612.1 | pvdj | BGC0001130 | NRP + Polyketide | 26.0 | 43.7 | 97.0 | 7.3e-20 |
| ATY37591.1 | BogD | BGC0001532 | NRP | 24.0 | 45.3 | 97.0 | 7.3e-20 |
| BBA20967.1 | nonribosomal\_peptide\_synthetase | BGC0001763 | NRP + Polyketide | 28.0 | 42.8 | 97.0 | 7.3e-20 |
| WP\_044618979.1 | non-ribosomal\_peptide\_synthetase | BGC0001791 | NRP | 32.0 | 29.7 | 97.0 | 7.3e-20 |
| ABS74206.1 | fengycin\_synthetase\_D | BGC0001095 | NRP | 31.0 | 29.4 | 97.0 | 9.5e-20 |
| ACZ55946.1 | non-ribosomal\_peptide\_synthetase | BGC0000302 | NRP | 24.0 | 45.2 | 97.0 | 1.2e-19 |
| CAB38518.1 | CDA\_peptide\_synthetase\_I\_(CdaPs1) | BGC0000315 | NRP:Ca+-dependent lipopeptide | 32.0 | 30.0 | 96.0 | 1.6e-19 |
| AEP18655.1 | WAPS2 | BGC0000461 | NRP | 30.0 | 31.1 | 96.0 | 1.6e-19 |
| BAX90000.1 | Non-ribosomal\_peptide\_synthetase | BGC0001628 | NRP | 33.0 | 27.9 | 96.0 | 2.1e-19 |
| CAE53350.1 | non-ribosomal\_peptide\_synthetase | BGC0000440 | NRP:Glycopeptide | 25.0 | 45.5 | 96.0 | 2.8e-19 |
| CAG15009.1 | peptide\_synthetase,\_module\_1-2 | BGC0000441 | NRP | 25.0 | 45.5 | 96.0 | 2.8e-19 |
| AEI58866.1 | peptide\_synthetase | BGC0000455 | NRP | 32.0 | 29.8 | 96.0 | 2.8e-19 |
| AEP18656.1 | WAPS1 | BGC0000461 | NRP | 33.0 | 28.6 | 96.0 | 2.8e-19 |
| AFR69334.1 | nonribosomal\_peptide\_synthetase\_SpiDE1 | BGC0001045 | NRP:Cyclic depsipeptide + Polyketide:Modular type I | 31.0 | 26.9 | 96.0 | 2.8e-19 |
| ABC94347.1 | vicibactin\_biosynthesis\_non-ribosomal\_peptide\_synthase\_protein | BGC0000457 | NRP | 31.0 | 32.1 | 95.0 | 3.6e-19 |
| WP\_064118559.1 | non-ribosomal\_peptide\_synthetase | BGC0001509 | NRP | 27.0 | 43.3 | 95.0 | 3.6e-19 |
| AXN93605.1 | PuwA | BGC0001953 | NRP | 28.0 | 28.0 | 95.0 | 3.6e-19 |
| AAK81826.1 | peptide\_synthetase | BGC0000326 | NRP | 31.0 | 28.2 | 95.0 | 4.7e-19 |
| AEA30272.1 | peptide\_synthetase | BGC0000429 | Polyketide + NRP:Cyclic depsipeptide | 25.0 | 47.5 | 95.0 | 4.7e-19 |
| EAW09117.1 | hybrid\_NRPS/PKS\_enzyme,\_putative | BGC0000983 | NRP + Polyketide:Iterative type I | 23.0 | 76.3 | 95.0 | 4.7e-19 |
| ATG32076.1 | nonribosomal\_peptide\_synthetase | BGC0001750 | NRP + Polyketide | 30.0 | 26.8 | 95.0 | 4.7e-19 |
| ctg1\_orf20 |  | BGC0001767 | NRP | 31.0 | 31.1 | 95.0 | 4.7e-19 |
| APU91750.1 | Non-Ribosomal\_Peptide\_Synthetase | BGC0001806 | NRP | 32.0 | 30.9 | 95.0 | 4.7e-19 |
| QED88055.1 | nonribosomal\_peptide\_synthetase | BGC0001967 | NRP | 31.0 | 31.2 | 95.0 | 4.7e-19 |
| AEW31020.1 | plipastatin\_synthetase | BGC0000407 | NRP | 24.0 | 46.9 | 94.0 | 8.1e-19 |
| AEE88284.1 | CurF | BGC0000976 | NRP + Polyketide:Modular type I | 29.0 | 33.2 | 94.0 | 8.1e-19 |
| AAT70101.1 | CurF | BGC0001165 | NRP + Polyketide:Modular type I | 29.0 | 33.2 | 94.0 | 8.1e-19 |
| BAC67536.1 | arthrofactin\_synthetase\_C | BGC0000305 | NRP:Lipopeptide | 33.0 | 28.7 | 94.0 | 1.1e-18 |
| AAC06346.1 | bacitracin\_synthetase\_1 | BGC0000310 | NRP | 28.0 | 30.1 | 94.0 | 1.1e-18 |
| ADH04660.1 | TugD | BGC0001342 | NRP + Polyketide | 27.0 | 36.2 | 94.0 | 1.1e-18 |
| EME52989.1 | amino\_acid\_adenylation\_protein | BGC0001460 | NRP:Glycopeptide | 32.0 | 29.8 | 94.0 | 1.1e-18 |
| CAD91212.1 | putative\_non-ribosomal\_peptide\_synthetase,\_modules\_4-6 | BGC0000289 | NRP:Glycopeptide + Saccharide:Hybrid/tailoring | 33.0 | 29.8 | 93.0 | 1.4e-18 |
| AAM80537.1 | StaC | BGC0000290 | NRP:Glycopeptide | 32.0 | 27.1 | 93.0 | 1.4e-18 |
| AAZ55900.1 | non-ribosomal\_peptide\_synthase:Amino\_acid\_adenylation | BGC0000359 | NRP | 35.0 | 27.1 | 93.0 | 1.4e-18 |
| BAH43869.1 | linear\_pentadecapeptide\_gramicidin\_synthetase\_LgrA | BGC0000367 | NRP | 27.0 | 29.8 | 93.0 | 1.4e-18 |
| CCJ67639.1 | TaaD | BGC0000447 | NRP:Lipopeptide | 29.0 | 43.1 | 93.0 | 1.4e-18 |
| CAJ96470.1 | non-ribosomal\_peptide\_synthetase | BGC0000330 | NRP:NRP siderophore | 32.0 | 28.6 | 93.0 | 1.8e-18 |
| QBC75022.1 | non-ribosomal\_peptide\_synthetase | BGC0001968 | NRP | 31.0 | 28.4 | 93.0 | 1.8e-18 |
| APZ78855.1 | nonribosomal\_peptide\_synthetase | BGC0001432 | NRP:Cyclic depsipeptide + Polyketide:Iterative type I | 23.0 | 49.8 | 92.0 | 2.3e-18 |
| QDF82255.1 | non-ribosomal\_peptide\_synthetase | BGC0001980 | NRP | 30.0 | 31.4 | 92.0 | 2.3e-18 |
| ARU08075.1 | mlcM | BGC0001448 | NRP:Ca+-dependent lipopeptide | 32.0 | 28.8 | 92.0 | 3.1e-18 |
| AFJ23825.1 | WLIP\_synthetase\_B | BGC0001838 | NRP | 31.0 | 35.5 | 92.0 | 3.1e-18 |
| AEC14348.1 | nonribosomal\_peptide\_synthetase | BGC0000377 | NRP | 27.0 | 30.0 | 92.0 | 4e-18 |
| AFH75328.1 | nonribosomal\_peptide\_synthetase | BGC0000398 | NRP:Cyclic depsipeptide | 31.0 | 29.6 | 92.0 | 4e-18 |
| AIG79242.1 | Non-ribosomal\_peptide\_synthetase | BGC0000419 | Saccharide + NRP:Glycopeptide | 29.0 | 30.5 | 92.0 | 4e-18 |
| CAQ34921.1 | nonribosomal\_peptide\_synthetase | BGC0000986 | NRP + Polyketide | 32.0 | 26.4 | 92.0 | 4e-18 |
| ARU08073.1 | mlcK | BGC0001448 | NRP:Ca+-dependent lipopeptide | 29.0 | 30.3 | 92.0 | 4e-18 |
| CAC48361.1 | peptide\_synthetase | BGC0000311 | NRP | 33.0 | 29.2 | 91.0 | 5.2e-18 |
| BAP16689.1 | nonribosomal\_peptide\_synthetase | BGC0000376 | NRP | 33.0 | 24.3 | 91.0 | 5.2e-18 |
| AIE77058.1 | peptide\_synthetase\_module\_3 | BGC0000418 | NRP | 29.0 | 30.5 | 91.0 | 5.2e-18 |
| ATQ39428.1 | cyclosporin\_C\_synthetase | BGC0001565 | NRP | 24.0 | 48.9 | 91.0 | 5.2e-18 |
| WP\_013310343.1 | non-ribosomal\_peptide\_synthetase | BGC0001993 | NRP | 29.0 | 30.7 | 91.0 | 5.2e-18 |
| CAE53353.1 | non-ribosomal\_peptide\_synthetase | BGC0000440 | NRP:Glycopeptide | 29.0 | 29.6 | 91.0 | 6.8e-18 |
| CAG15012.1 | peptide\_synthetase,\_module\_7 | BGC0000441 | NRP | 29.0 | 29.6 | 91.0 | 6.8e-18 |
| CCJ67640.1 | TaaE | BGC0000447 | NRP:Lipopeptide | 29.0 | 42.1 | 91.0 | 6.8e-18 |
| AJW76709.1 | DsaG | BGC0001196 | NRP | 29.0 | 31.4 | 91.0 | 6.8e-18 |
| KPN93063.1 | NupA | BGC0001416 | NRP | 32.0 | 28.6 | 91.0 | 6.8e-18 |
| QDF82259.1 | non-ribosomal\_peptide\_synthetase | BGC0001980 | NRP | 29.0 | 30.9 | 91.0 | 6.8e-18 |
| AAY93355.1 | non-ribosomal\_peptide\_synthetase\_PvdJ | BGC0000413 | NRP | 34.0 | 26.2 | 91.0 | 8.9e-18 |
| CAN89663.1 | putative\_non-ribosomal\_peptide\_synthetase | BGC0001070 | NRP + Polyketide:Modular type I + Polyketide:Trans-AT type I | 31.0 | 29.3 | 91.0 | 8.9e-18 |
| WP\_016638470.1 | non-ribosomal\_peptide\_synthetase | BGC0001519 | NRP + Polyketide | 24.0 | 46.7 | 91.0 | 8.9e-18 |
| WP\_013184322.1 | non-ribosomal\_peptide\_synthetase | BGC0001692 | NRP | 29.0 | 26.2 | 91.0 | 8.9e-18 |
| AWI62626.1 | nonribosomal\_peptide\_synthetase | BGC0001943 | NRP | 30.0 | 27.7 | 91.0 | 8.9e-18 |
| ABA73956.1 | putative\_non-ribosomal\_peptide\_synthetase | BGC0001842 | NRP:Lipopeptide | 26.0 | 43.7 | 91.0 | 8.9e-18 |
| AXN93592.1 | PuwA | BGC0001952 | NRP | 27.0 | 27.9 | 91.0 | 8.9e-18 |
| AEC14346.1 | nonribosomal\_peptide\_synthetase | BGC0000377 | NRP | 28.0 | 30.1 | 90.0 | 1.2e-17 |
| DAB41477.1 | nonribosomal\_peptide\_synthetase | BGC0001766 | NRP | 34.0 | 27.1 | 90.0 | 1.2e-17 |
| KFL51885.1 | amino\_acid\_adenylation\_protein | BGC0001711 | NRP + Polyketide | 31.0 | 28.9 | 90.0 | 1.2e-17 |
| CDG17982.1 | Non-ribosomal\_peptide\_synthetase | BGC0000464 | NRP:Cyclic depsipeptide | 30.0 | 33.3 | 90.0 | 1.5e-17 |
| AKC91856.1 | nonribosomal\_peptide\_synthetase | BGC0001414 | NRP | 33.0 | 26.1 | 90.0 | 1.5e-17 |
| WP\_019634550.1 | type\_I\_polyketide\_synthase | BGC0001443 | NRP + Polyketide | 32.0 | 21.8 | 89.0 | 2e-17 |
| RLV71193.1 | non-ribosomal\_peptide\_synthetase | BGC0001846 | NRP + Saccharide:Hybrid/tailoring | 31.0 | 30.6 | 89.0 | 2e-17 |
| AXN93601.1 | PuwE | BGC0001952 | NRP | 28.0 | 39.6 | 89.0 | 2e-17 |
| CAD91211.1 | putative\_non-ribosomal\_peptide\_synthetase,\_module\_7 | BGC0000289 | NRP:Glycopeptide + Saccharide:Hybrid/tailoring | 28.0 | 32.8 | 89.0 | 2.6e-17 |
| BAH43766.1 | tyrocidine\_synthetase\_III | BGC0000452 | NRP | 29.0 | 28.9 | 89.0 | 2.6e-17 |
| ALG65317.1 | Cal19 | BGC0001297 | NRP | 30.0 | 31.2 | 89.0 | 2.6e-17 |
| WP\_039806856.1 | non-ribosomal\_peptide\_synthetase | BGC0002001 | NRP + Polyketide | 31.0 | 28.4 | 89.0 | 2.6e-17 |
| AAC06348.1 | bacitracin\_synthetase\_3 | BGC0000310 | NRP | 29.0 | 27.1 | 89.0 | 3.4e-17 |
| AFH75320.1 | nonribosomal\_peptide\_synthetase | BGC0000425 | NRP:Cyclic depsipeptide | 30.0 | 30.6 | 89.0 | 3.4e-17 |
| CAJ76290.1 | putative\_non-ribosomal\_peptide\_synthase | BGC0000972 | NRP + Polyketide:Modular type I + Polyketide:Trans-AT type I | 30.0 | 32.1 | 89.0 | 3.4e-17 |
| AHD05627.1 | putative\_non-ribosomal\_peptide\_ligase\_domain\_protein | BGC0001033 | NRP + Polyketide | 30.0 | 27.8 | 89.0 | 3.4e-17 |
| WP\_008704344.1 | non-ribosomal\_peptide\_synthetase | BGC0001575 | NRP | 30.0 | 30.3 | 89.0 | 3.4e-17 |
| AXM42952.1 | non-ribosomal\_peptide\_synthetase | BGC0001941 | NRP + Polyketide | 33.0 | 28.4 | 89.0 | 3.4e-17 |
| AAZ23078.1 | peptide\_synthetase | BGC0000291 | NRP | 34.0 | 27.9 | 88.0 | 4.4e-17 |
| CCJ67636.1 | TaaA | BGC0000447 | NRP:Lipopeptide | 30.0 | 28.6 | 88.0 | 4.4e-17 |
| CAO91861.1 | PKS-NRPS\_hybrid | BGC0000968 | NRP + Polyketide:Iterative type I | 22.0 | 78.0 | 88.0 | 4.4e-17 |
| AHD05614.1 | putative\_non-ribosomal\_peptide\_ligase/\_polyketide\_synthase\_hybrid | BGC0001033 | NRP + Polyketide | 25.0 | 43.7 | 88.0 | 4.4e-17 |
| AGZ15459.1 | putative\_non-ribosomal\_peptide\_synthetase | BGC0001036 | NRP + Polyketide | 30.0 | 27.0 | 88.0 | 4.4e-17 |
| CBZ42146.1 | putative\_non-ribosomal\_peptide\_synthetase | BGC0001117 | NRP | 31.0 | 28.8 | 88.0 | 4.4e-17 |
| AGS77310.1 | NRPS\_module\_7 | BGC0001178 | NRP:Glycopeptide | 28.0 | 29.6 | 88.0 | 4.4e-17 |
| AJW76710.1 | DsaH | BGC0001196 | NRP | 32.0 | 26.0 | 88.0 | 4.4e-17 |
| AOC89001.1 | putative\_nonribosomal\_peptide\_synthetase | BGC0001652 | NRP | 25.0 | 51.6 | 88.0 | 4.4e-17 |
| KFL51886.1 | amino\_acid\_adenylation\_protein | BGC0001711 | NRP + Polyketide | 30.0 | 30.9 | 88.0 | 4.4e-17 |
| ACZ55943.1 | non-ribosomal\_peptide\_synthetase | BGC0000302 | NRP | 29.0 | 29.6 | 88.0 | 5.8e-17 |
| ABS75232.1 | DhbF | BGC0001185 | NRP:NRP siderophore | 33.0 | 25.9 | 88.0 | 5.8e-17 |
| AAY91420.2 | non-ribosomal\_peptide\_synthetase\_OfaB | BGC0000399 | NRP:Cyclic depsipeptide | 29.0 | 29.8 | 87.0 | 7.5e-17 |
| ABG94125.1 | non-ribosomal\_peptide\_synthetase | BGC0000417 | NRP | 32.0 | 28.3 | 87.0 | 7.5e-17 |
| AQZ26587.1 | obafluorin\_dimodular\_nonribosomal\_peptide\_synthetase | BGC0001710 | NRP | 34.0 | 26.2 | 87.0 | 7.5e-17 |
| BAV56270.1 |  | BGC0001657 | NRP | 32.0 | 25.7 | 87.0 | 7.5e-17 |
| EFL06867.1 | predicted\_protein | BGC0000300 | NRP | 30.0 | 31.2 | 87.0 | 9.8e-17 |
| AGS77307.1 | NRPS\_modules\_1-2 | BGC0001178 | NRP:Glycopeptide | 26.0 | 45.6 | 87.0 | 9.8e-17 |
| AOZ21320.1 | SulM | BGC0001790 | NRP | 24.0 | 45.3 | 87.0 | 9.8e-17 |
| AAZ55899.1 | amino\_acid\_adenylation | BGC0000359 | NRP | 28.0 | 29.8 | 87.0 | 1.3e-16 |
| ALV86867.1 | Tlo21 | BGC0001406 | NRP | 31.0 | 28.9 | 87.0 | 1.3e-16 |
| OLZ50885.1 | non-ribosomal\_peptide\_synthetase | BGC0001461 | NRP:Glycopeptide | 31.0 | 26.8 | 87.0 | 1.3e-16 |
| KFL51887.1 | amino\_acid\_adenylation\_protein | BGC0001711 | NRP + Polyketide | 33.0 | 25.7 | 87.0 | 1.3e-16 |
| AYA44686.1 | icosalide\_NRPS | BGC0001833 | NRP:Lipopeptide | 27.0 | 33.5 | 87.0 | 1.3e-16 |
| AYA22334.1 | KerC | BGC0001955 | Other | 31.0 | 26.8 | 87.0 | 1.3e-16 |
| CAM02313.1 | putative\_non-ribosomal\_peptide\_synthetase | BGC0000349 | NRP | 32.0 | 28.2 | 86.0 | 1.7e-16 |
| AFO85453.1 | non-ribosomal\_peptide\_synthetase | BGC0000391 | NRP | 33.0 | 22.3 | 86.0 | 1.7e-16 |
| AHD05616.1 | putative\_non-ribosomal\_peptide\_ligase\_domain\_protein | BGC0001033 | NRP + Polyketide | 29.0 | 29.6 | 86.0 | 1.7e-16 |
| CBJ89761.1 | Non-ribosomal\_peptide\_synthase\_involved\_in\_xenocoumacin\_synthesis | BGC0001054 | NRP + Polyketide:Modular type I | 29.0 | 28.9 | 86.0 | 1.7e-16 |
| ACM79806.1 | ZmaB | BGC0001059 | NRP + Polyketide | 23.0 | 43.9 | 86.0 | 1.7e-16 |
| WP\_013184463.1 | non-ribosomal\_peptide\_synthetase | BGC0001716 | NRP | 30.0 | 26.2 | 86.0 | 1.7e-16 |
| CAD91220.1 | putative\_non-ribosomal\_peptide\_synthetase,\_modules\_1-2 | BGC0000289 | NRP:Glycopeptide + Saccharide:Hybrid/tailoring | 29.0 | 30.9 | 86.0 | 2.2e-16 |
| BAE98155.1 | putative\_non-ribosomal\_peptide\_synthetase | BGC0000339 | NRP | 32.0 | 28.7 | 86.0 | 2.9e-16 |
| AGZ15458.1 | putative\_non-ribosomal\_peptide\_synthetase | BGC0001036 | NRP + Polyketide | 29.0 | 29.1 | 86.0 | 2.9e-16 |
| ANG60379.1 | nonribosomal\_peptide\_synthetase\_BudA | BGC0001698 | NRP | 26.0 | 39.4 | 86.0 | 2.9e-16 |
| AEA30273.1 | peptide\_synthetase | BGC0000429 | Polyketide + NRP:Cyclic depsipeptide | 33.0 | 28.4 | 85.0 | 3.7e-16 |
| CAQ71828.1 | non\_ribosomal\_peptide\_synthase,\_antibiotic\_synthesis;\_contains\_3\_condensation\_domains,\_2\_AMP-acid\_ligases\_II\_domains,\_2\_PP-binding,\_Phosphopantetheine\_attachment\_site | BGC0001189 | NRP | 30.0 | 29.2 | 85.0 | 3.7e-16 |
| AET13875.1 | epichloenin\_A\_synthetase | BGC0001250 | NRP | 28.0 | 31.9 | 85.0 | 3.7e-16 |
| AET13879.1 | epichloenin\_A\_synthetase | BGC0001251 | NRP | 28.0 | 31.9 | 85.0 | 3.7e-16 |
| CAJ96471.1 | non-ribosomal\_peptide\_synthetase | BGC0000330 | NRP:NRP siderophore | 30.0 | 30.2 | 85.0 | 4.9e-16 |
| CAE02630.1 | surfactin\_synthetase\_A | BGC0000433 | NRP:Lipopeptide | 27.0 | 28.7 | 85.0 | 4.9e-16 |
| AHD05617.1 | putative\_non-ribosomal\_peptide\_ligase\_domain\_protein | BGC0001033 | NRP + Polyketide | 27.0 | 30.1 | 85.0 | 4.9e-16 |
| ABL74945.1 | NRPS | BGC0001048 | NRP:Glycopeptide + Polyketide:Modular type I + Saccharide:Hybrid/tailoring | 30.0 | 25.2 | 85.0 | 4.9e-16 |
| ADH04641.1 | TgaC | BGC0001051 | NRP + Polyketide:Modular type I | 30.0 | 28.3 | 85.0 | 4.9e-16 |
| WP\_084702182.1 | non-ribosomal\_peptide\_synthetase | BGC0001211 | NRP | 32.0 | 28.0 | 85.0 | 4.9e-16 |
| AKC91849.1 | nonribosomal\_peptide\_synthetase | BGC0001414 | NRP | 33.0 | 26.1 | 85.0 | 4.9e-16 |
| WP\_050383084.1 | non-ribosomal\_peptide\_synthetase | BGC0001451 | NRP | 30.0 | 31.6 | 85.0 | 4.9e-16 |
| WP\_016638471.1 | non-ribosomal\_peptide\_synthetase | BGC0001519 | NRP + Polyketide | 31.0 | 25.9 | 85.0 | 4.9e-16 |
| AFJ23826.1 | WLIP\_synthetase\_C | BGC0001838 | NRP | 29.0 | 33.0 | 85.0 | 4.9e-16 |
| AAY93354.1 | non-ribosomal\_peptide\_synthetase\_PvdD | BGC0000413 | NRP | 30.0 | 25.7 | 84.0 | 6.4e-16 |
| CAN89638.1 | putative\_non-ribosomal\_peptide\_synthetase | BGC0001070 | NRP + Polyketide:Modular type I + Polyketide:Trans-AT type I | 29.0 | 31.2 | 84.0 | 6.4e-16 |
| WP\_039806850.1 | non-ribosomal\_peptide\_synthetase | BGC0002001 | NRP + Polyketide | 26.0 | 40.7 | 84.0 | 6.4e-16 |
| AIE77057.1 | peptide\_synthetase | BGC0000418 | NRP | 31.0 | 29.6 | 84.0 | 8.3e-16 |
| AHD05621.1 | non-ribosomal\_peptide\_ligase\_domain\_protein | BGC0001033 | NRP + Polyketide | 27.0 | 29.4 | 84.0 | 8.3e-16 |
| AAM80536.1 | StaD | BGC0000290 | NRP:Glycopeptide | 26.0 | 29.6 | 84.0 | 1.1e-15 |
| AJI44176.1 | nonribosomal\_peptide\_synthetase | BGC0001193 | NRP | 34.0 | 26.0 | 84.0 | 1.1e-15 |
| WP\_051700105.1 | non-ribosomal\_peptide\_synthetase | BGC0001368 | NRP | 28.0 | 27.5 | 84.0 | 1.1e-15 |
| EME52990.1 | amino\_acid\_adenylation\_protein | BGC0001460 | NRP:Glycopeptide | 25.0 | 45.6 | 84.0 | 1.1e-15 |
| AEI70245.1 | nonribosomal\_peptide\_synthetase\_NRPS | BGC0000401 | NRP | 31.0 | 25.6 | 83.0 | 1.4e-15 |
| CCJ67647.1 | JagC | BGC0001127 | NRP | 31.0 | 28.0 | 83.0 | 1.4e-15 |
| ATY37609.1 | BreD | BGC0001536 | NRP | 23.0 | 48.9 | 83.0 | 1.4e-15 |
| AWI62627.1 | nonribosomal\_peptide\_synthetase | BGC0001943 | NRP | 29.0 | 28.7 | 83.0 | 1.4e-15 |
| ABI22131.1 | putative\_non-ribosomal\_peptide\_synthetase | BGC0000422 | NRP | 32.0 | 24.8 | 83.0 | 1.9e-15 |
| AJV88376.1 | MfnD | BGC0001214 | NRP | 32.0 | 27.4 | 83.0 | 1.9e-15 |
| AXA20091.1 | hybrid\_trans-AT\_PKS/NRPS\_LgaB | BGC0001946 | NRP + Polyketide | 25.0 | 49.7 | 83.0 | 1.9e-15 |
| AYJ71721.1 | non-ribosomal\_peptide\_synthetase | BGC0001942 | NRP + Polyketide | 29.0 | 29.1 | 83.0 | 1.9e-15 |
| EFG10344.1 | Non-ribosomal\_peptide\_synthetase | BGC0000373 | NRP | 31.0 | 26.2 | 82.0 | 2.4e-15 |
| AAY93356.2 | non-ribosomal\_peptide\_synthetase\_PvdI | BGC0000413 | NRP | 30.0 | 26.2 | 82.0 | 2.4e-15 |
| AET98906.1 | putative\_non-ribosomal\_peptide\_synthetase | BGC0000415 | NRP | 29.0 | 28.3 | 82.0 | 2.4e-15 |
| AFH75321.1 | nonribosomal\_peptide\_synthetase | BGC0000425 | NRP:Cyclic depsipeptide | 27.0 | 29.1 | 82.0 | 2.4e-15 |
| BAI63283.1 | putative\_non-ribosomal\_peptide\_synthetase | BGC0000434 | NRP | 26.0 | 42.1 | 82.0 | 2.4e-15 |
| WP\_080679150.1 | non-ribosomal\_peptide\_synthetase | BGC0001228 | NRP:Cyclic depsipeptide | 31.0 | 28.3 | 82.0 | 2.4e-15 |
| WP\_047197779.1 | non-ribosomal\_peptide\_synthetase | BGC0002003 | NRP | 28.0 | 27.9 | 82.0 | 2.4e-15 |
| BAV56271.1 |  | BGC0001657 | NRP | 30.0 | 28.3 | 82.0 | 2.4e-15 |
| WP\_013184596.1 | non-ribosomal\_peptide\_synthetase | BGC0001825 | NRP | 29.0 | 29.1 | 82.0 | 3.2e-15 |
| ADL64235.1 | aureusimine\_non-ribosomal\_peptide\_synthetase | BGC0000308 | NRP | 28.0 | 26.4 | 82.0 | 4.1e-15 |
| ABA59547.1 | NRPS | BGC0000453 | NRP:Cyclic depsipeptide | 29.0 | 27.9 | 82.0 | 4.1e-15 |
| PHM49485.1 | Amino\_acid\_adenylation | BGC0001131 | NRP | 30.0 | 30.3 | 82.0 | 4.1e-15 |
| ALG65318.1 | Cal18 | BGC0001297 | NRP | 30.0 | 28.6 | 82.0 | 4.1e-15 |
| WP\_047197778.1 | non-ribosomal\_peptide\_synthetase | BGC0002003 | NRP | 26.0 | 30.2 | 82.0 | 4.1e-15 |
| BAW32334.1 | hybrid\_cis-AT\_polyketide\_synthase\_-\_nonribosomal\_peptide\_synthetase | BGC0001631 | NRP + Polyketide | 24.0 | 45.6 | 82.0 | 4.1e-15 |
| AAN32981.1 | BarG | BGC0000962 | NRP + Polyketide:Modular type I | 29.0 | 29.3 | 81.0 | 5.4e-15 |
| CBJ79916.1 | putative\_Ornithine\_racemase | BGC0001133 | NRP | 28.0 | 25.2 | 81.0 | 5.4e-15 |
| CAQ71829.1 | non\_ribosomal\_peptide\_synthase,\_antibiotic\_synthesis;\_contains\_4\_condensation\_domains,\_3\_AMP-acid\_ligases\_II\_domains,\_3\_PP-binding,\_Phosphopantetheine\_attachment\_site\_and\_a\_putative\_thioesterase\_domain | BGC0001189 | NRP | 31.0 | 26.4 | 81.0 | 5.4e-15 |
| AXA20090.1 | hybrid\_trans-AT\_PKS/NRPS\_LgaA | BGC0001946 | NRP + Polyketide | 26.0 | 38.3 | 81.0 | 5.4e-15 |
| AYA22335.1 | KerB | BGC0001955 | Other | 27.0 | 29.3 | 81.0 | 7.1e-15 |
| AIG79243.1 | Non-ribosomal\_peptide\_synthetase | BGC0000419 | Saccharide + NRP:Glycopeptide | 30.0 | 29.6 | 81.0 | 9.2e-15 |
| EHK22005.1 | putative\_non-ribosomal\_peptide\_synthetase\_GliP | BGC0001609 | NRP | 24.0 | 51.7 | 80.0 | 1.2e-14 |
| AQX14497.1 | monobactam\_NRPS\_scaffold\_4 | BGC0001672 | NRP | 27.0 | 28.2 | 80.0 | 1.2e-14 |
| EWS95124.1 | hypothetical\_protein | BGC0000306 | NRP:Lipopeptide | 24.0 | 45.5 | 80.0 | 1.6e-14 |
| BAI63288.1 | putative\_non-ribosomal\_peptide\_synthetase | BGC0000434 | NRP | 30.0 | 29.1 | 80.0 | 1.6e-14 |
| ABC39418.1 | dihydroaeruginoic\_acid\_synthetase | BGC0000964 | NRP:Cyclic depsipeptide + Polyketide:Trans-AT type I | 32.0 | 27.1 | 80.0 | 1.6e-14 |
| WP\_019032755.1 | non-ribosomal\_peptide\_synthetase | BGC0001331 | NRP:Cyclic depsipeptide + Polyketide:Modular type I | 30.0 | 26.8 | 80.0 | 1.6e-14 |
| AJD77023.1 | IkaA | BGC0001435 | NRP + Polyketide:Iterative type I | 27.0 | 31.0 | 80.0 | 1.6e-14 |
| ASX95241.1 | IlaS | BGC0001620 | Polyketide | 29.0 | 25.7 | 80.0 | 1.6e-14 |
| AQX14441.1 | EM5400\_NRPS\_scaffold | BGC0001671 | NRP | 23.0 | 49.6 | 80.0 | 1.6e-14 |
| QBC75023.1 | non-ribosomal\_peptide\_synthetase | BGC0001968 | NRP | 23.0 | 44.6 | 80.0 | 1.6e-14 |
| BAH04162.1 | trsJ | BGC0000450 | NRP | 30.0 | 28.4 | 79.0 | 2.1e-14 |
| BBA21071.1 | putative\_non-ribosomal\_peptide\_synthetase | BGC0001740 | NRP + Polyketide | 31.0 | 28.2 | 79.0 | 2.1e-14 |
| TXD00259.1 | amino\_acid\_adenylation\_domain-containing\_protein | BGC0001877 | Polyketide | 31.0 | 22.7 | 79.0 | 2.1e-14 |
| CAC48360.1 | peptide\_synthetase | BGC0000311 | NRP | 29.0 | 27.1 | 79.0 | 3.5e-14 |
| WP\_078857610.1 | non-ribosomal\_peptide\_synthetase | BGC0001368 | NRP | 29.0 | 28.3 | 79.0 | 3.5e-14 |
| AUD11994.1 | OrbI | BGC0001721 | NRP | 29.0 | 30.5 | 79.0 | 3.5e-14 |
| WP\_039806852.1 | non-ribosomal\_peptide\_synthetase | BGC0002001 | NRP + Polyketide | 29.0 | 28.0 | 79.0 | 3.5e-14 |
| AHB38497.1 | non-ribosomal\_peptide\_synthetase | BGC0000346 | NRP + Polyketide:Modular type I | 29.0 | 28.3 | 78.0 | 4.6e-14 |
| CAE52334.1 | non-ribosomal\_peptide\_synthase | BGC0001088 | NRP + Polyketide | 29.0 | 26.2 | 78.0 | 4.6e-14 |
| BAX64247.1 | NRPS | BGC0001623 | NRP + Polyketide | 30.0 | 32.3 | 78.0 | 4.6e-14 |
| CAC17499.1 | putative\_non-ribosomal\_peptide\_synthase | BGC0000324 | NRP | 29.0 | 27.7 | 78.0 | 6e-14 |
| AEF16059.1 | non-ribosomal\_peptide\_synthetase | BGC0000953 | Saccharide:Aminoglycoside | 28.0 | 32.3 | 78.0 | 6e-14 |
| AFK57219.1 | DidH | BGC0000985 | Polyketide + NRP:Cyclic depsipeptide | 24.0 | 48.5 | 78.0 | 6e-14 |
| AVV61987.1 | putative\_non-ribosomal\_peptide\_synthetase | BGC0001477 | NRP + Polyketide:Modular type I | 25.0 | 42.4 | 78.0 | 6e-14 |
| ADQ55475.1 | NRPS | BGC0000350 | NRP:Beta-lactam | 27.0 | 38.7 | 77.0 | 7.8e-14 |
| AGD80618.1 | non-ribosomal\_peptide\_synthetase | BGC0000394 | NRP | 26.0 | 30.5 | 77.0 | 7.8e-14 |
| CAJ14037.1 | peptide\_synthetase | BGC0000406 | NRP | 31.0 | 25.9 | 77.0 | 1e-13 |
| ABP57745.1 | DepA | BGC0000993 | NRP:Cyclic depsipeptide + Polyketide:Modular type I | 30.0 | 27.1 | 77.0 | 1e-13 |
| ABR67750.1 | CmnG | BGC0000316 | NRP | 28.0 | 31.5 | 76.0 | 1.7e-13 |
| EDY47118.1 | N-(5-amino-5-carboxypentanoyl)-L-cysteinyl-D-\_valine\_synthase | BGC0000319 | NRP:Beta-lactam | 27.0 | 31.9 | 76.0 | 1.7e-13 |
| CAF05649.1 | TubD\_protein | BGC0001053 | NRP + Polyketide | 26.0 | 26.1 | 76.0 | 1.7e-13 |
| AEF33078.1 | dimodular\_nonribosomal\_peptide\_synthetase | BGC0001039 | NRP + Polyketide | 28.0 | 41.7 | 76.0 | 2.3e-13 |
| ctg1\_orf13 |  | BGC0001329 | Polyketide + NRP:Cyclic depsipeptide | 27.0 | 27.9 | 76.0 | 3e-13 |
| AQX14493.1 | monobactam\_NRPS\_scaffold\_2 | BGC0001672 | NRP | 29.0 | 26.1 | 76.0 | 3e-13 |
| BBA21068.1 | putative\_non-ribosomal\_peptide\_synthetase | BGC0001740 | NRP + Polyketide | 27.0 | 42.4 | 75.0 | 5.1e-13 |
| BAP16693.1 | nonribosomal\_peptide\_synthetase | BGC0000376 | NRP | 30.0 | 26.5 | 74.0 | 6.6e-13 |
| BAG17643.1 | putative\_NRPS-type-I\_PKS\_fusion\_protein | BGC0001043 | NRP + Polyketide | 26.0 | 26.5 | 74.0 | 6.6e-13 |
| ADA82585.1 | hybrid\_trans-AT\_polyketide\_synthase\_-\_nonribosomal\_peptide\_synthetase | BGC0001110 | NRP + Polyketide:Trans-AT type I | 28.0 | 28.8 | 74.0 | 6.6e-13 |
| ADH04680.1 | hybrid\_polyketide\_synthase/non-ribosomal\_peptide\_synthetase | BGC0001344 | NRP + Polyketide | 26.0 | 26.2 | 74.0 | 6.6e-13 |
| AIW58892.1 | non-ribosomal\_peptide\_synthetase | BGC0001582 | NRP | 30.0 | 28.3 | 74.0 | 6.6e-13 |
| WP\_013184461.1 | non-ribosomal\_peptide\_synthetase | BGC0001716 | NRP | 28.0 | 27.3 | 74.0 | 6.6e-13 |
| ABI22133.1 | putative\_non-ribosomal\_peptide\_synthetase | BGC0000422 | NRP | 31.0 | 27.0 | 74.0 | 1.1e-12 |
| CCE88377.1 | non-ribosomal\_peptide\_synthetase/polyketide\_synthase | BGC0001034 | NRP + Polyketide:Modular type I | 34.0 | 20.6 | 74.0 | 1.1e-12 |
| BAB69699.1 | iturin\_A\_synthetase\_B | BGC0001098 | NRP + Polyketide | 28.0 | 26.2 | 74.0 | 1.1e-12 |
| ATD51280.1 | nonribosomal\_peptide\_synthase | BGC0001650 | NRP | 31.0 | 30.0 | 74.0 | 1.1e-12 |
| CBK62746.1 |  | BGC0001115 | NRP + Polyketide | 29.0 | 26.0 | 73.0 | 1.5e-12 |
| BAK26562.1 | PKS-NRPS\_hybrid | BGC0000977 | NRP + Polyketide | 21.0 | 52.1 | 73.0 | 1.9e-12 |
| ABU70377.1 | hypothetical\_protein | BGC0001890 | NRP | 27.0 | 28.3 | 73.0 | 1.9e-12 |
| CAD70195.1 | non-ribosomal\_peptide\_synthetase | BGC0001047 | NRP + Polyketide | 22.0 | 46.0 | 72.0 | 2.5e-12 |
| AAF00961.1 | mcyB | BGC0001017 | NRP + Polyketide:Modular type I | 29.0 | 33.7 | 72.0 | 3.3e-12 |
| CAG23957.2 | hybrid\_NRPS/PKS\_protein | BGC0001089 | Polyketide + NRP | 23.0 | 48.0 | 72.0 | 4.3e-12 |
| ANI24100.1 | nonribosomal\_peptide\_synthetase | BGC0001235 | NRP + Polyketide | 31.0 | 21.0 | 72.0 | 4.3e-12 |
| AWO77084.1 | hybrid\_non-ribosomal\_peptide\_synthetase/type\_I\_polyketide\_synthase | BGC0001556 | NRP + Polyketide | 25.0 | 28.7 | 72.0 | 4.3e-12 |
| AEO14743.1 | NdaA | BGC0000396 | NRP | 32.0 | 25.1 | 71.0 | 5.6e-12 |
| AGN74876.1 | nonribosomal\_peptide\_synthetase | BGC0000459 | NRP:Cyclic depsipeptide + Polyketide:Trans-AT type I | 28.0 | 28.4 | 71.0 | 5.6e-12 |
| CAL80821.1 | sylD-like\_NRPS/PKS | BGC0000997 | NRP + Polyketide | 23.0 | 46.9 | 71.0 | 5.6e-12 |
| AFR69331.1 | nonribosomal\_peptide\_synthetase\_SpiA | BGC0001045 | NRP:Cyclic depsipeptide + Polyketide:Modular type I | 31.0 | 20.5 | 71.0 | 5.6e-12 |
| ABW00331.1 | amino\_acid\_adenylation\_domain | BGC0000333 | NRP | 27.0 | 38.2 | 71.0 | 7.3e-12 |
| EAL88817.1 | nonribosomal\_peptide\_synthase\_GliP | BGC0000361 | NRP | 22.0 | 66.7 | 71.0 | 7.3e-12 |
| CAG23960.2 | hybrid\_NRPS/PKS\_protein | BGC0001089 | Polyketide + NRP | 26.0 | 29.1 | 71.0 | 7.3e-12 |
| CAL69889.1 | RhiB\_protein | BGC0001112 | NRP + Polyketide:Trans-AT type I | 29.0 | 20.2 | 71.0 | 7.3e-12 |
| AFA26384.1 | polyketide\_synthase\_A | BGC0001874 | NRP + Polyketide | 22.0 | 52.4 | 71.0 | 9.5e-12 |
| AFD30953.1 | CrmB | BGC0000966 | NRP + Polyketide | 30.0 | 25.7 | 69.0 | 2.1e-11 |
| ATD51278.1 | nonribosomal\_peptide\_synthase | BGC0001650 | NRP | 27.0 | 28.9 | 69.0 | 2.1e-11 |
| DAC76734.1 | type\_I\_polyketide\_synthase/non-ribosomal\_peptide\_synthetase | BGC0001885 | Polyketide | 33.0 | 21.0 | 69.0 | 2.1e-11 |
| ADQ55476.1 | NRPS | BGC0000350 | NRP:Beta-lactam | 27.0 | 25.5 | 69.0 | 2.8e-11 |
| CAH55654.1 | putative\_L-prolyl-AMP\_ligase | BGC0000259 | Polyketide | 28.0 | 30.3 | 69.0 | 3.6e-11 |
| CAJ76298.1 | putative\_hybrid\_polyketide-non-ribosomal\_peptide\_synthetase | BGC0000972 | NRP + Polyketide:Modular type I + Polyketide:Trans-AT type I | 29.0 | 28.3 | 69.0 | 3.6e-11 |
| AAN85506.1 | nonribosomal\_peptide\_synthetase | BGC0001101 | NRP + Polyketide:Modular type I + Polyketide:Trans-AT type I | 27.0 | 28.3 | 69.0 | 3.6e-11 |
| ABC35522.1 | thiotemplate\_mechanism\_natural\_product\_synthetase | BGC0000186 | NRP + Polyketide:Modular type I | 24.0 | 44.8 | 68.0 | 4.7e-11 |
| CAJ96472.1 | non-ribosomal\_peptide\_synthetase | BGC0000330 | NRP:NRP siderophore | 28.0 | 29.3 | 68.0 | 4.7e-11 |
| AAC32048.1 | FxbC | BGC0000351 | NRP | 30.0 | 28.3 | 68.0 | 4.7e-11 |
| AAV97877.1 | OnnI | BGC0001105 | NRP + Polyketide:Trans-AT type I | 26.0 | 28.4 | 68.0 | 4.7e-11 |
| AAS47564.1 | mixed\_type\_I\_polyketide\_synthase/nonribosomal\_peptide\_synthetase | BGC0001108 | Polyketide:Trans-AT type I | 28.0 | 30.2 | 68.0 | 4.7e-11 |
| ctg1\_orf6 |  | BGC0001109 | NRP + Polyketide | 28.0 | 30.2 | 68.0 | 4.7e-11 |
| AGN74898.1 | nonribosomal\_peptide\_synthetase | BGC0000459 | NRP:Cyclic depsipeptide + Polyketide:Trans-AT type I | 28.0 | 27.4 | 68.0 | 6.2e-11 |
| WP\_053065269.1 | non-ribosomal\_peptide\_synthetase | BGC0001330 | NRP:Cyclic depsipeptide + Polyketide:Modular type I | 27.0 | 28.9 | 67.0 | 8.1e-11 |
| AKP45395.1 | CysG | BGC0001413 | NRP | 27.0 | 25.9 | 67.0 | 8.1e-11 |
| AZF85940.1 | non-ribosomal\_peptide\_synthase | BGC0001963 | NRP + Polyketide | 27.0 | 27.9 | 67.0 | 1.1e-10 |
| ADQ74618.1 | amino\_acid\_adenylation\_protein | BGC0000921 | Polyketide + NRP + Other:Shikimate-derived | 29.0 | 28.3 | 66.0 | 1.8e-10 |
| CAJ76292.1 | putative\_non-ribosomal\_peptide\_synthase | BGC0000972 | NRP + Polyketide:Modular type I + Polyketide:Trans-AT type I | 23.0 | 53.9 | 66.0 | 3.1e-10 |
| ALG65314.1 | Cal22 | BGC0001297 | NRP | 29.0 | 20.6 | 65.0 | 4e-10 |
| CAL69597.1 | PKS-NRPS | BGC0001049 | NRP + Polyketide:Iterative type I | 28.0 | 21.9 | 65.0 | 5.2e-10 |
| CAJ34374.1 | NRPS\_protein | BGC0000445 | NRP:Cyclic depsipeptide | 27.0 | 29.1 | 64.0 | 6.8e-10 |
| AAG02364.1 | peptide\_synthetase\_NRPS2-1 | BGC0000963 | NRP:Glycopeptide + Polyketide:Modular type I + Saccharide:Hybrid/tailoring | 30.0 | 24.8 | 64.0 | 6.8e-10 |
| AQH32481.1 | hybrid\_polyketide\_synthase/peptide\_synthetase | BGC0001667 | NRP + Polyketide | 25.0 | 27.8 | 64.0 | 8.9e-10 |
| DAB41917.1 | ArzO\_-\_NRPS\_(Cy,\_A,\_Ox,\_PCP) | BGC0001884 | NRP + Polyketide | 29.0 | 23.2 | 64.0 | 8.9e-10 |
| QBE85649.1 | BuaA | BGC0001978 | NRP + Polyketide | 32.0 | 21.1 | 64.0 | 1.2e-09 |
| AGD80616.1 | amino\_acid\_adenylation\_domain-containing\_protein | BGC0000394 | NRP | 26.0 | 32.8 | 63.0 | 1.5e-09 |
| ABS74207.1 | fengycin\_synthetase\_C | BGC0001095 | NRP | 26.0 | 30.2 | 63.0 | 2e-09 |
| ABS74209.1 | fengycin\_synthetase\_A | BGC0001095 | NRP | 26.0 | 30.2 | 63.0 | 2e-09 |
| AKC54422.1 | fumosorinone\_biosynthesis\_polyketide\_synthase | BGC0001218 | NRP + Polyketide | 29.0 | 22.0 | 63.0 | 2e-09 |
| CCB53267.1 | non-ribosomal\_peptide\_synthetase | BGC0001393 | NRP | 24.0 | 40.7 | 63.0 | 2e-09 |
| CAD29798.1 | peptide\_synthetase | BGC0001015 | NRP + Polyketide | 28.0 | 30.5 | 62.0 | 3.4e-09 |
| AQH32485.1 | peptide\_synthetase | BGC0001667 | NRP + Polyketide | 26.0 | 30.2 | 61.0 | 5.8e-09 |
| BAL15726.1 | NRPS | BGC0000432 | NRP | 27.0 | 22.2 | 61.0 | 7.6e-09 |
| AAG02359.1 | peptide\_synthetase\_NRPS5-4-3 | BGC0000963 | NRP:Glycopeptide + Polyketide:Modular type I + Saccharide:Hybrid/tailoring | 28.0 | 23.7 | 61.0 | 7.6e-09 |
| AAO62587.1 | peptide\_sythetase | BGC0001016 | NRP + Polyketide | 25.0 | 30.3 | 61.0 | 7.6e-09 |
| EAL85113.2 | hybrid\_PKS-NRPS\_enzyme | BGC0001037 | NRP + Polyketide:Iterative type I | 27.0 | 22.5 | 61.0 | 9.9e-09 |
| ADN43685.1 | DmbS | BGC0001136 | NRP + Polyketide:Iterative type I | 29.0 | 21.5 | 60.0 | 1.3e-08 |
| EAU38971.1 | hypothetical\_protein | BGC0001122 | NRP + Polyketide:Iterative type I | 21.0 | 53.6 | 60.0 | 1.7e-08 |
| gene3 |  | BGC0002035 | NRP + Polyketide | 27.0 | 25.4 | 59.0 | 2.2e-08 |
| AAX31555.1 | acyl-CoA\_ligase | BGC0000336 | NRP | 31.0 | 21.9 | 59.0 | 2.9e-08 |
| AAO62582.1 | polyketide\_synthase\_peptide\_sythetase\_fusion\_protein | BGC0001016 | NRP + Polyketide | 24.0 | 27.8 | 59.0 | 2.9e-08 |
| AAF00958.1 | mcyE | BGC0001017 | NRP + Polyketide:Modular type I | 25.0 | 27.1 | 59.0 | 2.9e-08 |
| AAM12937.1 | MupU | BGC0000182 | Polyketide:Iterative type I + Polyketide:Trans-AT type I | 28.0 | 20.6 | 59.0 | 3.7e-08 |
| ABV56588.1 | KtzH | BGC0000378 | NRP | 32.0 | 21.3 | 59.0 | 3.7e-08 |
| CAJ76286.1 | putative\_non-ribosomal\_peptide\_synthetase | BGC0000972 | NRP + Polyketide:Modular type I + Polyketide:Trans-AT type I | 24.0 | 32.1 | 58.0 | 4.9e-08 |
| AGO86662.1 | equisetin\_synthetase | BGC0001255 | NRP + Polyketide | 21.0 | 32.9 | 58.0 | 4.9e-08 |
| AEO57481.1 | PKS-NRPSs | BGC0001449 | NRP + Alkaloid + Polyketide:Iterative type I | 21.0 | 45.3 | 58.0 | 4.9e-08 |
| EPS29069.1 | hypothetical\_protein | BGC0001724 | NRP + Polyketide | 23.0 | 35.0 | 58.0 | 6.4e-08 |
| CCE28989.1 | non-ribosomal\_peptide\_synthetase | BGC0001365 | NRP | 21.0 | 46.1 | 57.0 | 8.4e-08 |
| BBA21084.1 | putative\_5-aminolevulinate\_CoA-ligase | BGC0001740 | NRP + Polyketide | 28.0 | 22.2 | 57.0 | 8.4e-08 |
| AQX36215.1 | siderophore\_synthetase | BGC0001527 | Other | 21.0 | 39.8 | 57.0 | 1.1e-07 |
| ATY69590.1 | adenylation\_protein | BGC0001823 | NRP + Polyketide | 25.0 | 34.3 | 57.0 | 1.4e-07 |
| ADC79613.1 | BafX | BGC0000028 | Polyketide:Modular type I | 25.0 | 23.4 | 56.0 | 1.9e-07 |
| AIL50189.1 | putative\_5-aminolevulinate\_CoA-ligase | BGC0000213 | Polyketide:Type II | 24.0 | 22.2 | 56.0 | 3.2e-07 |
| AAV66110.2 | fusaridione\_A\_synthetase | BGC0000992 | NRP + Polyketide | 21.0 | 39.2 | 53.0 | 1.6e-06 |
| AGY30675.1 | Ann3 | BGC0001298 | Polyketide | 23.0 | 23.7 | 53.0 | 1.6e-06 |
| AAZ23074.1 | acyl-CoA\_ligase | BGC0000291 | NRP | 26.0 | 20.2 | 51.0 | 7.8e-06 |
