## Supplementary Results for "A draft genome of the ascomycotal fungal species *Pseudopithomyces maydicus* (family *Didymosphaeriaceae*)": input.path1.gene398_mibig_hits.html

| MIBiG Protein | Description | MIBiG Cluster | MiBiG Product | % ID | % Coverage | BLAST Score | E-value |
| --- | --- | --- | --- | --- | --- | --- | --- |
| CBF73453.1 | nonribosomal\_peptide\_synthase,\_putative\_(JCVI) | BGC0001515 | NRP | 32.0 | 101.6 | 697.0 | 5.1e-200 |
| CAP93139.1 | cyclic\_hydrophobic\_tetrapeptide | BGC0000357 | NRP:Cyclic depsipeptide | 33.0 | 89.8 | 652.0 | 1.4e-186 |
| CBF87869.1 | nonribosomal\_peptide\_synthase,\_putative\_(Eurofung) | BGC0001699 | Polyketide | 30.0 | 99.9 | 610.0 | 6.3e-174 |
| EAL89049.1 | nonribosomal\_peptide\_synthetase | BGC0000355 | NRP | 29.0 | 102.4 | 595.0 | 2.1e-169 |
| ADY16697.1 | TqaA | BGC0001142 | NRP | 29.0 | 104.8 | 551.0 | 4.5e-156 |
| QCC62999.1 | BII-rafflesfungin\_nonribosomal\_protein\_synthetase | BGC0001966 | NRP | 35.0 | 67.5 | 545.0 | 3.2e-154 |
| EXU96269.1 | nonribosomal\_peptide\_synthetase,\_serinocyclin\_synthetase\_NPS1 | BGC0001240 | NRP | 34.0 | 67.6 | 524.0 | 4.5e-148 |
| EAW16180.1 | nonribosomal\_peptide\_synthase,\_putative | BGC0000293 | NRP | 29.0 | 85.7 | 500.0 | 7e-141 |
| ACZ66258.1 | APS1 | BGC0000304 | NRP | 32.0 | 72.7 | 483.0 | 1.5e-135 |
| CBF87069.1 | nonribosomal\_peptide\_synthase,\_putative\_(Eurofung) | BGC0001290 | NRP | 34.0 | 57.4 | 441.0 | 6.6e-123 |
| XP\_001217690.1 | hypothetical\_protein | BGC0001517 | NRP | 35.0 | 57.1 | 432.0 | 2.3e-120 |
| ctg1\_orf000000 |  | BGC0000900 | Other | 34.0 | 43.6 | 336.0 | 3e-91 |
| ctg1\_orf003 |  | BGC0000334 | NRP | 33.0 | 58.7 | 420.0 | 1.2e-116 |
| ctg1\_orf00001 |  | BGC0000901 | Other | 32.0 | 58.0 | 418.0 | 6e-116 |
| AGQ43600.1 | HC-toxin\_synthetase | BGC0001166 | NRP | 32.0 | 60.7 | 410.0 | 9.5e-114 |
| OAQ83772.1 | nonribosomal\_peptide\_synthase | BGC0001358 | Polyketide | 34.0 | 58.5 | 409.0 | 2.1e-113 |
| MAA\_10043 | non-ribosomal\_peptide\_synthetase | BGC0000337 | NRP | 30.0 | 66.2 | 406.0 | 1.4e-112 |
| AUS29494.1 | non-ribosomal\_peptide\_synthetase | BGC0001030 | NRP + Polyketide | 32.0 | 57.8 | 396.0 | 1.9e-109 |
| AHB82071.1 | non\_ribosomal\_peptide\_synthetase | BGC0001231 | NRP + Polyketide:Modular type I | 27.0 | 96.5 | 392.0 | 3.5e-108 |
| CBF76038.1 | nonribosomal\_peptide\_synthase,\_putative\_(Eurofung) | BGC0001399 | NRP | 32.0 | 56.7 | 387.0 | 1.1e-106 |
| ABM91454.1 | lysergyl\_peptide\_synthetase\_LpsB | BGC0000348 | NRP | 31.0 | 59.1 | 386.0 | 2.5e-106 |
| BBC83957.1 | nonribosomal\_peptide\_synthetase | BGC0001636 | NRP | 30.0 | 64.3 | 381.0 | 4.7e-105 |
| XP\_011325838.1 | hypothetical\_protein | BGC0001545 | NRP | 32.0 | 58.3 | 377.0 | 8.9e-104 |
| XP\_003044554.1 | hypothetical\_protein | BGC0001768 | NRP | 31.0 | 66.1 | 376.0 | 2e-103 |
| AMM63162.1 | AniA | BGC0001371 | NRP | 31.0 | 59.5 | 374.0 | 9.9e-103 |
| XP\_002379984.1 | nonribosomal\_peptide\_synthase,\_putative | BGC0001621 | NRP | 32.0 | 59.3 | 372.0 | 3.8e-102 |
| ADM34138.1 | non-ribosomal\_peptide\_synthetase | BGC0001084 | NRP + Terpene + Alkaloid | 32.0 | 57.6 | 371.0 | 8.4e-102 |
| AHB82058.1 | non\_ribosomal\_peptide\_synthetase | BGC0001019 | NRP + Polyketide:Modular type I | 27.0 | 97.1 | 369.0 | 1.9e-101 |
| EPE34341.1 | non-ribosomal\_peptide\_synthetase | BGC0001035 | Polyketide + NRP + Other:Aminocoumarin | 30.0 | 59.4 | 369.0 | 2.4e-101 |
| BAN91483.1 | putative\_Aculeacin\_A\_nonribosomal\_peptide\_synthase | BGC0001220 | NRP | 30.0 | 60.0 | 366.0 | 2.1e-100 |
| CAP96445.1 | NRPS | BGC0000420 | NRP | 35.0 | 45.6 | 363.0 | 1.7e-99 |
| XP\_020057667.1 | hypothetical\_protein | BGC0001718 | NRP | 35.0 | 47.1 | 362.0 | 3.9e-99 |
| AGC83576.1 | NRPS | BGC0000818 | NRP | 38.0 | 42.2 | 361.0 | 8.7e-99 |
| CRG85572.1 | nonribosomal\_peptide\_synthase,\_putative | BGC0001402 | NRP | 29.0 | 61.0 | 359.0 | 1.9e-98 |
| WP\_043882190.1 | non-ribosomal\_peptide\_synthetase | BGC0001993 | NRP | 26.0 | 98.4 | 357.0 | 9.6e-98 |
| CBF77087.1 | conserved\_hypothetical\_protein | BGC0001679 | NRP | 34.0 | 47.7 | 356.0 | 1.6e-97 |
| CZT62785.1 | Non-ribosomal\_peptide\_synthase,\_involved\_in\_Hassallidin\_biosynthesis | BGC0001614 | NRP | 30.0 | 62.4 | 356.0 | 2.8e-97 |
| BBB04327.1 | nonribosomal\_peptide\_synthetase | BGC0001717 | Alkaloid | 35.0 | 46.1 | 351.0 | 5.3e-96 |
| BAH23995.1 | nonribosomal\_peptide\_synthetase | BGC0000356 | NRP + Alkaloid | 37.0 | 44.7 | 350.0 | 1.2e-95 |
| ACR78148.1 | BSLS | BGC0000312 | NRP | 32.0 | 50.2 | 347.0 | 7.6e-95 |
| CAA79245.2 | enniatin\_synthetase | BGC0000342 | NRP | 28.0 | 63.3 | 347.0 | 7.6e-95 |
| ANY57892.1 | PenN | BGC0001372 | Terpene | 31.0 | 55.3 | 347.0 | 9.9e-95 |
| ATO51563.1 | non-ribosomal\_peptide\_synthetase | BGC0001796 | NRP | 25.0 | 97.1 | 344.0 | 1.1e-93 |
| AQM58286.1 | non-ribosomal\_peptide\_synthase | BGC0001816 | NRP + Polyketide | 29.0 | 59.9 | 340.0 | 1.6e-92 |
| ACI30655.1 | BEAS\_beauvericin\_nonribosomal\_cyclodepsipeptide\_synthetase | BGC0000313 | NRP | 29.0 | 63.6 | 339.0 | 2.1e-92 |
| ATP76246.1 | SpuB | BGC0001748 | NRP + Polyketide | 28.0 | 64.3 | 339.0 | 2.1e-92 |
| AQM58288.1 | non-ribosomal\_peptide\_synthase | BGC0001816 | NRP + Polyketide | 35.0 | 44.5 | 337.0 | 1.3e-91 |
| ABX37382.1 | amino\_acid\_adenylation\_domain\_protein | BGC0000984 | NRP + Polyketide | 28.0 | 76.1 | 331.0 | 5.6e-90 |
| CBZ42146.1 | putative\_non-ribosomal\_peptide\_synthetase | BGC0001117 | NRP | 30.0 | 63.3 | 330.0 | 1.3e-89 |
| ACZ55942.1 | non-ribosomal\_peptide\_synthetase | BGC0000302 | NRP | 29.0 | 64.9 | 329.0 | 2.1e-89 |
| EAL89046.1 | nonribosomal\_peptide\_synthetase | BGC0000355 | NRP | 33.0 | 50.0 | 329.0 | 2.8e-89 |
| AAY93356.2 | non-ribosomal\_peptide\_synthetase\_PvdI | BGC0000413 | NRP | 31.0 | 62.5 | 329.0 | 3.6e-89 |
| AGA37267.1 | NRPS | BGC0000816 | NRP + Alkaloid | 31.0 | 58.1 | 329.0 | 3.6e-89 |
| AAF01762.1 | AM-toxin\_synthetase | BGC0001261 | NRP | 30.0 | 57.6 | 327.0 | 8.1e-89 |
| BAH43871.1 | truncated\_linear\_pentadecapeptide\_gramicidin\_synthetase\_LgrC | BGC0000367 | NRP | 28.0 | 63.8 | 325.0 | 4e-88 |
| AHF21228.1 | TriD | BGC0000449 | NRP | 28.0 | 68.9 | 324.0 | 9e-88 |
| AEA30274.1 | peptide\_synthetase | BGC0000429 | Polyketide + NRP:Cyclic depsipeptide | 31.0 | 63.6 | 323.0 | 1.5e-87 |
| PHM49485.1 | Amino\_acid\_adenylation | BGC0001131 | NRP | 30.0 | 67.1 | 323.0 | 1.5e-87 |
| ADY16689.1 | TqaB | BGC0001142 | NRP | 32.0 | 51.1 | 323.0 | 1.5e-87 |
| QDA77059.1 | polyketide\_synthase/nonribosomal\_peptide\_synthetase | BGC0002026 | NRP | 30.0 | 63.1 | 322.0 | 2.6e-87 |
| AFP87549.1 | NrpS | BGC0001135 | NRP | 29.0 | 67.9 | 322.0 | 4.5e-87 |
| AHZ20773.1 | non-ribosomal\_peptide\_synthase | BGC0000369 | NRP + Saccharide:Hybrid/tailoring | 28.0 | 65.5 | 321.0 | 9.9e-87 |
| EFE73313.1 | nonribosomal\_peptide\_synthetase | BGC0000431 | NRP:Cyclic depsipeptide | 29.0 | 66.1 | 320.0 | 1.3e-86 |
| WP\_015507661.1 | non-ribosomal\_peptide\_synthetase | BGC0001792 | NRP | 27.0 | 83.4 | 320.0 | 1.7e-86 |
| BAH43870.1 | putative\_linear\_pentadecapeptide\_gramicidin\_synthetase\_LgrB | BGC0000367 | NRP | 29.0 | 65.3 | 319.0 | 2.2e-86 |
| AFN69082.1 | nonribosomal\_peptide\_synthetase\_6 | BGC0001249 | NRP | 33.0 | 45.0 | 319.0 | 2.9e-86 |
| BAW32324.1 | nonribosomal\_peptide\_synthetase | BGC0001630 | NRP + Polyketide | 29.0 | 69.4 | 319.0 | 3.8e-86 |
| ABB90279.1 | non-ribosomal\_peptide\_synthetase | BGC0001057 | NRP + Polyketide | 32.0 | 49.0 | 318.0 | 6.4e-86 |
| ACJ04424.1 | aureobasidin\_A1\_biosynthesis\_complex | BGC0000307 | NRP | 31.0 | 51.2 | 316.0 | 1.9e-85 |
| AGM16412.1 | paenibacterin\_synthetase\_A | BGC0000400 | NRP | 28.0 | 67.0 | 314.0 | 7.1e-85 |
| AQH32484.1 | peptide\_synthetase | BGC0001667 | NRP + Polyketide | 28.0 | 66.4 | 314.0 | 7.1e-85 |
| ABD14711.1 | cesA | BGC0000320 | NRP:Cyclic depsipeptide | 27.0 | 69.4 | 314.0 | 9.3e-85 |
| CUX79061.1 | Octapeptin\_synthase\_subunit\_B | BGC0001715 | NRP | 27.0 | 64.3 | 313.0 | 2.1e-84 |
| ARB50207.1 | lysergyl\_peptide\_synthetase\_21 | BGC0001573 | Alkaloid | 31.0 | 46.6 | 312.0 | 3.5e-84 |
| AAF00960.1 | mcyA | BGC0001017 | NRP + Polyketide:Modular type I | 27.0 | 86.8 | 310.0 | 1.3e-83 |
| CAD29798.1 | peptide\_synthetase | BGC0001015 | NRP + Polyketide | 30.0 | 66.0 | 309.0 | 2.3e-83 |
| AHZ20784.1 | non-ribosomal\_peptide\_synthase | BGC0000369 | NRP + Saccharide:Hybrid/tailoring | 30.0 | 59.8 | 309.0 | 3.9e-83 |
| ACC81021.1 | non-ribosomal\_peptide\_synthetase | BGC0001479 | NRP | 30.0 | 52.0 | 309.0 | 3.9e-83 |
| AQM58287.1 | non-ribosomal\_peptide\_synthase | BGC0001816 | NRP + Polyketide | 28.0 | 56.9 | 308.0 | 5.1e-83 |
| WP\_050383082.1 | non-ribosomal\_peptide\_synthetase | BGC0001451 | NRP | 29.0 | 63.8 | 307.0 | 8.7e-83 |
| AXN93581.1 | PuwF-G | BGC0001950 | NRP | 28.0 | 70.1 | 307.0 | 8.7e-83 |
| CAM02313.1 | putative\_non-ribosomal\_peptide\_synthetase | BGC0000349 | NRP | 29.0 | 66.6 | 307.0 | 1.5e-82 |
| AAY93354.1 | non-ribosomal\_peptide\_synthetase\_PvdD | BGC0000413 | NRP | 31.0 | 58.4 | 307.0 | 1.5e-82 |
| ANZ15840.1 | non-ribosomal\_peptide\_synthase/amino\_acid\_adenylation\_enzyme | BGC0001569 | NRP | 27.0 | 63.0 | 307.0 | 1.5e-82 |
| BBA21073.1 | putative\_non-ribosomal\_peptide\_synthetase | BGC0001740 | NRP + Polyketide | 29.0 | 61.3 | 307.0 | 1.5e-82 |
| BAH22764.1 | nonribosomal\_peptide\_synthetase | BGC0001018 | NRP + Polyketide | 28.0 | 69.6 | 304.0 | 7.4e-82 |
| CCJ67640.1 | TaaE | BGC0000447 | NRP:Lipopeptide | 28.0 | 68.8 | 304.0 | 9.6e-82 |
| AIW82284.1 | PuwG | BGC0001125 | NRP + Polyketide | 28.0 | 69.8 | 304.0 | 1.3e-81 |
| AXN93590.1 | PuwF-G | BGC0001951 | NRP | 28.0 | 70.1 | 303.0 | 2.1e-81 |
| AQH32485.1 | peptide\_synthetase | BGC0001667 | NRP + Polyketide | 28.0 | 64.2 | 301.0 | 8.1e-81 |
| CAY48789.1 | putative\_non-ribosomal\_peptide\_synthetase | BGC0001312 | NRP | 30.0 | 63.4 | 300.0 | 1.4e-80 |
| AJF34464.1 | Txo2 | BGC0001207 | NRP | 28.0 | 67.5 | 299.0 | 2.4e-80 |
| CAE15637.1 |  | BGC0001128 | NRP | 29.0 | 57.1 | 298.0 | 5.3e-80 |
| AAO62587.1 | peptide\_sythetase | BGC0001016 | NRP + Polyketide | 28.0 | 67.9 | 297.0 | 1.2e-79 |
| AEH59100.1 | amino\_acid\_adenylation\_domain-containing\_protein/NRPS | BGC0000385 | NRP | 29.0 | 58.9 | 297.0 | 1.5e-79 |
| AJV88375.1 | MfnC | BGC0001214 | NRP | 29.0 | 66.7 | 296.0 | 2.6e-79 |
| AFH75329.1 | nonribosomal\_peptide\_synthetase | BGC0000398 | NRP:Cyclic depsipeptide | 30.0 | 57.9 | 295.0 | 5.8e-79 |
| AFH75320.1 | nonribosomal\_peptide\_synthetase | BGC0000425 | NRP:Cyclic depsipeptide | 30.0 | 57.9 | 294.0 | 7.6e-79 |
| ACA97580.1 | PmxE | BGC0000408 | NRP | 28.0 | 66.2 | 294.0 | 1e-78 |
| ABC36785.1 | peptide\_synthetase,\_putative | BGC0000964 | NRP:Cyclic depsipeptide + Polyketide:Trans-AT type I | 24.0 | 99.7 | 294.0 | 1.3e-78 |
| ALV82388.1 | CDA\_peptide\_synthetase\_III | BGC0001370 | NRP | 28.0 | 63.9 | 294.0 | 1.3e-78 |
| ANI24100.1 | nonribosomal\_peptide\_synthetase | BGC0001235 | NRP + Polyketide | 29.0 | 66.3 | 293.0 | 1.7e-78 |
| WP\_015507662.1 | non-ribosomal\_peptide\_synthetase | BGC0001792 | NRP | 26.0 | 84.4 | 293.0 | 2.2e-78 |
| AJW76710.1 | DsaH | BGC0001196 | NRP | 29.0 | 57.6 | 292.0 | 3.8e-78 |
| CAY48788.1 | putative\_non-ribosomal\_peptide\_synthetase | BGC0001312 | NRP | 29.0 | 68.3 | 292.0 | 3.8e-78 |
| ATY37591.1 | BogD | BGC0001532 | NRP | 27.0 | 61.8 | 292.0 | 4.9e-78 |
| BAH43764.1 | tyrocidine\_synthetase\_I | BGC0000452 | NRP | 29.0 | 52.6 | 291.0 | 6.5e-78 |
| AJW76709.1 | DsaG | BGC0001196 | NRP | 28.0 | 66.7 | 291.0 | 6.5e-78 |
| CCJ67637.1 | TaaB | BGC0000447 | NRP:Lipopeptide | 28.0 | 57.8 | 291.0 | 8.4e-78 |
| ATY37609.1 | BreD | BGC0001536 | NRP | 26.0 | 69.0 | 291.0 | 8.4e-78 |
| WP\_047195042.1 | non-ribosomal\_peptide\_synthetase | BGC0002003 | NRP | 29.0 | 65.5 | 291.0 | 8.4e-78 |
| AXN93582.1 | PuwH | BGC0001950 | NRP | 30.0 | 58.5 | 290.0 | 1.4e-77 |
| AXN93591.1 | PuwH | BGC0001951 | NRP | 30.0 | 58.5 | 290.0 | 1.4e-77 |
| AAY37647.1 | Amino\_acid\_adenylation | BGC0000437 | NRP | 28.0 | 61.3 | 289.0 | 2.5e-77 |
| CDG12864.1 | non-ribosomal\_peptide\_synthetase | BGC0001415 | NRP | 28.0 | 68.0 | 289.0 | 2.5e-77 |
| ARB50206.1 | lysergyl\_peptide\_synthetase\_2 | BGC0001573 | Alkaloid | 26.0 | 62.0 | 289.0 | 2.5e-77 |
| ABV79985.1 | ApnA | BGC0000301 | NRP | 26.0 | 65.3 | 289.0 | 3.2e-77 |
| AEU11006.1 | NpnB | BGC0001029 | NRP + Polyketide | 29.0 | 57.2 | 289.0 | 4.2e-77 |
| ALD82526.1 | non-ribosomal\_peptide\_synthase | BGC0001212 | NRP + Polyketide | 30.0 | 59.9 | 289.0 | 4.2e-77 |
| WP\_039806850.1 | non-ribosomal\_peptide\_synthetase | BGC0002001 | NRP + Polyketide | 31.0 | 56.4 | 289.0 | 4.2e-77 |
| AEZ51516.1 | pmxA | BGC0001153 | NRP:Lipopeptide | 28.0 | 59.8 | 288.0 | 5.5e-77 |
| WP\_011146892.1 | non-ribosomal\_peptide\_synthetase | BGC0001641 | NRP | 34.0 | 43.0 | 288.0 | 7.1e-77 |
| AHI59109.1 | locillomycin\_synthase\_B | BGC0001005 | NRP + Polyketide | 26.0 | 70.9 | 287.0 | 9.3e-77 |
| ANG60379.1 | nonribosomal\_peptide\_synthetase\_BudA | BGC0001698 | NRP | 27.0 | 58.3 | 287.0 | 9.3e-77 |
| AAT01807.1 | non-ribosomal\_peptide\_synthetase | BGC0000365 | NRP | 29.0 | 53.7 | 287.0 | 1.2e-76 |
| ABS74179.1 | bacillomycin\_D\_synthetase\_C | BGC0001090 | Polyketide + NRP:Lipopeptide | 28.0 | 60.8 | 287.0 | 1.2e-76 |
| ABS74180.1 | bacillomycin\_D\_synthetase\_B | BGC0001090 | Polyketide + NRP:Lipopeptide | 27.0 | 66.9 | 287.0 | 1.6e-76 |
| ABH06368.2 | MassB | BGC0000389 | NRP:Cyclic depsipeptide | 30.0 | 57.6 | 286.0 | 2.1e-76 |
| EFE73312.1 | nonribosomal\_peptide\_synthetase | BGC0000431 | NRP:Cyclic depsipeptide | 29.0 | 60.6 | 286.0 | 2.1e-76 |
| AAD44234.1 | PstB | BGC0000362 | NRP | 30.0 | 55.7 | 286.0 | 2.7e-76 |
| AYA22336.1 | KerA | BGC0001955 | Other | 28.0 | 62.7 | 285.0 | 3.5e-76 |
| ACZ55943.1 | non-ribosomal\_peptide\_synthetase | BGC0000302 | NRP | 29.0 | 60.6 | 285.0 | 6e-76 |
| AHZ34232.1 | CifA | BGC0000323 | NRP:Lipopeptide | 29.0 | 58.6 | 284.0 | 7.9e-76 |
| APU91750.1 | Non-Ribosomal\_Peptide\_Synthetase | BGC0001806 | NRP | 30.0 | 55.9 | 284.0 | 7.9e-76 |
| APZ78782.1 | nonribosomal\_peptide\_synthetase | BGC0001426 | NRP:Cyclic depsipeptide + Polyketide:Iterative type I | 28.0 | 67.6 | 284.0 | 1e-75 |
| BAE60013.1 |  | BGC0001518 | Terpene | 29.0 | 54.1 | 284.0 | 1e-75 |
| ATY37590.1 | BogC | BGC0001532 | NRP | 25.0 | 70.1 | 284.0 | 1e-75 |
| KUM80514.1 | hypothetical\_protein | BGC0001562 | NRP | 29.0 | 59.3 | 284.0 | 1e-75 |
| AAO23334.1 | NcpB | BGC0000397 | NRP | 27.0 | 66.8 | 284.0 | 1.3e-75 |
| AAO56328.1 | non-ribosomal\_peptide\_synthetase\_SyfA | BGC0000435 | NRP | 29.0 | 58.9 | 284.0 | 1.3e-75 |
| BS330\_28385 | non-ribosomal\_peptide\_synthetase | BGC0001461 | NRP:Glycopeptide | 27.0 | 62.7 | 284.0 | 1.3e-75 |
| AAT01806.1 | non-ribosomal\_peptide\_synthetase | BGC0000365 | NRP | 27.0 | 63.0 | 283.0 | 1.8e-75 |
| WP\_013184463.1 | non-ribosomal\_peptide\_synthetase | BGC0001716 | NRP | 27.0 | 76.4 | 283.0 | 1.8e-75 |
| AAO23333.1 | NcpA | BGC0000397 | NRP | 27.0 | 66.2 | 282.0 | 3e-75 |
| AAY37654.1 | Amino\_acid\_adenylation | BGC0000437 | NRP | 30.0 | 59.0 | 282.0 | 3e-75 |
| AED90002.1 | non-ribosomal\_peptide\_synthetase\_ThaA | BGC0000443 | NRP:Beta-lactam | 29.0 | 58.9 | 282.0 | 3.9e-75 |
| APZ78729.1 | nonribosomal\_peptide\_synthetase | BGC0001421 | NRP:Cyclic depsipeptide + Polyketide:Iterative type I | 28.0 | 66.8 | 282.0 | 3.9e-75 |
| APZ78769.1 | nonribosomal\_peptide\_synthetase | BGC0001425 | NRP:Cyclic depsipeptide + Polyketide:Iterative type I | 28.0 | 67.6 | 282.0 | 3.9e-75 |
| APZ78809.1 | nonribosomal\_peptide\_synthetase | BGC0001428 | NRP:Cyclic depsipeptide + Polyketide:Iterative type I | 28.0 | 67.9 | 282.0 | 3.9e-75 |
| RAT94091.1 | NRPS | BGC0001469 | NRP | 27.0 | 61.0 | 282.0 | 3.9e-75 |
| AIG79241.1 | Hypothetical\_protein | BGC0000419 | Saccharide + NRP:Glycopeptide | 28.0 | 62.2 | 282.0 | 5.1e-75 |
| KUM80513.1 | hypothetical\_protein | BGC0001562 | NRP | 29.0 | 63.8 | 282.0 | 5.1e-75 |
| CAE53350.1 | non-ribosomal\_peptide\_synthetase | BGC0000440 | NRP:Glycopeptide | 28.0 | 61.9 | 281.0 | 6.7e-75 |
| CCJ67636.1 | TaaA | BGC0000447 | NRP:Lipopeptide | 29.0 | 57.6 | 281.0 | 6.7e-75 |
| APZ78795.1 | nonribosomal\_peptide\_synthetase | BGC0001427 | NRP:Cyclic depsipeptide + Polyketide:Iterative type I | 28.0 | 67.6 | 281.0 | 6.7e-75 |
| AAZ03551.1 | McnB | BGC0000332 | NRP | 28.0 | 60.2 | 281.0 | 8.7e-75 |
| AEU11003.1 | NpnC | BGC0001029 | NRP + Polyketide | 28.0 | 62.0 | 281.0 | 8.7e-75 |
| KFL51886.1 | amino\_acid\_adenylation\_protein | BGC0001711 | NRP + Polyketide | 28.0 | 67.1 | 281.0 | 8.7e-75 |
| AAM80539.1 | StaA | BGC0000290 | NRP:Glycopeptide | 28.0 | 63.3 | 280.0 | 1.1e-74 |
| AAD44233.1 | PstA | BGC0000362 | NRP | 29.0 | 56.2 | 280.0 | 1.1e-74 |
| CAJ77716.1 | Mps2\_protein | BGC0000364 | NRP | 30.0 | 52.5 | 280.0 | 1.1e-74 |
| ABV79986.1 | ApnB | BGC0000301 | NRP | 29.0 | 60.0 | 280.0 | 1.5e-74 |
| ABA73955.1 | putative\_non-ribosomal\_peptide\_synthetase | BGC0001842 | NRP:Lipopeptide | 28.0 | 58.0 | 280.0 | 1.5e-74 |
| CAG15009.1 | peptide\_synthetase,\_module\_1-2 | BGC0000441 | NRP | 28.0 | 61.9 | 280.0 | 1.9e-74 |
| AJI44167.1 | long-chain-fatty-acid-CoA\_ligase | BGC0001193 | NRP | 29.0 | 68.1 | 280.0 | 1.9e-74 |
| ATY37608.1 | BreC | BGC0001536 | NRP | 27.0 | 63.2 | 280.0 | 1.9e-74 |
| AJV88376.1 | MfnD | BGC0001214 | NRP | 29.0 | 66.2 | 279.0 | 2.5e-74 |
| AHZ34240.1 | CipC | BGC0001389 | NRP | 29.0 | 57.6 | 279.0 | 2.5e-74 |
| AEC14347.1 | nonribosomal\_peptide\_synthetase | BGC0000377 | NRP | 26.0 | 64.0 | 279.0 | 3.3e-74 |
| BAH43765.1 | tyrocidine\_synthetase\_II | BGC0000452 | NRP | 27.0 | 65.6 | 279.0 | 3.3e-74 |
| ABW00331.1 | amino\_acid\_adenylation\_domain | BGC0000333 | NRP | 28.0 | 65.6 | 279.0 | 4.3e-74 |
| BAV56271.1 |  | BGC0001657 | NRP | 28.0 | 63.0 | 279.0 | 4.3e-74 |
| AHZ34233.1 | CifB | BGC0000323 | NRP:Lipopeptide | 30.0 | 64.4 | 278.0 | 5.7e-74 |
| CAG23957.2 | hybrid\_NRPS/PKS\_protein | BGC0001089 | Polyketide + NRP | 29.0 | 57.7 | 278.0 | 7.4e-74 |
| CAD17792.1 | probable\_non\_ribosomal\_peptide\_synthetase\_protein | BGC0001754 | NRP + Polyketide | 28.0 | 66.4 | 278.0 | 7.4e-74 |
| AIE77059.1 | peptide\_synthetase | BGC0000418 | NRP | 28.0 | 62.8 | 277.0 | 9.6e-74 |
| AHD05618.1 | putative\_non-ribosomal\_peptide\_ligase\_domain\_protein | BGC0001033 | NRP + Polyketide | 28.0 | 56.9 | 277.0 | 9.6e-74 |
| CBG75492.1 | putative\_NRPS/siderophore\_biosynthesis\_protein | BGC0000423 | NRP | 28.0 | 60.8 | 277.0 | 1.3e-73 |
| ANY58984.1 | non-ribosomal\_synthetase | BGC0001615 | NRP | 27.0 | 57.5 | 277.0 | 1.3e-73 |
| CAB38518.1 | CDA\_peptide\_synthetase\_I\_(CdaPs1) | BGC0000315 | NRP:Ca+-dependent lipopeptide | 28.0 | 64.5 | 277.0 | 1.6e-73 |
| AFH75330.1 | nonribosomal\_peptide\_synthetase | BGC0000398 | NRP:Cyclic depsipeptide | 28.0 | 60.1 | 277.0 | 1.6e-73 |
| AHD05614.1 | putative\_non-ribosomal\_peptide\_ligase/\_polyketide\_synthase\_hybrid | BGC0001033 | NRP + Polyketide | 27.0 | 59.5 | 277.0 | 1.6e-73 |
| ABQ96384.2 | fusaricidin\_synthetase | BGC0001152 | Polyketide + NRP:Lipopeptide | 31.0 | 49.2 | 277.0 | 1.6e-73 |
| KFL51887.1 | amino\_acid\_adenylation\_protein | BGC0001711 | NRP + Polyketide | 26.0 | 68.4 | 277.0 | 1.6e-73 |
| BAH22763.1 | nonribosomal\_peptide\_synthetase | BGC0001018 | NRP + Polyketide | 28.0 | 60.6 | 276.0 | 2.8e-73 |
| ALV82356.1 | CDA\_peptide\_synthetase\_I | BGC0001370 | NRP | 28.0 | 62.5 | 276.0 | 2.8e-73 |
| AKC91849.1 | nonribosomal\_peptide\_synthetase | BGC0001414 | NRP | 28.0 | 66.1 | 276.0 | 2.8e-73 |
| CAD91220.1 | putative\_non-ribosomal\_peptide\_synthetase,\_modules\_1-2 | BGC0000289 | NRP:Glycopeptide + Saccharide:Hybrid/tailoring | 28.0 | 63.8 | 275.0 | 3.7e-73 |
| AEH41793.1 | HrmO | BGC0000374 | NRP:Cyclic depsipeptide | 27.0 | 66.5 | 275.0 | 3.7e-73 |
| AAO72425.1 | syringopeptin\_synthetase\_C | BGC0000438 | NRP | 27.0 | 64.0 | 275.0 | 3.7e-73 |
| WP\_079164394.1 | non-ribosomal\_peptide\_synthetase | BGC0001567 | NRP | 30.0 | 57.5 | 275.0 | 3.7e-73 |
| WP\_003981346.1 | non-ribosomal\_peptide\_synthetase | BGC0001813 | NRP | 28.0 | 63.6 | 275.0 | 3.7e-73 |
| AAY37655.1 | Amino\_acid\_adenylation | BGC0000437 | NRP | 27.0 | 64.9 | 275.0 | 4.8e-73 |
| BAP05597.1 | calI | BGC0000967 | NRP + Polyketide:Trans-AT type I | 29.0 | 58.8 | 275.0 | 4.8e-73 |
| ERG36783.1 | phenylalanine\_racemase | BGC0000299 | NRP | 28.0 | 63.7 | 275.0 | 6.2e-73 |
| WP\_010369428.1 | non-ribosomal\_peptide\_synthase | BGC0000314 | Polyketide + NRP:Cyclic depsipeptide + Other:Aminocoumarin | 28.0 | 63.7 | 275.0 | 6.2e-73 |
| CAQ43084.1 | non\_ribosomal\_polypeptide\_synthetase | BGC0000970 | NRP + Polyketide:Modular type I | 29.0 | 57.4 | 275.0 | 6.2e-73 |
| APZ78704.1 | nonribosomal\_peptide\_synthetase | BGC0001419 | NRP:Cyclic depsipeptide + Polyketide:Iterative type I | 32.0 | 51.3 | 275.0 | 6.2e-73 |
| ACU36654.1 | amino\_acid\_adenylation\_domain\_protein | BGC0000392 | NRP | 29.0 | 62.1 | 274.0 | 8.2e-73 |
| APZ78716.1 | nonribosomal\_peptide\_synthetase | BGC0001420 | NRP:Cyclic depsipeptide + Polyketide:Iterative type I | 29.0 | 66.5 | 274.0 | 8.2e-73 |
| BAC67536.1 | arthrofactin\_synthetase\_C | BGC0000305 | NRP:Lipopeptide | 28.0 | 57.5 | 274.0 | 1.1e-72 |
| AQZ69228.1 | hypothetical\_protein | BGC0001635 | NRP + Polyketide | 29.0 | 56.5 | 274.0 | 1.1e-72 |
| ABA73956.1 | putative\_non-ribosomal\_peptide\_synthetase | BGC0001842 | NRP:Lipopeptide | 28.0 | 58.2 | 274.0 | 1.1e-72 |
| WP\_013428324.1 | non-ribosomal\_peptide\_synthetase | BGC0001758 | NRP | 28.0 | 62.6 | 273.0 | 2.4e-72 |
| AVI26393.1 | nonribosomal\_peptide\_synthase | BGC0001800 | NRP + Polyketide | 28.0 | 66.5 | 273.0 | 2.4e-72 |
| AXN93602.1 | PuwF-G | BGC0001952 | NRP | 27.0 | 64.4 | 273.0 | 2.4e-72 |
| CAJ14039.1 | peptide\_synthetase | BGC0000406 | NRP | 29.0 | 56.6 | 272.0 | 3.1e-72 |
| ACA97576.1 | PmxA | BGC0000408 | NRP | 27.0 | 65.8 | 272.0 | 3.1e-72 |
| AGE11899.1 | nonribosomal\_peptide\_synthetase | BGC0000366 | NRP | 28.0 | 62.8 | 272.0 | 4.1e-72 |
| AAU39359.1 | lichenysin\_synthase\_LchAA | BGC0000381 | NRP | 26.0 | 63.5 | 272.0 | 4.1e-72 |
| BAC67535.1 | arthrofactin\_synthetase\_B | BGC0000305 | NRP:Lipopeptide | 28.0 | 58.0 | 272.0 | 5.3e-72 |
| AFH75322.1 | nonribosomal\_peptide\_synthetase | BGC0000425 | NRP:Cyclic depsipeptide | 28.0 | 67.3 | 272.0 | 5.3e-72 |
| AGS77307.1 | NRPS\_modules\_1-2 | BGC0001178 | NRP:Glycopeptide | 28.0 | 62.8 | 272.0 | 5.3e-72 |
| APZ78692.1 | nonribosomal\_peptide\_synthetase | BGC0001418 | NRP:Cyclic depsipeptide + Polyketide:Iterative type I | 29.0 | 66.5 | 272.0 | 5.3e-72 |
| AWI62626.1 | nonribosomal\_peptide\_synthetase | BGC0001943 | NRP | 27.0 | 69.8 | 272.0 | 5.3e-72 |
| AZF85940.1 | non-ribosomal\_peptide\_synthase | BGC0001963 | NRP + Polyketide | 28.0 | 62.8 | 272.0 | 5.3e-72 |
| QED55422.1 | nonribosomal\_peptide\_synthetase | BGC0001984 | NRP | 29.0 | 64.5 | 272.0 | 5.3e-72 |
| AEH41794.1 | HrmP | BGC0000374 | NRP:Cyclic depsipeptide | 26.0 | 64.9 | 271.0 | 9e-72 |
| AEA30273.1 | peptide\_synthetase | BGC0000429 | Polyketide + NRP:Cyclic depsipeptide | 27.0 | 66.0 | 271.0 | 9e-72 |
| CBL93730.1 | NRPS | BGC0000360 | NRP | 30.0 | 59.2 | 270.0 | 1.2e-71 |
| APZ78846.1 | nonribosomal\_peptide\_synthetase | BGC0001431 | NRP:Cyclic depsipeptide + Polyketide:Iterative type I | 28.0 | 67.3 | 270.0 | 1.2e-71 |
| AXN93614.1 | PuwF | BGC0001953 | NRP | 28.0 | 59.6 | 270.0 | 1.2e-71 |
| BAB69700.1 | iturin\_A\_synthetase\_C | BGC0001098 | NRP + Polyketide | 27.0 | 60.2 | 270.0 | 1.5e-71 |
| APZ78834.1 | nonribosomal\_peptide\_synthetase | BGC0001430 | NRP:Cyclic depsipeptide + Polyketide:Iterative type I | 29.0 | 65.6 | 270.0 | 1.5e-71 |
| AXN93592.1 | PuwA | BGC0001952 | NRP | 28.0 | 59.5 | 270.0 | 2e-71 |
| AXN93601.1 | PuwE | BGC0001952 | NRP | 27.0 | 61.2 | 270.0 | 2e-71 |
| APZ78856.1 | nonribosomal\_peptide\_synthetase | BGC0001432 | NRP:Cyclic depsipeptide + Polyketide:Iterative type I | 28.0 | 69.4 | 269.0 | 2.6e-71 |
| AAY91419.3 | non-ribosomal\_peptide\_synthetase\_OfaA | BGC0000399 | NRP:Cyclic depsipeptide | 28.0 | 63.0 | 269.0 | 3.4e-71 |
| BBA20967.1 | nonribosomal\_peptide\_synthetase | BGC0001763 | NRP + Polyketide | 28.0 | 63.6 | 269.0 | 3.4e-71 |
| BAH43766.1 | tyrocidine\_synthetase\_III | BGC0000452 | NRP | 25.0 | 67.9 | 269.0 | 4.5e-71 |
| AAS98786.1 | nonribosomal\_peptide\_synthetase | BGC0001001 | NRP + Polyketide | 29.0 | 56.8 | 269.0 | 4.5e-71 |
| CAG29032.1 | nonribosomal\_peptide\_synthetase\_(modules\_3\_to\_6) | BGC0001023 | NRP + Polyketide:Modular type I | 31.0 | 51.2 | 269.0 | 4.5e-71 |
| ACM79810.1 | ZmaO | BGC0001059 | NRP + Polyketide | 29.0 | 49.2 | 269.0 | 4.5e-71 |
| AQM37584.1 | nonribosomal\_peptide\_synthetase | BGC0001424 | NRP:Cyclic depsipeptide + Polyketide:Iterative type I | 28.0 | 68.0 | 269.0 | 4.5e-71 |
| APZ78822.1 | nonribosomal\_peptide\_synthetase | BGC0001429 | NRP:Cyclic depsipeptide + Polyketide:Iterative type I | 31.0 | 51.2 | 269.0 | 4.5e-71 |
| ASX95241.1 | IlaS | BGC0001620 | Polyketide | 28.0 | 63.8 | 269.0 | 4.5e-71 |
| WP\_039806852.1 | non-ribosomal\_peptide\_synthetase | BGC0002001 | NRP + Polyketide | 29.0 | 56.3 | 269.0 | 4.5e-71 |
| AGM14933.1 | xantholysin\_synthetase\_B | BGC0000463 | NRP:Lipopeptide | 28.0 | 57.4 | 268.0 | 5.8e-71 |
| WP\_018540607.1 | non-ribosomal\_peptide\_synthetase | BGC0001332 | NRP + Polyketide | 29.0 | 63.2 | 268.0 | 5.8e-71 |
| WP\_050383094.1 | non-ribosomal\_peptide\_synthetase | BGC0001451 | NRP | 32.0 | 42.2 | 268.0 | 5.8e-71 |
| AEC14346.1 | nonribosomal\_peptide\_synthetase | BGC0000377 | NRP | 26.0 | 68.5 | 268.0 | 7.6e-71 |
| OLZ52457.1 | non-ribosomal\_peptide\_synthetase | BGC0001462 | NRP:Glycopeptide | 27.0 | 62.3 | 268.0 | 7.6e-71 |
| ctg1\_orf20 |  | BGC0001767 | NRP | 29.0 | 63.8 | 268.0 | 7.6e-71 |
| WP\_044618979.1 | non-ribosomal\_peptide\_synthetase | BGC0001791 | NRP | 26.0 | 65.1 | 268.0 | 7.6e-71 |
| CAD55498.1 | CDA\_peptide\_synthetase\_III\_(CdaPs3) | BGC0000315 | NRP:Ca+-dependent lipopeptide | 28.0 | 60.6 | 267.0 | 1e-70 |
| CAE02630.1 | surfactin\_synthetase\_A | BGC0000433 | NRP:Lipopeptide | 25.0 | 66.4 | 267.0 | 1e-70 |
| AGZ15459.1 | putative\_non-ribosomal\_peptide\_synthetase | BGC0001036 | NRP + Polyketide | 28.0 | 56.8 | 267.0 | 1e-70 |
| AEO14744.1 | NdaB | BGC0000396 | NRP | 28.0 | 58.7 | 267.0 | 1.3e-70 |
| ATU31794.1 | NRPS | BGC0001814 | NRP | 28.0 | 58.3 | 267.0 | 1.3e-70 |
| AFJ23825.1 | WLIP\_synthetase\_B | BGC0001838 | NRP | 27.0 | 65.6 | 267.0 | 1.3e-70 |
| QDF82255.1 | non-ribosomal\_peptide\_synthetase | BGC0001980 | NRP | 28.0 | 58.1 | 267.0 | 1.3e-70 |
| AAF08797.1 | MycC | BGC0001103 | NRP + Polyketide | 28.0 | 57.6 | 267.0 | 1.7e-70 |
| WP\_041826607.1 | non-ribosomal\_peptide\_synthetase | BGC0001792 | NRP | 28.0 | 58.8 | 267.0 | 1.7e-70 |
| AWI62627.1 | nonribosomal\_peptide\_synthetase | BGC0001943 | NRP | 28.0 | 65.9 | 267.0 | 1.7e-70 |
| AAO56329.1 | non-ribosomal\_peptide\_synthetase\_SyfB | BGC0000435 | NRP | 28.0 | 57.1 | 266.0 | 2.2e-70 |
| CAJ46692.1 | non-ribosomal\_peptide\_synthase | BGC0000969 | NRP:Cyclic depsipeptide + Polyketide:Modular type I | 28.0 | 59.1 | 266.0 | 2.2e-70 |
| APZ78680.1 | nonribosomal\_peptide\_synthetase | BGC0001417 | NRP:Cyclic depsipeptide + Polyketide:Iterative type I | 29.0 | 62.2 | 266.0 | 2.2e-70 |
| OTA20325.1 | peptide\_synthase | BGC0001824 | NRP | 27.0 | 65.4 | 266.0 | 2.2e-70 |
| AAC06346.1 | bacitracin\_synthetase\_1 | BGC0000310 | NRP | 25.0 | 63.6 | 266.0 | 2.9e-70 |
| CAM56770.1 |  | BGC0000354 | NRP | 28.0 | 64.6 | 266.0 | 2.9e-70 |
| AAG02355.1 | peptide\_synthetase\_NRPS9-8 | BGC0000963 | NRP:Glycopeptide + Polyketide:Modular type I + Saccharide:Hybrid/tailoring | 30.0 | 55.2 | 266.0 | 2.9e-70 |
| APZ78756.1 | nonribosomal\_peptide\_synthetase | BGC0001423 | NRP:Cyclic depsipeptide + Polyketide:Iterative type I | 28.0 | 64.0 | 266.0 | 2.9e-70 |
| WP\_050383088.1 | non-ribosomal\_peptide\_synthetase | BGC0001451 | NRP | 32.0 | 42.2 | 266.0 | 2.9e-70 |
| AXN93605.1 | PuwA | BGC0001953 | NRP | 28.0 | 59.5 | 266.0 | 2.9e-70 |
| CAJ77696.1 | MPS2\_protein | BGC0000363 | NRP | 30.0 | 48.5 | 265.0 | 3.8e-70 |
| AAT12283.1 | LtxA | BGC0000384 | NRP | 28.0 | 59.2 | 265.0 | 3.8e-70 |
| AEP18656.1 | WAPS1 | BGC0000461 | NRP | 29.0 | 63.9 | 265.0 | 3.8e-70 |
| AGM14934.1 | xantholysin\_synthetase\_C | BGC0000463 | NRP:Lipopeptide | 27.0 | 66.8 | 265.0 | 3.8e-70 |
| CDG17980.1 | Putative\_Ornithine\_racemase\_(fragment) | BGC0000464 | NRP:Cyclic depsipeptide | 28.0 | 60.4 | 265.0 | 3.8e-70 |
| ALV82384.1 | CDA\_peptide\_synthetase\_II | BGC0001370 | NRP | 28.0 | 64.3 | 265.0 | 3.8e-70 |
| APZ78744.1 | nonribosomal\_peptide\_synthetase | BGC0001422 | NRP:Cyclic depsipeptide + Polyketide:Iterative type I | 28.0 | 64.5 | 265.0 | 3.8e-70 |
| APU91751.1 | Non-Ribosomal\_Peptide\_Synthetase | BGC0001806 | NRP | 27.0 | 57.6 | 265.0 | 3.8e-70 |
| ACZ55945.1 | non-ribosomal\_peptide\_synthetase | BGC0000302 | NRP | 29.0 | 49.5 | 265.0 | 5e-70 |
| CAN89656.1 | putative\_non-ribosomal\_peptide\_synthetase | BGC0001070 | NRP + Polyketide:Modular type I + Polyketide:Trans-AT type I | 27.0 | 70.7 | 265.0 | 5e-70 |
| WP\_051206727.1 | non-ribosomal\_peptide\_synthetase | BGC0001467 | NRP:Cyclic depsipeptide + Polyketide:Modular type I | 29.0 | 48.9 | 265.0 | 5e-70 |
| AXN93575.1 | PuwA | BGC0001950 | NRP | 27.0 | 61.5 | 265.0 | 5e-70 |
| AXN93584.1 | PuwA | BGC0001951 | NRP | 27.0 | 61.5 | 265.0 | 5e-70 |
| AAC06348.1 | bacitracin\_synthetase\_3 | BGC0000310 | NRP | 25.0 | 68.5 | 265.0 | 6.5e-70 |
| AEG64698.1 | LpmD | BGC0000379 | NRP | 29.0 | 64.0 | 265.0 | 6.5e-70 |
| AHJ31215.1 | Long-chain-fatty-acid--CoA\_ligase | BGC0000430 | NRP + Polyketide:Modular type I | 28.0 | 62.0 | 265.0 | 6.5e-70 |
| BAX64247.1 | NRPS | BGC0001623 | NRP + Polyketide | 27.0 | 60.1 | 265.0 | 6.5e-70 |
| CAB38517.1 | CDA\_peptide\_synthetase\_II\_(CdaPs2) | BGC0000315 | NRP:Ca+-dependent lipopeptide | 27.0 | 65.1 | 264.0 | 8.4e-70 |
| ABI26077.1 | OciA | BGC0000331 | NRP | 27.0 | 61.2 | 264.0 | 8.4e-70 |
| CCA29203.1 | non-ribosomal\_peptide\_synthetase/polyketide\_synthase | BGC0000955 | NRP + Polyketide:Modular type I | 29.0 | 61.3 | 264.0 | 8.4e-70 |
| AEF33078.1 | dimodular\_nonribosomal\_peptide\_synthetase | BGC0001039 | NRP + Polyketide | 27.0 | 58.5 | 264.0 | 8.4e-70 |
| AIW82282.1 | PuwE | BGC0001125 | NRP + Polyketide | 29.0 | 57.8 | 264.0 | 8.4e-70 |
| AEH59099.1 | amino\_acid\_adenylation\_domain-containing\_protein/NRPS | BGC0000385 | NRP | 27.0 | 64.9 | 264.0 | 1.1e-69 |
| AEW31021.1 | plipastatin\_synthetase | BGC0000407 | NRP | 28.0 | 65.5 | 264.0 | 1.1e-69 |
| AHH53508.1 | non-ribosomal\_peptide\_synthetase | BGC0000439 | NRP:Ca+-dependent lipopeptide | 28.0 | 63.7 | 264.0 | 1.1e-69 |
| ABS74208.1 | fengycin\_synthetase\_B | BGC0001095 | NRP | 25.0 | 68.0 | 264.0 | 1.1e-69 |
| ABL74936.1 | NRPS | BGC0001048 | NRP:Glycopeptide + Polyketide:Modular type I + Saccharide:Hybrid/tailoring | 29.0 | 57.5 | 264.0 | 1.4e-69 |
| AGS77309.1 | NRPS\_modules\_4-6 | BGC0001178 | NRP:Glycopeptide | 28.0 | 63.4 | 264.0 | 1.4e-69 |
| CAD91212.1 | putative\_non-ribosomal\_peptide\_synthetase,\_modules\_4-6 | BGC0000289 | NRP:Glycopeptide + Saccharide:Hybrid/tailoring | 27.0 | 65.0 | 263.0 | 1.9e-69 |
| ABF87031.1 | non-ribosomal\_peptide\_synthetase/polyketide\_synthase | BGC0000393 | NRP + Polyketide:Modular type I | 29.0 | 58.4 | 263.0 | 1.9e-69 |
| ARU08073.1 | mlcK | BGC0001448 | NRP:Ca+-dependent lipopeptide | 26.0 | 85.1 | 263.0 | 2.5e-69 |
| AGA37269.1 | NRPS | BGC0000819 | NRP + Alkaloid | 26.0 | 55.3 | 262.0 | 3.2e-69 |
| AHZ34243.1 | CipF | BGC0001389 | NRP | 26.0 | 69.0 | 262.0 | 3.2e-69 |
| AQH32486.1 | peptide\_synthetase | BGC0001667 | NRP + Polyketide | 28.0 | 57.8 | 262.0 | 3.2e-69 |
| AIW82277.1 | PuwA | BGC0001125 | NRP + Polyketide | 27.0 | 61.3 | 262.0 | 4.2e-69 |
| WP\_084702182.1 | non-ribosomal\_peptide\_synthetase | BGC0001211 | NRP | 29.0 | 62.3 | 262.0 | 4.2e-69 |
| CAG23960.2 | hybrid\_NRPS/PKS\_protein | BGC0001089 | Polyketide + NRP | 27.0 | 59.3 | 262.0 | 5.5e-69 |
| QDF82259.1 | non-ribosomal\_peptide\_synthetase | BGC0001980 | NRP | 27.0 | 58.1 | 262.0 | 5.5e-69 |
| ABD65957.1 | nonribosomal\_peptide\_synthetase | BGC0000341 | NRP | 27.0 | 63.1 | 261.0 | 7.2e-69 |
| AQW44894.1 | non-ribosomal\_peptide\_synthetase | BGC0001737 | NRP + Polyketide | 28.0 | 60.6 | 261.0 | 7.2e-69 |
| AFR69334.1 | nonribosomal\_peptide\_synthetase\_SpiDE1 | BGC0001045 | NRP:Cyclic depsipeptide + Polyketide:Modular type I | 26.0 | 65.9 | 261.0 | 9.3e-69 |
| WP\_013428774.1 | non-ribosomal\_peptide\_synthetase | BGC0001844 | NRP:Lipopeptide | 27.0 | 65.4 | 261.0 | 9.3e-69 |
| QBG38784.1 | Atr23 | BGC0001975 | NRP | 27.0 | 65.9 | 261.0 | 9.3e-69 |
| AGM16414.1 | paenibacterin\_synthetase\_C | BGC0000400 | NRP | 27.0 | 59.9 | 260.0 | 1.2e-68 |
| CCM44336.1 | Nonribosomal\_peptide\_synthetase | BGC0001056 | NRP + Polyketide:Modular type I + Polyketide:PUFA synthase or related | 27.0 | 59.4 | 260.0 | 1.2e-68 |
| ATY37592.1 | BogE | BGC0001532 | NRP | 25.0 | 65.4 | 260.0 | 1.6e-68 |
| ADJ63842.1 | Serobactin\_synthetase | BGC0000424 | NRP:NRP siderophore | 29.0 | 57.6 | 259.0 | 2.7e-68 |
| CAC48361.1 | peptide\_synthetase | BGC0000311 | NRP | 27.0 | 62.8 | 259.0 | 3.5e-68 |
| CAQ71829.1 | non\_ribosomal\_peptide\_synthase,\_antibiotic\_synthesis;\_contains\_4\_condensation\_domains,\_3\_AMP-acid\_ligases\_II\_domains,\_3\_PP-binding,\_Phosphopantetheine\_attachment\_site\_and\_a\_putative\_thioesterase\_domain | BGC0001189 | NRP | 30.0 | 57.7 | 259.0 | 3.5e-68 |
| QED55423.1 | nonribosomal\_peptide\_synthetase | BGC0001984 | NRP | 30.0 | 45.1 | 259.0 | 3.5e-68 |
| WP\_010369430.1 | non-ribosomal\_peptide\_synthetase | BGC0000314 | Polyketide + NRP:Cyclic depsipeptide + Other:Aminocoumarin | 26.0 | 65.1 | 258.0 | 6.1e-68 |
| AID65225.1 | nonribosomal\_peptide\_synthetase | BGC0000335 | NRP | 26.0 | 62.4 | 258.0 | 6.1e-68 |
| ABS74207.1 | fengycin\_synthetase\_C | BGC0001095 | NRP | 27.0 | 66.9 | 258.0 | 6.1e-68 |
| CBJ90358.1 | putative\_Peptide\_synthetase | BGC0000465 | NRP | 30.0 | 46.3 | 258.0 | 7.9e-68 |
| CAD17793.1 | probable\_non\_ribosomal\_peptide\_synthetase\_protein | BGC0001754 | NRP + Polyketide | 27.0 | 64.5 | 258.0 | 7.9e-68 |
| WP\_013184461.1 | non-ribosomal\_peptide\_synthetase | BGC0001716 | NRP | 26.0 | 66.7 | 258.0 | 7.9e-68 |
| QED88054.1 | nonribosomal\_peptide\_synthetase | BGC0001967 | NRP | 26.0 | 69.6 | 258.0 | 7.9e-68 |
| AAU34202.1 | mannopeptimycin\_peptide\_synthetase\_MppA | BGC0000388 | NRP | 28.0 | 58.7 | 257.0 | 1e-67 |
| AXN93580.1 | PuwE | BGC0001950 | NRP | 28.0 | 58.9 | 257.0 | 1e-67 |
| AXN93589.1 | PuwE | BGC0001951 | NRP | 28.0 | 58.9 | 257.0 | 1e-67 |
| EFL06865.1 | hypothetical\_protein | BGC0000300 | NRP | 28.0 | 63.3 | 257.0 | 1.3e-67 |
| EWS95124.1 | hypothetical\_protein | BGC0000306 | NRP:Lipopeptide | 27.0 | 60.4 | 257.0 | 1.3e-67 |
| AAK81826.1 | peptide\_synthetase | BGC0000326 | NRP | 27.0 | 62.6 | 257.0 | 1.3e-67 |
| WP\_064118559.1 | non-ribosomal\_peptide\_synthetase | BGC0001509 | NRP | 27.0 | 57.6 | 257.0 | 1.8e-67 |
| KPN93065.1 | NunD | BGC0001416 | NRP | 27.0 | 65.1 | 256.0 | 2.3e-67 |
| AAZ55898.1 | amino\_acid\_adenylation | BGC0000359 | NRP | 28.0 | 58.2 | 256.0 | 3e-67 |
| AHI59108.1 | locillomycin\_synthase\_A | BGC0001005 | NRP + Polyketide | 24.0 | 69.1 | 256.0 | 3e-67 |
| OKA09423.1 | non-ribosomal\_peptide\_synthetase | BGC0001459 | NRP:Glycopeptide | 29.0 | 57.6 | 256.0 | 3e-67 |
| BAD55612.1 | non-ribosomal\_peptide\_synthetase | BGC0001027 | NRP + Polyketide | 29.0 | 63.2 | 255.0 | 3.9e-67 |
| AEZ51520.1 | pmxE | BGC0001153 | NRP:Lipopeptide | 26.0 | 67.6 | 255.0 | 3.9e-67 |
| WP\_019032755.1 | non-ribosomal\_peptide\_synthetase | BGC0001331 | NRP:Cyclic depsipeptide + Polyketide:Modular type I | 29.0 | 60.1 | 255.0 | 3.9e-67 |
| AGC45618.1 | non-ribosomal\_peptide\_synthetase | BGC0001394 | NRP + Polyketide | 28.0 | 58.3 | 255.0 | 3.9e-67 |
| KFL51883.1 | amino\_acid\_adenylation\_protein | BGC0001711 | NRP + Polyketide | 28.0 | 68.2 | 255.0 | 3.9e-67 |
| KPN90376.1 | NupC | BGC0001416 | NRP | 26.0 | 67.4 | 255.0 | 5.1e-67 |
| BAV56270.1 |  | BGC0001657 | NRP | 29.0 | 64.9 | 255.0 | 5.1e-67 |
| AAL15600.1 | SimH | BGC0000270 | Polyketide | 26.0 | 56.9 | 255.0 | 6.7e-67 |
| KFH48607.1 | N-like\_protein | BGC0000317 | NRP | 27.0 | 64.0 | 255.0 | 6.7e-67 |
| AEG64697.1 | LpmC | BGC0000379 | NRP | 28.0 | 64.1 | 255.0 | 6.7e-67 |
| AAG34184.1 | SimH | BGC0000835 | Polyketide | 26.0 | 56.9 | 255.0 | 6.7e-67 |
| WP\_039806856.1 | non-ribosomal\_peptide\_synthetase | BGC0002001 | NRP + Polyketide | 27.0 | 59.3 | 255.0 | 6.7e-67 |
| AAC06347.1 | bacitracin\_synthetase\_2 | BGC0000310 | NRP | 25.0 | 64.5 | 254.0 | 8.7e-67 |
| ALV86867.1 | Tlo21 | BGC0001406 | NRP | 27.0 | 63.6 | 254.0 | 8.7e-67 |
| WP\_015507663.1 | non-ribosomal\_peptide\_synthase | BGC0001792 | NRP | 29.0 | 63.4 | 254.0 | 8.7e-67 |
| EFL06867.1 | predicted\_protein | BGC0000300 | NRP | 29.0 | 62.3 | 254.0 | 1.1e-66 |
| AHB82069.1 | non\_ribosomal\_peptide\_synthetase | BGC0001231 | NRP + Polyketide:Modular type I | 28.0 | 58.8 | 254.0 | 1.1e-66 |
| CDF96614.1 | NRPS | BGC0001149 | NRP:Lipopeptide + Saccharide:Hybrid/tailoring | 30.0 | 50.1 | 253.0 | 1.5e-66 |
| ctg1\_orf1265 |  | BGC0001752 | NRP | 28.0 | 63.3 | 253.0 | 1.5e-66 |
| AWI62628.1 | nonribosomal\_peptide\_synthetase | BGC0001943 | NRP | 28.0 | 57.9 | 253.0 | 1.5e-66 |
| AJW76711.1 | DsaI | BGC0001196 | NRP | 27.0 | 63.4 | 253.0 | 1.9e-66 |
| KFL51885.1 | amino\_acid\_adenylation\_protein | BGC0001711 | NRP + Polyketide | 30.0 | 47.0 | 253.0 | 1.9e-66 |
| CAK15814.1 | putative\_non-ribosomal\_peptide\_synthetase,\_terminal\_component | BGC0000344 | NRP | 28.0 | 58.0 | 253.0 | 2.5e-66 |
| ADA69239.2 | trans-AT\_hybrid\_polyketide\_synthase-NRPS | BGC0001071 | NRP + Polyketide:Modular type I + Polyketide:Trans-AT type I | 28.0 | 58.0 | 253.0 | 2.5e-66 |
| AEG64696.1 | LpmB | BGC0000379 | NRP | 29.0 | 49.0 | 252.0 | 3.3e-66 |
| AAK06804.1 | Tyroxyl-AMP-forming\_enzyme | BGC0001072 | Saccharide + Polyketide:Modular type I + Polyketide:Type II + Other:Aminocoumarin | 26.0 | 56.9 | 252.0 | 3.3e-66 |
| AJM89738.1 | PmxE | BGC0001192 | NRP | 26.0 | 67.5 | 252.0 | 3.3e-66 |
| WP\_144411596.1 | amino\_acid\_adenylation\_domain-containing\_protein | BGC0002001 | NRP + Polyketide | 28.0 | 57.9 | 252.0 | 3.3e-66 |
| AAY93355.1 | non-ribosomal\_peptide\_synthetase\_PvdJ | BGC0000413 | NRP | 29.0 | 58.5 | 252.0 | 4.3e-66 |
| AAF15891.2 | nosA | BGC0001028 | Polyketide + NRP:Cyclic depsipeptide | 27.0 | 65.8 | 252.0 | 4.3e-66 |
| AAS47564.1 | mixed\_type\_I\_polyketide\_synthase/nonribosomal\_peptide\_synthetase | BGC0001108 | Polyketide:Trans-AT type I | 27.0 | 59.1 | 252.0 | 4.3e-66 |
| ctg1\_orf6 |  | BGC0001109 | NRP + Polyketide | 27.0 | 59.1 | 252.0 | 4.3e-66 |
| BAB69699.1 | iturin\_A\_synthetase\_B | BGC0001098 | NRP + Polyketide | 26.0 | 64.6 | 252.0 | 5.7e-66 |
| ARF06222.1 | non-ribosomal\_peptide\_synthetase | BGC0001593 | NRP | 28.0 | 59.8 | 252.0 | 5.7e-66 |
| WP\_047197779.1 | non-ribosomal\_peptide\_synthetase | BGC0002003 | NRP | 29.0 | 58.9 | 252.0 | 5.7e-66 |
| AYA44686.1 | icosalide\_NRPS | BGC0001833 | NRP:Lipopeptide | 30.0 | 43.5 | 252.0 | 5.7e-66 |
| ctg1\_orf1264 |  | BGC0001752 | NRP | 29.0 | 65.2 | 251.0 | 7.4e-66 |
| AAO62588.1 | peptide\_sythetase | BGC0001016 | NRP + Polyketide | 27.0 | 57.2 | 251.0 | 9.7e-66 |
| CZT62784.1 | Non-ribosomal\_peptide\_synthase,\_involved\_in\_Hassallidin\_biosynthesis | BGC0001614 | NRP | 27.0 | 57.4 | 251.0 | 9.7e-66 |
| ATY37589.1 | BogB | BGC0001532 | NRP | 25.0 | 67.2 | 250.0 | 1.6e-65 |
| BAX90000.1 | Non-ribosomal\_peptide\_synthetase | BGC0001628 | NRP | 28.0 | 64.9 | 250.0 | 1.6e-65 |
| QBC75022.1 | non-ribosomal\_peptide\_synthetase | BGC0001968 | NRP | 26.0 | 65.6 | 250.0 | 1.6e-65 |
| AJF34463.1 | Txo1 | BGC0001207 | NRP | 29.0 | 58.2 | 250.0 | 2.2e-65 |
| BAP05596.1 | calH | BGC0000967 | NRP + Polyketide:Trans-AT type I | 28.0 | 58.1 | 249.0 | 2.8e-65 |
| CBJ79915.1 | putative\_Phenylalanine\_racemase\_(ATP-hydrolyzing) | BGC0001133 | NRP | 26.0 | 64.8 | 249.0 | 3.7e-65 |
| ABW17376.1 | PsoB | BGC0000411 | NRP | 28.0 | 57.6 | 248.0 | 4.8e-65 |
| ABW17377.1 | PsoC | BGC0000411 | NRP | 28.0 | 58.3 | 248.0 | 4.8e-65 |
| CDG17987.1 | Putative\_Ornithine\_racemase\_(fragment) | BGC0000464 | NRP:Cyclic depsipeptide | 27.0 | 60.0 | 248.0 | 6.3e-65 |
| WP\_064118560.1 | non-ribosomal\_peptide\_synthetase | BGC0001509 | NRP | 28.0 | 59.1 | 248.0 | 6.3e-65 |
| AFJ23826.1 | WLIP\_synthetase\_C | BGC0001838 | NRP | 29.0 | 60.2 | 248.0 | 6.3e-65 |
| AHH53507.1 | non-ribosomal\_peptide\_synthetase | BGC0000439 | NRP:Ca+-dependent lipopeptide | 26.0 | 67.1 | 248.0 | 8.2e-65 |
| BAH22762.1 | nonribosomal\_peptide\_synthetase | BGC0001018 | NRP + Polyketide | 27.0 | 59.7 | 248.0 | 8.2e-65 |
| WP\_054234643.1 | non-ribosomal\_peptide\_synthetase | BGC0002014 | NRP + Polyketide | 27.0 | 66.8 | 248.0 | 8.2e-65 |
| CBJ90287.1 | peptide\_synthetase | BGC0000416 | NRP | 27.0 | 48.6 | 247.0 | 1.1e-64 |
| PHM26612.1 | pvdj | BGC0001130 | NRP + Polyketide | 25.0 | 70.3 | 247.0 | 1.1e-64 |
| ALG65318.1 | Cal18 | BGC0001297 | NRP | 26.0 | 65.1 | 247.0 | 1.1e-64 |
| WP\_013184317.1 | non-ribosomal\_peptide\_synthetase | BGC0001692 | NRP | 27.0 | 48.6 | 247.0 | 1.1e-64 |
| AGD80618.1 | non-ribosomal\_peptide\_synthetase | BGC0000394 | NRP | 28.0 | 61.6 | 247.0 | 1.4e-64 |
| ACA97577.1 | PmxB | BGC0000408 | NRP | 28.0 | 58.3 | 247.0 | 1.4e-64 |
| AIG79242.1 | Non-ribosomal\_peptide\_synthetase | BGC0000419 | Saccharide + NRP:Glycopeptide | 27.0 | 58.9 | 247.0 | 1.4e-64 |
| QDA77045.1 | polyketide\_synthase/nonribosomal\_peptide\_synthetase | BGC0002025 | NRP | 30.0 | 51.6 | 247.0 | 1.4e-64 |
| AHD05678.1 | nonribosomal\_peptide\_ligase\_subunit | BGC0000402 | NRP | 26.0 | 68.6 | 247.0 | 1.8e-64 |
| CBJ90289.1 | peptide\_synthetase | BGC0000416 | NRP | 29.0 | 46.5 | 247.0 | 1.8e-64 |
| ARU08074.1 | mlcL | BGC0001448 | NRP:Ca+-dependent lipopeptide | 28.0 | 64.1 | 247.0 | 1.8e-64 |
| WP\_013184319.1 | non-ribosomal\_peptide\_synthetase | BGC0001692 | NRP | 29.0 | 46.5 | 247.0 | 1.8e-64 |
| ABG94125.1 | non-ribosomal\_peptide\_synthetase | BGC0000417 | NRP | 27.0 | 58.0 | 246.0 | 2.4e-64 |
| WP\_053065269.1 | non-ribosomal\_peptide\_synthetase | BGC0001330 | NRP:Cyclic depsipeptide + Polyketide:Modular type I | 28.0 | 62.9 | 246.0 | 3.1e-64 |
| ACM68690.1 | AerG1 | BGC0000298 | NRP | 27.0 | 57.3 | 245.0 | 4.1e-64 |
| ABS75232.1 | DhbF | BGC0001185 | NRP:NRP siderophore | 28.0 | 48.9 | 245.0 | 4.1e-64 |
| BAH43869.1 | linear\_pentadecapeptide\_gramicidin\_synthetase\_LgrA | BGC0000367 | NRP | 28.0 | 46.8 | 245.0 | 5.3e-64 |
| ABP57749.1 | DepE | BGC0000993 | NRP:Cyclic depsipeptide + Polyketide:Modular type I | 27.0 | 64.5 | 245.0 | 5.3e-64 |
| AHZ34241.1 | CipD | BGC0001389 | NRP | 28.0 | 57.8 | 245.0 | 5.3e-64 |
| YP\_856988.1 | nonribosomal\_peptide\_synthetase | BGC0001502 | NRP | 28.0 | 62.3 | 245.0 | 5.3e-64 |
| ANZ15839.1 | peptide\_synthetase\_ScpsB | BGC0001569 | NRP | 27.0 | 73.0 | 245.0 | 5.3e-64 |
| AHH53506.1 | non-ribosomal\_peptide\_synthetase | BGC0000439 | NRP:Ca+-dependent lipopeptide | 30.0 | 47.5 | 245.0 | 6.9e-64 |
| CAK15815.1 | putative\_non\_ribosomal\_peptide\_synthetase | BGC0000344 | NRP | 28.0 | 58.2 | 244.0 | 9e-64 |
| ACY06285.1 | non-ribosomal\_peptide\_synthetase | BGC0001042 | NRP + Polyketide | 26.0 | 64.3 | 244.0 | 9e-64 |
| ABV56585.1 | KtzE | BGC0000378 | NRP | 30.0 | 47.2 | 244.0 | 1.2e-63 |
| AJY78094.1 | nonribosomal\_peptide\_synthetase | BGC0001902 | NRP + Polyketide | 27.0 | 59.1 | 244.0 | 1.2e-63 |
| QED55421.1 | nonribosomal\_peptide\_synthetase | BGC0001984 | NRP | 27.0 | 63.6 | 244.0 | 1.2e-63 |
| CAQ34921.1 | nonribosomal\_peptide\_synthetase | BGC0000986 | NRP + Polyketide | 29.0 | 57.8 | 243.0 | 1.5e-63 |
| ANG60380.1 | nonribosomal\_peptide\_synthetase\_BudB | BGC0001698 | NRP | 29.0 | 46.7 | 243.0 | 1.5e-63 |
| ATL73036.1 | amino\_acid\_adenylation\_protein | BGC0001807 | NRP + Polyketide | 25.0 | 65.3 | 243.0 | 2e-63 |
| ATU31795.1 | NRPS | BGC0001814 | NRP | 28.0 | 66.0 | 243.0 | 2e-63 |
| AGC09528.1 | NRPS | BGC0001183 | Polyketide | 25.0 | 65.6 | 243.0 | 2.6e-63 |
| KPN93064.1 | NupB | BGC0001416 | NRP | 27.0 | 58.3 | 243.0 | 2.6e-63 |
| ABP57748.1 | DepD | BGC0000993 | NRP:Cyclic depsipeptide + Polyketide:Modular type I | 27.0 | 63.9 | 242.0 | 3.4e-63 |
| AHD05627.1 | putative\_non-ribosomal\_peptide\_ligase\_domain\_protein | BGC0001033 | NRP + Polyketide | 28.0 | 47.1 | 242.0 | 3.4e-63 |
| ATD51278.1 | nonribosomal\_peptide\_synthase | BGC0001650 | NRP | 27.0 | 59.2 | 242.0 | 3.4e-63 |
| AID65224.1 | nonribosomal\_peptide\_synthetase | BGC0000335 | NRP | 25.0 | 61.2 | 241.0 | 7.7e-63 |
| ABS74209.1 | fengycin\_synthetase\_A | BGC0001095 | NRP | 26.0 | 66.2 | 241.0 | 7.7e-63 |
| AJM89734.1 | PmxB | BGC0001192 | NRP | 27.0 | 57.8 | 241.0 | 7.7e-63 |
| ABD65958.1 | nonribosomal\_peptide\_synthetase | BGC0000341 | NRP | 27.0 | 59.2 | 241.0 | 1e-62 |
| AET98906.1 | putative\_non-ribosomal\_peptide\_synthetase | BGC0000415 | NRP | 30.0 | 47.9 | 241.0 | 1e-62 |
| EME52974.1 | non-ribosomal\_peptide\_synthetase | BGC0001460 | NRP:Glycopeptide | 32.0 | 40.7 | 241.0 | 1e-62 |
| AAZ55900.1 | non-ribosomal\_peptide\_synthase:Amino\_acid\_adenylation | BGC0000359 | NRP | 31.0 | 48.0 | 240.0 | 1.7e-62 |
| ABF85931.1 | non-ribosomal\_peptide\_synthase/polyketide\_synthase\_Ta1 | BGC0001025 | NRP + Polyketide:Trans-AT type I | 28.0 | 59.8 | 240.0 | 1.7e-62 |
| WP\_018540603.1 | non-ribosomal\_peptide\_synthetase | BGC0001332 | NRP + Polyketide | 28.0 | 57.6 | 240.0 | 1.7e-62 |
| ABX37383.1 | amino\_acid\_adenylation\_domain\_protein | BGC0000984 | NRP + Polyketide | 28.0 | 52.9 | 240.0 | 2.2e-62 |
| CCJ67646.1 | JagB | BGC0001127 | NRP | 26.0 | 62.6 | 239.0 | 2.9e-62 |
| AEW31019.1 | plipastatin\_synthetase | BGC0000407 | NRP | 27.0 | 64.8 | 238.0 | 5e-62 |
| AAK57184.1 | MxaA | BGC0001022 | NRP + Polyketide | 28.0 | 58.8 | 238.0 | 5e-62 |
| ALG65317.1 | Cal19 | BGC0001297 | NRP | 28.0 | 58.1 | 238.0 | 5e-62 |
| AIE77058.1 | peptide\_synthetase\_module\_3 | BGC0000418 | NRP | 27.0 | 58.9 | 238.0 | 6.5e-62 |
| ABV56588.1 | KtzH | BGC0000378 | NRP | 31.0 | 46.0 | 238.0 | 8.5e-62 |
| CBJ90359.1 | putative\_Peptide\_synthetase | BGC0000465 | NRP | 28.0 | 45.8 | 238.0 | 8.5e-62 |
| DAB41477.1 | nonribosomal\_peptide\_synthetase | BGC0001766 | NRP | 27.0 | 64.5 | 237.0 | 1.1e-61 |
| WP\_016638471.1 | non-ribosomal\_peptide\_synthetase | BGC0001519 | NRP + Polyketide | 28.0 | 58.3 | 237.0 | 1.1e-61 |
| WP\_013184462.1 | non-ribosomal\_peptide\_synthetase | BGC0001716 | NRP | 27.0 | 58.3 | 237.0 | 1.1e-61 |
| AEZ51517.1 | pmxB | BGC0001153 | NRP:Lipopeptide | 26.0 | 63.0 | 237.0 | 1.4e-61 |
| AYJ71721.1 | non-ribosomal\_peptide\_synthetase | BGC0001942 | NRP + Polyketide | 27.0 | 63.8 | 237.0 | 1.9e-61 |
| AEI58879.1 | peptide\_synthetase | BGC0000455 | NRP | 32.0 | 40.7 | 236.0 | 2.5e-61 |
| AAY32966.1 | DszC | BGC0001093 | NRP + Polyketide | 34.0 | 34.9 | 236.0 | 2.5e-61 |
| AIG79224.1 | Non-ribosomal\_peptide\_synthetase/andenylation\_domain | BGC0000419 | Saccharide + NRP:Glycopeptide | 31.0 | 40.8 | 236.0 | 3.2e-61 |
| WP\_078857611.1 | non-ribosomal\_peptide\_synthetase | BGC0001368 | NRP | 27.0 | 57.8 | 236.0 | 3.2e-61 |
| AKP45399.1 | CysK | BGC0001413 | NRP | 24.0 | 70.2 | 236.0 | 3.2e-61 |
| AKC91856.1 | nonribosomal\_peptide\_synthetase | BGC0001414 | NRP | 28.0 | 51.9 | 236.0 | 3.2e-61 |
| ADG27358.1 | peptide\_synthetase | BGC0000296 | NRP | 30.0 | 48.4 | 235.0 | 4.2e-61 |
| AAO62586.1 | peptide\_sythetase | BGC0001016 | NRP + Polyketide | 30.0 | 43.2 | 235.0 | 4.2e-61 |
| BAO84861.1 | putative\_non-ribosomal\_peptide\_synthetase | BGC0000414 | NRP | 28.0 | 56.5 | 235.0 | 5.5e-61 |
| CAL80824.1 | NRPS\_module\_protein | BGC0000997 | NRP + Polyketide | 28.0 | 60.8 | 235.0 | 5.5e-61 |
| OLZ52442.1 | non-ribosomal\_peptide\_synthetase | BGC0001462 | NRP:Glycopeptide | 32.0 | 40.9 | 235.0 | 5.5e-61 |
| OKA09664.1 | non-ribosomal\_peptide\_synthetase | BGC0001459 | NRP:Glycopeptide | 32.0 | 40.9 | 235.0 | 7.2e-61 |
| AIE77076.1 | peptide\_synthetase | BGC0000418 | NRP | 32.0 | 40.9 | 234.0 | 9.4e-61 |
| ABX37385.1 | amino\_acid\_adenylation\_domain\_protein | BGC0000984 | NRP + Polyketide | 27.0 | 58.2 | 234.0 | 9.4e-61 |
| BAW27693.1 | NRPS(C-A-T-TE) | BGC0001764 | NRP | 26.0 | 58.9 | 234.0 | 9.4e-61 |
| AYA22318.1 | KerE | BGC0001955 | Other | 31.0 | 40.8 | 234.0 | 9.4e-61 |
| AAZ23078.1 | peptide\_synthetase | BGC0000291 | NRP | 26.0 | 64.3 | 234.0 | 1.2e-60 |
| BAI63283.1 | putative\_non-ribosomal\_peptide\_synthetase | BGC0000434 | NRP | 31.0 | 40.1 | 233.0 | 1.6e-60 |
| BAH04162.1 | trsJ | BGC0000450 | NRP | 29.0 | 49.2 | 233.0 | 1.6e-60 |
| ATP76244.1 | NdaB | BGC0001705 | NRP + Polyketide | 27.0 | 57.8 | 233.0 | 1.6e-60 |
| AAC32047.1 | FxbB | BGC0000351 | NRP | 30.0 | 49.0 | 233.0 | 2.1e-60 |
| CCJ67648.1 | JagD | BGC0001127 | NRP | 26.0 | 64.7 | 233.0 | 2.1e-60 |
| AAZ23077.1 | peptide\_synthetase | BGC0000291 | NRP | 27.0 | 69.2 | 233.0 | 2.7e-60 |
| BAC67534.2 | arthrofactin\_synthetase\_A | BGC0000305 | NRP:Lipopeptide | 28.0 | 52.6 | 233.0 | 2.7e-60 |
| CAC48369.1 | peptide\_synthetase | BGC0000311 | NRP | 31.0 | 40.9 | 232.0 | 3.6e-60 |
| AHZ34239.1 | CipB | BGC0001389 | NRP | 27.0 | 57.6 | 232.0 | 3.6e-60 |
| OLZ50899.1 | non-ribosomal\_peptide\_synthetase | BGC0001461 | NRP:Glycopeptide | 30.0 | 40.8 | 232.0 | 3.6e-60 |
| AAN85512.1 | nonribosomal\_peptide\_synthetase | BGC0001101 | NRP + Polyketide:Modular type I + Polyketide:Trans-AT type I | 27.0 | 60.3 | 232.0 | 4.6e-60 |
| AVR48535.1 | CusC | BGC0001564 | NRP + Polyketide | 26.0 | 67.5 | 232.0 | 4.6e-60 |
| BAP05591.1 | calC | BGC0000967 | NRP + Polyketide:Trans-AT type I | 30.0 | 44.6 | 232.0 | 6.1e-60 |
| ARU08075.1 | mlcM | BGC0001448 | NRP:Ca+-dependent lipopeptide | 27.0 | 65.4 | 232.0 | 6.1e-60 |
| WP\_008704344.1 | non-ribosomal\_peptide\_synthetase | BGC0001575 | NRP | 30.0 | 41.7 | 231.0 | 1e-59 |
| CBZ42143.1 | putative\_non-ribosomal\_peptide\_synthetase | BGC0001117 | NRP | 29.0 | 49.5 | 230.0 | 1.4e-59 |
| ALV86868.1 | Tlo22 | BGC0001406 | NRP | 26.0 | 63.4 | 230.0 | 1.4e-59 |
| BAO98812.1 | putative\_nonribosomal\_peptide\_synthetase | BGC0001002 | NRP + Polyketide | 28.0 | 50.0 | 230.0 | 1.8e-59 |
| PHM26613.1 | pyoverdine\_synthetase\_D | BGC0001130 | NRP + Polyketide | 26.0 | 62.0 | 230.0 | 2.3e-59 |
| WP\_051462298.1 | non-ribosomal\_peptide\_synthetase | BGC0001873 | NRP:Lipopeptide | 27.0 | 69.0 | 229.0 | 3e-59 |
| CAJ88610.1 | putative\_non-ribosomal\_peptide\_synthetase | BGC0000327 | NRP | 29.0 | 53.7 | 229.0 | 3.9e-59 |
| BAP05590.1 | calB | BGC0000967 | NRP + Polyketide:Trans-AT type I | 26.0 | 58.0 | 229.0 | 3.9e-59 |
| CCA89328.1 | mixed\_trans-AT\_type\_I\_polyketide\_synthase/nonribosomal\_peptide\_synthetase | BGC0001111 | NRP + Polyketide:Trans-AT type I | 32.0 | 42.4 | 229.0 | 3.9e-59 |
| ATP76243.1 | NdaA | BGC0001705 | NRP + Polyketide | 31.0 | 42.7 | 229.0 | 3.9e-59 |
| QBC75021.1 | non-ribosomal\_peptide\_synthetase | BGC0001968 | NRP | 27.0 | 65.8 | 229.0 | 3.9e-59 |
| ABW17375.1 | PsoA | BGC0000411 | NRP | 26.0 | 54.6 | 228.0 | 5.1e-59 |
| CAJ18237.2 | non-ribosomal\_peptide\_synthetase\_B | BGC0000354 | NRP | 27.0 | 64.0 | 227.0 | 1.1e-58 |
| CBJ90288.1 | Peptide\_synthetase | BGC0000416 | NRP | 27.0 | 46.2 | 227.0 | 1.1e-58 |
| ALG65313.1 | Cal23 | BGC0001297 | NRP | 31.0 | 41.7 | 227.0 | 1.1e-58 |
| WP\_013184318.1 | non-ribosomal\_peptide\_synthetase | BGC0001692 | NRP | 27.0 | 46.2 | 227.0 | 1.1e-58 |
| AVI26390.1 | polyketide\_synthase\_/\_nonribosomal\_peptide\_synthase\_hybrid | BGC0001800 | NRP + Polyketide | 27.0 | 64.1 | 227.0 | 1.1e-58 |
| Xekj\_RS17945 | non-ribosomal\_peptide\_synthetase | BGC0001826 | NRP | 25.0 | 65.8 | 227.0 | 1.5e-58 |
| BAX64246.1 | NRPS | BGC0001623 | NRP + Polyketide | 26.0 | 65.3 | 227.0 | 2e-58 |
| BAI63288.1 | putative\_non-ribosomal\_peptide\_synthetase | BGC0000434 | NRP | 29.0 | 50.5 | 226.0 | 2.5e-58 |
| AHD05617.1 | putative\_non-ribosomal\_peptide\_ligase\_domain\_protein | BGC0001033 | NRP + Polyketide | 29.0 | 47.7 | 226.0 | 2.5e-58 |
| CZT62792.1 | non-ribosomal\_peptide\_synthase\_involved\_in\_Hassallidin\_biosynthesis | BGC0001614 | NRP | 30.0 | 39.6 | 226.0 | 2.5e-58 |
| ABL86391.1 | hybrid\_polyketide\_synthase\_and\_nonribosomal\_peptide\_synthetase | BGC0000999 | NRP + Polyketide | 26.0 | 59.8 | 226.0 | 3.3e-58 |
| ACG60773.1 | NRPS(C/A/PCP) | BGC0001058 | NRP:Glycopeptide + Polyketide:Modular type I + Saccharide:Hybrid/tailoring | 31.0 | 41.8 | 226.0 | 3.3e-58 |
| BAW32323.1 | hybrid\_cis-AT\_polyketide\_synthase\_-\_nonribosomal\_peptide\_synthetase | BGC0001630 | NRP + Polyketide | 26.0 | 64.7 | 226.0 | 3.3e-58 |
| AHZ20781.1 | non-ribosomal\_peptide\_synthase | BGC0000369 | NRP + Saccharide:Hybrid/tailoring | 25.0 | 60.4 | 225.0 | 4.3e-58 |
| ABL74937.1 | NRPS | BGC0001048 | NRP:Glycopeptide + Polyketide:Modular type I + Saccharide:Hybrid/tailoring | 31.0 | 45.0 | 225.0 | 4.3e-58 |
| AEW31015.1 | plipastatin\_synthetase | BGC0000407 | NRP | 25.0 | 61.1 | 225.0 | 5.7e-58 |
| ABD65956.1 | nonribosomal\_peptide\_synthetase | BGC0000341 | NRP | 27.0 | 57.6 | 225.0 | 7.4e-58 |
| AXM43052.1 | non-ribosomal\_peptide\_synthetase | BGC0001945 | NRP | 26.0 | 68.1 | 225.0 | 7.4e-58 |
| AAY89051.1 | hybrid\_nonribosomal\_peptide\_synthetase/polyketide\_synthase | BGC0001069 | NRP + Polyketide:Trans-AT type I | 30.0 | 43.3 | 224.0 | 9.7e-58 |
| CAJ14037.1 | peptide\_synthetase | BGC0000406 | NRP | 28.0 | 61.4 | 224.0 | 1.3e-57 |
| AJW65406.1 | nonribosomal\_peptide\_synthetase | BGC0001195 | NRP + Polyketide | 25.0 | 71.1 | 223.0 | 2.8e-57 |
| QDF82254.1 | non-ribosomal\_peptide\_synthetase | BGC0001980 | NRP | 28.0 | 52.6 | 222.0 | 3.7e-57 |
| AEO14743.1 | NdaA | BGC0000396 | NRP | 31.0 | 38.7 | 222.0 | 4.8e-57 |
| AYJ71720.1 | non-ribosomal\_peptide\_synthetase | BGC0001942 | NRP + Polyketide | 29.0 | 47.6 | 221.0 | 1.1e-56 |
| EDY47118.1 | N-(5-amino-5-carboxypentanoyl)-L-cysteinyl-D-\_valine\_synthase | BGC0000319 | NRP:Beta-lactam | 27.0 | 58.7 | 220.0 | 1.4e-56 |
| WP\_078857610.1 | non-ribosomal\_peptide\_synthetase | BGC0001368 | NRP | 26.0 | 59.9 | 220.0 | 1.4e-56 |
| ABI22131.1 | putative\_non-ribosomal\_peptide\_synthetase | BGC0000422 | NRP | 28.0 | 45.4 | 220.0 | 1.8e-56 |
| CAJ46691.1 | non-ribosomal\_peptide\_synthase | BGC0000969 | NRP:Cyclic depsipeptide + Polyketide:Modular type I | 26.0 | 59.3 | 220.0 | 1.8e-56 |
| CAN89663.1 | putative\_non-ribosomal\_peptide\_synthetase | BGC0001070 | NRP + Polyketide:Modular type I + Polyketide:Trans-AT type I | 27.0 | 58.5 | 220.0 | 2.4e-56 |
| AKP45395.1 | CysG | BGC0001413 | NRP | 25.0 | 61.1 | 219.0 | 3.1e-56 |
| BAE98162.1 | putative\_non-ribosomal\_peptide\_synthetase | BGC0000339 | NRP | 31.0 | 40.8 | 219.0 | 4.1e-56 |
| AQX14497.1 | monobactam\_NRPS\_scaffold\_4 | BGC0001672 | NRP | 26.0 | 59.3 | 219.0 | 4.1e-56 |
| ADH04679.1 | non-ribosomal\_peptide\_synthetase | BGC0001344 | NRP + Polyketide | 27.0 | 68.3 | 218.0 | 5.3e-56 |
| ATQ39428.1 | cyclosporin\_C\_synthetase | BGC0001565 | NRP | 26.0 | 58.9 | 218.0 | 5.3e-56 |
| DAC76734.1 | type\_I\_polyketide\_synthase/non-ribosomal\_peptide\_synthetase | BGC0001885 | Polyketide | 28.0 | 49.3 | 218.0 | 9.1e-56 |
| WP\_054234617.1 | non-ribosomal\_peptide\_synthase | BGC0002014 | NRP + Polyketide | 28.0 | 47.7 | 217.0 | 1.2e-55 |
| BAJ19066.1 | L-lysine\_activating\_non-ribosomal\_peptide\_synthetase | BGC0000288 | NRP | 27.0 | 57.7 | 217.0 | 1.5e-55 |
| AOZ21320.1 | SulM | BGC0001790 | NRP | 26.0 | 59.3 | 217.0 | 1.5e-55 |
| QED88055.1 | nonribosomal\_peptide\_synthetase | BGC0001967 | NRP | 27.0 | 59.4 | 217.0 | 1.5e-55 |
| CAJ45639.1 | vanchrobactin\_non\_ribosomal\_peptide\_synthetase | BGC0000454 | NRP | 27.0 | 49.6 | 217.0 | 2e-55 |
| ACU71638.1 | peptide\_synthetase-like\_protein | BGC0001154 | Other | 30.0 | 41.0 | 217.0 | 2e-55 |
| AHI59110.1 | locillomycin\_synthase\_C | BGC0001005 | NRP + Polyketide | 26.0 | 51.2 | 216.0 | 3.4e-55 |
| CBJ79916.1 | putative\_Ornithine\_racemase | BGC0001133 | NRP | 25.0 | 64.9 | 215.0 | 4.5e-55 |
| WP\_004571779.1 | non-ribosomal\_peptide\_synthetase | BGC0001760 | NRP | 31.0 | 42.7 | 215.0 | 4.5e-55 |
| QBC75023.1 | non-ribosomal\_peptide\_synthetase | BGC0001968 | NRP | 28.0 | 49.7 | 215.0 | 4.5e-55 |
| WP\_013184596.1 | non-ribosomal\_peptide\_synthetase | BGC0001825 | NRP | 25.0 | 64.9 | 215.0 | 7.7e-55 |
| BAO84868.1 | putative\_non-ribosomal\_peptide\_synthetase | BGC0000414 | NRP | 32.0 | 35.3 | 214.0 | 1e-54 |
| AWI62629.1 | nonribosomal\_peptide\_synthetase | BGC0001943 | NRP | 28.0 | 49.2 | 214.0 | 1e-54 |
| AHH25592.1 | NRPS | BGC0000957 | NRP + Polyketide | 30.0 | 42.3 | 213.0 | 1.7e-54 |
| AZH23822.1 | MgiJ | BGC0001971 | NRP + Polyketide | 28.0 | 47.3 | 213.0 | 1.7e-54 |
| ABC35522.1 | thiotemplate\_mechanism\_natural\_product\_synthetase | BGC0000186 | NRP + Polyketide:Modular type I | 25.0 | 58.1 | 213.0 | 2.2e-54 |
| ABA70582.1 | alpha-aminoadypil-cysteinyl-valine\_synthetase | BGC0000404 | NRP | 26.0 | 51.2 | 213.0 | 2.2e-54 |
| ABR12615.1 | ACV\_synthetase | BGC0000405 | NRP:Beta-lactam | 26.0 | 51.2 | 213.0 | 2.2e-54 |
| AZH23792.1 | MgcJ | BGC0001970 | NRP + Polyketide | 28.0 | 47.1 | 213.0 | 2.2e-54 |
| ABV56604.1 | adenylation\_domain\_protein | BGC0000378 | NRP | 31.0 | 41.1 | 212.0 | 3.8e-54 |
| AKA54626.1 | NRPS | BGC0001216 | NRP + Polyketide | 29.0 | 50.3 | 212.0 | 3.8e-54 |
| AEP18655.1 | WAPS2 | BGC0000461 | NRP | 28.0 | 47.7 | 212.0 | 6.5e-54 |
| CCC55921.1 | non-ribosomal\_peptide\_synthetase/polyketide\_synthase\_hybrid\_protein | BGC0000973 | NRP + Polyketide:Modular type I | 30.0 | 47.6 | 211.0 | 1.1e-53 |
| ADL64235.1 | aureusimine\_non-ribosomal\_peptide\_synthetase | BGC0000308 | NRP | 27.0 | 52.0 | 210.0 | 1.4e-53 |
| AHB82072.1 | non\_ribosomal\_peptide\_synthetase/polyketide\_synthase | BGC0001231 | NRP + Polyketide:Modular type I | 26.0 | 58.2 | 210.0 | 1.4e-53 |
| WP\_053065270.1 | non-ribosomal\_peptide\_synthetase | BGC0001330 | NRP:Cyclic depsipeptide + Polyketide:Modular type I | 29.0 | 47.5 | 210.0 | 1.4e-53 |
| AFD30953.1 | CrmB | BGC0000966 | NRP + Polyketide | 25.0 | 59.0 | 210.0 | 1.9e-53 |
| AAK81824.1 | peptide\_synthetase | BGC0000326 | NRP | 27.0 | 53.9 | 210.0 | 2.5e-53 |
| ALD82525.1 | non-ribosomal\_peptide\_synthase | BGC0001212 | NRP + Polyketide | 29.0 | 47.5 | 210.0 | 2.5e-53 |
| WP\_019032753.1 | non-ribosomal\_peptide\_synthetase | BGC0001331 | NRP:Cyclic depsipeptide + Polyketide:Modular type I | 28.0 | 48.4 | 210.0 | 2.5e-53 |
| KPN90369.1 | NunE | BGC0001416 | NRP | 25.0 | 59.3 | 210.0 | 2.5e-53 |
| CRI73800.1 | loading\_module\_of\_NRPS-PKS | BGC0001215 | NRP | 31.0 | 35.5 | 209.0 | 4.2e-53 |
| AQX14499.1 | monobactam\_NRPS\_scaffold\_1 | BGC0001672 | NRP | 24.0 | 56.8 | 209.0 | 4.2e-53 |
| CBL93723.1 | NRPS\_didomain\_A-PCP | BGC0000360 | NRP | 30.0 | 41.5 | 208.0 | 9.4e-53 |
| APZ78703.1 | nonribosomal\_peptide\_synthetase | BGC0001419 | NRP:Cyclic depsipeptide + Polyketide:Iterative type I | 28.0 | 49.4 | 208.0 | 9.4e-53 |
| ALV86866.1 | Tlo20 | BGC0001406 | NRP | 26.0 | 64.3 | 207.0 | 1.2e-52 |
| ADG27359.1 | peptide\_synthetase | BGC0000296 | NRP | 30.0 | 47.3 | 207.0 | 1.6e-52 |
| CBL93720.1 | NRPS\_didomain\_PCP-A | BGC0000360 | NRP | 30.0 | 42.2 | 207.0 | 1.6e-52 |
| AAT09804.1 | NocA | BGC0000395 | NRP | 28.0 | 50.2 | 207.0 | 2.1e-52 |
| AAP92491.1 | nonribosomal\_peptide\_synthetase | BGC0000458 | NRP | 28.0 | 47.8 | 207.0 | 2.1e-52 |
| AED90003.1 | non-ribosomal\_peptide\_synthetase\_ThaB | BGC0000443 | NRP:Beta-lactam | 25.0 | 59.3 | 206.0 | 2.7e-52 |
| CAA60461.1 | pipecolate\_incorporating\_enzyme | BGC0001040 | NRP + Polyketide | 26.0 | 65.4 | 206.0 | 2.7e-52 |
| ABI22132.1 | putative\_non-ribosomal\_peptide\_synthetase | BGC0000422 | NRP | 26.0 | 59.7 | 206.0 | 3.6e-52 |
| AZF85944.1 | hypothetical\_protein | BGC0001963 | NRP + Polyketide | 29.0 | 45.3 | 205.0 | 4.7e-52 |
| ABS90470.1 | NRPS/PKS | BGC0001106 | NRP + Polyketide | 27.0 | 57.6 | 205.0 | 6.1e-52 |
| AZF85929.1 | non-ribosomal\_peptide\_synthase | BGC0001963 | NRP + Polyketide | 28.0 | 42.9 | 205.0 | 6.1e-52 |
| CAJ34382.1 | NRPS\_protein | BGC0000445 | NRP:Cyclic depsipeptide | 26.0 | 57.8 | 205.0 | 7.9e-52 |
| APZ78833.1 | nonribosomal\_peptide\_synthetase | BGC0001430 | NRP:Cyclic depsipeptide + Polyketide:Iterative type I | 28.0 | 46.8 | 205.0 | 7.9e-52 |
| ADC79642.1 | TamD | BGC0001052 | NRP + Polyketide:Modular type I | 26.0 | 58.0 | 204.0 | 1e-51 |
| APZ78715.1 | nonribosomal\_peptide\_synthetase | BGC0001420 | NRP:Cyclic depsipeptide + Polyketide:Iterative type I | 27.0 | 47.7 | 204.0 | 1e-51 |
| AGN74885.1 | nonribosomal\_peptide\_synthetase | BGC0000459 | NRP:Cyclic depsipeptide + Polyketide:Trans-AT type I | 28.0 | 49.2 | 204.0 | 1.4e-51 |
| ACG60776.1 | NRPS(AL/ACP/C/A/PCP/C/A) | BGC0001058 | NRP:Glycopeptide + Polyketide:Modular type I + Saccharide:Hybrid/tailoring | 28.0 | 48.9 | 203.0 | 1.8e-51 |
| APZ78691.1 | nonribosomal\_peptide\_synthetase | BGC0001418 | NRP:Cyclic depsipeptide + Polyketide:Iterative type I | 27.0 | 47.7 | 203.0 | 1.8e-51 |
| BBA21071.1 | putative\_non-ribosomal\_peptide\_synthetase | BGC0001740 | NRP + Polyketide | 29.0 | 43.8 | 203.0 | 1.8e-51 |
| BAP05589.1 | calA | BGC0000967 | NRP + Polyketide:Trans-AT type I | 31.0 | 32.3 | 203.0 | 3e-51 |
| CCC55922.1 | putative\_non-ribosomal\_peptide\_synthetase | BGC0000973 | NRP + Polyketide:Modular type I | 25.0 | 58.7 | 203.0 | 3e-51 |
| BAW27670.1 | NRPS(A-T) | BGC0001764 | NRP | 29.0 | 41.2 | 202.0 | 3.9e-51 |
| AEC14349.1 | nonribosomal\_peptide\_synthetase | BGC0000377 | NRP | 28.0 | 43.1 | 202.0 | 5.1e-51 |
| AAG02356.1 | peptide\_synthetase\_NRPS7 | BGC0000963 | NRP:Glycopeptide + Polyketide:Modular type I + Saccharide:Hybrid/tailoring | 25.0 | 56.9 | 202.0 | 5.1e-51 |
| ADH04680.1 | hybrid\_polyketide\_synthase/non-ribosomal\_peptide\_synthetase | BGC0001344 | NRP + Polyketide | 29.0 | 40.3 | 201.0 | 6.7e-51 |
| AJD77023.1 | IkaA | BGC0001435 | NRP + Polyketide:Iterative type I | 25.0 | 63.4 | 201.0 | 6.7e-51 |
| APZ78781.1 | nonribosomal\_peptide\_synthetase | BGC0001426 | NRP:Cyclic depsipeptide + Polyketide:Iterative type I | 28.0 | 49.4 | 201.0 | 1.1e-50 |
| BAG17643.1 | putative\_NRPS-type-I\_PKS\_fusion\_protein | BGC0001043 | NRP + Polyketide | 26.0 | 59.1 | 200.0 | 1.5e-50 |
| AJO72717.1 | adenylation\_domain-containing\_protein | BGC0001381 | Polyketide | 28.0 | 43.0 | 200.0 | 1.5e-50 |
| CUX96957.1 | TmcJ | BGC0001829 | NRP + Polyketide | 28.0 | 42.5 | 200.0 | 1.5e-50 |
| DAC76731.1 | methionyl-tRNA\_formyltransferase | BGC0001885 | Polyketide | 30.0 | 35.2 | 200.0 | 1.5e-50 |
| AEF16059.1 | non-ribosomal\_peptide\_synthetase | BGC0000953 | Saccharide:Aminoglycoside | 27.0 | 48.2 | 200.0 | 2e-50 |
| AFU82614.1 | mixed\_NRPS\_PKS | BGC0000998 | NRP + Polyketide | 29.0 | 42.4 | 200.0 | 2.6e-50 |
| APZ78855.1 | nonribosomal\_peptide\_synthetase | BGC0001432 | NRP:Cyclic depsipeptide + Polyketide:Iterative type I | 27.0 | 50.3 | 200.0 | 2.6e-50 |
| APZ78743.1 | nonribosomal\_peptide\_synthetase | BGC0001422 | NRP:Cyclic depsipeptide + Polyketide:Iterative type I | 28.0 | 50.0 | 199.0 | 3.3e-50 |
| APZ78808.1 | nonribosomal\_peptide\_synthetase | BGC0001428 | NRP:Cyclic depsipeptide + Polyketide:Iterative type I | 28.0 | 49.2 | 199.0 | 4.4e-50 |
| APZ78768.1 | nonribosomal\_peptide\_synthetase | BGC0001425 | NRP:Cyclic depsipeptide + Polyketide:Iterative type I | 28.0 | 49.4 | 198.0 | 5.7e-50 |
| APZ78794.1 | nonribosomal\_peptide\_synthetase | BGC0001427 | NRP:Cyclic depsipeptide + Polyketide:Iterative type I | 28.0 | 49.4 | 198.0 | 7.4e-50 |
| AAX31559.1 | peptide\_synthetase\_3 | BGC0000336 | NRP | 25.0 | 58.5 | 198.0 | 9.7e-50 |
| AAG02359.1 | peptide\_synthetase\_NRPS5-4-3 | BGC0000963 | NRP:Glycopeptide + Polyketide:Modular type I + Saccharide:Hybrid/tailoring | 26.0 | 64.7 | 198.0 | 9.7e-50 |
| AJM89735.1 | PmxA | BGC0001192 | NRP | 28.0 | 43.9 | 198.0 | 9.7e-50 |
| AZF85912.1 | non-ribosomal\_peptide\_synthase | BGC0001963 | NRP + Polyketide | 29.0 | 41.9 | 198.0 | 9.7e-50 |
| AQM37583.1 | nonribosomal\_peptide\_synthetase | BGC0001424 | NRP:Cyclic depsipeptide + Polyketide:Iterative type I | 28.0 | 49.2 | 197.0 | 1.3e-49 |
| OAL11475.1 | non-ribosomal\_peptide\_synthetase\_module | BGC0001570 | NRP | 30.0 | 41.2 | 197.0 | 1.3e-49 |
| TXC99989.1 | non-ribosomal\_peptide\_synthetase | BGC0001877 | Polyketide | 28.0 | 43.5 | 197.0 | 1.7e-49 |
| CAE14171.1 |  | BGC0000383 | NRP + Polyketide:Modular type I | 28.0 | 42.8 | 196.0 | 2.2e-49 |
| APZ78755.1 | nonribosomal\_peptide\_synthetase | BGC0001423 | NRP:Cyclic depsipeptide + Polyketide:Iterative type I | 28.0 | 48.0 | 196.0 | 2.2e-49 |
| CAE14173.1 | hypothetical\_protein | BGC0000383 | NRP + Polyketide:Modular type I | 25.0 | 57.0 | 196.0 | 2.8e-49 |
| APZ78728.1 | nonribosomal\_peptide\_synthetase | BGC0001421 | NRP:Cyclic depsipeptide + Polyketide:Iterative type I | 27.0 | 48.2 | 196.0 | 2.8e-49 |
| CAG29031.1 | nonribosomal\_peptide\_synthetase\_(modules\_1\_and\_2) | BGC0001023 | NRP + Polyketide:Modular type I | 27.0 | 48.2 | 196.0 | 3.7e-49 |
| APZ78821.1 | nonribosomal\_peptide\_synthetase | BGC0001429 | NRP:Cyclic depsipeptide + Polyketide:Iterative type I | 27.0 | 48.2 | 196.0 | 3.7e-49 |
| APZ78679.1 | nonribosomal\_peptide\_synthetase | BGC0001417 | NRP:Cyclic depsipeptide + Polyketide:Iterative type I | 27.0 | 47.9 | 195.0 | 4.8e-49 |
| AAF86395.1 | FkbP | BGC0000994 | NRP + Polyketide | 25.0 | 64.0 | 195.0 | 6.3e-49 |
| BAP27942.1 | nonribosomal\_peptide\_synthetase | BGC0001085 | NRP + Terpene | 36.0 | 30.6 | 195.0 | 6.3e-49 |
| QBF51786.1 | Nrps | BGC0001856 | Polyketide:Modular type I | 28.0 | 45.6 | 195.0 | 6.3e-49 |
| APZ78845.1 | nonribosomal\_peptide\_synthetase | BGC0001431 | NRP:Cyclic depsipeptide + Polyketide:Iterative type I | 27.0 | 48.6 | 195.0 | 8.2e-49 |
| AQX36215.1 | siderophore\_synthetase | BGC0001527 | Other | 28.0 | 43.9 | 195.0 | 8.2e-49 |
| AIW58892.1 | non-ribosomal\_peptide\_synthetase | BGC0001582 | NRP | 34.0 | 29.7 | 194.0 | 1.1e-48 |
| ABR67749.1 | CmnF | BGC0000316 | NRP | 27.0 | 49.8 | 194.0 | 1.4e-48 |
| AQI70\_32580 |  | BGC0001561 | NRP | 28.0 | 46.6 | 193.0 | 2.4e-48 |
| AUD11994.1 | OrbI | BGC0001721 | NRP | 30.0 | 37.6 | 193.0 | 2.4e-48 |
| CAL69889.1 | RhiB\_protein | BGC0001112 | NRP + Polyketide:Trans-AT type I | 28.0 | 47.2 | 193.0 | 3.1e-48 |
| WP\_019634550.1 | type\_I\_polyketide\_synthase | BGC0001443 | NRP + Polyketide | 30.0 | 39.6 | 192.0 | 4.1e-48 |
| ACZ55944.1 | non-ribosomal\_peptide\_synthetase | BGC0000302 | NRP | 25.0 | 49.9 | 191.0 | 9.1e-48 |
| AUD08663.1 | iPKS-NRPS | BGC0001553 | NRP + Polyketide | 26.0 | 61.7 | 189.0 | 4.5e-47 |
| AAP92505.1 | nonribosomal\_peptide\_synthetase | BGC0000458 | NRP | 28.0 | 43.2 | 188.0 | 1e-46 |
| ACG60772.1 | NRPS(C/A/PCP/Cy/A/PCP/Cy) | BGC0001058 | NRP:Glycopeptide + Polyketide:Modular type I + Saccharide:Hybrid/tailoring | 27.0 | 49.1 | 188.0 | 1e-46 |
| ALJ49920.1 | TtmI | BGC0001236 | Polyketide | 29.0 | 41.0 | 188.0 | 1e-46 |
| AHB38497.1 | non-ribosomal\_peptide\_synthetase | BGC0000346 | NRP + Polyketide:Modular type I | 33.0 | 30.0 | 186.0 | 2.2e-46 |
| AAW03326.1 | CtaC | BGC0000982 | NRP + Polyketide | 26.0 | 58.6 | 186.0 | 2.2e-46 |
| CAF05648.1 | TubC\_protein | BGC0001053 | NRP + Polyketide | 28.0 | 46.0 | 186.0 | 2.2e-46 |
| CAD70194.1 | non-ribosomal\_peptide\_synthetase | BGC0001047 | NRP + Polyketide | 25.0 | 48.1 | 186.0 | 2.9e-46 |
| ABL74940.1 | NRPS | BGC0001048 | NRP:Glycopeptide + Polyketide:Modular type I + Saccharide:Hybrid/tailoring | 26.0 | 52.9 | 186.0 | 2.9e-46 |
| AAZ23075.1 | peptide\_synthetase | BGC0000291 | NRP | 26.0 | 50.8 | 185.0 | 5e-46 |
| EJK79843.1 | amino\_acid\_adenylation\_enzyme/thioester\_reductase\_family\_protein | BGC0000436 | NRP | 25.0 | 57.6 | 185.0 | 5e-46 |
| CAJ76290.1 | putative\_non-ribosomal\_peptide\_synthase | BGC0000972 | NRP + Polyketide:Modular type I + Polyketide:Trans-AT type I | 25.0 | 67.0 | 185.0 | 6.5e-46 |
| CAQ34918.1 | nonribosomal\_peptide\_synthetase/\_polyketide\_synthase | BGC0000986 | NRP + Polyketide | 24.0 | 60.8 | 184.0 | 1.5e-45 |
| AQV04230.1 | SwnK | BGC0001794 | Polyketide | 28.0 | 39.5 | 183.0 | 3.2e-45 |
| BAP16693.1 | nonribosomal\_peptide\_synthetase | BGC0000376 | NRP | 25.0 | 58.8 | 182.0 | 4.2e-45 |
| ALG65314.1 | Cal22 | BGC0001297 | NRP | 27.0 | 46.6 | 181.0 | 7.2e-45 |
| EWS95122.1 | hypothetical\_protein | BGC0000306 | NRP:Lipopeptide | 32.0 | 34.7 | 181.0 | 1.2e-44 |
| CCB53264.1 | non-ribosomal\_peptide\_synthetase | BGC0001393 | NRP | 25.0 | 51.0 | 180.0 | 2.1e-44 |
| CAF05649.1 | TubD\_protein | BGC0001053 | NRP + Polyketide | 27.0 | 44.2 | 180.0 | 2.7e-44 |
| CBJ89761.1 | Non-ribosomal\_peptide\_synthase\_involved\_in\_xenocoumacin\_synthesis | BGC0001054 | NRP + Polyketide:Modular type I | 27.0 | 44.1 | 180.0 | 2.7e-44 |
| CAE52342.1 | non-ribosomal\_peptide\_synthase | BGC0001088 | NRP + Polyketide | 32.0 | 30.4 | 179.0 | 4.7e-44 |
| CCJ67647.1 | JagC | BGC0001127 | NRP | 27.0 | 53.1 | 179.0 | 4.7e-44 |
| EFL02193.1 | amino\_acid\_adenylation\_domain-containing\_protein | BGC0000996 | NRP + Polyketide:Iterative type I | 27.0 | 51.9 | 178.0 | 1e-43 |
| WP\_051700105.1 | non-ribosomal\_peptide\_synthetase | BGC0001368 | NRP | 27.0 | 47.6 | 178.0 | 1e-43 |
| AAF19811.1 | mtaC | BGC0001024 | NRP + Polyketide:Modular type I | 26.0 | 55.7 | 177.0 | 1.4e-43 |
| AWO77084.1 | hybrid\_non-ribosomal\_peptide\_synthetase/type\_I\_polyketide\_synthase | BGC0001556 | NRP + Polyketide | 24.0 | 60.7 | 177.0 | 1.4e-43 |
| ARR97038.1 | SphE | BGC0001780 | NRP | 29.0 | 42.1 | 175.0 | 8.8e-43 |
| AAN85506.1 | nonribosomal\_peptide\_synthetase | BGC0001101 | NRP + Polyketide:Modular type I + Polyketide:Trans-AT type I | 27.0 | 42.8 | 174.0 | 1.1e-42 |
| ARR97036.1 | SphC | BGC0001780 | NRP | 30.0 | 42.0 | 174.0 | 1.1e-42 |
| AHZ34242.1 | CipE | BGC0001389 | NRP | 27.0 | 44.1 | 173.0 | 2.6e-42 |
| CDE97356.1 | plipastatin\_synthase\_subunit\_C | BGC0001686 | NRP | 23.0 | 56.5 | 172.0 | 4.4e-42 |
| AGD80616.1 | amino\_acid\_adenylation\_domain-containing\_protein | BGC0000394 | NRP | 30.0 | 38.0 | 172.0 | 5.7e-42 |
| CAN89638.1 | putative\_non-ribosomal\_peptide\_synthetase | BGC0001070 | NRP + Polyketide:Modular type I + Polyketide:Trans-AT type I | 24.0 | 59.1 | 171.0 | 7.5e-42 |
| AVV61987.1 | putative\_non-ribosomal\_peptide\_synthetase | BGC0001477 | NRP + Polyketide:Modular type I | 25.0 | 62.3 | 171.0 | 7.5e-42 |
| AKP45396.1 | CysH | BGC0001413 | NRP | 33.0 | 30.0 | 171.0 | 9.7e-42 |
| BAL15726.1 | NRPS | BGC0000432 | NRP | 27.0 | 44.2 | 170.0 | 2.2e-41 |
| ADZ45339.1 | non-ribosomal\_peptide\_synthetase | BGC0001020 | NRP + Polyketide | 30.0 | 34.5 | 170.0 | 2.2e-41 |
| BAF50727.1 | hybrid\_polyketide\_synthase-non\_ribosomal\_peptide\_synthetase | BGC0001116 | NRP + Polyketide | 29.0 | 34.1 | 169.0 | 3.7e-41 |
| EAU38971.1 | hypothetical\_protein | BGC0001122 | NRP + Polyketide:Iterative type I | 25.0 | 60.4 | 168.0 | 6.3e-41 |
| EJK79842.1 | amino\_acid\_adenylation\_enzyme/thioester\_reductase\_family\_protein | BGC0000436 | NRP | 25.0 | 54.4 | 168.0 | 8.2e-41 |
| ADH04660.1 | TugD | BGC0001342 | NRP + Polyketide | 25.0 | 48.1 | 167.0 | 1.4e-40 |
| ADZ45324.1 | non-ribosomal\_peptide\_synthetase | BGC0001020 | NRP + Polyketide | 25.0 | 55.3 | 166.0 | 3.1e-40 |
| BBA21068.1 | putative\_non-ribosomal\_peptide\_synthetase | BGC0001740 | NRP + Polyketide | 25.0 | 57.2 | 166.0 | 3.1e-40 |
| ATG32071.1 | proline\_specific\_adenylation\_domain-containing\_protein | BGC0001750 | NRP + Polyketide | 27.0 | 35.0 | 166.0 | 3.1e-40 |
| AAW03327.1 | CtaD | BGC0000982 | NRP + Polyketide | 27.0 | 46.6 | 165.0 | 5.3e-40 |
| AJV88377.1 | MfnE | BGC0001214 | NRP | 27.0 | 48.0 | 165.0 | 7e-40 |
| AAQ90177.1 | putative\_acyl-CoA\_synthetase | BGC0000128 | Polyketide | 27.0 | 36.1 | 165.0 | 9.1e-40 |
| AGN74892.1 | nonribosomal\_peptide\_synthetase/polyketide\_synthase\_hybrid\_protein | BGC0000459 | NRP:Cyclic depsipeptide + Polyketide:Trans-AT type I | 28.0 | 34.6 | 165.0 | 9.1e-40 |
| WP\_024483797.1 | amino\_acid\_adenylation\_domain-containing\_protein | BGC0002002 | NRP | 26.0 | 40.3 | 165.0 | 9.1e-40 |
| CBL93718.1 | NRPS\_didomain\_PCP-C | BGC0000360 | NRP | 24.0 | 53.8 | 164.0 | 1.2e-39 |
| AGZ15476.1 | putative\_non\_ribosomal\_peptide\_synthetase | BGC0001036 | NRP + Polyketide | 27.0 | 42.8 | 164.0 | 1.6e-39 |
| BAI63289.1 | putative\_non-ribosomal\_peptide\_synthetase | BGC0000434 | NRP | 24.0 | 52.5 | 163.0 | 2e-39 |
| CAJ76292.1 | putative\_non-ribosomal\_peptide\_synthase | BGC0000972 | NRP + Polyketide:Modular type I + Polyketide:Trans-AT type I | 29.0 | 34.5 | 163.0 | 2.7e-39 |
| ADH04641.1 | TgaC | BGC0001051 | NRP + Polyketide:Modular type I | 29.0 | 31.9 | 163.0 | 2.7e-39 |
| QCE20608.1 | AsmM | BGC0001961 | NRP | 28.0 | 41.5 | 162.0 | 4.5e-39 |
| CAC17499.1 | putative\_non-ribosomal\_peptide\_synthase | BGC0000324 | NRP | 24.0 | 58.2 | 161.0 | 1e-38 |
| AET98905.1 | putative\_non-ribosomal\_peptide\_synthetase | BGC0000415 | NRP | 24.0 | 54.4 | 161.0 | 1e-38 |
| ATY69551.1 | hybrid\_nonribosomal\_peptide\_synthetase/type\_I\_polyketide\_synthase | BGC0001611 | NRP + Polyketide | 30.0 | 35.4 | 161.0 | 1.3e-38 |
| WP\_078620912.1 | non-ribosomal\_peptide\_synthetase | BGC0001519 | NRP + Polyketide | 24.0 | 43.9 | 160.0 | 2.2e-38 |
| BAH04161.1 | putative\_non-ribosomal\_peptide\_synthetase | BGC0000450 | NRP | 25.0 | 49.7 | 158.0 | 6.5e-38 |
| AGC65516.1 | NRPS/PKS\_hybrid | BGC0001050 | NRP:Lipopeptide + Polyketide:Trans-AT type I | 25.0 | 51.9 | 158.0 | 6.5e-38 |
| AET13875.1 | epichloenin\_A\_synthetase | BGC0001250 | NRP | 26.0 | 41.1 | 158.0 | 8.5e-38 |
| CCB53266.1 | non-ribosomal\_peptide\_synthetase | BGC0001393 | NRP | 22.0 | 50.8 | 158.0 | 8.5e-38 |
| AET13879.1 | epichloenin\_A\_synthetase | BGC0001251 | NRP | 25.0 | 43.0 | 158.0 | 1.1e-37 |
| AHA12086.1 | amino\_acid\_adenyltransferase | BGC0001172 | NRP + Polyketide:Modular type I | 27.0 | 35.6 | 156.0 | 2.5e-37 |
| AIT38302.1 | LstE | BGC0000382 | NRP | 33.0 | 27.1 | 155.0 | 5.5e-37 |
| ABY83146.1 | Azi7 | BGC0000960 | NRP + Polyketide | 26.0 | 42.9 | 155.0 | 5.5e-37 |
| AOC89000.1 | putative\_nonribosomal\_peptide\_synthetase | BGC0001652 | NRP | 27.0 | 42.0 | 153.0 | 2.7e-36 |
| BAW32333.1 | nonribosomal\_peptide\_synthetase | BGC0001631 | NRP + Polyketide | 24.0 | 59.8 | 152.0 | 4.7e-36 |
| ABW71852.1 | nonribosomal\_peptide\_synthetase | BGC0000303 | NRP | 29.0 | 34.1 | 152.0 | 6.1e-36 |
| BAU50944.1 | nonribosomal\_peptide\_synthetase | BGC0001379 | NRP | 27.0 | 42.0 | 152.0 | 6.1e-36 |
| CDG76959.1 | non-ribosomal\_peptide\_synthetase,\_terminal\_component | BGC0001805 | NRP | 23.0 | 56.5 | 151.0 | 8e-36 |
| ATJ04411.1 | NRPS,\_TomB\_binding | BGC0001637 | NRP | 23.0 | 57.6 | 151.0 | 8e-36 |
| CAQ18831.1 | nonribosomal\_polypetide\_synthetase | BGC0000954 | NRP + Polyketide:Modular type I | 25.0 | 47.1 | 150.0 | 1.8e-35 |
| CAD15508.1 | polyketide\_synthase/non-ribosomal\_peptide\_synthetase | BGC0001014 | NRP:NRP siderophore + Polyketide:Modular type I + Polyketide:Iterative type I | 29.0 | 31.4 | 150.0 | 2.3e-35 |
| WP\_028678148.1 | non-ribosomal\_peptide\_synthetase | BGC0001228 | NRP:Cyclic depsipeptide | 25.0 | 49.3 | 150.0 | 2.3e-35 |
| CAO91861.1 | PKS-NRPS\_hybrid | BGC0000968 | NRP + Polyketide:Iterative type I | 25.0 | 43.9 | 148.0 | 6.8e-35 |
| AFK57221.1 | DidJ | BGC0000985 | Polyketide + NRP:Cyclic depsipeptide | 26.0 | 45.1 | 147.0 | 2e-34 |
| BAO66533.1 | nonribosomal\_peptide\_synthase | BGC0000042 | Polyketide | 28.0 | 36.2 | 146.0 | 2.6e-34 |
| ALA09365.1 | AMP-dependent\_synthetase\_and\_ligase | BGC0001303 | Polyketide | 25.0 | 36.2 | 146.0 | 4.4e-34 |
| CCE28989.1 | non-ribosomal\_peptide\_synthetase | BGC0001365 | NRP | 26.0 | 42.8 | 145.0 | 5.7e-34 |
| CBJ89771.1 | Non-ribosomal\_peptide\_synthase\_involved\_in\_Xenocoumacin\_synthesis | BGC0001054 | NRP + Polyketide:Modular type I | 24.0 | 47.1 | 144.0 | 9.8e-34 |
| CAJ76289.1 | putative\_hybrid\_non-ribosomal\_peptide-polyketide\_synthetase | BGC0000972 | NRP + Polyketide:Modular type I + Polyketide:Trans-AT type I | 28.0 | 31.0 | 144.0 | 1.7e-33 |
| AGC65517.1 | NRPS | BGC0001050 | NRP:Lipopeptide + Polyketide:Trans-AT type I | 30.0 | 36.3 | 143.0 | 2.2e-33 |
| AAD24881.1 | putative\_acyl-CoA\_synthetase | BGC0000127 | Polyketide | 26.0 | 36.1 | 143.0 | 2.8e-33 |
| CBF80487.1 | hybrid\_PKS-NRPS\_(Eurofung) | BGC0000959 | NRP + Polyketide:Iterative type I | 25.0 | 42.4 | 143.0 | 2.8e-33 |
| ATY69590.1 | adenylation\_protein | BGC0001823 | NRP + Polyketide | 29.0 | 35.7 | 143.0 | 2.8e-33 |
| AKD43754.1 | HerJ | BGC0001349 | NRP + Polyketide | 28.0 | 33.8 | 142.0 | 4.8e-33 |
| BBC43184.1 | PKS-NRPS\_hybrid | BGC0001738 | NRP + Polyketide | 26.0 | 48.2 | 142.0 | 6.3e-33 |
| AXN93615.1 | PuwG | BGC0001953 | NRP | 30.0 | 26.6 | 140.0 | 1.8e-32 |
| AKA59436.1 | non-ribosomal\_peptide\_synthetase | BGC0001202 | NRP + Polyketide | 26.0 | 40.5 | 140.0 | 2.4e-32 |
| ACO94461.1 | putative\_AMP-dependent\_acyl-CoA\_synthetase/ligase | BGC0000029 | Polyketide:Modular type I | 27.0 | 33.8 | 139.0 | 4.1e-32 |
| ACO94489.1 | putative\_AMP-dependent\_acyl-CoA\_synthetase/ligase | BGC0000097 | Polyketide:Modular type I | 27.0 | 33.8 | 139.0 | 5.4e-32 |
| ABP55217.1 | AMP-dependent\_synthetase\_and\_ligase | BGC0000142 | Polyketide | 25.0 | 36.0 | 138.0 | 7e-32 |
| AFL68053.1 | amino\_acid\_adenylation\_enzyme/thioester\_reductase\_family\_protein | BGC0001524 | NRP + Polyketide | 23.0 | 58.1 | 137.0 | 1.6e-31 |
| ADB92568.1 | CcbC | BGC0001225 | NRP | 27.0 | 34.6 | 137.0 | 2e-31 |
| sipL1 | AMP-dependent\_synthetase\_and\_ligase | BGC0001452 | Polyketide | 25.0 | 36.4 | 136.0 | 4.5e-31 |
| AKC54422.1 | fumosorinone\_biosynthesis\_polyketide\_synthase | BGC0001218 | NRP + Polyketide | 25.0 | 43.2 | 132.0 | 5e-30 |
| gene3 |  | BGC0002035 | NRP + Polyketide | 24.0 | 60.8 | 132.0 | 6.5e-30 |
| AAC83656.1 | dihydroaeruginoic\_acid\_synthetase | BGC0000412 | NRP | 26.0 | 47.7 | 131.0 | 1.1e-29 |
| AXM42952.1 | non-ribosomal\_peptide\_synthetase | BGC0001941 | NRP + Polyketide | 30.0 | 27.2 | 130.0 | 1.9e-29 |
| AWR88409.1 | putative\_AMP-dependent\_synthetase\_and\_ligase | BGC0001522 | Polyketide | 25.0 | 36.3 | 129.0 | 3.2e-29 |
| CAD15513.1 | non-ribosomal\_peptide\_synthetase | BGC0001014 | NRP:NRP siderophore + Polyketide:Modular type I + Polyketide:Iterative type I | 28.0 | 31.4 | 126.0 | 3.6e-28 |
| AGC24265.1 | prlK | BGC0001038 | NRP + Polyketide:Modular type I | 26.0 | 34.7 | 125.0 | 6.1e-28 |
| BAP34707.1 | AMP-dependent\_synthetase\_and\_ligase | BGC0000078 | Polyketide | 25.0 | 36.5 | 125.0 | 8e-28 |
| ALU98443.1 | hypothetical\_protein | BGC0001397 | NRP + Polyketide | 28.0 | 34.7 | 124.0 | 1e-27 |
| CCG06115.1 | hybrid\_NRPS/PKS | BGC0001543 | Polyketide | 25.0 | 43.1 | 124.0 | 1.4e-27 |
| ABX00600.1 | LmbC | BGC0000907 | Other | 26.0 | 35.2 | 123.0 | 2.3e-27 |
| ACS68554.1 | hybrid\_PKS-NRPS\_protein | BGC0001026 | NRP + Polyketide | 24.0 | 43.1 | 123.0 | 2.3e-27 |
| WP\_032920652.1 | amino\_acid\_adenylation\_domain-containing\_protein | BGC0001813 | NRP | 25.0 | 36.8 | 121.0 | 8.8e-27 |
| AAL06681.1 | A-domain\_type\_II\_peptide\_synthetase | BGC0000965 | Polyketide:Iterative type I + Polyketide:Enediyne type I | 31.0 | 25.9 | 120.0 | 2.6e-26 |
| AAF19815.1 | mtaG | BGC0001024 | NRP + Polyketide:Modular type I | 26.0 | 27.2 | 120.0 | 2.6e-26 |
| BAL15732.1 | NRPS | BGC0000432 | NRP | 27.0 | 31.7 | 119.0 | 4.4e-26 |
| AAV66110.2 | fusaridione\_A\_synthetase | BGC0000992 | NRP + Polyketide | 22.0 | 57.7 | 117.0 | 1.7e-25 |
| BAO84864.1 | putative\_non-ribosomal\_peptide\_synthetase | BGC0000414 | NRP | 23.0 | 61.8 | 116.0 | 4.9e-25 |
| QCS37521.1 | PyiS | BGC0001982 | NRP + Polyketide | 24.0 | 44.6 | 116.0 | 4.9e-25 |
| EPS29069.1 | hypothetical\_protein | BGC0001724 | NRP + Polyketide | 23.0 | 66.6 | 114.0 | 1.8e-24 |
| ARS01470.1 | NcmB | BGC0001702 | NRP + Polyketide | 26.0 | 33.1 | 111.0 | 9.2e-24 |
| CAP20364.1 | equibactin\_equibactin\_nonribosomal\_peptide\_synthase\_protein | BGC0000347 | NRP | 22.0 | 46.0 | 111.0 | 1.6e-23 |
| AVI26392.1 | putative\_nonribosomal\_peptide\_synthase | BGC0001800 | NRP + Polyketide | 26.0 | 23.8 | 111.0 | 1.6e-23 |
| EAW09117.1 | hybrid\_NRPS/PKS\_enzyme,\_putative | BGC0000983 | NRP + Polyketide:Iterative type I | 22.0 | 58.8 | 110.0 | 2e-23 |
| AFP87519.1 | proline\_adenyltransferase | BGC0001159 | NRP + Polyketide:Modular type I | 27.0 | 25.9 | 110.0 | 2.7e-23 |
| WP\_020636845.1 | type\_I\_polyketide\_synthase | BGC0002011 | Polyketide | 27.0 | 34.5 | 109.0 | 4.5e-23 |
| ctg1\_orf4 |  | BGC0001329 | Polyketide + NRP:Cyclic depsipeptide | 26.0 | 25.2 | 99.0 | 6.1e-20 |
| AEO57481.1 | PKS-NRPSs | BGC0001449 | NRP + Alkaloid + Polyketide:Iterative type I | 22.0 | 59.3 | 98.0 | 8e-20 |
| Xekj\_RS17950 | non-ribosomal\_peptide\_synthetase | BGC0001826 | NRP | 27.0 | 21.1 | 97.0 | 2.3e-19 |
| AAF08796.1 | MycB | BGC0001103 | NRP + Polyketide | 25.0 | 20.4 | 76.0 | 5.6e-13 |
| DAB41661.1 | nonribosomal\_peptide\_synthetase | BGC0001585 | Alkaloid | 23.0 | 30.9 | 75.0 | 7.3e-13 |
| AAF81723.1 | putative\_acyl-CoA\_ligase\_EncH | BGC0000220 | Polyketide:Type II | 22.0 | 34.2 | 75.0 | 9.5e-13 |
| CCP45172.1 | Bifunctional\_enzyme\_MbtA:\_salicyl-AMP\_ligase\_(SAL-AMP\_ligase)\_+\_salicyl-S-ArCP\_synthetase | BGC0001021 | NRP + Polyketide | 23.0 | 36.8 | 72.0 | 4.7e-12 |
