## Supplementary Results for "A draft genome of the ascomycotal fungal species *Pseudopithomyces maydicus* (family *Didymosphaeriaceae*)": input.path1.gene401_mibig_hits.html

| MIBiG Protein | Description | MIBiG Cluster | MiBiG Product | % ID | % Coverage | BLAST Score | E-value |
| --- | --- | --- | --- | --- | --- | --- | --- |
| EHA28233.1 | hypothetical\_protein | BGC0001143 | Polyketide | 34.0 | 76.3 | 137.0 | 5.4e-32 |
| AMM63166.1 | AniI | BGC0001371 | NRP | 32.0 | 70.2 | 130.0 | 5.1e-30 |
| CBF87076.1 | conserved\_hypothetical\_protein | BGC0001290 | NRP | 29.0 | 74.7 | 123.0 | 1.1e-27 |
| QCC62990.1 | AMP\_dependent\_ligase | BGC0001966 | NRP | 30.0 | 71.4 | 113.0 | 6.4e-25 |
| EPE34349.1 | acyl-AMP\_ligase | BGC0001035 | Polyketide + NRP + Other:Aminocoumarin | 28.0 | 70.7 | 108.0 | 2.1e-23 |
| OAQ83766.1 | phenylacetyl-\_ligase\_protein | BGC0001358 | Polyketide | 30.0 | 64.8 | 102.0 | 1.5e-21 |
| CAA07759.1 | acyl\_CoA\_ligase | BGC0000246 | Polyketide | 28.0 | 71.7 | 101.0 | 3.3e-21 |
| CAK50779.1 | acyl\_CoA\_ligase | BGC0000247 | Polyketide:Type II + Saccharide:Oligosaccharide | 28.0 | 71.7 | 101.0 | 3.3e-21 |
| AKA59447.1 | non-ribosomal\_peptide\_synthetase | BGC0001203 | NRP + Polyketide | 27.0 | 81.4 | 90.0 | 7.6e-18 |
| EJK79843.1 | amino\_acid\_adenylation\_enzyme/thioester\_reductase\_family\_protein | BGC0000436 | NRP | 25.0 | 86.2 | 89.0 | 1e-17 |
| AEZ64574.1 | fatty-acid-CoA\_ligase | BGC0001065 | Polyketide | 26.0 | 86.7 | 89.0 | 1e-17 |
| AQV04230.1 | SwnK | BGC0001794 | Polyketide | 29.0 | 71.2 | 86.0 | 1.4e-16 |
| QCT05736.1 | Tri3 | BGC0001983 | Other | 27.0 | 84.2 | 85.0 | 2.5e-16 |
| ADZ24989.1 | prolin\_adenylation\_protein | BGC0000380 | NRP + Polyketide:Modular type I | 27.0 | 77.0 | 84.0 | 3.2e-16 |
| CAN89633.1 | putative\_hybrid\_non-ribosomal\_peptide\_synthetase/polyketide\_synthase | BGC0001070 | NRP + Polyketide:Modular type I + Polyketide:Trans-AT type I | 27.0 | 70.9 | 84.0 | 3.2e-16 |
| AVR48535.1 | CusC | BGC0001564 | NRP + Polyketide | 26.0 | 74.0 | 84.0 | 4.2e-16 |
| AEU11006.1 | NpnB | BGC0001029 | NRP + Polyketide | 26.0 | 70.9 | 79.0 | 1e-14 |
| AXL88825.1 | hypothetical\_protein | BGC0001895 | Polyketide | 23.0 | 67.3 | 79.0 | 1.8e-14 |
| AQV04224.1 | SwnK | BGC0001793 | Polyketide | 26.0 | 71.9 | 77.0 | 3.9e-14 |
| ORC16618.1 | hypothetical\_protein | BGC0001341 | NRP | 24.0 | 81.1 | 77.0 | 5.1e-14 |
| CAH55654.1 | putative\_L-prolyl-AMP\_ligase | BGC0000259 | Polyketide | 25.0 | 74.7 | 76.0 | 8.7e-14 |
| BAK64635.1 | putative\_CoA\_ligase | BGC0000135 | Polyketide | 23.0 | 85.7 | 76.0 | 1.1e-13 |
| CAJ34375.1 | NRPS | BGC0000445 | NRP:Cyclic depsipeptide | 27.0 | 75.5 | 74.0 | 3.3e-13 |
| BAI63289.1 | putative\_non-ribosomal\_peptide\_synthetase | BGC0000434 | NRP | 26.0 | 76.5 | 73.0 | 7.4e-13 |
| AGN74885.1 | nonribosomal\_peptide\_synthetase | BGC0000459 | NRP:Cyclic depsipeptide + Polyketide:Trans-AT type I | 28.0 | 77.8 | 72.0 | 1.6e-12 |
| CBJ89761.1 | Non-ribosomal\_peptide\_synthase\_involved\_in\_xenocoumacin\_synthesis | BGC0001054 | NRP + Polyketide:Modular type I | 23.0 | 73.5 | 72.0 | 1.6e-12 |
| AET98905.1 | putative\_non-ribosomal\_peptide\_synthetase | BGC0000415 | NRP | 27.0 | 74.2 | 72.0 | 2.1e-12 |
| BAH04161.1 | putative\_non-ribosomal\_peptide\_synthetase | BGC0000450 | NRP | 26.0 | 70.7 | 72.0 | 2.1e-12 |
| WP\_028678148.1 | non-ribosomal\_peptide\_synthetase | BGC0001228 | NRP:Cyclic depsipeptide | 27.0 | 74.5 | 71.0 | 2.8e-12 |
| AHB82071.1 | non\_ribosomal\_peptide\_synthetase | BGC0001231 | NRP + Polyketide:Modular type I | 27.0 | 72.4 | 71.0 | 2.8e-12 |
| CAJ76286.1 | putative\_non-ribosomal\_peptide\_synthetase | BGC0000972 | NRP + Polyketide:Modular type I + Polyketide:Trans-AT type I | 24.0 | 72.7 | 71.0 | 3.7e-12 |
| ABX37383.1 | amino\_acid\_adenylation\_domain\_protein | BGC0000984 | NRP + Polyketide | 25.0 | 77.8 | 70.0 | 6.3e-12 |
| CCA89326.1 | mixed\_trans-AT\_type\_I\_polyketide\_synthase/nonribosomal\_peptide\_synthetase | BGC0001111 | NRP + Polyketide:Trans-AT type I | 26.0 | 70.9 | 70.0 | 6.3e-12 |
| ADQ55476.1 | NRPS | BGC0000350 | NRP:Beta-lactam | 23.0 | 83.7 | 70.0 | 8.2e-12 |
| ADJ63842.1 | Serobactin\_synthetase | BGC0000424 | NRP:NRP siderophore | 25.0 | 74.7 | 70.0 | 8.2e-12 |
| BAE98156.1 | putative\_non-ribosomal\_peptide\_synthetase | BGC0000339 | NRP | 25.0 | 74.7 | 69.0 | 1.1e-11 |
| ASX95241.1 | IlaS | BGC0001620 | Polyketide | 27.0 | 70.4 | 69.0 | 1.8e-11 |
| BBA20967.1 | nonribosomal\_peptide\_synthetase | BGC0001763 | NRP + Polyketide | 27.0 | 70.4 | 69.0 | 1.8e-11 |
| ABP55169.1 | amino\_acid\_adenylation\_domain | BGC0000150 | NRP + Polyketide:Enediyne type I | 25.0 | 72.2 | 68.0 | 3.1e-11 |
| ABX37382.1 | amino\_acid\_adenylation\_domain\_protein | BGC0000984 | NRP + Polyketide | 25.0 | 78.6 | 68.0 | 3.1e-11 |
| AIW58892.1 | non-ribosomal\_peptide\_synthetase | BGC0001582 | NRP | 25.0 | 70.7 | 68.0 | 3.1e-11 |
| ABC34305.1 | peptide\_synthetase,\_putative | BGC0000961 | NRP + Polyketide | 26.0 | 74.0 | 66.0 | 9e-11 |
| PHM26613.1 | pyoverdine\_synthetase\_D | BGC0001130 | NRP + Polyketide | 25.0 | 76.0 | 66.0 | 1.2e-10 |
| AGZ15460.1 | putative\_non-ribosomal\_peptide\_synthetase | BGC0001036 | NRP + Polyketide | 24.0 | 77.8 | 64.0 | 4.5e-10 |
| AGZ15458.1 | putative\_non-ribosomal\_peptide\_synthetase | BGC0001036 | NRP + Polyketide | 23.0 | 74.5 | 63.0 | 7.6e-10 |
| AKJ75110.1 | Bmp4 | BGC0001464 | Other | 23.0 | 69.1 | 63.0 | 7.6e-10 |
| CAC17499.1 | putative\_non-ribosomal\_peptide\_synthase | BGC0000324 | NRP | 24.0 | 71.2 | 63.0 | 1e-09 |
| CCA53799.1 | iron\_aquisition\_yersiniabactin\_synthesis\_enzyme | BGC0001801 | NRP | 25.0 | 71.7 | 62.0 | 1.7e-09 |
| CAJ88192.1 | putative\_peptide\_synthetase\_NRPS5-4-3 | BGC0000151 | Polyketide:Modular type I + Saccharide:Hybrid/tailoring | 24.0 | 83.4 | 61.0 | 2.9e-09 |
| AAC06346.1 | bacitracin\_synthetase\_1 | BGC0000310 | NRP | 22.0 | 79.6 | 61.0 | 3.8e-09 |
| AAY37650.1 | Amino\_acid\_adenylation | BGC0000437 | NRP | 25.0 | 74.7 | 61.0 | 5e-09 |
| AET13875.1 | epichloenin\_A\_synthetase | BGC0001250 | NRP | 25.0 | 79.6 | 60.0 | 6.5e-09 |
