## Supplementary Results for "A draft genome of the ascomycotal fungal species *Pseudopithomyces maydicus* (family *Didymosphaeriaceae*)": input.path1.gene405_mibig_hits.html

| MIBiG Protein | Description | MIBiG Cluster | MiBiG Product | % ID | % Coverage | BLAST Score | E-value |
| --- | --- | --- | --- | --- | --- | --- | --- |
| EED57519.1 | MFS\_glucose\_transporter,\_putative | BGC0001446 | Polyketide:Iterative type I | 43.0 | 97.2 | 387.0 | 3.8e-107 |
| XP\_023094066.1 |  | BGC0001996 | Other | 28.0 | 87.4 | 161.0 | 2.6e-39 |
| AEZ53939.1 | putative\_glucose-6-phosphate\_1-dehydrogenase | BGC0000144 | Polyketide:Modular type I | 30.0 | 87.2 | 153.0 | 9.2e-37 |
| BAV32173.1 | putative\_MFS\_sugar\_transporter | BGC0001373 | Polyketide | 23.0 | 87.6 | 135.0 | 2e-31 |
| AAQ21378.1 | QbsN | BGC0000925 | Other | 25.0 | 59.3 | 72.0 | 1.6e-12 |
